# Urinary single-cell sequencing captures intrarenal injury and repair processes in human acute kidney injury

**DOI:** 10.1101/2022.02.15.479234

**Authors:** Jan Klocke, Seung Joon Kim, Christopher M. Skopnik, Christian Hinze, Anastasiya Boltengagen, Diana Metzke, Emil Grothgar, Luka Prskalo, Leonie Wagner, Paul Freund, Nina Görlich, Frédéric Muench, Kai M. Schmidt-Ott, Mir-Farzin Mashreghi, Christine Kocks, Kai-Uwe Eckardt, Nikolaus Rajewsky, Philipp Enghard

**Affiliations:** Department of Nephrology and Medical Intensive Care, Charité-Universitätsmedizin Berlin, Corporate Member of Freie Universität Berlin and Humboldt-Universität zu Berlin, Berlin, Germany; Deutsches Rheumaforschungszentrum, an Institute of the Leibniz Foundation, Berlin, Germany; Max-Delbrück-Center for Molecular Medicine in the Helmholtz Association (MDC), Berlin Institute for Medical Systems Biology (BIMSB), Systems Biology of Gene-regulatory Elements, Berlin, Germany; Max-Delbrück-Center for Molecular Medicine in the Helmholtz Association (MDC), Molecular and Translational Kidney Research, Berlin, Germany

## Abstract

Acute kidney injury (AKI) is a major health issue, the outcome of which depends primarily on damage and reparative processes of tubular epithelial cells (TEC). Mechanisms underlying AKI remain incompletely understood, specific therapies are lacking and monitoring the course of AKI in clinical routine is confined to measuring urine output and plasma levels of filtration markers.

Here we demonstrate feasibility and potential of a novel approach to assess the cellular and molecular dynamics of AKI by establishing a robust urine-to-single cell RNA sequencing (scRNAseq) pipeline for excreted kidney cells via flow cytometry sorting. We analyzed 42,608 single cell transcriptomes of 40 urine samples from 32 AKI patients and compared our data with reference material from human AKI post-mortem biopsies and published mouse data. We demonstrate that TEC transcriptomes mirror intrarenal pathology and reflect distinct injury and repair processes, including oxidative stress, inflammation, and tissue rearrangement. We also describe an AKI-specific abundant urinary excretion of progenitor-like cells.

In conclusion, single cell transcriptomics of kidney cells excreted in urine provides non-invasive, unprecedented insight into cellular processes underlying AKI, thereby opening novel opportunities for target identification, AKI sub-categorization and monitoring of natural disease course and interventions.

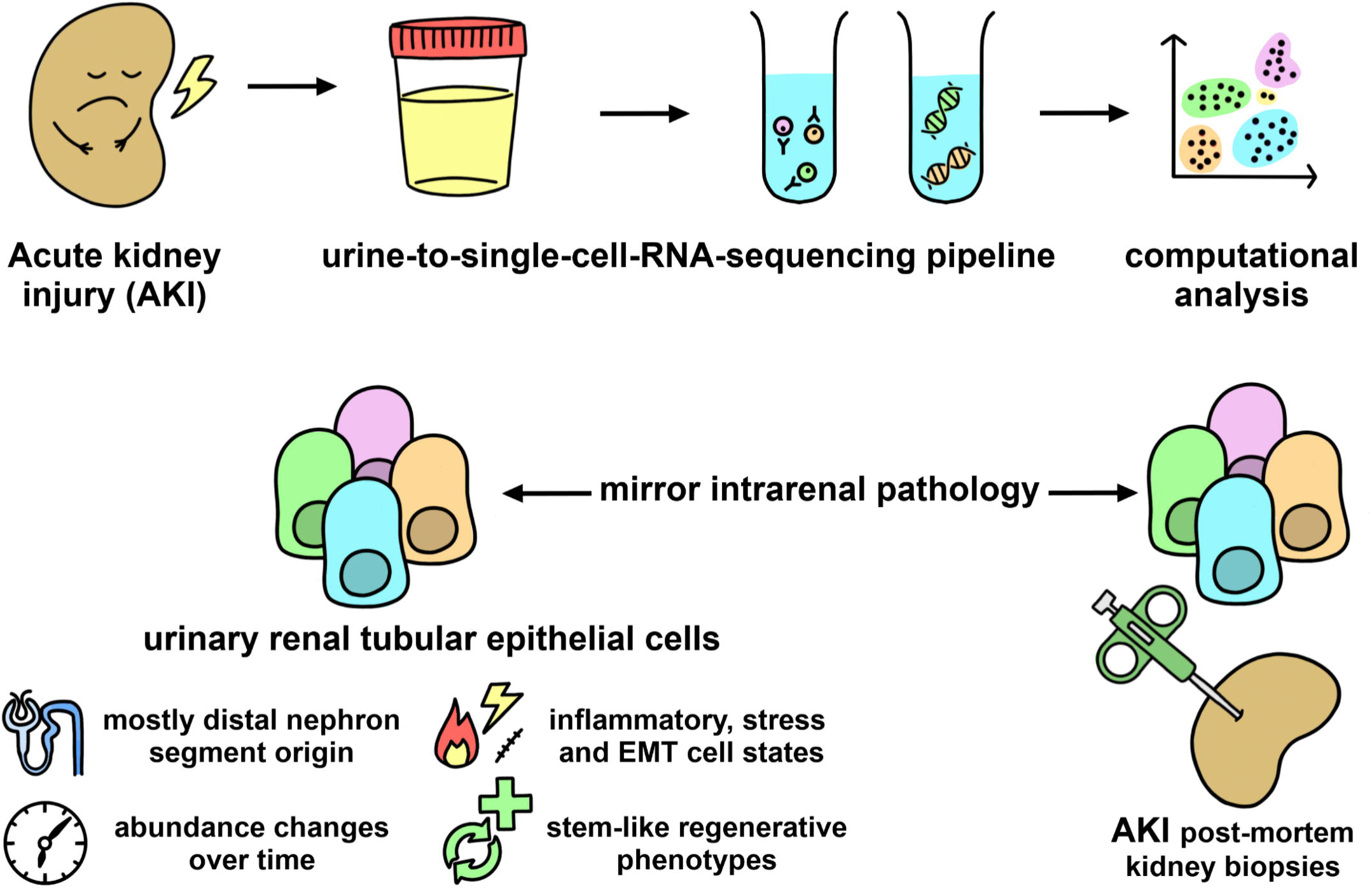

## Introduction

Acute kidney injury (AKI) is a major health concern associated with significant morbidity and mortality (1, 2). Causal therapies for preventing AKI or improving its recovery are still missing. Renal tubular epithelial cells (TEC) are the primarily affected cells in AKI and also a key component in the resulting inflammation and healing processes (3–6).

Research on AKI and on the role of TEC must overcome three major challenges: a) finding a suitable way to study human disease pathomechanisms or effectively translate findings from mouse models to the human setting, b) provide clinicians with meaningful biomarkers to predict outcomes and c) understanding TEC repair to enable strategies to enhance regeneration and reduce scarring(7). It is reasonable to believe that answers to all three challenges may lie in the urine.

Transcriptomic analyses in mouse AKI models on the single-cell level have rapidly enhanced and transformed our understanding of injured and recovering TEC states (5, 6, 8–10). Recently the first biopsy-based human single-cell studies of the kidney and AKI have been reported (8, 11, 12). However, the invasive nature of kidney biopsies limits scalability and represents a hurdle for translation into clinical routine. In contrast, analysis of kidney cells excreted in urine may provide non-invasive insights into altered cell physiology both in inflammation (13–16) and injury (17). The feasibility of urinary single-cell RNA sequencing (scRNAseq) has been proven in chronic kidney diseases such as diabetic nephropathy (DN) and focal-segmental glomerulosclerosis (FSGS) (18–20) and indicates that the urine has untapped potential as a source for the non-invasive study of renal epithelial cells. Moreover, urine-derived progenitor or stem cells, which were first isolated in pediatric patients (21), have since been studied extensively (22). They are now a common source for inducible pluripotent stem cells (23) and can be detected even in healthy individuals (20).

Therefore, we hypothesized that urinary TEC, which are excreted with the urine after kidney damage, may be an apt target to study inflammation and regeneration in AKI via urinary scRNAseq. By analyzing 40 urine samples from 32 individuals with AKI, we observed different injury-related cell states that mirror findings from mouse disease models and human AKI kidney tissue. Our data show that urinary progenitor-like cells are a prominent feature in AKI patients and reveal dynamic changes in urinary cell composition over time.

## Results

### Urine cellular composition is representative for major kidney and immune cell types

The first goal of our study was to understand what cell types are to be expected in the urine sediment after AKI. Therefore, we performed scRNAseq on sediment derived from 40 fresh urine samples of 32 patients (Suppl. Table 1, Suppl. Fig. 1). We collected samples at different timepoints 0 to 21 days after onset of AKI due to cardiac surgery (n=7), pneumonia (n=15, among them 14 cases of Coronavirus disease 2019 (COVID-19)), or other prerenal causes (n=10). After quality control filtering, we obtained a total of 42608 single cells with a median of 472 cells per sample as well as 1436 detected genes and 3991 unique transcripts per cell.

Data analysis (Uniform manifold approximation and projection (UMAP) Visualization, unbiased clustering and annotation based on known marker genes) revealed three cell type categories to be the main features of the AKI urine sediment (Fig. 1A+D, Suppl. Table 2): *Renal parenchymal cells* were comprised of a small, homogenous fraction of podocytes (“PDC”, *NPHS2, PODXL*) and many cells from the renal tubules (“TEC”, *CRYAB, EPCAM*) with various injury reactive traits, which are detailed further below and in Fig. 2. *Immune cells* included a dominant myeloid signal with multiple monocyte/macrophage subsets showing tissue residency (24) (“MO_kdnrs”, *C1QB, CD74*), pro-inflammatory (“MO_infl”, *IL1B, TIMP1*), pro-fibrotic (25, 26) (“MO_SPP1+”, *SPP1, ALOX5AP*), anti-oxidative (“MO_MT+”, various metallothioneine genes) or severely injured (“MO_dmg”) phenotypes. Granulocytes were excluded from the analysis via flow sort (see Suppl. Fig. 2) and are featured here only as a small residual cluster (“GRAN”, *CSF3R, MNDA*). Lymphocytes were also regularly featured, with T cells being more frequent than B cells. Finally, cells from the *urogenital tract* (“UGEC”) included epithelia from the reproductive system (*KRT13, PSCA*) and urothelial cells expressing uroplakins (*UPK2*).

**Figure 1:**
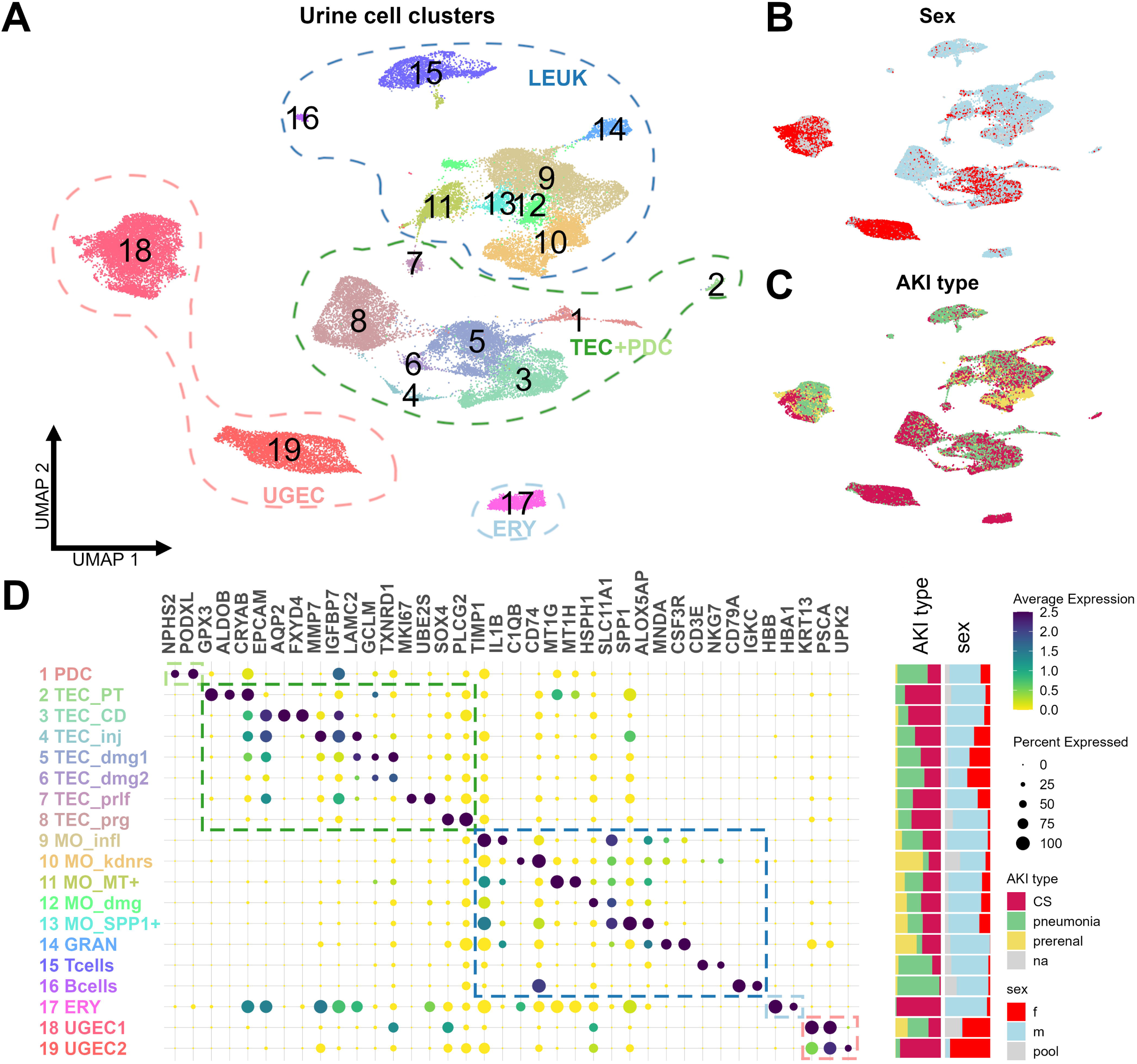
Cellular composition of urine in AKI is diverse and individual, but representative for major epithelial and immune cell types. ***A.*** *Uniform manifold approximation and projection (UMAP) of* 42608 *scRNA-seq urine cells from 32 individuals with AKI. Cellular composition is individually diverse but can be broadly grouped into kidney cells (PDC – podocytes, TEC – tubular epithelial cells), urogenital epithelial cells (UGEC) and leukocytes (LEUK) with sporadic contamination of erythrocytes (ERY). See Suppl Fig. 4 for detailed sample info. **B+C.** Distribution of cells in UMAP by sex (B) and etiology of AKI (C). Females (red) excrete more UGEC via urine (also Suppl. Fig. 5). Patients with (mild) prerenal causes of AKI (yellow) excrete mainly myeloid cells and UGEC and less TEC than cardiac surgery assoc. AKI (CS, red) or pneumonia (green), also note Suppl. Fig. 6A+C. **D.** Dotplot of marker gene expression for each cell type. PT – proximal tubule, CD – collecting duct, MO –* monocytes/macrophages*, GRAN – granulocytes, inj – injured, dmg – damaged, prlf – proliferating, prg – progenitor-like, infl – inflammatory, kdnrs – kidney resident*.

**Figure 2:**
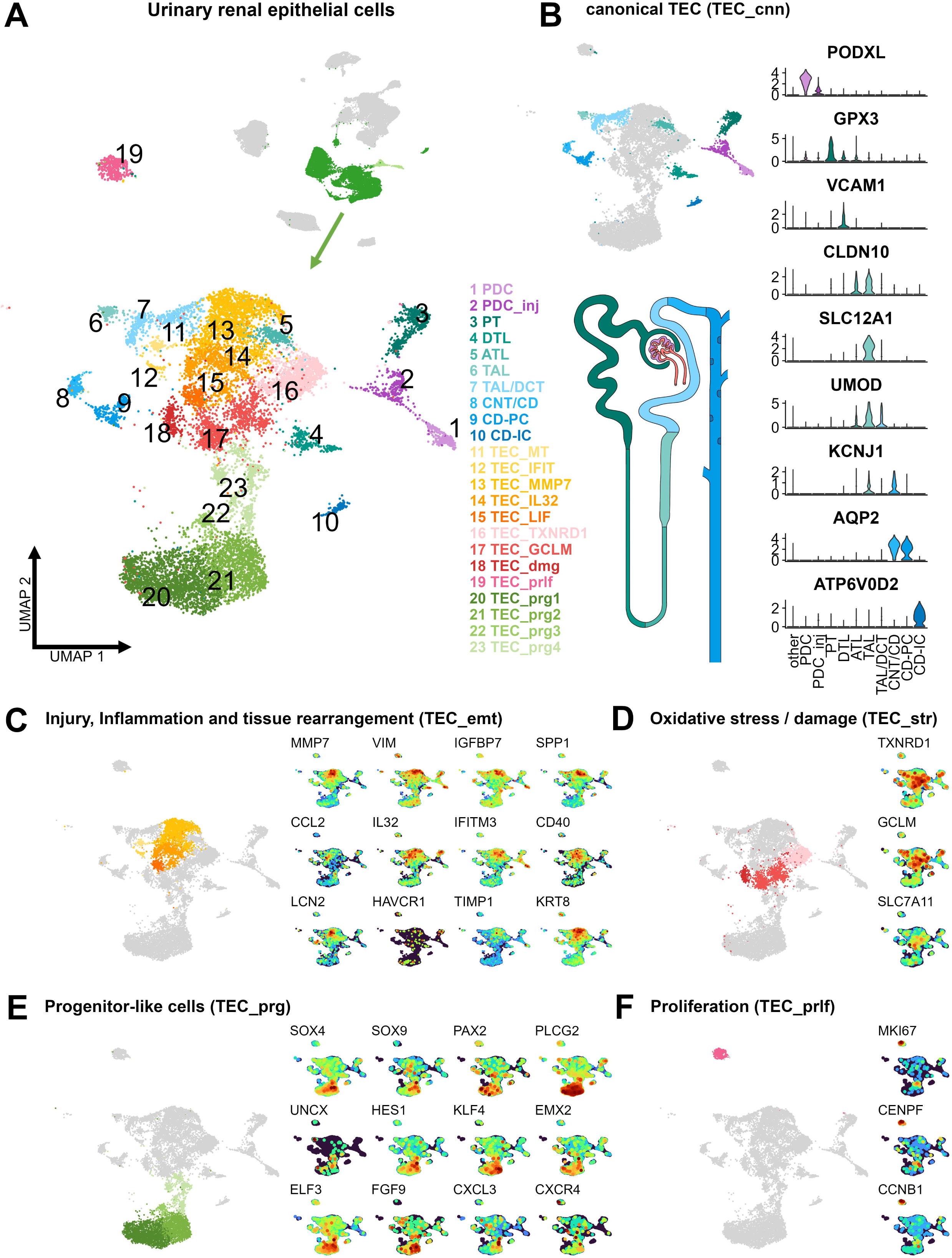
Urinary epithelial cells display different injury reactions. ***A.*** *UMAP of 12853 urinary renal parenchymal scRNAseq transcriptomes in 23 distinct clusters. Clusters can be grouped into five major groups: **B.** Canonical renal cells (TEC_cnn), including tubular cells from the proximal tubule (PT), descending thin limb (DTL), ascending thin limb (ATL), thick ascending limb (TAL), distal convoluted tubule (DCT), connecting tubule (CNT), collecting duct principal cells (CD-PC), collecting duct intercalated cells (CD-IC), as well as podocytes (PDC), see violin plot and Suppl. Fig. 8 for detailed marker genes. **C-F.** TEC clusters with markers for injury, inflammatory and epithelial to mesenchymal transition (EMT) (**C**), oxidative stress (**D**), progenitor / dedifferentiation markers (**E**) and proliferation markers (**F**)*.

Interestingly, all three major cell types were detectable across most samples (Suppl. Fig. 4D), despite the diversity of the overall quantity of captured high-quality single cell transcriptomes across samples, with a median of 472 (range 8 – 5,900) cells/sample (Suppl. Fig.4D). UGEC were featured more prominently in female patients (Fig. 1B+D, Suppl. Fig 5). Thus, we asked whether other factors like differing AKI etiology (Fig. 1C+D) or an infection with SARS-Cov-2 also influence the cellular urine composition in AKI.

Therefore, we examined the data of different AKI entities separately and compared their cellular composition: Individuals with cardiac-surgery (CS) or pneumonia / COVID-19 associated AKI showed a similar pattern of urinary composition with many renal cells (33 and 36% of all single-cell transcriptomes) and differed only in a more pronounced (B) lymphocyte signal in the pneumonia patient pool (Cluster 17 in Suppl. Fig. 6.1A, 6.2). The prerenal AKI pool contributed mostly to myeloid (clusters 10-15) and urogenital (clusters 19, 20) cells and had much less kidney parenchymal cells (7%, clusters 1-9), consistent with less severe tubular damage.

SARS-Cov-2 infection, however, had only marginal effects on the AKI urine sediment, despite being discussed as an active driver of kidney inflammation and fibrosis (27): Urinary cells from COVID-19 AKI patients showed no transcriptional activity for the SARS-Cov-2 genome. Although previously reported COVID-19 associated genes (27, 28) (*PLCG2, AFDN*) as well as inflammation (*CCL2, IFITM3*) and damage markers (*HSPA1A*) were differentially expressed in TEC of COVID-19 patients in comparison to the Non-Covid AKI group (Suppl. Fig. 7), the differences were marginal, none of these signatures were exclusive for COVID-19 patients and no distinct TEC population was detected.

Overall, urine samples from AKI patients regularly featured immune and most importantly epithelial cells from the kidney, potentially reflecting kidney damage type and severity.

### Urinary epithelial cells display different injury reactions

To use urinary cellular analysis as a proxy for AKI pathophysiology, it is crucial to know which kidney features are released into the urine and how detailed those cells can be characterized. For that reason, we re-analyzed the 12,853 AKI single cell transcriptomes which were identified as TEC or PDC before (Fig.2A, Suppl. Fig. 8, Suppl. Table 3): In this focused analysis on renal parenchymal cells, ten cell subsets resembled cells from different segments of the renal tubule system with expression of specific marker genes (Fig. 2B, Suppl. Fig. 9A). However, those “canonical” clusters (summarized as “TEC_cnn”) only composed a small fraction of urinary TEC, while most other transcriptomes showed one of several injury-related cell states (Fig. 2C-F): A large group of cells (“TEC_emt”, Fig. 2C) combined high expression of injury markers (*LCN2, IGFBP7, KRT8*) with proinflammatory cyto- and chemokines (*IL32, CD40, CCL2, IFIT1-3*) and *epithelial-to-mesenchymal transition* (EMT) markers (*VIM, MMP7*). *Oxidative stress* signatures (*TXNRD1, GCLM, SLC7A11*) were dominant in another group (“TEC_str”, Fig. 2D). Particularly interesting were potentially regenerating cells showing *proliferation* (*CENPF, MKI67*, “TEC_prlf”, Fig. 2F), and a large subset expressing transcription factors connected to kidney development and repair (*SOX4, SOX9, PAX2*) presumably reflecting (dedifferentiated) *nephron progenitors* (“TEC_prg”, Fig. 2E). The phenotype of these progenitor-like cells was further supported by automatic annotation (SingleR) using an expression atlas of human primary cells(29): Here, most urinary renal cells were unsurprisingly annotated as epithelial cells, while a substantial fraction of TEC_prg was annotated as tissue- or embryonic stem cells. (Suppl. Fig. 10AB). Additionally, high expression of markers for kidney development (30, 31) (*SOX4, CITED2*), stemness (32, 33) (*PLCG2, HES1, KLF4*) and differentiation processes (34–37) (*ELF3, IER2, DDX17*) in these subsets (Suppl. Fig. 9C) further underline their regenerative potential.

All injury related subsets weakly expressed either proximal tubule (PT) markers (*GPX3*), thick ascending limb (TAL) markers *SLC12A1/UMOD* or collecting duct (CD) markers *AQP2/FXYD4*, without co-expression of these markers in single cells (Suppl. Fig. 10AB), indicating a preserved injury reaction from all nephron segments, that is not limited to the much-investigated PT.

Next, we wanted to know whether any of these TEC phenotypes are specific for the damage introduced by AKI: We compared our data with recently published urinary scRNAseq data of chronic glomerular diseases DN (18) and FSGS (19) (Suppl. Fig. 6): TEC_emt and TEC_str were also the most prominently featured TEC in DN and FSGS (Suppl. Fig. 6B). Interestingly, TEC_prg were detectable in DN and FSGS urine only to a much lesser extent than in AKI (Suppl. Fig. 6.1+6.3), which may indicate a possible AKI-specific abundant excretion of TEC_prg. Taken together, the phenotype of urinary TEC seemed to be primarily shaped by injury-related dedifferentiation processes and only to a lesser extent by their tubule segment origin.

### Urinary cells from AKI patients mirror AKI post-mortem biopsies

It is important to investigate to what extent urinary cells indeed reflect kidney pathophysiology. Therefore, we used single nuclei RNA seq (snRNAseq) data from human post-mortem AKI kidney biopsies generated previously(12) (Fig. 3B, Suppl. Fig. 11A) and compared it to our urinary data. Briefly, data of human AKI kidney tissue contained 106,971 single nuclei with transcriptomes indicating tubular, leukocyte, endothelial and fibrocyte identities. In most nephron segments, Hinze et al. describe several injury-related altered tubular phenotypes, most notably inflammation, oxidative stress, and signs of EMT, here annotated as PT_New, TL_New, TAL_New, DCT_New, CNT_New, CD-PC_New and CD-IC_New (12).

**Figure 3.**
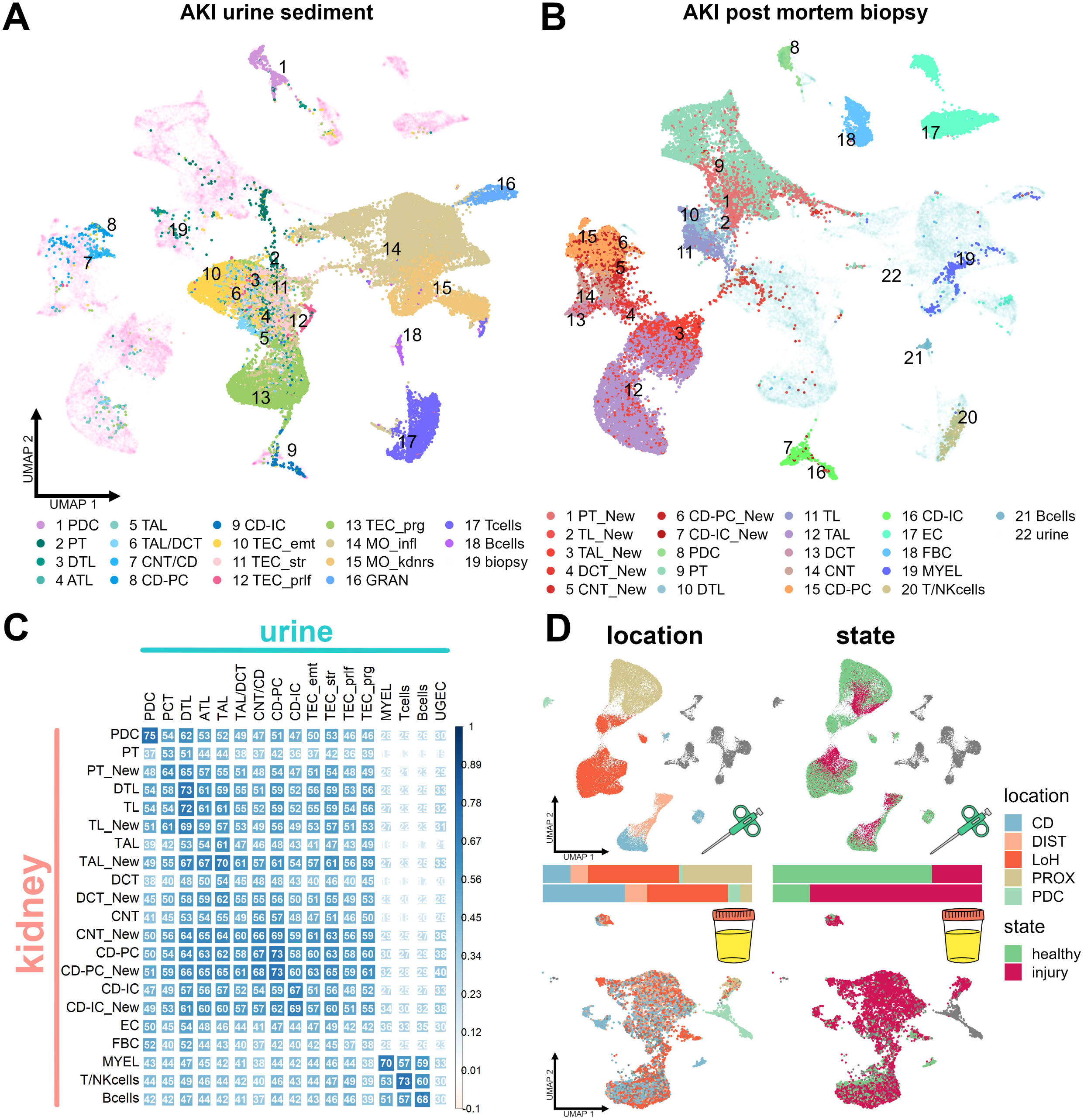
Urinary cells from AKI patients mirror AKI post-mortem biopsies. ***A+B.*** *Integrated AKI urinary scRNAseq (**A**) and AKI post-mortem biopsy snRNAseq* (12) *(**B**) datasets. Note how kidney tissue injury related cell types (“_New”, red shadings in B) partially overlap with injury related urine TEC (transparent turquois clusters in middle) and how few segment specific urine TEC (green and blue shadings in A) cluster with respective tissue cell types. **C**. Correlation plot for gene expression of each 3000 most variable expressed genes in urine (columns) and biopsy clusters (rows). Size and color represent Spearman’s rho (x100), all p <0.001. Most urine TEC subsets correlate more closely with injured (“_New”) tissue counterparts. **D**. UMAP Distribution of epithelial single cells in biopsy (above) and urine (below) by location along the nephron (PROX – proximal, LoH – loop of henle, DIST – distal, CD – collecting duct) and by state (healthy / injured). Note how urine excreted cells have a bias towards distal nephron segments and injured states. Endothelial and immune cells in grey. Annotation of urine cells done by SingleR (see Suppl. Fig. 12). _New – AKI reactive cell state, PDC – podocytes, PCT – proximal tubule, DTL – descending thin limb, ATL – ascending thin limb, TAL – thick ascending limb, DCT – distal convoluted tubule, CNT – connecting tubule, CD-PC – collecting duct principal cells, CD-IC – collecting duct intercalated cells, MYEL – myeloid cells, EC – endothelial cells, FBR – fibrocytes, MO – monocytes/macrophages, GRAN – granulocytes, emt – epithelial-mesenchymal transition, str – stressed, prlf – proliferating, prg – progenitor-like, infl – inflammatory, kdnrs – kidney resident*.

We compared the gene expression between urine and biopsy datasets. Correlations of expression profiles showed molecular similarity of corresponding urine and tissue cell types (Fig. 3C) and most urinary tubular subsets had a better expression correlation with injury-related “New” biopsy clusters than healthy counterparts.

Encouraged by these findings, we then integrated biopsy AKI data with urine data (Fig. 3AB). This data overlap of urine and tissue validated our prior findings with few urinary TEC clustering with healthy tissue TEC, while most injury related urinary subsets clustered separately, overlapping only partially with biopsy “New” clusters. Integration also revealed very low abundance of urinary fibrocytes and endothelial cells (Fig. 3A) that had previously been clustered with TEC and leukocytes in the non-integrated urine data (Fig. 1).

Wanting to validate our findings with another approach, automated cell annotation (SingleR) of our urine data using AKI biopsy as reference yielded similar results: Aside from small subsets of healthy, segment specific tubular cells, urinary TEC appeared to be of injured phenotypes and derived mostly from distal segments of the tubule (Suppl. Fig. 12). This urinary bias toward distal and injured nephron segments (Fig. 3D) indicates that the urinary renal cell signature is influenced by injured TEC being excreted following loss of tubular integrity and/or detachment from the basal membrane, with viable TAL and CD cells being more prone to final urinary excretion.

Working with urine derived cells always brings forth two major skeptic questions: “Are those cells really derived from the kidney and not from the urogenital tract?” and “To what extent does prolonged contact to urine alter the phenotype of these cells?”. We addressed these questions by comparison to two further references: In an integrated dataset with tumor adjacent healthy human kidney tissue (KPMP (38), Suppl. Fig. 13C) and healthy bladder tissue (39) (Suppl. Fig. 13D), urinary TEC clustered with kidney derived TEC, while urinary UGEC clustered with bladder cells (Suppl. Fig. 13), providing further evidence for the renal origin of urinary TEC. We then compared TEC from all tissues and urinary datasets at hand for the occurrence of gene signatures associated with salt-, fluid shear-, osmotic- or pH-stress. Urine samples uniformly had a higher mean expression of the salt-stress signature genes, but all other stress related gene sets were not enriched in the urinary datasets, suggesting that the impact of exposure to urine was confined. (Suppl. Fig. 14).

In conclusion, mostly injured and distal TEC were excreted with urine. Even though the urinary environment seemed to alter the transcriptional signature to a certain extent, information about tubular segment origin and pathophysiology was preserved in these cells.

### scRNAseq analysis of human urinary cells resembles central motifs of mouse AKI

The occurrence of multiple injury related TEC subsets with inflammatory, fibrotic and regenerative traits motivated us to ask whether our urinary data of human AKI are in accordance with recent findings in mouse AKI models describing “maladaptive” (6) or “failed-repair” (5, 9) TEC, which are suspected to drive fibrosis post AKI. We downloaded the mouse ischemia reperfusion injury dataset by Kirita et al (5), who initially described these cell states, and reconstructed the investigated PT clusters showing the emergence of injured, repairing and “failed repair” out of healthy PT cells after ischemia-reperfusion injury (Fig. 4A). The urinary TEC were compared with automated cell annotation (SingleR) using the mouse TEC as reference (Fig. 4B). Unsurprisingly, most urinary TEC showed a phenotype most reminiscent of injured PT cells. However, especially in the progenitor-like clusters, a fraction of cells correlated most with repairing and “failed-repair”-like PT phenotypes (Fig. 4CD). This indicated that TEC subsets with potentially beneficial or detrimental effects on the disease course may be detected via urine.

**Figure 4.**
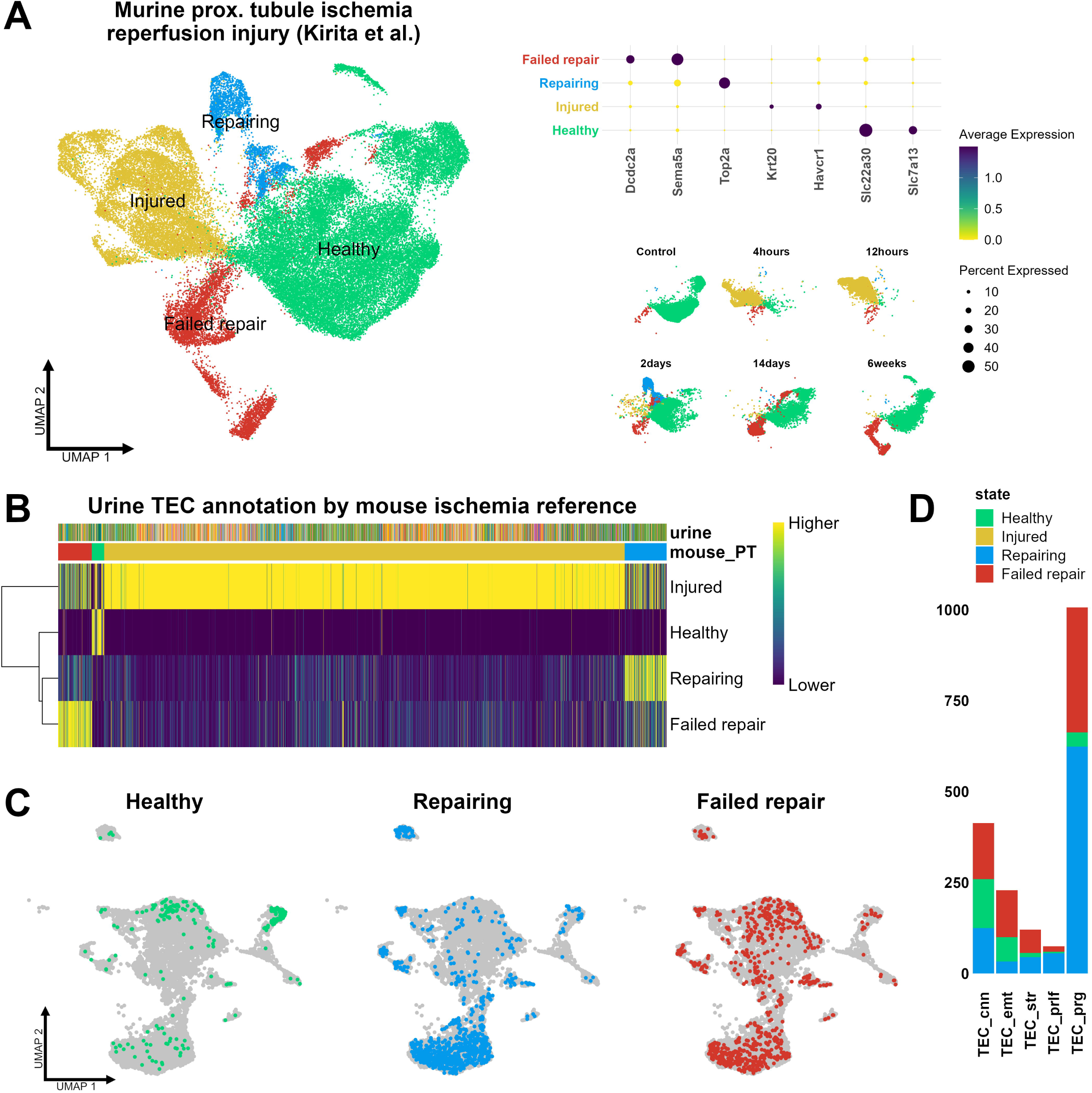
Urinary tubule cells partly reflect “repairing” and “failed-repair” mouse ischemia model phenotypes. ***A.*** *UMAP and marker genes for proximal tubule (PT) dynamic cell states after ischemia reperfusion injury in a mouse model*(5)*. **B.** Heatmap of automatic annotation scores (SingleR) of urinary cells with mouse model cell types as reference. Most, but not all cells are labeled as “Injured”. **C.** Urine TEC labeled as healthy, repairing or “failed-repair” by automatic annotation plotted in UMAP. Note how “repairing and “failed repair” labels are predominantly found in presumed progenitor cells (bottom middle clusters) and EMT clusters (top middle clusters) **D.** Relative abundance of healthy, repairing or “failed-repair” annotation labels in urinary TEC subsets (compare Fig. 2)*.

### Urinary progenitor-like cells show regenerative signatures

The subset TEC_prg remained the most elusive because no evidence for a tissue correlate was found, even though similar *SOX4+SOX9+* cells occur in the reference kidney tissue samples after AKI (Suppl. Fig. 11B-D). Several hints indicated that these cells might be a “snapshot” of (re-)differentiating tubular cells: These cells were *reminiscent of repairing and partially “failed-repair” cell states* in the mouse data. Furthermore, Phospholipase C gamma 2 (*PLCG2*) – together with proliferation markers *FOS* and *JUN* – were among the most highly expressed genes in these clusters (Fig. 5A). Although *PLCG2* is mostly associated with immune cells (40), recent evidence for *PLCG2*-dependent signaling in epithelia (33, 41) and in *in vitro* proliferation of hepatocytes (42) suggest a *role of PLCG2 in epithelial regeneration*, possibly via interactors in the fibroblast growth factor (FGF)/ERK pathway (43) or WNT pathways (44). Fittingly, activity for AKI regeneration associated pathways PI3K (45, 46), Estrogen (47) and WNT (44) (inferred by PROGENy) was more pronounced in the TEC_prg clusters (Fig. 5C). c) We noticed a divide between embryogenesis/developmental transcription factors *PAX2*, *SOX4* and *SOX9*, which were predominantly expressed in the presumed progenitor clusters and other known surface stem cell markers like *CD24*, *CD44* and *PROM1*, which were expressed most in the EMT clusters (Fig. 5B). Appropriately, gene signatures in *PAX2+* cells were similar to developmental terms found in the Gene Ontology (GO) database, many of them kidney specific like “metanephric tubule development” with additional stem cell terms like “somatic stem cell population maintenance”, indicating a *developmental stage* (Fig. 4D). *PROM1+* cell signatures yielded similar GO terms like “nephron tubule development” but in contrast also “extracellular matrix organization” and “regulation of TGF-ß production”, hinting at potential EMT (Suppl. Fig. 15). Distinct cellular crosstalk of TEC_prg with epithelial and immune cells in comparison to other TEC further emphasized these findings (Suppl. Fig. 16).

**Figure 5.**
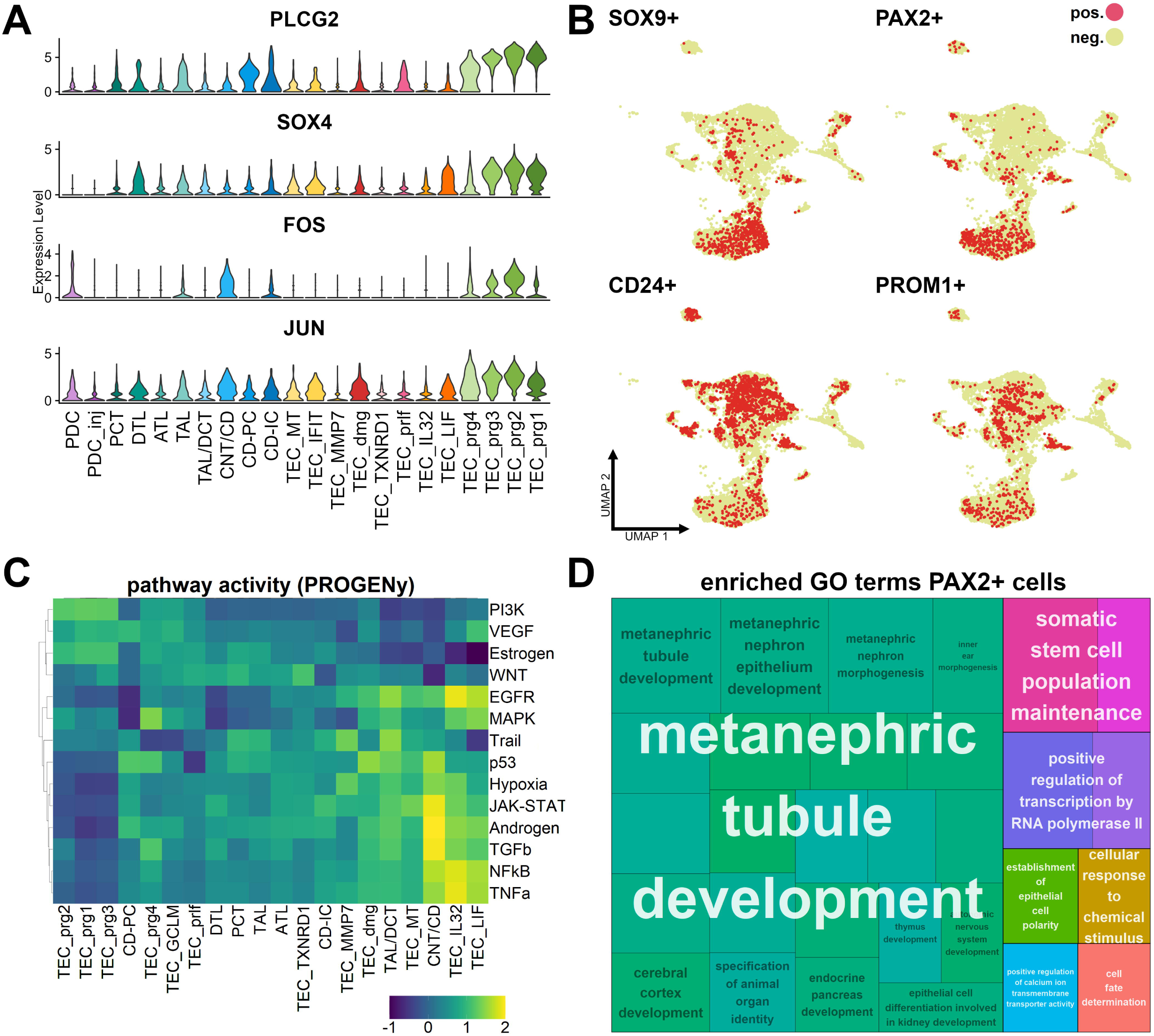
Urinary progenitor-like cells differentiate towards injury-related kidney cells. ***A.*** *Violin plot of cell differentiation marker expression across renal tubular cell (TEC) clusters. **B.** Heatmap differing pathway activity in TEC clusters (inferred from PROGENy). Note the divide between progenitor-like (TEC_prg) and other subsets concerning regeneration-associated pathways (WNT, PI3K, Estrogen). **C.** Distribution of stem cell marker positive cells across UMAP representation. **D.** Treemap plot of enriched gene ontology (GO) terms in PAX2+ cells. Each rectangle is a single GO term (black text), sized based on -log10(adj. p-value). The terms are joined into GO clusters by similarity, with the largest rectangle of the cluster providing the group name (white text). visualized with different colors. proximal tubule (PT), descending thin limb (DTL), ascending thin limb (ATL), thick ascending limb (TAL), distal convoluted tubule (DCT), connecting tubule (CNT), collecting duct principal cells (CD-PC), collecting duct intercalated cells (CD-IC), as well as podocytes (PDC), prg – progenitor-like, prlf – proliferating, dmg – damage*.

### Urinary tubular epithelial cell abundance and cell type proportions change over time after AKI

One of the major advantages of cellular urinalysis is its repeatability, allowing to evaluate temporal changes in the emergence of TEC after AKI onset more easily than biopsy-based approaches. Therefore, we compared absolute TEC counts in different phases (Fig. 6) after onset of AKI (early = up to five days after AKI onset, mid = day six to ten, late = after day ten). Overall, the highest urinary TEC excretion seemed to happen from day six onward, with total TEC counts being significantly increased in patients sampled between day six and ten (p < 0.018, Unpaired Two-Sample Wilcoxon Test). This rise in TEC counts seems to be driven mainly by urinary shedding of TEC_emt and TEC_str after AKI, which both significantly increased in the mid phase compared to early and late AKI (p < 0.005 and p < 0.0059 respectively). On the contrary, amounts of excreted TEC_prg remained unchanged in the analyzed timeframe. TEC_cnn seemed to be a prominent urinary feature mostly in the early course of AKI (Fig. 6B), although time-dependent differences did not reach statistical significance in this dataset. The interindividual variability in the AKI disease course is difficult to control for but these dynamic changes also observed in several repeatedly sampled patients (Fig. 6C) hint at a common underlying temporal shift in the urine cell signature after AKI.

**Figure 6.**
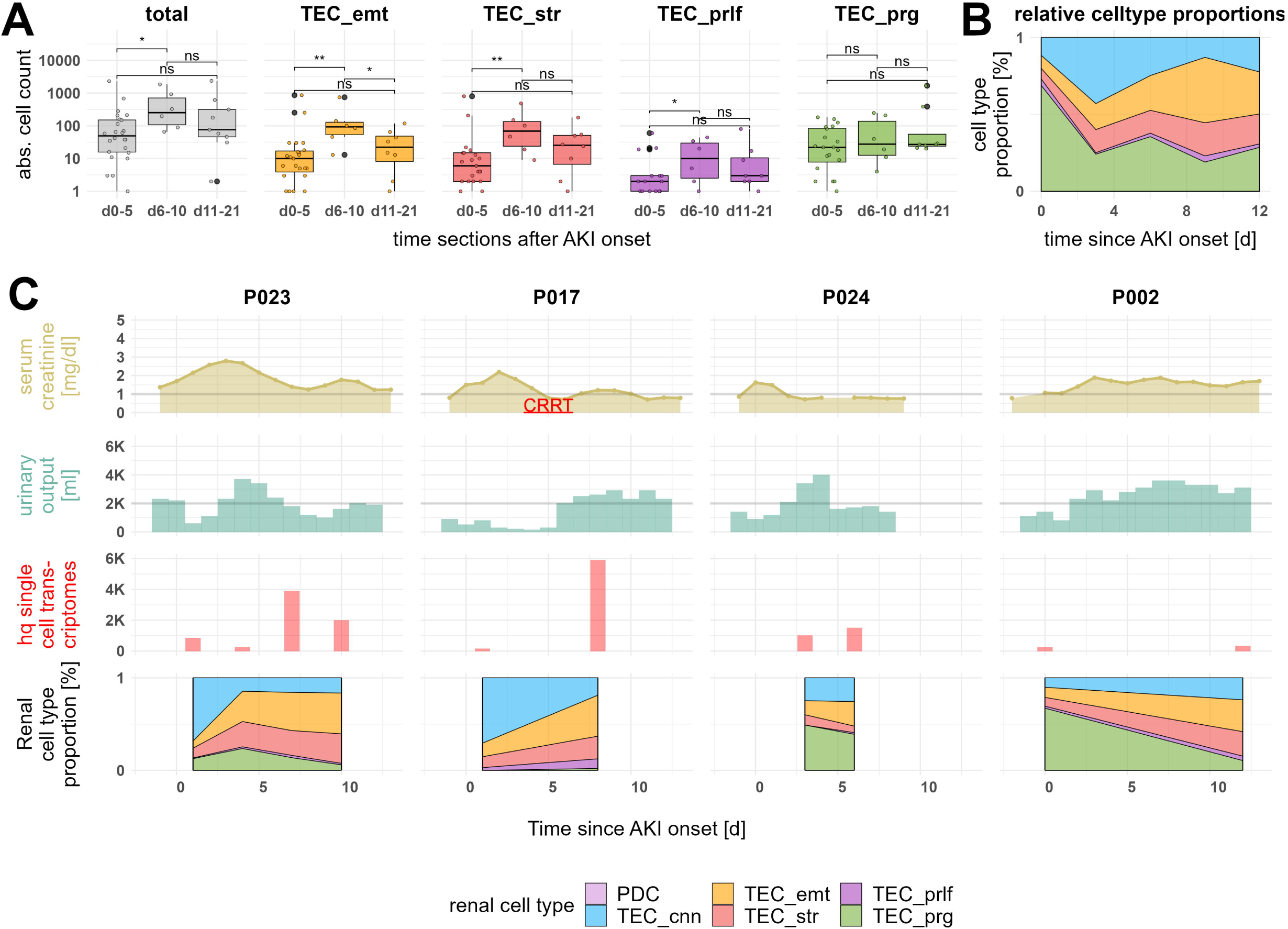
Urinary AKI tubular epithelial cell abundance and cell type proportions change over time. ***A.*** *Absolute amount of urinary renal tubular epithelial cells (TEC) and subsets by patient over time. Excretion of TEC peaks around d5-10, mainly via more injured cell types (TEC_emt, TEC_str). **B.** Relative TEC subset proportions over time (samples were binned by timepoint in three-day intervals. The mean proportion of cell type across samples in each bin was calculated). **C.** Individual AKI course over time with serum creatinine, urinary output, single cell transcriptome yield and relative amount of urinary renal parenchymal cell subsets. AKI disease course is individual diverse but temporal changes of cell type proportions are similar. cnn – canonical, emt – epithelial to mesenchymal transition, str – oxidative stress, prg – progenitor-like., prlf – proliferating*.

## Discussion

This study positions urinary scRNAseq as a new, powerful tool to study human AKI. Kidney parenchymal cells including TEC and podocytes can reliably be detected in urine of AKI patients. These urine-excreted kidney cells are mainly of distal tubular origin and show diverse injury reactive phenotypes mirroring findings in transcriptional analyses of human AKI tissue samples and mouse ischemia reperfusion injury models. Furthermore, urine TEC quantity and composition change over time, enabling temporally resolved insights of the disease course.

### Feasibility of urinary single cell RNA sequencing

Together with data from DN (18) and FSGS (19) patients, urinary scRNAseq has now been tested in three credible cohorts across different diseases. These studies revealed a high variability of the viable cell count across samples. This variability which likely relates to disease severity but is probably also influenced by varying sample sizes due to fluctuating urinary output (Suppl. Fig1B). Two strategies to overcome this issue are a) pooling of multiple samples and b) repeated sampling of the same patient over time, both of which were successfully applied here.

The urinary cellular signal – independent from disease type or severity – is dominated by leukocytes and urogenital epithelia. (18, 19). Therefore, most urinary cellular studies, single-cell (48) or otherwise (14, 16, 49), have focused on analyses of urinary immune cells. Here, we demonstrate that by flow cytometric pre-selection and gathering of sufficient viable cell counts, the focus of urinary cellular analysis can be shifted towards renal parenchymal cells.

### Comparison with human AKI tissue data

Urinary TEC transcriptomes correlate well with respective reference cells from tissue and mirror the damaged and injured TEC phenotypes in post-mortem AKI biopsy samples. This result is essential as it shows that the molecular makeup of TEC is sufficiently preserved in urine to infer data informative about molecular expression patterns in a patient’s kidney without biopsies. Moreover, together with the data by Hinze et al.^12^, our results challenge the proximal tubular centricity of (murine) AKI research, with other nephron segments like TAL and CD-PC seemingly contributing to injury related cell pools as much as PT cells. Urine appears to be especially suited to study changes in the more distal nephron segments, as those TEC seem to be preferentially excreted into urine.

### Injury related tubular cell states in the urine

Our current understanding of TEC injury reaction assumes a dedifferentiation and redifferentiation after successful repair, which has been demonstrated for PT cells in several mouse models (5, 8, 9) and cells in human AKI biopsies have been reported to mirror this process (8). A central question is how this repair takes place and why it can fail. The Humphreys group first described a “failed-repair” subset in mouse AKI (5), characterized by downregulation of differentiation markers of PT cells, expression of VCAM1 and persistence after AKI. These cells have since been reported in several other mouse models for AKI, indicating an inflammatory signature, which may fuel progression to CKD (50) as well as a potential recovery of these cells governed by ferroptotic stress (9).

The most prominent urinary TEC subset in our data, TEC_emt, is also the most reminiscent of the “failed repair” state. Although mostly VCAM1-negative, (VCAM1 expression in our data is mostly confined to fractions of the descending thin limb cluster), this subset has pro-inflammatory and EMT phenotypes similar to the failed state and shows a dynamic excretion after injury, potentially indicating wound healing processes.

### Urinary tubular progenitor-like cell signature

Though not unprecedented (21, 51), urinary tubular progenitor-like cells (TEC_prg) and their high abundance after AKI are a particularly interesting finding. They show markers of specific TEC subsets, especially CD-PC and TAL, and expression of dedifferentiation markers SOX4, SOX9 and PAX2 while not expressing urogenital markers. Similar cells in urine of FSGS and DN patients can be detected, albeit to a much lesser extent. SOX4+SOX9+ TEC can also be found in post-mortal kidney samples with increased frequency in TAL and CD clusters after AKI compared to control, suggesting an injury-related increased excretion of these cells.

The origin of the regenerative progenitors (dedifferentiation (5) vs. fixed stem cells (52)) cannot be delineated from this dataset. RNA velocity analysis was tested to infer (de-)differentiation trajectories in the urine TEC and suggested, among other findings, a MALAT1 / PRMT1 dependent kinetic potentially linked to progenitor self-renewal as shown in embryonic nephron progenitors (53). However, more research is needed to validate these findings, as robustness of velocity analysis in urine samples is still lacking.

Though speculative, based on the urine cell signature’s bias toward distal nephron segments, those progenitor-like cells – if not dedifferentiated distal TEC – may also be a separate entity of distal/medullary/papillary tip origin, like the recently characterized distal multipotent Aqp2+ progenitor cells in mice(54), and therefore harder to detect via biopsy.

### Study limitations

Although urinary scRNAseq enables safe and easily repeatable examination of kidney cells in AKI, it is limited to patients with preserved diuresis, therefore excluding a substantial part of severe, oligo-to-anuric AKI cases from analysis. On a similar note, because healthy donors usually excrete almost no tubular cells, urinary data cannot be compared to a healthy urinary control group. While our findings are overlapping quite well with reference tissue data, it is not yet evident how exact urinary analysis can reflect the individual disease course, as gathering of urine and concurrent biopsy tissue from the same patient was not possible. Lastly, our method currently requires immediate processing of fresh urine samples, thus limiting its use for prospective clinical and research applications.

## Conclusions

With the present dataset, we demonstrate feasibility of urinary scRNAseq in human AKI. The epithelial cells detected in the urine originate mainly from the distal nephron segments and reflect changes observed in kidney tissue of mice and men. With further standardization of sample collection and conservation, non-invasive and temporally resolved pathophysiological insights through urinary scRNAseq may facilitate kidney diagnostics and monitoring.

## Methods

### Patients

Between 2019 and 2021 we collected 40 urine samples of 32 patients with AKI as defined by Kidney Disease: Improving Global Outcomes (KDIGO) criteria (Suppl. Fig. 1, Suppl. Table 1). Patients were sampled at a variable timepoint within the first 21 days after AKI onset. Five patients were sampled at two separate timepoints and a single patient on four occasions.

Seven patients underwent cardiac surgery within max. 48 h prior to AKI onset, 15 patients were admitted to intensive care units (ICU) because of pneumonia (all fulfilling sepsis criteria) and developed AKI during the first five days of their ICU stay; the majority of these (14/15) suffered from COVID-19. An additional 10 patients had other, mostly prerenal causes of AKI, including gastrointestinal bleeding (1), diarrhea (2), exsiccosis (4) or decompensated heart failure (3). Children and patients with kidney transplants, active oncological disease, urinary tract infections or postrenal causes of AKI were excluded from the study. Additionally, all patients were screened via urinary dipstick prior to sample preparation, excluding samples with > 70 leukocytes/µl (one of three “+” in urinary dipstick) and patients with less than 50 ml/ 4 h urinary output.

### Sample preparation

Samples (median 150 ml, range 60-440 ml) were collected either as first morning void urine or via urinary catheter (using the pooled urine output of 4 hours) and stored on ice and transported to the lab. After exclusion of leukocyturia via urinary dipstick, samples were centrifuged at 600g / 4 °C, and resuspended with Annexin binding buffer (55) / 1 % bovine serum albumin (ABB/BSA). Incubation with Actinomycin D for 30 min on ice was applied to prevent subsequent alteration of transcriptomes(56) followed by blocking Fc receptors using human FcR blocking reagent (Miltenyi, Bergisch Gladbach, Germany). Cells were then incubated with fluorescent dyes and fluorochrome-conjugated monoclonal antibodies for fluorescence-activated cell sorting (FACS). The following antibodies and dyes were used for cell labeling: Calcein AM (BD Biosciences, San Jose, CA, USA, 564061), Hoechst 33342 (BD Biosciences, San Jose, CA, USA, 561908), AnnexinV-PE (Biolegend, San Diego, CA, USA, 640908), CD45-APC/Vio770 and CD66b-PE/Vio770 (both Miltenyi, 130-110-635 and 130-119-768). Cells were washed, filtered through a 70 µm cell strainer and labeled with To-Pro3 iodide (T3605, Life technologies, Eugene, Oregon) for dead cell discrimination immediately prior to sorting.

Samples were sorted using a Sony MA900 (Sony, Tokyo, Japan) with a 100µm nozzle into phosphate buffered saline (PBS) / 1 % BSA gating To-Pro3^-^AnnexinV^-^ Hoechst33342^+^CalceinAM^+^ singlets for obtaining single viable urinary cells (Suppl. Fig. 1). Urinary granulocytes, which make up a large fraction of the urinary cellular signal (median 58% in FACS gating, Suppl. Fig. 1) but for which a renal tissue origin is not verifiable, and which may interfere with downstream analysis, were excluded. In the initial cohort of patients (n=7) only CD45^-^ cells were obtained, excluding most leukocytes, to maximize the amount of analyzed renal parenchymal cells; for the subsequent patients (n=25) CD45^-^CD66b^-^ and CD45^+^CD66b^-^ were sorted, including lymphocytes and monocytes/macrophages in the measurements (Suppl. Fig. 2). To adjust for low total cell counts in urine (median 3455 viable cells /100ml), samples P008-016 and P048-119 (Suppl. Fig. 4D) were labelled with TotalSeq B Hashtag antibodies 1-4 (Biolegend, clone LNH-94) during the staining procedure described above and pooled in groups of four.

### Single cell sequencing

Relative single cell counts in sorted suspensions were counted using a MACSQuant Analyzer (Miltenyi). Cells were centrifuged, resuspended, and re-counted on a hemocytometer. Cell suspensions were subjected to single-cell sequencing following the 10x Genomics protocol for Chromium Next GEM Single Cell 3’ v3.1 chemistry targeting between 1000 - 10000 cells depending on sample cell count. In brief, cells were partitioned into a droplet with a barcoded gel bead using the 10x Chromium controller (10x Genomics, Pleasanton, California, USA) and lysed. RNA was reverse transcribed into complementary DNA (cDNA) within each droplet. cDNA was amplified and fragmented, followed by the addition of Illumina adapters using Single Cell 3′ Library & Gel Bead Kit v3.1 (10x Genomics). Libraries were sequenced on Illumina HiSeq 4000 (Illumina, San Diego, California, USA) sequencers.

### Data processing of scRNAseq libraries

Demultiplexing, barcode processing, read alignment and gene expression quantification was carried out using Cell Ranger software (v3.1.0, 10x Genomics). First, Cell Ranger mkfastq demultiplexed the sequencing by sample index. The quality of the data was checked using FastQC (v0.11.5) and all samples showed high quality RNAseq data with good median per-base quality across most of the read length. Cell Ranger count used STAR software to align sequenced reads to the reference genome (GRCh38.p12). Samples from COVID-19 patients included SARS-CoV1/2 genomic information to reference genome.

### Sample Pool Demultiplexing

Demultiplexing of unfiltered hashed cell sample pools was performed in Seurat v4(57) (Suppl. Fig. 3). Hashtag oligo (HTO) libraries were added to the Seurat object as an independent assay and normalized (“NormalizeData(SO, assay = “HTO”, normalization.method = “CLR”)”). Demultiplexing was performed with “HTODemux(SO, assay = “HTO”)” with “positive.quantile =” between 0.90-0.99. Cross sample doublets were excluded. HTO negative cells were further analyzed as “POOL” samples (Suppl. Fig. 3+4).

### scRNAseq data analysis

Initial quality control was done by removing cells with less than 200 or more than 4,000 genes detected as well as cells with > 10% mitochondrial (mt) reads. For renal cells, a cut-off of 20% mt reads was applied to adjust for higher mitochondrial abundance in the kidney (58). ScDblFinder (59) was used for removing multiplets. Seurat v4 was used for downstream analysis. We combined all AKI sample datasets. SCTransform (60) was used for normalization, scaling, and determining variable features. After principal component analysis (“RunPCA” function), integration and batch correction were done by Harmony (61) (“RunHarmony” function). Dimensional reduction by UMAP (62) (dims=1:45) and clustering was performed using reduction=”harmony”. Cluster resolution was determined manually. “FindAllMarkers()” was applied to determine differentially expressed genes (DEG) in all clusters. Cluster annotation and rearrangement was done based on review of specific lineage markers.

For further detailed analysis of kidney parenchymal cells in urine, cells of corresponding clusters were subsetted and re-analyzed using the same workflow as above. Cluster resolution and annotation was again performed manually based on DEG and known lineage markers. Clustering revealed 25 clusters, two of which were comprised of few remaining leukocytes and urogenital cells, which were excluded from further analysis.

### Reference datasets

For referencing of urinary cells, multiple datasets were used: AKI post-mortem biopsy data was retrieved as an annotated Seurat object from Hinze et al.; data were processed and analyzed as reported in the original publication (12). Healthy tumor adjacent kidney tissue data was downloaded from the Kidney Precision Medicine Project (KPMP) (38) database (participants 17-1606, 18-139, 18-162, 18-342, 18142-5). scRNAseq data of 3 human bladder samples (39) (GSE129845) was downloaded from the National Center for the Biotechnology Information Database (NCBI) / Gene expression omnibus (GEO) repository. Urine scRNAseq data including 16 samples from 5 patients with diabetic nephropathy (18) (GSE157640) and 23 samples from 12 patients focal segmental glomerulosclerosis (FSGS) (19) (GSE176465) were also downloaded from the NCBI/GEO repository. Data integration/analysis/dimensional reduction and clustering of these datasets was done exactly as with in urine AKI samples. Mouse ischemia reperfusion injury PT data(5) (GSE139107) was downloaded from NCBI/GEO database. Ischemia reperfusion injury datasets were subsetted for PT cells based on the provided metadata, merged, normalized and scaled with base Seurat functions (“FindVariableFeatures()”, “ScaleData()”, “RunPCA()”). UMAP dimensional reduction and clustering was followed by differentially expressed gene discovery through “FindAllMarkers()” function and cluster annotation based on marker genes provided by the original paper. Human orthologous genes were generated using Biomart (63).

For gene expression correlation, the top 3000 variable genes each per urine and reference datasets were used. Integration of urine and reference datasets was done by Harmony. SingleR (64) was used for automatic unbiased cell type annotation in the urine TEC dataset using Human primary cell atlas bulk data (29) as well as aforementioned datasets as reference. Briefly, SingleR computes Spearman’s rank correlation coefficient for the gene expression of each cell with set reference groups. Wilcoxon ranked sum test is used to identify the top markers used for each comparison. A score is defined for each cell based on the reference correlation in comparison to the other cells. Scores are determined across all reference groups. Cells are labeled according to their highest score.

### Ligand receptor interactions

We used CellPhoneDB (2.1.1) (65) to assess cellular crosstalk between different cell types. As input data, we calculated normalized data with scale factor 10,000. To identify putative cell-cell interactions via multi-subunit ligand-receptor complex pairs, label permutation was performed. Finally, we conducted statistical analyses by randomly permuting the cluster labels of each cell 10,000 times. Tests were conducted intraindividually.

### Gene set and pathway analysis

GO term enrichment analysis was performed with topGO (66) using all DEG with an average log-fold change of > 0.5 in *PAX2* and *PROM1* positive cells (“runTest(GOdata, statistic = “fisher”, algorithm = “elim”)”). GO set visualization was done with rrvgo (67). Rrvgo groups the provided GO terms based on their similarity. GO group representatives were selected by rrvgo based on adjusted p value. Pathway activity across TEC subsets was calculated with PROGENy (68).

### Statistics

All RNA-sequencing data were analyzed using R (69). Normalization, logarithmic transformation, identification of highly variable features, scaling, principal component analysis and extraction of differentially expressed features were done using Seurat. Significant differences between cell counts were tested using two-tailed Unpaired Two-Sample Wilcoxon Test.

### Study approval

The ethics committee of Charité University Hospital (Charité EA2/141/19) approved the study. Informed consent was obtained from all patients or next of kin before participation.

## Author contributions

JK, KUE, CK, NR and PE designed the study. JK, CMS, AB, DM, EG, LP, LW, PF, NG, FM, MFM and PE conducted experiments and acquired data. JK, SJK, CMS, CH, CK and PE analyzed data. JK, SJK, CMS, CH, KMSO, MFM, CK, KUE, NR and PE interpreted the results. JK, CK, KUE and PE wrote the manuscript.

## Acknowledgments

This work was supported by grants from the Berlin Institute of Health (BIH), the Jackstädt-Stiftung, the Deutsche Forschungsgemeinschaft (DFG, HI 2238/2-1) and Bundesministerium für Bildung und Forschung (BMBF).

JK was supported by a research scholarship of the Deutsche Gesellschaft für Nephrologie (DGfN). KMSO was supported by the Deutsche Forschungsgemeinschaft (DFG; SFB 1365, GRK 2318 and FOR 2841).

We thank the Flow Cytometry & Cell Sorting Facility (FCCF) and lab managers at DRFZ for technical support and helpful insights. We thank all staff at the Charité’s nephrology, cardiac surgery and intensive care wards for helping with sample collection and we thank all patients for participating.

## Supplementals

Supplemental Material – Supplemental Table 1 and Supplemental Figure 1-16

Supplemental Table 2 – Differentially expressed genes for clusters of all urine cells

Supplemental Table 3 – Differentially expressed genes for clusters of all kidney parenchymal cells

