## Supplemental Material for "Urinary single-cell sequencing captures intrarenal injury and repair processes in human acute kidney injury"

| patient | gender | age | AKI etiology | proteinuria mg/gCrea | CKD | outcome (90d) | RRT | COVID-19 infection | gating | sampling | repeated samples | Creatinine baseline (mg/dl) | Creatinine max. (mg/dl) | KDIGO AKI stage | AKI timepoint (d) | sc captured (post qc) |
|---|---|---|---|---|---|---|---|---|---|---|---|---|---|---|---|---|
| **P001** | f | 43 | pneumonia | 300 | no | baseline | no | yes | CD45- | single | 1 | 0.60 | 3.20 | 3 | 1 | 1595 |
| **P002** | m | 41 | pneumonia | 356 | no | baseline | no | yes | CD45- | single | 2 | 0.80 | 1.89 | 2 | 0; 12 | 229; 344 |
| **P003** | m | 55 | pneumonia | 3000 | no | baseline | no | yes | CD45- | single | 1 | 0.50 | 1.13 | 2 | 3 | 48 |
| **P005** | m | 68 | cardiac surgery | 150 | no | baseline | no | no | CD45- | single | 1 | 1.10 | 4.16 | 3 | 12 | 692 |
| **P006** | m | 76 | cardiac surgery | 300 | no | baseline | no | no | CD45- | single | 1 | 0.90 | 3.01 | 3 | 4 | 371 |
| **P007** | f | 70 | prerenal | 81 | no | progress CKD | no | no | CD45- | single | 1 | 0.90 | 2.82 | 3 | 4 | 1676 |
| **P008** | f | 83 | cardiac surgery | 102 | G3bA1 | progress CKD | no | no | CD66b- | pool | 1 | 1.40 | 2.51 | 2 | 1 | 8 |
| **P009** | m | 80 | prerenal | na | no | baseline | no | no | CD66b- | pool | 1 | 0.90 | 1.24 | 1 | 3 | 8 |
| **P010** | f | 69 | prerenal | na | no | baseline | no | no | CD66b- | pool | 1 | 0.30 | 2.40 | 3 | 14 | 801 |
| **P012** | m | 24 | prerenal | 215 | G4A2 | progress CKD | no | no | CD66b- | pool | 1 | 2.40 | 5.81 | 2 | 4 | 567 |
| **P013** | m | 78 | prerenal | 150 | G2A1 | progress CKD | no | no | CD66b- | pool | 1 | 1.20 | 4.42 | 3 | 5 | 242 |
| **P014** | m | 74 | prerenal | 1000 | G3A2 | baseline | no | no | CD66b- | pool | 1 | 1.80 | 3.46 | 1 | 5 | 186 |
| **P015** | f | 73 | prerenal | 410 | no | progress CKD | no | no | CD66b- | pool | 1 | 1.10 | 4.37 | 3 | 5 | 205 |
| **P016** | m | 58 | prerenal | 150 | no | progress CKD | no | no | CD66b- | pool | 1 | 0.70 | 2.10 | 3 | 5 | 240 |
| **P017** | f | 67 | cardiac surgery | na | no | baseline | CVVHD | no | CD45- | single | 2 | 0.74 | 2.19 | 2 | 1; 8 | 153; 5900 |
| **P018** | f | 71 | pneumonia | 150 | no | death | CVVHD | no | CD66b- | single | 2 | 0.68 | 3.09 | 3 | 4; 11 | 2899; 248 |
| **P019** | m | 56 | cardiac surgery | na | no | baseline | no | no | CD66b- | single | 2 | 1.50 | 4.83 | 3 | 6; 16 | 1004; 2585 |
| **P021** | m | 67 | prerenal | 200 | no | progress CKD | no | no | CD66b- | single | 1 | 1.20 | 2.48 | 2 | 4 | 817 |
| **P022** | m | 84 | prerenal | 3000 | G3bA3 | progress CKD | CVVHD | no | CD66b- | single | 1 | 1.80 | 2.84 | 1 | 4 | 64 |
| **P023** | m | 81 | cardiac surgery | na | no | baseline | no | no | CD66b- | single | 4 | 0.98 | 2.79 | 2 | 1; 4; 7; 10 | 843; 266; 3909; 2008 |
| **P024** | m | 51 | cardiac surgery | 0 | no | baseline | no | no | CD66b- | single | 2 | 0.80 | 1.62 | 2 | 3; 6 | 1010; 1500 |
| **P048** | f | 64 | pneumonia | 3608 | no | death | CVVHD | yes | CD66b- | pool | 1 | 0.80 | 1.84 | 3 | 12 | 376 |
| **P049** | m | 44 | pneumonia | 2318 | no | death | CVVHD | yes | CD66b- | pool | 1 | 1.00 | 3.16 | 3 | 2 | 92 |
| **P050** | m | 33 | pneumonia | 3790 | no | death | CVVHD | yes | CD66b- | pool | 1 | 1.00 | 4.93 | 3 | 7 | 928 |
| **P051** | m | 61 | pneumonia | 702 | no | baseline | no | yes | CD66b- | pool | 1 | 1.00 | 2.05 | 2 | 13 | 126 |
| **P052** | m | 78 | pneumonia | 1402 | G4A2 | death | CVVHD | yes | CD66b- | pool | 1 | 3.00 | 4.80 | 1 | 11 | 1781 |
| **P053** | m | 77 | pneumonia | 717 | no | baseline | no | yes | CD66b- | pool | 1 | 0.90 | 4.22 | 3 | 21 | 185 |
| **P058** | m | 55 | pneumonia | 863 | no | baseline | no | yes | CD66b- | pool | 1 | 0.60 | 1.10 | 1 | 0 | 1048 |
| **P059** | m | 62 | pneumonia | 651 | no | progress CKD | no | yes | CD66b- | pool | 1 | 0.70 | 1.75 | 2 | 5 | 81 |
| **P060** | m | 62 | pneumonia | 1459 | no | baseline | no | yes | CD66b- | pool | 1 | 0.50 | 1.58 | 3 | 0 | 163 |
| **P061** | m | 77 | pneumonia | 2839 | no | progress CKD | CVVHD | yes | CD66b- | pool | 1 | 1.20 | 6.60 | 3 | 9 | 1474 |
| **P119** | m | 73 | pneumonia | 2625 | no | baseline | CVVHD | yes | CD66b- | pool | 1 | 1.10 | 2.98 | 2 | 1 | 98 |

### Supplemental Table 1. Patient information.

### Supplemental information

A

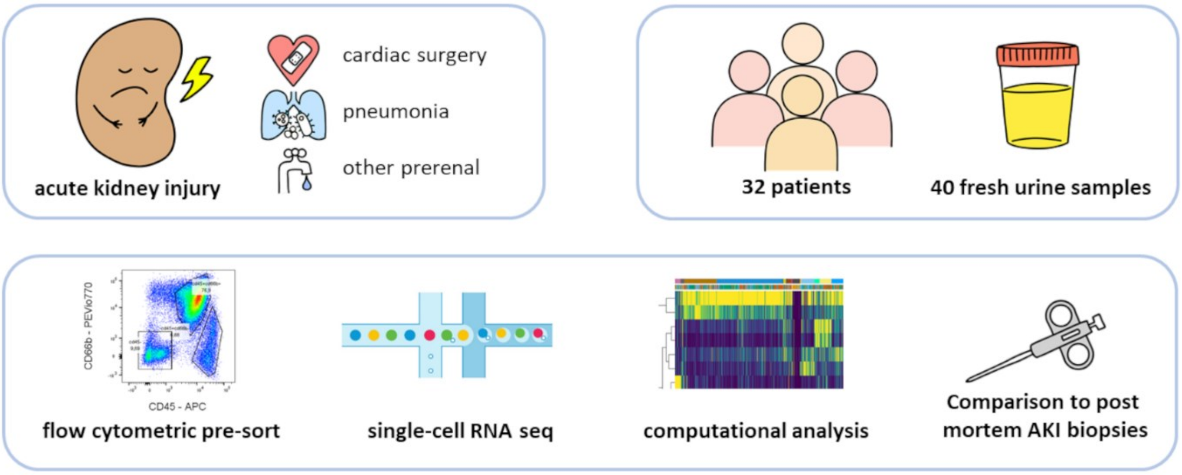


B

| AKI etiology | cardiac surgery | pneumonia | prerenal | total |
|---|---|---|---|---|
| patients (samples) | 7 (13) | 15 (17) | 10 (10) | 32 (40) |
| mean age (range) | 69 (51-83) | 60 (33-78) | 68 (24-84) | 64 (24-84) |
| sex (f/m) | 2/5 | 2/13 | 3/7 | 7/25 |
| KDIGO AKI stage (1/2/3) | 0/4/3 | 2/5/8 | 3/2/5 | 5/11/16 |
| Total post-QC sc transcriptomes | 20249 | 15582 | 6777 | 42608 |

**Supplemental Figure 1. Study design and patient characteristics.**

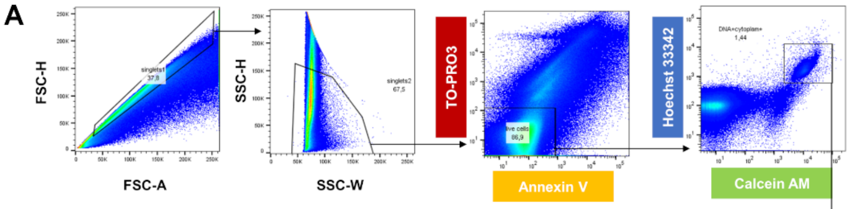


B

| celltype | Median cell count [/dl] | Median fraction |
|---|---|---|
| CD45- | 458 | 16% |
| CD45+CD66b- | 471 | 12% |
| CD45+CD66b+ | 2678 | 58% |

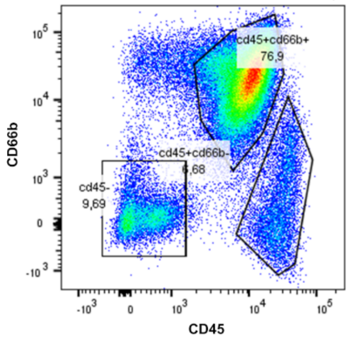


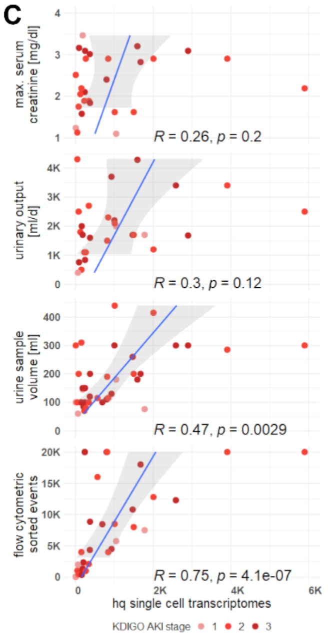


**Supplemental Figure 2. Flow cytometry can provide viable pre-sorted urine cell fractions**

***A.*** *Gating strategy for sorting single viable cells from urine. After singlet gating in forward and sideward-scatter, dead (TO-PRO3+) and apoptotic (Annexin V) cells are excluded, before viable (Calcein AM+) events with DNA (Hoechst 33342+) are singled out. For single-cell suspension, CD45-CD66b- and CD45+CD66b- cells were sorted.* ***B.*** *Table of urinary cell type fractions. Granulocytes (CD45+CD66b+) provide the largest fraction of viable urine cells.* ***C.*** *Correlation of captured high-quality single cell transcriptomes with patient metrics. The absolute number of captured cells is dependent on urine sample size.*

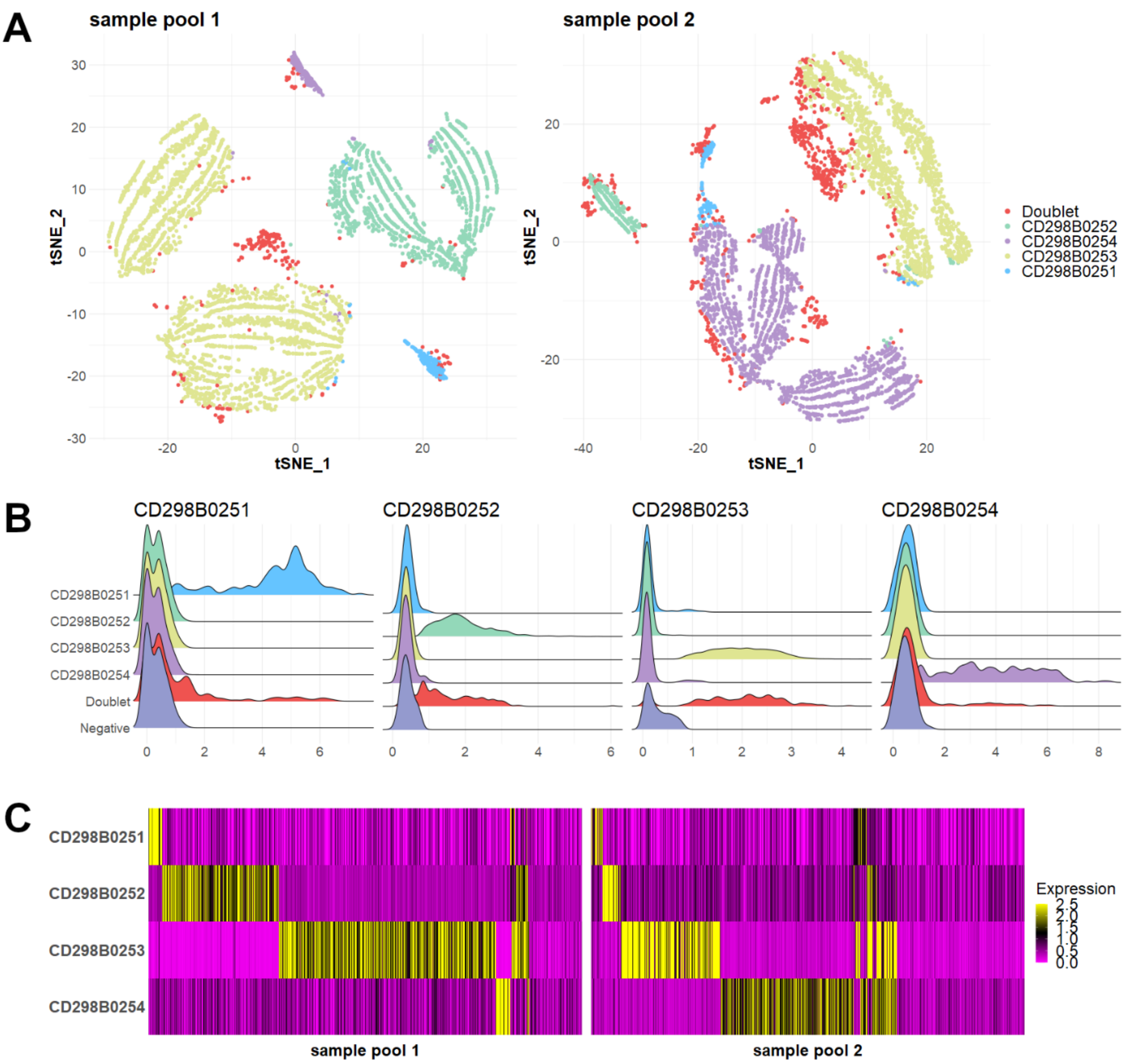


**Supplemental Figure 3. Urine single cell libraries can be demultiplexed after hashing and pooling of samples.**

***A.*** *t-Distributed Stochastic Neighbor Embedding (tSNE) dimensional reduction of pooled patient samples that were labeled by barcode cell surface antibodies (CD298B0251-4).* ***B.*** *Ridgeplots of barcode antibody detection per sample. Inter-sample doublets (red) can be easily identified.* ***C.*** *Heatmap of barcode antibody detection per cell. Doublets were excluded, cells without barcode detection were further analyzed as pool samples.*

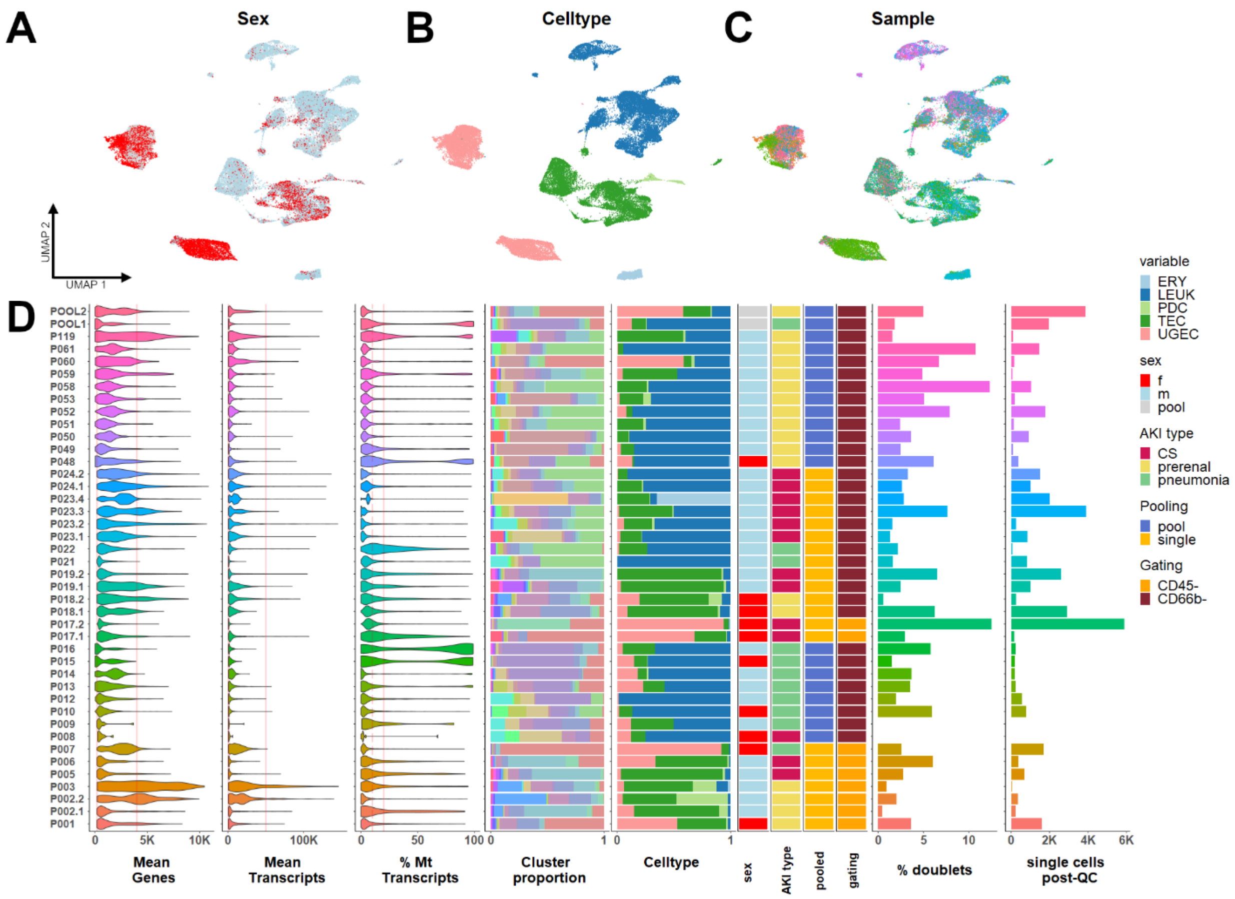


**Supplemental Figure 4. Amount of captured urinary cells is diverse.**

*Uniform manifold approximation and projection (UMAP) of* 42608 *scRNA-seq urine cells from 32 individuals with AKI.* ***A-C.*** *Distribution of cells in UMAP by sex (A), celltype (B) and etiology of AKI (C). Females (red) excrete more UGEC via urine (also Suppl. Fig. 5).* ***D.*** *Sample metrics by sample. Patients with multiple samples are numbered (1.1, 1.2 …). Amount of captured urinary cells is diverse (panel on the right), but major cell types (kidney cells (PDC – podocytes, TEC – tubular epithelial cells), urogenital epithelial cells (UGEC) and leukocytes (LEUK)) are regularly featured. (Color code – see Legend on the right)*

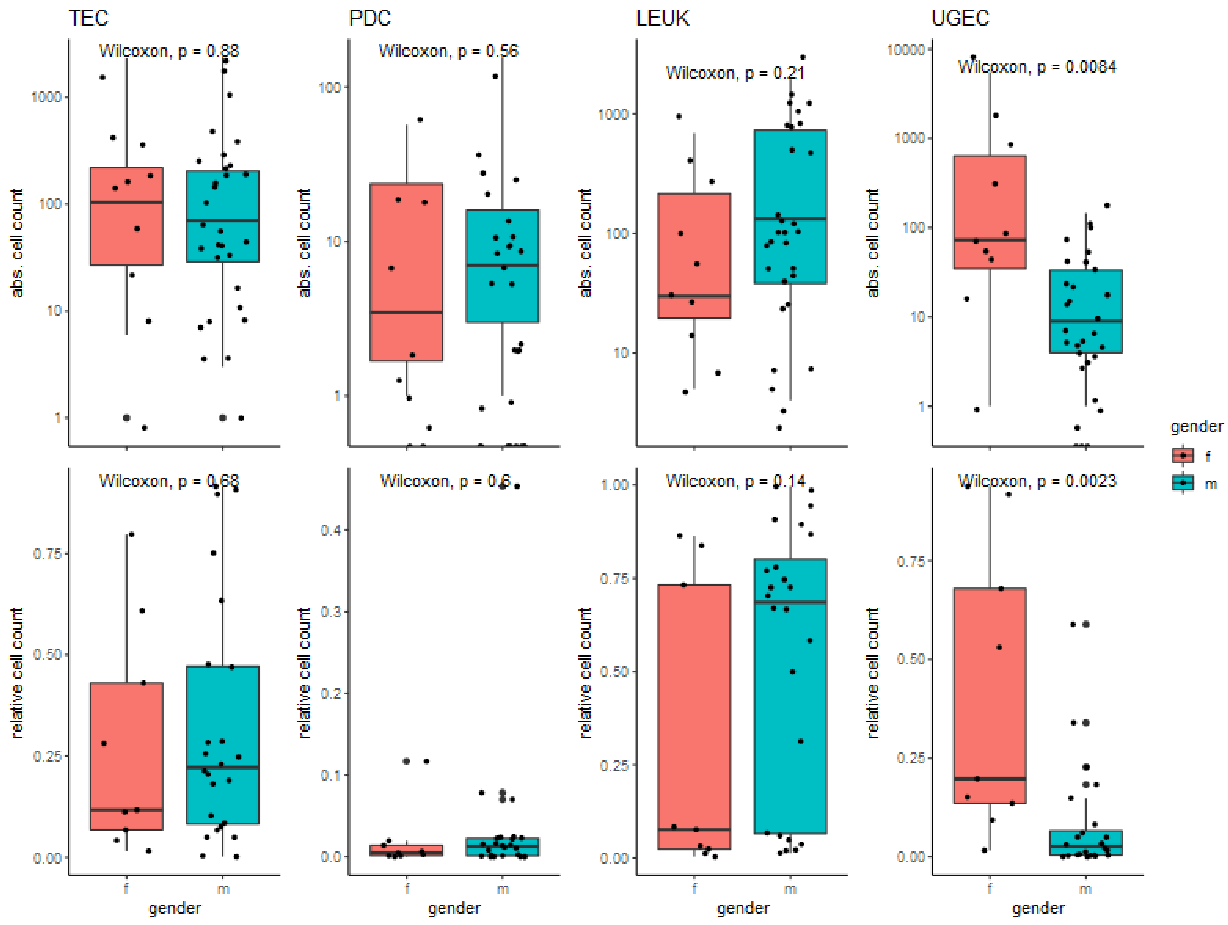


**Supplemental Figure 5. Urogenital cell abundance in urine is higher in females**

*Boxplots of absolute counts and relative proportions of urinary cells per sample in female (red) and male (green) patients. PDC – podocytes, TEC – tubular epithelial cells), UGEC – urogenital epithelial cells, LEUK – leukocytes.*

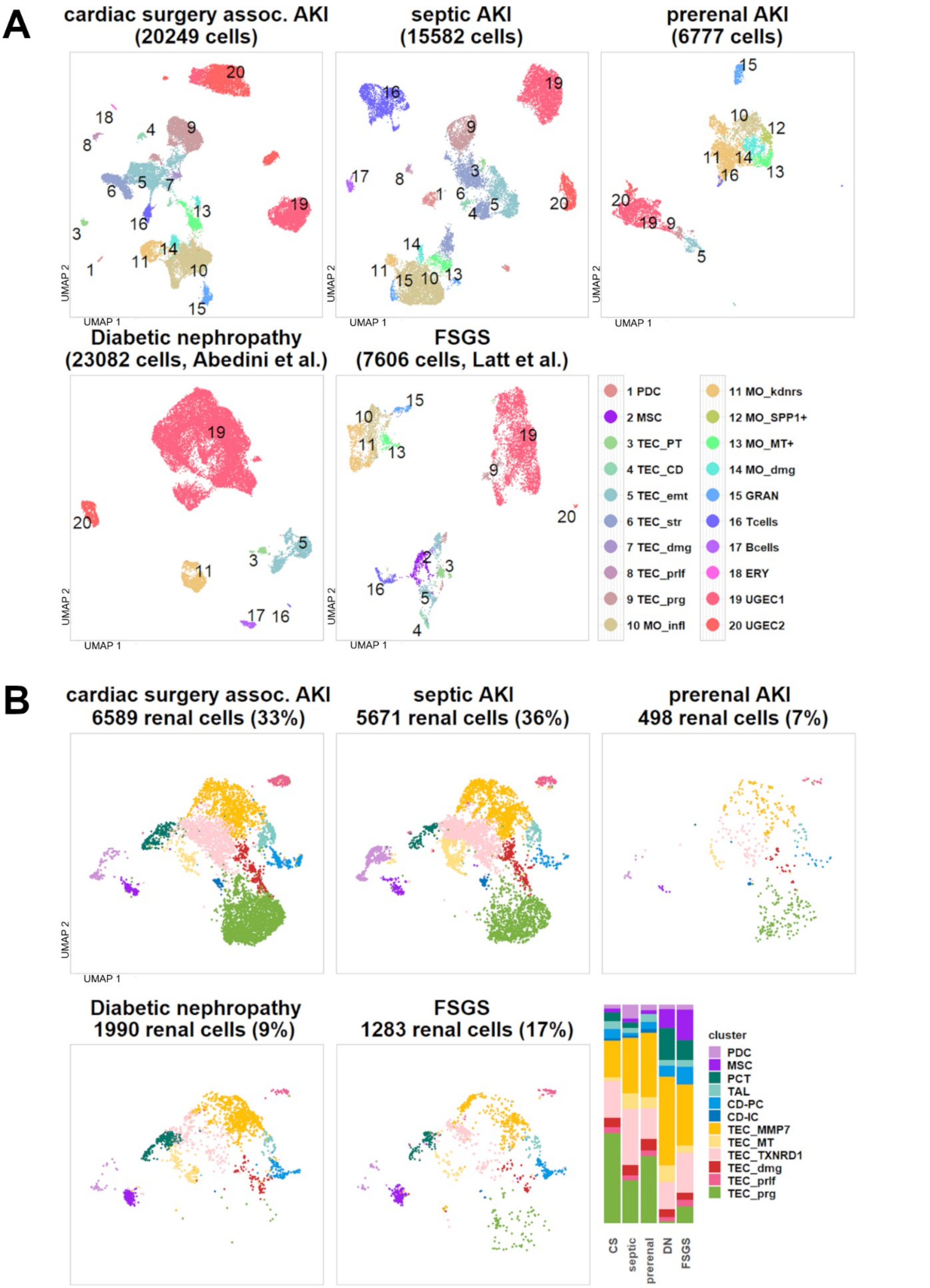


**Supplemental Figure 6.1. Urine cell abundance and proportions vary between diseases**
***A.*** *UMAP of urinary cell signatures by AKI etiology (panel 1-3) compared to diabetic nephropathy*(18) *(DN, panel 4) and focal segmental glomerulosclerosis*(19) *(FSGS, panel 5).* ***B****. Integrated UMAP of all renal parenchymal clusters split by disease type*

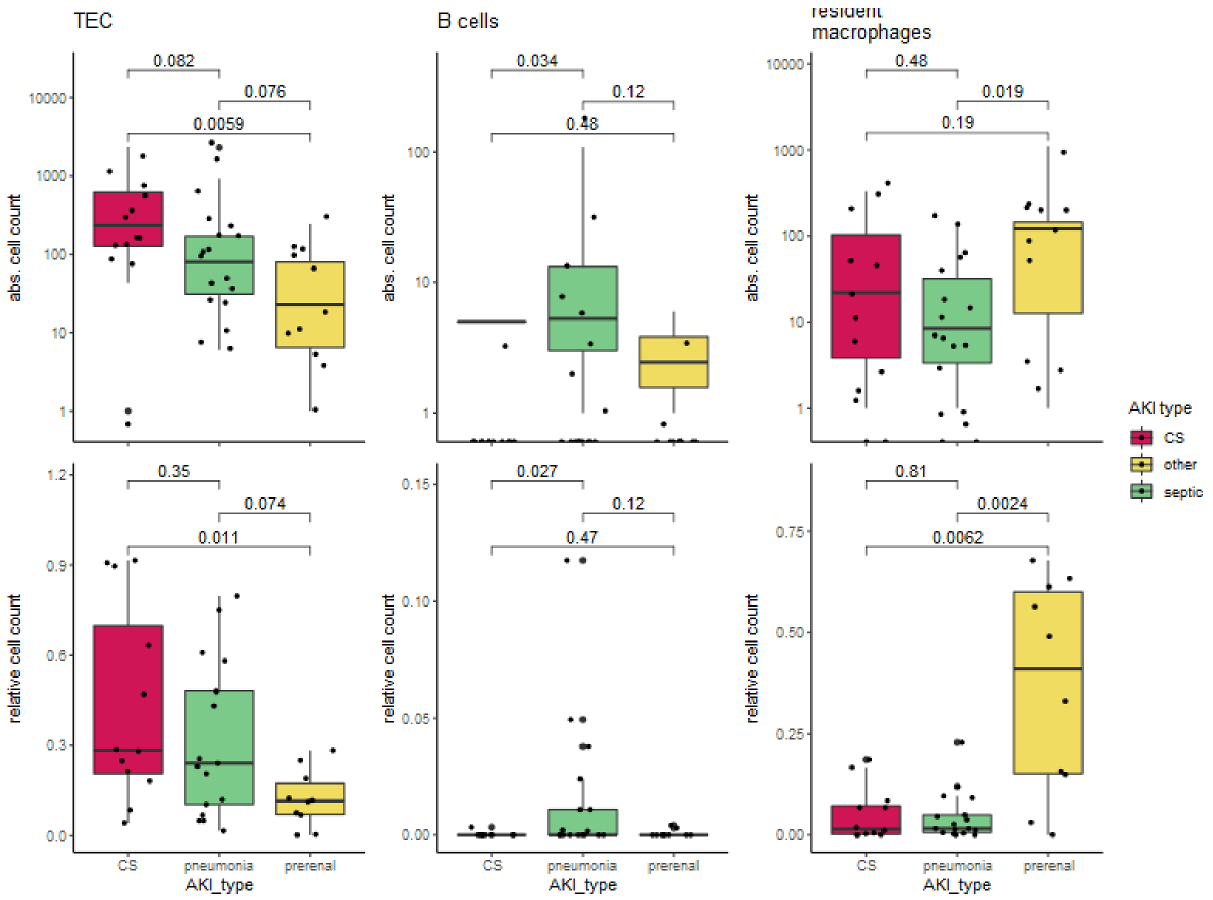


**Supplemental Figure 6.2. Urine cell abundance and proportions vary between diseases**
*Boxplots of absolute counts and relative proportions of urinary cells per sample in different AKI etiologies (CS – cardiac surgery (CS) – red, pneumonia – green, prerenal – yellow). B cells are more frequent in pneumonia patients, macrophages in prerenal patients.*

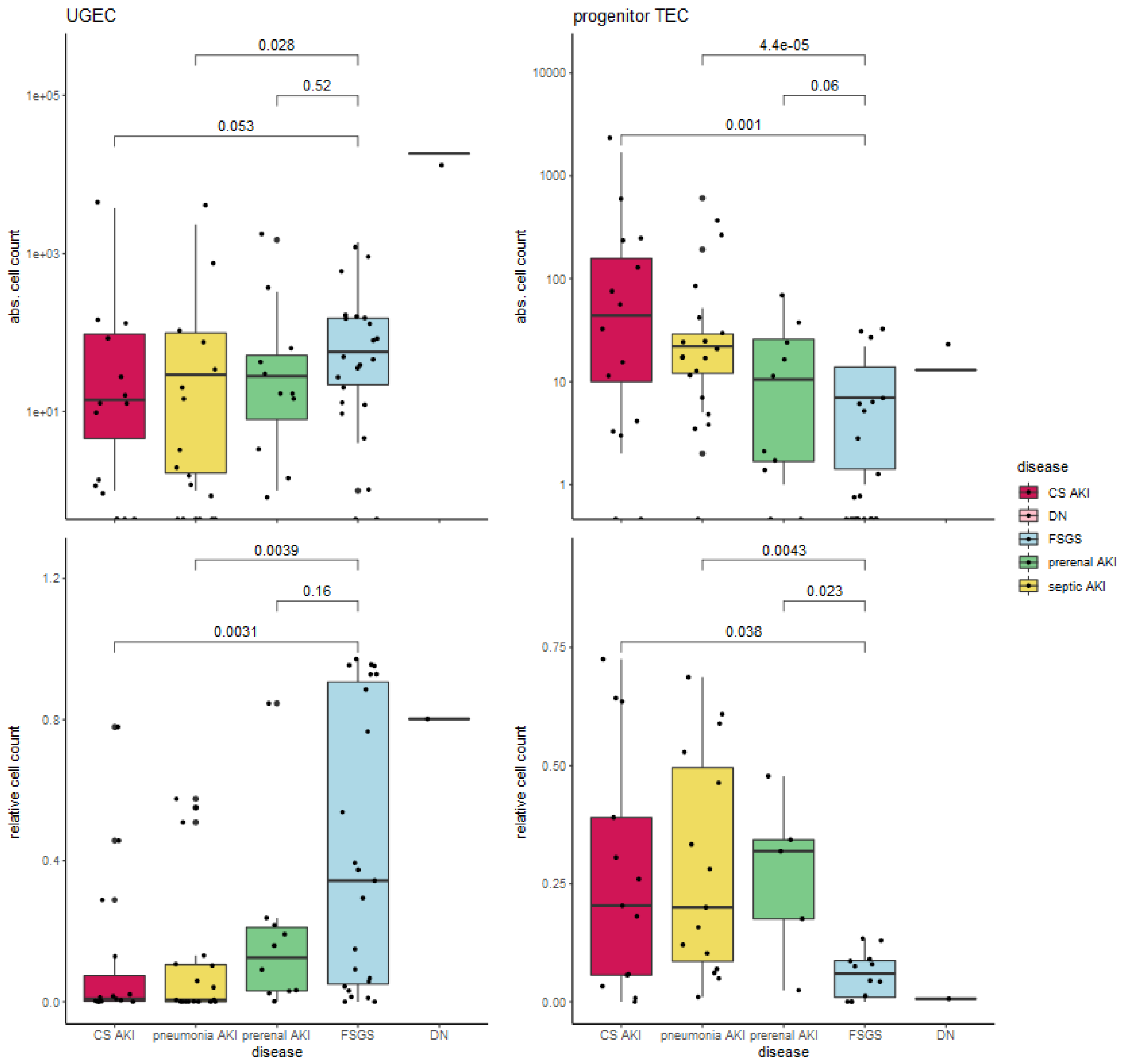


**Supplemental Figure 6.3. Urine cell abundance and proportions vary between diseases**
*Boxplots of absolute counts and relative proportions of urinary cells per sample in different AKI etiologies and diseases (FSGS – blue, DN – pink). Progenitor-like cells are more abundant in AKI urine.*

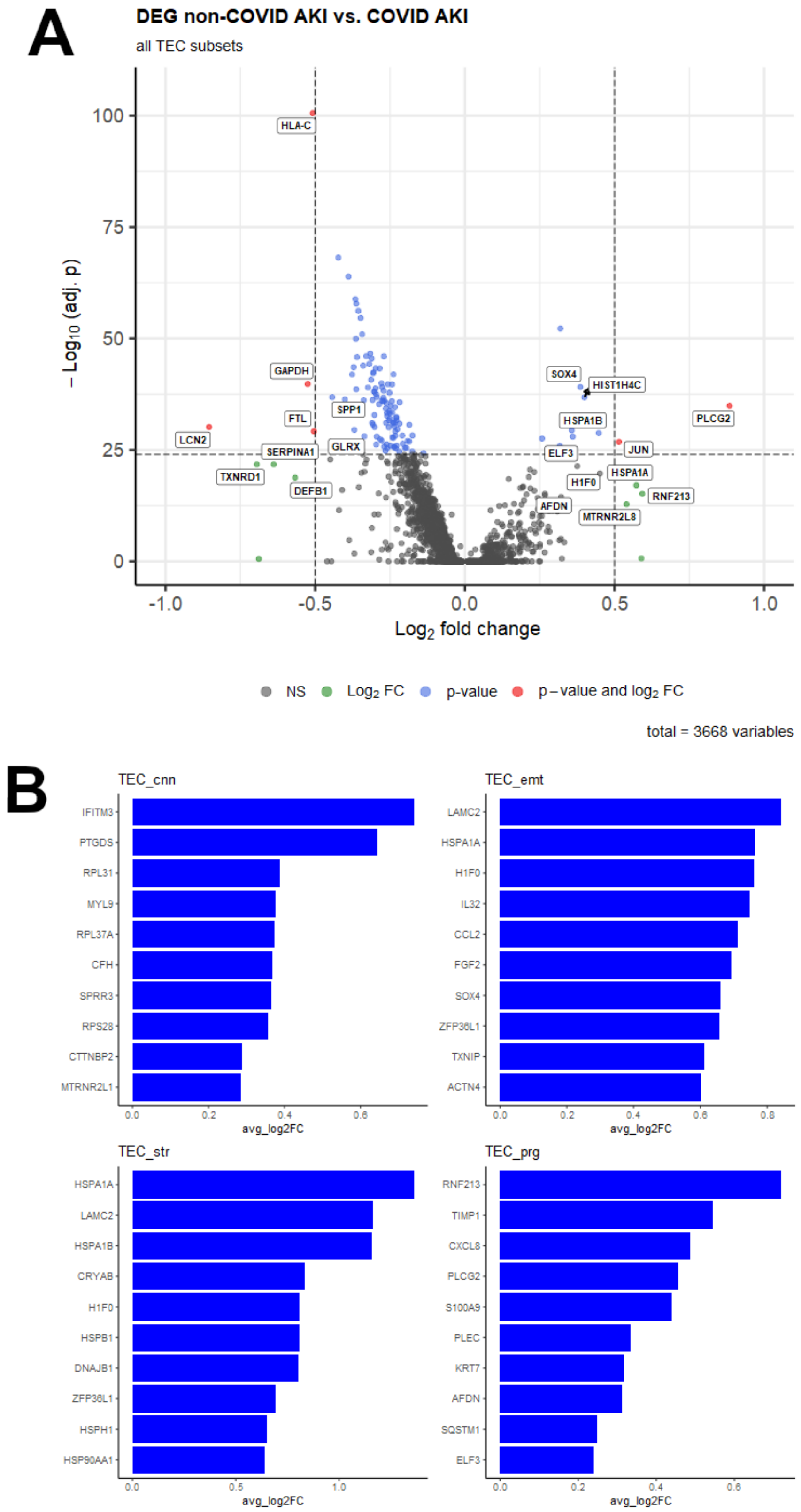


**Supplemental Figure 7. Urine cells show some COVID-19 associated transcriptional signatures.**

***A*** *Volcano plot indicating significance (y) and differential expression of genes (x) between all tubular epithelial cells (TEC) from COVID-19 AKI patients (right) and other AKI patients (left).* ***B*** *Bar plots with the top ten most differentially expressed genes per TEC subgroup. All adj. p values < 10e-5. emt – epithelial-mesenchymal transition, str – stressed, prg – progenitor-like.*

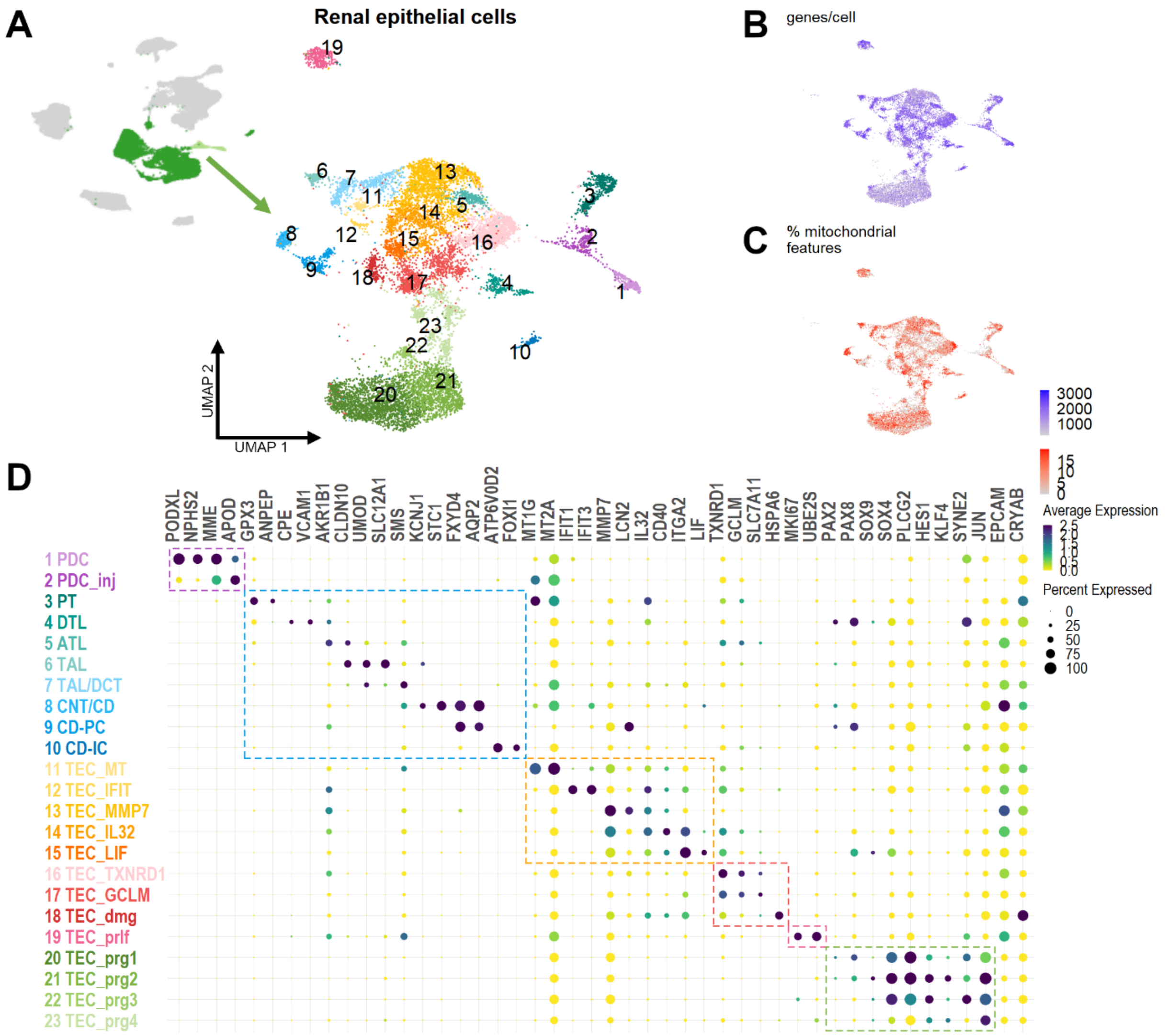


**Supplemental Figure 8. Diverse tubular cell reactions to AKI**

***A.*** *UMAP of 12853 urinary renal parenchymal scRNAseq transcriptomes in 23 distinct clusters.* ***B+C.*** *Quality metrics genes/cell (B) and percentage of mitochondrial genes per cell (C) plotted in UMAP.* ***D.*** *Dotplot of marker gene expression for each cell type. Next to podocytes (PDC, cluster 1-2) and segment specific tubular epithelial cells (TEC, clusters 3-10), several injury related subtypes of TEC (clusters 11-23) can be distinguished. PT – proximal tubule, DTL – descending thin limb, ATL – ascending thin limb, TAL – thick ascending limb, DT – distal tubule, CNT – connecting tubule, CD-PC – collecting duct principal cells, CD-IC – collecting duct intercalated cells, inj – injured, dmg – damaged, prlf – proliferating, prg – progenitor-like.*

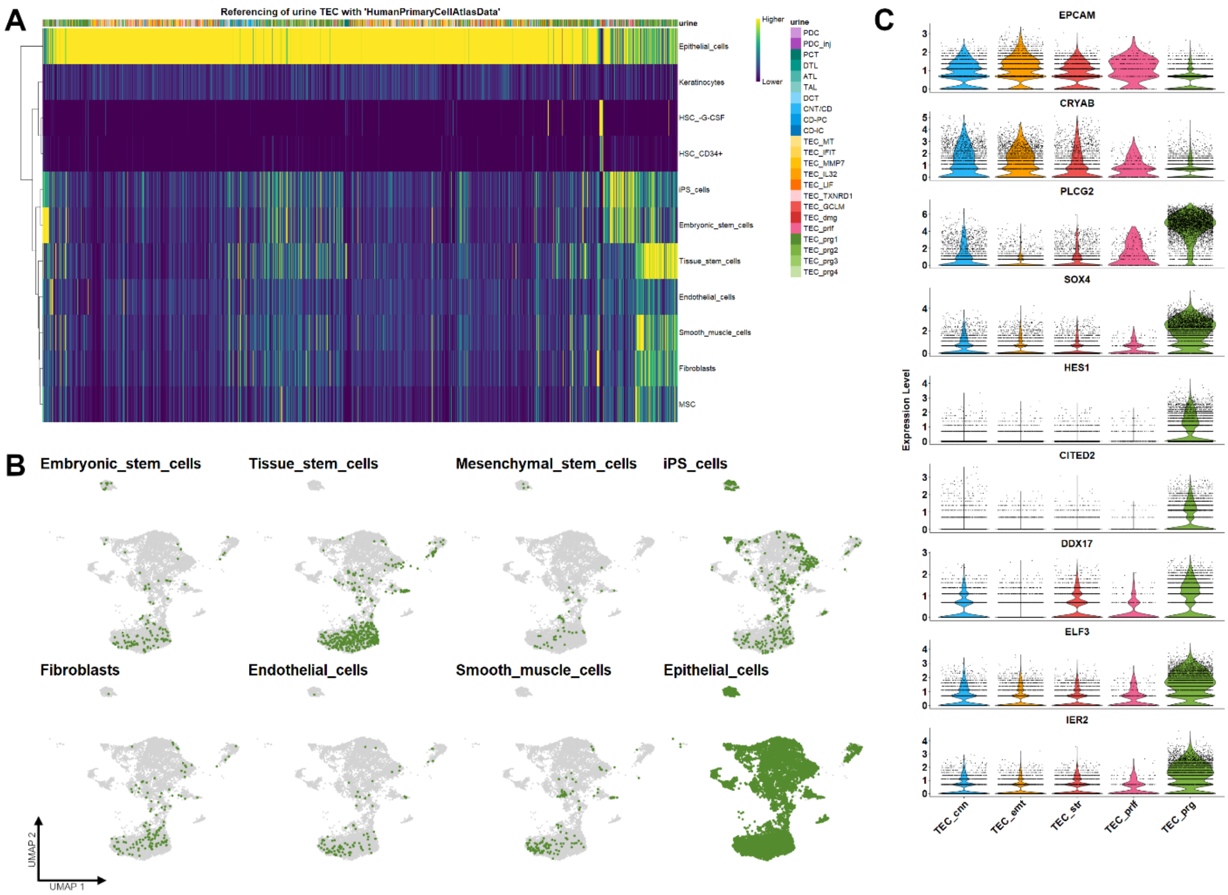


**Supplemental Figure 9. Urinary tubular epithelial cells partially resemble stem cells**
***A.*** *Heatmap of automatic annotation scores (SingleR) of urinary tubular epithelial cells (TEC) with a human primary cell atlas*(29). *Most cells are annotated as epithelial, while some are annotated as stem- or mesenchymal cells, pointing towards dedifferentiation.* ***B.*** *UMAP Distribution of automatic cell labels of A in urine TEC. Most stem labels occur in progenitor-like (TEC_prg) subsets (bottom middle clusters).* ***C.*** *Violin plot of marker gene expression across TEC subgroups. Tubular markers (EPCAM, CRYAB) are downregulated in TEC_prg, while stemness markers (PLCG2, HES1) are abundantly expressed.*

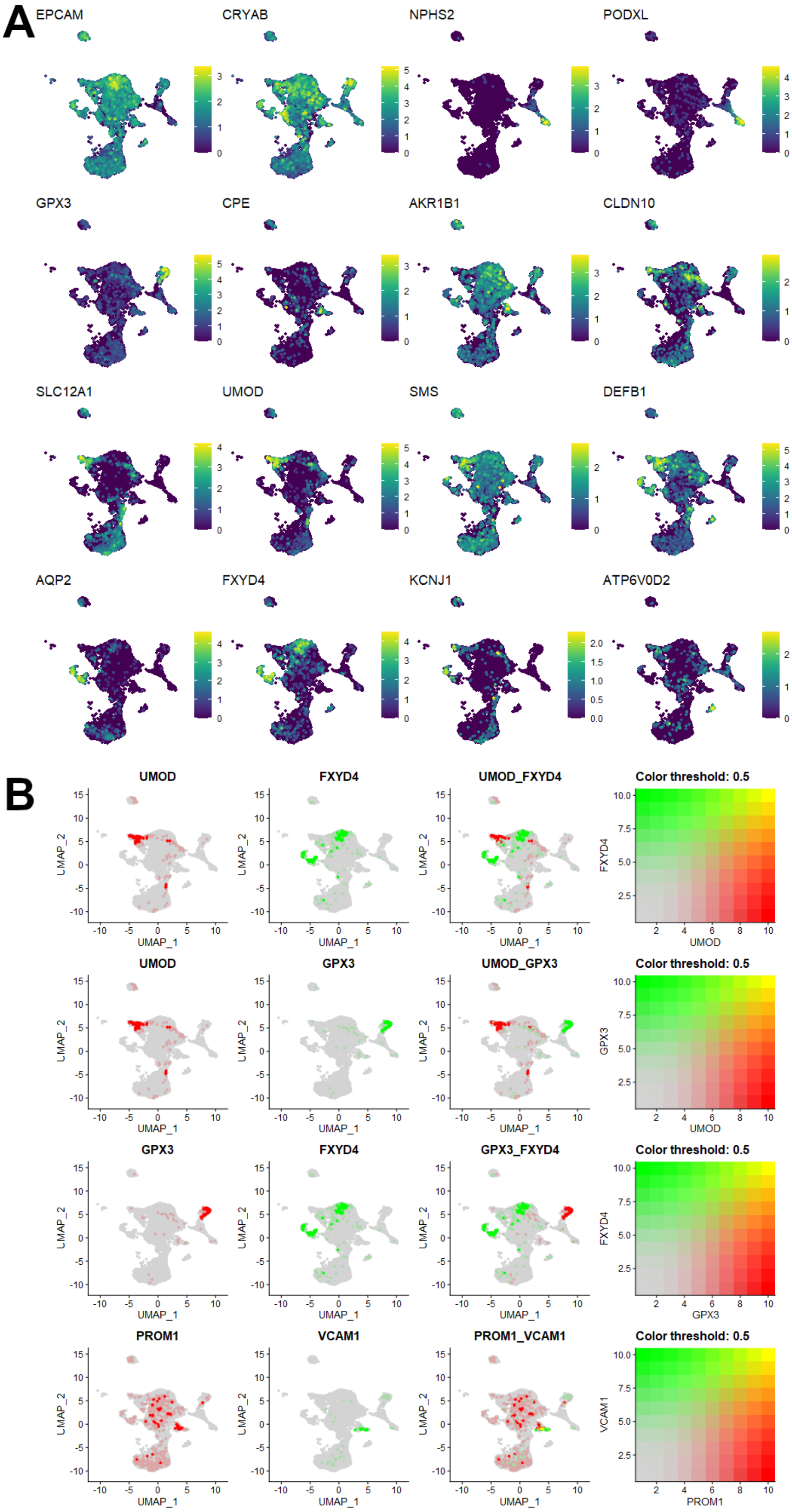


**Supplemental Figure 10. Injured cell states are of mixed tubular identity**

***A.*** *Tubular cell marker expression in urinary renal cells plotted in UMAP. Injured clusters (top to bottom middle) dimly express several segment specific marker genes.* ***B.*** *Co-Expression of select markers shows no overlap between segment specific markers. One cluster of descending thin limb (DTL) identity has co-expression of VCAM1 and PROM1, hinting at resident stem cells.*

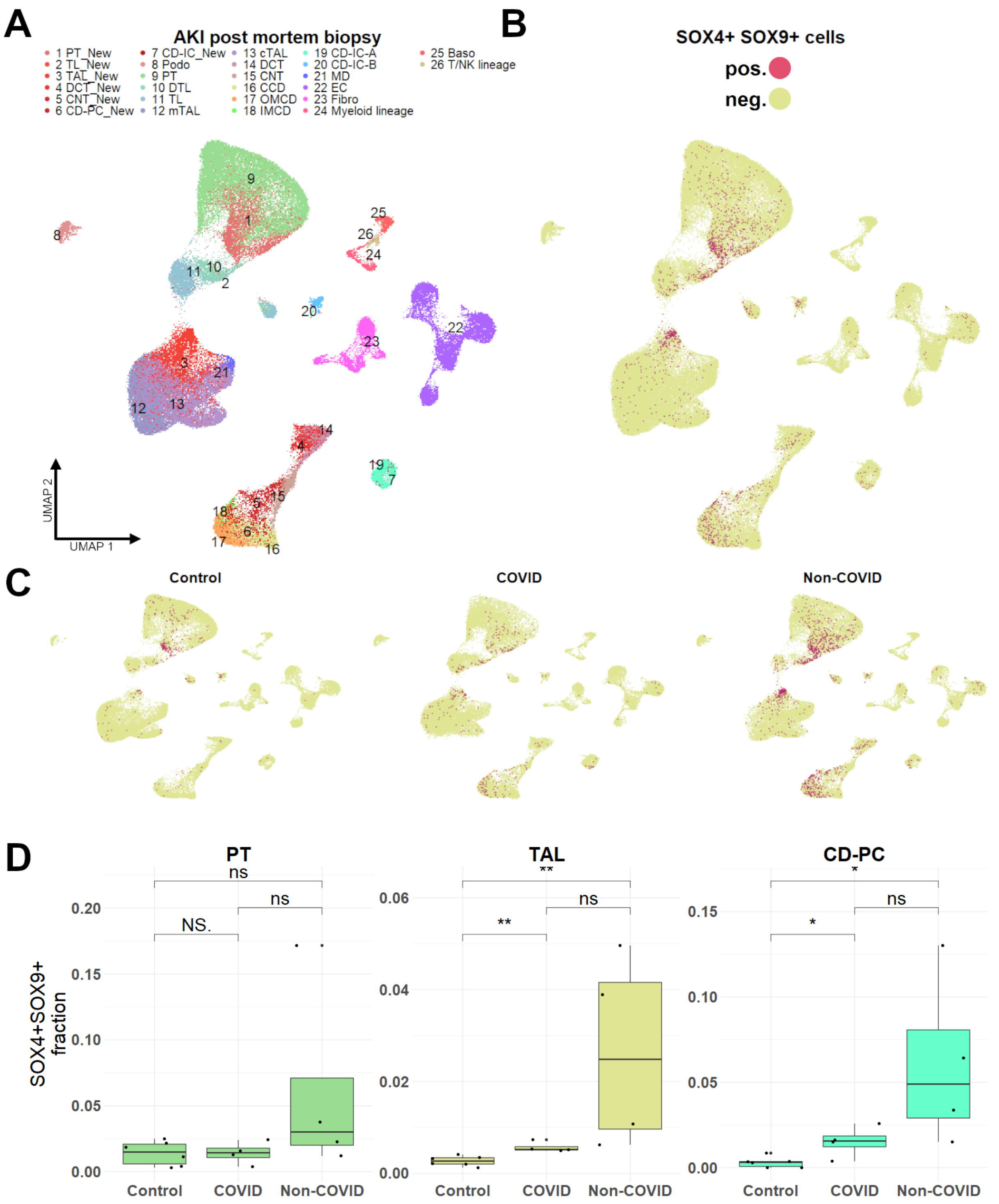


**Supplemental Figure 11. SOX4+SOX9+ progenitor-like cells occur in kidney AKI tissue.** ***A.*** *UMAP dimensional reduction of AKI post-mortem biopsy snRNAseq dataset. "_New" clusters in red shadings indicate injury reactive tubular cell states occurring in AKI. PDC – podocytes, PCT – proximal convoluted tubule, DTL – descending thin limb, ATL – ascending thin limb, TAL – thick ascending limb, DCT – distal convoluted tubule, CNT – connecting tubule, CD-PC – collecting duct principal cells, CD-IC – collecting duct intercalated cells, MYEL – myeloid cells, EC – endothelial cells, FBR – fibrocytes, MO – monocytes/macrophages, GRAN – granulocytes, emt – epithelial-mesenchymal transition, str – stressed, prlf – proliferating, prg – progenitor-like* ***B+C.*** *Localization of SOX4+SOX9+ cells in dimensional reduction of all samples (**B**) and by disease group (**C**).* ***D****. Fraction of SOX4+SOX9+ TEC in proximal tubule (PT), thick ascending limb (TAL) and collecting duct principal cells (CD-PC) divided by disease group. In both COVID-AKI and non-COVID-AKI the relative SOX4+SOX9+ cell amount increases compared to healthy tissue.*

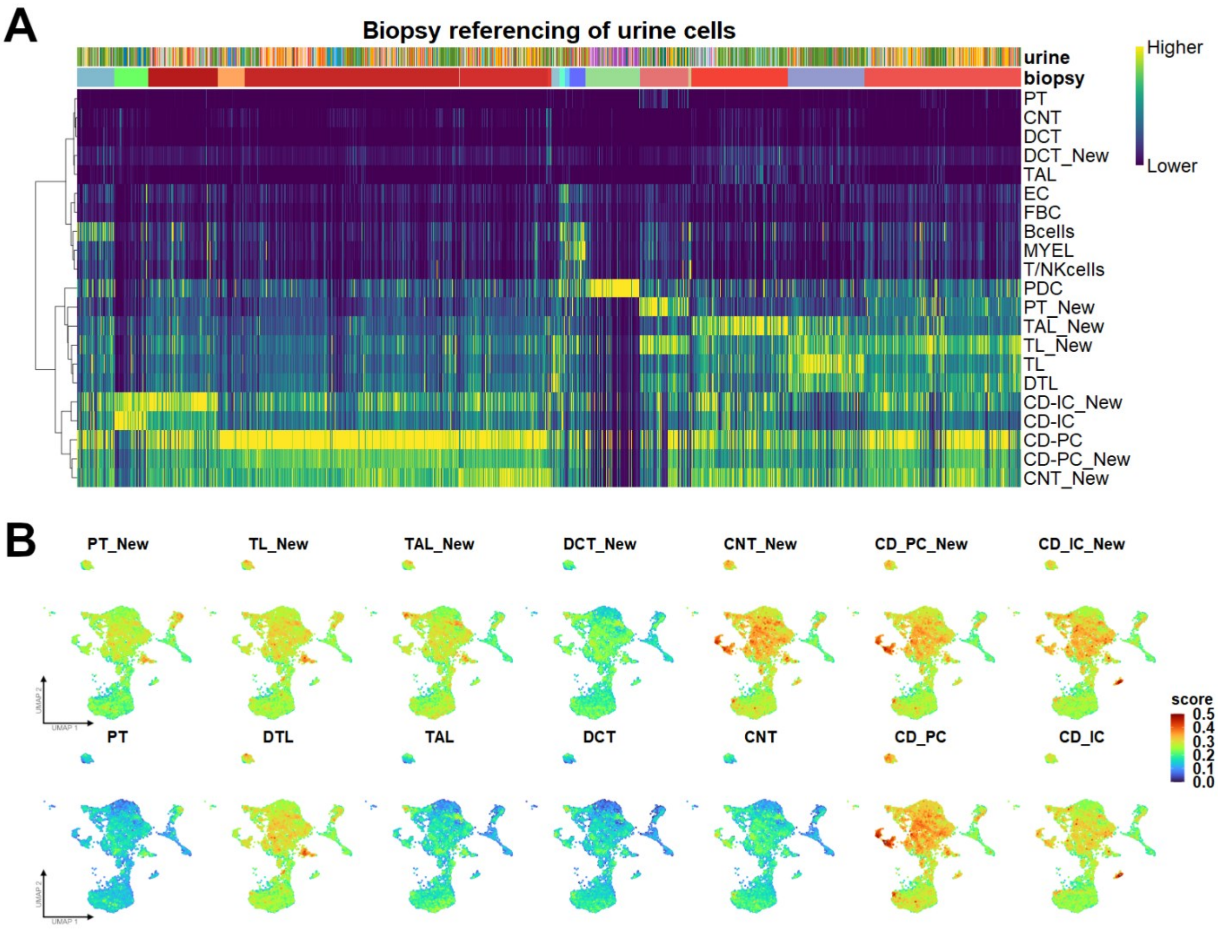


**Supplemental Figure 12. Urinary tubular cells resemble injured and distal tubules**

***A.*** *Heatmap of automatic annotation scores (SingleR) of urinary tubular epithelial cells (TEC) with human post-mortal biopsy AKI tissue*(12)*. Most cells are most similar to injury reactive cell states from AKI kidney tissue ("_New") and distal tubular segments like collecting duct.* ***B.*** *UMAP Distribution automatic annotation scores of A in urine TEC. PDC – podocytes, PCT – proximal convoluted tubule, DTL – descending thin limb, ATL – ascending thin limb, TAL – thick ascending limb, DCT – distal convoluted tubule, CNT – connecting tubule, CD-PC – collecting duct principal cells, CD-IC – collecting duct intercalated cells.*

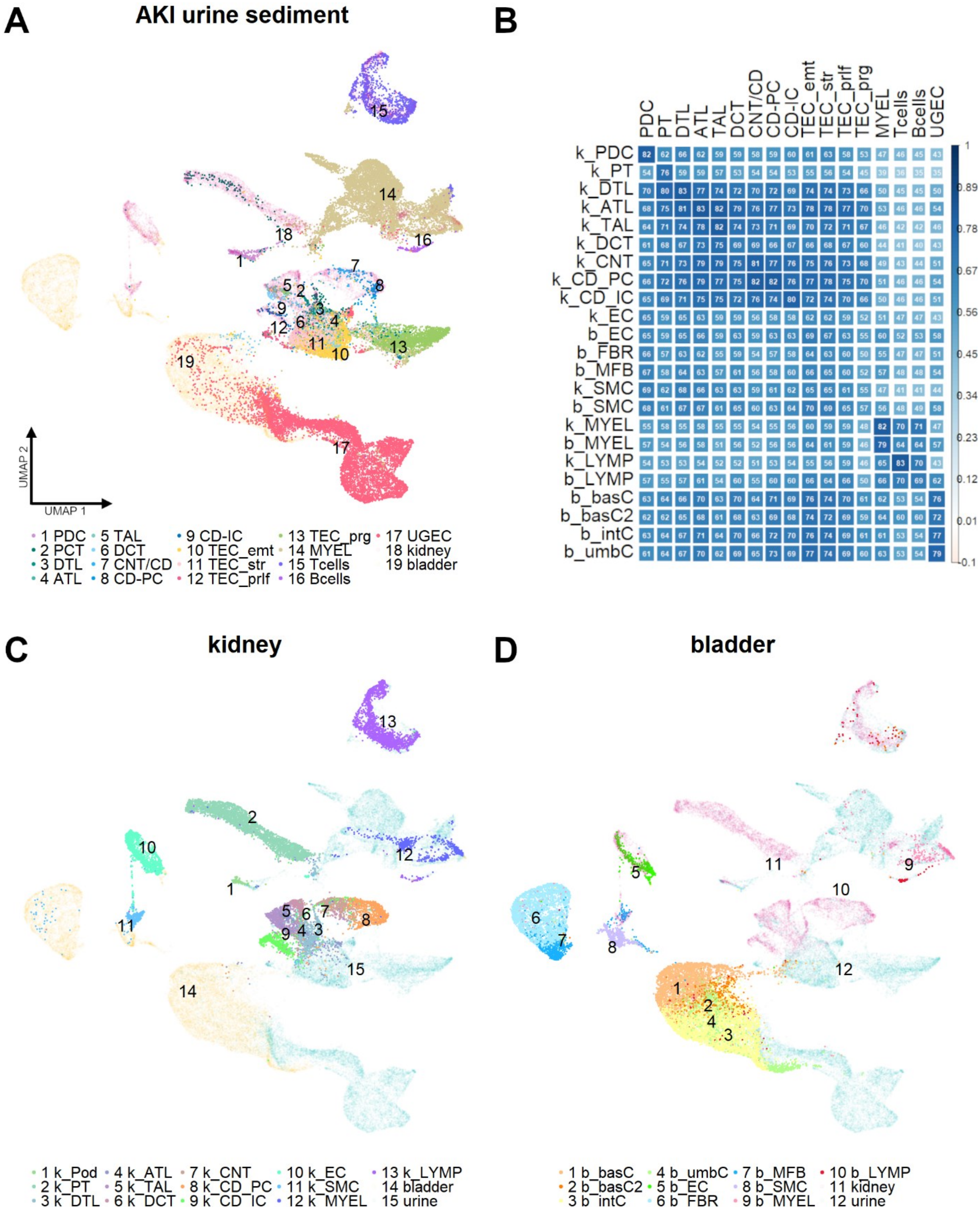


**Supplemental Figure 13. Urine tubular cells cluster with kidney, not bladder tissue**

*UMAP dimensional reduction of tumor-adjacent healthy kidney tissue scRNAseq*(38) *and healthy bladder tissue*(39) *dataset.* ***A+C-D.*** *Integrated healthy kidney (**C**), bladder (**D**) and AKI urinary scRNAseq (**A**) datasets. Urine tubular epithelial cells (TEC) cluster with kidney epithelial cells while urine urogenital cells cluster with bladder cells, confirming a kidney origin of urinary TEC.* ***B****. Correlation plot for gene expression urine clusters (columns) vs. kidney (k clusters) and bladder (b clusters) tissue clusters (rows). Size and color represent Spearman R (displayed as percentage), all p <0.001. PDC – podocytes, PT – proximal tubule, DTL – descending thin limb, ATL – ascending thin limb, TAL – thick ascending limb, DCT – distal convoluted tubule, CNT – connecting tubule, CD-PC – collecting duct* *principal cells, CD-IC – collecting duct intercalated cells, MYEL – myeloid cells, LYMP – lymphocytes, EC – endothelial cells, FBR – fibrocytes, MFB – myofibroblasts, SMC – smooth muscle cells, UGEC – urogenital epithelial cells, including basC - basal cells, intC intermediate cells, umbC – umbrella cells, emt – epithelial-mesenchymal transition, str – stressed, prlf – proliferating, prg – progenitor-like, infl – inflammatory, kdnrs – kidney resident.*

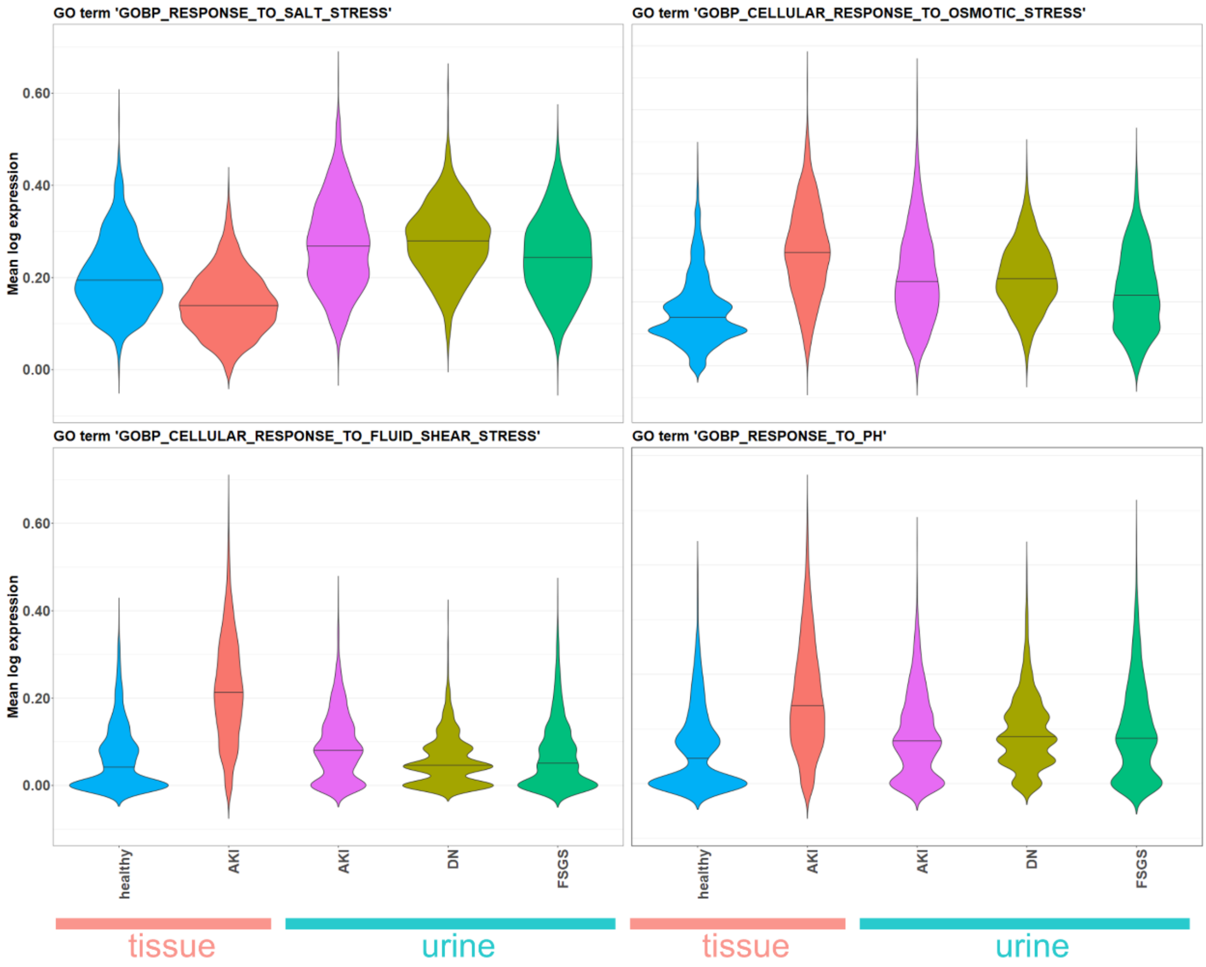


**Supplemental Figure 14. Stress genes are partially upregulated in urine cells**
*Violin plot of mean log normalized expression per cell of genes contained in named GO term.*

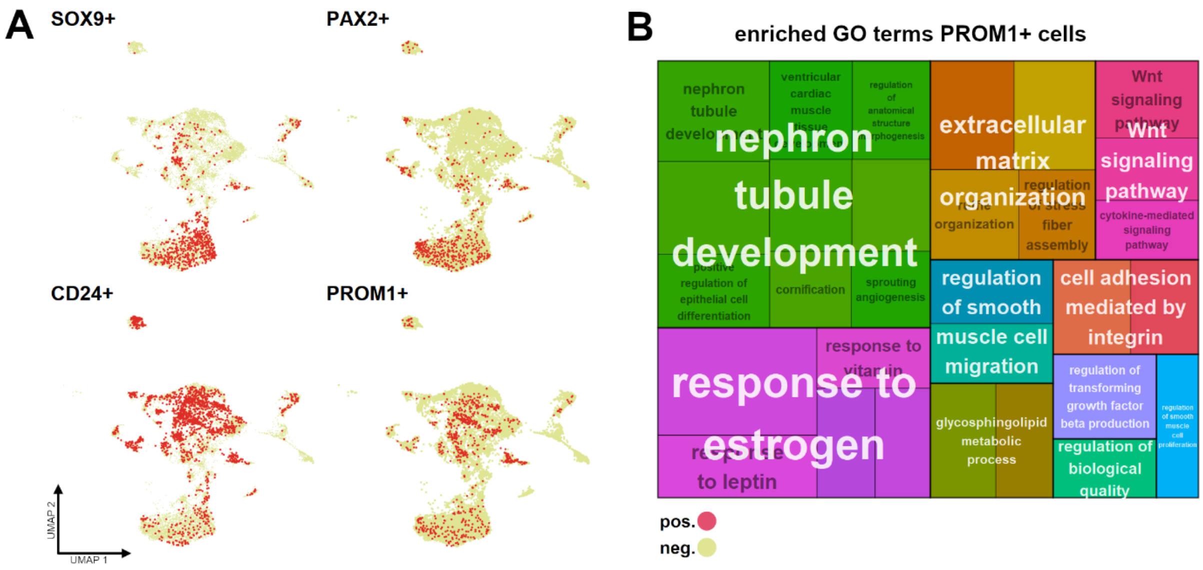


**Supplemental Figure 15. Distribution of stem cell markers**

***A***. *Distribution of stem cell marker positive cells across UMAP representation.* ***B***. *Treemap plot of enriched gene ontology (GO) terms in PROM1+ cells. Each rectangle is a single GO term (black text), sized based on -log10(adj. pvalue). The terms are joined into GO clusters by similarity, with the largest rectangle of the cluster providing the group name (white text). visualized with different colors.*

**Urinary tubular subsets show distinct interactions with epithelial and immune cells (Supplemental Figure 16)**

Emphasizing the difference between TEC and presumed progenitors we used the CellPhone database to search for distinct cellular crosstalk of these cells with other epithelia and leukocytes (Suppl. Fig. 16): FGF/FGF receptor (FGFR) crosstalk in epithelial cells was pronounced in TEC_prg (70). Adrenoreceptor Beta 2 (ADRB2) signaling with IL1B and VEGF which is not usually expressed in epithelia but in mesenchymal stem cells (71) and renal clear-cell carcinoma (72), was also expressed exclusively on TEC_prg. In contrast, other TEC contributed to wound healing via the CD44-osteopontin (SPP1) axis (73–75) and EGFR-signaling via various ligands including EGFR-TGFß1 transactivation, known for inducing EMT phenotypes and renal fibrosis (76–78). The epithelial-leukocyte interaction via plexin B (PLXNB) and semaphorin 4D (SEMA4D), which has implications in developmental ureteric branching(79) are redundantly featured in both TEC and progenitors.

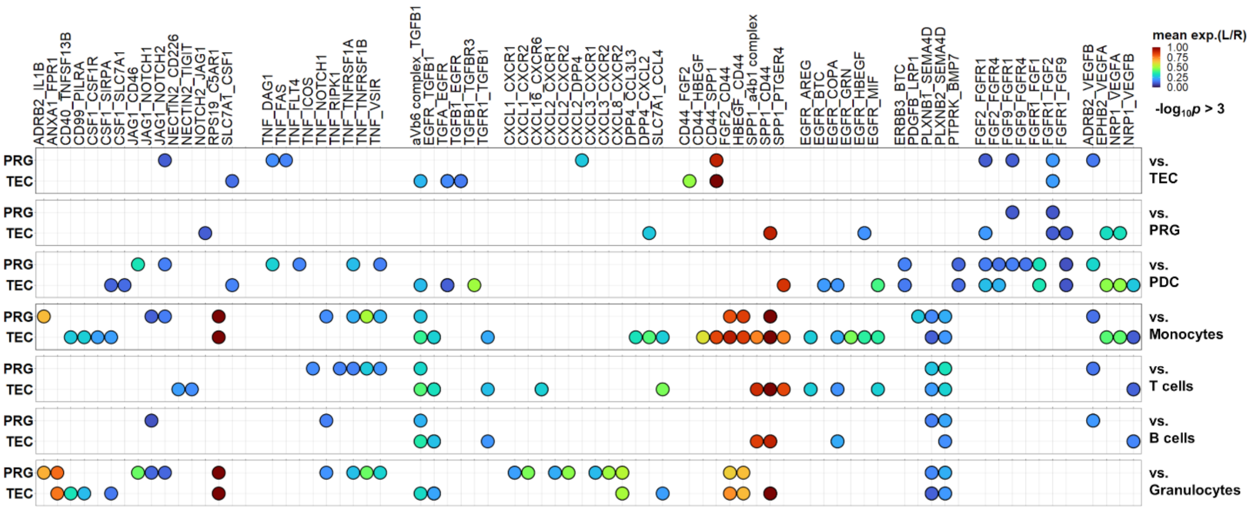


**Supplemental Figure 16. Subset specific cellular crosstalk of urinary tubular cells.**
*Cellphone DB cell crosstalk of progenitor-like cells (PRG) and tubular epithelial cells (TEC) (right, first interactor) with selected cell types (left, second interactor). Color indicates sum of receptor and ligand expression in respective cells, all interactions -log10(adj. p value) >3.*
