## Supplemental Table 2 for "Urinary single-cell sequencing captures intrarenal injury and repair processes in human acute kidney injury"

| p_val | avg_log2FC | pct.1 | pct.2 | p_val_adj | cluster | gene |
|---|---|---|---|---|---|---|
| 0 | 3.82687727 | 0.644 | 0.023 | 0 | PDC | PTGDS |
| 0 | 3.3062233 | 0.698 | 0.024 | 0 | PDC | PODXL |
| 0 | 2.92494961 | 0.883 | 0.127 | 0 | PDC | NUPR1 |
| 0 | 2.7860233 | 0.804 | 0.027 | 0 | PDC | MME |
| 0 | 2.72486732 | 0.644 | 0.014 | 0 | PDC | CHI3L1 |
| 0 | 2.5196939 | 0.851 | 0.065 | 0 | PDC | MYL9 |
| 0 | 2.41692537 | 0.572 | 0.044 | 0 | PDC | SERPINE2 |
| 0 | 2.3655583 | 0.737 | 0.042 | 0 | PDC | HTRA1 |
| 0 | 2.23330373 | 0.534 | 0.007 | 0 | PDC | NPHS2 |
| 0 | 2.22090988 | 0.604 | 0.006 | 0 | PDC | DCN |
| 0 | 2.16143665 | 0.68 | 0.012 | 0 | PDC | APOD |
| 0 | 2.00396448 | 0.685 | 0.013 | 0 | PDC | CLIC5 |
| 0 | 1.97095878 | 0.68 | 0.127 | 0 | PDC | UCHL1 |
| 0 | 1.96909865 | 0.421 | 0.003 | 0 | PDC | IGFBP1 |
| 0 | 1.89691344 | 0.638 | 0.015 | 0 | PDC | PTPRO |
| 0 | 1.8513816 | 0.635 | 0.016 | 0 | PDC | PLA2R1 |
| 0 | 1.72533456 | 0.387 | 0.012 | 0 | PDC | TPPP3 |
| 0 | 1.71789493 | 0.532 | 0.005 | 0 | PDC | ENPEP |
| 0 | 1.7042596 | 0.737 | 0.12 | 0 | PDC | BEX2 |
| 0 | 1.69047524 | 0.714 | 0.032 | 0 | PDC | TGFBR3 |
| 0 | 1.65543592 | 0.603 | 0.003 | 0 | PDC | TCF21 |
| 0 | 1.65016319 | 0.667 | 0.076 | 0 | PDC | IGFBP2 |
| 0 | 1.57495302 | 0.788 | 0.075 | 0 | PDC | CALD1 |
| 0 | 1.53894197 | 0.581 | 0.043 | 0 | PDC | NPNT |
| 0 | 1.41858864 | 0.703 | 0.03 | 0 | PDC | MAP1B |
| 0 | 1.4151281 | 0.754 | 0.122 | 0 | PDC | FOXC1 |
| 0 | 1.41424958 | 0.77 | 0.172 | 0 | PDC | DST |
| 0 | 1.41363839 | 0.61 | 0.03 | 0 | PDC | IGFBP4 |
| 0 | 1.40974252 | 0.878 | 0.237 | 0 | PDC | PTMS |
| 0 | 1.38778711 | 0.757 | 0.189 | 0 | PDC | CCPG1 |
| 0 | 1.34227911 | 0.527 | 0.01 | 0 | PDC | DACH1 |
| 0 | 1.2843003 | 0.419 | 0.008 | 0 | PDC | NES |
| 0 | 1.26836785 | 0.572 | 0.014 | 0 | PDC | SPOCK1 |
| 0 | 1.24264091 | 0.698 | 0.069 | 0 | PDC | NEBL |
| 0 | 1.21334705 | 0.696 | 0.123 | 0 | PDC | PALLD |
| 0 | 1.21028071 | 0.489 | 0.009 | 0 | PDC | FGF1 |
| 0 | 1.17425105 | 0.574 | 0.017 | 0 | PDC | NTNG1 |
| 0 | 1.15454408 | 0.455 | 0.001 | 0 | PDC | NDNF |
| 0 | 1.15184537 | 0.471 | 0.023 | 0 | PDC | SBSPON |
| 0 | 1.15160576 | 0.576 | 0.061 | 0 | PDC | THSD7A |
| 0 | 1.15038419 | 0.739 | 0.152 | 0 | PDC | BEX4 |
| 0 | 1.15028855 | 0.745 | 0.152 | 0 | PDC | SELENOM |
| 0 | 1.10136923 | 0.549 | 0.02 | 0 | PDC | F2R |
| 0 | 1.09569832 | 0.572 | 0.05 | 0 | PDC | SPOCK2 |
| 0 | 1.09497402 | 0.444 | 0.002 | 0 | PDC | WT1 |
| 0 | 1.09289815 | 0.585 | 0.073 | 0 | PDC | GOLIM4 |
| 0 | 1.09077304 | 0.653 | 0.062 | 0 | PDC | RASSF8 |
| 0 | 1.06702806 | 0.536 | 0.087 | 0 | PDC | GBE1 |
| 0 | 1.06216521 | 0.403 | 0.008 | 0 | PDC | GAS1 |

| | | | | | | |
|---|---|---|---|---|---|---|
| 0 | 1.04755061 | 0.629 | 0.073 | 0 | PDC | RETREG1 |
| 0 | 1.04456751 | 0.671 | 0.134 | 0 | PDC | BEX3 |
| 0 | 1.03682848 | 0.673 | 0.134 | 0 | PDC | NFIC |
| 0 | 1.03408505 | 0.518 | 0.004 | 0 | PDC | ARHGAP28 |
| 0 | 1.03004783 | 0.583 | 0.096 | 0 | PDC | IQGAP2 |
| 0 | 1.02960309 | 0.712 | 0.127 | 0 | PDC | ADH5 |
| 0 | 1.02559354 | 0.606 | 0.021 | 0 | PDC | MYLK |
| 0 | 1.01794805 | 0.529 | 0.018 | 0 | PDC | SERPINI1 |
| 0 | 0.9832185 | 0.513 | 0.029 | 0 | PDC | TCEAL2 |
| 0 | 0.96967741 | 0.629 | 0.09 | 0 | PDC | FERMT2 |
| 0 | 0.95799208 | 0.585 | 0.084 | 0 | PDC | CMYA5 |
| 0 | 0.95600639 | 0.522 | 0.032 | 0 | PDC | PPM1L |
| 0 | 0.94609462 | 0.477 | 0.005 | 0 | PDC | PTPRD |
| 0 | 0.94509013 | 0.372 | 0.011 | 0 | PDC | SPARC |
| 0 | 0.93559564 | 0.423 | 0.005 | 0 | PDC | MXRA8 |
| 0 | 0.88920025 | 0.522 | 0.054 | 0 | PDC | KLHL42 |
| 0 | 0.86160467 | 0.552 | 0.029 | 0 | PDC | PARD3B |
| 0 | 0.86053226 | 0.302 | 0.024 | 0 | PDC | PLTP |
| 0 | 0.85245001 | 0.471 | 0.033 | 0 | PDC | SEMA5A |
| 0 | 0.83680663 | 0.595 | 0.101 | 0 | PDC | CFAP36 |
| 0 | 0.83554276 | 0.504 | 0.069 | 0 | PDC | MGAT5 |
| 0 | 0.83257504 | 0.505 | 0.069 | 0 | PDC | TIMP3 |
| 0 | 0.82037314 | 0.457 | 0.019 | 0 | PDC | PCOLCE2 |
| 0 | 0.81951606 | 0.439 | 0.005 | 0 | PDC | CLGN |
| 0 | 0.80298894 | 0.529 | 0.03 | 0 | PDC | SNCA |
| 0 | 0.7979511 | 0.496 | 0.006 | 0 | PDC | LRRC2 |
| 0 | 0.79013619 | 0.441 | 0.003 | 0 | PDC | NECAB1 |
| 0 | 0.7782727 | 0.338 | 0.015 | 0 | PDC | C1R |
| 0 | 0.77343917 | 0.496 | 0.029 | 0 | PDC | PLCE1 |
| 0 | 0.76821237 | 0.433 | 0.026 | 0 | PDC | MAGI2 |
| 0 | 0.7660108 | 0.489 | 0.044 | 0 | PDC | EFEMP1 |
| 0 | 0.75076125 | 0.466 | 0.005 | 0 | PDC | ITIH5 |
| 0 | 0.75001674 | 0.173 | 0.002 | 0 | PDC | DDN |
| 0 | 0.74437573 | 0.399 | 0.018 | 0 | PDC | ARMH4 |
| 0 | 0.73953634 | 0.468 | 0.035 | 0 | PDC | FAM89A |
| 0 | 0.73267075 | 0.561 | 0.088 | 0 | PDC | HINT2 |
| 0 | 0.729725 | 0.424 | 0.006 | 0 | PDC | RERG |
| 0 | 0.72597737 | 0.489 | 0.046 | 0 | PDC | COL4A4 |
| 0 | 0.71819511 | 0.475 | 0.012 | 0 | PDC | DPYSL3 |
| 0 | 0.70685926 | 0.473 | 0.053 | 0 | PDC | SRGAP2C |
| 0 | 0.70012517 | 0.322 | 0.002 | 0 | PDC | MRGPRF |
| 0 | 0.6992009 | 0.396 | 0.002 | 0 | PDC | CCBE1 |
| 0 | 0.69429905 | 0.318 | 0.003 | 0 | PDC | NPHS1 |
| 0 | 0.6930468 | 0.426 | 0.007 | 0 | PDC | WIPF3 |
| 0 | 0.69302297 | 0.362 | 0.001 | 0 | PDC | SULF1 |
| 0 | 0.68987111 | 0.358 | 0.025 | 0 | PDC | COL4A3 |
| 0 | 0.67857509 | 0.164 | 0.003 | 0 | PDC | C1QL1 |
| 0 | 0.67407652 | 0.326 | 0.014 | 0 | PDC | TYRO3 |
| 0 | 0.66920151 | 0.275 | 0.01 | 0 | PDC | RAMP1 |
| 0 | 0.65728316 | 0.381 | 0.002 | 0 | PDC | PHYHIP |

| | | | | | | |
|---|---|---|---|---|---|---|
| 0 | 0.65492086 | 0.426 | 0.03 | 0 | PDC | KANK2 |
| 0 | 0.6452497 | 0.286 | 0.006 | 0 | PDC | ADAMTS5 |
| 0 | 0.64300944 | 0.335 | 0.013 | 0 | PDC | SYNPO |
| 0 | 0.64155396 | 0.406 | 0.016 | 0 | PDC | FHL1 |
| 0 | 0.63802739 | 0.353 | 0.01 | 0 | PDC | FXYD1 |
| 0 | 0.63074477 | 0.351 | 0.035 | 0 | PDC | LTBP4 |
| 0 | 0.62464883 | 0.369 | 0.032 | 0 | PDC | TMEM98 |
| 0 | 0.61482406 | 0.477 | 0.064 | 0 | PDC | C9orf3 |
| 0 | 0.58626718 | 0.414 | 0.008 | 0 | PDC | MSRB3 |
| 0 | 0.58434813 | 0.363 | 0.021 | 0 | PDC | FLRT3 |
| 0 | 0.58372258 | 0.358 | 0.018 | 0 | PDC | FOXD1 |
| 0 | 0.57644918 | 0.23 | 0.001 | 0 | PDC | ADCYAP1 |
| 0 | 0.57640722 | 0.39 | 0.009 | 0 | PDC | PTP4A3 |
| 0 | 0.56831566 | 0.335 | 0.005 | 0 | PDC | LINC00839 |
| 0 | 0.56797517 | 0.356 | 0.006 | 0 | PDC | ST6GALNAC3 |
| 0 | 0.55131422 | 0.381 | 0.042 | 0 | PDC | C1orf21 |
| 0 | 0.54352269 | 0.396 | 0.038 | 0 | PDC | RASSF8-AS1 |
| 0 | 0.52721171 | 0.304 | 0.005 | 0 | PDC | SAMD11 |
| 0 | 0.51998261 | 0.32 | 0.008 | 0 | PDC | TMEM150C |
| 0 | 0.50137685 | 0.27 | 0.002 | 0 | PDC | FAM180A |
| 0 | 0.50073274 | 0.32 | 0.009 | 0 | PDC | NPR3 |
| 0 | 0.49018221 | 0.218 | 0.004 | 0 | PDC | BGN |
| 0 | 0.48734356 | 0.351 | 0.035 | 0 | PDC | TGFB1I1 |
| 0 | 0.48061884 | 0.34 | 0.023 | 0 | PDC | CTTNBP2 |
| 0 | 0.47977151 | 0.32 | 0.001 | 0 | PDC | DPP6 |
| 0 | 0.47952186 | 0.302 | 0.007 | 0 | PDC | NR2F1 |
| 0 | 0.47723602 | 0.351 | 0.021 | 0 | PDC | NDN |
| 0 | 0.47180452 | 0.369 | 0.035 | 0 | PDC | AC084033.3 |
| 0 | 0.46787382 | 0.255 | 0.017 | 0 | PDC | SH3BGR |
| 0 | 0.46676369 | 0.336 | 0.03 | 0 | PDC | TMEM47 |
| 0 | 0.46634689 | 0.295 | 0.002 | 0 | PDC | TRIM54 |
| 0 | 0.46626749 | 0.335 | 0.021 | 0 | PDC | HOXA7 |
| 0 | 0.46129699 | 0.201 | 0.002 | 0 | PDC | MFAP4 |
| 0 | 0.45971833 | 0.349 | 0.033 | 0 | PDC | SGCE |
| 0 | 0.4573851 | 0.36 | 0.027 | 0 | PDC | UNG |
| 0 | 0.45728926 | 0.318 | 0.014 | 0 | PDC | HSD17B14 |
| 0 | 0.45638657 | 0.336 | 0.018 | 0 | PDC | BAG2 |
| 0 | 0.45147526 | 0.32 | 0.016 | 0 | PDC | PRKAR2B |
| 0 | 0.43402469 | 0.309 | 0.024 | 0 | PDC | ZNRF3 |
| 0 | 0.43174937 | 0.275 | 0.006 | 0 | PDC | ABLIM2 |
| 0 | 0.42485738 | 0.187 | 0.001 | 0 | PDC | AC092813.2 |
| 0 | 0.418709 | 0.196 | 0.001 | 0 | PDC | ROBO2 |
| 0 | 0.40805571 | 0.259 | 0.018 | 0 | PDC | FRMD3 |
| 0 | 0.40633563 | 0.295 | 0.016 | 0 | PDC | B9D1 |
| 0 | 0.40538337 | 0.302 | 0.026 | 0 | PDC | HOXD-AS2 |
| 0 | 0.39982435 | 0.259 | 0.011 | 0 | PDC | CORO2B |
| 0 | 0.39799417 | 0.241 | 0.005 | 0 | PDC | NPR1 |
| 0 | 0.3949458 | 0.214 | 0.008 | 0 | PDC | MIR503HG |
| 0 | 0.39444625 | 0.259 | 0.003 | 0 | PDC | FOXD2 |
| 0 | 0.39428481 | 0.212 | 0.009 | 0 | PDC | EDIL3 |

| | | | | | | |
|---|---|---|---|---|---|---|
| 0 | 0.39349047 | 0.288 | 0.012 | 0 | PDC | GSTA4 |
| 0 | 0.3933321 | 0.266 | 0.002 | 0 | PDC | LRRN4CL |
| 0 | 0.391531 | 0.176 | 0.002 | 0 | PDC | SEMA3G |
| 0 | 0.39070147 | 0.272 | 0.007 | 0 | PDC | GPC6 |
| 0 | 0.38935947 | 0.263 | 0.001 | 0 | PDC | LMX1B |
| 0 | 0.38101313 | 0.169 | 0.002 | 0 | PDC | PHACTR3 |
| 0 | 0.38027906 | 0.273 | 0.009 | 0 | PDC | B4GAT1 |
| 0 | 0.37927933 | 0.209 | 0.006 | 0 | PDC | CSDC2 |
| 0 | 0.37619373 | 0.23 | 0.014 | 0 | PDC | TMOD1 |
| 0 | 0.37434755 | 0.236 | 0.013 | 0 | PDC | KIRREL1 |
| 0 | 0.37316338 | 0.264 | 0.017 | 0 | PDC | FGF13 |
| 0 | 0.37198946 | 0.248 | 0.01 | 0 | PDC | TCF7L1 |
| 0 | 0.37192525 | 0.275 | 0.007 | 0 | PDC | RAB3IL1 |
| 0 | 0.36817869 | 0.191 | 0 | 0 | PDC | TCEAL6 |
| 0 | 0.36250839 | 0.216 | 0.009 | 0 | PDC | FGFR4 |
| 0 | 0.36015888 | 0.237 | 0.009 | 0 | PDC | TCEAL7 |
| 0 | 0.36001126 | 0.221 | 0.003 | 0 | PDC | CLDN11 |
| 0 | 0.35762735 | 0.27 | 0.015 | 0 | PDC | SMIM1 |
| 0 | 0.35722676 | 0.277 | 0.019 | 0 | PDC | SNTA1 |
| 0 | 0.35237979 | 0.232 | 0.01 | 0 | PDC | PALMD |
| 0 | 0.35089442 | 0.241 | 0.015 | 0 | PDC | AC007405.3 |
| 0 | 0.34749821 | 0.223 | 0.012 | 0 | PDC | PARD6G |
| 0 | 0.34304029 | 0.185 | 0.001 | 0 | PDC | TRPC6 |
| 0 | 0.34152736 | 0.207 | 0.01 | 0 | PDC | TNS2 |
| 0 | 0.34000229 | 0.227 | 0.007 | 0 | PDC | DIRAS1 |
| 0 | 0.33883792 | 0.165 | 0.002 | 0 | PDC | CRB2 |
| 0 | 0.32838852 | 0.225 | 0.006 | 0 | PDC | SCN9A |
| 0 | 0.32835277 | 0.218 | 0.011 | 0 | PDC | LOXL1 |
| 0 | 0.32117873 | 0.225 | 0.006 | 0 | PDC | MAP6 |
| 0 | 0.31550031 | 0.147 | 0.003 | 0 | PDC | XKR4 |
| 0 | 0.31051691 | 0.126 | 0.001 | 0 | PDC | CLDN5 |
| 0 | 0.31014033 | 0.218 | 0.007 | 0 | PDC | FIGN |
| 0 | 0.30914621 | 0.201 | 0.002 | 0 | PDC | FRMD1 |
| 0 | 0.30661384 | 0.14 | 0.003 | 0 | PDC | CFAP45 |
| 0 | 0.30410357 | 0.18 | 0.007 | 0 | PDC | BMP7 |
| 0 | 0.29699151 | 0.156 | 0.001 | 0 | PDC | OLFML2A |
| 0 | 0.29620202 | 0.225 | 0.01 | 0 | PDC | PDPN |
| 0 | 0.29254434 | 0.207 | 0.004 | 0 | PDC | LIMS2 |
| 0 | 0.28923497 | 0.191 | 0.006 | 0 | PDC | SMCO3 |
| 0 | 0.28838566 | 0.182 | 0.005 | 0 | PDC | CA10 |
| 0 | 0.28239087 | 0.14 | 0.001 | 0 | PDC | PTPRQ |
| 0 | 0.28143952 | 0.191 | 0.003 | 0 | PDC | NPAS3 |
| 0 | 0.27729403 | 0.171 | 0.004 | 0 | PDC | TENM4 |
| 0 | 0.2746152 | 0.203 | 0.005 | 0 | PDC | SMIM10 |
| 0 | 0.2726368 | 0.189 | 0.006 | 0 | PDC | ERICH2 |
| 0 | 0.27113775 | 0.183 | 0.006 | 0 | PDC | FIBIN |
| 0 | 0.27077134 | 0.196 | 0.002 | 0 | PDC | SCUBE1 |
| 0 | 0.27062399 | 0.165 | 0.002 | 0 | PDC | FOXC2 |
| 0 | 0.26881299 | 0.173 | 0.004 | 0 | PDC | ARHGAP20 |
| 0 | 0.26670187 | 0.171 | 0.004 | 0 | PDC | CLIP3 |

| | | | | | | |
|---|---|---|---|---|---|---|
| 0 | 0.26568076 | 0.167 | 0.004 | 0 | PDC | TMIE |
| 1.141E-303 | 0.89562692 | 0.635 | 0.124 | 2.696E-299 | PDC | C7orf50 |
| 2.702E-302 | 1.06886533 | 0.692 | 0.149 | 6.383E-298 | PDC | NFE2L1 |
| 2.373E-301 | 1.31658506 | 0.631 | 0.127 | 5.607E-297 | PDC | MAP3K20 |
| 1.018E-300 | 0.352733 | 0.223 | 0.015 | 2.404E-296 | PDC | C1S |
| 7.87E-299 | 0.30096858 | 0.183 | 0.01 | 1.859E-294 | PDC | ADAMTS9-AS1 |
| 1.653E-298 | 0.73241571 | 0.408 | 0.05 | 3.905E-294 | PDC | RARRES2 |
| 1.955E-297 | 0.81298506 | 0.612 | 0.113 | 4.618E-293 | PDC | PSIP1 |
| 4.414E-297 | 1.90392004 | 0.777 | 0.226 | 1.043E-292 | PDC | BST2 |
| 1.507E-290 | 1.32045263 | 0.818 | 0.245 | 3.56E-286 | PDC | EIF3M |
| 4.335E-288 | 0.99316424 | 0.621 | 0.124 | 1.024E-283 | PDC | NUCB2 |
| 1.347E-287 | 0.49744222 | 0.379 | 0.045 | 3.183E-283 | PDC | CTDSPL |
| 5.263E-287 | 0.74029003 | 0.349 | 0.039 | 1.243E-282 | PDC | PTH1R |
| 1.682E-282 | 0.29760687 | 0.225 | 0.016 | 3.974E-278 | PDC | CEP126 |
| 1.766E-282 | 1.32718443 | 0.576 | 0.112 | 4.171E-278 | PDC | GMDS |
| 1.228E-281 | 0.96703212 | 0.619 | 0.128 | 2.902E-277 | PDC | RNF187 |
| 1.408E-280 | 0.63661036 | 0.532 | 0.089 | 3.326E-276 | PDC | RTL8A |
| 5.676E-279 | 0.83269533 | 0.417 | 0.058 | 1.341E-274 | PDC | ARHGEF3 |
| 1.044E-277 | 0.40164247 | 0.302 | 0.03 | 2.467E-273 | PDC | SFMBT2 |
| 9.32E-277 | 0.25449415 | 0.192 | 0.012 | 2.202E-272 | PDC | WASF3 |
| 5.198E-276 | 0.70371953 | 0.478 | 0.074 | 1.228E-271 | PDC | PTPN14 |
| 5.595E-275 | 0.4946941 | 0.351 | 0.04 | 1.322E-270 | PDC | IFIT1 |
| 3.064E-274 | 0.91612917 | 0.57 | 0.107 | 7.237E-270 | PDC | MTSS1 |
| 7.489E-274 | 0.49103123 | 0.354 | 0.042 | 1.769E-269 | PDC | SRGAP2B |
| 1.567E-269 | 0.26139792 | 0.198 | 0.013 | 3.701E-265 | PDC | FAM171B |
| 3.014E-269 | 1.37125331 | 0.826 | 0.275 | 7.121E-265 | PDC | ISCU |
| 2.221E-267 | 0.32365746 | 0.248 | 0.021 | 5.246E-263 | PDC | BTC |
| 4.952E-264 | 0.77918886 | 0.595 | 0.12 | 1.17E-259 | PDC | ARL2 |
| 6.574E-263 | 0.69153274 | 0.486 | 0.078 | 1.553E-258 | PDC | NR2F2 |
| 1.713E-261 | 0.62769942 | 0.468 | 0.074 | 4.046E-257 | PDC | EPHX1 |
| 8.094E-261 | 0.54277534 | 0.252 | 0.022 | 1.912E-256 | PDC | TPM2 |
| 2.36E-260 | 0.37106236 | 0.291 | 0.029 | 5.574E-256 | PDC | HSPA12A |
| 3.536E-260 | 0.78037281 | 0.628 | 0.135 | 8.352E-256 | PDC | ARL3 |
| 3.352E-259 | 0.80941739 | 0.516 | 0.093 | 7.92E-255 | PDC | XPOT |
| 1.773E-257 | 0.46835622 | 0.381 | 0.05 | 4.189E-253 | PDC | TCEA2 |
| 2.38E-254 | 0.26041259 | 0.185 | 0.012 | 5.623E-250 | PDC | GRK5 |
| 3.366E-253 | 0.67970585 | 0.559 | 0.108 | 7.951E-249 | PDC | CHPT1 |
| 5.417E-252 | 1.02120115 | 0.637 | 0.143 | 1.28E-247 | PDC | PLS3 |
| 5.965E-252 | 0.37245252 | 0.299 | 0.032 | 1.409E-247 | PDC | C1orf115 |
| 5.758E-251 | 1.00726099 | 0.601 | 0.127 | 1.36E-246 | PDC | PDE4DIP |
| 1.138E-250 | 0.4719678 | 0.39 | 0.054 | 2.689E-246 | PDC | PSMG1 |
| 4.036E-249 | 0.38540478 | 0.299 | 0.032 | 9.535E-245 | PDC | DYNC2H1 |
| 3.949E-248 | 0.71252737 | 0.527 | 0.096 | 9.328E-244 | PDC | EPB41L5 |
| 3.022E-246 | 1.75777444 | 0.867 | 0.315 | 7.138E-242 | PDC | IFITM3 |
| 9.505E-245 | 0.51022869 | 0.396 | 0.056 | 2.245E-240 | PDC | PRKACB |
| 1.04E-243 | 0.92376358 | 0.579 | 0.12 | 2.456E-239 | PDC | SELENOP |
| 7.618E-242 | 0.91875027 | 0.572 | 0.122 | 1.8E-237 | PDC | DUSP23 |
| 1.405E-241 | 0.25307027 | 0.198 | 0.015 | 3.319E-237 | PDC | ROBO1 |
| 2.001E-241 | 0.66864804 | 0.331 | 0.041 | 4.728E-237 | PDC | SLC40A1 |
| 2.156E-240 | 0.63398091 | 0.36 | 0.049 | 5.094E-236 | PDC | CAMTA2 |

| | | | | | | |
|---|---|---|---|---|---|---|
| 1.31E-239 | 0.40643754 | 0.282 | 0.03 | 3.096E-235 | PDC | COX7A1 |
| 2.593E-239 | 0.89347423 | 0.59 | 0.128 | 6.126E-235 | PDC | SEPT11 |
| 6.378E-238 | 0.55249662 | 0.356 | 0.048 | 1.507E-233 | PDC | ST3GAL6 |
| 1.282E-236 | 0.64037783 | 0.326 | 0.041 | 3.029E-232 | PDC | TMEM91 |
| 1.674E-236 | 0.34924032 | 0.255 | 0.025 | 3.955E-232 | PDC | LAMP3 |
| 3.605E-236 | 0.88039977 | 0.689 | 0.179 | 8.516E-232 | PDC | NDUFAF3 |
| 7.374E-235 | 0.28027023 | 0.196 | 0.015 | 1.742E-230 | PDC | ADAMTS9-AS2 |
| 1.522E-234 | 0.73837378 | 0.5 | 0.094 | 3.596E-230 | PDC | HSPA13 |
| 4.227E-234 | 0.35342045 | 0.214 | 0.018 | 9.986E-230 | PDC | GJA1 |
| 1.178E-231 | 0.25588389 | 0.201 | 0.016 | 2.782E-227 | PDC | SPACA9 |
| 1.223E-231 | 0.67872484 | 0.576 | 0.12 | 2.888E-227 | PDC | PLA2G16 |
| 1.308E-231 | 0.87529201 | 0.676 | 0.173 | 3.091E-227 | PDC | SARS |
| 2.034E-231 | 0.85428015 | 0.612 | 0.138 | 4.805E-227 | PDC | CRIM1 |
| 3.989E-231 | 0.75637332 | 0.462 | 0.082 | 9.423E-227 | PDC | PDLIM2 |
| 9.975E-231 | 0.43959832 | 0.353 | 0.047 | 2.356E-226 | PDC | SSBP2 |
| 3.751E-229 | 0.40430228 | 0.272 | 0.029 | 8.861E-225 | PDC | TSLP |
| 9.227E-229 | 0.53949518 | 0.441 | 0.074 | 2.18E-224 | PDC | CDK4 |
| 1.339E-228 | 0.32615177 | 0.227 | 0.02 | 3.162E-224 | PDC | CA11 |
| 2.201E-228 | 0.50210813 | 0.428 | 0.069 | 5.2E-224 | PDC | ZNF667-AS1 |
| 1.109E-227 | 0.78632478 | 0.561 | 0.122 | 2.621E-223 | PDC | UQCC2 |
| 2.741E-226 | 0.62672012 | 0.534 | 0.108 | 6.475E-222 | PDC | DMAC1 |
| 3.631E-225 | 0.78598822 | 0.577 | 0.127 | 8.579E-221 | PDC | RFC1 |
| 9.947E-225 | 0.75514893 | 0.577 | 0.126 | 2.35E-220 | PDC | DNAJB9 |
| 1.441E-223 | 0.49901498 | 0.397 | 0.061 | 3.403E-219 | PDC | SLC48A1 |
| 7.531E-222 | 1.04000974 | 0.791 | 0.261 | 1.779E-217 | PDC | EID1 |
| 8.755E-221 | 0.33618221 | 0.266 | 0.029 | 2.068E-216 | PDC | WFS1 |
| 5.248E-219 | 0.53827544 | 0.39 | 0.061 | 1.24E-214 | PDC | FBXO2 |
| 7.056E-218 | 1.36400081 | 0.766 | 0.275 | 1.667E-213 | PDC | TCEA1 |
| 1.92E-217 | 0.90906132 | 0.586 | 0.134 | 4.535E-213 | PDC | GPNMB |
| 1.763E-216 | 0.79476046 | 0.565 | 0.127 | 4.164E-212 | PDC | RRAS |
| 2.366E-215 | 0.50781384 | 0.246 | 0.025 | 5.589E-211 | PDC | CES1 |
| 2.654E-213 | 0.25197497 | 0.196 | 0.016 | 6.269E-209 | PDC | ZBTB7C |
| 3.18E-213 | 0.43248526 | 0.342 | 0.048 | 7.513E-209 | PDC | SGCB |
| 3.607E-213 | 0.33607406 | 0.277 | 0.032 | 8.521E-209 | PDC | TUSC1 |
| 3.278E-212 | 0.71596789 | 0.646 | 0.166 | 7.743E-208 | PDC | STUB1 |
| 4.732E-212 | 0.50820539 | 0.423 | 0.072 | 1.118E-207 | PDC | PAM |
| 5.774E-211 | 1.27085362 | 0.728 | 0.224 | 1.364E-206 | PDC | TJP1 |
| 4.141E-210 | 0.86705083 | 0.653 | 0.175 | 9.782E-206 | PDC | REXO2 |
| 4.295E-210 | 0.98910465 | 0.77 | 0.249 | 1.015E-205 | PDC | VAMP2 |
| 1.847E-209 | 0.49280069 | 0.376 | 0.058 | 4.364E-205 | PDC | FRY |
| 1.951E-209 | 0.49096781 | 0.353 | 0.052 | 4.609E-205 | PDC | KANK1 |
| 3.039E-209 | 0.33226036 | 0.263 | 0.029 | 7.18E-205 | PDC | RBM43 |
| 4.145E-207 | 0.54461621 | 0.335 | 0.047 | 9.791E-203 | PDC | SLC7A8 |
| 7.355E-207 | 0.59833666 | 0.496 | 0.1 | 1.737E-202 | PDC | TPD52 |
| 2.411E-205 | 1.01811326 | 0.804 | 0.3 | 5.695E-201 | PDC | ANAPC16 |
| 4.696E-205 | 1.00068902 | 0.827 | 0.292 | 1.109E-200 | PDC | NUCKS1 |
| 9.643E-205 | 0.56603116 | 0.428 | 0.077 | 2.278E-200 | PDC | CLIP4 |
| 1.071E-204 | 0.32443544 | 0.185 | 0.015 | 2.529E-200 | PDC | SHISA2 |
| 1.128E-204 | 0.50820183 | 0.39 | 0.063 | 2.664E-200 | PDC | PLOD2 |
| 1.155E-204 | 0.54691806 | 0.498 | 0.1 | 2.728E-200 | PDC | MRPS26 |

| | | | | | | |
|---|---|---|---|---|---|---|
| 6.652E-204 | 0.43381596 | 0.376 | 0.059 | 1.572E-199 | PDC | ZNF32 |
| 5.757E-202 | 0.44847105 | 0.342 | 0.05 | 1.36E-197 | PDC | LAMB2 |
| 6.984E-201 | 1.39858942 | 0.946 | 0.62 | 1.65E-196 | PDC | DSTN |
| 6.995E-201 | 0.91662913 | 0.689 | 0.201 | 1.652E-196 | PDC | MAP4 |
| 2.118E-200 | 0.8375601 | 0.655 | 0.18 | 5.004E-196 | PDC | SEL1L |
| 2.912E-200 | 0.56355726 | 0.426 | 0.078 | 6.879E-196 | PDC | CERS6 |
| 3.792E-200 | 0.74383364 | 0.644 | 0.174 | 8.958E-196 | PDC | POLR2I |
| 1.523E-199 | 0.44664459 | 0.295 | 0.039 | 3.598E-195 | PDC | CDC14A |
| 2.216E-199 | 0.48388342 | 0.412 | 0.072 | 5.236E-195 | PDC | CFL2 |
| 6.253E-198 | 0.31606689 | 0.264 | 0.031 | 1.477E-193 | PDC | TCTN1 |
| 7.324E-198 | 1.01657985 | 0.883 | 0.342 | 1.73E-193 | PDC | PEBP1 |
| 3.481E-197 | 0.61444637 | 0.513 | 0.11 | 8.223E-193 | PDC | VEGFB |
| 3.5E-196 | 0.51244209 | 0.326 | 0.047 | 8.268E-192 | PDC | ST6GAL1 |
| 1.319E-195 | 0.37538156 | 0.327 | 0.047 | 3.116E-191 | PDC | THAP7 |
| 8.05E-195 | -2.2271925 | 0.57 | 0.944 | 1.902E-190 | PDC | SAT1 |
| 8.786E-195 | 0.78703154 | 0.617 | 0.167 | 2.076E-190 | PDC | ATRAID |
| 1.215E-194 | 0.82033085 | 0.594 | 0.148 | 2.87E-190 | PDC | ARHGEF12 |
| 4.574E-194 | 1.01349063 | 0.572 | 0.143 | 1.081E-189 | PDC | CDC42EP3 |
| 4.874E-194 | 0.415204 | 0.351 | 0.054 | 1.151E-189 | PDC | LZTS2 |
| 1.921E-193 | 0.4693316 | 0.347 | 0.054 | 4.537E-189 | PDC | PLPP1 |
| 2.206E-193 | 0.3748911 | 0.302 | 0.041 | 5.211E-189 | PDC | ITGA1 |
| 3.019E-192 | 0.32868492 | 0.263 | 0.032 | 7.132E-188 | PDC | LCA5 |
| 1.119E-191 | 0.9349917 | 0.727 | 0.232 | 2.644E-187 | PDC | C12orf57 |
| 2.948E-191 | 0.70788352 | 0.601 | 0.154 | 6.964E-187 | PDC | DUT |
| 5.565E-190 | 0.5806129 | 0.502 | 0.109 | 1.315E-185 | PDC | COPRS |
| 2.497E-188 | 0.69234747 | 0.59 | 0.151 | 5.898E-184 | PDC | HAX1 |
| 3.306E-188 | 0.88557032 | 0.685 | 0.208 | 7.809E-184 | PDC | LMAN1 |
| 8.398E-188 | 0.51030896 | 0.372 | 0.063 | 1.984E-183 | PDC | CHPF |
| 1.21E-187 | 0.35256447 | 0.284 | 0.038 | 2.858E-183 | PDC | ARMCX1 |
| 2.241E-186 | 0.3099103 | 0.266 | 0.033 | 5.294E-182 | PDC | FAM229B |
| 3.152E-186 | 0.58936384 | 0.522 | 0.118 | 7.446E-182 | PDC | DNAJC19 |
| 4.921E-186 | 0.41350352 | 0.324 | 0.049 | 1.162E-181 | PDC | ENDOD1 |
| 5.582E-184 | 0.36065831 | 0.282 | 0.038 | 1.319E-179 | PDC | HOXA9 |
| 1.226E-183 | 0.25452138 | 0.207 | 0.021 | 2.897E-179 | PDC | RILPL1 |
| 3.194E-183 | 0.59475624 | 0.369 | 0.064 | 7.546E-179 | PDC | DAG1 |
| 4.691E-183 | 0.63345704 | 0.493 | 0.109 | 1.108E-178 | PDC | UBE2L6 |
| 7.783E-183 | 0.51076128 | 0.401 | 0.073 | 1.839E-178 | PDC | LARP6 |
| 1.032E-181 | 0.41400058 | 0.322 | 0.049 | 2.437E-177 | PDC | MAP9 |
| 4.252E-181 | 0.51224328 | 0.383 | 0.067 | 1.004E-176 | PDC | PTGER4 |
| 7.356E-181 | 1.07346776 | 0.599 | 0.168 | 1.738E-176 | PDC | VKORC1 |
| 9.792E-180 | 0.43550275 | 0.371 | 0.064 | 2.313E-175 | PDC | HYI |
| 1.653E-179 | 0.84898384 | 0.716 | 0.211 | 3.905E-175 | PDC | ZBTB20 |
| 5.309E-179 | 1.85871263 | 0.879 | 0.329 | 1.254E-174 | PDC | IGFBP7 |
| 7.698E-179 | 0.64750337 | 0.583 | 0.153 | 1.818E-174 | PDC | ESD |
| 1.863E-178 | 1.03043134 | 0.84 | 0.363 | 4.4E-174 | PDC | PRDX6 |
| 4.636E-178 | 0.71626037 | 0.683 | 0.202 | 1.095E-173 | PDC | TCEAL4 |
| 4.774E-178 | 0.30479651 | 0.245 | 0.029 | 1.128E-173 | PDC | PIR |
| 1.928E-176 | 1.06485415 | 0.844 | 0.364 | 4.554E-172 | PDC | CIRBP |
| 4.693E-176 | 0.398366 | 0.336 | 0.054 | 1.109E-171 | PDC | UGGT2 |
| 5.813E-176 | 0.29987528 | 0.239 | 0.029 | 1.373E-171 | PDC | ME1 |

| | | | | | | |
|---|---|---|---|---|---|---|
| 1.275E-174 | 0.30871856 | 0.259 | 0.034 | 3.012E-170 | PDC | LANCL1 |
| 1.891E-174 | 0.40488729 | 0.333 | 0.054 | 4.468E-170 | PDC | FAM98A |
| 1.94E-174 | 0.93220006 | 0.683 | 0.221 | 4.583E-170 | PDC | PCMTD1 |
| 2.867E-174 | 0.32700814 | 0.268 | 0.036 | 6.774E-170 | PDC | EFEMP2 |
| 3.814E-174 | 0.58627395 | 0.815 | 0.241 | 9.011E-170 | PDC | CRYAB |
| 4.27E-174 | 0.48830659 | 0.468 | 0.1 | 1.009E-169 | PDC | GCSH |
| 2.863E-173 | 0.41662665 | 0.113 | 0.007 | 6.764E-169 | PDC | GSTA1 |
| 4.343E-173 | 0.3431874 | 0.243 | 0.03 | 1.026E-168 | PDC | MCC |
| 6.691E-173 | 0.87002059 | 0.642 | 0.198 | 1.581E-168 | PDC | C6orf48 |
| 2.151E-172 | 0.57628389 | 0.457 | 0.1 | 5.081E-168 | PDC | HDDC2 |
| 2.461E-172 | 0.43181736 | 0.381 | 0.07 | 5.815E-168 | PDC | MOCS2 |
| 7.33E-172 | 0.54218043 | 0.52 | 0.122 | 1.732E-167 | PDC | DNPH1 |
| 1.323E-171 | 0.37136 | 0.318 | 0.05 | 3.125E-167 | PDC | QDPR |
| 3.875E-171 | 0.27536218 | 0.209 | 0.023 | 9.153E-167 | PDC | GPX8 |
| 5.067E-171 | 1.61357431 | 0.89 | 0.469 | 1.197E-166 | PDC | CD81 |
| 1.359E-170 | 0.26124669 | 0.198 | 0.02 | 3.21E-166 | PDC | TRAM2 |
| 1.425E-170 | 0.50862454 | 0.442 | 0.093 | 3.365E-166 | PDC | DTD1 |
| 1.943E-170 | 0.54101932 | 0.468 | 0.102 | 4.59E-166 | PDC | CFAP97 |
| 2.096E-169 | 0.34617785 | 0.232 | 0.028 | 4.951E-165 | PDC | SLC2A13 |
| 6.108E-169 | 0.48987127 | 0.448 | 0.094 | 1.443E-164 | PDC | SLC25A4 |
| 6.856E-169 | 0.8470568 | 0.752 | 0.276 | 1.62E-164 | PDC | COX7A2L |
| 9.352E-169 | 1.55109536 | 0.707 | 0.26 | 2.209E-164 | PDC | DDIT3 |
| 3.87E-168 | 1.07221933 | 0.946 | 0.732 | 9.143E-164 | PDC | CCNI |
| 1.473E-167 | 0.95821883 | 0.612 | 0.188 | 3.48E-163 | PDC | SEC11C |
| 1.878E-167 | 0.29179386 | 0.236 | 0.029 | 4.437E-163 | PDC | SOX6 |
| 9.902E-166 | 0.29226235 | 0.183 | 0.018 | 2.339E-161 | PDC | ATP6V0D2 |
| 3.497E-165 | 0.39331777 | 0.322 | 0.053 | 8.262E-161 | PDC | PKIG |
| 5.5E-165 | 0.69087477 | 0.59 | 0.163 | 1.299E-160 | PDC | EPRS |
| 9.14E-165 | 0.25726669 | 0.203 | 0.022 | 2.159E-160 | PDC | TTLL7 |
| 3.021E-164 | 0.4447593 | 0.405 | 0.081 | 7.136E-160 | PDC | FRA10AC1 |
| 5.395E-164 | 0.78648226 | 0.419 | 0.091 | 1.274E-159 | PDC | SULF2 |
| 3.494E-163 | 0.41629229 | 0.34 | 0.059 | 8.253E-159 | PDC | PARVA |
| 1.084E-161 | 0.45820939 | 0.405 | 0.082 | 2.561E-157 | PDC | ESYT1 |
| 2.62E-161 | 0.68473153 | 0.475 | 0.11 | 6.189E-157 | PDC | NQO1 |
| 2.951E-161 | 0.38409733 | 0.212 | 0.025 | 6.971E-157 | PDC | UNC5B |
| 1.058E-160 | 0.32254114 | 0.27 | 0.039 | 2.499E-156 | PDC | HOXD8 |
| 1.458E-160 | 0.72242957 | 0.577 | 0.162 | 3.445E-156 | PDC | HMGN3 |
| 2.551E-160 | 0.33538053 | 0.237 | 0.031 | 6.026E-156 | PDC | TSHZ1 |
| 3.061E-159 | 0.30574352 | 0.246 | 0.033 | 7.23E-155 | PDC | CDC25B |
| 4.839E-159 | -5.3069134 | 0.149 | 0.735 | 1.143E-154 | PDC | S100A9 |
| 3.95E-158 | 0.28139975 | 0.239 | 0.031 | 9.331E-154 | PDC | FBXO17 |
| 4.895E-158 | 0.30504609 | 0.264 | 0.038 | 1.156E-153 | PDC | LINC00662 |
| 9.942E-158 | 0.56209465 | 0.486 | 0.118 | 2.349E-153 | PDC | MRPL40 |
| 2.308E-157 | 0.54338099 | 0.507 | 0.127 | 5.453E-153 | PDC | METTL5 |
| 6.02E-157 | 0.29667125 | 0.237 | 0.031 | 1.422E-152 | PDC | HSD17B8 |
| 7.124E-157 | 0.29650238 | 0.261 | 0.037 | 1.683E-152 | PDC | FAM228B |
| 1.379E-155 | 1.26417141 | 0.808 | 0.312 | 3.259E-151 | PDC | GADD45A |
| 2.114E-154 | 0.59377204 | 0.493 | 0.124 | 4.993E-150 | PDC | NDUFAF2 |
| 2.533E-154 | 0.51299482 | 0.441 | 0.099 | 5.985E-150 | PDC | IARS |
| 3.289E-154 | 0.2927435 | 0.255 | 0.036 | 7.769E-150 | PDC | WDSUB1 |

| | | | | | | |
|---|---|---|---|---|---|---|
| 3.991E-154 | 0.30024068 | 0.227 | 0.029 | 9.428E-150 | PDC | ZBED3 |
| 9.743E-154 | 0.55726674 | 0.536 | 0.143 | 2.302E-149 | PDC | PYURF |
| 1.006E-153 | 0.43788033 | 0.353 | 0.066 | 2.377E-149 | PDC | ITGB5 |
| 4.191E-153 | 0.79041464 | 0.781 | 0.274 | 9.901E-149 | PDC | MT1E |
| 1.073E-152 | 0.35984787 | 0.317 | 0.054 | 2.536E-148 | PDC | KIZ |
| 1.597E-152 | 0.9275567 | 0.8 | 0.351 | 3.773E-148 | PDC | TRMT112 |
| 2.917E-152 | 0.74650397 | 0.496 | 0.126 | 6.89E-148 | PDC | VAMP5 |
| 6.973E-152 | 0.55459879 | 0.423 | 0.096 | 1.647E-147 | PDC | EIF1AY |
| 1.453E-150 | 0.56481572 | 0.466 | 0.113 | 3.432E-146 | PDC | MAGED2 |
| 2.448E-150 | 0.63383962 | 0.421 | 0.093 | 5.784E-146 | PDC | SLC9A3R2 |
| 2.608E-150 | 0.53561126 | 0.48 | 0.118 | 6.161E-146 | PDC | SNRPN |
| 4.487E-150 | 0.56044825 | 0.549 | 0.151 | 1.06E-145 | PDC | COPS6 |
| 1.244E-149 | 0.60495264 | 0.43 | 0.098 | 2.94E-145 | PDC | TCEAL3 |
| 3.79E-149 | 0.50565653 | 0.45 | 0.107 | 8.954E-145 | PDC | TMEM126B |
| 2.854E-148 | 0.81033459 | 0.797 | 0.332 | 6.743E-144 | PDC | ARL6IP4 |
| 3.732E-148 | 0.79755487 | 0.538 | 0.155 | 8.816E-144 | PDC | GFPT1 |
| 1.343E-147 | 0.71631384 | 0.716 | 0.261 | 3.172E-143 | PDC | ERH |
| 4.414E-147 | 0.26379779 | 0.234 | 0.032 | 1.043E-142 | PDC | ZEB1 |
| 5.784E-147 | 0.35670536 | 0.338 | 0.063 | 1.366E-142 | PDC | NDUFAF4 |
| 2.518E-146 | 0.61982047 | 0.516 | 0.139 | 5.949E-142 | PDC | CLCN3 |
| 3.022E-146 | 0.26605734 | 0.227 | 0.03 | 7.138E-142 | PDC | SERGEF |
| 8.736E-146 | 0.31851322 | 0.288 | 0.047 | 2.064E-141 | PDC | RIC3 |
| 1.366E-145 | 0.41745236 | 0.371 | 0.075 | 3.226E-141 | PDC | A1BG |
| 1.397E-145 | 0.27425321 | 0.173 | 0.018 | 3.299E-141 | PDC | MSC-AS1 |
| 5.981E-145 | 0.37650295 | 0.327 | 0.06 | 1.413E-140 | PDC | C11orf1 |
| 6.586E-145 | 0.79376527 | 0.716 | 0.264 | 1.556E-140 | PDC | PHPT1 |
| 2.025E-144 | 0.29910396 | 0.263 | 0.04 | 4.783E-140 | PDC | DMWD |
| 4.784E-144 | 0.34955229 | 0.311 | 0.055 | 1.13E-139 | PDC | NOL3 |
| 5.122E-144 | 0.46051853 | 0.308 | 0.055 | 1.21E-139 | PDC | ALS2CL |
| 6.812E-144 | 0.39783445 | 0.36 | 0.071 | 1.609E-139 | PDC | PFN2 |
| 3.843E-143 | 0.68568857 | 0.629 | 0.209 | 9.077E-139 | PDC | RAD23A |
| 5.12E-143 | 0.47086315 | 0.441 | 0.104 | 1.21E-138 | PDC | MRPS18B |
| 6.024E-143 | 0.41691498 | 0.376 | 0.078 | 1.423E-138 | PDC | EBAG9 |
| 1.14E-142 | 0.95948973 | 0.682 | 0.262 | 2.693E-138 | PDC | HSPA9 |
| 2.791E-142 | 0.32475823 | 0.21 | 0.027 | 6.594E-138 | PDC | DDIT4L |
| 1.367E-141 | 0.31881062 | 0.277 | 0.045 | 3.228E-137 | PDC | PIGK |
| 1.441E-141 | 0.25504947 | 0.196 | 0.024 | 3.405E-137 | PDC | C16orf45 |
| 4.339E-141 | 0.39449345 | 0.329 | 0.063 | 1.025E-136 | PDC | WDR41 |
| 4.71E-141 | 0.25934526 | 0.227 | 0.031 | 1.113E-136 | PDC | ALG14 |
| 8.701E-141 | 0.32698316 | 0.29 | 0.049 | 2.055E-136 | PDC | PIP4P2 |
| 9.209E-141 | 0.45780354 | 0.41 | 0.093 | 2.176E-136 | PDC | PREPL |
| 1.436E-140 | 0.47291639 | 0.406 | 0.092 | 3.393E-136 | PDC | ARMC10 |
| 1.638E-140 | 0.49217616 | 0.45 | 0.108 | 3.869E-136 | PDC | FYN |
| 1.916E-140 | 0.4581103 | 0.371 | 0.078 | 4.527E-136 | PDC | PCYOX1 |
| 2.379E-140 | 0.32942347 | 0.284 | 0.048 | 5.62E-136 | PDC | C2orf74 |
| 2.583E-140 | 0.90486483 | 0.414 | 0.101 | 6.102E-136 | PDC | ZDHHC6 |
| 3.39E-140 | 0.64224753 | 0.538 | 0.152 | 8.008E-136 | PDC | SH3BP5 |
| 5.755E-139 | 0.28082052 | 0.245 | 0.036 | 1.359E-134 | PDC | HOXA10 |
| 8.806E-139 | 0.37254951 | 0.327 | 0.062 | 2.08E-134 | PDC | UFSP2 |
| 2.071E-138 | 0.59181726 | 0.563 | 0.168 | 4.893E-134 | PDC | ARMCX3 |

| | | | | | | |
|---|---|---|---|---|---|---|
| 3.119E-138 | 0.55105928 | 0.511 | 0.139 | 7.368E-134 | PDC | OTUD6B-AS1 |
| 1.075E-137 | 0.38646789 | 0.327 | 0.063 | 2.54E-133 | PDC | CMSS1 |
| 1.234E-137 | 0.34861283 | 0.309 | 0.057 | 2.916E-133 | PDC | WRB |
| 1.28E-137 | 0.32493968 | 0.254 | 0.039 | 3.024E-133 | PDC | ANKRD33B |
| 1.479E-137 | 0.26675114 | 0.218 | 0.03 | 3.493E-133 | PDC | KIAA1841 |
| 1.241E-136 | 0.35451607 | 0.329 | 0.064 | 2.931E-132 | PDC | THYN1 |
| 5.736E-136 | 0.27357493 | 0.23 | 0.033 | 1.355E-131 | PDC | APBB2 |
| 8.085E-136 | 0.55580882 | 0.496 | 0.135 | 1.91E-131 | PDC | PLD3 |
| 1.136E-135 | -1.5922355 | 0.881 | 0.966 | 2.684E-131 | PDC | S100A11 |
| 1.601E-135 | 0.26132682 | 0.133 | 0.012 | 3.783E-131 | PDC | POLA2 |
| 3.874E-135 | -4.2880609 | 0.131 | 0.678 | 9.151E-131 | PDC | S100A8 |
| 4.23E-135 | 0.49123269 | 0.45 | 0.112 | 9.992E-131 | PDC | CBX5 |
| 6.408E-135 | 0.26267193 | 0.227 | 0.032 | 1.514E-130 | PDC | MANEA |
| 9.252E-135 | 0.40050455 | 0.349 | 0.071 | 2.186E-130 | PDC | CKS1B |
| 1.535E-134 | 0.42353409 | 0.372 | 0.08 | 3.627E-130 | PDC | TUSC3 |
| 1.842E-134 | 1.17726095 | 0.381 | 0.089 | 4.35E-130 | PDC | BCAM |
| 2.25E-134 | 0.42383788 | 0.392 | 0.089 | 5.315E-130 | PDC | MRPL17 |
| 2.912E-134 | 0.39577411 | 0.336 | 0.068 | 6.879E-130 | PDC | HPF1 |
| 3.251E-134 | 0.40144346 | 0.344 | 0.069 | 7.681E-130 | PDC | KIF21A |
| 3.566E-134 | 0.49304336 | 0.46 | 0.118 | 8.425E-130 | PDC | LYRM2 |
| 3.778E-134 | 0.37915789 | 0.306 | 0.057 | 8.925E-130 | PDC | CPNE8 |
| 4.964E-134 | 0.59803405 | 0.559 | 0.169 | 1.173E-129 | PDC | RDX |
| 7.493E-134 | 0.56894647 | 0.543 | 0.162 | 1.77E-129 | PDC | TBCB |
| 2.174E-133 | 0.39096958 | 0.308 | 0.058 | 5.135E-129 | PDC | PCK2 |
| 2.452E-133 | 0.6624884 | 0.682 | 0.251 | 5.791E-129 | PDC | NDUFB10 |
| 4.519E-133 | 0.57432565 | 0.414 | 0.099 | 1.068E-128 | PDC | TRIB3 |
| 8.474E-133 | 0.34405252 | 0.317 | 0.061 | 2.002E-128 | PDC | UROS |
| 1.229E-132 | 0.59531716 | 0.549 | 0.167 | 2.903E-128 | PDC | MCFD2 |
| 1.93E-131 | -1.7133624 | 0.477 | 0.864 | 4.56E-127 | PDC | SH3BGRL3 |
| 2.254E-131 | 0.45310717 | 0.302 | 0.057 | 5.325E-127 | PDC | P3H2 |
| 2.806E-131 | 0.57082863 | 0.574 | 0.181 | 6.628E-127 | PDC | PCMT1 |
| 3.428E-131 | 0.41044605 | 0.381 | 0.085 | 8.099E-127 | PDC | CETN2 |
| 7.294E-131 | 0.71412589 | 0.728 | 0.29 | 1.723E-126 | PDC | RABAC1 |
| 1.239E-130 | 0.69338925 | 0.385 | 0.091 | 2.928E-126 | PDC | MPP5 |
| 1.341E-130 | 0.89184926 | 0.928 | 0.597 | 3.168E-126 | PDC | HINT1 |
| 2.747E-130 | 0.41565948 | 0.367 | 0.081 | 6.49E-126 | PDC | TMEM18 |
| 3.46E-130 | 0.5103146 | 0.475 | 0.127 | 8.173E-126 | PDC | DNAJC10 |
| 5.532E-130 | 0.28481434 | 0.255 | 0.042 | 1.307E-125 | PDC | NADK2 |
| 6.181E-130 | 0.64988675 | 0.692 | 0.255 | 1.46E-125 | PDC | CUTA |
| 1.956E-129 | 0.5567252 | 0.536 | 0.16 | 4.622E-125 | PDC | ZCRB1 |
| 3.364E-129 | 0.62566666 | 0.579 | 0.187 | 7.948E-125 | PDC | RHOQ |
| 1.247E-128 | 0.52086891 | 0.509 | 0.146 | 2.946E-124 | PDC | MRPL34 |
| 2.632E-128 | 0.32767674 | 0.299 | 0.056 | 6.218E-124 | PDC | TSEN15 |
| 4.488E-128 | 0.73930629 | 0.692 | 0.245 | 1.06E-123 | PDC | SYNE2 |
| 1.078E-127 | 0.37973375 | 0.257 | 0.043 | 2.546E-123 | PDC | BMP2 |
| 5.966E-127 | 0.75846699 | 0.705 | 0.282 | 1.409E-122 | PDC | NDUFC1 |
| 4.295E-126 | 0.69070421 | 0.635 | 0.23 | 1.015E-121 | PDC | PSMA3 |
| 4.584E-126 | 0.28796255 | 0.255 | 0.043 | 1.083E-121 | PDC | MED27 |
| 2.249E-125 | 0.2514995 | 0.207 | 0.029 | 5.312E-121 | PDC | CCDC34 |
| 2.294E-125 | 0.41296681 | 0.406 | 0.099 | 5.418E-121 | PDC | TSR3 |

| | | | | | | |
|---|---|---|---|---|---|---|
| 3.101E-125 | 0.31111832 | 0.27 | 0.047 | 7.326E-121 | PDC | CLUAP1 |
| 4.685E-125 | 0.37374253 | 0.333 | 0.07 | 1.107E-120 | PDC | ETFRF1 |
| 5.103E-125 | 0.38492981 | 0.369 | 0.083 | 1.205E-120 | PDC | CCDC115 |
| 5.564E-125 | -1.4273136 | 0.232 | 0.744 | 1.314E-120 | PDC | CTSB |
| 1.795E-124 | 0.40352242 | 0.293 | 0.056 | 4.24E-120 | PDC | NLRP1 |
| 2.094E-124 | 0.30385866 | 0.254 | 0.043 | 4.946E-120 | PDC | THRA |
| 3.57E-124 | 0.2694766 | 0.218 | 0.032 | 8.434E-120 | PDC | DUSP15 |
| 3.791E-124 | 0.30282418 | 0.27 | 0.048 | 8.956E-120 | PDC | SORBS3 |
| 1.115E-123 | 0.55389344 | 0.57 | 0.178 | 2.634E-119 | PDC | BNIP3 |
| 5.186E-123 | 0.31030572 | 0.281 | 0.052 | 1.225E-118 | PDC | GLT8D1 |
| 6.543E-123 | 0.42562923 | 0.417 | 0.105 | 1.546E-118 | PDC | WASHC3 |
| 1.325E-122 | 0.76461006 | 0.594 | 0.212 | 3.13E-118 | PDC | NUCB1 |
| 1.533E-122 | 0.45891748 | 0.419 | 0.107 | 3.621E-118 | PDC | CCDC91 |
| 3.788E-122 | -2.2862896 | 0.092 | 0.621 | 8.948E-118 | PDC | C15orf48 |
| 5.047E-122 | 0.2806406 | 0.252 | 0.043 | 1.192E-117 | PDC | C17orf75 |
| 5.47E-122 | 0.32171345 | 0.306 | 0.06 | 1.292E-117 | PDC | RDH14 |
| 6.73E-122 | 0.496051 | 0.498 | 0.143 | 1.59E-117 | PDC | SVIP |
| 1.22E-121 | 0.67136671 | 0.656 | 0.247 | 2.881E-117 | PDC | FIS1 |
| 1.553E-121 | 0.37921623 | 0.317 | 0.065 | 3.668E-117 | PDC | CDK5RAP2 |
| 4.668E-121 | 0.60519241 | 0.588 | 0.202 | 1.103E-116 | PDC | COX14 |
| 6.317E-121 | 0.4850169 | 0.394 | 0.097 | 1.492E-116 | PDC | TMEM245 |
| 9.436E-121 | 0.64060184 | 0.709 | 0.282 | 2.229E-116 | PDC | MRFAP1 |
| 1.093E-120 | 0.2782677 | 0.232 | 0.037 | 2.581E-116 | PDC | CBX7 |
| 2.319E-120 | 0.65003296 | 0.667 | 0.229 | 5.477E-116 | PDC | CDKN1C |
| 3.986E-119 | 0.45982288 | 0.469 | 0.133 | 9.417E-115 | PDC | ACTR10 |
| 4.225E-119 | 0.62233881 | 0.689 | 0.255 | 9.981E-115 | PDC | PRDX2 |
| 6.815E-119 | 0.54394643 | 0.52 | 0.161 | 1.61E-114 | PDC | NDUFAF8 |
| 6.952E-119 | 0.42620922 | 0.397 | 0.1 | 1.642E-114 | PDC | STX8 |
| 7.184E-119 | 0.72029955 | 0.806 | 0.369 | 1.697E-114 | PDC | PARK7 |
| 1.11E-118 | 0.26850358 | 0.228 | 0.037 | 2.623E-114 | PDC | LRFN4 |
| 1.304E-118 | 0.33373155 | 0.263 | 0.047 | 3.081E-114 | PDC | PSAT1 |
| 1.572E-118 | 0.40310681 | 0.399 | 0.099 | 3.713E-114 | PDC | ANAPC13 |
| 2.448E-118 | 0.46257052 | 0.504 | 0.148 | 5.782E-114 | PDC | SCOC |
| 3.277E-118 | 0.56362051 | 0.621 | 0.217 | 7.742E-114 | PDC | PPP2CB |
| 3.329E-118 | 0.55066383 | 0.477 | 0.132 | 7.864E-114 | PDC | ARHGAP29 |
| 8.383E-118 | 0.6145676 | 0.538 | 0.177 | 1.98E-113 | PDC | DNAJC1 |
| 9.943E-118 | 0.3820588 | 0.358 | 0.082 | 2.349E-113 | PDC | THAP5 |
| 2.913E-117 | 0.79615834 | 0.716 | 0.32 | 6.881E-113 | PDC | GHITM |
| 4.314E-117 | 0.38601825 | 0.326 | 0.07 | 1.019E-112 | PDC | JADE1 |
| 1.209E-116 | 0.41434292 | 0.371 | 0.089 | 2.856E-112 | PDC | PIGP |
| 2.573E-116 | 0.56821154 | 0.633 | 0.227 | 6.078E-112 | PDC | OPTN |
| 3.046E-116 | 0.35916259 | 0.322 | 0.069 | 7.194E-112 | PDC | TXNDC15 |
| 4.112E-116 | 0.28902932 | 0.261 | 0.047 | 9.714E-112 | PDC | CBY1 |
| 4.298E-116 | 0.45684007 | 0.453 | 0.127 | 1.015E-111 | PDC | SURF1 |
| 6.281E-116 | 0.36359435 | 0.342 | 0.076 | 1.484E-111 | PDC | SPAG16 |
| 1.364E-115 | 0.32240564 | 0.259 | 0.047 | 3.221E-111 | PDC | DBP |
| 5.073E-115 | 0.26533997 | 0.223 | 0.036 | 1.198E-110 | PDC | ARMCX2 |
| 8.881E-115 | 0.52191581 | 0.493 | 0.15 | 2.098E-110 | PDC | NDUFS4 |
| 9.414E-115 | 0.77928371 | 0.926 | 0.711 | 2.224E-110 | PDC | SKP1 |
| 2.99E-114 | 0.72997307 | 0.831 | 0.398 | 7.063E-110 | PDC | SOD1 |

| | | | | | | |
|---|---|---|---|---|---|---|
| 6.216E-114 | 0.37289974 | 0.369 | 0.088 | 1.468E-109 | PDC | MALSU1 |
| 1.156E-113 | 0.30517472 | 0.27 | 0.051 | 2.731E-109 | PDC | TMEM14A |
| 2.475E-113 | 0.38172948 | 0.326 | 0.072 | 5.848E-109 | PDC | ZBTB4 |
| 3.347E-113 | 0.26474577 | 0.246 | 0.043 | 7.906E-109 | PDC | STXBP1 |
| 7.251E-113 | 0.6032731 | 0.604 | 0.213 | 1.713E-108 | PDC | SPATS2L |
| 2.099E-112 | 0.75203884 | 0.586 | 0.208 | 4.958E-108 | PDC | BTG3 |
| 5.673E-112 | 0.37221651 | 0.315 | 0.069 | 1.34E-107 | PDC | P4HTM |
| 6.572E-112 | 0.39632112 | 0.378 | 0.094 | 1.553E-107 | PDC | ARL2BP |
| 8.11E-112 | 0.36038855 | 0.365 | 0.088 | 1.916E-107 | PDC | COPS8 |
| 8.259E-112 | 0.30698981 | 0.279 | 0.055 | 1.951E-107 | PDC | PPP1R16A |
| 9.388E-112 | 0.35865023 | 0.335 | 0.076 | 2.218E-107 | PDC | NFU1 |
| 1.774E-111 | 0.55797989 | 0.637 | 0.237 | 4.19E-107 | PDC | TMEM14C |
| 1.801E-111 | 0.38939409 | 0.327 | 0.074 | 4.254E-107 | PDC | SHMT2 |
| 2.229E-111 | 0.54791823 | 0.624 | 0.231 | 5.265E-107 | PDC | C1orf43 |
| 3.864E-111 | 0.39412365 | 0.399 | 0.104 | 9.128E-107 | PDC | SLC25A11 |
| 4.763E-111 | 0.39252384 | 0.401 | 0.103 | 1.125E-106 | PDC | CISD3 |
| 5.257E-111 | -1.4575245 | 0.117 | 0.619 | 1.242E-106 | PDC | CD44 |
| 9.049E-111 | 0.34692368 | 0.317 | 0.069 | 2.138E-106 | PDC | EXOSC7 |
| 1.526E-110 | 0.65611095 | 0.782 | 0.353 | 3.604E-106 | PDC | TOMM20 |
| 1.965E-110 | 0.55387115 | 0.563 | 0.19 | 4.642E-106 | PDC | RBPMS |
| 3.204E-110 | 0.43875251 | 0.415 | 0.113 | 7.57E-106 | PDC | EMC2 |
| 4.004E-110 | 0.57484625 | 0.617 | 0.224 | 9.458E-106 | PDC | PJA2 |
| 4.847E-110 | 0.53183783 | 0.55 | 0.186 | 1.145E-105 | PDC | ALKBH7 |
| 5.67E-110 | 0.28156238 | 0.255 | 0.047 | 1.34E-105 | PDC | SPRYD7 |
| 7.153E-110 | 0.68719168 | 0.791 | 0.38 | 1.69E-105 | PDC | CNBP |
| 1.448E-109 | 0.39610443 | 0.324 | 0.073 | 3.421E-105 | PDC | PYGB |
| 2.145E-109 | -1.3823737 | 0.291 | 0.733 | 5.066E-105 | PDC | CYBA |
| 2.431E-109 | 0.43517705 | 0.369 | 0.091 | 5.742E-105 | PDC | PON2 |
| 2.861E-109 | -1.5337803 | 0.146 | 0.648 | 6.759E-105 | PDC | IER3 |
| 4.01E-109 | 0.38042157 | 0.257 | 0.049 | 9.473E-105 | PDC | MAP2 |
| 5.773E-109 | 0.66705497 | 0.73 | 0.306 | 1.364E-104 | PDC | SBDS |
| 7.172E-109 | 0.25006988 | 0.227 | 0.039 | 1.694E-104 | PDC | NAGLU |
| 7.99E-108 | 0.33151786 | 0.291 | 0.061 | 1.888E-103 | PDC | C15orf61 |
| 8.375E-108 | 0.45665833 | 0.478 | 0.145 | 1.978E-103 | PDC | SEC63 |
| 8.5E-108 | 0.33042724 | 0.297 | 0.063 | 2.008E-103 | PDC | ZDHHC4 |
| 1.43E-107 | 0.47473714 | 0.46 | 0.139 | 3.378E-103 | PDC | TRAPPC2L |
| 1.498E-107 | 0.26172866 | 0.232 | 0.041 | 3.539E-103 | PDC | LRRC23 |
| 1.892E-107 | 0.40158119 | 0.392 | 0.103 | 4.469E-103 | PDC | UNC50 |
| 2.194E-107 | 0.59485775 | 0.606 | 0.228 | 5.182E-103 | PDC | PSMB8 |
| 2.228E-107 | 0.42814839 | 0.394 | 0.105 | 5.264E-103 | PDC | HIKESHI |
| 3.224E-107 | 0.38848593 | 0.383 | 0.098 | 7.617E-103 | PDC | USP1 |
| 4.331E-107 | 0.35855189 | 0.351 | 0.084 | 1.023E-102 | PDC | TMEM38B |
| 6.806E-107 | 0.33850541 | 0.327 | 0.075 | 1.608E-102 | PDC | PRKRA |
| 8.221E-107 | 0.28079026 | 0.275 | 0.055 | 1.942E-102 | PDC | NFKBIL1 |
| 1.537E-106 | 0.27040532 | 0.25 | 0.047 | 3.631E-102 | PDC | TMEM42 |
| 1.875E-106 | 0.47099677 | 0.513 | 0.164 | 4.429E-102 | PDC | METTL26 |
| 2.049E-106 | 0.31376518 | 0.281 | 0.057 | 4.84E-102 | PDC | TMEM140 |
| 2.267E-106 | 0.27653143 | 0.245 | 0.045 | 5.354E-102 | PDC | RPS6KA2 |
| 2.822E-106 | 0.34739468 | 0.318 | 0.072 | 6.666E-102 | PDC | TMEM109 |
| 3.386E-106 | 0.41169773 | 0.396 | 0.106 | 7.999E-102 | PDC | CRTAP |

| | | | | | | |
|---|---|---|---|---|---|---|
| 4.847E-106 | 0.77137848 | 0.732 | 0.344 | 1.145E-101 | PDC | FKBP2 |
| 4.868E-106 | 0.98414327 | 0.638 | 0.279 | 1.15E-101 | PDC | RPS4Y1 |
| 6.493E-106 | 0.52887131 | 0.629 | 0.237 | 1.534E-101 | PDC | NUDC |
| 8.113E-106 | -2.678493 | 0.086 | 0.577 | 1.917E-101 | PDC | SRGN |
| 1.024E-105 | 0.41099459 | 0.353 | 0.087 | 2.419E-101 | PDC | SRGAP2 |
| 3.171E-105 | 0.54179451 | 0.567 | 0.197 | 7.491E-101 | PDC | C19orf70 |
| 8.863E-105 | 0.50571093 | 0.505 | 0.163 | 2.094E-100 | PDC | DCXR |
| 2.256E-104 | 0.4717911 | 0.45 | 0.136 | 5.33E-100 | PDC | FEZ2 |
| 3.68E-104 | 0.32592288 | 0.272 | 0.055 | 8.694E-100 | PDC | LRP11 |
| 4.744E-104 | 0.47752303 | 0.523 | 0.173 | 1.121E-99 | PDC | ARL1 |
| 4.882E-104 | 0.5037188 | 0.597 | 0.212 | 1.153E-99 | PDC | MLEC |
| 6.575E-104 | 0.28639494 | 0.275 | 0.056 | 1.553E-99 | PDC | KIFAP3 |
| 7.246E-104 | 0.47804044 | 0.545 | 0.186 | 1.712E-99 | PDC | EMC7 |
| 8.188E-104 | -2.1626519 | 0.108 | 0.585 | 1.934E-99 | PDC | KRT19 |
| 1.364E-103 | 0.48902681 | 0.568 | 0.197 | 3.223E-99 | PDC | NDUFS8 |
| 1.462E-103 | 0.35632902 | 0.331 | 0.078 | 3.454E-99 | PDC | TMEM106C |
| 1.824E-103 | 0.55636106 | 0.635 | 0.246 | 4.31E-99 | PDC | NDUFAB1 |
| 1.861E-103 | 0.26013195 | 0.225 | 0.039 | 4.396E-99 | PDC | ZFYVE9 |
| 2.02E-103 | 0.59435036 | 0.766 | 0.347 | 4.772E-99 | PDC | SPCS1 |
| 2.924E-103 | 0.39762026 | 0.356 | 0.09 | 6.908E-99 | PDC | SLC2A4RG |
| 4.157E-103 | 0.55290365 | 0.653 | 0.261 | 9.819E-99 | PDC | RNH1 |
| 4.413E-103 | 0.74147858 | 0.824 | 0.435 | 1.0424E-98 | PDC | PDIA3 |
| 6.165E-103 | 0.34931192 | 0.304 | 0.068 | 1.4563E-98 | PDC | TACC2 |
| 6.769E-103 | 0.52712749 | 0.538 | 0.189 | 1.5989E-98 | PDC | MRPL54 |
| 7.763E-103 | 0.32684357 | 0.266 | 0.054 | 1.8339E-98 | PDC | SLC39A14 |
| 1.458E-102 | 0.52420795 | 0.469 | 0.146 | 3.4445E-98 | PDC | FAM129A |
| 1.883E-102 | 0.41224986 | 0.392 | 0.107 | 4.4487E-98 | PDC | ZNF428 |
| 1.917E-102 | 0.32974744 | 0.322 | 0.075 | 4.5287E-98 | PDC | SNAPC5 |
| 2.12E-102 | 0.58407319 | 0.594 | 0.225 | 5.009E-98 | PDC | MRPL57 |
| 2.197E-102 | 0.37212122 | 0.342 | 0.083 | 5.1907E-98 | PDC | MYO1D |
| 2.477E-102 | 0.44194637 | 0.435 | 0.128 | 5.8503E-98 | PDC | GANAB |
| 3.693E-102 | 0.30845994 | 0.266 | 0.054 | 8.7239E-98 | PDC | IGSF8 |
| 4.28E-102 | 0.35529676 | 0.27 | 0.055 | 1.0111E-97 | PDC | FAM43A |
| 5.731E-102 | 0.29683924 | 0.282 | 0.06 | 1.3538E-97 | PDC | FAM20B |
| 7.449E-102 | 0.46228543 | 0.451 | 0.135 | 1.7597E-97 | PDC | GULP1 |
| 8.242E-102 | 0.32967264 | 0.315 | 0.073 | 1.9469E-97 | PDC | TMED1 |
| 9.073E-102 | 0.30833308 | 0.272 | 0.056 | 2.1433E-97 | PDC | ACTR1B |
| 1.08E-101 | 0.28327288 | 0.281 | 0.059 | 2.5515E-97 | PDC | LINC00667 |
| 1.15E-101 | 0.3479374 | 0.326 | 0.078 | 2.7168E-97 | PDC | ZFYVE21 |
| 1.339E-101 | 0.4971779 | 0.536 | 0.186 | 3.1621E-97 | PDC | BTF3L4 |
| 1.351E-101 | 0.47366381 | 0.565 | 0.196 | 3.1924E-97 | PDC | PSMB5 |
| 1.999E-101 | 0.25846619 | 0.246 | 0.047 | 4.7222E-97 | PDC | DYRK4 |
| 2.28E-101 | -1.7495455 | 0.099 | 0.575 | 5.3864E-97 | PDC | TACSTD2 |
| 2.976E-101 | 0.58414936 | 0.338 | 0.086 | 7.0302E-97 | PDC | RIPOR1 |
| 3.847E-101 | 0.31923376 | 0.16 | 0.022 | 9.0886E-97 | PDC | PAPPA |
| 7.056E-101 | 0.28902614 | 0.248 | 0.048 | 1.6669E-96 | PDC | C3orf14 |
| 7.717E-101 | 0.3763999 | 0.354 | 0.089 | 1.823E-96 | PDC | PHYH |
| 8.052E-101 | 0.42356756 | 0.365 | 0.095 | 1.9022E-96 | PDC | MXRA7 |
| 8.94E-101 | 0.39340429 | 0.394 | 0.108 | 2.1118E-96 | PDC | ATXN10 |
| 1.159E-100 | 0.28438553 | 0.266 | 0.055 | 2.7382E-96 | PDC | FAM174A |

| | | | | | | |
|---|---|---|---|---|---|---|
| 1.385E-100 | 0.65739953 | 0.768 | 0.366 | 3.2719E-96 | PDC | RHEB |
| 1.397E-100 | -1.057235 | 0.081 | 0.557 | 3.3012E-96 | PDC | FAM107B |
| 1.451E-100 | 0.30905982 | 0.3 | 0.067 | 3.4276E-96 | PDC | ABHD14B |
| 3.283E-100 | 0.29297007 | 0.288 | 0.062 | 7.7545E-96 | PDC | IFT22 |
| 4.006E-100 | 0.53625293 | 0.43 | 0.129 | 9.4632E-96 | PDC | HYOU1 |
| 6.659E-100 | 0.54233619 | 0.692 | 0.286 | 1.573E-95 | PDC | NDUFS6 |
| 7.78E-100 | 0.33806292 | 0.295 | 0.066 | 1.8379E-95 | PDC | FAM20C |
| 3.117E-99 | 0.79497656 | 0.853 | 0.558 | 7.3633E-95 | PDC | TOMM7 |
| 3.627E-99 | 0.5469131 | 0.64 | 0.255 | 8.5686E-95 | PDC | OCIAD1 |
| 4.57E-99 | 0.35366163 | 0.308 | 0.072 | 1.0795E-94 | PDC | SLC35F6 |
| 4.669E-99 | 0.45718152 | 0.408 | 0.118 | 1.1029E-94 | PDC | PKN1 |
| 5.518E-99 | 0.40268054 | 0.421 | 0.122 | 1.3035E-94 | PDC | APMAP |
| 1.7964E-98 | 0.43937412 | 0.354 | 0.091 | 4.2435E-94 | PDC | PTPN13 |
| 2.2192E-98 | 0.43924886 | 0.36 | 0.095 | 5.2423E-94 | PDC | MIB1 |
| 6.6863E-98 | 0.4663518 | 0.464 | 0.146 | 1.5795E-93 | PDC | LARS |
| 1.7472E-97 | 0.33938741 | 0.327 | 0.08 | 4.1273E-93 | PDC | C1orf123 |
| 2.0554E-97 | -1.8571517 | 0.103 | 0.561 | 4.8554E-93 | PDC | ELF3 |
| 2.1109E-97 | 0.44980791 | 0.464 | 0.146 | 4.9866E-93 | PDC | RCN2 |
| 2.4953E-97 | 0.53490651 | 0.687 | 0.283 | 5.8945E-93 | PDC | NDUFB7 |
| 3.1212E-97 | 0.31232839 | 0.309 | 0.073 | 7.3732E-93 | PDC | PSMG4 |
| 3.5633E-97 | 0.26307874 | 0.241 | 0.047 | 8.4176E-93 | PDC | C6orf203 |
| 4.7481E-97 | 0.37233034 | 0.358 | 0.094 | 1.1216E-92 | PDC | MVB12A |
| 6.4097E-97 | 1.07104167 | 0.978 | 0.905 | 1.5142E-92 | PDC | ITM2B |
| 7.4603E-97 | 0.44799214 | 0.464 | 0.141 | 1.7623E-92 | PDC | SDC2 |
| 7.6466E-97 | 0.42925931 | 0.441 | 0.135 | 1.8064E-92 | PDC | TRAPPC4 |
| 1.1796E-96 | 0.43961852 | 0.403 | 0.116 | 2.7867E-92 | PDC | SLC1A5 |
| 1.4452E-96 | 0.46970785 | 0.315 | 0.076 | 3.414E-92 | PDC | VWA1 |
| 2.5365E-96 | 0.65717706 | 0.766 | 0.381 | 5.9919E-92 | PDC | DAD1 |
| 3.5226E-96 | 0.28241189 | 0.128 | 0.015 | 8.3214E-92 | PDC | TSPAN8 |
| 5.9242E-96 | 0.35612938 | 0.338 | 0.085 | 1.3995E-91 | PDC | PELO |
| 5.9261E-96 | 0.51713991 | 0.442 | 0.135 | 1.3999E-91 | PDC | MSI2 |
| 6.6068E-96 | 0.75382828 | 0.838 | 0.482 | 1.5607E-91 | PDC | CANX |
| 6.9121E-96 | 0.91851222 | 0.982 | 0.874 | 1.6328E-91 | PDC | RPL3 |
| 8.1941E-96 | 0.2587787 | 0.223 | 0.041 | 1.9357E-91 | PDC | PLA2G12A |
| 1.0678E-95 | 0.46554675 | 0.583 | 0.213 | 2.5225E-91 | PDC | NDUFA5 |
| 1.6659E-95 | -1.3093974 | 0.072 | 0.537 | 3.9353E-91 | PDC | CTSS |
| 2.0241E-95 | 0.36441324 | 0.322 | 0.079 | 4.7816E-91 | PDC | MYO1B |
| 2.0254E-95 | 0.32313815 | 0.315 | 0.076 | 4.7845E-91 | PDC | CFAP298 |
| 2.2813E-95 | 0.47949762 | 0.509 | 0.172 | 5.3892E-91 | PDC | PURA |
| 3.5006E-95 | 0.30809149 | 0.311 | 0.074 | 8.2694E-91 | PDC | FAM210B |
| 3.9886E-95 | 0.39516538 | 0.371 | 0.102 | 9.4222E-91 | PDC | YARS |
| 7.1031E-95 | 0.41915448 | 0.347 | 0.09 | 1.678E-90 | PDC | SPIRE1 |
| 7.7229E-95 | 0.55242006 | 0.712 | 0.306 | 1.8244E-90 | PDC | MZT2B |
| 7.7504E-95 | 0.40769057 | 0.475 | 0.149 | 1.8309E-90 | PDC | RCN1 |
| 8.1252E-95 | 0.46238897 | 0.46 | 0.148 | 1.9194E-90 | PDC | ECH1 |
| 1.2741E-94 | -1.7934284 | 0.094 | 0.551 | 3.0099E-90 | PDC | KRT7 |
| 1.2899E-94 | 0.28006133 | 0.277 | 0.061 | 3.0472E-90 | PDC | MRPL24 |
| 2.6619E-94 | 0.25132967 | 0.236 | 0.046 | 6.2883E-90 | PDC | AKIP1 |
| 2.7204E-94 | 0.55751409 | 0.514 | 0.179 | 6.4263E-90 | PDC | DAB2 |
| 3.3919E-94 | 0.47397787 | 0.5 | 0.17 | 8.0126E-90 | PDC | CYC1 |

| | | | | | | |
|---|---|---|---|---|---|---|
| 4.132E-94 | 0.39633814 | 0.41 | 0.121 | 9.761E-90 | PDC | MSRB2 |
| 6.8608E-94 | 0.56111118 | 0.694 | 0.298 | 1.6207E-89 | PDC | TMED9 |
| 1.0641E-93 | 0.3838963 | 0.405 | 0.118 | 2.5138E-89 | PDC | NDUFA8 |
| 1.2251E-93 | 0.48253341 | 0.527 | 0.185 | 2.894E-89 | PDC | NDUFS7 |
| 2.3838E-93 | 0.57405716 | 0.631 | 0.264 | 5.6312E-89 | PDC | NOL7 |
| 4.068E-93 | 0.31498432 | 0.302 | 0.072 | 9.6098E-89 | PDC | FMC1 |
| 6.5301E-93 | 0.31934341 | 0.288 | 0.066 | 1.5426E-88 | PDC | ARHGAP35 |
| 9.4863E-93 | 0.35879863 | 0.342 | 0.089 | 2.2409E-88 | PDC | WDR13 |
| 1.3226E-92 | 0.61639649 | 0.752 | 0.336 | 3.1243E-88 | PDC | PDIA6 |
| 1.7909E-92 | 0.53927117 | 0.594 | 0.236 | 4.2306E-88 | PDC | SMIM26 |
| 1.9129E-92 | 0.46055273 | 0.522 | 0.184 | 4.5187E-88 | PDC | ORMDL1 |
| 2.2585E-92 | 0.36340978 | 0.371 | 0.103 | 5.3353E-88 | PDC | SNX17 |
| 2.2912E-92 | 0.60844521 | 0.665 | 0.3 | 5.4125E-88 | PDC | JTB |
| 2.8796E-92 | 0.2772133 | 0.299 | 0.071 | 6.8026E-88 | PDC | GPX3 |
| 3.5807E-92 | 0.3513788 | 0.275 | 0.062 | 8.4588E-88 | PDC | P4HA2 |
| 1.0062E-91 | 0.39342709 | 0.433 | 0.133 | 2.3768E-87 | PDC | URI1 |
| 1.1873E-91 | 0.53352005 | 0.66 | 0.279 | 2.8048E-87 | PDC | PSMB6 |
| 1.4747E-91 | 0.5328769 | 0.647 | 0.275 | 3.4836E-87 | PDC | TMBIM4 |
| 1.653E-91 | 0.49864538 | 0.563 | 0.211 | 3.9049E-87 | PDC | RWDD1 |
| 1.9131E-91 | 0.28378329 | 0.264 | 0.058 | 4.5194E-87 | PDC | COX11 |
| 4.4758E-91 | 0.42309126 | 0.448 | 0.142 | 1.0573E-86 | PDC | COA4 |
| 4.7716E-91 | 0.46806333 | 0.459 | 0.15 | 1.1272E-86 | PDC | ZBTB38 |
| 8.0961E-91 | -2.4915997 | 0.094 | 0.542 | 1.9125E-86 | PDC | WFDC2 |
| 1.4337E-90 | 0.26880323 | 0.252 | 0.053 | 3.3867E-86 | PDC | DMAP1 |
| 1.5071E-90 | 0.3683121 | 0.423 | 0.127 | 3.5601E-86 | PDC | TMX4 |
| 2.019E-90 | 0.4683828 | 0.568 | 0.215 | 4.7694E-86 | PDC | PCNP |
| 2.5054E-90 | 0.62725069 | 0.646 | 0.289 | 5.9185E-86 | PDC | COX17 |
| 2.8408E-90 | 0.36745615 | 0.379 | 0.108 | 6.7108E-86 | PDC | MRPL3 |
| 2.9173E-90 | 0.28029121 | 0.268 | 0.06 | 6.8916E-86 | PDC | NUDT2 |
| 4.0094E-90 | 0.38412977 | 0.406 | 0.122 | 9.4713E-86 | PDC | CCDC107 |
| 4.4701E-90 | 0.73513586 | 0.908 | 0.685 | 1.056E-85 | PDC | CAST |
| 5.6412E-90 | 0.30842693 | 0.293 | 0.07 | 1.3326E-85 | PDC | PYCR2 |
| 7.1765E-90 | 0.4429846 | 0.559 | 0.206 | 1.6953E-85 | PDC | ANXA7 |
| 1.0367E-89 | 0.26926513 | 0.259 | 0.056 | 2.449E-85 | PDC | SDR39U1 |
| 1.1466E-89 | 0.2909628 | 0.241 | 0.05 | 2.7086E-85 | PDC | SUPV3L1 |
| 1.6155E-89 | 0.37966585 | 0.424 | 0.132 | 3.8162E-85 | PDC | PSMF1 |
| 2.1579E-89 | -1.5287775 | 0.101 | 0.544 | 5.0976E-85 | PDC | PLAUR |
| 2.3416E-89 | 0.35560891 | 0.313 | 0.078 | 5.5315E-85 | PDC | CEP290 |
| 2.7575E-89 | 0.4759713 | 0.603 | 0.235 | 6.514E-85 | PDC | MRPS21 |
| 4.8166E-89 | 0.58759762 | 0.775 | 0.378 | 1.1378E-84 | PDC | DYNLRB1 |
| 5.7441E-89 | 0.39114828 | 0.363 | 0.103 | 1.3569E-84 | PDC | SRM |
| 6.1453E-89 | 0.39488489 | 0.415 | 0.126 | 1.4517E-84 | PDC | AIG1 |
| 9.8979E-89 | -1.4827789 | 0.761 | 0.943 | 2.3382E-84 | PDC | NEAT1 |
| 1.6484E-88 | 0.43075657 | 0.468 | 0.158 | 3.8941E-84 | PDC | PPCS |
| 1.8241E-88 | 0.35173073 | 0.338 | 0.091 | 4.3091E-84 | PDC | AC027644.3 |
| 2.2794E-88 | 0.31777838 | 0.225 | 0.045 | 5.3846E-84 | PDC | EMX2OS |
| 2.8473E-88 | 0.3931008 | 0.437 | 0.138 | 6.7261E-84 | PDC | ERLEC1 |
| 4.1922E-88 | 0.55913671 | 0.705 | 0.322 | 9.9033E-84 | PDC | LAMP1 |
| 4.2073E-88 | 0.77352585 | 0.995 | 0.98 | 9.939E-84 | PDC | EEF1A1 |
| 4.2879E-88 | -1.7083531 | 0.104 | 0.545 | 1.0129E-83 | PDC | CLDN4 |

| | | | | | | |
|---|---|---|---|---|---|---|
| 5.1191E-88 | 0.40911315 | 0.435 | 0.14 | 1.2093E-83 | PDC | NELFE |
| 5.4092E-88 | 0.39623904 | 0.392 | 0.116 | 1.2778E-83 | PDC | NQO2 |
| 5.7668E-88 | 0.38305663 | 0.342 | 0.093 | 1.3623E-83 | PDC | TNS3 |
| 8.4883E-88 | 0.47938143 | 0.568 | 0.22 | 2.0052E-83 | PDC | NARS |
| 1.0176E-87 | 0.53428338 | 0.601 | 0.245 | 2.4039E-83 | PDC | MRPL20 |
| 1.734E-87 | 0.38352216 | 0.43 | 0.136 | 4.0963E-83 | PDC | PAIP1 |
| 4.7233E-87 | 0.29706874 | 0.295 | 0.072 | 1.1158E-82 | PDC | POLD2 |
| 5.4048E-87 | 0.31613322 | 0.187 | 0.033 | 1.2768E-82 | PDC | BEX1 |
| 8.2496E-87 | 0.52256059 | 0.482 | 0.17 | 1.9488E-82 | PDC | CSRP1 |
| 1.9741E-86 | 0.5027773 | 0.635 | 0.262 | 4.6634E-82 | PDC | GADD45GIP1 |
| 4.3927E-86 | 0.43999454 | 0.408 | 0.128 | 1.0377E-81 | PDC | CXXC5 |
| 5.2852E-86 | 0.44032232 | 0.489 | 0.167 | 1.2485E-81 | PDC | BBX |
| 6.7216E-86 | 0.50799942 | 0.655 | 0.283 | 1.5878E-81 | PDC | TMED10 |
| 7.0168E-86 | 0.36122897 | 0.363 | 0.104 | 1.6576E-81 | PDC | RTL8C |
| 9.4107E-86 | 0.27403666 | 0.263 | 0.06 | 2.2231E-81 | PDC | HABP4 |
| 9.6653E-86 | 0.28493568 | 0.243 | 0.053 | 2.2832E-81 | PDC | NLK |
| 1.6467E-85 | 0.52232791 | 0.642 | 0.27 | 3.8899E-81 | PDC | SSB |
| 1.8715E-85 | 0.38146556 | 0.299 | 0.075 | 4.4211E-81 | PDC | CBLB |
| 2.0348E-85 | 0.47956256 | 0.541 | 0.207 | 4.8068E-81 | PDC | PSMC5 |
| 4.9132E-85 | 0.25090774 | 0.191 | 0.035 | 1.1606E-80 | PDC | AL355338.1 |
| 6.1059E-85 | 0.61501354 | 0.815 | 0.421 | 1.4424E-80 | PDC | SLC3A2 |
| 7.4166E-85 | 0.41402678 | 0.46 | 0.155 | 1.752E-80 | PDC | FAM200B |
| 9.3905E-85 | 0.41107134 | 0.484 | 0.169 | 2.2183E-80 | PDC | PSMC3 |
| 9.9308E-85 | 0.30508573 | 0.291 | 0.072 | 2.346E-80 | PDC | DGCR6L |
| 1.2221E-84 | 0.78160255 | 0.637 | 0.279 | 2.887E-80 | PDC | CD151 |
| 1.442E-84 | 0.43016407 | 0.437 | 0.148 | 3.4065E-80 | PDC | MT1H |
| 3.1095E-84 | 0.40363459 | 0.46 | 0.156 | 7.3456E-80 | PDC | GLRX5 |
| 4.1988E-84 | 0.43226522 | 0.604 | 0.232 | 9.9188E-80 | PDC | PCBD1 |
| 4.7158E-84 | 0.4087525 | 0.43 | 0.14 | 1.114E-79 | PDC | SPART |
| 7.9507E-84 | 0.48520728 | 0.469 | 0.161 | 1.8782E-79 | PDC | EPB41L4A-AS1 |
| 1.3033E-83 | 0.52562639 | 0.586 | 0.239 | 3.0787E-79 | PDC | PDCD5 |
| 2.3401E-83 | 0.38130346 | 0.396 | 0.122 | 5.5279E-79 | PDC | EBNA1BP2 |
| 2.7729E-83 | 0.38520135 | 0.437 | 0.144 | 6.5503E-79 | PDC | MRPL43 |
| 3.33E-83 | 0.36835318 | 0.36 | 0.105 | 7.8664E-79 | PDC | EEF1E1 |
| 5.0299E-83 | 0.60794242 | 0.709 | 0.341 | 1.1882E-78 | PDC | CAPN2 |
| 9.0294E-83 | 0.41300352 | 0.426 | 0.14 | 2.133E-78 | PDC | PIN1 |
| 1.5554E-82 | 0.5599845 | 0.662 | 0.301 | 3.6743E-78 | PDC | NOP53 |
| 1.6998E-82 | 0.28978079 | 0.272 | 0.065 | 4.0155E-78 | PDC | AMZ2 |
| 4.7106E-82 | 0.36289668 | 0.388 | 0.119 | 1.1128E-77 | PDC | APPL1 |
| 4.7762E-82 | 0.35740835 | 0.374 | 0.111 | 1.1283E-77 | PDC | AC245297.3 |
| 7.9141E-82 | 0.34511305 | 0.351 | 0.1 | 1.8695E-77 | PDC | ILF3-DT |
| 8.0451E-82 | 0.39588439 | 0.453 | 0.154 | 1.9005E-77 | PDC | CNPY2 |
| 1.0321E-81 | -1.232052 | 0.115 | 0.546 | 2.438E-77 | PDC | CD24 |
| 1.2638E-81 | 0.33951072 | 0.338 | 0.095 | 2.9855E-77 | PDC | RPF2 |
| 2.3598E-81 | 0.54093447 | 0.718 | 0.326 | 5.5746E-77 | PDC | APP |
| 2.6337E-81 | 0.41958782 | 0.387 | 0.119 | 6.2216E-77 | PDC | ADD3 |
| 3.7005E-81 | 0.49873754 | 0.604 | 0.256 | 8.7416E-77 | PDC | SMDT1 |
| 5.0467E-81 | 0.419912 | 0.421 | 0.139 | 1.1922E-76 | PDC | NME3 |
| 6.3029E-81 | 0.76165938 | 0.968 | 0.816 | 1.4889E-76 | PDC | RPL5 |
| 6.4486E-81 | 0.34721359 | 0.318 | 0.086 | 1.5233E-76 | PDC | FAM114A1 |

| | | | | | | | |
|---|---|---|---|---|---|---|---|
| 6.9388E-81 | 0.36060909 | 0.353 | 0.102 | 1.6391E-76 | PDC | | GOT1 |
| 7.0207E-81 | 0.41641582 | 0.482 | 0.172 | 1.6585E-76 | PDC | | TMEM147 |
| 7.3399E-81 | 0.51515884 | 0.746 | 0.368 | 1.7339E-76 | PDC | | C11orf58 |
| 8.1966E-81 | 0.41184569 | 0.417 | 0.137 | 1.9363E-76 | PDC | | CEBPG |
| 9.4129E-81 | 0.31341048 | 0.333 | 0.092 | 2.2236E-76 | PDC | | ZNF770 |
| 1.1173E-80 | 0.39980871 | 0.381 | 0.115 | 2.6394E-76 | PDC | | FKBP5 |
| 1.8963E-80 | 0.32648026 | 0.308 | 0.082 | 4.4796E-76 | PDC | | SURF2 |
| 1.9463E-80 | 0.27135162 | 0.252 | 0.058 | 4.5978E-76 | PDC | | RANGRF |
| 4.9743E-80 | 0.30090917 | 0.318 | 0.086 | 1.1751E-75 | PDC | | HDGFL3 |
| 5.1384E-80 | 0.38448785 | 0.444 | 0.151 | 1.2139E-75 | PDC | | CHCHD5 |
| 6.0019E-80 | 0.82011316 | 0.959 | 0.772 | 1.4178E-75 | PDC | | CTSD |
| 9.4773E-80 | 0.37483419 | 0.412 | 0.134 | 2.2388E-75 | PDC | | DDRGK1 |
| 1.0558E-79 | 0.64899228 | 0.737 | 0.384 | 2.4942E-75 | PDC | | HMGN2 |
| 1.1161E-79 | 0.39321977 | 0.446 | 0.154 | 2.6366E-75 | PDC | | DCTN3 |
| 1.2897E-79 | 0.47574425 | 0.612 | 0.249 | 3.0467E-75 | PDC | | ODC1 |
| 1.3329E-79 | 0.26514813 | 0.232 | 0.051 | 3.1486E-75 | PDC | | LURAP1L |
| 1.3756E-79 | 0.31542952 | 0.347 | 0.1 | 3.2495E-75 | PDC | | AK6 |
| 1.4063E-79 | 0.33541564 | 0.326 | 0.091 | 3.3222E-75 | PDC | | PMVK |
| 1.4736E-79 | 0.44666929 | 0.237 | 0.053 | 3.4811E-75 | PDC | | PDK4 |
| 3.1721E-79 | 0.5272421 | 0.712 | 0.334 | 7.4935E-75 | PDC | | SNU13 |
| 3.8052E-79 | 0.53373463 | 0.712 | 0.33 | 8.989E-75 | PDC | | EIF4B |
| 3.8078E-79 | 0.31303044 | 0.284 | 0.072 | 8.9952E-75 | PDC | | FGFR1 |
| 6.4155E-79 | 0.29062759 | 0.29 | 0.075 | 1.5155E-74 | PDC | | ZFPM1 |
| 9.8932E-79 | 0.53846194 | 0.401 | 0.133 | 2.3371E-74 | PDC | | SRGAP1 |
| 1.0069E-78 | 0.28377327 | 0.299 | 0.078 | 2.3785E-74 | PDC | | ECI2 |
| 2.9261E-78 | 0.90470223 | 0.975 | 0.911 | 6.9124E-74 | PDC | | RPS8 |
| 3.1211E-78 | 0.30024779 | 0.286 | 0.074 | 7.3729E-74 | PDC | | POLR3GL |
| 3.5003E-78 | 0.4258075 | 0.529 | 0.207 | 8.2687E-74 | PDC | | UBE2M |
| 3.7198E-78 | 0.30150605 | 0.27 | 0.067 | 8.7874E-74 | PDC | | NUDT14 |
| 4.2329E-78 | 0.30548598 | 0.317 | 0.087 | 9.9995E-74 | PDC | | TYW3 |
| 4.8086E-78 | 0.29384744 | 0.291 | 0.075 | 1.1359E-73 | PDC | | ZNF37A |
| 5.4743E-78 | 0.54921452 | 0.689 | 0.322 | 1.2932E-73 | PDC | | EIF1B |
| 5.9086E-78 | 0.34544598 | 0.326 | 0.093 | 1.3958E-73 | PDC | | NSMCE1 |
| 8.3302E-78 | 0.33524114 | 0.354 | 0.106 | 1.9678E-73 | PDC | | FAM173A |
| 9.5106E-78 | 0.42236706 | 0.457 | 0.158 | 2.2467E-73 | PDC | | TMEM141 |
| 1.3931E-77 | 0.29345333 | 0.308 | 0.083 | 3.291E-73 | PDC | | BOD1 |
| 1.8692E-77 | 0.25297731 | 0.218 | 0.046 | 4.4157E-73 | PDC | | CCDC85C |
| 2.3828E-77 | 0.39631899 | 0.32 | 0.091 | 5.6289E-73 | PDC | | EXOSC6 |
| 3.1847E-77 | 0.49608846 | 0.48 | 0.179 | 7.5233E-73 | PDC | | PSMB9 |
| 3.4994E-77 | 0.36068237 | 0.399 | 0.128 | 8.2666E-73 | PDC | | PGRMC2 |
| 4.6598E-77 | 0.48820971 | 0.621 | 0.272 | 1.1008E-72 | PDC | | MRPL51 |
| 4.7995E-77 | 0.3246397 | 0.347 | 0.103 | 1.1338E-72 | PDC | | RRAGA |
| 1.0517E-76 | 0.37880433 | 0.313 | 0.088 | 2.4844E-72 | PDC | | FBXO21 |
| 1.0611E-76 | 0.33996328 | 0.326 | 0.093 | 2.5066E-72 | PDC | | CPQ |
| 1.3215E-76 | 0.37902848 | 0.388 | 0.126 | 3.1217E-72 | PDC | | BORCS7 |
| 1.9653E-76 | 0.46700348 | 0.595 | 0.253 | 4.6426E-72 | PDC | | RPS19BP1 |
| 2.016E-76 | 0.31591565 | 0.311 | 0.086 | 4.7623E-72 | PDC | | ZMYND11 |
| 2.1013E-76 | 0.32420267 | 0.326 | 0.093 | 4.9639E-72 | PDC | | DCTD |
| 2.3475E-76 | -1.0288252 | 0.822 | 0.952 | 5.5454E-72 | PDC | | MT-CO1 |
| 3.2642E-76 | 0.56442319 | 0.746 | 0.397 | 7.711E-72 | PDC | | CSDE1 |

| | | | | | | |
|---|---|---|---|---|---|---|
| 3.7452E-76 | 0.30423759 | 0.272 | 0.068 | 8.8473E-72 | PDC | MAOA |
| 4.2866E-76 | 0.46146469 | 0.513 | 0.201 | 1.0126E-71 | PDC | SLIRP |
| 5.5785E-76 | 0.56621181 | 0.763 | 0.405 | 1.3178E-71 | PDC | ATP5PF |
| 6.6236E-76 | -3.0289515 | 0.094 | 0.493 | 1.5647E-71 | PDC | CXCL8 |
| 6.715E-76 | 0.33779831 | 0.376 | 0.118 | 1.5863E-71 | PDC | HAGH |
| 7.8113E-76 | 0.30847504 | 0.29 | 0.077 | 1.8453E-71 | PDC | PMM1 |
| 8.0292E-76 | 0.399993 | 0.417 | 0.142 | 1.8967E-71 | PDC | ATF6 |
| 1.1426E-75 | 0.26469538 | 0.248 | 0.059 | 2.6992E-71 | PDC | ERI3 |
| 1.3107E-75 | 0.28678116 | 0.277 | 0.071 | 3.0962E-71 | PDC | RFNG |
| 1.8688E-75 | 0.38916108 | 0.439 | 0.153 | 4.4147E-71 | PDC | SMC3 |
| 2.0583E-75 | 0.46277502 | 0.665 | 0.296 | 4.8624E-71 | PDC | DDX24 |
| 2.7223E-75 | 0.64204994 | 0.682 | 0.321 | 6.4308E-71 | PDC | PPA1 |
| 3.8556E-75 | 0.28144322 | 0.275 | 0.071 | 9.108E-71 | PDC | UROD |
| 6.8296E-75 | 0.40417119 | 0.505 | 0.191 | 1.6134E-70 | PDC | C1QBP |
| 7.7236E-75 | 0.2679996 | 0.281 | 0.073 | 1.8245E-70 | PDC | ZFAND1 |
| 1.0737E-74 | 0.40817024 | 0.433 | 0.153 | 2.5365E-70 | PDC | FAM204A |
| 1.1248E-74 | 0.52938199 | 0.665 | 0.326 | 2.6571E-70 | PDC | UBXN1 |
| 1.1583E-74 | 0.47649141 | 0.423 | 0.149 | 2.7363E-70 | PDC | EIF4EBP1 |
| 2.0007E-74 | 0.49749159 | 0.626 | 0.283 | 4.7262E-70 | PDC | LAMTOR1 |
| 2.8762E-74 | 0.39778717 | 0.426 | 0.149 | 6.7944E-70 | PDC | DDOST |
| 2.9905E-74 | 0.4149476 | 0.482 | 0.179 | 7.0645E-70 | PDC | CALU |
| 3.329E-74 | 0.41252465 | 0.516 | 0.194 | 7.8641E-70 | PDC | TMEM106B |
| 4.178E-74 | 0.33462347 | 0.379 | 0.121 | 9.8696E-70 | PDC | IER3IP1 |
| 4.8437E-74 | 0.38231599 | 0.313 | 0.09 | 1.1442E-69 | PDC | INF2 |
| 6.2185E-74 | 0.43416808 | 0.545 | 0.222 | 1.469E-69 | PDC | PSMB2 |
| 1.0874E-73 | 0.49384909 | 0.603 | 0.267 | 2.5688E-69 | PDC | NDUFB8 |
| 1.1762E-73 | 0.32456101 | 0.631 | 0.277 | 2.7785E-69 | PDC | IFITM2 |
| 1.5587E-73 | 0.33164482 | 0.568 | 0.25 | 3.682E-69 | PDC | MT1G |
| 1.5871E-73 | 0.30706676 | 0.297 | 0.082 | 3.7492E-69 | PDC | GTPBP6 |
| 1.7214E-73 | 0.4400535 | 0.3 | 0.085 | 4.0665E-69 | PDC | CRIP2 |
| 2.3451E-73 | 0.2604576 | 0.259 | 0.065 | 5.5398E-69 | PDC | FLYWCH2 |
| 2.5232E-73 | 0.25884943 | 0.252 | 0.062 | 5.9605E-69 | PDC | DCAF8 |
| 2.818E-73 | 0.39190694 | 0.487 | 0.182 | 6.6571E-69 | PDC | DYNLT3 |
| 2.8414E-73 | 0.37335462 | 0.475 | 0.174 | 6.7123E-69 | PDC | REX1BD |
| 2.9527E-73 | 0.33831325 | 0.403 | 0.134 | 6.9752E-69 | PDC | MRPS7 |
| 3.7988E-73 | 0.2985091 | 0.308 | 0.087 | 8.9739E-69 | PDC | TSPAN31 |
| 3.8013E-73 | 0.45071939 | 0.601 | 0.264 | 8.9797E-69 | PDC | NDUFA12 |
| 4.0478E-73 | 0.36119812 | 0.367 | 0.116 | 9.5622E-69 | PDC | HRAS |
| 5.1979E-73 | 0.76084787 | 0.905 | 0.655 | 1.2279E-68 | PDC | CALM2 |
| 8.3824E-73 | 0.5767551 | 0.633 | 0.299 | 1.9802E-68 | PDC | BLVRB |
| 1.3585E-72 | 0.81594862 | 0.941 | 0.799 | 3.2091E-68 | PDC | RPS5 |
| 1.9874E-72 | 0.39605187 | 0.444 | 0.157 | 4.6948E-68 | PDC | NCKAP1 |
| 2.2157E-72 | 0.33295829 | 0.665 | 0.268 | 5.2342E-68 | PDC | MAL2 |
| 2.5242E-72 | 0.53730248 | 0.761 | 0.389 | 5.9629E-68 | PDC | NDUFB2 |
| 3.1177E-72 | 0.28797463 | 0.286 | 0.077 | 7.3649E-68 | PDC | MORN2 |
| 3.6947E-72 | 0.41020378 | 0.369 | 0.121 | 8.728E-68 | PDC | NKAP |
| 4.2642E-72 | 0.53054637 | 0.667 | 0.316 | 1.0073E-67 | PDC | KRT10 |
| 4.8043E-72 | 0.28175157 | 0.275 | 0.072 | 1.1349E-67 | PDC | PKP4 |
| 6.1632E-72 | -1.2169413 | 0.07 | 0.454 | 1.4559E-67 | PDC | MARCKS |
| 7.6507E-72 | 0.32052015 | 0.326 | 0.096 | 1.8073E-67 | PDC | SMAD5 |

| | | | | | | |
|---|---|---|---|---|---|---|
| 7.8477E-72 | 0.34627951 | 0.345 | 0.107 | 1.8539E-67 | PDC | TCEAL8 |
| 8.5161E-72 | -3.2122243 | 0.142 | 0.528 | 2.0118E-67 | PDC | LCN2 |
| 1.0329E-71 | 0.26609517 | 0.254 | 0.064 | 2.4401E-67 | PDC | PITPNA-AS1 |
| 1.1067E-71 | 0.51964003 | 0.66 | 0.316 | 2.6145E-67 | PDC | ERP29 |
| 1.2055E-71 | 0.32662461 | 0.363 | 0.116 | 2.8478E-67 | PDC | LSM2 |
| 1.434E-71 | 0.3840831 | 0.464 | 0.172 | 3.3875E-67 | PDC | SRP72 |
| 1.622E-71 | 0.62950275 | 0.871 | 0.669 | 3.8317E-67 | PDC | COX5B |
| 1.6406E-71 | 0.38615282 | 0.399 | 0.135 | 3.8756E-67 | PDC | MYO1E |
| 1.7901E-71 | 0.2706578 | 0.248 | 0.061 | 4.2287E-67 | PDC | DVL1 |
| 1.9249E-71 | -1.7679826 | 0.061 | 0.445 | 4.5472E-67 | PDC | TYROBP |
| 2.3104E-71 | 0.32981814 | 0.291 | 0.081 | 5.4579E-67 | PDC | WASHC2A |
| 3.1715E-71 | 0.40957243 | 0.505 | 0.2 | 7.492E-67 | PDC | TXNL4A |
| 3.5585E-71 | 0.3342953 | 0.367 | 0.118 | 8.4062E-67 | PDC | MRPL55 |
| 5.3064E-71 | -2.3143631 | 0.081 | 0.469 | 1.2535E-66 | PDC | LYZ |
| 5.5459E-71 | 0.43842331 | 0.612 | 0.271 | 1.3101E-66 | PDC | RSL24D1 |
| 6.632E-71 | 0.69666149 | 0.951 | 0.738 | 1.5667E-66 | PDC | PSAP |
| 7.1101E-71 | 0.30872149 | 0.356 | 0.112 | 1.6796E-66 | PDC | CLPP |
| 8.0935E-71 | 0.63202055 | 0.347 | 0.113 | 1.9119E-66 | PDC | IFI6 |
| 1.0829E-70 | 0.27538341 | 0.272 | 0.072 | 2.5581E-66 | PDC | LETMD1 |
| 1.3885E-70 | -1.7133031 | 0.103 | 0.496 | 3.2801E-66 | PDC | CD74 |
| 5.4548E-70 | 0.26009606 | 0.282 | 0.077 | 1.2886E-65 | PDC | NAE1 |
| 5.4818E-70 | 0.28205235 | 0.27 | 0.072 | 1.295E-65 | PDC | ANKRD40 |
| 6.7602E-70 | 0.26413967 | 0.273 | 0.073 | 1.597E-65 | PDC | ATIC |
| 7.9104E-70 | 0.26232313 | 0.25 | 0.063 | 1.8687E-65 | PDC | ZC2HC1A |
| 2.899E-69 | 0.38519142 | 0.426 | 0.153 | 6.8482E-65 | PDC | TROVE2 |
| 3.5497E-69 | 0.41969429 | 0.52 | 0.213 | 8.3855E-65 | PDC | COPZ1 |
| 5.2032E-69 | 0.33927772 | 0.392 | 0.134 | 1.2292E-64 | PDC | MRPL36 |
| 8.859E-69 | 0.77726962 | 0.982 | 0.918 | 2.0928E-64 | PDC | RPL9 |
| 9.5496E-69 | 0.41377716 | 0.514 | 0.208 | 2.2559E-64 | PDC | GTF3C6 |
| 9.9207E-69 | 0.33498834 | 0.358 | 0.116 | 2.3436E-64 | PDC | TIMMDC1 |
| 1.0116E-68 | 0.39682514 | 0.448 | 0.17 | 2.3897E-64 | PDC | MRPS18C |
| 1.1916E-68 | 0.30208016 | 0.326 | 0.099 | 2.8148E-64 | PDC | TSSC4 |
| 1.3593E-68 | 0.36428178 | 0.464 | 0.176 | 3.2111E-64 | PDC | THOC7 |
| 1.6626E-68 | 0.3235813 | 0.32 | 0.096 | 3.9276E-64 | PDC | TTC37 |
| 1.9522E-68 | 0.28223541 | 0.281 | 0.078 | 4.6117E-64 | PDC | MRPL2 |
| 2.0701E-68 | 0.34738831 | 0.189 | 0.04 | 4.8903E-64 | PDC | CHST2 |
| 2.2905E-68 | 0.25765206 | 0.279 | 0.076 | 5.4107E-64 | PDC | ENOPH1 |
| 2.5484E-68 | 0.27422765 | 0.272 | 0.074 | 6.0201E-64 | PDC | WASHC2C |
| 4.9144E-68 | 0.42667457 | 0.441 | 0.165 | 1.1609E-63 | PDC | GARS |
| 5.1293E-68 | 0.25783151 | 0.227 | 0.054 | 1.2117E-63 | PDC | TANC1 |
| 7.364E-68 | 0.5391671 | 0.649 | 0.317 | 1.7396E-63 | PDC | LEPROT |
| 7.3874E-68 | 0.39736599 | 0.493 | 0.195 | 1.7451E-63 | PDC | TMEM230 |
| 1.0639E-67 | 0.29633712 | 0.291 | 0.084 | 2.5133E-63 | PDC | SWI5 |
| 1.2178E-67 | 0.52574003 | 0.844 | 0.525 | 2.8767E-63 | PDC | EIF4A2 |
| 1.2586E-67 | 0.5105547 | 0.687 | 0.338 | 2.9731E-63 | PDC | BSG |
| 1.3145E-67 | 0.2750671 | 0.282 | 0.079 | 3.1052E-63 | PDC | EMG1 |
| 1.5607E-67 | 0.32065483 | 0.356 | 0.116 | 3.6868E-63 | PDC | FAM120AOS |
| 1.7652E-67 | 0.31002387 | 0.299 | 0.086 | 4.17E-63 | PDC | PLEKHA5 |
| 1.9032E-67 | 0.4050098 | 0.502 | 0.204 | 4.496E-63 | PDC | FAM50A |
| 2.1024E-67 | -1.3745095 | 0.094 | 0.462 | 4.9664E-63 | PDC | MXD1 |

| | | | | | | |
|---|---|---|---|---|---|---|
| 2.6427E-67 | 0.40377584 | 0.487 | 0.193 | 6.2427E-63 | PDC | TMED4 |
| 2.788E-67 | 0.32153649 | 0.344 | 0.11 | 6.5861E-63 | PDC | GRHPR |
| 2.9E-67 | 0.34993974 | 0.322 | 0.1 | 6.8508E-63 | PDC | MCUR1 |
| 2.9375E-67 | 0.34709232 | 0.392 | 0.135 | 6.9392E-63 | PDC | HEXA |
| 3.0708E-67 | 0.2563906 | 0.272 | 0.074 | 7.2541E-63 | PDC | COPS4 |
| 3.4865E-67 | 0.31776854 | 0.284 | 0.079 | 8.2362E-63 | PDC | CAVIN1 |
| 7.9639E-67 | 0.3781194 | 0.423 | 0.156 | 1.8813E-62 | PDC | DCTN2 |
| 8.3992E-67 | 0.96661614 | 0.982 | 0.933 | 1.9841E-62 | PDC | RPS12 |
| 8.6599E-67 | 0.96760174 | 0.908 | 0.765 | 2.0457E-62 | PDC | RPS26 |
| 1.0718E-66 | 0.43836657 | 0.606 | 0.275 | 2.532E-62 | PDC | MT1X |
| 2.8294E-66 | 0.5119976 | 0.583 | 0.254 | 6.684E-62 | PDC | TSC22D1 |
| 2.8938E-66 | 0.30125263 | 0.333 | 0.104 | 6.836E-62 | PDC | SMIM30 |
| 3.2805E-66 | 0.32156289 | 0.374 | 0.126 | 7.7495E-62 | PDC | PLEKHA3 |
| 3.3019E-66 | 0.29295537 | 0.277 | 0.078 | 7.8E-62 | PDC | AKT3 |
| 3.5284E-66 | 0.32968465 | 0.399 | 0.14 | 8.3351E-62 | PDC | MRPL27 |
| 4.1623E-66 | 0.33960087 | 0.354 | 0.117 | 9.8327E-62 | PDC | RRAGD |
| 4.9024E-66 | 0.33764015 | 0.464 | 0.175 | 1.1581E-61 | PDC | RANBP1 |
| 5.0772E-66 | 0.93770698 | 0.732 | 0.444 | 1.1994E-61 | PDC | ACTN4 |
| 7.1533E-66 | 0.32764156 | 0.378 | 0.129 | 1.6898E-61 | PDC | TM2D1 |
| 8.5288E-66 | -1.6482343 | 0.088 | 0.455 | 2.0148E-61 | PDC | SOX4 |
| 9.0831E-66 | 0.35019418 | 0.39 | 0.136 | 2.1457E-61 | PDC | ZNF593 |
| 1.0375E-65 | 0.38735945 | 0.529 | 0.216 | 2.4509E-61 | PDC | PCM1 |
| 1.1154E-65 | 0.30673979 | 0.36 | 0.118 | 2.6349E-61 | PDC | SMIM37 |
| 1.1609E-65 | 0.50525179 | 0.574 | 0.26 | 2.7424E-61 | PDC | GSN |
| 1.6709E-65 | 0.30294953 | 0.349 | 0.113 | 3.9471E-61 | PDC | MRPS23 |
| 2.578E-65 | 0.40154058 | 0.403 | 0.147 | 6.09E-61 | PDC | VGLL4 |
| 2.8442E-65 | 0.49884587 | 0.687 | 0.348 | 6.7189E-61 | PDC | EIF3E |
| 3.1619E-65 | 0.29141698 | 0.293 | 0.085 | 7.4693E-61 | PDC | RAB11FIP2 |
| 5.103E-65 | 0.35807602 | 0.426 | 0.158 | 1.2055E-60 | PDC | BLOC1S2 |
| 6.736E-65 | 0.38108449 | 0.46 | 0.177 | 1.5912E-60 | PDC | RBCK1 |
| 6.8213E-65 | 0.37663492 | 0.464 | 0.18 | 1.6114E-60 | PDC | DPP7 |
| 7.1661E-65 | 0.41333733 | 0.581 | 0.261 | 1.6928E-60 | PDC | TMEM219 |
| 1.287E-64 | 0.70976034 | 0.962 | 0.869 | 3.0403E-60 | PDC | HLA-C |
| 1.5985E-64 | 0.44799445 | 0.685 | 0.343 | 3.7762E-60 | PDC | PSMD8 |
| 2.0858E-64 | 0.34283773 | 0.471 | 0.182 | 4.9272E-60 | PDC | NHP2 |
| 2.1107E-64 | -0.9946391 | 0.236 | 0.576 | 4.9862E-60 | PDC | VMP1 |
| 2.2467E-64 | 0.63381842 | 0.912 | 0.69 | 5.3073E-60 | PDC | HSP90B1 |
| 2.384E-64 | 0.35066175 | 0.405 | 0.145 | 5.6317E-60 | PDC | TSPYL1 |
| 2.8613E-64 | 0.65847955 | 0.989 | 0.939 | 6.7592E-60 | PDC | RPS28 |
| 2.8966E-64 | 0.67875344 | 0.944 | 0.779 | 6.8427E-60 | PDC | RPL10A |
| 4.074E-64 | 0.41636872 | 0.484 | 0.194 | 9.624E-60 | PDC | TCEAL9 |
| 4.5777E-64 | 0.4290996 | 0.48 | 0.198 | 1.0814E-59 | PDC | KRTCAP2 |
| 4.877E-64 | -1.0202379 | 0.09 | 0.457 | 1.1521E-59 | PDC | CLDN7 |
| 6.2875E-64 | 0.30610241 | 0.313 | 0.098 | 1.4853E-59 | PDC | ALDH9A1 |
| 6.6298E-64 | 0.65949115 | 0.817 | 0.56 | 1.5662E-59 | PDC | SLC25A6 |
| 8.0044E-64 | 0.35408788 | 0.399 | 0.145 | 1.8909E-59 | PDC | USO1 |
| 1.1931E-63 | 0.71145421 | 0.987 | 0.947 | 2.8185E-59 | PDC | RPL10 |
| 4.7198E-63 | 0.37086256 | 0.417 | 0.154 | 1.1149E-58 | PDC | PRKDC |
| 6.1534E-63 | 0.28120046 | 0.293 | 0.088 | 1.4536E-58 | PDC | GLRX2 |
| 7.2012E-63 | 0.63677563 | 0.964 | 0.895 | 1.7011E-58 | PDC | RPL35A |

| | | | | | | |
|---|---|---|---|---|---|---|
| 8.9351E-63 | 0.43307254 | 0.516 | 0.221 | 2.1107E-58 | PDC | RSL1D1 |
| 9.0376E-63 | 0.7396402 | 0.978 | 0.898 | 2.1349E-58 | PDC | RPL21 |
| 1.1693E-62 | 0.27275895 | 0.237 | 0.063 | 2.7622E-58 | PDC | ZADH2 |
| 1.8542E-62 | 0.98095408 | 0.847 | 0.642 | 4.3803E-58 | PDC | HSPA5 |
| 1.8661E-62 | 0.36789926 | 0.439 | 0.169 | 4.4082E-58 | PDC | CPNE3 |
| 2.0144E-62 | 0.4423586 | 0.683 | 0.347 | 4.7585E-58 | PDC | SNX3 |
| 2.1408E-62 | 0.92848683 | 0.18 | 0.04 | 5.0571E-58 | PDC | MGP |
| 2.8001E-62 | 0.36094782 | 0.39 | 0.142 | 6.6147E-58 | PDC | TMEM256 |
| 3.0901E-62 | 0.49776191 | 0.658 | 0.328 | 7.2998E-58 | PDC | SNHG8 |
| 3.608E-62 | 0.44603005 | 0.66 | 0.327 | 8.5232E-58 | PDC | NDUFC2 |
| 3.6084E-62 | 0.73502081 | 0.975 | 0.926 | 8.524E-58 | PDC | RPL19 |
| 4.1734E-62 | 0.33995998 | 0.362 | 0.124 | 9.8588E-58 | PDC | RNF115 |
| 5.1361E-62 | 0.25535675 | 0.255 | 0.07 | 1.2133E-57 | PDC | ZNF302 |
| 5.5425E-62 | 0.66614353 | 0.969 | 0.897 | 1.3093E-57 | PDC | RPS25 |
| 5.7522E-62 | 0.31560837 | 0.371 | 0.129 | 1.3589E-57 | PDC | STOML2 |
| 6.2583E-62 | 0.38833646 | 0.421 | 0.16 | 1.4784E-57 | PDC | MTHFD2 |
| 7.1601E-62 | 0.2839598 | 0.302 | 0.093 | 1.6914E-57 | PDC | CHCHD3 |
| 7.5736E-62 | 0.37924443 | 0.549 | 0.23 | 1.7891E-57 | PDC | SNHG7 |
| 8.2624E-62 | 0.47806543 | 0.66 | 0.336 | 1.9518E-57 | PDC | ANP32B |
| 8.3147E-62 | 0.40889325 | 0.617 | 0.29 | 1.9642E-57 | PDC | EIF3G |
| 8.5914E-62 | 0.39888937 | 0.322 | 0.106 | 2.0295E-57 | PDC | PAK1 |
| 1.2654E-61 | 0.29565431 | 0.317 | 0.1 | 2.9894E-57 | PDC | MDFIC |
| 1.2921E-61 | 0.76238427 | 0.975 | 0.894 | 3.0523E-57 | PDC | RPL37A |
| 1.9916E-61 | 0.42864672 | 0.665 | 0.325 | 4.7046E-57 | PDC | AURKAIP1 |
| 2.7212E-61 | 0.31368888 | 0.365 | 0.127 | 6.4284E-57 | PDC | CERS2 |
| 2.7516E-61 | 0.31964032 | 0.358 | 0.124 | 6.5E-57 | PDC | PTS |
| 3.3848E-61 | 0.52974602 | 0.833 | 0.55 | 7.9959E-57 | PDC | MORF4L1 |
| 3.7589E-61 | -0.7552134 | 0.745 | 0.868 | 8.8795E-57 | PDC | ARPC3 |
| 4.6844E-61 | 0.50184855 | 0.883 | 0.655 | 1.1066E-56 | PDC | NDUFA4 |
| 5.3131E-61 | 0.25312772 | 0.232 | 0.061 | 1.2551E-56 | PDC | JMJD8 |
| 5.389E-61 | 0.31948094 | 0.322 | 0.105 | 1.273E-56 | PDC | OSBPL9 |
| 6.3842E-61 | 0.42067906 | 0.552 | 0.25 | 1.5082E-56 | PDC | RPN2 |
| 6.8013E-61 | 0.37060775 | 0.477 | 0.193 | 1.6067E-56 | PDC | ATP5MC1 |
| 7.7298E-61 | -1.1400185 | 0.149 | 0.508 | 1.826E-56 | PDC | DNAJB1 |
| 9.1645E-61 | 0.4049426 | 0.363 | 0.128 | 2.1649E-56 | PDC | SLC44A2 |
| 9.4357E-61 | 0.36832039 | 0.347 | 0.12 | 2.229E-56 | PDC | EVA1B |
| 9.8989E-61 | 0.39262922 | 0.457 | 0.185 | 2.3384E-56 | PDC | HADHB |
| 1.2536E-60 | 0.25460531 | 0.367 | 0.123 | 2.9614E-56 | PDC | SYNE1 |
| 1.6486E-60 | -0.8176937 | 0.088 | 0.432 | 3.8944E-56 | PDC | EFHD2 |
| 1.8913E-60 | 0.28010285 | 0.266 | 0.077 | 4.4678E-56 | PDC | AKAP11 |
| 2.8791E-60 | 0.7965618 | 0.973 | 0.886 | 6.8013E-56 | PDC | RPS6 |
| 3.7326E-60 | 0.3211339 | 0.432 | 0.163 | 8.8176E-56 | PDC | CCDC50 |
| 3.7606E-60 | 0.27889986 | 0.295 | 0.092 | 8.8837E-56 | PDC | CUEDC2 |
| 3.7938E-60 | 0.3959774 | 0.446 | 0.175 | 8.962E-56 | PDC | STAT1 |
| 4.1052E-60 | 0.26041216 | 0.173 | 0.038 | 9.6977E-56 | PDC | STON2 |
| 4.7959E-60 | 0.39416357 | 0.592 | 0.277 | 1.1329E-55 | PDC | ATP6V1E1 |
| 4.8995E-60 | 0.25266904 | 0.266 | 0.077 | 1.1574E-55 | PDC | MRPL1 |
| 5.6845E-60 | 0.29351534 | 0.39 | 0.141 | 1.3428E-55 | PDC | SUCLG1 |
| 5.691E-60 | 0.2666722 | 0.275 | 0.082 | 1.3444E-55 | PDC | POP5 |
| 6.6861E-60 | 0.32523979 | 0.295 | 0.092 | 1.5795E-55 | PDC | WASHC1 |

| | | | | | | |
|---|---|---|---|---|---|---|
| 9.0687E-60 | 0.367705 | 0.536 | 0.232 | 2.1423E-55 | PDC | ATP5F1A |
| 9.6019E-60 | 0.36196027 | 0.545 | 0.234 | 2.2683E-55 | PDC | TSTD1 |
| 1.3433E-59 | 0.47689095 | 0.689 | 0.368 | 3.1732E-55 | PDC | KIF5B |
| 1.4436E-59 | 0.31781738 | 0.304 | 0.097 | 3.4103E-55 | PDC | LONP1 |
| 1.6122E-59 | -1.2978582 | 0.383 | 0.708 | 3.8084E-55 | PDC | SOD2 |
| 1.7172E-59 | 0.27010929 | 0.309 | 0.098 | 4.0566E-55 | PDC | TMX2 |
| 1.7756E-59 | 0.26630689 | 0.268 | 0.079 | 4.1946E-55 | PDC | TMEM9 |
| 1.7816E-59 | 0.28385888 | 0.322 | 0.105 | 4.2086E-55 | PDC | DDX1 |
| 1.9389E-59 | -1.2991604 | 0.135 | 0.477 | 4.5802E-55 | PDC | ERO1A |
| 2.0465E-59 | 0.36411347 | 0.493 | 0.207 | 4.8345E-55 | PDC | STRAP |
| 2.1173E-59 | 0.25416851 | 0.295 | 0.091 | 5.0016E-55 | PDC | USE1 |
| 2.577E-59 | 0.3719168 | 0.466 | 0.183 | 6.0876E-55 | PDC | PAWR |
| 3.3021E-59 | -1.682675 | 0.05 | 0.39 | 7.8006E-55 | PDC | FCER1G |
| 4.2758E-59 | 0.33206533 | 0.318 | 0.105 | 1.0101E-54 | PDC | IGF1R |
| 4.7894E-59 | 0.34589869 | 0.414 | 0.16 | 1.1314E-54 | PDC | SMIM7 |
| 5.383E-59 | 0.50250115 | 0.876 | 0.644 | 1.2716E-54 | PDC | ATP5MC2 |
| 5.547E-59 | 0.60432686 | 0.906 | 0.683 | 1.3104E-54 | PDC | NPM1 |
| 6.5454E-59 | 0.35949045 | 0.397 | 0.148 | 1.5462E-54 | PDC | EEA1 |
| 6.7733E-59 | 0.26805359 | 0.277 | 0.083 | 1.6001E-54 | PDC | ZC3H14 |
| 8.5284E-59 | 0.26063443 | 0.279 | 0.084 | 2.0147E-54 | PDC | LYPLAL1 |
| 9.356E-59 | 0.36306917 | 0.394 | 0.148 | 2.2102E-54 | PDC | ECI1 |
| 1.0229E-58 | 0.68097846 | 0.982 | 0.943 | 2.4165E-54 | PDC | RPS27 |
| 1.2122E-58 | 0.29317581 | 0.349 | 0.117 | 2.8636E-54 | PDC | CTNNAL1 |
| 1.9569E-58 | 0.27363907 | 0.27 | 0.08 | 4.6228E-54 | PDC | CHID1 |
| 2.4448E-58 | 0.76241477 | 0.646 | 0.319 | 5.7754E-54 | PDC | VEGFA |
| 3.1034E-58 | 0.36318079 | 0.532 | 0.236 | 7.3313E-54 | PDC | TXNL1 |
| 4.1503E-58 | -2.486745 | 0.081 | 0.412 | 9.8043E-54 | PDC | S100P |
| 5.8675E-58 | 0.40276229 | 0.59 | 0.275 | 1.3861E-53 | PDC | MRPL41 |
| 6.8023E-58 | 0.26437106 | 0.3 | 0.095 | 1.6069E-53 | PDC | IFT57 |
| 7.2061E-58 | 0.25431046 | 0.223 | 0.059 | 1.7023E-53 | PDC | GAS6 |
| 7.895E-58 | 0.29978176 | 0.327 | 0.11 | 1.865E-53 | PDC | CISD1 |
| 1.0078E-57 | 0.28018562 | 0.322 | 0.107 | 2.3806E-53 | PDC | CCDC25 |
| 1.5653E-57 | 0.44013661 | 0.585 | 0.283 | 3.6977E-53 | PDC | NDUFA11 |
| 1.5877E-57 | 0.26599542 | 0.257 | 0.075 | 3.7505E-53 | PDC | CEP63 |
| 1.7084E-57 | 0.36662874 | 0.482 | 0.201 | 4.0358E-53 | PDC | NENF |
| 1.7905E-57 | 0.28048782 | 0.353 | 0.123 | 4.2297E-53 | PDC | NDUFS3 |
| 1.9635E-57 | 0.37626251 | 0.509 | 0.222 | 4.6383E-53 | PDC | COX20 |
| 2.5489E-57 | 0.47056902 | 0.703 | 0.36 | 6.0213E-53 | PDC | ATP5F1D |
| 2.6952E-57 | -0.8619209 | 0.201 | 0.543 | 6.3668E-53 | PDC | UPP1 |
| 2.9624E-57 | 0.36290492 | 0.48 | 0.204 | 6.9981E-53 | PDC | SIVA1 |
| 3.1399E-57 | 0.27862357 | 0.342 | 0.117 | 7.4173E-53 | PDC | MRPL42 |
| 3.414E-57 | 0.33027103 | 0.419 | 0.164 | 8.0649E-53 | PDC | EMC4 |
| 3.6177E-57 | 0.35994677 | 0.441 | 0.179 | 8.5462E-53 | PDC | EMD |
| 3.6303E-57 | 0.40395389 | 0.728 | 0.381 | 8.5757E-53 | PDC | TBCA |
| 4.1042E-57 | 0.56167971 | 0.433 | 0.182 | 9.6953E-53 | PDC | HIPK2 |
| 4.1456E-57 | 0.25500244 | 0.236 | 0.064 | 9.7931E-53 | PDC | PCLO |
| 4.5929E-57 | 0.34990465 | 0.489 | 0.208 | 1.085E-52 | PDC | RTF2 |
| 4.9645E-57 | 0.32881955 | 0.371 | 0.137 | 1.1728E-52 | PDC | RPA3 |
| 5.8254E-57 | -1.3569879 | 0.16 | 0.498 | 1.3761E-52 | PDC | KRT8 |
| 6.3305E-57 | 0.30946014 | 0.331 | 0.113 | 1.4955E-52 | PDC | BOLA3 |

| | | | | | | |
|---|---|---|---|---|---|---|
| 6.645E-57 | 0.30724082 | 0.406 | 0.154 | 1.5698E-52 | PDC | SUPT16H |
| 6.8661E-57 | 0.29997892 | 0.34 | 0.118 | 1.622E-52 | PDC | TUSC2 |
| 7.6823E-57 | 0.26279131 | 0.275 | 0.084 | 1.8148E-52 | PDC | TBC1D5 |
| 8.8329E-57 | 0.27058328 | 0.272 | 0.082 | 2.0866E-52 | PDC | TMCC3 |
| 8.9251E-57 | 0.30816203 | 0.397 | 0.15 | 2.1084E-52 | PDC | EIF2A |
| 9.6431E-57 | 0.30972518 | 0.327 | 0.112 | 2.278E-52 | PDC | SMIM19 |
| 1.6652E-56 | 0.58693023 | 0.933 | 0.753 | 3.9337E-52 | PDC | RPSA |
| 1.8556E-56 | -0.8241149 | 0.75 | 0.94 | 4.3836E-52 | PDC | MT-ATP6 |
| 2.2097E-56 | 0.30805731 | 0.405 | 0.154 | 5.2201E-52 | PDC | FUNDC2 |
| 2.2225E-56 | 0.31277139 | 0.399 | 0.152 | 5.2501E-52 | PDC | MRPS16 |
| 2.3516E-56 | 0.37713772 | 0.673 | 0.323 | 5.5553E-52 | PDC | LMNA |
| 2.4521E-56 | 0.38800816 | 0.554 | 0.26 | 5.7925E-52 | PDC | RTRAF |
| 2.7506E-56 | -0.7887696 | 0.149 | 0.49 | 6.4978E-52 | PDC | HMGA1 |
| 2.8449E-56 | -0.7781596 | 0.086 | 0.422 | 6.7205E-52 | PDC | JPT1 |
| 2.933E-56 | 0.31119542 | 0.329 | 0.112 | 6.9285E-52 | PDC | YAP1 |
| 3.1054E-56 | 0.46338787 | 0.658 | 0.346 | 7.3358E-52 | PDC | TMEM258 |
| 3.1364E-56 | 0.36601333 | 0.54 | 0.243 | 7.4092E-52 | PDC | CCT4 |
| 3.3583E-56 | 0.34835469 | 0.394 | 0.151 | 7.9333E-52 | PDC | TARS |
| 3.3644E-56 | 0.41229965 | 0.59 | 0.285 | 7.9478E-52 | PDC | NDUFB9 |
| 3.7333E-56 | 0.2952332 | 0.365 | 0.132 | 8.8193E-52 | PDC | ERGIC2 |
| 4.3282E-56 | 0.26338427 | 0.268 | 0.081 | 1.0225E-51 | PDC | ATN1 |
| 5.7485E-56 | 0.26348114 | 0.3 | 0.097 | 1.358E-51 | PDC | PLPBP |
| 6.3422E-56 | 0.27573322 | 0.293 | 0.094 | 1.4982E-51 | PDC | BLVRA |
| 6.7375E-56 | 0.2548608 | 0.257 | 0.076 | 1.5916E-51 | PDC | MUM1 |
| 7.9276E-56 | 0.31235471 | 0.365 | 0.134 | 1.8727E-51 | PDC | MRPL21 |
| 8.7047E-56 | 0.43812471 | 0.592 | 0.289 | 2.0563E-51 | PDC | HDLBP |
| 1.1435E-55 | -0.8951616 | 0.263 | 0.591 | 2.7012E-51 | PDC | KLF6 |
| 1.2543E-55 | 0.2962317 | 0.25 | 0.074 | 2.9631E-51 | PDC | TSPAN5 |
| 1.3204E-55 | 0.25999901 | 0.284 | 0.089 | 3.1192E-51 | PDC | SSRP1 |
| 1.4864E-55 | -2.4641929 | 0.122 | 0.457 | 3.5113E-51 | PDC | SPP1 |
| 1.5389E-55 | -0.624109 | 0.342 | 0.703 | 3.6354E-51 | PDC | JUNB |
| 1.6616E-55 | 0.4275507 | 0.523 | 0.24 | 3.9251E-51 | PDC | OSTC |
| 2.0211E-55 | 0.4163254 | 0.685 | 0.359 | 4.7746E-51 | PDC | C19orf53 |
| 2.3415E-55 | 0.3326346 | 0.39 | 0.151 | 5.5314E-51 | PDC | TMED7 |
| 2.3619E-55 | -1.2531282 | 0.018 | 0.339 | 5.5794E-51 | PDC | LAPTM5 |
| 3.532E-55 | 0.35566 | 0.354 | 0.13 | 8.3436E-51 | PDC | CCDC6 |
| 6.3177E-55 | -4.3649077 | 0.085 | 0.405 | 1.4924E-50 | PDC | SPRR3 |
| 6.4317E-55 | 0.25792881 | 0.281 | 0.088 | 1.5194E-50 | PDC | CREB3 |
| 6.5154E-55 | 0.28545896 | 0.288 | 0.093 | 1.5391E-50 | PDC | SDSL |
| 7.8094E-55 | 0.39829375 | 0.5 | 0.223 | 1.8448E-50 | PDC | MRPL52 |
| 8.0132E-55 | 0.3082414 | 0.293 | 0.096 | 1.893E-50 | PDC | DLGAP1-AS1 |
| 8.3629E-55 | 0.31511707 | 0.356 | 0.13 | 1.9756E-50 | PDC | UFL1 |
| 9.1135E-55 | 0.70530523 | 0.957 | 0.927 | 2.1529E-50 | PDC | RPL30 |
| 9.1436E-55 | 0.25553142 | 0.302 | 0.097 | 2.16E-50 | PDC | APOL1 |
| 9.6692E-55 | 0.25391991 | 0.306 | 0.101 | 2.2842E-50 | PDC | POLR2C |
| 1.1367E-54 | 0.26187738 | 0.29 | 0.092 | 2.6852E-50 | PDC | ZBTB10 |
| 1.1798E-54 | 0.42860379 | 0.372 | 0.145 | 2.787E-50 | PDC | SUN2 |
| 1.2105E-54 | 0.27171243 | 0.266 | 0.081 | 2.8596E-50 | PDC | SRSF8 |
| 1.8072E-54 | 0.27885157 | 0.317 | 0.108 | 4.2692E-50 | PDC | RETREG2 |
| 1.8968E-54 | 0.25814455 | 0.297 | 0.097 | 4.4808E-50 | PDC | IMP4 |

| | | | | | | |
|---|---|---|---|---|---|---|
| 2.1995E-54 | 0.25812032 | 0.275 | 0.085 | 5.196E-50 | PDC | ASNS |
| 2.9118E-54 | 0.29853662 | 0.333 | 0.118 | 6.8785E-50 | PDC | TIMM17B |
| 3.065E-54 | 0.26097788 | 0.272 | 0.085 | 7.2404E-50 | PDC | TIMM9 |
| 3.4076E-54 | 0.37577792 | 0.612 | 0.296 | 8.0498E-50 | PDC | SF3B5 |
| 3.5253E-54 | -1.8887754 | 0.074 | 0.401 | 8.3278E-50 | PDC | HLA-DRA |
| 3.9736E-54 | 0.26900477 | 0.311 | 0.105 | 9.3868E-50 | PDC | COMMD1 |
| 4.2147E-54 | 0.26927102 | 0.288 | 0.093 | 9.9565E-50 | PDC | ARL14EP |
| 5.2822E-54 | 0.31604188 | 0.428 | 0.173 | 1.2478E-49 | PDC | POLR2F |
| 5.7597E-54 | 0.44118018 | 0.845 | 0.579 | 1.3606E-49 | PDC | CLTA |
| 5.7802E-54 | 0.34474577 | 0.486 | 0.212 | 1.3654E-49 | PDC | NDUFB5 |
| 5.8277E-54 | 0.27232416 | 0.326 | 0.112 | 1.3767E-49 | PDC | MGMT |
| 6.2033E-54 | 0.37486483 | 0.462 | 0.196 | 1.4654E-49 | PDC | TTC3 |
| 9.0501E-54 | 0.2584242 | 0.264 | 0.081 | 2.1379E-49 | PDC | CDC42BPA |
| 9.2546E-54 | 0.35267971 | 0.345 | 0.126 | 2.1862E-49 | PDC | UTRN |
| 1.0461E-53 | 0.46137278 | 0.838 | 0.543 | 2.4711E-49 | PDC | PPIB |
| 1.1874E-53 | -0.7724148 | 0.014 | 0.329 | 2.8051E-49 | PDC | ARHGDIB |
| 1.3609E-53 | 0.30908681 | 0.345 | 0.124 | 3.2148E-49 | PDC | UPF3A |
| 1.6448E-53 | 0.43659674 | 0.367 | 0.139 | 3.8855E-49 | PDC | SERPING1 |
| 1.6774E-53 | 0.32354439 | 0.367 | 0.138 | 3.9624E-49 | PDC | CCDC90B |
| 1.8481E-53 | 0.28523652 | 0.356 | 0.13 | 4.3657E-49 | PDC | IAH1 |
| 1.9081E-53 | 0.39184508 | 0.577 | 0.278 | 4.5075E-49 | PDC | LRPAP1 |
| 1.9354E-53 | 0.53566893 | 0.835 | 0.608 | 4.5721E-49 | PDC | SEC61B |
| 2.5158E-53 | 0.42430901 | 0.701 | 0.385 | 5.943E-49 | PDC | SPCS2 |
| 2.8464E-53 | 0.37655386 | 0.486 | 0.218 | 6.724E-49 | PDC | TMEM14B |
| 2.8718E-53 | -0.6394246 | 0.608 | 0.834 | 6.7842E-49 | PDC | YWHAZ |
| 2.9215E-53 | 0.26375902 | 0.282 | 0.092 | 6.9014E-49 | PDC | STARD3NL |
| 3.0785E-53 | 0.26510542 | 0.304 | 0.102 | 7.2723E-49 | PDC | TSN |
| 4.0782E-53 | 0.33289714 | 0.415 | 0.168 | 9.634E-49 | PDC | PHB2 |
| 5.7102E-53 | 0.27785009 | 0.356 | 0.131 | 1.3489E-48 | PDC | UBA2 |
| 6.1044E-53 | 0.30525206 | 0.335 | 0.121 | 1.442E-48 | PDC | TMEM9B |
| 6.1513E-53 | -3.0450873 | 0.072 | 0.382 | 1.4531E-48 | PDC | KRT13 |
| 6.6449E-53 | -0.5964743 | 0.315 | 0.682 | 1.5697E-48 | PDC | IER2 |
| 7.3847E-53 | -1.5641031 | 0.027 | 0.337 | 1.7445E-48 | PDC | IL1RN |
| 8.2094E-53 | 0.36095996 | 0.417 | 0.172 | 1.9393E-48 | PDC | IFI27L2 |
| 8.6717E-53 | 0.25692781 | 0.263 | 0.081 | 2.0485E-48 | PDC | RNASEH2B |
| 1.1315E-52 | 0.26920709 | 0.333 | 0.118 | 2.6729E-48 | PDC | YIF1A |
| 1.1378E-52 | 0.2806603 | 0.365 | 0.135 | 2.6879E-48 | PDC | PRDX4 |
| 1.2626E-52 | -1.0970057 | 0.219 | 0.531 | 2.9827E-48 | PDC | ZFP36 |
| 1.3985E-52 | 0.60437774 | 0.986 | 0.923 | 3.3036E-48 | PDC | RPS27A |
| 1.6428E-52 | 0.30741679 | 0.39 | 0.152 | 3.8808E-48 | PDC | MEAF6 |
| 2.216E-52 | -1.0212821 | 0.493 | 0.787 | 5.2349E-48 | PDC | LGALS3 |
| 2.3009E-52 | 0.39405472 | 0.608 | 0.304 | 5.4354E-48 | PDC | SSBP1 |
| 2.4995E-52 | -0.6962257 | 0.052 | 0.371 | 5.9045E-48 | PDC | CTSH |
| 2.5687E-52 | 0.36719197 | 0.559 | 0.262 | 6.0681E-48 | PDC | XBP1 |
| 2.7706E-52 | 0.29116413 | 0.311 | 0.108 | 6.545E-48 | PDC | MID1IP1 |
| 2.8298E-52 | 0.33229311 | 0.286 | 0.094 | 6.6848E-48 | PDC | UACA |
| 3.5108E-52 | 0.41157116 | 0.759 | 0.45 | 8.2936E-48 | PDC | ST13 |
| 5.7213E-52 | -0.7737093 | 0.556 | 0.778 | 1.3515E-47 | PDC | TPI1 |
| 7.8777E-52 | 0.39619904 | 0.579 | 0.272 | 1.8609E-47 | PDC | SLK |
| 8.1613E-52 | 0.33350813 | 0.459 | 0.198 | 1.9279E-47 | PDC | VPS26A |

| | | | | | | |
|---|---|---|---|---|---|---|
| 8.2424E-52 | 0.35570274 | 0.66 | 0.326 | 1.9471E-47 | PDC | FAM96B |
| 8.3922E-52 | 0.38436705 | 0.95 | 0.831 | 1.9825E-47 | PDC | PPDPF |
| 8.973E-52 | 0.26436358 | 0.317 | 0.11 | 2.1197E-47 | PDC | GLOD4 |
| 1.139E-51 | 0.41908474 | 0.665 | 0.359 | 2.6908E-47 | PDC | PSMB1 |
| 1.2367E-51 | 0.36885926 | 0.423 | 0.176 | 2.9215E-47 | PDC | PHACTR4 |
| 1.4537E-51 | 0.26088336 | 0.288 | 0.096 | 3.434E-47 | PDC | TSR2 |
| 1.8983E-51 | 0.25944016 | 0.324 | 0.114 | 4.4843E-47 | PDC | ACOT13 |
| 1.9574E-51 | 0.35680065 | 0.523 | 0.242 | 4.624E-47 | PDC | VPS35 |
| 4.5993E-51 | 0.2703729 | 0.335 | 0.121 | 1.0865E-46 | PDC | CYHR1 |
| 6.2896E-51 | 0.25988967 | 0.308 | 0.106 | 1.4858E-46 | PDC | METTL23 |
| 7.5219E-51 | -1.0688134 | 0.049 | 0.355 | 1.7769E-46 | PDC | PHLDA2 |
| 7.5915E-51 | -1.4229367 | 0.092 | 0.409 | 1.7933E-46 | PDC | SLPI |
| 8.229E-51 | 0.31535669 | 0.459 | 0.196 | 1.9439E-46 | PDC | MDH1 |
| 1.0501E-50 | -1.6859584 | 0.067 | 0.373 | 2.4807E-46 | PDC | MAL |
| 1.2229E-50 | -0.6007184 | 0.532 | 0.773 | 2.8889E-46 | PDC | CDC42 |
| 1.2304E-50 | 0.29447086 | 0.406 | 0.163 | 2.9067E-46 | PDC | GLO1 |
| 1.2335E-50 | 0.2876272 | 0.304 | 0.105 | 2.9139E-46 | PDC | JPX |
| 1.3439E-50 | 0.25824265 | 0.297 | 0.101 | 3.1747E-46 | PDC | CEBPZOS |
| 1.4092E-50 | 0.30516784 | 0.381 | 0.15 | 3.329E-46 | PDC | SPAG7 |
| 1.4309E-50 | 0.40729729 | 0.563 | 0.274 | 3.3803E-46 | PDC | MYDGF |
| 1.831E-50 | -0.7689178 | 0.921 | 0.958 | 4.3253E-46 | PDC | H3F3A |
| 1.9112E-50 | 0.33784216 | 0.486 | 0.214 | 4.5149E-46 | PDC | CDK2AP2 |
| 1.9433E-50 | 0.28644606 | 0.315 | 0.112 | 4.5907E-46 | PDC | PSMC2 |
| 1.9819E-50 | 0.25586667 | 0.281 | 0.093 | 4.6819E-46 | PDC | TADA3 |
| 2.0652E-50 | 0.50385252 | 0.865 | 0.626 | 4.8785E-46 | PDC | YBX3 |
| 2.2018E-50 | 0.36440279 | 0.691 | 0.37 | 5.2014E-46 | PDC | LAMTOR5 |
| 2.9926E-50 | 0.41729488 | 0.845 | 0.597 | 7.0693E-46 | PDC | YWHAE |
| 4.0274E-50 | 0.26659676 | 0.284 | 0.096 | 9.5138E-46 | PDC | LZIC |
| 5.5846E-50 | 0.30162546 | 0.29 | 0.099 | 1.3192E-45 | PDC | ARG2 |
| 7.7159E-50 | -0.6583824 | 0.034 | 0.334 | 1.8227E-45 | PDC | RBM47 |
| 7.8894E-50 | 0.30966381 | 0.374 | 0.145 | 1.8637E-45 | PDC | SUCO |
| 9.2071E-50 | 0.30083699 | 0.363 | 0.141 | 2.175E-45 | PDC | SLC39A1 |
| 1.0978E-49 | 0.3664 | 0.475 | 0.211 | 2.5933E-45 | PDC | MKNK2 |
| 1.1176E-49 | 0.57197655 | 0.973 | 0.904 | 2.64E-45 | PDC | RPL7A |
| 1.1937E-49 | 0.31755426 | 0.394 | 0.159 | 2.8198E-45 | PDC | GLG1 |
| 1.7456E-49 | 0.29082945 | 0.379 | 0.149 | 4.1237E-45 | PDC | BCAS2 |
| 1.7837E-49 | 0.38336598 | 0.59 | 0.292 | 4.2136E-45 | PDC | KDELR1 |
| 1.8501E-49 | -0.7555445 | 0.219 | 0.517 | 4.3705E-45 | PDC | ANXA11 |
| 2.1833E-49 | 0.33177509 | 0.383 | 0.153 | 5.1576E-45 | PDC | LARP1 |
| 2.301E-49 | 0.64933923 | 0.977 | 0.922 | 5.4356E-45 | PDC | RPL8 |
| 2.4418E-49 | 0.25637777 | 0.243 | 0.075 | 5.7682E-45 | PDC | AARS |
| 2.6139E-49 | 0.27224789 | 0.417 | 0.169 | 6.1748E-45 | PDC | MPC2 |
| 2.7474E-49 | -0.8004353 | 0.317 | 0.606 | 6.4901E-45 | PDC | MCL1 |
| 4.0168E-49 | 0.34406831 | 0.511 | 0.237 | 9.4888E-45 | PDC | POLR2K |
| 4.665E-49 | -2.3302093 | 0.052 | 0.345 | 1.102E-44 | PDC | PSCA |
| 4.8926E-49 | 0.30640239 | 0.412 | 0.17 | 1.1558E-44 | PDC | RPL7L1 |
| 5.8218E-49 | 0.36279489 | 0.619 | 0.322 | 1.3753E-44 | PDC | SRP9 |
| 5.9029E-49 | 0.50594086 | 0.968 | 0.901 | 1.3944E-44 | PDC | RPL24 |
| 7.6534E-49 | 0.30456429 | 0.656 | 0.32 | 1.808E-44 | PDC | MGST3 |
| 7.7932E-49 | 0.43466955 | 0.806 | 0.529 | 1.841E-44 | PDC | COX6C |

| | | | | | | |
|---|---|---|---|---|---|---|
| 8.6643E-49 | 0.37476659 | 0.577 | 0.285 | 2.0468E-44 | PDC | APRT |
| 9.3211E-49 | 0.33900359 | 0.442 | 0.194 | 2.2019E-44 | PDC | VTI1B |
| 1.7931E-48 | 0.51250029 | 0.955 | 0.919 | 4.2358E-44 | PDC | NACA |
| 1.9046E-48 | 0.30845163 | 0.459 | 0.198 | 4.4993E-44 | PDC | PHB |
| 2.0207E-48 | 0.74233686 | 0.61 | 0.341 | 4.7736E-44 | PDC | GDF15 |
| 2.2773E-48 | 0.30670372 | 0.396 | 0.16 | 5.3797E-44 | PDC | CDKN1B |
| 2.5648E-48 | 0.35960254 | 0.486 | 0.226 | 6.0588E-44 | PDC | SAT2 |
| 3.0169E-48 | 0.27214146 | 0.338 | 0.127 | 7.1268E-44 | PDC | FBXO7 |
| 3.4378E-48 | 0.34470944 | 0.545 | 0.258 | 8.1212E-44 | PDC | MPHOSPH8 |
| 5.4446E-48 | 0.31839744 | 0.268 | 0.089 | 1.2862E-43 | PDC | SEMA3B |
| 1.111E-47 | 0.56876873 | 0.54 | 0.266 | 2.6246E-43 | PDC | TSC22D3 |
| 1.1588E-47 | 0.39008916 | 0.712 | 0.391 | 2.7375E-43 | PDC | COMMD6 |
| 1.1691E-47 | 0.25142097 | 0.261 | 0.085 | 2.7617E-43 | PDC | SRPRB |
| 1.5874E-47 | 0.3670534 | 0.543 | 0.27 | 3.7499E-43 | PDC | LAMP2 |
| 1.6662E-47 | 0.28447676 | 0.336 | 0.126 | 3.936E-43 | PDC | CBR1 |
| 2.0539E-47 | 0.27962137 | 0.293 | 0.103 | 4.8519E-43 | PDC | CREB3L2 |
| 3.0992E-47 | 0.28502679 | 0.338 | 0.129 | 7.3212E-43 | PDC | AP3S1 |
| 3.5205E-47 | 0.28013316 | 0.345 | 0.133 | 8.3165E-43 | PDC | MED4 |
| 3.579E-47 | 0.26147917 | 0.281 | 0.097 | 8.4546E-43 | PDC | MED21 |
| 4.3224E-47 | 0.30812474 | 0.381 | 0.157 | 1.0211E-42 | PDC | NDUFB6 |
| 4.767E-47 | -0.6708618 | 0.077 | 0.378 | 1.1261E-42 | PDC | METRNL |
| 4.9004E-47 | 0.33263839 | 0.498 | 0.233 | 1.1576E-42 | PDC | NDUFA13 |
| 5.648E-47 | 0.30180068 | 0.372 | 0.15 | 1.3342E-42 | PDC | PNKD |
| 5.8565E-47 | 0.25236923 | 0.279 | 0.096 | 1.3835E-42 | PDC | B3GAT3 |
| 6.2602E-47 | 0.33984049 | 0.446 | 0.191 | 1.4788E-42 | PDC | SDF2L1 |
| 6.6824E-47 | 0.30445072 | 0.369 | 0.146 | 1.5786E-42 | PDC | NUB1 |
| 8.159E-47 | 0.29370943 | 0.45 | 0.196 | 1.9274E-42 | PDC | TIMM13 |
| 8.6525E-47 | 0.31360491 | 0.525 | 0.246 | 2.044E-42 | PDC | DNAJC3 |
| 1.282E-46 | 0.28565381 | 0.399 | 0.164 | 3.0284E-42 | PDC | MRPS34 |
| 1.3034E-46 | 0.4258012 | 0.484 | 0.231 | 3.079E-42 | PDC | RBMS1 |
| 1.5757E-46 | -0.8549396 | 0.02 | 0.306 | 3.7222E-42 | PDC | SERPINA1 |
| 1.6325E-46 | -0.579921 | 0.295 | 0.641 | 3.8563E-42 | PDC | GADD45B |
| 1.9918E-46 | -0.9586581 | 0.013 | 0.295 | 4.7052E-42 | PDC | LCP1 |
| 2.3694E-46 | -0.9931342 | 0.311 | 0.605 | 5.5971E-42 | PDC | NFKBIA |
| 3.0554E-46 | 0.53700511 | 0.791 | 0.566 | 7.2178E-42 | PDC | RPS27L |
| 4.4537E-46 | -0.653961 | 0.36 | 0.628 | 1.0521E-41 | PDC | ACTR2 |
| 4.5611E-46 | -0.9770109 | 0.588 | 0.766 | 1.0775E-41 | PDC | BTG1 |
| 4.7266E-46 | -0.6595269 | 0.038 | 0.323 | 1.1166E-41 | PDC | SLC16A3 |
| 4.7334E-46 | 0.62580722 | 0.966 | 0.87 | 1.1182E-41 | PDC | RPL14 |
| 4.8612E-46 | 0.29743515 | 0.379 | 0.155 | 1.1484E-41 | PDC | DDAH2 |
| 5.9182E-46 | 0.43990283 | 0.928 | 0.76 | 1.3981E-41 | PDC | COX7C |
| 6.2357E-46 | -1.1233458 | 0.032 | 0.315 | 1.4731E-41 | PDC | EMP1 |
| 9.2239E-46 | 0.31440735 | 0.414 | 0.177 | 2.179E-41 | PDC | WARS |
| 9.6606E-46 | 0.30575494 | 0.435 | 0.193 | 2.2821E-41 | PDC | CHMP4A |
| 1.006E-45 | 0.31212574 | 0.379 | 0.158 | 2.3765E-41 | PDC | GINM1 |
| 1.0075E-45 | 0.25518682 | 0.308 | 0.113 | 2.38E-41 | PDC | SCFD1 |
| 1.0103E-45 | 0.26187974 | 0.3 | 0.109 | 2.3867E-41 | PDC | RFK |
| 1.1948E-45 | 0.39513929 | 0.73 | 0.423 | 2.8226E-41 | PDC | SEC62 |
| 1.2581E-45 | 0.34555551 | 0.52 | 0.25 | 2.972E-41 | PDC | PA2G4 |
| 1.2761E-45 | 0.28801003 | 0.403 | 0.17 | 3.0145E-41 | PDC | PSMD1 |

| | | | | | | |
|---|---|---|---|---|---|---|
| 1.7177E-45 | 0.31307041 | 0.414 | 0.179 | 4.0577E-41 | PDC | CCT7 |
| 1.7242E-45 | 0.27070219 | 0.183 | 0.05 | 4.073E-41 | PDC | CSF1 |
| 1.7658E-45 | -0.9232772 | 0.173 | 0.46 | 4.1714E-41 | PDC | RAB11FIP1 |
| 2.084E-45 | 0.60459326 | 0.966 | 0.888 | 4.9231E-41 | PDC | RPS23 |
| 2.4183E-45 | 0.38626386 | 0.68 | 0.371 | 5.7128E-41 | PDC | ATP5ME |
| 3.071E-45 | 0.33303737 | 0.647 | 0.341 | 7.2545E-41 | PDC | LAMTOR4 |
| 3.0854E-45 | -1.0191773 | 0.82 | 0.921 | 7.2886E-41 | PDC | S100A10 |
| 3.4527E-45 | 0.30684527 | 0.435 | 0.192 | 8.1564E-41 | PDC | SNRPD1 |
| 4.4008E-45 | -0.7189187 | 0.032 | 0.315 | 1.0396E-40 | PDC | CD68 |
| 4.4305E-45 | 0.36052541 | 0.667 | 0.363 | 1.0466E-40 | PDC | WDR83OS |
| 4.8281E-45 | 0.29272308 | 0.356 | 0.144 | 1.1405E-40 | PDC | RABGGTB |
| 5.5512E-45 | 0.27031077 | 0.354 | 0.141 | 1.3114E-40 | PDC | PFDN6 |
| 8.5413E-45 | -1.1150668 | 0.013 | 0.285 | 2.0177E-40 | PDC | PTPRC |
| 1.1515E-44 | 0.33729558 | 0.489 | 0.225 | 2.7201E-40 | PDC | CYB5A |
| 1.712E-44 | 0.27102197 | 0.329 | 0.128 | 4.0444E-40 | PDC | MEA1 |
| 1.7441E-44 | 0.25849312 | 0.257 | 0.087 | 4.1202E-40 | PDC | IRF2BPL |
| 2.3256E-44 | 0.36082123 | 0.381 | 0.159 | 5.4938E-40 | PDC | ALCAM |
| 2.3791E-44 | 0.34083439 | 0.581 | 0.297 | 5.6202E-40 | PDC | ATP5PO |
| 3.1933E-44 | -0.7112469 | 0.766 | 0.946 | 7.5434E-40 | PDC | MT-CO3 |
| 3.7284E-44 | 0.29241203 | 0.362 | 0.148 | 8.8076E-40 | PDC | TMEM179B |
| 3.7557E-44 | 0.2972863 | 0.362 | 0.148 | 8.8722E-40 | PDC | BCL7C |
| 4.1683E-44 | -1.4768154 | 0.415 | 0.66 | 9.8468E-40 | PDC | HSPA1A |
| 5.1222E-44 | 0.35647902 | 0.514 | 0.25 | 1.21E-39 | PDC | C1orf122 |
| 6.1661E-44 | 0.27129967 | 0.338 | 0.134 | 1.4566E-39 | PDC | YIPF6 |
| 6.1935E-44 | 0.30865835 | 0.324 | 0.127 | 1.4631E-39 | PDC | RAB34 |
| 6.9853E-44 | 0.33164589 | 0.61 | 0.317 | 1.6501E-39 | PDC | YWHAQ |
| 7.2882E-44 | 0.4826315 | 0.993 | 0.98 | 1.7217E-39 | PDC | RPL41 |
| 8.2719E-44 | 0.25462767 | 0.232 | 0.074 | 1.9541E-39 | PDC | CGNL1 |
| 8.8119E-44 | -1.1151886 | 0.04 | 0.316 | 2.0816E-39 | PDC | PHLDA1 |
| 9.2833E-44 | 0.57788864 | 0.989 | 0.945 | 2.193E-39 | PDC | PTMA |
| 9.3239E-44 | -0.639341 | 0.052 | 0.33 | 2.2026E-39 | PDC | CAPG |
| 9.9964E-44 | 0.28224986 | 0.338 | 0.135 | 2.3615E-39 | PDC | DPM3 |
| 1.2276E-43 | 0.34209245 | 0.543 | 0.277 | 2.8999E-39 | PDC | UFC1 |
| 1.2826E-43 | 0.34221203 | 0.552 | 0.279 | 3.0299E-39 | PDC | RTN3 |
| 1.3184E-43 | 0.27683939 | 0.536 | 0.254 | 3.1144E-39 | PDC | CCT3 |
| 1.3376E-43 | -1.275434 | 0.293 | 0.561 | 3.1599E-39 | PDC | KRT18 |
| 1.5667E-43 | 0.25094629 | 0.326 | 0.126 | 3.7009E-39 | PDC | DCTN6 |
| 1.6765E-43 | 0.33220868 | 0.495 | 0.238 | 3.9603E-39 | PDC | FXR1 |
| 1.7499E-43 | 0.25468004 | 0.299 | 0.111 | 4.1338E-39 | PDC | NUDCD2 |
| 2.1884E-43 | 0.28593564 | 0.354 | 0.141 | 5.1696E-39 | PDC | SVIL |
| 2.2679E-43 | 0.25095317 | 0.322 | 0.123 | 5.3574E-39 | PDC | CDC40 |
| 2.4369E-43 | 0.28164433 | 0.39 | 0.166 | 5.7566E-39 | PDC | TTC1 |
| 2.6142E-43 | 0.41187092 | 0.619 | 0.339 | 6.1755E-39 | PDC | ATP5IF1 |
| 2.7213E-43 | 0.25031842 | 0.295 | 0.109 | 6.4284E-39 | PDC | BAD |
| 2.9131E-43 | 0.35133312 | 0.662 | 0.363 | 6.8816E-39 | PDC | REEP5 |
| 2.9813E-43 | 0.41643666 | 0.84 | 0.64 | 7.0428E-39 | PDC | EDF1 |
| 3.8982E-43 | 0.43179956 | 0.603 | 0.33 | 9.2087E-39 | PDC | SSR3 |
| 4.6472E-43 | 0.36179288 | 0.96 | 0.863 | 1.0978E-38 | PDC | SRP14 |
| 5.2118E-43 | 0.26555758 | 0.286 | 0.105 | 1.2312E-38 | PDC | SMIM29 |
| 7.6081E-43 | 0.29547718 | 0.387 | 0.167 | 1.7973E-38 | PDC | MRPS15 |

| | | | | | | |
|---|---|---|---|---|---|---|
| 9.6704E-43 | 0.30508807 | 0.401 | 0.174 | 2.2844E-38 | PDC | SENP6 |
| 1.0012E-42 | 0.41250437 | 0.423 | 0.195 | 2.3651E-38 | PDC | ITGAV |
| 1.5556E-42 | -0.6579353 | 0.025 | 0.293 | 3.6747E-38 | PDC | SPINT1 |
| 1.6283E-42 | -0.684632 | 0.022 | 0.287 | 3.8465E-38 | PDC | MARCKSL1 |
| 1.6926E-42 | 0.28811172 | 0.282 | 0.103 | 3.9983E-38 | PDC | KIAA0040 |
| 1.7902E-42 | 0.305433 | 0.331 | 0.133 | 4.2289E-38 | PDC | DLGAP4 |
| 2.3641E-42 | 0.31126392 | 0.493 | 0.236 | 5.5848E-38 | PDC | C8orf59 |
| 2.5157E-42 | -0.9562019 | 0.937 | 0.97 | 5.9429E-38 | PDC | TMSB10 |
| 2.6096E-42 | 0.65031516 | 0.558 | 0.29 | 6.1646E-38 | PDC | CHCHD10 |
| 2.6329E-42 | 0.31147952 | 0.761 | 0.458 | 6.2198E-38 | PDC | SARAF |
| 2.8579E-42 | 0.3107784 | 0.363 | 0.152 | 6.7512E-38 | PDC | MPC1 |
| 3.6667E-42 | 0.34361396 | 0.529 | 0.263 | 8.6619E-38 | PDC | ATOX1 |
| 4.1281E-42 | 0.26012186 | 0.324 | 0.127 | 9.7518E-38 | PDC | LSM10 |
| 4.8659E-42 | 0.45133309 | 0.975 | 0.886 | 1.1495E-37 | PDC | RPS3 |
| 6.5036E-42 | 0.30005412 | 0.442 | 0.203 | 1.5363E-37 | PDC | SUGT1 |
| 7.7715E-42 | 0.2990424 | 0.293 | 0.11 | 1.8359E-37 | PDC | LGALS3BP |
| 1.0463E-41 | -0.7040697 | 0.013 | 0.271 | 2.4716E-37 | PDC | CD53 |
| 1.3611E-41 | 0.32694325 | 0.435 | 0.204 | 3.2154E-37 | PDC | CWC15 |
| 1.7053E-41 | 0.38353119 | 0.594 | 0.321 | 4.0285E-37 | PDC | SERINC1 |
| 1.9467E-41 | 0.26488842 | 0.344 | 0.14 | 4.5988E-37 | PDC | PRPF6 |
| 2.3037E-41 | 0.25105805 | 0.32 | 0.125 | 5.442E-37 | PDC | SH3GLB2 |
| 2.4218E-41 | 0.31785194 | 0.592 | 0.311 | 5.7211E-37 | PDC | RBX1 |
| 2.4257E-41 | 0.3647134 | 0.691 | 0.403 | 5.7303E-37 | PDC | NDUFB4 |
| 2.4322E-41 | 0.57486048 | 0.959 | 0.874 | 5.7456E-37 | PDC | RPL26 |
| 2.5286E-41 | 0.32139598 | 0.52 | 0.263 | 5.9734E-37 | PDC | VPS29 |
| 3.3791E-41 | -0.8476293 | 0.11 | 0.388 | 7.9825E-37 | PDC | TYMP |
| 3.4884E-41 | 0.29871357 | 0.469 | 0.219 | 8.2407E-37 | PDC | DDX18 |
| 3.7669E-41 | 0.30833833 | 0.475 | 0.227 | 8.8985E-37 | PDC | ENSA |
| 4.1161E-41 | 0.26458684 | 0.32 | 0.127 | 9.7234E-37 | PDC | APOPT1 |
| 5.1125E-41 | 0.26746784 | 0.333 | 0.136 | 1.2077E-36 | PDC | ASNA1 |
| 6.1209E-41 | 0.31474387 | 0.576 | 0.299 | 1.4459E-36 | PDC | ANAPC11 |
| 6.4945E-41 | 0.34671397 | 0.523 | 0.255 | 1.5342E-36 | PDC | WASL |
| 8.3995E-41 | 0.28538431 | 0.475 | 0.224 | 1.9842E-36 | PDC | DHX36 |
| 8.6894E-41 | 0.28514066 | 0.378 | 0.163 | 2.0527E-36 | PDC | CAMLG |
| 1.1009E-40 | 0.43206371 | 0.905 | 0.811 | 2.6006E-36 | PDC | RPL36AL |
| 1.1372E-40 | 0.54865494 | 0.975 | 0.915 | 2.6865E-36 | PDC | RPL32 |
| 1.4471E-40 | 0.25429991 | 0.327 | 0.131 | 3.4185E-36 | PDC | EI24 |
| 1.4538E-40 | 0.52567013 | 0.969 | 0.896 | 3.4342E-36 | PDC | RPS3A |
| 1.5731E-40 | 0.48236646 | 0.556 | 0.301 | 3.7162E-36 | PDC | RPL36A |
| 2.1664E-40 | -0.7258173 | 0.036 | 0.298 | 5.1177E-36 | PDC | GLRX |
| 3.3095E-40 | 0.29521218 | 0.297 | 0.114 | 7.8181E-36 | PDC | VASN |
| 3.6062E-40 | 0.27892529 | 0.466 | 0.222 | 8.5188E-36 | PDC | PSMC1 |
| 4.0778E-40 | 0.29291354 | 0.329 | 0.134 | 9.6329E-36 | PDC | CCDC85B |
| 4.5834E-40 | 0.34421104 | 0.509 | 0.26 | 1.0827E-35 | PDC | NDUFA6 |
| 5.9015E-40 | 0.27238793 | 0.378 | 0.164 | 1.3941E-35 | PDC | CCDC59 |
| 6.9341E-40 | 0.30812159 | 0.504 | 0.253 | 1.638E-35 | PDC | DNAJC8 |
| 8.0169E-40 | 0.49108296 | 0.959 | 0.847 | 1.8938E-35 | PDC | RPS4X |
| 9.4972E-40 | 0.3034419 | 0.504 | 0.251 | 2.2435E-35 | PDC | CCT6A |
| 1.1428E-39 | 0.29012768 | 0.426 | 0.196 | 2.6996E-35 | PDC | ETFB |
| 1.4244E-39 | 0.33042981 | 0.534 | 0.281 | 3.365E-35 | PDC | RNF181 |

| | | | | | | |
|---|---|---|---|---|---|---|
| 1.4884E-39 | 0.27782507 | 0.326 | 0.133 | 3.516E-35 | PDC | OSTM1 |
| 1.5624E-39 | 0.54481361 | 0.953 | 0.883 | 3.6909E-35 | PDC | RPS13 |
| 1.7394E-39 | 0.27242825 | 0.362 | 0.155 | 4.1091E-35 | PDC | TMEM205 |
| 1.8942E-39 | 0.31676904 | 0.559 | 0.292 | 4.4746E-35 | PDC | VPS28 |
| 2.2402E-39 | -2.4132228 | 0.085 | 0.342 | 5.292E-35 | PDC | MMP7 |
| 2.6235E-39 | 0.29589336 | 0.507 | 0.254 | 6.1975E-35 | PDC | MORF4L2 |
| 4.1099E-39 | 0.36931154 | 0.504 | 0.253 | 9.7087E-35 | PDC | PDLIM5 |
| 4.232E-39 | 0.42963563 | 0.885 | 0.702 | 9.9972E-35 | PDC | RPL4 |
| 4.7535E-39 | 0.2589943 | 0.412 | 0.186 | 1.1229E-34 | PDC | PRDX3 |
| 4.9544E-39 | 0.29272897 | 0.403 | 0.184 | 1.1704E-34 | PDC | GTF3A |
| 6.1109E-39 | 0.35388271 | 0.752 | 0.49 | 1.4436E-34 | PDC | H2AFJ |
| 6.9391E-39 | 0.42360508 | 0.275 | 0.107 | 1.6392E-34 | PDC | ITM2C |
| 7.043E-39 | -1.1262083 | 0.034 | 0.282 | 1.6638E-34 | PDC | DUSP5 |
| 7.5413E-39 | 0.26491999 | 0.394 | 0.176 | 1.7815E-34 | PDC | CALCOCO2 |
| 8.2135E-39 | -0.6620459 | 0.3 | 0.551 | 1.9403E-34 | PDC | TOP1 |
| 9.3095E-39 | -0.750445 | 0.007 | 0.249 | 2.1992E-34 | PDC | CSTA |
| 9.7956E-39 | 0.26159095 | 0.293 | 0.115 | 2.314E-34 | PDC | CLSTN1 |
| 1.0368E-38 | 0.32215271 | 0.308 | 0.121 | 2.4493E-34 | PDC | FAT1 |
| 1.0838E-38 | -0.5753336 | 0.034 | 0.283 | 2.5603E-34 | PDC | LYN |
| 1.3746E-38 | -0.6123143 | 0.665 | 0.801 | 3.2471E-34 | PDC | ARPC2 |
| 1.4112E-38 | 0.33248054 | 0.468 | 0.232 | 3.3336E-34 | PDC | BANF1 |
| 1.4481E-38 | 0.44670012 | 0.982 | 0.957 | 3.4209E-34 | PDC | RPL15 |
| 1.5225E-38 | -0.7846595 | 0.113 | 0.371 | 3.5965E-34 | PDC | COTL1 |
| 1.6803E-38 | 0.29690427 | 0.518 | 0.264 | 3.9694E-34 | PDC | KHDRBS1 |
| 2.7258E-38 | 0.28460884 | 0.439 | 0.209 | 6.4391E-34 | PDC | ARPC1A |
| 4.2523E-38 | 0.25564693 | 0.349 | 0.15 | 1.0045E-33 | PDC | TXNDC12 |
| 5.7093E-38 | 0.35973231 | 0.766 | 0.499 | 1.3487E-33 | PDC | LAPTM4A |
| 5.8389E-38 | 0.3117613 | 0.52 | 0.269 | 1.3793E-33 | PDC | UXT |
| 6.9418E-38 | 0.43113869 | 0.881 | 0.687 | 1.6399E-33 | PDC | EEF2 |
| 9.1519E-38 | 0.50753044 | 0.583 | 0.323 | 2.162E-33 | PDC | CEBPD |
| 1.0002E-37 | 0.32498624 | 0.59 | 0.324 | 2.3628E-33 | PDC | RNF7 |
| 1.0044E-37 | -0.8970266 | 0.604 | 0.754 | 2.3727E-33 | PDC | TPM4 |
| 1.0292E-37 | 0.25126172 | 0.282 | 0.109 | 2.4312E-33 | PDC | MRPS35 |
| 1.0347E-37 | 0.2830101 | 0.394 | 0.18 | 2.4442E-33 | PDC | POLE4 |
| 1.1508E-37 | -0.577756 | 0.05 | 0.299 | 2.7186E-33 | PDC | IFNGR2 |
| 1.406E-37 | 0.42094194 | 0.962 | 0.845 | 3.3215E-33 | PDC | RPL35 |
| 1.4534E-37 | -0.8498835 | 0.387 | 0.634 | 3.4334E-33 | PDC | ZFP36L1 |
| 1.4694E-37 | 0.53012788 | 0.973 | 0.946 | 3.4711E-33 | PDC | RPS16 |
| 1.6435E-37 | 0.39419563 | 0.727 | 0.475 | 3.8824E-33 | PDC | HSPD1 |
| 1.669E-37 | -0.6903057 | 0.052 | 0.302 | 3.9426E-33 | PDC | FAM129B |
| 1.6773E-37 | -1.306067 | 0.036 | 0.28 | 3.9623E-33 | PDC | HLA-DRB1 |
| 1.8708E-37 | -0.6237355 | 0.842 | 0.905 | 4.4194E-33 | PDC | CFL1 |
| 1.8809E-37 | 0.27331285 | 0.48 | 0.238 | 4.4433E-33 | PDC | PSMD4 |
| 1.9971E-37 | 0.26793388 | 0.356 | 0.156 | 4.7178E-33 | PDC | MTIF3 |
| 2.0281E-37 | 0.26431619 | 0.372 | 0.165 | 4.7909E-33 | PDC | CISD2 |
| 2.2639E-37 | 0.28966101 | 0.64 | 0.354 | 5.348E-33 | PDC | XRCC5 |
| 2.6696E-37 | 0.26476951 | 0.43 | 0.202 | 6.3065E-33 | PDC | CHURC1 |
| 2.8645E-37 | 0.29302496 | 0.464 | 0.224 | 6.7668E-33 | PDC | RGS10 |
| 3.0351E-37 | 0.25374351 | 0.367 | 0.158 | 7.1698E-33 | PDC | FARP1 |
| 3.9505E-37 | 0.28348048 | 0.471 | 0.231 | 9.3322E-33 | PDC | SCARB2 |

| | | | | | | |
|---|---|---|---|---|---|---|
| 4.1624E-37 | 0.28597063 | 0.509 | 0.26 | 9.8328E-33 | PDC | PSMA1 |
| 4.9326E-37 | 0.27300664 | 0.419 | 0.196 | 1.1652E-32 | PDC | SSR1 |
| 5.4028E-37 | 0.29484344 | 0.423 | 0.203 | 1.2763E-32 | PDC | CHMP3 |
| 5.6138E-37 | -0.6073061 | 0.34 | 0.629 | 1.3261E-32 | PDC | S100A4 |
| 6.8163E-37 | 0.39979074 | 0.671 | 0.411 | 1.6102E-32 | PDC | EIF3K |
| 6.9918E-37 | 0.25775401 | 0.331 | 0.142 | 1.6517E-32 | PDC | PPP1R11 |
| 7.3402E-37 | -1.1461097 | 0.426 | 0.646 | 1.734E-32 | PDC | GLUL |
| 7.7329E-37 | -0.9448435 | 0.009 | 0.241 | 1.8267E-32 | PDC | ADGRF1 |
| 9.0167E-37 | -0.7584077 | 0.504 | 0.701 | 2.13E-32 | PDC | PPP1R15A |
| 9.2712E-37 | 0.31628383 | 0.82 | 0.558 | 2.1901E-32 | PDC | SET |
| 1.0437E-36 | -0.7644216 | 0.099 | 0.35 | 2.4655E-32 | PDC | IER5 |
| 1.2072E-36 | 0.25563453 | 0.365 | 0.162 | 2.8519E-32 | PDC | SNRPC |
| 1.3373E-36 | 0.60436896 | 0.941 | 0.881 | 3.159E-32 | PDC | RPL12 |
| 1.5259E-36 | -0.7383639 | 0.027 | 0.263 | 3.6047E-32 | PDC | MUC1 |
| 1.6443E-36 | 0.2943499 | 0.531 | 0.279 | 3.8843E-32 | PDC | WAC |
| 1.7962E-36 | 0.27574984 | 0.387 | 0.179 | 4.2432E-32 | PDC | SNF8 |
| 2.1054E-36 | 0.62314132 | 0.773 | 0.655 | 4.9736E-32 | PDC | ZFAS1 |
| 2.1093E-36 | 0.28397048 | 0.408 | 0.19 | 4.9828E-32 | PDC | PGRMC1 |
| 2.4639E-36 | 0.2698806 | 0.264 | 0.101 | 5.8206E-32 | PDC | KLF9 |
| 2.4739E-36 | -1.1186162 | 0.268 | 0.508 | 5.8442E-32 | PDC | HSPA1B |
| 2.7085E-36 | 0.70524821 | 0.505 | 0.279 | 6.3982E-32 | PDC | MAFB |
| 2.935E-36 | 0.60899873 | 0.982 | 0.957 | 6.9334E-32 | PDC | RPS18 |
| 3.1658E-36 | 0.31683109 | 0.435 | 0.21 | 7.4786E-32 | PDC | RPL17 |
| 3.1734E-36 | 0.27459579 | 0.365 | 0.164 | 7.4965E-32 | PDC | GTF2H5 |
| 3.4108E-36 | -0.6469375 | 0.689 | 0.812 | 8.0574E-32 | PDC | DDX5 |
| 3.4232E-36 | -0.5311304 | 0.002 | 0.227 | 8.0867E-32 | PDC | KYNU |
| 3.6317E-36 | 0.35711998 | 0.777 | 0.534 | 8.5792E-32 | PDC | HMGN1 |
| 3.7661E-36 | 0.28418678 | 0.419 | 0.2 | 8.8966E-32 | PDC | NEU1 |
| 3.7904E-36 | 0.28589398 | 0.403 | 0.193 | 8.954E-32 | PDC | UFD1 |
| 4.2008E-36 | -0.6566784 | 0.013 | 0.245 | 9.9235E-32 | PDC | PRSS8 |
| 4.2584E-36 | 0.2945791 | 0.227 | 0.081 | 1.006E-31 | PDC | RNF220 |
| 4.6054E-36 | -1.6464286 | 0.007 | 0.235 | 1.0879E-31 | PDC | PRSS22 |
| 4.8619E-36 | 0.29900135 | 0.613 | 0.343 | 1.1485E-31 | PDC | SF3B6 |
| 1.0459E-35 | 0.296514 | 0.55 | 0.292 | 2.4707E-31 | PDC | SELENOS |
| 1.1685E-35 | 0.31501706 | 0.365 | 0.165 | 2.7604E-31 | PDC | CXADR |
| 1.2171E-35 | -0.5841762 | 0.029 | 0.262 | 2.8751E-31 | PDC | NABP1 |
| 2.105E-35 | -0.5906231 | 0.018 | 0.249 | 4.9727E-31 | PDC | GMFG |
| 2.1315E-35 | 0.32472451 | 0.691 | 0.404 | 5.0352E-31 | PDC | ATF4 |
| 2.3341E-35 | -0.8060563 | 0.018 | 0.249 | 5.5138E-31 | PDC | CST6 |
| 3.938E-35 | -1.0526143 | 0.059 | 0.295 | 9.3027E-31 | PDC | RND3 |
| 5.5928E-35 | -1.0899712 | 0.014 | 0.239 | 1.3212E-30 | PDC | BCL2A1 |
| 5.7674E-35 | 0.26630271 | 0.331 | 0.145 | 1.3624E-30 | PDC | PPP2R5E |
| 5.7869E-35 | 0.29255353 | 0.498 | 0.259 | 1.367E-30 | PDC | GTF2A2 |
| 7.3349E-35 | 0.25770424 | 0.353 | 0.157 | 1.7327E-30 | PDC | SLC25A36 |
| 8.2376E-35 | 0.39908296 | 0.906 | 0.782 | 1.946E-30 | PDC | EEF1D |
| 9.1674E-35 | 0.31908529 | 0.829 | 0.609 | 2.1656E-30 | PDC | PTGES3 |
| 9.1904E-35 | -0.6776037 | 0.013 | 0.237 | 2.1711E-30 | PDC | HCST |
| 1.282E-34 | -1.9928914 | 0.041 | 0.269 | 3.0285E-30 | PDC | S100A2 |
| 1.3778E-34 | 0.29570928 | 0.57 | 0.311 | 3.2547E-30 | PDC | EIF3F |
| 1.5206E-34 | 0.2636658 | 0.34 | 0.151 | 3.5922E-30 | PDC | DNAJB2 |

| | | | | | | |
|---|---|---|---|---|---|---|
| 2.2712E-34 | -0.4565046 | 0.022 | 0.247 | 5.3651E-30 | PDC | CD82 |
| 2.4969E-34 | -0.7150748 | 0.254 | 0.483 | 5.8983E-30 | PDC | RIOK3 |
| 2.4969E-34 | -0.6592852 | 0.572 | 0.73 | 5.8985E-30 | PDC | PKM |
| 2.5932E-34 | 0.34382257 | 0.604 | 0.351 | 6.1259E-30 | PDC | SEPT7 |
| 2.6899E-34 | 0.39023607 | 0.98 | 0.895 | 6.3544E-30 | PDC | HLA-A |
| 2.7442E-34 | -0.5734852 | 0.16 | 0.4 | 6.4826E-30 | PDC | H2AFY |
| 3.1875E-34 | -0.9042398 | 0.023 | 0.247 | 7.5299E-30 | PDC | SLC11A1 |
| 4.2187E-34 | 0.26995541 | 0.511 | 0.265 | 9.9658E-30 | PDC | BAG1 |
| 4.9776E-34 | -0.8010662 | 0.162 | 0.397 | 1.1759E-29 | PDC | HIF1A |
| 5.0792E-34 | 0.26390326 | 0.439 | 0.216 | 1.1999E-29 | PDC | MDH2 |
| 5.1708E-34 | 0.25958412 | 0.475 | 0.24 | 1.2215E-29 | PDC | NDUFA3 |
| 6.9977E-34 | 0.25637448 | 0.255 | 0.099 | 1.6531E-29 | PDC | RRM2B |
| 7.1684E-34 | -0.6151822 | 0.052 | 0.282 | 1.6934E-29 | PDC | ABHD5 |
| 8.967E-34 | -0.5941435 | 0.068 | 0.3 | 2.1183E-29 | PDC | REL |
| 9.1919E-34 | 0.2691642 | 0.333 | 0.149 | 2.1714E-29 | PDC | FGD5-AS1 |
| 1.0967E-33 | -0.5913605 | 0.887 | 0.93 | 2.5907E-29 | PDC | OAZ1 |
| 1.1535E-33 | -0.5294651 | 0.059 | 0.291 | 2.7249E-29 | PDC | RNF149 |
| 1.1607E-33 | 0.32708193 | 0.523 | 0.291 | 2.742E-29 | PDC | SEPT2 |
| 1.4168E-33 | 0.50980078 | 0.986 | 0.884 | 3.3468E-29 | PDC | RPS2 |
| 2.0281E-33 | -1.1115549 | 0.005 | 0.218 | 4.7909E-29 | PDC | S100A12 |
| 2.1443E-33 | 0.44890888 | 0.933 | 0.857 | 5.0655E-29 | PDC | RPL7 |
| 2.6207E-33 | 0.31366425 | 0.626 | 0.373 | 6.1908E-29 | PDC | EIF3H |
| 3.84E-33 | 0.29262734 | 0.347 | 0.159 | 9.0712E-29 | PDC | SORT1 |
| 3.8625E-33 | -1.0706979 | 0.009 | 0.222 | 9.1243E-29 | PDC | ALOX5AP |
| 3.901E-33 | 0.30990041 | 0.595 | 0.34 | 9.2154E-29 | PDC | SSR2 |
| 5.2041E-33 | -0.4781922 | 0.061 | 0.297 | 1.2294E-28 | PDC | FXYD5 |
| 6.5397E-33 | 0.32715327 | 0.646 | 0.383 | 1.5449E-28 | PDC | PSME1 |
| 6.9909E-33 | 0.26222101 | 0.482 | 0.25 | 1.6515E-28 | PDC | TMED2 |
| 7.532E-33 | 0.38146532 | 0.77 | 0.55 | 1.7793E-28 | PDC | SSR4 |
| 1.4734E-32 | -0.6137737 | 0.011 | 0.222 | 3.4806E-28 | PDC | ITGB2 |
| 1.5141E-32 | -0.4808566 | 0.817 | 0.947 | 3.5768E-28 | PDC | MT-CO2 |
| 1.6372E-32 | 0.2501382 | 0.473 | 0.243 | 3.8677E-28 | PDC | CCT8 |
| 2.2018E-32 | 0.34141138 | 0.403 | 0.201 | 5.2014E-28 | PDC | IGF2R |
| 2.3375E-32 | -0.9152222 | 0.045 | 0.267 | 5.5219E-28 | PDC | TSPAN1 |
| 3.3473E-32 | 0.28201888 | 0.558 | 0.312 | 7.9073E-28 | PDC | TAF10 |
| 3.9319E-32 | -0.5213965 | 0.009 | 0.217 | 9.2884E-28 | PDC | EHF |
| 4.2611E-32 | 0.29429667 | 0.441 | 0.223 | 1.0066E-27 | PDC | TPP1 |
| 4.6953E-32 | -0.4629293 | 0.061 | 0.289 | 1.1092E-27 | PDC | BAZ1A |
| 4.9282E-32 | 0.45631037 | 0.982 | 0.945 | 1.1642E-27 | PDC | RPL13A |
| 4.9629E-32 | 0.26253383 | 0.484 | 0.254 | 1.1724E-27 | PDC | SF3B2 |
| 7.0801E-32 | 0.50533526 | 0.993 | 0.964 | 1.6725E-27 | PDC | RPL13 |
| 9.9984E-32 | 0.26208439 | 0.331 | 0.151 | 2.3619E-27 | PDC | C19orf24 |
| 1.0727E-31 | 0.32602741 | 0.628 | 0.383 | 2.5341E-27 | PDC | RAB2A |
| 1.1084E-31 | -1.6819315 | 0.018 | 0.225 | 2.6184E-27 | PDC | VCAN |
| 1.1588E-31 | -0.5851303 | 0.556 | 0.705 | 2.7375E-27 | PDC | TPM3 |
| 1.9525E-31 | 0.26263746 | 0.565 | 0.304 | 4.6124E-27 | PDC | PFDN2 |
| 2.015E-31 | 0.25330466 | 0.27 | 0.11 | 4.7601E-27 | PDC | TFPI |
| 2.0957E-31 | 0.29684266 | 0.809 | 0.586 | 4.9507E-27 | PDC | SLC25A3 |
| 2.6423E-31 | 0.39049785 | 0.982 | 0.949 | 6.2418E-27 | PDC | RPL18 |
| 2.6881E-31 | -0.7205424 | 0.009 | 0.212 | 6.3502E-27 | PDC | TNFRSF1B |

| | | | | | | |
|---|---|---|---|---|---|---|
| 2.7532E-31 | -0.6225922 | 0.049 | 0.269 | 6.5039E-27 | PDC | ID2 |
| 3.05E-31 | -0.5420036 | 0.013 | 0.218 | 7.205E-27 | PDC | SPI1 |
| 3.1121E-31 | -0.6531754 | 0.002 | 0.201 | 7.3517E-27 | PDC | DUSP4 |
| 3.2817E-31 | -0.532809 | 0.5 | 0.712 | 7.7523E-27 | PDC | SDCBP |
| 4.9965E-31 | -0.5734671 | 0.736 | 0.87 | 1.1803E-26 | PDC | PABPC1 |
| 1.0948E-30 | 0.26628613 | 0.32 | 0.147 | 2.5862E-26 | PDC | PTPRA |
| 1.1741E-30 | 0.2975469 | 0.876 | 0.686 | 2.7736E-26 | PDC | PSMA7 |
| 1.3321E-30 | -0.5542234 | 0.092 | 0.314 | 3.1467E-26 | PDC | CYTOR |
| 1.4232E-30 | -0.4675819 | 0.007 | 0.205 | 3.362E-26 | PDC | LSP1 |
| 1.5585E-30 | 0.44995337 | 0.968 | 0.874 | 3.6816E-26 | PDC | RPL36 |
| 1.6133E-30 | -0.9606847 | 0.02 | 0.222 | 3.8111E-26 | PDC | CD14 |
| 1.6811E-30 | 0.2614752 | 0.387 | 0.188 | 3.9712E-26 | PDC | CHD9 |
| 1.948E-30 | 0.27049913 | 0.475 | 0.252 | 4.6018E-26 | PDC | RPL22L1 |
| 2.2453E-30 | -0.3987143 | 0.022 | 0.227 | 5.304E-26 | PDC | ST14 |
| 2.724E-30 | -0.5803711 | 0.018 | 0.22 | 6.4349E-26 | PDC | TGM2 |
| 3.3178E-30 | 0.26082585 | 0.383 | 0.185 | 7.8377E-26 | PDC | VCL |
| 4.7003E-30 | -0.6358098 | 0.013 | 0.211 | 1.1103E-25 | PDC | LST1 |
| 4.8207E-30 | 0.29077407 | 0.64 | 0.382 | 1.1388E-25 | PDC | NEDD8 |
| 4.8589E-30 | -0.5927904 | 0.104 | 0.325 | 1.1478E-25 | PDC | B4GALT1 |
| 5.7006E-30 | 0.26845086 | 0.466 | 0.251 | 1.3467E-25 | PDC | RAB14 |
| 7.1843E-30 | -0.550246 | 0.007 | 0.202 | 1.6972E-25 | PDC | ITGAX |
| 8.1939E-30 | -0.6854023 | 0.065 | 0.276 | 1.9356E-25 | PDC | TNFAIP2 |
| 1.0576E-29 | 0.28330997 | 0.423 | 0.212 | 2.4984E-25 | PDC | CNN3 |
| 1.1732E-29 | 0.27801619 | 0.613 | 0.357 | 2.7715E-25 | PDC | MTDH |
| 1.2248E-29 | 0.26527701 | 0.615 | 0.355 | 2.8933E-25 | PDC | SNRPD2 |
| 1.3683E-29 | 0.31279797 | 0.95 | 0.836 | 3.2323E-25 | PDC | SQSTM1 |
| 1.4057E-29 | -0.4105522 | 0.047 | 0.258 | 3.3207E-25 | PDC | MAP2K3 |
| 1.4994E-29 | -0.4897123 | 0.067 | 0.279 | 3.5421E-25 | PDC | VASP |
| 1.607E-29 | 0.30231283 | 0.45 | 0.24 | 3.7963E-25 | PDC | COMT |
| 2.0128E-29 | -1.6393609 | 0.009 | 0.201 | 4.7548E-25 | PDC | KRT17 |
| 2.0537E-29 | -1.4629825 | 0.007 | 0.199 | 4.8515E-25 | PDC | HOPX |
| 2.3154E-29 | 0.26745543 | 0.378 | 0.191 | 5.4696E-25 | PDC | NAPA |
| 2.363E-29 | -0.7247564 | 0.011 | 0.203 | 5.5822E-25 | PDC | CYBB |
| 2.7643E-29 | -0.5390156 | 0.011 | 0.203 | 6.5301E-25 | PDC | NCF2 |
| 2.7664E-29 | 0.33750673 | 0.869 | 0.758 | 6.535E-25 | PDC | UQCRB |
| 2.9945E-29 | 0.37850805 | 0.502 | 0.278 | 7.074E-25 | PDC | H1F0 |
| 3.8537E-29 | -1.1005791 | 0.874 | 0.883 | 9.1036E-25 | PDC | CSTB |
| 5.0215E-29 | 0.25534827 | 0.433 | 0.227 | 1.1862E-24 | PDC | RAB18 |
| 5.0898E-29 | -0.6116113 | 0.049 | 0.252 | 1.2024E-24 | PDC | KLF5 |
| 6.6311E-29 | -0.5398167 | 0.016 | 0.21 | 1.5665E-24 | PDC | MACC1 |
| 7.2634E-29 | -0.9286518 | 0.221 | 0.436 | 1.7158E-24 | PDC | HIST1H1C |
| 7.8974E-29 | -0.406879 | 0.065 | 0.276 | 1.8656E-24 | PDC | FAM49B |
| 8.0148E-29 | -0.4212768 | 0.016 | 0.21 | 1.8933E-24 | PDC | STK17B |
| 8.4472E-29 | -0.6208127 | 0.04 | 0.24 | 1.9955E-24 | PDC | TNFAIP3 |
| 8.7097E-29 | -0.9190702 | 0.004 | 0.19 | 2.0575E-24 | PDC | TMPRSS2 |
| 1.2524E-28 | -1.2436368 | 0.002 | 0.186 | 2.9586E-24 | PDC | DUOX2 |
| 1.2911E-28 | 0.2763371 | 0.475 | 0.262 | 3.05E-24 | PDC | PSMA4 |
| 1.2931E-28 | -0.5992463 | 0.113 | 0.329 | 3.0546E-24 | PDC | MAFF |
| 1.7662E-28 | 0.29211838 | 0.478 | 0.26 | 4.1723E-24 | PDC | TUBB |
| 1.7729E-28 | -0.3666497 | 0.002 | 0.186 | 4.1881E-24 | PDC | GNA15 |

| | | | | | | |
|---|---|---|---|---|---|---|
| 1.7982E-28 | -0.5979539 | 0.034 | 0.232 | 4.2479E-24 | PDC | PPIF |
| 2.7349E-28 | 0.3776458 | 0.89 | 0.745 | 6.4608E-24 | PDC | EEF1B2 |
| 2.9224E-28 | -0.3855143 | 0.005 | 0.19 | 6.9036E-24 | PDC | HCLS1 |
| 3.368E-28 | -0.6653656 | 0.059 | 0.261 | 7.9562E-24 | PDC | PMAIP1 |
| 3.9913E-28 | 0.40889093 | 0.363 | 0.186 | 9.4286E-24 | PDC | PMEPA1 |
| 4.4337E-28 | 0.32304214 | 0.689 | 0.449 | 1.0474E-23 | PDC | SEC61G |
| 4.5982E-28 | 0.28945284 | 0.264 | 0.116 | 1.0862E-23 | PDC | HCFC1R1 |
| 5.048E-28 | 0.35790161 | 0.962 | 0.892 | 1.1925E-23 | PDC | RPS15A |
| 5.317E-28 | -0.5623099 | 0.023 | 0.215 | 1.256E-23 | PDC | LAMB3 |
| 5.5922E-28 | 0.38928529 | 0.932 | 0.837 | 1.321E-23 | PDC | RACK1 |
| 5.7856E-28 | -0.703485 | 0.013 | 0.2 | 1.3667E-23 | PDC | CD52 |
| 6.7884E-28 | -0.3947352 | 0.007 | 0.191 | 1.6036E-23 | PDC | PYCARD |
| 8.4602E-28 | 0.25575333 | 0.522 | 0.299 | 1.9986E-23 | PDC | SCAND1 |
| 9.651E-28 | -0.4571236 | 0.079 | 0.287 | 2.2799E-23 | PDC | CXCL16 |
| 1.2431E-27 | 0.29509711 | 0.892 | 0.784 | 2.9367E-23 | PDC | BTF3 |
| 1.3095E-27 | 0.27688056 | 0.572 | 0.33 | 3.0935E-23 | PDC | GOLGA4 |
| 1.3892E-27 | 0.29229574 | 0.601 | 0.355 | 3.2817E-23 | PDC | MIF |
| 1.6739E-27 | -0.5112676 | 0.052 | 0.25 | 3.9542E-23 | PDC | SERINC2 |
| 1.8749E-27 | -0.5564273 | 0.009 | 0.19 | 4.429E-23 | PDC | FYB1 |
| 3.2794E-27 | -0.552406 | 0.3 | 0.498 | 7.7468E-23 | PDC | DDX3X |
| 3.3722E-27 | -0.674519 | 0.011 | 0.191 | 7.9661E-23 | PDC | EPS8L1 |
| 4.7903E-27 | -0.7129356 | 0.04 | 0.232 | 1.1316E-22 | PDC | ITGA2 |
| 7.1228E-27 | 0.32590863 | 0.964 | 0.933 | 1.6826E-22 | PDC | RPL34 |
| 7.769E-27 | -0.3485369 | 0.009 | 0.187 | 1.8353E-22 | PDC | CDCP1 |
| 8.1379E-27 | -0.6561935 | 0.016 | 0.198 | 1.9224E-22 | PDC | SFN |
| 1.062E-26 | -0.4776594 | 0.002 | 0.175 | 2.5088E-22 | PDC | SMIM25 |
| 1.0967E-26 | 0.28693374 | 0.527 | 0.31 | 2.5908E-22 | PDC | BLOC1S1 |
| 1.4354E-26 | -0.7475404 | 0.011 | 0.19 | 3.3909E-22 | PDC | AC020656.1 |
| 1.578E-26 | -1.0783421 | 0.971 | 0.962 | 3.7276E-22 | PDC | HSP90AA1 |
| 2.0879E-26 | -0.5368047 | 0.538 | 0.687 | 4.9322E-22 | PDC | BZW1 |
| 2.3952E-26 | 0.25282465 | 0.766 | 0.514 | 5.6583E-22 | PDC | NAP1L1 |
| 4.6198E-26 | 0.25014155 | 0.633 | 0.389 | 1.0913E-21 | PDC | UBXN4 |
| 4.9965E-26 | -0.5593929 | 0.302 | 0.496 | 1.1803E-21 | PDC | RSRP1 |
| 5.8074E-26 | -0.3514321 | 0.013 | 0.189 | 1.3719E-21 | PDC | GPSM3 |
| 5.9061E-26 | -0.6589165 | 0.004 | 0.174 | 1.3952E-21 | PDC | STEAP4 |
| 6.387E-26 | -0.4028849 | 0.014 | 0.19 | 1.5088E-21 | PDC | RAC2 |
| 6.5681E-26 | 0.35623166 | 0.977 | 0.922 | 1.5516E-21 | PDC | RPL11 |
| 6.8186E-26 | -0.6048568 | 0.005 | 0.177 | 1.6108E-21 | PDC | SPNS2 |
| 8.5806E-26 | -0.4150323 | 0.705 | 0.88 | 2.027E-21 | PDC | MT-ND3 |
| 9.6961E-26 | 0.26669605 | 0.888 | 0.793 | 2.2905E-21 | PDC | CHCHD2 |
| 9.7628E-26 | -0.5204169 | 0.011 | 0.184 | 2.3063E-21 | PDC | SAMSN1 |
| 1.1951E-25 | -0.3918453 | 0.099 | 0.3 | 2.8232E-21 | PDC | UBALD2 |
| 1.5123E-25 | -0.5744306 | 0.176 | 0.377 | 3.5725E-21 | PDC | CCNL1 |
| 1.6361E-25 | 0.30801228 | 0.703 | 0.511 | 3.8649E-21 | PDC | EIF5 |
| 1.688E-25 | -0.5940445 | 0.624 | 0.728 | 3.9876E-21 | PDC | ENO1 |
| 1.695E-25 | 0.27845114 | 0.358 | 0.188 | 4.004E-21 | PDC | CUX1 |
| 1.704E-25 | -0.3489549 | 0.005 | 0.174 | 4.0253E-21 | PDC | DAPP1 |
| 1.7464E-25 | 0.25719204 | 0.54 | 0.321 | 4.1256E-21 | PDC | NSA2 |
| 1.7528E-25 | -0.4668994 | 0.011 | 0.183 | 4.1406E-21 | PDC | S100A14 |
| 1.7605E-25 | -0.4410429 | 0.014 | 0.189 | 4.1589E-21 | PDC | PTPRE |

| | | | | | | |
|---|---|---|---|---|---|---|
| 1.8155E-25 | 0.31981763 | 0.746 | 0.544 | 4.2889E-21 | PDC | RPS10 |
| 1.9199E-25 | -1.2815742 | 0.002 | 0.168 | 4.5355E-21 | PDC | DUOXA2 |
| 2.0977E-25 | -0.567384 | 0.013 | 0.185 | 4.9553E-21 | PDC | IGSF6 |
| 2.3267E-25 | -0.6822201 | 0.011 | 0.182 | 5.4964E-21 | PDC | OLR1 |
| 3.0159E-25 | -0.5318483 | 0.005 | 0.172 | 7.1244E-21 | PDC | PRDM1 |
| 3.2902E-25 | 0.33194541 | 0.588 | 0.364 | 7.7724E-21 | PDC | MYH9 |
| 3.5805E-25 | 0.36184504 | 0.863 | 0.671 | 8.4583E-21 | PDC | CALR |
| 3.6055E-25 | 0.25196343 | 0.442 | 0.247 | 8.5173E-21 | PDC | CHD4 |
| 3.7608E-25 | -0.4680249 | 0.142 | 0.341 | 8.8841E-21 | PDC | GRB2 |
| 4.269E-25 | -0.3602422 | 0.784 | 0.932 | 1.0085E-20 | PDC | MT-CYB |
| 4.4975E-25 | 0.28768208 | 0.853 | 0.666 | 1.0624E-20 | PDC | RTN4 |
| 4.8054E-25 | -0.4973312 | 0.252 | 0.449 | 1.1352E-20 | PDC | DNAJB6 |
| 5.8408E-25 | -0.5687182 | 0.011 | 0.179 | 1.3798E-20 | PDC | ANPEP |
| 5.869E-25 | -0.5136113 | 0.009 | 0.177 | 1.3864E-20 | PDC | C6orf132 |
| 5.9907E-25 | -0.4646919 | 0.126 | 0.327 | 1.4152E-20 | PDC | FOSL2 |
| 6.318E-25 | -0.6306444 | 0.304 | 0.498 | 1.4925E-20 | PDC | CD9 |
| 6.6207E-25 | -0.5059089 | 0.054 | 0.239 | 1.564E-20 | PDC | NFKBIZ |
| 8.0738E-25 | 0.31847258 | 0.82 | 0.626 | 1.9073E-20 | PDC | OST4 |
| 8.4161E-25 | 0.2501535 | 0.613 | 0.386 | 1.9881E-20 | PDC | SEM1 |
| 8.6804E-25 | -0.4455049 | 0.005 | 0.17 | 2.0506E-20 | PDC | SLA |
| 1.1109E-24 | -0.6999404 | 0.054 | 0.237 | 2.6242E-20 | PDC | EPCAM |
| 1.1527E-24 | -0.3829805 | 0.083 | 0.276 | 2.723E-20 | PDC | ADAM10 |
| 1.188E-24 | -0.3965724 | 0.047 | 0.228 | 2.8064E-20 | PDC | PNP |
| 1.3678E-24 | -0.4915359 | 0.034 | 0.209 | 3.2312E-20 | PDC | SLC43A2 |
| 1.4086E-24 | -1.14677 | 0.002 | 0.163 | 3.3276E-20 | PDC | MUC21 |
| 1.7725E-24 | -0.3841891 | 0.007 | 0.17 | 4.1872E-20 | PDC | NECTIN4 |
| 1.8165E-24 | -0.6224441 | 0.007 | 0.17 | 4.291E-20 | PDC | FXYD3 |
| 1.9415E-24 | -0.7673733 | 0.067 | 0.248 | 4.5864E-20 | PDC | PLAU |
| 1.9499E-24 | -0.4650819 | 0.088 | 0.277 | 4.6062E-20 | PDC | PLIN3 |
| 2.0981E-24 | -1.0617527 | 0.002 | 0.162 | 4.9563E-20 | PDC | CEACAM5 |
| 2.1738E-24 | -0.7300844 | 0.011 | 0.177 | 5.1352E-20 | PDC | PLEK |
| 2.5269E-24 | -0.4467849 | 0.11 | 0.301 | 5.9694E-20 | PDC | TNIP1 |
| 3.0174E-24 | 0.34064971 | 0.975 | 0.923 | 7.128E-20 | PDC | RPL18A |
| 3.0785E-24 | -0.3347859 | 0.025 | 0.196 | 7.2724E-20 | PDC | KIFC3 |
| 3.3921E-24 | 0.35080573 | 0.962 | 0.897 | 8.0132E-20 | PDC | RPL29 |
| 3.7288E-24 | -0.8674659 | 0 | 0.158 | 8.8085E-20 | PDC | KRT15 |
| 3.7306E-24 | -0.7856464 | 0.009 | 0.172 | 8.8128E-20 | PDC | FCN1 |
| 4.5932E-24 | -0.3826727 | 0.014 | 0.181 | 1.0851E-19 | PDC | CD37 |
| 5.094E-24 | -0.5882318 | 0.02 | 0.186 | 1.2034E-19 | PDC | C5AR1 |
| 5.123E-24 | -0.5059045 | 0.02 | 0.188 | 1.2102E-19 | PDC | TNFSF10 |
| 5.6173E-24 | -0.5001177 | 0.018 | 0.184 | 1.327E-19 | PDC | FBP1 |
| 6.8422E-24 | -0.9536205 | 0.058 | 0.235 | 1.6163E-19 | PDC | SLC2A3 |
| 7.2979E-24 | 0.25467784 | 0.673 | 0.455 | 1.724E-19 | PDC | EIF2S2 |
| 8.5257E-24 | -0.7573122 | 0.221 | 0.407 | 2.014E-19 | PDC | RNF213 |
| 8.5576E-24 | 0.41585438 | 0.991 | 0.972 | 2.0216E-19 | PDC | RPLP1 |
| 8.5942E-24 | -0.6740817 | 0.268 | 0.435 | 2.0302E-19 | PDC | GSTO1 |
| 1.0582E-23 | -0.606896 | 0.013 | 0.174 | 2.4997E-19 | PDC | HLA-DQB1 |
| 1.1335E-23 | -0.4058435 | 0.018 | 0.183 | 2.6776E-19 | PDC | PFKFB3 |
| 1.2458E-23 | -0.3935075 | 0.013 | 0.175 | 2.9429E-19 | PDC | CORO1A |
| 1.2488E-23 | -0.4682768 | 0.005 | 0.163 | 2.95E-19 | PDC | AQP9 |

| | | | | | | |
|---|---|---|---|---|---|---|
| 1.3542E-23 | -0.6580375 | 0.002 | 0.157 | 3.199E-19 | PDC | MUC4 |
| 1.6417E-23 | -0.4501614 | 0.041 | 0.217 | 3.8782E-19 | PDC | ARL4C |
| 1.7043E-23 | -2.1969837 | 0.016 | 0.177 | 4.0262E-19 | PDC | IL1B |
| 1.7773E-23 | -0.4055538 | 0.036 | 0.206 | 4.1986E-19 | PDC | MAP3K8 |
| 1.8505E-23 | -0.3674098 | 0.072 | 0.255 | 4.3714E-19 | PDC | C4orf48 |
| 2.0774E-23 | -0.3812092 | 0.011 | 0.171 | 4.9074E-19 | PDC | CD48 |
| 2.1425E-23 | -0.5593811 | 0.09 | 0.271 | 5.0613E-19 | PDC | CCNG2 |
| 2.3816E-23 | -1.2043075 | 0.036 | 0.206 | 5.6261E-19 | PDC | HLA-DPA1 |
| 2.666E-23 | -0.4375498 | 0.027 | 0.192 | 6.298E-19 | PDC | ABLIM1 |
| 2.9284E-23 | -0.7577586 | 0.002 | 0.155 | 6.9178E-19 | PDC | SCEL |
| 2.9797E-23 | 0.26022285 | 0.505 | 0.299 | 7.039E-19 | PDC | PSME2 |
| 3.017E-23 | -0.3266395 | 0.022 | 0.185 | 7.1272E-19 | PDC | BID |
| 3.1128E-23 | -0.448965 | 0.164 | 0.357 | 7.3533E-19 | PDC | AKAP13 |
| 3.1438E-23 | -0.4188126 | 0.005 | 0.161 | 7.4266E-19 | PDC | SDC1 |
| 3.1491E-23 | -0.3139388 | 0.007 | 0.164 | 7.4392E-19 | PDC | FERMT3 |
| 3.2766E-23 | -0.4218842 | 0.009 | 0.166 | 7.7403E-19 | PDC | LCP2 |
| 3.6617E-23 | -0.497851 | 0.005 | 0.16 | 8.65E-19 | PDC | MALL |
| 3.8762E-23 | -0.981625 | 0.005 | 0.16 | 9.1568E-19 | PDC | PRSS27 |
| 5.3796E-23 | -0.29063 | 0.005 | 0.16 | 1.2708E-18 | PDC | ARHGAP30 |
| 5.5409E-23 | -0.5811978 | 0.016 | 0.174 | 1.3089E-18 | PDC | PPL |
| 5.6269E-23 | -0.5075483 | 0.002 | 0.153 | 1.3292E-18 | PDC | CLEC5A |
| 5.6417E-23 | -0.5069504 | 0.027 | 0.193 | 1.3327E-18 | PDC | PLK2 |
| 5.686E-23 | -0.9644056 | 0.004 | 0.156 | 1.3432E-18 | PDC | PITX1 |
| 5.7203E-23 | 0.30577891 | 0.944 | 0.876 | 1.3513E-18 | PDC | RPL23A |
| 6.2226E-23 | -0.3334264 | 0.009 | 0.165 | 1.47E-18 | PDC | MYO1F |
| 6.7257E-23 | -0.4673382 | 0.027 | 0.191 | 1.5888E-18 | PDC | DUSP6 |
| 6.9335E-23 | -0.3377017 | 0.02 | 0.181 | 1.6379E-18 | PDC | FMNL1 |
| 7.4551E-23 | -1.064722 | 0.549 | 0.683 | 1.7611E-18 | PDC | NAMPT |
| 7.584E-23 | -0.8443637 | 0.009 | 0.163 | 1.7916E-18 | PDC | APOBEC3A |
| 8.3021E-23 | -0.8351969 | 0.103 | 0.288 | 1.9612E-18 | PDC | HMOX1 |
| 9.8354E-23 | 0.25147223 | 0.556 | 0.34 | 2.3234E-18 | PDC | ATP5F1B |
| 1.0137E-22 | -0.4713155 | 0.487 | 0.617 | 2.3945E-18 | PDC | CAP1 |
| 1.0212E-22 | -0.4959806 | 0.014 | 0.172 | 2.4124E-18 | PDC | CXCR4 |
| 1.0958E-22 | -0.3624978 | 0.02 | 0.179 | 2.5887E-18 | PDC | GLIPR1 |
| 1.1466E-22 | 0.29147885 | 0.853 | 0.724 | 2.7086E-18 | PDC | ELOB |
| 1.1938E-22 | -0.3156862 | 0.009 | 0.163 | 2.8202E-18 | PDC | GALNT3 |
| 1.2041E-22 | -0.3965922 | 0.092 | 0.271 | 2.8445E-18 | PDC | UBE2D1 |
| 1.2396E-22 | -0.5532851 | 0.007 | 0.16 | 2.9283E-18 | PDC | CD300E |
| 1.3697E-22 | -0.8690446 | 0 | 0.148 | 3.2356E-18 | PDC | CEACAM6 |
| 1.4254E-22 | -0.4377846 | 0.09 | 0.273 | 3.3672E-18 | PDC | RAP2B |
| 1.5157E-22 | -0.2937313 | 0.022 | 0.183 | 3.5806E-18 | PDC | ELK3 |
| 1.5889E-22 | -0.7603285 | 0.002 | 0.15 | 3.7534E-18 | PDC | TMPRSS11D |
| 1.5962E-22 | -0.4170859 | 0 | 0.148 | 3.7708E-18 | PDC | VSIG2 |
| 1.7073E-22 | -0.4409621 | 0.005 | 0.156 | 4.0332E-18 | PDC | TMPRSS4 |
| 2.3417E-22 | -0.4387317 | 0.108 | 0.291 | 5.5317E-18 | PDC | MIR4435-2HG |
| 2.6474E-22 | -0.7761831 | 0 | 0.146 | 6.2539E-18 | PDC | TMPRSS11E |
| 3.3296E-22 | -0.6490538 | 0.002 | 0.148 | 7.8655E-18 | PDC | NCCRP1 |
| 4.7965E-22 | -0.3652132 | 0.029 | 0.189 | 1.1331E-17 | PDC | PIM1 |
| 4.8383E-22 | -0.8225363 | 0.002 | 0.148 | 1.143E-17 | PDC | CEACAM7 |
| 5.0066E-22 | -0.5134165 | 0.045 | 0.209 | 1.1827E-17 | PDC | GK |

| | | | | | | |
|---|---|---|---|---|---|---|
| 7.3926E-22 | -0.3314698 | 0.014 | 0.168 | 1.7464E-17 | PDC | CFD |
| 8.1982E-22 | -0.5209335 | 0.076 | 0.246 | 1.9367E-17 | PDC | KIAA1551 |
| 1.175E-21 | 0.25790154 | 0.764 | 0.548 | 2.7757E-17 | PDC | UQCR10 |
| 1.1808E-21 | -0.3769792 | 0.004 | 0.148 | 2.7894E-17 | PDC | TREM1 |
| 1.2457E-21 | -0.3989199 | 0.02 | 0.173 | 2.9426E-17 | PDC | SAMHD1 |
| 1.2562E-21 | -0.4472549 | 0.117 | 0.294 | 2.9675E-17 | PDC | SKIL |
| 1.3345E-21 | -0.3119969 | 0.063 | 0.234 | 3.1525E-17 | PDC | LASP1 |
| 1.5083E-21 | -1.3531104 | 0.007 | 0.154 | 3.5631E-17 | PDC | CCL20 |
| 1.5088E-21 | -0.587524 | 0.009 | 0.156 | 3.5643E-17 | PDC | CDC42EP5 |
| 1.5562E-21 | -0.5267054 | 0.02 | 0.172 | 3.6762E-17 | PDC | TGFBI |
| 1.5604E-21 | -0.3625418 | 0.007 | 0.153 | 3.6862E-17 | PDC | EVI2B |
| 1.8244E-21 | -0.4704743 | 0.147 | 0.327 | 4.3097E-17 | PDC | TMBIM1 |
| 1.9027E-21 | -0.310328 | 0.049 | 0.212 | 4.4947E-17 | PDC | CYTH1 |
| 1.9059E-21 | -0.3351587 | 0.041 | 0.203 | 4.5023E-17 | PDC | ZNF217 |
| 1.9653E-21 | 0.3500981 | 0.959 | 0.889 | 4.6426E-17 | PDC | HLA-B |
| 2.0252E-21 | -0.4685459 | 0.025 | 0.18 | 4.7842E-17 | PDC | EPHA2 |
| 2.4055E-21 | -0.7205672 | 0.104 | 0.283 | 5.6826E-17 | PDC | CLU |
| 3.416E-21 | 0.31829043 | 0.973 | 0.933 | 8.0696E-17 | PDC | RPS14 |
| 3.4275E-21 | -0.3651228 | 0.02 | 0.17 | 8.0968E-17 | PDC | TINAGL1 |
| 3.5547E-21 | -0.2926856 | 0.02 | 0.17 | 8.3973E-17 | PDC | VPS37B |
| 3.9314E-21 | -0.3314824 | 0.036 | 0.193 | 9.287E-17 | PDC | F11R |
| 4.5786E-21 | -0.458402 | 0.081 | 0.25 | 1.0816E-16 | PDC | GPCPD1 |
| 4.6106E-21 | -1.0308597 | 0.002 | 0.142 | 1.0892E-16 | PDC | CXCL6 |
| 4.9893E-21 | -0.3456223 | 0.027 | 0.182 | 1.1786E-16 | PDC | ANXA3 |
| 5.4543E-21 | -0.4080281 | 0.005 | 0.147 | 1.2885E-16 | PDC | MS4A7 |
| 5.5813E-21 | -0.3009589 | 0.04 | 0.199 | 1.3185E-16 | PDC | MYO9B |
| 6.8495E-21 | -0.3285095 | 0.023 | 0.173 | 1.6181E-16 | PDC | B3GNT5 |
| 7.8606E-21 | -0.5228743 | 0.432 | 0.574 | 1.8569E-16 | PDC | TSPO |
| 9.151E-21 | -0.2719188 | 0.016 | 0.162 | 2.1617E-16 | PDC | SH3BP2 |
| 9.2047E-21 | 0.28106489 | 0.923 | 0.83 | 2.1744E-16 | PDC | RPL22 |
| 9.3033E-21 | -1.6972294 | 0.005 | 0.145 | 2.1977E-16 | PDC | SPRR2A |
| 1.1111E-20 | -0.3965913 | 0 | 0.136 | 2.6247E-16 | PDC | BCAS1 |
| 1.2301E-20 | -0.493916 | 0.038 | 0.191 | 2.9059E-16 | PDC | FCGR2A |
| 1.2935E-20 | -0.4231968 | 0.007 | 0.147 | 3.0555E-16 | PDC | FPR1 |
| 1.3886E-20 | -0.3864179 | 0.067 | 0.23 | 3.2802E-16 | PDC | GALNT1 |
| 1.4955E-20 | -0.6133035 | 0.054 | 0.21 | 3.5329E-16 | PDC | CDKN2B |
| 1.8767E-20 | -0.3303443 | 0.027 | 0.178 | 4.4333E-16 | PDC | SMIM22 |
| 2.2979E-20 | -0.5009217 | 0.068 | 0.23 | 5.4284E-16 | PDC | IVNS1ABP |
| 2.4957E-20 | -0.3549245 | 0.007 | 0.145 | 5.8955E-16 | PDC | TLR2 |
| 3.3775E-20 | -0.9072867 | 0.002 | 0.136 | 7.9786E-16 | PDC | TMPRSS11B |
| 3.7019E-20 | -0.2562217 | 0.002 | 0.136 | 8.745E-16 | PDC | NCKAP1L |
| 3.9514E-20 | -0.3399487 | 0.016 | 0.158 | 9.3343E-16 | PDC | PELI1 |
| 3.9551E-20 | -0.9842178 | 0.029 | 0.175 | 9.3432E-16 | PDC | RDH10 |
| 4.1046E-20 | 0.31788192 | 0.89 | 0.785 | 9.6962E-16 | PDC | RPL31 |
| 4.7206E-20 | -0.4335313 | 0.004 | 0.138 | 1.1151E-15 | PDC | MCEMP1 |
| 4.7224E-20 | -0.6675657 | 0 | 0.132 | 1.1156E-15 | PDC | DHRS9 |
| 4.8601E-20 | -0.7050694 | 0.766 | 0.791 | 1.1481E-15 | PDC | PFN1 |
| 4.8809E-20 | -0.3007594 | 0.004 | 0.139 | 1.153E-15 | PDC | CLEC7A |
| 5.452E-20 | -0.7147454 | 0.032 | 0.184 | 1.2879E-15 | PDC | LAMC2 |
| 5.6325E-20 | -0.3825649 | 0.02 | 0.163 | 1.3306E-15 | PDC | SYTL2 |

| | | | | | | |
|---|---|---|---|---|---|---|
| 5.7531E-20 | -0.6433019 | 0.171 | 0.33 | 1.3591E-15 | PDC | MYO6 |
| 6.0657E-20 | -0.6286935 | 0.004 | 0.138 | 1.4329E-15 | PDC | CXCL17 |
| 6.0843E-20 | -0.3329947 | 0.061 | 0.22 | 1.4373E-15 | PDC | TUT7 |
| 6.3053E-20 | 0.28382533 | 0.815 | 0.702 | 1.4895E-15 | PDC | SUMO2 |
| 6.4853E-20 | -0.4859832 | 0.009 | 0.146 | 1.532E-15 | PDC | CD93 |
| 6.7393E-20 | -0.3253245 | 0.043 | 0.195 | 1.592E-15 | PDC | B4GALT5 |
| 7.0492E-20 | -0.4582883 | 0.011 | 0.148 | 1.6652E-15 | PDC | ALDH1A3 |
| 7.5335E-20 | -0.38977 | 0.018 | 0.159 | 1.7796E-15 | PDC | TTC9 |
| 8.1685E-20 | -0.2993557 | 0.023 | 0.167 | 1.9297E-15 | PDC | FLNB |
| 8.6101E-20 | -0.4286555 | 0.665 | 0.774 | 2.034E-15 | PDC | EIF4G2 |
| 8.918E-20 | -0.3131609 | 0.029 | 0.175 | 2.1067E-15 | PDC | TNFAIP8 |
| 8.9201E-20 | 0.25161386 | 0.41 | 0.243 | 2.1072E-15 | PDC | PEA15 |
| 9.0545E-20 | -0.4761809 | 0.047 | 0.199 | 2.139E-15 | PDC | BIRC3 |
| 1.0025E-19 | -0.6175509 | 0.065 | 0.223 | 2.3682E-15 | PDC | CCND1 |
| 1.115E-19 | -0.2812327 | 0.02 | 0.161 | 2.6339E-15 | PDC | DOCK8 |
| 1.1515E-19 | -0.6589779 | 0.002 | 0.133 | 2.7202E-15 | PDC | GABRP |
| 1.1966E-19 | 0.27311706 | 0.905 | 0.79 | 2.8268E-15 | PDC | RPS21 |
| 1.2783E-19 | -0.266342 | 0.007 | 0.142 | 3.0198E-15 | PDC | IL2RG |
| 1.2853E-19 | -0.2693663 | 0.005 | 0.139 | 3.0363E-15 | PDC | EVI2A |
| 1.3013E-19 | -0.2622877 | 0.002 | 0.132 | 3.074E-15 | PDC | APBB1IP |
| 1.3106E-19 | -0.3586461 | 0 | 0.129 | 3.096E-15 | PDC | LRG1 |
| 1.4637E-19 | -0.4445611 | 0.727 | 0.837 | 3.4578E-15 | PDC | HLA-E |
| 1.554E-19 | 0.33323943 | 0.622 | 0.4 | 3.6711E-15 | PDC | FLNA |
| 1.5677E-19 | -0.8049618 | 0.007 | 0.141 | 3.7033E-15 | PDC | CXCL1 |
| 1.7637E-19 | -0.2912527 | 0.011 | 0.146 | 4.1664E-15 | PDC | SLC44A4 |
| 1.8203E-19 | 0.28048953 | 0.977 | 0.895 | 4.3002E-15 | PDC | RPS9 |
| 1.8719E-19 | -1.5959797 | 0.004 | 0.134 | 4.4221E-15 | PDC | LYPD2 |
| 1.8991E-19 | -0.3834181 | 0.061 | 0.219 | 4.4862E-15 | PDC | DSP |
| 2.1169E-19 | -0.9064549 | 0.004 | 0.134 | 5.0008E-15 | PDC | RHCG |
| 2.5617E-19 | -0.2911623 | 0.004 | 0.134 | 6.0514E-15 | PDC | MPZL2 |
| 2.5642E-19 | -0.3303479 | 0.018 | 0.157 | 6.0575E-15 | PDC | ADGRE5 |
| 2.7476E-19 | 0.30192687 | 0.921 | 0.848 | 6.4907E-15 | PDC | RPL38 |
| 2.9947E-19 | -0.7220738 | 0.045 | 0.19 | 7.0744E-15 | PDC | ANKRD36C |
| 3.0276E-19 | -0.3661819 | 0.038 | 0.185 | 7.152E-15 | PDC | CDH1 |
| 3.3539E-19 | -1.1627087 | 0.004 | 0.133 | 7.923E-15 | PDC | PLAT |
| 3.5814E-19 | -0.3924432 | 0.011 | 0.144 | 8.4603E-15 | PDC | NCF1 |
| 4.2445E-19 | -1.0379302 | 0.005 | 0.135 | 1.0027E-14 | PDC | SERPINB2 |
| 4.8442E-19 | -0.8838946 | 0.308 | 0.454 | 1.1443E-14 | PDC | HSPH1 |
| 5.0322E-19 | -0.3840825 | 0.309 | 0.472 | 1.1888E-14 | PDC | RAP1B |
| 5.4793E-19 | -0.2705452 | 0.032 | 0.175 | 1.2944E-14 | PDC | ZDHHC20 |
| 5.7756E-19 | 0.34746682 | 0.935 | 0.859 | 1.3644E-14 | PDC | RPL23 |
| 6.267E-19 | -0.5871614 | 0.387 | 0.538 | 1.4805E-14 | PDC | SERPINB1 |
| 6.5786E-19 | -0.4249234 | 0.131 | 0.295 | 1.5541E-14 | PDC | TUBA1C |
| 6.6376E-19 | -0.3617452 | 0.023 | 0.162 | 1.568E-14 | PDC | SOCS1 |
| 7.7722E-19 | -0.3671948 | 0.013 | 0.145 | 1.836E-14 | PDC | ITGB6 |
| 8.6657E-19 | -0.3060846 | 0.018 | 0.153 | 2.0471E-14 | PDC | CPVL |
| 9.0226E-19 | 0.3223986 | 0.953 | 0.9 | 2.1314E-14 | PDC | RPS11 |
| 1.0135E-18 | -0.2635223 | 0.011 | 0.142 | 2.3942E-14 | PDC | RAB25 |
| 1.027E-18 | -0.2697003 | 0.007 | 0.136 | 2.426E-14 | PDC | FGR |
| 1.0971E-18 | -0.5044532 | 0.041 | 0.185 | 2.5917E-14 | PDC | SDCBP2 |

| | | | | | | |
|---|---|---|---|---|---|---|
| 1.1915E-18 | -0.6074517 | 0.013 | 0.144 | 2.8147E-14 | PDC | CDKN2A |
| 1.478E-18 | -0.2751601 | 0.009 | 0.138 | 3.4915E-14 | PDC | ITGA5 |
| 1.5492E-18 | -0.2876333 | 0.025 | 0.162 | 3.6597E-14 | PDC | WIPF1 |
| 1.5492E-18 | -0.3324081 | 0.022 | 0.156 | 3.6597E-14 | PDC | ITPKC |
| 1.6965E-18 | -0.8233806 | 0.047 | 0.195 | 4.0077E-14 | PDC | HSPA6 |
| 1.699E-18 | -0.3500203 | 0.092 | 0.249 | 4.0136E-14 | PDC | CLK1 |
| 1.7382E-18 | -0.5781888 | 0.187 | 0.347 | 4.1061E-14 | PDC | MIDN |
| 2.0984E-18 | -0.2793595 | 0.009 | 0.137 | 4.957E-14 | PDC | HCK |
| 2.3432E-18 | 0.28805957 | 0.903 | 0.819 | 5.5353E-14 | PDC | RPL27 |
| 2.5438E-18 | -0.9023579 | 0.009 | 0.135 | 6.0093E-14 | PDC | EREG |
| 2.5463E-18 | -0.4740124 | 0.126 | 0.045 | 6.0152E-14 | PDC | MT1A |
| 3.054E-18 | -0.372854 | 0.113 | 0.272 | 7.2144E-14 | PDC | QSOX1 |
| 3.3537E-18 | -0.2594502 | 0.014 | 0.144 | 7.9225E-14 | PDC | CPM |
| 3.6158E-18 | -0.3103419 | 0.106 | 0.268 | 8.5415E-14 | PDC | CDC42SE1 |
| 3.667E-18 | -0.3138047 | 0.016 | 0.147 | 8.6625E-14 | PDC | C1orf162 |
| 3.7067E-18 | -0.2714398 | 0.009 | 0.135 | 8.7564E-14 | PDC | CYTIP |
| 4.1769E-18 | -0.4231924 | 0.133 | 0.291 | 9.8671E-14 | PDC | CHMP1B |
| 4.6089E-18 | -0.3010333 | 0.027 | 0.163 | 1.0888E-13 | PDC | MET |
| 4.8183E-18 | -0.2503428 | 0.022 | 0.154 | 1.1382E-13 | PDC | EHBP1L1 |
| 4.9376E-18 | -0.3971379 | 0.45 | 0.585 | 1.1664E-13 | PDC | ACTR3 |
| 7.8835E-18 | -1.4957455 | 0.018 | 0.146 | 1.8623E-13 | PDC | CXCL3 |
| 8.8272E-18 | -0.406702 | 0.02 | 0.149 | 2.0852E-13 | PDC | RGS2 |
| 9.3034E-18 | -0.548734 | 0.004 | 0.123 | 2.1977E-13 | PDC | AGR2 |
| 1.0704E-17 | 0.31627795 | 0.966 | 0.923 | 2.5285E-13 | PDC | RPS24 |
| 1.1312E-17 | 0.25797335 | 0.543 | 0.362 | 2.6721E-13 | PDC | WASF2 |
| 1.2281E-17 | -0.2611007 | 0.009 | 0.132 | 2.9012E-13 | PDC | C1orf116 |
| 1.3867E-17 | -0.5790337 | 0.016 | 0.143 | 3.2759E-13 | PDC | SNCG |
| 1.4411E-17 | -0.2507095 | 0.018 | 0.146 | 3.4043E-13 | PDC | CCDC69 |
| 1.6803E-17 | -0.3012679 | 0.004 | 0.122 | 3.9694E-13 | PDC | LILRB2 |
| 1.8345E-17 | -0.3521252 | 0.007 | 0.128 | 4.3337E-13 | PDC | MARCO |
| 2.1158E-17 | -1.1528457 | 0.005 | 0.124 | 4.9981E-13 | PDC | CCL3 |
| 2.4338E-17 | -0.3111124 | 0.068 | 0.215 | 5.7494E-13 | PDC | RUNX1 |
| 2.5104E-17 | -0.2778131 | 0.072 | 0.22 | 5.9304E-13 | PDC | RIT1 |
| 2.9771E-17 | -0.3835681 | 0.113 | 0.266 | 7.0328E-13 | PDC | BACH1 |
| 3.0616E-17 | -0.2996554 | 0.005 | 0.123 | 7.2324E-13 | PDC | FOXQ1 |
| 3.3392E-17 | -0.397304 | 0.041 | 0.178 | 7.8883E-13 | PDC | FHL2 |
| 3.6239E-17 | -0.8009335 | 0.058 | 0.196 | 8.5608E-13 | PDC | C12orf75 |
| 3.6351E-17 | -0.5745348 | 0.128 | 0.277 | 8.5873E-13 | PDC | CAMK2N1 |
| 3.8987E-17 | -0.4600428 | 0 | 0.114 | 9.2098E-13 | PDC | TMPRSS11A |
| 3.9293E-17 | -0.2676344 | 0.011 | 0.132 | 9.2822E-13 | PDC | LILRB3 |
| 4.0035E-17 | -0.5954907 | 0 | 0.114 | 9.4575E-13 | PDC | KRT78 |
| 4.4136E-17 | -0.3310599 | 0.077 | 0.223 | 1.0426E-12 | PDC | ACTN1 |
| 4.935E-17 | -0.9998301 | 0.05 | 0.186 | 1.1658E-12 | PDC | HLA-DPB1 |
| 5.0774E-17 | -0.3694761 | 0.002 | 0.116 | 1.1994E-12 | PDC | CEACAM1 |
| 5.106E-17 | -0.2568723 | 0.045 | 0.183 | 1.2062E-12 | PDC | PFKP |
| 5.2719E-17 | -0.4566421 | 0.21 | 0.362 | 1.2454E-12 | PDC | CFLAR |
| 5.3518E-17 | -0.2691201 | 0.004 | 0.119 | 1.2643E-12 | PDC | CD86 |
| 5.722E-17 | -0.3715342 | 0.049 | 0.185 | 1.3517E-12 | PDC | ICAM1 |
| 5.7586E-17 | -0.2926221 | 0 | 0.112 | 1.3604E-12 | PDC | FUT3 |
| 5.7859E-17 | -0.4011788 | 0.491 | 0.624 | 1.3668E-12 | PDC | PRRC2C |

| | | | | | | |
|---|---|---|---|---|---|---|
| 8.2677E-17 | -0.4362554 | 0.192 | 0.342 | 1.9531E-12 | PDC | YTHDC1 |
| 8.5106E-17 | -1.279516 | 0.002 | 0.114 | 2.0105E-12 | PDC | SPRR1B |
| 8.5354E-17 | -0.3217297 | 0.011 | 0.129 | 2.0163E-12 | PDC | FAM84A |
| 9.2398E-17 | -0.3386766 | 0.128 | 0.28 | 2.1827E-12 | PDC | ADAM9 |
| 9.4404E-17 | -0.2671904 | 0.002 | 0.114 | 2.2301E-12 | PDC | PECAM1 |
| 1.0113E-16 | -0.3477488 | 0.004 | 0.117 | 2.389E-12 | PDC | CD163 |
| 1.0531E-16 | -0.4580436 | 0.085 | 0.227 | 2.4877E-12 | PDC | INTS6 |
| 1.1309E-16 | -0.3262492 | 0.036 | 0.165 | 2.6716E-12 | PDC | CLEC2B |
| 1.1702E-16 | -0.2980537 | 0.045 | 0.178 | 2.7643E-12 | PDC | ABCA1 |
| 1.2229E-16 | -0.4132668 | 0.128 | 0.276 | 2.8889E-12 | PDC | JUP |
| 1.3965E-16 | -0.2567247 | 0.031 | 0.16 | 3.2991E-12 | PDC | CRB3 |
| 1.8531E-16 | -0.4003567 | 0.067 | 0.205 | 4.3777E-12 | PDC | CTSC |
| 1.981E-16 | -0.2810997 | 0.02 | 0.141 | 4.6797E-12 | PDC | IFI30 |
| 2.5182E-16 | -0.3665555 | 0.077 | 0.217 | 5.9487E-12 | PDC | KDM6B |
| 2.6091E-16 | -0.2775184 | 0.02 | 0.14 | 6.1634E-12 | PDC | ZNRF1 |
| 2.6636E-16 | -0.4141966 | 0.065 | 0.204 | 6.2922E-12 | PDC | FOSB |
| 2.7378E-16 | 0.29279529 | 0.973 | 0.945 | 6.4676E-12 | PDC | RPS19 |
| 2.791E-16 | -0.4188834 | 0 | 0.108 | 6.5932E-12 | PDC | TMC5 |
| 3.0761E-16 | 0.30148738 | 0.951 | 0.903 | 7.2667E-12 | PDC | RPL37 |
| 3.1304E-16 | -0.3593093 | 0.007 | 0.12 | 7.395E-12 | PDC | FCGR3A |
| 3.2811E-16 | -0.667726 | 0.444 | 0.64 | 7.7509E-12 | PDC | MTRNR2L12 |
| 3.439E-16 | -0.2564176 | 0.025 | 0.148 | 8.1239E-12 | PDC | KCNK1 |
| 3.4517E-16 | -0.2591038 | 0.045 | 0.177 | 8.1539E-12 | PDC | TRAF4 |
| 3.9716E-16 | -0.3444222 | 0 | 0.107 | 9.3822E-12 | PDC | CSF3R |
| 4.0285E-16 | -0.2912989 | 0.004 | 0.113 | 9.5164E-12 | PDC | HIST1H2BG |
| 4.2768E-16 | -0.3274342 | 0 | 0.107 | 1.0103E-11 | PDC | A2ML1 |
| 4.6351E-16 | -1.1070552 | 0.007 | 0.118 | 1.0949E-11 | PDC | SPINK5 |
| 4.755E-16 | -0.2939414 | 0.025 | 0.146 | 1.1233E-11 | PDC | PDP1 |
| 4.7779E-16 | -0.2735676 | 0.016 | 0.133 | 1.1287E-11 | PDC | EPB41L3 |
| 4.8799E-16 | -0.2568709 | 0.018 | 0.135 | 1.1528E-11 | PDC | GPAT3 |
| 6.1083E-16 | -1.4311876 | 0.002 | 0.109 | 1.443E-11 | PDC | CNFN |
| 6.3299E-16 | -0.2551226 | 0.009 | 0.121 | 1.4953E-11 | PDC | CSF1R |
| 8.1364E-16 | -0.4532187 | 0.029 | 0.15 | 1.9221E-11 | PDC | RASD1 |
| 8.9909E-16 | -0.5562689 | 0.151 | 0.295 | 2.1239E-11 | PDC | ATP13A3 |
| 1.0014E-15 | -0.9757353 | 0.009 | 0.119 | 2.3655E-11 | PDC | CXCL2 |
| 1.2718E-15 | -0.4614508 | 0.047 | 0.173 | 3.0043E-11 | PDC | DNAJA4 |
| 1.3382E-15 | -0.3872836 | 0.022 | 0.137 | 3.1611E-11 | PDC | GCNT3 |
| 1.4363E-15 | -0.2701514 | 0.043 | 0.169 | 3.3929E-11 | PDC | GLIPR2 |
| 1.4964E-15 | -0.3239923 | 0.013 | 0.124 | 3.5349E-11 | PDC | KRT80 |
| 1.5093E-15 | -0.2997529 | 0.041 | 0.167 | 3.5655E-11 | PDC | JARID2 |
| 1.5387E-15 | -0.2974021 | 0.031 | 0.15 | 3.6348E-11 | PDC | BCAT1 |
| 1.6533E-15 | -0.3633985 | 0.011 | 0.121 | 3.9057E-11 | PDC | EGLN3 |
| 1.738E-15 | -0.7779312 | 0.002 | 0.106 | 4.1056E-11 | PDC | ZG16B |
| 1.8727E-15 | -0.6070708 | 0.088 | 0.223 | 4.4238E-11 | PDC | HES1 |
| 2.1353E-15 | -0.37488 | 0.101 | 0.239 | 5.0442E-11 | PDC | RNASET2 |
| 3.5881E-15 | -0.3621857 | 0.103 | 0.243 | 8.4761E-11 | PDC | BHLHE40 |
| 4.2441E-15 | -0.255134 | 0.052 | 0.178 | 1.0026E-10 | PDC | PPP1R18 |
| 4.5014E-15 | -0.5602175 | 0.011 | 0.118 | 1.0634E-10 | PDC | MNDA |
| 5.0451E-15 | -0.3825439 | 0.005 | 0.109 | 1.1918E-10 | PDC | KRT23 |
| 6.1286E-15 | -0.7445329 | 0.009 | 0.114 | 1.4478E-10 | PDC | IL7R |

| | | | | | | |
|---|---|---|---|---|---|---|
| 6.3886E-15 | -0.2823964 | 0.126 | 0.269 | 1.5092E-10 | PDC | DIAPH1 |
| 6.7282E-15 | -0.4444299 | 0.027 | 0.14 | 1.5894E-10 | PDC | PLAC8 |
| 7.3687E-15 | -0.2842984 | 0.005 | 0.108 | 1.7407E-10 | PDC | MS4A6A |
| 7.9858E-15 | -0.5084462 | 0.014 | 0.121 | 1.8865E-10 | PDC | TCIM |
| 8.341E-15 | -0.3408047 | 0.131 | 0.268 | 1.9704E-10 | PDC | USP15 |
| 9.0394E-15 | -0.4124648 | 0.079 | 0.206 | 2.1354E-10 | PDC | CSNK1E |
| 9.3714E-15 | -0.2502222 | 0.045 | 0.166 | 2.2138E-10 | PDC | SMAD3 |
| 9.4719E-15 | -0.4822595 | 0.002 | 0.101 | 2.2375E-10 | PDC | RETN |
| 1.0897E-14 | -0.5021951 | 0.451 | 0.58 | 2.5742E-10 | PDC | LITAF |
| 1.1002E-14 | -0.2966381 | 0.088 | 0.22 | 2.599E-10 | PDC | POLR2J3.1 |
| 1.1306E-14 | 0.26622045 | 0.953 | 0.898 | 2.6708E-10 | PDC | RPL27A |
| 1.2767E-14 | -1.0683259 | 0.007 | 0.109 | 3.0158E-10 | PDC | CCL4 |
| 1.3577E-14 | -0.2807233 | 0.022 | 0.132 | 3.2072E-10 | PDC | HIST1H2BC |
| 1.3799E-14 | -0.3036036 | 0.007 | 0.109 | 3.2597E-10 | PDC | FGL2 |
| 1.6644E-14 | -0.5637908 | 0.712 | 0.767 | 3.9317E-10 | PDC | CST3 |
| 1.9048E-14 | -1.5286039 | 0.007 | 0.108 | 4.4997E-10 | PDC | CRCT1 |
| 1.921E-14 | -0.3121968 | 0.122 | 0.258 | 4.5381E-10 | PDC | LYPLA1 |
| 2.1317E-14 | -0.4335681 | 0.025 | 0.136 | 5.0358E-10 | PDC | HLA-DRB5 |
| 2.5002E-14 | -0.3183676 | 0.07 | 0.195 | 5.9062E-10 | PDC | SNX10 |
| 2.5332E-14 | -0.4626299 | 0.806 | 0.831 | 5.9841E-10 | PDC | HNRNPA2B1 |
| 2.5623E-14 | 0.29733901 | 0.331 | 0.211 | 6.053E-10 | PDC | RNF145 |
| 2.8416E-14 | -0.4430668 | 0.049 | 0.165 | 6.7127E-10 | PDC | ANKRD37 |
| 2.9547E-14 | -0.2802181 | 0.032 | 0.146 | 6.9799E-10 | PDC | CLDN1 |
| 3.3518E-14 | -0.256972 | 0.023 | 0.132 | 7.9178E-10 | PDC | SCIN |
| 3.4889E-14 | -0.7850785 | 0.669 | 0.723 | 8.2418E-10 | PDC | HSPB1 |
| 3.7709E-14 | -0.4479033 | 0.408 | 0.532 | 8.9081E-10 | PDC | PTTG1IP |
| 4.4314E-14 | -0.3274306 | 0.086 | 0.211 | 1.0468E-09 | PDC | ARRB2 |
| 4.8467E-14 | -0.4606381 | 0.076 | 0.197 | 1.1449E-09 | PDC | SAMD9 |
| 7.3117E-14 | -0.3189378 | 0.016 | 0.119 | 1.7272E-09 | PDC | DUSP2 |
| 8.396E-14 | -0.2710612 | 0.031 | 0.139 | 1.9834E-09 | PDC | ETS1 |
| 9.3894E-14 | -0.2627035 | 0.007 | 0.104 | 2.2181E-09 | PDC | LILRB1 |
| 9.5576E-14 | -0.2926564 | 0.04 | 0.151 | 2.2578E-09 | PDC | DMXL2 |
| 1.1094E-13 | -0.3955237 | 0.14 | 0.268 | 2.6208E-09 | PDC | AFDN |
| 1.1595E-13 | -0.4049057 | 0.183 | 0.313 | 2.739E-09 | PDC | SMS |
| 1.1748E-13 | -0.3311783 | 0.164 | 0.302 | 2.7753E-09 | PDC | SERTAD1 |
| 1.1894E-13 | -0.3317244 | 0.013 | 0.111 | 2.8096E-09 | PDC | SMIM5 |
| 1.2977E-13 | -0.3131359 | 0.027 | 0.134 | 3.0655E-09 | PDC | RGCC |
| 1.3206E-13 | -0.3978138 | 0.48 | 0.63 | 3.1197E-09 | PDC | JUN |
| 1.6523E-13 | -0.3876097 | 0.011 | 0.108 | 3.9033E-09 | PDC | THBD |
| 1.7958E-13 | -0.2680444 | 0.022 | 0.124 | 4.2423E-09 | PDC | PHACTR1 |
| 2.2987E-13 | -0.3278736 | 0.18 | 0.313 | 5.4303E-09 | PDC | CTNNB1 |
| 2.5411E-13 | -0.5375061 | 0.509 | 0.607 | 6.0028E-09 | PDC | EZR |
| 2.8107E-13 | -0.527165 | 0.191 | 0.316 | 6.6397E-09 | PDC | HIST1H2AC |
| 3.4129E-13 | -0.7051657 | 0.178 | 0.293 | 8.0623E-09 | PDC | GPRC5A |
| 3.4692E-13 | -0.3008567 | 0.07 | 0.187 | 8.1952E-09 | PDC | CKLF |
| 3.6654E-13 | -0.5695918 | 0.009 | 0.102 | 8.6589E-09 | PDC | HLA-DQA1 |
| 3.9049E-13 | -0.2788511 | 0.009 | 0.102 | 9.2246E-09 | PDC | LRMP |
| 4.1109E-13 | -0.3028408 | 0.032 | 0.138 | 9.7111E-09 | PDC | HLA-DMA |
| 4.1243E-13 | -0.3132906 | 0.014 | 0.111 | 9.7428E-09 | PDC | TNF |
| 4.4203E-13 | -0.4332003 | 0.333 | 0.494 | 1.0442E-08 | PDC | MTRNR2L8 |

| | | | | | | |
|---|---|---|---|---|---|---|
| 4.9257E-13 | -0.2829027 | 0.065 | 0.181 | 1.1636E-08 | PDC | SLC20A1 |
| 5.0022E-13 | -0.4519043 | 0.304 | 0.415 | 1.1817E-08 | PDC | SFPQ |
| 6.3193E-13 | -0.2861458 | 0.137 | 0.263 | 1.4928E-08 | PDC | CHP1 |
| 6.4129E-13 | -0.4577788 | 0.975 | 0.965 | 1.5149E-08 | PDC | UBC |
| 7.4863E-13 | -0.3217754 | 0.031 | 0.132 | 1.7685E-08 | PDC | CD83 |
| 7.8623E-13 | -0.2791302 | 0.056 | 0.167 | 1.8573E-08 | PDC | CEP170 |
| 8.9064E-13 | -0.3047234 | 0.085 | 0.203 | 2.104E-08 | PDC | NFKB1 |
| 1.0503E-12 | -0.2817588 | 0.112 | 0.233 | 2.4812E-08 | PDC | BCL10 |
| 1.227E-12 | -0.4457683 | 0.137 | 0.26 | 2.8985E-08 | PDC | IDI1 |
| 1.602E-12 | -0.3199254 | 0.182 | 0.311 | 3.7844E-08 | PDC | OGA |
| 1.6508E-12 | -0.2886987 | 0.018 | 0.111 | 3.8996E-08 | PDC | STS |
| 1.6686E-12 | -0.3064317 | 0.05 | 0.157 | 3.9416E-08 | PDC | COBLL1 |
| 1.8169E-12 | -0.3652867 | 0.027 | 0.124 | 4.292E-08 | PDC | TMEM40 |
| 2.3767E-12 | -0.3418129 | 0.108 | 0.225 | 5.6145E-08 | PDC | STXBP2 |
| 2.4338E-12 | -0.4029242 | 0.547 | 0.641 | 5.7494E-08 | PDC | ARPC5 |
| 2.8122E-12 | -0.2991949 | 0.103 | 0.219 | 6.6432E-08 | PDC | STK4 |
| 3.0217E-12 | -0.3307389 | 0.254 | 0.386 | 7.1383E-08 | PDC | HNRNPH1 |
| 3.0438E-12 | -0.3006379 | 0.137 | 0.258 | 7.1904E-08 | PDC | CHD2 |
| 3.1261E-12 | -0.6906788 | 0.018 | 0.11 | 7.3847E-08 | PDC | APOC1 |
| 3.3911E-12 | -0.3301287 | 0.187 | 0.313 | 8.0108E-08 | PDC | TMEM165 |
| 3.8236E-12 | -0.9963854 | 0.076 | 0.182 | 9.0325E-08 | PDC | SPINK1 |
| 4.4817E-12 | -0.3726155 | 0.032 | 0.13 | 1.0587E-07 | PDC | AC023157.3 |
| 4.562E-12 | -0.2970763 | 0.121 | 0.238 | 1.0777E-07 | PDC | PNPLA2 |
| 4.9836E-12 | -0.2737054 | 0.973 | 0.993 | 1.1773E-07 | PDC | TPT1 |
| 5.2368E-12 | -0.3382251 | 0.104 | 0.219 | 1.2371E-07 | PDC | DSE |
| 5.2524E-12 | -0.5311776 | 0.031 | 0.13 | 1.2408E-07 | PDC | TFPI2 |
| 5.5834E-12 | -0.3523503 | 0.147 | 0.275 | 1.319E-07 | PDC | TNFRSF12A |
| 5.9671E-12 | -0.341268 | 0.183 | 0.309 | 1.4096E-07 | PDC | NUFIP2 |
| 6.6863E-12 | -0.2683278 | 0.07 | 0.179 | 1.5795E-07 | PDC | EHD1 |
| 6.7529E-12 | -0.3609104 | 0.52 | 0.6 | 1.5952E-07 | PDC | VAMP8 |
| 6.8051E-12 | -0.2695401 | 0.013 | 0.1 | 1.6076E-07 | PDC | LY6K |
| 9.3457E-12 | -0.4113338 | 0.063 | 0.167 | 2.2077E-07 | PDC | HIST3H2A |
| 9.6384E-12 | -0.264754 | 0.02 | 0.11 | 2.2769E-07 | PDC | YOD1 |
| 1.0107E-11 | -0.5083729 | 0.029 | 0.121 | 2.3875E-07 | PDC | HILPDA |
| 1.0983E-11 | -0.2557999 | 0.104 | 0.221 | 2.5946E-07 | PDC | TES |
| 1.2899E-11 | -0.3557036 | 0.246 | 0.36 | 3.0472E-07 | PDC | PLSCR1 |
| 1.4674E-11 | -0.2536822 | 0.076 | 0.184 | 3.4664E-07 | PDC | STK17A |
| 1.665E-11 | -0.4509854 | 0.574 | 0.648 | 3.9332E-07 | PDC | LDHA |
| 1.9485E-11 | -0.3264645 | 0.227 | 0.347 | 4.6028E-07 | PDC | ZNF207 |
| 2.0348E-11 | -0.3398551 | 0.498 | 0.585 | 4.8067E-07 | PDC | NOP10 |
| 2.3909E-11 | -0.6447817 | 0.018 | 0.105 | 5.6479E-07 | PDC | CCL5 |
| 2.6887E-11 | -0.3319484 | 0.183 | 0.297 | 6.3514E-07 | PDC | VPS4B |
| 3.0899E-11 | -0.3927345 | 0.737 | 0.781 | 7.2992E-07 | PDC | MAP1LC3B |
| 3.2037E-11 | -0.7377647 | 0.209 | 0.114 | 7.568E-07 | PDC | MT1M |
| 3.2853E-11 | -0.2812145 | 0.133 | 0.246 | 7.7609E-07 | PDC | PTPN12 |
| 4.3019E-11 | -0.9410672 | 0.095 | 0.201 | 1.0162E-06 | PDC | THBS1 |
| 4.9376E-11 | -0.3004371 | 0.219 | 0.332 | 1.1664E-06 | PDC | MOB1A |
| 5.6801E-11 | -0.2678814 | 0.072 | 0.175 | 1.3418E-06 | PDC | KCTD12 |
| 5.8174E-11 | -0.2830238 | 0.149 | 0.262 | 1.3742E-06 | PDC | MBP |
| 5.9786E-11 | -0.4627055 | 0.086 | 0.19 | 1.4123E-06 | PDC | HBEGF |

| | | | | | | |
|---|---|---|---|---|---|---|
| 6.1681E-11 | -0.2577229 | 0.14 | 0.257 | 1.4571E-06 | PDC | TACC1 |
| 7.3813E-11 | -0.3717641 | 0.058 | 0.156 | 1.7437E-06 | PDC | CYR61 |
| 7.774E-11 | -0.3682129 | 0.149 | 0.266 | 1.8364E-06 | PDC | MGST1 |
| 1.1136E-10 | -0.2917461 | 0.038 | 0.127 | 2.6306E-06 | PDC | POLB |
| 1.1308E-10 | -0.2695797 | 0.04 | 0.13 | 2.6713E-06 | PDC | GAN |
| 1.1607E-10 | -0.3204756 | 0.128 | 0.25 | 2.7419E-06 | PDC | EGR1 |
| 1.366E-10 | -0.2865061 | 0.088 | 0.192 | 3.227E-06 | PDC | LIMA1 |
| 1.5062E-10 | -0.4039909 | 0.086 | 0.187 | 3.558E-06 | PDC | ACSL1 |
| 1.5707E-10 | -1.091314 | 0.049 | 0.144 | 3.7105E-06 | PDC | CCL2 |
| 1.5837E-10 | -0.2887472 | 0.049 | 0.144 | 3.7411E-06 | PDC | PAX8 |
| 1.7187E-10 | -0.4652529 | 0.04 | 0.13 | 4.06E-06 | PDC | AKAP12 |
| 1.7586E-10 | -0.4510193 | 0.153 | 0.263 | 4.1544E-06 | PDC | HES4 |
| 1.9177E-10 | 0.39054845 | 0.243 | 0.391 | 4.5301E-06 | PDC | FOS |
| 1.926E-10 | -3.3022483 | 0.432 | 0.543 | 4.5499E-06 | PDC | PLCG2 |
| 1.9437E-10 | -0.433421 | 0.065 | 0.159 | 4.5916E-06 | PDC | BPGM |
| 1.9725E-10 | -0.3370111 | 0.049 | 0.142 | 4.6597E-06 | PDC | ID1 |
| 2.0416E-10 | -0.2951456 | 0.153 | 0.26 | 4.8229E-06 | PDC | ANKLE2 |
| 2.5302E-10 | -1.1269603 | 0.649 | 0.71 | 5.9771E-06 | PDC | ANXA1 |
| 2.7238E-10 | -0.2538991 | 0.149 | 0.26 | 6.4345E-06 | PDC | ERBIN |
| 3.2098E-10 | -0.3075349 | 0.21 | 0.319 | 7.5825E-06 | PDC | CMTM6 |
| 3.4399E-10 | -0.35758 | 0.023 | 0.105 | 8.1261E-06 | PDC | LTB |
| 4.4376E-10 | -0.3180011 | 0.124 | 0.227 | 1.0483E-05 | PDC | RAB9A |
| 4.7378E-10 | -0.3702837 | 0.09 | 0.188 | 1.1192E-05 | PDC | BAG3 |
| 5.1437E-10 | -0.3676702 | 0.067 | 0.163 | 1.2151E-05 | PDC | CSKMT |
| 5.1743E-10 | -0.2711414 | 0.128 | 0.232 | 1.2223E-05 | PDC | ATP6V1B2 |
| 5.189E-10 | -0.2991848 | 0.696 | 0.748 | 1.2258E-05 | PDC | ATP6V0E1 |
| 5.6422E-10 | -0.2611543 | 0.126 | 0.23 | 1.3329E-05 | PDC | PGD |
| 6.0182E-10 | -0.3097821 | 0.076 | 0.171 | 1.4217E-05 | PDC | FAM214A |
| 7.0307E-10 | -0.2635723 | 0.185 | 0.295 | 1.6609E-05 | PDC | SYAP1 |
| 8.7143E-10 | -0.7866292 | 0.049 | 0.138 | 2.0586E-05 | PDC | DEFB1 |
| 9.3448E-10 | -0.2736919 | 0.387 | 0.484 | 2.2075E-05 | PDC | TMEM50A |
| 9.7819E-10 | -0.2798337 | 0.167 | 0.27 | 2.3108E-05 | PDC | AGFG1 |
| 1.1285E-09 | -0.5120803 | 0.128 | 0.229 | 2.6658E-05 | PDC | IL32 |
| 1.3261E-09 | -0.3510269 | 0.027 | 0.104 | 3.1326E-05 | PDC | CLIC3 |
| 1.3272E-09 | -0.3265324 | 0.237 | 0.336 | 3.1353E-05 | PDC | RBM25 |
| 1.8837E-09 | -0.2946051 | 0.086 | 0.179 | 4.45E-05 | PDC | LTA4H |
| 2.2037E-09 | -0.2668987 | 0.129 | 0.229 | 5.2059E-05 | PDC | NUMB |
| 2.5883E-09 | -0.269575 | 0.04 | 0.12 | 6.1143E-05 | PDC | AC007952.4 |
| 3.2082E-09 | -0.4377996 | 0.063 | 0.149 | 7.5788E-05 | PDC | SMC4 |
| 3.7124E-09 | -0.8954787 | 0.187 | 0.264 | 8.7698E-05 | PDC | LMO7 |
| 3.7844E-09 | -0.3273227 | 0.149 | 0.243 | 8.9398E-05 | PDC | GIPC1 |
| 4.1121E-09 | -0.2765672 | 0.223 | 0.321 | 9.7141E-05 | PDC | AP2S1 |
| 4.5119E-09 | -0.2667385 | 0.236 | 0.337 | 0.00010658 | PDC | CAPZA1 |
| 5.2085E-09 | -0.2540049 | 0.086 | 0.177 | 0.00012304 | PDC | P4HA1 |
| 8.5441E-09 | -0.3930929 | 0.284 | 0.362 | 0.00020184 | PDC | GNAI2 |
| 9.7725E-09 | -0.2646329 | 0.173 | 0.269 | 0.00023086 | PDC | RAB5IF |
| 1.2446E-08 | -0.3855134 | 0.439 | 0.502 | 0.000294 | PDC | ASAH1 |
| 1.2727E-08 | 0.38455297 | 0.98 | 0.937 | 0.00030065 | PDC | TMSB4X |
| 1.5765E-08 | -0.284406 | 0.228 | 0.325 | 0.00037243 | PDC | ARL6IP1 |
| 1.7426E-08 | -0.2797824 | 0.115 | 0.205 | 0.00041165 | PDC | TIPARP |

| | | | | | | |
|---|---|---|---|---|---|---|
| 1.9517E-08 | -0.3994365 | 0.221 | 0.308 | 0.00046105 | PDC | ZNF292 |
| 2.0893E-08 | -0.3035549 | 0.155 | 0.245 | 0.00049355 | PDC | RAB31 |
| 2.2416E-08 | -0.2854281 | 0.038 | 0.111 | 0.00052954 | PDC | FXYD2 |
| 2.7331E-08 | -0.2997913 | 0.453 | 0.521 | 0.00064563 | PDC | ATP6V1F |
| 2.9477E-08 | 0.3347754 | 0.122 | 0.066 | 0.00069634 | PDC | CFH |
| 3.1732E-08 | -0.5199834 | 0.935 | 0.932 | 0.00074961 | PDC | GAPDH |
| 3.3647E-08 | -0.2677308 | 0.11 | 0.198 | 0.00079484 | PDC | KLF4 |
| 3.9411E-08 | -0.2697381 | 0.077 | 0.157 | 0.00093101 | PDC | TSPAN6 |
| 4.2474E-08 | -0.3345221 | 0.854 | 0.864 | 0.00100336 | PDC | CALM1 |
| 4.3872E-08 | -0.3619119 | 0.138 | 0.226 | 0.0010364 | PDC | PPP1R10 |
| 4.4873E-08 | -0.3570176 | 0.099 | 0.184 | 0.00106004 | PDC | INSIG1 |
| 4.5122E-08 | -0.3557106 | 0.147 | 0.235 | 0.00106592 | PDC | TOB1 |
| 4.971E-08 | -0.346818 | 0.072 | 0.152 | 0.00117431 | PDC | SLC7A11 |
| 5.246E-08 | -0.3194781 | 0.081 | 0.161 | 0.00123926 | PDC | SERPINB9 |
| 5.8625E-08 | -0.2943232 | 0.061 | 0.138 | 0.00138491 | PDC | DCDC2 |
| 6.9186E-08 | -0.2797331 | 0.041 | 0.113 | 0.00163437 | PDC | ANGPTL4 |
| 1.0426E-07 | -0.2707328 | 0.228 | 0.322 | 0.00246294 | PDC | YPEL3 |
| 1.0841E-07 | -0.3093827 | 0.466 | 0.508 | 0.00256092 | PDC | DBI |
| 1.3837E-07 | -0.254731 | 0.203 | 0.291 | 0.00326863 | PDC | CHMP2B |
| 1.6783E-07 | -0.9382791 | 0.059 | 0.132 | 0.00396461 | PDC | APOE |
| 1.7033E-07 | -0.4506186 | 0.153 | 0.237 | 0.00402362 | PDC | FABP5 |
| 1.8641E-07 | -0.2825921 | 0.987 | 0.98 | 0.0044035 | PDC | MALAT1 |
| 2.068E-07 | -0.510393 | 0.115 | 0.198 | 0.00488514 | PDC | TM4SF1 |
| 2.582E-07 | -0.3050148 | 0.113 | 0.194 | 0.00609957 | PDC | KATNBL1 |
| 2.6835E-07 | -0.3386101 | 0.349 | 0.417 | 0.00633922 | PDC | FUS |
| 3.4184E-07 | -0.2746474 | 0.138 | 0.223 | 0.00807534 | PDC | POLR2A |
| 3.8255E-07 | -0.4471538 | 0.491 | 0.523 | 0.00903688 | PDC | HEBP2 |
| 4.5413E-07 | -0.6206409 | 0.155 | 0.234 | 0.01072799 | PDC | PLIN2 |
| 7.0644E-07 | -0.2657733 | 0.257 | 0.332 | 0.01668828 | PDC | MSN |
| 8.6308E-07 | -0.3401091 | 0.309 | 0.378 | 0.02038864 | PDC | GCC2 |
| 1.1253E-06 | -0.4896131 | 0.16 | 0.228 | 0.02658268 | PDC | ZEB2 |
| 1.4786E-06 | -0.2898962 | 0.669 | 0.695 | 0.03492987 | PDC | LRRFIP1 |
| 1.8681E-06 | -0.2523207 | 0.7 | 0.723 | 0.04412986 | PDC | YWHAB |
| 2.1659E-06 | -0.6328509 | 0.192 | 0.263 | 0.05116463 | PDC | AREG |
| 2.463E-06 | -0.3653958 | 0.201 | 0.275 | 0.05818278 | PDC | ADM |
| 3.8427E-06 | 0.28066962 | 0.851 | 0.909 | 0.09077697 | PDC | MT-ND2 |
| 4.2746E-06 | -0.4492943 | 0.282 | 0.345 | 0.10097803 | PDC | DDIT4 |
| 5.3114E-06 | -0.2948965 | 0.532 | 0.575 | 0.12547036 | PDC | PPP1CB |
| 8.3406E-06 | -0.2652972 | 0.259 | 0.325 | 0.19703016 | PDC | YPEL5 |
| 1.2979E-05 | -0.3222912 | 0.309 | 0.367 | 0.30660719 | PDC | RALBP1 |
| 2.0592E-05 | -0.4824242 | 0.165 | 0.227 | 0.48644344 | PDC | HEXIM1 |
| 2.1812E-05 | -0.3525978 | 0.162 | 0.227 | 0.5152559 | PDC | IER5L |
| 2.4667E-05 | -0.2928393 | 0.146 | 0.214 | 0.5827086 | PDC | BASP1 |
| 3.302E-05 | -0.4200676 | 0.273 | 0.187 | 0.78002663 | PDC | RARRES3 |
| 4.1761E-05 | -0.2901372 | 0.372 | 0.416 | 0.98652704 | PDC | GRN |
| 0.0001131 | -0.3157141 | 0.311 | 0.35 | 1 | PDC | UBE2H |
| 0.00011371 | -0.2651042 | 0.059 | 0.107 | 1 | PDC | SLC16A10 |
| 0.00018779 | -0.2637979 | 0.061 | 0.108 | 1 | PDC | TCHP |
| 0.00024242 | -1.2375771 | 0.119 | 0.163 | 1 | PDC | ECM1 |
| 0.000331 | -0.2651322 | 0.257 | 0.304 | 1 | PDC | SRSF2 |

| | | | | | | |
|---|---|---|---|---|---|---|
| 0.00033722 | -0.372711 | 0.306 | 0.351 | 1 | PDC | GLS |
| 0.00037984 | -0.3876202 | 0.75 | 0.731 | 1 | PDC | HSPA8 |
| 0.00046663 | -0.2692826 | 0.241 | 0.289 | 1 | PDC | ATP1B3 |
| 0.00059902 | -0.2949587 | 0.689 | 0.688 | 1 | PDC | DNAJA1 |
| 0.00070039 | -0.3033087 | 0.351 | 0.386 | 1 | PDC | C9orf16 |
| 0.0008249 | -0.2571363 | 0.486 | 0.495 | 1 | PDC | SRSF3 |
| 0.00087293 | -0.3172976 | 0.085 | 0.125 | 1 | PDC | SRD5A3 |
| 0.00142152 | -0.3265258 | 0.612 | 0.597 | 1 | PDC | NPC2 |
| 0.00161125 | -0.3513872 | 0.299 | 0.323 | 1 | PDC | LIMS1 |
| 0.00197864 | 0.25950307 | 0.104 | 0.071 | 1 | PDC | TAGLN |
| 0.00274118 | -0.2941856 | 0.372 | 0.397 | 1 | PDC | ARGLU1 |
| 0.00284423 | -0.2606288 | 0.353 | 0.389 | 1 | PDC | TUBB4B |
| 0.00326066 | -0.8544988 | 0.192 | 0.223 | 1 | PDC | G0S2 |
| 0.0054112 | -0.2972689 | 0.291 | 0.316 | 1 | PDC | HMGB2 |
| 0.0056998 | -0.5637359 | 0.432 | 0.425 | 1 | PDC | EMP3 |
| 0.0058648 | 0.58373575 | 0.124 | 0.093 | 1 | PDC | CTGF |
| 0.00714347 | -0.2716451 | 0.374 | 0.391 | 1 | PDC | RSRC2 |
| 0 | 4.86288053 | 0.929 | 0.068 | 0 | TEC_PT | GPX3 |
| 0 | 4.41803764 | 0.687 | 0.002 | 0 | TEC_PT | ALDOB |
| 0 | 2.77254517 | 0.539 | 0.004 | 0 | TEC_PT | GSTA1 |
| 0 | 2.76936871 | 0.64 | 0.029 | 0 | TEC_PT | GATM |
| 0 | 2.57606767 | 0.589 | 0.004 | 0 | TEC_PT | MIOX |
| 0 | 2.57432551 | 0.418 | 0.001 | 0 | TEC_PT | FABP1 |
| 0 | 2.57134039 | 0.912 | 0.125 | 0 | TEC_PT | APOE |
| 0 | 2.51570313 | 0.808 | 0.041 | 0 | TEC_PT | CXCL14 |
| 0 | 2.17072517 | 0.38 | 0 | 0 | TEC_PT | GSTA2 |
| 0 | 1.98570119 | 0.623 | 0.04 | 0 | TEC_PT | CMBL |
| 0 | 1.97050361 | 0.599 | 0.005 | 0 | TEC_PT | RBP5 |
| 0 | 1.93902231 | 0.36 | 0.011 | 0 | TEC_PT | PCK1 |
| 0 | 1.78417498 | 0.519 | 0.004 | 0 | TEC_PT | HPD |
| 0 | 1.59479566 | 0.572 | 0.038 | 0 | TEC_PT | RIDA |
| 0 | 1.4956702 | 0.542 | 0.052 | 0 | TEC_PT | UGT2B7 |
| 0 | 1.47794683 | 0.461 | 0.034 | 0 | TEC_PT | BHMT |
| 0 | 1.41516276 | 0.401 | 0.004 | 0 | TEC_PT | BBOX1 |
| 0 | 1.31859284 | 0.508 | 0.017 | 0 | TEC_PT | PDZK1 |
| 0 | 1.31717053 | 0.434 | 0.001 | 0 | TEC_PT | NAT8 |
| 0 | 1.31473253 | 0.441 | 0.009 | 0 | TEC_PT | KHK |
| 0 | 1.27011897 | 0.569 | 0.038 | 0 | TEC_PT | BHMT2 |
| 0 | 1.26888303 | 0.374 | 0 | 0 | TEC_PT | ALB |
| 0 | 1.08779182 | 0.444 | 0.001 | 0 | TEC_PT | DPYS |
| 0 | 1.07689596 | 0.424 | 0.021 | 0 | TEC_PT | SMIM24 |
| 0 | 1.05852207 | 0.354 | 0.001 | 0 | TEC_PT | GLYAT |
| 0 | 1.03424949 | 0.377 | 0.007 | 0 | TEC_PT | AKR7A3 |
| 0 | 0.99355128 | 0.441 | 0.002 | 0 | TEC_PT | DPEP1 |
| 0 | 0.98827734 | 0.455 | 0.032 | 0 | TEC_PT | SHMT1 |
| 0 | 0.94029726 | 0.414 | 0.026 | 0 | TEC_PT | RAB11FIP3 |
| 0 | 0.92889447 | 0.364 | 0.023 | 0 | TEC_PT | ALDH4A1 |
| 0 | 0.85303072 | 0.343 | 0.005 | 0 | TEC_PT | AQP1 |
| 0 | 0.84962733 | 0.354 | 0.005 | 0 | TEC_PT | AGMAT |
| 0 | 0.83949787 | 0.31 | 0.001 | 0 | TEC_PT | PRODH2 |

| | | | | | | |
|---|---|---|---|---|---|---|
| 0 | 0.83419703 | 0.367 | 0.002 | 0 | TEC_PT | ACSM2B |
| 0 | 0.80671951 | 0.296 | 0.001 | 0 | TEC_PT | PAH |
| 0 | 0.78180373 | 0.35 | 0.001 | 0 | TEC_PT | DDC |
| 0 | 0.77311968 | 0.31 | 0.006 | 0 | TEC_PT | GPD1 |
| 0 | 0.71896242 | 0.269 | 0 | 0 | TEC_PT | AGXT2 |
| 0 | 0.71648968 | 0.316 | 0.004 | 0 | TEC_PT | LRP2 |
| 0 | 0.71389172 | 0.343 | 0.016 | 0 | TEC_PT | EHHADH |
| 0 | 0.70878869 | 0.333 | 0.013 | 0 | TEC_PT | APOM |
| 0 | 0.69097814 | 0.347 | 0.002 | 0 | TEC_PT | ACSM2A |
| 0 | 0.68804363 | 0.313 | 0.002 | 0 | TEC_PT | FTCD |
| 0 | 0.67974869 | 0.215 | 0.001 | 0 | TEC_PT | GLYATL1 |
| 0 | 0.63262636 | 0.246 | 0.005 | 0 | TEC_PT | CUBN |
| 0 | 0.62603139 | 0.3 | 0.003 | 0 | TEC_PT | ACMSD |
| 0 | 0.60249403 | 0.273 | 0 | 0 | TEC_PT | SLC22A12 |
| 0 | 0.5910984 | 0.29 | 0.005 | 0 | TEC_PT | SLC6A13 |
| 0 | 0.58440282 | 0.306 | 0.013 | 0 | TEC_PT | SLC22A2 |
| 0 | 0.5793468 | 0.253 | 0 | 0 | TEC_PT | PRAP1 |
| 0 | 0.56622983 | 0.29 | 0.002 | 0 | TEC_PT | SLC13A3 |
| 0 | 0.56588986 | 0.263 | 0.003 | 0 | TEC_PT | PIPOX |
| 0 | 0.56107497 | 0.293 | 0.01 | 0 | TEC_PT | HGD |
| 0 | 0.54752099 | 0.249 | 0.001 | 0 | TEC_PT | SLC17A3 |
| 0 | 0.53125062 | 0.215 | 0.002 | 0 | TEC_PT | ACY3 |
| 0 | 0.52680693 | 0.229 | 0 | 0 | TEC_PT | SLC22A8 |
| 0 | 0.52261429 | 0.226 | 0.001 | 0 | TEC_PT | SLC5A12 |
| 0 | 0.51998306 | 0.276 | 0.006 | 0 | TEC_PT | GIPC2 |
| 0 | 0.50438091 | 0.327 | 0.014 | 0 | TEC_PT | HPN |
| 0 | 0.49468493 | 0.276 | 0.003 | 0 | TEC_PT | USH1C |
| 0 | 0.48742488 | 0.222 | 0.003 | 0 | TEC_PT | AOX1 |
| 0 | 0.48577966 | 0.222 | 0 | 0 | TEC_PT | ASPDH |
| 0 | 0.47523565 | 0.232 | 0 | 0 | TEC_PT | SLC22A6 |
| 0 | 0.46530376 | 0.135 | 0.002 | 0 | TEC_PT | SERPINA6 |
| 0 | 0.45912209 | 0.263 | 0.011 | 0 | TEC_PT | NAPSA |
| 0 | 0.45391816 | 0.199 | 0 | 0 | TEC_PT | SMIM32 |
| 0 | 0.42848044 | 0.168 | 0.001 | 0 | TEC_PT | SLC13A1 |
| 0 | 0.41605576 | 0.172 | 0.004 | 0 | TEC_PT | AGXT |
| 0 | 0.40323885 | 0.222 | 0.003 | 0 | TEC_PT | AGT |
| 0 | 0.40316166 | 0.205 | 0.005 | 0 | TEC_PT | ALDH8A1 |
| 0 | 0.39329996 | 0.195 | 0.002 | 0 | TEC_PT | ANGPTL3 |
| 0 | 0.38896629 | 0.138 | 0 | 0 | TEC_PT | SLC34A1 |
| 0 | 0.38889811 | 0.229 | 0 | 0 | TEC_PT | SLC39A5 |
| 0 | 0.3713345 | 0.209 | 0.004 | 0 | TEC_PT | DNAJC12 |
| 0 | 0.3624556 | 0.185 | 0.002 | 0 | TEC_PT | SLC47A2 |
| 0 | 0.35902122 | 0.135 | 0 | 0 | TEC_PT | HAO2 |
| 0 | 0.35776044 | 0.192 | 0.001 | 0 | TEC_PT | RNF186 |
| 0 | 0.34754828 | 0.185 | 0 | 0 | TEC_PT | FAM151A |
| 0 | 0.32419199 | 0.172 | 0 | 0 | TEC_PT | A1CF |
| 0 | 0.32177355 | 0.155 | 0.003 | 0 | TEC_PT | UGT2A3 |
| 0 | 0.32167211 | 0.168 | 0.002 | 0 | TEC_PT | FMO1 |
| 0 | 0.30910599 | 0.168 | 0.002 | 0 | TEC_PT | NKAIN4 |
| 0 | 0.30134662 | 0.152 | 0 | 0 | TEC_PT | LINC01874 |

| | | | | | | |
|---|---|---|---|---|---|---|
| 0 | 0.29729715 | 0.141 | 0 | 0 | TEC_PT | DAO |
| 0 | 0.29610435 | 0.138 | 0.001 | 0 | TEC_PT | CYP17A1 |
| 0 | 0.28599885 | 0.141 | 0.002 | 0 | TEC_PT | NOX4 |
| 0 | 0.27583011 | 0.145 | 0.001 | 0 | TEC_PT | CLDN2 |
| 0 | 0.27236046 | 0.138 | 0.002 | 0 | TEC_PT | LINC00671 |
| 0 | 0.26365511 | 0.145 | 0.001 | 0 | TEC_PT | SLC28A1 |
| 0 | 0.2604599 | 0.125 | 0.001 | 0 | TEC_PT | PLG |
| 1.908E-307 | 0.40460739 | 0.185 | 0.006 | 4.508E-303 | TEC_PT | IL17RB |
| 4.36E-307 | 0.92724121 | 0.421 | 0.031 | 1.03E-302 | TEC_PT | SLC3A1 |
| 3.09E-289 | 0.40706164 | 0.199 | 0.007 | 7.3E-285 | TEC_PT | MAOB |
| 1.639E-285 | 0.33532831 | 0.205 | 0.008 | 3.871E-281 | TEC_PT | TINAG |
| 5.582E-278 | 0.80592995 | 0.32 | 0.02 | 1.319E-273 | TEC_PT | CLTRN |
| 7.561E-276 | 0.38460944 | 0.131 | 0.003 | 1.786E-271 | TEC_PT | SLC47A1 |
| 1.272E-273 | 0.57290741 | 0.286 | 0.016 | 3.004E-269 | TEC_PT | CHDH |
| 8.563E-270 | 0.78716201 | 0.38 | 0.029 | 2.023E-265 | TEC_PT | GAMT |
| 7.84E-260 | 0.88514127 | 0.323 | 0.022 | 1.852E-255 | TEC_PT | PBLD |
| 9.276E-258 | 0.27412176 | 0.145 | 0.004 | 2.191E-253 | TEC_PT | CDH6 |
| 2.346E-257 | 0.51811957 | 0.259 | 0.014 | 5.541E-253 | TEC_PT | SLC4A4 |
| 9.362E-257 | 0.40014895 | 0.148 | 0.004 | 2.211E-252 | TEC_PT | MCCD1 |
| 5.457E-247 | 1.50754931 | 0.488 | 0.054 | 1.289E-242 | TEC_PT | ALDH6A1 |
| 5.759E-244 | 0.29688687 | 0.111 | 0.002 | 1.36E-239 | TEC_PT | ABCC2 |
| 2.473E-238 | 0.25594532 | 0.168 | 0.006 | 5.842E-234 | TEC_PT | MAPT |
| 1.819E-236 | 1.11766369 | 0.34 | 0.026 | 4.296E-232 | TEC_PT | S100A1 |
| 5.191E-236 | 0.32947996 | 0.192 | 0.008 | 1.226E-231 | TEC_PT | GDA |
| 3.955E-227 | 1.13980043 | 0.455 | 0.049 | 9.343E-223 | TEC_PT | LGALS2 |
| 3.891E-226 | 1.57947495 | 0.495 | 0.06 | 9.192E-222 | TEC_PT | AK4 |
| 2.72E-223 | 0.33076317 | 0.192 | 0.009 | 6.426E-219 | TEC_PT | HAAO |
| 1.196E-215 | 0.74954908 | 0.38 | 0.035 | 2.825E-211 | TEC_PT | IGFBP4 |
| 1.065E-207 | 2.20132089 | 0.872 | 0.229 | 2.517E-203 | TEC_PT | PDZK1IP1 |
| 8.079E-207 | 1.22768098 | 0.552 | 0.079 | 1.908E-202 | TEC_PT | AKR1C3 |
| 4.113E-204 | 0.39100774 | 0.222 | 0.013 | 9.717E-200 | TEC_PT | SLC25A10 |
| 3.76E-202 | 0.55641102 | 0.182 | 0.009 | 8.882E-198 | TEC_PT | VCAM1 |
| 4.17E-198 | 1.52724146 | 0.633 | 0.107 | 9.851E-194 | TEC_PT | FXYD2 |
| 5.891E-196 | 0.57565905 | 0.249 | 0.017 | 1.392E-191 | TEC_PT | THY1 |
| 2.103E-192 | 0.55553137 | 0.296 | 0.024 | 4.969E-188 | TEC_PT | AZGP1 |
| 3.767E-192 | 1.19844322 | 0.535 | 0.08 | 8.898E-188 | TEC_PT | QPRT |
| 2.89E-191 | 0.60111956 | 0.148 | 0.006 | 6.827E-187 | TEC_PT | AKR1B10 |
| 1.731E-189 | 1.58251558 | 0.589 | 0.104 | 4.09E-185 | TEC_PT | PEPD |
| 1.268E-187 | 0.44158529 | 0.242 | 0.017 | 2.995E-183 | TEC_PT | HSD17B14 |
| 1.812E-187 | 0.6226984 | 0.323 | 0.029 | 4.281E-183 | TEC_PT | AC245595.1 |
| 4.061E-186 | 2.63250389 | 0.882 | 0.244 | 9.594E-182 | TEC_PT | CRYAB |
| 2.895E-180 | 0.74372023 | 0.374 | 0.041 | 6.839E-176 | TEC_PT | PTH1R |
| 1.131E-178 | 0.3925214 | 0.199 | 0.012 | 2.672E-174 | TEC_PT | RGN |
| 1.464E-176 | 1.19556436 | 0.471 | 0.068 | 3.459E-172 | TEC_PT | CRYL1 |
| 4.572E-175 | 1.05157034 | 0.529 | 0.085 | 1.08E-170 | TEC_PT | ISOC2 |
| 3.674E-171 | 1.96542578 | 0.646 | 0.141 | 8.679E-167 | TEC_PT | SUCLG1 |
| 1.808E-170 | 1.33788185 | 0.519 | 0.086 | 4.272E-166 | TEC_PT | ACAT1 |
| 3.753E-165 | 0.43735546 | 0.263 | 0.022 | 8.867E-161 | TEC_PT | NR1H4 |
| 1.073E-164 | 1.09844297 | 0.495 | 0.078 | 2.535E-160 | TEC_PT | PCSK1N |
| 1.65E-164 | 0.59359789 | 0.266 | 0.023 | 3.898E-160 | TEC_PT | ETNK2 |

| | | | | | | |
|---|---|---|---|---|---|---|
| 7.206E-162 | 1.8439081 | 0.848 | 0.279 | 1.702E-157 | TEC_PT | LDHB |
| 7.069E-160 | 0.86632115 | 0.431 | 0.06 | 1.67E-155 | TEC_PT | ALDH1A1 |
| 5.566E-158 | 1.05871194 | 0.481 | 0.077 | 1.315E-153 | TEC_PT | IMPA2 |
| 1.84E-157 | 0.5689638 | 0.212 | 0.015 | 4.346E-153 | TEC_PT | ANK2 |
| 8.598E-153 | 1.48479807 | 0.475 | 0.08 | 2.031E-148 | TEC_PT | C11orf54 |
| 1.069E-152 | 0.31628672 | 0.145 | 0.007 | 2.524E-148 | TEC_PT | AC087482.1 |
| 8.241E-152 | 0.75034891 | 0.384 | 0.051 | 1.947E-147 | TEC_PT | AIFM1 |
| 7.21E-151 | 0.77228263 | 0.125 | 0.005 | 1.703E-146 | TEC_PT | RBP4 |
| 1.084E-149 | 2.13128613 | 0.774 | 0.225 | 2.562E-145 | TEC_PT | CYB5A |
| 2.455E-147 | 0.43264494 | 0.249 | 0.022 | 5.798E-143 | TEC_PT | EPHX2 |
| 7.844E-147 | 0.91650933 | 0.444 | 0.07 | 1.853E-142 | TEC_PT | BDH2 |
| 1.212E-146 | 1.23292217 | 0.582 | 0.124 | 2.864E-142 | TEC_PT | DNPH1 |
| 1.249E-146 | 1.27530176 | 0.556 | 0.112 | 2.951E-142 | TEC_PT | SLC9A3R1 |
| 2.855E-145 | 1.57405474 | 0.677 | 0.177 | 6.745E-141 | TEC_PT | ADI1 |
| 3.685E-145 | 1.91637995 | 0.727 | 0.217 | 8.705E-141 | TEC_PT | ECHS1 |
| 1.534E-141 | 0.90668932 | 0.421 | 0.065 | 3.624E-137 | TEC_PT | MSRA |
| 3.488E-141 | 0.5173905 | 0.303 | 0.034 | 8.239E-137 | TEC_PT | MACROD1 |
| 1.558E-139 | 0.27356919 | 0.165 | 0.01 | 3.679E-135 | TEC_PT | DHDH |
| 1.875E-139 | 0.8984884 | 0.461 | 0.078 | 4.43E-135 | TEC_PT | ECI2 |
| 3.398E-139 | 0.31264039 | 0.165 | 0.01 | 8.026E-135 | TEC_PT | LDHD |
| 4.537E-139 | 0.30957516 | 0.175 | 0.012 | 1.072E-134 | TEC_PT | UGT3A1 |
| 7.025E-139 | 0.52005536 | 0.296 | 0.033 | 1.659E-134 | TEC_PT | TTC38 |
| 2.464E-138 | 0.43429442 | 0.219 | 0.018 | 5.82E-134 | TEC_PT | ACSF2 |
| 2.812E-137 | 0.29852497 | 0.178 | 0.012 | 6.644E-133 | TEC_PT | NPR3 |
| 7.071E-134 | 1.77224177 | 0.768 | 0.223 | 1.67E-129 | TEC_PT | IL32 |
| 1.369E-133 | 0.50475845 | 0.313 | 0.037 | 3.234E-129 | TEC_PT | AMN |
| 6.394E-132 | 0.42819135 | 0.239 | 0.022 | 1.51E-127 | TEC_PT | TMEM37 |
| 1.446E-129 | 2.22082808 | 0.838 | 0.345 | 3.416E-125 | TEC_PT | PEBP1 |
| 1.134E-128 | 0.30430206 | 0.185 | 0.014 | 2.679E-124 | TEC_PT | SPX |
| 2.032E-128 | 0.61362137 | 0.286 | 0.033 | 4.8E-124 | TEC_PT | PXMP2 |
| 8.65E-127 | 0.58351355 | 0.306 | 0.037 | 2.043E-122 | TEC_PT | FABP3 |
| 7.315E-124 | 1.46490434 | 0.529 | 0.115 | 1.728E-119 | TEC_PT | ASS1 |
| 8.839E-123 | 0.26254928 | 0.125 | 0.007 | 2.088E-118 | TEC_PT | AMBP |
| 1.174E-122 | 0.64450191 | 0.34 | 0.048 | 2.774E-118 | TEC_PT | ALDH1L1 |
| 1.626E-121 | 2.25959192 | 0.768 | 0.251 | 3.842E-117 | TEC_PT | MT1G |
| 5.181E-121 | 0.66868176 | 0.337 | 0.048 | 1.224E-116 | TEC_PT | ECHDC3 |
| 5.781E-118 | 0.67963122 | 0.33 | 0.047 | 1.366E-113 | TEC_PT | BPHL |
| 2.732E-116 | 1.0471521 | 0.552 | 0.128 | 6.453E-112 | TEC_PT | TMEM176A |
| 3.29E-116 | 0.69263584 | 0.343 | 0.052 | 7.771E-112 | TEC_PT | QDPR |
| 8.971E-115 | 0.81156702 | 0.276 | 0.034 | 2.119E-110 | TEC_PT | RNF152 |
| 1.572E-114 | 0.92111435 | 0.401 | 0.071 | 3.713E-110 | TEC_PT | DHRS4L2 |
| 7.003E-114 | 0.33816002 | 0.135 | 0.008 | 1.654E-109 | TEC_PT | SLC16A9 |
| 5.552E-113 | -2.6301953 | 0.549 | 0.941 | 1.312E-108 | TEC_PT | TMSB4X |
| 4.059E-112 | 0.46304307 | 0.229 | 0.024 | 9.588E-108 | TEC_PT | WDR72 |
| 1.104E-110 | 1.32232631 | 0.559 | 0.144 | 2.607E-106 | TEC_PT | ALDH2 |
| 5.617E-109 | 0.54390557 | 0.343 | 0.053 | 1.327E-104 | TEC_PT | ALDH7A1 |
| 1.264E-108 | 0.26719719 | 0.152 | 0.011 | 2.986E-104 | TEC_PT | ENPEP |
| 5.348E-108 | 1.03486748 | 0.481 | 0.108 | 1.263E-103 | TEC_PT | ACAA2 |
| 8.457E-108 | 0.28985401 | 0.175 | 0.015 | 1.998E-103 | TEC_PT | DECR2 |
| 6.52E-104 | 1.34772337 | 0.572 | 0.148 | 1.54E-99 | TEC_PT | MT1H |

| | | | | | | |
|---|---|---|---|---|---|---|
| 9.767E-104 | 1.73580296 | 0.576 | 0.164 | 2.307E-99 | TEC_PT | DCXR |
| 1.135E-103 | 1.03930042 | 0.515 | 0.124 | 2.681E-99 | TEC_PT | SELENOP |
| 4.664E-103 | 0.62869634 | 0.337 | 0.053 | 1.1017E-98 | TEC_PT | RARRES2 |
| 9.092E-103 | 1.03387261 | 0.556 | 0.149 | 2.1477E-98 | TEC_PT | ECH1 |
| 2.808E-102 | 0.26189374 | 0.158 | 0.013 | 6.6325E-98 | TEC_PT | SLC25A42 |
| 1.128E-101 | 0.69704128 | 0.418 | 0.082 | 2.6654E-97 | TEC_PT | FAHD1 |
| 1.128E-100 | 1.51263387 | 0.721 | 0.291 | 2.6656E-96 | TEC_PT | CHCHD10 |
| 3.496E-99 | 1.133932 | 0.576 | 0.169 | 8.2594E-95 | TEC_PT | MPC2 |
| 4.4229E-98 | 1.23878286 | 0.579 | 0.174 | 1.0448E-93 | TEC_PT | AKR1A1 |
| 1.5185E-97 | 0.26484225 | 0.152 | 0.012 | 3.587E-93 | TEC_PT | KL |
| 3.0667E-94 | 1.43717994 | 0.717 | 0.261 | 7.2444E-90 | TEC_PT | MGST1 |
| 2.046E-92 | 0.55035729 | 0.313 | 0.052 | 4.8333E-88 | TEC_PT | HADH |
| 1.0124E-91 | 0.48834051 | 0.279 | 0.042 | 2.3916E-87 | TEC_PT | ACOT7 |
| 1.1681E-91 | 0.7100858 | 0.397 | 0.082 | 2.7593E-87 | TEC_PT | LACTB2 |
| 2.9763E-91 | 0.25779593 | 0.141 | 0.011 | 7.031E-87 | TEC_PT | PRUNE2 |
| 3.2273E-91 | 0.7244382 | 0.407 | 0.082 | 7.6239E-87 | TEC_PT | CALD1 |
| 1.7887E-89 | 0.97701007 | 0.481 | 0.124 | 4.2256E-85 | TEC_PT | PLA2G16 |
| 2.7711E-89 | 0.46148772 | 0.296 | 0.048 | 6.5462E-85 | TEC_PT | SLC16A4 |
| 3.686E-89 | 0.53879228 | 0.229 | 0.03 | 8.7074E-85 | TEC_PT | ACADSB |
| 4.6334E-89 | 0.78030507 | 0.3 | 0.049 | 1.0946E-84 | TEC_PT | SLC7A8 |
| 8.3468E-88 | 0.31090063 | 0.182 | 0.019 | 1.9718E-83 | TEC_PT | SELENBP1 |
| 1.2121E-86 | -1.5271732 | 0.768 | 0.982 | 2.8634E-82 | TEC_PT | MALAT1 |
| 1.1933E-85 | 0.25496841 | 0.152 | 0.014 | 2.8189E-81 | TEC_PT | SAMD5 |
| 2.7909E-84 | 0.71756443 | 0.407 | 0.092 | 6.593E-80 | TEC_PT | AKR7A2 |
| 1.7297E-83 | -2.1739975 | 0.646 | 0.939 | 4.086E-79 | TEC_PT | S100A6 |
| 1.9538E-82 | 0.8504639 | 0.424 | 0.102 | 4.6156E-78 | TEC_PT | GCSH |
| 8.381E-82 | 0.86399624 | 0.468 | 0.125 | 1.9798E-77 | TEC_PT | DUSP23 |
| 1.1983E-81 | 0.88083142 | 0.916 | 0.45 | 2.8306E-77 | TEC_PT | SPP1 |
| 2.6759E-81 | 0.51654535 | 0.3 | 0.053 | 6.3212E-77 | TEC_PT | ACO1 |
| 3.4056E-81 | -1.3303501 | 0.633 | 0.943 | 8.0451E-77 | TEC_PT | NEAT1 |
| 9.7439E-81 | 0.96170783 | 0.29 | 0.049 | 2.3018E-76 | TEC_PT | SERPINE2 |
| 1.7075E-80 | 1.06308115 | 0.687 | 0.279 | 4.0336E-76 | TEC_PT | ISCU |
| 5.0559E-80 | 0.57507814 | 0.306 | 0.056 | 1.1944E-75 | TEC_PT | PPP1R16A |
| 5.3868E-80 | 0.40207592 | 0.266 | 0.042 | 1.2725E-75 | TEC_PT | FAH |
| 9.3496E-80 | 1.47240573 | 0.768 | 0.401 | 2.2087E-75 | TEC_PT | SOD1 |
| 1.603E-79 | 1.27721539 | 0.556 | 0.181 | 3.7867E-75 | TEC_PT | DAB2 |
| 2.0205E-79 | 0.34604759 | 0.219 | 0.03 | 4.773E-75 | TEC_PT | NUDT8 |
| 6.3171E-79 | 1.42372159 | 0.734 | 0.368 | 1.4923E-74 | TEC_PT | ATP5MC3 |
| 9.1864E-79 | 0.694715 | 0.34 | 0.068 | 2.1701E-74 | TEC_PT | MAOA |
| 3.0318E-78 | 0.84710935 | 0.448 | 0.118 | 7.1621E-74 | TEC_PT | NQO2 |
| 4.289E-78 | 0.45656802 | 0.266 | 0.043 | 1.0132E-73 | TEC_PT | CLDN10 |
| 5.4623E-77 | 0.30298169 | 0.242 | 0.036 | 1.2903E-72 | TEC_PT | MME |
| 9.4473E-76 | 0.51492642 | 0.343 | 0.071 | 2.2317E-71 | TEC_PT | CRYZ |
| 2.057E-75 | 1.12503912 | 0.556 | 0.197 | 4.8592E-71 | TEC_PT | ETFB |
| 4.8731E-75 | 1.10343654 | 0.532 | 0.173 | 1.1512E-70 | TEC_PT | ACAA1 |
| 8.1086E-75 | 0.32225377 | 0.236 | 0.035 | 1.9155E-70 | TEC_PT | TCEA3 |
| 2.9584E-74 | 0.42886751 | 0.273 | 0.047 | 6.9885E-70 | TEC_PT | PNPO |
| 4.2081E-74 | 0.31195131 | 0.192 | 0.025 | 9.9408E-70 | TEC_PT | DMGDH |
| 2.4621E-73 | 1.50561942 | 0.714 | 0.372 | 5.8162E-69 | TEC_PT | LAMTOR5 |
| 1.4018E-72 | 0.50898924 | 0.276 | 0.05 | 3.3114E-68 | TEC_PT | RAB17 |

| | | | | | | |
|---|---|---|---|---|---|---|
| 2.6588E-72 | 0.64840386 | 0.374 | 0.085 | 6.2809E-68 | TEC_PT | CA2 |
| 3.3373E-71 | 0.46617175 | 0.303 | 0.059 | 7.8837E-67 | TEC_PT | PCK2 |
| 1.6278E-70 | 1.64192086 | 0.98 | 0.987 | 3.8452E-66 | TEC_PT | FTL |
| 5.5326E-70 | 0.88274002 | 0.424 | 0.113 | 1.307E-65 | TEC_PT | NQO1 |
| 2.2635E-69 | 1.29017156 | 0.875 | 0.6 | 5.3471E-65 | TEC_PT | HINT1 |
| 4.1789E-69 | 0.85833996 | 0.539 | 0.176 | 9.8719E-65 | TEC_PT | TMEM176B |
| 1.0367E-68 | 0.25980704 | 0.192 | 0.026 | 2.4489E-64 | TEC_PT | PAIP2B |
| 1.5345E-68 | 0.43063976 | 0.215 | 0.032 | 3.625E-64 | TEC_PT | PPP1R1A |
| 8.7406E-68 | 0.35139295 | 0.219 | 0.034 | 2.0648E-63 | TEC_PT | CYS1 |
| 2.1936E-67 | 0.62786835 | 0.33 | 0.073 | 5.182E-63 | TEC_PT | DDAH1 |
| 5.6592E-67 | 0.72234198 | 0.421 | 0.117 | 1.3369E-62 | TEC_PT | NDUFV1 |
| 9.6653E-67 | 1.1665642 | 0.727 | 0.375 | 2.2832E-62 | TEC_PT | POLR2L |
| 1.0685E-66 | 1.37805088 | 0.808 | 0.505 | 2.524E-62 | TEC_PT | DBI |
| 6.4428E-66 | -1.4678792 | 0.32 | 0.755 | 1.522E-61 | TEC_PT | TPM4 |
| 1.1819E-65 | 0.36753065 | 0.226 | 0.037 | 2.792E-61 | TEC_PT | FOLR1 |
| 2.0284E-65 | 1.03074193 | 0.495 | 0.154 | 4.7916E-61 | TEC_PT | MT1F |
| 4.3126E-65 | 0.82705804 | 0.556 | 0.205 | 1.0188E-60 | TEC_PT | COX14 |
| 6.7251E-65 | 0.64805092 | 0.357 | 0.087 | 1.5887E-60 | TEC_PT | NIT2 |
| 7.4428E-65 | 1.23732096 | 0.923 | 0.761 | 1.7582E-60 | TEC_PT | COX7C |
| 8.2391E-65 | 0.51756032 | 0.259 | 0.049 | 1.9463E-60 | TEC_PT | PSAT1 |
| 1.292E-64 | 1.2844525 | 0.694 | 0.354 | 3.052E-60 | TEC_PT | UQCRQ |
| 2.4565E-63 | 0.92359246 | 0.626 | 0.258 | 5.8031E-59 | TEC_PT | PRDX2 |
| 2.8285E-63 | 0.45392165 | 0.259 | 0.049 | 6.6817E-59 | TEC_PT | NIPSNAP1 |
| 3.5837E-63 | 0.63417252 | 0.33 | 0.077 | 8.4658E-59 | TEC_PT | RAB29 |
| 1.743E-62 | 0.95781704 | 0.596 | 0.234 | 4.1176E-58 | TEC_PT | PCBD1 |
| 2.688E-62 | 0.82939994 | 0.535 | 0.174 | 6.35E-58 | TEC_PT | ANPEP |
| 4.2302E-62 | 1.03439348 | 0.616 | 0.264 | 9.9931E-58 | TEC_PT | ATOX1 |
| 1.7941E-61 | 0.30746869 | 0.158 | 0.02 | 4.2382E-57 | TEC_PT | SLC27A2 |
| 2.1744E-61 | -1.593811 | 0.411 | 0.786 | 5.1367E-57 | TEC_PT | LGALS3 |
| 2.4776E-61 | 0.25327029 | 0.195 | 0.029 | 5.8529E-57 | TEC_PT | SOD3 |
| 4.9576E-61 | 0.34316604 | 0.215 | 0.036 | 1.1711E-56 | TEC_PT | PDK2 |
| 5.2939E-61 | 0.49538182 | 0.33 | 0.077 | 1.2506E-56 | TEC_PT | EPHX1 |
| 5.7559E-61 | 1.11707705 | 0.556 | 0.227 | 1.3597E-56 | TEC_PT | SAT2 |
| 1.5597E-60 | 0.92863406 | 0.559 | 0.217 | 3.6846E-56 | TEC_PT | CNDP2 |
| 1.8447E-60 | 1.06118342 | 0.731 | 0.391 | 4.3577E-56 | TEC_PT | NDUFB2 |
| 2.7211E-60 | 0.75624034 | 0.424 | 0.127 | 6.428E-56 | TEC_PT | CBR1 |
| 3.3411E-60 | 0.80974591 | 0.542 | 0.199 | 7.8926E-56 | TEC_PT | PHB |
| 3.8319E-60 | 0.48234959 | 0.293 | 0.064 | 9.0522E-56 | TEC_PT | HIBCH |
| 8.6608E-60 | 1.09793405 | 0.67 | 0.34 | 2.0459E-55 | TEC_PT | ATP5F1B |
| 1.3501E-59 | 0.42895882 | 0.249 | 0.048 | 3.1894E-55 | TEC_PT | TSPAN33 |
| 1.5893E-59 | 0.46553533 | 0.313 | 0.071 | 3.7545E-55 | TEC_PT | BIN1 |
| 2.8266E-59 | -1.3897556 | 0.916 | 0.99 | 6.6773E-55 | TEC_PT | B2M |
| 4.7693E-59 | 0.31142103 | 0.219 | 0.038 | 1.1267E-54 | TEC_PT | TTC39A |
| 5.7822E-59 | 0.35818257 | 0.229 | 0.041 | 1.3659E-54 | TEC_PT | RGS14 |
| 1.3045E-57 | 0.47906223 | 0.323 | 0.078 | 3.0816E-53 | TEC_PT | TM7SF3 |
| 1.3657E-57 | 0.64453902 | 0.391 | 0.11 | 3.2262E-53 | TEC_PT | CISD1 |
| 1.5514E-57 | 1.17220685 | 0.902 | 0.707 | 3.6649E-53 | TEC_PT | GPX4 |
| 2.8405E-57 | 1.13747295 | 0.714 | 0.408 | 6.7101E-53 | TEC_PT | ATP5PF |
| 5.0375E-57 | -1.6197438 | 0.788 | 0.94 | 1.19E-52 | TEC_PT | SAT1 |
| 1.0898E-56 | 0.9285387 | 0.515 | 0.197 | 2.5744E-52 | TEC_PT | MDH1 |

| | | | | | | |
|---|---|---|---|---|---|---|
| 9.0314E-56 | -1.1986333 | 0.848 | 0.966 | 2.1335E-51 | TEC_PT | S100A11 |
| 9.0315E-56 | 0.82977945 | 0.519 | 0.195 | 2.1335E-51 | TEC_PT | ATP5MC1 |
| 1.0064E-55 | 0.25723896 | 0.118 | 0.013 | 2.3774E-51 | TEC_PT | CYP24A1 |
| 2.898E-55 | 0.59289172 | 0.333 | 0.086 | 6.8458E-51 | TEC_PT | MSRB1 |
| 5.0685E-55 | 1.05571439 | 0.66 | 0.322 | 1.1973E-50 | TEC_PT | MGST3 |
| 5.7931E-55 | 0.32446695 | 0.222 | 0.041 | 1.3685E-50 | TEC_PT | PCYT2 |
| 5.9516E-55 | 0.90558642 | 0.556 | 0.234 | 1.4059E-50 | TEC_PT | ATP5F1A |
| 8.4514E-55 | 1.01792297 | 0.616 | 0.285 | 1.9965E-50 | TEC_PT | NDUFC1 |
| 2.1242E-54 | 0.9342963 | 0.569 | 0.222 | 5.0181E-50 | TEC_PT | OCIAD2 |
| 2.3092E-54 | 0.28923839 | 0.199 | 0.034 | 5.455E-50 | TEC_PT | RTKN |
| 4.5774E-54 | 0.69677357 | 0.377 | 0.111 | 1.0813E-49 | TEC_PT | GRHPR |
| 6.1865E-54 | 1.11748261 | 0.808 | 0.522 | 1.4614E-49 | TEC_PT | HSPE1 |
| 6.6888E-54 | 0.52246403 | 0.343 | 0.093 | 1.5801E-49 | TEC_PT | HINT2 |
| 1.2939E-53 | 0.28337097 | 0.199 | 0.034 | 3.0565E-49 | TEC_PT | C1orf115 |
| 3.1805E-53 | -4.7736296 | 0.418 | 0.729 | 7.5133E-49 | TEC_PT | S100A9 |
| 3.544E-53 | 0.56539285 | 0.364 | 0.102 | 8.3721E-49 | TEC_PT | SUCLG2 |
| 4.3668E-53 | 0.53447247 | 0.354 | 0.098 | 1.0316E-48 | TEC_PT | MCRIP2 |
| 4.4359E-53 | 0.29879544 | 0.158 | 0.023 | 1.0479E-48 | TEC_PT | OGDHL |
| 5.5044E-53 | 0.83496809 | 0.525 | 0.195 | 1.3003E-48 | TEC_PT | PTGR1 |
| 7.9033E-53 | 0.45480256 | 0.306 | 0.076 | 1.867E-48 | TEC_PT | POLDIP2 |
| 9.0901E-53 | -1.1991356 | 0.855 | 0.954 | 2.1473E-48 | TEC_PT | H3F3B |
| 1.8443E-52 | 0.51199807 | 0.367 | 0.105 | 4.3567E-48 | TEC_PT | HMGCL |
| 3.1131E-52 | 0.48764404 | 0.269 | 0.061 | 7.3541E-48 | TEC_PT | GALM |
| 4.3178E-52 | 0.5833232 | 0.337 | 0.091 | 1.02E-47 | TEC_PT | PHYH |
| 1.268E-51 | 1.19869884 | 0.663 | 0.34 | 2.9953E-47 | TEC_PT | ATP5IF1 |
| 1.6755E-51 | 0.33145194 | 0.256 | 0.055 | 3.9581E-47 | TEC_PT | PKIG |
| 1.7132E-51 | 0.26576528 | 0.182 | 0.03 | 4.0472E-47 | TEC_PT | CCDC51 |
| 2.4318E-51 | 0.40810286 | 0.215 | 0.041 | 5.7446E-47 | TEC_PT | ASRGL1 |
| 8.7852E-51 | 1.05034112 | 0.657 | 0.362 | 2.0753E-46 | TEC_PT | ATP5F1D |
| 1.2459E-50 | 0.41073478 | 0.266 | 0.06 | 2.9431E-46 | TEC_PT | ABAT |
| 1.3465E-50 | 0.92211606 | 0.428 | 0.135 | 3.1809E-46 | TEC_PT | NUPR1 |
| 2.3675E-50 | 0.58183355 | 0.391 | 0.123 | 5.5928E-46 | TEC_PT | MSRB2 |
| 5.0815E-50 | 0.73832824 | 0.461 | 0.165 | 1.2004E-45 | TEC_PT | HMGN3 |
| 6.772E-50 | 0.66311541 | 0.323 | 0.087 | 1.5998E-45 | TEC_PT | MAF |
| 1.5146E-49 | 1.02066608 | 0.832 | 0.614 | 3.5779E-45 | TEC_PT | COX8A |
| 6.2329E-49 | 0.80857785 | 0.593 | 0.259 | 1.4724E-44 | TEC_PT | RHOC |
| 9.7323E-49 | 0.89321849 | 0.481 | 0.183 | 2.2991E-44 | TEC_PT | SEPHS2 |
| 1.2626E-48 | 0.43158802 | 0.343 | 0.096 | 2.9825E-44 | TEC_PT | ECHDC2 |
| 1.3356E-48 | 0.54213608 | 0.32 | 0.087 | 3.1551E-44 | TEC_PT | SLC7A7 |
| 3.0126E-48 | 0.97654603 | 0.892 | 0.723 | 7.1167E-44 | TEC_PT | COX6A1 |
| 5.4029E-48 | 1.04071364 | 0.663 | 0.373 | 1.2763E-43 | TEC_PT | ATP5ME |
| 5.7046E-48 | -1.2335543 | 0.687 | 0.891 | 1.3476E-43 | TEC_PT | HLA-B |
| 1.3917E-47 | 1.06518324 | 0.663 | 0.396 | 3.2876E-43 | TEC_PT | NDUFA1 |
| 1.8306E-47 | 0.61886265 | 0.421 | 0.145 | 4.3245E-43 | TEC_PT | AK2 |
| 2.1346E-47 | 0.37748693 | 0.242 | 0.053 | 5.0425E-43 | TEC_PT | CCDC198 |
| 6.2749E-47 | 0.51994602 | 0.347 | 0.103 | 1.4823E-42 | TEC_PT | COMMD8 |
| 8.4681E-47 | 0.84581829 | 0.993 | 0.98 | 2.0004E-42 | TEC_PT | EEF1A1 |
| 9.3465E-47 | 0.27784344 | 0.189 | 0.035 | 2.2079E-42 | TEC_PT | CFI |
| 1.4582E-46 | 0.41417369 | 0.226 | 0.048 | 3.4447E-42 | TEC_PT | CRYM |
| 2.3872E-46 | 0.62054269 | 0.438 | 0.159 | 5.6394E-42 | TEC_PT | SDHB |

| | | | | | | |
|---|---|---|---|---|---|---|
| 2.6132E-46 | 0.49842545 | 0.354 | 0.107 | 6.1733E-42 | TEC_PT | TPMT |
| 2.974E-46 | 0.9334109 | 0.835 | 0.67 | 7.0255E-42 | TEC_PT | COX5B |
| 3.5836E-46 | 0.40719663 | 0.273 | 0.067 | 8.4656E-42 | TEC_PT | CES2 |
| 4.5454E-46 | 0.48450194 | 0.502 | 0.186 | 1.0738E-41 | TEC_PT | RARRES3 |
| 9.5876E-46 | 0.75397676 | 0.475 | 0.181 | 2.2649E-41 | TEC_PT | BNIP3 |
| 1.2127E-45 | 0.40313718 | 0.34 | 0.097 | 2.8649E-41 | TEC_PT | SLC25A4 |
| 1.2247E-45 | 0.42964254 | 0.266 | 0.066 | 2.893E-41 | TEC_PT | MTHFS |
| 1.4748E-45 | 0.59860415 | 0.414 | 0.144 | 3.484E-41 | TEC_PT | COA3 |
| 1.576E-45 | 0.57387092 | 0.347 | 0.105 | 3.723E-41 | TEC_PT | TST |
| 3.9678E-45 | -1.9473073 | 0.273 | 0.627 | 9.3732E-41 | TEC_PT | S100A4 |
| 1.0729E-44 | 0.41529097 | 0.3 | 0.081 | 2.5345E-40 | TEC_PT | MRPL12 |
| 1.3999E-44 | 0.60528612 | 0.418 | 0.149 | 3.3071E-40 | TEC_PT | ECI1 |
| 1.6581E-44 | -1.3919302 | 0.923 | 0.973 | 3.9169E-40 | TEC_PT | ACTB |
| 2.398E-44 | 0.30169122 | 0.185 | 0.035 | 5.6648E-40 | TEC_PT | DHRS4 |
| 5.2049E-44 | 0.72374082 | 0.488 | 0.2 | 1.2296E-39 | TEC_PT | NDUFS8 |
| 5.3223E-44 | 0.8694295 | 0.818 | 0.587 | 1.2573E-39 | TEC_PT | SLC25A3 |
| 6.391E-44 | 0.30962222 | 0.199 | 0.04 | 1.5098E-39 | TEC_PT | ATP6V0E2 |
| 2.4959E-43 | -1.0115975 | 0.189 | 0.578 | 5.8961E-39 | TEC_PT | CD55 |
| 3.2078E-43 | 0.94358423 | 0.916 | 0.745 | 7.5778E-39 | TEC_PT | PRDX1 |
| 5.8162E-43 | 0.99427752 | 0.761 | 0.531 | 1.374E-38 | TEC_PT | COX6C |
| 8.628E-43 | 0.48589633 | 0.347 | 0.109 | 2.0382E-38 | TEC_PT | MTCH2 |
| 8.9168E-43 | -1.3223491 | 0.253 | 0.615 | 2.1064E-38 | TEC_PT | CD44 |
| 1.3668E-42 | 0.31226545 | 0.168 | 0.031 | 3.2288E-38 | TEC_PT | TKFC |
| 1.4948E-42 | 0.78527229 | 0.976 | 0.954 | 3.5311E-38 | TEC_PT | SERF2 |
| 1.7737E-42 | 0.7429772 | 0.552 | 0.243 | 4.19E-38 | TEC_PT | PTMS |
| 2.3415E-42 | 0.29181328 | 0.185 | 0.036 | 5.5313E-38 | TEC_PT | SCRN2 |
| 2.6384E-42 | -0.7964694 | 0.552 | 0.837 | 6.2326E-38 | TEC_PT | HLA-E |
| 3.2065E-42 | 0.8021871 | 0.522 | 0.241 | 7.5748E-38 | TEC_PT | NDUFA3 |
| 3.9047E-42 | 0.47646586 | 0.296 | 0.084 | 9.224E-38 | TEC_PT | FUOM |
| 5.0554E-42 | -3.9097121 | 0.387 | 0.673 | 1.1942E-37 | TEC_PT | S100A8 |
| 6.7189E-42 | 0.88506704 | 0.657 | 0.378 | 1.5872E-37 | TEC_PT | PRELID1 |
| 9.5853E-42 | 0.89344512 | 0.764 | 0.476 | 2.2643E-37 | TEC_PT | HSPD1 |
| 1.0682E-41 | 0.33582805 | 0.269 | 0.07 | 2.5235E-37 | TEC_PT | TSFM |
| 1.0887E-41 | 0.29345285 | 0.209 | 0.045 | 2.5718E-37 | TEC_PT | PGM1 |
| 1.9598E-41 | 0.77683203 | 0.579 | 0.287 | 4.6297E-37 | TEC_PT | NDUFB7 |
| 4.0938E-41 | 0.77020277 | 0.993 | 0.932 | 9.6708E-37 | TEC_PT | GAPDH |
| 5.3303E-41 | 0.50318118 | 0.313 | 0.092 | 1.2592E-36 | TEC_PT | SORD |
| 5.7758E-41 | 0.97623253 | 0.845 | 0.656 | 1.3644E-36 | TEC_PT | NDUFA4 |
| 1.4083E-40 | -0.8916055 | 0.431 | 0.76 | 3.3267E-36 | TEC_PT | ITGB1 |
| 1.9838E-40 | 0.69955434 | 0.62 | 0.327 | 4.6863E-36 | TEC_PT | AURKAIP1 |
| 2.6659E-40 | 0.31072 | 0.209 | 0.047 | 6.2977E-36 | TEC_PT | ACADS |
| 3.7365E-40 | 0.46182974 | 0.306 | 0.092 | 8.8268E-36 | TEC_PT | PMVK |
| 4.2715E-40 | 0.32925845 | 0.222 | 0.052 | 1.0091E-35 | TEC_PT | SLC22A18 |
| 5.2772E-40 | 0.63736761 | 0.333 | 0.11 | 1.2466E-35 | TEC_PT | ACO2 |
| 5.9059E-40 | -1.7191624 | 0.394 | 0.712 | 1.3952E-35 | TEC_PT | ANXA1 |
| 6.5916E-40 | 0.33209942 | 0.232 | 0.056 | 1.5571E-35 | TEC_PT | IVD |
| 7.761E-40 | 0.51549222 | 0.259 | 0.07 | 1.8334E-35 | TEC_PT | STK19 |
| 8.3531E-40 | 0.88159817 | 0.448 | 0.18 | 1.9732E-35 | TEC_PT | FBP1 |
| 8.8207E-40 | -1.6441174 | 0.384 | 0.683 | 2.0837E-35 | TEC_PT | NAMPT |
| 1.0158E-39 | 0.99044728 | 0.623 | 0.361 | 2.3996E-35 | TEC_PT | SLC25A5 |

| | | | | | | |
|---|---|---|---|---|---|---|
| 2.1846E-39 | -0.8914623 | 0.872 | 0.958 | 5.1606E-35 | TEC_PT | H3F3A |
| 3.3162E-39 | -2.0500584 | 0.3 | 0.616 | 7.8337E-35 | TEC_PT | C15orf48 |
| 1.0196E-38 | -1.2047886 | 0.168 | 0.533 | 2.4085E-34 | TEC_PT | CTSS |
| 1.1076E-38 | 1.02997353 | 0.673 | 0.448 | 2.6164E-34 | TEC_PT | ATP5MD |
| 3.042E-38 | 0.3061008 | 0.205 | 0.046 | 7.1862E-34 | TEC_PT | EMX2OS |
| 3.5949E-38 | 0.40377127 | 0.266 | 0.073 | 8.4923E-34 | TEC_PT | RASSF4 |
| 5.8249E-38 | 0.53085677 | 0.34 | 0.114 | 1.376E-33 | TEC_PT | BOLA3 |
| 1.1362E-37 | 0.64993124 | 0.451 | 0.188 | 2.684E-33 | TEC_PT | NDUFS7 |
| 1.1538E-37 | 0.50696762 | 0.34 | 0.115 | 2.7255E-33 | TEC_PT | MRPS23 |
| 1.6813E-37 | -1.1334909 | 0.875 | 0.955 | 3.9718E-33 | TEC_PT | ACTG1 |
| 1.6925E-37 | 0.71663037 | 0.519 | 0.25 | 3.9981E-33 | TEC_PT | FIS1 |
| 1.796E-37 | 0.37874817 | 0.242 | 0.064 | 4.2426E-33 | TEC_PT | HIBADH |
| 1.8353E-37 | 0.54604355 | 0.31 | 0.099 | 4.3355E-33 | TEC_PT | MPST |
| 1.9913E-37 | 0.70993193 | 0.512 | 0.243 | 4.7041E-33 | TEC_PT | UQCRFS1 |
| 2.2011E-37 | 0.47497589 | 0.323 | 0.103 | 5.1997E-33 | TEC_PT | AHCY |
| 3.1796E-37 | 0.60295267 | 0.475 | 0.203 | 7.5113E-33 | TEC_PT | SLIRP |
| 3.965E-37 | 0.52247056 | 0.357 | 0.125 | 9.3665E-33 | TEC_PT | AP1S1 |
| 4.4076E-37 | 0.55954787 | 0.367 | 0.134 | 1.0412E-32 | TEC_PT | ETFA |
| 5.7737E-37 | 0.79227236 | 0.545 | 0.268 | 1.3639E-32 | TEC_PT | PHPT1 |
| 6.178E-37 | 0.5967766 | 0.391 | 0.149 | 1.4594E-32 | TEC_PT | MRPL34 |
| 8.4147E-37 | 0.5412934 | 0.461 | 0.194 | 1.9878E-32 | TEC_PT | C1QBP |
| 9.2154E-37 | -1.6263529 | 0.266 | 0.571 | 2.177E-32 | TEC_PT | TACSTD2 |
| 2.0835E-36 | 0.5156383 | 0.367 | 0.133 | 4.922E-32 | TEC_PT | ADH5 |
| 2.7394E-36 | 0.25872862 | 0.192 | 0.043 | 6.4713E-32 | TEC_PT | PCCB |
| 3.2926E-36 | 0.69414327 | 0.946 | 0.831 | 7.7781E-32 | TEC_PT | UBB |
| 3.6559E-36 | 0.29378014 | 0.219 | 0.054 | 8.6365E-32 | TEC_PT | CDIP1 |
| 6.925E-36 | 0.59718213 | 0.34 | 0.12 | 1.6359E-31 | TEC_PT | HAGH |
| 7.3002E-36 | 0.4799401 | 0.374 | 0.135 | 1.7245E-31 | TEC_PT | GNPDA1 |
| 1.1627E-35 | 0.75180207 | 0.545 | 0.277 | 2.7466E-31 | TEC_PT | MRPL41 |
| 2.3822E-35 | 0.78686561 | 0.505 | 0.232 | 5.6276E-31 | TEC_PT | PLIN2 |
| 2.6655E-35 | -1.4485092 | 0.212 | 0.541 | 6.2967E-31 | TEC_PT | PLAUR |
| 2.9576E-35 | -2.0069827 | 0.296 | 0.581 | 6.9867E-31 | TEC_PT | KRT19 |
| 9.7337E-35 | 0.51589486 | 0.293 | 0.093 | 2.2994E-30 | TEC_PT | AP1M2 |
| 1.1443E-34 | 0.44378946 | 0.279 | 0.085 | 2.7032E-30 | TEC_PT | METRN |
| 1.7981E-34 | 0.47723227 | 0.29 | 0.092 | 4.2477E-30 | TEC_PT | SLC2A4RG |
| 1.9634E-34 | 0.52668009 | 0.401 | 0.158 | 4.6382E-30 | TEC_PT | GLRX5 |
| 3.3019E-34 | 0.62223902 | 0.431 | 0.187 | 7.8001E-30 | TEC_PT | PRDX3 |
| 3.4567E-34 | 0.82749972 | 0.785 | 0.567 | 8.1658E-30 | TEC_PT | RPS27L |
| 3.6392E-34 | 0.34877282 | 0.242 | 0.066 | 8.5969E-30 | TEC_PT | HOOK1 |
| 5.09E-34 | -1.0638538 | 0.892 | 0.971 | 1.2024E-29 | TEC_PT | TMSB10 |
| 5.1999E-34 | 0.52495119 | 0.333 | 0.118 | 1.2284E-29 | TEC_PT | CAT |
| 5.3174E-34 | 0.31519011 | 0.189 | 0.044 | 1.2561E-29 | TEC_PT | SLC40A1 |
| 5.5284E-34 | 0.40679375 | 0.296 | 0.095 | 1.306E-29 | TEC_PT | ALDH3A2 |
| 7.0974E-34 | 0.50191087 | 0.391 | 0.152 | 1.6766E-29 | TEC_PT | NDUFS2 |
| 1.0381E-33 | 1.04938467 | 0.68 | 0.51 | 2.4523E-29 | TEC_PT | COX7B |
| 1.415E-33 | 0.60178787 | 0.434 | 0.189 | 3.3426E-29 | TEC_PT | SDHC |
| 2.0418E-33 | 0.54253547 | 0.323 | 0.113 | 4.8234E-29 | TEC_PT | GCHFR |
| 2.4258E-33 | 0.4016799 | 0.3 | 0.099 | 5.7304E-29 | TEC_PT | ALDH9A1 |
| 1.0042E-32 | 0.55609021 | 0.374 | 0.145 | 2.3722E-28 | TEC_PT | COA4 |
| 1.7202E-32 | 0.5439562 | 0.414 | 0.173 | 4.0635E-28 | TEC_PT | CYC1 |

| | | | | | | |
|---|---|---|---|---|---|---|
| 1.8987E-32 | -2.4845493 | 0.283 | 0.572 | 4.4854E-28 | TEC_PT | SRGN |
| 2.0723E-32 | -0.9971079 | 0.172 | 0.498 | 4.8955E-28 | TEC_PT | CD9 |
| 2.1813E-32 | 0.38106115 | 0.296 | 0.098 | 5.153E-28 | TEC_PT | RNF5 |
| 2.9735E-32 | 0.42665615 | 0.31 | 0.106 | 7.0244E-28 | TEC_PT | CISD3 |
| 3.2713E-32 | -1.4507666 | 0.377 | 0.645 | 7.7279E-28 | TEC_PT | GLUL |
| 3.5672E-32 | 0.73564979 | 0.636 | 0.389 | 8.4267E-28 | TEC_PT | ATP5MPL |
| 3.6157E-32 | 0.74828646 | 0.771 | 0.553 | 8.5414E-28 | TEC_PT | NDUFS5 |
| 3.7548E-32 | 0.89564836 | 0.579 | 0.347 | 8.87E-28 | TEC_PT | DDT |
| 5.5847E-32 | 0.74574653 | 0.697 | 0.435 | 1.3193E-27 | TEC_PT | CYSTM1 |
| 5.795E-32 | 0.5163415 | 0.448 | 0.197 | 1.3689E-27 | TEC_PT | TIMM13 |
| 5.973E-32 | 0.25758367 | 0.175 | 0.04 | 1.411E-27 | TEC_PT | RENBP |
| 7.1459E-32 | 0.56818814 | 0.377 | 0.149 | 1.6881E-27 | TEC_PT | CLIC4 |
| 7.3991E-32 | -1.1392673 | 0.125 | 0.452 | 1.7479E-27 | TEC_PT | MARCKS |
| 8.2953E-32 | 0.4772179 | 0.387 | 0.156 | 1.9596E-27 | TEC_PT | GRPEL1 |
| 8.3678E-32 | 1.25862224 | 0.556 | 0.3 | 1.9767E-27 | TEC_PT | SERPINA1 |
| 1.0149E-31 | -0.8073111 | 0 | 0.327 | 2.3974E-27 | TEC_PT | ARHGDIB |
| 1.08E-31 | -1.6242813 | 0.266 | 0.547 | 2.5512E-27 | TEC_PT | KRT7 |
| 1.5568E-31 | 0.94988328 | 0.545 | 0.293 | 3.6775E-27 | TEC_PT | GLRX |
| 2.2072E-31 | 0.53632998 | 0.226 | 0.063 | 5.2141E-27 | TEC_PT | PIGR |
| 2.9371E-31 | 0.46614941 | 0.316 | 0.113 | 6.9383E-27 | TEC_PT | SIGIRR |
| 3.2862E-31 | 0.26012153 | 0.212 | 0.056 | 7.7631E-27 | TEC_PT | TRAP1 |
| 3.6E-31 | 0.49431127 | 0.357 | 0.132 | 8.5042E-27 | TEC_PT | ACADM |
| 4.2289E-31 | -0.7261315 | 0.02 | 0.35 | 9.9899E-27 | TEC_PT | TIMP2 |
| 7.0249E-31 | 0.69451038 | 0.589 | 0.336 | 1.6595E-26 | TEC_PT | ARL6IP4 |
| 1.1864E-30 | -1.2726966 | 0.017 | 0.338 | 2.8026E-26 | TEC_PT | LAPTM5 |
| 1.2984E-30 | 0.79500527 | 0.609 | 0.387 | 3.0672E-26 | TEC_PT | UQCR11 |
| 1.4823E-30 | 0.64980964 | 0.936 | 0.835 | 3.5017E-26 | TEC_PT | COX4I1 |
| 1.5385E-30 | 0.48739629 | 0.377 | 0.152 | 3.6344E-26 | TEC_PT | SLC25A39 |
| 1.587E-30 | 0.38611686 | 0.256 | 0.079 | 3.7489E-26 | TEC_PT | MORN2 |
| 1.9888E-30 | 0.55525838 | 0.441 | 0.2 | 4.6981E-26 | TEC_PT | C19orf70 |
| 2.1201E-30 | 0.70458875 | 0.912 | 0.765 | 5.0084E-26 | TEC_PT | RPS26 |
| 2.4157E-30 | 0.75123449 | 0.764 | 0.562 | 5.7066E-26 | TEC_PT | SLC25A6 |
| 2.4485E-30 | 0.3077702 | 0.242 | 0.073 | 5.7842E-26 | TEC_PT | GSS |
| 3.0446E-30 | -0.8837866 | 0.074 | 0.405 | 7.1922E-26 | TEC_PT | FLNA |
| 9.6272E-30 | 0.27498564 | 0.215 | 0.06 | 2.2742E-25 | TEC_PT | RAB40B |
| 1.4071E-29 | 0.81866869 | 0.643 | 0.432 | 3.324E-25 | TEC_PT | ATP5MF |
| 1.5794E-29 | 0.72768035 | 0.219 | 0.062 | 3.731E-25 | TEC_PT | AKR1C1 |
| 1.6138E-29 | 0.39081078 | 0.195 | 0.051 | 3.8123E-25 | TEC_PT | SGCB |
| 1.7955E-29 | -0.7128431 | 0.054 | 0.376 | 4.2416E-25 | TEC_PT | METRNL |
| 3.129E-29 | 0.62013702 | 0.566 | 0.309 | 7.3917E-25 | TEC_PT | MZT2B |
| 4.0624E-29 | -0.6782786 | 0.407 | 0.697 | 9.5965E-25 | TEC_PT | LRRFIP1 |
| 4.3821E-29 | 0.34148446 | 0.572 | 0.279 | 1.0352E-24 | TEC_PT | MT1E |
| 4.6943E-29 | 0.65020292 | 0.498 | 0.255 | 1.1089E-24 | TEC_PT | NDUFB10 |
| 4.8944E-29 | 0.55654621 | 0.458 | 0.213 | 1.1562E-24 | TEC_PT | CNN3 |
| 5.8328E-29 | -1.1800707 | 0.007 | 0.314 | 1.3779E-24 | TEC_PT | EMP1 |
| 6.4298E-29 | 0.36597477 | 0.337 | 0.124 | 1.5189E-24 | TEC_PT | FAM213A |
| 8.3075E-29 | 0.4484012 | 0.31 | 0.112 | 1.9625E-24 | TEC_PT | UGDH |
| 1.083E-28 | 0.74555618 | 0.838 | 0.713 | 2.5584E-24 | TEC_PT | ATP5MG |
| 1.1008E-28 | 0.38600109 | 0.306 | 0.11 | 2.6005E-24 | TEC_PT | MRPS35 |
| 1.3562E-28 | 0.31507944 | 0.222 | 0.064 | 3.2037E-24 | TEC_PT | IFT22 |

| | | | | | | |
|---|---|---|---|---|---|---|
| 1.4185E-28 | 0.41150135 | 0.313 | 0.115 | 3.3509E-24 | TEC_PT | ACOT13 |
| 1.4856E-28 | 0.61524311 | 0.512 | 0.271 | 3.5095E-24 | TEC_PT | MINOS1 |
| 1.8231E-28 | 0.72607719 | 0.549 | 0.323 | 4.3066E-24 | TEC_PT | GHITM |
| 1.8708E-28 | -0.7223541 | 0.673 | 0.869 | 4.4193E-24 | TEC_PT | ANXA2 |
| 2.5877E-28 | 0.34256504 | 0.246 | 0.078 | 6.1129E-24 | TEC_PT | ASL |
| 3.9529E-28 | 0.35729164 | 0.202 | 0.056 | 9.3379E-24 | TEC_PT | RGS3 |
| 4.1052E-28 | 0.59858681 | 0.522 | 0.285 | 9.6976E-24 | TEC_PT | NDUFA11 |
| 4.7486E-28 | 0.7278108 | 0.471 | 0.25 | 1.1218E-23 | TEC_PT | NDUFAB1 |
| 4.8378E-28 | -1.0447723 | 0.38 | 0.643 | 1.1428E-23 | TEC_PT | TXNIP |
| 6.4191E-28 | 0.79014898 | 0.889 | 0.784 | 1.5164E-23 | TEC_PT | TXN |
| 7.1839E-28 | 0.37774311 | 0.32 | 0.12 | 1.6971E-23 | TEC_PT | HSD17B10 |
| 1.5817E-27 | 0.26299781 | 0.185 | 0.049 | 3.7365E-23 | TEC_PT | TCN2 |
| 1.6768E-27 | 0.35843927 | 0.3 | 0.108 | 3.9612E-23 | TEC_PT | FAM173A |
| 3.3003E-27 | 0.28098968 | 0.259 | 0.085 | 7.7964E-23 | TEC_PT | LYPLAL1 |
| 3.7379E-27 | 0.51234363 | 0.404 | 0.184 | 8.8301E-23 | TEC_PT | NDUFAF3 |
| 4.6799E-27 | 0.27549364 | 0.219 | 0.065 | 1.1055E-22 | TEC_PT | MRPL46 |
| 5.6357E-27 | -0.8123601 | 0.397 | 0.679 | 1.3313E-22 | TEC_PT | IER2 |
| 7.6949E-27 | 0.26281912 | 0.189 | 0.051 | 1.8178E-22 | TEC_PT | TRIM47 |
| 8.0317E-27 | 0.42910311 | 0.35 | 0.142 | 1.8973E-22 | TEC_PT | MRPL27 |
| 9.3559E-27 | 0.76274278 | 0.795 | 0.648 | 2.2101E-22 | TEC_PT | UQCRH |
| 9.9704E-27 | 0.26366569 | 0.212 | 0.062 | 2.3553E-22 | TEC_PT | KIF9 |
| 1.3071E-26 | 0.5758307 | 0.461 | 0.237 | 3.0877E-22 | TEC_PT | TUFM |
| 1.3347E-26 | 0.58236911 | 0.98 | 0.939 | 3.1529E-22 | TEC_PT | RPS28 |
| 1.456E-26 | 0.53179687 | 0.441 | 0.218 | 3.4395E-22 | TEC_PT | MDH2 |
| 1.5654E-26 | 0.37507324 | 0.195 | 0.056 | 3.698E-22 | TEC_PT | SLC39A14 |
| 1.6382E-26 | 0.36679551 | 0.293 | 0.108 | 3.87E-22 | TEC_PT | FAM96A |
| 1.8243E-26 | 0.45752122 | 0.374 | 0.161 | 4.3095E-22 | TEC_PT | TMEM141 |
| 1.8789E-26 | 0.37553308 | 0.323 | 0.125 | 4.4386E-22 | TEC_PT | ARL2 |
| 1.9396E-26 | 0.25014159 | 0.185 | 0.05 | 4.5818E-22 | TEC_PT | MIR4458HG |
| 2.9602E-26 | -1.282338 | 0.195 | 0.475 | 6.9928E-22 | TEC_PT | ERO1A |
| 3.0701E-26 | 0.40490372 | 0.313 | 0.12 | 7.2526E-22 | TEC_PT | TUSC2 |
| 3.1451E-26 | 0.29641441 | 0.266 | 0.09 | 7.4298E-22 | TEC_PT | ENOSF1 |
| 3.3323E-26 | 0.27906848 | 0.236 | 0.075 | 7.8719E-22 | TEC_PT | MYL6B |
| 3.4714E-26 | 0.51442403 | 0.394 | 0.178 | 8.2004E-22 | TEC_PT | UQCRC1 |
| 3.6442E-26 | -1.0364651 | 0.01 | 0.294 | 8.6086E-22 | TEC_PT | GPRC5A |
| 3.9521E-26 | 0.67269395 | 0.613 | 0.384 | 9.3359E-22 | TEC_PT | TBCA |
| 4.3753E-26 | 0.49207724 | 0.343 | 0.144 | 1.0336E-21 | TEC_PT | TMEM256 |
| 4.6748E-26 | 0.68843246 | 0.502 | 0.288 | 1.1043E-21 | TEC_PT | NDUFB3 |
| 5.3076E-26 | 0.58087022 | 0.838 | 0.725 | 1.2538E-21 | TEC_PT | ELOB |
| 5.5832E-26 | -1.596075 | 0.165 | 0.442 | 1.3189E-21 | TEC_PT | TYROBP |
| 5.8759E-26 | -0.9482979 | 0.461 | 0.7 | 1.3881E-21 | TEC_PT | JUNB |
| 6.6428E-26 | -1.1312399 | 0.003 | 0.283 | 1.5692E-21 | TEC_PT | PTPRC |
| 7.5875E-26 | -0.6212892 | 0.003 | 0.282 | 1.7924E-21 | TEC_PT | LYN |
| 1.0025E-25 | -1.2015393 | 0.774 | 0.859 | 2.3681E-21 | TEC_PT | SH3BGRL3 |
| 1.1052E-25 | -0.9453145 | 0.01 | 0.293 | 2.6107E-21 | TEC_PT | LCP1 |
| 1.1569E-25 | -0.5905521 | 0.013 | 0.296 | 2.7329E-21 | TEC_PT | FXYD5 |
| 1.1627E-25 | 0.67804865 | 0.572 | 0.356 | 2.7467E-21 | TEC_PT | ATP5PD |
| 1.2573E-25 | -1.2736168 | 0 | 0.277 | 2.9701E-21 | TEC_PT | AIF1 |
| 1.5927E-25 | 0.68637805 | 0.798 | 0.7 | 3.7624E-21 | TEC_PT | COX6B1 |
| 1.6953E-25 | 0.71669266 | 0.869 | 0.758 | 4.0049E-21 | TEC_PT | UQCRB |

| | | | | | | |
|---|---|---|---|---|---|---|
| 1.6973E-25 | -0.6464739 | 0.034 | 0.321 | 4.0095E-21 | TEC_PT | SLC16A3 |
| 1.8018E-25 | -0.6310957 | 0.128 | 0.43 | 4.2564E-21 | TEC_PT | IQGAP1 |
| 1.8792E-25 | 0.44425244 | 0.431 | 0.203 | 4.4393E-21 | TEC_PT | NENF |
| 1.9597E-25 | -0.7585612 | 0.293 | 0.58 | 4.6294E-21 | TEC_PT | LITAF |
| 2.4945E-25 | -1.5167078 | 0.054 | 0.335 | 5.8928E-21 | TEC_PT | IL1RN |
| 2.7053E-25 | -0.5750417 | 0.623 | 0.832 | 6.3908E-21 | TEC_PT | YWHAZ |
| 2.9678E-25 | 0.56048942 | 0.502 | 0.27 | 7.0107E-21 | TEC_PT | NDUFB8 |
| 3.2548E-25 | -0.6216288 | 0.478 | 0.711 | 7.6887E-21 | TEC_PT | SDCBP |
| 3.9721E-25 | 0.6044126 | 0.778 | 0.574 | 9.3833E-21 | TEC_PT | PRDX5 |
| 4.6534E-25 | 0.62145818 | 0.606 | 0.368 | 1.0993E-20 | TEC_PT | PRDX6 |
| 5.2803E-25 | 0.39231731 | 0.357 | 0.153 | 1.2474E-20 | TEC_PT | NDUFS4 |
| 5.8612E-25 | -0.7070456 | 0.906 | 0.966 | 1.3846E-20 | TEC_PT | UBC |
| 6.5134E-25 | 0.3383946 | 0.215 | 0.067 | 1.5387E-20 | TEC_PT | NFIB |
| 1.1266E-24 | -0.8559807 | 0.562 | 0.765 | 2.6614E-20 | TEC_PT | BTG1 |
| 1.257E-24 | 0.45217548 | 0.303 | 0.12 | 2.9695E-20 | TEC_PT | SMIM37 |
| 1.2686E-24 | -0.9688519 | 0.185 | 0.458 | 2.9968E-20 | TEC_PT | RAB11FIP1 |
| 1.3007E-24 | 0.45874562 | 0.397 | 0.185 | 3.0726E-20 | TEC_PT | ILF2 |
| 1.8239E-24 | 0.35434645 | 0.276 | 0.102 | 4.3086E-20 | TEC_PT | HMOX2 |
| 1.9197E-24 | 0.62193452 | 0.461 | 0.252 | 4.5349E-20 | TEC_PT | SCP2 |
| 1.9903E-24 | 0.54644391 | 0.35 | 0.153 | 4.7017E-20 | TEC_PT | MPC1 |
| 2.0897E-24 | 0.58449969 | 0.512 | 0.29 | 4.9365E-20 | TEC_PT | NDUFS6 |
| 2.1457E-24 | -0.7241701 | 0.003 | 0.269 | 5.0688E-20 | TEC_PT | CD53 |
| 2.2847E-24 | -0.8226876 | 0.461 | 0.7 | 5.3972E-20 | TEC_PT | PPP1R15A |
| 2.6751E-24 | 0.31843582 | 0.286 | 0.107 | 6.3194E-20 | TEC_PT | MRPS18B |
| 2.8412E-24 | 0.47712282 | 0.451 | 0.224 | 6.7118E-20 | TEC_PT | COX20 |
| 3.0038E-24 | 0.26687498 | 0.175 | 0.049 | 7.0959E-20 | TEC_PT | CTDSPL |
| 3.1529E-24 | -0.6299458 | 0.024 | 0.299 | 7.4481E-20 | TEC_PT | REL |
| 3.2741E-24 | -0.5808457 | 0.694 | 0.87 | 7.7344E-20 | TEC_PT | PABPC1 |
| 3.2978E-24 | 0.35898828 | 0.33 | 0.136 | 7.7905E-20 | TEC_PT | MRPS7 |
| 4.1307E-24 | 0.63891959 | 0.801 | 0.678 | 9.7581E-20 | TEC_PT | COX7A2 |
| 4.9752E-24 | -0.6537632 | 0.684 | 0.872 | 1.1753E-19 | TEC_PT | HLA-C |
| 5.2765E-24 | -0.7835001 | 0 | 0.262 | 1.2465E-19 | TEC_PT | MUC1 |
| 5.4722E-24 | 0.61195068 | 0.953 | 0.892 | 1.2927E-19 | TEC_PT | ATP5F1E |
| 5.5852E-24 | 0.31371165 | 0.286 | 0.106 | 1.3194E-19 | TEC_PT | KIAA1191 |
| 5.834E-24 | 0.29473784 | 0.246 | 0.083 | 1.3782E-19 | TEC_PT | CSRP2 |
| 6.7569E-24 | -0.7704501 | 0.215 | 0.484 | 1.5962E-19 | TEC_PT | CD46 |
| 8.3906E-24 | 0.29950407 | 0.259 | 0.092 | 1.9821E-19 | TEC_PT | RRAS2 |
| 8.8533E-24 | 0.54977986 | 0.502 | 0.283 | 2.0914E-19 | TEC_PT | PSMB6 |
| 1.1366E-23 | 0.38641588 | 0.32 | 0.129 | 2.685E-19 | TEC_PT | AIG1 |
| 1.1953E-23 | 0.2779091 | 0.215 | 0.069 | 2.8236E-19 | TEC_PT | ABHD14B |
| 1.5301E-23 | -2.2848826 | 0.215 | 0.466 | 3.6145E-19 | TEC_PT | LYZ |
| 1.6958E-23 | 0.50440705 | 0.99 | 0.98 | 4.0061E-19 | TEC_PT | RPL41 |
| 2.7462E-23 | 0.28873124 | 0.259 | 0.093 | 6.4872E-19 | TEC_PT | TOMM5 |
| 3.4005E-23 | -0.5952736 | 0.364 | 0.626 | 8.033E-19 | TEC_PT | ACTR2 |
| 3.4133E-23 | -0.716431 | 0.034 | 0.3 | 8.0632E-19 | TEC_PT | FAM129B |
| 3.5342E-23 | -0.5349335 | 0.013 | 0.272 | 8.3487E-19 | TEC_PT | QSOX1 |
| 3.9375E-23 | 0.32257321 | 0.269 | 0.101 | 9.3015E-19 | TEC_PT | DERA |
| 4.0391E-23 | -1.205595 | 0.202 | 0.459 | 9.5415E-19 | TEC_PT | MXD1 |
| 8.0195E-23 | -1.0613981 | 0.781 | 0.884 | 1.8944E-18 | TEC_PT | CSTB |
| 8.0542E-23 | 0.55754705 | 0.438 | 0.228 | 1.9026E-18 | TEC_PT | MRPL57 |

| | | | | | | |
|---|---|---|---|---|---|---|
| 8.697E-23 | 0.37170079 | 0.269 | 0.102 | 2.0545E-18 | TEC_PT | SLC25A1 |
| 8.7782E-23 | 0.35050042 | 0.32 | 0.131 | 2.0737E-18 | TEC_PT | STOML2 |
| 8.8123E-23 | -0.6368397 | 0.623 | 0.812 | 2.0817E-18 | TEC_PT | DDX5 |
| 1.0811E-22 | 0.26093145 | 0.215 | 0.071 | 2.5539E-18 | TEC_PT | COQ9 |
| 1.1801E-22 | 0.47122002 | 0.354 | 0.161 | 2.7877E-18 | TEC_PT | DECR1 |
| 1.7848E-22 | -0.7624084 | 0 | 0.248 | 4.2162E-18 | TEC_PT | CSTA |
| 1.7895E-22 | 0.31657935 | 0.283 | 0.11 | 4.2272E-18 | TEC_PT | ZNF428 |
| 1.8143E-22 | -1.5412483 | 0.152 | 0.407 | 4.2859E-18 | TEC_PT | SLPI |
| 1.9077E-22 | 0.47709826 | 0.458 | 0.238 | 4.5065E-18 | TEC_PT | MRPS21 |
| 2.1741E-22 | -1.0668248 | 0.027 | 0.284 | 5.136E-18 | TEC_PT | IFITM2 |
| 2.1947E-22 | -1.099589 | 0.411 | 0.643 | 5.1845E-18 | TEC_PT | IER3 |
| 2.257E-22 | 0.48824432 | 0.485 | 0.266 | 5.3317E-18 | TEC_PT | GADD45GIP1 |
| 2.6669E-22 | -0.552527 | 0.081 | 0.36 | 6.3001E-18 | TEC_PT | PLSCR1 |
| 2.6841E-22 | 0.72578037 | 0.485 | 0.283 | 6.3407E-18 | TEC_PT | NDUFB1 |
| 3.7663E-22 | 0.54564801 | 0.811 | 0.645 | 8.8972E-18 | TEC_PT | ATP5MC2 |
| 3.9393E-22 | 0.35232408 | 0.266 | 0.102 | 9.3058E-18 | TEC_PT | MCUR1 |
| 4.5572E-22 | 0.60715529 | 0.401 | 0.211 | 1.0765E-17 | TEC_PT | SDHD |
| 4.5997E-22 | -0.5853292 | 0.626 | 0.8 | 1.0866E-17 | TEC_PT | ARPC2 |
| 5.1361E-22 | 0.31951261 | 0.273 | 0.104 | 1.2133E-17 | TEC_PT | GOT1 |
| 5.2714E-22 | 0.25299047 | 0.215 | 0.072 | 1.2453E-17 | TEC_PT | MRPL37 |
| 5.7697E-22 | 0.30903008 | 0.215 | 0.073 | 1.363E-17 | TEC_PT | MOCS2 |
| 7.8776E-22 | -0.6217876 | 0.003 | 0.247 | 1.8609E-17 | TEC_PT | GMFG |
| 8.6229E-22 | -0.8889151 | 0.212 | 0.454 | 2.037E-17 | TEC_PT | CLDN7 |
| 9.0263E-22 | -0.6279739 | 0.222 | 0.489 | 2.1323E-17 | TEC_PT | ANKRD12 |
| 9.267E-22 | 0.39240785 | 0.367 | 0.168 | 2.1891E-17 | TEC_PT | GNG12 |
| 9.6522E-22 | -0.661426 | 0.165 | 0.43 | 2.2801E-17 | TEC_PT | EFHD2 |
| 9.6622E-22 | 0.2700686 | 0.232 | 0.082 | 2.2825E-17 | TEC_PT | SUCLA2 |
| 1.029E-21 | -0.8131457 | 0.269 | 0.52 | 2.4308E-17 | TEC_PT | H1FX |
| 1.0296E-21 | -1.1649509 | 0.017 | 0.264 | 2.4321E-17 | TEC_PT | AREG |
| 1.0366E-21 | 0.25408139 | 0.104 | 0.022 | 2.4487E-17 | TEC_PT | CALML3 |
| 1.2642E-21 | 0.25073878 | 0.236 | 0.084 | 2.9864E-17 | TEC_PT | GTPBP6 |
| 1.3085E-21 | -0.9612789 | 0 | 0.24 | 3.0911E-17 | TEC_PT | ADGRF1 |
| 1.3417E-21 | 0.4027983 | 0.36 | 0.166 | 3.1694E-17 | TEC_PT | MRPS34 |
| 1.4363E-21 | 0.31248436 | 0.215 | 0.074 | 3.393E-17 | TEC_PT | FMC1 |
| 1.5326E-21 | 0.38423412 | 0.293 | 0.121 | 3.6205E-17 | TEC_PT | SNRPN |
| 1.603E-21 | 0.27839343 | 0.226 | 0.079 | 3.7868E-17 | TEC_PT | PMM1 |
| 1.6427E-21 | 0.56034926 | 0.455 | 0.257 | 3.8805E-17 | TEC_PT | LSM3 |
| 1.6463E-21 | 0.44479451 | 0.401 | 0.2 | 3.8891E-17 | TEC_PT | PSMB5 |
| 1.6595E-21 | 0.48715784 | 0.441 | 0.236 | 3.9202E-17 | TEC_PT | PSMB7 |
| 1.8149E-21 | 0.47057509 | 0.434 | 0.225 | 4.2873E-17 | TEC_PT | MRPL52 |
| 1.8969E-21 | -0.6763467 | 0.064 | 0.322 | 4.481E-17 | TEC_PT | PERP |
| 2.049E-21 | 0.33502106 | 0.296 | 0.123 | 4.8404E-17 | TEC_PT | CLPTM1L |
| 2.0833E-21 | -0.8786205 | 0.212 | 0.472 | 4.9213E-17 | TEC_PT | DUSP1 |
| 2.5803E-21 | 0.43244706 | 0.38 | 0.184 | 6.0954E-17 | TEC_PT | ANXA4 |
| 2.9948E-21 | -0.7479823 | 0.32 | 0.574 | 7.0746E-17 | TEC_PT | VMP1 |
| 3.1571E-21 | 0.32162729 | 0.263 | 0.102 | 7.4581E-17 | TEC_PT | ANAPC13 |
| 3.6222E-21 | 0.26977288 | 0.215 | 0.074 | 8.5568E-17 | TEC_PT | SLC39A4 |
| 3.8052E-21 | 0.5356798 | 0.973 | 0.922 | 8.9889E-17 | TEC_PT | RPL8 |
| 4.1132E-21 | -3.1800646 | 0.495 | 0.677 | 9.7166E-17 | TEC_PT | TIMP1 |
| 4.3957E-21 | -1.1070674 | 0 | 0.235 | 1.0384E-16 | TEC_PT | SLC2A3 |

| | | | | | | |
|---|---|---|---|---|---|---|
| 4.9525E-21 | -3.2747796 | 0.313 | 0.525 | 1.1699E-16 | TEC_PT | LCN2 |
| 5.2593E-21 | 0.525094 | 0.478 | 0.271 | 1.2424E-16 | TEC_PT | ZNHIT1 |
| 6.0301E-21 | -1.6636235 | 0 | 0.233 | 1.4245E-16 | TEC_PT | PRSS22 |
| 6.1354E-21 | 0.3448663 | 0.283 | 0.115 | 1.4494E-16 | TEC_PT | VEGFB |
| 6.1513E-21 | -1.5643044 | 0.148 | 0.387 | 1.4531E-16 | TEC_PT | FCER1G |
| 6.6785E-21 | -0.8698022 | 0.29 | 0.54 | 1.5777E-16 | TEC_PT | UPP1 |
| 6.9169E-21 | 0.34470411 | 0.303 | 0.129 | 1.634E-16 | TEC_PT | CERS2 |
| 7.9258E-21 | 0.62352253 | 0.569 | 0.367 | 1.8723E-16 | TEC_PT | TXNDC17 |
| 7.9287E-21 | 0.48271499 | 0.505 | 0.288 | 1.873E-16 | TEC_PT | NDUFB9 |
| 8.1053E-21 | 0.55728133 | 0.502 | 0.3 | 1.9147E-16 | TEC_PT | ATP5PO |
| 8.8143E-21 | -0.5465159 | 0.135 | 0.399 | 2.0822E-16 | TEC_PT | H2AFY |
| 9.0856E-21 | 0.3312297 | 0.229 | 0.083 | 2.1463E-16 | TEC_PT | ZSCAN16-AS1 |
| 9.8288E-21 | -1.0777157 | 0.003 | 0.237 | 2.3219E-16 | TEC_PT | BCL2A1 |
| 1.1248E-20 | 0.53937487 | 0.559 | 0.356 | 2.6572E-16 | TEC_PT | COX5A |
| 1.1482E-20 | -0.4914862 | 0.003 | 0.235 | 2.7123E-16 | TEC_PT | BST2 |
| 1.325E-20 | 0.62207715 | 0.886 | 0.725 | 3.13E-16 | TEC_PT | ENO1 |
| 1.4956E-20 | 0.36045297 | 0.175 | 0.054 | 3.533E-16 | TEC_PT | HKDC1 |
| 1.6715E-20 | -0.724287 | 0.296 | 0.537 | 3.9486E-16 | TEC_PT | SERPINB1 |
| 1.7649E-20 | 0.67594047 | 0.623 | 0.453 | 4.1693E-16 | TEC_PT | ST13 |
| 1.9468E-20 | -0.6115188 | 0.414 | 0.641 | 4.5989E-16 | TEC_PT | ARPC5 |
| 2.2583E-20 | -1.0878285 | 0.04 | 0.28 | 5.3349E-16 | TEC_PT | DUSP5 |
| 2.2704E-20 | -0.6179963 | 0.071 | 0.324 | 5.3633E-16 | TEC_PT | B4GALT1 |
| 2.2811E-20 | -1.1070813 | 0.054 | 0.294 | 5.3885E-16 | TEC_PT | RND3 |
| 2.3285E-20 | -0.5518463 | 0.013 | 0.249 | 5.5007E-16 | TEC_PT | GPCPD1 |
| 2.4564E-20 | 0.53607891 | 0.431 | 0.235 | 5.8028E-16 | TEC_PT | NDUFA13 |
| 2.8822E-20 | 0.29717862 | 0.249 | 0.096 | 6.8085E-16 | TEC_PT | MRPL19 |
| 3.016E-20 | 0.39443998 | 0.354 | 0.167 | 7.1247E-16 | TEC_PT | METTL26 |
| 3.4291E-20 | 0.25269273 | 0.239 | 0.089 | 8.1005E-16 | TEC_PT | VRK3 |
| 3.5531E-20 | 0.32923908 | 0.357 | 0.164 | 8.3936E-16 | TEC_PT | MRPL14 |
| 3.5667E-20 | 0.50619878 | 0.963 | 0.895 | 8.4257E-16 | TEC_PT | RPL37A |
| 3.7944E-20 | 0.44319261 | 0.401 | 0.205 | 8.9636E-16 | TEC_PT | NAA38 |
| 4.8031E-20 | -0.5994716 | 0.125 | 0.39 | 1.1346E-15 | TEC_PT | AHNAK |
| 4.8619E-20 | -2.0961991 | 0.343 | 0.537 | 1.1485E-15 | TEC_PT | WFDC2 |
| 5.2804E-20 | -0.5318499 | 0.101 | 0.356 | 1.2474E-15 | TEC_PT | AKAP13 |
| 5.4829E-20 | 0.58014175 | 0.552 | 0.348 | 1.2952E-15 | TEC_PT | FKBP2 |
| 6.1106E-20 | -0.8299691 | 0.017 | 0.247 | 1.4435E-15 | TEC_PT | CST6 |
| 7.3359E-20 | 0.29334672 | 0.266 | 0.106 | 1.733E-15 | TEC_PT | SLC25A11 |
| 7.8104E-20 | 0.31628009 | 0.286 | 0.121 | 1.8451E-15 | TEC_PT | NDUFA8 |
| 8.2003E-20 | 0.28976603 | 0.253 | 0.098 | 1.9371E-15 | TEC_PT | IMP4 |
| 8.7146E-20 | 0.57354065 | 0.758 | 0.642 | 2.0587E-15 | TEC_PT | EDF1 |
| 9.8864E-20 | 0.52883095 | 0.438 | 0.248 | 2.3355E-15 | TEC_PT | ATP5PB |
| 1.1027E-19 | -1.1462981 | 0.034 | 0.264 | 2.6049E-15 | TEC_PT | LMO7 |
| 1.1509E-19 | 0.4575784 | 0.394 | 0.205 | 2.7187E-15 | TEC_PT | TIMM8B |
| 1.193E-19 | 0.32945602 | 0.273 | 0.111 | 2.8182E-15 | TEC_PT | SETD3 |
| 1.2224E-19 | 0.28800138 | 0.229 | 0.086 | 2.8876E-15 | TEC_PT | TPRG1L |
| 1.2847E-19 | 0.29279519 | 0.296 | 0.124 | 3.0348E-15 | TEC_PT | EBNA1BP2 |
| 1.3309E-19 | 0.34430917 | 0.219 | 0.079 | 3.144E-15 | TEC_PT | RETREG1 |
| 1.3378E-19 | -0.4668092 | 0.017 | 0.246 | 3.1604E-15 | TEC_PT | CD82 |
| 1.3753E-19 | 0.57158949 | 0.579 | 0.379 | 3.2488E-15 | TEC_PT | EIF5A |
| 1.5452E-19 | 0.57930844 | 0.582 | 0.406 | 3.6503E-15 | TEC_PT | NDUFB4 |

| | | | | | | |
|---|---|---|---|---|---|---|
| 1.7216E-19 | 0.50119987 | 0.488 | 0.286 | 4.067E-15 | TEC_PT | LAMTOR1 |
| 1.7232E-19 | -0.533104 | 0.377 | 0.617 | 4.0707E-15 | TEC_PT | KTN1 |
| 1.872E-19 | 0.38624172 | 0.313 | 0.142 | 4.4222E-15 | TEC_PT | TRAPPC2L |
| 1.8844E-19 | -0.5023434 | 0.347 | 0.585 | 4.4514E-15 | TEC_PT | ACTR3 |
| 2.0095E-19 | 0.36513053 | 0.293 | 0.127 | 4.747E-15 | TEC_PT | SDHA |
| 2.1282E-19 | -0.460819 | 0.364 | 0.624 | 5.0274E-15 | TEC_PT | PRRC2C |
| 2.1377E-19 | 0.55821762 | 0.37 | 0.18 | 5.0499E-15 | TEC_PT | GCLM |
| 2.1644E-19 | -0.913363 | 0.02 | 0.246 | 5.113E-15 | TEC_PT | SLC11A1 |
| 2.3448E-19 | 0.31189254 | 0.31 | 0.137 | 5.5391E-15 | TEC_PT | ASNA1 |
| 2.6995E-19 | 0.47964978 | 0.38 | 0.201 | 6.3769E-15 | TEC_PT | MRPS36 |
| 3.0907E-19 | 0.48044995 | 0.976 | 0.927 | 7.3012E-15 | TEC_PT | RPL19 |
| 3.1293E-19 | 0.49677016 | 0.401 | 0.211 | 7.3924E-15 | TEC_PT | GTF3C6 |
| 3.2333E-19 | 0.30458952 | 0.259 | 0.104 | 7.6379E-15 | TEC_PT | MRPS26 |
| 3.5378E-19 | 0.27216412 | 0.189 | 0.063 | 8.3572E-15 | TEC_PT | KIF22 |
| 4.1347E-19 | 0.28665051 | 0.276 | 0.111 | 9.7674E-15 | TEC_PT | TFPI |
| 4.1479E-19 | -0.9324329 | 0.303 | 0.529 | 9.7986E-15 | TEC_PT | ZFP36 |
| 4.7254E-19 | 0.56495791 | 0.545 | 0.348 | 1.1163E-14 | TEC_PT | VDAC1 |
| 5.0142E-19 | 0.68440839 | 0.704 | 0.55 | 1.1845E-14 | TEC_PT | UQCR10 |
| 5.5085E-19 | -0.6943383 | 0.101 | 0.349 | 1.3013E-14 | TEC_PT | IER5 |
| 5.9868E-19 | 0.26125523 | 0.195 | 0.068 | 1.4143E-14 | TEC_PT | SNX4 |
| 6.352E-19 | 0.39564524 | 0.323 | 0.149 | 1.5005E-14 | TEC_PT | CENPX |
| 8.1145E-19 | -0.5678658 | 0.003 | 0.217 | 1.9169E-14 | TEC_PT | SPI1 |
| 8.9988E-19 | 0.37286545 | 0.313 | 0.144 | 2.1258E-14 | TEC_PT | DYNLL2 |
| 9.8522E-19 | -0.7162668 | 0.795 | 0.92 | 2.3274E-14 | TEC_PT | S100A10 |
| 9.9421E-19 | -0.9803767 | 0.007 | 0.221 | 2.3486E-14 | TEC_PT | CD14 |
| 1.2618E-18 | -1.0625767 | 0.007 | 0.221 | 2.9808E-14 | TEC_PT | ALOX5AP |
| 1.3575E-18 | -0.5271238 | 0.003 | 0.215 | 3.2069E-14 | TEC_PT | EHF |
| 1.3772E-18 | -0.5099146 | 0.751 | 0.876 | 3.2534E-14 | TEC_PT | MYL12A |
| 1.4224E-18 | -0.6799643 | 0.091 | 0.325 | 3.3601E-14 | TEC_PT | LIMS1 |
| 1.4336E-18 | -0.6145935 | 0.007 | 0.221 | 3.3866E-14 | TEC_PT | ITGB2 |
| 1.5349E-18 | 0.58813525 | 0.455 | 0.272 | 3.6258E-14 | TEC_PT | ATP5F1C |
| 2.176E-18 | -0.2617697 | 0.862 | 0.658 | 5.1405E-14 | TEC_PT | MT2A |
| 2.2285E-18 | -0.6055055 | 0.088 | 0.328 | 5.2643E-14 | TEC_PT | MAFF |
| 2.4713E-18 | 0.29829095 | 0.269 | 0.113 | 5.8379E-14 | TEC_PT | CHPT1 |
| 2.5138E-18 | -0.9173349 | 0.145 | 0.391 | 5.9384E-14 | TEC_PT | FOS |
| 2.5261E-18 | 0.32231876 | 0.394 | 0.2 | 5.9673E-14 | TEC_PT | MLF2 |
| 2.5873E-18 | -0.6718098 | 0.02 | 0.236 | 6.1121E-14 | TEC_PT | HCST |
| 2.8172E-18 | 0.33721236 | 0.283 | 0.125 | 6.6551E-14 | TEC_PT | NDUFS3 |
| 4.013E-18 | -0.4887296 | 0.027 | 0.245 | 9.4799E-14 | TEC_PT | RAB31 |
| 4.8704E-18 | -0.4303125 | 0.017 | 0.228 | 1.1505E-13 | TEC_PT | RGS10 |
| 4.9229E-18 | 0.39929519 | 0.343 | 0.166 | 1.1629E-13 | TEC_PT | LAP3 |
| 5.0428E-18 | -1.3094306 | 0.367 | 0.54 | 1.1913E-13 | TEC_PT | CLDN4 |
| 5.4808E-18 | 0.46666316 | 0.576 | 0.334 | 1.2947E-13 | TEC_PT | IGFBP7 |
| 5.8978E-18 | -0.6439269 | 0.003 | 0.209 | 1.3932E-13 | TEC_PT | LST1 |
| 6.2266E-18 | 0.31965244 | 0.273 | 0.118 | 1.4709E-13 | TEC_PT | TIMMDC1 |
| 7.4518E-18 | -0.6318469 | 0.37 | 0.604 | 1.7603E-13 | TEC_PT | MCL1 |
| 7.6733E-18 | 0.52390304 | 0.394 | 0.216 | 1.8127E-13 | TEC_PT | MLEC |
| 8.1721E-18 | 0.33651175 | 0.31 | 0.14 | 1.9305E-13 | TEC_PT | BEX3 |
| 8.5269E-18 | -0.7368156 | 0 | 0.202 | 2.0143E-13 | TEC_PT | CYBB |
| 8.734E-18 | -1.0882293 | 0.01 | 0.217 | 2.0632E-13 | TEC_PT | S100A12 |

| | | | | | | |
|---|---|---|---|---|---|---|
| 9.7683E-18 | -0.7301616 | 0.111 | 0.345 | 2.3076E-13 | TEC_PT | DDIT4 |
| 1.1255E-17 | -0.6032435 | 0.034 | 0.251 | 2.6588E-13 | TEC_PT | KLF5 |
| 1.2072E-17 | 0.50050193 | 1 | 0.993 | 2.8518E-13 | TEC_PT | TPT1 |
| 1.3442E-17 | -0.3931213 | 0.044 | 0.267 | 3.1754E-13 | TEC_PT | CDC42SE1 |
| 1.4967E-17 | -1.9495938 | 0.054 | 0.268 | 3.5356E-13 | TEC_PT | S100A2 |
| 1.5237E-17 | -0.9663245 | 0.202 | 0.426 | 3.5993E-13 | TEC_PT | EMP3 |
| 1.6194E-17 | -0.487526 | 0.064 | 0.29 | 3.8256E-13 | TEC_PT | RNF149 |
| 1.6495E-17 | -2.3662887 | 0.215 | 0.409 | 3.8966E-13 | TEC_PT | S100P |
| 1.7269E-17 | 0.50830809 | 0.438 | 0.256 | 4.0795E-13 | TEC_PT | RPS19BP1 |
| 1.7498E-17 | -0.5031323 | 0.081 | 0.317 | 4.1336E-13 | TEC_PT | HMGB2 |
| 1.8183E-17 | -0.5947663 | 0.01 | 0.214 | 4.2955E-13 | TEC_PT | BASP1 |
| 1.9011E-17 | 0.54532536 | 0.475 | 0.297 | 4.491E-13 | TEC_PT | ROMO1 |
| 1.9074E-17 | -1.0110115 | 0.091 | 0.314 | 4.5059E-13 | TEC_PT | PHLDA1 |
| 2.0993E-17 | -0.6450883 | 0.044 | 0.263 | 4.9592E-13 | TEC_PT | HES4 |
| 2.405E-17 | 0.39445997 | 0.323 | 0.158 | 5.6814E-13 | TEC_PT | NDUFB6 |
| 2.5468E-17 | 0.33371888 | 0.283 | 0.126 | 6.0162E-13 | TEC_PT | UQCC2 |
| 2.6075E-17 | -0.5054057 | 0.343 | 0.596 | 6.1596E-13 | TEC_PT | DDX17 |
| 2.8646E-17 | -0.7209483 | 0.01 | 0.211 | 6.767E-13 | TEC_PT | TNFRSF1B |
| 3.0701E-17 | -0.5398274 | 0.003 | 0.202 | 7.2525E-13 | TEC_PT | NCF2 |
| 3.0725E-17 | 0.37893562 | 0.421 | 0.228 | 7.2583E-13 | TEC_PT | HDGF |
| 3.8564E-17 | 0.26698671 | 0.165 | 0.055 | 9.1101E-13 | TEC_PT | PDK4 |
| 4.0842E-17 | -0.6476891 | 0.003 | 0.2 | 9.648E-13 | TEC_PT | DUSP4 |
| 4.2115E-17 | 0.40751189 | 0.387 | 0.21 | 9.9488E-13 | TEC_PT | PSMC5 |
| 4.3737E-17 | -0.7438862 | 0.003 | 0.199 | 1.0332E-12 | TEC_PT | CD52 |
| 4.6834E-17 | -0.4588043 | 0.242 | 0.495 | 1.1064E-12 | TEC_PT | RSRP1 |
| 4.8876E-17 | -2.5558374 | 0.276 | 0.49 | 1.1546E-12 | TEC_PT | CXCL8 |
| 5.1045E-17 | 0.30204226 | 0.269 | 0.117 | 1.2058E-12 | TEC_PT | HCFC1R1 |
| 5.2891E-17 | -0.8909685 | 0.064 | 0.283 | 1.2494E-12 | TEC_PT | MAFB |
| 5.6086E-17 | -0.4298821 | 0.236 | 0.485 | 1.3249E-12 | TEC_PT | TMEM50A |
| 5.6186E-17 | 0.27452281 | 0.246 | 0.102 | 1.3273E-12 | TEC_PT | TSR3 |
| 5.6384E-17 | -0.8266907 | 0.128 | 0.353 | 1.332E-12 | TEC_PT | PHLDA2 |
| 6.0888E-17 | -0.5037537 | 0.36 | 0.576 | 1.4384E-12 | TEC_PT | PPP1CB |
| 6.3972E-17 | -0.4088117 | 0.007 | 0.203 | 1.5112E-12 | TEC_PT | IFI16 |
| 6.4417E-17 | -0.4685704 | 0.088 | 0.318 | 1.5217E-12 | TEC_PT | BOD1L1 |
| 6.6951E-17 | -1.4680066 | 0.003 | 0.197 | 1.5816E-12 | TEC_PT | HOPX |
| 6.9248E-17 | -0.4781286 | 0.091 | 0.313 | 1.6358E-12 | TEC_PT | TMEM165 |
| 7.3899E-17 | 0.52928225 | 0.842 | 0.689 | 1.7457E-12 | TEC_PT | EEF2 |
| 7.8496E-17 | 0.49075036 | 0.939 | 0.8 | 1.8543E-12 | TEC_PT | RPS5 |
| 7.8872E-17 | 0.69252144 | 0.313 | 0.151 | 1.8632E-12 | TEC_PT | MT-ND6 |
| 8.0242E-17 | -0.3583759 | 0.013 | 0.215 | 1.8956E-12 | TEC_PT | BTG3 |
| 8.368E-17 | -0.578191 | 0.003 | 0.196 | 1.9768E-12 | TEC_PT | SAMD9 |
| 9.7517E-17 | -0.5561508 | 0.007 | 0.201 | 2.3037E-12 | TEC_PT | ITGAX |
| 1.0246E-16 | 0.40311261 | 0.327 | 0.164 | 2.4205E-12 | TEC_PT | NDUFAF8 |
| 1.133E-16 | 0.52850066 | 0.862 | 0.703 | 2.6764E-12 | TEC_PT | SOD2 |
| 1.1993E-16 | 0.5189623 | 0.471 | 0.287 | 2.8331E-12 | TEC_PT | APRT |
| 1.4739E-16 | 0.28876641 | 0.283 | 0.128 | 3.4817E-12 | TEC_PT | MRPS12 |
| 1.4891E-16 | 0.42659818 | 0.455 | 0.266 | 3.5176E-12 | TEC_PT | HSPA9 |
| 1.5669E-16 | -0.391247 | 0 | 0.188 | 3.7015E-12 | TEC_PT | HCLS1 |
| 1.6419E-16 | 0.33050193 | 0.31 | 0.147 | 3.8787E-12 | TEC_PT | SNX5 |
| 1.6984E-16 | -0.4622529 | 0.077 | 0.294 | 4.0122E-12 | TEC_PT | JMJD1C |

| | | | | | | |
|---|---|---|---|---|---|---|
| 1.7247E-16 | -0.4029117 | 0.017 | 0.215 | 4.0741E-12 | TEC_PT | RUNX1 |
| 1.8552E-16 | -0.3753791 | 0 | 0.188 | 4.3825E-12 | TEC_PT | GPSM3 |
| 1.8949E-16 | -0.5171846 | 0.118 | 0.344 | 4.4763E-12 | TEC_PT | WSB1 |
| 1.9252E-16 | 0.48978839 | 0.488 | 0.312 | 4.5478E-12 | TEC_PT | BLOC1S1 |
| 2.0021E-16 | -0.4395209 | 0.094 | 0.319 | 4.7296E-12 | TEC_PT | CMTM6 |
| 2.3242E-16 | 0.38178995 | 0.337 | 0.172 | 5.4906E-12 | TEC_PT | STUB1 |
| 2.3434E-16 | 0.3677961 | 0.333 | 0.167 | 5.5358E-12 | TEC_PT | MVP |
| 2.3691E-16 | 0.44632818 | 0.909 | 0.834 | 5.5964E-12 | TEC_PT | PPIA |
| 2.3788E-16 | -0.4504147 | 0.121 | 0.348 | 5.6194E-12 | TEC_PT | CAPN2 |
| 2.4643E-16 | 0.36144279 | 0.35 | 0.179 | 5.8213E-12 | TEC_PT | POLR2I |
| 2.6361E-16 | 0.45546443 | 0.421 | 0.244 | 6.2271E-12 | TEC_PT | METTL9 |
| 2.8375E-16 | 0.46361623 | 0.407 | 0.233 | 6.7031E-12 | TEC_PT | PSMA5 |
| 3.0573E-16 | 0.61042343 | 0.512 | 0.348 | 7.2222E-12 | TEC_PT | PRR13 |
| 3.1349E-16 | 0.5183744 | 0.549 | 0.373 | 7.4055E-12 | TEC_PT | PARK7 |
| 3.2799E-16 | 0.29812517 | 0.316 | 0.152 | 7.748E-12 | TEC_PT | PNKD |
| 3.3498E-16 | 0.25081521 | 0.192 | 0.071 | 7.9132E-12 | TEC_PT | METTL7A |
| 3.4589E-16 | -1.6201012 | 0.027 | 0.224 | 8.171E-12 | TEC_PT | VCAN |
| 3.5223E-16 | -1.2411261 | 0 | 0.185 | 8.3206E-12 | TEC_PT | DUOX2 |
| 3.5643E-16 | -0.3698749 | 0 | 0.185 | 8.4199E-12 | TEC_PT | GNA15 |
| 4.0571E-16 | 0.35998376 | 0.394 | 0.216 | 9.5842E-12 | TEC_PT | LAMTOR2 |
| 4.2403E-16 | -0.5380571 | 0.118 | 0.347 | 1.0017E-11 | TEC_PT | MIDN |
| 4.2806E-16 | 0.52361716 | 0.465 | 0.282 | 1.0112E-11 | TEC_PT | RPS4Y1 |
| 4.2999E-16 | 0.4902988 | 0.963 | 0.898 | 1.0158E-11 | TEC_PT | RPL21 |
| 4.6581E-16 | -0.7801949 | 0.003 | 0.189 | 1.1004E-11 | TEC_PT | AC020656.1 |
| 4.7555E-16 | 0.43879755 | 0.956 | 0.879 | 1.1234E-11 | TEC_PT | RPS29 |
| 4.9799E-16 | -0.4135948 | 0.054 | 0.265 | 1.1764E-11 | TEC_PT | GSN |
| 5.852E-16 | -2.1725458 | 0.152 | 0.343 | 1.3824E-11 | TEC_PT | PSCA |
| 6.6654E-16 | -0.6992104 | 0.067 | 0.275 | 1.5746E-11 | TEC_PT | MAL2 |
| 6.8162E-16 | -0.5053407 | 0.512 | 0.689 | 1.6102E-11 | TEC_PT | CAST |
| 7.5951E-16 | 0.49550275 | 0.502 | 0.329 | 1.7942E-11 | TEC_PT | FAM96B |
| 7.7485E-16 | 0.5042492 | 0.896 | 0.864 | 1.8304E-11 | TEC_PT | SRP14 |
| 7.8724E-16 | -0.4000433 | 0.067 | 0.274 | 1.8597E-11 | TEC_PT | FAM49B |
| 8.007E-16 | -0.5442732 | 0.044 | 0.246 | 1.8915E-11 | TEC_PT | KIAA1551 |
| 9.2377E-16 | -0.4650142 | 0.081 | 0.293 | 2.1822E-11 | TEC_PT | SKIL |
| 9.2956E-16 | -0.5641304 | 0.306 | 0.515 | 2.1959E-11 | TEC_PT | ANXA11 |
| 9.4256E-16 | -1.6177583 | 0.013 | 0.2 | 2.2266E-11 | TEC_PT | KRT17 |
| 9.6866E-16 | 0.33181012 | 0.246 | 0.106 | 2.2883E-11 | TEC_PT | CFAP36 |
| 1.0243E-15 | -0.4810026 | 0.141 | 0.369 | 2.4196E-11 | TEC_PT | MYH9 |
| 1.0959E-15 | -0.5057927 | 0.148 | 0.376 | 2.5888E-11 | TEC_PT | CCNL1 |
| 1.201E-15 | -0.7776903 | 0.007 | 0.19 | 2.8372E-11 | TEC_PT | ANKRD36C |
| 1.3573E-15 | 0.55290022 | 0.953 | 0.846 | 3.2063E-11 | TEC_PT | RPL35 |
| 1.4207E-15 | -0.4443366 | 0.017 | 0.204 | 3.3561E-11 | TEC_PT | LSP1 |
| 1.5534E-15 | -0.5857326 | 0.003 | 0.184 | 3.6696E-11 | TEC_PT | IGSF6 |
| 1.7701E-15 | 0.34480777 | 0.347 | 0.178 | 4.1816E-11 | TEC_PT | GPI |
| 1.8708E-15 | -0.8572205 | 0.051 | 0.247 | 4.4194E-11 | TEC_PT | PLAU |
| 1.9688E-15 | -0.4402151 | 0.771 | 0.867 | 4.6508E-11 | TEC_PT | ARPC3 |
| 2.1964E-15 | 0.43763225 | 0.566 | 0.384 | 5.1884E-11 | TEC_PT | NEDD8 |
| 2.2429E-15 | -0.5160385 | 0.189 | 0.417 | 5.2984E-11 | TEC_PT | FUS |
| 2.3703E-15 | 0.26690077 | 0.199 | 0.078 | 5.5993E-11 | TEC_PT | CDK4 |
| 2.501E-15 | -0.3873476 | 0.067 | 0.275 | 5.9082E-11 | TEC_PT | ADAM10 |

| | | | | | | |
|---|---|---|---|---|---|---|
| 2.5983E-15 | -0.7651396 | 0 | 0.176 | 6.1379E-11 | TEC_PT | PLEK |
| 2.6267E-15 | -0.705197 | 0.003 | 0.181 | 6.2051E-11 | TEC_PT | OLR1 |
| 2.6917E-15 | -0.7035157 | 0.037 | 0.23 | 6.3586E-11 | TEC_PT | ITGA2 |
| 2.7723E-15 | -0.6115876 | 0.007 | 0.185 | 6.549E-11 | TEC_PT | C5AR1 |
| 2.7946E-15 | -1.4772826 | 0.168 | 0.37 | 6.6018E-11 | TEC_PT | MAL |
| 3.0256E-15 | -0.4118068 | 0.003 | 0.18 | 7.1473E-11 | TEC_PT | CD37 |
| 3.0485E-15 | -0.6006983 | 0.269 | 0.476 | 7.2014E-11 | TEC_PT | CD81 |
| 3.05E-15 | 0.39845919 | 0.963 | 0.83 | 7.2049E-11 | TEC_PT | RPL22 |
| 3.1703E-15 | -0.3450478 | 0.007 | 0.186 | 7.4892E-11 | TEC_PT | CDCP1 |
| 3.2695E-15 | 0.25383503 | 0.229 | 0.097 | 7.7236E-11 | TEC_PT | MRPL23 |
| 3.2701E-15 | 0.48867904 | 0.468 | 0.301 | 7.725E-11 | TEC_PT | SCAND1 |
| 3.2859E-15 | 0.46980698 | 0.532 | 0.362 | 7.7623E-11 | TEC_PT | C19orf53 |
| 3.2933E-15 | -0.3824311 | 0.071 | 0.283 | 7.7798E-11 | TEC_PT | ATRX |
| 3.3622E-15 | -0.3250728 | 0.044 | 0.243 | 7.9425E-11 | TEC_PT | OSTF1 |
| 3.5491E-15 | -0.4506948 | 0.044 | 0.239 | 8.384E-11 | TEC_PT | SMCHD1 |
| 3.615E-15 | -0.4746503 | 0.027 | 0.216 | 8.5397E-11 | TEC_PT | ARL4C |
| 3.6283E-15 | -0.4223668 | 0 | 0.174 | 8.571E-11 | TEC_PT | CORO1A |
| 3.7496E-15 | -0.357353 | 0.003 | 0.18 | 8.8576E-11 | TEC_PT | FMNL1 |
| 4.0201E-15 | -0.4140098 | 0.024 | 0.209 | 9.4967E-11 | TEC_PT | STK17B |
| 4.2318E-15 | 0.34530034 | 0.323 | 0.166 | 9.9968E-11 | TEC_PT | CISD2 |
| 4.2352E-15 | -0.4116404 | 0.02 | 0.206 | 1.0005E-10 | TEC_PT | MAP3K8 |
| 4.2398E-15 | 0.31193127 | 0.293 | 0.137 | 1.0016E-10 | TEC_PT | MGLL |
| 4.3744E-15 | -0.4315826 | 0.108 | 0.326 | 1.0334E-10 | TEC_PT | FOSL2 |
| 4.3927E-15 | 0.38501508 | 0.276 | 0.133 | 1.0377E-10 | TEC_PT | MRPS33 |
| 4.5391E-15 | 0.31020232 | 0.303 | 0.147 | 1.0723E-10 | TEC_PT | NME1 |
| 4.7139E-15 | -0.3548625 | 0 | 0.173 | 1.1136E-10 | TEC_PT | DAPP1 |
| 4.7534E-15 | -0.4046175 | 0.01 | 0.189 | 1.1229E-10 | TEC_PT | RAC2 |
| 4.8145E-15 | -0.8981775 | 0.017 | 0.198 | 1.1373E-10 | TEC_PT | TM4SF1 |
| 5.1545E-15 | -0.5287212 | 0.007 | 0.183 | 1.2177E-10 | TEC_PT | SAMSN1 |
| 5.8437E-15 | 0.37796803 | 0.303 | 0.152 | 1.3805E-10 | TEC_PT | C19orf24 |
| 6.172E-15 | 0.33186053 | 0.313 | 0.158 | 1.458E-10 | TEC_PT | FDX1 |
| 6.5671E-15 | -0.5722236 | 0.01 | 0.186 | 1.5514E-10 | TEC_PT | KLF2 |
| 6.6083E-15 | 0.36524694 | 0.852 | 0.775 | 1.5611E-10 | TEC_PT | TPI1 |
| 6.7322E-15 | 0.31227539 | 0.266 | 0.122 | 1.5904E-10 | TEC_PT | ESRRA |
| 6.7415E-15 | -0.4480209 | 0.01 | 0.187 | 1.5925E-10 | TEC_PT | PTPRE |
| 6.9442E-15 | -0.5369239 | 0 | 0.171 | 1.6404E-10 | TEC_PT | PRDM1 |
| 7.1393E-15 | 0.43571993 | 0.933 | 0.848 | 1.6865E-10 | TEC_PT | RPL38 |
| 7.143E-15 | -0.4656512 | 0.148 | 0.361 | 1.6874E-10 | TEC_PT | TMOD3 |
| 7.7711E-15 | -0.3594357 | 0.03 | 0.217 | 1.8358E-10 | TEC_PT | TGFB1 |
| 7.9906E-15 | -0.8026271 | 0 | 0.171 | 1.8876E-10 | TEC_PT | FCN1 |
| 8.2887E-15 | -0.4101277 | 0 | 0.17 | 1.958E-10 | TEC_PT | CD48 |
| 9.3261E-15 | -0.3299672 | 0.017 | 0.198 | 2.2031E-10 | TEC_PT | MYO9B |
| 9.6192E-15 | -0.5310174 | 0.091 | 0.291 | 2.2723E-10 | TEC_PT | SPINT1 |
| 9.7764E-15 | 0.31042468 | 0.286 | 0.138 | 2.3095E-10 | TEC_PT | DUSP3 |
| 9.9215E-15 | -0.4110062 | 0.108 | 0.318 | 2.3438E-10 | TEC_PT | KMT2E |
| 1.0429E-14 | -0.7525938 | 0.33 | 0.526 | 2.4637E-10 | TEC_PT | CEBPB |
| 1.0752E-14 | -0.6530136 | 0.013 | 0.19 | 2.5398E-10 | TEC_PT | EPS8L1 |
| 1.0981E-14 | -0.3136398 | 0.013 | 0.189 | 2.5941E-10 | TEC_PT | CNN2 |
| 1.1205E-14 | 0.39782889 | 0.387 | 0.217 | 2.6471E-10 | TEC_PT | SPATS2L |
| 1.1412E-14 | 0.36598196 | 0.3 | 0.151 | 2.6959E-10 | TEC_PT | IDH2 |

| | | | | | | |
|---|---|---|---|---|---|---|
| 1.1545E-14 | -0.4208863 | 0.01 | 0.184 | 2.7274E-10 | TEC_PT | CDH1 |
| 1.1682E-14 | 0.57633567 | 0.374 | 0.216 | 2.7596E-10 | TEC_PT | CDK2AP2 |
| 1.1806E-14 | -0.5005752 | 0.512 | 0.686 | 2.789E-10 | TEC_PT | BZW1 |
| 1.185E-14 | 0.36188027 | 0.313 | 0.161 | 2.7994E-10 | TEC_PT | PPCS |
| 1.3396E-14 | 0.48258144 | 0.475 | 0.31 | 3.1646E-10 | TEC_PT | NDUFA2 |
| 1.351E-14 | -0.3417208 | 0.034 | 0.219 | 3.1915E-10 | TEC_PT | TUT7 |
| 1.3611E-14 | -0.4632006 | 0.003 | 0.174 | 3.2153E-10 | TEC_PT | SMIM25 |
| 1.4175E-14 | 0.27001216 | 0.266 | 0.123 | 3.3485E-10 | TEC_PT | EGLN1 |
| 1.4416E-14 | -0.6586019 | 0.003 | 0.173 | 3.4054E-10 | TEC_PT | STEAP4 |
| 1.5628E-14 | 0.40183184 | 0.377 | 0.217 | 3.6918E-10 | TEC_PT | NDUFA5 |
| 1.5799E-14 | -0.5869735 | 0.003 | 0.173 | 3.7323E-10 | TEC_PT | PPL |
| 1.6334E-14 | 0.4240006 | 0.96 | 0.901 | 3.8586E-10 | TEC_PT | RPS7 |
| 1.7201E-14 | 0.44942083 | 0.525 | 0.35 | 4.0633E-10 | TEC_PT | SNX3 |
| 1.7747E-14 | -1.2789662 | 0 | 0.167 | 4.1923E-10 | TEC_PT | DUOXA2 |
| 1.9263E-14 | 0.31861435 | 0.32 | 0.16 | 4.5504E-10 | TEC_PT | FARP1 |
| 2.0644E-14 | -0.4907184 | 0.152 | 0.363 | 4.8768E-10 | TEC_PT | GNAI2 |
| 2.4006E-14 | -0.8431273 | 0.01 | 0.181 | 5.6709E-10 | TEC_PT | F3 |
| 2.4261E-14 | -0.4444427 | 0.175 | 0.396 | 5.7312E-10 | TEC_PT | PNISR |
| 2.4353E-14 | -0.681352 | 0.047 | 0.228 | 5.7529E-10 | TEC_PT | ZEB2 |
| 2.4437E-14 | -0.4352093 | 0 | 0.165 | 5.7729E-10 | TEC_PT | LCP2 |
| 2.4989E-14 | -0.4808386 | 0.121 | 0.327 | 5.9032E-10 | TEC_PT | CAPG |
| 2.5197E-14 | 0.38589555 | 0.377 | 0.215 | 5.9523E-10 | TEC_PT | NDUFB5 |
| 2.6946E-14 | -0.3782414 | 0 | 0.165 | 6.3655E-10 | TEC_PT | CLEC2B |
| 2.9012E-14 | 0.35290203 | 0.3 | 0.154 | 6.8536E-10 | TEC_PT | NDUFV3 |
| 2.9801E-14 | -0.566037 | 0.013 | 0.184 | 7.0398E-10 | TEC_PT | SDCBP2 |
| 3.0257E-14 | -0.8863684 | 0.017 | 0.189 | 7.1477E-10 | TEC_PT | TMPRSS2 |
| 3.4643E-14 | -0.3903266 | 0.034 | 0.214 | 8.1836E-10 | TEC_PT | ZFYVE16 |
| 3.4812E-14 | -0.3697026 | 0.071 | 0.27 | 8.2237E-10 | TEC_PT | UBE2D1 |
| 3.7875E-14 | -0.6151694 | 0.003 | 0.169 | 8.9473E-10 | TEC_PT | FXYD3 |
| 3.9473E-14 | 0.27344982 | 0.259 | 0.12 | 9.3246E-10 | TEC_PT | MRPL55 |
| 3.9729E-14 | 0.26728403 | 0.249 | 0.114 | 9.3851E-10 | TEC_PT | COPRS |
| 4.0436E-14 | 0.41771731 | 0.939 | 0.838 | 9.5521E-10 | TEC_PT | RACK1 |
| 4.1457E-14 | 0.3131961 | 0.434 | 0.252 | 9.7934E-10 | TEC_PT | C1orf122 |
| 4.1855E-14 | -0.3326746 | 0.003 | 0.169 | 9.8875E-10 | TEC_PT | GLIPR2 |
| 4.1895E-14 | 0.41939976 | 0.97 | 0.817 | 9.8967E-10 | TEC_PT | RPL5 |
| 4.2107E-14 | -0.4510873 | 0.172 | 0.386 | 9.9468E-10 | TEC_PT | HNRNPH1 |
| 4.5927E-14 | 0.34272957 | 0.327 | 0.172 | 1.0849E-09 | TEC_PT | FAM162A |
| 4.9163E-14 | 0.34117234 | 0.333 | 0.178 | 1.1614E-09 | TEC_PT | UQCRC2 |
| 5.0057E-14 | 0.28406132 | 0.279 | 0.135 | 1.1825E-09 | TEC_PT | COMMD4 |
| 5.2951E-14 | -0.4125062 | 0.081 | 0.277 | 1.2509E-09 | TEC_PT | VASP |
| 5.7064E-14 | -1.2404433 | 0.03 | 0.205 | 1.348E-09 | TEC_PT | HLA-DPA1 |
| 6.3037E-14 | -0.4573273 | 0.054 | 0.238 | 1.4891E-09 | TEC_PT | RNASET2 |
| 7.0003E-14 | -0.5431997 | 0.007 | 0.171 | 1.6537E-09 | TEC_PT | TGFBI |
| 7.2124E-14 | 0.37223307 | 0.448 | 0.282 | 1.7038E-09 | TEC_PT | RAB5C |
| 7.4607E-14 | -0.4653771 | 0.111 | 0.312 | 1.7624E-09 | TEC_PT | CYTOR |
| 7.8426E-14 | -0.5330593 | 0.192 | 0.398 | 1.8527E-09 | TEC_PT | ARGLU1 |
| 7.8522E-14 | 0.42694841 | 0.933 | 0.819 | 1.8549E-09 | TEC_PT | RPL27 |
| 7.9587E-14 | -0.4175485 | 0.428 | 0.617 | 1.8801E-09 | TEC_PT | CAP1 |
| 8.1773E-14 | -0.6071451 | 0.071 | 0.26 | 1.9317E-09 | TEC_PT | PMAIP1 |
| 8.2069E-14 | 0.47631253 | 0.727 | 0.628 | 1.9387E-09 | TEC_PT | OST4 |

| | | | | | | |
|---|---|---|---|---|---|---|
| 8.2513E-14 | -0.5353881 | 0.02 | 0.189 | 1.9492E-09 | TEC_PT | FYB1 |
| 8.553E-14 | 0.39751703 | 0.946 | 0.875 | 2.0205E-09 | TEC_PT | RPL3 |
| 8.72E-14 | -0.6210005 | 0.01 | 0.177 | 2.0599E-09 | TEC_PT | IGFBP3 |
| 8.8106E-14 | -0.4855013 | 0.114 | 0.313 | 2.0813E-09 | TEC_PT | FNDC3B |
| 8.8811E-14 | -0.3474512 | 0.003 | 0.164 | 2.098E-09 | TEC_PT | MYO1F |
| 9.3804E-14 | -0.4464068 | 0.007 | 0.169 | 2.2159E-09 | TEC_PT | SLA |
| 9.9909E-14 | -0.4125413 | 0.36 | 0.562 | 2.3602E-09 | TEC_PT | CSNK1A1 |
| 1.0093E-13 | 0.27539914 | 0.259 | 0.124 | 2.3842E-09 | TEC_PT | IER3IP1 |
| 1.0499E-13 | -0.2905477 | 0.01 | 0.176 | 2.4801E-09 | TEC_PT | TCIRG1 |
| 1.1286E-13 | -0.4170704 | 0.094 | 0.288 | 2.666E-09 | TEC_PT | BAZ1A |
| 1.3655E-13 | 0.2690212 | 0.306 | 0.157 | 3.2257E-09 | TEC_PT | TXN2 |
| 1.4264E-13 | -1.139655 | 0.003 | 0.162 | 3.3695E-09 | TEC_PT | MUC21 |
| 1.5054E-13 | -0.4687558 | 0.003 | 0.162 | 3.5562E-09 | TEC_PT | AQP9 |
| 1.506E-13 | -0.8414353 | 0.003 | 0.162 | 3.5577E-09 | TEC_PT | APOBEC3A |
| 1.5441E-13 | -0.3534481 | 0.007 | 0.167 | 3.6476E-09 | TEC_PT | CFD |
| 1.6377E-13 | -0.6600027 | 0 | 0.156 | 3.8688E-09 | TEC_PT | MUC4 |
| 1.7652E-13 | -0.4860173 | 0.074 | 0.26 | 4.17E-09 | TEC_PT | NABP1 |
| 1.7689E-13 | -0.4680997 | 0.034 | 0.205 | 4.1787E-09 | TEC_PT | CSNK1E |
| 1.8067E-13 | 0.37919829 | 0.451 | 0.281 | 4.2679E-09 | TEC_PT | COX7A2L |
| 1.8112E-13 | -0.5026697 | 0.114 | 0.313 | 4.2786E-09 | TEC_PT | CD68 |
| 1.8286E-13 | -0.5163467 | 0.024 | 0.19 | 4.3197E-09 | TEC_PT | FCGR2A |
| 1.8896E-13 | -2.8206728 | 0.219 | 0.379 | 4.4639E-09 | TEC_PT | KRT13 |
| 1.9056E-13 | -0.5998894 | 0.104 | 0.3 | 4.5016E-09 | TEC_PT | BTG2 |
| 1.9349E-13 | 0.32366624 | 0.273 | 0.136 | 4.5708E-09 | TEC_PT | MRPL21 |
| 2.0423E-13 | -1.0403799 | 0.003 | 0.161 | 4.8246E-09 | TEC_PT | CEACAM5 |
| 2.1016E-13 | 0.45035072 | 0.424 | 0.257 | 4.9647E-09 | TEC_PT | CCT3 |
| 2.2498E-13 | -0.5106049 | 0.01 | 0.171 | 5.3148E-09 | TEC_PT | CXCR4 |
| 2.2798E-13 | -0.3893057 | 0.098 | 0.29 | 5.3856E-09 | TEC_PT | ARID4B |
| 2.557E-13 | -0.2947023 | 0.017 | 0.182 | 6.0404E-09 | TEC_PT | ELK3 |
| 2.5657E-13 | -0.4042539 | 0.114 | 0.316 | 6.0609E-09 | TEC_PT | AKAP9 |
| 2.5767E-13 | -0.4091668 | 0.094 | 0.286 | 6.087E-09 | TEC_PT | CXCL16 |
| 2.5949E-13 | -0.756749 | 0 | 0.154 | 6.1298E-09 | TEC_PT | SCEL |
| 2.7223E-13 | -0.3130432 | 0.003 | 0.16 | 6.4309E-09 | TEC_PT | MYADM |
| 2.9254E-13 | -0.3744368 | 0.037 | 0.21 | 6.9106E-09 | TEC_PT | ARRB2 |
| 2.9409E-13 | -0.3731159 | 0.01 | 0.169 | 6.9473E-09 | TEC_PT | NECTIN4 |
| 2.9459E-13 | -0.5562998 | 0.003 | 0.159 | 6.9592E-09 | TEC_PT | CD300E |
| 3.3296E-13 | 0.52638411 | 0.556 | 0.412 | 7.8655E-09 | TEC_PT | PCBP2 |
| 3.3812E-13 | 0.37123335 | 0.411 | 0.248 | 7.9873E-09 | TEC_PT | MRPL20 |
| 3.394E-13 | 0.36316777 | 0.337 | 0.19 | 8.0177E-09 | TEC_PT | ALKBH7 |
| 3.4103E-13 | -0.3765449 | 0.013 | 0.175 | 8.0562E-09 | TEC_PT | KCTD12 |
| 3.4268E-13 | -0.3735643 | 0 | 0.152 | 8.095E-09 | TEC_PT | EVI2B |
| 3.6322E-13 | 0.34429603 | 0.33 | 0.18 | 8.5804E-09 | TEC_PT | REXO2 |
| 3.6656E-13 | -0.4407475 | 0.731 | 0.832 | 8.6594E-09 | TEC_PT | HNRNPA2B1 |
| 3.7414E-13 | -0.3546374 | 0.027 | 0.193 | 8.8384E-09 | TEC_PT | ITPRIPL2 |
| 3.9403E-13 | -0.4934665 | 0.374 | 0.549 | 9.3083E-09 | TEC_PT | TOP1 |
| 4.0262E-13 | -0.3408314 | 0.013 | 0.172 | 9.5111E-09 | TEC_PT | B3GNT5 |
| 4.3082E-13 | 0.4741703 | 0.976 | 0.897 | 1.0177E-08 | TEC_PT | RPL29 |
| 4.3402E-13 | 0.27434558 | 0.29 | 0.147 | 1.0253E-08 | TEC_PT | MRPL43 |
| 4.3428E-13 | -0.4137188 | 0.013 | 0.172 | 1.0259E-08 | TEC_PT | SAMHD1 |
| 4.3461E-13 | 0.30058735 | 0.316 | 0.168 | 1.0267E-08 | TEC_PT | MRPS15 |

| | | | | | | |
|---|---|---|---|---|---|---|
| 4.4999E-13 | -0.3429423 | 0.057 | 0.238 | 1.063E-08 | TEC_PT | HOOK3 |
| 4.6543E-13 | -0.6207792 | 0.04 | 0.209 | 1.0995E-08 | TEC_PT | CDKN2B |
| 4.7337E-13 | -0.4741758 | 0.013 | 0.174 | 1.1182E-08 | TEC_PT | XIST |
| 4.7462E-13 | -0.7409734 | 0.064 | 0.237 | 1.1212E-08 | TEC_PT | FABP5 |
| 4.8022E-13 | -0.3314233 | 0.007 | 0.163 | 1.1344E-08 | TEC_PT | FERMT3 |
| 4.8047E-13 | -0.468955 | 0.175 | 0.378 | 1.135E-08 | TEC_PT | GCC2 |
| 4.8189E-13 | -0.3214668 | 0.017 | 0.179 | 1.1384E-08 | TEC_PT | PIK3IP1 |
| 5.0787E-13 | -0.5838649 | 0.013 | 0.173 | 1.1997E-08 | TEC_PT | HLA-DQB1 |
| 5.2224E-13 | -0.5078105 | 0.017 | 0.176 | 1.2337E-08 | TEC_PT | C6orf132 |
| 5.2809E-13 | -0.3672453 | 0.081 | 0.266 | 1.2475E-08 | TEC_PT | PLP2 |
| 5.3361E-13 | -0.4904842 | 0.01 | 0.166 | 1.2606E-08 | TEC_PT | HIST3H2A |
| 5.433E-13 | -0.3578123 | 0.118 | 0.318 | 1.2834E-08 | TEC_PT | PTBP3 |
| 5.6067E-13 | -0.6801137 | 0.468 | 0.632 | 1.3245E-08 | TEC_PT | ZFP36L1 |
| 5.7205E-13 | -0.9662266 | 0.003 | 0.156 | 1.3514E-08 | TEC_PT | PITX1 |
| 5.8195E-13 | -0.3082223 | 0.074 | 0.263 | 1.3747E-08 | TEC_PT | ELF1 |
| 6.0159E-13 | 0.31141928 | 0.32 | 0.172 | 1.4211E-08 | TEC_PT | NDUFA10 |
| 6.0179E-13 | -0.4446831 | 0.178 | 0.382 | 1.4216E-08 | TEC_PT | N4BP2L2 |
| 6.2501E-13 | -0.483158 | 0.387 | 0.574 | 1.4765E-08 | TEC_PT | TSPO |
| 6.3055E-13 | -0.3700875 | 0.047 | 0.218 | 1.4895E-08 | TEC_PT | STK4 |
| 6.3663E-13 | -0.5952522 | 0.003 | 0.155 | 1.5039E-08 | TEC_PT | CDC42EP5 |
| 6.4406E-13 | -0.7593096 | 0 | 0.15 | 1.5215E-08 | TEC_PT | TMPRSS11D |
| 6.4861E-13 | 0.3570981 | 0.36 | 0.21 | 1.5322E-08 | TEC_PT | ANXA7 |
| 6.6512E-13 | -0.4255766 | 0.007 | 0.161 | 1.5712E-08 | TEC_PT | SERPINB9 |
| 6.703E-13 | -0.5505523 | 0.067 | 0.244 | 1.5834E-08 | TEC_PT | ATP2B1 |
| 6.7152E-13 | -0.308417 | 0.04 | 0.211 | 1.5863E-08 | TEC_PT | CYTH1 |
| 6.7312E-13 | -0.4225571 | 0.088 | 0.272 | 1.5901E-08 | TEC_PT | RAP2B |
| 6.8761E-13 | -0.4377735 | 0.926 | 0.945 | 1.6243E-08 | TEC_PT | PTMA |
| 6.935E-13 | 0.36219696 | 0.29 | 0.153 | 1.6383E-08 | TEC_PT | PFKL |
| 6.9474E-13 | 0.28930559 | 0.269 | 0.134 | 1.6412E-08 | TEC_PT | TIMM17A |
| 7.3658E-13 | 0.37148296 | 0.377 | 0.22 | 1.74E-08 | TEC_PT | LSM4 |
| 7.4702E-13 | -1.5858045 | 0.01 | 0.163 | 1.7647E-08 | TEC_PT | ECM1 |
| 7.7509E-13 | -0.4300603 | 0.128 | 0.326 | 1.831E-08 | TEC_PT | TMBIM1 |
| 8.0286E-13 | 0.25599432 | 0.199 | 0.085 | 1.8966E-08 | TEC_PT | BCAR1 |
| 8.0673E-13 | 0.38004306 | 0.919 | 0.791 | 1.9057E-08 | TEC_PT | RPS21 |
| 8.7652E-13 | -1.0825514 | 0.448 | 0.556 | 2.0706E-08 | TEC_PT | ELF3 |
| 8.7781E-13 | 0.29624584 | 0.293 | 0.152 | 2.0736E-08 | TEC_PT | SCOC |
| 9.8384E-13 | -0.2851805 | 0.007 | 0.159 | 2.3241E-08 | TEC_PT | ARHGAP30 |
| 9.9516E-13 | -0.4040712 | 0.007 | 0.158 | 2.3509E-08 | TEC_PT | TTC9 |
| 1.0143E-12 | -0.4569982 | 0.04 | 0.207 | 2.3961E-08 | TEC_PT | SLC43A2 |
| 1.0219E-12 | -0.3801552 | 0 | 0.147 | 2.4141E-08 | TEC_PT | TREM1 |
| 1.0406E-12 | -0.5400593 | 0.064 | 0.239 | 2.4583E-08 | TEC_PT | TNFAIP3 |
| 1.046E-12 | -0.3815083 | 0.145 | 0.337 | 2.4709E-08 | TEC_PT | CAPZA1 |
| 1.0572E-12 | -0.2701827 | 0.003 | 0.153 | 2.4975E-08 | TEC_PT | EHBP1L1 |
| 1.1411E-12 | -0.8238944 | 0 | 0.147 | 2.6957E-08 | TEC_PT | CEACAM7 |
| 1.1695E-12 | -0.8537496 | 0.007 | 0.157 | 2.7628E-08 | TEC_PT | KRT15 |
| 1.1997E-12 | -0.4228904 | 0.094 | 0.276 | 2.834E-08 | TEC_PT | PLIN3 |
| 1.2357E-12 | -0.4162474 | 0 | 0.146 | 2.9191E-08 | TEC_PT | MS4A7 |
| 1.264E-12 | 0.42188674 | 0.697 | 0.561 | 2.9859E-08 | TEC_PT | TOMM7 |
| 1.3343E-12 | -0.9508155 | 0.03 | 0.194 | 3.152E-08 | TEC_PT | HSPA6 |
| 1.3361E-12 | 0.36392807 | 0.35 | 0.203 | 3.1562E-08 | TEC_PT | TXNL4A |

| | | | | | | |
|---|---|---|---|---|---|---|
| 1.3413E-12 | 0.43403851 | 0.438 | 0.283 | 3.1685E-08 | TEC_PT | RNF181 |
| 1.3519E-12 | -2.1615562 | 0.02 | 0.176 | 3.1937E-08 | TEC_PT | IL1B |
| 1.4691E-12 | -0.7725038 | 0 | 0.145 | 3.4704E-08 | TEC_PT | TMPRSS11E |
| 1.4767E-12 | -0.3201945 | 0.027 | 0.187 | 3.4885E-08 | TEC_PT | TET2 |
| 1.4845E-12 | -0.354879 | 0.007 | 0.156 | 3.5068E-08 | TEC_PT | ADGRE5 |
| 1.4895E-12 | -0.3964465 | 0.024 | 0.182 | 3.5186E-08 | TEC_PT | PFKFB3 |
| 1.4903E-12 | -0.5283241 | 0.047 | 0.214 | 3.5204E-08 | TEC_PT | LAMB3 |
| 1.5075E-12 | 0.43036955 | 0.572 | 0.413 | 3.5612E-08 | TEC_PT | EIF3K |
| 1.6333E-12 | 0.38239333 | 0.99 | 0.964 | 3.8584E-08 | TEC_PT | RPL13 |
| 1.7055E-12 | -2.1716298 | 0.168 | 0.34 | 4.029E-08 | TEC_PT | MMP7 |
| 1.8761E-12 | -1.6988888 | 0 | 0.144 | 4.4319E-08 | TEC_PT | SPRR2A |
| 1.8963E-12 | 0.28038166 | 0.249 | 0.122 | 4.4797E-08 | TEC_PT | MRPL40 |
| 1.9862E-12 | 0.25638498 | 0.239 | 0.114 | 4.6919E-08 | TEC_PT | CLPP |
| 2.0222E-12 | 0.35862521 | 0.32 | 0.181 | 4.7771E-08 | TEC_PT | SSNA1 |
| 2.0273E-12 | 0.29924757 | 0.259 | 0.128 | 4.7891E-08 | TEC_PT | BORCS7 |
| 2.0609E-12 | 0.28239449 | 0.34 | 0.188 | 4.8684E-08 | TEC_PT | HADHB |
| 2.1059E-12 | 0.39697564 | 0.424 | 0.259 | 4.9747E-08 | TEC_PT | CUTA |
| 2.1168E-12 | -0.4522913 | 0.098 | 0.275 | 5.0005E-08 | TEC_PT | JUP |
| 2.1511E-12 | -0.9682687 | 0.01 | 0.159 | 5.0815E-08 | TEC_PT | PRSS27 |
| 2.1645E-12 | -0.4083298 | 0.074 | 0.249 | 5.1132E-08 | TEC_PT | PKN2 |
| 2.4301E-12 | 0.2877154 | 0.327 | 0.18 | 5.7406E-08 | TEC_PT | POLR2E |
| 2.5389E-12 | -0.3412651 | 0.047 | 0.213 | 5.9976E-08 | TEC_PT | VPS13C |
| 2.6557E-12 | -0.3498642 | 0.03 | 0.189 | 6.2736E-08 | TEC_PT | PMEPA1 |
| 2.6825E-12 | -0.6435836 | 0.003 | 0.148 | 6.3368E-08 | TEC_PT | NCCRP1 |
| 2.7667E-12 | -0.2872008 | 0.044 | 0.21 | 6.5358E-08 | TEC_PT | NECTIN2 |
| 2.8917E-12 | -0.8601749 | 0.003 | 0.147 | 6.8312E-08 | TEC_PT | CEACAM6 |
| 2.9013E-12 | 0.30863207 | 0.316 | 0.173 | 6.8538E-08 | TEC_PT | VKORC1 |
| 2.9539E-12 | -0.3184855 | 0.007 | 0.153 | 6.9781E-08 | TEC_PT | HK2 |
| 3.1465E-12 | 0.31919643 | 0.259 | 0.127 | 7.4331E-08 | TEC_PT | BEX2 |
| 3.2402E-12 | -0.409992 | 0.003 | 0.147 | 7.6543E-08 | TEC_PT | VSIG2 |
| 3.2985E-12 | 0.52192453 | 0.704 | 0.631 | 7.7921E-08 | TEC_PT | SAP18 |
| 3.3044E-12 | 0.38415656 | 0.96 | 0.875 | 7.806E-08 | TEC_PT | RPL36 |
| 3.3288E-12 | -0.312692 | 0.013 | 0.162 | 7.8636E-08 | TEC_PT | GALNT3 |
| 3.4675E-12 | -0.3071342 | 0.017 | 0.168 | 8.1914E-08 | TEC_PT | CXADR |
| 3.5404E-12 | -0.4264711 | 0.003 | 0.146 | 8.3635E-08 | TEC_PT | FPR1 |
| 3.7071E-12 | -0.4248022 | 0.098 | 0.277 | 8.7573E-08 | TEC_PT | SLK |
| 3.7659E-12 | 0.40775329 | 0.394 | 0.244 | 8.8962E-08 | TEC_PT | FAM177A1 |
| 4.0501E-12 | -0.3188385 | 0.007 | 0.151 | 9.5676E-08 | TEC_PT | FAM129A |
| 4.205E-12 | -0.3678578 | 0.044 | 0.205 | 9.9335E-08 | TEC_PT | SLFN5 |
| 4.3977E-12 | -0.4612481 | 0.098 | 0.277 | 1.0389E-07 | TEC_PT | ETS2 |
| 4.4142E-12 | -0.4047252 | 0.024 | 0.176 | 1.0428E-07 | TEC_PT | NR4A1 |
| 4.5408E-12 | -0.5508433 | 0.101 | 0.282 | 1.0727E-07 | TEC_PT | H1F0 |
| 4.7001E-12 | 0.42165881 | 0.96 | 0.919 | 1.1103E-07 | TEC_PT | NACA |
| 4.9974E-12 | -0.3585199 | 0.003 | 0.145 | 1.1805E-07 | TEC_PT | TLR2 |
| 5.0017E-12 | -0.2843794 | 0.024 | 0.177 | 1.1815E-07 | TEC_PT | PPP1R18 |
| 5.0362E-12 | -0.3403602 | 0.017 | 0.166 | 1.1897E-07 | TEC_PT | JARID2 |
| 5.0699E-12 | -0.3691592 | 0.101 | 0.278 | 1.1977E-07 | TEC_PT | ROCK1 |
| 5.2713E-12 | -0.4888549 | 0.013 | 0.159 | 1.2452E-07 | TEC_PT | MALL |
| 5.5732E-12 | 0.29398866 | 0.276 | 0.143 | 1.3166E-07 | TEC_PT | PFDN6 |
| 6.3703E-12 | -0.2782635 | 0 | 0.138 | 1.5049E-07 | TEC_PT | EVI2A |

| | | | | | | |
|---|---|---|---|---|---|---|
| 6.4281E-12 | -0.3817052 | 0.098 | 0.285 | 1.5185E-07 | TEC_PT | MARCKSL1 |
| 6.4487E-12 | -0.2734677 | 0.771 | 0.897 | 1.5234E-07 | TEC_PT | HLA-A |
| 6.4646E-12 | -0.3350565 | 0.077 | 0.256 | 1.5271E-07 | TEC_PT | MGAT1 |
| 6.6894E-12 | -0.3833175 | 0.081 | 0.251 | 1.5802E-07 | TEC_PT | PLEC |
| 6.9635E-12 | -0.2769511 | 0.047 | 0.214 | 1.645E-07 | TEC_PT | RNF145 |
| 6.9725E-12 | 0.27075976 | 0.283 | 0.147 | 1.6471E-07 | TEC_PT | TSPYL1 |
| 7.2266E-12 | 0.44077112 | 0.468 | 0.317 | 1.7071E-07 | TEC_PT | CALM3 |
| 7.4898E-12 | -0.4364445 | 0 | 0.137 | 1.7693E-07 | TEC_PT | MCEMP1 |
| 7.5531E-12 | -1.0864193 | 0.034 | 0.186 | 1.7843E-07 | TEC_PT | HLA-DPB1 |
| 7.6125E-12 | 0.47867341 | 0.32 | 0.187 | 1.7983E-07 | TEC_PT | SH3YL1 |
| 7.7841E-12 | -0.2766982 | 0.024 | 0.174 | 1.8388E-07 | TEC_PT | ZDHHC20 |
| 8.0909E-12 | -0.3181607 | 0.037 | 0.194 | 1.9113E-07 | TEC_PT | ACSL4 |
| 8.2058E-12 | -0.2985327 | 0 | 0.137 | 1.9385E-07 | TEC_PT | HCK |
| 8.548E-12 | -0.4889531 | 0.098 | 0.276 | 2.0193E-07 | TEC_PT | ADM |
| 8.7965E-12 | -0.316274 | 0.064 | 0.231 | 2.078E-07 | TEC_PT | KDM5A |
| 8.8877E-12 | -0.4811691 | 0.051 | 0.209 | 2.0995E-07 | TEC_PT | MACC1 |
| 8.9124E-12 | 0.26484236 | 0.869 | 0.73 | 2.1054E-07 | TEC_PT | HSPA8 |
| 9.0797E-12 | -0.4480359 | 0.434 | 0.614 | 2.1449E-07 | TEC_PT | SRRM2 |
| 9.3202E-12 | -0.2781299 | 0.003 | 0.142 | 2.2017E-07 | TEC_PT | IL2RG |
| 9.741E-12 | -0.3230271 | 0.061 | 0.228 | 2.3011E-07 | TEC_PT | IFNGR1 |
| 1.0283E-11 | 0.48998118 | 0.502 | 0.357 | 2.4291E-07 | TEC_PT | MIF |
| 1.0374E-11 | -1.0292676 | 0.003 | 0.141 | 2.4506E-07 | TEC_PT | CXCL6 |
| 1.0969E-11 | -0.3469704 | 0.003 | 0.141 | 2.5911E-07 | TEC_PT | EMP2 |
| 1.115E-11 | -0.2811846 | 0 | 0.135 | 2.634E-07 | TEC_PT | FGR |
| 1.1266E-11 | 0.39586768 | 0.956 | 0.87 | 2.6614E-07 | TEC_PT | RPL14 |
| 1.138E-11 | -0.2573916 | 0 | 0.135 | 2.6883E-07 | TEC_PT | NCKAP1L |
| 1.202E-11 | -0.3621739 | 0.128 | 0.312 | 2.8394E-07 | TEC_PT | CTNNB1 |
| 1.2666E-11 | -0.2854993 | 0 | 0.135 | 2.9921E-07 | TEC_PT | CYTIP |
| 1.2753E-11 | -0.7019615 | 0.212 | 0.385 | 3.0126E-07 | TEC_PT | TYMP |
| 1.2825E-11 | -0.4808843 | 0.003 | 0.14 | 3.0296E-07 | TEC_PT | PLAC8 |
| 1.3087E-11 | -0.7978676 | 0.027 | 0.174 | 3.0915E-07 | TEC_PT | NDRG2 |
| 1.3218E-11 | -0.322737 | 0.192 | 0.392 | 3.1224E-07 | TEC_PT | RSRC2 |
| 1.3321E-11 | -0.3807756 | 0.138 | 0.322 | 3.1468E-07 | TEC_PT | YPEL3 |
| 1.4117E-11 | -0.4478279 | 0.593 | 0.743 | 3.3348E-07 | TEC_PT | APLP2 |
| 1.4165E-11 | -0.3153913 | 0.027 | 0.178 | 3.3462E-07 | TEC_PT | GLIPR1 |
| 1.4325E-11 | 0.37612188 | 0.418 | 0.267 | 3.3841E-07 | TEC_PT | BAG1 |
| 1.4561E-11 | -0.347585 | 0.03 | 0.181 | 3.4398E-07 | TEC_PT | ANXA3 |
| 1.4855E-11 | -0.3176163 | 0.077 | 0.246 | 3.5092E-07 | TEC_PT | PTPN12 |
| 1.4879E-11 | -0.2669143 | 0.013 | 0.156 | 3.5148E-07 | TEC_PT | SERINC5 |
| 1.5669E-11 | -0.4821986 | 0.007 | 0.145 | 3.7015E-07 | TEC_PT | CD93 |
| 1.6209E-11 | 0.32877968 | 0.97 | 0.888 | 3.8292E-07 | TEC_PT | RPS23 |
| 1.636E-11 | -0.2638124 | 0.027 | 0.176 | 3.8648E-07 | TEC_PT | NCOA3 |
| 1.6419E-11 | -0.3400718 | 0.081 | 0.248 | 3.8787E-07 | TEC_PT | CD47 |
| 1.6582E-11 | 0.53594539 | 0.522 | 0.385 | 3.9172E-07 | TEC_PT | DAD1 |
| 1.695E-11 | -0.3710301 | 0.367 | 0.555 | 4.0042E-07 | TEC_PT | HNRNPA3 |
| 1.7754E-11 | 0.26170203 | 0.273 | 0.141 | 4.1941E-07 | TEC_PT | NME3 |
| 1.8926E-11 | -0.3303106 | 0.104 | 0.285 | 4.4709E-07 | TEC_PT | TPR |
| 1.9444E-11 | -0.3046204 | 0.003 | 0.138 | 4.5932E-07 | TEC_PT | CLEC7A |
| 1.9564E-11 | -0.3572209 | 0.148 | 0.336 | 4.6216E-07 | TEC_PT | RBM25 |
| 2.0439E-11 | 0.4588323 | 0.778 | 0.688 | 4.8282E-07 | TEC_PT | PSMA7 |

| | | | | | | |
|---|---|---|---|---|---|---|
| 2.1045E-11 | -0.3939078 | 0.007 | 0.143 | 4.9714E-07 | TEC_PT | NCF1 |
| 2.1822E-11 | -0.6583401 | 0 | 0.132 | 5.155E-07 | TEC_PT | GABRP |
| 2.191E-11 | 0.30445847 | 0.29 | 0.158 | 5.1758E-07 | TEC_PT | TMEM160 |
| 2.2205E-11 | -0.3959194 | 0.064 | 0.226 | 5.2455E-07 | TEC_PT | KYNU |
| 2.2641E-11 | 0.48449064 | 0.276 | 0.149 | 5.3486E-07 | TEC_PT | LGMN |
| 2.3625E-11 | -0.2844059 | 0.037 | 0.189 | 5.5809E-07 | TEC_PT | UBA6 |
| 2.402E-11 | -0.2894384 | 0 | 0.131 | 5.6743E-07 | TEC_PT | LILRB3 |
| 2.4271E-11 | -0.6288186 | 0.003 | 0.137 | 5.7335E-07 | TEC_PT | CXCL17 |
| 2.5723E-11 | -0.4909015 | 0.077 | 0.243 | 6.0765E-07 | TEC_PT | PRSS8 |
| 2.6577E-11 | -0.3191427 | 0.04 | 0.192 | 6.2782E-07 | TEC_PT | F11R |
| 2.6592E-11 | -0.4686801 | 0.155 | 0.33 | 6.2819E-07 | TEC_PT | ASPH |
| 2.764E-11 | 0.28685861 | 0.219 | 0.103 | 6.5294E-07 | TEC_PT | DPP4 |
| 2.8017E-11 | 0.31749723 | 0.367 | 0.221 | 6.6185E-07 | TEC_PT | PPP2CB |
| 2.8257E-11 | 0.58957458 | 0.869 | 0.837 | 6.6752E-07 | TEC_PT | SQSTM1 |
| 2.8529E-11 | -0.3199034 | 0.084 | 0.254 | 6.7394E-07 | TEC_PT | NSD3 |
| 2.8694E-11 | -0.3335747 | 0.01 | 0.147 | 6.7783E-07 | TEC_PT | C1orf162 |
| 2.8808E-11 | -0.4286265 | 0.034 | 0.182 | 6.8053E-07 | TEC_PT | S100A14 |
| 2.9066E-11 | 0.33003268 | 0.35 | 0.207 | 6.8662E-07 | TEC_PT | GSTK1 |
| 2.911E-11 | -0.7247996 | 0.034 | 0.183 | 6.8768E-07 | TEC_PT | LAMC2 |
| 2.9326E-11 | -0.2966582 | 0.067 | 0.232 | 6.9277E-07 | TEC_PT | TAOK3 |
| 3.0158E-11 | -0.8150656 | 0.121 | 0.286 | 7.1243E-07 | TEC_PT | SLC25A37 |
| 3.0759E-11 | -0.4161257 | 0.316 | 0.497 | 7.2662E-07 | TEC_PT | DDX3X |
| 3.1414E-11 | 0.31060246 | 0.902 | 0.754 | 7.4208E-07 | TEC_PT | RPSA |
| 3.1794E-11 | -0.9008928 | 0.003 | 0.135 | 7.5108E-07 | TEC_PT | TMPRSS11B |
| 3.202E-11 | 0.26542317 | 0.266 | 0.139 | 7.5641E-07 | TEC_PT | ZNF593 |
| 3.2143E-11 | -0.2894016 | 0.01 | 0.146 | 7.5931E-07 | TEC_PT | CARD16 |
| 3.4343E-11 | -0.5517276 | 0.209 | 0.392 | 8.1127E-07 | TEC_PT | SRSF7 |
| 3.4769E-11 | -0.3191043 | 0.064 | 0.226 | 8.2135E-07 | TEC_PT | ST14 |
| 3.5318E-11 | 0.37036413 | 0.374 | 0.232 | 8.3432E-07 | TEC_PT | OPTN |
| 3.5537E-11 | -0.4417282 | 0.034 | 0.184 | 8.3949E-07 | TEC_PT | INSIG1 |
| 3.6264E-11 | -0.2870285 | 0.044 | 0.196 | 8.5666E-07 | TEC_PT | HLA-F |
| 3.6273E-11 | -0.3493643 | 0.492 | 0.661 | 8.5687E-07 | TEC_PT | RBM39 |
| 3.6419E-11 | -0.4222272 | 0.017 | 0.155 | 8.6034E-07 | TEC_PT | TMPRSS4 |
| 3.6715E-11 | -0.3569693 | 0.104 | 0.276 | 8.6732E-07 | TEC_PT | BPTF |
| 3.6921E-11 | -0.3659962 | 0.111 | 0.29 | 8.7219E-07 | TEC_PT | MIR4435-2HG |
| 3.7164E-11 | -0.3233582 | 0.03 | 0.177 | 8.7794E-07 | TEC_PT | SMIM22 |
| 3.8032E-11 | 0.42280717 | 0.414 | 0.266 | 8.9842E-07 | TEC_PT | EID1 |
| 3.8125E-11 | -0.3113222 | 0.013 | 0.151 | 9.0063E-07 | TEC_PT | SORL1 |
| 3.8527E-11 | -0.377585 | 0.071 | 0.229 | 9.1012E-07 | TEC_PT | GALNT1 |
| 4.1958E-11 | -0.2986937 | 0.108 | 0.281 | 9.9118E-07 | TEC_PT | TMBIM4 |
| 4.1997E-11 | -0.4548007 | 0.013 | 0.152 | 9.921E-07 | TEC_PT | CLEC5A |
| 4.2413E-11 | -0.3764874 | 0.01 | 0.145 | 1.0019E-06 | TEC_PT | ITGB6 |
| 4.3777E-11 | 0.28993988 | 0.32 | 0.178 | 1.0341E-06 | TEC_PT | MGAT4B |
| 4.6105E-11 | 0.30188924 | 0.229 | 0.112 | 1.0891E-06 | TEC_PT | WWC1 |
| 4.7435E-11 | -0.4549298 | 0.131 | 0.308 | 1.1206E-06 | TEC_PT | ZNF292 |
| 4.9094E-11 | -1.0764309 | 0.037 | 0.181 | 1.1597E-06 | TEC_PT | SPINK1 |
| 4.923E-11 | -0.3634035 | 0.098 | 0.266 | 1.163E-06 | TEC_PT | BACH1 |
| 4.9802E-11 | -0.2846598 | 0 | 0.128 | 1.1765E-06 | TEC_PT | IRAK3 |
| 4.9897E-11 | -0.2638201 | 0.007 | 0.139 | 1.1787E-06 | TEC_PT | EMILIN2 |
| 5.003E-11 | -0.3596268 | 0.168 | 0.342 | 1.1819E-06 | TEC_PT | NDFIP1 |

| | | | | | | |
|---|---|---|---|---|---|---|
| 5.0454E-11 | 0.29527373 | 0.354 | 0.21 | 1.1919E-06 | TEC_PT | STRAP |
| 5.1202E-11 | -0.3552826 | 0.003 | 0.133 | 1.2096E-06 | TEC_PT | RGCC |
| 5.2289E-11 | -0.3177639 | 0.121 | 0.302 | 1.2352E-06 | TEC_PT | SERTAD1 |
| 5.3557E-11 | 0.2818435 | 0.195 | 0.089 | 1.2652E-06 | TEC_PT | TYW3 |
| 5.8283E-11 | 0.42214702 | 0.468 | 0.328 | 1.3768E-06 | TEC_PT | COPS9 |
| 5.9255E-11 | 0.40577581 | 0.973 | 0.887 | 1.3998E-06 | TEC_PT | RPL6 |
| 6.7123E-11 | -0.6594197 | 0.003 | 0.132 | 1.5857E-06 | TEC_PT | DHRS9 |
| 6.7524E-11 | -0.4264829 | 0.034 | 0.177 | 1.5951E-06 | TEC_PT | FHL2 |
| 7.0919E-11 | 0.33792452 | 0.397 | 0.26 | 1.6753E-06 | TEC_PT | CAMTA1 |
| 7.1001E-11 | -0.2816645 | 0.094 | 0.266 | 1.6772E-06 | TEC_PT | KRAS |
| 7.1085E-11 | -0.4053953 | 0.246 | 0.429 | 1.6792E-06 | TEC_PT | SRSF5 |
| 7.4168E-11 | -0.29455 | 0.003 | 0.132 | 1.7521E-06 | TEC_PT | SCIN |
| 7.803E-11 | 0.50066186 | 0.502 | 0.355 | 1.8433E-06 | TEC_PT | TKT |
| 7.8187E-11 | -0.2677516 | 0.003 | 0.131 | 1.847E-06 | TEC_PT | C1orf116 |
| 7.8624E-11 | -0.2727331 | 0.01 | 0.142 | 1.8573E-06 | TEC_PT | LRP1 |
| 8.0562E-11 | 0.47466173 | 0.966 | 0.945 | 1.9031E-06 | TEC_PT | RPL13A |
| 8.5414E-11 | -4.0656698 | 0.273 | 0.402 | 2.0177E-06 | TEC_PT | SPRR3 |
| 8.9138E-11 | -0.3847328 | 0.007 | 0.135 | 2.1057E-06 | TEC_PT | BCAS1 |
| 8.9274E-11 | 0.38370185 | 0.505 | 0.355 | 2.1089E-06 | TEC_PT | TRMT112 |
| 9.6817E-11 | -0.4000906 | 0.067 | 0.218 | 2.2871E-06 | TEC_PT | DSE |
| 9.7876E-11 | -0.3649253 | 0.098 | 0.266 | 2.3121E-06 | TEC_PT | HECTD1 |
| 1.0784E-10 | -0.3009259 | 0.074 | 0.234 | 2.5475E-06 | TEC_PT | BAZ2B |
| 1.0898E-10 | -0.896656 | 0.007 | 0.135 | 2.5744E-06 | TEC_PT | EREG |
| 1.1013E-10 | -0.3841651 | 0.165 | 0.34 | 2.6017E-06 | TEC_PT | GPBP1 |
| 1.1029E-10 | -0.2891828 | 0.007 | 0.135 | 2.6055E-06 | TEC_PT | MAP3K20 |
| 1.1486E-10 | 0.33908463 | 0.367 | 0.228 | 2.7134E-06 | TEC_PT | DNAJC15 |
| 1.1733E-10 | -0.2992485 | 0.04 | 0.188 | 2.7717E-06 | TEC_PT | PIM1 |
| 1.3219E-10 | -1.5784935 | 0.007 | 0.133 | 3.1227E-06 | TEC_PT | LYPD2 |
| 1.3234E-10 | 0.30020346 | 0.3 | 0.169 | 3.1261E-06 | TEC_PT | PSMC4 |
| 1.3673E-10 | -0.4105796 | 0.525 | 0.662 | 3.2299E-06 | TEC_PT | HNRNPU |
| 1.3945E-10 | -0.8955677 | 0.007 | 0.133 | 3.2943E-06 | TEC_PT | RHCG |
| 1.431E-10 | -0.2789356 | 0.017 | 0.15 | 3.3804E-06 | TEC_PT | RALGDS |
| 1.4454E-10 | -0.6953839 | 0.296 | 0.454 | 3.4145E-06 | TEC_PT | HSPH1 |
| 1.5925E-10 | -0.418633 | 0.061 | 0.21 | 3.7621E-06 | TEC_PT | SOCS3 |
| 1.5998E-10 | -0.6089233 | 0.013 | 0.143 | 3.7792E-06 | TEC_PT | CDKN2A |
| 1.6302E-10 | -0.5207583 | 0.411 | 0.603 | 3.8509E-06 | TEC_PT | NFKBIA |
| 1.6547E-10 | -0.2716322 | 0.064 | 0.219 | 3.909E-06 | TEC_PT | ACAP2 |
| 1.7262E-10 | -0.3048124 | 0 | 0.121 | 4.0778E-06 | TEC_PT | LILRB2 |
| 1.7345E-10 | -0.3303742 | 0.037 | 0.178 | 4.0975E-06 | TEC_PT | CCDC88A |
| 1.8849E-10 | -0.4428179 | 0.017 | 0.148 | 4.4528E-06 | TEC_PT | ALDH1A3 |
| 1.8937E-10 | -0.2571878 | 0.024 | 0.161 | 4.4734E-06 | TEC_PT | AP1S2 |
| 1.9222E-10 | -0.2771969 | 0.01 | 0.137 | 4.5409E-06 | TEC_PT | ITGA5 |
| 1.9325E-10 | -0.9922668 | 0.054 | 0.2 | 4.5651E-06 | TEC_PT | THBS1 |
| 2.0016E-10 | -0.2764487 | 0.03 | 0.168 | 4.7284E-06 | TEC_PT | MYOF |
| 2.091E-10 | -0.2692664 | 0 | 0.12 | 4.9397E-06 | TEC_PT | CSF1R |
| 2.1136E-10 | -0.2754776 | 0.044 | 0.191 | 4.9929E-06 | TEC_PT | GBP2 |
| 2.2003E-10 | -0.3498659 | 0.141 | 0.31 | 5.1978E-06 | TEC_PT | OGA |
| 2.2378E-10 | -0.2633429 | 0.017 | 0.147 | 5.2863E-06 | TEC_PT | SPECC1 |
| 2.2931E-10 | -0.3806529 | 0 | 0.12 | 5.417E-06 | TEC_PT | FCGR3A |
| 2.3522E-10 | -0.359518 | 0.027 | 0.163 | 5.5565E-06 | TEC_PT | SYTL2 |

| | | | | | | |
|---|---|---|---|---|---|---|
| 2.3537E-10 | -0.2709497 | 0.017 | 0.146 | 5.5601E-06 | TEC_PT | UCP2 |
| 2.3697E-10 | 0.42873156 | 0.323 | 0.192 | 5.5979E-06 | TEC_PT | PGRMC1 |
| 2.443E-10 | 0.91281382 | 0.428 | 0.285 | 5.7712E-06 | TEC_PT | HMOX1 |
| 2.4494E-10 | -0.3156465 | 0.03 | 0.166 | 5.7863E-06 | TEC_PT | CEP170 |
| 2.5564E-10 | 0.26341402 | 0.313 | 0.181 | 6.0389E-06 | TEC_PT | SNF8 |
| 2.6179E-10 | 0.36980393 | 0.37 | 0.239 | 6.1842E-06 | TEC_PT | SMIM26 |
| 2.8448E-10 | -0.6896831 | 0.492 | 0.618 | 6.7203E-06 | TEC_PT | LGALS1 |
| 2.9443E-10 | -0.2529103 | 0.037 | 0.176 | 6.9552E-06 | TEC_PT | EHD4 |
| 3.0698E-10 | -0.5326909 | 0.323 | 0.481 | 7.2518E-06 | TEC_PT | RIOK3 |
| 3.217E-10 | -0.2500952 | 0.007 | 0.131 | 7.5995E-06 | TEC_PT | CELF2 |
| 3.2839E-10 | 0.27584332 | 0.162 | 0.07 | 7.7574E-06 | TEC_PT | TPD52L1 |
| 3.4339E-10 | -0.3839211 | 0.084 | 0.232 | 8.1118E-06 | TEC_PT | TJP1 |
| 3.4487E-10 | -0.3634951 | 0.165 | 0.341 | 8.1468E-06 | TEC_PT | YTHDC1 |
| 3.4983E-10 | -1.0221847 | 0.01 | 0.134 | 8.2641E-06 | TEC_PT | SERPINB2 |
| 3.502E-10 | -0.3418496 | 0.051 | 0.197 | 8.2727E-06 | TEC_PT | SFN |
| 3.507E-10 | -0.3972597 | 0.003 | 0.124 | 8.2845E-06 | TEC_PT | TMEM40 |
| 3.6232E-10 | -0.348647 | 0.111 | 0.268 | 8.5592E-06 | TEC_PT | USP15 |
| 3.6304E-10 | -0.4094697 | 0.047 | 0.19 | 8.5761E-06 | TEC_PT | DUSP6 |
| 3.7144E-10 | -1.1126917 | 0 | 0.117 | 8.7745E-06 | TEC_PT | SPINK5 |
| 3.7793E-10 | -0.5806484 | 0 | 0.117 | 8.9278E-06 | TEC_PT | MNDA |
| 3.9136E-10 | -0.3711003 | 0.071 | 0.227 | 9.2451E-06 | TEC_PT | IER5L |
| 3.9389E-10 | -0.3069485 | 0.034 | 0.17 | 9.305E-06 | TEC_PT | JMY |
| 4.1792E-10 | -0.4010667 | 0.057 | 0.203 | 9.8724E-06 | TEC_PT | FOSB |
| 4.1964E-10 | 0.39386886 | 0.431 | 0.286 | 9.9132E-06 | TEC_PT | MRFAP1 |
| 4.2271E-10 | 0.41562478 | 0.96 | 0.901 | 9.9857E-06 | TEC_PT | RPL24 |
| 4.2513E-10 | -1.1493885 | 0.545 | 0.658 | 1.0043E-05 | TEC_PT | HSPA1A |
| 4.321E-10 | -1.2121712 | 0.01 | 0.132 | 1.0207E-05 | TEC_PT | PLAT |
| 4.6592E-10 | 0.42060549 | 0.502 | 0.367 | 1.1006E-05 | TEC_PT | NDUFV2 |
| 4.9484E-10 | 0.34736784 | 0.391 | 0.253 | 1.169E-05 | TEC_PT | CCT6A |
| 4.9899E-10 | -0.3081166 | 0.071 | 0.222 | 1.1788E-05 | TEC_PT | ACTN1 |
| 5.0237E-10 | -0.3258685 | 0.192 | 0.365 | 1.1867E-05 | TEC_PT | WASF2 |
| 5.0472E-10 | -0.2980175 | 0.098 | 0.257 | 1.1923E-05 | TEC_PT | TACC1 |
| 5.1359E-10 | -0.2626124 | 0.034 | 0.169 | 1.2132E-05 | TEC_PT | TP53BP2 |
| 5.196E-10 | -0.2892172 | 0.02 | 0.15 | 1.2275E-05 | TEC_PT | BCAT1 |
| 5.201E-10 | -0.3052945 | 0.034 | 0.169 | 1.2286E-05 | TEC_PT | SLC38A1 |
| 5.2084E-10 | -0.3294927 | 0.051 | 0.189 | 1.2304E-05 | TEC_PT | PYCARD |
| 5.2319E-10 | -0.295498 | 0.034 | 0.17 | 1.2359E-05 | TEC_PT | ISG20 |
| 5.235E-10 | -0.4534259 | 0.044 | 0.179 | 1.2367E-05 | TEC_PT | EPHA2 |
| 5.3096E-10 | 0.34677159 | 0.387 | 0.247 | 1.2543E-05 | TEC_PT | RPS17 |
| 5.3542E-10 | -0.5905806 | 0.67 | 0.775 | 1.2648E-05 | TEC_PT | CTSD |
| 5.3931E-10 | -0.2594086 | 0.02 | 0.149 | 1.274E-05 | TEC_PT | CDC42EP3 |
| 5.4217E-10 | 0.70423506 | 0.933 | 0.906 | 1.2808E-05 | TEC_PT | ITM2B |
| 5.5333E-10 | -0.4509683 | 0.212 | 0.386 | 1.3071E-05 | TEC_PT | C9orf16 |
| 6.1392E-10 | -0.2644216 | 0.091 | 0.251 | 1.4503E-05 | TEC_PT | IDS |
| 6.1625E-10 | -0.3050801 | 0.088 | 0.239 | 1.4558E-05 | TEC_PT | ACSL3 |
| 6.2201E-10 | -0.2956921 | 0.091 | 0.245 | 1.4694E-05 | TEC_PT | JOSD1 |
| 6.4325E-10 | -0.4384462 | 0.024 | 0.152 | 1.5196E-05 | TEC_PT | SLC7A11 |
| 6.575E-10 | 0.36846556 | 0.32 | 0.197 | 1.5532E-05 | TEC_PT | VTI1B |
| 6.652E-10 | -0.8462809 | 0.36 | 0.506 | 1.5714E-05 | TEC_PT | HSPA1B |
| 6.7068E-10 | 0.3468741 | 0.404 | 0.273 | 1.5843E-05 | TEC_PT | AP2M1 |

| | | | | | | |
|---|---|---|---|---|---|---|
| 6.8736E-10 | -0.2907917 | 0.411 | 0.608 | 1.6238E-05 | TEC_PT | SON |
| 7.0503E-10 | -0.2929479 | 0.071 | 0.22 | 1.6655E-05 | TEC_PT | ZFC3H1 |
| 7.1868E-10 | -0.2559762 | 0.017 | 0.141 | 1.6977E-05 | TEC_PT | RAB25 |
| 7.2684E-10 | 0.26515043 | 0.269 | 0.149 | 1.717E-05 | TEC_PT | MRPL13 |
| 7.2773E-10 | -0.3042713 | 0.101 | 0.254 | 1.7191E-05 | TEC_PT | DHRS7 |
| 7.3947E-10 | 0.26786727 | 0.189 | 0.089 | 1.7468E-05 | TEC_PT | CETN2 |
| 7.56E-10 | -1.2769133 | 0 | 0.114 | 1.7859E-05 | TEC_PT | SPRR1B |
| 7.8485E-10 | 0.32487124 | 0.485 | 0.331 | 1.8541E-05 | TEC_PT | SNHG8 |
| 7.8698E-10 | -0.3189624 | 0.094 | 0.248 | 1.8591E-05 | TEC_PT | CLK1 |
| 7.9404E-10 | 0.26075246 | 0.279 | 0.159 | 1.8758E-05 | TEC_PT | TRAPPC3 |
| 7.9629E-10 | 0.39565591 | 0.384 | 0.263 | 1.8811E-05 | TEC_PT | NDUFA6 |
| 8.059E-10 | -1.8744576 | 0.219 | 0.345 | 1.9038E-05 | TEC_PT | HPGD |
| 8.0735E-10 | 0.39934138 | 0.859 | 0.786 | 1.9072E-05 | TEC_PT | RPL31 |
| 8.4986E-10 | -0.2971376 | 0.02 | 0.145 | 2.0076E-05 | TEC_PT | PDP1 |
| 8.6189E-10 | -0.9486341 | 0.303 | 0.452 | 2.036E-05 | TEC_PT | SOX4 |
| 8.7979E-10 | -0.592501 | 0 | 0.113 | 2.0783E-05 | TEC_PT | KRT78 |
| 8.8643E-10 | 0.25481316 | 0.249 | 0.134 | 2.094E-05 | TEC_PT | HSDL2 |
| 9.1146E-10 | 0.25646857 | 0.296 | 0.17 | 2.1531E-05 | TEC_PT | PHB2 |
| 9.2586E-10 | -0.361593 | 0.057 | 0.198 | 2.1872E-05 | TEC_PT | SERTAD2 |
| 9.3448E-10 | -0.3210137 | 0.205 | 0.386 | 2.2075E-05 | TEC_PT | DEK |
| 9.749E-10 | -0.3537789 | 0.131 | 0.289 | 2.303E-05 | TEC_PT | ATP1B3 |
| 9.8191E-10 | -0.6079513 | 0.101 | 0.249 | 2.3196E-05 | TEC_PT | EGR1 |
| 1.023E-09 | -0.2992192 | 0.145 | 0.31 | 2.4166E-05 | TEC_PT | ARPC1B |
| 1.0241E-09 | -0.3136283 | 0.111 | 0.268 | 2.4192E-05 | TEC_PT | SMG1 |
| 1.0548E-09 | -0.3306292 | 0.007 | 0.123 | 2.4917E-05 | TEC_PT | KRT80 |
| 1.0646E-09 | -0.2630878 | 0.003 | 0.118 | 2.5148E-05 | TEC_PT | CD86 |
| 1.079E-09 | -0.2618186 | 0.088 | 0.241 | 2.549E-05 | TEC_PT | CASP4 |
| 1.088E-09 | -0.2811797 | 0.034 | 0.163 | 2.5702E-05 | TEC_PT | PPP4R1 |
| 1.1039E-09 | 0.33209518 | 0.374 | 0.235 | 2.6078E-05 | TEC_PT | PSMA3 |
| 1.13E-09 | -0.3973011 | 0.024 | 0.149 | 2.6695E-05 | TEC_PT | RGS2 |
| 1.1843E-09 | -0.5317487 | 0.007 | 0.123 | 2.7977E-05 | TEC_PT | AGR2 |
| 1.2353E-09 | -0.2606115 | 0.027 | 0.154 | 2.9182E-05 | TEC_PT | VSIR |
| 1.2398E-09 | -1.0706016 | 0.007 | 0.123 | 2.9288E-05 | TEC_PT | CCL3 |
| 1.2737E-09 | -0.2779384 | 0.027 | 0.153 | 3.0089E-05 | TEC_PT | SNX18 |
| 1.3077E-09 | 0.26767561 | 0.279 | 0.157 | 3.0891E-05 | TEC_PT | MTIF3 |
| 1.3492E-09 | -0.566008 | 0.172 | 0.324 | 3.1872E-05 | TEC_PT | VEGFA |
| 1.3542E-09 | -0.3302432 | 0.054 | 0.192 | 3.1989E-05 | TEC_PT | LIMA1 |
| 1.3599E-09 | -0.3068918 | 0.01 | 0.129 | 3.2124E-05 | TEC_PT | FAM84A |
| 1.365E-09 | 0.25503227 | 0.242 | 0.13 | 3.2246E-05 | TEC_PT | TOLLIP |
| 1.4124E-09 | -0.3460308 | 0.01 | 0.127 | 3.3364E-05 | TEC_PT | MARCO |
| 1.4305E-09 | -0.2843107 | 0.168 | 0.339 | 3.3793E-05 | TEC_PT | TM9SF3 |
| 1.4385E-09 | -0.5348807 | 0.141 | 0.294 | 3.3981E-05 | TEC_PT | ATP13A3 |
| 1.4879E-09 | -0.4139857 | 0.091 | 0.237 | 3.5148E-05 | TEC_PT | NFKBIZ |
| 1.5334E-09 | -0.2917593 | 0.118 | 0.275 | 3.6224E-05 | TEC_PT | NIPBL |
| 1.555E-09 | 0.26273749 | 0.239 | 0.13 | 3.6734E-05 | TEC_PT | C7orf50 |
| 1.5933E-09 | -0.2868674 | 0.128 | 0.29 | 3.7639E-05 | TEC_PT | CLIP1 |
| 1.6144E-09 | -0.3058641 | 0.054 | 0.189 | 3.8137E-05 | TEC_PT | VCL |
| 1.6715E-09 | -0.3166817 | 0.138 | 0.296 | 3.9487E-05 | TEC_PT | ARF6 |
| 1.6799E-09 | -0.2559742 | 0.044 | 0.176 | 3.9683E-05 | TEC_PT | ADAM17 |
| 1.8171E-09 | 0.40058984 | 0.428 | 0.302 | 4.2926E-05 | TEC_PT | ANAPC11 |

| | | | | | | |
|---|---|---|---|---|---|---|
| 1.8683E-09 | -0.4162618 | 0.01 | 0.125 | 4.4136E-05 | TEC_PT | SRD5A3 |
| 1.8787E-09 | 0.36906012 | 0.687 | 0.546 | 4.4381E-05 | TEC_PT | RPS10 |
| 1.9976E-09 | 0.30142598 | 0.949 | 0.869 | 4.7189E-05 | TEC_PT | RPLP0 |
| 2.0789E-09 | -0.2974823 | 0.084 | 0.23 | 4.911E-05 | TEC_PT | PGD |
| 2.129E-09 | -0.2554576 | 0.037 | 0.166 | 5.0292E-05 | TEC_PT | DDR1 |
| 2.1309E-09 | 0.29192957 | 0.387 | 0.252 | 5.0339E-05 | TEC_PT | HADHA |
| 2.1328E-09 | -0.390883 | 0 | 0.108 | 5.0383E-05 | TEC_PT | KRT23 |
| 2.193E-09 | -0.756795 | 0.003 | 0.114 | 5.1805E-05 | TEC_PT | IL7R |
| 2.2358E-09 | -0.2824352 | 0.027 | 0.15 | 5.2816E-05 | TEC_PT | SMAP2 |
| 2.4169E-09 | -0.5397624 | 0.003 | 0.113 | 5.7094E-05 | TEC_PT | CYP1B1 |
| 2.4966E-09 | -0.4166562 | 0 | 0.107 | 5.8978E-05 | TEC_PT | TMC5 |
| 2.501E-09 | -0.2593996 | 0.064 | 0.203 | 5.9081E-05 | TEC_PT | PRDM2 |
| 2.5338E-09 | -0.2829578 | 0.175 | 0.341 | 5.9856E-05 | TEC_PT | ANP32B |
| 2.6104E-09 | -0.2905261 | 0 | 0.107 | 6.1667E-05 | TEC_PT | MS4A6A |
| 2.665E-09 | -1.538345 | 0 | 0.107 | 6.2954E-05 | TEC_PT | CRCT1 |
| 2.6703E-09 | -0.2537197 | 0.003 | 0.113 | 6.308E-05 | TEC_PT | PECAM1 |
| 2.6738E-09 | 0.27793638 | 0.316 | 0.193 | 6.3163E-05 | TEC_PT | MRPL54 |
| 2.7548E-09 | -0.4430555 | 0.003 | 0.113 | 6.5077E-05 | TEC_PT | TMPRSS11A |
| 2.7946E-09 | -0.3386822 | 0.007 | 0.118 | 6.6016E-05 | TEC_PT | DUSP2 |
| 2.8004E-09 | 0.31143802 | 0.953 | 0.905 | 6.6154E-05 | TEC_PT | RPL7A |
| 2.8194E-09 | 0.407427 | 0.62 | 0.52 | 6.6604E-05 | TEC_PT | ATP6V1F |
| 2.8453E-09 | 0.25461212 | 0.172 | 0.082 | 6.7215E-05 | TEC_PT | PXDC1 |
| 2.8862E-09 | -0.259781 | 0.088 | 0.235 | 6.818E-05 | TEC_PT | RBMS1 |
| 2.8949E-09 | -0.5958026 | 0.646 | 0.741 | 6.8386E-05 | TEC_PT | PSAP |
| 2.9225E-09 | 0.25627948 | 0.35 | 0.215 | 6.9039E-05 | TEC_PT | RWDD1 |
| 2.9303E-09 | -0.3324822 | 0.013 | 0.129 | 6.9223E-05 | TEC_PT | LRG1 |
| 3.0177E-09 | -0.3425454 | 0 | 0.106 | 7.1287E-05 | TEC_PT | CSF3R |
| 3.0273E-09 | -0.2784062 | 0.01 | 0.124 | 7.1514E-05 | TEC_PT | PHACTR1 |
| 3.0429E-09 | -0.29224 | 0.077 | 0.219 | 7.1882E-05 | TEC_PT | POLR2J3.1 |
| 3.0549E-09 | 0.27748627 | 0.249 | 0.136 | 7.2165E-05 | TEC_PT | MRPL36 |
| 3.0964E-09 | -0.2618702 | 0.121 | 0.28 | 7.3147E-05 | TEC_PT | TMF1 |
| 3.1275E-09 | -0.3503624 | 0.899 | 0.945 | 7.3881E-05 | TEC_PT | MT-CO2 |
| 3.1401E-09 | -0.3256399 | 0 | 0.106 | 7.418E-05 | TEC_PT | A2ML1 |
| 3.211E-09 | -0.3498873 | 0.074 | 0.213 | 7.5854E-05 | TEC_PT | TLE4 |
| 3.3505E-09 | 0.40772512 | 0.431 | 0.299 | 7.9149E-05 | TEC_PT | SF3B5 |
| 3.6977E-09 | -0.3508926 | 0.003 | 0.111 | 8.7352E-05 | TEC_PT | SMIM5 |
| 3.7969E-09 | -0.2567406 | 0.061 | 0.196 | 8.9694E-05 | TEC_PT | RAB8B |
| 3.8002E-09 | -0.7768487 | 0 | 0.105 | 8.9772E-05 | TEC_PT | ZG16B |
| 3.8698E-09 | -0.2694166 | 0.027 | 0.147 | 9.1415E-05 | TEC_PT | PLEKHO1 |
| 4.0159E-09 | -0.3537857 | 0.071 | 0.205 | 9.4869E-05 | TEC_PT | TIPARP |
| 4.1768E-09 | 0.58117507 | 0.549 | 0.428 | 9.8669E-05 | TEC_PT | SELENOK |
| 4.1888E-09 | -0.3342549 | 0.007 | 0.116 | 9.8953E-05 | TEC_PT | CD163 |
| 4.6434E-09 | 0.28014209 | 0.38 | 0.247 | 0.00010969 | TEC_PT | TRAPPC1 |
| 4.6976E-09 | -0.3078841 | 0.35 | 0.509 | 0.00011097 | TEC_PT | HNRNPDL |
| 4.8943E-09 | -0.2501441 | 0.003 | 0.109 | 0.00011562 | TEC_PT | GATA3 |
| 4.9034E-09 | -0.2598841 | 0.044 | 0.169 | 0.00011583 | TEC_PT | VPS37B |
| 5.0995E-09 | -0.2793877 | 0 | 0.103 | 0.00012047 | TEC_PT | LILRB1 |
| 5.2006E-09 | -0.2641907 | 0.003 | 0.109 | 0.00012285 | TEC_PT | PDE4B |
| 5.5023E-09 | 0.3095187 | 0.872 | 0.746 | 0.00012998 | TEC_PT | EEF1B2 |
| 5.847E-09 | 0.32779167 | 0.263 | 0.151 | 0.00013812 | TEC_PT | GLUD1 |

| | | | | | | |
|---|---|---|---|---|---|---|
| 5.8586E-09 | -0.2752157 | 0.064 | 0.199 | 0.0001384 | TEC_PT | ARHGAP18 |
| 5.9221E-09 | -0.3076114 | 0.03 | 0.151 | 0.0001399 | TEC_PT | LPIN1 |
| 6.0114E-09 | -0.392568 | 0.357 | 0.502 | 0.00014201 | TEC_PT | ASAH1 |
| 6.0649E-09 | 0.2564198 | 0.178 | 0.088 | 0.00014327 | TEC_PT | NPL |
| 6.1011E-09 | -0.2566243 | 0.061 | 0.192 | 0.00014413 | TEC_PT | FCHO2 |
| 6.4517E-09 | -0.2525302 | 0.03 | 0.15 | 0.00015241 | TEC_PT | RASGEF1B |
| 6.5016E-09 | -0.2686281 | 0.098 | 0.242 | 0.00015359 | TEC_PT | KMT2C |
| 6.546E-09 | -0.3438713 | 0.024 | 0.141 | 0.00015464 | TEC_PT | GPNMB |
| 6.5647E-09 | -0.2963407 | 0.094 | 0.235 | 0.00015508 | TEC_PT | FNBP4 |
| 6.5813E-09 | -1.0572925 | 0.003 | 0.108 | 0.00015547 | TEC_PT | CCL4 |
| 6.7043E-09 | 0.34410192 | 0.953 | 0.904 | 0.00015838 | TEC_PT | RPL37 |
| 7.2818E-09 | -0.2642891 | 0.114 | 0.267 | 0.00017202 | TEC_PT | TMEM123 |
| 7.3332E-09 | -0.2550916 | 0.098 | 0.241 | 0.00017323 | TEC_PT | ABI1 |
| 7.3549E-09 | -0.4184824 | 0.047 | 0.172 | 0.00017374 | TEC_PT | DNAJA4 |
| 7.3669E-09 | -0.4473651 | 0.125 | 0.271 | 0.00017403 | TEC_PT | TSC22D3 |
| 7.3681E-09 | 0.28452891 | 0.212 | 0.111 | 0.00017406 | TEC_PT | ADGRG1 |
| 7.5971E-09 | -0.4013726 | 0.003 | 0.107 | 0.00017947 | TEC_PT | THBD |
| 7.6252E-09 | -0.2515174 | 0.047 | 0.173 | 0.00018013 | TEC_PT | ABHD2 |
| 7.7569E-09 | -0.4204509 | 0.33 | 0.486 | 0.00018324 | TEC_PT | HMGA1 |
| 7.7721E-09 | 0.33172029 | 0.842 | 0.703 | 0.0001836 | TEC_PT | RPL4 |
| 8.0636E-09 | 0.25585879 | 0.333 | 0.203 | 0.00019049 | TEC_PT | C6orf48 |
| 8.1356E-09 | -0.2823185 | 0.051 | 0.177 | 0.00019219 | TEC_PT | ABCA1 |
| 8.3818E-09 | 0.29646402 | 0.976 | 0.922 | 0.000198 | TEC_PT | RPL11 |
| 8.3821E-09 | 0.28456815 | 0.283 | 0.167 | 0.00019801 | TEC_PT | FBXW5 |
| 8.8552E-09 | 0.37326177 | 0.455 | 0.334 | 0.00020919 | TEC_PT | EIF4B |
| 9.0065E-09 | -0.4849543 | 0 | 0.1 | 0.00021276 | TEC_PT | RETN |
| 9.2461E-09 | -0.2554144 | 0.013 | 0.123 | 0.00021842 | TEC_PT | TCP11L2 |
| 9.2708E-09 | -0.3586163 | 0.333 | 0.493 | 0.000219 | TEC_PT | TAGLN2 |
| 9.5421E-09 | -0.3688419 | 0.205 | 0.361 | 0.00022541 | TEC_PT | CFLAR |
| 9.8253E-09 | 0.2664404 | 0.279 | 0.166 | 0.0002321 | TEC_PT | GTF2H5 |
| 9.8288E-09 | -0.2683857 | 0.017 | 0.128 | 0.00023219 | TEC_PT | TP53INP2 |
| 9.984E-09 | -0.2609544 | 0.02 | 0.133 | 0.00023585 | TEC_PT | MPZL2 |
| 1.0412E-08 | 0.26144483 | 0.966 | 0.923 | 0.00024596 | TEC_PT | RPS27A |
| 1.0497E-08 | -0.2971296 | 0.037 | 0.157 | 0.00024796 | TEC_PT | PELI1 |
| 1.0767E-08 | -0.2555434 | 0.027 | 0.143 | 0.00025435 | TEC_PT | ARL4A |
| 1.1464E-08 | 0.38429122 | 0.68 | 0.611 | 0.00027082 | TEC_PT | SEC61B |
| 1.2348E-08 | -0.5328696 | 0.027 | 0.142 | 0.00029169 | TEC_PT | SNCG |
| 1.2761E-08 | 0.39729478 | 0.68 | 0.582 | 0.00030145 | TEC_PT | CLTA |
| 1.3113E-08 | -0.2554025 | 0.121 | 0.269 | 0.00030977 | TEC_PT | DIAPH1 |
| 1.3914E-08 | -0.3828575 | 0.003 | 0.104 | 0.00032869 | TEC_PT | CLIC3 |
| 1.4911E-08 | 0.38843674 | 0.761 | 0.687 | 0.00035224 | TEC_PT | BRI3 |
| 1.5389E-08 | 0.54896427 | 0.343 | 0.234 | 0.00036355 | TEC_PT | MT-ATP8 |
| 1.5682E-08 | -0.2644613 | 0.071 | 0.202 | 0.00037047 | TEC_PT | ZNF217 |
| 1.629E-08 | -0.2655341 | 0.071 | 0.201 | 0.00038481 | TEC_PT | ZNF91 |
| 1.6356E-08 | 0.38791837 | 0.485 | 0.362 | 0.00038637 | TEC_PT | PSMB1 |
| 1.6618E-08 | -0.2570065 | 0.067 | 0.198 | 0.00039256 | TEC_PT | CHD1 |
| 1.7228E-08 | -0.2624445 | 0.037 | 0.155 | 0.00040698 | TEC_PT | ITPKC |
| 1.7827E-08 | -0.4850598 | 0.064 | 0.189 | 0.00042113 | TEC_PT | HBEGF |
| 1.8412E-08 | 0.54348043 | 0.128 | 0.056 | 0.00043495 | TEC_PT | FSTL3 |
| 1.9002E-08 | -0.4268079 | 0.088 | 0.219 | 0.00044888 | TEC_PT | TGM2 |

| | | | | | | |
|---|---|---|---|---|---|---|
| 1.9014E-08 | -0.2775539 | 0.125 | 0.27 | 0.00044917 | TEC_PT | NPEPPS |
| 1.9908E-08 | 0.26866422 | 0.343 | 0.219 | 0.00047028 | TEC_PT | TMEM208 |
| 2.0424E-08 | -0.3911947 | 0.209 | 0.367 | 0.00048247 | TEC_PT | RALBP1 |
| 2.0514E-08 | -0.3191427 | 0.04 | 0.157 | 0.0004846 | TEC_PT | TSPAN6 |
| 2.0719E-08 | 0.30836485 | 0.838 | 0.794 | 0.00048944 | TEC_PT | CHCHD2 |
| 2.0808E-08 | 0.3979444 | 0.882 | 0.864 | 0.00049154 | TEC_PT | CALM1 |
| 2.1184E-08 | -0.2559218 | 0.286 | 0.448 | 0.00050044 | TEC_PT | DNAJB6 |
| 2.1207E-08 | -0.3809398 | 0.074 | 0.208 | 0.00050098 | TEC_PT | KCNQ1OT1 |
| 2.1433E-08 | -0.2552309 | 0.061 | 0.186 | 0.00050631 | TEC_PT | PHACTR2 |
| 2.1792E-08 | 0.29621853 | 0.448 | 0.325 | 0.0005148 | TEC_PT | SRP9 |
| 2.1975E-08 | 0.26248839 | 0.263 | 0.151 | 0.00051911 | TEC_PT | PLEKHA1 |
| 2.2145E-08 | -0.280154 | 0.111 | 0.251 | 0.00052312 | TEC_PT | UBL3 |
| 2.2239E-08 | -0.2560053 | 0.051 | 0.174 | 0.00052534 | TEC_PT | TNFAIP8 |
| 2.237E-08 | 0.29962189 | 0.98 | 0.895 | 0.00052844 | TEC_PT | RPS9 |
| 2.2946E-08 | 0.59363831 | 0.276 | 0.168 | 0.00054204 | TEC_PT | AKR1B1 |
| 2.3098E-08 | -0.2526635 | 0.034 | 0.149 | 0.00054565 | TEC_PT | RALGPS2 |
| 2.3116E-08 | 0.30632047 | 0.347 | 0.223 | 0.00054607 | TEC_PT | NARS |
| 2.3596E-08 | 0.39348414 | 0.99 | 0.957 | 0.0005574 | TEC_PT | RPS18 |
| 2.4103E-08 | 0.47803589 | 0.461 | 0.317 | 0.00056938 | TEC_PT | GADD45A |
| 2.5283E-08 | -0.2856781 | 0.114 | 0.253 | 0.00059727 | TEC_PT | C4orf48 |
| 2.6742E-08 | 0.3054175 | 0.367 | 0.242 | 0.00063172 | TEC_PT | TMEM14C |
| 2.766E-08 | 0.25135912 | 0.367 | 0.237 | 0.00065342 | TEC_PT | TSTD1 |
| 2.8084E-08 | 0.63698545 | 0.596 | 0.506 | 0.00066343 | TEC_PT | MT-ND4L |
| 2.8344E-08 | -0.3549409 | 0.189 | 0.334 | 0.00066957 | TEC_PT | TLN1 |
| 2.9125E-08 | -0.7087834 | 0.677 | 0.74 | 0.00068803 | TEC_PT | VIM |
| 2.9167E-08 | -0.2761375 | 0.104 | 0.24 | 0.00068901 | TEC_PT | SP100 |
| 2.945E-08 | 0.29869684 | 0.391 | 0.265 | 0.00069571 | TEC_PT | VPS29 |
| 2.9591E-08 | -0.2723362 | 0.128 | 0.27 | 0.00069902 | TEC_PT | AGFG1 |
| 3.0362E-08 | -0.2694113 | 0.088 | 0.22 | 0.00071723 | TEC_PT | TES |
| 3.0672E-08 | -0.2935496 | 0.01 | 0.111 | 0.00072457 | TEC_PT | STS |
| 3.192E-08 | -0.2889538 | 0.269 | 0.421 | 0.00075404 | TEC_PT | CAPZB |
| 3.276E-08 | -1.6259157 | 0.279 | 0.397 | 0.00077388 | TEC_PT | HLA-DRA |
| 3.2783E-08 | -0.3101062 | 0.148 | 0.297 | 0.00077444 | TEC_PT | IFNGR2 |
| 3.3981E-08 | -0.2751504 | 0.094 | 0.228 | 0.00080273 | TEC_PT | TRIM56 |
| 3.5155E-08 | -0.3137824 | 0.185 | 0.337 | 0.00083046 | TEC_PT | LUC7L3 |
| 3.6836E-08 | -0.3660492 | 0.131 | 0.267 | 0.00087019 | TEC_PT | AFDN |
| 3.8207E-08 | 0.27206801 | 0.263 | 0.154 | 0.00090257 | TEC_PT | CHCHD5 |
| 4.015E-08 | 0.34283915 | 0.414 | 0.289 | 0.00094845 | TEC_PT | HIGD2A |
| 4.0274E-08 | -0.2955914 | 0.024 | 0.129 | 0.0009514 | TEC_PT | GAN |
| 4.1049E-08 | -0.2550206 | 0.253 | 0.417 | 0.0009697 | TEC_PT | GRN |
| 4.1945E-08 | -0.3061213 | 0.357 | 0.507 | 0.00099087 | TEC_PT | JAK1 |
| 4.2621E-08 | -0.3054973 | 0.01 | 0.109 | 0.00100684 | TEC_PT | FGL2 |
| 4.2885E-08 | -0.3806778 | 0.202 | 0.347 | 0.00101306 | TEC_PT | GOLGB1 |
| 4.6881E-08 | 0.28427293 | 0.966 | 0.915 | 0.00110747 | TEC_PT | RPL32 |
| 4.7441E-08 | 0.32795859 | 0.404 | 0.286 | 0.00112071 | TEC_PT | ACADVL |
| 4.7751E-08 | -1.4088684 | 0.01 | 0.108 | 0.00112803 | TEC_PT | CNFN |
| 4.9951E-08 | -0.2613499 | 0.04 | 0.152 | 0.00117999 | TEC_PT | SLC44A1 |
| 5.0424E-08 | 0.29682198 | 0.303 | 0.188 | 0.00119117 | TEC_PT | ABCF1 |
| 5.2303E-08 | -0.3679904 | 0.209 | 0.35 | 0.00123555 | TEC_PT | UBE2H |
| 5.5635E-08 | -0.2538329 | 0.414 | 0.579 | 0.00131427 | TEC_PT | UBE2B |

| | | | | | | |
|---|---|---|---|---|---|---|
| 6.0757E-08 | 0.32900335 | 0.966 | 0.896 | 0.00143527 | TEC_PT | RPL35A |
| 6.2123E-08 | -0.2753083 | 0.195 | 0.346 | 0.00146752 | TEC_PT | SCAF11 |
| 6.818E-08 | -0.2768813 | 0.131 | 0.27 | 0.00161062 | TEC_PT | FAM133B |
| 6.9036E-08 | -0.2527517 | 0.034 | 0.141 | 0.00163084 | TEC_PT | AL133453.1 |
| 7.2712E-08 | 0.26141854 | 0.414 | 0.282 | 0.00171768 | TEC_PT | EIF6 |
| 7.339E-08 | 0.33700137 | 0.801 | 0.762 | 0.0017337 | TEC_PT | UBL5 |
| 7.4797E-08 | -0.2876881 | 0.04 | 0.15 | 0.00176693 | TEC_PT | DMXL2 |
| 7.5281E-08 | -0.2708619 | 0.084 | 0.213 | 0.00177835 | TEC_PT | SETX |
| 7.6908E-08 | 0.34950488 | 0.879 | 0.783 | 0.00181679 | TEC_PT | DYNLL1 |
| 9.1225E-08 | -0.7164247 | 0.374 | 0.504 | 0.002155 | TEC_PT | DNAJB1 |
| 9.3809E-08 | -0.2875902 | 0.067 | 0.186 | 0.00221605 | TEC_PT | CKLF |
| 9.4682E-08 | -0.3635415 | 0.067 | 0.189 | 0.00223667 | TEC_PT | SLC5A3 |
| 1.0451E-07 | -0.3109162 | 0.111 | 0.243 | 0.00246877 | TEC_PT | IFRD1 |
| 1.0452E-07 | -0.5171929 | 0.192 | 0.315 | 0.00246911 | TEC_PT | HIST1H2AC |
| 1.052E-07 | -0.3248261 | 0.168 | 0.308 | 0.00248512 | TEC_PT | NUFIP2 |
| 1.1172E-07 | -0.2670523 | 0.088 | 0.211 | 0.00263922 | TEC_PT | NFAT5 |
| 1.1618E-07 | -0.2851871 | 0.108 | 0.233 | 0.00274459 | TEC_PT | RYBP |
| 1.1917E-07 | -0.2527701 | 0.125 | 0.26 | 0.00281507 | TEC_PT | ANKLE2 |
| 1.2172E-07 | -0.2501458 | 0.202 | 0.355 | 0.00287536 | TEC_PT | SEPT7 |
| 1.2661E-07 | -0.2581953 | 0.148 | 0.29 | 0.00299081 | TEC_PT | CHMP2B |
| 1.3121E-07 | -0.2596096 | 0.074 | 0.195 | 0.00309959 | TEC_PT | IRS2 |
| 1.3531E-07 | -0.3007124 | 0.195 | 0.34 | 0.00319647 | TEC_PT | GRB2 |
| 1.4364E-07 | 0.26650273 | 0.263 | 0.156 | 0.00339316 | TEC_PT | FNIP2 |
| 1.4509E-07 | 0.36013836 | 0.343 | 0.232 | 0.00342752 | TEC_PT | CTSA |
| 1.465E-07 | -0.5779788 | 0.01 | 0.102 | 0.00346085 | TEC_PT | HLA-DQA1 |
| 1.738E-07 | 0.31944748 | 0.418 | 0.301 | 0.00410568 | TEC_PT | PGAM1 |
| 1.8393E-07 | 0.31247796 | 0.182 | 0.096 | 0.00434489 | TEC_PT | CITED4 |
| 1.9301E-07 | -0.4354244 | 0.034 | 0.136 | 0.00455953 | TEC_PT | HLA-DRB5 |
| 1.9324E-07 | -0.3412431 | 0.121 | 0.242 | 0.0045648 | TEC_PT | GIPC1 |
| 2.0991E-07 | 0.25606559 | 0.306 | 0.194 | 0.00495881 | TEC_PT | RBPMS |
| 2.12E-07 | -0.3019768 | 0.266 | 0.415 | 0.00500796 | TEC_PT | SFPQ |
| 2.1295E-07 | -0.2911532 | 0.027 | 0.126 | 0.00503052 | TEC_PT | POLB |
| 2.1562E-07 | 0.32329742 | 0.401 | 0.294 | 0.00509353 | TEC_PT | VPS28 |
| 2.1604E-07 | -0.3053236 | 0.074 | 0.194 | 0.0051036 | TEC_PT | KATNBL1 |
| 2.3153E-07 | -0.2682122 | 0.03 | 0.131 | 0.00546946 | TEC_PT | ID3 |
| 2.5824E-07 | -0.3763456 | 0.03 | 0.129 | 0.00610042 | TEC_PT | AC023157.3 |
| 2.7452E-07 | -0.2563221 | 0.067 | 0.183 | 0.00648502 | TEC_PT | SPTSSA |
| 2.756E-07 | -0.2849665 | 0.118 | 0.244 | 0.00651061 | TEC_PT | RBPJ |
| 2.7693E-07 | -0.3220464 | 0.02 | 0.115 | 0.00654194 | TEC_PT | CEACAM1 |
| 2.7825E-07 | 0.3554041 | 0.397 | 0.292 | 0.00657313 | TEC_PT | COX17 |
| 2.9005E-07 | -0.2834773 | 0.667 | 0.771 | 0.00685181 | TEC_PT | RHOA |
| 2.9933E-07 | -0.2623815 | 0.03 | 0.131 | 0.00707114 | TEC_PT | FOXC1 |
| 3.0836E-07 | -0.2740653 | 0.189 | 0.332 | 0.0072845 | TEC_PT | MSN |
| 3.1811E-07 | -0.3482461 | 0.125 | 0.248 | 0.00751462 | TEC_PT | SERINC2 |
| 3.2467E-07 | 0.26011451 | 0.983 | 0.972 | 0.00766979 | TEC_PT | RPLP1 |
| 3.3381E-07 | -0.3358413 | 0.135 | 0.261 | 0.00788571 | TEC_PT | ARRDC3 |
| 4.0179E-07 | -0.3447568 | 0.394 | 0.519 | 0.00949156 | TEC_PT | NAP1L1 |
| 4.1514E-07 | -0.4101666 | 0.111 | 0.229 | 0.00980686 | TEC_PT | IVNS1ABP |
| 4.2409E-07 | -0.3023888 | 0.209 | 0.351 | 0.01001836 | TEC_PT | PNN |
| 4.3793E-07 | 0.29881481 | 0.384 | 0.275 | 0.01034533 | TEC_PT | MRPL51 |

| | | | | | | |
|---|---|---|---|---|---|---|
| 4.5794E-07 | 0.27171022 | 0.354 | 0.243 | 0.01081789 | TEC_PT | CFDP1 |
| 4.5988E-07 | 0.3650195 | 0.424 | 0.319 | 0.01086381 | TEC_PT | ERP29 |
| 4.6601E-07 | -0.266117 | 0.209 | 0.347 | 0.01100849 | TEC_PT | ZNF207 |
| 4.7481E-07 | -0.2524306 | 0.02 | 0.112 | 0.01121634 | TEC_PT | FUT3 |
| 6.261E-07 | -0.252668 | 0.047 | 0.15 | 0.01479041 | TEC_PT | PLXDC2 |
| 6.3223E-07 | -0.4994014 | 0.084 | 0.192 | 0.01493525 | TEC_PT | FBXO32 |
| 6.4984E-07 | -0.3628299 | 0.973 | 0.978 | 0.01535118 | TEC_PT | RPL28 |
| 6.6806E-07 | -0.4674853 | 0.219 | 0.329 | 0.01578148 | TEC_PT | MYO6 |
| 6.9316E-07 | 0.27879653 | 0.852 | 0.77 | 0.01637456 | TEC_PT | TMA7 |
| 7.1042E-07 | 0.25799461 | 0.263 | 0.164 | 0.01678216 | TEC_PT | SNRPC |
| 7.5024E-07 | -0.3565869 | 0.04 | 0.136 | 0.01772293 | TEC_PT | GCNT3 |
| 7.7117E-07 | -0.4575864 | 0.508 | 0.638 | 0.01821727 | TEC_PT | GADD45B |
| 7.921E-07 | -0.3838445 | 0.606 | 0.722 | 0.01871173 | TEC_PT | JUND |
| 7.9618E-07 | 0.26800137 | 0.943 | 0.876 | 0.01880806 | TEC_PT | RPL23A |
| 8.08E-07 | -0.2893814 | 0.02 | 0.11 | 0.01908737 | TEC_PT | TNF |
| 8.3745E-07 | 0.26391281 | 0.414 | 0.304 | 0.01978304 | TEC_PT | JTB |
| 8.4463E-07 | -0.3312187 | 0.189 | 0.329 | 0.01995265 | TEC_PT | SLC38A2 |
| 8.9408E-07 | -0.2549885 | 0.145 | 0.268 | 0.02112088 | TEC_PT | TOR1AIP2 |
| 8.9784E-07 | -0.4236333 | 0.418 | 0.523 | 0.02120958 | TEC_PT | HEBP2 |
| 9.0451E-07 | -0.6006314 | 0.323 | 0.434 | 0.02136734 | TEC_PT | HIST1H1C |
| 9.1152E-07 | -0.3267806 | 0.02 | 0.108 | 0.02153274 | TEC_PT | TCHP |
| 9.85E-07 | 0.26256599 | 0.731 | 0.623 | 0.02326876 | TEC_PT | DSTN |
| 1.027E-06 | 0.31080521 | 0.492 | 0.385 | 0.0242614 | TEC_PT | PSME1 |
| 1.07E-06 | 0.2537519 | 0.687 | 0.6 | 0.025277 | TEC_PT | YWHAE |
| 1.085E-06 | -0.2567664 | 0.024 | 0.113 | 0.02563117 | TEC_PT | HIST1H2BG |
| 1.2066E-06 | -0.2754903 | 0.195 | 0.325 | 0.02850272 | TEC_PT | YPEL5 |
| 1.2102E-06 | -0.2549413 | 0.138 | 0.26 | 0.02858887 | TEC_PT | MBNL1 |
| 1.2652E-06 | -0.2774003 | 0.249 | 0.391 | 0.02988802 | TEC_PT | STAT3 |
| 1.2676E-06 | 0.31072572 | 0.364 | 0.259 | 0.02994499 | TEC_PT | SMDT1 |
| 1.3997E-06 | -0.2582909 | 0.158 | 0.282 | 0.03306543 | TEC_PT | TMEM30A |
| 1.4753E-06 | -0.3886162 | 0.131 | 0.252 | 0.03485102 | TEC_PT | SNHG12 |
| 1.5513E-06 | -0.2626664 | 0.259 | 0.396 | 0.03664589 | TEC_PT | CTNNA1 |
| 1.7115E-06 | 0.30941474 | 0.485 | 0.383 | 0.04043103 | TEC_PT | DYNLRB1 |
| 1.8732E-06 | -0.2686127 | 0.114 | 0.234 | 0.04424988 | TEC_PT | PARP14 |
| 1.9235E-06 | -0.331358 | 0.101 | 0.216 | 0.0454395 | TEC_PT | TUBA1A |
| 1.9814E-06 | 0.25500652 | 0.411 | 0.298 | 0.04680637 | TEC_PT | NUCKS1 |
| 2.0114E-06 | -0.250253 | 0.104 | 0.217 | 0.04751582 | TEC_PT | DSP |
| 2.0256E-06 | -0.3105814 | 0.606 | 0.704 | 0.04785063 | TEC_PT | TPM3 |
| 2.1574E-06 | 0.28574284 | 0.471 | 0.354 | 0.05096511 | TEC_PT | CYCS |
| 2.1648E-06 | -0.3628947 | 0.108 | 0.215 | 0.05114008 | TEC_PT | CREM |
| 2.2835E-06 | -0.8854242 | 0.03 | 0.118 | 0.05394241 | TEC_PT | CXCL2 |
| 2.3604E-06 | -0.2717858 | 0.29 | 0.418 | 0.05575962 | TEC_PT | JPT1 |
| 2.4896E-06 | 0.30685394 | 0.404 | 0.303 | 0.05881171 | TEC_PT | BLVRB |
| 2.62E-06 | -0.5203862 | 0.596 | 0.668 | 0.06189189 | TEC_PT | ANXA5 |
| 2.8094E-06 | -0.2705361 | 0.091 | 0.197 | 0.06636549 | TEC_PT | KLF4 |
| 2.9203E-06 | -0.380318 | 0.145 | 0.259 | 0.06898717 | TEC_PT | IDI1 |
| 3.3361E-06 | 0.25022178 | 0.283 | 0.19 | 0.07880825 | TEC_PT | CHCHD1 |
| 3.4439E-06 | -0.32743 | 0.074 | 0.171 | 0.08135606 | TEC_PT | FAM214A |
| 3.5184E-06 | -0.2597409 | 0.293 | 0.429 | 0.08311444 | TEC_PT | EIF3A |
| 3.5793E-06 | -0.6829708 | 0.027 | 0.109 | 0.0845528 | TEC_PT | APOC1 |

| | | | | | | |
|---|---|---|---|---|---|---|
| 3.6534E-06 | -0.3466287 | 0.535 | 0.63 | 0.08630367 | TEC_PT | TAX1BP1 |
| 3.7236E-06 | -0.2687961 | 0.135 | 0.248 | 0.08796335 | TEC_PT | CREBRF |
| 3.8092E-06 | -0.4007058 | 0.054 | 0.149 | 0.08998362 | TEC_PT | SMC4 |
| 3.9439E-06 | -0.2703861 | 0.865 | 0.905 | 0.093167 | TEC_PT | CFL1 |
| 3.9691E-06 | -0.6448701 | 0.047 | 0.14 | 0.09376212 | TEC_PT | CXCL1 |
| 4.04E-06 | 0.27660164 | 0.953 | 0.9 | 0.09543767 | TEC_PT | RPS11 |
| 5.3348E-06 | 0.67450957 | 0.195 | 0.114 | 0.12602437 | TEC_PT | RRAD |
| 5.5382E-06 | 0.2615416 | 0.374 | 0.265 | 0.13082782 | TEC_PT | TCP1 |
| 5.5472E-06 | 0.2560643 | 0.202 | 0.122 | 0.13104128 | TEC_PT | LYRM2 |
| 6.1766E-06 | -0.3825951 | 0.162 | 0.27 | 0.14590925 | TEC_PT | CCNG2 |
| 6.3368E-06 | 0.25276034 | 0.367 | 0.264 | 0.1496942 | TEC_PT | PSMA4 |
| 6.5099E-06 | 0.28318083 | 0.34 | 0.241 | 0.15378426 | TEC_PT | NUDC |
| 6.5854E-06 | 0.25143952 | 0.761 | 0.694 | 0.15556697 | TEC_PT | GUK1 |
| 6.8424E-06 | -0.4184109 | 0.135 | 0.244 | 0.16163754 | TEC_PT | ITGB8 |
| 7.2316E-06 | -0.2704014 | 0.131 | 0.242 | 0.17083099 | TEC_PT | BHLHE40 |
| 7.4243E-06 | 0.28616444 | 0.451 | 0.352 | 0.17538451 | TEC_PT | SPCS1 |
| 7.623E-06 | -0.2604668 | 0.239 | 0.369 | 0.18007782 | TEC_PT | ANKRD11 |
| 8.5694E-06 | -0.3047274 | 0.121 | 0.231 | 0.20243548 | TEC_PT | ISG15 |
| 8.5777E-06 | -0.2561767 | 0.168 | 0.282 | 0.20263018 | TEC_PT | PRPF4B |
| 9.0231E-06 | 0.28651869 | 0.879 | 0.814 | 0.21315372 | TEC_PT | YBX1 |
| 9.7714E-06 | 0.25667858 | 0.31 | 0.212 | 0.23082927 | TEC_PT | RPL17 |
| 1.0345E-05 | -0.2586202 | 0.148 | 0.258 | 0.2443711 | TEC_PT | PDLIM5 |
| 1.0559E-05 | -0.5857477 | 0.963 | 0.962 | 0.24943355 | TEC_PT | HSP90AA1 |
| 1.0826E-05 | 0.27313978 | 0.438 | 0.342 | 0.25573983 | TEC_PT | BSG |
| 1.1379E-05 | -0.2681425 | 0.182 | 0.289 | 0.26881775 | TEC_PT | CHMP1B |
| 1.2182E-05 | 0.34569566 | 0.916 | 0.858 | 0.28778468 | TEC_PT | RPL7 |
| 1.3171E-05 | 0.35474591 | 0.283 | 0.194 | 0.31114724 | TEC_PT | SDF2L1 |
| 1.4055E-05 | 0.37464834 | 0.468 | 0.394 | 0.33203137 | TEC_PT | NDUFB11 |
| 1.4176E-05 | 0.38550117 | 0.374 | 0.273 | 0.33487581 | TEC_PT | TNFRSF12A |
| 1.9815E-05 | -0.276256 | 0.088 | 0.179 | 0.46809891 | TEC_PT | LTA4H |
| 2.0094E-05 | -0.3428634 | 0.475 | 0.571 | 0.47467541 | TEC_PT | ZFAND5 |
| 2.3565E-05 | 0.27712973 | 0.431 | 0.327 | 0.55666618 | TEC_PT | LMNA |
| 2.4779E-05 | 0.31587412 | 0.232 | 0.152 | 0.58535543 | TEC_PT | CKS2 |
| 2.6215E-05 | 0.8572711 | 0.872 | 0.909 | 0.61927201 | TEC_PT | MT-ND2 |
| 2.6498E-05 | -0.4306692 | 0.239 | 0.347 | 0.62595476 | TEC_PT | ZFP36L2 |
| 2.6578E-05 | 0.26405471 | 0.488 | 0.394 | 0.62784649 | TEC_PT | COMMD6 |
| 2.7705E-05 | 0.27674301 | 0.384 | 0.304 | 0.65447452 | TEC_PT | RAB1A |
| 2.7938E-05 | 0.26887948 | 0.367 | 0.277 | 0.65998367 | TEC_PT | PLEKHB2 |
| 2.8215E-05 | -0.3625136 | 0.182 | 0.28 | 0.66652391 | TEC_PT | ABHD5 |
| 3.5054E-05 | 0.27953538 | 0.418 | 0.325 | 0.82808979 | TEC_PT | PPA1 |
| 3.5662E-05 | -0.2853681 | 0.034 | 0.107 | 0.84243697 | TEC_PT | SLC16A10 |
| 3.6279E-05 | -0.4478235 | 0.044 | 0.121 | 0.85701157 | TEC_PT | HILPDA |
| 3.822E-05 | 0.25367458 | 0.323 | 0.239 | 0.9028673 | TEC_PT | TXNL1 |
| 4.8032E-05 | -0.2616584 | 0.091 | 0.181 | 1 | TEC_PT | WARS |
| 4.8052E-05 | 0.33293772 | 0.475 | 0.399 | 1 | TEC_PT | CIB1 |
| 5.2692E-05 | 0.80922256 | 0.825 | 0.823 | 1 | TEC_PT | MT-ND5 |
| 5.4331E-05 | -0.3631902 | 0.044 | 0.121 | 1 | TEC_PT | TCIM |
| 5.6556E-05 | 0.27358639 | 0.36 | 0.281 | 1 | TEC_PT | ATP6V1E1 |
| 6.1491E-05 | 0.25152574 | 0.798 | 0.685 | 1 | TEC_PT | NPM1 |
| 7.4934E-05 | -0.2851145 | 0.067 | 0.145 | 1 | TEC_PT | SDC2 |

| | | | | | | |
|---|---|---|---|---|---|---|
| 7.5816E-05 | -0.6386244 | 0.576 | 0.646 | 1 | TEC_PT | HSPA5 |
| 9.0422E-05 | 0.25066565 | 0.343 | 0.261 | 1 | TEC_PT | EIF3J |
| 9.2604E-05 | 0.31262697 | 0.498 | 0.433 | 1 | TEC_PT | TRIR |
| 0.0001073 | 0.25454783 | 0.286 | 0.207 | 1 | TEC_PT | SIVA1 |
| 0.00010861 | -1.1694806 | 0.067 | 0.145 | 1 | TEC_PT | CXCL3 |
| 0.00011464 | 0.25057462 | 0.34 | 0.254 | 1 | TEC_PT | GABARAPL1 |
| 0.0001209 | 0.26041674 | 0.306 | 0.233 | 1 | TEC_PT | FKBP8 |
| 0.00012873 | -0.4699478 | 0.138 | 0.222 | 1 | TEC_PT | HES1 |
| 0.00013754 | -0.5661582 | 0.037 | 0.104 | 1 | TEC_PT | CCL5 |
| 0.00014784 | -0.3153381 | 0.138 | 0.225 | 1 | TEC_PT | INTS6 |
| 0.00014939 | 0.25960828 | 0.306 | 0.234 | 1 | TEC_PT | AHCYL1 |
| 0.00016736 | -0.6050954 | 0.108 | 0.184 | 1 | TEC_PT | AQP3 |
| 0.00017676 | 0.29241801 | 0.559 | 0.486 | 1 | TEC_PT | DYNLT1 |
| 0.00018442 | -0.2562771 | 0.098 | 0.18 | 1 | TEC_PT | DST |
| 0.00027178 | -0.3118115 | 0.542 | 0.628 | 1 | TEC_PT | JUN |
| 0.00035233 | -0.3192797 | 0.155 | 0.238 | 1 | TEC_PT | CITED2 |
| 0.00046368 | 0.27849333 | 0.36 | 0.285 | 1 | TEC_PT | ATP6V1D |
| 0.00047738 | 0.26209274 | 0.391 | 0.315 | 1 | TEC_PT | MMP24OS |
| 0.00051274 | -0.3615975 | 0.31 | 0.395 | 1 | TEC_PT | HIF1A |
| 0.00054047 | 0.27142406 | 0.438 | 0.375 | 1 | TEC_PT | BRK1 |
| 0.00058175 | -0.5001556 | 0.064 | 0.129 | 1 | TEC_PT | TFPI2 |
| 0.00058325 | 0.25904299 | 0.569 | 0.491 | 1 | TEC_PT | MTRNR2L8 |
| 0.00060765 | -0.8572081 | 0.205 | 0.277 | 1 | TEC_PT | HLA-DRB1 |
| 0.00064495 | -0.3880986 | 0.158 | 0.235 | 1 | TEC_PT | CDKN1C |
| 0.00076729 | -0.2698118 | 0.626 | 0.675 | 1 | TEC_PT | TALDO1 |
| 0.00077339 | 0.26500658 | 0.293 | 0.226 | 1 | TEC_PT | PSMB2 |
| 0.00077537 | 0.25690407 | 0.391 | 0.327 | 1 | TEC_PT | RNF7 |
| 0.00083662 | -0.787939 | 0.613 | 0.541 | 1 | TEC_PT | PLCG2 |
| 0.00087697 | -0.3073386 | 0.071 | 0.134 | 1 | TEC_PT | UCHL1 |
| 0.00093928 | -0.319805 | 0.343 | 0.409 | 1 | TEC_PT | CD59 |
| 0.0010287 | 0.25606989 | 0.801 | 0.785 | 1 | TEC_PT | BTF3 |
| 0.00121723 | 0.38658248 | 0.266 | 0.202 | 1 | TEC_PT | NEU1 |
| 0.00128004 | -0.2805878 | 0.084 | 0.148 | 1 | TEC_PT | RASD1 |
| 0.00147252 | -0.3716088 | 0.148 | 0.222 | 1 | TEC_PT | CCND1 |
| 0.00150136 | -0.3038613 | 0.101 | 0.164 | 1 | TEC_PT | ANKRD37 |
| 0.00217197 | -0.2697568 | 0.36 | 0.427 | 1 | TEC_PT | SPINT2 |
| 0.00217477 | -0.2916948 | 0.199 | 0.267 | 1 | TEC_PT | UGCG |
| 0.00314287 | -0.3793153 | 0.118 | 0.175 | 1 | TEC_PT | SPNS2 |
| 0.00315364 | -0.3467634 | 0.101 | 0.159 | 1 | TEC_PT | BPGM |
| 0.00322507 | 0.47807946 | 0.882 | 0.877 | 1 | TEC_PT | MT-ND3 |
| 0.00479927 | -0.2630908 | 0.172 | 0.235 | 1 | TEC_PT | TOB1 |
| 0.00524827 | 0.30347209 | 0.199 | 0.149 | 1 | TEC_PT | RRAGC |
| 0.00607822 | 0.25409447 | 0.411 | 0.364 | 1 | TEC_PT | ELOC |
| 0.00660788 | -0.7660712 | 0.088 | 0.137 | 1 | TEC_PT | DEFB1 |
| 0.00792426 | 0.54143791 | 0.32 | 0.266 | 1 | TEC_PT | DDIT3 |
| 0.00797933 | -0.295053 | 0.168 | 0.23 | 1 | TEC_PT | PPIF |
| 0.00848204 | 0.26701572 | 0.976 | 0.945 | 1 | TEC_PT | RPS19 |
| 0 | 4.01953169 | 0.877 | 0.013 | 0 | TEC_CD | AQP2 |
| 0 | 3.55402135 | 0.83 | 0.13 | 0 | TEC_CD | DEFB1 |
| 0 | 3.51417179 | 0.892 | 0.047 | 0 | TEC_CD | FXYD4 |

| | | | | | | |
|---|---|---|---|---|---|---|
| 0 | 3.41416914 | 0.411 | 0.016 | 0 | TEC_CD | CALB1 |
| 0 | 2.28365934 | 0.426 | 0.026 | 0 | TEC_CD | STC1 |
| 0 | 2.19463266 | 0.603 | 0.017 | 0 | TEC_CD | HSD11B2 |
| 0 | 1.95025143 | 0.488 | 0.051 | 0 | TEC_CD | PDK4 |
| 0 | 1.88659908 | 0.613 | 0.082 | 0 | TEC_CD | CA2 |
| 0 | 1.7777627 | 0.781 | 0.141 | 0 | TEC_CD | GSTM3 |
| 0 | 1.75579957 | 0.852 | 0.149 | 0 | TEC_CD | COBLL1 |
| 0 | 1.62856515 | 0.677 | 0.054 | 0 | TEC_CD | CDH16 |
| 0 | 1.29403477 | 0.53 | 0.021 | 0 | TEC_CD | PPARGC1A |
| 0 | 1.06351059 | 0.389 | 0.009 | 0 | TEC_CD | HMGCS2 |
| 0 | 1.00994589 | 0.325 | 0.024 | 0 | TEC_CD | TMEM52B |
| 0 | 0.99400786 | 0.441 | 0.019 | 0 | TEC_CD | TMEM213 |
| 0 | 0.97960446 | 0.32 | 0.006 | 0 | TEC_CD | PVALB |
| 0 | 0.91407527 | 0.387 | 0.013 | 0 | TEC_CD | AVPR2 |
| 0 | 0.87451888 | 0.527 | 0.057 | 0 | TEC_CD | MYH10 |
| 0 | 0.85939161 | 0.475 | 0.047 | 0 | TEC_CD | TBX2 |
| 0 | 0.85864772 | 0.451 | 0.034 | 0 | TEC_CD | FOLR1 |
| 0 | 0.85855809 | 0.485 | 0.047 | 0 | TEC_CD | ST6GAL1 |
| 0 | 0.84727038 | 0.505 | 0.028 | 0 | TEC_CD | AC019197.1 |
| 0 | 0.79487591 | 0.51 | 0.058 | 0 | TEC_CD | ALDH1A1 |
| 0 | 0.78793764 | 0.493 | 0.034 | 0 | TEC_CD | KCNJ16 |
| 0 | 0.77106372 | 0.426 | 0.034 | 0 | TEC_CD | NOV |
| 0 | 0.75524196 | 0.397 | 0.02 | 0 | TEC_CD | PAPPA |
| 0 | 0.69702298 | 0.483 | 0.045 | 0 | TEC_CD | SLC16A4 |
| 0 | 0.67539128 | 0.397 | 0.025 | 0 | TEC_CD | EMX1 |
| 0 | 0.6451985 | 0.246 | 0.009 | 0 | TEC_CD | KCNJ1 |
| 0 | 0.64415237 | 0.382 | 0.01 | 0 | TEC_CD | CLDN8 |
| 0 | 0.63634595 | 0.382 | 0.032 | 0 | TEC_CD | CYS1 |
| 0 | 0.62954839 | 0.438 | 0.035 | 0 | TEC_CD | NUPR2 |
| 0 | 0.59012999 | 0.335 | 0.021 | 0 | TEC_CD | OGDHL |
| 0 | 0.57748359 | 0.32 | 0.006 | 0 | TEC_CD | PTGER1 |
| 0 | 0.57186077 | 0.259 | 0.005 | 0 | TEC_CD | NR0B2 |
| 0 | 0.54501292 | 0.308 | 0.009 | 0 | TEC_CD | PIK3C2G |
| 0 | 0.5374544 | 0.342 | 0.02 | 0 | TEC_CD | GATA3-AS1 |
| 0 | 0.5353208 | 0.283 | 0.009 | 0 | TEC_CD | CLCNKB |
| 0 | 0.51270383 | 0.35 | 0.009 | 0 | TEC_CD | VEPH1 |
| 0 | 0.42175007 | 0.3 | 0.014 | 0 | TEC_CD | ELF5 |
| 0 | 0.41321378 | 0.296 | 0.019 | 0 | TEC_CD | GPRC5B |
| 0 | 0.39489289 | 0.227 | 0.011 | 0 | TEC_CD | PQLC2L |
| 0 | 0.3798849 | 0.224 | 0.009 | 0 | TEC_CD | LINC00982 |
| 0 | 0.37475156 | 0.217 | 0.001 | 0 | TEC_CD | ADH1C |
| 0 | 0.360418 | 0.207 | 0.001 | 0 | TEC_CD | TEX41 |
| 0 | 0.33856703 | 0.167 | 0.005 | 0 | TEC_CD | SLC14A2 |
| 0 | 0.33166507 | 0.234 | 0.01 | 0 | TEC_CD | FYB2 |
| 0 | 0.30689435 | 0.175 | 0.005 | 0 | TEC_CD | KCNK3 |
| 0 | 0.27199752 | 0.2 | 0.006 | 0 | TEC_CD | INPP5J |
| 1.408E-302 | 0.91350668 | 0.527 | 0.065 | 3.325E-298 | TEC_CD | L1CAM |
| 7.125E-297 | 1.28421808 | 0.495 | 0.06 | 1.683E-292 | TEC_CD | PIGR |
| 1.927E-294 | 0.34917034 | 0.229 | 0.012 | 4.551E-290 | TEC_CD | CKMT2 |
| 7.286E-294 | 0.55466023 | 0.369 | 0.032 | 1.721E-289 | TEC_CD | SPTBN2 |

| | | | | | | |
|---|---|---|---|---|---|---|
| 1.656E-293 | 0.96832472 | 0.611 | 0.088 | 3.912E-289 | TEC_CD | BCAM |
| 6.367E-292 | 0.86024238 | 0.438 | 0.047 | 1.504E-287 | TEC_CD | TOX3 |
| 1.065E-289 | 0.43978213 | 0.296 | 0.021 | 2.516E-285 | TEC_CD | NR1H4 |
| 5.096E-274 | 1.48070425 | 0.574 | 0.087 | 1.204E-269 | TEC_CD | CA12 |
| 3.016E-272 | 0.79739555 | 0.323 | 0.027 | 7.126E-268 | TEC_CD | FAM83F |
| 2.357E-271 | 0.87736432 | 0.544 | 0.077 | 5.569E-267 | TEC_CD | TWSG1 |
| 2.171E-269 | 0.30791039 | 0.207 | 0.011 | 5.13E-265 | TEC_CD | UGT3A1 |
| 5.185E-269 | 0.48417146 | 0.298 | 0.023 | 1.225E-264 | TEC_CD | AOC1 |
| 9.807E-266 | 0.26995064 | 0.187 | 0.009 | 2.317E-261 | TEC_CD | CLCNKA |
| 4.57E-265 | 0.54802123 | 0.217 | 0.012 | 1.079E-260 | TEC_CD | SERPINA5 |
| 5.217E-265 | 0.37470754 | 0.286 | 0.021 | 1.232E-260 | TEC_CD | PDE1A |
| 1.276E-262 | 2.31974282 | 0.901 | 0.275 | 3.013E-258 | TEC_CD | CLU |
| 1.666E-253 | 0.32726813 | 0.229 | 0.014 | 3.936E-249 | TEC_CD | SGK2 |
| 1.018E-251 | 0.27163165 | 0.2 | 0.011 | 2.405E-247 | TEC_CD | BMPR1B |
| 4.28E-250 | 1.05608041 | 0.672 | 0.125 | 1.011E-245 | TEC_CD | AIF1L |
| 1.731E-249 | 0.56306745 | 0.424 | 0.049 | 4.088E-245 | TEC_CD | BLNK |
| 7.773E-247 | 1.19653165 | 0.709 | 0.137 | 1.836E-242 | TEC_CD | PAX8 |
| 8.112E-243 | 0.56859125 | 0.394 | 0.044 | 1.916E-238 | TEC_CD | EMX2OS |
| 1.62E-240 | 0.26475454 | 0.163 | 0.007 | 3.827E-236 | TEC_CD | EYA4 |
| 4.952E-238 | 0.40400207 | 0.256 | 0.019 | 1.17E-233 | TEC_CD | MNS1 |
| 1.399E-237 | 0.25277404 | 0.165 | 0.008 | 3.304E-233 | TEC_CD | GABRA2 |
| 7.703E-235 | 0.60053591 | 0.409 | 0.048 | 1.82E-230 | TEC_CD | PAX2 |
| 5.351E-233 | 0.27183434 | 0.19 | 0.011 | 1.264E-228 | TEC_CD | LCN12 |
| 1.997E-230 | 0.65693486 | 0.421 | 0.052 | 4.718E-226 | TEC_CD | SULT1C2 |
| 2.462E-226 | 0.93193685 | 0.658 | 0.126 | 5.817E-222 | TEC_CD | SCIN |
| 3.989E-225 | 0.90338551 | 0.458 | 0.064 | 9.424E-221 | TEC_CD | BHLHE41 |
| 2.114E-218 | 1.50630918 | 0.857 | 0.229 | 4.993E-214 | TEC_CD | EPCAM |
| 5.96E-212 | 0.41856027 | 0.268 | 0.023 | 1.408E-207 | TEC_CD | ARHGAP6 |
| 1.161E-210 | 1.49270695 | 0.805 | 0.223 | 2.742E-206 | TEC_CD | CYB5A |
| 1.162E-209 | 0.51727725 | 0.347 | 0.039 | 2.746E-205 | TEC_CD | GATA2 |
| 4.686E-209 | 0.36739953 | 0.16 | 0.008 | 1.107E-204 | TEC_CD | MEST |
| 5.68E-209 | 0.29493331 | 0.202 | 0.013 | 1.342E-204 | TEC_CD | FRAS1 |
| 8.747E-207 | 1.88083123 | 0.746 | 0.178 | 2.066E-202 | TEC_CD | AQP3 |
| 1.886E-205 | 0.41628747 | 0.328 | 0.035 | 4.455E-201 | TEC_CD | PLLP |
| 7.681E-205 | 0.36958279 | 0.254 | 0.022 | 1.814E-200 | TEC_CD | SMIM24 |
| 2.783E-204 | 0.3908506 | 0.254 | 0.022 | 6.574E-200 | TEC_CD | PALM3 |
| 3.762E-204 | 0.34411063 | 0.209 | 0.015 | 8.887E-200 | TEC_CD | ATP6V0A4 |
| 1.135E-203 | 0.63126755 | 0.453 | 0.067 | 2.681E-199 | TEC_CD | TPD52L1 |
| 1.905E-201 | 0.35013697 | 0.202 | 0.014 | 4.501E-197 | TEC_CD | AC025263.1 |
| 9.247E-200 | 0.46586707 | 0.219 | 0.017 | 2.184E-195 | TEC_CD | AC129507.4 |
| 4.312E-195 | 0.25107001 | 0.182 | 0.012 | 1.019E-190 | TEC_CD | SLC9A4 |
| 1.004E-194 | 0.64373273 | 0.303 | 0.033 | 2.372E-190 | TEC_CD | AC092803.2 |
| 4.106E-194 | 0.34277149 | 0.229 | 0.019 | 9.701E-190 | TEC_CD | COLCA1 |
| 7.983E-194 | 0.77879263 | 0.453 | 0.07 | 1.886E-189 | TEC_CD | KITLG |
| 1.203E-191 | 0.9051687 | 0.557 | 0.108 | 2.842E-187 | TEC_CD | ADGRG1 |
| 1.719E-188 | 0.42173629 | 0.241 | 0.021 | 4.06E-184 | TEC_CD | MXRA5 |
| 3.087E-186 | 0.36913206 | 0.3 | 0.033 | 7.293E-182 | TEC_CD | SHMT1 |
| 7E-182 | 1.56052979 | 0.512 | 0.098 | 1.653E-177 | TEC_CD | RCAN1 |
| 2.517E-181 | 0.54368748 | 0.438 | 0.069 | 5.946E-177 | TEC_CD | CRYZ |
| 5.394E-177 | 1.14064501 | 0.392 | 0.059 | 1.274E-172 | TEC_CD | NEURL3 |

| | | | | | | |
|---|---|---|---|---|---|---|
| 6.561E-177 | 0.48112261 | 0.372 | 0.051 | 1.55E-172 | TEC_CD | CCDC198 |
| 1.589E-176 | 0.58581213 | 0.436 | 0.07 | 3.753E-172 | TEC_CD | SLC25A25 |
| 9.074E-174 | 0.48026369 | 0.382 | 0.055 | 2.144E-169 | TEC_CD | PRKAA2 |
| 1.254E-170 | 1.15137254 | 0.606 | 0.141 | 2.962E-166 | TEC_CD | NCOA7 |
| 3.121E-170 | 0.26777494 | 0.19 | 0.015 | 7.372E-166 | TEC_CD | IQUB |
| 8.854E-170 | 0.35454074 | 0.256 | 0.026 | 2.092E-165 | TEC_CD | SIM1 |
| 1.646E-168 | 0.53250641 | 0.357 | 0.049 | 3.889E-164 | TEC_CD | PTGER3 |
| 3.285E-168 | 0.78112957 | 0.446 | 0.077 | 7.76E-164 | TEC_CD | PCSK1N |
| 4.308E-168 | 0.4395254 | 0.239 | 0.023 | 1.018E-163 | TEC_CD | SCNN1G |
| 9.187E-168 | 0.93726784 | 0.603 | 0.14 | 2.17E-163 | TEC_CD | GPRC5C |
| 2.659E-167 | 0.71256138 | 0.466 | 0.083 | 6.281E-163 | TEC_CD | SLC25A29 |
| 1.87E-166 | 0.75154242 | 0.53 | 0.108 | 4.417E-162 | TEC_CD | RHOBTB3 |
| 2.021E-165 | 0.61028917 | 0.431 | 0.071 | 4.775E-161 | TEC_CD | EMX2 |
| 4.71E-165 | 0.5561638 | 0.303 | 0.038 | 1.113E-160 | TEC_CD | ABCB1 |
| 1.116E-164 | 1.76649651 | 0.857 | 0.361 | 2.636E-160 | TEC_CD | RALBP1 |
| 8.565E-163 | 0.62372309 | 0.431 | 0.073 | 2.023E-158 | TEC_CD | TFCP2L1 |
| 7.894E-161 | 0.50196677 | 0.291 | 0.035 | 1.865E-156 | TEC_CD | LHX1 |
| 3.268E-160 | 0.27026642 | 0.18 | 0.014 | 7.72E-156 | TEC_CD | SLC4A4 |
| 3.692E-160 | 0.51786542 | 0.399 | 0.063 | 8.722E-156 | TEC_CD | ANK3 |
| 1.77E-159 | 0.60297074 | 0.419 | 0.07 | 4.182E-155 | TEC_CD | BICC1 |
| 8.445E-159 | 0.71799154 | 0.382 | 0.06 | 1.995E-154 | TEC_CD | NUAK2 |
| 3.118E-158 | 0.67025435 | 0.564 | 0.122 | 7.365E-154 | TEC_CD | SELENOP |
| 6.66E-158 | 0.27785676 | 0.214 | 0.02 | 1.573E-153 | TEC_CD | HOXB-AS3 |
| 2.544E-157 | 0.37086552 | 0.286 | 0.034 | 6.011E-153 | TEC_CD | TCEA3 |
| 2.81E-157 | 0.29023021 | 0.224 | 0.022 | 6.639E-153 | TEC_CD | TBX2-AS1 |
| 4.146E-153 | 0.26250494 | 0.214 | 0.02 | 9.793E-149 | TEC_CD | MPPED2 |
| 1.374E-152 | 0.62686844 | 0.52 | 0.11 | 3.246E-148 | TEC_CD | CHPT1 |
| 3.57E-152 | 0.50907297 | 0.389 | 0.064 | 8.434E-148 | TEC_CD | SLC19A2 |
| 9.458E-152 | 0.51106055 | 0.315 | 0.043 | 2.234E-147 | TEC_CD | CADPS2 |
| 1.153E-151 | 0.97286905 | 0.567 | 0.137 | 2.724E-147 | TEC_CD | BEX3 |
| 2.547E-151 | 0.93306846 | 0.702 | 0.213 | 6.018E-147 | TEC_CD | NDUFA5 |
| 5.75E-151 | 0.48342373 | 0.298 | 0.039 | 1.358E-146 | TEC_CD | HOXD8 |
| 8.639E-151 | 0.25697792 | 0.204 | 0.019 | 2.041E-146 | TEC_CD | ACSM3 |
| 1.062E-150 | 1.7150662 | 0.933 | 0.432 | 2.508E-146 | TEC_CD | ATP1B1 |
| 1.211E-150 | 0.51469236 | 0.352 | 0.054 | 2.861E-146 | TEC_CD | PLEKHG3 |
| 1.479E-150 | 0.30334857 | 0.229 | 0.024 | 3.493E-146 | TEC_CD | ALDH4A1 |
| 2.024E-150 | 0.8116806 | 0.527 | 0.119 | 4.781E-146 | TEC_CD | PDHA1 |
| 1.585E-149 | 0.46028283 | 0.33 | 0.047 | 3.744E-145 | TEC_CD | GMNN |
| 7.395E-149 | 0.64074087 | 0.357 | 0.057 | 1.747E-144 | TEC_CD | HS6ST1 |
| 4.05E-148 | 0.65706956 | 0.473 | 0.094 | 9.567E-144 | TEC_CD | CAPS |
| 3.045E-147 | 0.75785156 | 0.502 | 0.105 | 7.194E-143 | TEC_CD | GATA3 |
| 4.086E-147 | 2.17639184 | 0.842 | 0.34 | 9.651E-143 | TEC_CD | GDF15 |
| 3.931E-146 | 0.31466408 | 0.204 | 0.02 | 9.287E-142 | TEC_CD | RASSF9 |
| 7.554E-146 | 0.26657512 | 0.217 | 0.022 | 1.785E-141 | TEC_CD | EPHX2 |
| 1.231E-145 | 0.31802752 | 0.187 | 0.017 | 2.907E-141 | TEC_CD | RORC |
| 3.207E-145 | 0.38950495 | 0.286 | 0.037 | 7.577E-141 | TEC_CD | FAM110B |
| 2.086E-144 | 0.66414109 | 0.53 | 0.118 | 4.928E-140 | TEC_CD | SNRPN |
| 3.577E-144 | 0.75285849 | 0.473 | 0.098 | 8.449E-140 | TEC_CD | ARG2 |
| 9.386E-144 | 0.3885744 | 0.187 | 0.017 | 2.217E-139 | TEC_CD | THY1 |
| 1.524E-143 | 0.31274136 | 0.224 | 0.024 | 3.601E-139 | TEC_CD | FHOD3 |

| | | | | | | |
|---|---|---|---|---|---|---|
| 1.153E-142 | 0.70708755 | 0.404 | 0.073 | 2.723E-138 | TEC_CD | ZNF503 |
| 1.197E-142 | 0.37355149 | 0.313 | 0.045 | 2.827E-138 | TEC_CD | SGPP1 |
| 2.048E-142 | 0.40634347 | 0.328 | 0.049 | 4.838E-138 | TEC_CD | RAB17 |
| 2.747E-141 | 0.77406536 | 0.631 | 0.164 | 6.489E-137 | TEC_CD | SLC38A1 |
| 7.361E-141 | 0.51642465 | 0.399 | 0.07 | 1.739E-136 | TEC_CD | MECOM |
| 1.352E-140 | 0.85689693 | 0.527 | 0.126 | 3.195E-136 | TEC_CD | SCPEP1 |
| 1.976E-140 | 0.60666836 | 0.5 | 0.109 | 4.667E-136 | TEC_CD | CISD1 |
| 2.504E-140 | 0.25573764 | 0.187 | 0.017 | 5.915E-136 | TEC_CD | CLYBL |
| 4.76E-140 | 0.40719948 | 0.195 | 0.019 | 1.124E-135 | TEC_CD | AC026250.1 |
| 7.191E-140 | 0.5279008 | 0.291 | 0.039 | 1.699E-135 | TEC_CD | MGP |
| 9.055E-140 | 0.47300098 | 0.357 | 0.058 | 2.139E-135 | TEC_CD | SHROOM3 |
| 1.259E-138 | 0.30957232 | 0.249 | 0.029 | 2.975E-134 | TEC_CD | SOX6 |
| 2.554E-137 | 0.46070274 | 0.379 | 0.065 | 6.032E-133 | TEC_CD | NFIB |
| 4.139E-137 | 0.35505924 | 0.298 | 0.042 | 9.777E-133 | TEC_CD | ZNF704 |
| 4.375E-137 | 0.41383535 | 0.342 | 0.055 | 1.034E-132 | TEC_CD | IVD |
| 2.686E-136 | 0.26650271 | 0.224 | 0.025 | 6.345E-132 | TEC_CD | MID1 |
| 1.911E-135 | 0.31761975 | 0.241 | 0.029 | 4.514E-131 | TEC_CD | DZIP3 |
| 1.209E-134 | 0.49883578 | 0.382 | 0.067 | 2.856E-130 | TEC_CD | GNG11 |
| 2.014E-134 | 0.27689575 | 0.19 | 0.018 | 4.757E-130 | TEC_CD | MUC15 |
| 2.798E-134 | 0.32772291 | 0.259 | 0.033 | 6.61E-130 | TEC_CD | CDKL2 |
| 4.411E-134 | 1.28082064 | 0.771 | 0.289 | 1.042E-129 | TEC_CD | CHCHD10 |
| 7.498E-133 | 0.48918873 | 0.382 | 0.069 | 1.771E-128 | TEC_CD | COQ9 |
| 2.864E-132 | 0.35163061 | 0.273 | 0.037 | 6.766E-128 | TEC_CD | MAP3K21 |
| 5.683E-132 | 0.36945022 | 0.234 | 0.028 | 1.343E-127 | TEC_CD | GLIS2 |
| 7.502E-131 | 0.82500952 | 0.564 | 0.151 | 1.772E-126 | TEC_CD | MPC1 |
| 2.889E-130 | 0.49190091 | 0.278 | 0.039 | 6.825E-126 | TEC_CD | SMIM6 |
| 5.556E-130 | 0.35767059 | 0.276 | 0.038 | 1.313E-125 | TEC_CD | CTSF |
| 1.35E-129 | 0.56080065 | 0.451 | 0.094 | 3.188E-125 | TEC_CD | FERMT2 |
| 8.647E-129 | 0.5358863 | 0.397 | 0.076 | 2.043E-124 | TEC_CD | SLC16A7 |
| 9.633E-129 | 0.36506267 | 0.244 | 0.031 | 2.276E-124 | TEC_CD | EDNRB |
| 1.288E-127 | 0.42587231 | 0.36 | 0.063 | 3.042E-123 | TEC_CD | KCNQ1 |
| 1.644E-127 | 0.28304108 | 0.227 | 0.027 | 3.883E-123 | TEC_CD | LINC02532 |
| 2.239E-127 | 0.26432002 | 0.167 | 0.015 | 5.289E-123 | TEC_CD | CREB3L1 |
| 3.145E-127 | 0.5766712 | 0.399 | 0.077 | 7.429E-123 | TEC_CD | RETREG1 |
| 5.646E-127 | 0.32844049 | 0.278 | 0.039 | 1.334E-122 | TEC_CD | BHMT2 |
| 5.912E-127 | 0.27938154 | 0.209 | 0.023 | 1.397E-122 | TEC_CD | PFKM |
| 8.183E-127 | 0.59647229 | 0.522 | 0.124 | 1.933E-122 | TEC_CD | HACD3 |
| 1.818E-126 | 0.70011645 | 0.576 | 0.155 | 4.295E-122 | TEC_CD | FDX1 |
| 1.855E-126 | 0.29294617 | 0.232 | 0.028 | 4.382E-122 | TEC_CD | GSTM4 |
| 5.047E-126 | 0.62834057 | 0.399 | 0.078 | 1.192E-121 | TEC_CD | RAPGEF3 |
| 6.185E-126 | 1.18803995 | 0.613 | 0.181 | 1.461E-121 | TEC_CD | SEPHS2 |
| 7.983E-126 | 0.25903457 | 0.212 | 0.024 | 1.886E-121 | TEC_CD | WDR72 |
| 1.575E-125 | 0.59467786 | 0.431 | 0.091 | 3.721E-121 | TEC_CD | SPIRE1 |
| 6.677E-125 | -5.3736012 | 0.111 | 0.733 | 1.577E-120 | TEC_CD | S100A9 |
| 7.783E-125 | 0.8905751 | 0.672 | 0.222 | 1.839E-120 | TEC_CD | COX20 |
| 2.363E-124 | 0.73479371 | 0.645 | 0.197 | 5.582E-120 | TEC_CD | SNX2 |
| 2.576E-124 | 0.53257733 | 0.49 | 0.113 | 6.084E-120 | TEC_CD | CBX5 |
| 2.606E-124 | 0.66829983 | 0.591 | 0.155 | 6.155E-120 | TEC_CD | SELENOM |
| 5.694E-123 | 0.65452962 | 0.645 | 0.178 | 1.345E-118 | TEC_CD | CDH1 |
| 1.606E-122 | 0.4258799 | 0.251 | 0.034 | 3.795E-118 | TEC_CD | LGALSL |

| | | | | | | |
|---|---|---|---|---|---|---|
| 8.558E-122 | 0.38316034 | 0.236 | 0.03 | 2.022E-117 | TEC_CD | AL139274.2 |
| 1.668E-121 | 0.38371965 | 0.32 | 0.052 | 3.94E-117 | TEC_CD | PPP1R9A |
| 1.486E-120 | 0.97328341 | 0.845 | 0.344 | 3.509E-116 | TEC_CD | PEBP1 |
| 3.456E-120 | 0.36867285 | 0.259 | 0.036 | 8.165E-116 | TEC_CD | HNF1B |
| 4.22E-120 | 1.01011678 | 0.845 | 0.345 | 9.969E-116 | TEC_CD | GLS |
| 1.159E-119 | 0.79648346 | 0.527 | 0.142 | 2.737E-115 | TEC_CD | COA3 |
| 1.322E-119 | 0.67732002 | 0.493 | 0.119 | 3.123E-115 | TEC_CD | RAB3IP |
| 3.115E-119 | 0.38766235 | 0.303 | 0.049 | 7.358E-115 | TEC_CD | DCPS |
| 1.639E-117 | 0.32789886 | 0.256 | 0.036 | 3.871E-113 | TEC_CD | TMEM136 |
| 8.127E-117 | 0.38211598 | 0.337 | 0.06 | 1.92E-112 | TEC_CD | ERMP1 |
| 1.578E-116 | 0.72077084 | 0.32 | 0.057 | 3.728E-112 | TEC_CD | PRPS2 |
| 4.316E-116 | 0.34506028 | 0.281 | 0.043 | 1.02E-111 | TEC_CD | THRA |
| 4.505E-116 | 0.42368837 | 0.372 | 0.072 | 1.064E-111 | TEC_CD | DDAH1 |
| 8.003E-116 | 0.53390494 | 0.406 | 0.086 | 1.891E-111 | TEC_CD | ACAT1 |
| 4.219E-115 | 0.26686202 | 0.175 | 0.018 | 9.966E-111 | TEC_CD | RASL11A |
| 8.703E-115 | 0.41664273 | 0.352 | 0.065 | 2.056E-110 | TEC_CD | ERBB2 |
| 1.167E-114 | 0.43716431 | 0.14 | 0.012 | 2.757E-110 | TEC_CD | LINC02154 |
| 1.44E-114 | 0.29435923 | 0.264 | 0.039 | 3.401E-110 | TEC_CD | TMEM116 |
| 2.734E-114 | 0.64026132 | 0.544 | 0.148 | 6.457E-110 | TEC_CD | ECH1 |
| 4.004E-114 | 0.90341327 | 0.766 | 0.295 | 9.459E-110 | TEC_CD | NUCKS1 |
| 5.212E-114 | 0.53142294 | 0.323 | 0.058 | 1.231E-109 | TEC_CD | KLF11 |
| 8.959E-114 | 0.25035696 | 0.108 | 0.007 | 2.116E-109 | TEC_CD | FIBIN |
| 8.781E-113 | 0.86857419 | 0.618 | 0.186 | 2.074E-108 | TEC_CD | LIMA1 |
| 1.878E-112 | 0.88344915 | 0.776 | 0.269 | 4.437E-108 | TEC_CD | MAL2 |
| 5.002E-111 | 0.64278246 | 0.202 | 0.024 | 1.182E-106 | TEC_CD | LTF |
| 6.189E-111 | 0.82132522 | 0.389 | 0.082 | 1.462E-106 | TEC_CD | IFIT3 |
| 2.995E-110 | 0.52003859 | 0.397 | 0.084 | 7.074E-106 | TEC_CD | CRIP2 |
| 5.581E-110 | -4.4623418 | 0.084 | 0.677 | 1.318E-105 | TEC_CD | S100A8 |
| 1.693E-109 | 0.35936337 | 0.296 | 0.049 | 4E-105 | TEC_CD | PKHD1 |
| 2.019E-109 | 0.31154798 | 0.268 | 0.042 | 4.769E-105 | TEC_CD | PCCB |
| 2.094E-109 | 0.54848783 | 0.392 | 0.085 | 4.947E-105 | TEC_CD | RAB11FIP2 |
| 2.279E-109 | 0.25133971 | 0.143 | 0.013 | 5.383E-105 | TEC_CD | DOK7 |
| 5.18E-109 | 0.32814881 | 0.219 | 0.029 | 1.224E-104 | TEC_CD | FGFR2 |
| 1.301E-107 | 0.37526996 | 0.256 | 0.039 | 3.073E-103 | TEC_CD | VAV2 |
| 1.596E-107 | 0.57185735 | 0.532 | 0.146 | 3.77E-103 | TEC_CD | SINHCAF |
| 3.283E-106 | 1.10627741 | 0.948 | 0.536 | 7.756E-102 | TEC_CD | CD24 |
| 4.387E-106 | 0.82555275 | 0.5 | 0.14 | 1.036E-101 | TEC_CD | AFF1 |
| 1.411E-105 | 0.26222703 | 0.2 | 0.025 | 3.333E-101 | TEC_CD | HACD1 |
| 1.792E-105 | 0.29253537 | 0.224 | 0.031 | 4.233E-101 | TEC_CD | SETBP1 |
| 6.852E-105 | 0.50153102 | 0.337 | 0.067 | 1.619E-100 | TEC_CD | BCAR3 |
| 9.421E-105 | -1.9604706 | 0.458 | 0.863 | 2.225E-100 | TEC_CD | SH3BGRL3 |
| 1.483E-104 | 0.28565159 | 0.246 | 0.037 | 3.504E-100 | TEC_CD | TSPAN12 |
| 8.13E-104 | 0.2907221 | 0.244 | 0.036 | 1.921E-99 | TEC_CD | ARMCX2 |
| 4.287E-103 | 0.45378783 | 0.355 | 0.072 | 1.0128E-98 | TEC_CD | PLPP3 |
| 1.14E-102 | 0.50156142 | 0.495 | 0.128 | 2.6926E-98 | TEC_CD | DMKN |
| 1.935E-102 | 0.62930993 | 0.51 | 0.142 | 4.5715E-98 | TEC_CD | LLGL2 |
| 2.321E-102 | 0.54443978 | 0.411 | 0.095 | 5.4832E-98 | TEC_CD | SLC25A4 |
| 4.658E-102 | 0.63663504 | 0.416 | 0.1 | 1.1004E-97 | TEC_CD | KIAA1522 |
| 1.09E-101 | 0.38231455 | 0.337 | 0.066 | 2.5739E-97 | TEC_CD | EPB41L1 |
| 1.216E-101 | 0.40511553 | 0.281 | 0.048 | 2.8734E-97 | TEC_CD | PRRG1 |

| | | | | | | |
|---|---|---|---|---|---|---|
| 1.386E-101 | 1.0474897 | 0.342 | 0.072 | 3.2747E-97 | TEC_CD | TIMP3 |
| 1.606E-101 | 0.28219788 | 0.234 | 0.035 | 3.7939E-97 | TEC_CD | FAM229B |
| 2.806E-101 | 0.55154579 | 0.35 | 0.073 | 6.6284E-97 | TEC_CD | MTMR10 |
| 3.791E-101 | 0.46255867 | 0.133 | 0.012 | 8.9547E-97 | TEC_CD | KNG1 |
| 7.292E-101 | 0.7387337 | 0.704 | 0.268 | 1.7227E-96 | TEC_CD | MINOS1 |
| 8.332E-101 | 0.79831706 | 0.603 | 0.193 | 1.9684E-96 | TEC_CD | PTGR1 |
| 1.231E-100 | 0.89778371 | 0.579 | 0.193 | 2.9083E-96 | TEC_CD | ATP5MC1 |
| 1.866E-100 | 0.26962299 | 0.224 | 0.032 | 4.4073E-96 | TEC_CD | SIAE |
| 2.259E-100 | 0.41280543 | 0.345 | 0.07 | 5.337E-96 | TEC_CD | BDH2 |
| 3.582E-99 | 0.54958331 | 0.525 | 0.145 | 8.4612E-95 | TEC_CD | SCCPDH |
| 3.595E-99 | 0.35585674 | 0.3 | 0.055 | 8.4929E-95 | TEC_CD | KIZ |
| 3.925E-99 | 0.52721159 | 0.404 | 0.093 | 9.2718E-95 | TEC_CD | EFNA1 |
| 9.352E-99 | 0.94967522 | 0.712 | 0.256 | 2.2092E-94 | TEC_CD | MUC1 |
| 1.318E-98 | 0.28125757 | 0.217 | 0.031 | 3.1135E-94 | TEC_CD | ZNF618 |
| 3.0036E-98 | 0.40625934 | 0.35 | 0.073 | 7.0955E-94 | TEC_CD | PFN2 |
| 5.7978E-98 | 0.52727816 | 0.493 | 0.133 | 1.3696E-93 | TEC_CD | SCNN1A |
| 9.0821E-98 | 0.53252632 | 0.515 | 0.144 | 2.1455E-93 | TEC_CD | SVIP |
| 1.1567E-97 | 0.52586276 | 0.453 | 0.119 | 2.7324E-93 | TEC_CD | NDUFA8 |
| 2.2352E-97 | 0.43004292 | 0.401 | 0.094 | 5.2801E-93 | TEC_CD | MRPL19 |
| 4.9109E-97 | 0.27886069 | 0.219 | 0.032 | 1.1601E-92 | TEC_CD | LINC01116 |
| 1.1745E-96 | -3.9168686 | 0.15 | 0.681 | 2.7744E-92 | TEC_CD | TIMP1 |
| 2.6119E-96 | 0.46635188 | 0.421 | 0.102 | 6.17E-92 | TEC_CD | GOT1 |
| 7.3189E-96 | 1.03512761 | 0.7 | 0.279 | 1.7289E-91 | TEC_CD | LDHB |
| 1.0382E-95 | 1.2194984 | 0.544 | 0.177 | 2.4526E-91 | TEC_CD | PFKFB3 |
| 1.0845E-95 | 0.58852022 | 0.522 | 0.156 | 2.562E-91 | TEC_CD | GLRX5 |
| 1.9283E-95 | 0.59419417 | 0.623 | 0.204 | 4.5551E-91 | TEC_CD | TCEAL4 |
| 2.308E-95 | 0.45332225 | 0.347 | 0.074 | 5.4522E-91 | TEC_CD | MTR |
| 2.5566E-95 | 0.4687502 | 0.431 | 0.109 | 6.0394E-91 | TEC_CD | ACO2 |
| 3.2497E-95 | 0.36570101 | 0.273 | 0.048 | 7.6767E-91 | TEC_CD | SLC16A5 |
| 6.275E-95 | 1.47203245 | 0.475 | 0.143 | 1.4824E-90 | TEC_CD | FAM102A |
| 1.2418E-94 | 0.47138503 | 0.414 | 0.102 | 2.9334E-90 | TEC_CD | AFG3L2 |
| 1.4064E-94 | 0.52035067 | 0.515 | 0.149 | 3.3223E-90 | TEC_CD | SCOC |
| 1.4163E-94 | 0.40125156 | 0.214 | 0.031 | 3.3457E-90 | TEC_CD | COX7A1 |
| 2.9596E-94 | 0.3421048 | 0.333 | 0.067 | 6.9914E-90 | TEC_CD | RASSF8 |
| 4.352E-94 | 0.66265083 | 0.579 | 0.183 | 1.0281E-89 | TEC_CD | PAWR |
| 1.2941E-93 | 0.48367979 | 0.325 | 0.066 | 3.0572E-89 | TEC_CD | THSD7A |
| 1.8501E-93 | 0.82516252 | 0.68 | 0.264 | 4.3705E-89 | TEC_CD | BAG1 |
| 4.4868E-93 | 0.68946699 | 0.717 | 0.283 | 1.0599E-88 | TEC_CD | NDUFC1 |
| 4.6545E-93 | 0.69508854 | 0.717 | 0.283 | 1.0995E-88 | TEC_CD | ACADVL |
| 6.7141E-93 | 0.51358888 | 0.389 | 0.093 | 1.5861E-88 | TEC_CD | RBBP8 |
| 2.2405E-92 | 2.34338214 | 0.451 | 0.134 | 5.2927E-88 | TEC_CD | NUPR1 |
| 2.7171E-92 | 0.34282147 | 0.222 | 0.034 | 6.4187E-88 | TEC_CD | TCEAL2 |
| 5.1356E-92 | -2.0489604 | 0.175 | 0.714 | 1.2132E-87 | TEC_CD | ANXA1 |
| 8.084E-92 | 1.02300618 | 0.687 | 0.262 | 1.9097E-87 | TEC_CD | DDIT3 |
| 8.8836E-92 | 0.25889431 | 0.121 | 0.011 | 2.0986E-87 | TEC_CD | AC007906.2 |
| 9.2198E-92 | 0.28931044 | 0.229 | 0.036 | 2.178E-87 | TEC_CD | SLC25A30 |
| 1.792E-91 | 0.49862604 | 0.483 | 0.131 | 4.2331E-87 | TEC_CD | COL18A1 |
| 2.5115E-91 | 0.55936685 | 0.461 | 0.125 | 5.9329E-87 | TEC_CD | BEX2 |
| 2.6525E-91 | 1.36735651 | 0.687 | 0.244 | 6.266E-87 | TEC_CD | CRYAB |
| 5.7444E-91 | 0.3180082 | 0.286 | 0.054 | 1.357E-86 | TEC_CD | PCCA |

| | | | | | | |
|---|---|---|---|---|---|---|
| 6.782E-91 | -1.6382529 | 0.865 | 0.971 | 1.6021E-86 | TEC_CD | TMSB10 |
| 2.0308E-90 | 0.578252 | 0.586 | 0.196 | 4.7975E-86 | TEC_CD | MDH1 |
| 5.9308E-90 | 0.29453807 | 0.182 | 0.024 | 1.401E-85 | TEC_CD | DEGS2 |
| 8.4428E-90 | 0.54453948 | 0.591 | 0.193 | 1.9944E-85 | TEC_CD | TMED4 |
| 2.0773E-89 | -1.488978 | 0.266 | 0.742 | 4.9072E-85 | TEC_CD | CTSB |
| 3.8976E-89 | 0.4413324 | 0.347 | 0.078 | 9.2073E-85 | TEC_CD | ECI2 |
| 1.8248E-88 | 1.0198777 | 0.818 | 0.433 | 4.3108E-84 | TEC_CD | CYSTM1 |
| 3.4372E-88 | 0.39112087 | 0.315 | 0.065 | 8.1198E-84 | TEC_CD | HOXB6 |
| 4.5413E-88 | 0.65189045 | 0.601 | 0.21 | 1.0728E-83 | TEC_CD | BTG3 |
| 8.9097E-88 | 0.39593896 | 0.268 | 0.05 | 2.1047E-83 | TEC_CD | COL4A4 |
| 1.3034E-87 | -2.8732197 | 0.047 | 0.575 | 3.079E-83 | TEC_CD | SRGN |
| 1.6109E-87 | 0.26456721 | 0.212 | 0.032 | 3.8055E-83 | TEC_CD | NAALADL2 |
| 1.9673E-87 | 0.57378652 | 0.473 | 0.138 | 4.6474E-83 | TEC_CD | NFIC |
| 2.5284E-87 | -2.2382264 | 0.094 | 0.619 | 5.9729E-83 | TEC_CD | C15orf48 |
| 3.1303E-87 | 0.54437704 | 0.426 | 0.116 | 7.3948E-83 | TEC_CD | NDUFV1 |
| 4.2317E-87 | 0.50423883 | 0.485 | 0.134 | 9.9965E-83 | TEC_CD | DCDC2 |
| 6.2633E-87 | 0.48044308 | 0.372 | 0.09 | 1.4796E-82 | TEC_CD | PHYH |
| 1.2538E-86 | 0.42073075 | 0.374 | 0.09 | 2.9618E-82 | TEC_CD | TBC1D9 |
| 1.9989E-86 | 0.27061335 | 0.204 | 0.031 | 4.722E-82 | TEC_CD | TDRP |
| 2.4521E-86 | 0.30189393 | 0.261 | 0.048 | 5.7925E-82 | TEC_CD | CTDSPL |
| 4.1878E-86 | 0.50170447 | 0.421 | 0.113 | 9.8929E-82 | TEC_CD | VEGFB |
| 7.0308E-86 | 0.44531317 | 0.365 | 0.087 | 1.6609E-81 | TEC_CD | HOXB2 |
| 8.7476E-86 | 0.25920922 | 0.222 | 0.036 | 2.0664E-81 | TEC_CD | C6orf136 |
| 1.1153E-85 | 0.69312799 | 0.69 | 0.278 | 2.6346E-81 | TEC_CD | ISCU |
| 1.2096E-85 | 0.42112241 | 0.389 | 0.096 | 2.8574E-81 | TEC_CD | TRIM2 |
| 6.3689E-85 | 0.2738783 | 0.236 | 0.04 | 1.5045E-80 | TEC_CD | RGL3 |
| 1.2011E-84 | 0.35627365 | 0.305 | 0.064 | 2.8374E-80 | TEC_CD | NDUFAF4 |
| 1.242E-84 | 0.92887107 | 0.749 | 0.351 | 2.9341E-80 | TEC_CD | CYCS |
| 1.4041E-84 | 0.45644349 | 0.34 | 0.078 | 3.3169E-80 | TEC_CD | STAM |
| 1.5249E-84 | 0.65929755 | 0.926 | 0.532 | 3.6022E-80 | TEC_CD | WFDC2 |
| 1.7944E-84 | 0.46824151 | 0.384 | 0.097 | 4.2388E-80 | TEC_CD | MCRIP2 |
| 3.6331E-84 | 0.5311968 | 0.483 | 0.143 | 8.5825E-80 | TEC_CD | ALDH2 |
| 3.3151E-83 | 0.92309696 | 0.357 | 0.092 | 7.8312E-79 | TEC_CD | PPP1R3B |
| 4.4206E-83 | 0.61545905 | 0.404 | 0.108 | 1.0443E-78 | TEC_CD | SLC2A1 |
| 5.0731E-83 | 0.35177031 | 0.3 | 0.063 | 1.1984E-78 | TEC_CD | HIBCH |
| 6.7678E-83 | 0.63468426 | 0.461 | 0.141 | 1.5988E-78 | TEC_CD | SUCLG1 |
| 2.0871E-82 | 0.32224517 | 0.291 | 0.06 | 4.9304E-78 | TEC_CD | NME7 |
| 3.1966E-82 | 0.25723894 | 0.202 | 0.031 | 7.5514E-78 | TEC_CD | TNFRSF14-AS1 |
| 3.7838E-82 | 0.57060521 | 0.463 | 0.137 | 8.9384E-78 | TEC_CD | LPIN2 |
| 4.2308E-82 | 0.50257416 | 0.426 | 0.117 | 9.9945E-78 | TEC_CD | MYO10 |
| 4.6184E-82 | -1.0880809 | 0.532 | 0.872 | 1.091E-77 | TEC_CD | PABPC1 |
| 6.9378E-82 | 0.34344321 | 0.298 | 0.063 | 1.6389E-77 | TEC_CD | HIBADH |
| 7.4061E-82 | 0.73979268 | 0.65 | 0.243 | 1.7496E-77 | TEC_CD | SPTBN1 |
| 1.0631E-81 | 0.41071516 | 0.352 | 0.084 | 2.5114E-77 | TEC_CD | SCRN1 |
| 1.7442E-81 | 1.19231283 | 0.291 | 0.063 | 4.1204E-77 | TEC_CD | RSAD2 |
| 1.8243E-81 | 0.25225879 | 0.212 | 0.034 | 4.3094E-77 | TEC_CD | RPP25 |
| 1.9506E-81 | 0.73559586 | 0.488 | 0.153 | 4.6079E-77 | TEC_CD | FNIP2 |
| 4.5353E-81 | 0.48923637 | 0.47 | 0.133 | 1.0714E-76 | TEC_CD | ARHGAP29 |
| 1.0713E-80 | 0.26334035 | 0.227 | 0.039 | 2.5307E-76 | TEC_CD | ADCY6 |
| 3.6611E-80 | 0.49670982 | 0.463 | 0.136 | 8.6485E-76 | TEC_CD | MSI2 |

| | | | | | | |
|---|---|---|---|---|---|---|
| 1.4371E-79 | 0.37419564 | 0.303 | 0.066 | 3.3949E-75 | TEC_CD | NNT |
| 2.1786E-79 | 0.40722641 | 0.387 | 0.1 | 5.1464E-75 | TEC_CD | TCEAL3 |
| 2.6731E-79 | 0.64366346 | 0.766 | 0.327 | 6.3147E-75 | TEC_CD | APP |
| 2.7493E-79 | -2.2140827 | 0.308 | 0.744 | 6.4947E-75 | TEC_CD | VIM |
| 2.7827E-79 | 0.95912525 | 0.675 | 0.269 | 6.5735E-75 | TEC_CD | TNFRSF12A |
| 2.8606E-79 | 0.94928324 | 0.621 | 0.257 | 6.7576E-75 | TEC_CD | PRDX2 |
| 5.7355E-79 | 0.25161054 | 0.207 | 0.034 | 1.3549E-74 | TEC_CD | PRR15 |
| 6.7432E-79 | 0.39556374 | 0.325 | 0.075 | 1.5929E-74 | TEC_CD | FAM241A |
| 7.85E-79 | 0.43814826 | 0.32 | 0.074 | 1.8544E-74 | TEC_CD | TRIM24 |
| 1.2982E-78 | 0.42532542 | 0.3 | 0.067 | 3.0668E-74 | TEC_CD | FAM20C |
| 2.5678E-78 | 0.25897056 | 0.108 | 0.01 | 6.066E-74 | TEC_CD | AJAP1 |
| 3.3658E-78 | 0.31424392 | 0.259 | 0.05 | 7.9511E-74 | TEC_CD | MAP9 |
| 5.3285E-78 | 0.57403987 | 0.495 | 0.159 | 1.2588E-73 | TEC_CD | TMEM141 |
| 5.5901E-78 | 0.5870777 | 0.333 | 0.08 | 1.3205E-73 | TEC_CD | SNAPC1 |
| 6.3843E-78 | 0.78362931 | 0.633 | 0.249 | 1.5082E-73 | TEC_CD | CD2AP |
| 7.9177E-78 | -1.1486265 | 0.803 | 0.959 | 1.8704E-73 | TEC_CD | H3F3A |
| 1.1592E-77 | 0.98153894 | 0.502 | 0.179 | 2.7383E-73 | TEC_CD | SPTSSA |
| 1.7298E-77 | 0.2882629 | 0.2 | 0.032 | 4.0863E-73 | TEC_CD | PCDH7 |
| 1.7321E-77 | 0.32369542 | 0.305 | 0.068 | 4.0917E-73 | TEC_CD | ABHD14B |
| 4.1432E-77 | 0.30119679 | 0.276 | 0.057 | 9.7875E-73 | TEC_CD | KIFAP3 |
| 5.7767E-77 | 0.45151967 | 0.394 | 0.108 | 1.3646E-72 | TEC_CD | ACAA2 |
| 8.1276E-77 | 0.29440961 | 0.261 | 0.052 | 1.92E-72 | TEC_CD | LAMB2 |
| 1.0265E-76 | 0.45757312 | 0.441 | 0.127 | 2.425E-72 | TEC_CD | NUCB2 |
| 1.2869E-76 | 0.25525564 | 0.19 | 0.03 | 3.0401E-72 | TEC_CD | REPS2 |
| 2.057E-76 | 0.87961134 | 0.682 | 0.303 | 4.8594E-72 | TEC_CD | ZNF292 |
| 2.5575E-76 | 0.41404851 | 0.416 | 0.116 | 6.0415E-72 | TEC_CD | CAT |
| 3.471E-76 | 0.27771187 | 0.256 | 0.05 | 8.1995E-72 | TEC_CD | DYNC2LI1 |
| 4.8577E-76 | 0.60572777 | 0.547 | 0.194 | 1.1475E-71 | TEC_CD | TCEAL9 |
| 5.0137E-76 | 0.40629389 | 0.276 | 0.059 | 1.1844E-71 | TEC_CD | RAB40B |
| 9.8617E-76 | 0.31247423 | 0.244 | 0.046 | 2.3296E-71 | TEC_CD | NIPAL1 |
| 1.1804E-75 | 0.40440223 | 0.384 | 0.103 | 2.7885E-71 | TEC_CD | TPD52 |
| 1.8984E-75 | 0.36284071 | 0.325 | 0.078 | 4.4847E-71 | TEC_CD | MORN2 |
| 2.177E-75 | 0.33378052 | 0.256 | 0.051 | 5.1428E-71 | TEC_CD | POU3F3 |
| 2.3591E-75 | 0.32566748 | 0.283 | 0.061 | 5.5729E-71 | TEC_CD | KIF9 |
| 2.3955E-75 | -1.4339808 | 0.145 | 0.617 | 5.6589E-71 | TEC_CD | CD44 |
| 2.904E-75 | 0.93144307 | 0.756 | 0.367 | 6.86E-71 | TEC_CD | ATP5MC3 |
| 7.5837E-75 | 0.26924337 | 0.249 | 0.048 | 1.7915E-70 | TEC_CD | CLUAP1 |
| 8.3719E-75 | 0.40371547 | 0.357 | 0.092 | 1.9777E-70 | TEC_CD | HINT2 |
| 9.3528E-75 | 0.41920111 | 0.436 | 0.127 | 2.2094E-70 | TEC_CD | AIG1 |
| 1.0494E-74 | 0.28398242 | 0.251 | 0.049 | 2.479E-70 | TEC_CD | MIR4458HG |
| 1.3346E-74 | 0.32782653 | 0.31 | 0.072 | 3.1528E-70 | TEC_CD | ZNF667-AS1 |
| 1.7675E-74 | 0.48110272 | 0.433 | 0.128 | 4.1754E-70 | TEC_CD | CXXC5 |
| 2.1057E-74 | -1.3810896 | 0.318 | 0.731 | 4.9743E-70 | TEC_CD | CYBA |
| 2.7516E-74 | 0.33339837 | 0.219 | 0.04 | 6.5002E-70 | TEC_CD | PINX1.1 |
| 2.9505E-74 | 0.40821663 | 0.315 | 0.075 | 6.97E-70 | TEC_CD | VAV3 |
| 3.0197E-74 | 0.49350549 | 0.431 | 0.123 | 7.1334E-70 | TEC_CD | SYNE1 |
| 4.1781E-74 | 0.2792062 | 0.266 | 0.055 | 9.8698E-70 | TEC_CD | ALDH6A1 |
| 2.4255E-73 | 0.9819949 | 0.768 | 0.408 | 5.7298E-69 | TEC_CD | CDV3 |
| 4.3781E-73 | 0.614387 | 0.438 | 0.131 | 1.0342E-68 | TEC_CD | HMGCS1 |
| 5.3459E-73 | 0.42159497 | 0.357 | 0.092 | 1.2629E-68 | TEC_CD | PTPN13 |

| | | | | | | |
|---|---|---|---|---|---|---|
| 5.357E-73 | 0.29368268 | 0.268 | 0.056 | 1.2655E-68 | TEC_CD | HOXB9 |
| 6.8314E-73 | 0.38027256 | 0.365 | 0.097 | 1.6138E-68 | TEC_CD | MTX2 |
| 8.0391E-73 | 0.39918704 | 0.374 | 0.102 | 1.8991E-68 | TEC_CD | SUCLG2 |
| 8.3571E-73 | 0.3032351 | 0.276 | 0.06 | 1.9742E-68 | TEC_CD | IMMP2L |
| 9.2571E-73 | 0.39006016 | 0.342 | 0.087 | 2.1868E-68 | TEC_CD | NUDT9 |
| 1.1196E-72 | 0.45680178 | 0.389 | 0.107 | 2.6447E-68 | TEC_CD | WWTR1 |
| 1.3669E-72 | 0.51803073 | 0.461 | 0.15 | 3.2289E-68 | TEC_CD | PNKD |
| 1.581E-72 | 0.3632683 | 0.328 | 0.08 | 3.7349E-68 | TEC_CD | MYO1B |
| 2.251E-72 | 0.35421922 | 0.296 | 0.068 | 5.3174E-68 | TEC_CD | C16orf87 |
| 3.0011E-72 | 0.87660777 | 0.389 | 0.112 | 7.0896E-68 | TEC_CD | RRAD |
| 3.0479E-72 | 0.44290324 | 0.419 | 0.123 | 7.2001E-68 | TEC_CD | TRIP6 |
| 5.1483E-72 | 0.35314693 | 0.342 | 0.087 | 1.2162E-67 | TEC_CD | DLD |
| 9.7279E-72 | 0.33517302 | 0.268 | 0.058 | 2.298E-67 | TEC_CD | MLF1 |
| 1.6332E-71 | 0.8302315 | 0.929 | 0.761 | 3.8581E-67 | TEC_CD | COX7C |
| 1.8604E-71 | 0.2862941 | 0.293 | 0.065 | 4.3949E-67 | TEC_CD | PLXNB1 |
| 2.3042E-71 | 0.71697679 | 0.707 | 0.321 | 5.4431E-67 | TEC_CD | MGST3 |
| 3.7688E-71 | 0.38002691 | 0.389 | 0.108 | 8.9031E-67 | TEC_CD | TMEM45B |
| 7.4067E-71 | 1.18779753 | 0.865 | 0.656 | 1.7497E-66 | TEC_CD | NDUFA4 |
| 9.969E-71 | -0.9974439 | 0.714 | 0.931 | 2.355E-66 | TEC_CD | OAZ1 |
| 1.5476E-70 | 0.37840176 | 0.357 | 0.095 | 3.656E-66 | TEC_CD | ECHDC2 |
| 1.9281E-70 | 0.32278501 | 0.3 | 0.071 | 4.5547E-66 | TEC_CD | TOM1L1 |
| 3.0727E-70 | 0.34303597 | 0.315 | 0.077 | 7.2587E-66 | TEC_CD | HOXB7 |
| 3.1328E-70 | 0.46754208 | 0.493 | 0.164 | 7.4007E-66 | TEC_CD | HMGN3 |
| 3.1331E-70 | 0.3144297 | 0.264 | 0.057 | 7.4013E-66 | TEC_CD | PKD2 |
| 4.475E-70 | 2.14309497 | 0.658 | 0.333 | 1.0571E-65 | TEC_CD | IGFBP7 |
| 6.4893E-70 | 0.29359295 | 0.266 | 0.057 | 1.533E-65 | TEC_CD | THAP4 |
| 8.3145E-70 | 0.39872702 | 0.158 | 0.023 | 1.9641E-65 | TEC_CD | RAET1L |
| 1.6934E-69 | 0.35971328 | 0.315 | 0.077 | 4.0004E-65 | TEC_CD | FRK |
| 1.7477E-69 | 0.52347169 | 0.451 | 0.148 | 4.1287E-65 | TEC_CD | MRPL34 |
| 1.8494E-69 | 0.2627252 | 0.241 | 0.049 | 4.3689E-65 | TEC_CD | NIPSNAP1 |
| 2.5431E-69 | 0.25883501 | 0.219 | 0.041 | 6.0075E-65 | TEC_CD | DNAL1 |
| 3.3741E-69 | 0.25795502 | 0.204 | 0.036 | 7.9706E-65 | TEC_CD | NAPEPLD |
| 3.6904E-69 | 0.64820861 | 0.7 | 0.297 | 8.7178E-65 | TEC_CD | DDX24 |
| 8.5765E-69 | 0.4170467 | 0.392 | 0.111 | 2.026E-64 | TEC_CD | AHI1 |
| 8.8883E-69 | 0.40592975 | 0.399 | 0.117 | 2.0997E-64 | TEC_CD | RRAGD |
| 9.7792E-69 | 0.60478781 | 0.475 | 0.153 | 2.3101E-64 | TEC_CD | CLDN3 |
| 1.0735E-68 | 0.52750618 | 0.569 | 0.202 | 2.536E-64 | TEC_CD | PARD6B |
| 1.1363E-68 | 0.46480277 | 0.377 | 0.105 | 2.6844E-64 | TEC_CD | WEE1 |
| 1.338E-68 | 0.32560704 | 0.172 | 0.027 | 3.1609E-64 | TEC_CD | KLHDC7B |
| 1.6731E-68 | 0.72485353 | 0.916 | 0.621 | 3.9524E-64 | TEC_CD | DSTN |
| 1.9224E-68 | 0.28900965 | 0.138 | 0.018 | 4.5412E-64 | TEC_CD | CXCL11 |
| 2.5425E-68 | 0.27778485 | 0.239 | 0.049 | 6.0061E-64 | TEC_CD | CLCN5 |
| 2.7116E-68 | 0.28691275 | 0.244 | 0.05 | 6.4056E-64 | TEC_CD | TLE2 |
| 3.5789E-68 | 0.75661108 | 0.732 | 0.355 | 8.4544E-64 | TEC_CD | ATP1A1 |
| 4.4685E-68 | 0.7492387 | 0.7 | 0.339 | 1.0556E-63 | TEC_CD | ATP5IF1 |
| 4.6835E-68 | 0.40628015 | 0.416 | 0.125 | 1.1064E-63 | TEC_CD | SH3GLB2 |
| 4.8227E-68 | 0.29001087 | 0.271 | 0.06 | 1.1393E-63 | TEC_CD | TFAP2A |
| 5.661E-68 | 0.46024496 | 0.392 | 0.115 | 1.3373E-63 | TEC_CD | HCFC1R1 |
| 6.8793E-68 | 0.39911925 | 0.372 | 0.104 | 1.6251E-63 | TEC_CD | ASAP2 |
| 1.2463E-67 | 0.68337448 | 0.574 | 0.233 | 2.9441E-63 | TEC_CD | ATP5F1A |

| | | | | | | |
|---|---|---|---|---|---|---|
| 1.4089E-67 | 0.58758508 | 0.49 | 0.176 | 3.3282E-63 | TEC_CD | UQCRC2 |
| 1.5508E-67 | 0.37674534 | 0.416 | 0.123 | 3.6635E-63 | TEC_CD | FAM213A |
| 1.7022E-67 | 0.69864213 | 0.732 | 0.351 | 4.021E-63 | TEC_CD | MRPL33 |
| 1.7057E-67 | 0.25373245 | 0.214 | 0.04 | 4.0294E-63 | TEC_CD | PIGQ |
| 2.3022E-67 | 0.63960393 | 0.744 | 0.37 | 5.4385E-63 | TEC_CD | LAMTOR5 |
| 2.4305E-67 | 0.53911993 | 0.488 | 0.171 | 5.7416E-63 | TEC_CD | CYC1 |
| 2.9009E-67 | 0.89980523 | 0.675 | 0.339 | 6.8529E-63 | TEC_CD | ATP5F1B |
| 3.9452E-67 | 0.27473426 | 0.2 | 0.036 | 9.3199E-63 | TEC_CD | SERTAD4-AS1 |
| 5.4532E-67 | 0.37046386 | 0.298 | 0.073 | 1.2882E-62 | TEC_CD | MCPH1 |
| 5.4547E-67 | 0.44142985 | 0.414 | 0.126 | 1.2886E-62 | TEC_CD | NUDT4 |
| 6.3648E-67 | 0.93681689 | 0.899 | 0.666 | 1.5035E-62 | TEC_CD | RTN4 |
| 6.517E-67 | 0.38198079 | 0.345 | 0.092 | 1.5395E-62 | TEC_CD | ZBTB10 |
| 1.0112E-66 | 0.34985374 | 0.335 | 0.088 | 2.3887E-62 | TEC_CD | ZSCAN18 |
| 1.6396E-66 | 0.34320825 | 0.293 | 0.071 | 3.8733E-62 | TEC_CD | AC118549.1 |
| 2.0891E-66 | 0.46819914 | 0.372 | 0.106 | 4.935E-62 | TEC_CD | ITM2C |
| 2.1626E-66 | 0.57381858 | 0.559 | 0.218 | 5.1086E-62 | TEC_CD | ECHS1 |
| 4.2989E-66 | 0.29180567 | 0.251 | 0.054 | 1.0155E-61 | TEC_CD | GAB1 |
| 4.3263E-66 | 0.70751912 | 0.534 | 0.211 | 1.022E-61 | TEC_CD | METAP2 |
| 1.2357E-65 | 0.41015152 | 0.392 | 0.117 | 2.919E-61 | TEC_CD | KIF13B |
| 1.675E-65 | 0.44178731 | 0.458 | 0.154 | 3.9568E-61 | TEC_CD | HSD17B12 |
| 2.2999E-65 | 1.31284874 | 0.82 | 0.584 | 5.4331E-61 | TEC_CD | KLF6 |
| 3.1873E-65 | 0.64353892 | 0.515 | 0.185 | 7.5293E-61 | TEC_CD | VCL |
| 3.2491E-65 | 0.91780496 | 0.576 | 0.242 | 7.6754E-61 | TEC_CD | PTMS |
| 4.6141E-65 | 0.3210407 | 0.323 | 0.083 | 1.09E-60 | TEC_CD | KANSL1L |
| 4.7316E-65 | 0.30919936 | 0.283 | 0.067 | 1.1177E-60 | TEC_CD | TNKS1BP1 |
| 4.8906E-65 | 0.28844068 | 0.268 | 0.061 | 1.1553E-60 | TEC_CD | ATPAF1 |
| 7.7199E-65 | 0.36120641 | 0.446 | 0.139 | 1.8237E-60 | TEC_CD | LAPTM4B |
| 8.5813E-65 | 0.2701613 | 0.266 | 0.06 | 2.0272E-60 | TEC_CD | GNA11 |
| 1.6417E-64 | 0.29149358 | 0.217 | 0.043 | 3.8781E-60 | TEC_CD | PPFIBP2 |
| 2.2133E-64 | 0.3789094 | 0.406 | 0.124 | 5.2285E-60 | TEC_CD | ARL2 |
| 2.2936E-64 | 0.27384092 | 0.241 | 0.051 | 5.4181E-60 | TEC_CD | CROT |
| 3.2303E-64 | 0.32793003 | 0.318 | 0.081 | 7.6308E-60 | TEC_CD | QPRT |
| 3.4105E-64 | 0.47191412 | 0.589 | 0.231 | 8.0566E-60 | TEC_CD | SYPL1 |
| 3.9546E-64 | 0.26150847 | 0.239 | 0.051 | 9.342E-60 | TEC_CD | TRIM47 |
| 7.5371E-64 | -1.7407044 | 0.323 | 0.684 | 1.7805E-59 | TEC_CD | NAMPT |
| 9.0095E-64 | 0.56870853 | 0.571 | 0.214 | 2.1283E-59 | TEC_CD | ZBTB20 |
| 1.175E-63 | 0.25209704 | 0.241 | 0.051 | 2.7757E-59 | TEC_CD | THEM4 |
| 1.2358E-63 | -1.7599797 | 0.209 | 0.629 | 2.9193E-59 | TEC_CD | S100A4 |
| 1.5773E-63 | 0.51213414 | 0.596 | 0.234 | 3.7261E-59 | TEC_CD | PCBD1 |
| 2.761E-63 | 0.26502741 | 0.239 | 0.051 | 6.5222E-59 | TEC_CD | IFT74 |
| 2.8944E-63 | 0.41342351 | 0.374 | 0.11 | 6.8374E-59 | TEC_CD | LGALS3BP |
| 2.9039E-63 | -2.5530394 | 0.034 | 0.468 | 6.8599E-59 | TEC_CD | LYZ |
| 6.4339E-63 | 0.30977254 | 0.296 | 0.074 | 1.5199E-58 | TEC_CD | MITF |
| 9.9098E-63 | 0.81272699 | 0.736 | 0.407 | 2.341E-58 | TEC_CD | ATP5PF |
| 1.5085E-62 | 0.40989756 | 0.685 | 0.271 | 3.5634E-58 | TEC_CD | CAMK2N1 |
| 1.8395E-62 | 0.28470204 | 0.305 | 0.077 | 4.3454E-58 | TEC_CD | SPAG16 |
| 2.0169E-62 | 0.35144809 | 0.355 | 0.101 | 4.7646E-58 | TEC_CD | TMEM251 |
| 2.6934E-62 | 0.31107957 | 0.308 | 0.079 | 6.3625E-58 | TEC_CD | HIP1R |
| 2.708E-62 | 0.27443519 | 0.202 | 0.039 | 6.3972E-58 | TEC_CD | FAM20A |
| 2.7685E-62 | 0.4305755 | 0.448 | 0.151 | 6.5401E-58 | TEC_CD | NDUFS4 |

| | | | | | | |
|---|---|---|---|---|---|---|
| 3.4144E-62 | 0.29026475 | 0.229 | 0.048 | 8.0658E-58 | TEC_CD | COL6A1 |
| 5.2772E-62 | 0.34692807 | 0.323 | 0.087 | 1.2466E-57 | TEC_CD | KIF3B |
| 5.8206E-62 | 0.39846722 | 0.397 | 0.124 | 1.375E-57 | TEC_CD | NDUFS3 |
| 6.1066E-62 | 0.34635591 | 0.323 | 0.087 | 1.4426E-57 | TEC_CD | PPIC |
| 7.0743E-62 | 0.46891858 | 0.404 | 0.126 | 1.6712E-57 | TEC_CD | KAZN |
| 8.0164E-62 | 0.61266619 | 0.719 | 0.339 | 1.8937E-57 | TEC_CD | BSG |
| 8.155E-62 | 0.27484325 | 0.234 | 0.05 | 1.9265E-57 | TEC_CD | SLC29A1 |
| 2.4446E-61 | 0.76637366 | 0.502 | 0.191 | 5.7749E-57 | TEC_CD | IRS2 |
| 2.6768E-61 | 0.35298287 | 0.362 | 0.105 | 6.3234E-57 | TEC_CD | CFAP97 |
| 3.5201E-61 | 0.42311762 | 0.473 | 0.163 | 8.3156E-57 | TEC_CD | GLO1 |
| 3.6184E-61 | 0.27904701 | 0.251 | 0.057 | 8.5478E-57 | TEC_CD | ARHGEF28 |
| 5.8645E-61 | 0.33333417 | 0.281 | 0.07 | 1.3854E-56 | TEC_CD | IFIT5 |
| 6.8497E-61 | 0.4500684 | 0.443 | 0.146 | 1.6181E-56 | TEC_CD | PLS3 |
| 9.0554E-61 | 0.38912789 | 0.365 | 0.109 | 2.1392E-56 | TEC_CD | NIPSNAP2 |
| 1.6425E-60 | 0.71163385 | 0.692 | 0.335 | 3.88E-56 | TEC_CD | NDRG1 |
| 1.8393E-60 | 0.37922267 | 0.347 | 0.101 | 4.3449E-56 | TEC_CD | TEX264 |
| 2.1456E-60 | 0.43916943 | 0.502 | 0.186 | 5.0685E-56 | TEC_CD | HADHB |
| 2.252E-60 | 0.29963992 | 0.256 | 0.06 | 5.3199E-56 | TEC_CD | KLHL8 |
| 3.8431E-60 | -1.8656157 | 0.025 | 0.444 | 9.0784E-56 | TEC_CD | TYROBP |
| 6.5615E-60 | 0.37558753 | 0.335 | 0.096 | 1.55E-55 | TEC_CD | ACBD5 |
| 1.3296E-59 | 0.33933599 | 0.296 | 0.076 | 3.1409E-55 | TEC_CD | PPP2R3A |
| 1.6908E-59 | 0.58865859 | 0.515 | 0.193 | 3.9941E-55 | TEC_CD | NEDD4L |
| 2.0432E-59 | 0.41363329 | 0.448 | 0.154 | 4.8266E-55 | TEC_CD | TMEM205 |
| 2.2532E-59 | 0.52011416 | 0.441 | 0.153 | 5.3228E-55 | TEC_CD | NFE2L1 |
| 2.4995E-59 | 0.48924676 | 0.626 | 0.264 | 5.9046E-55 | TEC_CD | EID1 |
| 6.1858E-59 | 0.29754839 | 0.256 | 0.061 | 1.4613E-54 | TEC_CD | RGP1 |
| 9.7147E-59 | 0.25111851 | 0.185 | 0.034 | 2.2949E-54 | TEC_CD | RNF152 |
| 4.3306E-58 | 0.37373112 | 0.35 | 0.103 | 1.023E-53 | TEC_CD | ABHD11 |
| 5.9729E-58 | 0.70951507 | 0.564 | 0.238 | 1.411E-53 | TEC_CD | BHLHE40 |
| 6.8036E-58 | 0.37653522 | 0.416 | 0.134 | 1.6072E-53 | TEC_CD | PATJ |
| 1.5977E-57 | 0.39676876 | 0.404 | 0.131 | 3.7741E-53 | TEC_CD | SEPT11 |
| 1.8997E-57 | 0.61490246 | 0.712 | 0.359 | 4.4876E-53 | TEC_CD | NFE2L2 |
| 2.6934E-57 | 0.43730647 | 0.429 | 0.148 | 6.3626E-53 | TEC_CD | ECI1 |
| 2.8805E-57 | 0.32156229 | 0.286 | 0.074 | 6.8047E-53 | TEC_CD | CYFIP2 |
| 2.9169E-57 | 0.35866494 | 0.365 | 0.112 | 6.8906E-53 | TEC_CD | COPRS |
| 4.1409E-57 | 0.29968744 | 0.273 | 0.069 | 9.7821E-53 | TEC_CD | TACC2 |
| 6.3113E-57 | 0.39205659 | 0.365 | 0.113 | 1.4909E-52 | TEC_CD | YAP1 |
| 9.3928E-57 | 0.45672964 | 0.429 | 0.151 | 2.2189E-52 | TEC_CD | NDUFS2 |
| 1.0077E-56 | 0.44666859 | 0.369 | 0.114 | 2.3805E-52 | TEC_CD | VASN |
| 1.1966E-56 | 0.2900363 | 0.165 | 0.029 | 2.8266E-52 | TEC_CD | HRK |
| 1.9813E-56 | 0.31941175 | 0.268 | 0.068 | 4.6805E-52 | TEC_CD | SAMD12 |
| 2.7765E-56 | 0.29990882 | 0.303 | 0.083 | 6.559E-52 | TEC_CD | TMEM263 |
| 2.8593E-55 | 0.42151108 | 0.397 | 0.134 | 6.7544E-51 | TEC_CD | GNPDA1 |
| 3.2064E-55 | 0.35389361 | 0.401 | 0.132 | 7.5744E-51 | TEC_CD | ADH5 |
| 3.3376E-55 | 0.29424074 | 0.276 | 0.072 | 7.8845E-51 | TEC_CD | JADE1 |
| 4.2689E-55 | 0.26722208 | 0.224 | 0.051 | 1.0084E-50 | TEC_CD | EFHD1 |
| 4.5497E-55 | 0.42100433 | 0.441 | 0.157 | 1.0748E-50 | TEC_CD | BEX4 |
| 4.9639E-55 | 0.3740363 | 0.458 | 0.162 | 1.1726E-50 | TEC_CD | DDR1 |
| 5.0068E-55 | 0.26893947 | 0.246 | 0.059 | 1.1827E-50 | TEC_CD | CDC42EP4 |
| 5.9892E-55 | 0.49849615 | 0.505 | 0.199 | 1.4148E-50 | TEC_CD | NDUFS8 |

| | | | | | | |
|---|---|---|---|---|---|---|
| 8.3776E-55 | 0.31577899 | 0.273 | 0.072 | 1.979E-50 | TEC_CD | MOCS2 |
| 8.9431E-55 | 0.27665156 | 0.273 | 0.07 | 2.1126E-50 | TEC_CD | METTL7A |
| 9.2636E-55 | 0.27288203 | 0.256 | 0.064 | 2.1883E-50 | TEC_CD | ZC2HC1A |
| 1.141E-54 | 0.52343197 | 0.586 | 0.26 | 2.6954E-50 | TEC_CD | NDUFA6 |
| 1.2418E-54 | 0.40253294 | 0.453 | 0.161 | 2.9336E-50 | TEC_CD | MINDY2 |
| 1.2486E-54 | 0.43583434 | 0.52 | 0.21 | 2.9495E-50 | TEC_CD | SDHD |
| 1.2671E-54 | -1.5551414 | 0.249 | 0.62 | 2.9932E-50 | TEC_CD | LGALS1 |
| 1.4106E-54 | 0.5436181 | 0.537 | 0.211 | 3.3323E-50 | TEC_CD | EHF |
| 1.4227E-54 | 0.51197253 | 0.559 | 0.235 | 3.3609E-50 | TEC_CD | TSTD1 |
| 2.1917E-54 | 0.56269672 | 0.672 | 0.318 | 5.1773E-50 | TEC_CD | KRT10 |
| 2.2773E-54 | 0.56271088 | 0.426 | 0.159 | 5.3797E-50 | TEC_CD | DECR1 |
| 2.3864E-54 | 0.41509855 | 0.458 | 0.17 | 5.6373E-50 | TEC_CD | NDUFA10 |
| 3.6721E-54 | 0.33622226 | 0.337 | 0.102 | 8.6746E-50 | TEC_CD | HDDC2 |
| 3.9793E-54 | 0.25240238 | 0.222 | 0.051 | 9.4002E-50 | TEC_CD | ZNF639 |
| 4.6912E-54 | 0.26833891 | 0.239 | 0.057 | 1.1082E-49 | TEC_CD | GNAI1 |
| 4.8961E-54 | 0.58078811 | 0.988 | 0.905 | 1.1566E-49 | TEC_CD | ITM2B |
| 6.1125E-54 | 0.28513226 | 0.209 | 0.045 | 1.444E-49 | TEC_CD | EGOT |
| 6.3898E-54 | 0.75679453 | 0.825 | 0.548 | 1.5095E-49 | TEC_CD | FAM107B |
| 9.4084E-54 | 0.4440614 | 0.466 | 0.178 | 2.2226E-49 | TEC_CD | ADI1 |
| 1.3189E-53 | 0.30988447 | 0.276 | 0.074 | 3.1156E-49 | TEC_CD | MYL6B |
| 2.0794E-53 | 0.41517655 | 0.337 | 0.103 | 4.9122E-49 | TEC_CD | BBC3 |
| 2.1549E-53 | 0.32652625 | 0.305 | 0.087 | 5.0906E-49 | TEC_CD | IRF2BPL |
| 4.0438E-53 | 0.60400435 | 0.522 | 0.212 | 9.5526E-49 | TEC_CD | CKB |
| 4.3778E-53 | 0.42182509 | 0.498 | 0.192 | 1.0342E-48 | TEC_CD | RBPMS |
| 5.6304E-53 | 0.25171735 | 0.246 | 0.061 | 1.3301E-48 | TEC_CD | MBIP |
| 5.9668E-53 | 0.31705797 | 0.288 | 0.08 | 1.4095E-48 | TEC_CD | TMEM106C |
| 6.2306E-53 | 0.65828291 | 0.722 | 0.395 | 1.4719E-48 | TEC_CD | NDUFA1 |
| 6.3234E-53 | 0.36710413 | 0.441 | 0.157 | 1.4938E-48 | TEC_CD | SLC25A36 |
| 7.6004E-53 | 0.27092676 | 0.212 | 0.048 | 1.7954E-48 | TEC_CD | SBF2-AS1 |
| 8.5683E-53 | 0.33613411 | 0.36 | 0.114 | 2.0241E-48 | TEC_CD | BOLA3 |
| 1.096E-52 | 0.77075358 | 0.313 | 0.094 | 2.5891E-48 | TEC_CD | IFIT2 |
| 1.2978E-52 | 0.58452188 | 0.584 | 0.255 | 3.0658E-48 | TEC_CD | TSC22D1 |
| 1.4685E-52 | 0.45223144 | 0.599 | 0.255 | 3.469E-48 | TEC_CD | ARHGAP5 |
| 2.0688E-52 | 0.2601871 | 0.232 | 0.056 | 4.887E-48 | TEC_CD | ADAP1 |
| 2.4619E-52 | 0.44521431 | 0.377 | 0.127 | 5.8158E-48 | TEC_CD | CBR1 |
| 2.5217E-52 | 0.35470019 | 0.266 | 0.07 | 5.957E-48 | TEC_CD | LAMB1 |
| 2.9952E-52 | 0.31015738 | 0.281 | 0.077 | 7.0755E-48 | TEC_CD | FBXO25 |
| 3.6518E-52 | 0.42619725 | 0.286 | 0.08 | 8.6267E-48 | TEC_CD | EFNB2 |
| 3.7212E-52 | 0.34379374 | 0.34 | 0.105 | 8.7907E-48 | TEC_CD | CFAP36 |
| 3.8023E-52 | 0.29261324 | 0.303 | 0.087 | 8.9822E-48 | TEC_CD | GOT2 |
| 4.1339E-52 | 0.25272501 | 0.234 | 0.057 | 9.7655E-48 | TEC_CD | GBA2 |
| 4.1955E-52 | 0.95912172 | 0.498 | 0.218 | 9.9109E-48 | TEC_CD | WNK1 |
| 4.3113E-52 | 0.35310742 | 0.266 | 0.072 | 1.0185E-47 | TEC_CD | SLC12A7 |
| 8.2809E-52 | 0.3368047 | 0.281 | 0.078 | 1.9562E-47 | TEC_CD | TM7SF3 |
| 8.6426E-52 | 0.35572025 | 0.431 | 0.155 | 2.0416E-47 | TEC_CD | TXN2 |
| 8.7786E-52 | 0.26152606 | 0.264 | 0.069 | 2.0738E-47 | TEC_CD | TMC4 |
| 1.1244E-51 | 0.4737316 | 0.631 | 0.261 | 2.6561E-47 | TEC_CD | TSPAN1 |
| 1.1973E-51 | 0.48789229 | 0.406 | 0.147 | 2.8284E-47 | TEC_CD | RRAGC |
| 1.5763E-51 | 0.28287455 | 0.271 | 0.073 | 3.7237E-47 | TEC_CD | REPIN1 |
| 1.8051E-51 | -0.7768776 | 0.663 | 0.868 | 4.2641E-47 | TEC_CD | ARPC3 |

| | | | | | | |
|---|---|---|---|---|---|---|
| 1.9621E-51 | -0.8798461 | 0.84 | 0.948 | 4.6351E-47 | TEC_CD | RPS16 |
| 2.0892E-51 | -2.7750265 | 0.101 | 0.492 | 4.9353E-47 | TEC_CD | CXCL8 |
| 2.6568E-51 | 0.32381208 | 0.278 | 0.077 | 6.2762E-47 | TEC_CD | C3orf52 |
| 3.9988E-51 | -0.5290515 | 0.823 | 0.539 | 9.4463E-47 | TEC_CD | PLCG2 |
| 5.9411E-51 | 0.32730926 | 0.352 | 0.111 | 1.4035E-46 | TEC_CD | WWC1 |
| 9.1635E-51 | 0.3523826 | 0.468 | 0.175 | 2.1647E-46 | TEC_CD | ARL1 |
| 1.0613E-50 | 0.34400936 | 0.36 | 0.117 | 2.507E-46 | TEC_CD | LRPPRC |
| 1.172E-50 | 0.28204188 | 0.246 | 0.063 | 2.7685E-46 | TEC_CD | DUSP8 |
| 1.1908E-50 | 0.49602559 | 0.502 | 0.213 | 2.813E-46 | TEC_CD | NDUFB5 |
| 1.2471E-50 | 0.28342393 | 0.293 | 0.084 | 2.9461E-46 | TEC_CD | HINT3 |
| 1.4223E-50 | 0.264898 | 0.217 | 0.051 | 3.36E-46 | TEC_CD | MMD |
| 3.8028E-50 | 0.47910138 | 0.441 | 0.17 | 8.9834E-46 | TEC_CD | MPC2 |
| 4.0033E-50 | -0.7304181 | 0.961 | 0.993 | 9.457E-46 | TEC_CD | TPT1 |
| 4.3313E-50 | 0.34803953 | 0.483 | 0.183 | 1.0232E-45 | TEC_CD | ANXA4 |
| 5.5138E-50 | 0.55756858 | 0.712 | 0.364 | 1.3025E-45 | TEC_CD | CTSH |
| 8.953E-50 | -2.5595644 | 0.042 | 0.411 | 2.115E-45 | TEC_CD | S100P |
| 9.0409E-50 | 0.26663898 | 0.33 | 0.1 | 2.1357E-45 | TEC_CD | EPB41L5 |
| 1.2227E-49 | 0.2575327 | 0.259 | 0.069 | 2.8884E-45 | TEC_CD | CRYL1 |
| 1.3097E-49 | 0.46455811 | 0.392 | 0.138 | 3.0939E-45 | TEC_CD | GPNMB |
| 1.8089E-49 | 0.52042839 | 0.751 | 0.39 | 4.2733E-45 | TEC_CD | NDUFB2 |
| 2.374E-49 | 0.31378679 | 0.261 | 0.07 | 5.6082E-45 | TEC_CD | ARL4D |
| 2.6953E-49 | 0.31595375 | 0.35 | 0.113 | 6.3672E-45 | TEC_CD | SPINT1-AS1 |
| 2.7189E-49 | 0.58836039 | 0.443 | 0.166 | 6.423E-45 | TEC_CD | AKR1B1 |
| 3.207E-49 | 0.26343029 | 0.268 | 0.073 | 7.5758E-45 | TEC_CD | PRKCA |
| 4.975E-49 | -1.7732958 | 0.025 | 0.389 | 1.1752E-44 | TEC_CD | FCER1G |
| 8.6402E-49 | 0.59560291 | 0.692 | 0.361 | 2.0411E-44 | TEC_CD | ATP5F1D |
| 9.6108E-49 | 0.3716106 | 0.318 | 0.099 | 2.2704E-44 | TEC_CD | RRM2B |
| 9.6747E-49 | -1.9699957 | 0.966 | 0.987 | 2.2855E-44 | TEC_CD | FTL |
| 1.0087E-48 | 0.42681524 | 0.466 | 0.186 | 2.3829E-44 | TEC_CD | PRDX3 |
| 1.81E-48 | 0.40704906 | 0.522 | 0.205 | 4.2757E-44 | TEC_CD | MACC1 |
| 1.9307E-48 | 0.37098272 | 0.345 | 0.108 | 4.5608E-44 | TEC_CD | SMIM5 |
| 1.9769E-48 | 0.30730841 | 0.286 | 0.083 | 4.6699E-44 | TEC_CD | SIAH1 |
| 2.0896E-48 | 0.31667915 | 0.291 | 0.085 | 4.9363E-44 | TEC_CD | SYNJ2BP |
| 2.3859E-48 | 0.40080913 | 0.34 | 0.113 | 5.6362E-44 | TEC_CD | CLN5 |
| 2.9872E-48 | 0.36750837 | 0.357 | 0.121 | 7.0567E-44 | TEC_CD | ESRRA |
| 3.1167E-48 | 0.45694677 | 0.727 | 0.363 | 7.3625E-44 | TEC_CD | REEP5 |
| 3.2761E-48 | 0.26919561 | 0.254 | 0.068 | 7.7391E-44 | TEC_CD | ACYP1 |
| 3.5345E-48 | 0.52296424 | 0.549 | 0.244 | 8.3496E-44 | TEC_CD | CREBRF |
| 4.4174E-48 | 0.32229398 | 0.305 | 0.093 | 1.0435E-43 | TEC_CD | GSE1 |
| 6.3524E-48 | 0.31334755 | 0.335 | 0.107 | 1.5006E-43 | TEC_CD | AK3 |
| 7.0069E-48 | 0.29386653 | 0.313 | 0.095 | 1.6552E-43 | TEC_CD | IFT57 |
| 7.8131E-48 | 0.46525726 | 0.525 | 0.22 | 1.8457E-43 | TEC_CD | MRPS6 |
| 1.0465E-47 | 0.2902969 | 0.291 | 0.085 | 2.4723E-43 | TEC_CD | TFDP2 |
| 1.9661E-47 | 0.60026123 | 0.732 | 0.4 | 4.6445E-43 | TEC_CD | SOD1 |
| 3.1422E-47 | 0.67316545 | 0.727 | 0.423 | 7.4229E-43 | TEC_CD | SLC3A2 |
| 3.2403E-47 | 0.34224989 | 0.424 | 0.158 | 7.6545E-43 | TEC_CD | GINM1 |
| 3.5635E-47 | 0.42271992 | 0.441 | 0.166 | 8.4181E-43 | TEC_CD | CDKN2AIP |
| 3.7461E-47 | 0.45217252 | 0.569 | 0.258 | 8.8494E-43 | TEC_CD | CUTA |
| 5.0786E-47 | -1.4481818 | 0.328 | 0.646 | 1.1997E-42 | TEC_CD | GLUL |
| 5.1318E-47 | 0.38886678 | 0.515 | 0.211 | 1.2123E-42 | TEC_CD | LMAN1 |

| | | | | | | |
|---|---|---|---|---|---|---|
| 1.16E-46 | 0.68036173 | 0.512 | 0.232 | 2.7403E-42 | TEC_CD | MT-ATP8 |
| 1.5565E-46 | 0.4583376 | 0.374 | 0.133 | 3.6768E-42 | TEC_CD | KLHL21 |
| 1.5615E-46 | 0.27290466 | 0.305 | 0.093 | 3.6887E-42 | TEC_CD | MPP5 |
| 1.7568E-46 | 0.93298109 | 0.367 | 0.139 | 4.1501E-42 | TEC_CD | OTUD1 |
| 1.9038E-46 | 0.29577465 | 0.335 | 0.109 | 4.4974E-42 | TEC_CD | MRPS35 |
| 2.0262E-46 | 0.57478266 | 0.695 | 0.372 | 4.7865E-42 | TEC_CD | ATP5ME |
| 3.3757E-46 | -1.1429127 | 0.872 | 0.94 | 7.9743E-42 | TEC_CD | SAT1 |
| 3.6639E-46 | 0.25151882 | 0.259 | 0.071 | 8.6552E-42 | TEC_CD | ZNF626 |
| 3.7526E-46 | 0.33489912 | 0.379 | 0.134 | 8.8647E-42 | TEC_CD | RDH11 |
| 4.0393E-46 | 0.27482431 | 0.268 | 0.076 | 9.5421E-42 | TEC_CD | SH3RF1 |
| 4.8454E-46 | 0.54291989 | 0.394 | 0.149 | 1.1446E-41 | TEC_CD | CPVL |
| 4.9339E-46 | 0.42917044 | 0.369 | 0.132 | 1.1655E-41 | TEC_CD | CDC42BPB |
| 5.8225E-46 | 0.4168854 | 0.377 | 0.136 | 1.3754E-41 | TEC_CD | KIAA1109 |
| 6.1175E-46 | 0.33371419 | 0.298 | 0.091 | 1.4451E-41 | TEC_CD | FAM84B |
| 7.2965E-46 | 0.4786745 | 0.586 | 0.279 | 1.7236E-41 | TEC_CD | LRPAP1 |
| 8.4328E-46 | -4.3767908 | 0.054 | 0.405 | 1.9921E-41 | TEC_CD | SPRR3 |
| 8.6895E-46 | 0.56823863 | 0.19 | 0.044 | 2.0527E-41 | TEC_CD | ARL14 |
| 9.0279E-46 | -0.7978747 | 0.18 | 0.539 | 2.1327E-41 | TEC_CD | SERPINB1 |
| 9.3556E-46 | 0.34498856 | 0.298 | 0.089 | 2.2101E-41 | TEC_CD | CMYA5 |
| 1.3038E-45 | -1.8922698 | 0.995 | 0.994 | 3.08E-41 | TEC_CD | FTH1 |
| 2.1768E-45 | 0.38639866 | 0.305 | 0.096 | 5.1422E-41 | TEC_CD | SH3BP4 |
| 2.1837E-45 | 0.32561104 | 0.305 | 0.095 | 5.1585E-41 | TEC_CD | GRAMD2B |
| 2.2954E-45 | 0.35244812 | 0.308 | 0.098 | 5.4225E-41 | TEC_CD | PTPRF |
| 2.3684E-45 | -3.0368256 | 0.034 | 0.381 | 5.5949E-41 | TEC_CD | KRT13 |
| 2.4865E-45 | 0.64097057 | 0.667 | 0.324 | 5.8739E-41 | TEC_CD | LMNA |
| 2.8852E-45 | 0.26549369 | 0.288 | 0.087 | 6.8156E-41 | TEC_CD | PRCP |
| 3.9671E-45 | 0.27894247 | 0.259 | 0.073 | 9.3715E-41 | TEC_CD | PKP4 |
| 7.0243E-45 | 0.4119319 | 0.35 | 0.122 | 1.6593E-40 | TEC_CD | TNFRSF10B |
| 9.3487E-45 | 0.4075782 | 0.286 | 0.089 | 2.2084E-40 | TEC_CD | ELMSAN1 |
| 1.0271E-44 | 0.40158085 | 0.365 | 0.128 | 2.4262E-40 | TEC_CD | NTN4 |
| 1.2284E-44 | 0.30378154 | 0.315 | 0.102 | 2.9019E-40 | TEC_CD | MOB1B |
| 1.4676E-44 | 0.40334688 | 0.409 | 0.155 | 3.4668E-40 | TEC_CD | CHD3 |
| 2.0459E-44 | 0.27830363 | 0.266 | 0.077 | 4.8329E-40 | TEC_CD | GLIS3 |
| 2.3452E-44 | 0.30804698 | 0.357 | 0.124 | 5.5401E-40 | TEC_CD | THUMPD1 |
| 2.5185E-44 | 0.38679159 | 0.451 | 0.177 | 5.9494E-40 | TEC_CD | FNBP1L |
| 2.7937E-44 | 0.25401455 | 0.251 | 0.07 | 6.5995E-40 | TEC_CD | TMEM30B |
| 2.8074E-44 | 0.27064528 | 0.261 | 0.074 | 6.6318E-40 | TEC_CD | CGNL1 |
| 3.5085E-44 | 0.32842594 | 0.438 | 0.172 | 8.2882E-40 | TEC_CD | SMIM15 |
| 3.529E-44 | 0.25463364 | 0.256 | 0.073 | 8.3366E-40 | TEC_CD | SLC39A4 |
| 4.0616E-44 | 0.25808782 | 0.266 | 0.077 | 9.5946E-40 | TEC_CD | MRPL1 |
| 5.4779E-44 | -1.2967303 | 0.002 | 0.338 | 1.294E-39 | TEC_CD | LAPTM5 |
| 6.2615E-44 | 0.30077478 | 0.2 | 0.049 | 1.4791E-39 | TEC_CD | AC004130.1 |
| 6.3743E-44 | 0.35311578 | 0.372 | 0.135 | 1.5058E-39 | TEC_CD | PACSIN2 |
| 6.8281E-44 | 0.2641714 | 0.271 | 0.08 | 1.613E-39 | TEC_CD | ZNF44 |
| 7.1748E-44 | 0.29555088 | 0.214 | 0.055 | 1.6949E-39 | TEC_CD | ZNF165 |
| 9.0643E-44 | 0.41282996 | 0.36 | 0.13 | 2.1413E-39 | TEC_CD | ROCK2 |
| 9.4676E-44 | 0.31484128 | 0.318 | 0.106 | 2.2365E-39 | TEC_CD | NDUFA9 |
| 1.0119E-43 | 0.35657865 | 0.335 | 0.115 | 2.3904E-39 | TEC_CD | MAGED2 |
| 1.4406E-43 | 0.47305738 | 0.256 | 0.074 | 3.4031E-39 | TEC_CD | FGF2 |
| 1.7734E-43 | 0.25337033 | 0.271 | 0.079 | 4.1893E-39 | TEC_CD | LIMCH1 |

| | | | | | | |
|---|---|---|---|---|---|---|
| 1.9992E-43 | 0.25188826 | 0.256 | 0.074 | 4.7227E-39 | TEC_CD | ZBTB4 |
| 2.3438E-43 | 0.36399227 | 0.564 | 0.257 | 5.5368E-39 | TEC_CD | OCIAD1 |
| 2.4645E-43 | -1.0285433 | 0.983 | 0.99 | 5.8219E-39 | TEC_CD | B2M |
| 2.7086E-43 | 0.36606642 | 0.382 | 0.141 | 6.3986E-39 | TEC_CD | SLC39A1 |
| 3.1558E-43 | 0.58652924 | 0.645 | 0.354 | 7.4549E-39 | TEC_CD | COX5A |
| 3.3576E-43 | 0.25005086 | 0.244 | 0.068 | 7.9317E-39 | TEC_CD | TSPAN15 |
| 4.8298E-43 | 0.70314967 | 0.768 | 0.53 | 1.1409E-38 | TEC_CD | COX6C |
| 5.292E-43 | 0.28051717 | 0.217 | 0.056 | 1.2501E-38 | TEC_CD | NAV2 |
| 5.4373E-43 | 0.60104834 | 0.741 | 0.446 | 1.2844E-38 | TEC_CD | ATP5MD |
| 7.4214E-43 | 0.31980029 | 0.362 | 0.131 | 1.7532E-38 | TEC_CD | CMC2 |
| 8.6856E-43 | 0.5876297 | 0.401 | 0.152 | 2.0518E-38 | TEC_CD | CYR61 |
| 1.7543E-42 | 0.27863008 | 0.249 | 0.071 | 4.1441E-38 | TEC_CD | KIF21A |
| 2.17E-42 | 0.56061891 | 0.655 | 0.354 | 5.1262E-38 | TEC_CD | UQCRQ |
| 2.3736E-42 | 0.80498835 | 0.293 | 0.099 | 5.6072E-38 | TEC_CD | SLC8A1 |
| 2.7739E-42 | 0.25039484 | 0.268 | 0.08 | 6.5527E-38 | TEC_CD | EBAG9 |
| 3.1852E-42 | 0.49572664 | 0.569 | 0.276 | 7.5245E-38 | TEC_CD | MRPL41 |
| 3.19E-42 | 0.41027383 | 0.411 | 0.159 | 7.5357E-38 | TEC_CD | MET |
| 3.3442E-42 | 0.35427543 | 0.389 | 0.148 | 7.9001E-38 | TEC_CD | VGLL4 |
| 3.3918E-42 | 0.40684866 | 0.594 | 0.286 | 8.0124E-38 | TEC_CD | NDUFB3 |
| 3.6235E-42 | -1.1102284 | 0.217 | 0.534 | 8.5598E-38 | TEC_CD | CTSS |
| 3.7119E-42 | 0.30794912 | 0.3 | 0.097 | 8.7686E-38 | TEC_CD | TTC37 |
| 4.4548E-42 | 0.87401193 | 0.613 | 0.316 | 1.0524E-37 | TEC_CD | GADD45A |
| 6.175E-42 | 0.51049684 | 0.655 | 0.346 | 1.4587E-37 | TEC_CD | FKBP2 |
| 8.4922E-42 | 0.5330036 | 0.621 | 0.309 | 2.0061E-37 | TEC_CD | SDC4 |
| 9.2703E-42 | 0.46936794 | 0.47 | 0.2 | 2.1899E-37 | TEC_CD | NEU1 |
| 1.138E-41 | 0.34246575 | 0.429 | 0.169 | 2.6882E-37 | TEC_CD | BBX |
| 2.3739E-41 | 0.32340737 | 0.35 | 0.126 | 5.6079E-37 | TEC_CD | NDUFS1 |
| 2.6336E-41 | 0.32577429 | 0.254 | 0.076 | 6.2214E-37 | TEC_CD | STIM2 |
| 3.2171E-41 | 0.40021575 | 0.288 | 0.094 | 7.5998E-37 | TEC_CD | NEDD9 |
| 3.5718E-41 | 0.27783799 | 0.283 | 0.089 | 8.4376E-37 | TEC_CD | RUFY3 |
| 4.6794E-41 | 0.66406893 | 0.889 | 0.758 | 1.1054E-36 | TEC_CD | UQCRB |
| 5.0221E-41 | 0.37550102 | 0.532 | 0.239 | 1.1864E-36 | TEC_CD | SRI |
| 6.0947E-41 | 0.25284404 | 0.224 | 0.062 | 1.4398E-36 | TEC_CD | RDH14 |
| 6.9478E-41 | 0.62814228 | 0.921 | 0.773 | 1.6413E-36 | TEC_CD | CTSD |
| 7.2363E-41 | -0.7878335 | 0.108 | 0.431 | 1.7094E-36 | TEC_CD | EFHD2 |
| 7.6088E-41 | -1.200351 | 0.101 | 0.428 | 1.7974E-36 | TEC_CD | EMP3 |
| 8.2693E-41 | 0.30033308 | 0.347 | 0.124 | 1.9535E-36 | TEC_CD | PTS |
| 8.568E-41 | 0.26946454 | 0.18 | 0.043 | 2.024E-36 | TEC_CD | IFIT1 |
| 9.4231E-41 | 0.27100198 | 0.273 | 0.085 | 2.226E-36 | TEC_CD | RCOR3 |
| 9.8192E-41 | 0.32496657 | 0.347 | 0.123 | 2.3196E-36 | TEC_CD | HOOK2 |
| 1.1164E-40 | 0.35718587 | 0.47 | 0.199 | 2.6372E-36 | TEC_CD | PHB |
| 1.3366E-40 | 0.31053845 | 0.382 | 0.144 | 3.1575E-36 | TEC_CD | AK2 |
| 1.5924E-40 | 0.27870332 | 0.308 | 0.103 | 3.7617E-36 | TEC_CD | MRPS26 |
| 1.6133E-40 | 0.33179711 | 0.397 | 0.152 | 3.8112E-36 | TEC_CD | ARHGEF12 |
| 1.8248E-40 | 0.31859767 | 0.19 | 0.047 | 4.3107E-36 | TEC_CD | PEG10 |
| 2.7408E-40 | 0.38078955 | 0.539 | 0.251 | 6.4745E-36 | TEC_CD | RAB14 |
| 3.0939E-40 | 0.26253054 | 0.266 | 0.082 | 7.3087E-36 | TEC_CD | SLC7A1 |
| 3.496E-40 | 0.36644552 | 0.416 | 0.168 | 8.2585E-36 | TEC_CD | MRPS5 |
| 3.6657E-40 | 0.50314381 | 0.525 | 0.252 | 8.6595E-36 | TEC_CD | GABARAPL1 |
| 4.7952E-40 | 0.37729176 | 0.387 | 0.149 | 1.1328E-35 | TEC_CD | SLC44A1 |

| | | | | | | |
|---|---|---|---|---|---|---|
| 5.4798E-40 | 0.52659088 | 0.66 | 0.34 | 1.2945E-35 | TEC_CD | WSB1 |
| 5.6916E-40 | 0.26639825 | 0.264 | 0.082 | 1.3445E-35 | TEC_CD | ATN1 |
| 6.1093E-40 | 0.35187657 | 0.369 | 0.14 | 1.4432E-35 | TEC_CD | BACE2 |
| 7.0313E-40 | 0.30791671 | 0.283 | 0.091 | 1.661E-35 | TEC_CD | ATP8B1 |
| 7.5014E-40 | 0.92081095 | 0.82 | 0.629 | 1.7721E-35 | TEC_CD | ZFP36L1 |
| 8.4061E-40 | 1.18946718 | 0.483 | 0.235 | 1.9858E-35 | TEC_CD | CITED2 |
| 1.0248E-39 | 0.55305735 | 0.399 | 0.169 | 2.4209E-35 | TEC_CD | POR |
| 1.5903E-39 | 0.31221903 | 0.352 | 0.129 | 3.7567E-35 | TEC_CD | PGRMC2 |
| 1.9479E-39 | -2.3147976 | 0.032 | 0.344 | 4.6016E-35 | TEC_CD | PSCA |
| 1.9683E-39 | 0.30119604 | 0.389 | 0.151 | 4.6497E-35 | TEC_CD | SPAG7 |
| 2.1291E-39 | 0.32389287 | 0.305 | 0.105 | 5.0295E-35 | TEC_CD | SLC39A6 |
| 2.4109E-39 | 0.26301144 | 0.261 | 0.081 | 5.6953E-35 | TEC_CD | SUCLA2 |
| 2.507E-39 | 0.26440873 | 0.254 | 0.077 | 5.9222E-35 | TEC_CD | BEND7 |
| 2.5766E-39 | 0.27042394 | 0.273 | 0.088 | 6.0868E-35 | TEC_CD | CETN2 |
| 2.8463E-39 | 0.450063 | 0.431 | 0.188 | 6.7238E-35 | TEC_CD | NDUFS7 |
| 3.4305E-39 | -0.7928084 | 0.813 | 0.918 | 8.1039E-35 | TEC_CD | UBA52 |
| 4.0473E-39 | 0.38360531 | 0.608 | 0.304 | 9.5608E-35 | TEC_CD | ANAPC16 |
| 5.6927E-39 | 0.6291878 | 0.522 | 0.258 | 1.3448E-34 | TEC_CD | MBP |
| 7.0837E-39 | 0.3542583 | 0.5 | 0.223 | 1.6734E-34 | TEC_CD | ARPP19 |
| 7.2103E-39 | 0.32803772 | 0.394 | 0.156 | 1.7033E-34 | TEC_CD | ESD |
| 8.7353E-39 | 0.25584909 | 0.278 | 0.09 | 2.0635E-34 | TEC_CD | MRPS25 |
| 9.7153E-39 | 0.34791991 | 0.249 | 0.076 | 2.2951E-34 | TEC_CD | AL121944.1 |
| 9.9003E-39 | 0.45817498 | 0.347 | 0.126 | 2.3387E-34 | TEC_CD | TFPI2 |
| 1.6583E-38 | 0.51285418 | 0.672 | 0.343 | 3.9174E-34 | TEC_CD | ZFP36L2 |
| 1.6778E-38 | 1.15993849 | 0.564 | 0.264 | 3.9635E-34 | TEC_CD | S100A2 |
| 1.8124E-38 | 0.31537955 | 0.281 | 0.092 | 4.2814E-34 | TEC_CD | LRBA |
| 2.1047E-38 | -0.7655536 | 0.406 | 0.677 | 4.9719E-34 | TEC_CD | TALDO1 |
| 2.796E-38 | 0.52358462 | 0.589 | 0.295 | 6.6049E-34 | TEC_CD | PIM3 |
| 2.8213E-38 | 0.33090158 | 0.416 | 0.172 | 6.6648E-34 | TEC_CD | DPY30 |
| 2.928E-38 | -0.7625511 | 0.362 | 0.642 | 6.9167E-34 | TEC_CD | ARPC5 |
| 3.0217E-38 | 0.33155726 | 0.401 | 0.158 | 7.1382E-34 | TEC_CD | NCKAP1 |
| 3.0984E-38 | 0.32847594 | 0.478 | 0.205 | 7.3193E-34 | TEC_CD | MAP4 |
| 3.152E-38 | 0.2645922 | 0.31 | 0.107 | 7.446E-34 | TEC_CD | MRPS18B |
| 3.3122E-38 | 0.52703193 | 0.532 | 0.27 | 7.8243E-34 | TEC_CD | ATP5F1C |
| 3.6156E-38 | 0.59589065 | 0.399 | 0.172 | 8.5411E-34 | TEC_CD | CSRP1 |
| 3.6684E-38 | 0.28079554 | 0.372 | 0.141 | 8.6658E-34 | TEC_CD | SPART |
| 3.9485E-38 | 0.36165491 | 0.542 | 0.256 | 9.3274E-34 | TEC_CD | TMCO1 |
| 4.3872E-38 | 0.39218411 | 0.631 | 0.312 | 1.0364E-33 | TEC_CD | AKAP9 |
| 4.6629E-38 | 0.2627502 | 0.251 | 0.078 | 1.1015E-33 | TEC_CD | TSC1 |
| 4.7342E-38 | 0.40500965 | 0.347 | 0.133 | 1.1184E-33 | TEC_CD | ESF1 |
| 4.7596E-38 | 0.42723141 | 0.616 | 0.317 | 1.1244E-33 | TEC_CD | ERP29 |
| 5.1407E-38 | 0.28385853 | 0.293 | 0.099 | 1.2144E-33 | TEC_CD | GGCT |
| 7.7004E-38 | 0.27662484 | 0.305 | 0.106 | 1.8191E-33 | TEC_CD | PLEKHF2 |
| 8.3932E-38 | 0.2798805 | 0.328 | 0.118 | 1.9827E-33 | TEC_CD | PPIL4 |
| 8.5988E-38 | 0.40505394 | 0.438 | 0.188 | 2.0313E-33 | TEC_CD | CHD9 |
| 9.6434E-38 | 0.38097376 | 0.517 | 0.243 | 2.2781E-33 | TEC_CD | METTL9 |
| 1.143E-37 | 0.40491135 | 0.633 | 0.321 | 2.7002E-33 | TEC_CD | TGOLN2 |
| 1.5083E-37 | 0.45544386 | 0.537 | 0.247 | 3.5631E-33 | TEC_CD | KLF5 |
| 1.7247E-37 | 0.27339307 | 0.315 | 0.111 | 4.0743E-33 | TEC_CD | DMAC1 |
| 1.7824E-37 | 0.26007717 | 0.256 | 0.081 | 4.2105E-33 | TEC_CD | MRPL12 |

| | | | | | | |
|---|---|---|---|---|---|---|
| 1.8861E-37 | 0.25739595 | 0.298 | 0.1 | 4.4555E-33 | TEC_CD | IQGAP2 |
| 2.0008E-37 | 0.26156972 | 0.264 | 0.085 | 4.7265E-33 | TEC_CD | DMXL1 |
| 2.4284E-37 | 0.28305856 | 0.271 | 0.088 | 5.7365E-33 | TEC_CD | CPEB2 |
| 2.533E-37 | 0.33449924 | 0.372 | 0.142 | 5.9837E-33 | TEC_CD | EPS8 |
| 2.577E-37 | 0.27865973 | 0.293 | 0.101 | 6.0876E-33 | TEC_CD | MCUR1 |
| 2.6275E-37 | 0.26574833 | 0.259 | 0.081 | 6.207E-33 | TEC_CD | BCL2 |
| 4.0852E-37 | 0.25248828 | 0.251 | 0.078 | 9.6504E-33 | TEC_CD | TRAK1 |
| 4.7054E-37 | 0.28438154 | 0.367 | 0.14 | 1.1116E-32 | TEC_CD | NME3 |
| 7.0408E-37 | 0.35679276 | 0.527 | 0.25 | 1.6632E-32 | TEC_CD | HADHA |
| 7.6993E-37 | 0.32008452 | 0.328 | 0.118 | 1.8188E-32 | TEC_CD | CTNNAL1 |
| 8.9785E-37 | 0.27076227 | 0.286 | 0.096 | 2.121E-32 | TEC_CD | CDC37L1 |
| 1.2661E-36 | 0.34130644 | 0.362 | 0.142 | 2.9909E-32 | TEC_CD | MAP1LC3A |
| 1.7516E-36 | 0.39530199 | 0.468 | 0.217 | 4.1379E-32 | TEC_CD | MDH2 |
| 1.7625E-36 | 0.30520663 | 0.328 | 0.121 | 4.1636E-32 | TEC_CD | TMEM51 |
| 1.7918E-36 | 0.31786416 | 0.217 | 0.063 | 4.2329E-32 | TEC_CD | AGPAT5 |
| 2.0742E-36 | -0.9272439 | 0.012 | 0.305 | 4.8998E-32 | TEC_CD | SERPINA1 |
| 2.4461E-36 | 0.69820751 | 0.798 | 0.67 | 5.7784E-32 | TEC_CD | COX5B |
| 3.3484E-36 | 0.35989841 | 0.268 | 0.09 | 7.9099E-32 | TEC_CD | SEMA3B |
| 3.3536E-36 | 0.53100835 | 0.783 | 0.549 | 7.9222E-32 | TEC_CD | UQCR10 |
| 3.3904E-36 | 0.44829718 | 0.512 | 0.254 | 8.0092E-32 | TEC_CD | NDUFB10 |
| 3.7576E-36 | 0.27929455 | 0.305 | 0.108 | 8.8765E-32 | TEC_CD | GPR160 |
| 7.2897E-36 | 0.26689061 | 0.296 | 0.103 | 1.722E-31 | TEC_CD | RHOD |
| 8.1294E-36 | 0.34775515 | 0.32 | 0.117 | 1.9204E-31 | TEC_CD | MUC20 |
| 1.1838E-35 | 0.31578332 | 0.387 | 0.159 | 2.7966E-31 | TEC_CD | SDHB |
| 1.3308E-35 | 0.29660209 | 0.379 | 0.15 | 3.1437E-31 | TEC_CD | PLEKHA1 |
| 1.6477E-35 | -1.1381343 | 0 | 0.284 | 3.8923E-31 | TEC_CD | PTPRC |
| 1.869E-35 | -1.2321961 | 0.17 | 0.46 | 4.4152E-31 | TEC_CD | MXD1 |
| 1.9721E-35 | 0.31986334 | 0.36 | 0.143 | 4.6587E-31 | TEC_CD | DYNLL2 |
| 3.3889E-35 | 0.25680531 | 0.288 | 0.101 | 8.0056E-31 | TEC_CD | ZMPSTE24 |
| 4.1515E-35 | 0.32947029 | 0.419 | 0.174 | 9.8072E-31 | TEC_CD | SMIM22 |
| 4.5705E-35 | 0.26484422 | 0.271 | 0.092 | 1.0797E-30 | TEC_CD | SLC2A4RG |
| 4.5798E-35 | 0.38279081 | 0.655 | 0.347 | 1.0819E-30 | TEC_CD | VDAC1 |
| 4.6398E-35 | 0.26866864 | 0.281 | 0.098 | 1.0961E-30 | TEC_CD | PIP4P1 |
| 5.476E-35 | 0.35840826 | 0.429 | 0.188 | 1.2936E-30 | TEC_CD | TBL1XR1 |
| 5.7764E-35 | 0.49712 | 0.48 | 0.226 | 1.3646E-30 | TEC_CD | IVNS1ABP |
| 6.8051E-35 | 0.26378189 | 0.278 | 0.095 | 1.6076E-30 | TEC_CD | KAT6B |
| 6.8465E-35 | 0.25867249 | 0.345 | 0.131 | 1.6173E-30 | TEC_CD | EI24 |
| 8.4562E-35 | -0.9055038 | 0.212 | 0.542 | 1.9976E-30 | TEC_CD | PLAUR |
| 9.7523E-35 | 0.33114964 | 0.387 | 0.161 | 2.3038E-30 | TEC_CD | TMEM87A |
| 9.8935E-35 | 0.25470388 | 0.268 | 0.091 | 2.3371E-30 | TEC_CD | PDHB |
| 1.0019E-34 | 0.29564958 | 0.323 | 0.122 | 2.3668E-30 | TEC_CD | DNAJC19 |
| 1.0845E-34 | 0.27311866 | 0.273 | 0.094 | 2.562E-30 | TEC_CD | MAP4K3 |
| 1.2249E-34 | -0.8589239 | 0.66 | 0.87 | 2.8937E-30 | TEC_CD | ANXA2 |
| 1.2371E-34 | 0.29050282 | 0.328 | 0.123 | 2.9223E-30 | TEC_CD | ZBTB44 |
| 1.3236E-34 | 0.29718041 | 0.296 | 0.106 | 3.1268E-30 | TEC_CD | EEF1E1 |
| 1.4754E-34 | 0.27073433 | 0.323 | 0.121 | 3.4854E-30 | TEC_CD | TMEM9B |
| 1.7648E-34 | 0.28102855 | 0.337 | 0.129 | 4.1689E-30 | TEC_CD | MXD4 |
| 1.8798E-34 | 0.25192372 | 0.222 | 0.068 | 4.4406E-30 | TEC_CD | ARHGAP35 |
| 2.3033E-34 | 0.25733009 | 0.271 | 0.093 | 5.4411E-30 | TEC_CD | OARD1 |
| 2.7561E-34 | 0.50304212 | 0.429 | 0.186 | 6.5106E-30 | TEC_CD | SLC5A3 |

| | | | | | | |
|---|---|---|---|---|---|---|
| 2.7674E-34 | 0.38021887 | 0.544 | 0.265 | 6.5374E-30 | TEC_CD | HSPA9 |
| 3.5844E-34 | 0.34459421 | 0.345 | 0.136 | 8.4675E-30 | TEC_CD | MYO1E |
| 3.8574E-34 | 0.39560595 | 0.389 | 0.166 | 9.1123E-30 | TEC_CD | NRIP1 |
| 3.8919E-34 | 0.29850199 | 0.328 | 0.126 | 9.1939E-30 | TEC_CD | DNPH1 |
| 3.8954E-34 | 0.31052845 | 0.337 | 0.132 | 9.202E-30 | TEC_CD | MRPS33 |
| 4.7261E-34 | -1.272237 | 0.002 | 0.277 | 1.1164E-29 | TEC_CD | AIF1 |
| 4.9406E-34 | 0.25935211 | 0.266 | 0.09 | 1.1671E-29 | TEC_CD | C1orf35 |
| 6.5665E-34 | 0.35630958 | 0.517 | 0.251 | 1.5512E-29 | TEC_CD | DHRS7 |
| 7.0592E-34 | 0.30278279 | 0.404 | 0.172 | 1.6676E-29 | TEC_CD | RNF114 |
| 1.1043E-33 | 0.31367929 | 0.485 | 0.214 | 2.6087E-29 | TEC_CD | DSP |
| 1.1649E-33 | 0.38723352 | 0.534 | 0.269 | 2.7518E-29 | TEC_CD | NDUFB8 |
| 1.3199E-33 | 0.41136276 | 0.655 | 0.368 | 3.118E-29 | TEC_CD | CIRBP |
| 1.3751E-33 | 0.43691652 | 0.717 | 0.425 | 3.2483E-29 | TEC_CD | SEC62 |
| 1.5278E-33 | 0.27673431 | 0.296 | 0.107 | 3.609E-29 | TEC_CD | WDR43 |
| 1.8994E-33 | 0.53250006 | 0.458 | 0.223 | 4.4869E-29 | TEC_CD | PNP |
| 2.0907E-33 | 0.41872142 | 0.493 | 0.241 | 4.9388E-29 | TEC_CD | NDUFA3 |
| 2.4486E-33 | -0.9645543 | 0.02 | 0.294 | 5.7844E-29 | TEC_CD | LCP1 |
| 2.4693E-33 | 0.294889 | 0.273 | 0.097 | 5.8333E-29 | TEC_CD | RNF24 |
| 2.8466E-33 | 0.26566313 | 0.251 | 0.084 | 6.7244E-29 | TEC_CD | BRIX1 |
| 3.6537E-33 | 0.8425237 | 0.53 | 0.287 | 8.6312E-29 | TEC_CD | MIR4435-2HG |
| 4.1969E-33 | 0.43135331 | 0.414 | 0.173 | 9.9143E-29 | TEC_CD | IGFBP3 |
| 4.496E-33 | 0.35385076 | 0.436 | 0.199 | 1.0621E-28 | TEC_CD | C19orf70 |
| 4.6821E-33 | 0.25084727 | 0.268 | 0.094 | 1.106E-28 | TEC_CD | PPA2 |
| 4.9713E-33 | 0.44423697 | 0.581 | 0.322 | 1.1744E-28 | TEC_CD | GHITM |
| 5.3826E-33 | -0.7269478 | 0.002 | 0.27 | 1.2715E-28 | TEC_CD | CD53 |
| 5.9004E-33 | 0.29240936 | 0.374 | 0.156 | 1.3938E-28 | TEC_CD | EAPP |
| 6.1284E-33 | 0.38421334 | 0.525 | 0.248 | 1.4477E-28 | TEC_CD | SYNE2 |
| 6.9871E-33 | 0.38185349 | 0.564 | 0.291 | 1.6506E-28 | TEC_CD | COX17 |
| 7.5442E-33 | 0.42095841 | 0.931 | 0.864 | 1.7822E-28 | TEC_CD | SRP14 |
| 8.258E-33 | 0.37166598 | 0.5 | 0.239 | 1.9508E-28 | TEC_CD | IFRD1 |
| 8.5628E-33 | 0.38595652 | 0.606 | 0.324 | 2.0228E-28 | TEC_CD | EIF1B |
| 9.5103E-33 | 0.30112314 | 0.377 | 0.158 | 2.2466E-28 | TEC_CD | DUT |
| 9.656E-33 | 0.35709413 | 0.443 | 0.205 | 2.281E-28 | TEC_CD | COX14 |
| 1.1594E-32 | 0.34404455 | 0.52 | 0.258 | 2.7388E-28 | TEC_CD | CAMTA1 |
| 1.2464E-32 | 0.27579548 | 0.278 | 0.1 | 2.9443E-28 | TEC_CD | PISD |
| 1.2517E-32 | 0.29210101 | 0.236 | 0.078 | 2.9569E-28 | TEC_CD | ITPR3 |
| 1.3816E-32 | 0.80630646 | 0.687 | 0.51 | 3.2637E-28 | TEC_CD | COX7B |
| 1.8627E-32 | 1.06333918 | 0.567 | 0.351 | 4.4003E-28 | TEC_CD | ATF3 |
| 2.0922E-32 | 0.25257281 | 0.187 | 0.053 | 4.9425E-28 | TEC_CD | SOCS2 |
| 2.3222E-32 | 0.45893445 | 0.554 | 0.298 | 5.4856E-28 | TEC_CD | ATP5PO |
| 2.914E-32 | 0.36615875 | 0.53 | 0.265 | 6.8838E-28 | TEC_CD | TOR1AIP2 |
| 3.2004E-32 | -0.964032 | 0.264 | 0.527 | 7.5602E-28 | TEC_CD | CEBPB |
| 3.4565E-32 | 0.32764281 | 0.436 | 0.198 | 8.1654E-28 | TEC_CD | NAA20 |
| 3.4664E-32 | 0.27629461 | 0.217 | 0.068 | 8.1886E-28 | TEC_CD | CNKSR3 |
| 3.7804E-32 | 0.28482051 | 0.303 | 0.115 | 8.9303E-28 | TEC_CD | MRPS23 |
| 5.3619E-32 | 0.36772937 | 0.433 | 0.199 | 1.2667E-27 | TEC_CD | ZNF217 |
| 5.5895E-32 | 0.32808312 | 0.335 | 0.134 | 1.3204E-27 | TEC_CD | MBNL2 |
| 6.3487E-32 | 0.34303766 | 0.399 | 0.177 | 1.4997E-27 | TEC_CD | GPI |
| 7.8069E-32 | 0.34736695 | 0.658 | 0.346 | 1.8442E-27 | TEC_CD | UBE2H |
| 9.6151E-32 | 0.26113313 | 0.214 | 0.067 | 2.2714E-27 | TEC_CD | C19orf48 |

| | | | | | | |
|---|---|---|---|---|---|---|
| 1.0856E-31 | 0.29339112 | 0.397 | 0.171 | 2.5644E-27 | TEC_CD | FAM162A |
| 1.1421E-31 | 0.33456821 | 0.502 | 0.247 | 2.6979E-27 | TEC_CD | ATP5PB |
| 1.1571E-31 | 0.37566309 | 0.411 | 0.185 | 2.7335E-27 | TEC_CD | HSBP1L1 |
| 1.2128E-31 | 0.32028133 | 0.384 | 0.166 | 2.8651E-27 | TEC_CD | EFCAB14 |
| 1.5932E-31 | 0.26567292 | 0.251 | 0.086 | 3.7637E-27 | TEC_CD | ZMAT3 |
| 1.728E-31 | 0.37112977 | 0.564 | 0.3 | 4.082E-27 | TEC_CD | PRKAR1A |
| 1.8438E-31 | 0.46845859 | 0.603 | 0.355 | 4.3556E-27 | TEC_CD | ATP5PD |
| 1.9087E-31 | 0.36390448 | 0.547 | 0.28 | 4.5089E-27 | TEC_CD | RTN3 |
| 1.9984E-31 | 0.25868125 | 0.246 | 0.083 | 4.7207E-27 | TEC_CD | IGFBP2 |
| 2.0823E-31 | 0.40590543 | 0.239 | 0.08 | 4.9189E-27 | TEC_CD | AKR1C3 |
| 2.5589E-31 | -0.6239715 | 0.367 | 0.627 | 6.0449E-27 | TEC_CD | ACTR2 |
| 2.5724E-31 | 0.39831066 | 0.495 | 0.249 | 6.0768E-27 | TEC_CD | NDUFAB1 |
| 2.9654E-31 | 0.26527345 | 0.268 | 0.097 | 7.0052E-27 | TEC_CD | RPF2 |
| 3.2138E-31 | 0.28270238 | 0.352 | 0.146 | 7.592E-27 | TEC_CD | MRPL43 |
| 4.3731E-31 | 0.30211162 | 0.431 | 0.195 | 1.033E-26 | TEC_CD | CCPG1 |
| 4.9631E-31 | 0.32752647 | 0.621 | 0.338 | 1.1724E-26 | TEC_CD | NDFIP1 |
| 5.1642E-31 | 0.25379031 | 0.342 | 0.138 | 1.2199E-26 | TEC_CD | ZNF593 |
| 5.2267E-31 | 0.31010664 | 0.357 | 0.149 | 1.2347E-26 | TEC_CD | ZNF83 |
| 5.3756E-31 | 0.38552932 | 0.663 | 0.384 | 1.2699E-26 | TEC_CD | DAD1 |
| 6.2303E-31 | 0.37320028 | 0.564 | 0.293 | 1.4718E-26 | TEC_CD | TRAM1 |
| 7.9546E-31 | -2.1095243 | 0.084 | 0.347 | 1.8791E-26 | TEC_CD | HPGD |
| 8.0139E-31 | 0.2869642 | 0.345 | 0.142 | 1.8931E-26 | TEC_CD | AGPAT2 |
| 1.0379E-30 | 0.29419365 | 0.281 | 0.104 | 2.4519E-26 | TEC_CD | MAP3K1 |
| 1.5163E-30 | 0.54128232 | 0.433 | 0.207 | 3.5819E-26 | TEC_CD | MDM2 |
| 1.8404E-30 | 0.29899747 | 0.333 | 0.137 | 4.3477E-26 | TEC_CD | HEXA |
| 1.9598E-30 | 0.33545253 | 0.456 | 0.212 | 4.6296E-26 | TEC_CD | CNN3 |
| 1.9842E-30 | 0.35991587 | 0.549 | 0.278 | 4.6873E-26 | TEC_CD | H1F0 |
| 1.9845E-30 | 0.28421385 | 0.31 | 0.123 | 4.688E-26 | TEC_CD | MSRB2 |
| 2.1135E-30 | 0.43566009 | 0.335 | 0.141 | 4.9928E-26 | TEC_CD | PHLDA3 |
| 2.9774E-30 | 0.27575896 | 0.335 | 0.137 | 7.0334E-26 | TEC_CD | PCGF5 |
| 3.3906E-30 | -0.7639372 | 0 | 0.248 | 8.0097E-26 | TEC_CD | CSTA |
| 3.4379E-30 | 0.3323918 | 0.547 | 0.285 | 8.1214E-26 | TEC_CD | MRFAP1 |
| 3.4478E-30 | -0.6376814 | 0 | 0.248 | 8.1446E-26 | TEC_CD | GMFG |
| 3.527E-30 | 1.05618408 | 0.643 | 0.505 | 8.3318E-26 | TEC_CD | MT-ND4L |
| 4.5347E-30 | 0.28267782 | 0.323 | 0.129 | 1.0712E-25 | TEC_CD | PPFIBP1 |
| 4.5557E-30 | 0.34076318 | 0.461 | 0.227 | 1.0762E-25 | TEC_CD | SAT2 |
| 4.8639E-30 | 0.31577489 | 0.392 | 0.177 | 1.149E-25 | TEC_CD | UQCRC1 |
| 5.072E-30 | 0.27604757 | 0.296 | 0.114 | 1.1982E-25 | TEC_CD | VEZT |
| 5.8169E-30 | 0.33567347 | 0.532 | 0.275 | 1.3741E-25 | TEC_CD | PLEKHB2 |
| 5.9288E-30 | 0.2755251 | 0.362 | 0.154 | 1.4006E-25 | TEC_CD | RPAIN |
| 6.2231E-30 | 0.50887816 | 0.845 | 0.723 | 1.4701E-25 | TEC_CD | COX6A1 |
| 7.6547E-30 | -0.7312331 | 0.973 | 0.978 | 1.8083E-25 | TEC_CD | RPL28 |
| 8.5995E-30 | 0.2547827 | 0.333 | 0.136 | 2.0315E-25 | TEC_CD | MRPS7 |
| 9.428E-30 | 0.27234313 | 0.286 | 0.109 | 2.2272E-25 | TEC_CD | POGZ |
| 9.4545E-30 | 0.267346 | 0.52 | 0.245 | 2.2334E-25 | TEC_CD | SERINC2 |
| 9.9392E-30 | -0.5969547 | 0.623 | 0.801 | 2.3479E-25 | TEC_CD | ARPC2 |
| 1.1178E-29 | 0.3132463 | 0.382 | 0.171 | 2.6405E-25 | TEC_CD | ATRAID |
| 1.2358E-29 | 0.3266639 | 0.379 | 0.169 | 2.9194E-25 | TEC_CD | SREK1 |
| 1.4375E-29 | 0.25114853 | 0.31 | 0.123 | 3.3957E-25 | TEC_CD | C12orf49 |
| 1.4551E-29 | 0.28157384 | 0.414 | 0.189 | 3.4375E-25 | TEC_CD | SDHC |

| | | | | | | |
|---|---|---|---|---|---|---|
| 1.5763E-29 | -0.9056618 | 0.182 | 0.452 | 3.7237E-25 | TEC_CD | MARCKS |
| 1.6237E-29 | 0.28768368 | 0.362 | 0.156 | 3.8357E-25 | TEC_CD | MTIF3 |
| 1.8726E-29 | 0.26475853 | 0.421 | 0.194 | 4.4235E-25 | TEC_CD | HIGD1A |
| 2.0538E-29 | -0.6198085 | 0.764 | 0.906 | 4.8516E-25 | TEC_CD | CFL1 |
| 2.4012E-29 | 0.38165069 | 0.667 | 0.382 | 5.6724E-25 | TEC_CD | TBCA |
| 2.8038E-29 | 0.31660727 | 0.739 | 0.424 | 6.6233E-25 | TEC_CD | SPINT2 |
| 2.9141E-29 | 0.42902439 | 0.631 | 0.369 | 6.884E-25 | TEC_CD | RHEB |
| 2.9455E-29 | 0.35324264 | 0.448 | 0.22 | 6.9581E-25 | TEC_CD | PPP2CB |
| 3.0961E-29 | 0.28719294 | 0.315 | 0.128 | 7.314E-25 | TEC_CD | GSR |
| 3.1006E-29 | 0.26681788 | 0.256 | 0.092 | 7.3247E-25 | TEC_CD | SCNN1B |
| 3.3146E-29 | 0.27211353 | 0.347 | 0.147 | 7.83E-25 | TEC_CD | NUB1 |
| 3.3971E-29 | 0.3117896 | 0.365 | 0.158 | 8.0249E-25 | TEC_CD | ADSS |
| 3.7539E-29 | 0.44834893 | 0.458 | 0.233 | 8.8679E-25 | TEC_CD | PLXNB2 |
| 4.6953E-29 | 0.30081762 | 0.399 | 0.183 | 1.1092E-24 | TEC_CD | NDUFAF3 |
| 5.314E-29 | 0.34472622 | 0.493 | 0.25 | 1.2553E-24 | TEC_CD | FIS1 |
| 5.4163E-29 | 0.45265008 | 0.586 | 0.346 | 1.2795E-24 | TEC_CD | DDT |
| 5.9206E-29 | 0.26555554 | 0.278 | 0.106 | 1.3986E-24 | TEC_CD | MAP4K5 |
| 5.9548E-29 | 0.33551729 | 0.401 | 0.182 | 1.4067E-24 | TEC_CD | EPS8L2 |
| 6.5398E-29 | 0.36870722 | 0.451 | 0.225 | 1.5449E-24 | TEC_CD | PCMTD1 |
| 7.8697E-29 | 0.2840385 | 0.411 | 0.19 | 1.8591E-24 | TEC_CD | ATP6AP1 |
| 8.0267E-29 | 0.30119491 | 0.537 | 0.272 | 1.8961E-24 | TEC_CD | SSB |
| 1.1852E-28 | 0.36561258 | 0.337 | 0.139 | 2.7998E-24 | TEC_CD | MSMO1 |
| 1.4988E-28 | 0.25491933 | 0.308 | 0.123 | 3.5407E-24 | TEC_CD | PTK2 |
| 1.5539E-28 | -0.9403596 | 0.135 | 0.386 | 3.6709E-24 | TEC_CD | TYMP |
| 1.9673E-28 | 0.28294735 | 0.345 | 0.146 | 4.6473E-24 | TEC_CD | SUCO |
| 2.0032E-28 | 0.3037246 | 0.296 | 0.12 | 4.7321E-24 | TEC_CD | PKN1 |
| 2.523E-28 | 0.34266436 | 0.229 | 0.081 | 5.96E-24 | TEC_CD | PXDC1 |
| 2.5504E-28 | 0.25318478 | 0.31 | 0.126 | 6.0247E-24 | TEC_CD | SDHA |
| 2.6908E-28 | 0.55583523 | 0.552 | 0.296 | 6.3564E-24 | TEC_CD | BTG2 |
| 2.8791E-28 | 0.2992752 | 0.278 | 0.109 | 6.8014E-24 | TEC_CD | KMT5A |
| 2.8989E-28 | 0.3120851 | 0.36 | 0.159 | 6.848E-24 | TEC_CD | PRRC2B |
| 2.9727E-28 | 0.28264456 | 0.32 | 0.131 | 7.0224E-24 | TEC_CD | RASSF7 |
| 2.9916E-28 | -0.647445 | 0.502 | 0.705 | 7.067E-24 | TEC_CD | TPM3 |
| 3.9637E-28 | 0.32167593 | 0.303 | 0.124 | 9.3635E-24 | TEC_CD | MAN1A1 |
| 4.0878E-28 | 0.30669967 | 0.419 | 0.197 | 9.6565E-24 | TEC_CD | TIMM13 |
| 4.3779E-28 | 0.3550689 | 0.505 | 0.26 | 1.0342E-23 | TEC_CD | MPHOSPH8 |
| 4.6529E-28 | 0.38483652 | 0.818 | 0.599 | 1.0992E-23 | TEC_CD | YWHAE |
| 5.5308E-28 | 0.27890635 | 0.281 | 0.109 | 1.3065E-23 | TEC_CD | ERC1 |
| 7.1143E-28 | 0.2702243 | 0.313 | 0.128 | 1.6806E-23 | TEC_CD | COMTD1 |
| 7.8155E-28 | -1.1054975 | 0.002 | 0.235 | 1.8463E-23 | TEC_CD | SLC2A3 |
| 8.894E-28 | -1.1110728 | 0.005 | 0.238 | 2.101E-23 | TEC_CD | BCL2A1 |
| 9.8803E-28 | 0.26267482 | 0.345 | 0.149 | 2.334E-23 | TEC_CD | FKBP3 |
| 1.0087E-27 | 0.28104131 | 0.286 | 0.114 | 2.3829E-23 | TEC_CD | PBDC1 |
| 1.0754E-27 | 0.25789239 | 0.335 | 0.141 | 2.5404E-23 | TEC_CD | MYO1C |
| 1.1068E-27 | 0.72801244 | 0.983 | 0.98 | 2.6147E-23 | TEC_CD | MALAT1 |
| 1.2967E-27 | 0.35496282 | 0.458 | 0.232 | 3.0632E-23 | TEC_CD | SCARB2 |
| 1.3564E-27 | 0.29892574 | 0.313 | 0.131 | 3.2042E-23 | TEC_CD | VEZF1 |
| 1.3909E-27 | 0.39328455 | 0.635 | 0.372 | 3.2857E-23 | TEC_CD | CCNL1 |
| 1.9256E-27 | 0.34081389 | 0.665 | 0.381 | 4.5489E-23 | TEC_CD | DYNLRB1 |
| 2.0778E-27 | 0.27195325 | 0.394 | 0.179 | 4.9083E-23 | TEC_CD | MEF2A |

| | | | | | | |
|---|---|---|---|---|---|---|
| 2.219E-27 | 0.25324442 | 0.379 | 0.168 | 5.2421E-23 | TEC_CD | GNG12 |
| 2.228E-27 | 0.32038405 | 0.397 | 0.184 | 5.2632E-23 | TEC_CD | DYNLT3 |
| 2.2377E-27 | 0.25816388 | 0.298 | 0.121 | 5.2861E-23 | TEC_CD | LYRM2 |
| 2.7946E-27 | -0.7350289 | 0.094 | 0.349 | 6.6016E-23 | TEC_CD | IER5 |
| 3.1667E-27 | 0.33494243 | 0.539 | 0.289 | 7.4806E-23 | TEC_CD | HNRNPH3 |
| 3.1985E-27 | 0.32203333 | 0.473 | 0.24 | 7.5558E-23 | TEC_CD | TMEM14C |
| 5.0432E-27 | -0.7553263 | 0.002 | 0.229 | 1.1914E-22 | TEC_CD | ZEB2 |
| 5.0482E-27 | 0.25227849 | 0.239 | 0.086 | 1.1925E-22 | TEC_CD | PDE4D |
| 5.1492E-27 | 0.341024 | 0.463 | 0.229 | 1.2164E-22 | TEC_CD | BCL10 |
| 5.4553E-27 | 0.27174939 | 0.34 | 0.147 | 1.2887E-22 | TEC_CD | LAMTOR3 |
| 5.9765E-27 | 0.37461103 | 0.53 | 0.281 | 1.4118E-22 | TEC_CD | RPS4Y1 |
| 6.1849E-27 | 0.33169724 | 0.515 | 0.267 | 1.4611E-22 | TEC_CD | PHPT1 |
| 6.3374E-27 | -1.4203898 | 0.094 | 0.335 | 1.4971E-22 | TEC_CD | IL1RN |
| 6.6967E-27 | 0.25866477 | 0.283 | 0.114 | 1.582E-22 | TEC_CD | OGDH |
| 7.5616E-27 | 0.33556828 | 0.219 | 0.077 | 1.7863E-22 | TEC_CD | CRABP2 |
| 9.6542E-27 | 0.29667269 | 0.488 | 0.251 | 2.2806E-22 | TEC_CD | SCP2 |
| 9.7983E-27 | 0.47118265 | 0.293 | 0.124 | 2.3146E-22 | TEC_CD | PLA2G16 |
| 1.107E-26 | 0.27228396 | 0.431 | 0.206 | 2.6151E-22 | TEC_CD | PPP2R1A |
| 1.1247E-26 | 0.25757223 | 0.288 | 0.116 | 2.657E-22 | TEC_CD | TULP4 |
| 1.1705E-26 | -0.5285872 | 0.002 | 0.227 | 2.765E-22 | TEC_CD | KYNU |
| 1.2409E-26 | 0.4714252 | 0.547 | 0.314 | 2.9315E-22 | TEC_CD | PTBP3 |
| 2.322E-26 | -1.7133579 | 0.002 | 0.225 | 5.4852E-22 | TEC_CD | VCAN |
| 2.3329E-26 | -1.0887131 | 0 | 0.221 | 5.5109E-22 | TEC_CD | ALOX5AP |
| 2.3805E-26 | -0.6353064 | 0 | 0.221 | 5.6234E-22 | TEC_CD | ITGB2 |
| 2.4007E-26 | -0.7959506 | 0.914 | 0.904 | 5.6712E-22 | TEC_CD | RPLP2 |
| 2.7924E-26 | -0.9313358 | 0.02 | 0.246 | 6.5965E-22 | TEC_CD | SLC11A1 |
| 2.8857E-26 | 0.25246943 | 0.337 | 0.146 | 6.817E-22 | TEC_CD | NUMA1 |
| 2.9544E-26 | -0.8394146 | 0.012 | 0.238 | 6.9792E-22 | TEC_CD | FABP5 |
| 2.9906E-26 | 0.26914267 | 0.345 | 0.152 | 7.0647E-22 | TEC_CD | PFKL |
| 3.3736E-26 | -0.7476361 | 0.869 | 0.924 | 7.9694E-22 | TEC_CD | RPL18A |
| 4.2466E-26 | -0.7819257 | 0.882 | 0.941 | 1.0032E-21 | TEC_CD | NEAT1 |
| 4.7709E-26 | 0.27206236 | 0.584 | 0.309 | 1.127E-21 | TEC_CD | SBDS |
| 6.0171E-26 | -0.9932168 | 0.002 | 0.222 | 1.4214E-21 | TEC_CD | CD14 |
| 7.6521E-26 | 0.29581939 | 0.498 | 0.259 | 1.8077E-21 | TEC_CD | EIF3J |
| 8.3309E-26 | 0.26646221 | 0.323 | 0.139 | 1.968E-21 | TEC_CD | ARL3 |
| 8.6278E-26 | 0.32915073 | 0.766 | 0.498 | 2.0381E-21 | TEC_CD | ASAH1 |
| 8.9483E-26 | 0.32507094 | 0.227 | 0.083 | 2.1138E-21 | TEC_CD | AC015912.3 |
| 1.0236E-25 | 0.32362916 | 0.475 | 0.248 | 2.418E-21 | TEC_CD | SH3BGRL |
| 1.0491E-25 | 0.29937568 | 0.308 | 0.132 | 2.4784E-21 | TEC_CD | RNF187 |
| 1.0702E-25 | 0.39037939 | 0.611 | 0.366 | 2.5281E-21 | TEC_CD | NDUFV2 |
| 1.1646E-25 | 0.26713534 | 0.352 | 0.16 | 2.7511E-21 | TEC_CD | SLBP |
| 1.6852E-25 | -1.1218934 | 0.234 | 0.475 | 3.9809E-21 | TEC_CD | ERO1A |
| 1.9083E-25 | 0.32867035 | 0.539 | 0.294 | 4.508E-21 | TEC_CD | EIF5B |
| 2.1444E-25 | 0.32293436 | 0.677 | 0.406 | 5.0656E-21 | TEC_CD | PTP4A2 |
| 2.3597E-25 | -1.1156207 | 0.002 | 0.217 | 5.5742E-21 | TEC_CD | S100A12 |
| 3.041E-25 | 0.25446492 | 0.31 | 0.133 | 7.1837E-21 | TEC_CD | ADNP |
| 3.089E-25 | -0.6869628 | 0.017 | 0.236 | 7.2972E-21 | TEC_CD | HCST |
| 3.1593E-25 | -0.5312683 | 0.399 | 0.617 | 7.4631E-21 | TEC_CD | CAP1 |
| 3.766E-25 | 0.38927913 | 0.803 | 0.6 | 8.8963E-21 | TEC_CD | HINT1 |
| 4.1817E-25 | 0.39871535 | 0.404 | 0.2 | 9.8784E-21 | TEC_CD | MUC20-OT1 |

| | | | | | | |
|---|---|---|---|---|---|---|
| 4.1859E-25 | -0.8100646 | 0.825 | 0.9 | 9.8884E-21 | TEC_CD | RPL39 |
| 4.2037E-25 | 0.34943182 | 0.416 | 0.208 | 9.9304E-21 | TEC_CD | NFAT5 |
| 4.3344E-25 | 0.25709321 | 0.404 | 0.193 | 1.0239E-20 | TEC_CD | CCT2 |
| 4.5422E-25 | -0.7418254 | 0 | 0.212 | 1.073E-20 | TEC_CD | TNFRSF1B |
| 4.5995E-25 | -0.9852833 | 0.057 | 0.288 | 1.0865E-20 | TEC_CD | HMOX1 |
| 5.0874E-25 | -0.4377459 | 0.012 | 0.229 | 1.2018E-20 | TEC_CD | RGS10 |
| 5.1026E-25 | 0.28731718 | 0.411 | 0.198 | 1.2054E-20 | TEC_CD | TTC3 |
| 5.7375E-25 | 0.48130614 | 0.318 | 0.15 | 1.3554E-20 | TEC_CD | IDH2 |
| 8.416E-25 | 0.28003702 | 0.608 | 0.337 | 1.9881E-20 | TEC_CD | SNU13 |
| 1.5123E-24 | -0.6131284 | 0.005 | 0.215 | 3.5725E-20 | TEC_CD | BASP1 |
| 1.5241E-24 | 0.30923714 | 0.631 | 0.364 | 3.6003E-20 | TEC_CD | AES |
| 1.8275E-24 | 0.29035698 | 0.259 | 0.104 | 4.3171E-20 | TEC_CD | RHOF |
| 1.8419E-24 | 0.26602902 | 0.414 | 0.198 | 4.3511E-20 | TEC_CD | ZNF91 |
| 1.9275E-24 | 0.49434298 | 0.729 | 0.505 | 4.5534E-20 | TEC_CD | DBI |
| 1.9742E-24 | -0.5632685 | 0.007 | 0.217 | 4.6636E-20 | TEC_CD | SPI1 |
| 3.375E-24 | -1.0681063 | 0.818 | 0.884 | 7.9727E-20 | TEC_CD | CSTB |
| 3.3972E-24 | -0.5395457 | 0.478 | 0.734 | 8.0253E-20 | TEC_CD | HSPA8 |
| 3.6181E-24 | 0.28280803 | 0.323 | 0.144 | 8.547E-20 | TEC_CD | USP12 |
| 3.688E-24 | -0.7765834 | 0.498 | 0.701 | 8.7121E-20 | TEC_CD | JUNB |
| 3.96E-24 | 0.27821204 | 0.325 | 0.146 | 9.3547E-20 | TEC_CD | DDX3Y |
| 5.4214E-24 | 0.28014894 | 0.365 | 0.171 | 1.2807E-19 | TEC_CD | CEBPZ |
| 6.4422E-24 | -0.8092937 | 0.399 | 0.66 | 1.5218E-19 | TEC_CD | HSPA1A |
| 9.0983E-24 | 0.27906903 | 0.436 | 0.222 | 2.1493E-19 | TEC_CD | NARS |
| 9.8786E-24 | 0.75085744 | 0.451 | 0.257 | 2.3336E-19 | TEC_CD | PMAIP1 |
| 1.0785E-23 | 0.28128549 | 0.397 | 0.195 | 2.5477E-19 | TEC_CD | GNS |
| 1.1342E-23 | -0.4755935 | 0.002 | 0.205 | 2.6792E-19 | TEC_CD | LSP1 |
| 1.1493E-23 | 0.27156928 | 0.394 | 0.194 | 2.7149E-19 | TEC_CD | FNIP1 |
| 1.1986E-23 | 0.25422903 | 0.387 | 0.189 | 2.8313E-19 | TEC_CD | ALKBH7 |
| 1.3141E-23 | 0.27364545 | 0.48 | 0.255 | 3.1043E-19 | TEC_CD | DYNC1I2 |
| 1.4669E-23 | 0.29879536 | 0.419 | 0.214 | 3.4653E-19 | TEC_CD | SMIM14 |
| 2.0386E-23 | -0.5372036 | 0.628 | 0.786 | 4.8157E-19 | TEC_CD | EEF1D |
| 2.3734E-23 | 0.38192915 | 0.495 | 0.277 | 5.6068E-19 | TEC_CD | PTP4A1 |
| 2.3914E-23 | -0.5303827 | 0.682 | 0.837 | 5.6493E-19 | TEC_CD | HLA-E |
| 2.4465E-23 | -0.7347355 | 0.002 | 0.202 | 5.7794E-19 | TEC_CD | CYBB |
| 2.4525E-23 | 0.30768998 | 0.67 | 0.409 | 5.7936E-19 | TEC_CD | VAPA |
| 2.4795E-23 | 0.27449532 | 0.485 | 0.258 | 5.8572E-19 | TEC_CD | SMDT1 |
| 3.4127E-23 | -0.5648008 | 0.002 | 0.202 | 8.0619E-19 | TEC_CD | ITGAX |
| 5.29E-23 | -0.7466944 | 0.002 | 0.2 | 1.2497E-18 | TEC_CD | CD52 |
| 5.7307E-23 | 0.28008898 | 0.626 | 0.37 | 1.3538E-18 | TEC_CD | C11orf58 |
| 6.3443E-23 | 0.30823644 | 0.507 | 0.287 | 1.4987E-18 | TEC_CD | NDUFB7 |
| 6.6188E-23 | -0.548449 | 0.005 | 0.203 | 1.5636E-18 | TEC_CD | NCF2 |
| 7.781E-23 | -0.8008622 | 0.818 | 0.92 | 1.8381E-18 | TEC_CD | S100A10 |
| 8.7053E-23 | 0.54605799 | 0.571 | 0.361 | 2.0565E-18 | TEC_CD | SLC25A5 |
| 9.8907E-23 | 0.33180202 | 0.638 | 0.388 | 2.3365E-18 | TEC_CD | RSRC2 |
| 1.097E-22 | -0.5679418 | 0.01 | 0.209 | 2.5914E-18 | TEC_CD | GK |
| 1.5971E-22 | 0.30687347 | 0.357 | 0.174 | 3.7728E-18 | TEC_CD | ATP6V1A |
| 1.6214E-22 | 0.27376225 | 0.557 | 0.318 | 3.8303E-18 | TEC_CD | ZNF706 |
| 1.7193E-22 | -0.8333031 | 0.653 | 0.768 | 4.0614E-18 | TEC_CD | CST3 |
| 1.7945E-22 | -1.3710082 | 0.086 | 0.297 | 4.239E-18 | TEC_CD | ADIRF |
| 1.9944E-22 | -0.9938588 | 0.099 | 0.314 | 4.7114E-18 | TEC_CD | EMP1 |

| | | | | | | |
|---|---|---|---|---|---|---|
| 2.1069E-22 | 0.33482174 | 0.505 | 0.283 | 4.9772E-18 | TEC_CD | YWHAH |
| 2.4048E-22 | 0.26526517 | 0.291 | 0.129 | 5.6809E-18 | TEC_CD | C7orf50 |
| 2.5793E-22 | 0.25111553 | 0.345 | 0.163 | 6.0931E-18 | TEC_CD | ZCRB1 |
| 3.0754E-22 | 0.29819629 | 0.515 | 0.287 | 7.265E-18 | TEC_CD | NDUFB9 |
| 3.1024E-22 | 0.3328393 | 0.436 | 0.237 | 7.3288E-18 | TEC_CD | TUFM |
| 3.372E-22 | -0.6308581 | 0.116 | 0.327 | 7.9657E-18 | TEC_CD | ARHGDIB |
| 3.4345E-22 | 0.43470914 | 0.911 | 0.837 | 8.1134E-18 | TEC_CD | SQSTM1 |
| 4.5129E-22 | -0.7866271 | 0 | 0.189 | 1.0661E-17 | TEC_CD | AC020656.1 |
| 4.854E-22 | 0.3983313 | 0.138 | 0.042 | 1.1467E-17 | TEC_CD | LIF |
| 5.0535E-22 | 0.31130846 | 0.544 | 0.313 | 1.1938E-17 | TEC_CD | MMP24OS |
| 6.6123E-22 | 0.38546249 | 0.515 | 0.291 | 1.562E-17 | TEC_CD | TUBA1C |
| 6.9517E-22 | -0.6280298 | 0.015 | 0.21 | 1.6422E-17 | TEC_CD | LST1 |
| 7.1604E-22 | 0.27124416 | 0.485 | 0.266 | 1.6915E-17 | TEC_CD | NDUFA12 |
| 9.6857E-22 | -0.4825451 | 0.616 | 0.774 | 2.2881E-17 | TEC_CD | EIF4G2 |
| 1.1321E-21 | 0.29101876 | 0.397 | 0.204 | 2.6744E-17 | TEC_CD | NAA38 |
| 1.1775E-21 | 0.31441369 | 0.271 | 0.118 | 2.7817E-17 | TEC_CD | AMOTL2 |
| 1.1901E-21 | -0.5725985 | 0.845 | 0.901 | 2.8113E-17 | TEC_CD | RPS11 |
| 1.1911E-21 | 0.29620512 | 0.268 | 0.113 | 2.8138E-17 | TEC_CD | NQO1 |
| 1.196E-21 | -0.6087755 | 0.835 | 0.9 | 2.8253E-17 | TEC_CD | RPL27A |
| 1.3893E-21 | -0.3885567 | 0.002 | 0.189 | 3.282E-17 | TEC_CD | HCLS1 |
| 1.651E-21 | 0.37424634 | 0.761 | 0.568 | 3.9E-17 | TEC_CD | ZFAND5 |
| 2.266E-21 | 0.26111873 | 0.443 | 0.237 | 5.3529E-17 | TEC_CD | RNMT |
| 2.7545E-21 | -0.8054014 | 0.906 | 0.885 | 6.507E-17 | TEC_CD | RPS2 |
| 3.7302E-21 | -1.2397109 | 0.002 | 0.186 | 8.812E-17 | TEC_CD | DUOX2 |
| 5.6617E-21 | 0.28248832 | 0.446 | 0.242 | 1.3375E-16 | TEC_CD | TERF2IP |
| 1.2174E-20 | 0.31779015 | 0.522 | 0.306 | 2.8759E-16 | TEC_CD | YY1 |
| 1.2559E-20 | -0.8204909 | 0.628 | 0.786 | 2.9667E-16 | TEC_CD | TXN |
| 1.3265E-20 | 0.2525189 | 0.399 | 0.206 | 3.1337E-16 | TEC_CD | GSTK1 |
| 1.3379E-20 | 0.56437375 | 0.374 | 0.195 | 3.1606E-16 | TEC_CD | BIRC3 |
| 1.5474E-20 | 0.26675558 | 0.379 | 0.197 | 3.6554E-16 | TEC_CD | TMEM230 |
| 1.6661E-20 | -0.4625742 | 0.007 | 0.188 | 3.9358E-16 | TEC_CD | PTPRE |
| 1.6805E-20 | -0.5895852 | 0.005 | 0.184 | 3.9697E-16 | TEC_CD | IGSF6 |
| 1.6886E-20 | -2.2263817 | 0 | 0.177 | 3.989E-16 | TEC_CD | IL1B |
| 1.8103E-20 | 0.26906802 | 0.569 | 0.329 | 4.2765E-16 | TEC_CD | PPP1R14B |
| 1.8943E-20 | 0.30795749 | 0.429 | 0.238 | 4.4749E-16 | TEC_CD | SMIM26 |
| 2.1136E-20 | 0.29655519 | 0.352 | 0.177 | 4.993E-16 | TEC_CD | ERICH1 |
| 2.7266E-20 | -0.5722468 | 0.16 | 0.363 | 6.441E-16 | TEC_CD | GNAI2 |
| 3.0821E-20 | -0.7370939 | 0.052 | 0.248 | 7.2808E-16 | TEC_CD | CST6 |
| 3.0878E-20 | 0.2547314 | 0.586 | 0.342 | 7.2942E-16 | TEC_CD | NORAD |
| 3.1074E-20 | 0.27626779 | 0.33 | 0.159 | 7.3407E-16 | TEC_CD | GFPT1 |
| 3.2596E-20 | -0.4233122 | 0 | 0.175 | 7.7002E-16 | TEC_CD | CORO1A |
| 3.4221E-20 | 0.31659082 | 0.729 | 0.518 | 8.084E-16 | TEC_CD | ATP6V1F |
| 3.4897E-20 | -0.5264736 | 0.01 | 0.19 | 8.2437E-16 | TEC_CD | FCGR2A |
| 3.9111E-20 | 0.28695504 | 0.456 | 0.251 | 9.2392E-16 | TEC_CD | NSD3 |
| 3.9625E-20 | 0.76809144 | 0.512 | 0.325 | 9.3607E-16 | TEC_CD | MAFF |
| 4.4422E-20 | -0.5151635 | 0.525 | 0.686 | 1.0494E-15 | TEC_CD | BZW1 |
| 4.9463E-20 | -0.3564674 | 0.007 | 0.185 | 1.1685E-15 | TEC_CD | GNA15 |
| 5.322E-20 | 0.27557973 | 0.567 | 0.336 | 1.2572E-15 | TEC_CD | GPBP1 |
| 6.0922E-20 | -0.7631033 | 0.002 | 0.176 | 1.4392E-15 | TEC_CD | PLEK |
| 7.14E-20 | 0.26028184 | 0.419 | 0.223 | 1.6867E-15 | TEC_CD | TMED5 |

| | | | | | | |
|---|---|---|---|---|---|---|
| 7.8124E-20 | -0.4126923 | 0.027 | 0.211 | 1.8455E-15 | TEC_CD | ARRB2 |
| 9.1523E-20 | -0.6375659 | 0.852 | 0.86 | 2.1621E-15 | TEC_CD | RPL23 |
| 9.7185E-20 | -0.4071823 | 0.012 | 0.189 | 2.2958E-15 | TEC_CD | RAC2 |
| 9.7392E-20 | -0.4752501 | 0.002 | 0.175 | 2.3007E-15 | TEC_CD | SMIM25 |
| 1.0066E-19 | -0.4110494 | 0 | 0.171 | 2.378E-15 | TEC_CD | CD48 |
| 1.0138E-19 | -0.5780782 | 0.005 | 0.178 | 2.395E-15 | TEC_CD | ANPEP |
| 1.0935E-19 | -1.4435079 | 0.02 | 0.198 | 2.5833E-15 | TEC_CD | HOPX |
| 1.1906E-19 | 0.47540734 | 0.766 | 0.648 | 2.8125E-15 | TEC_CD | UQCRH |
| 1.2446E-19 | 0.27494893 | 0.426 | 0.231 | 2.94E-15 | TEC_CD | RYBP |
| 1.338E-19 | -0.695087 | 0.007 | 0.182 | 3.1608E-15 | TEC_CD | OLR1 |
| 1.4189E-19 | -0.4571487 | 0 | 0.17 | 3.3518E-15 | TEC_CD | SLA |
| 1.6916E-19 | -0.5078897 | 0.067 | 0.261 | 3.996E-15 | TEC_CD | NABP1 |
| 2.0266E-19 | 0.4611236 | 0.741 | 0.615 | 4.7874E-15 | TEC_CD | COX8A |
| 2.2009E-19 | -0.5602231 | 0.002 | 0.172 | 5.1991E-15 | TEC_CD | TGFBI |
| 2.6531E-19 | 0.25078073 | 0.522 | 0.301 | 6.2673E-15 | TEC_CD | BLVRB |
| 2.725E-19 | 0.27148221 | 0.47 | 0.267 | 6.4372E-15 | TEC_CD | NOL7 |
| 3.4938E-19 | 0.30528029 | 0.239 | 0.104 | 8.2534E-15 | TEC_CD | LDLR |
| 4.3309E-19 | 0.26349445 | 0.431 | 0.239 | 1.0231E-14 | TEC_CD | POLR2K |
| 4.3343E-19 | -0.3585283 | 0.01 | 0.181 | 1.0239E-14 | TEC_CD | FMNL1 |
| 4.4024E-19 | -0.4361794 | 0 | 0.166 | 1.04E-14 | TEC_CD | LCP2 |
| 4.5059E-19 | -0.4033696 | 0.01 | 0.18 | 1.0644E-14 | TEC_CD | CD37 |
| 5.174E-19 | -0.5288138 | 0.687 | 0.816 | 1.2222E-14 | TEC_CD | YBX1 |
| 6.6057E-19 | 0.27548924 | 0.411 | 0.228 | 1.5605E-14 | TEC_CD | DNAJC15 |
| 7.0193E-19 | -0.92149 | 0.094 | 0.284 | 1.6582E-14 | TEC_CD | MAFB |
| 7.4171E-19 | -0.6672172 | 0.217 | 0.424 | 1.7521E-14 | TEC_CD | TXNRD1 |
| 7.4678E-19 | -0.6528168 | 0.217 | 0.405 | 1.7641E-14 | TEC_CD | FLNA |
| 7.9364E-19 | -0.7970876 | 0.005 | 0.171 | 1.8748E-14 | TEC_CD | FCN1 |
| 9.5413E-19 | -0.4334047 | 0.84 | 0.965 | 2.2539E-14 | TEC_CD | MYL6 |
| 9.9183E-19 | -1.2528916 | 0.002 | 0.167 | 2.343E-14 | TEC_CD | DUOXA2 |
| 1.1552E-18 | -0.65416 | 0.007 | 0.173 | 2.729E-14 | TEC_CD | STEAP4 |
| 1.1999E-18 | -0.4746721 | 0 | 0.162 | 2.8346E-14 | TEC_CD | AQP9 |
| 1.2225E-18 | -1.1465591 | 0 | 0.162 | 2.8878E-14 | TEC_CD | MUC21 |
| 1.2414E-18 | -0.3513861 | 0.017 | 0.188 | 2.9326E-14 | TEC_CD | GPSM3 |
| 1.6203E-18 | -0.5716651 | 0.611 | 0.759 | 3.8275E-14 | TEC_CD | ITGB1 |
| 1.7023E-18 | -0.3495587 | 0.002 | 0.165 | 4.0213E-14 | TEC_CD | MYO1F |
| 1.7641E-18 | -0.4458161 | 0.101 | 0.295 | 4.1672E-14 | TEC_CD | FXYD5 |
| 1.7961E-18 | -0.5834707 | 0.837 | 0.89 | 4.243E-14 | TEC_CD | HLA-B |
| 1.8279E-18 | -0.8298628 | 0.025 | 0.199 | 4.3181E-14 | TEC_CD | TM4SF1 |
| 2.0416E-18 | -0.5942891 | 0.017 | 0.186 | 4.8228E-14 | TEC_CD | C5AR1 |
| 2.3677E-18 | 0.25097602 | 0.495 | 0.285 | 5.5933E-14 | TEC_CD | CLINT1 |
| 2.6224E-18 | -0.5672371 | 0 | 0.159 | 6.1948E-14 | TEC_CD | CD300E |
| 3.0497E-18 | 0.25337619 | 0.608 | 0.372 | 7.2043E-14 | TEC_CD | PARK7 |
| 3.2096E-18 | 0.25433034 | 0.35 | 0.182 | 7.5821E-14 | TEC_CD | UBE2Z |
| 3.4834E-18 | 0.28327886 | 0.426 | 0.24 | 8.2289E-14 | TEC_CD | MAFG |
| 3.8953E-18 | -0.6377428 | 0.892 | 0.902 | 9.2018E-14 | TEC_CD | RPS7 |
| 4.3424E-18 | 0.3566554 | 0.81 | 0.713 | 1.0258E-13 | TEC_CD | SKP1 |
| 4.5232E-18 | -1.0581178 | 0.002 | 0.161 | 1.0685E-13 | TEC_CD | CEACAM5 |
| 4.584E-18 | 0.51178328 | 0.118 | 0.037 | 1.0829E-13 | TEC_CD | TAC1 |
| 5.5383E-18 | -1.5212312 | 0.03 | 0.2 | 1.3083E-13 | TEC_CD | KRT17 |
| 5.8397E-18 | -1.595721 | 0.005 | 0.164 | 1.3795E-13 | TEC_CD | ECM1 |

| | | | | | | |
|---|---|---|---|---|---|---|
| 7.0207E-18 | -0.9826149 | 0 | 0.156 | 1.6585E-13 | TEC_CD | PITX1 |
| 7.1026E-18 | -0.5716127 | 0.424 | 0.574 | 1.6778E-13 | TEC_CD | TSPO |
| 7.4423E-18 | -0.6069386 | 0.446 | 0.643 | 1.7581E-13 | TEC_CD | TXNIP |
| 8.1759E-18 | -0.8505308 | 0.005 | 0.163 | 1.9314E-13 | TEC_CD | APOBEC3A |
| 1.1056E-17 | -0.7582692 | 0 | 0.154 | 2.6118E-13 | TEC_CD | SCEL |
| 1.6157E-17 | -0.3744141 | 0 | 0.153 | 3.8167E-13 | TEC_CD | EVI2B |
| 1.7255E-17 | -0.3613104 | 0.007 | 0.165 | 4.0762E-13 | TEC_CD | CLEC2B |
| 1.7255E-17 | -0.5086366 | 0 | 0.153 | 4.0763E-13 | TEC_CD | CLEC5A |
| 1.8569E-17 | 0.33021974 | 0.591 | 0.386 | 4.3865E-13 | TEC_CD | UQCR11 |
| 2.225E-17 | -0.3409335 | 0.007 | 0.163 | 5.2561E-13 | TEC_CD | FERMT3 |
| 2.7082E-17 | 0.28234846 | 0.825 | 0.658 | 6.3976E-13 | TEC_CD | RBM39 |
| 3.1454E-17 | 0.32864007 | 0.86 | 0.725 | 7.4305E-13 | TEC_CD | ENO1 |
| 3.665E-17 | -0.4140669 | 0.015 | 0.173 | 8.6579E-13 | TEC_CD | SAMHD1 |
| 3.8207E-17 | -0.7608336 | 0 | 0.15 | 9.0257E-13 | TEC_CD | TMPRSS11D |
| 4.2887E-17 | -0.5167755 | 0.015 | 0.172 | 1.0131E-12 | TEC_CD | PRDM1 |
| 4.8739E-17 | -0.4202383 | 0.135 | 0.322 | 1.1514E-12 | TEC_CD | AP2S1 |
| 5.556E-17 | -0.5014798 | 0.165 | 0.349 | 1.3125E-12 | TEC_CD | TIMP2 |
| 6.3943E-17 | -0.6674064 | 0.303 | 0.505 | 1.5105E-12 | TEC_CD | DNAJB1 |
| 7.0663E-17 | -1.0136414 | 0.091 | 0.264 | 1.6693E-12 | TEC_CD | AREG |
| 7.1712E-17 | -0.3810181 | 0 | 0.148 | 1.694E-12 | TEC_CD | TREM1 |
| 7.262E-17 | -0.6609428 | 0.899 | 0.948 | 1.7155E-12 | TEC_CD | RPL10 |
| 8.0082E-17 | 0.29194619 | 0.825 | 0.656 | 1.8918E-12 | TEC_CD | CALM2 |
| 8.9141E-17 | -0.4323085 | 0 | 0.147 | 2.1058E-12 | TEC_CD | FPR1 |
| 9.2919E-17 | -0.417181 | 0 | 0.147 | 2.195E-12 | TEC_CD | MS4A7 |
| 1.1573E-16 | -0.5108022 | 0.03 | 0.19 | 2.7339E-12 | TEC_CD | FYB1 |
| 1.1802E-16 | -0.4143077 | 0.429 | 0.644 | 2.7879E-12 | TEC_CD | IER3 |
| 1.273E-16 | -0.5026894 | 0 | 0.145 | 3.0073E-12 | TEC_CD | CD93 |
| 1.2752E-16 | -0.3183473 | 0.015 | 0.169 | 3.0124E-12 | TEC_CD | GLIPR2 |
| 1.3525E-16 | 0.2901279 | 0.463 | 0.274 | 3.195E-12 | TEC_CD | ATP6AP2 |
| 1.3982E-16 | -0.5089197 | 0.017 | 0.172 | 3.3031E-12 | TEC_CD | CXCR4 |
| 1.5387E-16 | 0.34563208 | 0.788 | 0.678 | 3.6348E-12 | TEC_CD | COX7A2 |
| 1.6817E-16 | -0.9752709 | 0.01 | 0.159 | 3.9728E-12 | TEC_CD | PRSS27 |
| 2.3286E-16 | -0.8219464 | 0.002 | 0.147 | 5.5008E-12 | TEC_CD | CEACAM7 |
| 2.6887E-16 | -0.4660714 | 0.887 | 0.931 | 6.3515E-12 | TEC_CD | FAU |
| 3.0116E-16 | -0.3356234 | 0.005 | 0.15 | 7.1144E-12 | TEC_CD | BCAT1 |
| 3.1917E-16 | -0.2836709 | 0 | 0.142 | 7.5397E-12 | TEC_CD | IL2RG |
| 3.3016E-16 | -0.7704788 | 0.002 | 0.146 | 7.7993E-12 | TEC_CD | TMPRSS11E |
| 3.3185E-16 | -1.8773752 | 0.261 | 0.455 | 7.8393E-12 | TEC_CD | SPP1 |
| 3.5034E-16 | 0.28663314 | 0.374 | 0.212 | 8.276E-12 | TEC_CD | PHF3 |
| 5.0428E-16 | -0.3391647 | 0.017 | 0.167 | 1.1913E-11 | TEC_CD | CFD |
| 5.2096E-16 | -0.4868229 | 0 | 0.14 | 1.2307E-11 | TEC_CD | PLAC8 |
| 5.9066E-16 | -0.4058282 | 0.002 | 0.144 | 1.3953E-11 | TEC_CD | NCF1 |
| 6.0445E-16 | 0.25376492 | 0.219 | 0.099 | 1.4279E-11 | TEC_CD | FOSL1 |
| 6.1926E-16 | -0.4296453 | 0.01 | 0.155 | 1.4629E-11 | TEC_CD | TMPRSS4 |
| 6.2313E-16 | -0.8525033 | 0.005 | 0.148 | 1.472E-11 | TEC_CD | CEACAM6 |
| 6.5753E-16 | 0.58125094 | 0.328 | 0.185 | 1.5533E-11 | TEC_CD | PIM1 |
| 7.3231E-16 | -0.3418103 | 0.005 | 0.147 | 1.7299E-11 | TEC_CD | C1orf162 |
| 1.0843E-15 | -0.4374169 | 0 | 0.138 | 2.5614E-11 | TEC_CD | MCEMP1 |
| 1.1152E-15 | -0.4808844 | 0.148 | 0.321 | 2.6345E-11 | TEC_CD | SLC16A3 |
| 1.1714E-15 | -0.3734862 | 0.049 | 0.209 | 2.7672E-11 | TEC_CD | STK17B |

| | | | | | | |
|---|---|---|---|---|---|---|
| 1.1943E-15 | -0.4211066 | 0.424 | 0.585 | 2.8213E-11 | TEC_CD | ACTR3 |
| 1.228E-15 | -0.2992291 | 0 | 0.137 | 2.9009E-11 | TEC_CD | HCK |
| 1.2542E-15 | 0.28547957 | 0.783 | 0.537 | 2.9628E-11 | TEC_CD | CLDN4 |
| 1.2969E-15 | 0.253736 | 0.596 | 0.375 | 3.0637E-11 | TEC_CD | POLR2L |
| 1.4207E-15 | -0.6190874 | 0.012 | 0.156 | 3.356E-11 | TEC_CD | MUC4 |
| 1.6568E-15 | -0.2839699 | 0.015 | 0.159 | 3.9138E-11 | TEC_CD | ARHGAP30 |
| 1.6615E-15 | -0.6101302 | 0.943 | 0.973 | 3.9251E-11 | TEC_CD | RPLP1 |
| 1.7665E-15 | -0.4334272 | 0.362 | 0.535 | 4.1731E-11 | TEC_CD | P4HB |
| 1.8652E-15 | -0.2818443 | 0 | 0.135 | 4.4061E-11 | TEC_CD | FGR |
| 1.9128E-15 | -0.9074967 | 0 | 0.136 | 4.5186E-11 | TEC_CD | TMPRSS11B |
| 1.9178E-15 | -0.2580004 | 0 | 0.135 | 4.5303E-11 | TEC_CD | NCKAP1L |
| 2.0029E-15 | 0.30191997 | 0.167 | 0.068 | 4.7315E-11 | TEC_CD | TNFRSF11B |
| 2.0734E-15 | 0.2900161 | 0.85 | 0.793 | 4.898E-11 | TEC_CD | CHCHD2 |
| 2.1455E-15 | 0.32248109 | 0.712 | 0.574 | 5.0682E-11 | TEC_CD | PRDX5 |
| 2.3259E-15 | -1.0482736 | 0 | 0.135 | 5.4945E-11 | TEC_CD | SERPINB2 |
| 2.4128E-15 | -0.429617 | 0.133 | 0.307 | 5.6999E-11 | TEC_CD | BNIP3L |
| 2.4977E-15 | -0.584107 | 0.047 | 0.2 | 5.9004E-11 | TEC_CD | DUSP4 |
| 2.5743E-15 | -0.3956304 | 0.069 | 0.23 | 6.0812E-11 | TEC_CD | GALNT1 |
| 2.652E-15 | -0.2753397 | 0.002 | 0.138 | 6.2648E-11 | TEC_CD | EVI2A |
| 2.8112E-15 | -0.355445 | 0.069 | 0.23 | 6.6409E-11 | TEC_CD | PGD |
| 2.8567E-15 | -0.6714983 | 0.347 | 0.529 | 6.7484E-11 | TEC_CD | ZFP36 |
| 3.1116E-15 | -1.6003322 | 0 | 0.134 | 7.3504E-11 | TEC_CD | LYPD2 |
| 3.1609E-15 | -0.4108277 | 0.081 | 0.245 | 7.4671E-11 | TEC_CD | RAB31 |
| 3.4803E-15 | -0.2676758 | 0.007 | 0.145 | 8.2216E-11 | TEC_CD | CCDC69 |
| 3.564E-15 | -1.0218282 | 0.005 | 0.141 | 8.4192E-11 | TEC_CD | CXCL6 |
| 3.5936E-15 | 0.43374152 | 0.352 | 0.2 | 8.4892E-11 | TEC_CD | FOSB |
| 4.598E-15 | -0.4375454 | 0.111 | 0.276 | 1.0862E-10 | TEC_CD | PLIN3 |
| 4.8139E-15 | 0.36654951 | 0.32 | 0.179 | 1.1372E-10 | TEC_CD | GNA13 |
| 5.8365E-15 | -0.5103134 | 0.751 | 0.821 | 1.3788E-10 | TEC_CD | RPL27 |
| 5.8543E-15 | -0.8368456 | 0.017 | 0.157 | 1.383E-10 | TEC_CD | KRT15 |
| 6.1385E-15 | -0.6179443 | 0.007 | 0.143 | 1.4501E-10 | TEC_CD | CDKN2A |
| 6.242E-15 | -0.9093865 | 0.002 | 0.135 | 1.4745E-10 | TEC_CD | EREG |
| 7.7037E-15 | -0.2935336 | 0.02 | 0.162 | 1.8198E-10 | TEC_CD | WIPF1 |
| 7.8803E-15 | -0.303072 | 0.005 | 0.138 | 1.8616E-10 | TEC_CD | CLEC7A |
| 7.9482E-15 | 0.29058558 | 0.268 | 0.139 | 1.8776E-10 | TEC_CD | TNFRSF14 |
| 9.9839E-15 | -0.4705254 | 0.02 | 0.16 | 2.3585E-10 | TEC_CD | MALL |
| 1.1362E-14 | -0.3362123 | 0.032 | 0.178 | 2.6841E-10 | TEC_CD | GLIPR1 |
| 1.1864E-14 | -0.6214778 | 0.012 | 0.148 | 2.8026E-10 | TEC_CD | NCCRP1 |
| 1.3115E-14 | -0.9400059 | 0.126 | 0.28 | 3.0982E-10 | TEC_CD | DUSP5 |
| 1.3803E-14 | -0.4574584 | 0.872 | 0.918 | 3.2606E-10 | TEC_CD | RPS20 |
| 1.4059E-14 | -0.3980111 | 0.012 | 0.147 | 3.3212E-10 | TEC_CD | VSIG2 |
| 1.4152E-14 | -0.2995659 | 0.037 | 0.185 | 3.3431E-10 | TEC_CD | BID |
| 1.4655E-14 | -0.2604707 | 0.002 | 0.132 | 3.4619E-10 | TEC_CD | APBB1IP |
| 1.4729E-14 | 0.27878693 | 0.591 | 0.392 | 3.4795E-10 | TEC_CD | HIF1A |
| 1.5835E-14 | -0.2683522 | 0.007 | 0.14 | 3.7408E-10 | TEC_CD | EMILIN2 |
| 1.7344E-14 | -0.5651731 | 0.97 | 0.964 | 4.0973E-10 | TEC_CD | RPL13 |
| 1.9882E-14 | -0.2790226 | 0.005 | 0.135 | 4.6967E-10 | TEC_CD | CYTIP |
| 2.081E-14 | 0.49405341 | 0.328 | 0.182 | 4.9161E-10 | TEC_CD | SGK1 |
| 2.228E-14 | -0.4834079 | 0.067 | 0.219 | 5.2633E-10 | TEC_CD | TGM2 |
| 2.2419E-14 | -0.6511668 | 0.323 | 0.507 | 5.296E-10 | TEC_CD | HSPA1B |

| | | | | | | |
|---|---|---|---|---|---|---|
| 2.5359E-14 | 0.26560929 | 0.431 | 0.26 | 5.9906E-10 | TEC_CD | WTAP |
| 3.6485E-14 | -0.652572 | 0.005 | 0.132 | 8.6187E-10 | TEC_CD | GABRP |
| 3.8586E-14 | 0.46743057 | 0.675 | 0.494 | 9.1151E-10 | TEC_CD | CD9 |
| 4.1431E-14 | -0.4663606 | 0.039 | 0.183 | 9.7872E-10 | TEC_CD | SAMSN1 |
| 4.2109E-14 | -0.4977219 | 0.929 | 0.95 | 9.9473E-10 | TEC_CD | RPL18 |
| 4.2883E-14 | -0.2817497 | 0.002 | 0.128 | 1.013E-09 | TEC_CD | IRAK3 |
| 5.2904E-14 | -0.3578582 | 0.002 | 0.127 | 1.2497E-09 | TEC_CD | MARCO |
| 5.3697E-14 | -0.4049891 | 0.025 | 0.162 | 1.2685E-09 | TEC_CD | SERPINB9 |
| 6.8887E-14 | 0.69339992 | 0.463 | 0.31 | 1.6273E-09 | TEC_CD | CYTOR |
| 6.946E-14 | -0.2675867 | 0.03 | 0.168 | 1.6409E-09 | TEC_CD | SLC12A6 |
| 7.132E-14 | -0.2692558 | 0.02 | 0.154 | 1.6848E-09 | TEC_CD | VSIR |
| 7.5871E-14 | -0.2646082 | 0.03 | 0.168 | 1.7923E-09 | TEC_CD | SIRPA |
| 7.8008E-14 | -0.3055219 | 0 | 0.122 | 1.8428E-09 | TEC_CD | LILRB2 |
| 7.9245E-14 | 0.30898925 | 0.315 | 0.174 | 1.872E-09 | TEC_CD | NR4A1 |
| 8.098E-14 | -0.4043147 | 0.153 | 0.312 | 1.913E-09 | TEC_CD | CTNNB1 |
| 8.1213E-14 | -0.6962919 | 0.086 | 0.235 | 1.9185E-09 | TEC_CD | PDZK1IP1 |
| 8.1295E-14 | -0.2782063 | 0.01 | 0.137 | 1.9204E-09 | TEC_CD | ITGA5 |
| 9.3462E-14 | -0.3503742 | 0.005 | 0.129 | 2.2078E-09 | TEC_CD | LRG1 |
| 1.0156E-13 | 0.41846403 | 0.786 | 0.721 | 2.3991E-09 | TEC_CD | JUND |
| 1.0296E-13 | 0.28985474 | 0.502 | 0.323 | 2.4323E-09 | TEC_CD | FOSL2 |
| 1.0454E-13 | 0.28082424 | 0.397 | 0.24 | 2.4696E-09 | TEC_CD | KANSL1 |
| 1.0622E-13 | -1.0115041 | 0.074 | 0.224 | 2.5092E-09 | TEC_CD | G0S2 |
| 1.1486E-13 | -0.3815168 | 0 | 0.12 | 2.7133E-09 | TEC_CD | FCGR3A |
| 1.3126E-13 | -0.2527405 | 0.044 | 0.188 | 3.1007E-09 | TEC_CD | SQOR |
| 1.3636E-13 | -0.2794106 | 0.007 | 0.132 | 3.2212E-09 | TEC_CD | LILRB3 |
| 1.3644E-13 | -0.418524 | 0.648 | 0.772 | 3.2232E-09 | TEC_CD | RHOA |
| 1.3995E-13 | 0.29423669 | 0.424 | 0.263 | 3.306E-09 | TEC_CD | HECTD1 |
| 1.4285E-13 | 0.34749089 | 0.786 | 0.713 | 3.3745E-09 | TEC_CD | ATP5MG |
| 1.464E-13 | -0.4304666 | 0.244 | 0.398 | 3.4584E-09 | TEC_CD | H2AFY |
| 1.6892E-13 | -0.5007302 | 0.158 | 0.313 | 3.9905E-09 | TEC_CD | CD68 |
| 1.6977E-13 | 0.26297629 | 0.599 | 0.412 | 4.0104E-09 | TEC_CD | EIF3K |
| 1.8084E-13 | -0.2734687 | 0 | 0.118 | 4.272E-09 | TEC_CD | CD86 |
| 1.8987E-13 | 0.28408937 | 0.781 | 0.627 | 4.4854E-09 | TEC_CD | OST4 |
| 1.9093E-13 | -0.4582891 | 0.032 | 0.167 | 4.5104E-09 | TEC_CD | HIST3H2A |
| 1.9562E-13 | 0.25519372 | 0.397 | 0.243 | 4.6212E-09 | TEC_CD | UQCRFS1 |
| 2.0004E-13 | 0.43897126 | 0.167 | 0.073 | 4.7255E-09 | TEC_CD | RARRES1 |
| 2.1565E-13 | 0.31192351 | 0.818 | 0.746 | 5.0943E-09 | TEC_CD | ATP6V0E1 |
| 2.2684E-13 | -0.5818825 | 0 | 0.118 | 5.3585E-09 | TEC_CD | MNDA |
| 2.5233E-13 | -0.5315726 | 0.948 | 0.92 | 5.9607E-09 | TEC_CD | RPS15 |
| 2.63E-13 | -0.4301958 | 0.064 | 0.21 | 6.2129E-09 | TEC_CD | LY6E |
| 2.7163E-13 | -0.387131 | 0.436 | 0.585 | 6.4167E-09 | TEC_CD | NOP10 |
| 2.8437E-13 | -0.5940535 | 0.069 | 0.209 | 6.7176E-09 | TEC_CD | CDKN2B |
| 2.907E-13 | -1.1552675 | 0.01 | 0.133 | 6.8673E-09 | TEC_CD | PLAT |
| 3.0285E-13 | -0.2663233 | 0.002 | 0.121 | 7.1543E-09 | TEC_CD | CSF1R |
| 3.375E-13 | 0.30076663 | 0.249 | 0.13 | 7.9728E-09 | TEC_CD | PALLD |
| 3.3762E-13 | -0.644371 | 0.01 | 0.132 | 7.9757E-09 | TEC_CD | DHRS9 |
| 3.472E-13 | -0.3884925 | 0.14 | 0.296 | 8.2018E-09 | TEC_CD | CD99 |
| 3.9505E-13 | -1.1471512 | 0.005 | 0.124 | 9.3324E-09 | TEC_CD | CCL3 |
| 4.359E-13 | 0.78416336 | 0.736 | 0.824 | 1.0297E-08 | TEC_CD | MT-ND5 |
| 4.7503E-13 | -0.547136 | 0.882 | 0.928 | 1.1222E-08 | TEC_CD | RPL30 |

| | | | | | | |
|---|---|---|---|---|---|---|
| 5.9592E-13 | -0.4307619 | 0.86 | 0.896 | 1.4077E-08 | TEC_CD | HLA-A |
| 6.2284E-13 | -1.1111285 | 0.002 | 0.118 | 1.4713E-08 | TEC_CD | SPINK5 |
| 6.5357E-13 | -0.4640357 | 0.929 | 0.946 | 1.5439E-08 | TEC_CD | RPL13A |
| 6.7355E-13 | -0.5046458 | 0.665 | 0.747 | 1.5911E-08 | TEC_CD | EEF1B2 |
| 7.0313E-13 | -0.458642 | 0 | 0.113 | 1.661E-08 | TEC_CD | TMPRSS11A |
| 8.2149E-13 | -0.3740378 | 0.015 | 0.136 | 1.9406E-08 | TEC_CD | BCAS1 |
| 8.6917E-13 | -0.4126648 | 0.094 | 0.239 | 2.0532E-08 | TEC_CD | RNASET2 |
| 9.9383E-13 | -0.269563 | 0.032 | 0.161 | 2.3477E-08 | TEC_CD | AP1S2 |
| 1.0247E-12 | -0.4385897 | 0.049 | 0.186 | 2.4206E-08 | TEC_CD | KLF2 |
| 1.1841E-12 | 0.33215499 | 0.719 | 0.562 | 2.7973E-08 | TEC_CD | SLC25A6 |
| 1.5545E-12 | 0.27975677 | 0.547 | 0.356 | 3.6722E-08 | TEC_CD | MIF |
| 1.59E-12 | 0.29243493 | 0.766 | 0.664 | 3.756E-08 | TEC_CD | HNRNPC |
| 1.6421E-12 | -1.2755369 | 0.002 | 0.114 | 3.8791E-08 | TEC_CD | SPRR1B |
| 1.6977E-12 | -0.7596229 | 0.002 | 0.114 | 4.0104E-08 | TEC_CD | IL7R |
| 1.7956E-12 | -0.265359 | 0.002 | 0.114 | 4.2417E-08 | TEC_CD | PECAM1 |
| 2.0306E-12 | -0.5901838 | 0.002 | 0.113 | 4.7968E-08 | TEC_CD | KRT78 |
| 2.6667E-12 | -0.5759722 | 0.022 | 0.142 | 6.2995E-08 | TEC_CD | SNCG |
| 2.7659E-12 | -0.4119162 | 0.653 | 0.748 | 6.5339E-08 | TEC_CD | CLIC1 |
| 2.9007E-12 | -0.3846686 | 0.081 | 0.218 | 6.8523E-08 | TEC_CD | DSE |
| 2.957E-12 | -0.4165767 | 0.106 | 0.249 | 6.9854E-08 | TEC_CD | GPCPD1 |
| 2.9974E-12 | -0.2943328 | 0.076 | 0.217 | 7.0808E-08 | TEC_CD | TGFB1 |
| 3.1516E-12 | -0.2912056 | 0 | 0.107 | 7.445E-08 | TEC_CD | MS4A6A |
| 3.1561E-12 | -0.4071619 | 0 | 0.107 | 7.4556E-08 | TEC_CD | THBD |
| 3.5109E-12 | -0.6032306 | 0.325 | 0.434 | 8.2938E-08 | TEC_CD | GSTO1 |
| 3.7706E-12 | -0.3125372 | 0.067 | 0.202 | 8.9074E-08 | TEC_CD | IFI16 |
| 4.0527E-12 | -0.3263926 | 0 | 0.106 | 9.5736E-08 | TEC_CD | A2ML1 |
| 4.1857E-12 | -0.3511838 | 0.571 | 0.696 | 9.8879E-08 | TEC_CD | GUK1 |
| 4.4124E-12 | -0.5607886 | 0.865 | 0.894 | 1.0423E-07 | TEC_CD | RPS15A |
| 4.6223E-12 | 0.31117591 | 0.7 | 0.588 | 1.0919E-07 | TEC_CD | SLC25A3 |
| 4.8045E-12 | -0.5217486 | 0.722 | 0.793 | 1.135E-07 | TEC_CD | RPS21 |
| 5.4766E-12 | -0.5386701 | 0.005 | 0.113 | 1.2937E-07 | TEC_CD | CYP1B1 |
| 5.4838E-12 | -0.4527746 | 0.145 | 0.284 | 1.2954E-07 | TEC_CD | CTSZ |
| 6.2211E-12 | -0.3986177 | 0.946 | 0.945 | 1.4696E-07 | TEC_CD | RPS19 |
| 7.174E-12 | 0.26064024 | 0.438 | 0.283 | 1.6947E-07 | TEC_CD | NDUFB1 |
| 7.4818E-12 | -0.3495258 | 0.076 | 0.213 | 1.7674E-07 | TEC_CD | TLE4 |
| 7.5726E-12 | -0.2583535 | 0.017 | 0.132 | 1.7889E-07 | TEC_CD | VAMP5 |
| 7.8404E-12 | -0.2800436 | 0 | 0.104 | 1.8521E-07 | TEC_CD | LILRB1 |
| 8.2499E-12 | -0.4056282 | 0.14 | 0.281 | 1.9489E-07 | TEC_CD | LYN |
| 8.4718E-12 | -1.6421083 | 0.027 | 0.144 | 2.0013E-07 | TEC_CD | SPRR2A |
| 9.4701E-12 | -0.2501071 | 0.017 | 0.131 | 2.2371E-07 | TEC_CD | CELF2 |
| 1.1344E-11 | -0.3397567 | 0.002 | 0.107 | 2.6799E-07 | TEC_CD | CSF3R |
| 1.2421E-11 | -0.296142 | 0 | 0.102 | 2.9341E-07 | TEC_CD | LRMP |
| 1.3178E-11 | -0.75563 | 0.057 | 0.181 | 3.113E-07 | TEC_CD | F3 |
| 1.38E-11 | -0.2693397 | 0.032 | 0.153 | 3.26E-07 | TEC_CD | HK2 |
| 1.7001E-11 | -0.4860176 | 0 | 0.101 | 4.0162E-07 | TEC_CD | RETN |
| 1.734E-11 | 0.83972085 | 0.766 | 0.934 | 4.0963E-07 | TEC_CD | MT-ND4 |
| 1.7535E-11 | -0.2564644 | 0.052 | 0.177 | 4.1423E-07 | TEC_CD | PPP1R18 |
| 1.8165E-11 | -0.2526049 | 0.039 | 0.16 | 4.2911E-07 | TEC_CD | DOCK8 |
| 2.069E-11 | -0.3810678 | 0.005 | 0.108 | 4.8877E-07 | TEC_CD | KRT23 |
| 2.0935E-11 | -0.3850357 | 0.002 | 0.104 | 4.9455E-07 | TEC_CD | CLIC3 |

| | | | | | | |
|---|---|---|---|---|---|---|
| 2.3122E-11 | -0.3166775 | 0.02 | 0.131 | 5.4622E-07 | TEC_CD | FOXC1 |
| 2.3838E-11 | 0.80469412 | 0.764 | 0.91 | 5.6312E-07 | TEC_CD | MT-ND2 |
| 2.4013E-11 | -0.3566789 | 0.01 | 0.115 | 5.6725E-07 | TEC_CD | CEACAM1 |
| 2.5858E-11 | -1.5301299 | 0.005 | 0.107 | 6.1084E-07 | TEC_CD | CRCT1 |
| 2.7534E-11 | -0.5260386 | 0.931 | 0.912 | 6.5044E-07 | TEC_CD | RPS8 |
| 2.7562E-11 | -0.3972722 | 0.059 | 0.187 | 6.5111E-07 | TEC_CD | TNFSF10 |
| 2.9227E-11 | -0.4069989 | 0.081 | 0.21 | 6.9044E-07 | TEC_CD | SOCS3 |
| 2.9236E-11 | -0.4496198 | 0.424 | 0.573 | 6.9064E-07 | TEC_CD | VMP1 |
| 3.1195E-11 | -0.5033838 | 0.163 | 0.3 | 7.3691E-07 | TEC_CD | FAM129B |
| 3.1776E-11 | 0.25445888 | 0.466 | 0.315 | 7.5064E-07 | TEC_CD | KMT2E |
| 3.2181E-11 | 0.5024818 | 0.823 | 0.832 | 7.6022E-07 | TEC_CD | UBB |
| 3.381E-11 | -0.5902291 | 0.002 | 0.102 | 7.9869E-07 | TEC_CD | HLA-DQA1 |
| 4.1106E-11 | -0.4434877 | 0.138 | 0.275 | 9.7105E-07 | TEC_CD | TNFAIP2 |
| 4.2096E-11 | -0.2559448 | 0.032 | 0.147 | 9.9443E-07 | TEC_CD | PLEKHO1 |
| 4.3381E-11 | -0.4576306 | 0.071 | 0.196 | 1.0248E-06 | TEC_CD | SAMD9 |
| 4.3668E-11 | -0.7677184 | 0.005 | 0.105 | 1.0316E-06 | TEC_CD | ZG16B |
| 4.4558E-11 | -0.258996 | 0.007 | 0.109 | 1.0526E-06 | TEC_CD | PDE4B |
| 4.941E-11 | -0.3366797 | 0.012 | 0.117 | 1.1672E-06 | TEC_CD | CD163 |
| 5.0002E-11 | -0.5232852 | 0.357 | 0.481 | 1.1812E-06 | TEC_CD | RIOK3 |
| 5.1112E-11 | -0.2605374 | 0.081 | 0.211 | 1.2074E-06 | TEC_CD | CYTH1 |
| 5.2556E-11 | -0.3082416 | 0.022 | 0.133 | 1.2415E-06 | TEC_CD | RGCC |
| 5.4393E-11 | -0.3230162 | 0.121 | 0.256 | 1.2849E-06 | TEC_CD | MGAT1 |
| 5.7914E-11 | -1.0433502 | 0.007 | 0.109 | 1.3681E-06 | TEC_CD | CCL4 |
| 7.4614E-11 | -0.3869052 | 0.18 | 0.31 | 1.7626E-06 | TEC_CD | ARPC1B |
| 8.2254E-11 | -0.3901208 | 0.163 | 0.297 | 1.9431E-06 | TEC_CD | IFNGR2 |
| 1.159E-10 | -0.5279987 | 0.914 | 0.916 | 2.7379E-06 | TEC_CD | RPL32 |
| 1.2429E-10 | -0.4857343 | 0.872 | 0.881 | 2.9362E-06 | TEC_CD | RPL12 |
| 1.2957E-10 | -0.3149438 | 0.034 | 0.145 | 3.0609E-06 | TEC_CD | TLR2 |
| 1.4618E-10 | -0.3987117 | 0.562 | 0.692 | 3.4533E-06 | TEC_CD | PNRC1 |
| 1.4798E-10 | -0.383673 | 0.071 | 0.197 | 3.4958E-06 | TEC_CD | SFN |
| 1.5186E-10 | -0.3690582 | 0.02 | 0.124 | 3.5874E-06 | TEC_CD | TMEM40 |
| 1.6423E-10 | -0.6548903 | 0.594 | 0.668 | 3.8795E-06 | TEC_CD | ANXA5 |
| 1.6962E-10 | -0.2703482 | 0.012 | 0.112 | 4.0068E-06 | TEC_CD | FUT3 |
| 2.2916E-10 | -0.2604968 | 0.089 | 0.214 | 5.4134E-06 | TEC_CD | RNF145 |
| 2.7048E-10 | -0.3290548 | 0.204 | 0.337 | 6.3895E-06 | TEC_CD | CAPZA1 |
| 3.2476E-10 | 0.47525245 | 0.355 | 0.237 | 7.6719E-06 | TEC_CD | TNFAIP3 |
| 3.7383E-10 | -0.427878 | 0.84 | 0.849 | 8.831E-06 | TEC_CD | RPL38 |
| 4.2762E-10 | -0.7026734 | 0.259 | 0.143 | 1.0102E-05 | TEC_CD | CXCL3 |
| 4.7708E-10 | -0.2766134 | 0.052 | 0.166 | 1.127E-05 | TEC_CD | JARID2 |
| 6.215E-10 | -0.4732126 | 0.059 | 0.172 | 1.4682E-05 | TEC_CD | HSPB8 |
| 6.8719E-10 | -0.7111443 | 0.015 | 0.109 | 1.6233E-05 | TEC_CD | APOC1 |
| 8.1408E-10 | -0.2684559 | 0.069 | 0.184 | 1.9231E-05 | TEC_CD | PSMB9 |
| 8.4102E-10 | -0.6412873 | 0.076 | 0.189 | 1.9867E-05 | TEC_CD | ANKRD36C |
| 9.154E-10 | -0.3335915 | 0.429 | 0.554 | 2.1624E-05 | TEC_CD | SSR4 |
| 9.6424E-10 | 0.87004731 | 0.751 | 0.932 | 2.2778E-05 | TEC_CD | MT-CYB |
| 1.1575E-09 | -0.4382499 | 0.276 | 0.398 | 2.7343E-05 | TEC_CD | ARGLU1 |
| 1.1718E-09 | 0.31363979 | 0.746 | 0.7 | 2.7681E-05 | TEC_CD | COX6B1 |
| 1.1749E-09 | -0.3372359 | 0.3 | 0.421 | 2.7755E-05 | TEC_CD | CAPZB |
| 1.2916E-09 | -0.4245315 | 0.921 | 0.962 | 3.0512E-05 | TEC_CD | HSP90AA1 |
| 1.3727E-09 | -0.4649997 | 0.833 | 0.87 | 3.2428E-05 | TEC_CD | RPLP0 |

| | | | | | | |
|---|---|---|---|---|---|---|
| 1.8983E-09 | 0.75900345 | 0.749 | 0.916 | 4.4844E-05 | TEC_CD | MT-ND1 |
| 2.3685E-09 | 0.27242029 | 0.589 | 0.463 | 5.5952E-05 | TEC_CD | CDKN1A |
| 2.5892E-09 | -0.5078958 | 0.052 | 0.155 | 6.1166E-05 | TEC_CD | CDC42EP5 |
| 2.6516E-09 | -0.3943077 | 0.03 | 0.126 | 6.264E-05 | TEC_CD | SRD5A3 |
| 2.691E-09 | -0.3293402 | 0.525 | 0.679 | 6.3569E-05 | TEC_CD | IER2 |
| 2.8886E-09 | -0.3868967 | 0.094 | 0.205 | 6.8237E-05 | TEC_CD | CSNK1E |
| 3.014E-09 | -0.4120905 | 0.879 | 0.884 | 7.1199E-05 | TEC_CD | RPS13 |
| 3.1926E-09 | 0.25837762 | 0.291 | 0.177 | 7.5419E-05 | TEC_CD | STAT1 |
| 3.4836E-09 | -0.2818072 | 0.02 | 0.111 | 8.2292E-05 | TEC_CD | STS |
| 3.5392E-09 | -0.3975912 | 0.894 | 0.919 | 8.3606E-05 | TEC_CD | RPL9 |
| 3.9321E-09 | 0.87481771 | 0.727 | 0.879 | 9.2888E-05 | TEC_CD | MT-ND3 |
| 3.9339E-09 | -0.5632632 | 0.039 | 0.137 | 9.293E-05 | TEC_CD | CXCL17 |
| 3.9522E-09 | -0.4514556 | 0.921 | 0.933 | 9.3363E-05 | TEC_CD | RPS14 |
| 4.0748E-09 | -0.3718641 | 0.884 | 0.896 | 9.6258E-05 | TEC_CD | RPS9 |
| 4.1538E-09 | 0.37159285 | 0.328 | 0.211 | 9.8126E-05 | TEC_CD | SETX |
| 4.2765E-09 | 0.82691802 | 0.783 | 0.952 | 0.00010102 | TEC_CD | MT-CO1 |
| 4.5767E-09 | -0.3926044 | 0.049 | 0.152 | 0.00010812 | TEC_CD | SLC7A11 |
| 4.9972E-09 | -0.5282683 | 0.069 | 0.173 | 0.00011805 | TEC_CD | HLA-DQB1 |
| 5.6127E-09 | 0.38288073 | 0.461 | 0.325 | 0.00013259 | TEC_CD | CEBPD |
| 6.7813E-09 | -0.4718598 | 0.882 | 0.879 | 0.00016019 | TEC_CD | RPS29 |
| 6.8695E-09 | -0.285338 | 0.054 | 0.156 | 0.00016228 | TEC_CD | ADGRE5 |
| 7.7164E-09 | -0.9760775 | 0.049 | 0.153 | 0.00018228 | TEC_CD | CCL20 |
| 9.0601E-09 | -0.7753461 | 0.458 | 0.545 | 0.00021403 | TEC_CD | KRT7 |
| 1.0125E-08 | -0.3649009 | 0.48 | 0.581 | 0.00023919 | TEC_CD | PGK1 |
| 1.0244E-08 | -0.5962972 | 0.766 | 0.791 | 0.000242 | TEC_CD | PFN1 |
| 1.0446E-08 | -0.2878033 | 0.074 | 0.178 | 0.00024676 | TEC_CD | CCDC88A |
| 1.1817E-08 | -0.3417271 | 0.894 | 0.958 | 0.00027915 | TEC_CD | RPL15 |
| 1.1918E-08 | -0.3238406 | 0.182 | 0.3 | 0.00028155 | TEC_CD | TNIP1 |
| 1.3007E-08 | -0.2697123 | 0.15 | 0.267 | 0.00030726 | TEC_CD | CDC42SE1 |
| 1.4232E-08 | -0.291861 | 0.032 | 0.123 | 0.0003362 | TEC_CD | KRT80 |
| 1.4308E-08 | 0.42477007 | 0.323 | 0.206 | 0.00033799 | TEC_CD | KCNQ1OT1 |
| 1.521E-08 | -0.4867765 | 0.185 | 0.296 | 0.00035931 | TEC_CD | GLRX |
| 1.5698E-08 | -0.3926102 | 0.103 | 0.207 | 0.00037083 | TEC_CD | SLC43A2 |
| 1.8045E-08 | -0.5007028 | 0.081 | 0.194 | 0.00042627 | TEC_CD | HSPA6 |
| 1.9839E-08 | -0.2895946 | 0.195 | 0.308 | 0.00046865 | TEC_CD | SPCS3 |
| 2.1813E-08 | -0.4679748 | 0.51 | 0.603 | 0.00051529 | TEC_CD | MCL1 |
| 2.3586E-08 | 0.80975525 | 0.461 | 0.35 | 0.00055717 | TEC_CD | PHLDA2 |
| 2.4286E-08 | -0.4000539 | 0.48 | 0.576 | 0.00057372 | TEC_CD | CD55 |
| 2.4533E-08 | -0.2649558 | 0.18 | 0.298 | 0.00057955 | TEC_CD | UBALD2 |
| 2.5338E-08 | -0.326008 | 0.062 | 0.158 | 0.00059855 | TEC_CD | TTC9 |
| 2.5564E-08 | -0.3641021 | 0.172 | 0.277 | 0.00060391 | TEC_CD | VASP |
| 2.6903E-08 | -0.2724853 | 0.054 | 0.15 | 0.00063553 | TEC_CD | PLXDC2 |
| 2.693E-08 | -0.353516 | 0.232 | 0.34 | 0.00063616 | TEC_CD | GRB2 |
| 3.1961E-08 | -0.3578527 | 0.108 | 0.213 | 0.00075502 | TEC_CD | NINJ1 |
| 3.3845E-08 | -0.2882052 | 0.155 | 0.271 | 0.00079953 | TEC_CD | QSOX1 |
| 3.3906E-08 | -0.4966921 | 0.695 | 0.784 | 0.00080095 | TEC_CD | LGALS3 |
| 3.4045E-08 | -0.3609521 | 0.456 | 0.548 | 0.00080425 | TEC_CD | C4orf3 |
| 3.7116E-08 | -0.9522453 | 0.086 | 0.185 | 0.00087678 | TEC_CD | HLA-DPB1 |
| 4.1637E-08 | -0.3840711 | 0.946 | 0.924 | 0.00098358 | TEC_CD | RPS24 |
| 4.4799E-08 | -0.31677 | 0.118 | 0.225 | 0.00105829 | TEC_CD | STXBP2 |

| | | | | | | |
|---|---|---|---|---|---|---|
| 4.828E-08 | -0.2856304 | 0.749 | 0.811 | 0.00114052 | TEC_CD | RAC1 |
| 5.3802E-08 | -0.2624884 | 0.108 | 0.214 | 0.00127096 | TEC_CD | RUNX1 |
| 5.4204E-08 | -0.3313194 | 0.054 | 0.149 | 0.00128046 | TEC_CD | RGS2 |
| 6.0735E-08 | -0.417994 | 0.022 | 0.105 | 0.00143475 | TEC_CD | CCL5 |
| 6.4187E-08 | -0.4162172 | 0.877 | 0.898 | 0.00151628 | TEC_CD | RPL29 |
| 7.1708E-08 | 0.7997851 | 0.776 | 0.946 | 0.00169395 | TEC_CD | MT-CO3 |
| 8.6961E-08 | -0.5229472 | 0.153 | 0.254 | 0.00205429 | TEC_CD | CARHSP1 |
| 1.137E-07 | -0.5903133 | 0.687 | 0.764 | 0.00268599 | TEC_CD | BTG1 |
| 1.4147E-07 | -0.4262401 | 0.522 | 0.62 | 0.00334204 | TEC_CD | PCBP1 |
| 1.5425E-07 | -0.2859424 | 0.165 | 0.269 | 0.00364373 | TEC_CD | RAB5IF |
| 1.699E-07 | -0.3359959 | 0.392 | 0.497 | 0.00401355 | TEC_CD | DDX3X |
| 1.871E-07 | -0.3171881 | 0.182 | 0.286 | 0.00441998 | TEC_CD | CXCL16 |
| 2.6276E-07 | -0.2876804 | 0.241 | 0.342 | 0.00620712 | TEC_CD | ARPC4 |
| 2.9265E-07 | -0.2816371 | 0.044 | 0.126 | 0.00691332 | TEC_CD | POLB |
| 3.0231E-07 | -0.5180436 | 0.288 | 0.369 | 0.00714155 | TEC_CD | COTL1 |
| 3.3021E-07 | -0.3013005 | 0.217 | 0.319 | 0.00780065 | TEC_CD | CMTM6 |
| 3.6292E-07 | 0.3092025 | 0.702 | 0.627 | 0.00857317 | TEC_CD | JUN |
| 3.6764E-07 | -0.7486917 | 0.2 | 0.286 | 0.00868473 | TEC_CD | SLC25A37 |
| 4.1736E-07 | -0.3287371 | 0.643 | 0.71 | 0.00985924 | TEC_CD | SDCBP |
| 5.174E-07 | -0.3692695 | 0.096 | 0.189 | 0.01222245 | TEC_CD | HBEGF |
| 5.6516E-07 | -0.4646345 | 0.042 | 0.121 | 0.01335073 | TEC_CD | HILPDA |
| 6.8086E-07 | -0.2638238 | 0.039 | 0.118 | 0.01608388 | TEC_CD | DUSP2 |
| 7.2007E-07 | 0.69265914 | 0.638 | 0.638 | 0.01701016 | TEC_CD | MTRNR2L12 |
| 9.0881E-07 | -0.3947608 | 0.244 | 0.345 | 0.02146871 | TEC_CD | DDIT4 |
| 9.1081E-07 | -0.3686305 | 0.054 | 0.136 | 0.02151597 | TEC_CD | HLA-DRB5 |
| 1.1028E-06 | -0.3435252 | 0.672 | 0.704 | 0.02605175 | TEC_CD | RPL4 |
| 1.1063E-06 | -0.4823486 | 0.899 | 0.887 | 0.02613504 | TEC_CD | RPS3 |
| 1.1353E-06 | -0.5460691 | 0.133 | 0.231 | 0.02681835 | TEC_CD | ISG15 |
| 1.2412E-06 | -0.344603 | 0.185 | 0.284 | 0.02932168 | TEC_CD | MARCKSL1 |
| 1.5578E-06 | -0.4031385 | 0.97 | 0.939 | 0.0368008 | TEC_CD | RPS28 |
| 1.8568E-06 | -0.3515152 | 0.914 | 0.888 | 0.04386343 | TEC_CD | RPL6 |
| 2.4395E-06 | 0.62025002 | 0.768 | 0.939 | 0.05762843 | TEC_CD | MT-ATP6 |
| 2.4894E-06 | -0.2726273 | 0.727 | 0.801 | 0.05880701 | TEC_CD | GNG5 |
| 2.5672E-06 | 0.2570972 | 0.635 | 0.568 | 0.06064411 | TEC_CD | RPS27L |
| 2.5969E-06 | -0.3741057 | 0.951 | 0.957 | 0.06134774 | TEC_CD | RPS18 |
| 3.0574E-06 | -0.3264146 | 0.667 | 0.776 | 0.07222485 | TEC_CD | TPI1 |
| 3.2828E-06 | -0.4116187 | 0.103 | 0.183 | 0.07754992 | TEC_CD | SDCBP2 |
| 3.5826E-06 | -0.3851258 | 0.197 | 0.28 | 0.08463266 | TEC_CD | ABHD5 |
| 3.5961E-06 | -0.354897 | 0.621 | 0.674 | 0.08495127 | TEC_CD | CALR |
| 5.0341E-06 | -0.3281535 | 0.897 | 0.896 | 0.11892029 | TEC_CD | RPL35A |
| 5.6783E-06 | -0.8229222 | 0.064 | 0.144 | 0.13413786 | TEC_CD | CCL2 |
| 7.0401E-06 | -0.3725055 | 0.951 | 0.903 | 0.16630758 | TEC_CD | RPL37 |
| 7.1619E-06 | -1.3677465 | 0.042 | 0.108 | 0.16918489 | TEC_CD | CNFN |
| 7.9556E-06 | -0.3819793 | 0.084 | 0.162 | 0.18793512 | TEC_CD | CSKMT |
| 8.7406E-06 | -0.2923367 | 0.232 | 0.312 | 0.20647821 | TEC_CD | TMEM165 |
| 8.7501E-06 | -0.3273051 | 0.985 | 0.965 | 0.20670449 | TEC_CD | S100A11 |
| 1.2287E-05 | -0.2712438 | 0.436 | 0.52 | 0.29026539 | TEC_CD | H2AFZ |
| 1.2483E-05 | -0.4276825 | 0.796 | 0.801 | 0.29489022 | TEC_CD | RPS5 |
| 1.3977E-05 | -1.2288223 | 0.047 | 0.116 | 0.3301678 | TEC_CD | MT1M |
| 1.4608E-05 | -1.1056022 | 0.303 | 0.345 | 0.3450954 | TEC_CD | CTSL |

| | | | | | | |
|---|---|---|---|---|---|---|
| 1.5979E-05 | -0.3220047 | 0.217 | 0.298 | 0.37747256 | TEC_CD | REL |
| 1.7004E-05 | -1.0104654 | 0.064 | 0.131 | 0.40168432 | TEC_CD | APOE |
| 1.7998E-05 | -0.4083383 | 0.931 | 0.934 | 0.42517508 | TEC_CD | RPL34 |
| 3.5353E-05 | -0.2817969 | 0.163 | 0.239 | 0.83514469 | TEC_CD | SMCHD1 |
| 3.8608E-05 | -0.2536761 | 0.539 | 0.599 | 0.91203593 | TEC_CD | VAMP8 |
| 4.2012E-05 | -0.3187568 | 0.187 | 0.117 | 0.99243915 | TEC_CD | CXCL2 |
| 4.5525E-05 | -0.3977484 | 0.167 | 0.246 | 1 | TEC_CD | PLAU |
| 4.8873E-05 | -0.2796821 | 0.047 | 0.107 | 1 | TEC_CD | SLC16A10 |
| 4.9016E-05 | -0.3622612 | 0.943 | 0.922 | 1 | TEC_CD | RPL11 |
| 5.3214E-05 | -0.3353335 | 0.916 | 0.875 | 1 | TEC_CD | RPL36 |
| 7.344E-05 | -0.3842533 | 0.911 | 0.877 | 1 | TEC_CD | RPL23A |
| 7.8002E-05 | -0.2709329 | 0.116 | 0.189 | 1 | TEC_CD | DUSP6 |
| 8.4243E-05 | -0.341701 | 0.303 | 0.357 | 1 | TEC_CD | TKT |
| 8.6914E-05 | -1.2004297 | 0.081 | 0.152 | 1 | TEC_CD | MT1H |
| 0.00010531 | -1.4709397 | 0.35 | 0.397 | 1 | TEC_CD | HLA-DRA |
| 0.00010971 | -0.2601589 | 0.081 | 0.146 | 1 | TEC_CD | SDC2 |
| 0.00011689 | -0.3821002 | 0.475 | 0.523 | 1 | TEC_CD | HEBP2 |
| 0.00017093 | -0.4064713 | 0.182 | 0.243 | 1 | TEC_CD | ATP2B1 |
| 0.0001724 | -0.3148526 | 0.968 | 0.943 | 1 | TEC_CD | RPS27 |
| 0.00021264 | -0.3679566 | 0.453 | 0.519 | 1 | TEC_CD | H1FX |
| 0.00024762 | -0.2658505 | 0.493 | 0.547 | 1 | TEC_CD | RPS10 |
| 0.00027508 | -1.3417234 | 0.185 | 0.232 | 1 | TEC_CD | PRSS22 |
| 0.00028644 | -0.9988891 | 0.145 | 0.204 | 1 | TEC_CD | HLA-DPA1 |
| 0.00029932 | -0.3479079 | 0.195 | 0.262 | 1 | TEC_CD | HES4 |
| 0.00033426 | -0.4010896 | 0.207 | 0.265 | 1 | TEC_CD | MGST1 |
| 0.00040963 | -0.3307425 | 0.057 | 0.107 | 1 | TEC_CD | TMC5 |
| 0.00041742 | -0.2857312 | 0.2 | 0.257 | 1 | TEC_CD | PDLIM5 |
| 0.00049915 | -0.2555543 | 0.933 | 0.895 | 1 | TEC_CD | RPL37A |
| 0.00063373 | -0.3565375 | 0.123 | 0.175 | 1 | TEC_CD | C6orf132 |
| 0.00081215 | -0.3566585 | 0.214 | 0.27 | 1 | TEC_CD | TSC22D3 |
| 0.00086158 | -0.8191143 | 0.131 | 0.181 | 1 | TEC_CD | SPINK1 |
| 0.00089312 | 0.51836211 | 0.759 | 0.947 | 1 | TEC_CD | MT-CO2 |
| 0.00090703 | -0.3887469 | 0.879 | 0.933 | 1 | TEC_CD | GAPDH |
| 0.00098871 | -0.3019865 | 0.845 | 0.839 | 1 | TEC_CD | RACK1 |
| 0.00102065 | -0.2562994 | 0.059 | 0.108 | 1 | TEC_CD | TCHP |
| 0.00116053 | -0.2868525 | 0.86 | 0.858 | 1 | TEC_CD | RPL7 |
| 0.00118804 | -0.3765602 | 0.096 | 0.148 | 1 | TEC_CD | SMC4 |
| 0.00122672 | -0.307127 | 0.574 | 0.598 | 1 | TEC_CD | NPC2 |
| 0.00145061 | -0.3316885 | 0.51 | 0.513 | 1 | TEC_CD | ANXA11 |
| 0.00153712 | -0.3064153 | 0.924 | 0.897 | 1 | TEC_CD | RPS3A |
| 0.00186495 | -0.3631496 | 0.547 | 0.578 | 1 | TEC_CD | LITAF |
| 0.00216036 | -0.3999129 | 0.941 | 0.933 | 1 | TEC_CD | RPS12 |
| 0.00240865 | -0.3432327 | 0.901 | 0.846 | 1 | TEC_CD | RPL35 |
| 0.00242283 | -0.3861876 | 0.966 | 0.973 | 1 | TEC_CD | ACTB |
| 0.00290902 | -0.293183 | 0.882 | 0.875 | 1 | TEC_CD | RPL26 |
| 0.00337149 | -0.2870149 | 0.14 | 0.191 | 1 | TEC_CD | PLK2 |
| 0.00399913 | -0.4591162 | 0.384 | 0.281 | 1 | TEC_CD | IFITM2 |
| 0.00402353 | -0.417743 | 0.266 | 0.199 | 1 | TEC_CD | THBS1 |
| 0.00431448 | -0.831752 | 0.249 | 0.263 | 1 | TEC_CD | LMO7 |
| 0.00461816 | -0.303363 | 0.106 | 0.152 | 1 | TEC_CD | ZFAND2A |

| | | | | | | |
|---|---|---|---|---|---|---|
| 0.00487893 | -0.4263143 | 0.261 | 0.293 | 1 | TEC_CD | ATP13A3 |
| 0.00578157 | -0.2682991 | 0.288 | 0.325 | 1 | TEC_CD | TMBIM1 |
| 0.00590781 | -0.427006 | 0.153 | 0.189 | 1 | TEC_CD | EPS8L1 |
| 0.00900508 | -0.3038258 | 0.648 | 0.629 | 1 | TEC_CD | TAX1BP1 |
| 0 | 3.77206213 | 0.796 | 0.299 | 0 | TEC_inj | MMP7 |
| 0 | 2.70369231 | 0.724 | 0.226 | 0 | TEC_inj | S100A2 |
| 0 | 2.44783086 | 0.756 | 0.146 | 0 | TEC_inj | C12orf75 |
| 0 | 2.29917941 | 0.91 | 0.286 | 0 | TEC_inj | IGFBP7 |
| 0 | 2.12778421 | 0.561 | 0.09 | 0 | TEC_inj | TFPI2 |
| 0 | 2.12532572 | 0.808 | 0.422 | 0 | TEC_inj | SPP1 |
| 0 | 1.91147144 | 0.164 | 0.028 | 0 | TEC_inj | UMOD |
| 0 | 1.88982167 | 0.724 | 0.207 | 0 | TEC_inj | CRYAB |
| 0 | 1.87341846 | 0.654 | 0.383 | 0 | TEC_inj | SLPI |
| 0 | 1.85861442 | 0.468 | 0.108 | 0 | TEC_inj | DEFB1 |
| 0 | 1.82004213 | 0.755 | 0.221 | 0 | TEC_inj | TSPAN1 |
| 0 | 1.81267936 | 0.861 | 0.461 | 0 | TEC_inj | KRT8 |
| 0 | 1.71932741 | 0.926 | 0.862 | 0 | TEC_inj | ANXA2 |
| 0 | 1.64888266 | 0.979 | 0.933 | 0 | TEC_inj | S100A6 |
| 0 | 1.64806898 | 0.641 | 0.141 | 0 | TEC_inj | LAMC2 |
| 0 | 1.64167144 | 0.953 | 0.8 | 0 | TEC_inj | GSTP1 |
| 0 | 1.59572265 | 0.857 | 0.181 | 0 | TEC_inj | EPCAM |
| 0 | 1.47616559 | 0.697 | 0.244 | 0 | TEC_inj | CLU |
| 0 | 1.39844642 | 0.864 | 0.531 | 0 | TEC_inj | KRT18 |
| 0 | 1.3446081 | 0.93 | 0.73 | 0 | TEC_inj | PRDX1 |
| 0 | 1.31646434 | 0.658 | 0.183 | 0 | TEC_inj | CCND1 |
| 0 | 1.3100846 | 0.557 | 0.218 | 0 | TEC_inj | CST6 |
| 0 | 1.30161119 | 0.772 | 0.195 | 0 | TEC_inj | S100A13 |
| 0 | 1.27049211 | 0.614 | 0.214 | 0 | TEC_inj | PLAU |
| 0 | 1.24849229 | 0.739 | 0.295 | 0 | TEC_inj | PPP1R14B |
| 0 | 1.24179453 | 0.749 | 0.232 | 0 | TEC_inj | MAL2 |
| 0 | 1.23864641 | 0.907 | 0.508 | 0 | TEC_inj | CD24 |
| 0 | 1.2176197 | 0.871 | 0.399 | 0 | TEC_inj | CYSTM1 |
| 0 | 1.19577426 | 0.707 | 0.222 | 0 | TEC_inj | RHOC |
| 0 | 1.18992009 | 0.973 | 0.877 | 0 | TEC_inj | MYL12B |
| 0 | 1.12769746 | 0.95 | 0.738 | 0 | TEC_inj | RPSA |
| 0 | 1.12339148 | 0.911 | 0.636 | 0 | TEC_inj | NDUFA4 |
| 0 | 1.12145595 | 0.674 | 0.223 | 0 | TEC_inj | TPM1 |
| 0 | 1.11693997 | 0.975 | 0.86 | 0 | TEC_inj | RPLP0 |
| 0 | 1.11071506 | 0.657 | 0.2 | 0 | TEC_inj | S100A16 |
| 0 | 1.1063652 | 0.84 | 0.391 | 0 | TEC_inj | SPINT2 |
| 0 | 1.10386752 | 0.966 | 0.915 | 0 | TEC_inj | S100A10 |
| 0 | 1.09871304 | 0.736 | 0.286 | 0 | TEC_inj | IFITM3 |
| 0 | 1.08052687 | 0.977 | 0.9 | 0 | TEC_inj | ITM2B |
| 0 | 1.07073712 | 0.556 | 0.198 | 0 | TEC_inj | IL32 |
| 0 | 1.06230902 | 0.243 | 0.057 | 0 | TEC_inj | TAGLN |
| 0 | 1.05684884 | 0.677 | 0.184 | 0 | TEC_inj | OCIAD2 |
| 0 | 1.05253666 | 0.7 | 0.247 | 0 | TEC_inj | CD151 |
| 0 | 1.04651992 | 0.885 | 0.505 | 0 | TEC_inj | WFDC2 |
| 0 | 1.01918078 | 0.733 | 0.307 | 0 | TEC_inj | PDIA6 |
| 0 | 1.01209207 | 0.961 | 0.965 | 0 | TEC_inj | S100A11 |

| | | | | | | |
|---|---|---|---|---|---|---|
| 0 | 1.01167512 | 0.461 | 0.105 | 0 | TEC_inj | UCHL1 |
| 0 | 0.99397339 | 0.956 | 0.824 | 0 | TEC_inj | PPIA |
| 0 | 0.99256153 | 0.775 | 0.377 | 0 | TEC_inj | CD59 |
| 0 | 0.98809885 | 0.78 | 0.459 | 0 | TEC_inj | HMGA1 |
| 0 | 0.98609808 | 0.898 | 0.576 | 0 | TEC_inj | HINT1 |
| 0 | 0.97964301 | 0.404 | 0.085 | 0 | TEC_inj | FXYD2 |
| 0 | 0.96162497 | 0.925 | 0.711 | 0 | TEC_inj | PKM |
| 0 | 0.95890686 | 0.99 | 0.876 | 0 | TEC_inj | RPS2 |
| 0 | 0.95487538 | 0.634 | 0.242 | 0 | TEC_inj | TNFRSF12A |
| 0 | 0.9512616 | 0.987 | 0.866 | 0 | TEC_inj | RPL3 |
| 0 | 0.94678228 | 0.483 | 0.114 | 0 | TEC_inj | ITGB6 |
| 0 | 0.9330854 | 0.993 | 0.928 | 0 | TEC_inj | RPS12 |
| 0 | 0.93186446 | 0.935 | 0.782 | 0 | TEC_inj | CHCHD2 |
| 0 | 0.92094123 | 0.686 | 0.248 | 0 | TEC_inj | LDHB |
| 0 | 0.91804106 | 0.975 | 0.873 | 0 | TEC_inj | RPL12 |
| 0 | 0.91645479 | 0.512 | 0.139 | 0 | TEC_inj | AKR1B1 |
| 0 | 0.91253103 | 0.22 | 0.04 | 0 | TEC_inj | FXYD4 |
| 0 | 0.91033346 | 0.996 | 0.954 | 0 | TEC_inj | RPS18 |
| 0 | 0.90777675 | 0.96 | 0.787 | 0 | TEC_inj | RPS5 |
| 0 | 0.89646376 | 0.801 | 0.309 | 0 | TEC_inj | PEBP1 |
| 0 | 0.89523763 | 0.996 | 0.916 | 0 | TEC_inj | RPL8 |
| 0 | 0.89341776 | 0.73 | 0.309 | 0 | TEC_inj | BSG |
| 0 | 0.88584537 | 0.82 | 0.444 | 0 | TEC_inj | CD81 |
| 0 | 0.87787915 | 0.532 | 0.166 | 0 | TEC_inj | HIGD1A |
| 0 | 0.86637448 | 0.288 | 0.05 | 0 | TEC_inj | TNFRSF11B |
| 0 | 0.86481878 | 0.538 | 0.19 | 0 | TEC_inj | TGM2 |
| 0 | 0.86350711 | 0.519 | 0.204 | 0 | TEC_inj | ITGA2 |
| 0 | 0.85541135 | 0.713 | 0.298 | 0 | TEC_inj | APP |
| 0 | 0.84365357 | 0.569 | 0.105 | 0 | TEC_inj | LAPTM4B |
| 0 | 0.84081016 | 0.989 | 0.918 | 0 | TEC_inj | RPL18A |
| 0 | 0.83939439 | 0.79 | 0.526 | 0 | TEC_inj | PPIB |
| 0 | 0.83889224 | 0.707 | 0.291 | 0 | TEC_inj | MGST3 |
| 0 | 0.8331095 | 0.959 | 0.856 | 0 | TEC_inj | SRP14 |
| 0 | 0.82938452 | 0.994 | 0.941 | 0 | TEC_inj | RPS19 |
| 0 | 0.82248328 | 0.996 | 0.97 | 0 | TEC_inj | RPLP1 |
| 0 | 0.819624 | 0.999 | 0.979 | 0 | TEC_inj | EEF1A1 |
| 0 | 0.81454809 | 0.965 | 0.766 | 0 | TEC_inj | RPL10A |
| 0 | 0.81077758 | 0.56 | 0.146 | 0 | TEC_inj | ANXA3 |
| 0 | 0.80515077 | 0.498 | 0.153 | 0 | TEC_inj | S100A14 |
| 0 | 0.79598224 | 0.94 | 0.84 | 0 | TEC_inj | RPS4X |
| 0 | 0.79559848 | 0.81 | 0.468 | 0 | TEC_inj | CD9 |
| 0 | 0.79387044 | 0.534 | 0.118 | 0 | TEC_inj | ITGA3 |
| 0 | 0.78258987 | 0.639 | 0.283 | 0 | TEC_inj | SDC4 |
| 0 | 0.77445317 | 0.613 | 0.23 | 0 | TEC_inj | PRDX2 |
| 0 | 0.7650752 | 0.482 | 0.108 | 0 | TEC_inj | MGLL |
| 0 | 0.76386191 | 0.995 | 0.954 | 0 | TEC_inj | RPL15 |
| 0 | 0.75700196 | 0.983 | 0.888 | 0 | TEC_inj | RPS9 |
| 0 | 0.74736961 | 0.957 | 0.886 | 0 | TEC_inj | CD63 |
| 0 | 0.74487164 | 0.826 | 0.554 | 0 | TEC_inj | PRDX5 |
| 0 | 0.74441694 | 0.443 | 0.074 | 0 | TEC_inj | DPP4 |

| | | | | | | |
|---|---|---|---|---|---|---|
| 0 | 0.73787555 | 0.749 | 0.391 | 0 | TEC_inj | RAN |
| 0 | 0.73321341 | 0.669 | 0.322 | 0 | TEC_inj | GLS |
| 0 | 0.72872511 | 0.776 | 0.371 | 0 | TEC_inj | SOD1 |
| 0 | 0.72567599 | 0.978 | 0.835 | 0 | TEC_inj | RPL35 |
| 0 | 0.71656645 | 0.886 | 0.629 | 0 | TEC_inj | UQCRH |
| 0 | 0.71259531 | 0.457 | 0.109 | 0 | TEC_inj | DCDC2 |
| 0 | 0.70914685 | 0.982 | 0.866 | 0 | TEC_inj | RPL36 |
| 0 | 0.7087226 | 0.987 | 0.88 | 0 | TEC_inj | RPS23 |
| 0 | 0.7010923 | 0.983 | 0.898 | 0 | TEC_inj | RPL7A |
| 0 | 0.69178365 | 0.729 | 0.338 | 0 | TEC_inj | PRDX6 |
| 0 | 0.68528127 | 0.895 | 0.71 | 0 | TEC_inj | COX6A1 |
| 0 | 0.68466445 | 0.959 | 0.828 | 0 | TEC_inj | RACK1 |
| 0 | 0.68436626 | 0.991 | 0.891 | 0 | TEC_inj | RPL21 |
| 0 | 0.6815262 | 0.729 | 0.343 | 0 | TEC_inj | PARK7 |
| 0 | 0.67602484 | 0.855 | 0.652 | 0 | TEC_inj | RTN4 |
| 0 | 0.67602203 | 0.993 | 0.918 | 0 | TEC_inj | RPS27A |
| 0 | 0.67489928 | 0.457 | 0.133 | 0 | TEC_inj | SELENOM |
| 0 | 0.66175068 | 0.925 | 0.731 | 0 | TEC_inj | EEF1B2 |
| 0 | 0.65968492 | 0.647 | 0.333 | 0 | TEC_inj | ATP1A1 |
| 0 | 0.65494409 | 0.995 | 0.979 | 0 | TEC_inj | RPL41 |
| 0 | 0.65235141 | 0.604 | 0.205 | 0 | TEC_inj | PCBD1 |
| 0 | 0.64972274 | 0.496 | 0.099 | 0 | TEC_inj | DMKN |
| 0 | 0.646476 | 0.656 | 0.356 | 0 | TEC_inj | EIF5A |
| 0 | 0.64315474 | 0.991 | 0.912 | 0 | TEC_inj | RPL9 |
| 0 | 0.6409155 | 0.983 | 0.895 | 0 | TEC_inj | RPS7 |
| 0 | 0.63436454 | 0.992 | 0.889 | 0 | TEC_inj | RPS3A |
| 0 | 0.63298413 | 0.451 | 0.148 | 0 | TEC_inj | STMN1 |
| 0 | 0.63011086 | 0.491 | 0.19 | 0 | TEC_inj | CKB |
| 0 | 0.62862413 | 0.988 | 0.861 | 0 | TEC_inj | RPL14 |
| 0 | 0.62833018 | 0.354 | 0.066 | 0 | TEC_inj | DCBLD2 |
| 0 | 0.62813488 | 0.378 | 0.125 | 0 | TEC_inj | CD40 |
| 0 | 0.6250854 | 0.669 | 0.34 | 0 | TEC_inj | REEP5 |
| 0 | 0.62230278 | 0.974 | 0.848 | 0 | TEC_inj | RPL7 |
| 0 | 0.62141973 | 0.972 | 0.804 | 0 | TEC_inj | RPL5 |
| 0 | 0.61793901 | 0.991 | 0.916 | 0 | TEC_inj | RPL11 |
| 0 | 0.61776795 | 0.518 | 0.153 | 0 | TEC_inj | CDH1 |
| 0 | 0.61359061 | 0.628 | 0.3 | 0 | TEC_inj | CAPG |
| 0 | 0.6125492 | 0.887 | 0.601 | 0 | TEC_inj | DSTN |
| 0 | 0.60900221 | 0.998 | 0.961 | 0 | TEC_inj | RPL13 |
| 0 | 0.60152173 | 0.66 | 0.315 | 0 | TEC_inj | ATP5F1B |
| 0 | 0.59972758 | 0.573 | 0.243 | 0 | TEC_inj | PHPT1 |
| 0 | 0.59640441 | 0.69 | 0.36 | 0 | TEC_inj | DAD1 |
| 0 | 0.59540551 | 0.515 | 0.209 | 0 | TEC_inj | PDZK1IP1 |
| 0 | 0.59524614 | 0.39 | 0.136 | 0 | TEC_inj | CLDN3 |
| 0 | 0.59157104 | 0.617 | 0.284 | 0 | TEC_inj | MZT2B |
| 0 | 0.59090883 | 0.666 | 0.352 | 0 | TEC_inj | POLR2L |
| 0 | 0.58877007 | 0.637 | 0.321 | 0 | TEC_inj | CAPN2 |
| 0 | 0.58408317 | 0.994 | 0.921 | 0 | TEC_inj | RPL19 |
| 0 | 0.57447382 | 0.525 | 0.222 | 0 | TEC_inj | CD47 |
| 0 | 0.57430305 | 0.366 | 0.056 | 0 | TEC_inj | PCSK1N |

| | | | | | | |
|---|---|---|---|---|---|---|
| 0 | 0.57283656 | 0.655 | 0.328 | 0 | TEC_inj | MRPL33 |
| 0 | 0.56965527 | 0.491 | 0.193 | 0 | TEC_inj | CDK2AP2 |
| 0 | 0.56923944 | 0.587 | 0.268 | 0 | TEC_inj | CHCHD10 |
| 0 | 0.56876367 | 0.595 | 0.278 | 0 | TEC_inj | TMED9 |
| 0 | 0.56013801 | 0.304 | 0.059 | 0 | TEC_inj | AL353751.1 |
| 0 | 0.55732184 | 0.655 | 0.339 | 0 | TEC_inj | ATP5F1D |
| 0 | 0.55703339 | 0.254 | 0.06 | 0 | TEC_inj | FGF2 |
| 0 | 0.55410245 | 0.536 | 0.193 | 0 | TEC_inj | ECHS1 |
| 0 | 0.54957574 | 0.622 | 0.295 | 0 | TEC_inj | YWHAQ |
| 0 | 0.54950147 | 0.406 | 0.11 | 0 | TEC_inj | TMEM238 |
| 0 | 0.54351443 | 0.37 | 0.122 | 0 | TEC_inj | SERPING1 |
| 0 | 0.53965877 | 0.545 | 0.254 | 0 | TEC_inj | MYDGF |
| 0 | 0.53759724 | 0.99 | 0.879 | 0 | TEC_inj | RPS6 |
| 0 | 0.53586547 | 0.542 | 0.219 | 0 | TEC_inj | PTMS |
| 0 | 0.53253317 | 0.548 | 0.211 | 0 | TEC_inj | TSTD1 |
| 0 | 0.53140108 | 0.465 | 0.126 | 0 | TEC_inj | RCN1 |
| 0 | 0.53121844 | 0.296 | 0.06 | 0 | TEC_inj | CRABP2 |
| 0 | 0.53081842 | 0.537 | 0.236 | 0 | TEC_inj | CUTA |
| 0 | 0.52996605 | 0.58 | 0.271 | 0 | TEC_inj | KDELR1 |
| 0 | 0.52983854 | 0.371 | 0.069 | 0 | TEC_inj | CAV2 |
| 0 | 0.52821247 | 0.459 | 0.14 | 0 | TEC_inj | GLO1 |
| 0 | 0.52786424 | 0.148 | 0.015 | 0 | TEC_inj | SFRP1 |
| 0 | 0.52449769 | 0.289 | 0.066 | 0 | TEC_inj | IGFBP2 |
| 0 | 0.52319346 | 0.373 | 0.071 | 0 | TEC_inj | PON2 |
| 0 | 0.52032311 | 0.509 | 0.175 | 0 | TEC_inj | PHB |
| 0 | 0.51300243 | 0.306 | 0.064 | 0 | TEC_inj | QPRT |
| 0 | 0.50922822 | 0.542 | 0.216 | 0 | TEC_inj | TMEM14C |
| 0 | 0.50530851 | 0.339 | 0.071 | 0 | TEC_inj | BCAM |
| 0 | 0.50409909 | 0.539 | 0.201 | 0 | TEC_inj | CYB5A |
| 0 | 0.50280225 | 0.443 | 0.108 | 0 | TEC_inj | COL18A1 |
| 0 | 0.50139325 | 0.187 | 0.032 | 0 | TEC_inj | CLDN10 |
| 0 | 0.50080146 | 0.488 | 0.198 | 0 | TEC_inj | LSM4 |
| 0 | 0.50001486 | 0.561 | 0.264 | 0 | TEC_inj | TMED10 |
| 0 | 0.49780837 | 0.285 | 0.075 | 0 | TEC_inj | CA12 |
| 0 | 0.49593602 | 0.679 | 0.359 | 0 | TEC_inj | TBCA |
| 0 | 0.48528165 | 0.473 | 0.181 | 0 | TEC_inj | NENF |
| 0 | 0.48338468 | 0.625 | 0.299 | 0 | TEC_inj | PPA1 |
| 0 | 0.48205856 | 0.465 | 0.169 | 0 | TEC_inj | PGRMC1 |
| 0 | 0.48064661 | 0.35 | 0.117 | 0 | TEC_inj | ARHGAP29 |
| 0 | 0.4792214 | 0.391 | 0.116 | 0 | TEC_inj | PRDX4 |
| 0 | 0.47820169 | 0.246 | 0.029 | 0 | TEC_inj | GAL |
| 0 | 0.47671227 | 0.645 | 0.291 | 0 | TEC_inj | PERP |
| 0 | 0.47400788 | 0.49 | 0.184 | 0 | TEC_inj | PPP2R1A |
| 0 | 0.46970324 | 0.289 | 0.078 | 0 | TEC_inj | PDLIM4 |
| 0 | 0.46644791 | 0.41 | 0.14 | 0 | TEC_inj | MET |
| 0 | 0.46586239 | 0.428 | 0.122 | 0 | TEC_inj | GSTM3 |
| 0 | 0.45828071 | 0.437 | 0.159 | 0 | TEC_inj | PFKP |
| 0 | 0.45804455 | 0.457 | 0.171 | 0 | TEC_inj | SEC11C |
| 0 | 0.45665523 | 0.445 | 0.163 | 0 | TEC_inj | ANXA4 |
| 0 | 0.45306211 | 0.358 | 0.088 | 0 | TEC_inj | WWTR1 |

| | | | | | | |
|---|---|---|---|---|---|---|
| 0 | 0.45087787 | 0.431 | 0.167 | 0 | TEC_inj | PRDX3 |
| 0 | 0.44944151 | 0.282 | 0.052 | 0 | TEC_inj | GNG11 |
| 0 | 0.44267943 | 0.431 | 0.164 | 0 | TEC_inj | NHP2 |
| 0 | 0.43694376 | 0.486 | 0.196 | 0 | TEC_inj | MDH2 |
| 0 | 0.42967682 | 0.43 | 0.157 | 0 | TEC_inj | MGAT4B |
| 0 | 0.4293001 | 0.372 | 0.107 | 0 | TEC_inj | HACD3 |
| 0 | 0.42301519 | 0.376 | 0.128 | 0 | TEC_inj | NME1 |
| 0 | 0.4210817 | 0.41 | 0.141 | 0 | TEC_inj | TMEM141 |
| 0 | 0.42054927 | 0.269 | 0.042 | 0 | TEC_inj | CDH16 |
| 0 | 0.41486887 | 0.421 | 0.165 | 0 | TEC_inj | MZT2A |
| 0 | 0.41276675 | 0.354 | 0.1 | 0 | TEC_inj | CTNNAL1 |
| 0 | 0.41036686 | 0.385 | 0.13 | 0 | TEC_inj | RCN2 |
| 0 | 0.40969305 | 0.285 | 0.064 | 0 | TEC_inj | CAVIN1 |
| 0 | 0.40713502 | 0.283 | 0.076 | 0 | TEC_inj | ADAMTS9 |
| 0 | 0.40609104 | 0.388 | 0.133 | 0 | TEC_inj | SLC25A39 |
| 0 | 0.40085223 | 0.211 | 0.034 | 0 | TEC_inj | LAMA3 |
| 0 | 0.40011515 | 0.324 | 0.09 | 0 | TEC_inj | ITM2C |
| 0 | 0.39712037 | 0.34 | 0.092 | 0 | TEC_inj | ADGRG1 |
| 0 | 0.392716 | 0.289 | 0.062 | 0 | TEC_inj | LMTK3 |
| 0 | 0.39256304 | 0.352 | 0.108 | 0 | TEC_inj | DNPH1 |
| 0 | 0.38885017 | 0.396 | 0.131 | 0 | TEC_inj | SCOC |
| 0 | 0.38716279 | 0.4 | 0.142 | 0 | TEC_inj | NUTF2 |
| 0 | 0.38669897 | 0.357 | 0.125 | 0 | TEC_inj | SLC39A1 |
| 0 | 0.38542669 | 0.378 | 0.137 | 0 | TEC_inj | HSD17B12 |
| 0 | 0.38235381 | 0.594 | 0.247 | 0 | TEC_inj | CAMK2N1 |
| 0 | 0.38227838 | 0.33 | 0.094 | 0 | TEC_inj | LGALS3BP |
| 0 | 0.38219919 | 0.256 | 0.058 | 0 | TEC_inj | PRSS23 |
| 0 | 0.37744475 | 0.308 | 0.096 | 0 | TEC_inj | GOLM1 |
| 0 | 0.37463396 | 0.241 | 0.06 | 0 | TEC_inj | TIMP3 |
| 0 | 0.37378294 | 0.328 | 0.098 | 0 | TEC_inj | TRNP1 |
| 0 | 0.36628884 | 0.276 | 0.052 | 0 | TEC_inj | L1CAM |
| 0 | 0.36222902 | 0.293 | 0.071 | 0 | TEC_inj | DHCR24 |
| 0 | 0.35978159 | 0.312 | 0.076 | 0 | TEC_inj | AP1M2 |
| 0 | 0.35240695 | 0.329 | 0.109 | 0 | TEC_inj | ARL2 |
| 0 | 0.3474674 | 0.268 | 0.058 | 0 | TEC_inj | MYL9 |
| 0 | 0.34707241 | 0.255 | 0.046 | 0 | TEC_inj | LIPG |
| 0 | 0.33575347 | 0.275 | 0.057 | 0 | TEC_inj | DDAH1 |
| 0 | 0.33157226 | 0.253 | 0.047 | 0 | TEC_inj | SCD5 |
| 0 | 0.32825624 | 0.321 | 0.101 | 0 | TEC_inj | NDUFV1 |
| 0 | 0.32822527 | 0.3 | 0.087 | 0 | TEC_inj | GCSH |
| 0 | 0.32627523 | 0.312 | 0.095 | 0 | TEC_inj | CISD1 |
| 0 | 0.32421533 | 0.252 | 0.048 | 0 | TEC_inj | TMEM132A |
| 0 | 0.32321373 | 0.276 | 0.075 | 0 | TEC_inj | SEMA3B |
| 0 | 0.32218814 | 0.28 | 0.073 | 0 | TEC_inj | CETN2 |
| 0 | 0.31494245 | 0.277 | 0.073 | 0 | TEC_inj | DCTPP1 |
| 0 | 0.31237768 | 0.206 | 0.041 | 0 | TEC_inj | CAV1 |
| 0 | 0.31116866 | 0.343 | 0.111 | 0 | TEC_inj | AIF1L |
| 0 | 0.31004495 | 0.211 | 0.05 | 0 | TEC_inj | ALDH1A1 |
| 0 | 0.30709462 | 0.199 | 0.042 | 0 | TEC_inj | RBP1 |
| 0 | 0.30642761 | 0.275 | 0.067 | 0 | TEC_inj | TUSC3 |

| | | | | | | |
|---|---|---|---|---|---|---|
| 0 | 0.30422471 | 0.267 | 0.07 | 0 | TEC_inj | METRN |
| 0 | 0.30348441 | 0.158 | 0.023 | 0 | TEC_inj | PPP1R1A |
| 0 | 0.30084755 | 0.194 | 0.024 | 0 | TEC_inj | FOLR1 |
| 0 | 0.29598866 | 0.222 | 0.034 | 0 | TEC_inj | EFEMP1 |
| 0 | 0.27516698 | 0.248 | 0.063 | 0 | TEC_inj | LARP6 |
| 0 | 0.26254053 | 0.247 | 0.057 | 0 | TEC_inj | TOM1L1 |
| 0 | 0.26156801 | 0.227 | 0.052 | 0 | TEC_inj | CD320 |
| 0 | 0.25522387 | 0.192 | 0.04 | 0 | TEC_inj | EFHD1 |
| 0 | -0.9343315 | 0.837 | 0.993 | 0 | TEC_inj | MALAT1 |
| 0 | -1.0994892 | 0.384 | 0.726 | 0 | TEC_inj | JUNB |
| 0 | -1.1963486 | 0.183 | 0.557 | 0 | TEC_inj | ZFP36 |
| 0 | -1.3996722 | 0.11 | 0.488 | 0 | TEC_inj | MXD1 |
| 0 | -1.64332 | 0.265 | 0.677 | 0 | TEC_inj | GLUL |
| 0 | -1.7293182 | 0.114 | 0.469 | 0 | TEC_inj | TYROBP |
| 0 | -1.7405787 | 0.073 | 0.413 | 0 | TEC_inj | FCER1G |
| 0 | -1.7430298 | 0.266 | 0.644 | 0 | TEC_inj | C15orf48 |
| 0 | -1.83359 | 0.285 | 0.716 | 0 | TEC_inj | NAMPT |
| 0 | -1.9891004 | 0.63 | 0.968 | 0 | TEC_inj | NEAT1 |
| 0 | -2.2271114 | 0.148 | 0.492 | 0 | TEC_inj | LYZ |
| 0 | -2.4441448 | 0.089 | 0.436 | 0 | TEC_inj | S100P |
| 0 | -2.6824605 | 0.163 | 0.606 | 0 | TEC_inj | SRGN |
| 0 | -4.130217 | 0.194 | 0.713 | 0 | TEC_inj | S100A8 |
| 0 | -4.3486716 | 0.326 | 0.762 | 0 | TEC_inj | S100A9 |
| 5.309E-308 | -1.2480818 | 0.046 | 0.361 | 1.254E-303 | TEC_inj | LAPTM5 |
| 5.665E-308 | 0.92626336 | 0.623 | 0.328 | 1.338E-303 | TEC_inj | PHLDA2 |
| 6.01E-308 | 0.26385642 | 0.235 | 0.059 | 1.42E-303 | TEC_inj | DBN1 |
| 2.712E-307 | 0.29928593 | 0.23 | 0.058 | 6.407E-303 | TEC_inj | PROM1 |
| 5.393E-307 | 0.44132771 | 0.41 | 0.158 | 1.274E-302 | TEC_inj | LSR |
| 5.425E-307 | 0.64149585 | 0.985 | 0.89 | 1.281E-302 | TEC_inj | RPL29 |
| 2.22E-306 | 0.25383053 | 0.198 | 0.044 | 5.244E-302 | TEC_inj | FSTL3 |
| 1.099E-305 | 0.62153457 | 0.992 | 0.905 | 2.597E-301 | TEC_inj | RPS8 |
| 1.468E-305 | 0.39881749 | 0.417 | 0.164 | 3.468E-301 | TEC_inj | EMC10 |
| 6.598E-305 | 0.26069322 | 0.213 | 0.05 | 1.559E-300 | TEC_inj | COL4A2 |
| 7.501E-305 | 0.3470729 | 0.33 | 0.108 | 1.772E-300 | TEC_inj | FAM213A |
| 1.55E-304 | 0.43118011 | 0.365 | 0.129 | 3.663E-300 | TEC_inj | SCCPDH |
| 5.666E-304 | 0.30412379 | 0.281 | 0.083 | 1.338E-299 | TEC_inj | MXRA7 |
| 1.279E-303 | 0.28768 | 0.263 | 0.074 | 3.021E-299 | TEC_inj | PPIC |
| 4.249E-303 | 0.62660109 | 0.987 | 0.942 | 1.004E-298 | TEC_inj | RPL13A |
| 4.908E-303 | 0.67429554 | 0.984 | 0.977 | 1.159E-298 | TEC_inj | RPL28 |
| 6.401E-303 | -2.9862275 | 0.09 | 0.403 | 1.512E-298 | TEC_inj | KRT13 |
| 9.528E-303 | -4.2550715 | 0.111 | 0.427 | 2.251E-298 | TEC_inj | SPRR3 |
| 6.875E-302 | 0.39396977 | 0.443 | 0.18 | 1.624E-297 | TEC_inj | PSMB5 |
| 9.814E-302 | 0.56440681 | 0.974 | 0.819 | 2.318E-297 | TEC_inj | RPL22 |
| 1.449E-301 | 0.27213799 | 0.153 | 0.027 | 3.423E-297 | TEC_inj | NOV |
| 2.255E-301 | 0.343059 | 0.195 | 0.044 | 5.326E-297 | TEC_inj | SULT1C2 |
| 4.197E-301 | 0.61126178 | 0.985 | 0.879 | 9.915E-297 | TEC_inj | RPL6 |
| 9.872E-301 | 0.52304384 | 0.689 | 0.385 | 2.332E-296 | TEC_inj | ATP5PF |
| 5.596E-298 | 0.28427838 | 0.176 | 0.037 | 1.322E-293 | TEC_inj | TUBB2B |
| 4.37E-297 | -1.2989321 | 0.415 | 0.735 | 1.032E-292 | TEC_inj | ANXA1 |
| 4.571E-297 | 0.45807865 | 0.623 | 0.314 | 1.08E-292 | TEC_inj | SNU13 |

| | | | | | | |
|---|---|---|---|---|---|---|
| 1.475E-296 | 0.36223642 | 0.408 | 0.157 | 3.485E-292 | TEC_inj | ARL1 |
| 9.92E-296 | 0.40801645 | 0.554 | 0.258 | 2.343E-291 | TEC_inj | COX7A2L |
| 5.305E-295 | 0.53913348 | 0.936 | 0.827 | 1.253E-290 | TEC_inj | COX4I1 |
| 1.545E-294 | 0.31777176 | 0.248 | 0.068 | 3.651E-290 | TEC_inj | KHDRBS3 |
| 3.807E-294 | 0.34780245 | 0.352 | 0.124 | 8.993E-290 | TEC_inj | BACE2 |
| 5.56E-294 | -1.5674288 | 0.052 | 0.358 | 1.313E-289 | TEC_inj | IL1RN |
| 6.631E-294 | 0.38062403 | 0.328 | 0.112 | 1.566E-289 | TEC_inj | CBR1 |
| 1.129E-293 | 0.6149856 | 0.651 | 0.363 | 2.666E-289 | TEC_inj | PSME1 |
| 9.247E-293 | 0.32246598 | 0.317 | 0.105 | 2.184E-288 | TEC_inj | UCHL3 |
| 2.466E-292 | 0.54669593 | 0.507 | 0.232 | 5.826E-288 | TEC_inj | ODC1 |
| 8.235E-292 | 0.62338769 | 0.993 | 0.909 | 1.945E-287 | TEC_inj | RPL32 |
| 8.632E-292 | 0.77937655 | 0.979 | 0.928 | 2.039E-287 | TEC_inj | GAPDH |
| 1.75E-291 | 0.48555447 | 0.637 | 0.343 | 4.135E-287 | TEC_inj | WDR83OS |
| 2.018E-291 | 0.46900999 | 0.552 | 0.264 | 4.767E-287 | TEC_inj | ACADVL |
| 1.017E-288 | 0.33809098 | 0.308 | 0.099 | 2.403E-284 | TEC_inj | DSG2 |
| 4.707E-288 | 0.25758564 | 0.243 | 0.067 | 1.112E-283 | TEC_inj | TMEM9 |
| 6.927E-288 | 0.53534769 | 0.796 | 0.571 | 1.636E-283 | TEC_inj | SLC25A3 |
| 9.231E-288 | 0.62802934 | 0.883 | 0.713 | 2.181E-283 | TEC_inj | ENO1 |
| 2.091E-287 | -0.6758627 | 0.406 | 0.716 | 4.94E-283 | TEC_inj | PNRC1 |
| 1.532E-286 | 0.54109206 | 0.87 | 0.7 | 3.62E-282 | TEC_inj | SKP1 |
| 4.7E-286 | 0.94535645 | 0.85 | 0.749 | 1.11E-281 | TEC_inj | ITGB1 |
| 8.998E-286 | 0.51226955 | 0.899 | 0.687 | 2.126E-281 | TEC_inj | RPL4 |
| 9.445E-285 | 0.73717915 | 0.158 | 0.031 | 2.231E-280 | TEC_inj | MGP |
| 2.859E-284 | 0.45958973 | 0.39 | 0.156 | 6.754E-280 | TEC_inj | CSRP1 |
| 3.812E-284 | 0.2945996 | 0.162 | 0.032 | 9.005E-280 | TEC_inj | PTH1R |
| 5.126E-284 | -1.1694664 | 0.014 | 0.304 | 1.211E-279 | TEC_inj | PTPRC |
| 7.59E-284 | 0.93949095 | 0.79 | 0.523 | 1.793E-279 | TEC_inj | KRT7 |
| 1.468E-283 | 0.52193756 | 0.674 | 0.369 | 3.468E-279 | TEC_inj | NDUFB2 |
| 2.233E-283 | -2.2427254 | 0.067 | 0.365 | 5.275E-279 | TEC_inj | PSCA |
| 2.242E-283 | 0.49950762 | 0.658 | 0.359 | 5.297E-279 | TEC_inj | DYNLRB1 |
| 1.505E-282 | 0.43596948 | 0.493 | 0.222 | 3.555E-278 | TEC_inj | PDCD5 |
| 7.703E-282 | 0.53163212 | 0.633 | 0.334 | 1.82E-277 | TEC_inj | MIF |
| 9.822E-282 | 0.6115847 | 0.355 | 0.133 | 2.32E-277 | TEC_inj | MDK |
| 1.434E-281 | 0.42255778 | 0.525 | 0.247 | 3.388E-277 | TEC_inj | LMAN2 |
| 9.461E-281 | 0.41260772 | 0.48 | 0.216 | 2.235E-276 | TEC_inj | PSMB7 |
| 2.241E-279 | 0.52019991 | 0.712 | 0.426 | 5.295E-275 | TEC_inj | ATP5MD |
| 9.187E-279 | 0.555709 | 0.981 | 0.868 | 2.17E-274 | TEC_inj | RPL23A |
| 2.965E-278 | 0.49781288 | 0.921 | 0.749 | 7.003E-274 | TEC_inj | COX7C |
| 3.567E-278 | 0.37921208 | 0.349 | 0.126 | 8.425E-274 | TEC_inj | GPRC5C |
| 9.449E-277 | 0.26961449 | 0.208 | 0.052 | 2.232E-272 | TEC_inj | FBXO2 |
| 8.927E-276 | 0.60680557 | 0.938 | 0.871 | 2.109E-271 | TEC_inj | TMBIM6 |
| 1.258E-275 | 0.46752668 | 0.632 | 0.334 | 2.972E-271 | TEC_inj | SNRPD2 |
| 1.263E-275 | 0.49933364 | 0.498 | 0.232 | 2.983E-271 | TEC_inj | DHRS7 |
| 2.223E-275 | 0.46191162 | 0.342 | 0.124 | 5.251E-271 | TEC_inj | MSMO1 |
| 3.357E-275 | 0.25978435 | 0.244 | 0.069 | 7.93E-271 | TEC_inj | LAMC1 |
| 3.899E-275 | 0.36941759 | 0.343 | 0.126 | 9.21E-271 | TEC_inj | ETHE1 |
| 1.876E-274 | 0.51371358 | 0.432 | 0.175 | 4.431E-270 | TEC_inj | SFN |
| 5.85E-274 | 0.54420808 | 0.796 | 0.543 | 1.382E-269 | TEC_inj | SLC25A6 |
| 1.002E-273 | 0.44943506 | 0.589 | 0.297 | 2.366E-269 | TEC_inj | ERP29 |
| 1.021E-273 | 0.40014387 | 0.482 | 0.218 | 2.411E-269 | TEC_inj | TUFM |

| | | | | | | |
|---|---|---|---|---|---|---|
| 1.525E-273 | 0.44225261 | 0.523 | 0.251 | 3.603E-269 | TEC_inj | MINOS1 |
| 1.673E-273 | 0.4961897 | 0.644 | 0.363 | 3.952E-269 | TEC_inj | NEDD8 |
| 1.361E-272 | 0.54856103 | 0.821 | 0.627 | 3.215E-268 | TEC_inj | EDF1 |
| 1.346E-271 | 0.434769 | 0.623 | 0.325 | 3.179E-267 | TEC_inj | FKBP2 |
| 3.286E-271 | 0.47831785 | 0.985 | 0.894 | 7.763E-267 | TEC_inj | RPL24 |
| 1.126E-270 | 0.29949725 | 0.294 | 0.096 | 2.66E-266 | TEC_inj | MTURN |
| 1.188E-270 | 0.53617038 | 0.985 | 0.866 | 2.807E-266 | TEC_inj | RPL26 |
| 4.072E-270 | 0.45808882 | 0.581 | 0.303 | 9.619E-266 | TEC_inj | SRP9 |
| 6.494E-270 | 0.31293897 | 0.27 | 0.083 | 1.534E-265 | TEC_inj | SLC25A4 |
| 5.419E-269 | 0.30769273 | 0.129 | 0.022 | 1.28E-264 | TEC_inj | SOD3 |
| 7.374E-269 | -0.8415189 | 0.506 | 0.786 | 1.742E-264 | TEC_inj | BTG1 |
| 2.071E-268 | -0.7873142 | 0.34 | 0.625 | 4.892E-264 | TEC_inj | MCL1 |
| 1.759E-267 | 0.38485096 | 0.476 | 0.214 | 4.156E-263 | TEC_inj | BANF1 |
| 4.571E-267 | 0.46162202 | 0.633 | 0.351 | 1.08E-262 | TEC_inj | LAMTOR5 |
| 1.528E-266 | 0.32665086 | 0.356 | 0.134 | 3.608E-262 | TEC_inj | DDOST |
| 5.252E-266 | 0.53824282 | 0.741 | 0.481 | 1.241E-261 | TEC_inj | LAPTM4A |
| 1.375E-265 | 0.92937355 | 0.138 | 0.026 | 3.249E-261 | TEC_inj | SLC12A1 |
| 3.292E-264 | 0.44917992 | 0.531 | 0.261 | 7.776E-260 | TEC_inj | EIF6 |
| 2.285E-263 | 0.40442862 | 0.52 | 0.25 | 5.397E-259 | TEC_inj | NDUFB8 |
| 9.376E-263 | 0.25546112 | 0.251 | 0.075 | 2.215E-258 | TEC_inj | ACAT1 |
| 2.802E-262 | 0.55542434 | 0.982 | 0.892 | 6.618E-258 | TEC_inj | RPL27A |
| 9.291E-262 | 0.35559903 | 0.339 | 0.123 | 2.195E-257 | TEC_inj | BEX3 |
| 1.665E-261 | 0.71609932 | 0.774 | 0.648 | 3.934E-257 | TEC_inj | CALM2 |
| 3.219E-261 | 0.54766621 | 0.765 | 0.537 | 7.604E-257 | TEC_inj | NDUFS5 |
| 1.905E-260 | 0.30005922 | 0.339 | 0.124 | 4.5E-256 | TEC_inj | ARL3 |
| 2.07E-260 | 0.49790441 | 0.77 | 0.511 | 4.891E-256 | TEC_inj | COX6C |
| 6.046E-260 | 0.40657518 | 0.543 | 0.265 | 1.428E-255 | TEC_inj | NDUFC1 |
| 2.421E-259 | 0.37008259 | 0.451 | 0.197 | 5.718E-255 | TEC_inj | MLEC |
| 5.37E-259 | 0.25184313 | 0.122 | 0.02 | 1.269E-254 | TEC_inj | S100A1 |
| 1.684E-258 | 0.42444389 | 0.501 | 0.225 | 3.978E-254 | TEC_inj | SPTBN1 |
| 5.692E-258 | 0.35602323 | 0.422 | 0.18 | 1.345E-253 | TEC_inj | MDH1 |
| 6.173E-258 | 0.51553556 | 0.773 | 0.566 | 1.458E-253 | TEC_inj | CLTA |
| 9.507E-258 | 0.4120279 | 0.526 | 0.247 | 2.246E-253 | TEC_inj | FAM3C |
| 2.535E-257 | 0.27852641 | 0.27 | 0.087 | 5.988E-253 | TEC_inj | PLGRKT |
| 3.687E-257 | 0.29955445 | 0.246 | 0.073 | 8.71E-253 | TEC_inj | CA2 |
| 5.248E-257 | 0.45806455 | 0.488 | 0.232 | 1.24E-252 | TEC_inj | C4orf48 |
| 5.658E-256 | 0.36165374 | 0.403 | 0.169 | 1.337E-251 | TEC_inj | MFSD10 |
| 6.545E-256 | -0.4857103 | 0.746 | 0.971 | 1.546E-251 | TEC_inj | H3F3B |
| 9.383E-256 | 0.43514613 | 0.115 | 0.018 | 2.216E-251 | TEC_inj | LTF |
| 1.803E-255 | 0.53880654 | 0.909 | 0.773 | 4.259E-251 | TEC_inj | EEF1D |
| 5.791E-255 | 0.52517597 | 0.526 | 0.269 | 1.368E-250 | TEC_inj | NDUFB9 |
| 1.42E-254 | 0.4696296 | 0.542 | 0.277 | 3.355E-250 | TEC_inj | ROMO1 |
| 2.48E-254 | 0.32001581 | 0.3 | 0.105 | 5.859E-250 | TEC_inj | NBL1 |
| 3.867E-254 | 0.44212309 | 0.914 | 0.774 | 9.135E-250 | TEC_inj | BTF3 |
| 3.089E-252 | 0.35795766 | 0.433 | 0.189 | 7.298E-248 | TEC_inj | CNIH4 |
| 3.481E-252 | 0.25431659 | 0.185 | 0.045 | 8.223E-248 | TEC_inj | COL4A1 |
| 5.693E-252 | 0.64687017 | 0.747 | 0.536 | 1.345E-247 | TEC_inj | SSR4 |
| 1.923E-251 | 0.46650102 | 0.612 | 0.334 | 4.542E-247 | TEC_inj | UQCRQ |
| 3.787E-251 | 0.4088801 | 0.476 | 0.223 | 8.947E-247 | TEC_inj | OSTC |
| 1.61E-250 | 0.44007933 | 0.542 | 0.274 | 3.803E-246 | TEC_inj | TRAM1 |

| | | | | | | |
|---|---|---|---|---|---|---|
| 4.533E-250 | -0.7308575 | 0.022 | 0.289 | 1.071E-245 | TEC_inj | CD53 |
| 6.928E-250 | 0.2890679 | 0.304 | 0.106 | 1.637E-245 | TEC_inj | SNRPN |
| 8.658E-250 | 0.30219587 | 0.283 | 0.096 | 2.045E-245 | TEC_inj | ACAA2 |
| 9.273E-250 | 0.3256458 | 0.358 | 0.14 | 2.19E-245 | TEC_inj | TMEM205 |
| 4.828E-249 | 0.3867963 | 0.433 | 0.193 | 1.141E-244 | TEC_inj | GTF3C6 |
| 6.571E-249 | 0.77289588 | 0.837 | 0.68 | 1.552E-244 | TEC_inj | HSP90B1 |
| 9.111E-249 | 0.38939916 | 0.581 | 0.302 | 2.152E-244 | TEC_inj | GHITM |
| 7.037E-248 | 0.39449757 | 0.488 | 0.232 | 1.662E-243 | TEC_inj | FIS1 |
| 2.557E-247 | 0.44776224 | 0.538 | 0.275 | 6.041E-243 | TEC_inj | RABAC1 |
| 4.304E-247 | 0.49729034 | 0.983 | 0.875 | 1.017E-242 | TEC_inj | RPS13 |
| 6.622E-247 | 0.32050261 | 0.318 | 0.115 | 1.564E-242 | TEC_inj | SLC44A2 |
| 8.385E-247 | 0.51787428 | 0.671 | 0.413 | 1.981E-242 | TEC_inj | ATP5MF |
| 9.22E-247 | 0.33788202 | 0.385 | 0.158 | 2.178E-242 | TEC_inj | TMEM147 |
| 4.027E-246 | 0.33219692 | 0.45 | 0.194 | 9.512E-242 | TEC_inj | CNN3 |
| 6.399E-246 | 0.46520054 | 0.975 | 0.88 | 1.512E-241 | TEC_inj | RPS3 |
| 8.685E-246 | 0.33871411 | 0.386 | 0.16 | 2.052E-241 | TEC_inj | CYB5R3 |
| 8.895E-246 | 0.42270115 | 0.401 | 0.169 | 2.101E-241 | TEC_inj | VCL |
| 1.595E-245 | 0.31423874 | 0.29 | 0.101 | 3.767E-241 | TEC_inj | BOLA3 |
| 8.27E-245 | 0.57653177 | 0.949 | 0.948 | 1.954E-240 | TEC_inj | RPL10 |
| 2.348E-243 | 0.36274909 | 0.446 | 0.201 | 5.547E-239 | TEC_inj | TMEM208 |
| 5.718E-243 | 0.40973303 | 0.396 | 0.171 | 1.351E-238 | TEC_inj | PSMD2 |
| 2.096E-242 | 0.57494145 | 0.683 | 0.419 | 4.952E-238 | TEC_inj | PDIA3 |
| 6.869E-242 | -0.7844192 | 0.009 | 0.266 | 1.623E-237 | TEC_inj | CSTA |
| 1.108E-241 | 0.50337056 | 0.714 | 0.467 | 2.617E-237 | TEC_inj | DYNLT1 |
| 5.152E-241 | 0.43448354 | 0.602 | 0.331 | 1.217E-236 | TEC_inj | SPCS1 |
| 1.68E-240 | -0.9508836 | 0.048 | 0.312 | 3.969E-236 | TEC_inj | LCP1 |
| 2.977E-240 | 0.47851241 | 0.982 | 0.929 | 7.032E-236 | TEC_inj | RPS14 |
| 3.587E-240 | 0.34960671 | 0.418 | 0.182 | 8.473E-236 | TEC_inj | C19orf70 |
| 2.934E-239 | 0.4770039 | 0.556 | 0.299 | 6.931E-235 | TEC_inj | FKBP1A |
| 4.478E-239 | 0.48936963 | 0.491 | 0.245 | 1.058E-234 | TEC_inj | PLP2 |
| 2.749E-238 | 0.26312357 | 0.228 | 0.068 | 6.495E-234 | TEC_inj | PHGDH |
| 2.933E-238 | 0.35996521 | 0.49 | 0.235 | 6.928E-234 | TEC_inj | EIF3I |
| 3.72E-238 | 0.40946692 | 0.233 | 0.071 | 8.787E-234 | TEC_inj | CSRP2 |
| 3.879E-237 | 0.71092479 | 0.957 | 0.971 | 9.162E-233 | TEC_inj | TMSB10 |
| 4.154E-237 | 0.58873004 | 0.991 | 0.887 | 9.812E-233 | TEC_inj | RPL37A |
| 2.796E-236 | 0.44184051 | 0.545 | 0.292 | 6.606E-232 | TEC_inj | CAPNS1 |
| 3.155E-236 | 0.50423457 | 0.941 | 0.779 | 7.454E-232 | TEC_inj | RPS21 |
| 3.512E-236 | 0.3824238 | 0.19 | 0.051 | 8.295E-232 | TEC_inj | NEURL3 |
| 4.08E-236 | 0.56080342 | 0.311 | 0.113 | 9.638E-232 | TEC_inj | AKAP12 |
| 4.325E-236 | 0.30954492 | 0.376 | 0.15 | 1.022E-231 | TEC_inj | TINAGL1 |
| 8.351E-236 | 0.31432585 | 0.352 | 0.141 | 1.973E-231 | TEC_inj | FUNDC2 |
| 1.532E-235 | 0.39603986 | 0.39 | 0.168 | 3.619E-231 | TEC_inj | DYNLT3 |
| 6.003E-235 | 0.5122764 | 0.728 | 0.466 | 1.418E-230 | TEC_inj | CANX |
| 1.766E-234 | 0.37697752 | 0.434 | 0.195 | 4.171E-230 | TEC_inj | LMAN1 |
| 6.261E-234 | 0.28572825 | 0.325 | 0.123 | 1.479E-229 | TEC_inj | OLA1 |
| 2.827E-233 | 0.43806173 | 0.527 | 0.267 | 6.679E-229 | TEC_inj | APRT |
| 3.415E-233 | 0.28946954 | 0.365 | 0.143 | 8.066E-229 | TEC_inj | FARP1 |
| 6.225E-233 | -0.6077438 | 0.333 | 0.617 | 1.47E-228 | TEC_inj | DDX17 |
| 1.198E-231 | 0.30780899 | 0.317 | 0.118 | 2.83E-227 | TEC_inj | SEPT11 |
| 1.464E-231 | 0.32615225 | 0.35 | 0.142 | 3.457E-227 | TEC_inj | PRMT1 |

| | | | | | | |
|---|---|---|---|---|---|---|
| 7.938E-231 | -1.2442832 | 0.041 | 0.295 | 1.875E-226 | TEC_inj | AIF1 |
| 9.19E-231 | -0.678294 | 0.48 | 0.743 | 2.171E-226 | TEC_inj | JUND |
| 1.894E-229 | -0.6451379 | 0.722 | 0.882 | 4.474E-225 | TEC_inj | PABPC1 |
| 2.682E-229 | 0.27869066 | 0.256 | 0.084 | 6.335E-225 | TEC_inj | ITGA6 |
| 9.303E-229 | -0.6426412 | 0.017 | 0.266 | 2.198E-224 | TEC_inj | GMFG |
| 2.881E-228 | 0.34648384 | 0.395 | 0.172 | 6.805E-224 | TEC_inj | DAP |
| 2.428E-227 | 0.32711322 | 0.361 | 0.15 | 5.736E-223 | TEC_inj | TBCB |
| 3.131E-227 | 0.36755494 | 0.403 | 0.177 | 7.397E-223 | TEC_inj | C1QBP |
| 4.698E-227 | 0.51100777 | 0.991 | 0.918 | 1.11E-222 | TEC_inj | RPS24 |
| 4.72E-227 | -0.9173897 | 0.288 | 0.546 | 1.115E-222 | TEC_inj | CEBPB |
| 1.788E-226 | 0.41209105 | 0.571 | 0.309 | 4.224E-222 | TEC_inj | FAM96B |
| 1.833E-226 | 0.43680296 | 0.987 | 0.946 | 4.329E-222 | TEC_inj | RPL18 |
| 2.077E-226 | 0.30920053 | 0.372 | 0.155 | 4.906E-222 | TEC_inj | ATRAID |
| 3.118E-226 | 0.388584 | 0.514 | 0.264 | 7.365E-222 | TEC_inj | SELENOF |
| 4.454E-226 | 0.44939555 | 0.859 | 0.675 | 1.052E-221 | TEC_inj | EEF2 |
| 4.927E-225 | 0.42040601 | 0.476 | 0.234 | 1.164E-220 | TEC_inj | RPN2 |
| 5.386E-225 | 0.52766931 | 0.986 | 0.914 | 1.272E-220 | TEC_inj | RPS15 |
| 2.056E-224 | 0.26240727 | 0.224 | 0.069 | 4.857E-220 | TEC_inj | TMEM106C |
| 5.028E-224 | 0.61805748 | 0.816 | 0.684 | 1.188E-219 | TEC_inj | GUK1 |
| 5.418E-223 | 0.38792398 | 0.508 | 0.26 | 1.28E-218 | TEC_inj | UFC1 |
| 3.811E-222 | 0.61893718 | 0.926 | 0.805 | 9.002E-218 | TEC_inj | YBX1 |
| 8.578E-222 | 0.44826936 | 0.87 | 0.722 | 2.026E-217 | TEC_inj | GNAS |
| 1.026E-221 | 0.51973916 | 0.85 | 0.671 | 2.425E-217 | TEC_inj | NPM1 |
| 1.179E-221 | 0.32450961 | 0.407 | 0.182 | 2.784E-217 | TEC_inj | NAA20 |
| 2.061E-218 | 0.52312998 | 0.968 | 0.913 | 4.868E-214 | TEC_inj | RPS20 |
| 4.026E-218 | -0.9204834 | 0.292 | 0.551 | 9.511E-214 | TEC_inj | CTSS |
| 1.019E-217 | 0.78259789 | 0.541 | 0.299 | 2.406E-213 | TEC_inj | GADD45A |
| 1.474E-217 | 0.57071128 | 0.929 | 0.87 | 3.482E-213 | TEC_inj | MYL12A |
| 4.007E-217 | 0.56710999 | 0.981 | 0.897 | 9.466E-213 | TEC_inj | RPLP2 |
| 4.099E-217 | 0.45501678 | 0.426 | 0.194 | 9.683E-213 | TEC_inj | LAMB3 |
| 4.275E-217 | -1.137239 | 0.015 | 0.252 | 1.01E-212 | TEC_inj | SLC2A3 |
| 9.612E-217 | -0.7852554 | 0.011 | 0.246 | 2.271E-212 | TEC_inj | ZEB2 |
| 6.99E-216 | 0.27719542 | 0.291 | 0.108 | 1.651E-211 | TEC_inj | PDHA1 |
| 1.088E-215 | -0.5479119 | 0.009 | 0.243 | 2.571E-211 | TEC_inj | KYNU |
| 1.9E-215 | 0.30657611 | 0.419 | 0.186 | 4.489E-211 | TEC_inj | CTTN |
| 1.143E-213 | 0.37454003 | 0.496 | 0.253 | 2.7E-209 | TEC_inj | ATP5F1C |
| 4.841E-213 | 0.32014531 | 0.355 | 0.151 | 1.144E-208 | TEC_inj | EMC4 |
| 1.693E-212 | -0.6799905 | 0.383 | 0.64 | 4E-208 | TEC_inj | PCBP1 |
| 3.732E-212 | 0.62071208 | 0.968 | 0.963 | 8.817E-208 | TEC_inj | MYL6 |
| 1.152E-211 | 0.3106618 | 0.104 | 0.018 | 2.72E-207 | TEC_inj | IL1RL1 |
| 8.909E-211 | 0.30080766 | 0.249 | 0.085 | 2.105E-206 | TEC_inj | SH3BP4 |
| 1.846E-210 | -0.6613217 | 0.092 | 0.345 | 4.362E-206 | TEC_inj | ARHGDIB |
| 2.478E-210 | 0.31459289 | 0.375 | 0.164 | 5.853E-206 | TEC_inj | ADI1 |
| 2.579E-210 | 0.39724599 | 0.578 | 0.327 | 6.092E-206 | TEC_inj | PSMD8 |
| 2.796E-210 | 0.34781934 | 0.481 | 0.241 | 6.604E-206 | TEC_inj | SMDT1 |
| 3.022E-210 | -0.646711 | 0.425 | 0.658 | 7.139E-206 | TEC_inj | ARPC5 |
| 3.829E-210 | -0.8541492 | 0.385 | 0.646 | 9.046E-206 | TEC_inj | S100A4 |
| 1.543E-209 | -0.5706565 | 0.245 | 0.518 | 3.645E-205 | TEC_inj | DDX3X |
| 1.548E-209 | 0.40106093 | 0.601 | 0.333 | 3.656E-205 | TEC_inj | CYCS |
| 1.562E-208 | -0.6300984 | 0.396 | 0.663 | 3.69E-204 | TEC_inj | IER3 |

| | | | | | | |
|---|---|---|---|---|---|---|
| 6.879E-208 | 0.45980211 | 0.929 | 0.774 | 1.625E-203 | TEC_inj | RPL31 |
| 7.277E-208 | 0.51515848 | 0.796 | 0.614 | 1.719E-203 | TEC_inj | OST4 |
| 1.618E-207 | -0.6068067 | 0.069 | 0.317 | 3.822E-203 | TEC_inj | REL |
| 7.438E-207 | 0.3093093 | 0.226 | 0.072 | 1.757E-202 | TEC_inj | CALD1 |
| 7.523E-207 | -1.164527 | 0.008 | 0.233 | 1.777E-202 | TEC_inj | S100A12 |
| 1.937E-206 | -0.6546743 | 0.336 | 0.593 | 4.576E-202 | TEC_inj | VMP1 |
| 2.576E-206 | -0.8178889 | 0.231 | 0.491 | 6.085E-202 | TEC_inj | DUSP1 |
| 2.588E-206 | 0.47844021 | 0.592 | 0.343 | 6.114E-202 | TEC_inj | SLC25A5 |
| 6.774E-206 | 0.34685605 | 0.394 | 0.18 | 1.6E-201 | TEC_inj | ATP5MC1 |
| 1.532E-205 | 0.38031569 | 0.513 | 0.269 | 3.619E-201 | TEC_inj | NDUFB7 |
| 1.564E-205 | 0.27398446 | 0.292 | 0.112 | 3.696E-201 | TEC_inj | AP1S1 |
| 1.96E-205 | 0.35404367 | 0.368 | 0.159 | 4.629E-201 | TEC_inj | TMEM54 |
| 3.634E-205 | 0.35155285 | 0.411 | 0.195 | 8.583E-201 | TEC_inj | ARPC1A |
| 1.123E-204 | 0.35963037 | 0.55 | 0.291 | 2.653E-200 | TEC_inj | SBDS |
| 2.138E-204 | 0.26284596 | 0.187 | 0.053 | 5.052E-200 | TEC_inj | PIGR |
| 4.298E-204 | 0.41789648 | 0.381 | 0.172 | 1.015E-199 | TEC_inj | PDIA4 |
| 1.147E-203 | 0.48315484 | 0.456 | 0.22 | 2.71E-199 | TEC_inj | C12orf57 |
| 2.036E-203 | 0.38234812 | 0.501 | 0.239 | 4.808E-199 | TEC_inj | MUC1 |
| 4.795E-203 | -0.9518105 | 0.032 | 0.263 | 1.133E-198 | TEC_inj | SLC11A1 |
| 6.172E-203 | 0.31999751 | 0.427 | 0.203 | 1.458E-198 | TEC_inj | TMEM14B |
| 8.119E-203 | 0.3573882 | 0.613 | 0.346 | 1.918E-198 | TEC_inj | SEC11A |
| 3.616E-202 | -2.1629799 | 0.242 | 0.51 | 8.543E-198 | TEC_inj | CXCL8 |
| 5.583E-202 | 0.63679434 | 0.238 | 0.08 | 1.319E-197 | TEC_inj | SAA1 |
| 8.026E-202 | 0.45038435 | 0.139 | 0.033 | 1.896E-197 | TEC_inj | KRTAP2-3 |
| 1.193E-201 | -0.5055743 | 0.577 | 0.831 | 2.818E-197 | TEC_inj | DDX5 |
| 1.649E-201 | -0.8975522 | 0.222 | 0.469 | 3.896E-197 | TEC_inj | MARCKS |
| 2.09E-201 | 0.38262931 | 0.58 | 0.322 | 4.938E-197 | TEC_inj | ATP5IF1 |
| 2.363E-201 | 0.25032767 | 0.25 | 0.088 | 5.583E-197 | TEC_inj | GGCT |
| 3.775E-201 | 0.25667179 | 0.281 | 0.106 | 8.917E-197 | TEC_inj | HRAS |
| 9.37E-201 | 0.33308724 | 0.449 | 0.219 | 2.214E-196 | TEC_inj | SNRPE |
| 1.526E-200 | 0.39294175 | 0.537 | 0.287 | 3.605E-196 | TEC_inj | PFDN2 |
| 2.983E-200 | 0.31130972 | 0.398 | 0.182 | 7.046E-196 | TEC_inj | TIMM13 |
| 2.075E-199 | 0.32226466 | 0.375 | 0.169 | 4.902E-195 | TEC_inj | NDUFAF3 |
| 2.471E-199 | 0.38646415 | 0.553 | 0.312 | 5.836E-195 | TEC_inj | NDUFC2 |
| 2.696E-199 | 0.34886649 | 0.396 | 0.185 | 6.368E-195 | TEC_inj | NDUFS8 |
| 4.091E-199 | 0.29098592 | 0.381 | 0.17 | 9.664E-195 | TEC_inj | AP2B1 |
| 1.71E-197 | 0.30713519 | 0.254 | 0.091 | 4.04E-193 | TEC_inj | LINC00958 |
| 4.765E-197 | 0.25904446 | 0.307 | 0.124 | 1.126E-192 | TEC_inj | ASNA1 |
| 1.5E-196 | 0.42458682 | 0.632 | 0.377 | 3.543E-192 | TEC_inj | NDUFA1 |
| 3.824E-196 | 0.34200134 | 0.445 | 0.218 | 9.035E-192 | TEC_inj | ATP5F1A |
| 6.646E-196 | 0.28445114 | 0.325 | 0.136 | 1.57E-191 | TEC_inj | BCL7C |
| 9.391E-196 | 0.40530667 | 0.982 | 0.891 | 2.218E-191 | TEC_inj | RPS25 |
| 1.234E-195 | -0.7665412 | 0.01 | 0.227 | 2.915E-191 | TEC_inj | TNFRSF1B |
| 1.683E-195 | 0.4103228 | 0.559 | 0.314 | 3.976E-191 | TEC_inj | SSR3 |
| 1.91E-194 | 0.26203453 | 0.327 | 0.136 | 4.512E-190 | TEC_inj | BUD23 |
| 3.228E-194 | -0.6987826 | 0.029 | 0.252 | 7.625E-190 | TEC_inj | HCST |
| 4.802E-193 | 0.317833 | 0.104 | 0.02 | 1.134E-188 | TEC_inj | CLDN14 |
| 1.176E-191 | 0.39334557 | 0.862 | 0.701 | 2.777E-187 | TEC_inj | ATP5MG |
| 2.169E-191 | 0.42283736 | 0.269 | 0.101 | 5.123E-187 | TEC_inj | RRAD |
| 6.312E-191 | 0.29963876 | 0.33 | 0.142 | 1.491E-186 | TEC_inj | ITGB1BP1 |

| | | | | | | |
|---|---|---|---|---|---|---|
| 1.16E-190 | 0.33437347 | 0.494 | 0.257 | 2.74E-186 | TEC_inj | CCT5 |
| 4.243E-190 | 0.4171315 | 0.696 | 0.472 | 1.002E-185 | TEC_inj | HSBP1 |
| 5.128E-190 | 0.27716864 | 0.295 | 0.119 | 1.211E-185 | TEC_inj | RRAS |
| 1.294E-189 | 0.33179362 | 0.532 | 0.282 | 3.056E-185 | TEC_inj | PSME2 |
| 2.215E-189 | 0.30320949 | 0.441 | 0.216 | 5.233E-185 | TEC_inj | SYPL1 |
| 3.864E-189 | 0.27556116 | 0.314 | 0.127 | 9.128E-185 | TEC_inj | OCLN |
| 6.245E-189 | -0.5765577 | 0.018 | 0.233 | 1.475E-184 | TEC_inj | SPI1 |
| 7.043E-189 | 0.34978952 | 0.478 | 0.248 | 1.664E-184 | TEC_inj | ATOX1 |
| 9.5E-189 | 0.36078845 | 0.518 | 0.282 | 2.244E-184 | TEC_inj | ATP5PO |
| 1.205E-188 | 0.36699097 | 0.59 | 0.336 | 2.847E-184 | TEC_inj | TRMT112 |
| 6.056E-188 | 0.39482715 | 0.597 | 0.35 | 1.431E-183 | TEC_inj | ATP5MC3 |
| 7.195E-188 | 0.29106412 | 0.405 | 0.192 | 1.7E-183 | TEC_inj | ACP1 |
| 9.83E-188 | 0.48682481 | 0.231 | 0.081 | 2.322E-183 | TEC_inj | CTGF |
| 3.292E-187 | -0.4907392 | 0.01 | 0.22 | 7.777E-183 | TEC_inj | LSP1 |
| 2.886E-184 | 0.44621199 | 0.821 | 0.642 | 6.817E-180 | TEC_inj | ZFAS1 |
| 6.615E-184 | 0.28663176 | 0.364 | 0.166 | 1.563E-179 | TEC_inj | CCT7 |
| 3.667E-183 | 0.36022815 | 0.579 | 0.329 | 8.663E-179 | TEC_inj | VDAC1 |
| 1.62E-182 | 0.29803429 | 0.471 | 0.24 | 3.828E-178 | TEC_inj | CCT3 |
| 3.189E-182 | -0.4942397 | 0.391 | 0.642 | 7.534E-178 | TEC_inj | PRRC2C |
| 2.39E-181 | 0.26907799 | 0.263 | 0.1 | 5.646E-177 | TEC_inj | UGDH |
| 6.809E-180 | 0.47601096 | 0.768 | 0.603 | 1.608E-175 | TEC_inj | COX8A |
| 4.24E-179 | 0.30164186 | 0.565 | 0.307 | 1.002E-174 | TEC_inj | LMNA |
| 6.378E-179 | 0.32527923 | 0.453 | 0.235 | 1.507E-174 | TEC_inj | TMED2 |
| 1.099E-178 | 0.32737487 | 0.134 | 0.033 | 2.596E-174 | TEC_inj | PI3 |
| 1.48E-178 | 0.37242341 | 0.781 | 0.585 | 3.495E-174 | TEC_inj | YWHAE |
| 1.833E-178 | 0.30059917 | 0.195 | 0.063 | 4.331E-174 | TEC_inj | KITLG |
| 2.845E-178 | 0.52864971 | 0.803 | 0.662 | 6.72E-174 | TEC_inj | CALR |
| 1.412E-177 | -0.5300227 | 0.458 | 0.679 | 3.335E-173 | TEC_inj | HNRNPU |
| 1.513E-177 | 0.25649025 | 0.219 | 0.076 | 3.574E-173 | TEC_inj | ASNS |
| 1.953E-177 | -0.6219978 | 0.028 | 0.236 | 4.614E-173 | TEC_inj | ITGB2 |
| 2.338E-177 | 0.31385817 | 0.596 | 0.338 | 5.522E-173 | TEC_inj | TOMM20 |
| 3.448E-177 | -0.5380528 | 0.432 | 0.641 | 8.145E-173 | TEC_inj | ACTR2 |
| 5.486E-177 | 0.30797007 | 0.336 | 0.151 | 1.296E-172 | TEC_inj | NDUFAF8 |
| 5.822E-177 | -1.5041634 | 0.011 | 0.212 | 1.375E-172 | TEC_inj | HOPX |
| 5.9E-177 | -0.5836107 | 0.014 | 0.216 | 1.394E-172 | TEC_inj | ITGAX |
| 1.477E-176 | 0.38586264 | 0.82 | 0.666 | 3.49E-172 | TEC_inj | COX7A2 |
| 2.203E-176 | 0.32861275 | 0.287 | 0.118 | 5.205E-172 | TEC_inj | TMEM176A |
| 2.401E-176 | 0.54482325 | 0.94 | 0.902 | 5.671E-172 | TEC_inj | CFL1 |
| 2.78E-176 | 0.34501533 | 0.415 | 0.199 | 6.567E-172 | TEC_inj | DSP |
| 4.931E-176 | -0.4457397 | 0.021 | 0.225 | 1.165E-171 | TEC_inj | ARRB2 |
| 1.399E-175 | 0.41323998 | 0.6 | 0.366 | 3.306E-171 | TEC_inj | C9orf16 |
| 3.322E-175 | 0.2985332 | 0.447 | 0.229 | 7.847E-171 | TEC_inj | CCT8 |
| 1.234E-174 | 0.48135099 | 0.685 | 0.475 | 2.915E-170 | TEC_inj | TAGLN2 |
| 1.455E-174 | 0.45375929 | 0.513 | 0.292 | 3.437E-170 | TEC_inj | RAB13 |
| 2.853E-174 | -0.5230702 | 0.076 | 0.298 | 6.739E-170 | TEC_inj | LYN |
| 3.752E-174 | 0.27281644 | 0.296 | 0.122 | 8.864E-170 | TEC_inj | AGRN |
| 5.734E-174 | 0.32552755 | 0.492 | 0.254 | 1.355E-169 | TEC_inj | JUP |
| 3.811E-173 | 0.40032345 | 0.917 | 0.825 | 9.004E-169 | TEC_inj | UBB |
| 8.713E-173 | 0.49630576 | 0.471 | 0.264 | 2.058E-168 | TEC_inj | RBM3 |
| 9.422E-173 | 0.4493011 | 0.679 | 0.432 | 2.226E-168 | TEC_inj | CLDN7 |

| | | | | | | |
|---|---|---|---|---|---|---|
| 2.954E-172 | -0.5837011 | 0.022 | 0.223 | 6.978E-168 | TEC_inj | GK |
| 9.206E-172 | -1.7104456 | 0.036 | 0.239 | 2.175E-167 | TEC_inj | VCAN |
| 2.563E-171 | 0.40396454 | 0.992 | 0.935 | 6.055E-167 | TEC_inj | RPS28 |
| 3.79E-171 | 0.31390082 | 0.461 | 0.241 | 8.954E-167 | TEC_inj | TMCO1 |
| 6.14E-171 | 0.39647166 | 0.94 | 0.809 | 1.45E-166 | TEC_inj | RPL27 |
| 7.408E-171 | 0.45534703 | 0.406 | 0.201 | 1.75E-166 | TEC_inj | ZBTB20 |
| 1.563E-170 | -1.077183 | 0.032 | 0.236 | 3.692E-166 | TEC_inj | ALOX5AP |
| 8.341E-170 | 0.27346483 | 0.203 | 0.069 | 1.97E-165 | TEC_inj | RETREG1 |
| 1.12E-169 | -0.5335934 | 0.053 | 0.264 | 2.647E-165 | TEC_inj | GPCPD1 |
| 1.965E-169 | 0.35379576 | 0.539 | 0.311 | 4.641E-165 | TEC_inj | COPS9 |
| 1.112E-168 | 0.31015339 | 0.44 | 0.225 | 2.627E-164 | TEC_inj | COMT |
| 1.375E-168 | -0.4009796 | 0.01 | 0.203 | 3.247E-164 | TEC_inj | HCLS1 |
| 1.465E-168 | 0.45072272 | 0.976 | 0.894 | 3.46E-164 | TEC_inj | RPS11 |
| 1.658E-168 | -0.3814142 | 0.007 | 0.199 | 3.917E-164 | TEC_inj | GNA15 |
| 6.925E-168 | -0.9779851 | 0.035 | 0.236 | 1.636E-163 | TEC_inj | CD14 |
| 9.847E-168 | 0.44779104 | 0.237 | 0.09 | 2.326E-163 | TEC_inj | RCAN1 |
| 1.325E-167 | 0.28778201 | 0.395 | 0.195 | 3.13E-163 | TEC_inj | STRAP |
| 2.018E-167 | -0.553499 | 0.169 | 0.392 | 4.768E-163 | TEC_inj | METRNL |
| 3.404E-167 | 0.29897622 | 0.45 | 0.234 | 8.042E-163 | TEC_inj | NDUFAB1 |
| 7.001E-167 | 0.26280836 | 0.243 | 0.093 | 1.654E-162 | TEC_inj | RHOF |
| 8.1E-167 | 0.27529935 | 0.366 | 0.175 | 1.913E-162 | TEC_inj | SDHC |
| 8.859E-167 | -0.7385769 | 0.021 | 0.216 | 2.093E-162 | TEC_inj | CYBB |
| 1.952E-166 | 0.28994672 | 0.377 | 0.183 | 4.612E-162 | TEC_inj | TMEM230 |
| 2.879E-166 | -0.6473433 | 0.026 | 0.224 | 6.801E-162 | TEC_inj | LST1 |
| 5.545E-166 | -0.5632178 | 0.013 | 0.204 | 1.31E-161 | TEC_inj | FCGR2A |
| 7.186E-166 | 0.48262906 | 0.318 | 0.14 | 1.698E-161 | TEC_inj | CYR61 |
| 9.645E-166 | 0.31512821 | 0.454 | 0.242 | 2.279E-161 | TEC_inj | PGLS |
| 8.61E-165 | -1.7901444 | 0.197 | 0.414 | 2.034E-160 | TEC_inj | HLA-DRA |
| 1.468E-164 | 0.28596553 | 0.399 | 0.2 | 3.468E-160 | TEC_inj | NDUFB5 |
| 2.33E-164 | -0.5802125 | 0.013 | 0.203 | 5.504E-160 | TEC_inj | FYB1 |
| 2.94E-164 | -0.4801072 | 0.012 | 0.202 | 6.945E-160 | TEC_inj | PTPRE |
| 3.589E-164 | 0.35209897 | 0.477 | 0.267 | 8.479E-160 | TEC_inj | PSMB6 |
| 4.051E-164 | -0.595905 | 0.031 | 0.229 | 9.569E-160 | TEC_inj | BASP1 |
| 1.18E-163 | 0.30064619 | 0.48 | 0.258 | 2.788E-159 | TEC_inj | MRPL51 |
| 4.679E-163 | -0.4842206 | 0.4 | 0.632 | 1.105E-158 | TEC_inj | SRRM2 |
| 2.274E-162 | 0.27709466 | 0.339 | 0.157 | 5.371E-158 | TEC_inj | MCFD2 |
| 3.669E-161 | -1.2691277 | 0.013 | 0.199 | 8.668E-157 | TEC_inj | DUOX2 |
| 3.991E-160 | -0.6497171 | 0.023 | 0.214 | 9.429E-156 | TEC_inj | DUSP4 |
| 5.004E-160 | -0.945047 | 0.439 | 0.676 | 1.182E-155 | TEC_inj | HSPA1A |
| 1.103E-159 | 0.30145169 | 0.524 | 0.296 | 2.605E-155 | TEC_inj | EIF3F |
| 1.435E-159 | 0.34906504 | 0.566 | 0.339 | 3.39E-155 | TEC_inj | COX5A |
| 2.04E-159 | 0.28871214 | 0.982 | 0.885 | 4.82E-155 | TEC_inj | RPS15A |
| 4.066E-159 | 0.25113705 | 0.322 | 0.145 | 9.606E-155 | TEC_inj | GLRX5 |
| 4.872E-159 | 0.45986876 | 0.701 | 0.519 | 1.151E-154 | TEC_inj | P4HB |
| 6.039E-159 | 0.2865356 | 0.411 | 0.21 | 1.427E-154 | TEC_inj | ARPP19 |
| 1.101E-158 | 0.32730753 | 0.546 | 0.318 | 2.601E-154 | TEC_inj | CHMP5 |
| 1.745E-158 | 0.27045298 | 0.362 | 0.176 | 4.122E-154 | TEC_inj | EMC7 |
| 3.111E-158 | 0.29571115 | 0.279 | 0.118 | 7.349E-154 | TEC_inj | PPFIBP1 |
| 4.952E-158 | -0.9811152 | 0.193 | 0.406 | 1.17E-153 | TEC_inj | FOS |
| 9.348E-158 | 0.32854777 | 0.639 | 0.394 | 2.208E-153 | TEC_inj | EIF3K |

| | | | | | | |
|---|---|---|---|---|---|---|
| 9.79E-158 | 0.26200233 | 0.328 | 0.152 | 2.313E-153 | TEC_inj | MCRIP1 |
| 1.538E-157 | -0.6925513 | 0.006 | 0.186 | 3.633E-153 | TEC_inj | STEAP4 |
| 1.832E-157 | -0.5522342 | 0.013 | 0.196 | 4.327E-153 | TEC_inj | SAMSN1 |
| 1.165E-156 | 0.32892819 | 0.274 | 0.116 | 2.752E-152 | TEC_inj | BEX2 |
| 1.172E-156 | 0.25571664 | 0.337 | 0.157 | 2.769E-152 | TEC_inj | GLRX3 |
| 1.956E-156 | 0.26761582 | 0.37 | 0.179 | 4.621E-152 | TEC_inj | CCT2 |
| 3.41E-156 | 0.2551695 | 0.267 | 0.11 | 8.056E-152 | TEC_inj | RAB3IP |
| 7.821E-156 | 0.30043119 | 0.411 | 0.213 | 1.847E-151 | TEC_inj | HDGF |
| 8.689E-156 | 0.39387895 | 0.786 | 0.594 | 2.053E-151 | TEC_inj | HNRNPA1 |
| 1.018E-155 | -0.7925067 | 0.009 | 0.189 | 2.404E-151 | TEC_inj | PLEK |
| 1.118E-155 | -1.8485755 | 0.158 | 0.36 | 2.64E-151 | TEC_inj | HPGD |
| 1.119E-155 | -0.612269 | 0.923 | 0.96 | 2.644E-151 | TEC_inj | H3F3A |
| 2.323E-155 | 0.31614412 | 0.429 | 0.229 | 5.488E-151 | TEC_inj | UQCRFS1 |
| 3.84E-154 | -1.0430193 | 0.057 | 0.251 | 9.072E-150 | TEC_inj | BCL2A1 |
| 6.836E-154 | 0.27815653 | 0.408 | 0.21 | 1.615E-149 | TEC_inj | PSMB2 |
| 9.45E-154 | -0.7329443 | 0.014 | 0.195 | 2.232E-149 | TEC_inj | OLR1 |
| 2.822E-153 | -0.3756077 | 0.013 | 0.194 | 6.666E-149 | TEC_inj | FMNL1 |
| 6.711E-153 | -0.603105 | 0.016 | 0.197 | 1.585E-148 | TEC_inj | IGSF6 |
| 1.268E-152 | 0.53189029 | 0.17 | 0.056 | 2.995E-148 | TEC_inj | RSAD2 |
| 1.981E-152 | -0.7386611 | 0.028 | 0.213 | 4.68E-148 | TEC_inj | CD52 |
| 2.012E-152 | -0.6311809 | 0.018 | 0.199 | 4.754E-148 | TEC_inj | C5AR1 |
| 4.24E-152 | 0.27153161 | 0.336 | 0.158 | 1.002E-147 | TEC_inj | MPC2 |
| 1.272E-151 | 0.31119373 | 0.369 | 0.183 | 3.005E-147 | TEC_inj | ITGAV |
| 3.211E-151 | 0.27544108 | 0.369 | 0.182 | 7.584E-147 | TEC_inj | TMED4 |
| 3.8E-151 | -0.473254 | 0.007 | 0.182 | 8.977E-147 | TEC_inj | SLA |
| 5.914E-151 | 0.3032931 | 0.499 | 0.285 | 1.397E-146 | TEC_inj | ANAPC11 |
| 1.683E-150 | 0.32512216 | 0.188 | 0.065 | 3.976E-146 | TEC_inj | SAA2 |
| 2.279E-150 | 0.41956942 | 0.976 | 0.912 | 5.384E-146 | TEC_inj | UBA52 |
| 3.212E-150 | 0.27839441 | 0.396 | 0.202 | 7.587E-146 | TEC_inj | NDUFA5 |
| 3.658E-150 | 0.35249463 | 0.705 | 0.49 | 8.641E-146 | TEC_inj | DBI |
| 6.929E-150 | -0.4177792 | 0.015 | 0.193 | 1.637E-145 | TEC_inj | CD37 |
| 8.285E-150 | 0.25603248 | 0.235 | 0.094 | 1.957E-145 | TEC_inj | GOT1 |
| 9.42E-150 | 0.27923253 | 0.347 | 0.169 | 2.225E-145 | TEC_inj | CALU |
| 1.819E-149 | -1.3296088 | 0.007 | 0.18 | 4.296E-145 | TEC_inj | DUOXA2 |
| 2.336E-149 | -0.8371962 | 0.302 | 0.521 | 5.519E-145 | TEC_inj | DNAJB1 |
| 4.167E-149 | 0.31080605 | 0.478 | 0.273 | 9.843E-145 | TEC_inj | NDUFB3 |
| 4.575E-149 | 0.40170285 | 0.983 | 0.897 | 1.081E-144 | TEC_inj | RPL37 |
| 5.26E-149 | 0.31888732 | 0.342 | 0.164 | 1.242E-144 | TEC_inj | TMEM176B |
| 9.63E-149 | 0.2773489 | 0.33 | 0.155 | 2.275E-144 | TEC_inj | EPRS |
| 9.97E-149 | 0.29717994 | 0.139 | 0.04 | 2.355E-144 | TEC_inj | PEG10 |
| 2.488E-148 | -0.5640762 | 0.028 | 0.21 | 5.878E-144 | TEC_inj | SAMD9 |
| 3.455E-148 | 0.36163275 | 0.589 | 0.361 | 8.162E-144 | TEC_inj | PRELID1 |
| 2.133E-147 | 0.38732456 | 0.562 | 0.359 | 5.038E-143 | TEC_inj | BRK1 |
| 4.108E-147 | -0.4158973 | 0.023 | 0.202 | 9.703E-143 | TEC_inj | RAC2 |
| 4.348E-147 | 0.29076928 | 0.476 | 0.264 | 1.027E-142 | TEC_inj | LRPAP1 |
| 4.566E-147 | -0.4295387 | 0.013 | 0.187 | 1.079E-142 | TEC_inj | CORO1A |
| 1.117E-146 | -0.3784374 | 0.032 | 0.216 | 2.639E-142 | TEC_inj | IFI16 |
| 2.002E-146 | -0.4497784 | 0.007 | 0.178 | 4.73E-142 | TEC_inj | LCP2 |
| 5.781E-146 | -1.1885099 | 0.006 | 0.174 | 1.366E-141 | TEC_inj | MUC21 |
| 8.799E-146 | -0.4751363 | 0.106 | 0.308 | 2.079E-141 | TEC_inj | SKIL |

| | | | | | | |
|---|---|---|---|---|---|---|
| 9.167E-146 | 0.46541614 | 0.958 | 0.891 | 2.166E-141 | TEC_inj | HLA-A |
| 2.03E-145 | -0.5457945 | 0.012 | 0.184 | 4.796E-141 | TEC_inj | PRDM1 |
| 5.035E-145 | 0.36512389 | 0.646 | 0.435 | 1.189E-140 | TEC_inj | SEC61G |
| 1.333E-144 | -2.2778721 | 0.017 | 0.189 | 3.149E-140 | TEC_inj | IL1B |
| 2.064E-144 | 0.36379346 | 0.29 | 0.131 | 4.875E-140 | TEC_inj | CLDN1 |
| 2.677E-144 | -0.416127 | 0.012 | 0.183 | 6.325E-140 | TEC_inj | CD48 |
| 4.514E-144 | 0.26099642 | 0.298 | 0.137 | 1.066E-139 | TEC_inj | CENPX |
| 6.218E-144 | 0.31623224 | 0.558 | 0.34 | 1.469E-139 | TEC_inj | ATP5PD |
| 8.795E-144 | 0.30050056 | 0.476 | 0.271 | 2.078E-139 | TEC_inj | MRFAP1 |
| 1.407E-143 | 0.27433202 | 0.387 | 0.203 | 3.324E-139 | TEC_inj | COPZ1 |
| 9.791E-143 | -0.3193257 | 0.006 | 0.172 | 2.313E-138 | TEC_inj | DOCK8 |
| 1.554E-142 | 0.28818468 | 0.456 | 0.251 | 3.672E-138 | TEC_inj | GADD45GIP1 |
| 2.106E-142 | -1.0877359 | 0.007 | 0.173 | 4.976E-138 | TEC_inj | CEACAM5 |
| 2.83E-142 | 0.32562189 | 0.972 | 0.871 | 6.685E-138 | TEC_inj | RPS29 |
| 5.42E-142 | -0.7652536 | 0.145 | 0.339 | 1.28E-137 | TEC_inj | VEGFA |
| 7.035E-142 | -0.6104196 | 0.478 | 0.695 | 1.662E-137 | TEC_inj | IER2 |
| 1.484E-141 | 0.4208688 | 0.845 | 0.739 | 3.506E-137 | TEC_inj | CLIC1 |
| 1.628E-141 | 0.25561075 | 0.354 | 0.172 | 3.846E-137 | TEC_inj | PAWR |
| 1.51E-140 | 0.34541143 | 0.491 | 0.29 | 3.567E-136 | TEC_inj | BCAP31 |
| 2.694E-140 | -0.4904175 | 0.009 | 0.174 | 6.364E-136 | TEC_inj | AQP9 |
| 3.602E-140 | 0.29384257 | 0.539 | 0.32 | 8.51E-136 | TEC_inj | ARL6IP4 |
| 7.07E-140 | -1.0437347 | 0.089 | 0.278 | 1.67E-135 | TEC_inj | AREG |
| 1.577E-139 | -0.6142722 | 0.298 | 0.496 | 3.725E-135 | TEC_inj | RIOK3 |
| 1.626E-139 | -0.5209897 | 0.038 | 0.215 | 3.841E-135 | TEC_inj | NCF2 |
| 4.836E-139 | -0.3876018 | 0.397 | 0.625 | 1.142E-134 | TEC_inj | SON |
| 5.17E-139 | 0.36378081 | 0.822 | 0.69 | 1.221E-134 | TEC_inj | COX6B1 |
| 1.127E-138 | -0.3555327 | 0.027 | 0.2 | 2.661E-134 | TEC_inj | GPSM3 |
| 1.14E-138 | -0.3571891 | 0.011 | 0.175 | 2.692E-134 | TEC_inj | FERMT3 |
| 1.736E-138 | 0.35468985 | 0.584 | 0.371 | 4.101E-134 | TEC_inj | UQCR11 |
| 9.615E-138 | -1.0208395 | 0.007 | 0.167 | 2.271E-133 | TEC_inj | PITX1 |
| 1.443E-137 | -1.0201703 | 0.009 | 0.171 | 3.41E-133 | TEC_inj | PRSS27 |
| 3.45E-137 | 0.27258894 | 0.441 | 0.244 | 8.149E-133 | TEC_inj | OCIAD1 |
| 3.541E-137 | -0.859205 | 0.106 | 0.301 | 8.364E-133 | TEC_inj | HMOX1 |
| 4.223E-137 | -0.3097941 | 0.009 | 0.171 | 9.975E-133 | TEC_inj | ARHGAP30 |
| 4.343E-137 | 0.31067218 | 0.975 | 0.889 | 1.026E-132 | TEC_inj | RPL35A |
| 4.496E-137 | -0.8589553 | 0.011 | 0.174 | 1.062E-132 | TEC_inj | APOBEC3A |
| 5.14E-137 | 0.30655196 | 0.502 | 0.298 | 1.214E-132 | TEC_inj | RBX1 |
| 7.186E-137 | -0.5870214 | 0.009 | 0.171 | 1.698E-132 | TEC_inj | CD300E |
| 7.939E-137 | 0.2940197 | 0.426 | 0.233 | 1.875E-132 | TEC_inj | RPS17 |
| 8.538E-137 | 0.41068311 | 0.362 | 0.188 | 2.017E-132 | TEC_inj | MANF |
| 1.043E-136 | 0.34487665 | 0.216 | 0.085 | 2.463E-132 | TEC_inj | IFIT2 |
| 2.458E-136 | 0.34021477 | 0.863 | 0.754 | 5.808E-132 | TEC_inj | UBL5 |
| 2.542E-136 | 0.31227004 | 0.748 | 0.529 | 6.005E-132 | TEC_inj | RPS10 |
| 3.506E-136 | -0.4519465 | 0.068 | 0.253 | 8.282E-132 | TEC_inj | SMCHD1 |
| 7.824E-136 | -0.8173211 | 0.314 | 0.521 | 1.848E-131 | TEC_inj | HSPA1B |
| 9.127E-136 | 0.2657428 | 0.446 | 0.249 | 2.156E-131 | TEC_inj | TMEM219 |
| 1.061E-135 | -0.7942851 | 0.004 | 0.161 | 2.508E-131 | TEC_inj | TMPRSS11D |
| 1.35E-135 | -0.3538994 | 0.013 | 0.176 | 3.19E-131 | TEC_inj | MYO1F |
| 1.472E-135 | -0.4721449 | 0.421 | 0.661 | 3.477E-131 | TEC_inj | TXNIP |
| 2.333E-135 | -0.4695801 | 0.293 | 0.504 | 5.512E-131 | TEC_inj | ANKRD12 |

| | | | | | | |
|---|---|---|---|---|---|---|
| 3.234E-135 | 0.25697436 | 0.429 | 0.233 | 7.639E-131 | TEC_inj | ATP5PB |
| 4.201E-134 | 0.30538171 | 0.562 | 0.346 | 9.925E-130 | TEC_inj | C19orf53 |
| 5.378E-134 | 0.38744351 | 0.418 | 0.228 | 1.27E-129 | TEC_inj | ITGB8 |
| 1.074E-133 | 0.25884075 | 0.377 | 0.196 | 2.537E-129 | TEC_inj | NBDY |
| 1.308E-133 | 0.25943488 | 0.117 | 0.033 | 3.09E-129 | TEC_inj | FABP3 |
| 1.395E-133 | -0.4059664 | 0.046 | 0.222 | 3.296E-129 | TEC_inj | STK17B |
| 5.344E-133 | -0.8587179 | 0.004 | 0.158 | 1.262E-128 | TEC_inj | CEACAM7 |
| 7.853E-133 | -0.5627006 | 0.515 | 0.714 | 1.855E-128 | TEC_inj | PPP1R15A |
| 8.818E-133 | 0.27944443 | 0.713 | 0.514 | 2.083E-128 | TEC_inj | PTTG1IP |
| 7.475E-132 | -0.386665 | 0.008 | 0.164 | 1.766E-127 | TEC_inj | EVI2B |
| 1.717E-131 | 0.37178469 | 0.332 | 0.172 | 4.055E-127 | TEC_inj | FDPS |
| 1.814E-131 | 0.25680307 | 0.563 | 0.334 | 4.286E-127 | TEC_inj | EIF3E |
| 2.778E-131 | -0.3415567 | 0.598 | 0.349 | 6.563E-127 | TEC_inj | MAL |
| 9.696E-131 | -0.3725962 | 0.016 | 0.176 | 2.29E-126 | TEC_inj | CLEC2B |
| 2.12E-130 | 0.25648174 | 0.412 | 0.224 | 5.008E-126 | TEC_inj | MRPS21 |
| 2.761E-130 | 0.29566705 | 0.539 | 0.332 | 6.522E-126 | TEC_inj | DDT |
| 4.483E-130 | -0.5279375 | 0.004 | 0.156 | 1.059E-125 | TEC_inj | CD93 |
| 6.941E-130 | -0.4653 | 0.116 | 0.303 | 1.64E-125 | TEC_inj | RNF149 |
| 2.035E-129 | -0.4658433 | 0.024 | 0.186 | 4.806E-125 | TEC_inj | SMIM25 |
| 2.254E-129 | -0.5383725 | 0.216 | 0.413 | 5.325E-125 | TEC_inj | ARGLU1 |
| 1.046E-128 | 0.34059276 | 0.845 | 0.739 | 2.47E-124 | TEC_inj | ATP6V0E1 |
| 1.383E-128 | -0.8053292 | 0.022 | 0.182 | 3.267E-124 | TEC_inj | FCN1 |
| 6.411E-128 | 0.29844087 | 0.534 | 0.315 | 1.514E-123 | TEC_inj | SNHG8 |
| 7.53E-128 | 0.3348142 | 0.424 | 0.237 | 1.779E-123 | TEC_inj | CD2AP |
| 1.471E-127 | -1.6237485 | 0.019 | 0.175 | 3.474E-123 | TEC_inj | ECM1 |
| 2.281E-127 | -0.4492401 | 0.007 | 0.157 | 5.389E-123 | TEC_inj | FPR1 |
| 2.762E-127 | 0.28862086 | 0.513 | 0.313 | 6.525E-123 | TEC_inj | AURKAIP1 |
| 4.913E-127 | -0.296203 | 0.004 | 0.153 | 1.161E-122 | TEC_inj | IL2RG |
| 4.924E-127 | -0.8028308 | 0.006 | 0.157 | 1.163E-122 | TEC_inj | TMPRSS11E |
| 6.004E-127 | 0.29980137 | 0.963 | 0.851 | 1.418E-122 | TEC_inj | RPL23 |
| 1.243E-126 | -0.535169 | 0.259 | 0.442 | 2.936E-122 | TEC_inj | EFHD2 |
| 1.466E-126 | 0.31171312 | 0.478 | 0.286 | 3.462E-122 | TEC_inj | PGAM1 |
| 3.869E-126 | -0.490994 | 0.146 | 0.335 | 9.14E-122 | TEC_inj | SLC16A3 |
| 6.489E-126 | -0.5116614 | 0.004 | 0.151 | 1.533E-121 | TEC_inj | PLAC8 |
| 7.24E-125 | 0.49092523 | 0.892 | 0.782 | 1.71E-120 | TEC_inj | PFN1 |
| 8.118E-125 | 0.27654033 | 0.685 | 0.48 | 1.918E-120 | TEC_inj | VDAC2 |
| 8.432E-125 | 0.30989275 | 0.459 | 0.266 | 1.992E-120 | TEC_inj | ISCU |
| 1.705E-124 | -0.7388322 | 0.039 | 0.202 | 4.029E-120 | TEC_inj | ANKRD36C |
| 4.121E-124 | 0.26441558 | 0.438 | 0.251 | 9.735E-120 | TEC_inj | VPS29 |
| 4.79E-124 | -0.389657 | 0.009 | 0.158 | 1.132E-119 | TEC_inj | TREM1 |
| 1.179E-123 | 0.3428099 | 0.805 | 0.66 | 2.786E-119 | TEC_inj | COX5B |
| 1.289E-123 | 0.2910628 | 0.325 | 0.166 | 3.044E-119 | TEC_inj | RANBP1 |
| 2.59E-123 | 0.25013238 | 0.99 | 0.922 | 6.119E-119 | TEC_inj | RPL30 |
| 3.144E-123 | 0.40675556 | 0.98 | 0.952 | 7.427E-119 | TEC_inj | SERF2 |
| 3.68E-123 | 0.26959029 | 0.426 | 0.244 | 8.694E-119 | TEC_inj | LSM3 |
| 4.421E-123 | 0.25390404 | 0.394 | 0.216 | 1.044E-118 | TEC_inj | ENSA |
| 5.188E-123 | 0.27244581 | 0.463 | 0.272 | 1.226E-118 | TEC_inj | NDUFA11 |
| 6.289E-123 | 0.34562502 | 0.757 | 0.598 | 1.486E-118 | TEC_inj | SEC61B |
| 1.597E-122 | -0.5586066 | 0.031 | 0.189 | 3.773E-118 | TEC_inj | ANPEP |
| 1.985E-122 | -1.0285915 | 0.316 | 0.487 | 4.689E-118 | TEC_inj | ERO1A |

| | | | | | | |
|---|---|---|---|---|---|---|
| 5.907E-122 | 0.25439804 | 0.465 | 0.272 | 1.395E-117 | TEC_inj | LAMTOR1 |
| 8.467E-122 | -0.4197386 | 0.028 | 0.184 | 2E-117 | TEC_inj | SAMHD1 |
| 1.316E-121 | -0.4533735 | 0.17 | 0.353 | 3.109E-117 | TEC_inj | GRB2 |
| 1.708E-121 | 0.26001263 | 0.433 | 0.249 | 4.034E-117 | TEC_inj | RTRAF |
| 2.251E-121 | -0.3438421 | 0.061 | 0.231 | 5.318E-117 | TEC_inj | TUT7 |
| 4.803E-121 | -0.9493282 | 0.004 | 0.146 | 1.135E-116 | TEC_inj | TMPRSS11B |
| 4.979E-121 | 0.25551159 | 0.469 | 0.277 | 1.176E-116 | TEC_inj | MTCH1 |
| 5.758E-121 | -0.6636348 | 0.012 | 0.158 | 1.36E-116 | TEC_inj | NCCRP1 |
| 6.386E-121 | 0.27383678 | 0.553 | 0.348 | 1.509E-116 | TEC_inj | ELOC |
| 9.098E-121 | 0.26153416 | 0.408 | 0.227 | 2.149E-116 | TEC_inj | NUDC |
| 1.331E-120 | 0.35509501 | 0.545 | 0.356 | 3.144E-116 | TEC_inj | RHEB |
| 1.765E-120 | 0.60858217 | 0.425 | 0.252 | 4.17E-116 | TEC_inj | DDIT3 |
| 1.866E-120 | -0.3539588 | 0.024 | 0.177 | 4.408E-116 | TEC_inj | JARID2 |
| 2.159E-120 | -0.3129899 | 0.02 | 0.172 | 5.101E-116 | TEC_inj | WIPF1 |
| 4.13E-119 | -0.4261518 | 0.012 | 0.157 | 9.756E-115 | TEC_inj | MS4A7 |
| 6.114E-119 | -0.8774046 | 0.013 | 0.158 | 1.444E-114 | TEC_inj | CEACAM6 |
| 1.091E-118 | -0.4262497 | 0.085 | 0.258 | 2.577E-114 | TEC_inj | RAB31 |
| 1.267E-118 | 0.31272796 | 0.628 | 0.432 | 2.992E-114 | TEC_inj | ACTN4 |
| 1.591E-118 | -0.4176185 | 0.012 | 0.157 | 3.758E-114 | TEC_inj | VSIG2 |
| 2.877E-118 | 0.26687176 | 0.433 | 0.252 | 6.796E-114 | TEC_inj | KDELR2 |
| 9.159E-118 | 0.31697991 | 0.34 | 0.177 | 2.164E-113 | TEC_inj | LIMA1 |
| 1.106E-117 | -0.2634357 | 0.011 | 0.154 | 2.613E-113 | TEC_inj | RASSF5 |
| 1.159E-117 | 0.28472861 | 0.436 | 0.253 | 2.738E-113 | TEC_inj | EID1 |
| 1.539E-117 | -0.3675573 | 0.072 | 0.241 | 3.635E-113 | TEC_inj | NUMB |
| 1.587E-117 | -1.0854743 | 0.005 | 0.145 | 3.75E-113 | TEC_inj | SERPINB2 |
| 2.114E-117 | 0.28993573 | 0.247 | 0.113 | 4.993E-113 | TEC_inj | TNFRSF10B |
| 4.158E-117 | -0.5103899 | 0.017 | 0.163 | 9.823E-113 | TEC_inj | CLEC5A |
| 5.116E-117 | -0.6545211 | 0.02 | 0.167 | 1.209E-112 | TEC_inj | MUC4 |
| 7.314E-117 | 0.3800776 | 0.958 | 0.955 | 1.728E-112 | TEC_inj | ACTG1 |
| 1.118E-116 | 0.30253843 | 0.521 | 0.332 | 2.64E-112 | TEC_inj | SNRPB |
| 1.131E-116 | -0.4801792 | 0.1 | 0.273 | 2.671E-112 | TEC_inj | NABP1 |
| 1.655E-116 | -0.3082554 | 0.007 | 0.147 | 3.91E-112 | TEC_inj | HCK |
| 1.798E-116 | -0.4428451 | 0.7 | 0.847 | 4.247E-112 | TEC_inj | HLA-E |
| 3.626E-116 | -0.2662099 | 0.006 | 0.145 | 8.567E-112 | TEC_inj | NCKAP1L |
| 2.769E-115 | 0.25324491 | 0.46 | 0.27 | 6.541E-111 | TEC_inj | YWHAH |
| 3.686E-115 | -0.4414553 | 0.189 | 0.369 | 8.707E-111 | TEC_inj | AKAP13 |
| 5.954E-115 | -0.2821968 | 0.008 | 0.148 | 1.407E-110 | TEC_inj | EVI2A |
| 5.963E-115 | -0.8890726 | 0.274 | 0.438 | 1.409E-110 | TEC_inj | EMP3 |
| 6.402E-115 | -0.3661009 | 0.013 | 0.155 | 1.512E-110 | TEC_inj | TLR2 |
| 6.727E-115 | 0.72445593 | 0.882 | 0.949 | 1.589E-110 | TEC_inj | MT-CO3 |
| 1.355E-114 | 0.32107671 | 0.868 | 0.75 | 3.201E-110 | TEC_inj | UQCRB |
| 1.755E-114 | -0.3990796 | 0.007 | 0.146 | 4.145E-110 | TEC_inj | BCAS1 |
| 1.93E-114 | 0.25785492 | 0.426 | 0.249 | 4.559E-110 | TEC_inj | NDUFA6 |
| 3.579E-114 | -0.3764381 | 0.106 | 0.281 | 8.454E-110 | TEC_inj | USP15 |
| 4.089E-114 | -0.3446258 | 0.017 | 0.161 | 9.66E-110 | TEC_inj | DMXL2 |
| 4.867E-114 | -0.5143507 | 0.032 | 0.182 | 1.15E-109 | TEC_inj | CXCR4 |
| 7.722E-114 | -1.6498734 | 0.007 | 0.144 | 1.824E-109 | TEC_inj | LYPD2 |
| 6.497E-113 | -0.6876042 | 0.006 | 0.142 | 1.535E-108 | TEC_inj | GABRP |
| 6.743E-113 | -0.2879903 | 0.008 | 0.145 | 1.593E-108 | TEC_inj | FGR |
| 8.931E-113 | 0.2711102 | 0.52 | 0.328 | 2.11E-108 | TEC_inj | WDR1 |

| | | | | | | |
|---|---|---|---|---|---|---|
| 1.993E-112 | 0.35627983 | 0.456 | 0.278 | 4.709E-108 | TEC_inj | TUBA1C |
| 6.543E-112 | 0.25877119 | 0.483 | 0.29 | 1.546E-107 | TEC_inj | CLTB |
| 8.794E-112 | -0.5092894 | 0.12 | 0.289 | 2.077E-107 | TEC_inj | ETS2 |
| 1.85E-111 | -0.3437156 | 0.016 | 0.157 | 4.371E-107 | TEC_inj | C1orf162 |
| 5.098E-111 | -0.7259223 | 0.047 | 0.2 | 1.204E-106 | TEC_inj | AC020656.1 |
| 6.532E-111 | 0.27357984 | 0.565 | 0.359 | 1.543E-106 | TEC_inj | ATP5ME |
| 7.026E-111 | -0.270503 | 0.015 | 0.155 | 1.66E-106 | TEC_inj | CCDC69 |
| 9.504E-111 | -0.3226547 | 0.037 | 0.188 | 2.245E-106 | TEC_inj | ABCA1 |
| 1.835E-110 | -0.2966174 | 0.005 | 0.137 | 4.335E-106 | TEC_inj | IRAK3 |
| 2.316E-110 | -0.7574671 | 0.616 | 0.748 | 5.47E-106 | TEC_inj | CTSB |
| 4.657E-110 | -0.2690612 | 0.008 | 0.141 | 1.1E-105 | TEC_inj | APBB1IP |
| 5.021E-110 | -0.9441113 | 0.01 | 0.145 | 1.186E-105 | TEC_inj | EREG |
| 1.509E-109 | -0.4621956 | 0.39 | 0.561 | 3.564E-105 | TEC_inj | TOP1 |
| 1.515E-109 | 0.33570198 | 0.658 | 0.477 | 3.579E-105 | TEC_inj | SELENOW |
| 1.969E-109 | -0.8233554 | 0.127 | 0.295 | 4.651E-105 | TEC_inj | MAFB |
| 5.8E-109 | -0.3101211 | 0.012 | 0.148 | 1.37E-104 | TEC_inj | CLEC7A |
| 7.61E-109 | -0.2860688 | 0.022 | 0.164 | 1.798E-104 | TEC_inj | VSIR |
| 7.876E-109 | -0.3118561 | 0.055 | 0.21 | 1.86E-104 | TEC_inj | MYO9B |
| 9.979E-109 | -0.2881231 | 0.01 | 0.144 | 2.357E-104 | TEC_inj | CYTIP |
| 1.818E-108 | -0.2756633 | 0.013 | 0.149 | 4.295E-104 | TEC_inj | EMILIN2 |
| 1.988E-108 | -0.2963363 | 0.008 | 0.141 | 4.697E-104 | TEC_inj | LILRB3 |
| 3.089E-108 | -0.2944647 | 0.064 | 0.223 | 7.297E-104 | TEC_inj | CYTH1 |
| 3.354E-108 | -0.5553027 | 0.095 | 0.255 | 7.923E-104 | TEC_inj | ATP2B1 |
| 5.767E-108 | 0.26124173 | 0.454 | 0.276 | 1.362E-103 | TEC_inj | HIGD2A |
| 1.491E-107 | 0.25074268 | 0.486 | 0.303 | 3.523E-103 | TEC_inj | RER1 |
| 1.834E-107 | -0.4068876 | 0.017 | 0.153 | 4.334E-103 | TEC_inj | NCF1 |
| 2.329E-107 | -0.3248313 | 0.021 | 0.16 | 5.502E-103 | TEC_inj | BCAT1 |
| 2.53E-107 | 0.2527298 | 0.886 | 0.765 | 5.976E-103 | TEC_inj | CTSD |
| 3.647E-107 | -0.5681751 | 0.431 | 0.591 | 8.616E-103 | TEC_inj | LITAF |
| 7.099E-107 | -0.3128279 | 0.022 | 0.161 | 1.677E-102 | TEC_inj | SMAP2 |
| 1.02E-106 | -0.7457575 | 0.026 | 0.164 | 2.409E-102 | TEC_inj | SCEL |
| 2.576E-106 | 0.27805455 | 0.946 | 0.879 | 6.084E-102 | TEC_inj | HSP90AB1 |
| 2.63E-106 | -0.5955137 | 0.189 | 0.361 | 6.212E-102 | TEC_inj | IER5 |
| 2.846E-106 | -0.6330704 | 0.054 | 0.2 | 6.724E-102 | TEC_inj | EPS8L1 |
| 3.69E-106 | 0.25360491 | 0.432 | 0.26 | 8.717E-102 | TEC_inj | AP2M1 |
| 1.266E-105 | -0.2958395 | 0.055 | 0.207 | 2.99E-101 | TEC_inj | RAB8B |
| 1.375E-105 | -0.3861592 | 0.151 | 0.323 | 3.248E-101 | TEC_inj | QKI |
| 1.599E-105 | 0.26748575 | 0.455 | 0.277 | 3.778E-101 | TEC_inj | NDUFS6 |
| 3.488E-105 | -0.3255182 | 0.06 | 0.213 | 8.24E-101 | TEC_inj | NR3C1 |
| 7.047E-105 | -0.3595877 | 0.07 | 0.225 | 1.665E-100 | TEC_inj | ZFYVE16 |
| 7.644E-105 | -0.8573894 | 0.029 | 0.167 | 1.806E-100 | TEC_inj | KRT15 |
| 8.041E-105 | -0.3180721 | 0.004 | 0.131 | 1.899E-100 | TEC_inj | LILRB2 |
| 7.159E-104 | 0.33538936 | 0.577 | 0.393 | 1.691E-99 | TEC_inj | ATF4 |
| 1.209E-103 | 0.26077479 | 0.264 | 0.132 | 2.856E-99 | TEC_inj | CPM |
| 1.585E-103 | -0.8234978 | 0.043 | 0.184 | 3.743E-99 | TEC_inj | NDRG2 |
| 1.685E-103 | -0.6789578 | 0.012 | 0.141 | 3.98E-99 | TEC_inj | DHRS9 |
| 2.539E-103 | -0.3829673 | 0.415 | 0.618 | 5.997E-99 | TEC_inj | NFKBIA |
| 4.91E-103 | -0.4369583 | 0.016 | 0.147 | 1.1598E-98 | TEC_inj | MCEMP1 |
| 8.464E-103 | -1.0292557 | 0.018 | 0.151 | 1.9995E-98 | TEC_inj | CXCL6 |
| 1.537E-102 | -0.3484588 | 0.045 | 0.189 | 3.6311E-98 | TEC_inj | CCDC88A |

| | | | | | | |
|---|---|---|---|---|---|---|
| 2.962E-102 | -0.5684316 | 0.172 | 0.34 | 6.9978E-98 | TEC_inj | CEBPD |
| 3.711E-102 | -1.1646548 | 0.004 | 0.126 | 8.7676E-98 | TEC_inj | SPINK5 |
| 2.156E-101 | -0.4841672 | 0.001 | 0.122 | 5.0934E-97 | TEC_inj | TMPRSS11A |
| 3.498E-101 | -0.5761488 | 0.168 | 0.327 | 8.2639E-97 | TEC_inj | HIST1H2AC |
| 3.482E-100 | -0.3514098 | 0.158 | 0.331 | 8.2245E-96 | TEC_inj | KMT2E |
| 3.798E-100 | -0.6285177 | 0.018 | 0.146 | 8.9729E-96 | TEC_inj | CXCL17 |
| 1.464E-99 | -0.3977478 | 0.171 | 0.338 | 3.4587E-95 | TEC_inj | FOSL2 |
| 3.861E-99 | -0.2826143 | 0.024 | 0.156 | 9.122E-95 | TEC_inj | CARD16 |
| 5.344E-99 | -0.8566391 | 0.146 | 0.297 | 1.2625E-94 | TEC_inj | SLC25A37 |
| 6.105E-99 | 0.4568542 | 0.434 | 0.276 | 1.4423E-94 | TEC_inj | MIR4435-2HG |
| 1.0649E-98 | -0.3317676 | 0.027 | 0.16 | 2.5157E-94 | TEC_inj | PLXDC2 |
| 1.1345E-98 | -0.3240038 | 0.039 | 0.176 | 2.6801E-94 | TEC_inj | CEP170 |
| 1.531E-98 | 0.4180973 | 0.558 | 0.405 | 3.6166E-94 | TEC_inj | JPT1 |
| 4.3867E-98 | 0.25940314 | 0.831 | 0.698 | 1.0363E-93 | TEC_inj | GPX4 |
| 6.2119E-98 | 0.27202029 | 0.471 | 0.302 | 1.4674E-93 | TEC_inj | TAF10 |
| 1.2085E-97 | -0.9628812 | 0.13 | 0.274 | 2.855E-93 | TEC_inj | LMO7 |
| 2.2518E-97 | -0.6856813 | 0.466 | 0.642 | 5.3194E-93 | TEC_inj | JUN |
| 5.8496E-97 | -0.506077 | 0.055 | 0.197 | 1.3818E-92 | TEC_inj | KLF2 |
| 6.4503E-97 | -0.2752433 | 0.21 | 0.389 | 1.5238E-92 | TEC_inj | CCNL1 |
| 9.1914E-97 | 0.27262669 | 0.548 | 0.374 | 2.1713E-92 | TEC_inj | COPE |
| 2.4638E-96 | -0.2542072 | 0.006 | 0.125 | 5.8201E-92 | TEC_inj | ADGRE2 |
| 3.4375E-96 | -0.2786361 | 0.008 | 0.127 | 8.1203E-92 | TEC_inj | CD86 |
| 4.0615E-96 | -0.4442869 | 0.41 | 0.559 | 9.5944E-92 | TEC_inj | C4orf3 |
| 4.1125E-96 | -0.3605332 | 0.074 | 0.224 | 9.7151E-92 | TEC_inj | TLE4 |
| 2.8089E-95 | -0.6046197 | 0.008 | 0.126 | 6.6355E-91 | TEC_inj | MNDA |
| 8.3184E-95 | -0.3020252 | 0.005 | 0.12 | 1.9651E-90 | TEC_inj | FUT3 |
| 1.1651E-94 | -0.6184162 | 0.006 | 0.121 | 2.7523E-90 | TEC_inj | KRT78 |
| 1.152E-93 | -0.3897411 | 0.011 | 0.128 | 2.7215E-89 | TEC_inj | FCGR3A |
| 1.4069E-93 | -0.2505314 | 0.008 | 0.125 | 3.3236E-89 | TEC_inj | C3AR1 |
| 1.7317E-93 | -0.302727 | 0.046 | 0.182 | 4.0908E-89 | TEC_inj | B3GNT5 |
| 1.8515E-93 | 0.26284661 | 0.505 | 0.333 | 4.3738E-89 | TEC_inj | SF3B6 |
| 3.0458E-93 | -0.2803522 | 0.056 | 0.197 | 7.1952E-89 | TEC_inj | TET2 |
| 9.7561E-93 | -0.4993 | 0.461 | 0.645 | 2.3047E-88 | TEC_inj | ZFP36L1 |
| 1.682E-92 | -0.3411204 | 0.003 | 0.114 | 3.9734E-88 | TEC_inj | A2ML1 |
| 4.6863E-92 | 0.27861511 | 0.695 | 0.538 | 1.107E-87 | TEC_inj | UQCR10 |
| 1.5705E-91 | 0.33272328 | 0.697 | 0.561 | 3.71E-87 | TEC_inj | TSPO |
| 1.6119E-91 | -0.2839557 | 0.083 | 0.23 | 3.8079E-87 | TEC_inj | ACAP2 |
| 2.7884E-91 | 0.31389259 | 0.235 | 0.117 | 6.587E-87 | TEC_inj | PLA2G16 |
| 3.1318E-91 | -0.2534729 | 0.043 | 0.177 | 7.3982E-87 | TEC_inj | SLC12A6 |
| 7.3004E-91 | -0.3343573 | 0.085 | 0.229 | 1.7246E-86 | TEC_inj | STK4 |
| 8.1585E-91 | 0.29467393 | 0.944 | 0.896 | 1.9273E-86 | TEC_inj | RPL39 |
| 9.1205E-91 | -1.3950909 | 0.073 | 0.209 | 2.1545E-86 | TEC_inj | KRT17 |
| 2.1753E-90 | -0.4241805 | 0.399 | 0.564 | 5.1386E-86 | TEC_inj | FAM107B |
| 2.4183E-90 | -0.8178985 | 0.004 | 0.113 | 5.7128E-86 | TEC_inj | ZG16B |
| 7.2694E-90 | 0.29027939 | 0.466 | 0.298 | 1.7172E-85 | TEC_inj | SMS |
| 8.646E-90 | -0.2729205 | 0.009 | 0.122 | 2.0424E-85 | TEC_inj | PECAM1 |
| 1.0922E-89 | 0.26271144 | 0.426 | 0.263 | 2.5802E-85 | TEC_inj | ATP6AP2 |
| 2.1882E-89 | -0.3948324 | 0.34 | 0.507 | 5.1691E-85 | TEC_inj | RSRP1 |
| 3.0375E-89 | 0.2882994 | 0.45 | 0.294 | 7.1754E-85 | TEC_inj | BUD31 |
| 5.3303E-89 | -0.2932453 | 0.003 | 0.111 | 1.2592E-84 | TEC_inj | LILRB1 |

| | | | | | | |
|---|---|---|---|---|---|---|
| 1.0893E-88 | -0.4336855 | 0.006 | 0.115 | 2.5734E-84 | TEC_inj | TMC5 |
| 1.6723E-88 | 0.61763915 | 0.913 | 0.94 | 3.9504E-84 | TEC_inj | TMSB4X |
| 2.3194E-88 | -0.356851 | 0.005 | 0.114 | 5.4791E-84 | TEC_inj | CSF3R |
| 3.2317E-88 | -0.4524995 | 0.511 | 0.64 | 7.6343E-84 | TEC_inj | TAX1BP1 |
| 6.7548E-88 | -0.339577 | 0.144 | 0.299 | 1.5957E-83 | TEC_inj | BAZ1A |
| 9.0826E-88 | 0.55673946 | 0.703 | 0.568 | 2.1456E-83 | TEC_inj | KRT19 |
| 2.6116E-87 | -0.253175 | 0.021 | 0.139 | 6.1695E-83 | TEC_inj | CELF2 |
| 4.6346E-87 | -0.3067656 | 0.003 | 0.11 | 1.0948E-82 | TEC_inj | LRMP |
| 4.7346E-87 | -0.2623645 | 0.015 | 0.129 | 1.1184E-82 | TEC_inj | CSF1R |
| 5.1919E-87 | -1.1731857 | 0.017 | 0.132 | 1.2265E-82 | TEC_inj | CCL3 |
| 5.5243E-87 | 0.35159681 | 0.419 | 0.272 | 1.305E-82 | TEC_inj | RPS4Y1 |
| 6.1367E-87 | -0.3341212 | 0.023 | 0.142 | 1.4497E-82 | TEC_inj | RGCC |
| 6.5389E-87 | -0.4219392 | 0.142 | 0.288 | 1.5447E-82 | TEC_inj | SLK |
| 1.0271E-86 | -0.3736482 | 0.012 | 0.123 | 2.4263E-82 | TEC_inj | CEACAM1 |
| 2.9532E-86 | -0.2540015 | 0.048 | 0.177 | 6.9765E-82 | TEC_inj | SIRPA |
| 2.9654E-86 | -0.3241274 | 0.04 | 0.165 | 7.0053E-82 | TEC_inj | ADGRE5 |
| 3.1888E-86 | 0.34330785 | 0.915 | 0.866 | 7.5328E-82 | TEC_inj | HLA-C |
| 3.6779E-86 | -0.4335417 | 0.434 | 0.588 | 8.6882E-82 | TEC_inj | CD55 |
| 4.3689E-86 | -0.4352491 | 0.066 | 0.199 | 1.0321E-81 | TEC_inj | DUSP6 |
| 4.923E-86 | -0.3011727 | 0.047 | 0.176 | 1.163E-81 | TEC_inj | CFD |
| 9.5957E-86 | -0.7866896 | 0.012 | 0.122 | 2.2668E-81 | TEC_inj | IL7R |
| 5.6542E-85 | -0.4767748 | 0.053 | 0.181 | 1.3357E-80 | TEC_inj | TGFBI |
| 8.0049E-85 | 0.33149743 | 0.407 | 0.254 | 1.891E-80 | TEC_inj | UGCG |
| 1.3337E-84 | -0.3544322 | 0.014 | 0.125 | 3.1505E-80 | TEC_inj | CD163 |
| 1.3771E-84 | 0.25371182 | 0.495 | 0.331 | 3.2532E-80 | TEC_inj | EIF1AX |
| 1.5416E-84 | -0.4008063 | 0.006 | 0.112 | 3.6416E-80 | TEC_inj | CLIC3 |
| 2.3306E-84 | -1.07285 | 0.067 | 0.195 | 5.5055E-80 | TEC_inj | HLA-DPB1 |
| 2.9057E-84 | -0.4181293 | 0.077 | 0.213 | 6.8642E-80 | TEC_inj | FOSB |
| 4.7477E-84 | -0.4973575 | 0.136 | 0.281 | 1.1216E-79 | TEC_inj | TSC22D3 |
| 5.2414E-84 | -0.2676585 | 0.036 | 0.159 | 1.2382E-79 | TEC_inj | RALGDS |
| 1.8874E-83 | -0.3187591 | 0.031 | 0.149 | 4.4587E-79 | TEC_inj | EMP2 |
| 2.729E-83 | -0.3630183 | 0.161 | 0.308 | 6.4467E-79 | TEC_inj | VPS4B |
| 3.9692E-83 | -0.3458474 | 0.128 | 0.271 | 9.3765E-79 | TEC_inj | MBNL1 |
| 4.9895E-83 | -0.3071151 | 0.098 | 0.238 | 1.1787E-78 | TEC_inj | RGS10 |
| 5.0088E-83 | -0.4515751 | 0.179 | 0.323 | 1.1832E-78 | TEC_inj | FNDC3B |
| 7.7214E-83 | -0.2731126 | 0.031 | 0.149 | 1.824E-78 | TEC_inj | IFI30 |
| 9.9976E-83 | -0.2675181 | 0.068 | 0.2 | 2.3617E-78 | TEC_inj | N4BP1 |
| 1.269E-82 | -0.3051144 | 0.107 | 0.248 | 2.9978E-78 | TEC_inj | HOOK3 |
| 1.4354E-82 | -0.6747078 | 0.111 | 0.247 | 3.3909E-78 | TEC_inj | FABP5 |
| 1.5283E-82 | 0.38533601 | 0.712 | 0.556 | 3.6102E-78 | TEC_inj | TACSTD2 |
| 1.5423E-81 | -0.6761113 | 0.263 | 0.395 | 3.6433E-77 | TEC_inj | TYMP |
| 4.6476E-81 | -0.2951907 | 0.004 | 0.103 | 1.0979E-76 | TEC_inj | GPR183 |
| 5.9928E-81 | -0.8219811 | 0.186 | 0.323 | 1.4157E-76 | TEC_inj | PHLDA1 |
| 7.7345E-81 | -0.4137833 | 0.011 | 0.115 | 1.8271E-76 | TEC_inj | THBD |
| 1.0058E-80 | 0.39477909 | 0.442 | 0.3 | 2.376E-76 | TEC_inj | CYTOR |
| 1.2082E-80 | -0.4357159 | 0.065 | 0.192 | 2.8542E-76 | TEC_inj | FBP1 |
| 2.4627E-80 | -0.2916737 | 0.014 | 0.12 | 5.8177E-76 | TEC_inj | HIST1H2BG |
| 2.5681E-80 | 0.82543427 | 0.495 | 0.331 | 6.0666E-76 | TEC_inj | GDF15 |
| 3.2098E-80 | -0.9069205 | 0.029 | 0.141 | 7.5825E-76 | TEC_inj | RHCG |
| 3.4675E-80 | -0.3000567 | 0.14 | 0.284 | 8.1913E-76 | TEC_inj | FAM49B |

| | | | | | | |
|---|---|---|---|---|---|---|
| 6.1868E-80 | -0.3194436 | 0.092 | 0.226 | 1.4615E-75 | TEC_inj | EML4 |
| 7.5744E-80 | -0.2701784 | 0.042 | 0.162 | 1.7893E-75 | TEC_inj | SNX18 |
| 9.0147E-80 | -0.5448278 | 0.057 | 0.178 | 2.1295E-75 | TEC_inj | FXYD3 |
| 1.1253E-79 | -0.8326543 | 0.076 | 0.197 | 2.6583E-75 | TEC_inj | TMPRSS2 |
| 1.3742E-79 | -0.256486 | 0.06 | 0.186 | 3.2462E-75 | TEC_inj | PPP1R18 |
| 1.794E-79 | -0.3600668 | 0.197 | 0.342 | 4.2379E-75 | TEC_inj | TCF25 |
| 2.367E-79 | -0.3651723 | 0.136 | 0.276 | 5.5916E-75 | TEC_inj | BACH1 |
| 9.1265E-79 | 0.29885328 | 0.862 | 0.758 | 2.1559E-74 | TEC_inj | RPS26 |
| 1.4219E-78 | 0.30075405 | 0.363 | 0.223 | 3.359E-74 | TEC_inj | SNHG7 |
| 1.6358E-78 | -0.3805781 | 0.175 | 0.317 | 3.8643E-74 | TEC_inj | BNIP3L |
| 2.359E-78 | -0.3403399 | 0.019 | 0.126 | 5.5727E-74 | TEC_inj | DUSP2 |
| 2.3817E-78 | -0.2541379 | 0.004 | 0.102 | 5.6262E-74 | TEC_inj | B3GALT5 |
| 3.2953E-78 | -0.2557739 | 0.114 | 0.252 | 7.7844E-74 | TEC_inj | OSTF1 |
| 4.2915E-78 | -1.5794777 | 0.013 | 0.114 | 1.0138E-73 | TEC_inj | CRCT1 |
| 4.6356E-78 | 0.29805169 | 0.314 | 0.184 | 1.0951E-73 | TEC_inj | SDF2L1 |
| 4.8685E-78 | -1.0793232 | 0.013 | 0.116 | 1.1501E-73 | TEC_inj | CCL4 |
| 1.1087E-77 | -0.333058 | 0.027 | 0.137 | 2.6192E-73 | TEC_inj | LRG1 |
| 1.1517E-77 | -0.6291292 | 0.006 | 0.103 | 2.7205E-73 | TEC_inj | PTGS2 |
| 1.529E-77 | -0.754683 | 0.113 | 0.244 | 3.6119E-73 | TEC_inj | PLIN2 |
| 2.0493E-77 | -0.2726524 | 0.056 | 0.177 | 4.8412E-73 | TEC_inj | GLIPR2 |
| 3.1856E-77 | -2.8045432 | 0.386 | 0.555 | 7.5254E-73 | TEC_inj | PLCG2 |
| 4.4831E-77 | -0.5523623 | 0.296 | 0.445 | 1.059E-72 | TEC_inj | HIST1H1C |
| 5.0427E-77 | -0.2538739 | 0.033 | 0.145 | 1.1912E-72 | TEC_inj | ITGA5 |
| 7.4526E-77 | -0.3873624 | 0.024 | 0.131 | 1.7605E-72 | TEC_inj | TMEM40 |
| 2.7085E-76 | -0.2550656 | 0.014 | 0.117 | 6.3983E-72 | TEC_inj | PDE4B |
| 3.4692E-76 | 0.51061188 | 0.655 | 0.529 | 8.1953E-72 | TEC_inj | CLDN4 |
| 4.444E-76 | -0.2767728 | 0.096 | 0.227 | 1.0498E-71 | TEC_inj | TRIM38 |
| 7.2668E-76 | 0.29499405 | 0.249 | 0.136 | 1.7166E-71 | TEC_inj | NCOA7 |
| 8.5927E-76 | -1.1680604 | 0.032 | 0.14 | 2.0299E-71 | TEC_inj | PLAT |
| 9.6109E-76 | -0.4152621 | 0.091 | 0.216 | 2.2704E-71 | TEC_inj | SLC43A2 |
| 1.1535E-75 | -0.2677464 | 0.044 | 0.161 | 2.725E-71 | TEC_inj | HK2 |
| 1.6656E-75 | -0.4958248 | 0.181 | 0.32 | 3.9347E-71 | TEC_inj | IRF1 |
| 2.1552E-75 | -0.2881388 | 0.014 | 0.114 | 5.0912E-71 | TEC_inj | MS4A6A |
| 1.1111E-74 | -0.3780254 | 0.17 | 0.307 | 2.6247E-70 | TEC_inj | IFNGR2 |
| 1.5001E-74 | 0.39546878 | 0.876 | 0.857 | 3.5436E-70 | TEC_inj | SH3BGRL3 |
| 4.8585E-74 | -0.2644165 | 0.081 | 0.208 | 1.1477E-69 | TEC_inj | CHD1 |
| 5.4266E-74 | -0.292914 | 0.111 | 0.243 | 1.2819E-69 | TEC_inj | BAZ2B |
| 5.6821E-74 | -1.1611421 | 0.092 | 0.213 | 1.3423E-69 | TEC_inj | HLA-DPA1 |
| 7.1379E-74 | -0.2657345 | 0.109 | 0.241 | 1.6862E-69 | TEC_inj | TAOK3 |
| 8.7144E-74 | -0.4602356 | 0.19 | 0.322 | 2.0586E-69 | TEC_inj | CD68 |
| 2.4807E-73 | -0.3643251 | 0.119 | 0.255 | 5.8601E-69 | TEC_inj | KIAA1551 |
| 1.358E-72 | -0.3484834 | 0.065 | 0.185 | 3.208E-68 | TEC_inj | NR4A1 |
| 3.2677E-72 | -0.2639643 | 0.07 | 0.191 | 7.7194E-68 | TEC_inj | RILPL2 |
| 4.219E-72 | -0.2512916 | 0.043 | 0.155 | 9.9666E-68 | TEC_inj | PLEKHO1 |
| 4.6301E-72 | -0.3523207 | 0.062 | 0.179 | 1.0938E-67 | TEC_inj | FAM214A |
| 6.2454E-72 | -0.2540194 | 0.027 | 0.13 | 1.4754E-67 | TEC_inj | TCP11L2 |
| 9.3122E-72 | -0.3205297 | 0.106 | 0.233 | 2.1998E-67 | TEC_inj | GRINA |
| 1.3817E-70 | -0.5574318 | 0.1 | 0.218 | 3.264E-66 | TEC_inj | CDKN2B |
| 1.5573E-70 | -0.3091251 | 0.134 | 0.265 | 3.6788E-66 | TEC_inj | MGAT1 |
| 1.5673E-70 | -0.2820171 | 0.054 | 0.166 | 3.7024E-66 | TEC_inj | TRIP10 |

| | | | | | | |
|---|---|---|---|---|---|---|
| 1.6442E-70 | 0.54978295 | 0.452 | 0.319 | 3.8842E-66 | TEC_inj | HERPUD1 |
| 1.8727E-70 | -0.2504919 | 0.1 | 0.228 | 4.4239E-66 | TEC_inj | RIT1 |
| 8.3011E-70 | -0.4056852 | 0.12 | 0.246 | 1.961E-65 | TEC_inj | NFKBIZ |
| 1.4718E-69 | -0.2827004 | 0.112 | 0.24 | 3.4767E-65 | TEC_inj | PHF20L1 |
| 2.7918E-69 | -0.3008024 | 0.457 | 0.595 | 6.5952E-65 | TEC_inj | ACTR3 |
| 2.9172E-69 | -0.3192983 | 0.126 | 0.253 | 6.8912E-65 | TEC_inj | RBPJ |
| 6.3566E-69 | -0.2839042 | 0.155 | 0.289 | 1.5016E-64 | TEC_inj | TMF1 |
| 6.4277E-69 | -0.4134453 | 0.076 | 0.193 | 1.5184E-64 | TEC_inj | SGK1 |
| 9.1828E-69 | 0.30400262 | 0.385 | 0.253 | 2.1692E-64 | TEC_inj | TUBB |
| 1.1502E-68 | -0.4168544 | 0.109 | 0.235 | 2.7172E-64 | TEC_inj | INTS6 |
| 2.4093E-68 | -0.3076274 | 0.213 | 0.347 | 5.6915E-64 | TEC_inj | CAPZA1 |
| 2.0252E-67 | -0.4078018 | 0.033 | 0.133 | 4.7842E-63 | TEC_inj | SRD5A3 |
| 4.406E-67 | -0.312298 | 0.041 | 0.145 | 1.0408E-62 | TEC_inj | HLA-DMA |
| 6.575E-67 | -0.7103449 | 0.076 | 0.189 | 1.5532E-62 | TEC_inj | F3 |
| 7.3085E-67 | 0.27550365 | 0.965 | 0.974 | 1.7265E-62 | TEC_inj | ACTB |
| 1.0885E-66 | -0.537915 | 0.046 | 0.15 | 2.5713E-62 | TEC_inj | CDKN2A |
| 1.5007E-66 | -0.3811765 | 0.176 | 0.31 | 3.5451E-62 | TEC_inj | BTG2 |
| 2.9721E-66 | -0.3201617 | 0.296 | 0.439 | 7.0209E-62 | TEC_inj | SRSF5 |
| 3.1399E-66 | -0.333741 | 0.225 | 0.356 | 7.4174E-62 | TEC_inj | SCAF11 |
| 3.723E-66 | -0.2745621 | 0.014 | 0.105 | 8.7949E-62 | TEC_inj | RNF39 |
| 3.8972E-66 | -0.3805229 | 0.05 | 0.156 | 9.2063E-62 | TEC_inj | RGS2 |
| 1.6759E-65 | -0.2709586 | 0.125 | 0.251 | 3.9589E-61 | TEC_inj | KMT2C |
| 1.8243E-65 | -0.2938101 | 0.17 | 0.303 | 4.3095E-61 | TEC_inj | JMJD1C |
| 3.1432E-65 | -0.2832511 | 0.111 | 0.232 | 7.4252E-61 | TEC_inj | MAP3K2 |
| 4.2326E-65 | 0.40022617 | 0.286 | 0.166 | 9.9987E-61 | TEC_inj | IGFBP3 |
| 5.0605E-65 | -0.5031723 | 0.073 | 0.181 | 1.1954E-60 | TEC_inj | PPL |
| 6.4438E-65 | -0.2753466 | 0.145 | 0.271 | 1.5222E-60 | TEC_inj | UBE2R2 |
| 7.2896E-65 | -0.3370103 | 0.021 | 0.114 | 1.722E-60 | TEC_inj | SLC16A10 |
| 7.9109E-65 | 0.25436274 | 0.377 | 0.244 | 1.8688E-60 | TEC_inj | RPL22L1 |
| 7.9432E-65 | -0.5256865 | 0.025 | 0.12 | 1.8764E-60 | TEC_inj | CYP1B1 |
| 6.0799E-64 | -0.2628326 | 0.142 | 0.269 | 1.4362E-59 | TEC_inj | ERBIN |
| 1.8218E-63 | -0.323043 | 0.099 | 0.213 | 4.3037E-59 | TEC_inj | TIPARP |
| 1.9426E-63 | -0.8392371 | 0.176 | 0.287 | 4.5891E-59 | TEC_inj | DUSP5 |
| 2.2777E-63 | -0.3858379 | 0.1 | 0.213 | 5.3807E-59 | TEC_inj | CSNK1E |
| 2.2912E-63 | -0.2589976 | 0.111 | 0.231 | 5.4124E-59 | TEC_inj | ZBTB7A |
| 1.1893E-62 | -0.3129441 | 0.107 | 0.224 | 2.8094E-58 | TEC_inj | KDM6B |
| 1.2705E-62 | 0.84478802 | 0.592 | 0.534 | 3.0012E-58 | TEC_inj | UPP1 |
| 1.4312E-62 | 0.54111789 | 0.896 | 0.955 | 3.3808E-58 | TEC_inj | MT-CO1 |
| 8.3399E-62 | -0.5939413 | 0.02 | 0.108 | 1.9701E-57 | TEC_inj | HLA-DQA1 |
| 1.1947E-61 | -0.4203521 | 0.014 | 0.1 | 2.8223E-57 | TEC_inj | RNASE7 |
| 3.2416E-61 | 0.37476256 | 0.536 | 0.416 | 7.6575E-57 | TEC_inj | SLC3A2 |
| 4.3528E-61 | -0.4821566 | 0.02 | 0.107 | 1.0283E-56 | TEC_inj | RETN |
| 5.4139E-61 | -0.2551255 | 0.03 | 0.123 | 1.2789E-56 | TEC_inj | OAS1 |
| 7.5733E-61 | -0.338007 | 0.246 | 0.368 | 1.789E-56 | TEC_inj | PLSCR1 |
| 7.6208E-61 | -0.3168948 | 0.017 | 0.103 | 1.8003E-56 | TEC_inj | KLK10 |
| 1.1093E-60 | 0.31806598 | 0.607 | 0.482 | 2.6204E-56 | TEC_inj | MTRNR2L8 |
| 1.1716E-60 | -0.2833169 | 0.157 | 0.279 | 2.7676E-56 | TEC_inj | AGFG1 |
| 1.242E-60 | -0.3030309 | 0.166 | 0.292 | 2.934E-56 | TEC_inj | PRPF4B |
| 2.5758E-60 | -0.3354861 | 0.276 | 0.4 | 6.0848E-56 | TEC_inj | STAT3 |
| 6.0241E-60 | -0.3185721 | 0.039 | 0.134 | 1.4231E-55 | TEC_inj | MARCO |

| | | | | | | |
|---|---|---|---|---|---|---|
| 6.0364E-60 | -0.4695326 | 0.056 | 0.156 | 1.426E-55 | TEC_inj | SMC4 |
| 6.7549E-60 | -0.3918209 | 0.106 | 0.217 | 1.5957E-55 | TEC_inj | LY6E |
| 7.3783E-60 | -0.3270851 | 0.2 | 0.321 | 1.743E-55 | TEC_inj | TMEM165 |
| 7.3845E-60 | -0.5322467 | 0.077 | 0.181 | 1.7444E-55 | TEC_inj | HLA-DQB1 |
| 7.7193E-60 | -0.3160188 | 0.387 | 0.511 | 1.8235E-55 | TEC_inj | ASAH1 |
| 6.7993E-59 | -0.2737198 | 0.11 | 0.222 | 1.6062E-54 | TEC_inj | RUNX1 |
| 1.3223E-58 | -0.4341789 | 0.115 | 0.223 | 3.1236E-54 | TEC_inj | CREM |
| 2.2613E-58 | -0.2762135 | 0.063 | 0.165 | 5.3418E-54 | TEC_inj | PELI1 |
| 2.8569E-58 | -0.2510507 | 0.136 | 0.257 | 6.7489E-54 | TEC_inj | CLK1 |
| 6.4242E-58 | -0.3217671 | 0.25 | 0.369 | 1.5176E-53 | TEC_inj | TMOD3 |
| 8.31E-58 | -0.2605386 | 0.115 | 0.228 | 1.9631E-53 | TEC_inj | ZFC3H1 |
| 6.4075E-57 | -0.4229474 | 0.09 | 0.194 | 1.5136E-52 | TEC_inj | ACSL1 |
| 8.809E-57 | -0.2815283 | 0.127 | 0.24 | 2.0809E-52 | TEC_inj | ATP6V1B2 |
| 1.1155E-56 | -0.3871895 | 0.125 | 0.237 | 2.6351E-52 | TEC_inj | IVNS1ABP |
| 1.2593E-56 | -0.4893246 | 0.037 | 0.127 | 2.9749E-52 | TEC_inj | HILPDA |
| 8.539E-56 | -0.4098046 | 0.077 | 0.179 | 2.0172E-51 | TEC_inj | DNAJA4 |
| 1.6188E-55 | -0.3634534 | 0.111 | 0.217 | 3.8241E-51 | TEC_inj | SOCS3 |
| 1.704E-55 | -0.2803813 | 0.031 | 0.117 | 4.0254E-51 | TEC_inj | STS |
| 5.784E-55 | -0.3312227 | 0.029 | 0.114 | 1.3664E-50 | TEC_inj | TCHP |
| 9.9011E-55 | 0.45805325 | 0.883 | 0.95 | 2.3389E-50 | TEC_inj | MT-CO2 |
| 1.0972E-54 | -1.4393353 | 0.03 | 0.114 | 2.5919E-50 | TEC_inj | CNFN |
| 1.6708E-54 | -0.301031 | 0.086 | 0.189 | 3.947E-50 | TEC_inj | PFKFB3 |
| 7.3795E-54 | -0.3137644 | 0.284 | 0.404 | 1.7433E-49 | TEC_inj | PNISR |
| 8.6932E-54 | -0.3378947 | 0.452 | 0.581 | 2.0536E-49 | TEC_inj | ZFAND5 |
| 9.6294E-54 | -0.2511922 | 0.438 | 0.571 | 2.2748E-49 | TEC_inj | CSNK1A1 |
| 1.2078E-53 | -0.2901426 | 0.844 | 0.976 | 2.8532E-49 | TEC_inj | UBC |
| 1.3598E-53 | -0.2715676 | 0.13 | 0.237 | 3.2123E-49 | TEC_inj | PGD |
| 2.2272E-53 | -1.0261788 | 0.183 | 0.285 | 5.2614E-49 | TEC_inj | HLA-DRB1 |
| 3.2286E-53 | -0.3204273 | 0.201 | 0.316 | 7.627E-49 | TEC_inj | NUFIP2 |
| 6.3862E-53 | -0.2558535 | 0.149 | 0.262 | 1.5086E-48 | TEC_inj | NSD3 |
| 9.7968E-53 | -0.2686835 | 0.127 | 0.236 | 2.3143E-48 | TEC_inj | TRIM56 |
| 1.5257E-52 | -0.3202801 | 0.233 | 0.349 | 3.6042E-48 | TEC_inj | YTHDC1 |
| 2.1934E-52 | -0.3012218 | 0.191 | 0.307 | 5.1814E-48 | TEC_inj | PIM3 |
| 4.7752E-52 | -0.4459715 | 0.244 | 0.354 | 1.1281E-47 | TEC_inj | MIDN |
| 5.289E-52 | 0.46311967 | 0.386 | 0.274 | 1.2494E-47 | TEC_inj | MARCKSL1 |
| 1.7573E-51 | -0.2510174 | 0.064 | 0.158 | 4.1512E-47 | TEC_inj | SORL1 |
| 2.2187E-51 | -0.8387292 | 0.504 | 0.56 | 5.2413E-47 | TEC_inj | ELF3 |
| 3.154E-51 | -0.4433824 | 0.026 | 0.104 | 7.4508E-47 | TEC_inj | DHRS2 |
| 4.3892E-51 | 0.46655689 | 0.858 | 0.92 | 1.0369E-46 | TEC_inj | MT-ND1 |
| 1.0973E-50 | -0.9102493 | 0.089 | 0.181 | 2.5922E-46 | TEC_inj | RDH10 |
| 1.3312E-50 | -0.2802275 | 0.178 | 0.285 | 3.1446E-46 | TEC_inj | ROCK1 |
| 1.8415E-50 | -0.2628254 | 0.118 | 0.22 | 4.3502E-46 | TEC_inj | VPS13C |
| 1.879E-50 | -0.2554664 | 0.147 | 0.254 | 4.4386E-46 | TEC_inj | PTPN12 |
| 1.8879E-50 | 0.4107747 | 0.879 | 0.937 | 4.4598E-46 | TEC_inj | MT-ND4 |
| 2.0733E-50 | -0.3344885 | 0.279 | 0.389 | 4.8978E-46 | TEC_inj | N4BP2L2 |
| 4.6351E-50 | -0.2940444 | 0.134 | 0.236 | 1.095E-45 | TEC_inj | GALNT1 |
| 3.3385E-49 | -1.0802496 | 0.268 | 0.352 | 7.8864E-45 | TEC_inj | CTSL |
| 6.9485E-48 | -0.5095323 | 0.107 | 0.199 | 1.6414E-43 | TEC_inj | FBXO32 |
| 1.2694E-47 | -0.2821612 | 0.164 | 0.267 | 2.9988E-43 | TEC_inj | ABRACL |
| 1.2939E-47 | -0.2651274 | 0.112 | 0.212 | 3.0566E-43 | TEC_inj | MAP3K8 |

| | | | | | | |
|---|---|---|---|---|---|---|
| 2.1863E-47 | -0.2883311 | 0.05 | 0.132 | 5.1647E-43 | TEC_inj | POLB |
| 9.9801E-47 | -0.2655274 | 0.177 | 0.282 | 2.3576E-42 | TEC_inj | AAK1 |
| 1.2906E-46 | -0.2600037 | 0.242 | 0.355 | 3.0487E-42 | TEC_inj | ZNF207 |
| 2.2153E-46 | -0.2588774 | 0.173 | 0.278 | 5.2331E-42 | TEC_inj | FAM133B |
| 2.957E-46 | -0.474408 | 0.134 | 0.235 | 6.9854E-42 | TEC_inj | HEXIM1 |
| 7.5257E-46 | -0.2740982 | 0.037 | 0.114 | 1.7778E-41 | TEC_inj | FGL2 |
| 1.1333E-45 | -1.2132785 | 0.042 | 0.119 | 2.6772E-41 | TEC_inj | SPRR1B |
| 1.877E-45 | -0.2514854 | 0.074 | 0.161 | 4.4339E-41 | TEC_inj | HIST1H1E |
| 3.4371E-45 | -0.3284448 | 0.169 | 0.264 | 8.1194E-41 | TEC_inj | PDLIM5 |
| 5.3756E-45 | -0.2597807 | 0.104 | 0.196 | 1.2699E-40 | TEC_inj | PYCARD |
| 7.5429E-45 | -0.2991431 | 0.235 | 0.338 | 1.7819E-40 | TEC_inj | RBM47 |
| 2.5633E-44 | -0.4191235 | 0.235 | 0.336 | 6.0553E-40 | TEC_inj | SLC38A2 |
| 2.7023E-44 | -0.5096166 | 0.211 | 0.3 | 6.3836E-40 | TEC_inj | ATP13A3 |
| 7.0726E-44 | 0.45878226 | 0.849 | 0.914 | 1.6708E-39 | TEC_inj | MT-ND2 |
| 7.8435E-44 | -0.3437182 | 0.071 | 0.158 | 1.8529E-39 | TEC_inj | SLC7A11 |
| 1.1032E-43 | -0.3251702 | 0.126 | 0.219 | 2.6061E-39 | TEC_inj | NINJ1 |
| 1.4351E-43 | -0.3230762 | 0.081 | 0.167 | 3.3902E-39 | TEC_inj | SERPINB9 |
| 6.4134E-43 | -0.4110732 | 0.71 | 0.655 | 1.515E-38 | TEC_inj | MT2A |
| 8.3923E-43 | -0.6459901 | 0.163 | 0.245 | 1.9825E-38 | TEC_inj | ADGRF1 |
| 2.6728E-42 | -0.4984957 | 0.068 | 0.147 | 6.314E-38 | TEC_inj | SNCG |
| 3.716E-42 | 0.58312459 | 0.678 | 0.642 | 8.7782E-38 | TEC_inj | HSPA5 |
| 3.3893E-41 | 0.25739169 | 0.6 | 0.498 | 8.0066E-37 | TEC_inj | MT-ND4L |
| 3.7799E-41 | -0.2894429 | 0.043 | 0.116 | 8.9292E-37 | TEC_inj | SMIM5 |
| 5.7981E-41 | -0.6172632 | 0.108 | 0.2 | 1.3697E-36 | TEC_inj | HSPA6 |
| 2.5707E-40 | -0.300111 | 0.326 | 0.422 | 6.0727E-36 | TEC_inj | SFPQ |
| 1.3931E-39 | -0.2973946 | 0.042 | 0.113 | 3.2909E-35 | TEC_inj | KRT23 |
| 2.9215E-39 | -0.2946776 | 0.137 | 0.224 | 6.9014E-35 | TEC_inj | DSE |
| 8.8313E-39 | 0.48120834 | 0.543 | 0.458 | 2.0862E-34 | TEC_inj | CDKN1A |
| 2.2759E-38 | -0.2560617 | 0.214 | 0.308 | 5.3764E-34 | TEC_inj | PICALM |
| 3.06E-38 | -0.2621303 | 0.219 | 0.312 | 7.2287E-34 | TEC_inj | HP1BP3 |
| 3.3021E-38 | -0.2543725 | 0.098 | 0.181 | 7.8006E-34 | TEC_inj | KCTD12 |
| 5.5103E-38 | -0.2967121 | 0.196 | 0.283 | 1.3017E-33 | TEC_inj | VASP |
| 1.6914E-37 | -0.5455855 | 0.368 | 0.46 | 3.9957E-33 | TEC_inj | HSPH1 |
| 9.044E-37 | -0.2615385 | 0.063 | 0.137 | 2.1365E-32 | TEC_inj | CD83 |
| 1.7163E-36 | 0.25101437 | 0.81 | 0.782 | 4.0544E-32 | TEC_inj | DYNLL1 |
| 4.143E-36 | -0.2754068 | 0.168 | 0.255 | 9.7871E-32 | TEC_inj | PKN2 |
| 7.9583E-36 | -0.451956 | 0.171 | 0.255 | 1.88E-31 | TEC_inj | EGR1 |
| 8.815E-36 | -0.4712047 | 0.326 | 0.4 | 2.0824E-31 | TEC_inj | HIF1A |
| 9.9937E-36 | -0.4129949 | 0.114 | 0.189 | 2.3608E-31 | TEC_inj | SDCBP2 |
| 3.4626E-35 | -0.2842466 | 0.122 | 0.204 | 8.1797E-31 | TEC_inj | SERTAD2 |
| 7.1117E-35 | -0.4182714 | 0.269 | 0.353 | 1.68E-30 | TEC_inj | ZFP36L2 |
| 3.2953E-34 | 0.45743133 | 0.837 | 0.881 | 7.7845E-30 | TEC_inj | MT-ND3 |
| 7.0167E-34 | -0.28491 | 0.326 | 0.403 | 1.6575E-29 | TEC_inj | H2AFY |
| 9.6591E-34 | -0.3218756 | 0.267 | 0.339 | 2.2818E-29 | TEC_inj | TLN1 |
| 6.883E-33 | -0.3048573 | 0.159 | 0.241 | 1.626E-28 | TEC_inj | TOB1 |
| 1.1458E-32 | -0.8414244 | 0.057 | 0.123 | 2.7066E-28 | TEC_inj | CXCL2 |
| 1.6063E-32 | -0.2795189 | 0.151 | 0.23 | 3.7945E-28 | TEC_inj | STXBP2 |
| 1.8677E-32 | -0.3529451 | 0.251 | 0.328 | 4.4121E-28 | TEC_inj | B4GALT1 |
| 2.5054E-32 | -0.2916166 | 0.073 | 0.141 | 5.9186E-28 | TEC_inj | GCNT3 |
| 3.721E-32 | -0.471322 | 0.092 | 0.159 | 8.79E-28 | TEC_inj | CDC42EP5 |

| | | | | | | |
|---|---|---|---|---|---|---|
| 6.2717E-32 | -0.2550823 | 0.24 | 0.325 | 1.4816E-27 | TEC_inj | C6orf62 |
| 6.3677E-32 | 0.28407411 | 0.105 | 0.055 | 1.5043E-27 | TEC_inj | NEFM |
| 1.2192E-31 | -0.2778829 | 0.214 | 0.295 | 2.8801E-27 | TEC_inj | CHMP1B |
| 1.3291E-31 | -0.315848 | 0.153 | 0.233 | 3.1398E-27 | TEC_inj | IER5L |
| 2.0014E-31 | -0.2956041 | 0.286 | 0.366 | 4.728E-27 | TEC_inj | CFLAR |
| 2.4283E-31 | -0.3347443 | 0.097 | 0.17 | 5.7364E-27 | TEC_inj | ANKRD37 |
| 4.5262E-31 | -0.2840109 | 0.363 | 0.434 | 1.0692E-26 | TEC_inj | IQGAP1 |
| 5.5417E-31 | -0.2569891 | 0.066 | 0.134 | 1.3091E-26 | TEC_inj | AC023157.3 |
| 6.7414E-31 | -0.2758032 | 0.248 | 0.328 | 1.5925E-26 | TEC_inj | YPEL3 |
| 1.116E-30 | -0.4028711 | 0.233 | 0.304 | 2.6363E-26 | TEC_inj | FAM129B |
| 1.3862E-30 | -0.7628452 | 0.117 | 0.186 | 3.2747E-26 | TEC_inj | SPINK1 |
| 3.2726E-30 | 0.25862147 | 0.276 | 0.199 | 7.7308E-26 | TEC_inj | UBE2S |
| 6.8115E-30 | -0.3993236 | 0.264 | 0.328 | 1.6091E-25 | TEC_inj | LIMS1 |
| 7.1981E-30 | -0.2716805 | 0.656 | 0.714 | 1.7004E-25 | TEC_inj | SDCBP |
| 8.4539E-30 | -0.2531466 | 0.064 | 0.127 | 1.9971E-25 | TEC_inj | KRT80 |
| 9.2086E-30 | 0.34445343 | 0.874 | 0.935 | 2.1754E-25 | TEC_inj | MT-CYB |
| 1.1782E-29 | -0.2507972 | 0.198 | 0.277 | 2.7833E-25 | TEC_inj | OSBPL8 |
| 2.0776E-29 | -0.3998159 | 0.896 | 0.943 | 4.9079E-25 | TEC_inj | SAT1 |
| 6.9155E-29 | -0.2561848 | 0.637 | 0.762 | 1.6337E-24 | TEC_inj | TPM4 |
| 4.8349E-28 | -1.0637457 | 0.144 | 0.085 | 1.1421E-23 | TEC_inj | HBB |
| 5.5355E-28 | -0.5244016 | 0.288 | 0.192 | 1.3076E-23 | TEC_inj | THBS1 |
| 4.8296E-27 | -0.2704147 | 0.199 | 0.28 | 1.1409E-22 | TEC_inj | TNFAIP2 |
| 2.0677E-26 | -0.2542067 | 0.118 | 0.183 | 4.8845E-22 | TEC_inj | LTA4H |
| 2.7122E-26 | -0.3759639 | 0.212 | 0.274 | 6.4071E-22 | TEC_inj | CCNG2 |
| 3.7608E-26 | -0.2543756 | 0.798 | 0.834 | 8.8841E-22 | TEC_inj | HNRNPA2B1 |
| 1.9544E-25 | -0.2615441 | 0.174 | 0.243 | 4.6169E-21 | TEC_inj | RNASET2 |
| 6.0314E-25 | -0.2536864 | 0.63 | 0.74 | 1.4248E-20 | TEC_inj | HSPA8 |
| 7.5322E-25 | -0.2546948 | 0.355 | 0.421 | 1.7793E-20 | TEC_inj | FUS |
| 1.1849E-24 | -0.2781503 | 0.1 | 0.159 | 2.7991E-20 | TEC_inj | TMPRSS4 |
| 1.6992E-24 | -0.3224649 | 0.221 | 0.284 | 4.0141E-20 | TEC_inj | ABHD5 |
| 1.9928E-24 | -0.416581 | 0.162 | 0.227 | 4.7076E-20 | TEC_inj | HES1 |
| 2.5048E-24 | -0.2948968 | 0.992 | 0.989 | 5.9171E-20 | TEC_inj | B2M |
| 3.1254E-24 | -0.3545553 | 0.104 | 0.166 | 7.3832E-20 | TEC_inj | CSKMT |
| 7.7237E-24 | -0.6370703 | 0.264 | 0.316 | 1.8246E-19 | TEC_inj | EMP1 |
| 9.8394E-24 | -0.5272039 | 0.172 | 0.235 | 2.3244E-19 | TEC_inj | ISG15 |
| 1.1818E-23 | -0.3443852 | 0.172 | 0.235 | 2.7918E-19 | TEC_inj | PPIF |
| 1.8058E-23 | -0.305491 | 0.111 | 0.17 | 4.2659E-19 | TEC_inj | HIST3H2A |
| 9.6415E-23 | -0.2805143 | 0.212 | 0.28 | 2.2776E-18 | TEC_inj | ADM |
| 2.1585E-21 | -0.2620465 | 0.108 | 0.162 | 5.099E-17 | TEC_inj | TTC9 |
| 6.3079E-21 | -0.4079101 | 0.127 | 0.179 | 1.4901E-16 | TEC_inj | SPNS2 |
| 1.6258E-19 | -0.3048506 | 0.327 | 0.364 | 3.8405E-15 | TEC_inj | GNAI2 |
| 1.6536E-19 | -0.5399622 | 0.737 | 0.788 | 3.9062E-15 | TEC_inj | LGALS3 |
| 4.4069E-19 | -0.2679192 | 0.076 | 0.123 | 1.041E-14 | TEC_inj | EGLN3 |
| 1.2638E-18 | -0.289578 | 0.082 | 0.132 | 2.9855E-14 | TEC_inj | GBP1 |
| 2.1296E-18 | -1.1440691 | 0.194 | 0.235 | 5.0307E-14 | TEC_inj | PRSS22 |
| 3.3342E-18 | -0.3922122 | 0.127 | 0.175 | 7.8763E-14 | TEC_inj | HSPB8 |
| 5.529E-18 | -0.6848049 | 0.89 | 0.883 | 1.3061E-13 | TEC_inj | CSTB |
| 1.0405E-17 | 0.29639558 | 0.876 | 0.943 | 2.458E-13 | TEC_inj | MT-ATP6 |
| 1.0405E-17 | -0.2883601 | 0.097 | 0.145 | 2.458E-13 | TEC_inj | ID1 |
| 3.0749E-16 | -1.1567878 | 0.465 | 0.493 | 7.2638E-12 | TEC_inj | CD74 |

| | | | | | | |
|---|---|---|---|---|---|---|
| 1.6324E-15 | -0.5673049 | 0.264 | 0.294 | 3.8563E-11 | TEC_inj | RND3 |
| 2.8006E-15 | -0.9473321 | 0.997 | 0.993 | 6.6159E-11 | TEC_inj | FTH1 |
| 6.9233E-15 | -0.4609035 | 0.098 | 0.143 | 1.6355E-10 | TEC_inj | CXCL1 |
| 4.7931E-14 | -1.2215072 | 0.105 | 0.148 | 1.1323E-09 | TEC_inj | CXCL3 |
| 2.1354E-12 | -0.4962383 | 0.151 | 0.187 | 5.0445E-08 | TEC_inj | AQP3 |
| 5.2005E-12 | 0.29484259 | 0.408 | 0.355 | 1.2285E-07 | TEC_inj | TUBA1B |
| 1.292E-11 | -0.3021149 | 0.117 | 0.155 | 3.052E-07 | TEC_inj | ZFAND2A |
| 1.4668E-11 | -0.3436138 | 0.247 | 0.285 | 3.465E-07 | TEC_inj | RHOB |
| 4.4869E-11 | -0.6197648 | 0.419 | 0.453 | 1.0599E-06 | TEC_inj | SOX4 |
| 1.0618E-09 | -0.2578621 | 0.364 | 0.393 | 2.5083E-05 | TEC_inj | SRSF7 |
| 7.048E-08 | -1.5073637 | 0.124 | 0.154 | 0.00166496 | TEC_inj | MT1H |
| 1.8246E-07 | -0.3029985 | 0.376 | 0.407 | 0.00431029 | TEC_inj | RNF213 |
| 1.8468E-07 | -0.3972413 | 0.282 | 0.311 | 0.0043628 | TEC_inj | HIST1H4C |
| 2.431E-06 | -0.2716592 | 0.237 | 0.251 | 0.05742762 | TEC_inj | KLF5 |
| 3.3965E-06 | -0.538568 | 0.165 | 0.19 | 0.08023492 | TEC_inj | RARRES3 |
| 7.4742E-06 | -0.4770212 | 0.161 | 0.128 | 0.17656391 | TEC_inj | APOE |
| 7.7532E-06 | -0.3047448 | 0.116 | 0.137 | 0.18315347 | TEC_inj | HLA-DRB5 |
| 1.817E-05 | -0.6420473 | 0.991 | 0.986 | 0.42924037 | TEC_inj | FTL |
| 1.9596E-05 | -0.3385022 | 0.709 | 0.742 | 0.46290779 | TEC_inj | VIM |
| 2.0029E-05 | -0.2731524 | 0.146 | 0.114 | 0.47314515 | TEC_inj | HSPA2 |
| 3.6004E-05 | -1.2050309 | 0.096 | 0.117 | 0.85051525 | TEC_inj | MT1M |
| 0.0001114 | -0.7007396 | 0.299 | 0.295 | 1 | TEC_inj | ADIRF |
| 0.00017454 | -0.3670529 | 0.292 | 0.292 | 1 | TEC_inj | GPRC5A |
| 0.00035706 | 0.53587327 | 0.263 | 0.305 | 1 | TEC_inj | SERPINA1 |
| 0.0045559 | -0.5389608 | 0.103 | 0.085 | 1 | TEC_inj | SPRR2D |
| 0.00657506 | -0.2736327 | 0.583 | 0.535 | 1 | TEC_inj | PLAUR |
| 0 | 1.38171845 | 0.689 | 0.395 | 0 | TEC_dmg1 | TXNRD1 |
| 0 | 1.34439451 | 0.507 | 0.149 | 0 | TEC_dmg1 | LAMC2 |
| 0 | 1.29720608 | 0.816 | 0.628 | 0 | TEC_dmg1 | HSPA5 |
| 0 | 1.29251905 | 0.593 | 0.184 | 0 | TEC_dmg1 | CCND1 |
| 0 | 1.26741294 | 0.532 | 0.199 | 0 | TEC_dmg1 | ITGA2 |
| 0 | 1.2355969 | 0.593 | 0.214 | 0 | TEC_dmg1 | CRYAB |
| 0 | 1.16088907 | 0.62 | 0.177 | 0 | TEC_dmg1 | ZBTB20 |
| 0 | 1.10512795 | 0.601 | 0.225 | 0 | TEC_dmg1 | TPM1 |
| 0 | 1.08309502 | 0.716 | 0.187 | 0 | TEC_dmg1 | EPCAM |
| 0 | 1.07605153 | 0.496 | 0.217 | 0 | TEC_dmg1 | ITGB8 |
| 0 | 1.06704946 | 0.672 | 0.233 | 0 | TEC_dmg1 | MAL2 |
| 0 | 1.02414557 | 0.422 | 0.1 | 0 | TEC_dmg1 | AKAP12 |
| 0 | 0.99437896 | 0.651 | 0.32 | 0 | TEC_dmg1 | GLS |
| 0 | 0.96291698 | 0.831 | 0.511 | 0 | TEC_dmg1 | CD24 |
| 0 | 0.95088 | 0.595 | 0.233 | 0 | TEC_dmg1 | UGCG |
| 0 | 0.91698944 | 0.631 | 0.287 | 0 | TEC_dmg1 | GADD45A |
| 0 | 0.90848639 | 0.458 | 0.154 | 0 | TEC_dmg1 | GCLM |
| 0 | 0.89990177 | 0.64 | 0.237 | 0 | TEC_dmg1 | TNFRSF12A |
| 0 | 0.88985285 | 0.453 | 0.14 | 0 | TEC_dmg1 | AKR1B1 |
| 0 | 0.88110366 | 0.633 | 0.202 | 0 | TEC_dmg1 | S100A13 |
| 0 | 0.86206334 | 0.893 | 0.731 | 0 | TEC_dmg1 | PRDX1 |
| 0 | 0.85720397 | 0.714 | 0.462 | 0 | TEC_dmg1 | HMGA1 |
| 0 | 0.85564125 | 0.864 | 0.527 | 0 | TEC_dmg1 | KRT18 |
| 0 | 0.83053628 | 0.682 | 0.4 | 0 | TEC_dmg1 | SLC3A2 |

| | | | | | | |
|---|---|---|---|---|---|---|
| 0 | 0.81303142 | 0.44 | 0.106 | 0 | TEC_dmg1 | ARHGAP29 |
| 0 | 0.80974211 | 0.432 | 0.107 | 0 | TEC_dmg1 | DCDC2 |
| 0 | 0.78019478 | 0.582 | 0.188 | 0 | TEC_dmg1 | OCIAD2 |
| 0 | 0.77417802 | 0.458 | 0.188 | 0 | TEC_dmg1 | LAMB3 |
| 0 | 0.74828124 | 0.517 | 0.133 | 0 | TEC_dmg1 | SLC38A1 |
| 0 | 0.74621626 | 0.717 | 0.377 | 0 | TEC_dmg1 | ATF4 |
| 0 | 0.73607933 | 0.864 | 0.637 | 0 | TEC_dmg1 | CALM2 |
| 0 | 0.73189573 | 0.712 | 0.298 | 0 | TEC_dmg1 | IGFBP7 |
| 0 | 0.70008243 | 0.531 | 0.185 | 0 | TEC_dmg1 | DSP |
| 0 | 0.69719204 | 0.651 | 0.299 | 0 | TEC_dmg1 | APP |
| 0 | 0.68898132 | 0.553 | 0.15 | 0 | TEC_dmg1 | PAWR |
| 0 | 0.68547191 | 0.88 | 0.666 | 0 | TEC_dmg1 | NPM1 |
| 0 | 0.68092243 | 0.329 | 0.069 | 0 | TEC_dmg1 | ADAMTS9 |
| 0 | 0.67926095 | 0.86 | 0.649 | 0 | TEC_dmg1 | RTN4 |
| 0 | 0.67505992 | 0.456 | 0.121 | 0 | TEC_dmg1 | ITGA3 |
| 0 | 0.66911614 | 0.491 | 0.152 | 0 | TEC_dmg1 | CDH1 |
| 0 | 0.66567616 | 0.752 | 0.394 | 0 | TEC_dmg1 | SPINT2 |
| 0 | 0.66259707 | 0.451 | 0.183 | 0 | TEC_dmg1 | MACC1 |
| 0 | 0.65305112 | 0.367 | 0.133 | 0 | TEC_dmg1 | CYR61 |
| 0 | 0.64836285 | 0.546 | 0.217 | 0 | TEC_dmg1 | SPTBN1 |
| 0 | 0.64659934 | 0.466 | 0.131 | 0 | TEC_dmg1 | MET |
| 0 | 0.64483629 | 0.868 | 0.672 | 0 | TEC_dmg1 | EEF2 |
| 0 | 0.63109009 | 0.72 | 0.408 | 0 | TEC_dmg1 | CYSTM1 |
| 0 | 0.62906644 | 0.593 | 0.283 | 0 | TEC_dmg1 | SBDS |
| 0 | 0.62770636 | 0.534 | 0.207 | 0 | TEC_dmg1 | S100A16 |
| 0 | 0.62639842 | 0.559 | 0.266 | 0 | TEC_dmg1 | TUBA1C |
| 0 | 0.62605946 | 0.426 | 0.114 | 0 | TEC_dmg1 | PAX8 |
| 0 | 0.62028065 | 0.312 | 0.067 | 0 | TEC_dmg1 | DCBLD2 |
| 0 | 0.61909083 | 0.321 | 0.051 | 0 | TEC_dmg1 | FGF2 |
| 0 | 0.61588306 | 0.531 | 0.241 | 0 | TEC_dmg1 | HSPA9 |
| 0 | 0.6143647 | 0.417 | 0.13 | 0 | TEC_dmg1 | CLDN3 |
| 0 | 0.60069189 | 0.534 | 0.226 | 0 | TEC_dmg1 | ODC1 |
| 0 | 0.58340731 | 0.378 | 0.121 | 0 | TEC_dmg1 | CLDN1 |
| 0 | 0.58283849 | 0.356 | 0.079 | 0 | TEC_dmg1 | DPP4 |
| 0 | 0.5747586 | 0.562 | 0.231 | 0 | TEC_dmg1 | RHOC |
| 0 | 0.57187267 | 0.478 | 0.193 | 0 | TEC_dmg1 | TES |
| 0 | 0.56462018 | 0.368 | 0.11 | 0 | TEC_dmg1 | UCHL1 |
| 0 | 0.56207958 | 0.427 | 0.143 | 0 | TEC_dmg1 | GARS |
| 0 | 0.55850153 | 0.451 | 0.152 | 0 | TEC_dmg1 | ANXA3 |
| 0 | 0.55055762 | 0.403 | 0.135 | 0 | TEC_dmg1 | SELENOM |
| 0 | 0.54907395 | 0.444 | 0.141 | 0 | TEC_dmg1 | EPRS |
| 0 | 0.54790054 | 0.54 | 0.236 | 0 | TEC_dmg1 | TSPAN1 |
| 0 | 0.54746476 | 0.606 | 0.318 | 0 | TEC_dmg1 | NORAD |
| 0 | 0.5437953 | 0.294 | 0.049 | 0 | TEC_dmg1 | PROM1 |
| 0 | 0.53845977 | 0.447 | 0.138 | 0 | TEC_dmg1 | CCDC50 |
| 0 | 0.53591424 | 0.371 | 0.133 | 0 | TEC_dmg1 | SESN2 |
| 0 | 0.53518038 | 0.616 | 0.333 | 0 | TEC_dmg1 | TOMM20 |
| 0 | 0.53430911 | 0.262 | 0.043 | 0 | TEC_dmg1 | VGLL3 |
| 0 | 0.53223707 | 0.537 | 0.231 | 0 | TEC_dmg1 | ARHGAP5 |
| 0 | 0.5320927 | 0.512 | 0.226 | 0 | TEC_dmg1 | CD2AP |

| | | | | | | |
|---|---|---|---|---|---|---|
| 0 | 0.52951497 | 0.419 | 0.114 | 0 | TEC_dmg1 | OCLN |
| 0 | 0.52104504 | 0.41 | 0.135 | 0 | TEC_dmg1 | GFPT1 |
| 0 | 0.51769068 | 0.413 | 0.117 | 0 | TEC_dmg1 | CRIM1 |
| 0 | 0.51733098 | 0.442 | 0.165 | 0 | TEC_dmg1 | LIMA1 |
| 0 | 0.51274563 | 0.389 | 0.143 | 0 | TEC_dmg1 | EPB41L4A-AS1 |
| 0 | 0.51234314 | 0.541 | 0.271 | 0 | TEC_dmg1 | TRAM1 |
| 0 | 0.51130935 | 0.52 | 0.202 | 0 | TEC_dmg1 | TJP1 |
| 0 | 0.51114673 | 0.377 | 0.114 | 0 | TEC_dmg1 | MGLL |
| 0 | 0.50862391 | 0.283 | 0.064 | 0 | TEC_dmg1 | CALD1 |
| 0 | 0.50780433 | 0.385 | 0.094 | 0 | TEC_dmg1 | CTNNAL1 |
| 0 | 0.49967864 | 0.505 | 0.164 | 0 | TEC_dmg1 | C12orf75 |
| 0 | 0.4993483 | 0.437 | 0.133 | 0 | TEC_dmg1 | NCKAP1 |
| 0 | 0.49866264 | 0.33 | 0.076 | 0 | TEC_dmg1 | PHLDB2 |
| 0 | 0.49719185 | 0.458 | 0.189 | 0 | TEC_dmg1 | LMAN1 |
| 0 | 0.49475782 | 0.488 | 0.223 | 0 | TEC_dmg1 | CREBRF |
| 0 | 0.49120484 | 0.488 | 0.199 | 0 | TEC_dmg1 | ARPP19 |
| 0 | 0.48583373 | 0.36 | 0.078 | 0 | TEC_dmg1 | LINC00958 |
| 0 | 0.48560133 | 0.3 | 0.084 | 0 | TEC_dmg1 | TRIB3 |
| 0 | 0.48418947 | 0.642 | 0.319 | 0 | TEC_dmg1 | PEBP1 |
| 0 | 0.48139611 | 0.393 | 0.119 | 0 | TEC_dmg1 | GPRC5C |
| 0 | 0.47212299 | 0.404 | 0.157 | 0 | TEC_dmg1 | TEAD1 |
| 0 | 0.46591231 | 0.398 | 0.116 | 0 | TEC_dmg1 | LAPTM4B |
| 0 | 0.46180034 | 0.374 | 0.115 | 0 | TEC_dmg1 | GULP1 |
| 0 | 0.45992232 | 0.294 | 0.061 | 0 | TEC_dmg1 | CAVIN1 |
| 0 | 0.44768412 | 0.465 | 0.205 | 0 | TEC_dmg1 | IL6ST |
| 0 | 0.4363217 | 0.323 | 0.075 | 0 | TEC_dmg1 | GRAMD2B |
| 0 | 0.43171365 | 0.332 | 0.091 | 0 | TEC_dmg1 | WWC1 |
| 0 | 0.42942694 | 0.329 | 0.053 | 0 | TEC_dmg1 | BEND7 |
| 0 | 0.42890436 | 0.315 | 0.08 | 0 | TEC_dmg1 | EGFR |
| 0 | 0.42665193 | 0.45 | 0.183 | 0 | TEC_dmg1 | MAP4 |
| 0 | 0.41638124 | 0.411 | 0.157 | 0 | TEC_dmg1 | SARS |
| 0 | 0.41601276 | 0.346 | 0.1 | 0 | TEC_dmg1 | RAB3IP |
| 0 | 0.41419711 | 0.346 | 0.124 | 0 | TEC_dmg1 | SVIL |
| 0 | 0.41332801 | 0.399 | 0.137 | 0 | TEC_dmg1 | FARP1 |
| 0 | 0.41302553 | 0.206 | 0.026 | 0 | TEC_dmg1 | PXDN |
| 0 | 0.41239755 | 0.342 | 0.11 | 0 | TEC_dmg1 | PPFIBP1 |
| 0 | 0.40961871 | 0.417 | 0.173 | 0 | TEC_dmg1 | MED10 |
| 0 | 0.40875991 | 0.352 | 0.106 | 0 | TEC_dmg1 | BEX2 |
| 0 | 0.40640815 | 0.291 | 0.067 | 0 | TEC_dmg1 | ASNS |
| 0 | 0.38994643 | 0.32 | 0.089 | 0 | TEC_dmg1 | WWTR1 |
| 0 | 0.38881828 | 0.303 | 0.085 | 0 | TEC_dmg1 | GOT1 |
| 0 | 0.38830009 | 0.318 | 0.095 | 0 | TEC_dmg1 | DSG2 |
| 0 | 0.38739545 | 0.308 | 0.077 | 0 | TEC_dmg1 | SH3BP4 |
| 0 | 0.38701981 | 0.36 | 0.119 | 0 | TEC_dmg1 | CEBPG |
| 0 | 0.3862988 | 0.267 | 0.05 | 0 | TEC_dmg1 | RASSF8 |
| 0 | 0.38373642 | 0.395 | 0.147 | 0 | TEC_dmg1 | GNG12 |
| 0 | 0.38274515 | 0.231 | 0.033 | 0 | TEC_dmg1 | MAP2 |
| 0 | 0.38207368 | 0.28 | 0.06 | 0 | TEC_dmg1 | PHGDH |
| 0 | 0.37522445 | 0.275 | 0.053 | 0 | TEC_dmg1 | ARHGEF35 |
| 0 | 0.36163664 | 0.307 | 0.101 | 0 | TEC_dmg1 | AMOTL2 |

| | | | | | | |
|---|---|---|---|---|---|---|
| 0 | 0.35805005 | 0.278 | 0.064 | 0 | TEC_dmg1 | SLC7A1 |
| 0 | 0.35376674 | 0.249 | 0.064 | 0 | TEC_dmg1 | RAPGEF3 |
| 0 | 0.35349278 | 0.34 | 0.11 | 0 | TEC_dmg1 | DMKN |
| 0 | 0.34758037 | 0.278 | 0.07 | 0 | TEC_dmg1 | PGM3 |
| 0 | 0.3469285 | 0.25 | 0.058 | 0 | TEC_dmg1 | F2RL1 |
| 0 | 0.34561928 | 0.3 | 0.077 | 0 | TEC_dmg1 | FERMT2 |
| 0 | 0.33866452 | 0.257 | 0.055 | 0 | TEC_dmg1 | BICC1 |
| 0 | 0.33305832 | 0.291 | 0.079 | 0 | TEC_dmg1 | CHMP4C |
| 0 | 0.33191611 | 0.238 | 0.046 | 0 | TEC_dmg1 | SCD5 |
| 0 | 0.33095246 | 0.261 | 0.057 | 0 | TEC_dmg1 | SGPP2 |
| 0 | 0.32495368 | 0.234 | 0.063 | 0 | TEC_dmg1 | AL353751.1 |
| 0 | 0.32189584 | 0.275 | 0.074 | 0 | TEC_dmg1 | WWC2 |
| 0 | 0.31987456 | 0.288 | 0.085 | 0 | TEC_dmg1 | IARS |
| 0 | 0.31573835 | 0.256 | 0.061 | 0 | TEC_dmg1 | PTPN14 |
| 0 | 0.31293231 | 0.295 | 0.089 | 0 | TEC_dmg1 | YES1 |
| 0 | 0.31191524 | 0.259 | 0.056 | 0 | TEC_dmg1 | DDAH1 |
| 0 | 0.30741682 | 0.266 | 0.069 | 0 | TEC_dmg1 | SCRN1 |
| 0 | 0.30485649 | 0.266 | 0.076 | 0 | TEC_dmg1 | CAV2 |
| 0 | 0.30289679 | 0.238 | 0.053 | 0 | TEC_dmg1 | L1CAM |
| 0 | 0.29114601 | 0.252 | 0.06 | 0 | TEC_dmg1 | LARP6 |
| 0 | 0.29053489 | 0.265 | 0.075 | 0 | TEC_dmg1 | KIAA1217 |
| 0 | 0.28981813 | 0.256 | 0.061 | 0 | TEC_dmg1 | PPP2R3A |
| 0 | 0.28938668 | 0.267 | 0.076 | 0 | TEC_dmg1 | RRAS2 |
| 0 | 0.28262437 | 0.221 | 0.041 | 0 | TEC_dmg1 | PRKAA2 |
| 0 | 0.28037544 | 0.255 | 0.062 | 0 | TEC_dmg1 | MYEF2 |
| 0 | 0.27984861 | 0.235 | 0.048 | 0 | TEC_dmg1 | ARHGEF5 |
| 0 | 0.27648106 | 0.249 | 0.066 | 0 | TEC_dmg1 | LAMC1 |
| 0 | 0.27159345 | 0.245 | 0.055 | 0 | TEC_dmg1 | TOM1L1 |
| 0 | 0.27028233 | 0.25 | 0.071 | 0 | TEC_dmg1 | SYNJ2BP |
| 0 | 0.26675893 | 0.222 | 0.042 | 0 | TEC_dmg1 | HOXB9 |
| 0 | 0.26487852 | 0.236 | 0.056 | 0 | TEC_dmg1 | PVR |
| 0 | 0.26094051 | 0.239 | 0.063 | 0 | TEC_dmg1 | GLIS3 |
| 0 | 0.26047477 | 0.23 | 0.052 | 0 | TEC_dmg1 | CMTM4 |
| 0 | -0.7979391 | 0.026 | 0.355 | 0 | TEC_dmg1 | ARHGDIB |
| 0 | -0.9642738 | 0.975 | 0.991 | 0 | TEC_dmg1 | B2M |
| 0 | -1.0200953 | 0.44 | 0.725 | 0 | TEC_dmg1 | JUNB |
| 0 | -1.0272706 | 0.492 | 0.751 | 0 | TEC_dmg1 | CYBA |
| 0 | -1.0781146 | 0.246 | 0.559 | 0 | TEC_dmg1 | CTSS |
| 0 | -1.0990628 | 0.24 | 0.556 | 0 | TEC_dmg1 | ZFP36 |
| 0 | -1.1697485 | 0.011 | 0.308 | 0 | TEC_dmg1 | PTPRC |
| 0 | -1.2682224 | 0.029 | 0.366 | 0 | TEC_dmg1 | LAPTM5 |
| 0 | -1.4931468 | 0.358 | 0.672 | 0 | TEC_dmg1 | GLUL |
| 0 | -1.5377537 | 0.447 | 0.704 | 0 | TEC_dmg1 | NAMPT |
| 0 | -1.6698644 | 0.287 | 0.659 | 0 | TEC_dmg1 | S100A4 |
| 0 | -1.7259304 | 0.085 | 0.415 | 0 | TEC_dmg1 | FCER1G |
| 0 | -1.7516204 | 0.117 | 0.472 | 0 | TEC_dmg1 | TYROBP |
| 0 | -1.8095137 | 0.332 | 0.747 | 0 | TEC_dmg1 | ANXA1 |
| 0 | -1.8241431 | 0.287 | 0.646 | 0 | TEC_dmg1 | C15orf48 |
| 0 | -2.3369373 | 0.153 | 0.495 | 0 | TEC_dmg1 | LYZ |
| 0 | -2.6552945 | 0.192 | 0.608 | 0 | TEC_dmg1 | SRGN |

| | | | | | | |
|---|---|---|---|---|---|---|
| 0 | -4.061975 | 0.244 | 0.714 | 0 | TEC_dmg1 | S100A8 |
| 0 | -4.8148303 | 0.292 | 0.771 | 0 | TEC_dmg1 | S100A9 |
| 4.688E-308 | 0.39908684 | 0.396 | 0.159 | 1.107E-303 | TEC_dmg1 | RBCK1 |
| 9.145E-308 | 0.34484852 | 0.336 | 0.118 | 2.16E-303 | TEC_dmg1 | MYO1E |
| 1.225E-307 | 0.31122949 | 0.28 | 0.088 | 2.894E-303 | TEC_dmg1 | RHOD |
| 4.117E-307 | 0.9708818 | 0.856 | 0.676 | 9.726E-303 | TEC_dmg1 | HSP90B1 |
| 9.657E-307 | -1.9807348 | 0.127 | 0.423 | 2.281E-302 | TEC_dmg1 | HLA-DRA |
| 1.491E-306 | 0.31175177 | 0.269 | 0.082 | 3.523E-302 | TEC_dmg1 | HSPA13 |
| 2.584E-306 | 0.52972167 | 0.524 | 0.249 | 6.104E-302 | TEC_dmg1 | JUP |
| 2.252E-305 | 0.60833079 | 0.987 | 0.98 | 5.32E-301 | TEC_dmg1 | EEF1A1 |
| 4.376E-305 | 0.56157734 | 0.728 | 0.463 | 1.034E-300 | TEC_dmg1 | CANX |
| 6.218E-305 | 0.26417207 | 0.224 | 0.059 | 1.469E-300 | TEC_dmg1 | TCAF1 |
| 9.308E-305 | -2.3952119 | 0.151 | 0.433 | 2.199E-300 | TEC_dmg1 | S100P |
| 1.489E-304 | -0.7438923 | 0.012 | 0.293 | 3.516E-300 | TEC_dmg1 | CD53 |
| 1.204E-303 | 0.26175044 | 0.216 | 0.056 | 2.844E-299 | TEC_dmg1 | SAMD4A |
| 2.068E-303 | -1.5249368 | 0.069 | 0.36 | 4.886E-299 | TEC_dmg1 | IL1RN |
| 9.142E-302 | 0.37096503 | 0.376 | 0.145 | 2.16E-297 | TEC_dmg1 | SUN1 |
| 3.629E-301 | 0.37767028 | 0.36 | 0.136 | 8.574E-297 | TEC_dmg1 | NFE2L1 |
| 1.066E-300 | 0.57079849 | 0.666 | 0.383 | 2.518E-296 | TEC_dmg1 | CD59 |
| 7.347E-299 | 0.5284831 | 0.551 | 0.283 | 1.736E-294 | TEC_dmg1 | PFDN2 |
| 1.427E-298 | 0.39893477 | 0.367 | 0.135 | 3.371E-294 | TEC_dmg1 | COBLL1 |
| 1.852E-298 | -1.267022 | 0.021 | 0.3 | 4.375E-294 | TEC_dmg1 | AIF1 |
| 2.512E-298 | 0.25255581 | 0.203 | 0.051 | 5.934E-294 | TEC_dmg1 | TMEM132A |
| 3.244E-298 | 0.43317125 | 0.318 | 0.113 | 7.664E-294 | TEC_dmg1 | DNAJB9 |
| 2.126E-297 | 0.5011278 | 0.4 | 0.159 | 5.022E-293 | TEC_dmg1 | S100A14 |
| 2.699E-296 | 0.25883392 | 0.275 | 0.085 | 6.377E-292 | TEC_dmg1 | EPB41L5 |
| 5.988E-296 | 0.28880139 | 0.238 | 0.067 | 1.415E-291 | TEC_dmg1 | KHDRBS3 |
| 5.007E-294 | 0.31291479 | 0.279 | 0.089 | 1.183E-289 | TEC_dmg1 | SGMS2 |
| 4.082E-292 | 0.47168304 | 0.343 | 0.129 | 9.644E-288 | TEC_dmg1 | SCCPDH |
| 2.317E-291 | 0.3343674 | 0.202 | 0.051 | 5.474E-287 | TEC_dmg1 | RSAD2 |
| 1.413E-290 | 0.56810639 | 0.515 | 0.262 | 3.339E-286 | TEC_dmg1 | RB1CC1 |
| 1.846E-290 | 0.26201985 | 0.226 | 0.061 | 4.361E-286 | TEC_dmg1 | TFCP2L1 |
| 2.826E-290 | 0.33778447 | 0.348 | 0.129 | 6.675E-286 | TEC_dmg1 | SINHCAF |
| 1.263E-287 | -0.959039 | 0.04 | 0.316 | 2.983E-283 | TEC_dmg1 | LCP1 |
| 1.324E-286 | 0.86490984 | 0.567 | 0.306 | 3.127E-282 | TEC_dmg1 | HERPUD1 |
| 9.23E-284 | 0.53926922 | 0.512 | 0.254 | 2.181E-279 | TEC_dmg1 | MYDGF |
| 2.336E-283 | 0.37192717 | 0.369 | 0.147 | 5.519E-279 | TEC_dmg1 | STX3 |
| 2.037E-282 | 0.30001897 | 0.544 | 0.248 | 4.812E-278 | TEC_dmg1 | CAMK2N1 |
| 3.3E-282 | 0.53245468 | 0.561 | 0.287 | 7.795E-278 | TEC_dmg1 | SDC4 |
| 1.958E-281 | 0.28500591 | 0.287 | 0.094 | 4.625E-277 | TEC_dmg1 | RHOBTB3 |
| 5.807E-281 | 0.51271537 | 0.598 | 0.32 | 1.372E-276 | TEC_dmg1 | GOLGB1 |
| 6.067E-281 | 0.4271145 | 0.399 | 0.167 | 1.433E-276 | TEC_dmg1 | VCL |
| 1.797E-280 | 0.29206561 | 0.16 | 0.035 | 4.245E-276 | TEC_dmg1 | IQCD |
| 2.696E-280 | 0.49079699 | 0.783 | 0.464 | 6.368E-276 | TEC_dmg1 | KRT8 |
| 3.58E-280 | -0.6629276 | 0.008 | 0.27 | 8.458E-276 | TEC_dmg1 | GMFG |
| 4.79E-279 | 0.37007362 | 0.372 | 0.152 | 1.132E-274 | TEC_dmg1 | MCFD2 |
| 5.51E-279 | 0.50294483 | 0.972 | 0.876 | 1.302E-274 | TEC_dmg1 | HSP90AB1 |
| 1.807E-276 | 0.45382655 | 0.505 | 0.252 | 4.268E-272 | TEC_dmg1 | RSL24D1 |
| 5.547E-276 | 0.30154782 | 0.297 | 0.102 | 1.31E-271 | TEC_dmg1 | MYO10 |
| 1.966E-275 | 0.31946689 | 0.346 | 0.133 | 4.645E-271 | TEC_dmg1 | SCOC |

| | | | | | | |
|---|---|---|---|---|---|---|
| 2.409E-274 | -0.7813913 | 0.01 | 0.269 | 5.69E-270 | TEC_dmg1 | CSTA |
| 1.808E-273 | 0.37097648 | 0.389 | 0.163 | 4.27E-269 | TEC_dmg1 | UBE2Z |
| 4.159E-273 | 0.26329183 | 0.264 | 0.085 | 9.825E-269 | TEC_dmg1 | TMEM245 |
| 2.214E-272 | 0.28872935 | 0.279 | 0.093 | 5.23E-268 | TEC_dmg1 | PARD3 |
| 4.551E-272 | 0.54553061 | 0.459 | 0.204 | 1.075E-267 | TEC_dmg1 | IL32 |
| 1.086E-271 | -0.6892307 | 0.176 | 0.453 | 2.567E-267 | TEC_dmg1 | EFHD2 |
| 4.29E-271 | 0.52429515 | 0.592 | 0.316 | 1.013E-266 | TEC_dmg1 | PDIA6 |
| 4.638E-271 | -2.2362423 | 0.102 | 0.365 | 1.096E-266 | TEC_dmg1 | PSCA |
| 9.998E-269 | 0.54442141 | 0.559 | 0.311 | 2.362E-264 | TEC_dmg1 | SSR3 |
| 1.012E-267 | 0.75760653 | 0.318 | 0.119 | 2.389E-263 | TEC_dmg1 | NUPR1 |
| 4.134E-267 | 0.32120067 | 0.365 | 0.146 | 9.766E-263 | TEC_dmg1 | GLO1 |
| 3.413E-266 | 0.31321329 | 0.347 | 0.134 | 8.063E-262 | TEC_dmg1 | RCN1 |
| 5.481E-266 | 0.71358811 | 0.492 | 0.244 | 1.295E-261 | TEC_dmg1 | DDIT3 |
| 6.837E-266 | 0.58798927 | 0.817 | 0.511 | 1.615E-261 | TEC_dmg1 | CLDN4 |
| 7.562E-266 | 0.26349768 | 0.268 | 0.089 | 1.786E-261 | TEC_dmg1 | FAM168B |
| 6.457E-264 | 0.43042887 | 0.373 | 0.157 | 1.525E-259 | TEC_dmg1 | ZBTB43 |
| 1.093E-263 | 0.2761953 | 0.257 | 0.083 | 2.581E-259 | TEC_dmg1 | AC005261.1 |
| 2.174E-263 | -1.1703653 | 0.219 | 0.481 | 5.136E-259 | TEC_dmg1 | MXD1 |
| 8.756E-263 | 0.33008596 | 0.343 | 0.135 | 2.068E-258 | TEC_dmg1 | TARS |
| 9.431E-263 | 0.28012169 | 0.282 | 0.098 | 2.228E-258 | TEC_dmg1 | COPRS |
| 2.555E-262 | -2.8936263 | 0.144 | 0.402 | 6.035E-258 | TEC_dmg1 | KRT13 |
| 4.385E-261 | 0.26783788 | 0.198 | 0.053 | 1.036E-256 | TEC_dmg1 | PCLO |
| 2.064E-260 | 0.39661219 | 0.525 | 0.259 | 4.877E-256 | TEC_dmg1 | CD151 |
| 4.237E-260 | 0.36127822 | 0.457 | 0.212 | 1.001E-255 | TEC_dmg1 | SYPL1 |
| 4.542E-260 | 0.32839329 | 0.336 | 0.128 | 1.073E-255 | TEC_dmg1 | GSTM3 |
| 6.267E-260 | 0.37926956 | 0.334 | 0.132 | 1.48E-255 | TEC_dmg1 | CLIC4 |
| 2.781E-259 | 0.25794166 | 0.258 | 0.084 | 6.569E-255 | TEC_dmg1 | IPO5 |
| 1.058E-258 | 0.46769574 | 0.571 | 0.321 | 2.499E-254 | TEC_dmg1 | EIF1AX |
| 2.258E-258 | 0.29604724 | 0.314 | 0.116 | 5.334E-254 | TEC_dmg1 | CDC42BPB |
| 7.694E-257 | 0.41431942 | 0.449 | 0.216 | 1.818E-252 | TEC_dmg1 | PSMB7 |
| 9.589E-257 | -0.7295161 | 0.011 | 0.256 | 2.265E-252 | TEC_dmg1 | HCST |
| 7.941E-254 | -2.4514906 | 0.242 | 0.513 | 1.876E-249 | TEC_dmg1 | CXCL8 |
| 9.588E-254 | 0.41360439 | 0.397 | 0.174 | 2.265E-249 | TEC_dmg1 | IRS2 |
| 1.9E-251 | 0.49097513 | 0.756 | 0.565 | 4.488E-247 | TEC_dmg1 | CLTA |
| 2.657E-251 | 0.53287297 | 0.565 | 0.31 | 6.277E-247 | TEC_dmg1 | GOLGA4 |
| 1.985E-249 | 0.28952485 | 0.315 | 0.117 | 4.69E-245 | TEC_dmg1 | COL18A1 |
| 4.921E-249 | 0.3526269 | 0.358 | 0.147 | 1.163E-244 | TEC_dmg1 | MYOF |
| 8.26E-249 | 0.31358324 | 0.311 | 0.119 | 1.951E-244 | TEC_dmg1 | PPP2R5A |
| 1.079E-248 | 0.36959067 | 0.412 | 0.188 | 2.549E-244 | TEC_dmg1 | PPP2R1A |
| 1.507E-248 | 0.47761603 | 0.516 | 0.273 | 3.56E-244 | TEC_dmg1 | SELENOS |
| 1.568E-248 | -2.0912541 | 0.116 | 0.367 | 3.705E-244 | TEC_dmg1 | HPGD |
| 1.377E-247 | -0.9393126 | 0.022 | 0.267 | 3.254E-243 | TEC_dmg1 | SLC11A1 |
| 2.152E-246 | 0.34846033 | 0.586 | 0.302 | 5.084E-242 | TEC_dmg1 | LMNA |
| 3.144E-246 | 0.40640277 | 0.318 | 0.124 | 7.428E-242 | TEC_dmg1 | MSMO1 |
| 7.937E-246 | 0.28892959 | 0.29 | 0.107 | 1.875E-241 | TEC_dmg1 | ISCA1 |
| 9.102E-245 | 0.32936016 | 0.352 | 0.146 | 2.15E-240 | TEC_dmg1 | ZCRB1 |
| 1.117E-244 | -0.6960246 | 0.921 | 0.961 | 2.639E-240 | TEC_dmg1 | H3F3A |
| 4.914E-244 | -1.1736666 | 0.004 | 0.237 | 1.161E-239 | TEC_dmg1 | S100A12 |
| 8.522E-244 | 0.27175107 | 0.264 | 0.09 | 2.013E-239 | TEC_dmg1 | ASAP2 |
| 3.672E-243 | 0.28821345 | 0.261 | 0.09 | 8.675E-239 | TEC_dmg1 | RHOF |

| | | | | | | |
|---|---|---|---|---|---|---|
| 1.773E-242 | 0.27180682 | 0.23 | 0.073 | 4.187E-238 | TEC_dmg1 | MFHAS1 |
| 4.016E-242 | -0.5375273 | 0.011 | 0.246 | 9.487E-238 | TEC_dmg1 | KYNU |
| 4.985E-242 | 0.38042754 | 0.403 | 0.183 | 1.178E-237 | TEC_dmg1 | LRRFIP2 |
| 1.94E-241 | 0.31942653 | 0.222 | 0.069 | 4.583E-237 | TEC_dmg1 | QPRT |
| 1.11E-240 | 0.53181294 | 0.94 | 0.862 | 2.621E-236 | TEC_dmg1 | RPLP0 |
| 1.112E-240 | 0.26181049 | 0.25 | 0.083 | 2.627E-236 | TEC_dmg1 | SLC25A4 |
| 1.174E-240 | 0.33427158 | 0.412 | 0.184 | 2.774E-236 | TEC_dmg1 | CTTN |
| 2.785E-240 | -0.9062203 | 0.552 | 0.756 | 6.578E-236 | TEC_dmg1 | CTSB |
| 3.447E-240 | 0.46528403 | 0.437 | 0.213 | 8.142E-236 | TEC_dmg1 | LPP |
| 3.628E-240 | 0.26290828 | 0.232 | 0.074 | 8.571E-236 | TEC_dmg1 | DHCR24 |
| 1.771E-239 | 0.32069626 | 0.382 | 0.168 | 4.183E-235 | TEC_dmg1 | AP2B1 |
| 5.664E-239 | 0.43541295 | 0.485 | 0.254 | 1.338E-234 | TEC_dmg1 | UFM1 |
| 1.034E-238 | 0.26220541 | 0.183 | 0.05 | 2.443E-234 | TEC_dmg1 | NEURL3 |
| 1.89E-238 | 0.31041967 | 0.327 | 0.13 | 4.465E-234 | TEC_dmg1 | SUCO |
| 1.099E-237 | 0.49564748 | 0.243 | 0.081 | 2.596E-233 | TEC_dmg1 | IFIT2 |
| 1.879E-237 | 0.26723037 | 0.268 | 0.095 | 4.439E-233 | TEC_dmg1 | RFK |
| 1.417E-236 | -0.5966657 | 0.007 | 0.236 | 3.346E-232 | TEC_dmg1 | SPI1 |
| 1.563E-236 | -4.1745748 | 0.183 | 0.423 | 3.693E-232 | TEC_dmg1 | SPRR3 |
| 1.986E-236 | -1.1043199 | 0.02 | 0.254 | 4.692E-232 | TEC_dmg1 | SLC2A3 |
| 4.217E-236 | 0.30099664 | 0.318 | 0.125 | 9.962E-232 | TEC_dmg1 | BAG5 |
| 1.11E-235 | 0.32431331 | 0.358 | 0.152 | 2.622E-231 | TEC_dmg1 | CTBP2 |
| 1.617E-235 | 0.35112546 | 0.395 | 0.18 | 3.82E-231 | TEC_dmg1 | NAA20 |
| 2.363E-235 | 0.2716846 | 0.25 | 0.085 | 5.582E-231 | TEC_dmg1 | HSPG2 |
| 2.958E-235 | 0.28466824 | 0.21 | 0.064 | 6.988E-231 | TEC_dmg1 | OSGIN1 |
| 3.338E-235 | 0.26992908 | 0.313 | 0.119 | 7.885E-231 | TEC_dmg1 | PATJ |
| 7.222E-235 | 0.30444535 | 0.235 | 0.077 | 1.706E-230 | TEC_dmg1 | CDK6 |
| 4.265E-233 | 0.27363648 | 0.269 | 0.096 | 1.007E-228 | TEC_dmg1 | MTURN |
| 5.805E-233 | 0.31733747 | 0.302 | 0.114 | 1.371E-228 | TEC_dmg1 | SCIN |
| 6.079E-233 | 0.27412498 | 0.246 | 0.084 | 1.436E-228 | TEC_dmg1 | XPOT |
| 4.173E-232 | 0.28770572 | 0.288 | 0.108 | 9.857E-228 | TEC_dmg1 | TNFRSF10B |
| 6.535E-232 | -1.0212925 | 0.013 | 0.24 | 1.544E-227 | TEC_dmg1 | CD14 |
| 8.191E-232 | 0.42744384 | 0.564 | 0.317 | 1.935E-227 | TEC_dmg1 | SNU13 |
| 8.298E-232 | 0.36937791 | 0.377 | 0.16 | 1.96E-227 | TEC_dmg1 | DST |
| 1.617E-231 | 0.27791304 | 0.311 | 0.122 | 3.819E-227 | TEC_dmg1 | PAIP1 |
| 8.324E-231 | 0.29963278 | 0.297 | 0.111 | 1.966E-226 | TEC_dmg1 | TFPI2 |
| 9.46E-231 | 0.36227344 | 0.369 | 0.162 | 2.235E-226 | TEC_dmg1 | PFKP |
| 1.708E-230 | 0.58975252 | 0.516 | 0.286 | 4.034E-226 | TEC_dmg1 | ZNF292 |
| 1.921E-230 | 0.39093348 | 0.317 | 0.128 | 4.538E-226 | TEC_dmg1 | CD40 |
| 2.478E-230 | -2.9385186 | 0.499 | 0.694 | 5.853E-226 | TEC_dmg1 | TIMP1 |
| 3.299E-230 | 0.37777224 | 0.394 | 0.176 | 7.794E-226 | TEC_dmg1 | NEDD4L |
| 1.215E-229 | 0.27829878 | 0.269 | 0.096 | 2.869E-225 | TEC_dmg1 | ADGRG1 |
| 2.888E-229 | 0.27580171 | 0.29 | 0.109 | 6.823E-225 | TEC_dmg1 | THUMPD1 |
| 5.117E-229 | 0.31233993 | 0.333 | 0.138 | 1.209E-224 | TEC_dmg1 | RPAIN |
| 2.928E-228 | -0.6476163 | 0.014 | 0.24 | 6.917E-224 | TEC_dmg1 | ITGB2 |
| 4.915E-228 | 0.44363722 | 0.649 | 0.397 | 1.161E-223 | TEC_dmg1 | RAN |
| 9.946E-227 | 0.25137131 | 0.223 | 0.072 | 2.35E-222 | TEC_dmg1 | TLE1 |
| 3.164E-226 | 0.43494526 | 0.502 | 0.268 | 7.475E-222 | TEC_dmg1 | NDUFB9 |
| 1.09E-225 | 0.27483549 | 0.154 | 0.038 | 2.574E-221 | TEC_dmg1 | LAMA3 |
| 1.68E-225 | 0.30434218 | 0.297 | 0.115 | 3.968E-221 | TEC_dmg1 | SLC44A2 |
| 7.673E-225 | 0.41511896 | 0.478 | 0.228 | 1.813E-220 | TEC_dmg1 | SYNE2 |

| | | | | | | |
|---|---|---|---|---|---|---|
| 4.382E-224 | 0.32207513 | 0.405 | 0.19 | 1.035E-219 | TEC_dmg1 | ACP1 |
| 1.002E-223 | -0.7673572 | 0.01 | 0.23 | 2.367E-219 | TEC_dmg1 | TNFRSF1B |
| 1.64E-223 | 0.34484215 | 0.367 | 0.165 | 3.875E-219 | TEC_dmg1 | CALU |
| 3.913E-223 | 0.49132915 | 0.587 | 0.346 | 9.243E-219 | TEC_dmg1 | ANKRD11 |
| 5.522E-222 | 0.28819426 | 0.3 | 0.117 | 1.304E-217 | TEC_dmg1 | SEPT11 |
| 1.086E-221 | -1.7317948 | 0.021 | 0.243 | 2.566E-217 | TEC_dmg1 | VCAN |
| 2.204E-221 | 0.35333208 | 0.491 | 0.247 | 5.207E-217 | TEC_dmg1 | FAM3C |
| 3.971E-221 | 0.31059506 | 0.211 | 0.067 | 9.38E-217 | TEC_dmg1 | RETREG1 |
| 1.018E-220 | -0.4995643 | 0.007 | 0.223 | 2.404E-216 | TEC_dmg1 | LSP1 |
| 1.914E-220 | 0.25363552 | 0.192 | 0.056 | 4.523E-216 | TEC_dmg1 | THSD7A |
| 3.704E-220 | -1.0961725 | 0.031 | 0.256 | 8.75E-216 | TEC_dmg1 | BCL2A1 |
| 3.796E-219 | 0.36864097 | 0.509 | 0.275 | 8.966E-215 | TEC_dmg1 | KDELR1 |
| 4.421E-219 | 0.44380955 | 0.59 | 0.349 | 1.044E-214 | TEC_dmg1 | RHEB |
| 3.297E-218 | -0.7362638 | 0.024 | 0.248 | 7.788E-214 | TEC_dmg1 | ZEB2 |
| 4.133E-218 | 0.45700532 | 0.593 | 0.347 | 9.763E-214 | TEC_dmg1 | PRDX6 |
| 4.928E-218 | 0.2896823 | 0.298 | 0.119 | 1.164E-213 | TEC_dmg1 | TIMM17A |
| 1.486E-217 | 0.33965838 | 0.381 | 0.176 | 3.51E-213 | TEC_dmg1 | IPO7 |
| 2.139E-217 | -1.090533 | 0.019 | 0.239 | 5.053E-213 | TEC_dmg1 | ALOX5AP |
| 2.268E-217 | -0.6708666 | 0.012 | 0.228 | 5.357E-213 | TEC_dmg1 | LST1 |
| 2.27E-217 | 0.31856987 | 0.372 | 0.167 | 5.363E-213 | TEC_dmg1 | DYNLT3 |
| 6.932E-216 | 0.30324127 | 0.316 | 0.131 | 1.638E-211 | TEC_dmg1 | USP14 |
| 8.138E-216 | 0.3264927 | 0.418 | 0.194 | 1.922E-211 | TEC_dmg1 | CNN3 |
| 9.989E-216 | 0.44256463 | 0.532 | 0.3 | 2.36E-211 | TEC_dmg1 | KRT10 |
| 1.558E-215 | 0.32450212 | 0.405 | 0.192 | 3.68E-211 | TEC_dmg1 | RNPS1 |
| 2.109E-215 | 0.31250897 | 0.342 | 0.148 | 4.982E-211 | TEC_dmg1 | CAMLG |
| 3.713E-215 | 0.3554852 | 0.378 | 0.176 | 8.772E-211 | TEC_dmg1 | SEC61A1 |
| 1.153E-214 | -1.6204632 | 0.279 | 0.512 | 2.723E-210 | TEC_dmg1 | CD74 |
| 2.853E-214 | -0.7632639 | 0.008 | 0.22 | 6.74E-210 | TEC_dmg1 | CYBB |
| 4.511E-214 | -0.6489142 | 0.096 | 0.333 | 1.066E-209 | TEC_dmg1 | CD68 |
| 4.758E-214 | 0.36422809 | 0.446 | 0.227 | 1.124E-209 | TEC_dmg1 | CCT4 |
| 1.581E-213 | 0.25040692 | 0.232 | 0.079 | 3.736E-209 | TEC_dmg1 | ZNF844 |
| 1.859E-213 | -0.9204156 | 0.634 | 0.799 | 4.391E-209 | TEC_dmg1 | LGALS3 |
| 1.003E-212 | -1.0062424 | 0.211 | 0.446 | 2.371E-208 | TEC_dmg1 | EMP3 |
| 5.783E-211 | 0.30339895 | 0.407 | 0.189 | 1.366E-206 | TEC_dmg1 | TCEAL4 |
| 1.309E-210 | 0.4892532 | 0.568 | 0.33 | 3.093E-206 | TEC_dmg1 | TAF1D |
| 1.473E-210 | 0.31401163 | 0.357 | 0.159 | 3.481E-206 | TEC_dmg1 | ARL1 |
| 1.665E-210 | 0.41445849 | 0.498 | 0.266 | 3.933E-206 | TEC_dmg1 | MACF1 |
| 1.257E-208 | 0.42722656 | 0.554 | 0.325 | 2.97E-204 | TEC_dmg1 | CAPN2 |
| 1.646E-207 | -0.7821781 | 0.631 | 0.78 | 3.887E-203 | TEC_dmg1 | CST3 |
| 2.914E-207 | 0.3227001 | 0.359 | 0.16 | 6.885E-203 | TEC_dmg1 | LSR |
| 5.014E-207 | 0.43735436 | 0.394 | 0.194 | 1.185E-202 | TEC_dmg1 | DNTTIP2 |
| 5.347E-207 | 0.35252761 | 0.429 | 0.216 | 1.263E-202 | TEC_dmg1 | PSMA3 |
| 1.687E-206 | 0.29174965 | 0.311 | 0.131 | 3.984E-202 | TEC_dmg1 | PSMD14 |
| 2.212E-206 | -0.5727413 | 0.288 | 0.522 | 5.225E-202 | TEC_dmg1 | ASAH1 |
| 6.64E-206 | 0.33958503 | 0.44 | 0.217 | 1.569E-201 | TEC_dmg1 | PCBD1 |
| 8.295E-206 | 0.42541883 | 0.791 | 0.582 | 1.959E-201 | TEC_dmg1 | HINT1 |
| 3.215E-205 | 0.35037424 | 0.454 | 0.234 | 7.595E-201 | TEC_dmg1 | CCT6A |
| 3.777E-204 | 0.36942449 | 0.47 | 0.249 | 8.923E-200 | TEC_dmg1 | NOL7 |
| 4.848E-204 | -0.4478512 | 0.019 | 0.228 | 1.145E-199 | TEC_dmg1 | ARRB2 |
| 9.386E-204 | 0.52508754 | 0.593 | 0.368 | 2.217E-199 | TEC_dmg1 | AHNAK |

| | | | | | | |
|---|---|---|---|---|---|---|
| 1.45E-203 | 0.37104603 | 0.556 | 0.296 | 3.425E-199 | TEC_dmg1 | PERP |
| 1.879E-203 | -1.5069239 | 0.011 | 0.215 | 4.438E-199 | TEC_dmg1 | HOPX |
| 2.892E-203 | 0.37924628 | 0.53 | 0.3 | 6.832E-199 | TEC_dmg1 | YWHAQ |
| 1.305E-202 | 0.34444281 | 0.366 | 0.171 | 3.084E-198 | TEC_dmg1 | PSMD2 |
| 1.633E-202 | 0.28727202 | 0.304 | 0.127 | 3.857E-198 | TEC_dmg1 | MYO1C |
| 4.182E-202 | -0.6018156 | 0.022 | 0.232 | 9.88E-198 | TEC_dmg1 | BASP1 |
| 6.14E-202 | 0.26641705 | 0.287 | 0.116 | 1.45E-197 | TEC_dmg1 | ATG101 |
| 1.066E-201 | 0.3240763 | 0.334 | 0.147 | 2.519E-197 | TEC_dmg1 | MINDY2 |
| 1.388E-201 | 0.2801356 | 0.295 | 0.119 | 3.279E-197 | TEC_dmg1 | KLHL21 |
| 2.097E-201 | 0.33089494 | 0.451 | 0.225 | 4.955E-197 | TEC_dmg1 | PTMS |
| 2.483E-201 | 0.37503325 | 0.214 | 0.072 | 5.865E-197 | TEC_dmg1 | IFIT3 |
| 1.443E-200 | 0.47684568 | 0.934 | 0.806 | 3.409E-196 | TEC_dmg1 | RPL5 |
| 2.843E-200 | 0.33248495 | 0.443 | 0.229 | 6.716E-196 | TEC_dmg1 | EIF3D |
| 2.651E-199 | 0.27132813 | 0.3 | 0.125 | 6.262E-195 | TEC_dmg1 | ARL3 |
| 3.275E-199 | 0.37184827 | 0.481 | 0.262 | 7.737E-195 | TEC_dmg1 | COX7A2L |
| 1.406E-198 | -0.5589541 | 0.015 | 0.219 | 3.323E-194 | TEC_dmg1 | ITGAX |
| 7.19E-198 | 0.32533432 | 0.393 | 0.193 | 1.698E-193 | TEC_dmg1 | STRAP |
| 1.305E-197 | -0.7593338 | 0.015 | 0.216 | 3.083E-193 | TEC_dmg1 | CD52 |
| 2.203E-197 | 0.35459845 | 0.439 | 0.224 | 5.205E-193 | TEC_dmg1 | PDCD5 |
| 1.376E-196 | -0.8353199 | 0.447 | 0.661 | 3.251E-192 | TEC_dmg1 | IER3 |
| 9.873E-196 | -1.0148762 | 0.182 | 0.41 | 2.332E-191 | TEC_dmg1 | FOS |
| 7.86E-195 | -0.3815524 | 0.006 | 0.201 | 1.857E-190 | TEC_dmg1 | GNA15 |
| 2.314E-194 | -0.5022475 | 0.762 | 0.877 | 5.465E-190 | TEC_dmg1 | ARPC3 |
| 2.332E-194 | 0.44000652 | 0.436 | 0.231 | 5.51E-190 | TEC_dmg1 | DNAJC3 |
| 3.177E-194 | -0.5866232 | 0.01 | 0.206 | 7.505E-190 | TEC_dmg1 | FYB1 |
| 5.15E-194 | 0.40361374 | 0.895 | 0.741 | 1.217E-189 | TEC_dmg1 | RPSA |
| 1.645E-193 | 0.26136858 | 0.233 | 0.084 | 3.885E-189 | TEC_dmg1 | ITGA6 |
| 7.134E-193 | 0.26761286 | 0.266 | 0.105 | 1.685E-188 | TEC_dmg1 | SLC1A5 |
| 2.321E-192 | 0.27225062 | 0.298 | 0.125 | 5.482E-188 | TEC_dmg1 | NFIC |
| 8.854E-192 | -0.6192105 | 0.007 | 0.2 | 2.091E-187 | TEC_dmg1 | IGSF6 |
| 1.316E-191 | -1.26375 | 0.081 | 0.296 | 3.108E-187 | TEC_dmg1 | HLA-DRB1 |
| 1.537E-191 | 0.442982 | 0.419 | 0.216 | 3.632E-187 | TEC_dmg1 | SNHG7 |
| 3.196E-191 | -0.4355449 | 0.011 | 0.205 | 7.55E-187 | TEC_dmg1 | RAC2 |
| 7.342E-191 | -0.5605629 | 0.012 | 0.206 | 1.734E-186 | TEC_dmg1 | FCGR2A |
| 7.583E-189 | 0.25370224 | 0.279 | 0.115 | 1.791E-184 | TEC_dmg1 | PLEKHA3 |
| 1.508E-188 | 0.29132551 | 0.317 | 0.142 | 3.562E-184 | TEC_dmg1 | FUNDC2 |
| 5.374E-188 | 0.25344677 | 0.319 | 0.138 | 1.269E-183 | TEC_dmg1 | ARHGEF12 |
| 5.959E-188 | -0.5995858 | 0.928 | 0.955 | 1.408E-183 | TEC_dmg1 | H3F3B |
| 6.255E-188 | 0.28757341 | 0.25 | 0.097 | 1.478E-183 | TEC_dmg1 | NCEH1 |
| 3.499E-187 | -0.5909553 | 0.47 | 0.657 | 8.266E-183 | TEC_dmg1 | ARPC5 |
| 1.242E-186 | -0.7420965 | 0.008 | 0.197 | 2.933E-182 | TEC_dmg1 | OLR1 |
| 1.248E-186 | 0.32420827 | 0.431 | 0.223 | 2.949E-182 | TEC_dmg1 | NUDC |
| 2.326E-185 | 0.25297585 | 0.31 | 0.135 | 5.495E-181 | TEC_dmg1 | BUD23 |
| 6.915E-185 | 0.4035524 | 0.539 | 0.327 | 1.633E-180 | TEC_dmg1 | SF3B6 |
| 7.009E-185 | -0.3914656 | 0.013 | 0.205 | 1.656E-180 | TEC_dmg1 | HCLS1 |
| 7.776E-185 | 0.42640224 | 0.853 | 0.689 | 1.837E-180 | TEC_dmg1 | RPL4 |
| 1.006E-183 | -0.8031868 | 0.006 | 0.192 | 2.377E-179 | TEC_dmg1 | PLEK |
| 1.78E-183 | 0.40240655 | 0.516 | 0.306 | 4.204E-179 | TEC_dmg1 | GHITM |
| 5.423E-183 | 0.27560531 | 0.224 | 0.082 | 1.281E-178 | TEC_dmg1 | PDLIM4 |
| 1.52E-182 | -0.5508795 | 0.011 | 0.199 | 3.592E-178 | TEC_dmg1 | SAMSN1 |

| | | | | | | |
|---|---|---|---|---|---|---|
| 2.097E-182 | 0.25022215 | 0.579 | 0.303 | 4.954E-178 | TEC_dmg1 | MYO6 |
| 5.34E-182 | -0.4650853 | 0.906 | 0.959 | 1.261E-177 | TEC_dmg1 | SERF2 |
| 1.108E-181 | 0.39243525 | 0.528 | 0.304 | 2.618E-177 | TEC_dmg1 | MGST3 |
| 1.231E-181 | 0.25917349 | 0.305 | 0.132 | 2.907E-177 | TEC_dmg1 | SVIP |
| 1.818E-181 | 0.50179813 | 0.955 | 0.958 | 4.295E-177 | TEC_dmg1 | RPL15 |
| 4.508E-181 | 0.34429652 | 0.369 | 0.181 | 1.065E-176 | TEC_dmg1 | ITGAV |
| 6.078E-181 | 0.25218564 | 0.253 | 0.099 | 1.436E-176 | TEC_dmg1 | UGDH |
| 9.774E-181 | -0.5204111 | 0.862 | 0.936 | 2.309E-176 | TEC_dmg1 | OAZ1 |
| 1.08E-180 | 0.33608974 | 0.355 | 0.171 | 2.551E-176 | TEC_dmg1 | PSMD12 |
| 1.183E-180 | -0.4377129 | 0.006 | 0.19 | 2.793E-176 | TEC_dmg1 | CORO1A |
| 1.427E-180 | 0.2857434 | 0.349 | 0.165 | 3.37E-176 | TEC_dmg1 | FUBP1 |
| 2.28E-180 | 0.28801946 | 0.336 | 0.155 | 5.386E-176 | TEC_dmg1 | BBX |
| 2.297E-180 | 0.29722469 | 0.302 | 0.134 | 5.426E-176 | TEC_dmg1 | RRAGC |
| 2.752E-180 | 0.37948715 | 0.486 | 0.282 | 6.5E-176 | TEC_dmg1 | VCP |
| 5.975E-180 | 0.47820539 | 0.884 | 0.805 | 1.411E-175 | TEC_dmg1 | GSTP1 |
| 7.263E-180 | -0.4972333 | 0.007 | 0.19 | 1.716E-175 | TEC_dmg1 | SMIM25 |
| 7.836E-180 | -0.7945589 | 0.016 | 0.205 | 1.851E-175 | TEC_dmg1 | AC020656.1 |
| 8.076E-180 | -0.5648043 | 0.03 | 0.224 | 1.908E-175 | TEC_dmg1 | GK |
| 9.722E-180 | -0.4812595 | 0.004 | 0.184 | 2.297E-175 | TEC_dmg1 | SLA |
| 2.123E-179 | -0.6966893 | 0.271 | 0.49 | 5.016E-175 | TEC_dmg1 | DUSP1 |
| 4.58E-179 | 0.34725379 | 0.337 | 0.156 | 1.082E-174 | TEC_dmg1 | STMN1 |
| 1.076E-178 | 0.27217519 | 0.334 | 0.155 | 2.542E-174 | TEC_dmg1 | GLRX3 |
| 3.885E-178 | 0.4745768 | 0.182 | 0.061 | 9.177E-174 | TEC_dmg1 | TAGLN |
| 8.265E-178 | -0.3748592 | 0.017 | 0.203 | 1.953E-173 | TEC_dmg1 | GPSM3 |
| 1.033E-177 | -0.5322115 | 0.167 | 0.387 | 2.44E-173 | TEC_dmg1 | CTSH |
| 1.718E-177 | 0.3706785 | 0.382 | 0.197 | 4.057E-173 | TEC_dmg1 | OSER1 |
| 2.219E-177 | 0.305184 | 0.396 | 0.199 | 5.243E-173 | TEC_dmg1 | MLEC |
| 4.412E-177 | 0.29049055 | 0.366 | 0.178 | 1.042E-172 | TEC_dmg1 | CCT2 |
| 1.376E-176 | 0.3487374 | 0.538 | 0.315 | 3.251E-172 | TEC_dmg1 | EIF4B |
| 1.62E-176 | 0.48711428 | 0.787 | 0.547 | 3.826E-172 | TEC_dmg1 | TACSTD2 |
| 2.889E-176 | -1.2926645 | 0.03 | 0.221 | 6.825E-172 | TEC_dmg1 | HLA-DPA1 |
| 9.781E-176 | 0.25868787 | 0.318 | 0.144 | 2.311E-171 | TEC_dmg1 | SLC25A36 |
| 1.453E-175 | 0.31255778 | 0.479 | 0.263 | 3.432E-171 | TEC_dmg1 | DYNC1H1 |
| 4.08E-175 | -0.3680312 | 0.013 | 0.196 | 9.639E-171 | TEC_dmg1 | FMNL1 |
| 1.189E-174 | 0.37993528 | 0.568 | 0.354 | 2.81E-170 | TEC_dmg1 | LAMTOR5 |
| 2.073E-174 | 0.36731011 | 0.231 | 0.089 | 4.897E-170 | TEC_dmg1 | RCAN1 |
| 6.193E-174 | -0.4230552 | 0.007 | 0.185 | 1.463E-169 | TEC_dmg1 | CD48 |
| 9.491E-174 | -2.2728735 | 0.012 | 0.192 | 2.242E-169 | TEC_dmg1 | IL1B |
| 2.496E-173 | 0.27633131 | 0.283 | 0.121 | 5.897E-169 | TEC_dmg1 | AGRN |
| 2.82E-173 | -0.4163404 | 0.014 | 0.195 | 6.663E-169 | TEC_dmg1 | CD37 |
| 4.894E-173 | 0.30256941 | 0.37 | 0.18 | 1.156E-168 | TEC_dmg1 | TCEAL9 |
| 5.288E-173 | 0.39921399 | 0.826 | 0.703 | 1.249E-168 | TEC_dmg1 | SKP1 |
| 1.052E-172 | 0.50593351 | 0.677 | 0.497 | 2.485E-168 | TEC_dmg1 | EIF5 |
| 6.183E-172 | 0.27570148 | 0.318 | 0.146 | 1.461E-167 | TEC_dmg1 | BTBD7 |
| 6.504E-172 | 0.26266886 | 0.23 | 0.088 | 1.536E-167 | TEC_dmg1 | ARG2 |
| 1.941E-171 | 0.32236821 | 0.406 | 0.213 | 4.586E-167 | TEC_dmg1 | ARPC5L |
| 2.094E-171 | 0.36220069 | 0.49 | 0.285 | 4.946E-167 | TEC_dmg1 | TMED9 |
| 2.141E-171 | 0.29222708 | 0.297 | 0.13 | 5.057E-167 | TEC_dmg1 | NCOA7 |
| 2.985E-171 | 0.2952739 | 0.322 | 0.151 | 7.051E-167 | TEC_dmg1 | CCDC59 |
| 3.92E-171 | 0.35604931 | 0.265 | 0.112 | 9.259E-167 | TEC_dmg1 | PLA2G16 |

| | | | | | | |
|---|---|---|---|---|---|---|
| 1.268E-170 | 0.36637915 | 0.755 | 0.585 | 2.996E-166 | TEC_dmg1 | YWHAE |
| 2.364E-170 | -0.5006949 | 0.004 | 0.176 | 5.585E-166 | TEC_dmg1 | AQP9 |
| 1.915E-169 | 0.29818177 | 0.369 | 0.184 | 4.523E-165 | TEC_dmg1 | PSMB5 |
| 2.406E-169 | -0.8279737 | 0.01 | 0.185 | 5.684E-165 | TEC_dmg1 | FCN1 |
| 2.684E-169 | 0.44429111 | 0.856 | 0.788 | 6.341E-165 | TEC_dmg1 | CHCHD2 |
| 2.966E-169 | 0.31887732 | 0.398 | 0.209 | 7.006E-165 | TEC_dmg1 | PSMB2 |
| 3.414E-169 | -0.4621049 | 0.021 | 0.203 | 8.064E-165 | TEC_dmg1 | PTPRE |
| 7.278E-169 | 0.36881336 | 0.618 | 0.382 | 1.719E-164 | TEC_dmg1 | SOD1 |
| 1.262E-168 | 0.28902817 | 0.397 | 0.204 | 2.981E-164 | TEC_dmg1 | PPP2CB |
| 2.05E-168 | -0.6321484 | 0.031 | 0.215 | 4.843E-164 | TEC_dmg1 | DUSP4 |
| 2.842E-168 | -0.4537951 | 0.007 | 0.18 | 6.715E-164 | TEC_dmg1 | LCP2 |
| 8.464E-168 | -1.2165077 | 0.021 | 0.2 | 1.999E-163 | TEC_dmg1 | DUOX2 |
| 3.656E-167 | -0.6722901 | 0.013 | 0.188 | 8.636E-163 | TEC_dmg1 | STEAP4 |
| 4.63E-167 | 0.58940361 | 0.78 | 0.644 | 1.094E-162 | TEC_dmg1 | ZFAS1 |
| 4.643E-167 | 0.33807932 | 0.81 | 0.605 | 1.097E-162 | TEC_dmg1 | DSTN |
| 6.381E-167 | 0.44092897 | 0.915 | 0.863 | 1.507E-162 | TEC_dmg1 | ANXA2 |
| 1.567E-166 | 0.31176789 | 0.273 | 0.118 | 3.701E-162 | TEC_dmg1 | HYOU1 |
| 2.979E-166 | 0.33323794 | 0.473 | 0.268 | 7.037E-162 | TEC_dmg1 | MRFAP1 |
| 3.76E-166 | 0.31861345 | 0.351 | 0.174 | 8.881E-162 | TEC_dmg1 | TBL1XR1 |
| 3.8E-166 | 0.39545754 | 0.547 | 0.345 | 8.977E-162 | TEC_dmg1 | PSMB1 |
| 4.808E-166 | -0.6241726 | 0.022 | 0.2 | 1.136E-161 | TEC_dmg1 | C5AR1 |
| 1.017E-165 | 0.34050745 | 0.497 | 0.282 | 2.402E-161 | TEC_dmg1 | DDX24 |
| 1.576E-165 | 0.28580444 | 0.436 | 0.234 | 3.722E-161 | TEC_dmg1 | EIF3M |
| 8.74E-165 | -0.595417 | 0.005 | 0.173 | 2.065E-160 | TEC_dmg1 | CD300E |
| 3.53E-164 | -0.9854659 | 0.844 | 0.887 | 8.339E-160 | TEC_dmg1 | CSTB |
| 7.759E-164 | -1.3114548 | 0.011 | 0.181 | 1.833E-159 | TEC_dmg1 | DUOXA2 |
| 1.037E-163 | -1.0969836 | 0.007 | 0.175 | 2.449E-159 | TEC_dmg1 | CEACAM5 |
| 3.352E-163 | 0.59116286 | 0.417 | 0.229 | 7.919E-159 | TEC_dmg1 | PLAU |
| 3.748E-163 | -1.1895102 | 0.008 | 0.176 | 8.853E-159 | TEC_dmg1 | MUC21 |
| 3.903E-163 | -0.3595607 | 0.008 | 0.177 | 9.22E-159 | TEC_dmg1 | FERMT3 |
| 1.031E-160 | -0.8585879 | 0.009 | 0.177 | 2.435E-156 | TEC_dmg1 | APOBEC3A |
| 2.045E-160 | -0.7617027 | 0.064 | 0.253 | 4.83E-156 | TEC_dmg1 | FABP5 |
| 2.755E-160 | -1.0277778 | 0.007 | 0.173 | 6.507E-156 | TEC_dmg1 | PRSS27 |
| 3.247E-160 | 0.3843849 | 0.467 | 0.271 | 7.67E-156 | TEC_dmg1 | MIR4435-2HG |
| 7.18E-160 | 0.36927259 | 0.619 | 0.409 | 1.696E-155 | TEC_dmg1 | EIF3A |
| 1.936E-159 | -1.0285564 | 0.088 | 0.28 | 4.572E-155 | TEC_dmg1 | AREG |
| 3.031E-159 | 0.37770745 | 0.495 | 0.292 | 7.159E-155 | TEC_dmg1 | CYTOR |
| 6.139E-159 | 0.37195302 | 0.289 | 0.129 | 1.45E-154 | TEC_dmg1 | ITGB6 |
| 9.789E-159 | -0.3509268 | 0.011 | 0.178 | 2.312E-154 | TEC_dmg1 | MYO1F |
| 2.426E-158 | 0.36031585 | 0.739 | 0.543 | 5.73E-154 | TEC_dmg1 | SET |
| 3.107E-158 | 0.35669522 | 0.575 | 0.364 | 7.34E-154 | TEC_dmg1 | DYNLRB1 |
| 3.785E-158 | 0.44580207 | 0.548 | 0.358 | 8.942E-154 | TEC_dmg1 | ARF4 |
| 1.083E-157 | 0.40106514 | 0.379 | 0.201 | 2.558E-153 | TEC_dmg1 | CDK2AP2 |
| 2.345E-157 | 0.44219469 | 0.919 | 0.831 | 5.54E-153 | TEC_dmg1 | RACK1 |
| 2.81E-157 | 0.43046415 | 0.352 | 0.178 | 6.637E-153 | TEC_dmg1 | SDF2L1 |
| 2.975E-157 | 0.25933347 | 0.304 | 0.142 | 7.027E-153 | TEC_dmg1 | ITGB1BP1 |
| 8.393E-157 | 0.33768177 | 0.507 | 0.306 | 1.983E-152 | TEC_dmg1 | SERINC1 |
| 2.948E-156 | 0.32478768 | 0.68 | 0.429 | 6.965E-152 | TEC_dmg1 | CLDN7 |
| 3.406E-156 | 0.29903081 | 0.362 | 0.186 | 8.046E-152 | TEC_dmg1 | USP16 |
| 5.469E-156 | 0.30106867 | 0.403 | 0.215 | 1.292E-151 | TEC_dmg1 | OPTN |

| | | | | | | |
|---|---|---|---|---|---|---|
| 6.368E-156 | 0.43320162 | 0.623 | 0.441 | 1.504E-151 | TEC_dmg1 | EIF2S2 |
| 7.852E-156 | 0.29510052 | 0.393 | 0.208 | 1.855E-151 | TEC_dmg1 | RSL1D1 |
| 1.842E-155 | 0.28134588 | 0.373 | 0.194 | 4.351E-151 | TEC_dmg1 | ANXA7 |
| 2.578E-155 | 0.34272013 | 0.39 | 0.208 | 6.091E-151 | TEC_dmg1 | ST14 |
| 3.32E-155 | -0.3117026 | 0.01 | 0.174 | 7.844E-151 | TEC_dmg1 | DOCK8 |
| 4.047E-154 | 0.28891059 | 0.418 | 0.229 | 9.56E-150 | TEC_dmg1 | CCT8 |
| 5.183E-154 | -0.5619927 | 0.022 | 0.192 | 1.224E-149 | TEC_dmg1 | ANPEP |
| 6.312E-154 | 0.27354141 | 0.384 | 0.2 | 1.491E-149 | TEC_dmg1 | DYNC1LI2 |
| 6.87E-154 | 0.2987079 | 0.557 | 0.328 | 1.623E-149 | TEC_dmg1 | UBE2H |
| 4.181E-153 | 0.34104096 | 0.547 | 0.341 | 9.876E-149 | TEC_dmg1 | SRSF11 |
| 4.894E-153 | 0.25326715 | 0.287 | 0.132 | 1.156E-148 | TEC_dmg1 | RABGGTB |
| 5.094E-153 | -1.0129604 | 0.009 | 0.169 | 1.203E-148 | TEC_dmg1 | PITX1 |
| 1.011E-152 | -0.5293198 | 0.007 | 0.166 | 2.389E-148 | TEC_dmg1 | CLEC5A |
| 1.065E-152 | -0.8681035 | 0.003 | 0.16 | 2.515E-148 | TEC_dmg1 | CEACAM7 |
| 1.601E-152 | 0.2569291 | 0.317 | 0.154 | 3.782E-148 | TEC_dmg1 | TTC1 |
| 3.428E-152 | -0.5324487 | 0.003 | 0.158 | 8.099E-148 | TEC_dmg1 | CD93 |
| 8.425E-152 | -1.6133803 | 0.015 | 0.177 | 1.99E-147 | TEC_dmg1 | ECM1 |
| 1.059E-151 | 0.2500357 | 0.297 | 0.139 | 2.503E-147 | TEC_dmg1 | DNAJB2 |
| 1.222E-151 | -0.4262352 | 0.02 | 0.187 | 2.886E-147 | TEC_dmg1 | SAMHD1 |
| 3.689E-151 | -0.4555942 | 0.004 | 0.16 | 8.714E-147 | TEC_dmg1 | FPR1 |
| 8.165E-151 | 0.33937551 | 0.133 | 0.04 | 1.929E-146 | TEC_dmg1 | PEG10 |
| 8.364E-151 | 0.27712429 | 0.33 | 0.164 | 1.976E-146 | TEC_dmg1 | MGAT4B |
| 8.843E-151 | 0.32667216 | 0.442 | 0.248 | 2.089E-146 | TEC_dmg1 | XBP1 |
| 1.007E-150 | -0.5586805 | 0.846 | 0.897 | 2.378E-146 | TEC_dmg1 | ATP5F1E |
| 1.137E-150 | -0.3799845 | 0.007 | 0.166 | 2.687E-146 | TEC_dmg1 | EVI2B |
| 1.35E-150 | 0.32823147 | 0.328 | 0.166 | 3.188E-146 | TEC_dmg1 | ELK3 |
| 2.776E-150 | -0.3599958 | 0.015 | 0.178 | 6.558E-146 | TEC_dmg1 | CLEC2B |
| 1.469E-149 | 0.27024643 | 0.355 | 0.181 | 3.47E-145 | TEC_dmg1 | CCPG1 |
| 1.564E-149 | -0.6819746 | 0.006 | 0.161 | 3.695E-145 | TEC_dmg1 | NCCRP1 |
| 6.039E-149 | 0.32185539 | 0.466 | 0.276 | 1.427E-144 | TEC_dmg1 | HDLBP |
| 6.979E-149 | 0.28126573 | 0.456 | 0.256 | 1.649E-144 | TEC_dmg1 | SSB |
| 1.126E-148 | -0.5272177 | 0.021 | 0.185 | 2.66E-144 | TEC_dmg1 | PRDM1 |
| 2.127E-148 | -1.1318839 | 0.032 | 0.2 | 5.024E-144 | TEC_dmg1 | HLA-DPB1 |
| 2.781E-148 | -0.8581417 | 0.108 | 0.299 | 6.569E-144 | TEC_dmg1 | MAFB |
| 3.037E-148 | -0.7873647 | 0.007 | 0.163 | 7.175E-144 | TEC_dmg1 | TMPRSS11D |
| 5.01E-148 | 0.43041529 | 0.899 | 0.791 | 1.184E-143 | TEC_dmg1 | RPS5 |
| 5.19E-148 | -0.9051239 | 0.006 | 0.16 | 1.226E-143 | TEC_dmg1 | CEACAM6 |
| 5.434E-148 | 0.34714621 | 0.356 | 0.18 | 1.284E-143 | TEC_dmg1 | SFN |
| 9.128E-148 | 0.37291464 | 0.683 | 0.476 | 2.156E-143 | TEC_dmg1 | CD9 |
| 1.187E-147 | 0.34614702 | 0.334 | 0.171 | 2.803E-143 | TEC_dmg1 | SEL1L |
| 2.885E-147 | -0.4355132 | 0.006 | 0.159 | 6.816E-143 | TEC_dmg1 | MS4A7 |
| 1.48E-146 | -0.3973065 | 0.007 | 0.16 | 3.497E-142 | TEC_dmg1 | TREM1 |
| 1.626E-146 | 0.41295835 | 0.646 | 0.437 | 3.841E-142 | TEC_dmg1 | RAB11FIP1 |
| 1.745E-146 | 0.26374532 | 0.324 | 0.161 | 4.121E-142 | TEC_dmg1 | SRP72 |
| 3.306E-146 | 0.42014138 | 0.911 | 0.882 | 7.809E-142 | TEC_dmg1 | MYL12B |
| 7.361E-146 | -0.8102008 | 0.443 | 0.634 | 1.739E-141 | TEC_dmg1 | LGALS1 |
| 9.747E-146 | 0.35022604 | 0.545 | 0.342 | 2.303E-141 | TEC_dmg1 | MTDH |
| 2.106E-145 | -0.5167751 | 0.003 | 0.153 | 4.976E-141 | TEC_dmg1 | PLAC8 |
| 2.65E-145 | 0.27877949 | 0.486 | 0.28 | 6.259E-141 | TEC_dmg1 | NUCKS1 |
| 3.798E-145 | 0.3513797 | 0.403 | 0.226 | 8.972E-141 | TEC_dmg1 | MAFG |

| | | | | | | |
|---|---|---|---|---|---|---|
| 6.398E-145 | -0.5182046 | 0.023 | 0.185 | 1.511E-140 | TEC_dmg1 | CXCR4 |
| 1.426E-144 | -0.4300917 | 0.007 | 0.16 | 3.368E-140 | TEC_dmg1 | VSIG2 |
| 1.63E-144 | -0.347726 | 0.02 | 0.18 | 3.851E-140 | TEC_dmg1 | CFD |
| 1.936E-144 | -0.29904 | 0.004 | 0.154 | 4.574E-140 | TEC_dmg1 | IL2RG |
| 2.153E-144 | 0.29271868 | 0.407 | 0.229 | 5.087E-140 | TEC_dmg1 | UQCRFS1 |
| 3.817E-144 | 0.36651555 | 0.66 | 0.48 | 9.016E-140 | TEC_dmg1 | VDAC2 |
| 4.266E-144 | 0.27694704 | 0.426 | 0.24 | 1.008E-139 | TEC_dmg1 | SF3B2 |
| 4.614E-143 | -0.3565667 | 0.065 | 0.243 | 1.09E-138 | TEC_dmg1 | RGS10 |
| 9.123E-143 | 0.38069108 | 0.921 | 0.829 | 2.155E-138 | TEC_dmg1 | SQSTM1 |
| 1.069E-142 | 0.33620597 | 0.523 | 0.325 | 2.524E-138 | TEC_dmg1 | BSG |
| 1.344E-142 | 0.29217503 | 0.417 | 0.236 | 3.175E-138 | TEC_dmg1 | TMED2 |
| 2.148E-142 | -0.4940522 | 0.047 | 0.216 | 5.074E-138 | TEC_dmg1 | NCF2 |
| 2.705E-142 | 0.44304735 | 0.318 | 0.158 | 6.39E-138 | TEC_dmg1 | XIST |
| 2.797E-142 | -0.5343197 | 0.024 | 0.185 | 6.607E-138 | TEC_dmg1 | TGFBI |
| 6.98E-142 | 0.38102152 | 0.548 | 0.324 | 1.649E-137 | TEC_dmg1 | GDF15 |
| 1.899E-141 | -0.8024236 | 0.008 | 0.158 | 4.486E-137 | TEC_dmg1 | TMPRSS11E |
| 8.655E-141 | -0.4274781 | 0.081 | 0.26 | 2.045E-136 | TEC_dmg1 | RAB31 |
| 1.523E-140 | -0.7379438 | 0.218 | 0.401 | 3.598E-136 | TEC_dmg1 | TYMP |
| 2.216E-139 | -0.3578764 | 0.01 | 0.159 | 5.236E-135 | TEC_dmg1 | C1orf162 |
| 4.264E-139 | 0.3343431 | 0.6 | 0.392 | 1.007E-134 | TEC_dmg1 | CDV3 |
| 4.502E-139 | 0.31959251 | 0.5 | 0.306 | 1.064E-134 | TEC_dmg1 | ARL6IP1 |
| 5.442E-139 | -0.4568849 | 0.085 | 0.264 | 1.285E-134 | TEC_dmg1 | GPCPD1 |
| 1.323E-138 | -0.2910136 | 0.018 | 0.172 | 3.124E-134 | TEC_dmg1 | ARHGAP30 |
| 1.397E-138 | 0.44745507 | 0.609 | 0.431 | 3.301E-134 | TEC_dmg1 | ACTN4 |
| 1.955E-138 | 0.25440423 | 0.369 | 0.197 | 4.619E-134 | TEC_dmg1 | CSNK2A1 |
| 1.967E-138 | 0.37359524 | 0.517 | 0.32 | 4.646E-134 | TEC_dmg1 | NDRG1 |
| 2.867E-138 | 0.28776728 | 0.301 | 0.15 | 6.772E-134 | TEC_dmg1 | BCL2L1 |
| 2.928E-138 | -0.3247187 | 0.005 | 0.15 | 6.918E-134 | TEC_dmg1 | CLEC7A |
| 2.972E-138 | 0.37954208 | 0.685 | 0.501 | 7.02E-134 | TEC_dmg1 | NAP1L1 |
| 3.835E-138 | 0.27008987 | 0.384 | 0.209 | 9.06E-134 | TEC_dmg1 | GGNBP2 |
| 6.712E-138 | -0.3113456 | 0.004 | 0.149 | 1.586E-133 | TEC_dmg1 | HCK |
| 7.439E-138 | -0.4582704 | 0.005 | 0.15 | 1.757E-133 | TEC_dmg1 | MCEMP1 |
| 1.151E-137 | 0.28528659 | 0.332 | 0.173 | 2.72E-133 | TEC_dmg1 | SRP19 |
| 1.766E-137 | 0.29033493 | 0.586 | 0.376 | 4.172E-133 | TEC_dmg1 | CTNNA1 |
| 2.074E-137 | 0.25024217 | 0.324 | 0.165 | 4.9E-133 | TEC_dmg1 | ARCN1 |
| 2.205E-137 | 0.28698979 | 0.385 | 0.213 | 5.209E-133 | TEC_dmg1 | HDGF |
| 2.87E-137 | -0.4736569 | 0.149 | 0.337 | 6.779E-133 | TEC_dmg1 | SLC16A3 |
| 3.198E-137 | -0.4926873 | 0.036 | 0.196 | 7.554E-133 | TEC_dmg1 | FBP1 |
| 4.539E-137 | 0.276778 | 0.331 | 0.168 | 1.072E-132 | TEC_dmg1 | BNIP3 |
| 5.059E-137 | -0.4181505 | 0.009 | 0.156 | 1.195E-132 | TEC_dmg1 | NCF1 |
| 9.809E-137 | 0.32640979 | 0.204 | 0.082 | 2.317E-132 | TEC_dmg1 | CTGF |
| 3.643E-136 | -0.2855688 | 0.006 | 0.15 | 8.607E-132 | TEC_dmg1 | EVI2A |
| 5.853E-136 | -0.2989266 | 0.004 | 0.147 | 1.383E-131 | TEC_dmg1 | CYTIP |
| 6.538E-136 | 0.32304677 | 0.519 | 0.33 | 1.544E-131 | TEC_dmg1 | SNRPB |
| 3.541E-135 | 0.31879374 | 0.479 | 0.296 | 8.365E-131 | TEC_dmg1 | POLR1D |
| 9.375E-135 | 0.26336908 | 0.435 | 0.248 | 2.215E-130 | TEC_dmg1 | TMEM123 |
| 1.108E-134 | 0.28393378 | 0.348 | 0.181 | 2.617E-130 | TEC_dmg1 | USP53 |
| 1.412E-134 | 0.31478038 | 0.446 | 0.265 | 3.335E-130 | TEC_dmg1 | TCEA1 |
| 2.876E-134 | -0.4615752 | 0.387 | 0.563 | 6.794E-130 | TEC_dmg1 | C4orf3 |
| 3.145E-134 | 0.40457519 | 0.733 | 0.526 | 7.429E-130 | TEC_dmg1 | KRT7 |

| | | | | | | |
|---|---|---|---|---|---|---|
| 3.934E-134 | -1.0998512 | 0.005 | 0.146 | 9.293E-130 | TEC_dmg1 | SERPINB2 |
| 4.482E-134 | -0.2642896 | 0.005 | 0.147 | 1.059E-129 | TEC_dmg1 | NCKAP1L |
| 5.506E-134 | 0.29343159 | 0.284 | 0.139 | 1.301E-129 | TEC_dmg1 | EIF4EBP1 |
| 6.842E-134 | -1.6706698 | 0.005 | 0.145 | 1.616E-129 | TEC_dmg1 | LYPD2 |
| 7.615E-134 | 0.28345881 | 0.411 | 0.234 | 1.799E-129 | TEC_dmg1 | SEC31A |
| 8.713E-134 | -0.4607223 | 0.12 | 0.296 | 2.058E-129 | TEC_dmg1 | LYN |
| 1.097E-133 | -0.9393712 | 0.006 | 0.147 | 2.592E-129 | TEC_dmg1 | TMPRSS11B |
| 2.726E-133 | -0.6633024 | 0.007 | 0.149 | 6.44E-129 | TEC_dmg1 | CXCL17 |
| 7.63E-133 | -0.3633797 | 0.074 | 0.249 | 1.803E-128 | TEC_dmg1 | BST2 |
| 1.061E-132 | 0.47748612 | 0.825 | 0.751 | 2.507E-128 | TEC_dmg1 | ITGB1 |
| 1.138E-132 | 0.29496139 | 0.218 | 0.094 | 2.687E-128 | TEC_dmg1 | LDLR |
| 1.287E-132 | -0.2859321 | 0.006 | 0.147 | 3.04E-128 | TEC_dmg1 | FGR |
| 2.241E-132 | 0.4379499 | 0.533 | 0.351 | 5.294E-128 | TEC_dmg1 | MYH9 |
| 2.928E-132 | 0.30609388 | 0.546 | 0.355 | 6.917E-128 | TEC_dmg1 | C11orf58 |
| 7.233E-132 | 0.29603612 | 0.671 | 0.455 | 1.709E-127 | TEC_dmg1 | CD81 |
| 1.633E-131 | -0.8215922 | 0.034 | 0.187 | 3.858E-127 | TEC_dmg1 | NDRG2 |
| 2.152E-131 | 0.40231694 | 0.374 | 0.208 | 5.084E-127 | TEC_dmg1 | MRPS6 |
| 2.237E-131 | 0.31889092 | 0.295 | 0.15 | 5.285E-127 | TEC_dmg1 | MTHFD2 |
| 3.728E-131 | -0.7639477 | 0.02 | 0.166 | 8.808E-127 | TEC_dmg1 | SCEL |
| 1.22E-130 | 0.26219482 | 0.422 | 0.242 | 2.883E-126 | TEC_dmg1 | CCT3 |
| 1.341E-130 | -1.0664436 | 0.011 | 0.153 | 3.168E-126 | TEC_dmg1 | CXCL6 |
| 7.068E-130 | 0.45568363 | 0.41 | 0.235 | 1.67E-125 | TEC_dmg1 | SNHG12 |
| 8.224E-130 | 0.25198622 | 0.356 | 0.192 | 1.943E-125 | TEC_dmg1 | CWC15 |
| 9.712E-130 | 0.30544626 | 0.465 | 0.284 | 2.294E-125 | TEC_dmg1 | SF3B5 |
| 2.248E-129 | -0.3017562 | 0.015 | 0.158 | 5.311E-125 | TEC_dmg1 | CARD16 |
| 2.459E-129 | 0.30934624 | 0.408 | 0.236 | 5.81E-125 | TEC_dmg1 | BCLAF1 |
| 6.991E-129 | -0.9462348 | 0.008 | 0.146 | 1.651E-124 | TEC_dmg1 | EREG |
| 2.359E-128 | 0.26787904 | 0.431 | 0.252 | 5.573E-124 | TEC_dmg1 | BRD4 |
| 2.46E-128 | 0.26348316 | 0.398 | 0.226 | 5.811E-124 | TEC_dmg1 | FXR1 |
| 4.996E-128 | 0.34564074 | 0.943 | 0.869 | 1.18E-123 | TEC_dmg1 | RPL3 |
| 1.358E-127 | 0.25421395 | 0.349 | 0.184 | 3.208E-123 | TEC_dmg1 | TMEM106B |
| 1.441E-127 | 0.31756565 | 0.232 | 0.103 | 3.405E-123 | TEC_dmg1 | RRAD |
| 2.391E-127 | 0.30927791 | 0.418 | 0.241 | 5.648E-123 | TEC_dmg1 | CHD2 |
| 5.62E-127 | -0.5913798 | 0.033 | 0.182 | 1.328E-122 | TEC_dmg1 | FXYD3 |
| 6.729E-127 | -0.2797674 | 0.012 | 0.151 | 1.59E-122 | TEC_dmg1 | EMILIN2 |
| 7.286E-127 | -0.3976559 | 0.009 | 0.147 | 1.721E-122 | TEC_dmg1 | BCAS1 |
| 1.185E-126 | 0.42493268 | 0.443 | 0.265 | 2.8E-122 | TEC_dmg1 | H1F0 |
| 1.262E-126 | 0.28054259 | 0.507 | 0.313 | 2.982E-122 | TEC_dmg1 | PPP1R14B |
| 7.88E-126 | -0.296828 | 0.005 | 0.139 | 1.862E-121 | TEC_dmg1 | IRAK3 |
| 1.652E-125 | -0.3612293 | 0.016 | 0.156 | 3.902E-121 | TEC_dmg1 | TLR2 |
| 4.976E-125 | 0.27217714 | 0.438 | 0.26 | 1.175E-120 | TEC_dmg1 | MRPL51 |
| 6.179E-125 | -0.2956059 | 0.008 | 0.143 | 1.46E-120 | TEC_dmg1 | LILRB3 |
| 7.74E-125 | -0.676497 | 0.008 | 0.143 | 1.828E-120 | TEC_dmg1 | GABRP |
| 6.577E-124 | 0.25306711 | 0.369 | 0.206 | 1.554E-119 | TEC_dmg1 | TMEM14B |
| 2.81E-123 | 0.26005015 | 0.416 | 0.244 | 6.637E-119 | TEC_dmg1 | OCIAD1 |
| 2.982E-123 | 0.37430349 | 0.91 | 0.853 | 7.045E-119 | TEC_dmg1 | RPL7 |
| 4.807E-123 | 0.2905429 | 0.56 | 0.368 | 1.136E-118 | TEC_dmg1 | CNBP |
| 7.623E-123 | 0.30728664 | 0.699 | 0.54 | 1.801E-118 | TEC_dmg1 | MORF4L1 |
| 1.367E-122 | 0.32021027 | 0.719 | 0.576 | 3.228E-118 | TEC_dmg1 | SLC25A3 |
| 5.523E-122 | 0.34916771 | 0.483 | 0.311 | 1.305E-117 | TEC_dmg1 | EIF1B |

| | | | | | | |
|---|---|---|---|---|---|---|
| 8.045E-122 | -1.077735 | 0.64 | 0.75 | 1.901E-117 | TEC_dmg1 | VIM |
| 8.262E-122 | -0.3278911 | 0.057 | 0.216 | 1.952E-117 | TEC_dmg1 | IFI16 |
| 8.916E-122 | 0.37691242 | 0.788 | 0.686 | 2.106E-117 | TEC_dmg1 | LRRFIP1 |
| 1.711E-121 | 0.29767175 | 0.569 | 0.385 | 4.043E-117 | TEC_dmg1 | CSDE1 |
| 2.164E-121 | -0.3200843 | 0.004 | 0.132 | 5.113E-117 | TEC_dmg1 | LILRB2 |
| 2.976E-121 | -0.2908885 | 0.002 | 0.129 | 7.031E-117 | TEC_dmg1 | CD86 |
| 3.368E-121 | 0.27049333 | 0.527 | 0.335 | 7.956E-117 | TEC_dmg1 | EIF3E |
| 6.872E-121 | 0.27026722 | 0.357 | 0.202 | 1.623E-116 | TEC_dmg1 | SURF4 |
| 1.004E-120 | -0.4029936 | 0.003 | 0.131 | 2.372E-116 | TEC_dmg1 | FCGR3A |
| 1.773E-120 | -0.6785454 | 0.011 | 0.143 | 4.188E-116 | TEC_dmg1 | DHRS9 |
| 2.117E-120 | 0.25568248 | 0.385 | 0.22 | 5.001E-116 | TEC_dmg1 | ETF1 |
| 2.171E-120 | 0.25075062 | 0.34 | 0.185 | 5.13E-116 | TEC_dmg1 | PRNP |
| 3.873E-120 | 0.29866411 | 0.514 | 0.336 | 9.15E-116 | TEC_dmg1 | SYF2 |
| 5.153E-120 | -1.4476483 | 0.061 | 0.212 | 1.217E-115 | TEC_dmg1 | KRT17 |
| 8.232E-120 | 0.26113092 | 0.389 | 0.225 | 1.945E-115 | TEC_dmg1 | SMIM26 |
| 1.018E-119 | -0.2601303 | 0.019 | 0.156 | 2.405E-115 | TEC_dmg1 | CCDC69 |
| 1.201E-119 | -0.265831 | 0.002 | 0.127 | 2.836E-115 | TEC_dmg1 | C3AR1 |
| 1.689E-119 | 0.29492026 | 0.449 | 0.275 | 3.989E-115 | TEC_dmg1 | HNRNPH3 |
| 3.61E-119 | -0.3690169 | 0.01 | 0.14 | 8.528E-115 | TEC_dmg1 | LRG1 |
| 1.585E-118 | 0.27280686 | 0.412 | 0.246 | 3.743E-114 | TEC_dmg1 | PSMD7 |
| 1.599E-118 | 0.3126459 | 0.476 | 0.305 | 3.778E-114 | TEC_dmg1 | LEPROT |
| 2.03E-118 | 0.35917806 | 0.157 | 0.06 | 4.795E-114 | TEC_dmg1 | TNFRSF11B |
| 4.203E-118 | 0.3009658 | 0.754 | 0.591 | 9.929E-114 | TEC_dmg1 | EZR |
| 5.021E-118 | 0.26189652 | 0.355 | 0.197 | 1.186E-113 | TEC_dmg1 | CELF1 |
| 5.55E-118 | 0.26148115 | 0.343 | 0.19 | 1.311E-113 | TEC_dmg1 | UBE2J1 |
| 6.777E-118 | 0.32261288 | 0.111 | 0.034 | 1.601E-113 | TEC_dmg1 | KRTAP2-3 |
| 1.119E-117 | 0.30025614 | 0.534 | 0.356 | 2.644E-113 | TEC_dmg1 | KIF5B |
| 2.184E-117 | -0.3577577 | 0.013 | 0.144 | 5.158E-113 | TEC_dmg1 | RGCC |
| 6.184E-117 | 0.29816172 | 0.58 | 0.397 | 1.461E-112 | TEC_dmg1 | PCBP2 |
| 1.112E-116 | 0.28121407 | 0.413 | 0.245 | 2.628E-112 | TEC_dmg1 | CUTA |
| 1.26E-116 | 0.33835249 | 0.546 | 0.37 | 2.977E-112 | TEC_dmg1 | DAD1 |
| 1.293E-116 | -0.2798665 | 0.005 | 0.131 | 3.054E-112 | TEC_dmg1 | CSF1R |
| 2.416E-116 | 0.25808044 | 0.386 | 0.222 | 5.708E-112 | TEC_dmg1 | SNRPE |
| 3.453E-116 | -0.304142 | 0.038 | 0.18 | 8.158E-112 | TEC_dmg1 | GLIPR2 |
| 4.374E-116 | 0.35377589 | 0.675 | 0.514 | 1.033E-111 | TEC_dmg1 | EIF4A2 |
| 4.908E-116 | -0.2778134 | 0.027 | 0.165 | 1.16E-111 | TEC_dmg1 | VSIR |
| 6.572E-116 | 0.33601309 | 0.34 | 0.19 | 1.552E-111 | TEC_dmg1 | SLFN5 |
| 1.382E-115 | 0.27646149 | 0.461 | 0.28 | 3.266E-111 | TEC_dmg1 | EIF5B |
| 2.716E-115 | -1.2084589 | 0.008 | 0.134 | 6.415E-111 | TEC_dmg1 | CCL3 |
| 7.148E-115 | -0.8140821 | 0.799 | 0.865 | 1.689E-110 | TEC_dmg1 | SH3BGRL3 |
| 1.016E-114 | 0.25415777 | 0.18 | 0.075 | 2.4E-110 | TEC_dmg1 | CSRP2 |
| 8.974E-114 | 0.29559631 | 0.75 | 0.624 | 2.12E-109 | TEC_dmg1 | ARF1 |
| 1.053E-113 | 0.27441856 | 0.382 | 0.225 | 2.486E-109 | TEC_dmg1 | TXNL1 |
| 1.634E-113 | 0.43875151 | 0.334 | 0.189 | 3.859E-109 | TEC_dmg1 | MANF |
| 1.723E-113 | -0.4388225 | 0.069 | 0.222 | 4.071E-109 | TEC_dmg1 | LY6E |
| 2.085E-113 | 0.33717563 | 0.631 | 0.448 | 4.926E-109 | TEC_dmg1 | CDKN1A |
| 2.259E-113 | -0.6278157 | 0.003 | 0.123 | 5.337E-109 | TEC_dmg1 | KRT78 |
| 4.855E-113 | -0.2770224 | 0.095 | 0.256 | 1.147E-108 | TEC_dmg1 | OSTF1 |
| 9.973E-113 | -0.31332 | 0.026 | 0.161 | 2.356E-108 | TEC_dmg1 | BCAT1 |
| 3.191E-112 | -0.6070146 | 0.006 | 0.128 | 7.537E-108 | TEC_dmg1 | MNDA |

| | | | | | | |
|---|---|---|---|---|---|---|
| 3.264E-112 | 0.29573505 | 0.574 | 0.392 | 7.711E-108 | TEC_dmg1 | PTP4A2 |
| 3.668E-112 | -0.4387028 | 0.225 | 0.389 | 8.664E-108 | TEC_dmg1 | METRNL |
| 4.677E-112 | -0.3661551 | 0.005 | 0.127 | 1.105E-107 | TEC_dmg1 | CD163 |
| 8.258E-112 | -1.1624791 | 0.006 | 0.128 | 1.951E-107 | TEC_dmg1 | SPINK5 |
| 8.644E-112 | 0.25382597 | 0.239 | 0.115 | 2.042E-107 | TEC_dmg1 | ABL2 |
| 1.293E-111 | -0.3637398 | 0.012 | 0.137 | 3.055E-107 | TEC_dmg1 | MARCO |
| 1.685E-111 | 0.27406786 | 0.432 | 0.268 | 3.98E-107 | TEC_dmg1 | EIF6 |
| 1.897E-111 | -0.483244 | 0.004 | 0.123 | 4.48E-107 | TEC_dmg1 | TMPRSS11A |
| 1.921E-111 | -0.5766846 | 0.195 | 0.362 | 4.537E-107 | TEC_dmg1 | IER5 |
| 2.133E-111 | -0.846946 | 0.034 | 0.168 | 5.04E-107 | TEC_dmg1 | KRT15 |
| 3.385E-111 | -0.2764401 | 0.004 | 0.124 | 7.995E-107 | TEC_dmg1 | PECAM1 |
| 6.155E-111 | 0.25523893 | 0.534 | 0.35 | 1.454E-106 | TEC_dmg1 | REEP5 |
| 1.629E-110 | 0.25404185 | 0.315 | 0.17 | 3.848E-106 | TEC_dmg1 | LMO4 |
| 1.838E-110 | 0.28837297 | 0.25 | 0.122 | 4.341E-106 | TEC_dmg1 | HMGCS1 |
| 2.081E-110 | -0.2539232 | 0.006 | 0.126 | 4.915E-106 | TEC_dmg1 | ADGRE2 |
| 2.609E-110 | -0.3472147 | 0.088 | 0.242 | 6.164E-106 | TEC_dmg1 | FCGRT |
| 3.021E-110 | -0.2815837 | 0.036 | 0.173 | 7.136E-106 | TEC_dmg1 | WIPF1 |
| 1.6E-109 | 0.27736635 | 0.527 | 0.352 | 3.78E-105 | TEC_dmg1 | GNB1 |
| 1.955E-109 | 0.33778877 | 0.594 | 0.425 | 4.619E-105 | TEC_dmg1 | PDIA3 |
| 2.18E-109 | -0.5876151 | 0.033 | 0.167 | 5.149E-105 | TEC_dmg1 | MUC4 |
| 3.862E-109 | -0.4441139 | 0.152 | 0.312 | 9.124E-105 | TEC_dmg1 | REL |
| 4.56E-109 | -0.3497348 | 0.02 | 0.148 | 1.077E-104 | TEC_dmg1 | HLA-DMA |
| 1.119E-108 | -0.2513907 | 0.016 | 0.142 | 2.644E-104 | TEC_dmg1 | APBB1IP |
| 1.305E-108 | 0.31497191 | 0.713 | 0.548 | 3.083E-104 | TEC_dmg1 | SLC25A6 |
| 1.928E-108 | -1.1305516 | 0.156 | 0.309 | 4.555E-104 | TEC_dmg1 | ADIRF |
| 2.356E-108 | 0.25722166 | 0.361 | 0.212 | 5.566E-104 | TEC_dmg1 | PSMC1 |
| 2.449E-108 | 0.26873926 | 0.417 | 0.252 | 5.786E-104 | TEC_dmg1 | GADD45GIP1 |
| 3.033E-108 | -0.7999151 | 0.006 | 0.124 | 7.165E-104 | TEC_dmg1 | IL7R |
| 1.992E-107 | 0.31605718 | 0.405 | 0.249 | 4.706E-103 | TEC_dmg1 | TUBB |
| 1.997E-107 | 0.29026012 | 0.752 | 0.595 | 4.718E-103 | TEC_dmg1 | HNRNPA1 |
| 2.009E-107 | 0.29141894 | 0.538 | 0.371 | 4.745E-103 | TEC_dmg1 | RAB2A |
| 2.713E-107 | 0.32292962 | 0.84 | 0.78 | 6.408E-103 | TEC_dmg1 | BTF3 |
| 3.186E-107 | -0.4432783 | 0.014 | 0.136 | 7.526E-103 | TEC_dmg1 | SRD5A3 |
| 3.688E-107 | 0.28677487 | 0.754 | 0.617 | 8.713E-103 | TEC_dmg1 | YBX3 |
| 4.066E-107 | -0.3476285 | 0.002 | 0.116 | 9.604E-103 | TEC_dmg1 | A2ML1 |
| 4.084E-107 | -0.4864074 | 0.066 | 0.208 | 9.647E-103 | TEC_dmg1 | SAMD9 |
| 9.431E-107 | 0.25030234 | 0.361 | 0.21 | 2.228E-102 | TEC_dmg1 | NARS |
| 4.044E-106 | -0.4788096 | 0.057 | 0.198 | 9.553E-102 | TEC_dmg1 | KLF2 |
| 3.3E-105 | -1.134148 | 0.004 | 0.118 | 7.795E-101 | TEC_dmg1 | CCL4 |
| 3.441E-105 | 0.38372198 | 0.981 | 0.984 | 8.129E-101 | TEC_dmg1 | EIF1 |
| 4.039E-105 | 0.26050201 | 0.426 | 0.262 | 9.541E-101 | TEC_dmg1 | TAF7 |
| 4.086E-105 | 0.3250398 | 0.95 | 0.897 | 9.652E-101 | TEC_dmg1 | RPS7 |
| 4.226E-105 | 0.27132662 | 0.382 | 0.23 | 9.983E-101 | TEC_dmg1 | OSTC |
| 4.747E-105 | -0.2702555 | 0.024 | 0.151 | 1.121E-100 | TEC_dmg1 | IFI30 |
| 4.298E-104 | 0.39946972 | 0.718 | 0.6 | 1.015E-99 | TEC_dmg1 | SEC61B |
| 5.329E-104 | -0.3176913 | 0.032 | 0.161 | 1.259E-99 | TEC_dmg1 | PLXDC2 |
| 9.481E-104 | -0.2535179 | 0.021 | 0.143 | 2.24E-99 | TEC_dmg1 | PILRA |
| 3.533E-103 | 0.26803324 | 0.666 | 0.476 | 8.345E-99 | TEC_dmg1 | NCL |
| 4.023E-103 | 0.30851309 | 0.962 | 0.944 | 9.503E-99 | TEC_dmg1 | RPL13A |
| 4.389E-103 | 0.33046439 | 0.947 | 0.883 | 1.0368E-98 | TEC_dmg1 | RPS23 |

| | | | | | | |
|---|---|---|---|---|---|---|
| 6.501E-103 | -0.2955465 | 0.003 | 0.113 | 1.5357E-98 | TEC_dmg1 | LILRB1 |
| 1.039E-102 | -1.2140559 | 0.022 | 0.143 | 2.4547E-98 | TEC_dmg1 | PLAT |
| 2.535E-102 | 0.28725561 | 0.645 | 0.492 | 5.9896E-98 | TEC_dmg1 | JAK1 |
| 3.007E-102 | 0.32242974 | 0.936 | 0.864 | 7.1032E-98 | TEC_dmg1 | RPL14 |
| 3.526E-102 | -0.3236379 | 0.062 | 0.201 | 8.33E-98 | TEC_dmg1 | PYCARD |
| 4.326E-102 | 0.33389985 | 0.947 | 0.901 | 1.022E-97 | TEC_dmg1 | RPL7A |
| 6.716E-102 | -0.3340194 | 0.077 | 0.221 | 1.5865E-97 | TEC_dmg1 | STK17B |
| 6.752E-102 | -0.259919 | 0.027 | 0.152 | 1.5951E-97 | TEC_dmg1 | LRP1 |
| 6.933E-102 | -0.3011463 | 0.005 | 0.116 | 1.6378E-97 | TEC_dmg1 | MS4A6A |
| 1.095E-101 | 0.28546752 | 0.282 | 0.152 | 2.5878E-97 | TEC_dmg1 | MIR22HG |
| 1.405E-100 | 0.25045865 | 0.338 | 0.193 | 3.3183E-96 | TEC_dmg1 | RBBP6 |
| 9.914E-100 | 0.25703284 | 0.474 | 0.303 | 2.3421E-95 | TEC_dmg1 | BRD2 |
| 1.59E-99 | -0.355423 | 0.006 | 0.116 | 3.7561E-95 | TEC_dmg1 | CSF3R |
| 4.11E-99 | -0.81474 | 0.006 | 0.114 | 9.709E-95 | TEC_dmg1 | ZG16B |
| 7.692E-99 | -0.3750771 | 0.012 | 0.125 | 1.817E-94 | TEC_dmg1 | CEACAM1 |
| 2.187E-98 | -0.306045 | 0.004 | 0.111 | 5.1664E-94 | TEC_dmg1 | LRMP |
| 2.7975E-98 | 0.3714265 | 0.677 | 0.534 | 6.6085E-94 | TEC_dmg1 | PPIB |
| 3.5474E-98 | 0.25466971 | 0.42 | 0.263 | 8.3801E-94 | TEC_dmg1 | PLEKHB2 |
| 3.6185E-98 | -0.5880781 | 0.031 | 0.152 | 8.548E-94 | TEC_dmg1 | SNCG |
| 4.6702E-98 | -0.4054167 | 0.005 | 0.113 | 1.1032E-93 | TEC_dmg1 | CLIC3 |
| 6.2355E-98 | -0.3844071 | 0.035 | 0.159 | 1.473E-93 | TEC_dmg1 | RGS2 |
| 8.4642E-98 | -0.5174666 | 0.489 | 0.614 | 1.9995E-93 | TEC_dmg1 | MCL1 |
| 1.0754E-97 | -0.2869596 | 0.055 | 0.188 | 2.5404E-93 | TEC_dmg1 | ABCA1 |
| 1.5073E-97 | -0.2929264 | 0.011 | 0.121 | 3.5606E-93 | TEC_dmg1 | FUT3 |
| 4.3945E-97 | 0.29363323 | 0.642 | 0.488 | 1.0381E-92 | TEC_dmg1 | LAPTM4A |
| 6.054E-97 | 0.34699302 | 0.921 | 0.917 | 1.4301E-92 | TEC_dmg1 | RPS20 |
| 3.4441E-96 | 0.30935066 | 0.942 | 0.916 | 8.136E-92 | TEC_dmg1 | RPL9 |
| 5.3082E-96 | -0.4185124 | 0.008 | 0.116 | 1.254E-91 | TEC_dmg1 | THBD |
| 8.0256E-96 | 0.2716859 | 0.508 | 0.35 | 1.8959E-91 | TEC_dmg1 | ELOC |
| 1.0211E-95 | 0.32351628 | 0.513 | 0.356 | 2.4122E-91 | TEC_dmg1 | CIRBP |
| 1.2863E-95 | 0.30618033 | 0.503 | 0.342 | 3.0386E-91 | TEC_dmg1 | TRMT112 |
| 2.0991E-95 | 0.25051195 | 0.327 | 0.184 | 4.9586E-91 | TEC_dmg1 | PTGR1 |
| 2.6963E-95 | -0.9128065 | 0.027 | 0.143 | 6.3694E-91 | TEC_dmg1 | RHCG |
| 1.0697E-94 | -0.6235479 | 0.006 | 0.111 | 2.5269E-90 | TEC_dmg1 | HLA-DQA1 |
| 1.0809E-94 | -0.3258593 | 0.119 | 0.263 | 2.5534E-90 | TEC_dmg1 | SH3BGRL |
| 2.0682E-94 | -1.3089501 | 0.014 | 0.123 | 4.8857E-90 | TEC_dmg1 | SPRR1B |
| 2.5026E-94 | 0.25735469 | 0.328 | 0.191 | 5.9119E-90 | TEC_dmg1 | C6orf48 |
| 5.271E-94 | 0.30387127 | 0.834 | 0.717 | 1.2452E-89 | TEC_dmg1 | PKM |
| 1.1231E-93 | -0.4311868 | 0.064 | 0.195 | 2.6532E-89 | TEC_dmg1 | SGK1 |
| 2.447E-93 | 0.26279419 | 0.765 | 0.647 | 5.7805E-89 | TEC_dmg1 | NDUFA4 |
| 3.9374E-93 | -0.3913121 | 0.484 | 0.657 | 9.3014E-89 | TEC_dmg1 | TXNIP |
| 5.2332E-93 | 0.27550566 | 0.896 | 0.861 | 1.2362E-88 | TEC_dmg1 | SRP14 |
| 6.9247E-93 | 0.33671888 | 0.843 | 0.775 | 1.6358E-88 | TEC_dmg1 | MAP1LC3B |
| 7.4674E-93 | -1.581189 | 0.01 | 0.116 | 1.764E-88 | TEC_dmg1 | CRCT1 |
| 7.6827E-93 | -0.307207 | 0.051 | 0.176 | 1.8149E-88 | TEC_dmg1 | JARID2 |
| 1.418E-92 | -0.494704 | 0.006 | 0.109 | 3.3497E-88 | TEC_dmg1 | RETN |
| 2.6197E-92 | -0.3347433 | 0.017 | 0.128 | 6.1885E-88 | TEC_dmg1 | DUSP2 |
| 3.0846E-92 | -0.2726819 | 0.002 | 0.1 | 7.2867E-88 | TEC_dmg1 | KLK11 |
| 3.9469E-92 | 0.25836905 | 0.418 | 0.265 | 9.3237E-88 | TEC_dmg1 | PTP4A1 |
| 7.2771E-92 | -0.2942082 | 0.004 | 0.104 | 1.7191E-87 | TEC_dmg1 | GPR183 |

| | | | | | | |
|---|---|---|---|---|---|---|
| 7.0403E-91 | -0.4131924 | 0.011 | 0.116 | 1.6631E-86 | TEC_dmg1 | TMC5 |
| 1.3889E-90 | 0.29591953 | 0.481 | 0.307 | 3.281E-86 | TEC_dmg1 | IFITM3 |
| 8.8365E-90 | 0.26997162 | 0.106 | 0.038 | 2.0874E-85 | TEC_dmg1 | IFIT1 |
| 1.7218E-89 | 0.37049378 | 0.558 | 0.416 | 4.0673E-85 | TEC_dmg1 | SELENOK |
| 2.1436E-89 | -0.5659069 | 0.899 | 0.942 | 5.0639E-85 | TEC_dmg1 | TMSB4X |
| 2.4898E-89 | 0.32885156 | 0.96 | 0.957 | 5.8817E-85 | TEC_dmg1 | RPS18 |
| 5.2864E-89 | 0.33283632 | 0.503 | 0.338 | 1.2488E-84 | TEC_dmg1 | ATF3 |
| 6.0104E-89 | -0.4041335 | 0.082 | 0.214 | 1.4198E-84 | TEC_dmg1 | FOSB |
| 1.5732E-88 | 0.30039811 | 0.952 | 0.92 | 3.7163E-84 | TEC_dmg1 | RPL8 |
| 3.1082E-88 | -0.2543873 | 0.026 | 0.138 | 7.3425E-84 | TEC_dmg1 | PLBD1 |
| 1.72E-87 | -0.6179482 | 0.074 | 0.2 | 4.0631E-83 | TEC_dmg1 | ANKRD36C |
| 1.7536E-87 | 0.31767788 | 0.585 | 0.439 | 4.1425E-83 | TEC_dmg1 | SEC61G |
| 2.6264E-87 | -0.8317353 | 0.541 | 0.669 | 6.2044E-83 | TEC_dmg1 | HSPA1A |
| 6.5003E-87 | 0.26420367 | 0.484 | 0.334 | 1.5356E-82 | TEC_dmg1 | PSMD8 |
| 6.8115E-87 | 0.26717302 | 0.916 | 0.842 | 1.6091E-82 | TEC_dmg1 | RPS4X |
| 8.8248E-87 | 0.27517673 | 0.617 | 0.464 | 2.0847E-82 | TEC_dmg1 | HSPD1 |
| 1.4569E-86 | -0.2964445 | 0.043 | 0.16 | 3.4416E-82 | TEC_dmg1 | DMXL2 |
| 1.5001E-86 | -0.5537785 | 0.062 | 0.183 | 3.5438E-82 | TEC_dmg1 | HLA-DQB1 |
| 2.6816E-86 | 0.27195926 | 0.758 | 0.655 | 6.3348E-82 | TEC_dmg1 | HNRNPC |
| 2.8097E-86 | -1.4765005 | 0.015 | 0.117 | 6.6374E-82 | TEC_dmg1 | CNFN |
| 4.4781E-86 | -0.4668897 | 0.034 | 0.145 | 1.0579E-81 | TEC_dmg1 | HLA-DRB5 |
| 5.2076E-86 | 0.26040367 | 0.6 | 0.44 | 1.2302E-81 | TEC_dmg1 | ST13 |
| 7.5252E-86 | -0.2909144 | 0.064 | 0.189 | 1.7777E-81 | TEC_dmg1 | CCDC88A |
| 1.2819E-85 | -0.2708898 | 0.044 | 0.16 | 3.0282E-81 | TEC_dmg1 | SMAP2 |
| 2.0752E-85 | 0.30880222 | 0.338 | 0.206 | 4.9023E-81 | TEC_dmg1 | TGM2 |
| 2.3037E-85 | 0.31722511 | 0.933 | 0.876 | 5.442E-81 | TEC_dmg1 | RPL12 |
| 4.1563E-85 | -0.4167176 | 0.461 | 0.616 | 9.8185E-81 | TEC_dmg1 | NFKBIA |
| 6.0396E-85 | -0.3111959 | 0.016 | 0.117 | 1.4267E-80 | TEC_dmg1 | FGL2 |
| 1.6146E-84 | -0.771484 | 0.189 | 0.324 | 3.8141E-80 | TEC_dmg1 | PHLDA1 |
| 4.3382E-84 | 0.2992194 | 0.951 | 0.931 | 1.0248E-79 | TEC_dmg1 | RPS12 |
| 5.5589E-84 | -0.3668942 | 0.75 | 0.844 | 1.3132E-79 | TEC_dmg1 | HLA-E |
| 7.6084E-84 | 0.25649278 | 0.783 | 0.678 | 1.7973E-79 | TEC_dmg1 | CAST |
| 7.6977E-84 | -0.2789281 | 0.065 | 0.188 | 1.8184E-79 | TEC_dmg1 | GLIPR1 |
| 1.0535E-83 | -0.3770397 | 0.016 | 0.117 | 2.4886E-79 | TEC_dmg1 | KRT23 |
| 1.6496E-83 | -1.1128951 | 0.235 | 0.356 | 3.8969E-79 | TEC_dmg1 | CTSL |
| 6.648E-83 | -0.311852 | 0.135 | 0.271 | 1.5705E-78 | TEC_dmg1 | ABRACL |
| 1.3016E-82 | -0.2695375 | 0.161 | 0.316 | 3.0748E-78 | TEC_dmg1 | SERPINA1 |
| 2.6118E-82 | -0.380131 | 0.094 | 0.22 | 6.1698E-78 | TEC_dmg1 | SOCS3 |
| 1.1391E-81 | -0.2632724 | 0.085 | 0.213 | 2.6908E-77 | TEC_dmg1 | NR3C1 |
| 2.1152E-81 | -0.2687 | 0.076 | 0.198 | 4.9966E-77 | TEC_dmg1 | RNF130 |
| 7.5755E-81 | -0.5816954 | 0.01 | 0.104 | 1.7896E-76 | TEC_dmg1 | PTGS2 |
| 8.8312E-81 | -0.378466 | 0.029 | 0.132 | 2.0862E-76 | TEC_dmg1 | TMEM40 |
| 1.0294E-80 | -0.6888219 | 0.076 | 0.195 | 2.4317E-76 | TEC_dmg1 | AQP3 |
| 3.3007E-80 | 0.25968748 | 0.956 | 0.908 | 7.7972E-76 | TEC_dmg1 | RPS8 |
| 8.7452E-80 | -0.333291 | 0.011 | 0.105 | 2.0659E-75 | TEC_dmg1 | KLK10 |
| 1.2471E-79 | -0.6464536 | 0.377 | 0.518 | 2.9461E-75 | TEC_dmg1 | HSPA1B |
| 3.916E-79 | -0.3681121 | 0.528 | 0.634 | 9.2507E-75 | TEC_dmg1 | ACTR2 |
| 1.6611E-78 | -0.3686682 | 0.099 | 0.223 | 3.924E-74 | TEC_dmg1 | NINJ1 |
| 1.7191E-78 | 0.29522493 | 0.919 | 0.854 | 4.061E-74 | TEC_dmg1 | RPL23 |
| 3.7206E-78 | -0.4490616 | 0.01 | 0.101 | 8.7893E-74 | TEC_dmg1 | RNASE7 |

| | | | | | | |
|---|---|---|---|---|---|---|
| 4.4918E-78 | -0.2637817 | 0.622 | 0.737 | 1.0611E-73 | TEC_dmg1 | ELOB |
| 9.6153E-78 | 0.28082702 | 0.943 | 0.894 | 2.2714E-73 | TEC_dmg1 | RPL21 |
| 4.5848E-77 | 0.26278577 | 0.402 | 0.269 | 1.0831E-72 | TEC_dmg1 | RBM3 |
| 1.1225E-76 | -0.9102587 | 0.077 | 0.191 | 2.6516E-72 | TEC_dmg1 | SPINK1 |
| 3.6923E-76 | -0.3734249 | 0.056 | 0.164 | 8.7223E-72 | TEC_dmg1 | TMPRSS4 |
| 5.1259E-76 | 0.27454919 | 0.427 | 0.295 | 1.2109E-71 | TEC_dmg1 | BUD31 |
| 9.5635E-76 | 0.25300211 | 0.363 | 0.235 | 2.2592E-71 | TEC_dmg1 | CD47 |
| 9.7973E-76 | 0.28245707 | 0.788 | 0.729 | 2.3144E-71 | TEC_dmg1 | CCNI |
| 1.0694E-75 | -0.4002845 | 0.624 | 0.765 | 2.5263E-71 | TEC_dmg1 | TPM4 |
| 2.7251E-75 | 0.27603896 | 0.25 | 0.141 | 6.4375E-71 | TEC_dmg1 | MDK |
| 3.2298E-75 | -0.3571204 | 0.178 | 0.307 | 7.6298E-71 | TEC_dmg1 | IFNGR2 |
| 6.8114E-75 | 0.52492833 | 0.642 | 0.528 | 1.6091E-70 | TEC_dmg1 | UPP1 |
| 2.1053E-74 | -0.5522241 | 0.088 | 0.199 | 4.9733E-70 | TEC_dmg1 | EPS8L1 |
| 2.3378E-74 | 0.2801678 | 0.932 | 0.898 | 5.5226E-70 | TEC_dmg1 | RPL24 |
| 7.1512E-74 | -0.3353005 | 0.072 | 0.186 | 1.6893E-69 | TEC_dmg1 | NR4A1 |
| 1.3895E-73 | 0.2571681 | 0.511 | 0.377 | 3.2824E-69 | TEC_dmg1 | SPCS2 |
| 3.5215E-73 | 0.25198096 | 0.783 | 0.679 | 8.3187E-69 | TEC_dmg1 | PSMA7 |
| 4E-73 | -0.5923816 | 0.429 | 0.535 | 9.4492E-69 | TEC_dmg1 | CEBPB |
| 8.1324E-73 | -0.523201 | 0.059 | 0.163 | 1.9211E-68 | TEC_dmg1 | CDC42EP5 |
| 2.046E-72 | -0.2922833 | 0.058 | 0.165 | 4.8332E-68 | TEC_dmg1 | ADGRE5 |
| 2.1256E-72 | -0.4712191 | 0.017 | 0.106 | 5.0213E-68 | TEC_dmg1 | DHRS2 |
| 8.8354E-72 | -0.7840021 | 0.091 | 0.197 | 2.0872E-67 | TEC_dmg1 | TMPRSS2 |
| 2.0355E-71 | 0.3381307 | 0.296 | 0.177 | 4.8085E-67 | TEC_dmg1 | SLC5A3 |
| 7.5111E-71 | -0.3451087 | 0.478 | 0.59 | 1.7744E-66 | TEC_dmg1 | PGK1 |
| 1.898E-70 | -0.4083185 | 0.166 | 0.286 | 4.4835E-66 | TEC_dmg1 | ETS2 |
| 4.2086E-70 | -0.3125691 | 0.179 | 0.305 | 9.9419E-66 | TEC_dmg1 | FXYD5 |
| 6.8707E-70 | -0.7005243 | 0.082 | 0.19 | 1.6231E-65 | TEC_dmg1 | F3 |
| 1.2274E-69 | 0.27788655 | 0.735 | 0.633 | 2.8996E-65 | TEC_dmg1 | EDF1 |
| 2.6733E-69 | -0.3307818 | 0.145 | 0.27 | 6.315E-65 | TEC_dmg1 | NABP1 |
| 2.7186E-69 | -0.3182055 | 0.122 | 0.239 | 6.4221E-65 | TEC_dmg1 | GALNT1 |
| 5.8717E-69 | -0.4321744 | 0.512 | 0.606 | 1.3871E-64 | TEC_dmg1 | NPC2 |
| 7.167E-69 | -0.3214695 | 0.177 | 0.303 | 1.6931E-64 | TEC_dmg1 | SKIL |
| 9.7912E-69 | -1.4380224 | 0.052 | 0.154 | 2.313E-64 | TEC_dmg1 | CXCL3 |
| 2.776E-68 | -0.5311365 | 0.051 | 0.151 | 6.5577E-64 | TEC_dmg1 | CDKN2A |
| 5.3972E-68 | -0.6764722 | 0.048 | 0.148 | 1.275E-63 | TEC_dmg1 | CXCL1 |
| 2.2401E-67 | 0.26438987 | 0.46 | 0.32 | 5.2917E-63 | TEC_dmg1 | SNHG8 |
| 2.6367E-67 | 0.27403446 | 0.846 | 0.809 | 6.2287E-63 | TEC_dmg1 | RPL36AL |
| 3.9373E-67 | 0.28357157 | 0.947 | 0.891 | 9.301E-63 | TEC_dmg1 | RPS9 |
| 1.27E-66 | -0.7349155 | 0.663 | 0.728 | 3E-62 | TEC_dmg1 | HSPB1 |
| 1.3553E-66 | -0.2883123 | 0.029 | 0.118 | 3.2017E-62 | TEC_dmg1 | STS |
| 8.7242E-66 | -0.3526523 | 0.692 | 0.784 | 2.0609E-61 | TEC_dmg1 | TPI1 |
| 1.4428E-65 | -0.4632767 | 0.038 | 0.13 | 3.4082E-61 | TEC_dmg1 | AGR2 |
| 2.3177E-65 | -0.4033222 | 0.211 | 0.338 | 5.475E-61 | TEC_dmg1 | CEBPD |
| 3.7225E-65 | -0.4545416 | 0.191 | 0.31 | 8.7938E-61 | TEC_dmg1 | BTG2 |
| 4.5542E-65 | -0.3747718 | 0.108 | 0.216 | 1.0758E-60 | TEC_dmg1 | SLC43A2 |
| 6.1307E-65 | 0.47558433 | 0.986 | 0.98 | 1.4483E-60 | TEC_dmg1 | MALAT1 |
| 1.0296E-64 | -0.5164268 | 0.113 | 0.218 | 2.4322E-60 | TEC_dmg1 | CDKN2B |
| 1.2177E-64 | -0.3522198 | 0.068 | 0.169 | 2.8766E-60 | TEC_dmg1 | SERPINB9 |
| 2.4419E-64 | -0.3113169 | 0.462 | 0.583 | 5.7684E-60 | TEC_dmg1 | VMP1 |
| 1.8533E-63 | -0.2988365 | 0.123 | 0.233 | 4.3781E-59 | TEC_dmg1 | GRINA |

| | | | | | | |
|---|---|---|---|---|---|---|
| 2.7092E-63 | -0.5419071 | 0.931 | 0.94 | 6.4E-59 | TEC_dmg1 | SAT1 |
| 2.7156E-63 | -0.2729014 | 0.089 | 0.195 | 6.4151E-59 | TEC_dmg1 | CKLF |
| 5.0678E-63 | -0.3751304 | 0.093 | 0.198 | 1.1972E-58 | TEC_dmg1 | DUSP6 |
| 8.6497E-63 | -1.3436484 | 0.14 | 0.241 | 2.0433E-58 | TEC_dmg1 | PRSS22 |
| 4.546E-61 | -0.3097947 | 0.218 | 0.336 | 1.0739E-56 | TEC_dmg1 | FOSL2 |
| 6.1454E-61 | -0.2895145 | 0.047 | 0.139 | 1.4517E-56 | TEC_dmg1 | CD83 |
| 9.1734E-61 | -0.3015938 | 0.136 | 0.248 | 2.167E-56 | TEC_dmg1 | SMCHD1 |
| 3.5281E-60 | -0.2565907 | 0.024 | 0.105 | 8.3344E-56 | TEC_dmg1 | RNF39 |
| 4.5431E-60 | -0.3249041 | 0.106 | 0.213 | 1.0732E-55 | TEC_dmg1 | CTSC |
| 1.3998E-58 | -0.605491 | 0.401 | 0.514 | 3.3068E-54 | TEC_dmg1 | DNAJB1 |
| 1.7275E-58 | -0.257771 | 0.077 | 0.174 | 4.081E-54 | TEC_dmg1 | CEP170 |
| 2.6559E-58 | -0.3299382 | 0.189 | 0.298 | 6.274E-54 | TEC_dmg1 | RNF149 |
| 5.0417E-58 | -0.3207002 | 0.138 | 0.247 | 1.191E-53 | TEC_dmg1 | RNASET2 |
| 5.5663E-58 | -0.443258 | 0.148 | 0.252 | 1.3149E-53 | TEC_dmg1 | ATP2B1 |
| 5.7298E-58 | -0.4127223 | 0.17 | 0.28 | 1.3536E-53 | TEC_dmg1 | TSC22D3 |
| 6.2359E-58 | 0.3043043 | 0.279 | 0.18 | 1.4731E-53 | TEC_dmg1 | PDIA4 |
| 3.0209E-57 | -0.3639798 | 0.078 | 0.174 | 7.1363E-53 | TEC_dmg1 | HIST3H2A |
| 4.3295E-57 | -0.7855258 | 0.096 | 0.197 | 1.0228E-52 | TEC_dmg1 | RARRES3 |
| 1.2663E-56 | 0.25281884 | 0.937 | 0.923 | 2.9913E-52 | TEC_dmg1 | RPL18A |
| 2.4034E-56 | -0.3664968 | 0.074 | 0.167 | 5.6775E-52 | TEC_dmg1 | MALL |
| 5.4493E-56 | -0.8425645 | 0.183 | 0.271 | 1.2873E-51 | TEC_dmg1 | LMO7 |
| 6.3635E-56 | -0.9201693 | 0.041 | 0.125 | 1.5033E-51 | TEC_dmg1 | CXCL2 |
| 7.1731E-56 | -0.2628058 | 0.059 | 0.148 | 1.6945E-51 | TEC_dmg1 | EMP2 |
| 7.531E-56 | -0.2887802 | 0.073 | 0.164 | 1.7791E-51 | TEC_dmg1 | TSPAN6 |
| 1.8256E-55 | -0.624461 | 0.803 | 0.836 | 4.3126E-51 | TEC_dmg1 | PPDPF |
| 2.0236E-55 | -0.3977189 | 0.124 | 0.223 | 4.7803E-51 | TEC_dmg1 | CREM |
| 2.6402E-55 | 0.25314187 | 0.916 | 0.881 | 6.2369E-51 | TEC_dmg1 | RPS13 |
| 1.5005E-54 | -0.2989529 | 0.212 | 0.318 | 3.5445E-50 | TEC_dmg1 | ARPC1B |
| 6.7275E-54 | -1.0126948 | 0.068 | 0.16 | 1.5892E-49 | TEC_dmg1 | CCL20 |
| 7.0609E-54 | -0.3413589 | 0.465 | 0.556 | 1.668E-49 | TEC_dmg1 | TOP1 |
| 1.1721E-53 | -0.3119708 | 0.392 | 0.502 | 2.7688E-49 | TEC_dmg1 | SELENOW |
| 7.4446E-53 | -0.3092876 | 0.035 | 0.114 | 1.7586E-48 | TEC_dmg1 | SLC16A10 |
| 4.2071E-52 | -1.7641355 | 0.071 | 0.16 | 9.9384E-48 | TEC_dmg1 | MT1H |
| 5.0705E-52 | -0.4383054 | 0.599 | 0.685 | 1.1978E-47 | TEC_dmg1 | IER2 |
| 4.037E-50 | -0.3981353 | 0.103 | 0.194 | 9.5366E-46 | TEC_dmg1 | ACSL1 |
| 7.084E-50 | -0.3858379 | 0.141 | 0.237 | 1.6735E-45 | TEC_dmg1 | IVNS1ABP |
| 8.4574E-50 | -0.8704914 | 0.408 | 0.479 | 1.9979E-45 | TEC_dmg1 | ERO1A |
| 9.5168E-50 | -0.3338715 | 0.835 | 0.872 | 2.2482E-45 | TEC_dmg1 | PABPC1 |
| 1.3542E-49 | -0.2892478 | 0.266 | 0.368 | 3.1991E-45 | TEC_dmg1 | PLSCR1 |
| 2.6432E-49 | -0.5942757 | 0.039 | 0.116 | 6.2441E-45 | TEC_dmg1 | APOC1 |
| 7.8031E-49 | -0.3574361 | 0.066 | 0.149 | 1.8433E-44 | TEC_dmg1 | ID1 |
| 8.8711E-49 | -0.3500928 | 0.964 | 0.965 | 2.0956E-44 | TEC_dmg1 | S100A11 |
| 1.3771E-48 | 0.25425477 | 0.617 | 0.509 | 3.2531E-44 | TEC_dmg1 | H1FX |
| 5.9185E-48 | -0.2975627 | 0.055 | 0.136 | 1.3981E-43 | TEC_dmg1 | AC023157.3 |
| 1.1727E-47 | -0.6584029 | 0.106 | 0.202 | 2.7703E-43 | TEC_dmg1 | HSPA6 |
| 1.2356E-47 | -0.3946681 | 0.226 | 0.324 | 2.9188E-43 | TEC_dmg1 | HIST1H2AC |
| 2.2759E-47 | -0.5886939 | 0.381 | 0.456 | 5.3764E-43 | TEC_dmg1 | MARCKS |
| 4.7397E-47 | -0.2526796 | 0.394 | 0.269 | 1.1197E-42 | TEC_dmg1 | MT1E |
| 5.8984E-47 | -0.2892686 | 0.173 | 0.276 | 1.3934E-42 | TEC_dmg1 | ID2 |
| 9.5264E-46 | -0.3478928 | 0.092 | 0.179 | 2.2504E-41 | TEC_dmg1 | DNAJA4 |

| | | | | | | |
|---|---|---|---|---|---|---|
| 1.1273E-45 | -0.3334671 | 0.159 | 0.253 | 2.6631E-41 | TEC_dmg1 | KIAA1551 |
| 1.6175E-45 | -0.276082 | 0.742 | 0.805 | 3.821E-41 | TEC_dmg1 | ARPC2 |
| 5.9717E-45 | 0.31535513 | 0.357 | 0.262 | 1.4107E-40 | TEC_dmg1 | RAP2B |
| 7.6197E-45 | -0.2503356 | 0.127 | 0.22 | 1.8E-40 | TEC_dmg1 | TLE4 |
| 1.5997E-44 | -0.5216165 | 0.349 | 0.442 | 3.779E-40 | TEC_dmg1 | HIST1H1C |
| 2.0692E-44 | 0.36613589 | 0.93 | 0.945 | 4.888E-40 | TEC_dmg1 | MT-CO3 |
| 6.7038E-44 | -0.3388058 | 0.508 | 0.579 | 1.5836E-39 | TEC_dmg1 | TSPO |
| 8.3356E-44 | -0.7637263 | 0.752 | 0.65 | 1.9691E-39 | TEC_dmg1 | MT2A |
| 1.3491E-42 | -0.2906103 | 0.145 | 0.231 | 3.187E-38 | TEC_dmg1 | STXBP2 |
| 4.4268E-42 | 0.32574557 | 0.921 | 0.907 | 1.0457E-37 | TEC_dmg1 | MT-ND2 |
| 5.5297E-41 | -0.7031976 | 0.209 | 0.285 | 1.3063E-36 | TEC_dmg1 | DUSP5 |
| 2.4441E-40 | -0.4512101 | 0.05 | 0.119 | 5.7737E-36 | TEC_dmg1 | CYP1B1 |
| 1.064E-39 | -0.2867752 | 0.09 | 0.166 | 2.5136E-35 | TEC_dmg1 | SDC1 |
| 1.9068E-38 | -0.3793661 | 0.232 | 0.317 | 4.5043E-34 | TEC_dmg1 | IRF1 |
| 1.7196E-37 | -0.2648889 | 0.199 | 0.283 | 4.0623E-33 | TEC_dmg1 | VASP |
| 3.0975E-37 | -0.2624837 | 0.064 | 0.132 | 7.3172E-33 | TEC_dmg1 | POLB |
| 5.2365E-37 | -0.3214299 | 0.15 | 0.232 | 1.237E-32 | TEC_dmg1 | INTS6 |
| 8.4905E-37 | -0.2967687 | 0.049 | 0.113 | 2.0057E-32 | TEC_dmg1 | TCHP |
| 1.246E-36 | -0.3917197 | 0.559 | 0.625 | 2.9433E-32 | TEC_dmg1 | PCBP1 |
| 2.0341E-36 | -0.4298994 | 0.109 | 0.179 | 4.8051E-32 | TEC_dmg1 | PPL |
| 3.2642E-36 | -0.4771487 | 0.73 | 0.767 | 7.711E-32 | TEC_dmg1 | BTG1 |
| 3.8062E-36 | -0.3225844 | 0.097 | 0.171 | 8.9915E-32 | TEC_dmg1 | ANKRD37 |
| 2.4223E-35 | -0.4415987 | 0.151 | 0.229 | 5.7221E-31 | TEC_dmg1 | HES1 |
| 7.8313E-35 | -0.324812 | 0.153 | 0.233 | 1.85E-30 | TEC_dmg1 | IER5L |
| 9.2046E-35 | -0.4529946 | 0.956 | 0.975 | 2.1744E-30 | TEC_dmg1 | ACTB |
| 1.2668E-34 | -0.877119 | 0.113 | 0.179 | 2.9925E-30 | TEC_dmg1 | RDH10 |
| 2.1237E-34 | -0.2958404 | 0.151 | 0.233 | 5.0168E-30 | TEC_dmg1 | PPP1R10 |
| 3.0796E-33 | -0.4534887 | 0.156 | 0.234 | 7.275E-29 | TEC_dmg1 | HEXIM1 |
| 6.7557E-33 | -0.4123648 | 0.528 | 0.583 | 1.5959E-28 | TEC_dmg1 | LITAF |
| 1.1283E-32 | -0.2782491 | 0.275 | 0.345 | 2.6653E-28 | TEC_dmg1 | GRB2 |
| 1.4752E-32 | -0.4025687 | 0.063 | 0.126 | 3.485E-28 | TEC_dmg1 | HILPDA |
| 2.4815E-32 | -0.2695993 | 0.452 | 0.544 | 5.8619E-28 | TEC_dmg1 | SERPINB1 |
| 8.054E-32 | -0.2592916 | 0.664 | 0.714 | 1.9026E-27 | TEC_dmg1 | SDCBP |
| 1.2642E-31 | -0.2896671 | 0.548 | 0.639 | 2.9865E-27 | TEC_dmg1 | ZFP36L1 |
| 2.8915E-31 | 0.2537179 | 0.924 | 0.914 | 6.8306E-27 | TEC_dmg1 | MT-ND1 |
| 1.7231E-30 | -0.4676698 | 0.135 | 0.197 | 4.0704E-26 | TEC_dmg1 | FBXO32 |
| 1.9092E-30 | -2.3411363 | 0.492 | 0.526 | 4.5101E-26 | TEC_dmg1 | LCN2 |
| 4.5834E-30 | -0.2655421 | 0.058 | 0.115 | 1.0827E-25 | TEC_dmg1 | SMIM5 |
| 5.6033E-30 | -0.2808909 | 0.428 | 0.5 | 1.3237E-25 | TEC_dmg1 | H2AFJ |
| 1.8136E-29 | -0.4106254 | 0.114 | 0.177 | 4.2843E-25 | TEC_dmg1 | HSPB8 |
| 2.2667E-29 | -0.2720394 | 0.62 | 0.68 | 5.3546E-25 | TEC_dmg1 | TALDO1 |
| 2.9078E-29 | -0.2650121 | 0.051 | 0.109 | 6.869E-25 | TEC_dmg1 | CCL5 |
| 3.1652E-29 | -0.6827791 | 0.219 | 0.288 | 7.4773E-25 | TEC_dmg1 | IFITM2 |
| 3.3826E-28 | -0.2551717 | 0.112 | 0.176 | 7.9906E-24 | TEC_dmg1 | FAM214A |
| 7.6254E-28 | -0.3604533 | 0.104 | 0.167 | 1.8013E-23 | TEC_dmg1 | CSKMT |
| 1.2747E-27 | -0.3172087 | 0.211 | 0.281 | 3.0113E-23 | TEC_dmg1 | ADM |
| 2.6459E-27 | -0.7165588 | 0.563 | 0.554 | 6.2504E-23 | TEC_dmg1 | ELF3 |
| 1.4794E-26 | -0.4344741 | 0.27 | 0.328 | 3.4949E-22 | TEC_dmg1 | VEGFA |
| 1.5046E-25 | -0.2858357 | 0.227 | 0.288 | 3.5544E-21 | TEC_dmg1 | CTSZ |
| 3.6948E-25 | -0.6706942 | 0.526 | 0.446 | 8.7283E-21 | TEC_dmg1 | SPP1 |

| | | | | | | |
|---|---|---|---|---|---|---|
| 9.9878E-25 | -0.2690119 | 0.072 | 0.124 | 2.3594E-20 | TEC_dmg1 | EGLN3 |
| 2.068E-24 | -0.2666383 | 0.203 | 0.27 | 4.8852E-20 | TEC_dmg1 | MGST1 |
| 2.3368E-24 | -0.3360388 | 0.682 | 0.725 | 5.5203E-20 | TEC_dmg1 | JUND |
| 3.2155E-24 | -0.3987618 | 0.33 | 0.372 | 7.596E-20 | TEC_dmg1 | COTL1 |
| 4.0151E-24 | -0.3721649 | 0.177 | 0.235 | 9.4849E-20 | TEC_dmg1 | PPIF |
| 4.4084E-24 | -0.7139588 | 0.363 | 0.257 | 1.0414E-19 | TEC_dmg1 | S100A2 |
| 2.3526E-23 | -0.8077184 | 0.987 | 0.994 | 5.5575E-19 | TEC_dmg1 | FTH1 |
| 6.0423E-23 | -0.3146423 | 0.909 | 0.888 | 1.4274E-18 | TEC_dmg1 | HLA-B |
| 1.0767E-22 | -0.5832511 | 0.239 | 0.29 | 2.5435E-18 | TEC_dmg1 | SLC25A37 |
| 1.1584E-21 | -0.2952729 | 0.082 | 0.133 | 2.7366E-17 | TEC_dmg1 | GBP1 |
| 1.5354E-21 | -0.3872639 | 0.176 | 0.235 | 3.627E-17 | TEC_dmg1 | ISG15 |
| 1.8349E-21 | -1.3417152 | 0.098 | 0.148 | 4.3347E-17 | TEC_dmg1 | SPRR2A |
| 7.7549E-21 | -0.2766513 | 0.292 | 0.352 | 1.8319E-16 | TEC_dmg1 | ZFP36L2 |
| 2.4247E-20 | -0.2722808 | 0.325 | 0.365 | 5.7279E-16 | TEC_dmg1 | GNAI2 |
| 6.234E-20 | -0.3606831 | 0.228 | 0.291 | 1.4727E-15 | TEC_dmg1 | HMOX1 |
| 6.6466E-19 | -0.4688726 | 0.594 | 0.614 | 1.5701E-14 | TEC_dmg1 | CD44 |
| 7.2364E-19 | -0.3651536 | 0.104 | 0.152 | 1.7095E-14 | TEC_dmg1 | SMC4 |
| 1.5637E-18 | -0.2720291 | 0.543 | 0.578 | 3.6939E-14 | TEC_dmg1 | CD55 |
| 1.1665E-17 | -0.4803503 | 0.173 | 0.228 | 2.7555E-13 | TEC_dmg1 | G0S2 |
| 2.2486E-17 | -1.243685 | 0.075 | 0.119 | 5.3119E-13 | TEC_dmg1 | MT1M |
| 3.7822E-17 | -0.8968666 | 0.099 | 0.147 | 8.9347E-13 | TEC_dmg1 | CCL2 |
| 8.1238E-16 | -0.2529275 | 0.201 | 0.253 | 1.9191E-11 | TEC_dmg1 | EGR1 |
| 3.6382E-15 | -0.6106095 | 0.287 | 0.314 | 8.5945E-11 | TEC_dmg1 | EMP1 |
| 4.1563E-15 | -0.3353934 | 0.148 | 0.186 | 9.8183E-11 | TEC_dmg1 | SDCBP2 |
| 4.3405E-14 | -0.7714214 | 0.095 | 0.134 | 1.0254E-09 | TEC_dmg1 | APOE |
| 8.2116E-14 | -0.3223068 | 0.424 | 0.434 | 1.9398E-09 | TEC_dmg1 | GSTO1 |
| 2.9599E-13 | -0.5037237 | 0.216 | 0.24 | 6.9922E-09 | TEC_dmg1 | ADGRF1 |
| 1.9464E-12 | -0.478806 | 0.786 | 0.736 | 4.598E-08 | TEC_dmg1 | PSAP |
| 2.0916E-11 | -0.2758572 | 0.917 | 0.934 | 4.9409E-07 | TEC_dmg1 | GAPDH |
| 5.2116E-11 | -2.8883308 | 0.507 | 0.545 | 1.2311E-06 | TEC_dmg1 | PLCG2 |
| 6.9248E-11 | -0.4413585 | 0.28 | 0.312 | 1.6358E-06 | TEC_dmg1 | HIST1H4C |
| 1.1553E-10 | -0.2559243 | 0.313 | 0.348 | 2.7292E-06 | TEC_dmg1 | MIDN |
| 3.4074E-10 | -0.2585045 | 0.298 | 0.324 | 8.0492E-06 | TEC_dmg1 | B4GALT1 |
| 6.4044E-10 | -0.4998712 | 0.204 | 0.236 | 1.5129E-05 | TEC_dmg1 | PLIN2 |
| 1.0525E-09 | -1.256542 | 0.256 | 0.282 | 2.4862E-05 | TEC_dmg1 | MT1X |
| 2.7425E-08 | -0.3041187 | 0.134 | 0.161 | 0.00064787 | TEC_dmg1 | BPGM |
| 5.356E-08 | -0.4989655 | 0.537 | 0.539 | 0.00126524 | TEC_dmg1 | PLAUR |
| 1.3389E-05 | -0.3187314 | 0.666 | 0.708 | 0.31629259 | TEC_dmg1 | SOD2 |
| 2.3912E-05 | 0.40369664 | 0.313 | 0.293 | 0.56488284 | TEC_dmg1 | GLRX |
| 2.8906E-05 | -0.3884856 | 0.181 | 0.199 | 0.68283884 | TEC_dmg1 | TM4SF1 |
| 3.3403E-05 | -0.4354791 | 0.138 | 0.158 | 0.78907451 | TEC_dmg1 | MT1F |
| 5.024E-05 | -0.4087319 | 0.294 | 0.292 | 1 | TEC_dmg1 | RND3 |
| 8.0752E-05 | -0.3059849 | 0.401 | 0.393 | 1 | TEC_dmg1 | HIF1A |
| 0.00010078 | -1.6587348 | 0.111 | 0.087 | 1 | TEC_dmg1 | HBB |
| 0.00010917 | -0.4397525 | 0.621 | 0.639 | 1 | TEC_dmg1 | MTRNR2L12 |
| 0.00046312 | -0.3873548 | 0.441 | 0.452 | 1 | TEC_dmg1 | SOX4 |
| 0.00059369 | -0.7997419 | 0.429 | 0.363 | 1 | TEC_dmg1 | MAL |
| 0.00150599 | -1.8728103 | 0.248 | 0.255 | 1 | TEC_dmg1 | MT1G |
| 0.00182785 | -0.3403691 | 0.633 | 0.627 | 1 | TEC_dmg1 | JUN |
| 0.00224576 | -0.2544253 | 0.171 | 0.175 | 1 | TEC_dmg1 | C6orf132 |

| | | | | | | |
|---|---|---|---|---|---|---|
| 7.283E-228 | 1.82111494 | 1 | 0.937 | 1.72E-223 | TEC_dmg2 | MT-ATP6 |
| 2.937E-225 | 1.80924242 | 1 | 0.932 | 6.939E-221 | TEC_dmg2 | MT-ND4 |
| 4.958E-217 | 2.02156513 | 1 | 0.943 | 1.171E-212 | TEC_dmg2 | MT-CO3 |
| 4.15E-212 | 1.76104377 | 1 | 0.95 | 9.803E-208 | TEC_dmg2 | MT-CO1 |
| 2.85E-211 | 2.00617034 | 1 | 0.914 | 6.733E-207 | TEC_dmg2 | MT-ND1 |
| 8.931E-211 | 2.11813028 | 1 | 0.907 | 2.11E-206 | TEC_dmg2 | MT-ND2 |
| 3.851E-206 | 1.67577734 | 1 | 0.944 | 9.098E-202 | TEC_dmg2 | MT-CO2 |
| 2.295E-204 | 1.80510351 | 1 | 0.929 | 5.421E-200 | TEC_dmg2 | MT-CYB |
| 5.849E-158 | 1.63280562 | 1 | 0.876 | 1.382E-153 | TEC_dmg2 | MT-ND3 |
| 9.868E-154 | 1.57100978 | 0.998 | 0.821 | 2.331E-149 | TEC_dmg2 | MT-ND5 |
| 1.721E-104 | 1.60276416 | 0.875 | 0.488 | 4.066E-100 | TEC_dmg2 | MTRNR2L8 |
| 4.096E-99 | 1.19597348 | 0.861 | 0.503 | 9.6762E-95 | TEC_dmg2 | MT-ND4L |
| 1.0332E-82 | -1.4395092 | 0.935 | 0.99 | 2.4407E-78 | TEC_dmg2 | B2M |
| 7.9386E-73 | 0.85014265 | 0.573 | 0.232 | 1.8753E-68 | TEC_dmg2 | MT-ATP8 |
| 1.3474E-71 | -1.0620971 | 0.854 | 0.958 | 3.1829E-67 | TEC_dmg2 | H3F3A |
| 6.9549E-68 | 0.84876043 | 0.918 | 0.635 | 1.643E-63 | TEC_dmg2 | MTRNR2L12 |
| 1.0672E-65 | -4.110639 | 0.441 | 0.73 | 2.5211E-61 | TEC_dmg2 | S100A9 |
| 1.4252E-65 | -1.3890617 | 0.832 | 0.939 | 3.3668E-61 | TEC_dmg2 | TMSB4X |
| 1.0077E-58 | -1.071871 | 0.933 | 0.965 | 2.3804E-54 | TEC_dmg2 | S100A11 |
| 1.4567E-58 | -1.2043875 | 0.422 | 0.73 | 3.4411E-54 | TEC_dmg2 | CYBA |
| 2.0513E-58 | -1.1249626 | 0.902 | 0.971 | 4.8459E-54 | TEC_dmg2 | TMSB10 |
| 4.7964E-55 | 0.75990399 | 0.412 | 0.149 | 1.133E-50 | TEC_dmg2 | MT-ND6 |
| 9.3855E-54 | -1.3390421 | 0.676 | 0.861 | 2.2171E-49 | TEC_dmg2 | SH3BGRL3 |
| 9.4426E-54 | -1.0637391 | 0.451 | 0.755 | 2.2306E-49 | TEC_dmg2 | TPM4 |
| 9.5622E-54 | 0.75581633 | 0.516 | 0.215 | 2.2589E-49 | TEC_dmg2 | ZBTB20 |
| 1.8099E-53 | -0.7255628 | 0.635 | 0.869 | 4.2756E-49 | TEC_dmg2 | ARPC3 |
| 3.8344E-51 | -3.4086504 | 0.41 | 0.674 | 9.0581E-47 | TEC_dmg2 | S100A8 |
| 1.2278E-50 | -1.3075777 | 0.724 | 0.885 | 2.9005E-46 | TEC_dmg2 | CSTB |
| 3.7035E-47 | -0.6915912 | 0.835 | 0.956 | 8.7488E-43 | TEC_dmg2 | SERF2 |
| 2.4423E-44 | -0.7615529 | 0.729 | 0.894 | 5.7693E-40 | TEC_dmg2 | ATP5F1E |
| 3.7892E-43 | -1.7883922 | 0.343 | 0.616 | 8.9513E-39 | TEC_dmg2 | C15orf48 |
| 3.8657E-43 | -1.1458148 | 0.568 | 0.786 | 9.132E-39 | TEC_dmg2 | LGALS3 |
| 1.1064E-41 | -1.5254327 | 0.46 | 0.712 | 2.6138E-37 | TEC_dmg2 | ANXA1 |
| 2.913E-41 | -2.357193 | 0.293 | 0.573 | 6.8813E-37 | TEC_dmg2 | SRGN |
| 7.1468E-41 | -0.767904 | 0.012 | 0.328 | 1.6883E-36 | TEC_dmg2 | ARHGDIB |
| 1.3188E-39 | -0.6118838 | 0.928 | 0.994 | 3.1153E-35 | TEC_dmg2 | TPT1 |
| 1.3458E-38 | 0.64961691 | 0.408 | 0.179 | 3.1792E-34 | TEC_dmg2 | GCLM |
| 1.9236E-38 | -0.9672888 | 0.494 | 0.74 | 4.5442E-34 | TEC_dmg2 | CTSB |
| 3.4912E-38 | 0.46813683 | 0.396 | 0.166 | 8.2473E-34 | TEC_dmg2 | SLC38A1 |
| 5.1486E-38 | -1.0371002 | 0.235 | 0.534 | 1.2163E-33 | TEC_dmg2 | CTSS |
| 1.0748E-37 | -0.6666004 | 0.803 | 0.931 | 2.5389E-33 | TEC_dmg2 | OAZ1 |
| 1.0743E-36 | 0.40691694 | 0.237 | 0.074 | 2.5379E-32 | TEC_dmg2 | FGF2 |
| 1.1648E-36 | -0.6640157 | 0.866 | 0.94 | 2.7515E-32 | TEC_dmg2 | SAT1 |
| 1.5239E-35 | -0.9303206 | 0.65 | 0.835 | 3.5999E-31 | TEC_dmg2 | PPDPF |
| 3.1335E-35 | -1.4951468 | 0.386 | 0.627 | 7.4024E-31 | TEC_dmg2 | S100A4 |
| 3.3461E-35 | -1.0851918 | 0.046 | 0.338 | 7.9045E-31 | TEC_dmg2 | LAPTM5 |
| 9.1557E-35 | -1.4507671 | 0.957 | 0.994 | 2.1629E-30 | TEC_dmg2 | FTH1 |
| 2.1385E-34 | -1.5486543 | 0.173 | 0.443 | 5.0518E-30 | TEC_dmg2 | TYROBP |
| 4.8171E-34 | -0.6764893 | 0.835 | 0.918 | 1.1379E-29 | TEC_dmg2 | UBA52 |
| 5.9749E-34 | -1.3688645 | 0.482 | 0.683 | 1.4115E-29 | TEC_dmg2 | NAMPT |

| | | | | | | |
|---|---|---|---|---|---|---|
| 1.6599E-33 | -1.437513 | 0.055 | 0.336 | 3.9213E-29 | TEC_dmg2 | IL1RN |
| 2.1343E-33 | 0.64071753 | 0.6 | 0.366 | 5.0419E-29 | TEC_dmg2 | ANKRD11 |
| 4.7037E-33 | -1.2250221 | 0.424 | 0.646 | 1.1112E-28 | TEC_dmg2 | GLUL |
| 1.1778E-32 | -0.6021308 | 0.137 | 0.431 | 2.7824E-28 | TEC_dmg2 | EFHD2 |
| 1.2102E-31 | -0.9812631 | 0.269 | 0.53 | 2.8588E-27 | TEC_dmg2 | ZFP36 |
| 1.6317E-31 | -1.3110297 | 0.403 | 0.619 | 3.8547E-27 | TEC_dmg2 | LGALS1 |
| 1.9723E-31 | -1.5723679 | 0.129 | 0.388 | 4.6591E-27 | TEC_dmg2 | FCER1G |
| 2.3014E-31 | 0.31750128 | 0.276 | 0.102 | 5.4367E-27 | TEC_dmg2 | LINC00958 |
| 1.6096E-30 | -1.2048409 | 0.024 | 0.277 | 3.8023E-26 | TEC_dmg2 | AIF1 |
| 1.275E-29 | -3.0833041 | 0.499 | 0.678 | 3.0119E-25 | TEC_dmg2 | TIMP1 |
| 1.2758E-29 | -0.6278169 | 0.95 | 0.98 | 3.0138E-25 | TEC_dmg2 | RPL41 |
| 1.938E-29 | -0.623955 | 0.007 | 0.248 | 4.5781E-25 | TEC_dmg2 | GMFG |
| 2.2519E-29 | -0.9139016 | 0.863 | 0.933 | 5.3196E-25 | TEC_dmg2 | GAPDH |
| 2.3398E-29 | -0.6626689 | 0.022 | 0.27 | 5.5273E-25 | TEC_dmg2 | CD53 |
| 2.5732E-29 | -0.6855858 | 0.899 | 0.94 | 6.0786E-25 | TEC_dmg2 | RPS28 |
| 2.9518E-29 | -0.5699349 | 0.098 | 0.37 | 6.9731E-25 | TEC_dmg2 | CTSH |
| 3.8403E-29 | -0.4195439 | 0.477 | 0.729 | 9.0721E-25 | TEC_dmg2 | ELOB |
| 3.9153E-29 | -0.7331046 | 0.007 | 0.248 | 9.249E-25 | TEC_dmg2 | CSTA |
| 7.7702E-29 | -0.5020201 | 0.882 | 0.964 | 1.8356E-24 | TEC_dmg2 | MYL6 |
| 1.4068E-28 | -0.8074289 | 0.849 | 0.9 | 3.3234E-24 | TEC_dmg2 | RPL39 |
| 1.5418E-28 | -0.7735931 | 0.83 | 0.89 | 3.6422E-24 | TEC_dmg2 | HLA-B |
| 5.801E-28 | -1.2618781 | 0.036 | 0.279 | 1.3704E-23 | TEC_dmg2 | HLA-DRB1 |
| 6.1879E-28 | -0.8545453 | 0.048 | 0.293 | 1.4618E-23 | TEC_dmg2 | LCP1 |
| 7.2848E-28 | -0.8768485 | 0.945 | 0.973 | 1.7209E-23 | TEC_dmg2 | ACTB |
| 1.3996E-27 | 0.29574199 | 0.293 | 0.119 | 3.3062E-23 | TEC_dmg2 | CTNNAL1 |
| 1.4964E-27 | 0.42734626 | 0.367 | 0.175 | 3.5348E-23 | TEC_dmg2 | ZBTB43 |
| 2.3397E-27 | -2.1189191 | 0.237 | 0.466 | 5.5272E-23 | TEC_dmg2 | LYZ |
| 6.433E-27 | 0.81247588 | 0.99 | 0.98 | 1.5197E-22 | TEC_dmg2 | MALAT1 |
| 9.1993E-27 | -0.8177488 | 0.628 | 0.768 | 2.1732E-22 | TEC_dmg2 | CST3 |
| 1.2664E-26 | -0.5248335 | 0.106 | 0.361 | 2.9916E-22 | TEC_dmg2 | PLSCR1 |
| 1.3563E-26 | -0.6435495 | 0.806 | 0.907 | 3.204E-22 | TEC_dmg2 | ITM2B |
| 1.7749E-26 | -0.6741244 | 0.012 | 0.236 | 4.1927E-22 | TEC_dmg2 | HCST |
| 3.0231E-26 | -0.5956037 | 0.887 | 0.954 | 7.1415E-22 | TEC_dmg2 | H3F3B |
| 8.7135E-26 | -0.5282899 | 0.938 | 0.978 | 2.0584E-21 | TEC_dmg2 | RPL28 |
| 8.9237E-26 | -0.602831 | 0.326 | 0.549 | 2.108E-21 | TEC_dmg2 | C4orf3 |
| 1.4385E-25 | -0.737676 | 0.508 | 0.701 | 3.3982E-21 | TEC_dmg2 | JUNB |
| 1.4786E-25 | -1.8257382 | 0.168 | 0.399 | 3.4929E-21 | TEC_dmg2 | HLA-DRA |
| 3.9872E-25 | -0.4686387 | 0.247 | 0.503 | 9.419E-21 | TEC_dmg2 | ASAH1 |
| 4.2303E-25 | -0.5374949 | 0.643 | 0.801 | 9.9931E-21 | TEC_dmg2 | ARPC2 |
| 6.4884E-25 | 0.37361869 | 0.376 | 0.185 | 1.5327E-20 | TEC_dmg2 | PAWR |
| 6.7518E-25 | 0.60655834 | 0.542 | 0.344 | 1.595E-20 | TEC_dmg2 | GOLGB1 |
| 9.3868E-25 | -0.5183824 | 0.784 | 0.906 | 2.2174E-20 | TEC_dmg2 | CFL1 |
| 1.5711E-24 | -0.9490902 | 0.012 | 0.222 | 3.7114E-20 | TEC_dmg2 | CD14 |
| 1.5829E-24 | -0.554276 | 0.698 | 0.837 | 3.7393E-20 | TEC_dmg2 | HLA-E |
| 3.449E-24 | -0.4731589 | 0.624 | 0.802 | 8.1477E-20 | TEC_dmg2 | GNG5 |
| 3.7236E-24 | -0.6295297 | 0.089 | 0.314 | 8.7962E-20 | TEC_dmg2 | CD68 |
| 5.5736E-24 | -0.6647519 | 0.89 | 0.944 | 1.3166E-19 | TEC_dmg2 | RPS27 |
| 6.162E-24 | -0.4595995 | 0.065 | 0.296 | 1.4556E-19 | TEC_dmg2 | FXYD5 |
| 8.8028E-24 | 0.32160855 | 0.295 | 0.128 | 2.0795E-19 | TEC_dmg2 | AKAP12 |
| 1.4412E-23 | -0.46274 | 0.424 | 0.63 | 3.4044E-19 | TEC_dmg2 | OST4 |

| | | | | | | |
|---|---|---|---|---|---|---|
| 1.5709E-23 | -0.7879031 | 0.055 | 0.283 | 3.7108E-19 | TEC_dmg2 | PTPRC |
| 1.6504E-23 | -0.9544416 | 0.024 | 0.235 | 3.8986E-19 | TEC_dmg2 | SLC2A3 |
| 1.6733E-23 | -0.508197 | 0.691 | 0.817 | 3.9529E-19 | TEC_dmg2 | PFDN5 |
| 1.9859E-23 | -0.6853441 | 0.077 | 0.304 | 4.6913E-19 | TEC_dmg2 | SERPINA1 |
| 2.7202E-23 | -0.4642171 | 0.019 | 0.226 | 6.4258E-19 | TEC_dmg2 | KYNU |
| 4.9298E-23 | -0.7365622 | 0.005 | 0.2 | 1.1646E-18 | TEC_dmg2 | CD52 |
| 7.6285E-23 | -0.583128 | 0.753 | 0.871 | 1.8021E-18 | TEC_dmg2 | HLA-C |
| 9.047E-23 | 0.27620792 | 0.245 | 0.102 | 2.1372E-18 | TEC_dmg2 | IARS |
| 1.6011E-22 | 0.26485426 | 0.223 | 0.088 | 3.7823E-18 | TEC_dmg2 | PGM3 |
| 2.3263E-22 | -0.5261436 | 0.602 | 0.777 | 5.4955E-18 | TEC_dmg2 | TPI1 |
| 2.8898E-22 | -0.5424259 | 0.017 | 0.215 | 6.8265E-18 | TEC_dmg2 | BASP1 |
| 3.0013E-22 | 0.3180071 | 0.247 | 0.104 | 7.0899E-18 | TEC_dmg2 | LDLR |
| 3.3395E-22 | -1.5957365 | 0.293 | 0.493 | 7.889E-18 | TEC_dmg2 | CD74 |
| 3.9768E-22 | -0.6673635 | 0.823 | 0.88 | 9.3945E-18 | TEC_dmg2 | RPS29 |
| 4.0646E-22 | -0.550891 | 0.022 | 0.221 | 9.6018E-18 | TEC_dmg2 | ITGB2 |
| 6.2117E-22 | -0.5390584 | 0.489 | 0.676 | 1.4674E-17 | TEC_dmg2 | TALDO1 |
| 7.2139E-22 | -1.0370812 | 0.058 | 0.264 | 1.7041E-17 | TEC_dmg2 | AREG |
| 7.6192E-22 | -0.9817298 | 0.024 | 0.221 | 1.7999E-17 | TEC_dmg2 | ALOX5AP |
| 7.6391E-22 | 0.32387478 | 0.336 | 0.164 | 1.8046E-17 | TEC_dmg2 | CCDC50 |
| 1.0855E-21 | -0.6021998 | 0.41 | 0.599 | 2.5642E-17 | TEC_dmg2 | NPC2 |
| 1.0867E-21 | -0.3935633 | 0.017 | 0.211 | 2.5672E-17 | TEC_dmg2 | ARRB2 |
| 1.1806E-21 | -0.8981876 | 0.168 | 0.391 | 2.7888E-17 | TEC_dmg2 | FOS |
| 1.2268E-21 | -0.7187461 | 0.247 | 0.472 | 2.8982E-17 | TEC_dmg2 | DUSP1 |
| 1.2326E-21 | -0.4146459 | 0.005 | 0.189 | 2.9118E-17 | TEC_dmg2 | RAC2 |
| 1.9365E-21 | -0.9667071 | 0.038 | 0.238 | 4.5746E-17 | TEC_dmg2 | BCL2A1 |
| 2.2085E-21 | -1.0284512 | 0.024 | 0.217 | 5.2172E-17 | TEC_dmg2 | S100A12 |
| 3.0421E-21 | -0.4274362 | 0.084 | 0.296 | 7.1862E-17 | TEC_dmg2 | CD99 |
| 4.0267E-21 | -1.3633663 | 0.655 | 0.741 | 9.5122E-17 | TEC_dmg2 | VIM |
| 4.0967E-21 | -2.5461177 | 0.293 | 0.49 | 9.6776E-17 | TEC_dmg2 | CXCL8 |
| 4.1434E-21 | -0.5937243 | 0.933 | 0.973 | 9.788E-17 | TEC_dmg2 | RPLP1 |
| 4.5266E-21 | 0.49791672 | 0.444 | 0.266 | 1.0693E-16 | TEC_dmg2 | HSPA9 |
| 4.9263E-21 | -0.3965996 | 0.036 | 0.235 | 1.1637E-16 | TEC_dmg2 | BST2 |
| 6.0001E-21 | -0.3534747 | 0.005 | 0.185 | 1.4174E-16 | TEC_dmg2 | GNA15 |
| 6.9811E-21 | 0.41270605 | 0.288 | 0.137 | 1.6491E-16 | TEC_dmg2 | MGLL |
| 7.2752E-21 | -0.4091793 | 0.168 | 0.397 | 1.7186E-16 | TEC_dmg2 | NDUFB11 |
| 8.4228E-21 | 0.54407429 | 0.516 | 0.331 | 1.9897E-16 | TEC_dmg2 | GOLGA4 |
| 1.0099E-20 | -0.9300312 | 0.106 | 0.314 | 2.3857E-16 | TEC_dmg2 | PHLDA1 |
| 1.0873E-20 | -1.5256511 | 0.034 | 0.224 | 2.5686E-16 | TEC_dmg2 | VCAN |
| 1.1112E-20 | -1.3652075 | 0.014 | 0.198 | 2.6249E-16 | TEC_dmg2 | HOPX |
| 1.154E-20 | 0.4615951 | 0.379 | 0.211 | 2.726E-16 | TEC_dmg2 | DNTTIP2 |
| 1.4534E-20 | -0.5878141 | 0.329 | 0.538 | 3.4333E-16 | TEC_dmg2 | SERPINB1 |
| 1.6657E-20 | -1.9137538 | 0.156 | 0.346 | 3.9348E-16 | TEC_dmg2 | HPGD |
| 1.7284E-20 | -0.5071989 | 0.151 | 0.363 | 4.083E-16 | TEC_dmg2 | GNAI2 |
| 1.9819E-20 | -0.7422382 | 0.01 | 0.189 | 4.6818E-16 | TEC_dmg2 | AC020656.1 |
| 2.097E-20 | -0.4895086 | 0.029 | 0.217 | 4.9538E-16 | TEC_dmg2 | SPI1 |
| 2.5068E-20 | -0.7733581 | 0.048 | 0.238 | 5.9218E-16 | TEC_dmg2 | FABP5 |
| 2.5951E-20 | -0.4518276 | 0.108 | 0.321 | 6.1303E-16 | TEC_dmg2 | SLC16A3 |
| 2.9864E-20 | -0.4849018 | 0.386 | 0.582 | 7.0547E-16 | TEC_dmg2 | PGK1 |
| 3.2182E-20 | -0.520999 | 0.163 | 0.387 | 7.6024E-16 | TEC_dmg2 | C9orf16 |
| 3.3939E-20 | -0.6901304 | 0.168 | 0.386 | 8.0174E-16 | TEC_dmg2 | TYMP |

| | | | | | | |
|---|---|---|---|---|---|---|
| 3.7127E-20 | -0.4632099 | 0.271 | 0.494 | 8.7705E-16 | TEC_dmg2 | SELENOW |
| 7.2078E-20 | -0.3486927 | 0.012 | 0.188 | 1.7027E-15 | TEC_dmg2 | GPSM3 |
| 7.9495E-20 | 0.56268507 | 0.561 | 0.406 | 1.8779E-15 | TEC_dmg2 | ATF4 |
| 1.2103E-19 | -0.4212087 | 0.024 | 0.205 | 2.8592E-15 | TEC_dmg2 | LSP1 |
| 1.5391E-19 | 0.29721979 | 0.319 | 0.159 | 3.6358E-15 | TEC_dmg2 | NCKAP1 |
| 1.5909E-19 | -0.5893587 | 0.022 | 0.2 | 3.7583E-15 | TEC_dmg2 | DUSP4 |
| 2.0145E-19 | -0.3935147 | 0.01 | 0.181 | 4.7589E-15 | TEC_dmg2 | CD37 |
| 2.5468E-19 | -0.5656302 | 0.029 | 0.21 | 6.0163E-15 | TEC_dmg2 | LST1 |
| 2.7248E-19 | -0.3503865 | 0.058 | 0.252 | 6.4368E-15 | TEC_dmg2 | SH3BGRL |
| 2.8619E-19 | -1.0981361 | 0.014 | 0.186 | 6.7608E-15 | TEC_dmg2 | HLA-DPB1 |
| 2.9187E-19 | -0.6279016 | 0.024 | 0.202 | 6.8947E-15 | TEC_dmg2 | CYBB |
| 3.561E-19 | -0.7008792 | 0.727 | 0.792 | 8.4121E-15 | TEC_dmg2 | PFN1 |
| 4.1578E-19 | -0.6346542 | 0.031 | 0.212 | 9.8221E-15 | TEC_dmg2 | TNFRSF1B |
| 4.5085E-19 | 0.29770267 | 0.288 | 0.142 | 1.065E-14 | TEC_dmg2 | BAG5 |
| 6.2521E-19 | 0.80018874 | 0.573 | 0.42 | 1.4769E-14 | TEC_dmg2 | TXNRD1 |
| 7.143E-19 | -0.3954597 | 0.07 | 0.26 | 1.6874E-14 | TEC_dmg2 | ABRACL |
| 7.4334E-19 | -1.1897576 | 0.103 | 0.297 | 1.756E-14 | TEC_dmg2 | ADIRF |
| 7.5525E-19 | 0.27223644 | 0.319 | 0.16 | 1.7841E-14 | TEC_dmg2 | MET |
| 8.2222E-19 | -0.5085158 | 0.388 | 0.574 | 1.9423E-14 | TEC_dmg2 | TSPO |
| 1.0505E-18 | 0.47012516 | 0.516 | 0.342 | 2.4816E-14 | TEC_dmg2 | NORAD |
| 1.0958E-18 | -2.1224812 | 0.247 | 0.409 | 2.5886E-14 | TEC_dmg2 | S100P |
| 1.1017E-18 | -0.4967254 | 0.019 | 0.19 | 2.6025E-14 | TEC_dmg2 | FCGR2A |
| 1.2165E-18 | 0.36849524 | 0.381 | 0.211 | 2.8737E-14 | TEC_dmg2 | LAMB3 |
| 1.4958E-18 | -0.6008766 | 0.906 | 0.948 | 3.5335E-14 | TEC_dmg2 | RPL10 |
| 1.5707E-18 | -0.4438275 | 0.115 | 0.31 | 3.7105E-14 | TEC_dmg2 | ARPC1B |
| 1.6403E-18 | -1.1927725 | 0.031 | 0.205 | 3.875E-14 | TEC_dmg2 | HLA-DPA1 |
| 1.7956E-18 | -1.0480408 | 0.002 | 0.161 | 4.2419E-14 | TEC_dmg2 | CEACAM5 |
| 1.8898E-18 | -0.3644454 | 0.048 | 0.228 | 4.4642E-14 | TEC_dmg2 | RGS10 |
| 2.1751E-18 | -0.5623298 | 0.861 | 0.928 | 5.1382E-14 | TEC_dmg2 | RPL30 |
| 2.2535E-18 | 0.28355939 | 0.288 | 0.135 | 5.3235E-14 | TEC_dmg2 | DCDC2 |
| 2.7737E-18 | -1.9030371 | 0.173 | 0.343 | 6.5522E-14 | TEC_dmg2 | PSCA |
| 3.2228E-18 | -0.4513182 | 0.012 | 0.175 | 7.6132E-14 | TEC_dmg2 | SMIM25 |
| 3.3274E-18 | -0.3903781 | 0.01 | 0.171 | 7.8603E-14 | TEC_dmg2 | CD48 |
| 3.7034E-18 | -0.7598331 | 0.01 | 0.171 | 8.7485E-14 | TEC_dmg2 | FCN1 |
| 3.8067E-18 | -0.3923723 | 0.012 | 0.175 | 8.9925E-14 | TEC_dmg2 | CORO1A |
| 4.0993E-18 | -0.3701939 | 0.05 | 0.229 | 9.6839E-14 | TEC_dmg2 | FCGRT |
| 4.3032E-18 | -0.6305803 | 0.012 | 0.173 | 1.0165E-13 | TEC_dmg2 | STEAP4 |
| 4.5407E-18 | 0.49643965 | 0.429 | 0.265 | 1.0727E-13 | TEC_dmg2 | UGCG |
| 5.1952E-18 | -0.6287232 | 0.017 | 0.182 | 1.2273E-13 | TEC_dmg2 | OLR1 |
| 5.4155E-18 | -0.7324379 | 0.014 | 0.176 | 1.2793E-13 | TEC_dmg2 | PLEK |
| 6.4308E-18 | 0.321309 | 0.307 | 0.159 | 1.5191E-13 | TEC_dmg2 | GFPT1 |
| 6.6867E-18 | -0.4102074 | 0.089 | 0.286 | 1.5796E-13 | TEC_dmg2 | RPS4Y1 |
| 8.4075E-18 | -0.3084776 | 0.59 | 0.764 | 1.9861E-13 | TEC_dmg2 | UBL5 |
| 9.678E-18 | 0.32922278 | 0.341 | 0.185 | 2.2862E-13 | TEC_dmg2 | GPATCH2L |
| 1.0152E-17 | -0.5488902 | 0.022 | 0.184 | 2.3983E-13 | TEC_dmg2 | IGSF6 |
| 1.0242E-17 | 0.36958693 | 0.345 | 0.194 | 2.4194E-13 | TEC_dmg2 | MED10 |
| 1.2065E-17 | -0.492124 | 0.041 | 0.21 | 2.8501E-13 | TEC_dmg2 | LY6E |
| 1.291E-17 | -0.4574589 | 0.007 | 0.162 | 3.0497E-13 | TEC_dmg2 | AQP9 |
| 1.4216E-17 | 0.4108283 | 0.492 | 0.317 | 3.3583E-13 | TEC_dmg2 | BRD2 |
| 1.5552E-17 | -0.5857937 | 0.88 | 0.934 | 3.674E-13 | TEC_dmg2 | RPL34 |

| | | | | | | |
|---|---|---|---|---|---|---|
| 1.6799E-17 | 0.35983728 | 0.336 | 0.178 | 3.9684E-13 | TEC_dmg2 | DST |
| 1.9994E-17 | -0.3829009 | 0.065 | 0.246 | 4.7231E-13 | TEC_dmg2 | RAB31 |
| 2.1924E-17 | -0.4882497 | 0.492 | 0.641 | 5.1792E-13 | TEC_dmg2 | ARPC5 |
| 2.3001E-17 | -0.3494881 | 0.523 | 0.703 | 5.4335E-13 | TEC_dmg2 | COX6B1 |
| 2.4921E-17 | 0.36924664 | 0.417 | 0.25 | 5.8871E-13 | TEC_dmg2 | BCLAF1 |
| 2.6171E-17 | -0.471985 | 0.038 | 0.208 | 6.1824E-13 | TEC_dmg2 | GK |
| 2.6727E-17 | -1.1158202 | 0.01 | 0.162 | 6.3137E-13 | TEC_dmg2 | MUC21 |
| 2.6907E-17 | -2.1689056 | 0.019 | 0.177 | 6.3562E-13 | TEC_dmg2 | IL1B |
| 2.7148E-17 | -0.3153341 | 0.034 | 0.203 | 6.4133E-13 | TEC_dmg2 | IFI16 |
| 2.824E-17 | -0.4925085 | 0.026 | 0.19 | 6.6712E-13 | TEC_dmg2 | FYB1 |
| 2.8975E-17 | -0.4452785 | 0.758 | 0.87 | 6.8449E-13 | TEC_dmg2 | PABPC1 |
| 3.0588E-17 | -0.3330514 | 0.012 | 0.167 | 7.2259E-13 | TEC_dmg2 | CFD |
| 3.1666E-17 | -0.3042845 | 0.024 | 0.189 | 7.4806E-13 | TEC_dmg2 | HCLS1 |
| 4.4587E-17 | -0.538015 | 0.062 | 0.246 | 1.0533E-12 | TEC_dmg2 | SLC11A1 |
| 4.9775E-17 | -0.5140076 | 0.849 | 0.898 | 1.1758E-12 | TEC_dmg2 | RPL29 |
| 4.9934E-17 | 0.2871874 | 0.379 | 0.21 | 1.1796E-12 | TEC_dmg2 | BDP1 |
| 5.2222E-17 | -0.6013666 | 0.856 | 0.894 | 1.2336E-12 | TEC_dmg2 | RPS15A |
| 5.469E-17 | -0.4126378 | 0.014 | 0.169 | 1.292E-12 | TEC_dmg2 | SLA |
| 6.2036E-17 | -0.4013222 | 0.034 | 0.202 | 1.4655E-12 | TEC_dmg2 | ITGAX |
| 6.348E-17 | -0.4320456 | 0.113 | 0.297 | 1.4996E-12 | TEC_dmg2 | IFNGR2 |
| 6.3929E-17 | -1.2626972 | 0.484 | 0.58 | 1.5102E-12 | TEC_dmg2 | KRT19 |
| 7.0101E-17 | 0.25872688 | 0.23 | 0.105 | 1.656E-12 | TEC_dmg2 | SGMS2 |
| 7.2856E-17 | -0.6921907 | 0.269 | 0.451 | 1.7211E-12 | TEC_dmg2 | MARCKS |
| 1.04E-16 | -0.5172968 | 0.022 | 0.178 | 2.4569E-12 | TEC_dmg2 | ANPEP |
| 1.1534E-16 | -0.5765128 | 0.017 | 0.169 | 2.7247E-12 | TEC_dmg2 | FXYD3 |
| 1.3205E-16 | -0.4052958 | 0.029 | 0.188 | 3.1195E-12 | TEC_dmg2 | PTPRE |
| 1.5239E-16 | -0.3910105 | 0.019 | 0.173 | 3.5999E-12 | TEC_dmg2 | SAMHD1 |
| 1.5928E-16 | 0.61059118 | 0.372 | 0.22 | 3.7626E-12 | TEC_dmg2 | CCND1 |
| 1.7991E-16 | -0.4936031 | 0.019 | 0.172 | 4.2499E-12 | TEC_dmg2 | PRDM1 |
| 2.1456E-16 | 0.25225905 | 0.259 | 0.128 | 5.0687E-12 | TEC_dmg2 | GOLGA3 |
| 2.2486E-16 | -0.8564941 | 0.005 | 0.148 | 5.312E-12 | TEC_dmg2 | CEACAM6 |
| 2.4136E-16 | -0.5025325 | 0.019 | 0.172 | 5.7017E-12 | TEC_dmg2 | TGFBI |
| 3.0483E-16 | -0.9206181 | 0.655 | 0.723 | 7.201E-12 | TEC_dmg2 | HSPB1 |
| 3.2386E-16 | -0.4051679 | 0.101 | 0.282 | 7.6506E-12 | TEC_dmg2 | LYN |
| 3.4917E-16 | -0.4185316 | 0.005 | 0.147 | 8.2485E-12 | TEC_dmg2 | FPR1 |
| 3.5417E-16 | -0.4647446 | 0.849 | 0.899 | 8.3665E-12 | TEC_dmg2 | RPS25 |
| 3.9064E-16 | -0.4387949 | 0.026 | 0.183 | 9.228E-12 | TEC_dmg2 | SAMSN1 |
| 3.9228E-16 | -0.7603512 | 0.005 | 0.146 | 9.2668E-12 | TEC_dmg2 | TMPRSS11E |
| 4.0396E-16 | -0.7312902 | 0.014 | 0.163 | 9.5428E-12 | TEC_dmg2 | APOBEC3A |
| 4.3415E-16 | -0.4279297 | 0.329 | 0.509 | 1.0256E-11 | TEC_dmg2 | DBI |
| 4.5296E-16 | 0.50510382 | 0.463 | 0.305 | 1.07E-11 | TEC_dmg2 | ZNF292 |
| 4.6677E-16 | -1.0365775 | 0.031 | 0.185 | 1.1027E-11 | TEC_dmg2 | DUOX2 |
| 4.7612E-16 | 0.28591665 | 0.365 | 0.21 | 1.1247E-11 | TEC_dmg2 | RNPS1 |
| 5.3536E-16 | -0.5042336 | 0.058 | 0.229 | 1.2647E-11 | TEC_dmg2 | ZEB2 |
| 6.0387E-16 | -1.4385494 | 0.043 | 0.2 | 1.4265E-11 | TEC_dmg2 | KRT17 |
| 6.2121E-16 | -0.9384283 | 0.014 | 0.159 | 1.4675E-11 | TEC_dmg2 | PRSS27 |
| 6.7925E-16 | -0.4799789 | 0.002 | 0.14 | 1.6046E-11 | TEC_dmg2 | PLAC8 |
| 6.9408E-16 | -1.1328979 | 0.019 | 0.167 | 1.6396E-11 | TEC_dmg2 | DUOXA2 |
| 7.094E-16 | -0.3318685 | 0.034 | 0.19 | 1.6758E-11 | TEC_dmg2 | PYCARD |
| 8.7283E-16 | -0.4033941 | 0.345 | 0.522 | 2.0619E-11 | TEC_dmg2 | ATP6V1F |

| | | | | | | |
|---|---|---|---|---|---|---|
| 8.9723E-16 | -0.4406272 | 0.048 | 0.205 | 2.1195E-11 | TEC_dmg2 | CTSC |
| 1.1081E-15 | -0.7265973 | 0.01 | 0.15 | 2.6176E-11 | TEC_dmg2 | TMPRSS11D |
| 1.1462E-15 | -0.4550491 | 0.031 | 0.183 | 2.7077E-11 | TEC_dmg2 | FBP1 |
| 1.2158E-15 | -0.3020239 | 0.022 | 0.169 | 2.8721E-11 | TEC_dmg2 | GLIPR2 |
| 1.239E-15 | -0.3033314 | 0.002 | 0.138 | 2.9269E-11 | TEC_dmg2 | CLEC7A |
| 1.3861E-15 | 0.37729014 | 0.4 | 0.246 | 3.2743E-11 | TEC_dmg2 | CREBRF |
| 1.9381E-15 | -0.567212 | 0.026 | 0.174 | 4.5783E-11 | TEC_dmg2 | HLA-DQB1 |
| 2.0215E-15 | -0.3699887 | 0.141 | 0.322 | 4.7754E-11 | TEC_dmg2 | AP2S1 |
| 2.1407E-15 | 0.55006176 | 0.772 | 0.644 | 5.0571E-11 | TEC_dmg2 | HSPA5 |
| 2.3459E-15 | -0.3332532 | 0.012 | 0.153 | 5.5417E-11 | TEC_dmg2 | EVI2B |
| 2.4098E-15 | -0.395128 | 0.01 | 0.147 | 5.6927E-11 | TEC_dmg2 | VSIG2 |
| 2.4889E-15 | -0.4754249 | 0.904 | 0.924 | 5.8795E-11 | TEC_dmg2 | RPS24 |
| 2.962E-15 | -0.403082 | 0.885 | 0.947 | 6.997E-11 | TEC_dmg2 | RPS16 |
| 3.1412E-15 | -0.2874294 | 0.01 | 0.146 | 7.4203E-11 | TEC_dmg2 | CARD16 |
| 3.1866E-15 | 0.45226585 | 0.314 | 0.171 | 7.5278E-11 | TEC_dmg2 | XIST |
| 3.3425E-15 | -0.2951465 | 0.137 | 0.317 | 7.8961E-11 | TEC_dmg2 | TAF10 |
| 3.3582E-15 | -0.7610312 | 0.01 | 0.147 | 7.933E-11 | TEC_dmg2 | CEACAM7 |
| 3.5474E-15 | -0.3967964 | 0.055 | 0.213 | 8.3801E-11 | TEC_dmg2 | NINJ1 |
| 3.7103E-15 | -0.2685566 | 0.005 | 0.138 | 8.7649E-11 | TEC_dmg2 | EVI2A |
| 3.731E-15 | -0.5392799 | 0.259 | 0.435 | 8.8138E-11 | TEC_dmg2 | GSTO1 |
| 3.9755E-15 | 0.35088883 | 0.504 | 0.329 | 9.3914E-11 | TEC_dmg2 | APP |
| 4.2925E-15 | -0.5172429 | 0.887 | 0.916 | 1.014E-10 | TEC_dmg2 | RPL32 |
| 4.382E-15 | -0.4628835 | 0.026 | 0.172 | 1.0352E-10 | TEC_dmg2 | CXCR4 |
| 4.3915E-15 | -0.4236414 | 0.005 | 0.138 | 1.0374E-10 | TEC_dmg2 | MCEMP1 |
| 4.7375E-15 | -0.2835944 | 0.019 | 0.163 | 1.1191E-10 | TEC_dmg2 | FERMT3 |
| 5.2185E-15 | -0.2818267 | 0 | 0.129 | 1.2328E-10 | TEC_dmg2 | PLBD1 |
| 5.6364E-15 | 0.35853423 | 0.489 | 0.317 | 1.3315E-10 | TEC_dmg2 | GADD45A |
| 5.6751E-15 | 0.31190283 | 0.264 | 0.137 | 1.3406E-10 | TEC_dmg2 | GULP1 |
| 5.7893E-15 | -0.5414853 | 0.038 | 0.186 | 1.3676E-10 | TEC_dmg2 | C5AR1 |
| 6.2937E-15 | -0.5160293 | 0.019 | 0.159 | 1.4868E-10 | TEC_dmg2 | CD300E |
| 6.5746E-15 | -1.5155332 | 0.024 | 0.164 | 1.5531E-10 | TEC_dmg2 | ECM1 |
| 8.4493E-15 | -0.532594 | 0.911 | 0.904 | 1.996E-10 | TEC_dmg2 | RPLP2 |
| 9.4851E-15 | -0.3990639 | 0.192 | 0.375 | 2.2407E-10 | TEC_dmg2 | METRNL |
| 9.6684E-15 | -0.268295 | 0.012 | 0.147 | 2.284E-10 | TEC_dmg2 | UCP2 |
| 1.1579E-14 | -0.3629047 | 0.849 | 0.932 | 2.7352E-10 | TEC_dmg2 | FAU |
| 1.1608E-14 | -0.8865894 | 0.122 | 0.284 | 2.7421E-10 | TEC_dmg2 | IFITM2 |
| 1.3586E-14 | -0.6158412 | 0.007 | 0.137 | 3.2094E-10 | TEC_dmg2 | CXCL17 |
| 1.3619E-14 | -2.5254278 | 0.242 | 0.379 | 3.2172E-10 | TEC_dmg2 | KRT13 |
| 1.4662E-14 | -0.3366812 | 0.012 | 0.145 | 3.4636E-10 | TEC_dmg2 | TLR2 |
| 1.5946E-14 | -0.6019939 | 0.014 | 0.148 | 3.7669E-10 | TEC_dmg2 | NCCRP1 |
| 1.5968E-14 | -0.3977818 | 0.513 | 0.648 | 3.7721E-10 | TEC_dmg2 | ATP5MC2 |
| 1.6026E-14 | -0.5443685 | 0.199 | 0.37 | 3.7858E-10 | TEC_dmg2 | COTL1 |
| 1.9775E-14 | -0.301179 | 0.041 | 0.185 | 4.6714E-10 | TEC_dmg2 | BID |
| 2.0333E-14 | -0.3141641 | 0.055 | 0.209 | 4.8033E-10 | TEC_dmg2 | STK17B |
| 2.0469E-14 | -0.438909 | 0.892 | 0.927 | 4.8353E-10 | TEC_dmg2 | RPL19 |
| 2.0896E-14 | -0.9224333 | 0.314 | 0.459 | 4.9363E-10 | TEC_dmg2 | MXD1 |
| 2.2845E-14 | -1.0451169 | 0.962 | 0.987 | 5.3966E-10 | TEC_dmg2 | FTL |
| 2.7682E-14 | 0.35434325 | 0.403 | 0.255 | 6.5394E-10 | TEC_dmg2 | CHD2 |
| 2.7945E-14 | -1.0277584 | 0.007 | 0.135 | 6.6013E-10 | TEC_dmg2 | SERPINB2 |
| 2.8322E-14 | -0.3641453 | 0.165 | 0.346 | 6.6906E-10 | TEC_dmg2 | DDIT4 |

| | | | | | | |
|---|---|---|---|---|---|---|
| 2.8404E-14 | -0.3768605 | 0.755 | 0.865 | 6.7098E-10 | TEC_dmg2 | CALM1 |
| 2.9791E-14 | -0.314577 | 0.014 | 0.147 | 7.0375E-10 | TEC_dmg2 | C1orf162 |
| 3.3457E-14 | -0.9951014 | 0.012 | 0.141 | 7.9036E-10 | TEC_dmg2 | CXCL6 |
| 3.4315E-14 | -0.4073412 | 0.875 | 0.92 | 8.1061E-10 | TEC_dmg2 | RPS15 |
| 3.6284E-14 | 0.32742394 | 0.408 | 0.257 | 8.5714E-10 | TEC_dmg2 | ARHGAP5 |
| 3.6885E-14 | -0.3177461 | 0.026 | 0.165 | 8.7134E-10 | TEC_dmg2 | CLEC2B |
| 3.7693E-14 | -0.4264358 | 0.062 | 0.21 | 8.9043E-10 | TEC_dmg2 | SOCS3 |
| 3.9498E-14 | -1.4065528 | 0.084 | 0.233 | 9.3306E-10 | TEC_dmg2 | PRSS22 |
| 4.1655E-14 | -0.4574039 | 0.019 | 0.153 | 9.8402E-10 | TEC_dmg2 | CLEC5A |
| 4.2034E-14 | -0.4672812 | 0.921 | 0.955 | 9.9296E-10 | TEC_dmg2 | ACTG1 |
| 4.4544E-14 | -0.3277857 | 0.043 | 0.187 | 1.0523E-09 | TEC_dmg2 | CKLF |
| 4.5824E-14 | -0.2847446 | 0.062 | 0.217 | 1.0825E-09 | TEC_dmg2 | TGFB1 |
| 4.6962E-14 | -0.444742 | 0.014 | 0.145 | 1.1094E-09 | TEC_dmg2 | CD93 |
| 5.7848E-14 | -0.8733013 | 0.01 | 0.136 | 1.3666E-09 | TEC_dmg2 | TMPRSS11B |
| 6.013E-14 | -0.6416362 | 0.007 | 0.132 | 1.4204E-09 | TEC_dmg2 | DHRS9 |
| 6.0175E-14 | -0.6805018 | 0.022 | 0.154 | 1.4215E-09 | TEC_dmg2 | SCEL |
| 6.4094E-14 | -0.3229443 | 0.376 | 0.553 | 1.5141E-09 | TEC_dmg2 | UQCR10 |
| 6.5867E-14 | -0.2860439 | 0.549 | 0.705 | 1.556E-09 | TEC_dmg2 | SUMO2 |
| 7.8276E-14 | -0.3384963 | 0.211 | 0.39 | 1.8491E-09 | TEC_dmg2 | UQCR11 |
| 8.9754E-14 | -0.3297276 | 0.017 | 0.147 | 2.1203E-09 | TEC_dmg2 | TREM1 |
| 9.3514E-14 | -0.4297789 | 0.005 | 0.126 | 2.2091E-09 | TEC_dmg2 | SRD5A3 |
| 9.3527E-14 | -0.872352 | 0.024 | 0.156 | 2.2094E-09 | TEC_dmg2 | PITX1 |
| 9.9649E-14 | -1.5360892 | 0.01 | 0.134 | 2.354E-09 | TEC_dmg2 | LYPD2 |
| 1.1336E-13 | -0.2517667 | 0.01 | 0.135 | 2.6779E-09 | TEC_dmg2 | CYTIP |
| 1.1415E-13 | -0.348361 | 0.029 | 0.165 | 2.6966E-09 | TEC_dmg2 | LCP2 |
| 1.1708E-13 | 0.33125842 | 0.293 | 0.163 | 2.7657E-09 | TEC_dmg2 | MIR22HG |
| 1.3474E-13 | 0.30690909 | 0.326 | 0.197 | 3.183E-09 | TEC_dmg2 | MDM4 |
| 1.4042E-13 | -0.4746549 | 0.82 | 0.877 | 3.3172E-09 | TEC_dmg2 | RPL23A |
| 1.4634E-13 | 0.28518666 | 0.348 | 0.206 | 3.4571E-09 | TEC_dmg2 | MAP4 |
| 1.493E-13 | 0.32607755 | 0.254 | 0.133 | 3.527E-09 | TEC_dmg2 | UCHL1 |
| 1.7221E-13 | -0.5169573 | 0.024 | 0.155 | 4.0682E-09 | TEC_dmg2 | CDC42EP5 |
| 1.7653E-13 | -0.2899973 | 0.453 | 0.618 | 4.1703E-09 | TEC_dmg2 | COX8A |
| 2.0301E-13 | -0.4087438 | 0.765 | 0.82 | 4.7958E-09 | TEC_dmg2 | RPL27 |
| 2.0573E-13 | -0.313062 | 0.082 | 0.239 | 4.8601E-09 | TEC_dmg2 | RNASET2 |
| 2.7814E-13 | -0.277087 | 0.24 | 0.421 | 6.5706E-09 | TEC_dmg2 | CAPZB |
| 3.3741E-13 | -1.133851 | 0.201 | 0.346 | 7.9707E-09 | TEC_dmg2 | CTSL |
| 3.5864E-13 | -0.3402705 | 0.01 | 0.129 | 8.4722E-09 | TEC_dmg2 | LRG1 |
| 3.6268E-13 | -0.4733759 | 0.885 | 0.934 | 8.5675E-09 | TEC_dmg2 | RPS14 |
| 3.6806E-13 | -0.2817704 | 0.031 | 0.164 | 8.6947E-09 | TEC_dmg2 | MYO1F |
| 4.0878E-13 | -0.3957568 | 0.245 | 0.419 | 9.6565E-09 | TEC_dmg2 | JPT1 |
| 4.0977E-13 | 0.34219351 | 0.369 | 0.232 | 9.6799E-09 | TEC_dmg2 | LPP |
| 4.596E-13 | -0.3303499 | 0.002 | 0.117 | 1.0857E-08 | TEC_dmg2 | CD163 |
| 4.7314E-13 | -0.3953725 | 0.326 | 0.493 | 1.1177E-08 | TEC_dmg2 | TAGLN2 |
| 5.2919E-13 | -0.416281 | 0.062 | 0.202 | 1.2501E-08 | TEC_dmg2 | NCF2 |
| 6.4335E-13 | -0.3210577 | 0.674 | 0.785 | 1.5198E-08 | TEC_dmg2 | DYNLL1 |
| 7.0469E-13 | -3.891051 | 0.283 | 0.402 | 1.6647E-08 | TEC_dmg2 | SPRR3 |
| 7.1082E-13 | 0.37374894 | 0.489 | 0.351 | 1.6792E-08 | TEC_dmg2 | TAF1D |
| 7.3907E-13 | -0.3555508 | 0.233 | 0.397 | 1.7459E-08 | TEC_dmg2 | COMMD6 |
| 7.8475E-13 | -0.5348674 | 0.029 | 0.156 | 1.8538E-08 | TEC_dmg2 | MUC4 |
| 7.9091E-13 | -0.3340125 | 0.014 | 0.136 | 1.8684E-08 | TEC_dmg2 | BCAS1 |

| | | | | | | |
|---|---|---|---|---|---|---|
| 8.115E-13 | -0.5475152 | 0.019 | 0.142 | 1.917E-08 | TEC_dmg2 | SNCG |
| 8.3987E-13 | -0.4232184 | 0.036 | 0.167 | 1.984E-08 | TEC_dmg2 | HIST3H2A |
| 8.833E-13 | -0.3007454 | 0.029 | 0.156 | 2.0866E-08 | TEC_dmg2 | ADGRE5 |
| 8.8336E-13 | -0.7742973 | 0.031 | 0.157 | 2.0868E-08 | TEC_dmg2 | KRT15 |
| 8.9854E-13 | 0.36154244 | 0.348 | 0.218 | 2.1226E-08 | TEC_dmg2 | TES |
| 9.1501E-13 | -0.2780532 | 0.024 | 0.15 | 2.1615E-08 | TEC_dmg2 | BCAT1 |
| 9.9545E-13 | 0.25437889 | 0.362 | 0.224 | 2.3516E-08 | TEC_dmg2 | ARPP19 |
| 1.0054E-12 | -0.3425385 | 0.022 | 0.146 | 2.3751E-08 | TEC_dmg2 | MS4A7 |
| 1.1252E-12 | -0.3456147 | 0.933 | 0.945 | 2.6581E-08 | TEC_dmg2 | PTMA |
| 1.1336E-12 | -0.6152721 | 0.014 | 0.132 | 2.6779E-08 | TEC_dmg2 | GABRP |
| 1.1391E-12 | 0.25711023 | 0.4 | 0.24 | 2.6909E-08 | TEC_dmg2 | S100A13 |
| 1.1471E-12 | -0.8815514 | 0.336 | 0.474 | 2.7098E-08 | TEC_dmg2 | ERO1A |
| 1.1846E-12 | 0.33658498 | 0.376 | 0.241 | 2.7983E-08 | TEC_dmg2 | MAFG |
| 1.3836E-12 | -0.2546252 | 0.043 | 0.177 | 3.2684E-08 | TEC_dmg2 | ABCA1 |
| 1.4129E-12 | -0.2580358 | 0.017 | 0.137 | 3.3378E-08 | TEC_dmg2 | HCK |
| 1.5477E-12 | -0.7305349 | 0.046 | 0.174 | 3.6561E-08 | TEC_dmg2 | NDRG2 |
| 1.7757E-12 | 0.41373965 | 0.516 | 0.379 | 4.1947E-08 | TEC_dmg2 | N4BP2L2 |
| 1.8195E-12 | 0.48430733 | 0.338 | 0.205 | 4.2982E-08 | TEC_dmg2 | KCNQ1OT1 |
| 1.8841E-12 | -0.3437306 | 0.007 | 0.119 | 4.4508E-08 | TEC_dmg2 | DUSP2 |
| 1.9165E-12 | 0.26389392 | 0.288 | 0.164 | 4.5272E-08 | TEC_dmg2 | EPB41L4A-AS1 |
| 1.9308E-12 | 0.52856571 | 0.525 | 0.425 | 4.561E-08 | TEC_dmg2 | SLC3A2 |
| 1.9433E-12 | -0.3524518 | 0.446 | 0.6 | 4.5907E-08 | TEC_dmg2 | VAMP8 |
| 1.9642E-12 | -0.4296273 | 0.892 | 0.904 | 4.64E-08 | TEC_dmg2 | RPL37 |
| 2.0063E-12 | 0.37753943 | 0.482 | 0.345 | 4.7394E-08 | TEC_dmg2 | ZNF207 |
| 2.2725E-12 | -0.3314475 | 0.022 | 0.144 | 5.3683E-08 | TEC_dmg2 | NCF1 |
| 2.3128E-12 | -0.4051657 | 0.197 | 0.349 | 5.4634E-08 | TEC_dmg2 | TIMP2 |
| 2.4482E-12 | -0.3742458 | 0.201 | 0.359 | 5.7835E-08 | TEC_dmg2 | MIF |
| 2.4913E-12 | -0.3100998 | 0.201 | 0.37 | 5.8851E-08 | TEC_dmg2 | TXNDC17 |
| 2.5764E-12 | -1.0907684 | 0.007 | 0.118 | 6.0862E-08 | TEC_dmg2 | SPINK5 |
| 2.6657E-12 | 0.26861873 | 0.29 | 0.168 | 6.2973E-08 | TEC_dmg2 | GARS |
| 2.8466E-12 | -0.5646975 | 0.007 | 0.118 | 6.7246E-08 | TEC_dmg2 | MNDA |
| 2.8965E-12 | -0.3153695 | 0.837 | 0.901 | 6.8424E-08 | TEC_dmg2 | RPS11 |
| 3.2082E-12 | -0.5765735 | 0.005 | 0.113 | 7.5788E-08 | TEC_dmg2 | KRT78 |
| 3.3757E-12 | -0.4379849 | 0.005 | 0.113 | 7.9744E-08 | TEC_dmg2 | TMPRSS11A |
| 3.3881E-12 | -0.3197621 | 0.422 | 0.585 | 8.0037E-08 | TEC_dmg2 | NOP10 |
| 3.4032E-12 | -0.7939513 | 0.055 | 0.189 | 8.0393E-08 | TEC_dmg2 | RARRES3 |
| 3.5368E-12 | -0.4500923 | 0.132 | 0.277 | 8.3549E-08 | TEC_dmg2 | ETS2 |
| 3.6956E-12 | -0.2604727 | 0.113 | 0.267 | 8.7302E-08 | TEC_dmg2 | CDC42SE1 |
| 3.766E-12 | -0.3278074 | 0.252 | 0.416 | 8.8964E-08 | TEC_dmg2 | EIF3K |
| 4.1327E-12 | 0.32268559 | 0.767 | 0.667 | 9.7626E-08 | TEC_dmg2 | RTN4 |
| 4.3488E-12 | -0.2791666 | 0.156 | 0.315 | 1.0273E-07 | TEC_dmg2 | BLOC1S1 |
| 4.5787E-12 | -0.339208 | 0.405 | 0.555 | 1.0816E-07 | TEC_dmg2 | SSR4 |
| 4.6923E-12 | -0.3286518 | 0.134 | 0.29 | 1.1085E-07 | TEC_dmg2 | RNF149 |
| 4.7674E-12 | 0.32748366 | 0.312 | 0.187 | 1.1262E-07 | TEC_dmg2 | SLC5A3 |
| 5.6711E-12 | -0.3270624 | 0.014 | 0.127 | 1.3397E-07 | TEC_dmg2 | MARCO |
| 5.8498E-12 | -0.3604807 | 0.777 | 0.849 | 1.3819E-07 | TEC_dmg2 | RPL38 |
| 6.232E-12 | -0.8145168 | 0.019 | 0.133 | 1.4722E-07 | TEC_dmg2 | RHCG |
| 6.4464E-12 | -0.3195114 | 0.002 | 0.106 | 1.5228E-07 | TEC_dmg2 | A2ML1 |
| 6.9363E-12 | -0.4138813 | 0.055 | 0.187 | 1.6386E-07 | TEC_dmg2 | TNFSF10 |
| 7.1729E-12 | -0.7330913 | 0.055 | 0.181 | 1.6945E-07 | TEC_dmg2 | F3 |

| | | | | | | |
|---|---|---|---|---|---|---|
| 7.5669E-12 | -0.3798566 | 0.947 | 0.964 | 1.7875E-07 | TEC_dmg2 | RPL13 |
| 8.1336E-12 | -0.2516419 | 0.053 | 0.184 | 1.9214E-07 | TEC_dmg2 | PSMB9 |
| 8.4501E-12 | -0.3065569 | 0.755 | 0.876 | 1.9962E-07 | TEC_dmg2 | MYL12A |
| 9.8426E-12 | -0.3248862 | 0.127 | 0.269 | 2.3251E-07 | TEC_dmg2 | RAB5IF |
| 1.0377E-11 | -0.4106042 | 0.065 | 0.196 | 2.4514E-07 | TEC_dmg2 | SAMD9 |
| 1.1199E-11 | -0.3779794 | 0.005 | 0.108 | 2.6455E-07 | TEC_dmg2 | KRT23 |
| 1.1328E-11 | -0.5415778 | 0.887 | 0.887 | 2.676E-07 | TEC_dmg2 | RPS3 |
| 1.1424E-11 | -0.2662232 | 0.01 | 0.117 | 2.6986E-07 | TEC_dmg2 | OAS1 |
| 1.1506E-11 | -0.3701551 | 0.014 | 0.124 | 2.7181E-07 | TEC_dmg2 | TMEM40 |
| 1.167E-11 | 0.25567753 | 0.297 | 0.179 | 2.7569E-07 | TEC_dmg2 | SARS |
| 1.1695E-11 | -0.2508313 | 0.012 | 0.121 | 2.7627E-07 | TEC_dmg2 | LILRB2 |
| 1.194E-11 | -0.8418652 | 0.022 | 0.135 | 2.8205E-07 | TEC_dmg2 | EREG |
| 1.2193E-11 | -0.4788444 | 0.53 | 0.648 | 2.8804E-07 | TEC_dmg2 | LDHA |
| 1.2222E-11 | -0.7733786 | 0.065 | 0.189 | 2.8873E-07 | TEC_dmg2 | TMPRSS2 |
| 1.3133E-11 | -0.347153 | 0.012 | 0.12 | 3.1025E-07 | TEC_dmg2 | FCGR3A |
| 1.4438E-11 | -0.3095708 | 0.089 | 0.224 | 3.4107E-07 | TEC_dmg2 | GRINA |
| 1.5486E-11 | -0.2589251 | 0.731 | 0.837 | 3.6582E-07 | TEC_dmg2 | COX4I1 |
| 1.6067E-11 | 0.41610292 | 0.513 | 0.393 | 3.7955E-07 | TEC_dmg2 | PNISR |
| 1.7083E-11 | 0.26771994 | 0.336 | 0.21 | 4.0354E-07 | TEC_dmg2 | CELF1 |
| 1.7775E-11 | 0.2609455 | 0.4 | 0.265 | 4.199E-07 | TEC_dmg2 | SMG1 |
| 1.8287E-11 | -0.3787301 | 0.854 | 0.897 | 4.32E-07 | TEC_dmg2 | RPL35A |
| 1.9841E-11 | -0.3448089 | 0.333 | 0.483 | 4.687E-07 | TEC_dmg2 | GABARAPL2 |
| 2.3482E-11 | -0.2960916 | 0.024 | 0.137 | 5.5471E-07 | TEC_dmg2 | HLA-DMA |
| 2.3796E-11 | -0.3029644 | 0.022 | 0.133 | 5.6213E-07 | TEC_dmg2 | RGCC |
| 2.4116E-11 | -0.3459573 | 0.588 | 0.71 | 5.697E-07 | TEC_dmg2 | GPX4 |
| 2.7298E-11 | -0.4601119 | 0.177 | 0.328 | 6.4486E-07 | TEC_dmg2 | CEBPD |
| 2.7855E-11 | -1.1236839 | 0.017 | 0.124 | 6.5802E-07 | TEC_dmg2 | CCL3 |
| 2.9112E-11 | -1.4135369 | 0.007 | 0.108 | 6.8772E-07 | TEC_dmg2 | CNFN |
| 3.07E-11 | -1.4400195 | 0.031 | 0.145 | 7.2524E-07 | TEC_dmg2 | CXCL3 |
| 3.0963E-11 | -0.3265445 | 0.23 | 0.388 | 7.3144E-07 | TEC_dmg2 | PSME1 |
| 3.1717E-11 | -0.6845976 | 0.062 | 0.185 | 7.4926E-07 | TEC_dmg2 | AQP3 |
| 3.2883E-11 | -0.3350996 | 0.211 | 0.358 | 7.768E-07 | TEC_dmg2 | UQCRQ |
| 3.2941E-11 | -0.3154604 | 0.058 | 0.184 | 7.7816E-07 | TEC_dmg2 | ICAM1 |
| 3.6631E-11 | -0.2768261 | 0.053 | 0.174 | 8.6534E-07 | TEC_dmg2 | DHRS3 |
| 3.7573E-11 | -0.3433871 | 0.012 | 0.115 | 8.8758E-07 | TEC_dmg2 | CEACAM1 |
| 3.8756E-11 | -0.7464801 | 0.029 | 0.14 | 9.1554E-07 | TEC_dmg2 | CXCL1 |
| 3.8785E-11 | -0.3006979 | 0.213 | 0.37 | 9.1622E-07 | TEC_dmg2 | NDUFV2 |
| 4.0922E-11 | -0.6112543 | 0.189 | 0.324 | 9.667E-07 | TEC_dmg2 | IFITM3 |
| 4.1E-11 | -0.558117 | 0.031 | 0.143 | 9.6853E-07 | TEC_dmg2 | CDKN2A |
| 4.1508E-11 | -0.2773897 | 0.007 | 0.107 | 9.8055E-07 | TEC_dmg2 | MS4A6A |
| 4.4477E-11 | -1.1380267 | 0.024 | 0.133 | 1.0507E-06 | TEC_dmg2 | PLAT |
| 4.6918E-11 | -0.3929778 | 0.065 | 0.19 | 1.1083E-06 | TEC_dmg2 | DUSP6 |
| 5.1998E-11 | -0.4851236 | 0.904 | 0.933 | 1.2283E-06 | TEC_dmg2 | RPS12 |
| 6.1223E-11 | -0.4306701 | 0.026 | 0.136 | 1.4463E-06 | TEC_dmg2 | HLA-DRB5 |
| 6.3284E-11 | -0.3860717 | 0.88 | 0.924 | 1.4949E-06 | TEC_dmg2 | RPL18A |
| 6.6512E-11 | 0.25369171 | 0.329 | 0.205 | 1.5712E-06 | TEC_dmg2 | RBBP6 |
| 6.7191E-11 | -0.8885956 | 0.062 | 0.182 | 1.5873E-06 | TEC_dmg2 | SPINK1 |
| 6.8772E-11 | 0.31218005 | 0.415 | 0.286 | 1.6246E-06 | TEC_dmg2 | MACF1 |
| 7.5799E-11 | -0.4181941 | 0.098 | 0.23 | 1.7906E-06 | TEC_dmg2 | CYB5A |
| 8.0527E-11 | -0.4797193 | 0.46 | 0.579 | 1.9023E-06 | TEC_dmg2 | LITAF |

| | | | | | | |
|---|---|---|---|---|---|---|
| 8.3555E-11 | -1.229699 | 0.038 | 0.153 | 1.9738E-06 | TEC_dmg2 | CCL20 |
| 8.5325E-11 | -0.4722547 | 0.005 | 0.101 | 2.0156E-06 | TEC_dmg2 | RETN |
| 9.1947E-11 | -0.3608388 | 0.556 | 0.674 | 2.1721E-06 | TEC_dmg2 | CALR |
| 9.2294E-11 | 0.43290807 | 0.46 | 0.328 | 2.1803E-06 | TEC_dmg2 | HERPUD1 |
| 9.6523E-11 | -0.2964267 | 0.012 | 0.111 | 2.2802E-06 | TEC_dmg2 | STS |
| 9.8322E-11 | -0.397482 | 0.17 | 0.316 | 2.3227E-06 | TEC_dmg2 | HIST1H2AC |
| 1.0389E-10 | -0.3043493 | 0.177 | 0.321 | 2.4541E-06 | TEC_dmg2 | FKBP1A |
| 1.1741E-10 | -0.2538786 | 0.012 | 0.112 | 2.7736E-06 | TEC_dmg2 | FUT3 |
| 1.2487E-10 | -0.3452692 | 0.408 | 0.549 | 2.9498E-06 | TEC_dmg2 | TOP1 |
| 1.3421E-10 | -0.4183583 | 0.88 | 0.898 | 3.1705E-06 | TEC_dmg2 | RPS3A |
| 1.3499E-10 | 0.328011 | 0.544 | 0.414 | 3.1889E-06 | TEC_dmg2 | FUS |
| 1.3838E-10 | -0.4118113 | 0.134 | 0.27 | 3.269E-06 | TEC_dmg2 | CCNG2 |
| 1.4118E-10 | -0.2859104 | 0.043 | 0.157 | 3.3351E-06 | TEC_dmg2 | TSPAN6 |
| 1.5007E-10 | -0.3487713 | 0.904 | 0.895 | 3.5451E-06 | TEC_dmg2 | RPL37A |
| 1.6078E-10 | 0.33473566 | 0.405 | 0.284 | 3.798E-06 | TEC_dmg2 | RB1CC1 |
| 1.671E-10 | -0.2847059 | 0.635 | 0.772 | 3.9475E-06 | TEC_dmg2 | RHOA |
| 1.6726E-10 | -0.3152792 | 0.096 | 0.225 | 3.9513E-06 | TEC_dmg2 | STXBP2 |
| 1.6942E-10 | -0.566331 | 0.007 | 0.102 | 4.0022E-06 | TEC_dmg2 | HLA-DQA1 |
| 1.8293E-10 | -0.2754344 | 0.007 | 0.102 | 4.3213E-06 | TEC_dmg2 | LRMP |
| 1.8982E-10 | -0.7475707 | 0.01 | 0.105 | 4.4841E-06 | TEC_dmg2 | ZG16B |
| 2.1003E-10 | -0.4849822 | 0.89 | 0.888 | 4.9615E-06 | TEC_dmg2 | RPS6 |
| 2.2359E-10 | -0.3097338 | 0.161 | 0.299 | 5.2819E-06 | TEC_dmg2 | ARL6IP5 |
| 2.3695E-10 | 0.30724435 | 0.333 | 0.213 | 5.5974E-06 | TEC_dmg2 | NKTR |
| 2.4326E-10 | -0.2584465 | 0.041 | 0.151 | 5.7464E-06 | TEC_dmg2 | SMAP2 |
| 2.4372E-10 | -0.3398655 | 0.866 | 0.876 | 5.7574E-06 | TEC_dmg2 | RPL36 |
| 2.4427E-10 | 0.28176188 | 0.36 | 0.237 | 5.7704E-06 | TEC_dmg2 | RNMT |
| 2.635E-10 | 0.38126581 | 0.41 | 0.292 | 6.2247E-06 | TEC_dmg2 | TUBA1C |
| 2.7107E-10 | -0.4613027 | 0.022 | 0.123 | 6.4036E-06 | TEC_dmg2 | AGR2 |
| 2.786E-10 | -0.4450597 | 0.902 | 0.924 | 6.5813E-06 | TEC_dmg2 | RPS27A |
| 2.8756E-10 | -1.4999619 | 0.012 | 0.107 | 6.793E-06 | TEC_dmg2 | CRCT1 |
| 3.1918E-10 | -0.2593307 | 0.01 | 0.104 | 7.5399E-06 | TEC_dmg2 | LILRB1 |
| 3.3416E-10 | -0.2735117 | 0.688 | 0.772 | 7.8938E-06 | TEC_dmg2 | TMA7 |
| 3.5633E-10 | -0.4505625 | 0.07 | 0.184 | 8.4176E-06 | TEC_dmg2 | SDCBP2 |
| 3.5993E-10 | -0.2802879 | 0.484 | 0.617 | 8.5027E-06 | TEC_dmg2 | CAP1 |
| 3.9082E-10 | -0.6822629 | 0.302 | 0.426 | 9.2323E-06 | TEC_dmg2 | EMP3 |
| 4.1378E-10 | -0.3163995 | 0.115 | 0.249 | 9.7747E-06 | TEC_dmg2 | GPCPD1 |
| 4.2374E-10 | -0.3856755 | 0.882 | 0.905 | 1.001E-05 | TEC_dmg2 | RPL7A |
| 5.002E-10 | -0.2541464 | 0.225 | 0.374 | 1.1816E-05 | TEC_dmg2 | GNB2 |
| 5.1053E-10 | -0.3763613 | 0.05 | 0.159 | 1.206E-05 | TEC_dmg2 | MALL |
| 5.5725E-10 | -0.3674503 | 0.36 | 0.494 | 1.3164E-05 | TEC_dmg2 | H2AFJ |
| 5.9636E-10 | -0.3439488 | 0.041 | 0.149 | 1.4088E-05 | TEC_dmg2 | RGS2 |
| 6.4036E-10 | -0.2818871 | 0.228 | 0.379 | 1.5127E-05 | TEC_dmg2 | POLR2L |
| 6.8265E-10 | -0.2802472 | 0.206 | 0.346 | 1.6126E-05 | TEC_dmg2 | LAMTOR4 |
| 7.6268E-10 | 0.49692606 | 0.345 | 0.228 | 1.8017E-05 | TEC_dmg2 | ITGA2 |
| 7.6345E-10 | -0.4132161 | 0.118 | 0.244 | 1.8035E-05 | TEC_dmg2 | ATP2B1 |
| 8.2228E-10 | -0.3919075 | 0.894 | 0.912 | 1.9425E-05 | TEC_dmg2 | RPS8 |
| 8.3513E-10 | -1.5691581 | 0.041 | 0.144 | 1.9728E-05 | TEC_dmg2 | SPRR2A |
| 8.3767E-10 | -0.3728079 | 0.014 | 0.107 | 1.9788E-05 | TEC_dmg2 | THBD |
| 9.1082E-10 | -0.3406935 | 0.012 | 0.104 | 2.1516E-05 | TEC_dmg2 | CLIC3 |
| 9.1093E-10 | 0.32676301 | 0.393 | 0.274 | 2.1519E-05 | TEC_dmg2 | UFM1 |

| | | | | | | |
|---|---|---|---|---|---|---|
| 9.3373E-10 | -0.6598502 | 0.125 | 0.239 | 2.2058E-05 | TEC_dmg2 | ADGRF1 |
| 9.3794E-10 | -0.2651414 | 0.29 | 0.435 | 2.2157E-05 | TEC_dmg2 | ATP5MF |
| 9.4617E-10 | 0.36460748 | 0.499 | 0.387 | 2.2351E-05 | TEC_dmg2 | AHNAK |
| 9.5928E-10 | -0.6781461 | 0.54 | 0.643 | 2.2661E-05 | TEC_dmg2 | IER3 |
| 1.0267E-09 | -0.766296 | 0.436 | 0.539 | 2.4253E-05 | TEC_dmg2 | PLAUR |
| 1.1689E-09 | -0.3876622 | 0.468 | 0.576 | 2.7612E-05 | TEC_dmg2 | CD55 |
| 1.25E-09 | -0.2549156 | 0.177 | 0.319 | 2.9528E-05 | TEC_dmg2 | CMTM6 |
| 1.2985E-09 | -0.4186777 | 0.492 | 0.62 | 3.0675E-05 | TEC_dmg2 | PCBP1 |
| 1.2998E-09 | -0.2759994 | 0.235 | 0.377 | 3.0705E-05 | TEC_dmg2 | BRK1 |
| 1.3361E-09 | -0.2898426 | 0.017 | 0.109 | 3.1563E-05 | TEC_dmg2 | FGL2 |
| 1.3927E-09 | -0.2933984 | 0.029 | 0.127 | 3.29E-05 | TEC_dmg2 | POLB |
| 1.4607E-09 | 0.27421505 | 0.312 | 0.201 | 3.4506E-05 | TEC_dmg2 | MUC20-OT1 |
| 1.5546E-09 | 0.39480093 | 0.386 | 0.276 | 3.6724E-05 | TEC_dmg2 | TAF7 |
| 1.6417E-09 | -0.2753705 | 0.64 | 0.742 | 3.8782E-05 | TEC_dmg2 | SERP1 |
| 1.849E-09 | 0.35705275 | 0.417 | 0.308 | 4.3678E-05 | TEC_dmg2 | OGA |
| 1.849E-09 | -0.6139433 | 0.122 | 0.235 | 4.3678E-05 | TEC_dmg2 | PDZK1IP1 |
| 2.1544E-09 | -0.2990324 | 0.034 | 0.132 | 5.0893E-05 | TEC_dmg2 | CD83 |
| 2.2258E-09 | -0.2673919 | 0.07 | 0.18 | 5.2581E-05 | TEC_dmg2 | TMEM176B |
| 2.5285E-09 | -0.2618415 | 0.348 | 0.491 | 5.9731E-05 | TEC_dmg2 | HSBP1 |
| 2.5686E-09 | -0.2882402 | 0.161 | 0.293 | 6.0679E-05 | TEC_dmg2 | SKIL |
| 2.5726E-09 | -0.4180551 | 0.139 | 0.266 | 6.0774E-05 | TEC_dmg2 | MGST1 |
| 2.9227E-09 | 0.32369168 | 0.324 | 0.217 | 6.9043E-05 | TEC_dmg2 | POLR2J3.1 |
| 2.9249E-09 | 0.26483866 | 0.369 | 0.255 | 6.9094E-05 | TEC_dmg2 | VAMP2 |
| 3.2416E-09 | 0.36957174 | 0.743 | 0.689 | 7.6576E-05 | TEC_dmg2 | EEF2 |
| 3.2537E-09 | -0.4263435 | 0.918 | 0.885 | 7.6862E-05 | TEC_dmg2 | RPS2 |
| 3.3281E-09 | -0.6546525 | 0.019 | 0.109 | 7.862E-05 | TEC_dmg2 | APOC1 |
| 3.4698E-09 | -0.31886 | 0.811 | 0.847 | 8.1966E-05 | TEC_dmg2 | RPL35 |
| 3.9144E-09 | -0.2614988 | 0.041 | 0.141 | 9.247E-05 | TEC_dmg2 | EMP2 |
| 4.2012E-09 | 0.33977644 | 0.436 | 0.314 | 9.9244E-05 | TEC_dmg2 | AKAP9 |
| 4.25E-09 | -0.2975356 | 0.113 | 0.229 | 0.0001004 | TEC_dmg2 | GALNT1 |
| 4.3044E-09 | -0.7914022 | 0.156 | 0.264 | 0.00010168 | TEC_dmg2 | LMO7 |
| 4.5512E-09 | -0.4684752 | 0.153 | 0.283 | 0.00010751 | TEC_dmg2 | MAFB |
| 4.8843E-09 | -0.2550678 | 0.254 | 0.391 | 0.00011538 | TEC_dmg2 | CHMP2A |
| 5.2058E-09 | -0.3485962 | 0.528 | 0.623 | 0.00012298 | TEC_dmg2 | TMEM59 |
| 5.2342E-09 | -0.3218918 | 0.837 | 0.892 | 0.00012365 | TEC_dmg2 | CD63 |
| 5.2459E-09 | -1.1982518 | 0.024 | 0.114 | 0.00012392 | TEC_dmg2 | SPRR1B |
| 5.2873E-09 | -0.9976941 | 0.019 | 0.109 | 0.0001249 | TEC_dmg2 | CCL4 |
| 5.3694E-09 | -0.296358 | 0.259 | 0.395 | 0.00012684 | TEC_dmg2 | NDUFB2 |
| 5.5105E-09 | -0.5222661 | 0.415 | 0.526 | 0.00013018 | TEC_dmg2 | CEBPB |
| 6.0084E-09 | -0.2617289 | 0.573 | 0.687 | 0.00014194 | TEC_dmg2 | ATP6V0B |
| 7.2376E-09 | -0.2848257 | 0.511 | 0.626 | 0.00017097 | TEC_dmg2 | ACTR2 |
| 7.3962E-09 | -0.3019997 | 0.916 | 0.958 | 0.00017472 | TEC_dmg2 | RPS18 |
| 8.0256E-09 | 0.26476841 | 0.285 | 0.187 | 0.00018959 | TEC_dmg2 | PSMD12 |
| 8.1586E-09 | -0.3823987 | 0.813 | 0.832 | 0.00019273 | TEC_dmg2 | RPL22 |
| 8.6094E-09 | -0.2822653 | 0.741 | 0.835 | 0.00020338 | TEC_dmg2 | PPIA |
| 9.6328E-09 | -0.5067429 | 0.141 | 0.254 | 0.00022756 | TEC_dmg2 | CARHSP1 |
| 1.0038E-08 | -0.2852477 | 0.156 | 0.285 | 0.00023712 | TEC_dmg2 | MARCKSL1 |
| 1.0744E-08 | -0.3537661 | 0.079 | 0.186 | 0.0002538 | TEC_dmg2 | KLF2 |
| 1.3323E-08 | 0.28695756 | 0.429 | 0.315 | 0.00031472 | TEC_dmg2 | BOD1L1 |
| 1.4128E-08 | -0.27021 | 0.127 | 0.244 | 0.00033375 | TEC_dmg2 | RBPJ |

| | | | | | | |
|---|---|---|---|---|---|---|
| 1.444E-08 | -0.3045575 | 0.175 | 0.299 | 0.00034111 | TEC_dmg2 | PIM3 |
| 1.706E-08 | -0.3285545 | 0.043 | 0.136 | 0.000403 | TEC_dmg2 | GCNT3 |
| 1.7111E-08 | -1.7303322 | 0.055 | 0.152 | 0.00040421 | TEC_dmg2 | MT1H |
| 1.8981E-08 | 0.305282 | 0.484 | 0.367 | 0.00044838 | TEC_dmg2 | SPAG9 |
| 1.9303E-08 | -0.6690245 | 0.026 | 0.114 | 0.000456 | TEC_dmg2 | IL7R |
| 1.9332E-08 | -0.2977256 | 0.077 | 0.179 | 0.00045667 | TEC_dmg2 | LTA4H |
| 2.0495E-08 | -0.3480265 | 0.439 | 0.548 | 0.00048415 | TEC_dmg2 | RPS10 |
| 2.0576E-08 | -0.3015936 | 0.911 | 0.95 | 0.00048606 | TEC_dmg2 | RPL18 |
| 2.0596E-08 | -0.3759514 | 0.038 | 0.129 | 0.00048655 | TEC_dmg2 | GBP1 |
| 2.2356E-08 | -0.2886614 | 0.022 | 0.106 | 0.00052811 | TEC_dmg2 | CSF3R |
| 2.2743E-08 | -0.8611 | 0.118 | 0.224 | 0.00053727 | TEC_dmg2 | G0S2 |
| 2.4166E-08 | -0.9217691 | 0.031 | 0.118 | 0.00057087 | TEC_dmg2 | CXCL2 |
| 2.651E-08 | -0.5292097 | 0.882 | 0.92 | 0.00062624 | TEC_dmg2 | S100A10 |
| 2.6701E-08 | -0.3086508 | 0.026 | 0.111 | 0.00063076 | TEC_dmg2 | SMIM5 |
| 2.7789E-08 | -0.3475085 | 0.201 | 0.323 | 0.00065645 | TEC_dmg2 | B4GALT1 |
| 2.8593E-08 | -0.2640794 | 0.199 | 0.318 | 0.00067544 | TEC_dmg2 | RAB11A |
| 3.0764E-08 | -2.6714818 | 0.475 | 0.524 | 0.00072673 | TEC_dmg2 | LCN2 |
| 3.1395E-08 | -0.2589211 | 0.444 | 0.563 | 0.00074166 | TEC_dmg2 | TOMM7 |
| 3.2332E-08 | -0.8077836 | 0.18 | 0.279 | 0.00076377 | TEC_dmg2 | DUSP5 |
| 3.2475E-08 | -0.2819134 | 0.235 | 0.361 | 0.00076716 | TEC_dmg2 | TMOD3 |
| 3.7265E-08 | 0.3593769 | 0.403 | 0.311 | 0.00088032 | TEC_dmg2 | SBDS |
| 4.0469E-08 | -0.2925011 | 0.062 | 0.161 | 0.00095599 | TEC_dmg2 | SERPINB9 |
| 4.1492E-08 | -0.8656519 | 0.309 | 0.406 | 0.00098017 | TEC_dmg2 | SLPI |
| 4.4703E-08 | -0.2651268 | 0.024 | 0.107 | 0.00105603 | TEC_dmg2 | SLC16A10 |
| 4.602E-08 | 0.32559509 | 0.679 | 0.606 | 0.00108714 | TEC_dmg2 | SON |
| 4.6992E-08 | -0.3466254 | 0.909 | 0.922 | 0.00111009 | TEC_dmg2 | RPL11 |
| 5.0049E-08 | -0.3288009 | 0.053 | 0.148 | 0.00118231 | TEC_dmg2 | ALDH1A3 |
| 5.065E-08 | -0.2956453 | 0.101 | 0.207 | 0.0011965 | TEC_dmg2 | SLC43A2 |
| 6.2967E-08 | -1.5259715 | 0.175 | 0.281 | 0.00148746 | TEC_dmg2 | MT1X |
| 6.8509E-08 | -0.4009091 | 0.532 | 0.642 | 0.00161839 | TEC_dmg2 | TXNIP |
| 6.9504E-08 | -0.3284363 | 0.173 | 0.284 | 0.00164189 | TEC_dmg2 | CTSZ |
| 7.7809E-08 | -0.684335 | 0.089 | 0.194 | 0.00183809 | TEC_dmg2 | HSPA6 |
| 7.9553E-08 | -0.3429483 | 0.026 | 0.107 | 0.00187927 | TEC_dmg2 | TMC5 |
| 8.0334E-08 | -0.3751541 | 0.053 | 0.142 | 0.00189773 | TEC_dmg2 | ID1 |
| 8.1404E-08 | -0.2546332 | 0.237 | 0.365 | 0.001923 | TEC_dmg2 | SLC25A5 |
| 8.3502E-08 | -0.3453615 | 0.077 | 0.177 | 0.00197258 | TEC_dmg2 | IGFBP3 |
| 9.2468E-08 | -0.3188875 | 0.851 | 0.884 | 0.00218436 | TEC_dmg2 | RPS13 |
| 9.8781E-08 | -0.2595072 | 0.863 | 0.899 | 0.0023335 | TEC_dmg2 | RPL27A |
| 9.9701E-08 | -0.4951048 | 0.141 | 0.247 | 0.00235525 | TEC_dmg2 | CST6 |
| 1.0375E-07 | -0.3834902 | 0.767 | 0.782 | 0.00245078 | TEC_dmg2 | RPL10A |
| 1.2027E-07 | -0.6091796 | 0.098 | 0.198 | 0.0028412 | TEC_dmg2 | TM4SF1 |
| 1.2538E-07 | -0.2601725 | 0.043 | 0.129 | 0.00296177 | TEC_dmg2 | FAM84A |
| 1.2785E-07 | 0.29438559 | 0.348 | 0.249 | 0.00302016 | TEC_dmg2 | DNAJC3 |
| 1.3555E-07 | -0.2640684 | 0.436 | 0.552 | 0.00320199 | TEC_dmg2 | FAM107B |
| 1.3673E-07 | -0.3248263 | 0.084 | 0.184 | 0.00322996 | TEC_dmg2 | SGK1 |
| 1.3942E-07 | -0.3427808 | 0.825 | 0.86 | 0.00329348 | TEC_dmg2 | RPL23 |
| 1.4023E-07 | -0.2621463 | 0.189 | 0.305 | 0.00331267 | TEC_dmg2 | RPL36A |
| 1.4407E-07 | -0.2836804 | 0.667 | 0.748 | 0.00340348 | TEC_dmg2 | CLIC1 |
| 1.7344E-07 | -1.1791833 | 0.055 | 0.144 | 0.00409723 | TEC_dmg2 | CCL2 |
| 1.8186E-07 | 0.25249435 | 0.468 | 0.349 | 0.00429607 | TEC_dmg2 | GLS |

| | | | | | | |
|---|---|---|---|---|---|---|
| 2.0058E-07 | -0.3802115 | 0.199 | 0.31 | 0.00473833 | TEC_dmg2 | IRF1 |
| 2.046E-07 | -0.2545428 | 0.084 | 0.182 | 0.00483337 | TEC_dmg2 | PFKFB3 |
| 2.151E-07 | -0.5804498 | 0.026 | 0.105 | 0.00508139 | TEC_dmg2 | CCL5 |
| 2.4823E-07 | -0.6036203 | 0.18 | 0.287 | 0.00586383 | TEC_dmg2 | HMOX1 |
| 2.8005E-07 | -1.3165205 | 0.489 | 0.536 | 0.00661573 | TEC_dmg2 | WFDC2 |
| 2.8853E-07 | -0.2644614 | 0.213 | 0.327 | 0.00681583 | TEC_dmg2 | CAPG |
| 3.0118E-07 | -0.389801 | 0.07 | 0.159 | 0.00711484 | TEC_dmg2 | BPGM |
| 3.3468E-07 | -0.3497764 | 0.815 | 0.849 | 0.00790617 | TEC_dmg2 | RPS4X |
| 3.3533E-07 | -0.2648135 | 0.082 | 0.176 | 0.00792156 | TEC_dmg2 | NR4A1 |
| 3.5929E-07 | -0.3590703 | 0.386 | 0.483 | 0.00848761 | TEC_dmg2 | CD46 |
| 3.6224E-07 | -0.4981743 | 0.504 | 0.603 | 0.00855711 | TEC_dmg2 | NFKBIA |
| 3.7557E-07 | -0.3348369 | 0.113 | 0.209 | 0.00887208 | TEC_dmg2 | CDKN2B |
| 4.0097E-07 | -0.4182728 | 0.041 | 0.121 | 0.00947219 | TEC_dmg2 | HILPDA |
| 4.2326E-07 | 0.25252868 | 0.305 | 0.215 | 0.00999857 | TEC_dmg2 | DDX6 |
| 4.3174E-07 | -0.315256 | 0.607 | 0.691 | 0.01019892 | TEC_dmg2 | PNRC1 |
| 4.4093E-07 | -1.140488 | 0.283 | 0.37 | 0.01041612 | TEC_dmg2 | MAL |
| 5.0204E-07 | -0.280597 | 0.175 | 0.277 | 0.01185964 | TEC_dmg2 | VASP |
| 5.2133E-07 | -0.2995957 | 0.885 | 0.881 | 0.01231547 | TEC_dmg2 | RPL12 |
| 5.3175E-07 | -0.5213651 | 0.388 | 0.504 | 0.01256151 | TEC_dmg2 | DNAJB1 |
| 6.1163E-07 | -0.5443484 | 0.132 | 0.234 | 0.01444846 | TEC_dmg2 | PLIN2 |
| 6.2105E-07 | -0.5888605 | 0.338 | 0.434 | 0.0146711 | TEC_dmg2 | HIST1H1C |
| 6.3364E-07 | -0.3427247 | 0.098 | 0.189 | 0.01496843 | TEC_dmg2 | EPS8L1 |
| 7.1666E-07 | 0.25801948 | 0.403 | 0.306 | 0.01692969 | TEC_dmg2 | NUFIP2 |
| 8.1955E-07 | -0.2593181 | 0.866 | 0.902 | 0.01936025 | TEC_dmg2 | RPL24 |
| 8.248E-07 | -0.2569369 | 0.187 | 0.298 | 0.01948417 | TEC_dmg2 | REL |
| 8.3253E-07 | -0.3088436 | 0.871 | 0.896 | 0.0196668 | TEC_dmg2 | HLA-A |
| 9.0876E-07 | 0.32645549 | 0.444 | 0.37 | 0.02146758 | TEC_dmg2 | CIRBP |
| 9.3654E-07 | -0.4007207 | 0.65 | 0.732 | 0.02212393 | TEC_dmg2 | HSPA8 |
| 9.8937E-07 | -0.2642877 | 0.892 | 0.888 | 0.02337189 | TEC_dmg2 | RPL6 |
| 1.041E-06 | 0.26773522 | 0.302 | 0.207 | 0.02459213 | TEC_dmg2 | MACC1 |
| 1.1654E-06 | -0.2542104 | 0.647 | 0.731 | 0.02753051 | TEC_dmg2 | ATP6V1G1 |
| 1.1845E-06 | -0.8437161 | 0.091 | 0.174 | 0.02798035 | TEC_dmg2 | RDH10 |
| 1.4014E-06 | -0.4473427 | 0.302 | 0.395 | 0.03310535 | TEC_dmg2 | HIF1A |
| 1.4384E-06 | -0.2759794 | 0.225 | 0.324 | 0.0339788 | TEC_dmg2 | VEGFA |
| 1.4611E-06 | -0.3391473 | 0.962 | 0.937 | 0.03451498 | TEC_dmg2 | S100A6 |
| 1.4754E-06 | -0.3708196 | 0.796 | 0.792 | 0.03485224 | TEC_dmg2 | RPS21 |
| 1.8941E-06 | -0.2917759 | 0.153 | 0.242 | 0.04474491 | TEC_dmg2 | GIPC1 |
| 1.9154E-06 | -0.3344082 | 0.141 | 0.239 | 0.04524694 | TEC_dmg2 | TNFAIP3 |
| 1.9812E-06 | -0.7659952 | 0.535 | 0.555 | 0.04680264 | TEC_dmg2 | ELF3 |
| 1.9859E-06 | -0.3745057 | 0.089 | 0.173 | 0.04691211 | TEC_dmg2 | PPL |
| 2.2493E-06 | 0.26467746 | 0.381 | 0.3 | 0.05313556 | TEC_dmg2 | VCP |
| 2.4724E-06 | 0.27148924 | 0.71 | 0.659 | 0.05840605 | TEC_dmg2 | RBM39 |
| 2.6496E-06 | -0.7716481 | 0.597 | 0.658 | 0.06259144 | TEC_dmg2 | HSPA1A |
| 3.0438E-06 | -0.2530634 | 0.161 | 0.26 | 0.0719028 | TEC_dmg2 | NABP1 |
| 3.1592E-06 | -0.2999493 | 0.679 | 0.742 | 0.07462894 | TEC_dmg2 | APLP2 |
| 4.0095E-06 | -0.4580564 | 0.043 | 0.113 | 0.09471608 | TEC_dmg2 | CYP1B1 |
| 4.1169E-06 | 0.33165312 | 0.564 | 0.513 | 0.09725307 | TEC_dmg2 | EIF5 |
| 4.2372E-06 | -0.2687381 | 0.038 | 0.108 | 0.10009532 | TEC_dmg2 | TCHP |
| 4.5299E-06 | -0.6813975 | 0.058 | 0.131 | 0.10700997 | TEC_dmg2 | APOE |
| 4.8969E-06 | -1.2613277 | 0.046 | 0.116 | 0.11567943 | TEC_dmg2 | MT1M |

| | | | | | | |
|---|---|---|---|---|---|---|
| 5.1059E-06 | -0.3001393 | 0.036 | 0.105 | 0.12061783 | TEC_dmg2 | LTB |
| 5.8683E-06 | -0.3988708 | 0.094 | 0.172 | 0.13862668 | TEC_dmg2 | HSPB8 |
| 5.8928E-06 | -0.3695449 | 0.899 | 0.889 | 0.13920478 | TEC_dmg2 | RPS23 |
| 6.1727E-06 | 0.30958813 | 0.266 | 0.181 | 0.14581754 | TEC_dmg2 | LAMC2 |
| 6.9836E-06 | -0.4847457 | 0.211 | 0.293 | 0.16497408 | TEC_dmg2 | GPRC5A |
| 7.2557E-06 | 0.25509203 | 0.42 | 0.334 | 0.17140101 | TEC_dmg2 | RBM25 |
| 7.4291E-06 | -0.431988 | 0.216 | 0.31 | 0.1754982 | TEC_dmg2 | HIST1H4C |
| 7.9139E-06 | -0.3227199 | 0.194 | 0.296 | 0.18694928 | TEC_dmg2 | GLRX |
| 8.0835E-06 | -0.2745973 | 0.633 | 0.704 | 0.19095632 | TEC_dmg2 | TPM3 |
| 9.2198E-06 | -0.5501284 | 0.082 | 0.157 | 0.21780042 | TEC_dmg2 | MT1F |
| 9.2679E-06 | -0.2915643 | 0.199 | 0.3 | 0.21893528 | TEC_dmg2 | BTG2 |
| 9.5732E-06 | -0.3304732 | 0.106 | 0.186 | 0.22614758 | TEC_dmg2 | ACSL1 |
| 9.6353E-06 | -0.4157262 | 0.724 | 0.764 | 0.22761369 | TEC_dmg2 | BTG1 |
| 1.0811E-05 | -0.2706842 | 0.084 | 0.158 | 0.25537753 | TEC_dmg2 | TTC9 |
| 1.2052E-05 | -0.3369764 | 0.094 | 0.172 | 0.28471571 | TEC_dmg2 | DNAJA4 |
| 1.2196E-05 | -0.4551262 | 0.118 | 0.192 | 0.28811016 | TEC_dmg2 | FBXO32 |
| 1.2885E-05 | 0.30574081 | 0.506 | 0.447 | 0.30438053 | TEC_dmg2 | ACTN4 |
| 1.3139E-05 | -0.2753845 | 0.712 | 0.767 | 0.31038352 | TEC_dmg2 | RPS26 |
| 1.3325E-05 | -0.3564183 | 0.655 | 0.722 | 0.31477392 | TEC_dmg2 | JUND |
| 1.4161E-05 | -0.2712194 | 0.811 | 0.839 | 0.33453086 | TEC_dmg2 | RACK1 |
| 1.6075E-05 | 0.2774357 | 0.36 | 0.282 | 0.37973202 | TEC_dmg2 | DYNC1H1 |
| 1.6913E-05 | -0.2670288 | 0.177 | 0.271 | 0.3995258 | TEC_dmg2 | TSC22D3 |
| 2.0689E-05 | 0.2696055 | 0.592 | 0.528 | 0.48873115 | TEC_dmg2 | EIF4A2 |
| 2.3449E-05 | -0.292022 | 0.175 | 0.262 | 0.55394152 | TEC_dmg2 | HES4 |
| 2.4227E-05 | -0.4555491 | 0.724 | 0.785 | 0.57231513 | TEC_dmg2 | TXN |
| 2.5698E-05 | -0.2544054 | 0.892 | 0.902 | 0.60706878 | TEC_dmg2 | RPS7 |
| 2.977E-05 | -0.2701744 | 0.077 | 0.146 | 0.70326449 | TEC_dmg2 | SDC2 |
| 3.2526E-05 | -0.294656 | 0.866 | 0.875 | 0.76835891 | TEC_dmg2 | RPL26 |
| 3.3126E-05 | -0.3328202 | 0.139 | 0.215 | 0.78253699 | TEC_dmg2 | CREM |
| 3.3284E-05 | -0.2574203 | 0.192 | 0.275 | 0.78626957 | TEC_dmg2 | PLIN3 |
| 3.5741E-05 | 0.27192973 | 0.751 | 0.685 | 0.84431296 | TEC_dmg2 | NPM1 |
| 4.6202E-05 | -0.2663705 | 0.144 | 0.227 | 1 | TEC_dmg2 | IER5L |
| 5.8692E-05 | -0.3135164 | 0.18 | 0.261 | 1 | TEC_dmg2 | MUC1 |
| 6.1193E-05 | 0.25569443 | 0.64 | 0.593 | 1 | TEC_dmg2 | DDX17 |
| 6.2379E-05 | -1.407716 | 0.206 | 0.267 | 1 | TEC_dmg2 | S100A2 |
| 7.4336E-05 | -0.2996913 | 0.453 | 0.514 | 1 | TEC_dmg2 | ANXA11 |
| 7.599E-05 | -0.3127589 | 0.952 | 0.965 | 1 | TEC_dmg2 | UBC |
| 8.2301E-05 | -0.2723684 | 0.182 | 0.267 | 1 | TEC_dmg2 | ID2 |
| 8.5822E-05 | -0.4757101 | 0.415 | 0.506 | 1 | TEC_dmg2 | HSPA1B |
| 8.5965E-05 | -0.2589898 | 0.249 | 0.326 | 1 | TEC_dmg2 | PPA1 |
| 0.00010097 | -0.3036263 | 0.108 | 0.175 | 1 | TEC_dmg2 | C6orf132 |
| 0.00014621 | -0.286089 | 0.259 | 0.348 | 1 | TEC_dmg2 | IER5 |
| 0.00016314 | -0.877837 | 0.566 | 0.452 | 1 | TEC_dmg2 | SPP1 |
| 0.00018099 | -0.5261089 | 0.357 | 0.247 | 1 | TEC_dmg2 | CRYAB |
| 0.00027152 | -0.2596559 | 0.451 | 0.525 | 1 | TEC_dmg2 | HSPE1 |
| 0.00028991 | 0.26223194 | 0.353 | 0.28 | 1 | TEC_dmg2 | H1F0 |
| 0.00031077 | -0.3141461 | 0.177 | 0.249 | 1 | TEC_dmg2 | EGR1 |
| 0.00033138 | -0.3594136 | 0.158 | 0.231 | 1 | TEC_dmg2 | ISG15 |
| 0.00033888 | -0.5273564 | 0.237 | 0.292 | 1 | TEC_dmg2 | RND3 |
| 0.0003665 | -0.3430983 | 0.477 | 0.523 | 1 | TEC_dmg2 | HEBP2 |

| | | | | | | |
|---|---|---|---|---|---|---|
| 0.0005858 | -1.2374602 | 0.801 | 0.658 | 1 | TEC_dmg2 | MT2A |
| 0.00062201 | -0.4857533 | 0.595 | 0.612 | 1 | TEC_dmg2 | CD44 |
| 0.00071469 | -0.295647 | 0.165 | 0.23 | 1 | TEC_dmg2 | PPIF |
| 0.00116942 | -0.2689178 | 0.108 | 0.164 | 1 | TEC_dmg2 | ANKRD37 |
| 0.00119865 | -0.2844996 | 0.177 | 0.238 | 1 | TEC_dmg2 | CITED2 |
| 0.00136941 | -2.0516356 | 0.199 | 0.255 | 1 | TEC_dmg2 | MT1G |
| 0.00153358 | -0.3229633 | 0.163 | 0.222 | 1 | TEC_dmg2 | HES1 |
| 0.00162647 | -0.3558092 | 0.734 | 0.741 | 1 | TEC_dmg2 | PSAP |
| 0.00188242 | -0.3077917 | 0.77 | 0.775 | 1 | TEC_dmg2 | CTSD |
| 0.00229949 | -0.2670039 | 0.175 | 0.235 | 1 | TEC_dmg2 | CDKN1C |
| 0.00231451 | -0.4287317 | 0.228 | 0.286 | 1 | TEC_dmg2 | SLC25A37 |
| 0.00358271 | -0.299925 | 0.847 | 0.868 | 1 | TEC_dmg2 | ANXA2 |
| 0.0050151 | -0.3149311 | 0.129 | 0.175 | 1 | TEC_dmg2 | SPNS2 |
| 0.00530562 | -0.4706449 | 0.669 | 0.704 | 1 | TEC_dmg2 | SOD2 |
| 0.00564174 | -0.3486301 | 0.439 | 0.452 | 1 | TEC_dmg2 | CLDN7 |
| 0.00712473 | -0.2777185 | 0.106 | 0.148 | 1 | TEC_dmg2 | RASD1 |
| 0.00735255 | -0.3476735 | 0.59 | 0.631 | 1 | TEC_dmg2 | ZFP36L1 |
| 0.00742747 | -0.7660808 | 0.153 | 0.2 | 1 | TEC_dmg2 | THBS1 |
| 0.00857822 | -0.3282814 | 0.233 | 0.282 | 1 | TEC_dmg2 | RHOB |
| 0.00928546 | 0.2890805 | 0.249 | 0.208 | 1 | TEC_dmg2 | MDM2 |
| 0.00968535 | -0.4716271 | 0.412 | 0.451 | 1 | TEC_dmg2 | SOX4 |
| 0 | 2.92701791 | 0.86 | 0.02 | 0 | TEC_prlf | CENPF |
| 0 | 2.3833543 | 0.878 | 0.166 | 0 | TEC_prlf | STMN1 |
| 0 | 2.0747794 | 0.732 | 0.026 | 0 | TEC_prlf | CCNB1 |
| 0 | 2.05088287 | 0.7 | 0.054 | 0 | TEC_prlf | PTTG1 |
| 0 | 1.97860666 | 0.673 | 0.011 | 0 | TEC_prlf | MKI67 |
| 0 | 1.69906626 | 0.685 | 0.011 | 0 | TEC_prlf | HMMR |
| 0 | 1.60805186 | 0.624 | 0.029 | 0 | TEC_prlf | NUSAP1 |
| 0 | 1.58154909 | 0.659 | 0.01 | 0 | TEC_prlf | TPX2 |
| 0 | 1.5380074 | 0.534 | 0.011 | 0 | TEC_prlf | TOP2A |
| 0 | 1.44343989 | 0.636 | 0.012 | 0 | TEC_prlf | BIRC5 |
| 0 | 1.37940312 | 0.563 | 0.009 | 0 | TEC_prlf | CDC20 |
| 0 | 1.32836532 | 0.507 | 0.034 | 0 | TEC_prlf | CKAP2 |
| 0 | 1.2305955 | 0.469 | 0.008 | 0 | TEC_prlf | ASPM |
| 0 | 1.21705636 | 0.536 | 0.015 | 0 | TEC_prlf | CDKN3 |
| 0 | 1.16569568 | 0.484 | 0.015 | 0 | TEC_prlf | AURKA |
| 0 | 1.14087402 | 0.493 | 0.005 | 0 | TEC_prlf | GTSE1 |
| 0 | 1.05570667 | 0.507 | 0.006 | 0 | TEC_prlf | CEP55 |
| 0 | 1.01241248 | 0.312 | 0.016 | 0 | TEC_prlf | THY1 |
| 0 | 0.95776046 | 0.437 | 0.009 | 0 | TEC_prlf | CCNB2 |
| 0 | 0.95399297 | 0.478 | 0.003 | 0 | TEC_prlf | DLGAP5 |
| 0 | 0.94099138 | 0.466 | 0.033 | 0 | TEC_prlf | ECT2 |
| 0 | 0.92873763 | 0.513 | 0.035 | 0 | TEC_prlf | TACC3 |
| 0 | 0.86342132 | 0.405 | 0.003 | 0 | TEC_prlf | DEPDC1 |
| 0 | 0.86084068 | 0.446 | 0.038 | 0 | TEC_prlf | KIF20B |
| 0 | 0.83321275 | 0.397 | 0.01 | 0 | TEC_prlf | CENPE |
| 0 | 0.79495625 | 0.434 | 0.004 | 0 | TEC_prlf | PBK |
| 0 | 0.79407546 | 0.397 | 0.006 | 0 | TEC_prlf | CCNA2 |
| 0 | 0.76851106 | 0.414 | 0.008 | 0 | TEC_prlf | ANLN |
| 0 | 0.76183258 | 0.394 | 0.004 | 0 | TEC_prlf | NUF2 |

| | | | | | | |
|---|---|---|---|---|---|---|
| 0 | 0.75954943 | 0.379 | 0.007 | 0 | TEC_prlf | PLK1 |
| 0 | 0.75650112 | 0.373 | 0.011 | 0 | TEC_prlf | PRC1 |
| 0 | 0.75101798 | 0.443 | 0.032 | 0 | TEC_prlf | TK1 |
| 0 | 0.74379275 | 0.382 | 0.015 | 0 | TEC_prlf | MAD2L1 |
| 0 | 0.73545428 | 0.347 | 0.006 | 0 | TEC_prlf | CDCA8 |
| 0 | 0.72612779 | 0.385 | 0.012 | 0 | TEC_prlf | KIF23 |
| 0 | 0.72009131 | 0.376 | 0.022 | 0 | TEC_prlf | KNSTRN |
| 0 | 0.7171363 | 0.359 | 0.009 | 0 | TEC_prlf | ARHGAP11A |
| 0 | 0.66912254 | 0.303 | 0.004 | 0 | TEC_prlf | CENPA |
| 0 | 0.66049106 | 0.332 | 0.003 | 0 | TEC_prlf | KIF2C |
| 0 | 0.62789362 | 0.327 | 0.005 | 0 | TEC_prlf | TROAP |
| 0 | 0.61464313 | 0.376 | 0.017 | 0 | TEC_prlf | DIAPH3 |
| 0 | 0.60239776 | 0.347 | 0.002 | 0 | TEC_prlf | KIF4A |
| 0 | 0.60086541 | 0.324 | 0.017 | 0 | TEC_prlf | UBE2T |
| 0 | 0.58400476 | 0.335 | 0.021 | 0 | TEC_prlf | TYMS |
| 0 | 0.58324363 | 0.329 | 0.005 | 0 | TEC_prlf | BUB1 |
| 0 | 0.56963547 | 0.329 | 0.016 | 0 | TEC_prlf | RACGAP1 |
| 0 | 0.56229134 | 0.294 | 0.003 | 0 | TEC_prlf | HJURP |
| 0 | 0.55333362 | 0.294 | 0.009 | 0 | TEC_prlf | KNL1 |
| 0 | 0.55112482 | 0.274 | 0.002 | 0 | TEC_prlf | NEK2 |
| 0 | 0.54920729 | 0.248 | 0.009 | 0 | TEC_prlf | UBE2C |
| 0 | 0.52460747 | 0.309 | 0.004 | 0 | TEC_prlf | CKAP2L |
| 0 | 0.52336073 | 0.3 | 0.003 | 0 | TEC_prlf | KIF20A |
| 0 | 0.52060966 | 0.303 | 0.005 | 0 | TEC_prlf | NDC80 |
| 0 | 0.51800102 | 0.245 | 0.009 | 0 | TEC_prlf | CDK1 |
| 0 | 0.50884994 | 0.265 | 0.004 | 0 | TEC_prlf | KIF18A |
| 0 | 0.50045313 | 0.28 | 0.013 | 0 | TEC_prlf | CIT |
| 0 | 0.48574677 | 0.303 | 0.013 | 0 | TEC_prlf | ZWINT |
| 0 | 0.48112889 | 0.268 | 0.006 | 0 | TEC_prlf | KIFC1 |
| 0 | 0.47628341 | 0.268 | 0.005 | 0 | TEC_prlf | SGO1 |
| 0 | 0.448535 | 0.277 | 0.005 | 0 | TEC_prlf | KIF14 |
| 0 | 0.44102153 | 0.242 | 0.003 | 0 | TEC_prlf | TTK |
| 0 | 0.43524263 | 0.254 | 0.004 | 0 | TEC_prlf | CDCA3 |
| 0 | 0.43252742 | 0.245 | 0.006 | 0 | TEC_prlf | KIF11 |
| 0 | 0.43068333 | 0.236 | 0.005 | 0 | TEC_prlf | IQGAP3 |
| 0 | 0.42606077 | 0.251 | 0.004 | 0 | TEC_prlf | NCAPG |
| 0 | 0.41609313 | 0.213 | 0.002 | 0 | TEC_prlf | BUB1B |
| 0 | 0.38300435 | 0.224 | 0.006 | 0 | TEC_prlf | PSRC1 |
| 0 | 0.36759737 | 0.236 | 0.005 | 0 | TEC_prlf | CDCA2 |
| 0 | 0.35974154 | 0.224 | 0.006 | 0 | TEC_prlf | FOXM1 |
| 0 | 0.34330221 | 0.21 | 0.008 | 0 | TEC_prlf | PRR11 |
| 0 | 0.34204368 | 0.219 | 0.007 | 0 | TEC_prlf | PARPBP |
| 0 | 0.330874 | 0.239 | 0.01 | 0 | TEC_prlf | SPAG5 |
| 0 | 0.32942238 | 0.236 | 0.01 | 0 | TEC_prlf | FANCI |
| 0 | 0.31561857 | 0.21 | 0.004 | 0 | TEC_prlf | SKA3 |
| 0 | 0.31265937 | 0.184 | 0.004 | 0 | TEC_prlf | TMSB15A |
| 0 | 0.30738328 | 0.19 | 0.005 | 0 | TEC_prlf | AURKB |
| 0 | 0.28125155 | 0.175 | 0.005 | 0 | TEC_prlf | CDCA5 |
| 0 | 0.26939147 | 0.178 | 0.006 | 0 | TEC_prlf | DEPDC1B |
| 0 | 0.25116409 | 0.149 | 0.002 | 0 | TEC_prlf | PIMREG |

| | | | | | | |
|---|---|---|---|---|---|---|
| 7.185E-301 | 0.58337612 | 0.309 | 0.019 | 1.697E-296 | TEC_prlf | SGO2 |
| 1.249E-265 | 0.39142531 | 0.262 | 0.015 | 2.95E-261 | TEC_prlf | CENPK |
| 3.028E-257 | 0.57553639 | 0.35 | 0.029 | 7.152E-253 | TEC_prlf | CCDC34 |
| 2.555E-248 | 0.47070422 | 0.321 | 0.025 | 6.036E-244 | TEC_prlf | CENPU |
| 2.017E-236 | 0.30339002 | 0.125 | 0.004 | 4.764E-232 | TEC_prlf | HIST1H3G |
| 2.162E-225 | 2.2435877 | 0.796 | 0.201 | 5.108E-221 | TEC_prlf | UBE2S |
| 3.904E-218 | 0.69103216 | 0.397 | 0.043 | 9.222E-214 | TEC_prlf | SMC2 |
| 1.358E-213 | 0.27881325 | 0.195 | 0.011 | 3.208E-209 | TEC_prlf | CIP2A |
| 7.643E-207 | 0.25505389 | 0.169 | 0.008 | 1.806E-202 | TEC_prlf | RAD51AP1 |
| 1.947E-200 | 0.55475783 | 0.356 | 0.038 | 4.599E-196 | TEC_prlf | CENPW |
| 2.217E-195 | 0.40441516 | 0.257 | 0.02 | 5.237E-191 | TEC_prlf | CENPN |
| 1.812E-194 | 2.05596875 | 0.848 | 0.295 | 4.28E-190 | TEC_prlf | NUCKS1 |
| 8.678E-194 | 1.75271103 | 0.671 | 0.149 | 2.05E-189 | TEC_prlf | CKS2 |
| 3.687E-187 | 2.04708092 | 0.799 | 0.241 | 8.71E-183 | TEC_prlf | PTMS |
| 8.874E-185 | 0.31760666 | 0.175 | 0.01 | 2.096E-180 | TEC_prlf | RRM2 |
| 1.17E-184 | 0.65925142 | 0.318 | 0.033 | 2.764E-180 | TEC_prlf | CYS1 |
| 1.851E-184 | 0.48475797 | 0.315 | 0.032 | 4.374E-180 | TEC_prlf | SPDL1 |
| 1.232E-176 | 0.57132078 | 0.28 | 0.026 | 2.91E-172 | TEC_prlf | PCLAF |
| 1.432E-173 | 1.1578728 | 0.286 | 0.028 | 3.383E-169 | TEC_prlf | SOD3 |
| 5.071E-169 | 0.6589743 | 0.423 | 0.062 | 1.198E-164 | TEC_prlf | DTYMK |
| 1.329E-162 | 0.30371765 | 0.163 | 0.01 | 3.139E-158 | TEC_prlf | INCENP |
| 5.374E-159 | 0.54510496 | 0.367 | 0.049 | 1.269E-154 | TEC_prlf | PHF19 |
| 6.834E-159 | 0.45075854 | 0.3 | 0.033 | 1.614E-154 | TEC_prlf | GGH |
| 3.103E-154 | 0.43074497 | 0.294 | 0.033 | 7.33E-150 | TEC_prlf | NCAPD2 |
| 6.041E-150 | 0.25473436 | 0.175 | 0.012 | 1.427E-145 | TEC_prlf | ASF1B |
| 1.666E-146 | 1.84881155 | 0.848 | 0.385 | 3.936E-142 | TEC_prlf | HMGN2 |
| 2.076E-146 | 1.82817821 | 0.988 | 0.8 | 4.904E-142 | TEC_prlf | HMGB1 |
| 1.539E-144 | 0.80618447 | 0.286 | 0.033 | 3.635E-140 | TEC_prlf | SLC12A1 |
| 1.086E-138 | 0.25260085 | 0.163 | 0.011 | 2.566E-134 | TEC_prlf | MXD3 |
| 5.89E-137 | 1.11180177 | 0.327 | 0.044 | 1.392E-132 | TEC_prlf | CXCL14 |
| 5.968E-136 | 1.02602108 | 0.603 | 0.144 | 1.41E-131 | TEC_prlf | SMC4 |
| 7.223E-133 | 0.66264569 | 0.353 | 0.053 | 1.706E-128 | TEC_prlf | ISYNA1 |
| 6.994E-132 | 0.62099304 | 0.42 | 0.074 | 1.652E-127 | TEC_prlf | HMGB3 |
| 1.358E-130 | 0.76114597 | 0.408 | 0.072 | 3.208E-126 | TEC_prlf | CKS1B |
| 1.02E-127 | 0.80971204 | 0.367 | 0.059 | 2.409E-123 | TEC_prlf | MYH10 |
| 1.24E-127 | 1.69209096 | 0.904 | 0.516 | 2.929E-123 | TEC_prlf | H2AFZ |
| 2.037E-127 | 0.33318904 | 0.23 | 0.024 | 4.813E-123 | TEC_prlf | ORC6 |
| 8.104E-126 | 0.28173002 | 0.14 | 0.009 | 1.914E-121 | TEC_prlf | GPC3 |
| 3.161E-123 | 0.37379406 | 0.204 | 0.02 | 7.467E-119 | TEC_prlf | MNS1 |
| 4.824E-123 | 0.26655065 | 0.163 | 0.013 | 1.14E-118 | TEC_prlf | C21orf58 |
| 1.396E-122 | 1.98734575 | 0.755 | 0.312 | 3.297E-118 | TEC_prlf | HMGB2 |
| 3.979E-122 | 0.91299126 | 0.507 | 0.118 | 9.4E-118 | TEC_prlf | ANP32E |
| 5.203E-115 | 0.26002198 | 0.163 | 0.014 | 1.229E-110 | TEC_prlf | SCARA3 |
| 3.093E-114 | 0.54777167 | 0.327 | 0.052 | 7.307E-110 | TEC_prlf | SMARCA1 |
| 5.17E-114 | 0.35909665 | 0.268 | 0.036 | 1.221E-109 | TEC_prlf | LMNB2 |
| 8.566E-114 | 0.89699611 | 0.414 | 0.081 | 2.023E-109 | TEC_prlf | IGFBP2 |
| 1.483E-113 | 1.26730754 | 0.324 | 0.053 | 3.503E-109 | TEC_prlf | RARRES2 |
| 5.882E-112 | 0.54141035 | 0.312 | 0.049 | 1.39E-107 | TEC_prlf | CAVIN3 |
| 3.336E-111 | 0.84557963 | 0.481 | 0.113 | 7.881E-107 | TEC_prlf | CBX5 |
| 4.463E-110 | 0.26314135 | 0.149 | 0.012 | 1.054E-105 | TEC_prlf | CHL1 |

| | | | | | | |
|---|---|---|---|---|---|---|
| 8.172E-108 | 0.65989413 | 0.394 | 0.077 | 1.93E-103 | TEC_prlf | MYEF2 |
| 4.233E-107 | 0.84583459 | 0.426 | 0.09 | 9.999E-103 | TEC_prlf | BCAM |
| 8.905E-107 | 1.0856932 | 0.72 | 0.231 | 2.104E-102 | TEC_prlf | EPCAM |
| 5.196E-103 | 0.44110834 | 0.309 | 0.051 | 1.2274E-98 | TEC_prlf | CEP70 |
| 1.074E-102 | 1.27991494 | 0.726 | 0.308 | 2.5369E-98 | TEC_prlf | MZT2B |
| 1.264E-100 | 0.47946304 | 0.257 | 0.037 | 2.9871E-96 | TEC_prlf | TSPAN12 |
| 2.851E-100 | 0.37984481 | 0.251 | 0.035 | 6.7358E-96 | TEC_prlf | SPA17 |
| 3.992E-100 | 0.4672927 | 0.324 | 0.057 | 9.4314E-96 | TEC_prlf | CKAP5 |
| 1.1299E-98 | 0.55263836 | 0.327 | 0.058 | 2.6692E-94 | TEC_prlf | CDH16 |
| 3.6935E-98 | 0.55195985 | 0.367 | 0.071 | 8.7253E-94 | TEC_prlf | BICC1 |
| 6.1571E-98 | 1.34310968 | 0.802 | 0.332 | 1.4545E-93 | TEC_prlf | IGFBP7 |
| 1.9466E-97 | 0.4757737 | 0.3 | 0.05 | 4.5983E-93 | TEC_prlf | EFHD1 |
| 1.0306E-95 | 0.75681696 | 0.402 | 0.091 | 2.4347E-91 | TEC_prlf | BRD8 |
| 1.7477E-95 | 0.85655923 | 0.58 | 0.182 | 4.1286E-91 | TEC_prlf | MZT2A |
| 8.4672E-95 | 0.55119904 | 0.362 | 0.072 | 2.0002E-90 | TEC_prlf | PKP4 |
| 1.9991E-93 | 0.98933098 | 0.539 | 0.164 | 4.7224E-89 | TEC_prlf | HMGN3 |
| 3.5843E-93 | 0.5564809 | 0.394 | 0.087 | 8.4672E-89 | TEC_prlf | SKA2 |
| 7.0382E-91 | 0.85782548 | 0.627 | 0.191 | 1.6626E-86 | TEC_prlf | C12orf75 |
| 1.4005E-90 | 0.38953449 | 0.227 | 0.032 | 3.3085E-86 | TEC_prlf | HMGN5 |
| 3.0268E-89 | 0.34476573 | 0.143 | 0.013 | 7.1502E-85 | TEC_prlf | CPE |
| 6.9988E-89 | 0.44040772 | 0.294 | 0.052 | 1.6533E-84 | TEC_prlf | LMNB1 |
| 4.595E-88 | 0.87027152 | 0.484 | 0.139 | 1.0855E-83 | TEC_prlf | LSM5 |
| 5.4359E-87 | 0.32918861 | 0.23 | 0.034 | 1.2841E-82 | TEC_prlf | SHMT1 |
| 1.3277E-86 | 0.95774789 | 0.516 | 0.158 | 3.1365E-82 | TEC_prlf | TMPO |
| 7.913E-86 | 0.92316408 | 0.484 | 0.133 | 1.8693E-81 | TEC_prlf | ARHGAP29 |
| 1.5395E-85 | 1.00483525 | 0.685 | 0.298 | 3.6368E-81 | TEC_prlf | H2AFV |
| 2.4747E-85 | 0.50241884 | 0.297 | 0.054 | 5.8461E-81 | TEC_prlf | COL4A1 |
| 2.5201E-85 | 1.08536643 | 0.743 | 0.339 | 5.9532E-81 | TEC_prlf | ATP5IF1 |
| 6.1686E-83 | 0.34079565 | 0.236 | 0.037 | 1.4572E-78 | TEC_prlf | ODF2 |
| 1.2122E-82 | 0.52746668 | 0.292 | 0.055 | 2.8636E-78 | TEC_prlf | DBF4 |
| 6.7554E-81 | 0.30802517 | 0.216 | 0.032 | 1.5958E-76 | TEC_prlf | DHFR |
| 1.7275E-80 | 0.99704131 | 0.481 | 0.144 | 4.0809E-76 | TEC_prlf | GSTM3 |
| 4.8785E-80 | -1.905299 | 0.799 | 0.942 | 1.1524E-75 | TEC_prlf | NEAT1 |
| 8.7685E-80 | 1.22411359 | 0.679 | 0.301 | 2.0714E-75 | TEC_prlf | HP1BP3 |
| 3.6252E-78 | 0.3254696 | 0.222 | 0.034 | 8.5638E-74 | TEC_prlf | CDC25B |
| 5.3535E-78 | 0.3531939 | 0.233 | 0.038 | 1.2647E-73 | TEC_prlf | CCDC18 |
| 5.4683E-77 | 0.47099042 | 0.394 | 0.097 | 1.2918E-72 | TEC_prlf | RANGAP1 |
| 6.3337E-77 | 0.59060318 | 0.324 | 0.07 | 1.4962E-72 | TEC_prlf | METTL7A |
| 1.2009E-76 | 0.44331384 | 0.294 | 0.059 | 2.837E-72 | TEC_prlf | EMC9 |
| 5.6165E-76 | 0.47812289 | 0.312 | 0.065 | 1.3268E-71 | TEC_prlf | NSD2 |
| 1.3334E-75 | 0.76892229 | 0.466 | 0.131 | 3.1499E-71 | TEC_prlf | UCHL1 |
| 1.7948E-75 | 0.35128236 | 0.254 | 0.045 | 4.2398E-71 | TEC_prlf | CENPM |
| 2.9773E-75 | 1.02587693 | 0.671 | 0.28 | 7.0333E-71 | TEC_prlf | LDHB |
| 1.6778E-73 | 0.59829668 | 0.431 | 0.122 | 3.9635E-69 | TEC_prlf | DDX39A |
| 2.0538E-72 | 0.45407605 | 0.216 | 0.035 | 4.8517E-68 | TEC_prlf | CRNDE |
| 2.1554E-72 | 0.82872533 | 0.586 | 0.226 | 5.0918E-68 | TEC_prlf | HDGF |
| 3.0256E-72 | 1.43317997 | 0.968 | 0.811 | 7.1474E-68 | TEC_prlf | GSTP1 |
| 4.7975E-72 | 0.42492004 | 0.297 | 0.061 | 1.1333E-67 | TEC_prlf | COL4A2 |
| 6.9603E-71 | 0.46591728 | 0.28 | 0.057 | 1.6442E-66 | TEC_prlf | GNAI1 |
| 2.843E-70 | 1.13671157 | 0.942 | 0.691 | 6.716E-66 | TEC_prlf | HSP90B1 |

| | | | | | | |
|---|---|---|---|---|---|---|
| 4.0414E-70 | 1.14074772 | 0.892 | 0.537 | 9.547E-66 | TEC_prlf | CD24 |
| 2.5011E-69 | 0.8227079 | 0.717 | 0.324 | 5.9083E-65 | TEC_prlf | LMNA |
| 3.4977E-69 | 0.47087227 | 0.286 | 0.06 | 8.2627E-65 | TEC_prlf | NDE1 |
| 9.5484E-69 | 1.02620596 | 0.977 | 0.83 | 2.2556E-64 | TEC_prlf | HNRNPA2B1 |
| 1.9638E-68 | 0.99416084 | 0.883 | 0.599 | 4.6391E-64 | TEC_prlf | HINT1 |
| 3.7432E-68 | 0.26399758 | 0.157 | 0.02 | 8.8426E-64 | TEC_prlf | BRCA2 |
| 8.5339E-68 | 1.13651701 | 0.618 | 0.244 | 2.016E-63 | TEC_prlf | SPTBN1 |
| 2.7652E-67 | 0.92717723 | 0.895 | 0.607 | 6.5322E-63 | TEC_prlf | HNRNPA1 |
| 5.1927E-67 | 0.7054839 | 0.49 | 0.154 | 1.2267E-62 | TEC_prlf | CLDN3 |
| 5.561E-67 | 0.33214559 | 0.254 | 0.049 | 1.3137E-62 | TEC_prlf | RPL39L |
| 1.2654E-65 | 0.42129772 | 0.315 | 0.074 | 2.9892E-61 | TEC_prlf | MYL6B |
| 4.8775E-65 | 1.1249596 | 0.729 | 0.417 | 1.1522E-60 | TEC_prlf | RAN |
| 5.6386E-65 | 0.72881582 | 0.504 | 0.176 | 1.332E-60 | TEC_prlf | RANBP1 |
| 1.9416E-63 | 0.27868474 | 0.192 | 0.031 | 4.5867E-59 | TEC_prlf | RNF26 |
| 2.7553E-63 | 0.47093116 | 0.335 | 0.086 | 6.5088E-59 | TEC_prlf | MZT1 |
| 1.1959E-62 | 0.3159505 | 0.198 | 0.033 | 2.8251E-58 | TEC_prlf | CDKL2 |
| 2.9791E-62 | 0.3975803 | 0.303 | 0.07 | 7.0375E-58 | TEC_prlf | LAMB1 |
| 8.6869E-62 | 0.41467858 | 0.242 | 0.049 | 2.0521E-57 | TEC_prlf | PCNT |
| 1.5078E-61 | 0.28106048 | 0.222 | 0.041 | 3.5619E-57 | TEC_prlf | ASRGL1 |
| 2.0868E-61 | 0.25368513 | 0.19 | 0.031 | 4.9297E-57 | TEC_prlf | DNALI1 |
| 6.5593E-61 | 0.57989141 | 0.394 | 0.117 | 1.5495E-56 | TEC_prlf | PSIP1 |
| 1.7645E-60 | 0.30542 | 0.198 | 0.034 | 4.1683E-56 | TEC_prlf | TCEAL2 |
| 3.6016E-60 | -2.3182175 | 0.956 | 0.987 | 8.508E-56 | TEC_prlf | FTL |
| 5.4596E-60 | 0.79579279 | 0.621 | 0.286 | 1.2897E-55 | TEC_prlf | CBX3 |
| 1.4703E-59 | 0.53545169 | 0.364 | 0.103 | 3.4734E-55 | TEC_prlf | GOT1 |
| 1.167E-58 | 0.64513038 | 0.414 | 0.132 | 2.7567E-54 | TEC_prlf | TMEM238 |
| 1.1724E-58 | 0.91061044 | 0.971 | 0.831 | 2.7696E-54 | TEC_prlf | UBB |
| 1.309E-58 | 0.27206664 | 0.227 | 0.045 | 3.0922E-54 | TEC_prlf | BCL2L12 |
| 1.9891E-58 | 0.34395387 | 0.201 | 0.036 | 4.6989E-54 | TEC_prlf | GAS2L3 |
| 3.4321E-58 | 0.45869324 | 0.327 | 0.086 | 8.1076E-54 | TEC_prlf | ISOC2 |
| 6.9045E-58 | 0.31254874 | 0.198 | 0.035 | 1.631E-53 | TEC_prlf | TCEA3 |
| 6.9448E-58 | 0.4161323 | 0.303 | 0.076 | 1.6406E-53 | TEC_prlf | NUDT1 |
| 1.8302E-57 | 0.64067825 | 0.452 | 0.157 | 4.3235E-53 | TEC_prlf | GLRX5 |
| 5.9552E-57 | 0.26353935 | 0.192 | 0.034 | 1.4068E-52 | TEC_prlf | TMEM19 |
| 6.7937E-57 | 0.34227115 | 0.274 | 0.064 | 1.6049E-52 | TEC_prlf | IFT22 |
| 9.4297E-57 | 0.46709459 | 0.306 | 0.078 | 2.2276E-52 | TEC_prlf | IMPA2 |
| 2.2551E-56 | 0.5761056 | 0.379 | 0.115 | 5.3273E-52 | TEC_prlf | CBX1 |
| 5.9613E-56 | 0.3459579 | 0.245 | 0.052 | 1.4082E-51 | TEC_prlf | CCDC198 |
| 7.9968E-56 | 0.78557686 | 0.603 | 0.274 | 1.8891E-51 | TEC_prlf | MRPL51 |
| 1.3911E-55 | 0.5485136 | 0.364 | 0.107 | 3.2863E-51 | TEC_prlf | ITM2C |
| 2.1142E-55 | 0.30820826 | 0.204 | 0.039 | 4.9944E-51 | TEC_prlf | CETN3 |
| 2.2233E-55 | 1.2372847 | 0.668 | 0.357 | 5.252E-51 | TEC_prlf | TUBA1B |
| 5.1804E-55 | 0.9945612 | 0.601 | 0.257 | 1.2238E-50 | TEC_prlf | TPM1 |
| 5.9963E-55 | 0.43337367 | 0.341 | 0.096 | 1.4165E-50 | TEC_prlf | IFT57 |
| 8.7371E-55 | 0.6020994 | 0.423 | 0.143 | 2.064E-50 | TEC_prlf | COA3 |
| 1.4862E-54 | 1.24352845 | 0.673 | 0.346 | 3.5108E-50 | TEC_prlf | PEBP1 |
| 1.5105E-54 | 0.62783511 | 0.539 | 0.219 | 3.5683E-50 | TEC_prlf | LSM4 |
| 1.2942E-53 | 1.05152807 | 0.58 | 0.248 | 3.0572E-49 | TEC_prlf | SYNE2 |
| 2.84E-53 | 0.45268672 | 0.248 | 0.056 | 6.7089E-49 | TEC_prlf | NAV2 |
| 3.6804E-53 | 0.37300397 | 0.303 | 0.079 | 8.6943E-49 | TEC_prlf | PHGDH |

| | | | | | | |
|---|---|---|---|---|---|---|
| 5.4548E-53 | 0.38372395 | 0.303 | 0.08 | 1.2886E-48 | TEC_prlf | TMEM106C |
| 7.593E-53 | 0.3408959 | 0.262 | 0.062 | 1.7937E-48 | TEC_prlf | KIF22 |
| 1.0371E-52 | 0.3266542 | 0.242 | 0.054 | 2.4498E-48 | TEC_prlf | UGT2B7 |
| 1.1536E-52 | 0.88518248 | 0.743 | 0.424 | 2.7252E-48 | TEC_prlf | SPINT2 |
| 1.2786E-52 | 0.85144743 | 0.985 | 0.945 | 3.0205E-48 | TEC_prlf | PTMA |
| 8.9375E-52 | 0.73130228 | 0.589 | 0.239 | 2.1113E-47 | TEC_prlf | S100A13 |
| 3.134E-51 | 0.83026609 | 0.808 | 0.535 | 7.4034E-47 | TEC_prlf | HMGN1 |
| 8.6557E-51 | 0.74200987 | 0.965 | 0.814 | 2.0447E-46 | TEC_prlf | YBX1 |
| 9.4847E-51 | 0.90278245 | 0.936 | 0.783 | 2.2406E-46 | TEC_prlf | DYNLL1 |
| 1.4186E-50 | 0.49724298 | 0.391 | 0.126 | 3.3511E-46 | TEC_prlf | H2AFX |
| 2.8549E-50 | 0.89537742 | 0.921 | 0.726 | 6.744E-46 | TEC_prlf | PKM |
| 4.0244E-50 | 0.34940419 | 0.271 | 0.069 | 9.5068E-46 | TEC_prlf | RUVBL2 |
| 4.0907E-50 | -3.8548722 | 0.364 | 0.673 | 9.6635E-46 | TEC_prlf | S100A8 |
| 7.6694E-50 | 0.42249888 | 0.321 | 0.093 | 1.8117E-45 | TEC_prlf | HINT2 |
| 8.806E-50 | 0.38956329 | 0.219 | 0.047 | 2.0803E-45 | TEC_prlf | PEG10 |
| 1.1826E-49 | 0.53030275 | 0.257 | 0.063 | 2.7937E-45 | TEC_prlf | SFTA2 |
| 1.7021E-49 | 0.28182325 | 0.23 | 0.052 | 4.0208E-45 | TEC_prlf | RRM1 |
| 3.2377E-49 | 0.30666327 | 0.271 | 0.068 | 7.6485E-45 | TEC_prlf | ACTL6A |
| 1.0126E-48 | 0.82988073 | 0.577 | 0.282 | 2.392E-44 | TEC_prlf | RAD21 |
| 4.681E-48 | -4.432326 | 0.469 | 0.729 | 1.1058E-43 | TEC_prlf | S100A9 |
| 5.0783E-48 | -1.3383591 | 0.851 | 0.94 | 1.1996E-43 | TEC_prlf | SAT1 |
| 5.2677E-48 | 0.27645374 | 0.187 | 0.037 | 1.2444E-43 | TEC_prlf | FOLR1 |
| 1.0403E-47 | 0.56712142 | 0.399 | 0.139 | 2.4575E-43 | TEC_prlf | BEX3 |
| 2.732E-47 | 0.42630222 | 0.259 | 0.067 | 6.4537E-43 | TEC_prlf | CDK5RAP2 |
| 6.002E-47 | 0.30847037 | 0.224 | 0.051 | 1.4178E-42 | TEC_prlf | IFT74 |
| 1.4745E-46 | 0.56282906 | 0.431 | 0.159 | 3.4831E-42 | TEC_prlf | FARP1 |
| 3.051E-46 | 0.29575019 | 0.102 | 0.013 | 7.2074E-42 | TEC_prlf | CLDN16 |
| 6.7034E-46 | 0.33317609 | 0.236 | 0.057 | 1.5835E-41 | TEC_prlf | ATAD2 |
| 7.9494E-46 | 0.55955883 | 0.385 | 0.132 | 1.8779E-41 | TEC_prlf | CDC42BPB |
| 1.0113E-45 | 0.66268934 | 0.598 | 0.283 | 2.3891E-41 | TEC_prlf | YWHAH |
| 1.2831E-45 | 0.64769178 | 0.51 | 0.226 | 3.0311E-41 | TEC_prlf | PDAP1 |
| 1.1034E-44 | 0.62909112 | 0.563 | 0.258 | 2.6066E-40 | TEC_prlf | PRDX2 |
| 1.2102E-44 | 0.31947384 | 0.248 | 0.063 | 2.8588E-40 | TEC_prlf | DNAJC9 |
| 1.3069E-44 | 0.63916318 | 0.557 | 0.256 | 3.0873E-40 | TEC_prlf | CCT3 |
| 2.1069E-44 | 0.25019849 | 0.213 | 0.049 | 4.9771E-40 | TEC_prlf | R3HDM1 |
| 2.2852E-44 | 0.25144624 | 0.184 | 0.038 | 5.3984E-40 | TEC_prlf | KCNJ16 |
| 6.921E-44 | 0.93143214 | 0.499 | 0.221 | 1.635E-39 | TEC_prlf | MRPS6 |
| 1.4584E-43 | -1.6535403 | 0.411 | 0.683 | 3.4452E-39 | TEC_prlf | NAMPT |
| 1.4989E-43 | 0.85581666 | 0.767 | 0.491 | 3.5408E-39 | TEC_prlf | NCL |
| 1.4998E-43 | 0.32425341 | 0.143 | 0.025 | 3.5429E-39 | TEC_prlf | SFRP1 |
| 1.7226E-43 | 0.28754171 | 0.224 | 0.054 | 4.0694E-39 | TEC_prlf | GPSM2 |
| 3.2409E-43 | 0.29182807 | 0.219 | 0.051 | 7.6561E-39 | TEC_prlf | POU3F3 |
| 7.0793E-43 | 0.58172464 | 0.452 | 0.184 | 1.6724E-38 | TEC_prlf | ILF2 |
| 1.1689E-42 | 0.30926452 | 0.274 | 0.076 | 2.7614E-38 | TEC_prlf | ESRP1 |
| 1.4908E-42 | 0.72118831 | 0.913 | 0.725 | 3.5218E-38 | TEC_prlf | ENO1 |
| 4.3985E-42 | 0.56963587 | 0.397 | 0.141 | 1.0391E-37 | TEC_prlf | PAX8 |
| 4.5468E-42 | 0.92003547 | 0.426 | 0.167 | 1.0741E-37 | TEC_prlf | AKR1B1 |
| 5.0505E-42 | 0.40351443 | 0.338 | 0.111 | 1.1931E-37 | TEC_prlf | TIMM10 |
| 5.4304E-42 | 0.26897687 | 0.198 | 0.045 | 1.2828E-37 | TEC_prlf | TBC1D4 |
| 9.0457E-42 | 0.82118738 | 0.758 | 0.505 | 2.1369E-37 | TEC_prlf | DBI |

| | | | | | | |
|---|---|---|---|---|---|---|
| 1.8885E-41 | 0.33490473 | 0.283 | 0.082 | 4.4613E-37 | TEC_prlf | TUSC3 |
| 3.3392E-41 | 0.43096576 | 0.367 | 0.126 | 7.8882E-37 | TEC_prlf | HACD3 |
| 3.962E-41 | 0.28584307 | 0.23 | 0.059 | 9.3594E-37 | TEC_prlf | IMMP1L |
| 5.2297E-41 | 0.27871504 | 0.254 | 0.069 | 1.2354E-36 | TEC_prlf | RUVBL1 |
| 5.931E-41 | -1.9014013 | 0.315 | 0.616 | 1.4011E-36 | TEC_prlf | C15orf48 |
| 6.5699E-41 | 0.31730227 | 0.274 | 0.078 | 1.552E-36 | TEC_prlf | SPAG16 |
| 8.8409E-41 | 0.31838718 | 0.306 | 0.093 | 2.0885E-36 | TEC_prlf | PON2 |
| 1.9398E-40 | 0.84450712 | 0.615 | 0.329 | 4.5823E-36 | TEC_prlf | APP |
| 1.9659E-40 | 0.43794831 | 0.329 | 0.109 | 4.644E-36 | TEC_prlf | KMT5A |
| 2.7932E-40 | -0.8776506 | 0.889 | 0.958 | 6.5984E-36 | TEC_prlf | H3F3A |
| 3.2643E-40 | 0.68915707 | 0.793 | 0.551 | 7.7113E-36 | TEC_prlf | HNRNPA3 |
| 3.6707E-40 | 0.43579899 | 0.379 | 0.139 | 8.6713E-36 | TEC_prlf | VDAC3 |
| 5.3403E-40 | 0.59046554 | 0.356 | 0.128 | 1.2615E-35 | TEC_prlf | NUCB2 |
| 5.5623E-40 | 0.69674075 | 0.983 | 0.845 | 1.314E-35 | TEC_prlf | RPL35 |
| 5.8593E-40 | 0.52367768 | 0.466 | 0.2 | 1.3841E-35 | TEC_prlf | NDUFS8 |
| 6.3663E-40 | 0.955406 | 0.577 | 0.279 | 1.5039E-35 | TEC_prlf | CLU |
| 1.0458E-39 | 0.78492021 | 0.586 | 0.316 | 2.4705E-35 | TEC_prlf | CALM3 |
| 1.2478E-39 | -2.0497708 | 0.991 | 0.994 | 2.9477E-35 | TEC_prlf | FTH1 |
| 1.3351E-39 | 0.28762182 | 0.262 | 0.075 | 3.154E-35 | TEC_prlf | MAGOHB |
| 2.2461E-39 | 0.77434318 | 0.895 | 0.733 | 5.306E-35 | TEC_prlf | GNAS |
| 2.9664E-39 | 0.45209107 | 0.399 | 0.154 | 7.0075E-35 | TEC_prlf | BUB3 |
| 4.7572E-39 | 0.27968123 | 0.233 | 0.061 | 1.1238E-34 | TEC_prlf | ALDH1A1 |
| 5.1293E-39 | 0.34944067 | 0.248 | 0.069 | 1.2117E-34 | TEC_prlf | MAOA |
| 5.6477E-39 | 0.41697492 | 0.3 | 0.097 | 1.3342E-34 | TEC_prlf | RNF5 |
| 6.0252E-39 | 0.36416768 | 0.274 | 0.081 | 1.4233E-34 | TEC_prlf | CAVIN1 |
| 6.5819E-39 | 0.28987983 | 0.239 | 0.065 | 1.5548E-34 | TEC_prlf | CD320 |
| 7.7985E-39 | 0.30372393 | 0.268 | 0.079 | 1.8422E-34 | TEC_prlf | EXOSC8 |
| 1.4804E-38 | 0.93699374 | 0.886 | 0.555 | 3.4971E-34 | TEC_prlf | KRT18 |
| 2.0684E-38 | 0.4040588 | 0.344 | 0.121 | 4.8862E-34 | TEC_prlf | PDHA1 |
| 2.9905E-38 | 0.61749257 | 0.484 | 0.224 | 7.0644E-34 | TEC_prlf | COX20 |
| 3.6923E-38 | 0.2699301 | 0.216 | 0.055 | 8.7224E-34 | TEC_prlf | MYO5C |
| 5.7765E-38 | -2.5324377 | 0.283 | 0.572 | 1.3646E-33 | TEC_prlf | SRGN |
| 1.0558E-37 | 0.50828845 | 0.391 | 0.156 | 2.4941E-33 | TEC_prlf | DCTN3 |
| 1.3501E-37 | 0.38865394 | 0.315 | 0.106 | 3.1894E-33 | TEC_prlf | CFAP36 |
| 1.6091E-37 | 0.84115689 | 0.598 | 0.335 | 3.8011E-33 | TEC_prlf | SRRM1 |
| 1.9927E-37 | 0.70773807 | 0.945 | 0.833 | 4.7072E-33 | TEC_prlf | PPIA |
| 2.9058E-37 | -0.7145301 | 0.606 | 0.837 | 6.8643E-33 | TEC_prlf | HLA-E |
| 6.4328E-37 | 0.64028715 | 0.621 | 0.339 | 1.5196E-32 | TEC_prlf | PDIA6 |
| 8.299E-37 | 0.59434376 | 0.478 | 0.215 | 1.9605E-32 | TEC_prlf | CDK2AP2 |
| 1.2931E-36 | 0.53467807 | 0.455 | 0.202 | 3.0546E-32 | TEC_prlf | NENF |
| 1.6618E-36 | 0.31084134 | 0.219 | 0.059 | 3.9257E-32 | TEC_prlf | EZH2 |
| 1.6725E-36 | 0.54555114 | 0.353 | 0.133 | 3.951E-32 | TEC_prlf | SUDS3 |
| 1.8708E-36 | 0.72333973 | 0.603 | 0.338 | 4.4193E-32 | TEC_prlf | ANP32B |
| 2.4256E-36 | -1.6406626 | 0.446 | 0.711 | 5.73E-32 | TEC_prlf | ANXA1 |
| 2.5048E-36 | 0.65829029 | 0.898 | 0.684 | 5.9171E-32 | TEC_prlf | NPM1 |
| 2.8282E-36 | 0.76514275 | 0.63 | 0.383 | 6.6811E-32 | TEC_prlf | DEK |
| 3.3017E-36 | -1.0120729 | 0.411 | 0.701 | 7.7997E-32 | TEC_prlf | JUNB |
| 3.7578E-36 | 0.71358297 | 0.595 | 0.341 | 8.877E-32 | TEC_prlf | BSG |
| 3.7632E-36 | 0.30977178 | 0.294 | 0.095 | 8.8897E-32 | TEC_prlf | MRPL19 |
| 5.5838E-36 | 0.28802034 | 0.251 | 0.073 | 1.3191E-31 | TEC_prlf | DDAH1 |

| | | | | | | |
|---|---|---|---|---|---|---|
| 8.5187E-36 | 0.27326054 | 0.224 | 0.062 | 2.0124E-31 | TEC_prlf | SNRNP25 |
| 8.6082E-36 | 0.39184645 | 0.181 | 0.043 | 2.0335E-31 | TEC_prlf | CLDN10 |
| 1.0784E-35 | 0.27377668 | 0.122 | 0.022 | 2.5476E-31 | TEC_prlf | IRX3 |
| 1.1218E-35 | 0.573197 | 0.373 | 0.15 | 2.6501E-31 | TEC_prlf | LBR |
| 1.2073E-35 | 0.34035302 | 0.259 | 0.079 | 2.8519E-31 | TEC_prlf | KDM1A |
| 1.3834E-35 | 0.46995202 | 0.397 | 0.154 | 3.2679E-31 | TEC_prlf | COBLL1 |
| 1.6427E-35 | 0.38821458 | 0.364 | 0.138 | 3.8806E-31 | TEC_prlf | RPA3 |
| 2.5371E-35 | 0.57503927 | 0.49 | 0.237 | 5.9933E-31 | TEC_prlf | GSPT1 |
| 3.0018E-35 | 0.45271773 | 0.265 | 0.083 | 7.0913E-31 | TEC_prlf | CSRP2 |
| 4.1604E-35 | 0.60548097 | 0.397 | 0.17 | 9.8282E-31 | TEC_prlf | MPC2 |
| 5.5798E-35 | 0.4521519 | 0.388 | 0.156 | 1.3181E-30 | TEC_prlf | PRMT1 |
| 9.1046E-35 | 0.59054621 | 0.985 | 0.945 | 2.1508E-30 | TEC_prlf | RPL13A |
| 1.4059E-34 | 0.51664998 | 0.501 | 0.24 | 3.3212E-30 | TEC_prlf | NUDC |
| 2.7326E-34 | 0.42172799 | 0.324 | 0.119 | 6.4552E-30 | TEC_prlf | MAZ |
| 3.1881E-34 | 0.622464 | 0.988 | 0.875 | 7.5312E-30 | TEC_prlf | RPL3 |
| 3.1965E-34 | 0.73551033 | 0.615 | 0.37 | 7.551E-30 | TEC_prlf | KIF5B |
| 6.4856E-34 | 0.3206202 | 0.254 | 0.079 | 1.5321E-29 | TEC_prlf | MORN2 |
| 7.0995E-34 | 0.27277981 | 0.23 | 0.066 | 1.6771E-29 | TEC_prlf | EPB41L2 |
| 7.3607E-34 | -0.9812045 | 0.983 | 0.99 | 1.7388E-29 | TEC_prlf | B2M |
| 9.8112E-34 | 0.59548557 | 0.633 | 0.356 | 2.3177E-29 | TEC_prlf | SNRPD2 |
| 1.0417E-33 | 0.38354957 | 0.329 | 0.119 | 2.4609E-29 | TEC_prlf | CTNNAL1 |
| 1.4132E-33 | 0.57922554 | 0.574 | 0.319 | 3.3384E-29 | TEC_prlf | YWHAQ |
| 1.5818E-33 | 0.36181259 | 0.277 | 0.091 | 3.7366E-29 | TEC_prlf | KPNA2 |
| 1.6222E-33 | 0.50211865 | 0.536 | 0.274 | 3.8321E-29 | TEC_prlf | SNRPD3 |
| 1.9037E-33 | 0.39994119 | 0.347 | 0.128 | 4.4971E-29 | TEC_prlf | AIF1L |
| 2.0736E-33 | 0.66448917 | 0.507 | 0.262 | 4.8985E-29 | TEC_prlf | PLP2 |
| 2.4711E-33 | -1.2007242 | 0.023 | 0.338 | 5.8375E-29 | TEC_prlf | LAPTM5 |
| 2.9585E-33 | 0.42817133 | 0.408 | 0.172 | 6.9888E-29 | TEC_prlf | RDX |
| 4.9726E-33 | 0.5394156 | 0.569 | 0.305 | 1.1747E-28 | TEC_prlf | HNRNPM |
| 5.3696E-33 | 0.3905519 | 0.318 | 0.116 | 1.2685E-28 | TEC_prlf | HMG20B |
| 5.8951E-33 | 0.25861054 | 0.216 | 0.061 | 1.3926E-28 | TEC_prlf | SEPHS1 |
| 6.4807E-33 | 0.74569701 | 0.857 | 0.656 | 1.5309E-28 | TEC_prlf | CALM2 |
| 8.8055E-33 | 0.33750313 | 0.137 | 0.028 | 2.0801E-28 | TEC_prlf | DDIT4L |
| 1.0658E-32 | 0.26229792 | 0.227 | 0.066 | 2.5178E-28 | TEC_prlf | HOOK1 |
| 1.0938E-32 | 0.33939394 | 0.306 | 0.109 | 2.5838E-28 | TEC_prlf | BAD |
| 1.1398E-32 | 0.59533265 | 0.746 | 0.472 | 2.6925E-28 | TEC_prlf | CD81 |
| 1.2469E-32 | 0.4239997 | 0.391 | 0.159 | 2.9455E-28 | TEC_prlf | NCKAP1 |
| 1.4907E-32 | 0.25403378 | 0.224 | 0.065 | 3.5214E-28 | TEC_prlf | REEP4 |
| 3.1673E-32 | 0.33063445 | 0.306 | 0.106 | 7.4821E-28 | TEC_prlf | WEE1 |
| 3.3234E-32 | 0.48216875 | 0.306 | 0.109 | 7.851E-28 | TEC_prlf | FXYD2 |
| 4.8622E-32 | -1.0493558 | 0.242 | 0.533 | 1.1486E-27 | TEC_prlf | CTSS |
| 6.0638E-32 | 0.61643332 | 0.569 | 0.327 | 1.4325E-27 | TEC_prlf | AURKAIP1 |
| 7.4208E-32 | 0.45480222 | 0.42 | 0.184 | 1.753E-27 | TEC_prlf | ANXA4 |
| 1.1317E-31 | 0.34705488 | 0.332 | 0.126 | 2.6733E-27 | TEC_prlf | RBBP7 |
| 1.2508E-31 | 0.36266566 | 0.265 | 0.089 | 2.9548E-27 | TEC_prlf | DNMT1 |
| 1.4625E-31 | 0.55415139 | 0.746 | 0.476 | 3.4549E-27 | TEC_prlf | HSPD1 |
| 1.4725E-31 | 0.62846155 | 0.65 | 0.39 | 3.4784E-27 | TEC_prlf | SERBP1 |
| 1.9386E-31 | 0.59740608 | 0.799 | 0.599 | 4.5796E-27 | TEC_prlf | YWHAE |
| 2.5345E-31 | 0.28876799 | 0.239 | 0.074 | 5.9872E-27 | TEC_prlf | MITF |
| 2.6729E-31 | 0.45767092 | 0.42 | 0.19 | 6.3143E-27 | TEC_prlf | TECR |

| | | | | | | |
|---|---|---|---|---|---|---|
| 3.0914E-31 | 0.27042444 | 0.233 | 0.071 | 7.3028E-27 | TEC_prlf | CRYZ |
| 7.7604E-31 | 0.45966783 | 0.411 | 0.184 | 1.8332E-26 | TEC_prlf | NDUFAF3 |
| 1.019E-30 | 0.75396801 | 0.496 | 0.261 | 2.4072E-26 | TEC_prlf | TUBB |
| 1.3058E-30 | 0.4315384 | 0.429 | 0.194 | 3.0847E-26 | TEC_prlf | SNRPD1 |
| 1.3669E-30 | 0.36485948 | 0.329 | 0.122 | 3.229E-26 | TEC_prlf | FAT1 |
| 1.4092E-30 | 0.69532452 | 0.583 | 0.347 | 3.3289E-26 | TEC_prlf | FKBP2 |
| 2.1505E-30 | 0.35951626 | 0.3 | 0.112 | 5.08E-26 | TEC_prlf | NUDCD2 |
| 2.5066E-30 | 0.37611531 | 0.341 | 0.132 | 5.9213E-26 | TEC_prlf | SEPT11 |
| 3.0605E-30 | 0.51318402 | 0.583 | 0.318 | 7.2298E-26 | TEC_prlf | ERP29 |
| 3.1581E-30 | 0.53187343 | 0.475 | 0.236 | 7.4603E-26 | TEC_prlf | TSTD1 |
| 3.3779E-30 | 0.26249601 | 0.222 | 0.067 | 7.9796E-26 | TEC_prlf | HYI |
| 3.7072E-30 | 0.30005313 | 0.297 | 0.106 | 8.7575E-26 | TEC_prlf | YES1 |
| 6.0964E-30 | 0.27033559 | 0.222 | 0.068 | 1.4401E-25 | TEC_prlf | FOPNL |
| 1.3256E-29 | 0.57127088 | 0.776 | 0.559 | 3.1315E-25 | TEC_prlf | SET |
| 1.4561E-29 | 0.27442376 | 0.254 | 0.084 | 3.4399E-25 | TEC_prlf | GAR1 |
| 1.4635E-29 | -1.6376834 | 0.172 | 0.442 | 3.4573E-25 | TEC_prlf | TYROBP |
| 1.8994E-29 | 0.47080074 | 0.42 | 0.198 | 4.4869E-25 | TEC_prlf | MDH1 |
| 2.1261E-29 | 0.44363355 | 0.359 | 0.147 | 5.0224E-25 | TEC_prlf | SCCPDH |
| 2.3021E-29 | 0.51172482 | 0.321 | 0.124 | 5.4383E-25 | TEC_prlf | SYNE1 |
| 3.8563E-29 | 0.3192513 | 0.286 | 0.103 | 9.1097E-25 | TEC_prlf | MRPS26 |
| 4.8051E-29 | 0.49396436 | 0.56 | 0.309 | 1.1351E-24 | TEC_prlf | SMS |
| 6.0363E-29 | -1.0847034 | 0.262 | 0.529 | 1.426E-24 | TEC_prlf | ZFP36 |
| 6.6263E-29 | 0.64008733 | 0.647 | 0.412 | 1.5653E-24 | TEC_prlf | SFPQ |
| 1.2622E-28 | 0.38000365 | 0.356 | 0.147 | 2.9816E-24 | TEC_prlf | BAZ1B |
| 1.5869E-28 | 0.55911924 | 0.802 | 0.614 | 3.7486E-24 | TEC_prlf | COX8A |
| 1.5993E-28 | 0.41708779 | 0.327 | 0.129 | 3.7779E-24 | TEC_prlf | NTN4 |
| 1.6752E-28 | 0.46379213 | 0.522 | 0.274 | 3.9573E-24 | TEC_prlf | CCT5 |
| 1.8067E-28 | 0.60927511 | 0.612 | 0.373 | 4.2679E-24 | TEC_prlf | ATP5ME |
| 1.9718E-28 | -1.4893261 | 0.055 | 0.335 | 4.6579E-24 | TEC_prlf | IL1RN |
| 2.3789E-28 | 0.62903246 | 0.962 | 0.905 | 5.6197E-24 | TEC_prlf | ITM2B |
| 2.6582E-28 | 0.28010033 | 0.251 | 0.085 | 6.2794E-24 | TEC_prlf | EIF2AK4 |
| 2.7717E-28 | 0.25323336 | 0.207 | 0.062 | 6.5476E-24 | TEC_prlf | SCD5 |
| 2.8398E-28 | -0.9800203 | 0.105 | 0.386 | 6.7085E-24 | TEC_prlf | TYMP |
| 2.8919E-28 | 0.75165761 | 0.603 | 0.364 | 6.8315E-24 | TEC_prlf | RALBP1 |
| 3.0499E-28 | -1.1514693 | 0.717 | 0.86 | 7.2049E-24 | TEC_prlf | SH3BGRL3 |
| 3.1457E-28 | -1.0340457 | 0.563 | 0.739 | 7.4311E-24 | TEC_prlf | CTSB |
| 3.5181E-28 | 0.64172932 | 0.793 | 0.491 | 8.3108E-24 | TEC_prlf | KRT8 |
| 3.6276E-28 | 0.52532969 | 0.504 | 0.267 | 8.5695E-24 | TEC_prlf | NOL7 |
| 4.7472E-28 | -0.7360148 | 0.038 | 0.314 | 1.1214E-23 | TEC_prlf | CD68 |
| 9.2552E-28 | 0.59019612 | 0.65 | 0.401 | 2.1864E-23 | TEC_prlf | SOD1 |
| 9.6224E-28 | 0.38243499 | 0.338 | 0.139 | 2.2731E-23 | TEC_prlf | ARL3 |
| 9.7017E-28 | -1.2329366 | 0.009 | 0.277 | 2.2918E-23 | TEC_prlf | AIF1 |
| 9.9082E-28 | 0.29639204 | 0.219 | 0.069 | 2.3406E-23 | TEC_prlf | GNG11 |
| 1.1067E-27 | 0.40595225 | 0.353 | 0.151 | 2.6143E-23 | TEC_prlf | SCOC |
| 1.3436E-27 | 0.25552549 | 0.245 | 0.083 | 3.1739E-23 | TEC_prlf | TMEM18 |
| 1.4569E-27 | 0.54126454 | 0.86 | 0.622 | 3.4417E-23 | TEC_prlf | DSTN |
| 1.7435E-27 | 0.31308219 | 0.254 | 0.088 | 4.1186E-23 | TEC_prlf | HOXB2 |
| 1.9596E-27 | 0.41533481 | 0.37 | 0.163 | 4.6292E-23 | TEC_prlf | ZCRB1 |
| 2.0916E-27 | 0.56140028 | 0.834 | 0.656 | 4.9411E-23 | TEC_prlf | NDUFA4 |
| 2.303E-27 | 0.426758 | 0.324 | 0.129 | 5.4404E-23 | TEC_prlf | SCIN |

| | | | | | | |
|---|---|---|---|---|---|---|
| 2.314E-27 | 0.35163571 | 0.353 | 0.149 | 5.4664E-23 | TEC_prlf | FKBP3 |
| 2.6305E-27 | -1.3270165 | 0.443 | 0.645 | 6.2139E-23 | TEC_prlf | GLUL |
| 2.9293E-27 | 0.26236283 | 0.265 | 0.094 | 6.92E-23 | TEC_prlf | ANAPC15 |
| 3.773E-27 | 0.2670202 | 0.207 | 0.064 | 8.9129E-23 | TEC_prlf | DOCK1 |
| 4.6982E-27 | 0.45505305 | 0.303 | 0.121 | 1.1098E-22 | TEC_prlf | MIS18BP1 |
| 5.4546E-27 | 0.60323556 | 0.566 | 0.341 | 1.2885E-22 | TEC_prlf | ATP5F1B |
| 5.7105E-27 | 0.4550807 | 0.359 | 0.158 | 1.349E-22 | TEC_prlf | DUT |
| 5.7836E-27 | 0.29598635 | 0.329 | 0.133 | 1.3663E-22 | TEC_prlf | MRPS33 |
| 5.8592E-27 | 0.45022418 | 0.429 | 0.206 | 1.3841E-22 | TEC_prlf | RBMX |
| 7.2897E-27 | 0.26230936 | 0.257 | 0.091 | 1.722E-22 | TEC_prlf | SMARCB1 |
| 7.3176E-27 | -1.5955026 | 0.125 | 0.387 | 1.7286E-22 | TEC_prlf | FCER1G |
| 7.5271E-27 | -0.8704013 | 0.198 | 0.472 | 1.7781E-22 | TEC_prlf | DUSP1 |
| 8.0598E-27 | 0.4889168 | 0.481 | 0.257 | 1.904E-22 | TEC_prlf | LSM3 |
| 8.1414E-27 | 0.42577152 | 0.487 | 0.255 | 1.9232E-22 | TEC_prlf | MORF4L2 |
| 8.5054E-27 | 0.40998919 | 0.443 | 0.212 | 2.0092E-22 | TEC_prlf | LMAN1 |
| 8.5488E-27 | -0.7874593 | 0.915 | 0.966 | 2.0195E-22 | TEC_prlf | UBC |
| 1.0503E-26 | -0.6867254 | 0.155 | 0.43 | 2.4811E-22 | TEC_prlf | EFHD2 |
| 1.0903E-26 | -2.1362516 | 0.216 | 0.466 | 2.5757E-22 | TEC_prlf | LYZ |
| 1.1632E-26 | -1.2057789 | 0.507 | 0.705 | 2.7479E-22 | TEC_prlf | SOD2 |
| 1.3579E-26 | -1.1719062 | 0.195 | 0.46 | 3.2078E-22 | TEC_prlf | MXD1 |
| 2.3457E-26 | 0.3193236 | 0.347 | 0.146 | 5.5412E-22 | TEC_prlf | SNX5 |
| 2.7963E-26 | 0.33507297 | 0.306 | 0.121 | 6.6056E-22 | TEC_prlf | ADD3 |
| 3.071E-26 | 0.33141684 | 0.294 | 0.116 | 7.2545E-22 | TEC_prlf | CLSTN1 |
| 4.3625E-26 | 0.52548837 | 0.539 | 0.309 | 1.0305E-21 | TEC_prlf | NDUFA2 |
| 4.7877E-26 | -0.9371959 | 0.277 | 0.527 | 1.131E-21 | TEC_prlf | CEBPB |
| 5.2699E-26 | 0.4158554 | 0.347 | 0.151 | 1.2449E-21 | TEC_prlf | PNKD |
| 6.4572E-26 | 0.30719703 | 0.338 | 0.135 | 1.5254E-21 | TEC_prlf | DCDC2 |
| 6.6458E-26 | 0.38655596 | 0.356 | 0.157 | 1.5699E-21 | TEC_prlf | DDAH2 |
| 9.1359E-26 | 0.52761273 | 0.586 | 0.352 | 2.1582E-21 | TEC_prlf | SEPT7 |
| 1.1826E-25 | 0.41486676 | 0.414 | 0.2 | 2.7937E-21 | TEC_prlf | PHB |
| 1.3107E-25 | 0.33627826 | 0.306 | 0.124 | 3.0962E-21 | TEC_prlf | EBNA1BP2 |
| 1.4195E-25 | -0.8772739 | 0.032 | 0.293 | 3.3532E-21 | TEC_prlf | LCP1 |
| 1.6396E-25 | 0.5356499 | 0.507 | 0.287 | 3.8733E-21 | TEC_prlf | NDUFB7 |
| 2.2272E-25 | 0.40850214 | 0.455 | 0.235 | 5.2613E-21 | TEC_prlf | ATP5F1A |
| 2.2813E-25 | 0.27748934 | 0.245 | 0.086 | 5.3891E-21 | TEC_prlf | CRIP2 |
| 2.4931E-25 | 0.33883806 | 0.309 | 0.126 | 5.8894E-21 | TEC_prlf | UQCC2 |
| 2.8937E-25 | 0.51688599 | 0.977 | 0.876 | 6.8358E-21 | TEC_prlf | RPL23A |
| 3.1021E-25 | 0.66297026 | 0.609 | 0.386 | 7.3282E-21 | TEC_prlf | TUBB4B |
| 3.9026E-25 | 0.27983512 | 0.289 | 0.113 | 9.2192E-21 | TEC_prlf | COPRS |
| 4.9795E-25 | 0.37040174 | 0.341 | 0.149 | 1.1763E-20 | TEC_prlf | TBC1D1 |
| 4.9874E-25 | 0.32533066 | 0.23 | 0.079 | 1.1782E-20 | TEC_prlf | PCSK1N |
| 5.494E-25 | 0.64672918 | 0.577 | 0.369 | 1.2978E-20 | TEC_prlf | ATP5MC3 |
| 6.2447E-25 | 0.56191799 | 0.714 | 0.531 | 1.4752E-20 | TEC_prlf | COX6C |
| 6.4554E-25 | 0.36093973 | 0.394 | 0.187 | 1.525E-20 | TEC_prlf | PRDX3 |
| 6.7789E-25 | 0.31560575 | 0.289 | 0.115 | 1.6014E-20 | TEC_prlf | BOLA3 |
| 8.1399E-25 | 0.30555993 | 0.242 | 0.086 | 1.9229E-20 | TEC_prlf | CA2 |
| 8.165E-25 | 0.26088615 | 0.271 | 0.103 | 1.9288E-20 | TEC_prlf | CDK16 |
| 8.2205E-25 | 0.39198207 | 0.408 | 0.2 | 1.9419E-20 | TEC_prlf | MLF2 |
| 9.0494E-25 | 0.32733752 | 0.286 | 0.112 | 2.1378E-20 | TEC_prlf | GOLM1 |
| 9.4317E-25 | 0.3883979 | 0.388 | 0.178 | 2.2281E-20 | TEC_prlf | ANXA3 |

| | | | | | | |
|---|---|---|---|---|---|---|
| 9.5721E-25 | -0.6757265 | 0.02 | 0.27 | 2.2612E-20 | TEC_prlf | CD53 |
| 9.9012E-25 | 0.29548976 | 0.289 | 0.112 | 2.3389E-20 | TEC_prlf | WWC1 |
| 1.0357E-24 | 0.30527963 | 0.274 | 0.107 | 2.4466E-20 | TEC_prlf | EEF1E1 |
| 1.1279E-24 | 0.46173922 | 0.478 | 0.25 | 2.6644E-20 | TEC_prlf | CD2AP |
| 1.2218E-24 | 0.44134668 | 0.376 | 0.179 | 2.8862E-20 | TEC_prlf | ADI1 |
| 1.3154E-24 | 0.26816546 | 0.222 | 0.076 | 3.1075E-20 | TEC_prlf | VAV3 |
| 1.3747E-24 | 0.38456593 | 0.245 | 0.088 | 3.2474E-20 | TEC_prlf | DCBLD2 |
| 1.6417E-24 | -0.7515034 | 0.516 | 0.723 | 3.8781E-20 | TEC_prlf | JUND |
| 1.8176E-24 | 0.36525498 | 0.143 | 0.038 | 4.2938E-20 | TEC_prlf | UMOD |
| 1.9233E-24 | -0.9621523 | 0.029 | 0.283 | 4.5435E-20 | TEC_prlf | PTPRC |
| 2.324E-24 | 0.48666484 | 0.513 | 0.294 | 5.49E-20 | TEC_prlf | KDELR1 |
| 2.5066E-24 | 0.3908079 | 0.452 | 0.233 | 5.9214E-20 | TEC_prlf | NONO |
| 3.0289E-24 | 0.413154 | 0.429 | 0.218 | 7.1551E-20 | TEC_prlf | SNRPF |
| 3.1141E-24 | 0.46778576 | 0.44 | 0.234 | 7.3565E-20 | TEC_prlf | BANF1 |
| 3.5069E-24 | 0.44822806 | 0.481 | 0.265 | 8.2843E-20 | TEC_prlf | ERH |
| 3.8376E-24 | 0.47483777 | 0.983 | 0.922 | 9.0655E-20 | TEC_prlf | RPL8 |
| 4.402E-24 | 0.42164444 | 0.44 | 0.215 | 1.0399E-19 | TEC_prlf | DSP |
| 4.5975E-24 | -0.7379137 | 0.009 | 0.248 | 1.0861E-19 | TEC_prlf | CSTA |
| 7.1786E-24 | 0.34335445 | 0.356 | 0.163 | 1.6958E-19 | TEC_prlf | SNRPC |
| 7.2255E-24 | 0.68193622 | 0.697 | 0.504 | 1.7069E-19 | TEC_prlf | JAK1 |
| 1.1265E-23 | 0.47078897 | 0.452 | 0.242 | 2.6612E-19 | TEC_prlf | PDCD5 |
| 1.2475E-23 | 0.55658831 | 0.662 | 0.453 | 2.947E-19 | TEC_prlf | ST13 |
| 1.2572E-23 | 0.53969843 | 0.574 | 0.358 | 2.9698E-19 | TEC_prlf | SRSF11 |
| 1.4349E-23 | 0.2827674 | 0.213 | 0.073 | 3.3897E-19 | TEC_prlf | KITLG |
| 1.5981E-23 | -1.3231124 | 0.423 | 0.626 | 3.7753E-19 | TEC_prlf | S100A4 |
| 1.6489E-23 | -1.0574253 | 0.411 | 0.643 | 3.8953E-19 | TEC_prlf | IER3 |
| 1.7584E-23 | 0.49555616 | 0.671 | 0.435 | 4.154E-19 | TEC_prlf | CYSTM1 |
| 1.7659E-23 | 0.29120261 | 0.315 | 0.13 | 4.1717E-19 | TEC_prlf | DMKN |
| 1.9461E-23 | 0.26552411 | 0.259 | 0.099 | 4.5973E-19 | TEC_prlf | TPGS2 |
| 2.7147E-23 | 0.27839686 | 0.262 | 0.102 | 6.4129E-19 | TEC_prlf | AAMDC |
| 2.7303E-23 | -2.6226627 | 0.257 | 0.49 | 6.4497E-19 | TEC_prlf | CXCL8 |
| 3.299E-23 | -1.099489 | 0.006 | 0.235 | 7.7933E-19 | TEC_prlf | SLC2A3 |
| 3.5688E-23 | -1.0337468 | 0.668 | 0.785 | 8.4306E-19 | TEC_prlf | LGALS3 |
| 3.7199E-23 | 0.37596749 | 0.42 | 0.212 | 8.7875E-19 | TEC_prlf | NAP1L4 |
| 3.9255E-23 | 0.37038942 | 0.359 | 0.168 | 9.2731E-19 | TEC_prlf | MRPS15 |
| 4.3225E-23 | 0.54612759 | 0.531 | 0.323 | 1.0211E-18 | TEC_prlf | MGST3 |
| 4.3501E-23 | 0.32866531 | 0.335 | 0.149 | 1.0276E-18 | TEC_prlf | RCN2 |
| 4.8689E-23 | -0.5745666 | 0.073 | 0.321 | 1.1502E-18 | TEC_prlf | SLC16A3 |
| 5.0895E-23 | 0.47766486 | 0.469 | 0.261 | 1.2023E-18 | TEC_prlf | HNRNPR |
| 5.7994E-23 | 0.33633516 | 0.356 | 0.164 | 1.37E-18 | TEC_prlf | GLO1 |
| 5.9307E-23 | 0.49414907 | 0.493 | 0.285 | 1.401E-18 | TEC_prlf | MRFAP1 |
| 6.043E-23 | 0.40996531 | 0.423 | 0.216 | 1.4275E-18 | TEC_prlf | MLEC |
| 7.8405E-23 | 0.51124331 | 0.484 | 0.272 | 1.8522E-18 | TEC_prlf | SLTM |
| 7.9801E-23 | 0.28864911 | 0.297 | 0.124 | 1.8851E-18 | TEC_prlf | RPL26L1 |
| 8.0586E-23 | 0.34197491 | 0.309 | 0.135 | 1.9037E-18 | TEC_prlf | COMMD4 |
| 8.7944E-23 | 0.4707187 | 0.773 | 0.562 | 2.0775E-18 | TEC_prlf | SLC25A6 |
| 9.0456E-23 | 0.47787006 | 0.799 | 0.611 | 2.1368E-18 | TEC_prlf | PTGES3 |
| 1.1027E-22 | 0.38843325 | 0.414 | 0.208 | 2.6049E-18 | TEC_prlf | ILF3 |
| 1.1384E-22 | -0.4936484 | 0.994 | 0.993 | 2.6892E-18 | TEC_prlf | TPT1 |
| 1.2365E-22 | 0.5234505 | 0.595 | 0.383 | 2.921E-18 | TEC_prlf | TBCA |

| | | | | | | |
|---|---|---|---|---|---|---|
| 1.2806E-22 | 0.26286305 | 0.294 | 0.121 | 3.025E-18 | TEC_prlf | PGP |
| 1.4499E-22 | 0.39477989 | 0.391 | 0.193 | 3.425E-18 | TEC_prlf | CCT2 |
| 1.5085E-22 | 0.38637255 | 0.429 | 0.219 | 3.5635E-18 | TEC_prlf | PCM1 |
| 1.6138E-22 | 0.44408873 | 0.825 | 0.648 | 3.8122E-18 | TEC_prlf | UQCRH |
| 2.0566E-22 | 0.27495904 | 0.239 | 0.089 | 4.8583E-18 | TEC_prlf | HERC2 |
| 2.1239E-22 | 0.4499951 | 0.461 | 0.235 | 5.0173E-18 | TEC_prlf | S100A16 |
| 2.6915E-22 | 0.25368614 | 0.262 | 0.103 | 6.3581E-18 | TEC_prlf | HDDC2 |
| 4.4876E-22 | 0.26241091 | 0.271 | 0.109 | 1.0601E-17 | TEC_prlf | ZNF428 |
| 4.8735E-22 | 0.41325478 | 0.466 | 0.254 | 1.1513E-17 | TEC_prlf | DNAJC8 |
| 4.9803E-22 | 0.251767 | 0.187 | 0.061 | 1.1765E-17 | TEC_prlf | ERMP1 |
| 5.0693E-22 | 0.254757 | 0.262 | 0.104 | 1.1975E-17 | TEC_prlf | IARS2 |
| 5.8255E-22 | -0.6632818 | 0.35 | 0.577 | 1.3762E-17 | TEC_prlf | CD55 |
| 6.4682E-22 | 0.31536469 | 0.262 | 0.105 | 1.528E-17 | TEC_prlf | ASAP2 |
| 8.8788E-22 | 0.53588295 | 0.571 | 0.37 | 2.0974E-17 | TEC_prlf | RHEB |
| 9.4725E-22 | 0.45499458 | 0.475 | 0.271 | 2.2377E-17 | TEC_prlf | MINOS1 |
| 1.0241E-21 | 0.36127089 | 0.42 | 0.216 | 2.4191E-17 | TEC_prlf | NDUFA5 |
| 1.0357E-21 | 0.274953 | 0.242 | 0.093 | 2.4467E-17 | TEC_prlf | JPT2 |
| 1.0449E-21 | -1.0877769 | 0.017 | 0.238 | 2.4684E-17 | TEC_prlf | BCL2A1 |
| 1.1911E-21 | 0.29413187 | 0.227 | 0.083 | 2.8136E-17 | TEC_prlf | CALD1 |
| 1.305E-21 | -0.9872877 | 0.006 | 0.221 | 3.0828E-17 | TEC_prlf | CD14 |
| 1.3771E-21 | 0.4124209 | 0.977 | 0.87 | 3.2532E-17 | TEC_prlf | RPL14 |
| 1.5948E-21 | 0.43540203 | 0.554 | 0.336 | 3.7673E-17 | TEC_prlf | ARL6IP4 |
| 1.8886E-21 | -0.728728 | 0.012 | 0.229 | 4.4615E-17 | TEC_prlf | ZEB2 |
| 1.9449E-21 | 0.25157795 | 0.245 | 0.095 | 4.5945E-17 | TEC_prlf | CHCHD3 |
| 2.0194E-21 | -0.6488694 | 0.015 | 0.236 | 4.7704E-17 | TEC_prlf | HCST |
| 2.0362E-21 | 0.48416487 | 0.924 | 0.754 | 4.8101E-17 | TEC_prlf | RPSA |
| 2.1983E-21 | -0.872004 | 0.023 | 0.246 | 5.193E-17 | TEC_prlf | SLC11A1 |
| 2.4401E-21 | 0.45482828 | 0.863 | 0.687 | 5.7643E-17 | TEC_prlf | PSMA7 |
| 2.4872E-21 | 0.2850532 | 0.315 | 0.137 | 5.8755E-17 | TEC_prlf | MGLL |
| 3.3838E-21 | 0.32466211 | 0.324 | 0.147 | 7.9936E-17 | TEC_prlf | USP14 |
| 3.6683E-21 | -0.5673934 | 0.137 | 0.376 | 8.6656E-17 | TEC_prlf | METRNL |
| 4.4145E-21 | 0.64929476 | 0.577 | 0.356 | 1.0428E-16 | TEC_prlf | MIF |
| 4.4553E-21 | 0.40863108 | 0.458 | 0.255 | 1.0525E-16 | TEC_prlf | NDUFB10 |
| 4.5759E-21 | 0.30243066 | 0.292 | 0.126 | 1.081E-16 | TEC_prlf | SMC1A |
| 4.6668E-21 | -1.1054139 | 0.006 | 0.217 | 1.1024E-16 | TEC_prlf | S100A12 |
| 4.8965E-21 | -1.1696577 | 0.472 | 0.659 | 1.1567E-16 | TEC_prlf | HSPA1A |
| 5.1541E-21 | -1.6862011 | 0.012 | 0.224 | 1.2176E-16 | TEC_prlf | VCAN |
| 5.5805E-21 | 0.35365894 | 0.274 | 0.117 | 1.3183E-16 | TEC_prlf | HCFC1R1 |
| 5.6358E-21 | 0.26194934 | 0.254 | 0.102 | 1.3313E-16 | TEC_prlf | ITGAE |
| 5.861E-21 | 0.41748334 | 0.81 | 0.604 | 1.3845E-16 | TEC_prlf | EZR |
| 5.8766E-21 | 0.49345716 | 0.936 | 0.884 | 1.3882E-16 | TEC_prlf | MYL12B |
| 5.984E-21 | -0.9820035 | 0.082 | 0.314 | 1.4136E-16 | TEC_prlf | PHLDA1 |
| 6.6304E-21 | 0.25431905 | 0.23 | 0.088 | 1.5663E-16 | TEC_prlf | CETN2 |
| 6.9477E-21 | 0.30410782 | 0.265 | 0.108 | 1.6412E-16 | TEC_prlf | WWTR1 |
| 9.7996E-21 | 0.28254574 | 0.265 | 0.11 | 2.315E-16 | TEC_prlf | ACAA2 |
| 1.021E-20 | 0.35407046 | 0.335 | 0.158 | 2.4119E-16 | TEC_prlf | TMEM160 |
| 1.0475E-20 | 0.39148006 | 0.37 | 0.186 | 2.4746E-16 | TEC_prlf | AP2B1 |
| 1.1213E-20 | 0.41775416 | 0.388 | 0.2 | 2.6489E-16 | TEC_prlf | STIP1 |
| 1.2563E-20 | -0.5427729 | 0.026 | 0.247 | 2.9677E-16 | TEC_prlf | GMFG |
| 1.5894E-20 | 0.27488893 | 0.254 | 0.103 | 3.7547E-16 | TEC_prlf | PARP1 |

| | | | | | | |
|---|---|---|---|---|---|---|
| 1.6168E-20 | 0.42407665 | 0.569 | 0.352 | 3.8194E-16 | TEC_prlf | MRPL33 |
| 1.7048E-20 | 0.47579635 | 0.574 | 0.356 | 4.0272E-16 | TEC_prlf | ATP1A1 |
| 2.0498E-20 | 0.43060436 | 0.574 | 0.362 | 4.8423E-16 | TEC_prlf | ATP5F1D |
| 2.3289E-20 | 0.26886281 | 0.122 | 0.033 | 5.5015E-16 | TEC_prlf | PPP1R1A |
| 2.7895E-20 | 0.5039175 | 0.3 | 0.136 | 6.5896E-16 | TEC_prlf | NUPR1 |
| 3.1829E-20 | 0.33984593 | 0.344 | 0.165 | 7.519E-16 | TEC_prlf | GTF2H5 |
| 3.818E-20 | 0.46822637 | 0.615 | 0.407 | 9.0193E-16 | TEC_prlf | CD59 |
| 4.0551E-20 | 0.2526245 | 0.207 | 0.075 | 9.5793E-16 | TEC_prlf | TFCP2L1 |
| 4.0901E-20 | -0.8442224 | 0.968 | 0.98 | 9.662E-16 | TEC_prlf | MALAT1 |
| 4.6112E-20 | -0.493917 | 0.017 | 0.226 | 1.0893E-15 | TEC_prlf | KYNU |
| 4.744E-20 | 0.47418088 | 0.402 | 0.212 | 1.1207E-15 | TEC_prlf | METAP2 |
| 4.9258E-20 | -0.5480964 | 0.012 | 0.217 | 1.1636E-15 | TEC_prlf | SPI1 |
| 5.0133E-20 | 0.30006796 | 0.286 | 0.125 | 1.1843E-15 | TEC_prlf | HDAC1 |
| 5.225E-20 | 0.42206258 | 0.493 | 0.289 | 1.2343E-15 | TEC_prlf | HNRNPH3 |
| 5.8843E-20 | 0.25129817 | 0.213 | 0.08 | 1.3901E-15 | TEC_prlf | TWSG1 |
| 5.9118E-20 | 0.30053818 | 0.265 | 0.113 | 1.3965E-15 | TEC_prlf | RSRC1 |
| 6.0464E-20 | -1.1740849 | 0.741 | 0.54 | 1.4283E-15 | TEC_prlf | PLCG2 |
| 6.7041E-20 | -0.7199798 | 0.009 | 0.211 | 1.5837E-15 | TEC_prlf | TNFRSF1B |
| 6.7252E-20 | 0.30835464 | 0.292 | 0.13 | 1.5887E-15 | TEC_prlf | SMARCA4 |
| 7.1727E-20 | -0.3981854 | 0.02 | 0.228 | 1.6944E-15 | TEC_prlf | TPP1 |
| 9.2028E-20 | 0.26965595 | 0.312 | 0.143 | 2.174E-15 | TEC_prlf | PIN1 |
| 1.0026E-19 | 0.40016014 | 0.362 | 0.188 | 2.3684E-15 | TEC_prlf | PSMD2 |
| 1.0863E-19 | -0.6760317 | 0.399 | 0.604 | 2.5662E-15 | TEC_prlf | MCL1 |
| 1.2784E-19 | 0.33162852 | 0.324 | 0.154 | 3.02E-15 | TEC_prlf | CHCHD5 |
| 1.3813E-19 | 0.38677704 | 0.452 | 0.253 | 3.2631E-15 | TEC_prlf | CCT6A |
| 1.5641E-19 | 0.26049964 | 0.245 | 0.101 | 3.6948E-15 | TEC_prlf | CDC123 |
| 1.6627E-19 | 0.29658184 | 0.268 | 0.115 | 3.9278E-15 | TEC_prlf | ASXL1 |
| 1.8097E-19 | 0.40355398 | 0.475 | 0.264 | 4.2751E-15 | TEC_prlf | AFDN |
| 2.1291E-19 | 0.48765485 | 0.458 | 0.273 | 5.0296E-15 | TEC_prlf | PDXK |
| 2.5049E-19 | 0.42012734 | 0.98 | 0.958 | 5.9174E-15 | TEC_prlf | RPL15 |
| 2.5553E-19 | 0.25896221 | 0.286 | 0.126 | 6.0364E-15 | TEC_prlf | UPF3A |
| 2.7438E-19 | 0.49536826 | 0.566 | 0.378 | 6.4817E-15 | TEC_prlf | EIF5A |
| 2.8702E-19 | -0.6380798 | 0.012 | 0.21 | 6.7802E-15 | TEC_prlf | LST1 |
| 3.095E-19 | 0.30073246 | 0.292 | 0.131 | 7.3114E-15 | TEC_prlf | ROCK2 |
| 3.6681E-19 | 0.41910096 | 0.499 | 0.299 | 8.6651E-15 | TEC_prlf | ATP5PO |
| 3.7259E-19 | -0.7635556 | 0.746 | 0.838 | 8.8016E-15 | TEC_prlf | SQSTM1 |
| 4.0684E-19 | 0.26767943 | 0.271 | 0.118 | 9.6107E-15 | TEC_prlf | LSM2 |
| 4.7032E-19 | -1.0255813 | 0.02 | 0.221 | 1.111E-14 | TEC_prlf | ALOX5AP |
| 4.7183E-19 | 0.57887429 | 0.571 | 0.367 | 1.1146E-14 | TEC_prlf | ANKRD11 |
| 4.9894E-19 | -0.7483847 | 0.636 | 0.765 | 1.1786E-14 | TEC_prlf | BTG1 |
| 5.0989E-19 | -0.9255658 | 0.07 | 0.287 | 1.2045E-14 | TEC_prlf | HMOX1 |
| 5.346E-19 | 0.37845352 | 0.458 | 0.263 | 1.2629E-14 | TEC_prlf | PSMA4 |
| 5.5251E-19 | -1.2018184 | 0.067 | 0.278 | 1.3052E-14 | TEC_prlf | HLA-DRB1 |
| 5.6497E-19 | 0.42692028 | 0.382 | 0.202 | 1.3346E-14 | TEC_prlf | CAPRIN1 |
| 5.789E-19 | 0.39876712 | 0.977 | 0.917 | 1.3675E-14 | TEC_prlf | RPS20 |
| 6.4299E-19 | 0.40647117 | 0.402 | 0.215 | 1.5189E-14 | TEC_prlf | RTF1 |
| 6.8668E-19 | -0.4876558 | 0.102 | 0.327 | 1.6221E-14 | TEC_prlf | FOSL2 |
| 7.3927E-19 | 0.38146565 | 0.382 | 0.205 | 1.7464E-14 | TEC_prlf | NAA38 |
| 8.46E-19 | -0.5707405 | 0.793 | 0.869 | 1.9985E-14 | TEC_prlf | PABPC1 |
| 8.5243E-19 | 0.30181669 | 0.274 | 0.122 | 2.0137E-14 | TEC_prlf | DNAJC19 |

| | | | | | | |
|---|---|---|---|---|---|---|
| 9.3561E-19 | 0.2737327 | 0.268 | 0.118 | 2.2102E-14 | TEC_prlf | VBP1 |
| 9.6378E-19 | -0.5316766 | 0.102 | 0.327 | 2.2767E-14 | TEC_prlf | ARHGDIB |
| 9.7312E-19 | 0.35309237 | 0.431 | 0.235 | 2.2988E-14 | TEC_prlf | PCBD1 |
| 1.0511E-18 | 0.39619016 | 0.522 | 0.324 | 2.4831E-14 | TEC_prlf | SNRPG |
| 1.4102E-18 | 0.31913992 | 0.385 | 0.202 | 3.3314E-14 | TEC_prlf | TXNL4A |
| 1.8144E-18 | -0.6223119 | 0.528 | 0.699 | 4.2862E-14 | TEC_prlf | PPP1R15A |
| 1.8638E-18 | -0.6122995 | 0.362 | 0.574 | 4.4029E-14 | TEC_prlf | VMP1 |
| 1.8788E-18 | -0.7164623 | 0.012 | 0.202 | 4.4383E-14 | TEC_prlf | CYBB |
| 1.9413E-18 | 0.34253474 | 0.344 | 0.172 | 4.5859E-14 | TEC_prlf | PSMC3 |
| 1.9622E-18 | 0.46658794 | 0.475 | 0.292 | 4.6352E-14 | TEC_prlf | HDLBP |
| 2.0346E-18 | 0.34793261 | 0.367 | 0.181 | 4.8064E-14 | TEC_prlf | CDH1 |
| 2.113E-18 | -0.4724376 | 0.044 | 0.245 | 4.9915E-14 | TEC_prlf | RAB31 |
| 2.3495E-18 | 0.25663235 | 0.259 | 0.112 | 5.5503E-14 | TEC_prlf | DMAC1 |
| 2.4097E-18 | -0.5502386 | 0.012 | 0.201 | 5.6925E-14 | TEC_prlf | ITGAX |
| 2.8005E-18 | 0.26876702 | 0.286 | 0.129 | 6.6157E-14 | TEC_prlf | TMCO3 |
| 2.8924E-18 | -0.4819281 | 0.07 | 0.282 | 6.8326E-14 | TEC_prlf | LYN |
| 2.9565E-18 | -0.5737254 | 0.02 | 0.214 | 6.9841E-14 | TEC_prlf | BASP1 |
| 3.1247E-18 | 0.25394916 | 0.187 | 0.069 | 7.3816E-14 | TEC_prlf | TSPAN15 |
| 3.2778E-18 | 0.39282952 | 0.49 | 0.29 | 7.7433E-14 | TEC_prlf | HNRNPAB |
| 3.4931E-18 | -0.5715688 | 0.061 | 0.271 | 8.2517E-14 | TEC_prlf | TSC22D3 |
| 3.8631E-18 | 0.25744122 | 0.227 | 0.093 | 9.1257E-14 | TEC_prlf | UBAC1 |
| 4.2355E-18 | -0.5765169 | 0.026 | 0.221 | 1.0005E-13 | TEC_prlf | ITGB2 |
| 4.8315E-18 | 0.29316067 | 0.312 | 0.15 | 1.1413E-13 | TEC_prlf | IDH2 |
| 5.1789E-18 | 0.28505573 | 0.268 | 0.12 | 1.2234E-13 | TEC_prlf | SUMF2 |
| 5.2544E-18 | -0.4414524 | 0.586 | 0.742 | 1.2412E-13 | TEC_prlf | SERP1 |
| 5.6025E-18 | 0.37759965 | 0.411 | 0.229 | 1.3235E-13 | TEC_prlf | ARPC5L |
| 6.2163E-18 | -0.7772956 | 0.006 | 0.189 | 1.4685E-13 | TEC_prlf | AC020656.1 |
| 6.4565E-18 | 0.74663116 | 0.776 | 0.616 | 1.5252E-13 | TEC_prlf | LGALS1 |
| 7.0456E-18 | -0.5296205 | 0.006 | 0.19 | 1.6644E-13 | TEC_prlf | FCGR2A |
| 7.6569E-18 | 0.46696 | 0.659 | 0.493 | 1.8088E-13 | TEC_prlf | SRSF3 |
| 8.1923E-18 | 0.38826838 | 0.452 | 0.266 | 1.9353E-13 | TEC_prlf | KHDRBS1 |
| 9.7377E-18 | 0.26979416 | 0.268 | 0.12 | 2.3003E-13 | TEC_prlf | HAGH |
| 1.0014E-17 | -0.5515521 | 0.02 | 0.208 | 2.3657E-13 | TEC_prlf | GK |
| 1.1557E-17 | -1.0721876 | 0.277 | 0.475 | 2.7301E-13 | TEC_prlf | ERO1A |
| 1.1851E-17 | 0.37805848 | 0.356 | 0.188 | 2.7996E-13 | TEC_prlf | ABCF1 |
| 1.3805E-17 | 0.25818615 | 0.248 | 0.108 | 3.2613E-13 | TEC_prlf | ZDHHC12 |
| 1.4579E-17 | -0.9200808 | 0.837 | 0.884 | 3.444E-13 | TEC_prlf | CSTB |
| 1.6229E-17 | -0.6092338 | 0.006 | 0.186 | 3.8337E-13 | TEC_prlf | C5AR1 |
| 1.6477E-17 | -1.7325822 | 0.198 | 0.398 | 3.8923E-13 | TEC_prlf | HLA-DRA |
| 1.7998E-17 | -0.3617894 | 0.006 | 0.185 | 4.2516E-13 | TEC_prlf | GNA15 |
| 2.9429E-17 | 0.25131846 | 0.233 | 0.1 | 6.9519E-13 | TEC_prlf | TMX2 |
| 3.2305E-17 | -0.6196953 | 0.017 | 0.2 | 7.6314E-13 | TEC_prlf | DUSP4 |
| 3.3943E-17 | 0.36481425 | 0.341 | 0.178 | 8.0183E-13 | TEC_prlf | THOC7 |
| 3.7556E-17 | -0.3873486 | 0.58 | 0.748 | 8.8719E-13 | TEC_prlf | ATP6V0E1 |
| 3.8648E-17 | 0.38983712 | 0.988 | 0.895 | 9.1299E-13 | TEC_prlf | RPS9 |
| 4.7528E-17 | -0.9733827 | 0.236 | 0.426 | 1.1228E-12 | TEC_prlf | EMP3 |
| 4.7965E-17 | -0.6987243 | 0.006 | 0.181 | 1.1331E-12 | TEC_prlf | OLR1 |
| 4.8651E-17 | -0.5995383 | 0.131 | 0.347 | 1.1493E-12 | TEC_prlf | MIDN |
| 4.8906E-17 | 0.4413144 | 0.985 | 0.919 | 1.1553E-12 | TEC_prlf | RPS15 |
| 4.9352E-17 | -1.0486928 | 0.373 | 0.54 | 1.1658E-12 | TEC_prlf | PLAUR |

| | | | | | | |
|---|---|---|---|---|---|---|
| 5.7357E-17 | 0.38234431 | 0.475 | 0.286 | 1.355E-12 | TEC_prlf | NDUFC1 |
| 5.7516E-17 | 0.27481465 | 0.306 | 0.15 | 1.3587E-12 | TEC_prlf | BCL7C |
| 6.5072E-17 | 0.30964631 | 0.321 | 0.16 | 1.5372E-12 | TEC_prlf | NASP |
| 6.9031E-17 | 0.31978138 | 0.977 | 0.932 | 1.6307E-12 | TEC_prlf | GAPDH |
| 7.955E-17 | 0.4497026 | 0.51 | 0.332 | 1.8792E-12 | TEC_prlf | HNRNPD |
| 8.4592E-17 | 0.25553841 | 0.236 | 0.102 | 1.9983E-12 | TEC_prlf | AK6 |
| 9.0055E-17 | 0.35052964 | 0.848 | 0.713 | 2.1274E-12 | TEC_prlf | ATP5MG |
| 9.1459E-17 | -0.3547285 | 0.012 | 0.188 | 2.1605E-12 | TEC_prlf | GPSM3 |
| 9.2793E-17 | 0.332894 | 0.417 | 0.223 | 2.192E-12 | TEC_prlf | OCIAD2 |
| 9.5667E-17 | 0.26151245 | 0.227 | 0.098 | 2.26E-12 | TEC_prlf | TXLNA |
| 1.023E-16 | 0.36849775 | 0.466 | 0.283 | 2.4167E-12 | TEC_prlf | PSMB6 |
| 1.0451E-16 | -0.9703609 | 0.085 | 0.28 | 2.4688E-12 | TEC_prlf | DUSP5 |
| 1.0564E-16 | 0.31462083 | 0.312 | 0.156 | 2.4956E-12 | TEC_prlf | SUPT16H |
| 1.0764E-16 | -0.8460201 | 0.082 | 0.283 | 2.5429E-12 | TEC_prlf | MAFB |
| 1.0976E-16 | 0.40215296 | 0.525 | 0.329 | 2.5929E-12 | TEC_prlf | PPP1R14B |
| 1.1341E-16 | 0.32740652 | 0.942 | 0.78 | 2.679E-12 | TEC_prlf | RPL10A |
| 1.211E-16 | 0.29886659 | 0.306 | 0.152 | 2.8606E-12 | TEC_prlf | CCAR1 |
| 1.2325E-16 | -0.5621611 | 0.012 | 0.187 | 2.9114E-12 | TEC_prlf | KLF2 |
| 1.2684E-16 | 0.36892673 | 0.883 | 0.793 | 2.9964E-12 | TEC_prlf | CHCHD2 |
| 1.4339E-16 | -0.8491554 | 0.178 | 0.391 | 3.3874E-12 | TEC_prlf | FOS |
| 1.527E-16 | -0.895954 | 0.093 | 0.287 | 3.6073E-12 | TEC_prlf | SLC25A37 |
| 1.818E-16 | 0.31492555 | 0.394 | 0.213 | 4.2947E-12 | TEC_prlf | CNN3 |
| 1.9206E-16 | 0.28640175 | 0.128 | 0.04 | 4.5371E-12 | TEC_prlf | AKR1C2 |
| 2.2421E-16 | -0.5666868 | 0.012 | 0.184 | 5.2965E-12 | TEC_prlf | IGSF6 |
| 2.4679E-16 | 0.36119881 | 0.169 | 0.062 | 5.83E-12 | TEC_prlf | AKR1C1 |
| 2.7401E-16 | -0.4655792 | 0.006 | 0.174 | 6.473E-12 | TEC_prlf | SMIM25 |
| 2.8136E-16 | 0.41180176 | 0.475 | 0.296 | 6.6465E-12 | TEC_prlf | UBALD2 |
| 2.8674E-16 | 0.34179971 | 0.385 | 0.215 | 6.7737E-12 | TEC_prlf | U2SURP |
| 4.8806E-16 | 0.36355057 | 0.569 | 0.365 | 1.1529E-11 | TEC_prlf | AES |
| 5.3364E-16 | 0.36893535 | 0.417 | 0.25 | 1.2606E-11 | TEC_prlf | XRCC6 |
| 5.4683E-16 | 0.49656082 | 0.569 | 0.409 | 1.2918E-11 | TEC_prlf | ATP5PF |
| 5.5357E-16 | -0.7948701 | 0.006 | 0.171 | 1.3077E-11 | TEC_prlf | FCN1 |
| 5.6006E-16 | 0.3238131 | 0.297 | 0.148 | 1.323E-11 | TEC_prlf | TAF15 |
| 5.6815E-16 | -0.4740691 | 0 | 0.162 | 1.3421E-11 | TEC_prlf | AQP9 |
| 5.9453E-16 | 0.28827107 | 0.341 | 0.179 | 1.4045E-11 | TEC_prlf | LARP7 |
| 6.1521E-16 | 0.42930269 | 0.563 | 0.385 | 1.4533E-11 | TEC_prlf | PSME1 |
| 6.4476E-16 | 0.33718159 | 0.496 | 0.3 | 1.5231E-11 | TEC_prlf | PSME2 |
| 6.4828E-16 | -0.6523975 | 0.149 | 0.348 | 1.5314E-11 | TEC_prlf | IER5 |
| 6.7806E-16 | 0.32530035 | 0.446 | 0.265 | 1.6018E-11 | TEC_prlf | SYNCRIP |
| 7.1889E-16 | -0.5263502 | 0.303 | 0.495 | 1.6982E-11 | TEC_prlf | RSRP1 |
| 7.6108E-16 | 0.28628878 | 0.257 | 0.118 | 1.7979E-11 | TEC_prlf | DAPK1 |
| 8.4748E-16 | 0.40013768 | 0.464 | 0.288 | 2.002E-11 | TEC_prlf | NDUFB3 |
| 8.5632E-16 | 0.33848238 | 0.373 | 0.207 | 2.0229E-11 | TEC_prlf | CNIH4 |
| 8.5665E-16 | 0.42933469 | 0.51 | 0.346 | 2.0237E-11 | TEC_prlf | SNRPB |
| 9.5835E-16 | 0.37227063 | 0.822 | 0.713 | 2.2639E-11 | TEC_prlf | SKP1 |
| 1.0083E-15 | -2.2037745 | 0.012 | 0.177 | 2.3819E-11 | TEC_prlf | IL1B |
| 1.0098E-15 | 0.29652403 | 0.469 | 0.28 | 2.3855E-11 | TEC_prlf | TCEA1 |
| 1.1547E-15 | 0.40432397 | 0.464 | 0.29 | 2.7278E-11 | TEC_prlf | NDUFS6 |
| 1.1874E-15 | 0.3112775 | 0.44 | 0.262 | 2.805E-11 | TEC_prlf | RTRAF |
| 1.1915E-15 | -0.4692539 | 0.064 | 0.249 | 2.8146E-11 | TEC_prlf | GPCPD1 |

| | | | | | | |
|---|---|---|---|---|---|---|
| 1.2119E-15 | -0.3928918 | 0.035 | 0.21 | 2.8628E-11 | TEC_prlf | ARRB2 |
| 1.217E-15 | 0.32155818 | 0.994 | 0.911 | 2.875E-11 | TEC_prlf | RPS8 |
| 1.2262E-15 | 0.4247964 | 0.466 | 0.292 | 2.8966E-11 | TEC_prlf | COX17 |
| 1.2277E-15 | -0.660806 | 0.429 | 0.579 | 2.9002E-11 | TEC_prlf | LITAF |
| 1.2745E-15 | -1.2714731 | 0.006 | 0.167 | 3.0108E-11 | TEC_prlf | DUOXA2 |
| 1.3602E-15 | -0.4554217 | 0.105 | 0.297 | 3.2133E-11 | TEC_prlf | IFNGR2 |
| 1.4105E-15 | 0.32866352 | 0.265 | 0.128 | 3.3321E-11 | TEC_prlf | CBR1 |
| 1.508E-15 | -0.5408697 | 0.417 | 0.603 | 3.5625E-11 | TEC_prlf | NFKBIA |
| 1.5177E-15 | -0.7406468 | 0.012 | 0.176 | 3.5852E-11 | TEC_prlf | PLEK |
| 1.6324E-15 | 0.30638598 | 0.297 | 0.15 | 3.8562E-11 | TEC_prlf | MRPL34 |
| 1.6552E-15 | 0.35949038 | 0.501 | 0.324 | 3.91E-11 | TEC_prlf | SRP9 |
| 1.7098E-15 | 0.39493852 | 0.551 | 0.374 | 4.0391E-11 | TEC_prlf | SRSF9 |
| 1.7891E-15 | 0.36408171 | 0.697 | 0.494 | 4.2263E-11 | TEC_prlf | CD9 |
| 1.8605E-15 | -0.5137802 | 0.017 | 0.183 | 4.3951E-11 | TEC_prlf | SAMSN1 |
| 1.8698E-15 | 0.30989297 | 0.338 | 0.179 | 4.4171E-11 | TEC_prlf | POLR2I |
| 1.91E-15 | -0.3523437 | 0.079 | 0.267 | 4.512E-11 | TEC_prlf | CDC42SE1 |
| 1.9728E-15 | 0.30900792 | 0.434 | 0.256 | 4.6604E-11 | TEC_prlf | RPS19BP1 |
| 2.1711E-15 | 0.25817128 | 0.402 | 0.224 | 5.1287E-11 | TEC_prlf | ARPP19 |
| 2.2639E-15 | 0.28111642 | 0.294 | 0.146 | 5.3481E-11 | TEC_prlf | MRPL43 |
| 2.3768E-15 | -0.5932353 | 0.026 | 0.199 | 5.6147E-11 | TEC_prlf | CD52 |
| 2.8852E-15 | 0.33245402 | 0.446 | 0.266 | 6.8158E-11 | TEC_prlf | GADD45GIP1 |
| 3.0059E-15 | -1.3839142 | 0.029 | 0.197 | 7.1009E-11 | TEC_prlf | HOPX |
| 3.0383E-15 | 0.42104152 | 0.466 | 0.272 | 7.1774E-11 | TEC_prlf | MAL2 |
| 3.2185E-15 | -0.5623148 | 0.003 | 0.159 | 7.603E-11 | TEC_prlf | CD300E |
| 3.5995E-15 | 0.38261742 | 0.962 | 0.869 | 8.5031E-11 | TEC_prlf | RPLP0 |
| 3.6963E-15 | 0.38647249 | 0.472 | 0.295 | 8.7317E-11 | TEC_prlf | EIF5B |
| 3.8436E-15 | -0.6223765 | 0.012 | 0.173 | 9.0798E-11 | TEC_prlf | STEAP4 |
| 3.922E-15 | 0.35787408 | 0.461 | 0.286 | 9.2648E-11 | TEC_prlf | ACADVL |
| 3.9597E-15 | 0.29249104 | 0.726 | 0.522 | 9.354E-11 | TEC_prlf | HSPE1 |
| 4.1799E-15 | -0.4003126 | 0.032 | 0.204 | 9.8742E-11 | TEC_prlf | LSP1 |
| 4.1987E-15 | 0.29990214 | 0.353 | 0.193 | 9.9186E-11 | TEC_prlf | BRD7 |
| 4.3066E-15 | -0.4694064 | 0.65 | 0.782 | 1.0173E-10 | TEC_prlf | MAP1LC3B |
| 4.7102E-15 | 0.3440738 | 0.895 | 0.762 | 1.1127E-10 | TEC_prlf | COX7C |
| 5.2036E-15 | -0.636508 | 0.041 | 0.209 | 1.2292E-10 | TEC_prlf | CDKN2B |
| 5.2352E-15 | -0.4777371 | 0.37 | 0.548 | 1.2367E-10 | TEC_prlf | C4orf3 |
| 6.1957E-15 | 0.32680459 | 0.265 | 0.127 | 1.4636E-10 | TEC_prlf | UTRN |
| 6.3373E-15 | -0.422979 | 0.009 | 0.165 | 1.4971E-10 | TEC_prlf | LCP2 |
| 6.8022E-15 | -0.439738 | 0.012 | 0.169 | 1.6069E-10 | TEC_prlf | SLA |
| 6.8444E-15 | -0.38119 | 0.012 | 0.17 | 1.6168E-10 | TEC_prlf | CD48 |
| 7.2711E-15 | -0.3416972 | 0.023 | 0.188 | 1.7176E-10 | TEC_prlf | HCLS1 |
| 7.2834E-15 | 0.26634344 | 0.289 | 0.144 | 1.7206E-10 | TEC_prlf | TUBB6 |
| 7.5434E-15 | 0.3190538 | 0.391 | 0.227 | 1.782E-10 | TEC_prlf | THRAP3 |
| 7.9029E-15 | -0.4495584 | 0.12 | 0.307 | 1.8669E-10 | TEC_prlf | BNIP3L |
| 8.0254E-15 | -0.6978922 | 0.14 | 0.325 | 1.8958E-10 | TEC_prlf | VEGFA |
| 8.0873E-15 | 0.28394988 | 0.429 | 0.251 | 1.9105E-10 | TEC_prlf | TMED2 |
| 8.3759E-15 | -0.4887504 | 0.035 | 0.202 | 1.9786E-10 | TEC_prlf | NCF2 |
| 9.6048E-15 | -0.3387049 | 0.017 | 0.177 | 2.2689E-10 | TEC_prlf | ABCA1 |
| 1.06E-14 | -0.759952 | 0 | 0.15 | 2.504E-10 | TEC_prlf | TMPRSS11D |
| 1.162E-14 | -0.4350794 | 0.026 | 0.188 | 2.7451E-10 | TEC_prlf | PTPRE |
| 1.2177E-14 | 0.3794867 | 0.56 | 0.382 | 2.8766E-10 | TEC_prlf | DYNLRB1 |

| | | | | | | |
|---|---|---|---|---|---|---|
| 1.3475E-14 | 0.45070757 | 0.615 | 0.446 | 3.1832E-10 | TEC_prlf | ACTN4 |
| 1.3813E-14 | -0.4479126 | 0.079 | 0.26 | 3.2631E-10 | TEC_prlf | NABP1 |
| 1.3889E-14 | 0.26209086 | 0.411 | 0.234 | 3.281E-10 | TEC_prlf | PSMA3 |
| 1.5187E-14 | -0.3535413 | 0.052 | 0.229 | 3.5876E-10 | TEC_prlf | TRIM56 |
| 1.5255E-14 | 0.41527262 | 0.805 | 0.672 | 3.6038E-10 | TEC_prlf | CALR |
| 1.5895E-14 | -0.5037736 | 0.003 | 0.152 | 3.755E-10 | TEC_prlf | CLEC5A |
| 1.6382E-14 | -0.8232806 | 0.009 | 0.163 | 3.8699E-10 | TEC_prlf | APOBEC3A |
| 1.7348E-14 | 0.27664481 | 0.327 | 0.175 | 4.0981E-10 | TEC_prlf | POLR2F |
| 1.8195E-14 | 0.30267599 | 0.283 | 0.141 | 4.2981E-10 | TEC_prlf | LAPTM4B |
| 2.0986E-14 | 0.37265067 | 0.86 | 0.707 | 4.9575E-10 | TEC_prlf | GPX4 |
| 2.1507E-14 | -0.5641836 | 0.513 | 0.679 | 5.0807E-10 | TEC_prlf | IER2 |
| 2.2767E-14 | 0.35911053 | 0.883 | 0.745 | 5.3782E-10 | TEC_prlf | PRDX1 |
| 2.3787E-14 | 0.53497762 | 0.469 | 0.323 | 5.6191E-10 | TEC_prlf | ARL6IP1 |
| 2.4407E-14 | -0.6042611 | 0.464 | 0.632 | 5.7657E-10 | TEC_prlf | ZFP36L1 |
| 2.4603E-14 | -0.8467283 | 0.029 | 0.189 | 5.8119E-10 | TEC_prlf | TMPRSS2 |
| 2.5895E-14 | 0.42568052 | 0.464 | 0.306 | 6.1171E-10 | TEC_prlf | YY1 |
| 2.9329E-14 | 0.31064204 | 0.44 | 0.273 | 6.9283E-10 | TEC_prlf | AP2M1 |
| 3.274E-14 | 0.33567529 | 0.402 | 0.238 | 7.7341E-10 | TEC_prlf | C12orf57 |
| 3.3424E-14 | -0.5081008 | 0.017 | 0.171 | 7.8958E-10 | TEC_prlf | PRDM1 |
| 3.4794E-14 | -0.3794786 | 0.023 | 0.18 | 8.2193E-10 | TEC_prlf | CD37 |
| 3.5589E-14 | -1.5894484 | 0.245 | 0.409 | 8.4073E-10 | TEC_prlf | S100P |
| 3.7823E-14 | 0.44303979 | 0.545 | 0.376 | 8.935E-10 | TEC_prlf | POLR2L |
| 3.8917E-14 | -1.0996689 | 0.012 | 0.162 | 9.1934E-10 | TEC_prlf | MUC21 |
| 4.8133E-14 | 0.26585529 | 0.318 | 0.17 | 1.1371E-09 | TEC_prlf | PPP1CC |
| 5.003E-14 | -0.6381713 | 0.035 | 0.193 | 1.1819E-09 | TEC_prlf | FBXO32 |
| 5.0843E-14 | 0.27673255 | 0.239 | 0.114 | 1.2011E-09 | TEC_prlf | FRMD4B |
| 5.37E-14 | -0.3762927 | 0.003 | 0.147 | 1.2686E-09 | TEC_prlf | TREM1 |
| 5.4586E-14 | 0.29661019 | 0.382 | 0.221 | 1.2895E-09 | TEC_prlf | PPP2CB |
| 5.5604E-14 | 0.40297469 | 0.458 | 0.297 | 1.3135E-09 | TEC_prlf | GTF2I |
| 5.8095E-14 | 0.37084277 | 0.758 | 0.615 | 1.3724E-09 | TEC_prlf | KTN1 |
| 6.1983E-14 | 0.34519822 | 0.408 | 0.248 | 1.4642E-09 | TEC_prlf | CHD4 |
| 6.4089E-14 | -0.4275263 | 0.003 | 0.147 | 1.514E-09 | TEC_prlf | FPR1 |
| 6.9284E-14 | 0.27188263 | 0.283 | 0.145 | 1.6367E-09 | TEC_prlf | COA4 |
| 7.4049E-14 | 0.36069424 | 0.417 | 0.265 | 1.7493E-09 | TEC_prlf | VPS29 |
| 7.4158E-14 | 0.36228829 | 0.723 | 0.546 | 1.7518E-09 | TEC_prlf | PPIB |
| 7.5723E-14 | 0.46293256 | 0.513 | 0.362 | 1.7888E-09 | TEC_prlf | SLC25A5 |
| 8.287E-14 | -1.193486 | 0.181 | 0.346 | 1.9576E-09 | TEC_prlf | CTSL |
| 9.8134E-14 | 0.3292145 | 0.37 | 0.216 | 2.3182E-09 | TEC_prlf | PPM1G |
| 1.0732E-13 | 0.32447884 | 0.481 | 0.313 | 2.5352E-09 | TEC_prlf | EIF3F |
| 1.0983E-13 | 0.28756081 | 0.385 | 0.213 | 2.5945E-09 | TEC_prlf | EHF |
| 1.1121E-13 | 0.35101926 | 0.735 | 0.595 | 2.6272E-09 | TEC_prlf | HNRNPK |
| 1.2071E-13 | 0.38507401 | 0.688 | 0.55 | 2.8514E-09 | TEC_prlf | UQCR10 |
| 1.2183E-13 | 0.28013268 | 0.286 | 0.149 | 2.8781E-09 | TEC_prlf | CENPX |
| 1.2233E-13 | 0.28292927 | 0.971 | 0.816 | 2.8898E-09 | TEC_prlf | RPL5 |
| 1.3523E-13 | 0.37505773 | 0.548 | 0.373 | 3.1945E-09 | TEC_prlf | PARK7 |
| 1.3886E-13 | 0.3041931 | 0.315 | 0.173 | 3.2804E-09 | TEC_prlf | CYC1 |
| 1.4445E-13 | -0.9605872 | 0.012 | 0.156 | 3.4124E-09 | TEC_prlf | PITX1 |
| 1.4845E-13 | 0.28398837 | 0.443 | 0.273 | 3.5068E-09 | TEC_prlf | SSB |
| 1.5375E-13 | -0.3410388 | 0.061 | 0.228 | 3.6321E-09 | TEC_prlf | RGS10 |
| 1.5574E-13 | -1.1521214 | 0.047 | 0.205 | 3.679E-09 | TEC_prlf | HLA-DPA1 |

| | | | | | | |
|---|---|---|---|---|---|---|
| 1.5772E-13 | -0.3075272 | 0.009 | 0.153 | 3.7257E-09 | TEC_prlf | HK2 |
| 1.6117E-13 | -0.352912 | 0.009 | 0.153 | 3.8073E-09 | TEC_prlf | EVI2B |
| 1.9322E-13 | 0.44482969 | 0.63 | 0.435 | 4.5644E-09 | TEC_prlf | ATP1B1 |
| 1.9735E-13 | 0.32864313 | 0.449 | 0.282 | 4.6621E-09 | TEC_prlf | TPR |
| 2.0012E-13 | -0.5252221 | 0.102 | 0.276 | 4.7275E-09 | TEC_prlf | ADM |
| 2.0538E-13 | -1.8265722 | 0.187 | 0.342 | 4.8517E-09 | TEC_prlf | PSCA |
| 2.1634E-13 | 0.3212751 | 0.95 | 0.799 | 5.1106E-09 | TEC_prlf | RPS5 |
| 2.1778E-13 | -0.4040007 | 0.006 | 0.146 | 5.1447E-09 | TEC_prlf | MS4A7 |
| 2.262E-13 | 0.25676283 | 0.312 | 0.167 | 5.3436E-09 | TEC_prlf | METTL26 |
| 2.2965E-13 | 0.30506029 | 0.991 | 0.945 | 5.4251E-09 | TEC_prlf | RPS19 |
| 2.514E-13 | -0.3272357 | 0.082 | 0.251 | 5.9389E-09 | TEC_prlf | IDS |
| 2.5441E-13 | -0.3191912 | 0.017 | 0.164 | 6.01E-09 | TEC_prlf | MYO1F |
| 2.6207E-13 | 0.25472344 | 0.297 | 0.158 | 6.191E-09 | TEC_prlf | NDUFB6 |
| 2.7487E-13 | 0.29074391 | 0.259 | 0.13 | 6.4932E-09 | TEC_prlf | TMEM176A |
| 2.76E-13 | -0.563234 | 0.137 | 0.316 | 6.5199E-09 | TEC_prlf | HIST1H2AC |
| 3.0642E-13 | 0.27029279 | 0.289 | 0.149 | 7.2387E-09 | TEC_prlf | ATF7IP |
| 3.2276E-13 | 0.26089797 | 0.259 | 0.132 | 7.6247E-09 | TEC_prlf | TRIOBP |
| 3.3053E-13 | -0.3300584 | 0.02 | 0.167 | 7.8082E-09 | TEC_prlf | CFD |
| 3.4236E-13 | -1.0471626 | 0 | 0.135 | 8.0877E-09 | TEC_prlf | SERPINB2 |
| 3.4503E-13 | 0.257298 | 0.259 | 0.127 | 8.1507E-09 | TEC_prlf | TFPI2 |
| 3.531E-13 | 0.29208117 | 0.338 | 0.189 | 8.3414E-09 | TEC_prlf | TBL1XR1 |
| 3.718E-13 | 0.26563096 | 0.324 | 0.181 | 8.783E-09 | TEC_prlf | SSNA1 |
| 3.8581E-13 | -0.3362543 | 0.02 | 0.166 | 9.114E-09 | TEC_prlf | JARID2 |
| 3.9154E-13 | -0.2871985 | 0.032 | 0.189 | 9.2493E-09 | TEC_prlf | RAC2 |
| 3.9324E-13 | -0.313562 | 0.017 | 0.163 | 9.2895E-09 | TEC_prlf | FERMT3 |
| 4.0282E-13 | -0.9945254 | 0.017 | 0.161 | 9.5159E-09 | TEC_prlf | CEACAM5 |
| 4.2318E-13 | -0.4994898 | 0.038 | 0.189 | 9.9968E-09 | TEC_prlf | FYB1 |
| 4.7902E-13 | -0.4278857 | 0.122 | 0.299 | 1.1316E-08 | TEC_prlf | REL |
| 5.2638E-13 | -0.4326291 | 0.003 | 0.137 | 1.2435E-08 | TEC_prlf | MCEMP1 |
| 5.36E-13 | -0.4302816 | 0.052 | 0.21 | 1.2662E-08 | TEC_prlf | SOCS3 |
| 5.395E-13 | 0.37007979 | 0.985 | 0.978 | 1.2745E-08 | TEC_prlf | RPL28 |
| 6.009E-13 | -0.2945985 | 0.003 | 0.137 | 1.4195E-08 | TEC_prlf | HCK |
| 6.3921E-13 | -0.3658865 | 0.05 | 0.206 | 1.51E-08 | TEC_prlf | MAP3K8 |
| 6.8815E-13 | -0.2897238 | 0 | 0.131 | 1.6256E-08 | TEC_prlf | LILRB3 |
| 7.5335E-13 | -0.485233 | 0.009 | 0.145 | 1.7796E-08 | TEC_prlf | CD93 |
| 7.569E-13 | 0.29576517 | 0.983 | 0.905 | 1.788E-08 | TEC_prlf | RPL7A |
| 7.7296E-13 | -0.4943071 | 0.029 | 0.178 | 1.826E-08 | TEC_prlf | ANPEP |
| 7.9099E-13 | 0.29874608 | 0.802 | 0.664 | 1.8686E-08 | TEC_prlf | HNRNPC |
| 7.9418E-13 | -0.3511023 | 0.009 | 0.145 | 1.8761E-08 | TEC_prlf | TLR2 |
| 8.0472E-13 | -0.3212985 | 0.012 | 0.15 | 1.901E-08 | TEC_prlf | BCAT1 |
| 8.5282E-13 | -0.4790679 | 0.557 | 0.692 | 2.0146E-08 | TEC_prlf | PNRC1 |
| 9.5001E-13 | 0.29296651 | 0.455 | 0.292 | 2.2442E-08 | TEC_prlf | SEPT2 |
| 9.5045E-13 | -0.2910423 | 0.044 | 0.196 | 2.2453E-08 | TEC_prlf | RAB8B |
| 9.7907E-13 | -0.5635852 | 0.321 | 0.481 | 2.3129E-08 | TEC_prlf | RIOK3 |
| 9.8474E-13 | 0.28079471 | 0.327 | 0.185 | 2.3263E-08 | TEC_prlf | PCMT1 |
| 9.8694E-13 | 0.278071 | 0.414 | 0.253 | 2.3314E-08 | TEC_prlf | RPN2 |
| 1.0851E-12 | -1.0747757 | 0.038 | 0.185 | 2.5634E-08 | TEC_prlf | DUOX2 |
| 1.1282E-12 | -0.4302003 | 0.586 | 0.711 | 2.665E-08 | TEC_prlf | SDCBP |
| 1.1533E-12 | -0.5566249 | 0.029 | 0.172 | 2.7245E-08 | TEC_prlf | HSPB8 |
| 1.2006E-12 | 0.28662052 | 0.397 | 0.244 | 2.8362E-08 | TEC_prlf | UQCRFS1 |

| | | | | | | |
|---|---|---|---|---|---|---|
| 1.2515E-12 | -0.8299399 | 0.35 | 0.506 | 2.9565E-08 | TEC_prlf | HSPA1B |
| 1.3402E-12 | -0.3013576 | 0.006 | 0.138 | 3.166E-08 | TEC_prlf | CLEC7A |
| 1.4019E-12 | 0.34133167 | 0.452 | 0.289 | 3.3118E-08 | TEC_prlf | ZC3H15 |
| 1.4541E-12 | 0.42522568 | 0.286 | 0.157 | 3.435E-08 | TEC_prlf | CHD3 |
| 1.5077E-12 | -0.7829251 | 0.012 | 0.147 | 3.5617E-08 | TEC_prlf | CEACAM7 |
| 1.5115E-12 | 0.35426132 | 0.519 | 0.356 | 3.5705E-08 | TEC_prlf | ATP5PD |
| 1.5573E-12 | -0.6025612 | 0.017 | 0.156 | 3.6788E-08 | TEC_prlf | MUC4 |
| 1.5701E-12 | 0.42799059 | 0.638 | 0.51 | 3.709E-08 | TEC_prlf | COX7B |
| 1.5882E-12 | -0.3010138 | 0.026 | 0.169 | 3.7518E-08 | TEC_prlf | GLIPR2 |
| 1.5951E-12 | -0.2849409 | 0 | 0.128 | 3.768E-08 | TEC_prlf | IRAK3 |
| 1.697E-12 | 1.51467669 | 0.743 | 0.637 | 4.0088E-08 | TEC_prlf | MTRNR2L12 |
| 1.7087E-12 | 0.28611467 | 0.397 | 0.242 | 4.0365E-08 | TEC_prlf | LSM7 |
| 1.8864E-12 | -0.2848653 | 0.076 | 0.241 | 4.4562E-08 | TEC_prlf | CASP4 |
| 1.9021E-12 | -0.4919308 | 0.114 | 0.277 | 4.4932E-08 | TEC_prlf | ETS2 |
| 2.0592E-12 | 0.2502889 | 0.268 | 0.142 | 4.8645E-08 | TEC_prlf | WBP11 |
| 2.0936E-12 | -0.2723796 | 0.041 | 0.188 | 4.9458E-08 | TEC_prlf | ETV6 |
| 2.1126E-12 | 0.35810601 | 0.464 | 0.306 | 4.9906E-08 | TEC_prlf | ANAPC16 |
| 2.2968E-12 | -0.2611644 | 0.012 | 0.145 | 5.4257E-08 | TEC_prlf | CCDC69 |
| 2.3877E-12 | 0.25647454 | 0.294 | 0.159 | 5.6404E-08 | TEC_prlf | SAFB2 |
| 2.4581E-12 | -0.6286927 | 0.93 | 0.938 | 5.8067E-08 | TEC_prlf | TMSB4X |
| 2.5734E-12 | 0.25079938 | 0.289 | 0.156 | 6.0792E-08 | TEC_prlf | ITGB1BP1 |
| 2.8312E-12 | -0.8993271 | 0.006 | 0.135 | 6.688E-08 | TEC_prlf | EREG |
| 2.8544E-12 | -0.5656034 | 0.032 | 0.173 | 6.7429E-08 | TEC_prlf | HLA-DQB1 |
| 2.9409E-12 | -0.4610422 | 0.009 | 0.14 | 6.9472E-08 | TEC_prlf | PLAC8 |
| 3.2126E-12 | 0.37866515 | 0.507 | 0.356 | 7.5892E-08 | TEC_prlf | XRCC5 |
| 3.4062E-12 | 0.36772568 | 0.764 | 0.646 | 8.0465E-08 | TEC_prlf | LDHA |
| 3.4564E-12 | 0.26296949 | 0.356 | 0.211 | 8.1651E-08 | TEC_prlf | GTF3C6 |
| 3.4781E-12 | -0.2604435 | 0.006 | 0.135 | 8.2164E-08 | TEC_prlf | FGR |
| 3.5685E-12 | -0.2894414 | 0.05 | 0.202 | 8.4298E-08 | TEC_prlf | IFI16 |
| 3.6823E-12 | -0.4044307 | 0.19 | 0.36 | 8.6987E-08 | TEC_prlf | PLSCR1 |
| 3.6956E-12 | -1.5779551 | 0.006 | 0.134 | 8.7301E-08 | TEC_prlf | LYPD2 |
| 3.7521E-12 | -0.6691407 | 0.691 | 0.775 | 8.8636E-08 | TEC_prlf | CTSD |
| 4.0834E-12 | -0.3274785 | 0.015 | 0.147 | 9.6462E-08 | TEC_prlf | C1orf162 |
| 4.0921E-12 | -0.3902913 | 0.067 | 0.218 | 9.6667E-08 | TEC_prlf | DSE |
| 4.1043E-12 | 0.27689127 | 0.356 | 0.206 | 9.6957E-08 | TEC_prlf | MAP4 |
| 4.3099E-12 | 0.39563221 | 0.481 | 0.33 | 1.0181E-07 | TEC_prlf | NDUFC2 |
| 4.3215E-12 | -0.7621848 | 0.499 | 0.613 | 1.0209E-07 | TEC_prlf | CD44 |
| 4.5153E-12 | -0.2616992 | 0.009 | 0.138 | 1.0667E-07 | TEC_prlf | EVI2A |
| 4.6149E-12 | 0.29610583 | 0.344 | 0.206 | 1.0902E-07 | TEC_prlf | SIVA1 |
| 4.7026E-12 | -0.4957638 | 0.041 | 0.184 | 1.1109E-07 | TEC_prlf | SDCBP2 |
| 5.091E-12 | 0.30151531 | 0.344 | 0.203 | 1.2027E-07 | TEC_prlf | C6orf48 |
| 5.1484E-12 | -0.3176215 | 0.035 | 0.178 | 1.2162E-07 | TEC_prlf | GLIPR1 |
| 5.6272E-12 | 0.33196768 | 0.452 | 0.301 | 1.3293E-07 | TEC_prlf | ANAPC11 |
| 5.7733E-12 | -0.7233045 | 0.015 | 0.145 | 1.3638E-07 | TEC_prlf | TMPRSS11E |
| 6.2756E-12 | -0.6296176 | 0.006 | 0.132 | 1.4825E-07 | TEC_prlf | GABRP |
| 6.5701E-12 | 0.39320297 | 0.574 | 0.433 | 1.5521E-07 | TEC_prlf | ATP5MF |
| 7.2802E-12 | -0.3466037 | 0.099 | 0.266 | 1.7198E-07 | TEC_prlf | BACH1 |
| 7.3356E-12 | 0.25735589 | 0.289 | 0.16 | 1.7329E-07 | TEC_prlf | RAB8A |
| 7.9189E-12 | -1.5114543 | 0.029 | 0.163 | 1.8707E-07 | TEC_prlf | ECM1 |
| 8.0994E-12 | 0.36560501 | 0.528 | 0.384 | 1.9133E-07 | TEC_prlf | NEDD8 |

| | | | | | | |
|---|---|---|---|---|---|---|
| 8.8245E-12 | -1.345763 | 0.02 | 0.153 | 2.0846E-07 | TEC_prlf | CCL20 |
| 9.2832E-12 | -0.2576602 | 0.012 | 0.139 | 2.193E-07 | TEC_prlf | EMILIN2 |
| 9.3552E-12 | -0.4775805 | 0.052 | 0.196 | 2.21E-07 | TEC_prlf | SAMD9 |
| 9.8099E-12 | -0.2977751 | 0.047 | 0.191 | 2.3174E-07 | TEC_prlf | GBP2 |
| 1.0153E-11 | 0.33138911 | 0.455 | 0.307 | 2.3985E-07 | TEC_prlf | SSBP1 |
| 1.0689E-11 | -0.2963173 | 0.026 | 0.165 | 2.525E-07 | TEC_prlf | CLEC2B |
| 1.0818E-11 | 0.32367749 | 0.813 | 0.724 | 2.5556E-07 | TEC_prlf | COX6A1 |
| 1.089E-11 | 0.30243875 | 0.714 | 0.553 | 2.5726E-07 | TEC_prlf | MORF4L1 |
| 1.1192E-11 | 0.3198982 | 0.942 | 0.863 | 2.6438E-07 | TEC_prlf | CALM1 |
| 1.139E-11 | -0.707537 | 0.023 | 0.154 | 2.6907E-07 | TEC_prlf | SCEL |
| 1.1482E-11 | -0.3898193 | 0.192 | 0.356 | 2.7123E-07 | TEC_prlf | AKAP13 |
| 1.1602E-11 | -0.7916235 | 0.017 | 0.147 | 2.7408E-07 | TEC_prlf | CEACAM6 |
| 1.2465E-11 | 0.29709402 | 0.525 | 0.366 | 2.9447E-07 | TEC_prlf | SEC11A |
| 1.2603E-11 | -0.6139564 | 0.012 | 0.137 | 2.9773E-07 | TEC_prlf | CXCL17 |
| 1.2642E-11 | 0.32162751 | 0.869 | 0.785 | 2.9864E-07 | TEC_prlf | BTF3 |
| 1.2664E-11 | 0.27702984 | 0.994 | 0.887 | 2.9916E-07 | TEC_prlf | RPL6 |
| 1.3782E-11 | -0.4215497 | 0.184 | 0.349 | 3.2557E-07 | TEC_prlf | TIMP2 |
| 1.4E-11 | 0.44331225 | 0.455 | 0.308 | 3.3073E-07 | TEC_prlf | RAB13 |
| 1.4304E-11 | -0.5458681 | 0.172 | 0.324 | 3.3791E-07 | TEC_prlf | LIMS1 |
| 1.5839E-11 | 0.31225564 | 0.297 | 0.169 | 3.7416E-07 | TEC_prlf | AP3D1 |
| 1.604E-11 | 0.25465294 | 0.213 | 0.105 | 3.7891E-07 | TEC_prlf | KIF1C |
| 1.6429E-11 | 0.34446659 | 0.501 | 0.351 | 3.8811E-07 | TEC_prlf | SPCS1 |
| 1.6594E-11 | -0.3691259 | 0.026 | 0.158 | 3.9199E-07 | TEC_prlf | TTC9 |
| 1.752E-11 | -0.4705975 | 0.146 | 0.31 | 4.1388E-07 | TEC_prlf | IRF1 |
| 1.7999E-11 | -0.8951719 | 0.111 | 0.264 | 4.2519E-07 | TEC_prlf | AREG |
| 1.9373E-11 | -0.3132545 | 0.07 | 0.218 | 4.5766E-07 | TEC_prlf | STK4 |
| 1.9486E-11 | 0.30073208 | 0.443 | 0.294 | 4.6032E-07 | TEC_prlf | EIF3G |
| 1.9858E-11 | -0.3008838 | 0.003 | 0.121 | 4.6911E-07 | TEC_prlf | LILRB2 |
| 2.0723E-11 | -0.2549483 | 0.026 | 0.159 | 4.8954E-07 | TEC_prlf | ARHGAP30 |
| 2.0731E-11 | -0.2863162 | 0.035 | 0.172 | 4.8972E-07 | TEC_prlf | B3GNT5 |
| 2.1468E-11 | -0.2855433 | 0.035 | 0.174 | 5.0714E-07 | TEC_prlf | CORO1A |
| 2.2263E-11 | -0.4628547 | 0.163 | 0.323 | 5.2591E-07 | TEC_prlf | B4GALT1 |
| 2.2982E-11 | -0.3587679 | 0.079 | 0.224 | 5.4291E-07 | TEC_prlf | GRINA |
| 2.4528E-11 | -0.9143162 | 0.029 | 0.159 | 5.7942E-07 | TEC_prlf | PRSS27 |
| 2.46E-11 | -0.406263 | 0.058 | 0.203 | 5.8112E-07 | TEC_prlf | FOSB |
| 2.4734E-11 | -0.3795453 | 0.017 | 0.143 | 5.8428E-07 | TEC_prlf | NCF1 |
| 2.4927E-11 | -0.2653054 | 0.003 | 0.12 | 5.8885E-07 | TEC_prlf | CSF1R |
| 2.495E-11 | -0.5179449 | 0.096 | 0.244 | 5.8938E-07 | TEC_prlf | ATP2B1 |
| 2.6259E-11 | 0.30359256 | 0.464 | 0.313 | 6.2031E-07 | TEC_prlf | RBX1 |
| 2.7084E-11 | -0.5828995 | 0.02 | 0.148 | 6.398E-07 | TEC_prlf | NCCRP1 |
| 3.2358E-11 | -0.3139425 | 0.125 | 0.288 | 7.644E-07 | TEC_prlf | BAZ1A |
| 3.3329E-11 | -0.3894212 | 0.181 | 0.345 | 7.8734E-07 | TEC_prlf | CPEB4 |
| 3.4323E-11 | -1.1482127 | 0.006 | 0.123 | 8.1081E-07 | TEC_prlf | CCL3 |
| 3.5206E-11 | 0.31194591 | 0.708 | 0.588 | 8.3168E-07 | TEC_prlf | SLC25A3 |
| 3.5828E-11 | 0.25128857 | 0.35 | 0.207 | 8.4637E-07 | TEC_prlf | TCEAL4 |
| 3.6577E-11 | -0.2710494 | 0.015 | 0.137 | 8.6405E-07 | TEC_prlf | ITGA5 |
| 3.7892E-11 | -0.2685706 | 0 | 0.114 | 8.9511E-07 | TEC_prlf | PECAM1 |
| 3.8217E-11 | -1.3979036 | 0.061 | 0.199 | 9.0281E-07 | TEC_prlf | KRT17 |
| 4.0772E-11 | -1.4314803 | 0.093 | 0.233 | 9.6316E-07 | TEC_prlf | PRSS22 |
| 4.1241E-11 | -0.2543683 | 0.029 | 0.16 | 9.7424E-07 | TEC_prlf | DOCK8 |

| | | | | | | |
|---|---|---|---|---|---|---|
| 4.2092E-11 | -0.2516751 | 0.029 | 0.159 | 9.9435E-07 | TEC_prlf | SUSD6 |
| 4.2688E-11 | -0.7940695 | 0.029 | 0.157 | 1.0084E-06 | TEC_prlf | KRT15 |
| 4.3075E-11 | -1.6761827 | 0.207 | 0.345 | 1.0176E-06 | TEC_prlf | HPGD |
| 4.8845E-11 | -0.8689616 | 0.015 | 0.135 | 1.1539E-06 | TEC_prlf | TMPRSS11B |
| 5.191E-11 | -0.2666842 | 0.038 | 0.174 | 1.2263E-06 | TEC_prlf | DHRS3 |
| 5.3047E-11 | -0.6110646 | 0.134 | 0.283 | 1.2531E-06 | TEC_prlf | RHOB |
| 5.4701E-11 | -0.2912867 | 0 | 0.112 | 1.2922E-06 | TEC_prlf | FUT3 |
| 5.5577E-11 | 0.31514478 | 0.878 | 0.758 | 1.3129E-06 | TEC_prlf | UQCRB |
| 5.5717E-11 | 0.27675466 | 0.364 | 0.229 | 1.3162E-06 | TEC_prlf | ENSA |
| 5.6085E-11 | 0.3243626 | 0.379 | 0.241 | 1.3249E-06 | TEC_prlf | TMEM14C |
| 5.6972E-11 | -0.3497768 | 0.003 | 0.117 | 1.3458E-06 | TEC_prlf | CD163 |
| 5.7654E-11 | -0.2583821 | 0.026 | 0.154 | 1.362E-06 | TEC_prlf | VSIR |
| 5.8807E-11 | 0.34585423 | 0.51 | 0.362 | 1.3892E-06 | TEC_prlf | C19orf53 |
| 6.4441E-11 | -0.3020517 | 0.047 | 0.18 | 1.5223E-06 | TEC_prlf | FMNL1 |
| 7.2234E-11 | 0.25877032 | 0.338 | 0.204 | 1.7064E-06 | TEC_prlf | SLIRP |
| 7.5303E-11 | -0.8815827 | 0.015 | 0.133 | 1.7789E-06 | TEC_prlf | RHCG |
| 7.6649E-11 | -0.2903807 | 0.061 | 0.202 | 1.8107E-06 | TEC_prlf | NR3C1 |
| 7.9684E-11 | -0.3449208 | 0.015 | 0.135 | 1.8824E-06 | TEC_prlf | BCAS1 |
| 8.1493E-11 | 0.27716834 | 0.405 | 0.261 | 1.9251E-06 | TEC_prlf | GTF2A2 |
| 9.9808E-11 | -0.4950367 | 0.388 | 0.537 | 2.3578E-06 | TEC_prlf | SERPINB1 |
| 1.0539E-10 | 0.28842713 | 0.356 | 0.223 | 2.4897E-06 | TEC_prlf | NARS |
| 1.0592E-10 | 0.25289299 | 0.362 | 0.217 | 2.5021E-06 | TEC_prlf | SPATS2L |
| 1.0902E-10 | -0.3592856 | 0.137 | 0.293 | 2.5754E-06 | TEC_prlf | SKIL |
| 1.1119E-10 | -2.3848602 | 0.254 | 0.379 | 2.6267E-06 | TEC_prlf | KRT13 |
| 1.1226E-10 | -0.3004029 | 0.157 | 0.317 | 2.652E-06 | TEC_prlf | MMP24OS |
| 1.1282E-10 | -0.9688951 | 0.02 | 0.141 | 2.6652E-06 | TEC_prlf | CXCL6 |
| 1.1823E-10 | -0.2646531 | 0.006 | 0.118 | 2.793E-06 | TEC_prlf | CD86 |
| 1.195E-10 | -0.2625462 | 0.023 | 0.146 | 2.8229E-06 | TEC_prlf | CARD16 |
| 1.2183E-10 | 0.25911996 | 0.283 | 0.162 | 2.8781E-06 | TEC_prlf | DENR |
| 1.2221E-10 | -0.4538314 | 0.003 | 0.113 | 2.8869E-06 | TEC_prlf | TMPRSS11A |
| 1.2268E-10 | 0.3792901 | 0.469 | 0.322 | 2.8981E-06 | TEC_prlf | TGOLN2 |
| 1.2373E-10 | -0.5888072 | 0.003 | 0.113 | 2.9228E-06 | TEC_prlf | KRT78 |
| 1.2831E-10 | 0.29562746 | 0.851 | 0.703 | 3.0311E-06 | TEC_prlf | RPL4 |
| 1.3339E-10 | 0.2531853 | 0.362 | 0.225 | 3.151E-06 | TEC_prlf | PSMB2 |
| 1.3704E-10 | -0.3275763 | 0.085 | 0.229 | 3.2374E-06 | TEC_prlf | GALNT1 |
| 1.4372E-10 | -0.3273753 | 0.07 | 0.209 | 3.395E-06 | TEC_prlf | STK17B |
| 1.4885E-10 | -0.3289918 | 0.023 | 0.147 | 3.5163E-06 | TEC_prlf | VSIG2 |
| 1.4908E-10 | -0.2583162 | 0.026 | 0.15 | 3.5217E-06 | TEC_prlf | RALGDS |
| 1.5207E-10 | -0.5926886 | 0.023 | 0.143 | 3.5922E-06 | TEC_prlf | CDKN2A |
| 1.8202E-10 | -0.4285381 | 0.052 | 0.184 | 4.2998E-06 | TEC_prlf | SGK1 |
| 1.8925E-10 | -0.3259572 | 0 | 0.106 | 4.4706E-06 | TEC_prlf | A2ML1 |
| 1.9439E-10 | -0.2827982 | 0.12 | 0.274 | 4.592E-06 | TEC_prlf | FAM49B |
| 2.0914E-10 | -0.3806338 | 0.114 | 0.258 | 4.9405E-06 | TEC_prlf | PDLIM5 |
| 2.228E-10 | -0.4969438 | 0.125 | 0.275 | 5.2632E-06 | TEC_prlf | TNFAIP2 |
| 2.3415E-10 | -0.3641851 | 0.009 | 0.12 | 5.5312E-06 | TEC_prlf | FCGR3A |
| 2.3577E-10 | -0.7775023 | 0 | 0.105 | 5.5696E-06 | TEC_prlf | ZG16B |
| 2.3699E-10 | 0.27345859 | 0.51 | 0.359 | 5.5985E-06 | TEC_prlf | RBM8A |
| 2.4166E-10 | -0.3353274 | 0.017 | 0.133 | 5.7087E-06 | TEC_prlf | RGCC |
| 2.4372E-10 | 0.36480564 | 0.496 | 0.366 | 5.7574E-06 | TEC_prlf | PNRC2 |
| 2.4924E-10 | -0.388845 | 0.07 | 0.205 | 5.8878E-06 | TEC_prlf | CSNK1E |

| | | | | | | |
|---|---|---|---|---|---|---|
| 2.5015E-10 | -0.9924385 | 0.055 | 0.186 | 5.9094E-06 | TEC_prlf | HLA-DPB1 |
| 2.5317E-10 | -0.3132885 | 0.09 | 0.232 | 5.9806E-06 | TEC_prlf | ATP6V1B2 |
| 2.9655E-10 | -0.4321115 | 0.041 | 0.166 | 7.0053E-06 | TEC_prlf | HIST3H2A |
| 3.14E-10 | -0.2981164 | 0.029 | 0.15 | 7.4176E-06 | TEC_prlf | PLXDC2 |
| 3.2828E-10 | -0.2904294 | 0.108 | 0.255 | 7.755E-06 | TEC_prlf | GABARAPL1 |
| 3.3086E-10 | -0.2796641 | 0 | 0.104 | 7.8159E-06 | TEC_prlf | LILRB1 |
| 3.6318E-10 | 0.30948185 | 0.872 | 0.77 | 8.5794E-06 | TEC_prlf | TMA7 |
| 3.632E-10 | 0.2878033 | 0.402 | 0.27 | 8.5799E-06 | TEC_prlf | NDUFB8 |
| 3.7149E-10 | -0.4811997 | 0.306 | 0.466 | 8.7758E-06 | TEC_prlf | CDKN1A |
| 3.7651E-10 | 0.32391436 | 0.248 | 0.139 | 8.8943E-06 | TEC_prlf | LPIN2 |
| 3.7784E-10 | -1.0884224 | 0.009 | 0.117 | 8.9257E-06 | TEC_prlf | SPINK5 |
| 3.9838E-10 | 0.25861054 | 0.338 | 0.209 | 9.4109E-06 | TEC_prlf | RBM17 |
| 4.0223E-10 | -0.4051975 | 0.163 | 0.313 | 9.5018E-06 | TEC_prlf | FNDC3B |
| 4.0442E-10 | 0.41625144 | 0.965 | 0.954 | 9.5536E-06 | TEC_prlf | SERF2 |
| 4.1879E-10 | 0.294019 | 0.743 | 0.635 | 9.893E-06 | TEC_prlf | ARF1 |
| 4.3015E-10 | -0.4440187 | 0.02 | 0.136 | 1.0161E-05 | TEC_prlf | HLA-DRB5 |
| 4.5043E-10 | 0.2798497 | 0.397 | 0.259 | 1.0641E-05 | TEC_prlf | OCIAD1 |
| 4.5355E-10 | -0.6511391 | 0.513 | 0.638 | 1.0714E-05 | TEC_prlf | GADD45B |
| 4.5402E-10 | -0.3726804 | 0.052 | 0.183 | 1.0725E-05 | TEC_prlf | FBP1 |
| 4.5566E-10 | 0.26063336 | 0.968 | 0.901 | 1.0764E-05 | TEC_prlf | RPL24 |
| 4.5716E-10 | 0.25108843 | 0.983 | 0.898 | 1.0799E-05 | TEC_prlf | RPL21 |
| 4.6639E-10 | 0.28426788 | 0.516 | 0.371 | 1.1017E-05 | TEC_prlf | C11orf58 |
| 4.7344E-10 | -2.954387 | 0.638 | 0.676 | 1.1184E-05 | TEC_prlf | TIMP1 |
| 4.7441E-10 | -0.3668821 | 0.461 | 0.579 | 1.1207E-05 | TEC_prlf | UBE2B |
| 4.7609E-10 | -0.5312843 | 0.009 | 0.117 | 1.1247E-05 | TEC_prlf | MNDA |
| 4.9592E-10 | 0.25111292 | 0.306 | 0.184 | 1.1715E-05 | TEC_prlf | FDPS |
| 5.3421E-10 | -0.3386503 | 0.003 | 0.106 | 1.262E-05 | TEC_prlf | CSF3R |
| 5.4196E-10 | -0.3018882 | 0.044 | 0.166 | 1.2803E-05 | TEC_prlf | CEP170 |
| 5.509E-10 | -0.262441 | 0.029 | 0.147 | 1.3014E-05 | TEC_prlf | PLEKHO1 |
| 5.7465E-10 | 0.25754327 | 0.364 | 0.236 | 1.3575E-05 | TEC_prlf | SNRPE |
| 6.3132E-10 | -0.3536085 | 0.009 | 0.115 | 1.4914E-05 | TEC_prlf | CEACAM1 |
| 6.4387E-10 | 0.28423304 | 0.391 | 0.257 | 1.521E-05 | TEC_prlf | DYNC1I2 |
| 7.1187E-10 | -0.3125337 | 0.05 | 0.175 | 1.6817E-05 | TEC_prlf | KCTD12 |
| 7.2352E-10 | -0.2880266 | 0.032 | 0.15 | 1.7092E-05 | TEC_prlf | DMXL2 |
| 7.2554E-10 | -0.3524569 | 0.449 | 0.585 | 1.7139E-05 | TEC_prlf | NOP10 |
| 7.6445E-10 | -0.4067914 | 0.047 | 0.171 | 1.8059E-05 | TEC_prlf | CXCR4 |
| 7.9129E-10 | -0.3732977 | 0.023 | 0.136 | 1.8693E-05 | TEC_prlf | GCNT3 |
| 8.4693E-10 | -0.2588978 | 0.015 | 0.123 | 2.0007E-05 | TEC_prlf | TCP11L2 |
| 8.6394E-10 | -0.386002 | 0.105 | 0.245 | 2.0409E-05 | TEC_prlf | KIAA1551 |
| 9.0795E-10 | 0.34901423 | 0.767 | 0.678 | 2.1448E-05 | TEC_prlf | COX7A2 |
| 9.4552E-10 | 0.27092204 | 0.315 | 0.188 | 2.2336E-05 | TEC_prlf | PDIA4 |
| 9.5853E-10 | -0.2504459 | 0.052 | 0.177 | 2.2643E-05 | TEC_prlf | PPP1R18 |
| 9.599E-10 | -0.2657934 | 0.085 | 0.219 | 2.2676E-05 | TEC_prlf | RIT1 |
| 9.6185E-10 | 0.26148012 | 0.318 | 0.19 | 2.2722E-05 | TEC_prlf | LIMA1 |
| 9.7438E-10 | 0.27522664 | 0.405 | 0.274 | 2.3018E-05 | TEC_prlf | SNX6 |
| 1.0035E-09 | 0.32750884 | 0.461 | 0.323 | 2.3707E-05 | TEC_prlf | NSA2 |
| 1.0143E-09 | 0.3370879 | 0.589 | 0.457 | 2.3962E-05 | TEC_prlf | EIF2S2 |
| 1.0637E-09 | -0.6807832 | 0.096 | 0.231 | 2.5128E-05 | TEC_prlf | ISG15 |
| 1.1721E-09 | -0.2634359 | 0.026 | 0.14 | 2.7688E-05 | TEC_prlf | IFI30 |
| 1.1795E-09 | -0.6433103 | 0.061 | 0.194 | 2.7863E-05 | TEC_prlf | HSPA6 |

| | | | | | | |
|---|---|---|---|---|---|---|
| 1.1904E-09 | 0.3296395 | 0.764 | 0.66 | 2.8122E-05 | TEC_prlf | HNRNPU |
| 1.2246E-09 | -0.2635297 | 0.082 | 0.219 | 2.893E-05 | TEC_prlf | TUT7 |
| 1.2916E-09 | 0.28027544 | 0.898 | 0.765 | 3.0511E-05 | TEC_prlf | RPS26 |
| 1.3503E-09 | -0.3283443 | 0.05 | 0.172 | 3.1898E-05 | TEC_prlf | SAMHD1 |
| 1.3861E-09 | -0.3348986 | 0.076 | 0.205 | 3.2745E-05 | TEC_prlf | TIPARP |
| 1.39E-09 | 0.32629404 | 0.484 | 0.354 | 3.2837E-05 | TEC_prlf | CYCS |
| 1.4187E-09 | -1.5060178 | 0.006 | 0.107 | 3.3513E-05 | TEC_prlf | CRCT1 |
| 1.4277E-09 | 0.26753208 | 0.329 | 0.206 | 3.3728E-05 | TEC_prlf | CWC15 |
| 1.5503E-09 | 0.28299419 | 0.93 | 0.838 | 3.6622E-05 | TEC_prlf | RACK1 |
| 1.5618E-09 | -0.3936253 | 0.137 | 0.28 | 3.6894E-05 | TEC_prlf | ABHD5 |
| 1.8522E-09 | -0.4811781 | 0.003 | 0.101 | 4.3755E-05 | TEC_prlf | RETN |
| 1.9057E-09 | 0.25995045 | 0.268 | 0.153 | 4.5018E-05 | TEC_prlf | ARHGEF12 |
| 1.9276E-09 | -0.4298128 | 0.05 | 0.171 | 4.5536E-05 | TEC_prlf | TGFBI |
| 1.9342E-09 | -0.300888 | 0.114 | 0.251 | 4.5691E-05 | TEC_prlf | UBL3 |
| 1.9543E-09 | -0.7305253 | 0.055 | 0.174 | 4.6167E-05 | TEC_prlf | NDRG2 |
| 2.1133E-09 | -0.6068529 | 0.023 | 0.131 | 4.9923E-05 | TEC_prlf | DHRS9 |
| 2.1713E-09 | 0.31833272 | 0.37 | 0.246 | 5.1292E-05 | TEC_prlf | CD47 |
| 2.3066E-09 | -3.7853076 | 0.294 | 0.402 | 5.4488E-05 | TEC_prlf | SPRR3 |
| 2.4039E-09 | -0.3316249 | 0.02 | 0.127 | 5.6787E-05 | TEC_prlf | MARCO |
| 2.4595E-09 | -0.3546456 | 0.181 | 0.322 | 5.8102E-05 | TEC_prlf | YPEL3 |
| 2.4755E-09 | -0.2883825 | 0.085 | 0.214 | 5.8479E-05 | TEC_prlf | RUNX1 |
| 2.4941E-09 | -0.7751056 | 0.163 | 0.293 | 5.8918E-05 | TEC_prlf | RND3 |
| 2.573E-09 | 0.28050099 | 0.42 | 0.288 | 6.0781E-05 | TEC_prlf | NDUFB9 |
| 2.6016E-09 | -0.2705107 | 0.05 | 0.17 | 6.1458E-05 | TEC_prlf | ISG20 |
| 2.6287E-09 | -1.4166046 | 0.009 | 0.108 | 6.2097E-05 | TEC_prlf | CNFN |
| 2.6361E-09 | -0.4363628 | 0.102 | 0.239 | 6.2273E-05 | TEC_prlf | TNFAIP3 |
| 2.81E-09 | -0.3201266 | 0.015 | 0.118 | 6.6379E-05 | TEC_prlf | DUSP2 |
| 2.9012E-09 | 0.28534822 | 0.548 | 0.412 | 6.8536E-05 | TEC_prlf | PCBP2 |
| 3.1282E-09 | 0.54786986 | 0.49 | 0.366 | 7.3897E-05 | TEC_prlf | MYH9 |
| 3.1749E-09 | -0.8408041 | 0.067 | 0.189 | 7.5E-05 | TEC_prlf | RARRES3 |
| 3.182E-09 | 0.27219942 | 0.889 | 0.745 | 7.5169E-05 | TEC_prlf | EEF1B2 |
| 3.2167E-09 | 0.33087542 | 0.402 | 0.28 | 7.5988E-05 | TEC_prlf | LRPAP1 |
| 3.3227E-09 | 0.26346234 | 0.98 | 0.901 | 7.8492E-05 | TEC_prlf | RPS7 |
| 3.5918E-09 | -0.2781653 | 0.009 | 0.107 | 8.4849E-05 | TEC_prlf | MS4A6A |
| 3.6175E-09 | 0.31715784 | 0.417 | 0.272 | 8.5456E-05 | TEC_prlf | TNFRSF12A |
| 3.8632E-09 | -0.3856256 | 0.009 | 0.107 | 9.126E-05 | TEC_prlf | THBD |
| 4.0628E-09 | -0.5762574 | 0.006 | 0.102 | 9.5975E-05 | TEC_prlf | HLA-DQA1 |
| 4.2329E-09 | -0.3853302 | 0.638 | 0.731 | 9.9994E-05 | TEC_prlf | ATP6V1G1 |
| 4.3434E-09 | -0.3431339 | 0.149 | 0.289 | 0.0001026 | TEC_prlf | RNF149 |
| 4.365E-09 | 0.34787741 | 0.414 | 0.295 | 0.00010312 | TEC_prlf | RABAC1 |
| 4.4555E-09 | 0.33472921 | 0.519 | 0.391 | 0.00010525 | TEC_prlf | UBXN4 |
| 4.7611E-09 | 0.26275668 | 0.318 | 0.2 | 0.00011247 | TEC_prlf | KRTCAP2 |
| 4.8309E-09 | -2.767071 | 0.443 | 0.524 | 0.00011412 | TEC_prlf | LCN2 |
| 5.1235E-09 | -0.3373998 | 0.09 | 0.216 | 0.00012103 | TEC_prlf | KDM6B |
| 5.6309E-09 | 0.30404819 | 0.315 | 0.203 | 0.00013302 | TEC_prlf | LRRFIP2 |
| 5.7042E-09 | 0.25260106 | 0.443 | 0.306 | 0.00013475 | TEC_prlf | PFDN2 |
| 6.0342E-09 | -0.8632313 | 0.099 | 0.224 | 0.00014255 | TEC_prlf | G0S2 |
| 6.1748E-09 | -0.3033502 | 0.268 | 0.414 | 0.00014587 | TEC_prlf | DAZAP2 |
| 6.3926E-09 | 0.25456607 | 0.423 | 0.291 | 0.00015101 | TEC_prlf | PPP1CA |
| 6.6778E-09 | -0.4022003 | 0.07 | 0.19 | 0.00015775 | TEC_prlf | DUSP6 |

| | | | | | | |
|---|---|---|---|---|---|---|
| 6.9944E-09 | -0.51997 | 0.015 | 0.113 | 0.00016523 | TEC_prlf | CYP1B1 |
| 7.2497E-09 | 0.27278277 | 0.394 | 0.272 | 0.00017126 | TEC_prlf | ATP5F1C |
| 7.5293E-09 | -0.3702467 | 0.012 | 0.108 | 0.00017787 | TEC_prlf | KRT23 |
| 8.3534E-09 | 0.26514755 | 0.98 | 0.923 | 0.00019733 | TEC_prlf | RPL18A |
| 9.7546E-09 | -0.362066 | 0.09 | 0.213 | 0.00023043 | TEC_prlf | NINJ1 |
| 1.0665E-08 | -0.3748315 | 0.041 | 0.149 | 0.00025193 | TEC_prlf | RGS2 |
| 1.0836E-08 | -0.2851155 | 0.195 | 0.337 | 0.00025598 | TEC_prlf | CAPZA1 |
| 1.1148E-08 | -0.2844324 | 0.933 | 0.964 | 0.00026334 | TEC_prlf | MYL6 |
| 1.1817E-08 | -0.371228 | 0.012 | 0.107 | 0.00027915 | TEC_prlf | TMC5 |
| 1.2185E-08 | 0.31243892 | 0.522 | 0.406 | 0.00028785 | TEC_prlf | NDUFB4 |
| 1.2619E-08 | -0.2607479 | 0.026 | 0.128 | 0.00029809 | TEC_prlf | TP53INP2 |
| 1.3315E-08 | -0.2828756 | 0.055 | 0.169 | 0.00031455 | TEC_prlf | NECTIN4 |
| 1.3581E-08 | 0.2980379 | 0.711 | 0.612 | 0.00032082 | TEC_prlf | SRRM2 |
| 1.4077E-08 | 0.35069845 | 0.362 | 0.244 | 0.00033255 | TEC_prlf | SPTAN1 |
| 1.4569E-08 | -0.3533763 | 0.023 | 0.124 | 0.00034417 | TEC_prlf | TMEM40 |
| 1.4935E-08 | -0.920643 | 0.064 | 0.174 | 0.00035282 | TEC_prlf | RDH10 |
| 1.5751E-08 | -0.3743259 | 0.087 | 0.207 | 0.00037208 | TEC_prlf | SLC43A2 |
| 1.7072E-08 | -1.2118347 | 0.017 | 0.114 | 0.00040329 | TEC_prlf | SPRR1B |
| 1.7186E-08 | 0.28476376 | 0.329 | 0.216 | 0.00040599 | TEC_prlf | DYNC1LI2 |
| 1.7327E-08 | -0.3569501 | 0.192 | 0.326 | 0.00040933 | TEC_prlf | TMBIM1 |
| 1.88E-08 | -0.4352045 | 0.169 | 0.3 | 0.00044411 | TEC_prlf | FAM129B |
| 1.8967E-08 | -0.3851608 | 0.114 | 0.237 | 0.00044806 | TEC_prlf | NFKBIZ |
| 2.0136E-08 | -1.6140244 | 0.041 | 0.144 | 0.00047566 | TEC_prlf | SPRR2A |
| 2.2214E-08 | -1.0317645 | 0.015 | 0.108 | 0.00052477 | TEC_prlf | CCL4 |
| 2.2431E-08 | 0.26240775 | 0.49 | 0.355 | 0.00052989 | TEC_prlf | UQCRQ |
| 2.2632E-08 | -0.3596017 | 0.531 | 0.625 | 0.00053464 | TEC_prlf | ACTR2 |
| 2.2854E-08 | -0.6446307 | 0.017 | 0.114 | 0.00053988 | TEC_prlf | IL7R |
| 2.3841E-08 | 0.26825817 | 0.35 | 0.233 | 0.0005632 | TEC_prlf | SYPL1 |
| 2.4062E-08 | 0.25448179 | 0.429 | 0.295 | 0.00056843 | TEC_prlf | TRAM1 |
| 2.4455E-08 | -0.3604801 | 0.044 | 0.152 | 0.0005777 | TEC_prlf | SLC7A11 |
| 2.5589E-08 | 0.28161961 | 0.487 | 0.36 | 0.00060449 | TEC_prlf | MTDH |
| 2.6087E-08 | -0.2596482 | 0.05 | 0.158 | 0.00061625 | TEC_prlf | TRIP10 |
| 2.6917E-08 | 0.2881936 | 0.528 | 0.393 | 0.00063586 | TEC_prlf | NDUFB2 |
| 2.7937E-08 | -0.4273423 | 0.067 | 0.176 | 0.00065995 | TEC_prlf | C6orf132 |
| 2.8256E-08 | -0.2533955 | 0.076 | 0.193 | 0.0006675 | TEC_prlf | ITPRIPL2 |
| 2.8276E-08 | 0.25564147 | 0.624 | 0.491 | 0.00066797 | TEC_prlf | TAGLN2 |
| 3.156E-08 | -0.3987529 | 0.052 | 0.159 | 0.00074555 | TEC_prlf | MALL |
| 3.1804E-08 | 0.28433393 | 0.776 | 0.687 | 0.0007513 | TEC_prlf | CAST |
| 3.2207E-08 | -0.2797306 | 0.128 | 0.256 | 0.00076082 | TEC_prlf | MGAT1 |
| 3.2478E-08 | 0.35188235 | 0.825 | 0.725 | 0.00076722 | TEC_prlf | ELOB |
| 3.3863E-08 | -0.3174546 | 0.341 | 0.471 | 0.00079994 | TEC_prlf | RAP1B |
| 3.5853E-08 | -0.4821173 | 0.248 | 0.369 | 0.00084696 | TEC_prlf | COTL1 |
| 3.6432E-08 | -0.7306634 | 0.682 | 0.74 | 0.00086064 | TEC_prlf | VIM |
| 4.0012E-08 | -0.2887121 | 0.802 | 0.867 | 0.00094521 | TEC_prlf | ARPC3 |
| 4.5263E-08 | -0.2629759 | 0.032 | 0.132 | 0.00106926 | TEC_prlf | IL18 |
| 4.6401E-08 | 0.39983885 | 0.638 | 0.49 | 0.00109612 | TEC_prlf | MTRNR2L8 |
| 4.7031E-08 | -0.4301231 | 0.195 | 0.327 | 0.00111102 | TEC_prlf | CEBPD |
| 4.727E-08 | -0.6845865 | 0.117 | 0.234 | 0.00111666 | TEC_prlf | PLIN2 |
| 4.8018E-08 | 0.30232465 | 0.525 | 0.397 | 0.00113433 | TEC_prlf | NDUFA1 |
| 4.9149E-08 | -0.4452968 | 0.117 | 0.236 | 0.00116105 | TEC_prlf | CDKN1C |

| | | | | | | |
|---|---|---|---|---|---|---|
| 5.1017E-08 | -0.2547626 | 0.464 | 0.587 | 0.00120518 | TEC_prlf | KLF6 |
| 5.1174E-08 | -0.3411297 | 0.169 | 0.299 | 0.00120887 | TEC_prlf | PIM3 |
| 5.5114E-08 | -1.111954 | 0.035 | 0.132 | 0.00130196 | TEC_prlf | PLAT |
| 6.0161E-08 | -0.3881264 | 0.195 | 0.328 | 0.00142118 | TEC_prlf | MAFF |
| 6.0853E-08 | -0.3090806 | 0.233 | 0.376 | 0.00143753 | TEC_prlf | CCNL1 |
| 6.1482E-08 | -0.2610721 | 0.076 | 0.189 | 0.00145238 | TEC_prlf | PYCARD |
| 6.3701E-08 | 0.38243317 | 0.423 | 0.312 | 0.0015048 | TEC_prlf | BLOC1S1 |
| 6.6587E-08 | -0.5124711 | 0.437 | 0.539 | 0.00157299 | TEC_prlf | UPP1 |
| 6.9781E-08 | -0.3390743 | 0.219 | 0.34 | 0.00164844 | TEC_prlf | GRB2 |
| 7.2337E-08 | -0.3545549 | 0.032 | 0.129 | 0.00170883 | TEC_prlf | AC023157.3 |
| 7.3841E-08 | -0.3152314 | 0.484 | 0.595 | 0.00174433 | TEC_prlf | DDX17 |
| 7.4883E-08 | -0.3302028 | 0.163 | 0.29 | 0.00176896 | TEC_prlf | CHMP1B |
| 7.6923E-08 | -0.2786785 | 0.052 | 0.156 | 0.00181714 | TEC_prlf | ADGRE5 |
| 7.8687E-08 | 0.28537846 | 0.569 | 0.448 | 0.00185883 | TEC_prlf | ATP5MD |
| 8.2588E-08 | -0.2705575 | 0.015 | 0.102 | 0.00195099 | TEC_prlf | LRMP |
| 8.2723E-08 | -0.3108273 | 0.085 | 0.198 | 0.00195416 | TEC_prlf | SERTAD2 |
| 9.3617E-08 | -0.2956614 | 0.064 | 0.171 | 0.00221151 | TEC_prlf | FAM214A |
| 1.0053E-07 | 0.29441453 | 0.411 | 0.293 | 0.00237473 | TEC_prlf | CHCHD10 |
| 1.1412E-07 | 0.26233786 | 0.499 | 0.387 | 0.00269595 | TEC_prlf | COPE |
| 1.1678E-07 | 0.28147986 | 0.636 | 0.517 | 0.00275869 | TEC_prlf | NAP1L1 |
| 1.1749E-07 | -0.2652545 | 0.163 | 0.295 | 0.00277549 | TEC_prlf | FXYD5 |
| 1.2818E-07 | -0.5868464 | 0.02 | 0.109 | 0.00302793 | TEC_prlf | APOC1 |
| 1.4474E-07 | -0.2703749 | 0.085 | 0.197 | 0.00341908 | TEC_prlf | KLF4 |
| 1.4999E-07 | 0.26809447 | 0.394 | 0.273 | 0.00354323 | TEC_prlf | BPTF |
| 1.643E-07 | 0.27089545 | 0.924 | 0.876 | 0.00388131 | TEC_prlf | TMBIM6 |
| 1.6809E-07 | -0.3943427 | 0.219 | 0.345 | 0.00397091 | TEC_prlf | DDIT4 |
| 1.7533E-07 | -0.4308119 | 0.131 | 0.243 | 0.00414188 | TEC_prlf | PRSS8 |
| 1.8204E-07 | -0.4673912 | 0.055 | 0.154 | 0.00430033 | TEC_prlf | CDC42EP5 |
| 1.9317E-07 | 0.60529714 | 0.915 | 0.944 | 0.00456324 | TEC_prlf | MT-CO3 |
| 2.1975E-07 | -0.3602679 | 0.128 | 0.238 | 0.00519111 | TEC_prlf | RNASET2 |
| 2.2893E-07 | -0.2516329 | 0.149 | 0.27 | 0.0054081 | TEC_prlf | UBE2D1 |
| 2.7533E-07 | 0.28247508 | 0.37 | 0.263 | 0.00650416 | TEC_prlf | IK |
| 2.793E-07 | -0.6781577 | 0.079 | 0.181 | 0.00659797 | TEC_prlf | F3 |
| 2.8031E-07 | -0.4067453 | 0.172 | 0.293 | 0.00662186 | TEC_prlf | GPRC5A |
| 3.0374E-07 | -0.3284908 | 0.061 | 0.161 | 0.00717514 | TEC_prlf | SERPINB9 |
| 3.1315E-07 | -0.3546318 | 0.02 | 0.104 | 0.00739743 | TEC_prlf | CLIC3 |
| 3.1667E-07 | -0.253537 | 0.076 | 0.184 | 0.00748077 | TEC_prlf | ICAM1 |
| 3.4776E-07 | -0.64115 | 0.137 | 0.239 | 0.00821506 | TEC_prlf | ADGRF1 |
| 3.646E-07 | 0.33761201 | 0.595 | 0.501 | 0.00861299 | TEC_prlf | LAPTM4A |
| 3.7238E-07 | -0.2732036 | 0.102 | 0.213 | 0.00879675 | TEC_prlf | TLE4 |
| 3.9543E-07 | 0.27986597 | 0.808 | 0.741 | 0.00934128 | TEC_prlf | APLP2 |
| 4.1911E-07 | 0.26742299 | 0.347 | 0.245 | 0.00990067 | TEC_prlf | CCT8 |
| 4.6387E-07 | -0.2636831 | 0.032 | 0.119 | 0.01095801 | TEC_prlf | ASS1 |
| 5.0137E-07 | -0.2785401 | 0.035 | 0.123 | 0.01184376 | TEC_prlf | KRT80 |
| 5.4807E-07 | -0.3739429 | 0.038 | 0.125 | 0.01294711 | TEC_prlf | SRD5A3 |
| 5.5972E-07 | -0.3592795 | 0.204 | 0.329 | 0.0132223 | TEC_prlf | SLC38A2 |
| 5.6696E-07 | -0.2780436 | 0.038 | 0.126 | 0.0133933 | TEC_prlf | POLB |
| 5.7285E-07 | 0.25215354 | 0.767 | 0.671 | 0.01353242 | TEC_prlf | COX5B |
| 6.2994E-07 | -0.3338744 | 0.134 | 0.249 | 0.01488102 | TEC_prlf | EGR1 |
| 6.4444E-07 | -0.4090535 | 0.087 | 0.186 | 0.01522363 | TEC_prlf | ACSL1 |

| | | | | | | |
|---|---|---|---|---|---|---|
| 6.4579E-07 | -0.2500734 | 0.184 | 0.302 | 0.01525553 | TEC_prlf | SERTAD1 |
| 6.5372E-07 | 0.27565755 | 0.469 | 0.362 | 0.01544277 | TEC_prlf | PSMB1 |
| 6.6158E-07 | -0.3919944 | 0.073 | 0.172 | 0.0156285 | TEC_prlf | DNAJA4 |
| 6.9995E-07 | 0.28193498 | 0.397 | 0.282 | 0.01653492 | TEC_prlf | CD151 |
| 8.5795E-07 | -0.2635651 | 0.155 | 0.268 | 0.02026725 | TEC_prlf | USP15 |
| 8.6453E-07 | -0.3368911 | 0.213 | 0.339 | 0.02042279 | TEC_prlf | NDRG1 |
| 9.3025E-07 | 0.29551579 | 0.781 | 0.703 | 0.0219753 | TEC_prlf | TPM3 |
| 9.4409E-07 | -0.4036171 | 0.122 | 0.227 | 0.02230213 | TEC_prlf | IER5L |
| 1.0375E-06 | 0.2518549 | 0.411 | 0.306 | 0.02450867 | TEC_prlf | BUD31 |
| 1.0531E-06 | -0.2970359 | 0.624 | 0.686 | 0.02487762 | TEC_prlf | ATP6V0B |
| 1.1054E-06 | -0.2761522 | 0.128 | 0.23 | 0.02611209 | TEC_prlf | PGD |
| 1.491E-06 | -0.2836333 | 0.079 | 0.176 | 0.03522235 | TEC_prlf | NR4A1 |
| 1.5153E-06 | -0.4499246 | 0.093 | 0.189 | 0.03579544 | TEC_prlf | EPS8L1 |
| 1.5619E-06 | -0.398211 | 0.91 | 0.889 | 0.03689741 | TEC_prlf | HLA-B |
| 1.5859E-06 | -0.3173722 | 0.064 | 0.155 | 0.0374638 | TEC_prlf | TMPRSS4 |
| 1.6484E-06 | -0.2792062 | 0.166 | 0.277 | 0.0389404 | TEC_prlf | VASP |
| 1.6745E-06 | -0.4453066 | 0.052 | 0.142 | 0.03955673 | TEC_prlf | SNCG |
| 1.7242E-06 | -1.2296005 | 0.055 | 0.143 | 0.04073107 | TEC_prlf | CCL2 |
| 1.7308E-06 | -0.397618 | 0.067 | 0.159 | 0.04088662 | TEC_prlf | BPGM |
| 1.8113E-06 | -0.3356366 | 0.108 | 0.209 | 0.0427888 | TEC_prlf | LY6E |
| 1.8252E-06 | 0.25519274 | 0.831 | 0.747 | 0.04311778 | TEC_prlf | CLIC1 |
| 1.9621E-06 | -0.3636546 | 0.087 | 0.183 | 0.04635007 | TEC_prlf | INSIG1 |
| 2.4168E-06 | -0.2737204 | 0.187 | 0.296 | 0.05709211 | TEC_prlf | ZFAND6 |
| 2.5345E-06 | -0.2676677 | 0.067 | 0.155 | 0.05987293 | TEC_prlf | ITPKC |
| 2.7398E-06 | 0.56316936 | 0.956 | 0.95 | 0.06472165 | TEC_prlf | MT-CO1 |
| 2.7516E-06 | -0.3169224 | 0.058 | 0.147 | 0.06500096 | TEC_prlf | ALDH1A3 |
| 2.7981E-06 | -0.315141 | 0.481 | 0.575 | 0.06609838 | TEC_prlf | PPP1CB |
| 2.8338E-06 | -0.28587 | 0.032 | 0.111 | 0.06694376 | TEC_prlf | SMIM5 |
| 2.8853E-06 | -0.4428283 | 0.085 | 0.175 | 0.06815878 | TEC_prlf | SPNS2 |
| 3.0855E-06 | 0.25438062 | 0.816 | 0.762 | 0.07288805 | TEC_prlf | UBL5 |
| 3.1391E-06 | -0.2549942 | 0.14 | 0.244 | 0.07415588 | TEC_prlf | RBPJ |
| 3.3625E-06 | -0.4291782 | 0.079 | 0.169 | 0.07943165 | TEC_prlf | FXYD3 |
| 3.7618E-06 | -1.3151266 | 0.058 | 0.145 | 0.08886519 | TEC_prlf | CXCL3 |
| 4.3144E-06 | -0.2667706 | 0.047 | 0.128 | 0.10191817 | TEC_prlf | LRG1 |
| 4.3288E-06 | -0.5795725 | 0.37 | 0.45 | 0.10225816 | TEC_prlf | MARCKS |
| 4.6778E-06 | -0.2894669 | 0.035 | 0.112 | 0.11050449 | TEC_prlf | ANGPTL4 |
| 4.6935E-06 | -0.3930288 | 0.122 | 0.215 | 0.11087545 | TEC_prlf | CREM |
| 5.0537E-06 | 0.28938591 | 0.496 | 0.388 | 0.11938465 | TEC_prlf | AHNAK |
| 5.071E-06 | -0.2644218 | 0.05 | 0.132 | 0.11979146 | TEC_prlf | CD83 |
| 5.0738E-06 | -0.255166 | 0.035 | 0.111 | 0.11985837 | TEC_prlf | STS |
| 5.7184E-06 | -0.4694246 | 0.029 | 0.104 | 0.13508476 | TEC_prlf | CCL5 |
| 5.7673E-06 | -0.2895493 | 0.251 | 0.361 | 0.13624068 | TEC_prlf | CFLAR |
| 6.6702E-06 | -0.4770162 | 0.706 | 0.727 | 0.15756975 | TEC_prlf | CYBA |
| 7.1939E-06 | -0.2546487 | 0.184 | 0.29 | 0.16994162 | TEC_prlf | MIR4435-2HG |
| 8.1467E-06 | -0.3491581 | 0.05 | 0.129 | 0.19245004 | TEC_prlf | GBP1 |
| 8.5429E-06 | -0.3886704 | 0.099 | 0.189 | 0.20180854 | TEC_prlf | HBEGF |
| 8.8751E-06 | -0.3315277 | 0.044 | 0.121 | 0.20965544 | TEC_prlf | HILPDA |
| 1.0632E-05 | -0.2563831 | 0.09 | 0.181 | 0.25115647 | TEC_prlf | WARS |
| 1.0717E-05 | -0.294444 | 0.035 | 0.107 | 0.25316824 | TEC_prlf | SLC16A10 |
| 1.1063E-05 | -0.2647254 | 0.991 | 0.984 | 0.26132947 | TEC_prlf | EIF1 |

| | | | | | | |
|---|---|---|---|---|---|---|
| 1.6468E-05 | -0.3573101 | 0.082 | 0.165 | 0.38903527 | TEC_prlf | ANKRD37 |
| 1.6925E-05 | 0.46622711 | 0.845 | 0.823 | 0.39980807 | TEC_prlf | MT-ND5 |
| 1.9622E-05 | -0.3391826 | 0.58 | 0.64 | 0.46352174 | TEC_prlf | ARPC5 |
| 2.0256E-05 | 0.4198854 | 0.589 | 0.506 | 0.47851675 | TEC_prlf | MT-ND4L |
| 2.0677E-05 | -0.7497413 | 0.534 | 0.555 | 0.48845291 | TEC_prlf | ELF3 |
| 2.3851E-05 | -0.2650275 | 0.079 | 0.161 | 0.56343998 | TEC_prlf | SOCS1 |
| 2.6541E-05 | -0.6876379 | 0.236 | 0.312 | 0.62697642 | TEC_prlf | EMP1 |
| 2.6634E-05 | 0.26086871 | 0.437 | 0.357 | 0.6291714 | TEC_prlf | COX5A |
| 2.7659E-05 | -0.495318 | 0.475 | 0.338 | 0.65339394 | TEC_prlf | MMP7 |
| 2.8009E-05 | -0.3276179 | 0.665 | 0.688 | 0.66165774 | TEC_prlf | BRI3 |
| 2.8847E-05 | 0.25087883 | 0.324 | 0.234 | 0.68144785 | TEC_prlf | MT-ATP8 |
| 2.9867E-05 | -0.2559626 | 0.134 | 0.224 | 0.70554556 | TEC_prlf | STXBP2 |
| 3.0568E-05 | -0.3459484 | 0.991 | 0.965 | 0.7221006 | TEC_prlf | S100A11 |
| 3.198E-05 | -0.6202313 | 0.431 | 0.504 | 0.7554693 | TEC_prlf | DNAJB1 |
| 3.6375E-05 | -0.7732644 | 0.047 | 0.118 | 0.85928895 | TEC_prlf | CXCL2 |
| 3.9483E-05 | -0.3388009 | 0.137 | 0.225 | 0.93270926 | TEC_prlf | INTS6 |
| 4.6494E-05 | -0.4695131 | 0.152 | 0.237 | 1 | TEC_prlf | FABP5 |
| 4.7758E-05 | -0.3012896 | 0.327 | 0.427 | 1 | TEC_prlf | SLC3A2 |
| 5.0017E-05 | -0.549361 | 0.105 | 0.185 | 1 | TEC_prlf | AQP3 |
| 5.0236E-05 | -0.2786498 | 0.417 | 0.502 | 1 | TEC_prlf | ASAH1 |
| 6.2738E-05 | 0.43565151 | 0.913 | 0.933 | 1 | TEC_prlf | MT-ND4 |
| 6.3728E-05 | -0.2589875 | 0.093 | 0.173 | 1 | TEC_prlf | XIST |
| 6.7064E-05 | -0.2771746 | 0.539 | 0.628 | 1 | TEC_prlf | JUN |
| 7.5098E-05 | -0.6183236 | 0.102 | 0.181 | 1 | TEC_prlf | SPINK1 |
| 7.5643E-05 | -0.3434294 | 0.501 | 0.571 | 1 | TEC_prlf | ZFAND5 |
| 8.3571E-05 | -0.3890882 | 0.259 | 0.352 | 1 | TEC_prlf | PHLDA2 |
| 0.00011815 | 0.38872656 | 0.898 | 0.93 | 1 | TEC_prlf | MT-CYB |
| 0.00013197 | -0.3550513 | 0.082 | 0.152 | 1 | TEC_prlf | ZFAND2A |
| 0.00017364 | -0.3142544 | 0.259 | 0.347 | 1 | TEC_prlf | ZFP36L2 |
| 0.00018931 | -0.3241462 | 0.108 | 0.179 | 1 | TEC_prlf | EPHA2 |
| 0.00024079 | -0.5306421 | 0.12 | 0.189 | 1 | TEC_prlf | ANKRD36C |
| 0.00028858 | -0.2808233 | 0.265 | 0.343 | 1 | TEC_prlf | WSB1 |
| 0.00029171 | -0.2520729 | 0.3 | 0.381 | 1 | TEC_prlf | N4BP2L2 |
| 0.00036389 | -0.2542963 | 0.773 | 0.811 | 1 | TEC_prlf | DDX5 |
| 0.00038318 | -0.3282442 | 0.093 | 0.162 | 1 | TEC_prlf | CSKMT |
| 0.00040788 | -1.3156015 | 0.085 | 0.152 | 1 | TEC_prlf | MT1H |
| 0.00044915 | -0.2572644 | 0.122 | 0.191 | 1 | TEC_prlf | ABLIM1 |
| 0.0004572 | -1.2286298 | 0.458 | 0.491 | 1 | TEC_prlf | CD74 |
| 0.00047354 | -0.2507655 | 0.061 | 0.122 | 1 | TEC_prlf | AGR2 |
| 0.00053732 | -0.3725009 | 0.44 | 0.482 | 1 | TEC_prlf | CD46 |
| 0.00055027 | -0.3161401 | 0.085 | 0.148 | 1 | TEC_prlf | RASD1 |
| 0.00059935 | -0.2828275 | 0.187 | 0.265 | 1 | TEC_prlf | MGST1 |
| 0.00065933 | -0.2807718 | 0.163 | 0.238 | 1 | TEC_prlf | CITED2 |
| 0.00070173 | -0.4654842 | 0.364 | 0.434 | 1 | TEC_prlf | HIST1H1C |
| 0.00076614 | -0.2598883 | 0.19 | 0.267 | 1 | TEC_prlf | ID2 |
| 0.00111497 | -0.5367126 | 0.082 | 0.14 | 1 | TEC_prlf | CXCL1 |
| 0.0011168 | -0.253389 | 0.187 | 0.25 | 1 | TEC_prlf | KLF5 |
| 0.00114294 | -0.3649571 | 0.224 | 0.296 | 1 | TEC_prlf | GLRX |
| 0.00125941 | -0.3496775 | 0.58 | 0.619 | 1 | TEC_prlf | PCBP1 |
| 0.00128889 | -0.4150226 | 0.236 | 0.294 | 1 | TEC_prlf | ATP13A3 |

| | | | | | | |
|---|---|---|---|---|---|---|
| 0.00149665 | -0.2710053 | 0.213 | 0.284 | 1 | TEC_prlf | MARCKSL1 |
| 0.00156344 | -0.3855426 | 0.948 | 0.899 | 1 | TEC_prlf | RPL39 |
| 0.00167246 | -0.2909976 | 0.478 | 0.514 | 1 | TEC_prlf | ANXA11 |
| 0.00177213 | -0.7573233 | 0.294 | 0.345 | 1 | TEC_prlf | GDF15 |
| 0.00198065 | -0.3487206 | 0.391 | 0.262 | 1 | TEC_prlf | LMO7 |
| 0.0022342 | -0.3220224 | 0.169 | 0.23 | 1 | TEC_prlf | PPIF |
| 0.00259437 | 0.27477299 | 0.35 | 0.292 | 1 | TEC_prlf | TUBA1C |
| 0.00291357 | -0.3369825 | 0.353 | 0.422 | 1 | TEC_prlf | TXNRD1 |
| 0.00296123 | 0.25017782 | 0.207 | 0.152 | 1 | TEC_prlf | MT-ND6 |
| 0.00298099 | -0.8283705 | 0.219 | 0.28 | 1 | TEC_prlf | MT1X |
| 0.00364605 | -0.3645566 | 0.122 | 0.173 | 1 | TEC_prlf | PPL |
| 0.00495771 | -0.7837215 | 0.414 | 0.451 | 1 | TEC_prlf | SOX4 |
| 0.00508228 | -1.5765166 | 0.198 | 0.255 | 1 | TEC_prlf | MT1G |
| 0.00616417 | 0.26187442 | 0.324 | 0.275 | 1 | TEC_prlf | CHMP4B |
| 0.0088461 | -0.2827059 | 0.137 | 0.188 | 1 | TEC_prlf | BAG3 |
| 0 | 6.10497449 | 0.998 | 0.489 | 0 | TEC_prg | PLCG2 |
| 0 | 2.35810663 | 0.855 | 0.404 | 0 | TEC_prg | SOX4 |
| 0 | 2.33222761 | 0.901 | 0.607 | 0 | TEC_prg | MTRNR2L12 |
| 0 | 2.13308676 | 0.701 | 0.264 | 0 | TEC_prg | HIST1H4C |
| 0 | 2.07988725 | 0.605 | 0.06 | 0 | TEC_prg | HSPA2 |
| 0 | 1.98357384 | 0.668 | 0.176 | 0 | TEC_prg | HEXIM1 |
| 0 | 1.93934872 | 0.914 | 0.595 | 0 | TEC_prg | JUN |
| 0 | 1.92283345 | 0.636 | 0.241 | 0 | TEC_prg | MAFB |
| 0 | 1.78681377 | 0.583 | 0.113 | 0 | TEC_prg | CSKMT |
| 0 | 1.7556387 | 0.522 | 0.074 | 0 | TEC_prg | TCIM |
| 0 | 1.62743319 | 0.587 | 0.179 | 0 | TEC_prg | HES1 |
| 0 | 1.56254731 | 0.556 | 0.101 | 0 | TEC_prg | RASD1 |
| 0 | 1.51271852 | 0.557 | 0.149 | 0 | TEC_prg | PLK2 |
| 0 | 1.45821359 | 0.582 | 0.184 | 0 | TEC_prg | PPP1R10 |
| 0 | 1.44820945 | 0.573 | 0.248 | 0 | TEC_prg | RHOB |
| 0 | 1.44421156 | 0.636 | 0.293 | 0 | TEC_prg | SLC38A2 |
| 0 | 1.38008616 | 0.863 | 0.656 | 0 | TEC_prg | IER2 |
| 0 | 1.35362362 | 0.464 | 0.035 | 0 | TEC_prg | PIK3R3 |
| 0 | 1.3218932 | 0.396 | 0.05 | 0 | TEC_prg | ADAMTS1 |
| 0 | 1.31633289 | 0.63 | 0.363 | 0 | TEC_prg | SRSF7 |
| 0 | 1.3023835 | 0.893 | 0.702 | 0 | TEC_prg | JUND |
| 0 | 1.26741418 | 0.576 | 0.213 | 0 | TEC_prg | SYNE2 |
| 0 | 1.25788756 | 0.51 | 0.192 | 0 | TEC_prg | INTS6 |
| 0 | 1.24800164 | 0.529 | 0.191 | 0 | TEC_prg | IER5L |
| 0 | 1.24059104 | 0.461 | 0.087 | 0 | TEC_prg | ID4 |
| 0 | 1.18934103 | 0.462 | 0.126 | 0 | TEC_prg | SOCS1 |
| 0 | 1.14759217 | 0.833 | 0.614 | 0 | TEC_prg | GADD45B |
| 0 | 1.14396088 | 0.904 | 0.515 | 0 | TEC_prg | ELF3 |
| 0 | 1.13390952 | 0.485 | 0.192 | 0 | TEC_prg | POLR2A |
| 0 | 1.11127318 | 0.408 | 0.086 | 0 | TEC_prg | EGLN3 |
| 0 | 1.05704229 | 0.527 | 0.295 | 0 | TEC_prg | BRD2 |
| 0 | 1.05074826 | 0.861 | 0.616 | 0 | TEC_prg | IER3 |
| 0 | 1.04111058 | 0.466 | 0.223 | 0 | TEC_prg | EGR1 |
| 0 | 1.03700124 | 0.178 | 0.038 | 0 | TEC_prg | CCDC80 |
| 0 | 1.03579312 | 0.472 | 0.15 | 0 | TEC_prg | SGK1 |

| | | | | | | |
|---|---|---|---|---|---|---|
| 0 | 1.03074983 | 0.279 | 0.027 | 0 | TEC_prg | COL7A1 |
| 0 | 1.02127996 | 0.494 | 0.232 | 0 | TEC_prg | TPM1 |
| 0 | 1.01666995 | 0.493 | 0.223 | 0 | TEC_prg | SNHG12 |
| 0 | 0.99945383 | 1 | 0.978 | 0 | TEC_prg | MALAT1 |
| 0 | 0.99690126 | 0.775 | 0.594 | 0 | TEC_prg | SRRM2 |
| 0 | 0.97810132 | 0.429 | 0.11 | 0 | TEC_prg | PAX8 |
| 0 | 0.95005681 | 0.43 | 0.175 | 0 | TEC_prg | AMD1 |
| 0 | 0.94669736 | 0.536 | 0.251 | 0 | TEC_prg | CLU |
| 0 | 0.93351812 | 0.465 | 0.211 | 0 | TEC_prg | CITED2 |
| 0 | 0.92100813 | 0.361 | 0.099 | 0 | TEC_prg | SYNE1 |
| 0 | 0.92065936 | 0.204 | 0.042 | 0 | TEC_prg | NEFM |
| 0 | 0.86749071 | 0.442 | 0.211 | 0 | TEC_prg | CDKN1C |
| 0 | 0.85994883 | 0.264 | 0.04 | 0 | TEC_prg | AP001160.1 |
| 0 | 0.85939418 | 0.33 | 0.07 | 0 | TEC_prg | AC025164.1 |
| 0 | 0.85216232 | 0.406 | 0.189 | 0 | TEC_prg | TLE4 |
| 0 | 0.84320333 | 0.449 | 0.212 | 0 | TEC_prg | NFKBIZ |
| 0 | 0.83779363 | 0.329 | 0.056 | 0 | TEC_prg | HIST2H2BF |
| 0 | 0.81874665 | 0.327 | 0.105 | 0 | TEC_prg | H2AFX |
| 0 | 0.81728555 | 0.365 | 0.091 | 0 | TEC_prg | AC007952.4 |
| 0 | 0.78347252 | 0.353 | 0.102 | 0 | TEC_prg | ERN1 |
| 0 | 0.76785914 | 0.232 | 0.007 | 0 | TEC_prg | PCDH10 |
| 0 | 0.73530525 | 0.291 | 0.027 | 0 | TEC_prg | HIST4H4 |
| 0 | 0.72670953 | 0.284 | 0.047 | 0 | TEC_prg | HOMER1 |
| 0 | 0.72623304 | 0.36 | 0.147 | 0 | TEC_prg | CDKN2AIP |
| 0 | 0.71340869 | 0.286 | 0.052 | 0 | TEC_prg | CSRNP2 |
| 0 | 0.70979254 | 0.274 | 0.037 | 0 | TEC_prg | AL355075.4 |
| 0 | 0.70864227 | 0.208 | 0.035 | 0 | TEC_prg | ARC |
| 0 | 0.68986007 | 0.35 | 0.132 | 0 | TEC_prg | CYR61 |
| 0 | 0.67590889 | 0.277 | 0.043 | 0 | TEC_prg | AL135925.1 |
| 0 | 0.66164396 | 0.243 | 0.023 | 0 | TEC_prg | PCDH9 |
| 0 | 0.65516547 | 0.256 | 0.018 | 0 | TEC_prg | MAFA |
| 0 | 0.65419615 | 0.26 | 0.016 | 0 | TEC_prg | AL137077.2 |
| 0 | 0.64599171 | 0.248 | 0.009 | 0 | TEC_prg | HIST1H3A |
| 0 | 0.64363353 | 0.182 | 0.012 | 0 | TEC_prg | HIST1H1A |
| 0 | 0.64352709 | 0.311 | 0.114 | 0 | TEC_prg | COL18A1 |
| 0 | 0.63985821 | 0.186 | 0.031 | 0 | TEC_prg | HIST1H2AG |
| 0 | 0.61966986 | 0.28 | 0.09 | 0 | TEC_prg | AL118516.1 |
| 0 | 0.61300518 | 0.363 | 0.154 | 0 | TEC_prg | RERE |
| 0 | 0.60808073 | 0.209 | 0.033 | 0 | TEC_prg | AL021453.1 |
| 0 | 0.60289165 | 0.238 | 0.03 | 0 | TEC_prg | Z93241.1 |
| 0 | 0.59975837 | 0.246 | 0.036 | 0 | TEC_prg | AC012306.2 |
| 0 | 0.59405953 | 0.253 | 0.048 | 0 | TEC_prg | PLEKHH2 |
| 0 | 0.57498909 | 0.288 | 0.094 | 0 | TEC_prg | MYO9A |
| 0 | 0.56745596 | 0.208 | 0.028 | 0 | TEC_prg | EGOT |
| 0 | 0.56711875 | 0.188 | 0.017 | 0 | TEC_prg | POU3F1 |
| 0 | 0.56184722 | 0.257 | 0.027 | 0 | TEC_prg | PKHD1 |
| 0 | 0.54194121 | 0.248 | 0.021 | 0 | TEC_prg | LINC00472 |
| 0 | 0.54175633 | 0.292 | 0.099 | 0 | TEC_prg | RNPC3 |
| 0 | 0.5165542 | 0.246 | 0.073 | 0 | TEC_prg | MEPCE |
| 0 | 0.49353319 | 0.202 | 0.022 | 0 | TEC_prg | TAF4B |

| | | | | | | |
|---|---|---|---|---|---|---|
| 0 | 0.48867906 | 0.204 | 0.036 | 0 | TEC_prg | NRARP |
| 0 | 0.4730636 | 0.214 | 0.022 | 0 | TEC_prg | KIF12 |
| 0 | 0.47036806 | 0.171 | 0.04 | 0 | TEC_prg | PARD6G-AS1 |
| 0 | 0.46966626 | 0.215 | 0.051 | 0 | TEC_prg | THSD7A |
| 0 | 0.45983469 | 0.212 | 0.04 | 0 | TEC_prg | AC087239.1 |
| 0 | 0.45583514 | 0.247 | 0.072 | 0 | TEC_prg | HERC2 |
| 0 | 0.45444376 | 0.234 | 0.048 | 0 | TEC_prg | PLXNB1 |
| 0 | 0.44851666 | 0.164 | 0.011 | 0 | TEC_prg | AC005837.1 |
| 0 | 0.4442666 | 0.233 | 0.07 | 0 | TEC_prg | WDR60 |
| 0 | 0.44099831 | 0.152 | 0.021 | 0 | TEC_prg | SLC12A1 |
| 0 | 0.44036102 | 0.193 | 0.011 | 0 | TEC_prg | AC069224.1 |
| 0 | 0.43810197 | 0.21 | 0.058 | 0 | TEC_prg | DALRD3 |
| 0 | 0.43785164 | 0.229 | 0.057 | 0 | TEC_prg | EMX2 |
| 0 | 0.41515387 | 0.152 | 0.019 | 0 | TEC_prg | AC069185.1 |
| 0 | 0.413605 | 0.18 | 0.037 | 0 | TEC_prg | FDXR |
| 0 | 0.39999062 | 0.149 | 0.025 | 0 | TEC_prg | ADRB2 |
| 0 | 0.39408484 | 0.136 | 0.006 | 0 | TEC_prg | LINC02432 |
| 0 | 0.39287612 | 0.206 | 0.048 | 0 | TEC_prg | GOLGA8A |
| 0 | 0.38003044 | 0.164 | 0.034 | 0 | TEC_prg | PDGFB |
| 0 | 0.37935203 | 0.162 | 0.024 | 0 | TEC_prg | FZD8 |
| 0 | 0.36567002 | 0.144 | 0.022 | 0 | TEC_prg | C2CD4B |
| 0 | 0.35651959 | 0.163 | 0.008 | 0 | TEC_prg | AL022069.1 |
| 0 | 0.34142794 | 0.138 | 0.004 | 0 | TEC_prg | HIST2H4B |
| 0 | 0.34046518 | 0.145 | 0.007 | 0 | TEC_prg | HIST1H4B |
| 0 | 0.3360558 | 0.129 | 0.022 | 0 | TEC_prg | AC019197.1 |
| 0 | 0.32961505 | 0.166 | 0.019 | 0 | TEC_prg | MST1 |
| 0 | 0.32792286 | 0.141 | 0.012 | 0 | TEC_prg | AL136040.1 |
| 0 | 0.32079804 | 0.184 | 0.043 | 0 | TEC_prg | GOLGA8B |
| 0 | 0.30353367 | 0.141 | 0.017 | 0 | TEC_prg | COL27A1 |
| 0 | 0.3033942 | 0.127 | 0.007 | 0 | TEC_prg | HIST1H4A |
| 0 | 0.29609225 | 0.116 | 0.002 | 0 | TEC_prg | HIST1H3C |
| 0 | 0.28171309 | 0.135 | 0.017 | 0 | TEC_prg | DDIT4L |
| 0 | 0.27986541 | 0.115 | 0.009 | 0 | TEC_prg | HIST1H2AH |
| 0 | 0.27837686 | 0.145 | 0.027 | 0 | TEC_prg | INTU |
| 0 | 0.27741482 | 0.122 | 0.013 | 0 | TEC_prg | NANOS1 |
| 0 | 0.27627768 | 0.136 | 0.022 | 0 | TEC_prg | CCSER1 |
| 0 | 0.26845317 | 0.11 | 0.007 | 0 | TEC_prg | AC011498.1 |
| 0 | 0.25781603 | 0.126 | 0.02 | 0 | TEC_prg | EVC |
| 0 | 0.25528641 | 0.123 | 0.018 | 0 | TEC_prg | SIM2 |
| 0 | -0.4641213 | 0.087 | 0.397 | 0 | TEC_prg | ELOC |
| 0 | -0.4663481 | 0.372 | 0.666 | 0 | TEC_prg | ARF1 |
| 0 | -0.4673376 | 0.092 | 0.405 | 0 | TEC_prg | GNB2 |
| 0 | -0.4742843 | 0.081 | 0.383 | 0 | TEC_prg | SNX3 |
| 0 | -0.4749311 | 0.067 | 0.36 | 0 | TEC_prg | COPS9 |
| 0 | -0.4843556 | 0.074 | 0.379 | 0 | TEC_prg | SNRPB |
| 0 | -0.4869508 | 0.059 | 0.357 | 0 | TEC_prg | SRP9 |
| 0 | -0.4905552 | 0.048 | 0.336 | 0 | TEC_prg | BCAP31 |
| 0 | -0.4922026 | 0.113 | 0.441 | 0 | TEC_prg | NDUFB4 |
| 0 | -0.493028 | 0.046 | 0.349 | 0 | TEC_prg | RER1 |
| 0 | -0.4937796 | 0.074 | 0.374 | 0 | TEC_prg | SSR2 |

| | | | | | | |
|---|---|---|---|---|---|---|
| 0 | -0.49588 | 0.089 | 0.409 | 0 | TEC_prg | EIF3H |
| 0 | -0.5042328 | 0.047 | 0.331 | 0 | TEC_prg | PGAM1 |
| 0 | -0.5059954 | 0.094 | 0.42 | 0 | TEC_prg | RAB2A |
| 0 | -0.5064651 | 0.05 | 0.352 | 0 | TEC_prg | LEPROT |
| 0 | -0.5077222 | 0.058 | 0.368 | 0 | TEC_prg | CHMP5 |
| 0 | -0.5084227 | 0.093 | 0.398 | 0 | TEC_prg | REEP5 |
| 0 | -0.5107383 | 0.101 | 0.424 | 0 | TEC_prg | ATP5MPL |
| 0 | -0.5109962 | 0.101 | 0.436 | 0 | TEC_prg | CSDE1 |
| 0 | -0.5150025 | 0.059 | 0.368 | 0 | TEC_prg | CAPZA1 |
| 0 | -0.5165679 | 0.075 | 0.39 | 0 | TEC_prg | ATP5PD |
| 0 | -0.5201573 | 0.06 | 0.346 | 0 | TEC_prg | RAB11A |
| 0 | -0.5207873 | 0.071 | 0.399 | 0 | TEC_prg | RAB7A |
| 0 | -0.5219371 | 0.067 | 0.377 | 0 | TEC_prg | MTPN |
| 0 | -0.5257105 | 0.07 | 0.39 | 0 | TEC_prg | COX5A |
| 0 | -0.5269703 | 0.045 | 0.364 | 0 | TEC_prg | NDUFC2 |
| 0 | -0.5273462 | 0.065 | 0.373 | 0 | TEC_prg | NDFIP1 |
| 0 | -0.5277349 | 0.02 | 0.286 | 0 | TEC_prg | ABRACL |
| 0 | -0.5335206 | 0.074 | 0.397 | 0 | TEC_prg | PSMB1 |
| 0 | -0.543026 | 0.079 | 0.426 | 0 | TEC_prg | CHMP2A |
| 0 | -0.5436319 | 0.068 | 0.382 | 0 | TEC_prg | VDAC1 |
| 0 | -0.5445702 | 0.088 | 0.42 | 0 | TEC_prg | NEDD8 |
| 0 | -0.5482207 | 0.09 | 0.419 | 0 | TEC_prg | CNBP |
| 0 | -0.5486868 | 0.143 | 0.468 | 0 | TEC_prg | ATP5MF |
| 0 | -0.5492588 | 0.507 | 0.75 | 0 | TEC_prg | COX6A1 |
| 0 | -0.551586 | 0.086 | 0.401 | 0 | TEC_prg | TXNDC17 |
| 0 | -0.5519969 | 0.097 | 0.402 | 0 | TEC_prg | ATP5MC3 |
| 0 | -0.5525349 | 0.429 | 0.707 | 0 | TEC_prg | COX7A2 |
| 0 | -0.5530999 | 0.038 | 0.324 | 0 | TEC_prg | CD99 |
| 0 | -0.5554508 | 0.681 | 0.848 | 0 | TEC_prg | YWHAZ |
| 0 | -0.5557145 | 0.331 | 0.637 | 0 | TEC_prg | SUB1 |
| 0 | -0.5568302 | 0.05 | 0.351 | 0 | TEC_prg | AP2S1 |
| 0 | -0.5568411 | 0.083 | 0.396 | 0 | TEC_prg | SLC25A5 |
| 0 | -0.557589 | 0.033 | 0.314 | 0 | TEC_prg | CXCL16 |
| 0 | -0.5581531 | 0.043 | 0.364 | 0 | TEC_prg | PSMB3 |
| 0 | -0.5594765 | 0.083 | 0.401 | 0 | TEC_prg | NDUFV2 |
| 0 | -0.5662421 | 0.085 | 0.424 | 0 | TEC_prg | SPCS2 |
| 0 | -0.5690618 | 0.068 | 0.361 | 0 | TEC_prg | MSN |
| 0 | -0.5817337 | 0.076 | 0.431 | 0 | TEC_prg | NDUFB11 |
| 0 | -0.5834663 | 0.054 | 0.366 | 0 | TEC_prg | SSR3 |
| 0 | -0.5837924 | 0.518 | 0.767 | 0 | TEC_prg | SERP1 |
| 0 | -0.5846536 | 0.065 | 0.403 | 0 | TEC_prg | SEC11A |
| 0 | -0.58763 | 0.032 | 0.353 | 0 | TEC_prg | FKBP1A |
| 0 | -0.5907669 | 0.454 | 0.723 | 0 | TEC_prg | GUK1 |
| 0 | -0.5917301 | 0.095 | 0.432 | 0 | TEC_prg | H2AFY |
| 0 | -0.5922276 | 0.381 | 0.657 | 0 | TEC_prg | OST4 |
| 0 | -0.5925148 | 0.091 | 0.42 | 0 | TEC_prg | PSME1 |
| 0 | -0.5936644 | 0.102 | 0.429 | 0 | TEC_prg | COMMD6 |
| 0 | -0.5977882 | 0.037 | 0.325 | 0 | TEC_prg | IFNGR2 |
| 0 | -0.5987823 | 0.034 | 0.376 | 0 | TEC_prg | ARPC4 |
| 0 | -0.599402 | 0.438 | 0.734 | 0 | TEC_prg | SUMO2 |

| | | | | | | |
|---|---|---|---|---|---|---|
| 0 | -0.5999278 | 0.039 | 0.34 | 0 | TEC_prg | ARPC1B |
| 0 | -0.599943 | 0.496 | 0.748 | 0 | TEC_prg | YWHAB |
| 0 | -0.6004403 | 0.908 | 0.96 | 0 | TEC_prg | SERF2 |
| 0 | -0.6010009 | 0.088 | 0.408 | 0 | TEC_prg | PARK7 |
| 0 | -0.60313 | 0.15 | 0.521 | 0 | TEC_prg | TMEM50A |
| 0 | -0.6079209 | 0.297 | 0.606 | 0 | TEC_prg | PPP1CB |
| 0 | -0.6092293 | 0.029 | 0.351 | 0 | TEC_prg | CMTM6 |
| 0 | -0.6095444 | 0.325 | 0.65 | 0 | TEC_prg | COX8A |
| 0 | -0.6098479 | 0.482 | 0.741 | 0 | TEC_prg | ATP5MG |
| 0 | -0.6104053 | 0.417 | 0.673 | 0 | TEC_prg | ATP5MC2 |
| 0 | -0.6155251 | 0.375 | 0.653 | 0 | TEC_prg | ACTR2 |
| 0 | -0.615594 | 0.054 | 0.356 | 0 | TEC_prg | TMBIM1 |
| 0 | -0.6191482 | 0.105 | 0.45 | 0 | TEC_prg | EIF3K |
| 0 | -0.6193811 | 0.287 | 0.618 | 0 | TEC_prg | ACTR3 |
| 0 | -0.6205917 | 0.07 | 0.423 | 0 | TEC_prg | DAD1 |
| 0 | -0.6207509 | 0.146 | 0.507 | 0 | TEC_prg | RAP1B |
| 0 | -0.6207535 | 0.136 | 0.462 | 0 | TEC_prg | SELENOK |
| 0 | -0.6268782 | 0.081 | 0.414 | 0 | TEC_prg | EIF5A |
| 0 | -0.6280972 | 0.187 | 0.521 | 0 | TEC_prg | DYNLT1 |
| 0 | -0.6284061 | 0.055 | 0.387 | 0 | TEC_prg | EIF3E |
| 0 | -0.6353039 | 0.47 | 0.71 | 0 | TEC_prg | BZW1 |
| 0 | -0.6366352 | 0.086 | 0.446 | 0 | TEC_prg | PTP4A2 |
| 0 | -0.6367617 | 0.075 | 0.403 | 0 | TEC_prg | PRDX6 |
| 0 | -0.6369643 | 0.012 | 0.326 | 0 | TEC_prg | FXYD5 |
| 0 | -0.6380411 | 0.219 | 0.555 | 0 | TEC_prg | ATP6V1F |
| 0 | -0.6456721 | 0.117 | 0.45 | 0 | TEC_prg | GRN |
| 0 | -0.6458321 | 0.411 | 0.677 | 0 | TEC_prg | UQCRH |
| 0 | -0.6489439 | 0.039 | 0.414 | 0 | TEC_prg | BRK1 |
| 0 | -0.6512856 | 0.071 | 0.415 | 0 | TEC_prg | PRELID1 |
| 0 | -0.6576827 | 0.075 | 0.394 | 0 | TEC_prg | GNAI2 |
| 0 | -0.6596975 | 0.189 | 0.521 | 0 | TEC_prg | CANX |
| 0 | -0.6611268 | 0.602 | 0.764 | 0 | TEC_prg | CLIC1 |
| 0 | -0.6635025 | 0.175 | 0.54 | 0 | TEC_prg | LAPTM4A |
| 0 | -0.6658717 | 0.533 | 0.753 | 0 | TEC_prg | ATP6V1G1 |
| 0 | -0.6662796 | 0.172 | 0.517 | 0 | TEC_prg | GABARAPL2 |
| 0 | -0.6669197 | 0.054 | 0.353 | 0 | TEC_prg | B4GALT1 |
| 0 | -0.6674519 | 0.731 | 0.846 | 0 | TEC_prg | PPIA |
| 0 | -0.6729901 | 0.211 | 0.524 | 0 | TEC_prg | TAGLN2 |
| 0 | -0.6735221 | 0.601 | 0.807 | 0 | TEC_prg | BTF3 |
| 0 | -0.6775746 | 0.337 | 0.643 | 0 | TEC_prg | SEC61B |
| 0 | -0.6786121 | 0.272 | 0.62 | 0 | TEC_prg | NOP10 |
| 0 | -0.6816712 | 0.204 | 0.535 | 0 | TEC_prg | ASAH1 |
| 0 | -0.6819316 | 0.002 | 0.274 | 0 | TEC_prg | GMFG |
| 0 | -0.6824654 | 0.145 | 0.499 | 0 | TEC_prg | SARAF |
| 0 | -0.684114 | 0.137 | 0.489 | 0 | TEC_prg | SEC61G |
| 0 | -0.6843724 | 0.522 | 0.798 | 0 | TEC_prg | CDC42 |
| 0 | -0.6861041 | 0.061 | 0.438 | 0 | TEC_prg | CIB1 |
| 0 | -0.6881232 | 0.969 | 0.979 | 0 | TEC_prg | RPL28 |
| 0 | -0.6881735 | 0.06 | 0.461 | 0 | TEC_prg | CAPZB |
| 0 | -0.6919719 | 0.222 | 0.58 | 0 | TEC_prg | POMP |

| | | | | | | |
|---|---|---|---|---|---|---|
| 0 | -0.6929887 | 0.26 | 0.587 | 0 | TEC_prg | SSR4 |
| 0 | -0.6998372 | 0.681 | 0.853 | 0 | TEC_prg | HLA-E |
| 0 | -0.7034005 | 0.33 | 0.63 | 0 | TEC_prg | VAMP8 |
| 0 | -0.7110914 | 0.683 | 0.854 | 0 | TEC_prg | COX4I1 |
| 0 | -0.7129551 | 0.714 | 0.895 | 0 | TEC_prg | TMBIM6 |
| 0 | -0.7130672 | 0.443 | 0.716 | 0 | TEC_prg | PSMA7 |
| 0 | -0.7143243 | 0.062 | 0.374 | 0 | TEC_prg | PDIA6 |
| 0 | -0.7151629 | 0.427 | 0.716 | 0 | TEC_prg | ATP6V0B |
| 0 | -0.7194864 | 0.867 | 0.903 | 0 | TEC_prg | RPL27A |
| 0 | -0.7195064 | 0.064 | 0.353 | 0 | TEC_prg | LIMS1 |
| 0 | -0.7197775 | 0.306 | 0.651 | 0 | TEC_prg | CAP1 |
| 0 | -0.7207365 | 0.86 | 0.9 | 0 | TEC_prg | RPL35A |
| 0 | -0.7234917 | 0.535 | 0.798 | 0 | TEC_prg | RHOA |
| 0 | -0.723732 | 0.36 | 0.652 | 0 | TEC_prg | TMEM59 |
| 0 | -0.7285438 | 0.608 | 0.815 | 0 | TEC_prg | CHCHD2 |
| 0 | -0.7370762 | 0.485 | 0.777 | 0 | TEC_prg | ATP6V0E1 |
| 0 | -0.7432186 | 0.269 | 0.579 | 0 | TEC_prg | PPIB |
| 0 | -0.7466872 | 0.92 | 0.923 | 0 | TEC_prg | RPL8 |
| 0 | -0.7537207 | 0.112 | 0.478 | 0 | TEC_prg | PDIA3 |
| 0 | -0.7601293 | 0.026 | 0.292 | 0 | TEC_prg | MGST1 |
| 0 | -0.7659885 | 0.87 | 0.904 | 0 | TEC_prg | RPS11 |
| 0 | -0.7674812 | 0.005 | 0.298 | 0 | TEC_prg | CD53 |
| 0 | -0.7686189 | 0.586 | 0.825 | 0 | TEC_prg | GNG5 |
| 0 | -0.7697401 | 0.396 | 0.668 | 0 | TEC_prg | ARPC5 |
| 0 | -0.7705995 | 0.453 | 0.739 | 0 | TEC_prg | SDCBP |
| 0 | -0.7762063 | 0.283 | 0.565 | 0 | TEC_prg | SERPINB1 |
| 0 | -0.7778032 | 0.594 | 0.835 | 0 | TEC_prg | RAC1 |
| 0 | -0.7797941 | 0.251 | 0.618 | 0 | TEC_prg | PGK1 |
| 0 | -0.7797971 | 0.23 | 0.584 | 0 | TEC_prg | C4orf3 |
| 0 | -0.7826248 | 0.107 | 0.465 | 0 | TEC_prg | EFHD2 |
| 0 | -0.7992369 | 0.319 | 0.605 | 0 | TEC_prg | CD55 |
| 0 | -0.8024708 | 0.787 | 0.918 | 0 | TEC_prg | CFL1 |
| 0 | -0.803279 | 0.006 | 0.273 | 0 | TEC_prg | CSTA |
| 0 | -0.8042958 | 0.019 | 0.346 | 0 | TEC_prg | CD68 |
| 0 | -0.8074284 | 0.468 | 0.713 | 0 | TEC_prg | BRI3 |
| 0 | -0.8104351 | 0.737 | 0.891 | 0 | TEC_prg | MYL12A |
| 0 | -0.8132291 | 0.857 | 0.907 | 0 | TEC_prg | RPL24 |
| 0 | -0.822293 | 0.747 | 0.9 | 0 | TEC_prg | MYL12B |
| 0 | -0.823747 | 0.089 | 0.4 | 0 | TEC_prg | COTL1 |
| 0 | -0.8267497 | 0.925 | 0.968 | 0 | TEC_prg | MYL6 |
| 0 | -0.829655 | 0.868 | 0.878 | 0 | TEC_prg | RPL23A |
| 0 | -0.8298739 | 0.716 | 0.832 | 0 | TEC_prg | RPL27 |
| 0 | -0.8331436 | 0.012 | 0.361 | 0 | TEC_prg | ARHGDIB |
| 0 | -0.8355634 | 0.766 | 0.882 | 0 | TEC_prg | HLA-C |
| 0 | -0.8372224 | 0.882 | 0.9 | 0 | TEC_prg | RPL29 |
| 0 | -0.8384092 | 0.885 | 0.936 | 0 | TEC_prg | FAU |
| 0 | -0.8512977 | 0.572 | 0.825 | 0 | TEC_prg | ARPC2 |
| 0 | -0.8543794 | 0.267 | 0.608 | 0 | TEC_prg | TSPO |
| 0 | -0.8575119 | 0.833 | 0.903 | 0 | TEC_prg | HLA-A |
| 0 | -0.857551 | 0.82 | 0.916 | 0 | TEC_prg | ITM2B |

| | | | | | | |
|---|---|---|---|---|---|---|
| 0 | -0.8610974 | 0.315 | 0.63 | 0 | TEC_prg | NPC2 |
| 0 | -0.8663784 | 0.991 | 0.979 | 0 | TEC_prg | EEF1A1 |
| 0 | -0.8667671 | 0.037 | 0.332 | 0 | TEC_prg | SERPINA1 |
| 0 | -0.8699831 | 0.868 | 0.886 | 0 | TEC_prg | HSP90AB1 |
| 0 | -0.8752562 | 0.303 | 0.61 | 0 | TEC_prg | LITAF |
| 0 | -0.8802527 | 0.419 | 0.704 | 0 | TEC_prg | TALDO1 |
| 0 | -0.8866915 | 0.848 | 0.904 | 0 | TEC_prg | RPS25 |
| 0 | -0.8868352 | 0.898 | 0.906 | 0 | TEC_prg | RPL7A |
| 0 | -0.8878301 | 0.783 | 0.845 | 0 | TEC_prg | RACK1 |
| 0 | -0.8953368 | 0.916 | 0.925 | 0 | TEC_prg | RPL18A |
| 0 | -0.9001085 | 0.843 | 0.929 | 0 | TEC_prg | NACA |
| 0 | -0.9008866 | 0.131 | 0.468 | 0 | TEC_prg | GSTO1 |
| 0 | -0.9029645 | 0.868 | 0.876 | 0 | TEC_prg | RPL26 |
| 0 | -0.9036402 | 0.866 | 0.896 | 0 | TEC_prg | RPS15A |
| 0 | -0.9056577 | 0.903 | 0.955 | 0 | TEC_prg | RPL18 |
| 0 | -0.9105731 | 0.796 | 0.82 | 0 | TEC_prg | RPL5 |
| 0 | -0.9260774 | 0.915 | 0.928 | 0 | TEC_prg | RPL19 |
| 0 | -0.935381 | 0.814 | 0.901 | 0 | TEC_prg | ATP5F1E |
| 0 | -0.9436971 | 0.922 | 0.922 | 0 | TEC_prg | RPL11 |
| 0 | -0.9443765 | 0.892 | 0.953 | 0 | TEC_prg | RPS16 |
| 0 | -0.9479515 | 0.648 | 0.78 | 0 | TEC_prg | CST3 |
| 0 | -0.9577079 | 0.926 | 0.924 | 0 | TEC_prg | RPS24 |
| 0 | -0.9727598 | 0.816 | 0.943 | 0 | TEC_prg | OAZ1 |
| 0 | -0.9785309 | 0.915 | 0.916 | 0 | TEC_prg | RPL32 |
| 0 | -0.9819987 | 0.894 | 0.899 | 0 | TEC_prg | RPL21 |
| 0 | -0.9903024 | 0.969 | 0.981 | 0 | TEC_prg | RPL41 |
| 0 | -1.0030972 | 0.49 | 0.688 | 0 | TEC_prg | ANXA5 |
| 0 | -1.0053784 | 0.801 | 0.867 | 0 | TEC_prg | RPL23 |
| 0 | -1.0203216 | 0.944 | 0.964 | 0 | TEC_prg | HSP90AA1 |
| 0 | -1.0327753 | 0.897 | 0.905 | 0 | TEC_prg | RPLP2 |
| 0 | -1.0354959 | 0.595 | 0.754 | 0 | TEC_prg | CTSB |
| 0 | -1.0389279 | 0.958 | 0.971 | 0 | TEC_prg | TMSB10 |
| 0 | -1.0416615 | 0.009 | 0.324 | 0 | TEC_prg | LCP1 |
| 0 | -1.042202 | 0.908 | 0.93 | 0 | TEC_prg | RPL30 |
| 0 | -1.049045 | 0.91 | 0.886 | 0 | TEC_prg | RPS23 |
| 0 | -1.0514887 | 0.795 | 0.865 | 0 | TEC_prg | RPL7 |
| 0 | -1.0556596 | 0.891 | 0.938 | 0 | TEC_prg | RPL34 |
| 0 | -1.0609223 | 0.036 | 0.314 | 0 | TEC_prg | HMOX1 |
| 0 | -1.0684598 | 0.615 | 0.768 | 0 | TEC_prg | TPM4 |
| 0 | -1.0698938 | 0.673 | 0.786 | 0 | TEC_prg | CTSD |
| 0 | -1.0712761 | 0.907 | 0.964 | 0 | TEC_prg | RPL15 |
| 0 | -1.0804985 | 0.832 | 0.927 | 0 | TEC_prg | UBA52 |
| 0 | -1.0808364 | 0.914 | 0.942 | 0 | TEC_prg | RPS28 |
| 0 | -1.0834025 | 0.679 | 0.804 | 0 | TEC_prg | PFN1 |
| 0 | -1.0885682 | 0.635 | 0.893 | 0 | TEC_prg | ARPC3 |
| 0 | -1.0965636 | 0.955 | 0.931 | 0 | TEC_prg | RPS12 |
| 0 | -1.118074 | 0.872 | 0.924 | 0 | TEC_prg | RPL9 |
| 0 | -1.1373286 | 0.851 | 0.888 | 0 | TEC_prg | RPS13 |
| 0 | -1.1406376 | 0.581 | 0.744 | 0 | TEC_prg | CYBA |
| 0 | -1.1441964 | 0.863 | 0.89 | 0 | TEC_prg | RPS3 |

| | | | | | | |
|---|---|---|---|---|---|---|
| 0 | -1.1552616 | 0.979 | 0.972 | 0 | TEC_prg | RPLP1 |
| 0 | -1.1633072 | 0.244 | 0.564 | 0 | TEC_prg | CTSS |
| 0 | -1.1730814 | 0.833 | 0.91 | 0 | TEC_prg | RPS7 |
| 0 | -1.1749286 | 0.01 | 0.312 | 0 | TEC_prg | PTPRC |
| 0 | -1.1916484 | 0.912 | 0.935 | 0 | TEC_prg | RPS14 |
| 0 | -1.1942417 | 0.831 | 0.896 | 0 | TEC_prg | HLA-B |
| 0 | -1.2402104 | 0.752 | 0.881 | 0 | TEC_prg | ANXA2 |
| 0 | -1.242808 | 0.649 | 0.8 | 0 | TEC_prg | TXN |
| 0 | -1.2598432 | 0.866 | 0.903 | 0 | TEC_prg | RPL39 |
| 0 | -1.2622601 | 0.886 | 0.899 | 0 | TEC_prg | RPS3A |
| 0 | -1.2782643 | 0.899 | 0.927 | 0 | TEC_prg | RPS27A |
| 0 | -1.2992042 | 0.752 | 0.908 | 0 | TEC_prg | CD63 |
| 0 | -1.3301547 | 0.895 | 0.887 | 0 | TEC_prg | RPS6 |
| 0 | -1.331053 | 0.008 | 0.305 | 0 | TEC_prg | AIF1 |
| 0 | -1.3373005 | 0.017 | 0.372 | 0 | TEC_prg | LAPTM5 |
| 0 | -1.3448781 | 0.789 | 0.855 | 0 | TEC_prg | RPS4X |
| 0 | -1.3487737 | 0.881 | 0.938 | 0 | TEC_prg | GAPDH |
| 0 | -1.3627699 | 0.985 | 0.994 | 0 | TEC_prg | TPT1 |
| 0 | -1.4034207 | 0.02 | 0.306 | 0 | TEC_prg | HLA-DRB1 |
| 0 | -1.411001 | 0.858 | 0.969 | 0 | TEC_prg | H3F3A |
| 0 | -1.413391 | 0.635 | 0.801 | 0 | TEC_prg | LGALS3 |
| 0 | -1.4336983 | 0.934 | 0.957 | 0 | TEC_prg | ACTG1 |
| 0 | -1.5086656 | 0.855 | 0.927 | 0 | TEC_prg | S100A10 |
| 0 | -1.5166055 | 0.569 | 0.694 | 0 | TEC_prg | NAMPT |
| 0 | -1.5193489 | 0.786 | 0.894 | 0 | TEC_prg | CSTB |
| 0 | -1.5599859 | 0.922 | 0.951 | 0 | TEC_prg | RPL10 |
| 0 | -1.5848262 | 0.066 | 0.364 | 0 | TEC_prg | IL1RN |
| 0 | -1.6728368 | 0.152 | 0.412 | 0 | TEC_prg | FCER1G |
| 0 | -1.7294436 | 0.929 | 0.969 | 0 | TEC_prg | S100A11 |
| 0 | -1.8052218 | 0.739 | 0.872 | 0 | TEC_prg | SH3BGRL3 |
| 0 | -1.8633027 | 0.595 | 0.723 | 0 | TEC_prg | ANXA1 |
| 0 | -1.9499495 | 0.911 | 0.941 | 0 | TEC_prg | TMSB4X |
| 0 | -1.9593536 | 0.954 | 0.994 | 0 | TEC_prg | B2M |
| 0 | -2.1294044 | 0.919 | 0.979 | 0 | TEC_prg | ACTB |
| 0 | -2.7743581 | 0.966 | 0.989 | 0 | TEC_prg | FTL |
| 0 | -2.9612733 | 0.981 | 0.995 | 0 | TEC_prg | FTH1 |
| 0 | -4.4060966 | 0.755 | 0.724 | 0 | TEC_prg | S100A9 |
| 5.54E-308 | -0.465446 | 0.06 | 0.348 | 1.309E-303 | TEC_prg | CALM3 |
| 5.836E-308 | -0.5688107 | 0.034 | 0.308 | 1.379E-303 | TEC_prg | LYN |
| 7.445E-308 | -0.7541382 | 0.005 | 0.26 | 1.759E-303 | TEC_prg | HCST |
| 1.534E-307 | -0.5762568 | 0.101 | 0.405 | 3.625E-303 | TEC_prg | METRNL |
| 3.387E-307 | -0.5200009 | 0.572 | 0.781 | 8E-303 | TEC_prg | UQCRB |
| 1.319E-306 | -0.4635521 | 0.069 | 0.361 | 3.116E-302 | TEC_prg | MOB1A |
| 1.883E-306 | -0.5481962 | 0.053 | 0.335 | 4.448E-302 | TEC_prg | BNIP3L |
| 2.959E-306 | 0.67512437 | 0.328 | 0.134 | 6.989E-302 | TEC_prg | HIST1H1E |
| 6.205E-306 | -0.4614105 | 0.108 | 0.421 | 1.466E-301 | TEC_prg | SEM1 |
| 5.329E-305 | -1.1685072 | 0.007 | 0.262 | 1.259E-300 | TEC_prg | BCL2A1 |
| 1.428E-304 | 0.72290986 | 0.309 | 0.12 | 3.374E-300 | TEC_prg | GADD45G |
| 2.676E-304 | -0.446269 | 0.1 | 0.407 | 6.321E-300 | TEC_prg | SRSF9 |
| 3.158E-304 | -0.4633661 | 0.065 | 0.354 | 7.461E-300 | TEC_prg | SERINC1 |

| | | | | | | |
|---|---|---|---|---|---|---|
| 3.373E-304 | -0.6063023 | 0.769 | 0.88 | 7.968E-300 | TEC_prg | PABPC1 |
| 4.736E-304 | -0.4893394 | 0.079 | 0.375 | 1.119E-299 | TEC_prg | EIF1AX |
| 1.757E-303 | -0.4530503 | 0.027 | 0.294 | 4.151E-299 | TEC_prg | KDELR2 |
| 5.111E-303 | 0.53723622 | 0.271 | 0.096 | 1.207E-298 | TEC_prg | AHI1 |
| 8.753E-303 | -0.5415563 | 0.036 | 0.305 | 2.068E-298 | TEC_prg | MYDGF |
| 9.989E-303 | 0.31593744 | 0.167 | 0.038 | 2.36E-298 | TEC_prg | PAX2 |
| 1.051E-302 | -0.4232359 | 0.038 | 0.311 | 2.483E-298 | TEC_prg | RAB5C |
| 2.826E-302 | -0.4779674 | 0.207 | 0.523 | 6.676E-298 | TEC_prg | HSBP1 |
| 4.143E-302 | -0.815718 | 0.867 | 0.881 | 9.787E-298 | TEC_prg | RPS29 |
| 2.229E-301 | -0.4617948 | 0.067 | 0.354 | 5.265E-297 | TEC_prg | GHITM |
| 1.52E-300 | -0.4272116 | 0.108 | 0.42 | 3.591E-296 | TEC_prg | COPE |
| 4.342E-300 | -0.4588871 | 0.035 | 0.304 | 1.026E-295 | TEC_prg | ATP6AP2 |
| 5.053E-300 | -1.0933239 | 0.181 | 0.453 | 1.194E-295 | TEC_prg | EMP3 |
| 5.534E-300 | 0.26996252 | 0.113 | 0.018 | 1.307E-295 | TEC_prg | AC083973.1 |
| 6.322E-299 | -0.5519996 | 0.23 | 0.527 | 1.493E-294 | TEC_prg | VDAC2 |
| 1.07E-298 | -0.4536702 | 0.043 | 0.316 | 2.528E-294 | TEC_prg | TMED10 |
| 1.218E-298 | -0.4868283 | 0.111 | 0.42 | 2.876E-294 | TEC_prg | UQCR11 |
| 1.872E-298 | -0.6543411 | 0.909 | 0.894 | 4.422E-294 | TEC_prg | RPS9 |
| 1.993E-298 | -0.5858549 | 0.044 | 0.316 | 4.707E-294 | TEC_prg | ATP1B3 |
| 3.126E-298 | -0.4400601 | 0.043 | 0.316 | 7.384E-294 | TEC_prg | LAMTOR1 |
| 1.449E-297 | -0.5869852 | 0.073 | 0.353 | 3.423E-293 | TEC_prg | MGST3 |
| 3.082E-297 | -0.4775458 | 0.088 | 0.383 | 7.28E-293 | TEC_prg | SPCS1 |
| 4.425E-297 | -0.467641 | 0.071 | 0.357 | 1.045E-292 | TEC_prg | LAMP1 |
| 1.425E-296 | -0.4489422 | 0.05 | 0.327 | 3.367E-292 | TEC_prg | TMEM167A |
| 1.927E-296 | -0.4499483 | 0.082 | 0.377 | 4.553E-292 | TEC_prg | SF3B6 |
| 1.29E-295 | -1.0508008 | 0.067 | 0.34 | 3.047E-291 | TEC_prg | EMP1 |
| 1.836E-295 | -0.5002434 | 0.032 | 0.295 | 4.338E-291 | TEC_prg | RAB5IF |
| 3.009E-295 | -0.496795 | 0.132 | 0.442 | 7.108E-291 | TEC_prg | ATP5PF |
| 6.332E-295 | 0.75454852 | 0.393 | 0.185 | 1.496E-290 | TEC_prg | KCNQ1OT1 |
| 1.073E-293 | -0.4143216 | 0.049 | 0.322 | 2.535E-289 | TEC_prg | SUMO1 |
| 1.168E-293 | 0.36265835 | 0.166 | 0.039 | 2.759E-289 | TEC_prg | PTGER3 |
| 1.434E-293 | -0.4681767 | 0.086 | 0.379 | 3.388E-289 | TEC_prg | DDT |
| 1.926E-293 | 0.36079993 | 0.162 | 0.037 | 4.55E-289 | TEC_prg | MYCL |
| 7.158E-293 | 0.33046518 | 0.169 | 0.041 | 1.691E-288 | TEC_prg | CCDC198 |
| 7.906E-293 | -0.5008514 | 0.105 | 0.402 | 1.868E-288 | TEC_prg | RHEB |
| 8.885E-293 | -0.5538231 | 0.1 | 0.398 | 2.099E-288 | TEC_prg | CTSH |
| 1.659E-292 | -1.2497795 | 0.337 | 0.562 | 3.92E-288 | TEC_prg | PLAUR |
| 2.119E-292 | -0.4784602 | 0.743 | 0.878 | 5.005E-288 | TEC_prg | SRP14 |
| 3.656E-292 | -0.5726362 | 0.074 | 0.355 | 8.635E-288 | TEC_prg | CAPG |
| 1.413E-291 | -1.646797 | 0.235 | 0.464 | 3.337E-287 | TEC_prg | TYROBP |
| 2.592E-291 | -0.5425577 | 0.017 | 0.27 | 6.123E-287 | TEC_prg | RAB31 |
| 3.626E-291 | -0.5768806 | 0.075 | 0.354 | 8.565E-287 | TEC_prg | PPA1 |
| 3.775E-291 | -0.8372825 | 0.265 | 0.522 | 8.918E-287 | TEC_prg | CD9 |
| 8.567E-291 | -0.7157528 | 0.919 | 0.902 | 2.024E-286 | TEC_prg | RPL37 |
| 1.87E-290 | -0.4307069 | 0.072 | 0.357 | 4.418E-286 | TEC_prg | RNF7 |
| 2.08E-290 | -1.1408122 | 0.011 | 0.259 | 4.913E-286 | TEC_prg | SLC2A3 |
| 2.529E-290 | -0.9450944 | 0.698 | 0.762 | 5.975E-286 | TEC_prg | RPSA |
| 4.232E-290 | -0.4108924 | 0.016 | 0.267 | 9.998E-286 | TEC_prg | OSTF1 |
| 1.846E-289 | -0.4734473 | 0.085 | 0.375 | 4.361E-285 | TEC_prg | LAMTOR4 |
| 1.925E-289 | -1.068849 | 0.046 | 0.309 | 4.547E-285 | TEC_prg | IFITM2 |

| | | | | | | |
|---|---|---|---|---|---|---|
| 4.769E-289 | -0.5480655 | 0.125 | 0.425 | 1.127E-284 | TEC_prg | NDUFB2 |
| 8.221E-289 | -0.6039928 | 0.72 | 0.826 | 1.942E-284 | TEC_prg | PFDN5 |
| 2.112E-288 | -0.6790524 | 0.003 | 0.244 | 4.989E-284 | TEC_prg | ITGB2 |
| 3.018E-288 | -1.2089362 | 0.245 | 0.499 | 7.13E-284 | TEC_prg | ERO1A |
| 9.355E-288 | -0.5139077 | 0.095 | 0.387 | 2.21E-283 | TEC_prg | UQCRQ |
| 3.933E-287 | -0.4764788 | 0.107 | 0.405 | 9.292E-283 | TEC_prg | LAMTOR5 |
| 1.053E-285 | -3.7037553 | 0.683 | 0.67 | 2.487E-281 | TEC_prg | S100A8 |
| 1.573E-285 | -0.417705 | 0.067 | 0.347 | 3.716E-281 | TEC_prg | CAPZA2 |
| 4.522E-285 | 0.84387586 | 0.49 | 0.277 | 1.068E-280 | TEC_prg | BTG2 |
| 4.838E-285 | -0.4709113 | 0.009 | 0.252 | 1.143E-280 | TEC_prg | RGS10 |
| 1.951E-284 | -0.4817168 | 0.287 | 0.581 | 4.61E-280 | TEC_prg | UQCR10 |
| 2.928E-284 | -0.4358093 | 0.065 | 0.341 | 6.917E-280 | TEC_prg | CAPNS1 |
| 9.44E-284 | -0.6583588 | 0.271 | 0.541 | 2.23E-279 | TEC_prg | ANXA11 |
| 1.089E-282 | -1.4508442 | 0.486 | 0.632 | 2.571E-278 | TEC_prg | LGALS1 |
| 2.182E-282 | -0.4638014 | 0.105 | 0.4 | 5.155E-278 | TEC_prg | PAK2 |
| 8.335E-282 | 0.3273284 | 0.151 | 0.034 | 1.969E-277 | TEC_prg | FAM117A |
| 7.427E-280 | -0.4896128 | 0.118 | 0.414 | 1.755E-275 | TEC_prg | DYNLRB1 |
| 7.886E-280 | -0.5004882 | 0.058 | 0.325 | 1.863E-275 | TEC_prg | ARL6IP5 |
| 1.744E-279 | -1.0432074 | 0.007 | 0.244 | 4.119E-275 | TEC_prg | CD14 |
| 5.459E-279 | -0.4329338 | 0.061 | 0.33 | 1.289E-274 | TEC_prg | ANAPC11 |
| 6.358E-279 | -1.1406805 | 0.007 | 0.244 | 1.502E-274 | TEC_prg | ALOX5AP |
| 6.948E-279 | -0.6906491 | 0.162 | 0.437 | 1.641E-274 | TEC_prg | CD59 |
| 9.767E-279 | 0.9868529 | 0.936 | 0.81 | 2.307E-274 | TEC_prg | MT-ND5 |
| 1.996E-278 | -0.7411367 | 0.457 | 0.669 | 4.715E-274 | TEC_prg | LDHA |
| 3.116E-278 | -0.8439201 | 0.025 | 0.271 | 7.361E-274 | TEC_prg | CST6 |
| 9.47E-278 | -1.346004 | 0.111 | 0.372 | 2.237E-273 | TEC_prg | CTSL |
| 1.327E-277 | -0.4588069 | 0.063 | 0.331 | 3.134E-273 | TEC_prg | TMED9 |
| 2.019E-277 | -0.6031587 | 0.305 | 0.575 | 4.77E-273 | TEC_prg | RPS10 |
| 2.545E-277 | -0.5772136 | 0.262 | 0.547 | 6.012E-273 | TEC_prg | NAP1L1 |
| 3.63E-276 | -0.6045216 | 0.005 | 0.24 | 8.575E-272 | TEC_prg | SPI1 |
| 6.227E-276 | -0.4576987 | 0.668 | 0.829 | 1.471E-271 | TEC_prg | RPL36AL |
| 9.183E-276 | -1.6337676 | 0.513 | 0.638 | 2.169E-271 | TEC_prg | S100A4 |
| 9.563E-276 | -0.4434258 | 0.238 | 0.543 | 2.259E-271 | TEC_prg | COX7B |
| 1.947E-275 | -0.4642392 | 0.054 | 0.315 | 4.601E-271 | TEC_prg | RALA |
| 3.281E-275 | 0.78810262 | 0.528 | 0.33 | 7.752E-271 | TEC_prg | GLS |
| 3.892E-275 | -0.4355786 | 0.035 | 0.287 | 9.195E-271 | TEC_prg | CHP1 |
| 4.724E-275 | -0.4314711 | 0.106 | 0.399 | 1.116E-270 | TEC_prg | GNB1 |
| 4.053E-274 | -0.6230202 | 0.94 | 0.946 | 9.574E-270 | TEC_prg | RPL13A |
| 4.51E-274 | -0.4413296 | 0.095 | 0.379 | 1.065E-269 | TEC_prg | TMEM258 |
| 3.574E-273 | -0.4917514 | 0.017 | 0.258 | 8.444E-269 | TEC_prg | BST2 |
| 1.706E-272 | -1.1719106 | 0.007 | 0.239 | 4.03E-268 | TEC_prg | S100A12 |
| 2.704E-272 | -0.4891434 | 0.182 | 0.48 | 6.387E-268 | TEC_prg | ATP5MD |
| 2.957E-272 | -0.4912057 | 0.102 | 0.383 | 6.985E-268 | TEC_prg | SEPT7 |
| 3.388E-272 | -0.4110084 | 0.064 | 0.332 | 8.003E-268 | TEC_prg | RAB1A |
| 6.366E-271 | -0.7689297 | 0.015 | 0.252 | 1.504E-266 | TEC_prg | ZEB2 |
| 6.767E-271 | 0.83676751 | 0.527 | 0.326 | 1.599E-266 | TEC_prg | ZFP36L2 |
| 1.372E-270 | 0.39407247 | 0.2 | 0.059 | 3.24E-266 | TEC_prg | TM7SF2 |
| 1.593E-269 | -0.938258 | 0.028 | 0.269 | 3.764E-265 | TEC_prg | SLC11A1 |
| 6.257E-269 | -0.4327861 | 0.074 | 0.343 | 1.478E-264 | TEC_prg | EIF3F |
| 7.704E-269 | -0.5100473 | 0.046 | 0.301 | 1.82E-264 | TEC_prg | PLIN3 |

| | | | | | | |
|---|---|---|---|---|---|---|
| 2.059E-268 | -0.5317707 | 0.014 | 0.249 | 4.864E-264 | TEC_prg | KYNU |
| 2.454E-266 | 0.74369647 | 0.448 | 0.246 | 5.797E-262 | TEC_prg | ID2 |
| 8.398E-266 | -0.4843525 | 0.061 | 0.322 | 1.984E-261 | TEC_prg | ZFAND6 |
| 1.144E-265 | -0.4802153 | 0.328 | 0.607 | 2.702E-261 | TEC_prg | UBE2B |
| 5.407E-265 | -0.5387174 | 0.107 | 0.385 | 1.277E-260 | TEC_prg | TKT |
| 6.814E-265 | -0.45193 | 0.112 | 0.397 | 1.61E-260 | TEC_prg | RAB10 |
| 8.157E-264 | 0.8309419 | 0.463 | 0.285 | 1.927E-259 | TEC_prg | SRSF2 |
| 8.478E-263 | -0.4024635 | 0.054 | 0.311 | 2.003E-258 | TEC_prg | RNF181 |
| 9.88E-263 | -0.4523417 | 0.093 | 0.368 | 2.334E-258 | TEC_prg | SNU13 |
| 2.106E-262 | 0.77132716 | 0.348 | 0.166 | 4.974E-258 | TEC_prg | SEPHS2 |
| 2.848E-262 | -0.7770736 | 0.007 | 0.233 | 6.729E-258 | TEC_prg | TNFRSF1B |
| 3.33E-262 | -0.3929648 | 0.042 | 0.291 | 7.867E-258 | TEC_prg | TMEM219 |
| 5.932E-262 | -0.4443491 | 0.108 | 0.388 | 1.401E-257 | TEC_prg | TOMM20 |
| 2.891E-261 | -0.9474726 | 0.647 | 0.751 | 6.829E-257 | TEC_prg | PSAP |
| 6.175E-261 | -0.8048391 | 0.09 | 0.349 | 1.459E-256 | TEC_prg | IFITM3 |
| 7.408E-261 | -0.4260881 | 0.071 | 0.334 | 1.75E-256 | TEC_prg | SPCS3 |
| 1.452E-260 | -0.4669842 | 0.266 | 0.557 | 3.429E-256 | TEC_prg | CD164 |
| 1.064E-259 | -0.477982 | 0.109 | 0.388 | 2.513E-255 | TEC_prg | TMOD3 |
| 2.029E-259 | -0.5885432 | 0.553 | 0.721 | 4.792E-255 | TEC_prg | RPL4 |
| 4.313E-259 | 0.47843455 | 0.169 | 0.045 | 1.019E-254 | TEC_prg | AC093323.1 |
| 5.953E-259 | -0.4578187 | 0.044 | 0.29 | 1.406E-254 | TEC_prg | PLP2 |
| 6.579E-259 | -0.3849537 | 0.04 | 0.285 | 1.554E-254 | TEC_prg | RAP1A |
| 8.367E-259 | -0.8005304 | 0.001 | 0.221 | 1.977E-254 | TEC_prg | CD52 |
| 2.497E-258 | -0.3905545 | 0.062 | 0.319 | 5.899E-254 | TEC_prg | PPP1CA |
| 3.692E-258 | -0.4784274 | 0.02 | 0.252 | 8.721E-254 | TEC_prg | GALNT1 |
| 8.21E-258 | -0.400983 | 0.036 | 0.278 | 1.939E-253 | TEC_prg | SCP2 |
| 6.989E-257 | -0.4086005 | 0.036 | 0.276 | 1.651E-252 | TEC_prg | NDUFAB1 |
| 5.318E-256 | -0.5928668 | 0.137 | 0.414 | 1.256E-251 | TEC_prg | C9orf16 |
| 5.871E-256 | -0.415333 | 0.58 | 0.784 | 1.387E-251 | TEC_prg | UBL5 |
| 1.779E-255 | -0.5043577 | 0.005 | 0.226 | 4.204E-251 | TEC_prg | LSP1 |
| 3.564E-255 | -0.4032068 | 0.089 | 0.358 | 8.42E-251 | TEC_prg | HNRNPF |
| 1.009E-254 | -0.8831318 | 0.222 | 0.476 | 2.383E-250 | TEC_prg | MARCKS |
| 1.358E-254 | 0.42162636 | 0.207 | 0.067 | 3.207E-250 | TEC_prg | HIP1R |
| 1.011E-253 | -0.8142911 | 0.149 | 0.411 | 2.389E-249 | TEC_prg | TYMP |
| 1.661E-253 | -0.6655819 | 0.602 | 0.758 | 3.923E-249 | TEC_prg | APLP2 |
| 2.447E-253 | 0.6883286 | 0.365 | 0.185 | 5.78E-249 | TEC_prg | MAT2A |
| 3.465E-253 | -0.6708476 | 0.593 | 0.742 | 8.185E-249 | TEC_prg | ENO1 |
| 7.728E-253 | -0.4859561 | 0.171 | 0.458 | 1.826E-248 | TEC_prg | IQGAP1 |
| 8.872E-253 | -0.4553829 | 0.049 | 0.295 | 2.096E-248 | TEC_prg | QSOX1 |
| 3.418E-252 | -0.779541 | 0.005 | 0.223 | 8.074E-248 | TEC_prg | CYBB |
| 1.989E-251 | -0.5569932 | 0.093 | 0.356 | 4.7E-247 | TEC_prg | ASPH |
| 2.068E-251 | 0.65043368 | 0.321 | 0.149 | 4.886E-247 | TEC_prg | CHCHD7 |
| 7.603E-251 | -0.6288043 | 0.014 | 0.236 | 1.796E-246 | TEC_prg | BASP1 |
| 1.4E-250 | -0.3791915 | 0.052 | 0.3 | 3.308E-246 | TEC_prg | LAMP2 |
| 2.22E-250 | 0.80942443 | 0.447 | 0.274 | 5.245E-246 | TEC_prg | HNRNPAB |
| 2.939E-250 | -0.3850594 | 0.05 | 0.295 | 6.942E-246 | TEC_prg | LMAN2 |
| 4.4E-250 | -0.5099915 | 0.328 | 0.589 | 1.039E-245 | TEC_prg | TOMM7 |
| 4.736E-250 | -0.4094245 | 0.059 | 0.308 | 1.119E-245 | TEC_prg | COX7A2L |
| 2.087E-249 | 0.52823932 | 0.267 | 0.104 | 4.931E-245 | TEC_prg | TSPYL2 |
| 3.099E-249 | -0.4402269 | 0.101 | 0.37 | 7.32E-245 | TEC_prg | ATP5F1B |

| | | | | | | |
|---|---|---|---|---|---|---|
| 1.488E-248 | 0.36041481 | 0.148 | 0.035 | 3.516E-244 | TEC_prg | CXCL14 |
| 6.115E-248 | 0.44293235 | 0.225 | 0.078 | 1.445E-243 | TEC_prg | ING1 |
| 2.461E-247 | -0.7208833 | 0.662 | 0.756 | 5.813E-243 | TEC_prg | EEF1B2 |
| 3.798E-247 | -0.3843419 | 0.044 | 0.284 | 8.973E-243 | TEC_prg | ATP6V0D1 |
| 2.412E-246 | 0.53220865 | 0.281 | 0.112 | 5.699E-242 | TEC_prg | FOXC1 |
| 2.58E-246 | -0.6350091 | 0.401 | 0.644 | 6.094E-242 | TEC_prg | PCBP1 |
| 6.986E-246 | 0.48173342 | 0.273 | 0.106 | 1.65E-241 | TEC_prg | FAT1 |
| 7.291E-246 | 0.80013139 | 0.539 | 0.38 | 1.722E-241 | TEC_prg | ARGLU1 |
| 7.616E-246 | -0.5775711 | 0.008 | 0.223 | 1.799E-241 | TEC_prg | NCF2 |
| 1.653E-245 | -0.3928212 | 0.065 | 0.316 | 3.905E-241 | TEC_prg | HIGD2A |
| 3.918E-245 | -0.4376772 | 0.04 | 0.274 | 9.255E-241 | TEC_prg | SH3BGRL |
| 3.929E-245 | -0.4194011 | 0.491 | 0.725 | 9.281E-241 | TEC_prg | COX6B1 |
| 6.091E-245 | -1.967698 | 0.196 | 0.419 | 1.439E-240 | TEC_prg | HLA-DRA |
| 6.444E-245 | -0.6392833 | 0.201 | 0.464 | 1.522E-240 | TEC_prg | CYSTM1 |
| 1.068E-244 | -0.5480079 | 0.064 | 0.308 | 2.524E-240 | TEC_prg | CTSZ |
| 1.848E-244 | -0.4703631 | 0.353 | 0.609 | 4.366E-240 | TEC_prg | CLTA |
| 2.612E-244 | -0.4372064 | 0.103 | 0.368 | 6.17E-240 | TEC_prg | ANP32B |
| 1.757E-243 | -0.8297294 | 0.032 | 0.257 | 4.149E-239 | TEC_prg | PDZK1IP1 |
| 3.233E-243 | -0.4029464 | 0.071 | 0.322 | 7.637E-239 | TEC_prg | KDELR1 |
| 7.146E-243 | -0.4720153 | 0.383 | 0.639 | 1.688E-238 | TEC_prg | PTGES3 |
| 1.063E-242 | 0.69933032 | 0.346 | 0.16 | 2.512E-238 | TEC_prg | DST |
| 2.027E-242 | -0.6075912 | 0.478 | 0.678 | 4.789E-238 | TEC_prg | NDUFA4 |
| 3.948E-242 | -0.4527083 | 0.015 | 0.231 | 9.327E-238 | TEC_prg | ARRB2 |
| 5.125E-242 | -1.3602539 | 0.012 | 0.226 | 1.211E-237 | TEC_prg | HLA-DPA1 |
| 5.39E-242 | -0.5933326 | 0.009 | 0.222 | 1.273E-237 | TEC_prg | ITGAX |
| 9.385E-242 | 0.46005887 | 0.152 | 0.039 | 2.217E-237 | TEC_prg | PGGHG |
| 1.151E-241 | -0.6446612 | 0.908 | 0.886 | 2.719E-237 | TEC_prg | RPL6 |
| 1.311E-241 | -0.4298144 | 0.046 | 0.281 | 3.098E-237 | TEC_prg | LYPLA1 |
| 3.381E-241 | -0.3942515 | 0.071 | 0.322 | 7.988E-237 | TEC_prg | RABAC1 |
| 4.981E-241 | -0.4925152 | 0.109 | 0.375 | 1.177E-236 | TEC_prg | TIMP2 |
| 6.699E-241 | -1.7182704 | 0.027 | 0.245 | 1.583E-236 | TEC_prg | VCAN |
| 7.692E-241 | -0.3530885 | 0.043 | 0.279 | 1.817E-236 | TEC_prg | PSMB4 |
| 4.545E-240 | -0.3599095 | 0.061 | 0.307 | 1.074E-235 | TEC_prg | ATP6V1E1 |
| 8.2E-240 | 0.27974663 | 0.151 | 0.039 | 1.937E-235 | TEC_prg | CAPRIN2 |
| 9.496E-240 | -0.3991025 | 0.085 | 0.34 | 2.243E-235 | TEC_prg | ARHGDIA |
| 5.651E-239 | -0.7949976 | 0.9 | 0.873 | 1.335E-234 | TEC_prg | RPL3 |
| 6.241E-239 | -0.3658377 | 0.074 | 0.324 | 1.474E-234 | TEC_prg | UBE2L3 |
| 8.82E-239 | -0.4309289 | 0.068 | 0.314 | 2.084E-234 | TEC_prg | APRT |
| 1.071E-238 | 0.31848702 | 0.141 | 0.034 | 2.53E-234 | TEC_prg | HIST1H2BN |
| 3.297E-238 | -0.3740519 | 0.057 | 0.297 | 7.788E-234 | TEC_prg | UXT |
| 3.298E-238 | -0.3904363 | 0.039 | 0.267 | 7.79E-234 | TEC_prg | OSTC |
| 4.787E-238 | -0.6373371 | 0.621 | 0.773 | 1.131E-233 | TEC_prg | ITGB1 |
| 7.068E-238 | -0.558053 | 0.9 | 0.919 | 1.67E-233 | TEC_prg | RPS20 |
| 1.172E-237 | -0.6646827 | 0.821 | 0.833 | 2.769E-233 | TEC_prg | RPL22 |
| 1.278E-237 | -0.3812108 | 0.061 | 0.303 | 3.018E-233 | TEC_prg | GDI2 |
| 1.311E-237 | -0.3642753 | 0.053 | 0.291 | 3.097E-233 | TEC_prg | VPS29 |
| 1.929E-237 | -0.4336628 | 0.116 | 0.386 | 4.557E-233 | TEC_prg | PLSCR1 |
| 2.731E-237 | -0.3578423 | 0.019 | 0.234 | 6.452E-233 | TEC_prg | ARPC1A |
| 3.025E-237 | -0.4548748 | 0.003 | 0.209 | 7.145E-233 | TEC_prg | RAC2 |
| 5.865E-237 | -0.5120487 | 0.057 | 0.294 | 1.385E-232 | TEC_prg | FAM3C |

| | | | | | | |
|---|---|---|---|---|---|---|
| 6.207E-236 | -0.5851724 | 0.004 | 0.21 | 1.466E-231 | TEC_prg | FCGR2A |
| 6.463E-236 | -1.6985613 | 0.521 | 0.624 | 1.527E-231 | TEC_prg | C15orf48 |
| 2.535E-235 | -0.6544032 | 0.017 | 0.23 | 5.988E-231 | TEC_prg | LST1 |
| 4.475E-235 | -0.4069479 | 0.074 | 0.321 | 1.057E-230 | TEC_prg | TRAM1 |
| 4.824E-235 | -0.6370678 | 0.977 | 0.963 | 1.14E-230 | TEC_prg | RPL13 |
| 7.694E-235 | -0.648713 | 0.964 | 0.956 | 1.818E-230 | TEC_prg | RPS18 |
| 1.28E-234 | -0.3753428 | 0.059 | 0.298 | 3.023E-230 | TEC_prg | ARL8B |
| 1.743E-234 | 0.47236919 | 0.279 | 0.117 | 4.117E-230 | TEC_prg | PAXBP1 |
| 3.123E-234 | 0.5750852 | 0.297 | 0.123 | 7.378E-230 | TEC_prg | ID1 |
| 3.736E-234 | -0.3787218 | 0.095 | 0.352 | 8.826E-230 | TEC_prg | SNRPG |
| 3.889E-234 | 0.39709091 | 0.211 | 0.072 | 9.188E-230 | TEC_prg | SLC25A29 |
| 8.839E-234 | 0.3299143 | 0.174 | 0.051 | 2.088E-229 | TEC_prg | MICAL3 |
| 1.511E-233 | -0.3758206 | 0.081 | 0.33 | 3.569E-229 | TEC_prg | JTB |
| 2.112E-233 | -0.4063269 | 0.354 | 0.624 | 4.99E-229 | TEC_prg | HNRNPK |
| 5.746E-233 | -0.4974936 | 0.267 | 0.535 | 1.357E-228 | TEC_prg | DBI |
| 1.728E-232 | -0.9402446 | 0.062 | 0.287 | 4.081E-228 | TEC_prg | TSPAN1 |
| 4.629E-232 | -0.36915 | 0.107 | 0.371 | 1.094E-227 | TEC_prg | WDR1 |
| 5.14E-232 | -0.4214386 | 0.072 | 0.315 | 1.214E-227 | TEC_prg | CHMP2B |
| 5.442E-232 | 0.33024052 | 0.182 | 0.055 | 1.286E-227 | TEC_prg | LAMA5 |
| 1.348E-231 | -0.3600737 | 0.043 | 0.272 | 3.185E-227 | TEC_prg | TRAPPC1 |
| 5.278E-231 | -0.4034652 | 0.063 | 0.303 | 1.247E-226 | TEC_prg | ADAM9 |
| 5.503E-231 | -0.3977474 | 0.005 | 0.207 | 1.3E-226 | TEC_prg | GPSM3 |
| 7.535E-231 | -0.3616038 | 0.085 | 0.337 | 1.78E-226 | TEC_prg | NDUFA2 |
| 3.368E-230 | -0.3674952 | 0.083 | 0.333 | 7.956E-226 | TEC_prg | BUD31 |
| 8.942E-230 | -0.5033282 | 0.038 | 0.26 | 2.112E-225 | TEC_prg | RNASET2 |
| 9.808E-230 | -0.390341 | 0.1 | 0.357 | 2.317E-225 | TEC_prg | FAM96B |
| 1.167E-229 | -0.3582472 | 0.045 | 0.273 | 2.757E-225 | TEC_prg | ATP5PB |
| 1.676E-229 | -0.8290413 | 0.006 | 0.208 | 3.96E-225 | TEC_prg | AC020656.1 |
| 3.769E-229 | -0.3966772 | 0.049 | 0.277 | 8.905E-225 | TEC_prg | DHRS7 |
| 8.16E-229 | -0.3765539 | 0.049 | 0.278 | 1.928E-224 | TEC_prg | RPN2 |
| 9.067E-229 | -2.4325914 | 0.479 | 0.581 | 2.142E-224 | TEC_prg | SRGN |
| 9.857E-229 | -0.3965913 | 0.146 | 0.427 | 2.329E-224 | TEC_prg | NDUFA1 |
| 1.975E-228 | -0.386008 | 0.067 | 0.307 | 4.666E-224 | TEC_prg | RTN3 |
| 2.958E-228 | 0.7497344 | 0.61 | 0.48 | 6.988E-224 | TEC_prg | RSRP1 |
| 2.96E-228 | -0.9038561 | 0.452 | 0.631 | 6.992E-224 | TEC_prg | CD44 |
| 4.309E-228 | -0.3811251 | 0.081 | 0.326 | 1.018E-223 | TEC_prg | ATP5PO |
| 7.948E-228 | -0.3943452 | 0.447 | 0.69 | 1.878E-223 | TEC_prg | HNRNPC |
| 1.482E-227 | -0.6347607 | 0.004 | 0.203 | 3.5E-223 | TEC_prg | IGSF6 |
| 3.201E-227 | -0.4676448 | 0.066 | 0.3 | 7.561E-223 | TEC_prg | VASP |
| 3.879E-227 | -0.3647914 | 0.113 | 0.376 | 9.164E-223 | TEC_prg | PRR13 |
| 6.482E-227 | -0.3509731 | 0.051 | 0.28 | 1.531E-222 | TEC_prg | EIF3I |
| 2.402E-226 | -0.3768636 | 0.054 | 0.285 | 5.673E-222 | TEC_prg | UBE2R2 |
| 2.536E-226 | -0.3579749 | 0.08 | 0.327 | 5.99E-222 | TEC_prg | H2AFV |
| 6.114E-226 | -0.3671078 | 0.051 | 0.279 | 1.444E-221 | TEC_prg | EIF3L |
| 7.166E-226 | -0.3794278 | 0.065 | 0.302 | 1.693E-221 | TEC_prg | PLEKHB2 |
| 9.982E-226 | 0.66088581 | 0.701 | 0.582 | 2.358E-221 | TEC_prg | DDX17 |
| 1.31E-225 | -0.4023538 | 0.033 | 0.251 | 3.094E-221 | TEC_prg | PGD |
| 1.696E-225 | -0.5537681 | 0.021 | 0.23 | 4.005E-221 | TEC_prg | LY6E |
| 3.582E-225 | -0.3593498 | 0.05 | 0.276 | 8.462E-221 | TEC_prg | TMED2 |
| 3.832E-225 | -0.3709488 | 0.041 | 0.263 | 9.053E-221 | TEC_prg | CASP4 |

| | | | | | | |
|---|---|---|---|---|---|---|
| 5.119E-225 | -0.400277 | 0.133 | 0.403 | 1.209E-220 | TEC_prg | ARF4 |
| 5.378E-225 | -0.3566175 | 0.068 | 0.305 | 1.27E-220 | TEC_prg | UFC1 |
| 5.416E-225 | 0.45964737 | 0.251 | 0.1 | 1.28E-220 | TEC_prg | SNHG25 |
| 6.491E-225 | -0.3886161 | 0.005 | 0.204 | 1.533E-220 | TEC_prg | GNA15 |
| 1.01E-224 | 0.42656965 | 0.225 | 0.084 | 2.387E-220 | TEC_prg | SNX29 |
| 1.191E-224 | -0.4422564 | 0.111 | 0.365 | 2.813E-220 | TEC_prg | GRB2 |
| 1.778E-224 | -0.6046702 | 0.418 | 0.623 | 4.201E-220 | TEC_prg | HINT1 |
| 2.501E-224 | -0.3781883 | 0.127 | 0.395 | 5.907E-220 | TEC_prg | WDR83OS |
| 2.538E-224 | -0.3658305 | 0.09 | 0.34 | 5.996E-220 | TEC_prg | RBX1 |
| 2.708E-224 | -0.3732316 | 0.305 | 0.583 | 6.396E-220 | TEC_prg | MORF4L1 |
| 3.221E-224 | -0.5926522 | 0.261 | 0.511 | 7.609E-220 | TEC_prg | HMGA1 |
| 3.815E-224 | -0.4054047 | 0.008 | 0.208 | 9.012E-220 | TEC_prg | HCLS1 |
| 4.842E-224 | -0.4102748 | 0.096 | 0.346 | 1.144E-219 | TEC_prg | ERP29 |
| 6.907E-224 | -0.3783362 | 0.121 | 0.384 | 1.632E-219 | TEC_prg | TRMT112 |
| 8.189E-224 | -0.6571716 | 0.007 | 0.205 | 1.934E-219 | TEC_prg | C5AR1 |
| 2.247E-223 | -0.6561374 | 0.884 | 0.87 | 5.308E-219 | TEC_prg | RPL14 |
| 2.823E-223 | -0.4330127 | 0.2 | 0.475 | 6.67E-219 | TEC_prg | DNAJB6 |
| 4.042E-223 | -0.3373104 | 0.008 | 0.207 | 9.549E-219 | TEC_prg | SQOR |
| 7.771E-223 | -0.6777715 | 0.082 | 0.32 | 1.836E-218 | TEC_prg | GLRX |
| 1.293E-222 | 0.2662244 | 0.155 | 0.043 | 3.053E-218 | TEC_prg | TRIM73 |
| 3.32E-222 | -0.4968179 | 0.047 | 0.27 | 7.844E-218 | TEC_prg | GPCPD1 |
| 2.151E-221 | -0.6690645 | 0.084 | 0.317 | 5.081E-217 | TEC_prg | ATP13A3 |
| 2.703E-221 | -0.4885912 | 0.009 | 0.207 | 6.385E-217 | TEC_prg | PTPRE |
| 3.375E-221 | -0.5621871 | 0.312 | 0.555 | 7.972E-217 | TEC_prg | PTTG1IP |
| 6.467E-221 | 0.71609273 | 0.359 | 0.184 | 1.528E-216 | TEC_prg | FOSB |
| 6.698E-221 | -0.4147784 | 0.088 | 0.331 | 1.582E-216 | TEC_prg | CLTB |
| 8.328E-221 | -0.3423525 | 0.027 | 0.238 | 1.967E-216 | TEC_prg | ADIPOR1 |
| 9.515E-221 | -0.4109143 | 0.084 | 0.325 | 2.248E-216 | TEC_prg | PICALM |
| 3.15E-220 | -0.3622683 | 0.064 | 0.297 | 7.44E-216 | TEC_prg | ATP5F1C |
| 3.603E-220 | -0.7583372 | 0.006 | 0.2 | 8.512E-216 | TEC_prg | OLR1 |
| 5.26E-220 | 0.58800934 | 0.924 | 0.82 | 1.243E-215 | TEC_prg | HNRNPA2B1 |
| 8.738E-220 | -0.4071631 | 0.077 | 0.314 | 2.064E-215 | TEC_prg | NDUFB9 |
| 6.857E-219 | -0.4626478 | 0.056 | 0.28 | 1.62E-214 | TEC_prg | PDLIM5 |
| 7.121E-219 | -0.3615696 | 0.02 | 0.224 | 1.682E-214 | TEC_prg | RHOG |
| 7.695E-219 | -0.3282447 | 0.029 | 0.239 | 1.818E-214 | TEC_prg | COPZ1 |
| 1.037E-218 | 0.30294938 | 0.147 | 0.04 | 2.45E-214 | TEC_prg | HOXD1 |
| 1.642E-218 | -0.3408957 | 0.061 | 0.29 | 3.878E-214 | TEC_prg | CSNK2B |
| 2.315E-218 | -0.3332342 | 0.055 | 0.282 | 5.468E-214 | TEC_prg | LSM3 |
| 2.317E-218 | -0.4168859 | 0.102 | 0.35 | 5.472E-214 | TEC_prg | YPEL5 |
| 2.38E-218 | -0.3846026 | 0.13 | 0.391 | 5.623E-214 | TEC_prg | WASF2 |
| 1.06E-217 | -0.3616337 | 0.075 | 0.312 | 2.504E-213 | TEC_prg | HM13 |
| 1.015E-216 | -0.3249015 | 0.055 | 0.279 | 2.398E-212 | TEC_prg | POLR2J |
| 1.51E-216 | -0.402413 | 0.147 | 0.413 | 3.568E-212 | TEC_prg | TBCA |
| 3.126E-216 | -0.5635953 | 0.008 | 0.201 | 7.385E-212 | TEC_prg | SAMSN1 |
| 5.219E-216 | -0.3691601 | 0.067 | 0.296 | 1.233E-211 | TEC_prg | MINOS1 |
| 1.048E-215 | -0.452386 | 0.003 | 0.193 | 2.475E-211 | TEC_prg | CORO1A |
| 1.196E-215 | -0.4100997 | 0.296 | 0.559 | 2.826E-211 | TEC_prg | COX6C |
| 2.003E-215 | -0.6598902 | 0.943 | 0.908 | 4.732E-211 | TEC_prg | RPS8 |
| 2.257E-215 | -0.3520583 | 0.06 | 0.288 | 5.331E-211 | TEC_prg | PSMA4 |
| 2.853E-215 | -0.4818433 | 0.638 | 0.777 | 6.74E-211 | TEC_prg | COX7C |

| | | | | | | |
|---|---|---|---|---|---|---|
| 3.226E-215 | -0.3945568 | 0.055 | 0.278 | 7.622E-211 | TEC_prg | MGAT1 |
| 3.531E-215 | -0.4012613 | 0.052 | 0.273 | 8.341E-211 | TEC_prg | DNAJC3 |
| 4.589E-215 | -0.4455494 | 0.001 | 0.188 | 1.084E-210 | TEC_prg | CD48 |
| 7.74E-215 | -0.343827 | 0.048 | 0.268 | 1.828E-210 | TEC_prg | UQCRFS1 |
| 1.613E-214 | -0.5106554 | 0.003 | 0.193 | 3.81E-210 | TEC_prg | SMIM25 |
| 1.84E-214 | -0.3859129 | 0.007 | 0.199 | 4.347E-210 | TEC_prg | FMNL1 |
| 1.853E-214 | -0.3397134 | 0.058 | 0.284 | 4.377E-210 | TEC_prg | SMDT1 |
| 2.052E-214 | -0.3457294 | 0.075 | 0.31 | 4.847E-210 | TEC_prg | ATP6V1D |
| 2.912E-214 | -0.5748468 | 0.032 | 0.239 | 6.88E-210 | TEC_prg | TGM2 |
| 8.773E-214 | -0.33015 | 0.065 | 0.295 | 2.072E-209 | TEC_prg | STMP1 |
| 1.01E-213 | 0.62608801 | 0.383 | 0.216 | 2.386E-209 | TEC_prg | KMT2A |
| 5.914E-213 | -0.5789202 | 0.257 | 0.5 | 1.397E-208 | TEC_prg | CD81 |
| 1.152E-212 | -0.4120443 | 0.012 | 0.205 | 2.721E-208 | TEC_prg | CKLF |
| 1.551E-212 | -0.3900277 | 0.056 | 0.278 | 3.665E-208 | TEC_prg | MAP2K3 |
| 1.835E-212 | 0.32971237 | 0.182 | 0.059 | 4.335E-208 | TEC_prg | TCTA |
| 1.852E-212 | 0.39445948 | 0.219 | 0.082 | 4.374E-208 | TEC_prg | KMT2E-AS1 |
| 2.386E-212 | -0.3359686 | 0.061 | 0.287 | 5.636E-208 | TEC_prg | PSMA1 |
| 3.354E-212 | -0.3701464 | 0.068 | 0.296 | 7.922E-208 | TEC_prg | FAM49B |
| 3.949E-212 | -0.3436638 | 0.072 | 0.304 | 9.329E-208 | TEC_prg | TMBIM4 |
| 4.398E-212 | -0.5505172 | 0.445 | 0.651 | 1.039E-207 | TEC_prg | TAX1BP1 |
| 5.605E-212 | -0.3697761 | 0.056 | 0.275 | 1.324E-207 | TEC_prg | EIF3M |
| 6.986E-212 | -0.3352262 | 0.096 | 0.341 | 1.65E-207 | TEC_prg | TAF10 |
| 6.987E-212 | -0.3641503 | 0.051 | 0.268 | 1.651E-207 | TEC_prg | METTL9 |
| 7.648E-212 | -0.4026426 | 0.096 | 0.337 | 1.807E-207 | TEC_prg | SBDS |
| 9.868E-212 | 0.27688874 | 0.143 | 0.039 | 2.331E-207 | TEC_prg | ZNF821 |
| 2.307E-211 | -0.3391953 | 0.038 | 0.248 | 5.451E-207 | TEC_prg | PSMB2 |
| 5.945E-211 | -0.3875856 | 0.038 | 0.249 | 1.404E-206 | TEC_prg | IFNGR1 |
| 1.063E-210 | -0.8528724 | 0.355 | 0.559 | 2.511E-206 | TEC_prg | UPP1 |
| 1.215E-210 | -0.322547 | 0.164 | 0.441 | 2.869E-206 | TEC_prg | DAZAP2 |
| 1.563E-210 | -0.8103132 | 0.006 | 0.194 | 3.693E-206 | TEC_prg | PLEK |
| 3.572E-210 | -0.5039127 | 0.081 | 0.307 | 8.437E-206 | TEC_prg | RPS4Y1 |
| 9.147E-210 | -0.5209548 | 0.5 | 0.688 | 2.161E-205 | TEC_prg | RTN4 |
| 1.392E-209 | -0.3346155 | 0.065 | 0.291 | 3.289E-205 | TEC_prg | NDUFA12 |
| 1.595E-209 | -0.3448283 | 0.095 | 0.338 | 3.767E-205 | TEC_prg | BLOC1S1 |
| 1.698E-209 | -0.4143263 | 0.566 | 0.754 | 4.012E-205 | TEC_prg | CCNI |
| 3.725E-209 | -0.3681368 | 0.084 | 0.318 | 8.799E-205 | TEC_prg | SYAP1 |
| 2.18E-208 | -0.3851853 | 0.023 | 0.222 | 5.149E-204 | TEC_prg | IFI16 |
| 3.655E-208 | -0.9392127 | 0.022 | 0.217 | 8.633E-204 | TEC_prg | TM4SF1 |
| 6.043E-208 | -1.1859476 | 0.014 | 0.204 | 1.428E-203 | TEC_prg | HLA-DPB1 |
| 2.087E-207 | -0.542838 | 0.012 | 0.201 | 4.93E-203 | TEC_prg | FBP1 |
| 3.193E-207 | -0.3489525 | 0.166 | 0.44 | 7.544E-203 | TEC_prg | VAPA |
| 3.444E-207 | -0.404283 | 0.009 | 0.196 | 8.136E-203 | TEC_prg | GLIPR1 |
| 4.347E-207 | -0.3127062 | 0.031 | 0.235 | 1.027E-202 | TEC_prg | SFT2D1 |
| 1.179E-206 | -0.331501 | 0.079 | 0.311 | 2.785E-202 | TEC_prg | PPP4C |
| 1.472E-206 | 0.76343639 | 0.554 | 0.37 | 3.476E-202 | TEC_prg | FOS |
| 1.735E-206 | -0.3690233 | 0.218 | 0.486 | 4.1E-202 | TEC_prg | EIF2S2 |
| 2.172E-206 | -0.3264345 | 0.066 | 0.29 | 5.132E-202 | TEC_prg | ERH |
| 3.016E-206 | 0.31718289 | 0.194 | 0.067 | 7.125E-202 | TEC_prg | MYEF2 |
| 3.132E-206 | -1.4375193 | 0.022 | 0.216 | 7.398E-202 | TEC_prg | HOPX |
| 3.251E-206 | -0.3465773 | 0.061 | 0.282 | 7.679E-202 | TEC_prg | PGLS |

| | | | | | | |
|---|---|---|---|---|---|---|
| 3.436E-206 | 0.41314044 | 0.259 | 0.109 | 8.118E-202 | TEC_prg | HOOK2 |
| 6.746E-206 | -0.4074859 | 0.078 | 0.304 | 1.594E-201 | TEC_prg | RBM3 |
| 1.467E-205 | -0.3165895 | 0.034 | 0.237 | 3.465E-201 | TEC_prg | NDUFB5 |
| 2.324E-205 | -0.3437651 | 0.094 | 0.332 | 5.49E-201 | TEC_prg | ANAPC16 |
| 2.74E-205 | -0.4849109 | 0.004 | 0.187 | 6.473E-201 | TEC_prg | SLA |
| 4.135E-205 | -0.3529091 | 0.066 | 0.287 | 9.768E-201 | TEC_prg | KRAS |
| 5.227E-205 | -0.8452914 | 0.048 | 0.255 | 1.235E-200 | TEC_prg | PLIN2 |
| 7.344E-205 | -0.3364969 | 0.064 | 0.285 | 1.735E-200 | TEC_prg | GTF2A2 |
| 1.221E-204 | -0.8497268 | 0.005 | 0.188 | 2.885E-200 | TEC_prg | FCN1 |
| 1.32E-204 | 0.37317014 | 0.195 | 0.067 | 3.117E-200 | TEC_prg | SOX9 |
| 1.396E-204 | -0.3437926 | 0.078 | 0.308 | 3.298E-200 | TEC_prg | NDUFB1 |
| 5.374E-204 | -0.306981 | 0.032 | 0.234 | 1.269E-199 | TEC_prg | SDHD |
| 5.985E-204 | -0.3403413 | 0.053 | 0.267 | 1.414E-199 | TEC_prg | PEA15 |
| 8.335E-204 | 0.39076436 | 0.208 | 0.077 | 1.969E-199 | TEC_prg | C12orf65 |
| 2.033E-203 | -0.3104212 | 0.026 | 0.223 | 4.803E-199 | TEC_prg | HNRNPH2 |
| 2.468E-203 | -0.3968319 | 0.042 | 0.249 | 5.83E-199 | TEC_prg | FCGRT |
| 5.479E-203 | -0.5442386 | 0.029 | 0.227 | 1.294E-198 | TEC_prg | GK |
| 8.866E-203 | -0.3171097 | 0.109 | 0.354 | 2.094E-198 | TEC_prg | PAIP2 |
| 1.05E-202 | -0.3151798 | 0.019 | 0.211 | 2.48E-198 | TEC_prg | AGTRAP |
| 1.736E-202 | -0.3533843 | 0.014 | 0.203 | 4.102E-198 | TEC_prg | BID |
| 3.042E-202 | -0.3590416 | 0.517 | 0.734 | 7.185E-198 | TEC_prg | UBE2D3 |
| 5.27E-202 | -0.6588774 | 0.008 | 0.191 | 1.245E-197 | TEC_prg | HLA-DQB1 |
| 5.519E-202 | 0.55486573 | 0.333 | 0.174 | 1.304E-197 | TEC_prg | CHD9 |
| 6.089E-202 | -0.3450357 | 0.089 | 0.32 | 1.438E-197 | TEC_prg | KPNB1 |
| 1.256E-201 | -0.4651761 | 0.003 | 0.183 | 2.968E-197 | TEC_prg | LCP2 |
| 1.64E-201 | 0.27728852 | 0.148 | 0.043 | 3.874E-197 | TEC_prg | KLHL15 |
| 1.435E-200 | -0.3282442 | 0.082 | 0.311 | 3.389E-196 | TEC_prg | NDUFA11 |
| 2.797E-200 | -0.4071597 | 0.095 | 0.327 | 6.607E-196 | TEC_prg | BLVRB |
| 5.325E-200 | -0.6921416 | 0.009 | 0.19 | 1.258E-195 | TEC_prg | STEAP4 |
| 8.267E-200 | -0.401903 | 0.004 | 0.182 | 1.953E-195 | TEC_prg | CLEC2B |
| 3.977E-199 | -0.3327306 | 0.04 | 0.242 | 9.394E-195 | TEC_prg | LSM4 |
| 4.079E-199 | -0.3919211 | 0.33 | 0.581 | 9.636E-195 | TEC_prg | NDUFS5 |
| 4.676E-199 | -0.6938763 | 0.787 | 0.802 | 1.105E-194 | TEC_prg | RPS5 |
| 4.998E-199 | -0.3329738 | 0.081 | 0.308 | 1.181E-194 | TEC_prg | PSMB6 |
| 1.017E-198 | -0.3944852 | 0.02 | 0.208 | 2.403E-194 | TEC_prg | PYCARD |
| 1.026E-198 | -0.5624783 | 0.02 | 0.207 | 2.423E-194 | TEC_prg | FYB1 |
| 1.461E-198 | -0.3503875 | 0.186 | 0.455 | 3.451E-194 | TEC_prg | SEC62 |
| 1.677E-198 | -0.563694 | 0.266 | 0.504 | 3.961E-194 | TEC_prg | RIOK3 |
| 6.528E-198 | 0.28745639 | 0.163 | 0.051 | 1.542E-193 | TEC_prg | ANKRD26 |
| 7.695E-198 | -0.3296274 | 0.074 | 0.294 | 1.818E-193 | TEC_prg | NDUFB8 |
| 8.474E-198 | -3.3089724 | 0.653 | 0.679 | 2.002E-193 | TEC_prg | TIMP1 |
| 1.344E-197 | -0.689105 | 0.084 | 0.295 | 3.174E-193 | TEC_prg | MAL2 |
| 1.394E-197 | -0.5422969 | 0.916 | 0.893 | 3.292E-193 | TEC_prg | RPL37A |
| 2.222E-197 | -0.3487091 | 0.097 | 0.326 | 5.249E-193 | TEC_prg | PRKAR1A |
| 2.321E-197 | -0.3206703 | 0.069 | 0.286 | 5.484E-193 | TEC_prg | RTRAF |
| 6.631E-197 | -0.4076488 | 0.086 | 0.308 | 1.566E-192 | TEC_prg | LRP10 |
| 6.966E-197 | -0.5079571 | 0.003 | 0.179 | 1.646E-192 | TEC_prg | AQP9 |
| 1.364E-196 | -0.4491833 | 0.01 | 0.19 | 3.222E-192 | TEC_prg | SAMHD1 |
| 1.727E-196 | -0.5867037 | 0.009 | 0.189 | 4.08E-192 | TEC_prg | TGFBI |
| 2.138E-196 | -0.7367848 | 0.66 | 0.756 | 5.051E-192 | TEC_prg | PRDX1 |

| | | | | | | |
|---|---|---|---|---|---|---|
| 3.455E-196 | -0.3832882 | 0.083 | 0.307 | 8.162E-192 | TEC_prg | PDCD6 |
| 5.805E-196 | -0.4271827 | 0.098 | 0.326 | 1.371E-191 | TEC_prg | PSME2 |
| 6.18E-196 | -0.6245624 | 0.028 | 0.218 | 1.46E-191 | TEC_prg | DUSP4 |
| 2.055E-195 | -1.2110531 | 0.004 | 0.179 | 4.855E-191 | TEC_prg | MUC21 |
| 2.784E-195 | -0.3569842 | 0.022 | 0.21 | 6.578E-191 | TEC_prg | GBP2 |
| 4.533E-195 | -0.4856267 | 0.339 | 0.571 | 1.071E-190 | TEC_prg | TOP1 |
| 5.061E-195 | 0.46450281 | 0.299 | 0.14 | 1.196E-190 | TEC_prg | COBLL1 |
| 5.375E-195 | -0.297323 | 0.038 | 0.235 | 1.27E-190 | TEC_prg | SUPT4H1 |
| 1.075E-194 | -0.3291045 | 0.049 | 0.254 | 2.539E-190 | TEC_prg | M6PR |
| 1.729E-194 | -0.3274923 | 0.053 | 0.258 | 4.085E-190 | TEC_prg | PSMB7 |
| 2.633E-194 | -0.3152793 | 0.127 | 0.373 | 6.22E-190 | TEC_prg | PSMD8 |
| 3.338E-194 | -0.3271697 | 0.083 | 0.308 | 7.886E-190 | TEC_prg | RAD21 |
| 4.726E-194 | -0.3032924 | 0.067 | 0.283 | 1.117E-189 | TEC_prg | CAMTA1 |
| 1.279E-193 | -0.3210863 | 0.089 | 0.315 | 3.021E-189 | TEC_prg | EIF4H |
| 1.696E-193 | -0.3190071 | 0.069 | 0.284 | 4.007E-189 | TEC_prg | PSMD7 |
| 2.034E-193 | -0.303453 | 0.04 | 0.239 | 4.805E-189 | TEC_prg | SELENOT |
| 6.712E-193 | -0.3694072 | 0.005 | 0.18 | 1.586E-188 | TEC_prg | FERMT3 |
| 8.05E-193 | -0.3152178 | 0.031 | 0.224 | 1.902E-188 | TEC_prg | UBE2J1 |
| 9.062E-193 | 0.39059244 | 0.222 | 0.088 | 2.141E-188 | TEC_prg | EPB41L5 |
| 1.415E-192 | -0.3500623 | 0.009 | 0.186 | 3.344E-188 | TEC_prg | GLIPR2 |
| 1.416E-192 | -0.4880654 | 0.383 | 0.598 | 3.344E-188 | TEC_prg | PRDX5 |
| 1.701E-192 | -0.6346999 | 0.183 | 0.419 | 4.018E-188 | TEC_prg | HIF1A |
| 1.468E-191 | -0.3419597 | 0.138 | 0.384 | 3.469E-187 | TEC_prg | SNRPD2 |
| 2.026E-191 | -0.3875476 | 0.128 | 0.367 | 4.786E-187 | TEC_prg | BSG |
| 2.324E-191 | -0.328446 | 0.026 | 0.215 | 5.489E-187 | TEC_prg | RAB8B |
| 5.391E-191 | 0.54823959 | 0.355 | 0.195 | 1.274E-186 | TEC_prg | BDP1 |
| 6.822E-191 | 1.09236034 | 0.63 | 0.476 | 1.611E-186 | TEC_prg | MTRNR2L8 |
| 1.596E-190 | -0.5841618 | 0.016 | 0.195 | 3.771E-186 | TEC_prg | ANPEP |
| 1.944E-190 | -1.4537447 | 0.24 | 0.424 | 4.593E-186 | TEC_prg | SLPI |
| 3.614E-190 | 0.36646978 | 0.24 | 0.101 | 8.538E-186 | TEC_prg | CREBZF |
| 3.872E-190 | -0.3285303 | 0.097 | 0.324 | 9.148E-186 | TEC_prg | SF3B5 |
| 4.393E-190 | -0.3770748 | 0.059 | 0.265 | 1.038E-185 | TEC_prg | CD82 |
| 5.983E-190 | -0.5440179 | 0.101 | 0.321 | 1.413E-185 | TEC_prg | FAM129B |
| 1.441E-189 | -0.3318875 | 0.08 | 0.297 | 3.405E-185 | TEC_prg | UFM1 |
| 2.265E-189 | -0.3400235 | 0.052 | 0.253 | 5.35E-185 | TEC_prg | RYBP |
| 2.333E-189 | -0.5464598 | 0.769 | 0.82 | 5.511E-185 | TEC_prg | YBX1 |
| 2.496E-189 | -0.7303505 | 0.057 | 0.257 | 5.897E-185 | TEC_prg | FABP5 |
| 2.971E-189 | -0.3007514 | 0.033 | 0.225 | 7.02E-185 | TEC_prg | RPN1 |
| 3.97E-189 | -0.4684294 | 0.04 | 0.232 | 9.379E-185 | TEC_prg | NINJ1 |
| 5.149E-189 | -0.3262185 | 0.03 | 0.218 | 1.216E-184 | TEC_prg | PRNP |
| 7.304E-189 | 0.42191904 | 0.243 | 0.105 | 1.726E-184 | TEC_prg | DAPK1 |
| 9.848E-189 | -0.3014864 | 0.03 | 0.22 | 2.326E-184 | TEC_prg | NANS |
| 1.034E-188 | -1.1062616 | 0.006 | 0.177 | 2.442E-184 | TEC_prg | CEACAM5 |
| 1.091E-188 | 0.49124085 | 0.273 | 0.121 | 2.578E-184 | TEC_prg | DCDC2 |
| 2.132E-188 | -0.4022763 | 0.028 | 0.213 | 5.037E-184 | TEC_prg | SNX10 |
| 6.159E-188 | -1.1744152 | 0.022 | 0.203 | 1.455E-183 | TEC_prg | DUOX2 |
| 6.703E-188 | -0.2865105 | 0.038 | 0.232 | 1.584E-183 | TEC_prg | RTF2 |
| 7.651E-188 | 0.6369276 | 0.418 | 0.266 | 1.807E-183 | TEC_prg | PRPF4B |
| 7.832E-188 | -0.6197326 | 0.969 | 0.933 | 1.85E-183 | TEC_prg | S100A6 |
| 1.272E-187 | -1.6629461 | 0.009 | 0.18 | 3.004E-183 | TEC_prg | ECM1 |

| | | | | | | |
|---|---|---|---|---|---|---|
| 2.371E-187 | -0.3101229 | 0.09 | 0.312 | 5.601E-183 | TEC_prg | NDUFB3 |
| 2.679E-187 | -1.5695668 | 0.033 | 0.217 | 6.329E-183 | TEC_prg | KRT17 |
| 3.137E-187 | -0.3888588 | 0.079 | 0.293 | 7.411E-183 | TEC_prg | OSBPL8 |
| 5.989E-187 | -0.8803154 | 0.007 | 0.179 | 1.415E-182 | TEC_prg | APOBEC3A |
| 6.779E-187 | -0.7945729 | 0.226 | 0.444 | 1.602E-182 | TEC_prg | TXNRD1 |
| 9.744E-187 | -0.3028234 | 0.08 | 0.296 | 2.302E-182 | TEC_prg | AP2M1 |
| 1.545E-186 | -0.3047512 | 0.024 | 0.207 | 3.651E-182 | TEC_prg | CNN2 |
| 4.863E-186 | -0.3599409 | 0.105 | 0.331 | 1.149E-181 | TEC_prg | PFDN2 |
| 5.983E-186 | -0.5590373 | 0.063 | 0.263 | 1.413E-181 | TEC_prg | ATP2B1 |
| 6.028E-186 | -0.4017988 | 0.002 | 0.169 | 1.424E-181 | TEC_prg | EVI2B |
| 1.926E-185 | -0.4065222 | 0.988 | 0.984 | 4.549E-181 | TEC_prg | EIF1 |
| 2.818E-185 | -0.2984985 | 0.025 | 0.207 | 6.657E-181 | TEC_prg | PRDX3 |
| 3.074E-185 | 0.27114584 | 0.159 | 0.051 | 7.261E-181 | TEC_prg | MAPK8IP3 |
| 3.12E-185 | -0.5948998 | 0.006 | 0.175 | 7.37E-181 | TEC_prg | CD300E |
| 3.546E-185 | -0.3514157 | 0.032 | 0.218 | 8.376E-181 | TEC_prg | CREG1 |
| 5.405E-185 | -0.3057968 | 0.065 | 0.272 | 1.277E-180 | TEC_prg | STAU1 |
| 6.661E-185 | -0.5714663 | 0.269 | 0.502 | 1.574E-180 | TEC_prg | HSPD1 |
| 7.721E-185 | -0.335603 | 0.06 | 0.263 | 1.824E-180 | TEC_prg | SRI |
| 1.438E-184 | -0.4209076 | 0.05 | 0.246 | 3.396E-180 | TEC_prg | RAB9A |
| 3.474E-184 | -0.2837893 | 0.075 | 0.288 | 8.206E-180 | TEC_prg | SSU72 |
| 3.806E-184 | -0.3058056 | 0.054 | 0.254 | 8.99E-180 | TEC_prg | PSMB8 |
| 6.275E-184 | -0.3569296 | 0.029 | 0.212 | 1.482E-179 | TEC_prg | ACSL4 |
| 3.499E-183 | 0.48212982 | 0.245 | 0.11 | 8.265E-179 | TEC_prg | MED29 |
| 5.382E-183 | -0.5319583 | 0.015 | 0.188 | 1.271E-178 | TEC_prg | PRDM1 |
| 9.391E-183 | -2.0181723 | 0.185 | 0.362 | 2.218E-178 | TEC_prg | HPGD |
| 1.18E-182 | -0.2839979 | 0.056 | 0.257 | 2.788E-178 | TEC_prg | C1orf43 |
| 2.237E-182 | 0.70426091 | 0.419 | 0.272 | 5.284E-178 | TEC_prg | MACF1 |
| 2.835E-182 | -0.2963962 | 0.049 | 0.245 | 6.698E-178 | TEC_prg | EMC3 |
| 3.314E-182 | -0.2896569 | 0.047 | 0.241 | 7.829E-178 | TEC_prg | TMEM14B |
| 6.942E-182 | -0.3569547 | 0.011 | 0.181 | 1.64E-177 | TEC_prg | MYO1F |
| 8.545E-182 | -0.3295725 | 0.089 | 0.305 | 2.019E-177 | TEC_prg | EIF6 |
| 1.248E-181 | -0.5422864 | 0.004 | 0.168 | 2.949E-177 | TEC_prg | CLEC5A |
| 1.28E-181 | -0.2929189 | 0.051 | 0.249 | 3.024E-177 | TEC_prg | WBP2 |
| 1.99E-181 | -0.3152415 | 0.084 | 0.298 | 4.701E-177 | TEC_prg | MRPL51 |
| 3.532E-181 | -0.3291619 | 0.024 | 0.203 | 8.343E-177 | TEC_prg | CDCP1 |
| 9.85E-181 | -0.3573857 | 0.048 | 0.241 | 2.327E-176 | TEC_prg | ECHS1 |
| 1.748E-180 | -0.6356339 | 0.075 | 0.274 | 4.129E-176 | TEC_prg | CARHSP1 |
| 1.792E-180 | -0.2753034 | 0.194 | 0.461 | 4.233E-176 | TEC_prg | TRIR |
| 2.721E-180 | -0.3548594 | 0.1 | 0.318 | 6.428E-176 | TEC_prg | SELENOS |
| 5.24E-180 | -0.8103476 | 0.003 | 0.165 | 1.238E-175 | TEC_prg | TMPRSS11D |
| 5.953E-180 | -0.3565311 | 0.111 | 0.334 | 1.406E-175 | TEC_prg | TMEM165 |
| 5.978E-180 | -0.529081 | 0.009 | 0.176 | 1.412E-175 | TEC_prg | MALL |
| 7.19E-180 | -0.2872042 | 0.051 | 0.248 | 1.698E-175 | TEC_prg | CMPK1 |
| 1.147E-179 | -0.4673688 | 0.001 | 0.162 | 2.709E-175 | TEC_prg | FPR1 |
| 1.25E-179 | 0.30639591 | 0.191 | 0.071 | 2.953E-175 | TEC_prg | HECTD4 |
| 1.304E-179 | -0.323382 | 0.086 | 0.298 | 3.081E-175 | TEC_prg | CCT5 |
| 1.403E-179 | -0.3497278 | 0.08 | 0.287 | 3.313E-175 | TEC_prg | TMEM123 |
| 2.567E-179 | 0.85108742 | 0.347 | 0.2 | 6.063E-175 | TEC_prg | TUBA1A |
| 3.537E-179 | -0.3049924 | 0.15 | 0.398 | 8.355E-175 | TEC_prg | KIF5B |
| 4.716E-179 | -0.3281281 | 0.044 | 0.235 | 1.114E-174 | TEC_prg | TGFB1 |

| | | | | | | |
|---|---|---|---|---|---|---|
| 7.453E-179 | -0.3147607 | 0.033 | 0.214 | 1.761E-174 | TEC_prg | C1QBP |
| 7.794E-179 | -0.3551174 | 0.013 | 0.183 | 1.841E-174 | TEC_prg | CFD |
| 8.857E-179 | -0.35806 | 0.142 | 0.379 | 2.092E-174 | TEC_prg | MRPL33 |
| 1.365E-178 | -0.3825323 | 0.644 | 0.786 | 3.223E-174 | TEC_prg | TMA7 |
| 2.487E-178 | -0.2871272 | 0.076 | 0.285 | 5.874E-174 | TEC_prg | NDUFA6 |
| 3.09E-178 | -1.1869018 | 0.114 | 0.316 | 7.3E-174 | TEC_prg | ADIRF |
| 3.292E-178 | -2.2619431 | 0.022 | 0.193 | 7.777E-174 | TEC_prg | IL1B |
| 3.504E-178 | -0.7096968 | 0.791 | 0.78 | 8.278E-174 | TEC_prg | RPL10A |
| 9.656E-178 | -0.4393783 | 0.114 | 0.334 | 2.281E-173 | TEC_prg | FNDC3B |
| 1.001E-177 | -0.4482576 | 0.002 | 0.162 | 2.365E-173 | TEC_prg | MS4A7 |
| 1.381E-177 | -0.3297463 | 0.011 | 0.177 | 3.262E-173 | TEC_prg | WIPF1 |
| 1.524E-177 | -1.2654742 | 0.015 | 0.183 | 3.601E-173 | TEC_prg | DUOXA2 |
| 2.033E-177 | -0.3405592 | 0.033 | 0.215 | 4.803E-173 | TEC_prg | HIGD1A |
| 2.327E-177 | -0.3886217 | 0.022 | 0.197 | 5.496E-173 | TEC_prg | CD37 |
| 2.45E-176 | -0.8285073 | 0.002 | 0.161 | 5.787E-172 | TEC_prg | TMPRSS11E |
| 2.464E-176 | -1.0171399 | 0.008 | 0.171 | 5.822E-172 | TEC_prg | PITX1 |
| 3.85E-176 | 0.42110739 | 0.268 | 0.121 | 9.095E-172 | TEC_prg | ARHGAP29 |
| 5.532E-176 | -0.2976613 | 0.04 | 0.226 | 1.307E-171 | TEC_prg | SNAP23 |
| 5.943E-176 | -0.5558064 | 0.233 | 0.449 | 1.404E-171 | TEC_prg | SPINT2 |
| 1.047E-175 | -0.515666 | 0.048 | 0.233 | 2.473E-171 | TEC_prg | CREM |
| 2.655E-175 | -0.55965 | 0.027 | 0.201 | 6.272E-171 | TEC_prg | SDCBP2 |
| 3.301E-175 | -1.0169595 | 0.011 | 0.174 | 7.797E-171 | TEC_prg | PRSS27 |
| 4.896E-175 | -0.4043259 | 0.003 | 0.163 | 1.156E-170 | TEC_prg | TREM1 |
| 1.224E-174 | -0.4571023 | 0.012 | 0.177 | 2.89E-170 | TEC_prg | SERPINB9 |
| 1.369E-174 | -0.6300988 | 0.018 | 0.185 | 3.234E-170 | TEC_prg | FXYD3 |
| 1.681E-174 | -0.2768557 | 0.035 | 0.216 | 3.972E-170 | TEC_prg | ERP44 |
| 1.923E-174 | -0.2915391 | 0.063 | 0.261 | 4.542E-170 | TEC_prg | POLR2K |
| 2.018E-174 | -0.3513262 | 0.057 | 0.25 | 4.768E-170 | TEC_prg | ATP6V1B2 |
| 2.365E-174 | -0.2976022 | 0.033 | 0.214 | 5.588E-170 | TEC_prg | GLTP |
| 2.722E-174 | -0.2789618 | 0.046 | 0.234 | 6.431E-170 | TEC_prg | SERINC3 |
| 3.549E-174 | -0.3356956 | 0.019 | 0.189 | 8.385E-170 | TEC_prg | DAPP1 |
| 1.004E-173 | -0.3168764 | 0.047 | 0.235 | 2.372E-169 | TEC_prg | PPP2R5C |
| 1.035E-173 | -0.3005674 | 0.011 | 0.175 | 2.444E-169 | TEC_prg | ARHGAP30 |
| 1.209E-173 | -0.3897049 | 0.363 | 0.586 | 2.857E-169 | TEC_prg | SLC25A6 |
| 2.025E-173 | 0.5546599 | 0.837 | 0.683 | 4.785E-169 | TEC_prg | JUNB |
| 7.04E-173 | -0.2880795 | 0.023 | 0.196 | 1.663E-168 | TEC_prg | IL13RA1 |
| 9.178E-173 | -0.3566383 | 0.006 | 0.165 | 2.168E-168 | TEC_prg | BCAT1 |
| 9.45E-173 | -0.3316626 | 0.027 | 0.2 | 2.232E-168 | TEC_prg | PSMB9 |
| 1.706E-172 | -0.3210688 | 0.004 | 0.161 | 4.031E-168 | TEC_prg | CARD16 |
| 1.727E-172 | -0.401153 | 0.034 | 0.213 | 4.079E-168 | TEC_prg | SDF2L1 |
| 2.555E-172 | -0.688924 | 0.005 | 0.163 | 6.036E-168 | TEC_prg | NCCRP1 |
| 6.165E-172 | 0.34479952 | 0.165 | 0.057 | 1.456E-167 | TEC_prg | SLC12A2 |
| 9.371E-172 | -0.2770618 | 0.068 | 0.266 | 2.214E-167 | TEC_prg | SNRPB2 |
| 1.46E-171 | -0.3381092 | 0.021 | 0.19 | 3.45E-167 | TEC_prg | DHRS3 |
| 1.713E-171 | -0.5355591 | 0.004 | 0.16 | 4.048E-167 | TEC_prg | CD93 |
| 2.733E-171 | -0.7821916 | 0.01 | 0.169 | 6.457E-167 | TEC_prg | SCEL |
| 3.208E-171 | -0.29059 | 0.076 | 0.279 | 7.579E-167 | TEC_prg | TMCO1 |
| 3.241E-171 | -0.3660024 | 0.065 | 0.257 | 7.656E-167 | TEC_prg | PCBD1 |
| 5.017E-171 | -0.4661229 | 0.358 | 0.573 | 1.185E-166 | TEC_prg | FAM107B |
| 5.631E-171 | -0.3369644 | 0.033 | 0.209 | 1.33E-166 | TEC_prg | F11R |

| | | | | | | |
|---|---|---|---|---|---|---|
| 5.782E-171 | -2.2149784 | 0.343 | 0.478 | 1.366E-166 | TEC_prg | LYZ |
| 6.326E-171 | -0.3089854 | 0.009 | 0.169 | 1.494E-166 | TEC_prg | VSIR |
| 9.085E-171 | -0.3043214 | 0.002 | 0.157 | 2.146E-166 | TEC_prg | IL2RG |
| 9.646E-171 | -0.3406471 | 0.069 | 0.266 | 2.279E-166 | TEC_prg | SNX9 |
| 3.975E-170 | -0.3984918 | 0.152 | 0.382 | 9.39E-166 | TEC_prg | MIF |
| 4.474E-170 | -0.3649209 | 0.026 | 0.196 | 1.057E-165 | TEC_prg | TMEM176B |
| 6.713E-170 | -0.3140227 | 0.089 | 0.296 | 1.586E-165 | TEC_prg | SSB |
| 1.986E-169 | -0.2967592 | 0.124 | 0.348 | 4.692E-165 | TEC_prg | DRAP1 |
| 2.103E-169 | -0.3104972 | 0.091 | 0.297 | 4.968E-165 | TEC_prg | RSL24D1 |
| 2.115E-169 | -0.9555965 | 0.033 | 0.206 | 4.995E-165 | TEC_prg | RARRES3 |
| 2.221E-169 | 0.2966684 | 0.162 | 0.055 | 5.246E-165 | TEC_prg | PCLO |
| 2.225E-169 | -0.3212237 | 0.08 | 0.281 | 5.256E-165 | TEC_prg | CUTA |
| 2.235E-169 | 0.31120424 | 0.198 | 0.078 | 5.28E-165 | TEC_prg | SHPRH |
| 2.575E-169 | -0.2660723 | 0.026 | 0.197 | 6.083E-165 | TEC_prg | SUMO3 |
| 2.607E-169 | -0.3582789 | 0.115 | 0.332 | 6.158E-165 | TEC_prg | RAB13 |
| 3.5E-169 | -0.292623 | 0.052 | 0.238 | 8.269E-165 | TEC_prg | MDH2 |
| 4.063E-169 | -0.294545 | 0.101 | 0.315 | 9.599E-165 | TEC_prg | COX17 |
| 4.541E-169 | -0.2993804 | 0.049 | 0.236 | 1.073E-164 | TEC_prg | SMIM14 |
| 5.519E-169 | -0.4801768 | 0.569 | 0.719 | 1.304E-164 | TEC_prg | TPM3 |
| 6.828E-169 | -0.2977214 | 0.037 | 0.215 | 1.613E-164 | TEC_prg | GNS |
| 1.202E-168 | 0.41085939 | 0.275 | 0.135 | 2.841E-164 | TEC_prg | CCNL2 |
| 1.418E-168 | -0.2811597 | 0.06 | 0.253 | 3.349E-164 | TEC_prg | CTSA |
| 2.392E-168 | -0.3333786 | 0.065 | 0.258 | 5.651E-164 | TEC_prg | TSTD1 |
| 2.879E-168 | -0.3277632 | 0.032 | 0.205 | 6.801E-164 | TEC_prg | RNF130 |
| 2.894E-168 | -0.3643888 | 0.006 | 0.162 | 6.836E-164 | TEC_prg | C1orf162 |
| 3.222E-168 | -0.2862397 | 0.075 | 0.274 | 7.611E-164 | TEC_prg | RAB14 |
| 4.741E-168 | -0.3209414 | 0.037 | 0.212 | 1.12E-163 | TEC_prg | SEC11C |
| 5.305E-168 | -0.5234633 | 0.002 | 0.155 | 1.253E-163 | TEC_prg | PLAC8 |
| 6.078E-168 | 0.59391191 | 0.325 | 0.181 | 1.436E-163 | TEC_prg | KLF4 |
| 1.037E-167 | -0.5369897 | 0.063 | 0.249 | 2.449E-163 | TEC_prg | PPIF |
| 1.162E-167 | -0.2699546 | 0.066 | 0.261 | 2.746E-163 | TEC_prg | PSMD4 |
| 2.026E-167 | -0.2888212 | 0.058 | 0.248 | 4.785E-163 | TEC_prg | SYNGR2 |
| 2.532E-167 | -0.2638376 | 0.041 | 0.22 | 5.982E-163 | TEC_prg | VPS26A |
| 2.938E-167 | -0.355445 | 0.021 | 0.187 | 6.941E-163 | TEC_prg | ISG20 |
| 5.399E-167 | -0.331302 | 0.038 | 0.214 | 1.275E-162 | TEC_prg | TSPAN3 |
| 2.3E-166 | -0.2993984 | 0.057 | 0.245 | 5.433E-162 | TEC_prg | TMED5 |
| 2.523E-166 | 0.40291103 | 0.217 | 0.093 | 5.96E-162 | TEC_prg | SLC39A10 |
| 2.758E-166 | -0.2974106 | 0.107 | 0.321 | 6.515E-162 | TEC_prg | ROMO1 |
| 3.059E-166 | -0.4517112 | 0.046 | 0.225 | 7.227E-162 | TEC_prg | SLC43A2 |
| 5.391E-166 | -0.3193112 | 0.085 | 0.287 | 1.274E-161 | TEC_prg | ATOX1 |
| 5.828E-166 | -0.6729251 | 0.062 | 0.249 | 1.377E-161 | TEC_prg | ISG15 |
| 7.145E-166 | -0.3186799 | 0.004 | 0.156 | 1.688E-161 | TEC_prg | AL133453.1 |
| 7.443E-166 | -0.3823084 | 0.006 | 0.16 | 1.758E-161 | TEC_prg | TLR2 |
| 7.621E-166 | 0.34369308 | 0.229 | 0.096 | 1.8E-161 | TEC_prg | FXYD2 |
| 1.269E-165 | -0.2895401 | 0.038 | 0.213 | 2.997E-161 | TEC_prg | NCOA4 |
| 1.377E-165 | -0.26698 | 0.047 | 0.23 | 3.252E-161 | TEC_prg | STRAP |
| 1.722E-165 | -0.2844303 | 0.073 | 0.268 | 4.068E-161 | TEC_prg | EIF3D |
| 2.024E-165 | -0.2680571 | 0.038 | 0.214 | 4.781E-161 | TEC_prg | UBE2N |
| 2.24E-165 | -0.3119793 | 0.048 | 0.229 | 5.291E-161 | TEC_prg | NBDY |
| 2.621E-165 | -0.899727 | 0.008 | 0.162 | 6.192E-161 | TEC_prg | CEACAM6 |

| | | | | | | |
|---|---|---|---|---|---|---|
| 2.868E-165 | -0.2819516 | 0.073 | 0.27 | 6.775E-161 | TEC_prg | MIEN1 |
| 3.606E-165 | -0.2995201 | 0.002 | 0.152 | 8.518E-161 | TEC_prg | EVI2A |
| 7.513E-165 | -0.2677897 | 0.047 | 0.229 | 1.775E-160 | TEC_prg | MAPK1IP1L |
| 7.732E-165 | -0.3510067 | 0.064 | 0.251 | 1.827E-160 | TEC_prg | CACYBP |
| 8.055E-165 | -0.2806097 | 0.081 | 0.281 | 1.903E-160 | TEC_prg | OCIAD1 |
| 1.183E-164 | -1.7763299 | 0.006 | 0.159 | 2.796E-160 | TEC_prg | SPRR2A |
| 1.314E-164 | -0.2797119 | 0.008 | 0.163 | 3.104E-160 | TEC_prg | SH3KBP1 |
| 2.126E-164 | -0.3219046 | 0.002 | 0.151 | 5.021E-160 | TEC_prg | HCK |
| 2.976E-164 | -0.2967183 | 0.13 | 0.353 | 7.029E-160 | TEC_prg | SH3GLB1 |
| 3.274E-164 | -0.8561655 | 0.008 | 0.162 | 7.735E-160 | TEC_prg | CEACAM7 |
| 3.775E-164 | -0.3662933 | 0.058 | 0.242 | 8.917E-160 | TEC_prg | GRINA |
| 4.8E-164 | -0.2872696 | 0.088 | 0.291 | 1.134E-159 | TEC_prg | PDCD6IP |
| 6.223E-164 | 0.36572296 | 0.184 | 0.07 | 1.47E-159 | TEC_prg | EFNB2 |
| 6.412E-164 | -0.2897553 | 0.053 | 0.237 | 1.515E-159 | TEC_prg | RIT1 |
| 1.208E-163 | -0.2627406 | 0.04 | 0.217 | 2.854E-159 | TEC_prg | TMEM230 |
| 1.719E-163 | -0.2520278 | 0.035 | 0.209 | 4.06E-159 | TEC_prg | EMC7 |
| 3.563E-163 | -0.3044731 | 0.101 | 0.309 | 8.417E-159 | TEC_prg | NDUFC1 |
| 4.438E-163 | -0.5238754 | 0.075 | 0.261 | 1.048E-158 | TEC_prg | PRSS8 |
| 4.503E-163 | -0.3073426 | 0.05 | 0.231 | 1.064E-158 | TEC_prg | RNF13 |
| 5.144E-163 | -0.4682707 | 0.003 | 0.152 | 1.215E-158 | TEC_prg | MCEMP1 |
| 6.364E-163 | -0.5225444 | 0.072 | 0.255 | 1.503E-158 | TEC_prg | S100A16 |
| 9.738E-163 | -1.0925928 | 0.005 | 0.155 | 2.3E-158 | TEC_prg | CXCL6 |
| 1.072E-162 | -0.313034 | 0.06 | 0.245 | 2.532E-158 | TEC_prg | TPP1 |
| 1.616E-162 | -0.2666001 | 0.153 | 0.387 | 3.817E-158 | TEC_prg | C19orf53 |
| 2.393E-162 | -0.3269966 | 0.039 | 0.211 | 5.654E-158 | TEC_prg | B4GALT5 |
| 3.933E-162 | -0.3629332 | 0.014 | 0.171 | 9.292E-158 | TEC_prg | ADGRE5 |
| 4.158E-162 | -0.4411907 | 0.046 | 0.222 | 9.822E-158 | TEC_prg | CTSC |
| 4.76E-162 | -0.9687674 | 0.002 | 0.15 | 1.124E-157 | TEC_prg | TMPRSS11B |
| 5.295E-162 | -0.3180517 | 0.015 | 0.173 | 1.251E-157 | TEC_prg | SH3BP5 |
| 7.143E-162 | -0.3262474 | 0.012 | 0.168 | 1.687E-157 | TEC_prg | HK2 |
| 7.922E-162 | -0.964771 | 0.294 | 0.476 | 1.871E-157 | TEC_prg | MXD1 |
| 8.894E-162 | -0.2782383 | 0.002 | 0.149 | 2.101E-157 | TEC_prg | NCKAP1L |
| 1.144E-161 | -0.2730748 | 0.09 | 0.293 | 2.703E-157 | TEC_prg | PPP1R2 |
| 1.294E-161 | -0.27912 | 0.093 | 0.297 | 3.056E-157 | TEC_prg | SNRPD3 |
| 1.3E-161 | -0.2590974 | 0.059 | 0.245 | 3.07E-157 | TEC_prg | PSMC1 |
| 1.367E-161 | -0.3152578 | 0.137 | 0.358 | 3.229E-157 | TEC_prg | EIF4B |
| 4.404E-161 | -0.3127445 | 0.121 | 0.338 | 1.04E-156 | TEC_prg | MMP24OS |
| 9.478E-161 | -0.5643112 | 0.756 | 0.796 | 2.239E-156 | TEC_prg | RPS21 |
| 9.926E-161 | -0.5403025 | 0.585 | 0.697 | 2.345E-156 | TEC_prg | NPM1 |
| 1.022E-160 | -0.2541886 | 0.021 | 0.182 | 2.414E-156 | TEC_prg | CNIH1 |
| 1.148E-160 | -0.2815446 | 0.008 | 0.16 | 2.712E-156 | TEC_prg | CCDC69 |
| 1.989E-160 | -0.2515382 | 0.044 | 0.22 | 4.7E-156 | TEC_prg | ARF5 |
| 2.696E-160 | -0.276521 | 0.073 | 0.266 | 6.369E-156 | TEC_prg | CCT8 |
| 3.318E-160 | -0.3490436 | 0.556 | 0.732 | 7.838E-156 | TEC_prg | SKP1 |
| 4.235E-160 | -0.3056049 | 0.002 | 0.149 | 1E-155 | TEC_prg | CYTIP |
| 5.298E-160 | -0.2663189 | 0.032 | 0.2 | 1.251E-155 | TEC_prg | POLE4 |
| 1.811E-159 | -0.5146951 | 0.493 | 0.639 | 4.279E-155 | TEC_prg | DSTN |
| 3.068E-159 | -0.3133228 | 0.09 | 0.289 | 7.248E-155 | TEC_prg | UBE2D1 |
| 4.787E-159 | -0.2778547 | 0.067 | 0.255 | 1.131E-154 | TEC_prg | PSMA3 |
| 5.047E-159 | -0.3309257 | 0.206 | 0.444 | 1.192E-154 | TEC_prg | RAN |

| | | | | | | |
|---|---|---|---|---|---|---|
| 5.145E-159 | -0.2820767 | 0.089 | 0.288 | 1.215E-154 | TEC_prg | DIAPH1 |
| 6.474E-159 | -0.2570424 | 0.036 | 0.205 | 1.529E-154 | TEC_prg | DGUOK |
| 8.111E-159 | -0.2761502 | 0.018 | 0.175 | 1.916E-154 | TEC_prg | PPT1 |
| 9.261E-159 | -0.2984214 | 0.003 | 0.149 | 2.188E-154 | TEC_prg | FGR |
| 1.004E-158 | -0.270582 | 0.048 | 0.226 | 2.373E-154 | TEC_prg | EIF4EBP2 |
| 1.183E-158 | 0.52523583 | 0.31 | 0.169 | 2.794E-154 | TEC_prg | EPS8L2 |
| 1.635E-158 | -0.2582845 | 0.105 | 0.314 | 3.862E-154 | TEC_prg | MTCH1 |
| 1.716E-158 | -0.5301533 | 0.004 | 0.15 | 4.054E-154 | TEC_prg | HLA-DRB5 |
| 4.166E-158 | -0.2705133 | 0.043 | 0.217 | 9.842E-154 | TEC_prg | SSR1 |
| 5.185E-158 | -0.3160239 | 0.184 | 0.419 | 1.225E-153 | TEC_prg | CTNNA1 |
| 7.468E-158 | -0.2666365 | 0.07 | 0.259 | 1.764E-153 | TEC_prg | TXNL1 |
| 1.282E-157 | -0.3023245 | 0.07 | 0.256 | 3.028E-153 | TEC_prg | ATP5F1A |
| 1.891E-157 | -0.3031803 | 0.11 | 0.316 | 4.466E-153 | TEC_prg | ARF6 |
| 1.895E-157 | -0.2630265 | 0.154 | 0.384 | 4.478E-153 | TEC_prg | RBM8A |
| 1.898E-157 | -0.4208832 | 0.009 | 0.158 | 4.484E-153 | TEC_prg | NCF1 |
| 2.019E-157 | -0.2788823 | 0.093 | 0.293 | 4.769E-153 | TEC_prg | GNAI3 |
| 2.146E-157 | 0.2612471 | 0.142 | 0.046 | 5.068E-153 | TEC_prg | COL4A1 |
| 2.148E-157 | -0.3242207 | 0.005 | 0.152 | 5.073E-153 | TEC_prg | CLEC7A |
| 6.375E-157 | -0.2754718 | 0.07 | 0.258 | 1.506E-152 | TEC_prg | NAA50 |
| 7.17E-157 | -0.307733 | 0.033 | 0.198 | 1.694E-152 | TEC_prg | RILPL2 |
| 7.791E-157 | -0.3230413 | 0.163 | 0.39 | 1.84E-152 | TEC_prg | AES |
| 8.006E-157 | -0.2802702 | 0.078 | 0.269 | 1.891E-152 | TEC_prg | MRPL20 |
| 1.218E-156 | -0.277295 | 0.023 | 0.183 | 2.878E-152 | TEC_prg | SIRPA |
| 1.256E-156 | -0.2705596 | 0.053 | 0.23 | 2.966E-152 | TEC_prg | GTF3C6 |
| 1.72E-156 | -0.4500178 | 0.092 | 0.281 | 4.063E-152 | TEC_prg | RHOC |
| 6.865E-156 | -0.3511393 | 0.123 | 0.333 | 1.622E-151 | TEC_prg | SMS |
| 1.396E-155 | -0.2551437 | 0.033 | 0.199 | 3.297E-151 | TEC_prg | CCT7 |
| 2.991E-155 | -0.6745302 | 0.006 | 0.151 | 7.067E-151 | TEC_prg | CXCL17 |
| 3.515E-155 | -0.2615836 | 0.08 | 0.271 | 8.304E-151 | TEC_prg | XRCC6 |
| 4.208E-155 | -0.2783833 | 0.07 | 0.254 | 9.941E-151 | TEC_prg | BANF1 |
| 4.667E-155 | -0.3499435 | 0.122 | 0.333 | 1.103E-150 | TEC_prg | CYTOR |
| 6.561E-155 | 0.49326642 | 0.244 | 0.118 | 1.55E-150 | TEC_prg | TMX4 |
| 8.21E-155 | -0.2836038 | 0.114 | 0.322 | 1.94E-150 | TEC_prg | TAPBP |
| 8.272E-155 | 0.62780526 | 0.395 | 0.243 | 1.954E-150 | TEC_prg | PMAIP1 |
| 9.36E-155 | -0.4169707 | 0.012 | 0.161 | 2.211E-150 | TEC_prg | VSIG2 |
| 1.041E-154 | -0.2731476 | 0.169 | 0.402 | 2.459E-150 | TEC_prg | PAPOLA |
| 1.428E-154 | -0.3215384 | 0.102 | 0.302 | 3.374E-150 | TEC_prg | TMEM30A |
| 1.933E-154 | -0.3876056 | 0.077 | 0.26 | 4.565E-150 | TEC_prg | GIPC1 |
| 2.602E-154 | -0.3732171 | 0.019 | 0.172 | 6.147E-150 | TEC_prg | TSPAN6 |
| 3.07E-154 | -0.4461168 | 0.828 | 0.851 | 7.252E-150 | TEC_prg | RPL38 |
| 3.159E-154 | -0.2702333 | 0.035 | 0.2 | 7.462E-150 | TEC_prg | CALU |
| 3.467E-154 | -0.2643581 | 0.065 | 0.248 | 8.19E-150 | TEC_prg | RAB18 |
| 3.49E-154 | 0.70063441 | 0.67 | 0.599 | 8.245E-150 | TEC_prg | SON |
| 5.069E-154 | -0.2506105 | 0.04 | 0.208 | 1.197E-149 | TEC_prg | SDHC |
| 7.435E-154 | -0.2986632 | 0.049 | 0.221 | 1.756E-149 | TEC_prg | ANP32A |
| 7.668E-154 | -1.4626186 | 0.214 | 0.387 | 1.811E-149 | TEC_prg | MAL |
| 1.097E-153 | -0.276806 | 0.079 | 0.269 | 2.591E-149 | TEC_prg | IDS |
| 1.58E-153 | -0.2877634 | 0.086 | 0.278 | 3.732E-149 | TEC_prg | CCT3 |
| 1.736E-153 | 0.41322564 | 0.248 | 0.117 | 4.102E-149 | TEC_prg | SCIN |
| 2.74E-153 | -0.2737189 | 0.112 | 0.319 | 6.472E-149 | TEC_prg | UBALD2 |

| | | | | | | |
|---|---|---|---|---|---|---|
| 1.537E-152 | -0.2556113 | 0.054 | 0.229 | 3.632E-148 | TEC_prg | ANXA7 |
| 2.316E-152 | -0.2548204 | 0.05 | 0.223 | 5.471E-148 | TEC_prg | CHURC1 |
| 2.946E-152 | -0.2646925 | 0.03 | 0.191 | 6.958E-148 | TEC_prg | VKORC1 |
| 2.986E-152 | -1.639006 | 0.005 | 0.147 | 7.055E-148 | TEC_prg | LYPD2 |
| 3.057E-152 | -0.2693067 | 0.061 | 0.238 | 7.221E-148 | TEC_prg | SNRPF |
| 4.351E-152 | -0.2777862 | 0.127 | 0.338 | 1.028E-147 | TEC_prg | CLTC |
| 5.115E-152 | -0.2896004 | 0.03 | 0.189 | 1.208E-147 | TEC_prg | ABHD2 |
| 5.565E-152 | -0.2578539 | 0.06 | 0.239 | 1.315E-147 | TEC_prg | TMEM208 |
| 8.148E-152 | -0.5855181 | 0.019 | 0.169 | 1.925E-147 | TEC_prg | CDC42EP5 |
| 1.183E-151 | -0.3134307 | 0.179 | 0.409 | 2.795E-147 | TEC_prg | DEK |
| 1.243E-151 | -0.2910657 | 0.078 | 0.266 | 2.937E-147 | TEC_prg | CD47 |
| 1.55E-151 | -0.2599482 | 0.052 | 0.225 | 3.662E-147 | TEC_prg | SIVA1 |
| 2.436E-151 | -0.3015488 | 0.57 | 0.744 | 5.755E-147 | TEC_prg | ELOB |
| 2.527E-151 | -0.3652283 | 0.064 | 0.242 | 5.969E-147 | TEC_prg | STXBP2 |
| 2.824E-151 | -0.2639861 | 0.155 | 0.381 | 6.672E-147 | TEC_prg | XRCC5 |
| 3.138E-151 | -0.2607722 | 0.025 | 0.18 | 7.413E-147 | TEC_prg | CNPY3 |
| 8.024E-151 | 0.64775306 | 0.459 | 0.337 | 1.895E-146 | TEC_prg | PNN |
| 8.214E-151 | -0.2542733 | 0.048 | 0.219 | 1.94E-146 | TEC_prg | KRTCAP2 |
| 8.942E-151 | -0.3189475 | 0.016 | 0.165 | 2.112E-146 | TEC_prg | FAM129A |
| 1.216E-150 | -0.2666493 | 0.049 | 0.219 | 2.872E-146 | TEC_prg | PSMB5 |
| 4.98E-150 | 0.3536706 | 0.186 | 0.075 | 1.176E-145 | TEC_prg | DEPP1 |
| 7.458E-150 | -0.2655322 | 0.032 | 0.193 | 1.762E-145 | TEC_prg | TMEM147 |
| 9.225E-150 | -0.2503474 | 0.106 | 0.308 | 2.179E-145 | TEC_prg | MRFAP1 |
| 1.084E-149 | -0.3494869 | 0.054 | 0.225 | 2.56E-145 | TEC_prg | STK17B |
| 1.552E-149 | -0.333247 | 0.078 | 0.262 | 3.666E-145 | TEC_prg | RBPJ |
| 2.053E-149 | -0.3334514 | 0.134 | 0.343 | 4.85E-145 | TEC_prg | YPEL3 |
| 2.663E-149 | -0.40359 | 0.03 | 0.189 | 6.292E-145 | TEC_prg | XIST |
| 3.606E-149 | 0.79902466 | 0.455 | 0.334 | 8.517E-145 | TEC_prg | IER5 |
| 7.331E-149 | -0.3099305 | 0.06 | 0.234 | 1.732E-144 | TEC_prg | MRPL18 |
| 7.993E-149 | -0.2513766 | 0.037 | 0.201 | 1.888E-144 | TEC_prg | MAP2K1 |
| 1.872E-148 | -0.2766068 | 0.006 | 0.145 | 4.422E-144 | TEC_prg | APBB1IP |
| 2.117E-148 | -0.2975399 | 0.009 | 0.151 | 5.001E-144 | TEC_prg | ITGA5 |
| 2.291E-148 | -0.2594971 | 0.04 | 0.204 | 5.411E-144 | TEC_prg | GTF3A |
| 2.391E-148 | -0.3018446 | 0.003 | 0.141 | 5.649E-144 | TEC_prg | IRAK3 |
| 3.515E-148 | -0.5335418 | 0.358 | 0.542 | 8.303E-144 | TEC_prg | HEBP2 |
| 3.689E-148 | -0.2590146 | 0.071 | 0.253 | 8.714E-144 | TEC_prg | PSMA5 |
| 5.49E-148 | -0.3594777 | 0.053 | 0.222 | 1.297E-143 | TEC_prg | MAP3K8 |
| 5.87E-148 | 0.70194325 | 0.45 | 0.312 | 1.387E-143 | TEC_prg | CEBPD |
| 7.788E-148 | -1.6146401 | 0.359 | 0.506 | 1.84E-143 | TEC_prg | CD74 |
| 8.196E-148 | -0.2501268 | 0.055 | 0.227 | 1.936E-143 | TEC_prg | ACP1 |
| 1.012E-147 | -0.3611893 | 0.038 | 0.2 | 2.39E-143 | TEC_prg | ICAM1 |
| 1.152E-147 | -0.2913735 | 0.495 | 0.692 | 2.72E-143 | TEC_prg | COX5B |
| 1.898E-147 | -0.3073399 | 0.219 | 0.452 | 4.485E-143 | TEC_prg | EIF3A |
| 2.19E-147 | -0.2972434 | 0.034 | 0.193 | 5.173E-143 | TEC_prg | TMEM159 |
| 2.654E-147 | -0.2672724 | 0.111 | 0.313 | 6.269E-143 | TEC_prg | NDUFS6 |
| 3.062E-147 | -0.389679 | 0.119 | 0.318 | 7.234E-143 | TEC_prg | REL |
| 3.419E-147 | -0.6881972 | 0.006 | 0.145 | 8.077E-143 | TEC_prg | DHRS9 |
| 4.228E-147 | -0.3122397 | 0.084 | 0.269 | 9.987E-143 | TEC_prg | UBL3 |
| 4.399E-147 | -0.2567463 | 0.034 | 0.192 | 1.039E-142 | TEC_prg | ATP6V1A |
| 5.648E-147 | 0.35117634 | 0.168 | 0.065 | 1.334E-142 | TEC_prg | FADS3 |

| | | | | | | |
|---|---|---|---|---|---|---|
| 6.735E-147 | -0.2680571 | 0.046 | 0.212 | 1.591E-142 | TEC_prg | PHC2 |
| 1.175E-146 | -0.3283034 | 0.063 | 0.235 | 2.775E-142 | TEC_prg | STK4 |
| 1.197E-146 | -0.2595481 | 0.075 | 0.258 | 2.827E-142 | TEC_prg | TUFM |
| 2.952E-146 | -0.2850912 | 0.024 | 0.175 | 6.975E-142 | TEC_prg | AP1S2 |
| 3.911E-146 | -1.0986274 | 0.008 | 0.148 | 9.239E-142 | TEC_prg | SERPINB2 |
| 7.009E-146 | -0.2649437 | 0.056 | 0.226 | 1.656E-141 | TEC_prg | CNIH4 |
| 8.293E-146 | -0.2696759 | 0.023 | 0.175 | 1.959E-141 | TEC_prg | DOCK8 |
| 1.01E-145 | 0.27391025 | 0.161 | 0.061 | 2.387E-141 | TEC_prg | AL450998.2 |
| 1.033E-145 | -0.4313641 | 0.944 | 0.917 | 2.439E-141 | TEC_prg | RPS15 |
| 1.863E-145 | -0.3315757 | 0.001 | 0.134 | 4.402E-141 | TEC_prg | LILRB2 |
| 2.293E-145 | -1.4779428 | 0.778 | 0.735 | 5.416E-141 | TEC_prg | VIM |
| 2.644E-145 | -0.4155758 | 0.055 | 0.221 | 6.247E-141 | TEC_prg | CSNK1E |
| 4.435E-145 | -0.2703864 | 0.016 | 0.16 | 1.048E-140 | TEC_prg | LEPROTL1 |
| 5.206E-145 | -0.4563353 | 0.049 | 0.212 | 1.23E-140 | TEC_prg | SAMD9 |
| 7.432E-145 | -0.381058 | 0.004 | 0.14 | 1.756E-140 | TEC_prg | MARCO |
| 9.796E-145 | -0.2600926 | 0.1 | 0.292 | 2.314E-140 | TEC_prg | RAB21 |
| 1.092E-144 | 0.61587675 | 0.429 | 0.302 | 2.581E-140 | TEC_prg | AKAP9 |
| 1.273E-144 | -0.3541078 | 0.053 | 0.219 | 3.008E-140 | TEC_prg | MANF |
| 1.467E-144 | -0.2554308 | 0.059 | 0.231 | 3.466E-140 | TEC_prg | OSER1 |
| 1.749E-144 | -0.27536 | 0.071 | 0.25 | 4.131E-140 | TEC_prg | BCL10 |
| 1.907E-144 | -0.3119761 | 0.095 | 0.284 | 4.504E-140 | TEC_prg | GSN |
| 4.526E-144 | -0.3879816 | 0.096 | 0.278 | 1.069E-139 | TEC_prg | WASL |
| 7.529E-144 | -0.2507839 | 0.076 | 0.256 | 1.779E-139 | TEC_prg | TM9SF2 |
| 1.284E-143 | -0.7714315 | 0.824 | 0.834 | 3.034E-139 | TEC_prg | PPDPF |
| 1.575E-143 | -0.2915194 | 0.001 | 0.133 | 3.72E-139 | TEC_prg | CSF1R |
| 2.073E-143 | 0.2552046 | 0.144 | 0.051 | 4.896E-139 | TEC_prg | DNAJC3-DT |
| 2.33E-143 | 0.62960404 | 0.509 | 0.403 | 5.505E-139 | TEC_prg | SFPQ |
| 2.799E-143 | 0.27382084 | 0.182 | 0.074 | 6.613E-139 | TEC_prg | KANSL1L |
| 5.271E-143 | -0.270122 | 0.126 | 0.329 | 1.245E-138 | TEC_prg | SSBP1 |
| 1.062E-142 | -0.3041801 | 0.298 | 0.53 | 2.508E-138 | TEC_prg | JAK1 |
| 1.156E-142 | -0.2888819 | 0.132 | 0.338 | 2.73E-138 | TEC_prg | PTBP3 |
| 1.581E-142 | -0.9471588 | 0.01 | 0.148 | 3.736E-138 | TEC_prg | EREG |
| 2.327E-142 | -0.819388 | 0.025 | 0.171 | 5.497E-138 | TEC_prg | KRT15 |
| 4.65E-142 | -0.4074715 | 0.15 | 0.352 | 1.099E-137 | TEC_prg | PPP1R14B |
| 4.67E-142 | -0.2925229 | 0.008 | 0.144 | 1.103E-137 | TEC_prg | LILRB3 |
| 1.094E-141 | -0.2971796 | 0.001 | 0.131 | 2.584E-137 | TEC_prg | CD86 |
| 1.573E-141 | -0.4192425 | 0.165 | 0.37 | 3.716E-137 | TEC_prg | PEBP1 |
| 1.952E-141 | -0.2604548 | 0.09 | 0.276 | 4.612E-137 | TEC_prg | MORF4L2 |
| 3.209E-141 | -0.2844711 | 0.082 | 0.261 | 7.581E-137 | TEC_prg | COMT |
| 4.772E-141 | -0.5178296 | 0.646 | 0.737 | 1.127E-136 | TEC_prg | PKM |
| 7.218E-141 | 0.60307215 | 0.44 | 0.3 | 1.705E-136 | TEC_prg | HIST1H2AC |
| 9.554E-141 | 1.21827888 | 0.496 | 0.394 | 2.257E-136 | TEC_prg | RNF213 |
| 1.13E-140 | -0.2624532 | 0.082 | 0.262 | 2.669E-136 | TEC_prg | NDUFA3 |
| 1.285E-140 | -0.3266322 | 0.118 | 0.312 | 3.036E-136 | TEC_prg | SKIL |
| 1.543E-140 | -1.3634947 | 0.022 | 0.167 | 3.646E-136 | TEC_prg | CCL20 |
| 2.025E-140 | 0.55452415 | 0.381 | 0.244 | 4.784E-136 | TEC_prg | TSC22D1 |
| 3.271E-140 | -0.2508449 | 0.094 | 0.28 | 7.726E-136 | TEC_prg | YWHAG |
| 3.689E-140 | -0.2683673 | 0.09 | 0.273 | 8.715E-136 | TEC_prg | CCT6A |
| 7.115E-140 | -2.5041761 | 0.105 | 0.271 | 1.681E-135 | TEC_prg | MT1G |
| 1.203E-139 | -0.2556642 | 0.079 | 0.256 | 2.842E-135 | TEC_prg | SNRPE |

| | | | | | | |
|---|---|---|---|---|---|---|
| 1.333E-139 | -0.2540195 | 0.014 | 0.152 | 3.149E-135 | TEC_prg | MGST2 |
| 1.494E-139 | -0.2667792 | 0.076 | 0.252 | 3.53E-135 | TEC_prg | SYPL1 |
| 1.681E-139 | -0.2754391 | 0.089 | 0.271 | 3.971E-135 | TEC_prg | PSMC6 |
| 1.975E-139 | -0.2539771 | 0.084 | 0.264 | 4.665E-135 | TEC_prg | VPS35 |
| 2.284E-139 | -0.7791764 | 0.039 | 0.189 | 5.394E-135 | TEC_prg | NDRG2 |
| 2.345E-139 | 0.26410804 | 0.148 | 0.053 | 5.54E-135 | TEC_prg | COL4A2 |
| 2.465E-139 | -0.2647663 | 0.01 | 0.146 | 5.822E-135 | TEC_prg | PILRA |
| 2.732E-139 | -0.2544029 | 0.052 | 0.215 | 6.453E-135 | TEC_prg | TWF1 |
| 6.375E-139 | 0.43788211 | 0.274 | 0.146 | 1.506E-134 | TEC_prg | TUT4 |
| 8.355E-139 | -0.2553031 | 0.059 | 0.226 | 1.974E-134 | TEC_prg | NECTIN2 |
| 8.701E-139 | -0.2559546 | 0.076 | 0.252 | 2.055E-134 | TEC_prg | SCARB2 |
| 9.337E-139 | -0.676701 | 0.01 | 0.145 | 2.206E-134 | TEC_prg | GABRP |
| 9.35E-139 | -0.3107288 | 0.023 | 0.166 | 2.209E-134 | TEC_prg | CPVL |
| 1.56E-138 | -0.4055708 | 0.003 | 0.132 | 3.685E-134 | TEC_prg | FCGR3A |
| 4.68E-138 | -0.2522152 | 0.037 | 0.19 | 1.105E-133 | TEC_prg | ERGIC1 |
| 4.977E-138 | -0.3598479 | 0.011 | 0.146 | 1.176E-133 | TEC_prg | RGCC |
| 5.768E-138 | -0.4051034 | 0.117 | 0.302 | 1.363E-133 | TEC_prg | LDHB |
| 7.492E-138 | -1.1006773 | 0.046 | 0.196 | 1.77E-133 | TEC_prg | SPINK1 |
| 8.719E-138 | -0.3026824 | 0.022 | 0.164 | 2.06E-133 | TEC_prg | SMAP2 |
| 9.012E-138 | 0.26263634 | 0.164 | 0.063 | 2.129E-133 | TEC_prg | KITLG |
| 9.445E-138 | -0.4298014 | 0.006 | 0.136 | 2.231E-133 | TEC_prg | TMEM40 |
| 1.237E-137 | -0.5586606 | 0.885 | 0.867 | 2.921E-133 | TEC_prg | RPLP0 |
| 1.404E-137 | -0.2566632 | 0.037 | 0.189 | 3.317E-133 | TEC_prg | ZDHHC20 |
| 2.134E-137 | -0.2679803 | 0.001 | 0.128 | 5.04E-133 | TEC_prg | C3AR1 |
| 8.482E-137 | -0.2571239 | 0.005 | 0.135 | 2.004E-132 | TEC_prg | PTAFR |
| 1.522E-136 | -0.617913 | 0.018 | 0.156 | 3.595E-132 | TEC_prg | CDKN2A |
| 2.181E-136 | 0.58041499 | 0.718 | 0.588 | 5.152E-132 | TEC_prg | NFKBIA |
| 3.545E-136 | 0.4115421 | 0.242 | 0.121 | 8.375E-132 | TEC_prg | ASCC3 |
| 3.879E-136 | -0.7465475 | 0.048 | 0.2 | 9.164E-132 | TEC_prg | AQP3 |
| 4.719E-136 | -0.3045931 | 0.123 | 0.316 | 1.115E-131 | TEC_prg | VPS4B |
| 5.823E-136 | -0.2585619 | 0.071 | 0.242 | 1.376E-131 | TEC_prg | RSL1D1 |
| 6.911E-136 | -0.6234617 | 0.003 | 0.13 | 1.633E-131 | TEC_prg | MNDA |
| 9.396E-136 | 0.66510468 | 0.806 | 0.686 | 2.22E-131 | TEC_prg | PPP1R15A |
| 1.973E-135 | -0.2615483 | 0.057 | 0.218 | 4.66E-131 | TEC_prg | PHB |
| 2.46E-135 | -0.4655731 | 0.047 | 0.201 | 5.812E-131 | TEC_prg | ACSL1 |
| 2.79E-135 | -0.3802633 | 0.002 | 0.129 | 6.59E-131 | TEC_prg | CD163 |
| 3.025E-135 | -0.3290458 | 0.151 | 0.351 | 7.145E-131 | TEC_prg | RBM47 |
| 4.408E-135 | -0.3788423 | 0.029 | 0.172 | 1.041E-130 | TEC_prg | TTC9 |
| 4.904E-135 | -0.2743055 | 0.011 | 0.144 | 1.158E-130 | TEC_prg | CELF2 |
| 5.484E-135 | -0.2785194 | 0.12 | 0.311 | 1.296E-130 | TEC_prg | STK24 |
| 2.096E-134 | -0.2911874 | 0.001 | 0.125 | 4.95E-130 | TEC_prg | PECAM1 |
| 2.513E-134 | -0.4540732 | 0.008 | 0.138 | 5.936E-130 | TEC_prg | SRD5A3 |
| 3.657E-134 | -1.1790898 | 0.004 | 0.13 | 8.638E-130 | TEC_prg | SPINK5 |
| 4.161E-134 | 0.4242785 | 0.87 | 0.751 | 9.83E-130 | TEC_prg | BTG1 |
| 5.959E-134 | -0.2512126 | 0.007 | 0.135 | 1.408E-129 | TEC_prg | DOCK10 |
| 9.347E-134 | -0.2924899 | 0.037 | 0.187 | 2.208E-129 | TEC_prg | B3GNT5 |
| 9.52E-134 | -0.4949717 | 0.001 | 0.125 | 2.249E-129 | TEC_prg | TMPRSS11A |
| 1.295E-133 | -0.2879878 | 0.041 | 0.191 | 3.059E-129 | TEC_prg | TMEM54 |
| 3.202E-133 | -0.3526781 | 0.016 | 0.15 | 7.563E-129 | TEC_prg | HLA-DMA |
| 3.737E-133 | -0.3269163 | 0.036 | 0.184 | 8.827E-129 | TEC_prg | NECTIN4 |

| | | | | | | |
|---|---|---|---|---|---|---|
| 4.136E-133 | -0.9264587 | 0.014 | 0.146 | 9.77E-129 | TEC_prg | RHCG |
| 9.398E-133 | 0.42516479 | 0.22 | 0.102 | 2.22E-128 | TEC_prg | RRAD |
| 3.59E-132 | -1.2069235 | 0.008 | 0.136 | 8.481E-128 | TEC_prg | CCL3 |
| 4.014E-132 | -1.2575972 | 0.014 | 0.145 | 9.482E-128 | TEC_prg | PLAT |
| 4.858E-132 | -0.2854275 | 0.136 | 0.331 | 1.148E-127 | TEC_prg | MZT2B |
| 5.76E-132 | 0.61021556 | 0.304 | 0.175 | 1.361E-127 | TEC_prg | SLC5A3 |
| 1.246E-131 | -0.2523593 | 0.059 | 0.219 | 2.944E-127 | TEC_prg | TANK |
| 1.424E-131 | -0.6016142 | 0.076 | 0.238 | 3.365E-127 | TEC_prg | CCND1 |
| 4.478E-131 | -0.7658891 | 0.054 | 0.209 | 1.058E-126 | TEC_prg | HSPA6 |
| 4.621E-131 | -0.2511683 | 0.054 | 0.21 | 1.092E-126 | TEC_prg | GNAQ |
| 5.516E-131 | -0.2543559 | 0.057 | 0.216 | 1.303E-126 | TEC_prg | NT5C2 |
| 2.565E-130 | -0.2543615 | 0.096 | 0.275 | 6.06E-126 | TEC_prg | VAMP2 |
| 2.612E-130 | -0.2616436 | 0.005 | 0.128 | 6.171E-126 | TEC_prg | ADGRE2 |
| 2.763E-130 | -0.3600983 | 0.057 | 0.213 | 6.526E-126 | TEC_prg | SERTAD2 |
| 3.147E-130 | -0.4540094 | 0.043 | 0.19 | 7.435E-126 | TEC_prg | C6orf132 |
| 6.434E-130 | -0.4852616 | 0.701 | 0.774 | 1.52E-125 | TEC_prg | RPS26 |
| 8.148E-130 | -0.2771106 | 0.012 | 0.141 | 1.925E-125 | TEC_prg | PLBD1 |
| 1.166E-129 | -0.2941936 | 0.051 | 0.205 | 2.755E-125 | TEC_prg | PDIA4 |
| 1.695E-129 | 0.44887385 | 0.28 | 0.15 | 4.004E-125 | TEC_prg | ANKRD37 |
| 1.727E-129 | 0.2879472 | 0.154 | 0.06 | 4.081E-125 | TEC_prg | AC091271.1 |
| 4.127E-129 | -0.3235475 | 0.123 | 0.308 | 9.75E-125 | TEC_prg | CHMP1B |
| 5.549E-129 | -0.3052539 | 0.12 | 0.308 | 1.311E-124 | TEC_prg | RNF149 |
| 1.217E-128 | 0.271044 | 0.167 | 0.068 | 2.874E-124 | TEC_prg | GLIS3 |
| 1.442E-128 | -0.2597772 | 0.031 | 0.173 | 3.407E-124 | TEC_prg | MYADM |
| 1.753E-128 | -0.8118083 | 0.004 | 0.125 | 4.14E-124 | TEC_prg | IL7R |
| 3.025E-128 | -0.2764408 | 0.024 | 0.16 | 7.146E-124 | TEC_prg | PLEKHO1 |
| 1.359E-127 | -0.6302007 | 0.004 | 0.125 | 3.21E-123 | TEC_prg | KRT78 |
| 1.811E-127 | 0.27457547 | 0.174 | 0.073 | 4.279E-123 | TEC_prg | NR2F2 |
| 4.27E-127 | -1.391134 | 0.094 | 0.248 | 1.009E-122 | TEC_prg | PRSS22 |
| 1.034E-126 | -1.3462738 | 0.005 | 0.125 | 2.443E-122 | TEC_prg | SPRR1B |
| 1.301E-126 | -1.9242895 | 0.029 | 0.166 | 3.073E-122 | TEC_prg | MT1H |
| 2.558E-126 | -0.2564924 | 0.105 | 0.284 | 6.043E-122 | TEC_prg | CDC42SE1 |
| 2.883E-126 | -0.2809505 | 0.036 | 0.177 | 6.811E-122 | TEC_prg | NDFIP2 |
| 8.157E-126 | 0.2859726 | 0.185 | 0.082 | 1.927E-121 | TEC_prg | GPS2 |
| 8.222E-126 | 0.59733487 | 0.433 | 0.325 | 1.942E-121 | TEC_prg | LUC7L3 |
| 1.504E-125 | -0.2619166 | 0.043 | 0.189 | 3.553E-121 | TEC_prg | FAM162A |
| 4.836E-125 | -0.8076233 | 0.059 | 0.202 | 1.142E-120 | TEC_prg | TMPRSS2 |
| 6.469E-125 | -0.3054037 | 0.052 | 0.202 | 1.528E-120 | TEC_prg | PIM1 |
| 8.265E-125 | -0.3158538 | 0.028 | 0.163 | 1.952E-120 | TEC_prg | PLXDC2 |
| 6.495E-124 | -0.3220529 | 0.182 | 0.387 | 1.534E-119 | TEC_prg | RALBP1 |
| 8.122E-124 | -0.3018465 | 0.046 | 0.191 | 1.919E-119 | TEC_prg | SMIM22 |
| 9.872E-124 | 0.39496017 | 0.283 | 0.162 | 2.332E-119 | TEC_prg | CDK12 |
| 1.058E-123 | 0.35247581 | 0.242 | 0.125 | 2.499E-119 | TEC_prg | PATJ |
| 2.642E-123 | -0.3804027 | 0.007 | 0.127 | 6.242E-119 | TEC_prg | CEACAM1 |
| 2.838E-123 | -0.2538284 | 0.09 | 0.259 | 6.705E-119 | TEC_prg | MAFG |
| 3.509E-123 | -0.2544552 | 0.032 | 0.17 | 8.289E-119 | TEC_prg | CSGALNACT2 |
| 4.44E-123 | -0.2722456 | 0.004 | 0.121 | 1.049E-118 | TEC_prg | ACP5 |
| 7.803E-123 | -0.311781 | 0.002 | 0.118 | 1.843E-118 | TEC_prg | MS4A6A |
| 1.159E-122 | -0.6319262 | 0.963 | 0.876 | 2.739E-118 | TEC_prg | RPS2 |
| 1.254E-122 | 0.26851461 | 0.122 | 0.042 | 2.962E-118 | TEC_prg | SLC7A8 |

| | | | | | | |
|---|---|---|---|---|---|---|
| 1.279E-122 | -0.3325898 | 0.005 | 0.122 | 3.021E-118 | TEC_prg | STS |
| 1.503E-121 | -0.34369 | 0.077 | 0.233 | 3.552E-117 | TEC_prg | DSE |
| 1.898E-121 | -0.372907 | 0.133 | 0.308 | 4.485E-117 | TEC_prg | SPINT1 |
| 2.071E-121 | -0.2699042 | 0.018 | 0.144 | 4.892E-117 | TEC_prg | VAMP5 |
| 6.339E-121 | -0.2539311 | 0.094 | 0.261 | 1.497E-116 | TEC_prg | PDCD5 |
| 1.052E-120 | 0.25928612 | 0.143 | 0.056 | 2.484E-116 | TEC_prg | DDB2 |
| 1.303E-120 | -0.5636702 | 0.139 | 0.309 | 3.078E-116 | TEC_prg | GPRC5A |
| 1.593E-120 | -0.5774913 | 0.489 | 0.619 | 3.763E-116 | TEC_prg | EZR |
| 3.254E-120 | -0.3673854 | 0.1 | 0.264 | 7.687E-116 | TEC_prg | SERINC2 |
| 4.176E-120 | -0.4149177 | 0.058 | 0.204 | 9.864E-116 | TEC_prg | DUSP6 |
| 5.382E-120 | -0.2763384 | 0.138 | 0.323 | 1.271E-115 | TEC_prg | RPL36A |
| 5.526E-120 | -0.4567344 | 0.032 | 0.165 | 1.305E-115 | TEC_prg | ZFAND2A |
| 8.918E-120 | 0.51393217 | 0.428 | 0.276 | 2.107E-115 | TEC_prg | RND3 |
| 9.584E-120 | -0.3007426 | 0.002 | 0.114 | 2.264E-115 | TEC_prg | LILRB1 |
| 1.26E-119 | -0.255659 | 0.073 | 0.23 | 2.977E-115 | TEC_prg | LMAN1 |
| 1.285E-119 | 0.68967767 | 0.584 | 0.497 | 3.035E-115 | TEC_prg | MT-ND4L |
| 1.296E-119 | -0.3639676 | 0.003 | 0.117 | 3.061E-115 | TEC_prg | CSF3R |
| 1.777E-119 | -0.3472881 | 0.051 | 0.195 | 4.197E-115 | TEC_prg | WARS |
| 1.817E-119 | 0.79186382 | 0.447 | 0.349 | 4.292E-115 | TEC_prg | TUBA1B |
| 4.45E-119 | -0.337479 | 0.005 | 0.12 | 1.051E-114 | TEC_prg | FGL2 |
| 4.689E-119 | -1.620483 | 0.004 | 0.118 | 1.108E-114 | TEC_prg | CRCT1 |
| 4.748E-119 | 0.51302976 | 0.378 | 0.253 | 1.122E-114 | TEC_prg | AFDN |
| 7.572E-119 | -0.3027641 | 0.05 | 0.192 | 1.789E-114 | TEC_prg | CCDC88A |
| 1.435E-118 | -0.3757153 | 0.039 | 0.173 | 3.39E-114 | TEC_prg | SDC1 |
| 1.452E-118 | -0.5571646 | 0.014 | 0.134 | 3.431E-114 | TEC_prg | AGR2 |
| 1.743E-118 | -0.2566936 | 0.056 | 0.203 | 4.118E-114 | TEC_prg | HSBP1L1 |
| 2.149E-118 | 0.30792958 | 0.258 | 0.135 | 5.077E-114 | TEC_prg | RGS2 |
| 2.712E-118 | -0.3424508 | 0.004 | 0.117 | 6.407E-114 | TEC_prg | A2ML1 |
| 2.955E-118 | -0.282375 | 0.132 | 0.312 | 6.981E-114 | TEC_prg | CHCHD10 |
| 3.774E-118 | -0.2526535 | 0.181 | 0.382 | 8.916E-114 | TEC_prg | MTDH |
| 4.228E-118 | -0.2615413 | 0.037 | 0.173 | 9.988E-114 | TEC_prg | CRB3 |
| 4.335E-118 | 0.43317313 | 0.289 | 0.171 | 1.024E-113 | TEC_prg | LMO4 |
| 4.849E-118 | -0.4083257 | 0.118 | 0.287 | 1.146E-113 | TEC_prg | CCNG2 |
| 5.407E-118 | 0.3234635 | 0.181 | 0.082 | 1.277E-113 | TEC_prg | SLC9A1 |
| 1.553E-117 | -0.3226099 | 0.051 | 0.193 | 3.669E-113 | TEC_prg | LTA4H |
| 1.618E-117 | 0.28215545 | 0.19 | 0.088 | 3.822E-113 | TEC_prg | SMG6 |
| 3.009E-117 | -0.2990961 | 0.11 | 0.278 | 7.109E-113 | TEC_prg | PRDX2 |
| 3.945E-117 | 0.64114009 | 0.428 | 0.321 | 9.319E-113 | TEC_prg | SNHG8 |
| 4.146E-117 | -0.4132946 | 0.018 | 0.141 | 9.795E-113 | TEC_prg | GBP1 |
| 4.979E-117 | -0.2763602 | 0.006 | 0.12 | 1.176E-112 | TEC_prg | PDE4B |
| 6.069E-117 | -0.6304952 | 0.037 | 0.17 | 1.434E-112 | TEC_prg | MT1F |
| 8.897E-117 | 0.26277034 | 0.185 | 0.083 | 2.102E-112 | TEC_prg | LRBA |
| 1.048E-116 | 0.25853745 | 0.146 | 0.058 | 2.476E-112 | TEC_prg | RND1 |
| 1.542E-116 | -0.4228194 | 0.035 | 0.169 | 3.642E-112 | TEC_prg | MUC4 |
| 6.48E-116 | 0.49589687 | 0.72 | 0.654 | 1.531E-111 | TEC_prg | HNRNPU |
| 6.498E-116 | 0.27979916 | 0.193 | 0.09 | 1.535E-111 | TEC_prg | CHD6 |
| 1.458E-115 | -0.2604981 | 0.032 | 0.163 | 3.445E-111 | TEC_prg | PLS3 |
| 2.585E-115 | 0.3188713 | 0.118 | 0.041 | 6.107E-111 | TEC_prg | FILIP1L |
| 2.968E-115 | -0.3932232 | 0.025 | 0.148 | 7.011E-111 | TEC_prg | GCNT3 |
| 3.641E-115 | -0.4265331 | 0.006 | 0.118 | 8.601E-111 | TEC_prg | THBD |

| | | | | | | |
|---|---|---|---|---|---|---|
| 6.294E-115 | -0.2836527 | 0.011 | 0.128 | 1.487E-110 | TEC_prg | OAS1 |
| 8.125E-115 | -0.6337395 | 0.003 | 0.113 | 1.919E-110 | TEC_prg | HLA-DQA1 |
| 1.406E-114 | -0.2644171 | 0.127 | 0.304 | 3.322E-110 | TEC_prg | RB1CC1 |
| 1.663E-114 | -0.8223935 | 0.005 | 0.116 | 3.929E-110 | TEC_prg | ZG16B |
| 2.292E-114 | -0.4492076 | 0.049 | 0.184 | 5.415E-110 | TEC_prg | CXCR4 |
| 2.86E-114 | -0.2676816 | 0.035 | 0.165 | 6.756E-110 | TEC_prg | SLC44A1 |
| 4.574E-114 | -0.3859607 | 0.052 | 0.191 | 1.081E-109 | TEC_prg | FHL2 |
| 4.734E-114 | -0.2676041 | 0.023 | 0.146 | 1.118E-109 | TEC_prg | MAP3K20 |
| 1.036E-113 | -0.2674821 | 0.031 | 0.159 | 2.447E-109 | TEC_prg | CD40 |
| 1.854E-113 | -2.5376034 | 0.391 | 0.499 | 4.38E-109 | TEC_prg | CXCL8 |
| 2.75E-113 | -0.2504123 | 0.002 | 0.108 | 6.497E-109 | TEC_prg | IKZF1 |
| 4.453E-113 | -0.2681992 | 0.022 | 0.144 | 1.052E-108 | TEC_prg | EPB41L3 |
| 5.848E-113 | 0.36972355 | 0.23 | 0.119 | 1.381E-108 | TEC_prg | ID3 |
| 2.404E-112 | -0.3526479 | 0.037 | 0.167 | 5.68E-108 | TEC_prg | TMPRSS4 |
| 2.601E-112 | -0.3738915 | 0.118 | 0.287 | 6.143E-108 | TEC_prg | TSC22D3 |
| 3.815E-112 | 0.33226251 | 0.21 | 0.106 | 9.012E-108 | TEC_prg | ASXL1 |
| 3.947E-112 | -1.4989485 | 0.008 | 0.119 | 9.324E-108 | TEC_prg | CNFN |
| 7.745E-112 | -0.3043101 | 0.001 | 0.106 | 1.83E-107 | TEC_prg | GPR183 |
| 1.988E-111 | -0.2668458 | 0.055 | 0.196 | 4.696E-107 | TEC_prg | SPTSSA |
| 2.316E-111 | -0.3220199 | 0.561 | 0.702 | 5.471E-107 | TEC_prg | CAST |
| 2.387E-111 | -0.322773 | 0.02 | 0.138 | 5.64E-107 | TEC_prg | POLB |
| 5.905E-111 | -0.3425346 | 0.057 | 0.196 | 1.395E-106 | TEC_prg | GCLM |
| 1.074E-110 | -0.2804186 | 0.045 | 0.179 | 2.538E-106 | TEC_prg | JARID2 |
| 1.287E-110 | -1.9809578 | 0.242 | 0.353 | 3.04E-106 | TEC_prg | PSCA |
| 2.183E-110 | -0.3569236 | 0.692 | 0.791 | 5.158E-106 | TEC_prg | MAP1LC3B |
| 3.031E-110 | -0.3930203 | 0.122 | 0.283 | 7.16E-106 | TEC_prg | UGCG |
| 4.302E-110 | -0.5127637 | 0.004 | 0.111 | 1.016E-105 | TEC_prg | RETN |
| 5.692E-110 | -0.2771435 | 0.069 | 0.214 | 1.345E-105 | TEC_prg | TMEM106B |
| 7.145E-110 | -0.3582808 | 0.002 | 0.107 | 1.688E-105 | TEC_prg | KLK10 |
| 8.325E-110 | -0.4184214 | 0.062 | 0.202 | 1.967E-105 | TEC_prg | BAG3 |
| 9.927E-110 | 0.44536486 | 0.297 | 0.185 | 2.345E-105 | TEC_prg | UPF2 |
| 1.129E-109 | -0.3942506 | 0.006 | 0.114 | 2.668E-105 | TEC_prg | CLIC3 |
| 1.763E-109 | -0.3133332 | 0.021 | 0.14 | 4.165E-105 | TEC_prg | LRG1 |
| 1.993E-109 | -0.2700012 | 0.131 | 0.303 | 4.707E-105 | TEC_prg | YWHAH |
| 1.238E-107 | -0.4872582 | 0.07 | 0.21 | 2.925E-103 | TEC_prg | SFN |
| 1.66E-107 | -0.6511842 | 0.162 | 0.329 | 3.922E-103 | TEC_prg | PHLDA1 |
| 2.061E-107 | -1.1260409 | 0.01 | 0.119 | 4.868E-103 | TEC_prg | CCL4 |
| 2.72E-107 | -0.2833263 | 0.012 | 0.122 | 6.426E-103 | TEC_prg | FUT3 |
| 8.538E-107 | -0.4013982 | 0.058 | 0.192 | 2.017E-102 | TEC_prg | EPHA2 |
| 1.463E-106 | 0.2641536 | 0.147 | 0.062 | 3.456E-102 | TEC_prg | HIST1H4H |
| 5.157E-106 | -0.4188322 | 0.42 | 0.589 | 1.218E-101 | TEC_prg | VMP1 |
| 5.497E-106 | -0.2987118 | 0.007 | 0.112 | 1.299E-101 | TEC_prg | LRMP |
| 6.918E-106 | -0.2769541 | 0.154 | 0.334 | 1.634E-101 | TEC_prg | HMGB2 |
| 6.928E-106 | -0.3914785 | 0.062 | 0.199 | 1.637E-101 | TEC_prg | KLF2 |
| 1.101E-105 | -0.2954928 | 0.031 | 0.152 | 2.6E-101 | TEC_prg | EMP2 |
| 4.144E-105 | -0.3643337 | 0.612 | 0.72 | 9.79E-101 | TEC_prg | GPX4 |
| 7.648E-105 | -0.2718267 | 0.061 | 0.197 | 1.807E-100 | TEC_prg | BNIP3 |
| 2.583E-104 | 0.422567 | 0.885 | 0.802 | 6.102E-100 | TEC_prg | DDX5 |
| 4.001E-104 | -0.4197173 | 0.098 | 0.243 | 9.451E-100 | TEC_prg | CYB5A |
| 9.249E-104 | -0.3820082 | 0.012 | 0.118 | 2.185E-99 | TEC_prg | KRT23 |

| | | | | | | |
|---|---|---|---|---|---|---|
| 1.228E-103 | 0.43934651 | 0.224 | 0.117 | 2.9E-99 | TEC_prg | AC023157.3 |
| 2.065E-102 | -0.7413191 | 0.01 | 0.115 | 4.8773E-98 | TEC_prg | CCL5 |
| 5.717E-102 | -0.3220261 | 0.571 | 0.709 | 1.3505E-97 | TEC_prg | LRRFIP1 |
| 5.792E-102 | -0.5394727 | 0.035 | 0.153 | 1.3683E-97 | TEC_prg | SNCG |
| 1.534E-101 | -0.3923989 | 0.085 | 0.227 | 3.6236E-97 | TEC_prg | LAMB3 |
| 2.51E-101 | -1.3809135 | 0.017 | 0.126 | 5.9298E-97 | TEC_prg | MT1M |
| 4.202E-101 | -0.2537278 | 0.044 | 0.169 | 9.9273E-97 | TEC_prg | OGFRL1 |
| 4.233E-101 | -0.2685472 | 0.144 | 0.31 | 9.999E-97 | TEC_prg | TUBA1C |
| 1.323E-100 | 0.45918349 | 0.309 | 0.201 | 3.125E-96 | TEC_prg | VPS13C |
| 1.1216E-98 | -0.2632436 | 0.005 | 0.101 | 2.6496E-94 | TEC_prg | KLK11 |
| 1.8495E-98 | 0.32616435 | 0.162 | 0.076 | 4.369E-94 | TEC_prg | C6orf120 |
| 2.0933E-98 | -0.3140238 | 0.032 | 0.146 | 4.9451E-94 | TEC_prg | BCAS1 |
| 2.2197E-98 | -0.261638 | 0.106 | 0.257 | 5.2435E-94 | TEC_prg | BHLHE40 |
| 3.3863E-98 | -0.3815069 | 0.373 | 0.541 | 7.9994E-94 | TEC_prg | HSPE1 |
| 3.675E-98 | -0.3593553 | 0.449 | 0.602 | 8.6814E-94 | TEC_prg | KLF6 |
| 8.0556E-98 | -0.7503285 | 0.016 | 0.119 | 1.903E-93 | TEC_prg | APOC1 |
| 1.0449E-97 | -0.3419664 | 0.185 | 0.364 | 2.4683E-93 | TEC_prg | MIDN |
| 1.1123E-97 | 0.3617546 | 0.923 | 0.828 | 2.6275E-93 | TEC_prg | SQSTM1 |
| 1.4753E-97 | -0.2697842 | 0.042 | 0.162 | 3.485E-93 | TEC_prg | DMXL2 |
| 2.7425E-97 | -0.6041505 | 0.116 | 0.261 | 6.4785E-93 | TEC_prg | PLAU |
| 5.7625E-97 | -0.2656267 | 0.054 | 0.178 | 1.3613E-92 | TEC_prg | CEP170 |
| 7.6597E-97 | -0.2859572 | 0.032 | 0.146 | 1.8095E-92 | TEC_prg | ACADM |
| 8.5937E-97 | -0.3230915 | 0.135 | 0.292 | 2.0301E-92 | TEC_prg | ETS2 |
| 8.6695E-96 | -0.2734729 | 0.03 | 0.143 | 2.048E-91 | TEC_prg | IL18 |
| 1.2158E-95 | -0.4738502 | 0.932 | 0.875 | 2.8722E-91 | TEC_prg | RPL12 |
| 2.0264E-95 | -0.2842507 | 0.182 | 0.351 | 4.787E-91 | TEC_prg | TLN1 |
| 4.2786E-95 | -0.3824194 | 0.044 | 0.163 | 1.0107E-90 | TEC_prg | SLC7A11 |
| 5.7855E-95 | -0.3529058 | 0.577 | 0.701 | 1.3667E-90 | TEC_prg | DNAJA1 |
| 6.3587E-95 | -0.3531921 | 0.328 | 0.5 | 1.5021E-90 | TEC_prg | CD46 |
| 6.5303E-95 | -0.2567527 | 0.013 | 0.112 | 1.5426E-90 | TEC_prg | DUSP14 |
| 6.8951E-95 | 0.40620398 | 0.303 | 0.199 | 1.6288E-90 | TEC_prg | TRIM44 |
| 2.5088E-94 | -0.2802713 | 0.137 | 0.292 | 5.9265E-90 | TEC_prg | SLK |
| 3.8046E-94 | 0.29198939 | 0.234 | 0.126 | 8.9876E-90 | TEC_prg | NUPR1 |
| 5.7064E-93 | -0.2525443 | 0.198 | 0.379 | 1.348E-88 | TEC_prg | CFLAR |
| 5.8376E-93 | 0.26944373 | 0.172 | 0.084 | 1.379E-88 | TEC_prg | PHYH |
| 1.1131E-92 | -0.6282083 | 0.01 | 0.105 | 2.6295E-88 | TEC_prg | PTGS2 |
| 3.6538E-92 | 0.25070556 | 0.172 | 0.084 | 8.6313E-88 | TEC_prg | MEX3C |
| 5.2075E-92 | -0.4750018 | 0.972 | 0.94 | 1.2302E-87 | TEC_prg | RPS27 |
| 7.6015E-92 | -0.7712591 | 0.137 | 0.277 | 1.7957E-87 | TEC_prg | AREG |
| 1.1486E-91 | 0.28701053 | 0.196 | 0.103 | 2.7132E-87 | TEC_prg | BRD1 |
| 1.6402E-91 | -0.2645584 | 0.814 | 0.869 | 3.8748E-87 | TEC_prg | CALM1 |
| 5.9847E-91 | -0.4389937 | 0.045 | 0.16 | 1.4138E-86 | TEC_prg | SMC4 |
| 3.7201E-90 | -0.2526178 | 0.031 | 0.139 | 8.7881E-86 | TEC_prg | TP53INP2 |
| 4.054E-90 | 0.27398731 | 0.173 | 0.084 | 9.5767E-86 | TEC_prg | CTGF |
| 1.0487E-89 | 0.49958135 | 0.331 | 0.231 | 2.4774E-85 | TEC_prg | IFRD1 |
| 1.6082E-89 | -2.1782234 | 0.355 | 0.414 | 3.799E-85 | TEC_prg | S100P |
| 2.6931E-89 | 0.46666834 | 0.339 | 0.237 | 6.362E-85 | TEC_prg | PKN2 |
| 4.7316E-89 | -0.4411167 | 0.01 | 0.102 | 1.1177E-84 | TEC_prg | RNASE7 |
| 9.3888E-89 | 0.63036178 | 0.971 | 0.901 | 2.2179E-84 | TEC_prg | MT-ND2 |
| 3.5511E-88 | 0.39213915 | 0.171 | 0.084 | 8.3888E-84 | TEC_prg | ADAMTS9 |

| | | | | | | |
|---|---|---|---|---|---|---|
| 7.0629E-88 | -0.3277192 | 0.022 | 0.12 | 1.6685E-83 | TEC_prg | SMIM5 |
| 2.2697E-87 | 0.64047581 | 0.355 | 0.27 | 5.3617E-83 | TEC_prg | YME1L1 |
| 6.2875E-87 | 0.31173308 | 0.142 | 0.065 | 1.4853E-82 | TEC_prg | FSCN1 |
| 1.1733E-86 | 0.35953389 | 0.17 | 0.086 | 2.7718E-82 | TEC_prg | AKT1S1 |
| 1.2576E-86 | 0.39629785 | 0.298 | 0.199 | 2.9709E-82 | TEC_prg | ACIN1 |
| 6.6852E-86 | -0.3957636 | 0.065 | 0.183 | 1.5792E-81 | TEC_prg | DNAJA4 |
| 6.9909E-86 | -0.6516834 | 0.321 | 0.468 | 1.6515E-81 | TEC_prg | HSPH1 |
| 9.2937E-86 | 0.55281905 | 0.391 | 0.31 | 2.1954E-81 | TEC_prg | C6orf62 |
| 9.6552E-86 | 0.54277898 | 0.397 | 0.323 | 2.2809E-81 | TEC_prg | TCF25 |
| 3.6246E-85 | 0.47576283 | 0.328 | 0.235 | 8.5624E-81 | TEC_prg | SPTAN1 |
| 1.3988E-84 | -0.3182892 | 0.047 | 0.156 | 3.3043E-80 | TEC_prg | SDC2 |
| 2.7798E-84 | 0.26809261 | 0.14 | 0.064 | 6.5667E-80 | TEC_prg | ARL4D |
| 3.074E-84 | 0.26753623 | 0.187 | 0.099 | 7.2616E-80 | TEC_prg | EPM2AIP1 |
| 3.7595E-84 | 0.39604335 | 0.266 | 0.168 | 8.8812E-80 | TEC_prg | ERICH1 |
| 4.8141E-84 | -0.3009525 | 0.749 | 0.791 | 1.1372E-79 | TEC_prg | RPL31 |
| 6.0358E-84 | -0.6825213 | 0.076 | 0.192 | 1.4258E-79 | TEC_prg | F3 |
| 1.0071E-83 | 0.61404163 | 0.465 | 0.41 | 2.379E-79 | TEC_prg | FUS |
| 1.1142E-83 | 0.33848709 | 0.166 | 0.084 | 2.6321E-79 | TEC_prg | KPNA2 |
| 9.905E-83 | -0.2910141 | 0.249 | 0.421 | 2.3399E-78 | TEC_prg | FLNA |
| 1.4458E-82 | 0.31771295 | 0.184 | 0.099 | 3.4155E-78 | TEC_prg | KIAA1191 |
| 1.3155E-81 | 0.3358266 | 0.319 | 0.206 | 3.1076E-77 | TEC_prg | ZBTB20 |
| 5.0909E-81 | -0.2800665 | 0.099 | 0.229 | 1.2026E-76 | TEC_prg | ARL4C |
| 6.6051E-81 | 0.2600624 | 0.181 | 0.095 | 1.5603E-76 | TEC_prg | XPC |
| 9.1582E-81 | -0.2502535 | 0.204 | 0.366 | 2.1634E-76 | TEC_prg | UBE2H |
| 1.4566E-80 | 0.28600768 | 0.172 | 0.089 | 3.441E-76 | TEC_prg | SLC8B1 |
| 1.0815E-79 | 0.40235396 | 0.303 | 0.209 | 2.5548E-75 | TEC_prg | ZFC3H1 |
| 1.095E-79 | -0.28154 | 0.03 | 0.128 | 2.5867E-75 | TEC_prg | DUSP2 |
| 1.4438E-79 | 0.31772088 | 0.246 | 0.152 | 3.4107E-75 | TEC_prg | RBM6 |
| 3.9654E-79 | 0.25178911 | 0.139 | 0.066 | 9.3675E-75 | TEC_prg | CYB561A3 |
| 4.419E-78 | -0.2671039 | 0.343 | 0.51 | 1.0439E-73 | TEC_prg | H2AFJ |
| 2.912E-77 | 0.43753283 | 0.691 | 0.656 | 6.8789E-73 | TEC_prg | RBM39 |
| 4.3893E-77 | 0.55425615 | 0.968 | 0.867 | 1.0369E-72 | TEC_prg | MT-ND3 |
| 1.8979E-76 | 0.27479897 | 0.196 | 0.111 | 4.4834E-72 | TEC_prg | PHF14 |
| 2.0212E-76 | -0.3324993 | 0.025 | 0.116 | 4.7747E-72 | TEC_prg | TMC5 |
| 3.1411E-76 | -0.454454 | 0.016 | 0.101 | 7.4201E-72 | TEC_prg | SAA1 |
| 2.4446E-75 | 0.5144718 | 0.566 | 0.524 | 5.775E-71 | TEC_prg | EIF4A2 |
| 4.4957E-75 | -0.6853939 | 0.165 | 0.292 | 1.062E-70 | TEC_prg | DUSP5 |
| 9.2602E-75 | -0.7529652 | 0.097 | 0.206 | 2.1875E-70 | TEC_prg | C12orf75 |
| 1.795E-74 | 0.25945643 | 0.197 | 0.112 | 4.2403E-70 | TEC_prg | GON4L |
| 2.5977E-74 | -0.4641556 | 0.092 | 0.203 | 6.1365E-70 | TEC_prg | FBXO32 |
| 3.1637E-74 | -0.4061784 | 0.064 | 0.169 | 7.4736E-70 | TEC_prg | BPGM |
| 4.9848E-74 | 0.25335799 | 0.186 | 0.103 | 1.1776E-69 | TEC_prg | SUGP2 |
| 5.2684E-74 | 0.30910118 | 0.166 | 0.088 | 1.2446E-69 | TEC_prg | HIST2H2AC |
| 6.7826E-74 | -0.3257033 | 0.128 | 0.258 | 1.6023E-69 | TEC_prg | KIAA1551 |
| 9.6025E-73 | -0.2519469 | 0.599 | 0.7 | 2.2684E-68 | TEC_prg | EEF2 |
| 1.6563E-72 | -0.5944209 | 0.17 | 0.299 | 3.9126E-68 | TEC_prg | SLC25A37 |
| 6.0848E-72 | -0.5963127 | 0.653 | 0.697 | 1.4374E-67 | TEC_prg | HSP90B1 |
| 1.1634E-71 | 0.4148375 | 0.319 | 0.234 | 2.7483E-67 | TEC_prg | ZRANB2 |
| 2.056E-71 | 0.30469412 | 0.209 | 0.125 | 4.8568E-67 | TEC_prg | NXF1 |
| 2.4E-71 | -0.4224563 | 0.079 | 0.185 | 5.6695E-67 | TEC_prg | SPNS2 |

| | | | | | | |
|---|---|---|---|---|---|---|
| 4.1064E-71 | 0.25929808 | 0.157 | 0.082 | 9.7005E-67 | TEC_prg | FRAT2 |
| 5.022E-71 | 0.49754534 | 0.437 | 0.374 | 1.1863E-66 | TEC_prg | N4BP2L2 |
| 2.8176E-70 | -0.3392104 | 0.885 | 0.842 | 6.656E-66 | TEC_prg | RPL35 |
| 3.0395E-70 | -0.3313695 | 0.119 | 0.236 | 7.1802E-66 | TEC_prg | OCIAD2 |
| 3.1918E-70 | 0.39276692 | 0.803 | 0.723 | 7.54E-66 | TEC_prg | HSPA8 |
| 3.4264E-70 | -0.3230437 | 0.07 | 0.176 | 8.0942E-66 | TEC_prg | HIST3H2A |
| 1.0705E-69 | -0.3327285 | 0.125 | 0.251 | 2.5288E-65 | TEC_prg | TNFAIP3 |
| 1.2433E-69 | -0.2874391 | 0.975 | 0.964 | 2.937E-65 | TEC_prg | UBC |
| 1.2699E-69 | 0.4891598 | 0.415 | 0.338 | 2.9998E-65 | TEC_prg | GOLGB1 |
| 2.9031E-68 | -0.3377335 | 0.028 | 0.113 | 6.858E-64 | TEC_prg | LTB |
| 2.9729E-68 | 0.28031562 | 0.172 | 0.095 | 7.0228E-64 | TEC_prg | C3orf58 |
| 1.8806E-67 | 0.43836644 | 0.24 | 0.153 | 4.4424E-63 | TEC_prg | SYTL2 |
| 5.5786E-67 | -0.3037347 | 0.918 | 0.871 | 1.3178E-62 | TEC_prg | RPL36 |
| 1.3916E-66 | 0.2872174 | 0.415 | 0.312 | 3.2873E-62 | TEC_prg | VEGFA |
| 5.3423E-66 | 0.34376405 | 0.231 | 0.149 | 1.262E-61 | TEC_prg | CHD3 |
| 1.411E-65 | -0.8532595 | 0.123 | 0.234 | 3.3332E-61 | TEC_prg | G0S2 |
| 9.1394E-65 | 0.29712932 | 0.22 | 0.136 | 2.159E-60 | TEC_prg | EPS8 |
| 1.8461E-64 | 0.49842597 | 0.97 | 0.909 | 4.3609E-60 | TEC_prg | MT-ND1 |
| 2.5983E-64 | 0.3135351 | 0.226 | 0.142 | 6.1379E-60 | TEC_prg | ZNF83 |
| 2.607E-64 | 0.44369711 | 0.285 | 0.2 | 6.1586E-60 | TEC_prg | MDM2 |
| 2.4548E-63 | 0.49762378 | 0.439 | 0.389 | 5.7991E-59 | TEC_prg | PNISR |
| 3.2057E-63 | -0.2856703 | 0.057 | 0.15 | 7.5729E-59 | TEC_prg | GPNMB |
| 3.7525E-63 | 0.48687466 | 0.399 | 0.331 | 8.8646E-59 | TEC_prg | NDRG1 |
| 6.802E-63 | 0.31294804 | 0.3 | 0.198 | 1.6068E-58 | TEC_prg | CDKN2B |
| 1.36E-62 | -0.3476742 | 0.137 | 0.255 | 3.2127E-58 | TEC_prg | ITGB8 |
| 1.8268E-62 | 0.45494029 | 0.176 | 0.102 | 4.3155E-58 | TEC_prg | TNF |
| 5.0361E-62 | -0.5340183 | 0.09 | 0.192 | 1.1897E-57 | TEC_prg | LAMC2 |
| 5.7131E-62 | -0.2524546 | 0.044 | 0.131 | 1.3496E-57 | TEC_prg | KRT80 |
| 8.1466E-62 | -0.4152459 | 0.028 | 0.106 | 1.9245E-57 | TEC_prg | DHRS2 |
| 4.0578E-61 | -0.2734109 | 0.079 | 0.179 | 9.5857E-57 | TEC_prg | AKR1B1 |
| 6.5763E-61 | -0.7061179 | 0.174 | 0.273 | 1.5535E-56 | TEC_prg | LMO7 |
| 1.2702E-60 | -0.4343842 | 0.365 | 0.462 | 3.0006E-56 | TEC_prg | CLDN7 |
| 1.6786E-60 | 0.39976475 | 0.342 | 0.267 | 3.9653E-56 | TEC_prg | BPTF |
| 2.8312E-60 | 0.42293819 | 0.54 | 0.504 | 6.6883E-56 | TEC_prg | HNRNPDL |
| 1.4218E-59 | -0.3617212 | 0.097 | 0.199 | 3.3587E-55 | TEC_prg | HBEGF |
| 2.6944E-59 | 0.51095222 | 0.807 | 0.772 | 6.365E-55 | TEC_prg | TPI1 |
| 2.8343E-59 | 0.41174281 | 0.303 | 0.228 | 6.6954E-55 | TEC_prg | PNPLA2 |
| 5.1699E-59 | 0.30952114 | 0.207 | 0.131 | 1.2213E-54 | TEC_prg | NARF |
| 1.4988E-57 | 0.27353588 | 0.153 | 0.085 | 3.5407E-53 | TEC_prg | SLC6A8 |
| 2.5065E-57 | 0.2805233 | 0.204 | 0.128 | 5.9212E-53 | TEC_prg | SCNN1A |
| 4.6289E-56 | 0.43691999 | 0.298 | 0.228 | 1.0935E-51 | TEC_prg | PLXNB2 |
| 2.6689E-55 | 0.25137283 | 0.206 | 0.13 | 6.3047E-51 | TEC_prg | KIAA1109 |
| 8.449E-55 | -2.9799158 | 0.552 | 0.52 | 1.9959E-50 | TEC_prg | LCN2 |
| 1.956E-54 | -2.5972682 | 0.348 | 0.382 | 4.6206E-50 | TEC_prg | KRT13 |
| 4.5979E-54 | -0.7450144 | 0.609 | 0.649 | 1.0862E-49 | TEC_prg | HSPA5 |
| 9.4785E-54 | 0.35434053 | 0.269 | 0.193 | 2.2391E-49 | TEC_prg | ZNF91 |
| 2.0239E-53 | 0.44463353 | 0.362 | 0.3 | 4.781E-49 | TEC_prg | NUFIP2 |
| 3.2154E-53 | 0.26129162 | 0.447 | 0.333 | 7.5958E-49 | TEC_prg | GDF15 |
| 4.8945E-53 | -0.5181685 | 0.138 | 0.238 | 1.1562E-48 | TEC_prg | IL32 |
| 9.5498E-53 | -0.3746286 | 0.135 | 0.24 | 2.2559E-48 | TEC_prg | ITGA2 |

| | | | | | | |
|---|---|---|---|---|---|---|
| 1.4765E-52 | -1.2419798 | 0.653 | 0.57 | 3.4878E-48 | TEC_prg | KRT19 |
| 4.2849E-52 | -0.4382326 | 0.827 | 0.811 | 1.0122E-47 | TEC_prg | GSTP1 |
| 2.0992E-51 | -1.0284243 | 0.191 | 0.291 | 4.9589E-47 | TEC_prg | MT1E |
| 1.133E-50 | -0.4655009 | 0.153 | 0.248 | 2.6765E-46 | TEC_prg | ADGRF1 |
| 2.2326E-50 | 0.40905805 | 0.405 | 0.337 | 5.274E-46 | TEC_prg | DDIT4 |
| 2.7717E-50 | 0.41427158 | 0.401 | 0.346 | 6.5475E-46 | TEC_prg | TAF1D |
| 1.4219E-49 | 0.33606281 | 0.303 | 0.234 | 3.3591E-45 | TEC_prg | KANSL1 |
| 2.014E-48 | -0.336357 | 0.053 | 0.128 | 4.7578E-44 | TEC_prg | HILPDA |
| 5.4296E-48 | 0.32575824 | 0.282 | 0.213 | 1.2826E-43 | TEC_prg | PCM1 |
| 1.661E-47 | -0.2555525 | 0.585 | 0.666 | 3.9239E-43 | TEC_prg | CALM2 |
| 2.4519E-47 | 0.34861508 | 0.309 | 0.246 | 5.7921E-43 | TEC_prg | NSD3 |
| 3.6274E-47 | 0.40174715 | 0.292 | 0.23 | 8.5689E-43 | TEC_prg | PABPC4 |
| 4.0412E-47 | -0.3527178 | 0.097 | 0.181 | 9.5464E-43 | TEC_prg | PPL |
| 8.4161E-47 | -0.2890119 | 0.168 | 0.271 | 1.9881E-42 | TEC_prg | MUC1 |
| 1.4509E-46 | -1.0715598 | 0.066 | 0.138 | 3.4275E-42 | TEC_prg | APOE |
| 5.7454E-46 | 0.34862757 | 0.321 | 0.26 | 1.3572E-41 | TEC_prg | SMG1 |
| 8.6046E-46 | 0.35672637 | 0.367 | 0.302 | 2.0327E-41 | TEC_prg | IRF1 |
| 1.347E-45 | 0.44293039 | 0.982 | 0.927 | 3.1821E-41 | TEC_prg | MT-ND4 |
| 1.8707E-45 | -0.4637156 | 0.161 | 0.244 | 4.4192E-41 | TEC_prg | EPCAM |
| 1.9781E-45 | -1.1329443 | 0.073 | 0.151 | 4.6728E-41 | TEC_prg | CCL2 |
| 2.2315E-45 | 0.33816677 | 0.244 | 0.174 | 5.2715E-41 | TEC_prg | PFKFB3 |
| 1.3642E-44 | -0.2776386 | 0.221 | 0.342 | 3.2226E-40 | TEC_prg | HERPUD1 |
| 3.3606E-42 | -0.296948 | 0.36 | 0.467 | 7.9386E-38 | TEC_prg | RAB11FIP1 |
| 1.1409E-41 | 0.25312875 | 0.204 | 0.141 | 2.695E-37 | TEC_prg | NUMA1 |
| 2.0989E-41 | 0.3634135 | 0.313 | 0.252 | 4.9582E-37 | TEC_prg | IDI1 |
| 4.4261E-41 | -2.2344884 | 0.731 | 0.651 | 1.0456E-36 | TEC_prg | MT2A |
| 4.6641E-41 | 0.37708517 | 0.337 | 0.277 | 1.1018E-36 | TEC_prg | MARCKSL1 |
| 5.628E-41 | 0.55512965 | 0.414 | 0.385 | 1.3295E-36 | TEC_prg | TUBB4B |
| 1.4858E-40 | 0.39051967 | 0.358 | 0.311 | 3.51E-36 | TEC_prg | BOD1L1 |
| 3.6029E-40 | 0.2512583 | 0.212 | 0.149 | 8.5112E-36 | TEC_prg | RNF169 |
| 5.283E-40 | 0.26403848 | 0.222 | 0.16 | 1.248E-35 | TEC_prg | NSD1 |
| 4.4597E-39 | 0.26560737 | 0.211 | 0.149 | 1.0535E-34 | TEC_prg | OFD1 |
| 5.863E-39 | 0.2542173 | 0.246 | 0.18 | 1.385E-34 | TEC_prg | PAWR |
| 8.9338E-39 | -0.255043 | 0.05 | 0.114 | 2.1104E-34 | TEC_prg | TCHP |
| 1.4603E-38 | -1.450317 | 0.206 | 0.288 | 3.4497E-34 | TEC_prg | MT1X |
| 1.908E-38 | -0.4237362 | 0.054 | 0.119 | 4.5073E-34 | TEC_prg | CYP1B1 |
| 2.9538E-38 | 0.38301914 | 0.287 | 0.229 | 6.9777E-34 | TEC_prg | MT-ATP8 |
| 3.6926E-38 | 0.30289684 | 0.265 | 0.206 | 8.723E-34 | TEC_prg | USP34 |
| 8.7255E-38 | 0.36245657 | 0.287 | 0.229 | 2.0612E-33 | TEC_prg | SNHG7 |
| 4.0055E-37 | -0.3074765 | 0.103 | 0.179 | 9.4622E-33 | TEC_prg | HSPB8 |
| 5.553E-37 | 0.2542914 | 0.202 | 0.143 | 1.3118E-32 | TEC_prg | ATXN7L3B |
| 1.5612E-36 | 0.43376802 | 0.405 | 0.371 | 3.688E-32 | TEC_prg | CCNL1 |
| 1.9074E-36 | -0.3063804 | 0.192 | 0.285 | 4.5058E-32 | TEC_prg | CAMK2N1 |
| 5.3073E-36 | 0.26583675 | 0.23 | 0.172 | 1.2538E-31 | TEC_prg | SRSF1 |
| 2.5658E-35 | 0.28900905 | 0.638 | 0.598 | 6.0612E-31 | TEC_prg | MCL1 |
| 1.7278E-34 | 0.33896174 | 0.304 | 0.257 | 4.0816E-30 | TEC_prg | MPHOSPH8 |
| 5.1386E-33 | 0.34906638 | 0.78 | 0.771 | 1.2139E-28 | TEC_prg | EIF4G2 |
| 6.0629E-33 | 0.38451002 | 0.979 | 0.924 | 1.4322E-28 | TEC_prg | MT-CYB |
| 1.8325E-32 | 0.34699154 | 0.375 | 0.339 | 4.3289E-28 | TEC_prg | WSB1 |
| 3.5819E-32 | 0.27172239 | 0.378 | 0.312 | 8.4616E-28 | TEC_prg | GADD45A |

| | | | | | | |
|---|---|---|---|---|---|---|
| 9.0497E-32 | 0.26081831 | 0.264 | 0.208 | 2.1378E-27 | TEC_prg | NKTR |
| 1.2135E-31 | 0.34691639 | 0.337 | 0.301 | 2.8667E-27 | TEC_prg | HP1BP3 |
| 1.0884E-30 | 0.31822832 | 0.185 | 0.135 | 2.5712E-26 | TEC_prg | IRF7 |
| 1.8285E-30 | -1.9828575 | 0.436 | 0.455 | 4.3194E-26 | TEC_prg | SPP1 |
| 2.8151E-30 | 0.448337 | 0.316 | 0.268 | 6.6501E-26 | TEC_prg | TNFAIP2 |
| 2.934E-28 | 0.35477674 | 0.372 | 0.34 | 6.9309E-24 | TEC_prg | CPEB4 |
| 3.885E-28 | 0.30589887 | 0.155 | 0.107 | 9.1775E-24 | TEC_prg | HIST1H2BG |
| 5.3486E-28 | 0.34109303 | 0.315 | 0.279 | 1.2635E-23 | TEC_prg | DYNC1H1 |
| 6.1248E-28 | -0.3937498 | 0.629 | 0.643 | 1.4469E-23 | TEC_prg | TXNIP |
| 3.1873E-26 | -0.2888863 | 0.118 | 0.182 | 7.5293E-22 | TEC_prg | IGFBP3 |
| 3.8803E-26 | 0.27155591 | 0.257 | 0.212 | 9.1664E-22 | TEC_prg | U2SURP |
| 8.066E-26 | 0.3045059 | 0.277 | 0.237 | 1.9054E-21 | TEC_prg | KMT2C |
| 2.2141E-25 | 0.30853879 | 0.614 | 0.623 | 5.2304E-21 | TEC_prg | PRRC2C |
| 6.4686E-25 | 0.32588517 | 0.298 | 0.263 | 1.5281E-20 | TEC_prg | DDX46 |
| 1.4249E-24 | 0.34526374 | 0.99 | 0.939 | 3.3659E-20 | TEC_prg | MT-CO3 |
| 5.8959E-24 | 0.29946289 | 0.64 | 0.628 | 1.3928E-19 | TEC_prg | YBX3 |
| 3.4681E-23 | 0.2735184 | 0.243 | 0.204 | 8.1928E-19 | TEC_prg | RBMX |
| 4.1497E-22 | 0.45751382 | 0.375 | 0.368 | 9.8029E-18 | TEC_prg | ANKRD11 |
| 5.1219E-22 | 0.44989874 | 0.483 | 0.497 | 1.21E-17 | TEC_prg | DDX3X |
| 7.1118E-22 | -1.7478636 | 0.309 | 0.343 | 1.68E-17 | TEC_prg | MMP7 |
| 2.5329E-21 | 0.37267644 | 0.508 | 0.514 | 5.9834E-17 | TEC_prg | EIF5 |
| 6.6818E-21 | -0.3055406 | 0.383 | 0.443 | 1.5785E-16 | TEC_prg | ATP1B1 |
| 2.3739E-20 | 0.29521743 | 0.263 | 0.227 | 5.6078E-16 | TEC_prg | THUMPD3-AS1 |
| 5.447E-20 | 0.35037736 | 0.409 | 0.407 | 1.2867E-15 | TEC_prg | ATF4 |
| 8.0094E-20 | 0.39016858 | 0.991 | 0.946 | 1.8921E-15 | TEC_prg | MT-CO1 |
| 4.4053E-19 | 0.38853732 | 0.312 | 0.299 | 1.0407E-14 | TEC_prg | EIF4A1 |
| 1.4563E-18 | 0.32204683 | 0.303 | 0.279 | 3.4402E-14 | TEC_prg | ATRX |
| 2.5015E-18 | -0.7259854 | 0.15 | 0.205 | 5.9094E-14 | TEC_prg | THBS1 |
| 3.5469E-18 | 0.37295134 | 0.361 | 0.358 | 8.3789E-14 | TEC_prg | ATP1A1 |
| 5.7682E-18 | -0.9371124 | 0.711 | 0.651 | 1.3626E-13 | TEC_prg | HSPA1A |
| 6.7017E-18 | 0.26475043 | 0.188 | 0.148 | 1.5831E-13 | TEC_prg | MT-ND6 |
| 1.5524E-17 | 0.40034713 | 0.502 | 0.521 | 3.6672E-13 | TEC_prg | H2AFZ |
| 2.5522E-16 | 0.31649997 | 0.316 | 0.3 | 6.0291E-12 | TEC_prg | DDX24 |
| 4.6692E-16 | 0.31005688 | 0.987 | 0.931 | 1.103E-11 | TEC_prg | MT-ATP6 |
| 7.1537E-16 | -0.4302098 | 0.091 | 0.133 | 1.6899E-11 | TEC_prg | TFPI2 |
| 9.199E-16 | 0.34079326 | 0.369 | 0.368 | 2.1731E-11 | TEC_prg | SPAG9 |
| 1.1235E-15 | 0.31255288 | 0.3 | 0.285 | 2.654E-11 | TEC_prg | ACADVL |
| 2.2142E-15 | 0.42568545 | 0.41 | 0.418 | 5.2306E-11 | TEC_prg | JPT1 |
| 2.4342E-15 | 0.27281034 | 0.162 | 0.129 | 5.7502E-11 | TEC_prg | SQLE |
| 4.7935E-15 | -0.870482 | 0.224 | 0.271 | 1.1324E-10 | TEC_prg | S100A2 |
| 1.6142E-14 | -0.7157601 | 0.103 | 0.141 | 3.8131E-10 | TEC_prg | DEFB1 |
| 2.8495E-14 | 0.38003146 | 0.302 | 0.278 | 6.7313E-10 | TEC_prg | H1F0 |
| 3.8542E-14 | -0.7076312 | 0.607 | 0.537 | 9.1047E-10 | TEC_prg | KRT7 |
| 4.89E-14 | 0.39252548 | 0.558 | 0.57 | 1.1552E-09 | TEC_prg | RPS27L |
| 1.012E-13 | 0.33953633 | 0.414 | 0.429 | 2.3907E-09 | TEC_prg | SRSF5 |
| 1.6044E-13 | -0.4337901 | 0.103 | 0.143 | 3.7901E-09 | TEC_prg | CXCL1 |
| 1.7327E-13 | 0.31985423 | 0.347 | 0.344 | 4.0932E-09 | TEC_prg | NORAD |
| 3.546E-13 | -0.4898252 | 0.714 | 0.54 | 8.3767E-09 | TEC_prg | KRT18 |
| 2.2164E-12 | 0.27721093 | 0.221 | 0.196 | 5.2359E-08 | TEC_prg | MDM4 |
| 2.3043E-11 | -3.9452885 | 0.46 | 0.394 | 5.4435E-07 | TEC_prg | SPRR3 |

| | | | | | | |
|---|---|---|---|---|---|---|
| 7.216E-11 | 0.27539565 | 0.316 | 0.315 | 1.7046E-06 | TEC_prg | AFF4 |
| 2.4085E-10 | -0.3293439 | 0.5 | 0.53 | 5.6895E-06 | TEC_prg | ZFP36 |
| 2.6736E-10 | 0.27141618 | 0.29 | 0.283 | 6.3158E-06 | TEC_prg | TPR |
| 3.7362E-10 | -0.5714968 | 0.724 | 0.702 | 8.826E-06 | TEC_prg | SOD2 |
| 5.4765E-09 | -0.6858705 | 0.526 | 0.49 | 0.00012937 | TEC_prg | KRT8 |
| 4.7132E-06 | 0.25426971 | 0.504 | 0.52 | 0.11134072 | TEC_prg | H1FX |
| 1.3512E-05 | 0.3568306 | 0.266 | 0.279 | 0.31920466 | TEC_prg | UGP2 |
| 2.8186E-05 | -1.421278 | 0.631 | 0.525 | 0.6658453 | TEC_prg | WFDC2 |
| 4.2863E-05 | 0.28796234 | 0.358 | 0.394 | 1 | TEC_prg | STAT3 |
| 0.00011022 | -0.7362979 | 0.239 | 0.249 | 1 | TEC_prg | CRYAB |
| 0.0002041 | -0.5736506 | 0.159 | 0.175 | 1 | TEC_prg | RDH10 |
| 0.00071541 | 0.28448711 | 0.298 | 0.308 | 1 | TEC_prg | ZNF292 |
| 0.00091124 | 0.28380316 | 0.449 | 0.466 | 1 | TEC_prg | CDKN1A |
| 0.00119279 | 0.26184514 | 0.35 | 0.38 | 1 | TEC_prg | GCC2 |
| 0.00402317 | 0.26395396 | 0.371 | 0.407 | 1 | TEC_prg | SOD1 |
| 0 | 3.77290562 | 0.985 | 0.628 | 0 | MO_infl | TIMP1 |
| 0 | 3.59978433 | 0.64 | 0.103 | 0 | MO_infl | IL1B |
| 0 | 3.13137558 | 0.412 | 0.102 | 0 | MO_infl | CXCL3 |
| 0 | 3.12602088 | 0.712 | 0.453 | 0 | MO_infl | CXCL8 |
| 0 | 3.12080822 | 0.877 | 0.12 | 0 | MO_infl | VCAN |
| 0 | 2.86003004 | 0.998 | 0.503 | 0 | MO_infl | SRGN |
| 0 | 2.53784703 | 0.9 | 0.396 | 0 | MO_infl | LYZ |
| 0 | 2.44645767 | 0.247 | 0.045 | 0 | MO_infl | CXCL5 |
| 0 | 2.42629788 | 0.309 | 0.117 | 0 | MO_infl | CCL2 |
| 0 | 2.41922516 | 0.399 | 0.113 | 0 | MO_infl | CCL20 |
| 0 | 2.33738863 | 0.417 | 0.076 | 0 | MO_infl | CCL3 |
| 0 | 2.33505414 | 0.546 | 0.069 | 0 | MO_infl | EREG |
| 0 | 2.31705147 | 0.351 | 0.081 | 0 | MO_infl | CXCL2 |
| 0 | 2.28679794 | 0.287 | 0.109 | 0 | MO_infl | SERPINB2 |
| 0 | 2.25614923 | 0.413 | 0.166 | 0 | MO_infl | THBS1 |
| 0 | 2.24724179 | 0.97 | 0.294 | 0 | MO_infl | FCER1G |
| 0 | 2.06560097 | 0.814 | 0.145 | 0 | MO_infl | BCL2A1 |
| 0 | 2.05989002 | 0.714 | 0.287 | 0 | MO_infl | CTSL |
| 0 | 1.96307434 | 0.975 | 0.356 | 0 | MO_infl | TYROBP |
| 0 | 1.95757032 | 0.786 | 0.146 | 0 | MO_infl | SLC2A3 |
| 0 | 1.95637587 | 0.993 | 0.7 | 0 | MO_infl | VIM |
| 0 | 1.94089472 | 0.652 | 0.094 | 0 | MO_infl | FCN1 |
| 0 | 1.92942952 | 0.887 | 0.484 | 0 | MO_infl | PLAUR |
| 0 | 1.87648832 | 0.741 | 0.138 | 0 | MO_infl | CD14 |
| 0 | 1.86971698 | 1 | 0.993 | 0 | MO_infl | FTH1 |
| 0 | 1.80618264 | 0.582 | 0.158 | 0 | MO_infl | S100A12 |
| 0 | 1.77028873 | 0.243 | 0.028 | 0 | MO_infl | CCL3L1 |
| 0 | 1.74989372 | 0.896 | 0.582 | 0 | MO_infl | S100A4 |
| 0 | 1.74722826 | 0.374 | 0.051 | 0 | MO_infl | PTGS2 |
| 0 | 1.73308688 | 0.964 | 0.557 | 0 | MO_infl | CD44 |
| 0 | 1.69682624 | 0.838 | 0.151 | 0 | MO_infl | SLC11A1 |
| 0 | 1.67012027 | 0.926 | 0.243 | 0 | MO_infl | LAPTM5 |
| 0 | 1.65851425 | 0.594 | 0.124 | 0 | MO_infl | AC020656.1 |
| 0 | 1.64146914 | 0.786 | 0.207 | 0 | MO_infl | SLC25A37 |
| 0 | 1.61491013 | 0.68 | 0.076 | 0 | MO_infl | CD300E |

| | | | | | | |
|---|---|---|---|---|---|---|
| 0 | 1.61368423 | 0.831 | 0.188 | 0 | MO_infl | AIF1 |
| 0 | 1.60372898 | 0.849 | 0.481 | 0 | MO_infl | CTSS |
| 0 | 1.58649005 | 0.727 | 0.129 | 0 | MO_infl | TNFRSF1B |
| 0 | 1.57531312 | 0.697 | 0.058 | 0 | MO_infl | CD93 |
| 0 | 1.56886981 | 0.951 | 0.565 | 0 | MO_infl | LGALS1 |
| 0 | 1.56760651 | 0.879 | 0.65 | 0 | MO_infl | NAMPT |
| 0 | 1.55876957 | 0.246 | 0.086 | 0 | MO_infl | CCL4 |
| 0 | 1.55187861 | 0.828 | 0.684 | 0 | MO_infl | SOD2 |
| 0 | 1.54148204 | 0.894 | 0.351 | 0 | MO_infl | EMP3 |
| 0 | 1.53359746 | 0.994 | 0.838 | 0 | MO_infl | SH3BGRL3 |
| 0 | 1.50614671 | 0.595 | 0.16 | 0 | MO_infl | ALOX5AP |
| 0 | 1.50553865 | 0.771 | 0.142 | 0 | MO_infl | ZEB2 |
| 0 | 1.49054884 | 0.573 | 0.18 | 0 | MO_infl | PLIN2 |
| 0 | 1.48756827 | 0.842 | 0.205 | 0 | MO_infl | LCP1 |
| 0 | 1.47991283 | 0.679 | 0.098 | 0 | MO_infl | ANPEP |
| 0 | 1.47816717 | 0.408 | 0.193 | 0 | MO_infl | G0S2 |
| 0 | 1.46680048 | 0.672 | 0.108 | 0 | MO_infl | C5AR1 |
| 0 | 1.43621511 | 0.95 | 0.623 | 0 | MO_infl | ANXA5 |
| 0 | 1.43472928 | 0.566 | 0.113 | 0 | MO_infl | PLEK |
| 0 | 1.42278688 | 0.589 | 0.116 | 0 | MO_infl | OLR1 |
| 0 | 1.39460764 | 0.733 | 0.224 | 0 | MO_infl | ATP13A3 |
| 0 | 1.34611776 | 0.46 | 0.051 | 0 | MO_infl | THBD |
| 0 | 1.33976668 | 0.781 | 0.334 | 0 | MO_infl | HIF1A |
| 0 | 1.32707707 | 0.346 | 0.076 | 0 | MO_infl | CYP1B1 |
| 0 | 1.30949358 | 0.856 | 0.473 | 0 | MO_infl | CEBPB |
| 0 | 1.29101411 | 0.519 | 0.137 | 0 | MO_infl | HBEGF |
| 0 | 1.29002153 | 0.638 | 0.132 | 0 | MO_infl | CYBB |
| 0 | 1.27740502 | 0.819 | 0.181 | 0 | MO_infl | CD53 |
| 0 | 1.24013096 | 0.678 | 0.08 | 0 | MO_infl | AQP9 |
| 0 | 1.21023764 | 0.61 | 0.114 | 0 | MO_infl | SAMSN1 |
| 0 | 1.20863821 | 0.631 | 0.182 | 0 | MO_infl | ATP2B1 |
| 0 | 1.20157862 | 1 | 0.985 | 0 | MO_infl | FTL |
| 0 | 1.1980605 | 0.676 | 0.133 | 0 | MO_infl | SLC43A2 |
| 0 | 1.18340274 | 0.156 | 0.016 | 0 | MO_infl | CCL7 |
| 0 | 1.17816846 | 0.986 | 0.909 | 0 | MO_infl | S100A10 |
| 0 | 1.17197427 | 0.933 | 0.707 | 0 | MO_infl | CTSB |
| 0 | 1.17170476 | 0.994 | 0.97 | 0 | MO_infl | ACTB |
| 0 | 1.17115557 | 0.35 | 0.08 | 0 | MO_infl | MNDA |
| 0 | 1.16581668 | 0.568 | 0.069 | 0 | MO_infl | MCEMP1 |
| 0 | 1.14866476 | 0.611 | 0.143 | 0 | MO_infl | GK |
| 0 | 1.14274499 | 0.958 | 0.691 | 0 | MO_infl | CYBA |
| 0 | 1.1355373 | 0.818 | 0.338 | 0 | MO_infl | FLNA |
| 0 | 1.13149868 | 0.561 | 0.098 | 0 | MO_infl | SERPINB9 |
| 0 | 1.13054623 | 0.777 | 0.304 | 0 | MO_infl | COTL1 |
| 0 | 1.12847247 | 0.527 | 0.132 | 0 | MO_infl | ACSL1 |
| 0 | 1.12810841 | 0.66 | 0.27 | 0 | MO_infl | VEGFA |
| 0 | 1.12741532 | 0.994 | 0.923 | 0 | MO_infl | GAPDH |
| 0 | 1.12353947 | 0.845 | 0.369 | 0 | MO_infl | GSTO1 |
| 0 | 1.12087216 | 0.351 | 0.06 | 0 | MO_infl | RETN |
| 0 | 1.11875303 | 0.745 | 0.257 | 0 | MO_infl | LIMS1 |

| | | | | | | |
|---|---|---|---|---|---|---|
| 0 | 1.11624307 | 0.542 | 0.181 | 0 | MO_infl | PPIF |
| 0 | 1.11188449 | 0.544 | 0.09 | 0 | MO_infl | CLEC5A |
| 0 | 1.09867088 | 0.554 | 0.131 | 0 | MO_infl | DUSP6 |
| 0 | 1.09252915 | 0.619 | 0.072 | 0 | MO_infl | TREM1 |
| 0 | 1.08577711 | 0.672 | 0.126 | 0 | MO_infl | ITGAX |
| 0 | 1.07594758 | 0.531 | 0.114 | 0 | MO_infl | TGFBI |
| 0 | 1.06480055 | 0.567 | 0.079 | 0 | MO_infl | FPR1 |
| 0 | 1.06367361 | 0.769 | 0.205 | 0 | MO_infl | PTPRC |
| 0 | 1.05690933 | 0.797 | 0.224 | 0 | MO_infl | SERPINA1 |
| 0 | 1.05461102 | 0.848 | 0.564 | 0 | MO_infl | MCL1 |
| 0 | 1.04570895 | 0.577 | 0.104 | 0 | MO_infl | SLA |
| 0 | 1.04453479 | 0.613 | 0.07 | 0 | MO_infl | TLR2 |
| 0 | 1.03827811 | 0.639 | 0.115 | 0 | MO_infl | PTPRE |
| 0 | 1.03711472 | 0.654 | 0.116 | 0 | MO_infl | FCGR2A |
| 0 | 1.03320627 | 0.561 | 0.102 | 0 | MO_infl | LCP2 |
| 0 | 1.02496159 | 0.578 | 0.174 | 0 | MO_infl | IVNS1ABP |
| 0 | 1.01894613 | 0.957 | 0.707 | 0 | MO_infl | PSAP |
| 0 | 1.01208608 | 0.935 | 0.712 | 0 | MO_infl | APLP2 |
| 0 | 1.00976021 | 0.799 | 0.319 | 0 | MO_infl | TYMP |
| 0 | 0.99072584 | 0.699 | 0.14 | 0 | MO_infl | SPI1 |
| 0 | 0.98802651 | 0.898 | 0.599 | 0 | MO_infl | ARPC5 |
| 0 | 0.98076864 | 0.219 | 0.017 | 0 | MO_infl | TNIP3 |
| 0 | 0.97872806 | 0.711 | 0.173 | 0 | MO_infl | GMFG |
| 0 | 0.97853496 | 0.301 | 0.029 | 0 | MO_infl | MMP19 |
| 0 | 0.97808275 | 0.642 | 0.218 | 0 | MO_infl | ETS2 |
| 0 | 0.97450251 | 0.633 | 0.152 | 0 | MO_infl | DSE |
| 0 | 0.97069634 | 0.42 | 0.068 | 0 | MO_infl | CD163 |
| 0 | 0.96942214 | 0.618 | 0.157 | 0 | MO_infl | ITGB2 |
| 0 | 0.96031623 | 0.584 | 0.109 | 0 | MO_infl | SMIM25 |
| 0 | 0.95417401 | 0.625 | 0.143 | 0 | MO_infl | LST1 |
| 0 | 0.94770024 | 0.633 | 0.133 | 0 | MO_infl | NCF2 |
| 0 | 0.93046349 | 0.351 | 0.068 | 0 | MO_infl | SLC16A10 |
| 0 | 0.92556266 | 0.668 | 0.256 | 0 | MO_infl | FNDC3B |
| 0 | 0.92259503 | 0.423 | 0.101 | 0 | MO_infl | SDC2 |
| 0 | 0.9037283 | 0.753 | 0.243 | 0 | MO_infl | CD68 |
| 0 | 0.90140585 | 0.42 | 0.053 | 0 | MO_infl | LILRB1 |
| 0 | 0.90139616 | 0.65 | 0.18 | 0 | MO_infl | RAB31 |
| 0 | 0.90133757 | 0.78 | 0.373 | 0 | MO_infl | EFHD2 |
| 0 | 0.90089533 | 0.984 | 0.921 | 0 | MO_infl | OAZ1 |
| 0 | 0.89685005 | 0.786 | 0.619 | 0 | MO_infl | IER3 |
| 0 | 0.89209403 | 0.401 | 0.148 | 0 | MO_infl | INSIG1 |
| 0 | 0.88959566 | 0.542 | 0.088 | 0 | MO_infl | DMXL2 |
| 0 | 0.88684966 | 0.488 | 0.063 | 0 | MO_infl | LILRB2 |
| 0 | 0.87845326 | 0.552 | 0.125 | 0 | MO_infl | IGSF6 |
| 0 | 0.875568 | 0.568 | 0.156 | 0 | MO_infl | NINJ1 |
| 0 | 0.87287045 | 0.309 | 0.044 | 0 | MO_infl | SLC39A8 |
| 0 | 0.86961382 | 0.734 | 0.581 | 0 | MO_infl | NFKBIA |
| 0 | 0.86452135 | 0.501 | 0.09 | 0 | MO_infl | MS4A7 |
| 0 | 0.86255618 | 0.579 | 0.096 | 0 | MO_infl | FERMT3 |
| 0 | 0.85974429 | 0.581 | 0.168 | 0 | MO_infl | STXBP2 |

| | | | | | | |
|---|---|---|---|---|---|---|
| 0 | 0.85119061 | 0.725 | 0.272 | 0 | MO_infl | TLN1 |
| 0 | 0.84963543 | 0.902 | 0.672 | 0 | MO_infl | TPM3 |
| 0 | 0.84941696 | 0.962 | 0.764 | 0 | MO_infl | PFN1 |
| 0 | 0.83785231 | 0.736 | 0.302 | 0 | MO_infl | GNAI2 |
| 0 | 0.82803149 | 0.513 | 0.149 | 0 | MO_infl | CD52 |
| 0 | 0.82501981 | 0.537 | 0.062 | 0 | MO_infl | IRAK3 |
| 0 | 0.80637328 | 0.656 | 0.216 | 0 | MO_infl | VASP |
| 0 | 0.80636557 | 0.864 | 0.587 | 0 | MO_infl | ACTR2 |
| 0 | 0.79787495 | 0.479 | 0.116 | 0 | MO_infl | JARID2 |
| 0 | 0.79755353 | 0.707 | 0.281 | 0 | MO_infl | GRB2 |
| 0 | 0.78927006 | 0.64 | 0.17 | 0 | MO_infl | HCST |
| 0 | 0.77861395 | 0.418 | 0.08 | 0 | MO_infl | MARCO |
| 0 | 0.77679809 | 0.49 | 0.169 | 0 | MO_infl | BASP1 |
| 0 | 0.77626819 | 0.626 | 0.144 | 0 | MO_infl | ARRB2 |
| 0 | 0.77549207 | 0.506 | 0.169 | 0 | MO_infl | CREM |
| 0 | 0.7687691 | 0.445 | 0.136 | 0 | MO_infl | LTA4H |
| 0 | 0.76438593 | 0.447 | 0.06 | 0 | MO_infl | PECAM1 |
| 0 | 0.75879982 | 0.633 | 0.243 | 0 | MO_infl | IFNGR2 |
| 0 | 0.75605525 | 0.47 | 0.144 | 0 | MO_infl | FYB1 |
| 0 | 0.75220898 | 0.886 | 0.744 | 0 | MO_infl | BTG1 |
| 0 | 0.74870061 | 0.514 | 0.075 | 0 | MO_infl | FGR |
| 0 | 0.73895526 | 0.514 | 0.08 | 0 | MO_infl | EMILIN2 |
| 0 | 0.73824689 | 0.523 | 0.109 | 0 | MO_infl | CEP170 |
| 0 | 0.73792193 | 0.59 | 0.213 | 0 | MO_infl | BACH1 |
| 0 | 0.7375367 | 0.615 | 0.237 | 0 | MO_infl | ATP1B3 |
| 0 | 0.73192404 | 0.373 | 0.106 | 0 | MO_infl | NCF1 |
| 0 | 0.72931103 | 0.533 | 0.121 | 0 | MO_infl | CCDC88A |
| 0 | 0.72878699 | 0.438 | 0.206 | 0 | MO_infl | TNFAIP3 |
| 0 | 0.72458541 | 0.495 | 0.079 | 0 | MO_infl | HCK |
| 0 | 0.71716896 | 0.469 | 0.081 | 0 | MO_infl | MAP3K20 |
| 0 | 0.71625163 | 0.493 | 0.08 | 0 | MO_infl | ITGA5 |
| 0 | 0.71601013 | 0.995 | 0.951 | 0 | MO_infl | H3F3A |
| 0 | 0.71562358 | 0.356 | 0.039 | 0 | MO_infl | NLRP3 |
| 0 | 0.71477459 | 0.579 | 0.172 | 0 | MO_infl | NUMB |
| 0 | 0.71353691 | 0.538 | 0.109 | 0 | MO_infl | GLIPR2 |
| 0 | 0.70957748 | 0.307 | 0.033 | 0 | MO_infl | FLT1 |
| 0 | 0.70869712 | 0.531 | 0.106 | 0 | MO_infl | MYO1F |
| 0 | 0.70712092 | 0.916 | 0.748 | 0 | MO_infl | RHOA |
| 0 | 0.70706144 | 0.482 | 0.075 | 0 | MO_infl | LILRB3 |
| 0 | 0.7068308 | 0.785 | 0.625 | 0 | MO_infl | LDHA |
| 0 | 0.7031219 | 0.563 | 0.128 | 0 | MO_infl | HCLS1 |
| 0 | 0.70305892 | 0.461 | 0.061 | 0 | MO_infl | ADGRE2 |
| 0 | 0.70193063 | 0.445 | 0.129 | 0 | MO_infl | SAMHD1 |
| 0 | 0.70144155 | 0.746 | 0.378 | 0 | MO_infl | IQGAP1 |
| 0 | 0.69883057 | 0.994 | 0.873 | 0 | MO_infl | HLA-B |
| 0 | 0.69714369 | 0.456 | 0.085 | 0 | MO_infl | DDX60L |
| 0 | 0.69529543 | 0.421 | 0.036 | 0 | MO_infl | CFP |
| 0 | 0.69465386 | 0.572 | 0.168 | 0 | MO_infl | GRINA |
| 0 | 0.69433371 | 0.461 | 0.096 | 0 | MO_infl | C1orf162 |
| 0 | 0.69317065 | 0.405 | 0.053 | 0 | MO_infl | LUCAT1 |

| | | | | | | |
|---|---|---|---|---|---|---|
| 0 | 0.68643428 | 0.541 | 0.175 | 0 | MO_infl | KYNU |
| 0 | 0.68492204 | 0.283 | 0.036 | 0 | MO_infl | TMEM158 |
| 0 | 0.68322068 | 0.442 | 0.172 | 0 | MO_infl | SOCS3 |
| 0 | 0.68047834 | 0.956 | 0.793 | 0 | MO_infl | PFDN5 |
| 0 | 0.67911417 | 0.502 | 0.17 | 0 | MO_infl | KDM6B |
| 0 | 0.67892998 | 0.397 | 0.029 | 0 | MO_infl | AC245128.3 |
| 0 | 0.67771649 | 0.553 | 0.121 | 0 | MO_infl | FMNL1 |
| 0 | 0.67354229 | 0.743 | 0.545 | 0 | MO_infl | VMP1 |
| 0 | 0.66456793 | 0.546 | 0.178 | 0 | MO_infl | FCGRT |
| 0 | 0.66388011 | 0.402 | 0.089 | 0 | MO_infl | EPB41L3 |
| 0 | 0.66027444 | 0.842 | 0.667 | 0 | MO_infl | PNRC1 |
| 0 | 0.66009059 | 0.438 | 0.065 | 0 | MO_infl | C3AR1 |
| 0 | 0.65610034 | 0.889 | 0.682 | 0 | MO_infl | SDCBP |
| 0 | 0.65564711 | 0.537 | 0.133 | 0 | MO_infl | RAC2 |
| 0 | 0.6548663 | 0.35 | 0.098 | 0 | MO_infl | RGCC |
| 0 | 0.65027699 | 0.471 | 0.089 | 0 | MO_infl | LRP1 |
| 0 | 0.64828294 | 0.58 | 0.171 | 0 | MO_infl | TPP1 |
| 0 | 0.64418762 | 0.516 | 0.128 | 0 | MO_infl | RILPL2 |
| 0 | 0.64219272 | 0.445 | 0.066 | 0 | MO_infl | MYO1G |
| 0 | 0.64119334 | 0.917 | 0.721 | 0 | MO_infl | CLIC1 |
| 0 | 0.64100944 | 0.401 | 0.116 | 0 | MO_infl | ADGRE5 |
| 0 | 0.64039974 | 0.404 | 0.06 | 0 | MO_infl | PIK3AP1 |
| 0 | 0.63824878 | 0.633 | 0.258 | 0 | MO_infl | ARPC1B |
| 0 | 0.63816284 | 0.979 | 0.878 | 0 | MO_infl | ATP5F1E |
| 0 | 0.63690744 | 0.877 | 0.749 | 0 | MO_infl | CST3 |
| 0 | 0.63675192 | 0.388 | 0.081 | 0 | MO_infl | PHACTR1 |
| 0 | 0.6363909 | 0.233 | 0.024 | 0 | MO_infl | TNFAIP6 |
| 0 | 0.63570565 | 0.393 | 0.056 | 0 | MO_infl | CLEC4E |
| 0 | 0.63158398 | 0.942 | 0.777 | 0 | MO_infl | ARPC2 |
| 0 | 0.63112935 | 0.447 | 0.135 | 0 | MO_infl | GLIPR1 |
| 0 | 0.62824101 | 0.385 | 0.059 | 0 | MO_infl | CCR1 |
| 0 | 0.62467903 | 0.465 | 0.107 | 0 | MO_infl | CSGALNACT2 |
| 0 | 0.62043323 | 0.662 | 0.279 | 0 | MO_infl | MSN |
| 0 | 0.61768062 | 0.157 | 0.037 | 0 | MO_infl | IL1A |
| 0 | 0.61589545 | 0.401 | 0.112 | 0 | MO_infl | EVI2B |
| 0 | 0.61476451 | 0.476 | 0.149 | 0 | MO_infl | ACSL4 |
| 0 | 0.6145011 | 0.55 | 0.267 | 0 | MO_infl | BNIP3L |
| 0 | 0.61309406 | 0.542 | 0.182 | 0 | MO_infl | ATP6V1B2 |
| 0 | 0.61304581 | 0.431 | 0.07 | 0 | MO_infl | MPP1 |
| 0 | 0.61216742 | 0.44 | 0.09 | 0 | MO_infl | CLEC7A |
| 0 | 0.61074254 | 0.408 | 0.15 | 0 | MO_infl | KLF2 |
| 0 | 0.61031852 | 0.346 | 0.043 | 0 | MO_infl | SEMA6B |
| 0 | 0.60933702 | 0.443 | 0.163 | 0 | MO_infl | NFKB1 |
| 0 | 0.6093294 | 0.307 | 0.047 | 0 | MO_infl | MPEG1 |
| 0 | 0.60855769 | 0.456 | 0.084 | 0 | MO_infl | NCKAP1L |
| 0 | 0.60784955 | 0.547 | 0.195 | 0 | MO_infl | RBPJ |
| 0 | 0.60596216 | 0.373 | 0.062 | 0 | MO_infl | HPSE |
| 0 | 0.60472581 | 0.247 | 0.035 | 0 | MO_infl | CD36 |
| 0 | 0.60312534 | 0.438 | 0.112 | 0 | MO_infl | SH3BP5 |
| 0 | 0.60272633 | 0.441 | 0.103 | 0 | MO_infl | PLXDC2 |

| | | | | | | |
|---|---|---|---|---|---|---|
| 0 | 0.59678075 | 0.573 | 0.237 | 0 | MO_infl | CTSZ |
| 0 | 0.59474129 | 0.534 | 0.172 | 0 | MO_infl | ACTN1 |
| 0 | 0.59443519 | 0.349 | 0.067 | 0 | MO_infl | ENG |
| 0 | 0.59407725 | 0.327 | 0.03 | 0 | MO_infl | LILRA5 |
| 0 | 0.59248133 | 0.55 | 0.203 | 0 | MO_infl | SH3BGRL |
| 0 | 0.59163233 | 0.539 | 0.199 | 0 | MO_infl | PTPN12 |
| 0 | 0.59157834 | 0.615 | 0.273 | 0 | MO_infl | SLC16A3 |
| 0 | 0.58727223 | 0.571 | 0.244 | 0 | MO_infl | RNF149 |
| 0 | 0.58507391 | 0.281 | 0.081 | 0 | MO_infl | FGL2 |
| 0 | 0.58423517 | 0.494 | 0.139 | 0 | MO_infl | RNF130 |
| 0 | 0.58398036 | 0.726 | 0.455 | 0 | MO_infl | TAGLN2 |
| 0 | 0.58186324 | 0.497 | 0.126 | 0 | MO_infl | PPP1R18 |
| 0 | 0.58022965 | 0.454 | 0.114 | 0 | MO_infl | AP1S2 |
| 0 | 0.58004902 | 0.511 | 0.135 | 0 | MO_infl | GPSM3 |
| 0 | 0.5772805 | 0.501 | 0.168 | 0 | MO_infl | RUNX1 |
| 0 | 0.57646347 | 0.429 | 0.116 | 0 | MO_infl | MYADM |
| 0 | 0.57414454 | 0.427 | 0.125 | 0 | MO_infl | CFD |
| 0 | 0.57289703 | 0.398 | 0.143 | 0 | MO_infl | EHD1 |
| 0 | 0.57221887 | 0.482 | 0.159 | 0 | MO_infl | LSP1 |
| 0 | 0.56854714 | 0.428 | 0.157 | 0 | MO_infl | SNX10 |
| 0 | 0.56766851 | 0.387 | 0.115 | 0 | MO_infl | SLC44A1 |
| 0 | 0.56478972 | 0.547 | 0.238 | 0 | MO_infl | LYN |
| 0 | 0.56460566 | 0.444 | 0.148 | 0 | MO_infl | PYCARD |
| 0 | 0.56357037 | 0.799 | 0.566 | 0 | MO_infl | NPC2 |
| 0 | 0.56140186 | 0.362 | 0.079 | 0 | MO_infl | CD86 |
| 0 | 0.55891393 | 0.39 | 0.114 | 0 | MO_infl | CPVL |
| 0 | 0.55812898 | 0.226 | 0.047 | 0 | MO_infl | IL1R2 |
| 0 | 0.555567 | 0.574 | 0.23 | 0 | MO_infl | ROCK1 |
| 0 | 0.55382789 | 0.636 | 0.3 | 0 | MO_infl | MTPN |
| 0 | 0.55329255 | 0.494 | 0.197 | 0 | MO_infl | RNASET2 |
| 0 | 0.55071573 | 0.855 | 0.661 | 0 | MO_infl | BRI3 |
| 0 | 0.54728054 | 0.438 | 0.143 | 0 | MO_infl | BID |
| 0 | 0.54675428 | 0.41 | 0.157 | 0 | MO_infl | FNDC3A |
| 0 | 0.54514789 | 0.39 | 0.066 | 0 | MO_infl | LAIR1 |
| 0 | 0.54305466 | 0.577 | 0.236 | 0 | MO_infl | MAFB |
| 0 | 0.54147996 | 0.285 | 0.032 | 0 | MO_infl | PID1 |
| 0 | 0.53915518 | 0.885 | 0.701 | 0 | MO_infl | ENO1 |
| 0 | 0.53848358 | 0.324 | 0.072 | 0 | MO_infl | MS4A6A |
| 0 | 0.53829719 | 0.423 | 0.087 | 0 | MO_infl | PILRA |
| 0 | 0.53633436 | 0.397 | 0.071 | 0 | MO_infl | ITGAM |
| 0 | 0.53527985 | 0.409 | 0.086 | 0 | MO_infl | IL6R |
| 0 | 0.5347528 | 0.447 | 0.133 | 0 | MO_infl | ATXN1 |
| 0 | 0.53450328 | 0.405 | 0.104 | 0 | MO_infl | CARD16 |
| 0 | 0.53216447 | 0.396 | 0.174 | 0 | MO_infl | MAP3K8 |
| 0 | 0.53186931 | 0.338 | 0.048 | 0 | MO_infl | ANTXR2 |
| 0 | 0.53151748 | 0.438 | 0.131 | 0 | MO_infl | CORO1A |
| 0 | 0.52820775 | 0.398 | 0.081 | 0 | MO_infl | DPYD |
| 0 | 0.52774651 | 0.466 | 0.187 | 0 | MO_infl | PNP |
| 0 | 0.52755779 | 0.448 | 0.113 | 0 | MO_infl | ARHGAP30 |
| 0 | 0.52618697 | 0.885 | 0.637 | 0 | MO_infl | S100A8 |

| | | | | | | |
|---|---|---|---|---|---|---|
| 0 | 0.52528649 | 0.474 | 0.238 | 0 | MO_infl | TSC22D3 |
| 0 | 0.52422955 | 0.402 | 0.146 | 0 | MO_infl | PFKFB3 |
| 0 | 0.52225263 | 0.996 | 0.932 | 0 | MO_infl | NEAT1 |
| 0 | 0.52200936 | 0.397 | 0.084 | 0 | MO_infl | ADAM8 |
| 0 | 0.52162336 | 0.756 | 0.552 | 0 | MO_infl | PGK1 |
| 0 | 0.51873972 | 0.599 | 0.263 | 0 | MO_infl | QKI |
| 0 | 0.51805402 | 0.355 | 0.105 | 0 | MO_infl | GCA |
| 0 | 0.51406772 | 0.984 | 0.906 | 0 | MO_infl | UBA52 |
| 0 | 0.51362501 | 0.415 | 0.097 | 0 | MO_infl | ARHGAP26 |
| 0 | 0.51299861 | 0.395 | 0.096 | 0 | MO_infl | EVI2A |
| 0 | 0.51260133 | 0.416 | 0.136 | 0 | MO_infl | KCTD12 |
| 0 | 0.51157769 | 0.331 | 0.086 | 0 | MO_infl | ATP2B1-AS1 |
| 0 | 0.51146398 | 0.477 | 0.153 | 0 | MO_infl | MYO9B |
| 0 | 0.51141879 | 0.499 | 0.184 | 0 | MO_infl | SKAP2 |
| 0 | 0.51106767 | 0.385 | 0.151 | 0 | MO_infl | ICAM1 |
| 0 | 0.50876039 | 0.565 | 0.282 | 0 | MO_infl | AP2S1 |
| 0 | 0.50840898 | 0.307 | 0.034 | 0 | MO_infl | FPR2 |
| 0 | 0.50808966 | 0.27 | 0.085 | 0 | MO_infl | TNF |
| 0 | 0.50778634 | 0.606 | 0.272 | 0 | MO_infl | CMTM6 |
| 0 | 0.5077737 | 0.532 | 0.257 | 0 | MO_infl | CD99 |
| 0 | 0.50748762 | 0.527 | 0.247 | 0 | MO_infl | CXCL16 |
| 0 | 0.50543387 | 0.405 | 0.109 | 0 | MO_infl | BCAT1 |
| 0 | 0.50420786 | 0.381 | 0.062 | 0 | MO_infl | CSF3R |
| 0 | 0.50377637 | 0.352 | 0.068 | 0 | MO_infl | HAVCR2 |
| 0 | 0.50229189 | 0.75 | 0.341 | 0 | MO_infl | HLA-DRA |
| 0 | 0.50167723 | 0.479 | 0.173 | 0 | MO_infl | ZYX |
| 0 | 0.5002864 | 0.489 | 0.169 | 0 | MO_infl | RNF13 |
| 0 | 0.49954408 | 0.408 | 0.14 | 0 | MO_infl | ABCA1 |
| 0 | 0.4992673 | 0.525 | 0.255 | 0 | MO_infl | SKIL |
| 0 | 0.4964708 | 0.992 | 0.949 | 0 | MO_infl | SERF2 |
| 0 | 0.49425582 | 0.773 | 0.554 | 0 | MO_infl | ACTR3 |
| 0 | 0.49213661 | 0.36 | 0.085 | 0 | MO_infl | DOCK10 |
| 0 | 0.49191532 | 0.322 | 0.1 | 0 | MO_infl | IL1R1 |
| 0 | 0.48996692 | 0.459 | 0.135 | 0 | MO_infl | CD37 |
| 0 | 0.48970345 | 0.474 | 0.151 | 0 | MO_infl | RAB8B |
| 0 | 0.48637496 | 0.375 | 0.105 | 0 | MO_infl | NUP214 |
| 0 | 0.48614867 | 0.52 | 0.227 | 0 | MO_infl | USP15 |
| 0 | 0.48178387 | 0.317 | 0.054 | 0 | MO_infl | RNASE6 |
| 0 | 0.48167112 | 0.391 | 0.148 | 0 | MO_infl | ELL2 |
| 0 | 0.48099239 | 0.388 | 0.101 | 0 | MO_infl | IFI30 |
| 0 | 0.48065285 | 0.433 | 0.135 | 0 | MO_infl | ADAM17 |
| 0 | 0.47807136 | 0.4 | 0.098 | 0 | MO_infl | PTPN6 |
| 0 | 0.47557414 | 0.389 | 0.106 | 0 | MO_infl | PCNX1 |
| 0 | 0.47384704 | 0.532 | 0.262 | 0 | MO_infl | TMEM167A |
| 0 | 0.47237848 | 0.385 | 0.08 | 0 | MO_infl | DENND5A |
| 0 | 0.46889201 | 0.343 | 0.086 | 0 | MO_infl | TRPS1 |
| 0 | 0.4681275 | 0.568 | 0.271 | 0 | MO_infl | CTNNB1 |
| 0 | 0.46770791 | 0.609 | 0.293 | 0 | MO_infl | CAPZA1 |
| 0 | 0.46506171 | 0.959 | 0.896 | 0 | MO_infl | CFL1 |
| 0 | 0.46434901 | 0.279 | 0.055 | 0 | MO_infl | RFLNB |

| | | | | | | |
|---|---|---|---|---|---|---|
| 0 | 0.463651 | 0.492 | 0.199 | 0 | MO_infl | SP100 |
| 0 | 0.46218447 | 0.439 | 0.133 | 0 | MO_infl | TCIRG1 |
| 0 | 0.46207333 | 0.658 | 0.374 | 0 | MO_infl | DAZAP2 |
| 0 | 0.46061812 | 0.422 | 0.127 | 0 | MO_infl | SIRPA |
| 0 | 0.45929295 | 0.396 | 0.111 | 0 | MO_infl | ASAP1 |
| 0 | 0.4583152 | 0.369 | 0.145 | 0 | MO_infl | P4HA1 |
| 0 | 0.45771353 | 0.459 | 0.163 | 0 | MO_infl | RHOG |
| 0 | 0.45639242 | 0.336 | 0.087 | 0 | MO_infl | PTAFR |
| 0 | 0.45497943 | 0.896 | 0.702 | 0 | MO_infl | PKM |
| 0 | 0.45486507 | 0.488 | 0.206 | 0 | MO_infl | CD82 |
| 0 | 0.45478684 | 0.573 | 0.286 | 0 | MO_infl | ARHGDIB |
| 0 | 0.45449044 | 0.307 | 0.077 | 0 | MO_infl | PDE4B |
| 0 | 0.45375718 | 0.371 | 0.086 | 0 | MO_infl | SYK |
| 0 | 0.45271691 | 0.393 | 0.134 | 0 | MO_infl | CD48 |
| 0 | 0.45169051 | 0.355 | 0.08 | 0 | MO_infl | CASP1 |
| 0 | 0.44965384 | 0.949 | 0.854 | 0 | MO_infl | ARPC3 |
| 0 | 0.44900021 | 0.43 | 0.147 | 0 | MO_infl | TET2 |
| 0 | 0.44711318 | 0.436 | 0.216 | 0 | MO_infl | METTL9 |
| 0 | 0.44680235 | 0.232 | 0.062 | 0 | MO_infl | POU2F2 |
| 0 | 0.44677113 | 0.529 | 0.233 | 0 | MO_infl | FAM49B |
| 0 | 0.44627143 | 0.815 | 0.495 | 0 | MO_infl | UPP1 |
| 0 | 0.44602543 | 0.393 | 0.114 | 0 | MO_infl | EHBP1L1 |
| 0 | 0.44598381 | 0.31 | 0.076 | 0 | MO_infl | IL1RAP |
| 0 | 0.4429792 | 0.343 | 0.069 | 0 | MO_infl | PRKCB |
| 0 | 0.44234211 | 0.345 | 0.07 | 0 | MO_infl | CSF2RA |
| 0 | 0.44199795 | 0.503 | 0.232 | 0 | MO_infl | AGFG1 |
| 0 | 0.44151591 | 0.356 | 0.074 | 0 | MO_infl | SLC43A3 |
| 0 | 0.43893569 | 0.392 | 0.109 | 0 | MO_infl | SH3KBP1 |
| 0 | 0.43702169 | 0.622 | 0.348 | 0 | MO_infl | DEK |
| 0 | 0.43677689 | 0.453 | 0.169 | 0 | MO_infl | STK17B |
| 0 | 0.43583432 | 0.361 | 0.082 | 0 | MO_infl | WAS |
| 0 | 0.43489546 | 0.321 | 0.053 | 0 | MO_infl | CHST15 |
| 0 | 0.43271571 | 0.473 | 0.193 | 0 | MO_infl | FLOT1 |
| 0 | 0.43216236 | 0.606 | 0.332 | 0 | MO_infl | PAK2 |
| 0 | 0.43204117 | 0.374 | 0.097 | 0 | MO_infl | SLC6A6 |
| 0 | 0.43158571 | 0.3 | 0.043 | 0 | MO_infl | TLR4 |
| 0 | 0.43104397 | 0.333 | 0.06 | 0 | MO_infl | TBXAS1 |
| 0 | 0.43083982 | 0.241 | 0.026 | 0 | MO_infl | CR1 |
| 0 | 0.42905064 | 0.385 | 0.105 | 0 | MO_infl | RXRA |
| 0 | 0.42872556 | 0.234 | 0.055 | 0 | MO_infl | RIPOR2 |
| 0 | 0.4286826 | 0.182 | 0.03 | 0 | MO_infl | OSM |
| 0 | 0.42578876 | 0.323 | 0.044 | 0 | MO_infl | PIK3R5 |
| 0 | 0.42503075 | 0.442 | 0.188 | 0 | MO_infl | MAP3K2 |
| 0 | 0.42484879 | 0.571 | 0.293 | 0 | MO_infl | MOB1A |
| 0 | 0.4237831 | 0.397 | 0.123 | 0 | MO_infl | WIPF1 |
| 0 | 0.42343667 | 0.224 | 0.019 | 0 | MO_infl | RNASE2 |
| 0 | 0.4221772 | 0.469 | 0.189 | 0 | MO_infl | RGS10 |
| 0 | 0.42165296 | 0.565 | 0.251 | 0 | MO_infl | FXYD5 |
| 0 | 0.4207822 | 0.389 | 0.154 | 0 | MO_infl | HIPK2 |
| 0 | 0.41935661 | 0.335 | 0.083 | 0 | MO_infl | MTF1 |

| | | | | | | |
|---|---|---|---|---|---|---|
| 0 | 0.41935272 | 0.317 | 0.062 | 0 | MO_infl | DOCK4 |
| 0 | 0.41828686 | 0.582 | 0.303 | 0 | MO_infl | ARPC4 |
| 0 | 0.41748223 | 0.33 | 0.059 | 0 | MO_infl | LAT2 |
| 0 | 0.41609584 | 0.231 | 0.066 | 0 | MO_infl | CRIP1 |
| 0 | 0.41484914 | 0.564 | 0.286 | 0 | MO_infl | TGOLN2 |
| 0 | 0.4139582 | 0.271 | 0.095 | 0 | MO_infl | FCGR3A |
| 0 | 0.41352358 | 0.428 | 0.16 | 0 | MO_infl | GNS |
| 0 | 0.41278731 | 0.609 | 0.225 | 0 | MO_infl | HLA-DRB1 |
| 0 | 0.41242682 | 0.398 | 0.132 | 0 | MO_infl | NUP58 |
| 0 | 0.41240719 | 0.374 | 0.131 | 0 | MO_infl | ZNF267 |
| 0 | 0.4107312 | 0.395 | 0.155 | 0 | MO_infl | AGO2 |
| 0 | 0.40707547 | 0.176 | 0.03 | 0 | MO_infl | CLEC12A |
| 0 | 0.40543792 | 0.353 | 0.096 | 0 | MO_infl | APBB1IP |
| 0 | 0.40486907 | 0.576 | 0.32 | 0 | MO_infl | AKAP13 |
| 0 | 0.40402197 | 0.383 | 0.154 | 0 | MO_infl | KLF10 |
| 0 | 0.40319935 | 0.497 | 0.228 | 0 | MO_infl | PLP2 |
| 0 | 0.40318826 | 0.438 | 0.197 | 0 | MO_infl | PHF20L1 |
| 0 | 0.40253816 | 0.451 | 0.19 | 0 | MO_infl | TMED5 |
| 0 | 0.40069819 | 0.426 | 0.166 | 0 | MO_infl | NR3C1 |
| 0 | 0.3998924 | 0.444 | 0.18 | 0 | MO_infl | TGFB1 |
| 0 | 0.39888985 | 0.304 | 0.059 | 0 | MO_infl | PLXNC1 |
| 0 | 0.39813154 | 0.265 | 0.023 | 0 | MO_infl | LILRA1 |
| 0 | 0.39177065 | 0.311 | 0.083 | 0 | MO_infl | IL17RA |
| 0 | 0.39051862 | 0.357 | 0.122 | 0 | MO_infl | CKAP4 |
| 0 | 0.39007587 | 0.308 | 0.068 | 0 | MO_infl | TFEC |
| 0 | 0.3896854 | 0.338 | 0.085 | 0 | MO_infl | CSF1R |
| 0 | 0.38965258 | 0.417 | 0.169 | 0 | MO_infl | TANK |
| 0 | 0.38911212 | 0.527 | 0.267 | 0 | MO_infl | PRKAR1A |
| 0 | 0.38907087 | 0.395 | 0.139 | 0 | MO_infl | ZDHHC20 |
| 0 | 0.38904853 | 0.377 | 0.127 | 0 | MO_infl | SH3BP2 |
| 0 | 0.38871958 | 0.312 | 0.063 | 0 | MO_infl | CYTH4 |
| 0 | 0.38837138 | 0.235 | 0.065 | 0 | MO_infl | LRRK2 |
| 0 | 0.38622064 | 0.465 | 0.213 | 0 | MO_infl | GPCPD1 |
| 0 | 0.38489243 | 0.336 | 0.114 | 0 | MO_infl | SIPA1L1 |
| 0 | 0.38475315 | 0.285 | 0.065 | 0 | MO_infl | GK5 |
| 0 | 0.38417912 | 0.529 | 0.273 | 0 | MO_infl | SPCS3 |
| 0 | 0.38366612 | 0.337 | 0.077 | 0 | MO_infl | PREX1 |
| 0 | 0.38152429 | 0.317 | 0.093 | 0 | MO_infl | HPCAL1 |
| 0 | 0.38002232 | 0.399 | 0.163 | 0 | MO_infl | PHC2 |
| 0 | 0.37970964 | 0.264 | 0.071 | 0 | MO_infl | LHFPL2 |
| 0 | 0.37811393 | 0.351 | 0.115 | 0 | MO_infl | DDX3Y |
| 0 | 0.37678674 | 0.267 | 0.046 | 0 | MO_infl | STAB1 |
| 0 | 0.37670233 | 0.363 | 0.127 | 0 | MO_infl | DOCK8 |
| 0 | 0.37532054 | 0.308 | 0.092 | 0 | MO_infl | LPCAT1 |
| 0 | 0.37487278 | 0.366 | 0.141 | 0 | MO_infl | B3GNT5 |
| 0 | 0.37351388 | 0.548 | 0.284 | 0 | MO_infl | FKBP1A |
| 0 | 0.37265713 | 0.396 | 0.154 | 0 | MO_infl | RAPGEF1 |
| 0 | 0.37036422 | 0.366 | 0.144 | 0 | MO_infl | ERGIC1 |
| 0 | 0.3700209 | 0.347 | 0.127 | 0 | MO_infl | OGFRL1 |
| 0 | 0.36898675 | 0.379 | 0.138 | 0 | MO_infl | LACTB |

| | | | | | | |
|---|---|---|---|---|---|---|
| 0 | 0.36860169 | 0.317 | 0.079 | 0 | MO_infl | LY96 |
| 0 | 0.36848696 | 0.383 | 0.17 | 0 | MO_infl | PRNP |
| 0 | 0.36847936 | 0.33 | 0.112 | 0 | MO_infl | FGD4 |
| 0 | 0.36669379 | 0.268 | 0.041 | 0 | MO_infl | MCTP1 |
| 0 | 0.3663174 | 0.457 | 0.223 | 0 | MO_infl | MGAT1 |
| 0 | 0.36523704 | 0.227 | 0.055 | 0 | MO_infl | MARCH1 |
| 0 | 0.36483847 | 0.42 | 0.177 | 0 | MO_infl | TAOK1 |
| 0 | 0.36434932 | 0.431 | 0.195 | 0 | MO_infl | IFNGR1 |
| 0 | 0.36424049 | 0.416 | 0.181 | 0 | MO_infl | ZFYVE16 |
| 0 | 0.36399805 | 0.356 | 0.129 | 0 | MO_infl | CPD |
| 0 | 0.36395359 | 0.342 | 0.119 | 0 | MO_infl | SMAP2 |
| 0 | 0.36382394 | 0.252 | 0.024 | 0 | MO_infl | LILRA6 |
| 0 | 0.36226078 | 0.253 | 0.051 | 0 | MO_infl | NAIP |
| 0 | 0.36149515 | 0.338 | 0.108 | 0 | MO_infl | KCTD20 |
| 0 | 0.36018337 | 0.269 | 0.051 | 0 | MO_infl | LRRC25 |
| 0 | 0.35840258 | 0.312 | 0.089 | 0 | MO_infl | P2RX4 |
| 0 | 0.35825472 | 0.194 | 0.017 | 0 | MO_infl | TNFSF8 |
| 0 | 0.3580812 | 0.279 | 0.064 | 0 | MO_infl | LILRB4 |
| 0 | 0.35767616 | 0.488 | 0.24 | 0 | MO_infl | ADAM10 |
| 0 | 0.35629039 | 0.27 | 0.049 | 0 | MO_infl | RASSF2 |
| 0 | 0.35515328 | 0.311 | 0.084 | 0 | MO_infl | MCUB |
| 0 | 0.35464131 | 0.371 | 0.162 | 0 | MO_infl | PTPN2 |
| 0 | 0.35391924 | 0.344 | 0.111 | 0 | MO_infl | CBL |
| 0 | 0.35331207 | 0.271 | 0.049 | 0 | MO_infl | APOBR |
| 0 | 0.35314133 | 0.439 | 0.201 | 0 | MO_infl | M6PR |
| 0 | 0.35167938 | 0.276 | 0.045 | 0 | MO_infl | OSCAR |
| 0 | 0.35160822 | 0.145 | 0.018 | 0 | MO_infl | MIR3945HG |
| 0 | 0.3514549 | 0.29 | 0.06 | 0 | MO_infl | PARVG |
| 0 | 0.35066461 | 0.246 | 0.058 | 0 | MO_infl | PLA2G7 |
| 0 | 0.34974456 | 0.301 | 0.081 | 0 | MO_infl | ALOX5 |
| 0 | 0.3483949 | 0.261 | 0.062 | 0 | MO_infl | NFKBID |
| 0 | 0.34800975 | 0.298 | 0.068 | 0 | MO_infl | DOCK2 |
| 0 | 0.34799284 | 0.466 | 0.227 | 0 | MO_infl | MBNL1 |
| 0 | 0.34708536 | 0.387 | 0.159 | 0 | MO_infl | CUX1 |
| 0 | 0.3467324 | 0.321 | 0.1 | 0 | MO_infl | PTPRJ |
| 0 | 0.34645141 | 0.382 | 0.169 | 0 | MO_infl | CHD1 |
| 0 | 0.345891 | 0.418 | 0.187 | 0 | MO_infl | ACAP2 |
| 0 | 0.34568586 | 0.266 | 0.065 | 0 | MO_infl | STARD4 |
| 0 | 0.34504791 | 0.303 | 0.073 | 0 | MO_infl | CARD8 |
| 0 | 0.34331698 | 0.295 | 0.081 | 0 | MO_infl | C9orf72 |
| 0 | 0.34108072 | 0.302 | 0.092 | 0 | MO_infl | SERPINB8 |
| 0 | 0.33903935 | 0.389 | 0.174 | 0 | MO_infl | USP3 |
| 0 | 0.3389344 | 0.507 | 0.247 | 0 | MO_infl | IFITM2 |
| 0 | 0.3385811 | 0.26 | 0.061 | 0 | MO_infl | HIP1 |
| 0 | 0.3360506 | 0.314 | 0.095 | 0 | MO_infl | FKBP15 |
| 0 | 0.33487291 | 0.291 | 0.068 | 0 | MO_infl | FAM49A |
| 0 | 0.33459992 | 0.283 | 0.063 | 0 | MO_infl | LPXN |
| 0 | 0.333184 | 0.303 | 0.084 | 0 | MO_infl | SLC36A4 |
| 0 | 0.33099957 | 0.174 | 0.037 | 0 | MO_infl | EGR3 |
| 0 | 0.33056136 | 0.318 | 0.102 | 0 | MO_infl | THEMIS2 |

| | | | | | | |
|---|---|---|---|---|---|---|
| 0 | 0.33054614 | 0.159 | 0.018 | 0 | MO_infl | C5AR2 |
| 0 | 0.33029858 | 0.258 | 0.059 | 0 | MO_infl | GSAP |
| 0 | 0.32937356 | 0.376 | 0.163 | 0 | MO_infl | DICER1 |
| 0 | 0.32819206 | 0.2 | 0.025 | 0 | MO_infl | GPR84 |
| 0 | 0.3266967 | 0.215 | 0.025 | 0 | MO_infl | AC005280.2 |
| 0 | 0.326138 | 0.323 | 0.117 | 0 | MO_infl | ZMIZ1 |
| 0 | 0.32586066 | 0.364 | 0.156 | 0 | MO_infl | RPS6KA3 |
| 0 | 0.32548711 | 0.33 | 0.111 | 0 | MO_infl | CD58 |
| 0 | 0.32531699 | 0.33 | 0.111 | 0 | MO_infl | PITPNA |
| 0 | 0.324225 | 0.234 | 0.039 | 0 | MO_infl | MLKL |
| 0 | 0.32264565 | 0.305 | 0.107 | 0 | MO_infl | CYTIP |
| 0 | 0.32206273 | 0.197 | 0.041 | 0 | MO_infl | SELL |
| 0 | 0.32202343 | 0.23 | 0.028 | 0 | MO_infl | LILRA2 |
| 0 | 0.32133831 | 0.213 | 0.043 | 0 | MO_infl | FCGR1A |
| 0 | 0.31993154 | 0.267 | 0.075 | 0 | MO_infl | SLC8A1 |
| 0 | 0.31814782 | 0.262 | 0.081 | 0 | MO_infl | GPRIN3 |
| 0 | 0.3177821 | 0.304 | 0.098 | 0 | MO_infl | HSD17B11 |
| 0 | 0.31773756 | 0.233 | 0.035 | 0 | MO_infl | BST1 |
| 0 | 0.3168103 | 0.269 | 0.067 | 0 | MO_infl | RGS19 |
| 0 | 0.31544129 | 0.278 | 0.069 | 0 | MO_infl | BIN2 |
| 0 | 0.31508133 | 0.299 | 0.099 | 0 | MO_infl | IL4R |
| 0 | 0.31495544 | 0.31 | 0.118 | 0 | MO_infl | IRAK1 |
| 0 | 0.31358973 | 0.243 | 0.048 | 0 | MO_infl | ST8SIA4 |
| 0 | 0.3134393 | 0.306 | 0.093 | 0 | MO_infl | STK10 |
| 0 | 0.31058817 | 0.247 | 0.061 | 0 | MO_infl | CDA |
| 0 | 0.30523082 | 0.253 | 0.072 | 0 | MO_infl | DENND3 |
| 0 | 0.30495638 | 0.204 | 0.039 | 0 | MO_infl | SLC2A6 |
| 0 | 0.30493022 | 0.244 | 0.062 | 0 | MO_infl | DOK2 |
| 0 | 0.30475264 | 0.248 | 0.058 | 0 | MO_infl | PDE4A |
| 0 | 0.30383149 | 0.28 | 0.092 | 0 | MO_infl | EDEM3 |
| 0 | 0.30202266 | 0.186 | 0.039 | 0 | MO_infl | SLC16A6 |
| 0 | 0.30040513 | 0.237 | 0.052 | 0 | MO_infl | FAM102B |
| 0 | 0.30020783 | 0.239 | 0.058 | 0 | MO_infl | AC004687.1 |
| 0 | 0.29976264 | 0.236 | 0.037 | 0 | MO_infl | DOK3 |
| 0 | 0.29953682 | 0.221 | 0.06 | 0 | MO_infl | AC020916.1 |
| 0 | 0.29934988 | 0.272 | 0.091 | 0 | MO_infl | SPHK1 |
| 0 | 0.29881285 | 0.257 | 0.064 | 0 | MO_infl | RARA |
| 0 | 0.29792413 | 0.192 | 0.033 | 0 | MO_infl | ADAM19 |
| 0 | 0.2955778 | 0.249 | 0.069 | 0 | MO_infl | DOT1L |
| 0 | 0.29207966 | 0.261 | 0.072 | 0 | MO_infl | PLEKHO2 |
| 0 | 0.29191307 | 0.253 | 0.062 | 0 | MO_infl | NPL |
| 0 | 0.29082184 | 0.264 | 0.06 | 0 | MO_infl | SLC15A3 |
| 0 | 0.29036405 | 0.272 | 0.086 | 0 | MO_infl | COLGALT1 |
| 0 | 0.28929541 | 0.213 | 0.027 | 0 | MO_infl | SIRPB1 |
| 0 | 0.2892904 | 0.189 | 0.026 | 0 | MO_infl | HRH2 |
| 0 | 0.28923884 | 0.216 | 0.054 | 0 | MO_infl | MEF2C |
| 0 | 0.28890544 | 0.236 | 0.061 | 0 | MO_infl | LFNG |
| 0 | 0.28770011 | 0.294 | 0.099 | 0 | MO_infl | MFSD1 |
| 0 | 0.28768417 | 0.243 | 0.064 | 0 | MO_infl | SLC7A7 |
| 0 | 0.2872992 | 0.256 | 0.07 | 0 | MO_infl | EMB |

| | | | | | | |
|---|---|---|---|---|---|---|
| 0 | 0.28497957 | 0.269 | 0.078 | 0 | MO_infl | DOCK5 |
| 0 | 0.28343473 | 0.3 | 0.105 | 0 | MO_infl | ADPGK |
| 0 | 0.28212928 | 0.288 | 0.108 | 0 | MO_infl | CARD19 |
| 0 | 0.28126906 | 0.208 | 0.045 | 0 | MO_infl | AOAH |
| 0 | 0.27890854 | 0.263 | 0.089 | 0 | MO_infl | RAB27A |
| 0 | 0.27830225 | 0.187 | 0.035 | 0 | MO_infl | CD302 |
| 0 | 0.27813753 | 0.245 | 0.06 | 0 | MO_infl | MOB3A |
| 0 | 0.27567002 | 0.268 | 0.085 | 0 | MO_infl | IL10RA |
| 0 | 0.27458603 | 0.186 | 0.032 | 0 | MO_infl | GAS7 |
| 0 | 0.27258937 | 0.289 | 0.105 | 0 | MO_infl | KDM7A |
| 0 | 0.27065621 | 0.259 | 0.078 | 0 | MO_infl | DIAPH2 |
| 0 | 0.26979826 | 0.212 | 0.044 | 0 | MO_infl | SLFN11 |
| 0 | 0.26751009 | 0.192 | 0.039 | 0 | MO_infl | CCRL2 |
| 0 | 0.26698253 | 0.222 | 0.045 | 0 | MO_infl | AC008105.3 |
| 0 | 0.26608478 | 0.291 | 0.103 | 0 | MO_infl | SHKBP1 |
| 0 | 0.26581253 | 0.224 | 0.055 | 0 | MO_infl | PGS1 |
| 0 | 0.26406835 | 0.204 | 0.044 | 0 | MO_infl | CHSY1 |
| 0 | 0.26360854 | 0.195 | 0.037 | 0 | MO_infl | CLEC4A |
| 0 | 0.2630977 | 0.253 | 0.085 | 0 | MO_infl | STX6 |
| 0 | 0.26263064 | 0.242 | 0.068 | 0 | MO_infl | GNB4 |
| 0 | 0.26217012 | 0.249 | 0.078 | 0 | MO_infl | ACER3 |
| 0 | 0.26172653 | 0.106 | 0.01 | 0 | MO_infl | IL10 |
| 0 | 0.26164373 | 0.272 | 0.091 | 0 | MO_infl | ARAP1 |
| 0 | 0.25918337 | 0.114 | 0.015 | 0 | MO_infl | FFAR2 |
| 0 | 0.25803471 | 0.219 | 0.054 | 0 | MO_infl | PTPN7 |
| 0 | 0.25696037 | 0.162 | 0.026 | 0 | MO_infl | FAM198B |
| 0 | 0.25617587 | 0.208 | 0.054 | 0 | MO_infl | CSF2RB |
| 0 | 0.25445158 | 0.242 | 0.081 | 0 | MO_infl | MTMR6 |
| 0 | 0.25388261 | 0.208 | 0.04 | 0 | MO_infl | SASH3 |
| 0 | 0.25219467 | 0.244 | 0.079 | 0 | MO_infl | BMP2K |
| 0 | 0.2512371 | 0.219 | 0.059 | 0 | MO_infl | NCF4 |
| 0 | 0.25075744 | 0.217 | 0.067 | 0 | MO_infl | IRAK2 |
| 0 | 0.25072791 | 0.254 | 0.081 | 0 | MO_infl | USP4 |
| 0 | -0.414661 | 0.004 | 0.225 | 0 | MO_infl | RBPMS |
| 0 | -0.4360142 | 0.005 | 0.237 | 0 | MO_infl | CTTN |
| 0 | -0.4408741 | 0.003 | 0.215 | 0 | MO_infl | PAWR |
| 0 | -0.4827396 | 0.004 | 0.211 | 0 | MO_infl | CDH1 |
| 0 | -0.4968126 | 0.004 | 0.248 | 0 | MO_infl | CNN3 |
| 0 | -0.5244655 | 0.053 | 0.283 | 0 | MO_infl | CD2AP |
| 0 | -0.5306994 | 0.002 | 0.219 | 0 | MO_infl | ABLIM1 |
| 0 | -0.5702791 | 0.004 | 0.266 | 0 | MO_infl | TJP1 |
| 0 | -0.576161 | 0.003 | 0.237 | 0 | MO_infl | PARD6B |
| 0 | -0.5788423 | 0.012 | 0.246 | 0 | MO_infl | CKB |
| 0 | -0.5795466 | 0.005 | 0.25 | 0 | MO_infl | DSP |
| 0 | -0.5991794 | 0.002 | 0.247 | 0 | MO_infl | EHF |
| 0 | -0.5996914 | 0.011 | 0.277 | 0 | MO_infl | GIPC1 |
| 0 | -0.6299459 | 0.008 | 0.239 | 0 | MO_infl | MACC1 |
| 0 | -0.6393383 | 0.081 | 0.315 | 0 | MO_infl | LDHB |
| 0 | -0.6483165 | 0.004 | 0.211 | 0 | MO_infl | SDCBP2 |
| 0 | -0.6559981 | 0.026 | 0.304 | 0 | MO_infl | AFDN |

| | | | | | | |
|---|---|---|---|---|---|---|
| 0 | -0.6561442 | 0.019 | 0.283 | 0 | MO_infl | SPTBN1 |
| 0 | -0.6674145 | 0.014 | 0.257 | 0 | MO_infl | OCIAD2 |
| 0 | -0.693267 | 0.344 | 0.612 | 0 | MO_infl | PRDX5 |
| 0 | -0.6935464 | 0.01 | 0.315 | 0 | MO_infl | JUP |
| 0 | -0.7091118 | 0.407 | 0.632 | 0 | MO_infl | HINT1 |
| 0 | -0.7272395 | 0.04 | 0.329 | 0 | MO_infl | SPINT1 |
| 0 | -0.7349413 | 0.006 | 0.272 | 0 | MO_infl | S100A16 |
| 0 | -0.7494067 | 0.008 | 0.217 | 0 | MO_infl | EPS8L1 |
| 0 | -0.7495646 | 0.309 | 0.556 | 0 | MO_infl | HEBP2 |
| 0 | -0.7669146 | 0.007 | 0.225 | 0 | MO_infl | SFN |
| 0 | -0.7674626 | 0.007 | 0.221 | 0 | MO_infl | FBXO32 |
| 0 | -0.7706494 | 0.008 | 0.288 | 0 | MO_infl | KLF5 |
| 0 | -0.7742283 | 0.003 | 0.279 | 0 | MO_infl | PRSS8 |
| 0 | -0.7760325 | 0.041 | 0.265 | 0 | MO_infl | CDKN1C |
| 0 | -0.7840978 | 0.143 | 0.451 | 0 | MO_infl | CD59 |
| 0 | -0.8033282 | 0.017 | 0.277 | 0 | MO_infl | S100A13 |
| 0 | -0.8064642 | 0.477 | 0.718 | 0 | MO_infl | NPM1 |
| 0 | -0.8508891 | 0.011 | 0.299 | 0 | MO_infl | MUC1 |
| 0 | -0.8588272 | 0.009 | 0.254 | 0 | MO_infl | CCND1 |
| 0 | -0.8595401 | 0.012 | 0.263 | 0 | MO_infl | ITGA2 |
| 0 | -0.866944 | 0.006 | 0.289 | 0 | MO_infl | SYNE2 |
| 0 | -0.8732226 | 0.011 | 0.368 | 0 | MO_infl | PERP |
| 0 | -0.8779051 | 0.033 | 0.279 | 0 | MO_infl | CST6 |
| 0 | -0.9118337 | 0.111 | 0.351 | 0 | MO_infl | GADD45A |
| 0 | -0.937379 | 0.003 | 0.318 | 0 | MO_infl | CAMK2N1 |
| 0 | -0.9407004 | 0.011 | 0.27 | 0 | MO_infl | EPCAM |
| 0 | -0.943204 | 0.192 | 0.543 | 0 | MO_infl | CD9 |
| 0 | -0.9451247 | 0.008 | 0.315 | 0 | MO_infl | MAL2 |
| 0 | -0.9464856 | 0.61 | 0.778 | 0 | MO_infl | RPSA |
| 0 | -0.9583495 | 0.004 | 0.27 | 0 | MO_infl | PDZK1IP1 |
| 0 | -0.9706945 | 0.048 | 0.396 | 0 | MO_infl | PEBP1 |
| 0 | -0.9739205 | 0.057 | 0.291 | 0 | MO_infl | TPM1 |
| 0 | -0.9886867 | 0.207 | 0.495 | 0 | MO_infl | RAB11FIP1 |
| 0 | -0.99613 | 0.018 | 0.26 | 0 | MO_infl | IL32 |
| 0 | -1.0099515 | 0.004 | 0.216 | 0 | MO_infl | TMPRSS2 |
| 0 | -1.0159332 | 0.852 | 0.88 | 0 | MO_infl | RPL3 |
| 0 | -1.0184898 | 0.076 | 0.482 | 0 | MO_infl | SPINT2 |
| 0 | -1.0247943 | 0.714 | 0.894 | 0 | MO_infl | RPLP0 |
| 0 | -1.0265069 | 0.935 | 0.961 | 0 | MO_infl | RPS18 |
| 0 | -1.0326441 | 0.013 | 0.223 | 0 | MO_infl | C12orf75 |
| 0 | -1.0515975 | 0.005 | 0.379 | 0 | MO_infl | MYO6 |
| 0 | -1.058138 | 0.002 | 0.275 | 0 | MO_infl | ADGRF1 |
| 0 | -1.1092309 | 0.025 | 0.334 | 0 | MO_infl | GPRC5A |
| 0 | -1.1225479 | 0.223 | 0.687 | 0 | MO_infl | DSTN |
| 0 | -1.1269394 | 0.597 | 0.846 | 0 | MO_infl | GSTP1 |
| 0 | -1.1318727 | 0.009 | 0.304 | 0 | MO_infl | TSPAN1 |
| 0 | -1.2771852 | 0.05 | 0.515 | 0 | MO_infl | CLDN7 |
| 0 | -1.2880997 | 0.045 | 0.498 | 0 | MO_infl | ATP1B1 |
| 0 | -1.3271865 | 0.008 | 0.303 | 0 | MO_infl | LMO7 |
| 0 | -1.3273302 | 0.003 | 0.337 | 0 | MO_infl | RND3 |

| | | | | | | |
|---|---|---|---|---|---|---|
| 0 | -1.350994 | 0.005 | 0.212 | 0 | MO_infl | DUOX2 |
| 0 | -1.4940568 | 0.529 | 0.88 | 0 | MO_infl | PPDPF |
| 0 | -1.5930589 | 0.006 | 0.226 | 0 | MO_infl | HOPX |
| 0 | -1.6528382 | 0.309 | 0.787 | 0 | MO_infl | HSPB1 |
| 0 | -1.6775768 | 0.107 | 0.372 | 0 | MO_infl | IGFBP7 |
| 0 | -1.6818184 | 0.012 | 0.339 | 0 | MO_infl | ADIRF |
| 0 | -1.682608 | 0.03 | 0.394 | 0 | MO_infl | GDF15 |
| 0 | -1.7151886 | 0.082 | 0.612 | 0 | MO_infl | CD24 |
| 0 | -1.764413 | 0.115 | 0.503 | 0 | MO_infl | SOX4 |
| 0 | -1.7661775 | 0.009 | 0.228 | 0 | MO_infl | KRT17 |
| 0 | -1.7855933 | 0.008 | 0.267 | 0 | MO_infl | PRSS22 |
| 0 | -1.8038367 | 0.077 | 0.559 | 0 | MO_infl | KRT8 |
| 0 | -1.8044447 | 0.058 | 0.46 | 0 | MO_infl | SLPI |
| 0 | -1.8219354 | 0.026 | 0.283 | 0 | MO_infl | CRYAB |
| 0 | -1.8796972 | 0.04 | 0.42 | 0 | MO_infl | MAL |
| 0 | -1.9923212 | 0.1 | 0.642 | 0 | MO_infl | TACSTD2 |
| 0 | -2.0377283 | 0.088 | 0.616 | 0 | MO_infl | KRT7 |
| 0 | -2.0515706 | 0.091 | 0.609 | 0 | MO_infl | CLDN4 |
| 0 | -2.0911605 | 0.084 | 0.629 | 0 | MO_infl | ELF3 |
| 0 | -2.1154954 | 0.098 | 0.63 | 0 | MO_infl | KRT18 |
| 0 | -2.1599927 | 0.033 | 0.303 | 0 | MO_infl | S100A2 |
| 0 | -2.3410937 | 0.054 | 0.389 | 0 | MO_infl | HPGD |
| 0 | -2.4311276 | 0.12 | 0.65 | 0 | MO_infl | KRT19 |
| 0 | -2.4604705 | 0.172 | 0.445 | 0 | MO_infl | S100P |
| 0 | -2.4661323 | 0.061 | 0.385 | 0 | MO_infl | PSCA |
| 0 | -2.6270172 | 0.092 | 0.378 | 0 | MO_infl | MMP7 |
| 0 | -2.7525538 | 0.101 | 0.604 | 0 | MO_infl | WFDC2 |
| 0 | -3.1993717 | 0.079 | 0.425 | 0 | MO_infl | KRT13 |
| 0 | -3.803597 | 0.122 | 0.586 | 0 | MO_infl | LCN2 |
| 0 | -4.5345411 | 0.101 | 0.448 | 0 | MO_infl | SPRR3 |
| 4.704E-308 | -0.4371574 | 0.005 | 0.212 | 1.111E-303 | MO_infl | EPS8L2 |
| 1.234E-307 | 0.38094541 | 0.352 | 0.152 | 2.916E-303 | MO_infl | TMEM176B |
| 1.823E-307 | 0.43087509 | 0.469 | 0.24 | 4.306E-303 | MO_infl | OSBPL8 |
| 1.986E-307 | -0.7372062 | 0.685 | 0.861 | 4.692E-303 | MO_infl | SQSTM1 |
| 2.082E-307 | 0.35112405 | 0.405 | 0.187 | 4.917E-303 | MO_infl | TRIM38 |
| 2.82E-307 | -0.6414497 | 0.06 | 0.293 | 6.662E-303 | MO_infl | HES4 |
| 7.841E-306 | 0.2912857 | 0.339 | 0.14 | 1.852E-301 | MO_infl | CORO1C |
| 1.105E-305 | -0.6687845 | 0.2 | 0.435 | 2.61E-301 | MO_infl | SOD1 |
| 2.085E-305 | -0.432759 | 0.003 | 0.207 | 4.926E-301 | MO_infl | TEAD1 |
| 2.572E-305 | -0.5552219 | 0.115 | 0.358 | 6.075E-301 | MO_infl | PPA1 |
| 6.269E-305 | 0.29611696 | 0.34 | 0.142 | 1.481E-300 | MO_infl | HEXB |
| 8.8E-305 | -0.6685338 | 0.32 | 0.544 | 2.079E-300 | MO_infl | ANXA11 |
| 1.786E-304 | 1.22662865 | 0.156 | 0.038 | 4.22E-300 | MO_infl | CCL4L2 |
| 2.764E-304 | -0.5475467 | 0.004 | 0.208 | 6.529E-300 | MO_infl | S100A14 |
| 3.898E-304 | -0.629091 | 0.068 | 0.297 | 9.208E-300 | MO_infl | UGCG |
| 1.838E-303 | -1.3288168 | 0.447 | 0.668 | 4.342E-299 | MO_infl | MTRNR2L12 |
| 9.181E-303 | -0.8622355 | 0.01 | 0.216 | 2.169E-298 | MO_infl | ANKRD36C |
| 2.125E-302 | 0.39630888 | 0.283 | 0.107 | 5.02E-298 | MO_infl | CD83 |
| 2.791E-302 | 0.41465139 | 0.674 | 0.438 | 6.593E-298 | MO_infl | RAP1B |
| 8.155E-302 | -0.3892738 | 0.003 | 0.205 | 1.926E-297 | MO_infl | LSR |

| | | | | | | |
|---|---|---|---|---|---|---|
| 1.893E-301 | 0.30805429 | 0.224 | 0.072 | 4.473E-297 | MO_infl | TNFSF13B |
| 4.321E-300 | -0.4402225 | 0.003 | 0.203 | 1.021E-295 | MO_infl | SMIM22 |
| 1.747E-299 | -0.5802288 | 0.004 | 0.206 | 4.126E-295 | MO_infl | EPHA2 |
| 2.402E-299 | 0.46778052 | 0.615 | 0.384 | 5.675E-295 | MO_infl | GRN |
| 1.034E-298 | 0.41675484 | 0.591 | 0.34 | 2.444E-294 | MO_infl | METRNL |
| 4.738E-298 | -0.5974756 | 0.002 | 0.202 | 1.119E-293 | MO_infl | C6orf132 |
| 5.326E-298 | -0.6860014 | 0.002 | 0.201 | 1.258E-293 | MO_infl | SPNS2 |
| 2.705E-297 | -0.5746952 | 0.056 | 0.277 | 6.389E-293 | MO_infl | SERINC2 |
| 3.485E-296 | 0.33346413 | 0.245 | 0.086 | 8.233E-292 | MO_infl | MEGF9 |
| 1.825E-295 | -0.4072079 | 0.006 | 0.206 | 4.312E-291 | MO_infl | FNBP1L |
| 2.758E-295 | -0.9434173 | 0.002 | 0.2 | 6.516E-291 | MO_infl | NDRG2 |
| 1.03E-294 | 0.34373596 | 0.328 | 0.139 | 2.432E-290 | MO_infl | TLE3 |
| 1.058E-293 | -0.5408666 | 0.005 | 0.203 | 2.5E-289 | MO_infl | FHL2 |
| 1.079E-293 | 0.27642217 | 0.308 | 0.123 | 2.55E-289 | MO_infl | ZSWIM6 |
| 1.444E-293 | -0.7498533 | 0.082 | 0.303 | 3.412E-289 | MO_infl | TNFRSF12A |
| 5.006E-293 | 0.2660959 | 0.411 | 0.187 | 1.183E-288 | MO_infl | STK4 |
| 6.966E-293 | 0.33409307 | 0.411 | 0.197 | 1.646E-288 | MO_infl | XRN2 |
| 9.771E-293 | -0.7473479 | 0.005 | 0.202 | 2.308E-288 | MO_infl | IGFBP3 |
| 1.475E-292 | 0.25349203 | 0.243 | 0.084 | 3.486E-288 | MO_infl | ARID5A |
| 1.093E-291 | 0.27476094 | 0.249 | 0.087 | 2.582E-287 | MO_infl | PAG1 |
| 3.619E-291 | -0.6693271 | 0.002 | 0.197 | 8.55E-287 | MO_infl | HSPB8 |
| 1.305E-290 | 0.29583216 | 0.347 | 0.15 | 3.082E-286 | MO_infl | MBOAT7 |
| 2.831E-290 | -0.5207622 | 0.113 | 0.343 | 6.688E-286 | MO_infl | SBDS |
| 2.397E-289 | -0.3895566 | 0.006 | 0.202 | 5.661E-285 | MO_infl | TMEM54 |
| 5.139E-289 | -0.8774862 | 0.01 | 0.208 | 1.214E-284 | MO_infl | LAMC2 |
| 1.517E-288 | -0.9297219 | 0.012 | 0.211 | 3.584E-284 | MO_infl | AQP3 |
| 3.36E-288 | -0.4869947 | 0.744 | 0.906 | 7.938E-284 | MO_infl | HSP90AB1 |
| 9.022E-288 | -0.9720348 | 0.021 | 0.224 | 2.131E-283 | MO_infl | TM4SF1 |
| 2.335E-287 | -0.5580214 | 0.069 | 0.288 | 5.515E-283 | MO_infl | WASL |
| 4.742E-287 | 0.36644809 | 0.637 | 0.376 | 1.12E-282 | MO_infl | CDV3 |
| 5.132E-287 | 0.31455699 | 0.347 | 0.151 | 1.212E-282 | MO_infl | PURB |
| 1.126E-286 | -0.3830941 | 0.001 | 0.193 | 2.66E-282 | MO_infl | CXADR |
| 1.601E-286 | 0.28826712 | 0.32 | 0.133 | 3.781E-282 | MO_infl | PPT1 |
| 1.686E-286 | 0.34754209 | 0.375 | 0.171 | 3.982E-282 | MO_infl | ARHGAP18 |
| 7.629E-286 | 0.34466807 | 0.582 | 0.335 | 1.802E-281 | MO_infl | GNB1 |
| 1.626E-285 | -0.3968092 | 0.015 | 0.215 | 3.841E-281 | MO_infl | SH3YL1 |
| 2.98E-285 | 0.26506176 | 0.186 | 0.055 | 7.041E-281 | MO_infl | NRIP3 |
| 1.427E-283 | 0.44127939 | 0.563 | 0.328 | 3.371E-279 | MO_infl | CFLAR |
| 3.23E-283 | 0.29313993 | 0.378 | 0.173 | 7.631E-279 | MO_infl | DCP2 |
| 3.853E-283 | -0.7863256 | 0.599 | 0.769 | 9.103E-279 | MO_infl | PRDX1 |
| 4.101E-283 | 0.33675442 | 0.449 | 0.229 | 9.689E-279 | MO_infl | ATP6V0D1 |
| 7.542E-283 | -0.6626659 | 0.006 | 0.198 | 1.782E-278 | MO_infl | PPL |
| 1.396E-281 | -0.6972987 | 0.003 | 0.194 | 3.299E-277 | MO_infl | FXYD3 |
| 1.661E-281 | 0.67307475 | 0.705 | 0.558 | 3.923E-277 | MO_infl | LITAF |
| 3.79E-281 | -0.43412 | 0.011 | 0.206 | 8.952E-277 | MO_infl | ANXA3 |
| 7.138E-281 | -0.550647 | 0.068 | 0.288 | 1.686E-276 | MO_infl | TSC22D1 |
| 4.53E-280 | 0.29529646 | 0.377 | 0.175 | 1.07E-275 | MO_infl | BNIP2 |
| 1.093E-279 | 0.26079138 | 0.28 | 0.106 | 2.582E-275 | MO_infl | CELF2 |
| 1.948E-279 | 0.65890981 | 0.159 | 0.043 | 4.601E-275 | MO_infl | RNASE1 |
| 5.071E-279 | 0.35492279 | 0.384 | 0.182 | 1.198E-274 | MO_infl | PNPLA8 |

| | | | | | | |
|---|---|---|---|---|---|---|
| 9.843E-279 | 0.4018075 | 0.334 | 0.148 | 2.325E-274 | MO_infl | TNFAIP8 |
| 6.492E-278 | 0.48352831 | 0.953 | 0.882 | 1.534E-273 | MO_infl | CD63 |
| 7.477E-278 | -1.3928533 | 0.003 | 0.191 | 1.766E-273 | MO_infl | DUOXA2 |
| 8.088E-278 | 0.31913895 | 0.45 | 0.23 | 1.911E-273 | MO_infl | RAP1A |
| 2.267E-277 | 0.29064659 | 0.322 | 0.137 | 5.355E-273 | MO_infl | NCOR2 |
| 1.171E-276 | 0.29026433 | 0.325 | 0.139 | 2.766E-272 | MO_infl | CNPY3 |
| 3.561E-276 | 0.3708601 | 0.555 | 0.309 | 8.413E-272 | MO_infl | WSB1 |
| 8.208E-276 | 0.34950886 | 0.44 | 0.22 | 1.939E-271 | MO_infl | PLEC |
| 1.053E-275 | 0.33381047 | 0.505 | 0.265 | 2.487E-271 | MO_infl | REL |
| 1.083E-275 | 0.2744543 | 0.316 | 0.133 | 2.558E-271 | MO_infl | BLOC1S6 |
| 1.119E-275 | -0.4326594 | 0.005 | 0.194 | 2.643E-271 | MO_infl | NECTIN4 |
| 1.565E-275 | 0.26251359 | 0.159 | 0.042 | 3.698E-271 | MO_infl | IL3RA |
| 5.462E-275 | -0.3471996 | 0.007 | 0.195 | 1.29E-270 | MO_infl | GNG12 |
| 1.877E-274 | -0.5830312 | 0.013 | 0.206 | 4.434E-270 | MO_infl | DST |
| 1.963E-274 | 0.30161516 | 0.301 | 0.122 | 4.638E-270 | MO_infl | PLEKHO1 |
| 2.787E-274 | 0.35970952 | 0.352 | 0.159 | 6.583E-270 | MO_infl | CKLF |
| 7.002E-274 | 0.32466469 | 0.472 | 0.253 | 1.654E-269 | MO_infl | RAB5C |
| 1.289E-273 | -0.6301394 | 0.05 | 0.256 | 3.046E-269 | MO_infl | CYB5A |
| 2.869E-273 | -1.1657828 | 0.001 | 0.185 | 6.778E-269 | MO_infl | CEACAM5 |
| 3.461E-273 | -0.770327 | 0.844 | 0.875 | 8.176E-269 | MO_infl | RPL14 |
| 1.281E-272 | 0.31157878 | 0.299 | 0.123 | 3.026E-268 | MO_infl | LYST |
| 8.297E-272 | 0.41643214 | 0.986 | 0.96 | 1.96E-267 | MO_infl | MYL6 |
| 2.168E-271 | -0.37452 | 0.003 | 0.188 | 5.122E-267 | MO_infl | NDFIP2 |
| 2.657E-271 | -1.2538676 | 0.002 | 0.186 | 6.277E-267 | MO_infl | MUC21 |
| 5.716E-271 | -0.3654417 | 0.001 | 0.183 | 1.35E-266 | MO_infl | CRB3 |
| 6.076E-271 | -1.1189033 | 0.01 | 0.199 | 1.435E-266 | MO_infl | RDH10 |
| 8.529E-270 | 0.36148997 | 0.544 | 0.314 | 2.015E-265 | MO_infl | LAMTOR4 |
| 2.984E-269 | 0.32674706 | 0.399 | 0.193 | 7.049E-265 | MO_infl | ZBTB7A |
| 4.94E-268 | 0.26989841 | 0.273 | 0.107 | 1.167E-263 | MO_infl | FBXL5 |
| 1.047E-267 | -0.4303459 | 0.008 | 0.193 | 2.474E-263 | MO_infl | TINAGL1 |
| 1.698E-267 | -0.5674663 | 0.002 | 0.183 | 4.012E-263 | MO_infl | MALL |
| 2.342E-267 | -0.3501908 | 0.003 | 0.185 | 5.533E-263 | MO_infl | NCKAP1 |
| 5.474E-267 | -0.6427862 | 0.416 | 0.635 | 1.293E-262 | MO_infl | EZR |
| 1.391E-266 | -0.5046428 | 0.079 | 0.289 | 3.287E-262 | MO_infl | PRDX2 |
| 1.484E-266 | -0.4849785 | 0.002 | 0.183 | 3.505E-262 | MO_infl | SDC1 |
| 1.009E-265 | 0.27402796 | 0.186 | 0.057 | 2.382E-261 | MO_infl | PSTPIP2 |
| 1.71E-263 | -0.4331628 | 0.001 | 0.18 | 4.038E-259 | MO_infl | TSPAN6 |
| 2.544E-263 | -0.4674561 | 0.002 | 0.182 | 6.009E-259 | MO_infl | TTC9 |
| 2.576E-263 | -0.4063001 | 0.009 | 0.193 | 6.085E-259 | MO_infl | SLC38A1 |
| 5.194E-263 | -0.7682477 | 0.71 | 0.793 | 1.227E-258 | MO_infl | RPL10A |
| 8.998E-263 | 0.35135903 | 0.501 | 0.277 | 2.126E-258 | MO_infl | HNRNPM |
| 1.757E-262 | 0.31033813 | 0.295 | 0.122 | 4.151E-258 | MO_infl | UCP2 |
| 1.655E-261 | 0.2718487 | 0.294 | 0.123 | 3.909E-257 | MO_infl | AZI2 |
| 1.843E-261 | 0.45750915 | 0.478 | 0.271 | 4.353E-257 | MO_infl | TNIP1 |
| 3.04E-261 | 0.42637501 | 0.739 | 0.559 | 7.181E-257 | MO_infl | NOP10 |
| 8.485E-261 | -0.3459244 | 0.008 | 0.189 | 2.004E-256 | MO_infl | DDR1 |
| 4.504E-260 | 0.59798719 | 0.2 | 0.066 | 1.064E-255 | MO_infl | GBP5 |
| 7.009E-260 | 0.34251978 | 0.919 | 0.822 | 1.656E-255 | MO_infl | HLA-E |
| 1.117E-259 | 0.75198551 | 0.63 | 0.445 | 2.639E-255 | MO_infl | DUSP1 |
| 1.604E-259 | 0.27434764 | 0.246 | 0.092 | 3.788E-255 | MO_infl | MTSS1 |

| | | | | | | |
|---|---|---|---|---|---|---|
| 2.507E-259 | -1.0839513 | 0.102 | 0.306 | 5.922E-255 | MO_infl | DUSP5 |
| 5.172E-259 | -0.7319442 | 0.002 | 0.179 | 1.222E-254 | MO_infl | MUC4 |
| 7.172E-259 | -1.0823453 | 0.004 | 0.182 | 1.694E-254 | MO_infl | PRSS27 |
| 1.254E-258 | 0.30109468 | 0.348 | 0.161 | 2.962E-254 | MO_infl | SQOR |
| 2.229E-258 | -0.9509358 | 0.003 | 0.179 | 5.266E-254 | MO_infl | KRT15 |
| 2.375E-258 | -0.4773985 | 0.013 | 0.197 | 5.61E-254 | MO_infl | STMN1 |
| 2.592E-258 | 0.33133534 | 0.408 | 0.205 | 6.122E-254 | MO_infl | ZNF106 |
| 6.002E-258 | -0.4984776 | 0.002 | 0.178 | 1.418E-253 | MO_infl | TMPRSS4 |
| 6.514E-258 | -0.480974 | 0.003 | 0.18 | 1.539E-253 | MO_infl | CLDN3 |
| 2.455E-257 | -0.4619224 | 0.007 | 0.186 | 5.799E-253 | MO_infl | SYTL2 |
| 2.676E-257 | 0.42817096 | 0.622 | 0.397 | 6.322E-253 | MO_infl | SEC62 |
| 2.686E-257 | -0.3751823 | 0.003 | 0.179 | 6.346E-253 | MO_infl | ERBB3 |
| 4.938E-257 | -0.3756261 | 0.024 | 0.213 | 1.166E-252 | MO_infl | HSBP1L1 |
| 1.743E-256 | -1.0774107 | 0.003 | 0.178 | 4.118E-252 | MO_infl | PITX1 |
| 5.204E-256 | 0.32588829 | 0.389 | 0.188 | 1.229E-251 | MO_infl | EML4 |
| 8.934E-256 | -0.4613121 | 0.079 | 0.287 | 2.11E-251 | MO_infl | ARHGAP5 |
| 2.547E-255 | -0.8381199 | 0.002 | 0.176 | 6.016E-251 | MO_infl | SCEL |
| 3.862E-255 | 0.26613979 | 0.319 | 0.14 | 9.122E-251 | MO_infl | TNFRSF1A |
| 1.109E-254 | -1.7205525 | 0.009 | 0.186 | 2.619E-250 | MO_infl | ECM1 |
| 1.173E-254 | -0.4192111 | 0.033 | 0.225 | 2.772E-250 | MO_infl | TMEM106B |
| 2.264E-254 | -0.4354669 | 0.004 | 0.18 | 5.349E-250 | MO_infl | COBLL1 |
| 8.705E-254 | -0.656835 | 0.305 | 0.51 | 2.056E-249 | MO_infl | CD46 |
| 1.051E-253 | 0.25276264 | 0.268 | 0.107 | 2.482E-249 | MO_infl | AGO3 |
| 1.928E-253 | -0.6649445 | 0.003 | 0.177 | 4.555E-249 | MO_infl | CDC42EP5 |
| 2.868E-252 | 0.48553778 | 0.294 | 0.125 | 6.776E-248 | MO_infl | RGS2 |
| 3.195E-251 | 0.2698586 | 0.143 | 0.038 | 7.548E-247 | MO_infl | LGALS2 |
| 8.731E-251 | -0.4456712 | 0.029 | 0.216 | 2.062E-246 | MO_infl | LIMA1 |
| 6.025E-250 | -0.528445 | 0.093 | 0.297 | 1.423E-245 | MO_infl | FAM3C |
| 8.148E-250 | -0.5408542 | 0.005 | 0.178 | 1.925E-245 | MO_infl | CYR61 |
| 1.901E-249 | -0.5436158 | 0.574 | 0.755 | 4.491E-245 | MO_infl | ATP6V1G1 |
| 2.355E-249 | 0.26437831 | 0.252 | 0.099 | 5.563E-245 | MO_infl | RIPK2 |
| 2.865E-249 | 0.32093796 | 0.871 | 0.754 | 6.768E-245 | MO_infl | CDC42 |
| 7.944E-249 | 0.28691382 | 0.299 | 0.127 | 1.877E-244 | MO_infl | SORL1 |
| 1.071E-248 | 0.37202119 | 0.361 | 0.174 | 2.529E-244 | MO_infl | CREG1 |
| 1.164E-248 | -0.4558857 | 0.341 | 0.568 | 2.75E-244 | MO_infl | HMGN1 |
| 1.039E-247 | -0.3448618 | 0.002 | 0.172 | 2.454E-243 | MO_infl | PLS3 |
| 3.199E-247 | 0.57227311 | 0.703 | 0.552 | 7.557E-243 | MO_infl | TSPO |
| 4.654E-247 | -0.4053918 | 0.065 | 0.265 | 1.099E-242 | MO_infl | TSTD1 |
| 6.062E-247 | -0.702784 | 0.045 | 0.234 | 1.432E-242 | MO_infl | CDKN2B |
| 1.401E-246 | 0.2587029 | 0.316 | 0.14 | 3.309E-242 | MO_infl | SLC25A24 |
| 2.416E-246 | -0.9136821 | 0 | 0.168 | 5.707E-242 | MO_infl | CEACAM7 |
| 2.67E-246 | -0.3342438 | 0.001 | 0.169 | 6.306E-242 | MO_infl | KCNK1 |
| 5.885E-246 | -0.3458358 | 0.019 | 0.199 | 1.39E-241 | MO_infl | TRAF4 |
| 1.088E-245 | 0.37239236 | 0.983 | 0.931 | 2.569E-241 | MO_infl | TMSB4X |
| 1.1E-245 | -0.8400791 | 0.002 | 0.171 | 2.599E-241 | MO_infl | TMPRSS11D |
| 1.213E-245 | -0.533432 | 0.001 | 0.169 | 2.866E-241 | MO_infl | ALDH1A3 |
| 6.269E-245 | -0.7212718 | 0.001 | 0.169 | 1.481E-240 | MO_infl | NCCRP1 |
| 6.52E-245 | -0.4660492 | 0.001 | 0.168 | 1.54E-240 | MO_infl | VSIG2 |
| 7.724E-245 | 0.26737856 | 0.383 | 0.185 | 1.825E-240 | MO_infl | CELF1 |
| 2.979E-244 | -0.9564048 | 0.001 | 0.169 | 7.036E-240 | MO_infl | CEACAM6 |

| | | | | | | |
|---|---|---|---|---|---|---|
| 3.945E-242 | 0.31900914 | 0.546 | 0.315 | 9.319E-238 | MO_infl | ZNF207 |
| 1.072E-241 | -0.4219025 | 0.005 | 0.174 | 2.532E-237 | MO_infl | MDK |
| 2.279E-241 | -0.8565323 | 0.001 | 0.167 | 5.384E-237 | MO_infl | TMPRSS11E |
| 2.548E-241 | -0.7696066 | 0.739 | 0.81 | 6.02E-237 | MO_infl | RPS5 |
| 9.237E-241 | 0.26190654 | 0.359 | 0.171 | 2.182E-236 | MO_infl | MAPK1 |
| 2.543E-240 | 0.28714094 | 0.353 | 0.168 | 6.008E-236 | MO_infl | GNAQ |
| 2.12E-239 | 0.3542688 | 0.553 | 0.328 | 5.008E-235 | MO_infl | PLSCR1 |
| 9.726E-239 | 0.26208066 | 0.333 | 0.153 | 2.298E-234 | MO_infl | UBR4 |
| 1.52E-237 | -1.2044226 | 0.027 | 0.204 | 3.59E-233 | MO_infl | SPINK1 |
| 5.316E-237 | 0.3197034 | 0.519 | 0.301 | 1.256E-232 | MO_infl | PSMB3 |
| 6.13E-237 | 0.25062347 | 0.298 | 0.129 | 1.448E-232 | MO_infl | LBR |
| 7.428E-237 | -0.4068379 | 0.007 | 0.174 | 1.755E-232 | MO_infl | ITGA3 |
| 2.231E-236 | -0.3419274 | 0.003 | 0.167 | 5.27E-232 | MO_infl | SLC44A4 |
| 3.909E-236 | -0.434376 | 0.002 | 0.166 | 9.235E-232 | MO_infl | ITGB6 |
| 1.305E-235 | -0.3351207 | 0.002 | 0.165 | 3.083E-231 | MO_infl | PHLDA3 |
| 6.12E-235 | -0.6608464 | 0.088 | 0.285 | 1.446E-230 | MO_infl | PMAIP1 |
| 7.352E-233 | 0.35741879 | 0.59 | 0.367 | 1.737E-228 | MO_infl | H2AFY |
| 1.09E-232 | -0.5416446 | 0.357 | 0.556 | 2.575E-228 | MO_infl | EIF4A2 |
| 1.789E-232 | -0.2900635 | 0.003 | 0.164 | 4.227E-228 | MO_infl | BACE2 |
| 2.696E-232 | 0.30317646 | 0.45 | 0.246 | 6.369E-228 | MO_infl | ARL8B |
| 4.557E-232 | -0.5752482 | 0.294 | 0.503 | 1.076E-227 | MO_infl | CD81 |
| 7.804E-232 | -0.3150032 | 0.002 | 0.162 | 1.844E-227 | MO_infl | RAB25 |
| 2.932E-231 | 0.25039589 | 0.285 | 0.123 | 6.926E-227 | MO_infl | STX4 |
| 1.322E-230 | 0.27334981 | 0.414 | 0.215 | 3.123E-226 | MO_infl | DNAJC7 |
| 3.111E-230 | -0.3238095 | 0.029 | 0.205 | 7.349E-226 | MO_infl | REXO2 |
| 4.769E-230 | 0.51641402 | 0.839 | 0.765 | 1.127E-225 | MO_infl | TPI1 |
| 2.392E-229 | -0.3988054 | 0.005 | 0.166 | 5.651E-225 | MO_infl | CLDN1 |
| 5.292E-228 | -0.5460663 | 0.563 | 0.758 | 1.25E-223 | MO_infl | HSPA8 |
| 5.47E-228 | 0.37328113 | 0.81 | 0.667 | 1.292E-223 | MO_infl | ATP6V0B |
| 1.183E-227 | -0.5983516 | 0.059 | 0.243 | 2.795E-223 | MO_infl | ZBTB20 |
| 2.607E-227 | 0.31289374 | 0.998 | 0.96 | 6.158E-223 | MO_infl | S100A11 |
| 4.568E-227 | -0.4651433 | 0.042 | 0.222 | 1.079E-222 | MO_infl | PTGR1 |
| 5.584E-227 | -0.340639 | 0.007 | 0.168 | 1.319E-222 | MO_infl | FAM102A |
| 6.694E-227 | -0.3118433 | 0.002 | 0.16 | 1.581E-222 | MO_infl | GULP1 |
| 9.849E-227 | 0.26919192 | 0.415 | 0.219 | 2.327E-222 | MO_infl | OS9 |
| 1.432E-225 | 0.3075964 | 0.536 | 0.315 | 3.384E-221 | MO_infl | SCAF11 |
| 4.2E-225 | 0.39623135 | 0.689 | 0.512 | 9.922E-221 | MO_infl | SERPINB1 |
| 4.646E-225 | 0.31283139 | 0.21 | 0.077 | 1.097E-220 | MO_infl | GPR183 |
| 6.109E-225 | 0.30306871 | 0.227 | 0.089 | 1.443E-220 | MO_infl | NBN |
| 9.966E-225 | 0.27833785 | 0.928 | 0.816 | 2.354E-220 | MO_infl | HNRNPA2B1 |
| 1.537E-224 | 0.30924126 | 0.496 | 0.285 | 3.63E-220 | MO_infl | ARHGDIA |
| 5.699E-224 | -0.3148198 | 0.007 | 0.166 | 1.346E-219 | MO_infl | GPRC5C |
| 2.107E-223 | 0.43864932 | 0.41 | 0.229 | 4.977E-219 | MO_infl | C1orf122 |
| 3.438E-223 | -0.4689299 | 0.002 | 0.157 | 8.122E-219 | MO_infl | GCNT3 |
| 4.829E-223 | -0.6911259 | 0.005 | 0.162 | 1.141E-218 | MO_infl | SNCG |
| 1.263E-222 | -0.7093719 | 0.002 | 0.157 | 2.983E-218 | MO_infl | CXCL17 |
| 1.431E-222 | -1.8262364 | 0.007 | 0.165 | 3.379E-218 | MO_infl | SPRR2A |
| 3.793E-222 | -0.3002063 | 0.003 | 0.158 | 8.961E-218 | MO_infl | PATJ |
| 1.091E-221 | -0.3554609 | 0.003 | 0.159 | 2.578E-217 | MO_infl | MTUS1 |
| 3.239E-221 | -0.4423245 | 0.002 | 0.155 | 7.651E-217 | MO_infl | BCAS1 |

| | | | | | | |
|---|---|---|---|---|---|---|
| 6.849E-221 | -0.9991148 | 0.002 | 0.155 | 1.618E-216 | MO_infl | TMPRSS11B |
| 1.11E-220 | -0.463317 | 0.067 | 0.251 | 2.622E-216 | MO_infl | RAB9A |
| 1.33E-220 | 0.29080586 | 0.395 | 0.208 | 3.143E-216 | MO_infl | CTSA |
| 1.649E-220 | -0.3898599 | 0.044 | 0.22 | 3.896E-216 | MO_infl | NEDD4L |
| 5.703E-220 | 0.31329595 | 0.548 | 0.336 | 1.347E-215 | MO_infl | RAB7A |
| 6.088E-220 | -0.5249929 | 0.468 | 0.646 | 1.438E-215 | MO_infl | TMEM59 |
| 9.975E-220 | -0.7094887 | 0.093 | 0.278 | 2.356E-215 | MO_infl | CARHSP1 |
| 1.229E-219 | -0.3269481 | 0.007 | 0.163 | 2.903E-215 | MO_infl | OCLN |
| 3.072E-219 | -0.4549992 | 0.004 | 0.158 | 7.257E-215 | MO_infl | DCDC2 |
| 3.404E-219 | 0.27011047 | 0.335 | 0.163 | 8.041E-215 | MO_infl | ATG3 |
| 1.018E-218 | 1.26531766 | 0.49 | 0.308 | 2.406E-214 | MO_infl | IL1RN |
| 3.51E-218 | 0.29599962 | 0.265 | 0.115 | 8.291E-214 | MO_infl | RNF19B |
| 1.008E-217 | -0.4971178 | 0.026 | 0.196 | 2.382E-213 | MO_infl | XIST |
| 2.008E-217 | 0.29367082 | 0.343 | 0.171 | 4.744E-213 | MO_infl | NCOA4 |
| 2.587E-217 | -0.4531021 | 0.298 | 0.506 | 6.11E-213 | MO_infl | HSPD1 |
| 4.205E-217 | -0.7943008 | 0.246 | 0.449 | 9.933E-213 | MO_infl | TXNRD1 |
| 4.561E-217 | -0.4887413 | 0.329 | 0.523 | 1.077E-212 | MO_infl | VDAC2 |
| 1.11E-216 | -0.2872146 | 0.01 | 0.167 | 2.622E-212 | MO_infl | LLGL2 |
| 2.58E-216 | 0.26770672 | 0.296 | 0.135 | 6.096E-212 | MO_infl | HNMT |
| 5.913E-216 | 0.33484944 | 0.273 | 0.121 | 1.397E-211 | MO_infl | OTUD1 |
| 1.707E-215 | -1.729907 | 0.002 | 0.153 | 4.032E-211 | MO_infl | LYPD2 |
| 1.966E-215 | 0.25636209 | 0.329 | 0.157 | 4.644E-211 | MO_infl | GNA13 |
| 2.52E-215 | -0.6983197 | 0.008 | 0.163 | 5.953E-211 | MO_infl | CDKN2A |
| 4.088E-215 | -0.5619519 | 0.187 | 0.374 | 9.656E-211 | MO_infl | UBE2H |
| 2.146E-214 | 0.30597929 | 0.461 | 0.261 | 5.07E-210 | MO_infl | HM13 |
| 4.779E-214 | -1.0024548 | 0.003 | 0.153 | 1.129E-209 | MO_infl | RHCG |
| 7.024E-214 | -0.7689213 | 0.005 | 0.157 | 1.659E-209 | MO_infl | NUPR1 |
| 7.119E-214 | -0.3070863 | 0.001 | 0.15 | 1.682E-209 | MO_infl | C1orf116 |
| 2.457E-213 | -0.294782 | 0.005 | 0.157 | 5.803E-209 | MO_infl | SCNN1A |
| 2.766E-213 | -0.5103151 | 0.004 | 0.154 | 6.533E-209 | MO_infl | UCHL1 |
| 3.776E-213 | -0.4463343 | 0.028 | 0.192 | 8.919E-209 | MO_infl | FAM214A |
| 8.029E-213 | -0.2935353 | 0.002 | 0.151 | 1.897E-208 | MO_infl | DMKN |
| 1.272E-212 | -0.4286052 | 0.005 | 0.157 | 3.006E-208 | MO_infl | ARHGAP29 |
| 2.861E-212 | -0.3045516 | 0.002 | 0.151 | 6.759E-208 | MO_infl | NTN4 |
| 6.555E-212 | -0.3628488 | 0.002 | 0.15 | 1.548E-207 | MO_infl | AIF1L |
| 7.394E-212 | 0.28488417 | 0.187 | 0.066 | 1.747E-207 | MO_infl | JAML |
| 1.182E-211 | 0.2849468 | 0.441 | 0.246 | 2.792E-207 | MO_infl | GNAI3 |
| 1.537E-211 | -0.4827002 | 0.345 | 0.552 | 3.631E-207 | MO_infl | HSPE1 |
| 5.758E-211 | -0.531767 | 0.208 | 0.403 | 1.36E-206 | MO_infl | GCC2 |
| 6.568E-211 | -0.7276783 | 0.003 | 0.151 | 1.552E-206 | MO_infl | GABRP |
| 1.592E-210 | -0.3836637 | 0.001 | 0.148 | 3.761E-206 | MO_infl | FAM84A |
| 1.192E-209 | -0.5698276 | 0.255 | 0.442 | 2.815E-205 | MO_infl | JPT1 |
| 4.631E-209 | -0.5320875 | 0.334 | 0.516 | 1.094E-204 | MO_infl | SELENOW |
| 5.913E-209 | -0.538117 | 0.408 | 0.594 | 1.397E-204 | MO_infl | RPS27L |
| 1.345E-208 | -0.3753186 | 0.019 | 0.176 | 3.178E-204 | MO_infl | ITPKC |
| 2.599E-208 | -0.3333743 | 0.023 | 0.183 | 6.14E-204 | MO_infl | GALNT3 |
| 3.494E-208 | 0.27941592 | 0.273 | 0.122 | 8.255E-204 | MO_infl | TRIM25 |
| 3.566E-208 | -1.3464453 | 0.004 | 0.152 | 8.424E-204 | MO_infl | PLAT |
| 3.451E-207 | -0.3265187 | 0.005 | 0.152 | 8.153E-203 | MO_infl | MPZL2 |
| 6.779E-207 | -0.4784388 | 0.202 | 0.392 | 1.601E-202 | MO_infl | AES |

| | | | | | | |
|---|---|---|---|---|---|---|
| 8.349E-207 | 0.28111727 | 0.561 | 0.346 | 1.972E-202 | MO_infl | BRK1 |
| 1.114E-205 | -0.3124603 | 0.003 | 0.149 | 2.632E-201 | MO_infl | KAZN |
| 1.445E-205 | -0.3352159 | 0.004 | 0.151 | 3.413E-201 | MO_infl | SCIN |
| 6.339E-205 | -0.2977794 | 0.019 | 0.175 | 1.497E-200 | MO_infl | ARHGEF12 |
| 3.339E-204 | -0.2685657 | 0.003 | 0.147 | 7.889E-200 | MO_infl | MYO5B |
| 3.843E-204 | -0.385386 | 0.004 | 0.15 | 9.078E-200 | MO_infl | FOXC1 |
| 1.503E-203 | 0.27440458 | 0.94 | 0.859 | 3.549E-199 | MO_infl | HLA-C |
| 1.675E-203 | -1.1000911 | 0.011 | 0.16 | 3.958E-199 | MO_infl | CXCL6 |
| 1.885E-203 | 0.32330001 | 0.999 | 0.992 | 4.452E-199 | MO_infl | TPT1 |
| 2.02E-203 | 0.26840334 | 0.335 | 0.168 | 4.772E-199 | MO_infl | AGTRAP |
| 3.455E-203 | 0.26047416 | 0.994 | 0.966 | 8.162E-199 | MO_infl | TMSB10 |
| 7.315E-203 | -0.81897 | 0.093 | 0.27 | 1.728E-198 | MO_infl | PLAU |
| 3.713E-202 | -0.3840745 | 0.054 | 0.223 | 8.772E-198 | MO_infl | ZNF91 |
| 5.306E-202 | -0.3752323 | 0.011 | 0.16 | 1.254E-197 | MO_infl | EMP2 |
| 5.554E-202 | -0.33194 | 0.001 | 0.143 | 1.312E-197 | MO_infl | FAT1 |
| 7.019E-202 | -0.2987164 | 0.003 | 0.146 | 1.658E-197 | MO_infl | SELENOP |
| 7.273E-202 | -0.4340595 | 0.002 | 0.145 | 1.718E-197 | MO_infl | ID4 |
| 3.305E-201 | -0.3317266 | 0.008 | 0.155 | 7.808E-197 | MO_infl | COL18A1 |
| 3.519E-200 | 0.34874727 | 0.667 | 0.48 | 8.312E-196 | MO_infl | JAK1 |
| 5.144E-200 | -0.3830239 | 0.001 | 0.141 | 1.215E-195 | MO_infl | KRT80 |
| 5.283E-200 | -0.6640535 | 0.189 | 0.378 | 1.248E-195 | MO_infl | ATF3 |
| 3.419E-199 | -0.2717147 | 0.003 | 0.145 | 8.078E-195 | MO_infl | FAM213A |
| 3.914E-199 | -0.6096378 | 0.229 | 0.41 | 9.247E-195 | MO_infl | C9orf16 |
| 4.655E-199 | -0.4556572 | 0.002 | 0.142 | 1.1E-194 | MO_infl | TMEM40 |
| 6.723E-199 | -0.2884393 | 0.01 | 0.156 | 1.588E-194 | MO_infl | AGRN |
| 7.091E-198 | 0.33830173 | 0.559 | 0.352 | 1.675E-193 | MO_infl | PRELID1 |
| 1.164E-196 | -0.3106233 | 0.016 | 0.164 | 2.75E-192 | MO_infl | CRIM1 |
| 1.2E-196 | -0.2871983 | 0.002 | 0.141 | 2.834E-192 | MO_infl | PTPN3 |
| 5.215E-196 | -0.4464587 | 0.015 | 0.163 | 1.232E-191 | MO_infl | PAX8 |
| 1.858E-195 | -0.2727657 | 0.018 | 0.168 | 4.39E-191 | MO_infl | SVIP |
| 2.937E-195 | -0.848367 | 0.925 | 0.882 | 6.938E-191 | MO_infl | RPS6 |
| 4.431E-195 | 0.29414858 | 0.598 | 0.392 | 1.047E-190 | MO_infl | CAPZB |
| 1.541E-194 | -0.6129363 | 0.003 | 0.141 | 3.64E-190 | MO_infl | AGR2 |
| 1.811E-194 | -0.2712701 | 0.002 | 0.14 | 4.278E-190 | MO_infl | TC2N |
| 3.084E-194 | 0.25630181 | 0.441 | 0.25 | 7.286E-190 | MO_infl | ATP6AP2 |
| 4.95E-194 | -0.3235979 | 0.007 | 0.147 | 1.169E-189 | MO_infl | BEX2 |
| 7.355E-194 | -0.7337822 | 0.97 | 0.941 | 1.737E-189 | MO_infl | RPS19 |
| 4.648E-193 | -0.2678997 | 0.002 | 0.138 | 1.098E-188 | MO_infl | AMOTL2 |
| 5.235E-193 | -0.6842353 | 0.007 | 0.147 | 1.237E-188 | MO_infl | TFPI2 |
| 8.377E-193 | -0.4132107 | 0.479 | 0.656 | 1.979E-188 | MO_infl | SAP18 |
| 2.011E-192 | -0.278851 | 0.011 | 0.154 | 4.75E-188 | MO_infl | TMEM238 |
| 8.864E-192 | -0.4121147 | 0.425 | 0.626 | 2.094E-187 | MO_infl | VAMP8 |
| 9.331E-192 | -0.2620787 | 0.001 | 0.137 | 2.204E-187 | MO_infl | GRB7 |
| 1.105E-191 | -0.6138082 | 0.653 | 0.805 | 2.609E-187 | MO_infl | TXN |
| 1.419E-191 | 0.32586814 | 0.448 | 0.261 | 3.352E-187 | MO_infl | BAZ1A |
| 1.426E-191 | -0.3066387 | 0.016 | 0.162 | 3.369E-187 | MO_infl | LAPTM4B |
| 5.723E-191 | -0.5206708 | 0.088 | 0.257 | 1.352E-186 | MO_infl | TOB1 |
| 2.143E-190 | -0.2832413 | 0.003 | 0.139 | 5.061E-186 | MO_infl | BICDL2 |
| 2.984E-190 | -0.2880817 | 0.002 | 0.136 | 7.05E-186 | MO_infl | C19orf33 |
| 4.879E-190 | 0.28095774 | 0.468 | 0.275 | 1.152E-185 | MO_infl | PGAM1 |

| | | | | | | |
|---|---|---|---|---|---|---|
| 6.779E-190 | -1.2221658 | 0.001 | 0.135 | 1.601E-185 | MO_infl | SPINK5 |
| 8.962E-190 | -0.4500595 | 0.069 | 0.238 | 2.117E-185 | MO_infl | ARL4C |
| 9.665E-190 | -0.2515879 | 0.369 | 0.178 | 2.283E-185 | MO_infl | HLA-DPA1 |
| 1.029E-189 | -0.3514602 | 0.002 | 0.136 | 2.431E-185 | MO_infl | ASS1 |
| 1.534E-189 | -0.4933178 | 0.066 | 0.231 | 3.624E-185 | MO_infl | LY6E |
| 4.252E-189 | 0.25630894 | 0.486 | 0.292 | 1.005E-184 | MO_infl | CAPZA2 |
| 5.68E-189 | -0.3794121 | 0.029 | 0.18 | 1.342E-184 | MO_infl | SELENOM |
| 5.835E-189 | -0.5876344 | 0.003 | 0.138 | 1.378E-184 | MO_infl | TCIM |
| 6.373E-189 | 0.31552554 | 0.746 | 0.583 | 1.505E-184 | MO_infl | SUB1 |
| 4.316E-188 | -0.3590543 | 0.036 | 0.19 | 1.02E-183 | MO_infl | JMY |
| 7.717E-188 | -0.5461041 | 0.029 | 0.178 | 1.823E-183 | MO_infl | BPGM |
| 1.205E-186 | -0.6047723 | 0.94 | 0.946 | 2.847E-182 | MO_infl | RPL13A |
| 5.163E-186 | -0.3404669 | 0.051 | 0.21 | 1.22E-181 | MO_infl | MPRIP |
| 5.185E-186 | 0.32643311 | 0.696 | 0.519 | 1.225E-181 | MO_infl | POMP |
| 8.735E-186 | 0.29495162 | 0.494 | 0.305 | 2.063E-181 | MO_infl | SH3GLB1 |
| 9.989E-186 | 0.31772048 | 0.912 | 0.8 | 2.36E-181 | MO_infl | YBX1 |
| 1.539E-185 | -0.2793801 | 0.002 | 0.134 | 3.636E-181 | MO_infl | VASN |
| 8.139E-185 | 0.25025243 | 0.388 | 0.21 | 1.923E-180 | MO_infl | RBMS1 |
| 9E-185 | 0.28335673 | 0.54 | 0.34 | 2.126E-180 | MO_infl | RAB10 |
| 1.624E-184 | -0.3665654 | 0.075 | 0.241 | 3.837E-180 | MO_infl | SPATS2L |
| 4.747E-184 | -0.2721762 | 0.002 | 0.133 | 1.121E-179 | MO_infl | DSG2 |
| 7.589E-184 | -0.2921259 | 0.004 | 0.137 | 1.793E-179 | MO_infl | MUC20 |
| 3.852E-183 | -0.2603232 | 0.003 | 0.135 | 9.099E-179 | MO_infl | TRNP1 |
| 1.167E-182 | -1.3959701 | 0.001 | 0.13 | 2.757E-178 | MO_infl | SPRR1B |
| 2.35E-182 | -0.5136122 | 0.001 | 0.13 | 5.552E-178 | MO_infl | TMPRSS11A |
| 2.807E-182 | -0.3918904 | 0.093 | 0.26 | 6.63E-178 | MO_infl | PCBD1 |
| 3.014E-182 | -0.2556892 | 0.002 | 0.132 | 7.121E-178 | MO_infl | MYH14 |
| 6.053E-182 | -0.3286669 | 0.032 | 0.182 | 1.43E-177 | MO_infl | MET |
| 2.214E-181 | -0.329052 | 0 | 0.128 | 5.231E-177 | MO_infl | FUT3 |
| 3.735E-181 | -0.6607181 | 0.001 | 0.129 | 8.823E-177 | MO_infl | KRT78 |
| 4.534E-181 | -0.4180085 | 0.005 | 0.137 | 1.071E-176 | MO_infl | EGLN3 |
| 7.938E-181 | 0.60291074 | 0.725 | 0.616 | 1.875E-176 | MO_infl | ZFP36L1 |
| 2.31E-179 | 0.30803163 | 0.538 | 0.341 | 5.458E-175 | MO_infl | SPAG9 |
| 9.572E-179 | 0.25558976 | 0.482 | 0.29 | 2.261E-174 | MO_infl | CLTC |
| 1.136E-178 | -0.2556728 | 0.002 | 0.13 | 2.682E-174 | MO_infl | WWC1 |
| 1.916E-178 | -0.6128382 | 0.822 | 0.85 | 4.527E-174 | MO_infl | RPL35 |
| 2.146E-178 | -0.5332569 | 0.509 | 0.648 | 5.069E-174 | MO_infl | TAX1BP1 |
| 1.452E-177 | -0.3048346 | 0.011 | 0.144 | 3.43E-173 | MO_infl | MAST4 |
| 2.158E-177 | 0.256658 | 0.361 | 0.195 | 5.099E-173 | MO_infl | RIT1 |
| 2.918E-177 | -0.4049525 | 0.137 | 0.309 | 6.894E-173 | MO_infl | LRP10 |
| 4.111E-177 | -0.2689362 | 0.001 | 0.126 | 9.711E-173 | MO_infl | PLS1 |
| 4.476E-176 | -0.287026 | 0.003 | 0.129 | 1.057E-171 | MO_infl | DENND2C |
| 6.009E-176 | 0.2565559 | 0.367 | 0.199 | 1.419E-171 | MO_infl | MAP4K4 |
| 2.009E-175 | -0.4758063 | 0.147 | 0.317 | 4.745E-171 | MO_infl | CHCHD10 |
| 5.8E-175 | -0.3982154 | 0.002 | 0.127 | 1.37E-170 | MO_infl | SMIM5 |
| 2.218E-174 | -0.4844658 | 0.299 | 0.471 | 5.24E-170 | MO_infl | ACTN4 |
| 1.057E-173 | 0.30339926 | 0.74 | 0.596 | 2.498E-169 | MO_infl | CAP1 |
| 7.023E-173 | -0.4077343 | 0.005 | 0.131 | 1.659E-168 | MO_infl | CEACAM1 |
| 1.883E-172 | -0.2854342 | 0.003 | 0.128 | 4.447E-168 | MO_infl | GRHL1 |
| 5.173E-172 | -0.4136629 | 0.16 | 0.329 | 1.222E-167 | MO_infl | CLTB |

| | | | | | | |
|---|---|---|---|---|---|---|
| 6.948E-172 | -0.2856605 | 0.002 | 0.125 | 1.641E-167 | MO_infl | GATA3 |
| 7.035E-172 | 0.29954798 | 0.658 | 0.476 | 1.662E-167 | MO_infl | ASAH1 |
| 7.461E-172 | 0.2643191 | 0.589 | 0.38 | 1.763E-167 | MO_infl | PTP4A2 |
| 7.715E-172 | -0.4680307 | 0.001 | 0.123 | 1.823E-167 | MO_infl | TMC5 |
| 1.158E-171 | -0.3153411 | 0.073 | 0.232 | 2.735E-167 | MO_infl | NBDY |
| 1.041E-169 | 0.33501798 | 0.343 | 0.182 | 2.46E-165 | MO_infl | CTSC |
| 1.181E-169 | -0.3668919 | 0.001 | 0.122 | 2.791E-165 | MO_infl | A2ML1 |
| 4.246E-169 | -0.4615926 | 0.167 | 0.338 | 1.003E-164 | MO_infl | AKAP9 |
| 4.547E-169 | -0.4676632 | 0.043 | 0.189 | 1.074E-164 | MO_infl | AKR1B1 |
| 5.605E-169 | 0.25708968 | 0.415 | 0.241 | 1.324E-164 | MO_infl | KRAS |
| 4.887E-167 | -0.3001207 | 0.038 | 0.18 | 1.154E-162 | MO_infl | FARP1 |
| 9.65E-167 | 0.2930174 | 0.119 | 0.037 | 2.28E-162 | MO_infl | NT5E |
| 1.832E-166 | -0.2566197 | 0.003 | 0.124 | 4.328E-162 | MO_infl | WEE1 |
| 2.202E-166 | 0.26780557 | 0.424 | 0.252 | 5.202E-162 | MO_infl | PDXK |
| 2.228E-166 | -0.3949632 | 0.005 | 0.127 | 5.262E-162 | MO_infl | FXYD2 |
| 4.004E-166 | 0.68010007 | 0.69 | 0.602 | 9.458E-162 | MO_infl | C15orf48 |
| 2.908E-165 | 0.28958607 | 0.27 | 0.134 | 6.869E-161 | MO_infl | VSIR |
| 3.511E-165 | -1.1026035 | 0.023 | 0.155 | 8.294E-161 | MO_infl | DEFB1 |
| 5.119E-165 | -0.6411084 | 0.795 | 0.821 | 1.209E-160 | MO_infl | RPL5 |
| 1.205E-164 | 0.2918687 | 0.45 | 0.274 | 2.847E-160 | MO_infl | ARL6IP5 |
| 2.626E-164 | -0.2776568 | 0.034 | 0.172 | 6.204E-160 | MO_infl | RCN1 |
| 2.708E-164 | -0.4316742 | 0.004 | 0.124 | 6.398E-160 | MO_infl | KRT23 |
| 5.456E-164 | 0.29092971 | 0.36 | 0.203 | 1.289E-159 | MO_infl | ARFGAP3 |
| 8.343E-164 | -0.5363018 | 0.356 | 0.514 | 1.971E-159 | MO_infl | H2AFJ |
| 1.651E-163 | -0.4345737 | 0.002 | 0.119 | 3.9E-159 | MO_infl | CLIC3 |
| 3.207E-163 | -0.3114826 | 0.052 | 0.197 | 7.576E-159 | MO_infl | PIK3IP1 |
| 1.038E-162 | -1.6655989 | 0.004 | 0.122 | 2.452E-158 | MO_infl | CRCT1 |
| 2.524E-162 | -0.2662978 | 0.029 | 0.165 | 5.962E-158 | MO_infl | NR1D2 |
| 3.735E-162 | -0.3821725 | 0.005 | 0.125 | 8.823E-158 | MO_infl | NR1D1 |
| 7.698E-161 | 0.29295909 | 0.41 | 0.235 | 1.819E-156 | MO_infl | NABP1 |
| 3.261E-160 | -0.260467 | 0.001 | 0.115 | 7.703E-156 | MO_infl | RNF223 |
| 2.108E-159 | -0.2711098 | 0.003 | 0.12 | 4.98E-155 | MO_infl | DPP4 |
| 2.396E-158 | 0.31889793 | 0.314 | 0.171 | 5.66E-154 | MO_infl | GBP2 |
| 9.504E-158 | -0.2575237 | 0.001 | 0.114 | 2.245E-153 | MO_infl | LIPH |
| 1.241E-157 | -0.8544638 | 0.004 | 0.12 | 2.932E-153 | MO_infl | ZG16B |
| 1.964E-157 | -0.3201421 | 0 | 0.113 | 4.639E-153 | MO_infl | RNF39 |
| 6.142E-157 | -0.3404415 | 0.008 | 0.126 | 1.451E-152 | MO_infl | STS |
| 2.901E-156 | -0.4984867 | 0.934 | 0.952 | 6.853E-152 | MO_infl | RPL18 |
| 6.559E-156 | -0.3753719 | 0.934 | 0.966 | 1.549E-151 | MO_infl | HSP90AA1 |
| 6.807E-156 | -0.5610801 | 0.673 | 0.758 | 1.608E-151 | MO_infl | EEF1B2 |
| 1.001E-155 | -0.6426397 | 0.909 | 0.877 | 2.364E-151 | MO_infl | RPL12 |
| 1.187E-155 | 0.31417219 | 0.401 | 0.24 | 2.803E-151 | MO_infl | WTAP |
| 4.726E-155 | -0.5240685 | 0.135 | 0.29 | 1.116E-150 | MO_infl | CCNG2 |
| 5.192E-155 | -0.2985255 | 0.027 | 0.155 | 1.226E-150 | MO_infl | ZNRF1 |
| 1.003E-154 | -0.2711368 | 0.029 | 0.159 | 2.369E-150 | MO_infl | SERPING1 |
| 1.772E-153 | -0.2720693 | 0.061 | 0.205 | 4.186E-149 | MO_infl | MZT2A |
| 3.953E-153 | -0.256341 | 0 | 0.11 | 9.337E-149 | MO_infl | C4orf19 |
| 8.176E-153 | 0.25582272 | 0.525 | 0.349 | 1.931E-148 | MO_infl | GNB2 |
| 8.98E-153 | -0.2930368 | 0.035 | 0.166 | 2.121E-148 | MO_infl | RALGPS2 |
| 1.153E-152 | -0.3180795 | 0.071 | 0.215 | 2.723E-148 | MO_infl | TSPAN3 |

| | | | | | | |
|---|---|---|---|---|---|---|
| 1.275E-152 | -1.5417332 | 0.008 | 0.123 | 3.011E-148 | MO_infl | CNFN |
| 3.608E-152 | -0.5523871 | 0.109 | 0.264 | 8.523E-148 | MO_infl | ITGB8 |
| 4.612E-152 | -0.5115853 | 0.068 | 0.21 | 1.089E-147 | MO_infl | PLK2 |
| 5.785E-152 | -0.2879925 | 0.037 | 0.168 | 1.367E-147 | MO_infl | ZNF83 |
| 1.023E-151 | -0.7144118 | 0.866 | 0.846 | 2.417E-147 | MO_infl | RPS4X |
| 1.039E-151 | -0.2810813 | 0 | 0.109 | 2.455E-147 | MO_infl | B3GALT5 |
| 1.479E-151 | 0.41042957 | 0.23 | 0.111 | 3.493E-147 | MO_infl | PLBD1 |
| 1.671E-150 | 0.25210482 | 0.484 | 0.3 | 3.947E-146 | MO_infl | FOSL2 |
| 2.106E-150 | -0.3739158 | 0.002 | 0.111 | 4.974E-146 | MO_infl | KLK10 |
| 4.004E-149 | -0.3136593 | 0.015 | 0.132 | 9.458E-145 | MO_infl | ST3GAL4 |
| 6.51E-149 | 0.38644946 | 0.696 | 0.556 | 1.538E-144 | MO_infl | CD55 |
| 6.525E-149 | -0.9318511 | 0.067 | 0.207 | 1.541E-144 | MO_infl | RARRES3 |
| 9.536E-149 | -0.2944215 | 0.057 | 0.195 | 2.253E-144 | MO_infl | TMEM159 |
| 2.818E-148 | -0.5427541 | 0.004 | 0.113 | 6.657E-144 | MO_infl | DHRS2 |
| 5.234E-148 | -0.3203371 | 0.067 | 0.207 | 1.237E-143 | MO_infl | VCL |
| 1.727E-147 | 0.25176746 | 0.885 | 0.798 | 4.079E-143 | MO_infl | RAC1 |
| 1.031E-146 | -0.4475447 | 0.117 | 0.265 | 2.435E-142 | MO_infl | PTMS |
| 1.167E-146 | -0.491655 | 0.001 | 0.107 | 2.757E-142 | MO_infl | RNASE7 |
| 3.294E-146 | -0.2548066 | 0.05 | 0.184 | 7.781E-142 | MO_infl | SUN1 |
| 5.393E-146 | -0.6165387 | 0.917 | 0.903 | 1.274E-141 | MO_infl | RPL7A |
| 1.385E-145 | -0.3283166 | 0.037 | 0.164 | 3.271E-141 | MO_infl | GSTM3 |
| 2.419E-145 | 0.29319044 | 0.986 | 0.932 | 5.714E-141 | MO_infl | SAT1 |
| 2.504E-145 | 0.36391162 | 0.709 | 0.567 | 5.915E-141 | MO_infl | KLF6 |
| 3.613E-145 | -0.2503008 | 0.025 | 0.147 | 8.536E-141 | MO_infl | PPFIBP1 |
| 8.063E-145 | -0.2872416 | 0.001 | 0.105 | 1.905E-140 | MO_infl | KLK11 |
| 2.036E-144 | 0.25069675 | 0.808 | 0.709 | 4.811E-140 | MO_infl | YWHAB |
| 2.935E-144 | -0.4801003 | 0.051 | 0.182 | 6.932E-140 | MO_infl | ANKRD37 |
| 2.257E-143 | -0.2620813 | 0.044 | 0.173 | 5.331E-139 | MO_infl | OFD1 |
| 4.513E-143 | 0.36781957 | 0.321 | 0.179 | 1.066E-138 | MO_infl | DUSP4 |
| 4.758E-143 | -0.3275805 | 0.002 | 0.107 | 1.124E-138 | MO_infl | CTGF |
| 1.993E-142 | -0.4904455 | 0.448 | 0.589 | 4.708E-138 | MO_infl | ZFAND5 |
| 5.344E-142 | -0.4797447 | 0.235 | 0.387 | 1.262E-137 | MO_infl | RALBP1 |
| 5.663E-142 | -0.3042828 | 0.071 | 0.21 | 1.338E-137 | MO_infl | F11R |
| 8.367E-142 | -0.2956381 | 0.075 | 0.215 | 1.977E-137 | MO_infl | USP53 |
| 3.489E-141 | -0.324995 | 0 | 0.102 | 8.242E-137 | MO_infl | SLC6A14 |
| 1.704E-140 | -0.331056 | 0.055 | 0.186 | 4.025E-136 | MO_infl | CDKN2AIP |
| 1.979E-140 | 0.326527 | 0.784 | 0.681 | 4.676E-136 | MO_infl | LRRFIP1 |
| 2.635E-140 | -0.5035176 | 0.219 | 0.365 | 6.224E-136 | MO_infl | GOLGB1 |
| 3.45E-139 | -0.2583135 | 0.003 | 0.107 | 8.15E-135 | MO_infl | BCAM |
| 4.545E-139 | -0.5617016 | 0.003 | 0.106 | 1.074E-134 | MO_infl | SAA1 |
| 6.901E-139 | -0.3045798 | 0.004 | 0.107 | 1.63E-134 | MO_infl | ADAMTS9 |
| 1.134E-138 | -0.2807198 | 0.009 | 0.117 | 2.678E-134 | MO_infl | DUSP14 |
| 1.632E-138 | -0.2858291 | 0.033 | 0.155 | 3.855E-134 | MO_infl | MGLL |
| 2.536E-138 | -0.331102 | 0.001 | 0.101 | 5.991E-134 | MO_infl | FAM3D |
| 5.711E-138 | 0.25424622 | 0.156 | 0.063 | 1.349E-133 | MO_infl | FCGR2B |
| 6.597E-137 | -0.3936997 | 0.789 | 0.876 | 1.558E-132 | MO_infl | SRP14 |
| 2.246E-136 | -0.3830002 | 0.128 | 0.273 | 5.306E-132 | MO_infl | ODC1 |
| 6.697E-136 | -0.9395238 | 0.001 | 0.1 | 1.582E-131 | MO_infl | SPRR2D |
| 3.292E-135 | 0.4082967 | 0.296 | 0.162 | 7.777E-131 | MO_infl | WARS |
| 3.522E-135 | -0.2976948 | 0.032 | 0.151 | 8.321E-131 | MO_infl | GCLC |

| | | | | | | |
|---|---|---|---|---|---|---|
| 3.931E-135 | -0.3400205 | 0.026 | 0.141 | 9.287E-131 | MO_infl | POLB |
| 1.645E-133 | 0.2513144 | 0.165 | 0.071 | 3.887E-129 | MO_infl | RIN2 |
| 2.333E-133 | -0.4854323 | 0.356 | 0.505 | 5.512E-129 | MO_infl | HMGA1 |
| 6.431E-132 | 0.25331046 | 0.792 | 0.672 | 1.519E-127 | MO_infl | PSMA7 |
| 8.249E-131 | -0.2763324 | 0.062 | 0.19 | 1.949E-126 | MO_infl | GNL3 |
| 4.398E-130 | -0.5609818 | 0.883 | 0.876 | 1.039E-125 | MO_infl | RPL23A |
| 1.536E-129 | -0.420439 | 0.09 | 0.222 | 3.629E-125 | MO_infl | CSNK1E |
| 1.855E-129 | -0.3322914 | 0.123 | 0.264 | 4.381E-125 | MO_infl | SPTAN1 |
| 3.897E-128 | -0.461824 | 0.212 | 0.351 | 9.206E-124 | MO_infl | SNHG8 |
| 4.366E-128 | -0.3795791 | 0.215 | 0.363 | 1.031E-123 | MO_infl | ATP5IF1 |
| 9.404E-127 | -0.2942255 | 0.038 | 0.154 | 2.222E-122 | MO_infl | ETS1 |
| 9.785E-127 | -0.2809648 | 0.04 | 0.157 | 2.311E-122 | MO_infl | MSMO1 |
| 3.947E-126 | -0.4375799 | 0.03 | 0.14 | 9.324E-122 | MO_infl | SRD5A3 |
| 4.085E-126 | 0.35399796 | 0.487 | 0.336 | 9.651E-122 | MO_infl | TKT |
| 7.276E-126 | -0.3545566 | 0.607 | 0.731 | 1.719E-121 | MO_infl | SKP1 |
| 1.415E-125 | 0.65687113 | 0.527 | 0.368 | 3.343E-121 | MO_infl | FOS |
| 3.264E-125 | 0.74339682 | 0.55 | 0.434 | 7.709E-121 | MO_infl | MARCKS |
| 8.65E-125 | -0.6704958 | 0.035 | 0.146 | 2.043E-120 | MO_infl | DHRS9 |
| 1.993E-124 | 0.29052316 | 0.402 | 0.255 | 4.709E-120 | MO_infl | DDX21 |
| 2.152E-123 | -0.5985856 | 0.152 | 0.284 | 5.084E-119 | MO_infl | DDIT3 |
| 2.544E-123 | -0.3559348 | 0.163 | 0.303 | 6.011E-119 | MO_infl | PDCD6 |
| 2.859E-123 | 0.2687254 | 0.501 | 0.34 | 6.755E-119 | MO_infl | NFE2L2 |
| 1.999E-122 | -0.7442111 | 0.957 | 0.929 | 4.723E-118 | MO_infl | RPS12 |
| 2.291E-122 | -0.262257 | 0.12 | 0.26 | 5.413E-118 | MO_infl | POLR2K |
| 2.222E-121 | -0.2704693 | 0.028 | 0.136 | 5.249E-117 | MO_infl | FOXQ1 |
| 2.688E-121 | -0.5765248 | 0.215 | 0.347 | 6.349E-117 | MO_infl | HERPUD1 |
| 1.032E-120 | -1.1202823 | 0.036 | 0.146 | 2.439E-116 | MO_infl | APOE |
| 1.582E-120 | -0.2564219 | 0.062 | 0.182 | 3.737E-116 | MO_infl | CCDC50 |
| 4.166E-120 | -0.4592773 | 0.198 | 0.33 | 9.84E-116 | MO_infl | SDC4 |
| 2.127E-117 | -0.2617023 | 0.031 | 0.136 | 5.024E-113 | MO_infl | TCP11L2 |
| 2.784E-117 | -0.3043453 | 0.109 | 0.239 | 6.577E-113 | MO_infl | ECHS1 |
| 1.18E-116 | -0.2878926 | 0.057 | 0.172 | 2.788E-112 | MO_infl | TRIP10 |
| 1.826E-116 | -0.3218437 | 0.107 | 0.234 | 4.314E-112 | MO_infl | CDK2AP2 |
| 1.958E-116 | -0.2563624 | 0.092 | 0.219 | 4.624E-112 | MO_infl | PHB |
| 4.421E-116 | -0.6461786 | 0.997 | 0.978 | 1.044E-111 | MO_infl | MALAT1 |
| 1.335E-115 | -0.4675572 | 0.104 | 0.23 | 3.154E-111 | MO_infl | LAMB3 |
| 3.331E-115 | -0.5057278 | 0.14 | 0.268 | 7.868E-111 | MO_infl | SNHG12 |
| 3.219E-114 | -0.3476804 | 0.472 | 0.595 | 7.605E-110 | MO_infl | UBE2B |
| 7.069E-114 | -0.6543407 | 0.946 | 0.875 | 1.67E-109 | MO_infl | RPS2 |
| 7.142E-114 | -0.5516601 | 0.029 | 0.13 | 1.687E-109 | MO_infl | HSPA2 |
| 1.17E-113 | -0.6561832 | 0.947 | 0.927 | 2.763E-109 | MO_infl | MT-CYB |
| 2.803E-111 | 0.41682179 | 0.83 | 0.766 | 6.622E-107 | MO_infl | CTSD |
| 2.779E-109 | -0.3587342 | 0.139 | 0.265 | 6.565E-105 | MO_infl | PKN2 |
| 1.591E-107 | -0.3264719 | 0.227 | 0.359 | 3.758E-103 | MO_infl | NDFIP1 |
| 3.065E-107 | 0.28034596 | 0.633 | 0.503 | 7.242E-103 | MO_infl | ATP6V1F |
| 3.378E-107 | -0.4238165 | 0.05 | 0.155 | 7.979E-103 | MO_infl | ID1 |
| 4.734E-107 | -0.3478974 | 0.206 | 0.334 | 1.118E-102 | MO_infl | RAB11A |
| 6.117E-107 | -0.4725292 | 0.097 | 0.211 | 1.445E-102 | MO_infl | SAMD9 |
| 1.391E-106 | -0.2620661 | 0.052 | 0.16 | 3.287E-102 | MO_infl | NCOA7 |
| 1.576E-105 | -0.4506519 | 0.947 | 0.959 | 3.722E-101 | MO_infl | RPL15 |

| | | | | | | |
|---|---|---|---|---|---|---|
| 5.23E-104 | -0.267871 | 0.128 | 0.256 | 1.236E-99 | MO_infl | C12orf57 |
| 2.518E-103 | -0.534014 | 0.901 | 0.872 | 5.947E-99 | MO_infl | RPL36 |
| 1.773E-102 | -0.3104016 | 0.554 | 0.692 | 4.1875E-98 | MO_infl | CALR |
| 3.712E-102 | -0.4151058 | 0.075 | 0.185 | 8.7685E-98 | MO_infl | PRDM1 |
| 5.783E-102 | -0.2834659 | 0.13 | 0.256 | 1.3661E-97 | MO_infl | FKBP4 |
| 3.035E-100 | -0.3078612 | 0.184 | 0.314 | 7.1684E-96 | MO_infl | EIF5B |
| 4.344E-99 | -0.7034805 | 0.811 | 0.825 | 1.0262E-94 | MO_infl | MT-ND5 |
| 4.385E-99 | -0.332588 | 0.201 | 0.322 | 1.0359E-94 | MO_infl | NOP53 |
| 5.85E-99 | -0.6307867 | 0.318 | 0.451 | 1.382E-94 | MO_infl | HIST1H1C |
| 8.432E-99 | 0.3324756 | 0.384 | 0.248 | 1.9918E-94 | MO_infl | ID2 |
| 2.5654E-98 | -0.5083221 | 0.25 | 0.367 | 6.0602E-94 | MO_infl | PHLDA2 |
| 3.9886E-98 | -0.4991452 | 0.239 | 0.364 | 9.4223E-94 | MO_infl | ZFP36L2 |
| 6.1519E-98 | -0.2953512 | 0.049 | 0.147 | 1.4533E-93 | MO_infl | ACADM |
| 1.9483E-97 | -0.3082688 | 0.175 | 0.299 | 4.6024E-93 | MO_infl | ISCU |
| 3.6752E-97 | -0.547683 | 0.941 | 0.957 | 8.6818E-93 | MO_infl | ACTG1 |
| 9.7208E-97 | -0.3646912 | 0.304 | 0.476 | 2.2963E-92 | MO_infl | SPP1 |
| 1.1011E-96 | -0.250509 | 0.096 | 0.209 | 2.601E-92 | MO_infl | SEC11C |
| 5.0045E-96 | -0.305452 | 0.213 | 0.337 | 1.1822E-91 | MO_infl | ERP29 |
| 9.189E-96 | -0.2740421 | 0.083 | 0.191 | 2.1707E-91 | MO_infl | RERE |
| 1.6153E-95 | -1.4990833 | 0.018 | 0.101 | 3.8159E-91 | MO_infl | HBB |
| 3.8131E-95 | -0.3705313 | 0.559 | 0.671 | 9.0076E-91 | MO_infl | ZFAS1 |
| 3.869E-95 | -0.3433796 | 0.097 | 0.208 | 9.1396E-91 | MO_infl | KATNBL1 |
| 3.7122E-94 | -0.5968858 | 0.916 | 0.883 | 8.7693E-90 | MO_infl | RPS3 |
| 5.0079E-94 | -0.5267595 | 0.537 | 0.662 | 1.183E-89 | MO_infl | HSPA5 |
| 2.0641E-93 | -0.3528797 | 0.074 | 0.18 | 4.876E-89 | MO_infl | HIST3H2A |
| 7.6292E-92 | -0.4474087 | 0.853 | 0.914 | 1.8022E-87 | MO_infl | ITM2B |
| 8.5442E-91 | -0.3773267 | 0.702 | 0.772 | 2.0184E-86 | MO_infl | COX7C |
| 1.763E-90 | -0.2506974 | 0.048 | 0.143 | 4.1648E-86 | MO_infl | ID3 |
| 8.2647E-90 | -0.2867537 | 0.636 | 0.749 | 1.9524E-85 | MO_infl | GNAS |
| 3.2286E-89 | -0.5006266 | 0.898 | 0.872 | 7.627E-85 | MO_infl | RPL26 |
| 7.2588E-88 | -0.4196737 | 0.235 | 0.346 | 1.7147E-83 | MO_infl | PPP1R14B |
| 1.5297E-87 | -0.3189656 | 0.406 | 0.517 | 3.6136E-83 | MO_infl | LAPTM4A |
| 9.8062E-87 | -0.2729425 | 0.035 | 0.12 | 2.3165E-82 | MO_infl | YOD1 |
| 3.1222E-86 | 0.35043625 | 0.143 | 0.069 | 7.3755E-82 | MO_infl | MRC1 |
| 3.5555E-86 | -0.2637437 | 0.062 | 0.157 | 8.3992E-82 | MO_infl | PDP1 |
| 4.9283E-86 | -0.5683996 | 0.132 | 0.245 | 1.1642E-81 | MO_infl | ISG15 |
| 2.1946E-85 | -0.3747453 | 0.184 | 0.291 | 5.1844E-81 | MO_infl | SLK |
| 3.971E-85 | -0.3150249 | 0.215 | 0.326 | 9.3807E-81 | MO_infl | MZT2B |
| 4.3723E-85 | -0.5648318 | 0.266 | 0.363 | 1.0329E-80 | MO_infl | GLS |
| 4.5129E-85 | -0.7118171 | 0.032 | 0.115 | 1.0661E-80 | MO_infl | CCL5 |
| 9.5441E-85 | -0.7127847 | 0.19 | 0.295 | 2.2546E-80 | MO_infl | CLU |
| 1.1148E-83 | -0.3065172 | 0.219 | 0.337 | 2.6334E-79 | MO_infl | YPEL3 |
| 1.755E-83 | -0.4307273 | 0.165 | 0.273 | 4.1457E-79 | MO_infl | IDI1 |
| 2.4288E-83 | -0.3756439 | 0.292 | 0.168 | 5.7376E-79 | MO_infl | HLA-DPB1 |
| 2.7219E-83 | 0.29435503 | 0.383 | 0.256 | 6.4299E-79 | MO_infl | TNFAIP2 |
| 3.661E-83 | -0.346366 | 0.172 | 0.279 | 8.6485E-79 | MO_infl | HECTD1 |
| 9.7408E-83 | -0.2604133 | 0.152 | 0.263 | 2.3011E-78 | MO_infl | CCDC186 |
| 1.9687E-82 | -0.4830207 | 0.955 | 0.918 | 4.6507E-78 | MO_infl | RPL8 |
| 4.529E-82 | -0.4661989 | 0.982 | 0.961 | 1.0699E-77 | MO_infl | RPL13 |
| 1.5656E-81 | -0.5184223 | 0.131 | 0.236 | 3.6984E-77 | MO_infl | HES1 |

| | | | | | | |
|---|---|---|---|---|---|---|
| 1.7085E-81 | -0.2927658 | 0.05 | 0.138 | 4.0359E-77 | MO_infl | SYNE1 |
| 4.3732E-81 | -0.3874406 | 0.263 | 0.366 | 1.0331E-76 | MO_infl | TAF1D |
| 3.2089E-78 | -0.4100872 | 0.035 | 0.115 | 7.5804E-74 | MO_infl | LTB |
| 4.722E-78 | -0.6440998 | 0.039 | 0.119 | 1.1155E-73 | MO_infl | APOC1 |
| 5.4062E-78 | -0.463243 | 0.929 | 0.923 | 1.2771E-73 | MO_infl | RPL18A |
| 6.5002E-78 | -0.2605855 | 0.168 | 0.275 | 1.5355E-73 | MO_infl | CUTA |
| 2.551E-77 | -0.3356376 | 0.169 | 0.276 | 6.0262E-73 | MO_infl | RHOC |
| 1.6147E-76 | -0.5166055 | 0.936 | 0.891 | 3.8145E-72 | MO_infl | RPS3A |
| 2.1967E-76 | 0.28121432 | 0.494 | 0.372 | 5.1893E-72 | MO_infl | AHNAK |
| 5.6369E-76 | -0.3527515 | 0.732 | 0.804 | 1.3316E-71 | MO_infl | CHCHD2 |
| 1.0199E-75 | -0.4180509 | 0.242 | 0.34 | 2.4092E-71 | MO_infl | MAFF |
| 3.558E-75 | -0.419457 | 0.141 | 0.239 | 8.4051E-71 | MO_infl | IER5L |
| 5.3694E-75 | -0.267162 | 0.154 | 0.257 | 1.2684E-70 | MO_infl | CFDP1 |
| 2.2655E-74 | -0.2638086 | 0.178 | 0.282 | 5.3518E-70 | MO_infl | BAG1 |
| 2.3589E-74 | -0.3772448 | 0.787 | 0.842 | 5.5723E-70 | MO_infl | PPIA |
| 5.6534E-74 | -0.2692477 | 0.15 | 0.25 | 1.3355E-69 | MO_infl | PNPLA2 |
| 1.1174E-73 | 0.62424723 | 0.197 | 0.117 | 2.6397E-69 | MO_infl | GBP1 |
| 8.7186E-73 | -0.2790599 | 0.202 | 0.303 | 2.0596E-68 | MO_infl | CHMP2B |
| 1.3782E-71 | -0.3777988 | 0.073 | 0.16 | 3.2558E-67 | MO_infl | RASD1 |
| 2.9682E-70 | -0.4014993 | 0.599 | 0.707 | 7.0117E-66 | MO_infl | HSP90B1 |
| 3.2545E-70 | -0.3168794 | 0.897 | 0.923 | 7.6881E-66 | MO_infl | NACA |
| 4.6378E-70 | -0.2655942 | 0.037 | 0.111 | 1.0956E-65 | MO_infl | LRMP |
| 3.2316E-69 | -0.277192 | 0.265 | 0.366 | 7.634E-65 | MO_infl | SPCS1 |
| 1.4727E-68 | -0.3496201 | 0.189 | 0.295 | 3.4789E-64 | MO_infl | H1F0 |
| 4.7029E-68 | 0.2811845 | 0.422 | 0.307 | 1.111E-63 | MO_infl | IFITM3 |
| 1.3328E-66 | -0.8451109 | 0.404 | 0.484 | 3.1485E-62 | MO_infl | ERO1A |
| 4.7555E-66 | -0.3040059 | 0.969 | 0.965 | 1.1234E-61 | MO_infl | UBC |
| 8.2411E-66 | -0.253825 | 0.203 | 0.301 | 1.9468E-61 | MO_infl | NDUFC1 |
| 1.4872E-65 | -0.3046797 | 0.109 | 0.197 | 3.5133E-61 | MO_infl | SEPHS2 |
| 6.2833E-65 | -0.3211729 | 0.278 | 0.366 | 1.4843E-60 | MO_infl | MRPL33 |
| 9.2153E-65 | -0.2628341 | 0.511 | 0.594 | 2.1769E-60 | MO_infl | CLTA |
| 9.4011E-65 | -0.3291523 | 0.194 | 0.292 | 2.2208E-60 | MO_infl | ABHD5 |
| 9.8107E-65 | -0.3491764 | 0.248 | 0.337 | 2.3176E-60 | MO_infl | TMBIM1 |
| 1.5239E-64 | -1.5492649 | 0.08 | 0.163 | 3.6E-60 | MO_infl | MT1H |
| 3.2872E-63 | -0.4631944 | 0.285 | 0.371 | 7.7654E-59 | MO_infl | TUBA1B |
| 1.5087E-62 | -0.2804311 | 0.207 | 0.299 | 3.564E-58 | MO_infl | CLINT1 |
| 1.1161E-61 | 0.48295863 | 0.216 | 0.141 | 2.6366E-57 | MO_infl | SLC7A11 |
| 2.6723E-61 | -0.2587356 | 0.408 | 0.499 | 6.3128E-57 | MO_infl | DYNLT1 |
| 2.9184E-61 | -0.2939352 | 0.247 | 0.336 | 6.8941E-57 | MO_infl | YPEL5 |
| 2.9237E-61 | -0.2628426 | 0.571 | 0.654 | 6.9067E-57 | MO_infl | EDF1 |
| 6.4503E-61 | -0.2946292 | 0.159 | 0.246 | 1.5237E-56 | MO_infl | SNHG7 |
| 1.2228E-60 | -0.3034322 | 0.556 | 0.618 | 2.8886E-56 | MO_infl | HNRNPA1 |
| 1.4434E-60 | -0.3168948 | 0.222 | 0.311 | 3.4097E-56 | MO_infl | NUCKS1 |
| 4.5045E-59 | -0.2517725 | 0.041 | 0.108 | 1.0641E-54 | MO_infl | LY6K |
| 7.6337E-59 | -0.2829086 | 0.191 | 0.279 | 1.8033E-54 | MO_infl | HSPA9 |
| 4.0386E-58 | -0.2984349 | 0.417 | 0.492 | 9.5403E-54 | MO_infl | GABARAPL2 |
| 1.0849E-57 | -0.2939966 | 0.652 | 0.712 | 2.5628E-53 | MO_infl | RPL4 |
| 2.4538E-56 | 0.8980977 | 0.201 | 0.129 | 5.7967E-52 | MO_infl | CXCL1 |
| 5.6726E-56 | -0.4959821 | 0.963 | 0.918 | 1.34E-51 | MO_infl | RPS27A |
| 8.8673E-56 | -0.3947042 | 0.648 | 0.733 | 2.0947E-51 | MO_infl | JUND |

| | | | | | | |
|---|---|---|---|---|---|---|
| 5.4232E-55 | -0.306998 | 0.554 | 0.625 | 1.2811E-50 | MO_infl | KTN1 |
| 8.3024E-55 | -0.2536666 | 0.194 | 0.279 | 1.9613E-50 | MO_infl | TOR1AIP2 |
| 7.9349E-54 | -0.2569268 | 0.662 | 0.735 | 1.8745E-49 | MO_infl | COX6A1 |
| 1.241E-53 | -0.4224109 | 0.962 | 0.916 | 2.9317E-49 | MO_infl | RPL11 |
| 1.4504E-53 | -0.4872038 | 0.957 | 0.934 | 3.4264E-49 | MO_infl | MT-ATP6 |
| 9.7828E-53 | -0.4891518 | 0.941 | 0.881 | 2.311E-48 | MO_infl | RPS23 |
| 1.57E-52 | -0.3437269 | 0.923 | 0.898 | 3.7089E-48 | MO_infl | RPL24 |
| 2.2502E-52 | -0.2946893 | 0.104 | 0.182 | 5.3157E-48 | MO_infl | DNAJA4 |
| 4.0436E-52 | -0.2788014 | 0.165 | 0.248 | 9.5523E-48 | MO_infl | ARID5B |
| 5.2445E-52 | -0.2636514 | 0.301 | 0.382 | 1.2389E-47 | MO_infl | RHEB |
| 2.1657E-51 | -0.2677768 | 0.497 | 0.571 | 5.116E-47 | MO_infl | SET |
| 3.1646E-51 | -0.2696539 | 0.727 | 0.795 | 7.4757E-47 | MO_infl | BTF3 |
| 4.7822E-51 | -0.39872 | 0.927 | 0.893 | 1.1297E-46 | MO_infl | RPL29 |
| 1.1542E-50 | -0.3219817 | 0.295 | 0.369 | 2.7267E-46 | MO_infl | RRBP1 |
| 6.9602E-50 | -0.4064323 | 0.112 | 0.19 | 1.6442E-45 | MO_infl | F3 |
| 7.5821E-50 | -0.3099347 | 0.053 | 0.116 | 1.7911E-45 | MO_infl | TCHP |
| 8.2888E-50 | -0.4415348 | 0.088 | 0.157 | 1.9581E-45 | MO_infl | SMC4 |
| 1.2027E-49 | -0.9903667 | 0.214 | 0.291 | 2.8412E-45 | MO_infl | MT1E |
| 2.9288E-49 | -1.8941451 | 0.187 | 0.265 | 6.9188E-45 | MO_infl | MT1G |
| 9.6335E-49 | -0.2681382 | 0.476 | 0.541 | 2.2757E-44 | MO_infl | COX6C |
| 6.7296E-48 | -0.2944359 | 0.395 | 0.473 | 1.5897E-43 | MO_infl | SARAF |
| 7.2223E-47 | -0.2642009 | 0.099 | 0.17 | 1.7061E-42 | MO_infl | SOCS1 |
| 2.4362E-45 | -0.3365732 | 0.995 | 0.978 | 5.755E-41 | MO_infl | EEF1A1 |
| 2.2067E-44 | -0.278707 | 0.12 | 0.196 | 5.2128E-40 | MO_infl | TNFSF10 |
| 1.1026E-43 | -0.3322951 | 0.274 | 0.34 | 2.6047E-39 | MO_infl | APP |
| 1.8105E-43 | -0.3529475 | 0.95 | 0.914 | 4.2769E-39 | MO_infl | RPL9 |
| 1.9757E-43 | -0.3304197 | 0.612 | 0.665 | 4.6671E-39 | MO_infl | NDUFA4 |
| 4.206E-43 | -0.2900485 | 0.606 | 0.656 | 9.9358E-39 | MO_infl | UQCRH |
| 1.4626E-42 | -0.2807419 | 0.729 | 0.792 | 3.4552E-38 | MO_infl | DYNLL1 |
| 3.3947E-42 | -0.3667836 | 0.932 | 0.897 | 8.0192E-38 | MO_infl | RPS7 |
| 1.5718E-41 | -0.3194874 | 0.329 | 0.397 | 3.713E-37 | MO_infl | TUBB4B |
| 4.6973E-41 | -0.2965332 | 0.225 | 0.292 | 1.1096E-36 | MO_infl | CD151 |
| 2.6829E-40 | -0.275179 | 0.627 | 0.709 | 6.3377E-36 | MO_infl | PPP1R15A |
| 4.2956E-40 | -0.2593484 | 0.266 | 0.33 | 1.0148E-35 | MO_infl | KRT10 |
| 6.0992E-40 | -0.3574908 | 0.179 | 0.247 | 1.4408E-35 | MO_infl | CITED2 |
| 6.4302E-40 | -0.393129 | 0.952 | 0.906 | 1.519E-35 | MO_infl | RPS8 |
| 1.5417E-39 | -0.2980793 | 0.507 | 0.553 | 3.642E-35 | MO_infl | RPS10 |
| 1.0604E-38 | -0.2863513 | 0.461 | 0.528 | 2.505E-34 | MO_infl | H2AFZ |
| 1.3223E-38 | -0.5488845 | 0.8 | 0.695 | 3.1237E-34 | MO_infl | ANXA1 |
| 4.3954E-38 | -0.2538227 | 0.243 | 0.31 | 1.0383E-33 | MO_infl | DDX24 |
| 2.4692E-37 | -0.2909912 | 0.616 | 0.664 | 5.8331E-33 | MO_infl | CALM2 |
| 5.4745E-37 | -0.3288643 | 0.452 | 0.5 | 1.2932E-32 | MO_infl | NCL |
| 1.5544E-35 | -0.3999697 | 0.951 | 0.91 | 3.6719E-31 | MO_infl | RPL32 |
| 3.2359E-35 | -0.2648042 | 0.144 | 0.205 | 7.6442E-31 | MO_infl | SERTAD2 |
| 9.9296E-35 | -0.2583278 | 0.353 | 0.402 | 2.3457E-30 | MO_infl | CTNNA1 |
| 1.5979E-34 | -0.341303 | 0.386 | 0.432 | 3.7748E-30 | MO_infl | SLC3A2 |
| 3.8756E-34 | -0.3928568 | 0.411 | 0.441 | 9.1552E-30 | MO_infl | CYSTM1 |
| 4.0262E-34 | -0.2720622 | 0.206 | 0.265 | 9.5112E-30 | MO_infl | PDLIM5 |
| 1.072E-33 | -0.3239031 | 0.987 | 0.979 | 2.5323E-29 | MO_infl | RPL41 |
| 1.1342E-33 | -0.3200493 | 0.297 | 0.349 | 2.6793E-29 | MO_infl | PDIA6 |

| | | | | | | |
|---|---|---|---|---|---|---|
| 1.6231E-33 | -0.2899217 | 0.384 | 0.425 | 3.8342E-29 | MO_infl | RAN |
| 2.8204E-33 | -0.3760726 | 0.349 | 0.398 | 6.6627E-29 | MO_infl | SRSF7 |
| 6.1423E-33 | -0.6281642 | 0.45 | 0.498 | 1.451E-28 | MO_infl | MTRNR2L8 |
| 2.7708E-32 | -0.250753 | 0.34 | 0.39 | 6.5455E-28 | MO_infl | DYNLRB1 |
| 2.8864E-32 | -0.3363243 | 0.346 | 0.413 | 6.8186E-28 | MO_infl | RNF213 |
| 6.2177E-32 | 0.39445882 | 0.295 | 0.227 | 1.4688E-27 | MO_infl | FABP5 |
| 3.1002E-31 | -0.2851649 | 0.288 | 0.34 | 7.3235E-27 | MO_infl | GOLGA4 |
| 3.1735E-30 | -0.2697551 | 0.123 | 0.179 | 7.4967E-26 | MO_infl | STEAP4 |
| 4.0521E-30 | -0.3126155 | 0.854 | 0.828 | 9.5722E-26 | MO_infl | RPL22 |
| 7.6403E-30 | -0.3034276 | 0.309 | 0.356 | 1.8049E-25 | MO_infl | PNN |
| 1.8789E-29 | -1.2292406 | 0.072 | 0.122 | 4.4385E-25 | MO_infl | MT1M |
| 4.4113E-28 | -0.321567 | 0.933 | 0.893 | 1.0421E-23 | MO_infl | RPS25 |
| 4.6687E-28 | -0.3751257 | 0.916 | 0.89 | 1.1029E-23 | MO_infl | RPS15A |
| 8.366E-28 | -3.6475609 | 0.521 | 0.545 | 1.9763E-23 | MO_infl | PLCG2 |
| 1.2837E-27 | -0.3554761 | 0.474 | 0.511 | 3.0324E-23 | MO_infl | MT-ND4L |
| 3.1317E-27 | 0.25742182 | 0.27 | 0.21 | 7.3981E-23 | MO_infl | TGM2 |
| 6.9855E-27 | -0.2843351 | 0.492 | 0.536 | 1.6502E-22 | MO_infl | PTTG1IP |
| 4.0818E-25 | -0.2981571 | 0.927 | 0.895 | 9.6424E-21 | MO_infl | RPL27A |
| 1.7402E-24 | -0.4561852 | 0.925 | 0.906 | 4.1108E-20 | MO_infl | MT-ND2 |
| 1.1359E-23 | -0.2752796 | 0.782 | 0.787 | 2.6833E-19 | MO_infl | RPL31 |
| 2.0699E-21 | -0.2876872 | 0.987 | 0.929 | 4.8896E-17 | MO_infl | S100A6 |
| 3.53E-20 | 0.28847743 | 0.145 | 0.106 | 8.339E-16 | MO_infl | ANGPTL4 |
| 8.851E-20 | -0.3183225 | 0.086 | 0.125 | 2.0909E-15 | MO_infl | HILPDA |
| 1.1251E-19 | -0.3095789 | 0.767 | 0.766 | 2.6577E-15 | MO_infl | RPS26 |
| 1.4424E-19 | -0.4215488 | 0.191 | 0.232 | 3.4074E-15 | MO_infl | HEXIM1 |
| 1.673E-19 | -0.297583 | 0.287 | 0.324 | 3.9522E-15 | MO_infl | BRD2 |
| 3.9048E-19 | -0.2970484 | 0.879 | 0.885 | 9.2244E-15 | MO_infl | MYL12B |
| 9.1976E-18 | -0.2804102 | 0.967 | 0.921 | 2.1727E-13 | MO_infl | RPL19 |
| 3.2771E-16 | -0.5523593 | 0.287 | 0.312 | 7.7416E-12 | MO_infl | HIST1H4C |
| 1.1549E-15 | -0.2881208 | 0.961 | 0.929 | 2.7283E-11 | MO_infl | RPS14 |
| 4.6596E-15 | -0.3311978 | 0.815 | 0.788 | 1.1007E-10 | MO_infl | RPS21 |
| 6.8786E-14 | -0.2524533 | 0.967 | 0.913 | 1.6249E-09 | MO_infl | RPS15 |
| 1.8298E-13 | -0.2980025 | 0.958 | 0.93 | 4.3226E-09 | MO_infl | RPL34 |
| 1.2193E-12 | -0.2703785 | 0.882 | 0.832 | 2.8804E-08 | MO_infl | RACK1 |
| 2.9143E-12 | -0.345597 | 0.138 | 0.165 | 6.8846E-08 | MO_infl | CSKMT |
| 3.4275E-11 | -0.2985913 | 0.956 | 0.89 | 8.0968E-07 | MO_infl | RPL21 |
| 6.852E-11 | -0.3054494 | 0.953 | 0.929 | 1.6186E-06 | MO_infl | MT-ND4 |
| 2.6951E-10 | -0.2700315 | 0.964 | 0.942 | 6.3667E-06 | MO_infl | MT-CO2 |
| 8.326E-10 | -0.253218 | 0.189 | 0.212 | 1.9669E-05 | MO_infl | MDM2 |
| 7.6464E-09 | -0.439867 | 0.789 | 0.783 | 0.00018063 | MO_infl | LGALS3 |
| 1.8042E-08 | -0.3196855 | 0.975 | 0.938 | 0.0004262 | MO_infl | RPS27 |
| 1.2596E-07 | -0.4212194 | 0.579 | 0.672 | 0.00297562 | MO_infl | MT2A |
| 1.6962E-07 | -0.2659546 | 0.402 | 0.396 | 0.00400686 | MO_infl | ARGLU1 |
| 1.9589E-07 | -0.294508 | 0.585 | 0.634 | 0.00462761 | MO_infl | JUN |
| 7.0508E-07 | -0.3244707 | 0.958 | 0.942 | 0.01665614 | MO_infl | MT-CO3 |
| 2.244E-05 | -0.2902469 | 0.982 | 0.943 | 0.53008856 | MO_infl | RPL10 |
| 3.5116E-05 | 0.4264142 | 0.281 | 0.259 | 0.8295339 | MO_infl | AREG |
| 0.00015433 | 0.26158871 | 0.598 | 0.666 | 1 | MO_infl | HSPA1A |
| 0.00022431 | -0.2918117 | 0.222 | 0.226 | 1 | MO_infl | PPP1R10 |
| 0.00991455 | -0.3478003 | 0.095 | 0.102 | 1 | MO_infl | HLA-DQA1 |

| | | | | | | |
|---|---|---|---|---|---|---|
| 0 | 3.4731496 | 0.991 | 0.343 | 0 | MO_kdnrs | HLA-DRA |
| 0 | 3.3847854 | 0.993 | 0.446 | 0 | MO_kdnrs | CD74 |
| 0 | 3.38055854 | 0.949 | 0.137 | 0 | MO_kdnrs | HLA-DPA1 |
| 0 | 3.23433254 | 0.922 | 0.119 | 0 | MO_kdnrs | HLA-DPB1 |
| 0 | 2.85931589 | 0.919 | 0.219 | 0 | MO_kdnrs | HLA-DRB1 |
| 0 | 2.44523346 | 0.767 | 0.042 | 0 | MO_kdnrs | HLA-DQA1 |
| 0 | 2.23596641 | 0.567 | 0.092 | 0 | MO_kdnrs | APOE |
| 0 | 2.12093673 | 0.823 | 0.114 | 0 | MO_kdnrs | HLA-DQB1 |
| 0 | 2.04606632 | 0.488 | 0.036 | 0 | MO_kdnrs | C1QB |
| 0 | 1.9959162 | 0.536 | 0.036 | 0 | MO_kdnrs | C1QA |
| 0 | 1.9319956 | 0.548 | 0.069 | 0 | MO_kdnrs | APOC1 |
| 0 | 1.86549639 | 0.963 | 0.419 | 0 | MO_kdnrs | LYZ |
| 0 | 1.83609645 | 0.647 | 0.089 | 0 | MO_kdnrs | HLA-DRB5 |
| 0 | 1.67529744 | 0.887 | 0.22 | 0 | MO_kdnrs | AIF1 |
| 0 | 1.67063974 | 0.412 | 0.025 | 0 | MO_kdnrs | C1QC |
| 0 | 1.61114027 | 0.89 | 0.339 | 0 | MO_kdnrs | TYMP |
| 0 | 1.53384074 | 0.98 | 0.392 | 0 | MO_kdnrs | TYROBP |
| 0 | 1.51723804 | 0.43 | 0.01 | 0 | MO_kdnrs | CXCL9 |
| 0 | 1.47201023 | 0.979 | 0.748 | 0 | MO_kdnrs | CST3 |
| 0 | 1.45584577 | 0.729 | 0.083 | 0 | MO_kdnrs | HLA-DMA |
| 0 | 1.36862572 | 0.965 | 0.717 | 0 | MO_kdnrs | CTSB |
| 0 | 1.32499675 | 0.999 | 0.967 | 0 | MO_kdnrs | TMSB10 |
| 0 | 1.3225 | 0.973 | 0.72 | 0 | MO_kdnrs | PSAP |
| 0 | 1.30423133 | 1 | 0.986 | 0 | MO_kdnrs | FTL |
| 0 | 1.28880489 | 0.999 | 0.932 | 0 | MO_kdnrs | TMSB4X |
| 0 | 1.21631489 | 0.945 | 0.566 | 0 | MO_kdnrs | NPC2 |
| 0 | 1.20927941 | 0.946 | 0.335 | 0 | MO_kdnrs | FCER1G |
| 0 | 1.20776 | 0.795 | 0.168 | 0 | MO_kdnrs | ITGB2 |
| 0 | 1.19737316 | 0.721 | 0.157 | 0 | MO_kdnrs | CTSC |
| 0 | 1.17519832 | 0.59 | 0.077 | 0 | MO_kdnrs | FCGR3A |
| 0 | 1.16630686 | 0.763 | 0.187 | 0 | MO_kdnrs | HCST |
| 0 | 1.14582529 | 0.894 | 0.285 | 0 | MO_kdnrs | LAPTM5 |
| 0 | 1.13198066 | 0.796 | 0.268 | 0 | MO_kdnrs | CD68 |
| 0 | 1.12402541 | 0.834 | 0.388 | 0 | MO_kdnrs | EMP3 |
| 0 | 1.11783932 | 0.769 | 0.239 | 0 | MO_kdnrs | CTSZ |
| 0 | 1.11162096 | 0.392 | 0.044 | 0 | MO_kdnrs | FN1 |
| 0 | 1.10692384 | 0.418 | 0.048 | 0 | MO_kdnrs | MRC1 |
| 0 | 1.10253837 | 0.743 | 0.16 | 0 | MO_kdnrs | LST1 |
| 0 | 1.0975337 | 0.726 | 0.154 | 0 | MO_kdnrs | CYBB |
| 0 | 1.08867263 | 0.694 | 0.137 | 0 | MO_kdnrs | IGSF6 |
| 0 | 1.08663127 | 0.664 | 0.126 | 0 | MO_kdnrs | TGFBI |
| 0 | 1.05457514 | 0.828 | 0.327 | 0 | MO_kdnrs | COTL1 |
| 0 | 1.04772287 | 0.996 | 0.971 | 0 | MO_kdnrs | ACTB |
| 0 | 1.0208679 | 0.329 | 0.017 | 0 | MO_kdnrs | CXCL10 |
| 0 | 1.0154581 | 0.976 | 0.705 | 0 | MO_kdnrs | CYBA |
| 0 | 1.00429922 | 0.655 | 0.128 | 0 | MO_kdnrs | SAMHD1 |
| 0 | 1.00403731 | 0.575 | 0.066 | 0 | MO_kdnrs | FGL2 |
| 0 | 1.00262307 | 1 | 0.989 | 0 | MO_kdnrs | B2M |
| 0 | 0.99742537 | 0.75 | 0.168 | 0 | MO_kdnrs | SPI1 |
| 0 | 0.99558038 | 0.56 | 0.038 | 0 | MO_kdnrs | HLA-DMB |

| | | | | | | |
|---|---|---|---|---|---|---|
| 0 | 0.98413202 | 0.525 | 0.069 | 0 | MO_kdnrs | MS4A6A |
| 0 | 0.97532793 | 0.653 | 0.146 | 0 | MO_kdnrs | AC020656.1 |
| 0 | 0.97271384 | 0.968 | 0.775 | 0 | MO_kdnrs | PFN1 |
| 0 | 0.96347639 | 0.894 | 0.601 | 0 | MO_kdnrs | S100A4 |
| 0 | 0.96340208 | 0.319 | 0.038 | 0 | MO_kdnrs | RGS1 |
| 0 | 0.9447183 | 0.567 | 0.079 | 0 | MO_kdnrs | CSF1R |
| 0 | 0.93872629 | 0.454 | 0.063 | 0 | MO_kdnrs | GPR183 |
| 0 | 0.93810923 | 0.565 | 0.165 | 0 | MO_kdnrs | CD52 |
| 0 | 0.92840598 | 0.995 | 0.891 | 0 | MO_kdnrs | RPL39 |
| 0 | 0.92776825 | 0.991 | 0.883 | 0 | MO_kdnrs | ATP5F1E |
| 0 | 0.92181565 | 0.662 | 0.137 | 0 | MO_kdnrs | OLR1 |
| 0 | 0.91986004 | 0.399 | 0.008 | 0 | MO_kdnrs | HLA-DQA2 |
| 0 | 0.90996527 | 0.655 | 0.181 | 0 | MO_kdnrs | CD14 |
| 0 | 0.905115 | 0.997 | 0.88 | 0 | MO_kdnrs | HLA-B |
| 0 | 0.90150933 | 0.911 | 0.497 | 0 | MO_kdnrs | CTSS |
| 0 | 0.89365573 | 0.589 | 0.107 | 0 | MO_kdnrs | PLEKHO1 |
| 0 | 0.89323348 | 0.657 | 0.27 | 0 | MO_kdnrs | PSME2 |
| 0 | 0.87562908 | 0.362 | 0.042 | 0 | MO_kdnrs | CST7 |
| 0 | 0.86905922 | 0.513 | 0.112 | 0 | MO_kdnrs | MS4A7 |
| 0 | 0.86608324 | 0.552 | 0.147 | 0 | MO_kdnrs | WARS |
| 0 | 0.86483121 | 0.996 | 0.896 | 0 | MO_kdnrs | RPLP2 |
| 0 | 0.86340091 | 0.599 | 0.203 | 0 | MO_kdnrs | FABP5 |
| 0 | 0.86301177 | 0.667 | 0.254 | 0 | MO_kdnrs | ATP1B3 |
| 0 | 0.86131484 | 0.791 | 0.323 | 0 | MO_kdnrs | GNAI2 |
| 0 | 0.85560302 | 0.993 | 0.928 | 0 | MO_kdnrs | RPL34 |
| 0 | 0.85446766 | 0.653 | 0.159 | 0 | MO_kdnrs | ITGAX |
| 0 | 0.85408201 | 0.282 | 0.02 | 0 | MO_kdnrs | A2M |
| 0 | 0.84203774 | 0.563 | 0.108 | 0 | MO_kdnrs | C1orf162 |
| 0 | 0.83635999 | 0.647 | 0.189 | 0 | MO_kdnrs | RGS10 |
| 0 | 0.83483858 | 0.369 | 0.12 | 0 | MO_kdnrs | GPNMB |
| 0 | 0.83482221 | 0.833 | 0.521 | 0 | MO_kdnrs | RPS10 |
| 0 | 0.82988097 | 0.738 | 0.27 | 0 | MO_kdnrs | ARPC1B |
| 0 | 0.82926075 | 0.939 | 0.588 | 0 | MO_kdnrs | LGALS1 |
| 0 | 0.8154696 | 0.998 | 0.934 | 0 | MO_kdnrs | RPS28 |
| 0 | 0.80861494 | 0.9 | 0.763 | 0 | MO_kdnrs | CTSD |
| 0 | 0.8033526 | 0.797 | 0.246 | 0 | MO_kdnrs | LCP1 |
| 0 | 0.79726344 | 0.564 | 0.135 | 0 | MO_kdnrs | CXCR4 |
| 0 | 0.79064985 | 0.994 | 0.896 | 0 | MO_kdnrs | RPL37 |
| 0 | 0.78640665 | 0.529 | 0.115 | 0 | MO_kdnrs | PLXDC2 |
| 0 | 0.78326449 | 0.778 | 0.383 | 0 | MO_kdnrs | GRN |
| 0 | 0.7822429 | 0.999 | 0.943 | 0 | MO_kdnrs | RPL10 |
| 0 | 0.77793623 | 0.628 | 0.165 | 0 | MO_kdnrs | LSP1 |
| 0 | 0.77520343 | 0.979 | 0.848 | 0 | MO_kdnrs | SH3BGRL3 |
| 0 | 0.77435138 | 0.991 | 0.717 | 0 | MO_kdnrs | VIM |
| 0 | 0.77396911 | 0.44 | 0.103 | 0 | MO_kdnrs | CD83 |
| 0 | 0.77210969 | 0.661 | 0.272 | 0 | MO_kdnrs | RPL36A |
| 0 | 0.77059989 | 0.997 | 0.917 | 0 | MO_kdnrs | RPS24 |
| 0 | 0.77046579 | 0.68 | 0.259 | 0 | MO_kdnrs | FXYD5 |
| 0 | 0.76035058 | 0.361 | 0.027 | 0 | MO_kdnrs | MSR1 |
| 0 | 0.75695731 | 0.989 | 0.87 | 0 | MO_kdnrs | RPS29 |

| | | | | | | |
|---|---|---|---|---|---|---|
| 0 | 0.75537954 | 0.588 | 0.152 | 0 | MO_kdnrs | PYCARD |
| 0 | 0.74944792 | 0.48 | 0.058 | 0 | MO_kdnrs | TNFSF13B |
| 0 | 0.74638337 | 0.275 | 0.025 | 0 | MO_kdnrs | TREM2 |
| 0 | 0.74613014 | 0.984 | 0.836 | 0 | MO_kdnrs | RPL38 |
| 0 | 0.74484119 | 0.416 | 0.081 | 0 | MO_kdnrs | ACP5 |
| 0 | 0.73458399 | 0.741 | 0.334 | 0 | MO_kdnrs | CTSH |
| 0 | 0.73054609 | 0.72 | 0.29 | 0 | MO_kdnrs | ARHGDIB |
| 0 | 0.72748615 | 0.386 | 0.079 | 0 | MO_kdnrs | LTB |
| 0 | 0.72419227 | 0.542 | 0.141 | 0 | MO_kdnrs | KCTD12 |
| 0 | 0.72405672 | 0.988 | 0.885 | 0 | MO_kdnrs | RPS15A |
| 0 | 0.71958251 | 0.573 | 0.154 | 0 | MO_kdnrs | FCGR2A |
| 0 | 0.71146625 | 0.99 | 0.909 | 0 | MO_kdnrs | RPL32 |
| 0 | 0.7112158 | 0.32 | 0.018 | 0 | MO_kdnrs | SDS |
| 0 | 0.7096623 | 0.992 | 0.922 | 0 | MO_kdnrs | RPL30 |
| 0 | 0.70434678 | 0.683 | 0.207 | 0 | MO_kdnrs | GMFG |
| 0 | 0.70319039 | 0.992 | 0.925 | 0 | MO_kdnrs | FAU |
| 0 | 0.70279465 | 0.593 | 0.195 | 0 | MO_kdnrs | FCGRT |
| 0 | 0.70096995 | 0.491 | 0.158 | 0 | MO_kdnrs | KLF2 |
| 0 | 0.69949598 | 0.989 | 0.888 | 0 | MO_kdnrs | HLA-A |
| 0 | 0.69946645 | 0.649 | 0.252 | 0 | MO_kdnrs | CXCL16 |
| 0 | 0.69579555 | 0.603 | 0.151 | 0 | MO_kdnrs | FYB1 |
| 0 | 0.69492095 | 0.49 | 0.084 | 0 | MO_kdnrs | CD86 |
| 0 | 0.69444173 | 0.719 | 0.327 | 0 | MO_kdnrs | TUBA1B |
| 0 | 0.69327066 | 0.445 | 0.103 | 0 | MO_kdnrs | VAMP5 |
| 0 | 0.6926174 | 0.555 | 0.143 | 0 | MO_kdnrs | CCDC88A |
| 0 | 0.69206855 | 0.421 | 0.085 | 0 | MO_kdnrs | LIPA |
| 0 | 0.69131327 | 0.525 | 0.187 | 0 | MO_kdnrs | CREM |
| 0 | 0.69016836 | 0.642 | 0.215 | 0 | MO_kdnrs | SH3BGRL |
| 0 | 0.68905102 | 0.288 | 0.033 | 0 | MO_kdnrs | INHBA |
| 0 | 0.68787491 | 0.524 | 0.102 | 0 | MO_kdnrs | CLEC7A |
| 0 | 0.6772125 | 0.58 | 0.207 | 0 | MO_kdnrs | RNASET2 |
| 0 | 0.67397635 | 0.963 | 0.777 | 0 | MO_kdnrs | RPS21 |
| 0 | 0.6714973 | 0.402 | 0.034 | 0 | MO_kdnrs | FPR3 |
| 0 | 0.6696579 | 0.216 | 0.044 | 0 | MO_kdnrs | RNASE1 |
| 0 | 0.66726537 | 0.993 | 0.879 | 0 | MO_kdnrs | RPS23 |
| 0 | 0.66688943 | 0.707 | 0.312 | 0 | MO_kdnrs | CTSL |
| 0 | 0.6628281 | 0.605 | 0.189 | 0 | MO_kdnrs | GRINA |
| 0 | 0.65857092 | 0.476 | 0.151 | 0 | MO_kdnrs | STAT1 |
| 0 | 0.65832354 | 0.695 | 0.29 | 0 | MO_kdnrs | LIMS1 |
| 0 | 0.65795172 | 0.995 | 0.939 | 0 | MO_kdnrs | RPS27 |
| 0 | 0.6556533 | 0.545 | 0.198 | 0 | MO_kdnrs | IFNGR1 |
| 0 | 0.65463069 | 0.723 | 0.227 | 0 | MO_kdnrs | CD53 |
| 0 | 0.6540732 | 0.592 | 0.258 | 0 | MO_kdnrs | YWHAH |
| 0 | 0.6536499 | 0.346 | 0.051 | 0 | MO_kdnrs | FCGR2B |
| 0 | 0.65163637 | 0.968 | 0.807 | 0 | MO_kdnrs | RPL27 |
| 0 | 0.65019885 | 0.991 | 0.89 | 0 | MO_kdnrs | RPS25 |
| 0 | 0.64892847 | 0.52 | 0.136 | 0 | MO_kdnrs | LAP3 |
| 0 | 0.64631497 | 0.993 | 0.928 | 0 | MO_kdnrs | RPS12 |
| 0 | 0.64501784 | 0.647 | 0.19 | 0 | MO_kdnrs | ZEB2 |
| 0 | 0.64441309 | 0.991 | 0.91 | 0 | MO_kdnrs | UBA52 |

| | | | | | | |
|---|---|---|---|---|---|---|
| 0 | 0.64350259 | 0.559 | 0.151 | 0 | MO_kdnrs | C5AR1 |
| 0 | 0.64180861 | 0.37 | 0.058 | 0 | MO_kdnrs | IL4I1 |
| 0 | 0.6417675 | 0.999 | 0.976 | 0 | MO_kdnrs | RPL28 |
| 0 | 0.63856748 | 0.491 | 0.151 | 0 | MO_kdnrs | TMEM176B |
| 0 | 0.63819919 | 0.543 | 0.14 | 0 | MO_kdnrs | CORO1A |
| 0 | 0.6376218 | 0.998 | 0.941 | 0 | MO_kdnrs | PTMA |
| 0 | 0.63716161 | 0.399 | 0.087 | 0 | MO_kdnrs | IL7R |
| 0 | 0.63627524 | 0.452 | 0.125 | 0 | MO_kdnrs | CPVL |
| 0 | 0.63596177 | 0.658 | 0.29 | 0 | MO_kdnrs | AP2S1 |
| 0 | 0.63590788 | 0.984 | 0.533 | 0 | MO_kdnrs | SRGN |
| 0 | 0.63330232 | 0.495 | 0.14 | 0 | MO_kdnrs | FCN1 |
| 0 | 0.63321918 | 0.323 | 0.049 | 0 | MO_kdnrs | VSIG4 |
| 0 | 0.62937089 | 0.549 | 0.154 | 0 | MO_kdnrs | RNF130 |
| 0 | 0.62585251 | 0.797 | 0.496 | 0 | MO_kdnrs | ATP6V1F |
| 0 | 0.625637 | 0.995 | 0.916 | 0 | MO_kdnrs | RPL11 |
| 0 | 0.62315471 | 0.255 | 0.043 | 0 | MO_kdnrs | VMO1 |
| 0 | 0.62304543 | 0.993 | 0.951 | 0 | MO_kdnrs | SERF2 |
| 0 | 0.62255836 | 0.514 | 0.127 | 0 | MO_kdnrs | PPT1 |
| 0 | 0.6224127 | 0.43 | 0.101 | 0 | MO_kdnrs | GBP1 |
| 0 | 0.62127467 | 0.971 | 0.802 | 0 | MO_kdnrs | PFDN5 |
| 0 | 0.61846262 | 0.422 | 0.133 | 0 | MO_kdnrs | OGFRL1 |
| 0 | 0.61824963 | 0.538 | 0.239 | 0 | MO_kdnrs | GSN |
| 0 | 0.61824825 | 0.535 | 0.147 | 0 | MO_kdnrs | CD37 |
| 0 | 0.61683236 | 0.486 | 0.115 | 0 | MO_kdnrs | UCP2 |
| 0 | 0.61661395 | 0.902 | 0.616 | 0 | MO_kdnrs | ARPC5 |
| 0 | 0.61477143 | 0.263 | 0.042 | 0 | MO_kdnrs | C3 |
| 0 | 0.6140427 | 1 | 0.993 | 0 | MO_kdnrs | FTH1 |
| 0 | 0.61394372 | 0.616 | 0.3 | 0 | MO_kdnrs | CAPG |
| 0 | 0.61346163 | 0.785 | 0.236 | 0 | MO_kdnrs | PTPRC |
| 0 | 0.6134282 | 0.365 | 0.072 | 0 | MO_kdnrs | ZNF331 |
| 0 | 0.61179887 | 0.997 | 0.979 | 0 | MO_kdnrs | RPL41 |
| 0 | 0.60926809 | 0.434 | 0.071 | 0 | MO_kdnrs | SLC8A1 |
| 0 | 0.60736798 | 0.538 | 0.136 | 0 | MO_kdnrs | CD48 |
| 0 | 0.60704111 | 0.299 | 0.009 | 0 | MO_kdnrs | CLEC10A |
| 0 | 0.60545835 | 0.501 | 0.154 | 0 | MO_kdnrs | PSMB9 |
| 0 | 0.60191295 | 0.998 | 0.94 | 0 | MO_kdnrs | RPS19 |
| 0 | 0.60156587 | 0.98 | 0.818 | 0 | MO_kdnrs | RPL22 |
| 0 | 0.60146887 | 0.436 | 0.122 | 0 | MO_kdnrs | RGS2 |
| 0 | 0.6002865 | 0.787 | 0.402 | 0 | MO_kdnrs | GSTO1 |
| 0 | 0.59791178 | 0.481 | 0.117 | 0 | MO_kdnrs | ALDH2 |
| 0 | 0.59547781 | 0.552 | 0.155 | 0 | MO_kdnrs | HCLS1 |
| 0 | 0.59176267 | 0.327 | 0.054 | 0 | MO_kdnrs | MEF2C |
| 0 | 0.59159853 | 0.478 | 0.157 | 0 | MO_kdnrs | SGK1 |
| 0 | 0.5903448 | 0.997 | 0.875 | 0 | MO_kdnrs | RPS2 |
| 0 | 0.5878285 | 0.999 | 0.97 | 0 | MO_kdnrs | RPLP1 |
| 0 | 0.58735741 | 0.567 | 0.168 | 0 | MO_kdnrs | NCF2 |
| 0 | 0.58502355 | 0.995 | 0.921 | 0 | MO_kdnrs | RPL19 |
| 0 | 0.58460163 | 0.363 | 0.05 | 0 | MO_kdnrs | CD84 |
| 0 | 0.5831672 | 0.586 | 0.225 | 0 | MO_kdnrs | MGAT1 |
| 0 | 0.58153168 | 0.385 | 0.035 | 0 | MO_kdnrs | CD4 |

| | | | | | | |
|---|---|---|---|---|---|---|
| 0 | 0.5802506 | 0.99 | 0.872 | 0 | MO_kdnrs | RPL12 |
| 0 | 0.57967746 | 0.432 | 0.159 | 0 | MO_kdnrs | FBP1 |
| 0 | 0.57780064 | 0.983 | 0.866 | 0 | MO_kdnrs | RPL26 |
| 0 | 0.57771772 | 0.929 | 0.689 | 0 | MO_kdnrs | GPX4 |
| 0 | 0.57692353 | 0.984 | 0.888 | 0 | MO_kdnrs | RPL35A |
| 0 | 0.57516364 | 0.468 | 0.11 | 0 | MO_kdnrs | IFI30 |
| 0 | 0.57482381 | 0.987 | 0.834 | 0 | MO_kdnrs | RPL35 |
| 0 | 0.57270443 | 0.994 | 0.928 | 0 | MO_kdnrs | RPS14 |
| 0 | 0.57190307 | 0.597 | 0.137 | 0 | MO_kdnrs | PLEK |
| 0 | 0.56740541 | 0.552 | 0.154 | 0 | MO_kdnrs | GPSM3 |
| 0 | 0.56089307 | 0.318 | 0.018 | 0 | MO_kdnrs | GNLY |
| 0 | 0.55964636 | 0.99 | 0.918 | 0 | MO_kdnrs | RPS27A |
| 0 | 0.55915584 | 0.477 | 0.115 | 0 | MO_kdnrs | CARD16 |
| 0 | 0.55718242 | 0.409 | 0.144 | 0 | MO_kdnrs | CFD |
| 0 | 0.55596147 | 0.996 | 0.913 | 0 | MO_kdnrs | RPS15 |
| 0 | 0.55300934 | 0.533 | 0.191 | 0 | MO_kdnrs | ALOX5AP |
| 0 | 0.55106814 | 0.416 | 0.064 | 0 | MO_kdnrs | LILRB4 |
| 0 | 0.54561893 | 0.385 | 0.083 | 0 | MO_kdnrs | CSF2RA |
| 0 | 0.54217559 | 0.536 | 0.155 | 0 | MO_kdnrs | PTPRE |
| 0 | 0.53840079 | 0.985 | 0.875 | 0 | MO_kdnrs | RPS13 |
| 0 | 0.53819061 | 0.992 | 0.889 | 0 | MO_kdnrs | RPS3A |
| 0 | 0.5354881 | 0.383 | 0.095 | 0 | MO_kdnrs | ATF5 |
| 0 | 0.53450787 | 0.879 | 0.626 | 0 | MO_kdnrs | ATP5MC2 |
| 0 | 0.53341734 | 0.459 | 0.124 | 0 | MO_kdnrs | EVI2B |
| 0 | 0.5320414 | 0.716 | 0.366 | 0 | MO_kdnrs | COMMD6 |
| 0 | 0.53037313 | 0.29 | 0.031 | 0 | MO_kdnrs | ENTPD1 |
| 0 | 0.5287111 | 0.471 | 0.16 | 0 | MO_kdnrs | CKLF |
| 0 | 0.52864734 | 0.963 | 0.802 | 0 | MO_kdnrs | YBX1 |
| 0 | 0.52748382 | 0.494 | 0.083 | 0 | MO_kdnrs | MNDA |
| 0 | 0.52744977 | 0.235 | 0.01 | 0 | MO_kdnrs | IGKC |
| 0 | 0.52660479 | 0.351 | 0.067 | 0 | MO_kdnrs | RNASE6 |
| 0 | 0.52659728 | 0.491 | 0.139 | 0 | MO_kdnrs | GLIPR2 |
| 0 | 0.5263517 | 0.545 | 0.182 | 0 | MO_kdnrs | NINJ1 |
| 0 | 0.52558127 | 0.995 | 0.886 | 0 | MO_kdnrs | RPL37A |
| 0 | 0.52541479 | 0.624 | 0.253 | 0 | MO_kdnrs | RPS4Y1 |
| 0 | 0.52444173 | 0.32 | 0.043 | 0 | MO_kdnrs | FCGR1A |
| 0 | 0.52280718 | 0.329 | 0.028 | 0 | MO_kdnrs | CIITA |
| 0 | 0.52248095 | 0.317 | 0.063 | 0 | MO_kdnrs | PLA2G7 |
| 0 | 0.52144663 | 0.51 | 0.132 | 0 | MO_kdnrs | CLEC2B |
| 0 | 0.52056919 | 0.391 | 0.091 | 0 | MO_kdnrs | CD163 |
| 0 | 0.51962999 | 0.488 | 0.176 | 0 | MO_kdnrs | IFI16 |
| 0 | 0.51709205 | 0.536 | 0.242 | 0 | MO_kdnrs | ATOX1 |
| 0 | 0.51541855 | 0.554 | 0.178 | 0 | MO_kdnrs | ARRB2 |
| 0 | 0.51260698 | 0.791 | 0.516 | 0 | MO_kdnrs | PLAUR |
| 0 | 0.51200363 | 0.414 | 0.094 | 0 | MO_kdnrs | LILRB2 |
| 0 | 0.50975643 | 0.625 | 0.21 | 0 | MO_kdnrs | SLC11A1 |
| 0 | 0.50913275 | 0.456 | 0.148 | 0 | MO_kdnrs | SMIM25 |
| 0 | 0.50735067 | 0.44 | 0.105 | 0 | MO_kdnrs | PILRA |
| 0 | 0.50688054 | 0.991 | 0.865 | 0 | MO_kdnrs | RPL36 |
| 0 | 0.5005141 | 0.353 | 0.054 | 0 | MO_kdnrs | MARCH1 |

| | | | | | | |
|---|---|---|---|---|---|---|
| 0 | 0.50022632 | 0.679 | 0.362 | 0 | MO_kdnrs | UQCR11 |
| 0 | 0.50015655 | 0.54 | 0.216 | 0 | MO_kdnrs | RBPJ |
| 0 | 0.50012408 | 0.435 | 0.112 | 0 | MO_kdnrs | PTPN6 |
| 0 | 0.49907397 | 0.419 | 0.093 | 0 | MO_kdnrs | LIMD2 |
| 0 | 0.49821763 | 0.597 | 0.2 | 0 | MO_kdnrs | SLC2A3 |
| 0 | 0.4961283 | 0.505 | 0.176 | 0 | MO_kdnrs | RHOG |
| 0 | 0.49589308 | 0.427 | 0.103 | 0 | MO_kdnrs | CELF2 |
| 0 | 0.49399611 | 0.882 | 0.62 | 0 | MO_kdnrs | TXNIP |
| 0 | 0.49272747 | 0.396 | 0.08 | 0 | MO_kdnrs | HAVCR2 |
| 0 | 0.49013493 | 0.339 | 0.09 | 0 | MO_kdnrs | NR4A2 |
| 0 | 0.48984997 | 0.999 | 0.961 | 0 | MO_kdnrs | RPL13 |
| 0 | 0.48892354 | 0.39 | 0.085 | 0 | MO_kdnrs | LAIR1 |
| 0 | 0.48818039 | 0.602 | 0.175 | 0 | MO_kdnrs | TNFRSF1B |
| 0 | 0.48810971 | 0.396 | 0.086 | 0 | MO_kdnrs | LY96 |
| 0 | 0.48730976 | 0.473 | 0.132 | 0 | MO_kdnrs | WIPF1 |
| 0 | 0.48729738 | 0.308 | 0.036 | 0 | MO_kdnrs | LY86 |
| 0 | 0.48514656 | 0.978 | 0.861 | 0 | MO_kdnrs | HLA-C |
| 0 | 0.48301154 | 0.373 | 0.069 | 0 | MO_kdnrs | ADA2 |
| 0 | 0.48264863 | 0.496 | 0.153 | 0 | MO_kdnrs | SAMSN1 |
| 0 | 0.48257064 | 0.255 | 0.052 | 0 | MO_kdnrs | NR4A3 |
| 0 | 0.48207253 | 0.443 | 0.154 | 0 | MO_kdnrs | LTA4H |
| 0 | 0.48164698 | 0.521 | 0.205 | 0 | MO_kdnrs | ATP6V1B2 |
| 0 | 0.48132699 | 0.305 | 0.044 | 0 | MO_kdnrs | MS4A4A |
| 0 | 0.48026226 | 0.182 | 0.018 | 0 | MO_kdnrs | IDO1 |
| 0 | 0.47890531 | 0.992 | 0.879 | 0 | MO_kdnrs | RPL6 |
| 0 | 0.47884556 | 0.434 | 0.13 | 0 | MO_kdnrs | ADGRE5 |
| 0 | 0.47863709 | 0.522 | 0.207 | 0 | MO_kdnrs | M6PR |
| 0 | 0.47848055 | 0.458 | 0.187 | 0 | MO_kdnrs | SOCS3 |
| 0 | 0.47440036 | 0.492 | 0.151 | 0 | MO_kdnrs | FMNL1 |
| 0 | 0.47255918 | 0.329 | 0.045 | 0 | MO_kdnrs | NKG7 |
| 0 | 0.47194506 | 0.51 | 0.189 | 0 | MO_kdnrs | TGFB1 |
| 0 | 0.47093498 | 0.643 | 0.318 | 0 | MO_kdnrs | LAMTOR4 |
| 0 | 0.46835721 | 0.391 | 0.084 | 0 | MO_kdnrs | IL10RA |
| 0 | 0.4671109 | 0.307 | 0.055 | 0 | MO_kdnrs | STAB1 |
| 0 | 0.46644107 | 0.359 | 0.093 | 0 | MO_kdnrs | C3AR1 |
| 0 | 0.46640087 | 0.346 | 0.061 | 0 | MO_kdnrs | GBP5 |
| 0 | 0.46435712 | 0.44 | 0.111 | 0 | MO_kdnrs | EMILIN2 |
| 0 | 0.46415381 | 0.528 | 0.199 | 0 | MO_kdnrs | TPP1 |
| 0 | 0.46411203 | 0.323 | 0.05 | 0 | MO_kdnrs | ADAP2 |
| 0 | 0.46288381 | 0.635 | 0.312 | 0 | MO_kdnrs | GRB2 |
| 0 | 0.46151792 | 0.585 | 0.272 | 0 | MO_kdnrs | ARL6IP5 |
| 0 | 0.46007308 | 0.356 | 0.109 | 0 | MO_kdnrs | C1orf56 |
| 0 | 0.45709394 | 0.472 | 0.19 | 0 | MO_kdnrs | ZFYVE16 |
| 0 | 0.45650154 | 0.288 | 0.017 | 0 | MO_kdnrs | CALHM6 |
| 0 | 0.45618607 | 0.623 | 0.315 | 0 | MO_kdnrs | ARPC4 |
| 0 | 0.45492247 | 0.407 | 0.127 | 0 | MO_kdnrs | SMAP2 |
| 0 | 0.45110529 | 0.514 | 0.186 | 0 | MO_kdnrs | BASP1 |
| 0 | 0.45080471 | 0.429 | 0.136 | 0 | MO_kdnrs | AP1S2 |
| 0 | 0.45002076 | 0.529 | 0.206 | 0 | MO_kdnrs | BST2 |
| 0 | 0.44630961 | 0.397 | 0.107 | 0 | MO_kdnrs | EPB41L3 |

| | | | | | | |
|---|---|---|---|---|---|---|
| 0 | 0.44600001 | 0.434 | 0.14 | 0 | MO_kdnrs | CNPY3 |
| 0 | 0.44534745 | 0.357 | 0.063 | 0 | MO_kdnrs | SLC15A3 |
| 0 | 0.4448418 | 0.306 | 0.053 | 0 | MO_kdnrs | RASSF4 |
| 0 | 0.44463983 | 0.346 | 0.113 | 0 | MO_kdnrs | RGCC |
| 0 | 0.44451289 | 0.351 | 0.112 | 0 | MO_kdnrs | TMEM176A |
| 0 | 0.44109518 | 0.343 | 0.097 | 0 | MO_kdnrs | DUSP2 |
| 0 | 0.44102543 | 0.342 | 0.07 | 0 | MO_kdnrs | GNB4 |
| 0 | 0.43941397 | 0.53 | 0.218 | 0 | MO_kdnrs | RAB31 |
| 0 | 0.43718868 | 0.396 | 0.11 | 0 | MO_kdnrs | CYTIP |
| 0 | 0.437039 | 0.397 | 0.107 | 0 | MO_kdnrs | THEMIS2 |
| 0 | 0.4358044 | 0.45 | 0.177 | 0 | MO_kdnrs | CREG1 |
| 0 | 0.43486609 | 0.421 | 0.14 | 0 | MO_kdnrs | MTHFD2 |
| 0 | 0.43362298 | 0.309 | 0.063 | 0 | MO_kdnrs | JAML |
| 0 | 0.43296602 | 0.411 | 0.128 | 0 | MO_kdnrs | NAGK |
| 0 | 0.43181355 | 0.395 | 0.096 | 0 | MO_kdnrs | WAS |
| 0 | 0.43169516 | 0.324 | 0.063 | 0 | MO_kdnrs | GIMAP4 |
| 0 | 0.42994606 | 0.41 | 0.109 | 0 | MO_kdnrs | NCKAP1L |
| 0 | 0.429257 | 0.265 | 0.033 | 0 | MO_kdnrs | LGALS2 |
| 0 | 0.42883122 | 0.396 | 0.114 | 0 | MO_kdnrs | LGALS9 |
| 0 | 0.42776585 | 0.368 | 0.099 | 0 | MO_kdnrs | PTAFR |
| 0 | 0.42732739 | 0.325 | 0.06 | 0 | MO_kdnrs | MPEG1 |
| 0 | 0.4247675 | 0.45 | 0.172 | 0 | MO_kdnrs | HLA-F |
| 0 | 0.42460464 | 0.424 | 0.146 | 0 | MO_kdnrs | HEXB |
| 0 | 0.42417367 | 0.456 | 0.137 | 0 | MO_kdnrs | MYO1F |
| 0 | 0.42388343 | 0.491 | 0.193 | 0 | MO_kdnrs | DSE |
| 0 | 0.42277855 | 0.23 | 0.014 | 0 | MO_kdnrs | TNFRSF4 |
| 0 | 0.42225534 | 0.359 | 0.078 | 0 | MO_kdnrs | TFEC |
| 0 | 0.42023518 | 0.341 | 0.084 | 0 | MO_kdnrs | GPRIN3 |
| 0 | 0.41960521 | 0.304 | 0.051 | 0 | MO_kdnrs | GBP4 |
| 0 | 0.41942069 | 0.442 | 0.161 | 0 | MO_kdnrs | DPP7 |
| 0 | 0.41338853 | 0.261 | 0.048 | 0 | MO_kdnrs | LPAR6 |
| 0 | 0.41279642 | 0.389 | 0.118 | 0 | MO_kdnrs | LRP1 |
| 0 | 0.41216193 | 0.414 | 0.156 | 0 | MO_kdnrs | GLIPR1 |
| 0 | 0.41138441 | 0.403 | 0.112 | 0 | MO_kdnrs | HCK |
| 0 | 0.40848521 | 0.399 | 0.113 | 0 | MO_kdnrs | EVI2A |
| 0 | 0.40648796 | 0.307 | 0.083 | 0 | MO_kdnrs | BMP2K |
| 0 | 0.39694471 | 0.402 | 0.106 | 0 | MO_kdnrs | LILRB3 |
| 0 | 0.39648788 | 0.241 | 0.008 | 0 | MO_kdnrs | HLA-DOA |
| 0 | 0.39583611 | 0.253 | 0.043 | 0 | MO_kdnrs | SLAMF7 |
| 0 | 0.3950301 | 0.436 | 0.137 | 0 | MO_kdnrs | FERMT3 |
| 0 | 0.39303185 | 0.349 | 0.081 | 0 | MO_kdnrs | LILRB1 |
| 0 | 0.39278346 | 0.265 | 0.047 | 0 | MO_kdnrs | CEBPA |
| 0 | 0.39185742 | 0.277 | 0.027 | 0 | MO_kdnrs | VASH1 |
| 0 | 0.39112686 | 0.304 | 0.066 | 0 | MO_kdnrs | DPYSL2 |
| 0 | 0.38847748 | 0.416 | 0.144 | 0 | MO_kdnrs | SIRPA |
| 0 | 0.38714566 | 0.41 | 0.141 | 0 | MO_kdnrs | TWF2 |
| 0 | 0.3863118 | 0.309 | 0.079 | 0 | MO_kdnrs | FAM49A |
| 0 | 0.38616161 | 0.325 | 0.067 | 0 | MO_kdnrs | SLC7A7 |
| 0 | 0.3838085 | 0.359 | 0.11 | 0 | MO_kdnrs | APBB1IP |
| 0 | 0.38070184 | 0.309 | 0.073 | 0 | MO_kdnrs | TTYH3 |

| | | | | | | |
|---|---|---|---|---|---|---|
| 0 | 0.37925093 | 0.231 | 0.014 | 0 | MO_kdnrs | SLCO2B1 |
| 0 | 0.37904084 | 0.254 | 0.018 | 0 | MO_kdnrs | SLAMF8 |
| 0 | 0.37772111 | 0.313 | 0.079 | 0 | MO_kdnrs | LPCAT2 |
| 0 | 0.37592993 | 0.291 | 0.052 | 0 | MO_kdnrs | GPR65 |
| 0 | 0.37558588 | 0.327 | 0.092 | 0 | MO_kdnrs | GM2A |
| 0 | 0.37492793 | 0.373 | 0.119 | 0 | MO_kdnrs | TNFRSF14 |
| 0 | 0.3745628 | 0.408 | 0.156 | 0 | MO_kdnrs | AKR1A1 |
| 0 | 0.37190224 | 0.304 | 0.083 | 0 | MO_kdnrs | GYPC |
| 0 | 0.36699974 | 0.349 | 0.097 | 0 | MO_kdnrs | CASP1 |
| 0 | 0.36698933 | 0.378 | 0.132 | 0 | MO_kdnrs | USF2 |
| 0 | 0.36695051 | 0.257 | 0.023 | 0 | MO_kdnrs | FGD2 |
| 0 | 0.36619228 | 0.348 | 0.111 | 0 | MO_kdnrs | FOXN3 |
| 0 | 0.36551712 | 0.148 | 0.008 | 0 | MO_kdnrs | GPR34 |
| 0 | 0.36379468 | 0.164 | 0.017 | 0 | MO_kdnrs | F13A1 |
| 0 | 0.36288349 | 0.398 | 0.144 | 0 | MO_kdnrs | CEP170 |
| 0 | 0.36234002 | 0.345 | 0.106 | 0 | MO_kdnrs | MFSD1 |
| 0 | 0.3603109 | 0.413 | 0.157 | 0 | MO_kdnrs | REX1BD |
| 0 | 0.35849135 | 0.312 | 0.088 | 0 | MO_kdnrs | SMCO4 |
| 0 | 0.35817637 | 0.311 | 0.072 | 0 | MO_kdnrs | PARVG |
| 0 | 0.35607725 | 0.222 | 0.047 | 0 | MO_kdnrs | PMP22 |
| 0 | 0.35547746 | 0.402 | 0.129 | 0 | MO_kdnrs | CLEC5A |
| 0 | 0.35415386 | 0.278 | 0.049 | 0 | MO_kdnrs | PLXND1 |
| 0 | 0.3531489 | 0.634 | 0.272 | 0 | MO_kdnrs | SERPINA1 |
| 0 | 0.35071518 | 0.422 | 0.154 | 0 | MO_kdnrs | PPP1R18 |
| 0 | 0.34900234 | 0.471 | 0.183 | 0 | MO_kdnrs | SLC43A2 |
| 0 | 0.34847543 | 0.308 | 0.079 | 0 | MO_kdnrs | TBXAS1 |
| 0 | 0.34380246 | 0.23 | 0.032 | 0 | MO_kdnrs | GRASP |
| 0 | 0.34351144 | 0.307 | 0.077 | 0 | MO_kdnrs | LAT2 |
| 0 | 0.34342653 | 0.431 | 0.14 | 0 | MO_kdnrs | LCP2 |
| 0 | 0.34342074 | 0.38 | 0.121 | 0 | MO_kdnrs | NCF1 |
| 0 | 0.34154087 | 0.321 | 0.086 | 0 | MO_kdnrs | ENG |
| 0 | 0.34135877 | 0.48 | 0.182 | 0 | MO_kdnrs | GK |
| 0 | 0.34042677 | 0.335 | 0.095 | 0 | MO_kdnrs | ITGAM |
| 0 | 0.33932004 | 0.324 | 0.092 | 0 | MO_kdnrs | ALOX5 |
| 0 | 0.33896415 | 0.335 | 0.113 | 0 | MO_kdnrs | MFSD12 |
| 0 | 0.33700638 | 0.284 | 0.069 | 0 | MO_kdnrs | DOK2 |
| 0 | 0.33600542 | 0.306 | 0.08 | 0 | MO_kdnrs | KIAA0930 |
| 0 | 0.33532244 | 0.294 | 0.076 | 0 | MO_kdnrs | RGS19 |
| 0 | 0.33248305 | 0.295 | 0.066 | 0 | MO_kdnrs | FUOM |
| 0 | 0.33187125 | 0.362 | 0.114 | 0 | MO_kdnrs | FGR |
| 0 | 0.33036824 | 0.295 | 0.074 | 0 | MO_kdnrs | PLXNC1 |
| 0 | 0.32976031 | 0.107 | 0.011 | 0 | MO_kdnrs | CX3CR1 |
| 0 | 0.32626207 | 0.336 | 0.103 | 0 | MO_kdnrs | DOCK10 |
| 0 | 0.32499891 | 0.175 | 0.025 | 0 | MO_kdnrs | C1orf54 |
| 0 | 0.32261354 | 0.299 | 0.085 | 0 | MO_kdnrs | IQGAP2 |
| 0 | 0.32207767 | 0.281 | 0.071 | 0 | MO_kdnrs | NPL |
| 0 | 0.32188695 | 0.315 | 0.094 | 0 | MO_kdnrs | PREX1 |
| 0 | 0.32115925 | 0.295 | 0.086 | 0 | MO_kdnrs | CSK |
| 0 | 0.31794257 | 0.218 | 0.042 | 0 | MO_kdnrs | TCF4 |
| 0 | 0.31362431 | 0.286 | 0.071 | 0 | MO_kdnrs | EVL |

| | | | | | | |
|---|---|---|---|---|---|---|
| 0 | 0.3134943 | 0.266 | 0.057 | 0 | MO_kdnrs | SH2B3 |
| 0 | 0.31309634 | 0.403 | 0.122 | 0 | MO_kdnrs | FPR1 |
| 0 | 0.31292627 | 0.249 | 0.063 | 0 | MO_kdnrs | RUNX3 |
| 0 | 0.31198368 | 0.297 | 0.08 | 0 | MO_kdnrs | PLEKHO2 |
| 0 | 0.31181142 | 0.23 | 0.025 | 0 | MO_kdnrs | SCIMP |
| 0 | 0.31131034 | 0.282 | 0.067 | 0 | MO_kdnrs | ARHGAP45 |
| 0 | 0.30914421 | 0.258 | 0.049 | 0 | MO_kdnrs | TNFSF12 |
| 0 | 0.30739134 | 0.243 | 0.052 | 0 | MO_kdnrs | IRF8 |
| 0 | 0.3069485 | 0.285 | 0.08 | 0 | MO_kdnrs | DOCK4 |
| 0 | 0.30544533 | 0.165 | 0.019 | 0 | MO_kdnrs | SLC1A3 |
| 0 | 0.30487746 | 0.26 | 0.06 | 0 | MO_kdnrs | RAB20 |
| 0 | 0.30480478 | 0.306 | 0.094 | 0 | MO_kdnrs | COLGALT1 |
| 0 | 0.30479361 | 0.2 | 0.044 | 0 | MO_kdnrs | NRP2 |
| 0 | 0.30168365 | 0.297 | 0.082 | 0 | MO_kdnrs | DOCK2 |
| 0 | 0.30167449 | 0.244 | 0.059 | 0 | MO_kdnrs | PSTPIP2 |
| 0 | 0.30109918 | 0.267 | 0.058 | 0 | MO_kdnrs | ARHGAP4 |
| 0 | 0.29814453 | 0.267 | 0.067 | 0 | MO_kdnrs | LYSMD2 |
| 0 | 0.29789278 | 0.297 | 0.079 | 0 | MO_kdnrs | CYTH4 |
| 0 | 0.29611776 | 0.212 | 0.044 | 0 | MO_kdnrs | IL3RA |
| 0 | 0.28940103 | 0.246 | 0.064 | 0 | MO_kdnrs | CHST11 |
| 0 | 0.28936989 | 0.226 | 0.044 | 0 | MO_kdnrs | CLEC4A |
| 0 | 0.28783732 | 0.233 | 0.041 | 0 | MO_kdnrs | LYL1 |
| 0 | 0.28705022 | 0.245 | 0.058 | 0 | MO_kdnrs | EPSTI1 |
| 0 | 0.28494432 | 0.241 | 0.049 | 0 | MO_kdnrs | ADCY7 |
| 0 | 0.28209977 | 0.217 | 0.041 | 0 | MO_kdnrs | CD302 |
| 0 | 0.2770396 | 0.181 | 0.017 | 0 | MO_kdnrs | ABI3 |
| 0 | 0.27499757 | 0.237 | 0.06 | 0 | MO_kdnrs | ST8SIA4 |
| 0 | 0.27430112 | 0.229 | 0.052 | 0 | MO_kdnrs | AOAH |
| 0 | 0.27376005 | 0.255 | 0.06 | 0 | MO_kdnrs | GRK3 |
| 0 | 0.27313653 | 0.23 | 0.051 | 0 | MO_kdnrs | CD300A |
| 0 | 0.2721754 | 0.204 | 0.033 | 0 | MO_kdnrs | CD300LF |
| 0 | 0.26962478 | 0.212 | 0.041 | 0 | MO_kdnrs | P2RX7 |
| 0 | 0.26584516 | 0.243 | 0.065 | 0 | MO_kdnrs | A1BG |
| 0 | 0.26550519 | 0.21 | 0.035 | 0 | MO_kdnrs | CLEC12A |
| 0 | 0.26335345 | 0.246 | 0.064 | 0 | MO_kdnrs | PSMB10 |
| 0 | 0.26331869 | 0.208 | 0.037 | 0 | MO_kdnrs | SPN |
| 0 | 0.26311322 | 0.222 | 0.048 | 0 | MO_kdnrs | ASGR1 |
| 0 | 0.26306906 | 0.246 | 0.053 | 0 | MO_kdnrs | GIMAP7 |
| 0 | 0.26065126 | 0.289 | 0.08 | 0 | MO_kdnrs | IKZF1 |
| 0 | 0.26061837 | 0.123 | 0.015 | 0 | MO_kdnrs | PADI2 |
| 0 | 0.25577885 | 0.25 | 0.066 | 0 | MO_kdnrs | NCF4 |
| 0 | 0.25295442 | 0.4 | 0.136 | 0 | MO_kdnrs | CD300E |
| 0 | 0.25160414 | 0.154 | 0.017 | 0 | MO_kdnrs | RAB7B |
| 0 | 0.25105181 | 0.219 | 0.05 | 0 | MO_kdnrs | KCNAB2 |
| 0 | 0.25063226 | 0.204 | 0.04 | 0 | MO_kdnrs | OTULINL |
| 0 | -0.6555242 | 0.396 | 0.13 | 0 | MO_kdnrs | MT1H |
| 0 | -0.7027849 | 0.9 | 0.656 | 0 | MO_kdnrs | TIMP1 |
| 0 | -0.8316289 | 0.015 | 0.347 | 0 | MO_kdnrs | PERP |
| 0 | -0.9173951 | 0.746 | 0.954 | 0 | MO_kdnrs | MT-ATP6 |
| 0 | -0.9320939 | 0.313 | 0.652 | 0 | MO_kdnrs | DSTN |

| | | | | | | |
|---|---|---|---|---|---|---|
| 0 | -1.0046584 | 0.005 | 0.357 | 0 | MO_kdnrs | MYO6 |
| 0 | -1.0052279 | 0.676 | 0.929 | 0 | MO_kdnrs | MT-ND2 |
| 0 | -1.1585839 | 0.741 | 0.947 | 0 | MO_kdnrs | MT-CYB |
| 0 | -1.1808459 | 0.083 | 0.485 | 0 | MO_kdnrs | CLDN7 |
| 0 | -1.2658033 | 0.007 | 0.317 | 0 | MO_kdnrs | RND3 |
| 0 | -1.5226053 | 0.327 | 0.665 | 0 | MO_kdnrs | MTRNR2L12 |
| 0 | -1.5741626 | 0.058 | 0.37 | 0 | MO_kdnrs | GDF15 |
| 0 | -1.6313842 | 0.1 | 0.579 | 0 | MO_kdnrs | CD24 |
| 0 | -1.6360695 | 0.009 | 0.321 | 0 | MO_kdnrs | ADIRF |
| 0 | -1.6978826 | 0.134 | 0.479 | 0 | MO_kdnrs | SOX4 |
| 0 | -1.7286408 | 0.095 | 0.529 | 0 | MO_kdnrs | KRT8 |
| 0 | -1.7374991 | 0.08 | 0.434 | 0 | MO_kdnrs | SLPI |
| 0 | -1.7406432 | 0.08 | 0.395 | 0 | MO_kdnrs | MAL |
| 0 | -1.8894176 | 0.117 | 0.609 | 0 | MO_kdnrs | TACSTD2 |
| 0 | -1.9099199 | 0.113 | 0.583 | 0 | MO_kdnrs | KRT7 |
| 0 | -1.9579711 | 0.111 | 0.578 | 0 | MO_kdnrs | CLDN4 |
| 0 | -1.9846222 | 0.11 | 0.595 | 0 | MO_kdnrs | ELF3 |
| 0 | -2.0232776 | 0.12 | 0.597 | 0 | MO_kdnrs | KRT18 |
| 0 | -2.3445215 | 0.133 | 0.618 | 0 | MO_kdnrs | KRT19 |
| 0 | -2.6184032 | 0.122 | 0.573 | 0 | MO_kdnrs | WFDC2 |
| 0 | -3.6348721 | 0.145 | 0.557 | 0 | MO_kdnrs | LCN2 |
| 1.876E-307 | 0.47354021 | 0.995 | 0.905 | 4.432E-303 | MO_kdnrs | RPS8 |
| 1.618E-306 | 0.42615004 | 0.39 | 0.14 | 3.822E-302 | MO_kdnrs | SERPINB9 |
| 2.828E-306 | 0.41044673 | 0.411 | 0.16 | 6.68E-302 | MO_kdnrs | GNA13 |
| 1.075E-305 | 0.28422813 | 0.281 | 0.083 | 2.539E-301 | MO_kdnrs | UNC93B1 |
| 2.42E-305 | -0.7684921 | 0.435 | 0.152 | 5.716E-301 | MO_kdnrs | IL1B |
| 2.594E-305 | -0.8629332 | 0.775 | 0.966 | 6.127E-301 | MO_kdnrs | MT-CO1 |
| 9.56E-305 | 0.41599319 | 0.53 | 0.235 | 2.258E-300 | MO_kdnrs | MBNL1 |
| 1.43E-304 | -0.8376987 | 0.538 | 0.229 | 3.379E-300 | MO_kdnrs | MT1G |
| 9.883E-304 | 0.36211733 | 0.248 | 0.068 | 2.335E-299 | MO_kdnrs | CYP27A1 |
| 5.174E-303 | -0.9355801 | 0.468 | 0.719 | 1.222E-298 | MO_kdnrs | PPP1R15A |
| 1.125E-302 | 0.28492361 | 0.263 | 0.075 | 2.657E-298 | MO_kdnrs | DNMT1 |
| 2.354E-301 | -0.846661 | 0.125 | 0.434 | 5.561E-297 | MO_kdnrs | CD59 |
| 2.678E-301 | 0.46588477 | 0.995 | 0.942 | 6.326E-297 | MO_kdnrs | RPS16 |
| 7.355E-301 | 0.48325497 | 0.685 | 0.371 | 1.737E-296 | MO_kdnrs | H2AFY |
| 2.855E-299 | 0.46791291 | 0.621 | 0.296 | 6.744E-295 | MO_kdnrs | IFITM3 |
| 9.859E-299 | -3.0509509 | 0.103 | 0.403 | 2.329E-294 | MO_kdnrs | KRT13 |
| 5.211E-298 | 0.33383911 | 0.352 | 0.122 | 1.231E-293 | MO_kdnrs | IL2RG |
| 6.691E-298 | -1.0601234 | 0.017 | 0.316 | 1.581E-293 | MO_kdnrs | GPRC5A |
| 6.755E-298 | 0.30866309 | 0.279 | 0.084 | 1.596E-293 | MO_kdnrs | DNASE2 |
| 7.32E-297 | -0.9395375 | 0.823 | 0.955 | 1.729E-292 | MO_kdnrs | MT-CO3 |
| 9.536E-297 | 0.37547441 | 0.461 | 0.192 | 2.253E-292 | MO_kdnrs | ZYX |
| 2.792E-296 | 0.38136069 | 0.365 | 0.131 | 6.594E-292 | MO_kdnrs | SORL1 |
| 3.798E-296 | 0.37496978 | 0.424 | 0.172 | 8.972E-292 | MO_kdnrs | ATP6AP1 |
| 1.712E-295 | 0.28268788 | 0.272 | 0.081 | 4.044E-291 | MO_kdnrs | GAA |
| 5.919E-295 | -0.8736053 | 0.697 | 0.934 | 1.398E-290 | MO_kdnrs | MT-ND1 |
| 1.823E-294 | 0.4947412 | 0.81 | 0.557 | 4.307E-290 | MO_kdnrs | LITAF |
| 2.487E-293 | -0.9105503 | 0.006 | 0.297 | 5.876E-289 | MO_kdnrs | MAL2 |
| 5.618E-293 | -0.3880617 | 0.361 | 0.123 | 1.327E-288 | MO_kdnrs | CCL2 |
| 9.749E-293 | 0.45332102 | 0.989 | 0.878 | 2.303E-288 | MO_kdnrs | RPS3 |

| | | | | | | |
|---|---|---|---|---|---|---|
| 4.235E-290 | 0.48705482 | 0.989 | 0.893 | 1.001E-285 | MO_kdnrs | RPS11 |
| 4.58E-290 | 0.39603329 | 0.496 | 0.221 | 1.082E-285 | MO_kdnrs | NDUFA3 |
| 5.84E-290 | 0.48116246 | 0.943 | 0.729 | 1.38E-285 | MO_kdnrs | EEF1B2 |
| 7.885E-290 | 0.45846557 | 0.985 | 0.924 | 1.863E-285 | MO_kdnrs | OAZ1 |
| 2.795E-289 | 0.31857781 | 0.353 | 0.126 | 6.602E-285 | MO_kdnrs | LEPROTL1 |
| 3.732E-289 | 0.38960215 | 0.437 | 0.177 | 8.815E-285 | MO_kdnrs | ARHGAP18 |
| 3.777E-289 | 0.45346478 | 0.599 | 0.307 | 8.923E-285 | MO_kdnrs | PSMB3 |
| 2.998E-288 | 0.33040319 | 0.43 | 0.166 | 7.083E-284 | MO_kdnrs | RAC2 |
| 4.698E-288 | -0.9636216 | 0.556 | 0.847 | 1.11E-283 | MO_kdnrs | MT-ND5 |
| 1.264E-287 | 0.49323279 | 0.989 | 0.894 | 2.987E-283 | MO_kdnrs | RPS7 |
| 1.395E-287 | 0.46908004 | 0.616 | 0.3 | 3.296E-283 | MO_kdnrs | CEBPD |
| 1.457E-287 | 0.29323716 | 0.305 | 0.099 | 3.442E-283 | MO_kdnrs | ARAP1 |
| 2.871E-287 | 0.44461357 | 0.577 | 0.29 | 6.782E-283 | MO_kdnrs | BLOC1S1 |
| 2.324E-286 | 0.33590136 | 0.367 | 0.134 | 5.49E-282 | MO_kdnrs | VSIR |
| 3.707E-286 | 0.47786199 | 0.585 | 0.294 | 8.757E-282 | MO_kdnrs | CMTM6 |
| 2.257E-285 | -0.8846073 | 0.012 | 0.299 | 5.331E-281 | MO_kdnrs | CAMK2N1 |
| 2.309E-283 | 0.4127326 | 0.47 | 0.207 | 5.455E-279 | MO_kdnrs | SYNGR2 |
| 8.479E-283 | 0.35736962 | 0.297 | 0.097 | 2.003E-278 | MO_kdnrs | SATB1 |
| 2.447E-282 | -1.096193 | 0.004 | 0.287 | 5.781E-278 | MO_kdnrs | TSPAN1 |
| 4.27E-282 | 0.31201109 | 0.317 | 0.107 | 1.009E-277 | MO_kdnrs | SYK |
| 1.596E-280 | -4.3468869 | 0.136 | 0.425 | 3.771E-276 | MO_kdnrs | SPRR3 |
| 5.9E-280 | 0.25903575 | 0.566 | 0.26 | 1.394E-275 | MO_kdnrs | HMOX1 |
| 2.607E-278 | 0.29375748 | 0.289 | 0.092 | 6.157E-274 | MO_kdnrs | PAK1 |
| 2.927E-278 | -2.3436016 | 0.079 | 0.365 | 6.914E-274 | MO_kdnrs | PSCA |
| 2.969E-278 | 0.26236097 | 0.257 | 0.076 | 7.014E-274 | MO_kdnrs | ZNF385A |
| 4.069E-278 | 0.32586219 | 0.374 | 0.14 | 9.613E-274 | MO_kdnrs | DOCK8 |
| 7.187E-278 | 0.31140606 | 0.323 | 0.112 | 1.698E-273 | MO_kdnrs | SCPEP1 |
| 1.156E-277 | 0.45333983 | 0.816 | 0.539 | 2.732E-273 | MO_kdnrs | TOMM7 |
| 1.53E-277 | 0.47513675 | 0.994 | 0.945 | 3.614E-273 | MO_kdnrs | RPL18 |
| 1.762E-277 | -2.5114655 | 0.154 | 0.43 | 4.162E-273 | MO_kdnrs | S100P |
| 5.697E-277 | -2.2235963 | 0.083 | 0.367 | 1.346E-272 | MO_kdnrs | HPGD |
| 1.068E-276 | 0.39256021 | 0.623 | 0.305 | 2.522E-272 | MO_kdnrs | MSN |
| 3.171E-276 | 0.37501271 | 0.56 | 0.264 | 7.491E-272 | MO_kdnrs | APRT |
| 3.895E-276 | 0.25180404 | 0.121 | 0.019 | 9.201E-272 | MO_kdnrs | PLTP |
| 8.575E-276 | 0.40392382 | 0.336 | 0.122 | 2.026E-271 | MO_kdnrs | PLD3 |
| 3.607E-275 | 0.45935421 | 0.69 | 0.389 | 8.521E-271 | MO_kdnrs | EIF3K |
| 5.281E-275 | -1.0075765 | 0.177 | 0.46 | 1.248E-270 | MO_kdnrs | ATP1B1 |
| 5.584E-274 | 0.63228762 | 0.273 | 0.084 | 1.319E-269 | MO_kdnrs | RETN |
| 1.453E-273 | -2.5734385 | 0.078 | 0.362 | 3.433E-269 | MO_kdnrs | MMP7 |
| 7.441E-273 | 0.29072093 | 0.269 | 0.082 | 1.758E-268 | MO_kdnrs | LHFPL2 |
| 4.359E-272 | 0.51917036 | 0.77 | 0.484 | 1.03E-267 | MO_kdnrs | DBI |
| 1.584E-271 | 0.29032269 | 0.296 | 0.098 | 3.741E-267 | MO_kdnrs | UBE2L6 |
| 7.707E-271 | 0.33803928 | 0.476 | 0.208 | 1.821E-266 | MO_kdnrs | PPIF |
| 1.194E-270 | 0.3101416 | 0.297 | 0.096 | 2.822E-266 | MO_kdnrs | PECAM1 |
| 2.713E-270 | 0.32456637 | 0.309 | 0.106 | 6.41E-266 | MO_kdnrs | SOAT1 |
| 3.851E-270 | 0.34045582 | 0.429 | 0.176 | 9.098E-266 | MO_kdnrs | MYO9B |
| 4.298E-270 | 0.42368033 | 0.659 | 0.35 | 1.015E-265 | MO_kdnrs | ATP5ME |
| 3.104E-269 | -0.645668 | 0.019 | 0.296 | 7.332E-265 | MO_kdnrs | JUP |
| 5.41E-269 | 0.34734927 | 0.448 | 0.186 | 1.278E-264 | MO_kdnrs | STK17B |
| 1.016E-268 | 0.62082331 | 0.909 | 0.754 | 2.4E-264 | MO_kdnrs | RPS26 |

| | | | | | | |
|---|---|---|---|---|---|---|
| 1.086E-268 | 0.33587834 | 0.46 | 0.193 | 2.566E-264 | MO_kdnrs | TUBA1A |
| 1.793E-268 | 0.26461262 | 0.193 | 0.048 | 4.235E-264 | MO_kdnrs | SPRED1 |
| 1.883E-268 | 0.38741957 | 0.545 | 0.257 | 4.447E-264 | MO_kdnrs | RBM3 |
| 1.889E-268 | 0.25866332 | 0.28 | 0.087 | 4.461E-264 | MO_kdnrs | CCR1 |
| 8.733E-268 | -1.2599178 | 0.011 | 0.285 | 2.063E-263 | MO_kdnrs | LMO7 |
| 2.19E-267 | 0.2612856 | 0.241 | 0.07 | 5.173E-263 | MO_kdnrs | PDE4A |
| 2.511E-267 | 0.40147111 | 0.615 | 0.308 | 5.932E-263 | MO_kdnrs | TLN1 |
| 2.758E-267 | -0.808054 | 0.009 | 0.282 | 6.516E-263 | MO_kdnrs | MUC1 |
| 1.404E-266 | 0.43836706 | 0.994 | 0.912 | 3.317E-262 | MO_kdnrs | RPL9 |
| 6.772E-266 | 0.32632251 | 0.363 | 0.138 | 1.6E-261 | MO_kdnrs | HNMT |
| 6.807E-266 | 0.29586603 | 0.371 | 0.139 | 1.608E-261 | MO_kdnrs | ARHGAP30 |
| 7.261E-266 | 0.31989738 | 0.343 | 0.127 | 1.715E-261 | MO_kdnrs | STX4 |
| 1.4E-265 | 0.43299306 | 1 | 0.979 | 3.307E-261 | MO_kdnrs | EEF1A1 |
| 4.317E-265 | 0.27668499 | 0.28 | 0.09 | 1.02E-260 | MO_kdnrs | ARID5A |
| 1.635E-264 | 0.27157441 | 0.268 | 0.084 | 3.862E-260 | MO_kdnrs | ANKRD44 |
| 1.839E-264 | 0.26971154 | 0.261 | 0.081 | 4.345E-260 | MO_kdnrs | CMTM3 |
| 8.712E-264 | -0.7418446 | 0.003 | 0.272 | 2.058E-259 | MO_kdnrs | KLF5 |
| 9.229E-264 | 0.45990029 | 0.751 | 0.45 | 2.18E-259 | MO_kdnrs | CD81 |
| 9.884E-263 | -1.6578962 | 0.083 | 0.359 | 2.335E-258 | MO_kdnrs | IGFBP7 |
| 2.513E-262 | 0.43381106 | 0.997 | 0.887 | 5.936E-258 | MO_kdnrs | RPS9 |
| 6.94E-261 | 0.47901913 | 0.799 | 0.552 | 1.639E-256 | MO_kdnrs | TSPO |
| 7.118E-261 | 0.35476767 | 0.504 | 0.23 | 1.681E-256 | MO_kdnrs | C4orf48 |
| 1.568E-259 | 0.41429972 | 0.993 | 0.918 | 3.704E-255 | MO_kdnrs | RPL18A |
| 1.714E-259 | -0.8017193 | 0.278 | 0.545 | 4.05E-255 | MO_kdnrs | HEBP2 |
| 6.579E-259 | 0.31251857 | 0.343 | 0.128 | 1.554E-254 | MO_kdnrs | ZMIZ1 |
| 6.866E-259 | 0.42703334 | 0.989 | 0.891 | 1.622E-254 | MO_kdnrs | RPL21 |
| 1.657E-258 | 0.37403427 | 0.513 | 0.241 | 3.914E-254 | MO_kdnrs | TUBB |
| 2.995E-258 | 0.2826783 | 0.299 | 0.101 | 7.075E-254 | MO_kdnrs | MYO1G |
| 4.444E-258 | 0.29924206 | 0.355 | 0.131 | 1.05E-253 | MO_kdnrs | DMXL2 |
| 7.631E-257 | -0.9346303 | 0.707 | 0.849 | 1.803E-252 | MO_kdnrs | SQSTM1 |
| 3.489E-256 | 0.29282426 | 0.327 | 0.119 | 8.242E-252 | MO_kdnrs | CCDC85B |
| 9.675E-256 | 0.41213009 | 0.519 | 0.254 | 2.286E-251 | MO_kdnrs | ATP6AP2 |
| 1.554E-255 | -2.1143585 | 0.022 | 0.288 | 3.67E-251 | MO_kdnrs | S100A2 |
| 9.299E-254 | -0.7343087 | 0.003 | 0.263 | 2.197E-249 | MO_kdnrs | PRSS8 |
| 2.085E-253 | 0.4237575 | 0.96 | 0.858 | 4.925E-249 | MO_kdnrs | ARPC3 |
| 3.642E-253 | 0.2859927 | 0.289 | 0.098 | 8.603E-249 | MO_kdnrs | IL17RA |
| 5.012E-252 | 0.27527542 | 0.299 | 0.103 | 1.184E-247 | MO_kdnrs | MPP1 |
| 4.445E-251 | -1.0102737 | 0.002 | 0.259 | 1.05E-246 | MO_kdnrs | ADGRF1 |
| 4.056E-250 | 0.41455193 | 0.976 | 0.898 | 9.581E-246 | MO_kdnrs | CFL1 |
| 5.295E-250 | -0.3004517 | 0.501 | 0.198 | 1.251E-245 | MO_kdnrs | VCAN |
| 1.027E-249 | -0.7536959 | 0.8 | 0.958 | 2.426E-245 | MO_kdnrs | MT-CO2 |
| 2.022E-249 | 0.29301573 | 0.29 | 0.1 | 4.776E-245 | MO_kdnrs | VEGFB |
| 4.905E-249 | 0.33958573 | 0.441 | 0.192 | 1.159E-244 | MO_kdnrs | RNF13 |
| 1.275E-248 | 0.29916585 | 0.397 | 0.161 | 3.013E-244 | MO_kdnrs | RILPL2 |
| 5.059E-247 | -0.7072 | 0.003 | 0.257 | 1.195E-242 | MO_kdnrs | S100A16 |
| 2.01E-246 | 0.35080401 | 0.4 | 0.168 | 4.749E-242 | MO_kdnrs | RAPGEF1 |
| 2.499E-245 | 0.3782988 | 0.411 | 0.174 | 5.903E-241 | MO_kdnrs | SNX10 |
| 4.713E-245 | 0.38258408 | 0.525 | 0.253 | 1.113E-240 | MO_kdnrs | VASP |
| 8.975E-243 | 0.40279709 | 0.579 | 0.309 | 2.12E-238 | MO_kdnrs | MOB1A |
| 1.039E-242 | -0.9165891 | 0.003 | 0.254 | 2.455E-238 | MO_kdnrs | PDZK1IP1 |

| | | | | | | |
|---|---|---|---|---|---|---|
| 3.596E-242 | -0.9128103 | 0.005 | 0.256 | 8.496E-238 | MO_kdnrs | EPCAM |
| 1.104E-241 | -0.5460518 | 0.002 | 0.251 | 2.609E-237 | MO_kdnrs | TJP1 |
| 3.606E-241 | 0.31361266 | 0.38 | 0.157 | 8.517E-237 | MO_kdnrs | IFI27L2 |
| 4.015E-241 | 0.40787391 | 0.671 | 0.369 | 9.485E-237 | MO_kdnrs | NDUFB2 |
| 6.067E-241 | 0.34450688 | 0.955 | 0.738 | 1.433E-236 | MO_kdnrs | RPSA |
| 2.683E-240 | 0.3155598 | 0.397 | 0.167 | 6.338E-236 | MO_kdnrs | ATG3 |
| 7.752E-240 | 0.58277471 | 0.544 | 0.275 | 1.831E-235 | MO_kdnrs | REL |
| 2.152E-239 | -0.669144 | 0.048 | 0.311 | 5.084E-235 | MO_kdnrs | SPINT1 |
| 2.243E-239 | 0.31000351 | 0.38 | 0.156 | 5.299E-235 | MO_kdnrs | TCIRG1 |
| 9.3E-239 | -0.9600901 | 0.22 | 0.477 | 2.197E-234 | MO_kdnrs | RAB11FIP1 |
| 2.003E-238 | 0.38368031 | 0.378 | 0.161 | 4.731E-234 | MO_kdnrs | IL13RA1 |
| 2.025E-238 | 0.32617 | 0.379 | 0.155 | 4.785E-234 | MO_kdnrs | TNFAIP8 |
| 4.224E-238 | 0.4233337 | 0.981 | 0.837 | 9.979E-234 | MO_kdnrs | RPS4X |
| 4.389E-235 | 0.26196096 | 0.49 | 0.22 | 1.037E-230 | MO_kdnrs | ATP2B1 |
| 5.234E-235 | 0.29934429 | 0.373 | 0.152 | 1.236E-230 | MO_kdnrs | LACTB |
| 3.696E-234 | -1.1296377 | 0.238 | 0.514 | 8.73E-230 | MO_kdnrs | MTRNR2L8 |
| 1.736E-233 | 0.34663392 | 0.675 | 0.369 | 4.101E-229 | MO_kdnrs | HIF1A |
| 4.582E-233 | 0.40408468 | 0.882 | 0.699 | 1.082E-228 | MO_kdnrs | ATP5MG |
| 1.034E-232 | 0.26137976 | 0.194 | 0.052 | 2.444E-228 | MO_kdnrs | CD36 |
| 4.086E-232 | 0.40533973 | 0.943 | 0.772 | 9.653E-228 | MO_kdnrs | RPL31 |
| 6.445E-231 | 0.36184955 | 0.509 | 0.252 | 1.523E-226 | MO_kdnrs | FAM49B |
| 8.806E-231 | 0.39115789 | 0.983 | 0.867 | 2.08E-226 | MO_kdnrs | RPL23A |
| 1.018E-230 | -1.7029188 | 0.008 | 0.252 | 2.405E-226 | MO_kdnrs | PRSS22 |
| 1.9E-230 | 0.25790777 | 0.276 | 0.094 | 4.488E-226 | MO_kdnrs | PAG1 |
| 2.528E-230 | 0.26662982 | 0.309 | 0.115 | 5.973E-226 | MO_kdnrs | C7orf50 |
| 5.846E-230 | 0.28728417 | 0.291 | 0.103 | 1.381E-225 | MO_kdnrs | FKBP5 |
| 1.155E-229 | 0.39453595 | 0.988 | 0.89 | 2.729E-225 | MO_kdnrs | RPL29 |
| 2.464E-229 | 0.35325591 | 0.529 | 0.268 | 5.82E-225 | MO_kdnrs | HIGD2A |
| 3.961E-229 | 0.27838928 | 0.331 | 0.128 | 9.356E-225 | MO_kdnrs | GRK2 |
| 5.61E-229 | 0.31964833 | 0.425 | 0.189 | 1.325E-224 | MO_kdnrs | GSTK1 |
| 9.499E-229 | 0.27938415 | 0.343 | 0.135 | 2.244E-224 | MO_kdnrs | EHBP1L1 |
| 1.594E-227 | -0.8317078 | 0.009 | 0.249 | 3.765E-223 | MO_kdnrs | ITGA2 |
| 1.494E-225 | 0.511838 | 0.161 | 0.039 | 3.53E-221 | MO_kdnrs | MMP9 |
| 1.399E-224 | -1.019358 | 0.051 | 0.302 | 3.305E-220 | MO_kdnrs | CLU |
| 1.469E-224 | 0.25915091 | 0.247 | 0.082 | 3.47E-220 | MO_kdnrs | SDSL |
| 1.573E-224 | 0.35279657 | 0.494 | 0.246 | 3.716E-220 | MO_kdnrs | CSNK2B |
| 4.398E-224 | 0.30877362 | 0.408 | 0.18 | 1.039E-219 | MO_kdnrs | SSR1 |
| 8.559E-224 | 0.2571064 | 0.276 | 0.097 | 2.022E-219 | MO_kdnrs | SLC43A3 |
| 7.44E-223 | 0.36236014 | 0.572 | 0.297 | 1.758E-218 | MO_kdnrs | SLC16A3 |
| 3.228E-222 | 0.33910855 | 0.494 | 0.245 | 7.626E-218 | MO_kdnrs | RNH1 |
| 3.327E-222 | 0.30595603 | 0.513 | 0.248 | 7.859E-218 | MO_kdnrs | TSC22D3 |
| 6.622E-221 | -1.7249068 | 0.026 | 0.268 | 1.564E-216 | MO_kdnrs | CRYAB |
| 9.446E-221 | -0.5542717 | 0.003 | 0.236 | 2.231E-216 | MO_kdnrs | DSP |
| 1.854E-220 | 0.35082125 | 0.507 | 0.257 | 4.381E-216 | MO_kdnrs | GDI2 |
| 1.21E-219 | 0.26366945 | 0.305 | 0.116 | 2.859E-215 | MO_kdnrs | ADPGK |
| 7.316E-219 | 0.32501223 | 0.506 | 0.252 | 1.728E-214 | MO_kdnrs | MINOS1 |
| 1.249E-217 | 0.37386673 | 0.904 | 0.686 | 2.951E-213 | MO_kdnrs | RPL4 |
| 2.886E-217 | -0.4700046 | 0.004 | 0.233 | 6.819E-213 | MO_kdnrs | CNN3 |
| 8.29E-217 | 0.33078194 | 0.4 | 0.178 | 1.958E-212 | MO_kdnrs | GNS |
| 4.305E-216 | 0.29920373 | 0.279 | 0.101 | 1.017E-211 | MO_kdnrs | IFI6 |

| | | | | | | |
|---|---|---|---|---|---|---|
| 2.638E-215 | -0.5621757 | 0.004 | 0.233 | 6.231E-211 | MO_kdnrs | EHF |
| 3.771E-215 | -0.9062478 | 0.09 | 0.339 | 8.909E-211 | MO_kdnrs | GADD45A |
| 3.68E-214 | 0.32332771 | 0.47 | 0.229 | 8.693E-210 | MO_kdnrs | RPS17 |
| 4.63E-214 | -0.7388874 | 0.025 | 0.262 | 1.094E-209 | MO_kdnrs | ITGB8 |
| 4.732E-212 | 0.27333637 | 0.283 | 0.105 | 1.118E-207 | MO_kdnrs | ATP6V0A1 |
| 2.197E-211 | 0.36227003 | 0.335 | 0.141 | 5.19E-207 | MO_kdnrs | PTPN1 |
| 3.424E-211 | 0.35905311 | 0.987 | 0.848 | 8.088E-207 | MO_kdnrs | RPL23 |
| 5.055E-210 | 0.36719898 | 0.991 | 0.891 | 1.194E-205 | MO_kdnrs | RPL27A |
| 5.801E-210 | 0.28447593 | 0.54 | 0.272 | 1.37E-205 | MO_kdnrs | CHCHD10 |
| 1.221E-209 | 0.39290471 | 0.921 | 0.788 | 2.884E-205 | MO_kdnrs | ARPC2 |
| 5.751E-209 | -0.6017839 | 0.033 | 0.266 | 1.359E-204 | MO_kdnrs | SPTBN1 |
| 7.166E-209 | 0.34359384 | 0.611 | 0.336 | 1.693E-204 | MO_kdnrs | SNRPD2 |
| 5.679E-208 | 0.28046039 | 0.316 | 0.125 | 1.342E-203 | MO_kdnrs | MGAT4A |
| 4.401E-207 | 0.31185907 | 0.424 | 0.199 | 1.04E-202 | MO_kdnrs | LAMTOR2 |
| 3.194E-206 | 0.36155401 | 0.628 | 0.358 | 7.546E-202 | MO_kdnrs | EIF5A |
| 5.249E-206 | -0.5484731 | 0.004 | 0.224 | 1.24E-201 | MO_kdnrs | PARD6B |
| 7.441E-206 | 0.33576896 | 0.988 | 0.911 | 1.758E-201 | MO_kdnrs | RPS20 |
| 2.402E-205 | 0.3660998 | 0.8 | 0.542 | 5.675E-201 | MO_kdnrs | SLC25A6 |
| 1.318E-204 | 0.26597972 | 0.328 | 0.135 | 3.114E-200 | MO_kdnrs | VMA21 |
| 1.412E-204 | 0.30699838 | 0.378 | 0.17 | 3.337E-200 | MO_kdnrs | SQOR |
| 7.789E-204 | 0.33835097 | 0.69 | 0.404 | 1.84E-199 | MO_kdnrs | EFHD2 |
| 1.549E-203 | 0.37942533 | 0.919 | 0.758 | 3.659E-199 | MO_kdnrs | TMA7 |
| 1.561E-203 | 0.2754309 | 0.286 | 0.108 | 3.687E-199 | MO_kdnrs | MIS18BP1 |
| 5.353E-203 | 0.32143627 | 0.443 | 0.214 | 1.265E-198 | MO_kdnrs | PSMB8 |
| 7.633E-202 | 0.35625907 | 0.633 | 0.364 | 1.803E-197 | MO_kdnrs | PSME1 |
| 1.621E-201 | 0.34307065 | 0.555 | 0.304 | 3.83E-197 | MO_kdnrs | DRAP1 |
| 2.071E-201 | 0.39664178 | 0.868 | 0.671 | 4.893E-197 | MO_kdnrs | BRI3 |
| 2.38E-200 | 0.25501795 | 0.287 | 0.11 | 5.623E-196 | MO_kdnrs | FKBP15 |
| 3.664E-200 | 0.31968827 | 0.441 | 0.215 | 8.655E-196 | MO_kdnrs | CTSA |
| 4.517E-200 | 0.26552021 | 0.323 | 0.131 | 1.067E-195 | MO_kdnrs | SH3KBP1 |
| 4.672E-200 | 0.30544481 | 0.388 | 0.175 | 1.104E-195 | MO_kdnrs | RHOQ |
| 4.721E-200 | -0.4103142 | 0.007 | 0.223 | 1.115E-195 | MO_kdnrs | CTTN |
| 8.761E-200 | 0.36779303 | 0.898 | 0.747 | 2.07E-195 | MO_kdnrs | UQCRB |
| 9.043E-200 | 0.29902717 | 0.261 | 0.093 | 2.136E-195 | MO_kdnrs | SLC16A10 |
| 7.325E-199 | 0.29414872 | 0.426 | 0.201 | 1.73E-194 | MO_kdnrs | SNRPF |
| 7.35E-199 | -0.62447 | 0.74 | 0.95 | 1.736E-194 | MO_kdnrs | MT-ND4 |
| 3.574E-197 | 0.29749517 | 0.414 | 0.195 | 8.442E-193 | MO_kdnrs | RPL17 |
| 4.785E-197 | -0.7246041 | 0.036 | 0.26 | 1.13E-192 | MO_kdnrs | S100A13 |
| 8.527E-197 | 0.36487853 | 0.991 | 0.897 | 2.014E-192 | MO_kdnrs | RPL7A |
| 4.564E-196 | 0.33736526 | 0.452 | 0.226 | 1.078E-191 | MO_kdnrs | PEA15 |
| 6.822E-196 | -0.3957529 | 0.002 | 0.212 | 1.612E-191 | MO_kdnrs | RBPMS |
| 3.706E-195 | 0.35782217 | 0.477 | 0.247 | 8.755E-191 | MO_kdnrs | CDC42SE1 |
| 7.203E-195 | -0.7381267 | 0.003 | 0.213 | 1.702E-190 | MO_kdnrs | SFN |
| 1.227E-194 | 0.34291441 | 0.34 | 0.146 | 2.897E-190 | MO_kdnrs | ALCAM |
| 1.511E-194 | 0.29695439 | 0.373 | 0.166 | 3.569E-190 | MO_kdnrs | BID |
| 1.76E-194 | 0.26995298 | 0.313 | 0.127 | 4.157E-190 | MO_kdnrs | RASSF5 |
| 3.079E-194 | -0.986587 | 0.004 | 0.214 | 7.274E-190 | MO_kdnrs | TM4SF1 |
| 1.467E-193 | 0.36055655 | 0.972 | 0.804 | 3.466E-189 | MO_kdnrs | RPL5 |
| 3.757E-193 | -0.6326579 | 0.135 | 0.368 | 8.874E-189 | MO_kdnrs | UBE2H |
| 8.254E-193 | 0.26089328 | 0.225 | 0.076 | 1.95E-188 | MO_kdnrs | MAF |

| | | | | | | |
|---|---|---|---|---|---|---|
| 1.413E-191 | 0.35094413 | 0.534 | 0.294 | 3.339E-187 | MO_kdnrs | ARHGDIA |
| 1.637E-191 | 0.34430362 | 0.748 | 0.505 | 3.868E-187 | MO_kdnrs | CEBPB |
| 9.718E-191 | 0.42576379 | 0.284 | 0.113 | 2.296E-186 | MO_kdnrs | TFRC |
| 1.157E-190 | 0.35823257 | 0.981 | 0.914 | 2.733E-186 | MO_kdnrs | NACA |
| 7.379E-190 | 0.30308125 | 0.963 | 0.86 | 1.743E-185 | MO_kdnrs | PABPC1 |
| 2.642E-189 | 0.2749542 | 0.362 | 0.163 | 6.242E-185 | MO_kdnrs | SUMO3 |
| 6.967E-189 | 0.29674951 | 0.321 | 0.136 | 1.646E-184 | MO_kdnrs | RB1 |
| 2.592E-188 | -0.701011 | 0.28 | 0.5 | 6.123E-184 | MO_kdnrs | CD46 |
| 2.75E-187 | 0.37819244 | 0.761 | 0.523 | 6.496E-183 | MO_kdnrs | POMP |
| 3.167E-187 | 0.36695982 | 0.857 | 0.671 | 7.482E-183 | MO_kdnrs | ATP6V0B |
| 1.327E-186 | -0.5615058 | 0.046 | 0.265 | 3.135E-182 | MO_kdnrs | SERINC2 |
| 2.076E-186 | -0.53818 | 0.021 | 0.232 | 4.903E-182 | MO_kdnrs | CKB |
| 2.292E-186 | 0.29501272 | 0.449 | 0.224 | 5.415E-182 | MO_kdnrs | LSM7 |
| 4.509E-186 | 0.25041375 | 0.312 | 0.13 | 1.065E-181 | MO_kdnrs | TMEM256 |
| 8.373E-186 | 0.29954178 | 0.431 | 0.209 | 1.978E-181 | MO_kdnrs | SKAP2 |
| 4.122E-185 | 0.27039969 | 0.3 | 0.124 | 9.736E-181 | MO_kdnrs | HEXA |
| 6.997E-185 | -0.557885 | 0.057 | 0.277 | 1.653E-180 | MO_kdnrs | WASL |
| 7.345E-185 | 0.34479887 | 0.67 | 0.413 | 1.735E-180 | MO_kdnrs | ATP5MF |
| 1.146E-184 | 0.28714025 | 0.439 | 0.214 | 2.708E-180 | MO_kdnrs | PARP14 |
| 3.295E-184 | 0.34978762 | 0.833 | 0.61 | 7.785E-180 | MO_kdnrs | OST4 |
| 7.28E-184 | 0.29815872 | 0.349 | 0.154 | 1.72E-179 | MO_kdnrs | FNBP1 |
| 1.372E-183 | -0.9106721 | 0.146 | 0.37 | 3.24E-179 | MO_kdnrs | PHLDA2 |
| 1.551E-183 | 0.3554131 | 0.591 | 0.338 | 3.663E-179 | MO_kdnrs | PLSCR1 |
| 2.279E-183 | -0.9596902 | 0.004 | 0.204 | 5.384E-179 | MO_kdnrs | TMPRSS2 |
| 4.893E-183 | -0.5518386 | 0.063 | 0.284 | 1.156E-178 | MO_kdnrs | AFDN |
| 1.954E-182 | -0.4181832 | 0.004 | 0.203 | 4.616E-178 | MO_kdnrs | PAWR |
| 6.966E-182 | 0.40609743 | 0.49 | 0.27 | 1.646E-177 | MO_kdnrs | RALA |
| 1.588E-181 | 0.26930961 | 0.391 | 0.184 | 3.752E-177 | MO_kdnrs | KRTCAP2 |
| 3.517E-181 | 0.33532578 | 0.596 | 0.34 | 8.309E-177 | MO_kdnrs | MTDH |
| 5.767E-181 | 0.33277159 | 0.956 | 0.766 | 1.362E-176 | MO_kdnrs | RPL10A |
| 1.993E-180 | -0.7136564 | 0.021 | 0.225 | 4.708E-176 | MO_kdnrs | CDKN2B |
| 3.992E-180 | -0.6405983 | 0.118 | 0.345 | 9.431E-176 | MO_kdnrs | MAFF |
| 7.091E-180 | -1.2982955 | 0.004 | 0.2 | 1.675E-175 | MO_kdnrs | DUOX2 |
| 9.555E-180 | -1.6533127 | 0.014 | 0.215 | 2.257E-175 | MO_kdnrs | KRT17 |
| 1.686E-179 | 0.27252707 | 0.357 | 0.162 | 3.983E-175 | MO_kdnrs | STX7 |
| 5.996E-179 | -0.5253842 | 0.002 | 0.197 | 1.417E-174 | MO_kdnrs | S100A14 |
| 9.003E-179 | 0.32573142 | 0.649 | 0.381 | 2.127E-174 | MO_kdnrs | FLNA |
| 5.037E-178 | -0.413759 | 0.002 | 0.195 | 1.19E-173 | MO_kdnrs | TEAD1 |
| 2.391E-177 | 0.3835825 | 0.792 | 0.589 | 5.648E-173 | MO_kdnrs | SUB1 |
| 5.026E-177 | -0.9542202 | 0.224 | 0.452 | 1.187E-172 | MO_kdnrs | HIST1H1C |
| 5.053E-177 | -0.4575677 | 0.004 | 0.199 | 1.194E-172 | MO_kdnrs | CDH1 |
| 6.019E-177 | -0.613397 | 0.004 | 0.199 | 1.422E-172 | MO_kdnrs | SDCBP2 |
| 8.066E-177 | 0.33340263 | 0.565 | 0.32 | 1.906E-172 | MO_kdnrs | ANP32B |
| 1.317E-176 | -0.7096354 | 0.009 | 0.205 | 3.11E-172 | MO_kdnrs | EPS8L1 |
| 8.707E-175 | 0.31431723 | 0.985 | 0.861 | 2.057E-170 | MO_kdnrs | RPL14 |
| 1.553E-174 | 0.3181211 | 0.212 | 0.074 | 3.669E-170 | MO_kdnrs | POU2F2 |
| 2.272E-174 | -3.9711537 | 0.355 | 0.558 | 5.368E-170 | MO_kdnrs | PLCG2 |
| 2.792E-174 | 0.32628982 | 0.625 | 0.369 | 6.596E-170 | MO_kdnrs | ATP5MPL |
| 1.18E-173 | 0.30854066 | 0.492 | 0.266 | 2.788E-169 | MO_kdnrs | NDUFB1 |
| 1.237E-173 | -0.5543073 | 0.003 | 0.194 | 2.922E-169 | MO_kdnrs | EPHA2 |

| | | | | | | |
|---|---|---|---|---|---|---|
| 1.374E-173 | 0.26669664 | 0.367 | 0.171 | 3.247E-169 | MO_kdnrs | GTF3A |
| 2.548E-173 | 0.27829419 | 0.567 | 0.308 | 6.018E-169 | MO_kdnrs | HERPUD1 |
| 4.245E-173 | -0.4171071 | 0.002 | 0.192 | 1.003E-168 | MO_kdnrs | SMIM22 |
| 5.595E-173 | 0.27487748 | 0.317 | 0.139 | 1.322E-168 | MO_kdnrs | TAP1 |
| 8.644E-173 | 0.44520706 | 0.728 | 0.499 | 2.042E-168 | MO_kdnrs | NAP1L1 |
| 1.057E-172 | 0.28196717 | 0.328 | 0.146 | 2.496E-168 | MO_kdnrs | RAB32 |
| 4.087E-172 | -0.8543862 | 0.006 | 0.197 | 9.654E-168 | MO_kdnrs | LAMC2 |
| 6.815E-172 | -0.5088012 | 0.051 | 0.258 | 1.61E-167 | MO_kdnrs | GIPC1 |
| 3.644E-171 | 0.2673382 | 0.297 | 0.125 | 8.608E-167 | MO_kdnrs | ITPR2 |
| 7.517E-171 | 0.27898262 | 0.355 | 0.167 | 1.776E-166 | MO_kdnrs | EMD |
| 1.172E-170 | 0.31149048 | 0.528 | 0.295 | 2.769E-166 | MO_kdnrs | RBX1 |
| 3.24E-170 | 0.29484879 | 0.428 | 0.219 | 7.655E-166 | MO_kdnrs | NDUFA13 |
| 3.57E-170 | -0.5638648 | 0.003 | 0.19 | 8.433E-166 | MO_kdnrs | C6orf132 |
| 4.197E-170 | -0.5155622 | 0.004 | 0.192 | 9.913E-166 | MO_kdnrs | FHL2 |
| 5.046E-170 | 0.29921138 | 0.409 | 0.2 | 1.192E-165 | MO_kdnrs | STK4 |
| 9.303E-170 | -0.4213761 | 0.006 | 0.195 | 2.198E-165 | MO_kdnrs | ANXA3 |
| 2.9E-169 | -0.3722443 | 0.004 | 0.191 | 6.852E-165 | MO_kdnrs | TMEM54 |
| 6.786E-169 | 0.31256182 | 0.775 | 0.533 | 1.603E-164 | MO_kdnrs | SSR4 |
| 7.132E-169 | 0.28375087 | 0.491 | 0.257 | 1.685E-164 | MO_kdnrs | ROCK1 |
| 1.003E-168 | -0.638539 | 0.001 | 0.186 | 2.369E-164 | MO_kdnrs | HSPB8 |
| 1.139E-168 | 0.3241947 | 0.627 | 0.378 | 2.691E-164 | MO_kdnrs | NDUFA1 |
| 5.425E-168 | 0.3036815 | 0.913 | 0.749 | 1.281E-163 | MO_kdnrs | COX7C |
| 6.229E-168 | -0.6414254 | 0.004 | 0.19 | 1.472E-163 | MO_kdnrs | SPNS2 |
| 1.061E-167 | 0.32816721 | 0.968 | 0.827 | 2.507E-163 | MO_kdnrs | RACK1 |
| 1.208E-167 | 0.29267592 | 0.492 | 0.269 | 2.854E-163 | MO_kdnrs | NDUFA11 |
| 1.241E-167 | 0.28313273 | 0.444 | 0.231 | 2.931E-163 | MO_kdnrs | TRAPPC1 |
| 3.485E-167 | -0.7150124 | 0.017 | 0.207 | 8.232E-163 | MO_kdnrs | FBXO32 |
| 1.813E-166 | 0.27174231 | 0.394 | 0.194 | 4.284E-162 | MO_kdnrs | CYTH1 |
| 1.056E-165 | 0.29996853 | 0.579 | 0.338 | 2.495E-161 | MO_kdnrs | ATP5PD |
| 1.961E-165 | 0.28222972 | 0.365 | 0.174 | 4.633E-161 | MO_kdnrs | PTPN2 |
| 2.564E-165 | 0.30825624 | 0.891 | 0.76 | 6.057E-161 | MO_kdnrs | RHOA |
| 3.708E-164 | -0.6312331 | 0.005 | 0.187 | 8.76E-160 | MO_kdnrs | PPL |
| 4.23E-164 | -0.3337665 | 0.003 | 0.185 | 9.992E-160 | MO_kdnrs | GNG12 |
| 1.421E-163 | 0.32875985 | 0.85 | 0.687 | 3.357E-159 | MO_kdnrs | COX6B1 |
| 1.635E-163 | -0.4805098 | 0.017 | 0.205 | 3.862E-159 | MO_kdnrs | ABLIM1 |
| 2.687E-163 | -0.5963467 | 0.033 | 0.229 | 6.347E-159 | MO_kdnrs | LAMB3 |
| 3.614E-163 | 0.30996797 | 0.585 | 0.343 | 8.537E-159 | MO_kdnrs | SLC25A5 |
| 7.647E-163 | -0.400615 | 0.013 | 0.199 | 1.806E-158 | MO_kdnrs | EPS8L2 |
| 1.991E-162 | -0.6652389 | 0.003 | 0.183 | 4.702E-158 | MO_kdnrs | FXYD3 |
| 2.16E-162 | 0.28698751 | 0.412 | 0.207 | 5.103E-158 | MO_kdnrs | STXBP2 |
| 1.155E-161 | 0.30459904 | 0.863 | 0.696 | 2.729E-157 | MO_kdnrs | SDCBP |
| 2.581E-161 | -0.7047123 | 0.009 | 0.19 | 6.096E-157 | MO_kdnrs | IGFBP3 |
| 5.048E-160 | 0.25668486 | 0.422 | 0.214 | 1.193E-155 | MO_kdnrs | FLOT1 |
| 1.003E-159 | 0.28309757 | 0.928 | 0.827 | 2.37E-155 | MO_kdnrs | HLA-E |
| 2.038E-159 | 0.30133646 | 0.569 | 0.33 | 4.814E-155 | MO_kdnrs | TMEM258 |
| 1.214E-158 | 0.41166903 | 0.702 | 0.483 | 2.868E-154 | MO_kdnrs | ASAH1 |
| 2.174E-158 | 0.26091442 | 0.224 | 0.086 | 5.135E-154 | MO_kdnrs | GPR137B |
| 2.949E-158 | -0.5451467 | 0.033 | 0.223 | 6.967E-154 | MO_kdnrs | MACC1 |
| 4.368E-158 | -0.6810908 | 0.074 | 0.275 | 1.032E-153 | MO_kdnrs | PMAIP1 |
| 1.128E-157 | -0.3554467 | 0.005 | 0.182 | 2.665E-153 | MO_kdnrs | CXADR |

| | | | | | | |
|---|---|---|---|---|---|---|
| 1.133E-157 | -0.7227468 | 0.07 | 0.267 | 2.677E-153 | MO_kdnrs | SYNE2 |
| 2.062E-157 | -1.332716 | 0.004 | 0.18 | 4.871E-153 | MO_kdnrs | DUOXA2 |
| 4.24E-157 | 0.33929973 | 0.629 | 0.401 | 1.002E-152 | MO_kdnrs | CAPZB |
| 4.948E-157 | 0.37511078 | 0.271 | 0.113 | 1.169E-152 | MO_kdnrs | MARCO |
| 7.412E-156 | 0.29233144 | 0.512 | 0.287 | 1.751E-151 | MO_kdnrs | NOP53 |
| 7.754E-156 | -0.4116221 | 0.007 | 0.183 | 1.832E-151 | MO_kdnrs | TINAGL1 |
| 2.17E-155 | -0.4647668 | 0.151 | 0.363 | 5.126E-151 | MO_kdnrs | CAPN2 |
| 5.792E-155 | 0.26749443 | 0.592 | 0.344 | 1.368E-150 | MO_kdnrs | ATP5F1D |
| 1.492E-154 | -0.3327853 | 0.003 | 0.175 | 3.524E-150 | MO_kdnrs | NCKAP1 |
| 5.293E-154 | 0.3062424 | 0.519 | 0.292 | 1.25E-149 | MO_kdnrs | CTNNB1 |
| 5.683E-154 | -0.8971595 | 0.081 | 0.275 | 1.343E-149 | MO_kdnrs | TPM1 |
| 9.176E-154 | -0.3513852 | 0.014 | 0.193 | 2.168E-149 | MO_kdnrs | LSR |
| 1.068E-153 | -0.3445561 | 0.002 | 0.173 | 2.522E-149 | MO_kdnrs | CRB3 |
| 1.376E-153 | 0.29717981 | 0.989 | 0.917 | 3.251E-149 | MO_kdnrs | RPL8 |
| 2.603E-153 | -0.4067796 | 0.008 | 0.183 | 6.149E-149 | MO_kdnrs | NECTIN4 |
| 7.079E-153 | -0.5392216 | 0.537 | 0.698 | 1.672E-148 | MO_kdnrs | BZW1 |
| 1.077E-152 | -1.1057778 | 0.003 | 0.174 | 2.545E-148 | MO_kdnrs | CEACAM5 |
| 1.235E-152 | -0.5367625 | 0.002 | 0.172 | 2.919E-148 | MO_kdnrs | MALL |
| 1.326E-152 | -0.3501954 | 0.005 | 0.177 | 3.132E-148 | MO_kdnrs | NDFIP2 |
| 1.833E-152 | -0.5838515 | 0.665 | 0.791 | 4.329E-148 | MO_kdnrs | MAP1LC3B |
| 4.531E-152 | -0.4090077 | 0.001 | 0.17 | 1.07E-147 | MO_kdnrs | TSPAN6 |
| 8.038E-152 | -0.3297518 | 0.007 | 0.179 | 1.899E-147 | MO_kdnrs | DDR1 |
| 1.184E-151 | -0.7056458 | 0.096 | 0.289 | 2.796E-147 | MO_kdnrs | TNFRSF12A |
| 2.32E-151 | -0.4576671 | 0.003 | 0.173 | 5.48E-147 | MO_kdnrs | SDC1 |
| 1.317E-149 | -1.1175516 | 0.306 | 0.488 | 3.111E-145 | MO_kdnrs | ERO1A |
| 1.491E-149 | 0.25155324 | 0.996 | 0.941 | 3.522E-145 | MO_kdnrs | RPL13A |
| 1.611E-149 | -0.4585813 | 0.003 | 0.17 | 3.806E-145 | MO_kdnrs | CLDN3 |
| 1.828E-149 | -1.1831002 | 0.005 | 0.175 | 4.318E-145 | MO_kdnrs | MUC21 |
| 2.439E-149 | -0.4167986 | 0.003 | 0.17 | 5.761E-145 | MO_kdnrs | COBLL1 |
| 4.103E-149 | -0.3677524 | 0.018 | 0.194 | 9.692E-145 | MO_kdnrs | FNBP1L |
| 1.754E-148 | -1.0309882 | 0.002 | 0.168 | 4.143E-144 | MO_kdnrs | PITX1 |
| 1.998E-148 | -0.6353876 | 0.001 | 0.167 | 4.719E-144 | MO_kdnrs | CDC42EP5 |
| 3.692E-148 | -0.5194929 | 0.002 | 0.168 | 8.722E-144 | MO_kdnrs | CYR61 |
| 3.817E-148 | 0.32947952 | 0.588 | 0.361 | 9.018E-144 | MO_kdnrs | PRELID1 |
| 7.381E-148 | -0.4744462 | 0.002 | 0.167 | 1.744E-143 | MO_kdnrs | TMPRSS4 |
| 1.269E-147 | 0.27570062 | 0.537 | 0.311 | 2.998E-143 | MO_kdnrs | COPS9 |
| 4.357E-147 | 0.28970501 | 0.537 | 0.318 | 1.029E-142 | MO_kdnrs | SELENOH |
| 4.371E-147 | 0.28090871 | 0.607 | 0.365 | 1.032E-142 | MO_kdnrs | DEK |
| 1.075E-146 | -0.7989465 | 0.002 | 0.166 | 2.54E-142 | MO_kdnrs | SCEL |
| 1.203E-146 | -1.0223452 | 0.005 | 0.171 | 2.841E-142 | MO_kdnrs | PRSS27 |
| 1.339E-146 | -0.7573141 | 0.255 | 0.102 | 3.163E-142 | MO_kdnrs | MT1M |
| 3.084E-146 | 0.25990271 | 0.464 | 0.254 | 7.285E-142 | MO_kdnrs | NDUFB8 |
| 4.087E-146 | -1.0573851 | 0.016 | 0.187 | 9.654E-142 | MO_kdnrs | RDH10 |
| 4.362E-146 | -0.9027644 | 0.004 | 0.169 | 1.03E-141 | MO_kdnrs | KRT15 |
| 6.648E-146 | 0.27604371 | 0.523 | 0.301 | 1.57E-141 | MO_kdnrs | FKBP1A |
| 1.846E-145 | -0.3539007 | 0.004 | 0.169 | 4.361E-141 | MO_kdnrs | ERBB3 |
| 1.939E-145 | 0.25343495 | 0.43 | 0.23 | 4.579E-141 | MO_kdnrs | BAX |
| 7.672E-145 | -0.6871073 | 0.004 | 0.169 | 1.812E-140 | MO_kdnrs | MUC4 |
| 9.862E-145 | -0.5372587 | 0.697 | 0.893 | 2.33E-140 | MO_kdnrs | MT-ND3 |
| 1.409E-144 | -0.5690628 | 0.057 | 0.239 | 3.328E-140 | MO_kdnrs | OCIAD2 |

| | | | | | | |
|---|---|---|---|---|---|---|
| 1.687E-144 | 0.29098575 | 0.663 | 0.42 | 3.984E-140 | MO_kdnrs | PDIA3 |
| 1.962E-144 | -0.4345049 | 0.006 | 0.171 | 4.636E-140 | MO_kdnrs | TTC9 |
| 2.275E-144 | -0.3906045 | 0.033 | 0.211 | 5.374E-140 | MO_kdnrs | NEDD4L |
| 3.583E-144 | 0.28246276 | 0.983 | 0.894 | 8.465E-140 | MO_kdnrs | RPL24 |
| 1.376E-143 | -0.7217642 | 0.053 | 0.236 | 3.25E-139 | MO_kdnrs | CCND1 |
| 9.613E-143 | -0.8778917 | 0.023 | 0.194 | 2.271E-138 | MO_kdnrs | F3 |
| 1.824E-142 | -0.9808133 | 0.136 | 0.328 | 4.31E-138 | MO_kdnrs | PHLDA1 |
| 3.304E-142 | 0.25735197 | 0.439 | 0.239 | 7.805E-138 | MO_kdnrs | EIF3L |
| 7.01E-142 | 0.28834161 | 0.659 | 0.445 | 1.656E-137 | MO_kdnrs | SARAF |
| 9.004E-142 | -0.8011952 | 0.002 | 0.162 | 2.127E-137 | MO_kdnrs | TMPRSS11D |
| 1.141E-141 | -1.0738025 | 0.969 | 0.938 | 2.696E-137 | MO_kdnrs | NEAT1 |
| 1.494E-141 | 0.28330316 | 0.47 | 0.268 | 3.528E-137 | MO_kdnrs | RNF181 |
| 1.464E-140 | 0.28640254 | 0.738 | 0.514 | 3.458E-136 | MO_kdnrs | COX6C |
| 3.475E-140 | 0.25087194 | 0.444 | 0.244 | 8.21E-136 | MO_kdnrs | RAP1A |
| 4.001E-140 | -0.6862203 | 0.002 | 0.159 | 9.451E-136 | MO_kdnrs | NCCRP1 |
| 7.317E-140 | 0.28587861 | 0.339 | 0.167 | 1.729E-135 | MO_kdnrs | MEF2A |
| 1.085E-139 | -0.3152362 | 0.002 | 0.159 | 2.563E-135 | MO_kdnrs | KCNK1 |
| 1.099E-139 | -0.440994 | 0.001 | 0.159 | 2.595E-135 | MO_kdnrs | VSIG2 |
| 1.471E-139 | -0.3289668 | 0.001 | 0.157 | 3.476E-135 | MO_kdnrs | SLC44A4 |
| 2.048E-139 | 0.26496897 | 0.504 | 0.29 | 4.839E-135 | MO_kdnrs | SPCS3 |
| 7.972E-139 | -0.3658283 | 0.006 | 0.165 | 1.883E-134 | MO_kdnrs | ITGA3 |
| 2.004E-138 | -0.4175028 | 0.001 | 0.156 | 4.734E-134 | MO_kdnrs | ITGB6 |
| 6.307E-138 | -0.5006834 | 0.003 | 0.16 | 1.49E-133 | MO_kdnrs | ALDH1A3 |
| 1.202E-137 | -0.9074096 | 0.003 | 0.159 | 2.84E-133 | MO_kdnrs | CEACAM6 |
| 9.638E-137 | -0.8661588 | 0.003 | 0.158 | 2.277E-132 | MO_kdnrs | CEACAM7 |
| 2.538E-136 | 0.28698751 | 0.587 | 0.355 | 5.996E-132 | MO_kdnrs | METRNL |
| 4.106E-136 | -0.4924743 | 0.099 | 0.285 | 9.7E-132 | MO_kdnrs | FAM3C |
| 4.57E-136 | -0.3531815 | 0.013 | 0.175 | 1.08E-131 | MO_kdnrs | MET |
| 4.851E-136 | -0.5351394 | 0.136 | 0.328 | 1.146E-131 | MO_kdnrs | SDC4 |
| 1.166E-135 | -0.8098317 | 0.003 | 0.157 | 2.755E-131 | MO_kdnrs | TMPRSS11E |
| 1.456E-135 | -0.3175042 | 0.006 | 0.162 | 3.439E-131 | MO_kdnrs | PLS3 |
| 1.62E-135 | -0.5638906 | 0.035 | 0.205 | 3.826E-131 | MO_kdnrs | PLK2 |
| 2.478E-134 | 0.28092525 | 0.859 | 0.689 | 5.854E-130 | MO_kdnrs | TPM3 |
| 7.399E-134 | -1.6375891 | 0.017 | 0.175 | 1.748E-129 | MO_kdnrs | ECM1 |
| 1.267E-133 | -0.2991365 | 0.001 | 0.151 | 2.993E-129 | MO_kdnrs | GULP1 |
| 1.645E-133 | -0.2986789 | 0.001 | 0.152 | 3.885E-129 | MO_kdnrs | RAB25 |
| 1.85E-133 | -0.9211153 | 0.144 | 0.327 | 4.37E-129 | MO_kdnrs | EMP1 |
| 3.309E-133 | -0.425533 | 0.016 | 0.175 | 7.817E-129 | MO_kdnrs | SYTL2 |
| 3.83E-133 | -0.2741507 | 0.003 | 0.154 | 9.047E-129 | MO_kdnrs | BACE2 |
| 5.442E-133 | 0.25141038 | 0.491 | 0.284 | 1.286E-128 | MO_kdnrs | H2AFV |
| 8.854E-133 | -0.6525638 | 0.071 | 0.249 | 2.092E-128 | MO_kdnrs | CDKN1C |
| 9.069E-133 | 0.26187716 | 0.412 | 0.221 | 2.142E-128 | MO_kdnrs | PABPC4 |
| 1.564E-132 | -1.170769 | 0.03 | 0.194 | 3.694E-128 | MO_kdnrs | SPINK1 |
| 1.763E-132 | -0.5532009 | 0.158 | 0.344 | 4.164E-128 | MO_kdnrs | ASPH |
| 2.295E-132 | -0.5405406 | 0.488 | 0.634 | 5.421E-128 | MO_kdnrs | TMEM59 |
| 4.908E-132 | 0.29278637 | 0.385 | 0.2 | 1.16E-127 | MO_kdnrs | ARL4C |
| 5.7E-132 | 0.26908926 | 0.165 | 0.059 | 1.347E-127 | MO_kdnrs | EPB41L2 |
| 6.05E-132 | -1.0898681 | 0.002 | 0.152 | 1.429E-127 | MO_kdnrs | CXCL6 |
| 1.567E-131 | -0.7809335 | 0.037 | 0.202 | 3.701E-127 | MO_kdnrs | ANKRD36C |
| 2.874E-131 | -0.6627107 | 0.003 | 0.153 | 6.79E-127 | MO_kdnrs | SNCG |

| | | | | | | |
|---|---|---|---|---|---|---|
| 3.834E-131 | 0.25854555 | 0.557 | 0.334 | 9.058E-127 | MO_kdnrs | EIF3E |
| 2.339E-130 | -0.313141 | 0.005 | 0.155 | 5.525E-126 | MO_kdnrs | OCLN |
| 6.259E-130 | 0.25169815 | 0.436 | 0.243 | 1.479E-125 | MO_kdnrs | ATP6V0D1 |
| 1.051E-129 | 0.25372284 | 0.534 | 0.318 | 2.484E-125 | MO_kdnrs | CAPZA1 |
| 1.669E-128 | -1.7519035 | 0.006 | 0.155 | 3.942E-124 | MO_kdnrs | SPRR2A |
| 2.102E-127 | -0.4428906 | 0.002 | 0.147 | 4.965E-123 | MO_kdnrs | GCNT3 |
| 3.507E-127 | 0.28385079 | 0.78 | 0.583 | 8.285E-123 | MO_kdnrs | VAMP8 |
| 9.159E-127 | -0.7817624 | 0.996 | 0.979 | 2.164E-122 | MO_kdnrs | MALAT1 |
| 1.156E-126 | -0.4109527 | 0.003 | 0.148 | 2.73E-122 | MO_kdnrs | ARHGAP29 |
| 1.206E-126 | -0.523533 | 0.113 | 0.294 | 2.848E-122 | MO_kdnrs | ABHD5 |
| 1.495E-126 | 0.25084285 | 0.515 | 0.308 | 3.532E-122 | MO_kdnrs | SNRPG |
| 1.817E-126 | -0.43276 | 0.003 | 0.149 | 4.293E-122 | MO_kdnrs | DCDC2 |
| 3.235E-126 | -0.9546024 | 0.002 | 0.146 | 7.643E-122 | MO_kdnrs | TMPRSS11B |
| 4.591E-126 | -0.721607 | 0.066 | 0.235 | 1.085E-121 | MO_kdnrs | HES1 |
| 8.192E-126 | 0.33900133 | 0.818 | 0.66 | 1.935E-121 | MO_kdnrs | CALR |
| 8.725E-126 | 0.3058341 | 0.794 | 0.609 | 2.061E-121 | MO_kdnrs | ACTR2 |
| 1.24E-123 | -0.697475 | 0.001 | 0.143 | 2.929E-119 | MO_kdnrs | GABRP |
| 1.244E-123 | -0.6533724 | 0.004 | 0.148 | 2.938E-119 | MO_kdnrs | CXCL17 |
| 1.246E-123 | -0.413217 | 0.003 | 0.146 | 2.943E-119 | MO_kdnrs | BCAS1 |
| 1.289E-123 | 0.29602133 | 0.495 | 0.302 | 3.045E-119 | MO_kdnrs | CALM3 |
| 1.497E-123 | 0.25568355 | 0.508 | 0.308 | 3.536E-119 | MO_kdnrs | CDC37 |
| 1.894E-123 | -0.9702916 | 0.121 | 0.293 | 4.474E-119 | MO_kdnrs | DUSP5 |
| 3.924E-123 | -0.9595777 | 0.003 | 0.144 | 9.27E-119 | MO_kdnrs | RHCG |
| 6.529E-123 | -0.2786066 | 0.005 | 0.148 | 1.542E-118 | MO_kdnrs | SCNN1A |
| 7.782E-123 | -0.3050494 | 0.009 | 0.155 | 1.838E-118 | MO_kdnrs | PHLDA3 |
| 1.041E-122 | 0.27154664 | 0.359 | 0.187 | 2.458E-118 | MO_kdnrs | NFKB1 |
| 1.533E-122 | -0.2897168 | 0.001 | 0.141 | 3.621E-118 | MO_kdnrs | C1orf116 |
| 2.31E-122 | -0.2780717 | 0.002 | 0.143 | 5.456E-118 | MO_kdnrs | DMKN |
| 4.632E-122 | -0.2794178 | 0.006 | 0.149 | 1.094E-117 | MO_kdnrs | PATJ |
| 6.302E-122 | 0.2531708 | 0.8 | 0.592 | 1.489E-117 | MO_kdnrs | HNRNPA1 |
| 7.684E-122 | -0.3432454 | 0.002 | 0.141 | 1.815E-117 | MO_kdnrs | AIF1L |
| 7.746E-122 | -0.289004 | 0.002 | 0.142 | 1.83E-117 | MO_kdnrs | NTN4 |
| 1.691E-121 | -0.3008035 | 0.001 | 0.14 | 3.995E-117 | MO_kdnrs | KAZN |
| 1.396E-120 | -0.333201 | 0.041 | 0.2 | 3.298E-116 | MO_kdnrs | HSBP1L1 |
| 1.755E-120 | -0.3622199 | 0.001 | 0.139 | 4.147E-116 | MO_kdnrs | FAM84A |
| 1.197E-119 | -0.5605949 | 0.375 | 0.526 | 2.828E-115 | MO_kdnrs | ANXA11 |
| 1.7E-119 | -0.5964026 | 0.279 | 0.44 | 4.016E-115 | MO_kdnrs | SPINT2 |
| 4.457E-119 | -0.3169145 | 0.024 | 0.174 | 1.053E-114 | MO_kdnrs | GALNT3 |
| 6.399E-119 | -0.4150724 | 0.001 | 0.137 | 1.512E-114 | MO_kdnrs | ID4 |
| 7.397E-119 | -0.3812095 | 0.017 | 0.163 | 1.747E-114 | MO_kdnrs | MDK |
| 1.644E-118 | -0.6780137 | 0.003 | 0.14 | 3.884E-114 | MO_kdnrs | TFPI2 |
| 2.113E-118 | -0.2861025 | 0.013 | 0.156 | 4.992E-114 | MO_kdnrs | GPRC5C |
| 4.515E-118 | -0.3534716 | 0.02 | 0.166 | 1.067E-113 | MO_kdnrs | ITPKC |
| 7.012E-118 | -0.3163199 | 0 | 0.135 | 1.656E-113 | MO_kdnrs | FAT1 |
| 1.433E-117 | -0.8422935 | 0.035 | 0.185 | 3.385E-113 | MO_kdnrs | NDRG2 |
| 1.578E-117 | -0.331572 | 0.044 | 0.201 | 3.727E-113 | MO_kdnrs | SH3YL1 |
| 2.233E-117 | -1.6337855 | 0.006 | 0.144 | 5.275E-113 | MO_kdnrs | LYPD2 |
| 5.75E-117 | -0.4793653 | 0.007 | 0.145 | 1.358E-112 | MO_kdnrs | UCHL1 |
| 6.395E-117 | -0.2518452 | 0.003 | 0.139 | 1.511E-112 | MO_kdnrs | MYO5B |
| 1.213E-116 | 0.26227327 | 0.69 | 0.495 | 2.865E-112 | MO_kdnrs | COX7B |

| | | | | | | |
|---|---|---|---|---|---|---|
| 2.253E-116 | -0.465035 | 0.1 | 0.272 | 5.321E-112 | MO_kdnrs | TSC22D1 |
| 4.196E-116 | -0.6746614 | 0.103 | 0.267 | 9.911E-112 | MO_kdnrs | CARHSP1 |
| 5.733E-116 | -0.4914806 | 0.013 | 0.153 | 1.354E-111 | MO_kdnrs | ID1 |
| 1.187E-115 | -1.0124592 | 0.666 | 0.727 | 2.805E-111 | MO_kdnrs | HSPB1 |
| 1.213E-115 | -0.2741567 | 0.001 | 0.133 | 2.866E-111 | MO_kdnrs | PTPN3 |
| 2.223E-115 | -0.4298118 | 0.014 | 0.154 | 5.251E-111 | MO_kdnrs | PAX8 |
| 3.017E-115 | -0.4267324 | 0.053 | 0.21 | 7.127E-111 | MO_kdnrs | PTGR1 |
| 4.585E-115 | -0.2942779 | 0.013 | 0.154 | 1.083E-110 | MO_kdnrs | LAPTM4B |
| 5.993E-115 | -0.3618275 | 0.006 | 0.141 | 1.416E-110 | MO_kdnrs | FOXC1 |
| 8.168E-115 | -0.4335229 | 0.001 | 0.134 | 1.93E-110 | MO_kdnrs | TMEM40 |
| 1.448E-114 | -0.5301895 | 0.025 | 0.17 | 3.42E-110 | MO_kdnrs | BPGM |
| 2.232E-114 | -0.2731516 | 0 | 0.131 | 5.272E-110 | MO_kdnrs | BICDL2 |
| 6.098E-114 | -0.5855232 | 0.001 | 0.133 | 1.441E-109 | MO_kdnrs | AGR2 |
| 8.953E-114 | -0.5446245 | 0.524 | 0.639 | 2.115E-109 | MO_kdnrs | TAX1BP1 |
| 5.314E-113 | -0.3601156 | 0.002 | 0.133 | 1.255E-108 | MO_kdnrs | KRT80 |
| 1.363E-112 | -0.2547336 | 0.001 | 0.13 | 3.22E-108 | MO_kdnrs | AMOTL2 |
| 8.941E-112 | -1.019782 | 0.942 | 0.936 | 2.112E-107 | MO_kdnrs | S100A6 |
| 2.497E-111 | 0.25034875 | 0.472 | 0.286 | 5.898E-107 | MO_kdnrs | TAPBP |
| 6.739E-111 | -0.5600869 | 0.002 | 0.131 | 1.592E-106 | MO_kdnrs | TCIM |
| 1.066E-110 | -1.2819077 | 0.01 | 0.142 | 2.517E-106 | MO_kdnrs | PLAT |
| 1.273E-110 | 0.27540613 | 0.273 | 0.135 | 3.007E-106 | MO_kdnrs | MALT1 |
| 2.929E-110 | -1.1739325 | 0 | 0.127 | 6.919E-106 | MO_kdnrs | SPINK5 |
| 7.454E-110 | -0.6445844 | 0.017 | 0.153 | 1.761E-105 | MO_kdnrs | CDKN2A |
| 1.384E-109 | -0.2983753 | 0.01 | 0.143 | 3.271E-105 | MO_kdnrs | MPZL2 |
| 2.41E-109 | -0.4109128 | 0.037 | 0.182 | 5.693E-105 | MO_kdnrs | FAM214A |
| 5.751E-109 | -0.3322059 | 0.002 | 0.129 | 1.358E-104 | MO_kdnrs | ASS1 |
| 6.572E-109 | 0.25287778 | 0.591 | 0.382 | 1.552E-104 | MO_kdnrs | CIB1 |
| 6.61E-109 | -0.2794061 | 0.003 | 0.129 | 1.561E-104 | MO_kdnrs | MUC20 |
| 1.65E-108 | 0.25251736 | 0.57 | 0.372 | 3.898E-104 | MO_kdnrs | SEM1 |
| 3.15E-108 | -0.3098491 | 0.013 | 0.146 | 7.441E-104 | MO_kdnrs | COL18A1 |
| 2.023E-107 | -0.2695992 | 0.003 | 0.129 | 4.779E-103 | MO_kdnrs | C19orf33 |
| 3.548E-107 | -0.2637136 | 0.002 | 0.127 | 8.381E-103 | MO_kdnrs | VASN |
| 7.619E-107 | 0.25511978 | 0.824 | 0.712 | 1.8E-102 | MO_kdnrs | JUND |
| 1.227E-106 | -0.3443609 | 0.019 | 0.155 | 2.9E-102 | MO_kdnrs | CLDN1 |
| 3.342E-105 | -0.4884857 | 0.001 | 0.122 | 7.896E-101 | MO_kdnrs | TMPRSS11A |
| 4.742E-104 | -0.3115177 | 0.001 | 0.121 | 1.12E-99 | MO_kdnrs | FUT3 |
| 5.018E-104 | -0.5117189 | 0.158 | 0.329 | 1.185E-99 | MO_kdnrs | HIST1H2AC |
| 1.422E-103 | -0.337571 | 0.024 | 0.158 | 3.36E-99 | MO_kdnrs | GSTM3 |
| 5.297E-103 | -0.6238841 | 0.002 | 0.122 | 1.2514E-98 | MO_kdnrs | KRT78 |
| 5.402E-103 | -0.4884519 | 0.025 | 0.159 | 1.276E-98 | MO_kdnrs | RASD1 |
| 1.621E-102 | -0.4068765 | 0.109 | 0.265 | 3.8291E-98 | MO_kdnrs | CD2AP |
| 3.608E-102 | -0.3882373 | 0.007 | 0.129 | 8.5229E-98 | MO_kdnrs | EGLN3 |
| 4.765E-102 | -0.3490046 | 0.471 | 0.264 | 1.1256E-97 | MO_kdnrs | MT1E |
| 5.971E-102 | -1.3322901 | 0.003 | 0.123 | 1.4106E-97 | MO_kdnrs | SPRR1B |
| 2.834E-101 | -0.4524748 | 0.133 | 0.289 | 6.6956E-97 | MO_kdnrs | SLK |
| 6.108E-101 | -0.2712591 | 0.003 | 0.122 | 1.4428E-96 | MO_kdnrs | DENND2C |
| 8.471E-101 | -0.5902503 | 0.352 | 0.491 | 2.0012E-96 | MO_kdnrs | RIOK3 |
| 1.621E-100 | -0.2625186 | 0.002 | 0.119 | 3.8294E-96 | MO_kdnrs | PTGES |
| 4.379E-100 | -0.2815214 | 0.038 | 0.176 | 1.0345E-95 | MO_kdnrs | SMAD3 |
| 1.045E-99 | -0.2525924 | 0.002 | 0.119 | 2.4684E-95 | MO_kdnrs | PLS1 |

| | | | | | | |
|---|---|---|---|---|---|---|
| 1.6296E-98 | -0.7923052 | 0.285 | 0.434 | 3.8496E-94 | MO_kdnrs | TXNRD1 |
| 1.8129E-98 | -0.7281975 | 0.963 | 0.962 | 4.2827E-94 | MO_kdnrs | HSP90AA1 |
| 3.855E-98 | -0.4364809 | 0.071 | 0.216 | 9.1067E-94 | MO_kdnrs | CSNK1E |
| 5.9387E-98 | -0.2906222 | 0.027 | 0.157 | 1.4029E-93 | MO_kdnrs | FAM102A |
| 1.0254E-97 | -0.3475303 | 0.001 | 0.115 | 2.4223E-93 | MO_kdnrs | A2ML1 |
| 1.3405E-97 | -0.6014705 | 0.047 | 0.183 | 3.1667E-93 | MO_kdnrs | STEAP4 |
| 1.4951E-97 | -0.4433298 | 0.002 | 0.116 | 3.5318E-93 | MO_kdnrs | TMC5 |
| 2.8847E-97 | -0.5357545 | 0.086 | 0.23 | 6.8144E-93 | MO_kdnrs | ZBTB20 |
| 8.5108E-97 | -0.4124789 | 0.003 | 0.117 | 2.0105E-92 | MO_kdnrs | KRT23 |
| 9.2322E-97 | -0.3742989 | 0.004 | 0.12 | 2.1809E-92 | MO_kdnrs | SMIM5 |
| 1.8802E-96 | -0.3846968 | 0.007 | 0.124 | 4.4417E-92 | MO_kdnrs | CEACAM1 |
| 8.7621E-96 | -0.3826772 | 0.144 | 0.298 | 2.0699E-91 | MO_kdnrs | LRP10 |
| 1.6128E-95 | -0.342271 | 0.068 | 0.21 | 3.8098E-91 | MO_kdnrs | TMEM106B |
| 5.4163E-95 | -0.2570236 | 0.034 | 0.165 | 1.2795E-90 | MO_kdnrs | ARHGEF12 |
| 6.1619E-95 | -0.2651777 | 0.006 | 0.121 | 1.4556E-90 | MO_kdnrs | GRHL1 |
| 1.1401E-94 | -0.4698315 | 0.191 | 0.344 | 2.6932E-90 | MO_kdnrs | APP |
| 6.7833E-94 | -0.3622121 | 0.063 | 0.202 | 1.6024E-89 | MO_kdnrs | LIMA1 |
| 2.0534E-93 | -0.2847447 | 0.031 | 0.159 | 4.8508E-89 | MO_kdnrs | RALGPS2 |
| 3.8632E-93 | -0.5385617 | 0.359 | 0.496 | 9.1259E-89 | MO_kdnrs | HMGA1 |
| 1.7792E-92 | -1.6040058 | 0.005 | 0.115 | 4.203E-88 | MO_kdnrs | CRCT1 |
| 2.1833E-92 | -0.2953583 | 0.014 | 0.131 | 5.1576E-88 | MO_kdnrs | FOXQ1 |
| 3.1366E-92 | -0.80898 | 0.063 | 0.195 | 7.4096E-88 | MO_kdnrs | AQP3 |
| 1.3535E-91 | -0.3234731 | 0.007 | 0.119 | 3.1974E-87 | MO_kdnrs | STS |
| 1.9508E-91 | -0.3692436 | 0.007 | 0.119 | 4.6083E-87 | MO_kdnrs | FXYD2 |
| 2.0573E-91 | -0.3590583 | 0.007 | 0.118 | 4.86E-87 | MO_kdnrs | NR1D1 |
| 1.3638E-90 | -0.5169298 | 0.02 | 0.139 | 3.2217E-86 | MO_kdnrs | AKAP12 |
| 5.7551E-89 | -0.3001063 | 0.001 | 0.106 | 1.3595E-84 | MO_kdnrs | RNF39 |
| 4.8837E-88 | -0.3038108 | 0.064 | 0.199 | 1.1537E-83 | MO_kdnrs | VCL |
| 1.018E-87 | -0.5582451 | 0.014 | 0.126 | 2.4047E-83 | MO_kdnrs | HSPA2 |
| 1.1101E-87 | -0.5169235 | 0.002 | 0.106 | 2.6224E-83 | MO_kdnrs | DHRS2 |
| 3.8256E-87 | -0.7787619 | 0.791 | 0.814 | 9.0372E-83 | MO_kdnrs | GSTP1 |
| 7.4057E-87 | -0.4870555 | 0.36 | 0.519 | 1.7494E-82 | MO_kdnrs | MT-ND4L |
| 1.0171E-86 | -0.5850649 | 0.327 | 0.446 | 2.4027E-82 | MO_kdnrs | CYSTM1 |
| 1.2596E-86 | -0.2891562 | 0.028 | 0.147 | 2.9756E-82 | MO_kdnrs | MTUS1 |
| 4.3702E-86 | -0.2741287 | 0.021 | 0.136 | 1.0324E-81 | MO_kdnrs | MAST4 |
| 9.0464E-86 | -0.2640551 | 0.001 | 0.102 | 2.137E-81 | MO_kdnrs | B3GALT5 |
| 1.3637E-85 | -0.359453 | 0.679 | 0.78 | 3.2214E-81 | MO_kdnrs | EIF4G2 |
| 1.6519E-85 | -0.4075884 | 0.333 | 0.175 | 3.9023E-81 | MO_kdnrs | RARRES3 |
| 4.8733E-85 | -0.3517023 | 0.003 | 0.105 | 1.1512E-80 | MO_kdnrs | KLK10 |
| 5.1499E-85 | -0.4794685 | 0.481 | 0.584 | 1.2166E-80 | MO_kdnrs | PRDX5 |
| 3.5926E-84 | -0.2920547 | 0.001 | 0.101 | 8.4869E-80 | MO_kdnrs | ADAMTS9 |
| 1.6273E-83 | -0.5326611 | 0.001 | 0.101 | 3.8441E-79 | MO_kdnrs | SAA1 |
| 3.0174E-83 | -0.3221799 | 0.12 | 0.259 | 7.128E-79 | MO_kdnrs | CCDC186 |
| 4.3824E-83 | -0.2595172 | 0.035 | 0.154 | 1.0353E-78 | MO_kdnrs | CRIM1 |
| 6.6757E-83 | -0.4638585 | 0.002 | 0.101 | 1.577E-78 | MO_kdnrs | RNASE7 |
| 9.5562E-83 | -0.410189 | 0.194 | 0.339 | 2.2575E-78 | MO_kdnrs | LMNA |
| 1.1471E-82 | -0.3078598 | 0.002 | 0.101 | 2.7097E-78 | MO_kdnrs | CTGF |
| 1.3169E-82 | -0.3057402 | 0.054 | 0.178 | 3.1109E-78 | MO_kdnrs | SLC38A1 |
| 1.5161E-82 | -0.8045093 | 0.009 | 0.113 | 3.5815E-78 | MO_kdnrs | ZG16B |
| 2.2059E-82 | -0.396379 | 0.271 | 0.406 | 5.211E-78 | MO_kdnrs | CTNNA1 |

| | | | | | | |
|---|---|---|---|---|---|---|
| 2.4501E-82 | -0.4081161 | 0.381 | 0.507 | 5.7878E-78 | MO_kdnrs | VDAC2 |
| 3.6697E-82 | -0.323754 | 0.08 | 0.211 | 8.669E-78 | MO_kdnrs | ZNF91 |
| 6.3556E-82 | -0.5203979 | 0.687 | 0.764 | 1.5014E-77 | MO_kdnrs | ITGB1 |
| 1.041E-81 | -0.4209172 | 0.513 | 0.625 | 2.4591E-77 | MO_kdnrs | KTN1 |
| 1.4879E-81 | 0.30259898 | 0.143 | 0.061 | 3.5149E-77 | MO_kdnrs | BHLHE41 |
| 2.2872E-81 | -0.8749828 | 0.082 | 0.205 | 5.403E-77 | MO_kdnrs | C12orf75 |
| 1.4886E-80 | -0.3312547 | 0.054 | 0.183 | 3.5164E-76 | MO_kdnrs | XIST |
| 1.5771E-80 | -0.4634313 | 0.418 | 0.54 | 3.7256E-76 | MO_kdnrs | PTTG1IP |
| 1.7106E-80 | -1.3608538 | 0.085 | 0.206 | 4.0409E-76 | MO_kdnrs | HOPX |
| 1.9618E-80 | -0.6589116 | 0.126 | 0.256 | 4.6343E-76 | MO_kdnrs | CST6 |
| 4.9803E-80 | -0.354584 | 0.015 | 0.12 | 1.1765E-75 | MO_kdnrs | ANGPTL4 |
| 8.3433E-80 | -0.2915408 | 0.019 | 0.125 | 1.9709E-75 | MO_kdnrs | ST3GAL4 |
| 1.6364E-79 | -0.7079048 | 0.54 | 0.645 | 3.8656E-75 | MO_kdnrs | GADD45B |
| 6.8825E-78 | -0.493336 | 0.118 | 0.248 | 1.6259E-73 | MO_kdnrs | CITED2 |
| 7.1834E-78 | -0.3155013 | 0.114 | 0.247 | 1.6969E-73 | MO_kdnrs | PNPLA2 |
| 8.1708E-78 | -0.2880777 | 0.073 | 0.198 | 1.9302E-73 | MO_kdnrs | MPRIP |
| 1.29E-77 | -0.4536592 | 0.445 | 0.56 | 3.0474E-73 | MO_kdnrs | FAM107B |
| 3.011E-77 | -0.2605003 | 0.024 | 0.131 | 7.1128E-73 | MO_kdnrs | TCP11L2 |
| 5.6078E-77 | -0.2977893 | 0.009 | 0.107 | 1.3247E-72 | MO_kdnrs | LY6K |
| 1.8464E-76 | -0.3247081 | 0.029 | 0.137 | 4.3618E-72 | MO_kdnrs | LRG1 |
| 8.2931E-75 | -0.2607764 | 0.054 | 0.171 | 1.9591E-70 | MO_kdnrs | FARP1 |
| 1.2499E-74 | -0.4885139 | 0.114 | 0.239 | 2.9527E-70 | MO_kdnrs | CYB5A |
| 3.8706E-74 | -1.4739717 | 0.017 | 0.115 | 9.1434E-70 | MO_kdnrs | CNFN |
| 1.0411E-73 | -0.7309 | 0.132 | 0.256 | 2.4594E-69 | MO_kdnrs | PLAU |
| 2.723E-73 | -0.2813854 | 0.083 | 0.206 | 6.4325E-69 | MO_kdnrs | USP53 |
| 1.2313E-72 | -0.367876 | 0.192 | 0.322 | 2.9087E-68 | MO_kdnrs | SBDS |
| 2.6772E-72 | -0.2729251 | 0.079 | 0.201 | 6.3243E-68 | MO_kdnrs | F11R |
| 3.297E-72 | -0.3838923 | 0.023 | 0.123 | 7.7884E-68 | MO_kdnrs | RRAD |
| 6.5152E-72 | -0.3270056 | 0.092 | 0.214 | 1.5391E-67 | MO_kdnrs | TIPARP |
| 9.7073E-72 | -0.2563503 | 0.015 | 0.11 | 2.2932E-67 | MO_kdnrs | DUSP14 |
| 7.3974E-71 | -1.0153809 | 0.041 | 0.145 | 1.7475E-66 | MO_kdnrs | DEFB1 |
| 2.0649E-70 | -0.4398047 | 0.395 | 0.5 | 4.8779E-66 | MO_kdnrs | SELENOW |
| 5.9984E-70 | -0.6319818 | 0.117 | 0.236 | 1.417E-65 | MO_kdnrs | HEXIM1 |
| 6.1282E-68 | -0.2658898 | 0.043 | 0.148 | 1.4477E-63 | MO_kdnrs | EMP2 |
| 6.4962E-68 | -0.3255514 | 0.771 | 0.836 | 1.5346E-63 | MO_kdnrs | YWHAZ |
| 1.041E-67 | -0.4790363 | 0.282 | 0.398 | 2.4592E-63 | MO_kdnrs | TUBB4B |
| 6.5817E-67 | -0.6152143 | 0.835 | 0.87 | 1.5548E-62 | MO_kdnrs | ANXA2 |
| 1.0425E-66 | -0.2962466 | 0.109 | 0.228 | 2.4627E-62 | MO_kdnrs | SPATS2L |
| 1.2911E-66 | -0.2723197 | 0.037 | 0.139 | 3.0501E-62 | MO_kdnrs | HIST1H2BC |
| 2.3304E-66 | -0.2749746 | 0.058 | 0.166 | 5.505E-62 | MO_kdnrs | TRIP10 |
| 2.5596E-66 | -0.7626129 | 0.8 | 0.836 | 6.0464E-62 | MO_kdnrs | PPDPF |
| 2.601E-65 | -0.3573356 | 0.153 | 0.274 | 6.1444E-61 | MO_kdnrs | HECTD1 |
| 3.3557E-65 | -0.3083969 | 0.145 | 0.269 | 7.9272E-61 | MO_kdnrs | ARHGAP5 |
| 2.1908E-64 | 0.35070896 | 0.46 | 0.313 | 5.1754E-60 | MO_kdnrs | PPA1 |
| 2.2175E-64 | -0.5266345 | 0.159 | 0.274 | 5.2385E-60 | MO_kdnrs | MGST1 |
| 2.4962E-64 | -0.2933683 | 0.155 | 0.279 | 5.8968E-60 | MO_kdnrs | NPEPPS |
| 6.0983E-64 | -0.3810743 | 0.622 | 0.693 | 1.4406E-59 | MO_kdnrs | CAST |
| 6.6023E-63 | -0.3493893 | 0.547 | 0.636 | 1.5597E-58 | MO_kdnrs | YBX3 |
| 1.3616E-62 | -1.0105491 | 0.044 | 0.142 | 3.2166E-58 | MO_kdnrs | SERPINB2 |
| 7.1785E-62 | -0.3436593 | 0.142 | 0.257 | 1.6958E-57 | MO_kdnrs | PKN2 |

| | | | | | | |
|---|---|---|---|---|---|---|
| 6.819E-61 | -0.478663 | 0.17 | 0.284 | 1.6108E-56 | MO_kdnrs | ADM |
| 2.7034E-60 | -0.334829 | 0.122 | 0.235 | 6.3863E-56 | MO_kdnrs | RAB9A |
| 3.733E-60 | -0.3313246 | 0.649 | 0.72 | 8.8184E-56 | MO_kdnrs | SKP1 |
| 8.2232E-60 | -0.4203513 | 0.282 | 0.385 | 1.9426E-55 | MO_kdnrs | GCC2 |
| 1.0107E-59 | -0.4011931 | 0.35 | 0.456 | 2.3877E-55 | MO_kdnrs | ACTN4 |
| 5.0357E-58 | -0.5717586 | 0.607 | 0.683 | 1.1896E-53 | MO_kdnrs | IER2 |
| 2.7164E-57 | -0.4036872 | 0.077 | 0.179 | 6.417E-53 | MO_kdnrs | DNAJA4 |
| 6.6563E-57 | -0.2774249 | 0.183 | 0.301 | 1.5724E-52 | MO_kdnrs | STK24 |
| 2.4182E-56 | -0.3213442 | 0.221 | 0.333 | 5.7125E-52 | MO_kdnrs | YPEL5 |
| 5.1066E-56 | -0.5469095 | 0.536 | 0.612 | 1.2063E-51 | MO_kdnrs | EZR |
| 6.9538E-56 | -0.2739829 | 0.045 | 0.136 | 1.6427E-51 | MO_kdnrs | GAN |
| 1.294E-55 | -0.6150243 | 0.049 | 0.138 | 3.0568E-51 | MO_kdnrs | DHRS9 |
| 1.3932E-55 | -0.5623943 | 0.209 | 0.109 | 3.2911E-51 | MO_kdnrs | CXCL2 |
| 5.6267E-54 | -0.2772027 | 0.156 | 0.266 | 1.3292E-49 | MO_kdnrs | LYPLA1 |
| 8.0088E-54 | -0.783954 | 0.578 | 0.647 | 1.8919E-49 | MO_kdnrs | IER3 |
| 1.3934E-53 | -0.4620167 | 0.203 | 0.307 | 3.2915E-49 | MO_kdnrs | FAM129B |
| 1.8243E-53 | -0.4390605 | 0.41 | 0.5 | 4.3095E-49 | MO_kdnrs | H2AFJ |
| 2.6008E-53 | -0.3629691 | 0.243 | 0.348 | 6.1439E-49 | MO_kdnrs | YTHDC1 |
| 1.1922E-52 | -0.2501811 | 0.102 | 0.204 | 2.8164E-48 | MO_kdnrs | HIGD1A |
| 1.3589E-52 | -0.2882242 | 0.168 | 0.276 | 3.2101E-48 | MO_kdnrs | TOR1AIP2 |
| 1.6237E-52 | -0.2773252 | 0.08 | 0.177 | 3.8357E-48 | MO_kdnrs | CDKN2AIP |
| 2.8387E-52 | -0.2554123 | 0.184 | 0.295 | 6.7059E-48 | MO_kdnrs | ATP6V1D |
| 3.0886E-52 | -0.2597843 | 0.033 | 0.115 | 7.2962E-48 | MO_kdnrs | YOD1 |
| 1.3724E-51 | -0.386324 | 0.09 | 0.188 | 3.2421E-47 | MO_kdnrs | DST |
| 3.4162E-51 | -0.4348165 | 0.247 | 0.346 | 8.0702E-47 | MO_kdnrs | NDRG1 |
| 7.4313E-51 | -0.4037701 | 0.174 | 0.278 | 1.7555E-46 | MO_kdnrs | CCNG2 |
| 8.9286E-51 | -0.4098188 | 0.144 | 0.242 | 2.1092E-46 | MO_kdnrs | TOB1 |
| 1.0872E-50 | -0.4219492 | 0.258 | 0.353 | 2.5683E-46 | MO_kdnrs | GOLGB1 |
| 2.0193E-50 | -0.5005538 | 0.044 | 0.127 | 4.7703E-46 | MO_kdnrs | HILPDA |
| 5.697E-50 | -0.2918846 | 0.195 | 0.298 | 1.3458E-45 | MO_kdnrs | CHMP2B |
| 1.0505E-49 | -0.5459397 | 0.158 | 0.256 | 2.4815E-45 | MO_kdnrs | EGR1 |
| 1.3393E-49 | -0.3651984 | 0.079 | 0.173 | 3.1639E-45 | MO_kdnrs | HIST3H2A |
| 1.4784E-49 | -0.3672579 | 0.234 | 0.333 | 3.4924E-45 | MO_kdnrs | TMBIM1 |
| 4.6646E-49 | -0.4178447 | 0.217 | 0.315 | 1.1019E-44 | MO_kdnrs | ZNF292 |
| 4.8498E-49 | -0.4793718 | 0.703 | 0.734 | 1.1457E-44 | MO_kdnrs | HSPA8 |
| 8.5237E-48 | -0.3280618 | 0.172 | 0.269 | 2.0136E-43 | MO_kdnrs | PRDX2 |
| 1.0007E-47 | -0.3057631 | 0.226 | 0.325 | 2.3639E-43 | MO_kdnrs | RAB11A |
| 1.029E-47 | -0.5840198 | 0.977 | 0.953 | 2.4309E-43 | MO_kdnrs | ACTG1 |
| 7.2757E-47 | -0.3186742 | 0.287 | 0.383 | 1.7187E-42 | MO_kdnrs | ARF4 |
| 2.3027E-46 | -0.3287576 | 0.113 | 0.052 | 5.4396E-42 | MO_kdnrs | CCL3L1 |
| 4.5198E-46 | -0.3907773 | 0.177 | 0.274 | 1.0677E-41 | MO_kdnrs | UGCG |
| 6.7846E-46 | -0.3460856 | 0.273 | 0.361 | 1.6027E-41 | MO_kdnrs | MRPL33 |
| 7.3746E-45 | -0.3753721 | 0.259 | 0.351 | 1.7421E-40 | MO_kdnrs | CPEB4 |
| 2.1098E-44 | -0.4194524 | 0.874 | 0.885 | 4.9839E-40 | MO_kdnrs | MYL12B |
| 6.8434E-44 | -0.2827754 | 0.054 | 0.132 | 1.6166E-39 | MO_kdnrs | POLB |
| 8.4014E-44 | -0.4937954 | 0.061 | 0.144 | 1.9847E-39 | MO_kdnrs | NUPR1 |
| 1.1552E-43 | -0.2543851 | 0.09 | 0.176 | 2.729E-39 | MO_kdnrs | JMY |
| 3.4426E-43 | -0.3672402 | 0.466 | 0.534 | 8.1325E-39 | MO_kdnrs | EIF4A2 |
| 9.0902E-43 | 0.36720716 | 0.378 | 0.252 | 2.1474E-38 | MO_kdnrs | AREG |
| 9.3703E-43 | -0.2573045 | 0.157 | 0.252 | 2.2136E-38 | MO_kdnrs | SPTAN1 |

| | | | | | | |
|---|---|---|---|---|---|---|
| 2.1781E-42 | -0.4076831 | 0.507 | 0.581 | 5.1453E-38 | MO_kdnrs | CD55 |
| 2.2934E-42 | -0.5574367 | 0.913 | 0.882 | 5.4176E-38 | MO_kdnrs | HSP90AB1 |
| 3.3341E-42 | -0.2527561 | 0.17 | 0.267 | 7.876E-38 | MO_kdnrs | ANKLE2 |
| 1.1274E-41 | -0.3138179 | 0.174 | 0.264 | 2.6633E-37 | MO_kdnrs | PDLIM5 |
| 1.5556E-41 | -0.2910315 | 0.201 | 0.293 | 3.6748E-37 | MO_kdnrs | RB1CC1 |
| 2.7922E-41 | -0.3461343 | 0.058 | 0.135 | 6.596E-37 | MO_kdnrs | AC023157.3 |
| 4.0708E-41 | -0.370484 | 0.637 | 0.693 | 9.6165E-37 | MO_kdnrs | DNAJA1 |
| 6.8856E-39 | -0.4107955 | 0.122 | 0.202 | 1.6266E-34 | MO_kdnrs | SAMD9 |
| 1.7312E-37 | -0.3437796 | 0.042 | 0.109 | 4.0896E-33 | MO_kdnrs | CLIC3 |
| 2.7022E-37 | -0.2749006 | 0.516 | 0.584 | 6.3834E-33 | MO_kdnrs | UBE2B |
| 2.7481E-37 | -0.2864084 | 0.444 | 0.297 | 6.4918E-33 | MO_kdnrs | HIST1H4C |
| 1.3207E-36 | -0.2957208 | 0.287 | 0.366 | 3.1198E-32 | MO_kdnrs | TMOD3 |
| 1.9129E-35 | -0.6950958 | 0.074 | 0.145 | 4.5189E-31 | MO_kdnrs | CXCL1 |
| 7.9938E-35 | -0.743564 | 0.593 | 0.631 | 1.8884E-30 | MO_kdnrs | JUN |
| 1.086E-33 | -0.2733475 | 0.215 | 0.293 | 2.5654E-29 | MO_kdnrs | CLINT1 |
| 3.4743E-33 | -0.5042735 | 0.294 | 0.355 | 8.2074E-29 | MO_kdnrs | GLS |
| 1.0005E-32 | -0.2758301 | 0.17 | 0.248 | 2.3635E-28 | MO_kdnrs | IFRD1 |
| 1.5496E-32 | -0.3813527 | 0.1 | 0.17 | 3.6606E-28 | MO_kdnrs | ANKRD37 |
| 1.5542E-32 | -0.3743031 | 0.307 | 0.372 | 3.6714E-28 | MO_kdnrs | RALBP1 |
| 1.8485E-32 | -0.2653715 | 0.298 | 0.373 | 4.3666E-28 | MO_kdnrs | PNRC2 |
| 2.4306E-32 | -0.6339158 | 0.398 | 0.457 | 5.7419E-28 | MO_kdnrs | HSPH1 |
| 2.5476E-32 | -0.5356227 | 0.439 | 0.509 | 6.0182E-28 | MO_kdnrs | DNAJB1 |
| 2.5506E-32 | -0.2778797 | 0.238 | 0.312 | 6.0252E-28 | MO_kdnrs | CLTB |
| 3.8496E-32 | -0.2544058 | 0.252 | 0.327 | 9.094E-28 | MO_kdnrs | YWHAQ |
| 3.8831E-32 | -0.303634 | 0.119 | 0.194 | 9.1729E-28 | MO_kdnrs | SLC5A3 |
| 8.5793E-32 | -0.3897976 | 0.19 | 0.264 | 2.0267E-27 | MO_kdnrs | IDI1 |
| 6.2801E-31 | -0.3056448 | 0.126 | 0.199 | 1.4835E-26 | MO_kdnrs | KATNBL1 |
| 9.2223E-31 | -0.2669992 | 0.117 | 0.19 | 2.1786E-26 | MO_kdnrs | ERRFI1 |
| 1.4413E-30 | -0.2637293 | 0.226 | 0.301 | 3.4047E-26 | MO_kdnrs | ZFAND6 |
| 1.6058E-30 | -0.3249202 | 0.22 | 0.289 | 3.7934E-26 | MO_kdnrs | CD151 |
| 2.5237E-27 | -0.2848981 | 0.261 | 0.327 | 5.9617E-23 | MO_kdnrs | YPEL3 |
| 6.7589E-27 | -0.3438116 | 0.072 | 0.129 | 1.5967E-22 | MO_kdnrs | SRD5A3 |
| 3.9198E-26 | -0.2704049 | 0.121 | 0.187 | 9.2597E-22 | MO_kdnrs | GCLM |
| 5.0269E-26 | -0.3264524 | 0.127 | 0.192 | 1.1875E-21 | MO_kdnrs | BAG3 |
| 5.0808E-26 | -1.0532157 | 0.215 | 0.138 | 1.2002E-21 | MO_kdnrs | CXCL3 |
| 5.3417E-26 | -0.2662912 | 0.136 | 0.203 | 1.2619E-21 | MO_kdnrs | SERTAD2 |
| 5.8203E-26 | -0.2891972 | 0.251 | 0.317 | 1.3749E-21 | MO_kdnrs | SMS |
| 1.6267E-24 | -0.7411416 | 0.436 | 0.459 | 3.8427E-20 | MO_kdnrs | MXD1 |
| 1.8637E-24 | -0.3575943 | 0.169 | 0.23 | 4.4026E-20 | MO_kdnrs | INTS6 |
| 3.0179E-24 | -0.3392201 | 0.263 | 0.324 | 7.1291E-20 | MO_kdnrs | BRD2 |
| 4.4598E-24 | -0.3189294 | 0.149 | 0.09 | 1.0535E-19 | MO_kdnrs | PTGS2 |
| 4.1547E-23 | -0.3191726 | 0.303 | 0.354 | 9.8147E-19 | MO_kdnrs | PNN |
| 8.275E-23 | -0.2829411 | 0.061 | 0.111 | 1.9548E-18 | MO_kdnrs | TCHP |
| 2.2554E-21 | -0.2813322 | 0.154 | 0.214 | 5.3279E-17 | MO_kdnrs | MDM2 |
| 3.2242E-21 | -0.2802117 | 0.708 | 0.732 | 7.6165E-17 | MO_kdnrs | ATP6V1G1 |
| 5.4066E-21 | -0.3969172 | 0.302 | 0.349 | 1.2772E-16 | MO_kdnrs | MIDN |
| 3.1757E-20 | -0.2509344 | 0.542 | 0.577 | 7.5019E-16 | MO_kdnrs | PPP1CB |
| 1.0435E-18 | -0.3136427 | 0.159 | 0.211 | 2.4651E-14 | MO_kdnrs | KCNQ1OT1 |
| 2.5906E-18 | -0.3369035 | 0.106 | 0.156 | 6.1198E-14 | MO_kdnrs | ZFAND2A |
| 8.5085E-18 | -0.2801911 | 0.323 | 0.362 | 2.01E-13 | MO_kdnrs | RRBP1 |

| | | | | | | |
|---|---|---|---|---|---|---|
| 1.4083E-16 | -0.2807349 | 0.476 | 0.497 | 3.3269E-12 | MO_kdnrs | DDX3X |
| 1.9321E-16 | -0.3201671 | 0.559 | 0.571 | 4.5642E-12 | MO_kdnrs | ZFAND5 |
| 3.1677E-16 | -0.2664698 | 0.276 | 0.318 | 7.483E-12 | MO_kdnrs | AKAP9 |
| 3.3061E-16 | -0.3203861 | 0.369 | 0.399 | 7.8101E-12 | MO_kdnrs | ARGLU1 |
| 2.7813E-15 | -0.3957559 | 0.325 | 0.355 | 6.5702E-11 | MO_kdnrs | ATF3 |
| 3.1321E-15 | -0.282147 | 0.474 | 0.497 | 7.399E-11 | MO_kdnrs | CD9 |
| 3.2798E-15 | -0.2668273 | 0.312 | 0.347 | 7.7479E-11 | MO_kdnrs | NORAD |
| 3.4665E-15 | -0.4421856 | 0.77 | 0.75 | 8.1889E-11 | MO_kdnrs | TPM4 |
| 3.938E-15 | -0.2722071 | 0.534 | 0.549 | 9.3028E-11 | MO_kdnrs | TOP1 |
| 6.8418E-15 | -0.2678287 | 0.182 | 0.226 | 1.6162E-10 | MO_kdnrs | POLR2A |
| 3.2241E-14 | -0.2933619 | 0.35 | 0.377 | 7.6163E-10 | MO_kdnrs | CCNL1 |
| 1.351E-13 | -0.347743 | 0.731 | 0.65 | 3.1916E-09 | MO_kdnrs | HSPA1A |
| 4.7112E-13 | -0.3061013 | 0.52 | 0.537 | 1.1129E-08 | MO_kdnrs | SERPINB1 |
| 2.8053E-12 | -0.2535198 | 0.366 | 0.393 | 6.6268E-08 | MO_kdnrs | RSRC2 |
| 1.2892E-11 | -0.3374971 | 0.195 | 0.228 | 3.0454E-07 | MO_kdnrs | PPP1R10 |
| 4.012E-11 | -0.4010359 | 0.351 | 0.349 | 9.4776E-07 | MO_kdnrs | PEBP1 |
| 9.9531E-11 | -0.3135449 | 0.168 | 0.201 | 2.3512E-06 | MO_kdnrs | DUSP4 |
| 1.9655E-09 | -0.3088842 | 0.228 | 0.253 | 4.643E-05 | MO_kdnrs | SNHG12 |
| 3.3261E-09 | -0.3671942 | 0.314 | 0.329 | 7.8572E-05 | MO_kdnrs | SLC38A2 |
| 4.4464E-09 | -0.2846832 | 0.234 | 0.264 | 0.00010504 | MO_kdnrs | HES4 |
| 7.1218E-08 | -0.2830079 | 0.82 | 0.781 | 0.00168238 | MO_kdnrs | DYNLL1 |
| 1.2249E-07 | -2.7398047 | 0.799 | 0.721 | 0.00289369 | MO_kdnrs | S100A9 |
| 1.5155E-07 | -0.4089375 | 0.256 | 0.267 | 0.00358001 | MO_kdnrs | DDIT3 |
| 2.1936E-07 | -0.7419267 | 0.202 | 0.217 | 0.005182 | MO_kdnrs | S100A12 |
| 1.1073E-06 | -1.6684838 | 0.118 | 0.087 | 0.026158 | MO_kdnrs | HBB |
| 1.2202E-06 | -0.3152728 | 0.21 | 0.228 | 0.02882509 | MO_kdnrs | IER5L |
| 2.434E-05 | -0.2722817 | 0.155 | 0.171 | 0.57497511 | MO_kdnrs | PRDM1 |
| 4.189E-05 | -0.2696298 | 0.094 | 0.114 | 0.98957372 | MO_kdnrs | CYP1B1 |
| 4.2371E-05 | -0.3649326 | 0.348 | 0.28 | 1 | MO_kdnrs | SLC25A37 |
| 4.2688E-05 | -0.3492666 | 0.132 | 0.149 | 1 | MO_kdnrs | SMC4 |
| 4.5289E-05 | -1.578801 | 0.534 | 0.484 | 1 | MO_kdnrs | CXCL8 |
| 5.3647E-05 | -0.2541726 | 0.373 | 0.368 | 1 | MO_kdnrs | ANKRD11 |
| 5.7781E-05 | -0.9043952 | 0.136 | 0.153 | 1 | MO_kdnrs | CCL20 |
| 0.00041003 | -1.0744362 | 0.781 | 0.703 | 1 | MO_kdnrs | ANXA1 |
| 0.00042101 | -0.324572 | 0.62 | 0.531 | 1 | MO_kdnrs | UPP1 |
| 0.0004224 | -0.4493393 | 0.399 | 0.327 | 1 | MO_kdnrs | IL1RN |
| 0.00059826 | -1.6285397 | 0.741 | 0.665 | 1 | MO_kdnrs | S100A8 |
| 0.00654327 | -0.3269486 | 0.155 | 0.162 | 1 | MO_kdnrs | CSKMT |
| 0.00821785 | -0.2503929 | 0.162 | 0.17 | 1 | MO_kdnrs | AKR1B1 |
| 0 | 5.51871721 | 0.965 | 0.225 | 0 | MO_MT+ | MT1G |
| 0 | 5.23480524 | 0.809 | 0.124 | 0 | MO_MT+ | MT1H |
| 0 | 4.91475787 | 0.752 | 0.088 | 0 | MO_MT+ | MT1M |
| 0 | 4.63800751 | 0.983 | 0.251 | 0 | MO_MT+ | MT1X |
| 0 | 4.47488096 | 1 | 0.645 | 0 | MO_MT+ | MT2A |
| 0 | 4.12850413 | 0.498 | 0.028 | 0 | MO_MT+ | MT1A |
| 0 | 3.87015384 | 0.922 | 0.254 | 0 | MO_MT+ | MT1E |
| 0 | 3.11422392 | 0.84 | 0.128 | 0 | MO_MT+ | MT1F |
| 0 | 2.1335087 | 0.126 | 0.009 | 0 | MO_MT+ | MT1B |
| 0 | 1.1388852 | 0.573 | 0.118 | 0 | MO_MT+ | SLC30A1 |
| 0 | 0.73695045 | 0.464 | 0.136 | 0 | MO_MT+ | BCAT1 |

| | | | | | | |
|---|---|---|---|---|---|---|
| 2E-286 | 2.94347213 | 0.289 | 0.063 | 4.724E-282 | MO_MT+ | CXCL5 |
| 3.926E-278 | 0.41973078 | 0.309 | 0.071 | 9.274E-274 | MO_MT+ | LRRC25 |
| 5.409E-252 | 0.83661734 | 0.517 | 0.199 | 1.278E-247 | MO_MT+ | NINJ1 |
| 3.708E-250 | 1.13554187 | 0.55 | 0.22 | 8.759E-246 | MO_MT+ | PLIN2 |
| 4.834E-249 | 0.30597324 | 0.116 | 0.012 | 1.142E-244 | MO_MT+ | SLAMF9 |
| 6.683E-246 | 0.61575983 | 0.454 | 0.148 | 1.579E-241 | MO_MT+ | AQP9 |
| 6.583E-243 | 1.03984097 | 0.882 | 0.658 | 1.555E-238 | MO_MT+ | ANXA5 |
| 1.095E-241 | 0.94063695 | 0.561 | 0.222 | 2.586E-237 | MO_MT+ | BCL2A1 |
| 1.248E-240 | 0.86019164 | 0.423 | 0.14 | 2.948E-236 | MO_MT+ | CLEC5A |
| 3.34E-233 | 0.57901624 | 0.395 | 0.125 | 7.889E-229 | MO_MT+ | ITGA5 |
| 1.289E-232 | 1.12081855 | 0.999 | 0.993 | 3.046E-228 | MO_MT+ | FTH1 |
| 1.102E-229 | 1.04308031 | 0.998 | 0.986 | 2.603E-225 | MO_MT+ | FTL |
| 4.732E-227 | 0.78673796 | 0.647 | 0.31 | 1.118E-222 | MO_MT+ | LIMS1 |
| 7.194E-223 | 0.73380511 | 0.702 | 0.32 | 1.7E-218 | MO_MT+ | LAPTM5 |
| 1.165E-221 | 1.04620998 | 0.517 | 0.207 | 2.752E-217 | MO_MT+ | ALOX5AP |
| 8.036E-221 | 1.05097263 | 0.702 | 0.372 | 1.898E-216 | MO_MT+ | FCER1G |
| 8.091E-218 | 0.25341177 | 0.178 | 0.031 | 1.911E-213 | MO_MT+ | ASCL2 |
| 1.11E-207 | -1.496408 | 0.195 | 0.57 | 2.622E-203 | MO_MT+ | ELF3 |
| 3.29E-207 | 1.15810295 | 0.634 | 0.333 | 7.772E-203 | MO_MT+ | CTSL |
| 4.039E-206 | -1.5631596 | 0.226 | 0.593 | 9.54E-202 | MO_MT+ | KRT19 |
| 2.2E-198 | 1.06948305 | 0.887 | 0.734 | 5.196E-194 | MO_MT+ | VIM |
| 1.458E-197 | -1.0145857 | 0.798 | 0.961 | 3.444E-193 | MO_MT+ | ACTG1 |
| 4.455E-197 | 0.49590089 | 0.355 | 0.113 | 1.052E-192 | MO_MT+ | PHACTR1 |
| 8.083E-195 | 0.53521204 | 0.44 | 0.162 | 1.91E-190 | MO_MT+ | SMIM25 |
| 9.122E-195 | 0.53726814 | 0.239 | 0.059 | 2.155E-190 | MO_MT+ | PID1 |
| 4.323E-191 | 0.77137318 | 0.745 | 0.412 | 1.021E-186 | MO_MT+ | EMP3 |
| 1.842E-190 | 0.55710745 | 0.484 | 0.189 | 4.351E-186 | MO_MT+ | NCF2 |
| 2.222E-188 | 0.44059968 | 0.3 | 0.089 | 5.25E-184 | MO_MT+ | LHFPL2 |
| 4.465E-184 | 0.70090853 | 0.727 | 0.421 | 1.055E-179 | MO_MT+ | GSTO1 |
| 1.09E-183 | 0.37144431 | 0.306 | 0.092 | 2.575E-179 | MO_MT+ | TFEC |
| 1.47E-183 | 0.29771927 | 0.221 | 0.053 | 3.473E-179 | MO_MT+ | CCRL2 |
| 9.35E-183 | 0.4485067 | 0.36 | 0.123 | 2.209E-178 | MO_MT+ | PILRA |
| 1.955E-182 | 0.41380633 | 0.34 | 0.112 | 4.619E-178 | MO_MT+ | DENND5A |
| 2.122E-182 | 0.81335662 | 0.811 | 0.56 | 5.013E-178 | MO_MT+ | SRGN |
| 6.066E-182 | 1.53600298 | 0.383 | 0.142 | 1.433E-177 | MO_MT+ | CCL20 |
| 8.186E-182 | 1.28267788 | 0.832 | 0.67 | 1.934E-177 | MO_MT+ | TIMP1 |
| 6.239E-180 | 0.81466104 | 0.576 | 0.281 | 1.474E-175 | MO_MT+ | ATP13A3 |
| 1.134E-176 | -1.9860765 | 0.208 | 0.55 | 2.68E-172 | MO_MT+ | WFDC2 |
| 2.011E-173 | 0.75594583 | 0.515 | 0.221 | 4.751E-169 | MO_MT+ | SLC2A3 |
| 3.469E-171 | -3.0409622 | 0.203 | 0.537 | 8.195E-167 | MO_MT+ | LCN2 |
| 9.657E-171 | 0.64690884 | 0.853 | 0.602 | 2.281E-166 | MO_MT+ | CD44 |
| 3.107E-170 | 0.35770781 | 0.24 | 0.065 | 7.339E-166 | MO_MT+ | NBPF19 |
| 4.296E-170 | 0.80728832 | 0.376 | 0.141 | 1.015E-165 | MO_MT+ | SLC7A11 |
| 7.28E-169 | 0.52585325 | 0.371 | 0.133 | 1.72E-164 | MO_MT+ | NCF1 |
| 1.624E-168 | 0.3599909 | 0.29 | 0.089 | 3.836E-164 | MO_MT+ | DOCK4 |
| 2.551E-167 | 0.50861894 | 0.545 | 0.232 | 6.026E-163 | MO_MT+ | SLC11A1 |
| 1.922E-165 | 0.32857128 | 0.233 | 0.062 | 4.54E-161 | MO_MT+ | TNS1 |
| 2.902E-165 | 0.46089328 | 0.404 | 0.155 | 6.855E-161 | MO_MT+ | CEP170 |
| 3.073E-165 | 0.46867783 | 0.378 | 0.137 | 7.258E-161 | MO_MT+ | TREM1 |
| 5.879E-164 | 0.74466221 | 0.728 | 0.428 | 1.389E-159 | MO_MT+ | TYROBP |

| | | | | | | |
|---|---|---|---|---|---|---|
| 5.694E-161 | 0.53231991 | 0.505 | 0.216 | 1.345E-156 | MO_MT+ | ZEB2 |
| 1.053E-159 | 0.72759835 | 0.884 | 0.72 | 2.488E-155 | MO_MT+ | ENO1 |
| 1.333E-159 | 0.89619912 | 0.817 | 0.609 | 3.149E-155 | MO_MT+ | LGALS1 |
| 1.307E-158 | -1.1656177 | 0.226 | 0.558 | 3.088E-154 | MO_MT+ | KRT7 |
| 1.364E-158 | 0.3852943 | 0.26 | 0.077 | 3.222E-154 | MO_MT+ | SEMA6B |
| 1.54E-158 | 0.80054084 | 0.877 | 0.735 | 3.638E-154 | MO_MT+ | PSAP |
| 1.648E-158 | 0.43816729 | 0.614 | 0.278 | 3.892E-154 | MO_MT+ | LCP1 |
| 6.272E-158 | 0.47449501 | 0.432 | 0.177 | 1.482E-153 | MO_MT+ | RAC2 |
| 3.017E-156 | 0.47789103 | 0.562 | 0.255 | 7.126E-152 | MO_MT+ | CD53 |
| 6.479E-156 | 0.4012111 | 0.336 | 0.118 | 1.53E-151 | MO_MT+ | IRAK3 |
| 7.084E-156 | -1.0649702 | 0.257 | 0.581 | 1.673E-151 | MO_MT+ | TACSTD2 |
| 8.077E-155 | 0.43486065 | 0.419 | 0.171 | 1.908E-150 | MO_MT+ | RILPL2 |
| 1.276E-154 | 1.15942691 | 0.411 | 0.166 | 3.013E-150 | MO_MT+ | IL1B |
| 5.464E-153 | 0.53185064 | 0.416 | 0.167 | 1.291E-148 | MO_MT+ | ANPEP |
| 1.317E-152 | 0.60506369 | 0.581 | 0.301 | 3.111E-148 | MO_MT+ | CD68 |
| 5.897E-152 | 0.63656991 | 0.461 | 0.199 | 1.393E-147 | MO_MT+ | TNFRSF1B |
| 5.058E-151 | 0.53008587 | 0.424 | 0.17 | 1.195E-146 | MO_MT+ | OLR1 |
| 3.157E-149 | 0.25653383 | 0.173 | 0.039 | 7.458E-145 | MO_MT+ | TNFSF14 |
| 6.948E-148 | 1.10121782 | 0.766 | 0.607 | 1.641E-143 | MO_MT+ | C15orf48 |
| 2.669E-146 | 0.73317132 | 0.549 | 0.274 | 6.305E-142 | MO_MT+ | SLC25A37 |
| 2.677E-146 | 0.43158909 | 0.478 | 0.204 | 6.323E-142 | MO_MT+ | SPI1 |
| 1.301E-145 | -0.8773134 | 0.225 | 0.553 | 3.073E-141 | MO_MT+ | CD24 |
| 1.429E-145 | 0.40012668 | 0.457 | 0.189 | 3.375E-141 | MO_MT+ | ITGAX |
| 1.216E-144 | 0.47371288 | 0.47 | 0.213 | 2.874E-140 | MO_MT+ | GRINA |
| 6.388E-143 | 0.45101791 | 0.453 | 0.196 | 1.509E-138 | MO_MT+ | SLC43A2 |
| 6.418E-142 | 0.8471902 | 0.973 | 0.93 | 1.516E-137 | MO_MT+ | GAPDH |
| 6.466E-142 | 0.38305474 | 0.372 | 0.147 | 1.528E-137 | MO_MT+ | CSGALNACT2 |
| 1.349E-140 | 0.48078218 | 0.45 | 0.19 | 3.186E-136 | MO_MT+ | CYBB |
| 1.817E-140 | 0.38158587 | 0.336 | 0.127 | 4.292E-136 | MO_MT+ | SLC6A6 |
| 2.522E-139 | 0.40947573 | 0.228 | 0.068 | 5.958E-135 | MO_MT+ | NSMAF |
| 1.413E-138 | -1.0330275 | 0.244 | 0.551 | 3.338E-134 | MO_MT+ | CLDN4 |
| 5.011E-138 | 0.37491052 | 0.244 | 0.076 | 1.184E-133 | MO_MT+ | CYP27A1 |
| 6.428E-137 | 0.38517433 | 0.32 | 0.12 | 1.519E-132 | MO_MT+ | SHKBP1 |
| 1.231E-136 | -4.2098929 | 0.127 | 0.413 | 2.908E-132 | MO_MT+ | SPRR3 |
| 1.089E-135 | 0.3111419 | 0.284 | 0.096 | 2.572E-131 | MO_MT+ | CCR1 |
| 2.599E-135 | 0.35320416 | 0.382 | 0.153 | 6.14E-131 | MO_MT+ | FERMT3 |
| 1.272E-134 | 0.63540213 | 0.467 | 0.228 | 3.005E-130 | MO_MT+ | RNASET2 |
| 1.962E-134 | 0.50857177 | 0.445 | 0.203 | 4.634E-130 | MO_MT+ | ZFYVE16 |
| 2.152E-134 | 0.37865919 | 0.328 | 0.123 | 5.083E-130 | MO_MT+ | EPB41L3 |
| 2.541E-132 | -0.8016487 | 0.163 | 0.464 | 6.002E-128 | MO_MT+ | CLDN7 |
| 2.54E-131 | 0.31574758 | 0.304 | 0.111 | 6E-127 | MO_MT+ | P2RX4 |
| 3.365E-130 | -2.1124062 | 0.08 | 0.355 | 7.95E-126 | MO_MT+ | HPGD |
| 6.14E-129 | -2.1799858 | 0.078 | 0.352 | 1.451E-124 | MO_MT+ | PSCA |
| 2.029E-128 | -2.8883338 | 0.116 | 0.389 | 4.792E-124 | MO_MT+ | KRT13 |
| 1.576E-127 | 0.72325021 | 0.473 | 0.233 | 3.723E-123 | MO_MT+ | ATP2B1 |
| 4.463E-126 | 0.46749947 | 0.382 | 0.168 | 1.054E-121 | MO_MT+ | ABCA1 |
| 1.588E-125 | 1.56401981 | 0.302 | 0.115 | 3.751E-121 | MO_MT+ | CCL3 |
| 7.026E-125 | -1.0026487 | 0.274 | 0.569 | 1.66E-120 | MO_MT+ | KRT18 |
| 2.1E-124 | 0.27892652 | 0.208 | 0.06 | 4.962E-120 | MO_MT+ | MMP19 |
| 8.208E-124 | 0.4183778 | 0.434 | 0.203 | 1.939E-119 | MO_MT+ | RNF13 |

| | | | | | | |
|---|---|---|---|---|---|---|
| 3.027E-123 | 0.35322718 | 0.396 | 0.17 | 7.151E-119 | MO_MT+ | FMNL1 |
| 5.241E-123 | 0.58746836 | 0.959 | 0.918 | 1.238E-118 | MO_MT+ | S100A10 |
| 6.126E-123 | 0.45511615 | 0.398 | 0.176 | 1.447E-118 | MO_MT+ | KLF10 |
| 2.972E-122 | 0.4684244 | 0.642 | 0.374 | 7.021E-118 | MO_MT+ | CNBP |
| 4.666E-122 | 0.31318118 | 0.33 | 0.128 | 1.102E-117 | MO_MT+ | HCK |
| 1.362E-121 | 0.50139439 | 0.556 | 0.301 | 3.217E-117 | MO_MT+ | FNDC3B |
| 1.848E-121 | 0.27866415 | 0.217 | 0.067 | 4.365E-117 | MO_MT+ | NRIP3 |
| 1.87E-121 | -0.8356649 | 0.164 | 0.448 | 4.417E-117 | MO_MT+ | ATP1B1 |
| 1.807E-120 | 0.30238376 | 0.291 | 0.108 | 4.268E-116 | MO_MT+ | SPHK1 |
| 2.939E-120 | 0.28861413 | 0.286 | 0.105 | 6.943E-116 | MO_MT+ | SLC43A3 |
| 4.871E-120 | 0.33113668 | 0.346 | 0.142 | 1.151E-115 | MO_MT+ | ASAP1 |
| 1.663E-119 | 0.38561655 | 0.406 | 0.177 | 3.928E-115 | MO_MT+ | PTPRE |
| 1.122E-118 | -1.0014361 | 0.225 | 0.505 | 2.65E-114 | MO_MT+ | KRT8 |
| 6.811E-118 | 0.43234447 | 0.378 | 0.162 | 1.609E-113 | MO_MT+ | CXCR4 |
| 7.473E-117 | 0.66931278 | 0.174 | 0.048 | 1.765E-112 | MO_MT+ | TNFAIP6 |
| 2.828E-116 | -1.3246956 | 0.195 | 0.461 | 6.68E-112 | MO_MT+ | SOX4 |
| 1.298E-115 | 0.34338662 | 0.338 | 0.136 | 3.066E-111 | MO_MT+ | TLR2 |
| 7.153E-115 | 0.32308185 | 0.323 | 0.129 | 1.69E-110 | MO_MT+ | EVI2A |
| 7.784E-115 | 0.29973811 | 0.274 | 0.1 | 1.839E-110 | MO_MT+ | PIK3AP1 |
| 1.663E-114 | 0.41026472 | 0.452 | 0.221 | 3.929E-110 | MO_MT+ | ATP6V1B2 |
| 4.359E-113 | 0.39105468 | 0.412 | 0.192 | 1.03E-108 | MO_MT+ | NR3C1 |
| 3.408E-112 | -1.8635313 | 0.165 | 0.418 | 8.052E-108 | MO_MT+ | S100P |
| 3.828E-112 | 0.36189734 | 0.448 | 0.217 | 9.043E-108 | MO_MT+ | TPP1 |
| 4.608E-112 | 0.57733499 | 0.828 | 0.682 | 1.089E-107 | MO_MT+ | BRI3 |
| 5.447E-112 | 0.52120782 | 0.328 | 0.137 | 1.287E-107 | MO_MT+ | SDC2 |
| 1.411E-110 | 0.36416834 | 0.311 | 0.126 | 3.333E-106 | MO_MT+ | PDE4DIP |
| 1.781E-110 | 0.35306313 | 0.187 | 0.056 | 4.208E-106 | MO_MT+ | SERPINE1 |
| 2.47E-110 | 0.51193114 | 0.445 | 0.221 | 5.836E-106 | MO_MT+ | PPIF |
| 3.94E-109 | -1.5138626 | 0.121 | 0.379 | 9.308E-105 | MO_MT+ | MAL |
| 9.217E-109 | 0.39374454 | 0.32 | 0.129 | 2.177E-104 | MO_MT+ | MCEMP1 |
| 1.16E-108 | 0.55203962 | 0.418 | 0.204 | 2.74E-104 | MO_MT+ | BASP1 |
| 1.536E-108 | 0.41993394 | 0.459 | 0.236 | 3.628E-104 | MO_MT+ | PTPN12 |
| 2.474E-108 | 0.42064951 | 0.443 | 0.222 | 5.844E-104 | MO_MT+ | FLOT1 |
| 1.971E-107 | 0.28099098 | 0.366 | 0.156 | 4.656E-103 | MO_MT+ | LCP2 |
| 3.056E-106 | 0.47866025 | 0.432 | 0.216 | 7.22E-102 | MO_MT+ | KYNU |
| 3.153E-106 | 0.54841595 | 0.373 | 0.166 | 7.449E-102 | MO_MT+ | PLEK |
| 6.285E-106 | 0.44072062 | 0.657 | 0.392 | 1.485E-101 | MO_MT+ | FLNA |
| 1.408E-105 | 0.27481347 | 0.232 | 0.081 | 3.327E-101 | MO_MT+ | IRAK2 |
| 1.463E-105 | 0.3420636 | 0.408 | 0.194 | 3.456E-101 | MO_MT+ | TANK |
| 1.863E-105 | 0.29130888 | 0.377 | 0.168 | 4.4E-101 | MO_MT+ | PPP1R18 |
| 3.668E-105 | 0.27342396 | 0.258 | 0.094 | 8.664E-101 | MO_MT+ | LUCAT1 |
| 1.012E-104 | 0.45071911 | 0.365 | 0.168 | 2.39E-100 | MO_MT+ | P4HA1 |
| 1.103E-104 | 0.52346601 | 0.942 | 0.855 | 2.605E-100 | MO_MT+ | SH3BGRL3 |
| 1.805E-104 | 0.37750473 | 0.34 | 0.149 | 4.265E-100 | MO_MT+ | SH3BP5 |
| 2.917E-104 | -0.7588463 | 0.091 | 0.338 | 6.891E-100 | MO_MT+ | MYO6 |
| 6.684E-104 | 0.43631275 | 0.337 | 0.149 | 1.579E-99 | MO_MT+ | FNIP2 |
| 8.658E-104 | -1.1003471 | 0.154 | 0.416 | 2.045E-99 | MO_MT+ | SLPI |
| 3.203E-103 | 0.48370671 | 0.632 | 0.384 | 7.567E-99 | MO_MT+ | HIF1A |
| 1.081E-102 | -0.8942054 | 0.666 | 0.84 | 2.5544E-98 | MO_MT+ | PPDPF |
| 1.648E-102 | 0.2582958 | 0.158 | 0.044 | 3.8942E-98 | MO_MT+ | HIF1A-AS2 |

| | | | | | | |
|---|---|---|---|---|---|---|
| 3.251E-102 | 0.42851742 | 0.302 | 0.124 | 7.6795E-98 | MO_MT+ | CD83 |
| 7.869E-102 | 0.29465456 | 0.335 | 0.142 | 1.8589E-97 | MO_MT+ | DMXL2 |
| 1.788E-101 | 0.31011288 | 0.326 | 0.14 | 4.2228E-97 | MO_MT+ | SH3KBP1 |
| 1.007E-100 | 0.2996281 | 0.345 | 0.151 | 2.3784E-96 | MO_MT+ | MYADM |
| 4.413E-100 | 0.52350669 | 0.736 | 0.53 | 1.0424E-95 | MO_MT+ | PLAUR |
| 2.125E-99 | 0.33433264 | 0.342 | 0.152 | 5.0197E-95 | MO_MT+ | AP1S2 |
| 7.547E-99 | 0.5040367 | 0.483 | 0.262 | 1.7829E-94 | MO_MT+ | OSBPL8 |
| 3.572E-98 | 0.28880049 | 0.227 | 0.081 | 8.438E-94 | MO_MT+ | ANTXR2 |
| 4.8826E-98 | 0.40275816 | 0.323 | 0.137 | 1.1534E-93 | MO_MT+ | CD93 |
| 1.0284E-97 | 0.46473073 | 0.517 | 0.297 | 2.4293E-93 | MO_MT+ | BNIP3L |
| 2.5734E-97 | 0.50995662 | 0.287 | 0.119 | 6.0791E-93 | MO_MT+ | MARCO |
| 6.471E-97 | 0.50622868 | 0.307 | 0.133 | 1.5286E-92 | MO_MT+ | GPNMB |
| 3.2875E-95 | 1.01428094 | 0.419 | 0.214 | 7.766E-91 | MO_MT+ | G0S2 |
| 7.6764E-95 | -0.8340607 | 0.924 | 0.93 | 1.8134E-90 | MO_MT+ | MT-CYB |
| 1.2398E-94 | 0.45990758 | 0.382 | 0.191 | 2.9288E-90 | MO_MT+ | CREG1 |
| 2.3016E-94 | 0.33286706 | 0.356 | 0.161 | 5.437E-90 | MO_MT+ | CD48 |
| 2.7665E-94 | -0.7064098 | 0.511 | 0.731 | 6.5352E-90 | MO_MT+ | HSPB1 |
| 4.0695E-94 | 0.41670802 | 0.562 | 0.324 | 9.6133E-90 | MO_MT+ | TLN1 |
| 4.5566E-92 | 1.0027768 | 0.486 | 0.277 | 1.0764E-87 | MO_MT+ | HMOX1 |
| 5.8391E-92 | 0.8375487 | 0.168 | 0.052 | 1.3794E-87 | MO_MT+ | CCL3L1 |
| 1.0356E-91 | 0.4122617 | 0.375 | 0.175 | 2.4463E-87 | MO_MT+ | IGSF6 |
| 2.0874E-91 | 0.44966709 | 0.435 | 0.214 | 4.9311E-87 | MO_MT+ | VCAN |
| 2.2562E-91 | 0.47284285 | 0.424 | 0.211 | 5.3298E-87 | MO_MT+ | CD14 |
| 2.5597E-91 | -1.211116 | 0.075 | 0.304 | 6.0467E-87 | MO_MT+ | ADIRF |
| 3.4971E-90 | -0.6590099 | 0.267 | 0.491 | 8.2612E-86 | MO_MT+ | CD46 |
| 2.9356E-89 | 0.31322634 | 0.356 | 0.168 | 6.9349E-85 | MO_MT+ | ATXN1 |
| 3.6545E-89 | -0.9812582 | 0.08 | 0.301 | 8.6331E-85 | MO_MT+ | RND3 |
| 4.0142E-88 | 0.27055414 | 0.229 | 0.087 | 9.4828E-84 | MO_MT+ | PLXNC1 |
| 1.7038E-87 | 0.3329003 | 0.429 | 0.225 | 4.0248E-83 | MO_MT+ | M6PR |
| 1.9698E-87 | 0.3009545 | 0.249 | 0.099 | 4.6534E-83 | MO_MT+ | ENG |
| 4.314E-87 | 0.28588577 | 0.315 | 0.138 | 1.0191E-82 | MO_MT+ | MS4A7 |
| 1.0872E-86 | -0.8441377 | 0.039 | 0.246 | 2.5684E-82 | MO_MT+ | ADGRF1 |
| 2.3301E-86 | 0.28931249 | 0.296 | 0.129 | 5.5045E-82 | MO_MT+ | DDX60L |
| 2.4379E-86 | 0.25052278 | 0.229 | 0.087 | 5.759E-82 | MO_MT+ | LILRB4 |
| 2.4594E-86 | 0.30831901 | 0.335 | 0.157 | 5.8098E-82 | MO_MT+ | ZNF267 |
| 2.7237E-86 | -1.0443265 | 0.061 | 0.271 | 6.4343E-82 | MO_MT+ | LMO7 |
| 2.8669E-86 | 0.26522117 | 0.337 | 0.15 | 6.7725E-82 | MO_MT+ | CD300E |
| 5.7285E-85 | 0.32235478 | 0.48 | 0.266 | 1.3532E-80 | MO_MT+ | PDXK |
| 8.1536E-85 | 0.36349135 | 0.464 | 0.256 | 1.9261E-80 | MO_MT+ | PLP2 |
| 9.5071E-85 | 1.74773643 | 0.619 | 0.483 | 2.2459E-80 | MO_MT+ | CXCL8 |
| 1.2861E-84 | 0.34147357 | 0.379 | 0.19 | 3.0381E-80 | MO_MT+ | TCEAL9 |
| 2.0766E-84 | 0.37849464 | 0.438 | 0.229 | 4.9055E-80 | MO_MT+ | TNFAIP3 |
| 8.5267E-83 | 0.26811767 | 0.278 | 0.12 | 2.0143E-78 | MO_MT+ | ADAM8 |
| 1.2011E-82 | 0.82368219 | 0.778 | 0.701 | 2.8373E-78 | MO_MT+ | SOD2 |
| 1.2786E-82 | 0.3535308 | 0.339 | 0.161 | 3.0205E-78 | MO_MT+ | SLA |
| 1.8462E-82 | 0.39258184 | 0.496 | 0.283 | 4.3612E-78 | MO_MT+ | SKIL |
| 2.1248E-82 | 0.310083 | 0.298 | 0.136 | 5.0194E-78 | MO_MT+ | PRKAG2 |
| 2.8287E-82 | -0.6177548 | 0.323 | 0.531 | 6.6823E-78 | MO_MT+ | HEBP2 |
| 5.8891E-82 | 0.66616764 | 0.859 | 0.771 | 1.3912E-77 | MO_MT+ | CTSD |
| 1.1212E-81 | 0.86120404 | 0.156 | 0.05 | 2.6487E-77 | MO_MT+ | CCL4L2 |

| | | | | | | |
|---|---|---|---|---|---|---|
| 1.4484E-81 | 0.2744483 | 0.25 | 0.103 | 3.4215E-77 | MO_MT+ | PAG1 |
| 1.7027E-80 | -0.5846088 | 0.651 | 0.819 | 4.0222E-76 | MO_MT+ | GSTP1 |
| 1.896E-80 | -0.6445344 | 0.064 | 0.268 | 4.4789E-76 | MO_MT+ | MUC1 |
| 1.9757E-80 | -1.5627326 | 0.044 | 0.239 | 4.6672E-76 | MO_MT+ | PRSS22 |
| 3.8931E-80 | -1.1933356 | 0.008 | 0.191 | 9.1968E-76 | MO_MT+ | DUOX2 |
| 5.6203E-80 | 0.42460816 | 0.819 | 0.705 | 1.3277E-75 | MO_MT+ | SDCBP |
| 9.3904E-80 | 0.31704996 | 0.285 | 0.126 | 2.2183E-75 | MO_MT+ | RGCC |
| 1.1231E-79 | 0.26808298 | 0.374 | 0.181 | 2.6531E-75 | MO_MT+ | FCGR2A |
| 1.3303E-79 | 0.55541619 | 0.697 | 0.518 | 3.1426E-75 | MO_MT+ | CEBPB |
| 3.201E-79 | 0.27757757 | 0.26 | 0.112 | 7.5618E-75 | MO_MT+ | CASP1 |
| 3.3352E-79 | 0.37672268 | 0.477 | 0.267 | 7.8787E-75 | MO_MT+ | ETS2 |
| 1.5002E-78 | 0.36245799 | 0.145 | 0.045 | 3.5439E-74 | MO_MT+ | MMP9 |
| 1.7374E-78 | 0.32656677 | 0.405 | 0.214 | 4.1043E-74 | MO_MT+ | MAP3K2 |
| 1.999E-78 | 0.28130045 | 0.362 | 0.174 | 4.7222E-74 | MO_MT+ | SAMSN1 |
| 3.6679E-78 | 0.27312409 | 0.298 | 0.138 | 8.6646E-74 | MO_MT+ | IRAK1 |
| 2.0544E-77 | -0.4861098 | 0.431 | 0.632 | 4.8532E-73 | MO_MT+ | DSTN |
| 1.4878E-76 | 0.28046017 | 0.36 | 0.18 | 3.5147E-72 | MO_MT+ | RNF130 |
| 1.4997E-76 | 0.32090361 | 0.432 | 0.235 | 3.5428E-72 | MO_MT+ | RBPJ |
| 4.2708E-76 | -1.6434384 | 0.14 | 0.347 | 1.0089E-71 | MO_MT+ | MMP7 |
| 1.1623E-75 | 0.31562321 | 0.434 | 0.226 | 2.7457E-71 | MO_MT+ | HCST |
| 1.8444E-75 | -0.7012912 | 0.604 | 0.758 | 4.357E-71 | MO_MT+ | TPM4 |
| 2.7104E-75 | 1.30497754 | 0.287 | 0.137 | 6.4029E-71 | MO_MT+ | CCL2 |
| 7.6742E-75 | -0.4756407 | 0.209 | 0.436 | 1.8129E-70 | MO_MT+ | SPINT2 |
| 1.8266E-74 | 0.30359316 | 0.543 | 0.322 | 4.3151E-70 | MO_MT+ | MSN |
| 3.0254E-74 | 0.30218 | 0.399 | 0.21 | 7.1469E-70 | MO_MT+ | DSE |
| 3.1584E-74 | 0.27130322 | 0.291 | 0.134 | 7.461E-70 | MO_MT+ | LRP1 |
| 5.3237E-74 | 0.40610452 | 0.84 | 0.723 | 1.2576E-69 | MO_MT+ | PKM |
| 6.8653E-74 | 0.39911206 | 0.687 | 0.513 | 1.6218E-69 | MO_MT+ | ATP6V1F |
| 1.253E-73 | -0.5126387 | 0.097 | 0.297 | 2.9599E-69 | MO_MT+ | SPINT1 |
| 2.2809E-73 | 0.45558228 | 0.837 | 0.723 | 5.3881E-69 | MO_MT+ | CYBA |
| 3.5918E-73 | 0.25314532 | 0.235 | 0.099 | 8.485E-69 | MO_MT+ | HPSE |
| 4.4055E-73 | 0.25009048 | 0.297 | 0.139 | 1.0407E-68 | MO_MT+ | ZMIZ1 |
| 5.0404E-73 | 0.30847551 | 0.398 | 0.213 | 1.1907E-68 | MO_MT+ | ACTN1 |
| 1.6957E-72 | -0.6393896 | 0.26 | 0.464 | 4.0058E-68 | MO_MT+ | RAB11FIP1 |
| 1.7325E-72 | 0.26633971 | 0.196 | 0.075 | 4.0926E-68 | MO_MT+ | SLC39A8 |
| 2.0029E-72 | -0.6573616 | 0.097 | 0.3 | 4.7315E-68 | MO_MT+ | GPRC5A |
| 3.6815E-72 | -1.2827734 | 0.006 | 0.172 | 8.6968E-68 | MO_MT+ | DUOXA2 |
| 3.9165E-72 | 0.28391371 | 0.265 | 0.121 | 9.252E-68 | MO_MT+ | DRAM1 |
| 6.5663E-72 | -0.7130248 | 0.054 | 0.241 | 1.5512E-67 | MO_MT+ | PDZK1IP1 |
| 9.8405E-72 | 0.29304278 | 0.322 | 0.157 | 2.3246E-67 | MO_MT+ | CLEC2B |
| 1.6711E-71 | -1.134452 | 0.004 | 0.167 | 3.9476E-67 | MO_MT+ | MUC21 |
| 1.7476E-71 | 0.31125292 | 0.377 | 0.196 | 4.1283E-67 | MO_MT+ | LSP1 |
| 3.8358E-71 | 0.35518317 | 0.416 | 0.232 | 9.0614E-67 | MO_MT+ | FKBP4 |
| 8.4346E-71 | 0.32625998 | 0.45 | 0.257 | 1.9925E-66 | MO_MT+ | BACH1 |
| 1.7088E-70 | 0.25173922 | 0.254 | 0.112 | 4.0367E-66 | MO_MT+ | CD86 |
| 2.1323E-70 | 0.26402692 | 0.36 | 0.186 | 5.0371E-66 | MO_MT+ | ACSL4 |
| 2.4257E-70 | -0.5547942 | 0.063 | 0.249 | 5.7303E-66 | MO_MT+ | PRSS8 |
| 4.198E-70 | -0.5419138 | 0.331 | 0.521 | 9.9169E-66 | MO_MT+ | ANXA11 |
| 9.3037E-70 | 0.33379424 | 0.431 | 0.247 | 2.1978E-65 | MO_MT+ | MGAT1 |
| 1.4034E-69 | 0.2927818 | 0.49 | 0.294 | 3.3153E-65 | MO_MT+ | H2AFV |

| | | | | | | |
|---|---|---|---|---|---|---|
| 1.5949E-69 | 0.36822875 | 0.475 | 0.287 | 3.7677E-65 | MO_MT+ | CD99 |
| 2.0495E-69 | 0.30091422 | 0.334 | 0.172 | 4.8415E-65 | MO_MT+ | GPI |
| 4.5939E-69 | -0.5308903 | 0.825 | 0.909 | 1.0852E-64 | MO_MT+ | ITM2B |
| 6.9212E-69 | 0.25322779 | 0.363 | 0.188 | 1.635E-64 | MO_MT+ | RAB8B |
| 2.8509E-68 | -0.8376034 | 0.017 | 0.18 | 6.7347E-64 | MO_MT+ | NDRG2 |
| 3.3014E-68 | 0.32940726 | 0.502 | 0.293 | 7.7988E-64 | MO_MT+ | SERPINA1 |
| 4.9395E-68 | 0.33406275 | 0.307 | 0.154 | 1.1669E-63 | MO_MT+ | CPD |
| 5.6588E-68 | -1.4127958 | 0.034 | 0.203 | 1.3368E-63 | MO_MT+ | HOPX |
| 8.1578E-68 | 0.30463195 | 0.347 | 0.179 | 1.9271E-63 | MO_MT+ | ACSL1 |
| 3.2941E-67 | -1.0214578 | 0.008 | 0.166 | 7.7816E-63 | MO_MT+ | CEACAM5 |
| 6.7859E-67 | -0.8610971 | 0.029 | 0.194 | 1.603E-62 | MO_MT+ | TMPRSS2 |
| 8.7077E-67 | 0.59894826 | 0.768 | 0.642 | 2.057E-62 | MO_MT+ | LDHA |
| 1.4518E-66 | -1.569687 | 0.011 | 0.169 | 3.4295E-62 | MO_MT+ | ECM1 |
| 1.9323E-66 | -0.986034 | 0.008 | 0.164 | 4.5646E-62 | MO_MT+ | PRSS27 |
| 5.743E-66 | 0.29545292 | 0.46 | 0.269 | 1.3567E-61 | MO_MT+ | HLA-DRB1 |
| 6.1569E-66 | -0.7706513 | 0.003 | 0.154 | 1.4545E-61 | MO_MT+ | TMPRSS11D |
| 6.4831E-66 | 0.2505626 | 0.254 | 0.116 | 1.5315E-61 | MO_MT+ | LPCAT1 |
| 1.3622E-65 | -0.6163379 | 0.031 | 0.195 | 3.2179E-61 | MO_MT+ | EPS8L1 |
| 1.4635E-65 | -0.9383812 | 0.034 | 0.205 | 3.4573E-61 | MO_MT+ | KRT17 |
| 1.7649E-65 | 0.48006374 | 0.506 | 0.315 | 4.1692E-61 | MO_MT+ | VEGFA |
| 2.8992E-65 | -0.9747913 | 0.008 | 0.161 | 6.8489E-61 | MO_MT+ | PITX1 |
| 3.279E-65 | 0.27157545 | 0.358 | 0.19 | 7.7459E-61 | MO_MT+ | MYO9B |
| 4.0022E-65 | 1.04916734 | 0.229 | 0.103 | 9.4544E-61 | MO_MT+ | CCL4 |
| 6.19E-65 | -0.7568134 | 0.007 | 0.159 | 1.4623E-60 | MO_MT+ | SCEL |
| 1.8374E-64 | 0.29618758 | 0.457 | 0.27 | 4.3405E-60 | MO_MT+ | PLEKHB2 |
| 2.035E-64 | 0.25847555 | 0.745 | 0.53 | 4.8072E-60 | MO_MT+ | UPP1 |
| 2.1475E-64 | -0.8313163 | 0.003 | 0.152 | 5.073E-60 | MO_MT+ | CEACAM7 |
| 2.4045E-64 | -0.4669836 | 0.455 | 0.629 | 5.6801E-60 | MO_MT+ | TMEM59 |
| 3.3458E-64 | 0.2750067 | 0.34 | 0.177 | 7.9039E-60 | MO_MT+ | DAB2 |
| 7.0921E-64 | -0.6479616 | 0.004 | 0.152 | 1.6754E-59 | MO_MT+ | NCCRP1 |
| 8.9995E-64 | -0.7166756 | 0.087 | 0.271 | 2.1259E-59 | MO_MT+ | TSPAN1 |
| 1.056E-63 | 0.30615069 | 0.478 | 0.266 | 2.4947E-59 | MO_MT+ | AIF1 |
| 1.9557E-63 | -0.5570772 | 0.206 | 0.393 | 4.62E-59 | MO_MT+ | C9orf16 |
| 6.001E-63 | -0.592998 | 0.019 | 0.174 | 1.4176E-58 | MO_MT+ | FXYD3 |
| 1.3197E-62 | -0.850238 | 0.012 | 0.161 | 3.1176E-58 | MO_MT+ | KRT15 |
| 2.0283E-62 | 0.2582413 | 0.411 | 0.232 | 4.7915E-58 | MO_MT+ | SP100 |
| 2.0868E-62 | 0.25240019 | 0.462 | 0.268 | 4.9297E-58 | MO_MT+ | VASP |
| 2.2447E-62 | -0.6143194 | 0.945 | 0.945 | 5.3026E-58 | MO_MT+ | MT-CO2 |
| 5.5823E-62 | 0.35889318 | 0.345 | 0.187 | 1.3187E-57 | MO_MT+ | SNX10 |
| 1.3757E-61 | 0.25794846 | 0.313 | 0.162 | 3.2499E-57 | MO_MT+ | NUP58 |
| 1.4466E-61 | -1.3102972 | 0.096 | 0.273 | 3.4174E-57 | MO_MT+ | S100A2 |
| 2.2546E-61 | -0.7306716 | 0.745 | 0.826 | 5.3261E-57 | MO_MT+ | MT-ND5 |
| 2.7803E-61 | 0.35006125 | 0.189 | 0.078 | 6.5679E-57 | MO_MT+ | SNAPC1 |
| 8.6748E-61 | -0.4208273 | 0.077 | 0.248 | 2.0493E-56 | MO_MT+ | GIPC1 |
| 1.1895E-60 | 0.30841791 | 0.46 | 0.275 | 2.8099E-56 | MO_MT+ | CTSZ |
| 1.3838E-60 | 0.28200132 | 0.492 | 0.301 | 3.269E-56 | MO_MT+ | QKI |
| 1.5962E-60 | -0.7711667 | 0.007 | 0.15 | 3.7706E-56 | MO_MT+ | TMPRSS11E |
| 1.7826E-60 | 0.29424842 | 0.409 | 0.237 | 4.211E-56 | MO_MT+ | CD82 |
| 3.2057E-60 | 0.28751019 | 0.586 | 0.376 | 7.5729E-56 | MO_MT+ | TYMP |
| 3.6526E-60 | -0.9259571 | 0.001 | 0.14 | 8.6284E-56 | MO_MT+ | TMPRSS11B |

| | | | | | | |
|---|---|---|---|---|---|---|
| 5.1895E-60 | 0.38548043 | 0.817 | 0.704 | 1.2259E-55 | MO_MT+ | GPX4 |
| 7.3029E-60 | -0.5363085 | 0.028 | 0.181 | 1.7252E-55 | MO_MT+ | SPNS2 |
| 8.018E-60 | 0.47181006 | 0.14 | 0.05 | 1.8941E-55 | MO_MT+ | INHBA |
| 1.1245E-59 | -0.4928167 | 0.029 | 0.181 | 2.6563E-55 | MO_MT+ | C6orf132 |
| 1.851E-59 | 0.76546401 | 0.238 | 0.112 | 4.3725E-55 | MO_MT+ | CXCL2 |
| 7.4267E-59 | -0.398875 | 0.14 | 0.327 | 1.7544E-54 | MO_MT+ | PERP |
| 1.1563E-58 | -0.56331 | 0.029 | 0.178 | 2.7316E-54 | MO_MT+ | PPL |
| 1.9841E-58 | 0.25180668 | 0.319 | 0.168 | 4.687E-54 | MO_MT+ | TCIRG1 |
| 2.2737E-58 | 0.65258891 | 0.924 | 0.881 | 5.3712E-54 | MO_MT+ | CSTB |
| 2.4995E-58 | -0.393265 | 0.101 | 0.281 | 5.9046E-54 | MO_MT+ | JUP |
| 3.4374E-58 | 0.27634504 | 0.257 | 0.125 | 8.1201E-54 | MO_MT+ | IL1R1 |
| 3.6382E-58 | 0.26873181 | 0.244 | 0.118 | 8.5944E-54 | MO_MT+ | SOAT1 |
| 4.8231E-58 | 0.34509285 | 0.537 | 0.35 | 1.1394E-53 | MO_MT+ | MIF |
| 6.341E-58 | -0.8248794 | 0.011 | 0.152 | 1.4979E-53 | MO_MT+ | CEACAM6 |
| 6.3791E-58 | -0.4426285 | 0.087 | 0.254 | 1.5069E-53 | MO_MT+ | SERINC2 |
| 6.6425E-58 | 0.52156348 | 0.786 | 0.735 | 1.5692E-53 | MO_MT+ | CTSB |
| 1.0227E-57 | 0.57631579 | 0.676 | 0.576 | 2.4159E-53 | MO_MT+ | PGK1 |
| 1.4191E-57 | 0.36972601 | 0.328 | 0.177 | 3.3522E-53 | MO_MT+ | INSIG1 |
| 1.5383E-57 | -0.5535703 | 0.029 | 0.177 | 3.6339E-53 | MO_MT+ | HSPB8 |
| 1.8615E-57 | -0.3421244 | 0.032 | 0.182 | 4.3975E-53 | MO_MT+ | SMIM22 |
| 2.3924E-57 | -0.3662005 | 0.019 | 0.162 | 5.6516E-53 | MO_MT+ | TSPAN6 |
| 2.6168E-57 | -0.4205425 | 0.017 | 0.159 | 6.1817E-53 | MO_MT+ | TMPRSS4 |
| 1.025E-56 | -1.5980586 | 0.004 | 0.138 | 2.4213E-52 | MO_MT+ | LYPD2 |
| 1.2419E-56 | -0.5988755 | 0.018 | 0.161 | 2.9337E-52 | MO_MT+ | MUC4 |
| 1.3111E-56 | 0.31102335 | 0.467 | 0.294 | 3.0973E-52 | MO_MT+ | PGAM1 |
| 1.5104E-56 | 0.31631925 | 0.863 | 0.767 | 3.5679E-52 | MO_MT+ | RHOA |
| 5.3406E-56 | -0.39759 | 0.014 | 0.151 | 1.2616E-51 | MO_MT+ | VSIG2 |
| 6.4357E-56 | 0.27890147 | 0.616 | 0.42 | 1.5203E-51 | MO_MT+ | IQGAP1 |
| 7.3306E-56 | -0.6383461 | 0.004 | 0.136 | 1.7317E-51 | MO_MT+ | GABRP |
| 1.2642E-55 | 0.26935269 | 0.565 | 0.389 | 2.9865E-51 | MO_MT+ | HLA-DRA |
| 1.334E-55 | 0.27529807 | 0.519 | 0.331 | 3.1513E-51 | MO_MT+ | GRB2 |
| 1.5721E-55 | -0.6126701 | 0.168 | 0.352 | 3.7139E-51 | MO_MT+ | GDF15 |
| 2.5468E-55 | -0.6411022 | 0.936 | 0.937 | 6.0164E-51 | MO_MT+ | MT-ATP6 |
| 2.72E-55 | -0.5810244 | 0.032 | 0.177 | 6.4255E-51 | MO_MT+ | STEAP4 |
| 8.9216E-55 | 0.48090869 | 0.799 | 0.739 | 2.1075E-50 | MO_MT+ | APLP2 |
| 9.2652E-55 | -0.6875812 | 0.046 | 0.194 | 2.1887E-50 | MO_MT+ | ANKRD36C |
| 1.0431E-54 | -0.4530648 | 0.733 | 0.875 | 2.4641E-50 | MO_MT+ | RPLP0 |
| 1.311E-54 | -0.4132999 | 0.064 | 0.22 | 3.0969E-50 | MO_MT+ | EHF |
| 1.9076E-54 | -0.430839 | 0.09 | 0.256 | 4.5062E-50 | MO_MT+ | KLF5 |
| 2.1267E-54 | -0.53581 | 0.111 | 0.282 | 5.0239E-50 | MO_MT+ | CAMK2N1 |
| 2.3076E-54 | -0.5787134 | 0.063 | 0.214 | 5.4513E-50 | MO_MT+ | CDKN2B |
| 3.4055E-54 | -0.3792403 | 0.008 | 0.14 | 8.0449E-50 | MO_MT+ | BCAS1 |
| 4.2082E-54 | -0.4384022 | 0.419 | 0.582 | 9.9411E-50 | MO_MT+ | PRDX5 |
| 8.3165E-54 | -0.9029121 | 0.127 | 0.285 | 1.9646E-49 | MO_MT+ | DUSP5 |
| 9.6999E-54 | -1.6399132 | 0.015 | 0.149 | 2.2914E-49 | MO_MT+ | SPRR2A |
| 1.0062E-53 | -0.9815259 | 0.037 | 0.179 | 2.377E-49 | MO_MT+ | RDH10 |
| 2.0375E-53 | -1.0231171 | 0.014 | 0.145 | 4.8132E-49 | MO_MT+ | CXCL6 |
| 3.5031E-53 | 0.33248811 | 0.416 | 0.255 | 8.2753E-49 | MO_MT+ | WTAP |
| 4.2696E-53 | -0.4824087 | 0.043 | 0.189 | 1.0086E-48 | MO_MT+ | SDCBP2 |
| 4.4E-53 | 0.29589125 | 0.375 | 0.217 | 1.0394E-48 | MO_MT+ | STXBP2 |

| | | | | | | |
|---|---|---|---|---|---|---|
| 6.143E-53 | 0.30252168 | 0.46 | 0.291 | 1.4512E-48 | MO_MT+ | ARL6IP5 |
| 9.1718E-53 | -1.2284975 | 0.009 | 0.137 | 2.1667E-48 | MO_MT+ | PLAT |
| 2.3212E-52 | -0.4293993 | 0.027 | 0.164 | 5.4833E-48 | MO_MT+ | MALL |
| 2.9183E-52 | 0.30708632 | 0.454 | 0.289 | 6.8939E-48 | MO_MT+ | IFNGR2 |
| 7.3004E-52 | -0.3640665 | 0.425 | 0.606 | 1.7246E-47 | MO_MT+ | VAMP8 |
| 2.0069E-51 | -0.9071381 | 0.011 | 0.137 | 4.741E-47 | MO_MT+ | RHCG |
| 4.459E-51 | -0.4067193 | 0.014 | 0.141 | 1.0533E-46 | MO_MT+ | GCNT3 |
| 7.9764E-51 | 0.29367351 | 0.442 | 0.282 | 1.8843E-46 | MO_MT+ | RALA |
| 8.2123E-51 | 0.35810028 | 0.946 | 0.89 | 1.94E-46 | MO_MT+ | ATP5F1E |
| 1.891E-50 | 0.37738077 | 0.711 | 0.593 | 4.4671E-46 | MO_MT+ | NPC2 |
| 2.4822E-50 | 0.59292458 | 0.265 | 0.139 | 5.8638E-46 | MO_MT+ | CXCL3 |
| 4.2088E-50 | 0.3507954 | 0.1 | 0.033 | 9.9425E-46 | MO_MT+ | CCL7 |
| 5.227E-50 | 0.28946178 | 0.319 | 0.18 | 1.2348E-45 | MO_MT+ | CKLF |
| 5.2566E-50 | -1.1328939 | 0.003 | 0.121 | 1.2418E-45 | MO_MT+ | SPINK5 |
| 6.5037E-50 | -0.4109235 | 0.115 | 0.272 | 1.5364E-45 | MO_MT+ | AFDN |
| 1.0347E-49 | -0.5800997 | 0.057 | 0.197 | 2.4444E-45 | MO_MT+ | FBXO32 |
| 1.4814E-49 | 0.53730406 | 0.198 | 0.091 | 3.4994E-45 | MO_MT+ | PTGS2 |
| 1.8319E-49 | -0.3699525 | 0.03 | 0.163 | 4.3275E-45 | MO_MT+ | TTC9 |
| 1.8938E-49 | 0.2728225 | 0.343 | 0.195 | 4.4737E-45 | MO_MT+ | NFKB1 |
| 1.9882E-49 | 0.25583911 | 0.428 | 0.267 | 4.6967E-45 | MO_MT+ | ARL8B |
| 2.9901E-49 | 0.29891556 | 0.753 | 0.635 | 7.0636E-45 | MO_MT+ | ARPC5 |
| 3.0528E-49 | 0.26912277 | 0.348 | 0.198 | 7.2115E-45 | MO_MT+ | MAP3K8 |
| 3.1245E-49 | -0.6072623 | 0.001 | 0.117 | 7.381E-45 | MO_MT+ | KRT78 |
| 4.7908E-49 | 0.30472815 | 0.618 | 0.458 | 1.1317E-44 | MO_MT+ | LYZ |
| 8.9047E-49 | -0.4674652 | 0.002 | 0.117 | 2.1036E-44 | MO_MT+ | TMPRSS11A |
| 9.565E-49 | -0.5720618 | 0.956 | 0.95 | 2.2595E-44 | MO_MT+ | MT-CO1 |
| 1.086E-48 | -0.6416896 | 0.991 | 0.98 | 2.5654E-44 | MO_MT+ | MALAT1 |
| 1.1854E-48 | -0.4720013 | 0.899 | 0.96 | 2.8004E-44 | MO_MT+ | RPS18 |
| 1.525E-48 | -0.5580029 | 0.02 | 0.146 | 3.6024E-44 | MO_MT+ | SNCG |
| 3.1326E-48 | -0.2875295 | 0.047 | 0.185 | 7.4002E-44 | MO_MT+ | FNBP1L |
| 1.139E-47 | -1.283247 | 0.004 | 0.117 | 2.6906E-43 | MO_MT+ | SPRR1B |
| 1.706E-47 | -0.7124504 | 0.116 | 0.268 | 4.0301E-43 | MO_MT+ | AREG |
| 2.825E-47 | 0.30357422 | 0.229 | 0.115 | 6.6736E-43 | MO_MT+ | ATP2B1-AS1 |
| 3.4048E-47 | -0.5978204 | 0.942 | 0.944 | 8.0432E-43 | MO_MT+ | MT-CO3 |
| 1.1818E-46 | 0.25481247 | 0.559 | 0.378 | 2.7918E-42 | MO_MT+ | DEK |
| 2.0249E-46 | -0.3798557 | 0.076 | 0.22 | 4.7833E-42 | MO_MT+ | CKB |
| 2.4597E-46 | -0.3684368 | 0.07 | 0.211 | 5.8105E-42 | MO_MT+ | PARD6B |
| 3.7953E-46 | -0.4051982 | 0.117 | 0.265 | 8.9657E-42 | MO_MT+ | WASL |
| 4.0643E-46 | -0.4611434 | 0.455 | 0.612 | 9.6012E-42 | MO_MT+ | EZR |
| 6.3704E-46 | -0.5368212 | 0.02 | 0.141 | 1.5049E-41 | MO_MT+ | CXCL17 |
| 7.361E-46 | -0.577934 | 0.025 | 0.147 | 1.7389E-41 | MO_MT+ | CDKN2A |
| 7.8968E-46 | -0.2903293 | 0.005 | 0.115 | 1.8655E-41 | MO_MT+ | FUT3 |
| 9.2008E-46 | 0.31571932 | 0.47 | 0.313 | 2.1735E-41 | MO_MT+ | SLC16A3 |
| 9.6954E-46 | 0.3091968 | 0.296 | 0.168 | 2.2903E-41 | MO_MT+ | FAM162A |
| 1.8946E-45 | -0.5260269 | 0.034 | 0.159 | 4.4756E-41 | MO_MT+ | CDC42EP5 |
| 1.943E-45 | -0.5773002 | 0.055 | 0.189 | 4.5899E-41 | MO_MT+ | AQP3 |
| 2.976E-45 | -0.3597253 | 0.037 | 0.164 | 7.0301E-41 | MO_MT+ | SDC1 |
| 3.3131E-45 | -0.3200829 | 0.09 | 0.237 | 7.8265E-41 | MO_MT+ | TJP1 |
| 4.6517E-45 | -0.2868728 | 0.077 | 0.22 | 1.0989E-40 | MO_MT+ | CNN3 |
| 7.4584E-45 | -0.2772181 | 0.035 | 0.16 | 1.7619E-40 | MO_MT+ | ERBB3 |

| | | | | | | |
|---|---|---|---|---|---|---|
| 1.2479E-44 | -0.2757671 | 0.041 | 0.168 | 2.9479E-40 | MO_MT+ | NDFIP2 |
| 1.2624E-44 | -0.3137731 | 0.006 | 0.115 | 2.9823E-40 | MO_MT+ | STS |
| 1.3462E-44 | -1.561708 | 0.004 | 0.111 | 3.1801E-40 | MO_MT+ | CRCT1 |
| 2.8625E-44 | -0.3955641 | 0.124 | 0.28 | 6.7622E-40 | MO_MT+ | MAL2 |
| 2.9007E-44 | -0.3877904 | 0.014 | 0.127 | 6.8523E-40 | MO_MT+ | TMEM40 |
| 4.1379E-44 | -0.3285497 | 0.004 | 0.11 | 9.7749E-40 | MO_MT+ | A2ML1 |
| 6.0743E-44 | 0.26616929 | 0.224 | 0.115 | 1.4349E-39 | MO_MT+ | ATF5 |
| 9.9787E-44 | -1.4468238 | 0.005 | 0.112 | 2.3573E-39 | MO_MT+ | CNFN |
| 3.2109E-43 | -0.5231914 | 0.099 | 0.24 | 7.5852E-39 | MO_MT+ | CDKN1C |
| 5.0591E-43 | -0.4839202 | 0.091 | 0.235 | 1.1951E-38 | MO_MT+ | ITGA2 |
| 5.3864E-43 | 0.33884907 | 0.838 | 0.761 | 1.2724E-38 | MO_MT+ | BTG1 |
| 5.6259E-43 | -0.3908462 | 0.342 | 0.502 | 1.329E-38 | MO_MT+ | CD9 |
| 7.7235E-43 | -0.6216065 | 0.022 | 0.135 | 1.8245E-38 | MO_MT+ | DHRS9 |
| 7.7824E-43 | 0.32661274 | 0.545 | 0.391 | 1.8384E-38 | MO_MT+ | H2AFY |
| 1.1556E-42 | -0.3805223 | 0.032 | 0.151 | 2.73E-38 | MO_MT+ | ALDH1A3 |
| 1.3038E-42 | -0.3681091 | 0.065 | 0.195 | 3.08E-38 | MO_MT+ | ABLIM1 |
| 1.9018E-42 | -0.3640654 | 0.787 | 0.879 | 4.4926E-38 | MO_MT+ | MYL12A |
| 2.4981E-42 | 0.26499411 | 0.35 | 0.209 | 5.9013E-38 | MO_MT+ | KDM6B |
| 2.7498E-42 | -0.2900554 | 0.047 | 0.173 | 6.4958E-38 | MO_MT+ | NECTIN4 |
| 4.5178E-42 | -0.3068416 | 0.02 | 0.132 | 1.0672E-37 | MO_MT+ | FAM84A |
| 2.5699E-41 | -0.417071 | 0.058 | 0.183 | 6.0709E-37 | MO_MT+ | EPHA2 |
| 2.5081E-40 | -0.279952 | 0.062 | 0.189 | 5.925E-36 | MO_MT+ | EPS8L2 |
| 3.0025E-40 | -0.3600352 | 0.113 | 0.253 | 7.0928E-36 | MO_MT+ | SPTBN1 |
| 3.3919E-40 | -0.3447471 | 0.014 | 0.118 | 8.0126E-36 | MO_MT+ | CEACAM1 |
| 4.487E-40 | -0.5791752 | 0.127 | 0.258 | 1.06E-35 | MO_MT+ | CARHSP1 |
| 5.4944E-40 | -0.3977055 | 0.009 | 0.111 | 1.2979E-35 | MO_MT+ | TMC5 |
| 1.0289E-39 | -0.491824 | 0.005 | 0.102 | 2.4305E-35 | MO_MT+ | DHRS2 |
| 1.8452E-39 | -0.3934312 | 0.912 | 0.947 | 4.3589E-35 | MO_MT+ | RPL13A |
| 3.3112E-39 | 0.47946852 | 0.709 | 0.597 | 7.822E-35 | MO_MT+ | NFKBIA |
| 4.3821E-39 | -0.2530129 | 0.073 | 0.2 | 1.0352E-34 | MO_MT+ | RBPMS |
| 5.6208E-39 | -0.3643889 | 0.011 | 0.111 | 1.3278E-34 | MO_MT+ | KRT23 |
| 5.8827E-39 | -0.4010271 | 0.277 | 0.423 | 1.3897E-34 | MO_MT+ | JPT1 |
| 7.0206E-39 | -0.4685017 | 0.048 | 0.163 | 1.6585E-34 | MO_MT+ | BPGM |
| 8.071E-39 | -0.3201636 | 0.555 | 0.693 | 1.9066E-34 | MO_MT+ | CAST |
| 2.6855E-38 | -0.2611588 | 0.032 | 0.142 | 6.3441E-34 | MO_MT+ | MTUS1 |
| 4.1385E-38 | -0.3255937 | 0.228 | 0.372 | 9.7763E-34 | MO_MT+ | AES |
| 8.786E-38 | -0.4026978 | 0.069 | 0.191 | 2.0755E-33 | MO_MT+ | TNFSF10 |
| 1.1575E-37 | -0.4604866 | 0.121 | 0.256 | 2.7344E-33 | MO_MT+ | SYNE2 |
| 1.3766E-37 | -0.4358822 | 0.06 | 0.18 | 3.252E-33 | MO_MT+ | IGFBP3 |
| 1.5052E-37 | 0.27600909 | 0.381 | 0.248 | 3.5558E-33 | MO_MT+ | C1orf122 |
| 1.5518E-37 | -0.3142446 | 0.006 | 0.1 | 3.6657E-33 | MO_MT+ | KLK10 |
| 2.3759E-37 | -0.2853666 | 0.035 | 0.144 | 5.6126E-33 | MO_MT+ | EMP2 |
| 3.4004E-37 | -0.9251869 | 0.509 | 0.643 | 8.0328E-33 | MO_MT+ | MTRNR2L12 |
| 3.67E-37 | -0.4344762 | 0.577 | 0.704 | 8.6697E-33 | MO_MT+ | JUNB |
| 4.089E-37 | 0.78304445 | 0.553 | 0.449 | 9.6595E-33 | MO_MT+ | SPP1 |
| 4.4885E-37 | -0.4860151 | 0.077 | 0.202 | 1.0603E-32 | MO_MT+ | TM4SF1 |
| 4.5656E-37 | -0.3022455 | 0.024 | 0.126 | 1.0785E-32 | MO_MT+ | KRT80 |
| 7.6799E-37 | -0.3229678 | 0.134 | 0.266 | 1.8142E-32 | MO_MT+ | PRDX2 |
| 8.4548E-37 | 0.27648543 | 0.828 | 0.773 | 1.9973E-32 | MO_MT+ | TPI1 |
| 9.5282E-37 | -0.2929023 | 0.029 | 0.134 | 2.2508E-32 | MO_MT+ | FOXC1 |

| | | | | | | |
|---|---|---|---|---|---|---|
| 1.4885E-36 | 0.25304021 | 0.579 | 0.421 | 3.5164E-32 | MO_MT+ | SEC62 |
| 1.595E-36 | -0.3967183 | 0.027 | 0.129 | 3.768E-32 | MO_MT+ | SRD5A3 |
| 2.6005E-36 | 0.26976259 | 0.694 | 0.579 | 6.1432E-32 | MO_MT+ | NOP10 |
| 3.3635E-36 | 0.42498884 | 0.763 | 0.688 | 7.9456E-32 | MO_MT+ | PNRC1 |
| 3.4392E-36 | -0.7644717 | 0.014 | 0.108 | 8.1245E-32 | MO_MT+ | ZG16B |
| 3.4419E-36 | -0.3451622 | 0.087 | 0.213 | 8.1307E-32 | MO_MT+ | MACC1 |
| 3.516E-36 | -0.3140359 | 0.04 | 0.148 | 8.3058E-32 | MO_MT+ | ITGB6 |
| 4.0552E-36 | -0.3996315 | 0.114 | 0.239 | 9.5797E-32 | MO_MT+ | TOB1 |
| 4.6861E-36 | -0.2676203 | 0.066 | 0.184 | 1.107E-31 | MO_MT+ | TEAD1 |
| 1.0514E-35 | -0.6406887 | 0.12 | 0.254 | 2.4838E-31 | MO_MT+ | CRYAB |
| 1.995E-35 | -0.5285407 | 0.124 | 0.251 | 4.7127E-31 | MO_MT+ | CST6 |
| 3.672E-35 | -0.3153434 | 0.161 | 0.291 | 8.6743E-31 | MO_MT+ | LRP10 |
| 2.8159E-34 | 0.45331516 | 0.289 | 0.177 | 6.652E-30 | MO_MT+ | FBP1 |
| 4.9988E-34 | -0.3375851 | 0.067 | 0.181 | 1.1809E-29 | MO_MT+ | FHL2 |
| 7.4403E-34 | -0.5817833 | 0.071 | 0.184 | 1.7576E-29 | MO_MT+ | F3 |
| 7.7896E-34 | -0.3533955 | 0.15 | 0.275 | 1.8401E-29 | MO_MT+ | FAM3C |
| 1.2208E-33 | -0.2723197 | 0.535 | 0.679 | 2.8838E-29 | MO_MT+ | CALR |
| 1.667E-33 | -0.3042881 | 0.222 | 0.354 | 3.9379E-29 | MO_MT+ | PEBP1 |
| 1.8268E-33 | -0.3157444 | 0.195 | 0.322 | 4.3154E-29 | MO_MT+ | RAB11A |
| 1.8578E-33 | -0.3880274 | 0.373 | 0.498 | 4.3888E-29 | MO_MT+ | H2AFJ |
| 3.8184E-33 | 0.33494234 | 0.533 | 0.411 | 9.0203E-29 | MO_MT+ | GRN |
| 3.8419E-33 | -0.3923743 | 0.041 | 0.143 | 9.0757E-29 | MO_MT+ | PLAC8 |
| 4.0974E-33 | -0.4458324 | 0.842 | 0.877 | 9.6793E-29 | MO_MT+ | RPL3 |
| 5.8151E-33 | -0.2550082 | 0.013 | 0.101 | 1.3737E-28 | MO_MT+ | RNF39 |
| 6.9391E-33 | -0.4257372 | 0.894 | 0.906 | 1.6392E-28 | MO_MT+ | RPL7A |
| 7.3913E-33 | -0.3031691 | 0.133 | 0.257 | 1.7461E-28 | MO_MT+ | CD2AP |
| 1.5037E-32 | -0.3959567 | 0.162 | 0.284 | 3.5523E-28 | MO_MT+ | ABHD5 |
| 2.7881E-32 | -0.7618624 | 0.075 | 0.185 | 6.5863E-28 | MO_MT+ | SPINK1 |
| 4.0725E-32 | -0.4115206 | 0.643 | 0.76 | 9.6205E-28 | MO_MT+ | RPSA |
| 5.1567E-32 | -0.3685304 | 0.231 | 0.354 | 1.2182E-27 | MO_MT+ | UBE2H |
| 6.2132E-32 | -0.401905 | 0.552 | 0.633 | 1.4678E-27 | MO_MT+ | TAX1BP1 |
| 1.1261E-31 | -0.3753332 | 0.031 | 0.126 | 2.6602E-27 | MO_MT+ | AGR2 |
| 1.3779E-31 | -0.3129766 | 0.024 | 0.114 | 3.255E-27 | MO_MT+ | SMIM5 |
| 1.8386E-31 | 0.28047694 | 0.751 | 0.678 | 4.3433E-27 | MO_MT+ | NAMPT |
| 2.1566E-31 | -0.2726035 | 0.102 | 0.221 | 5.0946E-27 | MO_MT+ | DSP |
| 2.3896E-31 | -0.3255292 | 0.406 | 0.534 | 5.645E-27 | MO_MT+ | EIF4A2 |
| 2.469E-31 | -0.2641787 | 0.092 | 0.205 | 5.8325E-27 | MO_MT+ | ZNF91 |
| 3.7021E-31 | -0.3490673 | 0.255 | 0.382 | 8.7454E-27 | MO_MT+ | GCC2 |
| 3.8264E-31 | -0.3086464 | 0.063 | 0.166 | 9.039E-27 | MO_MT+ | SYTL2 |
| 6.5597E-31 | -0.476668 | 0.075 | 0.186 | 1.5496E-26 | MO_MT+ | LAMC2 |
| 1.4121E-30 | 0.26816547 | 0.932 | 0.89 | 3.3357E-26 | MO_MT+ | CD63 |
| 2.0757E-30 | -0.4284387 | 0.123 | 0.233 | 4.9035E-26 | MO_MT+ | CYB5A |
| 3.4932E-30 | -0.310692 | 0.379 | 0.501 | 8.2519E-26 | MO_MT+ | VDAC2 |
| 6.2226E-30 | -0.4366246 | 0.23 | 0.351 | 1.47E-25 | MO_MT+ | ZFP36L2 |
| 1.0783E-29 | -0.3195903 | 0.072 | 0.174 | 2.5472E-25 | MO_MT+ | FAM214A |
| 2.6653E-29 | -0.2632779 | 0.121 | 0.241 | 6.2961E-25 | MO_MT+ | S100A16 |
| 3.1218E-29 | -0.3591869 | 0.248 | 0.363 | 7.3745E-25 | MO_MT+ | RRBP1 |
| 3.1799E-29 | -0.2637504 | 0.021 | 0.104 | 7.5119E-25 | MO_MT+ | LRMP |
| 3.4833E-29 | -0.3748303 | 0.392 | 0.496 | 8.2286E-25 | MO_MT+ | SELENOW |
| 3.7597E-29 | -0.4073353 | 0.096 | 0.199 | 8.8815E-25 | MO_MT+ | SAMD9 |

| | | | | | | |
|---|---|---|---|---|---|---|
| 5.0877E-29 | -0.5246467 | 0.891 | 0.887 | 1.2019E-24 | MO_MT+ | RPS6 |
| 6.6064E-29 | -0.3128731 | 0.141 | 0.252 | 1.5606E-24 | MO_MT+ | PKN2 |
| 7.1803E-29 | -0.3247726 | 0.035 | 0.123 | 1.6962E-24 | MO_MT+ | EGLN3 |
| 1.1311E-28 | 0.25016086 | 0.763 | 0.683 | 2.6719E-24 | MO_MT+ | ATP6V0B |
| 1.8128E-28 | -0.3488797 | 0.081 | 0.184 | 4.2824E-24 | MO_MT+ | DST |
| 2.381E-28 | -0.3542012 | 0.69 | 0.785 | 5.6246E-24 | MO_MT+ | RPL10A |
| 2.6655E-28 | -0.3331507 | 0.024 | 0.106 | 6.2968E-24 | MO_MT+ | CLIC3 |
| 2.8451E-28 | -0.3420823 | 0.104 | 0.208 | 6.7209E-24 | MO_MT+ | CSNK1E |
| 4.8836E-28 | -0.2567092 | 0.203 | 0.325 | 1.1537E-23 | MO_MT+ | ERP29 |
| 6.3992E-28 | -0.2881156 | 0.041 | 0.129 | 1.5117E-23 | MO_MT+ | POLB |
| 6.4795E-28 | -0.296106 | 0.33 | 0.452 | 1.5306E-23 | MO_MT+ | ACTN4 |
| 1.9387E-27 | -0.2913988 | 0.093 | 0.2 | 4.5798E-23 | MO_MT+ | SFN |
| 2.1287E-27 | -0.4046453 | 0.416 | 0.532 | 5.0285E-23 | MO_MT+ | ZFP36 |
| 2.8439E-27 | -0.3240082 | 0.293 | 0.414 | 6.7181E-23 | MO_MT+ | CD59 |
| 3.3582E-27 | -0.2533261 | 0.063 | 0.158 | 7.933E-23 | MO_MT+ | ITPKC |
| 1.0386E-26 | -0.2948799 | 0.211 | 0.33 | 2.4535E-22 | MO_MT+ | PPA1 |
| 1.9582E-26 | -0.2715827 | 0.155 | 0.261 | 4.626E-22 | MO_MT+ | LYPLA1 |
| 3.7088E-26 | -0.408874 | 0.892 | 0.909 | 8.7612E-22 | MO_MT+ | MT-ND2 |
| 3.9808E-26 | -0.3058921 | 0.044 | 0.129 | 9.404E-22 | MO_MT+ | ID4 |
| 6.8903E-26 | -0.3853034 | 0.096 | 0.195 | 1.6277E-21 | MO_MT+ | PLK2 |
| 1.4885E-25 | -0.3059713 | 0.543 | 0.683 | 3.5162E-21 | MO_MT+ | IER2 |
| 3.7009E-25 | -0.3676962 | 0.081 | 0.174 | 8.7426E-21 | MO_MT+ | PRDM1 |
| 8.159E-25 | -0.2918249 | 0.809 | 0.871 | 1.9274E-20 | MO_MT+ | PABPC1 |
| 1.1093E-24 | -0.5392496 | 0.074 | 0.165 | 2.6206E-20 | MO_MT+ | APOBEC3A |
| 1.426E-24 | -0.3744761 | 0.041 | 0.123 | 3.3687E-20 | MO_MT+ | TCIM |
| 1.8796E-24 | -0.2920771 | 0.208 | 0.319 | 4.4402E-20 | MO_MT+ | AKAP9 |
| 3.133E-24 | -0.3415705 | 0.734 | 0.803 | 7.401E-20 | MO_MT+ | RPS5 |
| 4.8605E-24 | 0.29148307 | 0.172 | 0.097 | 1.1482E-19 | MO_MT+ | RETN |
| 4.9161E-24 | -0.3629678 | 0.268 | 0.37 | 1.1613E-19 | MO_MT+ | RALBP1 |
| 7.7302E-24 | -1.6541787 | 0.593 | 0.674 | 1.8261E-19 | MO_MT+ | S100A8 |
| 7.8736E-24 | 0.26262483 | 0.979 | 0.97 | 1.86E-19 | MO_MT+ | TMSB10 |
| 9.9404E-24 | -0.380862 | 0.937 | 0.932 | 2.3482E-19 | MO_MT+ | MT-ND4 |
| 2.5497E-23 | -0.2977491 | 0.304 | 0.399 | 6.0232E-19 | MO_MT+ | CTNNA1 |
| 2.6517E-23 | -0.2867437 | 0.395 | 0.491 | 6.2642E-19 | MO_MT+ | DYNLT1 |
| 4.5587E-23 | -0.3776967 | 0.176 | 0.273 | 1.0769E-18 | MO_MT+ | CCNG2 |
| 8.4832E-23 | -0.2635177 | 0.19 | 0.291 | 2.004E-18 | MO_MT+ | CLINT1 |
| 8.7993E-23 | -0.3159566 | 0.937 | 0.959 | 2.0787E-18 | MO_MT+ | RPL15 |
| 9.5884E-23 | -0.3309503 | 0.668 | 0.75 | 2.2651E-18 | MO_MT+ | EEF1B2 |
| 1.236E-22 | -0.5994609 | 0.108 | 0.198 | 2.9198E-18 | MO_MT+ | C12orf75 |
| 1.5894E-22 | -0.2534089 | 0.057 | 0.137 | 3.7547E-18 | MO_MT+ | ACADM |
| 1.7242E-22 | -0.2987268 | 0.136 | 0.229 | 4.0732E-18 | MO_MT+ | RAB9A |
| 1.9147E-22 | -0.2718413 | 0.189 | 0.288 | 4.523E-18 | MO_MT+ | PDCD6 |
| 3.5629E-22 | -0.7287471 | 0.245 | 0.34 | 8.4167E-18 | MO_MT+ | IGFBP7 |
| 5.0341E-22 | -0.4213012 | 0.325 | 0.438 | 1.1892E-17 | MO_MT+ | HIST1H1C |
| 5.5223E-22 | -0.3159012 | 0.846 | 0.872 | 1.3045E-17 | MO_MT+ | RPL14 |
| 6.7552E-22 | -0.2889124 | 0.124 | 0.219 | 1.5958E-17 | MO_MT+ | ARL4C |
| 9.1652E-22 | -0.3701684 | 0.141 | 0.239 | 2.1651E-17 | MO_MT+ | EPCAM |
| 9.8242E-22 | -0.3682567 | 0.084 | 0.167 | 2.3208E-17 | MO_MT+ | ANKRD37 |
| 1.3551E-21 | -0.2666461 | 0.929 | 0.95 | 3.2012E-17 | MO_MT+ | RPL18 |
| 1.5699E-21 | -0.4828208 | 0.154 | 0.25 | 3.7085E-17 | MO_MT+ | PLAU |

| | | | | | | |
|---|---|---|---|---|---|---|
| 2.7161E-21 | -0.3632251 | 0.167 | 0.262 | 6.4163E-17 | MO_MT+ | PMAIP1 |
| 3.1999E-21 | -2.1468288 | 0.681 | 0.729 | 7.5591E-17 | MO_MT+ | S100A9 |
| 1.5568E-20 | -0.3313467 | 0.22 | 0.323 | 3.6776E-16 | MO_MT+ | GADD45A |
| 2.8033E-20 | -3.0507641 | 0.467 | 0.545 | 6.6222E-16 | MO_MT+ | PLCG2 |
| 3.6102E-20 | -0.3201113 | 0.237 | 0.33 | 8.5283E-16 | MO_MT+ | MAFF |
| 4.718E-20 | -0.7135558 | 0.106 | 0.191 | 1.1145E-15 | MO_MT+ | RARRES3 |
| 5.9694E-20 | -0.3605141 | 0.172 | 0.262 | 1.4102E-15 | MO_MT+ | IDI1 |
| 8.1121E-20 | -0.6498404 | 0.235 | 0.315 | 1.9163E-15 | MO_MT+ | EMP1 |
| 1.0379E-19 | -0.2916416 | 0.19 | 0.28 | 2.4518E-15 | MO_MT+ | SLK |
| 1.1614E-19 | -0.4727892 | 0.174 | 0.263 | 2.7435E-15 | MO_MT+ | TPM1 |
| 2.4665E-19 | -0.2919361 | 0.042 | 0.11 | 5.8266E-15 | MO_MT+ | TCHP |
| 3.937E-19 | -0.3103006 | 0.074 | 0.151 | 9.3004E-15 | MO_MT+ | RASD1 |
| 1.173E-18 | -0.5384961 | 0.558 | 0.647 | 2.7711E-14 | MO_MT+ | GLUL |
| 2.0299E-18 | -0.2953532 | 0.45 | 0.534 | 4.7953E-14 | MO_MT+ | PTTG1IP |
| 2.316E-18 | -0.3626161 | 0.842 | 0.849 | 5.4711E-14 | MO_MT+ | RPS4X |
| 3.0972E-17 | -0.3158282 | 0.904 | 0.898 | 7.3166E-13 | MO_MT+ | RPL29 |
| 3.5673E-17 | -0.2571979 | 0.118 | 0.196 | 8.427E-13 | MO_MT+ | KATNBL1 |
| 5.5338E-17 | -0.3088081 | 0.968 | 0.964 | 1.3072E-12 | MO_MT+ | RPL13 |
| 7.8287E-17 | -0.3708068 | 0.093 | 0.164 | 1.8494E-12 | MO_MT+ | CSKMT |
| 1.1907E-16 | -0.4235821 | 0.055 | 0.119 | 2.8128E-12 | MO_MT+ | HSPA2 |
| 1.2032E-16 | -0.3040691 | 0.642 | 0.725 | 2.8424E-12 | MO_MT+ | JUND |
| 1.2232E-16 | -0.3447989 | 0.93 | 0.933 | 2.8895E-12 | MO_MT+ | RPS12 |
| 1.4794E-16 | -0.2648263 | 0.133 | 0.212 | 3.4948E-12 | MO_MT+ | LY6E |
| 1.6872E-16 | 0.28138645 | 0.171 | 0.107 | 3.9857E-12 | MO_MT+ | TNF |
| 1.9514E-16 | -0.2607718 | 0.243 | 0.324 | 4.6098E-12 | MO_MT+ | YPEL3 |
| 3.0456E-16 | -0.2928362 | 0.866 | 0.877 | 7.1946E-12 | MO_MT+ | RPL23A |
| 4.5734E-16 | -0.3527082 | 0.913 | 0.884 | 1.0804E-11 | MO_MT+ | RPS2 |
| 8.1931E-16 | -0.2758927 | 0.503 | 0.573 | 1.9355E-11 | MO_MT+ | ZFAND5 |
| 2.1557E-15 | -0.2936872 | 0.799 | 0.818 | 5.0924E-11 | MO_MT+ | RPL5 |
| 2.5788E-15 | -0.3355162 | 0.913 | 0.897 | 6.0919E-11 | MO_MT+ | RPS3A |
| 3.0594E-15 | -0.2636356 | 0.885 | 0.9 | 7.2271E-11 | MO_MT+ | RPL27A |
| 3.6226E-15 | -0.2549745 | 0.143 | 0.221 | 8.5576E-11 | MO_MT+ | ZBTB20 |
| 4.141E-15 | -0.4035775 | 0.202 | 0.284 | 9.7823E-11 | MO_MT+ | CLU |
| 4.6845E-15 | -0.2888949 | 0.918 | 0.887 | 1.1066E-10 | MO_MT+ | RPL6 |
| 6.8283E-15 | -0.4344763 | 0.157 | 0.229 | 1.613E-10 | MO_MT+ | HEXIM1 |
| 1.776E-14 | -0.3251721 | 0.437 | 0.509 | 4.1956E-10 | MO_MT+ | MT-ND4L |
| 1.8492E-14 | -0.2626918 | 0.19 | 0.26 | 4.3683E-10 | MO_MT+ | PDLIM5 |
| 7.1746E-14 | 0.26652014 | 0.313 | 0.233 | 1.6949E-09 | MO_MT+ | FABP5 |
| 2.0724E-13 | -0.3013832 | 0.159 | 0.229 | 4.8956E-09 | MO_MT+ | IER5L |
| 3.0113E-13 | -0.2514954 | 0.85 | 0.886 | 7.1136E-09 | MO_MT+ | MYL12B |
| 6.4528E-13 | -0.2673982 | 0.494 | 0.55 | 1.5243E-08 | MO_MT+ | TOP1 |
| 9.4838E-13 | -0.2796207 | 0.282 | 0.348 | 2.2404E-08 | MO_MT+ | GOLGB1 |
| 1.0388E-12 | -0.318834 | 0.265 | 0.324 | 2.4539E-08 | MO_MT+ | B4GALT1 |
| 1.1661E-12 | -0.4273057 | 0.342 | 0.407 | 2.7547E-08 | MO_MT+ | RNF213 |
| 1.241E-12 | -0.2968935 | 0.95 | 0.945 | 2.9316E-08 | MO_MT+ | RPS19 |
| 1.3866E-12 | -0.2536527 | 0.264 | 0.327 | 3.2755E-08 | MO_MT+ | TMBIM1 |
| 1.5046E-12 | -0.2507397 | 0.856 | 0.858 | 3.5544E-08 | MO_MT+ | RPL7 |
| 2.0233E-12 | -0.2553613 | 0.906 | 0.925 | 4.7796E-08 | MO_MT+ | RPL18A |
| 8.1387E-12 | -0.6292749 | 0.422 | 0.459 | 1.9226E-07 | MO_MT+ | MXD1 |
| 9.5961E-12 | -0.3631816 | 0.245 | 0.301 | 2.2669E-07 | MO_MT+ | FAM129B |

| | | | | | | |
|---|---|---|---|---|---|---|
| 1.0401E-11 | -0.2562137 | 0.29 | 0.352 | 2.4569E-07 | MO_MT+ | PNN |
| 1.1542E-11 | -0.2686066 | 0.93 | 0.922 | 2.7266E-07 | MO_MT+ | RPL8 |
| 1.1696E-11 | -0.2501578 | 0.932 | 0.933 | 2.7629E-07 | MO_MT+ | RPS14 |
| 1.8186E-11 | 0.3715951 | 0.112 | 0.071 | 4.2961E-07 | MO_MT+ | FN1 |
| 4.1805E-11 | -0.3700293 | 0.247 | 0.301 | 9.8757E-07 | MO_MT+ | BTG2 |
| 4.4188E-11 | 0.36254467 | 0.157 | 0.11 | 1.0439E-06 | MO_MT+ | ANGPTL4 |
| 5.2959E-11 | -0.2825731 | 0.478 | 0.538 | 1.251E-06 | MO_MT+ | SERPINB1 |
| 6.167E-11 | -0.3037776 | 0.163 | 0.224 | 1.4568E-06 | MO_MT+ | HES1 |
| 8.191E-11 | -0.2815678 | 0.945 | 0.921 | 1.935E-06 | MO_MT+ | RPL11 |
| 1.2624E-10 | -0.3735043 | 0.301 | 0.352 | 2.9823E-06 | MO_MT+ | GLS |
| 1.7605E-10 | -0.2681495 | 0.901 | 0.887 | 4.1589E-06 | MO_MT+ | RPS3 |
| 6.4264E-10 | -0.2874624 | 0.355 | 0.398 | 1.5181E-05 | MO_MT+ | ARGLU1 |
| 9.0492E-10 | -0.3689641 | 0.101 | 0.15 | 2.1377E-05 | MO_MT+ | SMC4 |
| 1.8813E-09 | -0.2594261 | 0.914 | 0.915 | 4.4442E-05 | MO_MT+ | MT-ND1 |
| 3.9385E-09 | -0.424022 | 0.906 | 0.876 | 9.304E-05 | MO_MT+ | MT-ND3 |
| 5.3579E-09 | -0.347723 | 0.091 | 0.139 | 0.00012657 | MO_MT+ | DEFB1 |
| 1.3411E-08 | -0.3654749 | 0.179 | 0.232 | 0.00031681 | MO_MT+ | ISG15 |
| 3.0616E-08 | -0.4624029 | 0.15 | 0.202 | 0.00072323 | MO_MT+ | THBS1 |
| 1.419E-07 | 0.25558296 | 0.417 | 0.507 | 0.00335203 | MO_MT+ | DNAJB1 |
| 5.0234E-07 | -0.3690742 | 0.21 | 0.247 | 0.01186687 | MO_MT+ | CSTA |
| 1.7217E-06 | -0.3505399 | 0.968 | 0.936 | 0.04067175 | MO_MT+ | S100A6 |
| 2.4515E-06 | 0.40440253 | 0.559 | 0.661 | 0.0579117 | MO_MT+ | HSPA1A |
| 5.6491E-06 | -0.7913193 | 0.701 | 0.71 | 0.13344754 | MO_MT+ | ANXA1 |
| 5.8568E-06 | -0.5575101 | 0.098 | 0.132 | 0.13835465 | MO_MT+ | APOE |
| 0.00011522 | -0.2551485 | 0.598 | 0.62 | 1 | MO_MT+ | PCBP1 |
| 0.00030347 | -0.2528144 | 0.326 | 0.346 | 1 | MO_MT+ | MIDN |
| 0.00082646 | 0.54471064 | 0.157 | 0.133 | 1 | MO_MT+ | SERPINB2 |
| 0.0028428 | -0.3161243 | 0.955 | 0.94 | 1 | MO_MT+ | NEAT1 |
| 0.00340294 | -0.4014227 | 0.293 | 0.31 | 1 | MO_MT+ | HIST1H4C |
| 0.00439588 | -0.2573799 | 0.182 | 0.199 | 1 | MO_MT+ | DUSP4 |
| 2.661E-245 | 1.30363994 | 0.638 | 0.233 | 6.287E-241 | MO_dmg | SLC11A1 |
| 4.075E-188 | -1.3054774 | 0.855 | 0.958 | 9.627E-184 | MO_dmg | ACTG1 |
| 1.539E-140 | 0.95619332 | 0.934 | 0.734 | 3.635E-136 | MO_dmg | VIM |
| 1.372E-135 | 1.53855832 | 0.997 | 0.986 | 3.24E-131 | MO_dmg | FTL |
| 9.2E-135 | 0.67890408 | 0.476 | 0.193 | 2.173E-130 | MO_dmg | NCF2 |
| 4.825E-115 | 1.08691721 | 0.486 | 0.226 | 1.14E-110 | MO_dmg | PLIN2 |
| 1.473E-114 | -1.4340369 | 0.297 | 0.576 | 3.481E-110 | MO_dmg | TACSTD2 |
| 8.233E-109 | 0.72306537 | 0.849 | 0.706 | 1.945E-104 | MO_dmg | SDCBP |
| 2.388E-108 | 0.58457378 | 0.504 | 0.215 | 5.641E-104 | MO_dmg | VCAN |
| 1.026E-107 | -1.4202372 | 0.289 | 0.562 | 2.425E-103 | MO_dmg | ELF3 |
| 9.212E-106 | -1.1407074 | 0.257 | 0.548 | 2.176E-101 | MO_dmg | CD24 |
| 1.328E-103 | -1.733975 | 0.346 | 0.585 | 3.136E-99 | MO_dmg | KRT19 |
| 7.38E-102 | -0.9346982 | 0.155 | 0.46 | 1.7434E-97 | MO_dmg | CLDN7 |
| 1.087E-101 | 0.71431705 | 0.426 | 0.195 | 2.5671E-97 | MO_dmg | CHORDC1 |
| 3.14E-101 | -1.4171247 | 0.291 | 0.552 | 7.4174E-97 | MO_dmg | KRT7 |
| 2.469E-99 | 0.65797607 | 0.837 | 0.606 | 5.8331E-95 | MO_dmg | CD44 |
| 3.601E-99 | 1.11170379 | 0.796 | 0.637 | 8.5075E-95 | MO_dmg | TXNIP |
| 4.734E-95 | -1.4396383 | 0.313 | 0.564 | 1.1183E-90 | MO_dmg | KRT18 |
| 6.7162E-95 | 0.47316107 | 0.761 | 0.431 | 1.5866E-90 | MO_dmg | TYROBP |
| 7.0613E-92 | -0.6254956 | 0.892 | 0.966 | 1.6681E-87 | MO_dmg | MYL6 |

| | | | | | | |
|---|---|---|---|---|---|---|
| 8.497E-92 | 0.50147605 | 0.317 | 0.124 | 2.0073E-87 | MO_dmg | ARRDC4 |
| 2.7638E-91 | 0.81472741 | 0.528 | 0.309 | 6.529E-87 | MO_dmg | HMGB2 |
| 4.2459E-91 | -0.8932379 | 0.15 | 0.445 | 1.003E-86 | MO_dmg | ATP1B1 |
| 6.8568E-91 | 0.43416216 | 0.352 | 0.143 | 1.6198E-86 | MO_dmg | BCAT1 |
| 5.0581E-89 | -1.2882486 | 0.295 | 0.546 | 1.1949E-84 | MO_dmg | CLDN4 |
| 1.455E-88 | 0.53025411 | 0.631 | 0.327 | 3.4371E-84 | MO_dmg | LAPTM5 |
| 1.0853E-87 | -1.2970297 | 0.243 | 0.5 | 2.5639E-83 | MO_dmg | KRT8 |
| 1.3352E-87 | 0.67041923 | 0.896 | 0.736 | 3.1542E-83 | MO_dmg | PSAP |
| 5.899E-86 | -0.7350159 | 0.15 | 0.434 | 1.3935E-81 | MO_dmg | SPINT2 |
| 8.7408E-86 | 0.67510146 | 0.386 | 0.179 | 2.0648E-81 | MO_dmg | KLF10 |
| 9.5921E-86 | -0.6241651 | 0.722 | 0.879 | 2.2659E-81 | MO_dmg | MYL12A |
| 8.8434E-84 | 0.610046 | 0.529 | 0.303 | 2.0891E-79 | MO_dmg | OGA |
| 1.4491E-82 | -2.0412854 | 0.316 | 0.542 | 3.4233E-78 | MO_dmg | WFDC2 |
| 1.2334E-81 | 0.65292934 | 0.836 | 0.611 | 2.9138E-77 | MO_dmg | LGALS1 |
| 2.5835E-81 | 1.09077056 | 0.985 | 0.939 | 6.103E-77 | MO_dmg | NEAT1 |
| 7.9544E-81 | 0.41006509 | 0.348 | 0.15 | 1.8791E-76 | MO_dmg | CSGALNACT2 |
| 1.2044E-80 | 1.80889569 | 0.622 | 0.448 | 2.8453E-76 | MO_dmg | HSPH1 |
| 3.6037E-80 | -1.3577428 | 0.041 | 0.302 | 8.513E-76 | MO_dmg | ADIRF |
| 4.6849E-80 | 0.73624292 | 0.996 | 0.993 | 1.1067E-75 | MO_dmg | FTH1 |
| 1.1517E-78 | 0.50387184 | 0.296 | 0.118 | 2.7206E-74 | MO_dmg | PHACTR1 |
| 1.6015E-77 | 0.54234172 | 0.415 | 0.194 | 3.7831E-73 | MO_dmg | ITGAX |
| 3.0977E-76 | 0.40966362 | 0.325 | 0.14 | 7.3177E-72 | MO_dmg | TDP2 |
| 2.7988E-75 | -0.7211443 | 0.073 | 0.335 | 6.6117E-71 | MO_dmg | MYO6 |
| 7.6919E-74 | -1.55579 | 0.128 | 0.375 | 1.817E-69 | MO_dmg | MAL |
| 1.7801E-72 | -0.6040456 | 0.073 | 0.327 | 4.205E-68 | MO_dmg | PERP |
| 2.0203E-72 | -0.8526443 | 0.668 | 0.816 | 4.7725E-68 | MO_dmg | GSTP1 |
| 1.8249E-71 | 2.30139016 | 0.762 | 0.654 | 4.311E-67 | MO_dmg | HSPA1A |
| 2.3482E-71 | 0.45190099 | 0.365 | 0.17 | 5.5471E-67 | MO_dmg | P4HA1 |
| 2.7439E-69 | 0.63593423 | 0.683 | 0.509 | 6.4819E-65 | MO_dmg | EIF5 |
| 6.2373E-69 | 0.5247745 | 0.442 | 0.221 | 1.4734E-64 | MO_dmg | ZEB2 |
| 9.8801E-67 | -1.088735 | 0.042 | 0.269 | 2.334E-62 | MO_dmg | LMO7 |
| 1.0043E-66 | -1.3182256 | 0.226 | 0.457 | 2.3725E-62 | MO_dmg | SOX4 |
| 1.5935E-66 | -0.9147776 | 0.06 | 0.298 | 3.7644E-62 | MO_dmg | RND3 |
| 1.7952E-66 | -0.6120038 | 0.431 | 0.629 | 4.2409E-62 | MO_dmg | DSTN |
| 2.827E-66 | -0.8018701 | 0.063 | 0.298 | 6.6783E-62 | MO_dmg | GPRC5A |
| 2.0539E-65 | 0.48971048 | 0.41 | 0.217 | 4.852E-61 | MO_dmg | MAP3K2 |
| 2.5668E-65 | 0.40791202 | 0.343 | 0.16 | 6.0637E-61 | MO_dmg | CEP170 |
| 5.3962E-65 | -1.4101706 | 0.188 | 0.411 | 1.2747E-60 | MO_dmg | SLPI |
| 5.4722E-65 | -0.8922447 | 0.026 | 0.244 | 1.2927E-60 | MO_dmg | ADGRF1 |
| 1.7582E-64 | -2.6130701 | 0.336 | 0.528 | 4.1533E-60 | MO_dmg | LCN2 |
| 4.9088E-64 | -0.7179372 | 0.053 | 0.28 | 1.1596E-59 | MO_dmg | MAL2 |
| 6.1961E-64 | 0.45419043 | 0.561 | 0.339 | 1.4637E-59 | MO_dmg | CTSL |
| 1.5036E-63 | -1.9673302 | 0.133 | 0.35 | 3.552E-59 | MO_dmg | HPGD |
| 2.63E-63 | -0.6728919 | 0.045 | 0.266 | 6.2128E-59 | MO_dmg | MUC1 |
| 3.272E-63 | 0.54025576 | 0.512 | 0.306 | 7.7295E-59 | MO_dmg | CD68 |
| 5.2373E-63 | 0.50413529 | 0.414 | 0.228 | 1.2372E-58 | MO_dmg | RYBP |
| 3.9009E-62 | -0.7006927 | 0.04 | 0.256 | 9.2151E-58 | MO_dmg | SYNE2 |
| 6.6392E-62 | -0.5717161 | 0.37 | 0.581 | 1.5684E-57 | MO_dmg | PRDX5 |
| 1.4282E-61 | -0.4933849 | 0.034 | 0.247 | 3.3738E-57 | MO_dmg | GIPC1 |
| 2.4525E-61 | 0.45276149 | 0.837 | 0.663 | 5.7936E-57 | MO_dmg | ANXA5 |

| | | | | | | |
|---|---|---|---|---|---|---|
| 1.1609E-60 | -0.4874917 | 0.057 | 0.28 | 2.7423E-56 | MO_dmg | JUP |
| 1.1663E-60 | -0.742159 | 0.909 | 0.931 | 2.7552E-56 | MO_dmg | MT-CYB |
| 2.2583E-60 | 0.77394743 | 0.485 | 0.28 | 5.3347E-56 | MO_dmg | SLC25A37 |
| 5.5938E-60 | 0.28907233 | 0.241 | 0.097 | 1.3214E-55 | MO_dmg | TFEC |
| 6.3776E-60 | 1.82493436 | 0.636 | 0.5 | 1.5066E-55 | MO_dmg | DNAJB1 |
| 8.7783E-60 | -0.8578341 | 0.054 | 0.27 | 2.0737E-55 | MO_dmg | TSPAN1 |
| 1.249E-59 | -1.2245751 | 0.128 | 0.351 | 2.9505E-55 | MO_dmg | GDF15 |
| 1.5069E-59 | -1.0129258 | 0.957 | 0.936 | 3.5599E-55 | MO_dmg | S100A6 |
| 2.8995E-59 | -0.6492522 | 0.061 | 0.281 | 6.8494E-55 | MO_dmg | CAMK2N1 |
| 1.5809E-58 | -0.6033379 | 0.186 | 0.415 | 3.7345E-54 | MO_dmg | CD59 |
| 2.2983E-58 | -0.7821122 | 0.035 | 0.239 | 5.4293E-54 | MO_dmg | PDZK1IP1 |
| 3.5019E-58 | 0.38808229 | 0.356 | 0.182 | 8.2726E-54 | MO_dmg | FGFR1OP2 |
| 5.5252E-58 | -0.5618101 | 0.035 | 0.242 | 1.3052E-53 | MO_dmg | S100A16 |
| 7.2388E-58 | -0.4812243 | 0.393 | 0.604 | 1.71E-53 | MO_dmg | VAMP8 |
| 1.3298E-57 | 0.37441723 | 0.304 | 0.143 | 3.1414E-53 | MO_dmg | FAR1 |
| 1.8859E-57 | 0.58821002 | 0.613 | 0.428 | 4.4551E-53 | MO_dmg | GSTO1 |
| 2.4436E-57 | 0.38535308 | 0.392 | 0.195 | 5.7725E-53 | MO_dmg | CYBB |
| 3.6659E-57 | -0.5902306 | 0.042 | 0.247 | 8.66E-53 | MO_dmg | PRSS8 |
| 9.3929E-57 | 0.58415736 | 0.6 | 0.421 | 2.2189E-52 | MO_dmg | SLC3A2 |
| 1.256E-56 | 0.36189066 | 0.351 | 0.176 | 2.9672E-52 | MO_dmg | RILPL2 |
| 1.408E-56 | 0.67405686 | 0.631 | 0.419 | 3.3261E-52 | MO_dmg | EMP3 |
| 5.5835E-56 | -0.4636434 | 0.059 | 0.266 | 1.319E-51 | MO_dmg | PRDX2 |
| 6.98E-56 | 0.38085852 | 0.364 | 0.19 | 1.6489E-51 | MO_dmg | RAB8B |
| 8.0835E-56 | -0.5354707 | 0.082 | 0.295 | 1.9096E-51 | MO_dmg | SPINT1 |
| 1.1201E-55 | -0.6606756 | 0.139 | 0.355 | 2.646E-51 | MO_dmg | PEBP1 |
| 4.1577E-55 | 0.29495485 | 0.412 | 0.214 | 9.8218E-51 | MO_dmg | ALOX5AP |
| 1.0178E-54 | 0.73462981 | 0.388 | 0.227 | 2.4043E-50 | MO_dmg | CACYBP |
| 2.6853E-54 | -1.3226851 | 0.14 | 0.341 | 6.3434E-50 | MO_dmg | IGFBP7 |
| 3.1192E-54 | 0.39630816 | 0.398 | 0.221 | 7.3685E-50 | MO_dmg | XPO1 |
| 3.8619E-54 | 1.75821524 | 0.619 | 0.502 | 9.1229E-50 | MO_dmg | HSPA1B |
| 3.9764E-54 | 0.54103955 | 0.803 | 0.684 | 9.3934E-50 | MO_dmg | BRI3 |
| 4.0312E-54 | -1.3926642 | 0.041 | 0.237 | 9.5228E-50 | MO_dmg | PRSS22 |
| 6.1277E-54 | 0.46628531 | 0.424 | 0.243 | 1.4475E-49 | MO_dmg | GPCPD1 |
| 8.7402E-54 | 0.55018336 | 0.774 | 0.684 | 2.0647E-49 | MO_dmg | ATP6V0B |
| 2.3535E-53 | 0.59246813 | 0.422 | 0.228 | 5.5596E-49 | MO_dmg | SLC2A3 |
| 3.0601E-53 | -0.4962031 | 0.054 | 0.252 | 7.2289E-49 | MO_dmg | SPTBN1 |
| 8.1585E-53 | 0.43405414 | 0.448 | 0.273 | 1.9273E-48 | MO_dmg | PLEKHB2 |
| 8.9889E-53 | -0.6204702 | 0.289 | 0.501 | 2.1235E-48 | MO_dmg | CD9 |
| 1.4671E-52 | 0.4452357 | 0.447 | 0.259 | 3.4658E-48 | MO_dmg | BACH1 |
| 4.9756E-52 | 0.36796471 | 0.341 | 0.173 | 1.1754E-47 | MO_dmg | CCDC88A |
| 5.114E-52 | -1.7382647 | 0.076 | 0.272 | 1.2081E-47 | MO_dmg | S100A2 |
| 5.5195E-52 | -0.4263338 | 0.044 | 0.236 | 1.3039E-47 | MO_dmg | TJP1 |
| 6.1602E-52 | 0.46765909 | 0.839 | 0.739 | 1.4552E-47 | MO_dmg | APLP2 |
| 1.3787E-51 | 0.26219484 | 0.166 | 0.059 | 3.257E-47 | MO_dmg | HSPA4L |
| 1.8201E-51 | 0.3101886 | 0.356 | 0.182 | 4.2997E-47 | MO_dmg | RNF130 |
| 4.1869E-51 | -0.4320028 | 0.152 | 0.372 | 9.8908E-47 | MO_dmg | AES |
| 8.2914E-51 | 0.40691467 | 0.351 | 0.186 | 1.9587E-46 | MO_dmg | FNDC3A |
| 1.9475E-50 | -0.6907087 | 0.64 | 0.755 | 4.6005E-46 | MO_dmg | TPM4 |
| 2.7529E-50 | -0.7586848 | 0.044 | 0.232 | 6.5031E-46 | MO_dmg | IL32 |
| 3.7343E-50 | 0.57560289 | 0.653 | 0.514 | 8.8216E-46 | MO_dmg | NAP1L1 |

| | | | | | | |
|---|---|---|---|---|---|---|
| 3.7935E-50 | -2.0407534 | 0.169 | 0.346 | 8.9614E-46 | MO_dmg | PSCA |
| 3.81E-50 | 0.30828718 | 0.327 | 0.166 | 9.0004E-46 | MO_dmg | LACTB |
| 9.3941E-50 | 1.48715541 | 0.973 | 0.962 | 2.2192E-45 | MO_dmg | HSP90AA1 |
| 1.167E-49 | 0.49720128 | 0.439 | 0.265 | 2.7567E-45 | MO_dmg | TSC22D3 |
| 5.3246E-49 | 0.28434832 | 0.737 | 0.533 | 1.2578E-44 | MO_dmg | UPP1 |
| 1.2691E-48 | -1.2523396 | 0.025 | 0.201 | 2.9979E-44 | MO_dmg | HOPX |
| 1.559E-48 | -0.3676195 | 0.038 | 0.219 | 3.6828E-44 | MO_dmg | CNN3 |
| 1.5655E-48 | 0.4563716 | 0.379 | 0.208 | 3.6981E-44 | MO_dmg | BASP1 |
| 2.1944E-48 | 0.30986784 | 0.296 | 0.141 | 5.1838E-44 | MO_dmg | MS4A7 |
| 2.3083E-48 | -0.5688931 | 0.042 | 0.226 | 5.4529E-44 | MO_dmg | CCND1 |
| 4.6846E-48 | -0.439046 | 0.024 | 0.195 | 1.1066E-43 | MO_dmg | ABLIM1 |
| 5.0734E-48 | 0.49304625 | 0.483 | 0.307 | 1.1985E-43 | MO_dmg | FNDC3B |
| 1.0313E-47 | 0.55421299 | 0.807 | 0.706 | 2.4362E-43 | MO_dmg | GPX4 |
| 1.6459E-47 | -0.7747813 | 0.02 | 0.188 | 3.8882E-43 | MO_dmg | AQP3 |
| 3.3577E-47 | -0.4823361 | 0.037 | 0.212 | 7.9319E-43 | MO_dmg | MACC1 |
| 4.2811E-47 | -0.4988078 | 0.049 | 0.229 | 1.0113E-42 | MO_dmg | OCIAD2 |
| 4.6518E-47 | 2.03904189 | 0.326 | 0.189 | 1.0989E-42 | MO_dmg | HSPA6 |
| 6.2082E-47 | 0.93294173 | 0.938 | 0.883 | 1.4666E-42 | MO_dmg | HSP90AB1 |
| 7.7805E-47 | -0.695488 | 0.067 | 0.251 | 1.838E-42 | MO_dmg | CST6 |
| 9.3493E-47 | 0.46159601 | 0.343 | 0.184 | 2.2086E-42 | MO_dmg | SLC5A3 |
| 9.6421E-47 | -0.521418 | 0.068 | 0.255 | 2.2777E-42 | MO_dmg | KLF5 |
| 1.0997E-46 | 0.53259191 | 0.736 | 0.582 | 2.5978E-42 | MO_dmg | KLF6 |
| 3.1923E-46 | -0.3675037 | 0.017 | 0.181 | 7.5411E-42 | MO_dmg | SMIM22 |
| 3.3741E-46 | -0.6819512 | 0.061 | 0.24 | 7.9707E-42 | MO_dmg | EPCAM |
| 3.9995E-46 | 0.33440204 | 0.191 | 0.076 | 9.448E-42 | MO_dmg | MRC1 |
| 4.6248E-46 | 0.35819839 | 0.292 | 0.145 | 1.0925E-41 | MO_dmg | DMXL2 |
| 5.371E-46 | -0.4399456 | 0.082 | 0.271 | 1.2688E-41 | MO_dmg | AFDN |
| 1.2285E-45 | -0.5148442 | 0.102 | 0.288 | 2.902E-41 | MO_dmg | LDHB |
| 1.3389E-45 | -1.2893242 | 0.033 | 0.203 | 3.1629E-41 | MO_dmg | KRT17 |
| 1.7065E-45 | -0.8065064 | 0.026 | 0.192 | 4.0314E-41 | MO_dmg | TMPRSS2 |
| 5.7022E-45 | 0.56778224 | 0.288 | 0.147 | 1.347E-40 | MO_dmg | SLC7A11 |
| 8.4968E-45 | 0.28615282 | 0.725 | 0.533 | 2.0072E-40 | MO_dmg | PLAUR |
| 2.6934E-44 | 0.28598252 | 0.26 | 0.123 | 6.3625E-40 | MO_dmg | IRAK3 |
| 2.7563E-44 | -0.5921298 | 0.016 | 0.172 | 6.5112E-40 | MO_dmg | FXYD3 |
| 2.7914E-44 | 0.27462101 | 0.297 | 0.147 | 6.5941E-40 | MO_dmg | EVI2B |
| 2.852E-44 | -0.5712824 | 0.844 | 0.907 | 6.7373E-40 | MO_dmg | ITM2B |
| 3.7098E-44 | 0.32836961 | 0.31 | 0.164 | 8.7637E-40 | MO_dmg | NUP58 |
| 4.0192E-44 | -0.3723383 | 0.047 | 0.221 | 9.4946E-40 | MO_dmg | DSP |
| 4.6997E-44 | -0.7671082 | 0.035 | 0.201 | 1.1102E-39 | MO_dmg | TM4SF1 |
| 4.799E-44 | -2.4392409 | 0.22 | 0.383 | 1.1337E-39 | MO_dmg | KRT13 |
| 5.441E-44 | -0.3179457 | 0.034 | 0.199 | 1.2853E-39 | MO_dmg | RBPMS |
| 5.9315E-44 | -0.430597 | 0.047 | 0.219 | 1.4012E-39 | MO_dmg | CKB |
| 6.2612E-44 | -0.3960424 | 0.079 | 0.257 | 1.4791E-39 | MO_dmg | CD2AP |
| 9.1501E-44 | -0.5803076 | 0.034 | 0.2 | 2.1615E-39 | MO_dmg | SFN |
| 1.0288E-43 | 0.31772422 | 0.413 | 0.231 | 2.4304E-39 | MO_dmg | BCL2A1 |
| 1.1428E-43 | -2.1476046 | 0.173 | 0.344 | 2.6997E-39 | MO_dmg | MMP7 |
| 1.3055E-43 | -1.0579031 | 0.027 | 0.188 | 3.084E-39 | MO_dmg | DUOX2 |
| 1.6579E-43 | -1.2349876 | 0.017 | 0.17 | 3.9164E-39 | MO_dmg | DUOXA2 |
| 1.9367E-43 | -0.4864569 | 0.332 | 0.518 | 4.5751E-39 | MO_dmg | ANXA11 |
| 2.1088E-43 | 0.33393743 | 0.255 | 0.123 | 4.9817E-39 | MO_dmg | ABL2 |

| | | | | | | |
|---|---|---|---|---|---|---|
| 3.1961E-43 | -0.3455224 | 0.028 | 0.188 | 7.5501E-39 | MO_dmg | EPS8L2 |
| 4.6163E-43 | 0.65927019 | 0.757 | 0.653 | 1.0905E-38 | MO_dmg | ZFAS1 |
| 4.9585E-43 | -0.5585799 | 0.021 | 0.176 | 1.1713E-38 | MO_dmg | PPL |
| 5.6076E-43 | -0.6022368 | 0.961 | 0.973 | 1.3247E-38 | MO_dmg | ACTB |
| 1.0615E-42 | 0.25106054 | 0.255 | 0.122 | 2.5076E-38 | MO_dmg | NFIL3 |
| 1.4144E-42 | -0.3149756 | 0.024 | 0.18 | 3.3413E-38 | MO_dmg | TMEM54 |
| 1.5421E-42 | -0.5404587 | 0.059 | 0.234 | 3.6429E-38 | MO_dmg | ITGA2 |
| 1.7076E-42 | 0.26870172 | 0.387 | 0.211 | 4.0338E-38 | MO_dmg | SPI1 |
| 3.5805E-42 | -0.5533407 | 0.357 | 0.527 | 8.4583E-38 | MO_dmg | HEBP2 |
| 4.5156E-42 | -0.4658746 | 0.024 | 0.179 | 1.0667E-37 | MO_dmg | C6orf132 |
| 5.049E-42 | -0.9654416 | 0.014 | 0.164 | 1.1927E-37 | MO_dmg | CEACAM5 |
| 6.41E-42 | -0.5083625 | 0.073 | 0.248 | 1.5142E-37 | MO_dmg | ITGB8 |
| 8.5066E-42 | 0.29432364 | 0.269 | 0.133 | 2.0095E-37 | MO_dmg | EVI2A |
| 9.3396E-42 | -0.3641676 | 0.029 | 0.187 | 2.2063E-37 | MO_dmg | CDH1 |
| 1.0157E-41 | -0.8429691 | 0.039 | 0.199 | 2.3994E-37 | MO_dmg | C12orf75 |
| 1.1295E-41 | 0.3403276 | 0.316 | 0.171 | 2.6682E-37 | MO_dmg | ATXN1 |
| 1.1555E-41 | -0.426292 | 0.029 | 0.185 | 2.7297E-37 | MO_dmg | S100A14 |
| 1.571E-41 | 0.35896534 | 0.32 | 0.166 | 3.7112E-37 | MO_dmg | CXCR4 |
| 2.2594E-41 | 0.25460229 | 0.229 | 0.106 | 5.3373E-37 | MO_dmg | PAG1 |
| 3.4255E-41 | -0.4720122 | 0.051 | 0.213 | 8.092E-37 | MO_dmg | LY6E |
| 3.8056E-41 | -0.4337936 | 0.145 | 0.33 | 8.9901E-37 | MO_dmg | PPA1 |
| 4.521E-41 | -0.4316329 | 0.027 | 0.18 | 1.068E-36 | MO_dmg | FHL2 |
| 4.5695E-41 | -0.4440368 | 0.028 | 0.183 | 1.0795E-36 | MO_dmg | EPHA2 |
| 6.0685E-41 | -0.5444429 | 0.035 | 0.193 | 1.4336E-36 | MO_dmg | EPS8L1 |
| 8.5359E-41 | -0.3749363 | 0.052 | 0.218 | 2.0164E-36 | MO_dmg | EHF |
| 8.6007E-41 | 0.42305541 | 0.39 | 0.238 | 2.0318E-36 | MO_dmg | MAFG |
| 8.9239E-41 | -0.489868 | 0.031 | 0.187 | 2.1081E-36 | MO_dmg | SDCBP2 |
| 1.2104E-40 | 0.92982219 | 0.316 | 0.184 | 2.8594E-36 | MO_dmg | BAG3 |
| 1.329E-40 | -0.3610102 | 0.015 | 0.16 | 3.1395E-36 | MO_dmg | TSPAN6 |
| 1.3536E-40 | -0.4042634 | 0.017 | 0.163 | 3.1977E-36 | MO_dmg | SDC1 |
| 1.5432E-40 | -0.6235822 | 0.009 | 0.15 | 3.6456E-36 | MO_dmg | NCCRP1 |
| 1.8385E-40 | 0.45185013 | 0.829 | 0.735 | 4.343E-36 | MO_dmg | CTSB |
| 2.2155E-40 | 0.47675127 | 0.554 | 0.39 | 5.2336E-36 | MO_dmg | HIF1A |
| 2.342E-40 | -0.3939529 | 0.047 | 0.21 | 5.5325E-36 | MO_dmg | PARD6B |
| 2.7289E-40 | 0.71502164 | 0.987 | 0.965 | 6.4464E-36 | MO_dmg | UBC |
| 2.8245E-40 | -0.3036009 | 0.048 | 0.209 | 6.6723E-36 | MO_dmg | CTTN |
| 4.1866E-40 | -1.0534715 | 0.019 | 0.165 | 9.8899E-36 | MO_dmg | MUC21 |
| 5.0473E-40 | 0.48533661 | 0.384 | 0.235 | 1.1923E-35 | MO_dmg | FKBP4 |
| 5.7392E-40 | -0.5662472 | 0.081 | 0.246 | 1.3558E-35 | MO_dmg | S100A13 |
| 6.8387E-40 | -0.5417501 | 0.028 | 0.18 | 1.6155E-35 | MO_dmg | IGFBP3 |
| 1.0557E-39 | -0.302686 | 0.024 | 0.171 | 2.4938E-35 | MO_dmg | CXADR |
| 1.2036E-39 | -0.6122345 | 0.007 | 0.145 | 2.8433E-35 | MO_dmg | SNCG |
| 1.3299E-39 | 0.43341026 | 0.456 | 0.289 | 3.1415E-35 | MO_dmg | ATP13A3 |
| 1.4113E-39 | -0.4765141 | 0.105 | 0.278 | 3.3339E-35 | MO_dmg | TNFRSF12A |
| 1.4434E-39 | 0.25687858 | 0.185 | 0.08 | 3.4098E-35 | MO_dmg | CYP27A1 |
| 1.5846E-39 | -0.8402433 | 0.114 | 0.283 | 3.7434E-35 | MO_dmg | DUSP5 |
| 1.7645E-39 | -0.3829179 | 0.017 | 0.161 | 4.1684E-35 | MO_dmg | TTC9 |
| 1.943E-39 | -0.3298283 | 0.032 | 0.183 | 4.59E-35 | MO_dmg | TEAD1 |
| 2.1278E-39 | 0.4102122 | 0.403 | 0.25 | 5.0264E-35 | MO_dmg | MGAT1 |
| 2.2311E-39 | -0.3774147 | 0.461 | 0.634 | 5.2705E-35 | MO_dmg | YBX3 |

| | | | | | | |
|---|---|---|---|---|---|---|
| 2.3407E-39 | -0.5355596 | 0.027 | 0.175 | 5.5293E-35 | MO_dmg | HSPB8 |
| 2.5688E-39 | 0.32139281 | 0.341 | 0.195 | 6.0684E-35 | MO_dmg | PRNP |
| 3.4439E-39 | 0.36240152 | 0.399 | 0.238 | 8.1355E-35 | MO_dmg | ATP2B1 |
| 3.8131E-39 | -0.8078944 | 0.017 | 0.159 | 9.0076E-35 | MO_dmg | KRT15 |
| 4.4496E-39 | 0.37605167 | 0.392 | 0.239 | 1.0511E-34 | MO_dmg | RBPJ |
| 4.9668E-39 | 0.31999843 | 0.323 | 0.18 | 1.1733E-34 | MO_dmg | DAB2 |
| 5.2974E-39 | 0.25046493 | 0.206 | 0.094 | 1.2514E-34 | MO_dmg | DOCK4 |
| 5.8043E-39 | -0.4763961 | 0.02 | 0.162 | 1.3711E-34 | MO_dmg | MALL |
| 7.5423E-39 | 0.44939161 | 0.499 | 0.348 | 1.7817E-34 | MO_dmg | TAF1D |
| 7.5868E-39 | -0.3070558 | 0.017 | 0.159 | 1.7922E-34 | MO_dmg | ERBB3 |
| 8.4137E-39 | -0.6963034 | 0.084 | 0.25 | 1.9876E-34 | MO_dmg | PLAU |
| 1.0598E-38 | 0.27157867 | 0.264 | 0.133 | 2.5035E-34 | MO_dmg | ITGA5 |
| 1.0967E-38 | -0.7096817 | 0.017 | 0.157 | 2.5907E-34 | MO_dmg | SCEL |
| 1.2178E-38 | -0.7613106 | 0.035 | 0.184 | 2.8768E-34 | MO_dmg | F3 |
| 1.7785E-38 | -0.9192316 | 0.018 | 0.158 | 4.2014E-34 | MO_dmg | PITX1 |
| 1.838E-38 | -0.3320261 | 0.034 | 0.184 | 4.3419E-34 | MO_dmg | ANXA3 |
| 2.9077E-38 | -0.3156431 | 0.026 | 0.172 | 6.8689E-34 | MO_dmg | NECTIN4 |
| 3.7911E-38 | -0.3092984 | 0.042 | 0.192 | 8.9558E-34 | MO_dmg | SH3YL1 |
| 3.8103E-38 | -0.2673968 | 0.028 | 0.174 | 9.001E-34 | MO_dmg | GNG12 |
| 4.2857E-38 | -0.7373223 | 0.031 | 0.177 | 1.0124E-33 | MO_dmg | NDRG2 |
| 4.7389E-38 | 0.30821983 | 0.333 | 0.19 | 1.1195E-33 | MO_dmg | SNX10 |
| 7.3982E-38 | -0.2935057 | 0.035 | 0.182 | 1.7477E-33 | MO_dmg | LSR |
| 7.5414E-38 | 0.52485627 | 0.622 | 0.498 | 1.7815E-33 | MO_dmg | ASAH1 |
| 1.0248E-37 | -0.5211002 | 0.032 | 0.178 | 2.4208E-33 | MO_dmg | SPNS2 |
| 1.6731E-37 | -0.3685417 | 0.313 | 0.492 | 3.9523E-33 | MO_dmg | DYNLT1 |
| 2.3497E-37 | -0.4980355 | 0.316 | 0.486 | 5.5507E-33 | MO_dmg | CD46 |
| 4.8833E-37 | -0.7580934 | 0.015 | 0.149 | 1.1536E-32 | MO_dmg | CEACAM7 |
| 5.7582E-37 | -0.3790777 | 0.015 | 0.149 | 1.3603E-32 | MO_dmg | VSIG2 |
| 6.4477E-37 | 0.42892146 | 0.387 | 0.243 | 1.5231E-32 | MO_dmg | CREBRF |
| 8.0987E-37 | 0.48899092 | 0.531 | 0.393 | 1.9132E-32 | MO_dmg | H2AFY |
| 1.1638E-36 | -0.868979 | 0.024 | 0.161 | 2.7492E-32 | MO_dmg | PRSS27 |
| 1.2326E-36 | 0.2633927 | 0.235 | 0.116 | 2.9118E-32 | MO_dmg | RIPK2 |
| 1.4906E-36 | -1.9501357 | 0.281 | 0.411 | 3.5213E-32 | MO_dmg | S100P |
| 1.6137E-36 | -0.55006 | 0.021 | 0.159 | 3.8121E-32 | MO_dmg | MUC4 |
| 1.9735E-36 | -0.3963632 | 0.024 | 0.16 | 4.6619E-32 | MO_dmg | CLDN3 |
| 2.6511E-36 | -0.2852086 | 0.027 | 0.166 | 6.2628E-32 | MO_dmg | NDFIP2 |
| 2.6891E-36 | -0.2603236 | 0.028 | 0.169 | 6.3525E-32 | MO_dmg | DDR1 |
| 3.3856E-36 | 0.31877078 | 0.39 | 0.241 | 7.9979E-32 | MO_dmg | PTPN12 |
| 3.8525E-36 | -0.5374457 | 0.022 | 0.157 | 9.1007E-32 | MO_dmg | CDC42EP5 |
| 4.3078E-36 | 1.13582027 | 0.591 | 0.475 | 1.0176E-31 | MO_dmg | HSPD1 |
| 4.5082E-36 | 0.7704937 | 0.481 | 0.343 | 1.065E-31 | MO_dmg | IER5 |
| 5.9215E-36 | -0.7293341 | 0.017 | 0.148 | 1.3988E-31 | MO_dmg | TMPRSS11E |
| 6.5827E-36 | 0.28369276 | 0.267 | 0.142 | 1.555E-31 | MO_dmg | WASHC4 |
| 8.8381E-36 | -0.3685656 | 0.028 | 0.165 | 2.0878E-31 | MO_dmg | SYTL2 |
| 9.4865E-36 | -0.3001839 | 0.04 | 0.183 | 2.241E-31 | MO_dmg | FNBP1L |
| 1.2644E-35 | 0.28885539 | 0.283 | 0.148 | 2.9869E-31 | MO_dmg | CLEC5A |
| 1.4367E-35 | -0.4006068 | 0.011 | 0.139 | 3.394E-31 | MO_dmg | GCNT3 |
| 1.6427E-35 | -1.4248217 | 0.03 | 0.166 | 3.8806E-31 | MO_dmg | ECM1 |
| 1.7461E-35 | 0.28934089 | 0.33 | 0.192 | 4.1248E-31 | MO_dmg | FNIP1 |
| 2.2132E-35 | -0.8735284 | 0.01 | 0.136 | 5.2282E-31 | MO_dmg | RHCG |

| | | | | | | |
|---|---|---|---|---|---|---|
| 2.5553E-35 | 0.36111479 | 0.328 | 0.194 | 6.0365E-31 | MO_dmg | MYO9B |
| 2.587E-35 | -0.378503 | 0.011 | 0.138 | 6.1113E-31 | MO_dmg | BCAS1 |
| 3.0115E-35 | -0.3053209 | 0.045 | 0.19 | 7.1141E-31 | MO_dmg | PAWR |
| 3.1331E-35 | -0.9424942 | 0.038 | 0.177 | 7.4013E-31 | MO_dmg | RDH10 |
| 4.365E-35 | -0.3296508 | 0.049 | 0.194 | 1.0311E-30 | MO_dmg | LIMA1 |
| 4.3763E-35 | 0.64066043 | 0.272 | 0.148 | 1.0338E-30 | MO_dmg | ZFAND2A |
| 5.383E-35 | -0.5588479 | 0.043 | 0.185 | 1.2716E-30 | MO_dmg | LAMC2 |
| 5.6958E-35 | -0.6830021 | 0.021 | 0.152 | 1.3455E-30 | MO_dmg | TMPRSS11D |
| 5.849E-35 | -0.715933 | 0.018 | 0.15 | 1.3817E-30 | MO_dmg | CEACAM6 |
| 6.2187E-35 | 0.29081111 | 0.337 | 0.198 | 1.469E-30 | MO_dmg | TANK |
| 6.4329E-35 | 0.6808865 | 0.844 | 0.773 | 1.5197E-30 | MO_dmg | CTSD |
| 7.8042E-35 | -0.8411824 | 0.049 | 0.192 | 1.8436E-30 | MO_dmg | RARRES3 |
| 1.0532E-34 | -0.2752557 | 0.018 | 0.148 | 2.488E-30 | MO_dmg | SLC44A4 |
| 1.0689E-34 | 0.34412535 | 0.334 | 0.2 | 2.5252E-30 | MO_dmg | C6orf48 |
| 1.1146E-34 | -0.2768255 | 0.028 | 0.162 | 2.633E-30 | MO_dmg | CRB3 |
| 1.2109E-34 | 0.29743645 | 0.291 | 0.154 | 2.8606E-30 | MO_dmg | CD300E |
| 1.2177E-34 | 0.39317782 | 0.214 | 0.107 | 2.8765E-30 | MO_dmg | TMEM70 |
| 1.2269E-34 | -0.5312617 | 0.037 | 0.175 | 2.8984E-30 | MO_dmg | STEAP4 |
| 1.4528E-34 | 0.36810299 | 0.348 | 0.209 | 3.4319E-30 | MO_dmg | ZFYVE16 |
| 1.8196E-34 | -0.4280841 | 0.02 | 0.15 | 4.2985E-30 | MO_dmg | ALDH1A3 |
| 1.8215E-34 | -0.3348311 | 0.023 | 0.154 | 4.3028E-30 | MO_dmg | MDK |
| 1.8906E-34 | 0.31934773 | 0.349 | 0.202 | 4.4661E-30 | MO_dmg | SLC43A2 |
| 2.3632E-34 | -0.3101526 | 0.034 | 0.172 | 5.5826E-30 | MO_dmg | TINAGL1 |
| 2.8185E-34 | -0.428051 | 0.816 | 0.886 | 6.6581E-30 | MO_dmg | MYL12B |
| 3.0538E-34 | 0.25008256 | 0.244 | 0.127 | 7.214E-30 | MO_dmg | KDM7A |
| 3.6455E-34 | 0.39243316 | 0.491 | 0.342 | 8.6118E-30 | MO_dmg | ZNF207 |
| 4.0135E-34 | 0.32755116 | 0.401 | 0.255 | 9.481E-30 | MO_dmg | MBNL1 |
| 4.3089E-34 | -0.3283017 | 0.126 | 0.29 | 1.0179E-29 | MO_dmg | LRP10 |
| 5.4581E-34 | -0.6211229 | 0.011 | 0.134 | 1.2894E-29 | MO_dmg | GABRP |
| 6.4262E-34 | 0.30900383 | 0.351 | 0.215 | 1.5181E-29 | MO_dmg | ACAP2 |
| 6.9947E-34 | -0.8654118 | 0.014 | 0.138 | 1.6524E-29 | MO_dmg | TMPRSS11B |
| 9.0486E-34 | 0.35134511 | 0.936 | 0.857 | 2.1376E-29 | MO_dmg | SH3BGRL3 |
| 1.0086E-33 | -0.3231685 | 0.02 | 0.148 | 2.3827E-29 | MO_dmg | CLDN1 |
| 1.2186E-33 | -0.25097 | 0.017 | 0.143 | 2.8787E-29 | MO_dmg | RAB25 |
| 1.354E-33 | 0.2663966 | 0.251 | 0.131 | 3.1987E-29 | MO_dmg | FGR |
| 1.4321E-33 | -3.3439545 | 0.279 | 0.405 | 3.3831E-29 | MO_dmg | SPRR3 |
| 1.9952E-33 | -0.3358476 | 0.146 | 0.31 | 4.7134E-29 | MO_dmg | CLTB |
| 2.1816E-33 | -0.4065367 | 0.1 | 0.251 | 5.1535E-29 | MO_dmg | SERINC2 |
| 3.0934E-33 | 0.72924553 | 0.289 | 0.168 | 7.3075E-29 | MO_dmg | DNAJA4 |
| 3.3212E-33 | -0.4499633 | 0.051 | 0.19 | 7.8457E-29 | MO_dmg | TNFSF10 |
| 4.1015E-33 | 0.46527023 | 0.598 | 0.484 | 9.689E-29 | MO_dmg | CANX |
| 5.8738E-33 | -0.3716631 | 0.028 | 0.157 | 1.3876E-28 | MO_dmg | TMPRSS4 |
| 6.7628E-33 | 0.25569703 | 0.289 | 0.16 | 1.5976E-28 | MO_dmg | MYO1F |
| 1.3627E-32 | 0.316014 | 0.342 | 0.209 | 3.2191E-28 | MO_dmg | RNF13 |
| 1.5439E-32 | 0.30349196 | 0.302 | 0.169 | 3.6471E-28 | MO_dmg | SMIM25 |
| 1.6372E-32 | 0.33555908 | 0.325 | 0.195 | 3.8676E-28 | MO_dmg | CREG1 |
| 1.7785E-32 | -0.2796472 | 0.017 | 0.138 | 4.2014E-28 | MO_dmg | COL18A1 |
| 2.0172E-32 | 0.31682447 | 0.299 | 0.173 | 4.7653E-28 | MO_dmg | ZBTB43 |
| 2.3838E-32 | -0.3847017 | 0.13 | 0.287 | 5.6314E-28 | MO_dmg | CD151 |
| 2.4173E-32 | 0.29068536 | 0.301 | 0.173 | 5.7103E-28 | MO_dmg | ABCA1 |

| | | | | | | |
|---|---|---|---|---|---|---|
| 2.5339E-32 | -0.547113 | 0.059 | 0.195 | 5.9859E-28 | MO_dmg | FBXO32 |
| 2.6206E-32 | -0.3036168 | 0.014 | 0.133 | 6.1907E-28 | MO_dmg | AIF1L |
| 2.8677E-32 | -0.3474068 | 0.023 | 0.147 | 6.7743E-28 | MO_dmg | ITGB6 |
| 2.928E-32 | -1.5797318 | 0.023 | 0.147 | 6.9168E-28 | MO_dmg | SPRR2A |
| 3.3022E-32 | -0.843055 | 0.02 | 0.143 | 7.8007E-28 | MO_dmg | CXCL6 |
| 5.5875E-32 | -0.3697258 | 0.071 | 0.217 | 1.3199E-27 | MO_dmg | LAMB3 |
| 6.8409E-32 | 0.41446086 | 0.482 | 0.339 | 1.616E-27 | MO_dmg | WSB1 |
| 1.1322E-31 | -0.6241366 | 0.318 | 0.46 | 2.6745E-27 | MO_dmg | RAB11FIP1 |
| 1.2922E-31 | -0.2631452 | 0.028 | 0.153 | 3.0525E-27 | MO_dmg | PLS3 |
| 1.822E-31 | -0.3909138 | 0.115 | 0.263 | 4.3042E-27 | MO_dmg | WASL |
| 1.8698E-31 | -0.604036 | 0.128 | 0.285 | 4.417E-27 | MO_dmg | CLU |
| 1.9296E-31 | -0.5911509 | 0.02 | 0.139 | 4.5582E-27 | MO_dmg | CXCL17 |
| 2.1135E-31 | 0.35198981 | 0.412 | 0.275 | 4.9928E-27 | MO_dmg | PTP4A1 |
| 4.2066E-31 | -0.566017 | 0.024 | 0.145 | 9.9373E-27 | MO_dmg | CDKN2A |
| 5.5491E-31 | 0.30479643 | 0.358 | 0.227 | 1.3109E-26 | MO_dmg | TAOK3 |
| 5.6153E-31 | 0.33964032 | 0.361 | 0.227 | 1.3265E-26 | MO_dmg | ATP6V1B2 |
| 5.8564E-31 | -0.4361472 | 0.023 | 0.142 | 1.3835E-26 | MO_dmg | PLAC8 |
| 6.0034E-31 | -0.3028909 | 0.031 | 0.155 | 1.4182E-26 | MO_dmg | ITGA3 |
| 7.648E-31 | -1.316956 | 0.018 | 0.136 | 1.8067E-26 | MO_dmg | LYPD2 |
| 8.4552E-31 | 0.26917073 | 0.264 | 0.147 | 1.9974E-26 | MO_dmg | ASAP1 |
| 9.5464E-31 | 0.26151265 | 0.359 | 0.223 | 2.2552E-26 | MO_dmg | TPP1 |
| 9.6089E-31 | 0.53713261 | 0.465 | 0.319 | 2.2699E-26 | MO_dmg | VEGFA |
| 1.1319E-30 | -0.3749712 | 0.159 | 0.316 | 2.674E-26 | MO_dmg | SDC4 |
| 1.3401E-30 | 0.27802563 | 0.267 | 0.148 | 3.1657E-26 | MO_dmg | SLC44A1 |
| 1.4503E-30 | -0.4109373 | 0.493 | 0.625 | 3.4261E-26 | MO_dmg | TMEM59 |
| 1.5486E-30 | -0.7938326 | 0.119 | 0.266 | 3.6582E-26 | MO_dmg | AREG |
| 1.573E-30 | 0.27523317 | 0.253 | 0.136 | 3.716E-26 | MO_dmg | IFI30 |
| 1.7273E-30 | 0.28686129 | 0.119 | 0.047 | 4.0805E-26 | MO_dmg | HIF1A-AS2 |
| 2.598E-30 | -1.085654 | 0.01 | 0.12 | 6.1372E-26 | MO_dmg | SPINK5 |
| 2.6542E-30 | 0.3579363 | 0.389 | 0.256 | 6.2701E-26 | MO_dmg | TRA2B |
| 3.5494E-30 | -0.3536106 | 0.057 | 0.192 | 8.3847E-26 | MO_dmg | ANKRD36C |
| 3.9343E-30 | -0.3849014 | 0.014 | 0.126 | 9.2939E-26 | MO_dmg | TMEM40 |
| 4.0574E-30 | 0.26091522 | 0.277 | 0.157 | 9.5849E-26 | MO_dmg | CPD |
| 5.8098E-30 | 0.28455184 | 0.24 | 0.128 | 1.3724E-25 | MO_dmg | TENT5A |
| 6.0501E-30 | -0.3606471 | 0.288 | 0.452 | 1.4292E-25 | MO_dmg | ACTN4 |
| 6.0612E-30 | 0.28634357 | 0.24 | 0.128 | 1.4318E-25 | MO_dmg | CD83 |
| 6.5911E-30 | -0.3350479 | 0.023 | 0.14 | 1.557E-25 | MO_dmg | DCDC2 |
| 8.9771E-30 | -0.3171988 | 0.014 | 0.125 | 2.1207E-25 | MO_dmg | KRT80 |
| 9.384E-30 | 0.28703024 | 0.276 | 0.16 | 2.2168E-25 | MO_dmg | MTHFD2 |
| 9.6743E-30 | -0.2517366 | 0.018 | 0.132 | 2.2853E-25 | MO_dmg | KAZN |
| 9.7529E-30 | 0.3161884 | 0.329 | 0.202 | 2.3039E-25 | MO_dmg | UBE2S |
| 1.0551E-29 | -0.4298253 | 0.149 | 0.303 | 2.4924E-25 | MO_dmg | BTG2 |
| 1.0595E-29 | -1.0474962 | 0.02 | 0.135 | 2.503E-25 | MO_dmg | PLAT |
| 1.1971E-29 | 0.25362881 | 0.273 | 0.153 | 2.8279E-25 | MO_dmg | OGFRL1 |
| 1.2326E-29 | 0.26662464 | 0.197 | 0.098 | 2.9118E-25 | MO_dmg | LUCAT1 |
| 1.2546E-29 | 0.40415408 | 0.37 | 0.243 | 2.9638E-25 | MO_dmg | CLK1 |
| 1.4808E-29 | -0.5462304 | 0.116 | 0.257 | 3.498E-25 | MO_dmg | CARHSP1 |
| 1.7206E-29 | -0.3025678 | 0.026 | 0.143 | 4.0647E-25 | MO_dmg | EMP2 |
| 2.1282E-29 | -0.3372509 | 0.007 | 0.113 | 5.0275E-25 | MO_dmg | FXYD2 |
| 2.4345E-29 | -0.2514502 | 0.028 | 0.145 | 5.7509E-25 | MO_dmg | OCLN |

| | | | | | | |
|---|---|---|---|---|---|---|
| 2.6511E-29 | -0.2790636 | 0.045 | 0.171 | 6.2627E-25 | MO_dmg | SLC38A1 |
| 2.8895E-29 | 0.28917967 | 0.296 | 0.177 | 6.8258E-25 | MO_dmg | GNA13 |
| 3.4935E-29 | -0.2541909 | 0.041 | 0.165 | 8.2527E-25 | MO_dmg | GALNT3 |
| 8.3728E-29 | 0.28283802 | 0.541 | 0.38 | 1.9779E-24 | MO_dmg | TYMP |
| 8.9308E-29 | -0.5283008 | 0.49 | 0.609 | 2.1097E-24 | MO_dmg | EZR |
| 1.1652E-28 | -0.273354 | 0.013 | 0.121 | 2.7527E-24 | MO_dmg | ASS1 |
| 1.2525E-28 | 0.27032498 | 0.274 | 0.159 | 2.9588E-24 | MO_dmg | SLBP |
| 2.3224E-28 | -0.4126819 | 0.544 | 0.661 | 5.4863E-24 | MO_dmg | CALM2 |
| 3.3071E-28 | 0.29487413 | 0.955 | 0.888 | 7.8124E-24 | MO_dmg | HLA-B |
| 3.9337E-28 | 0.27998269 | 0.364 | 0.238 | 9.2926E-24 | MO_dmg | FXR1 |
| 5.5734E-28 | -0.4741665 | 0.081 | 0.212 | 1.3166E-23 | MO_dmg | CDKN2B |
| 6.1352E-28 | -0.2644768 | 0.01 | 0.114 | 1.4493E-23 | MO_dmg | FUT3 |
| 6.245E-28 | 0.33405641 | 0.328 | 0.208 | 1.4753E-23 | MO_dmg | CELF1 |
| 6.9104E-28 | -0.2794307 | 0.103 | 0.241 | 1.6324E-23 | MO_dmg | TSTD1 |
| 7.2926E-28 | -1.4855055 | 0.008 | 0.109 | 1.7227E-23 | MO_dmg | CRCT1 |
| 7.8807E-28 | -0.3761306 | 0.009 | 0.11 | 1.8617E-23 | MO_dmg | KRT23 |
| 9.385E-28 | -0.3164352 | 0.042 | 0.159 | 2.217E-23 | MO_dmg | COBLL1 |
| 9.4283E-28 | -0.2944014 | 0.01 | 0.113 | 2.2272E-23 | MO_dmg | STS |
| 1.1588E-27 | -0.2777161 | 0.073 | 0.202 | 2.7375E-23 | MO_dmg | TMEM106B |
| 1.1675E-27 | -0.8739285 | 0.061 | 0.184 | 2.758E-23 | MO_dmg | SPINK1 |
| 1.2278E-27 | -0.570632 | 0.012 | 0.115 | 2.9004E-23 | MO_dmg | KRT78 |
| 1.2471E-27 | -0.3263611 | 0.027 | 0.139 | 2.946E-23 | MO_dmg | ARHGAP29 |
| 1.3373E-27 | -0.3005152 | 0.142 | 0.288 | 3.1592E-23 | MO_dmg | PDCD6 |
| 1.735E-27 | 0.35575009 | 0.301 | 0.188 | 4.0986E-23 | MO_dmg | AHSA1 |
| 1.9013E-27 | 0.30147178 | 0.337 | 0.214 | 4.4915E-23 | MO_dmg | TRIM38 |
| 2.1423E-27 | -0.2648552 | 0.02 | 0.127 | 5.0607E-23 | MO_dmg | FAT1 |
| 2.2072E-27 | -0.3258573 | 0.032 | 0.146 | 5.214E-23 | MO_dmg | PAX8 |
| 2.2812E-27 | 0.261724 | 0.32 | 0.198 | 5.3889E-23 | MO_dmg | NR3C1 |
| 2.3408E-27 | 0.32169235 | 0.275 | 0.16 | 5.5297E-23 | MO_dmg | CLEC2B |
| 2.551E-27 | 0.360466 | 0.537 | 0.399 | 6.0263E-23 | MO_dmg | FLNA |
| 3.3749E-27 | -0.2568613 | 0.066 | 0.192 | 7.9725E-23 | MO_dmg | MPRIP |
| 4.6261E-27 | -0.2837898 | 0.023 | 0.131 | 1.0928E-22 | MO_dmg | FAM84A |
| 6.4703E-27 | -0.3055608 | 0.179 | 0.324 | 1.5285E-22 | MO_dmg | ERP29 |
| 6.4992E-27 | -0.3341187 | 0.012 | 0.113 | 1.5353E-22 | MO_dmg | SMIM5 |
| 7.284E-27 | -0.2853837 | 0.042 | 0.158 | 1.7207E-22 | MO_dmg | ITPKC |
| 7.9415E-27 | -0.4325412 | 0.014 | 0.115 | 1.876E-22 | MO_dmg | TMPRSS11A |
| 8.1633E-27 | 0.31684007 | 0.462 | 0.331 | 1.9284E-22 | MO_dmg | RBM25 |
| 8.1644E-27 | -0.4998288 | 0.954 | 0.944 | 1.9287E-22 | MO_dmg | MT-CO3 |
| 8.3095E-27 | -0.3469812 | 0.018 | 0.122 | 1.9629E-22 | MO_dmg | EGLN3 |
| 8.6249E-27 | 0.28779435 | 0.326 | 0.207 | 2.0375E-22 | MO_dmg | TAOK1 |
| 9.1843E-27 | -0.2798948 | 0.024 | 0.133 | 2.1696E-22 | MO_dmg | FOXC1 |
| 9.293E-27 | 0.30431563 | 0.392 | 0.27 | 2.1953E-22 | MO_dmg | ARL8B |
| 1.0209E-26 | 0.26819679 | 0.318 | 0.2 | 2.4116E-22 | MO_dmg | USP3 |
| 1.1652E-26 | -0.273612 | 0.144 | 0.286 | 2.7524E-22 | MO_dmg | EIF6 |
| 1.3531E-26 | 0.31039779 | 0.254 | 0.145 | 3.1965E-22 | MO_dmg | CDC42EP3 |
| 1.3721E-26 | -0.4011318 | 0.062 | 0.183 | 3.2413E-22 | MO_dmg | DST |
| 1.4884E-26 | -0.3834273 | 0.073 | 0.199 | 3.5161E-22 | MO_dmg | SAMD9 |
| 1.728E-26 | -0.3385892 | 0.016 | 0.117 | 4.0821E-22 | MO_dmg | CEACAM1 |
| 1.9885E-26 | -0.3111652 | 0.043 | 0.158 | 4.6973E-22 | MO_dmg | CYR61 |
| 1.9907E-26 | 0.30616919 | 0.219 | 0.12 | 4.7027E-22 | MO_dmg | MIS18BP1 |

| | | | | | | |
|---|---|---|---|---|---|---|
| 2.1791E-26 | -0.5223157 | 0.184 | 0.326 | 5.1477E-22 | MO_dmg | IFITM3 |
| 2.591E-26 | 0.33287777 | 0.31 | 0.199 | 6.1207E-22 | MO_dmg | STIP1 |
| 3.2076E-26 | 0.31611763 | 0.459 | 0.319 | 7.5774E-22 | MO_dmg | LIMS1 |
| 3.346E-26 | -0.3758962 | 0.029 | 0.137 | 7.9043E-22 | MO_dmg | UCHL1 |
| 3.6919E-26 | -0.2580042 | 0.102 | 0.239 | 8.7215E-22 | MO_dmg | PLXNB2 |
| 3.7587E-26 | 0.25796013 | 0.313 | 0.194 | 8.8792E-22 | MO_dmg | CHD1 |
| 4.0466E-26 | -0.5425798 | 0.893 | 0.875 | 9.5592E-22 | MO_dmg | RPL3 |
| 5.1064E-26 | -0.6102846 | 0.031 | 0.14 | 1.2063E-21 | MO_dmg | NUPR1 |
| 6.3671E-26 | 0.27359293 | 0.303 | 0.183 | 1.5041E-21 | MO_dmg | PTPRE |
| 7.0872E-26 | -0.2722588 | 0.075 | 0.2 | 1.6742E-21 | MO_dmg | NEDD4L |
| 7.4548E-26 | -0.3806656 | 0.185 | 0.33 | 1.761E-21 | MO_dmg | MAFF |
| 1.1709E-25 | -2.6587397 | 0.76 | 0.726 | 2.7659E-21 | MO_dmg | S100A9 |
| 1.1766E-25 | 0.34036873 | 0.394 | 0.279 | 2.7794E-21 | MO_dmg | RTN3 |
| 1.7973E-25 | 0.45064901 | 0.495 | 0.388 | 4.2457E-21 | MO_dmg | RSRC2 |
| 1.8054E-25 | -0.2512723 | 0.17 | 0.316 | 4.265E-21 | MO_dmg | CAPNS1 |
| 1.9552E-25 | 0.29968795 | 0.26 | 0.154 | 4.6188E-21 | MO_dmg | FNIP2 |
| 2.2381E-25 | 0.29417015 | 0.693 | 0.595 | 5.2871E-21 | MO_dmg | NPC2 |
| 2.2853E-25 | -0.2997031 | 0.012 | 0.108 | 5.3985E-21 | MO_dmg | A2ML1 |
| 2.4551E-25 | -0.5360024 | 0.027 | 0.131 | 5.7997E-21 | MO_dmg | TFPI2 |
| 2.6214E-25 | -1.4064118 | 0.014 | 0.11 | 6.1926E-21 | MO_dmg | CNFN |
| 3.3748E-25 | 0.36629358 | 0.385 | 0.268 | 7.9723E-21 | MO_dmg | OSBPL8 |
| 4.2462E-25 | -0.4272663 | 0.947 | 0.957 | 1.0031E-20 | MO_dmg | RPS18 |
| 4.6683E-25 | 0.36117991 | 0.426 | 0.304 | 1.1028E-20 | MO_dmg | NUFIP2 |
| 5.983E-25 | -0.3859062 | 0.014 | 0.109 | 1.4134E-20 | MO_dmg | TMC5 |
| 6.1645E-25 | -0.2765261 | 0.796 | 0.866 | 1.4562E-20 | MO_dmg | SRP14 |
| 6.4431E-25 | 0.29760701 | 0.462 | 0.335 | 1.5221E-20 | MO_dmg | GRB2 |
| 7.4914E-25 | -0.4049019 | 0.162 | 0.302 | 1.7697E-20 | MO_dmg | FAM129B |
| 7.7491E-25 | -0.4619384 | 0.009 | 0.1 | 1.8306E-20 | MO_dmg | DHRS2 |
| 8.9892E-25 | 0.32110771 | 0.767 | 0.688 | 2.1235E-20 | MO_dmg | PNRC1 |
| 1.0886E-24 | 0.29737907 | 0.304 | 0.194 | 2.5717E-20 | MO_dmg | CCPG1 |
| 1.2052E-24 | 0.40791829 | 0.748 | 0.686 | 2.8471E-20 | MO_dmg | PSMA7 |
| 1.2886E-24 | 0.25691332 | 0.288 | 0.176 | 3.044E-20 | MO_dmg | FMNL1 |
| 1.4106E-24 | -1.111982 | 0.018 | 0.115 | 3.3323E-20 | MO_dmg | SPRR1B |
| 1.4237E-24 | -0.4646756 | 0.024 | 0.125 | 3.3632E-20 | MO_dmg | AGR2 |
| 3.423E-24 | -0.3840346 | 0.028 | 0.127 | 8.0861E-20 | MO_dmg | SRD5A3 |
| 3.5688E-24 | 0.34046938 | 0.458 | 0.342 | 8.4307E-20 | MO_dmg | SCAF11 |
| 5.0448E-24 | 0.32767725 | 0.376 | 0.257 | 1.1917E-19 | MO_dmg | ARRDC3 |
| 6.1803E-24 | -0.2590565 | 0.103 | 0.228 | 1.46E-19 | MO_dmg | ST14 |
| 6.7949E-24 | 0.3546097 | 0.375 | 0.263 | 1.6052E-19 | MO_dmg | TCP1 |
| 7.4216E-24 | -0.3602928 | 0.014 | 0.106 | 1.7532E-19 | MO_dmg | CLIC3 |
| 7.6865E-24 | 0.54267598 | 0.746 | 0.687 | 1.8158E-19 | MO_dmg | DNAJA1 |
| 7.8054E-24 | 0.35636607 | 0.605 | 0.505 | 1.8439E-19 | MO_dmg | HNRNPDL |
| 7.889E-24 | 0.36228074 | 0.51 | 0.391 | 1.8636E-19 | MO_dmg | PNISR |
| 1.0513E-23 | 0.25416657 | 0.319 | 0.206 | 2.4836E-19 | MO_dmg | PNPLA8 |
| 1.1649E-23 | 0.2714177 | 0.287 | 0.178 | 2.7518E-19 | MO_dmg | ELL2 |
| 1.1893E-23 | 0.32923948 | 0.934 | 0.891 | 2.8094E-19 | MO_dmg | ATP5F1E |
| 1.8488E-23 | -0.3487359 | 0.04 | 0.144 | 4.3675E-19 | MO_dmg | ID1 |
| 1.9633E-23 | -0.3665486 | 0.199 | 0.334 | 4.6379E-19 | MO_dmg | PPP1R14B |
| 3.5065E-23 | -0.2830422 | 0.186 | 0.32 | 8.2835E-19 | MO_dmg | RAB11A |
| 4.0235E-23 | -0.3705716 | 0.562 | 0.66 | 9.5048E-19 | MO_dmg | NDUFA4 |

| | | | | | | |
|---|---|---|---|---|---|---|
| 4.9673E-23 | 0.29931723 | 0.904 | 0.829 | 1.1734E-18 | MO_dmg | HNRNPA2B1 |
| 5.1228E-23 | -0.6718921 | 0.04 | 0.142 | 1.2101E-18 | MO_dmg | CXCL1 |
| 5.2866E-23 | 0.4852097 | 0.291 | 0.182 | 1.2488E-18 | MO_dmg | KLF2 |
| 7.9447E-23 | 0.32627094 | 0.81 | 0.77 | 1.8768E-18 | MO_dmg | RHOA |
| 8.102E-23 | 0.30743556 | 0.408 | 0.285 | 1.9139E-18 | MO_dmg | ATP1B3 |
| 8.4271E-23 | 0.28535347 | 0.2 | 0.11 | 1.9907E-18 | MO_dmg | DNAJB4 |
| 8.708E-23 | -0.3608013 | 0.284 | 0.421 | 2.0571E-18 | MO_dmg | JPT1 |
| 1.002E-22 | -0.2600073 | 0.025 | 0.119 | 2.3671E-18 | MO_dmg | ST3GAL4 |
| 1.0107E-22 | -0.351187 | 0.142 | 0.265 | 2.3877E-18 | MO_dmg | RHOC |
| 1.0132E-22 | -0.4113411 | 0.26 | 0.389 | 2.3936E-18 | MO_dmg | C9orf16 |
| 1.1064E-22 | 0.27781957 | 0.33 | 0.222 | 2.6137E-18 | MO_dmg | TMED5 |
| 1.1325E-22 | 0.74399591 | 0.764 | 0.691 | 2.6754E-18 | MO_dmg | HSP90B1 |
| 1.3735E-22 | -0.3094807 | 0.062 | 0.172 | 3.2447E-18 | MO_dmg | AKR1B1 |
| 1.7325E-22 | 0.27896484 | 0.314 | 0.207 | 4.0928E-18 | MO_dmg | CYTH1 |
| 2.0076E-22 | -0.2535956 | 0.013 | 0.1 | 4.7424E-18 | MO_dmg | RNF39 |
| 3.0523E-22 | -0.6504426 | 0.059 | 0.164 | 7.2105E-18 | MO_dmg | APOBEC3A |
| 3.1593E-22 | -0.3154929 | 0.032 | 0.128 | 7.4633E-18 | MO_dmg | ID4 |
| 3.5432E-22 | -0.4439532 | 0.666 | 0.748 | 8.37E-18 | MO_dmg | PRDX1 |
| 3.6965E-22 | 0.26338746 | 0.335 | 0.224 | 8.7323E-18 | MO_dmg | IFNGR1 |
| 3.8037E-22 | 0.80222105 | 0.726 | 0.643 | 8.9855E-18 | MO_dmg | HSPA5 |
| 3.9432E-22 | 0.2521619 | 0.341 | 0.226 | 9.3151E-18 | MO_dmg | PHF20L1 |
| 4.0782E-22 | 0.32432401 | 0.587 | 0.491 | 9.634E-18 | MO_dmg | RSRP1 |
| 4.1946E-22 | -0.6827631 | 0.018 | 0.107 | 9.9089E-18 | MO_dmg | ZG16B |
| 4.5069E-22 | 0.25074539 | 0.346 | 0.236 | 1.0647E-17 | MO_dmg | NAA50 |
| 4.6488E-22 | -0.2730045 | 0.039 | 0.136 | 1.0982E-17 | MO_dmg | ACADM |
| 4.7711E-22 | -0.299432 | 0.236 | 0.372 | 1.1271E-17 | MO_dmg | TXNDC17 |
| 4.8168E-22 | -0.4122223 | 0.088 | 0.201 | 1.1379E-17 | MO_dmg | DUSP4 |
| 4.934E-22 | -0.3955401 | 0.362 | 0.478 | 1.1656E-17 | MO_dmg | CD81 |
| 5.0562E-22 | 0.26525918 | 0.389 | 0.273 | 1.1944E-17 | MO_dmg | ROCK1 |
| 5.4615E-22 | -0.2853156 | 0.402 | 0.536 | 1.2902E-17 | MO_dmg | COX6C |
| 5.7809E-22 | 0.32886962 | 0.406 | 0.289 | 1.3656E-17 | MO_dmg | SKIL |
| 7.3803E-22 | 0.26999227 | 0.336 | 0.221 | 1.7435E-17 | MO_dmg | KYNU |
| 7.807E-22 | 0.39702601 | 0.639 | 0.522 | 1.8443E-17 | MO_dmg | CEBPB |
| 8.0974E-22 | -0.6777822 | 0.197 | 0.315 | 1.9128E-17 | MO_dmg | EMP1 |
| 1.8384E-21 | -0.5662904 | 0.039 | 0.133 | 4.3428E-17 | MO_dmg | DHRS9 |
| 2.271E-21 | -0.4350815 | 0.031 | 0.122 | 5.3647E-17 | MO_dmg | TCIM |
| 2.4086E-21 | -0.2622546 | 0.035 | 0.13 | 5.6897E-17 | MO_dmg | LRG1 |
| 3.1481E-21 | -0.2512849 | 0.563 | 0.682 | 7.4367E-17 | MO_dmg | COX7A2 |
| 4.4314E-21 | -0.3898008 | 0.955 | 0.945 | 1.0468E-16 | MO_dmg | MT-CO2 |
| 6.253E-21 | 0.30941309 | 0.944 | 0.929 | 1.4771E-16 | MO_dmg | OAZ1 |
| 6.297E-21 | 0.39547622 | 0.454 | 0.354 | 1.4875E-16 | MO_dmg | TKT |
| 9.4996E-21 | -0.373866 | 0.378 | 0.496 | 2.2441E-16 | MO_dmg | H2AFJ |
| 9.7705E-21 | -0.2654149 | 0.142 | 0.262 | 2.3081E-16 | MO_dmg | ABRACL |
| 2.6975E-20 | -0.8880904 | 0.046 | 0.139 | 6.3724E-16 | MO_dmg | DEFB1 |
| 2.7798E-20 | -0.421277 | 0.063 | 0.161 | 6.5667E-16 | MO_dmg | BPGM |
| 6.4294E-20 | -0.4235513 | 0.04 | 0.132 | 1.5188E-15 | MO_dmg | AKAP12 |
| 7.9392E-20 | -0.2814723 | 0.249 | 0.372 | 1.8755E-15 | MO_dmg | NDUFV2 |
| 9.2231E-20 | -0.2968107 | 0.716 | 0.796 | 2.1788E-15 | MO_dmg | CHCHD2 |
| 1.071E-19 | -0.257261 | 0.554 | 0.681 | 2.5301E-15 | MO_dmg | IER2 |
| 1.1133E-19 | 0.28058621 | 0.186 | 0.104 | 2.63E-15 | MO_dmg | SLC16A10 |

| | | | | | | |
|---|---|---|---|---|---|---|
| 1.5587E-19 | 0.26988097 | 0.417 | 0.306 | 3.682E-15 | MO_dmg | QKI |
| 1.7726E-19 | -0.2562061 | 0.443 | 0.558 | 4.1874E-15 | MO_dmg | NDUFS5 |
| 2.095E-19 | 0.34891361 | 0.404 | 0.303 | 4.9491E-15 | MO_dmg | BNIP3L |
| 2.3278E-19 | -0.3526531 | 0.146 | 0.262 | 5.499E-15 | MO_dmg | PMAIP1 |
| 2.3594E-19 | 0.27868072 | 0.483 | 0.372 | 5.5736E-15 | MO_dmg | CCNL1 |
| 3.7316E-19 | -0.2544399 | 0.194 | 0.315 | 8.8152E-15 | MO_dmg | SBDS |
| 4.2367E-19 | -0.2519216 | 0.148 | 0.26 | 1.0008E-14 | MO_dmg | LYPLA1 |
| 4.5412E-19 | -0.2523266 | 0.205 | 0.331 | 1.0728E-14 | MO_dmg | LMNA |
| 6.7135E-19 | 0.30049351 | 0.384 | 0.286 | 1.5859E-14 | MO_dmg | RALA |
| 6.8004E-19 | -2.0653239 | 0.672 | 0.671 | 1.6065E-14 | MO_dmg | S100A8 |
| 1.041E-18 | 0.25800799 | 0.37 | 0.264 | 2.4591E-14 | MO_dmg | SMG1 |
| 1.092E-18 | -0.367003 | 0.076 | 0.173 | 2.5797E-14 | MO_dmg | PRDM1 |
| 1.1176E-18 | -0.2693959 | 0.075 | 0.175 | 2.6402E-14 | MO_dmg | STMN1 |
| 1.139E-18 | -0.3012389 | 0.373 | 0.485 | 2.6906E-14 | MO_dmg | GABARAPL2 |
| 1.2019E-18 | -0.3538487 | 0.218 | 0.328 | 2.8393E-14 | MO_dmg | ARHGDIB |
| 1.4443E-18 | -0.271695 | 0.11 | 0.218 | 3.412E-14 | MO_dmg | ARL4C |
| 1.6004E-18 | 0.40212961 | 0.348 | 0.248 | 3.7807E-14 | MO_dmg | SNHG12 |
| 1.7028E-18 | -0.2819509 | 0.102 | 0.207 | 4.0225E-14 | MO_dmg | CSNK1E |
| 3.4124E-18 | 0.81588116 | 0.585 | 0.522 | 8.0612E-14 | MO_dmg | HSPE1 |
| 3.4248E-18 | -0.2905233 | 0.245 | 0.357 | 8.0904E-14 | MO_dmg | MRPL33 |
| 3.5461E-18 | 0.27596193 | 0.377 | 0.274 | 8.3769E-14 | MO_dmg | TAF7 |
| 3.802E-18 | 0.29635086 | 0.483 | 0.382 | 8.9816E-14 | MO_dmg | HNRNPH1 |
| 4.5985E-18 | -0.3938277 | 0.13 | 0.231 | 1.0863E-13 | MO_dmg | CYB5A |
| 4.6418E-18 | 0.29296076 | 0.466 | 0.365 | 1.0965E-13 | MO_dmg | SPAG9 |
| 7.1738E-18 | -0.3830922 | 0.281 | 0.408 | 1.6947E-13 | MO_dmg | RNF213 |
| 8.0088E-18 | 0.26872485 | 0.369 | 0.271 | 1.8919E-13 | MO_dmg | SLTM |
| 9.9066E-18 | -1.0851315 | 0.178 | 0.283 | 2.3402E-13 | MO_dmg | MT1E |
| 1.0529E-17 | -0.2521155 | 0.384 | 0.505 | 2.4872E-13 | MO_dmg | LAPTM4A |
| 1.1681E-17 | -0.2781731 | 0.212 | 0.328 | 2.7595E-13 | MO_dmg | TMBIM1 |
| 1.1883E-17 | -0.6375523 | 0.206 | 0.315 | 2.807E-13 | MO_dmg | PHLDA1 |
| 1.2639E-17 | -0.2914497 | 0.141 | 0.248 | 2.9858E-13 | MO_dmg | PTMS |
| 1.661E-17 | -0.3247496 | 0.074 | 0.168 | 3.9237E-13 | MO_dmg | HIST3H2A |
| 1.75E-17 | 0.30772866 | 0.415 | 0.325 | 4.1341E-13 | MO_dmg | LAMP1 |
| 2.4527E-17 | 0.34732644 | 0.953 | 0.898 | 5.7941E-13 | MO_dmg | RPL39 |
| 2.4976E-17 | 0.46437125 | 0.915 | 0.882 | 5.9001E-13 | MO_dmg | CSTB |
| 2.9835E-17 | 0.3070313 | 0.574 | 0.485 | 7.0479E-13 | MO_dmg | ANKRD12 |
| 3.0603E-17 | 0.25699475 | 0.251 | 0.163 | 7.2293E-13 | MO_dmg | JARID2 |
| 3.616E-17 | 0.69437415 | 0.38 | 0.283 | 8.542E-13 | MO_dmg | HMOX1 |
| 3.7383E-17 | 0.29628983 | 0.409 | 0.308 | 8.831E-13 | MO_dmg | CTNNB1 |
| 4.4304E-17 | 0.3110029 | 0.338 | 0.248 | 1.0466E-12 | MO_dmg | DNAJC3 |
| 4.4498E-17 | -0.6279008 | 0.167 | 0.262 | 1.0512E-12 | MO_dmg | TPM1 |
| 5.9571E-17 | 0.31296352 | 0.509 | 0.426 | 1.4073E-12 | MO_dmg | IQGAP1 |
| 6.3453E-17 | -0.250715 | 0.279 | 0.398 | 1.4989E-12 | MO_dmg | CTNNA1 |
| 7.5573E-17 | -0.3483516 | 0.097 | 0.193 | 1.7853E-12 | MO_dmg | PLK2 |
| 1.1861E-16 | -0.3877065 | 0.342 | 0.452 | 2.8019E-12 | MO_dmg | MARCKS |
| 1.3124E-16 | 0.27601046 | 0.427 | 0.338 | 3.1002E-12 | MO_dmg | ANP32B |
| 1.5924E-16 | -0.3599127 | 0.379 | 0.488 | 3.7618E-12 | MO_dmg | HMGA1 |
| 1.6606E-16 | 0.31887048 | 0.51 | 0.413 | 3.9228E-12 | MO_dmg | FUS |
| 1.9045E-16 | 0.26870765 | 0.343 | 0.252 | 4.4991E-12 | MO_dmg | RPL22L1 |
| 2.0102E-16 | -0.6581428 | 0.031 | 0.106 | 4.7487E-12 | MO_dmg | CCL5 |

| | | | | | | |
|---|---|---|---|---|---|---|
| 2.0762E-16 | -0.2874628 | 0.717 | 0.764 | 4.9047E-12 | MO_dmg | COX7C |
| 2.6629E-16 | -0.3778757 | 0.032 | 0.106 | 6.2906E-12 | MO_dmg | LTB |
| 2.69E-16 | -0.28099 | 0.131 | 0.228 | 6.3546E-12 | MO_dmg | RAB9A |
| 2.7586E-16 | 0.35893088 | 0.582 | 0.519 | 6.5167E-12 | MO_dmg | ATP6V1F |
| 4.1935E-16 | -0.3140178 | 0.122 | 0.22 | 9.9063E-12 | MO_dmg | ZBTB20 |
| 4.5246E-16 | 0.30701242 | 0.3 | 0.213 | 1.0688E-11 | MO_dmg | MRPL18 |
| 7.2948E-16 | 0.27652765 | 0.471 | 0.382 | 1.7233E-11 | MO_dmg | DEK |
| 8.7025E-16 | -0.6580208 | 0.184 | 0.285 | 2.0558E-11 | MO_dmg | IFITM2 |
| 9.8857E-16 | -0.257705 | 0.05 | 0.128 | 2.3353E-11 | MO_dmg | POLB |
| 2.3797E-15 | -0.4537815 | 0.753 | 0.755 | 5.6216E-11 | MO_dmg | RPSA |
| 3.4895E-15 | 0.259564 | 0.437 | 0.349 | 8.2433E-11 | MO_dmg | SNX3 |
| 6.0861E-15 | -0.3999536 | 0.959 | 0.937 | 1.4377E-10 | MO_dmg | MT-ATP6 |
| 8.1299E-15 | 0.34718838 | 0.99 | 0.984 | 1.9205E-10 | MO_dmg | EIF1 |
| 1.014E-14 | -0.2559092 | 0.269 | 0.38 | 2.3955E-10 | MO_dmg | GCC2 |
| 1.3535E-14 | 0.25835063 | 0.302 | 0.215 | 3.1975E-10 | MO_dmg | DSE |
| 1.7546E-14 | 0.26664076 | 0.748 | 0.693 | 4.1448E-10 | MO_dmg | LRRFIP1 |
| 1.8324E-14 | -0.2598831 | 0.205 | 0.305 | 4.3286E-10 | MO_dmg | PSME2 |
| 2.009E-14 | -0.391647 | 0.252 | 0.349 | 4.7459E-10 | MO_dmg | ZFP36L2 |
| 2.0188E-14 | 0.30287861 | 0.45 | 0.367 | 4.7689E-10 | MO_dmg | PRDX6 |
| 2.0333E-14 | -0.4096697 | 0.552 | 0.644 | 4.8034E-10 | MO_dmg | IER3 |
| 2.4377E-14 | 0.27929464 | 0.518 | 0.453 | 5.7586E-10 | MO_dmg | ST13 |
| 2.6439E-14 | -0.2535073 | 0.278 | 0.377 | 6.2458E-10 | MO_dmg | PARK7 |
| 2.86E-14 | -0.3728453 | 0.794 | 0.781 | 6.7562E-10 | MO_dmg | RPL10A |
| 3.8292E-14 | -0.3484085 | 0.849 | 0.87 | 9.0458E-10 | MO_dmg | RPLP0 |
| 4.4821E-14 | -0.2722752 | 0.185 | 0.279 | 1.0588E-09 | MO_dmg | SLK |
| 6.8396E-14 | -0.2920511 | 0.183 | 0.272 | 1.6157E-09 | MO_dmg | FAM3C |
| 9.0483E-14 | -0.310511 | 0.166 | 0.261 | 2.1375E-09 | MO_dmg | IDI1 |
| 1.276E-13 | 0.25590807 | 0.538 | 0.468 | 3.0143E-09 | MO_dmg | RAP1B |
| 1.7431E-13 | -0.2534322 | 0.114 | 0.2 | 4.1176E-09 | MO_dmg | PTGR1 |
| 2.3134E-13 | -0.3058749 | 0.538 | 0.603 | 5.465E-09 | MO_dmg | HINT1 |
| 3.3228E-13 | 0.26475233 | 0.364 | 0.273 | 7.8495E-09 | MO_dmg | ETS2 |
| 4.3265E-13 | -0.5516694 | 0.223 | 0.311 | 1.022E-08 | MO_dmg | HIST1H4C |
| 4.3353E-13 | 0.25269453 | 0.83 | 0.762 | 1.0241E-08 | MO_dmg | BTG1 |
| 5.2193E-13 | -0.316699 | 0.639 | 0.7 | 1.233E-08 | MO_dmg | JUNB |
| 6.6307E-13 | -0.3400639 | 0.917 | 0.905 | 1.5664E-08 | MO_dmg | RPL7A |
| 7.3116E-13 | 0.29340835 | 0.431 | 0.363 | 1.7272E-08 | MO_dmg | ELOC |
| 8.7475E-13 | -0.3273713 | 0.302 | 0.201 | 2.0664E-08 | MO_dmg | HLA-DPA1 |
| 8.9659E-13 | 0.34145687 | 0.149 | 0.091 | 2.118E-08 | MO_dmg | SERPINH1 |
| 9.4339E-13 | 0.26319724 | 0.504 | 0.425 | 2.2286E-08 | MO_dmg | SEC62 |
| 1.2649E-12 | -0.6923053 | 0.064 | 0.135 | 2.9881E-08 | MO_dmg | SERPINB2 |
| 1.331E-12 | 0.32142885 | 0.962 | 0.931 | 3.1441E-08 | MO_dmg | GAPDH |
| 2.2783E-12 | -0.299408 | 0.896 | 0.876 | 5.382E-08 | MO_dmg | RPL23A |
| 2.3857E-12 | 0.29073957 | 0.175 | 0.111 | 5.6358E-08 | MO_dmg | RECQL |
| 3.6369E-12 | -0.2600556 | 0.06 | 0.13 | 8.5915E-08 | MO_dmg | GBP1 |
| 4.1954E-12 | -0.3203752 | 0.233 | 0.321 | 9.9107E-08 | MO_dmg | GADD45A |
| 5.3837E-12 | 0.31206603 | 0.995 | 0.98 | 1.2718E-07 | MO_dmg | MALAT1 |
| 6.6583E-12 | -0.4500468 | 0.816 | 0.833 | 1.5729E-07 | MO_dmg | PPDPF |
| 7.9801E-12 | -0.262424 | 0.803 | 0.835 | 1.8851E-07 | MO_dmg | PPIA |
| 9.8839E-12 | -0.3258553 | 0.455 | 0.538 | 2.3349E-07 | MO_dmg | SERPINB1 |
| 1.634E-11 | -0.3303667 | 0.366 | 0.439 | 3.86E-07 | MO_dmg | CYSTM1 |

| | | | | | | |
|---|---|---|---|---|---|---|
| 3.6052E-11 | -1.1926298 | 0.053 | 0.117 | 8.5165E-07 | MO_dmg | MT1M |
| 4.3214E-11 | -0.393822 | 0.056 | 0.118 | 1.0209E-06 | MO_dmg | HSPA2 |
| 4.5525E-11 | -0.2694342 | 0.452 | 0.532 | 1.0754E-06 | MO_dmg | PTTG1IP |
| 1.8953E-10 | 0.51542959 | 0.556 | 0.518 | 4.4774E-06 | MO_dmg | H1FX |
| 2.3701E-10 | -0.2750175 | 0.271 | 0.354 | 5.5988E-06 | MO_dmg | PHLDA2 |
| 3.0189E-10 | 0.25524629 | 0.656 | 0.61 | 7.1316E-06 | MO_dmg | SEC61B |
| 4.1121E-10 | -0.2828333 | 0.955 | 0.945 | 9.7139E-06 | MO_dmg | RPL13A |
| 4.1469E-10 | 0.73082755 | 0.847 | 0.832 | 9.7962E-06 | MO_dmg | UBB |
| 4.4283E-10 | -0.629857 | 0.148 | 0.217 | 1.0461E-05 | MO_dmg | S100A12 |
| 6.5319E-10 | -0.3378052 | 0.948 | 0.923 | 1.543E-05 | MO_dmg | RPS27A |
| 2.8313E-09 | -0.292141 | 0.899 | 0.87 | 6.6883E-05 | MO_dmg | RPL14 |
| 3.0133E-09 | -0.4178601 | 0.828 | 0.823 | 7.1184E-05 | MO_dmg | MT-ND5 |
| 5.4647E-09 | 0.32449997 | 0.25 | 0.193 | 0.00012909 | MO_dmg | SDF2L1 |
| 6.1178E-09 | -0.2637643 | 0.971 | 0.95 | 0.00014452 | MO_dmg | MT-CO1 |
| 6.5031E-09 | 0.25438296 | 0.437 | 0.387 | 0.00015362 | MO_dmg | HMGN2 |
| 1.3399E-08 | 0.28185207 | 0.355 | 0.299 | 0.00031653 | MO_dmg | SERTAD1 |
| 3.7716E-08 | -0.3272974 | 0.885 | 0.847 | 0.00089097 | MO_dmg | RPS4X |
| 3.8606E-08 | -0.28739 | 0.108 | 0.165 | 0.00091199 | MO_dmg | ANKRD37 |
| 4.0879E-08 | -0.2891789 | 0.809 | 0.8 | 0.00096567 | MO_dmg | RPS5 |
| 4.1225E-08 | 0.27529762 | 0.689 | 0.66 | 0.00097386 | MO_dmg | HNRNPU |
| 4.3826E-08 | 0.28179218 | 0.482 | 0.439 | 0.0010353 | MO_dmg | PDIA3 |
| 4.9127E-08 | -0.3053284 | 0.47 | 0.529 | 0.00116052 | MO_dmg | ZFP36 |
| 6.2585E-08 | -0.317725 | 0.166 | 0.232 | 0.00147846 | MO_dmg | ISG15 |
| 6.4392E-08 | -0.3091209 | 0.256 | 0.183 | 0.00152113 | MO_dmg | HLA-DPB1 |
| 1.3308E-07 | -0.7767899 | 0.06 | 0.109 | 0.00314381 | MO_dmg | CCL4 |
| 3.5088E-07 | -0.6780251 | 0.083 | 0.132 | 0.00828893 | MO_dmg | APOE |
| 4.0699E-07 | 0.38048952 | 0.375 | 0.328 | 0.00961436 | MO_dmg | HERPUD1 |
| 8.7687E-07 | -0.3626676 | 0.272 | 0.335 | 0.02071419 | MO_dmg | IL1RN |
| 1.091E-06 | -0.3543358 | 0.914 | 0.876 | 0.02577323 | MO_dmg | MT-ND3 |
| 1.7935E-06 | 0.28796829 | 0.403 | 0.359 | 0.0423682 | MO_dmg | CFLAR |
| 1.9398E-06 | -0.3612671 | 0.942 | 0.886 | 0.04582386 | MO_dmg | RPS6 |
| 2.4822E-06 | -1.129436 | 0.226 | 0.281 | 0.05863716 | MO_dmg | MT1X |
| 3.7824E-06 | -0.2546248 | 0.932 | 0.896 | 0.08935184 | MO_dmg | RPS3A |
| 4.9937E-06 | -0.2821829 | 0.168 | 0.121 | 0.11796542 | MO_dmg | CCL3 |
| 6.1702E-06 | -0.2755562 | 0.861 | 0.816 | 0.14575771 | MO_dmg | RPL5 |
| 3.1525E-05 | -0.263323 | 0.385 | 0.435 | 0.74472556 | MO_dmg | HIST1H1C |
| 3.3514E-05 | -0.4308739 | 0.115 | 0.157 | 0.79169631 | MO_dmg | MT1F |
| 5.8722E-05 | -0.2750453 | 0.927 | 0.886 | 1 | MO_dmg | RPS3 |
| 0.00010045 | -0.6928571 | 0.103 | 0.144 | 1 | MO_dmg | CCL2 |
| 0.00019748 | 0.27362218 | 0.464 | 0.446 | 1 | MO_dmg | DNAJB6 |
| 0.0002361 | -0.2502784 | 0.151 | 0.112 | 1 | MO_dmg | IL7R |
| 0.00039173 | 0.50271489 | 0.733 | 0.731 | 1 | MO_dmg | HSPA8 |
| 0.00043497 | -1.7071476 | 0.22 | 0.255 | 1 | MO_dmg | MT1G |
| 0.00065445 | -0.3097547 | 0.116 | 0.149 | 1 | MO_dmg | SMC4 |
| 0.00089267 | -0.3147935 | 0.881 | 0.867 | 1 | MO_dmg | ANXA2 |
| 0.00241275 | 0.27164042 | 0.655 | 0.627 | 1 | MO_dmg | JUN |
| 0.00474518 | -0.3085496 | 0.688 | 0.612 | 1 | MO_dmg | C15orf48 |
| 0.0054756 | 0.36986393 | 0.586 | 0.572 | 1 | MO_dmg | VMP1 |
| 0.00601473 | -1.0763867 | 0.654 | 0.659 | 1 | MO_dmg | MT2A |
| 0 | 4.03237524 | 1 | 0.444 | 0 | MO_SPP1+ | SPP1 |

| | | | | | | |
|---|---|---|---|---|---|---|
| 0 | 2.10645299 | 0.793 | 0.21 | 0 | MO_SPP1+ | ALOX5AP |
| 0 | 1.40017301 | 0.693 | 0.143 | 0 | MO_SPP1+ | CLEC5A |
| 5.655E-273 | 1.3679722 | 0.798 | 0.235 | 1.336E-268 | MO_SPP1+ | SLC11A1 |
| 1.559E-272 | 1.20388872 | 0.582 | 0.129 | 3.682E-268 | MO_SPP1+ | MCEMP1 |
| 1.881E-250 | 1.37098186 | 0.418 | 0.073 | 4.443E-246 | MO_SPP1+ | MRC1 |
| 1.505E-234 | 1.3051197 | 0.861 | 0.327 | 3.556E-230 | MO_SPP1+ | LAPTM5 |
| 1.575E-227 | 1.01663319 | 0.625 | 0.166 | 3.72E-223 | MO_SPP1+ | SMIM25 |
| 5.628E-227 | 0.76616757 | 0.399 | 0.07 | 1.329E-222 | MO_SPP1+ | FCGR2B |
| 1.269E-222 | 1.01265835 | 0.666 | 0.193 | 2.997E-218 | MO_SPP1+ | NCF2 |
| 8.695E-219 | 1.5645096 | 0.987 | 0.736 | 2.054E-214 | MO_SPP1+ | VIM |
| 3.875E-218 | 1.07894839 | 0.639 | 0.173 | 9.154E-214 | MO_SPP1+ | OLR1 |
| 1.922E-213 | 0.64130442 | 0.378 | 0.067 | 4.54E-209 | MO_SPP1+ | VSIG4 |
| 7.166E-213 | 1.30087595 | 0.909 | 0.377 | 1.693E-208 | MO_SPP1+ | FCER1G |
| 6.238E-207 | 1.47709028 | 0.594 | 0.175 | 1.474E-202 | MO_SPP1+ | FBP1 |
| 7.032E-206 | 1.28791159 | 0.939 | 0.432 | 1.661E-201 | MO_SPP1+ | TYROBP |
| 1.62E-205 | 1.23737483 | 0.645 | 0.193 | 3.826E-201 | MO_SPP1+ | CYBB |
| 1.001E-197 | 1.23456081 | 0.703 | 0.215 | 2.364E-193 | MO_SPP1+ | VCAN |
| 9.587E-193 | 0.7930917 | 0.541 | 0.143 | 2.265E-188 | MO_SPP1+ | PLXDC2 |
| 1.462E-191 | 0.97251381 | 0.487 | 0.12 | 3.455E-187 | MO_SPP1+ | MARCO |
| 4.361E-186 | 1.40025478 | 0.946 | 0.612 | 1.03E-181 | MO_SPP1+ | LGALS1 |
| 8.963E-185 | 0.98761992 | 0.703 | 0.228 | 2.117E-180 | MO_SPP1+ | BCL2A1 |
| 8.434E-183 | 1.29901649 | 0.984 | 0.857 | 1.992E-178 | MO_SPP1+ | SH3BGRL3 |
| 4.076E-177 | 1.10775666 | 0.859 | 0.426 | 9.629E-173 | MO_SPP1+ | GSTO1 |
| 4.518E-174 | 1.42178108 | 0.969 | 0.671 | 1.067E-169 | MO_SPP1+ | TIMP1 |
| 1.097E-168 | 1.24487658 | 0.963 | 0.734 | 2.591E-164 | MO_SPP1+ | CTSB |
| 6.248E-165 | 0.86606946 | 0.715 | 0.261 | 1.476E-160 | MO_SPP1+ | CD53 |
| 9.156E-161 | 1.0137912 | 0.969 | 0.564 | 2.163E-156 | MO_SPP1+ | SRGN |
| 4.21E-156 | 0.5779448 | 0.33 | 0.066 | 9.945E-152 | MO_SPP1+ | FLT1 |
| 1.594E-154 | 0.95920598 | 0.571 | 0.182 | 3.766E-150 | MO_SPP1+ | FCGR2A |
| 1.641E-152 | 1.70198264 | 0.325 | 0.069 | 3.877E-148 | MO_SPP1+ | FN1 |
| 1.307E-151 | 0.59725715 | 0.442 | 0.114 | 3.088E-147 | MO_SPP1+ | CSF1R |
| 2.166E-150 | 0.959727 | 0.561 | 0.177 | 5.118E-146 | MO_SPP1+ | IGSF6 |
| 1.459E-148 | 0.83335517 | 0.739 | 0.267 | 3.447E-144 | MO_SPP1+ | AIF1 |
| 4.463E-147 | 1.0268171 | 0.933 | 0.607 | 1.054E-142 | MO_SPP1+ | CD44 |
| 1.416E-146 | 0.49672969 | 0.429 | 0.11 | 3.346E-142 | MO_SPP1+ | ITGAM |
| 2.76E-146 | 0.91710088 | 0.967 | 0.663 | 6.52E-142 | MO_SPP1+ | ANXA5 |
| 2.574E-145 | 0.72178073 | 0.517 | 0.16 | 6.082E-141 | MO_SPP1+ | CEP170 |
| 1.359E-144 | 1.25205963 | 0.385 | 0.095 | 3.21E-140 | MO_SPP1+ | RETN |
| 1.477E-144 | 0.86869425 | 0.64 | 0.228 | 3.489E-140 | MO_SPP1+ | HCST |
| 3.358E-141 | 1.04813493 | 0.969 | 0.737 | 7.933E-137 | MO_SPP1+ | PSAP |
| 2.641E-140 | 1.05638826 | 0.834 | 0.418 | 6.239E-136 | MO_SPP1+ | EMP3 |
| 1.961E-138 | 0.70523721 | 0.612 | 0.209 | 4.633E-134 | MO_SPP1+ | SPI1 |
| 6.343E-137 | 0.96299562 | 0.993 | 0.918 | 1.498E-132 | MO_SPP1+ | S100A10 |
| 8.419E-137 | 0.75068185 | 0.739 | 0.284 | 1.989E-132 | MO_SPP1+ | LCP1 |
| 1.712E-135 | 0.70401057 | 0.597 | 0.202 | 4.044E-131 | MO_SPP1+ | LST1 |
| 1.023E-133 | 0.63177383 | 0.478 | 0.14 | 2.416E-129 | MO_SPP1+ | MS4A7 |
| 1.591E-132 | 0.63023765 | 0.892 | 0.457 | 3.758E-128 | MO_SPP1+ | LYZ |
| 1.298E-131 | 0.8548778 | 0.706 | 0.305 | 3.067E-127 | MO_SPP1+ | CD68 |
| 8.012E-130 | 0.69278084 | 0.606 | 0.213 | 1.893E-125 | MO_SPP1+ | ITGB2 |
| 1.788E-127 | 0.78355579 | 0.7 | 0.295 | 4.224E-123 | MO_SPP1+ | SERPINA1 |

| | | | | | | |
|---|---|---|---|---|---|---|
| 1.349E-125 | 0.94724383 | 0.877 | 0.593 | 3.187E-121 | MO_SPP1+ | NPC2 |
| 7.234E-124 | 0.47842591 | 0.51 | 0.155 | 1.709E-119 | MO_SPP1+ | AQP9 |
| 2.681E-123 | 0.65588518 | 0.799 | 0.39 | 6.333E-119 | MO_SPP1+ | HLA-DRA |
| 1.544E-121 | 0.54237486 | 0.154 | 0.02 | 3.648E-117 | MO_SPP1+ | LPL |
| 1.371E-120 | 1.18730493 | 1 | 0.986 | 3.238E-116 | MO_SPP1+ | FTL |
| 4.802E-120 | 0.56790288 | 0.465 | 0.144 | 1.134E-115 | MO_SPP1+ | BCAT1 |
| 2.725E-119 | 0.96455782 | 0.999 | 0.993 | 6.437E-115 | MO_SPP1+ | FTH1 |
| 4.015E-119 | 0.74414666 | 0.402 | 0.114 | 9.484E-115 | MO_SPP1+ | FCGR3A |
| 8.985E-118 | 1.01381386 | 0.937 | 0.724 | 2.122E-113 | MO_SPP1+ | CYBA |
| 5.963E-117 | 0.51306021 | 0.4 | 0.113 | 1.409E-112 | MO_SPP1+ | CD86 |
| 2.228E-114 | 0.7705049 | 0.645 | 0.271 | 5.263E-110 | MO_SPP1+ | HLA-DRB1 |
| 3.577E-114 | 0.60165244 | 0.504 | 0.172 | 8.449E-110 | MO_SPP1+ | CCDC88A |
| 6.189E-113 | 0.47034704 | 0.372 | 0.101 | 1.462E-108 | MO_SPP1+ | ENG |
| 5.112E-112 | 0.53498219 | 0.454 | 0.141 | 1.208E-107 | MO_SPP1+ | TREM1 |
| 1.879E-111 | 0.77841173 | 0.696 | 0.317 | 4.439E-107 | MO_SPP1+ | LIMS1 |
| 3.855E-110 | 0.81822561 | 0.997 | 0.931 | 9.106E-106 | MO_SPP1+ | GAPDH |
| 1.452E-109 | 0.7338166 | 0.496 | 0.181 | 3.43E-105 | MO_SPP1+ | CKLF |
| 1.823E-109 | 0.59624163 | 0.508 | 0.171 | 4.306E-105 | MO_SPP1+ | ANPEP |
| 9.488E-107 | 0.93735646 | 0.564 | 0.214 | 2.241E-102 | MO_SPP1+ | CD14 |
| 1.001E-106 | 0.72468961 | 0.576 | 0.232 | 2.366E-102 | MO_SPP1+ | RNASET2 |
| 6.885E-105 | 0.62169144 | 0.607 | 0.24 | 1.626E-100 | MO_SPP1+ | GMFG |
| 8.144E-105 | 0.77090483 | 0.924 | 0.723 | 1.924E-100 | MO_SPP1+ | ENO1 |
| 1.067E-104 | 0.70184874 | 0.583 | 0.222 | 2.52E-100 | MO_SPP1+ | ZEB2 |
| 7.779E-99 | 0.93545927 | 0.576 | 0.228 | 1.8377E-94 | MO_SPP1+ | SLC2A3 |
| 4.6452E-97 | -1.8079815 | 0.18 | 0.561 | 1.0973E-92 | MO_SPP1+ | ELF3 |
| 2.344E-96 | 0.7056569 | 0.876 | 0.533 | 5.5373E-92 | MO_SPP1+ | PLAUR |
| 5.3707E-95 | -2.0892353 | 0.223 | 0.584 | 1.2687E-90 | MO_SPP1+ | KRT19 |
| 8.7901E-95 | 0.42230264 | 0.355 | 0.107 | 2.0765E-90 | MO_SPP1+ | COLGALT1 |
| 2.0205E-93 | 0.4837743 | 0.433 | 0.147 | 4.773E-89 | MO_SPP1+ | EVI2B |
| 2.5592E-93 | 0.56992943 | 0.217 | 0.046 | 6.0456E-89 | MO_SPP1+ | MMP9 |
| 5.9778E-93 | -1.6474012 | 0.208 | 0.574 | 1.4121E-88 | MO_SPP1+ | TACSTD2 |
| 1.3476E-92 | 0.75553145 | 0.979 | 0.891 | 3.1835E-88 | MO_SPP1+ | ATP5F1E |
| 3.9449E-92 | 0.6463468 | 0.558 | 0.238 | 9.319E-88 | MO_SPP1+ | RBPJ |
| 6.6717E-92 | 0.96637645 | 0.889 | 0.609 | 1.5761E-87 | MO_SPP1+ | C15orf48 |
| 1.3262E-91 | 0.44536285 | 0.312 | 0.086 | 3.1329E-87 | MO_SPP1+ | RNASE6 |
| 5.463E-91 | 0.44343429 | 0.373 | 0.118 | 1.2905E-86 | MO_SPP1+ | DOCK10 |
| 2.2018E-90 | -1.749462 | 0.192 | 0.564 | 5.2012E-86 | MO_SPP1+ | KRT18 |
| 5.8899E-90 | 0.55192879 | 0.495 | 0.194 | 1.3914E-85 | MO_SPP1+ | ARHGAP18 |
| 2.644E-89 | 0.41639001 | 0.252 | 0.062 | 6.2459E-85 | MO_SPP1+ | FPR3 |
| 6.6082E-89 | 0.42691476 | 0.339 | 0.101 | 1.561E-84 | MO_SPP1+ | GPRIN3 |
| 2.9778E-88 | 0.52773825 | 0.531 | 0.216 | 7.0343E-84 | MO_SPP1+ | ACTN1 |
| 4.2332E-88 | 0.98610378 | 0.534 | 0.229 | 1E-83 | MO_SPP1+ | PLIN2 |
| 9.5496E-88 | 0.70059308 | 0.922 | 0.739 | 2.2559E-83 | MO_SPP1+ | APLP2 |
| 9.6315E-88 | 0.74238716 | 0.958 | 0.813 | 2.2752E-83 | MO_SPP1+ | YBX1 |
| 5.1837E-87 | 0.73391166 | 0.96 | 0.788 | 1.2246E-82 | MO_SPP1+ | PFN1 |
| 1.8741E-86 | -1.3625402 | 0.177 | 0.546 | 4.4272E-82 | MO_SPP1+ | CD24 |
| 2.1633E-86 | 0.49784723 | 0.357 | 0.112 | 5.1105E-82 | MO_SPP1+ | CD163 |
| 5.1823E-86 | -1.6732142 | 0.192 | 0.55 | 1.2242E-81 | MO_SPP1+ | KRT7 |
| 1.657E-85 | -1.7514025 | 0.198 | 0.545 | 3.9142E-81 | MO_SPP1+ | CLDN4 |
| 1.3932E-84 | 0.33928723 | 0.162 | 0.03 | 3.2912E-80 | MO_SPP1+ | MATK |

| | | | | | | |
|---|---|---|---|---|---|---|
| 2.3659E-84 | 0.53380174 | 0.436 | 0.163 | 5.5891E-80 | MO_SPP1+ | GLIPR2 |
| 4.6838E-84 | 0.45060969 | 0.798 | 0.486 | 1.1065E-79 | MO_SPP1+ | CD74 |
| 6.6E-84 | 0.75238525 | 0.994 | 0.937 | 1.5591E-79 | MO_SPP1+ | TMSB4X |
| 1.7646E-83 | 0.54443965 | 0.45 | 0.171 | 4.1685E-79 | MO_SPP1+ | P4HA1 |
| 3.7667E-83 | 0.60014999 | 0.45 | 0.166 | 8.8981E-79 | MO_SPP1+ | TGFBI |
| 4.4335E-83 | 0.51925329 | 0.501 | 0.198 | 1.0473E-78 | MO_SPP1+ | LSP1 |
| 1.1709E-82 | 0.40927083 | 0.324 | 0.098 | 2.766E-78 | MO_SPP1+ | SLC8A1 |
| 1.2966E-82 | 0.52192231 | 0.45 | 0.17 | 3.0629E-78 | MO_SPP1+ | KCTD12 |
| 3.7259E-82 | 0.40838585 | 0.393 | 0.133 | 8.8017E-78 | MO_SPP1+ | CLEC7A |
| 3.7776E-82 | 0.94405196 | 0.663 | 0.34 | 8.9238E-78 | MO_SPP1+ | CTSL |
| 6.8959E-82 | 0.99885174 | 0.886 | 0.621 | 1.629E-77 | MO_SPP1+ | S100A4 |
| 2.8448E-81 | 0.57252862 | 0.448 | 0.174 | 6.7202E-77 | MO_SPP1+ | LTA4H |
| 8.1836E-81 | 0.47554676 | 0.402 | 0.14 | 1.9332E-76 | MO_SPP1+ | TLR2 |
| 9.661E-81 | 0.39202035 | 0.391 | 0.133 | 2.2822E-76 | MO_SPP1+ | EVI2A |
| 6.896E-80 | 0.36787047 | 0.349 | 0.112 | 1.629E-75 | MO_SPP1+ | ADGRE2 |
| 1.2181E-79 | 0.59263607 | 0.711 | 0.362 | 2.8776E-75 | MO_SPP1+ | COTL1 |
| 3.3203E-79 | 0.64162238 | 0.375 | 0.128 | 7.8436E-75 | MO_SPP1+ | RGCC |
| 2.9567E-78 | 0.57113941 | 0.414 | 0.158 | 6.9845E-74 | MO_SPP1+ | ALCAM |
| 5.3368E-78 | 0.56567017 | 0.988 | 0.888 | 1.2607E-73 | MO_SPP1+ | HLA-B |
| 2.0875E-77 | 0.60574953 | 0.927 | 0.725 | 4.9312E-73 | MO_SPP1+ | PKM |
| 3.4631E-77 | 0.51681783 | 0.436 | 0.164 | 8.1809E-73 | MO_SPP1+ | SLA |
| 3.8804E-77 | 0.52576908 | 0.466 | 0.189 | 9.1666E-73 | MO_SPP1+ | SNX10 |
| 9.3676E-77 | 0.6891163 | 0.904 | 0.705 | 2.2129E-72 | MO_SPP1+ | GPX4 |
| 1.6885E-76 | 0.74798684 | 0.871 | 0.643 | 3.9888E-72 | MO_SPP1+ | LDHA |
| 1.9193E-75 | 0.4984045 | 0.456 | 0.184 | 4.534E-71 | MO_SPP1+ | PYCARD |
| 2.6898E-75 | 0.26765775 | 0.216 | 0.053 | 6.3542E-71 | MO_SPP1+ | PARVB |
| 5.1439E-75 | 0.28156634 | 0.226 | 0.058 | 1.2151E-70 | MO_SPP1+ | MOB3B |
| 9.1823E-75 | 0.56007048 | 1 | 0.964 | 2.1691E-70 | MO_SPP1+ | S100A11 |
| 1.4274E-74 | 0.63826066 | 0.604 | 0.315 | 3.3719E-70 | MO_SPP1+ | AP2S1 |
| 3.0909E-74 | 0.35546106 | 0.31 | 0.097 | 7.3017E-70 | MO_SPP1+ | TFEC |
| 3.0914E-74 | 0.29015201 | 0.226 | 0.058 | 7.3028E-70 | MO_SPP1+ | NEK6 |
| 4.5345E-74 | 0.47123613 | 0.492 | 0.195 | 1.0712E-69 | MO_SPP1+ | ITGAX |
| 9.332E-74 | 0.48850034 | 0.511 | 0.218 | 2.2045E-69 | MO_SPP1+ | GRINA |
| 1.2904E-73 | -2.2337267 | 0.217 | 0.541 | 3.0484E-69 | MO_SPP1+ | WFDC2 |
| 1.3898E-73 | -1.0461722 | 0.115 | 0.457 | 3.283E-69 | MO_SPP1+ | CLDN7 |
| 1.6505E-73 | 0.46828481 | 0.462 | 0.182 | 3.899E-69 | MO_SPP1+ | PTPRE |
| 1.7449E-73 | 0.55569968 | 0.988 | 0.954 | 4.1219E-69 | MO_SPP1+ | SERF2 |
| 2.1041E-73 | 0.4451497 | 0.436 | 0.167 | 4.9704E-69 | MO_SPP1+ | SAMHD1 |
| 4.9512E-73 | 0.60462331 | 0.861 | 0.636 | 1.1696E-68 | MO_SPP1+ | ARPC5 |
| 5.3989E-73 | 0.43984599 | 0.46 | 0.183 | 1.2754E-68 | MO_SPP1+ | RAC2 |
| 1.2972E-72 | 0.58838978 | 0.928 | 0.797 | 3.0644E-68 | MO_SPP1+ | ARPC2 |
| 1.9124E-72 | -1.0826477 | 0.099 | 0.442 | 4.5176E-68 | MO_SPP1+ | ATP1B1 |
| 2.6638E-72 | 0.40184366 | 0.373 | 0.132 | 6.2926E-68 | MO_SPP1+ | HCK |
| 3.4483E-72 | 0.95839869 | 0.505 | 0.232 | 8.146E-68 | MO_SPP1+ | FABP5 |
| 1.6293E-71 | 0.55784642 | 0.382 | 0.141 | 3.8489E-67 | MO_SPP1+ | SDC2 |
| 3.2261E-71 | 0.56592631 | 0.471 | 0.199 | 7.6209E-67 | MO_SPP1+ | CTSC |
| 4.0739E-71 | 0.80814361 | 0.577 | 0.281 | 9.6238E-67 | MO_SPP1+ | SLC25A37 |
| 2.629E-70 | 0.4764738 | 0.379 | 0.144 | 6.2104E-66 | MO_SPP1+ | NME1 |
| 3.2776E-70 | 0.57864043 | 0.987 | 0.916 | 7.7428E-66 | MO_SPP1+ | UBA52 |
| 4.0307E-70 | -3.5058032 | 0.217 | 0.528 | 9.5217E-66 | MO_SPP1+ | LCN2 |

| | | | | | | |
|---|---|---|---|---|---|---|
| 4.2498E-70 | -1.49607 | 0.178 | 0.498 | 1.0039E-65 | MO_SPP1+ | KRT8 |
| 2.7654E-69 | 0.33764986 | 0.324 | 0.109 | 6.5328E-65 | MO_SPP1+ | SLC43A3 |
| 3.8938E-69 | 0.45046809 | 0.832 | 0.534 | 9.1984E-65 | MO_SPP1+ | UPP1 |
| 6.2736E-69 | 0.38935428 | 0.451 | 0.183 | 1.482E-64 | MO_SPP1+ | RNF130 |
| 1.1933E-67 | 0.43378349 | 0.159 | 0.033 | 2.819E-63 | MO_SPP1+ | CCL7 |
| 3.3501E-67 | 0.35607432 | 0.375 | 0.137 | 7.9139E-63 | MO_SPP1+ | LRP1 |
| 4.2521E-67 | 0.43080481 | 0.391 | 0.153 | 1.0045E-62 | MO_SPP1+ | HNMT |
| 7.4246E-67 | 0.42471377 | 0.405 | 0.158 | 1.7539E-62 | MO_SPP1+ | FERMT3 |
| 1.2358E-66 | 0.33924487 | 0.342 | 0.12 | 2.9193E-62 | MO_SPP1+ | DPYD |
| 1.8451E-66 | 0.58725289 | 0.466 | 0.194 | 4.3588E-62 | MO_SPP1+ | CD52 |
| 5.2719E-66 | 0.54571119 | 0.7 | 0.379 | 1.2454E-61 | MO_SPP1+ | TYMP |
| 5.5898E-66 | 0.49031421 | 0.477 | 0.211 | 1.3205E-61 | MO_SPP1+ | EML4 |
| 7.2591E-66 | 0.63893603 | 0.766 | 0.516 | 1.7148E-61 | MO_SPP1+ | ATP6V1F |
| 7.75E-66 | 0.65951096 | 0.871 | 0.685 | 1.8308E-61 | MO_SPP1+ | BRI3 |
| 1.6132E-65 | 0.3921952 | 0.361 | 0.133 | 3.8108E-61 | MO_SPP1+ | ITGA5 |
| 2.8655E-65 | 0.48753975 | 0.52 | 0.239 | 6.7691E-61 | MO_SPP1+ | RAB31 |
| 4.0276E-65 | 0.47352087 | 0.489 | 0.222 | 9.5144E-61 | MO_SPP1+ | TPP1 |
| 3.2604E-64 | 0.53249724 | 0.901 | 0.769 | 7.702E-60 | MO_SPP1+ | RHOA |
| 5.9996E-64 | 0.60507286 | 0.922 | 0.764 | 1.4173E-59 | MO_SPP1+ | CST3 |
| 8.1772E-64 | 0.59863495 | 0.595 | 0.307 | 1.9317E-59 | MO_SPP1+ | FNDC3B |
| 8.7627E-63 | 0.41129571 | 0.495 | 0.222 | 2.07E-58 | MO_SPP1+ | RGS10 |
| 9.3154E-63 | 0.53738621 | 0.651 | 0.356 | 2.2006E-58 | MO_SPP1+ | GNAI2 |
| 1.545E-62 | 0.51065587 | 0.538 | 0.271 | 3.6498E-58 | MO_SPP1+ | PDXK |
| 2.5218E-62 | 0.44709198 | 0.39 | 0.156 | 5.9572E-58 | MO_SPP1+ | AP1S2 |
| 3.1832E-62 | 0.56646714 | 0.67 | 0.392 | 7.5197E-58 | MO_SPP1+ | H2AFY |
| 5.461E-62 | 0.57019957 | 0.696 | 0.398 | 1.29E-57 | MO_SPP1+ | FLNA |
| 7.7507E-62 | 0.61155394 | 0.901 | 0.773 | 1.8309E-57 | MO_SPP1+ | TPI1 |
| 9.4223E-62 | 0.42253966 | 0.426 | 0.18 | 2.2258E-57 | MO_SPP1+ | DAB2 |
| 1.5349E-61 | 0.37398954 | 0.402 | 0.163 | 3.6259E-57 | MO_SPP1+ | SIRPA |
| 2.7628E-61 | 0.59350586 | 0.639 | 0.353 | 6.5265E-57 | MO_SPP1+ | MIF |
| 8.1768E-61 | 0.28659036 | 0.282 | 0.093 | 1.9316E-56 | MO_SPP1+ | LAT2 |
| 1.3222E-60 | 0.32669739 | 0.42 | 0.165 | 3.1235E-56 | MO_SPP1+ | FCN1 |
| 4.9351E-60 | 0.39714438 | 0.382 | 0.155 | 1.1658E-55 | MO_SPP1+ | PPT1 |
| 5.4864E-60 | 0.3052695 | 0.204 | 0.056 | 1.2961E-55 | MO_SPP1+ | IL3RA |
| 8.1295E-60 | 0.50936404 | 0.595 | 0.315 | 1.9204E-55 | MO_SPP1+ | SLC16A3 |
| 1.0594E-59 | -0.8785586 | 0.03 | 0.333 | 2.5026E-55 | MO_SPP1+ | MYO6 |
| 1.9974E-59 | 0.64852003 | 0.771 | 0.521 | 4.7184E-55 | MO_SPP1+ | CEBPB |
| 2.2227E-59 | 0.63909571 | 0.982 | 0.89 | 5.2507E-55 | MO_SPP1+ | CD63 |
| 3.1176E-59 | 0.54091927 | 0.549 | 0.279 | 7.3647E-55 | MO_SPP1+ | CTSZ |
| 5.0803E-59 | -0.7506378 | 0.024 | 0.325 | 1.2001E-54 | MO_SPP1+ | PERP |
| 2.8683E-58 | 0.57951976 | 0.957 | 0.929 | 6.7757E-54 | MO_SPP1+ | OAZ1 |
| 3.3187E-58 | 0.35577747 | 0.42 | 0.175 | 7.8398E-54 | MO_SPP1+ | FMNL1 |
| 1.8609E-57 | 0.47252614 | 0.54 | 0.272 | 4.3961E-53 | MO_SPP1+ | VASP |
| 2.1167E-57 | -0.8098014 | 0.133 | 0.432 | 5.0004E-53 | MO_SPP1+ | SPINT2 |
| 4.3321E-57 | 0.35614163 | 0.367 | 0.146 | 1.0234E-52 | MO_SPP1+ | ASAP1 |
| 2.7832E-56 | 0.39797976 | 0.328 | 0.126 | 6.5748E-52 | MO_SPP1+ | ARRDC4 |
| 3.7642E-56 | 0.71349469 | 0.84 | 0.638 | 8.8921E-52 | MO_SPP1+ | TXNIP |
| 4.3269E-56 | 0.53407899 | 0.994 | 0.944 | 1.0221E-51 | MO_SPP1+ | PTMA |
| 8.2196E-56 | 0.55617909 | 0.796 | 0.58 | 1.9417E-51 | MO_SPP1+ | NOP10 |
| 1.3002E-55 | 0.46906348 | 0.471 | 0.227 | 3.0714E-51 | MO_SPP1+ | ATP6V1B2 |

| | | | | | | |
|---|---|---|---|---|---|---|
| 1.3686E-55 | 0.34082055 | 0.318 | 0.118 | 3.233E-51 | MO_SPP1+ | PTAFR |
| 2.0947E-55 | 0.32779768 | 0.342 | 0.131 | 4.9484E-51 | MO_SPP1+ | FGR |
| 3.9228E-55 | -1.1751132 | 0.015 | 0.296 | 9.2669E-51 | MO_SPP1+ | RND3 |
| 4.2176E-55 | 0.41886826 | 0.474 | 0.224 | 9.9632E-51 | MO_SPP1+ | FCGRT |
| 6.9446E-55 | 0.36997666 | 0.361 | 0.142 | 1.6405E-50 | MO_SPP1+ | FPR1 |
| 1.0045E-54 | -1.5450597 | 0.021 | 0.3 | 2.373E-50 | MO_SPP1+ | ADIRF |
| 1.8757E-54 | 0.42772945 | 0.553 | 0.284 | 4.4309E-50 | MO_SPP1+ | ATP1B3 |
| 6.5947E-54 | -1.0173536 | 0.019 | 0.296 | 1.5579E-49 | MO_SPP1+ | GPRC5A |
| 6.9474E-54 | 0.50046892 | 0.999 | 0.97 | 1.6412E-49 | MO_SPP1+ | TMSB10 |
| 7.2702E-54 | 0.54881702 | 0.618 | 0.363 | 1.7174E-49 | MO_SPP1+ | CTSH |
| 8.3524E-54 | -0.7128342 | 0.375 | 0.628 | 1.9731E-49 | MO_SPP1+ | DSTN |
| 1.0166E-53 | 0.42929736 | 0.459 | 0.213 | 2.4016E-49 | MO_SPP1+ | DSE |
| 1.1899E-53 | -1.6613316 | 0.093 | 0.373 | 2.8109E-49 | MO_SPP1+ | MAL |
| 3.445E-53 | 0.40896823 | 0.289 | 0.104 | 8.1381E-49 | MO_SPP1+ | SLC16A10 |
| 4.6171E-53 | 0.26514458 | 0.237 | 0.077 | 1.0907E-48 | MO_SPP1+ | CHST11 |
| 5.0518E-53 | -1.5776028 | 0.135 | 0.41 | 1.1934E-48 | MO_SPP1+ | SLPI |
| 1.2152E-52 | 0.42990939 | 0.465 | 0.229 | 2.8707E-48 | MO_SPP1+ | M6PR |
| 2.8176E-52 | 0.284606 | 0.361 | 0.141 | 6.656E-48 | MO_SPP1+ | CD93 |
| 2.9569E-52 | 0.32658817 | 0.315 | 0.122 | 6.9852E-48 | MO_SPP1+ | MFSD1 |
| 4.3232E-52 | 0.50351324 | 0.747 | 0.504 | 1.0213E-47 | MO_SPP1+ | DBI |
| 5.4496E-52 | 0.47419493 | 0.439 | 0.203 | 1.2874E-47 | MO_SPP1+ | SLC43A2 |
| 2.6141E-51 | 0.49891988 | 0.456 | 0.22 | 6.1753E-47 | MO_SPP1+ | STXBP2 |
| 2.9552E-51 | 0.36994131 | 0.333 | 0.132 | 6.9811E-47 | MO_SPP1+ | DDX60L |
| 3.1419E-51 | 0.35493574 | 0.364 | 0.153 | 7.4221E-47 | MO_SPP1+ | OGFRL1 |
| 3.5118E-51 | 0.26236849 | 0.195 | 0.057 | 8.2959E-47 | MO_SPP1+ | SPRED1 |
| 6.0189E-51 | -0.6140244 | 0.013 | 0.278 | 1.4218E-46 | MO_SPP1+ | JUP |
| 6.2525E-51 | 0.46107058 | 0.826 | 0.621 | 1.477E-46 | MO_SPP1+ | ACTR2 |
| 1.3792E-50 | 0.46971652 | 0.549 | 0.292 | 3.2581E-46 | MO_SPP1+ | IFNGR2 |
| 2.537E-50 | 0.32136281 | 0.321 | 0.124 | 5.9931E-46 | MO_SPP1+ | IRAK3 |
| 2.5604E-50 | 0.39060123 | 0.346 | 0.142 | 6.0484E-46 | MO_SPP1+ | C1orf162 |
| 1.0834E-49 | 0.32044836 | 0.31 | 0.122 | 2.5593E-45 | MO_SPP1+ | SYK |
| 1.6019E-49 | 0.64427942 | 0.999 | 0.973 | 3.7841E-45 | MO_SPP1+ | ACTB |
| 2.5764E-49 | -1.3323091 | 0.078 | 0.349 | 6.0862E-45 | MO_SPP1+ | GDF15 |
| 3.2853E-49 | 0.31408269 | 0.376 | 0.16 | 7.7608E-45 | MO_SPP1+ | MYO1F |
| 4.2519E-49 | 0.87516667 | 0.892 | 0.773 | 1.0044E-44 | MO_SPP1+ | CTSD |
| 9.0725E-49 | 0.2503059 | 0.255 | 0.089 | 2.1432E-44 | MO_SPP1+ | GNB4 |
| 1.0099E-48 | 0.30964902 | 0.784 | 0.527 | 2.3857E-44 | MO_SPP1+ | CTSS |
| 1.36E-48 | -1.2018143 | 0.013 | 0.267 | 3.2127E-44 | MO_SPP1+ | LMO7 |
| 1.5476E-48 | 0.50285748 | 0.781 | 0.577 | 3.656E-44 | MO_SPP1+ | PGK1 |
| 1.9811E-48 | 0.25437215 | 0.19 | 0.057 | 4.68E-44 | MO_SPP1+ | CLEC4A |
| 3.924E-48 | -0.7628598 | 0.012 | 0.264 | 9.2697E-44 | MO_SPP1+ | MUC1 |
| 1.6585E-47 | -1.4801454 | 0.201 | 0.455 | 3.9179E-43 | MO_SPP1+ | SOX4 |
| 1.7267E-47 | -0.7114383 | 0.142 | 0.413 | 4.079E-43 | MO_SPP1+ | CD59 |
| 1.7959E-47 | -2.4272043 | 0.169 | 0.411 | 4.2424E-43 | MO_SPP1+ | S100P |
| 1.975E-47 | -0.8005411 | 0.025 | 0.279 | 4.6656E-43 | MO_SPP1+ | CAMK2N1 |
| 9.1818E-47 | 0.3104929 | 0.409 | 0.184 | 2.169E-42 | MO_SPP1+ | HCLS1 |
| 2.473E-46 | -0.9964993 | 0.021 | 0.268 | 5.8421E-42 | MO_SPP1+ | TSPAN1 |
| 4.4562E-46 | 0.44439447 | 0.949 | 0.813 | 1.0527E-41 | MO_SPP1+ | PFDN5 |
| 4.9699E-46 | 0.51117086 | 0.849 | 0.707 | 1.174E-41 | MO_SPP1+ | SDCBP |
| 5.3323E-46 | 0.28606385 | 0.291 | 0.112 | 1.2596E-41 | MO_SPP1+ | C3AR1 |

| | | | | | | |
|---|---|---|---|---|---|---|
| 6.763E-46 | 0.31148699 | 0.321 | 0.134 | 1.5976E-41 | MO_SPP1+ | LSM6 |
| 8.4932E-46 | 0.45138986 | 0.874 | 0.701 | 2.0063E-41 | MO_SPP1+ | TPM3 |
| 9.389E-46 | 1.14926422 | 0.324 | 0.14 | 2.218E-41 | MO_SPP1+ | CCL2 |
| 9.875E-46 | -0.806829 | 0.03 | 0.277 | 2.3328E-41 | MO_SPP1+ | MAL2 |
| 1.0087E-45 | 0.35768772 | 0.409 | 0.184 | 2.383E-41 | MO_SPP1+ | AC020656.1 |
| 1.3999E-45 | 0.44316665 | 0.534 | 0.298 | 3.307E-41 | MO_SPP1+ | PGAM1 |
| 1.4493E-45 | 0.40939798 | 0.586 | 0.327 | 3.4237E-41 | MO_SPP1+ | MSN |
| 2.1028E-45 | 0.51367336 | 0.63 | 0.39 | 4.9674E-41 | MO_SPP1+ | NDUFB2 |
| 2.6386E-45 | 0.28975442 | 0.268 | 0.101 | 6.2331E-41 | MO_SPP1+ | CCR1 |
| 4.7259E-45 | 0.48484764 | 0.918 | 0.798 | 1.1164E-40 | MO_SPP1+ | GNG5 |
| 7.1682E-45 | 0.27451821 | 0.409 | 0.181 | 1.6933E-40 | MO_SPP1+ | C5AR1 |
| 7.8623E-45 | -0.9501147 | 0.006 | 0.242 | 1.8573E-40 | MO_SPP1+ | ADGRF1 |
| 9.632E-45 | 0.34275909 | 0.363 | 0.166 | 2.2754E-40 | MO_SPP1+ | HEXB |
| 1.0461E-44 | 0.53395616 | 0.832 | 0.686 | 2.4712E-40 | MO_SPP1+ | PSMA7 |
| 1.4184E-44 | 0.26141641 | 0.237 | 0.084 | 3.3506E-40 | MO_SPP1+ | GSAP |
| 1.6273E-44 | 0.37530658 | 0.387 | 0.183 | 3.8441E-40 | MO_SPP1+ | SQOR |
| 1.9481E-44 | 0.44001941 | 0.582 | 0.335 | 4.6021E-40 | MO_SPP1+ | GRB2 |
| 2.406E-44 | 0.50712676 | 0.633 | 0.39 | 5.6836E-40 | MO_SPP1+ | HIF1A |
| 3.0022E-44 | 0.30560194 | 0.337 | 0.148 | 7.0921E-40 | MO_SPP1+ | VMA21 |
| 3.8711E-44 | 0.34231416 | 0.282 | 0.11 | 9.1446E-40 | MO_SPP1+ | PECAM1 |
| 4.5928E-44 | -2.1141884 | 0.109 | 0.348 | 1.085E-39 | MO_SPP1+ | HPGD |
| 5.7019E-44 | -0.6849692 | 0.012 | 0.246 | 1.3469E-39 | MO_SPP1+ | PRSS8 |
| 7.7849E-44 | 0.36011038 | 0.273 | 0.106 | 1.839E-39 | MO_SPP1+ | ACP5 |
| 8.7628E-44 | 0.28484967 | 0.325 | 0.135 | 2.07E-39 | MO_SPP1+ | EMILIN2 |
| 9.0169E-44 | 0.26486235 | 0.255 | 0.094 | 2.1301E-39 | MO_SPP1+ | DOCK4 |
| 1.0575E-43 | 0.48641227 | 0.834 | 0.684 | 2.4981E-39 | MO_SPP1+ | ATP6V0B |
| 1.2072E-43 | 0.44547518 | 0.969 | 0.858 | 2.8519E-39 | MO_SPP1+ | RPL23 |
| 2.2175E-43 | 0.26934352 | 0.204 | 0.067 | 5.2384E-39 | MO_SPP1+ | TMEM158 |
| 3.2268E-43 | -0.6814557 | 0.021 | 0.253 | 7.6226E-39 | MO_SPP1+ | KLF5 |
| 3.8003E-43 | 0.4346593 | 0.558 | 0.322 | 8.9774E-39 | MO_SPP1+ | CAPG |
| 4.5529E-43 | 0.44432348 | 0.579 | 0.346 | 1.0755E-38 | MO_SPP1+ | VDAC1 |
| 5.4265E-43 | 0.3065862 | 0.358 | 0.155 | 1.2819E-38 | MO_SPP1+ | CD300E |
| 6.2953E-43 | -0.6914386 | 0.019 | 0.254 | 1.4871E-38 | MO_SPP1+ | SYNE2 |
| 2.7962E-42 | 0.36313104 | 0.531 | 0.29 | 6.6055E-38 | MO_SPP1+ | FXYD5 |
| 2.9185E-42 | 0.30348936 | 0.357 | 0.161 | 6.8943E-38 | MO_SPP1+ | CNPY3 |
| 5.9129E-42 | 0.41680045 | 0.538 | 0.289 | 1.3968E-37 | MO_SPP1+ | ATP13A3 |
| 6.4991E-42 | -2.2324222 | 0.117 | 0.345 | 1.5353E-37 | MO_SPP1+ | PSCA |
| 1.7081E-41 | -1.6358143 | 0.012 | 0.235 | 4.0352E-37 | MO_SPP1+ | PRSS22 |
| 2.084E-41 | 0.30256649 | 0.337 | 0.146 | 4.9231E-37 | MO_SPP1+ | DMXL2 |
| 4.0864E-41 | 0.26096461 | 0.211 | 0.074 | 9.6533E-37 | MO_SPP1+ | CD84 |
| 5.0857E-41 | -2.9199524 | 0.159 | 0.382 | 1.2014E-36 | MO_SPP1+ | KRT13 |
| 5.2459E-41 | -0.83269 | 0.928 | 0.955 | 1.2392E-36 | MO_SPP1+ | ACTG1 |
| 6.6011E-41 | -1.0475343 | 0.049 | 0.282 | 1.5594E-36 | MO_SPP1+ | DUSP5 |
| 1.0164E-40 | 0.27883122 | 0.291 | 0.121 | 2.401E-36 | MO_SPP1+ | HPCAL1 |
| 1.0367E-40 | -0.7689647 | 0.394 | 0.609 | 2.449E-36 | MO_SPP1+ | EZR |
| 2.3963E-40 | 0.25960452 | 0.232 | 0.087 | 5.6609E-36 | MO_SPP1+ | TCF7L2 |
| 5.833E-40 | 0.29826348 | 0.355 | 0.161 | 1.3779E-35 | MO_SPP1+ | LCP2 |
| 7.4598E-40 | 0.44404396 | 0.648 | 0.424 | 1.7622E-35 | MO_SPP1+ | EFHD2 |
| 9.429E-40 | -0.4795299 | 0.015 | 0.234 | 2.2274E-35 | MO_SPP1+ | TJP1 |
| 9.9761E-40 | 0.29995239 | 0.421 | 0.206 | 2.3566E-35 | MO_SPP1+ | ARRB2 |

| | | | | | | |
|---|---|---|---|---|---|---|
| 1.2654E-39 | -0.5234128 | 0.004 | 0.217 | 2.9894E-35 | MO_SPP1+ | EHF |
| 2.6162E-39 | 0.4218027 | 0.546 | 0.328 | 6.1802E-35 | MO_SPP1+ | PSMB3 |
| 3.2685E-39 | -0.5088879 | 0.025 | 0.245 | 7.7212E-35 | MO_SPP1+ | GIPC1 |
| 3.5281E-39 | 0.39214748 | 0.382 | 0.188 | 8.3344E-35 | MO_SPP1+ | FNDC3A |
| 4.3204E-39 | -1.0111842 | 0.565 | 0.725 | 1.0206E-34 | MO_SPP1+ | HSPB1 |
| 1.0767E-38 | -0.6119425 | 0.022 | 0.24 | 2.5434E-34 | MO_SPP1+ | S100A16 |
| 1.3238E-38 | 0.26617689 | 0.217 | 0.08 | 3.1272E-34 | MO_SPP1+ | NRP1 |
| 2.545E-38 | -0.5483435 | 0.048 | 0.269 | 6.0119E-34 | MO_SPP1+ | AFDN |
| 2.7753E-38 | -4.2327162 | 0.192 | 0.404 | 6.5562E-34 | MO_SPP1+ | SPRR3 |
| 4.621E-38 | 0.30234807 | 0.373 | 0.18 | 1.0916E-33 | MO_SPP1+ | BID |
| 5.2122E-38 | 0.2583939 | 0.189 | 0.064 | 1.2313E-33 | MO_SPP1+ | PID1 |
| 5.5916E-38 | 0.33378722 | 0.388 | 0.19 | 1.3209E-33 | MO_SPP1+ | ACSL4 |
| 6.0531E-38 | 0.41554387 | 0.726 | 0.54 | 1.4299E-33 | MO_SPP1+ | POMP |
| 7.8932E-38 | 0.31369156 | 0.165 | 0.053 | 1.8646E-33 | MO_SPP1+ | MSR1 |
| 9.9192E-38 | -0.7518526 | 0.021 | 0.232 | 2.3432E-33 | MO_SPP1+ | ITGA2 |
| 1.5072E-37 | -0.8993995 | 0.063 | 0.284 | 3.5603E-33 | MO_SPP1+ | CLU |
| 1.5098E-37 | -0.6764513 | 0.045 | 0.265 | 3.5666E-33 | MO_SPP1+ | HES4 |
| 1.6259E-37 | -0.6886656 | 0.318 | 0.526 | 3.8408E-33 | MO_SPP1+ | HEBP2 |
| 2.0405E-37 | 0.2632357 | 0.258 | 0.105 | 4.8203E-33 | MO_SPP1+ | TGFBR1 |
| 2.0747E-37 | -0.859885 | 0.019 | 0.231 | 4.9011E-33 | MO_SPP1+ | IL32 |
| 2.5386E-37 | 0.26856465 | 0.262 | 0.107 | 5.9969E-33 | MO_SPP1+ | PAG1 |
| 2.6992E-37 | 0.39258475 | 0.42 | 0.223 | 6.3762E-33 | MO_SPP1+ | MRPL52 |
| 2.9224E-37 | -1.4809875 | 0 | 0.199 | 6.9037E-33 | MO_SPP1+ | HOPX |
| 3.0812E-37 | -0.7916116 | 0.027 | 0.238 | 7.2788E-33 | MO_SPP1+ | EPCAM |
| 3.8361E-37 | -0.7100531 | 0.027 | 0.238 | 9.062E-33 | MO_SPP1+ | CDKN1C |
| 7.6462E-37 | -0.6956754 | 0.048 | 0.262 | 1.8063E-32 | MO_SPP1+ | PMAIP1 |
| 8.35E-37 | -0.4881474 | 0.015 | 0.22 | 1.9725E-32 | MO_SPP1+ | DSP |
| 2.0256E-36 | -1.6535856 | 0.004 | 0.201 | 4.7851E-32 | MO_SPP1+ | KRT17 |
| 2.0822E-36 | -1.8972617 | 0.057 | 0.27 | 4.9189E-32 | MO_SPP1+ | S100A2 |
| 2.2879E-36 | 0.42381501 | 0.576 | 0.357 | 5.4048E-32 | MO_SPP1+ | MTDH |
| 2.436E-36 | 0.34039915 | 0.445 | 0.242 | 5.7545E-32 | MO_SPP1+ | PTPN12 |
| 3.842E-36 | 0.2749573 | 0.271 | 0.116 | 9.076E-32 | MO_SPP1+ | P2RX4 |
| 3.8846E-36 | 0.41014512 | 0.919 | 0.8 | 9.1766E-32 | MO_SPP1+ | HMGB1 |
| 5.0936E-36 | -0.5044603 | 0.01 | 0.209 | 1.2033E-31 | MO_SPP1+ | PARD6B |
| 5.2045E-36 | -0.4215145 | 0.016 | 0.218 | 1.2295E-31 | MO_SPP1+ | CNN3 |
| 5.6548E-36 | -0.7874325 | 0.03 | 0.237 | 1.3358E-31 | MO_SPP1+ | PDZK1IP1 |
| 1.5243E-35 | 0.32302665 | 0.457 | 0.247 | 3.6009E-31 | MO_SPP1+ | SH3BGRL |
| 2.4782E-35 | 0.37747296 | 0.991 | 0.903 | 5.8543E-31 | MO_SPP1+ | RPLP2 |
| 6.0582E-35 | 0.31890204 | 0.4 | 0.209 | 1.4311E-30 | MO_SPP1+ | RNF13 |
| 6.7075E-35 | 0.46828683 | 0.505 | 0.306 | 1.5845E-30 | MO_SPP1+ | RAB13 |
| 8.3162E-35 | 0.36119423 | 0.466 | 0.26 | 1.9645E-30 | MO_SPP1+ | TUBB |
| 1.0361E-34 | 0.25308931 | 0.303 | 0.138 | 2.4476E-30 | MO_SPP1+ | CD58 |
| 1.1955E-34 | 0.26339584 | 0.244 | 0.099 | 2.8242E-30 | MO_SPP1+ | CLEC4E |
| 2.137E-34 | -0.5418046 | 0.027 | 0.227 | 5.0482E-30 | MO_SPP1+ | OCIAD2 |
| 2.4641E-34 | 0.28617832 | 0.372 | 0.184 | 5.821E-30 | MO_SPP1+ | RAPGEF1 |
| 4.7435E-34 | 0.25380502 | 0.301 | 0.136 | 1.1206E-29 | MO_SPP1+ | PTPN6 |
| 5.0683E-34 | 0.29347901 | 0.342 | 0.168 | 1.1973E-29 | MO_SPP1+ | LACTB |
| 8.258E-34 | 0.27209477 | 0.319 | 0.149 | 1.9508E-29 | MO_SPP1+ | SLC44A1 |
| 8.8213E-34 | 0.39121913 | 0.645 | 0.425 | 2.0839E-29 | MO_SPP1+ | IQGAP1 |
| 9.635E-34 | -1.2441022 | 0.003 | 0.187 | 2.2761E-29 | MO_SPP1+ | DUOX2 |

| | | | | | | |
|---|---|---|---|---|---|---|
| 1.2071E-33 | 0.41070486 | 0.367 | 0.186 | 2.8515E-29 | MO_SPP1+ | DUSP6 |
| 1.2656E-33 | -0.9172753 | 0.012 | 0.2 | 2.9898E-29 | MO_SPP1+ | TM4SF1 |
| 1.8301E-33 | 0.26413157 | 0.277 | 0.123 | 4.3232E-29 | MO_SPP1+ | HSD17B11 |
| 2.7911E-33 | -0.5099275 | 0.049 | 0.25 | 6.5934E-29 | MO_SPP1+ | SERINC2 |
| 3.3708E-33 | -1.2758081 | 0.049 | 0.251 | 7.9629E-29 | MO_SPP1+ | CRYAB |
| 3.3845E-33 | -0.3536837 | 0.018 | 0.208 | 7.9952E-29 | MO_SPP1+ | CTTN |
| 3.4011E-33 | 0.39677567 | 0.807 | 0.644 | 8.0343E-29 | MO_SPP1+ | ATP5MC2 |
| 3.9674E-33 | -0.4668843 | 0.009 | 0.193 | 9.3721E-29 | MO_SPP1+ | ABLIM1 |
| 4.4938E-33 | 0.39844202 | 0.993 | 0.923 | 1.0616E-28 | MO_SPP1+ | RPS24 |
| 5.3494E-33 | -0.5831245 | 0.004 | 0.186 | 1.2637E-28 | MO_SPP1+ | SDCBP2 |
| 5.6113E-33 | 0.27516061 | 0.286 | 0.129 | 1.3256E-28 | MO_SPP1+ | EPB41L3 |
| 5.8465E-33 | -0.9044416 | 0.007 | 0.19 | 1.3811E-28 | MO_SPP1+ | TMPRSS2 |
| 6.4091E-33 | 0.33335684 | 0.304 | 0.14 | 1.514E-28 | MO_SPP1+ | NCF1 |
| 6.5379E-33 | -0.4901913 | 0.048 | 0.25 | 1.5444E-28 | MO_SPP1+ | SPTBN1 |
| 1.2615E-32 | 0.25364634 | 0.274 | 0.123 | 2.9801E-28 | MO_SPP1+ | SDCCAG8 |
| 1.3297E-32 | 0.25056556 | 0.207 | 0.081 | 3.1411E-28 | MO_SPP1+ | CYP27A1 |
| 1.4018E-32 | 0.28874581 | 0.351 | 0.176 | 3.3115E-28 | MO_SPP1+ | GPI |
| 1.4528E-32 | -0.3956869 | 0.001 | 0.179 | 3.432E-28 | MO_SPP1+ | SMIM22 |
| 1.6177E-32 | -0.5113715 | 0.063 | 0.262 | 3.8216E-28 | MO_SPP1+ | WASL |
| 1.7916E-32 | -0.6763943 | 0.012 | 0.194 | 4.2323E-28 | MO_SPP1+ | FBXO32 |
| 5.3442E-32 | 0.35366038 | 0.945 | 0.865 | 1.2625E-27 | MO_SPP1+ | ARPC3 |
| 5.6354E-32 | -0.4770101 | 0.028 | 0.218 | 1.3313E-27 | MO_SPP1+ | CKB |
| 6.227E-32 | 0.29181131 | 0.319 | 0.152 | 1.471E-27 | MO_SPP1+ | ADGRE5 |
| 6.2867E-32 | -0.6699008 | 0.012 | 0.191 | 1.4851E-27 | MO_SPP1+ | EPS8L1 |
| 6.7011E-32 | -0.5156656 | 0.024 | 0.211 | 1.583E-27 | MO_SPP1+ | MACC1 |
| 1.0911E-31 | -0.6036894 | 0.001 | 0.175 | 2.5776E-27 | MO_SPP1+ | PPL |
| 1.7996E-31 | 0.3257379 | 0.477 | 0.278 | 4.2512E-27 | MO_SPP1+ | RBM3 |
| 1.9872E-31 | -0.5631925 | 0.121 | 0.33 | 4.6945E-27 | MO_SPP1+ | MAFF |
| 2.1609E-31 | 0.35609782 | 0.477 | 0.285 | 5.1047E-27 | MO_SPP1+ | HM13 |
| 2.2878E-31 | 0.41257101 | 0.615 | 0.412 | 5.4045E-27 | MO_SPP1+ | GRN |
| 2.6636E-31 | -0.5342101 | 0.004 | 0.177 | 6.2922E-27 | MO_SPP1+ | C6orf132 |
| 2.8285E-31 | -0.6811183 | 0.154 | 0.352 | 6.6819E-27 | MO_SPP1+ | PEBP1 |
| 3.0609E-31 | -0.3809607 | 0.01 | 0.187 | 7.2307E-27 | MO_SPP1+ | EPS8L2 |
| 5.286E-31 | -0.7485985 | 0.121 | 0.322 | 1.2487E-26 | MO_SPP1+ | GADD45A |
| 6.3265E-31 | 0.31493037 | 0.346 | 0.18 | 1.4945E-26 | MO_SPP1+ | POLE4 |
| 7.9488E-31 | -0.6546147 | 0.019 | 0.198 | 1.8777E-26 | MO_SPP1+ | SFN |
| 8.2937E-31 | 0.28716916 | 0.366 | 0.186 | 1.9592E-26 | MO_SPP1+ | HBEGF |
| 8.6517E-31 | 0.46090125 | 0.985 | 0.898 | 2.0438E-26 | MO_SPP1+ | RPL39 |
| 8.763E-31 | 0.27315028 | 0.204 | 0.082 | 2.0701E-26 | MO_SPP1+ | PLA2G7 |
| 9.5626E-31 | 0.33433597 | 0.409 | 0.224 | 2.259E-26 | MO_SPP1+ | NUMB |
| 1.0811E-30 | 0.38090828 | 0.478 | 0.282 | 2.5538E-26 | MO_SPP1+ | CXCL16 |
| 1.2508E-30 | 0.32073335 | 0.963 | 0.904 | 2.9548E-26 | MO_SPP1+ | CFL1 |
| 1.8216E-30 | -0.5622315 | 0.1 | 0.293 | 4.3031E-26 | MO_SPP1+ | SPINT1 |
| 2.3424E-30 | -1.2842539 | 0.001 | 0.168 | 5.5335E-26 | MO_SPP1+ | DUOXA2 |
| 2.5286E-30 | 0.26461151 | 0.313 | 0.153 | 5.9734E-26 | MO_SPP1+ | CSGALNACT2 |
| 2.8388E-30 | -0.6255151 | 0.003 | 0.171 | 6.7061E-26 | MO_SPP1+ | FXYD3 |
| 3.7135E-30 | 0.36916531 | 0.456 | 0.273 | 8.7725E-26 | MO_SPP1+ | ATP6AP2 |
| 3.8122E-30 | 0.42950218 | 0.424 | 0.251 | 9.0056E-26 | MO_SPP1+ | C1orf122 |
| 4.5685E-30 | -0.3222132 | 0.019 | 0.198 | 1.0792E-25 | MO_SPP1+ | RBPMS |
| 4.7706E-30 | 0.32546082 | 0.405 | 0.224 | 1.127E-25 | MO_SPP1+ | IFNGR1 |

| | | | | | | |
|---|---|---|---|---|---|---|
| 5.2833E-30 | 0.43199367 | 0.727 | 0.57 | 1.2481E-25 | MO_SPP1+ | TSPO |
| 5.3284E-30 | -0.7609709 | 0.018 | 0.191 | 1.2587E-25 | MO_SPP1+ | ANKRD36C |
| 5.4298E-30 | -0.8435662 | 0.007 | 0.176 | 1.2827E-25 | MO_SPP1+ | NDRG2 |
| 5.8238E-30 | -0.5250987 | 0.177 | 0.38 | 1.3758E-25 | MO_SPP1+ | GCC2 |
| 6.3041E-30 | 0.35773043 | 0.531 | 0.338 | 1.4892E-25 | MO_SPP1+ | ARPC4 |
| 6.9E-30 | -0.6593325 | 0.009 | 0.178 | 1.63E-25 | MO_SPP1+ | IGFBP3 |
| 7.0736E-30 | -0.7846331 | 0.279 | 0.459 | 1.671E-25 | MO_SPP1+ | RAB11FIP1 |
| 8.042E-30 | 0.28386426 | 0.298 | 0.145 | 1.8998E-25 | MO_SPP1+ | SH3KBP1 |
| 8.3705E-30 | 0.33320515 | 0.697 | 0.488 | 1.9774E-25 | MO_SPP1+ | TAGLN2 |
| 8.4472E-30 | 0.27146853 | 0.325 | 0.165 | 1.9955E-25 | MO_SPP1+ | CORO1C |
| 8.7004E-30 | -0.6044304 | 0.009 | 0.177 | 2.0553E-25 | MO_SPP1+ | SPNS2 |
| 1.0293E-29 | -0.4020327 | 0.013 | 0.185 | 2.4315E-25 | MO_SPP1+ | CDH1 |
| 1.0488E-29 | -0.5091897 | 0.012 | 0.181 | 2.4777E-25 | MO_SPP1+ | EPHA2 |
| 1.059E-29 | 0.3637187 | 0.76 | 0.614 | 2.5017E-25 | MO_SPP1+ | COX8A |
| 1.1608E-29 | 0.31194992 | 0.367 | 0.196 | 2.7423E-25 | MO_SPP1+ | CREG1 |
| 1.1801E-29 | 0.35043699 | 0.408 | 0.225 | 2.7878E-25 | MO_SPP1+ | IVNS1ABP |
| 1.2599E-29 | -0.4384985 | 0.16 | 0.37 | 2.9764E-25 | MO_SPP1+ | AES |
| 1.33E-29 | 0.30188026 | 0.421 | 0.236 | 3.1418E-25 | MO_SPP1+ | SP100 |
| 1.3365E-29 | -0.6510468 | 0.034 | 0.211 | 3.1571E-25 | MO_SPP1+ | CDKN2B |
| 1.8466E-29 | 0.32034998 | 0.543 | 0.33 | 4.3621E-25 | MO_SPP1+ | TLN1 |
| 1.9083E-29 | 0.27788155 | 0.402 | 0.219 | 4.5079E-25 | MO_SPP1+ | MAP3K2 |
| 2.005E-29 | -0.4645229 | 0.013 | 0.183 | 4.7365E-25 | MO_SPP1+ | S100A14 |
| 2.0086E-29 | -0.3842664 | 0.013 | 0.182 | 4.7448E-25 | MO_SPP1+ | ANXA3 |
| 2.187E-29 | 0.8213345 | 0.157 | 0.057 | 5.1664E-25 | MO_SPP1+ | RNASE1 |
| 2.3002E-29 | -0.5059136 | 0.052 | 0.237 | 5.4339E-25 | MO_SPP1+ | TOB1 |
| 2.5675E-29 | -1.1492964 | 0.001 | 0.163 | 6.0651E-25 | MO_SPP1+ | MUC21 |
| 2.61E-29 | -0.380304 | 0.006 | 0.171 | 6.1656E-25 | MO_SPP1+ | NECTIN4 |
| 2.9496E-29 | 0.26348658 | 0.339 | 0.173 | 6.9679E-25 | MO_SPP1+ | ATXN1 |
| 2.9981E-29 | -0.2615306 | 0.351 | 0.173 | 7.0823E-25 | MO_SPP1+ | IL1B |
| 3.9224E-29 | -0.3426495 | 0.013 | 0.182 | 9.2659E-25 | MO_SPP1+ | FNBP1L |
| 4.0008E-29 | -0.4683883 | 0.012 | 0.179 | 9.4511E-25 | MO_SPP1+ | FHL2 |
| 5.0346E-29 | 0.31050387 | 0.987 | 0.916 | 1.1893E-24 | MO_SPP1+ | RPS20 |
| 5.101E-29 | 0.30063491 | 0.4 | 0.223 | 1.205E-24 | MO_SPP1+ | XPO1 |
| 5.8033E-29 | 0.29247226 | 0.459 | 0.27 | 1.3709E-24 | MO_SPP1+ | FAM49B |
| 6.7566E-29 | -0.3244579 | 0.001 | 0.161 | 1.5961E-24 | MO_SPP1+ | CRB3 |
| 1.1329E-28 | -0.9917546 | 0.001 | 0.16 | 2.6764E-24 | MO_SPP1+ | PRSS27 |
| 1.2033E-28 | 0.31167952 | 0.568 | 0.372 | 2.8426E-24 | MO_SPP1+ | BRK1 |
| 1.3464E-28 | -0.3290884 | 0.015 | 0.181 | 3.1805E-24 | MO_SPP1+ | LSR |
| 1.3937E-28 | -1.0119472 | 0.012 | 0.176 | 3.2923E-24 | MO_SPP1+ | RDH10 |
| 1.493E-28 | -0.5845173 | 0.01 | 0.173 | 3.527E-24 | MO_SPP1+ | HSPB8 |
| 1.6415E-28 | -0.9870427 | 0 | 0.157 | 3.8778E-24 | MO_SPP1+ | PITX1 |
| 1.6642E-28 | -0.4087756 | 0.001 | 0.16 | 3.9313E-24 | MO_SPP1+ | TTC9 |
| 1.8305E-28 | 0.30205814 | 0.925 | 0.833 | 4.3242E-24 | MO_SPP1+ | PPIA |
| 2.1481E-28 | -0.6073708 | 0.543 | 0.701 | 5.0745E-24 | MO_SPP1+ | PPP1R15A |
| 2.3937E-28 | -0.8667187 | 0.027 | 0.197 | 5.6546E-24 | MO_SPP1+ | C12orf75 |
| 3.0207E-28 | -1.2141712 | 0.153 | 0.339 | 7.1359E-24 | MO_SPP1+ | IGFBP7 |
| 3.329E-28 | -0.3544573 | 0.021 | 0.189 | 7.8641E-24 | MO_SPP1+ | PAWR |
| 3.4418E-28 | -1.0554959 | 0.004 | 0.162 | 8.1305E-24 | MO_SPP1+ | CEACAM5 |
| 3.7652E-28 | 0.27925162 | 0.439 | 0.249 | 8.8945E-24 | MO_SPP1+ | C4orf48 |
| 4.1058E-28 | -0.7742102 | 0.021 | 0.186 | 9.699E-24 | MO_SPP1+ | AQP3 |

| | | | | | | |
|---|---|---|---|---|---|---|
| 4.3384E-28 | -0.5220765 | 0.162 | 0.352 | 1.0249E-23 | MO_SPP1+ | UBE2H |
| 5.463E-28 | -0.3622822 | 0.018 | 0.182 | 1.2905E-23 | MO_SPP1+ | TEAD1 |
| 6.7128E-28 | -1.5980117 | 0.007 | 0.165 | 1.5858E-23 | MO_SPP1+ | ECM1 |
| 6.9283E-28 | -0.5315436 | 0.042 | 0.215 | 1.6367E-23 | MO_SPP1+ | LAMB3 |
| 6.9965E-28 | -0.6022648 | 0.001 | 0.156 | 1.6528E-23 | MO_SPP1+ | CDC42EP5 |
| 7.2282E-28 | -0.503303 | 0.004 | 0.161 | 1.7075E-23 | MO_SPP1+ | MALL |
| 9.785E-28 | -0.759753 | 0.001 | 0.155 | 2.3115E-23 | MO_SPP1+ | SCEL |
| 1.1865E-27 | 0.53245058 | 0.957 | 0.882 | 2.8028E-23 | MO_SPP1+ | CSTB |
| 1.2995E-27 | -0.6348069 | 0.046 | 0.224 | 3.0699E-23 | MO_SPP1+ | CCND1 |
| 1.3255E-27 | 0.36807924 | 0.61 | 0.431 | 3.1313E-23 | MO_SPP1+ | ATP5MF |
| 1.4343E-27 | 0.26421622 | 0.385 | 0.212 | 3.3883E-23 | MO_SPP1+ | ZYX |
| 1.5494E-27 | -0.6182522 | 0.067 | 0.246 | 3.6603E-23 | MO_SPP1+ | ITGB8 |
| 2.3672E-27 | -0.3261896 | 0.012 | 0.17 | 5.5921E-23 | MO_SPP1+ | CXADR |
| 2.5311E-27 | -0.3780383 | 0.004 | 0.158 | 5.9792E-23 | MO_SPP1+ | TSPAN6 |
| 2.7607E-27 | -0.7859428 | 0.022 | 0.184 | 6.5216E-23 | MO_SPP1+ | LAMC2 |
| 2.9879E-27 | -0.4146564 | 0.073 | 0.255 | 7.0582E-23 | MO_SPP1+ | CD2AP |
| 3.0437E-27 | -0.3005469 | 0.01 | 0.167 | 7.19E-23 | MO_SPP1+ | DDR1 |
| 3.0687E-27 | -0.3323834 | 0.025 | 0.19 | 7.2492E-23 | MO_SPP1+ | HSBP1L1 |
| 3.0692E-27 | 0.30502711 | 0.423 | 0.241 | 7.2503E-23 | MO_SPP1+ | CD82 |
| 3.454E-27 | 0.2686232 | 0.508 | 0.305 | 8.1594E-23 | MO_SPP1+ | ARPC1B |
| 3.9042E-27 | -0.7923952 | 0.021 | 0.182 | 9.2229E-23 | MO_SPP1+ | F3 |
| 4.4033E-27 | -2.2534349 | 0.171 | 0.342 | 1.0402E-22 | MO_SPP1+ | MMP7 |
| 4.9282E-27 | -0.9293598 | 0.027 | 0.19 | 1.1642E-22 | MO_SPP1+ | RARRES3 |
| 5.3261E-27 | -0.4432715 | 0.004 | 0.156 | 1.2582E-22 | MO_SPP1+ | TMPRSS4 |
| 6.5755E-27 | 0.30488838 | 0.411 | 0.24 | 1.5533E-22 | MO_SPP1+ | LSM7 |
| 7.1137E-27 | -0.2958715 | 0.015 | 0.172 | 1.6805E-22 | MO_SPP1+ | GNG12 |
| 7.8677E-27 | -0.7601486 | 0.001 | 0.151 | 1.8586E-22 | MO_SPP1+ | TMPRSS11D |
| 7.8819E-27 | 0.29943165 | 0.532 | 0.322 | 1.8619E-22 | MO_SPP1+ | ARHGDIB |
| 8.197E-27 | -0.3288203 | 0.019 | 0.178 | 1.9364E-22 | MO_SPP1+ | TMEM54 |
| 9.2149E-27 | -0.4180442 | 0 | 0.148 | 2.1768E-22 | MO_SPP1+ | VSIG2 |
| 9.399E-27 | 0.34715102 | 0.454 | 0.273 | 2.2203E-22 | MO_SPP1+ | ETS2 |
| 9.5325E-27 | -0.3283258 | 0.006 | 0.158 | 2.2519E-22 | MO_SPP1+ | ERBB3 |
| 9.6009E-27 | -0.8294213 | 0 | 0.148 | 2.268E-22 | MO_SPP1+ | CEACAM7 |
| 1.0266E-26 | -0.3168968 | 0.01 | 0.165 | 2.4252E-22 | MO_SPP1+ | NDFIP2 |
| 1.5118E-26 | 0.26967175 | 0.237 | 0.11 | 3.5712E-22 | MO_SPP1+ | LIPA |
| 1.5597E-26 | -0.3074946 | 0.003 | 0.152 | 3.6845E-22 | MO_SPP1+ | PLS3 |
| 1.7278E-26 | -0.5719973 | 0.07 | 0.244 | 4.0817E-22 | MO_SPP1+ | S100A13 |
| 2.1262E-26 | -0.6537847 | 0.007 | 0.157 | 5.0228E-22 | MO_SPP1+ | MUC4 |
| 2.1377E-26 | -0.8496843 | 0.007 | 0.158 | 5.05E-22 | MO_SPP1+ | KRT15 |
| 2.1565E-26 | -0.8685911 | 0.001 | 0.148 | 5.0943E-22 | MO_SPP1+ | CEACAM6 |
| 2.234E-26 | 0.36301444 | 0.277 | 0.138 | 5.2773E-22 | MO_SPP1+ | GPNMB |
| 2.7094E-26 | 0.31649327 | 0.444 | 0.262 | 6.4005E-22 | MO_SPP1+ | BACH1 |
| 2.7364E-26 | 0.26991058 | 0.46 | 0.274 | 6.4642E-22 | MO_SPP1+ | ROCK1 |
| 3.4332E-26 | 0.30281628 | 0.472 | 0.296 | 8.1102E-22 | MO_SPP1+ | TMEM167A |
| 3.6117E-26 | -0.4174579 | 0.018 | 0.172 | 8.5318E-22 | MO_SPP1+ | FAM214A |
| 3.8377E-26 | 0.27490579 | 0.445 | 0.262 | 9.0657E-22 | MO_SPP1+ | PLP2 |
| 4.0327E-26 | -1.114787 | 0.462 | 0.64 | 9.5264E-22 | MO_SPP1+ | MTRNR2L12 |
| 4.7653E-26 | -0.7755896 | 0.001 | 0.147 | 1.1257E-21 | MO_SPP1+ | TMPRSS11E |
| 4.8802E-26 | -0.5834427 | 0.163 | 0.356 | 1.1529E-21 | MO_SPP1+ | ATF3 |
| 5.085E-26 | 0.30154151 | 0.997 | 0.939 | 1.2012E-21 | MO_SPP1+ | RPS28 |

| | | | | | | |
|---|---|---|---|---|---|---|
| 5.7406E-26 | -0.6443416 | 0.003 | 0.149 | 1.3561E-21 | MO_SPP1+ | NCCRP1 |
| 6.7701E-26 | 0.44076488 | 0.445 | 0.283 | 1.5993E-21 | MO_SPP1+ | YWHAH |
| 6.9464E-26 | 0.4700108 | 0.423 | 0.261 | 1.6409E-21 | MO_SPP1+ | GSN |
| 8.6813E-26 | 0.25536481 | 0.328 | 0.174 | 2.0508E-21 | MO_SPP1+ | PPP1R18 |
| 1.2932E-25 | 0.32860095 | 0.52 | 0.337 | 3.055E-21 | MO_SPP1+ | ANP32B |
| 1.4647E-25 | -0.4736257 | 0.004 | 0.149 | 3.4601E-21 | MO_SPP1+ | ALDH1A3 |
| 1.5355E-25 | -0.460286 | 0.025 | 0.182 | 3.6274E-21 | MO_SPP1+ | DST |
| 1.5625E-25 | -0.303676 | 0.003 | 0.147 | 3.691E-21 | MO_SPP1+ | SLC44A4 |
| 1.6072E-25 | -0.7954543 | 0.729 | 0.814 | 3.7968E-21 | MO_SPP1+ | GSTP1 |
| 1.75E-25 | -0.5518051 | 0.1 | 0.276 | 4.134E-21 | MO_SPP1+ | TNFRSF12A |
| 1.8216E-25 | -0.3065129 | 0.013 | 0.164 | 4.3031E-21 | MO_SPP1+ | GALNT3 |
| 2.0109E-25 | 0.26359384 | 0.232 | 0.104 | 4.7504E-21 | MO_SPP1+ | THBD |
| 2.1442E-25 | -0.3640684 | 0.042 | 0.203 | 5.0652E-21 | MO_SPP1+ | ZNF91 |
| 2.9888E-25 | 0.32072414 | 0.573 | 0.387 | 7.0605E-21 | MO_SPP1+ | ATP5MPL |
| 3.0574E-25 | 0.34058397 | 0.42 | 0.252 | 7.2225E-21 | MO_SPP1+ | MGAT1 |
| 3.2271E-25 | -0.4141597 | 0.013 | 0.161 | 7.6235E-21 | MO_SPP1+ | SDC1 |
| 3.5001E-25 | -0.3237319 | 0.033 | 0.191 | 8.2682E-21 | MO_SPP1+ | SH3YL1 |
| 3.5873E-25 | -0.3679463 | 0.108 | 0.288 | 8.4742E-21 | MO_SPP1+ | LRP10 |
| 4.3399E-25 | -0.4141397 | 0.09 | 0.264 | 1.0252E-20 | MO_SPP1+ | PRDX2 |
| 5.3186E-25 | -0.3724471 | 0.012 | 0.158 | 1.2564E-20 | MO_SPP1+ | COBLL1 |
| 6.2517E-25 | -0.6274482 | 0.003 | 0.143 | 1.4769E-20 | MO_SPP1+ | SNCG |
| 6.5579E-25 | 0.26575464 | 0.225 | 0.101 | 1.5492E-20 | MO_SPP1+ | LILRB1 |
| 6.5897E-25 | -1.7011631 | 0.004 | 0.145 | 1.5567E-20 | MO_SPP1+ | SPRR2A |
| 8.6367E-25 | 0.26283293 | 0.277 | 0.142 | 2.0402E-20 | MO_SPP1+ | IRAK1 |
| 9.2656E-25 | -0.4219265 | 0 | 0.138 | 2.1888E-20 | MO_SPP1+ | GCNT3 |
| 1.2E-24 | 0.30130117 | 0.324 | 0.174 | 2.8346E-20 | MO_SPP1+ | ABCA1 |
| 1.4318E-24 | 0.29976517 | 0.574 | 0.382 | 3.3824E-20 | MO_SPP1+ | CNBP |
| 1.4721E-24 | -0.4060936 | 0.136 | 0.315 | 3.4775E-20 | MO_SPP1+ | SBDS |
| 1.8022E-24 | -0.2875921 | 0.007 | 0.148 | 4.2574E-20 | MO_SPP1+ | KCNK1 |
| 2.5014E-24 | 0.34191986 | 0.292 | 0.15 | 5.9091E-20 | MO_SPP1+ | CCL20 |
| 2.7331E-24 | 0.26558956 | 0.336 | 0.182 | 6.4564E-20 | MO_SPP1+ | KLF10 |
| 2.9465E-24 | 0.30041367 | 0.388 | 0.229 | 6.9604E-20 | MO_SPP1+ | FLOT1 |
| 3.2417E-24 | -0.2758879 | 0.004 | 0.142 | 7.6579E-20 | MO_SPP1+ | RAB25 |
| 3.7994E-24 | -0.4060877 | 0.015 | 0.159 | 8.9754E-20 | MO_SPP1+ | CLDN3 |
| 3.8762E-24 | 0.26894536 | 0.355 | 0.204 | 9.1567E-20 | MO_SPP1+ | TIMM8B |
| 4.8663E-24 | -0.9073263 | 0.001 | 0.136 | 1.1496E-19 | MO_SPP1+ | TMPRSS11B |
| 5.3585E-24 | 0.26528666 | 0.373 | 0.213 | 1.2658E-19 | MO_SPP1+ | TGFB1 |
| 5.4963E-24 | 0.29828015 | 0.544 | 0.353 | 1.2984E-19 | MO_SPP1+ | TKT |
| 6.4904E-24 | -0.6365492 | 0.003 | 0.138 | 1.5332E-19 | MO_SPP1+ | CXCL17 |
| 6.6923E-24 | 0.33006891 | 0.633 | 0.45 | 1.5809E-19 | MO_SPP1+ | SEC61G |
| 7.0907E-24 | -0.2969393 | 0.009 | 0.148 | 1.675E-19 | MO_SPP1+ | FAM102A |
| 7.141E-24 | 0.26016272 | 0.267 | 0.133 | 1.6869E-19 | MO_SPP1+ | HLA-DRB5 |
| 8.9992E-24 | -0.7140548 | 0.234 | 0.407 | 2.1259E-19 | MO_SPP1+ | RNF213 |
| 9.2565E-24 | -0.3154286 | 0.022 | 0.17 | 2.1867E-19 | MO_SPP1+ | SLC38A1 |
| 9.4286E-24 | -0.3774951 | 0.019 | 0.164 | 2.2273E-19 | MO_SPP1+ | SYTL2 |
| 9.8222E-24 | -0.3458647 | 0.006 | 0.142 | 2.3203E-19 | MO_SPP1+ | EMP2 |
| 1.0509E-23 | -0.2748191 | 0 | 0.132 | 2.4826E-19 | MO_SPP1+ | C1orf116 |
| 1.0673E-23 | -0.6799045 | 0.066 | 0.224 | 2.5212E-19 | MO_SPP1+ | HES1 |
| 1.1063E-23 | 0.2513746 | 0.384 | 0.222 | 2.6134E-19 | MO_SPP1+ | RSL1D1 |
| 1.2252E-23 | 0.30301816 | 0.513 | 0.333 | 2.8943E-19 | MO_SPP1+ | CAPZA1 |

| | | | | | | |
|---|---|---|---|---|---|---|
| 1.3032E-23 | 0.29265303 | 0.871 | 0.745 | 3.0785E-19 | MO_SPP1+ | CLIC1 |
| 1.3901E-23 | -0.3909561 | 0.003 | 0.137 | 3.2838E-19 | MO_SPP1+ | BCAS1 |
| 1.5723E-23 | -1.24146 | 0.001 | 0.134 | 3.7142E-19 | MO_SPP1+ | PLAT |
| 1.6957E-23 | 0.36947333 | 0.907 | 0.863 | 4.0058E-19 | MO_SPP1+ | CALM1 |
| 2.1275E-23 | 0.30427109 | 0.558 | 0.386 | 5.0259E-19 | MO_SPP1+ | HMGN2 |
| 2.1754E-23 | 0.29836723 | 0.883 | 0.769 | 5.139E-19 | MO_SPP1+ | TMA7 |
| 2.407E-23 | -1.0254526 | 0.007 | 0.142 | 5.686E-19 | MO_SPP1+ | CXCL6 |
| 2.6397E-23 | 0.33023142 | 0.694 | 0.515 | 6.2358E-19 | MO_SPP1+ | NAP1L1 |
| 2.7272E-23 | -0.657819 | 0.994 | 0.98 | 6.4424E-19 | MO_SPP1+ | MALAT1 |
| 3.1419E-23 | 0.32543095 | 0.369 | 0.21 | 7.4221E-19 | MO_SPP1+ | ZFYVE16 |
| 3.1514E-23 | -0.4643831 | 0.097 | 0.261 | 7.4446E-19 | MO_SPP1+ | TSC22D1 |
| 3.2157E-23 | -0.9117081 | 0.003 | 0.134 | 7.5964E-19 | MO_SPP1+ | RHCG |
| 4.4916E-23 | 0.27092813 | 0.387 | 0.231 | 1.061E-18 | MO_SPP1+ | CTSA |
| 4.8704E-23 | -0.6029673 | 0.094 | 0.256 | 1.1505E-18 | MO_SPP1+ | CARHSP1 |
| 5.179E-23 | 0.30023681 | 0.46 | 0.29 | 1.2234E-18 | MO_SPP1+ | TUBA1C |
| 5.1875E-23 | -0.6118861 | 0.546 | 0.679 | 1.2255E-18 | MO_SPP1+ | IER2 |
| 5.5891E-23 | -0.537088 | 0.841 | 0.907 | 1.3203E-18 | MO_SPP1+ | ITM2B |
| 5.6269E-23 | 0.26289976 | 0.982 | 0.93 | 1.3293E-18 | MO_SPP1+ | FAU |
| 6.0123E-23 | -0.6564563 | 0.003 | 0.133 | 1.4203E-18 | MO_SPP1+ | GABRP |
| 7.2192E-23 | -0.6250736 | 0.01 | 0.144 | 1.7054E-18 | MO_SPP1+ | CDKN2A |
| 8.0911E-23 | 0.29938314 | 0.727 | 0.552 | 1.9114E-18 | MO_SPP1+ | NDUFS5 |
| 8.1166E-23 | -1.5976655 | 0.004 | 0.135 | 1.9174E-18 | MO_SPP1+ | LYPD2 |
| 9.4393E-23 | 0.26706678 | 0.991 | 0.957 | 2.2298E-18 | MO_SPP1+ | H3F3A |
| 9.4578E-23 | -0.3711657 | 0.058 | 0.218 | 2.2342E-18 | MO_SPP1+ | ARL4C |
| 9.8757E-23 | 0.34839135 | 0.426 | 0.268 | 2.3329E-18 | MO_SPP1+ | OSBPL8 |
| 1.26E-22 | 0.28336027 | 0.366 | 0.211 | 2.9765E-18 | MO_SPP1+ | RUNX1 |
| 1.3885E-22 | -0.332516 | 0.016 | 0.154 | 3.2801E-18 | MO_SPP1+ | ITGA3 |
| 1.6457E-22 | -0.5744065 | 0.48 | 0.639 | 3.8875E-18 | MO_SPP1+ | GADD45B |
| 1.7546E-22 | -0.5015108 | 0.022 | 0.16 | 4.1449E-18 | MO_SPP1+ | BPGM |
| 2.6211E-22 | -0.3389896 | 0.003 | 0.13 | 6.1919E-18 | MO_SPP1+ | FAM84A |
| 2.7618E-22 | -0.4101188 | 0 | 0.125 | 6.5242E-18 | MO_SPP1+ | TMEM40 |
| 3.0183E-22 | 0.26218339 | 0.531 | 0.351 | 7.13E-18 | MO_SPP1+ | SEPT7 |
| 3.0451E-22 | 0.30926525 | 0.522 | 0.357 | 7.1935E-18 | MO_SPP1+ | RBM8A |
| 3.0872E-22 | -0.3562151 | 0.018 | 0.153 | 7.2928E-18 | MO_SPP1+ | MDK |
| 3.2335E-22 | -0.5732298 | 0.924 | 0.93 | 7.6385E-18 | MO_SPP1+ | MT-CYB |
| 3.6046E-22 | -0.2883693 | 0.006 | 0.134 | 8.5152E-18 | MO_SPP1+ | MPZL2 |
| 3.6797E-22 | 1.02199949 | 0.481 | 0.331 | 8.6926E-18 | MO_SPP1+ | IL1RN |
| 3.9479E-22 | -0.4147095 | 0.019 | 0.157 | 9.3262E-18 | MO_SPP1+ | CYR61 |
| 3.9712E-22 | -0.6790935 | 0.676 | 0.753 | 9.3811E-18 | MO_SPP1+ | TPM4 |
| 4.1723E-22 | 0.27678852 | 0.54 | 0.367 | 9.8563E-18 | MO_SPP1+ | PAK2 |
| 4.2867E-22 | -0.5812066 | 0.033 | 0.174 | 1.0126E-17 | MO_SPP1+ | STEAP4 |
| 4.9429E-22 | 0.29455474 | 0.462 | 0.3 | 1.1677E-17 | MO_SPP1+ | ANAPC11 |
| 6.1397E-22 | 0.30577571 | 0.433 | 0.273 | 1.4504E-17 | MO_SPP1+ | DDX21 |
| 6.7756E-22 | -0.4014143 | 0.052 | 0.2 | 1.6006E-17 | MO_SPP1+ | PTGR1 |
| 7.6187E-22 | 0.28818519 | 0.867 | 0.765 | 1.7998E-17 | MO_SPP1+ | RPS26 |
| 7.7198E-22 | 0.33676706 | 0.547 | 0.377 | 1.8236E-17 | MO_SPP1+ | EIF5A |
| 7.9245E-22 | -0.3229498 | 0.03 | 0.17 | 1.872E-17 | MO_SPP1+ | TINAGL1 |
| 8.5447E-22 | -0.2754642 | 0.013 | 0.144 | 2.0185E-17 | MO_SPP1+ | OCLN |
| 9.5934E-22 | -0.2759182 | 0.013 | 0.144 | 2.2662E-17 | MO_SPP1+ | SERPING1 |
| 1.0414E-21 | -0.4558575 | 0.438 | 0.578 | 2.4601E-17 | MO_SPP1+ | PRDX5 |

| | | | | | | |
|---|---|---|---|---|---|---|
| 1.1104E-21 | -0.3530425 | 0.048 | 0.193 | 2.6232E-17 | MO_SPP1+ | LIMA1 |
| 1.2159E-21 | -0.5508116 | 0.001 | 0.124 | 2.8723E-17 | MO_SPP1+ | AGR2 |
| 1.445E-21 | 0.32268294 | 0.472 | 0.308 | 3.4135E-17 | MO_SPP1+ | CTNNB1 |
| 1.4937E-21 | -0.3867183 | 0.154 | 0.324 | 3.5286E-17 | MO_SPP1+ | YPEL3 |
| 1.6363E-21 | -0.2713645 | 0.006 | 0.131 | 3.8654E-17 | MO_SPP1+ | KAZN |
| 1.7157E-21 | 0.30472513 | 0.532 | 0.368 | 4.053E-17 | MO_SPP1+ | ATP5MC3 |
| 1.7257E-21 | -0.2765246 | 0.015 | 0.145 | 4.0765E-17 | MO_SPP1+ | PHLDA3 |
| 1.7846E-21 | 0.31728833 | 0.489 | 0.325 | 4.2158E-17 | MO_SPP1+ | LAMP1 |
| 1.8882E-21 | -0.3530645 | 0.015 | 0.146 | 4.4605E-17 | MO_SPP1+ | ITGB6 |
| 2.2144E-21 | 0.28425188 | 0.687 | 0.525 | 5.2312E-17 | MO_SPP1+ | CD164 |
| 2.4168E-21 | -0.319192 | 0.033 | 0.171 | 5.7091E-17 | MO_SPP1+ | JMY |
| 2.4288E-21 | 0.27574793 | 0.363 | 0.209 | 5.7375E-17 | MO_SPP1+ | NINJ1 |
| 2.8834E-21 | -0.2612936 | 0.027 | 0.163 | 6.8115E-17 | MO_SPP1+ | NCKAP1 |
| 2.8853E-21 | -0.2507048 | 0.022 | 0.156 | 6.8159E-17 | MO_SPP1+ | ARHGEF12 |
| 3.0807E-21 | -0.3386642 | 0.003 | 0.124 | 7.2776E-17 | MO_SPP1+ | KRT80 |
| 3.2777E-21 | -0.7245904 | 0.778 | 0.834 | 7.7428E-17 | MO_SPP1+ | PPDPF |
| 3.5688E-21 | -0.2874499 | 0.027 | 0.164 | 8.4306E-17 | MO_SPP1+ | MET |
| 4.2635E-21 | -0.2509834 | 0.012 | 0.138 | 1.0072E-16 | MO_SPP1+ | SCNN1A |
| 4.5999E-21 | 0.27297576 | 0.163 | 0.069 | 1.0866E-16 | MO_SPP1+ | IL1R2 |
| 5.3476E-21 | -0.7483408 | 0.028 | 0.163 | 1.2633E-16 | MO_SPP1+ | APOBEC3A |
| 5.6688E-21 | -0.2565611 | 0.001 | 0.12 | 1.3391E-16 | MO_SPP1+ | C19orf33 |
| 5.7729E-21 | -0.3136101 | 0.001 | 0.12 | 1.3637E-16 | MO_SPP1+ | ASS1 |
| 6.3678E-21 | -0.3373947 | 0.009 | 0.132 | 1.5043E-16 | MO_SPP1+ | FOXC1 |
| 7.3645E-21 | 0.27074226 | 0.918 | 0.818 | 1.7397E-16 | MO_SPP1+ | RPL27 |
| 7.7626E-21 | -0.3905878 | 0.558 | 0.69 | 1.8337E-16 | MO_SPP1+ | CAST |
| 9.9311E-21 | 0.29017071 | 0.906 | 0.808 | 2.346E-16 | MO_SPP1+ | RAC1 |
| 1.0021E-20 | -0.3314924 | 0.057 | 0.199 | 2.3672E-16 | MO_SPP1+ | NEDD4L |
| 1.0328E-20 | 0.25948384 | 0.951 | 0.856 | 2.4397E-16 | MO_SPP1+ | RPL7 |
| 1.0668E-20 | -0.3703656 | 0.108 | 0.261 | 2.5201E-16 | MO_SPP1+ | ARHGAP5 |
| 1.1464E-20 | -1.1173401 | 0.001 | 0.118 | 2.7082E-16 | MO_SPP1+ | SPINK5 |
| 1.1886E-20 | -0.2615443 | 0.003 | 0.121 | 2.8078E-16 | MO_SPP1+ | MUC20 |
| 1.2248E-20 | 0.2621737 | 0.427 | 0.271 | 2.8934E-16 | MO_SPP1+ | ADAM10 |
| 1.3762E-20 | -0.2775653 | 0.007 | 0.128 | 3.2509E-16 | MO_SPP1+ | MAST4 |
| 1.4978E-20 | 0.25260892 | 0.858 | 0.768 | 3.5383E-16 | MO_SPP1+ | CDC42 |
| 1.5845E-20 | -0.3527318 | 0.013 | 0.138 | 3.743E-16 | MO_SPP1+ | ARHGAP29 |
| 1.6764E-20 | -0.2671469 | 0.033 | 0.167 | 3.9601E-16 | MO_SPP1+ | SMAD3 |
| 2.0507E-20 | -0.5278254 | 0.343 | 0.484 | 4.8444E-16 | MO_SPP1+ | CD46 |
| 2.2458E-20 | 0.28619491 | 0.511 | 0.342 | 5.3052E-16 | MO_SPP1+ | LAMTOR4 |
| 2.3841E-20 | -0.3810852 | 0.015 | 0.139 | 5.632E-16 | MO_SPP1+ | DCDC2 |
| 2.4275E-20 | 0.28304683 | 0.459 | 0.295 | 5.7346E-16 | MO_SPP1+ | ARL6IP5 |
| 2.4432E-20 | -0.4781682 | 0.054 | 0.193 | 5.7716E-16 | MO_SPP1+ | PLK2 |
| 3.3157E-20 | -0.2595091 | 0.009 | 0.128 | 7.8327E-16 | MO_SPP1+ | SELENOP |
| 3.4562E-20 | -0.7581577 | 0.117 | 0.262 | 8.1646E-16 | MO_SPP1+ | TPM1 |
| 4.3231E-20 | -0.2933229 | 0 | 0.113 | 1.0212E-15 | MO_SPP1+ | FUT3 |
| 4.3891E-20 | -0.2735368 | 0.007 | 0.126 | 1.0368E-15 | MO_SPP1+ | FAT1 |
| 4.5174E-20 | -0.4958367 | 0.379 | 0.515 | 1.0671E-15 | MO_SPP1+ | ANXA11 |
| 4.7125E-20 | 0.27600069 | 0.409 | 0.257 | 1.1132E-15 | MO_SPP1+ | MBNL1 |
| 5.1392E-20 | -0.2806474 | 0.012 | 0.133 | 1.214E-15 | MO_SPP1+ | SCIN |
| 5.6583E-20 | -0.3646968 | 0.081 | 0.228 | 1.3367E-15 | MO_SPP1+ | RAB9A |
| 5.7549E-20 | 0.26176294 | 0.418 | 0.269 | 1.3595E-15 | MO_SPP1+ | STMP1 |

| | | | | | | |
|---|---|---|---|---|---|---|
| 6.3083E-20 | -0.5221091 | 0.006 | 0.122 | 1.4902E-15 | MO_SPP1+ | TCIM |
| 7.5861E-20 | 0.29045795 | 0.478 | 0.321 | 1.7921E-15 | MO_SPP1+ | TGOLN2 |
| 7.7213E-20 | -0.4588986 | 0.001 | 0.114 | 1.824E-15 | MO_SPP1+ | TMPRSS11A |
| 1.0424E-19 | -0.9608902 | 0.048 | 0.183 | 2.4624E-15 | MO_SPP1+ | SPINK1 |
| 1.0788E-19 | -0.4727666 | 0.078 | 0.22 | 2.5484E-15 | MO_SPP1+ | ZBTB20 |
| 1.6527E-19 | -0.365739 | 0.01 | 0.127 | 3.9041E-15 | MO_SPP1+ | ID4 |
| 1.7405E-19 | 0.26604666 | 0.457 | 0.306 | 4.1115E-15 | MO_SPP1+ | SSBP1 |
| 1.7648E-19 | -0.3113793 | 0.022 | 0.146 | 4.1691E-15 | MO_SPP1+ | CLDN1 |
| 1.8671E-19 | -0.3563594 | 0.001 | 0.112 | 4.4107E-15 | MO_SPP1+ | SMIM5 |
| 1.881E-19 | -0.3019232 | 0.013 | 0.132 | 4.4435E-15 | MO_SPP1+ | AIF1L |
| 1.954E-19 | 0.31952687 | 0.465 | 0.313 | 4.6159E-15 | MO_SPP1+ | HMGB2 |
| 2.021E-19 | -0.3939005 | 0 | 0.109 | 4.7742E-15 | MO_SPP1+ | KRT23 |
| 2.227E-19 | -0.5924164 | 0.003 | 0.114 | 5.2608E-15 | MO_SPP1+ | KRT78 |
| 2.4607E-19 | -0.3157954 | 0.063 | 0.201 | 5.813E-15 | MO_SPP1+ | TMEM106B |
| 3.1177E-19 | 0.30984642 | 0.531 | 0.377 | 7.365E-15 | MO_SPP1+ | PRELID1 |
| 3.4432E-19 | -0.415736 | 0.171 | 0.33 | 8.1339E-15 | MO_SPP1+ | LMNA |
| 3.7653E-19 | -0.4923397 | 0.091 | 0.231 | 8.8949E-15 | MO_SPP1+ | CYB5A |
| 3.9463E-19 | 0.26370611 | 0.85 | 0.761 | 9.3223E-15 | MO_SPP1+ | UBL5 |
| 3.9763E-19 | 0.26098883 | 0.499 | 0.343 | 9.3931E-15 | MO_SPP1+ | MTPN |
| 4.3323E-19 | -0.662705 | 0.019 | 0.139 | 1.0234E-14 | MO_SPP1+ | NUPR1 |
| 4.4831E-19 | -1.2756865 | 0.004 | 0.115 | 1.059E-14 | MO_SPP1+ | SPRR1B |
| 4.9969E-19 | -0.277392 | 0.019 | 0.14 | 1.1804E-14 | MO_SPP1+ | MTUS1 |
| 5.1661E-19 | -0.311065 | 0.003 | 0.112 | 1.2204E-14 | MO_SPP1+ | STS |
| 5.5918E-19 | -0.6387084 | 0.016 | 0.132 | 1.321E-14 | MO_SPP1+ | DHRS9 |
| 6.739E-19 | -0.3712945 | 0.045 | 0.174 | 1.592E-14 | MO_SPP1+ | STMN1 |
| 7.8787E-19 | -0.3598837 | 0.006 | 0.116 | 1.8612E-14 | MO_SPP1+ | CEACAM1 |
| 8.23E-19 | -0.4176582 | 0.001 | 0.108 | 1.9442E-14 | MO_SPP1+ | TMC5 |
| 1.006E-18 | 0.29998722 | 0.562 | 0.392 | 2.3764E-14 | MO_SPP1+ | COMMD6 |
| 1.1316E-18 | 0.3585556 | 0.474 | 0.321 | 2.6733E-14 | MO_SPP1+ | VEGFA |
| 1.2438E-18 | -0.3907293 | 0 | 0.105 | 2.9381E-14 | MO_SPP1+ | CLIC3 |
| 1.3548E-18 | -0.2800889 | 0.057 | 0.19 | 3.2004E-14 | MO_SPP1+ | MPRIP |
| 1.3781E-18 | -0.2874261 | 0.06 | 0.193 | 3.2554E-14 | MO_SPP1+ | F11R |
| 1.6486E-18 | -0.2890478 | 0.16 | 0.071 | 3.8944E-14 | MO_SPP1+ | CXCL5 |
| 1.746E-18 | -0.479823 | 0.067 | 0.197 | 4.1245E-14 | MO_SPP1+ | SAMD9 |
| 1.7715E-18 | -0.2572188 | 0.028 | 0.15 | 4.1848E-14 | MO_SPP1+ | RALGPS2 |
| 1.7929E-18 | 0.27094815 | 0.375 | 0.227 | 4.2353E-14 | MO_SPP1+ | PPIF |
| 2.0958E-18 | -0.4043375 | 0.064 | 0.199 | 4.951E-14 | MO_SPP1+ | BIRC3 |
| 2.3363E-18 | -0.3607871 | 0.01 | 0.121 | 5.519E-14 | MO_SPP1+ | EGLN3 |
| 2.3956E-18 | 0.2640941 | 0.717 | 0.582 | 5.6591E-14 | MO_SPP1+ | ACTR3 |
| 2.4305E-18 | -0.6185282 | 0.016 | 0.13 | 5.7415E-14 | MO_SPP1+ | TFPI2 |
| 2.6024E-18 | -1.5423235 | 0.003 | 0.108 | 6.1476E-14 | MO_SPP1+ | CRCT1 |
| 2.6571E-18 | -0.286132 | 0.036 | 0.159 | 6.2769E-14 | MO_SPP1+ | TRIP10 |
| 3.9417E-18 | -0.323835 | 0.003 | 0.107 | 9.3115E-14 | MO_SPP1+ | A2ML1 |
| 4.2563E-18 | -0.2700348 | 0.024 | 0.14 | 1.0055E-13 | MO_SPP1+ | ZNRF1 |
| 4.6138E-18 | -0.2899549 | 0.045 | 0.171 | 1.0899E-13 | MO_SPP1+ | ISG20 |
| 4.8525E-18 | -0.310631 | 0.036 | 0.156 | 1.1463E-13 | MO_SPP1+ | ITPKC |
| 5.8979E-18 | -0.7777787 | 0.003 | 0.106 | 1.3933E-13 | MO_SPP1+ | ZG16B |
| 6.4256E-18 | -0.4545549 | 0.525 | 0.631 | 1.5179E-13 | MO_SPP1+ | TAX1BP1 |
| 6.6426E-18 | -0.4292407 | 0.027 | 0.143 | 1.5692E-13 | MO_SPP1+ | ID1 |
| 6.8018E-18 | 0.28564924 | 0.429 | 0.286 | 1.6068E-13 | MO_SPP1+ | APRT |

| | | | | | | |
|---|---|---|---|---|---|---|
| 7.9511E-18 | -0.2672603 | 0.018 | 0.13 | 1.8783E-13 | MO_SPP1+ | BEX2 |
| 8.3105E-18 | -0.4163786 | 0.306 | 0.45 | 1.9632E-13 | MO_SPP1+ | ACTN4 |
| 9.2177E-18 | 0.27215212 | 0.412 | 0.274 | 2.1775E-13 | MO_SPP1+ | CCT5 |
| 1.1166E-17 | 0.3090471 | 0.937 | 0.847 | 2.6377E-13 | MO_SPP1+ | RPL38 |
| 1.9472E-17 | 0.25912504 | 0.387 | 0.251 | 4.6E-13 | MO_SPP1+ | PA2G4 |
| 2.2769E-17 | 0.26760621 | 0.546 | 0.386 | 5.3786E-13 | MO_SPP1+ | UQCR11 |
| 2.8909E-17 | -0.2515969 | 0.024 | 0.136 | 6.8293E-13 | MO_SPP1+ | COL18A1 |
| 5.2987E-17 | 0.26357412 | 0.942 | 0.845 | 1.2517E-12 | MO_SPP1+ | RPL35 |
| 6.1569E-17 | -0.5260905 | 0.631 | 0.723 | 1.4544E-12 | MO_SPP1+ | JUND |
| 6.5619E-17 | -0.2605488 | 0.045 | 0.163 | 1.5501E-12 | MO_SPP1+ | FARP1 |
| 7.0744E-17 | -0.3982056 | 0.028 | 0.141 | 1.6712E-12 | MO_SPP1+ | PLAC8 |
| 7.297E-17 | -0.3264461 | 0.009 | 0.11 | 1.7238E-12 | MO_SPP1+ | NR1D1 |
| 7.5232E-17 | -0.5438463 | 0.942 | 0.939 | 1.7772E-12 | MO_SPP1+ | SAT1 |
| 9.1531E-17 | -0.3422666 | 0.186 | 0.326 | 2.1622E-12 | MO_SPP1+ | YPEL5 |
| 1.3708E-16 | -0.2544887 | 0.006 | 0.104 | 3.2382E-12 | MO_SPP1+ | DUSP14 |
| 1.6756E-16 | -0.2678077 | 0.022 | 0.129 | 3.9582E-12 | MO_SPP1+ | TP53INP2 |
| 2.5741E-16 | -0.3528745 | 0.15 | 0.291 | 6.0807E-12 | MO_SPP1+ | CHMP1B |
| 2.6448E-16 | -1.4167447 | 0.01 | 0.109 | 6.2478E-12 | MO_SPP1+ | CNFN |
| 2.7416E-16 | -0.2792422 | 0.058 | 0.178 | 6.4765E-12 | MO_SPP1+ | RERE |
| 2.7954E-16 | 0.3134883 | 0.708 | 0.594 | 6.6035E-12 | MO_SPP1+ | HNRNPK |
| 3.8114E-16 | -0.6385032 | 0.103 | 0.232 | 9.0038E-12 | MO_SPP1+ | ISG15 |
| 4.1537E-16 | 0.25272603 | 0.723 | 0.603 | 9.8123E-12 | MO_SPP1+ | SUB1 |
| 4.5795E-16 | 0.25588805 | 0.123 | 0.053 | 1.0818E-11 | MO_SPP1+ | INHBA |
| 6.0237E-16 | -0.3965223 | 0.085 | 0.21 | 1.423E-11 | MO_SPP1+ | LY6E |
| 6.2164E-16 | -0.3119814 | 0.024 | 0.127 | 1.4685E-11 | MO_SPP1+ | POLB |
| 8.181E-16 | 0.31924341 | 0.828 | 0.763 | 1.9326E-11 | MO_SPP1+ | BTG1 |
| 8.1868E-16 | -0.2765246 | 0.018 | 0.118 | 1.934E-11 | MO_SPP1+ | ST3GAL4 |
| 8.2925E-16 | -0.3460819 | 0.121 | 0.25 | 1.9589E-11 | MO_SPP1+ | PKN2 |
| 1.1787E-15 | 0.31173478 | 0.436 | 0.304 | 2.7844E-11 | MO_SPP1+ | BNIP3L |
| 1.3077E-15 | -0.2634028 | 0.076 | 0.198 | 3.0891E-11 | MO_SPP1+ | TSPAN3 |
| 1.315E-15 | 0.26110268 | 0.555 | 0.412 | 3.1065E-11 | MO_SPP1+ | EIF3K |
| 1.3939E-15 | -0.3964302 | 0.522 | 0.623 | 3.2927E-11 | MO_SPP1+ | TMEM59 |
| 1.5189E-15 | -0.3521343 | 0.264 | 0.397 | 3.5881E-11 | MO_SPP1+ | CTNNA1 |
| 1.7002E-15 | -0.5934087 | 0.225 | 0.349 | 4.0163E-11 | MO_SPP1+ | ZFP36L2 |
| 1.7384E-15 | -0.356399 | 0.037 | 0.144 | 4.1066E-11 | MO_SPP1+ | PAX8 |
| 2.09E-15 | 0.260498 | 0.978 | 0.894 | 4.9372E-11 | MO_SPP1+ | RPL37A |
| 2.2155E-15 | -0.6850874 | 0.126 | 0.248 | 5.2337E-11 | MO_SPP1+ | PLAU |
| 4.0827E-15 | -0.6236068 | 0.415 | 0.529 | 9.6445E-11 | MO_SPP1+ | ZFP36 |
| 5.3809E-15 | -0.4049484 | 0.219 | 0.348 | 1.2711E-10 | MO_SPP1+ | GOLGB1 |
| 5.6941E-15 | -0.3640789 | 0.087 | 0.206 | 1.3451E-10 | MO_SPP1+ | CSNK1E |
| 6.6978E-15 | -0.3196885 | 0.178 | 0.308 | 1.5822E-10 | MO_SPP1+ | CLTB |
| 7.2961E-15 | -0.2592496 | 0.066 | 0.179 | 1.7236E-10 | MO_SPP1+ | PIK3IP1 |
| 7.8736E-15 | -0.5369151 | 0.612 | 0.7 | 1.86E-10 | MO_SPP1+ | JUNB |
| 9.4458E-15 | -0.2623812 | 0.073 | 0.19 | 2.2314E-10 | MO_SPP1+ | VCL |
| 1.0137E-14 | -0.5054224 | 0.789 | 0.838 | 2.3946E-10 | MO_SPP1+ | SQSTM1 |
| 1.0181E-14 | -0.5336443 | 0.232 | 0.352 | 2.405E-10 | MO_SPP1+ | GLS |
| 1.1078E-14 | 0.28829915 | 0.4 | 0.268 | 2.6169E-10 | MO_SPP1+ | TSC22D3 |
| 1.5285E-14 | -0.4763435 | 0.318 | 0.451 | 3.6108E-10 | MO_SPP1+ | MARCKS |
| 1.6635E-14 | -0.3058632 | 0.088 | 0.206 | 3.9297E-10 | MO_SPP1+ | TIPARP |
| 1.9291E-14 | -0.92996 | 0.037 | 0.138 | 4.5571E-10 | MO_SPP1+ | DEFB1 |

| | | | | | | |
|---|---|---|---|---|---|---|
| 2.1253E-14 | 0.27042943 | 0.342 | 0.221 | 5.0206E-10 | MO_SPP1+ | MRPS6 |
| 2.5019E-14 | -0.2780226 | 0.054 | 0.161 | 5.9104E-10 | MO_SPP1+ | SELENOM |
| 2.8249E-14 | 0.25814125 | 0.489 | 0.363 | 6.6732E-10 | MO_SPP1+ | ELOC |
| 2.9067E-14 | -0.2760581 | 0.013 | 0.102 | 6.8666E-10 | MO_SPP1+ | LRMP |
| 3.4733E-14 | -0.3927054 | 0.075 | 0.188 | 8.205E-10 | MO_SPP1+ | TNFSF10 |
| 3.9361E-14 | -0.6715822 | 0.039 | 0.141 | 9.2984E-10 | MO_SPP1+ | CXCL1 |
| 4.0386E-14 | -0.7132332 | 0.144 | 0.264 | 9.5404E-10 | MO_SPP1+ | AREG |
| 4.9411E-14 | -0.4151857 | 0.066 | 0.172 | 1.1672E-09 | MO_SPP1+ | PRDM1 |
| 5.7598E-14 | -0.3433034 | 0.199 | 0.327 | 1.3606E-09 | MO_SPP1+ | PPA1 |
| 6.3247E-14 | -0.2607225 | 0.04 | 0.14 | 1.4941E-09 | MO_SPP1+ | ETS1 |
| 6.8757E-14 | -0.4504299 | 0.144 | 0.26 | 1.6243E-09 | MO_SPP1+ | IDI1 |
| 7.0413E-14 | -0.3481486 | 0.156 | 0.278 | 1.6634E-09 | MO_SPP1+ | SLK |
| 7.246E-14 | 0.29831044 | 0.25 | 0.149 | 1.7117E-09 | MO_SPP1+ | SLC7A11 |
| 7.3238E-14 | -0.4198155 | 0.478 | 0.572 | 1.7301E-09 | MO_SPP1+ | ZFAND5 |
| 7.8416E-14 | -0.4848774 | 0.025 | 0.118 | 1.8524E-09 | MO_SPP1+ | HSPA2 |
| 8.3849E-14 | -0.3716064 | 0.189 | 0.314 | 1.9808E-09 | MO_SPP1+ | SDC4 |
| 8.7642E-14 | -0.2675981 | 0.013 | 0.1 | 2.0704E-09 | MO_SPP1+ | LY6K |
| 1.0226E-13 | -0.2660889 | 0.1 | 0.216 | 2.4158E-09 | MO_SPP1+ | MKNK2 |
| 1.218E-13 | -0.2658828 | 0.118 | 0.238 | 2.8772E-09 | MO_SPP1+ | PNPLA2 |
| 1.3471E-13 | -0.3322461 | 0.516 | 0.617 | 3.1824E-09 | MO_SPP1+ | KTN1 |
| 1.7861E-13 | -0.3212962 | 0.12 | 0.239 | 4.2192E-09 | MO_SPP1+ | ARID5B |
| 2.8204E-13 | -0.4173069 | 0.064 | 0.165 | 6.6625E-09 | MO_SPP1+ | ANKRD37 |
| 3.85E-13 | -0.3584293 | 0.094 | 0.203 | 9.0948E-09 | MO_SPP1+ | FOSB |
| 3.9788E-13 | -0.3901678 | 0.051 | 0.149 | 9.3991E-09 | MO_SPP1+ | RASD1 |
| 5.0598E-13 | -0.2923373 | 0.471 | 0.58 | 1.1953E-08 | MO_SPP1+ | UBE2B |
| 6.3762E-13 | -0.3368318 | 0.156 | 0.271 | 1.5063E-08 | MO_SPP1+ | FAM3C |
| 6.865E-13 | -3.5884529 | 0.451 | 0.543 | 1.6217E-08 | MO_SPP1+ | PLCG2 |
| 7.0992E-13 | -0.3581477 | 0.183 | 0.3 | 1.6771E-08 | MO_SPP1+ | PIM3 |
| 8.1943E-13 | -0.2636428 | 0.13 | 0.246 | 1.9357E-08 | MO_SPP1+ | SPTAN1 |
| 8.7812E-13 | -0.2567984 | 0.051 | 0.149 | 2.0744E-08 | MO_SPP1+ | GSTM3 |
| 1.0171E-12 | -0.2855104 | 0.225 | 0.348 | 2.4028E-08 | MO_SPP1+ | CAPN2 |
| 1.0956E-12 | -0.6847446 | 0.04 | 0.135 | 2.5881E-08 | MO_SPP1+ | SERPINB2 |
| 1.2191E-12 | -0.302425 | 0.025 | 0.112 | 2.8799E-08 | MO_SPP1+ | FXYD2 |
| 1.5023E-12 | -0.252568 | 0.051 | 0.146 | 3.5488E-08 | MO_SPP1+ | PDP1 |
| 1.6096E-12 | 0.44875156 | 0.621 | 0.517 | 3.8025E-08 | MO_SPP1+ | H1FX |
| 1.9992E-12 | -0.3413736 | 0.199 | 0.317 | 4.7227E-08 | MO_SPP1+ | AKAP9 |
| 2.3971E-12 | -0.3042171 | 0.148 | 0.268 | 5.6627E-08 | MO_SPP1+ | UGCG |
| 2.947E-12 | -0.2905865 | 0.697 | 0.782 | 6.9616E-08 | MO_SPP1+ | MAP1LC3B |
| 7.4631E-12 | -0.3278523 | 0.045 | 0.135 | 1.763E-07 | MO_SPP1+ | UCHL1 |
| 8.0728E-12 | -0.3207268 | 0.031 | 0.116 | 1.907E-07 | MO_SPP1+ | RRAD |
| 1.011E-11 | -0.2768909 | 0.04 | 0.127 | 2.3883E-07 | MO_SPP1+ | SYNE1 |
| 1.2788E-11 | -0.3194963 | 0.12 | 0.224 | 3.0208E-07 | MO_SPP1+ | POLR2A |
| 1.5403E-11 | 0.3409981 | 0.7 | 0.63 | 3.6386E-07 | MO_SPP1+ | ZFP36L1 |
| 1.6724E-11 | -0.4333553 | 0.19 | 0.301 | 3.9507E-07 | MO_SPP1+ | BTG2 |
| 1.7863E-11 | -0.2689062 | 0.196 | 0.307 | 4.2197E-07 | MO_SPP1+ | NOP53 |
| 2.7014E-11 | -0.295282 | 0.229 | 0.341 | 6.3816E-07 | MO_SPP1+ | YTHDC1 |
| 3.322E-11 | -0.3053582 | 0.319 | 0.43 | 7.8477E-07 | MO_SPP1+ | SELENOK |
| 5.4385E-11 | -0.3444088 | 0.073 | 0.167 | 1.2847E-06 | MO_SPP1+ | HIST3H2A |
| 5.7847E-11 | -0.2628653 | 0.121 | 0.22 | 1.3665E-06 | MO_SPP1+ | SPATS2L |
| 7.5301E-11 | -0.3108023 | 0.045 | 0.13 | 1.7788E-06 | MO_SPP1+ | AKAP12 |

| | | | | | | |
|---|---|---|---|---|---|---|
| 7.6646E-11 | -0.3965406 | 0.283 | 0.387 | 1.8106E-06 | MO_SPP1+ | C9orf16 |
| 7.714E-11 | -0.3536246 | 0.079 | 0.173 | 1.8223E-06 | MO_SPP1+ | DNAJA4 |
| 7.8828E-11 | -0.4283285 | 0.411 | 0.494 | 1.8621E-06 | MO_SPP1+ | H2AFJ |
| 1.1924E-10 | -0.491961 | 0.318 | 0.423 | 2.8167E-06 | MO_SPP1+ | TXNRD1 |
| 1.408E-10 | -0.3705588 | 0.046 | 0.126 | 3.3261E-06 | MO_SPP1+ | SRD5A3 |
| 1.4366E-10 | 0.29758264 | 0.279 | 0.184 | 3.3936E-06 | MO_SPP1+ | KLF2 |
| 1.5469E-10 | -0.2793181 | 0.273 | 0.377 | 3.6542E-06 | MO_SPP1+ | ARF4 |
| 1.663E-10 | -0.3161657 | 0.397 | 0.493 | 3.9285E-06 | MO_SPP1+ | SELENOW |
| 2.1596E-10 | -0.3028572 | 0.166 | 0.266 | 5.1016E-06 | MO_SPP1+ | HECTD1 |
| 2.9717E-10 | -0.2779599 | 0.238 | 0.34 | 7.0201E-06 | MO_SPP1+ | GPBP1 |
| 3.5854E-10 | -0.5384529 | 0.136 | 0.228 | 8.4698E-06 | MO_SPP1+ | HEXIM1 |
| 4.5388E-10 | -0.3345565 | 0.22 | 0.323 | 1.0722E-05 | MO_SPP1+ | B4GALT1 |
| 4.8349E-10 | -0.2510101 | 0.051 | 0.13 | 1.1422E-05 | MO_SPP1+ | GAN |
| 6.8402E-10 | -0.3161707 | 0.18 | 0.28 | 1.6159E-05 | MO_SPP1+ | ABHD5 |
| 8.5554E-10 | -0.3667958 | 0.96 | 0.95 | 2.0211E-05 | MO_SPP1+ | MT-CO1 |
| 9.7354E-10 | -0.3191371 | 0.244 | 0.345 | 2.2998E-05 | MO_SPP1+ | CPEB4 |
| 1.0433E-09 | -0.3981725 | 0.222 | 0.324 | 2.4646E-05 | MO_SPP1+ | IFITM3 |
| 1.1257E-09 | -0.3038228 | 0.051 | 0.13 | 2.6592E-05 | MO_SPP1+ | AC023157.3 |
| 1.5327E-09 | -0.3175798 | 0.189 | 0.285 | 3.6207E-05 | MO_SPP1+ | LDHB |
| 1.6675E-09 | 0.26264218 | 0.186 | 0.111 | 3.9391E-05 | MO_SPP1+ | CYP1B1 |
| 2.0419E-09 | -0.2651763 | 0.264 | 0.36 | 4.8235E-05 | MO_SPP1+ | RRBP1 |
| 2.2846E-09 | -0.2860867 | 0.394 | 0.483 | 5.397E-05 | MO_SPP1+ | GABARAPL2 |
| 4.6975E-09 | -0.566646 | 0.109 | 0.201 | 0.00011097 | MO_SPP1+ | THBS1 |
| 6.5187E-09 | -0.2534225 | 0.231 | 0.322 | 0.00015399 | MO_SPP1+ | ERP29 |
| 7.6157E-09 | -0.3395253 | 0.414 | 0.497 | 0.0001799 | MO_SPP1+ | CD9 |
| 7.7222E-09 | -0.2919953 | 0.235 | 0.33 | 0.00018242 | MO_SPP1+ | ASPH |
| 1.5236E-08 | 0.33090401 | 0.178 | 0.107 | 0.00035993 | MO_SPP1+ | APOC1 |
| 2.2948E-08 | -0.3076 | 0.945 | 0.944 | 0.00054211 | MO_SPP1+ | MT-CO3 |
| 3.0883E-08 | -1.8554488 | 0.915 | 0.724 | 0.00072954 | MO_SPP1+ | S100A9 |
| 3.2574E-08 | -0.2840296 | 0.175 | 0.258 | 0.0007695 | MO_SPP1+ | PDLIM5 |
| 3.456E-08 | -0.3434471 | 0.156 | 0.239 | 0.00081642 | MO_SPP1+ | CITED2 |
| 3.5519E-08 | -0.2567851 | 0.268 | 0.356 | 0.00083907 | MO_SPP1+ | MRPL33 |
| 7.8614E-08 | -0.7332621 | 0.348 | 0.253 | 0.0018571 | MO_SPP1+ | MT1G |
| 1.3237E-07 | -0.2718058 | 0.496 | 0.577 | 0.00312702 | MO_SPP1+ | CD55 |
| 1.3841E-07 | -0.3418328 | 0.148 | 0.227 | 0.00326958 | MO_SPP1+ | IER5L |
| 2.0292E-07 | -0.285371 | 0.169 | 0.247 | 0.00479355 | MO_SPP1+ | PTMS |
| 2.924E-07 | -0.2664987 | 0.418 | 0.497 | 0.00690725 | MO_SPP1+ | DDX3X |
| 3.083E-07 | -0.281275 | 0.111 | 0.188 | 0.00728305 | MO_SPP1+ | BAG3 |
| 3.1347E-07 | -0.2828895 | 0.457 | 0.53 | 0.00740509 | MO_SPP1+ | EIF4A2 |
| 3.2491E-07 | -0.2937173 | 0.252 | 0.326 | 0.00767541 | MO_SPP1+ | TMBIM1 |
| 5.1736E-07 | -0.3683311 | 0.048 | 0.105 | 0.01222161 | MO_SPP1+ | LTB |
| 6.4725E-07 | -0.2605066 | 0.162 | 0.237 | 0.01528998 | MO_SPP1+ | NFKBIZ |
| 7.3064E-07 | -0.2546405 | 0.184 | 0.263 | 0.01725999 | MO_SPP1+ | RHOC |
| 8.2541E-07 | -0.6151632 | 0.048 | 0.105 | 0.01949861 | MO_SPP1+ | CCL5 |
| 9.2214E-07 | -0.264028 | 0.39 | 0.464 | 0.02178375 | MO_SPP1+ | SARAF |
| 9.4041E-07 | -0.4401507 | 0.288 | 0.352 | 0.02221524 | MO_SPP1+ | PHLDA2 |
| 1.0912E-06 | -0.3108715 | 0.498 | 0.552 | 0.02577631 | MO_SPP1+ | FAM107B |
| 1.1343E-06 | -0.263383 | 0.118 | 0.186 | 0.02679608 | MO_SPP1+ | SEPHS2 |
| 1.138E-06 | -0.3364402 | 0.127 | 0.199 | 0.02688284 | MO_SPP1+ | DUSP4 |
| 1.7564E-06 | -0.3206159 | 0.238 | 0.31 | 0.04149229 | MO_SPP1+ | IRF1 |

| | | | | | | |
|---|---|---|---|---|---|---|
| 1.7932E-06 | -0.2946182 | 0.351 | 0.418 | 0.04236191 | MO_SPP1+ | JPT1 |
| 1.8844E-06 | -0.2693483 | 0.052 | 0.108 | 0.04451426 | MO_SPP1+ | TCHP |
| 2.2065E-06 | -0.2662668 | 0.126 | 0.194 | 0.0521236 | MO_SPP1+ | KATNBL1 |
| 2.5262E-06 | -0.2813577 | 0.495 | 0.548 | 0.0596768 | MO_SPP1+ | TOP1 |
| 3.0496E-06 | -0.2828763 | 0.297 | 0.367 | 0.07203968 | MO_SPP1+ | RALBP1 |
| 3.2915E-06 | -0.2966101 | 0.07 | 0.129 | 0.07775397 | MO_SPP1+ | GBP1 |
| 3.3913E-06 | -0.4129595 | 0.829 | 0.823 | 0.08011279 | MO_SPP1+ | MT-ND5 |
| 4.3748E-06 | -0.2926816 | 0.249 | 0.32 | 0.10334619 | MO_SPP1+ | BRD2 |
| 7.5807E-06 | -0.3241335 | 0.207 | 0.27 | 0.17908 | MO_SPP1+ | CCNG2 |
| 7.7742E-06 | -0.2646811 | 0.346 | 0.404 | 0.1836491 | MO_SPP1+ | SOD1 |
| 8.6575E-06 | -0.2527309 | 0.102 | 0.161 | 0.20451657 | MO_SPP1+ | SOCS1 |
| 9.8965E-06 | -0.6154525 | 0.261 | 0.312 | 0.23378464 | MO_SPP1+ | EMP1 |
| 1.3443E-05 | -0.6631022 | 0.222 | 0.283 | 0.31756891 | MO_SPP1+ | IFITM2 |
| 1.6561E-05 | -0.3455492 | 0.211 | 0.274 | 0.39122192 | MO_SPP1+ | TNFAIP2 |
| 2.2457E-05 | -0.3088398 | 0.165 | 0.226 | 0.53050717 | MO_SPP1+ | PPP1R10 |
| 2.8506E-05 | -0.270869 | 0.247 | 0.308 | 0.67339913 | MO_SPP1+ | ZNF292 |
| 5.1396E-05 | -0.3374454 | 0.493 | 0.536 | 1 | MO_SPP1+ | SERPINB1 |
| 6.1971E-05 | -0.6576616 | 0.165 | 0.216 | 1 | MO_SPP1+ | S100A12 |
| 6.7694E-05 | -0.3087582 | 0.1 | 0.152 | 1 | MO_SPP1+ | ZFAND2A |
| 0.00010015 | -0.2843376 | 0.354 | 0.397 | 1 | MO_SPP1+ | ARGLU1 |
| 0.00010444 | -0.2716825 | 0.589 | 0.602 | 1 | MO_SPP1+ | HINT1 |
| 0.00021521 | -0.2764927 | 0.289 | 0.346 | 1 | MO_SPP1+ | MIDN |
| 0.00030565 | -0.3127669 | 0.412 | 0.437 | 1 | MO_SPP1+ | CYSTM1 |
| 0.00034541 | -0.3121511 | 0.447 | 0.504 | 1 | MO_SPP1+ | DNAJB1 |
| 0.00094948 | -0.2653567 | 0.45 | 0.48 | 1 | MO_SPP1+ | RIOK3 |
| 0.00105465 | -0.2989844 | 0.256 | 0.299 | 1 | MO_SPP1+ | FAM129B |
| 0.00149795 | -0.2791656 | 0.184 | 0.225 | 1 | MO_SPP1+ | INTS6 |
| 0.00154858 | -0.4653589 | 0.598 | 0.628 | 1 | MO_SPP1+ | JUN |
| 0.00196745 | -0.3408982 | 0.111 | 0.149 | 1 | MO_SPP1+ | SMC4 |
| 0.00229261 | -0.3327396 | 0.205 | 0.246 | 1 | MO_SPP1+ | CST6 |
| 0.00409984 | -0.5953844 | 0.454 | 0.458 | 1 | MO_SPP1+ | MXD1 |
| 0.00450166 | -0.2643948 | 0.207 | 0.249 | 1 | MO_SPP1+ | EGR1 |
| 0.00665184 | -0.5426767 | 0.289 | 0.312 | 1 | MO_SPP1+ | PHLDA1 |
| 0.00818339 | -0.2880009 | 0.237 | 0.267 | 1 | MO_SPP1+ | DDIT3 |
| 0 | 4.3591681 | 0.95 | 0.272 | 0 | GRAN | IFITM2 |
| 0 | 2.93240923 | 0.874 | 0.094 | 0 | GRAN | CSF3R |
| 0 | 2.85558461 | 1 | 0.676 | 0 | GRAN | NAMPT |
| 0 | 2.74462062 | 0.737 | 0.107 | 0 | GRAN | MNDA |
| 0 | 2.63114606 | 0.776 | 0.007 | 0 | GRAN | FCGR3B |
| 0 | 2.29963748 | 0.976 | 0.572 | 0 | GRAN | LITAF |
| 0 | 2.01814077 | 0.865 | 0.272 | 0 | GRAN | PTPRC |
| 0 | 1.91868472 | 0.731 | 0.2 | 0 | GRAN | LST1 |
| 0 | 1.83045612 | 0.716 | 0.18 | 0 | GRAN | FYB1 |
| 0 | 1.77449058 | 0.689 | 0.137 | 0 | GRAN | FPR1 |
| 0 | 1.77038462 | 0.615 | 0.071 | 0 | GRAN | RIPOR2 |
| 0 | 1.67056382 | 0.529 | 0.006 | 0 | GRAN | CMTM2 |
| 0 | 1.65051993 | 0.442 | 0.022 | 0 | GRAN | FFAR2 |
| 0 | 1.63286047 | 0.618 | 0.132 | 0 | GRAN | GCA |
| 0 | 1.61779431 | 0.468 | 0.065 | 0 | GRAN | IL1R2 |
| 0 | 1.54508806 | 0.531 | 0.004 | 0 | GRAN | CXCR2 |

| | | | | | | |
|---|---|---|---|---|---|---|
| 0 | 1.3886098 | 0.411 | 0.016 | 0 | GRAN | PROK2 |
| 0 | 1.30872307 | 0.502 | 0.082 | 0 | GRAN | LRRK2 |
| 0 | 1.27259115 | 0.513 | 0.074 | 0 | GRAN | TMEM154 |
| 0 | 1.14648577 | 0.44 | 0.057 | 0 | GRAN | SELL |
| 0 | 1.07835306 | 0.405 | 0.029 | 0 | GRAN | VNN2 |
| 0 | 0.81842253 | 0.247 | 0.012 | 0 | GRAN | HCAR3 |
| 0 | 0.74911499 | 0.315 | 0.02 | 0 | GRAN | ADGRG3 |
| 0 | 0.74039567 | 0.279 | 0.002 | 0 | GRAN | ALPL |
| 0 | 0.70212847 | 0.261 | 0.022 | 0 | GRAN | P2RY13 |
| 0 | 0.65718641 | 0.248 | 0.001 | 0 | GRAN | CXCR1 |
| 0 | 0.63870556 | 0.247 | 0.02 | 0 | GRAN | MGAM |
| 0 | 0.54813998 | 0.215 | 0.012 | 0 | GRAN | MMP25 |
| 0 | 0.47851102 | 0.192 | 0.011 | 0 | GRAN | TNFRSF10C |
| 0 | 0.45283919 | 0.187 | 0.006 | 0 | GRAN | CEACAM3 |
| 0 | 0.40711809 | 0.124 | 0.003 | 0 | GRAN | LINC01506 |
| 5.011E-301 | 1.59815144 | 0.611 | 0.144 | 1.184E-296 | GRAN | SORL1 |
| 7.809E-286 | 2.07319362 | 0.797 | 0.278 | 1.845E-281 | GRAN | SLC25A37 |
| 1.737E-279 | 2.06749598 | 1 | 0.564 | 4.104E-275 | GRAN | SRGN |
| 2.396E-277 | 0.72644757 | 0.244 | 0.022 | 5.661E-273 | GRAN | KCNJ2 |
| 1.112E-276 | 0.93294271 | 0.327 | 0.04 | 2.628E-272 | GRAN | IFITM1 |
| 1.925E-268 | 2.51120996 | 1 | 0.666 | 4.547E-264 | GRAN | S100A8 |
| 7.173E-267 | 1.53006594 | 0.661 | 0.182 | 1.694E-262 | GRAN | FCGR2A |
| 9.939E-265 | 2.03997191 | 1 | 0.94 | 2.348E-260 | GRAN | NEAT1 |
| 3.8E-255 | 1.58399074 | 0.571 | 0.144 | 8.978E-251 | GRAN | FAM129A |
| 4.859E-254 | 1.68792407 | 0.998 | 0.833 | 1.148E-249 | GRAN | HLA-E |
| 1.088E-253 | 1.01884525 | 0.402 | 0.066 | 2.57E-249 | GRAN | FPR2 |
| 3.345E-250 | 0.72404599 | 0.263 | 0.028 | 7.901E-246 | GRAN | CYP4F3 |
| 2.716E-249 | 0.71354777 | 0.289 | 0.034 | 6.416E-245 | GRAN | MME |
| 9.497E-247 | 2.28419288 | 0.998 | 0.7 | 2.243E-242 | GRAN | SOD2 |
| 3.187E-241 | 1.84148534 | 1 | 0.904 | 7.529E-237 | GRAN | ITM2B |
| 6.952E-236 | 1.07917928 | 0.397 | 0.07 | 1.642E-231 | GRAN | CSF2RB |
| 4.349E-235 | 1.64711126 | 0.676 | 0.232 | 1.027E-230 | GRAN | SMCHD1 |
| 3.504E-232 | 2.28485831 | 0.608 | 0.168 | 8.278E-228 | GRAN | PLEK |
| 8.048E-231 | 1.01429021 | 0.419 | 0.078 | 1.901E-226 | GRAN | JAML |
| 2.383E-225 | 1.77358059 | 0.705 | 0.282 | 5.629E-221 | GRAN | RNF149 |
| 1.731E-223 | 1.71143506 | 0.645 | 0.206 | 4.09E-219 | GRAN | BASP1 |
| 1.994E-212 | 1.39880999 | 0.568 | 0.155 | 4.711E-208 | GRAN | AQP9 |
| 2.558E-212 | 0.3945383 | 0.139 | 0.009 | 6.043E-208 | GRAN | HCAR2 |
| 2.843E-212 | 1.71678814 | 1 | 0.957 | 6.716E-208 | GRAN | H3F3A |
| 7.757E-211 | 1.41771394 | 1 | 0.989 | 1.832E-206 | GRAN | B2M |
| 4.754E-210 | 0.83968658 | 0.326 | 0.051 | 1.123E-205 | GRAN | CR1 |
| 8.339E-209 | 1.3514951 | 0.529 | 0.137 | 1.97E-204 | GRAN | NCF1 |
| 7.239E-208 | -2.0563389 | 0.99 | 0.98 | 1.71E-203 | GRAN | EEF1A1 |
| 2.504E-201 | -2.4665918 | 0.994 | 0.95 | 5.915E-197 | GRAN | MT-CO1 |
| 2.678E-194 | 1.54022269 | 0.584 | 0.198 | 6.326E-190 | GRAN | ANP32A |
| 1.35E-187 | 1.49245555 | 0.577 | 0.177 | 3.188E-183 | GRAN | IGSF6 |
| 2.868E-185 | 1.17242544 | 1 | 0.723 | 6.776E-181 | GRAN | S100A9 |
| 2.328E-184 | -2.0482126 | 0.99 | 0.964 | 5.5E-180 | GRAN | RPL13 |
| 4.713E-178 | 1.26783231 | 0.944 | 0.433 | 1.113E-173 | GRAN | TYROBP |
| 7.013E-177 | 1.30998125 | 0.589 | 0.195 | 1.657E-172 | GRAN | NCF2 |

| | | | | | | |
|---|---|---|---|---|---|---|
| 3.166E-175 | 1.15529054 | 0.468 | 0.125 | 7.48E-171 | GRAN | LILRB3 |
| 2.391E-171 | -1.9659907 | 0.979 | 0.957 | 5.648E-167 | GRAN | RPS18 |
| 8.132E-170 | 1.18485166 | 0.515 | 0.158 | 1.921E-165 | GRAN | MYO1F |
| 3.313E-169 | -1.8422485 | 0.981 | 0.944 | 7.827E-165 | GRAN | RPS19 |
| 8.346E-168 | 1.40912836 | 0.621 | 0.24 | 1.972E-163 | GRAN | GMFG |
| 6.04E-162 | 1.27624708 | 1 | 0.888 | 1.427E-157 | GRAN | HLA-B |
| 2.02E-161 | 1.18317163 | 0.492 | 0.146 | 4.771E-157 | GRAN | EVI2B |
| 6.767E-161 | 2.1253972 | 0.6 | 0.217 | 1.598E-156 | GRAN | G0S2 |
| 6.779E-156 | 1.23719488 | 0.869 | 0.451 | 1.601E-151 | GRAN | MXD1 |
| 5.395E-154 | 0.48758379 | 0.205 | 0.027 | 1.275E-149 | GRAN | MEFV |
| 1.71E-153 | 0.60053713 | 0.227 | 0.033 | 4.04E-149 | GRAN | RGS18 |
| 4.818E-153 | -1.4776242 | 0.977 | 0.957 | 1.138E-148 | GRAN | RPL15 |
| 2.881E-150 | -1.8309006 | 0.997 | 0.972 | 6.805E-146 | GRAN | RPLP1 |
| 4.134E-150 | 1.39312806 | 0.668 | 0.285 | 9.766E-146 | GRAN | LCP1 |
| 4.294E-147 | 0.97332704 | 0.276 | 0.049 | 1.014E-142 | GRAN | TNFAIP6 |
| 6.584E-147 | 1.42752401 | 0.997 | 0.868 | 1.555E-142 | GRAN | HLA-C |
| 1.222E-145 | 1.37883025 | 0.524 | 0.18 | 2.888E-141 | GRAN | ACSL1 |
| 2.457E-145 | -1.9024866 | 0.989 | 0.947 | 5.805E-141 | GRAN | RPL10 |
| 3.292E-145 | 1.55800309 | 0.608 | 0.23 | 7.776E-141 | GRAN | BCL2A1 |
| 4.19E-145 | 1.38561782 | 0.613 | 0.275 | 9.898E-141 | GRAN | LYN |
| 2.144E-144 | -1.4020225 | 0.979 | 0.945 | 5.066E-140 | GRAN | PTMA |
| 7.508E-144 | 1.18578458 | 1 | 0.973 | 1.774E-139 | GRAN | ACTB |
| 2.194E-142 | 1.57036777 | 0.944 | 0.626 | 5.182E-138 | GRAN | ZFP36L1 |
| 4.015E-140 | -1.3391716 | 0.976 | 0.949 | 9.485E-136 | GRAN | RPL18 |
| 2.655E-139 | 1.4245606 | 0.947 | 0.637 | 6.271E-135 | GRAN | TXNIP |
| 1.194E-137 | -2.0330173 | 0.985 | 0.932 | 2.821E-133 | GRAN | RPS12 |
| 5.692E-137 | 0.5853349 | 0.24 | 0.041 | 1.345E-132 | GRAN | MCTP2 |
| 1.263E-135 | 0.26886989 | 0.111 | 0.009 | 2.983E-131 | GRAN | NFE2 |
| 4.535E-133 | 0.71664641 | 0.237 | 0.041 | 1.071E-128 | GRAN | TAGAP |
| 5.847E-133 | 1.23454179 | 0.881 | 0.525 | 1.381E-128 | GRAN | CTSS |
| 6.083E-133 | 1.22061182 | 0.918 | 0.635 | 1.437E-128 | GRAN | ARPC5 |
| 1.457E-132 | 1.16873476 | 1 | 0.952 | 3.443E-128 | GRAN | H3F3B |
| 3.709E-132 | 1.39169069 | 0.577 | 0.214 | 8.763E-128 | GRAN | ALOX5AP |
| 1.343E-130 | -1.6988932 | 0.974 | 0.911 | 3.174E-126 | GRAN | RPS8 |
| 1.397E-130 | 0.32265366 | 0.137 | 0.014 | 3.301E-126 | GRAN | RGL4 |
| 1.836E-130 | 1.16871727 | 0.965 | 0.706 | 4.337E-126 | GRAN | SDCBP |
| 3.027E-130 | -1.4542663 | 0.982 | 0.945 | 7.15E-126 | GRAN | RPL13A |
| 1.362E-129 | 1.46433048 | 0.577 | 0.254 | 3.219E-125 | GRAN | NABP1 |
| 2.077E-126 | 1.40285412 | 0.444 | 0.143 | 4.906E-122 | GRAN | RGS2 |
| 6.114E-124 | -1.5029404 | 0.987 | 0.943 | 1.444E-119 | GRAN | RPS27 |
| 7.031E-124 | 0.54778356 | 0.213 | 0.035 | 1.661E-119 | GRAN | AL034397.3 |
| 1.228E-122 | 1.66774668 | 0.773 | 0.445 | 2.901E-118 | GRAN | MARCKS |
| 2.471E-121 | 0.83711591 | 0.308 | 0.073 | 5.837E-117 | GRAN | LIMK2 |
| 1.831E-120 | 1.86654079 | 0.603 | 0.305 | 4.326E-116 | GRAN | IRF1 |
| 4.047E-120 | -1.5186686 | 0.974 | 0.923 | 9.561E-116 | GRAN | RPL18A |
| 7.706E-118 | -1.7376865 | 0.994 | 0.936 | 1.82E-113 | GRAN | MT-ATP6 |
| 7.727E-118 | -1.5024872 | 0.984 | 0.923 | 1.825E-113 | GRAN | RPS24 |
| 7.798E-118 | 1.03961876 | 0.465 | 0.172 | 1.842E-113 | GRAN | MBOAT7 |
| 1.056E-115 | -1.501323 | 0.981 | 0.921 | 2.496E-111 | GRAN | RPL11 |
| 8.529E-114 | -1.4427512 | 0.981 | 0.926 | 2.015E-109 | GRAN | RPL19 |

| | | | | | | |
|---|---|---|---|---|---|---|
| 1.552E-113 | -1.1664483 | 0.965 | 0.917 | 3.666E-109 | GRAN | RPS20 |
| 9.594E-113 | -1.5636232 | 0.992 | 0.944 | 2.266E-108 | GRAN | MT-CO2 |
| 4.037E-112 | -1.3796182 | 0.979 | 0.922 | 9.537E-108 | GRAN | RPL8 |
| 1.515E-111 | 1.20798878 | 0.613 | 0.297 | 3.579E-107 | GRAN | SERPINA1 |
| 2.204E-110 | 1.13548976 | 0.497 | 0.171 | 5.207E-106 | GRAN | IL1B |
| 1.679E-109 | 0.70742055 | 0.366 | 0.1 | 3.966E-105 | GRAN | LTB |
| 8.792E-109 | -1.198557 | 0.99 | 0.977 | 2.077E-104 | GRAN | RPL28 |
| 1.445E-107 | -1.096703 | 0.982 | 0.946 | 3.413E-103 | GRAN | RPS16 |
| 5.216E-106 | -1.3220938 | 0.977 | 0.919 | 1.232E-101 | GRAN | RPS15 |
| 2.677E-105 | 1.35538064 | 0.352 | 0.105 | 6.323E-101 | GRAN | PDE4B |
| 4.591E-105 | -1.3146509 | 0.982 | 0.939 | 1.085E-100 | GRAN | RPS28 |
| 9.755E-105 | 0.75079507 | 0.319 | 0.088 | 2.304E-100 | GRAN | ICAM3 |
| 1.391E-104 | -1.4504715 | 0.974 | 0.904 | 3.286E-100 | GRAN | RPL7A |
| 1.56E-104 | 1.41402593 | 0.611 | 0.356 | 3.685E-100 | GRAN | CFLAR |
| 2.114E-104 | -1.4183569 | 0.985 | 0.932 | 4.993E-100 | GRAN | RPS14 |
| 3.646E-104 | -1.5197922 | 0.981 | 0.915 | 8.614E-100 | GRAN | RPL32 |
| 1.091E-103 | -1.2858691 | 0.979 | 0.918 | 2.577E-99 | GRAN | RPL9 |
| 5.006E-103 | -1.5999582 | 0.992 | 0.943 | 1.1827E-98 | GRAN | MT-CO3 |
| 1.465E-102 | -1.4353588 | 0.995 | 0.97 | 3.4604E-98 | GRAN | TMSB10 |
| 6.142E-102 | -1.314762 | 0.976 | 0.894 | 1.451E-97 | GRAN | RPL37A |
| 1.205E-101 | -1.3610546 | 0.981 | 0.903 | 2.8471E-97 | GRAN | RPLP2 |
| 1.4E-101 | -1.5662994 | 0.994 | 0.929 | 3.3074E-97 | GRAN | MT-CYB |
| 3.326E-100 | 1.51392975 | 0.506 | 0.224 | 7.8559E-96 | GRAN | IVNS1ABP |
| 2.168E-99 | -1.5799531 | 0.971 | 0.919 | 5.1216E-95 | GRAN | S100A10 |
| 2.286E-99 | -1.6015635 | 0.984 | 0.884 | 5.4009E-95 | GRAN | RPS2 |
| 1.0473E-98 | -1.5838365 | 0.976 | 0.888 | 2.474E-94 | GRAN | RPS23 |
| 7.2376E-98 | 0.78813432 | 0.34 | 0.103 | 1.7097E-93 | GRAN | MX2 |
| 3.7729E-97 | -1.3383604 | 0.977 | 0.897 | 8.9127E-93 | GRAN | RPL29 |
| 7.1846E-97 | 1.16255816 | 0.865 | 0.571 | 1.6972E-92 | GRAN | CD55 |
| 3.5229E-96 | -1.333715 | 0.969 | 0.875 | 8.3221E-92 | GRAN | RPL23A |
| 4.7575E-96 | 0.92729516 | 0.505 | 0.211 | 1.1239E-91 | GRAN | SPI1 |
| 6.3305E-96 | 0.68235628 | 0.289 | 0.078 | 1.4954E-91 | GRAN | NCF4 |
| 7.4061E-96 | -1.4505241 | 0.99 | 0.932 | 1.7495E-91 | GRAN | MT-ND4 |
| 1.0019E-94 | -1.2257374 | 0.974 | 0.887 | 2.3669E-90 | GRAN | RPL6 |
| 1.6516E-93 | 0.9615594 | 0.887 | 0.598 | 3.9016E-89 | GRAN | MCL1 |
| 2.8447E-93 | 1.01189693 | 0.545 | 0.228 | 6.7199E-89 | GRAN | SLC2A3 |
| 3.8294E-93 | 0.73481273 | 0.31 | 0.089 | 9.0461E-89 | GRAN | PLXNC1 |
| 4.1402E-93 | 1.14620704 | 0.535 | 0.272 | 9.7803E-89 | GRAN | VASP |
| 8.064E-93 | 0.33383302 | 0.148 | 0.023 | 1.905E-88 | GRAN | GLT1D1 |
| 2.5522E-92 | -1.1122662 | 0.973 | 0.9 | 6.029E-88 | GRAN | RPL24 |
| 1.4321E-91 | 0.52603947 | 0.235 | 0.054 | 3.383E-87 | GRAN | TLR1 |
| 1.4886E-91 | 0.59196296 | 0.25 | 0.06 | 3.5166E-87 | GRAN | KCNJ15 |
| 3.0595E-91 | 1.14693892 | 0.932 | 0.687 | 7.2276E-87 | GRAN | PNRC1 |
| 8.9764E-91 | -1.1115426 | 0.997 | 0.98 | 2.1205E-86 | GRAN | RPL41 |
| 9.7052E-91 | -1.3780757 | 0.981 | 0.923 | 2.2926E-86 | GRAN | RPS27A |
| 1.4182E-90 | 1.290238 | 0.508 | 0.24 | 3.3503E-86 | GRAN | KIAA1551 |
| 3.7186E-90 | 1.26008146 | 0.861 | 0.483 | 8.7845E-86 | GRAN | CXCL8 |
| 5.0226E-90 | -1.3955306 | 0.973 | 0.88 | 1.1865E-85 | GRAN | RPL12 |
| 5.2802E-89 | -1.3085869 | 0.953 | 0.868 | 1.2473E-84 | GRAN | RPLP0 |
| 1.716E-88 | 0.76718083 | 0.31 | 0.094 | 4.0538E-84 | GRAN | CYTH4 |

| | | | | | | |
|---|---|---|---|---|---|---|
| 1.8549E-88 | -1.3332153 | 0.968 | 0.883 | 4.3818E-84 | GRAN | HSP90AB1 |
| 5.2793E-87 | -1.5182315 | 0.977 | 0.886 | 1.2471E-82 | GRAN | RPS6 |
| 1.1486E-86 | -1.2419866 | 0.968 | 0.874 | 2.7133E-82 | GRAN | RPL36 |
| 1.3144E-86 | -1.2724191 | 0.94 | 0.866 | 3.1049E-82 | GRAN | ANXA2 |
| 6.7044E-85 | -1.4145726 | 0.965 | 0.875 | 1.5838E-80 | GRAN | RPL3 |
| 1.0684E-84 | 1.09549517 | 0.961 | 0.62 | 2.5239E-80 | GRAN | S100A4 |
| 3.2297E-84 | -1.1026472 | 0.966 | 0.895 | 7.6294E-80 | GRAN | RPL35A |
| 1.1317E-83 | 1.36685591 | 0.489 | 0.211 | 2.6735E-79 | GRAN | S100A12 |
| 9.5477E-83 | 0.64287061 | 0.285 | 0.085 | 2.2554E-78 | GRAN | MSRB1 |
| 3.069E-82 | -1.3863247 | 0.982 | 0.896 | 7.2499E-78 | GRAN | RPS3A |
| 1.2868E-81 | -1.2324007 | 0.976 | 0.883 | 3.0397E-77 | GRAN | RPS13 |
| 3.5182E-81 | -1.2382684 | 0.969 | 0.87 | 8.311E-77 | GRAN | RPL14 |
| 4.1592E-81 | -0.7093131 | 0.005 | 0.394 | 9.8253E-77 | GRAN | SEM1 |
| 6.1476E-81 | 0.95402259 | 0.435 | 0.168 | 1.4522E-76 | GRAN | STEAP4 |
| 7.795E-81 | 1.03949548 | 0.406 | 0.16 | 1.8414E-76 | GRAN | LCP2 |
| 1.9536E-80 | -1.1545553 | 0.971 | 0.901 | 4.6149E-76 | GRAN | RPS7 |
| 1.5227E-79 | 0.29674823 | 0.106 | 0.014 | 3.5971E-75 | GRAN | CEP19 |
| 2.0664E-79 | 0.86620479 | 0.858 | 0.568 | 4.8813E-75 | GRAN | VMP1 |
| 5.2042E-79 | 1.07829861 | 0.573 | 0.27 | 1.2294E-74 | GRAN | AIF1 |
| 1.2356E-78 | -1.1791055 | 0.937 | 0.83 | 2.9187E-74 | GRAN | RPL22 |
| 1.4044E-78 | -1.2740596 | 0.974 | 0.874 | 3.3176E-74 | GRAN | RPL26 |
| 1.5055E-78 | 0.78493804 | 0.352 | 0.127 | 3.5564E-74 | GRAN | CELF2 |
| 2.4276E-78 | 0.86395701 | 0.452 | 0.205 | 5.7347E-74 | GRAN | ARRB2 |
| 2.9033E-78 | 0.84472592 | 1 | 0.993 | 6.8584E-74 | GRAN | FTH1 |
| 3.5069E-78 | -1.1829509 | 0.942 | 0.845 | 8.2844E-74 | GRAN | RPL35 |
| 5.192E-78 | -1.209021 | 0.979 | 0.898 | 1.2265E-73 | GRAN | RPL21 |
| 8.3918E-78 | 0.90790148 | 0.353 | 0.131 | 1.9824E-73 | GRAN | BCL6 |
| 3.4053E-77 | 0.74495841 | 0.319 | 0.108 | 8.0443E-73 | GRAN | LY96 |
| 8.4443E-77 | -1.4426866 | 0.961 | 0.847 | 1.9948E-72 | GRAN | RPS4X |
| 2.1605E-75 | 0.78708311 | 0.389 | 0.157 | 5.1038E-71 | GRAN | WIPF1 |
| 2.2351E-75 | -0.8809939 | 0.016 | 0.394 | 5.2799E-71 | GRAN | AHNAK |
| 4.1131E-75 | -0.8023338 | 0.021 | 0.399 | 9.7163E-71 | GRAN | NDUFB2 |
| 8.7527E-75 | 0.94214606 | 0.803 | 0.521 | 2.0677E-70 | GRAN | CEBPB |
| 1.1959E-74 | 1.28099334 | 0.445 | 0.203 | 2.8251E-70 | GRAN | GK |
| 2.2644E-74 | -1.1102859 | 0.955 | 0.857 | 5.3492E-70 | GRAN | RPL7 |
| 7.0395E-74 | 0.32484195 | 0.142 | 0.025 | 1.6629E-69 | GRAN | DYSF |
| 2.1693E-73 | 0.3810923 | 0.171 | 0.036 | 5.1245E-69 | GRAN | LINC01001 |
| 2.7117E-73 | -1.2953147 | 0.935 | 0.816 | 6.4058E-69 | GRAN | RPL5 |
| 2.7464E-73 | 0.80519925 | 0.416 | 0.183 | 6.4878E-69 | GRAN | GPSM3 |
| 1.1899E-72 | -0.6801079 | 1 | 0.993 | 2.8109E-68 | GRAN | TPT1 |
| 1.3258E-72 | -1.0886362 | 0.937 | 0.837 | 3.1318E-68 | GRAN | RACK1 |
| 1.4014E-72 | 0.86783038 | 0.371 | 0.153 | 3.3104E-68 | GRAN | R3HDM4 |
| 4.3504E-72 | 0.60101788 | 0.255 | 0.076 | 1.0277E-67 | GRAN | ABTB1 |
| 5.0838E-72 | 0.64062809 | 0.244 | 0.066 | 1.201E-67 | GRAN | CST7 |
| 8.1059E-72 | -0.8759163 | 0.047 | 0.409 | 1.9149E-67 | GRAN | SOD1 |
| 1.2608E-71 | -0.733474 | 0.013 | 0.376 | 2.9784E-67 | GRAN | ATP5MC3 |
| 4.8566E-71 | -0.7128306 | 0.011 | 0.372 | 1.1473E-66 | GRAN | AES |
| 5.2889E-71 | -1.2305608 | 0.979 | 0.886 | 1.2494E-66 | GRAN | RPS3 |
| 7.5317E-71 | -0.9776126 | 0.976 | 0.897 | 1.7792E-66 | GRAN | RPS25 |
| 9.3774E-71 | -1.1299223 | 0.974 | 0.892 | 2.2152E-66 | GRAN | RPS15A |

| | | | | | | |
|---|---|---|---|---|---|---|
| 9.5011E-71 | -0.6829426 | 0.008 | 0.364 | 2.2444E-66 | GRAN | SNRPD2 |
| 1.552E-70 | -1.0943803 | 0.989 | 0.933 | 3.6662E-66 | GRAN | RPL34 |
| 1.9956E-70 | -0.6365225 | 0.029 | 0.4 | 4.7143E-66 | GRAN | NDUFB11 |
| 2.1304E-70 | -0.7039027 | 0.011 | 0.368 | 5.0327E-66 | GRAN | SLC25A5 |
| 2.6805E-70 | -0.7377828 | 0.006 | 0.36 | 6.3321E-66 | GRAN | CYCS |
| 4.769E-70 | -0.7840479 | 0.065 | 0.425 | 1.1266E-65 | GRAN | RAN |
| 4.8454E-69 | -0.6899871 | 0.008 | 0.358 | 1.1446E-64 | GRAN | EIF3E |
| 7.5151E-68 | -0.9242537 | 0.008 | 0.354 | 1.7753E-63 | GRAN | PEBP1 |
| 8.7121E-68 | -0.6540752 | 0.006 | 0.351 | 2.058E-63 | GRAN | CAPN2 |
| 1.2564E-67 | 0.86659797 | 0.926 | 0.7 | 2.9679E-63 | GRAN | TPM3 |
| 2.3217E-67 | 0.77523836 | 0.344 | 0.133 | 5.4846E-63 | GRAN | HCK |
| 2.4244E-67 | -0.6807943 | 0.018 | 0.369 | 5.7272E-63 | GRAN | ATP5F1D |
| 8.7238E-67 | -2.3048018 | 0.931 | 0.536 | 2.0608E-62 | GRAN | PLCG2 |
| 2.3194E-66 | 0.8593227 | 0.398 | 0.17 | 5.479E-62 | GRAN | SMIM25 |
| 3.435E-66 | -0.6275959 | 0.037 | 0.398 | 8.1146E-62 | GRAN | SERBP1 |
| 4.7602E-66 | -0.6531764 | 0.013 | 0.354 | 1.1245E-61 | GRAN | VDAC1 |
| 4.8582E-66 | -0.6512371 | 0.026 | 0.376 | 1.1477E-61 | GRAN | RHEB |
| 1.0416E-65 | -0.8053206 | 0.039 | 0.396 | 2.4605E-61 | GRAN | SRSF7 |
| 1.6952E-65 | -0.7272075 | 0.024 | 0.372 | 4.0045E-61 | GRAN | CTSH |
| 3.4552E-65 | -0.9750396 | 0.016 | 0.358 | 8.1623E-61 | GRAN | ATF3 |
| 4.8497E-65 | 0.80292244 | 0.352 | 0.144 | 1.1456E-60 | GRAN | LYST |
| 9.6942E-65 | 1.04742134 | 0.869 | 0.598 | 2.2901E-60 | GRAN | NFKBIA |
| 1.3056E-64 | -0.559683 | 0.031 | 0.381 | 3.0842E-60 | GRAN | EIF3H |
| 1.4181E-64 | -0.7791623 | 0.976 | 0.919 | 3.3499E-60 | GRAN | NACA |
| 1.4501E-64 | -1.3092065 | 0.937 | 0.831 | 3.4256E-60 | GRAN | PPDPF |
| 2.9947E-64 | 0.40842619 | 0.195 | 0.05 | 7.0744E-60 | GRAN | SIRPB1 |
| 3.7042E-64 | -0.7754483 | 0.006 | 0.337 | 8.7504E-60 | GRAN | SNHG8 |
| 4.0562E-64 | -0.9459184 | 0.003 | 0.333 | 9.5821E-60 | GRAN | MYO6 |
| 5.448E-64 | 0.82209511 | 0.426 | 0.2 | 1.287E-59 | GRAN | LSP1 |
| 6.4631E-64 | -0.6048928 | 0.023 | 0.362 | 1.5268E-59 | GRAN | COX5A |
| 6.6273E-64 | -0.6615222 | 0.031 | 0.38 | 1.5656E-59 | GRAN | PARK7 |
| 7.1467E-64 | -0.9121781 | 0.979 | 0.898 | 1.6883E-59 | GRAN | RPL27A |
| 7.2769E-64 | -0.6075909 | 0.023 | 0.364 | 1.719E-59 | GRAN | TOMM20 |
| 8.6124E-64 | -0.5776563 | 0.06 | 0.412 | 2.0345E-59 | GRAN | NDUFB4 |
| 6.1504E-63 | 0.8993349 | 0.371 | 0.16 | 1.4529E-58 | GRAN | CLEC2B |
| 7.5933E-63 | 0.76978255 | 0.339 | 0.134 | 1.7938E-58 | GRAN | CLEC7A |
| 1.0511E-62 | -0.5877537 | 0.027 | 0.368 | 2.483E-58 | GRAN | C19orf53 |
| 1.722E-62 | -0.7842291 | 0.006 | 0.332 | 4.0679E-58 | GRAN | LMNA |
| 2.815E-62 | 0.79427552 | 0.795 | 0.581 | 6.65E-58 | GRAN | ACTR3 |
| 2.8286E-62 | -0.6310965 | 0.052 | 0.4 | 6.682E-58 | GRAN | CTNNA1 |
| 4.0699E-62 | -0.6266862 | 0.027 | 0.366 | 9.6142E-58 | GRAN | MTDH |
| 4.7532E-62 | -0.7643482 | 0.016 | 0.346 | 1.1228E-57 | GRAN | PDIA6 |
| 5.8042E-62 | -0.5859659 | 0.018 | 0.349 | 1.3711E-57 | GRAN | EIF1AX |
| 7.1212E-62 | -0.7052387 | 0.008 | 0.33 | 1.6822E-57 | GRAN | PPA1 |
| 1.0105E-61 | -0.6098166 | 0.042 | 0.388 | 2.3871E-57 | GRAN | DYNLRB1 |
| 1.1889E-61 | -0.8020737 | 0.005 | 0.325 | 2.8086E-57 | GRAN | PERP |
| 1.2792E-61 | 0.46946112 | 0.195 | 0.051 | 3.0219E-57 | GRAN | LMNB1 |
| 1.7617E-61 | -0.7455256 | 0.047 | 0.393 | 4.1616E-57 | GRAN | TUBB4B |
| 4.6307E-61 | -0.6878466 | 0.008 | 0.329 | 1.0939E-56 | GRAN | MGST3 |
| 5.8278E-61 | -0.628835 | 0.015 | 0.338 | 1.3767E-56 | GRAN | SSR3 |

| | | | | | | |
|---|---|---|---|---|---|---|
| 7.3606E-61 | -0.8407475 | 0.916 | 0.833 | 1.7388E-56 | GRAN | PPIA |
| 9.0642E-61 | 0.73937309 | 0.415 | 0.181 | 2.1412E-56 | GRAN | C5AR1 |
| 1.0393E-60 | -1.1985108 | 0.002 | 0.316 | 2.4552E-56 | GRAN | EMP1 |
| 1.7039E-60 | -0.5771626 | 0.027 | 0.357 | 4.0251E-56 | GRAN | SPCS1 |
| 2.4393E-60 | 0.82240517 | 0.816 | 0.523 | 5.7624E-56 | GRAN | ZFP36 |
| 5.4378E-60 | 0.91700548 | 0.99 | 0.891 | 1.2846E-55 | GRAN | ATP5F1E |
| 7.1457E-60 | 0.81508551 | 0.935 | 0.673 | 1.688E-55 | GRAN | IER2 |
| 1.3676E-59 | 0.84388301 | 1 | 0.938 | 3.2308E-55 | GRAN | SAT1 |
| 1.4538E-59 | 0.82756672 | 0.6 | 0.331 | 3.4343E-55 | GRAN | LAPTM5 |
| 1.7023E-59 | -0.5939386 | 0.01 | 0.325 | 4.0213E-55 | GRAN | ERP29 |
| 1.9321E-59 | -0.6979573 | 0.016 | 0.336 | 4.5643E-55 | GRAN | APP |
| 2.7584E-59 | -0.6649208 | 0.013 | 0.331 | 6.5161E-55 | GRAN | CAPG |
| 4.4499E-59 | 0.85354843 | 1 | 0.965 | 1.0512E-54 | GRAN | S100A11 |
| 8.8368E-59 | 0.97585518 | 0.463 | 0.264 | 2.0875E-54 | GRAN | USP15 |
| 1.0128E-58 | -0.5770041 | 0.013 | 0.326 | 2.3924E-54 | GRAN | YWHAQ |
| 1.8187E-58 | -0.9296098 | 0.96 | 0.858 | 4.2962E-54 | GRAN | RPL23 |
| 2.6849E-58 | -0.9879021 | 0.985 | 0.927 | 6.3425E-54 | GRAN | RPL30 |
| 3.8437E-58 | -0.5678227 | 0.026 | 0.347 | 9.08E-54 | GRAN | ATP5F1B |
| 4.8312E-58 | -0.7971287 | 0 | 0.303 | 1.1413E-53 | GRAN | FAM129B |
| 6.0721E-58 | -0.6992634 | 0.008 | 0.316 | 1.4344E-53 | GRAN | SDC4 |
| 1.0804E-57 | -0.6012551 | 0.015 | 0.326 | 2.5522E-53 | GRAN | KRT10 |
| 1.2157E-57 | -0.6022312 | 0.079 | 0.415 | 2.8719E-53 | GRAN | ATP5PF |
| 1.4619E-57 | 0.6411417 | 0.263 | 0.09 | 3.4534E-53 | GRAN | TNFSF13B |
| 1.9983E-57 | -0.6346762 | 0.037 | 0.363 | 4.7205E-53 | GRAN | ATP1A1 |
| 2.3791E-57 | -0.5280356 | 0.024 | 0.341 | 5.62E-53 | GRAN | SELENOH |
| 2.644E-57 | -0.600928 | 0.048 | 0.382 | 6.246E-53 | GRAN | POLR2L |
| 3.239E-57 | -1.1221232 | 0.99 | 0.962 | 7.6515E-53 | GRAN | HSP90AA1 |
| 3.3511E-57 | -0.6128883 | 0 | 0.299 | 7.9164E-53 | GRAN | CD99 |
| 3.5118E-57 | -0.6085753 | 0.008 | 0.314 | 8.296E-53 | GRAN | RAB13 |
| 4.0501E-57 | 0.47102063 | 0.735 | 0.38 | 9.5676E-53 | GRAN | FCER1G |
| 4.0819E-57 | -0.5983054 | 0.006 | 0.31 | 9.6426E-53 | GRAN | NOP53 |
| 5.0139E-57 | -1.1587085 | 0.926 | 0.799 | 1.1844E-52 | GRAN | RPS5 |
| 9.7511E-57 | -0.5897386 | 0.052 | 0.385 | 2.3035E-52 | GRAN | EIF5A |
| 1.3113E-56 | -0.520062 | 0.023 | 0.336 | 3.0978E-52 | GRAN | NDUFC2 |
| 1.9009E-56 | -0.544883 | 0.019 | 0.329 | 4.4904E-52 | GRAN | NSA2 |
| 2.6699E-56 | -1.1621116 | 0.013 | 0.316 | 6.3072E-52 | GRAN | PHLDA1 |
| 5.9506E-56 | 0.6050775 | 0.268 | 0.097 | 1.4057E-51 | GRAN | XPO6 |
| 1.3852E-55 | 0.77114718 | 0.844 | 0.621 | 3.2723E-51 | GRAN | ACTR2 |
| 1.4357E-55 | -1.211995 | 0.002 | 0.296 | 3.3915E-51 | GRAN | RND3 |
| 1.816E-55 | -0.5997336 | 0.015 | 0.316 | 4.2899E-51 | GRAN | SBDS |
| 2.2044E-55 | -0.6070382 | 0.01 | 0.308 | 5.2076E-51 | GRAN | RPL36A |
| 2.6498E-55 | 0.406333 | 0.194 | 0.055 | 6.2597E-51 | GRAN | ARHGAP9 |
| 4.4198E-55 | 0.77093043 | 0.374 | 0.176 | 1.0441E-50 | GRAN | FMNL1 |
| 7.0165E-55 | -0.5449 | 0.032 | 0.345 | 1.6575E-50 | GRAN | ANP32B |
| 8.1292E-55 | -0.5829515 | 0.016 | 0.315 | 1.9204E-50 | GRAN | MZT2B |
| 1.0187E-54 | -0.4927048 | 0.035 | 0.352 | 2.4064E-50 | GRAN | PSMD8 |
| 1.053E-54 | -0.5443017 | 0.037 | 0.354 | 2.4874E-50 | GRAN | FKBP2 |
| 1.073E-54 | 0.36182742 | 0.16 | 0.039 | 2.5348E-50 | GRAN | ARHGAP25 |
| 1.3542E-54 | 0.56432054 | 0.232 | 0.076 | 3.199E-50 | GRAN | TLR4 |
| 1.6457E-54 | 0.87587961 | 0.413 | 0.212 | 3.8877E-50 | GRAN | KDM6B |

| | | | | | | |
|---|---|---|---|---|---|---|
| 2.2152E-54 | 0.462752 | 0.197 | 0.058 | 5.2331E-50 | GRAN | ZDHHC18 |
| 2.6246E-54 | -1.1688665 | 0.923 | 0.78 | 6.2002E-50 | GRAN | RPL10A |
| 3.4585E-54 | -0.5140789 | 0.015 | 0.312 | 8.1701E-50 | GRAN | SSBP1 |
| 3.643E-54 | -0.5377175 | 0.031 | 0.34 | 8.6058E-50 | GRAN | EIF4B |
| 5.1893E-54 | -0.5864593 | 0.052 | 0.374 | 1.2259E-49 | GRAN | PRDX6 |
| 5.8452E-54 | -1.0383312 | 0.006 | 0.296 | 1.3808E-49 | GRAN | GPRC5A |
| 5.997E-54 | 0.57652956 | 0.261 | 0.094 | 1.4167E-49 | GRAN | DENND3 |
| 7.4439E-54 | -0.5247178 | 0.013 | 0.308 | 1.7585E-49 | GRAN | TMED9 |
| 7.6007E-54 | -0.4725135 | 0.061 | 0.394 | 1.7955E-49 | GRAN | SPCS2 |
| 9.6404E-54 | -0.4964196 | 0.034 | 0.345 | 2.2774E-49 | GRAN | NDFIP1 |
| 1.1541E-53 | -0.7032276 | 0.002 | 0.287 | 2.7263E-49 | GRAN | LDHB |
| 1.3778E-53 | -1.1895279 | 0.926 | 0.811 | 3.2549E-49 | GRAN | GSTP1 |
| 1.6319E-53 | -0.4477211 | 0.053 | 0.377 | 3.855E-49 | GRAN | C11orf58 |
| 1.7401E-53 | -0.5144852 | 0.061 | 0.39 | 4.1107E-49 | GRAN | TBCA |
| 1.8696E-53 | -0.5019721 | 0.019 | 0.317 | 4.4167E-49 | GRAN | POLR1D |
| 2.1926E-53 | -0.5227559 | 0.069 | 0.404 | 5.1795E-49 | GRAN | CIB1 |
| 2.2332E-53 | -0.603866 | 0.013 | 0.306 | 5.2755E-49 | GRAN | DDX24 |
| 2.5241E-53 | -0.504895 | 0.029 | 0.334 | 5.9627E-49 | GRAN | COPS9 |
| 2.5513E-53 | -0.5107565 | 0.024 | 0.324 | 6.027E-49 | GRAN | ZNF706 |
| 3.3997E-53 | -0.5608431 | 0.006 | 0.292 | 8.0311E-49 | GRAN | APRT |
| 3.631E-53 | 0.90623616 | 0.332 | 0.147 | 8.5775E-49 | GRAN | SMAP2 |
| 6.4659E-53 | -0.6104183 | 0.013 | 0.303 | 1.5274E-48 | GRAN | NUCKS1 |
| 1.2841E-52 | -0.6058509 | 0.029 | 0.329 | 3.0335E-48 | GRAN | TMBIM1 |
| 1.3326E-52 | -0.6432162 | 0.005 | 0.287 | 3.148E-48 | GRAN | CD151 |
| 2.3986E-52 | -0.5998128 | 0.04 | 0.349 | 5.6661E-48 | GRAN | NORAD |
| 2.7847E-52 | -0.5267166 | 0.047 | 0.361 | 6.5783E-48 | GRAN | UQCRQ |
| 3.3831E-52 | -0.4935515 | 0.032 | 0.335 | 7.9918E-48 | GRAN | FAM96B |
| 4.0114E-52 | -1.0156108 | 0.984 | 0.898 | 9.4762E-48 | GRAN | RPL39 |
| 4.9273E-52 | 1.24042634 | 0.306 | 0.126 | 1.164E-47 | GRAN | GBP1 |
| 6.5598E-52 | -0.4920077 | 0.024 | 0.319 | 1.5496E-47 | GRAN | EIF3F |
| 8.3869E-52 | -0.6652227 | 0.01 | 0.294 | 1.9812E-47 | GRAN | SPINT1 |
| 1.114E-51 | -0.8528938 | 0.002 | 0.279 | 2.6315E-47 | GRAN | CAMK2N1 |
| 1.1577E-51 | -0.5236848 | 0.01 | 0.293 | 2.7349E-47 | GRAN | RALA |
| 1.3786E-51 | -1.016467 | 0.063 | 0.356 | 3.2566E-47 | GRAN | PHLDA2 |
| 1.4745E-51 | -0.5799517 | 0.115 | 0.432 | 3.4832E-47 | GRAN | EIF3A |
| 1.4789E-51 | -0.785189 | 0.005 | 0.285 | 3.4936E-47 | GRAN | H1F0 |
| 1.878E-51 | -0.5602606 | 0.015 | 0.301 | 4.4365E-47 | GRAN | EIF5B |
| 2.4064E-51 | -0.5968401 | 0.052 | 0.362 | 5.6847E-47 | GRAN | MIF |
| 2.7916E-51 | -0.4788361 | 0.015 | 0.299 | 6.5946E-47 | GRAN | EIF3G |
| 5.0474E-51 | -0.5533511 | 0.148 | 0.459 | 1.1924E-46 | GRAN | ST13 |
| 6.9505E-51 | -0.501324 | 0.006 | 0.283 | 1.6419E-46 | GRAN | MRPL41 |
| 7.1156E-51 | -0.4713683 | 0.045 | 0.352 | 1.6809E-46 | GRAN | ENY2 |
| 7.9218E-51 | -0.4773148 | 0.045 | 0.352 | 1.8714E-46 | GRAN | SNRPB |
| 8.6524E-51 | -0.6803009 | 0.026 | 0.316 | 2.0439E-46 | GRAN | CD68 |
| 9.7747E-51 | -1.4918702 | 0.018 | 0.299 | 2.3091E-46 | GRAN | ADIRF |
| 1.0376E-50 | -0.4505335 | 0.034 | 0.332 | 2.451E-46 | GRAN | RNF7 |
| 1.2517E-50 | -0.610093 | 0.039 | 0.337 | 2.9568E-46 | GRAN | GOLGA4 |
| 1.4749E-50 | -0.7256743 | 0.098 | 0.408 | 3.4841E-46 | GRAN | FLNA |
| 1.5397E-50 | -0.5057925 | 0.058 | 0.371 | 3.6372E-46 | GRAN | REEP5 |
| 1.8538E-50 | -0.6202526 | 0.016 | 0.298 | 4.3792E-46 | GRAN | CHCHD10 |

| | | | | | | |
|---|---|---|---|---|---|---|
| 1.9499E-50 | -0.826442 | 0.924 | 0.813 | 4.6063E-46 | GRAN | YBX1 |
| 1.9886E-50 | -0.5664856 | 0.021 | 0.307 | 4.6976E-46 | GRAN | BLVRB |
| 3.0923E-50 | -0.5140767 | 0.042 | 0.344 | 7.3049E-46 | GRAN | SNU13 |
| 3.9948E-50 | -0.4590109 | 0.058 | 0.371 | 9.4369E-46 | GRAN | PNRC2 |
| 5.7878E-50 | -0.5064208 | 0.056 | 0.364 | 1.3673E-45 | GRAN | SRSF11 |
| 6.6961E-50 | -0.5333079 | 0.052 | 0.359 | 1.5818E-45 | GRAN | MRPL33 |
| 6.8226E-50 | -0.5739362 | 0.056 | 0.363 | 1.6117E-45 | GRAN | RRBP1 |
| 7.8343E-50 | -0.4542609 | 0.035 | 0.33 | 1.8507E-45 | GRAN | SRP9 |
| 1.4531E-49 | -1.053358 | 0 | 0.268 | 3.4326E-45 | GRAN | TSPAN1 |
| 1.5265E-49 | -0.4885002 | 0.023 | 0.305 | 3.606E-45 | GRAN | ATP5PO |
| 1.5734E-49 | -0.4680518 | 0.029 | 0.318 | 3.7168E-45 | GRAN | RBX1 |
| 1.7958E-49 | -0.4624305 | 0.029 | 0.316 | 4.2422E-45 | GRAN | CAPNS1 |
| 1.9526E-49 | -0.8879332 | 0.027 | 0.313 | 4.6127E-45 | GRAN | HIST1H4C |
| 2.0424E-49 | -0.5169803 | 0.015 | 0.291 | 4.8247E-45 | GRAN | ACADVL |
| 2.0469E-49 | -0.8293481 | 0.934 | 0.818 | 4.8354E-45 | GRAN | RPL27 |
| 2.3783E-49 | -1.0287878 | 0.011 | 0.285 | 5.6182E-45 | GRAN | CLU |
| 2.4709E-49 | -0.618105 | 0.006 | 0.278 | 5.837E-45 | GRAN | JUP |
| 2.9996E-49 | -0.5183709 | 0.016 | 0.293 | 7.086E-45 | GRAN | NDUFB9 |
| 3.3295E-49 | -0.7894618 | 0.006 | 0.277 | 7.8653E-45 | GRAN | TNFRSF12A |
| 3.5879E-49 | -0.4563394 | 0.058 | 0.368 | 8.4757E-45 | GRAN | PSMB1 |
| 3.7959E-49 | -0.465048 | 0.019 | 0.3 | 8.967E-45 | GRAN | KDELR1 |
| 4.809E-49 | -0.6217997 | 0.147 | 0.445 | 1.136E-44 | GRAN | PDIA3 |
| 5.4844E-49 | -0.5567369 | 0.01 | 0.281 | 1.2956E-44 | GRAN | MYDGF |
| 5.8711E-49 | -0.4862657 | 0.063 | 0.373 | 1.3869E-44 | GRAN | NDUFV2 |
| 6.1021E-49 | -0.4677276 | 0.006 | 0.275 | 1.4415E-44 | GRAN | NDUFB8 |
| 7.5281E-49 | 0.63819186 | 0.263 | 0.099 | 1.7784E-44 | GRAN | CLEC4E |
| 9.1511E-49 | -0.5131986 | 0.035 | 0.324 | 2.1618E-44 | GRAN | AP2S1 |
| 1.0067E-48 | -0.4862983 | 0.021 | 0.3 | 2.3781E-44 | GRAN | TRAM1 |
| 1.2736E-48 | -0.5140239 | 0.105 | 0.419 | 3.0087E-44 | GRAN | EIF3K |
| 1.388E-48 | -0.4887016 | 0.005 | 0.271 | 3.279E-44 | GRAN | GADD45GIP1 |
| 1.4749E-48 | -0.6023054 | 0.006 | 0.274 | 3.4841E-44 | GRAN | FAM3C |
| 1.5045E-48 | -0.5798486 | 0.019 | 0.297 | 3.5541E-44 | GRAN | TUBA1C |
| 1.5298E-48 | -0.4564516 | 0.005 | 0.27 | 3.6139E-44 | GRAN | RNH1 |
| 1.7536E-48 | -0.4754505 | 0.065 | 0.375 | 4.1425E-44 | GRAN | CIRBP |
| 1.8606E-48 | -0.6182481 | 0 | 0.263 | 4.3953E-44 | GRAN | WASL |
| 1.9387E-48 | -0.4423927 | 0.045 | 0.342 | 4.5798E-44 | GRAN | ARL6IP4 |
| 1.9413E-48 | -0.7621648 | 0.044 | 0.334 | 4.5859E-44 | GRAN | HERPUD1 |
| 2.0784E-48 | -0.4588162 | 0.039 | 0.33 | 4.9097E-44 | GRAN | SNRPG |
| 2.1957E-48 | -0.5365004 | 0.006 | 0.274 | 5.1868E-44 | GRAN | PHPT1 |
| 2.3777E-48 | -0.5406262 | 0.031 | 0.315 | 5.6168E-44 | GRAN | SMS |
| 2.5715E-48 | -0.9285994 | 0.977 | 0.878 | 6.0746E-44 | GRAN | RPS29 |
| 2.7243E-48 | 0.63579759 | 0.258 | 0.093 | 6.4356E-44 | GRAN | PTGS2 |
| 2.947E-48 | 0.74087141 | 0.771 | 0.532 | 6.9618E-44 | GRAN | SERPINB1 |
| 3.1626E-48 | 0.77902239 | 0.808 | 0.613 | 7.4711E-44 | GRAN | CAP1 |
| 4.4947E-48 | -0.4815549 | 0.081 | 0.4 | 1.0618E-43 | GRAN | COMMD6 |
| 4.8754E-48 | -0.5144414 | 0.029 | 0.311 | 1.1517E-43 | GRAN | PFDN2 |
| 5.628E-48 | -0.8235829 | 0.01 | 0.277 | 1.3295E-43 | GRAN | MAL2 |
| 5.7927E-48 | 0.34977496 | 0.14 | 0.034 | 1.3684E-43 | GRAN | MIR3945HG |
| 5.837E-48 | -0.4525401 | 0.063 | 0.369 | 1.3789E-43 | GRAN | WASF2 |
| 7.6694E-48 | -0.4905793 | 0.026 | 0.304 | 1.8117E-43 | GRAN | EIF4A1 |

| | | | | | | |
|---|---|---|---|---|---|---|
| 7.6825E-48 | 0.7429965 | 0.334 | 0.152 | 1.8148E-43 | GRAN | ADGRE5 |
| 8.4552E-48 | 0.6538975 | 0.908 | 0.671 | 1.9974E-43 | GRAN | TALDO1 |
| 1.0481E-47 | -0.5176811 | 0.052 | 0.347 | 2.476E-43 | GRAN | ATP5IF1 |
| 1.3479E-47 | -1.2122666 | 0.005 | 0.267 | 3.1842E-43 | GRAN | LMO7 |
| 1.3675E-47 | -0.4851929 | 0.011 | 0.278 | 3.2305E-43 | GRAN | SSB |
| 1.4715E-47 | -0.4725198 | 0.06 | 0.361 | 3.476E-43 | GRAN | TRMT112 |
| 1.5206E-47 | 0.66883143 | 0.276 | 0.113 | 3.5922E-43 | GRAN | PYGL |
| 1.8469E-47 | -1.0377673 | 0.019 | 0.29 | 4.363E-43 | GRAN | HMOX1 |
| 2.2349E-47 | 0.73661593 | 1 | 0.894 | 5.2795E-43 | GRAN | HLA-A |
| 3.6636E-47 | -0.5338709 | 0.097 | 0.419 | 8.6544E-43 | GRAN | SFPQ |
| 3.6747E-47 | -0.7494615 | 0.006 | 0.268 | 8.6807E-43 | GRAN | MGST1 |
| 3.9234E-47 | -1.1084411 | 0.016 | 0.282 | 9.2683E-43 | GRAN | DUSP5 |
| 4.0611E-47 | -0.534743 | 0.003 | 0.262 | 9.5935E-43 | GRAN | ARHGAP5 |
| 4.4523E-47 | -0.4360096 | 0.055 | 0.353 | 1.0518E-42 | GRAN | DDT |
| 5.2523E-47 | -0.441656 | 0.016 | 0.284 | 1.2408E-42 | GRAN | UFC1 |
| 5.4419E-47 | -0.3917994 | 0.071 | 0.38 | 1.2855E-42 | GRAN | BRK1 |
| 6.5991E-47 | -0.4539744 | 0.019 | 0.288 | 1.5589E-42 | GRAN | PSMB6 |
| 1.2998E-46 | -0.4632366 | 0.079 | 0.391 | 3.0705E-42 | GRAN | DAD1 |
| 1.2999E-46 | -0.432241 | 0.006 | 0.264 | 3.0707E-42 | GRAN | CAMTA1 |
| 1.3128E-46 | -2.0176116 | 0.01 | 0.27 | 3.1013E-42 | GRAN | S100A2 |
| 1.3991E-46 | -0.6196801 | 0.048 | 0.335 | 3.3052E-42 | GRAN | PPP1R14B |
| 1.4745E-46 | -0.4177271 | 0.066 | 0.372 | 3.4832E-42 | GRAN | SEC11A |
| 1.5645E-46 | -0.5554082 | 0.098 | 0.412 | 3.6959E-42 | GRAN | ATF4 |
| 1.6838E-46 | -0.4607078 | 0.005 | 0.261 | 3.9777E-42 | GRAN | DYNC1I2 |
| 2.6988E-46 | -0.5470305 | 0.006 | 0.265 | 6.3754E-42 | GRAN | PRDX2 |
| 3.1273E-46 | -0.4203824 | 0.071 | 0.377 | 7.3877E-42 | GRAN | KIF5B |
| 3.9889E-46 | -0.4288777 | 0.011 | 0.272 | 9.4231E-42 | GRAN | NDUFA12 |
| 4.8178E-46 | 0.829326 | 0.369 | 0.198 | 1.1381E-41 | GRAN | SEC14L1 |
| 6.3443E-46 | -0.5273647 | 0.011 | 0.271 | 1.4987E-41 | GRAN | HSPA9 |
| 6.3504E-46 | 0.68173224 | 0.323 | 0.143 | 1.5001E-41 | GRAN | TREM1 |
| 7.0148E-46 | -0.4447911 | 0.019 | 0.286 | 1.6571E-41 | GRAN | COX7A2L |
| 7.5661E-46 | 0.56573446 | 0.248 | 0.096 | 1.7873E-41 | GRAN | RNF24 |
| 8.2752E-46 | -0.4525786 | 0.026 | 0.296 | 1.9548E-41 | GRAN | NDUFS6 |
| 8.3133E-46 | -0.6107074 | 0 | 0.251 | 1.9639E-41 | GRAN | SERINC2 |
| 8.5749E-46 | -0.5932785 | 0.011 | 0.27 | 2.0256E-41 | GRAN | AFDN |
| 9.4422E-46 | -0.4330259 | 0.008 | 0.264 | 2.2305E-41 | GRAN | SMDT1 |
| 1.1012E-45 | 0.91383827 | 0.432 | 0.27 | 2.6014E-41 | GRAN | FAM49B |
| 1.539E-45 | -0.4733934 | 0.081 | 0.389 | 3.6355E-41 | GRAN | HNRNPH1 |
| 1.7148E-45 | -0.656047 | 0.069 | 0.363 | 4.0509E-41 | GRAN | TUBA1B |
| 1.9745E-45 | -0.4262512 | 0.06 | 0.354 | 4.6644E-41 | GRAN | TMEM258 |
| 2.1923E-45 | -0.4445727 | 0.027 | 0.297 | 5.179E-41 | GRAN | HDLBP |
| 3.2838E-45 | -0.75565 | 0.01 | 0.264 | 7.7573E-41 | GRAN | MUC1 |
| 4.0883E-45 | -0.5958472 | 0.839 | 0.401 | 9.6579E-41 | GRAN | S100P |
| 4.0962E-45 | 0.59610554 | 0.969 | 0.724 | 9.6764E-41 | GRAN | CYBA |
| 4.2136E-45 | -0.581649 | 0.023 | 0.288 | 9.9538E-41 | GRAN | RPS4Y1 |
| 4.3532E-45 | 0.68772778 | 0.41 | 0.207 | 1.0284E-40 | GRAN | TNFRSF1B |
| 4.6699E-45 | -0.4857189 | 0.058 | 0.347 | 1.1032E-40 | GRAN | BSG |
| 5.5049E-45 | 0.95176784 | 0.4 | 0.238 | 1.3004E-40 | GRAN | CASP4 |
| 5.7376E-45 | -0.4340063 | 0.032 | 0.304 | 1.3554E-40 | GRAN | SF3B5 |
| 5.9523E-45 | -0.4439433 | 0.006 | 0.257 | 1.4061E-40 | GRAN | SCP2 |

| | | | | | | |
|---|---|---|---|---|---|---|
| 7.0605E-45 | -0.4664586 | 0.015 | 0.271 | 1.6679E-40 | GRAN | EID1 |
| 7.648E-45 | -0.5368424 | 0.029 | 0.295 | 1.8067E-40 | GRAN | HNRNPAB |
| 1.0201E-44 | -0.8438047 | 0.985 | 0.903 | 2.4098E-40 | GRAN | RPL37 |
| 1.065E-44 | -0.4740886 | 0.011 | 0.264 | 2.5159E-40 | GRAN | CUTA |
| 1.1708E-44 | -0.443052 | 0.003 | 0.251 | 2.7657E-40 | GRAN | CCT4 |
| 1.178E-44 | -0.4659925 | 0.023 | 0.286 | 2.7827E-40 | GRAN | DYNC1H1 |
| 1.6547E-44 | 1.72553455 | 0.539 | 0.33 | 3.9089E-40 | GRAN | IL1RN |
| 1.9353E-44 | -0.44512 | 0.01 | 0.262 | 4.5718E-40 | GRAN | CCT3 |
| 2.0611E-44 | -0.4576173 | 0.021 | 0.282 | 4.8688E-40 | GRAN | ADAM9 |
| 2.3589E-44 | -0.8209336 | 0.048 | 0.322 | 5.5725E-40 | GRAN | GADD45A |
| 2.5869E-44 | -0.5840587 | 0.01 | 0.262 | 6.1111E-40 | GRAN | TSC22D1 |
| 2.7413E-44 | -0.4389398 | 0.019 | 0.277 | 6.4757E-40 | GRAN | ATP5F1C |
| 3.0133E-44 | -0.6473466 | 0.11 | 0.389 | 7.1183E-40 | GRAN | C9orf16 |
| 3.2954E-44 | -0.4031642 | 0.035 | 0.306 | 7.7848E-40 | GRAN | ANAPC11 |
| 3.8327E-44 | -0.432633 | 0.003 | 0.248 | 9.0541E-40 | GRAN | PEA15 |
| 4.4503E-44 | -0.4498928 | 0.048 | 0.328 | 1.0513E-39 | GRAN | ARL6IP1 |
| 4.8174E-44 | -0.6006395 | 0.013 | 0.265 | 1.138E-39 | GRAN | RHOC |
| 5.4062E-44 | -0.4884999 | 0.148 | 0.438 | 1.2771E-39 | GRAN | ATP5MF |
| 5.6289E-44 | -0.4421706 | 0.008 | 0.256 | 1.3297E-39 | GRAN | EIF3M |
| 7.3378E-44 | -0.4142277 | 0.015 | 0.267 | 1.7334E-39 | GRAN | NDUFA6 |
| 8.1471E-44 | -0.4301074 | 0.013 | 0.265 | 1.9246E-39 | GRAN | YWHAG |
| 1.0922E-43 | -0.424906 | 0.05 | 0.329 | 2.58E-39 | GRAN | CDC37 |
| 1.1346E-43 | 0.60070605 | 0.26 | 0.105 | 2.6803E-39 | GRAN | CREB5 |
| 1.1667E-43 | -0.4211359 | 0.021 | 0.279 | 2.756E-39 | GRAN | UFM1 |
| 1.3547E-43 | -0.4176461 | 0.005 | 0.248 | 3.2002E-39 | GRAN | UQCRFS1 |
| 1.4101E-43 | -0.4419497 | 0.026 | 0.286 | 3.3311E-39 | GRAN | EIF6 |
| 1.416E-43 | -0.4356002 | 0.023 | 0.28 | 3.3449E-39 | GRAN | MRPL51 |
| 1.4827E-43 | -0.4799479 | 0.005 | 0.249 | 3.5025E-39 | GRAN | SNX9 |
| 1.5021E-43 | -0.4825502 | 0.029 | 0.291 | 3.5485E-39 | GRAN | CLINT1 |
| 1.5433E-43 | -0.8502043 | 0.005 | 0.249 | 3.6459E-39 | GRAN | CST6 |
| 1.7329E-43 | -0.4417001 | 0.01 | 0.257 | 4.0937E-39 | GRAN | PA2G4 |
| 1.9432E-43 | 0.57375541 | 0.265 | 0.11 | 4.5904E-39 | GRAN | PREX1 |
| 2.6136E-43 | -0.4121236 | 0.029 | 0.292 | 6.1741E-39 | GRAN | TMED10 |
| 2.8528E-43 | -0.4043991 | 0.076 | 0.373 | 6.7391E-39 | GRAN | TXNDC17 |
| 2.9108E-43 | -0.7436833 | 0.003 | 0.245 | 6.8761E-39 | GRAN | S100A13 |
| 2.9525E-43 | -0.6875267 | 0.003 | 0.245 | 6.9748E-39 | GRAN | PRSS8 |
| 3.0448E-43 | -0.4215339 | 0.006 | 0.25 | 7.1926E-39 | GRAN | CCT8 |
| 3.5838E-43 | -0.5229313 | 0.01 | 0.256 | 8.4659E-39 | GRAN | CD2AP |
| 3.5919E-43 | -0.4051141 | 0.023 | 0.279 | 8.4852E-39 | GRAN | SNRPD3 |
| 3.7981E-43 | -1.0163312 | 0.984 | 0.907 | 8.9722E-39 | GRAN | MT-ND2 |
| 3.8936E-43 | 0.75925681 | 0.39 | 0.218 | 9.1979E-39 | GRAN | ACTN1 |
| 3.8984E-43 | -0.5414559 | 0.003 | 0.245 | 9.2091E-39 | GRAN | GIPC1 |
| 3.9877E-43 | -0.4448971 | 0.005 | 0.248 | 9.4201E-39 | GRAN | PDCD5 |
| 4.1589E-43 | -0.4280396 | 0.011 | 0.258 | 9.8245E-39 | GRAN | CCT6A |
| 4.1786E-43 | -0.7121723 | 0.25 | 0.497 | 9.871E-39 | GRAN | NCL |
| 5.3871E-43 | -0.4596778 | 0.027 | 0.285 | 1.2726E-38 | GRAN | TCEA1 |
| 5.4117E-43 | -0.954991 | 0.002 | 0.241 | 1.2784E-38 | GRAN | ADGRF1 |
| 6.0854E-43 | -0.5426121 | 0.011 | 0.257 | 1.4376E-38 | GRAN | ODC1 |
| 6.0898E-43 | 0.4144733 | 0.935 | 0.639 | 1.4386E-38 | GRAN | GLUL |
| 6.8447E-43 | -0.4210354 | 0.01 | 0.255 | 1.6169E-38 | GRAN | NDUFAB1 |

| | | | | | | |
|---|---|---|---|---|---|---|
| 7.2902E-43 | -0.4025619 | 0.071 | 0.362 | 1.7222E-38 | GRAN | ATP5PD |
| 8.0613E-43 | -0.55511 | 0.052 | 0.326 | 1.9043E-38 | GRAN | B4GALT1 |
| 8.7441E-43 | 0.64000972 | 0.313 | 0.141 | 2.0656E-38 | GRAN | TLR2 |
| 8.9253E-43 | -0.4506269 | 0.035 | 0.299 | 2.1084E-38 | GRAN | SELENOS |
| 1.0072E-42 | -0.419252 | 0.031 | 0.292 | 2.3793E-38 | GRAN | NDUFB7 |
| 1.2803E-42 | -0.5344036 | 0.032 | 0.292 | 3.0244E-38 | GRAN | ATP1B3 |
| 1.43E-42 | -0.4459681 | 0.006 | 0.246 | 3.3782E-38 | GRAN | COMT |
| 1.5485E-42 | -0.4231932 | 0.082 | 0.38 | 3.6581E-38 | GRAN | ARF4 |
| 1.7652E-42 | -0.3835505 | 0.04 | 0.308 | 4.1698E-38 | GRAN | UBE2I |
| 1.803E-42 | 0.89798734 | 0.579 | 0.425 | 4.2592E-38 | GRAN | EFHD2 |
| 1.9326E-42 | -0.3974865 | 0.019 | 0.268 | 4.5654E-38 | GRAN | TMEM219 |
| 2.0745E-42 | -0.5246467 | 0.002 | 0.238 | 4.9005E-38 | GRAN | SNHG7 |
| 2.14E-42 | -0.4130187 | 0.019 | 0.27 | 5.0554E-38 | GRAN | KDELR2 |
| 2.3824E-42 | -1.0871479 | 0.956 | 0.821 | 5.6278E-38 | GRAN | MT-ND5 |
| 3.1952E-42 | -0.5091236 | 0.005 | 0.242 | 7.5479E-38 | GRAN | C12orf57 |
| 3.3467E-42 | -1.4392361 | 0.103 | 0.348 | 7.9059E-38 | GRAN | GDF15 |
| 3.4406E-42 | -0.5178919 | 0 | 0.234 | 8.1276E-38 | GRAN | TJP1 |
| 3.7091E-42 | -0.4853649 | 0.069 | 0.354 | 8.762E-38 | GRAN | PNN |
| 4.1162E-42 | -0.4792158 | 0.085 | 0.378 | 9.7237E-38 | GRAN | METRNL |
| 5.0371E-42 | -0.4222543 | 0.002 | 0.236 | 1.1899E-37 | GRAN | OPTN |
| 5.5161E-42 | -0.5013603 | 0.019 | 0.268 | 1.3031E-37 | GRAN | HECTD1 |
| 5.5644E-42 | -0.3831694 | 0.082 | 0.378 | 1.3145E-37 | GRAN | LAMTOR5 |
| 6.6774E-42 | 0.711539 | 0.884 | 0.692 | 1.5774E-37 | GRAN | LRRFIP1 |
| 7.8951E-42 | -0.6512771 | 0.011 | 0.253 | 1.8651E-37 | GRAN | KLF5 |
| 8.122E-42 | -0.7268477 | 0.008 | 0.246 | 1.9187E-37 | GRAN | ITGB8 |
| 8.3029E-42 | -0.705121 | 0.953 | 0.847 | 1.9614E-37 | GRAN | RPL38 |
| 9.6654E-42 | -0.4799102 | 0.005 | 0.24 | 2.2832E-37 | GRAN | PCBD1 |
| 1.0475E-41 | 0.93704692 | 0.466 | 0.319 | 2.4745E-37 | GRAN | YPEL3 |
| 1.0838E-41 | -0.424459 | 0.023 | 0.272 | 2.5603E-37 | GRAN | BAG1 |
| 1.1798E-41 | -0.4065781 | 0.002 | 0.234 | 2.787E-37 | GRAN | ARPC5L |
| 1.1957E-41 | 0.41929417 | 0.184 | 0.06 | 2.8247E-37 | GRAN | ATG16L2 |
| 1.2142E-41 | -0.4542707 | 0.032 | 0.288 | 2.8683E-37 | GRAN | PDCD6 |
| 1.2356E-41 | -0.435184 | 0.034 | 0.291 | 2.9188E-37 | GRAN | NDUFC1 |
| 1.3129E-41 | -0.67555 | 0.076 | 0.354 | 3.1015E-37 | GRAN | GLS |
| 1.4337E-41 | -0.6963988 | 0.981 | 0.9 | 3.3869E-37 | GRAN | RPS11 |
| 1.5356E-41 | -0.4202479 | 0.023 | 0.27 | 3.6276E-37 | GRAN | SYNCRIP |
| 1.6027E-41 | -0.4038177 | 0.061 | 0.337 | 3.7862E-37 | GRAN | HNRNPD |
| 1.6059E-41 | -0.4682838 | 0.168 | 0.453 | 3.7936E-37 | GRAN | ATP5MD |
| 1.9631E-41 | -0.3483294 | 0.04 | 0.302 | 4.6375E-37 | GRAN | UBE2L3 |
| 2.0781E-41 | -0.4731938 | 0.006 | 0.242 | 4.909E-37 | GRAN | FKBP4 |
| 2.1662E-41 | -1.7196065 | 0.015 | 0.252 | 5.1172E-37 | GRAN | CRYAB |
| 2.2168E-41 | -0.5704478 | 0.011 | 0.251 | 5.2367E-37 | GRAN | SPTBN1 |
| 2.3268E-41 | -0.4639876 | 0.005 | 0.238 | 5.4966E-37 | GRAN | PLXNB2 |
| 2.698E-41 | -0.3914851 | 0.016 | 0.259 | 6.3735E-37 | GRAN | EIF3L |
| 3.3823E-41 | -1.601781 | 0.035 | 0.284 | 7.99E-37 | GRAN | MT1E |
| 4.0609E-41 | -0.4406729 | 0.006 | 0.24 | 9.5931E-37 | GRAN | PABPC4 |
| 4.3287E-41 | -0.8904835 | 0.021 | 0.263 | 1.0226E-36 | GRAN | TPM1 |
| 4.6964E-41 | -0.4022761 | 0.01 | 0.246 | 1.1094E-36 | GRAN | TMEM14C |
| 5.1802E-41 | -0.8566729 | 0.005 | 0.237 | 1.2237E-36 | GRAN | PDZK1IP1 |
| 5.5746E-41 | -1.1353939 | 0.974 | 0.882 | 1.3169E-36 | GRAN | CSTB |

| | | | | | | |
|---|---|---|---|---|---|---|
| 5.6112E-41 | -0.3882553 | 0.037 | 0.294 | 1.3255E-36 | GRAN | HIGD2A |
| 6.5515E-41 | -0.6477567 | 0.006 | 0.24 | 1.5477E-36 | GRAN | S100A16 |
| 7.2728E-41 | -0.8523785 | 0.006 | 0.238 | 1.718E-36 | GRAN | EPCAM |
| 1.0229E-40 | 1.00644537 | 0.221 | 0.082 | 2.4164E-36 | GRAN | GBP5 |
| 1.0342E-40 | -0.8144069 | 0.882 | 0.782 | 2.4431E-36 | GRAN | DYNLL1 |
| 1.2299E-40 | -1.0485777 | 0.024 | 0.266 | 2.9055E-36 | GRAN | AREG |
| 1.2944E-40 | 0.33342913 | 0.155 | 0.045 | 3.0579E-36 | GRAN | DGAT2 |
| 1.2994E-40 | -0.403736 | 0.011 | 0.247 | 3.0695E-36 | GRAN | OSTC |
| 1.7309E-40 | -0.4022761 | 0.011 | 0.246 | 4.089E-36 | GRAN | LSM7 |
| 1.7464E-40 | -0.3887207 | 0.01 | 0.243 | 4.1256E-36 | GRAN | C8orf59 |
| 2.123E-40 | -0.6488698 | 0.016 | 0.254 | 5.0152E-36 | GRAN | SNHG12 |
| 2.229E-40 | -0.9228726 | 0.037 | 0.285 | 5.2655E-36 | GRAN | MAFB |
| 2.3169E-40 | -0.4129699 | 0.008 | 0.24 | 5.4731E-36 | GRAN | ATP5F1A |
| 3.1234E-40 | 0.52766095 | 0.956 | 0.695 | 7.3785E-36 | GRAN | JUNB |
| 3.8604E-40 | -0.3952022 | 0.005 | 0.233 | 9.1195E-36 | GRAN | DNAJC15 |
| 4.1472E-40 | -0.407739 | 0.016 | 0.252 | 9.797E-36 | GRAN | RPS17 |
| 4.1582E-40 | -0.4909011 | 0.077 | 0.356 | 9.823E-36 | GRAN | TAF1D |
| 4.4739E-40 | -0.3379511 | 0.061 | 0.334 | 1.0569E-35 | GRAN | UBXN1 |
| 4.7571E-40 | 0.4479011 | 0.21 | 0.077 | 1.1238E-35 | GRAN | RASSF2 |
| 4.8669E-40 | 0.32772692 | 0.144 | 0.04 | 1.1497E-35 | GRAN | TLR6 |
| 4.9508E-40 | -0.6191196 | 0.002 | 0.227 | 1.1695E-35 | GRAN | OCIAD2 |
| 5.8171E-40 | -0.4497444 | 0.019 | 0.258 | 1.3742E-35 | GRAN | RPL22L1 |
| 6.9953E-40 | -0.7422115 | 0.008 | 0.238 | 1.6525E-35 | GRAN | CDKN1C |
| 7.5279E-40 | 0.44197125 | 0.163 | 0.049 | 1.7783E-35 | GRAN | OSM |
| 7.6115E-40 | -0.5497006 | 0.203 | 0.456 | 1.7981E-35 | GRAN | SEC61G |
| 7.7599E-40 | -0.3930118 | 0.027 | 0.27 | 1.8331E-35 | GRAN | ERH |
| 7.8527E-40 | -0.3259149 | 0.084 | 0.369 | 1.855E-35 | GRAN | ELOC |
| 8.2329E-40 | -0.4525557 | 0.023 | 0.262 | 1.9449E-35 | GRAN | ABRACL |
| 8.6978E-40 | -0.4158726 | 0.011 | 0.243 | 2.0547E-35 | GRAN | GSPT1 |
| 9.1611E-40 | -0.374575 | 0.06 | 0.329 | 2.1641E-35 | GRAN | DRAP1 |
| 9.9431E-40 | -0.8266384 | 0.994 | 0.954 | 2.3489E-35 | GRAN | ACTG1 |
| 1.1076E-39 | -0.5754921 | 0.024 | 0.265 | 2.6164E-35 | GRAN | HES4 |
| 1.1098E-39 | -0.382751 | 0.011 | 0.243 | 2.6216E-35 | GRAN | MRPS21 |
| 1.2373E-39 | -0.3879954 | 0.018 | 0.253 | 2.923E-35 | GRAN | SEC31A |
| 1.2986E-39 | -0.3960608 | 0.013 | 0.245 | 3.0677E-35 | GRAN | NUDC |
| 1.4029E-39 | -1.6319901 | 0.008 | 0.235 | 3.3141E-35 | GRAN | PRSS22 |
| 1.5544E-39 | -0.4238707 | 0.05 | 0.31 | 3.6719E-35 | GRAN | CLTB |
| 1.7992E-39 | -0.3916862 | 0.011 | 0.24 | 4.2503E-35 | GRAN | PSMB7 |
| 2.967E-39 | -0.3266221 | 0.098 | 0.392 | 7.0089E-35 | GRAN | COPE |
| 3.3294E-39 | -0.3739579 | 0.006 | 0.231 | 7.8649E-35 | GRAN | CMPK1 |
| 4.0396E-39 | -0.4028943 | 0.005 | 0.228 | 9.5428E-35 | GRAN | RSL1D1 |
| 4.1139E-39 | -0.3728055 | 0.019 | 0.255 | 9.7182E-35 | GRAN | XRCC6 |
| 4.561E-39 | -0.9305658 | 0.006 | 0.231 | 1.0774E-34 | GRAN | IL32 |
| 4.7544E-39 | 0.31846006 | 0.132 | 0.036 | 1.1231E-34 | GRAN | TECPR2 |
| 4.9897E-39 | -0.886439 | 0.923 | 0.79 | 1.1787E-34 | GRAN | RPS21 |
| 5.2161E-39 | -0.3087974 | 0.09 | 0.38 | 1.2322E-34 | GRAN | SRSF9 |
| 5.2743E-39 | -0.4001322 | 0.008 | 0.233 | 1.246E-34 | GRAN | MRPL57 |
| 5.5548E-39 | -0.3842452 | 0.011 | 0.239 | 1.3122E-34 | GRAN | BANF1 |
| 5.8113E-39 | -0.4318824 | 0.005 | 0.228 | 1.3728E-34 | GRAN | ST14 |
| 6.5625E-39 | -0.4589347 | 0.01 | 0.236 | 1.5503E-34 | GRAN | KMT2A |

| | | | | | | |
|---|---|---|---|---|---|---|
| 7.1796E-39 | -0.444541 | 0.002 | 0.221 | 1.696E-34 | GRAN | SPATS2L |
| 8.7032E-39 | -0.3119364 | 0.106 | 0.406 | 2.056E-34 | GRAN | CSDE1 |
| 9.2047E-39 | -0.5495768 | 0 | 0.218 | 2.1744E-34 | GRAN | ARL4C |
| 9.2243E-39 | -0.390002 | 0.013 | 0.241 | 2.1791E-34 | GRAN | SNRPE |
| 1.0091E-38 | -0.3946833 | 0.027 | 0.266 | 2.3838E-34 | GRAN | HNRNPR |
| 1.2108E-38 | 0.56865792 | 0.294 | 0.138 | 2.8603E-34 | GRAN | ARHGAP26 |
| 1.2603E-38 | -0.3758912 | 0.034 | 0.279 | 2.9773E-34 | GRAN | PDXK |
| 1.3519E-38 | -0.4323737 | 0.058 | 0.319 | 3.1936E-34 | GRAN | AKAP9 |
| 1.409E-38 | 0.38688907 | 1 | 0.937 | 3.3285E-34 | GRAN | TMSB4X |
| 1.5368E-38 | -0.6119877 | 0.008 | 0.232 | 3.6305E-34 | GRAN | CYB5A |
| 1.7062E-38 | -0.5402978 | 0 | 0.217 | 4.0306E-34 | GRAN | EHF |
| 1.7783E-38 | -0.4592292 | 0.052 | 0.306 | 4.2009E-34 | GRAN | PSME2 |
| 2.0142E-38 | -0.3476905 | 0.034 | 0.277 | 4.7581E-34 | GRAN | AP2M1 |
| 2.226E-38 | -0.3702782 | 0.027 | 0.265 | 5.2584E-34 | GRAN | PSMD7 |
| 2.3962E-38 | -0.380344 | 0.035 | 0.28 | 5.6606E-34 | GRAN | CCT5 |
| 2.8205E-38 | 0.4056559 | 0.179 | 0.06 | 6.6628E-34 | GRAN | SELPLG |
| 3.1635E-38 | -0.3972655 | 0.027 | 0.265 | 7.4731E-34 | GRAN | EIF3J |
| 3.5196E-38 | -0.758713 | 0.01 | 0.232 | 8.3142E-34 | GRAN | ITGA2 |
| 3.5232E-38 | -0.3793423 | 0.016 | 0.244 | 8.3229E-34 | GRAN | POLR2K |
| 3.6314E-38 | -0.4302559 | 0.013 | 0.238 | 8.5785E-34 | GRAN | FNBP4 |
| 3.6758E-38 | -0.3810894 | 0.005 | 0.223 | 8.6832E-34 | GRAN | SNRPF |
| 3.8285E-38 | -0.3815819 | 0.013 | 0.237 | 9.044E-34 | GRAN | SCARB2 |
| 3.8414E-38 | -0.8167985 | 0.015 | 0.239 | 9.0746E-34 | GRAN | FABP5 |
| 4.0304E-38 | -0.4193868 | 0.042 | 0.289 | 9.5209E-34 | GRAN | YWHAH |
| 4.3096E-38 | -0.3799296 | 0.031 | 0.27 | 1.0181E-33 | GRAN | TCP1 |
| 4.3889E-38 | -0.5363523 | 0.019 | 0.249 | 1.0368E-33 | GRAN | PTMS |
| 4.4663E-38 | -0.3923439 | 0.06 | 0.319 | 1.0551E-33 | GRAN | MMP24OS |
| 4.5089E-38 | -0.4467037 | 0.002 | 0.218 | 1.0651E-33 | GRAN | CNN3 |
| 5.1482E-38 | -0.3488563 | 0.029 | 0.267 | 1.2162E-33 | GRAN | PSMA1 |
| 5.1649E-38 | 0.89432349 | 0.45 | 0.316 | 1.2201E-33 | GRAN | CMTM6 |
| 5.6087E-38 | -0.3274176 | 0.106 | 0.401 | 1.3249E-33 | GRAN | H2AFY |
| 5.6814E-38 | -0.383155 | 0.024 | 0.257 | 1.3421E-33 | GRAN | RPN2 |
| 6.1024E-38 | 0.679606 | 0.956 | 0.797 | 1.4416E-33 | GRAN | ARPC2 |
| 6.1755E-38 | -0.5192532 | 0.003 | 0.22 | 1.4588E-33 | GRAN | DSP |
| 6.5286E-38 | -0.3837699 | 0.037 | 0.279 | 1.5423E-33 | GRAN | RSL24D1 |
| 7.6768E-38 | 0.64218703 | 0.237 | 0.099 | 1.8135E-33 | GRAN | FAM126B |
| 7.9172E-38 | 1.13483175 | 0.427 | 0.271 | 1.8703E-33 | GRAN | TNFAIP2 |
| 8.4793E-38 | -0.337264 | 0.073 | 0.342 | 2.0031E-33 | GRAN | TM9SF3 |
| 8.8329E-38 | -0.7316054 | 0.895 | 0.785 | 2.0866E-33 | GRAN | RPL31 |
| 9.2624E-38 | -0.3450826 | 0.042 | 0.289 | 2.1881E-33 | GRAN | ATP6V1D |
| 9.3764E-38 | -0.3528867 | 0.053 | 0.306 | 2.215E-33 | GRAN | SCAND1 |
| 1.0409E-37 | -0.7699857 | 0.006 | 0.224 | 2.4588E-33 | GRAN | CCND1 |
| 1.0654E-37 | -0.3618352 | 0.055 | 0.311 | 2.5168E-33 | GRAN | HNRNPM |
| 1.1692E-37 | -1.0690519 | 0.905 | 0.753 | 2.7619E-33 | GRAN | RPSA |
| 1.2948E-37 | -0.5183941 | 0.016 | 0.24 | 3.0588E-33 | GRAN | ARID5B |
| 1.3265E-37 | -0.4199094 | 0.006 | 0.224 | 3.1337E-33 | GRAN | ECHS1 |
| 1.6336E-37 | -0.3658293 | 0.023 | 0.251 | 3.8592E-33 | GRAN | EIF3D |
| 1.6594E-37 | -0.3917063 | 0.005 | 0.22 | 3.92E-33 | GRAN | MLEC |
| 1.7177E-37 | -0.3824476 | 0.013 | 0.234 | 4.0578E-33 | GRAN | HNRNPA0 |
| 2.0131E-37 | -0.3755117 | 0.105 | 0.395 | 4.7555E-33 | GRAN | RSRC2 |

| | | | | | | |
|---|---|---|---|---|---|---|
| 2.2057E-37 | -0.3823451 | 0.003 | 0.217 | 5.2105E-33 | GRAN | METAP2 |
| 2.2175E-37 | -0.6817622 | 0.837 | 0.762 | 5.2385E-33 | GRAN | COX7C |
| 2.257E-37 | -0.4156727 | 0.003 | 0.217 | 5.3317E-33 | GRAN | LMAN1 |
| 2.4002E-37 | 0.582393 | 0.242 | 0.103 | 5.6699E-33 | GRAN | SP110 |
| 3.1661E-37 | -0.3716706 | 0.015 | 0.237 | 7.4792E-33 | GRAN | SYPL1 |
| 3.3217E-37 | -0.3767489 | 0.008 | 0.225 | 7.8469E-33 | GRAN | LSM4 |
| 3.3809E-37 | -0.4073412 | 0.042 | 0.284 | 7.9868E-33 | GRAN | RBM3 |
| 3.733E-37 | -1.4400216 | 0.047 | 0.283 | 8.8186E-33 | GRAN | MT1X |
| 4.1677E-37 | -0.3637085 | 0.013 | 0.233 | 9.8453E-33 | GRAN | SAT2 |
| 4.2878E-37 | 0.62480227 | 0.345 | 0.185 | 1.0129E-32 | GRAN | HCLS1 |
| 4.6567E-37 | -1.1852363 | 0.924 | 0.781 | 1.1E-32 | GRAN | LGALS3 |
| 4.8274E-37 | -0.3692671 | 0.006 | 0.221 | 1.1404E-32 | GRAN | NDUFA5 |
| 5.433E-37 | 0.59112932 | 0.215 | 0.084 | 1.2834E-32 | GRAN | CDA |
| 5.6849E-37 | -0.362943 | 0.003 | 0.215 | 1.3429E-32 | GRAN | GTF3C6 |
| 6.0704E-37 | -0.4263837 | 0.055 | 0.304 | 1.434E-32 | GRAN | SERTAD1 |
| 6.9074E-37 | 0.5865854 | 0.258 | 0.113 | 1.6317E-32 | GRAN | ADGRE2 |
| 7.4434E-37 | -0.358942 | 0.037 | 0.276 | 1.7584E-32 | GRAN | MINOS1 |
| 7.6713E-37 | -0.3458082 | 0.047 | 0.294 | 1.8122E-32 | GRAN | ZC3H15 |
| 7.6924E-37 | -0.5282607 | 0.005 | 0.218 | 1.8172E-32 | GRAN | CKB |
| 8.204E-37 | -0.3963559 | 0.003 | 0.213 | 1.938E-32 | GRAN | NBDY |
| 8.8938E-37 | -0.3168921 | 0.108 | 0.397 | 2.101E-32 | GRAN | UBXN4 |
| 9.65E-37 | -0.3964412 | 0 | 0.208 | 2.2796E-32 | GRAN | CTTN |
| 9.8643E-37 | -0.4105313 | 0.021 | 0.247 | 2.3302E-32 | GRAN | CFDP1 |
| 1.1075E-36 | -0.7056447 | 0.002 | 0.211 | 2.6162E-32 | GRAN | CDKN2B |
| 1.1388E-36 | -0.472867 | 0.039 | 0.274 | 2.6903E-32 | GRAN | RAP2B |
| 1.2389E-36 | 0.64849739 | 0.876 | 0.709 | 2.9266E-32 | GRAN | UBE2D3 |
| 1.274E-36 | -0.3832168 | 0.005 | 0.216 | 3.0096E-32 | GRAN | BTG3 |
| 1.3304E-36 | -0.3594618 | 0.008 | 0.222 | 3.1427E-32 | GRAN | MDH2 |
| 1.3593E-36 | -0.3639788 | 0.029 | 0.26 | 3.211E-32 | GRAN | MORF4L2 |
| 1.4087E-36 | -0.3593872 | 0.076 | 0.341 | 3.3279E-32 | GRAN | SRRM1 |
| 1.5167E-36 | -0.5699237 | 0.002 | 0.211 | 3.5829E-32 | GRAN | MACC1 |
| 1.8786E-36 | -0.3459876 | 0.029 | 0.259 | 4.4378E-32 | GRAN | EIF3I |
| 2.1317E-36 | -0.6103799 | 0.073 | 0.327 | 5.0357E-32 | GRAN | VEGFA |
| 2.1705E-36 | -0.3725892 | 0.013 | 0.229 | 5.1274E-32 | GRAN | MRPL52 |
| 2.1797E-36 | -0.3529376 | 0.027 | 0.256 | 5.1491E-32 | GRAN | HADHA |
| 2.2232E-36 | -0.5955443 | 0.853 | 0.784 | 5.2518E-32 | GRAN | BTF3 |
| 2.4067E-36 | -0.34372 | 0.031 | 0.262 | 5.6852E-32 | GRAN | LSM3 |
| 2.5395E-36 | -0.349447 | 0.048 | 0.29 | 5.999E-32 | GRAN | NDUFA11 |
| 2.9144E-36 | 0.63329222 | 0.269 | 0.127 | 6.8847E-32 | GRAN | USP10 |
| 2.9686E-36 | -0.3273981 | 0.068 | 0.325 | 7.0128E-32 | GRAN | LEPROT |
| 3.1992E-36 | 1.01495965 | 0.137 | 0.04 | 7.5575E-32 | GRAN | PI3 |
| 3.3873E-36 | -0.3518117 | 0.013 | 0.229 | 8.0019E-32 | GRAN | PSMB2 |
| 3.5464E-36 | 0.27410988 | 0.103 | 0.025 | 8.3776E-32 | GRAN | SCARF1 |
| 3.5777E-36 | 0.67704696 | 0.74 | 0.576 | 8.4517E-32 | GRAN | UBE2B |
| 3.7611E-36 | 0.64797609 | 0.294 | 0.143 | 8.8848E-32 | GRAN | CARD16 |
| 4.3618E-36 | -0.371033 | 0.053 | 0.298 | 1.0304E-31 | GRAN | ARF6 |
| 4.5033E-36 | -0.3858582 | 0.006 | 0.215 | 1.0638E-31 | GRAN | CTNND1 |
| 4.5052E-36 | -0.3978525 | 0.005 | 0.213 | 1.0643E-31 | GRAN | TRIM44 |
| 4.7549E-36 | -0.3803351 | 0.077 | 0.34 | 1.1232E-31 | GRAN | LUC7L3 |
| 4.8158E-36 | -0.3890345 | 0.063 | 0.315 | 1.1376E-31 | GRAN | CTNNB1 |

| | | | | | | |
|---|---|---|---|---|---|---|
| 5.1023E-36 | -0.4112145 | 0.015 | 0.23 | 1.2053E-31 | GRAN | RGS10 |
| 5.2999E-36 | -0.3374135 | 0.052 | 0.295 | 1.252E-31 | GRAN | EIF4H |
| 5.3378E-36 | -0.3320762 | 0.008 | 0.219 | 1.2609E-31 | GRAN | NDUFB5 |
| 5.9302E-36 | 0.65735638 | 0.368 | 0.197 | 1.4009E-31 | GRAN | ITGAX |
| 7.7086E-36 | -0.3832963 | 0.021 | 0.242 | 1.821E-31 | GRAN | RNMT |
| 7.9312E-36 | -0.3500818 | 0.035 | 0.267 | 1.8736E-31 | GRAN | RTRAF |
| 8.1205E-36 | -0.4015014 | 0.005 | 0.211 | 1.9183E-31 | GRAN | TCEAL4 |
| 8.6798E-36 | -0.3940978 | 0.044 | 0.277 | 2.0504E-31 | GRAN | AAK1 |
| 8.735E-36 | -0.348092 | 0.019 | 0.239 | 2.0635E-31 | GRAN | PSMA3 |
| 9.3152E-36 | -1.1550118 | 0.048 | 0.28 | 2.2005E-31 | GRAN | HLA-DRB1 |
| 9.3673E-36 | -0.5513464 | 0.01 | 0.221 | 2.2128E-31 | GRAN | TGM2 |
| 1.0992E-35 | -0.3726734 | 0.016 | 0.232 | 2.5965E-31 | GRAN | HDGF |
| 1.2944E-35 | 0.60427107 | 0.229 | 0.094 | 3.0577E-31 | GRAN | IFIT2 |
| 1.3756E-35 | -0.2997981 | 0.068 | 0.321 | 3.2495E-31 | GRAN | RER1 |
| 1.4106E-35 | -0.357061 | 0.005 | 0.211 | 3.3323E-31 | GRAN | ATP2A2 |
| 1.466E-35 | -0.3702366 | 0.015 | 0.229 | 3.4631E-31 | GRAN | COX20 |
| 1.5758E-35 | 0.42299766 | 0.19 | 0.069 | 3.7225E-31 | GRAN | LILRA5 |
| 1.6496E-35 | -0.4293773 | 0.089 | 0.353 | 3.8968E-31 | GRAN | UBE2H |
| 1.6606E-35 | -0.3662012 | 0.052 | 0.294 | 3.9229E-31 | GRAN | STK24 |
| 1.6942E-35 | -0.3577632 | 0 | 0.202 | 4.0023E-31 | GRAN | TIMM13 |
| 1.7089E-35 | -0.4143076 | 0.037 | 0.267 | 4.0369E-31 | GRAN | TUBB |
| 1.878E-35 | -0.4010607 | 0.026 | 0.248 | 4.4364E-31 | GRAN | SPTAN1 |
| 1.9147E-35 | -0.4088481 | 0.015 | 0.228 | 4.523E-31 | GRAN | PNP |
| 1.9357E-35 | -0.4768023 | 0.015 | 0.228 | 4.5727E-31 | GRAN | KYNU |
| 1.9932E-35 | -0.6327644 | 0.292 | 0.494 | 4.7085E-31 | GRAN | SELENOW |
| 2.0523E-35 | -0.3443406 | 0.023 | 0.242 | 4.8481E-31 | GRAN | TUFM |
| 2.0659E-35 | -0.3227704 | 0.055 | 0.299 | 4.8802E-31 | GRAN | VPS28 |
| 2.1936E-35 | -0.3201617 | 0.042 | 0.276 | 5.182E-31 | GRAN | UXT |
| 2.3354E-35 | -0.3401732 | 0.024 | 0.245 | 5.5168E-31 | GRAN | DNAJC7 |
| 2.3827E-35 | -0.3520354 | 0.003 | 0.207 | 5.6286E-31 | GRAN | NENF |
| 2.4786E-35 | -0.3466842 | 0.013 | 0.225 | 5.8552E-31 | GRAN | PPP2CB |
| 2.7901E-35 | -0.3824306 | 0.039 | 0.27 | 6.5912E-31 | GRAN | DDX46 |
| 3.2018E-35 | -0.3514478 | 0.005 | 0.209 | 7.5636E-31 | GRAN | SUGT1 |
| 3.3548E-35 | -0.3373512 | 0.029 | 0.252 | 7.9251E-31 | GRAN | ATP5PB |
| 3.3555E-35 | -0.5527105 | 0.006 | 0.211 | 7.9266E-31 | GRAN | LY6E |
| 3.55E-35 | -0.399388 | 0 | 0.201 | 8.3862E-31 | GRAN | TCEAL9 |
| 3.5757E-35 | -0.3320318 | 0.037 | 0.265 | 8.4469E-31 | GRAN | GTF2A2 |
| 3.9202E-35 | -0.5603426 | 0.016 | 0.229 | 9.2607E-31 | GRAN | IER5L |
| 4.1675E-35 | 0.90327696 | 0.632 | 0.489 | 9.845E-31 | GRAN | TAGLN2 |
| 4.7695E-35 | -0.3551868 | 0 | 0.2 | 1.1267E-30 | GRAN | SWAP70 |
| 5.2187E-35 | 0.66097334 | 0.305 | 0.158 | 1.2328E-30 | GRAN | CPD |
| 5.2883E-35 | -0.5048325 | 0.005 | 0.209 | 1.2492E-30 | GRAN | PARD6B |
| 5.6932E-35 | -0.7301724 | 0.29 | 0.477 | 1.3449E-30 | GRAN | CD81 |
| 6.1634E-35 | -0.3314904 | 0.035 | 0.263 | 1.456E-30 | GRAN | OCIAD1 |
| 6.5694E-35 | -1.0694435 | 0.985 | 0.876 | 1.5519E-30 | GRAN | MT-ND3 |
| 6.9512E-35 | -0.395059 | 0.023 | 0.239 | 1.6421E-30 | GRAN | PNPLA2 |
| 7.1762E-35 | 0.71297822 | 0.374 | 0.225 | 1.6952E-30 | GRAN | NUMB |
| 8.0139E-35 | -0.4198599 | 0.031 | 0.253 | 1.8931E-30 | GRAN | PLEC |
| 8.5222E-35 | -0.3497236 | 0 | 0.198 | 2.0132E-30 | GRAN | SNRPD1 |
| 8.6753E-35 | -0.3276549 | 0.044 | 0.277 | 2.0494E-30 | GRAN | ARL8B |

| | | | | | | |
|---|---|---|---|---|---|---|
| 8.9128E-35 | -0.5212895 | 0.289 | 0.511 | 2.1055E-30 | GRAN | HNRNPDL |
| 9.0187E-35 | -0.311275 | 0.034 | 0.26 | 2.1305E-30 | GRAN | RPS19BP1 |
| 9.3328E-35 | -0.3622788 | 0.003 | 0.205 | 2.2047E-30 | GRAN | PHB |
| 9.8103E-35 | -0.3548367 | 0.003 | 0.204 | 2.3175E-30 | GRAN | C19orf70 |
| 1.0181E-34 | -1.4799508 | 0 | 0.199 | 2.405E-30 | GRAN | HOPX |
| 1.021E-34 | 0.41194576 | 0.168 | 0.057 | 2.4119E-30 | GRAN | CLEC4A |
| 1.1141E-34 | -0.3843164 | 0.026 | 0.245 | 2.6318E-30 | GRAN | MAFG |
| 1.1173E-34 | -0.3242414 | 0.021 | 0.237 | 2.6394E-30 | GRAN | PSMA5 |
| 1.1184E-34 | -0.3521774 | 0.008 | 0.213 | 2.6419E-30 | GRAN | ILF3 |
| 1.1736E-34 | -0.3367884 | 0.003 | 0.203 | 2.7724E-30 | GRAN | AIMP1 |
| 1.3015E-34 | -0.6612731 | 0.99 | 0.965 | 3.0745E-30 | GRAN | UBC |
| 1.42E-34 | -0.3480769 | 0.016 | 0.227 | 3.3544E-30 | GRAN | NARS |
| 1.8049E-34 | -0.3150767 | 0.044 | 0.275 | 4.2638E-30 | GRAN | STMP1 |
| 1.8362E-34 | -0.3364629 | 0.003 | 0.203 | 4.3376E-30 | GRAN | NAA20 |
| 2.078E-34 | -0.3584304 | 0.005 | 0.205 | 4.9087E-30 | GRAN | NDUFS8 |
| 2.1008E-34 | -0.413523 | 0.156 | 0.431 | 4.9626E-30 | GRAN | SRSF5 |
| 2.6338E-34 | -0.7653851 | 0.281 | 0.481 | 6.2219E-30 | GRAN | HSPD1 |
| 2.8816E-34 | -0.5654599 | 0.026 | 0.24 | 6.8073E-30 | GRAN | CITED2 |
| 2.9218E-34 | -0.3293554 | 0.06 | 0.302 | 6.9021E-30 | GRAN | ROMO1 |
| 2.926E-34 | -0.3136309 | 0.076 | 0.328 | 6.9121E-30 | GRAN | GHITM |
| 2.9449E-34 | 0.62508281 | 0.273 | 0.126 | 6.9568E-30 | GRAN | LRG1 |
| 2.9583E-34 | -0.3096753 | 0.029 | 0.249 | 6.9884E-30 | GRAN | SNRPB2 |
| 3.0662E-34 | -0.5943228 | 0.095 | 0.347 | 7.2433E-30 | GRAN | DDIT4 |
| 3.1197E-34 | -0.4688197 | 0.087 | 0.342 | 7.3697E-30 | GRAN | NDRG1 |
| 3.197E-34 | 0.67265913 | 0.335 | 0.183 | 7.5523E-30 | GRAN | CKLF |
| 3.3514E-34 | -0.3908819 | 0.013 | 0.219 | 7.9171E-30 | GRAN | MRPL18 |
| 3.7612E-34 | -0.3928858 | 0.065 | 0.307 | 8.885E-30 | GRAN | SRSF2 |
| 3.8125E-34 | -0.3314737 | 0.01 | 0.212 | 9.0064E-30 | GRAN | ACP1 |
| 4.2233E-34 | -0.341678 | 0.005 | 0.204 | 9.9768E-30 | GRAN | PSMB5 |
| 4.5443E-34 | -0.3639883 | 0.052 | 0.285 | 1.0735E-29 | GRAN | ISCU |
| 4.7019E-34 | -0.6425674 | 0.003 | 0.201 | 1.1107E-29 | GRAN | DUSP4 |
| 5.0077E-34 | -0.3205023 | 0.106 | 0.379 | 1.183E-29 | GRAN | ATP5ME |
| 5.1106E-34 | -0.34453 | 0.044 | 0.273 | 1.2073E-29 | GRAN | NOL7 |
| 5.1362E-34 | -0.3708914 | 0.002 | 0.198 | 1.2133E-29 | GRAN | RBPMS |
| 5.387E-34 | -0.3248974 | 0.029 | 0.248 | 1.2726E-29 | GRAN | G3BP2 |
| 5.4146E-34 | -0.374584 | 0.006 | 0.207 | 1.2791E-29 | GRAN | C6orf48 |
| 5.58E-34 | -0.3492701 | 0.002 | 0.198 | 1.3182E-29 | GRAN | CCT2 |
| 5.6857E-34 | -0.3814969 | 0.029 | 0.246 | 1.3431E-29 | GRAN | ZRANB2 |
| 6.0149E-34 | -0.3439769 | 0.021 | 0.232 | 1.4209E-29 | GRAN | SYNGR2 |
| 6.5091E-34 | -0.3105184 | 0.061 | 0.304 | 1.5376E-29 | GRAN | VCP |
| 7.2329E-34 | -0.3706162 | 0.11 | 0.381 | 1.7086E-29 | GRAN | GCC2 |
| 7.5821E-34 | -0.3599427 | 0.003 | 0.2 | 1.7911E-29 | GRAN | ATP5MC1 |
| 7.9811E-34 | -0.7038927 | 0 | 0.194 | 1.8854E-29 | GRAN | FBXO32 |
| 8.0878E-34 | 1.04391396 | 0.442 | 0.297 | 1.9106E-29 | GRAN | BTG2 |
| 8.2383E-34 | -0.9327139 | 0.003 | 0.2 | 1.9461E-29 | GRAN | TM4SF1 |
| 8.3164E-34 | -0.4524441 | 0.039 | 0.26 | 1.9646E-29 | GRAN | PDLIM5 |
| 1.0744E-33 | -0.3496897 | 0.011 | 0.214 | 2.538E-29 | GRAN | G3BP1 |
| 1.1474E-33 | -0.35364 | 0.013 | 0.216 | 2.7106E-29 | GRAN | RPL17 |
| 1.2206E-33 | -0.2659473 | 0.097 | 0.364 | 2.8834E-29 | GRAN | RBM8A |
| 1.2209E-33 | -0.3501201 | 0.003 | 0.198 | 2.8841E-29 | GRAN | C1QBP |

| | | | | | | |
|---|---|---|---|---|---|---|
| 1.2413E-33 | -0.3688167 | 0.044 | 0.27 | 2.9324E-29 | GRAN | TOR1AIP2 |
| 1.3288E-33 | -0.4162962 | 0.213 | 0.461 | 3.1389E-29 | GRAN | EIF2S2 |
| 1.3825E-33 | -0.3292648 | 0.035 | 0.256 | 3.2659E-29 | GRAN | TMED2 |
| 1.4658E-33 | -0.4547748 | 0.021 | 0.229 | 3.4627E-29 | GRAN | RAB9A |
| 1.4939E-33 | -0.433282 | 0.094 | 0.349 | 3.5291E-29 | GRAN | GOLGB1 |
| 1.6298E-33 | 0.6244327 | 0.342 | 0.184 | 3.8501E-29 | GRAN | PTPRE |
| 1.6727E-33 | -0.3324667 | 0.015 | 0.218 | 3.9513E-29 | GRAN | RAD23A |
| 1.9463E-33 | -0.4141942 | 0.055 | 0.285 | 4.5977E-29 | GRAN | PRPF4B |
| 1.9844E-33 | -0.4623272 | 0.055 | 0.287 | 4.6877E-29 | GRAN | MARCKSL1 |
| 2.5131E-33 | -0.3729373 | 0.003 | 0.197 | 5.9367E-29 | GRAN | SEC11C |
| 2.6268E-33 | -0.3953529 | 0.061 | 0.297 | 6.2053E-29 | GRAN | FXYD5 |
| 2.9882E-33 | -1.2511596 | 0.134 | 0.348 | 7.0591E-29 | GRAN | CTSL |
| 3.1599E-33 | -0.3358844 | 0.006 | 0.202 | 7.4647E-29 | GRAN | MDH1 |
| 3.5064E-33 | -0.404155 | 0.005 | 0.199 | 8.2832E-29 | GRAN | NEDD4L |
| 3.5665E-33 | -0.4053154 | 0 | 0.191 | 8.4251E-29 | GRAN | PMEPA1 |
| 3.8209E-33 | -0.3684545 | 0.047 | 0.273 | 9.0261E-29 | GRAN | QSOX1 |
| 4.2001E-33 | -0.3343555 | 0.011 | 0.211 | 9.9218E-29 | GRAN | PPP2R1A |
| 4.2566E-33 | -0.5128945 | 0.027 | 0.237 | 1.0055E-28 | GRAN | TOB1 |
| 5.1864E-33 | -0.2931874 | 0.06 | 0.296 | 1.2252E-28 | GRAN | MTCH1 |
| 5.5312E-33 | -0.4279762 | 0.008 | 0.205 | 1.3066E-28 | GRAN | MANF |
| 5.5982E-33 | -0.4747993 | 0.002 | 0.193 | 1.3225E-28 | GRAN | ABLIM1 |
| 6.0444E-33 | -0.3229578 | 0.042 | 0.265 | 1.4279E-28 | GRAN | CHP1 |
| 6.4429E-33 | -0.4894821 | 0.01 | 0.207 | 1.522E-28 | GRAN | CSNK1E |
| 6.9692E-33 | -0.4757628 | 0.006 | 0.2 | 1.6463E-28 | GRAN | PTGR1 |
| 7.0522E-33 | -0.4054261 | 0.018 | 0.22 | 1.666E-28 | GRAN | CDK2AP2 |
| 7.1257E-33 | -0.4066449 | 0.008 | 0.203 | 1.6833E-28 | GRAN | ZNF91 |
| 7.3134E-33 | -0.3991765 | 0.015 | 0.215 | 1.7276E-28 | GRAN | BDP1 |
| 7.8357E-33 | -0.329118 | 0.045 | 0.27 | 1.851E-28 | GRAN | ATOX1 |
| 9.1621E-33 | -0.32532 | 0.015 | 0.215 | 2.1644E-28 | GRAN | CSNK2A1 |
| 9.9453E-33 | -0.3429342 | 0.013 | 0.211 | 2.3494E-28 | GRAN | RBMX |
| 9.9976E-33 | -0.8455747 | 0.873 | 0.745 | 2.3617E-28 | GRAN | EEF1B2 |
| 1.0534E-32 | -0.6789745 | 0.227 | 0.43 | 2.4885E-28 | GRAN | SPINT2 |
| 1.1358E-32 | -1.6197781 | 0.008 | 0.201 | 2.6831E-28 | GRAN | KRT17 |
| 1.1655E-32 | -0.378578 | 0.002 | 0.191 | 2.7533E-28 | GRAN | MPRIP |
| 1.1759E-32 | -0.3513786 | 0.008 | 0.203 | 2.7778E-28 | GRAN | TTC3 |
| 1.439E-32 | -0.3698919 | 0.013 | 0.21 | 3.3994E-28 | GRAN | MAP4 |
| 1.6792E-32 | -0.4341726 | 0.003 | 0.193 | 3.9667E-28 | GRAN | LIMA1 |
| 1.8182E-32 | -0.2644769 | 0.105 | 0.371 | 4.2951E-28 | GRAN | WDR83OS |
| 1.853E-32 | -0.6211651 | 0.042 | 0.256 | 4.3773E-28 | GRAN | CARHSP1 |
| 1.8785E-32 | -0.3681235 | 0.002 | 0.19 | 4.4377E-28 | GRAN | HSBP1L1 |
| 1.9392E-32 | -0.9183083 | 0.002 | 0.19 | 4.581E-28 | GRAN | TMPRSS2 |
| 1.9838E-32 | -0.2970077 | 0.066 | 0.302 | 4.6864E-28 | GRAN | TMEM167A |
| 2.0998E-32 | 0.7152296 | 0.353 | 0.216 | 4.9604E-28 | GRAN | PTEN |
| 2.2287E-32 | -0.5482593 | 0.002 | 0.19 | 5.2649E-28 | GRAN | BAG3 |
| 2.2943E-32 | -0.9417361 | 0.006 | 0.197 | 5.4199E-28 | GRAN | C12orf75 |
| 2.4589E-32 | -0.3420615 | 0.039 | 0.255 | 5.8087E-28 | GRAN | SRSF10 |
| 2.4849E-32 | -0.4721836 | 0.085 | 0.326 | 5.87E-28 | GRAN | LIMS1 |
| 2.509E-32 | -0.3157936 | 0.06 | 0.291 | 5.927E-28 | GRAN | MRFAP1 |
| 2.5348E-32 | -0.3314389 | 0.039 | 0.256 | 5.9878E-28 | GRAN | PSMC6 |
| 2.5416E-32 | -0.322849 | 0.008 | 0.201 | 6.0041E-28 | GRAN | MESD |

| | | | | | | |
|---|---|---|---|---|---|---|
| 2.7963E-32 | -0.3284007 | 0 | 0.186 | 6.6058E-28 | GRAN | CALU |
| 2.8128E-32 | -0.3524896 | 0.044 | 0.263 | 6.6447E-28 | GRAN | TRA2B |
| 2.8536E-32 | -1.2475151 | 0 | 0.187 | 6.7411E-28 | GRAN | DUOX2 |
| 3.1913E-32 | -0.3631878 | 0.008 | 0.201 | 7.5387E-28 | GRAN | ITGAV |
| 3.3784E-32 | -0.3249026 | 0.042 | 0.259 | 7.9808E-28 | GRAN | DNAJC8 |
| 3.4263E-32 | -0.7220482 | 0.029 | 0.236 | 8.0941E-28 | GRAN | PLIN2 |
| 3.4291E-32 | -0.3170178 | 0.021 | 0.223 | 8.1005E-28 | GRAN | TMEM208 |
| 3.9702E-32 | -0.3414341 | 0.016 | 0.215 | 9.3788E-28 | GRAN | SREK1IP1 |
| 4.0208E-32 | -0.3337619 | 0.029 | 0.236 | 9.4982E-28 | GRAN | M6PR |
| 4.2364E-32 | -0.330245 | 0.04 | 0.255 | 1.0008E-27 | GRAN | FIS1 |
| 4.4713E-32 | -0.2980943 | 0.063 | 0.297 | 1.0563E-27 | GRAN | COX17 |
| 4.5275E-32 | -0.5512828 | 0.198 | 0.429 | 1.0695E-27 | GRAN | SLC3A2 |
| 4.6442E-32 | -0.5891255 | 0 | 0.185 | 1.0971E-27 | GRAN | SDCBP2 |
| 4.818E-32 | -0.3736145 | 0.027 | 0.235 | 1.1382E-27 | GRAN | CACYBP |
| 5.0823E-32 | -0.3674944 | 0.003 | 0.191 | 1.2006E-27 | GRAN | SH3YL1 |
| 5.3057E-32 | -0.3297047 | 0.021 | 0.222 | 1.2534E-27 | GRAN | CNDP2 |
| 5.3936E-32 | 0.44201066 | 0.195 | 0.078 | 1.2741E-27 | GRAN | APOBR |
| 5.6699E-32 | -0.2893223 | 0.019 | 0.22 | 1.3394E-27 | GRAN | COPZ1 |
| 5.8554E-32 | -0.2537057 | 0.163 | 0.437 | 1.3832E-27 | GRAN | TRIR |
| 6.5464E-32 | -0.3168601 | 0.019 | 0.219 | 1.5465E-27 | GRAN | RWDD1 |
| 8.0686E-32 | -0.6685814 | 0.008 | 0.198 | 1.9061E-27 | GRAN | SFN |
| 8.9885E-32 | -0.393282 | 0.002 | 0.187 | 2.1233E-27 | GRAN | EPS8L2 |
| 9.1741E-32 | -0.3126959 | 0.003 | 0.189 | 2.1672E-27 | GRAN | UBE3A |
| 9.3165E-32 | -0.3118786 | 0.003 | 0.189 | 2.2008E-27 | GRAN | ILF2 |
| 9.947E-32 | 0.62952534 | 0.318 | 0.166 | 2.3498E-27 | GRAN | SLA |
| 1.0489E-31 | -0.2705829 | 0.073 | 0.31 | 2.4779E-27 | GRAN | ANAPC16 |
| 1.0588E-31 | -0.3152226 | 0.037 | 0.249 | 2.5013E-27 | GRAN | BAX |
| 1.1455E-31 | -0.3310862 | 0.003 | 0.188 | 2.7061E-27 | GRAN | NHP2 |
| 1.3197E-31 | -0.3050072 | 0.021 | 0.221 | 3.1175E-27 | GRAN | PPM1G |
| 1.418E-31 | -0.6699843 | 0.005 | 0.191 | 3.3497E-27 | GRAN | EPS8L1 |
| 1.5604E-31 | -0.3217533 | 0.008 | 0.197 | 3.686E-27 | GRAN | IPO7 |
| 1.6655E-31 | -0.7644394 | 0.005 | 0.191 | 3.9344E-27 | GRAN | ANKRD36C |
| 1.7576E-31 | -0.3289389 | 0.003 | 0.188 | 4.1519E-27 | GRAN | MZT2A |
| 1.8206E-31 | -0.3230853 | 0.003 | 0.188 | 4.3007E-27 | GRAN | DNAJC21 |
| 1.9495E-31 | -0.287041 | 0.055 | 0.279 | 4.6054E-27 | GRAN | ATP6AP2 |
| 2.0157E-31 | -0.3726215 | 0 | 0.182 | 4.7617E-27 | GRAN | FNBP1L |
| 2.1314E-31 | 0.51850127 | 0.242 | 0.112 | 5.035E-27 | GRAN | IL17RA |
| 2.681E-31 | -0.2704278 | 0.065 | 0.297 | 6.3334E-27 | GRAN | SEPT2 |
| 2.7255E-31 | -0.3121896 | 0.016 | 0.21 | 6.4383E-27 | GRAN | COX14 |
| 2.7864E-31 | -0.8550729 | 0.003 | 0.187 | 6.5823E-27 | GRAN | AQP3 |
| 2.8064E-31 | -0.3391873 | 0.002 | 0.184 | 6.6295E-27 | GRAN | PFKP |
| 2.9233E-31 | -0.369983 | 0.01 | 0.199 | 6.9057E-27 | GRAN | USP53 |
| 2.9713E-31 | -0.3118897 | 0.01 | 0.198 | 7.0192E-27 | GRAN | MED10 |
| 3.009E-31 | -0.2983406 | 0.015 | 0.208 | 7.1082E-27 | GRAN | SLIRP |
| 3.1204E-31 | -0.3428405 | 0.008 | 0.196 | 7.3713E-27 | GRAN | PGRMC1 |
| 3.1757E-31 | -0.4144013 | 0.084 | 0.322 | 7.5021E-27 | GRAN | BRD2 |
| 3.1994E-31 | -0.329435 | 0.002 | 0.184 | 7.5579E-27 | GRAN | REXO2 |
| 3.2164E-31 | -0.273959 | 0.073 | 0.309 | 7.5981E-27 | GRAN | BCAP31 |
| 3.975E-31 | -0.377814 | 0.003 | 0.186 | 9.3901E-27 | GRAN | DAB2 |
| 4.1496E-31 | -1.0936332 | 0.013 | 0.202 | 9.8026E-27 | GRAN | THBS1 |

| | | | | | | |
|---|---|---|---|---|---|---|
| 4.1878E-31 | -0.3946097 | 0.026 | 0.226 | 9.8929E-27 | GRAN | MRPS6 |
| 4.2913E-31 | -0.4916019 | 0.002 | 0.183 | 1.0137E-26 | GRAN | S100A14 |
| 4.3217E-31 | -0.3806311 | 0.005 | 0.189 | 1.0209E-26 | GRAN | PAWR |
| 4.525E-31 | 0.27986595 | 0.111 | 0.031 | 1.0689E-26 | GRAN | IL18R1 |
| 4.5491E-31 | -0.2874828 | 0.021 | 0.219 | 1.0746E-26 | GRAN | SURF4 |
| 4.7167E-31 | -0.3033512 | 0.04 | 0.253 | 1.1142E-26 | GRAN | MIEN1 |
| 5.3474E-31 | 0.68302711 | 0.419 | 0.265 | 1.2632E-26 | GRAN | CD53 |
| 5.6408E-31 | -0.4201287 | 0.01 | 0.197 | 1.3325E-26 | GRAN | SDF2L1 |
| 5.7614E-31 | -0.3898811 | 0.002 | 0.182 | 1.361E-26 | GRAN | TEAD1 |
| 5.9533E-31 | 0.69272795 | 0.655 | 0.467 | 1.4064E-26 | GRAN | DUSP1 |
| 6.1328E-31 | -0.3369009 | 0.026 | 0.225 | 1.4487E-26 | GRAN | DDX18 |
| 6.3036E-31 | 0.27583387 | 0.931 | 0.637 | 1.4891E-26 | GRAN | IER3 |
| 7.1627E-31 | -0.3098554 | 0.073 | 0.305 | 1.692E-26 | GRAN | PGAM1 |
| 7.8755E-31 | -0.5940923 | 0.926 | 0.831 | 1.8604E-26 | GRAN | UBB |
| 8.0502E-31 | -0.3118852 | 0.042 | 0.252 | 1.9017E-26 | GRAN | MRPL20 |
| 8.198E-31 | -0.4063884 | 0.011 | 0.199 | 1.9366E-26 | GRAN | KLF4 |
| 9.2713E-31 | -0.2996076 | 0.003 | 0.184 | 2.1902E-26 | GRAN | CCT7 |
| 9.3583E-31 | -0.3225304 | 0.013 | 0.201 | 2.2107E-26 | GRAN | CAPN1 |
| 9.4333E-31 | -0.520179 | 0.003 | 0.184 | 2.2284E-26 | GRAN | FBP1 |
| 9.7934E-31 | -0.3433635 | 0.005 | 0.187 | 2.3135E-26 | GRAN | CDCP1 |
| 1.0602E-30 | -0.2934477 | 0.019 | 0.214 | 2.5045E-26 | GRAN | STRAP |
| 1.1171E-30 | -0.300169 | 0.018 | 0.21 | 2.639E-26 | GRAN | SIVA1 |
| 1.1314E-30 | -0.5235512 | 0.002 | 0.181 | 2.6728E-26 | GRAN | EPHA2 |
| 1.1732E-30 | -0.3160048 | 0.013 | 0.202 | 2.7715E-26 | GRAN | PRNP |
| 1.1735E-30 | -0.3007253 | 0.14 | 0.415 | 2.7721E-26 | GRAN | CDV3 |
| 1.2914E-30 | -0.2818654 | 0.035 | 0.243 | 3.0507E-26 | GRAN | TXNL1 |
| 1.4686E-30 | -0.3509497 | 0.045 | 0.257 | 3.4694E-26 | GRAN | C1orf122 |
| 1.7143E-30 | -1.2251363 | 0.018 | 0.206 | 4.0496E-26 | GRAN | HLA-DPA1 |
| 1.7475E-30 | -0.5402505 | 0 | 0.177 | 4.128E-26 | GRAN | C6orf132 |
| 1.7778E-30 | -0.3298313 | 0.047 | 0.26 | 4.1997E-26 | GRAN | LYPLA1 |
| 1.833E-30 | -0.3766988 | 0.008 | 0.191 | 4.33E-26 | GRAN | PDIA4 |
| 1.8952E-30 | -0.3712485 | 0.016 | 0.207 | 4.4771E-26 | GRAN | SLFN5 |
| 1.9759E-30 | -0.365733 | 0.01 | 0.195 | 4.6677E-26 | GRAN | ITPRIPL2 |
| 2.0019E-30 | -0.3398336 | 0.048 | 0.262 | 4.7292E-26 | GRAN | ANKLE2 |
| 2.0082E-30 | -0.6212444 | 0 | 0.177 | 4.744E-26 | GRAN | SPNS2 |
| 2.0943E-30 | -0.3202777 | 0.006 | 0.188 | 4.9474E-26 | GRAN | ANXA4 |
| 2.113E-30 | -0.4219754 | 0.005 | 0.185 | 4.9916E-26 | GRAN | CDH1 |
| 2.1581E-30 | -0.5497348 | 0.01 | 0.194 | 5.098E-26 | GRAN | PLK2 |
| 2.2681E-30 | 0.5478797 | 0.258 | 0.127 | 5.358E-26 | GRAN | FBXL5 |
| 2.2742E-30 | -0.3153764 | 0.003 | 0.182 | 5.3724E-26 | GRAN | RANBP1 |
| 2.2811E-30 | -0.3056727 | 0.006 | 0.188 | 5.3886E-26 | GRAN | NDUFAF3 |
| 2.3566E-30 | 0.36234056 | 0.163 | 0.059 | 5.5671E-26 | GRAN | JAK3 |
| 2.3909E-30 | -0.3259418 | 0.002 | 0.179 | 5.6481E-26 | GRAN | PURA |
| 2.4113E-30 | -0.4226532 | 0.018 | 0.208 | 5.6962E-26 | GRAN | UBE2S |
| 2.5261E-30 | -0.2853051 | 0.048 | 0.261 | 5.9673E-26 | GRAN | TMCO1 |
| 2.6513E-30 | -0.2928407 | 0.008 | 0.191 | 6.2633E-26 | GRAN | AUP1 |
| 2.9884E-30 | -0.4870937 | 0.002 | 0.179 | 7.0595E-26 | GRAN | FHL2 |
| 3.0273E-30 | -0.3195976 | 0.01 | 0.193 | 7.1513E-26 | GRAN | AHSA1 |
| 3.169E-30 | -0.3083687 | 0.019 | 0.211 | 7.4862E-26 | GRAN | NECTIN2 |
| 3.322E-30 | -0.3904428 | 0.002 | 0.179 | 7.8477E-26 | GRAN | SMIM22 |

| | | | | | | |
|---|---|---|---|---|---|---|
| 3.3789E-30 | -0.3270174 | 0.006 | 0.187 | 7.9819E-26 | GRAN | FDPS |
| 3.4608E-30 | -0.3533643 | 0.013 | 0.198 | 8.1755E-26 | GRAN | HIGD1A |
| 3.7047E-30 | -0.2832479 | 0.021 | 0.214 | 8.7515E-26 | GRAN | PSMC5 |
| 3.7072E-30 | -0.8601925 | 0 | 0.176 | 8.7574E-26 | GRAN | NDRG2 |
| 3.908E-30 | -0.2605775 | 0.073 | 0.305 | 9.2318E-26 | GRAN | H2AFV |
| 3.9969E-30 | -0.2957668 | 0.015 | 0.202 | 9.442E-26 | GRAN | ETFB |
| 4.1828E-30 | -0.349619 | 0.002 | 0.178 | 9.881E-26 | GRAN | TMEM54 |
| 4.3024E-30 | 0.47532672 | 0.226 | 0.102 | 1.0164E-25 | GRAN | DOCK5 |
| 4.4346E-30 | -0.3470788 | 0 | 0.175 | 1.0476E-25 | GRAN | GNL3 |
| 4.5859E-30 | -0.2938247 | 0.019 | 0.21 | 1.0833E-25 | GRAN | SDF4 |
| 4.6107E-30 | -0.3508298 | 0.013 | 0.198 | 1.0892E-25 | GRAN | TSPAN3 |
| 4.6493E-30 | -0.3432473 | 0.003 | 0.181 | 1.0983E-25 | GRAN | LSR |
| 4.6576E-30 | -0.2972829 | 0.006 | 0.187 | 1.1003E-25 | GRAN | DPP7 |
| 4.8119E-30 | -0.308083 | 0.016 | 0.204 | 1.1367E-25 | GRAN | STIP1 |
| 4.99E-30 | -0.2982116 | 0.056 | 0.273 | 1.1788E-25 | GRAN | PDCD6IP |
| 5.2237E-30 | -0.4517521 | 0 | 0.175 | 1.234E-25 | GRAN | STMN1 |
| 5.9507E-30 | -0.3049884 | 0.003 | 0.18 | 1.4057E-25 | GRAN | ARL1 |
| 6.1064E-30 | -0.693271 | 0.031 | 0.224 | 1.4425E-25 | GRAN | HES1 |
| 6.4225E-30 | -0.3807327 | 0.016 | 0.201 | 1.5172E-25 | GRAN | TMEM106B |
| 6.7087E-30 | -0.2942495 | 0.042 | 0.249 | 1.5848E-25 | GRAN | SMARCA5 |
| 7.9105E-30 | -0.2576543 | 0.092 | 0.334 | 1.8687E-25 | GRAN | HNRNPF |
| 8.3048E-30 | 0.30101515 | 0.14 | 0.047 | 1.9618E-25 | GRAN | HRH2 |
| 9.6605E-30 | -0.5509575 | 0.382 | 0.534 | 2.2821E-25 | GRAN | COX6C |
| 9.7506E-30 | -0.3147525 | 0.005 | 0.182 | 2.3034E-25 | GRAN | MGAT4B |
| 1.0964E-29 | -0.3112202 | 0.003 | 0.179 | 2.59E-25 | GRAN | TMEM147 |
| 1.1334E-29 | -0.3003503 | 0.011 | 0.193 | 2.6775E-25 | GRAN | NDUFS7 |
| 1.1639E-29 | 0.56688794 | 0.232 | 0.11 | 2.7494E-25 | GRAN | CPPED1 |
| 1.1733E-29 | -0.3016035 | 0.008 | 0.187 | 2.7716E-25 | GRAN | EMC10 |
| 1.1848E-29 | -0.2728136 | 0.058 | 0.275 | 2.7989E-25 | GRAN | ZNHIT1 |
| 1.2799E-29 | -0.3194521 | 0 | 0.172 | 3.0235E-25 | GRAN | GNG12 |
| 1.3518E-29 | -0.2864939 | 0.011 | 0.194 | 3.1934E-25 | GRAN | ALKBH7 |
| 1.3751E-29 | -0.2758876 | 0.055 | 0.269 | 3.2485E-25 | GRAN | VPS29 |
| 1.4933E-29 | -0.3803184 | 0.008 | 0.186 | 3.5276E-25 | GRAN | BNIP3 |
| 1.5234E-29 | -0.2983365 | 0.002 | 0.174 | 3.5987E-25 | GRAN | MCFD2 |
| 1.53E-29 | -0.5279843 | 0.002 | 0.175 | 3.6142E-25 | GRAN | XIST |
| 1.5671E-29 | -0.2910507 | 0.019 | 0.207 | 3.7019E-25 | GRAN | TXNL4A |
| 1.6229E-29 | 0.37118605 | 0.145 | 0.05 | 3.8337E-25 | GRAN | RHOH |
| 1.6647E-29 | -0.5589643 | 0.35 | 0.52 | 3.9326E-25 | GRAN | NAP1L1 |
| 1.9163E-29 | -0.3301236 | 0 | 0.172 | 4.5268E-25 | GRAN | EPRS |
| 1.9448E-29 | -0.3000847 | 0.018 | 0.204 | 4.5942E-25 | GRAN | HSPA4 |
| 2.0226E-29 | -0.5982878 | 0.002 | 0.175 | 4.7781E-25 | GRAN | PPL |
| 2.0821E-29 | -0.6755267 | 0.018 | 0.201 | 4.9185E-25 | GRAN | CD52 |
| 2.106E-29 | 0.41260901 | 0.187 | 0.077 | 4.9751E-25 | GRAN | STAT5B |
| 2.1628E-29 | -0.3363765 | 0.029 | 0.223 | 5.1091E-25 | GRAN | PCM1 |
| 2.3539E-29 | -0.3005456 | 0.015 | 0.197 | 5.5606E-25 | GRAN | SEC61A1 |
| 2.4805E-29 | -0.2659431 | 0.031 | 0.227 | 5.8597E-25 | GRAN | EMC3 |
| 2.8996E-29 | -0.3951616 | 0 | 0.171 | 6.8497E-25 | GRAN | NECTIN4 |
| 2.9108E-29 | -0.5188654 | 0.006 | 0.182 | 6.8763E-25 | GRAN | DST |
| 3.177E-29 | -0.2766687 | 0.016 | 0.2 | 7.5051E-25 | GRAN | VTI1B |
| 3.3499E-29 | -0.2828727 | 0.019 | 0.206 | 7.9134E-25 | GRAN | CAPRIN1 |

| | | | | | | |
|---|---|---|---|---|---|---|
| 3.4248E-29 | -0.280484 | 0.003 | 0.176 | 8.0905E-25 | GRAN | PSMC3 |
| 3.6219E-29 | -0.2859938 | 0.006 | 0.182 | 8.5559E-25 | GRAN | CCNC |
| 3.6916E-29 | -0.2888176 | 0.024 | 0.214 | 8.7206E-25 | GRAN | RNPS1 |
| 3.7041E-29 | -0.2889282 | 0.019 | 0.206 | 8.7503E-25 | GRAN | REEP3 |
| 4.1854E-29 | -0.2848693 | 0.003 | 0.175 | 9.8872E-25 | GRAN | PFDN1 |
| 4.2759E-29 | -0.3200559 | 0.006 | 0.182 | 1.0101E-24 | GRAN | PHACTR4 |
| 4.2875E-29 | -1.0228686 | 0.003 | 0.176 | 1.0128E-24 | GRAN | RDH10 |
| 4.6195E-29 | -0.2981846 | 0.008 | 0.184 | 1.0913E-24 | GRAN | FUBP1 |
| 5.1369E-29 | -0.3094958 | 0.006 | 0.181 | 1.2135E-24 | GRAN | GPI |
| 5.6639E-29 | -0.2818491 | 0.002 | 0.172 | 1.338E-24 | GRAN | MRPS15 |
| 6.0348E-29 | -0.3269566 | 0.005 | 0.177 | 1.4256E-24 | GRAN | TRAF4 |
| 6.1728E-29 | -0.3092254 | 0.048 | 0.253 | 1.4582E-24 | GRAN | UBL3 |
| 6.2933E-29 | -0.4231795 | 0.01 | 0.187 | 1.4867E-24 | GRAN | ERRFI1 |
| 6.6287E-29 | -0.3443257 | 0.003 | 0.175 | 1.5659E-24 | GRAN | DHRS3 |
| 6.6348E-29 | -0.3249163 | 0.019 | 0.205 | 1.5673E-24 | GRAN | MUC20-OT1 |
| 7.0516E-29 | -0.7460348 | 0.994 | 0.931 | 1.6658E-24 | GRAN | GAPDH |
| 7.2087E-29 | -0.3287837 | 0 | 0.169 | 1.7029E-24 | GRAN | CCDC50 |
| 7.614E-29 | -0.2884448 | 0.016 | 0.196 | 1.7986E-24 | GRAN | MRPL54 |
| 7.7127E-29 | -0.497854 | 0.002 | 0.171 | 1.822E-24 | GRAN | AKR1B1 |
| 7.7802E-29 | -0.2888886 | 0.011 | 0.188 | 1.8379E-24 | GRAN | COA6 |
| 9.185E-29 | -0.3308226 | 0.018 | 0.2 | 2.1698E-24 | GRAN | SECISBP2L |
| 9.3183E-29 | -0.3339996 | 0 | 0.168 | 2.2013E-24 | GRAN | EPB41L4A-AS1 |
| 9.4089E-29 | -0.2861469 | 0.024 | 0.212 | 2.2227E-24 | GRAN | SERPINB6 |
| 9.9215E-29 | -0.5088659 | 0.295 | 0.49 | 2.3437E-24 | GRAN | CANX |
| 1.0189E-28 | -0.2885437 | 0.039 | 0.237 | 2.4069E-24 | GRAN | AHCYL1 |
| 1.0676E-28 | -0.3946374 | 0.002 | 0.171 | 2.5221E-24 | GRAN | TINAGL1 |
| 1.0815E-28 | -0.5962901 | 0.003 | 0.173 | 2.5549E-24 | GRAN | HSPB8 |
| 1.1259E-28 | -0.2659019 | 0.039 | 0.237 | 2.6596E-24 | GRAN | NONO |
| 1.1861E-28 | -0.6289646 | 0.002 | 0.17 | 2.802E-24 | GRAN | FXYD3 |
| 1.1913E-28 | -0.31981 | 0 | 0.167 | 2.8143E-24 | GRAN | DDR1 |
| 1.1962E-28 | 0.63508258 | 0.265 | 0.134 | 2.8258E-24 | GRAN | DDX60L |
| 1.2581E-28 | -0.2917601 | 0.006 | 0.179 | 2.9719E-24 | GRAN | AKR1A1 |
| 1.2846E-28 | 0.74771219 | 0.323 | 0.179 | 3.0347E-24 | GRAN | SAMSN1 |
| 1.337E-28 | -0.2520808 | 0.056 | 0.268 | 3.1585E-24 | GRAN | PSMA4 |
| 1.3891E-28 | -0.2924728 | 0.005 | 0.175 | 3.2814E-24 | GRAN | RPL7L1 |
| 1.4093E-28 | -0.278796 | 0.026 | 0.214 | 3.3292E-24 | GRAN | ADRM1 |
| 1.4343E-28 | -0.3224017 | 0.005 | 0.176 | 3.3883E-24 | GRAN | FAM162A |
| 1.5418E-28 | -0.3309477 | 0.006 | 0.179 | 3.6423E-24 | GRAN | TMEM159 |
| 1.589E-28 | -0.3547563 | 0.01 | 0.184 | 3.7537E-24 | GRAN | SPTSSA |
| 1.6091E-28 | -0.6592597 | 0.006 | 0.178 | 3.8011E-24 | GRAN | IGFBP3 |
| 1.6285E-28 | -0.2779434 | 0.027 | 0.217 | 3.8469E-24 | GRAN | NAP1L4 |
| 1.6845E-28 | -0.9062669 | 0.866 | 0.744 | 3.9792E-24 | GRAN | PRDX1 |
| 1.7935E-28 | -0.311289 | 0.047 | 0.25 | 4.2367E-24 | GRAN | EIF4A3 |
| 1.842E-28 | -0.2822951 | 0.024 | 0.21 | 4.3513E-24 | GRAN | FAM50A |
| 1.9018E-28 | -0.3444077 | 0.011 | 0.186 | 4.4926E-24 | GRAN | GNA15 |
| 1.9021E-28 | -0.4441892 | 0.006 | 0.178 | 4.4933E-24 | GRAN | NR4A1 |
| 2.107E-28 | -0.2837667 | 0.005 | 0.175 | 4.9774E-24 | GRAN | ATRAID |
| 2.1137E-28 | -0.3594631 | 0.124 | 0.372 | 4.9933E-24 | GRAN | ANKRD11 |
| 2.2409E-28 | -0.3452579 | 0.027 | 0.215 | 5.2937E-24 | GRAN | VPS13C |
| 2.2647E-28 | -0.2740601 | 0.032 | 0.224 | 5.3499E-24 | GRAN | TMEM14B |

| | | | | | | |
|---|---|---|---|---|---|---|
| 2.2763E-28 | -0.2608953 | 0.008 | 0.181 | 5.3773E-24 | GRAN | PSMG2 |
| 2.3125E-28 | -0.2830991 | 0.021 | 0.204 | 5.4629E-24 | GRAN | PTBP1 |
| 2.6576E-28 | -0.2992196 | 0.002 | 0.168 | 6.2781E-24 | GRAN | GLO1 |
| 2.7346E-28 | -0.347793 | 0.04 | 0.236 | 6.46E-24 | GRAN | BST2 |
| 2.7418E-28 | -0.2817092 | 0.073 | 0.293 | 6.4769E-24 | GRAN | CHMP2B |
| 2.7808E-28 | -0.3014088 | 0.006 | 0.177 | 6.569E-24 | GRAN | RDX |
| 2.902E-28 | -0.2972082 | 0.002 | 0.168 | 6.8554E-24 | GRAN | SUN1 |
| 2.971E-28 | -0.2860338 | 0.003 | 0.171 | 7.0183E-24 | GRAN | METTL26 |
| 3.0274E-28 | -0.2994912 | 0.015 | 0.191 | 7.1516E-24 | GRAN | PSMD2 |
| 3.0389E-28 | -0.3128856 | 0.135 | 0.392 | 7.1788E-24 | GRAN | HMGN2 |
| 3.0565E-28 | -0.520894 | 0.032 | 0.22 | 7.2203E-24 | GRAN | ZBTB20 |
| 3.2337E-28 | -1.276142 | 0.002 | 0.168 | 7.639E-24 | GRAN | DUOXA2 |
| 3.286E-28 | -0.2888561 | 0.002 | 0.168 | 7.7624E-24 | GRAN | NDUFAF8 |
| 3.2861E-28 | -0.2786724 | 0.018 | 0.197 | 7.7627E-24 | GRAN | BRD7 |
| 3.6041E-28 | -0.2799408 | 0.006 | 0.177 | 8.5141E-24 | GRAN | CYC1 |
| 3.6279E-28 | -0.3718879 | 0.008 | 0.18 | 8.5701E-24 | GRAN | CCDC88A |
| 3.8207E-28 | -0.5084864 | 0.002 | 0.168 | 9.0257E-24 | GRAN | HIST3H2A |
| 3.8911E-28 | -0.7823687 | 0.011 | 0.184 | 9.1918E-24 | GRAN | LAMC2 |
| 4.5612E-28 | -0.2706739 | 0.045 | 0.246 | 1.0775E-23 | GRAN | NDUFA3 |
| 4.6181E-28 | -0.2645145 | 0.037 | 0.232 | 1.0909E-23 | GRAN | PDAP1 |
| 5.1943E-28 | -0.4373759 | 0.015 | 0.19 | 1.2271E-23 | GRAN | SLC5A3 |
| 5.2051E-28 | -0.3362198 | 0.003 | 0.17 | 1.2296E-23 | GRAN | CXADR |
| 5.9369E-28 | -0.6350378 | 0.773 | 0.703 | 1.4025E-23 | GRAN | RPL4 |
| 6.0338E-28 | -0.4094587 | 0.063 | 0.269 | 1.4254E-23 | GRAN | UGCG |
| 7.9184E-28 | 0.7839649 | 0.482 | 0.366 | 1.8706E-23 | GRAN | COTL1 |
| 8.1412E-28 | -0.3259621 | 0.016 | 0.193 | 1.9232E-23 | GRAN | CHD9 |
| 8.1447E-28 | -0.277227 | 0.003 | 0.169 | 1.924E-23 | GRAN | GTF2H5 |
| 8.3187E-28 | -0.2708428 | 0.005 | 0.172 | 1.9651E-23 | GRAN | ERGIC3 |
| 8.3769E-28 | -0.2712083 | 0.013 | 0.187 | 1.9789E-23 | GRAN | PSMD11 |
| 9.0646E-28 | 0.86079547 | 0.511 | 0.427 | 2.1413E-23 | GRAN | IQGAP1 |
| 9.3255E-28 | 0.55237646 | 0.221 | 0.101 | 2.203E-23 | GRAN | CCR1 |
| 9.4579E-28 | -1.1505851 | 0 | 0.163 | 2.2342E-23 | GRAN | MUC21 |
| 9.8187E-28 | 0.80202631 | 0.352 | 0.228 | 2.3195E-23 | GRAN | PHF20L1 |
| 1.06E-27 | -0.3182808 | 0.05 | 0.25 | 2.504E-23 | GRAN | CD47 |
| 1.2627E-27 | -0.4240932 | 0.992 | 0.954 | 2.9828E-23 | GRAN | SERF2 |
| 1.2703E-27 | -0.3335993 | 0.002 | 0.165 | 3.0009E-23 | GRAN | NDFIP2 |
| 1.2791E-27 | -0.286151 | 0.094 | 0.32 | 3.0216E-23 | GRAN | RAB11A |
| 1.2929E-27 | -0.2770846 | 0.045 | 0.242 | 3.0543E-23 | GRAN | NAA50 |
| 1.3345E-27 | -1.6030878 | 0.002 | 0.165 | 3.1526E-23 | GRAN | ECM1 |
| 1.3409E-27 | -0.2726478 | 0.011 | 0.182 | 3.1677E-23 | GRAN | CORO1B |
| 1.3521E-27 | -0.2687064 | 0.018 | 0.194 | 3.1942E-23 | GRAN | CHCHD1 |
| 1.3949E-27 | -0.284629 | 0.039 | 0.23 | 3.2952E-23 | GRAN | DHX36 |
| 1.4173E-27 | -3.6405137 | 0.824 | 0.395 | 3.3481E-23 | GRAN | SPRR3 |
| 1.4548E-27 | 0.50915851 | 0.218 | 0.099 | 3.4368E-23 | GRAN | LUCAT1 |
| 1.4669E-27 | -1.0655085 | 0 | 0.162 | 3.4653E-23 | GRAN | CEACAM5 |
| 1.5056E-27 | -0.2717155 | 0.005 | 0.17 | 3.5568E-23 | GRAN | PLEKHJ1 |
| 1.6873E-27 | -0.3014618 | 0.005 | 0.17 | 3.9859E-23 | GRAN | HMGN3 |
| 1.7214E-27 | 0.71983627 | 0.366 | 0.235 | 4.0664E-23 | GRAN | RNASET2 |
| 1.8412E-27 | -0.3107524 | 0.003 | 0.167 | 4.3495E-23 | GRAN | SMAD3 |
| 1.8471E-27 | -0.8238619 | 0.013 | 0.182 | 4.3634E-23 | GRAN | F3 |

| | | | | | | |
|---|---|---|---|---|---|---|
| 1.8505E-27 | -0.2537005 | 0.061 | 0.269 | 4.3714E-23 | GRAN | CSNK2B |
| 2.019E-27 | -0.5405448 | 0.006 | 0.173 | 4.7696E-23 | GRAN | TGFBI |
| 2.1264E-27 | -1.416824 | 0.913 | 0.72 | 5.0233E-23 | GRAN | HSPB1 |
| 2.148E-27 | -0.2636519 | 0.011 | 0.182 | 5.0743E-23 | GRAN | SUMO3 |
| 2.1895E-27 | -0.3508627 | 0.002 | 0.164 | 5.1723E-23 | GRAN | MET |
| 2.2871E-27 | -0.2618708 | 0.008 | 0.175 | 5.4028E-23 | GRAN | SAR1B |
| 2.2936E-27 | -0.5025487 | 0.032 | 0.215 | 5.4183E-23 | GRAN | LAMB3 |
| 2.2995E-27 | -0.2627309 | 0.024 | 0.204 | 5.4321E-23 | GRAN | MRPS36 |
| 2.4189E-27 | -0.2623381 | 0.031 | 0.216 | 5.7141E-23 | GRAN | LSM8 |
| 2.5713E-27 | -0.2768678 | 0.013 | 0.184 | 6.0741E-23 | GRAN | DNAJC1 |
| 2.7902E-27 | -0.2637444 | 0.019 | 0.196 | 6.5913E-23 | GRAN | ORMDL2 |
| 2.8973E-27 | -0.2671631 | 0.01 | 0.178 | 6.8443E-23 | GRAN | SRP72 |
| 2.9089E-27 | -0.362276 | 0.006 | 0.172 | 6.8717E-23 | GRAN | JMY |
| 2.9502E-27 | -0.2605301 | 0.021 | 0.198 | 6.9693E-23 | GRAN | UBE2N |
| 2.9558E-27 | -0.3121599 | 0.002 | 0.163 | 6.9824E-23 | GRAN | NCKAP1 |
| 2.9807E-27 | -0.2604864 | 0.034 | 0.22 | 7.0414E-23 | GRAN | LAMTOR2 |
| 3.3735E-27 | -0.2749411 | 0.048 | 0.245 | 7.9692E-23 | GRAN | FXR1 |
| 3.4065E-27 | -0.2943949 | 0.144 | 0.398 | 8.0471E-23 | GRAN | PNISR |
| 3.505E-27 | -0.4552661 | 0.026 | 0.204 | 8.2799E-23 | GRAN | FOSB |
| 3.6397E-27 | -0.2906382 | 0.013 | 0.183 | 8.5981E-23 | GRAN | ADI1 |
| 3.7093E-27 | -0.2662394 | 0.003 | 0.165 | 8.7624E-23 | GRAN | NUTF2 |
| 3.8271E-27 | 0.55019162 | 0.86 | 0.608 | 9.0409E-23 | GRAN | CD44 |
| 4.3078E-27 | -0.2947593 | 0.002 | 0.162 | 1.0176E-22 | GRAN | BEX4 |
| 4.4619E-27 | 0.46386412 | 0.184 | 0.077 | 1.054E-22 | GRAN | NAIP |
| 4.5419E-27 | -0.2788803 | 0.023 | 0.199 | 1.0729E-22 | GRAN | UHMK1 |
| 4.5455E-27 | -0.2682658 | 0.045 | 0.239 | 1.0738E-22 | GRAN | NDUFA13 |
| 4.7113E-27 | -0.4405516 | 0.927 | 0.835 | 1.113E-22 | GRAN | COX4I1 |
| 5.0746E-27 | -0.3100355 | 0.042 | 0.234 | 1.1988E-22 | GRAN | THUMPD3-AS1 |
| 5.1102E-27 | -0.2887585 | 0.01 | 0.177 | 1.2072E-22 | GRAN | CSRP1 |
| 5.5894E-27 | -0.4591102 | 0.376 | 0.54 | 1.3204E-22 | GRAN | HMGN1 |
| 5.7286E-27 | -0.272663 | 0 | 0.158 | 1.3533E-22 | GRAN | RPAIN |
| 6.1504E-27 | -1.392876 | 0.044 | 0.225 | 1.4529E-22 | GRAN | VCAN |
| 6.2608E-27 | -0.2588717 | 0.04 | 0.232 | 1.479E-22 | GRAN | IL6ST |
| 6.3293E-27 | 0.26282721 | 0.124 | 0.041 | 1.4952E-22 | GRAN | TMEM71 |
| 6.3856E-27 | -0.2754951 | 0.018 | 0.19 | 1.5085E-22 | GRAN | MFSD10 |
| 7.7057E-27 | -0.2711136 | 0.002 | 0.161 | 1.8203E-22 | GRAN | PRMT1 |
| 8.1031E-27 | -0.2775784 | 0.034 | 0.219 | 1.9142E-22 | GRAN | SMIM14 |
| 8.1105E-27 | -0.4104161 | 0.003 | 0.164 | 1.916E-22 | GRAN | SYTL2 |
| 8.1548E-27 | -0.3192733 | 0.002 | 0.161 | 1.9264E-22 | GRAN | CRB3 |
| 8.1679E-27 | -0.2772957 | 0.018 | 0.189 | 1.9295E-22 | GRAN | GTF3A |
| 8.6489E-27 | -0.3388978 | 0 | 0.158 | 2.0431E-22 | GRAN | ERBB3 |
| 8.958E-27 | -0.5045066 | 0.002 | 0.161 | 2.1161E-22 | GRAN | MALL |
| 8.9797E-27 | -0.2561912 | 0.029 | 0.21 | 2.1213E-22 | GRAN | CWC15 |
| 9.375E-27 | -0.2593889 | 0.006 | 0.169 | 2.2146E-22 | GRAN | EMC4 |
| 9.4606E-27 | -0.2979699 | 0.027 | 0.206 | 2.2349E-22 | GRAN | TRIP11 |
| 1.0435E-26 | -0.2657559 | 0.002 | 0.16 | 2.465E-22 | GRAN | CNPY2 |
| 1.1301E-26 | -0.2664067 | 0.011 | 0.177 | 2.6697E-22 | GRAN | SMIM15 |
| 1.1308E-26 | -0.3033728 | 0.006 | 0.169 | 2.6713E-22 | GRAN | MYOF |
| 1.1492E-26 | -0.2976123 | 0.003 | 0.163 | 2.7148E-22 | GRAN | TMPO |
| 1.1774E-26 | -0.2661153 | 0.035 | 0.221 | 2.7814E-22 | GRAN | EIF4G1 |

| | | | | | | |
|---|---|---|---|---|---|---|
| 1.2669E-26 | -0.2658754 | 0.032 | 0.214 | 2.9929E-22 | GRAN | ACBD3 |
| 1.3118E-26 | -0.4167423 | 0.002 | 0.16 | 3.0988E-22 | GRAN | TTC9 |
| 1.331E-26 | 0.70362504 | 0.326 | 0.199 | 3.1442E-22 | GRAN | IFI16 |
| 1.3531E-26 | -0.2546526 | 0.111 | 0.347 | 3.1965E-22 | GRAN | SSR2 |
| 1.3675E-26 | -0.9862423 | 0 | 0.157 | 3.2304E-22 | GRAN | PITX1 |
| 1.3843E-26 | -0.2765799 | 0.003 | 0.162 | 3.2701E-22 | GRAN | SLC25A36 |
| 1.3938E-26 | -0.2820707 | 0.011 | 0.176 | 3.2926E-22 | GRAN | VKORC1 |
| 1.4161E-26 | -0.2840928 | 0.018 | 0.189 | 3.3454E-22 | GRAN | DYNLT3 |
| 1.4361E-26 | -0.2698415 | 0.026 | 0.203 | 3.3925E-22 | GRAN | NANS |
| 1.5006E-26 | 0.45004668 | 0.221 | 0.104 | 3.5448E-22 | GRAN | PRKCB |
| 1.5158E-26 | -0.501239 | 0.334 | 0.499 | 3.5807E-22 | GRAN | VDAC2 |
| 1.5201E-26 | -0.2761479 | 0.024 | 0.201 | 3.5909E-22 | GRAN | TWF1 |
| 1.5519E-26 | -0.2642004 | 0.01 | 0.174 | 3.666E-22 | GRAN | PHB2 |
| 1.6113E-26 | -0.2521684 | 0.04 | 0.228 | 3.8064E-22 | GRAN | MAP2K2 |
| 1.6836E-26 | -0.2587529 | 0.016 | 0.186 | 3.9772E-22 | GRAN | MAP2K1 |
| 1.686E-26 | -0.3432696 | 0.002 | 0.159 | 3.9828E-22 | GRAN | TRIP10 |
| 1.6898E-26 | -0.3161363 | 0.013 | 0.18 | 3.9917E-22 | GRAN | PIK3IP1 |
| 1.6947E-26 | -0.2679074 | 0.016 | 0.185 | 4.0034E-22 | GRAN | POLE4 |
| 1.7524E-26 | -0.3373786 | 0.027 | 0.204 | 4.1397E-22 | GRAN | CHORDC1 |
| 1.7688E-26 | -0.5448959 | 0.455 | 0.565 | 4.1785E-22 | GRAN | SLC25A6 |
| 1.7713E-26 | -0.4493887 | 0 | 0.156 | 4.1844E-22 | GRAN | TMPRSS4 |
| 1.7859E-26 | -0.3150329 | 0.006 | 0.168 | 4.2189E-22 | GRAN | FLNB |
| 1.8664E-26 | -0.3536578 | 0.008 | 0.17 | 4.4089E-22 | GRAN | SLC38A1 |
| 1.8826E-26 | -0.3046928 | 0.073 | 0.281 | 4.4473E-22 | GRAN | YME1L1 |
| 1.8837E-26 | -0.2609231 | 0.06 | 0.26 | 4.4499E-22 | GRAN | NDUFB10 |
| 1.9237E-26 | -0.2599934 | 0.021 | 0.193 | 4.5443E-22 | GRAN | SDHC |
| 1.9605E-26 | -0.6038959 | 0 | 0.156 | 4.6312E-22 | GRAN | CDC42EP5 |
| 1.9748E-26 | -0.2888772 | 0.015 | 0.182 | 4.6651E-22 | GRAN | SARS |
| 2.029E-26 | -0.2988192 | 0 | 0.155 | 4.7931E-22 | GRAN | RCN1 |
| 2.0445E-26 | -0.2574432 | 0.01 | 0.174 | 4.8298E-22 | GRAN | GLRX3 |
| 2.0648E-26 | -0.2611934 | 0.003 | 0.161 | 4.8776E-22 | GRAN | FDX1 |
| 2.143E-26 | -0.2943227 | 0.008 | 0.17 | 5.0624E-22 | GRAN | NRIP1 |
| 2.1443E-26 | -0.2520098 | 0.037 | 0.223 | 5.0655E-22 | GRAN | PCNP |
| 2.1762E-26 | 0.46762631 | 0.239 | 0.118 | 5.1408E-22 | GRAN | WAS |
| 2.2423E-26 | -0.2616831 | 0.003 | 0.161 | 5.297E-22 | GRAN | ESD |
| 2.2854E-26 | -0.2509545 | 0.034 | 0.215 | 5.3988E-22 | GRAN | SDHD |
| 2.3315E-26 | -0.4258998 | 0.003 | 0.161 | 5.5077E-22 | GRAN | SDC1 |
| 2.4304E-26 | -0.2880708 | 0.024 | 0.198 | 5.7414E-22 | GRAN | HUWE1 |
| 2.4899E-26 | -0.2560427 | 0.006 | 0.167 | 5.882E-22 | GRAN | SNRPC |
| 2.5544E-26 | -0.3815197 | 0.002 | 0.158 | 6.0343E-22 | GRAN | TSPAN6 |
| 2.6241E-26 | -0.8637753 | 0.002 | 0.158 | 6.1989E-22 | GRAN | KRT15 |
| 2.6468E-26 | -0.2893054 | 0.015 | 0.182 | 6.2526E-22 | GRAN | FDFT1 |
| 2.7371E-26 | -0.7612719 | 0 | 0.155 | 6.4658E-22 | GRAN | SCEL |
| 2.7941E-26 | -0.2938895 | 0.006 | 0.167 | 6.6004E-22 | GRAN | CDYL |
| 2.9129E-26 | 1.08886985 | 0.361 | 0.236 | 6.8811E-22 | GRAN | TNFAIP3 |
| 2.9431E-26 | -0.3588688 | 0.048 | 0.238 | 6.9525E-22 | GRAN | MT-ATP8 |
| 3.3442E-26 | -0.2653528 | 0.013 | 0.177 | 7.9001E-22 | GRAN | IFI27L2 |
| 3.4468E-26 | -0.4946303 | 0.006 | 0.166 | 8.1423E-22 | GRAN | ANKRD37 |
| 3.5215E-26 | 0.59739383 | 0.318 | 0.186 | 8.3188E-22 | GRAN | RAC2 |
| 3.8848E-26 | -0.2587151 | 0.002 | 0.156 | 9.1771E-22 | GRAN | NDUFS4 |

| | | | | | | |
|---|---|---|---|---|---|---|
| 4.0682E-26 | -0.2777011 | 0.01 | 0.172 | 9.6102E-22 | GRAN | ZNF644 |
| 4.224E-26 | -0.4883394 | 0.002 | 0.157 | 9.9784E-22 | GRAN | CYR61 |
| 4.4889E-26 | -0.3064935 | 0.005 | 0.163 | 1.0604E-21 | GRAN | ZFHX3 |
| 4.6683E-26 | -0.5833942 | 0.25 | 0.436 | 1.1028E-21 | GRAN | GSTO1 |
| 5.0136E-26 | -0.2785504 | 0.003 | 0.159 | 1.1844E-21 | GRAN | ZNF326 |
| 5.3349E-26 | -0.3715311 | 0.005 | 0.162 | 1.2603E-21 | GRAN | SELENOM |
| 5.6768E-26 | -0.2606652 | 0.016 | 0.183 | 1.341E-21 | GRAN | POLR2I |
| 5.9717E-26 | -0.2579694 | 0.002 | 0.155 | 1.4107E-21 | GRAN | EIF2A |
| 6.1398E-26 | -0.254038 | 0.008 | 0.167 | 1.4504E-21 | GRAN | CNIH1 |
| 6.4733E-26 | -0.3030427 | 0.019 | 0.188 | 1.5292E-21 | GRAN | HIPK2 |
| 6.9076E-26 | -0.4266106 | 0.003 | 0.159 | 1.6318E-21 | GRAN | CLDN3 |
| 7.0309E-26 | -0.2803717 | 0.003 | 0.159 | 1.6609E-21 | GRAN | HNMT |
| 7.1632E-26 | -0.3865629 | 0.003 | 0.158 | 1.6922E-21 | GRAN | COBLL1 |
| 7.4662E-26 | -0.2564528 | 0.018 | 0.184 | 1.7637E-21 | GRAN | SSNA1 |
| 7.6561E-26 | -0.260485 | 0.008 | 0.167 | 1.8086E-21 | GRAN | MRPL14 |
| 7.8043E-26 | -0.2803652 | 0.021 | 0.19 | 1.8436E-21 | GRAN | EIF4E |
| 8.0343E-26 | -0.2590818 | 0.021 | 0.19 | 1.898E-21 | GRAN | ORMDL1 |
| 8.1607E-26 | -0.2683195 | 0.011 | 0.173 | 1.9278E-21 | GRAN | CLDND1 |
| 8.8126E-26 | -0.3123129 | 0.006 | 0.163 | 2.0818E-21 | GRAN | FARP1 |
| 8.9118E-26 | -1.0567939 | 0.021 | 0.187 | 2.1052E-21 | GRAN | HLA-DPB1 |
| 9.0288E-26 | -0.2570825 | 0.026 | 0.198 | 2.1329E-21 | GRAN | CHMP4A |
| 9.4359E-26 | -0.2825808 | 0.092 | 0.308 | 2.229E-21 | GRAN | HP1BP3 |
| 9.5012E-26 | -0.4871646 | 0.371 | 0.522 | 2.2445E-21 | GRAN | ATP6V1F |
| 9.7216E-26 | -0.2620599 | 0.002 | 0.155 | 2.2965E-21 | GRAN | SCOC |
| 9.911E-26 | -0.3005286 | 0.015 | 0.179 | 2.3413E-21 | GRAN | ZBTB43 |
| 1.0304E-25 | -0.9744945 | 0.005 | 0.16 | 2.4342E-21 | GRAN | PRSS27 |
| 1.0736E-25 | -0.2589403 | 0.003 | 0.158 | 2.5361E-21 | GRAN | POLR2H |
| 1.0897E-25 | -0.2714336 | 0.039 | 0.219 | 2.5742E-21 | GRAN | U2SURP |
| 1.1274E-25 | -0.2510865 | 0.011 | 0.173 | 2.6634E-21 | GRAN | PSMC4 |
| 1.1318E-25 | -0.2849096 | 0.008 | 0.166 | 2.6737E-21 | GRAN | SMC5 |
| 1.1507E-25 | -0.27549 | 0.085 | 0.298 | 2.7182E-21 | GRAN | ZFAND6 |
| 1.1601E-25 | -0.6500468 | 0.003 | 0.157 | 2.7405E-21 | GRAN | MUC4 |
| 1.2424E-25 | -0.6813727 | 0.845 | 0.756 | 2.935E-21 | GRAN | ITGB1 |
| 1.248E-25 | -0.3840044 | 0.027 | 0.2 | 2.9482E-21 | GRAN | SERTAD2 |
| 1.3677E-25 | -0.2597962 | 0.044 | 0.228 | 3.2309E-21 | GRAN | XPO1 |
| 1.4053E-25 | -0.2557742 | 0.013 | 0.175 | 3.3197E-21 | GRAN | PSMD1 |
| 1.4332E-25 | -0.2886675 | 0.003 | 0.157 | 3.3856E-21 | GRAN | ARHGEF12 |
| 1.435E-25 | -0.272365 | 0.015 | 0.178 | 3.39E-21 | GRAN | TRIM28 |
| 1.661E-25 | -0.8333232 | 0.295 | 0.439 | 3.9239E-21 | GRAN | ATP1B1 |
| 1.6832E-25 | -0.3523024 | 0.003 | 0.157 | 3.9762E-21 | GRAN | ITPKC |
| 1.6917E-25 | -0.2665638 | 0.018 | 0.183 | 3.9963E-21 | GRAN | ELK3 |
| 1.8201E-25 | -0.7638441 | 0 | 0.151 | 4.2996E-21 | GRAN | TMPRSS11D |
| 1.8348E-25 | -0.2617401 | 0.003 | 0.156 | 4.3344E-21 | GRAN | SLC25A39 |
| 1.9133E-25 | -0.2830295 | 0.006 | 0.163 | 4.5198E-21 | GRAN | GFPT1 |
| 1.9135E-25 | -0.258285 | 0.006 | 0.161 | 4.5202E-21 | GRAN | PTPN11 |
| 1.9541E-25 | -0.2585713 | 0.002 | 0.153 | 4.6162E-21 | GRAN | ECI1 |
| 2.1205E-25 | -0.284721 | 0.061 | 0.257 | 5.0093E-21 | GRAN | MGAT1 |
| 2.1595E-25 | -1.4498476 | 0.174 | 0.338 | 5.1015E-21 | GRAN | IGFBP7 |
| 2.1631E-25 | -0.29253 | 0.005 | 0.159 | 5.1099E-21 | GRAN | CTTNBP2NL |
| 2.1947E-25 | -0.2753724 | 0.008 | 0.165 | 5.1845E-21 | GRAN | TMEM141 |

| | | | | | | |
|---|---|---|---|---|---|---|
| 2.3217E-25 | -0.2685854 | 0.002 | 0.153 | 5.4845E-21 | GRAN | SHTN1 |
| 2.4736E-25 | -0.2624256 | 0.016 | 0.179 | 5.8433E-21 | GRAN | RBBP4 |
| 2.7005E-25 | -0.2582562 | 0 | 0.149 | 6.3794E-21 | GRAN | PSMD14 |
| 2.7978E-25 | -0.252639 | 0.003 | 0.155 | 6.6094E-21 | GRAN | ARHGAP21 |
| 2.8585E-25 | -0.2653696 | 0.013 | 0.173 | 6.7526E-21 | GRAN | MRPS5 |
| 2.884E-25 | -0.2685686 | 0.016 | 0.179 | 6.8128E-21 | GRAN | SPSB3 |
| 3.2516E-25 | -0.3314098 | 0.018 | 0.181 | 7.6812E-21 | GRAN | TMEM176B |
| 3.4131E-25 | -0.2531462 | 0.003 | 0.155 | 8.0628E-21 | GRAN | PNKD |
| 3.4346E-25 | -0.2620106 | 0.011 | 0.17 | 8.1134E-21 | GRAN | MRPS34 |
| 3.5019E-25 | 0.58640491 | 0.297 | 0.175 | 8.2725E-21 | GRAN | PPP1R18 |
| 3.7468E-25 | -0.5019127 | 0.015 | 0.173 | 8.8511E-21 | GRAN | DNAJA4 |
| 4.0591E-25 | -0.2623419 | 0.029 | 0.2 | 9.5889E-21 | GRAN | WDR45B |
| 4.1198E-25 | -0.2618218 | 0.037 | 0.215 | 9.7322E-21 | GRAN | REST |
| 4.1223E-25 | -0.6524671 | 0 | 0.149 | 9.7382E-21 | GRAN | NCCRP1 |
| 4.2139E-25 | -0.3068538 | 0.002 | 0.151 | 9.9544E-21 | GRAN | PLS3 |
| 4.2404E-25 | -0.2592696 | 0.024 | 0.191 | 1.0017E-20 | GRAN | PRDX3 |
| 4.4428E-25 | -0.7899745 | 0.837 | 0.73 | 1.0495E-20 | GRAN | HSPA8 |
| 4.5868E-25 | -0.2710409 | 0 | 0.148 | 1.0835E-20 | GRAN | LLGL2 |
| 4.6644E-25 | -0.2574152 | 0.01 | 0.166 | 1.1019E-20 | GRAN | NPC1 |
| 4.7763E-25 | -0.717805 | 0.789 | 0.459 | 1.1283E-20 | GRAN | LYZ |
| 4.7816E-25 | -0.3659156 | 0.003 | 0.154 | 1.1296E-20 | GRAN | ITGA3 |
| 4.8635E-25 | -0.2765209 | 0.003 | 0.154 | 1.1489E-20 | GRAN | PLEKHA1 |
| 5.1867E-25 | -0.2658492 | 0 | 0.148 | 1.2253E-20 | GRAN | TUBB6 |
| 5.8571E-25 | -0.261461 | 0.027 | 0.197 | 1.3836E-20 | GRAN | GLTP |
| 5.993E-25 | -0.828715 | 0 | 0.148 | 1.4157E-20 | GRAN | CEACAM7 |
| 6.1089E-25 | -0.2534383 | 0.039 | 0.217 | 1.4431E-20 | GRAN | PUM1 |
| 6.116E-25 | -0.2688199 | 0.008 | 0.162 | 1.4448E-20 | GRAN | SSBP3 |
| 6.322E-25 | -0.3546314 | 0.034 | 0.207 | 1.4934E-20 | GRAN | TIPARP |
| 6.4976E-25 | -0.2572935 | 0.008 | 0.162 | 1.5349E-20 | GRAN | GLRX5 |
| 7.7392E-25 | -0.4982987 | 0.002 | 0.15 | 1.8282E-20 | GRAN | RASD1 |
| 7.7828E-25 | -0.4781307 | 0.39 | 0.535 | 1.8385E-20 | GRAN | P4HB |
| 7.8005E-25 | -0.3003056 | 0.005 | 0.156 | 1.8427E-20 | GRAN | AHR |
| 7.9012E-25 | -0.8670191 | 0.026 | 0.19 | 1.8665E-20 | GRAN | RARRES3 |
| 8.0122E-25 | -0.3744525 | 0.003 | 0.153 | 1.8927E-20 | GRAN | MDK |
| 8.0297E-25 | -0.2778244 | 0.002 | 0.15 | 1.8968E-20 | GRAN | NME1 |
| 8.7797E-25 | -0.3211232 | 0.006 | 0.159 | 2.074E-20 | GRAN | FNIP2 |
| 9.4154E-25 | -0.568943 | 0.016 | 0.174 | 2.2242E-20 | GRAN | HLA-DQB1 |
| 9.4482E-25 | -0.2604526 | 0.003 | 0.152 | 2.232E-20 | GRAN | RCN2 |
| 1.0142E-24 | -0.2654999 | 0.003 | 0.152 | 2.3959E-20 | GRAN | LARS |
| 1.0837E-24 | -0.2750356 | 0.034 | 0.206 | 2.56E-20 | GRAN | LRRFIP2 |
| 1.0974E-24 | -0.3471519 | 0.002 | 0.149 | 2.5923E-20 | GRAN | GSTM3 |
| 1.1688E-24 | -0.2803156 | 0.003 | 0.152 | 2.7611E-20 | GRAN | LGMN |
| 1.2131E-24 | -0.2823157 | 0.013 | 0.169 | 2.8657E-20 | GRAN | DCXR |
| 1.2575E-24 | -0.4750211 | 0.002 | 0.149 | 2.9705E-20 | GRAN | ALDH1A3 |
| 1.3007E-24 | -0.2727845 | 0.005 | 0.155 | 3.0726E-20 | GRAN | EIF4EBP1 |
| 1.4651E-24 | -0.324117 | 0.035 | 0.206 | 3.461E-20 | GRAN | MAT2A |
| 1.5427E-24 | -0.2502864 | 0.002 | 0.148 | 3.6443E-20 | GRAN | COA4 |
| 1.6478E-24 | -0.3243463 | 0.006 | 0.157 | 3.8925E-20 | GRAN | SESN2 |
| 1.6503E-24 | -0.4687633 | 0.345 | 0.51 | 3.8984E-20 | GRAN | DBI |
| 1.6633E-24 | -0.2935023 | 0.002 | 0.148 | 3.9292E-20 | GRAN | KCNK1 |

| | | | | | | |
|---|---|---|---|---|---|---|
| 1.6698E-24 | -0.2653918 | 0.053 | 0.24 | 3.9445E-20 | GRAN | HOOK3 |
| 1.6753E-24 | -0.3093435 | 0.002 | 0.148 | 3.9577E-20 | GRAN | FAM102A |
| 1.7179E-24 | -0.2923776 | 0.024 | 0.187 | 4.0582E-20 | GRAN | PHACTR2 |
| 1.733E-24 | -1.0893301 | 0.024 | 0.183 | 4.0939E-20 | GRAN | SPINK1 |
| 1.8053E-24 | -0.2613408 | 0.003 | 0.151 | 4.2646E-20 | GRAN | BAZ1B |
| 1.8072E-24 | -0.3212872 | 0.005 | 0.154 | 4.2691E-20 | GRAN | CPVL |
| 1.8387E-24 | -0.2782669 | 0.015 | 0.171 | 4.3436E-20 | GRAN | GARS |
| 1.9412E-24 | -0.259589 | 0.003 | 0.151 | 4.5858E-20 | GRAN | NOP56 |
| 2.1697E-24 | -0.5180822 | 0.087 | 0.284 | 5.1256E-20 | GRAN | RHOB |
| 2.1975E-24 | -0.3341833 | 0.015 | 0.171 | 5.1912E-20 | GRAN | CDKN2AIP |
| 2.2737E-24 | -0.313388 | 0.003 | 0.15 | 5.3712E-20 | GRAN | SCCPDH |
| 2.3278E-24 | -0.2600731 | 0.026 | 0.19 | 5.499E-20 | GRAN | PSMD12 |
| 2.4777E-24 | -0.2615275 | 0 | 0.144 | 5.853E-20 | GRAN | BACE2 |
| 2.5152E-24 | -0.2769533 | 0.011 | 0.165 | 5.9417E-20 | GRAN | MTHFD2 |
| 2.595E-24 | -0.2830784 | 0.015 | 0.17 | 6.1302E-20 | GRAN | TP53BP2 |
| 2.781E-24 | -0.2621785 | 0.013 | 0.167 | 6.5695E-20 | GRAN | ZCRB1 |
| 3.0157E-24 | -0.7724114 | 0.002 | 0.147 | 7.124E-20 | GRAN | TMPRSS11E |
| 3.0465E-24 | -0.4140454 | 0.947 | 0.868 | 7.1967E-20 | GRAN | PABPC1 |
| 3.3429E-24 | -0.3086492 | 0.079 | 0.278 | 7.8969E-20 | GRAN | PLIN3 |
| 3.4319E-24 | -0.2937885 | 0.026 | 0.19 | 8.1073E-20 | GRAN | VCL |
| 3.4686E-24 | -0.2871169 | 0.002 | 0.146 | 8.1939E-20 | GRAN | GPRC5C |
| 3.6394E-24 | -0.6020295 | 0.008 | 0.158 | 8.5974E-20 | GRAN | MT1F |
| 3.6963E-24 | -0.3600715 | 0.002 | 0.146 | 8.7317E-20 | GRAN | CLDN1 |
| 4.1521E-24 | -0.2581164 | 0.011 | 0.163 | 9.8086E-20 | GRAN | KRR1 |
| 4.4276E-24 | -0.8583573 | 0.003 | 0.148 | 1.0459E-19 | GRAN | CEACAM6 |
| 4.4794E-24 | -0.2829286 | 0.002 | 0.146 | 1.0582E-19 | GRAN | FUCA1 |
| 4.9339E-24 | -0.2746047 | 0.003 | 0.149 | 1.1655E-19 | GRAN | NR1D2 |
| 4.9914E-24 | 0.60206475 | 0.256 | 0.141 | 1.1791E-19 | GRAN | TRIM25 |
| 5.4231E-24 | -0.2758541 | 0.031 | 0.196 | 1.2811E-19 | GRAN | KIFC3 |
| 5.549E-24 | -0.2504073 | 0.008 | 0.156 | 1.3108E-19 | GRAN | PFKL |
| 5.9868E-24 | -0.4082572 | 0.003 | 0.148 | 1.4143E-19 | GRAN | VSIG2 |
| 6.5073E-24 | -0.295231 | 0.002 | 0.145 | 1.5372E-19 | GRAN | PHLDA3 |
| 6.952E-24 | 0.39332062 | 0.168 | 0.071 | 1.6423E-19 | GRAN | GPR65 |
| 7.7057E-24 | 0.48022474 | 0.245 | 0.129 | 1.8203E-19 | GRAN | APBB1IP |
| 7.9035E-24 | 0.54874893 | 0.247 | 0.136 | 1.867E-19 | GRAN | RALB |
| 8.8825E-24 | -0.4587357 | 0.074 | 0.261 | 2.0983E-19 | GRAN | PMAIP1 |
| 8.9014E-24 | -0.6371798 | 0.002 | 0.144 | 2.1028E-19 | GRAN | CDKN2A |
| 9.7091E-24 | -0.5051741 | 0.011 | 0.16 | 2.2936E-19 | GRAN | BPGM |
| 1.1179E-23 | -0.3214845 | 0.003 | 0.147 | 2.6409E-19 | GRAN | PDP1 |
| 1.1726E-23 | -0.2938922 | 0.003 | 0.146 | 2.77E-19 | GRAN | CRIM1 |
| 1.211E-23 | -0.2786073 | 0.013 | 0.164 | 2.8608E-19 | GRAN | ALCAM |
| 1.2868E-23 | -0.3417897 | 0.006 | 0.152 | 3.0398E-19 | GRAN | LPIN1 |
| 1.4118E-23 | -0.5112776 | 0.547 | 0.61 | 3.3351E-19 | GRAN | HNRNPA1 |
| 1.4189E-23 | -0.2898518 | 0.044 | 0.216 | 3.3518E-19 | GRAN | RUNX1 |
| 1.584E-23 | -0.277797 | 0.045 | 0.217 | 3.7419E-19 | GRAN | ARID1B |
| 1.6101E-23 | -0.2609363 | 0.047 | 0.222 | 3.8035E-19 | GRAN | TES |
| 1.6273E-23 | -1.6925905 | 0.003 | 0.145 | 3.8442E-19 | GRAN | SPRR2A |
| 1.6605E-23 | -0.2630343 | 0.006 | 0.151 | 3.9226E-19 | GRAN | ATXN7L3B |
| 1.6847E-23 | -0.3223678 | 0 | 0.14 | 3.9798E-19 | GRAN | MTUS1 |
| 1.7953E-23 | -0.3796463 | 0.003 | 0.146 | 4.2411E-19 | GRAN | ITGB6 |

| | | | | | | |
|---|---|---|---|---|---|---|
| 1.9007E-23 | -1.0088387 | 0.323 | 0.454 | 4.4901E-19 | GRAN | HSPH1 |
| 1.9402E-23 | -0.4082807 | 0.337 | 0.504 | 4.5833E-19 | GRAN | LAPTM4A |
| 2.0643E-23 | -0.2718949 | 0 | 0.139 | 4.8764E-19 | GRAN | PATJ |
| 2.1511E-23 | -0.2774571 | 0.002 | 0.142 | 5.0815E-19 | GRAN | RAB25 |
| 2.153E-23 | -0.3969757 | 0.002 | 0.142 | 5.0861E-19 | GRAN | GPNMB |
| 2.3428E-23 | -0.2762129 | 0.003 | 0.144 | 5.5344E-19 | GRAN | CPM |
| 2.3856E-23 | -0.7072427 | 0 | 0.139 | 5.6355E-19 | GRAN | NUPR1 |
| 2.5222E-23 | -0.3035026 | 0.005 | 0.148 | 5.9583E-19 | GRAN | CD40 |
| 2.5284E-23 | -0.3472018 | 0.002 | 0.142 | 5.9728E-19 | GRAN | EMP2 |
| 2.6206E-23 | -0.3294287 | 0.01 | 0.156 | 6.1906E-19 | GRAN | HIST1H1E |
| 2.7967E-23 | -0.29183 | 0.003 | 0.144 | 6.6067E-19 | GRAN | OCLN |
| 2.9678E-23 | -0.2660206 | 0 | 0.138 | 7.0108E-19 | GRAN | SRGAP1 |
| 3.0816E-23 | -0.388677 | 0.005 | 0.147 | 7.2798E-19 | GRAN | SDC2 |
| 3.1217E-23 | -0.2614989 | 0 | 0.138 | 7.3743E-19 | GRAN | MBNL2 |
| 3.2005E-23 | -0.2517065 | 0.01 | 0.156 | 7.5605E-19 | GRAN | ZBTB38 |
| 3.2793E-23 | -0.2869798 | 0.003 | 0.144 | 7.7468E-19 | GRAN | LAPTM4B |
| 3.41E-23 | -0.2781133 | 0.035 | 0.201 | 8.0554E-19 | GRAN | ARHGAP18 |
| 3.7288E-23 | -0.2766374 | 0.002 | 0.141 | 8.8084E-19 | GRAN | GULP1 |
| 4.138E-23 | -0.6195404 | 0.003 | 0.143 | 9.7752E-19 | GRAN | SNCG |
| 4.4506E-23 | 0.30517389 | 0.135 | 0.051 | 1.0514E-18 | GRAN | ENTPD1 |
| 4.5102E-23 | -0.3093181 | 0.002 | 0.14 | 1.0654E-18 | GRAN | MGLL |
| 4.5765E-23 | -0.4705454 | 0.003 | 0.143 | 1.0811E-18 | GRAN | ID1 |
| 4.9301E-23 | -0.2593727 | 0.002 | 0.14 | 1.1646E-18 | GRAN | MYO1E |
| 4.9532E-23 | -0.7570289 | 0.421 | 0.52 | 1.1701E-18 | GRAN | H1FX |
| 5.2464E-23 | -0.6892321 | 0.424 | 0.525 | 1.2394E-18 | GRAN | HSPE1 |
| 5.2911E-23 | -0.2882504 | 0.01 | 0.155 | 1.2499E-18 | GRAN | CKS2 |
| 5.3297E-23 | -0.2994605 | 0.002 | 0.14 | 1.259E-18 | GRAN | ZNRF1 |
| 5.3632E-23 | -0.282317 | 0.003 | 0.143 | 1.2669E-18 | GRAN | BEX3 |
| 5.6142E-23 | -0.5097867 | 0.977 | 0.89 | 1.3262E-18 | GRAN | CD63 |
| 5.8365E-23 | -0.5033017 | 0.084 | 0.269 | 1.3788E-18 | GRAN | DDIT3 |
| 6.3204E-23 | -0.2506394 | 0.016 | 0.166 | 1.4931E-18 | GRAN | THOC2 |
| 7.1759E-23 | -0.9109256 | 0 | 0.136 | 1.6952E-18 | GRAN | TMPRSS11B |
| 7.427E-23 | -0.4678422 | 0.01 | 0.154 | 1.7545E-18 | GRAN | ZFAND2A |
| 7.6432E-23 | -0.2907258 | 0.01 | 0.154 | 1.8055E-18 | GRAN | SLC44A1 |
| 7.8285E-23 | -0.462257 | 0.01 | 0.153 | 1.8493E-18 | GRAN | CLEC5A |
| 7.9047E-23 | -0.4903203 | 0.016 | 0.163 | 1.8673E-18 | GRAN | CSKMT |
| 8.0184E-23 | -0.4631876 | 0 | 0.136 | 1.8942E-18 | GRAN | UCHL1 |
| 8.1987E-23 | 0.79503978 | 0.495 | 0.393 | 1.9368E-18 | GRAN | HIF1A |
| 8.2241E-23 | -0.4061137 | 0.002 | 0.139 | 1.9428E-18 | GRAN | DCDC2 |
| 8.6454E-23 | -0.4033907 | 0.005 | 0.145 | 2.0423E-18 | GRAN | PAX8 |
| 9.159E-23 | -0.3600694 | 0.002 | 0.138 | 2.1636E-18 | GRAN | HLA-DMA |
| 9.7018E-23 | -0.5235264 | 0.458 | 0.555 | 2.2919E-18 | GRAN | SSR4 |
| 1.0254E-22 | -0.5188111 | 0.015 | 0.16 | 2.4223E-18 | GRAN | CD300E |
| 1.078E-22 | -0.265597 | 0.002 | 0.138 | 2.5465E-18 | GRAN | SCNN1A |
| 1.0877E-22 | 0.51496531 | 0.842 | 0.684 | 2.5695E-18 | GRAN | ATP6V0B |
| 1.1572E-22 | -0.2669354 | 0.002 | 0.138 | 2.7336E-18 | GRAN | AGRN |
| 1.2298E-22 | -0.3607211 | 0.021 | 0.172 | 2.9052E-18 | GRAN | FAM214A |
| 1.3171E-22 | 0.46458517 | 0.213 | 0.107 | 3.1114E-18 | GRAN | PAK1 |
| 1.4382E-22 | -0.4657329 | 0.384 | 0.531 | 3.3975E-18 | GRAN | EIF4A2 |
| 1.4433E-22 | -0.414476 | 0.002 | 0.138 | 3.4094E-18 | GRAN | GCNT3 |

| | | | | | | |
|---|---|---|---|---|---|---|
| 1.5018E-22 | -0.7321632 | 0.994 | 0.914 | 3.5477E-18 | GRAN | MT-ND1 |
| 1.5158E-22 | -0.2561092 | 0.013 | 0.158 | 3.5809E-18 | GRAN | NFE2L1 |
| 1.5897E-22 | 0.48824032 | 0.274 | 0.155 | 3.7554E-18 | GRAN | PELI1 |
| 1.6415E-22 | -0.3032499 | 0.002 | 0.137 | 3.8778E-18 | GRAN | SASH1 |
| 1.6735E-22 | -0.2571373 | 0.002 | 0.137 | 3.9533E-18 | GRAN | ESF1 |
| 1.6821E-22 | -0.2505584 | 0.008 | 0.149 | 3.9735E-18 | GRAN | MALT1 |
| 1.7004E-22 | -0.3151707 | 0.002 | 0.137 | 4.0169E-18 | GRAN | GCLC |
| 1.7905E-22 | 0.50203179 | 0.266 | 0.151 | 4.2297E-18 | GRAN | VSIR |
| 1.8103E-22 | -0.2729251 | 0.005 | 0.143 | 4.2764E-18 | GRAN | LRP1 |
| 1.8172E-22 | -0.2580598 | 0.008 | 0.149 | 4.2929E-18 | GRAN | YPEL2 |
| 2.2231E-22 | -1.2427026 | 0 | 0.134 | 5.2516E-18 | GRAN | PLAT |
| 2.3702E-22 | -0.2600878 | 0.002 | 0.136 | 5.5991E-18 | GRAN | TMEM238 |
| 2.4525E-22 | -0.2608789 | 0.01 | 0.151 | 5.7935E-18 | GRAN | RASGEF1B |
| 2.793E-22 | -0.6624007 | 0 | 0.133 | 6.5978E-18 | GRAN | GABRP |
| 2.8096E-22 | -0.2743401 | 0 | 0.133 | 6.6372E-18 | GRAN | PALLD |
| 3.4797E-22 | 0.33926639 | 0.168 | 0.073 | 8.2201E-18 | GRAN | ST8SIA4 |
| 3.5733E-22 | -0.6284821 | 0.003 | 0.138 | 8.4412E-18 | GRAN | CXCL17 |
| 3.6952E-22 | -0.4464961 | 0.034 | 0.191 | 8.7292E-18 | GRAN | HBEGF |
| 3.7597E-22 | -0.2842444 | 0.008 | 0.147 | 8.8816E-18 | GRAN | SLC44A4 |
| 3.9949E-22 | -0.274539 | 0 | 0.132 | 9.4373E-18 | GRAN | C1orf116 |
| 4.2201E-22 | -0.350128 | 0 | 0.132 | 9.9692E-18 | GRAN | FOXC1 |
| 4.3661E-22 | -1.6005828 | 0.002 | 0.134 | 1.0314E-17 | GRAN | LYPD2 |
| 4.4965E-22 | -0.2816894 | 0.002 | 0.135 | 1.0622E-17 | GRAN | SQLE |
| 4.6705E-22 | -0.2616491 | 0.003 | 0.138 | 1.1033E-17 | GRAN | MACO1 |
| 4.7874E-22 | -0.3005426 | 0.006 | 0.143 | 1.1309E-17 | GRAN | MSMO1 |
| 5.1595E-22 | -0.9082861 | 0.002 | 0.134 | 1.2188E-17 | GRAN | RHCG |
| 5.3047E-22 | -0.256828 | 0.198 | 0.419 | 1.2531E-17 | GRAN | GRN |
| 5.5665E-22 | 0.5487328 | 0.237 | 0.125 | 1.315E-17 | GRAN | IRAK3 |
| 5.5729E-22 | -1.0165506 | 0.006 | 0.142 | 1.3165E-17 | GRAN | CXCL6 |
| 6.258E-22 | -0.2594552 | 0.016 | 0.16 | 1.4783E-17 | GRAN | OXR1 |
| 6.6863E-22 | -0.3877376 | 0.003 | 0.136 | 1.5795E-17 | GRAN | BCAS1 |
| 6.7314E-22 | -0.577851 | 0.365 | 0.495 | 1.5902E-17 | GRAN | H2AFJ |
| 7.0203E-22 | -0.2525271 | 0.002 | 0.134 | 1.6584E-17 | GRAN | SLCO4A1 |
| 7.9054E-22 | -0.2994615 | 0 | 0.13 | 1.8675E-17 | GRAN | BEX2 |
| 8.1458E-22 | -0.3289697 | 0.003 | 0.136 | 1.9243E-17 | GRAN | ACADM |
| 8.1956E-22 | -0.2524126 | 0 | 0.13 | 1.9361E-17 | GRAN | HACD3 |
| 8.3072E-22 | 0.52627683 | 0.269 | 0.157 | 1.9624E-17 | GRAN | DOCK8 |
| 8.3849E-22 | -0.5329908 | 0.308 | 0.467 | 1.9808E-17 | GRAN | CDKN1A |
| 8.4692E-22 | -0.652489 | 0 | 0.13 | 2.0007E-17 | GRAN | TFPI2 |
| 8.9969E-22 | -0.4340366 | 0.013 | 0.153 | 2.1253E-17 | GRAN | SLC7A11 |
| 9.7736E-22 | -0.2833318 | 0.002 | 0.133 | 2.3088E-17 | GRAN | EPB41L3 |
| 9.9536E-22 | -0.3062299 | 0 | 0.13 | 2.3513E-17 | GRAN | H2AFX |
| 1.0788E-21 | -0.3771855 | 0.005 | 0.138 | 2.5485E-17 | GRAN | ARHGAP29 |
| 1.0953E-21 | -0.2672341 | 0.002 | 0.133 | 2.5874E-17 | GRAN | NTN4 |
| 1.3132E-21 | 0.53049333 | 0.276 | 0.166 | 3.1021E-17 | GRAN | STX3 |
| 1.5484E-21 | -0.3184509 | 0.002 | 0.132 | 3.6578E-17 | GRAN | AIF1L |
| 1.5874E-21 | 0.49248865 | 0.292 | 0.171 | 3.7498E-17 | GRAN | CORO1A |
| 1.6697E-21 | -0.2615509 | 0.002 | 0.131 | 3.9444E-17 | GRAN | INSR |
| 1.7203E-21 | -0.2723217 | 0 | 0.128 | 4.0638E-17 | GRAN | SELENOP |
| 1.8129E-21 | -0.2508422 | 0.016 | 0.158 | 4.2827E-17 | GRAN | OFD1 |

| | | | | | | |
|---|---|---|---|---|---|---|
| 1.9212E-21 | -0.4797669 | 0.005 | 0.137 | 4.5384E-17 | GRAN | HLA-DRB5 |
| 2.0491E-21 | -0.276113 | 0.052 | 0.216 | 4.8406E-17 | GRAN | NKTR |
| 2.4514E-21 | -0.3203578 | 0.947 | 0.863 | 5.7909E-17 | GRAN | SRP14 |
| 2.462E-21 | -0.3452801 | 0.085 | 0.269 | 5.816E-17 | GRAN | ID2 |
| 2.4878E-21 | -0.3913943 | 0 | 0.127 | 5.877E-17 | GRAN | ID4 |
| 2.4996E-21 | -0.5098932 | 0.729 | 0.689 | 5.9049E-17 | GRAN | EEF2 |
| 2.5675E-21 | 0.55608968 | 0.295 | 0.184 | 6.0652E-17 | GRAN | TET2 |
| 2.7018E-21 | -0.4930433 | 0.013 | 0.15 | 6.3825E-17 | GRAN | SMC4 |
| 2.9098E-21 | -0.3050088 | 0.097 | 0.286 | 6.8737E-17 | GRAN | CTSZ |
| 3.1604E-21 | -0.340669 | 0.002 | 0.13 | 7.4659E-17 | GRAN | FAM84A |
| 3.3845E-21 | -0.2946698 | 0.003 | 0.133 | 7.9952E-17 | GRAN | SCIN |
| 3.4154E-21 | -1.0288168 | 0.005 | 0.135 | 8.0681E-17 | GRAN | SERPINB2 |
| 3.5055E-21 | 0.42336917 | 0.179 | 0.084 | 8.2811E-17 | GRAN | MAPK14 |
| 3.6162E-21 | -0.4444011 | 0.45 | 0.557 | 8.5426E-17 | GRAN | NDUFS5 |
| 4.1476E-21 | 0.40358655 | 0.182 | 0.087 | 9.7979E-17 | GRAN | FLOT2 |
| 4.5013E-21 | 0.41312989 | 0.203 | 0.103 | 1.0633E-16 | GRAN | RASSF3 |
| 4.7101E-21 | -0.2510557 | 0.024 | 0.169 | 1.1127E-16 | GRAN | CHCHD7 |
| 4.8326E-21 | -0.2531238 | 0.218 | 0.432 | 1.1416E-16 | GRAN | SELENOK |
| 5.0508E-21 | -0.2595309 | 0.015 | 0.153 | 1.1932E-16 | GRAN | ZNF83 |
| 5.6039E-21 | 0.4543249 | 0.235 | 0.128 | 1.3238E-16 | GRAN | IL6R |
| 5.7067E-21 | 0.47805614 | 0.213 | 0.112 | 1.3481E-16 | GRAN | NMI |
| 5.7573E-21 | -0.6812514 | 0.787 | 0.684 | 1.36E-16 | GRAN | NPM1 |
| 5.8191E-21 | 0.54462406 | 0.271 | 0.163 | 1.3746E-16 | GRAN | TLE3 |
| 6.8073E-21 | -0.266423 | 0.002 | 0.128 | 1.6081E-16 | GRAN | PLA2G16 |
| 8.3295E-21 | -0.4097191 | 0 | 0.125 | 1.9677E-16 | GRAN | TMEM40 |
| 8.6268E-21 | -0.2581988 | 0 | 0.124 | 2.0379E-16 | GRAN | PTPN3 |
| 8.6844E-21 | -0.2537753 | 0.005 | 0.133 | 2.0515E-16 | GRAN | PPFIBP1 |
| 9.3292E-21 | 0.46643974 | 0.221 | 0.117 | 2.2038E-16 | GRAN | SLCO3A1 |
| 9.4705E-21 | 0.57797352 | 0.279 | 0.174 | 2.2372E-16 | GRAN | NRBF2 |
| 1.1587E-20 | -0.554662 | 0 | 0.124 | 2.7373E-16 | GRAN | AGR2 |
| 1.2856E-20 | -0.5859455 | 0.365 | 0.487 | 3.0369E-16 | GRAN | HMGA1 |
| 1.3701E-20 | -0.4000923 | 0.052 | 0.209 | 3.2365E-16 | GRAN | KCNQ1OT1 |
| 1.4701E-20 | -0.2527238 | 0.011 | 0.144 | 3.4729E-16 | GRAN | ARL4A |
| 1.5784E-20 | -0.2728278 | 0.008 | 0.137 | 3.7286E-16 | GRAN | IDH1 |
| 1.8721E-20 | 0.42706775 | 0.889 | 0.684 | 4.4224E-16 | GRAN | BRI3 |
| 1.9166E-20 | -0.2900945 | 0.002 | 0.125 | 4.5276E-16 | GRAN | FAT1 |
| 1.9361E-20 | -0.5629353 | 0.005 | 0.131 | 4.5737E-16 | GRAN | AKAP12 |
| 2.108E-20 | -0.3560184 | 0.003 | 0.128 | 4.9797E-16 | GRAN | MARCO |
| 2.1922E-20 | -0.2832992 | 0.01 | 0.14 | 5.1787E-16 | GRAN | ETS1 |
| 2.4281E-20 | 0.57485146 | 0.323 | 0.216 | 5.7358E-16 | GRAN | STK4 |
| 2.5129E-20 | -0.2511888 | 0.003 | 0.128 | 5.9363E-16 | GRAN | LRRC75A |
| 2.6203E-20 | 0.3895591 | 0.189 | 0.093 | 6.1899E-16 | GRAN | NADK |
| 2.9518E-20 | -0.2578522 | 0.174 | 0.4 | 6.973E-16 | GRAN | ARGLU1 |
| 3.128E-20 | 0.36010604 | 0.215 | 0.107 | 7.3892E-16 | GRAN | FGL2 |
| 3.2172E-20 | -0.4341022 | 0.369 | 0.516 | 7.6E-16 | GRAN | EIF5 |
| 3.2581E-20 | -0.2871801 | 0.008 | 0.136 | 7.6967E-16 | GRAN | HMGCS1 |
| 3.3769E-20 | -0.431513 | 0.003 | 0.126 | 7.9773E-16 | GRAN | SRD5A3 |
| 3.3917E-20 | 0.50492091 | 0.205 | 0.106 | 8.0123E-16 | GRAN | NBN |
| 3.4704E-20 | -0.3379955 | 0.002 | 0.124 | 8.1982E-16 | GRAN | KRT80 |
| 3.4847E-20 | -0.3821439 | 0 | 0.121 | 8.2319E-16 | GRAN | EGLN3 |

| | | | | | | |
|---|---|---|---|---|---|---|
| 3.7883E-20 | -0.3346467 | 0.053 | 0.211 | 8.949E-16 | GRAN | MDM2 |
| 3.8276E-20 | -0.2627529 | 0.095 | 0.279 | 9.042E-16 | GRAN | SLK |
| 4.0756E-20 | -0.2674847 | 0.01 | 0.138 | 9.6278E-16 | GRAN | SLC30A1 |
| 4.8611E-20 | -0.2998337 | 0.006 | 0.132 | 1.1483E-15 | GRAN | ID3 |
| 5.0601E-20 | -1.0173609 | 0.011 | 0.139 | 1.1954E-15 | GRAN | DEFB1 |
| 5.4422E-20 | 0.25442566 | 0.129 | 0.051 | 1.2856E-15 | GRAN | FGD3 |
| 5.729E-20 | -0.2787563 | 0.002 | 0.123 | 1.3534E-15 | GRAN | TSPYL2 |
| 6.4561E-20 | -0.7811647 | 0.373 | 0.453 | 1.5251E-15 | GRAN | CLDN7 |
| 6.5101E-20 | -0.2996822 | 0.019 | 0.154 | 1.5379E-15 | GRAN | MT-ND6 |
| 6.5622E-20 | -2.5314145 | 0.106 | 0.256 | 1.5502E-15 | GRAN | MT1G |
| 6.6599E-20 | -0.2757632 | 0.008 | 0.134 | 1.5733E-15 | GRAN | MPZL2 |
| 7.2E-20 | -0.2869312 | 0.016 | 0.147 | 1.7009E-15 | GRAN | NCOA7 |
| 7.8893E-20 | -1.3372217 | 0.021 | 0.154 | 1.8637E-15 | GRAN | CCL20 |
| 8.0703E-20 | -0.2553938 | 0.008 | 0.133 | 1.9065E-15 | GRAN | TMEM176A |
| 8.1134E-20 | -0.5254684 | 0.002 | 0.122 | 1.9166E-15 | GRAN | TCIM |
| 8.2536E-20 | -0.3658846 | 0.016 | 0.147 | 1.9498E-15 | GRAN | MS4A7 |
| 8.9208E-20 | -1.8549583 | 0.021 | 0.153 | 2.1074E-15 | GRAN | MT1H |
| 9.725E-20 | -0.2516913 | 0 | 0.118 | 2.2973E-15 | GRAN | VASN |
| 1.0499E-19 | -0.5464512 | 0 | 0.118 | 2.4802E-15 | GRAN | HSPA2 |
| 1.0808E-19 | -0.2632955 | 0.002 | 0.121 | 2.5532E-15 | GRAN | MUC20 |
| 1.1467E-19 | -0.3020747 | 0.034 | 0.176 | 2.7088E-15 | GRAN | KCTD12 |
| 1.382E-19 | -0.3350966 | 0.095 | 0.272 | 3.2648E-15 | GRAN | CCNG2 |
| 1.484E-19 | 0.54571698 | 0.253 | 0.15 | 3.5056E-15 | GRAN | LBR |
| 1.7897E-19 | 0.35992162 | 0.227 | 0.117 | 4.2278E-15 | GRAN | FCGR3A |
| 1.8754E-19 | -0.2590616 | 0.015 | 0.143 | 4.4302E-15 | GRAN | OTUD1 |
| 1.9012E-19 | -0.2569054 | 0 | 0.117 | 4.4912E-15 | GRAN | NQO1 |
| 2.2707E-19 | -0.327719 | 0.01 | 0.134 | 5.364E-15 | GRAN | RGCC |
| 2.3048E-19 | 0.39550846 | 0.982 | 0.788 | 5.4447E-15 | GRAN | PFN1 |
| 2.3143E-19 | -0.5410031 | 0.003 | 0.122 | 5.4671E-15 | GRAN | HILPDA |
| 2.3948E-19 | -0.3082967 | 0.008 | 0.13 | 5.6571E-15 | GRAN | GAN |
| 2.6222E-19 | 0.55340824 | 0.502 | 0.382 | 6.1944E-15 | GRAN | TYMP |
| 2.9582E-19 | -0.271437 | 0.998 | 0.984 | 6.9882E-15 | GRAN | EIF1 |
| 3.1012E-19 | -1.111634 | 0.002 | 0.118 | 7.3259E-15 | GRAN | SPINK5 |
| 3.1193E-19 | 0.7234442 | 0.392 | 0.315 | 7.3688E-15 | GRAN | BOD1L1 |
| 3.1216E-19 | -0.3095405 | 0.01 | 0.133 | 7.3742E-15 | GRAN | IL18 |
| 3.6638E-19 | -0.4633908 | 0.248 | 0.411 | 8.6551E-15 | GRAN | CD59 |
| 3.85E-19 | -0.2616608 | 0.245 | 0.451 | 9.0949E-15 | GRAN | ACTN4 |
| 4.3367E-19 | -0.5388574 | 0.073 | 0.232 | 1.0245E-14 | GRAN | ISG15 |
| 4.4371E-19 | -1.2834168 | 0 | 0.115 | 1.0482E-14 | GRAN | SPRR1B |
| 4.8291E-19 | -0.3835762 | 0.926 | 0.814 | 1.1408E-14 | GRAN | PFDN5 |
| 5.2443E-19 | -0.2531047 | 0 | 0.114 | 1.2389E-14 | GRAN | DNAJB4 |
| 5.5384E-19 | 0.40688578 | 0.948 | 0.744 | 1.3083E-14 | GRAN | CLIC1 |
| 5.9024E-19 | -0.4606431 | 0 | 0.114 | 1.3943E-14 | GRAN | TMPRSS11A |
| 6.0127E-19 | 0.74714766 | 0.498 | 0.388 | 1.4204E-14 | GRAN | FOS |
| 6.0797E-19 | -0.5962341 | 0 | 0.114 | 1.4362E-14 | GRAN | KRT78 |
| 6.7434E-19 | -0.2517183 | 0.069 | 0.23 | 1.593E-14 | GRAN | GALNT1 |
| 6.7953E-19 | -0.2669945 | 0.008 | 0.128 | 1.6053E-14 | GRAN | MAST4 |
| 6.9359E-19 | -0.3618063 | 0 | 0.113 | 1.6385E-14 | GRAN | ANGPTL4 |
| 7.2416E-19 | -0.3983514 | 0.002 | 0.116 | 1.7107E-14 | GRAN | RRAD |
| 7.5975E-19 | 0.46748497 | 0.202 | 0.107 | 1.7947E-14 | GRAN | MEGF9 |

| | | | | | | |
|---|---|---|---|---|---|---|
| 8.6418E-19 | -0.4267686 | 0.016 | 0.141 | 2.0415E-14 | GRAN | PLAC8 |
| 8.6922E-19 | 0.80467958 | 0.415 | 0.342 | 2.0534E-14 | GRAN | WSB1 |
| 9.1192E-19 | -0.2930258 | 0 | 0.113 | 2.1542E-14 | GRAN | FUT3 |
| 1.0156E-18 | -0.4908118 | 0.568 | 0.599 | 2.3992E-14 | GRAN | VAMP8 |
| 1.0645E-18 | 0.39997221 | 0.206 | 0.111 | 2.5148E-14 | GRAN | RAB27A |
| 1.1116E-18 | -0.2628553 | 0.018 | 0.144 | 2.6259E-14 | GRAN | PIK3R1 |
| 1.1241E-18 | -0.3201051 | 0.005 | 0.121 | 2.6553E-14 | GRAN | AC007952.4 |
| 1.1383E-18 | -0.5085014 | 0.66 | 0.649 | 2.689E-14 | GRAN | UQCRH |
| 1.265E-18 | -0.3600405 | 0 | 0.112 | 2.9883E-14 | GRAN | FXYD2 |
| 1.2659E-18 | -0.3151226 | 0 | 0.112 | 2.9904E-14 | GRAN | STS |
| 1.3201E-18 | -0.3581921 | 0 | 0.112 | 3.1185E-14 | GRAN | SMIM5 |
| 1.4337E-18 | -0.2868824 | 0.006 | 0.123 | 3.3867E-14 | GRAN | FOXQ1 |
| 1.4406E-18 | 0.31386868 | 0.163 | 0.077 | 3.4032E-14 | GRAN | PGS1 |
| 1.687E-18 | 0.65255805 | 0.287 | 0.189 | 3.9851E-14 | GRAN | GBP2 |
| 1.8057E-18 | -0.8589113 | 0.015 | 0.136 | 4.2656E-14 | GRAN | EREG |
| 1.8519E-18 | 0.43389456 | 0.202 | 0.106 | 4.3747E-14 | GRAN | IL1RAP |
| 1.9055E-18 | -0.2967643 | 0.005 | 0.12 | 4.5014E-14 | GRAN | ASS1 |
| 2.0851E-18 | -0.2569054 | 0 | 0.11 | 4.9256E-14 | GRAN | MTAP |
| 2.0946E-18 | -0.2540982 | 0.002 | 0.114 | 4.948E-14 | GRAN | DENND2C |
| 2.5951E-18 | -0.2541843 | 0.002 | 0.113 | 6.1305E-14 | GRAN | GRHL1 |
| 2.6282E-18 | -0.260673 | 0.005 | 0.119 | 6.2087E-14 | GRAN | CD86 |
| 2.8287E-18 | 0.50963777 | 0.577 | 0.494 | 6.6822E-14 | GRAN | DDX3X |
| 2.9115E-18 | 0.42886898 | 0.21 | 0.111 | 6.8779E-14 | GRAN | PECAM1 |
| 3.2404E-18 | 0.47383097 | 0.71 | 0.592 | 7.6548E-14 | GRAN | DDX17 |
| 3.2549E-18 | -1.2770476 | 0.021 | 0.145 | 7.689E-14 | GRAN | CCL2 |
| 3.9568E-18 | -1.4351771 | 0 | 0.109 | 9.3471E-14 | GRAN | CNFN |
| 4.3198E-18 | -1.1529772 | 0.015 | 0.132 | 1.0205E-13 | GRAN | APOE |
| 4.4909E-18 | -0.7109475 | 0.918 | 0.764 | 1.0609E-13 | GRAN | CST3 |
| 5.1755E-18 | -0.6175993 | 0.015 | 0.132 | 1.2226E-13 | GRAN | DHRS9 |
| 5.3011E-18 | -0.4194374 | 0 | 0.108 | 1.2523E-13 | GRAN | TMC5 |
| 5.5679E-18 | 0.41479067 | 0.219 | 0.124 | 1.3153E-13 | GRAN | HSD17B11 |
| 5.9666E-18 | -0.3875732 | 0.082 | 0.236 | 1.4095E-13 | GRAN | HCST |
| 6.0699E-18 | -1.545604 | 0 | 0.108 | 1.4339E-13 | GRAN | CRCT1 |
| 7.2256E-18 | 0.49402442 | 0.173 | 0.084 | 1.7069E-13 | GRAN | IFIT3 |
| 7.4186E-18 | -0.2521152 | 0.002 | 0.11 | 1.7525E-13 | GRAN | GATA3 |
| 9.2393E-18 | -0.5437333 | 0.79 | 0.707 | 2.1826E-13 | GRAN | GPX4 |
| 9.5844E-18 | -0.4813577 | 0.576 | 0.616 | 2.2641E-13 | GRAN | KTN1 |
| 9.829E-18 | -0.2795251 | 0.318 | 0.497 | 2.3219E-13 | GRAN | SRSF3 |
| 1.0981E-17 | 0.50786634 | 0.294 | 0.192 | 2.5939E-13 | GRAN | SNX10 |
| 1.2677E-17 | -0.7814622 | 0 | 0.106 | 2.9946E-13 | GRAN | ZG16B |
| 1.2739E-17 | 0.47535569 | 0.213 | 0.12 | 3.0093E-13 | GRAN | NIN |
| 1.4277E-17 | -0.3511746 | 0.002 | 0.109 | 3.3727E-13 | GRAN | TCHP |
| 1.503E-17 | -0.5483791 | 0.513 | 0.569 | 3.5506E-13 | GRAN | RPS27L |
| 1.8624E-17 | -0.3475512 | 0.002 | 0.108 | 4.3995E-13 | GRAN | SLC16A10 |
| 2.0178E-17 | 0.46334692 | 0.334 | 0.217 | 4.7666E-13 | GRAN | ITGB2 |
| 2.0691E-17 | -0.3903461 | 0 | 0.105 | 4.8879E-13 | GRAN | CLIC3 |
| 2.6616E-17 | -0.3231795 | 0.002 | 0.107 | 6.2874E-13 | GRAN | A2ML1 |
| 2.9016E-17 | 0.4192549 | 0.921 | 0.768 | 6.8545E-13 | GRAN | RHOA |
| 2.9087E-17 | -0.2872934 | 0 | 0.104 | 6.8712E-13 | GRAN | RCAN1 |
| 3.1246E-17 | -0.26296 | 0 | 0.104 | 7.3812E-13 | GRAN | DUSP14 |

| | | | | | | |
|---|---|---|---|---|---|---|
| 3.3512E-17 | -0.3652053 | 0.161 | 0.349 | 7.9165E-13 | GRAN | ZFP36L2 |
| 3.8362E-17 | -0.742924 | 0.006 | 0.114 | 9.0623E-13 | GRAN | IL7R |
| 3.9967E-17 | 0.33700167 | 0.766 | 0.584 | 9.4414E-13 | GRAN | KLF6 |
| 4.9594E-17 | 0.47495851 | 0.221 | 0.129 | 1.1715E-12 | GRAN | MSL1 |
| 5.8842E-17 | -0.589957 | 0.847 | 0.726 | 1.39E-12 | GRAN | PKM |
| 6.9408E-17 | 0.75660289 | 0.429 | 0.374 | 1.6396E-12 | GRAN | CCNL1 |
| 7.3614E-17 | -0.2952749 | 0.023 | 0.141 | 1.739E-12 | GRAN | GADD45G |
| 7.7603E-17 | -0.4166881 | 0.887 | 0.793 | 1.8332E-12 | GRAN | CHCHD2 |
| 8.2489E-17 | -0.4982978 | 0.477 | 0.548 | 1.9486E-12 | GRAN | PPIB |
| 9.3186E-17 | -0.4641515 | 0.094 | 0.248 | 2.2013E-12 | GRAN | PLAU |
| 1.1113E-16 | -0.4128443 | 0.487 | 0.563 | 2.6253E-12 | GRAN | TOMM7 |
| 1.142E-16 | 0.31272222 | 0.161 | 0.08 | 2.6977E-12 | GRAN | PHF21A |
| 1.1878E-16 | -0.2933622 | 0 | 0.1 | 2.8059E-12 | GRAN | LY6K |
| 1.2247E-16 | 0.34782769 | 0.185 | 0.097 | 2.8931E-12 | GRAN | ANKRD44 |
| 1.5358E-16 | -0.7202972 | 0.003 | 0.105 | 3.628E-12 | GRAN | CCL5 |
| 1.6842E-16 | -0.4913455 | 0.052 | 0.182 | 3.9786E-12 | GRAN | OLR1 |
| 1.7668E-16 | -0.6439279 | 1 | 0.98 | 4.1738E-12 | GRAN | MALAT1 |
| 1.845E-16 | -0.6971464 | 0.445 | 0.496 | 4.3584E-12 | GRAN | CD9 |
| 2.3171E-16 | -0.2859183 | 0.037 | 0.162 | 5.4736E-12 | GRAN | SOCS1 |
| 2.3867E-16 | 0.55977149 | 0.215 | 0.121 | 5.638E-12 | GRAN | PHACTR1 |
| 2.5685E-16 | -0.4506128 | 0.623 | 0.629 | 6.0677E-12 | GRAN | YBX3 |
| 2.8651E-16 | 0.37603969 | 0.182 | 0.097 | 6.7682E-12 | GRAN | LPCAT2 |
| 2.9981E-16 | -1.3894596 | 0.029 | 0.146 | 7.0825E-12 | GRAN | CXCL3 |
| 3.2654E-16 | 0.55257031 | 0.689 | 0.582 | 7.7138E-12 | GRAN | NOP10 |
| 3.6447E-16 | 0.41346032 | 0.224 | 0.134 | 8.6099E-12 | GRAN | SLC6A6 |
| 3.8763E-16 | 0.44394571 | 0.926 | 0.767 | 9.157E-12 | GRAN | CDC42 |
| 3.97E-16 | 0.45370067 | 0.25 | 0.155 | 9.3783E-12 | GRAN | OGFRL1 |
| 4.2173E-16 | 0.91607501 | 0.4 | 0.322 | 9.9626E-12 | GRAN | IFITM3 |
| 4.6053E-16 | -0.2544993 | 0.011 | 0.117 | 1.0879E-11 | GRAN | OAS1 |
| 5.0062E-16 | 0.26760797 | 0.137 | 0.063 | 1.1826E-11 | GRAN | DOK3 |
| 5.0237E-16 | -0.6032632 | 0.752 | 0.672 | 1.1867E-11 | GRAN | CALR |
| 5.0925E-16 | -0.3051772 | 0.011 | 0.117 | 1.203E-11 | GRAN | CD163 |
| 6.6341E-16 | -0.3531342 | 0.468 | 0.562 | 1.5672E-11 | GRAN | SET |
| 6.8859E-16 | -0.8509568 | 0.879 | 0.735 | 1.6267E-11 | GRAN | CTSB |
| 9.8356E-16 | 0.2658414 | 0.127 | 0.057 | 2.3235E-11 | GRAN | ALPK1 |
| 1.0317E-15 | -0.280406 | 0.948 | 0.862 | 2.4371E-11 | GRAN | CALM1 |
| 1.4487E-15 | -1.3262809 | 0.013 | 0.116 | 3.4223E-11 | GRAN | MT1M |
| 1.4506E-15 | 0.48748109 | 0.589 | 0.48 | 3.4268E-11 | GRAN | CD46 |
| 1.5873E-15 | -0.6982679 | 0.01 | 0.11 | 3.7497E-11 | GRAN | APOC1 |
| 2.0589E-15 | -0.3105097 | 0.01 | 0.11 | 4.8638E-11 | GRAN | NR1D1 |
| 3.1677E-15 | -0.2650369 | 0.131 | 0.295 | 7.4831E-11 | GRAN | ATP13A3 |
| 3.5569E-15 | -0.7216162 | 0.927 | 0.772 | 8.4024E-11 | GRAN | CTSD |
| 4.4422E-15 | 0.52671971 | 0.231 | 0.144 | 1.0494E-10 | GRAN | TDP2 |
| 4.5227E-15 | 0.44473549 | 0.994 | 0.928 | 1.0684E-10 | GRAN | OAZ1 |
| 6.2354E-15 | 0.48125884 | 0.981 | 0.865 | 1.473E-10 | GRAN | ARPC3 |
| 6.3031E-15 | 0.53687082 | 0.285 | 0.202 | 1.489E-10 | GRAN | RHOG |
| 8.4291E-15 | 0.39062662 | 0.206 | 0.12 | 1.9912E-10 | GRAN | PTAFR |
| 9.1851E-15 | 0.28313858 | 0.152 | 0.077 | 2.1698E-10 | GRAN | USP32 |
| 1.0198E-14 | 0.45014926 | 0.211 | 0.128 | 2.409E-10 | GRAN | SSH2 |
| 1.2958E-14 | -0.3577787 | 0.103 | 0.248 | 3.061E-10 | GRAN | CSTA |

| | | | | | | |
|---|---|---|---|---|---|---|
| 1.345E-14 | 0.41217727 | 0.963 | 0.829 | 3.1773E-10 | GRAN | YWHAZ |
| 1.4442E-14 | -0.4510732 | 0.485 | 0.548 | 3.4116E-10 | GRAN | RPS10 |
| 1.6663E-14 | -0.5158024 | 0.887 | 0.765 | 3.9362E-10 | GRAN | RPS26 |
| 1.9194E-14 | 0.34455947 | 0.195 | 0.113 | 4.5342E-10 | GRAN | WDFY3 |
| 1.9454E-14 | -0.3042378 | 0.06 | 0.184 | 4.5956E-10 | GRAN | INSIG1 |
| 2.5226E-14 | 0.42590861 | 0.216 | 0.131 | 5.9591E-10 | GRAN | PILRA |
| 2.7358E-14 | -0.3305033 | 0.524 | 0.59 | 6.4627E-10 | GRAN | SLC25A3 |
| 4.4686E-14 | 0.38456197 | 0.219 | 0.133 | 1.0556E-09 | GRAN | FGR |
| 4.6947E-14 | -0.2568023 | 0.015 | 0.11 | 1.109E-09 | GRAN | YOD1 |
| 5.9828E-14 | 0.55792254 | 0.263 | 0.181 | 1.4133E-09 | GRAN | PSMB9 |
| 7.674E-14 | -0.3344903 | 0.055 | 0.172 | 1.8128E-09 | GRAN | PRDM1 |
| 7.8145E-14 | 0.40576106 | 0.203 | 0.124 | 1.846E-09 | GRAN | CTBS |
| 7.8227E-14 | 0.2582813 | 0.137 | 0.067 | 1.8479E-09 | GRAN | ST20 |
| 9.002E-14 | -0.5409404 | 0.721 | 0.657 | 2.1265E-09 | GRAN | NDUFA4 |
| 9.0422E-14 | -0.4966392 | 0.405 | 0.508 | 2.136E-09 | GRAN | MT-ND4L |
| 9.909E-14 | 0.4255313 | 0.231 | 0.147 | 2.3408E-09 | GRAN | CDC42EP3 |
| 1.0098E-13 | -0.4447183 | 0.958 | 0.836 | 2.3855E-09 | GRAN | SQSTM1 |
| 1.466E-13 | -0.4041873 | 0.095 | 0.229 | 3.4632E-09 | GRAN | HEXIM1 |
| 2.0406E-13 | -0.5523 | 0.013 | 0.103 | 4.8205E-09 | GRAN | HLA-DQA1 |
| 2.4038E-13 | 0.68898268 | 0.337 | 0.281 | 5.6786E-09 | GRAN | RTN3 |
| 3.3124E-13 | -1.5517559 | 0.923 | 0.706 | 7.8248E-09 | GRAN | ANXA1 |
| 4.2346E-13 | 0.78834653 | 0.377 | 0.344 | 1.0003E-08 | GRAN | LAMTOR4 |
| 4.6066E-13 | -0.4124957 | 0.013 | 0.101 | 1.0882E-08 | GRAN | RETN |
| 4.7476E-13 | -0.3099301 | 0.835 | 0.758 | 1.1215E-08 | GRAN | UQCRB |
| 5.4078E-13 | 0.65248396 | 0.415 | 0.39 | 1.2775E-08 | GRAN | STAT3 |
| 1.099E-12 | -0.5395295 | 0.697 | 0.623 | 2.5961E-08 | GRAN | DSTN |
| 1.1149E-12 | -0.3714502 | 0.645 | 0.642 | 2.6337E-08 | GRAN | EDF1 |
| 1.2335E-12 | 0.55920103 | 0.298 | 0.226 | 2.9138E-08 | GRAN | SKAP2 |
| 1.3047E-12 | -0.4051289 | 0.679 | 0.646 | 3.082E-08 | GRAN | ATP5MC2 |
| 2.0594E-12 | -0.3427981 | 0.542 | 0.584 | 4.865E-08 | GRAN | CLTA |
| 2.2643E-12 | 0.40510013 | 0.218 | 0.139 | 5.349E-08 | GRAN | IL2RG |
| 2.5962E-12 | 0.48774182 | 0.284 | 0.211 | 6.133E-08 | GRAN | RNF13 |
| 2.8609E-12 | -0.5098198 | 0.624 | 0.601 | 6.7582E-08 | GRAN | HINT1 |
| 3.0136E-12 | -0.7563861 | 0.355 | 0.435 | 7.119E-08 | GRAN | HIST1H1C |
| 3.0646E-12 | 0.57448515 | 0.323 | 0.26 | 7.2395E-08 | GRAN | MBP |
| 5.377E-12 | 0.39347944 | 0.198 | 0.125 | 1.2702E-07 | GRAN | IL4R |
| 6.1378E-12 | -0.7810554 | 0.826 | 0.691 | 1.4499E-07 | GRAN | HSP90B1 |
| 6.9568E-12 | 0.43433352 | 0.658 | 0.545 | 1.6434E-07 | GRAN | C4orf3 |
| 7.8324E-12 | 0.49908688 | 0.255 | 0.18 | 1.8503E-07 | GRAN | RILPL2 |
| 8.9617E-12 | 0.6578559 | 0.3 | 0.241 | 2.117E-07 | GRAN | IFNAR1 |
| 1.0076E-11 | -0.325825 | 0.579 | 0.613 | 2.3802E-07 | GRAN | PTGES3 |
| 1.035E-11 | -0.369877 | 0.995 | 0.894 | 2.4449E-07 | GRAN | RPS9 |
| 1.1028E-11 | 0.62023563 | 0.311 | 0.246 | 2.6051E-07 | GRAN | CREBRF |
| 1.2494E-11 | 0.50935207 | 0.244 | 0.17 | 2.9515E-07 | GRAN | DAPP1 |
| 1.4278E-11 | 0.52570677 | 0.258 | 0.189 | 3.373E-07 | GRAN | N4BP1 |
| 1.8037E-11 | -0.4198929 | 0.027 | 0.114 | 4.2608E-07 | GRAN | CYP1B1 |
| 3.3904E-11 | 0.37501564 | 0.2 | 0.13 | 8.0092E-07 | GRAN | RICTOR |
| 3.5025E-11 | 0.30970627 | 0.165 | 0.096 | 8.274E-07 | GRAN | BIN2 |
| 3.7042E-11 | -0.3482727 | 0.871 | 0.774 | 8.7505E-07 | GRAN | TPI1 |
| 3.9011E-11 | 0.41944817 | 0.708 | 0.57 | 9.2156E-07 | GRAN | TSPO |

| | | | | | | |
|---|---|---|---|---|---|---|
| 4.4032E-11 | -0.8133566 | 0.032 | 0.119 | 1.0402E-06 | GRAN | CXCL2 |
| 5.0913E-11 | 0.25039094 | 0.129 | 0.068 | 1.2027E-06 | GRAN | AGTPBP1 |
| 5.2059E-11 | 0.5090212 | 0.281 | 0.217 | 1.2298E-06 | GRAN | ACAP2 |
| 5.6251E-11 | 0.53088821 | 0.268 | 0.203 | 1.3288E-06 | GRAN | IGF2R |
| 5.7623E-11 | 0.27370997 | 0.148 | 0.084 | 1.3612E-06 | GRAN | MOB3A |
| 6.9789E-11 | -0.2889048 | 0.556 | 0.601 | 1.6486E-06 | GRAN | YWHAE |
| 9.56E-11 | 0.54077844 | 0.247 | 0.177 | 2.2584E-06 | GRAN | EHD1 |
| 1.1789E-10 | -0.6553023 | 0.666 | 0.605 | 2.7849E-06 | GRAN | EZR |
| 1.1957E-10 | 0.41596882 | 0.927 | 0.769 | 2.8247E-06 | GRAN | TMA7 |
| 1.274E-10 | 0.68641721 | 0.382 | 0.362 | 3.0095E-06 | GRAN | NFE2L2 |
| 1.949E-10 | 0.3218227 | 0.177 | 0.11 | 4.6041E-06 | GRAN | ALOX5 |
| 2.0661E-10 | 0.32965587 | 0.166 | 0.1 | 4.8807E-06 | GRAN | ECE1 |
| 2.811E-10 | 0.38206756 | 0.21 | 0.141 | 6.6404E-06 | GRAN | NUP214 |
| 3.2808E-10 | -2.3772661 | 0.677 | 0.374 | 7.7501E-06 | GRAN | KRT13 |
| 5.356E-10 | 0.33290271 | 0.179 | 0.116 | 1.2653E-05 | GRAN | OAZ2 |
| 7.1302E-10 | 0.31068625 | 0.334 | 0.243 | 1.6844E-05 | GRAN | SLC11A1 |
| 7.8673E-10 | 0.4422823 | 0.218 | 0.152 | 1.8585E-05 | GRAN | TAP1 |
| 8.0124E-10 | 0.59166955 | 0.327 | 0.285 | 1.8928E-05 | GRAN | LRP10 |
| 8.5979E-10 | 0.59875089 | 0.369 | 0.324 | 2.0311E-05 | GRAN | ARHGDIB |
| 8.8056E-10 | 0.35033457 | 0.171 | 0.106 | 2.0801E-05 | GRAN | PIK3AP1 |
| 1.1501E-09 | 0.35464992 | 0.203 | 0.138 | 2.717E-05 | GRAN | PTPN6 |
| 1.2622E-09 | 0.30327493 | 0.155 | 0.093 | 2.9816E-05 | GRAN | PPP1R3B |
| 1.37E-09 | 0.40857235 | 0.224 | 0.163 | 3.2364E-05 | GRAN | TNFRSF1A |
| 1.3857E-09 | 0.39032875 | 0.229 | 0.165 | 3.2734E-05 | GRAN | SLC12A6 |
| 1.5053E-09 | -1.3899977 | 0.323 | 0.37 | 3.5559E-05 | GRAN | MAL |
| 3.7297E-09 | 0.65001688 | 0.377 | 0.369 | 8.8108E-05 | GRAN | PAK2 |
| 3.7713E-09 | 0.45358935 | 0.195 | 0.135 | 8.909E-05 | GRAN | RNF19B |
| 4.3445E-09 | -1.2742189 | 0.452 | 0.451 | 0.00010263 | GRAN | SOX4 |
| 4.3573E-09 | -0.250264 | 0.613 | 0.632 | 0.00010293 | GRAN | SAP18 |
| 4.7681E-09 | 0.40960622 | 0.245 | 0.187 | 0.00011264 | GRAN | CNN2 |
| 5.1204E-09 | 0.34016925 | 0.2 | 0.136 | 0.00012096 | GRAN | EVI2A |
| 5.2106E-09 | 0.33627604 | 0.192 | 0.13 | 0.00012309 | GRAN | PTK2B |
| 6.0838E-09 | 0.26363176 | 0.15 | 0.092 | 0.00014372 | GRAN | IQSEC1 |
| 9.9956E-09 | 0.3038701 | 0.187 | 0.125 | 0.00023613 | GRAN | ADAM8 |
| 1.0539E-08 | -0.2698621 | 0.635 | 0.628 | 0.00024897 | GRAN | OST4 |
| 1.2623E-08 | -0.4880348 | 0.084 | 0.171 | 0.00029818 | GRAN | FCN1 |
| 1.3285E-08 | 0.35976893 | 0.185 | 0.127 | 0.00031382 | GRAN | SHKBP1 |
| 1.4379E-08 | 0.31415774 | 0.174 | 0.115 | 0.00033967 | GRAN | BCL3 |
| 1.5357E-08 | -0.3523078 | 0.408 | 0.481 | 0.00036278 | GRAN | RIOK3 |
| 1.655E-08 | 0.58823662 | 0.369 | 0.325 | 0.00039096 | GRAN | CEBPD |
| 2.9013E-08 | 0.33886745 | 0.166 | 0.108 | 0.00068536 | GRAN | PAG1 |
| 2.9942E-08 | 0.32040945 | 0.203 | 0.135 | 0.00070732 | GRAN | MCEMP1 |
| 3.0853E-08 | 0.49825417 | 0.273 | 0.214 | 0.00072885 | GRAN | TUBA1A |
| 3.282E-08 | 0.39333632 | 0.198 | 0.142 | 0.0007753 | GRAN | RASSF5 |
| 4.0898E-08 | 0.40289001 | 0.19 | 0.128 | 0.00096612 | GRAN | AC023157.3 |
| 4.4354E-08 | 0.69909541 | 0.282 | 0.229 | 0.00104777 | GRAN | PPIF |
| 4.7443E-08 | 0.2582924 | 0.129 | 0.078 | 0.00112074 | GRAN | SLC15A4 |
| 5.8295E-08 | 0.29596879 | 0.147 | 0.094 | 0.0013771 | GRAN | GRAMD1A |
| 6.5286E-08 | 0.29694868 | 0.177 | 0.12 | 0.00154225 | GRAN | DENND5A |
| 8.7305E-08 | 0.45232931 | 0.252 | 0.205 | 0.0020624 | GRAN | ADAR |

| | | | | | | |
|---|---|---|---|---|---|---|
| 9.4498E-08 | 0.46367571 | 0.247 | 0.196 | 0.00223232 | GRAN | CCPG1 |
| 1.0973E-07 | -0.474415 | 0.731 | 0.645 | 0.00259207 | GRAN | LDHA |
| 1.4302E-07 | 0.51690409 | 0.377 | 0.361 | 0.00337846 | GRAN | GNAI2 |
| 2.0378E-07 | 0.58762771 | 0.51 | 0.487 | 0.0048138 | GRAN | ANKRD12 |
| 2.0806E-07 | 0.40568959 | 0.242 | 0.187 | 0.004915 | GRAN | PYCARD |
| 2.4013E-07 | 0.2722529 | 0.161 | 0.109 | 0.0056726 | GRAN | C9orf72 |
| 2.4811E-07 | -0.9305531 | 0.748 | 0.626 | 0.00586101 | GRAN | JUN |
| 2.5334E-07 | 0.62274075 | 0.363 | 0.373 | 0.00598472 | GRAN | GNB2 |
| 2.5606E-07 | 0.47872451 | 0.292 | 0.254 | 0.0060489 | GRAN | MAP2K3 |
| 2.7496E-07 | 0.59881794 | 0.39 | 0.413 | 0.00649527 | GRAN | DAZAP2 |
| 3.2735E-07 | 0.43431672 | 0.258 | 0.214 | 0.00773291 | GRAN | ZYX |
| 4.4223E-07 | 0.25046065 | 0.153 | 0.101 | 0.01044671 | GRAN | RNF144B |
| 4.5312E-07 | 0.4439235 | 0.261 | 0.217 | 0.01070403 | GRAN | TUT7 |
| 5.2957E-07 | 0.34923241 | 0.176 | 0.125 | 0.01250995 | GRAN | NFIL3 |
| 5.3544E-07 | 0.35615586 | 0.189 | 0.138 | 0.0126488 | GRAN | IRF2 |
| 5.9806E-07 | 0.68198648 | 0.335 | 0.33 | 0.01412802 | GRAN | SH3GLB1 |
| 6.628E-07 | 0.54367219 | 0.302 | 0.27 | 0.01565744 | GRAN | OSBPL8 |
| 8.5361E-07 | -0.2502497 | 0.794 | 0.713 | 0.02016488 | GRAN | SKP1 |
| 9.4748E-07 | 0.27264607 | 0.998 | 0.963 | 0.02238221 | GRAN | MYL6 |
| 1.0309E-06 | -0.3590059 | 0.861 | 0.724 | 0.02435324 | GRAN | ENO1 |
| 1.1627E-06 | -0.2591515 | 0.119 | 0.202 | 0.02746742 | GRAN | CYBB |
| 1.3714E-06 | 0.29340139 | 0.139 | 0.091 | 0.0323957 | GRAN | DSC2 |
| 2.1512E-06 | 0.2665357 | 0.147 | 0.101 | 0.05081711 | GRAN | RPS6KA1 |
| 2.5385E-06 | -0.8901941 | 0.666 | 0.538 | 0.05996798 | GRAN | CD24 |
| 2.6383E-06 | 0.25538012 | 0.99 | 0.903 | 0.0623251 | GRAN | CFL1 |
| 2.9582E-06 | -0.6597233 | 0.498 | 0.503 | 0.06988106 | GRAN | DNAJB1 |
| 3.2798E-06 | 0.42701344 | 0.273 | 0.238 | 0.07747952 | GRAN | SP100 |
| 3.5347E-06 | 0.32569053 | 0.187 | 0.142 | 0.08350028 | GRAN | KAT6A |
| 3.6623E-06 | -0.5883391 | 0.806 | 0.665 | 0.08651525 | GRAN | ANXA5 |
| 3.9147E-06 | 0.41078383 | 0.229 | 0.182 | 0.09247677 | GRAN | ICAM1 |
| 4.0138E-06 | 0.2584168 | 0.134 | 0.089 | 0.09481902 | GRAN | FRAT2 |
| 4.4793E-06 | 0.58593315 | 0.276 | 0.236 | 0.10581339 | GRAN | NFKBIZ |
| 4.8771E-06 | 0.3424996 | 0.189 | 0.143 | 0.11521237 | GRAN | PCNX1 |
| 5.5951E-06 | 0.31974431 | 0.161 | 0.118 | 0.13217248 | GRAN | CDKN2D |
| 6.1178E-06 | 0.32403287 | 0.213 | 0.167 | 0.14451969 | GRAN | GLIPR2 |
| 6.6986E-06 | 0.44375188 | 0.231 | 0.194 | 0.15824142 | GRAN | NCOA4 |
| 7.1856E-06 | 0.25041858 | 0.163 | 0.117 | 0.16974592 | GRAN | CASP1 |
| 7.9466E-06 | 0.30664416 | 0.171 | 0.128 | 0.18772195 | GRAN | CYB5R4 |
| 8.0825E-06 | 0.27789085 | 0.155 | 0.111 | 0.19093241 | GRAN | PXN |
| 9.6516E-06 | -0.3402804 | 0.115 | 0.189 | 0.22799964 | GRAN | AC020656.1 |
| 1.0035E-05 | 0.48538579 | 0.332 | 0.319 | 0.23704968 | GRAN | SLC16A3 |
| 1.0091E-05 | 0.2842363 | 0.232 | 0.185 | 0.23838414 | GRAN | TNFSF10 |
| 1.222E-05 | 0.29282551 | 0.932 | 0.808 | 0.28867229 | GRAN | RAC1 |
| 1.5276E-05 | 0.46372421 | 0.26 | 0.231 | 0.3608589 | GRAN | FLOT1 |
| 1.8688E-05 | 0.26094907 | 0.235 | 0.184 | 0.4414756 | GRAN | KLF2 |
| 1.8956E-05 | 0.28567539 | 0.147 | 0.106 | 0.44779166 | GRAN | AHCTF1 |
| 2.196E-05 | 0.44437659 | 0.226 | 0.193 | 0.51876947 | GRAN | CIR1 |
| 2.3054E-05 | 0.55112764 | 0.229 | 0.193 | 0.54459934 | GRAN | KATNBL1 |
| 3.721E-05 | -1.0796561 | 0.592 | 0.492 | 0.87901534 | GRAN | KRT8 |
| 4.1131E-05 | -0.2583605 | 0.827 | 0.729 | 0.97164123 | GRAN | ATP6V1G1 |

| | | | | | | |
|---|---|---|---|---|---|---|
| 4.7527E-05 | -1.9171598 | 0.968 | 0.672 | 1 | GRAN | TIMP1 |
| 5.8541E-05 | -2.8896885 | 0.881 | 0.518 | 1 | GRAN | LCN2 |
| 5.8829E-05 | 0.31029237 | 0.189 | 0.151 | 1 | GRAN | EHBP1L1 |
| 8.5702E-05 | 0.34128483 | 0.168 | 0.13 | 1 | GRAN | TRIB1 |
| 8.8555E-05 | 0.38282875 | 0.206 | 0.169 | 1 | GRAN | ISG20 |
| 0.00011179 | -0.5553878 | 0.395 | 0.422 | 1 | GRAN | TXNRD1 |
| 0.00011809 | 0.25684331 | 0.145 | 0.109 | 1 | GRAN | ELOVL5 |
| 0.00016773 | 0.47422246 | 0.277 | 0.266 | 1 | GRAN | CDC42SE1 |
| 0.00018963 | -1.1038982 | 0.797 | 0.565 | 1 | GRAN | TACSTD2 |
| 0.00019446 | -1.229543 | 0.768 | 0.552 | 1 | GRAN | ELF3 |
| 0.00019841 | 0.28971671 | 0.181 | 0.146 | 1 | GRAN | SH3KBP1 |
| 0.00021262 | -1.1244148 | 0.821 | 0.614 | 1 | GRAN | LGALS1 |
| 0.00021364 | 0.28146032 | 0.215 | 0.178 | 1 | GRAN | CD37 |
| 0.00022814 | 0.29169435 | 0.163 | 0.129 | 1 | GRAN | PTPRJ |
| 0.00023078 | 0.415087 | 0.305 | 0.3 | 1 | GRAN | PICALM |
| 0.00023124 | -0.3244835 | 0.737 | 0.655 | 1 | GRAN | ZFAS1 |
| 0.00023402 | 0.53598487 | 0.237 | 0.209 | 1 | GRAN | SOCS3 |
| 0.00033119 | 0.32648979 | 0.221 | 0.195 | 1 | GRAN | RAB8B |
| 0.00033427 | -0.6793551 | 0.074 | 0.123 | 1 | GRAN | CCL3 |
| 0.00034364 | 0.29726494 | 0.174 | 0.142 | 1 | GRAN | RXRA |
| 0.00036378 | 0.25421 | 0.645 | 0.596 | 1 | GRAN | HNRNPK |
| 0.00046728 | 0.42738433 | 0.329 | 0.333 | 1 | GRAN | TLN1 |
| 0.00049836 | 0.3941443 | 0.245 | 0.223 | 1 | GRAN | GRINA |
| 0.00052497 | 0.44647304 | 0.263 | 0.253 | 1 | GRAN | DHRS7 |
| 0.00055482 | -0.4177742 | 0.397 | 0.425 | 1 | GRAN | EMP3 |
| 0.00055876 | 0.39071121 | 0.248 | 0.23 | 1 | GRAN | ATP6V1B2 |
| 0.00069417 | 0.552049 | 0.324 | 0.349 | 1 | GRAN | PRR13 |
| 0.0007967 | 0.44413246 | 0.258 | 0.242 | 1 | GRAN | IFRD1 |
| 0.00110638 | 0.3313047 | 0.197 | 0.174 | 1 | GRAN | SHOC2 |
| 0.00113622 | 0.26301954 | 0.185 | 0.158 | 1 | GRAN | ARHGAP30 |
| 0.00122876 | 0.47319967 | 0.284 | 0.286 | 1 | GRAN | BAZ1A |
| 0.00143341 | 0.33914518 | 0.461 | 0.436 | 1 | GRAN | CYSTM1 |
| 0.00154212 | -0.7270944 | 0.553 | 0.504 | 1 | GRAN | HSPA1B |
| 0.00166862 | 0.43245387 | 0.253 | 0.249 | 1 | GRAN | IDS |
| 0.00166915 | 0.29591888 | 0.197 | 0.174 | 1 | GRAN | ERGIC1 |
| 0.00178214 | 0.48761376 | 0.239 | 0.223 | 1 | GRAN | STXBP2 |
| 0.00188201 | 0.41557277 | 0.237 | 0.222 | 1 | GRAN | MAP3K2 |
| 0.00191559 | -1.8820043 | 0.669 | 0.45 | 1 | GRAN | SPP1 |
| 0.00195188 | -1.1915952 | 0.748 | 0.536 | 1 | GRAN | CLDN4 |
| 0.00196155 | 0.28887307 | 0.184 | 0.158 | 1 | GRAN | MYADM |
| 0.00206191 | 0.34087954 | 0.534 | 0.5 | 1 | GRAN | ASAH1 |
| 0.00240306 | -0.3554023 | 0.84 | 0.72 | 1 | GRAN | JUND |
| 0.00261561 | -1.2415896 | 0.774 | 0.554 | 1 | GRAN | KRT18 |
| 0.00311946 | 0.37015901 | 0.332 | 0.345 | 1 | GRAN | MIDN |
| 0.00315432 | -2.0518319 | 0.358 | 0.339 | 1 | GRAN | MMP7 |
| 0.00358967 | 0.28724249 | 0.208 | 0.186 | 1 | GRAN | RNF130 |
| 0.00377312 | 0.40101282 | 0.342 | 0.368 | 1 | GRAN | MYH9 |
| 0.00387387 | 0.46037661 | 0.313 | 0.33 | 1 | GRAN | RBM47 |
| 0.00388283 | -0.3985929 | 0.998 | 0.936 | 1 | GRAN | S100A6 |
| 0.00401476 | -1.5199842 | 0.868 | 0.574 | 1 | GRAN | KRT19 |

| | | | | | | |
|---|---|---|---|---|---|---|
| 0.00448691 | 0.46531134 | 0.287 | 0.301 | 1 | GRAN | TAPBP |
| 0.00481158 | 0.25479331 | 0.147 | 0.123 | 1 | GRAN | EGLN1 |
| 0.00487119 | -1.177942 | 0.771 | 0.541 | 1 | GRAN | KRT7 |
| 0.0055169 | -0.2580573 | 0.658 | 0.597 | 1 | GRAN | NPC2 |
| 0.00559521 | -0.5378689 | 0.145 | 0.194 | 1 | GRAN | HSPA6 |
| 0.00647217 | 0.41094601 | 0.315 | 0.339 | 1 | GRAN | GRB2 |
| 0.00653432 | 0.32472908 | 0.268 | 0.264 | 1 | GRAN | BACH1 |
| 0.00823241 | 0.35014335 | 0.21 | 0.197 | 1 | GRAN | CHD1 |
| 0 | 2.83515242 | 0.633 | 0.076 | 0 | Tcells | CCL5 |
| 0 | 2.80154854 | 0.733 | 0.08 | 0 | Tcells | IL7R |
| 0 | 2.4465793 | 0.21 | 0.033 | 0 | Tcells | GNLY |
| 0 | 2.09061358 | 0.95 | 0.245 | 0 | Tcells | PTPRC |
| 0 | 1.96090586 | 0.468 | 0.047 | 0 | Tcells | NKG7 |
| 0 | 1.94817032 | 0.748 | 0.017 | 0 | Tcells | CD2 |
| 0 | 1.94264881 | 0.424 | 0.091 | 0 | Tcells | CCL4 |
| 0 | 1.93010969 | 0.767 | 0.025 | 0 | Tcells | CD3D |
| 0 | 1.85993467 | 0.745 | 0.034 | 0 | Tcells | TRBC2 |
| 0 | 1.83385469 | 0.779 | 0.198 | 0 | Tcells | IL32 |
| 0 | 1.71343087 | 0.691 | 0.022 | 0 | Tcells | CD7 |
| 0 | 1.66822509 | 0.319 | 0.016 | 0 | Tcells | GZMB |
| 0 | 1.64411564 | 0.996 | 0.881 | 0 | Tcells | RPS3 |
| 0 | 1.64152535 | 0.998 | 0.94 | 0 | Tcells | RPS27 |
| 0 | 1.61420974 | 0.992 | 0.873 | 0 | Tcells | RPS29 |
| 0 | 1.58496807 | 0.994 | 0.888 | 0 | Tcells | RPS15A |
| 0 | 1.55047721 | 0.742 | 0.021 | 0 | Tcells | CD3E |
| 0 | 1.52186219 | 0.508 | 0.021 | 0 | Tcells | TRBC1 |
| 0 | 1.5167483 | 0.665 | 0.022 | 0 | Tcells | TRAC |
| 0 | 1.51294325 | 0.848 | 0.32 | 0 | Tcells | ZFP36L2 |
| 0 | 1.49912424 | 0.721 | 0.141 | 0 | Tcells | CXCR4 |
| 0 | 1.48383002 | 0.702 | 0.171 | 0 | Tcells | CD52 |
| 0 | 1.46165122 | 0.996 | 0.924 | 0 | Tcells | RPL30 |
| 0 | 1.44043626 | 0.654 | 0.013 | 0 | Tcells | CD3G |
| 0 | 1.42618369 | 0.489 | 0.084 | 0 | Tcells | LTB |
| 0 | 1.42483057 | 0.378 | 0.016 | 0 | Tcells | GZMA |
| 0 | 1.41633821 | 0.859 | 0.441 | 0 | Tcells | SARAF |
| 0 | 1.40648024 | 0.498 | 0.01 | 0 | Tcells | CD8A |
| 0 | 1.37772135 | 0.616 | 0.039 | 0 | Tcells | GIMAP7 |
| 0 | 1.37742824 | 0.998 | 0.92 | 0 | Tcells | RPS27A |
| 0 | 1.36901055 | 0.5 | 0.036 | 0 | Tcells | CD69 |
| 0 | 1.31498862 | 0.96 | 0.753 | 0 | Tcells | BTG1 |
| 0 | 1.31190974 | 0.703 | 0.065 | 0 | Tcells | IKZF1 |
| 0 | 1.31006057 | 0.998 | 0.945 | 0 | Tcells | RPL10 |
| 0 | 1.29298343 | 0.558 | 0.047 | 0 | Tcells | SYTL3 |
| 0 | 1.29275924 | 0.703 | 0.247 | 0 | Tcells | TSC22D3 |
| 0 | 1.2726815 | 0.995 | 0.93 | 0 | Tcells | RPL34 |
| 0 | 1.26565316 | 0.997 | 0.93 | 0 | Tcells | RPS12 |
| 0 | 1.24234948 | 0.69 | 0.109 | 0 | Tcells | ETS1 |
| 0 | 1.22852992 | 0.996 | 0.93 | 0 | Tcells | RPS14 |
| 0 | 1.2064449 | 0.995 | 0.911 | 0 | Tcells | RPL32 |
| 0 | 1.19171034 | 0.625 | 0.037 | 0 | Tcells | CLEC2D |

| | | | | | | |
|---|---|---|---|---|---|---|
| 0 | 1.17270275 | 0.379 | 0.014 | 0 | Tcells | KLRD1 |
| 0 | 1.17105109 | 0.995 | 0.892 | 0 | Tcells | RPS3A |
| 0 | 1.16618155 | 0.633 | 0.085 | 0 | Tcells | FYN |
| 0 | 1.1635699 | 0.994 | 0.894 | 0 | Tcells | RPL39 |
| 0 | 1.16008184 | 0.999 | 0.962 | 0 | Tcells | RPL13 |
| 0 | 1.15841744 | 0.965 | 0.783 | 0 | Tcells | RPS21 |
| 0 | 1.15837868 | 0.603 | 0.117 | 0 | Tcells | PIK3R1 |
| 0 | 1.14357396 | 0.986 | 0.841 | 0 | Tcells | RPS4X |
| 0 | 1.13610714 | 0.692 | 0.141 | 0 | Tcells | CD48 |
| 0 | 1.10670793 | 0.998 | 0.935 | 0 | Tcells | TMSB4X |
| 0 | 1.10636834 | 0.676 | 0.156 | 0 | Tcells | STK17A |
| 0 | 1.10586725 | 0.994 | 0.883 | 0 | Tcells | RPS23 |
| 0 | 1.09455826 | 0.449 | 0.054 | 0 | Tcells | HIST1H1D |
| 0 | 1.09409804 | 1 | 0.942 | 0 | Tcells | RPS19 |
| 0 | 1.09153621 | 0.641 | 0.212 | 0 | Tcells | HCST |
| 0 | 1.08831643 | 0.636 | 0.13 | 0 | Tcells | PDCD4 |
| 0 | 1.08323624 | 0.974 | 0.744 | 0 | Tcells | RPSA |
| 0 | 1.07969336 | 0.983 | 0.809 | 0 | Tcells | RPL5 |
| 0 | 1.07535723 | 0.73 | 0.19 | 0 | Tcells | STK4 |
| 0 | 1.07482521 | 0.601 | 0.062 | 0 | Tcells | EVL |
| 0 | 1.06650609 | 0.996 | 0.923 | 0 | Tcells | RPL19 |
| 0 | 1.06143353 | 0.645 | 0.148 | 0 | Tcells | CORO1A |
| 0 | 1.057742 | 0.998 | 0.971 | 0 | Tcells | RPLP1 |
| 0 | 1.05620474 | 0.694 | 0.181 | 0 | Tcells | STK17B |
| 0 | 1.05450271 | 0.477 | 0.047 | 0 | Tcells | CST7 |
| 0 | 1.04468885 | 0.998 | 0.918 | 0 | Tcells | RPL11 |
| 0 | 1.04204163 | 0.998 | 0.936 | 0 | Tcells | RPS28 |
| 0 | 1.04166272 | 0.707 | 0.22 | 0 | Tcells | KIAA1551 |
| 0 | 1.03397954 | 0.986 | 0.865 | 0 | Tcells | RPL14 |
| 0 | 1.02859437 | 0.993 | 0.894 | 0 | Tcells | RPL21 |
| 0 | 1.02654541 | 0.987 | 0.871 | 0 | Tcells | RPL23A |
| 0 | 1.02608459 | 0.99 | 0.891 | 0 | Tcells | RPL35A |
| 0 | 1.01148505 | 0.645 | 0.193 | 0 | Tcells | PPP2R5C |
| 0 | 1.00829503 | 0.777 | 0.301 | 0 | Tcells | ARHGDIB |
| 0 | 1.00771709 | 0.993 | 0.899 | 0 | Tcells | RPLP2 |
| 0 | 1.00744439 | 0.611 | 0.12 | 0 | Tcells | LEPROTL1 |
| 0 | 0.99350255 | 0.552 | 0.093 | 0 | Tcells | CNOT6L |
| 0 | 0.99098987 | 0.309 | 0.013 | 0 | Tcells | GZMH |
| 0 | 0.98989959 | 0.986 | 0.894 | 0 | Tcells | RPS25 |
| 0 | 0.98348162 | 0.998 | 0.92 | 0 | Tcells | RPL18A |
| 0 | 0.97997393 | 0.541 | 0.031 | 0 | Tcells | SPOCK2 |
| 0 | 0.97912642 | 0.267 | 0.014 | 0 | Tcells | KLRB1 |
| 0 | 0.97523665 | 0.425 | 0.014 | 0 | Tcells | CTSW |
| 0 | 0.97320258 | 0.996 | 0.916 | 0 | Tcells | RPS15 |
| 0 | 0.97274883 | 0.991 | 0.87 | 0 | Tcells | RPL3 |
| 0 | 0.97151634 | 0.349 | 0.045 | 0 | Tcells | RGS1 |
| 0 | 0.96838217 | 0.341 | 0.009 | 0 | Tcells | PRF1 |
| 0 | 0.96161615 | 0.425 | 0.008 | 0 | Tcells | GPR171 |
| 0 | 0.95745923 | 0.985 | 0.879 | 0 | Tcells | RPS13 |
| 0 | 0.95714416 | 0.516 | 0.061 | 0 | Tcells | GIMAP4 |

| | | | | | | |
|---|---|---|---|---|---|---|
| 0 | 0.95014677 | 0.389 | 0.007 | 0 | Tcells | CD8B |
| 0 | 0.94911673 | 0.998 | 0.979 | 0 | Tcells | RPL41 |
| 0 | 0.94847936 | 0.994 | 0.899 | 0 | Tcells | RPL37 |
| 0 | 0.94684752 | 1 | 0.989 | 0 | Tcells | B2M |
| 0 | 0.94249387 | 0.515 | 0.01 | 0 | Tcells | CD247 |
| 0 | 0.92921999 | 0.55 | 0.071 | 0 | Tcells | EMB |
| 0 | 0.92813736 | 0.755 | 0.333 | 0 | Tcells | SEPT7 |
| 0 | 0.91894924 | 0.998 | 0.907 | 0 | Tcells | RPS8 |
| 0 | 0.91489362 | 0.687 | 0.262 | 0 | Tcells | RPS4Y1 |
| 0 | 0.90624147 | 0.997 | 0.879 | 0 | Tcells | RPS2 |
| 0 | 0.90562935 | 0.996 | 0.947 | 0 | Tcells | RPL18 |
| 0 | 0.90250978 | 0.647 | 0.164 | 0 | Tcells | FYB1 |
| 0 | 0.89817046 | 0.554 | 0.05 | 0 | Tcells | ACAP1 |
| 0 | 0.89379737 | 0.636 | 0.193 | 0 | Tcells | ARL4C |
| 0 | 0.87843439 | 0.615 | 0.165 | 0 | Tcells | RAC2 |
| 0 | 0.87473945 | 0.821 | 0.548 | 0 | Tcells | TOMM7 |
| 0 | 0.87035435 | 0.675 | 0.237 | 0 | Tcells | MBNL1 |
| 0 | 0.86836825 | 0.486 | 0.057 | 0 | Tcells | RUNX3 |
| 0 | 0.86796716 | 0.998 | 0.977 | 0 | Tcells | RPL28 |
| 0 | 0.86713096 | 0.991 | 0.893 | 0 | Tcells | RPL29 |
| 0 | 0.86511454 | 0.924 | 0.737 | 0 | Tcells | EEF1B2 |
| 0 | 0.86127068 | 0.991 | 0.928 | 0 | Tcells | FAU |
| 0 | 0.86073442 | 0.597 | 0.109 | 0 | Tcells | CYTIP |
| 0 | 0.85676382 | 0.616 | 0.141 | 0 | Tcells | CDC42SE2 |
| 0 | 0.85552 | 0.649 | 0.229 | 0 | Tcells | SYNE2 |
| 0 | 0.84864506 | 0.457 | 0.012 | 0 | Tcells | GZMM |
| 0 | 0.84751843 | 0.995 | 0.944 | 0 | Tcells | RPS16 |
| 0 | 0.84518687 | 0.99 | 0.897 | 0 | Tcells | RPS7 |
| 0 | 0.84338151 | 0.97 | 0.792 | 0 | Tcells | RPS5 |
| 0 | 0.84252905 | 0.458 | 0.01 | 0 | Tcells | CD96 |
| 0 | 0.84055262 | 0.996 | 0.9 | 0 | Tcells | RPL7A |
| 0 | 0.8362883 | 0.477 | 0.012 | 0 | Tcells | LCK |
| 0 | 0.83309473 | 0.991 | 0.882 | 0 | Tcells | RPL6 |
| 0 | 0.8218037 | 0.479 | 0.072 | 0 | Tcells | RORA |
| 0 | 0.82125304 | 0.694 | 0.285 | 0 | Tcells | NOP53 |
| 0 | 0.81978676 | 0.508 | 0.105 | 0 | Tcells | ATM |
| 0 | 0.81721086 | 0.58 | 0.148 | 0 | Tcells | ISG20 |
| 0 | 0.81057723 | 0.466 | 0.037 | 0 | Tcells | TRAF3IP3 |
| 0 | 0.80378086 | 0.609 | 0.218 | 0 | Tcells | SMCHD1 |
| 0 | 0.79014081 | 0.51 | 0.099 | 0 | Tcells | LIMD2 |
| 0 | 0.7897228 | 0.769 | 0.36 | 0 | Tcells | N4BP2L2 |
| 0 | 0.78870874 | 0.989 | 0.884 | 0 | Tcells | HLA-B |
| 0 | 0.78827372 | 0.411 | 0.014 | 0 | Tcells | IKZF3 |
| 0 | 0.78283112 | 0.451 | 0.058 | 0 | Tcells | PARP8 |
| 0 | 0.78188773 | 0.629 | 0.155 | 0 | Tcells | CD37 |
| 0 | 0.77699814 | 0.516 | 0.077 | 0 | Tcells | SEPT6 |
| 0 | 0.77426715 | 0.501 | 0.08 | 0 | Tcells | GYPC |
| 0 | 0.77325418 | 0.585 | 0.194 | 0 | Tcells | NKTR |
| 0 | 0.77192789 | 0.686 | 0.281 | 0 | Tcells | DDX24 |
| 0 | 0.7593148 | 0.513 | 0.093 | 0 | Tcells | AC245297.3 |

| | | | | | | |
|---|---|---|---|---|---|---|
| 0 | 0.73737136 | 0.391 | 0.026 | 0 | Tcells | PDE3B |
| 0 | 0.71752527 | 0.716 | 0.268 | 0 | Tcells | LCP1 |
| 0 | 0.71676484 | 0.329 | 0.009 | 0 | Tcells | LINC01871 |
| 0 | 0.70070662 | 0.521 | 0.187 | 0 | Tcells | SLFN5 |
| 0 | 0.69800249 | 0.758 | 0.371 | 0 | Tcells | SRSF7 |
| 0 | 0.69595055 | 0.459 | 0.109 | 0 | Tcells | TRIM22 |
| 0 | 0.69138687 | 0.535 | 0.129 | 0 | Tcells | SMAP2 |
| 0 | 0.67280381 | 0.517 | 0.121 | 0 | Tcells | IL2RG |
| 0 | 0.67125292 | 0.533 | 0.151 | 0 | Tcells | FNBP1 |
| 0 | 0.67036088 | 0.308 | 0.005 | 0 | Tcells | LINC00861 |
| 0 | 0.6667625 | 0.378 | 0.031 | 0 | Tcells | PTPN22 |
| 0 | 0.66568561 | 0.406 | 0.053 | 0 | Tcells | CCND2 |
| 0 | 0.66410349 | 0.374 | 0.038 | 0 | Tcells | ITGA4 |
| 0 | 0.65648173 | 0.468 | 0.112 | 0 | Tcells | CELF2 |
| 0 | 0.65597228 | 0.37 | 0.008 | 0 | Tcells | BCL11B |
| 0 | 0.6370267 | 0.43 | 0.102 | 0 | Tcells | FKBP5 |
| 0 | 0.63567826 | 0.632 | 0.225 | 0 | Tcells | GMFG |
| 0 | 0.63429538 | 0.504 | 0.16 | 0 | Tcells | PIK3IP1 |
| 0 | 0.63295334 | 0.392 | 0.033 | 0 | Tcells | RASGRP1 |
| 0 | 0.63023335 | 0.513 | 0.141 | 0 | Tcells | WIPF1 |
| 0 | 0.62959229 | 0.474 | 0.147 | 0 | Tcells | CLEC2B |
| 0 | 0.62578773 | 0.477 | 0.161 | 0 | Tcells | FOXP1 |
| 0 | 0.62476586 | 0.321 | 0.039 | 0 | Tcells | APOBEC3G |
| 0 | 0.62054567 | 0.528 | 0.164 | 0 | Tcells | PSMB9 |
| 0 | 0.61752854 | 0.457 | 0.101 | 0 | Tcells | PSIP1 |
| 0 | 0.61511733 | 0.503 | 0.179 | 0 | Tcells | TNRC6B |
| 0 | 0.6136514 | 0.373 | 0.025 | 0 | Tcells | OXNAD1 |
| 0 | 0.60790829 | 0.475 | 0.112 | 0 | Tcells | APBB1IP |
| 0 | 0.60679605 | 0.382 | 0.069 | 0 | Tcells | AKNA |
| 0 | 0.60533413 | 0.339 | 0.005 | 0 | Tcells | CAMK4 |
| 0 | 0.6028979 | 0.38 | 0.033 | 0 | Tcells | GIMAP1 |
| 0 | 0.60277808 | 0.435 | 0.129 | 0 | Tcells | GABPB1-AS1 |
| 0 | 0.58851645 | 0.163 | 0.001 | 0 | Tcells | XCL1 |
| 0 | 0.58714214 | 0.322 | 0.01 | 0 | Tcells | ITM2A |
| 0 | 0.57883669 | 0.263 | 0.033 | 0 | Tcells | ITGA1 |
| 0 | 0.57847788 | 0.43 | 0.095 | 0 | Tcells | EPB41 |
| 0 | 0.57734143 | 0.513 | 0.185 | 0 | Tcells | IFI16 |
| 0 | 0.57194903 | 0.142 | 0.007 | 0 | Tcells | GZMK |
| 0 | 0.56871179 | 0.413 | 0.086 | 0 | Tcells | IQGAP2 |
| 0 | 0.56838015 | 0.673 | 0.29 | 0 | Tcells | HIST1H4C |
| 0 | 0.56396971 | 0.353 | 0.074 | 0 | Tcells | PDE4D |
| 0 | 0.56084071 | 0.403 | 0.074 | 0 | Tcells | LBH |
| 0 | 0.55570088 | 0.506 | 0.15 | 0 | Tcells | SLC38A1 |
| 0 | 0.55563975 | 0.462 | 0.15 | 0 | Tcells | MYCBP2 |
| 0 | 0.55171196 | 0.397 | 0.081 | 0 | Tcells | BIN2 |
| 0 | 0.5483074 | 0.29 | 0.003 | 0 | Tcells | PYHIN1 |
| 0 | 0.54749654 | 0.372 | 0.053 | 0 | Tcells | SKAP1 |
| 0 | 0.54334937 | 0.457 | 0.144 | 0 | Tcells | TUT4 |
| 0 | 0.54047747 | 0.425 | 0.135 | 0 | Tcells | ABHD17A |
| 0 | 0.54021886 | 0.335 | 0.07 | 0 | Tcells | BCL2 |

| | | | | | | |
|---|---|---|---|---|---|---|
| 0 | 0.53765859 | 0.362 | 0.036 | 0 | Tcells | SEPT1 |
| 0 | 0.53729917 | 0.22 | 0.004 | 0 | Tcells | CXCR6 |
| 0 | 0.53573175 | 0.332 | 0.024 | 0 | Tcells | RNF125 |
| 0 | 0.53406943 | 0.186 | 0.013 | 0 | Tcells | CCR7 |
| 0 | 0.53352593 | 0.344 | 0.021 | 0 | Tcells | TBC1D10C |
| 0 | 0.53333408 | 0.357 | 0.079 | 0 | Tcells | RCAN3 |
| 0 | 0.53169968 | 0.383 | 0.083 | 0 | Tcells | ARHGEF1 |
| 0 | 0.52925473 | 0.356 | 0.074 | 0 | Tcells | CRIP1 |
| 0 | 0.52179752 | 0.3 | 0.005 | 0 | Tcells | SLFN12L |
| 0 | 0.51862936 | 0.361 | 0.062 | 0 | Tcells | GNG2 |
| 0 | 0.51759569 | 0.274 | 0.006 | 0 | Tcells | IL2RB |
| 0 | 0.51730859 | 0.387 | 0.083 | 0 | Tcells | ANKRD44 |
| 0 | 0.51579483 | 0.31 | 0.007 | 0 | Tcells | CD6 |
| 0 | 0.51260988 | 0.384 | 0.093 | 0 | Tcells | PIP4K2A |
| 0 | 0.51209182 | 0.326 | 0.02 | 0 | Tcells | STAT4 |
| 0 | 0.50972275 | 0.401 | 0.106 | 0 | Tcells | RBL2 |
| 0 | 0.50784339 | 0.416 | 0.113 | 0 | Tcells | UTRN |
| 0 | 0.50493196 | 0.3 | 0.005 | 0 | Tcells | ITK |
| 0 | 0.50452551 | 0.309 | 0.038 | 0 | Tcells | SPN |
| 0 | 0.50164328 | 0.222 | 0.012 | 0 | Tcells | TRGC2 |
| 0 | 0.49584538 | 0.342 | 0.035 | 0 | Tcells | RHOH |
| 0 | 0.4888384 | 0.101 | 0.006 | 0 | Tcells | FGFBP2 |
| 0 | 0.48749505 | 0.333 | 0.063 | 0 | Tcells | PTPN7 |
| 0 | 0.48705727 | 0.369 | 0.094 | 0 | Tcells | PDE4B |
| 0 | 0.48665212 | 0.361 | 0.077 | 0 | Tcells | PBXIP1 |
| 0 | 0.48384406 | 0.65 | 0.263 | 0 | Tcells | IFITM2 |
| 0 | 0.4807192 | 0.311 | 0.03 | 0 | Tcells | IFITM1 |
| 0 | 0.47447716 | 0.229 | 0.015 | 0 | Tcells | MYBL1 |
| 0 | 0.47384302 | 0.409 | 0.122 | 0 | Tcells | CCDC85B |
| 0 | 0.47231447 | 0.329 | 0.069 | 0 | Tcells | AC004687.1 |
| 0 | 0.4680098 | 0.331 | 0.066 | 0 | Tcells | FKBP11 |
| 0 | 0.46551416 | 0.139 | 0.002 | 0 | Tcells | XCL2 |
| 0 | 0.46143387 | 0.306 | 0.067 | 0 | Tcells | RIPOR2 |
| 0 | 0.46013538 | 0.3 | 0.066 | 0 | Tcells | CBLB |
| 0 | 0.45565131 | 0.356 | 0.084 | 0 | Tcells | CCND3 |
| 0 | 0.45154756 | 0.268 | 0.049 | 0 | Tcells | TCF7 |
| 0 | 0.45100269 | 0.122 | 0.002 | 0 | Tcells | KLRC1 |
| 0 | 0.448191 | 0.307 | 0.032 | 0 | Tcells | LAT |
| 0 | 0.44805029 | 0.374 | 0.094 | 0 | Tcells | KIF2A |
| 0 | 0.44388398 | 0.289 | 0.06 | 0 | Tcells | PTGER4 |
| 0 | 0.44360612 | 0.251 | 0.002 | 0 | Tcells | THEMIS |
| 0 | 0.44359991 | 0.314 | 0.049 | 0 | Tcells | TTC39C |
| 0 | 0.43543924 | 0.348 | 0.086 | 0 | Tcells | CASP8 |
| 0 | 0.43390647 | 0.344 | 0.083 | 0 | Tcells | SEMA4D |
| 0 | 0.42897438 | 0.348 | 0.078 | 0 | Tcells | ICAM3 |
| 0 | 0.42833598 | 0.255 | 0.011 | 0 | Tcells | SH2D2A |
| 0 | 0.42366414 | 0.322 | 0.062 | 0 | Tcells | PRKCH |
| 0 | 0.41757783 | 0.299 | 0.06 | 0 | Tcells | GPR65 |
| 0 | 0.41715824 | 0.286 | 0.052 | 0 | Tcells | BTN3A2 |
| 0 | 0.41560476 | 0.273 | 0.037 | 0 | Tcells | S1PR4 |

| | | | | | | |
|---|---|---|---|---|---|---|
| 0 | 0.40776871 | 0.269 | 0.039 | 0 | Tcells | PTPN4 |
| 0 | 0.40705264 | 0.287 | 0.05 | 0 | Tcells | SELPLG |
| 0 | 0.40149609 | 0.232 | 0.018 | 0 | Tcells | CD226 |
| 0 | 0.39960391 | 0.294 | 0.062 | 0 | Tcells | PDE7A |
| 0 | 0.39078322 | 0.235 | 0.003 | 0 | Tcells | GRAP2 |
| 0 | 0.38576587 | 0.154 | 0.002 | 0 | Tcells | LINC02446 |
| 0 | 0.38370967 | 0.231 | 0.038 | 0 | Tcells | AUTS2 |
| 0 | 0.37913676 | 0.297 | 0.065 | 0 | Tcells | CYFIP2 |
| 0 | 0.36729206 | 0.27 | 0.05 | 0 | Tcells | PRKACB |
| 0 | 0.36577103 | 0.172 | 0.005 | 0 | Tcells | TIGIT |
| 0 | 0.36423238 | 0.192 | 0.003 | 0 | Tcells | SAMD3 |
| 0 | 0.3637014 | 0.199 | 0.006 | 0 | Tcells | LAG3 |
| 0 | 0.36249818 | 0.223 | 0.004 | 0 | Tcells | AC243960.1 |
| 0 | 0.36203686 | 0.185 | 0.004 | 0 | Tcells | CD27 |
| 0 | 0.36035024 | 0.194 | 0.012 | 0 | Tcells | MIAT |
| 0 | 0.35983547 | 0.251 | 0.045 | 0 | Tcells | ITGAL |
| 0 | 0.35647919 | 0.196 | 0.004 | 0 | Tcells | TRAT1 |
| 0 | 0.35308399 | 0.227 | 0.031 | 0 | Tcells | CHST12 |
| 0 | 0.34795004 | 0.226 | 0.007 | 0 | Tcells | P2RY10 |
| 0 | 0.34662577 | 0.186 | 0.003 | 0 | Tcells | SLA2 |
| 0 | 0.34582464 | 0.2 | 0.002 | 0 | Tcells | SCML4 |
| 0 | 0.34378857 | 0.242 | 0.03 | 0 | Tcells | ARHGAP15 |
| 0 | 0.34338325 | 0.237 | 0.014 | 0 | Tcells | ANXA2R |
| 0 | 0.33990047 | 0.234 | 0.027 | 0 | Tcells | RASAL3 |
| 0 | 0.33335873 | 0.242 | 0.043 | 0 | Tcells | CCDC88C |
| 0 | 0.33274484 | 0.2 | 0.002 | 0 | Tcells | GPR174 |
| 0 | 0.33253405 | 0.221 | 0.006 | 0 | Tcells | ZAP70 |
| 0 | 0.33128226 | 0.236 | 0.029 | 0 | Tcells | GIMAP2 |
| 0 | 0.32502242 | 0.244 | 0.046 | 0 | Tcells | LINC00623 |
| 0 | 0.32149963 | 0.249 | 0.048 | 0 | Tcells | PSMB8-AS1 |
| 0 | 0.31453726 | 0.18 | 0.013 | 0 | Tcells | TRG-AS1 |
| 0 | 0.31229388 | 0.153 | 0.004 | 0 | Tcells | LEF1 |
| 0 | 0.30865236 | 0.221 | 0.04 | 0 | Tcells | HELB |
| 0 | 0.30440927 | 0.18 | 0.003 | 0 | Tcells | SH2D1A |
| 0 | 0.3020379 | 0.161 | 0.007 | 0 | Tcells | TXK |
| 0 | 0.3017066 | 0.207 | 0.021 | 0 | Tcells | SP140 |
| 0 | 0.30091015 | 0.235 | 0.043 | 0 | Tcells | FGD3 |
| 0 | 0.29337562 | 0.199 | 0.018 | 0 | Tcells | FCMR |
| 0 | 0.29185304 | 0.211 | 0.031 | 0 | Tcells | IL16 |
| 0 | 0.28065527 | 0.146 | 0.014 | 0 | Tcells | PASK |
| 0 | 0.27849344 | 0.181 | 0.017 | 0 | Tcells | NFATC2 |
| 0 | 0.27810494 | 0.178 | 0.024 | 0 | Tcells | GPR155 |
| 0 | 0.27432058 | 0.17 | 0.012 | 0 | Tcells | TNFRSF25 |
| 0 | 0.27374545 | 0.178 | 0.015 | 0 | Tcells | IPCEF1 |
| 0 | 0.2675243 | 0.116 | 0.003 | 0 | Tcells | ZNF683 |
| 0 | 0.26275504 | 0.176 | 0.019 | 0 | Tcells | F2R |
| 0 | 0.25742393 | 0.167 | 0.008 | 0 | Tcells | RGL4 |
| 0 | 0.25496011 | 0.168 | 0.01 | 0 | Tcells | PCED1B |
| 0 | 0.25345593 | 0.156 | 0.003 | 0 | Tcells | SIT1 |
| 0 | 0.25299628 | 0.136 | 0.013 | 0 | Tcells | AL357060.1 |

| | | | | | | |
|---|---|---|---|---|---|---|
| 0 | -0.8588843 | 0.446 | 0.829 | 0 | Tcells | RAC1 |
| 0 | -0.9720955 | 0.314 | 0.707 | 0 | Tcells | BRI3 |
| 0 | -1.1287306 | 0.307 | 0.719 | 0 | Tcells | PPP1R15A |
| 0 | -1.1450473 | 0.602 | 0.907 | 0 | Tcells | CD63 |
| 0 | -1.2035979 | 0.257 | 0.768 | 0 | Tcells | APLP2 |
| 0 | -1.2061391 | 0.046 | 0.474 | 0 | Tcells | CLDN7 |
| 0 | -1.2110127 | 0.367 | 0.76 | 0 | Tcells | PSAP |
| 0 | -1.3062638 | 0.414 | 0.794 | 0 | Tcells | CTSD |
| 0 | -1.3323879 | 0.529 | 0.854 | 0 | Tcells | SQSTM1 |
| 0 | -1.4090852 | 0.423 | 0.803 | 0 | Tcells | LGALS3 |
| 0 | -1.410702 | 0.249 | 0.763 | 0 | Tcells | CTSB |
| 0 | -1.5245837 | 0.187 | 0.798 | 0 | Tcells | CST3 |
| 0 | -1.5514866 | 0.488 | 0.888 | 0 | Tcells | ANXA2 |
| 0 | -1.6075279 | 0.245 | 0.665 | 0 | Tcells | GLUL |
| 0 | -1.6142868 | 0.089 | 0.564 | 0 | Tcells | CD24 |
| 0 | -1.6249478 | 0.659 | 0.895 | 0 | Tcells | CSTB |
| 0 | -1.6283091 | 0.309 | 0.701 | 0 | Tcells | NAMPT |
| 0 | -1.6552955 | 0.308 | 0.725 | 0 | Tcells | SOD2 |
| 0 | -1.7065324 | 0.168 | 0.667 | 0 | Tcells | IER3 |
| 0 | -1.7233591 | 0.068 | 0.516 | 0 | Tcells | KRT8 |
| 0 | -1.8685226 | 0.112 | 0.593 | 0 | Tcells | TACSTD2 |
| 0 | -1.8709445 | 0.352 | 0.742 | 0 | Tcells | HSPB1 |
| 0 | -1.9055449 | 0.11 | 0.568 | 0 | Tcells | KRT7 |
| 0 | -1.9056454 | 0.992 | 0.994 | 0 | Tcells | FTH1 |
| 0 | -1.9232294 | 0.104 | 0.562 | 0 | Tcells | CLDN4 |
| 0 | -1.9593789 | 0.101 | 0.579 | 0 | Tcells | ELF3 |
| 0 | -1.9886298 | 0.114 | 0.581 | 0 | Tcells | KRT18 |
| 0 | -2.0560426 | 0.781 | 0.975 | 0 | Tcells | S100A11 |
| 0 | -2.1164187 | 0.169 | 0.637 | 0 | Tcells | C15orf48 |
| 0 | -2.1474432 | 0.926 | 0.937 | 0 | Tcells | S100A6 |
| 0 | -2.2965696 | 0.137 | 0.602 | 0 | Tcells | KRT19 |
| 0 | -2.3038763 | 0.606 | 0.957 | 0 | Tcells | SAT1 |
| 0 | -2.538644 | 0.13 | 0.558 | 0 | Tcells | WFDC2 |
| 0 | -3.7647119 | 0.301 | 0.691 | 0 | Tcells | S100A8 |
| 0 | -4.6112525 | 0.364 | 0.746 | 0 | Tcells | S100A9 |
| 5.306E-307 | 0.51278413 | 0.49 | 0.17 | 1.253E-302 | Tcells | GPSM3 |
| 3.131E-306 | -2.2028919 | 0.982 | 0.987 | 7.396E-302 | Tcells | FTL |
| 6.282E-306 | 0.49617666 | 0.445 | 0.148 | 1.484E-301 | Tcells | KLF13 |
| 1.505E-304 | 0.26671706 | 0.191 | 0.031 | 3.554E-300 | Tcells | RASGRP2 |
| 3.316E-304 | 0.27063389 | 0.168 | 0.024 | 7.833E-300 | Tcells | MATK |
| 3.439E-301 | -1.2634389 | 0.052 | 0.471 | 8.124E-297 | Tcells | MARCKS |
| 5.032E-301 | 0.35918345 | 0.3 | 0.073 | 1.189E-296 | Tcells | ARHGAP45 |
| 1.515E-300 | 0.46302701 | 0.377 | 0.112 | 3.579E-296 | Tcells | PAXX |
| 2.278E-299 | 0.33569399 | 0.26 | 0.056 | 5.382E-295 | Tcells | RCSD1 |
| 3.627E-299 | 0.49208257 | 0.419 | 0.134 | 8.568E-295 | Tcells | LNPEP |
| 5.663E-299 | 0.40049965 | 0.297 | 0.071 | 1.338E-294 | Tcells | JAML |
| 1.3E-297 | 0.59054952 | 0.554 | 0.222 | 3.071E-293 | Tcells | C12orf57 |
| 1.308E-297 | 0.32120058 | 0.234 | 0.047 | 3.089E-293 | Tcells | ARHGAP9 |
| 1.449E-297 | 0.80320207 | 0.957 | 0.772 | 3.423E-293 | Tcells | RPL10A |
| 1.489E-297 | -1.4361394 | 0.141 | 0.56 | 3.518E-293 | Tcells | PLAUR |

| | | | | | | |
|---|---|---|---|---|---|---|
| 6.464E-297 | 0.68849222 | 0.612 | 0.277 | 1.527E-292 | Tcells | CD99 |
| 9.533E-297 | -1.6818899 | 0.906 | 0.942 | 2.252E-292 | Tcells | NEAT1 |
| 2.564E-296 | 0.29604258 | 0.203 | 0.036 | 6.056E-292 | Tcells | PTGER2 |
| 2.745E-295 | 0.26794849 | 0.21 | 0.038 | 6.485E-291 | Tcells | DEF6 |
| 1.223E-291 | 0.35423548 | 0.276 | 0.065 | 2.889E-287 | Tcells | PITPNC1 |
| 1.526E-291 | -1.7850427 | 0.064 | 0.471 | 3.605E-287 | Tcells | SOX4 |
| 1.11E-290 | -0.800122 | 0.327 | 0.705 | 2.622E-286 | Tcells | ATP6V0B |
| 1.501E-290 | 0.69812441 | 1 | 0.979 | 3.545E-286 | Tcells | EEF1A1 |
| 1.549E-287 | -0.9678571 | 0.395 | 0.746 | 3.659E-283 | Tcells | PKM |
| 1.915E-286 | 0.72277279 | 0.956 | 0.859 | 4.523E-282 | Tcells | CALM1 |
| 3.958E-283 | 0.34855725 | 0.209 | 0.04 | 9.349E-279 | Tcells | TENT5C |
| 1.35E-280 | 0.72334765 | 0.984 | 0.87 | 3.189E-276 | Tcells | RPL26 |
| 3.121E-280 | 0.6972364 | 0.997 | 0.943 | 7.372E-276 | Tcells | RPL13A |
| 1.793E-279 | 0.35071141 | 0.283 | 0.07 | 4.237E-275 | Tcells | CCDC66 |
| 1.903E-278 | -0.7430918 | 0.239 | 0.65 | 4.495E-274 | Tcells | YBX3 |
| 2.219E-278 | 0.43709305 | 0.486 | 0.171 | 5.242E-274 | Tcells | HCLS1 |
| 2.437E-277 | 0.5043294 | 0.453 | 0.164 | 5.757E-273 | Tcells | C9orf78 |
| 1.475E-275 | 0.34855787 | 0.261 | 0.061 | 3.485E-271 | Tcells | DENND2D |
| 3.052E-275 | -1.0337804 | 0.91 | 0.968 | 7.209E-271 | Tcells | UBC |
| 3.506E-275 | 0.57110876 | 0.463 | 0.172 | 8.283E-271 | Tcells | PRMT2 |
| 3.33E-274 | -0.901053 | 0.229 | 0.617 | 7.866E-270 | Tcells | NPC2 |
| 5.64E-274 | 0.46938134 | 0.351 | 0.105 | 1.332E-269 | Tcells | PCSK7 |
| 1.573E-272 | 0.52605461 | 0.476 | 0.18 | 3.716E-268 | Tcells | HLA-F |
| 5.489E-272 | 0.59174228 | 0.508 | 0.199 | 1.297E-267 | Tcells | EML4 |
| 1.625E-271 | 0.35891542 | 0.292 | 0.076 | 3.84E-267 | Tcells | NSMCE3 |
| 1.348E-270 | 0.42932951 | 0.33 | 0.093 | 3.185E-266 | Tcells | GPRIN3 |
| 1.776E-269 | 0.45703158 | 0.416 | 0.142 | 4.195E-265 | Tcells | BUB3 |
| 2.527E-267 | 0.76135377 | 0.954 | 0.866 | 5.969E-263 | Tcells | HLA-C |
| 2.863E-267 | 0.42785624 | 0.361 | 0.111 | 6.762E-263 | Tcells | TTC14 |
| 1.155E-266 | 0.61678366 | 0.61 | 0.281 | 2.728E-262 | Tcells | ARL6IP5 |
| 3.902E-266 | 0.33215185 | 0.259 | 0.062 | 9.217E-262 | Tcells | RNF166 |
| 5.332E-266 | -3.4831024 | 0.168 | 0.542 | 1.26E-261 | Tcells | LCN2 |
| 1.088E-265 | 0.74598479 | 0.992 | 0.915 | 2.57E-261 | Tcells | RPL9 |
| 4.561E-265 | 0.36400933 | 0.282 | 0.072 | 1.077E-260 | Tcells | PPM1K |
| 1.234E-263 | 0.28565494 | 0.232 | 0.05 | 2.914E-259 | Tcells | AC092683.1 |
| 2.568E-263 | -0.934126 | 0.494 | 0.799 | 6.066E-259 | Tcells | DYNLL1 |
| 8.894E-263 | 0.71249194 | 0.955 | 0.843 | 2.101E-258 | Tcells | RPL38 |
| 1.919E-262 | 0.45055249 | 0.368 | 0.117 | 4.532E-258 | Tcells | CCSER2 |
| 2.216E-262 | 0.64953595 | 0.929 | 0.776 | 5.235E-258 | Tcells | EEF1D |
| 3.695E-262 | 0.46207335 | 0.428 | 0.149 | 8.728E-258 | Tcells | CDKN1B |
| 7.77E-261 | 0.6415605 | 0.972 | 0.84 | 1.835E-256 | Tcells | RPL35 |
| 2.119E-260 | 0.4063036 | 0.357 | 0.109 | 5.005E-256 | Tcells | DOCK10 |
| 1.923E-259 | 0.71069161 | 0.792 | 0.534 | 4.543E-255 | Tcells | RPS10 |
| 5.521E-259 | -0.9983474 | 0.099 | 0.484 | 1.304E-254 | Tcells | CDKN1A |
| 1.761E-258 | 0.54365515 | 0.529 | 0.221 | 4.161E-254 | Tcells | PRPF38B |
| 1.808E-258 | 0.40375135 | 0.332 | 0.097 | 4.272E-254 | Tcells | IL10RA |
| 1.922E-258 | -0.8484431 | 0.346 | 0.692 | 4.54E-254 | Tcells | TALDO1 |
| 7.516E-257 | 0.44320091 | 0.401 | 0.136 | 1.775E-252 | Tcells | ZC3HAV1 |
| 1.271E-255 | 0.5571884 | 0.562 | 0.246 | 3.003E-251 | Tcells | MPHOSPH8 |
| 1.594E-254 | 0.70014526 | 0.344 | 0.105 | 3.765E-250 | Tcells | DUSP2 |

| | | | | | | |
|---|---|---|---|---|---|---|
| 1.85E-254 | -0.8194444 | 0.339 | 0.686 | 4.371E-250 | Tcells | RTN4 |
| 3.748E-254 | -1.2430298 | 0.529 | 0.798 | 8.853E-250 | Tcells | TXN |
| 4.88E-253 | 0.40300866 | 0.348 | 0.107 | 1.153E-248 | Tcells | CENPC |
| 1.354E-251 | 0.30530327 | 0.256 | 0.063 | 3.2E-247 | Tcells | BIN1 |
| 4.434E-251 | 0.44539949 | 0.416 | 0.145 | 1.047E-246 | Tcells | DOCK8 |
| 2.557E-249 | -0.7735938 | 0.382 | 0.727 | 6.04E-245 | Tcells | SDCBP |
| 8.439E-249 | -0.8406417 | 0.178 | 0.549 | 1.994E-244 | Tcells | PTTG1IP |
| 2.906E-248 | 0.42727862 | 0.334 | 0.101 | 6.866E-244 | Tcells | SATB1 |
| 1.12E-247 | 0.4342006 | 0.614 | 0.249 | 2.646E-243 | Tcells | CD53 |
| 4.148E-246 | 0.44016505 | 0.335 | 0.103 | 9.8E-242 | Tcells | CRYBG1 |
| 5.004E-245 | 0.44870016 | 0.41 | 0.145 | 1.182E-240 | Tcells | ARHGAP30 |
| 1.117E-244 | 0.69008486 | 0.996 | 0.955 | 2.639E-240 | Tcells | RPS18 |
| 8.849E-244 | 0.72302721 | 0.962 | 0.824 | 2.09E-239 | Tcells | RPL22 |
| 3.831E-243 | -1.7231034 | 0.061 | 0.424 | 9.049E-239 | Tcells | SLPI |
| 1.651E-241 | 0.59006026 | 0.467 | 0.187 | 3.9E-237 | Tcells | NR3C1 |
| 5.085E-241 | 0.50302946 | 0.549 | 0.241 | 1.201E-236 | Tcells | VAMP2 |
| 7.357E-241 | 0.5599236 | 0.565 | 0.258 | 1.738E-236 | Tcells | AAK1 |
| 1.011E-240 | 0.65380617 | 0.948 | 0.804 | 2.387E-236 | Tcells | DDX5 |
| 7.961E-240 | 0.62779799 | 0.977 | 0.917 | 1.881E-235 | Tcells | NACA |
| 1.441E-239 | -1.1787975 | 0.589 | 0.824 | 3.405E-235 | Tcells | GSTP1 |
| 1.891E-239 | 0.25897529 | 0.197 | 0.041 | 4.468E-235 | Tcells | TNRC6C |
| 3.018E-239 | 0.61817465 | 0.997 | 0.943 | 7.13E-235 | Tcells | PTMA |
| 9.309E-238 | 0.53249109 | 0.48 | 0.2 | 2.199E-233 | Tcells | NAP1L4 |
| 1.643E-237 | 0.64754166 | 0.984 | 0.87 | 3.881E-233 | Tcells | RPL36 |
| 7.363E-237 | -0.712175 | 0.525 | 0.815 | 1.739E-232 | Tcells | GNG5 |
| 2.367E-236 | 0.56938807 | 0.593 | 0.278 | 5.591E-232 | Tcells | FXYD5 |
| 2.383E-235 | -0.8025918 | 0.16 | 0.519 | 5.63E-231 | Tcells | ASAH1 |
| 1.514E-234 | 0.38786533 | 0.37 | 0.124 | 3.576E-230 | Tcells | URI1 |
| 5.35E-234 | 0.45795049 | 0.432 | 0.154 | 1.264E-229 | Tcells | SLA |
| 6.31E-234 | 0.51532062 | 0.565 | 0.258 | 1.491E-229 | Tcells | SF1 |
| 3.494E-233 | -0.8126113 | 0.51 | 0.795 | 8.254E-229 | Tcells | MAP1LC3B |
| 6.853E-233 | 0.50946287 | 0.225 | 0.054 | 1.619E-228 | Tcells | SELL |
| 2.346E-232 | -1.1036443 | 0.104 | 0.455 | 5.542E-228 | Tcells | ATP1B1 |
| 3.123E-230 | 0.35463274 | 0.309 | 0.093 | 7.378E-226 | Tcells | PARP1 |
| 3.81E-230 | 0.61157709 | 0.69 | 0.379 | 9E-226 | Tcells | COMMD6 |
| 7.656E-229 | 0.27575972 | 0.182 | 0.037 | 1.809E-224 | Tcells | AP3M2 |
| 6.73E-228 | -0.9133052 | 0.095 | 0.445 | 1.59E-223 | Tcells | SPINT2 |
| 1.178E-227 | 0.38498875 | 0.284 | 0.082 | 2.783E-223 | Tcells | LYAR |
| 3.594E-227 | 0.5741796 | 0.559 | 0.259 | 8.49E-223 | Tcells | BPTF |
| 6.389E-226 | 0.49349254 | 0.432 | 0.168 | 1.509E-221 | Tcells | CYLD |
| 2.768E-225 | 0.30409322 | 0.191 | 0.041 | 6.538E-221 | Tcells | LDLRAD4 |
| 4.173E-225 | -0.9824514 | 0.167 | 0.513 | 9.857E-221 | Tcells | CD9 |
| 9.417E-225 | -0.7453176 | 0.077 | 0.433 | 2.225E-220 | Tcells | GRN |
| 1.382E-224 | 0.28523274 | 0.217 | 0.051 | 3.264E-220 | Tcells | DDHD1 |
| 1.007E-223 | 0.49628968 | 0.393 | 0.142 | 2.378E-219 | Tcells | HIST1H1E |
| 1.106E-220 | -1.6066153 | 0.025 | 0.362 | 2.612E-216 | Tcells | GDF15 |
| 1.667E-220 | 0.63525527 | 0.938 | 0.814 | 3.939E-216 | Tcells | RPL27 |
| 2.101E-220 | 0.28254917 | 0.222 | 0.054 | 4.964E-216 | Tcells | INPP5D |
| 4.271E-220 | 0.44061907 | 0.382 | 0.138 | 1.009E-215 | Tcells | ATF7IP |
| 2.425E-219 | 0.53267551 | 0.536 | 0.246 | 5.728E-215 | Tcells | MBP |

| | | | | | | |
|---|---|---|---|---|---|---|
| 3.421E-219 | -1.1466628 | 0.031 | 0.368 | 8.081E-215 | Tcells | PHLDA2 |
| 4.963E-219 | -0.8177693 | 0.25 | 0.587 | 1.172E-214 | Tcells | ZFAND5 |
| 1.529E-217 | 0.25217187 | 0.194 | 0.043 | 3.612E-213 | Tcells | CEP85L |
| 1.442E-216 | 0.49634646 | 0.538 | 0.247 | 3.406E-212 | Tcells | ELF1 |
| 6.905E-216 | -0.7049342 | 0.015 | 0.347 | 1.631E-211 | Tcells | RBM47 |
| 1.304E-215 | 0.5503892 | 0.563 | 0.267 | 3.08E-211 | Tcells | ATRX |
| 1.931E-215 | -1.3126681 | 0.139 | 0.475 | 4.561E-211 | Tcells | MXD1 |
| 3.823E-215 | 0.44753068 | 0.434 | 0.174 | 9.032E-211 | Tcells | GTF3A |
| 4.327E-214 | 0.58729907 | 0.968 | 0.832 | 1.022E-209 | Tcells | RACK1 |
| 5.157E-214 | 0.51633024 | 0.402 | 0.158 | 1.218E-209 | Tcells | CLDND1 |
| 5.373E-214 | 0.41769516 | 0.475 | 0.188 | 1.269E-209 | Tcells | LSP1 |
| 6.507E-214 | 0.49246634 | 0.51 | 0.225 | 1.537E-209 | Tcells | SP100 |
| 1.208E-213 | 0.35130867 | 0.259 | 0.072 | 2.853E-209 | Tcells | RPAP2 |
| 1.267E-213 | -0.69841 | 0.071 | 0.412 | 2.993E-209 | Tcells | CTNNA1 |
| 2.12E-212 | 0.57808278 | 0.979 | 0.897 | 5.009E-208 | Tcells | RPL24 |
| 2.429E-211 | 0.40485878 | 0.348 | 0.121 | 5.738E-207 | Tcells | SMARCA2 |
| 6.911E-210 | 0.94798275 | 0.827 | 0.631 | 1.633E-205 | Tcells | TXNIP |
| 4.141E-209 | -0.7087604 | 0.434 | 0.775 | 9.782E-205 | Tcells | ITGB1 |
| 1.003E-208 | -2.6813472 | 0.141 | 0.47 | 2.37E-204 | Tcells | SPP1 |
| 5.44E-208 | -1.367552 | 0.36 | 0.673 | 1.285E-203 | Tcells | HSPA1A |
| 1.799E-207 | 0.53605142 | 0.688 | 0.378 | 4.249E-203 | Tcells | PNISR |
| 8.371E-206 | 0.40853088 | 0.432 | 0.175 | 1.977E-201 | Tcells | ORMDL1 |
| 1.369E-205 | -2.1084776 | 0.142 | 0.481 | 3.234E-201 | Tcells | LYZ |
| 4.427E-205 | 0.62305284 | 0.994 | 0.882 | 1.046E-200 | Tcells | RPS6 |
| 1.436E-204 | -0.9804828 | 0.044 | 0.369 | 3.393E-200 | Tcells | ATF3 |
| 1.583E-204 | 0.46162382 | 0.458 | 0.195 | 3.741E-200 | Tcells | GSTK1 |
| 4.195E-201 | 0.31627757 | 0.299 | 0.095 | 9.909E-197 | Tcells | SP110 |
| 5.453E-199 | 0.87711745 | 0.451 | 0.202 | 1.288E-194 | Tcells | CREM |
| 1.035E-198 | 0.29355479 | 0.233 | 0.063 | 2.445E-194 | Tcells | DGKA |
| 2.134E-197 | 0.51530078 | 0.595 | 0.302 | 5.04E-193 | Tcells | KMT2E |
| 2.012E-196 | 0.3727008 | 0.367 | 0.135 | 4.752E-192 | Tcells | SH3KBP1 |
| 2.323E-196 | -1.194927 | 0.017 | 0.327 | 5.488E-192 | Tcells | EMP1 |
| 3.304E-196 | 0.28179466 | 0.242 | 0.068 | 7.804E-192 | Tcells | POLR3GL |
| 5.655E-196 | 0.47453737 | 0.491 | 0.219 | 1.336E-191 | Tcells | KMT2A |
| 1.034E-195 | -0.6411961 | 0.016 | 0.325 | 2.442E-191 | Tcells | RAB13 |
| 1.311E-195 | -0.9873786 | 0.516 | 0.758 | 3.098E-191 | Tcells | PRDX1 |
| 1.401E-194 | 0.59844365 | 0.978 | 0.895 | 3.309E-190 | Tcells | RPL27A |
| 1.975E-194 | -2.5199051 | 0.119 | 0.423 | 4.666E-190 | Tcells | S100P |
| 2.289E-194 | -0.7170836 | 0.048 | 0.364 | 5.408E-190 | Tcells | TIMP2 |
| 3.474E-194 | 0.43227621 | 0.414 | 0.164 | 8.206E-190 | Tcells | GLIPR1 |
| 3.738E-194 | 0.25531764 | 0.203 | 0.05 | 8.829E-190 | Tcells | GOLGA8B |
| 6.935E-193 | -1.2379574 | 0.004 | 0.307 | 1.638E-188 | Tcells | RND3 |
| 1.651E-192 | 0.53943394 | 0.924 | 0.831 | 3.901E-188 | Tcells | HLA-E |
| 3.192E-191 | 0.31920456 | 0.267 | 0.082 | 7.54E-187 | Tcells | ANXA6 |
| 7.442E-191 | -0.8172486 | 0.189 | 0.501 | 1.758E-186 | Tcells | HMGA1 |
| 5.84E-190 | 0.5696222 | 0.987 | 0.891 | 1.38E-185 | Tcells | RPL37A |
| 1.014E-189 | -0.9885742 | 0.522 | 0.764 | 2.396E-185 | Tcells | TPM4 |
| 1.533E-189 | 0.38134756 | 0.36 | 0.136 | 3.62E-185 | Tcells | SYNRG |
| 1.674E-189 | -0.6988106 | 0.065 | 0.383 | 3.954E-185 | Tcells | CTSH |
| 2.596E-189 | 0.46884615 | 0.557 | 0.269 | 6.133E-185 | Tcells | LDHB |

| | | | | | | |
|---|---|---|---|---|---|---|
| 3.622E-189 | 0.41025192 | 0.276 | 0.085 | 8.555E-185 | Tcells | GPR183 |
| 5.121E-189 | 0.36424837 | 0.351 | 0.129 | 1.21E-184 | Tcells | LPIN2 |
| 5.791E-189 | 0.29506929 | 0.265 | 0.081 | 1.368E-184 | Tcells | DNMT1 |
| 2.213E-188 | -1.0700458 | 0.008 | 0.307 | 5.228E-184 | Tcells | GPRC5A |
| 2.333E-188 | -1.5870899 | 0.011 | 0.31 | 5.511E-184 | Tcells | ADIRF |
| 2.874E-188 | 0.37007585 | 0.387 | 0.15 | 6.788E-184 | Tcells | RSBN1L |
| 5.978E-188 | -1.5321834 | 0.042 | 0.349 | 1.412E-183 | Tcells | IL1RN |
| 1.403E-186 | -1.1990849 | 0.861 | 0.923 | 3.315E-182 | Tcells | S100A10 |
| 2.38E-186 | 0.39640444 | 0.348 | 0.13 | 5.623E-182 | Tcells | MGAT4A |
| 3.326E-186 | -0.7749819 | 0.195 | 0.509 | 7.856E-182 | Tcells | H2AFJ |
| 1.127E-184 | -1.0129005 | 0.355 | 0.652 | 2.663E-180 | Tcells | GADD45B |
| 2.772E-184 | 0.3417042 | 0.302 | 0.103 | 6.548E-180 | Tcells | RAB27A |
| 7.124E-184 | -0.7117976 | 0.027 | 0.327 | 1.683E-179 | Tcells | SDC4 |
| 1.065E-183 | -3.1937996 | 0.426 | 0.689 | 2.517E-179 | Tcells | TIMP1 |
| 1.764E-182 | -0.6320476 | 0.207 | 0.518 | 4.168E-178 | Tcells | LAPTM4A |
| 6.685E-182 | -0.7065077 | 0.046 | 0.346 | 1.579E-177 | Tcells | APP |
| 1.939E-181 | -2.330486 | 0.063 | 0.356 | 4.582E-177 | Tcells | PSCA |
| 2.738E-181 | 0.56010107 | 0.909 | 0.78 | 6.467E-177 | Tcells | RPL31 |
| 3.815E-181 | -0.8983288 | 0.046 | 0.343 | 9.012E-177 | Tcells | MYO6 |
| 2.711E-180 | -0.7786798 | 0.02 | 0.313 | 6.404E-176 | Tcells | FAM129B |
| 3.939E-180 | -0.7163134 | 0.045 | 0.344 | 9.304E-176 | Tcells | ASPH |
| 4.249E-180 | -1.6516206 | 0.056 | 0.351 | 1.004E-175 | Tcells | IGFBP7 |
| 5.718E-180 | 0.30898429 | 0.374 | 0.139 | 1.351E-175 | Tcells | EVI2B |
| 8.748E-180 | 0.4228884 | 0.465 | 0.209 | 2.067E-175 | Tcells | DDX18 |
| 2.143E-178 | 0.55025541 | 0.915 | 0.795 | 5.063E-174 | Tcells | HMGB1 |
| 5.148E-178 | -0.8510801 | 0.155 | 0.452 | 1.216E-173 | Tcells | CYSTM1 |
| 7.524E-178 | 0.38389149 | 0.329 | 0.12 | 1.777E-173 | Tcells | ODF2L |
| 8.489E-177 | 0.3181081 | 0.298 | 0.102 | 2.005E-172 | Tcells | PREX1 |
| 5.509E-176 | 0.44012776 | 0.449 | 0.201 | 1.301E-171 | Tcells | PHF3 |
| 1.492E-174 | 0.61167101 | 0.721 | 0.475 | 3.524E-170 | Tcells | ANKRD12 |
| 2.783E-174 | -0.7536956 | 0.369 | 0.638 | 6.575E-170 | Tcells | DSTN |
| 8.63E-174 | -0.6495117 | 0.007 | 0.288 | 2.039E-169 | Tcells | JUP |
| 7.354E-173 | -0.8116164 | 0.132 | 0.424 | 1.737E-168 | Tcells | CD59 |
| 3.467E-171 | 0.30531252 | 0.273 | 0.09 | 8.19E-167 | Tcells | DOCK2 |
| 1.179E-170 | -2.5543431 | 0.069 | 0.353 | 2.786E-166 | Tcells | MMP7 |
| 2.351E-170 | -0.7039631 | 0.023 | 0.304 | 5.553E-166 | Tcells | SPINT1 |
| 1.43E-169 | -0.8764121 | 0.011 | 0.287 | 3.378E-165 | Tcells | MAL2 |
| 3.1E-169 | 0.50902697 | 0.991 | 0.919 | 7.323E-165 | Tcells | RPL8 |
| 1.544E-168 | 0.34112092 | 0.395 | 0.161 | 3.647E-164 | Tcells | TNFAIP8 |
| 3.426E-168 | 0.27376569 | 0.243 | 0.075 | 8.093E-164 | Tcells | RNASEH2B |
| 1.058E-167 | -0.6944209 | 0.543 | 0.788 | 2.498E-163 | Tcells | TPI1 |
| 1.25E-166 | -0.7196516 | 0.04 | 0.326 | 2.953E-162 | Tcells | CD68 |
| 3.767E-166 | 0.28734532 | 0.299 | 0.105 | 8.9E-162 | Tcells | PDCD7 |
| 3.875E-166 | 0.42991725 | 0.515 | 0.253 | 9.155E-162 | Tcells | DDX46 |
| 4.06E-166 | 0.46403003 | 0.445 | 0.203 | 9.591E-162 | Tcells | TGFB1 |
| 8.451E-166 | -0.9211904 | 0.055 | 0.337 | 1.996E-161 | Tcells | VEGFA |
| 1.376E-165 | -0.786931 | 0.008 | 0.278 | 3.251E-161 | Tcells | MGST1 |
| 1.486E-165 | -1.0603203 | 0.007 | 0.278 | 3.51E-161 | Tcells | TSPAN1 |
| 3.415E-165 | 0.41260991 | 0.424 | 0.188 | 8.066E-161 | Tcells | TTC3 |
| 5.872E-165 | 0.2947811 | 0.31 | 0.112 | 1.387E-160 | Tcells | SNRPN |

| | | | | | | |
|---|---|---|---|---|---|---|
| 8.268E-165 | 0.48038265 | 0.884 | 0.765 | 1.953E-160 | Tcells | TMA7 |
| 8.955E-165 | -0.9697 | 0.189 | 0.471 | 2.115E-160 | Tcells | RAB11FIP1 |
| 1.544E-164 | -2.9911533 | 0.117 | 0.392 | 3.647E-160 | Tcells | KRT13 |
| 1.958E-164 | 0.40129605 | 0.576 | 0.297 | 4.626E-160 | Tcells | MZT2B |
| 4.816E-164 | -0.8920487 | 0.038 | 0.316 | 1.138E-159 | Tcells | SERPINA1 |
| 8.64E-164 | 0.411627 | 0.543 | 0.275 | 2.041E-159 | Tcells | APRT |
| 1.525E-163 | 0.30366946 | 0.281 | 0.096 | 3.604E-159 | Tcells | SLC39A10 |
| 4.82E-163 | -1.0914088 | 0.024 | 0.296 | 1.139E-158 | Tcells | MAFB |
| 5.535E-163 | -0.7710437 | 0.017 | 0.288 | 1.308E-158 | Tcells | ADM |
| 3.931E-162 | -0.7837144 | 0.262 | 0.537 | 9.286E-158 | Tcells | HEBP2 |
| 4.083E-162 | 0.35715383 | 0.404 | 0.174 | 9.646E-158 | Tcells | MZT2A |
| 2.852E-161 | 0.47826459 | 0.589 | 0.317 | 6.738E-157 | Tcells | MSN |
| 3.189E-161 | -0.5223942 | 0.022 | 0.292 | 7.532E-157 | Tcells | ADAM9 |
| 1.913E-160 | 0.34968851 | 0.316 | 0.119 | 4.52E-156 | Tcells | ARAP2 |
| 3.041E-160 | -0.9531817 | 0.93 | 0.956 | 7.184E-156 | Tcells | ACTG1 |
| 2.659E-159 | -1.3356161 | 0.167 | 0.455 | 6.282E-155 | Tcells | TYROBP |
| 4.027E-159 | -0.722877 | 0.063 | 0.34 | 9.512E-155 | Tcells | MAFF |
| 6.832E-159 | -0.5544224 | 0.216 | 0.505 | 1.614E-154 | Tcells | HSBP1 |
| 1.112E-157 | 0.25789663 | 0.206 | 0.059 | 2.627E-153 | Tcells | AOAH |
| 1.211E-157 | 0.39514072 | 1 | 0.979 | 2.86E-153 | Tcells | MALAT1 |
| 4.568E-157 | -0.7884074 | 0.013 | 0.273 | 1.079E-152 | Tcells | MUC1 |
| 7.881E-157 | 0.42166061 | 0.489 | 0.24 | 1.862E-152 | Tcells | NSD3 |
| 1.282E-156 | 0.40543898 | 0.448 | 0.208 | 3.027E-152 | Tcells | PCM1 |
| 1.53E-156 | 0.39936383 | 0.49 | 0.241 | 3.615E-152 | Tcells | PA2G4 |
| 1.552E-156 | 0.43035616 | 0.529 | 0.268 | 3.667E-152 | Tcells | PRPF4B |
| 1.775E-156 | 0.59930382 | 0.951 | 0.893 | 4.192E-152 | Tcells | HLA-A |
| 3.623E-155 | -0.6715076 | 0.081 | 0.357 | 8.558E-151 | Tcells | CPEB4 |
| 5.455E-154 | 0.29853548 | 0.408 | 0.169 | 1.289E-149 | Tcells | SAMSN1 |
| 5.936E-154 | -0.718479 | 0.007 | 0.263 | 1.402E-149 | Tcells | KLF5 |
| 6.334E-154 | 0.344867 | 0.357 | 0.144 | 1.496E-149 | Tcells | ADGRE5 |
| 9.879E-154 | 0.37809999 | 0.67 | 0.39 | 2.334E-149 | Tcells | RNF213 |
| 1.533E-153 | -0.7757861 | 0.03 | 0.294 | 3.622E-149 | Tcells | H1F0 |
| 1.558E-153 | 0.26925457 | 0.259 | 0.087 | 3.679E-149 | Tcells | HIST2H2AC |
| 2.549E-153 | 0.31332581 | 0.303 | 0.112 | 6.021E-149 | Tcells | TC2N |
| 3.266E-153 | 0.48049354 | 0.739 | 0.481 | 7.716E-149 | Tcells | NCL |
| 1.064E-152 | -0.9483616 | 0.458 | 0.678 | 2.513E-148 | Tcells | ANXA5 |
| 1.26E-152 | 0.33679931 | 0.339 | 0.137 | 2.977E-148 | Tcells | SUN2 |
| 5.836E-152 | -0.6163759 | 0.007 | 0.26 | 1.379E-147 | Tcells | SERINC2 |
| 2.049E-151 | -1.081361 | 0.037 | 0.299 | 4.841E-147 | Tcells | HMOX1 |
| 2.735E-151 | 0.46325823 | 0.631 | 0.373 | 6.46E-147 | Tcells | PSME1 |
| 4.627E-151 | 0.42715798 | 0.534 | 0.283 | 1.093E-146 | Tcells | RABAC1 |
| 1.607E-150 | -1.0337792 | 0.236 | 0.519 | 3.797E-146 | Tcells | HSPA1B |
| 1.67E-150 | 0.48189117 | 0.602 | 0.354 | 3.945E-146 | Tcells | AES |
| 1.715E-150 | -0.7587955 | 0.028 | 0.286 | 4.051E-146 | Tcells | TNFAIP2 |
| 3.628E-150 | -0.8830519 | 0.008 | 0.258 | 8.571E-146 | Tcells | CST6 |
| 2.347E-149 | 0.44370243 | 0.984 | 0.913 | 5.544E-145 | Tcells | UBA52 |
| 4.43E-149 | -0.7870131 | 0.031 | 0.286 | 1.047E-144 | Tcells | TNFRSF12A |
| 8.977E-149 | 0.75550449 | 0.379 | 0.175 | 2.121E-144 | Tcells | KLF2 |
| 1.33E-148 | 0.30729233 | 0.266 | 0.095 | 3.141E-144 | Tcells | MDFIC |
| 1.644E-147 | -0.9821957 | 0.005 | 0.25 | 3.884E-143 | Tcells | ADGRF1 |

| | | | | | | |
|---|---|---|---|---|---|---|
| 3.048E-147 | -0.7076737 | 0.007 | 0.254 | 7.199E-143 | Tcells | PRSS8 |
| 4.816E-147 | 0.31379349 | 0.346 | 0.141 | 1.138E-142 | Tcells | LBR |
| 1.479E-146 | -1.4065349 | 0.13 | 0.399 | 3.493E-142 | Tcells | FCER1G |
| 3.031E-146 | -0.7607049 | 0.009 | 0.255 | 7.16E-142 | Tcells | ITGB8 |
| 7.697E-146 | 0.48333575 | 0.734 | 0.527 | 1.818E-141 | Tcells | HMGN1 |
| 7.851E-146 | 0.29815762 | 0.364 | 0.15 | 1.855E-141 | Tcells | SOCS1 |
| 1E-145 | -0.5971269 | 0.052 | 0.309 | 2.363E-141 | Tcells | IFNGR2 |
| 1.319E-145 | 0.28669772 | 0.275 | 0.099 | 3.116E-141 | Tcells | CWF19L2 |
| 1.329E-145 | 0.43972274 | 0.515 | 0.271 | 3.14E-141 | Tcells | TPR |
| 3.133E-145 | -1.205016 | 0.229 | 0.486 | 7.4E-141 | Tcells | ERO1A |
| 1.016E-144 | 0.43662217 | 0.834 | 0.614 | 2.401E-140 | Tcells | S100A4 |
| 1.442E-144 | 0.47962059 | 0.628 | 0.372 | 3.405E-140 | Tcells | DEK |
| 1.58E-144 | -0.5287664 | 0.002 | 0.243 | 3.733E-140 | Tcells | TJP1 |
| 1.7E-144 | 0.40438644 | 0.497 | 0.26 | 4.015E-140 | Tcells | UXT |
| 3.075E-144 | -0.6812422 | 0.006 | 0.249 | 7.263E-140 | Tcells | S100A16 |
| 4.058E-144 | 0.46014037 | 0.62 | 0.364 | 9.585E-140 | Tcells | GCC2 |
| 2.537E-143 | 0.26765589 | 0.233 | 0.077 | 5.993E-139 | Tcells | CNTRL |
| 3.174E-143 | 0.26183554 | 0.226 | 0.074 | 7.497E-139 | Tcells | MTERF4 |
| 4.15E-143 | 0.29409694 | 0.275 | 0.1 | 9.803E-139 | Tcells | PAG1 |
| 6.21E-143 | 0.39944098 | 0.426 | 0.205 | 1.467E-138 | Tcells | DDX6 |
| 9.057E-143 | 0.259597 | 0.238 | 0.08 | 2.14E-138 | Tcells | AC016831.7 |
| 1.056E-142 | -2.0580652 | 0.032 | 0.279 | 2.494E-138 | Tcells | S100A2 |
| 1.199E-142 | -4.1736002 | 0.158 | 0.414 | 2.834E-138 | Tcells | SPRR3 |
| 1.284E-142 | 0.41133028 | 0.577 | 0.322 | 3.033E-138 | Tcells | EIF4B |
| 2.394E-142 | -0.6197829 | 0.95 | 0.964 | 5.655E-138 | Tcells | MYL6 |
| 4.17E-142 | -0.755924 | 0.013 | 0.254 | 9.851E-138 | Tcells | S100A13 |
| 5.916E-142 | -0.8832099 | 0.006 | 0.246 | 1.397E-137 | Tcells | PDZK1IP1 |
| 9.611E-142 | 0.31287106 | 0.312 | 0.124 | 2.27E-137 | Tcells | RFC1 |
| 1.351E-141 | 0.32284901 | 0.292 | 0.112 | 3.19E-137 | Tcells | ANP32E |
| 2.277E-141 | -0.5377786 | 0.044 | 0.297 | 5.379E-137 | Tcells | CXCL16 |
| 4.37E-141 | -0.8768836 | 0.077 | 0.336 | 1.032E-136 | Tcells | IFITM3 |
| 9.168E-141 | -2.7413656 | 0.245 | 0.501 | 2.166E-136 | Tcells | CXCL8 |
| 1.51E-140 | -0.9659043 | 0.017 | 0.258 | 3.567E-136 | Tcells | PLAU |
| 1.548E-140 | -1.614088 | 0.128 | 0.382 | 3.656E-136 | Tcells | MAL |
| 2.618E-140 | -0.8784677 | 0.009 | 0.247 | 6.185E-136 | Tcells | EPCAM |
| 3.271E-140 | -0.9494157 | 0.289 | 0.543 | 7.727E-136 | Tcells | CTSS |
| 6.256E-140 | 0.39474068 | 0.441 | 0.216 | 1.478E-135 | Tcells | DHX36 |
| 9.367E-140 | 0.2562099 | 0.218 | 0.07 | 2.213E-135 | Tcells | APOBEC3C |
| 1.098E-139 | -1.0903277 | 0.072 | 0.325 | 2.594E-135 | Tcells | PHLDA1 |
| 3.178E-139 | 0.33007066 | 0.387 | 0.176 | 7.508E-135 | Tcells | FGFR1OP2 |
| 3.914E-139 | 0.25297736 | 0.225 | 0.074 | 9.245E-135 | Tcells | SACM1L |
| 9.292E-139 | 0.40216661 | 0.542 | 0.291 | 2.195E-134 | Tcells | RPL36A |
| 1.546E-138 | 0.49982618 | 0.981 | 0.914 | 3.651E-134 | Tcells | RPS20 |
| 3.101E-138 | 0.44668906 | 0.902 | 0.808 | 7.325E-134 | Tcells | RPL36AL |
| 3.651E-138 | -0.972029 | 0.045 | 0.293 | 8.626E-134 | Tcells | CLU |
| 6.102E-138 | -0.7511881 | 0.019 | 0.258 | 1.441E-133 | Tcells | CSTA |
| 7.808E-138 | 0.28218576 | 0.283 | 0.106 | 1.844E-133 | Tcells | CREBZF |
| 7.857E-138 | 0.34474202 | 0.354 | 0.152 | 1.856E-133 | Tcells | SETD2 |
| 9.029E-138 | -1.7657444 | 0.023 | 0.26 | 2.133E-133 | Tcells | CRYAB |
| 5.869E-137 | 0.37151719 | 0.453 | 0.226 | 1.386E-132 | Tcells | NCOR1 |

| | | | | | | |
|---|---|---|---|---|---|---|
| 7.333E-137 | -0.4984666 | 0.033 | 0.276 | 1.732E-132 | Tcells | GSN |
| 1.47E-136 | 0.42470053 | 0.801 | 0.591 | 3.472E-132 | Tcells | HINT1 |
| 4.266E-136 | 0.46261615 | 0.59 | 0.342 | 1.008E-131 | Tcells | AKAP13 |
| 4.885E-136 | 0.33202698 | 0.342 | 0.147 | 1.154E-131 | Tcells | SMC3 |
| 3.043E-135 | -0.5559636 | 0.046 | 0.292 | 7.189E-131 | Tcells | LYN |
| 3.986E-135 | 0.31261187 | 0.31 | 0.126 | 9.415E-131 | Tcells | TGFBR2 |
| 1.357E-134 | 0.28030301 | 0.317 | 0.127 | 3.205E-130 | Tcells | EVI2A |
| 3.083E-134 | -0.5258232 | 0.02 | 0.256 | 7.284E-130 | Tcells | RAB31 |
| 4.12E-134 | -0.9922163 | 0.319 | 0.55 | 9.733E-130 | Tcells | UPP1 |
| 4.277E-134 | 0.28748144 | 0.242 | 0.085 | 1.01E-129 | Tcells | DIP2A |
| 2.346E-133 | 0.26110927 | 0.243 | 0.087 | 5.541E-129 | Tcells | ORAI1 |
| 3.518E-133 | -0.6159581 | 0.075 | 0.324 | 8.31E-129 | Tcells | FNDC3B |
| 8.575E-133 | -0.7758927 | 0.016 | 0.246 | 2.026E-128 | Tcells | CDKN1C |
| 1.361E-132 | -0.5552178 | 0.001 | 0.225 | 3.215E-128 | Tcells | EHF |
| 1.586E-132 | -0.4489576 | 0.009 | 0.236 | 3.746E-128 | Tcells | ST14 |
| 1.773E-132 | 0.29603158 | 0.287 | 0.113 | 4.188E-128 | Tcells | STK10 |
| 3.226E-132 | 0.42960609 | 0.623 | 0.376 | 7.62E-128 | Tcells | HMGN2 |
| 4.628E-132 | -1.2667118 | 0.105 | 0.358 | 1.093E-127 | Tcells | CTSL |
| 5.521E-132 | -0.7307669 | 0.036 | 0.274 | 1.304E-127 | Tcells | HES4 |
| 7.583E-131 | -0.5227871 | 0.35 | 0.595 | 1.791E-126 | Tcells | CLTA |
| 8.872E-131 | -1.6679988 | 0.016 | 0.243 | 2.096E-126 | Tcells | PRSS22 |
| 1.16E-130 | -0.5518052 | 0.117 | 0.371 | 2.741E-126 | Tcells | PLSCR1 |
| 3.19E-130 | -0.603013 | 0.086 | 0.34 | 7.536E-126 | Tcells | LMNA |
| 3.701E-130 | -0.8188921 | 0.009 | 0.233 | 8.744E-126 | Tcells | HES1 |
| 5.315E-130 | 0.30086744 | 0.288 | 0.114 | 1.256E-125 | Tcells | ADD3 |
| 5.35E-130 | -0.5315075 | 0.005 | 0.228 | 1.264E-125 | Tcells | DSP |
| 6.071E-130 | -2.0729217 | 0.122 | 0.356 | 1.434E-125 | Tcells | HPGD |
| 7.06E-130 | 0.40092235 | 0.528 | 0.295 | 1.668E-125 | Tcells | ANAPC16 |
| 7.82E-130 | 0.33156679 | 0.33 | 0.141 | 1.847E-125 | Tcells | OGT |
| 2.07E-129 | -0.5460537 | 0.073 | 0.316 | 4.891E-125 | Tcells | BLVRB |
| 2.683E-129 | 0.45350842 | 0.542 | 0.303 | 6.338E-125 | Tcells | AKAP9 |
| 3.499E-129 | 0.34029764 | 0.436 | 0.217 | 8.267E-125 | Tcells | THRAP3 |
| 3.564E-129 | 0.2974047 | 0.347 | 0.151 | 8.419E-125 | Tcells | TMPO |
| 3.695E-129 | -0.4621017 | 0.019 | 0.247 | 8.73E-125 | Tcells | PLXNB2 |
| 4.756E-129 | -1.5535418 | 0.15 | 0.409 | 1.124E-124 | Tcells | HLA-DRA |
| 5.04E-129 | 0.38460169 | 0.571 | 0.323 | 1.191E-124 | Tcells | LUC7L3 |
| 9.881E-129 | 0.39514673 | 0.589 | 0.34 | 2.334E-124 | Tcells | EIF3E |
| 1.219E-128 | 0.32690039 | 0.34 | 0.148 | 2.879E-124 | Tcells | PRKDC |
| 3.063E-128 | -0.5651803 | 0.043 | 0.278 | 7.236E-124 | Tcells | AFDN |
| 5.734E-128 | 0.37061494 | 0.415 | 0.201 | 1.355E-123 | Tcells | VPS13C |
| 3.951E-127 | -0.6610886 | 0.049 | 0.281 | 9.333E-123 | Tcells | CCNG2 |
| 4.99E-127 | 0.39680273 | 0.592 | 0.346 | 1.179E-122 | Tcells | SNRPD2 |
| 5.305E-127 | 0.44091941 | 0.986 | 0.896 | 1.253E-122 | Tcells | RPS11 |
| 5.425E-127 | 0.35579664 | 0.418 | 0.201 | 1.281E-122 | Tcells | BDP1 |
| 7.027E-127 | 0.33147294 | 0.439 | 0.215 | 1.66E-122 | Tcells | RGS10 |
| 2.08E-126 | 0.36349037 | 0.325 | 0.141 | 4.914E-122 | Tcells | FAM129A |
| 2.203E-126 | -0.7936449 | 0.012 | 0.232 | 5.204E-122 | Tcells | CCND1 |
| 2.799E-126 | -0.7890145 | 0.018 | 0.24 | 6.612E-122 | Tcells | ITGA2 |
| 3.128E-126 | -0.7081396 | 0.25 | 0.492 | 7.39E-122 | Tcells | RIOK3 |
| 3.903E-126 | 0.25802118 | 0.262 | 0.099 | 9.22E-122 | Tcells | CFAP97 |

| | | | | | | |
|---|---|---|---|---|---|---|
| 5E-126 | 0.28231114 | 0.273 | 0.107 | 1.181E-121 | Tcells | SLC9A3R1 |
| 1.727E-125 | -0.6208533 | 0.01 | 0.229 | 4.079E-121 | Tcells | TGM2 |
| 7.803E-125 | 0.42997033 | 0.992 | 0.92 | 1.843E-120 | Tcells | RPS24 |
| 1.609E-124 | 0.27242636 | 0.298 | 0.122 | 3.8E-120 | Tcells | TMX4 |
| 2.088E-124 | 0.34748139 | 0.332 | 0.146 | 4.931E-120 | Tcells | RIF1 |
| 4.613E-124 | 0.39465323 | 0.258 | 0.101 | 1.09E-119 | Tcells | ITM2C |
| 7.771E-124 | -0.4025872 | 0.003 | 0.216 | 1.836E-119 | Tcells | CTTN |
| 9.035E-124 | 0.42931568 | 0.778 | 0.601 | 2.134E-119 | Tcells | HNRNPA1 |
| 1.333E-123 | -0.6225381 | 0.008 | 0.224 | 3.15E-119 | Tcells | LAMB3 |
| 1.507E-123 | 0.30090226 | 0.324 | 0.141 | 3.56E-119 | Tcells | PHF20 |
| 2.005E-123 | -0.4501237 | 0.01 | 0.225 | 4.737E-119 | Tcells | CNN3 |
| 1.11E-122 | -0.4850726 | 0.051 | 0.281 | 2.622E-118 | Tcells | QSOX1 |
| 1.486E-122 | 0.26167104 | 0.283 | 0.113 | 3.511E-118 | Tcells | RSBN1 |
| 2.618E-122 | 0.2894785 | 0.319 | 0.137 | 6.184E-118 | Tcells | TLK1 |
| 3.63E-122 | -0.5135767 | 0.018 | 0.235 | 8.576E-118 | Tcells | KYNU |
| 4.446E-122 | 0.32299842 | 0.456 | 0.23 | 1.05E-117 | Tcells | KANSL1 |
| 1.014E-121 | 0.33332287 | 0.399 | 0.195 | 2.395E-117 | Tcells | CHURC1 |
| 1.032E-121 | 0.3429929 | 0.452 | 0.232 | 2.439E-117 | Tcells | ZRANB2 |
| 1.203E-121 | 0.37640082 | 0.994 | 0.956 | 2.843E-117 | Tcells | RPL15 |
| 2.102E-121 | 0.35162936 | 0.645 | 0.384 | 4.966E-117 | Tcells | ARGLU1 |
| 3.596E-121 | -0.5827991 | 0.007 | 0.218 | 8.495E-117 | Tcells | MACC1 |
| 4.698E-121 | -0.5223377 | 0.057 | 0.286 | 1.11E-116 | Tcells | PLIN3 |
| 8.892E-121 | 0.51637977 | 0.477 | 0.255 | 2.101E-116 | Tcells | ID2 |
| 1.317E-120 | 0.39160543 | 0.444 | 0.234 | 3.112E-116 | Tcells | G3BP2 |
| 3.627E-120 | 0.34213439 | 0.187 | 0.061 | 8.569E-116 | Tcells | SESN3 |
| 3.754E-120 | 0.3186023 | 0.299 | 0.127 | 8.869E-116 | Tcells | ZBTB1 |
| 8.148E-120 | 0.3895477 | 0.487 | 0.265 | 1.925E-115 | Tcells | ROCK1 |
| 1.016E-119 | -0.6139172 | 0.064 | 0.29 | 2.4E-115 | Tcells | ABHD5 |
| 1.045E-119 | 0.3306396 | 0.267 | 0.107 | 2.469E-115 | Tcells | GNPTAB |
| 5.51E-119 | 0.3903625 | 0.642 | 0.402 | 1.302E-114 | Tcells | EIF3K |
| 2.392E-118 | -0.7633186 | 0.063 | 0.287 | 5.65E-114 | Tcells | CAMK2N1 |
| 2.794E-118 | -1.1535111 | 0.057 | 0.274 | 6.599E-114 | Tcells | LMO7 |
| 9.117E-118 | 0.41099144 | 0.594 | 0.355 | 2.154E-113 | Tcells | MYH9 |
| 3.914E-117 | -0.4859751 | 0.38 | 0.612 | 9.245E-113 | Tcells | YWHAE |
| 5.21E-117 | -1.1165567 | 0.017 | 0.226 | 1.231E-112 | Tcells | S100A12 |
| 6.235E-117 | 0.32480339 | 0.367 | 0.174 | 1.473E-112 | Tcells | RPS6KA3 |
| 3.635E-116 | -0.5359364 | 0.017 | 0.225 | 8.586E-112 | Tcells | CKB |
| 4.708E-116 | 0.28431264 | 0.299 | 0.128 | 1.112E-111 | Tcells | MIA3 |
| 4.873E-116 | 0.38055879 | 0.555 | 0.325 | 1.151E-111 | Tcells | SRRM1 |
| 5.622E-116 | 0.28391625 | 0.308 | 0.134 | 1.328E-111 | Tcells | KAT6A |
| 6.612E-116 | 0.45034256 | 0.835 | 0.697 | 1.562E-111 | Tcells | RPL4 |
| 2.231E-115 | -0.4961045 | 0.006 | 0.208 | 5.27E-111 | Tcells | PTGR1 |
| 2.692E-115 | -0.706108 | 0.013 | 0.218 | 6.359E-111 | Tcells | CDKN2B |
| 3.83E-115 | 0.27698816 | 0.285 | 0.12 | 9.048E-111 | Tcells | R3HDM2 |
| 1.001E-114 | -0.936826 | 0.249 | 0.505 | 2.364E-110 | Tcells | MTRNR2L8 |
| 1.396E-114 | 0.30421032 | 0.352 | 0.163 | 3.299E-110 | Tcells | CEBPZ |
| 1.488E-114 | 0.28054712 | 0.297 | 0.127 | 3.516E-110 | Tcells | CSNK1G2 |
| 2.429E-114 | 0.2781583 | 0.285 | 0.12 | 5.737E-110 | Tcells | UBTF |
| 2.835E-114 | -0.379053 | 0.005 | 0.205 | 6.698E-110 | Tcells | RBPMS |
| 3.48E-114 | 0.40986642 | 0.558 | 0.334 | 8.221E-110 | Tcells | SCAF11 |

| | | | | | | |
|---|---|---|---|---|---|---|
| 6.224E-114 | -0.5249513 | 0.311 | 0.545 | 1.47E-109 | Tcells | P4HB |
| 1.133E-113 | 0.30451098 | 0.343 | 0.159 | 2.678E-109 | Tcells | DNAJB14 |
| 2.367E-113 | 0.45040472 | 0.695 | 0.496 | 5.591E-109 | Tcells | JAK1 |
| 2.488E-113 | 0.39854911 | 0.492 | 0.278 | 5.877E-109 | Tcells | ARID4B |
| 3.325E-113 | -0.419438 | 0.007 | 0.206 | 7.856E-109 | Tcells | NEDD4L |
| 3.718E-113 | 0.25510551 | 0.277 | 0.114 | 8.783E-109 | Tcells | CPNE1 |
| 4.332E-113 | 0.25122761 | 0.252 | 0.098 | 1.023E-108 | Tcells | PRKCB |
| 6.277E-113 | 0.26865262 | 0.309 | 0.134 | 1.483E-108 | Tcells | RASSF5 |
| 7.077E-113 | 0.39575719 | 0.982 | 0.876 | 1.672E-108 | Tcells | RPL12 |
| 1.807E-112 | 0.31205883 | 0.535 | 0.297 | 4.268E-108 | Tcells | IRF1 |
| 1.874E-112 | 0.35936204 | 0.895 | 0.718 | 4.427E-108 | Tcells | CYBA |
| 9.572E-112 | -0.6139381 | 0.542 | 0.717 | 2.261E-107 | Tcells | GPX4 |
| 1.999E-111 | -0.7004957 | 0.11 | 0.338 | 4.722E-107 | Tcells | CEBPD |
| 3.344E-111 | 0.28606358 | 0.306 | 0.133 | 7.899E-107 | Tcells | TUBA4A |
| 3.495E-111 | -0.9564051 | 0.01 | 0.207 | 8.255E-107 | Tcells | TM4SF1 |
| 5.605E-111 | -0.8599219 | 0.111 | 0.33 | 1.324E-106 | Tcells | GADD45A |
| 6.489E-111 | 0.29666277 | 0.307 | 0.135 | 1.533E-106 | Tcells | OTUD6B-AS1 |
| 7.172E-111 | -0.6033746 | 0.071 | 0.286 | 1.694E-106 | Tcells | ETS2 |
| 1.722E-110 | -0.7085445 | 0.009 | 0.206 | 4.067E-106 | Tcells | SFN |
| 2.746E-110 | -0.6015798 | 0.021 | 0.223 | 6.487E-106 | Tcells | BASP1 |
| 3.542E-110 | -0.3558405 | 0.018 | 0.219 | 8.368E-106 | Tcells | NECTIN2 |
| 1.5E-109 | 0.27757762 | 0.285 | 0.122 | 3.544E-105 | Tcells | NUCB2 |
| 2.337E-109 | 0.26020944 | 0.237 | 0.092 | 5.521E-105 | Tcells | OTULIN |
| 2.464E-109 | 0.26408211 | 0.363 | 0.169 | 5.821E-105 | Tcells | FMNL1 |
| 5.386E-109 | -0.9414137 | 0.005 | 0.197 | 1.272E-104 | Tcells | TMPRSS2 |
| 1.619E-108 | 0.28622657 | 0.311 | 0.14 | 3.825E-104 | Tcells | PTPRA |
| 2.396E-108 | 0.30922957 | 0.362 | 0.176 | 5.66E-104 | Tcells | PSMA3-AS1 |
| 2.809E-108 | 0.35851976 | 0.877 | 0.78 | 6.636E-104 | Tcells | BTF3 |
| 3.028E-108 | -0.8097746 | 0.427 | 0.641 | 7.154E-104 | Tcells | ZFP36L1 |
| 4.516E-108 | -0.511773 | 0.001 | 0.19 | 1.067E-103 | Tcells | S100A14 |
| 7.586E-108 | 0.34470354 | 0.858 | 0.676 | 1.792E-103 | Tcells | NPM1 |
| 9.268E-108 | 0.36622042 | 0.773 | 0.598 | 2.189E-103 | Tcells | SON |
| 1.452E-107 | -0.3897564 | 0.013 | 0.207 | 3.43E-103 | Tcells | TCEAL9 |
| 3.494E-107 | 0.30451008 | 0.366 | 0.178 | 8.255E-103 | Tcells | EPC1 |
| 3.657E-107 | 0.28354882 | 0.335 | 0.157 | 8.639E-103 | Tcells | SNRNP200 |
| 4.132E-107 | 0.28468027 | 0.311 | 0.141 | 9.76E-103 | Tcells | TAF15 |
| 4.585E-107 | -0.370519 | 0.023 | 0.223 | 1.083E-102 | Tcells | CTNND1 |
| 5.494E-107 | -0.5109293 | 0.863 | 0.554 | 1.298E-102 | Tcells | SRGN |
| 6.185E-107 | 0.41554302 | 0.387 | 0.198 | 1.461E-102 | Tcells | RNF19A |
| 9.088E-107 | -0.6157314 | 0.312 | 0.524 | 2.147E-102 | Tcells | ANXA11 |
| 1.208E-106 | 0.34682262 | 0.436 | 0.235 | 2.854E-102 | Tcells | PPP1R12A |
| 1.297E-106 | -1.6649421 | 0.016 | 0.208 | 3.064E-102 | Tcells | KRT17 |
| 3.188E-106 | -0.6931007 | 0.008 | 0.198 | 7.532E-102 | Tcells | EPS8L1 |
| 4.79E-106 | -0.8783599 | 0.96 | 0.962 | 1.132E-101 | Tcells | HSP90AA1 |
| 1.434E-105 | -0.380159 | 0.003 | 0.189 | 3.387E-101 | Tcells | FNBP1L |
| 2.326E-105 | -0.4429162 | 0.005 | 0.192 | 5.495E-101 | Tcells | CDH1 |
| 7.616E-105 | 0.25560293 | 0.256 | 0.106 | 1.799E-100 | Tcells | PLEKHA2 |
| 8.627E-105 | 0.33112708 | 0.532 | 0.309 | 2.038E-100 | Tcells | ERP29 |
| 9.66E-105 | 0.28034248 | 0.308 | 0.139 | 2.282E-100 | Tcells | SVIP |
| 4.596E-104 | 0.36364059 | 0.785 | 0.62 | 1.086E-99 | Tcells | OST4 |

| | | | | | | |
|---|---|---|---|---|---|---|
| 5.427E-104 | -0.4071308 | 0.002 | 0.185 | 1.282E-99 | Tcells | SMIM22 |
| 7.573E-104 | -0.4357881 | 0.016 | 0.206 | 1.789E-99 | Tcells | KLF4 |
| 8.284E-104 | -0.3609864 | 0.008 | 0.194 | 1.957E-99 | Tcells | CDCP1 |
| 1.272E-103 | -0.7637244 | 0.055 | 0.259 | 3.004E-99 | Tcells | EGR1 |
| 1.602E-103 | -0.3644274 | 0.002 | 0.185 | 3.785E-99 | Tcells | TMEM54 |
| 1.639E-103 | -0.7688698 | 0.389 | 0.613 | 3.872E-99 | Tcells | NFKBIA |
| 1.689E-103 | -0.3947943 | 0.005 | 0.189 | 3.99E-99 | Tcells | TEAD1 |
| 2.241E-103 | 0.36400054 | 0.547 | 0.324 | 5.294E-99 | Tcells | RBM25 |
| 2.574E-103 | 0.34869911 | 0.72 | 0.496 | 6.081E-99 | Tcells | HNRNPDL |
| 3.094E-103 | -0.6160119 | 0.135 | 0.349 | 7.309E-99 | Tcells | NDRG1 |
| 3.15E-103 | -1.0341426 | 0.084 | 0.289 | 7.442E-99 | Tcells | DUSP5 |
| 3.509E-103 | -0.4100694 | 0.005 | 0.189 | 8.29E-99 | Tcells | ANXA3 |
| 3.544E-103 | 0.25754354 | 0.309 | 0.139 | 8.372E-99 | Tcells | DDX3Y |
| 4.429E-103 | 0.32851533 | 0.431 | 0.234 | 1.0462E-98 | Tcells | TERF2IP |
| 5.346E-103 | 0.31428683 | 0.459 | 0.252 | 1.2629E-98 | Tcells | HNRNPR |
| 5.875E-103 | 0.25268682 | 0.239 | 0.096 | 1.3879E-98 | Tcells | ITGAE |
| 6.233E-103 | 0.36583632 | 0.524 | 0.314 | 1.4725E-98 | Tcells | NSA2 |
| 6.66E-103 | 0.34446888 | 0.918 | 0.81 | 1.5733E-98 | Tcells | PFDN5 |
| 6.832E-103 | -0.5359381 | 0.005 | 0.188 | 1.614E-98 | Tcells | EPHA2 |
| 1.05E-102 | 0.28646037 | 0.299 | 0.135 | 2.4796E-98 | Tcells | RASA2 |
| 1.334E-102 | -0.7652313 | 0.321 | 0.536 | 3.152E-98 | Tcells | CEBPB |
| 1.535E-102 | 0.27804758 | 0.348 | 0.169 | 3.6252E-98 | Tcells | REX1BD |
| 2.224E-102 | -0.3958906 | 0.011 | 0.196 | 5.2532E-98 | Tcells | PAWR |
| 2.286E-102 | 0.34520054 | 0.506 | 0.294 | 5.3994E-98 | Tcells | HP1BP3 |
| 3.129E-102 | 0.40655452 | 0.773 | 0.614 | 7.392E-98 | Tcells | PRRC2C |
| 3.941E-102 | -0.8972132 | 0.221 | 0.433 | 9.3092E-98 | Tcells | TXNRD1 |
| 5.37E-102 | -0.5251855 | 0.086 | 0.294 | 1.2687E-97 | Tcells | CD151 |
| 9.367E-102 | -0.5291879 | 0.116 | 0.33 | 2.2127E-97 | Tcells | SLC16A3 |
| 2.135E-101 | 0.34508106 | 0.512 | 0.304 | 5.0428E-97 | Tcells | EIF3F |
| 2.872E-101 | 0.44961452 | 0.42 | 0.228 | 6.784E-97 | Tcells | TNFAIP3 |
| 2.89E-101 | -1.2564492 | 0.011 | 0.193 | 6.8276E-97 | Tcells | DUOX2 |
| 3.411E-101 | -0.3288897 | 0.001 | 0.179 | 8.0581E-97 | Tcells | GNG12 |
| 5.913E-101 | 0.33477454 | 0.491 | 0.284 | 1.3967E-96 | Tcells | EIF3G |
| 7.571E-101 | -0.5908224 | 0.01 | 0.192 | 1.7885E-96 | Tcells | SDCBP2 |
| 1.148E-100 | -0.5218276 | 0.068 | 0.269 | 2.711E-96 | Tcells | WASL |
| 1.498E-100 | -0.6945584 | 0.084 | 0.292 | 3.5398E-96 | Tcells | RHOB |
| 1.522E-100 | 0.33482305 | 0.548 | 0.332 | 3.5958E-96 | Tcells | SSR2 |
| 1.857E-100 | -0.549637 | 0.005 | 0.184 | 4.3863E-96 | Tcells | C6orf132 |
| 3.728E-100 | -0.949247 | 0.038 | 0.229 | 8.8064E-96 | Tcells | CD14 |
| 5.114E-100 | 0.256556 | 0.194 | 0.07 | 1.2081E-95 | Tcells | XAF1 |
| 7.918E-100 | -0.5289363 | 0.023 | 0.21 | 1.8704E-95 | Tcells | NCF2 |
| 1.473E-99 | -0.6173943 | 0.003 | 0.18 | 3.4804E-95 | Tcells | HSPB8 |
| 4.279E-99 | -0.8236955 | 0.01 | 0.191 | 1.0109E-94 | Tcells | LAMC2 |
| 7.233E-99 | 0.27358098 | 0.271 | 0.118 | 1.7087E-94 | Tcells | RALGAPA1 |
| 1.1283E-98 | -0.6278603 | 0.006 | 0.183 | 2.6654E-94 | Tcells | SPNS2 |
| 1.2566E-98 | -1.215742 | 0.011 | 0.19 | 2.9684E-94 | Tcells | SPINK1 |
| 1.4354E-98 | -0.8757402 | 0.01 | 0.189 | 3.3908E-94 | Tcells | F3 |
| 2.3215E-98 | -0.4033518 | 0.002 | 0.177 | 5.4841E-94 | Tcells | NECTIN4 |
| 2.4571E-98 | -0.5140244 | 0.068 | 0.268 | 5.8044E-94 | Tcells | TSC22D1 |
| 2.5168E-98 | 0.39602529 | 0.697 | 0.508 | 5.9455E-94 | Tcells | NAP1L1 |

| | | | | | | |
|---|---|---|---|---|---|---|
| 3.6348E-98 | 0.27781591 | 0.387 | 0.199 | 8.5866E-94 | Tcells | RBMX |
| 4.3058E-98 | -0.4869585 | 0.068 | 0.267 | 1.0172E-93 | Tcells | PDLIM5 |
| 6.9082E-98 | -0.4902317 | 0.029 | 0.215 | 1.6319E-93 | Tcells | PARD6B |
| 1.3114E-97 | -0.3173163 | 0.009 | 0.187 | 3.0979E-93 | Tcells | IL13RA1 |
| 1.869E-97 | 0.29305466 | 0.327 | 0.159 | 4.4152E-93 | Tcells | HNRNPUL1 |
| 1.8961E-97 | 0.29162324 | 0.567 | 0.338 | 4.4792E-93 | Tcells | PNN |
| 1.9811E-97 | -0.5040886 | 0.099 | 0.303 | 4.68E-93 | Tcells | TUBA1C |
| 2.1648E-97 | -0.3258055 | 0.021 | 0.206 | 5.1139E-93 | Tcells | SWAP70 |
| 2.533E-97 | -0.4916228 | 0.008 | 0.185 | 5.9837E-93 | Tcells | FHL2 |
| 2.9681E-97 | -0.6759829 | 0.006 | 0.18 | 7.0114E-93 | Tcells | STEAP4 |
| 3.5906E-97 | -0.3875158 | 0.07 | 0.272 | 8.4821E-93 | Tcells | CHP1 |
| 5.4302E-97 | 0.3480037 | 0.538 | 0.33 | 1.2828E-92 | Tcells | ANP32B |
| 9.5243E-97 | -0.8603026 | 0.062 | 0.254 | 2.2499E-92 | Tcells | SLC11A1 |
| 1.1241E-96 | -0.6115408 | 0.006 | 0.181 | 2.6553E-92 | Tcells | PPL |
| 2.1466E-96 | 0.32994753 | 0.527 | 0.32 | 5.0709E-92 | Tcells | TCF25 |
| 3.8197E-96 | 0.30697082 | 0.264 | 0.113 | 9.0233E-92 | Tcells | TSPYL2 |
| 3.9574E-96 | 0.31762125 | 0.634 | 0.404 | 9.3487E-92 | Tcells | FUS |
| 4.413E-96 | -0.3636881 | 0.058 | 0.255 | 1.0425E-91 | Tcells | PEA15 |
| 4.9726E-96 | -1.2727061 | 0.045 | 0.232 | 1.1747E-91 | Tcells | G0S2 |
| 6.4044E-96 | 0.26975195 | 0.342 | 0.168 | 1.5129E-91 | Tcells | PURA |
| 1.4631E-95 | 0.29146371 | 0.318 | 0.151 | 3.4562E-91 | Tcells | DUT |
| 2.2303E-95 | 0.36349939 | 0.69 | 0.483 | 5.2687E-91 | Tcells | RSRP1 |
| 2.5472E-95 | -0.3925537 | 0.047 | 0.238 | 6.0172E-91 | Tcells | PGD |
| 4.3068E-95 | 0.30251882 | 0.352 | 0.177 | 1.0174E-90 | Tcells | CEP350 |
| 5.3023E-95 | 0.25893586 | 0.309 | 0.145 | 1.2526E-90 | Tcells | WDR82 |
| 7.6132E-95 | -0.6433245 | 0.006 | 0.177 | 1.7985E-90 | Tcells | FXYD3 |
| 2.6773E-94 | 0.29789645 | 0.438 | 0.242 | 6.3247E-90 | Tcells | SRSF10 |
| 1.1022E-93 | 0.37375253 | 0.725 | 0.544 | 2.6037E-89 | Tcells | HNRNPA3 |
| 1.7818E-93 | -0.5603011 | 0.022 | 0.2 | 4.2091E-89 | Tcells | PLK2 |
| 2.3604E-93 | -0.3669461 | 0.017 | 0.192 | 5.576E-89 | Tcells | DAB2 |
| 3.8725E-93 | -0.3625992 | 0.003 | 0.17 | 9.1479E-89 | Tcells | MET |
| 4.9026E-93 | -0.3365527 | 0.023 | 0.203 | 1.1582E-88 | Tcells | KIFC3 |
| 5.6373E-93 | -0.4090774 | 0.081 | 0.279 | 1.3317E-88 | Tcells | UBE2D1 |
| 5.8845E-93 | 0.25883549 | 0.29 | 0.133 | 1.3901E-88 | Tcells | RABGAP1L |
| 6.1262E-93 | -0.3622269 | 0.022 | 0.2 | 1.4472E-88 | Tcells | F11R |
| 7.6164E-93 | -0.3993388 | 0.007 | 0.177 | 1.7992E-88 | Tcells | TINAGL1 |
| 1.2024E-92 | 0.27918303 | 0.427 | 0.233 | 2.8404E-88 | Tcells | LSM7 |
| 1.3555E-92 | -0.3245819 | 0.006 | 0.175 | 3.2021E-88 | Tcells | MYOF |
| 2.849E-92 | -0.4952873 | 0.12 | 0.321 | 6.7301E-88 | Tcells | SMS |
| 5.5855E-92 | 0.3107995 | 0.444 | 0.254 | 1.3195E-87 | Tcells | IK |
| 5.9359E-92 | -0.4476997 | 0.002 | 0.167 | 1.4022E-87 | Tcells | SDC1 |
| 6.4549E-92 | -0.3402338 | 0.007 | 0.176 | 1.5249E-87 | Tcells | CXADR |
| 6.5016E-92 | 0.31170429 | 0.574 | 0.359 | 1.5359E-87 | Tcells | CIRBP |
| 8.5973E-92 | 0.28366643 | 0.383 | 0.204 | 2.0309E-87 | Tcells | LSM8 |
| 1.4139E-91 | 0.31257692 | 0.531 | 0.323 | 3.3401E-87 | Tcells | HNRNPD |
| 1.5033E-91 | -0.5236339 | 0.002 | 0.167 | 3.5513E-87 | Tcells | MALL |
| 2.0891E-91 | -0.5001708 | 0.041 | 0.225 | 4.9351E-87 | Tcells | SPI1 |
| 2.7973E-91 | -0.5337496 | 0.017 | 0.188 | 6.6081E-87 | Tcells | DST |
| 3.928E-91 | -1.0862866 | 0.004 | 0.168 | 9.2791E-87 | Tcells | CEACAM5 |
| 1.2443E-90 | -1.2950088 | 0.008 | 0.174 | 2.9393E-86 | Tcells | DUOXA2 |

| | | | | | | |
|---|---|---|---|---|---|---|
| 1.2989E-90 | 0.25519009 | 0.387 | 0.203 | 3.0683E-86 | Tcells | RPL17 |
| 2.2432E-90 | 0.25978813 | 0.257 | 0.115 | 5.299E-86 | Tcells | CCNH |
| 2.272E-90 | -0.3806521 | 0.017 | 0.187 | 5.3673E-86 | Tcells | TMEM176B |
| 2.3859E-90 | 0.35913591 | 0.961 | 0.852 | 5.6362E-86 | Tcells | RPL7 |
| 2.8514E-90 | 0.36805392 | 0.763 | 0.585 | 6.7359E-86 | Tcells | DDX17 |
| 4.3576E-90 | -0.5131776 | 0.166 | 0.369 | 1.0294E-85 | Tcells | RRBP1 |
| 4.7411E-90 | 0.34077939 | 0.984 | 0.891 | 1.12E-85 | Tcells | RPS9 |
| 5.1778E-90 | -0.4048522 | 0.002 | 0.164 | 1.2232E-85 | Tcells | COBLL1 |
| 5.5981E-90 | -0.3167587 | 0.008 | 0.173 | 1.3224E-85 | Tcells | DDR1 |
| 1.2749E-89 | -0.4446431 | 0.003 | 0.164 | 3.0117E-85 | Tcells | CLDN3 |
| 3.9707E-89 | -0.845973 | 0.083 | 0.269 | 9.3801E-85 | Tcells | TPM1 |
| 4.8652E-89 | -0.522503 | 0.025 | 0.197 | 1.1493E-84 | Tcells | FCGR2A |
| 7.2162E-89 | 0.28424332 | 0.364 | 0.189 | 1.7047E-84 | Tcells | CHD1 |
| 1.5285E-88 | -0.3958685 | 0.004 | 0.164 | 3.6108E-84 | Tcells | TSPAN6 |
| 2.2272E-88 | -0.3382032 | 0.026 | 0.2 | 5.2613E-84 | Tcells | FCHO2 |
| 2.3137E-88 | -0.675183 | 0.034 | 0.209 | 5.4657E-84 | Tcells | CYBB |
| 2.4585E-88 | 0.25521586 | 0.638 | 0.391 | 5.8077E-84 | Tcells | SOD1 |
| 2.7234E-88 | -1.1673973 | 0.007 | 0.169 | 6.4334E-84 | Tcells | MUC21 |
| 3.0033E-88 | -0.3526603 | 0.014 | 0.18 | 7.0948E-84 | Tcells | B3GNT5 |
| 4.1067E-88 | -0.676513 | 0.19 | 0.394 | 9.7012E-84 | Tcells | TYMP |
| 5.9134E-88 | -0.4984706 | 0.138 | 0.335 | 1.3969E-83 | Tcells | TMBIM1 |
| 1.4136E-87 | -0.3036588 | 0.008 | 0.169 | 3.3392E-83 | Tcells | SORT1 |
| 1.4636E-87 | 0.3313678 | 0.509 | 0.306 | 3.4575E-83 | Tcells | BOD1L1 |
| 1.8604E-87 | -0.8532117 | 0.017 | 0.181 | 4.3949E-83 | Tcells | NDRG2 |
| 1.8906E-87 | 0.30703808 | 0.538 | 0.322 | 4.4661E-83 | Tcells | TLN1 |
| 1.9162E-87 | 0.33514382 | 0.551 | 0.349 | 4.5266E-83 | Tcells | SRSF11 |
| 2.9339E-87 | 0.27066595 | 0.31 | 0.153 | 6.9307E-83 | Tcells | XRN1 |
| 4.4336E-87 | -0.4577214 | 0.004 | 0.162 | 1.0473E-82 | Tcells | TMPRSS4 |
| 4.7808E-87 | -0.7787215 | 0.003 | 0.161 | 1.1294E-82 | Tcells | SCEL |
| 5.3828E-87 | -1.621452 | 0.056 | 0.231 | 1.2716E-82 | Tcells | VCAN |
| 1.7034E-86 | -0.4997921 | 0.005 | 0.163 | 4.0239E-82 | Tcells | CYR61 |
| 3.1261E-86 | -0.5808083 | 0.028 | 0.197 | 7.3847E-82 | Tcells | HBEGF |
| 4.6421E-86 | -1.6177565 | 0.011 | 0.17 | 1.0966E-81 | Tcells | ECM1 |
| 1.0585E-85 | 0.28688327 | 0.381 | 0.207 | 2.5004E-81 | Tcells | RWDD1 |
| 1.2458E-85 | -0.3234667 | 0.008 | 0.167 | 2.9428E-81 | Tcells | CRB3 |
| 1.276E-85 | 0.33711378 | 0.611 | 0.424 | 3.0142E-81 | Tcells | TRIR |
| 1.3769E-85 | 0.27520611 | 0.365 | 0.193 | 3.2527E-81 | Tcells | STAG2 |
| 1.803E-85 | -0.6121444 | 0.005 | 0.162 | 4.2593E-81 | Tcells | CDC42EP5 |
| 1.9E-85 | -0.4385075 | 0.748 | 0.883 | 4.4884E-81 | Tcells | TMBIM6 |
| 3.28E-85 | -0.4937312 | 0.001 | 0.154 | 7.7483E-81 | Tcells | ALDH1A3 |
| 6.8994E-85 | 0.30550523 | 0.857 | 0.758 | 1.6299E-80 | Tcells | COX7C |
| 7.6554E-85 | 0.25706868 | 0.292 | 0.141 | 1.8084E-80 | Tcells | BAZ1B |
| 8.5584E-85 | 0.34183299 | 0.787 | 0.653 | 2.0217E-80 | Tcells | RBM39 |
| 1.1149E-84 | -0.6703549 | 0.007 | 0.163 | 2.6338E-80 | Tcells | MUC4 |
| 1.2614E-84 | -0.8792963 | 0.007 | 0.163 | 2.9799E-80 | Tcells | KRT15 |
| 3.039E-84 | -0.6587425 | 0.267 | 0.442 | 7.1791E-80 | Tcells | GSTO1 |
| 3.2945E-84 | -1.0016779 | 0.009 | 0.165 | 7.7826E-80 | Tcells | PRSS27 |
| 3.5593E-84 | -0.3412529 | 0.02 | 0.184 | 8.4081E-80 | Tcells | ABCA1 |
| 3.6366E-84 | -0.5346313 | 0.172 | 0.359 | 8.5907E-80 | Tcells | UBE2H |
| 5.3618E-84 | -0.9980578 | 0.007 | 0.162 | 1.2666E-79 | Tcells | PITX1 |

| | | | | | | |
|---|---|---|---|---|---|---|
| 1.1733E-83 | -0.2920478 | 0.014 | 0.175 | 2.7718E-79 | Tcells | SIRPA |
| 1.2829E-83 | 0.38698105 | 0.501 | 0.305 | 3.0307E-79 | Tcells | HMGB2 |
| 1.2984E-83 | -0.274052 | 0.011 | 0.169 | 3.0672E-79 | Tcells | RAB32 |
| 2.1047E-83 | 0.26816706 | 0.309 | 0.154 | 4.9719E-79 | Tcells | NASP |
| 2.8077E-83 | -0.8801103 | 0.112 | 0.294 | 6.6326E-79 | Tcells | SLC25A37 |
| 2.9583E-83 | -0.3131014 | 0.004 | 0.157 | 6.9885E-79 | Tcells | PLS3 |
| 3.9861E-83 | 0.27282511 | 0.367 | 0.197 | 9.4165E-79 | Tcells | ADAR |
| 4.852E-83 | 0.27262842 | 0.323 | 0.164 | 1.1462E-78 | Tcells | BBX |
| 5.383E-83 | -0.889121 | 0.003 | 0.154 | 1.2716E-78 | Tcells | CEACAM6 |
| 7.465E-83 | 0.30879671 | 0.439 | 0.256 | 1.7635E-78 | Tcells | TMEM123 |
| 7.9449E-83 | -0.6674117 | 0.003 | 0.154 | 1.8768E-78 | Tcells | NCCRP1 |
| 8.036E-83 | -0.4175431 | 0.605 | 0.781 | 1.8983E-78 | Tcells | EIF4G2 |
| 1.0386E-82 | -0.3021193 | 0.003 | 0.154 | 2.4536E-78 | Tcells | KCNK1 |
| 1.2247E-82 | -0.4264687 | 0.003 | 0.153 | 2.893E-78 | Tcells | VSIG2 |
| 1.5004E-82 | -0.5103345 | 0.045 | 0.215 | 3.5445E-78 | Tcells | GK |
| 2.669E-82 | -0.5657784 | 0.112 | 0.293 | 6.3049E-78 | Tcells | MARCKSL1 |
| 2.7848E-82 | -0.3155866 | 0.002 | 0.152 | 6.5786E-78 | Tcells | SLC44A4 |
| 3.7353E-82 | -0.297625 | 0.002 | 0.152 | 8.8239E-78 | Tcells | GPRC5C |
| 4.0303E-82 | 0.30870583 | 0.27 | 0.125 | 9.5208E-78 | Tcells | RGCC |
| 9.5477E-82 | -0.8448774 | 0.004 | 0.153 | 2.2554E-77 | Tcells | CEACAM7 |
| 1.0106E-81 | -0.3530551 | 0.034 | 0.2 | 2.3873E-77 | Tcells | ITPRIPL2 |
| 1.178E-81 | -0.3661162 | 0 | 0.147 | 2.7828E-77 | Tcells | EMP2 |
| 1.226E-81 | -0.771749 | 0.006 | 0.156 | 2.8963E-77 | Tcells | TMPRSS11D |
| 1.6787E-81 | -0.2861175 | 0.011 | 0.164 | 3.9655E-77 | Tcells | HNMT |
| 2.0748E-81 | -0.7942083 | 0.003 | 0.152 | 4.9014E-77 | Tcells | TMPRSS11E |
| 2.1355E-81 | 0.29793353 | 0.397 | 0.225 | 5.0448E-77 | Tcells | PSMB8 |
| 4.3966E-81 | -0.3713843 | 0.003 | 0.152 | 1.0386E-76 | Tcells | CLDN1 |
| 9.3009E-81 | -0.9440177 | 0.491 | 0.653 | 2.1972E-76 | Tcells | HSPA5 |
| 1.1557E-80 | -0.4548148 | 0.097 | 0.279 | 2.7302E-76 | Tcells | FAM3C |
| 1.2257E-80 | -0.4004581 | 0.003 | 0.151 | 2.8954E-76 | Tcells | ITGB6 |
| 1.24E-80 | 0.34841105 | 0.998 | 0.993 | 2.9294E-76 | Tcells | TPT1 |
| 1.2607E-80 | -0.4114488 | 0.004 | 0.152 | 2.9781E-76 | Tcells | SDC2 |
| 1.9119E-80 | -0.2982975 | 0.012 | 0.164 | 4.5165E-76 | Tcells | CTTNBP2NL |
| 2.2916E-80 | 0.29020052 | 0.377 | 0.206 | 5.4135E-76 | Tcells | ARID1B |
| 2.4217E-80 | -1.0792679 | 0.042 | 0.208 | 5.7207E-76 | Tcells | THBS1 |
| 2.8442E-80 | -0.3339647 | 0.04 | 0.206 | 6.7189E-76 | Tcells | ITGAV |
| 2.958E-80 | -0.4491197 | 0.023 | 0.181 | 6.9877E-76 | Tcells | SMIM25 |
| 4.1318E-80 | 0.31163361 | 0.403 | 0.23 | 9.7606E-76 | Tcells | GSPT1 |
| 4.2647E-80 | -0.264993 | 0.003 | 0.149 | 1.0074E-75 | Tcells | BACE2 |
| 6.574E-80 | -0.2694836 | 0.008 | 0.158 | 1.553E-75 | Tcells | SHTN1 |
| 7.8381E-80 | -0.8347787 | 0.279 | 0.462 | 1.8516E-75 | Tcells | HSPH1 |
| 9.5437E-80 | 0.29410103 | 0.813 | 0.684 | 2.2545E-75 | Tcells | PNRC1 |
| 1.6649E-79 | -0.3279919 | 0.03 | 0.191 | 3.933E-75 | Tcells | GNA15 |
| 1.8466E-79 | -0.5971119 | 0.361 | 0.545 | 4.3622E-75 | Tcells | SERPINB1 |
| 3.09E-79 | -0.3782141 | 0.009 | 0.158 | 7.2996E-75 | Tcells | MDK |
| 3.9382E-79 | -0.4633157 | 0.048 | 0.215 | 9.3033E-75 | Tcells | SLC43A2 |
| 4.9445E-79 | -0.5693219 | 0.032 | 0.192 | 1.168E-74 | Tcells | C5AR1 |
| 6.3759E-79 | -0.4239513 | 0.032 | 0.192 | 1.5062E-74 | Tcells | ERRFI1 |
| 7.7147E-79 | -0.3295284 | 0.013 | 0.163 | 1.8225E-74 | Tcells | ERBB3 |
| 8.8104E-79 | 0.26813963 | 0.958 | 0.865 | 2.0813E-74 | Tcells | RPLP0 |

| | | | | | | |
|---|---|---|---|---|---|---|
| 1.2963E-78 | -0.4462553 | 0.079 | 0.25 | 3.0623E-74 | Tcells | GIPC1 |
| 1.5033E-78 | -0.5231441 | 0.42 | 0.584 | 3.5513E-74 | Tcells | PRDX5 |
| 1.9908E-78 | 0.28714243 | 0.922 | 0.826 | 4.7028E-74 | Tcells | HNRNPA2B1 |
| 2.1947E-78 | -0.2869694 | 0.003 | 0.147 | 5.1844E-74 | Tcells | RAB25 |
| 2.5426E-78 | -0.3786677 | 0.026 | 0.182 | 6.0063E-74 | Tcells | KCTD12 |
| 3.2904E-78 | -0.6429146 | 0.004 | 0.148 | 7.7729E-74 | Tcells | SNCG |
| 4.3541E-78 | -0.2535261 | 0.019 | 0.174 | 1.0286E-73 | Tcells | CYFIP1 |
| 6.4757E-78 | 0.27680906 | 0.438 | 0.258 | 1.5297E-73 | Tcells | EID1 |
| 7.7579E-78 | -0.4239418 | 0.002 | 0.144 | 1.8327E-73 | Tcells | DCDC2 |
| 8.8107E-78 | 0.30038111 | 0.409 | 0.238 | 2.0814E-73 | Tcells | SMARCA5 |
| 9.1392E-78 | 0.37962682 | 0.348 | 0.189 | 2.159E-73 | Tcells | BIRC3 |
| 9.4848E-78 | 0.28621018 | 0.619 | 0.417 | 2.2406E-73 | Tcells | SRSF5 |
| 9.5131E-78 | -0.4907832 | 0.032 | 0.19 | 2.2473E-73 | Tcells | FBP1 |
| 1.6487E-77 | -0.3790082 | 0.144 | 0.33 | 3.8948E-73 | Tcells | LEPROT |
| 2.4868E-77 | 0.26609299 | 0.4 | 0.226 | 5.8745E-73 | Tcells | FNBP4 |
| 2.5821E-77 | -0.428587 | 0.006 | 0.15 | 6.0996E-73 | Tcells | PAX8 |
| 2.7825E-77 | 0.27624046 | 0.457 | 0.275 | 6.5732E-73 | Tcells | RAD21 |
| 5.5396E-77 | 0.32119804 | 0.571 | 0.383 | 1.3086E-72 | Tcells | SERBP1 |
| 6.2671E-77 | -0.4570165 | 0.018 | 0.168 | 1.4805E-72 | Tcells | AQP9 |
| 7.3274E-77 | -0.3121957 | 0.057 | 0.225 | 1.7309E-72 | Tcells | SMIM14 |
| 7.429E-77 | -0.7352669 | 0.038 | 0.195 | 1.755E-72 | Tcells | AC020656.1 |
| 7.5417E-77 | 0.28479501 | 0.483 | 0.304 | 1.7816E-72 | Tcells | UBE2D2 |
| 9.6594E-77 | -0.4081453 | 0.001 | 0.142 | 2.2818E-72 | Tcells | BCAS1 |
| 1.2054E-76 | 0.29914451 | 0.5 | 0.319 | 2.8475E-72 | Tcells | PAIP2 |
| 1.2361E-76 | 0.25585889 | 0.344 | 0.185 | 2.92E-72 | Tcells | CGGBP1 |
| 1.2474E-76 | -0.428585 | 0.348 | 0.529 | 2.9466E-72 | Tcells | ATP6V1F |
| 1.4778E-76 | -0.2604229 | 0.008 | 0.153 | 3.4909E-72 | Tcells | TUBB6 |
| 1.6779E-76 | 0.29810902 | 0.4 | 0.229 | 3.9637E-72 | Tcells | ASH1L |
| 4.1321E-76 | -0.6084732 | 0.174 | 0.356 | 9.7614E-72 | Tcells | IER5 |
| 4.9079E-76 | -0.5419939 | 0.034 | 0.19 | 1.1594E-71 | Tcells | IGSF6 |
| 5.0687E-76 | 0.25012679 | 0.1 | 0.028 | 1.1974E-71 | Tcells | TNFRSF4 |
| 5.4305E-76 | -0.2833409 | 0.005 | 0.146 | 1.2828E-71 | Tcells | GULP1 |
| 5.9029E-76 | -0.6363142 | 0.129 | 0.302 | 1.3944E-71 | Tcells | ATP13A3 |
| 6.5636E-76 | -0.2955456 | 0.018 | 0.168 | 1.5505E-71 | Tcells | ZFHX3 |
| 1.0626E-75 | -1.058067 | 0.006 | 0.147 | 2.5103E-71 | Tcells | CXCL6 |
| 1.1458E-75 | -0.567797 | 0.961 | 0.957 | 2.7067E-71 | Tcells | H3F3A |
| 1.2441E-75 | -0.646216 | 0.003 | 0.143 | 2.9389E-71 | Tcells | CXCL17 |
| 1.6486E-75 | -1.0097593 | 0.029 | 0.181 | 3.8946E-71 | Tcells | RDH10 |
| 1.7118E-75 | 0.53433596 | 0.142 | 0.05 | 4.0438E-71 | Tcells | CCL4L2 |
| 2.5966E-75 | -0.3972331 | 0.004 | 0.143 | 6.1339E-71 | Tcells | ARHGAP29 |
| 2.8876E-75 | 0.26686111 | 0.374 | 0.208 | 6.8214E-71 | Tcells | U2SURP |
| 3.0399E-75 | -0.3617802 | 0.063 | 0.229 | 7.1812E-71 | Tcells | ACTN1 |
| 3.1731E-75 | -0.3255175 | 0.013 | 0.159 | 7.4957E-71 | Tcells | CPVL |
| 3.5833E-75 | -0.5235355 | 0.602 | 0.733 | 8.4649E-71 | Tcells | ENO1 |
| 4.8895E-75 | -0.5028482 | 0.039 | 0.195 | 1.155E-70 | Tcells | BAG3 |
| 5.1804E-75 | -0.2805387 | 0.013 | 0.157 | 1.2238E-70 | Tcells | LGMN |
| 8.9128E-75 | -0.7165997 | 0.005 | 0.144 | 2.1055E-70 | Tcells | NUPR1 |
| 8.9303E-75 | -0.9301488 | 0.003 | 0.141 | 2.1096E-70 | Tcells | TMPRSS11B |
| 1.343E-74 | -0.3344495 | 0.047 | 0.207 | 3.1727E-70 | Tcells | CREG1 |
| 1.4669E-74 | -0.5235818 | 0.069 | 0.234 | 3.4653E-70 | Tcells | IER5L |

| | | | | | | |
|---|---|---|---|---|---|---|
| 1.5116E-74 | 0.2737162 | 0.423 | 0.252 | 3.5709E-70 | Tcells | SMDT1 |
| 2.0195E-74 | -0.9373586 | 0.002 | 0.139 | 4.7706E-70 | Tcells | RHCG |
| 2.0816E-74 | -0.423932 | 0.194 | 0.378 | 4.9174E-70 | Tcells | TXNDC17 |
| 2.2941E-74 | -0.2590106 | 0.013 | 0.158 | 5.4193E-70 | Tcells | CLIC4 |
| 2.3554E-74 | -0.6546068 | 0.034 | 0.188 | 5.5642E-70 | Tcells | OLR1 |
| 2.6554E-74 | -0.3767281 | 0.011 | 0.153 | 6.2727E-70 | Tcells | TREM1 |
| 3.3053E-74 | -0.3114708 | 0.004 | 0.142 | 7.808E-70 | Tcells | SASH1 |
| 4.499E-74 | -0.5874934 | 0.097 | 0.267 | 1.0628E-69 | Tcells | PMAIP1 |
| 5.5569E-74 | -0.4343126 | 0.204 | 0.385 | 1.3127E-69 | Tcells | ARF4 |
| 6.888E-74 | 0.3139681 | 0.406 | 0.241 | 1.6272E-69 | Tcells | IDS |
| 7.3857E-74 | -0.345128 | 0.017 | 0.162 | 1.7447E-69 | Tcells | ITPKC |
| 7.9237E-74 | 0.37215406 | 0.812 | 0.718 | 1.8718E-69 | Tcells | YWHAB |
| 1.0302E-73 | -0.3279042 | 0.019 | 0.164 | 2.4335E-69 | Tcells | TRIP10 |
| 1.1129E-73 | -0.3330272 | 0.001 | 0.137 | 2.629E-69 | Tcells | AIF1L |
| 1.1799E-73 | -0.427685 | 0.005 | 0.142 | 2.7873E-69 | Tcells | GCNT3 |
| 1.2927E-73 | -0.4862896 | 0.009 | 0.148 | 3.0538E-69 | Tcells | ID1 |
| 1.5747E-73 | -0.2776936 | 0.009 | 0.149 | 3.72E-69 | Tcells | CPM |
| 2.2E-73 | -0.2812017 | 0.002 | 0.137 | 5.197E-69 | Tcells | C1orf116 |
| 2.4518E-73 | -0.3590901 | 0.002 | 0.137 | 5.792E-69 | Tcells | FOXC1 |
| 2.6155E-73 | -0.4701546 | 0.015 | 0.158 | 6.1786E-69 | Tcells | SLC7A11 |
| 4.469E-73 | 0.28966024 | 0.831 | 0.755 | 1.0557E-68 | Tcells | UQCRB |
| 5.9556E-73 | -0.4691335 | 0.005 | 0.141 | 1.4069E-68 | Tcells | UCHL1 |
| 6.597E-73 | -0.3756283 | 0.068 | 0.231 | 1.5584E-68 | Tcells | GRINA |
| 6.8912E-73 | -0.2698697 | 0.006 | 0.143 | 1.6279E-68 | Tcells | SCNN1A |
| 8.2188E-73 | -0.3681542 | 0.071 | 0.236 | 1.9415E-68 | Tcells | FCGRT |
| 9.4844E-73 | -0.3581084 | 0.016 | 0.159 | 2.2405E-68 | Tcells | ITGA3 |
| 1.5462E-72 | 0.31619249 | 0.715 | 0.553 | 3.6526E-68 | Tcells | SET |
| 1.5734E-72 | -0.2894755 | 0.01 | 0.149 | 3.7168E-68 | Tcells | SERPING1 |
| 1.6513E-72 | -1.064753 | 0.007 | 0.144 | 3.9009E-68 | Tcells | DEFB1 |
| 1.706E-72 | -0.2903749 | 0.022 | 0.168 | 4.0301E-68 | Tcells | NCKAP1 |
| 2.469E-72 | -0.3053805 | 0.023 | 0.169 | 5.8325E-68 | Tcells | ALCAM |
| 3.1426E-72 | -0.6033086 | 0.54 | 0.707 | 7.4238E-68 | Tcells | JUNB |
| 7.5836E-72 | -0.3510763 | 0.002 | 0.134 | 1.7915E-67 | Tcells | FAM84A |
| 7.9536E-72 | -0.2698619 | 0.007 | 0.143 | 1.8789E-67 | Tcells | AGRN |
| 9.0837E-72 | -0.6762037 | 0.004 | 0.138 | 2.1458E-67 | Tcells | GABRP |
| 1.7938E-71 | -0.309868 | 0.004 | 0.138 | 4.2374E-67 | Tcells | SCIN |
| 1.9635E-71 | -0.6809479 | 0.004 | 0.137 | 4.6383E-67 | Tcells | DHRS9 |
| 2.6763E-71 | -0.6674341 | 0.003 | 0.135 | 6.3223E-67 | Tcells | TFPI2 |
| 3.081E-71 | 0.2937651 | 0.382 | 0.222 | 7.2783E-67 | Tcells | KDM5A |
| 3.6543E-71 | -0.6523912 | 0.11 | 0.275 | 8.6326E-67 | Tcells | DDIT3 |
| 4.6249E-71 | -1.7204118 | 0.013 | 0.15 | 1.0926E-66 | Tcells | SPRR2A |
| 5.8681E-71 | -0.2979408 | 0.076 | 0.242 | 1.3862E-66 | Tcells | SCARB2 |
| 8.8933E-71 | -1.6085453 | 0.006 | 0.139 | 2.1009E-66 | Tcells | LYPD2 |
| 1.6305E-70 | -0.3497596 | 0.013 | 0.151 | 3.8517E-66 | Tcells | TLR2 |
| 1.8307E-70 | -0.6990027 | 0.897 | 0.934 | 4.3246E-66 | Tcells | GAPDH |
| 2.7984E-70 | -0.4605232 | 0.123 | 0.291 | 6.6108E-66 | Tcells | CTSZ |
| 3.0982E-70 | 0.26919942 | 0.53 | 0.344 | 7.3188E-66 | Tcells | SF3B1 |
| 4.8469E-70 | -0.365577 | 0.004 | 0.134 | 1.145E-65 | Tcells | LRG1 |
| 7.0999E-70 | 0.26517413 | 0.413 | 0.243 | 1.6772E-65 | Tcells | BCLAF1 |
| 8.1264E-70 | -0.2869488 | 0.005 | 0.136 | 1.9197E-65 | Tcells | KAZN |

| | | | | | | |
|---|---|---|---|---|---|---|
| 1.0573E-69 | -0.5189628 | 0.037 | 0.184 | 2.4976E-65 | Tcells | ANPEP |
| 1.2368E-69 | 0.30219782 | 0.326 | 0.178 | 2.9218E-65 | Tcells | CKLF |
| 2.0459E-69 | -0.3261952 | 0.018 | 0.156 | 4.8331E-65 | Tcells | BCAT1 |
| 2.1237E-69 | -0.3030707 | 0.015 | 0.153 | 5.0169E-65 | Tcells | CD40 |
| 2.3244E-69 | -0.3045056 | 0.002 | 0.13 | 5.4909E-65 | Tcells | FAT1 |
| 4.0064E-69 | -0.4183254 | 0.001 | 0.129 | 9.4643E-65 | Tcells | TMEM40 |
| 4.7586E-69 | -0.3804943 | 0.041 | 0.191 | 1.1241E-64 | Tcells | ICAM1 |
| 6.0874E-69 | -0.3156176 | 0.001 | 0.128 | 1.438E-64 | Tcells | FOXQ1 |
| 6.7503E-69 | -0.4151695 | 0.208 | 0.38 | 1.5946E-64 | Tcells | RHEB |
| 7.4112E-69 | -0.3315859 | 0.018 | 0.156 | 1.7508E-64 | Tcells | PLXDC2 |
| 8.0738E-69 | 0.25986538 | 0.368 | 0.211 | 1.9073E-64 | Tcells | MIER1 |
| 8.1891E-69 | -0.3521338 | 0.001 | 0.128 | 1.9345E-64 | Tcells | KRT80 |
| 8.4352E-69 | -0.4531917 | 0.535 | 0.693 | 1.9927E-64 | Tcells | BZW1 |
| 1.0237E-68 | -0.5192575 | 0.224 | 0.397 | 2.4182E-64 | Tcells | TUBB4B |
| 1.0592E-68 | -0.3440588 | 0.069 | 0.226 | 2.5021E-64 | Tcells | SPATS2L |
| 1.0798E-68 | -0.3987821 | 0.004 | 0.132 | 2.5509E-64 | Tcells | ID4 |
| 1.1131E-68 | -0.272081 | 0.017 | 0.154 | 2.6294E-64 | Tcells | ALDH2 |
| 1.3006E-68 | -0.2643478 | 0.002 | 0.129 | 3.0723E-64 | Tcells | PTPN3 |
| 1.3996E-68 | -0.4031199 | 0.016 | 0.152 | 3.3064E-64 | Tcells | MS4A7 |
| 2.1386E-68 | -0.2718962 | 0.007 | 0.137 | 5.052E-64 | Tcells | NTN4 |
| 2.4788E-68 | -0.2618829 | 0.006 | 0.136 | 5.8557E-64 | Tcells | INSR |
| 3.0417E-68 | -0.417377 | 0.017 | 0.152 | 7.1855E-64 | Tcells | FPR1 |
| 3.2789E-68 | -0.4611263 | 0.047 | 0.196 | 7.7457E-64 | Tcells | DUSP6 |
| 3.4934E-68 | -0.3258658 | 0.124 | 0.294 | 8.2524E-64 | Tcells | ATP6V1D |
| 3.7153E-68 | -0.2585237 | 0.011 | 0.143 | 8.7766E-64 | Tcells | SRGAP1 |
| 3.743E-68 | -0.8244734 | 0.013 | 0.146 | 8.8421E-64 | Tcells | CXCL1 |
| 3.767E-68 | -0.3037358 | 0.02 | 0.159 | 8.8988E-64 | Tcells | HK2 |
| 3.9949E-68 | -0.2740396 | 0.005 | 0.133 | 9.4372E-64 | Tcells | SELENOP |
| 7.4235E-68 | -0.2858745 | 0.015 | 0.149 | 1.7536E-63 | Tcells | OCLN |
| 9.2153E-68 | -0.2611921 | 0.022 | 0.161 | 2.1769E-63 | Tcells | CKAP4 |
| 1.0814E-67 | -0.4377806 | 0.341 | 0.505 | 2.5546E-63 | Tcells | VDAC2 |
| 1.7474E-67 | -0.8159967 | 0.029 | 0.168 | 4.128E-63 | Tcells | APOBEC3A |
| 2.5358E-67 | -0.2745154 | 0.052 | 0.204 | 5.9903E-63 | Tcells | GNS |
| 7.509E-67 | -0.5671836 | 0.003 | 0.128 | 1.7739E-62 | Tcells | AGR2 |
| 1.7451E-66 | 0.2915027 | 0.433 | 0.267 | 4.1225E-62 | Tcells | PPIG |
| 2.4377E-66 | -0.3931491 | 0.015 | 0.147 | 5.7586E-62 | Tcells | GPNMB |
| 3.3905E-66 | -0.3127869 | 0.141 | 0.313 | 8.0094E-62 | Tcells | RAB1A |
| 3.4871E-66 | -0.25972 | 0.003 | 0.127 | 8.2376E-62 | Tcells | BICDL2 |
| 4.1366E-66 | -0.395277 | 0.128 | 0.298 | 9.772E-62 | Tcells | MIR4435-2HG |
| 4.3345E-66 | -0.4839196 | 0.023 | 0.158 | 1.0239E-61 | Tcells | CLEC5A |
| 4.7584E-66 | -0.5786851 | 0.008 | 0.136 | 1.1241E-61 | Tcells | AKAP12 |
| 6.0127E-66 | -0.3899584 | 0.002 | 0.125 | 1.4204E-61 | Tcells | EGLN3 |
| 6.7739E-66 | -0.4480171 | 0.057 | 0.208 | 1.6002E-61 | Tcells | ITGAX |
| 7.7845E-66 | -0.3227086 | 0.002 | 0.124 | 1.8389E-61 | Tcells | ASS1 |
| 8.0674E-66 | -0.334505 | 0.169 | 0.345 | 1.9058E-61 | Tcells | CHMP5 |
| 1.2141E-65 | 0.27574219 | 0.808 | 0.709 | 2.868E-61 | Tcells | ATP5MG |
| 1.3605E-65 | -1.2704569 | 0.129 | 0.289 | 3.2139E-61 | Tcells | MT1E |
| 1.5862E-65 | -0.2729129 | 0.016 | 0.147 | 3.747E-61 | Tcells | LRP1 |
| 1.6574E-65 | -0.3428778 | 0.082 | 0.238 | 3.9153E-61 | Tcells | ATP6V1B2 |
| 1.6995E-65 | -0.7461202 | 0.037 | 0.176 | 4.0146E-61 | Tcells | FCN1 |

| | | | | | | |
|---|---|---|---|---|---|---|
| 1.9058E-65 | -0.4550651 | 0.286 | 0.48 | 4.5021E-61 | Tcells | DUSP1 |
| 2.3876E-65 | -0.4301695 | 0.014 | 0.143 | 5.6403E-61 | Tcells | MCEMP1 |
| 2.3925E-65 | -0.3217261 | 0.033 | 0.173 | 5.6519E-61 | Tcells | CFD |
| 2.3952E-65 | 0.25853801 | 0.335 | 0.189 | 5.6582E-61 | Tcells | ZC3H13 |
| 3.287E-65 | -0.2599088 | 0.009 | 0.135 | 7.7649E-61 | Tcells | ARRDC4 |
| 4.3963E-65 | -0.2692748 | 0.011 | 0.138 | 1.0385E-60 | Tcells | TMEM176A |
| 5.5224E-65 | 0.26547811 | 0.541 | 0.36 | 1.3045E-60 | Tcells | PAK2 |
| 6.3733E-65 | -0.2875187 | 0.011 | 0.138 | 1.5056E-60 | Tcells | MPZL2 |
| 6.9709E-65 | -0.3063084 | 0.015 | 0.144 | 1.6467E-60 | Tcells | MTUS1 |
| 7.9273E-65 | -0.2678524 | 0.003 | 0.125 | 1.8727E-60 | Tcells | MUC20 |
| 8.4459E-65 | -0.4728912 | 0.022 | 0.155 | 1.9952E-60 | Tcells | RASD1 |
| 9.728E-65 | -0.3126649 | 0.026 | 0.162 | 2.298E-60 | Tcells | SESN2 |
| 1.021E-64 | -0.5044787 | 0.167 | 0.328 | 2.4119E-60 | Tcells | PERP |
| 1.2057E-64 | -0.5416883 | 0.004 | 0.126 | 2.8483E-60 | Tcells | TCIM |
| 2.0925E-64 | -0.2781817 | 0.009 | 0.134 | 4.9432E-60 | Tcells | PLBD1 |
| 2.1295E-64 | -0.2618552 | 0.003 | 0.124 | 5.0304E-60 | Tcells | C19orf33 |
| 2.6493E-64 | -0.2555415 | 0.002 | 0.123 | 6.2585E-60 | Tcells | VASN |
| 3.0632E-64 | -0.2650347 | 0.011 | 0.137 | 7.2362E-60 | Tcells | PALLD |
| 3.3532E-64 | -0.2877066 | 0.013 | 0.14 | 7.9214E-60 | Tcells | MAP3K20 |
| 3.455E-64 | -0.8532589 | 0.054 | 0.2 | 8.1618E-60 | Tcells | HSPA6 |
| 4.958E-64 | -0.3443849 | 0.012 | 0.138 | 1.1712E-59 | Tcells | IL18 |
| 5.3895E-64 | -0.3691695 | 0.144 | 0.309 | 1.2732E-59 | Tcells | PICALM |
| 5.4173E-64 | -1.1437721 | 0.003 | 0.123 | 1.2797E-59 | Tcells | SPINK5 |
| 5.6447E-64 | 0.25659834 | 0.523 | 0.349 | 1.3335E-59 | Tcells | XRCC5 |
| 7.3487E-64 | -0.251319 | 0.012 | 0.138 | 1.736E-59 | Tcells | DMKN |
| 7.3919E-64 | -0.2973778 | 0.016 | 0.145 | 1.7462E-59 | Tcells | MGLL |
| 7.5282E-64 | 0.27585597 | 0.455 | 0.293 | 1.7784E-59 | Tcells | H2AFV |
| 8.4642E-64 | 0.2795894 | 0.281 | 0.149 | 1.9995E-59 | Tcells | CEMIP2 |
| 1.0432E-63 | -0.4649348 | 0.598 | 0.737 | 2.4643E-59 | Tcells | ATP6V1G1 |
| 4.866E-63 | -1.3119001 | 0.001 | 0.119 | 1.1495E-58 | Tcells | SPRR1B |
| 1.0274E-62 | -0.274718 | 0.077 | 0.23 | 2.4271E-58 | Tcells | PPP2CB |
| 1.4529E-62 | -0.5412185 | 0.071 | 0.215 | 3.4323E-58 | Tcells | LST1 |
| 1.5377E-62 | -0.5166988 | 0.089 | 0.236 | 3.6324E-58 | Tcells | CYB5A |
| 1.5731E-62 | -0.6111881 | 0.001 | 0.118 | 3.7162E-58 | Tcells | KRT78 |
| 1.9407E-62 | -0.2826163 | 0.013 | 0.137 | 4.5844E-58 | Tcells | EPB41L3 |
| 2.1782E-62 | -0.3016572 | 0.001 | 0.117 | 5.1455E-58 | Tcells | FUT3 |
| 2.8126E-62 | -0.2919664 | 0.045 | 0.183 | 6.6442E-58 | Tcells | TMEM159 |
| 3.374E-62 | -0.2797558 | 0.016 | 0.142 | 7.9704E-58 | Tcells | HCK |
| 5.6363E-62 | -0.4234354 | 0.041 | 0.18 | 1.3315E-57 | Tcells | XIST |
| 1.4272E-61 | -0.30925 | 0.123 | 0.283 | 3.3714E-57 | Tcells | PDXK |
| 3.3501E-61 | -0.270567 | 0.022 | 0.149 | 7.914E-57 | Tcells | PHLDA3 |
| 4.0937E-61 | -0.2904644 | 0.123 | 0.282 | 9.6706E-57 | Tcells | LAMP2 |
| 4.1655E-61 | -0.2636639 | 0.002 | 0.117 | 9.8402E-57 | Tcells | GRHL1 |
| 6.1843E-61 | -0.4791382 | 0.024 | 0.15 | 1.4609E-56 | Tcells | CD93 |
| 9.7662E-61 | 0.32893904 | 0.895 | 0.785 | 2.3071E-56 | Tcells | PFN1 |
| 1.0572E-60 | -0.4685938 | 0.003 | 0.118 | 2.4975E-56 | Tcells | TMPRSS11A |
| 1.1496E-60 | -0.4961297 | 0.343 | 0.512 | 2.7157E-56 | Tcells | DNAJB1 |
| 1.3533E-60 | -0.4399689 | 0.226 | 0.398 | 3.1969E-56 | Tcells | FOS |
| 1.5706E-60 | -0.3105811 | 0.027 | 0.156 | 3.7101E-56 | Tcells | DMXL2 |
| 2.4517E-60 | -0.3270182 | 0.097 | 0.25 | 5.7916E-56 | Tcells | MAFG |

| | | | | | | |
|---|---|---|---|---|---|---|
| 4.2146E-60 | -1.0461988 | 0.018 | 0.14 | 9.9561E-56 | Tcells | SERPINB2 |
| 4.4711E-60 | -0.255077 | 0.002 | 0.115 | 1.0562E-55 | Tcells | PTGES |
| 6.9838E-60 | -0.4644343 | 0.055 | 0.192 | 1.6498E-55 | Tcells | ACSL1 |
| 8.5796E-60 | -0.3431414 | 0.096 | 0.245 | 2.0268E-55 | Tcells | PCBD1 |
| 9.3732E-60 | -0.3874973 | 0.243 | 0.405 | 2.2142E-55 | Tcells | H2AFY |
| 1.0973E-59 | -2.5815826 | 0.118 | 0.261 | 2.5922E-55 | Tcells | MT1G |
| 1.1633E-59 | -0.5114409 | 0.175 | 0.331 | 2.7481E-55 | Tcells | LIMS1 |
| 1.6626E-59 | -0.2745089 | 0.065 | 0.207 | 3.9276E-55 | Tcells | NT5C2 |
| 2.4439E-59 | -0.3368754 | 0 | 0.111 | 5.7732E-55 | Tcells | A2ML1 |
| 2.5113E-59 | -0.4062611 | 0.006 | 0.12 | 5.9323E-55 | Tcells | RRAD |
| 2.8951E-59 | -0.2838375 | 0.04 | 0.172 | 6.8391E-55 | Tcells | FLNB |
| 5.3216E-59 | -0.2814568 | 0.038 | 0.169 | 1.2571E-54 | Tcells | NDFIP2 |
| 8.4598E-59 | -0.2682425 | 0.024 | 0.148 | 1.9985E-54 | Tcells | LAPTM4B |
| 9.1117E-59 | -0.9429712 | 0.104 | 0.243 | 2.1525E-54 | Tcells | BCL2A1 |
| 1.3256E-58 | -0.3062766 | 0.476 | 0.644 | 3.1314E-54 | Tcells | ARF1 |
| 1.85E-58 | -0.5466703 | 0.422 | 0.58 | 4.3702E-54 | Tcells | VMP1 |
| 1.9901E-58 | -0.3604495 | 0.004 | 0.116 | 4.7013E-54 | Tcells | SMIM5 |
| 3.9537E-58 | -0.2914585 | 0.115 | 0.267 | 9.3398E-54 | Tcells | ATP6V0D1 |
| 4.1558E-58 | -0.3761665 | 0.153 | 0.306 | 9.8171E-54 | Tcells | TNIP1 |
| 5.1229E-58 | -1.7651301 | 0.541 | 0.666 | 1.2102E-53 | Tcells | MT2A |
| 7.6596E-58 | -0.4016575 | 0.003 | 0.113 | 1.8094E-53 | Tcells | KRT23 |
| 9.2662E-58 | -0.3509854 | 0.015 | 0.132 | 2.189E-53 | Tcells | MARCO |
| 1.3181E-57 | -0.2722205 | 0.039 | 0.168 | 3.1137E-53 | Tcells | FARP1 |
| 1.4698E-57 | -0.3172077 | 0.056 | 0.191 | 3.472E-53 | Tcells | EPS8L2 |
| 1.4844E-57 | -0.3634781 | 0.006 | 0.117 | 3.5067E-53 | Tcells | ANGPTL4 |
| 2.9712E-57 | -0.2527847 | 0.033 | 0.16 | 7.019E-53 | Tcells | RCN1 |
| 3.0769E-57 | -0.260623 | 0.048 | 0.182 | 7.2686E-53 | Tcells | TRAF4 |
| 3.5449E-57 | -0.3782359 | 0.067 | 0.204 | 8.374E-53 | Tcells | SERTAD2 |
| 4.2043E-57 | -0.2768294 | 0.022 | 0.143 | 9.9319E-53 | Tcells | CLEC7A |
| 5.6515E-57 | -0.5076603 | 0.038 | 0.164 | 1.3351E-52 | Tcells | CD300E |
| 8.3378E-57 | -0.4255467 | 0.004 | 0.112 | 1.9696E-52 | Tcells | TMC5 |
| 1.0944E-56 | -0.3678282 | 0.038 | 0.164 | 2.5853E-52 | Tcells | TTC9 |
| 1.1537E-56 | -0.3427995 | 0.216 | 0.373 | 2.7254E-52 | Tcells | RAB7A |
| 1.332E-56 | -0.2805377 | 0.025 | 0.146 | 3.1466E-52 | Tcells | IFI30 |
| 1.8453E-56 | -0.4651296 | 0.179 | 0.329 | 4.3592E-52 | Tcells | B4GALT1 |
| 2.3751E-56 | -0.4779909 | 0.047 | 0.177 | 5.6108E-52 | Tcells | TGFBI |
| 3.4806E-56 | -0.3668978 | 0.443 | 0.591 | 8.2223E-52 | Tcells | NOP10 |
| 3.6048E-56 | -0.3130119 | 0.113 | 0.263 | 8.5155E-52 | Tcells | MAP2K3 |
| 3.6249E-56 | -0.2895273 | 0.129 | 0.281 | 8.5632E-52 | Tcells | ARL8B |
| 4.4919E-56 | -0.5034731 | 0.481 | 0.626 | 1.0611E-51 | Tcells | PCBP1 |
| 6.829E-56 | -0.7945492 | 0.003 | 0.11 | 1.6132E-51 | Tcells | ZG16B |
| 1.8756E-55 | -0.2992325 | 0.096 | 0.239 | 4.4307E-51 | Tcells | BCL10 |
| 3.1486E-55 | -0.4311146 | 0.058 | 0.19 | 7.438E-51 | Tcells | SGK1 |
| 3.4658E-55 | -0.5449944 | 0.012 | 0.122 | 8.1873E-51 | Tcells | HSPA2 |
| 4.2653E-55 | -0.3391701 | 0.156 | 0.308 | 1.0076E-50 | Tcells | SERTAD1 |
| 6.2764E-55 | -0.3140044 | 0.008 | 0.116 | 1.4827E-50 | Tcells | STS |
| 6.6021E-55 | -0.2538438 | 0.009 | 0.117 | 1.5596E-50 | Tcells | DENND2C |
| 7.6879E-55 | -0.3588929 | 0.008 | 0.116 | 1.8161E-50 | Tcells | FXYD2 |
| 1.0553E-54 | -0.3674424 | 0.271 | 0.436 | 2.4929E-50 | Tcells | EFHD2 |
| 1.7106E-54 | -0.4681896 | 0.275 | 0.415 | 4.041E-50 | Tcells | ATF4 |

| | | | | | | |
|---|---|---|---|---|---|---|
| 1.7562E-54 | -0.3116338 | 0.067 | 0.201 | 4.1488E-50 | Tcells | IRS2 |
| 1.792E-54 | -0.8944917 | 0.024 | 0.14 | 4.2332E-50 | Tcells | EREG |
| 1.8029E-54 | -0.2801306 | 0.019 | 0.133 | 4.2591E-50 | Tcells | TP53INP2 |
| 1.8061E-54 | -0.5801671 | 0.055 | 0.182 | 4.2665E-50 | Tcells | IGFBP3 |
| 2.0096E-54 | -0.5359184 | 0.207 | 0.353 | 4.7473E-50 | Tcells | MIDN |
| 3.2715E-54 | -0.3075979 | 0.061 | 0.191 | 7.7283E-50 | Tcells | KLF10 |
| 3.2958E-54 | -0.2646689 | 0.019 | 0.132 | 7.7856E-50 | Tcells | IRAK3 |
| 3.667E-54 | -0.4211614 | 0.018 | 0.13 | 8.6625E-50 | Tcells | SRD5A3 |
| 4.013E-54 | 0.26157661 | 0.408 | 0.265 | 9.48E-50 | Tcells | PPP1R2 |
| 5.2321E-54 | -0.3627019 | 0.178 | 0.334 | 1.236E-49 | Tcells | CAPG |
| 5.5525E-54 | 0.25144654 | 0.373 | 0.234 | 1.3117E-49 | Tcells | RSF1 |
| 7.2721E-54 | -0.3006092 | 0.002 | 0.104 | 1.7179E-49 | Tcells | LY6K |
| 8.0935E-54 | -0.3732426 | 0.095 | 0.233 | 1.9119E-49 | Tcells | RAB9A |
| 8.994E-54 | -0.5432847 | 0.01 | 0.118 | 2.1247E-49 | Tcells | CYP1B1 |
| 1.3035E-53 | -0.9212206 | 0.151 | 0.281 | 3.0794E-49 | Tcells | AIF1 |
| 1.3433E-53 | -0.3405783 | 0.152 | 0.297 | 3.1733E-49 | Tcells | CHMP2B |
| 1.438E-53 | -0.2924354 | 0.001 | 0.102 | 3.3971E-49 | Tcells | RNF39 |
| 2.3808E-53 | -0.5946749 | 0.262 | 0.401 | 5.6241E-49 | Tcells | HIF1A |
| 2.9111E-53 | -0.2507377 | 0.015 | 0.125 | 6.877E-49 | Tcells | CSF1R |
| 4.9545E-53 | -0.4757531 | 0.135 | 0.273 | 1.1704E-48 | Tcells | UGCG |
| 5.5554E-53 | -0.2843562 | 0.064 | 0.195 | 1.3124E-48 | Tcells | VCL |
| 8.4662E-53 | -0.2872703 | 0.041 | 0.163 | 2E-48 | Tcells | FNIP2 |
| 1.4538E-52 | -0.4854869 | 0.1 | 0.236 | 3.4343E-48 | Tcells | PPIF |
| 2.0588E-52 | -0.2607579 | 0.023 | 0.136 | 4.8636E-48 | Tcells | LILRB3 |
| 3.8251E-52 | -0.3614885 | 0.013 | 0.12 | 9.036E-48 | Tcells | CEACAM1 |
| 4.4565E-52 | -0.5024461 | 0.003 | 0.103 | 1.0528E-47 | Tcells | DHRS2 |
| 5.7795E-52 | -0.2853779 | 0.019 | 0.128 | 1.3653E-47 | Tcells | PHACTR1 |
| 7.6873E-52 | -0.6989083 | 0.941 | 0.913 | 1.816E-47 | Tcells | MT-ND1 |
| 7.8764E-52 | -0.5380948 | 0.018 | 0.126 | 1.8607E-47 | Tcells | HILPDA |
| 1.5022E-51 | -0.3996237 | 0.485 | 0.605 | 3.5485E-47 | Tcells | VAMP8 |
| 1.9539E-51 | -0.336492 | 0.628 | 0.753 | 4.6158E-47 | Tcells | ATP6V0E1 |
| 2.2941E-51 | -0.3781782 | 0.17 | 0.313 | 5.4193E-47 | Tcells | BNIP3L |
| 2.4899E-51 | -0.4219666 | 0.498 | 0.629 | 5.8818E-47 | Tcells | TMEM59 |
| 2.9264E-51 | -0.3164948 | 0.112 | 0.251 | 6.913E-47 | Tcells | CD82 |
| 4.1876E-51 | -0.3399968 | 0.228 | 0.375 | 9.8925E-47 | Tcells | RAB10 |
| 4.313E-51 | -0.3868935 | 0.562 | 0.695 | 1.0189E-46 | Tcells | DNAJA1 |
| 4.9541E-51 | 0.26931396 | 0.711 | 0.6 | 1.1703E-46 | Tcells | SUB1 |
| 6.118E-51 | -0.3367213 | 0.206 | 0.353 | 1.4453E-46 | Tcells | CAPN2 |
| 6.4794E-51 | -1.0895191 | 0.355 | 0.498 | 1.5306E-46 | Tcells | CD74 |
| 1.4109E-50 | -0.2693209 | 0.054 | 0.178 | 3.3331E-46 | Tcells | DAPP1 |
| 3.45E-50 | 0.2506998 | 0.768 | 0.655 | 8.15E-46 | Tcells | HNRNPU |
| 4.7346E-50 | -0.3442391 | 0.092 | 0.224 | 1.1185E-45 | Tcells | DSE |
| 5.0387E-50 | -0.3361964 | 0.004 | 0.101 | 1.1903E-45 | Tcells | KLK10 |
| 5.1299E-50 | -0.2742254 | 0.05 | 0.17 | 1.2118E-45 | Tcells | MIR22HG |
| 9.0677E-50 | -0.2849076 | 0.017 | 0.122 | 2.1421E-45 | Tcells | ST3GAL4 |
| 1.5002E-49 | -0.4403677 | 0.454 | 0.579 | 3.5438E-45 | Tcells | TSPO |
| 2.0841E-49 | -0.3076652 | 0.136 | 0.276 | 4.9234E-45 | Tcells | AGFG1 |
| 2.391E-49 | -0.4477279 | 0.052 | 0.17 | 5.6483E-45 | Tcells | ANKRD37 |
| 6.8515E-49 | -0.5634353 | 0.018 | 0.122 | 1.6185E-44 | Tcells | MNDA |
| 8.1975E-49 | -0.3989296 | 0.012 | 0.111 | 1.9365E-44 | Tcells | THBD |

| | | | | | | |
|---|---|---|---|---|---|---|
| 1.1081E-48 | -0.2510111 | 0.019 | 0.122 | 2.6176E-44 | Tcells | CD86 |
| 1.1906E-48 | -0.279647 | 0.114 | 0.252 | 2.8127E-44 | Tcells | PTPN12 |
| 1.1911E-48 | -0.3238961 | 0.421 | 0.583 | 2.8138E-44 | Tcells | PPP1CB |
| 1.3495E-48 | -0.2698495 | 0.129 | 0.268 | 3.1879E-44 | Tcells | YWHAG |
| 1.4533E-48 | -0.3374478 | 0.178 | 0.318 | 3.4332E-44 | Tcells | CTNNB1 |
| 1.4993E-48 | -0.3626092 | 0.034 | 0.145 | 3.5418E-44 | Tcells | GADD45G |
| 2.2089E-48 | -0.2977746 | 0.231 | 0.38 | 5.2182E-44 | Tcells | GNB2 |
| 3.1449E-48 | -0.4615432 | 0.435 | 0.586 | 7.4291E-44 | Tcells | LITAF |
| 4.4534E-48 | -1.2129585 | 0.031 | 0.137 | 1.052E-43 | Tcells | PLAT |
| 7.775E-48 | -0.3388966 | 0.014 | 0.114 | 1.8367E-43 | Tcells | NR1D1 |
| 7.8815E-48 | -0.3319547 | 0.559 | 0.702 | 1.8619E-43 | Tcells | LRRFIP1 |
| 1.5063E-47 | -1.5534744 | 0.013 | 0.111 | 3.5583E-43 | Tcells | CRCT1 |
| 1.792E-47 | -0.4246411 | 0.186 | 0.321 | 4.2332E-43 | Tcells | HIST1H2AC |
| 1.9752E-47 | -0.3457705 | 0.353 | 0.494 | 4.6661E-43 | Tcells | DYNLT1 |
| 1.996E-47 | -0.290474 | 0.11 | 0.243 | 4.7152E-43 | Tcells | PNPLA2 |
| 2.1133E-47 | -0.3305464 | 0.061 | 0.181 | 4.9922E-43 | Tcells | NR4A1 |
| 2.7802E-47 | -0.3330122 | 0.388 | 0.535 | 6.5677E-43 | Tcells | CD164 |
| 3.3641E-47 | -0.432931 | 0.06 | 0.177 | 7.947E-43 | Tcells | DNAJA4 |
| 3.4772E-47 | -0.575483 | 0.08 | 0.198 | 8.2143E-43 | Tcells | FBXO32 |
| 5.0952E-47 | -0.467643 | 0.051 | 0.164 | 1.2036E-42 | Tcells | BPGM |
| 5.691E-47 | -0.2553901 | 0.065 | 0.184 | 1.3444E-42 | Tcells | LSR |
| 6.8064E-47 | -0.4079767 | 0.548 | 0.68 | 1.6079E-42 | Tcells | CALR |
| 7.0963E-47 | -0.2903441 | 0.068 | 0.188 | 1.6763E-42 | Tcells | SPTSSA |
| 3.6318E-46 | -0.3905411 | 0.089 | 0.21 | 8.5794E-42 | Tcells | CSNK1E |
| 5.0082E-46 | -0.2936403 | 0.106 | 0.234 | 1.1831E-41 | Tcells | NUMB |
| 7.0077E-46 | -0.4136508 | 0.174 | 0.305 | 1.6554E-41 | Tcells | PIM3 |
| 8.1666E-46 | -0.2590302 | 0.095 | 0.22 | 1.9292E-41 | Tcells | OSER1 |
| 9.2045E-46 | -0.313347 | 0.161 | 0.294 | 2.1744E-41 | Tcells | ACADVL |
| 1.2438E-45 | -0.2645227 | 0.024 | 0.126 | 2.9382E-41 | Tcells | LILRB2 |
| 1.3575E-45 | -1.4402128 | 0.016 | 0.112 | 3.2069E-41 | Tcells | CNFN |
| 1.7807E-45 | 0.4017914 | 0.885 | 0.868 | 4.2065E-41 | Tcells | PABPC1 |
| 6.1806E-45 | -0.3267187 | 0.022 | 0.121 | 1.46E-40 | Tcells | CD163 |
| 7.3788E-45 | -0.3196917 | 0.068 | 0.187 | 1.7431E-40 | Tcells | WARS |
| 8.1063E-45 | -0.3451141 | 0.801 | 0.87 | 1.915E-40 | Tcells | ARPC3 |
| 1.305E-44 | -0.2541466 | 0.075 | 0.193 | 3.0827E-40 | Tcells | HSBP1L1 |
| 2.4484E-44 | -0.3564726 | 0.164 | 0.295 | 5.7838E-40 | Tcells | CHMP1B |
| 2.5103E-44 | -0.4083499 | 0.515 | 0.636 | 5.9301E-40 | Tcells | TAX1BP1 |
| 4.0031E-44 | -0.2924484 | 0.118 | 0.245 | 9.4565E-40 | Tcells | FKBP4 |
| 6.5039E-44 | -0.3157425 | 0.418 | 0.549 | 1.5364E-39 | Tcells | POMP |
| 7.0824E-44 | -0.3137815 | 0.079 | 0.196 | 1.6731E-39 | Tcells | LIMA1 |
| 7.2184E-44 | -0.2641934 | 0.116 | 0.245 | 1.7052E-39 | Tcells | ACSL3 |
| 1.0441E-43 | -0.4809929 | 0.013 | 0.104 | 2.4666E-39 | Tcells | RETN |
| 1.1465E-43 | -0.5785408 | 0.044 | 0.147 | 2.7083E-39 | Tcells | CDKN2A |
| 1.5648E-43 | -0.2923845 | 0.144 | 0.273 | 3.6966E-39 | Tcells | ATOX1 |
| 2.0726E-43 | -0.2973455 | 0.431 | 0.567 | 4.896E-39 | Tcells | CSNK1A1 |
| 4.8655E-43 | -0.3641304 | 0.422 | 0.554 | 1.1494E-38 | Tcells | TOP1 |
| 5.5875E-43 | -0.2918568 | 0.235 | 0.372 | 1.3199E-38 | Tcells | ELOC |
| 8.9673E-43 | -0.3104468 | 0.191 | 0.322 | 2.1183E-38 | Tcells | MMP24OS |
| 1.145E-42 | -0.3289734 | 0.109 | 0.23 | 2.7048E-38 | Tcells | STXBP2 |
| 3.6867E-42 | -1.1109235 | 0.037 | 0.136 | 8.709E-38 | Tcells | APOE |

| | | | | | | |
|---|---|---|---|---|---|---|
| 4.6533E-42 | -0.2554059 | 0.083 | 0.199 | 1.0992E-37 | Tcells | B4GALT5 |
| 7.9033E-42 | -0.2660084 | 0.086 | 0.202 | 1.867E-37 | Tcells | HIGD1A |
| 1.229E-41 | -0.3043626 | 0.268 | 0.396 | 2.9032E-37 | Tcells | CHMP2A |
| 1.3071E-41 | -0.2794973 | 0.048 | 0.152 | 3.0877E-37 | Tcells | GSTM3 |
| 1.5354E-41 | -1.0007225 | 0.163 | 0.283 | 3.6271E-37 | Tcells | HLA-DRB1 |
| 1.9028E-41 | -0.2878262 | 0.147 | 0.273 | 4.495E-37 | Tcells | TOR1AIP2 |
| 3.2373E-41 | -0.2863222 | 0.112 | 0.231 | 7.6476E-37 | Tcells | PNP |
| 4.2387E-41 | -0.2764049 | 0.138 | 0.263 | 1.0013E-36 | Tcells | LYPLA1 |
| 4.9983E-41 | -0.2758095 | 0.072 | 0.183 | 1.1808E-36 | Tcells | CCDC88A |
| 8.6371E-41 | -0.2608276 | 0.104 | 0.223 | 2.0403E-36 | Tcells | FOXO3 |
| 9.514E-41 | -1.8695553 | 0.055 | 0.157 | 2.2475E-36 | Tcells | MT1H |
| 1.7131E-40 | -0.3513846 | 0.103 | 0.218 | 4.0468E-36 | Tcells | NINJ1 |
| 3.1456E-40 | -0.3691726 | 0.441 | 0.582 | 7.431E-36 | Tcells | CD55 |
| 6.2632E-40 | -0.3071787 | 0.19 | 0.315 | 1.4796E-35 | Tcells | QKI |
| 1.1108E-39 | -0.3179862 | 0.201 | 0.323 | 2.6241E-35 | Tcells | RAB11A |
| 2.2942E-39 | -0.259847 | 0.022 | 0.111 | 5.4196E-35 | Tcells | MS4A6A |
| 2.3516E-39 | -0.3224492 | 0.192 | 0.312 | 5.5552E-35 | Tcells | CLTB |
| 4.688E-39 | -0.4586857 | 0.563 | 0.683 | 1.1074E-34 | Tcells | IER2 |
| 5.1066E-39 | -0.2510088 | 0.09 | 0.201 | 1.2063E-34 | Tcells | TSPAN3 |
| 1.181E-38 | -0.2542762 | 0.09 | 0.202 | 2.7899E-34 | Tcells | USP53 |
| 1.2595E-38 | 0.27160376 | 0.885 | 0.874 | 2.9753E-34 | Tcells | MYL12A |
| 2.104E-38 | -0.2846995 | 0.09 | 0.199 | 4.9703E-34 | Tcells | SNX10 |
| 9.4702E-38 | -0.3765303 | 0.102 | 0.212 | 2.2371E-33 | Tcells | KCNQ1OT1 |
| 1.347E-37 | -0.3098117 | 0.152 | 0.27 | 3.1819E-33 | Tcells | BACH1 |
| 3.7861E-37 | -1.2734411 | 0.052 | 0.148 | 8.944E-33 | Tcells | CCL2 |
| 7.588E-37 | -0.5431539 | 0.88 | 0.885 | 1.7925E-32 | Tcells | HSP90AB1 |
| 1.0583E-36 | -0.2967025 | 0.133 | 0.248 | 2.5E-32 | Tcells | IFRD1 |
| 2.0841E-36 | -0.305711 | 0.025 | 0.11 | 4.9234E-32 | Tcells | CSF3R |
| 2.1481E-36 | -0.2827115 | 0.145 | 0.259 | 5.0743E-32 | Tcells | C1orf122 |
| 2.5854E-36 | -0.9521821 | 0.034 | 0.122 | 6.1076E-32 | Tcells | CXCL2 |
| 2.6361E-36 | -3.607152 | 0.441 | 0.547 | 6.2274E-32 | Tcells | PLCG2 |
| 2.9769E-36 | -1.2840695 | 0.062 | 0.157 | 7.0324E-32 | Tcells | CCL20 |
| 3.4935E-36 | -0.2708819 | 0.493 | 0.622 | 8.2527E-32 | Tcells | CAP1 |
| 4.0866E-36 | -0.2755248 | 0.173 | 0.288 | 9.6538E-32 | Tcells | EIF6 |
| 5.1566E-36 | -0.5541455 | 0.193 | 0.3 | 1.2182E-31 | Tcells | GLRX |
| 7.2778E-36 | -0.3664482 | 0.043 | 0.133 | 1.7192E-31 | Tcells | AC023157.3 |
| 1.2799E-35 | -0.3806725 | 0.258 | 0.362 | 3.0235E-31 | Tcells | TKT |
| 1.803E-35 | -0.3142859 | 0.674 | 0.776 | 4.2593E-31 | Tcells | RHOA |
| 4.6858E-35 | -0.2824112 | 0.524 | 0.638 | 1.1069E-30 | Tcells | SAP18 |
| 1.5977E-34 | -0.3012472 | 0.148 | 0.258 | 3.7743E-30 | Tcells | CD2AP |
| 1.7165E-34 | -0.7168877 | 0.136 | 0.239 | 4.0548E-30 | Tcells | PLIN2 |
| 3.7962E-34 | -0.2605545 | 0.066 | 0.161 | 8.9677E-30 | Tcells | PELI1 |
| 4.7127E-34 | -0.6737213 | 0.031 | 0.113 | 1.1133E-29 | Tcells | APOC1 |
| 7.5605E-34 | -0.2564634 | 0.026 | 0.106 | 1.786E-29 | Tcells | RCAN1 |
| 7.7428E-34 | -2.0595818 | 0.086 | 0.18 | 1.8291E-29 | Tcells | IL1B |
| 9.8289E-34 | -0.250657 | 0.111 | 0.218 | 2.3219E-29 | Tcells | ZFYVE16 |
| 1.916E-33 | -0.3659578 | 0.049 | 0.14 | 4.5262E-29 | Tcells | HLA-DRB5 |
| 5.4392E-33 | -0.2645313 | 0.56 | 0.694 | 1.2849E-28 | Tcells | CAST |
| 1.0315E-32 | -0.4369552 | 0.394 | 0.487 | 2.4367E-28 | Tcells | CD46 |
| 1.1279E-32 | -1.4467403 | 0.06 | 0.149 | 2.6645E-28 | Tcells | CXCL3 |

| | | | | | | |
|---|---|---|---|---|---|---|
| 1.1334E-32 | -0.2809128 | 0.182 | 0.291 | 2.6774E-28 | Tcells | LRP10 |
| 1.1569E-32 | -0.4988244 | 0.97 | 0.949 | 2.733E-28 | Tcells | MT-CO1 |
| 1.6099E-32 | -0.3908949 | 0.138 | 0.243 | 3.803E-28 | Tcells | CITED2 |
| 2.7782E-32 | -0.2889354 | 0.475 | 0.584 | 6.5629E-28 | Tcells | UBE2B |
| 1.6379E-31 | -0.3188284 | 0.338 | 0.453 | 3.8692E-27 | Tcells | ACTN4 |
| 2.2144E-31 | -0.626235 | 0.626 | 0.696 | 5.2311E-27 | Tcells | HSP90B1 |
| 6.9575E-31 | -0.3240095 | 0.174 | 0.276 | 1.6436E-26 | Tcells | RAP2B |
| 1.0326E-30 | -0.2525738 | 0.087 | 0.18 | 2.4393E-26 | Tcells | P4HA1 |
| 1.8326E-30 | -0.5836353 | 0.146 | 0.241 | 4.3292E-26 | Tcells | FABP5 |
| 2.349E-30 | -0.3547926 | 0.167 | 0.266 | 5.549E-26 | Tcells | RHOC |
| 2.5012E-30 | -0.2857073 | 0.052 | 0.135 | 5.9086E-26 | Tcells | CD83 |
| 2.7785E-30 | 0.26321891 | 0.337 | 0.24 | 6.5637E-26 | Tcells | FAM177A1 |
| 2.887E-30 | -0.326251 | 0.916 | 0.93 | 6.82E-26 | Tcells | OAZ1 |
| 3.1574E-30 | -0.3220729 | 0.393 | 0.492 | 7.4587E-26 | Tcells | CANX |
| 7.133E-30 | -0.2613172 | 0.626 | 0.731 | 1.685E-25 | Tcells | ELOB |
| 1.1703E-29 | -0.3285587 | 0.27 | 0.373 | 2.7647E-25 | Tcells | SPAG9 |
| 4.7761E-29 | -0.2848104 | 0.226 | 0.329 | 1.1283E-24 | Tcells | ARL6IP1 |
| 6.7736E-29 | -0.4621671 | 0.088 | 0.177 | 1.6001E-24 | Tcells | HLA-DQB1 |
| 7.2025E-29 | -0.3196969 | 0.394 | 0.486 | 1.7014E-24 | Tcells | GABARAPL2 |
| 1.2218E-28 | -0.3062977 | 0.036 | 0.11 | 2.8861E-24 | Tcells | SLC16A10 |
| 1.9391E-28 | -0.5506999 | 0.14 | 0.231 | 4.5807E-24 | Tcells | HEXIM1 |
| 1.1739E-27 | -0.2563372 | 0.115 | 0.209 | 2.7732E-23 | Tcells | MAP3K8 |
| 1.2178E-27 | -0.3053418 | 0.903 | 0.731 | 2.8768E-23 | Tcells | VIM |
| 1.7424E-27 | -0.3818641 | 0.073 | 0.156 | 4.116E-23 | Tcells | ZFAND2A |
| 2.3676E-27 | -0.3702782 | 0.1 | 0.187 | 5.5929E-23 | Tcells | INSIG1 |
| 2.4678E-27 | -0.3023724 | 0.252 | 0.348 | 5.8296E-23 | Tcells | BSG |
| 4.5725E-27 | -0.2873458 | 0.995 | 0.983 | 1.0802E-22 | Tcells | EIF1 |
| 1.3194E-26 | -0.2624545 | 0.582 | 0.676 | 3.1168E-22 | Tcells | COX5B |
| 1.5058E-26 | -0.6600434 | 0.314 | 0.19 | 3.5572E-22 | Tcells | HOPX |
| 2.9532E-26 | -0.2571741 | 0.629 | 0.73 | 6.9764E-22 | Tcells | COX6A1 |
| 4.4701E-25 | -0.3940588 | 0.533 | 0.609 | 1.056E-20 | Tcells | EZR |
| 1.0284E-24 | -1.3120771 | 0.048 | 0.118 | 2.4293E-20 | Tcells | MT1M |
| 1.6125E-24 | -0.3539282 | 0.339 | 0.421 | 3.8092E-20 | Tcells | JPT1 |
| 9.0345E-24 | -0.3182149 | 0.473 | 0.551 | 2.1342E-19 | Tcells | C4orf3 |
| 5.4031E-23 | -0.5022268 | 0.181 | 0.257 | 1.2764E-18 | Tcells | CARHSP1 |
| 1.018E-22 | -0.2713376 | 0.297 | 0.378 | 2.4049E-18 | Tcells | LAMTOR5 |
| 1.6907E-22 | -1.3494149 | 0.21 | 0.283 | 3.9939E-18 | Tcells | MT1X |
| 5.009E-22 | -0.6488924 | 0.555 | 0.62 | 1.1833E-17 | Tcells | LGALS1 |
| 5.1193E-22 | -0.2505153 | 0.092 | 0.169 | 1.2093E-17 | Tcells | HIST3H2A |
| 5.8563E-22 | -0.3839713 | 0.357 | 0.43 | 1.3834E-17 | Tcells | SLC3A2 |
| 1.2602E-21 | -0.2699456 | 0.132 | 0.213 | 2.977E-17 | Tcells | SOCS3 |
| 5.7521E-21 | -0.2928645 | 0.284 | 0.358 | 1.3588E-16 | Tcells | MRPL33 |
| 1.6522E-20 | -0.4498953 | 0.981 | 0.973 | 3.903E-16 | Tcells | ACTB |
| 4.2001E-20 | -0.3317639 | 0.125 | 0.202 | 9.9218E-16 | Tcells | DUSP4 |
| 8.4725E-20 | -0.2804562 | 0.391 | 0.456 | 2.0015E-15 | Tcells | SEC61G |
| 2.8423E-19 | -0.3092369 | 0.165 | 0.24 | 6.7143E-15 | Tcells | NFKBIZ |
| 5.1831E-19 | -0.2514433 | 0.112 | 0.181 | 1.2244E-14 | Tcells | LTA4H |
| 8.1433E-19 | -0.3253773 | 0.347 | 0.227 | 1.9237E-14 | Tcells | SLC2A3 |
| 7.968E-18 | -1.0062046 | 0.594 | 0.64 | 1.8823E-13 | Tcells | MTRNR2L12 |
| 7.4444E-17 | -0.2735136 | 0.964 | 0.954 | 1.7586E-12 | Tcells | SERF2 |

| | | | | | | |
|---|---|---|---|---|---|---|
| 3.3069E-16 | -0.3282352 | 0.824 | 0.814 | 7.812E-12 | Tcells | YBX1 |
| 3.4355E-16 | -0.293583 | 0.187 | 0.25 | 8.1156E-12 | Tcells | SPTBN1 |
| 4.4316E-15 | -0.4881516 | 0.116 | 0.178 | 1.0469E-10 | Tcells | PLEK |
| 4.5494E-15 | -0.8602073 | 0.674 | 0.711 | 1.0747E-10 | Tcells | ANXA1 |
| 9.7653E-15 | -0.6279589 | 0.135 | 0.186 | 2.3069E-10 | Tcells | AQP3 |
| 1.3456E-14 | -0.3672302 | 0.196 | 0.254 | 3.1787E-10 | Tcells | SNHG12 |
| 1.8522E-14 | -0.3896304 | 0.468 | 0.53 | 4.3754E-10 | Tcells | ZFP36 |
| 3.1453E-14 | -0.2587376 | 0.3 | 0.361 | 7.43E-10 | Tcells | ATP1A1 |
| 6.147E-14 | -0.2717402 | 0.521 | 0.571 | 1.4521E-09 | Tcells | RPS27L |
| 1.8215E-13 | -0.3009315 | 0.313 | 0.369 | 4.303E-09 | Tcells | RALBP1 |
| 2.7036E-13 | -0.3145477 | 0.861 | 0.886 | 6.3867E-09 | Tcells | MYL12B |
| 3.2976E-13 | -0.2590823 | 0.275 | 0.334 | 7.79E-09 | Tcells | PPP1R14B |
| 4.2296E-13 | -0.4922652 | 0.942 | 0.906 | 9.9915E-09 | Tcells | MT-ND2 |
| 5.0856E-13 | -0.3233768 | 0.364 | 0.405 | 1.2014E-08 | Tcells | FLNA |
| 7.1443E-13 | -0.3291775 | 0.12 | 0.172 | 1.6877E-08 | Tcells | AKR1B1 |
| 1.4475E-12 | -0.2805511 | 0.791 | 0.834 | 3.4194E-08 | Tcells | UBB |
| 1.7942E-12 | -0.2871063 | 0.607 | 0.659 | 4.2384E-08 | Tcells | ZFAS1 |
| 2.7672E-12 | -0.2531274 | 0.752 | 0.796 | 6.537E-08 | Tcells | CHCHD2 |
| 1.6829E-11 | -0.6189979 | 0.861 | 0.831 | 3.9755E-07 | Tcells | PPDPF |
| 1.8266E-11 | -0.2929155 | 0.642 | 0.659 | 4.315E-07 | Tcells | CALM2 |
| 2.3384E-11 | -0.4511345 | 0.061 | 0.103 | 5.5241E-07 | Tcells | HLA-DQA1 |
| 4.7507E-11 | -0.2668408 | 0.161 | 0.211 | 1.1223E-06 | Tcells | MDM2 |
| 7.7741E-11 | -0.2666973 | 0.556 | 0.605 | 1.8365E-06 | Tcells | MCL1 |
| 2.1212E-08 | -0.4877653 | 0.61 | 0.612 | 0.0005011 | Tcells | CD44 |
| 5.4248E-08 | -0.3542279 | 0.913 | 0.905 | 0.00128149 | Tcells | ITM2B |
| 1.1542E-07 | -0.2652648 | 0.315 | 0.343 | 0.00272665 | Tcells | PDIA6 |
| 8.9043E-07 | -0.2844521 | 0.154 | 0.188 | 0.02103464 | Tcells | TNFSF10 |
| 4.4871E-05 | -0.273097 | 0.219 | 0.244 | 1 | Tcells | ATP2B1 |
| 6.6795E-05 | -0.4523643 | 0.131 | 0.158 | 1 | Tcells | MT1F |
| 9.7944E-05 | -0.4429959 | 0.857 | 0.821 | 1 | Tcells | MT-ND5 |
| 0.00049231 | -0.7167802 | 0.102 | 0.124 | 1 | Tcells | CCL3 |
| 0.00066467 | -0.2926712 | 0.509 | 0.506 | 1 | Tcells | MT-ND4L |
| 0.00138335 | -0.285387 | 0.987 | 0.969 | 1 | Tcells | TMSB10 |
| 0.00148427 | -0.3462248 | 0.97 | 0.943 | 1 | Tcells | MT-CO3 |
| 0.00170402 | -0.3756366 | 0.276 | 0.228 | 1 | Tcells | ISG15 |
| 0.00649685 | -0.6228067 | 0.222 | 0.183 | 1 | Tcells | HLA-DPB1 |
| 0.0081037 | -0.2849823 | 0.961 | 0.931 | 1 | Tcells | MT-ND4 |
| 0 | 5.87572175 | 0.684 | 0.026 | 0 | Bcells | IGKC |
| 0 | 5.35964048 | 0.347 | 0.007 | 0 | Bcells | IGLC2 |
| 0 | 5.23501372 | 0.305 | 0.002 | 0 | Bcells | IGLC3 |
| 0 | 4.50297569 | 0.8 | 0.005 | 0 | Bcells | IGHM |
| 0 | 2.81378127 | 0.263 | 0.003 | 0 | Bcells | JCHAIN |
| 0 | 2.7770758 | 0.942 | 0.002 | 0 | Bcells | CD79A |
| 0 | 2.55470755 | 0.811 | 0.003 | 0 | Bcells | MS4A1 |
| 0 | 2.48638101 | 0.321 | 0.008 | 0 | Bcells | IGHA1 |
| 0 | 1.75352712 | 0.721 | 0.017 | 0 | Bcells | BANK1 |
| 0 | 1.75113019 | 0.721 | 0.006 | 0 | Bcells | CD79B |
| 0 | 1.4624125 | 0.653 | 0.003 | 0 | Bcells | LINC02397 |
| 0 | 1.40162255 | 0.774 | 0.064 | 0 | Bcells | RCSD1 |
| 0 | 1.37238683 | 0.495 | 0 | 0 | Bcells | IGHD |

| | | | | | | |
|---|---|---|---|---|---|---|
| 0 | 1.3396099 | 0.632 | 0.004 | 0 | Bcells | TNFRSF13C |
| 0 | 1.14831209 | 0.589 | 0.004 | 0 | Bcells | LINC00926 |
| 0 | 1.02571696 | 0.532 | 0.026 | 0 | Bcells | BCL11A |
| 0 | 0.97648886 | 0.553 | 0.02 | 0 | Bcells | CCR7 |
| 0 | 0.88933795 | 0.274 | 0 | 0 | Bcells | TCL1A |
| 0 | 0.8811054 | 0.484 | 0.007 | 0 | Bcells | SPIB |
| 0 | 0.87566783 | 0.526 | 0.032 | 0 | Bcells | IKZF3 |
| 0 | 0.8511083 | 0.516 | 0.025 | 0 | Bcells | FCMR |
| 0 | 0.81662368 | 0.432 | 0.003 | 0 | Bcells | AFF3 |
| 0 | 0.77908327 | 0.437 | 0.005 | 0 | Bcells | P2RX5 |
| 0 | 0.7448726 | 0.121 | 0 | 0 | Bcells | IGHA2 |
| 0 | 0.74483866 | 0.379 | 0.002 | 0 | Bcells | VPREB3 |
| 0 | 0.73780335 | 0.421 | 0 | 0 | Bcells | BLK |
| 0 | 0.70572472 | 0.316 | 0 | 0 | Bcells | TNFRSF13B |
| 0 | 0.68260577 | 0.395 | 0 | 0 | Bcells | PAX5 |
| 0 | 0.66790306 | 0.353 | 0 | 0 | Bcells | FCRL2 |
| 0 | 0.55898533 | 0.205 | 0 | 0 | Bcells | FCRL5 |
| 0 | 0.54446975 | 0.289 | 0.006 | 0 | Bcells | CD22 |
| 0 | 0.53469499 | 0.295 | 0.002 | 0 | Bcells | CPNE5 |
| 0 | 0.51188535 | 0.316 | 0 | 0 | Bcells | FCRL1 |
| 0 | 0.50546426 | 0.268 | 0.001 | 0 | Bcells | POU2AF1 |
| 0 | 0.48314986 | 0.263 | 0.002 | 0 | Bcells | FAM129C |
| 0 | 0.3953846 | 0.226 | 0 | 0 | Bcells | FAM30A |
| 0 | 0.38961639 | 0.226 | 0.004 | 0 | Bcells | HLA-DOB |
| 0 | 0.36589831 | 0.242 | 0.001 | 0 | Bcells | CD19 |
| 0 | 0.35308748 | 0.189 | 0.001 | 0 | Bcells | FCRL3 |
| 0 | 0.34870156 | 0.147 | 0 | 0 | Bcells | IGHG2 |
| 0 | 0.32153109 | 0.184 | 0.002 | 0 | Bcells | STAP1 |
| 0 | 0.31640694 | 0.2 | 0.002 | 0 | Bcells | FCRLA |
| 0 | 0.29673878 | 0.168 | 0.002 | 0 | Bcells | CXCR5 |
| 0 | 0.27886959 | 0.147 | 0.002 | 0 | Bcells | EBF1 |
| 5.568E-301 | 0.33051954 | 0.189 | 0.004 | 1.315E-296 | Bcells | SNX22 |
| 2.674E-293 | 0.40283538 | 0.226 | 0.006 | 6.317E-289 | Bcells | FCER2 |
| 9.608E-275 | 2.25473943 | 0.853 | 0.101 | 2.27E-270 | Bcells | LTB |
| 2.601E-271 | 0.51051189 | 0.353 | 0.016 | 6.144E-267 | Bcells | P2RY10 |
| 3.149E-239 | 0.77670984 | 0.5 | 0.037 | 7.438E-235 | Bcells | RASGRP2 |
| 7.037E-229 | 1.34654955 | 0.721 | 0.082 | 1.662E-224 | Bcells | POU2F2 |
| 3.438E-225 | 2.56797758 | 0.937 | 0.175 | 8.121E-221 | Bcells | CD37 |
| 6.363E-207 | 1.24123027 | 0.653 | 0.074 | 1.503E-202 | Bcells | MEF2C |
| 4.741E-204 | 1.05565158 | 0.558 | 0.054 | 1.12E-199 | Bcells | TCF4 |
| 1.8E-193 | 1.27369616 | 0.789 | 0.117 | 4.252E-189 | Bcells | LIMD2 |
| 1.042E-190 | 0.25972114 | 0.168 | 0.005 | 2.462E-186 | Bcells | NCR3 |
| 8.392E-186 | 1.35881562 | 0.737 | 0.098 | 1.982E-181 | Bcells | HLA-DQA1 |
| 2.193E-176 | 0.57763967 | 0.358 | 0.026 | 5.182E-172 | Bcells | RUBCNL |
| 3.797E-176 | 0.61073387 | 0.379 | 0.029 | 8.97E-172 | Bcells | SP140 |
| 7.014E-176 | 1.10887726 | 0.142 | 0.004 | 1.657E-171 | Bcells | MZB1 |
| 6.801E-164 | 0.7501864 | 0.584 | 0.068 | 1.607E-159 | Bcells | TRBC2 |
| 4.478E-163 | 0.28521729 | 0.205 | 0.009 | 1.058E-158 | Bcells | S1PR1 |
| 2.565E-160 | 0.31542264 | 0.105 | 0.002 | 6.059E-156 | Bcells | LINC01781 |
| 8.028E-159 | 1.06940552 | 0.6 | 0.077 | 1.896E-154 | Bcells | RIPOR2 |

| | | | | | | |
|---|---|---|---|---|---|---|
| 7.82E-152 | 0.95661605 | 0.589 | 0.08 | 1.847E-147 | Bcells | AC004687.1 |
| 3.549E-151 | 0.44511466 | 0.284 | 0.019 | 8.385E-147 | Bcells | GNG7 |
| 1.018E-149 | 0.34927114 | 0.211 | 0.01 | 2.406E-145 | Bcells | PARP15 |
| 2.42E-149 | 0.28992318 | 0.211 | 0.01 | 5.718E-145 | Bcells | LINC01215 |
| 1.306E-148 | 0.88168939 | 0.516 | 0.06 | 3.084E-144 | Bcells | SELL |
| 1.722E-145 | 0.30134285 | 0.211 | 0.011 | 4.067E-141 | Bcells | TMEM156 |
| 1.812E-140 | 0.33831959 | 0.205 | 0.011 | 4.28E-136 | Bcells | SHISAL2A |
| 3.262E-140 | 0.71766811 | 0.384 | 0.037 | 7.705E-136 | Bcells | ADAM28 |
| 1.361E-135 | 0.40478295 | 0.184 | 0.009 | 3.216E-131 | Bcells | LINC01857 |
| 2.745E-135 | 0.81744532 | 0.579 | 0.084 | 6.484E-131 | Bcells | ORAI2 |
| 1.716E-133 | 3.3094173 | 0.132 | 0.005 | 4.053E-129 | Bcells | IGHG1 |
| 4.096E-132 | 1.34481212 | 0.853 | 0.182 | 9.675E-128 | Bcells | HLA-DPB1 |
| 5.096E-132 | 1.31998373 | 0.816 | 0.169 | 1.204E-127 | Bcells | HLA-DQB1 |
| 1.149E-129 | 1.50726231 | 0.789 | 0.167 | 2.715E-125 | Bcells | CXCR4 |
| 1.581E-129 | 0.66419205 | 0.358 | 0.035 | 3.734E-125 | Bcells | HVCN1 |
| 4.851E-129 | 1.67890573 | 0.832 | 0.194 | 1.146E-124 | Bcells | BIRC3 |
| 9.848E-127 | 0.34988779 | 0.242 | 0.016 | 2.326E-122 | Bcells | C16orf74 |
| 6.891E-124 | 1.20670655 | 0.779 | 0.167 | 1.628E-119 | Bcells | ISG20 |
| 5.882E-123 | 0.68434506 | 0.511 | 0.071 | 1.39E-118 | Bcells | STRBP |
| 1.139E-121 | 0.84579805 | 0.484 | 0.065 | 2.69E-117 | Bcells | CLEC2D |
| 8.256E-119 | 0.34431167 | 0.179 | 0.01 | 1.95E-114 | Bcells | AIM2 |
| 7.729E-118 | 0.32649854 | 0.216 | 0.014 | 1.826E-113 | Bcells | CDCA7L |
| 2.22E-117 | 0.78589966 | 0.442 | 0.057 | 5.245E-113 | Bcells | ARHGAP24 |
| 9.749E-116 | 0.70892909 | 0.511 | 0.074 | 2.303E-111 | Bcells | ACAP1 |
| 6.976E-113 | 0.8218526 | 0.521 | 0.079 | 1.648E-108 | Bcells | HLA-DMB |
| 2.884E-112 | 0.45457072 | 0.247 | 0.019 | 6.812E-108 | Bcells | RAB30 |
| 2.162E-111 | 0.56636715 | 0.405 | 0.049 | 5.108E-107 | Bcells | ST6GAL1 |
| 1.396E-110 | 1.77161913 | 0.637 | 0.129 | 3.298E-106 | Bcells | CD83 |
| 6.541E-110 | 1.80603872 | 0.779 | 0.195 | 1.545E-105 | Bcells | CD52 |
| 1.152E-106 | 0.72268055 | 0.568 | 0.097 | 2.721E-102 | Bcells | SEPT6 |
| 6.455E-105 | 1.09723401 | 0.821 | 0.215 | 1.525E-100 | Bcells | STK4 |
| 3.651E-104 | 1.23713341 | 0.663 | 0.146 | 8.625E-100 | Bcells | RALGPS2 |
| 4.138E-104 | 0.80057982 | 0.505 | 0.081 | 9.776E-100 | Bcells | BCL2 |
| 9.205E-102 | 1.0617762 | 0.679 | 0.147 | 2.1745E-97 | Bcells | SMAP2 |
| 5.743E-101 | 1.52752738 | 0.884 | 0.274 | 1.3566E-96 | Bcells | HLA-DRB1 |
| 1.139E-100 | 0.2503593 | 0.179 | 0.011 | 2.6908E-96 | Bcells | PPP1R16B |
| 5.281E-100 | 0.25193779 | 0.168 | 0.01 | 1.2476E-95 | Bcells | WNT10A |
| 1.632E-99 | 2.00828493 | 0.995 | 0.489 | 3.8551E-95 | Bcells | CD74 |
| 9.558E-99 | 0.97763354 | 0.821 | 0.201 | 2.2579E-94 | Bcells | HLA-DPA1 |
| 8.126E-97 | 0.38844636 | 0.305 | 0.032 | 1.9196E-92 | Bcells | HSH2D |
| 1.5011E-94 | 0.45913866 | 0.295 | 0.031 | 3.5461E-90 | Bcells | BACH2 |
| 3.4897E-94 | 0.98221713 | 0.668 | 0.153 | 8.2438E-90 | Bcells | PDCD4 |
| 5.7099E-94 | -2.8050305 | 0.432 | 0.967 | 1.3488E-89 | Bcells | S100A11 |
| 3.429E-93 | 0.75573255 | 0.532 | 0.097 | 8.1003E-89 | Bcells | ANKRD44 |
| 9.9131E-93 | 0.55934378 | 0.458 | 0.072 | 2.3418E-88 | Bcells | PDE7A |
| 2.0537E-90 | 0.88020995 | 0.589 | 0.123 | 4.8516E-86 | Bcells | ATM |
| 1.2784E-85 | 0.31407134 | 0.153 | 0.01 | 3.0199E-81 | Bcells | CD70 |
| 3.4261E-85 | 0.72821736 | 0.516 | 0.095 | 8.0935E-81 | Bcells | IKZF1 |
| 3.7529E-85 | 0.45453767 | 0.363 | 0.05 | 8.8655E-81 | Bcells | RHOH |
| 1.1683E-84 | 0.55967474 | 0.4 | 0.061 | 2.7598E-80 | Bcells | INPP5D |

| | | | | | | |
|---|---|---|---|---|---|---|
| 2.4358E-84 | 1.12352779 | 0.679 | 0.18 | 5.754E-80 | Bcells | STK17A |
| 7.5712E-84 | 0.78991254 | 0.526 | 0.104 | 1.7885E-79 | Bcells | PRKCB |
| 1.0207E-82 | 0.72300339 | 0.516 | 0.102 | 2.4111E-78 | Bcells | PARP1 |
| 4.52E-82 | 1.64130675 | 0.995 | 0.943 | 1.0678E-77 | Bcells | RPS27 |
| 1.1497E-80 | 1.55054235 | 1 | 0.924 | 2.7159E-76 | Bcells | RPL18A |
| 2.0677E-80 | 0.63558575 | 0.405 | 0.066 | 4.8845E-76 | Bcells | IRF8 |
| 4.0215E-80 | 1.04362504 | 0.784 | 0.236 | 9.5001E-76 | Bcells | SMCHD1 |
| 1.1163E-79 | 0.5023896 | 0.179 | 0.014 | 2.6371E-75 | Bcells | TTN |
| 1.4554E-79 | -3.1669379 | 0.553 | 0.939 | 3.4381E-75 | Bcells | S100A6 |
| 2.4056E-79 | 0.89572865 | 0.563 | 0.125 | 5.6828E-75 | Bcells | TRIM22 |
| 2.5519E-79 | 0.78251272 | 0.605 | 0.133 | 6.0283E-75 | Bcells | HLA-DRB5 |
| 1.5627E-78 | 0.34847654 | 0.174 | 0.013 | 3.6916E-74 | Bcells | LY9 |
| 2.2769E-78 | 0.40555425 | 0.3 | 0.038 | 5.3786E-74 | Bcells | TRAF5 |
| 1.1481E-77 | 0.49139639 | 0.353 | 0.051 | 2.7121E-73 | Bcells | CIITA |
| 1.3074E-77 | 0.68481564 | 0.511 | 0.103 | 3.0884E-73 | Bcells | SP110 |
| 2.2617E-77 | 0.42591463 | 0.263 | 0.03 | 5.3429E-73 | Bcells | METTL8 |
| 2.4381E-77 | 0.33882551 | 0.189 | 0.016 | 5.7595E-73 | Bcells | CD72 |
| 3.1889E-77 | 0.84516978 | 0.679 | 0.171 | 7.5332E-73 | Bcells | CORO1A |
| 7.6761E-77 | 0.39122532 | 0.305 | 0.039 | 1.8133E-72 | Bcells | ARHGAP15 |
| 1.0589E-76 | 0.75312467 | 0.432 | 0.077 | 2.5014E-72 | Bcells | MARCH1 |
| 1.0953E-76 | 1.39330733 | 1 | 0.893 | 2.5874E-72 | Bcells | RPS15A |
| 7.5864E-76 | 0.95499044 | 0.595 | 0.142 | 1.7921E-71 | Bcells | GABPB1-AS1 |
| 1.8227E-75 | 1.42291698 | 0.921 | 0.394 | 4.3058E-71 | Bcells | HLA-DRA |
| 1.0353E-74 | 0.38766928 | 0.3 | 0.039 | 2.4457E-70 | Bcells | CAMK1D |
| 2.4023E-74 | 0.5504892 | 0.363 | 0.057 | 5.6749E-70 | Bcells | SEL1L3 |
| 8.0624E-74 | 1.06618438 | 0.826 | 0.303 | 1.9046E-69 | Bcells | NOP53 |
| 1.7649E-73 | 1.43087426 | 1 | 0.888 | 4.1693E-69 | Bcells | RPS23 |
| 2.1156E-72 | 0.39363024 | 0.284 | 0.036 | 4.9976E-68 | Bcells | TBC1D10C |
| 4.9461E-72 | 1.41602656 | 0.979 | 0.899 | 1.1684E-67 | Bcells | RPL21 |
| 6.9743E-72 | 1.28031567 | 0.989 | 0.9 | 1.6475E-67 | Bcells | RPS11 |
| 1.1543E-71 | 1.4143713 | 0.984 | 0.899 | 2.7268E-67 | Bcells | RPL39 |
| 1.293E-71 | 0.67239852 | 0.584 | 0.134 | 3.0545E-67 | Bcells | HLA-DMA |
| 1.8265E-71 | 1.37313263 | 0.995 | 0.933 | 4.3148E-67 | Bcells | RPL34 |
| 7.8084E-71 | 0.61952213 | 0.5 | 0.107 | 1.8446E-66 | Bcells | GGA2 |
| 1.3938E-70 | -1.7897197 | 0.342 | 0.894 | 3.2926E-66 | Bcells | CD63 |
| 1.4556E-70 | 1.03778554 | 0.721 | 0.237 | 3.4385E-66 | Bcells | SP100 |
| 1.2293E-69 | 1.28091608 | 0.995 | 0.912 | 2.9041E-65 | Bcells | RPS8 |
| 1.2774E-69 | 1.36649424 | 0.989 | 0.923 | 3.0177E-65 | Bcells | RPS27A |
| 4.69E-69 | 1.2860239 | 1 | 0.945 | 1.1079E-64 | Bcells | RPS19 |
| 3.5598E-68 | 0.77263117 | 0.616 | 0.163 | 8.4092E-64 | Bcells | MYCBP2 |
| 3.9247E-68 | 0.55795608 | 0.468 | 0.097 | 9.2714E-64 | Bcells | WDR11 |
| 5.8743E-68 | 0.37497379 | 0.311 | 0.045 | 1.3877E-63 | Bcells | BTK |
| 6.4898E-68 | 0.58428862 | 0.568 | 0.132 | 1.5331E-63 | Bcells | CYTIP |
| 6.5965E-68 | -2.0121593 | 0.121 | 0.787 | 1.5583E-63 | Bcells | LGALS3 |
| 1.0406E-67 | 1.2747296 | 0.989 | 0.903 | 2.4583E-63 | Bcells | RPL37 |
| 3.0312E-67 | 1.21468316 | 1 | 0.922 | 7.1607E-63 | Bcells | RPL11 |
| 7.9432E-67 | 1.1603277 | 1 | 0.964 | 1.8764E-62 | Bcells | RPL13 |
| 8.4326E-67 | 0.3647813 | 0.284 | 0.039 | 1.992E-62 | Bcells | IL16 |
| 1.4721E-66 | 0.26310596 | 0.137 | 0.01 | 3.4775E-62 | Bcells | CCR6 |
| 1.9967E-66 | 1.52077571 | 0.768 | 0.295 | 4.7168E-62 | Bcells | REL |

| | | | | | | |
|---|---|---|---|---|---|---|
| 2.0768E-66 | 1.19536569 | 0.989 | 0.915 | 4.906E-62 | Bcells | RPL32 |
| 6.2381E-66 | 1.31733804 | 0.968 | 0.879 | 1.4736E-61 | Bcells | RPS29 |
| 1.6203E-65 | 1.31433341 | 0.984 | 0.927 | 3.8277E-61 | Bcells | RPL30 |
| 1.9074E-65 | 0.73777215 | 0.495 | 0.113 | 4.5059E-61 | Bcells | AC245297.3 |
| 2.8503E-65 | 0.79004324 | 0.874 | 0.278 | 6.7333E-61 | Bcells | PTPRC |
| 4.8924E-65 | 0.85200338 | 0.347 | 0.058 | 1.1557E-60 | Bcells | CD69 |
| 9.468E-65 | 0.81322226 | 0.595 | 0.158 | 2.2366E-60 | Bcells | TUT4 |
| 1.1607E-64 | 0.44036878 | 0.337 | 0.054 | 2.742E-60 | Bcells | ITGA4 |
| 1.8856E-64 | 1.15238185 | 0.989 | 0.931 | 4.4544E-60 | Bcells | FAU |
| 2.7746E-64 | 1.17075294 | 0.989 | 0.904 | 6.5545E-60 | Bcells | RPLP2 |
| 2.0857E-63 | -2.1979161 | 0.689 | 0.94 | 4.9271E-59 | Bcells | SAT1 |
| 2.4662E-63 | 1.17786608 | 1 | 0.972 | 5.826E-59 | Bcells | RPLP1 |
| 9.886E-62 | 0.30992186 | 0.232 | 0.029 | 2.3354E-57 | Bcells | WDFY4 |
| 1.1134E-61 | 1.17137676 | 0.989 | 0.817 | 2.6301E-57 | Bcells | RPL5 |
| 2.9451E-61 | 0.52428662 | 0.363 | 0.066 | 6.9571E-57 | Bcells | SESN3 |
| 4.3635E-61 | 1.01795944 | 1 | 0.949 | 1.0308E-56 | Bcells | RPL18 |
| 8.6488E-61 | 1.18799794 | 0.995 | 0.897 | 2.0431E-56 | Bcells | RPS3A |
| 8.8391E-61 | 0.80207025 | 0.611 | 0.167 | 2.0881E-56 | Bcells | CD48 |
| 1.9233E-60 | 1.05445027 | 0.995 | 0.887 | 4.5435E-56 | Bcells | RPL6 |
| 1.9712E-60 | -2.0348376 | 0.521 | 0.921 | 4.6565E-56 | Bcells | S100A10 |
| 5.9983E-60 | 0.91848487 | 0.7 | 0.231 | 1.417E-55 | Bcells | PARP14 |
| 7.3339E-60 | 1.08904098 | 1 | 0.939 | 1.7325E-55 | Bcells | RPS28 |
| 7.6106E-60 | 0.47410599 | 0.326 | 0.056 | 1.7978E-55 | Bcells | DGKD |
| 1.0957E-59 | 0.32450479 | 0.247 | 0.033 | 2.5883E-55 | Bcells | PLEKHG1 |
| 1.4219E-59 | 1.06922738 | 0.995 | 0.945 | 3.3589E-55 | Bcells | RPL13A |
| 1.625E-59 | -1.5568382 | 0.126 | 0.77 | 3.8387E-55 | Bcells | CST3 |
| 2.0442E-59 | 1.13767077 | 0.979 | 0.754 | 4.8291E-55 | Bcells | RPSA |
| 2.17E-59 | 1.20003732 | 0.974 | 0.791 | 5.1261E-55 | Bcells | RPS21 |
| 2.6469E-59 | 0.96557487 | 0.905 | 0.333 | 6.2528E-55 | Bcells | LAPTM5 |
| 2.8331E-59 | -1.6962718 | 0.147 | 0.777 | 6.6927E-55 | Bcells | CTSD |
| 4.3079E-59 | 0.79358302 | 0.6 | 0.176 | 1.0177E-54 | Bcells | STX7 |
| 5.7203E-59 | 0.37918354 | 0.316 | 0.052 | 1.3513E-54 | Bcells | APOBEC3G |
| 6.239E-59 | 0.44103453 | 0.337 | 0.059 | 1.4738E-54 | Bcells | FCHSD2 |
| 1.417E-58 | 1.164731 | 0.984 | 0.887 | 3.3474E-54 | Bcells | RPS3 |
| 7.3239E-58 | 1.08090564 | 0.995 | 0.897 | 1.7301E-53 | Bcells | RPL29 |
| 8.1128E-58 | 1.11379381 | 0.979 | 0.8 | 1.9165E-53 | Bcells | RPS5 |
| 1.0863E-57 | 1.0305803 | 0.805 | 0.297 | 2.5661E-53 | Bcells | BTG2 |
| 1.401E-57 | 1.03811119 | 0.989 | 0.946 | 3.3095E-53 | Bcells | RPS16 |
| 1.7912E-57 | 1.08193561 | 0.995 | 0.927 | 4.2313E-53 | Bcells | RPL19 |
| 9.764E-57 | 0.25014828 | 0.163 | 0.016 | 2.3066E-52 | Bcells | CYSLTR1 |
| 2.2478E-56 | 0.56428029 | 0.437 | 0.098 | 5.31E-52 | Bcells | EIF1AY |
| 2.9027E-56 | 0.35577313 | 0.289 | 0.046 | 6.8571E-52 | Bcells | MARCH9 |
| 3.0211E-56 | 0.5904106 | 0.374 | 0.073 | 7.1367E-52 | Bcells | HIST1H1D |
| 4.7895E-56 | 0.63182398 | 0.453 | 0.107 | 1.1314E-51 | Bcells | TMEM243 |
| 8.1273E-56 | 1.13594652 | 0.979 | 0.884 | 1.9199E-51 | Bcells | RPS13 |
| 3.0356E-55 | 0.54966765 | 0.389 | 0.081 | 7.1711E-51 | Bcells | PPM1K |
| 1.1863E-54 | 0.48512076 | 0.232 | 0.032 | 2.8024E-50 | Bcells | PIM2 |
| 6.4015E-54 | 1.04248809 | 0.995 | 0.922 | 1.5122E-49 | Bcells | RPL8 |
| 6.5478E-54 | 0.43091115 | 0.321 | 0.058 | 1.5468E-49 | Bcells | DDHD1 |
| 9.7635E-54 | 0.39311065 | 0.316 | 0.056 | 2.3064E-49 | Bcells | ATP2A3 |

| | | | | | | |
|---|---|---|---|---|---|---|
| 1.0746E-53 | 0.40075223 | 0.289 | 0.048 | 2.5385E-49 | Bcells | TSPAN33 |
| 2.7908E-53 | 0.42899531 | 0.321 | 0.058 | 6.5928E-49 | Bcells | TRAF3IP3 |
| 2.9661E-53 | 0.85314837 | 0.568 | 0.175 | 7.0068E-49 | Bcells | FOXP1 |
| 1.047E-52 | 0.51229944 | 0.389 | 0.084 | 2.4734E-48 | Bcells | CNTRL |
| 1.1021E-52 | 1.03256038 | 0.984 | 0.917 | 2.6036E-48 | Bcells | RPS20 |
| 1.9896E-52 | 0.67109257 | 0.568 | 0.168 | 4.7E-48 | Bcells | FNBP1 |
| 1.9948E-52 | 1.0051027 | 0.989 | 0.896 | 4.7122E-48 | Bcells | RPL35A |
| 2.4235E-52 | 0.48201886 | 0.542 | 0.141 | 5.725E-48 | Bcells | NCF1 |
| 5.0977E-52 | -2.3600631 | 0.126 | 0.712 | 1.2042E-47 | Bcells | ANXA1 |
| 1.2436E-51 | 1.11274223 | 0.995 | 0.933 | 2.9377E-47 | Bcells | RPS12 |
| 2.6143E-51 | 1.01774137 | 0.995 | 0.848 | 6.1757E-47 | Bcells | RPS4X |
| 5.9669E-51 | 0.36129503 | 0.221 | 0.031 | 1.4096E-46 | Bcells | ZNF107 |
| 8.764E-51 | 1.01076022 | 0.995 | 0.948 | 2.0703E-46 | Bcells | RPL10 |
| 9.5192E-51 | 0.42625744 | 0.326 | 0.063 | 2.2487E-46 | Bcells | RBM38 |
| 9.9958E-51 | 0.85767867 | 0.863 | 0.392 | 2.3613E-46 | Bcells | PNISR |
| 1.0638E-50 | 0.49928809 | 0.463 | 0.116 | 2.5131E-46 | Bcells | EBLN3P |
| 1.1642E-50 | 0.83733007 | 0.668 | 0.241 | 2.7503E-46 | Bcells | LSM7 |
| 2.3263E-50 | 0.78675774 | 0.732 | 0.268 | 5.4955E-46 | Bcells | TSC22D3 |
| 2.68E-50 | -1.5908144 | 0.447 | 0.869 | 6.3308E-46 | Bcells | ANXA2 |
| 4.0464E-50 | 0.46392509 | 0.353 | 0.072 | 9.5588E-46 | Bcells | CYB561A3 |
| 1.5919E-49 | 0.38283951 | 0.279 | 0.048 | 3.7605E-45 | Bcells | BCAS4 |
| 4.5304E-49 | -2.5140467 | 0.984 | 0.994 | 1.0702E-44 | Bcells | FTH1 |
| 6.1606E-49 | 0.39211802 | 0.295 | 0.053 | 1.4553E-44 | Bcells | ZBTB24 |
| 2.732E-48 | 0.35167516 | 0.289 | 0.052 | 6.4537E-44 | Bcells | SMC6 |
| 1.4501E-47 | -1.8278909 | 0.511 | 0.885 | 3.4256E-43 | Bcells | CSTB |
| 1.8162E-47 | 0.46307587 | 0.411 | 0.098 | 4.2904E-43 | Bcells | DOCK2 |
| 2.4793E-47 | 0.36570033 | 0.247 | 0.04 | 5.8569E-43 | Bcells | ARHGAP25 |
| 2.646E-47 | 0.64533654 | 0.637 | 0.21 | 6.2507E-43 | Bcells | BDP1 |
| 8.2958E-47 | 0.54607086 | 0.437 | 0.112 | 1.9597E-42 | Bcells | PLEKHA2 |
| 9.2876E-47 | 0.8863019 | 0.995 | 0.92 | 2.194E-42 | Bcells | RPS15 |
| 1.0368E-46 | -4.9752196 | 0.232 | 0.729 | 2.4492E-42 | Bcells | S100A9 |
| 1.1613E-46 | 0.88085202 | 1 | 0.905 | 2.7433E-42 | Bcells | RPL7A |
| 2.3997E-46 | 0.91766024 | 0.979 | 0.918 | 5.6688E-42 | Bcells | RPL9 |
| 1.107E-45 | 0.92938873 | 1 | 0.933 | 2.615E-41 | Bcells | RPS14 |
| 1.4435E-45 | 0.4303103 | 0.342 | 0.074 | 3.4099E-41 | Bcells | ZCCHC7 |
| 1.4989E-45 | 0.9046823 | 0.989 | 0.876 | 3.5409E-41 | Bcells | RPL23A |
| 2.7543E-45 | 0.2574128 | 0.211 | 0.031 | 6.5066E-41 | Bcells | MAP4K2 |
| 1.0476E-44 | 0.84755034 | 1 | 0.887 | 2.4747E-40 | Bcells | RPS6 |
| 1.4139E-44 | 0.73699844 | 0.579 | 0.2 | 3.3401E-40 | Bcells | PRDM2 |
| 1.4175E-44 | 0.74955516 | 0.695 | 0.272 | 3.3487E-40 | Bcells | SF1 |
| 2.303E-44 | 0.86728164 | 0.684 | 0.273 | 5.4405E-40 | Bcells | BPTF |
| 2.5761E-44 | 0.88129893 | 0.995 | 0.87 | 6.0855E-40 | Bcells | RPL14 |
| 4.6651E-44 | 0.66055616 | 0.442 | 0.121 | 1.102E-39 | Bcells | MIS18BP1 |
| 6.6324E-44 | 0.87903462 | 0.995 | 0.895 | 1.5668E-39 | Bcells | RPL37A |
| 7.9195E-44 | 0.88889731 | 0.984 | 0.831 | 1.8708E-39 | Bcells | RPL22 |
| 2.7899E-43 | -1.8762543 | 0.195 | 0.706 | 6.5907E-39 | Bcells | SOD2 |
| 3.5404E-43 | 0.46591449 | 0.395 | 0.097 | 8.3636E-39 | Bcells | SNX29 |
| 4.6964E-43 | 0.49645115 | 0.5 | 0.144 | 1.1094E-38 | Bcells | CD40 |
| 7.9333E-43 | 0.81697502 | 1 | 0.846 | 1.8741E-38 | Bcells | RPL35 |
| 9.8836E-43 | 0.52296281 | 0.547 | 0.171 | 2.3348E-38 | Bcells | TNFAIP8 |

| | | | | | | |
|---|---|---|---|---|---|---|
| 1.3492E-42 | 0.39754724 | 0.347 | 0.079 | 3.1873E-38 | Bcells | UVRAG |
| 1.965E-42 | -3.9541941 | 0.158 | 0.673 | 4.642E-38 | Bcells | S100A8 |
| 2.342E-42 | 0.84786965 | 0.989 | 0.901 | 5.5325E-38 | Bcells | RPS7 |
| 2.5267E-42 | -2.3708703 | 0.053 | 0.581 | 5.9689E-38 | Bcells | KRT19 |
| 2.7288E-42 | 0.95085232 | 0.953 | 0.746 | 6.4462E-38 | Bcells | EEF1B2 |
| 3.5832E-42 | 0.42709766 | 0.421 | 0.107 | 8.4646E-38 | Bcells | PDE4B |
| 5.2093E-42 | -1.5341948 | 0.337 | 0.786 | 1.2306E-37 | Bcells | TXN |
| 6.0867E-42 | 0.47977461 | 0.368 | 0.091 | 1.4379E-37 | Bcells | SMARCB1 |
| 9.5374E-42 | 0.59954067 | 0.563 | 0.185 | 2.253E-37 | Bcells | GPSM3 |
| 1.6563E-41 | 0.61946427 | 0.568 | 0.186 | 3.9127E-37 | Bcells | RAC2 |
| 1.6962E-41 | 0.82350115 | 0.963 | 0.819 | 4.0069E-37 | Bcells | RPL27 |
| 2.9818E-41 | 0.83413028 | 0.984 | 0.898 | 7.044E-37 | Bcells | RPS25 |
| 3.8249E-41 | -2.0595533 | 0.037 | 0.56 | 9.0356E-37 | Bcells | KRT18 |
| 4.0633E-41 | 0.51802821 | 0.479 | 0.139 | 9.5987E-37 | Bcells | IL2RG |
| 4.518E-41 | 0.75568996 | 1 | 0.98 | 1.0673E-36 | Bcells | EEF1A1 |
| 4.7959E-41 | -1.7542592 | 0.132 | 0.646 | 1.1329E-36 | Bcells | GLUL |
| 5.1475E-41 | 0.84832212 | 0.989 | 0.838 | 1.216E-36 | Bcells | RACK1 |
| 6.7395E-41 | 0.33515159 | 0.268 | 0.052 | 1.5921E-36 | Bcells | BLNK |
| 7.7664E-41 | 0.55857043 | 0.537 | 0.166 | 1.8346E-36 | Bcells | SLC38A1 |
| 1.1424E-40 | -2.003007 | 0.037 | 0.557 | 2.6987E-36 | Bcells | ELF3 |
| 1.1602E-40 | 0.85201009 | 0.989 | 0.875 | 2.7408E-36 | Bcells | RPL3 |
| 1.2284E-40 | -1.5558181 | 0.121 | 0.644 | 2.9019E-36 | Bcells | IER3 |
| 1.6345E-40 | 0.42463988 | 0.4 | 0.104 | 3.8613E-36 | Bcells | TPD52 |
| 1.8643E-40 | -1.3264332 | 0.258 | 0.739 | 4.404E-36 | Bcells | CTSB |
| 2.4264E-40 | 0.32458241 | 0.268 | 0.052 | 5.732E-36 | Bcells | TCF3 |
| 3.0511E-40 | -1.9065504 | 0.063 | 0.571 | 7.2075E-36 | Bcells | TACSTD2 |
| 3.7094E-40 | 0.78499897 | 0.995 | 0.899 | 8.7626E-36 | Bcells | RPL27A |
| 1.1488E-39 | -1.9915841 | 0.026 | 0.542 | 2.7139E-35 | Bcells | CLDN4 |
| 1.8755E-39 | -1.5248175 | 0.716 | 0.933 | 4.4304E-35 | Bcells | GAPDH |
| 1.8795E-39 | 0.77744928 | 0.995 | 0.958 | 4.44E-35 | Bcells | RPL15 |
| 1.9224E-39 | 0.39355415 | 0.289 | 0.061 | 4.5413E-35 | Bcells | ZNF92 |
| 3.6359E-39 | 0.67483312 | 0.7 | 0.28 | 8.5891E-35 | Bcells | ATRX |
| 3.6488E-39 | -0.9569133 | 0.416 | 0.812 | 8.6195E-35 | Bcells | RAC1 |
| 5.4081E-39 | 0.48794152 | 0.516 | 0.164 | 1.2776E-34 | Bcells | CDC42SE2 |
| 5.5043E-39 | 0.37382302 | 0.363 | 0.088 | 1.3003E-34 | Bcells | EVL |
| 6.2001E-39 | 0.35497689 | 0.279 | 0.057 | 1.4647E-34 | Bcells | LY86 |
| 6.4811E-39 | -2.2021512 | 0.121 | 0.616 | 1.531E-34 | Bcells | C15orf48 |
| 8.6685E-39 | 0.4048085 | 0.379 | 0.095 | 2.0478E-34 | Bcells | AC025164.1 |
| 1.4042E-38 | -1.1681361 | 0.368 | 0.786 | 3.3173E-34 | Bcells | DYNLL1 |
| 2.2487E-38 | 0.75627039 | 0.747 | 0.352 | 5.3122E-34 | Bcells | SEPT7 |
| 2.7077E-38 | 0.59652456 | 0.558 | 0.194 | 6.3965E-34 | Bcells | TNRC6B |
| 3.7032E-38 | 0.4625692 | 0.474 | 0.145 | 8.748E-34 | Bcells | TLK1 |
| 3.9821E-38 | 1.0946372 | 0.926 | 0.763 | 9.4068E-34 | Bcells | BTG1 |
| 4.1384E-38 | -1.9449463 | 0.053 | 0.547 | 9.7762E-34 | Bcells | KRT7 |
| 5.6448E-38 | 0.27634942 | 0.232 | 0.042 | 1.3335E-33 | Bcells | FGD2 |
| 5.7473E-38 | 0.36048735 | 0.342 | 0.083 | 1.3577E-33 | Bcells | RNASEH2B |
| 6.0604E-38 | 0.44409006 | 0.395 | 0.106 | 1.4316E-33 | Bcells | EPM2AIP1 |
| 6.5833E-38 | -1.8063518 | 0.221 | 0.683 | 1.5552E-33 | Bcells | NAMPT |
| 7.428E-38 | 0.53213007 | 0.568 | 0.195 | 1.7547E-33 | Bcells | UPF2 |
| 7.755E-38 | 0.90488728 | 0.532 | 0.184 | 1.832E-33 | Bcells | KLF2 |

| | | | | | | |
|---|---|---|---|---|---|---|
| 3.0014E-37 | 0.696561 | 0.689 | 0.282 | 7.0902E-33 | Bcells | RPS4Y1 |
| 4.2746E-37 | 0.71782339 | 0.721 | 0.315 | 1.0098E-32 | Bcells | KMT2E |
| 4.5758E-37 | -1.1158946 | 0.158 | 0.626 | 1.0809E-32 | Bcells | DSTN |
| 4.6969E-37 | 0.45303977 | 0.353 | 0.09 | 1.1095E-32 | Bcells | SP140L |
| 5.2893E-37 | 0.49069774 | 0.595 | 0.206 | 1.2495E-32 | Bcells | STK17B |
| 6.4611E-37 | 0.47648225 | 0.347 | 0.088 | 1.5263E-32 | Bcells | NFKBID |
| 1.1092E-36 | 0.55736578 | 0.484 | 0.153 | 2.6202E-32 | Bcells | HIST1H1E |
| 1.156E-36 | 0.69383684 | 0.732 | 0.314 | 2.7309E-32 | Bcells | BOD1L1 |
| 1.5883E-36 | 0.32597053 | 0.268 | 0.056 | 3.7522E-32 | Bcells | LYL1 |
| 2.0873E-36 | 0.59972182 | 0.579 | 0.213 | 4.9309E-32 | Bcells | ARID1B |
| 5.7891E-36 | -2.649198 | 0.063 | 0.538 | 1.3676E-31 | Bcells | WFDC2 |
| 6.3285E-36 | -1.0430444 | 0.316 | 0.759 | 1.495E-31 | Bcells | ITGB1 |
| 1.0298E-35 | 0.34815247 | 0.205 | 0.036 | 2.4327E-31 | Bcells | OSBPL10 |
| 1.4392E-35 | 0.80712632 | 1 | 0.945 | 3.3999E-31 | Bcells | PTMA |
| 1.4953E-35 | -1.23531 | 0.895 | 0.965 | 3.5323E-31 | Bcells | UBC |
| 2.861E-35 | -1.7555914 | 0.016 | 0.496 | 6.7586E-31 | Bcells | KRT8 |
| 3.4113E-35 | 0.77857959 | 0.984 | 0.875 | 8.0585E-31 | Bcells | RPL26 |
| 6.7465E-35 | -1.0323891 | 0.258 | 0.689 | 1.5937E-30 | Bcells | BRI3 |
| 9.0049E-35 | 0.54982751 | 0.516 | 0.18 | 2.1272E-30 | Bcells | CYLD |
| 9.785E-35 | -1.3691163 | 0.068 | 0.54 | 2.3115E-30 | Bcells | UPP1 |
| 1.2633E-34 | 0.8116031 | 0.863 | 0.545 | 2.9842E-30 | Bcells | RPS10 |
| 1.3705E-34 | 0.35609812 | 0.332 | 0.084 | 3.2375E-30 | Bcells | ARHGAP45 |
| 1.4773E-34 | -1.1054272 | 0.316 | 0.73 | 3.4899E-30 | Bcells | PKM |
| 1.6039E-34 | 0.59645922 | 0.574 | 0.21 | 3.7889E-30 | Bcells | VPS13C |
| 1.701E-34 | 0.40435715 | 0.311 | 0.075 | 4.0183E-30 | Bcells | GRK3 |
| 1.7723E-34 | 0.44687188 | 0.411 | 0.121 | 4.1867E-30 | Bcells | APPL1 |
| 1.975E-34 | -1.0196486 | 0.384 | 0.777 | 4.6656E-30 | Bcells | TPI1 |
| 2.3532E-34 | 0.79080662 | 0.995 | 0.885 | 5.5591E-30 | Bcells | RPS2 |
| 3.0054E-34 | 0.57454313 | 0.537 | 0.195 | 7.0996E-30 | Bcells | SWAP70 |
| 3.646E-34 | 0.37975789 | 0.4 | 0.114 | 8.613E-30 | Bcells | PDCD7 |
| 8.1784E-34 | 0.70255275 | 0.979 | 0.98 | 1.932E-29 | Bcells | RPL41 |
| 1.7049E-33 | 0.57829862 | 0.658 | 0.279 | 4.0276E-29 | Bcells | TCEA1 |
| 1.7234E-33 | 0.58614468 | 0.563 | 0.212 | 4.0712E-29 | Bcells | NKTR |
| 1.8648E-33 | 0.41171659 | 0.411 | 0.122 | 4.4053E-29 | Bcells | ZNF33A |
| 2.2021E-33 | -1.3319828 | 0.826 | 0.955 | 5.2019E-29 | Bcells | ACTG1 |
| 3.3129E-33 | 0.36120557 | 0.311 | 0.078 | 7.826E-29 | Bcells | RAB29 |
| 3.3876E-33 | 0.38061568 | 0.389 | 0.11 | 8.0025E-29 | Bcells | EPB41 |
| 3.9976E-33 | 0.33821669 | 0.311 | 0.077 | 9.4435E-29 | Bcells | APOBEC3C |
| 4.1926E-33 | 0.39629874 | 0.474 | 0.15 | 9.9043E-29 | Bcells | EVI2B |
| 5.5111E-33 | 0.26279051 | 0.2 | 0.036 | 1.3019E-28 | Bcells | GAPT |
| 6.0031E-33 | 0.54000112 | 0.447 | 0.15 | 1.4181E-28 | Bcells | RB1 |
| 7.0631E-33 | 0.41465209 | 0.316 | 0.08 | 1.6685E-28 | Bcells | TMEM154 |
| 1.1771E-32 | 0.47921446 | 0.447 | 0.144 | 2.7807E-28 | Bcells | UCP2 |
| 1.36E-32 | -1.5988637 | 0.316 | 0.724 | 3.2127E-28 | Bcells | HSPB1 |
| 1.419E-32 | 0.26474007 | 0.137 | 0.019 | 3.352E-28 | Bcells | DERL3 |
| 1.8881E-32 | 0.28461083 | 0.189 | 0.034 | 4.4603E-28 | Bcells | ZEB1 |
| 1.9795E-32 | 0.40328545 | 0.232 | 0.048 | 4.6762E-28 | Bcells | TENT5C |
| 2.695E-32 | -3.5032596 | 0.263 | 0.678 | 6.3663E-28 | Bcells | TIMP1 |
| 4.7439E-32 | 0.49742458 | 0.442 | 0.147 | 1.1206E-27 | Bcells | LNPEP |
| 4.8067E-32 | 0.46235049 | 0.474 | 0.16 | 1.1355E-27 | Bcells | RBM6 |

| | | | | | | |
|---|---|---|---|---|---|---|
| 6.4853E-32 | 0.36965233 | 0.353 | 0.097 | 1.532E-27 | Bcells | CHD7 |
| 6.6688E-32 | -1.1800662 | 0.3 | 0.7 | 1.5754E-27 | Bcells | PPP1R15A |
| 6.7596E-32 | 0.40208236 | 0.316 | 0.082 | 1.5968E-27 | Bcells | MIR29B2CHG |
| 7.1699E-32 | -2.0806306 | 0.2 | 0.661 | 1.6938E-27 | Bcells | MT2A |
| 7.7581E-32 | -1.0200853 | 0.053 | 0.495 | 1.8327E-27 | Bcells | H2AFJ |
| 1.2252E-31 | 0.28782331 | 0.242 | 0.052 | 2.8944E-27 | Bcells | SEPT1 |
| 1.5855E-31 | -0.9505804 | 0.089 | 0.532 | 3.7454E-27 | Bcells | PTTG1IP |
| 1.7845E-31 | 0.70135003 | 0.963 | 0.726 | 4.2156E-27 | Bcells | CYBA |
| 2.1227E-31 | 0.45158735 | 0.432 | 0.137 | 5.0144E-27 | Bcells | ETS1 |
| 2.3298E-31 | 0.51361716 | 0.547 | 0.211 | 5.5036E-27 | Bcells | SETX |
| 2.5404E-31 | 0.56158871 | 0.732 | 0.343 | 6.0011E-27 | Bcells | SCAF11 |
| 2.5597E-31 | 0.57992742 | 0.558 | 0.215 | 6.0467E-27 | Bcells | TRIM38 |
| 3.1668E-31 | 0.3301281 | 0.268 | 0.063 | 7.481E-27 | Bcells | DCK |
| 3.4398E-31 | 0.27285379 | 0.242 | 0.053 | 8.1258E-27 | Bcells | ZNF224 |
| 4.9201E-31 | 0.32222474 | 0.311 | 0.081 | 1.1623E-26 | Bcells | MTERF4 |
| 5.1277E-31 | 0.40119831 | 0.342 | 0.096 | 1.2113E-26 | Bcells | IFT57 |
| 5.1499E-31 | -2.2642037 | 0.984 | 0.987 | 1.2166E-26 | Bcells | FTL |
| 5.322E-31 | -1.1555029 | 0.058 | 0.497 | 1.2572E-26 | Bcells | CD9 |
| 5.9996E-31 | 0.48490919 | 0.505 | 0.181 | 1.4173E-26 | Bcells | PSMB9 |
| 8.1816E-31 | 0.25615549 | 0.221 | 0.045 | 1.9327E-26 | Bcells | ZNF441 |
| 9.9604E-31 | 0.51659724 | 0.579 | 0.232 | 2.3529E-26 | Bcells | KMT2A |
| 1.1921E-30 | -0.8913149 | 0.305 | 0.711 | 2.8161E-26 | Bcells | SDCBP |
| 1.1989E-30 | 0.49777652 | 0.5 | 0.178 | 2.8322E-26 | Bcells | FMNL1 |
| 1.2513E-30 | 0.3303388 | 0.342 | 0.094 | 2.9559E-26 | Bcells | ARL14EP |
| 1.5754E-30 | 0.34412676 | 0.226 | 0.048 | 3.7216E-26 | Bcells | S1PR4 |
| 1.6263E-30 | -3.6707799 | 0.1 | 0.525 | 3.8419E-26 | Bcells | LCN2 |
| 1.7629E-30 | 0.55715512 | 0.605 | 0.243 | 4.1645E-26 | Bcells | KIAA1551 |
| 1.785E-30 | -1.0766414 | 0.679 | 0.907 | 4.2168E-26 | Bcells | ITM2B |
| 2.1151E-30 | -1.461767 | 0.095 | 0.54 | 4.9964E-26 | Bcells | PLAUR |
| 2.47E-30 | 0.38149129 | 0.389 | 0.118 | 5.835E-26 | Bcells | PSIP1 |
| 3.7157E-30 | -1.3470029 | 0.237 | 0.639 | 8.7776E-26 | Bcells | GADD45B |
| 3.8195E-30 | 0.61637465 | 0.763 | 0.379 | 9.0228E-26 | Bcells | N4BP2L2 |
| 4.1274E-30 | 0.40509641 | 0.374 | 0.113 | 9.7501E-26 | Bcells | CHPT1 |
| 4.619E-30 | 0.32979066 | 0.332 | 0.09 | 1.0911E-25 | Bcells | LBH |
| 7.2175E-30 | -1.1947901 | 0.026 | 0.454 | 1.705E-25 | Bcells | CLDN7 |
| 1.2171E-29 | 0.35428943 | 0.305 | 0.081 | 2.8752E-25 | Bcells | ZNF652 |
| 1.2241E-29 | 0.66593578 | 0.979 | 0.848 | 2.8916E-25 | Bcells | RPL38 |
| 1.4607E-29 | 0.43128661 | 0.5 | 0.175 | 3.4506E-25 | Bcells | GLIPR1 |
| 1.688E-29 | 0.51511216 | 0.732 | 0.334 | 3.9875E-25 | Bcells | LUC7L3 |
| 1.7043E-29 | 0.25740528 | 0.189 | 0.036 | 4.026E-25 | Bcells | QRSL1 |
| 2.2635E-29 | -0.9298773 | 0.3 | 0.67 | 5.347E-25 | Bcells | RTN4 |
| 2.7693E-29 | -1.211559 | 0.016 | 0.439 | 6.542E-25 | Bcells | ATP1B1 |
| 4.3161E-29 | -1.4476118 | 0.037 | 0.459 | 1.0196E-24 | Bcells | MXD1 |
| 4.6171E-29 | 0.80973579 | 0.784 | 0.486 | 1.0907E-24 | Bcells | ANKRD12 |
| 4.6715E-29 | 0.27396458 | 0.258 | 0.061 | 1.1036E-24 | Bcells | NDE1 |
| 5.0879E-29 | 0.30874045 | 0.289 | 0.075 | 1.2019E-24 | Bcells | PRKD3 |
| 7.3831E-29 | 0.60699727 | 0.774 | 0.393 | 1.7441E-24 | Bcells | COMMD6 |
| 8.2323E-29 | 0.53793706 | 0.511 | 0.2 | 1.9447E-24 | Bcells | SNX2 |
| 1.3351E-28 | -1.7284505 | 0.032 | 0.452 | 3.1539E-24 | Bcells | SOX4 |
| 1.418E-28 | 0.64488814 | 0.916 | 0.703 | 3.3498E-24 | Bcells | RPL4 |

| | | | | | | |
|---|---|---|---|---|---|---|
| 1.6556E-28 | 0.36560851 | 0.384 | 0.12 | 3.911E-24 | Bcells | RBM26 |
| 1.7371E-28 | -0.8919344 | 0.358 | 0.744 | 4.1037E-24 | Bcells | APLP2 |
| 1.9621E-28 | 0.28708684 | 0.258 | 0.062 | 4.6351E-24 | Bcells | DOCK11 |
| 2.2197E-28 | 0.38424183 | 0.274 | 0.071 | 5.2435E-24 | Bcells | DENND4A |
| 2.6648E-28 | 0.34974984 | 0.258 | 0.063 | 6.295E-24 | Bcells | AGPAT5 |
| 3.9674E-28 | 0.3843074 | 0.632 | 0.244 | 9.3722E-24 | Bcells | GMFG |
| 6.4689E-28 | 0.31542977 | 0.321 | 0.091 | 1.5282E-23 | Bcells | ZNF22 |
| 9.827E-28 | 0.34160077 | 0.284 | 0.075 | 2.3214E-23 | Bcells | PHF6 |
| 1.0024E-27 | 0.63014766 | 0.953 | 0.786 | 2.3681E-23 | Bcells | RPL31 |
| 1.3525E-27 | 0.34393403 | 0.311 | 0.088 | 3.195E-23 | Bcells | PWP1 |
| 1.6971E-27 | -0.9692795 | 0.395 | 0.728 | 4.009E-23 | Bcells | ENO1 |
| 1.7767E-27 | 0.31375941 | 0.332 | 0.095 | 4.197E-23 | Bcells | EMB |
| 2.7751E-27 | 0.47161649 | 0.437 | 0.155 | 6.5556E-23 | Bcells | PTPN1 |
| 3.6422E-27 | 0.34840094 | 0.295 | 0.081 | 8.604E-23 | Bcells | PIKFYVE |
| 3.8306E-27 | 0.37060995 | 0.374 | 0.118 | 9.0489E-23 | Bcells | CENPC |
| 5.1348E-27 | 0.40154161 | 0.326 | 0.094 | 1.213E-22 | Bcells | GPR183 |
| 6.1305E-27 | 0.27499227 | 0.226 | 0.052 | 1.4482E-22 | Bcells | CCDC88C |
| 6.9763E-27 | 0.61861537 | 0.847 | 0.56 | 1.648E-22 | Bcells | TOMM7 |
| 7.0028E-27 | 0.61442835 | 0.989 | 0.919 | 1.6543E-22 | Bcells | NACA |
| 1.0175E-26 | 0.59433975 | 0.674 | 0.344 | 2.4037E-22 | Bcells | MTPN |
| 1.1289E-26 | 0.33008951 | 0.263 | 0.068 | 2.6669E-22 | Bcells | AC008105.3 |
| 1.3733E-26 | -1.0988942 | 0.405 | 0.742 | 3.2441E-22 | Bcells | PSAP |
| 1.4737E-26 | 0.44692054 | 0.516 | 0.2 | 3.4813E-22 | Bcells | IFI16 |
| 1.6343E-26 | 0.57733996 | 1 | 0.977 | 3.8608E-22 | Bcells | RPL28 |
| 1.6692E-26 | 0.43010619 | 0.532 | 0.211 | 3.9431E-22 | Bcells | ITSN2 |
| 1.7046E-26 | 0.44619525 | 0.537 | 0.217 | 4.0268E-22 | Bcells | POLR2J3.1 |
| 2.0597E-26 | 0.38560891 | 0.395 | 0.132 | 4.8657E-22 | Bcells | ELK4 |
| 2.1214E-26 | 0.33307655 | 0.284 | 0.078 | 5.0114E-22 | Bcells | RASSF2 |
| 2.275E-26 | 0.44206072 | 0.505 | 0.197 | 5.3743E-22 | Bcells | MDM4 |
| 2.4193E-26 | 0.27478821 | 0.379 | 0.118 | 5.7151E-22 | Bcells | ATP2B1-AS1 |
| 2.9112E-26 | 0.41659861 | 0.4 | 0.136 | 6.8772E-22 | Bcells | DDX27 |
| 4.5455E-26 | 0.36496487 | 0.311 | 0.091 | 1.0738E-21 | Bcells | KDM4B |
| 4.9438E-26 | 0.50214225 | 0.537 | 0.227 | 1.1679E-21 | Bcells | THRAP3 |
| 5.0647E-26 | 0.36506737 | 0.5 | 0.186 | 1.1964E-21 | Bcells | HCLS1 |
| 5.6562E-26 | -0.8750993 | 0.463 | 0.782 | 1.3362E-21 | Bcells | MAP1LC3B |
| 6.1661E-26 | 0.66226118 | 0.984 | 0.859 | 1.4566E-21 | Bcells | RPL23 |
| 7.0835E-26 | 0.39522984 | 0.384 | 0.127 | 1.6733E-21 | Bcells | UTRN |
| 7.9204E-26 | 0.5217333 | 0.695 | 0.323 | 1.871E-21 | Bcells | ARHGDIB |
| 9.9839E-26 | 0.3592008 | 0.342 | 0.106 | 2.3585E-21 | Bcells | EHMT1 |
| 1.1065E-25 | 0.32054249 | 0.342 | 0.105 | 2.6138E-21 | Bcells | SLC39A10 |
| 1.8553E-25 | 0.36640837 | 0.384 | 0.131 | 4.3828E-21 | Bcells | AIDA |
| 1.9889E-25 | 0.3947181 | 0.353 | 0.112 | 4.6984E-21 | Bcells | MTSS1 |
| 2.1726E-25 | 0.31229036 | 0.295 | 0.085 | 5.1323E-21 | Bcells | VPS51 |
| 2.2797E-25 | -0.9014288 | 0.2 | 0.572 | 5.3854E-21 | Bcells | ZFAND5 |
| 2.3047E-25 | 0.61067348 | 0.974 | 0.916 | 5.4443E-21 | Bcells | UBA52 |
| 2.521E-25 | 0.33194228 | 0.342 | 0.108 | 5.9553E-21 | Bcells | FAM96A |
| 2.6127E-25 | -2.1188978 | 0.068 | 0.466 | 6.1719E-21 | Bcells | LYZ |
| 3.2162E-25 | 0.4755214 | 0.442 | 0.168 | 7.5975E-21 | Bcells | STAT6 |
| 4.8747E-25 | 0.34923107 | 0.237 | 0.06 | 1.1515E-20 | Bcells | PRKX |
| 4.9733E-25 | 0.50308901 | 1 | 0.98 | 1.1748E-20 | Bcells | MALAT1 |

| | | | | | | |
|---|---|---|---|---|---|---|
| 6.53E-25 | 0.37621415 | 0.442 | 0.159 | 1.5426E-20 | Bcells | WIPF1 |
| 6.5483E-25 | -1.1626988 | 0.611 | 0.838 | 1.5469E-20 | Bcells | SQSTM1 |
| 6.5929E-25 | 0.49758953 | 0.705 | 0.348 | 1.5574E-20 | Bcells | PNN |
| 6.7086E-25 | 0.39096354 | 0.368 | 0.125 | 1.5848E-20 | Bcells | CDC40 |
| 6.7822E-25 | -1.7516639 | 0.032 | 0.407 | 1.6022E-20 | Bcells | SLPI |
| 8.1191E-25 | -1.3896445 | 0.195 | 0.542 | 1.918E-20 | Bcells | CD24 |
| 1.0994E-24 | -1.0797743 | 0.142 | 0.527 | 2.597E-20 | Bcells | CEBPB |
| 1.579E-24 | 0.28963162 | 0.258 | 0.069 | 3.7301E-20 | Bcells | RUBCN |
| 2.0457E-24 | 0.28062041 | 0.668 | 0.307 | 4.8326E-20 | Bcells | HIST1H4C |
| 2.0763E-24 | 0.34438406 | 0.332 | 0.105 | 4.9048E-20 | Bcells | MAP3K1 |
| 2.334E-24 | 0.37722593 | 0.405 | 0.143 | 5.5137E-20 | Bcells | SIPA1L1 |
| 2.3597E-24 | 0.33747531 | 0.305 | 0.092 | 5.5743E-20 | Bcells | GPS2 |
| 2.7675E-24 | 0.30154064 | 0.279 | 0.078 | 6.5376E-20 | Bcells | RUNX3 |
| 4.245E-24 | 0.3339899 | 0.368 | 0.123 | 1.0028E-19 | Bcells | AKAP17A |
| 4.3519E-24 | 0.59766893 | 0.953 | 0.783 | 1.028E-19 | Bcells | EEF1D |
| 4.5825E-24 | 0.64596667 | 0.795 | 0.536 | 1.0825E-19 | Bcells | HMGN1 |
| 6.5888E-24 | -1.3674732 | 0.105 | 0.475 | 1.5565E-19 | Bcells | ERO1A |
| 6.7585E-24 | 0.37818963 | 0.358 | 0.122 | 1.5966E-19 | Bcells | CPNE1 |
| 7.9774E-24 | 0.51441801 | 0.579 | 0.271 | 1.8845E-19 | Bcells | UXT |
| 1.0996E-23 | -0.7537048 | 0.374 | 0.687 | 2.5976E-19 | Bcells | ATP6V0B |
| 1.245E-23 | 0.37237954 | 0.458 | 0.173 | 2.941E-19 | Bcells | CDK12 |
| 1.2568E-23 | 0.42580656 | 0.447 | 0.173 | 2.969E-19 | Bcells | ADD1 |
| 1.3803E-23 | 0.53183832 | 0.647 | 0.313 | 3.2607E-19 | Bcells | AKAP9 |
| 1.5744E-23 | -1.588349 | 0.068 | 0.442 | 3.7192E-19 | Bcells | TYROBP |
| 1.6078E-23 | -1.7890085 | 0.011 | 0.37 | 3.798E-19 | Bcells | MAL |
| 1.6574E-23 | -2.7124272 | 0.089 | 0.455 | 3.9152E-19 | Bcells | SPP1 |
| 1.6612E-23 | 0.36970936 | 0.421 | 0.156 | 3.9243E-19 | Bcells | SUPT16H |
| 1.9704E-23 | 0.31304963 | 0.405 | 0.145 | 4.6546E-19 | Bcells | WASHC4 |
| 2.0071E-23 | 0.37248848 | 0.321 | 0.104 | 4.7413E-19 | Bcells | CSNK1G3 |
| 2.2028E-23 | 0.39557291 | 0.489 | 0.196 | 5.2037E-19 | Bcells | CHD1 |
| 2.632E-23 | -2.5915827 | 0.053 | 0.409 | 6.2175E-19 | Bcells | S100P |
| 3.6404E-23 | 0.39694777 | 0.389 | 0.142 | 8.5998E-19 | Bcells | MBD4 |
| 4.286E-23 | 0.38421677 | 0.468 | 0.187 | 1.0125E-18 | Bcells | ORMDL1 |
| 4.3672E-23 | 0.51304102 | 0.521 | 0.227 | 1.0317E-18 | Bcells | SYNGR2 |
| 5.5302E-23 | 0.46844114 | 0.558 | 0.254 | 1.3064E-18 | Bcells | EIF3L |
| 5.589E-23 | -1.1699779 | 0 | 0.353 | 1.3203E-18 | Bcells | PHLDA2 |
| 6.0805E-23 | 0.26543633 | 0.3 | 0.09 | 1.4364E-18 | Bcells | ICAM3 |
| 6.4606E-23 | 0.37506668 | 0.363 | 0.128 | 1.5262E-18 | Bcells | PHKB |
| 7.4566E-23 | 0.45823746 | 0.668 | 0.334 | 1.7615E-18 | Bcells | EIF4B |
| 1.1657E-22 | -0.9369783 | 0.884 | 0.953 | 2.7538E-18 | Bcells | H3F3B |
| 2.1844E-22 | -0.9626744 | 0.116 | 0.466 | 5.1602E-18 | Bcells | CDKN1A |
| 2.3521E-22 | 0.33894845 | 0.305 | 0.097 | 5.5565E-18 | Bcells | ARHGEF1 |
| 3.2488E-22 | 0.48305579 | 0.558 | 0.258 | 7.6746E-18 | Bcells | MBNL1 |
| 3.4303E-22 | 0.26432904 | 0.284 | 0.085 | 8.1033E-18 | Bcells | UHRF2 |
| 3.5992E-22 | 0.31586765 | 0.358 | 0.122 | 8.5024E-18 | Bcells | PHACTR1 |
| 3.7365E-22 | 0.31544328 | 0.326 | 0.107 | 8.8267E-18 | Bcells | MITD1 |
| 4.5304E-22 | 0.5488419 | 1 | 0.895 | 1.0702E-17 | Bcells | RPS9 |
| 4.7678E-22 | 0.33434574 | 0.289 | 0.091 | 1.1263E-17 | Bcells | ANXA6 |
| 5.0879E-22 | 0.62485137 | 0.958 | 0.781 | 1.2019E-17 | Bcells | RPL10A |
| 5.2884E-22 | -0.923413 | 0.221 | 0.573 | 1.2493E-17 | Bcells | VMP1 |

| | | | | | | |
|---|---|---|---|---|---|---|
| 5.4303E-22 | 0.54969307 | 0.995 | 0.957 | 1.2828E-17 | Bcells | RPS18 |
| 6.0937E-22 | 0.25151747 | 0.237 | 0.064 | 1.4395E-17 | Bcells | NPAT |
| 6.2457E-22 | -3.086913 | 0.037 | 0.38 | 1.4754E-17 | Bcells | KRT13 |
| 7.129E-22 | -0.7718658 | 0.342 | 0.676 | 1.6841E-17 | Bcells | TALDO1 |
| 8.5737E-22 | 0.27881828 | 0.295 | 0.092 | 2.0254E-17 | Bcells | IQSEC1 |
| 9.5273E-22 | -1.0889674 | 0.521 | 0.814 | 2.2506E-17 | Bcells | GSTP1 |
| 1.0705E-21 | 0.50347076 | 0.542 | 0.251 | 2.5287E-17 | Bcells | NSD3 |
| 1.195E-21 | 0.42166564 | 0.411 | 0.159 | 2.823E-17 | Bcells | TMPO |
| 1.3414E-21 | -1.2943983 | 0.337 | 0.646 | 3.1688E-17 | Bcells | HSPA5 |
| 1.3961E-21 | 0.42013523 | 0.389 | 0.146 | 3.298E-17 | Bcells | LYST |
| 1.4888E-21 | 0.30516114 | 0.263 | 0.078 | 3.5171E-17 | Bcells | ZDHHC21 |
| 1.8076E-21 | -1.6004898 | 0.011 | 0.346 | 4.2702E-17 | Bcells | GDF15 |
| 1.9064E-21 | 0.53431254 | 0.868 | 0.6 | 4.5034E-17 | Bcells | HINT1 |
| 1.9131E-21 | 0.52269423 | 0.658 | 0.328 | 4.5193E-17 | Bcells | HERPUD1 |
| 1.9522E-21 | 0.30113677 | 0.295 | 0.093 | 4.6116E-17 | Bcells | RGS19 |
| 2.0459E-21 | 0.31036656 | 0.374 | 0.133 | 4.8331E-17 | Bcells | NCKAP1L |
| 2.1055E-21 | 0.25560677 | 0.205 | 0.052 | 4.9737E-17 | Bcells | TMED8 |
| 2.8618E-21 | -0.7639509 | 0.158 | 0.502 | 6.7604E-17 | Bcells | ASAH1 |
| 3.1539E-21 | 0.53291538 | 0.974 | 0.857 | 7.4505E-17 | Bcells | RPL7 |
| 3.3297E-21 | 0.2572953 | 0.232 | 0.063 | 7.8658E-17 | Bcells | MICAL3 |
| 3.3612E-21 | 0.42552604 | 0.484 | 0.207 | 7.9401E-17 | Bcells | RBMX |
| 3.3954E-21 | 0.72477334 | 0.837 | 0.604 | 8.021E-17 | Bcells | EZR |
| 3.8771E-21 | -0.7013611 | 0.547 | 0.801 | 9.1588E-17 | Bcells | GNG5 |
| 4.3581E-21 | 0.35219336 | 0.374 | 0.139 | 1.0295E-16 | Bcells | RPA3 |
| 4.4293E-21 | -0.7672487 | 0.932 | 0.964 | 1.0463E-16 | Bcells | MYL6 |
| 4.5772E-21 | 0.27643238 | 0.258 | 0.076 | 1.0813E-16 | Bcells | CYFIP2 |
| 4.7543E-21 | 0.53644826 | 0.747 | 0.461 | 1.1231E-16 | Bcells | SARAF |
| 5.555E-21 | 0.36790514 | 0.358 | 0.13 | 1.3122E-16 | Bcells | CXXC5 |
| 5.7671E-21 | 0.29715963 | 0.363 | 0.13 | 1.3624E-16 | Bcells | C7orf50 |
| 6.3524E-21 | 0.27404424 | 0.253 | 0.074 | 1.5006E-16 | Bcells | MGAT5 |
| 9.2638E-21 | 0.25184914 | 0.247 | 0.071 | 2.1884E-16 | Bcells | DGKA |
| 1.1316E-20 | 0.32800883 | 0.347 | 0.124 | 2.6732E-16 | Bcells | EPS15 |
| 1.1501E-20 | -1.2400718 | 0.316 | 0.659 | 2.7169E-16 | Bcells | HSPA1A |
| 1.4201E-20 | -0.9498166 | 0.005 | 0.329 | 3.3547E-16 | Bcells | MYO6 |
| 2.172E-20 | 0.34925566 | 0.384 | 0.148 | 5.131E-16 | Bcells | NUB1 |
| 2.8425E-20 | -2.3684628 | 0.021 | 0.343 | 6.7147E-16 | Bcells | PSCA |
| 3.1027E-20 | 0.34701998 | 0.384 | 0.149 | 7.3296E-16 | Bcells | PHF20 |
| 3.3343E-20 | 0.33965934 | 0.258 | 0.079 | 7.8767E-16 | Bcells | CAMK2D |
| 3.6905E-20 | 0.29776622 | 0.189 | 0.048 | 8.718E-16 | Bcells | GRASP |
| 4.5692E-20 | 0.29507158 | 0.3 | 0.1 | 1.0794E-15 | Bcells | DMTF1 |
| 5.8276E-20 | 0.26753683 | 0.342 | 0.12 | 1.3766E-15 | Bcells | ZNF791 |
| 5.8486E-20 | -2.2129949 | 0.026 | 0.346 | 1.3816E-15 | Bcells | HPGD |
| 5.8516E-20 | -0.6469481 | 0.184 | 0.535 | 1.3823E-15 | Bcells | P4HB |
| 6.1595E-20 | -1.4933821 | 0.942 | 0.941 | 1.4551E-15 | Bcells | NEAT1 |
| 7.18E-20 | 0.29063786 | 0.211 | 0.058 | 1.6961E-15 | Bcells | BCL7A |
| 7.8534E-20 | -0.7440768 | 0 | 0.313 | 1.8552E-15 | Bcells | SDC4 |
| 8.0008E-20 | 0.52917484 | 0.789 | 0.516 | 1.89E-15 | Bcells | NAP1L1 |
| 9.0947E-20 | -0.7352424 | 0.053 | 0.375 | 2.1485E-15 | Bcells | METRNL |
| 1.0301E-19 | 0.42733258 | 0.589 | 0.3 | 2.4333E-15 | Bcells | TAPBP |
| 1.0799E-19 | 0.27574648 | 0.279 | 0.09 | 2.551E-15 | Bcells | FAM13B |

| | | | | | | |
|---|---|---|---|---|---|---|
| 1.2087E-19 | 0.45448432 | 0.989 | 0.869 | 2.8553E-15 | Bcells | RPLP0 |
| 1.2903E-19 | 0.25750664 | 0.247 | 0.074 | 3.048E-15 | Bcells | MCPH1 |
| 1.3317E-19 | 0.26197848 | 0.321 | 0.112 | 3.1459E-15 | Bcells | RECQL |
| 1.3992E-19 | -0.9825288 | 0.037 | 0.354 | 3.3053E-15 | Bcells | ATF3 |
| 1.408E-19 | -1.591442 | 0.063 | 0.387 | 3.326E-15 | Bcells | FCER1G |
| 1.4728E-19 | 0.26194003 | 0.263 | 0.081 | 3.4792E-15 | Bcells | SAMD9L |
| 1.5541E-19 | 0.32432634 | 0.326 | 0.117 | 3.6713E-15 | Bcells | PCSK7 |
| 2.0122E-19 | 0.37392526 | 0.379 | 0.145 | 4.7535E-15 | Bcells | PLEKHO1 |
| 2.9825E-19 | -1.0074619 | 0.521 | 0.753 | 7.0454E-15 | Bcells | TPM4 |
| 3.0632E-19 | 0.52075393 | 0.874 | 0.689 | 7.2361E-15 | Bcells | EEF2 |
| 3.7486E-19 | -0.9881006 | 0.147 | 0.471 | 8.8552E-15 | Bcells | DUSP1 |
| 3.8573E-19 | -1.0300738 | 0.4 | 0.668 | 9.1122E-15 | Bcells | ANXA5 |
| 4.0659E-19 | 0.26861377 | 0.342 | 0.124 | 9.6049E-15 | Bcells | PPHLN1 |
| 4.1748E-19 | 0.30161546 | 0.358 | 0.135 | 9.8622E-15 | Bcells | CSNK1G2 |
| 5.5657E-19 | 0.41247468 | 0.411 | 0.174 | 1.3148E-14 | Bcells | NCOA3 |
| 5.8427E-19 | 0.37847459 | 0.353 | 0.135 | 1.3802E-14 | Bcells | ZBTB1 |
| 5.8568E-19 | 0.43216457 | 0.858 | 0.612 | 1.3835E-14 | Bcells | SRRM2 |
| 5.9355E-19 | 0.34876503 | 0.305 | 0.107 | 1.4021E-14 | Bcells | CWF19L2 |
| 6.9443E-19 | 0.39204039 | 0.474 | 0.212 | 1.6404E-14 | Bcells | PHF3 |
| 7.1936E-19 | -1.5397658 | 0.026 | 0.334 | 1.6993E-14 | Bcells | IL1RN |
| 7.3249E-19 | 0.27118225 | 0.279 | 0.092 | 1.7304E-14 | Bcells | HCG18 |
| 8.2003E-19 | -0.6400362 | 0.289 | 0.584 | 1.9371E-14 | Bcells | CLTA |
| 8.4912E-19 | 0.41613976 | 0.663 | 0.351 | 2.0059E-14 | Bcells | EIF3E |
| 9.9066E-19 | -0.7603289 | 0.021 | 0.329 | 2.3402E-14 | Bcells | LMNA |
| 1.1133E-18 | 0.37462633 | 0.468 | 0.202 | 2.6299E-14 | Bcells | LSP1 |
| 1.1178E-18 | 0.40360129 | 0.474 | 0.215 | 2.6405E-14 | Bcells | DDX6 |
| 1.2741E-18 | 0.54855461 | 0.653 | 0.365 | 3.0098E-14 | Bcells | AES |
| 1.805E-18 | 0.2752056 | 0.305 | 0.107 | 4.264E-14 | Bcells | RAB30-AS1 |
| 1.9522E-18 | 0.39577318 | 0.605 | 0.303 | 4.6117E-14 | Bcells | HP1BP3 |
| 2.0613E-18 | 0.26481546 | 0.605 | 0.266 | 4.8695E-14 | Bcells | CD53 |
| 2.1013E-18 | 0.47898173 | 0.984 | 0.875 | 4.9639E-14 | Bcells | RPL36 |
| 2.1725E-18 | -0.8257888 | 0.189 | 0.487 | 5.132E-14 | Bcells | HMGA1 |
| 2.2343E-18 | 0.29593826 | 0.495 | 0.212 | 5.278E-14 | Bcells | BASP1 |
| 2.2927E-18 | -0.7721814 | 0.021 | 0.321 | 5.4161E-14 | Bcells | PERP |
| 2.3364E-18 | 0.45085793 | 0.605 | 0.323 | 5.5193E-14 | Bcells | NSA2 |
| 2.3642E-18 | 0.48508219 | 0.947 | 0.815 | 5.585E-14 | Bcells | PFDN5 |
| 2.43E-18 | 0.35768412 | 0.395 | 0.162 | 5.7403E-14 | Bcells | SETD2 |
| 2.5295E-18 | 0.30416108 | 0.353 | 0.134 | 5.9754E-14 | Bcells | TGFBR2 |
| 2.7735E-18 | -0.8328647 | 0.316 | 0.604 | 6.5518E-14 | Bcells | MCL1 |
| 3.3546E-18 | -0.7328412 | 0.032 | 0.33 | 7.9245E-14 | Bcells | ASPH |
| 3.6362E-18 | -0.654759 | 0.179 | 0.488 | 8.5898E-14 | Bcells | DYNLT1 |
| 3.8414E-18 | -0.6551079 | 0.032 | 0.331 | 9.0744E-14 | Bcells | RBM47 |
| 4.1599E-18 | 0.33092799 | 0.395 | 0.162 | 9.827E-14 | Bcells | KLF13 |
| 4.7355E-18 | 0.44783504 | 0.389 | 0.165 | 1.1187E-13 | Bcells | CCDC50 |
| 5.3538E-18 | -4.2934773 | 0.1 | 0.403 | 1.2647E-13 | Bcells | SPRR3 |
| 6.1591E-18 | 0.40215888 | 0.568 | 0.287 | 1.455E-13 | Bcells | ARID4B |
| 6.5151E-18 | -0.7686514 | 0.032 | 0.328 | 1.5391E-13 | Bcells | MAFF |
| 7.394E-18 | 0.29061541 | 0.3 | 0.107 | 1.7467E-13 | Bcells | PLEKHF2 |
| 7.4995E-18 | -0.7224889 | 0.347 | 0.623 | 1.7716E-13 | Bcells | TMEM59 |
| 9.4634E-18 | 0.255334 | 0.279 | 0.095 | 2.2355E-13 | Bcells | ZNF770 |

| | | | | | | |
|---|---|---|---|---|---|---|
| 9.5372E-18 | -1.2053672 | 0.005 | 0.293 | 2.253E-13 | Bcells | RND3 |
| 9.6083E-18 | -1.0463517 | 0.005 | 0.293 | 2.2698E-13 | Bcells | GPRC5A |
| 9.9962E-18 | 0.30399713 | 0.347 | 0.133 | 2.3614E-13 | Bcells | NT5C |
| 1.0741E-17 | 0.40101106 | 0.584 | 0.296 | 2.5374E-13 | Bcells | ARL6IP5 |
| 1.1915E-17 | -0.7682465 | 0.011 | 0.3 | 2.8146E-13 | Bcells | FAM129B |
| 1.2306E-17 | -1.6506442 | 0.047 | 0.337 | 2.907E-13 | Bcells | IGFBP7 |
| 1.3717E-17 | 0.39732586 | 0.663 | 0.367 | 3.2403E-13 | Bcells | ANKRD11 |
| 1.5198E-17 | 0.27942038 | 0.305 | 0.11 | 3.5901E-13 | Bcells | MARF1 |
| 1.5376E-17 | -1.55981 | 0.011 | 0.297 | 3.6323E-13 | Bcells | ADIRF |
| 1.5512E-17 | 0.27486014 | 0.284 | 0.099 | 3.6644E-13 | Bcells | CASP8 |
| 1.5899E-17 | 0.25334081 | 0.316 | 0.115 | 3.7559E-13 | Bcells | CNOT6L |
| 1.6176E-17 | 0.39055366 | 0.542 | 0.265 | 3.8212E-13 | Bcells | DDX46 |
| 1.6183E-17 | 0.28158736 | 0.289 | 0.102 | 3.823E-13 | Bcells | CSK |
| 1.6471E-17 | -1.0097586 | 0.032 | 0.32 | 3.891E-13 | Bcells | GADD45A |
| 1.7125E-17 | -0.6815603 | 0.053 | 0.349 | 4.0454E-13 | Bcells | TIMP2 |
| 1.7932E-17 | 0.43272319 | 0.437 | 0.199 | 4.236E-13 | Bcells | TTC3 |
| 1.8798E-17 | 0.37336896 | 0.605 | 0.313 | 4.4407E-13 | Bcells | EIF3F |
| 2.126E-17 | 0.40679428 | 0.579 | 0.306 | 5.0223E-13 | Bcells | ANAPC16 |
| 2.5717E-17 | 0.25788445 | 0.263 | 0.089 | 6.075E-13 | Bcells | RNF216 |
| 2.5792E-17 | -0.8197228 | 0.121 | 0.41 | 6.0928E-13 | Bcells | CD59 |
| 2.6617E-17 | 0.28089474 | 0.268 | 0.092 | 6.2877E-13 | Bcells | LPXN |
| 2.6882E-17 | -0.9413179 | 0.032 | 0.324 | 6.3504E-13 | Bcells | IFITM3 |
| 3.2786E-17 | 0.2704845 | 0.279 | 0.097 | 7.7451E-13 | Bcells | CCND3 |
| 3.6369E-17 | -1.369263 | 0.053 | 0.346 | 8.5915E-13 | Bcells | CTSL |
| 3.866E-17 | 0.42423323 | 0.621 | 0.334 | 9.1327E-13 | Bcells | RBM25 |
| 3.9739E-17 | -1.2486197 | 0.1 | 0.391 | 9.3875E-13 | Bcells | FOS |
| 4.0098E-17 | 0.48014687 | 0.658 | 0.387 | 9.4724E-13 | Bcells | HMGN2 |
| 4.0722E-17 | 0.27714307 | 0.342 | 0.131 | 9.6197E-13 | Bcells | SMARCA2 |
| 4.5667E-17 | 0.26239746 | 0.211 | 0.063 | 1.0788E-12 | Bcells | RPS6KA5 |
| 4.8503E-17 | -1.1267574 | 0.205 | 0.493 | 1.1458E-12 | Bcells | MTRNR2L8 |
| 5.6998E-17 | -0.8520321 | 0.037 | 0.327 | 1.3465E-12 | Bcells | CEBPD |
| 5.8332E-17 | 0.26848244 | 0.247 | 0.082 | 1.378E-12 | Bcells | PACS1 |
| 6.9E-17 | 0.27272477 | 0.253 | 0.085 | 1.63E-12 | Bcells | INO80 |
| 7.0088E-17 | 0.27052665 | 0.758 | 0.423 | 1.6557E-12 | Bcells | EMP3 |
| 7.0876E-17 | 0.43440037 | 0.621 | 0.337 | 1.6743E-12 | Bcells | GPBP1 |
| 7.9835E-17 | -0.8007452 | 0.005 | 0.282 | 1.8859E-12 | Bcells | H1F0 |
| 8.1504E-17 | 0.42867201 | 0.647 | 0.345 | 1.9254E-12 | Bcells | ZFP36L2 |
| 1.0143E-16 | 0.30277772 | 0.3 | 0.113 | 2.3962E-12 | Bcells | DRAM2 |
| 1.0911E-16 | 0.43702882 | 0.863 | 0.593 | 2.5775E-12 | Bcells | DDX17 |
| 1.221E-16 | -0.9074076 | 0.026 | 0.303 | 2.8844E-12 | Bcells | SERPINA1 |
| 1.283E-16 | 0.28407692 | 0.279 | 0.099 | 3.0309E-12 | Bcells | MSL3 |
| 1.4584E-16 | 0.29314865 | 0.363 | 0.146 | 3.4451E-12 | Bcells | DDX3Y |
| 1.6106E-16 | 0.30318111 | 0.5 | 0.233 | 3.8048E-12 | Bcells | FNBP4 |
| 1.8565E-16 | 0.2711227 | 0.321 | 0.122 | 4.3856E-12 | Bcells | TTC14 |
| 2.0342E-16 | 0.26703105 | 0.284 | 0.103 | 4.8053E-12 | Bcells | ITGAE |
| 2.1573E-16 | -0.732378 | 0.058 | 0.339 | 5.0961E-12 | Bcells | NDRG1 |
| 2.1842E-16 | 0.40152227 | 0.605 | 0.332 | 5.1596E-12 | Bcells | HNRNPD |
| 2.2141E-16 | -0.9249277 | 0.047 | 0.324 | 5.2303E-12 | Bcells | VEGFA |
| 2.2724E-16 | 0.31612773 | 0.595 | 0.3 | 5.368E-12 | Bcells | DDX24 |
| 2.3621E-16 | 0.32914514 | 0.389 | 0.165 | 5.5799E-12 | Bcells | NSD1 |

| | | | | | | |
|---|---|---|---|---|---|---|
| 2.5842E-16 | 0.2760004 | 0.295 | 0.109 | 6.1046E-12 | Bcells | RABEP1 |
| 2.6812E-16 | 0.36556619 | 0.521 | 0.26 | 6.3337E-12 | Bcells | ELF1 |
| 2.696E-16 | 0.31531196 | 1 | 0.99 | 6.3688E-12 | Bcells | B2M |
| 2.7447E-16 | -2.5071022 | 0.063 | 0.34 | 6.4838E-12 | Bcells | MMP7 |
| 2.873E-16 | 0.35122412 | 0.516 | 0.251 | 6.787E-12 | Bcells | SRSF10 |
| 3.3185E-16 | 0.30524036 | 0.495 | 0.23 | 7.8393E-12 | Bcells | THUMPD3-AS1 |
| 3.3286E-16 | -0.7505669 | 0.253 | 0.514 | 7.8631E-12 | Bcells | ANXA11 |
| 3.5149E-16 | 0.37121547 | 0.347 | 0.138 | 8.3032E-12 | Bcells | PLAC8 |
| 4.6589E-16 | 0.41955644 | 0.289 | 0.108 | 1.1006E-11 | Bcells | ITM2C |
| 4.8853E-16 | 0.32691383 | 0.595 | 0.308 | 1.1541E-11 | Bcells | IRF1 |
| 5.1907E-16 | 0.26287893 | 0.326 | 0.127 | 1.2262E-11 | Bcells | ZNF800 |
| 5.4052E-16 | -1.0445646 | 0.016 | 0.282 | 1.2769E-11 | Bcells | CLU |
| 5.5696E-16 | 0.34876512 | 0.416 | 0.182 | 1.3157E-11 | Bcells | CDK13 |
| 5.7783E-16 | 0.31111125 | 0.684 | 0.39 | 1.365E-11 | Bcells | SRSF7 |
| 6.3459E-16 | 0.27171115 | 0.326 | 0.129 | 1.4991E-11 | Bcells | LSM10 |
| 6.4118E-16 | -0.8524942 | 0.011 | 0.275 | 1.5147E-11 | Bcells | MAL2 |
| 6.8552E-16 | 0.32672176 | 0.453 | 0.212 | 1.6194E-11 | Bcells | RPL17 |
| 6.9352E-16 | 0.28835245 | 0.4 | 0.17 | 1.6383E-11 | Bcells | DAPP1 |
| 6.9463E-16 | 0.28767741 | 0.316 | 0.122 | 1.6409E-11 | Bcells | ADD3 |
| 6.9992E-16 | -1.1275419 | 0.042 | 0.313 | 1.6534E-11 | Bcells | PHLDA1 |
| 8.0153E-16 | 0.25426085 | 0.332 | 0.131 | 1.8934E-11 | Bcells | ARIH2 |
| 8.1696E-16 | 0.41207183 | 0.468 | 0.225 | 1.9299E-11 | Bcells | XPO1 |
| 8.6362E-16 | -1.1040298 | 0.042 | 0.313 | 2.0401E-11 | Bcells | EMP1 |
| 9.2343E-16 | -0.5170569 | 0.016 | 0.279 | 2.1814E-11 | Bcells | ADAM9 |
| 1.0787E-15 | 0.40538168 | 0.511 | 0.261 | 2.5481E-11 | Bcells | MPHOSPH8 |
| 1.1003E-15 | -0.5852664 | 0.042 | 0.311 | 2.5992E-11 | Bcells | RAB13 |
| 1.1125E-15 | 0.29824557 | 0.247 | 0.086 | 2.628E-11 | Bcells | DMXL1 |
| 1.5245E-15 | -0.6577117 | 0.363 | 0.6 | 3.6012E-11 | Bcells | VAMP8 |
| 1.6403E-15 | -1.0641591 | 0.021 | 0.283 | 3.8749E-11 | Bcells | MAFB |
| 1.6864E-15 | 0.33615682 | 0.647 | 0.351 | 3.9838E-11 | Bcells | TAF1D |
| 1.7894E-15 | 0.35622406 | 0.547 | 0.279 | 4.2271E-11 | Bcells | LYN |
| 1.9317E-15 | -0.79064 | 0.016 | 0.275 | 4.5633E-11 | Bcells | TNFRSF12A |
| 1.9811E-15 | 0.35884686 | 0.558 | 0.295 | 4.6799E-11 | Bcells | RABAC1 |
| 2.0028E-15 | 0.43862453 | 0.642 | 0.385 | 4.7312E-11 | Bcells | PSME1 |
| 2.0081E-15 | 0.4161958 | 0.989 | 0.889 | 4.7436E-11 | Bcells | HLA-B |
| 2.0638E-15 | -2.0425869 | 0.316 | 0.571 | 4.8752E-11 | Bcells | SRGN |
| 2.0945E-15 | -0.6574811 | 0.153 | 0.429 | 4.9478E-11 | Bcells | EFHD2 |
| 2.1505E-15 | 0.27264479 | 0.326 | 0.129 | 5.0802E-11 | Bcells | CELF2 |
| 2.2138E-15 | 0.29548911 | 0.463 | 0.214 | 5.2297E-11 | Bcells | EML4 |
| 2.5599E-15 | 0.29288053 | 0.521 | 0.257 | 6.0473E-11 | Bcells | CCT3 |
| 2.8405E-15 | 0.275346 | 0.326 | 0.131 | 6.7101E-11 | Bcells | BRAF |
| 2.9165E-15 | -0.7593135 | 0.005 | 0.261 | 6.8897E-11 | Bcells | MUC1 |
| 3.2091E-15 | -0.672021 | 0.032 | 0.291 | 7.5808E-11 | Bcells | SPINT1 |
| 3.2425E-15 | 0.29846721 | 0.358 | 0.153 | 7.6598E-11 | Bcells | CCAR1 |
| 3.595E-15 | -0.9452523 | 0.195 | 0.457 | 8.4925E-11 | Bcells | RAB11FIP1 |
| 3.6935E-15 | 0.28227351 | 0.326 | 0.132 | 8.7252E-11 | Bcells | PPP3CA |
| 3.9711E-15 | -0.5601828 | 0.053 | 0.326 | 9.3809E-11 | Bcells | FOSL2 |
| 3.9937E-15 | 0.56313832 | 0.516 | 0.291 | 9.4342E-11 | Bcells | SKIL |
| 4.0803E-15 | 0.31158736 | 0.305 | 0.121 | 9.6389E-11 | Bcells | NIN |
| 4.1933E-15 | -0.5986074 | 0.032 | 0.29 | 9.9059E-11 | Bcells | MIR4435-2HG |

| | | | | | | |
|---|---|---|---|---|---|---|
| 4.2686E-15 | 0.5255085 | 0.879 | 0.721 | 1.0084E-10 | Bcells | JUND |
| 4.3641E-15 | -0.595569 | 0.121 | 0.396 | 1.0309E-10 | Bcells | CTNNA1 |
| 4.4906E-15 | 0.25075277 | 0.4 | 0.175 | 1.0608E-10 | Bcells | RBBP4 |
| 4.5033E-15 | -0.6758663 | 0.332 | 0.599 | 1.0638E-10 | Bcells | NPC2 |
| 5.5926E-15 | 0.4227995 | 0.389 | 0.175 | 1.3211E-10 | Bcells | RERE |
| 5.9394E-15 | 0.25184765 | 0.242 | 0.084 | 1.4031E-10 | Bcells | LFNG |
| 6.0185E-15 | -0.7649886 | 0.274 | 0.524 | 1.4218E-10 | Bcells | HEBP2 |
| 6.1057E-15 | 0.28298017 | 0.374 | 0.161 | 1.4423E-10 | Bcells | RSBN1L |
| 6.2188E-15 | 0.39501931 | 0.958 | 0.83 | 1.4691E-10 | Bcells | HNRNPA2B1 |
| 6.2931E-15 | -0.740677 | 0.021 | 0.275 | 1.4866E-10 | Bcells | ADM |
| 6.7578E-15 | -0.7528464 | 0.011 | 0.263 | 1.5964E-10 | Bcells | HES4 |
| 6.8891E-15 | -0.8668179 | 0 | 0.247 | 1.6274E-10 | Bcells | CST6 |
| 7.2387E-15 | 0.25711458 | 0.326 | 0.13 | 1.71E-10 | Bcells | APBB1IP |
| 7.5322E-15 | 0.2998324 | 0.437 | 0.205 | 1.7793E-10 | Bcells | PPP4R3A |
| 7.7712E-15 | -0.704148 | 0.053 | 0.313 | 1.8358E-10 | Bcells | CD68 |
| 7.93E-15 | -0.7781123 | 0.284 | 0.537 | 1.8733E-10 | Bcells | SERPINB1 |
| 8.0772E-15 | 0.36282066 | 0.484 | 0.247 | 1.9081E-10 | Bcells | RPS17 |
| 8.0919E-15 | 0.25273293 | 0.205 | 0.067 | 1.9115E-10 | Bcells | GNB5 |
| 9.0544E-15 | 0.31230829 | 0.258 | 0.095 | 2.1389E-10 | Bcells | HIVEP1 |
| 9.1104E-15 | -0.7485921 | 0.016 | 0.265 | 2.1521E-10 | Bcells | MGST1 |
| 9.2924E-15 | 0.33704685 | 0.326 | 0.138 | 2.1951E-10 | Bcells | PCGF5 |
| 9.4193E-15 | 0.34332391 | 0.563 | 0.292 | 2.2251E-10 | Bcells | FXYD5 |
| 1.0043E-14 | 0.28808183 | 0.368 | 0.161 | 2.3724E-10 | Bcells | RAB8A |
| 1.0107E-14 | 0.26337456 | 0.389 | 0.171 | 2.3875E-10 | Bcells | BBX |
| 1.0875E-14 | 0.30704616 | 0.495 | 0.242 | 2.5691E-10 | Bcells | ZRANB2 |
| 1.2095E-14 | 0.46412653 | 0.779 | 0.56 | 2.8571E-10 | Bcells | SET |
| 1.3091E-14 | 0.30343854 | 0.337 | 0.142 | 3.0924E-10 | Bcells | WBP11 |
| 1.4591E-14 | 0.39974568 | 0.353 | 0.158 | 3.4467E-10 | Bcells | NFKB2 |
| 1.478E-14 | 0.36258902 | 0.526 | 0.275 | 3.4914E-10 | Bcells | ROCK1 |
| 1.5812E-14 | -0.5980555 | 0.005 | 0.248 | 3.7353E-10 | Bcells | SERINC2 |
| 1.6138E-14 | -0.6570028 | 0.026 | 0.276 | 3.8124E-10 | Bcells | ETS2 |
| 1.746E-14 | -0.7986812 | 0.026 | 0.276 | 4.1247E-10 | Bcells | CAMK2N1 |
| 1.838E-14 | 0.28107267 | 0.411 | 0.186 | 4.3418E-10 | Bcells | EPC1 |
| 1.8516E-14 | 0.2666901 | 0.311 | 0.124 | 4.374E-10 | Bcells | SYK |
| 1.8527E-14 | -0.6217162 | 0.221 | 0.493 | 4.3766E-10 | Bcells | SELENOW |
| 1.8704E-14 | 0.2502986 | 0.305 | 0.122 | 4.4185E-10 | Bcells | TFAM |
| 2.0179E-14 | 0.307528 | 0.347 | 0.148 | 4.7668E-10 | Bcells | ZC3HAV1 |
| 2.205E-14 | -0.6542347 | 0.489 | 0.709 | 5.2089E-10 | Bcells | GPX4 |
| 2.2219E-14 | -1.015637 | 0.021 | 0.265 | 5.2487E-10 | Bcells | TSPAN1 |
| 2.2361E-14 | -0.5339413 | 0.084 | 0.347 | 5.2823E-10 | Bcells | CAPN2 |
| 2.2615E-14 | 0.28162587 | 0.289 | 0.113 | 5.3423E-10 | Bcells | AHI1 |
| 2.3143E-14 | 0.25552377 | 0.326 | 0.134 | 5.4671E-10 | Bcells | KIAA2026 |
| 2.5256E-14 | 0.28307447 | 0.295 | 0.118 | 5.9663E-10 | Bcells | CDKN2D |
| 3.3631E-14 | -0.58767 | 0.274 | 0.521 | 7.9446E-10 | Bcells | ATP6V1F |
| 3.4668E-14 | 0.40174593 | 0.684 | 0.426 | 8.1897E-10 | Bcells | SEC62 |
| 3.4906E-14 | 0.25605227 | 0.395 | 0.178 | 8.2459E-10 | Bcells | ATG12 |
| 3.5902E-14 | -0.5953288 | 0.105 | 0.359 | 8.4811E-10 | Bcells | PLSCR1 |
| 3.8152E-14 | -0.7777153 | 0 | 0.236 | 9.0126E-10 | Bcells | CDKN1C |
| 4.0068E-14 | -0.7694336 | 0.184 | 0.428 | 9.4652E-10 | Bcells | SPINT2 |
| 4.0287E-14 | 0.31505403 | 0.437 | 0.209 | 9.5169E-10 | Bcells | TRIM44 |

| | | | | | | |
|---|---|---|---|---|---|---|
| 4.2689E-14 | -0.5868247 | 0.258 | 0.503 | 1.0084E-09 | Bcells | LAPTM4A |
| 4.386E-14 | -0.5610463 | 0.111 | 0.37 | 1.0361E-09 | Bcells | TXNDC17 |
| 4.4781E-14 | -0.8709524 | 0 | 0.235 | 1.0579E-09 | Bcells | PDZK1IP1 |
| 4.4839E-14 | 0.29537192 | 0.363 | 0.158 | 1.0592E-09 | Bcells | DOCK8 |
| 4.6522E-14 | 0.34532985 | 0.458 | 0.229 | 1.099E-09 | Bcells | KDM5A |
| 4.7193E-14 | -0.5547791 | 0.053 | 0.304 | 1.1148E-09 | Bcells | BLVRB |
| 4.8337E-14 | -0.7952805 | 0.189 | 0.434 | 1.1419E-09 | Bcells | GSTO1 |
| 4.9013E-14 | -0.7359112 | 0.411 | 0.648 | 1.1578E-09 | Bcells | LDHA |
| 5.5575E-14 | -0.9257031 | 0.232 | 0.506 | 1.3128E-09 | Bcells | HSPA1B |
| 5.6447E-14 | 0.37853124 | 0.579 | 0.329 | 1.3335E-09 | Bcells | UBXN1 |
| 5.7594E-14 | 0.25680832 | 0.316 | 0.13 | 1.3605E-09 | Bcells | LMBRD1 |
| 5.9061E-14 | -0.6468353 | 0.037 | 0.28 | 1.3952E-09 | Bcells | ABHD5 |
| 5.9676E-14 | -0.9483132 | 0.011 | 0.247 | 1.4097E-09 | Bcells | PLAU |
| 6.5521E-14 | -0.5134742 | 0 | 0.232 | 1.5478E-09 | Bcells | TJP1 |
| 7.242E-14 | 0.30605359 | 0.311 | 0.131 | 1.7108E-09 | Bcells | DNAJC10 |
| 8.4228E-14 | 0.38840569 | 0.974 | 0.901 | 1.9897E-09 | Bcells | RPL24 |
| 8.8861E-14 | 0.33642397 | 0.574 | 0.32 | 2.0992E-09 | Bcells | YPEL3 |
| 8.994E-14 | 0.31754999 | 0.305 | 0.128 | 2.1246E-09 | Bcells | UBTF |
| 9.1864E-14 | -0.7963811 | 0 | 0.23 | 2.1701E-09 | Bcells | ITGA2 |
| 9.9037E-14 | -0.6075867 | 0.458 | 0.686 | 2.3395E-09 | Bcells | BZW1 |
| 1.0151E-13 | 0.31721954 | 0.337 | 0.149 | 2.398E-09 | Bcells | SINHCAF |
| 1.0154E-13 | -0.7215013 | 0.032 | 0.274 | 2.3988E-09 | Bcells | TNFAIP2 |
| 1.0207E-13 | -2.0365499 | 0.032 | 0.267 | 2.4112E-09 | Bcells | S100A2 |
| 1.0289E-13 | 0.26486695 | 0.279 | 0.11 | 2.4305E-09 | Bcells | USP48 |
| 1.0571E-13 | -0.8613879 | 0.005 | 0.236 | 2.4972E-09 | Bcells | EPCAM |
| 1.1069E-13 | -0.6830654 | 0.011 | 0.243 | 2.6148E-09 | Bcells | PRSS8 |
| 1.1119E-13 | -0.7365648 | 0.011 | 0.243 | 2.6266E-09 | Bcells | S100A13 |
| 1.1781E-13 | 0.27004246 | 0.305 | 0.126 | 2.7831E-09 | Bcells | UPF3A |
| 1.2443E-13 | -0.5379658 | 0.011 | 0.242 | 2.9394E-09 | Bcells | GIPC1 |
| 1.3403E-13 | 0.33964087 | 0.432 | 0.211 | 3.1662E-09 | Bcells | USP34 |
| 1.4459E-13 | 0.25158734 | 0.242 | 0.089 | 3.4157E-09 | Bcells | UPF3B |
| 1.4556E-13 | 0.25474345 | 0.321 | 0.135 | 3.4385E-09 | Bcells | MIA3 |
| 1.5481E-13 | -0.9365808 | 0.026 | 0.26 | 3.657E-09 | Bcells | TPM1 |
| 1.6623E-13 | 0.31840545 | 0.337 | 0.15 | 3.9268E-09 | Bcells | LPGAT1 |
| 1.6694E-13 | -0.493479 | 0.032 | 0.271 | 3.9436E-09 | Bcells | QSOX1 |
| 1.6853E-13 | -0.5428402 | 0.321 | 0.575 | 3.9813E-09 | Bcells | PPP1CB |
| 1.7317E-13 | 0.25322814 | 0.284 | 0.114 | 4.0907E-09 | Bcells | MCUB |
| 1.9049E-13 | -0.9443658 | 0.011 | 0.239 | 4.5001E-09 | Bcells | ADGRF1 |
| 1.9983E-13 | -0.6090278 | 0.084 | 0.332 | 4.7207E-09 | Bcells | APP |
| 2.0585E-13 | 0.36421083 | 0.6 | 0.35 | 4.8627E-09 | Bcells | SNX3 |
| 2.1138E-13 | -0.5203668 | 0.074 | 0.316 | 4.9935E-09 | Bcells | MMP24OS |
| 2.243E-13 | -0.4836393 | 0.037 | 0.275 | 5.2987E-09 | Bcells | AAK1 |
| 2.3055E-13 | 0.28646321 | 0.432 | 0.212 | 5.4463E-09 | Bcells | LSM8 |
| 2.4154E-13 | 0.35198572 | 0.463 | 0.234 | 5.7058E-09 | Bcells | SNHG7 |
| 2.4395E-13 | -0.7233387 | 0.016 | 0.247 | 5.7629E-09 | Bcells | CSTA |
| 2.5663E-13 | 0.41561542 | 0.653 | 0.432 | 6.0624E-09 | Bcells | TRIR |
| 2.6103E-13 | -0.6752458 | 0.021 | 0.251 | 6.1664E-09 | Bcells | KLF5 |
| 2.6184E-13 | -0.652751 | 0.011 | 0.237 | 6.1853E-09 | Bcells | S100A16 |
| 3.2441E-13 | -0.6588283 | 0.342 | 0.573 | 7.6635E-09 | Bcells | TSPO |
| 3.3843E-13 | 0.32983137 | 0.689 | 0.414 | 7.9946E-09 | Bcells | FUS |

| | | | | | | |
|---|---|---|---|---|---|---|
| 3.5917E-13 | -0.51134 | 0.547 | 0.748 | 8.4848E-09 | Bcells | ATP6V0E1 |
| 3.6073E-13 | 0.28822394 | 0.447 | 0.219 | 8.5214E-09 | Bcells | PCM1 |
| 3.6939E-13 | -0.4594332 | 0.389 | 0.637 | 8.7261E-09 | Bcells | ARF1 |
| 3.804E-13 | -0.7134899 | 0.016 | 0.244 | 8.9862E-09 | Bcells | ITGB8 |
| 3.81E-13 | -0.5609509 | 0.047 | 0.284 | 9.0003E-09 | Bcells | CD151 |
| 3.8998E-13 | 0.37649291 | 0.511 | 0.281 | 9.2124E-09 | Bcells | PRPF4B |
| 4.1555E-13 | 0.33934041 | 0.484 | 0.248 | 9.8165E-09 | Bcells | CHD4 |
| 4.7511E-13 | 0.25063143 | 0.3 | 0.124 | 1.1224E-08 | Bcells | ZBTB44 |
| 4.7673E-13 | -1.1724635 | 0.037 | 0.264 | 1.1262E-08 | Bcells | LMO7 |
| 5.6508E-13 | -1.0622835 | 0.732 | 0.859 | 1.3349E-08 | Bcells | SH3BGRL3 |
| 5.9642E-13 | -1.6310306 | 0.011 | 0.233 | 1.4089E-08 | Bcells | PRSS22 |
| 6.5251E-13 | 0.32526402 | 0.458 | 0.237 | 1.5414E-08 | Bcells | ASH1L |
| 7.2183E-13 | -0.5290827 | 0.063 | 0.302 | 1.7052E-08 | Bcells | SERTAD1 |
| 7.4387E-13 | 0.25532298 | 0.432 | 0.211 | 1.7572E-08 | Bcells | RNPS1 |
| 8.4667E-13 | -0.5690746 | 0.068 | 0.312 | 2.0001E-08 | Bcells | FNDC3B |
| 8.5681E-13 | -0.6895776 | 0.037 | 0.268 | 2.0241E-08 | Bcells | ID2 |
| 8.9374E-13 | -0.5357221 | 0 | 0.215 | 2.1113E-08 | Bcells | EHF |
| 9.1983E-13 | 0.29945577 | 0.284 | 0.12 | 2.1729E-08 | Bcells | ANP32E |
| 1.389E-12 | 0.31160631 | 0.489 | 0.261 | 3.2813E-08 | Bcells | HNRNPR |
| 1.4169E-12 | -0.6157585 | 0.174 | 0.418 | 3.3471E-08 | Bcells | JPT1 |
| 1.4173E-12 | -0.445152 | 0.032 | 0.255 | 3.3482E-08 | Bcells | GABARAPL1 |
| 1.4257E-12 | -0.437721 | 0.005 | 0.219 | 3.368E-08 | Bcells | SPATS2L |
| 1.5945E-12 | 0.36269386 | 0.505 | 0.282 | 3.7668E-08 | Bcells | TPR |
| 1.6134E-12 | -0.6789379 | 0.342 | 0.579 | 3.8114E-08 | Bcells | LITAF |
| 1.8058E-12 | -0.5165358 | 0.005 | 0.218 | 4.2658E-08 | Bcells | DSP |
| 1.9687E-12 | 0.29075364 | 0.384 | 0.186 | 4.6507E-08 | Bcells | GTF3A |
| 2.0049E-12 | 0.30795296 | 0.542 | 0.303 | 4.7362E-08 | Bcells | RPL36A |
| 2.3096E-12 | 0.259124 | 0.605 | 0.332 | 5.456E-08 | Bcells | TLN1 |
| 2.5485E-12 | 0.29233567 | 0.463 | 0.24 | 6.0203E-08 | Bcells | KANSL1 |
| 3.2228E-12 | 0.37856751 | 0.732 | 0.505 | 7.6132E-08 | Bcells | JAK1 |
| 3.3298E-12 | -0.5054079 | 0.079 | 0.307 | 7.8659E-08 | Bcells | CLTB |
| 3.5049E-12 | 0.31841036 | 0.505 | 0.272 | 8.2795E-08 | Bcells | SLTM |
| 3.6096E-12 | -0.7939049 | 0.263 | 0.504 | 8.527E-08 | Bcells | DNAJB1 |
| 3.9103E-12 | 0.25872232 | 0.284 | 0.121 | 9.2373E-08 | Bcells | CHD8 |
| 4.0869E-12 | -0.6472148 | 0.253 | 0.52 | 9.6545E-08 | Bcells | H1FX |
| 4.0881E-12 | -0.6329137 | 0.484 | 0.674 | 9.6574E-08 | Bcells | CALR |
| 4.5327E-12 | -0.5338626 | 0.142 | 0.376 | 1.0708E-07 | Bcells | ARF4 |
| 4.6661E-12 | 0.33293605 | 0.768 | 0.552 | 1.1023E-07 | Bcells | HNRNPA3 |
| 4.9869E-12 | -0.9963278 | 0.063 | 0.287 | 1.1781E-07 | Bcells | HMOX1 |
| 5.0826E-12 | 0.2829598 | 0.668 | 0.395 | 1.2007E-07 | Bcells | ARGLU1 |
| 5.2584E-12 | -1.702426 | 0.037 | 0.249 | 1.2422E-07 | Bcells | CRYAB |
| 6.0092E-12 | -0.3641358 | 0.005 | 0.209 | 1.4196E-07 | Bcells | NECTIN2 |
| 6.1533E-12 | -0.5952208 | 0.221 | 0.449 | 1.4536E-07 | Bcells | ACTN4 |
| 6.4907E-12 | -0.6970412 | 0.005 | 0.209 | 1.5333E-07 | Bcells | CDKN2B |
| 6.5658E-12 | 0.31877079 | 0.4 | 0.201 | 1.551E-07 | Bcells | STAG2 |
| 6.6266E-12 | -0.8522299 | 0.505 | 0.7 | 1.5654E-07 | Bcells | JUNB |
| 6.8729E-12 | -0.4211872 | 0.079 | 0.306 | 1.6236E-07 | Bcells | RAB1A |
| 7.8448E-12 | -0.7644079 | 0.016 | 0.222 | 1.8532E-07 | Bcells | CCND1 |
| 8.8352E-12 | -0.504915 | 0.495 | 0.696 | 2.0871E-07 | Bcells | GUK1 |
| 9.1503E-12 | -0.549827 | 0.058 | 0.275 | 2.1616E-07 | Bcells | JUP |

| | | | | | | |
|---|---|---|---|---|---|---|
| 9.8071E-12 | -0.6017877 | 0.011 | 0.214 | 2.3167E-07 | Bcells | LAMB3 |
| 1.006E-11 | -0.4878978 | 0 | 0.198 | 2.3765E-07 | Bcells | PTGR1 |
| 1.0325E-11 | -0.5058284 | 0.042 | 0.258 | 2.4391E-07 | Bcells | PDLIM5 |
| 1.1105E-11 | -0.5374679 | 0.074 | 0.294 | 2.6233E-07 | Bcells | TUBA1C |
| 1.1983E-11 | -0.560152 | 0.047 | 0.262 | 2.8307E-07 | Bcells | RHOC |
| 1.2179E-11 | 0.26614938 | 0.353 | 0.168 | 2.877E-07 | Bcells | NECAP2 |
| 1.3312E-11 | 0.25081906 | 0.453 | 0.232 | 3.1447E-07 | Bcells | BST2 |
| 1.3413E-11 | -0.5946258 | 0.016 | 0.219 | 3.1686E-07 | Bcells | TGM2 |
| 1.3578E-11 | 0.38087759 | 0.284 | 0.125 | 3.2074E-07 | Bcells | IL4R |
| 1.3698E-11 | 0.42396297 | 0.711 | 0.491 | 3.2359E-07 | Bcells | TAGLN2 |
| 1.3868E-11 | -0.5908626 | 0.021 | 0.227 | 3.2761E-07 | Bcells | IER5L |
| 1.3935E-11 | -0.3722347 | 0 | 0.196 | 3.2918E-07 | Bcells | RBPMS |
| 1.6481E-11 | -0.6992035 | 0.216 | 0.427 | 3.8934E-07 | Bcells | SLC3A2 |
| 1.7491E-11 | -1.092567 | 0.016 | 0.216 | 4.1319E-07 | Bcells | S100A12 |
| 2.1245E-11 | -0.5271253 | 0.368 | 0.581 | 5.0188E-07 | Bcells | PGK1 |
| 2.1696E-11 | -0.5196517 | 0.016 | 0.216 | 5.1253E-07 | Bcells | CKB |
| 2.4043E-11 | -1.0595545 | 0.063 | 0.276 | 5.6798E-07 | Bcells | AIF1 |
| 2.5639E-11 | 0.26440356 | 0.342 | 0.162 | 6.0566E-07 | Bcells | BIRC6 |
| 2.5889E-11 | 0.27226946 | 0.595 | 0.339 | 6.1157E-07 | Bcells | ANP32B |
| 2.7058E-11 | -0.5220811 | 0.153 | 0.372 | 6.3918E-07 | Bcells | RHEB |
| 2.731E-11 | -0.4253927 | 0.021 | 0.222 | 6.4514E-07 | Bcells | ACTN1 |
| 2.9523E-11 | -0.5381071 | 0.089 | 0.312 | 6.9743E-07 | Bcells | CYTOR |
| 2.953E-11 | 0.31498639 | 0.611 | 0.377 | 6.9759E-07 | Bcells | PAPOLA |
| 2.9649E-11 | 0.3263337 | 0.537 | 0.308 | 7.004E-07 | Bcells | OGA |
| 3.0803E-11 | -0.3777643 | 0.011 | 0.206 | 7.2765E-07 | Bcells | CTTN |
| 3.2914E-11 | -0.9320699 | 0.005 | 0.198 | 7.7753E-07 | Bcells | TM4SF1 |
| 3.4452E-11 | 0.54096847 | 0.768 | 0.641 | 8.1385E-07 | Bcells | TXNIP |
| 3.5466E-11 | 0.31713712 | 0.426 | 0.227 | 8.3782E-07 | Bcells | DHX36 |
| 3.6832E-11 | 0.26993641 | 0.4 | 0.201 | 8.7008E-07 | Bcells | TANK |
| 3.7147E-11 | -2.5783628 | 0.274 | 0.489 | 8.7753E-07 | Bcells | CXCL8 |
| 3.7566E-11 | -0.4542569 | 0.579 | 0.764 | 8.8741E-07 | Bcells | UBL5 |
| 4.0141E-11 | -0.6901378 | 0.005 | 0.196 | 9.4826E-07 | Bcells | SFN |
| 4.035E-11 | 0.25853813 | 0.437 | 0.231 | 9.5319E-07 | Bcells | EIF2S3 |
| 4.3861E-11 | -0.611483 | 0.774 | 0.885 | 1.0361E-06 | Bcells | MYL12B |
| 5.3721E-11 | -0.5410152 | 0.053 | 0.259 | 1.269E-06 | Bcells | TSC22D1 |
| 5.6153E-11 | -0.5312609 | 0.1 | 0.312 | 1.3265E-06 | Bcells | SMS |
| 5.8191E-11 | 0.35319711 | 0.921 | 0.801 | 1.3746E-06 | Bcells | HMGB1 |
| 6.2453E-11 | -0.4755451 | 0.126 | 0.36 | 1.4753E-06 | Bcells | RRBP1 |
| 7.1525E-11 | -1.6338312 | 0.011 | 0.199 | 1.6896E-06 | Bcells | KRT17 |
| 7.6323E-11 | 0.29690804 | 0.6 | 0.372 | 1.803E-06 | Bcells | C11orf58 |
| 7.6809E-11 | -0.4238421 | 0.037 | 0.236 | 1.8145E-06 | Bcells | PLXNB2 |
| 8.3325E-11 | -0.434395 | 0.005 | 0.191 | 1.9684E-06 | Bcells | LIMA1 |
| 8.5272E-11 | -0.5842141 | 0 | 0.184 | 2.0144E-06 | Bcells | SDCBP2 |
| 9.065E-11 | 0.33465387 | 0.963 | 0.929 | 2.1414E-06 | Bcells | OAZ1 |
| 9.2146E-11 | -0.3806885 | 0.011 | 0.198 | 2.1768E-06 | Bcells | TCEAL9 |
| 9.4941E-11 | -0.4752373 | 0.395 | 0.602 | 2.2428E-06 | Bcells | YWHAE |
| 1.0066E-10 | 0.30860971 | 0.521 | 0.307 | 2.3778E-06 | Bcells | YY1 |
| 1.0072E-10 | -0.8176286 | 0 | 0.182 | 2.3793E-06 | Bcells | LAMC2 |
| 1.0875E-10 | 0.26245711 | 0.221 | 0.088 | 2.569E-06 | Bcells | CRIP1 |
| 1.0887E-10 | 0.25357914 | 0.263 | 0.115 | 2.5718E-06 | Bcells | CBX5 |

| | | | | | | |
|---|---|---|---|---|---|---|
| 1.0958E-10 | -0.4290628 | 0.011 | 0.197 | 2.5887E-06 | Bcells | KLF4 |
| 1.1707E-10 | -0.5281819 | 0.047 | 0.246 | 2.7656E-06 | Bcells | PTMS |
| 1.2913E-10 | -0.3769123 | 0.068 | 0.275 | 3.0504E-06 | Bcells | LAMP2 |
| 1.292E-10 | -0.5396593 | 0.126 | 0.344 | 3.052E-06 | Bcells | CPEB4 |
| 1.3071E-10 | -0.3887467 | 0 | 0.18 | 3.0877E-06 | Bcells | TEAD1 |
| 1.3198E-10 | -0.542183 | 0 | 0.18 | 3.1179E-06 | Bcells | DST |
| 1.4293E-10 | -0.5418418 | 0.147 | 0.359 | 3.3765E-06 | Bcells | ATP1A1 |
| 1.4325E-10 | -0.3923531 | 0 | 0.18 | 3.384E-06 | Bcells | TMEM176B |
| 1.5046E-10 | -1.3834378 | 0.074 | 0.281 | 3.5542E-06 | Bcells | MT1E |
| 1.5209E-10 | 0.27457037 | 0.611 | 0.375 | 3.5929E-06 | Bcells | EIF3H |
| 1.6887E-10 | 0.25569649 | 0.289 | 0.133 | 3.9893E-06 | Bcells | RFC1 |
| 1.7605E-10 | -0.4320845 | 0.126 | 0.337 | 4.1588E-06 | Bcells | CHMP5 |
| 1.8154E-10 | -0.8568199 | 0.937 | 0.962 | 4.2884E-06 | Bcells | HSP90AA1 |
| 1.8632E-10 | -1.1423769 | 0.074 | 0.281 | 4.4015E-06 | Bcells | MT1X |
| 1.9434E-10 | -0.5480487 | 0.042 | 0.235 | 4.591E-06 | Bcells | TOB1 |
| 1.967E-10 | -0.524519 | 0.063 | 0.26 | 4.6466E-06 | Bcells | WASL |
| 2.0582E-10 | 0.25425484 | 0.568 | 0.335 | 4.862E-06 | Bcells | CAPZA1 |
| 2.1192E-10 | 0.28742789 | 0.626 | 0.391 | 5.0063E-06 | Bcells | SERBP1 |
| 2.3216E-10 | -0.4241725 | 0.063 | 0.265 | 5.4843E-06 | Bcells | GSN |
| 2.366E-10 | -0.5690845 | 0.011 | 0.192 | 5.5893E-06 | Bcells | PLK2 |
| 2.563E-10 | -0.4269146 | 0.005 | 0.183 | 6.0546E-06 | Bcells | CDH1 |
| 2.5716E-10 | -0.6383446 | 0.184 | 0.389 | 6.0749E-06 | Bcells | TUBB4B |
| 2.6227E-10 | -0.8856674 | 0.216 | 0.423 | 6.1956E-06 | Bcells | TXNRD1 |
| 2.6299E-10 | 0.27007562 | 0.742 | 0.493 | 6.2127E-06 | Bcells | NCL |
| 2.7224E-10 | -0.6161182 | 0 | 0.175 | 6.4311E-06 | Bcells | SPNS2 |
| 2.7928E-10 | 0.27480505 | 0.337 | 0.17 | 6.5975E-06 | Bcells | PPP1CC |
| 2.852E-10 | 0.26496923 | 0.374 | 0.194 | 6.7372E-06 | Bcells | HLA-F |
| 3.07E-10 | -0.6702421 | 0.011 | 0.189 | 7.2523E-06 | Bcells | EPS8L1 |
| 3.1868E-10 | -0.3824149 | 0.063 | 0.262 | 7.5283E-06 | Bcells | CHP1 |
| 3.3418E-10 | -0.4867613 | 0.005 | 0.182 | 7.8943E-06 | Bcells | S100A14 |
| 3.3574E-10 | -0.5947847 | 0.011 | 0.189 | 7.9311E-06 | Bcells | HBEGF |
| 3.4522E-10 | 0.40802105 | 0.905 | 0.836 | 8.1551E-06 | Bcells | COX4I1 |
| 3.5407E-10 | -0.903243 | 0.011 | 0.188 | 8.3643E-06 | Bcells | TMPRSS2 |
| 3.5833E-10 | 0.27450162 | 0.368 | 0.187 | 8.4647E-06 | Bcells | PMEPA1 |
| 3.7266E-10 | -0.8903043 | 0.032 | 0.22 | 8.8032E-06 | Bcells | CD14 |
| 3.7436E-10 | -0.6002992 | 0 | 0.173 | 8.8436E-06 | Bcells | PPL |
| 3.845E-10 | -0.4390035 | 0.279 | 0.491 | 9.083E-06 | Bcells | HSBP1 |
| 3.8811E-10 | -0.3965479 | 0.005 | 0.18 | 9.1684E-06 | Bcells | ANXA3 |
| 3.9869E-10 | -0.4722021 | 0.437 | 0.617 | 9.4182E-06 | Bcells | COX8A |
| 3.9927E-10 | 0.33256133 | 0.542 | 0.336 | 9.4321E-06 | Bcells | SRRM1 |
| 4.1344E-10 | 0.31328335 | 0.474 | 0.274 | 9.7667E-06 | Bcells | PPIG |
| 4.4682E-10 | -0.6006357 | 0 | 0.172 | 1.0555E-05 | Bcells | HSPB8 |
| 5.0846E-10 | 0.26290229 | 0.453 | 0.255 | 1.2011E-05 | Bcells | DNAJC8 |
| 5.1523E-10 | -0.3260554 | 0.016 | 0.194 | 1.2171E-05 | Bcells | GNAQ |
| 5.4626E-10 | 0.27413672 | 0.574 | 0.355 | 1.2904E-05 | Bcells | TRMT112 |
| 5.603E-10 | -0.6755708 | 0.016 | 0.192 | 1.3236E-05 | Bcells | FBXO32 |
| 6.1683E-10 | 0.26204496 | 0.616 | 0.376 | 1.4571E-05 | Bcells | GCC2 |
| 6.1841E-10 | -0.4814691 | 0.026 | 0.207 | 1.4609E-05 | Bcells | PARD6B |
| 6.2022E-10 | -0.3916532 | 0 | 0.169 | 1.4651E-05 | Bcells | NECTIN4 |
| 6.2891E-10 | 0.26698609 | 0.347 | 0.175 | 1.4857E-05 | Bcells | NSRP1 |

| | | | | | | |
|---|---|---|---|---|---|---|
| 6.3073E-10 | -0.3863596 | 0.005 | 0.177 | 1.49E-05 | Bcells | SMIM22 |
| 6.308E-10 | -0.4799608 | 0.005 | 0.177 | 1.4901E-05 | Bcells | FHL2 |
| 6.319E-10 | 0.25926923 | 0.347 | 0.176 | 1.4927E-05 | Bcells | SENP6 |
| 6.5016E-10 | 0.31001059 | 0.468 | 0.27 | 1.5359E-05 | Bcells | OSBPL8 |
| 6.6543E-10 | -0.7720883 | 0.063 | 0.249 | 1.5719E-05 | Bcells | EGR1 |
| 6.7137E-10 | -0.6284473 | 0 | 0.169 | 1.586E-05 | Bcells | FXYD3 |
| 6.9015E-10 | -0.3458567 | 0.005 | 0.176 | 1.6303E-05 | Bcells | TMEM54 |
| 7.1503E-10 | -0.3401649 | 0 | 0.168 | 1.6891E-05 | Bcells | CXADR |
| 7.1853E-10 | 0.31840052 | 0.384 | 0.209 | 1.6974E-05 | Bcells | RBM17 |
| 7.218E-10 | -0.5995028 | 0.121 | 0.324 | 1.7051E-05 | Bcells | LIMS1 |
| 7.3845E-10 | 0.28256212 | 0.542 | 0.33 | 1.7444E-05 | Bcells | MOB1A |
| 7.4081E-10 | 0.26359154 | 0.368 | 0.193 | 1.75E-05 | Bcells | CGGBP1 |
| 7.6583E-10 | -0.6618847 | 0.005 | 0.176 | 1.8091E-05 | Bcells | IGFBP3 |
| 7.6774E-10 | -0.5280785 | 0.005 | 0.175 | 1.8136E-05 | Bcells | C6orf132 |
| 7.7274E-10 | 0.25397969 | 0.458 | 0.255 | 1.8254E-05 | Bcells | VAMP2 |
| 7.9263E-10 | -0.35752 | 0.021 | 0.199 | 1.8724E-05 | Bcells | ITGAV |
| 8.2931E-10 | -0.9035188 | 0.095 | 0.286 | 1.9591E-05 | Bcells | SLC25A37 |
| 8.5687E-10 | -0.7109299 | 0.037 | 0.228 | 2.0242E-05 | Bcells | IL32 |
| 8.9671E-10 | -0.4220267 | 0.121 | 0.322 | 2.1183E-05 | Bcells | LEPROT |
| 9.1395E-10 | -1.2768251 | 0 | 0.166 | 2.159E-05 | Bcells | DUOXA2 |
| 9.4229E-10 | -1.024065 | 0.005 | 0.174 | 2.226E-05 | Bcells | RDH10 |
| 9.5341E-10 | 0.25372422 | 0.421 | 0.228 | 2.2522E-05 | Bcells | PJA2 |
| 9.5471E-10 | -0.3843594 | 0.021 | 0.197 | 2.2553E-05 | Bcells | NEDD4L |
| 9.8786E-10 | -1.1731884 | 0.011 | 0.181 | 2.3336E-05 | Bcells | SPINK1 |
| 9.9072E-10 | -0.3817488 | 0.089 | 0.287 | 2.3404E-05 | Bcells | ATP6V1D |
| 9.9797E-10 | 0.25757628 | 0.274 | 0.128 | 2.3575E-05 | Bcells | TRABD |
| 1.1183E-09 | -0.5043619 | 0.211 | 0.416 | 2.6418E-05 | Bcells | GRN |
| 1.1407E-09 | -0.4965672 | 0.758 | 0.877 | 2.6947E-05 | Bcells | TMBIM6 |
| 1.1653E-09 | -0.6545359 | 0.005 | 0.172 | 2.7527E-05 | Bcells | STEAP4 |
| 1.2366E-09 | -0.4569821 | 0.379 | 0.561 | 2.9213E-05 | Bcells | CSNK1A1 |
| 1.335E-09 | -0.3352398 | 0 | 0.163 | 3.1536E-05 | Bcells | NDFIP2 |
| 1.3705E-09 | -0.5220927 | 0.032 | 0.209 | 3.2375E-05 | Bcells | MACC1 |
| 1.5198E-09 | -0.3089437 | 0.005 | 0.171 | 3.5902E-05 | Bcells | GNG12 |
| 1.558E-09 | 0.25851679 | 0.579 | 0.359 | 3.6804E-05 | Bcells | SRSF11 |
| 1.5889E-09 | -0.3029617 | 0 | 0.162 | 3.7534E-05 | Bcells | SORT1 |
| 1.6289E-09 | -0.2817995 | 0 | 0.162 | 3.8479E-05 | Bcells | RAB32 |
| 1.6578E-09 | -0.7177449 | 0.042 | 0.222 | 3.9163E-05 | Bcells | HES1 |
| 1.6757E-09 | 0.29123175 | 0.363 | 0.195 | 3.9586E-05 | Bcells | ZC3H13 |
| 1.7005E-09 | 0.29122545 | 0.963 | 0.881 | 4.0171E-05 | Bcells | RPL12 |
| 1.7269E-09 | -0.3139748 | 0 | 0.161 | 4.0795E-05 | Bcells | NCKAP1 |
| 1.7292E-09 | -1.1867278 | 0.016 | 0.185 | 4.0848E-05 | Bcells | DUOX2 |
| 1.7394E-09 | -0.4548545 | 0.095 | 0.285 | 4.109E-05 | Bcells | CXCL16 |
| 1.7571E-09 | 0.2601875 | 0.332 | 0.169 | 4.1508E-05 | Bcells | ZNF644 |
| 1.7869E-09 | -0.4427361 | 0.016 | 0.185 | 4.2212E-05 | Bcells | ERRFI1 |
| 1.8182E-09 | -1.1425254 | 0 | 0.161 | 4.2952E-05 | Bcells | MUC21 |
| 1.8652E-09 | -0.5059792 | 0.205 | 0.398 | 4.4061E-05 | Bcells | H2AFY |
| 1.945E-09 | -0.3620384 | 0.026 | 0.2 | 4.5946E-05 | Bcells | CREG1 |
| 2.0364E-09 | -0.3610707 | 0.016 | 0.184 | 4.8106E-05 | Bcells | DAB2 |
| 2.0421E-09 | -1.1831063 | 0.042 | 0.223 | 4.824E-05 | Bcells | G0S2 |
| 2.0548E-09 | 0.28903456 | 0.774 | 0.562 | 4.854E-05 | Bcells | SLC25A6 |

| | | | | | | |
|---|---|---|---|---|---|---|
| 2.0754E-09 | -0.4820683 | 0.084 | 0.272 | 4.9028E-05 | Bcells | RAP2B |
| 2.0821E-09 | -1.0578512 | 0 | 0.16 | 4.9186E-05 | Bcells | CEACAM5 |
| 2.2696E-09 | 0.25000808 | 0.811 | 0.606 | 5.3615E-05 | Bcells | SON |
| 2.2879E-09 | -0.3212166 | 0.016 | 0.185 | 5.4046E-05 | Bcells | CDCP1 |
| 2.3253E-09 | -0.3233559 | 0 | 0.159 | 5.493E-05 | Bcells | CRB3 |
| 2.3795E-09 | 0.26455295 | 0.547 | 0.335 | 5.6211E-05 | Bcells | SELENOH |
| 2.447E-09 | 0.25533738 | 0.426 | 0.237 | 5.7807E-05 | Bcells | NCOR1 |
| 2.4627E-09 | -0.5071703 | 0 | 0.159 | 5.8175E-05 | Bcells | MALL |
| 2.4902E-09 | 0.39613372 | 0.453 | 0.265 | 5.8826E-05 | Bcells | TMEM123 |
| 2.7398E-09 | -0.3989599 | 0.184 | 0.391 | 6.4722E-05 | Bcells | CHMP2A |
| 2.8125E-09 | -0.5011996 | 0.005 | 0.166 | 6.644E-05 | Bcells | HIST3H2A |
| 2.8152E-09 | -0.3086023 | 0.005 | 0.167 | 6.6503E-05 | Bcells | MYOF |
| 2.8363E-09 | -0.4972175 | 0.016 | 0.182 | 6.7002E-05 | Bcells | FBP1 |
| 2.9243E-09 | -0.8430037 | 0.016 | 0.18 | 6.908E-05 | Bcells | F3 |
| 3.0505E-09 | -0.3704497 | 0.042 | 0.218 | 7.2061E-05 | Bcells | FOXO3 |
| 3.1073E-09 | 0.26418858 | 0.411 | 0.227 | 7.3404E-05 | Bcells | USP8 |
| 3.1505E-09 | -0.3446185 | 0.011 | 0.173 | 7.4424E-05 | Bcells | DHRS3 |
| 3.347E-09 | -0.3851052 | 0 | 0.157 | 7.9066E-05 | Bcells | TSPAN6 |
| 3.4712E-09 | 0.26589994 | 0.463 | 0.268 | 8.2001E-05 | Bcells | FAM133B |
| 3.5332E-09 | 0.29629785 | 0.526 | 0.327 | 8.3466E-05 | Bcells | PAIP2 |
| 3.5712E-09 | -0.4783834 | 0.384 | 0.556 | 8.4362E-05 | Bcells | NDUFS5 |
| 3.5781E-09 | -0.6818906 | 0.084 | 0.267 | 8.4526E-05 | Bcells | DDIT3 |
| 3.6018E-09 | -0.3358328 | 0 | 0.156 | 8.5086E-05 | Bcells | ERBB3 |
| 3.6117E-09 | -0.8618297 | 0 | 0.156 | 8.532E-05 | Bcells | KRT15 |
| 3.6984E-09 | 0.2730355 | 0.658 | 0.426 | 8.7368E-05 | Bcells | SRSF5 |
| 3.7201E-09 | -0.3985593 | 0.132 | 0.328 | 8.7881E-05 | Bcells | LAMP1 |
| 3.7362E-09 | -0.3442739 | 0.011 | 0.172 | 8.8261E-05 | Bcells | B3GNT5 |
| 4.0059E-09 | 0.25164536 | 0.937 | 0.762 | 9.463E-05 | Bcells | COX7C |
| 4.0362E-09 | -0.4887949 | 0 | 0.155 | 9.5346E-05 | Bcells | CYR61 |
| 4.2301E-09 | -0.3306023 | 0.026 | 0.194 | 9.9927E-05 | Bcells | KIFC3 |
| 4.3801E-09 | -0.4936023 | 0.016 | 0.179 | 0.00010347 | Bcells | EPHA2 |
| 4.6859E-09 | -0.413133 | 0.005 | 0.162 | 0.00011069 | Bcells | SYTL2 |
| 5.0521E-09 | 0.25784819 | 0.511 | 0.312 | 0.00011935 | Bcells | UBE2D2 |
| 5.142E-09 | -0.7552628 | 0 | 0.154 | 0.00012147 | Bcells | SCEL |
| 5.1504E-09 | 0.25380319 | 0.432 | 0.246 | 0.00012167 | Bcells | CD47 |
| 5.272E-09 | -0.4700035 | 0.126 | 0.32 | 0.00012454 | Bcells | SLC16A3 |
| 5.3272E-09 | -0.8959034 | 0.747 | 0.833 | 0.00012584 | Bcells | PPDPF |
| 5.3382E-09 | -0.3403992 | 0.058 | 0.235 | 0.0001261 | Bcells | SCARB2 |
| 5.4256E-09 | 0.29187336 | 0.905 | 0.81 | 0.00012817 | Bcells | DDX5 |
| 5.4818E-09 | -0.3642897 | 0.021 | 0.185 | 0.0001295 | Bcells | EPS8L2 |
| 5.6289E-09 | -0.6778882 | 0.279 | 0.437 | 0.00013297 | Bcells | CYSTM1 |
| 5.6448E-09 | -0.3806368 | 0.011 | 0.169 | 0.00013335 | Bcells | TINAGL1 |
| 5.6949E-09 | -0.3650587 | 0.053 | 0.23 | 0.00013453 | Bcells | PGD |
| 5.8208E-09 | -1.5810255 | 0.053 | 0.223 | 0.0001375 | Bcells | VCAN |
| 6.0504E-09 | -0.3388891 | 0.021 | 0.184 | 0.00014293 | Bcells | GNA15 |
| 6.2157E-09 | -0.624661 | 0.158 | 0.346 | 0.00014683 | Bcells | MIDN |
| 6.6254E-09 | -0.5018151 | 0.037 | 0.207 | 0.00015651 | Bcells | GK |
| 6.6722E-09 | -0.4261846 | 0.005 | 0.16 | 0.00015762 | Bcells | SDC1 |
| 7.0942E-09 | -0.8309513 | 0.016 | 0.174 | 0.00016759 | Bcells | NDRG2 |
| 7.0965E-09 | -0.3400947 | 0.026 | 0.192 | 0.00016764 | Bcells | F11R |

| | | | | | | |
|---|---|---|---|---|---|---|
| 7.7645E-09 | -0.9782426 | 0.005 | 0.158 | 0.00018342 | Bcells | PRSS27 |
| 8.2369E-09 | -0.310618 | 0.021 | 0.182 | 0.00019458 | Bcells | PFKP |
| 8.365E-09 | -0.4850377 | 0.026 | 0.189 | 0.00019761 | Bcells | DUSP6 |
| 8.8078E-09 | -0.4488088 | 0.095 | 0.275 | 0.00020807 | Bcells | PLIN3 |
| 9.4561E-09 | -0.2818159 | 0.005 | 0.157 | 0.00022338 | Bcells | HNMT |
| 9.7066E-09 | -0.334208 | 0.026 | 0.188 | 0.0002293 | Bcells | HSBP1L1 |
| 1.0023E-08 | 0.26630404 | 0.447 | 0.26 | 0.00023676 | Bcells | MBP |
| 1.1005E-08 | -1.5827673 | 0.011 | 0.163 | 0.00025997 | Bcells | ECM1 |
| 1.117E-08 | -0.8371228 | 0.484 | 0.613 | 0.00026386 | Bcells | CD44 |
| 1.1624E-08 | -0.4755737 | 0 | 0.147 | 0.00027458 | Bcells | ALDH1A3 |
| 1.1894E-08 | -0.6471366 | 0 | 0.147 | 0.00028098 | Bcells | NCCRP1 |
| 1.3197E-08 | -0.4139572 | 0 | 0.146 | 0.00031176 | Bcells | VSIG2 |
| 1.3381E-08 | -0.5912894 | 0.005 | 0.154 | 0.0003161 | Bcells | CDC42EP5 |
| 1.4373E-08 | -0.3372045 | 0.011 | 0.162 | 0.00033954 | Bcells | MET |
| 1.4497E-08 | -0.4014714 | 0 | 0.146 | 0.00034245 | Bcells | SDC2 |
| 1.5498E-08 | -0.2891386 | 0 | 0.145 | 0.00036612 | Bcells | GPRC5C |
| 1.5715E-08 | -0.7709942 | 0 | 0.145 | 0.00037125 | Bcells | TMPRSS11E |
| 1.6819E-08 | -0.4116366 | 0.184 | 0.366 | 0.00039732 | Bcells | ELOC |
| 1.6877E-08 | -0.3578811 | 0.026 | 0.184 | 0.0003987 | Bcells | BNIP3 |
| 1.7325E-08 | -0.3902007 | 0 | 0.144 | 0.00040927 | Bcells | ITGB6 |
| 1.7819E-08 | 0.33311267 | 0.716 | 0.55 | 0.00042093 | Bcells | FAM107B |
| 1.8008E-08 | -0.5429257 | 0.068 | 0.238 | 0.00042541 | Bcells | CITED2 |
| 1.8592E-08 | -0.4424557 | 0.011 | 0.161 | 0.0004392 | Bcells | AQP9 |
| 1.9467E-08 | -0.4206808 | 0 | 0.143 | 0.00045988 | Bcells | PAX8 |
| 1.9759E-08 | -0.4424112 | 0.111 | 0.289 | 0.00046676 | Bcells | RNF149 |
| 2.0777E-08 | -0.2631612 | 0.005 | 0.151 | 0.00049081 | Bcells | SHTN1 |
| 2.082E-08 | -0.8079317 | 0.079 | 0.245 | 0.00049182 | Bcells | SLC11A1 |
| 2.2167E-08 | -0.3852664 | 0.095 | 0.269 | 0.00052366 | Bcells | UBE2D1 |
| 2.3449E-08 | -0.3396928 | 0.016 | 0.166 | 0.00055393 | Bcells | CFD |
| 2.4344E-08 | -0.6260472 | 0 | 0.142 | 0.00057508 | Bcells | SNCG |
| 2.4627E-08 | -0.4417422 | 0.163 | 0.343 | 0.00058177 | Bcells | BSG |
| 2.4759E-08 | -0.3011115 | 0.005 | 0.15 | 0.00058488 | Bcells | PLS3 |
| 2.7354E-08 | -0.2795596 | 0 | 0.141 | 0.00064617 | Bcells | RAB25 |
| 2.7466E-08 | -0.4170133 | 0.011 | 0.157 | 0.00064882 | Bcells | CLDN3 |
| 2.7684E-08 | -0.3307962 | 0.011 | 0.157 | 0.00065397 | Bcells | FNIP2 |
| 2.9124E-08 | -0.413516 | 0.368 | 0.544 | 0.00068799 | Bcells | POMP |
| 2.9223E-08 | -0.8128523 | 0.032 | 0.185 | 0.00069035 | Bcells | AQP3 |
| 2.9247E-08 | 0.26241324 | 0.8 | 0.69 | 0.00069091 | Bcells | PNRC1 |
| 2.9372E-08 | -0.288291 | 0.037 | 0.195 | 0.00069385 | Bcells | RNF11 |
| 2.9999E-08 | -0.643516 | 0.011 | 0.156 | 0.00070866 | Bcells | MUC4 |
| 3.1436E-08 | -1.0598405 | 0.042 | 0.2 | 0.00074261 | Bcells | THBS1 |
| 3.1438E-08 | -0.9638402 | 0.011 | 0.155 | 0.00074266 | Bcells | PITX1 |
| 3.3223E-08 | -0.3410836 | 0.005 | 0.148 | 0.00078483 | Bcells | GSTM3 |
| 3.481E-08 | -0.2711665 | 0.021 | 0.172 | 0.00082231 | Bcells | CTBP2 |
| 3.581E-08 | -0.3037058 | 0 | 0.139 | 0.00084593 | Bcells | ZNRF1 |
| 3.6287E-08 | 0.28237897 | 0.379 | 0.218 | 0.0008572 | Bcells | ZFC3H1 |
| 3.7074E-08 | -0.3717126 | 0.005 | 0.147 | 0.0008758 | Bcells | TREM1 |
| 3.7279E-08 | -0.4303122 | 0.011 | 0.154 | 0.00088065 | Bcells | TMPRSS4 |
| 3.7508E-08 | -0.3194363 | 0 | 0.138 | 0.00088605 | Bcells | MTUS1 |
| 3.7856E-08 | -0.2878701 | 0.005 | 0.147 | 0.00089428 | Bcells | KCNK1 |

| | | | | | | |
|---|---|---|---|---|---|---|
| 4.16E-08 | -0.4095086 | 0 | 0.138 | 0.00098271 | Bcells | DCDC2 |
| 4.4699E-08 | -0.2635558 | 0 | 0.137 | 0.00105593 | Bcells | SRGAP1 |
| 4.585E-08 | -0.3877592 | 0 | 0.137 | 0.00108311 | Bcells | ARHGAP29 |
| 4.5908E-08 | -0.2994904 | 0.005 | 0.145 | 0.00108449 | Bcells | SLC44A4 |
| 4.69E-08 | -0.6372428 | 0 | 0.137 | 0.00110793 | Bcells | CXCL17 |
| 4.7878E-08 | -0.4174015 | 0.595 | 0.727 | 0.00113102 | Bcells | ELOB |
| 4.7926E-08 | -0.3539005 | 0.005 | 0.145 | 0.00113215 | Bcells | CLDN1 |
| 4.9218E-08 | -0.288064 | 0.005 | 0.145 | 0.00116267 | Bcells | SVIL |
| 4.9716E-08 | -0.4178068 | 0 | 0.136 | 0.00117444 | Bcells | GCNT3 |
| 4.9762E-08 | -0.2771003 | 0.037 | 0.191 | 0.00117553 | Bcells | AGTRAP |
| 5.0526E-08 | -0.3567861 | 0.011 | 0.152 | 0.00119358 | Bcells | ITGA3 |
| 5.1531E-08 | -0.4131604 | 0.526 | 0.672 | 0.00121733 | Bcells | COX5B |
| 5.317E-08 | -0.4912975 | 0.011 | 0.152 | 0.00125604 | Bcells | CLEC5A |
| 5.3832E-08 | -0.3294312 | 0.1 | 0.276 | 0.00127167 | Bcells | PDXK |
| 5.5329E-08 | -1.6812404 | 0.005 | 0.144 | 0.00130704 | Bcells | SPRR2A |
| 5.6661E-08 | -0.2919105 | 0.005 | 0.144 | 0.0013385 | Bcells | PHLDA3 |
| 5.7046E-08 | -0.361523 | 0.047 | 0.205 | 0.0013476 | Bcells | SLFN5 |
| 5.8012E-08 | -0.4655456 | 0.011 | 0.151 | 0.00137042 | Bcells | SLC7A11 |
| 5.8764E-08 | -0.9040626 | 0 | 0.135 | 0.00138819 | Bcells | TMPRSS11B |
| 5.9479E-08 | -0.4391504 | 0.026 | 0.174 | 0.00140508 | Bcells | SMIM25 |
| 6.0213E-08 | -0.9767944 | 0.121 | 0.279 | 0.00142242 | Bcells | DUSP5 |
| 6.0822E-08 | -0.4591657 | 0 | 0.135 | 0.00143679 | Bcells | UCHL1 |
| 6.1915E-08 | -0.3119372 | 0.011 | 0.152 | 0.00146261 | Bcells | CPVL |
| 6.3716E-08 | -0.2883688 | 0.005 | 0.143 | 0.00150517 | Bcells | LAPTM4B |
| 6.4042E-08 | -0.3168249 | 0.047 | 0.203 | 0.00151286 | Bcells | NEU1 |
| 6.6333E-08 | -0.2870384 | 0.005 | 0.142 | 0.00156698 | Bcells | SERPING1 |
| 6.9784E-08 | -0.4728111 | 0.005 | 0.142 | 0.0016485 | Bcells | ID1 |
| 7.1417E-08 | -0.6392904 | 0.032 | 0.181 | 0.00168709 | Bcells | OLR1 |
| 7.3096E-08 | -0.5909833 | 0.326 | 0.48 | 0.00172674 | Bcells | RIOK3 |
| 7.3558E-08 | -0.7299724 | 0.079 | 0.237 | 0.00173766 | Bcells | FABP5 |
| 7.3837E-08 | -0.7351578 | 0.011 | 0.149 | 0.00174424 | Bcells | TMPRSS11D |
| 7.475E-08 | -0.2867328 | 0.021 | 0.166 | 0.00176582 | Bcells | DDR1 |
| 7.5378E-08 | -0.3358451 | 0.011 | 0.149 | 0.00178066 | Bcells | PLXDC2 |
| 7.5415E-08 | -0.2814262 | 0.005 | 0.141 | 0.00178152 | Bcells | BEX3 |
| 7.593E-08 | -0.908418 | 0 | 0.133 | 0.00179371 | Bcells | RHCG |
| 7.9446E-08 | -0.2940847 | 0 | 0.132 | 0.00187675 | Bcells | MPZL2 |
| 8.0389E-08 | 0.33030518 | 0.405 | 0.243 | 0.00189904 | Bcells | CD82 |
| 8.392E-08 | -0.3424322 | 0.026 | 0.175 | 0.00198244 | Bcells | KCTD12 |
| 8.4595E-08 | -0.3434588 | 0.005 | 0.14 | 0.00199839 | Bcells | EMP2 |
| 8.5356E-08 | -0.26194 | 0 | 0.132 | 0.00201636 | Bcells | DMKN |
| 8.6508E-08 | -0.9616453 | 0.1 | 0.263 | 0.00204357 | Bcells | AREG |
| 8.6656E-08 | -0.3820275 | 0.079 | 0.238 | 0.00204706 | Bcells | PCBD1 |
| 8.9553E-08 | -0.6570061 | 0 | 0.132 | 0.0021155 | Bcells | GABRP |
| 8.9718E-08 | -0.2718053 | 0 | 0.131 | 0.00211941 | Bcells | PALLD |
| 9.1627E-08 | -0.271749 | 0 | 0.131 | 0.00216449 | Bcells | NTN4 |
| 9.4624E-08 | -0.9947125 | 0.005 | 0.141 | 0.00223529 | Bcells | CXCL6 |
| 9.8442E-08 | 0.28073265 | 0.516 | 0.332 | 0.0023255 | Bcells | GOLGA4 |
| 1.0007E-07 | -0.2720025 | 0 | 0.131 | 0.00236385 | Bcells | C1orf116 |
| 1.0145E-07 | -0.8071648 | 0.011 | 0.146 | 0.00239667 | Bcells | CEACAM7 |
| 1.0178E-07 | -0.3469731 | 0 | 0.131 | 0.00240439 | Bcells | FOXC1 |

| | | | | | | |
|---|---|---|---|---|---|---|
| 1.0268E-07 | -0.3225402 | 0 | 0.13 | 0.00242567 | Bcells | AIF1L |
| 1.031E-07 | -0.3058816 | 0.005 | 0.139 | 0.00243564 | Bcells | MGLL |
| 1.1266E-07 | -0.6226164 | 0.132 | 0.294 | 0.00266126 | Bcells | ATP13A3 |
| 1.143E-07 | -0.2635839 | 0.005 | 0.139 | 0.00270004 | Bcells | GULP1 |
| 1.1712E-07 | 0.25174431 | 0.295 | 0.162 | 0.00276666 | Bcells | DENR |
| 1.2157E-07 | -0.6939701 | 0.005 | 0.137 | 0.00287183 | Bcells | NUPR1 |
| 1.2247E-07 | -0.295553 | 0.021 | 0.162 | 0.00289314 | Bcells | FARP1 |
| 1.2408E-07 | -0.5689174 | 0.332 | 0.483 | 0.00293122 | Bcells | CD46 |
| 1.3583E-07 | -0.3399231 | 0 | 0.128 | 0.0032087 | Bcells | FAM84A |
| 1.3701E-07 | -0.2937425 | 0.021 | 0.162 | 0.00323658 | Bcells | GALNT3 |
| 1.3799E-07 | -0.260198 | 0.005 | 0.137 | 0.00325969 | Bcells | SCNN1A |
| 1.4062E-07 | -0.4747308 | 0.042 | 0.189 | 0.00332196 | Bcells | FCGR2A |
| 1.4634E-07 | 0.25428252 | 0.4 | 0.24 | 0.0034569 | Bcells | RSF1 |
| 1.4639E-07 | -0.2713045 | 0.021 | 0.161 | 0.00345808 | Bcells | CPD |
| 1.574E-07 | -0.2698038 | 0 | 0.127 | 0.00371824 | Bcells | SELENOP |
| 1.5747E-07 | -0.3692953 | 0.053 | 0.21 | 0.00371996 | Bcells | SOCS3 |
| 1.6995E-07 | -0.2714022 | 0.026 | 0.167 | 0.00401461 | Bcells | SIRPA |
| 1.71E-07 | -0.2969915 | 0.068 | 0.229 | 0.00403949 | Bcells | IL6ST |
| 1.7423E-07 | -0.2725086 | 0.047 | 0.197 | 0.00411577 | Bcells | USP53 |
| 1.79E-07 | 0.33408555 | 0.553 | 0.384 | 0.00422863 | Bcells | DEK |
| 1.8477E-07 | -0.4937734 | 0.479 | 0.63 | 0.00436475 | Bcells | TAX1BP1 |
| 1.8568E-07 | -0.3552385 | 0.121 | 0.283 | 0.00438642 | Bcells | RTN3 |
| 1.9978E-07 | -0.6427992 | 0.189 | 0.344 | 0.00471949 | Bcells | DDIT4 |
| 2.0335E-07 | -0.4721919 | 0.042 | 0.189 | 0.00480365 | Bcells | FYB1 |
| 2.071E-07 | -1.5878157 | 0.005 | 0.133 | 0.00489232 | Bcells | LYPD2 |
| 2.1335E-07 | 0.27744233 | 0.258 | 0.129 | 0.00504001 | Bcells | ID3 |
| 2.1674E-07 | -0.4907975 | 0.032 | 0.171 | 0.00512013 | Bcells | PRDM1 |
| 2.2483E-07 | -0.2944175 | 0 | 0.124 | 0.0053112 | Bcells | FAT1 |
| 2.2855E-07 | -0.4783335 | 0.016 | 0.149 | 0.00539915 | Bcells | RASD1 |
| 2.3319E-07 | -0.281433 | 0.058 | 0.209 | 0.00550862 | Bcells | ATP2A2 |
| 2.3445E-07 | -1.2266454 | 0.005 | 0.132 | 0.00553843 | Bcells | PLAT |
| 2.5027E-07 | -0.4217447 | 0.011 | 0.14 | 0.00591209 | Bcells | GADD45G |
| 2.5242E-07 | -0.8407434 | 0.016 | 0.147 | 0.00596303 | Bcells | CEACAM6 |
| 2.5638E-07 | -0.2662871 | 0.005 | 0.132 | 0.00605637 | Bcells | TMEM176A |
| 2.5648E-07 | -0.38937 | 0.179 | 0.342 | 0.00605882 | Bcells | NDFIP1 |
| 2.5684E-07 | -0.4060989 | 0 | 0.123 | 0.00606741 | Bcells | TMEM40 |
| 2.6857E-07 | -0.6553524 | 0.005 | 0.131 | 0.00634444 | Bcells | DHRS9 |
| 2.6907E-07 | -0.2959948 | 0.005 | 0.131 | 0.00635621 | Bcells | SCIN |
| 2.7697E-07 | -0.3396006 | 0 | 0.123 | 0.00654295 | Bcells | KRT80 |
| 2.7704E-07 | 0.26104174 | 0.368 | 0.216 | 0.00654463 | Bcells | SMIM14 |
| 2.8457E-07 | -0.5499864 | 0 | 0.122 | 0.00672251 | Bcells | AGR2 |
| 2.8555E-07 | -0.3305956 | 0.021 | 0.155 | 0.00674546 | Bcells | ITPKC |
| 2.9188E-07 | -0.7196626 | 0.032 | 0.17 | 0.00689517 | Bcells | FCN1 |
| 2.944E-07 | -0.286481 | 0.063 | 0.215 | 0.00695467 | Bcells | ZYX |
| 3.0079E-07 | -0.5085337 | 0.037 | 0.177 | 0.00710565 | Bcells | ANPEP |
| 3.018E-07 | -0.2612993 | 0.005 | 0.131 | 0.00712954 | Bcells | IL1R1 |
| 3.1538E-07 | -0.2769773 | 0.068 | 0.223 | 0.00745016 | Bcells | PPP2CB |
| 3.1819E-07 | -0.5171609 | 0.042 | 0.185 | 0.0075165 | Bcells | C5AR1 |
| 3.2615E-07 | -0.2931222 | 0.063 | 0.214 | 0.00770471 | Bcells | BTG3 |
| 3.4122E-07 | -0.254033 | 0 | 0.121 | 0.00806063 | Bcells | BICDL2 |

| | | | | | | |
|---|---|---|---|---|---|---|
| 3.4306E-07 | -0.7801732 | 0.026 | 0.162 | 0.00810412 | Bcells | APOBEC3A |
| 3.4881E-07 | -0.2659125 | 0.021 | 0.154 | 0.00823989 | Bcells | RCN1 |
| 3.5996E-07 | -0.5592962 | 0.005 | 0.13 | 0.00850337 | Bcells | AKAP12 |
| 3.6011E-07 | -0.5279826 | 0 | 0.121 | 0.0085069 | Bcells | TCIM |
| 3.6405E-07 | -0.3865499 | 0.142 | 0.301 | 0.0086 | Bcells | PICALM |
| 3.6794E-07 | -0.6395634 | 0.005 | 0.129 | 0.0086918 | Bcells | TFPI2 |
| 3.7769E-07 | -0.4984559 | 0.042 | 0.183 | 0.00892217 | Bcells | IGSF6 |
| 3.8997E-07 | -0.2663848 | 0.005 | 0.129 | 0.00921238 | Bcells | KAZN |
| 3.902E-07 | -0.2778216 | 0.016 | 0.145 | 0.00921773 | Bcells | CRIM1 |
| 3.9674E-07 | -0.3150838 | 0.053 | 0.196 | 0.00937229 | Bcells | HIGD1A |
| 4.003E-07 | -0.4836354 | 0.047 | 0.188 | 0.00945624 | Bcells | BAG3 |
| 4.0065E-07 | -0.3787366 | 0 | 0.12 | 0.00946461 | Bcells | EGLN3 |
| 4.1372E-07 | -0.2539632 | 0.011 | 0.136 | 0.00977325 | Bcells | AGRN |
| 4.242E-07 | -0.297316 | 0.021 | 0.152 | 0.01002081 | Bcells | HK2 |
| 4.4019E-07 | -0.3481155 | 0.111 | 0.267 | 0.01039872 | Bcells | ATOX1 |
| 4.4266E-07 | -0.4838677 | 0.047 | 0.187 | 0.01045691 | Bcells | TNFSF10 |
| 4.4729E-07 | -0.256085 | 0 | 0.119 | 0.01056627 | Bcells | C19orf33 |
| 4.4775E-07 | -0.3846578 | 0.637 | 0.773 | 0.01057719 | Bcells | EIF4G2 |
| 4.5158E-07 | -0.3126061 | 0 | 0.119 | 0.01066763 | Bcells | ASS1 |
| 4.6477E-07 | -0.3784377 | 0.011 | 0.135 | 0.01097923 | Bcells | BCAS1 |
| 4.7643E-07 | -0.2912968 | 0.011 | 0.135 | 0.01125474 | Bcells | COL18A1 |
| 5.0827E-07 | -0.7894623 | 0.395 | 0.528 | 0.01200683 | Bcells | ZFP36 |
| 5.102E-07 | -0.394371 | 0.584 | 0.715 | 0.01205246 | Bcells | SKP1 |
| 5.1078E-07 | -0.282454 | 0.011 | 0.136 | 0.01206625 | Bcells | SASH1 |
| 5.2477E-07 | -1.0293346 | 0.011 | 0.134 | 0.01239674 | Bcells | SERPINB2 |
| 5.2563E-07 | -0.6595688 | 0.047 | 0.188 | 0.01241701 | Bcells | AC020656.1 |
| 5.3396E-07 | -2.6222076 | 0.111 | 0.255 | 0.01261385 | Bcells | MT1G |
| 5.539E-07 | -0.3511293 | 0.084 | 0.231 | 0.01308485 | Bcells | ATP6V1B2 |
| 5.9971E-07 | -0.3228026 | 0.053 | 0.192 | 0.0141669 | Bcells | ITPRIPL2 |
| 6.191E-07 | -0.3114492 | 0.021 | 0.15 | 0.01462508 | Bcells | DMXL2 |
| 6.2204E-07 | -0.378674 | 0.063 | 0.205 | 0.01469438 | Bcells | TIPARP |
| 6.2856E-07 | -0.6119203 | 0.595 | 0.732 | 0.01484855 | Bcells | HSPA8 |
| 6.3675E-07 | -0.3863954 | 0.495 | 0.632 | 0.01504196 | Bcells | SAP18 |
| 6.5827E-07 | -0.286011 | 0.053 | 0.194 | 0.01555035 | Bcells | PGRMC1 |
| 7.6806E-07 | -0.2795048 | 0.011 | 0.132 | 0.01814395 | Bcells | EPB41L3 |
| 8.5157E-07 | -0.6426558 | 0.237 | 0.385 | 0.02011653 | Bcells | TYMP |
| 8.5255E-07 | -0.4926941 | 0.058 | 0.199 | 0.02013978 | Bcells | DUSP4 |
| 8.648E-07 | -0.2975097 | 0.005 | 0.122 | 0.02042916 | Bcells | FOXQ1 |
| 8.8389E-07 | -1.2747743 | 0 | 0.113 | 0.02088015 | Bcells | SPRR1B |
| 9.6598E-07 | -0.45664 | 0 | 0.113 | 0.02281931 | Bcells | TMPRSS11A |
| 9.6786E-07 | -0.4703729 | 0.021 | 0.145 | 0.02286376 | Bcells | CD93 |
| 9.7492E-07 | -0.5912748 | 0 | 0.113 | 0.02303056 | Bcells | KRT78 |
| 9.9524E-07 | -0.2802973 | 0.063 | 0.204 | 0.02351045 | Bcells | LRRFIP2 |
| 9.9947E-07 | -0.7551326 | 0.458 | 0.602 | 0.02361052 | Bcells | NFKBIA |
| 1.016E-06 | -0.4167158 | 0.453 | 0.579 | 0.02400174 | Bcells | UBE2B |
| 1.0423E-06 | -0.4987109 | 0.211 | 0.35 | 0.02462173 | Bcells | UBE2H |
| 1.0583E-06 | -0.2677584 | 0.111 | 0.267 | 0.02500134 | Bcells | KDELR2 |
| 1.0865E-06 | -0.2561314 | 0.037 | 0.168 | 0.02566746 | Bcells | STX3 |
| 1.1217E-06 | -0.3908821 | 0.095 | 0.242 | 0.02649716 | Bcells | BHLHE40 |
| 1.1634E-06 | -0.435426 | 0.211 | 0.355 | 0.0274841 | Bcells | MRPL33 |

| | | | | | | |
|---|---|---|---|---|---|---|
| 1.199E-06 | -0.3584772 | 0.332 | 0.484 | 0.02832501 | Bcells | TMEM50A |
| 1.2073E-06 | -0.3177893 | 0.126 | 0.274 | 0.02852084 | Bcells | ARL8B |
| 1.2142E-06 | -0.3022827 | 0.047 | 0.18 | 0.0286828 | Bcells | FNBP1L |
| 1.225E-06 | -0.3122501 | 0 | 0.111 | 0.02893807 | Bcells | STS |
| 1.2475E-06 | -0.4978709 | 0.032 | 0.158 | 0.02946855 | Bcells | CD300E |
| 1.289E-06 | -0.2565098 | 0.016 | 0.136 | 0.03044925 | Bcells | KLHL21 |
| 1.3106E-06 | 0.28899248 | 0.316 | 0.186 | 0.03096117 | Bcells | GPATCH2L |
| 1.3194E-06 | -0.4073222 | 0.074 | 0.212 | 0.03116744 | Bcells | NINJ1 |
| 1.3709E-06 | -0.3349003 | 0.147 | 0.297 | 0.03238551 | Bcells | KDELR1 |
| 1.3963E-06 | -0.3151498 | 0.132 | 0.29 | 0.03298575 | Bcells | CLIP1 |
| 1.3975E-06 | -0.41983 | 0.363 | 0.497 | 0.03301359 | Bcells | VDAC2 |
| 1.4061E-06 | -0.466182 | 0.032 | 0.159 | 0.03321619 | Bcells | BPGM |
| 1.4223E-06 | -0.5438549 | 0.5 | 0.62 | 0.03359987 | Bcells | PCBP1 |
| 1.4264E-06 | -0.2504828 | 0.021 | 0.143 | 0.03369528 | Bcells | CPM |
| 1.4545E-06 | -0.4757959 | 0.174 | 0.322 | 0.034359 | Bcells | B4GALT1 |
| 1.465E-06 | -0.3727563 | 0.011 | 0.126 | 0.03460791 | Bcells | ID4 |
| 1.5759E-06 | -1.103145 | 0.005 | 0.117 | 0.03722691 | Bcells | SPINK5 |
| 1.6136E-06 | -0.2863518 | 0.074 | 0.213 | 0.0381185 | Bcells | CTNND1 |
| 1.6437E-06 | -0.534236 | 0.005 | 0.117 | 0.03882829 | Bcells | HSPA2 |
| 1.6818E-06 | -0.2845486 | 0.026 | 0.149 | 0.03972841 | Bcells | SCCPDH |
| 1.7043E-06 | -0.4417375 | 0.211 | 0.357 | 0.0402599 | Bcells | TKT |
| 1.7073E-06 | -0.2994423 | 0.142 | 0.294 | 0.04033247 | Bcells | HDLBP |
| 1.7927E-06 | -0.7875906 | 0.021 | 0.14 | 0.04235007 | Bcells | CXCL1 |
| 1.8432E-06 | -0.264066 | 0.089 | 0.234 | 0.04354103 | Bcells | CTSA |
| 1.942E-06 | -0.3257188 | 0.105 | 0.251 | 0.04587671 | Bcells | DNAJC3 |
| 1.9536E-06 | -0.7993951 | 0.063 | 0.195 | 0.04615092 | Bcells | C12orf75 |
| 1.9569E-06 | -0.3254676 | 0.016 | 0.132 | 0.04622867 | Bcells | IL18 |
| 1.9994E-06 | -0.3433563 | 0.211 | 0.366 | 0.04723185 | Bcells | RAB7A |
| 2.1199E-06 | -0.3063662 | 0.084 | 0.225 | 0.05007946 | Bcells | ST14 |
| 2.1212E-06 | -0.3616628 | 0.005 | 0.115 | 0.05010908 | Bcells | CEACAM1 |
| 2.209E-06 | -0.2687922 | 0.032 | 0.154 | 0.0521829 | Bcells | AHR |
| 2.2168E-06 | -0.3928041 | 0.026 | 0.146 | 0.05236842 | Bcells | FPR1 |
| 2.2199E-06 | -0.3249044 | 0 | 0.106 | 0.0524408 | Bcells | A2ML1 |
| 2.3147E-06 | -0.2660918 | 0.016 | 0.131 | 0.05467963 | Bcells | LILRB3 |
| 2.3291E-06 | 0.26985773 | 0.732 | 0.622 | 0.0550199 | Bcells | PRRC2C |
| 2.3384E-06 | -1.0166581 | 0.021 | 0.137 | 0.05524019 | Bcells | DEFB1 |
| 2.5116E-06 | -0.7753327 | 0 | 0.105 | 0.05933076 | Bcells | ZG16B |
| 2.6327E-06 | -1.2986323 | 0.068 | 0.197 | 0.06219111 | Bcells | HOPX |
| 2.9864E-06 | -0.5990245 | 0.026 | 0.142 | 0.0705472 | Bcells | CDKN2A |
| 3.0444E-06 | -0.4559277 | 0.058 | 0.186 | 0.07191798 | Bcells | ACSL1 |
| 3.0851E-06 | -0.3332224 | 0.016 | 0.128 | 0.07287841 | Bcells | LRG1 |
| 3.1212E-06 | -0.3778887 | 0.489 | 0.616 | 0.07373115 | Bcells | CAP1 |
| 3.1682E-06 | -0.2833006 | 0.058 | 0.187 | 0.07484287 | Bcells | RNF130 |
| 3.2255E-06 | -0.282734 | 0.005 | 0.111 | 0.07619537 | Bcells | FUT3 |
| 3.3941E-06 | -0.4356932 | 0.721 | 0.794 | 0.0801795 | Bcells | CHCHD2 |
| 3.4111E-06 | -0.3358777 | 0.005 | 0.112 | 0.08058105 | Bcells | ANGPTL4 |
| 3.4373E-06 | -0.2577706 | 0.016 | 0.128 | 0.08119898 | Bcells | PLBD1 |
| 3.5065E-06 | -0.4002015 | 0.589 | 0.693 | 0.08283517 | Bcells | HSP90B1 |
| 3.6215E-06 | -0.284378 | 0.063 | 0.192 | 0.08555105 | Bcells | FCHO2 |
| 3.724E-06 | -0.4178048 | 0.068 | 0.201 | 0.0879712 | Bcells | ITGAX |

| | | | | | | |
|---|---|---|---|---|---|---|
| 3.812E-06 | -0.6046689 | 0.163 | 0.296 | 0.09005001 | Bcells | GLRX |
| 3.8396E-06 | -0.3590812 | 0.105 | 0.242 | 0.09070371 | Bcells | IFRD1 |
| 3.964E-06 | -0.296197 | 0.032 | 0.149 | 0.09364249 | Bcells | BCAT1 |
| 4.0731E-06 | -0.4362192 | 0.074 | 0.201 | 0.09621871 | Bcells | NCF2 |
| 4.1124E-06 | -0.2778758 | 0.042 | 0.165 | 0.09714629 | Bcells | MIR22HG |
| 4.3751E-06 | -0.8100694 | 0.479 | 0.628 | 0.10335355 | Bcells | JUN |
| 4.3981E-06 | -0.4120374 | 0.016 | 0.125 | 0.10389726 | Bcells | SRD5A3 |
| 4.8214E-06 | -0.6131273 | 0.653 | 0.747 | 0.1138963 | Bcells | PRDX1 |
| 4.9353E-06 | -0.3561634 | 0.168 | 0.312 | 0.11658684 | Bcells | SBDS |
| 4.9812E-06 | -0.3824198 | 0.005 | 0.108 | 0.11767195 | Bcells | KRT23 |
| 5.2923E-06 | -0.2739739 | 0.068 | 0.199 | 0.1250207 | Bcells | ARHGAP18 |
| 5.3525E-06 | -0.2848491 | 0.037 | 0.156 | 0.12644203 | Bcells | SESN2 |
| 5.4921E-06 | -0.4081443 | 0.005 | 0.107 | 0.12973876 | Bcells | TMC5 |
| 5.4972E-06 | -0.3248724 | 0.542 | 0.696 | 0.12986113 | Bcells | LRRFIP1 |
| 5.7529E-06 | -0.3977554 | 0.005 | 0.107 | 0.1359008 | Bcells | THBD |
| 5.8435E-06 | -0.2781918 | 0.053 | 0.177 | 0.1380413 | Bcells | ABCA1 |
| 5.8509E-06 | -0.3866477 | 0.011 | 0.115 | 0.13821541 | Bcells | RRAD |
| 6.3974E-06 | -0.2874233 | 0.063 | 0.189 | 0.15112693 | Bcells | VCL |
| 7.0756E-06 | -0.3175722 | 0.032 | 0.144 | 0.16714735 | Bcells | TLR2 |
| 7.1414E-06 | -0.3102939 | 0.068 | 0.195 | 0.16870222 | Bcells | IRS2 |
| 7.5509E-06 | -0.2655919 | 0.1 | 0.235 | 0.17837557 | Bcells | RBMS1 |
| 7.6041E-06 | -0.2835508 | 0.068 | 0.194 | 0.17963117 | Bcells | B4GALT5 |
| 7.6233E-06 | -0.5283368 | 0.011 | 0.113 | 0.18008496 | Bcells | CYP1B1 |
| 7.6661E-06 | -0.4575728 | 0.053 | 0.172 | 0.18109684 | Bcells | DNAJA4 |
| 7.7408E-06 | -0.2637755 | 0.026 | 0.137 | 0.18286025 | Bcells | CLEC7A |
| 8.1817E-06 | -0.4139191 | 0.232 | 0.368 | 0.1932767 | Bcells | SPAG9 |
| 8.4794E-06 | -0.2679681 | 0.058 | 0.179 | 0.20030834 | Bcells | LSR |
| 8.5999E-06 | -0.4986177 | 0.084 | 0.209 | 0.20315624 | Bcells | LST1 |
| 8.8239E-06 | -0.4372896 | 0.063 | 0.183 | 0.20844615 | Bcells | INSIG1 |
| 8.9449E-06 | -0.4300521 | 0.063 | 0.184 | 0.2113052 | Bcells | SGK1 |
| 9.1113E-06 | -0.2534407 | 0.032 | 0.143 | 0.21523645 | Bcells | OCLN |
| 9.2216E-06 | -0.3456034 | 0.047 | 0.165 | 0.21784232 | Bcells | LCP2 |
| 9.5912E-06 | -0.8022637 | 0.068 | 0.194 | 0.22657359 | Bcells | HSPA6 |
| 9.8149E-06 | -0.2638871 | 0.021 | 0.128 | 0.23185652 | Bcells | TP53INP2 |
| 1.0008E-05 | -0.3398131 | 0.011 | 0.111 | 0.23641406 | Bcells | SMIM5 |
| 1.002E-05 | -0.3416469 | 0.011 | 0.111 | 0.23669425 | Bcells | FXYD2 |
| 1.0607E-05 | -0.4179532 | 0.342 | 0.453 | 0.25056947 | Bcells | SEC61G |
| 1.0729E-05 | -0.2606266 | 0.026 | 0.134 | 0.25344337 | Bcells | MAP3K20 |
| 1.0736E-05 | -0.312318 | 0.179 | 0.317 | 0.25360633 | Bcells | CLTC |
| 1.1127E-05 | -0.7612492 | 0.111 | 0.234 | 0.26285555 | Bcells | PLIN2 |
| 1.1183E-05 | -0.2551664 | 0.063 | 0.186 | 0.26416775 | Bcells | PHACTR2 |
| 1.1231E-05 | -0.3554322 | 0.032 | 0.141 | 0.2653155 | Bcells | GPNMB |
| 1.1988E-05 | -0.331679 | 0.011 | 0.109 | 0.28319894 | Bcells | NR1D1 |
| 1.2885E-05 | -0.2768271 | 0.016 | 0.117 | 0.30437585 | Bcells | ST3GAL4 |
| 1.4021E-05 | -0.3301894 | 0.016 | 0.116 | 0.33122738 | Bcells | CD163 |
| 1.4062E-05 | -0.3448365 | 0.053 | 0.173 | 0.33219217 | Bcells | XIST |
| 1.4965E-05 | -0.3395737 | 0.042 | 0.157 | 0.35351266 | Bcells | MT1F |
| 1.5131E-05 | -0.3827826 | 0.137 | 0.267 | 0.35743704 | Bcells | AFDN |
| 1.5418E-05 | -0.2908528 | 0.089 | 0.214 | 0.36420823 | Bcells | RUNX1 |
| 1.6758E-05 | -0.390489 | 0.032 | 0.137 | 0.39586768 | Bcells | MCEMP1 |

| | | | | | | |
|---|---|---|---|---|---|---|
| 1.7297E-05 | -0.8972597 | 0.5 | 0.625 | 0.40859887 | Bcells | S100A4 |
| 1.7651E-05 | -0.5329391 | 0.295 | 0.404 | 0.41696091 | Bcells | FLNA |
| 1.815E-05 | -0.3266155 | 0.011 | 0.106 | 0.42875045 | Bcells | CSF3R |
| 1.8667E-05 | -0.320992 | 0.242 | 0.373 | 0.44095962 | Bcells | GNB2 |
| 1.9027E-05 | -0.4191116 | 0.084 | 0.202 | 0.44948534 | Bcells | FOSB |
| 1.9305E-05 | -0.3118254 | 0.158 | 0.29 | 0.45604319 | Bcells | CHMP2B |
| 2.0621E-05 | -0.6732764 | 0.963 | 0.973 | 0.48713966 | Bcells | ACTB |
| 2.0689E-05 | -0.2596175 | 0.105 | 0.234 | 0.4887278 | Bcells | LPP |
| 2.0781E-05 | -0.4062447 | 0.137 | 0.267 | 0.49091813 | Bcells | UGCG |
| 2.0918E-05 | -0.2550966 | 0.089 | 0.215 | 0.49415229 | Bcells | CNN3 |
| 2.1259E-05 | -0.2681162 | 0.042 | 0.151 | 0.50221137 | Bcells | FAM129A |
| 2.1422E-05 | -0.3890413 | 0.105 | 0.235 | 0.50604482 | Bcells | HCST |
| 2.3519E-05 | -0.3021412 | 0.189 | 0.324 | 0.55559132 | Bcells | ARL6IP1 |
| 2.5279E-05 | -0.3543563 | 0.105 | 0.224 | 0.59716038 | Bcells | STXBP2 |
| 2.535E-05 | -0.2622436 | 0.032 | 0.135 | 0.5988394 | Bcells | GCLC |
| 2.6903E-05 | -0.3326654 | 0.068 | 0.183 | 0.63552369 | Bcells | ICAM1 |
| 2.9483E-05 | -0.2861187 | 0.121 | 0.243 | 0.69648162 | Bcells | COMT |
| 2.9684E-05 | -1.8635851 | 0.047 | 0.152 | 0.70123394 | Bcells | MT1H |
| 3.1128E-05 | -0.4692755 | 0.111 | 0.23 | 0.7353451 | Bcells | PPIF |
| 3.2914E-05 | -1.118449 | 0.032 | 0.131 | 0.77751702 | Bcells | APOE |
| 3.4359E-05 | -0.8442651 | 0.121 | 0.236 | 0.81167217 | Bcells | BCL2A1 |
| 3.4998E-05 | -0.3217385 | 0.068 | 0.181 | 0.82676501 | Bcells | PFKFB3 |
| 3.8034E-05 | -0.3155112 | 0.242 | 0.368 | 0.89847234 | Bcells | RAB10 |
| 3.8693E-05 | -0.3379141 | 0.163 | 0.289 | 0.91405064 | Bcells | CHMP1B |
| 3.919E-05 | -0.3726112 | 0.153 | 0.277 | 0.92579214 | Bcells | SLK |
| 3.9322E-05 | -1.5133201 | 0.016 | 0.107 | 0.928913 | Bcells | CRCT1 |
| 4.0089E-05 | -0.3157295 | 0.047 | 0.151 | 0.94703019 | Bcells | MDK |
| 4.0761E-05 | -0.3614562 | 0.363 | 0.488 | 0.96290698 | Bcells | CANX |
| 4.2568E-05 | -0.3287239 | 0.016 | 0.107 | 1 | Bcells | SLC16A10 |
| 4.2785E-05 | -2.0154505 | 0.068 | 0.176 | 1 | Bcells | IL1B |
| 4.4555E-05 | -0.4941698 | 0.479 | 0.587 | 1 | Bcells | KLF6 |
| 4.4762E-05 | -0.3757857 | 0.058 | 0.171 | 1 | Bcells | TGFBI |
| 4.6516E-05 | -0.435695 | 0.426 | 0.519 | 1 | Bcells | H2AFZ |
| 4.7266E-05 | -0.3474036 | 0.089 | 0.204 | 1 | Bcells | CTSC |
| 4.7849E-05 | -0.3577151 | 0.111 | 0.225 | 1 | Bcells | KYNU |
| 4.9981E-05 | -0.2739959 | 0.153 | 0.278 | 1 | Bcells | PLEKHB2 |
| 5.4501E-05 | -0.3267965 | 0.153 | 0.268 | 1 | Bcells | RAB5IF |
| 5.5759E-05 | -0.4844718 | 0.137 | 0.251 | 1 | Bcells | SNHG12 |
| 6.1216E-05 | -0.3057273 | 0.137 | 0.254 | 1 | Bcells | C1orf122 |
| 6.1616E-05 | -0.6375006 | 0.484 | 0.617 | 1 | Bcells | LGALS1 |
| 6.223E-05 | -0.3695957 | 0.205 | 0.326 | 1 | Bcells | CAPG |
| 6.2279E-05 | -0.3372851 | 0.632 | 0.725 | 1 | Bcells | COX6A1 |
| 6.4016E-05 | -0.3147917 | 0.032 | 0.126 | 1 | Bcells | MARCO |
| 6.4027E-05 | -0.2689045 | 0.142 | 0.261 | 1 | Bcells | YWHAG |
| 7.6448E-05 | -0.3538049 | 0.116 | 0.226 | 1 | Bcells | RAB9A |
| 7.7426E-05 | -0.3194999 | 0.184 | 0.3 | 1 | Bcells | EIF4A1 |
| 7.7903E-05 | -0.3023406 | 0.226 | 0.344 | 1 | Bcells | WDR1 |
| 7.8413E-05 | -0.2738114 | 0.168 | 0.288 | 1 | Bcells | MRFAP1 |
| 7.9528E-05 | -0.2723357 | 0.111 | 0.226 | 1 | Bcells | PNP |
| 8.1246E-05 | -0.2961539 | 0.2 | 0.319 | 1 | Bcells | CALM3 |

| | | | | | | |
|---|---|---|---|---|---|---|
| 8.8776E-05 | -1.2581753 | 0.053 | 0.152 | 1 | Bcells | CCL20 |
| 9.1547E-05 | -0.3082296 | 0.195 | 0.317 | 1 | Bcells | RAB11A |
| 9.1884E-05 | -0.2932929 | 0.116 | 0.228 | 1 | Bcells | FCGRT |
| 9.4154E-05 | -0.4612406 | 0.016 | 0.1 | 1 | Bcells | RETN |
| 9.7814E-05 | -0.311877 | 0.174 | 0.287 | 1 | Bcells | ACADVL |
| 9.7828E-05 | -0.3779857 | 0.1 | 0.207 | 1 | Bcells | SLC43A2 |
| 0.00010262 | -0.5642787 | 0.174 | 0.282 | 1 | Bcells | RHOB |
| 0.00011689 | -0.2868003 | 0.142 | 0.255 | 1 | Bcells | MAP2K3 |
| 0.00011795 | -0.2510014 | 0.2 | 0.318 | 1 | Bcells | RER1 |
| 0.00012565 | -0.2590996 | 0.032 | 0.121 | 1 | Bcells | LILRB2 |
| 0.0001262 | -0.4933696 | 0.032 | 0.121 | 1 | Bcells | HILPDA |
| 0.00013753 | -0.3635632 | 0.447 | 0.281 | 1 | Bcells | IFITM2 |
| 0.00013754 | -0.3422474 | 0.095 | 0.198 | 1 | Bcells | SERTAD2 |
| 0.00014186 | -0.2969925 | 0.179 | 0.291 | 1 | Bcells | STK24 |
| 0.0001434 | -0.3293483 | 0.079 | 0.182 | 1 | Bcells | GCLM |
| 0.00014599 | -0.3908477 | 0.142 | 0.248 | 1 | Bcells | SPTBN1 |
| 0.00015027 | -0.4077257 | 0.584 | 0.658 | 1 | Bcells | CALM2 |
| 0.00015712 | -0.3772482 | 0.453 | 0.547 | 1 | Bcells | C4orf3 |
| 0.00017262 | -0.2715193 | 0.095 | 0.199 | 1 | Bcells | TMEM106B |
| 0.00017697 | -0.3117978 | 0.142 | 0.251 | 1 | Bcells | PLEC |
| 0.00017805 | -0.2500752 | 0.058 | 0.153 | 1 | Bcells | CKS2 |
| 0.00019084 | -0.2983251 | 0.032 | 0.119 | 1 | Bcells | FCGR3A |
| 0.00019829 | -0.2559184 | 0.053 | 0.145 | 1 | Bcells | PDP1 |
| 0.00020268 | -0.2705993 | 0.105 | 0.209 | 1 | Bcells | ARRB2 |
| 0.00020303 | -0.3071914 | 0.153 | 0.259 | 1 | Bcells | ABRACL |
| 0.00021632 | -0.2746489 | 0.074 | 0.17 | 1 | Bcells | JMY |
| 0.00021679 | -0.2729231 | 0.179 | 0.296 | 1 | Bcells | SELENOS |
| 0.00022531 | -0.2621411 | 0.126 | 0.233 | 1 | Bcells | RYBP |
| 0.00022919 | -0.2783596 | 0.074 | 0.172 | 1 | Bcells | SAMHD1 |
| 0.0002353 | -0.2611723 | 0.121 | 0.227 | 1 | Bcells | RGS10 |
| 0.00024725 | -0.2529719 | 0.147 | 0.254 | 1 | Bcells | SCP2 |
| 0.00025777 | -0.3263325 | 0.089 | 0.187 | 1 | Bcells | PTPRE |
| 0.00025848 | -0.2893773 | 0.426 | 0.528 | 1 | Bcells | CD164 |
| 0.00027879 | -0.398515 | 0.137 | 0.237 | 1 | Bcells | NFKBIZ |
| 0.000336 | -0.2578899 | 0.137 | 0.239 | 1 | Bcells | ACSL3 |
| 0.00038655 | -0.2890859 | 0.4 | 0.512 | 1 | Bcells | COX7B |
| 0.0003955 | -0.4456627 | 0.289 | 0.386 | 1 | Bcells | C9orf16 |
| 0.00043785 | -0.6771648 | 0.032 | 0.109 | 1 | Bcells | APOC1 |
| 0.0004403 | -0.3242685 | 0.111 | 0.206 | 1 | Bcells | UBE2S |
| 0.0004625 | -0.3489061 | 0.226 | 0.326 | 1 | Bcells | PPA1 |
| 0.00048141 | -0.4201216 | 0.174 | 0.27 | 1 | Bcells | CCNG2 |
| 0.00050043 | -0.3402352 | 0.979 | 0.984 | 1 | Bcells | EIF1 |
| 0.00050255 | -0.2735809 | 0.147 | 0.247 | 1 | Bcells | EIF4A3 |
| 0.00059412 | -1.0159647 | 0.042 | 0.123 | 1 | Bcells | CCL3 |
| 0.00059475 | -0.3710385 | 0.347 | 0.429 | 1 | Bcells | SELENOK |
| 0.00063832 | -0.3988905 | 0.489 | 0.569 | 1 | Bcells | RPS27L |
| 0.00069942 | -0.3441043 | 0.847 | 0.905 | 1 | Bcells | CFL1 |
| 0.00072964 | -0.4991468 | 0.274 | 0.36 | 1 | Bcells | TUBA1B |
| 0.00078343 | -0.3515821 | 0.232 | 0.325 | 1 | Bcells | TMBIM1 |
| 0.00079818 | -0.2709408 | 0.489 | 0.59 | 1 | Bcells | SLC25A3 |

| | | | | | | |
|---|---|---|---|---|---|---|
| 0.00080147 | -0.3954783 | 0.068 | 0.152 | 1 | Bcells | ZFAND2A |
| 0.00083167 | -0.2760966 | 0.295 | 0.389 | 1 | Bcells | SEM1 |
| 0.00086157 | -0.2753176 | 0.174 | 0.267 | 1 | Bcells | TOR1AIP2 |
| 0.00092118 | -0.25512 | 0.095 | 0.182 | 1 | Bcells | SPTSSA |
| 0.00098925 | -0.339369 | 0.195 | 0.289 | 1 | Bcells | ATP1B3 |
| 0.00103059 | -0.8281096 | 0.058 | 0.134 | 1 | Bcells | EREG |
| 0.00104449 | -1.2100444 | 0.063 | 0.143 | 1 | Bcells | CCL2 |
| 0.00113626 | -0.4454099 | 0.416 | 0.475 | 1 | Bcells | CD81 |
| 0.0011824 | -0.3347875 | 0.053 | 0.129 | 1 | Bcells | AC023157.3 |
| 0.00122396 | -0.2666445 | 0.226 | 0.32 | 1 | Bcells | FKBP1A |
| 0.00124034 | -0.3717029 | 0.211 | 0.298 | 1 | Bcells | PIM3 |
| 0.00126554 | -0.6219567 | 0.142 | 0.23 | 1 | Bcells | ISG15 |
| 0.00128341 | -1.2602828 | 0.042 | 0.115 | 1 | Bcells | MT1M |
| 0.00135727 | -0.9578394 | 0.037 | 0.108 | 1 | Bcells | CCL4 |
| 0.00138443 | -0.4878118 | 0.126 | 0.21 | 1 | Bcells | TNFRSF1B |
| 0.00159911 | -0.298282 | 0.068 | 0.146 | 1 | Bcells | MS4A7 |
| 0.00169985 | -0.304624 | 0.216 | 0.306 | 1 | Bcells | BNIP3L |
| 0.0017462 | -0.6553961 | 0.958 | 0.908 | 1 | Bcells | MT-ND2 |
| 0.00197396 | -0.2573973 | 0.168 | 0.255 | 1 | Bcells | MGAT1 |
| 0.00204587 | -0.3333685 | 0.174 | 0.259 | 1 | Bcells | NABP1 |
| 0.00219965 | -0.2980579 | 0.779 | 0.875 | 1 | Bcells | MYL12A |
| 0.00229554 | -0.3077589 | 0.311 | 0.384 | 1 | Bcells | DYNLRB1 |
| 0.00231302 | -0.3002178 | 0.595 | 0.643 | 1 | Bcells | EDF1 |
| 0.00247339 | 0.28525768 | 0.289 | 0.193 | 1 | Bcells | SEC11C |
| 0.00252473 | -0.4120253 | 0.153 | 0.229 | 1 | Bcells | CYB5A |
| 0.00268178 | -0.4860114 | 0.616 | 0.677 | 1 | Bcells | IER2 |
| 0.0026995 | -0.2736908 | 0.226 | 0.311 | 1 | Bcells | CTNNB1 |
| 0.00293587 | -0.6225432 | 0.979 | 0.915 | 1 | Bcells | MT-ND1 |
| 0.00298318 | -0.5575017 | 0.153 | 0.227 | 1 | Bcells | HEXIM1 |
| 0.00310117 | -1.3782681 | 0.074 | 0.145 | 1 | Bcells | CXCL3 |
| 0.00319576 | -0.3173214 | 0.468 | 0.548 | 1 | Bcells | TOP1 |
| 0.00337425 | -0.25541 | 0.147 | 0.228 | 1 | Bcells | GALNT1 |
| 0.00392242 | -0.2955387 | 0.658 | 0.688 | 1 | Bcells | PSMA7 |
| 0.00415376 | -0.7651741 | 0.147 | 0.22 | 1 | Bcells | ALOX5AP |
| 0.00579991 | -0.3558011 | 0.3 | 0.366 | 1 | Bcells | RALBP1 |
| 0.00616033 | -0.3115585 | 0.668 | 0.731 | 1 | Bcells | ATP6V1G1 |
| 0.00650815 | -0.2810859 | 0.174 | 0.244 | 1 | Bcells | RAB31 |
| 0.00747222 | -0.2699403 | 0.542 | 0.584 | 1 | Bcells | NOP10 |
| 0.00755471 | -0.3440935 | 0.537 | 0.576 | 1 | Bcells | PRDX5 |
| 0.00915138 | -0.3517265 | 0.311 | 0.358 | 1 | Bcells | MIF |
| 0 | 5.77067206 | 0.998 | 0.058 | 0 | ERY | HBB |
| 0 | 3.41844559 | 0.813 | 0.016 | 0 | ERY | HBA2 |
| 0 | 2.87515333 | 0.523 | 0.008 | 0 | ERY | HBA1 |
| 0 | 2.82639354 | 0.914 | 0.104 | 0 | ERY | APOE |
| 0 | 2.31989144 | 0.91 | 0.081 | 0 | ERY | APOC1 |
| 0 | 2.09846472 | 0.91 | 0.11 | 0 | ERY | DEFB1 |
| 0 | 1.86980549 | 0.927 | 0.225 | 0 | ERY | CRYAB |
| 0 | 1.83732571 | 0.974 | 0.523 | 0 | ERY | UPP1 |
| 0 | 1.80835545 | 0.972 | 0.317 | 0 | ERY | MMP7 |
| 0 | 1.66986106 | 0.983 | 0.543 | 0 | ERY | KRT18 |

| | | | | | | |
|---|---|---|---|---|---|---|
| 0 | 1.3153832 | 0.987 | 0.873 | 0 | ERY | MT-ND3 |
| 0 | 1.31048855 | 0.907 | 0.17 | 0 | ERY | C12orf75 |
| 0 | 1.271734 | 0.975 | 0.477 | 0 | ERY | KRT8 |
| 0 | 1.26910026 | 0.944 | 0.243 | 0 | ERY | S100A2 |
| 0 | 1.26045058 | 0.952 | 0.315 | 0 | ERY | IGFBP7 |
| 0 | 1.2356777 | 0.904 | 0.244 | 0 | ERY | DDIT3 |
| 0 | 1.22648723 | 0.892 | 0.201 | 0 | ERY | OCIAD2 |
| 0 | 1.21293489 | 0.91 | 0.212 | 0 | ERY | EPCAM |
| 0 | 1.19105571 | 0.893 | 0.219 | 0 | ERY | S100A13 |
| 0 | 1.18697947 | 0.999 | 0.931 | 0 | ERY | RPS12 |
| 0 | 1.17605941 | 0.982 | 0.565 | 0 | ERY | KRT19 |
| 0 | 1.17428655 | 0.989 | 0.935 | 0 | ERY | S100A6 |
| 0 | 1.14477204 | 0.856 | 0.066 | 0 | ERY | SAA1 |
| 0 | 1.1405154 | 0.988 | 0.885 | 0 | ERY | RPS23 |
| 0 | 1.13762536 | 0.86 | 0.157 | 0 | ERY | S100A14 |
| 0 | 1.12521871 | 0.99 | 0.872 | 0 | ERY | RPL36 |
| 0 | 1.07624757 | 0.845 | 0.172 | 0 | ERY | SDF2L1 |
| 0 | 1.07595743 | 0.976 | 0.435 | 0 | ERY | SPP1 |
| 0 | 1.07239982 | 0.9 | 0.31 | 0 | ERY | HERPUD1 |
| 0 | 1.0479072 | 1 | 0.986 | 0 | ERY | FTL |
| 0 | 1.04635781 | 0.893 | 0.252 | 0 | ERY | TNFRSF12A |
| 0 | 1.00705774 | 0.991 | 0.902 | 0 | ERY | RPL7A |
| 0 | 0.99943722 | 0.985 | 0.777 | 0 | ERY | DYNLL1 |
| 0 | 0.98588357 | 0.914 | 0.259 | 0 | ERY | MT1E |
| 0 | 0.98515541 | 0.871 | 0.214 | 0 | ERY | S100A16 |
| 0 | 0.97809713 | 0.82 | 0.18 | 0 | ERY | MANF |
| 0 | 0.97325188 | 0.988 | 0.881 | 0 | ERY | RPS2 |
| 0 | 0.97319235 | 0.809 | 0.134 | 0 | ERY | CLDN3 |
| 0 | 0.96888748 | 0.788 | 0.138 | 0 | ERY | SELENOM |
| 0 | 0.96604018 | 0.712 | 0.056 | 0 | ERY | AL353751.1 |
| 0 | 0.95641504 | 0.978 | 0.663 | 0 | ERY | CALR |
| 0 | 0.9539748 | 0.99 | 0.943 | 0 | ERY | RPS19 |
| 0 | 0.94174806 | 0.888 | 0.351 | 0 | ERY | PRDX6 |
| 0 | 0.94169539 | 0.995 | 0.972 | 0 | ERY | RPLP1 |
| 0 | 0.94057417 | 0.964 | 0.475 | 0 | ERY | MTRNR2L8 |
| 0 | 0.93616795 | 0.985 | 0.785 | 0 | ERY | RPS21 |
| 0 | 0.93252815 | 0.988 | 0.957 | 0 | ERY | RPL15 |
| 0 | 0.93178585 | 0.946 | 0.458 | 0 | ERY | CD81 |
| 0 | 0.93147196 | 0.987 | 0.872 | 0 | ERY | RPL3 |
| 0 | 0.9268306 | 0.988 | 0.871 | 0 | ERY | RPL26 |
| 0 | 0.92277475 | 0.894 | 0.322 | 0 | ERY | PDIA6 |
| 0 | 0.91424217 | 0.862 | 0.306 | 0 | ERY | MGST3 |
| 0 | 0.90643794 | 0.938 | 0.419 | 0 | ERY | CYSTM1 |
| 0 | 0.90501369 | 0.989 | 0.969 | 0 | ERY | TMSB10 |
| 0 | 0.90255989 | 0.988 | 0.899 | 0 | ERY | RPS7 |
| 0 | 0.90125663 | 0.91 | 0.332 | 0 | ERY | PHLDA2 |
| 0 | 0.89945841 | 0.987 | 0.826 | 0 | ERY | RPL22 |
| 0 | 0.89747984 | 0.831 | 0.258 | 0 | ERY | MYDGF |
| 0 | 0.89396808 | 0.99 | 0.92 | 0 | ERY | RPL8 |
| 0 | 0.89166807 | 0.992 | 0.921 | 0 | ERY | RPS27A |

| | | | | | | |
|---|---|---|---|---|---|---|
| 0 | 0.88673547 | 0.979 | 0.633 | 0 | ERY | HSPA5 |
| 0 | 0.88538411 | 0.918 | 0.436 | 0 | ERY | SEC61G |
| 0 | 0.87990355 | 0.987 | 0.767 | 0 | ERY | CTSD |
| 0 | 0.87459229 | 0.969 | 0.589 | 0 | ERY | HINT1 |
| 0 | 0.87354704 | 0.871 | 0.357 | 0 | ERY | PARK7 |
| 0 | 0.87313304 | 0.991 | 0.92 | 0 | ERY | RPL11 |
| 0 | 0.8669246 | 0.99 | 0.946 | 0 | ERY | RPL10 |
| 0 | 0.86609149 | 0.903 | 0.199 | 0 | ERY | G0S2 |
| 0 | 0.85877597 | 0.987 | 0.833 | 0 | ERY | RACK1 |
| 0 | 0.8563053 | 0.914 | 0.409 | 0 | ERY | SLC3A2 |
| 0 | 0.85006507 | 0.99 | 0.938 | 0 | ERY | RPS28 |
| 0 | 0.84920352 | 0.976 | 0.647 | 0 | ERY | NDUFA4 |
| 0 | 0.84709363 | 0.84 | 0.241 | 0 | ERY | RHOC |
| 0 | 0.83571281 | 0.987 | 0.841 | 0 | ERY | RPL35 |
| 0 | 0.83211808 | 0.968 | 0.386 | 0 | ERY | SLPI |
| 0 | 0.82829343 | 0.847 | 0.264 | 0 | ERY | RPS4Y1 |
| 0 | 0.81920763 | 0.67 | 0.089 | 0 | ERY | CRELD2 |
| 0 | 0.81429839 | 0.988 | 0.948 | 0 | ERY | RPL18 |
| 0 | 0.81369794 | 0.979 | 0.524 | 0 | ERY | CLDN4 |
| 0 | 0.81084239 | 0.638 | 0.052 | 0 | ERY | AC092069.1 |
| 0 | 0.80903363 | 0.848 | 0.337 | 0 | ERY | MRPL33 |
| 0 | 0.80711982 | 0.747 | 0.119 | 0 | ERY | GPNMB |
| 0 | 0.79478637 | 0.99 | 0.844 | 0 | ERY | RPS4X |
| 0 | 0.79332476 | 0.866 | 0.36 | 0 | ERY | POLR2L |
| 0 | 0.7928475 | 0.864 | 0.34 | 0 | ERY | MIF |
| 0 | 0.79249543 | 0.989 | 0.894 | 0 | ERY | RPS3A |
| 0 | 0.78793406 | 0.843 | 0.333 | 0 | ERY | TMEM258 |
| 0 | 0.78724355 | 0.988 | 0.898 | 0 | ERY | RPL24 |
| 0 | 0.78710612 | 0.99 | 0.956 | 0 | ERY | RPS18 |
| 0 | 0.78670607 | 0.974 | 0.645 | 0 | ERY | ZFAS1 |
| 0 | 0.78303739 | 0.822 | 0.289 | 0 | ERY | PFDN2 |
| 0 | 0.77225959 | 0.877 | 0.377 | 0 | ERY | NDUFB2 |
| 0 | 0.77155048 | 0.859 | 0.312 | 0 | ERY | PPP1R14B |
| 0 | 0.76581013 | 0.978 | 0.777 | 0 | ERY | EEF1D |
| 0 | 0.76240196 | 0.987 | 0.867 | 0 | ERY | RPL14 |
| 0 | 0.76178034 | 0.967 | 0.647 | 0 | ERY | CALM2 |
| 0 | 0.76087276 | 0.839 | 0.23 | 0 | ERY | SNHG12 |
| 0 | 0.76022949 | 0.95 | 0.632 | 0 | ERY | EDF1 |
| 0 | 0.75650346 | 0.819 | 0.312 | 0 | ERY | COPS9 |
| 0 | 0.75375454 | 0.876 | 0.253 | 0 | ERY | MAL2 |
| 0 | 0.75319763 | 0.98 | 0.788 | 0 | ERY | CHCHD2 |
| 0 | 0.75144957 | 0.984 | 0.738 | 0 | ERY | PRDX1 |
| 0 | 0.74898049 | 0.99 | 0.963 | 0 | ERY | RPL13 |
| 0 | 0.74493056 | 0.785 | 0.215 | 0 | ERY | SNHG7 |
| 0 | 0.7439063 | 0.739 | 0.17 | 0 | ERY | PDIA4 |
| 0 | 0.73905638 | 0.816 | 0.265 | 0 | ERY | CD151 |
| 0 | 0.73797962 | 0.946 | 0.539 | 0 | ERY | SSR4 |
| 0 | 0.73283239 | 0.98 | 0.755 | 0 | ERY | COX7C |
| 0 | 0.72820509 | 0.809 | 0.314 | 0 | ERY | FAM96B |
| 0 | 0.72782381 | 0.985 | 0.806 | 0 | ERY | GSTP1 |

| | | | | | | |
|---|---|---|---|---|---|---|
| 0 | 0.72193134 | 0.979 | 0.675 | 0 | ERY | NPM1 |
| 0 | 0.71660065 | 0.648 | 0.108 | 0 | ERY | LRRC75A |
| 0 | 0.71435203 | 0.994 | 0.984 | 0 | ERY | EIF1 |
| 0 | 0.71259041 | 0.987 | 0.888 | 0 | ERY | CD63 |
| 0 | 0.71251724 | 0.753 | 0.186 | 0 | ERY | UBE2S |
| 0 | 0.71007175 | 0.994 | 0.826 | 0 | ERY | UBB |
| 0 | 0.70890645 | 0.683 | 0.09 | 0 | ERY | FXYD2 |
| 0 | 0.70796389 | 0.913 | 0.41 | 0 | ERY | SPINT2 |
| 0 | 0.70396444 | 0.856 | 0.379 | 0 | ERY | COMMD6 |
| 0 | 0.70371672 | 0.892 | 0.242 | 0 | ERY | TSPAN1 |
| 0 | 0.70217248 | 0.664 | 0.128 | 0 | ERY | CD40 |
| 0 | 0.70139175 | 0.938 | 0.542 | 0 | ERY | NDUFS5 |
| 0 | 0.69716681 | 0.799 | 0.304 | 0 | ERY | KRT10 |
| 0 | 0.69006567 | 0.79 | 0.278 | 0 | ERY | SELENOS |
| 0 | 0.68894648 | 0.842 | 0.369 | 0 | ERY | TBCA |
| 0 | 0.68734772 | 0.827 | 0.352 | 0 | ERY | SEC11A |
| 0 | 0.68616539 | 0.771 | 0.236 | 0 | ERY | ODC1 |
| 0 | 0.67475767 | 0.85 | 0.314 | 0 | ERY | SNHG8 |
| 0 | 0.67162909 | 0.738 | 0.236 | 0 | ERY | C4orf48 |
| 0 | 0.66798616 | 0.775 | 0.291 | 0 | ERY | BUD31 |
| 0 | 0.65803479 | 0.814 | 0.317 | 0 | ERY | SSR3 |
| 0 | 0.65486666 | 0.731 | 0.23 | 0 | ERY | CD47 |
| 0 | 0.6459106 | 0.67 | 0.177 | 0 | ERY | SEC11C |
| 0 | 0.63609275 | 0.75 | 0.266 | 0 | ERY | COX7A2L |
| 0 | 0.62778424 | 0.608 | 0.115 | 0 | ERY | DMKN |
| 0 | 0.62635911 | 0.729 | 0.22 | 0 | ERY | PCBD1 |
| 0 | 0.62374643 | 0.776 | 0.287 | 0 | ERY | BLVRB |
| 0 | 0.6166731 | 0.78 | 0.295 | 0 | ERY | MZT2B |
| 0 | 0.61441372 | 0.707 | 0.221 | 0 | ERY | TSTD1 |
| 0 | 0.61238185 | 0.838 | 0.34 | 0 | ERY | TKT |
| 0 | 0.60804823 | 0.65 | 0.186 | 0 | ERY | PHB |
| 0 | 0.60777495 | 0.773 | 0.199 | 0 | ERY | TGM2 |
| 0 | 0.60415972 | 0.687 | 0.229 | 0 | ERY | PDCD5 |
| 0 | 0.5988338 | 0.718 | 0.261 | 0 | ERY | MRPL51 |
| 0 | 0.59464845 | 0.856 | 0.225 | 0 | ERY | PLAU |
| 0 | 0.59401409 | 0.687 | 0.236 | 0 | ERY | NDUFAB1 |
| 0 | 0.59055345 | 0.694 | 0.244 | 0 | ERY | PGLS |
| 0 | 0.58483804 | 0.564 | 0.087 | 0 | ERY | TRIB3 |
| 0 | 0.57798995 | 0.741 | 0.272 | 0 | ERY | NDUFC1 |
| 0 | 0.57193528 | 0.663 | 0.205 | 0 | ERY | TMEM208 |
| 0 | 0.56971933 | 0.645 | 0.135 | 0 | ERY | ITGA3 |
| 0 | 0.56967112 | 0.736 | 0.253 | 0 | ERY | PHPT1 |
| 0 | 0.56761028 | 0.66 | 0.225 | 0 | ERY | C8orf59 |
| 0 | 0.56080024 | 0.852 | 0.331 | 0 | ERY | PEBP1 |
| 0 | 0.55980513 | 0.861 | 0.304 | 0 | ERY | IFITM3 |
| 0 | 0.55765599 | 0.983 | 0.648 | 0 | ERY | MT2A |
| 0 | 0.55372452 | 0.45 | 0.009 | 0 | ERY | CCL18 |
| 0 | 0.55313751 | 0.751 | 0.107 | 0 | ERY | TFPI2 |
| 0 | 0.54896397 | 0.708 | 0.198 | 0 | ERY | CKB |
| 0 | 0.54814844 | 0.641 | 0.054 | 0 | ERY | C1QB |

| | | | | | | |
|---|---|---|---|---|---|---|
| 0 | 0.54740101 | 0.523 | 0.069 | 0 | ERY | CSRP2 |
| 0 | 0.54514089 | 0.683 | 0.201 | 0 | ERY | CDK2AP2 |
| 0 | 0.54058464 | 0.809 | 0.16 | 0 | ERY | LAMC2 |
| 0 | 0.5389233 | 0.443 | 0.015 | 0 | ERY | NNMT |
| 0 | 0.53363191 | 0.572 | 0.113 | 0 | ERY | BEX2 |
| 0 | 0.52909753 | 0.869 | 0.299 | 0 | ERY | GADD45A |
| 0 | 0.52364987 | 0.505 | 0.081 | 0 | ERY | PDLIM4 |
| 0 | 0.5208749 | 0.536 | 0.094 | 0 | ERY | ACP5 |
| 0 | 0.51701932 | 0.49 | 0.09 | 0 | ERY | AAMDC |
| 0 | 0.51372529 | 0.478 | 0.056 | 0 | ERY | GNG11 |
| 0 | 0.51117916 | 0.82 | 0.226 | 0 | ERY | CST6 |
| 0 | 0.50444108 | 0.576 | 0.151 | 0 | ERY | EPB41L4A-AS1 |
| 0 | 0.48986162 | 0.458 | 0.027 | 0 | ERY | KRTAP2-3 |
| 0 | 0.48382413 | 0.573 | 0.138 | 0 | ERY | CKS2 |
| 0 | 0.47419649 | 0.603 | 0.165 | 0 | ERY | ANXA3 |
| 0 | 0.47162771 | 0.545 | 0.13 | 0 | ERY | GPRC5C |
| 0 | 0.46946116 | 0.638 | 0.116 | 0 | ERY | UCHL1 |
| 0 | 0.46715623 | 0.448 | 0.075 | 0 | ERY | ISOC2 |
| 0 | 0.46691518 | 0.507 | 0.116 | 0 | ERY | C1orf56 |
| 0 | 0.45533485 | 0.599 | 0.059 | 0 | ERY | C1QA |
| 0 | 0.45360968 | 0.49 | 0.126 | 0 | ERY | PRDX4 |
| 0 | 0.45220459 | 0.445 | 0.083 | 0 | ERY | SDSL |
| 0 | 0.45067391 | 0.517 | 0.14 | 0 | ERY | DNAJB2 |
| 0 | 0.4488489 | 0.487 | 0.122 | 0 | ERY | MRPS33 |
| 0 | 0.44869251 | 0.415 | 0.075 | 0 | ERY | METRN |
| 0 | 0.44481667 | 0.592 | 0.098 | 0 | ERY | RRAD |
| 0 | 0.44415759 | 0.41 | 0.041 | 0 | ERY | PKIB |
| 0 | 0.42210054 | 0.506 | 0.057 | 0 | ERY | TAGLN |
| 0 | 0.41049118 | 0.457 | 0.092 | 0 | ERY | DPP4 |
| 0 | 0.40109264 | 0.425 | 0.094 | 0 | ERY | GOT1 |
| 0 | 0.39983379 | 0.788 | 0.215 | 0 | ERY | PDZK1IP1 |
| 0 | 0.39867545 | 0.406 | 0.076 | 0 | ERY | ASNS |
| 0 | 0.39320915 | 0.401 | 0.091 | 0 | ERY | PLGRKT |
| 0 | 0.39318279 | 0.45 | 0.055 | 0 | ERY | TNFRSF11B |
| 0 | 0.38234233 | 0.392 | 0.088 | 0 | ERY | DTD1 |
| 0 | 0.3810556 | 0.698 | 0.117 | 0 | ERY | NUPR1 |
| 0 | 0.37903046 | 0.42 | 0.09 | 0 | ERY | ARG2 |
| 0 | 0.37839306 | 0.348 | 0.036 | 0 | ERY | GAL |
| 0 | 0.3765863 | 0.317 | 0.022 | 0 | ERY | TSLP |
| 0 | 0.37400923 | 0.392 | 0.029 | 0 | ERY | MGP |
| 0 | 0.37138667 | 0.685 | 0.179 | 0 | ERY | SFN |
| 0 | 0.36948936 | 0.374 | 0.068 | 0 | ERY | AL121944.1 |
| 0 | 0.36705638 | 0.347 | 0.051 | 0 | ERY | EMC9 |
| 0 | 0.36442444 | 0.354 | 0.057 | 0 | ERY | NPDC1 |
| 0 | 0.34531942 | 0.327 | 0.038 | 0 | ERY | TUBB2B |
| 0 | 0.34366559 | 0.313 | 0.037 | 0 | ERY | SYCE1L |
| 0 | 0.34311993 | 0.327 | 0.044 | 0 | ERY | LURAP1L |
| 0 | 0.34201048 | 0.485 | 0.113 | 0 | ERY | MARCO |
| 0 | 0.34183828 | 0.831 | 0.262 | 0 | ERY | CLU |
| 0 | 0.34181558 | 0.38 | 0.07 | 0 | ERY | PCSK1N |

| | | | | | | |
|---|---|---|---|---|---|---|
| 0 | 0.33857206 | 0.347 | 0.06 | 0 | ERY | L1CAM |
| 0 | 0.3369234 | 0.32 | 0.057 | 0 | ERY | PITPNA-AS1 |
| 0 | 0.33502812 | 0.575 | 0.14 | 0 | ERY | CYR61 |
| 0 | 0.33401006 | 0.303 | 0.042 | 0 | ERY | RPL39L |
| 0 | 0.33269851 | 0.313 | 0.033 | 0 | ERY | PTH1R |
| 0 | 0.33234679 | 0.446 | 0.081 | 0 | ERY | CTGF |
| 0 | 0.32649493 | 0.314 | 0.045 | 0 | ERY | RBP1 |
| 0 | 0.32580611 | 0.311 | 0.029 | 0 | ERY | TAC1 |
| 0 | 0.31961127 | 0.317 | 0.056 | 0 | ERY | TMEM132A |
| 0 | 0.31643547 | 0.382 | 0.079 | 0 | ERY | DCBLD2 |
| 0 | 0.31446113 | 0.299 | 0.042 | 0 | ERY | CAVIN3 |
| 0 | 0.30990207 | 0.791 | 0.241 | 0 | ERY | TPM1 |
| 0 | 0.30853054 | 0.315 | 0.057 | 0 | ERY | MYC |
| 0 | 0.30491264 | 0.273 | 0.035 | 0 | ERY | BEX5 |
| 0 | 0.30108661 | 0.266 | 0.03 | 0 | ERY | DOK5 |
| 0 | 0.30106524 | 0.234 | 0.009 | 0 | ERY | CTSK |
| 0 | 0.30102921 | 0.285 | 0.047 | 0 | ERY | TMEM120B |
| 0 | 0.29196537 | 0.309 | 0.036 | 0 | ERY | TREM2 |
| 0 | 0.29182261 | 0.295 | 0.047 | 0 | ERY | UGT2B7 |
| 0 | 0.28952139 | 0.415 | 0.044 | 0 | ERY | C1QC |
| 0 | 0.27739835 | 0.687 | 0.205 | 0 | ERY | CCND1 |
| 0 | 0.27708243 | 0.231 | 0.015 | 0 | ERY | CCK |
| 0 | 0.27311994 | 0.271 | 0.031 | 0 | ERY | FABP3 |
| 0 | 0.27227735 | 0.701 | 0.213 | 0 | ERY | ITGA2 |
| 0 | 0.26949125 | 0.455 | 0.061 | 0 | ERY | SAA2 |
| 0 | 0.26632266 | 0.383 | 0.028 | 0 | ERY | MMP1 |
| 0 | 0.2659702 | 0.401 | 0.085 | 0 | ERY | IFIT2 |
| 0 | 0.26451393 | 0.28 | 0.036 | 0 | ERY | CLDN10 |
| 0 | 0.26232475 | 0.251 | 0.018 | 0 | ERY | LTF |
| 0 | 0.2600101 | 0.405 | 0.018 | 0 | ERY | PTGDS |
| 0 | 0.25731491 | 0.238 | 0.023 | 0 | ERY | HRK |
| 0 | 0.25708756 | 0.205 | 0.012 | 0 | ERY | AC034213.1 |
| 0 | 0.25337454 | 0.229 | 0.031 | 0 | ERY | SERTAD4-AS1 |
| 0 | 0.25333175 | 0.224 | 0.028 | 0 | ERY | DUSP15 |
| 0 | -0.3269112 | 0.701 | 0.094 | 0 | ERY | MT1M |
| 0 | -0.3310803 | 0.671 | 0.134 | 0 | ERY | CCL20 |
| 4.492E-307 | 0.61161778 | 0.8 | 0.323 | 1.061E-302 | ERY | SNU13 |
| 5.717E-307 | 0.56627544 | 0.979 | 0.683 | 1.351E-302 | ERY | HSP90B1 |
| 2.18E-303 | 0.51849249 | 0.605 | 0.195 | 5.149E-299 | ERY | CNIH4 |
| 2.753E-303 | 0.29281813 | 0.285 | 0.05 | 6.503E-299 | ERY | RGS3 |
| 1.62E-302 | 0.7315555 | 0.992 | 0.913 | 3.827E-298 | ERY | RPL32 |
| 2.025E-302 | 0.33889772 | 0.337 | 0.068 | 4.785E-298 | ERY | SHMT2 |
| 3.416E-302 | 0.68050907 | 0.992 | 0.98 | 8.069E-298 | ERY | RPL41 |
| 5.687E-302 | 0.52426406 | 0.665 | 0.229 | 1.343E-297 | ERY | OSTC |
| 6.634E-302 | 0.31919898 | 0.351 | 0.072 | 1.567E-297 | ERY | RAPGEF3 |
| 2.181E-299 | 0.27867151 | 0.328 | 0.063 | 5.152E-295 | ERY | VSIG4 |
| 4.215E-299 | 0.63934895 | 0.988 | 0.881 | 9.958E-295 | ERY | MYL12B |
| 7.565E-299 | 0.60743069 | 0.801 | 0.334 | 1.787E-294 | ERY | FKBP2 |
| 1.008E-297 | 0.56960755 | 0.738 | 0.285 | 2.381E-293 | ERY | SF3B5 |
| 8.587E-296 | 0.73658598 | 0.988 | 0.794 | 2.029E-291 | ERY | RPS5 |

| | | | | | | |
|---|---|---|---|---|---|---|
| 1.073E-295 | 0.55827503 | 0.763 | 0.294 | 2.534E-291 | ERY | RAB13 |
| 1.184E-295 | 0.50487389 | 0.584 | 0.186 | 2.796E-291 | ERY | TIMM13 |
| 1.39E-295 | 0.50454438 | 0.755 | 0.278 | 3.284E-291 | ERY | CHCHD10 |
| 3.553E-294 | 0.28833194 | 0.297 | 0.054 | 8.394E-290 | ERY | LIPG |
| 3.729E-294 | 0.35350799 | 0.453 | 0.115 | 8.809E-290 | ERY | PLA2G16 |
| 8.157E-294 | 0.55632211 | 0.748 | 0.288 | 1.927E-289 | ERY | TMED9 |
| 1.518E-292 | 0.64539155 | 0.873 | 0.434 | 3.587E-288 | ERY | ATP5MD |
| 1.019E-291 | 0.26193466 | 0.264 | 0.044 | 2.407E-287 | ERY | MAP2 |
| 7.053E-291 | 0.65548956 | 0.98 | 0.807 | 1.666E-286 | ERY | RPL36AL |
| 1.433E-289 | 0.46066601 | 0.619 | 0.202 | 3.385E-285 | ERY | MRPL18 |
| 1.552E-289 | 0.4153017 | 0.533 | 0.153 | 3.666E-285 | ERY | CFD |
| 2.025E-289 | 0.60765111 | 0.866 | 0.387 | 4.784E-285 | ERY | SOD1 |
| 3.295E-289 | 0.707743 | 0.982 | 0.738 | 7.783E-285 | ERY | EEF1B2 |
| 1.129E-288 | 0.65135432 | 0.967 | 0.678 | 2.666E-284 | ERY | BRI3 |
| 5.417E-288 | 0.3014264 | 0.297 | 0.056 | 1.28E-283 | ERY | TSPAN4 |
| 2.734E-287 | 0.37336509 | 0.487 | 0.13 | 6.458E-283 | ERY | LAPTM4B |
| 9.988E-287 | 0.45844451 | 0.841 | 0.326 | 2.359E-282 | ERY | DDIT4 |
| 5.366E-286 | 0.67936279 | 0.988 | 0.893 | 1.268E-281 | ERY | RPS9 |
| 1.59E-285 | 0.55926167 | 0.934 | 0.51 | 3.757E-281 | ERY | HSPE1 |
| 3.985E-285 | 0.61205475 | 0.941 | 0.556 | 9.415E-281 | ERY | RPS27L |
| 7.285E-282 | 0.57163926 | 0.759 | 0.305 | 1.721E-277 | ERY | ERP29 |
| 1.057E-281 | 0.34117219 | 0.551 | 0.159 | 2.496E-277 | ERY | STMN1 |
| 1.44E-280 | 0.53242892 | 0.664 | 0.243 | 3.401E-276 | ERY | RPS19BP1 |
| 1.569E-280 | 0.61573735 | 0.938 | 0.563 | 3.706E-276 | ERY | PRDX5 |
| 2.156E-280 | 0.33043403 | 0.35 | 0.077 | 5.093E-276 | ERY | SWI5 |
| 2.221E-280 | 0.64319524 | 0.903 | 0.494 | 5.246E-276 | ERY | DBI |
| 2.873E-280 | 0.39634619 | 0.428 | 0.11 | 6.788E-276 | ERY | NBL1 |
| 6.513E-280 | 0.25752991 | 0.269 | 0.047 | 1.539E-275 | ERY | CAV1 |
| 1.949E-279 | 0.67812184 | 0.992 | 0.917 | 4.605E-275 | ERY | RPS15 |
| 4.929E-279 | 0.64006499 | 0.972 | 0.699 | 1.164E-274 | ERY | GPX4 |
| 8.538E-279 | 0.46530101 | 0.553 | 0.173 | 2.017E-274 | ERY | MZT2A |
| 3.156E-278 | 0.45431221 | 0.508 | 0.15 | 7.455E-274 | ERY | TMEM141 |
| 1.083E-277 | 0.62440332 | 0.917 | 0.52 | 2.558E-273 | ERY | P4HB |
| 1.22E-277 | 0.7117326 | 0.988 | 0.812 | 2.882E-273 | ERY | RPL5 |
| 1.94E-277 | 0.29252103 | 0.283 | 0.053 | 4.584E-273 | ERY | FAIM |
| 2.821E-277 | 0.30610366 | 0.343 | 0.074 | 6.664E-273 | ERY | CYP27A1 |
| 1.163E-276 | 0.57081741 | 0.945 | 0.48 | 2.747E-272 | ERY | CD9 |
| 2.497E-274 | 0.61755205 | 0.966 | 0.756 | 5.898E-270 | ERY | UBL5 |
| 3.182E-274 | 0.35399757 | 0.392 | 0.095 | 7.517E-270 | ERY | MRPS26 |
| 1.55E-273 | 0.29728437 | 0.292 | 0.057 | 3.66E-269 | ERY | DTYMK |
| 1.123E-272 | 0.51198644 | 0.778 | 0.31 | 2.653E-268 | ERY | PPA1 |
| 1.357E-272 | 0.48362353 | 0.578 | 0.192 | 3.205E-268 | ERY | SLIRP |
| 2.388E-272 | 0.36062485 | 0.376 | 0.09 | 5.641E-268 | ERY | MTX2 |
| 5.652E-272 | 0.67563644 | 0.99 | 0.922 | 1.335E-267 | ERY | RPS24 |
| 3.271E-271 | 0.64766788 | 0.992 | 0.893 | 7.726E-267 | ERY | RPL35A |
| 1.121E-270 | 0.5162131 | 0.695 | 0.265 | 2.647E-266 | ERY | MRPL41 |
| 2.944E-270 | 0.52573045 | 0.673 | 0.256 | 6.955E-266 | ERY | LMAN2 |
| 3.328E-270 | 0.44590276 | 0.676 | 0.239 | 7.861E-266 | ERY | C1orf122 |
| 4.445E-270 | 0.57948638 | 0.809 | 0.356 | 1.05E-265 | ERY | RHEB |
| 1.296E-269 | 0.64812196 | 0.987 | 0.892 | 3.061E-265 | ERY | RPL37A |

| | | | | | | |
|---|---|---|---|---|---|---|
| 4.472E-269 | 0.25024543 | 0.224 | 0.035 | 1.056E-264 | ERY | TMEM125 |
| 1.075E-268 | 0.6587482 | 0.987 | 0.916 | 2.541E-264 | ERY | RPL9 |
| 9.046E-268 | 0.47158576 | 0.687 | 0.246 | 2.137E-263 | ERY | PRDX2 |
| 1.602E-267 | 0.58064855 | 0.93 | 0.435 | 3.784E-263 | ERY | CLDN7 |
| 4.234E-267 | 0.2701529 | 0.318 | 0.065 | 1E-262 | ERY | KITLG |
| 1.718E-266 | 0.60578104 | 0.943 | 0.669 | 4.059E-262 | ERY | COX7A2 |
| 1.83E-265 | 0.3290827 | 0.334 | 0.074 | 4.323E-261 | ERY | CHID1 |
| 2.054E-265 | 0.26960357 | 0.267 | 0.049 | 4.852E-261 | ERY | PKIG |
| 7.736E-265 | 0.51491931 | 0.675 | 0.253 | 1.827E-260 | ERY | GADD45GIP1 |
| 8.964E-265 | 0.34493345 | 0.461 | 0.123 | 2.118E-260 | ERY | COL18A1 |
| 2.767E-264 | 0.66026665 | 0.99 | 0.925 | 6.536E-260 | ERY | RPL19 |
| 1.395E-261 | 0.34861699 | 0.373 | 0.091 | 3.296E-257 | ERY | MCRIP2 |
| 2.854E-261 | 0.6804513 | 0.985 | 0.775 | 6.742E-257 | ERY | RPL10A |
| 1.905E-260 | 0.44183393 | 0.545 | 0.176 | 4.501E-256 | ERY | MFSD10 |
| 2.005E-260 | 0.42653132 | 0.495 | 0.149 | 4.737E-256 | ERY | RAB32 |
| 7.174E-260 | 0.39164989 | 0.427 | 0.116 | 1.695E-255 | ERY | AP1S1 |
| 1.124E-259 | 0.41742101 | 0.486 | 0.145 | 2.655E-255 | ERY | HAX1 |
| 4.734E-259 | 0.40540914 | 0.497 | 0.148 | 1.118E-254 | ERY | DUT |
| 7.168E-259 | 0.47550157 | 0.678 | 0.252 | 1.693E-254 | ERY | ATOX1 |
| 7.741E-259 | 0.57725169 | 0.822 | 0.383 | 1.829E-254 | ERY | NDUFA1 |
| 3.497E-258 | -0.8767521 | 0.981 | 0.666 | 8.261E-254 | ERY | TIMP1 |
| 1.182E-257 | 0.57316983 | 0.855 | 0.405 | 2.792E-253 | ERY | RAN |
| 3.456E-257 | 0.63040853 | 0.987 | 0.865 | 8.164E-253 | ERY | RPLP0 |
| 4.24E-257 | 0.37521295 | 0.42 | 0.113 | 1.002E-252 | ERY | MRPL40 |
| 5.145E-257 | 0.29336166 | 0.29 | 0.059 | 1.215E-252 | ERY | CD320 |
| 1.014E-255 | 0.6260749 | 0.973 | 0.739 | 2.396E-251 | ERY | CLIC1 |
| 1.166E-254 | 0.52925484 | 0.887 | 0.417 | 2.755E-250 | ERY | GSTO1 |
| 5.799E-253 | 0.57004799 | 0.805 | 0.372 | 1.37E-248 | ERY | DAD1 |
| 6.491E-253 | 0.57953373 | 0.845 | 0.42 | 1.533E-248 | ERY | ATP5MF |
| 1.007E-252 | 0.61683132 | 0.99 | 0.922 | 2.378E-248 | ERY | RPL18A |
| 1.57E-252 | 0.28357816 | 0.619 | 0.199 | 3.71E-248 | ERY | LAMB3 |
| 2.104E-250 | 0.49712091 | 0.692 | 0.275 | 4.97E-246 | ERY | NDUFB3 |
| 2.124E-250 | 0.5952098 | 0.917 | 0.571 | 5.018E-246 | ERY | CLTA |
| 3.156E-250 | 0.48269716 | 0.619 | 0.224 | 7.456E-246 | ERY | SNRPE |
| 1.285E-249 | 0.29334093 | 0.324 | 0.072 | 3.036E-245 | ERY | PHGDH |
| 1.65E-249 | -1.1863237 | 0.313 | 0.064 | 3.898E-245 | ERY | CXCL5 |
| 1.894E-249 | 0.49220709 | 0.899 | 0.471 | 4.474E-245 | ERY | HMGA1 |
| 5.037E-249 | 0.29572126 | 0.33 | 0.074 | 1.19E-244 | ERY | KHDRBS3 |
| 9.346E-249 | 0.51917338 | 0.962 | 0.525 | 2.208E-244 | ERY | CD24 |
| 1.483E-248 | 0.57290401 | 0.907 | 0.539 | 3.504E-244 | ERY | UQCR10 |
| 2.1E-248 | 0.55697248 | 0.81 | 0.376 | 4.961E-244 | ERY | ATP5MPL |
| 6.096E-247 | 0.30292643 | 0.335 | 0.077 | 1.44E-242 | ERY | TLE1 |
| 8.527E-247 | 0.39788245 | 0.475 | 0.141 | 2.014E-242 | ERY | EIF4EBP1 |
| 8.586E-247 | 0.38567297 | 0.436 | 0.123 | 2.028E-242 | ERY | TMEM50B |
| 1.261E-246 | 0.28225697 | 0.737 | 0.259 | 2.979E-242 | ERY | CAMK2N1 |
| 4.991E-246 | 0.57948709 | 0.963 | 0.679 | 1.179E-241 | ERY | PSMA7 |
| 7.901E-246 | 0.62718384 | 0.989 | 0.878 | 1.867E-241 | ERY | RPL12 |
| 1.111E-245 | 0.42043926 | 0.623 | 0.219 | 2.623E-241 | ERY | BST2 |
| 2.846E-245 | 0.5701329 | 0.904 | 0.519 | 6.723E-241 | ERY | COX6C |
| 4.179E-245 | 0.50365582 | 0.77 | 0.328 | 9.872E-241 | ERY | ATP5IF1 |

| | | | | | | |
|---|---|---|---|---|---|---|
| 6.519E-245 | 0.46684817 | 0.587 | 0.209 | 1.54E-240 | ERY | LSM4 |
| 1.148E-244 | 0.35166939 | 0.392 | 0.103 | 2.712E-240 | ERY | CISD1 |
| 2.957E-244 | 0.38284978 | 0.423 | 0.119 | 6.986E-240 | ERY | MRPS12 |
| 3.42E-243 | 0.5118471 | 0.717 | 0.294 | 8.078E-239 | ERY | SSBP1 |
| 5.553E-243 | 0.54947575 | 0.926 | 0.534 | 1.312E-238 | ERY | RPS10 |
| 2.923E-242 | 0.38904271 | 0.448 | 0.13 | 6.904E-238 | ERY | ARL3 |
| 3.242E-242 | 0.27444207 | 0.292 | 0.062 | 7.658E-238 | ERY | OAF |
| 4.528E-242 | -0.5560235 | 0.949 | 0.447 | 1.07E-237 | ERY | LYZ |
| 8.543E-242 | 0.63135768 | 0.993 | 0.931 | 2.018E-237 | ERY | RPS14 |
| 1.66E-241 | 0.53289033 | 0.989 | 0.942 | 3.922E-237 | ERY | RPS27 |
| 2.268E-241 | 0.4293745 | 0.71 | 0.274 | 5.358E-237 | ERY | MIR4435-2HG |
| 5.048E-241 | 0.43618589 | 0.59 | 0.208 | 1.192E-236 | ERY | ECHS1 |
| 9.209E-241 | 0.5930233 | 0.987 | 0.884 | 2.175E-236 | ERY | RPS3 |
| 9.779E-241 | 0.40852429 | 0.506 | 0.161 | 2.31E-236 | ERY | MPC2 |
| 1.164E-240 | 0.36063775 | 0.423 | 0.118 | 2.75E-236 | ERY | NDUFAF2 |
| 1.536E-239 | 0.28526644 | 0.292 | 0.063 | 3.629E-235 | ERY | PRR34-AS1 |
| 6.937E-239 | 0.37285713 | 0.465 | 0.139 | 1.639E-234 | ERY | LGMN |
| 8.214E-239 | 0.31055437 | 0.338 | 0.081 | 1.94E-234 | ERY | PAFAH1B3 |
| 1.568E-238 | 0.58649053 | 0.989 | 0.896 | 3.705E-234 | ERY | RPL39 |
| 2.167E-238 | 0.49303436 | 0.687 | 0.278 | 5.119E-234 | ERY | NDUFS6 |
| 2.706E-238 | 0.48414002 | 0.662 | 0.259 | 6.392E-234 | ERY | MINOS1 |
| 4.053E-238 | 0.43066229 | 0.973 | 0.53 | 9.575E-234 | ERY | KRT7 |
| 7.941E-238 | 0.29455107 | 0.318 | 0.073 | 1.876E-233 | ERY | SMOX |
| 6.848E-237 | 0.40101588 | 0.49 | 0.155 | 1.618E-232 | ERY | MCRIP1 |
| 3.177E-236 | 0.26750611 | 0.272 | 0.056 | 7.505E-232 | ERY | KIF9 |
| 6.987E-236 | 0.26087361 | 0.262 | 0.052 | 1.651E-231 | ERY | TEX30 |
| 5.575E-235 | 0.29935655 | 0.313 | 0.072 | 1.317E-230 | ERY | PMM1 |
| 6.147E-235 | 0.52036971 | 0.771 | 0.349 | 1.452E-230 | ERY | PSMB1 |
| 1.564E-234 | 0.26304135 | 0.326 | 0.075 | 3.695E-230 | ERY | QPRT |
| 2.63E-234 | 0.34343593 | 0.393 | 0.107 | 6.212E-230 | ERY | MRPS23 |
| 8.1E-234 | 0.37800267 | 0.424 | 0.123 | 1.913E-229 | ERY | CMC1 |
| 1.451E-233 | 0.31389676 | 0.349 | 0.086 | 3.428E-229 | ERY | PON2 |
| 2.547E-233 | 0.52091944 | 0.764 | 0.342 | 6.016E-229 | ERY | TRMT112 |
| 2.822E-233 | 0.27259554 | 0.285 | 0.061 | 6.666E-229 | ERY | C19orf48 |
| 2.856E-233 | 0.46532076 | 0.631 | 0.239 | 6.746E-229 | ERY | FIS1 |
| 8.176E-233 | 0.31536937 | 0.39 | 0.102 | 1.931E-228 | ERY | ADGRG1 |
| 1.641E-232 | 0.48161104 | 0.764 | 0.328 | 3.876E-228 | ERY | BSG |
| 1.493E-231 | 0.33819575 | 0.508 | 0.156 | 3.527E-227 | ERY | TINAGL1 |
| 1.655E-231 | 0.42435945 | 0.745 | 0.297 | 3.909E-227 | ERY | SDC4 |
| 9.684E-231 | 0.42081999 | 0.542 | 0.185 | 2.288E-226 | ERY | ATP5MC1 |
| 1.753E-230 | 0.54971497 | 0.924 | 0.6 | 4.14E-226 | ERY | SEC61B |
| 8.037E-230 | 0.40109881 | 0.463 | 0.144 | 1.899E-225 | ERY | CHCHD5 |
| 1.578E-229 | 0.25340021 | 0.346 | 0.083 | 3.729E-225 | ERY | CA12 |
| 2.612E-229 | 0.35450431 | 0.402 | 0.113 | 6.17E-225 | ERY | EVA1B |
| 9.523E-229 | 0.46983557 | 0.601 | 0.228 | 2.25E-224 | ERY | SMIM26 |
| 2.456E-227 | 0.41416231 | 0.538 | 0.184 | 5.801E-223 | ERY | C1QBP |
| 3.662E-227 | -2.0228719 | 0.989 | 0.939 | 8.65E-223 | ERY | NEAT1 |
| 2.099E-226 | 0.25423428 | 0.271 | 0.056 | 4.959E-222 | ERY | SCD5 |
| 6.641E-226 | 0.28212595 | 0.294 | 0.066 | 1.569E-221 | ERY | ZNF667-AS1 |
| 7.611E-226 | 0.52389023 | 0.988 | 0.942 | 1.798E-221 | ERY | MT-CO3 |

| | | | | | | |
|---|---|---|---|---|---|---|
| 1.199E-225 | 0.36582856 | 0.417 | 0.122 | 2.832E-221 | ERY | METTL5 |
| 5.005E-225 | 0.56449571 | 0.998 | 0.895 | 1.182E-220 | ERY | RPL21 |
| 1.593E-224 | 0.32028192 | 0.348 | 0.09 | 3.763E-220 | ERY | B3GAT3 |
| 1.971E-224 | 0.52343925 | 0.986 | 0.912 | 4.655E-220 | ERY | MT-ND1 |
| 3.892E-224 | 0.35703849 | 0.4 | 0.114 | 9.195E-220 | ERY | ISCA1 |
| 5.059E-224 | 0.29777162 | 0.47 | 0.14 | 1.195E-219 | ERY | MDK |
| 5.694E-224 | 0.48416149 | 0.801 | 0.361 | 1.345E-219 | ERY | ATP5ME |
| 8.635E-223 | 0.45405039 | 0.75 | 0.311 | 2.04E-218 | ERY | CAPG |
| 1.583E-222 | 0.37663336 | 0.462 | 0.145 | 3.74E-218 | ERY | POLR2H |
| 3.043E-222 | 0.5959684 | 0.989 | 0.935 | 7.189E-218 | ERY | MT-ATP6 |
| 3.899E-222 | 0.34009375 | 0.372 | 0.102 | 9.212E-218 | ERY | MRPL22 |
| 1.091E-221 | 0.27592525 | 0.31 | 0.073 | 2.578E-217 | ERY | LMTK3 |
| 1.332E-221 | 0.37283414 | 0.408 | 0.12 | 3.146E-217 | ERY | RAB34 |
| 5.625E-221 | 0.39490776 | 0.501 | 0.166 | 1.329E-216 | ERY | ARL1 |
| 2.676E-220 | 0.39611747 | 0.506 | 0.169 | 6.321E-216 | ERY | RBCK1 |
| 3.5E-220 | 0.43745941 | 0.599 | 0.226 | 8.268E-216 | ERY | TUFM |
| 1.043E-219 | 0.57138508 | 0.99 | 0.895 | 2.463E-215 | ERY | RPL29 |
| 8.53E-219 | -0.8555197 | 0.217 | 0.636 | 2.015E-214 | ERY | PRRC2C |
| 1.206E-218 | 0.38007835 | 0.495 | 0.163 | 2.85E-214 | ERY | CSRP1 |
| 9.925E-218 | 0.3198873 | 0.385 | 0.107 | 2.345E-213 | ERY | BOLA3 |
| 1.082E-217 | 0.32830203 | 0.378 | 0.104 | 2.556E-213 | ERY | TP53I3 |
| 2.43E-217 | 0.41374073 | 0.538 | 0.19 | 5.739E-213 | ERY | C19orf70 |
| 5.25E-217 | 0.44176928 | 0.636 | 0.247 | 1.24E-212 | ERY | CUTA |
| 1.784E-216 | 0.45646825 | 0.635 | 0.255 | 4.215E-212 | ERY | NDUFA12 |
| 2.076E-216 | 0.51755896 | 0.99 | 0.932 | 4.904E-212 | ERY | RPL34 |
| 2.199E-216 | 0.55637543 | 0.985 | 0.844 | 5.195E-212 | ERY | RPL38 |
| 5.422E-216 | 0.49254989 | 0.957 | 0.717 | 1.281E-211 | ERY | COX6A1 |
| 7.842E-216 | 0.35657287 | 0.429 | 0.13 | 1.852E-211 | ERY | PLD3 |
| 1.842E-215 | 0.54411253 | 0.991 | 0.929 | 4.352E-211 | ERY | FAU |
| 5.324E-215 | 0.43194761 | 0.569 | 0.213 | 1.258E-210 | ERY | MAP2K2 |
| 8.401E-215 | 0.53607592 | 0.983 | 0.747 | 1.985E-210 | ERY | RPSA |
| 7.016E-214 | 0.31113317 | 0.319 | 0.08 | 1.657E-209 | ERY | SRPRB |
| 1.898E-213 | 0.38215495 | 0.479 | 0.159 | 4.484E-209 | ERY | MRPS15 |
| 2.892E-213 | 0.32385699 | 0.453 | 0.138 | 6.833E-209 | ERY | SCCPDH |
| 1.145E-212 | 0.34289348 | 0.392 | 0.113 | 2.705E-208 | ERY | UCHL3 |
| 1.201E-212 | 0.2558232 | 0.272 | 0.06 | 2.836E-208 | ERY | CHPF |
| 1.672E-212 | 0.5747062 | 0.986 | 0.853 | 3.949E-208 | ERY | RPL7 |
| 2.021E-212 | 0.52133093 | 0.992 | 0.925 | 4.773E-208 | ERY | RPL30 |
| 2.142E-212 | 0.47193324 | 0.766 | 0.338 | 5.061E-208 | ERY | EIF3E |
| 2.507E-212 | 0.29640809 | 0.642 | 0.232 | 5.922E-208 | ERY | PTMS |
| 2.815E-212 | -0.7300031 | 0.463 | 0.132 | 6.649E-208 | ERY | CCL2 |
| 3.24E-211 | 0.5326256 | 0.991 | 0.917 | 7.653E-207 | ERY | NACA |
| 5.134E-211 | 0.36078316 | 0.974 | 0.604 | 1.213E-206 | ERY | LGALS1 |
| 9.51E-211 | 0.52136643 | 0.918 | 0.549 | 2.247E-206 | ERY | TOMM7 |
| 1.089E-210 | 0.47394741 | 0.787 | 0.366 | 2.573E-206 | ERY | EIF5A |
| 1.777E-210 | 0.43115926 | 0.659 | 0.268 | 4.197E-206 | ERY | LRPAP1 |
| 2.04E-210 | 0.43477183 | 0.585 | 0.225 | 4.819E-206 | ERY | PSMB7 |
| 7.98E-210 | 0.26480694 | 0.301 | 0.072 | 1.885E-205 | ERY | FKBP11 |
| 2.159E-209 | 0.34348902 | 0.457 | 0.143 | 5.099E-205 | ERY | RCN1 |
| 3.293E-209 | 0.52939862 | 0.99 | 0.901 | 7.779E-205 | ERY | RPL37 |

| | | | | | | |
|---|---|---|---|---|---|---|
| 5.606E-209 | 0.48648403 | 0.753 | 0.35 | 1.324E-204 | ERY | C19orf53 |
| 7.59E-209 | 0.48045937 | 0.769 | 0.36 | 1.793E-204 | ERY | LAMTOR5 |
| 3.812E-208 | 0.41444246 | 0.988 | 0.906 | 9.004E-204 | ERY | MT-ND2 |
| 5.174E-208 | 0.41260641 | 0.513 | 0.182 | 1.222E-203 | ERY | MRPL54 |
| 2.018E-207 | 0.42943206 | 0.841 | 0.402 | 4.767E-203 | ERY | JPT1 |
| 2.128E-207 | 0.52951105 | 0.992 | 0.98 | 5.028E-203 | ERY | EEF1A1 |
| 3.333E-207 | 0.27839382 | 0.29 | 0.069 | 7.873E-203 | ERY | MICA |
| 6.135E-207 | 0.30127223 | 0.342 | 0.09 | 1.449E-202 | ERY | SLC25A4 |
| 1.021E-206 | 0.4809005 | 0.98 | 0.86 | 2.412E-202 | ERY | SRP14 |
| 2.628E-206 | 0.38069135 | 0.467 | 0.155 | 6.209E-202 | ERY | NDUFAF8 |
| 5.267E-205 | 0.26705754 | 0.327 | 0.083 | 1.244E-200 | ERY | SEMA3B |
| 6.387E-205 | 0.54637493 | 0.987 | 0.873 | 1.509E-200 | ERY | RPL23A |
| 7.513E-205 | 0.51256629 | 0.989 | 0.884 | 1.775E-200 | ERY | RPS6 |
| 4.28E-204 | 0.2752207 | 0.33 | 0.085 | 1.011E-199 | ERY | CAV2 |
| 4.328E-204 | 0.52129437 | 0.99 | 0.977 | 1.022E-199 | ERY | RPL28 |
| 1.23E-202 | 0.38113768 | 0.497 | 0.172 | 2.906E-198 | ERY | POLE4 |
| 1.804E-202 | 0.51589351 | 0.979 | 0.831 | 4.261E-198 | ERY | COX4I1 |
| 3.362E-202 | 0.30009875 | 0.327 | 0.086 | 7.943E-198 | ERY | AKR7A2 |
| 3.616E-202 | 0.29086763 | 0.331 | 0.087 | 8.543E-198 | ERY | AP1M2 |
| 3.729E-202 | 0.33473757 | 0.383 | 0.112 | 8.809E-198 | ERY | SMIM37 |
| 5.138E-202 | 0.45200781 | 0.766 | 0.349 | 1.214E-197 | ERY | SLC25A5 |
| 2.154E-200 | -0.2500076 | 0.556 | 0.18 | 5.089E-196 | ERY | HSPA6 |
| 2.73E-200 | 0.49536526 | 0.967 | 0.764 | 6.45E-196 | ERY | TMA7 |
| 4.179E-200 | 0.34586366 | 0.417 | 0.13 | 9.871E-196 | ERY | ZNF593 |
| 9.579E-200 | 0.25932327 | 0.272 | 0.063 | 2.263E-195 | ERY | ACYP1 |
| 1.245E-199 | 0.25113634 | 0.608 | 0.215 | 2.942E-195 | ERY | CYB5A |
| 1.523E-199 | 0.28772916 | 0.317 | 0.083 | 3.599E-195 | ERY | GLRX2 |
| 2.776E-199 | 0.50143341 | 0.95 | 0.706 | 6.558E-195 | ERY | ATP5MG |
| 8.439E-199 | 0.31102365 | 0.348 | 0.096 | 1.994E-194 | ERY | GCSH |
| 3.286E-198 | 0.40533083 | 0.569 | 0.218 | 7.762E-194 | ERY | DNAJC15 |
| 4.262E-198 | 0.48687859 | 0.838 | 0.445 | 1.007E-193 | ERY | EIF2S2 |
| 8.468E-198 | 0.45956547 | 0.703 | 0.316 | 2E-193 | ERY | AURKAIP1 |
| 2.357E-197 | 0.37159728 | 0.693 | 0.279 | 5.567E-193 | ERY | TUBA1C |
| 6.885E-196 | -0.8889286 | 0.385 | 0.714 | 1.626E-191 | ERY | TPM3 |
| 1.658E-195 | 0.3933766 | 0.531 | 0.196 | 3.917E-191 | ERY | COX14 |
| 2.247E-195 | -1.2882024 | 0.14 | 0.541 | 5.308E-191 | ERY | ZFP36 |
| 3.996E-195 | 0.48892685 | 0.897 | 0.53 | 9.44E-191 | ERY | POMP |
| 5.691E-195 | 0.3725336 | 0.506 | 0.18 | 1.344E-190 | ERY | AHSA1 |
| 1.658E-194 | 0.47975418 | 0.942 | 0.677 | 3.916E-190 | ERY | ATP6V0B |
| 2.338E-194 | 0.38623664 | 0.484 | 0.171 | 5.524E-190 | ERY | CCT7 |
| 3.07E-194 | 0.32776593 | 0.398 | 0.122 | 7.253E-190 | ERY | ATG101 |
| 3.287E-194 | 0.44846256 | 0.727 | 0.334 | 7.764E-190 | ERY | PSMD8 |
| 3.351E-194 | 0.4502165 | 0.986 | 0.876 | 7.917E-190 | ERY | RPS29 |
| 1.652E-193 | 0.43380415 | 0.623 | 0.259 | 3.904E-189 | ERY | NDUFB8 |
| 1.249E-192 | 0.3175573 | 0.373 | 0.111 | 2.951E-188 | ERY | MPG |
| 1.357E-191 | 0.51136677 | 0.986 | 0.855 | 3.205E-187 | ERY | RPL23 |
| 3.616E-191 | 0.28556022 | 0.314 | 0.083 | 8.543E-187 | ERY | SLC52A2 |
| 2.424E-190 | 0.46623125 | 0.959 | 0.752 | 5.727E-186 | ERY | UQCRB |
| 4.232E-190 | 0.45010313 | 0.773 | 0.366 | 9.998E-186 | ERY | PRELID1 |
| 3.098E-189 | 0.35772741 | 0.49 | 0.174 | 7.319E-185 | ERY | FDPS |

| | | | | | | |
|---|---|---|---|---|---|---|
| 6.03E-189 | 0.40047054 | 0.879 | 0.506 | 1.425E-184 | ERY | H2AFZ |
| 1.257E-188 | 0.31849408 | 0.357 | 0.105 | 2.97E-184 | ERY | NUDCD2 |
| 2.854E-188 | 0.42559751 | 0.629 | 0.263 | 6.743E-184 | ERY | RSL24D1 |
| 1.488E-187 | 0.44658118 | 0.801 | 0.396 | 3.514E-183 | ERY | ATP5PF |
| 3.476E-187 | 0.2777222 | 0.299 | 0.078 | 8.211E-183 | ERY | SURF2 |
| 5.574E-187 | 0.46483945 | 0.936 | 0.639 | 1.317E-182 | ERY | UQCRH |
| 9.736E-187 | 0.32184979 | 0.375 | 0.115 | 2.3E-182 | ERY | ZNHIT3 |
| 9.899E-187 | 0.35253777 | 0.427 | 0.143 | 2.338E-182 | ERY | DDOST |
| 1.47E-186 | 0.25616406 | 0.277 | 0.068 | 3.472E-182 | ERY | CKS1B |
| 2.762E-186 | 0.39776784 | 0.643 | 0.272 | 6.524E-182 | ERY | NDUFB1 |
| 3.368E-186 | 0.3409568 | 0.483 | 0.167 | 7.956E-182 | ERY | LSR |
| 4.751E-186 | 0.30117001 | 0.352 | 0.102 | 1.122E-181 | ERY | MID1IP1 |
| 7.133E-186 | 0.28071078 | 0.317 | 0.086 | 1.685E-181 | ERY | TOMM5 |
| 4.002E-185 | 0.40217908 | 0.582 | 0.234 | 9.453E-181 | ERY | CCT8 |
| 6.182E-185 | 0.38421235 | 0.527 | 0.2 | 1.46E-180 | ERY | STRAP |
| 2.087E-184 | 0.41474015 | 0.651 | 0.281 | 4.93E-180 | ERY | COX17 |
| 2.351E-184 | 0.38139384 | 0.507 | 0.189 | 5.554E-180 | ERY | NAA20 |
| 3.024E-184 | -0.8603137 | 0.271 | 0.642 | 7.144E-180 | ERY | TAX1BP1 |
| 3.349E-184 | 0.27321051 | 0.31 | 0.083 | 7.911E-180 | ERY | TRIAP1 |
| 1.51E-183 | 0.3252397 | 0.412 | 0.134 | 3.567E-179 | ERY | MRPL27 |
| 3.481E-183 | 0.37334788 | 0.501 | 0.185 | 8.223E-179 | ERY | SNRPD1 |
| 5.714E-183 | 0.41278311 | 0.651 | 0.282 | 1.35E-178 | ERY | EIF3G |
| 1.619E-182 | -1.6710617 | 0.403 | 0.691 | 3.825E-178 | ERY | NAMPT |
| 5.197E-182 | 0.27547811 | 0.303 | 0.081 | 1.228E-177 | ERY | PDRG1 |
| 1.164E-181 | 0.40027147 | 0.599 | 0.248 | 2.751E-177 | ERY | SMDT1 |
| 1.234E-181 | 0.32198519 | 0.375 | 0.116 | 2.914E-177 | ERY | RPL26L1 |
| 1.289E-181 | 0.26854844 | 0.306 | 0.082 | 3.045E-177 | ERY | DCTPP1 |
| 2.176E-181 | 0.35734832 | 0.979 | 0.554 | 5.14E-177 | ERY | TACSTD2 |
| 3.18E-181 | 0.50361993 | 0.998 | 0.914 | 7.511E-177 | ERY | UBA52 |
| 4.003E-181 | 0.49062941 | 0.991 | 0.901 | 9.457E-177 | ERY | RPLP2 |
| 5.019E-181 | 0.41878124 | 0.657 | 0.29 | 1.186E-176 | ERY | ANAPC11 |
| 6.117E-181 | 0.39763473 | 0.653 | 0.276 | 1.445E-176 | ERY | APRT |
| 1.082E-180 | 0.3622145 | 0.448 | 0.157 | 2.556E-176 | ERY | GTF2H5 |
| 1.367E-180 | 0.33667696 | 0.429 | 0.145 | 3.228E-176 | ERY | NDUFS4 |
| 2.219E-180 | 0.31035561 | 0.987 | 0.863 | 5.241E-176 | ERY | ANXA2 |
| 2.387E-180 | 0.2853171 | 0.306 | 0.083 | 5.638E-176 | ERY | CCDC167 |
| 2.872E-180 | 0.30929099 | 0.348 | 0.103 | 6.784E-176 | ERY | SMIM20 |
| 4.895E-180 | 0.25904384 | 0.277 | 0.07 | 1.156E-175 | ERY | AARS |
| 5.794E-180 | 0.34402161 | 0.433 | 0.147 | 1.369E-175 | ERY | ITGB1BP1 |
| 1.069E-179 | 0.32517083 | 0.383 | 0.121 | 2.525E-175 | ERY | APOPT1 |
| 1.682E-179 | 0.28232144 | 0.327 | 0.092 | 3.974E-175 | ERY | EIF1AY |
| 2.199E-179 | 0.36620705 | 0.552 | 0.215 | 5.194E-175 | ERY | MRPL52 |
| 1.24E-178 | 0.41977053 | 0.911 | 0.534 | 2.93E-174 | ERY | PPIB |
| 1.544E-178 | -0.844721 | 0.266 | 0.625 | 3.648E-174 | ERY | SRRM2 |
| 2.489E-178 | 0.40660892 | 0.641 | 0.276 | 5.879E-174 | ERY | NDUFB7 |
| 4.697E-178 | 0.28875243 | 0.345 | 0.102 | 1.109E-173 | ERY | MRPL11 |
| 8.555E-178 | 0.26263166 | 0.297 | 0.079 | 2.021E-173 | ERY | LCMT1 |
| 1.163E-177 | 0.29606439 | 0.35 | 0.105 | 2.747E-173 | ERY | TIMM10 |
| 1.568E-177 | 0.25235341 | 0.775 | 0.335 | 3.705E-173 | ERY | GLS |
| 5.922E-177 | 0.37340475 | 0.457 | 0.165 | 1.399E-172 | ERY | IFI27L2 |

| | | | | | | |
|---|---|---|---|---|---|---|
| 7.085E-177 | 0.40729644 | 0.661 | 0.295 | 1.674E-172 | ERY | ANAPC16 |
| 7.296E-177 | 0.31130714 | 0.36 | 0.111 | 1.724E-172 | ERY | LSM2 |
| 9.07E-177 | 0.34627139 | 0.678 | 0.287 | 2.143E-172 | ERY | SERTAD1 |
| 1.889E-176 | 0.38230735 | 0.575 | 0.231 | 4.461E-172 | ERY | COMT |
| 2.464E-176 | 0.36297893 | 0.507 | 0.191 | 5.82E-172 | ERY | PSMB5 |
| 2.904E-176 | 0.31634456 | 0.386 | 0.123 | 6.859E-172 | ERY | STOML2 |
| 4.544E-176 | 0.31908308 | 0.454 | 0.157 | 1.073E-171 | ERY | HMGN3 |
| 5.621E-176 | 0.42664316 | 0.741 | 0.354 | 1.328E-171 | ERY | REEP5 |
| 3.951E-175 | 0.26273898 | 0.408 | 0.129 | 9.334E-171 | ERY | MGLL |
| 5.38E-174 | 0.36056339 | 0.594 | 0.243 | 1.271E-169 | ERY | RPL22L1 |
| 7.248E-174 | 0.43878327 | 0.956 | 0.74 | 1.712E-169 | ERY | ATP6V0E1 |
| 1.509E-173 | 0.47994637 | 0.987 | 0.943 | 3.565E-169 | ERY | MT-CO2 |
| 2.447E-173 | 0.33718842 | 0.429 | 0.148 | 5.782E-169 | ERY | CNPY2 |
| 2.456E-173 | 0.39757755 | 0.599 | 0.254 | 5.802E-169 | ERY | RNH1 |
| 5.21E-173 | 0.31029122 | 0.369 | 0.117 | 1.231E-168 | ERY | MSRB2 |
| 6.069E-173 | 0.30089997 | 0.352 | 0.107 | 1.434E-168 | ERY | TATDN1 |
| 8.242E-173 | 0.30640027 | 0.361 | 0.112 | 1.947E-168 | ERY | YIF1A |
| 5.43E-172 | 0.2639164 | 0.288 | 0.077 | 1.283E-167 | ERY | FAM207A |
| 5.815E-172 | 0.29812944 | 0.387 | 0.124 | 1.374E-167 | ERY | HYOU1 |
| 4.289E-171 | 0.32710988 | 0.413 | 0.141 | 1.013E-166 | ERY | MRPL13 |
| 4.743E-171 | 0.32491126 | 0.402 | 0.134 | 1.12E-166 | ERY | AGPAT2 |
| 5.279E-171 | 0.32132534 | 0.415 | 0.141 | 1.247E-166 | ERY | CENPX |
| 9.045E-171 | -0.801999 | 0.334 | 0.672 | 2.137E-166 | ERY | HNRNPU |
| 1.78E-170 | 0.26790323 | 0.31 | 0.088 | 4.204E-166 | ERY | MAD2L2 |
| 1.857E-170 | 0.29027885 | 0.376 | 0.119 | 4.388E-166 | ERY | DNPH1 |
| 6.529E-170 | -0.7374704 | 0.628 | 0.838 | 1.542E-165 | ERY | YWHAZ |
| 6.758E-169 | 0.39534773 | 0.566 | 0.237 | 1.596E-164 | ERY | EIF3D |
| 1.266E-168 | 0.28591082 | 0.317 | 0.092 | 2.99E-164 | ERY | MRPL15 |
| 3.259E-168 | 0.34371744 | 0.452 | 0.164 | 7.698E-164 | ERY | DPY30 |
| 5.695E-168 | 0.38637005 | 0.592 | 0.253 | 1.345E-163 | ERY | TMEM219 |
| 1.681E-167 | 0.29329163 | 0.334 | 0.101 | 3.972E-163 | ERY | HIKESHI |
| 2.895E-167 | 0.30193808 | 0.369 | 0.118 | 6.839E-163 | ERY | SNRPA1 |
| 3.411E-167 | 0.26377653 | 0.459 | 0.159 | 8.058E-163 | ERY | ISG20 |
| 6.946E-167 | 0.31319442 | 0.471 | 0.171 | 1.641E-162 | ERY | REXO2 |
| 3.55E-166 | 0.29740019 | 0.908 | 0.492 | 8.386E-162 | ERY | MT-ND4L |
| 3.828E-165 | 0.39987136 | 0.706 | 0.336 | 9.042E-161 | ERY | DDT |
| 4.899E-165 | 0.2964666 | 0.695 | 0.302 | 1.157E-160 | ERY | HMGB2 |
| 5.066E-165 | 0.42125368 | 0.954 | 0.733 | 1.197E-160 | ERY | SERP1 |
| 1.279E-164 | 0.38932557 | 0.571 | 0.245 | 3.021E-160 | ERY | POLR2J |
| 2.526E-164 | 0.32132992 | 0.423 | 0.148 | 5.968E-160 | ERY | TMEM205 |
| 3.712E-164 | 0.28588906 | 0.371 | 0.119 | 8.769E-160 | ERY | UQCC2 |
| 7.7E-164 | 0.30243159 | 0.401 | 0.136 | 1.819E-159 | ERY | TMEM256 |
| 1.848E-163 | 0.38339306 | 0.624 | 0.275 | 4.366E-159 | ERY | MRFAP1 |
| 2.715E-163 | 0.31884475 | 0.411 | 0.142 | 6.414E-159 | ERY | ECI1 |
| 6.838E-163 | 0.35259415 | 0.503 | 0.197 | 1.615E-158 | ERY | SIVA1 |
| 8.46E-163 | 0.32614218 | 0.462 | 0.169 | 1.999E-158 | ERY | MGAT4B |
| 1.346E-162 | 0.38852101 | 0.764 | 0.375 | 3.18E-158 | ERY | UQCR11 |
| 1.544E-162 | 0.30400629 | 0.391 | 0.131 | 3.649E-158 | ERY | PFDN4 |
| 2.567E-162 | 0.39752991 | 0.99 | 0.89 | 6.064E-158 | ERY | RPS15A |
| 8.45E-162 | 0.29412373 | 0.366 | 0.118 | 1.996E-157 | ERY | ARL2 |

| | | | | | | |
|---|---|---|---|---|---|---|
| 1.832E-160 | 0.38894009 | 0.672 | 0.313 | 4.328E-156 | ERY | SNRPG |
| 3.88E-160 | 0.26646278 | 0.348 | 0.108 | 9.165E-156 | ERY | TUBB2A |
| 6.151E-160 | 0.2664516 | 0.47 | 0.169 | 1.453E-155 | ERY | TMEM176B |
| 3.918E-159 | 0.36846707 | 0.575 | 0.248 | 9.255E-155 | ERY | LSM3 |
| 4.279E-159 | 0.33343348 | 0.457 | 0.171 | 1.011E-154 | ERY | POLR2I |
| 1.296E-158 | 0.36462352 | 0.564 | 0.238 | 3.062E-154 | ERY | MRPL20 |
| 2.72E-157 | 0.3973936 | 0.945 | 0.686 | 6.427E-153 | ERY | GUK1 |
| 3.001E-157 | 0.35060722 | 0.485 | 0.192 | 7.09E-153 | ERY | ARF5 |
| 3.857E-157 | 0.31356092 | 0.38 | 0.13 | 9.112E-153 | ERY | LSM1 |
| 7.343E-157 | 0.35324024 | 0.535 | 0.219 | 1.735E-152 | ERY | MRPL57 |
| 3.438E-156 | 0.37097145 | 0.667 | 0.313 | 8.122E-152 | ERY | GHITM |
| 5.388E-156 | 0.34926814 | 0.486 | 0.192 | 1.273E-151 | ERY | KRTCAP2 |
| 7.111E-156 | 0.35149758 | 0.512 | 0.207 | 1.68E-151 | ERY | LAMTOR2 |
| 9.717E-156 | 0.32295184 | 0.41 | 0.147 | 2.295E-151 | ERY | MRPS16 |
| 4.077E-155 | 0.2553902 | 0.341 | 0.105 | 9.631E-151 | ERY | UGDH |
| 8.6E-155 | -1.340905 | 0.122 | 0.469 | 2.032E-150 | ERY | MXD1 |
| 1.209E-154 | 0.3918584 | 0.99 | 0.931 | 2.855E-150 | ERY | MT-ND4 |
| 1.971E-154 | 0.35304174 | 0.513 | 0.209 | 4.655E-150 | ERY | MDH2 |
| 2.845E-154 | 0.26018294 | 0.317 | 0.096 | 6.721E-150 | ERY | TMEM251 |
| 8.677E-154 | -0.7760526 | 0.41 | 0.694 | 2.05E-149 | ERY | BZW1 |
| 1.068E-153 | -0.7500127 | 0.319 | 0.635 | 2.522E-149 | ERY | ACTR2 |
| 1.608E-153 | 0.3302715 | 0.443 | 0.167 | 3.798E-149 | ERY | POLR2F |
| 2.418E-153 | 0.41083564 | 0.783 | 0.401 | 5.711E-149 | ERY | EIF3K |
| 3.576E-153 | 0.28286568 | 0.765 | 0.372 | 8.448E-149 | ERY | C9orf16 |
| 6.858E-153 | 0.37528482 | 0.669 | 0.315 | 1.62E-148 | ERY | RNF7 |
| 1.313E-152 | 0.29591247 | 0.344 | 0.112 | 3.101E-148 | ERY | NTMT1 |
| 2.53E-152 | 0.3024167 | 0.599 | 0.252 | 5.976E-148 | ERY | TUBB |
| 2.422E-151 | 0.33419453 | 0.656 | 0.293 | 5.722E-147 | ERY | NOP53 |
| 4.556E-151 | 0.28024539 | 0.357 | 0.117 | 1.076E-146 | ERY | EBNA1BP2 |
| 9.21E-151 | 0.37411394 | 0.72 | 0.346 | 2.176E-146 | ERY | SNRPD2 |
| 1.277E-150 | 0.35146348 | 0.571 | 0.246 | 3.016E-146 | ERY | NDUFB10 |
| 1.868E-150 | 0.32297022 | 0.474 | 0.185 | 4.413E-146 | ERY | SEC61A1 |
| 2.378E-150 | -0.435487 | 0.503 | 0.173 | 5.617E-146 | ERY | HLA-DPB1 |
| 3.177E-150 | 0.25304447 | 0.303 | 0.09 | 7.506E-146 | ERY | IFT57 |
| 4.639E-150 | 0.39280594 | 0.683 | 0.331 | 1.096E-145 | ERY | SSR2 |
| 4.821E-150 | 0.38773657 | 0.89 | 0.573 | 1.139E-145 | ERY | NOP10 |
| 5.545E-149 | 0.36773443 | 0.69 | 0.335 | 1.31E-144 | ERY | SF3B6 |
| 9.597E-149 | 0.27465817 | 0.363 | 0.121 | 2.267E-144 | ERY | CBR1 |
| 1.172E-148 | 0.26214163 | 0.322 | 0.101 | 2.769E-144 | ERY | EEF1E1 |
| 1.377E-148 | 0.3346676 | 0.512 | 0.21 | 3.253E-144 | ERY | SNRPF |
| 4.123E-148 | 0.37176427 | 0.626 | 0.29 | 9.739E-144 | ERY | ATP5PO |
| 4.938E-148 | 0.30960267 | 0.445 | 0.169 | 1.167E-143 | ERY | REX1BD |
| 8.671E-148 | 0.25560977 | 0.303 | 0.092 | 2.048E-143 | ERY | IMP4 |
| 3.339E-147 | 0.3502795 | 0.572 | 0.251 | 7.887E-143 | ERY | GTF2A2 |
| 7.249E-147 | 0.327296 | 0.608 | 0.271 | 1.713E-142 | ERY | ISCU |
| 9.213E-147 | 0.27066875 | 0.332 | 0.108 | 2.176E-142 | ERY | CLPP |
| 1.982E-146 | 0.32817894 | 0.452 | 0.177 | 4.682E-142 | ERY | PCMT1 |
| 2.107E-146 | 0.38774631 | 0.831 | 0.475 | 4.978E-142 | ERY | DYNLT1 |
| 2.322E-146 | 0.36618062 | 0.715 | 0.347 | 5.485E-142 | ERY | TOMM20 |
| 6.049E-146 | 0.29663359 | 0.487 | 0.193 | 1.429E-141 | ERY | NEU1 |

| | | | | | | |
|---|---|---|---|---|---|---|
| 8.792E-146 | 0.41128004 | 0.985 | 0.81 | 2.077E-141 | ERY | PFDN5 |
| 1.306E-145 | 0.26582982 | 0.313 | 0.098 | 3.086E-141 | ERY | TSTA3 |
| 1.561E-145 | 0.36642301 | 0.848 | 0.5 | 3.688E-141 | ERY | COX7B |
| 3.924E-145 | 0.38345687 | 0.727 | 0.373 | 9.269E-141 | ERY | NEDD8 |
| 4.783E-145 | 0.38544923 | 0.893 | 0.552 | 1.13E-140 | ERY | SLC25A6 |
| 3.059E-144 | 0.29379699 | 0.409 | 0.15 | 7.227E-140 | ERY | PRMT1 |
| 3.337E-144 | 0.28538065 | 0.852 | 0.48 | 7.884E-140 | ERY | H2AFJ |
| 6.251E-144 | 0.29684355 | 0.655 | 0.292 | 1.477E-139 | ERY | RPL36A |
| 1.152E-143 | -1.3224891 | 0.994 | 0.98 | 2.721E-139 | ERY | MALAT1 |
| 2.713E-143 | 0.28812612 | 0.369 | 0.129 | 6.409E-139 | ERY | MRPL36 |
| 2.728E-143 | 0.31554904 | 0.66 | 0.296 | 6.445E-139 | ERY | ARPC1B |
| 2.807E-143 | 0.28974376 | 0.357 | 0.123 | 6.632E-139 | ERY | SURF1 |
| 3.693E-143 | 0.41018061 | 0.921 | 0.637 | 8.725E-139 | ERY | ATP5MC2 |
| 7.812E-142 | 0.3608708 | 0.671 | 0.325 | 1.845E-137 | ERY | SELENOH |
| 1.236E-141 | -0.8579396 | 0.841 | 0.381 | 2.919E-137 | ERY | HLA-DRA |
| 1.305E-141 | -4.0763774 | 0.076 | 0.412 | 3.083E-137 | ERY | SPRR3 |
| 1.314E-141 | 0.30205443 | 0.392 | 0.144 | 3.103E-137 | ERY | DERL2 |
| 1.812E-141 | 0.3250832 | 0.541 | 0.231 | 4.281E-137 | ERY | NUDC |
| 1.052E-140 | -0.7162286 | 0.425 | 0.697 | 2.485E-136 | ERY | CAST |
| 1.141E-140 | 0.31844221 | 0.61 | 0.276 | 2.695E-136 | ERY | ACADVL |
| 3.873E-140 | 0.36035668 | 0.637 | 0.306 | 9.15E-136 | ERY | RER1 |
| 2.809E-139 | 0.32812777 | 0.462 | 0.186 | 6.636E-135 | ERY | MED10 |
| 2.998E-139 | 0.30870536 | 0.457 | 0.182 | 7.082E-135 | ERY | ALKBH7 |
| 3.848E-139 | 0.2717821 | 0.39 | 0.139 | 9.091E-135 | ERY | NME1 |
| 5.529E-139 | 0.31441709 | 0.417 | 0.159 | 1.306E-134 | ERY | PLEKHJ1 |
| 1.625E-138 | 0.28123365 | 0.378 | 0.135 | 3.84E-134 | ERY | ETHE1 |
| 3.192E-138 | 0.25866697 | 0.368 | 0.128 | 7.54E-134 | ERY | CCDC85B |
| 5.034E-138 | 0.35928073 | 0.652 | 0.319 | 1.189E-133 | ERY | UBXN1 |
| 1.515E-137 | 0.28861843 | 0.399 | 0.148 | 3.578E-133 | ERY | CDC34 |
| 6.012E-137 | 0.27712565 | 0.355 | 0.124 | 1.42E-132 | ERY | VPS25 |
| 9.087E-137 | 0.2754211 | 0.357 | 0.125 | 2.147E-132 | ERY | RRAS |
| 1.107E-136 | 0.32655077 | 0.535 | 0.232 | 2.614E-132 | ERY | TMEM14C |
| 1.909E-136 | 0.28017124 | 0.473 | 0.188 | 4.509E-132 | ERY | TCEAL9 |
| 3.164E-136 | 0.33766707 | 0.719 | 0.358 | 7.474E-132 | ERY | ATP5MC3 |
| 1.403E-135 | 0.26638829 | 0.326 | 0.109 | 3.313E-131 | ERY | COA5 |
| 1.493E-135 | 0.31516096 | 0.468 | 0.191 | 3.527E-131 | ERY | NANS |
| 1.566E-135 | 0.32544177 | 0.571 | 0.256 | 3.698E-131 | ERY | KDELR2 |
| 3.68E-135 | 0.34517053 | 0.595 | 0.277 | 8.694E-131 | ERY | LAMTOR1 |
| 5.712E-135 | -0.7837914 | 0.441 | 0.699 | 1.349E-130 | ERY | PNRC1 |
| 8.521E-135 | -1.4592346 | 0.422 | 0.651 | 2.013E-130 | ERY | GLUL |
| 2.802E-134 | 0.31282574 | 0.657 | 0.308 | 6.62E-130 | ERY | AP2S1 |
| 4.349E-134 | 0.31051906 | 0.536 | 0.232 | 1.027E-129 | ERY | SRI |
| 2.056E-133 | 0.33473826 | 0.554 | 0.248 | 4.858E-129 | ERY | CCT3 |
| 2.497E-133 | 0.29853925 | 0.711 | 0.344 | 5.898E-129 | ERY | UQCRQ |
| 2.757E-133 | 0.33426908 | 0.398 | 0.151 | 6.513E-129 | ERY | GLRX5 |
| 3.103E-132 | 0.32174046 | 0.459 | 0.191 | 7.33E-128 | ERY | EIF4E2 |
| 3.295E-132 | 0.26602442 | 0.536 | 0.229 | 7.784E-128 | ERY | FKBP4 |
| 1.286E-131 | 0.3234047 | 0.564 | 0.256 | 3.038E-127 | ERY | VPS29 |
| 1.557E-131 | 0.26096817 | 0.327 | 0.111 | 3.679E-127 | ERY | NUDT5 |
| 5.418E-131 | 0.32917308 | 0.536 | 0.238 | 1.28E-126 | ERY | RPS17 |

| | | | | | | |
|---|---|---|---|---|---|---|
| 1.974E-130 | 0.27275522 | 0.336 | 0.117 | 4.664E-126 | ERY | IER3IP1 |
| 2.753E-130 | 0.25074113 | 0.593 | 0.258 | 6.504E-126 | ERY | FAM3C |
| 2.165E-129 | -0.7314662 | 0.32 | 0.603 | 5.114E-125 | ERY | DDX17 |
| 4.179E-129 | 0.28261194 | 0.4 | 0.154 | 9.872E-125 | ERY | MPLKIP |
| 1.534E-128 | 0.35651816 | 0.741 | 0.396 | 3.623E-124 | ERY | NDUFB4 |
| 2.268E-128 | 0.2598173 | 0.384 | 0.143 | 5.359E-124 | ERY | MRPL34 |
| 2.504E-128 | 0.3192904 | 0.569 | 0.259 | 5.916E-124 | ERY | NOL7 |
| 5.631E-128 | 0.27828017 | 0.38 | 0.143 | 1.33E-123 | ERY | BCL7C |
| 7.178E-128 | 0.36332651 | 0.987 | 0.896 | 1.696E-123 | ERY | RPL27A |
| 1.193E-127 | 0.25529476 | 0.319 | 0.109 | 2.818E-123 | ERY | MED8 |
| 1.288E-127 | 0.32850616 | 0.569 | 0.263 | 3.043E-123 | ERY | ATP5F1C |
| 4.799E-127 | 0.31208563 | 0.441 | 0.183 | 1.134E-122 | ERY | EMC6 |
| 5.357E-127 | 0.31663574 | 0.479 | 0.207 | 1.265E-122 | ERY | NDUFB5 |
| 9.893E-127 | 0.31790754 | 0.567 | 0.262 | 2.337E-122 | ERY | UXT |
| 1.308E-126 | 0.34084587 | 0.739 | 0.388 | 3.09E-122 | ERY | CIB1 |
| 7.246E-126 | 0.26646013 | 0.356 | 0.13 | 1.712E-121 | ERY | DDRGK1 |
| 1.191E-125 | -0.6982073 | 0.195 | 0.506 | 2.814E-121 | ERY | DDX3X |
| 1.401E-125 | 0.30896894 | 0.705 | 0.357 | 3.31E-121 | ERY | TXNDC17 |
| 1.539E-125 | 0.29458922 | 0.424 | 0.171 | 3.635E-121 | ERY | THOC7 |
| 1.828E-125 | -2.8129154 | 0.08 | 0.389 | 4.319E-121 | ERY | KRT13 |
| 2.058E-125 | 0.33039265 | 0.68 | 0.338 | 4.862E-121 | ERY | VDAC1 |
| 2.746E-125 | 0.30758123 | 0.462 | 0.195 | 6.488E-121 | ERY | NENF |
| 6.702E-125 | -0.7681473 | 0.693 | 0.815 | 1.583E-120 | ERY | DDX5 |
| 4.885E-124 | 0.32859419 | 0.716 | 0.372 | 1.154E-119 | ERY | DYNLRB1 |
| 5.469E-124 | 0.29666434 | 0.474 | 0.203 | 1.292E-119 | ERY | ARPC1A |
| 1.033E-123 | 0.2824439 | 0.971 | 0.719 | 2.44E-119 | ERY | PKM |
| 1.204E-123 | 0.32584106 | 0.997 | 0.888 | 2.845E-119 | ERY | ATP5F1E |
| 2.113E-123 | -0.714204 | 0.191 | 0.497 | 4.992E-119 | ERY | ANKRD12 |
| 2.438E-123 | 0.34426782 | 0.955 | 0.695 | 5.758E-119 | ERY | RPL4 |
| 2.854E-123 | 0.31066243 | 0.454 | 0.193 | 6.741E-119 | ERY | MRPS36 |
| 3.598E-123 | 0.27945669 | 0.45 | 0.184 | 8.499E-119 | ERY | PGRMC1 |
| 3.957E-123 | 0.32864228 | 0.643 | 0.32 | 9.348E-119 | ERY | NDUFC2 |
| 4.148E-123 | 0.28686252 | 0.364 | 0.137 | 9.799E-119 | ERY | HEBP1 |
| 7.366E-123 | 0.31426094 | 0.555 | 0.257 | 1.74E-118 | ERY | ERH |
| 3.01E-122 | 0.25829796 | 0.353 | 0.13 | 7.11E-118 | ERY | MRPS7 |
| 4.025E-122 | 0.30382338 | 0.52 | 0.233 | 9.509E-118 | ERY | LSM7 |
| 9.288E-122 | 0.30914288 | 0.508 | 0.227 | 2.194E-117 | ERY | NDUFA13 |
| 2.56E-121 | 0.25631814 | 0.342 | 0.124 | 6.048E-117 | ERY | MXD4 |
| 7.668E-121 | 0.29841531 | 0.604 | 0.289 | 1.811E-116 | ERY | EIF4A1 |
| 2.889E-120 | -0.7828941 | 0.19 | 0.49 | 6.826E-116 | ERY | RIOK3 |
| 5.625E-120 | 0.26032296 | 0.354 | 0.132 | 1.329E-115 | ERY | TRAPPC4 |
| 8.074E-119 | 0.32263267 | 0.643 | 0.325 | 1.907E-114 | ERY | CHMP5 |
| 1.742E-118 | 0.27040926 | 0.392 | 0.155 | 4.115E-114 | ERY | SMIM7 |
| 2.387E-118 | 0.29114559 | 0.468 | 0.202 | 5.64E-114 | ERY | PSMC5 |
| 2.657E-117 | 0.25127871 | 0.383 | 0.148 | 6.276E-113 | ERY | RPAIN |
| 3.737E-117 | 0.30476542 | 0.591 | 0.285 | 8.828E-113 | ERY | RABAC1 |
| 3.749E-117 | 0.25925669 | 0.335 | 0.123 | 8.857E-113 | ERY | MCTS1 |
| 7.333E-117 | 0.30216796 | 0.629 | 0.309 | 1.732E-112 | ERY | ZNF706 |
| 1.381E-116 | 0.2782776 | 0.403 | 0.164 | 3.262E-112 | ERY | HTATIP2 |
| 7.276E-116 | -1.1173716 | 0.551 | 0.704 | 1.719E-111 | ERY | JUNB |

| | | | | | | |
|---|---|---|---|---|---|---|
| 1.095E-115 | 0.32635138 | 0.819 | 0.491 | 2.588E-111 | ERY | LAPTM4A |
| 1.754E-115 | 0.26096302 | 0.41 | 0.166 | 4.144E-111 | ERY | CYC1 |
| 1.91E-115 | 0.27494661 | 0.459 | 0.197 | 4.513E-111 | ERY | NAA38 |
| 2.154E-115 | 0.29283865 | 0.499 | 0.225 | 5.087E-111 | ERY | PSMA5 |
| 3.907E-115 | 0.30648185 | 0.985 | 0.809 | 9.229E-111 | ERY | YBX1 |
| 7.923E-115 | 0.31124134 | 0.538 | 0.254 | 1.872E-110 | ERY | RTRAF |
| 1.068E-114 | 0.28405104 | 0.431 | 0.183 | 2.523E-110 | ERY | CHCHD1 |
| 2.322E-114 | 0.2620042 | 0.383 | 0.152 | 5.486E-110 | ERY | RNASEH2C |
| 2.724E-114 | 0.27057251 | 0.688 | 0.343 | 6.434E-110 | ERY | CYCS |
| 6.325E-114 | 0.27801451 | 0.544 | 0.254 | 1.494E-109 | ERY | NDUFA6 |
| 1.717E-113 | 0.27277684 | 0.402 | 0.165 | 4.056E-109 | ERY | STUB1 |
| 1.738E-113 | 0.35482884 | 0.976 | 0.814 | 4.105E-109 | ERY | RPL27 |
| 3.94E-113 | 0.30712787 | 0.48 | 0.217 | 9.308E-109 | ERY | PSMB2 |
| 5.861E-113 | 0.30665028 | 0.916 | 0.618 | 1.385E-108 | ERY | OST4 |
| 2.608E-111 | 0.31197413 | 0.932 | 0.719 | 6.161E-107 | ERY | ELOB |
| 1.084E-110 | 0.28398268 | 0.451 | 0.198 | 2.56E-106 | ERY | TIMM8B |
| 1.255E-110 | 0.28660557 | 0.513 | 0.238 | 2.965E-106 | ERY | CCT4 |
| 7.064E-110 | 0.25155687 | 0.378 | 0.151 | 1.669E-105 | ERY | DDAH2 |
| 7.236E-110 | 0.2801417 | 0.452 | 0.199 | 1.709E-105 | ERY | CWC15 |
| 1.704E-109 | 0.25691188 | 0.364 | 0.144 | 4.025E-105 | ERY | COPS5 |
| 4.557E-109 | -1.9492848 | 0.067 | 0.351 | 1.076E-104 | ERY | PSCA |
| 9.286E-109 | -1.8736649 | 0.13 | 0.417 | 2.194E-104 | ERY | S100P |
| 2.115E-108 | 0.27159945 | 0.439 | 0.191 | 4.997E-104 | ERY | MDH1 |
| 2.264E-108 | 0.27701467 | 0.422 | 0.183 | 5.348E-104 | ERY | EMC7 |
| 3.491E-108 | 0.25273602 | 0.364 | 0.145 | 8.246E-104 | ERY | MAF1 |
| 4.649E-108 | 0.2793252 | 0.415 | 0.178 | 1.098E-103 | ERY | COA6 |
| 1.515E-107 | 0.29912524 | 0.903 | 0.663 | 3.578E-103 | ERY | COX5B |
| 3.065E-107 | 0.28583332 | 0.645 | 0.327 | 7.241E-103 | ERY | ARL6IP4 |
| 3.291E-107 | 0.28578839 | 0.649 | 0.334 | 7.774E-103 | ERY | LAMTOR4 |
| 4.758E-107 | -0.8290947 | 0.383 | 0.61 | 1.124E-102 | ERY | MCL1 |
| 6.209E-107 | 0.27905645 | 0.428 | 0.188 | 1.467E-102 | ERY | CHMP4A |
| 1.645E-106 | -0.7186479 | 0.305 | 0.559 | 3.885E-102 | ERY | FAM107B |
| 2.499E-106 | 0.25241648 | 0.375 | 0.152 | 5.904E-102 | ERY | TMEM160 |
| 5.224E-106 | 0.25262096 | 0.881 | 0.562 | 1.234E-101 | ERY | TSPO |
| 3.47E-105 | 0.30416081 | 0.996 | 0.953 | 8.196E-101 | ERY | SERF2 |
| 1.13E-104 | 0.25824781 | 0.99 | 0.964 | 2.67E-100 | ERY | S100A11 |
| 1.815E-104 | 0.30240561 | 0.638 | 0.337 | 4.289E-100 | ERY | SNRPB |
| 3.194E-104 | 0.25027998 | 0.362 | 0.146 | 7.544E-100 | ERY | C19orf24 |
| 3.974E-104 | 0.28563966 | 0.493 | 0.231 | 9.388E-100 | ERY | MRPS21 |
| 6.226E-104 | 0.25949972 | 0.39 | 0.164 | 1.471E-99 | ERY | GLRX3 |
| 1.557E-103 | 0.28425922 | 0.652 | 0.342 | 3.679E-99 | ERY | SPCS1 |
| 2.701E-103 | 0.27629731 | 0.606 | 0.304 | 6.38E-99 | ERY | RBX1 |
| 1.919E-101 | -0.5850415 | 0.621 | 0.777 | 4.5332E-97 | ERY | EIF4G2 |
| 1.952E-101 | 0.25724587 | 0.381 | 0.16 | 4.6105E-97 | ERY | CISD2 |
| 2.492E-101 | 0.28427588 | 0.511 | 0.247 | 5.8874E-97 | ERY | EIF3I |
| 1.909E-100 | -0.6630458 | 0.117 | 0.386 | 4.5085E-96 | ERY | GCC2 |
| 5.7478E-98 | 0.25938062 | 0.434 | 0.196 | 1.3578E-93 | ERY | TXNL4A |
| 9.0696E-98 | 0.29531308 | 0.683 | 0.379 | 2.1425E-93 | ERY | SPCS2 |
| 1.1859E-97 | -0.5778044 | 0.257 | 0.514 | 2.8016E-93 | ERY | JAK1 |
| 1.8876E-97 | 0.29980636 | 0.664 | 0.366 | 4.4591E-93 | ERY | SRSF9 |

| | | | | | | |
|---|---|---|---|---|---|---|
| 4.2902E-97 | -0.9660325 | 0.208 | 0.458 | 1.0135E-92 | ERY | MARCKS |
| 8.569E-97 | 0.26465962 | 0.61 | 0.315 | 2.0243E-92 | ERY | DRAP1 |
| 1.6831E-95 | 0.26596902 | 0.482 | 0.233 | 3.9759E-91 | ERY | PSMD4 |
| 2.2771E-95 | 0.25970451 | 0.439 | 0.201 | 5.3792E-91 | ERY | ACP1 |
| 2.3518E-95 | -0.6111951 | 0.051 | 0.306 | 5.5557E-91 | ERY | REL |
| 2.4396E-95 | 0.2579473 | 0.435 | 0.198 | 5.763E-91 | ERY | SUGT1 |
| 8.7187E-95 | 0.28297045 | 0.935 | 0.681 | 2.0596E-90 | ERY | EEF2 |
| 2.9477E-94 | -1.8518183 | 0.097 | 0.353 | 6.9634E-90 | ERY | HPGD |
| 3.1406E-94 | 0.28048972 | 0.523 | 0.263 | 7.4191E-90 | ERY | ZNHIT1 |
| 8.1207E-94 | 0.26457286 | 0.581 | 0.302 | 1.9183E-89 | ERY | NDUFA2 |
| 1.0481E-93 | 0.28173821 | 0.646 | 0.35 | 2.4758E-89 | ERY | RBM8A |
| 4.7249E-93 | 0.25726751 | 0.488 | 0.236 | 1.1162E-88 | ERY | UQCRFS1 |
| 5.9298E-93 | 0.26881356 | 0.557 | 0.286 | 1.4008E-88 | ERY | VPS28 |
| 4.2156E-92 | 0.25880576 | 0.549 | 0.278 | 9.9586E-88 | ERY | NDUFA11 |
| 6.1812E-92 | -0.6623897 | 0.13 | 0.383 | 1.4602E-87 | ERY | CCNL1 |
| 1.2402E-91 | -1.0719728 | 0.05 | 0.289 | 2.9297E-87 | ERY | PTPRC |
| 1.5026E-91 | 0.26820534 | 0.551 | 0.283 | 3.5495E-87 | ERY | MTCH1 |
| 3.9515E-91 | 0.25186325 | 0.464 | 0.221 | 9.3347E-87 | ERY | SAT2 |
| 4.3715E-91 | -0.6252713 | 0.402 | 0.614 | 1.0327E-86 | ERY | SON |
| 1.2739E-90 | 0.25582134 | 0.564 | 0.289 | 3.0093E-86 | ERY | ROMO1 |
| 4.3442E-90 | -0.7168323 | 0.369 | 0.582 | 1.0262E-85 | ERY | CD55 |
| 5.1072E-90 | 0.25047812 | 0.389 | 0.175 | 1.2065E-85 | ERY | SNF8 |
| 3.1648E-89 | -0.6036548 | 0.192 | 0.436 | 7.4763E-85 | ERY | IQGAP1 |
| 1.8041E-88 | -0.6419864 | 0.329 | 0.555 | 4.2619E-84 | ERY | TOP1 |
| 2.5909E-88 | -0.5520609 | 0.118 | 0.368 | 6.1204E-84 | ERY | TMOD3 |
| 2.3209E-87 | 0.25500156 | 0.79 | 0.486 | 5.4826E-83 | ERY | VDAC2 |
| 3.3574E-87 | 0.26083407 | 0.49 | 0.247 | 7.9312E-83 | ERY | PSMB4 |
| 1.0237E-86 | -1.4272199 | 0.106 | 0.341 | 2.4183E-82 | ERY | IL1RN |
| 2.0983E-86 | 0.27528672 | 0.538 | 0.286 | 4.9569E-82 | ERY | SUMO1 |
| 2.9757E-85 | 0.26765116 | 0.769 | 0.48 | 7.0296E-81 | ERY | HSBP1 |
| 1.9239E-84 | 0.25001028 | 0.989 | 0.895 | 4.5449E-80 | ERY | RPS25 |
| 1.8133E-83 | -0.5751924 | 0.16 | 0.398 | 4.2836E-79 | ERY | STAT3 |
| 3.6341E-83 | -0.943527 | 0.691 | 0.754 | 8.5847E-79 | ERY | TPM4 |
| 1.7047E-82 | -1.2421172 | 0.485 | 0.633 | 4.0269E-78 | ERY | JUN |
| 2.1236E-82 | 0.26385031 | 0.661 | 0.379 | 5.0165E-78 | ERY | SEM1 |
| 4.2773E-82 | -0.4881139 | 0.068 | 0.3 | 1.0104E-77 | ERY | JMJD1C |
| 5.983E-82 | -0.5918005 | 0.141 | 0.376 | 1.4134E-77 | ERY | SPAG9 |
| 1.2405E-80 | -0.597469 | 0.207 | 0.435 | 2.9304E-76 | ERY | SRSF5 |
| 1.3557E-80 | -0.5575139 | 0.427 | 0.622 | 3.2026E-76 | ERY | KTN1 |
| 1.1008E-79 | 0.25083396 | 0.336 | 0.147 | 2.6004E-75 | ERY | SLC25A39 |
| 1.7852E-79 | -0.4872388 | 0.633 | 0.774 | 4.2172E-75 | ERY | CDC42 |
| 2.0497E-79 | -0.8148562 | 0.615 | 0.725 | 4.842E-75 | ERY | JUND |
| 1.1065E-78 | -0.5224309 | 0.431 | 0.622 | 2.6138E-74 | ERY | CAP1 |
| 3.8095E-78 | -1.0720661 | 0.052 | 0.27 | 8.9992E-74 | ERY | LMO7 |
| 7.9653E-77 | -2.2911865 | 0.918 | 0.51 | 1.8816E-72 | ERY | LCN2 |
| 1.2974E-74 | -0.4848756 | 0.084 | 0.303 | 3.0648E-70 | ERY | VPS4B |
| 7.0397E-74 | -0.5368074 | 0.125 | 0.347 | 1.663E-69 | ERY | YTHDC1 |
| 1.0737E-73 | -0.4963409 | 0.154 | 0.377 | 2.5364E-69 | ERY | PAK2 |
| 1.737E-72 | -0.5355483 | 0.146 | 0.362 | 4.1034E-68 | ERY | AKAP13 |
| 4.1482E-72 | -0.5046339 | 0.069 | 0.283 | 9.7993E-68 | ERY | SLK |

| | | | | | | |
|---|---|---|---|---|---|---|
| 6.3411E-72 | -0.5794679 | 0.131 | 0.353 | 1.498E-67 | ERY | GOLGB1 |
| 3.1887E-71 | -0.298826 | 0.437 | 0.189 | 7.5326E-67 | ERY | TM4SF1 |
| 4.2746E-71 | -0.6197848 | 0.851 | 0.869 | 1.0098E-66 | ERY | PABPC1 |
| 1.147E-70 | -0.4751622 | 0.126 | 0.344 | 2.7095E-66 | ERY | SRRM1 |
| 5.9132E-70 | -1.2038725 | 0.548 | 0.645 | 1.3969E-65 | ERY | IER3 |
| 1.9588E-69 | -0.6803169 | 0.445 | 0.195 | 4.6274E-65 | ERY | HLA-DPA1 |
| 2.3264E-69 | -0.6058423 | 0.146 | 0.043 | 5.4957E-65 | ERY | MT1A |
| 4.0499E-69 | -0.4648774 | 0.109 | 0.322 | 9.5672E-65 | ERY | AFF4 |
| 7.356E-69 | -0.5217069 | 0.505 | 0.665 | 1.7377E-64 | ERY | RBM39 |
| 1.1076E-68 | -0.4388881 | 0.069 | 0.274 | 2.6164E-64 | ERY | USP15 |
| 2.4392E-68 | -0.4749427 | 0.113 | 0.324 | 5.7621E-64 | ERY | KMT2E |
| 6.5824E-68 | -0.4619566 | 0.046 | 0.245 | 1.555E-63 | ERY | SMCHD1 |
| 1.4823E-67 | -1.1200489 | 0.295 | 0.479 | 3.5016E-63 | ERY | ERO1A |
| 1.5032E-67 | -1.2194853 | 0.008 | 0.19 | 3.5511E-63 | ERY | DUOX2 |
| 3.0994E-67 | -0.5701434 | 0.054 | 0.251 | 7.3216E-63 | ERY | KIAA1551 |
| 2.068E-66 | -0.3960844 | 0.052 | 0.247 | 4.8852E-62 | ERY | KMT2C |
| 4.4252E-66 | -1.0469302 | 0.206 | 0.396 | 1.0454E-61 | ERY | FOS |
| 9.3575E-66 | -0.5454588 | 0.018 | 0.201 | 2.2105E-61 | ERY | SAMD9 |
| 2.6906E-65 | -0.3887868 | 0.05 | 0.244 | 6.3561E-61 | ERY | ASH1L |
| 1.9539E-64 | -0.5965003 | 0.506 | 0.644 | 4.6157E-60 | ERY | ARPC5 |
| 7.9178E-64 | -0.4584297 | 0.615 | 0.739 | 1.8704E-59 | ERY | CCNI |
| 3.0176E-63 | -0.5739067 | 0.838 | 0.426 | 7.1284E-59 | ERY | TYROBP |
| 3.0362E-63 | -0.5610624 | 0.045 | 0.231 | 7.1723E-59 | ERY | PPP1R10 |
| 4.2047E-63 | -0.6534947 | 0.006 | 0.177 | 9.9328E-59 | ERY | STEAP4 |
| 4.3718E-63 | -0.40518 | 0.095 | 0.296 | 1.0328E-58 | ERY | CLIP1 |
| 7.8358E-63 | -0.6071659 | 0.142 | 0.331 | 1.851E-58 | ERY | ARHGDIB |
| 1.3129E-62 | -0.5330004 | 0.146 | 0.34 | 3.1014E-58 | ERY | TLN1 |
| 2.0536E-62 | -0.7573489 | 0.019 | 0.194 | 4.8512E-58 | ERY | ANKRD36C |
| 2.0633E-62 | -0.3954687 | 0.084 | 0.28 | 4.8742E-58 | ERY | NIPBL |
| 4.0351E-62 | -0.4675306 | 0.08 | 0.271 | 9.532E-58 | ERY | BACH1 |
| 1.5144E-61 | -0.4525591 | 0.587 | 0.264 | 3.5775E-57 | ERY | AIF1 |
| 2.6469E-61 | -0.4115515 | 0.094 | 0.29 | 6.2528E-57 | ERY | TPR |
| 3.0024E-61 | -0.498997 | 0.067 | 0.254 | 7.0925E-57 | ERY | GPCPD1 |
| 1.0764E-60 | -0.4732904 | 0.106 | 0.298 | 2.5428E-56 | ERY | SKIL |
| 1.2223E-60 | -1.3727732 | 0.027 | 0.202 | 2.8874E-56 | ERY | HOPX |
| 1.3019E-60 | -0.4208946 | 0.088 | 0.281 | 3.0756E-56 | ERY | BPTF |
| 7.0978E-60 | -0.4565219 | 0.157 | 0.352 | 1.6767E-55 | ERY | SCAF11 |
| 1.1518E-59 | -0.6164511 | 0.319 | 0.488 | 2.721E-55 | ERY | CD46 |
| 1.3552E-59 | -0.6102762 | 0.031 | 0.204 | 3.2013E-55 | ERY | DUSP4 |
| 5.2167E-59 | -1.1356755 | 0.006 | 0.166 | 1.2323E-54 | ERY | MUC21 |
| 6.6058E-59 | -0.4740101 | 0.132 | 0.323 | 1.5605E-54 | ERY | BOD1L1 |
| 1.3193E-58 | -0.8334766 | 0.123 | 0.297 | 3.1166E-54 | ERY | LCP1 |
| 2.5162E-58 | -0.2694191 | 0.776 | 0.413 | 5.9439E-54 | ERY | EMP3 |
| 7.1363E-58 | -0.3752547 | 0.059 | 0.239 | 1.6858E-53 | ERY | BAZ2B |
| 1.2006E-57 | -0.9857834 | 0.006 | 0.163 | 2.8361E-53 | ERY | PRSS27 |
| 2.2381E-57 | -1.2380421 | 0.011 | 0.171 | 5.287E-53 | ERY | DUOXA2 |
| 3.5274E-57 | -0.4519903 | 0.055 | 0.23 | 8.3327E-53 | ERY | KYNU |
| 1.0513E-56 | -0.4630452 | 0.158 | 0.345 | 2.4835E-52 | ERY | GRB2 |
| 1.9613E-56 | -0.4508094 | 0.436 | 0.589 | 4.6331E-52 | ERY | ACTR3 |
| 2.6386E-56 | -0.4248868 | 0.403 | 0.566 | 6.2332E-52 | ERY | CSNK1A1 |

| | | | | | | |
|---|---|---|---|---|---|---|
| 2.7739E-56 | -0.476036 | 0.105 | 0.286 | 6.5528E-52 | ERY | LYN |
| 4.1089E-56 | -1.0318256 | 0.009 | 0.165 | 9.7065E-52 | ERY | CEACAM5 |
| 8.5733E-56 | -0.3751678 | 0.099 | 0.285 | 2.0253E-51 | ERY | TMF1 |
| 2.3334E-55 | -0.8293169 | 0.02 | 0.178 | 5.5122E-51 | ERY | NDRG2 |
| 2.5662E-55 | -0.662338 | 0.126 | 0.305 | 6.0621E-51 | ERY | BTG2 |
| 5.4614E-55 | -0.5405465 | 0.227 | 0.4 | 1.2901E-50 | ERY | PNISR |
| 6.1367E-55 | -0.767314 | 0.65 | 0.7 | 1.4497E-50 | ERY | PPP1R15A |
| 1.0626E-54 | -0.6494109 | 0.003 | 0.152 | 2.5101E-50 | ERY | NCCRP1 |
| 1.1889E-54 | -1.5904208 | 0.014 | 0.167 | 2.8086E-50 | ERY | ECM1 |
| 2.0049E-54 | -0.964729 | 0.008 | 0.16 | 4.7362E-50 | ERY | PITX1 |
| 2.5692E-54 | -0.7524133 | 0.008 | 0.158 | 6.0693E-50 | ERY | SCEL |
| 2.7018E-54 | -0.411854 | 0.049 | 0.217 | 6.3825E-50 | ERY | TLE4 |
| 3.3994E-54 | -0.4914612 | 0.042 | 0.207 | 8.0305E-50 | ERY | FOSB |
| 3.5293E-54 | -0.5950895 | 0.45 | 0.574 | 8.3372E-50 | ERY | ZFAND5 |
| 4.1931E-54 | -0.4250969 | 0.078 | 0.254 | 9.9054E-50 | ERY | PKN2 |
| 7.8387E-54 | -0.3545413 | 0.069 | 0.243 | 1.8517E-49 | ERY | HOOK3 |
| 2.3591E-53 | -0.4126179 | 0.108 | 0.288 | 5.573E-49 | ERY | ATRX |
| 3.2417E-53 | -0.4714251 | 0.383 | 0.173 | 7.6578E-49 | ERY | SPINK1 |
| 4.7628E-53 | -0.7428797 | 0.006 | 0.153 | 1.1251E-48 | ERY | TMPRSS11D |
| 1.8744E-52 | -0.3370951 | 0.055 | 0.223 | 4.4278E-48 | ERY | TUT7 |
| 4.9517E-52 | -0.8350835 | 0.017 | 0.166 | 1.1697E-47 | ERY | APOBEC3A |
| 6.7114E-52 | -0.4278537 | 0.139 | 0.315 | 1.5854E-47 | ERY | QKI |
| 1.1793E-51 | -0.5014268 | 0.023 | 0.175 | 2.7859E-47 | ERY | PRDM1 |
| 1.8016E-51 | -0.6531344 | 0.456 | 0.582 | 4.2559E-47 | ERY | LITAF |
| 2.8328E-51 | -0.7552844 | 0.006 | 0.149 | 6.692E-47 | ERY | TMPRSS11E |
| 9.9365E-51 | -0.4889744 | 0.164 | 0.339 | 2.3473E-46 | ERY | GOLGA4 |
| 1.4897E-50 | -0.4031238 | 0.05 | 0.209 | 3.5191E-46 | ERY | TIPARP |
| 1.8757E-50 | -0.4415067 | 0.059 | 0.219 | 4.4311E-46 | ERY | EHF |
| 2.4634E-50 | -0.4517538 | 0.173 | 0.344 | 5.8192E-46 | ERY | GPBP1 |
| 3.3874E-50 | -0.3315702 | 0.08 | 0.247 | 8.0022E-46 | ERY | IFNAR1 |
| 3.5147E-50 | -0.6354182 | 0.092 | 0.256 | 8.3028E-46 | ERY | SYNE2 |
| 4.7553E-50 | -0.4314979 | 0.117 | 0.287 | 1.1234E-45 | ERY | PRPF4B |
| 5.5708E-50 | -0.3841522 | 0.165 | 0.342 | 1.316E-45 | ERY | CAPZA1 |
| 5.5829E-50 | -0.5329442 | 0.262 | 0.421 | 1.3188E-45 | ERY | FUS |
| 9.207E-50 | -0.3642095 | 0.092 | 0.261 | 2.175E-45 | ERY | TACC1 |
| 9.724E-50 | -0.6318178 | 0.016 | 0.16 | 2.2971E-45 | ERY | MUC4 |
| 2.977E-49 | -0.3051438 | 0.041 | 0.195 | 7.0326E-45 | ERY | N4BP1 |
| 5.1934E-49 | -0.6643139 | 0.061 | 0.215 | 1.2268E-44 | ERY | TNFRSF1B |
| 5.8544E-49 | -0.7825685 | 0.01 | 0.15 | 1.383E-44 | ERY | CEACAM7 |
| 6.2062E-49 | -0.4351546 | 0.141 | 0.313 | 1.4661E-44 | ERY | NUFIP2 |
| 8.1786E-49 | -0.6034723 | 0.058 | 0.213 | 1.932E-44 | ERY | CDKN2B |
| 1.1209E-48 | -1.0288779 | 0.008 | 0.144 | 2.6478E-44 | ERY | CXCL6 |
| 1.5837E-48 | -0.4189276 | 0.027 | 0.178 | 3.7412E-44 | ERY | XIST |
| 2.1141E-48 | -0.4765749 | 0.152 | 0.321 | 4.9942E-44 | ERY | AKAP9 |
| 2.5604E-48 | -0.3179094 | 0.038 | 0.188 | 6.0485E-44 | ERY | GNA15 |
| 6.5483E-48 | -1.6079441 | 0.004 | 0.137 | 1.5469E-43 | ERY | LYPD2 |
| 7.3024E-48 | -0.3689655 | 0.057 | 0.213 | 1.7251E-43 | ERY | STK17B |
| 1.0813E-47 | -0.5055514 | 0.059 | 0.212 | 2.5545E-43 | ERY | GK |
| 1.2236E-47 | -0.6030962 | 0.392 | 0.193 | 2.8906E-43 | ERY | THBS1 |
| 1.2556E-47 | -0.3508965 | 0.094 | 0.258 | 2.9661E-43 | ERY | NSD3 |

| | | | | | | |
|---|---|---|---|---|---|---|
| 1.6277E-47 | -0.5200698 | 0.228 | 0.386 | 3.8451E-43 | ERY | N4BP2L2 |
| 2.0217E-47 | -0.8432738 | 0.014 | 0.151 | 4.7758E-43 | ERY | CEACAM6 |
| 2.9702E-47 | -0.6604916 | 0.004 | 0.135 | 7.0165E-43 | ERY | GABRP |
| 4.8398E-47 | -0.3873937 | 0.188 | 0.359 | 1.1433E-42 | ERY | SF3B1 |
| 5.9321E-47 | -0.3266726 | 0.101 | 0.267 | 1.4013E-42 | ERY | ELF1 |
| 5.9794E-47 | -0.3877207 | 0.03 | 0.175 | 1.4125E-42 | ERY | FAM214A |
| 7.783E-47 | -0.4003058 | 0.113 | 0.276 | 1.8386E-42 | ERY | OSBPL8 |
| 8.8167E-47 | -0.3921283 | 0.104 | 0.265 | 2.0828E-42 | ERY | MBNL1 |
| 8.8969E-47 | -0.3136875 | 0.092 | 0.256 | 2.1017E-42 | ERY | TRIP12 |
| 9.3148E-47 | -0.3054334 | 0.042 | 0.191 | 2.2004E-42 | ERY | TET2 |
| 1.0783E-46 | -0.3398704 | 0.02 | 0.159 | 2.5472E-42 | ERY | HIST1H1E |
| 1.0848E-46 | -0.4158647 | 0.176 | 0.34 | 2.5626E-42 | ERY | RBM25 |
| 3.4869E-46 | -0.3394209 | 0.976 | 0.714 | 8.237E-42 | ERY | HSPB1 |
| 5.0318E-46 | -0.3973163 | 0.206 | 0.37 | 1.1887E-41 | ERY | WASF2 |
| 7.9164E-46 | -0.3705644 | 0.173 | 0.339 | 1.8701E-41 | ERY | HNRNPD |
| 1.7995E-45 | -0.361836 | 0.113 | 0.274 | 4.2509E-41 | ERY | AGFG1 |
| 2.5888E-45 | -0.2993203 | 0.068 | 0.221 | 6.1155E-41 | ERY | DDX6 |
| 2.8568E-45 | -1.0249942 | 0.074 | 0.22 | 6.7487E-41 | ERY | S100A12 |
| 3.5395E-45 | -0.3251184 | 0.057 | 0.206 | 8.3614E-41 | ERY | NR3C1 |
| 4.7812E-45 | -0.5186612 | 0.271 | 0.419 | 1.1295E-40 | ERY | SFPQ |
| 5.0904E-45 | -0.4589635 | 0.613 | 0.698 | 1.2025E-40 | ERY | LRRFIP1 |
| 6.9626E-45 | -0.3127362 | 0.066 | 0.219 | 1.6448E-40 | ERY | SPEN |
| 2.0313E-44 | -0.4617227 | 0.203 | 0.364 | 4.7986E-40 | ERY | RRBP1 |
| 2.4024E-44 | -0.3212595 | 0.073 | 0.225 | 5.6752E-40 | ERY | PCM1 |
| 3.1676E-44 | -0.5153682 | 0.22 | 0.374 | 7.4827E-40 | ERY | ANKRD11 |
| 3.6781E-44 | -0.6052249 | 0.938 | 0.614 | 8.6889E-40 | ERY | S100A4 |
| 6.8574E-44 | -0.3718393 | 0.081 | 0.233 | 1.6199E-39 | ERY | GALNT1 |
| 7.3355E-44 | -0.2669679 | 0.045 | 0.191 | 1.7329E-39 | ERY | CEP350 |
| 1.2091E-43 | -0.6624618 | 0.01 | 0.135 | 2.8562E-39 | ERY | DHRS9 |
| 2.0965E-43 | -0.3287859 | 0.129 | 0.29 | 4.9527E-39 | ERY | RAD21 |
| 2.1615E-43 | -0.5173667 | 0.13 | 0.28 | 5.1061E-39 | ERY | ETS2 |
| 3.9484E-43 | -0.3806475 | 0.072 | 0.22 | 9.3274E-39 | ERY | KDM6B |
| 4.0669E-43 | -0.6121568 | 0.103 | 0.251 | 9.6072E-39 | ERY | CSTA |
| 4.7263E-43 | -0.3343649 | 0.099 | 0.253 | 1.1165E-38 | ERY | CCDC186 |
| 5.0791E-43 | -0.3490282 | 0.141 | 0.299 | 1.1998E-38 | ERY | SYAP1 |
| 5.6871E-43 | -0.5781865 | 0.563 | 0.286 | 1.3435E-38 | ERY | ADIRF |
| 6.9154E-43 | -0.8627587 | 0.013 | 0.139 | 1.6336E-38 | ERY | TMPRSS11B |
| 7.7753E-43 | -0.4041531 | 0.194 | 0.351 | 1.8368E-38 | ERY | ZNF207 |
| 1.1641E-42 | -1.123661 | 0.002 | 0.121 | 2.7499E-38 | ERY | SPINK5 |
| 1.1933E-42 | -0.781739 | 0.1 | 0.243 | 2.8188E-38 | ERY | ADGRF1 |
| 3.1436E-42 | -0.4376155 | 0.216 | 0.367 | 7.4261E-38 | ERY | NFE2L2 |
| 4.5403E-42 | -0.5367254 | 0.101 | 0.247 | 1.0726E-37 | ERY | ATP2B1 |
| 4.6078E-42 | -0.8355145 | 0.056 | 0.192 | 1.0885E-37 | ERY | TMPRSS2 |
| 6.0108E-42 | -0.3710471 | 0.135 | 0.291 | 1.4199E-37 | ERY | RB1CC1 |
| 8.9251E-42 | -1.003873 | 0.787 | 0.481 | 2.1084E-37 | ERY | CD74 |
| 1.0916E-41 | -0.3129064 | 0.115 | 0.27 | 2.5786E-37 | ERY | KRAS |
| 1.4833E-41 | -0.3586696 | 0.138 | 0.294 | 3.5039E-37 | ERY | ARID4B |
| 1.5613E-41 | -0.3273032 | 0.104 | 0.257 | 3.6882E-37 | ERY | BCLAF1 |
| 3.7279E-41 | -0.5254208 | 0.856 | 0.835 | 8.8063E-37 | ERY | HLA-E |
| 5.0558E-41 | -0.9718517 | 0.045 | 0.178 | 1.1943E-36 | ERY | RDH10 |

| | | | | | | |
|---|---|---|---|---|---|---|
| 5.117E-41 | -0.4266907 | 0.243 | 0.39 | 1.2088E-36 | ERY | DEK |
| 5.728E-41 | -0.3622847 | 0.131 | 0.282 | 1.3531E-36 | ERY | ROCK1 |
| 6.376E-41 | -0.2998812 | 0.077 | 0.222 | 1.5062E-36 | ERY | U2SURP |
| 7.2868E-41 | -0.5986031 | 0.002 | 0.116 | 1.7214E-36 | ERY | KRT78 |
| 8.3814E-41 | -0.6059691 | 0.017 | 0.14 | 1.9799E-36 | ERY | CXCL17 |
| 8.8089E-41 | -0.3902532 | 0.22 | 0.372 | 2.0809E-36 | ERY | PNRC2 |
| 9.9403E-41 | -0.4374336 | 0.118 | 0.264 | 2.3482E-36 | ERY | WASL |
| 1.3079E-40 | -0.2998506 | 0.087 | 0.235 | 3.0896E-36 | ERY | KDM5A |
| 1.6874E-40 | -0.3560326 | 0.016 | 0.139 | 3.9861E-36 | ERY | BCAS1 |
| 2.16E-40 | -0.461307 | 0.02 | 0.143 | 5.1025E-36 | ERY | PLAC8 |
| 2.5189E-40 | -1.2867997 | 0.003 | 0.117 | 5.9504E-36 | ERY | SPRR1B |
| 3.5753E-40 | -0.3380696 | 0.076 | 0.218 | 8.446E-36 | ERY | ZFYVE16 |
| 3.5797E-40 | -0.2923747 | 0.097 | 0.246 | 8.4564E-36 | ERY | RSF1 |
| 4.4626E-40 | -0.395254 | 0.094 | 0.238 | 1.0542E-35 | ERY | PARP14 |
| 5.0434E-40 | -0.3333624 | 0.088 | 0.232 | 1.1914E-35 | ERY | NUMB |
| 5.3935E-40 | -0.3724357 | 0.159 | 0.309 | 1.2741E-35 | ERY | HP1BP3 |
| 5.8575E-40 | -0.3367887 | 0.076 | 0.217 | 1.3837E-35 | ERY | BDP1 |
| 6.1879E-40 | -0.3892686 | 0.167 | 0.316 | 1.4618E-35 | ERY | TMEM165 |
| 6.3318E-40 | -0.2737903 | 0.075 | 0.218 | 1.4958E-35 | ERY | ITSN2 |
| 7.9132E-40 | -0.4519217 | 0.003 | 0.116 | 1.8693E-35 | ERY | TMPRSS11A |
| 8.3139E-40 | -0.491429 | 0.369 | 0.498 | 1.964E-35 | ERY | NCL |
| 9.1322E-40 | -0.2785369 | 0.018 | 0.139 | 2.1573E-35 | ERY | SASH1 |
| 1.1195E-39 | -0.2971466 | 0.044 | 0.176 | 2.6446E-35 | ERY | B3GNT5 |
| 1.1236E-39 | -0.3065306 | 0.08 | 0.223 | 2.6542E-35 | ERY | POLR2J3.1 |
| 1.3881E-39 | -0.3707757 | 0.006 | 0.118 | 3.279E-35 | ERY | CEACAM1 |
| 2.0388E-39 | -0.5366049 | 0.048 | 0.176 | 4.8164E-35 | ERY | PPL |
| 2.424E-39 | -0.2980614 | 0.093 | 0.238 | 5.7261E-35 | ERY | ZNF106 |
| 4.2398E-39 | -0.3849568 | 0.159 | 0.305 | 1.0016E-34 | ERY | PICALM |
| 5.9527E-39 | -0.6299492 | 0.203 | 0.332 | 1.4062E-34 | ERY | MYO6 |
| 9.1745E-39 | -0.4734292 | 0.027 | 0.148 | 2.1673E-34 | ERY | CD93 |
| 1.753E-38 | -0.7799188 | 0.228 | 0.103 | 4.141E-34 | ERY | CCL4 |
| 1.7976E-38 | -0.2973112 | 0.997 | 0.993 | 4.2466E-34 | ERY | TPT1 |
| 2.5375E-38 | -0.394356 | 0.109 | 0.249 | 5.9943E-34 | ERY | RAB31 |
| 2.5581E-38 | -0.7789982 | 0.056 | 0.184 | 6.043E-34 | ERY | F3 |
| 2.6173E-38 | -0.5491073 | 0.705 | 0.743 | 6.1829E-34 | ERY | APLP2 |
| 2.8086E-38 | -0.3081839 | 0.132 | 0.28 | 6.6348E-34 | ERY | CHMP4B |
| 2.9502E-38 | -0.3086763 | 0.117 | 0.263 | 6.9692E-34 | ERY | ERBIN |
| 3.2309E-38 | -0.2502312 | 0.035 | 0.16 | 7.6325E-34 | ERY | FBXO34 |
| 3.3659E-38 | -0.3140101 | 0.097 | 0.239 | 7.9512E-34 | ERY | FNBP4 |
| 4.9558E-38 | -0.3629232 | 0.25 | 0.397 | 1.1707E-33 | ERY | UBXN4 |
| 5.892E-38 | -0.8050844 | 0.038 | 0.16 | 1.3919E-33 | ERY | KRT15 |
| 7.3674E-38 | -0.5902889 | 0.066 | 0.196 | 1.7404E-33 | ERY | FBXO32 |
| 7.5927E-38 | -0.3767246 | 0.075 | 0.207 | 1.7936E-33 | ERY | LSP1 |
| 7.8638E-38 | -0.3030925 | 0.087 | 0.226 | 1.8577E-33 | ERY | ZBTB7A |
| 9.2119E-38 | -0.4183221 | 0.003 | 0.11 | 2.1761E-33 | ERY | TMC5 |
| 9.9618E-38 | -0.327431 | 0.003 | 0.109 | 2.3533E-33 | ERY | A2ML1 |
| 1.2474E-37 | -0.4165355 | 0.104 | 0.241 | 2.9468E-33 | ERY | ARID5B |
| 1.7206E-37 | -0.485083 | 0.064 | 0.192 | 4.0646E-33 | ERY | FYB1 |
| 1.8032E-37 | -0.2796733 | 0.006 | 0.115 | 4.2597E-33 | ERY | FUT3 |
| 2.6683E-37 | -0.2868764 | 0.095 | 0.235 | 6.3034E-33 | ERY | TAOK3 |

| | | | | | | |
|---|---|---|---|---|---|---|
| 2.9019E-37 | -0.4908092 | 0.882 | 0.863 | 6.8551E-33 | ERY | CALM1 |
| 4.2344E-37 | -0.310511 | 0.096 | 0.234 | 1.0003E-32 | ERY | PHF20L1 |
| 4.3043E-37 | -0.4791131 | 0.094 | 0.229 | 1.0168E-32 | ERY | INTS6 |
| 4.4232E-37 | -0.3435119 | 0.217 | 0.362 | 1.0449E-32 | ERY | XRCC5 |
| 4.4847E-37 | -0.3240831 | 0.084 | 0.219 | 1.0594E-32 | ERY | EML4 |
| 5.4288E-37 | -0.3426944 | 0.099 | 0.238 | 1.2824E-32 | ERY | KMT2A |
| 6.1172E-37 | -1.0232224 | 0.022 | 0.137 | 1.4451E-32 | ERY | SERPINB2 |
| 6.2587E-37 | -0.4071712 | 0.153 | 0.293 | 1.4785E-32 | ERY | RNF149 |
| 7.2531E-37 | -0.2527277 | 0.04 | 0.163 | 1.7134E-32 | ERY | DOCK8 |
| 1.5649E-36 | -0.310446 | 0.045 | 0.169 | 3.6968E-32 | ERY | JARID2 |
| 4.5209E-36 | -0.4654056 | 0.199 | 0.333 | 1.068E-31 | ERY | ASPH |
| 4.9045E-36 | -0.7840489 | 0.005 | 0.108 | 1.1586E-31 | ERY | ZG16B |
| 7.8119E-36 | -0.2904372 | 0.112 | 0.25 | 1.8454E-31 | ERY | PPP1R12A |
| 1.1101E-35 | -0.2616547 | 0.074 | 0.206 | 2.6223E-31 | ERY | PRDM2 |
| 1.2626E-35 | -0.6209182 | 0.106 | 0.231 | 2.9827E-31 | ERY | ZEB2 |
| 1.8644E-35 | -0.3747603 | 0.048 | 0.168 | 4.4044E-31 | ERY | LCP2 |
| 1.9164E-35 | -0.2706793 | 0.096 | 0.233 | 4.5272E-31 | ERY | USP8 |
| 1.9343E-35 | -0.6953778 | 0.057 | 0.179 | 4.5694E-31 | ERY | PLEK |
| 2.1648E-35 | -0.3158393 | 0.077 | 0.205 | 5.1139E-31 | ERY | IFI16 |
| 3.5418E-35 | -0.3242802 | 0.033 | 0.148 | 8.3668E-31 | ERY | TLR2 |
| 3.8402E-35 | -0.2787707 | 0.073 | 0.202 | 9.0718E-31 | ERY | CHD1 |
| 4.8391E-35 | -0.5131976 | 0.045 | 0.162 | 1.1431E-30 | ERY | CD300E |
| 5.157E-35 | -0.3188157 | 0.109 | 0.243 | 1.2182E-30 | ERY | KLF3 |
| 5.3037E-35 | -0.2987594 | 0.091 | 0.223 | 1.2529E-30 | ERY | ZFC3H1 |
| 7.1935E-35 | -0.259302 | 0.085 | 0.217 | 1.6993E-30 | ERY | MARCH7 |
| 9.4833E-35 | -0.3849633 | 0.019 | 0.126 | 2.2402E-30 | ERY | TMEM40 |
| 1.1916E-34 | -0.3720861 | 0.133 | 0.269 | 2.815E-30 | ERY | HECTD1 |
| 1.3258E-34 | -0.5453704 | 0.54 | 0.622 | 3.1319E-30 | ERY | PCBP1 |
| 2.0678E-34 | -0.5874331 | 0.034 | 0.146 | 4.8847E-30 | ERY | CDKN2A |
| 2.0973E-34 | -0.5876563 | 0.073 | 0.193 | 4.9545E-30 | ERY | EPS8L1 |
| 2.5928E-34 | -0.3382839 | 0.212 | 0.35 | 6.125E-30 | ERY | MTPN |
| 2.9218E-34 | -0.3031685 | 0.109 | 0.243 | 6.9021E-30 | ERY | RNMT |
| 3.4228E-34 | -0.3803417 | 0.069 | 0.19 | 8.0857E-30 | ERY | PTPRE |
| 3.6791E-34 | -0.3563334 | 0.147 | 0.283 | 8.6911E-30 | ERY | YME1L1 |
| 4.4026E-34 | -0.3164647 | 0.022 | 0.129 | 1.04E-29 | ERY | POLB |
| 4.7573E-34 | -0.319326 | 0.093 | 0.222 | 1.1238E-29 | ERY | STK4 |
| 5.2654E-34 | -0.3149387 | 0.12 | 0.255 | 1.2438E-29 | ERY | UBL3 |
| 6.9972E-34 | -0.8849534 | 0.2 | 0.315 | 1.653E-29 | ERY | EMP1 |
| 7.0744E-34 | -0.3049156 | 0.088 | 0.216 | 1.6712E-29 | ERY | VPS13C |
| 1.1082E-33 | -0.2622172 | 0.092 | 0.223 | 2.6178E-29 | ERY | ACAP2 |
| 1.4261E-33 | -0.2862774 | 0.006 | 0.104 | 3.3688E-29 | ERY | LRMP |
| 1.4625E-33 | -0.2579145 | 0.078 | 0.208 | 3.4548E-29 | ERY | TRIP11 |
| 1.6803E-33 | -0.3029225 | 0.045 | 0.16 | 3.9695E-29 | ERY | PELI1 |
| 2.1137E-33 | -0.3343462 | 0.036 | 0.149 | 4.9933E-29 | ERY | VSIG2 |
| 2.4872E-33 | -0.4287244 | 0.224 | 0.354 | 5.8756E-29 | ERY | PNN |
| 2.7882E-33 | -0.2507269 | 0.02 | 0.126 | 6.5866E-29 | ERY | TCP11L2 |
| 3.8595E-33 | -0.4235751 | 0.135 | 0.263 | 9.1173E-29 | ERY | NABP1 |
| 5.5601E-33 | -0.3453977 | 0.648 | 0.714 | 1.3135E-28 | ERY | UBE2D3 |
| 8.6067E-33 | -0.2808253 | 0.137 | 0.272 | 2.0332E-28 | ERY | DIAPH1 |
| 1.5813E-32 | -0.2763019 | 0.086 | 0.212 | 3.7355E-28 | ERY | RNF19A |

| | | | | | | |
|---|---|---|---|---|---|---|
| 1.5872E-32 | -0.4524479 | 0.666 | 0.711 | 3.7494E-28 | ERY | SDCBP |
| 2.0563E-32 | -0.2583481 | 0.1 | 0.228 | 4.8576E-28 | ERY | KDM2A |
| 2.9398E-32 | -0.4434393 | 0.222 | 0.348 | 6.9447E-28 | ERY | CPEB4 |
| 3.1365E-32 | -0.4684737 | 0.284 | 0.401 | 7.4093E-28 | ERY | ARGLU1 |
| 3.6072E-32 | -1.1658299 | 0.276 | 0.14 | 8.5212E-28 | ERY | CXCL3 |
| 4.8818E-32 | -0.3167554 | 0.122 | 0.25 | 1.1532E-27 | ERY | SNX9 |
| 4.9686E-32 | -0.3256547 | 0.155 | 0.286 | 1.1737E-27 | ERY | TMEM30A |
| 5.6426E-32 | -0.2712734 | 0.083 | 0.206 | 1.333E-27 | ERY | MUC20-OT1 |
| 6.0223E-32 | -0.2585796 | 0.078 | 0.201 | 1.4226E-27 | ERY | PPP1R15B |
| 7.1068E-32 | -0.2869005 | 0.055 | 0.169 | 1.6788E-27 | ERY | CEP170 |
| 1.2282E-31 | -0.4081671 | 0.026 | 0.128 | 2.9014E-27 | ERY | SRD5A3 |
| 1.3374E-31 | -0.3033733 | 0.117 | 0.243 | 3.1593E-27 | ERY | SP100 |
| 1.6993E-31 | -0.2643911 | 0.076 | 0.195 | 4.0143E-27 | ERY | FCHO2 |
| 3.3379E-31 | -0.2666616 | 0.093 | 0.216 | 7.8852E-27 | ERY | USP34 |
| 4.164E-31 | -0.2858258 | 0.016 | 0.113 | 9.8367E-27 | ERY | STS |
| 4.9664E-31 | -0.264096 | 0.094 | 0.217 | 1.1732E-26 | ERY | PHF3 |
| 7.8339E-31 | -0.4264498 | 0.117 | 0.238 | 1.8506E-26 | ERY | TOB1 |
| 1.0226E-30 | -0.2683175 | 0.124 | 0.251 | 2.4158E-26 | ERY | SMARCA5 |
| 1.1087E-30 | -1.4187978 | 0.757 | 0.708 | 2.619E-26 | ERY | ANXA1 |
| 1.1439E-30 | -0.2902181 | 0.12 | 0.245 | 2.7023E-26 | ERY | KANSL1 |
| 1.2732E-30 | -0.2595497 | 0.081 | 0.201 | 3.0078E-26 | ERY | SECISBP2L |
| 1.8077E-30 | -0.3069778 | 0.109 | 0.231 | 4.2703E-26 | ERY | TRIM56 |
| 2.411E-30 | -0.2613141 | 0.04 | 0.146 | 5.6955E-26 | ERY | PIK3R1 |
| 3.3E-30 | -1.1831771 | 0.034 | 0.135 | 7.7956E-26 | ERY | PLAT |
| 3.6986E-30 | -0.362767 | 0.238 | 0.358 | 8.7372E-26 | ERY | SEPT7 |
| 4.4967E-30 | -0.4011324 | 0.15 | 0.27 | 1.0623E-25 | ERY | AFDN |
| 4.7881E-30 | -0.2533449 | 0.082 | 0.199 | 1.1311E-25 | ERY | RAB8B |
| 6.5157E-30 | -0.6367039 | 0.984 | 0.952 | 1.5392E-25 | ERY | H3F3B |
| 6.8141E-30 | -0.5708291 | 0.201 | 0.313 | 1.6097E-25 | ERY | IRF1 |
| 1.377E-29 | -0.2767481 | 0.039 | 0.143 | 3.2529E-25 | ERY | EMP2 |
| 1.8181E-29 | -0.3492453 | 0.08 | 0.194 | 4.295E-25 | ERY | ABLIM1 |
| 2.0035E-29 | -0.4354005 | 0.478 | 0.549 | 4.7329E-25 | ERY | C4orf3 |
| 2.3455E-29 | -0.9334124 | 0.13 | 0.237 | 5.5407E-25 | ERY | SLC2A3 |
| 2.5866E-29 | -0.3720691 | 0.142 | 0.261 | 6.1103E-25 | ERY | PDLIM5 |
| 2.9656E-29 | -0.6190512 | 0.116 | 0.23 | 7.0057E-25 | ERY | HEXIM1 |
| 3.3311E-29 | -0.4120989 | 0.047 | 0.15 | 7.869E-25 | ERY | ALDH1A3 |
| 6.532E-29 | -0.4325674 | 0.125 | 0.24 | 1.543E-24 | ERY | NFKBIZ |
| 8.4411E-29 | -0.42653 | 0.275 | 0.377 | 1.994E-24 | ERY | METRNL |
| 9.9056E-29 | -0.2923031 | 0.09 | 0.204 | 2.34E-24 | ERY | ZNF91 |
| 1.0269E-28 | -0.3881324 | 0.402 | 0.228 | 2.4258E-24 | ERY | PLIN2 |
| 1.0676E-28 | -0.2989156 | 0.099 | 0.216 | 2.5219E-24 | ERY | SETX |
| 1.3993E-28 | -0.3374665 | 0.185 | 0.31 | 3.3055E-24 | ERY | BNIP3L |
| 1.9214E-28 | -0.3190309 | 0.169 | 0.291 | 4.5389E-24 | ERY | CLINT1 |
| 2.1249E-28 | -0.3514598 | 0.217 | 0.334 | 5.0197E-24 | ERY | TCF25 |
| 2.7211E-28 | -0.2589015 | 0.05 | 0.153 | 6.4281E-24 | ERY | SMAP2 |
| 3.672E-28 | -0.6300796 | 0.307 | 0.397 | 8.6744E-24 | ERY | HIF1A |
| 3.8746E-28 | -0.2727557 | 0.108 | 0.226 | 9.153E-24 | ERY | MAP3K2 |
| 5.493E-28 | -0.2914553 | 0.157 | 0.278 | 1.2976E-23 | ERY | SF1 |
| 7.0546E-28 | -0.4732197 | 0.066 | 0.171 | 1.6665E-23 | ERY | FXYD3 |
| 9.3813E-28 | -0.364278 | 0.017 | 0.106 | 2.2161E-23 | ERY | CLIC3 |

| | | | | | | |
|---|---|---|---|---|---|---|
| 9.922E-28 | -1.0693582 | 0.424 | 0.225 | 2.3439E-23 | ERY | PRSS22 |
| 1.354E-27 | -0.258127 | 0.042 | 0.141 | 3.1986E-23 | ERY | MTUS1 |
| 1.6219E-27 | -0.5401655 | 0.246 | 0.349 | 3.8315E-23 | ERY | MIDN |
| 2.3098E-27 | -0.448394 | 0.198 | 0.31 | 5.4565E-23 | ERY | ZNF292 |
| 3.1168E-27 | -0.3150967 | 0.157 | 0.273 | 7.3627E-23 | ERY | FAM133B |
| 3.4402E-27 | -0.4509186 | 0.122 | 0.232 | 8.1267E-23 | ERY | IVNS1ABP |
| 3.5328E-27 | -0.2637216 | 0.102 | 0.217 | 8.3456E-23 | ERY | DNTTIP2 |
| 4.9041E-27 | -1.5258549 | 0.022 | 0.109 | 1.1585E-22 | ERY | CRCT1 |
| 6.6291E-27 | -0.2738978 | 0.086 | 0.195 | 1.566E-22 | ERY | FNDC3A |
| 1.1848E-26 | -0.3452276 | 0.093 | 0.201 | 2.7987E-22 | ERY | SERTAD2 |
| 1.3038E-26 | -0.3083684 | 0.173 | 0.29 | 3.08E-22 | ERY | BAZ1A |
| 1.9593E-26 | -0.2592025 | 0.086 | 0.194 | 4.6284E-22 | ERY | CHD9 |
| 2.4222E-26 | -0.3123093 | 0.15 | 0.263 | 5.722E-22 | ERY | TRA2B |
| 2.656E-26 | -0.2600354 | 0.016 | 0.1 | 6.2743E-22 | ERY | RNF39 |
| 2.748E-26 | -0.8614667 | 0.16 | 0.266 | 6.4916E-22 | ERY | AREG |
| 2.7901E-26 | -0.3904926 | 0.065 | 0.164 | 6.591E-22 | ERY | AQP9 |
| 2.9742E-26 | -0.3035377 | 0.255 | 0.369 | 7.0259E-22 | ERY | RAB7A |
| 4.2135E-26 | -0.3005488 | 0.159 | 0.273 | 9.9536E-22 | ERY | UBE2D1 |
| 5.1709E-26 | -0.4240179 | 0.083 | 0.185 | 1.2215E-21 | ERY | SAMSN1 |
| 6.1535E-26 | -0.4237788 | 0.264 | 0.363 | 1.4537E-21 | ERY | CFLAR |
| 7.7172E-26 | -0.2613283 | 0.103 | 0.212 | 1.823E-21 | ERY | PCF11 |
| 7.8494E-26 | -0.2594994 | 0.076 | 0.18 | 1.8543E-21 | ERY | RERE |
| 1.4859E-25 | -0.3017748 | 0.934 | 0.73 | 3.5101E-21 | ERY | CTSB |
| 1.672E-25 | -0.2512214 | 0.11 | 0.221 | 3.9497E-21 | ERY | TRIM38 |
| 2.1464E-25 | -0.2927371 | 0.975 | 0.828 | 5.0703E-21 | ERY | PPDPF |
| 2.732E-25 | -0.255616 | 0.157 | 0.272 | 6.4538E-21 | ERY | BRD4 |
| 3.194E-25 | -0.377939 | 0.183 | 0.291 | 7.5453E-21 | ERY | MACF1 |
| 3.4536E-25 | -0.2828031 | 0.11 | 0.218 | 8.1585E-21 | ERY | NKTR |
| 3.6857E-25 | -0.3589375 | 0.198 | 0.305 | 8.7067E-21 | ERY | DDX24 |
| 6.747E-25 | -0.2728798 | 0.073 | 0.172 | 1.5939E-20 | ERY | JMY |
| 8.3981E-25 | -0.367649 | 0.041 | 0.131 | 1.9839E-20 | ERY | AC023157.3 |
| 9.0097E-25 | -0.2657346 | 0.02 | 0.101 | 2.1284E-20 | ERY | LY6K |
| 9.7665E-25 | -0.8514927 | 0.046 | 0.135 | 2.3071E-20 | ERY | RHCG |
| 9.8176E-25 | -0.8750438 | 0.19 | 0.282 | 2.3192E-20 | ERY | DUSP5 |
| 1.3982E-24 | -0.2517418 | 0.159 | 0.271 | 3.303E-20 | ERY | KHDRBS1 |
| 1.7883E-24 | -0.3328311 | 0.079 | 0.179 | 4.2245E-20 | ERY | NR4A1 |
| 2.3692E-24 | -0.8584637 | 0.048 | 0.137 | 5.5968E-20 | ERY | EREG |
| 2.652E-24 | -0.563538 | 0.382 | 0.214 | 6.2647E-20 | ERY | ALOX5AP |
| 2.8583E-24 | -0.3078893 | 0.155 | 0.262 | 6.7522E-20 | ERY | ARHGAP5 |
| 3.0178E-24 | -0.2799578 | 0.15 | 0.258 | 7.129E-20 | ERY | GABARAPL1 |
| 3.2172E-24 | -0.2990063 | 0.211 | 0.32 | 7.6E-20 | ERY | PTBP3 |
| 4.3786E-24 | -0.3619988 | 0.266 | 0.361 | 1.0344E-19 | ERY | PLSCR1 |
| 6.9076E-24 | -0.3695295 | 0.229 | 0.328 | 1.6318E-19 | ERY | FOSL2 |
| 1.0055E-23 | -0.2518493 | 0.786 | 0.516 | 2.3753E-19 | ERY | CEBPB |
| 1.9103E-23 | -0.3139917 | 0.081 | 0.176 | 4.5128E-19 | ERY | CORO1A |
| 4.1157E-23 | -0.419345 | 0.218 | 0.315 | 9.7224E-19 | ERY | FNDC3B |
| 5.957E-23 | -0.2583979 | 0.093 | 0.19 | 1.4072E-18 | ERY | HCLS1 |
| 8.405E-23 | -0.3907979 | 0.299 | 0.165 | 1.9855E-18 | ERY | FCN1 |
| 1.7129E-22 | -0.2869117 | 0.19 | 0.293 | 4.0464E-18 | ERY | CHMP2B |
| 1.8655E-22 | -0.4474722 | 0.062 | 0.151 | 4.4069E-18 | ERY | SMC4 |

| | | | | | | |
|---|---|---|---|---|---|---|
| 1.9321E-22 | -0.2966538 | 0.097 | 0.191 | 4.5642E-18 | ERY | RAC2 |
| 2.1539E-22 | -0.2518915 | 0.087 | 0.182 | 5.0882E-18 | ERY | FMNL1 |
| 2.3275E-22 | -1.4076756 | 0.031 | 0.11 | 5.4982E-18 | ERY | CNFN |
| 2.6646E-22 | -0.4029589 | 0.113 | 0.21 | 6.2946E-18 | ERY | KCNQ1OT1 |
| 2.9123E-22 | -0.2511663 | 0.097 | 0.195 | 6.8796E-18 | ERY | ITPRIPL2 |
| 9.2768E-22 | -0.2818104 | 0.169 | 0.27 | 2.1915E-17 | ERY | SMG1 |
| 9.6559E-22 | -0.5707563 | 0.548 | 0.573 | 2.281E-17 | ERY | VMP1 |
| 1.228E-21 | -0.2513777 | 0.049 | 0.133 | 2.9009E-17 | ERY | HIST1H2BC |
| 1.4805E-21 | -0.3165194 | 0.092 | 0.184 | 3.4975E-17 | ERY | PFKFB3 |
| 2.6602E-21 | -1.5219768 | 0.676 | 0.611 | 6.2843E-17 | ERY | C15orf48 |
| 3.6035E-21 | -0.6222849 | 0.41 | 0.458 | 8.5125E-17 | ERY | RAB11FIP1 |
| 3.7961E-21 | -0.2966057 | 0.163 | 0.26 | 8.9674E-17 | ERY | CHD2 |
| 6.1044E-21 | -0.2853989 | 0.073 | 0.159 | 1.442E-16 | ERY | TSPAN6 |
| 6.5171E-21 | -0.3450177 | 0.32 | 0.398 | 1.5395E-16 | ERY | CTNNA1 |
| 8.2926E-21 | -0.2833553 | 0.069 | 0.154 | 1.959E-16 | ERY | EVI2B |
| 8.3642E-21 | -0.3626369 | 0.352 | 0.43 | 1.9759E-16 | ERY | EIF3A |
| 8.9061E-21 | -0.3428881 | 0.197 | 0.292 | 2.1039E-16 | ERY | CHMP1B |
| 1.2035E-20 | -0.2556768 | 0.148 | 0.246 | 2.843E-16 | ERY | ZRANB2 |
| 1.4885E-20 | -0.2972584 | 0.234 | 0.327 | 3.5164E-16 | ERY | TGOLN2 |
| 1.771E-20 | -0.3461515 | 0.244 | 0.333 | 4.1837E-16 | ERY | RBM47 |
| 2.6E-20 | -0.2846594 | 0.05 | 0.13 | 6.1419E-16 | ERY | LRG1 |
| 2.8481E-20 | -0.2711423 | 0.174 | 0.27 | 6.7281E-16 | ERY | TOR1AIP2 |
| 3.1563E-20 | -0.580469 | 0.099 | 0.183 | 7.4561E-16 | ERY | OLR1 |
| 3.3874E-20 | -0.6354998 | 0.254 | 0.325 | 8.002E-16 | ERY | VEGFA |
| 3.7317E-20 | -0.4280187 | 0.106 | 0.194 | 8.8153E-16 | ERY | PLK2 |
| 7.2314E-20 | -0.457013 | 0.988 | 0.956 | 1.7083E-15 | ERY | H3F3A |
| 7.2534E-20 | -0.6824634 | 0.719 | 0.676 | 1.7135E-15 | ERY | IER2 |
| 8.7988E-20 | -0.3453495 | 0.067 | 0.148 | 2.0785E-15 | ERY | FPR1 |
| 1.1267E-19 | -0.2823502 | 0.172 | 0.264 | 2.6615E-15 | ERY | MBP |
| 1.2276E-19 | -0.2624414 | 0.093 | 0.18 | 2.9E-15 | ERY | GLIPR1 |
| 1.6246E-19 | -0.7166813 | 0.653 | 0.63 | 3.8377E-15 | ERY | ZFP36L1 |
| 2.0426E-19 | -0.3764873 | 0.315 | 0.393 | 4.8253E-15 | ERY | RSRC2 |
| 2.1569E-19 | -0.5369072 | 0.217 | 0.296 | 5.0952E-15 | ERY | ATP13A3 |
| 2.318E-19 | -0.3075014 | 0.036 | 0.11 | 5.4757E-15 | ERY | TCHP |
| 2.633E-19 | -0.4241122 | 0.377 | 0.429 | 6.2201E-15 | ERY | EFHD2 |
| 4.4206E-19 | -0.4435153 | 0.171 | 0.252 | 1.0443E-14 | ERY | KLF5 |
| 5.6328E-19 | -0.2625939 | 0.095 | 0.18 | 1.3306E-14 | ERY | CCDC88A |
| 6.2004E-19 | -0.2500149 | 0.114 | 0.201 | 1.4647E-14 | ERY | ARHGAP18 |
| 6.5941E-19 | -0.2820027 | 0.171 | 0.262 | 1.5577E-14 | ERY | ABRACL |
| 1.0371E-18 | -0.313432 | 0.134 | 0.22 | 2.4499E-14 | ERY | DSE |
| 1.0775E-18 | -0.3028817 | 0.24 | 0.327 | 2.5453E-14 | ERY | YPEL5 |
| 1.9946E-18 | -0.3114186 | 0.341 | 0.415 | 4.7118E-14 | ERY | DAZAP2 |
| 2.0412E-18 | -0.3328708 | 0.078 | 0.156 | 4.8218E-14 | ERY | TMPRSS4 |
| 3.5045E-18 | -0.4044585 | 0.873 | 0.829 | 8.2787E-14 | ERY | HNRNPA2B1 |
| 4.0056E-18 | -0.4713306 | 0.587 | 0.327 | 9.4625E-14 | ERY | LAPTM5 |
| 4.39E-18 | -0.4997039 | 0.443 | 0.277 | 1.0371E-13 | ERY | IFITM2 |
| 5.351E-18 | -0.2731467 | 0.118 | 0.204 | 1.2641E-13 | ERY | AMD1 |
| 7.7269E-18 | -0.4587755 | 0.099 | 0.177 | 1.8253E-13 | ERY | SPNS2 |
| 8.1446E-18 | -0.2593075 | 0.222 | 0.311 | 1.924E-13 | ERY | YY1 |
| 1.0197E-17 | -0.6071018 | 0.874 | 0.544 | 2.4087E-13 | ERY | ELF3 |

| | | | | | | |
|---|---|---|---|---|---|---|
| 1.2358E-17 | -0.2580438 | 0.206 | 0.294 | 2.9193E-13 | ERY | STK24 |
| 1.3964E-17 | -0.4775831 | 0.11 | 0.187 | 3.2986E-13 | ERY | C5AR1 |
| 1.5631E-17 | -0.3375519 | 0.124 | 0.207 | 3.6925E-13 | ERY | CSNK1E |
| 1.7997E-17 | -0.3799596 | 0.112 | 0.191 | 4.2513E-13 | ERY | DUSP6 |
| 2.4316E-17 | -0.6676222 | 0.455 | 0.471 | 5.7442E-13 | ERY | DUSP1 |
| 2.8516E-17 | -0.3739603 | 0.103 | 0.182 | 6.7364E-13 | ERY | DST |
| 3.7663E-17 | -0.3011967 | 0.051 | 0.122 | 8.897E-13 | ERY | EGLN3 |
| 4.0849E-17 | -0.3266689 | 0.094 | 0.171 | 9.6498E-13 | ERY | SLA |
| 5.8277E-17 | -0.3486259 | 0.837 | 0.798 | 1.3767E-12 | ERY | ARPC2 |
| 8.3566E-17 | -0.2706993 | 0.303 | 0.381 | 1.9741E-12 | ERY | PAPOLA |
| 1.0081E-16 | -0.4081884 | 0.099 | 0.173 | 2.3813E-12 | ERY | CXCR4 |
| 1.8948E-16 | -0.302649 | 0.219 | 0.298 | 4.476E-12 | ERY | ZFAND6 |
| 2.7057E-16 | -0.2801775 | 0.127 | 0.207 | 6.3918E-12 | ERY | MAP3K8 |
| 3.1508E-16 | -0.2765329 | 0.206 | 0.285 | 7.4431E-12 | ERY | DYNC1H1 |
| 4.0326E-16 | -0.3442553 | 0.554 | 0.581 | 9.5263E-12 | ERY | PGK1 |
| 4.6866E-16 | -0.2680895 | 0.352 | 0.422 | 1.1071E-11 | ERY | CAPZB |
| 5.2056E-16 | -0.4629472 | 0.263 | 0.325 | 1.2297E-11 | ERY | LIMS1 |
| 5.4164E-16 | -0.3999225 | 0.463 | 0.495 | 1.2795E-11 | ERY | RSRP1 |
| 7.5943E-16 | -0.5368245 | 0.614 | 0.605 | 1.794E-11 | ERY | EZR |
| 1.0268E-15 | -0.8251478 | 0.118 | 0.19 | 2.4256E-11 | ERY | RARRES3 |
| 1.035E-15 | -0.3733489 | 0.09 | 0.161 | 2.4449E-11 | ERY | MALL |
| 1.0497E-15 | -0.8862237 | 0.207 | 0.12 | 2.4796E-11 | ERY | CCL3 |
| 1.2767E-15 | -0.3704588 | 0.947 | 0.903 | 3.016E-11 | ERY | CFL1 |
| 2.6847E-15 | -0.8297836 | 0.246 | 0.293 | 6.3421E-11 | ERY | RND3 |
| 8.1566E-15 | -3.745228 | 0.545 | 0.542 | 1.9268E-10 | ERY | PLCG2 |
| 1.9293E-14 | -0.2523003 | 0.06 | 0.124 | 4.5575E-10 | ERY | KRT80 |
| 1.9855E-14 | -0.3074954 | 0.271 | 0.333 | 4.6902E-10 | ERY | MSN |
| 2.7252E-14 | -0.7168728 | 0.909 | 0.697 | 6.4377E-10 | ERY | SOD2 |
| 4.1072E-14 | -0.6659987 | 0.844 | 0.761 | 9.7024E-10 | ERY | BTG1 |
| 9.1949E-14 | -0.5939641 | 0.754 | 0.496 | 2.1721E-09 | ERY | HSPA1B |
| 9.4841E-14 | -0.3792268 | 0.138 | 0.202 | 2.2404E-09 | ERY | ITGAX |
| 1.0113E-13 | -0.4168359 | 0.15 | 0.215 | 2.389E-09 | ERY | BASP1 |
| 1.2688E-13 | -0.325945 | 0.085 | 0.15 | 2.9973E-09 | ERY | RGS2 |
| 1.3332E-13 | -0.3177198 | 0.194 | 0.263 | 3.1494E-09 | ERY | ARRDC3 |
| 4.4155E-13 | -0.3107703 | 0.098 | 0.162 | 1.0431E-08 | ERY | SERPINB9 |
| 8.9573E-13 | -0.4997097 | 0.061 | 0.118 | 2.116E-08 | ERY | MNDA |
| 1.0487E-12 | -0.2682435 | 0.256 | 0.32 | 2.4774E-08 | ERY | CMTM6 |
| 1.2856E-12 | -0.4042086 | 0.124 | 0.185 | 3.0371E-08 | ERY | SDCBP2 |
| 2.4153E-12 | -0.2741148 | 0.052 | 0.107 | 5.7058E-08 | ERY | CSF3R |
| 2.469E-12 | -0.4768914 | 0.28 | 0.33 | 5.8325E-08 | ERY | SLC38A2 |
| 3.1219E-12 | -0.4360248 | 0.067 | 0.124 | 7.3749E-08 | ERY | AGR2 |
| 3.8075E-12 | -0.2779767 | 0.099 | 0.159 | 8.9944E-08 | ERY | TTC9 |
| 4.3567E-12 | -0.3343398 | 0.054 | 0.108 | 1.0292E-07 | ERY | THBD |
| 1.0321E-11 | -0.2680632 | 0.058 | 0.112 | 2.4382E-07 | ERY | SMIM5 |
| 1.2072E-11 | -0.2787266 | 0.262 | 0.32 | 2.8518E-07 | ERY | C6orf62 |
| 1.2493E-11 | -0.3663429 | 0.127 | 0.185 | 2.9513E-07 | ERY | SGK1 |
| 1.4972E-11 | -3.0814977 | 0.878 | 0.664 | 3.5368E-07 | ERY | S100A8 |
| 2.4722E-11 | -0.3654669 | 0.121 | 0.177 | 5.8402E-07 | ERY | C6orf132 |
| 2.5435E-11 | -0.3819213 | 0.731 | 0.524 | 6.0085E-07 | ERY | CTSS |
| 4.6025E-11 | -0.3672637 | 0.129 | 0.187 | 1.0873E-06 | ERY | ACSL1 |

| | | | | | | |
|---|---|---|---|---|---|---|
| 4.6417E-11 | -0.7771851 | 0.247 | 0.287 | 1.0965E-06 | ERY | HMOX1 |
| 6.6921E-11 | -0.3184322 | 0.195 | 0.249 | 1.5809E-06 | ERY | SERINC2 |
| 1.0455E-10 | -0.3771429 | 0.162 | 0.216 | 2.4697E-06 | ERY | CREM |
| 1.1799E-10 | -0.3417559 | 0.077 | 0.13 | 2.7872E-06 | ERY | GBP1 |
| 2.602E-10 | -0.4567852 | 0.234 | 0.276 | 6.1467E-06 | ERY | ADM |
| 2.614E-10 | -0.5601106 | 0.964 | 0.882 | 6.1752E-06 | ERY | HSP90AB1 |
| 2.6956E-10 | -0.5184252 | 0.648 | 0.446 | 6.3677E-06 | ERY | HSPH1 |
| 3.3891E-10 | -0.3029594 | 0.563 | 0.575 | 8.0061E-06 | ERY | PPP1CB |
| 3.6622E-10 | -0.3554668 | 0.138 | 0.19 | 8.6513E-06 | ERY | FCGR2A |
| 4.9075E-10 | -0.2698475 | 0.12 | 0.173 | 1.1593E-05 | ERY | SAMHD1 |
| 5.4574E-10 | -0.2515057 | 0.223 | 0.277 | 1.2892E-05 | ERY | DDX21 |
| 6.9838E-10 | -0.4542657 | 0.066 | 0.114 | 1.6498E-05 | ERY | CYP1B1 |
| 1.0696E-09 | -0.5095114 | 0.703 | 0.496 | 2.5266E-05 | ERY | DNAJB1 |
| 1.1903E-09 | -0.9446934 | 0.605 | 0.378 | 2.8117E-05 | ERY | FCER1G |
| 1.2243E-09 | -0.550102 | 0.178 | 0.223 | 2.8922E-05 | ERY | HES1 |
| 2.5812E-09 | -0.9011168 | 0.844 | 0.631 | 6.0976E-05 | ERY | MTRNR2L12 |
| 6.863E-09 | -0.3342164 | 0.175 | 0.217 | 0.00016212 | ERY | SPI1 |
| 7.4419E-09 | -0.8984502 | 0.664 | 0.443 | 0.0001758 | ERY | SOX4 |
| 7.4773E-09 | -0.3688564 | 0.285 | 0.32 | 0.00017664 | ERY | BRD2 |
| 1.3477E-08 | -0.4304251 | 0.329 | 0.227 | 0.00031837 | ERY | ISG15 |
| 1.4733E-08 | -0.3302913 | 0.164 | 0.21 | 0.00034804 | ERY | LY6E |
| 1.4921E-08 | -0.2849997 | 0.376 | 0.397 | 0.00035247 | ERY | H2AFY |
| 2.5105E-08 | -0.4040884 | 0.439 | 0.287 | 0.00059305 | ERY | GPRC5A |
| 3.2351E-08 | -0.3220807 | 0.361 | 0.244 | 0.00076424 | ERY | EGR1 |
| 3.3026E-08 | -0.2773972 | 0.102 | 0.147 | 0.00078017 | ERY | MS4A7 |
| 6.2717E-08 | -1.6808235 | 0.836 | 0.561 | 0.00148157 | ERY | SRGN |
| 7.3103E-08 | -0.6706068 | 0.074 | 0.114 | 0.00172691 | ERY | IL7R |
| 1.2666E-07 | -0.5528478 | 0.346 | 0.347 | 0.002992 | ERY | ZFP36L2 |
| 2.8122E-07 | -0.2983931 | 0.51 | 0.501 | 0.00664336 | ERY | ASAH1 |
| 4.5038E-07 | -0.8854976 | 0.081 | 0.119 | 0.01063927 | ERY | CXCL2 |
| 5.583E-07 | -0.7401008 | 0.269 | 0.282 | 0.0131887 | ERY | MAFB |
| 8.1918E-07 | -0.3355736 | 0.967 | 0.833 | 0.01935146 | ERY | SQSTM1 |
| 1.076E-06 | -0.2928201 | 0.857 | 0.8 | 0.02541718 | ERY | HMGB1 |
| 1.1247E-06 | -0.2519586 | 0.308 | 0.337 | 0.02656971 | ERY | LUC7L3 |
| 1.1983E-06 | -0.2708258 | 0.171 | 0.207 | 0.02830742 | ERY | PARD6B |
| 1.6961E-06 | -0.3539557 | 0.227 | 0.259 | 0.04006753 | ERY | IDI1 |
| 1.863E-06 | -0.2548714 | 0.26 | 0.293 | 0.04400987 | ERY | HNRNPAB |
| 2.4957E-06 | -0.4043901 | 0.085 | 0.121 | 0.05895613 | ERY | HILPDA |
| 3.8391E-06 | -0.4497504 | 0.308 | 0.315 | 0.09069216 | ERY | HIST1H2AC |
| 4.0122E-06 | -0.2518115 | 0.073 | 0.108 | 0.09477983 | ERY | SLC16A10 |
| 4.0276E-06 | -0.2888681 | 0.994 | 0.99 | 0.09514357 | ERY | B2M |
| 4.7871E-06 | -0.3157382 | 0.148 | 0.184 | 0.11308673 | ERY | INSIG1 |
| 7.6781E-06 | -0.4579535 | 0.345 | 0.347 | 0.18137965 | ERY | IER5 |
| 7.9847E-06 | -0.4380901 | 0.845 | 0.634 | 0.18862252 | ERY | TXNIP |
| 1.1071E-05 | -0.6654861 | 0.109 | 0.14 | 0.26153964 | ERY | CXCL1 |
| 1.6911E-05 | -0.2995446 | 0.783 | 0.729 | 0.39949287 | ERY | ATP6V1G1 |
| 3.507E-05 | -0.3510317 | 0.253 | 0.27 | 0.82846191 | ERY | CCNG2 |
| 4.299E-05 | -0.3044333 | 0.338 | 0.343 | 1 | ERY | WSB1 |
| 4.611E-05 | -0.6984761 | 0.686 | 0.599 | 1 | ERY | NFKBIA |
| 4.8039E-05 | -0.3784076 | 0.152 | 0.177 | 1 | ERY | ANPEP |

| | | | | | | |
|---|---|---|---|---|---|---|
| 5.7036E-05 | -0.3045856 | 0.122 | 0.152 | 1 | ERY | SLC7A11 |
| 6.5869E-05 | -0.3079216 | 0.554 | 0.529 | 1 | ERY | PTTG1IP |
| 0.00011018 | -0.2596648 | 0.283 | 0.205 | 1 | ERY | LST1 |
| 0.00011094 | -0.5547545 | 0.281 | 0.285 | 1 | ERY | SLC25A37 |
| 0.00012657 | -0.409268 | 0.131 | 0.154 | 1 | ERY | CDC42EP5 |
| 0.00018189 | -1.4007886 | 0.199 | 0.141 | 1 | ERY | SPRR2A |
| 0.00021846 | -0.3571921 | 0.267 | 0.268 | 1 | ERY | CD53 |
| 0.0002266 | -0.4447563 | 0.993 | 0.972 | 1 | ERY | ACTB |
| 0.00023022 | -0.3703503 | 0.294 | 0.299 | 1 | ERY | FAM129B |
| 0.00023331 | -0.3151294 | 0.237 | 0.246 | 1 | ERY | GMFG |
| 0.00024539 | -0.3237944 | 0.565 | 0.512 | 1 | ERY | ANXA11 |
| 0.00030836 | -0.2547962 | 0.112 | 0.136 | 1 | ERY | GCNT3 |
| 0.00036567 | -0.3049927 | 0.185 | 0.201 | 1 | ERY | NCF2 |
| 0.00036776 | -0.5548177 | 0.436 | 0.403 | 1 | ERY | RNF213 |
| 0.00053155 | -0.2858869 | 0.378 | 0.367 | 1 | ERY | MYH9 |
| 0.00060805 | -0.4201114 | 0.094 | 0.117 | 1 | ERY | HSPA2 |
| 0.00061864 | -0.2809674 | 0.19 | 0.207 | 1 | ERY | SLC43A2 |
| 0.00090888 | -0.5659117 | 0.452 | 0.421 | 1 | ERY | TXNRD1 |
| 0.00114314 | -0.2589473 | 0.379 | 0.367 | 1 | ERY | CTSH |
| 0.00159477 | -0.3079103 | 0.152 | 0.172 | 1 | ERY | DNAJA4 |
| 0.00176846 | -2.28783 | 0.661 | 0.482 | 1 | ERY | CXCL8 |
| 0.00206153 | -0.322319 | 0.222 | 0.17 | 1 | ERY | HLA-DQB1 |
| 0.00210198 | -0.3509639 | 0.988 | 0.909 | 1 | ERY | RPS8 |
| 0.00241395 | -0.3569381 | 0.424 | 0.387 | 1 | ERY | AHNAK |
| 0.00293972 | -0.2824501 | 0.378 | 0.366 | 1 | ERY | RALBP1 |
| 0.00376949 | -0.261834 | 0.197 | 0.209 | 1 | ERY | SOCS3 |
| 0.00825837 | -0.2505116 | 0.196 | 0.209 | 1 | ERY | MDM2 |
| 0 | 5.79743995 | 0.994 | 0.279 | 0 | UGEC1 | SPRR3 |
| 0 | 4.63288443 | 0.999 | 0.425 | 0 | UGEC1 | LCN2 |
| 0 | 4.26303376 | 0.99 | 0.252 | 0 | UGEC1 | KRT13 |
| 0 | 3.78025251 | 0.992 | 0.673 | 0 | UGEC1 | S100A9 |
| 0 | 3.54943797 | 0.674 | 0.021 | 0 | UGEC1 | LYPD2 |
| 0 | 3.50544322 | 0.573 | 0.055 | 0 | UGEC1 | SPRR2A |
| 0 | 3.50128186 | 0.786 | 0.034 | 0 | UGEC1 | ECM1 |
| 0 | 3.44994324 | 0.504 | 0.024 | 0 | UGEC1 | CRCT1 |
| 0 | 3.34472742 | 0.506 | 0.025 | 0 | UGEC1 | CNFN |
| 0 | 3.29455406 | 0.837 | 0.107 | 0 | UGEC1 | PRSS22 |
| 0 | 3.12036417 | 0.578 | 0.017 | 0 | UGEC1 | SPRR1B |
| 0 | 3.00609766 | 0.811 | 0.033 | 0 | UGEC1 | DUOXA2 |
| 0 | 3.00208553 | 0.914 | 0.227 | 0 | UGEC1 | HPGD |
| 0 | 2.95418142 | 0.977 | 0.608 | 0 | UGEC1 | S100A8 |
| 0 | 2.95257351 | 0.732 | 0.086 | 0 | UGEC1 | HOPX |
| 0 | 2.94491132 | 0.604 | 0.034 | 0 | UGEC1 | PLAT |
| 0 | 2.91222154 | 0.852 | 0.269 | 0 | UGEC1 | MAL |
| 0 | 2.8650322 | 0.6 | 0.017 | 0 | UGEC1 | SPINK5 |
| 0 | 2.85571814 | 0.819 | 0.025 | 0 | UGEC1 | MUC21 |
| 0 | 2.85503489 | 0.865 | 0.043 | 0 | UGEC1 | DUOX2 |
| 0 | 2.77524008 | 0.961 | 0.213 | 0 | UGEC1 | PSCA |
| 0 | 2.72072384 | 0.163 | 0.008 | 0 | UGEC1 | S100A7 |
| 0 | 2.63360066 | 0.67 | 0.031 | 0 | UGEC1 | CXCL6 |

| | | | | | | |
|---|---|---|---|---|---|---|
| 0 | 2.62299526 | 0.776 | 0.032 | 0 | UGEC1 | CEACAM5 |
| 0 | 2.59861893 | 0.66 | 0.103 | 0 | UGEC1 | KRT17 |
| 0 | 2.58990033 | 0.838 | 0.474 | 0 | UGEC1 | WFDC2 |
| 0 | 2.58873211 | 0.785 | 0.025 | 0 | UGEC1 | PITX1 |
| 0 | 2.58246705 | 0.79 | 0.027 | 0 | UGEC1 | PRSS27 |
| 0 | 2.53765553 | 0.932 | 0.663 | 0 | UGEC1 | ANXA1 |
| 0 | 2.53291019 | 0.96 | 0.294 | 0 | UGEC1 | S100P |
| 0 | 2.45520317 | 0.632 | 0.029 | 0 | UGEC1 | RHCG |
| 0 | 2.42920708 | 0.621 | 0.081 | 0 | UGEC1 | RDH10 |
| 0 | 2.41638438 | 0.656 | 0.027 | 0 | UGEC1 | TMPRSS11B |
| 0 | 2.35507357 | 0.918 | 0.381 | 0 | UGEC1 | ERO1A |
| 0 | 2.33531043 | 0.363 | 0.03 | 0 | UGEC1 | SPRR2D |
| 0 | 2.32773955 | 0.72 | 0.028 | 0 | UGEC1 | CEACAM6 |
| 0 | 2.30186654 | 0.552 | 0.012 | 0 | UGEC1 | ZG16B |
| 0 | 2.29945235 | 0.871 | 0.137 | 0 | UGEC1 | LMO7 |
| 0 | 2.29757094 | 0.787 | 0.064 | 0 | UGEC1 | TMPRSS2 |
| 0 | 2.29523066 | 0.722 | 0.039 | 0 | UGEC1 | KRT15 |
| 0 | 2.27848956 | 0.731 | 0.025 | 0 | UGEC1 | CEACAM7 |
| 0 | 2.2487831 | 0.829 | 0.165 | 0 | UGEC1 | DUSP5 |
| 0 | 2.24058005 | 0.768 | 0.051 | 0 | UGEC1 | NDRG2 |
| 0 | 2.23681267 | 0.752 | 0.019 | 0 | UGEC1 | TMPRSS11E |
| 0 | 2.20071568 | 0.772 | 0.02 | 0 | UGEC1 | TMPRSS11D |
| 0 | 2.18250845 | 0.349 | 0.006 | 0 | UGEC1 | SERPINB3 |
| 0 | 2.17237621 | 0.771 | 0.025 | 0 | UGEC1 | SCEL |
| 0 | 2.06003642 | 0.413 | 0.141 | 0 | UGEC1 | RARRES3 |
| 0 | 2.05648209 | 0.911 | 0.364 | 0 | UGEC1 | MXD1 |
| 0 | 2.01833402 | 0.688 | 0.016 | 0 | UGEC1 | GABRP |
| 0 | 1.96881419 | 0.283 | 0.017 | 0 | UGEC1 | FLG |
| 0 | 1.95546451 | 0.75 | 0.022 | 0 | UGEC1 | NCCRP1 |
| 0 | 1.94458898 | 0.614 | 0.031 | 0 | UGEC1 | DHRS9 |
| 0 | 1.93652845 | 0.798 | 0.123 | 0 | UGEC1 | ADGRF1 |
| 0 | 1.93284168 | 0.784 | 0.214 | 0 | UGEC1 | EMP1 |
| 0 | 1.92948345 | 0.71 | 0.23 | 0 | UGEC1 | PHLDA1 |
| 0 | 1.91393932 | 0.599 | 0.012 | 0 | UGEC1 | KRT78 |
| 0 | 1.89346741 | 0.623 | 0.088 | 0 | UGEC1 | F3 |
| 0 | 1.88609871 | 0.661 | 0.028 | 0 | UGEC1 | CXCL17 |
| 0 | 1.88134759 | 0.345 | 0.026 | 0 | UGEC1 | IFI27 |
| 0 | 1.87476536 | 0.663 | 0.035 | 0 | UGEC1 | CDKN2A |
| 0 | 1.84892499 | 0.749 | 0.033 | 0 | UGEC1 | MUC4 |
| 0 | 1.84700563 | 0.987 | 0.801 | 0 | UGEC1 | PPDPF |
| 0 | 1.82422406 | 0.278 | 0.02 | 0 | UGEC1 | OLFM4 |
| 0 | 1.80216259 | 0.309 | 0.007 | 0 | UGEC1 | CRISP3 |
| 0 | 1.7943444 | 0.74 | 0.079 | 0 | UGEC1 | FBXO32 |
| 0 | 1.76224429 | 0.789 | 0.189 | 0 | UGEC1 | RND3 |
| 0 | 1.7442602 | 0.766 | 0.069 | 0 | UGEC1 | EPS8L1 |
| 0 | 1.70709352 | 0.879 | 0.595 | 0 | UGEC1 | GLUL |
| 0 | 1.70614665 | 0.976 | 0.468 | 0 | UGEC1 | ELF3 |
| 0 | 1.70385607 | 0.783 | 0.361 | 0 | UGEC1 | HIST1H1C |
| 0 | 1.69424024 | 0.691 | 0.163 | 0 | UGEC1 | CARHSP1 |
| 0 | 1.69182478 | 0.963 | 0.747 | 0 | UGEC1 | LGALS3 |

| | | | | | | |
|---|---|---|---|---|---|---|
| 0 | 1.67843216 | 0.319 | 0.033 | 0 | UGEC1 | KLK6 |
| 0 | 1.67308961 | 0.686 | 0.086 | 0 | UGEC1 | ANKRD36C |
| 0 | 1.64412429 | 0.704 | 0.062 | 0 | UGEC1 | PPL |
| 0 | 1.64144108 | 0.688 | 0.069 | 0 | UGEC1 | SPNS2 |
| 0 | 1.64063872 | 0.716 | 0.059 | 0 | UGEC1 | STEAP4 |
| 0 | 1.62348869 | 0.604 | 0.061 | 0 | UGEC1 | CDC42EP5 |
| 0 | 1.61798718 | 0.342 | 0.003 | 0 | UGEC1 | SERPINB13 |
| 0 | 1.61228699 | 0.919 | 0.551 | 0 | UGEC1 | C15orf48 |
| 0 | 1.60997367 | 0.397 | 0.113 | 0 | UGEC1 | APOBEC3A |
| 0 | 1.59701056 | 0.507 | 0.133 | 0 | UGEC1 | TM4SF1 |
| 0 | 1.58414159 | 0.601 | 0.011 | 0 | UGEC1 | TMPRSS11A |
| 0 | 1.58294907 | 0.648 | 0.072 | 0 | UGEC1 | HSPB8 |
| 0 | 1.57676512 | 0.946 | 0.676 | 0 | UGEC1 | HSPB1 |
| 0 | 1.57468637 | 0.485 | 0.177 | 0 | UGEC1 | ISG15 |
| 0 | 1.52982262 | 0.709 | 0.074 | 0 | UGEC1 | SDCBP2 |
| 0 | 1.52975107 | 0.487 | 0.011 | 0 | UGEC1 | RNASE7 |
| 0 | 1.52543675 | 0.721 | 0.102 | 0 | UGEC1 | CDKN2B |
| 0 | 1.52479732 | 0.658 | 0.161 | 0 | UGEC1 | CSTA |
| 0 | 1.52323223 | 0.32 | 0.016 | 0 | UGEC1 | CRNN |
| 0 | 1.48256017 | 0.678 | 0.071 | 0 | UGEC1 | C6orf132 |
| 0 | 1.44311852 | 0.556 | 0.014 | 0 | UGEC1 | TMC5 |
| 0 | 1.43736692 | 0.637 | 0.151 | 0 | UGEC1 | PDZK1IP1 |
| 0 | 1.43511571 | 0.746 | 0.138 | 0 | UGEC1 | PRSS8 |
| 0 | 1.43501516 | 0.876 | 0.455 | 0 | UGEC1 | ZFP36 |
| 0 | 1.42697022 | 0.967 | 0.499 | 0 | UGEC1 | KRT19 |
| 0 | 1.42509815 | 0.557 | 0.035 | 0 | UGEC1 | SRD5A3 |
| 0 | 1.42257289 | 0.389 | 0.015 | 0 | UGEC1 | KRT16 |
| 0 | 1.40597612 | 0.387 | 0.019 | 0 | UGEC1 | IVL |
| 0 | 1.39874179 | 0.659 | 0.1 | 0 | UGEC1 | SAMD9 |
| 0 | 1.38133229 | 0.718 | 0.212 | 0 | UGEC1 | FAM129B |
| 0 | 1.3654477 | 0.589 | 0.026 | 0 | UGEC1 | TMEM40 |
| 0 | 1.36012502 | 0.421 | 0.092 | 0 | UGEC1 | SMC4 |
| 0 | 1.35574704 | 0.493 | 0.089 | 0 | UGEC1 | BPGM |
| 0 | 1.34623029 | 0.627 | 0.086 | 0 | UGEC1 | EPHA2 |
| 0 | 1.33807257 | 0.741 | 0.199 | 0 | UGEC1 | GPRC5A |
| 0 | 1.33629296 | 0.593 | 0.083 | 0 | UGEC1 | PRDM1 |
| 0 | 1.33511556 | 0.702 | 0.18 | 0 | UGEC1 | CCNG2 |
| 0 | 1.33302762 | 0.871 | 0.451 | 0 | UGEC1 | HEBP2 |
| 0 | 1.31954399 | 0.999 | 0.927 | 0 | UGEC1 | SAT1 |
| 0 | 1.31806967 | 0.59 | 0.042 | 0 | UGEC1 | GCNT3 |
| 0 | 1.31599089 | 0.973 | 0.706 | 0 | UGEC1 | TPM4 |
| 0 | 1.29469492 | 0.57 | 0.02 | 0 | UGEC1 | CEACAM1 |
| 0 | 1.29453512 | 0.603 | 0.067 | 0 | UGEC1 | MALL |
| 0 | 1.29147741 | 0.789 | 0.233 | 0 | UGEC1 | MYO6 |
| 0 | 1.28086104 | 0.838 | 0.406 | 0 | UGEC1 | RIOK3 |
| 0 | 1.27863355 | 0.828 | 0.318 | 0 | UGEC1 | SLPI |
| 0 | 1.27026074 | 0.996 | 0.946 | 0 | UGEC1 | ACTG1 |
| 0 | 1.26863929 | 0.361 | 0.019 | 0 | UGEC1 | CLCA4 |
| 0 | 1.26195944 | 0.517 | 0.01 | 0 | UGEC1 | KLK10 |
| 0 | 1.25471275 | 0.239 | 0.006 | 0 | UGEC1 | ALDH3A1 |

| | | | | | | |
|---|---|---|---|---|---|---|
| 0 | 1.24909572 | 0.631 | 0.055 | 0 | UGEC1 | TMPRSS4 |
| 0 | 1.24816364 | 0.58 | 0.008 | 0 | UGEC1 | A2ML1 |
| 0 | 1.24188769 | 0.688 | 0.172 | 0 | UGEC1 | MUC1 |
| 0 | 1.24186218 | 0.302 | 0.009 | 0 | UGEC1 | FAM25A |
| 0 | 1.24171503 | 0.64 | 0.058 | 0 | UGEC1 | TTC9 |
| 0 | 1.23992564 | 0.652 | 0.112 | 0 | UGEC1 | CSNK1E |
| 0 | 1.23512978 | 0.446 | 0.032 | 0 | UGEC1 | CLIC3 |
| 0 | 1.23506051 | 0.859 | 0.57 | 0 | UGEC1 | PCBP1 |
| 0 | 1.19657718 | 0.885 | 0.437 | 0 | UGEC1 | ANXA11 |
| 0 | 1.19246577 | 0.828 | 0.41 | 0 | UGEC1 | CD46 |
| 0 | 1.17484977 | 0.497 | 0.097 | 0 | UGEC1 | HIST3H2A |
| 0 | 1.17318175 | 0.555 | 0.125 | 0 | UGEC1 | DUSP4 |
| 0 | 1.17186978 | 0.541 | 0.065 | 0 | UGEC1 | ALDH1A3 |
| 0 | 1.15574969 | 0.372 | 0.07 | 0 | UGEC1 | AGR2 |
| 0 | 1.1525576 | 0.834 | 0.355 | 0 | UGEC1 | ATP1B1 |
| 0 | 1.15101393 | 0.444 | 0.011 | 0 | UGEC1 | HCG22 |
| 0 | 1.15009714 | 0.896 | 0.501 | 0 | UGEC1 | TACSTD2 |
| 0 | 1.14928027 | 0.433 | 0.04 | 0 | UGEC1 | KRT23 |
| 0 | 1.1271664 | 0.549 | 0.138 | 0 | UGEC1 | LY6E |
| 0 | 1.11234873 | 0.519 | 0.045 | 0 | UGEC1 | POLB |
| 0 | 1.11173322 | 0.472 | 0.009 | 0 | UGEC1 | SLC6A14 |
| 0 | 1.1067267 | 0.756 | 0.246 | 0 | UGEC1 | IL1RN |
| 0 | 1.10155262 | 0.435 | 0.078 | 0 | UGEC1 | PLAC8 |
| 0 | 1.10152674 | 0.526 | 0.024 | 0 | UGEC1 | STS |
| 0 | 1.09987894 | 0.286 | 0.003 | 0 | UGEC1 | KRT6A |
| 0 | 1.09977861 | 0.586 | 0.013 | 0 | UGEC1 | FUT3 |
| 0 | 1.09118507 | 0.688 | 0.17 | 0 | UGEC1 | WASL |
| 0 | 1.08871499 | 0.441 | 0.042 | 0 | UGEC1 | SMIM5 |
| 0 | 1.08620425 | 0.577 | 0.086 | 0 | UGEC1 | FAM214A |
| 0 | 1.0831711 | 0.362 | 0.054 | 0 | UGEC1 | TCHP |
| 0 | 1.08282624 | 0.435 | 0.016 | 0 | UGEC1 | FAM3D |
| 0 | 1.07634971 | 0.518 | 0.04 | 0 | UGEC1 | KRT80 |
| 0 | 1.0747063 | 0.675 | 0.152 | 0 | UGEC1 | GIPC1 |
| 0 | 1.07422167 | 0.831 | 0.374 | 0 | UGEC1 | CLDN7 |
| 0 | 1.07384325 | 0.915 | 0.625 | 0 | UGEC1 | TALDO1 |
| 0 | 1.07048829 | 0.679 | 0.17 | 0 | UGEC1 | PDLIM5 |
| 0 | 1.06762339 | 0.699 | 0.205 | 0 | UGEC1 | SPINT1 |
| 0 | 1.06626817 | 0.562 | 0.046 | 0 | UGEC1 | BCAS1 |
| 0 | 1.05657273 | 0.62 | 0.289 | 0 | UGEC1 | MIDN |
| 0 | 1.05562654 | 0.704 | 0.246 | 0 | UGEC1 | TMBIM1 |
| 0 | 1.0509593 | 0.918 | 0.57 | 0 | UGEC1 | TAX1BP1 |
| 0 | 1.0489302 | 0.511 | 0.005 | 0 | UGEC1 | KLK11 |
| 0 | 1.0438689 | 0.598 | 0.052 | 0 | UGEC1 | VSIG2 |
| 0 | 1.04141336 | 0.664 | 0.161 | 0 | UGEC1 | SERINC2 |
| 0 | 1.04068834 | 0.487 | 0.112 | 0 | UGEC1 | FHL2 |
| 0 | 1.0370159 | 0.548 | 0.125 | 0 | UGEC1 | SERTAD2 |
| 0 | 1.03525775 | 0.782 | 0.433 | 0 | UGEC1 | H2AFJ |
| 0 | 1.03383201 | 0.91 | 0.655 | 0 | UGEC1 | JUNB |
| 0 | 1.03279929 | 0.594 | 0.066 | 0 | UGEC1 | TSPAN6 |
| 0 | 1.03124833 | 0.482 | 0.018 | 0 | UGEC1 | RNF39 |

| | | | | | | |
|---|---|---|---|---|---|---|
| 0 | 1.03032484 | 0.577 | 0.11 | 0 | UGEC1 | ABLIM1 |
| 0 | 1.02815776 | 0.367 | 0.092 | 0 | UGEC1 | CXCL1 |
| 0 | 1.01772681 | 0.75 | 0.396 | 0 | UGEC1 | RAB11FIP1 |
| 0 | 1.01645293 | 0.244 | 0.029 | 0 | UGEC1 | DKK1 |
| 0 | 1.0133313 | 0.23 | 0.018 | 0 | UGEC1 | TFF1 |
| 0 | 1.01242026 | 0.582 | 0.152 | 0 | UGEC1 | RAB9A |
| 0 | 1.01231079 | 0.661 | 0.2 | 0 | UGEC1 | ABHD5 |
| 0 | 1.01007871 | 0.623 | 0.173 | 0 | UGEC1 | KLF5 |
| 0 | 1.00272395 | 0.456 | 0.112 | 0 | UGEC1 | DNAJA4 |
| 0 | 1.00083947 | 0.615 | 0.261 | 0 | UGEC1 | B4GALT1 |
| 0 | 0.99836223 | 0.469 | 0.004 | 0 | UGEC1 | C5orf66-AS1 |
| 0 | 0.99312823 | 0.641 | 0.126 | 0 | UGEC1 | EHF |
| 0 | 0.99291815 | 0.711 | 0.274 | 0 | UGEC1 | UBE2H |
| 0 | 0.99164579 | 0.88 | 0.569 | 0 | UGEC1 | TMEM59 |
| 0 | 0.98252442 | 0.527 | 0.207 | 0 | UGEC1 | HES4 |
| 0 | 0.97909653 | 0.601 | 0.207 | 0 | UGEC1 | PLIN3 |
| 0 | 0.97415456 | 0.612 | 0.338 | 0 | UGEC1 | C9orf16 |
| 0 | 0.96150624 | 0.753 | 0.484 | 0 | UGEC1 | PTTG1IP |
| 0 | 0.95894641 | 0.59 | 0.067 | 0 | UGEC1 | TRIP10 |
| 0 | 0.94840407 | 0.591 | 0.174 | 0 | UGEC1 | CST6 |
| 0 | 0.94544698 | 0.658 | 0.258 | 0 | UGEC1 | MAFF |
| 0 | 0.9432975 | 0.885 | 0.698 | 0 | UGEC1 | ATP6V1G1 |
| 0 | 0.94049549 | 0.415 | 0.139 | 0 | UGEC1 | TNFSF10 |
| 0 | 0.94048183 | 0.524 | 0.236 | 0 | UGEC1 | HMOX1 |
| 0 | 0.94028868 | 0.838 | 0.487 | 0 | UGEC1 | TOP1 |
| 0 | 0.939442 | 0.568 | 0.131 | 0 | UGEC1 | PARD6B |
| 0 | 0.93827907 | 0.299 | 0.121 | 0 | UGEC1 | ZFAND2A |
| 0 | 0.93718834 | 0.479 | 0.015 | 0 | UGEC1 | B3GALT5 |
| 0 | 0.93225173 | 0.996 | 0.959 | 0 | UGEC1 | UBC |
| 0 | 0.93175036 | 0.401 | 0.039 | 0 | UGEC1 | LRMP |
| 0 | 0.92287267 | 0.466 | 0.058 | 0 | UGEC1 | LRG1 |
| 0 | 0.91937895 | 0.431 | 0.013 | 0 | UGEC1 | C2orf54 |
| 0 | 0.9112367 | 0.774 | 0.421 | 0 | UGEC1 | GABARAPL2 |
| 0 | 0.91119193 | 0.972 | 0.96 | 0 | UGEC1 | HSP90AA1 |
| 0 | 0.90896985 | 0.809 | 0.521 | 0 | UGEC1 | ZFAND5 |
| 0 | 0.9074227 | 0.472 | 0.056 | 0 | UGEC1 | TP53INP2 |
| 0 | 0.90592136 | 0.18 | 0.053 | 0 | UGEC1 | SAA2 |
| 0 | 0.90288372 | 0.23 | 0.042 | 0 | UGEC1 | RARRES1 |
| 0 | 0.90119488 | 0.253 | 0.011 | 0 | UGEC1 | PIP |
| 0 | 0.90085166 | 0.271 | 0.003 | 0 | UGEC1 | MUC22 |
| 0 | 0.8994106 | 0.637 | 0.224 | 0 | UGEC1 | ZFAND6 |
| 0 | 0.89877374 | 0.29 | 0.085 | 0 | UGEC1 | HILPDA |
| 0 | 0.89812216 | 0.564 | 0.087 | 0 | UGEC1 | NECTIN4 |
| 0 | 0.89101175 | 0.376 | 0.054 | 0 | UGEC1 | NR1D1 |
| 0 | 0.88821241 | 0.527 | 0.058 | 0 | UGEC1 | ZNRF1 |
| 0 | 0.8844114 | 0.638 | 0.288 | 0 | UGEC1 | TIMP2 |
| 0 | 0.86675304 | 0.448 | 0.015 | 0 | UGEC1 | TTC22 |
| 0 | 0.86641673 | 0.131 | 0.004 | 0 | UGEC1 | SERPINB4 |
| 0 | 0.86632238 | 0.115 | 0.003 | 0 | UGEC1 | SPINK7 |
| 0 | 0.85625141 | 0.218 | 0.025 | 0 | UGEC1 | FCGBP |

| | | | | | | |
|---|---|---|---|---|---|---|
| 0 | 0.85099616 | 0.653 | 0.21 | 0 | UGEC1 | LRP10 |
| 0 | 0.84918394 | 0.359 | 0.124 | 0 | UGEC1 | ANKRD37 |
| 0 | 0.84271541 | 0.506 | 0.062 | 0 | UGEC1 | MTUS1 |
| 0 | 0.84258172 | 0.457 | 0.02 | 0 | UGEC1 | C4orf19 |
| 0 | 0.84172612 | 0.479 | 0.048 | 0 | UGEC1 | TCP11L2 |
| 0 | 0.84083824 | 0.674 | 0.47 | 0 | UGEC1 | HSPA1B |
| 0 | 0.84006344 | 0.418 | 0.054 | 0 | UGEC1 | ST3GAL4 |
| 0 | 0.83839458 | 0.953 | 0.814 | 0 | UGEC1 | SQSTM1 |
| 0 | 0.83833683 | 0.822 | 0.495 | 0 | UGEC1 | FAM107B |
| 0 | 0.83776759 | 0.234 | 0.007 | 0 | UGEC1 | KLK7 |
| 0 | 0.834245 | 0.64 | 0.225 | 0 | UGEC1 | VPS4B |
| 0 | 0.83341187 | 0.46 | 0.079 | 0 | UGEC1 | PDP1 |
| 0 | 0.83237542 | 0.533 | 0.136 | 0 | UGEC1 | TIPARP |
| 0 | 0.83183138 | 0.507 | 0.178 | 0 | UGEC1 | TOB1 |
| 0 | 0.82612307 | 0.603 | 0.209 | 0 | UGEC1 | SLK |
| 0 | 0.82246537 | 0.552 | 0.161 | 0 | UGEC1 | GALNT1 |
| 0 | 0.81855781 | 0.489 | 0.103 | 0 | UGEC1 | JMY |
| 0 | 0.81783327 | 0.579 | 0.204 | 0 | UGEC1 | RAB5IF |
| 0 | 0.81765884 | 0.316 | 0.006 | 0 | UGEC1 | CYSRT1 |
| 0 | 0.81147314 | 0.604 | 0.224 | 0 | UGEC1 | CHMP1B |
| 0 | 0.80403677 | 0.661 | 0.255 | 0 | UGEC1 | YPEL5 |
| 0 | 0.80107398 | 0.451 | 0.218 | 0 | UGEC1 | IDI1 |
| 0 | 0.79934278 | 0.352 | 0.05 | 0 | UGEC1 | DUSP14 |
| 0 | 0.79398613 | 0.788 | 0.381 | 0 | UGEC1 | SOX4 |
| 0 | 0.7936407 | 0.418 | 0.068 | 0 | UGEC1 | FAM84A |
| 0 | 0.79058664 | 0.453 | 0.022 | 0 | UGEC1 | TJP3 |
| 0 | 0.78962835 | 0.205 | 0.004 | 0 | UGEC1 | IL19 |
| 0 | 0.78941377 | 0.391 | 0.022 | 0 | UGEC1 | GALNT5 |
| 0 | 0.78161198 | 0.453 | 0.095 | 0 | UGEC1 | PELI1 |
| 0 | 0.77786803 | 0.358 | 0.009 | 0 | UGEC1 | FAM83A |
| 0 | 0.77685484 | 0.788 | 0.497 | 0 | UGEC1 | C4orf3 |
| 0 | 0.77378636 | 0.256 | 0.001 | 0 | UGEC1 | KLK12 |
| 0 | 0.77221697 | 0.662 | 0.297 | 0 | UGEC1 | TMOD3 |
| 0 | 0.77138633 | 0.94 | 0.748 | 0 | UGEC1 | MAP1LC3B |
| 0 | 0.77022604 | 0.471 | 0.186 | 0 | UGEC1 | CDKN1C |
| 0 | 0.76871321 | 0.642 | 0.255 | 0 | UGEC1 | YPEL3 |
| 0 | 0.76660692 | 0.337 | 0.092 | 0 | UGEC1 | ACADM |
| 0 | 0.76507792 | 0.372 | 0.149 | 0 | UGEC1 | BAG3 |
| 0 | 0.75992345 | 0.324 | 0.031 | 0 | UGEC1 | NTHL1 |
| 0 | 0.75981899 | 0.469 | 0.072 | 0 | UGEC1 | EMP2 |
| 0 | 0.75973368 | 0.692 | 0.504 | 0 | UGEC1 | SERPINB1 |
| 0 | 0.75823767 | 0.519 | 0.089 | 0 | UGEC1 | NDFIP2 |
| 0 | 0.75533505 | 0.316 | 0.066 | 0 | UGEC1 | YOD1 |
| 0 | 0.75274721 | 0.314 | 0.054 | 0 | UGEC1 | LY6K |
| 0 | 0.75223669 | 0.881 | 0.644 | 0 | UGEC1 | BZW1 |
| 0 | 0.74961015 | 0.447 | 0.071 | 0 | UGEC1 | SASH1 |
| 0 | 0.73698277 | 0.546 | 0.23 | 0 | UGEC1 | PDCD6 |
| 0 | 0.73692 | 0.51 | 0.116 | 0 | UGEC1 | GNA15 |
| 0 | 0.73643465 | 0.626 | 0.25 | 0 | UGEC1 | HIST1H2AC |
| 0 | 0.73583145 | 0.538 | 0.102 | 0 | UGEC1 | SMIM22 |

| | | | | | | |
|---|---|---|---|---|---|---|
| 0 | 0.73493601 | 0.529 | 0.203 | 0 | UGEC1 | ABRACL |
| 0 | 0.73410762 | 0.409 | 0.036 | 0 | UGEC1 | RNF223 |
| 0 | 0.73247689 | 0.34 | 0.142 | 0 | UGEC1 | IGFBP3 |
| 0 | 0.73120547 | 0.708 | 0.363 | 0 | UGEC1 | TXNRD1 |
| 0 | 0.72797463 | 0.533 | 0.209 | 0 | UGEC1 | HECTD1 |
| 0 | 0.72756343 | 0.359 | 0.015 | 0 | UGEC1 | PADI1 |
| 0 | 0.72739155 | 0.557 | 0.187 | 0 | UGEC1 | UBL3 |
| 0 | 0.72685339 | 0.446 | 0.092 | 0 | UGEC1 | SNX18 |
| 0 | 0.72548613 | 0.516 | 0.153 | 0 | UGEC1 | S100A12 |
| 0 | 0.72076388 | 0.594 | 0.247 | 0 | UGEC1 | CLTB |
| 0 | 0.7190638 | 0.416 | 0.071 | 0 | UGEC1 | HIST1H2BC |
| 0 | 0.71701168 | 0.432 | 0.045 | 0 | UGEC1 | GRHL1 |
| 0 | 0.71671863 | 0.774 | 0.533 | 0 | UGEC1 | PPP1CB |
| 0 | 0.71605129 | 0.423 | 0.116 | 0 | UGEC1 | RMND5A |
| 0 | 0.71594536 | 0.319 | 0.093 | 0 | UGEC1 | IL18 |
| 0 | 0.71593341 | 0.198 | 0.036 | 0 | UGEC1 | GRPEL2 |
| 0 | 0.71492776 | 0.281 | 0.027 | 0 | UGEC1 | HEG1 |
| 0 | 0.71165677 | 0.33 | 0.087 | 0 | UGEC1 | AC023157.3 |
| 0 | 0.71113753 | 0.403 | 0.15 | 0 | UGEC1 | KATNBL1 |
| 0 | 0.71008104 | 0.487 | 0.095 | 0 | UGEC1 | PPP4R1 |
| 0 | 0.70900213 | 0.421 | 0.083 | 0 | UGEC1 | AL133453.1 |
| 0 | 0.70782163 | 0.472 | 0.234 | 0 | UGEC1 | ADM |
| 0 | 0.70494093 | 0.571 | 0.161 | 0 | UGEC1 | HBP1 |
| 0 | 0.70417177 | 0.537 | 0.12 | 0 | UGEC1 | FCHO2 |
| 0 | 0.70408144 | 0.491 | 0.138 | 0 | UGEC1 | TMEM106B |
| 0 | 0.70203634 | 0.186 | 0.003 | 0 | UGEC1 | KRT4 |
| 0 | 0.70188007 | 0.406 | 0.017 | 0 | UGEC1 | TINCR |
| 0 | 0.6986251 | 0.468 | 0.112 | 0 | UGEC1 | DHRS3 |
| 0 | 0.69825196 | 0.432 | 0.066 | 0 | UGEC1 | KAZN |
| 0 | 0.69452983 | 0.503 | 0.122 | 0 | UGEC1 | HSBP1L1 |
| 0 | 0.69447982 | 0.408 | 0.034 | 0 | UGEC1 | SLC9A3R2 |
| 0 | 0.69383638 | 0.281 | 0.039 | 0 | UGEC1 | CHAC1 |
| 0 | 0.69170835 | 0.486 | 0.102 | 0 | UGEC1 | FXYD3 |
| 0 | 0.68466191 | 0.41 | 0.047 | 0 | UGEC1 | PLS1 |
| 0 | 0.68351516 | 0.438 | 0.096 | 0 | UGEC1 | ITPKC |
| 0 | 0.68341849 | 0.49 | 0.112 | 0 | UGEC1 | TMEM159 |
| 0 | 0.68319591 | 0.413 | 0.047 | 0 | UGEC1 | PTGES |
| 0 | 0.68262361 | 0.457 | 0.097 | 0 | UGEC1 | CRB3 |
| 0 | 0.68184822 | 0.688 | 0.334 | 0 | UGEC1 | CTNNA1 |
| 0 | 0.67739923 | 0.468 | 0.08 | 0 | UGEC1 | KCNK1 |
| 0 | 0.67024744 | 0.611 | 0.254 | 0 | UGEC1 | MMP24OS |
| 0 | 0.6695269 | 0.249 | 0.005 | 0 | UGEC1 | HAS2 |
| 0 | 0.6694381 | 0.37 | 0.091 | 0 | UGEC1 | SNRK |
| 0 | 0.66897335 | 0.51 | 0.254 | 0 | UGEC1 | PIM3 |
| 0 | 0.66855528 | 0.481 | 0.088 | 0 | UGEC1 | ERBB3 |
| 0 | 0.66700741 | 0.593 | 0.199 | 0 | UGEC1 | AFDN |
| 0 | 0.66625985 | 0.995 | 0.949 | 0 | UGEC1 | H3F3A |
| 0 | 0.66577186 | 0.395 | 0.062 | 0 | UGEC1 | ZNF117 |
| 0 | 0.66511389 | 0.253 | 0.006 | 0 | UGEC1 | AC116407.1 |
| 0 | 0.6620801 | 0.919 | 0.653 | 0 | UGEC1 | PPP1R15A |

| | | | | | | |
|---|---|---|---|---|---|---|
| 0 | 0.66131038 | 0.455 | 0.058 | 0 | UGEC1 | USP6NL |
| 0 | 0.66127806 | 0.632 | 0.303 | 0 | UGEC1 | RRBP1 |
| 0 | 0.6586891 | 0.46 | 0.05 | 0 | UGEC1 | BICDL2 |
| 0 | 0.65531439 | 0.46 | 0.204 | 0 | UGEC1 | GPCPD1 |
| 0 | 0.65462748 | 0.466 | 0.229 | 0 | UGEC1 | FAM3C |
| 0 | 0.65184397 | 0.125 | 0.001 | 0 | UGEC1 | FMO2 |
| 0 | 0.64998889 | 0.474 | 0.097 | 0 | UGEC1 | SORT1 |
| 0 | 0.64931535 | 0.359 | 0.009 | 0 | UGEC1 | VSIG10L |
| 0 | 0.64854032 | 0.83 | 0.526 | 0 | UGEC1 | UBE2B |
| 0 | 0.64826899 | 0.412 | 0.079 | 0 | UGEC1 | MACO1 |
| 0 | 0.64523699 | 0.592 | 0.322 | 0 | UGEC1 | NDUFV2 |
| 0 | 0.64424743 | 0.365 | 0.044 | 0 | UGEC1 | LPCAT4 |
| 0 | 0.6437967 | 0.344 | 0.018 | 0 | UGEC1 | ATG9B |
| 0 | 0.64361372 | 0.463 | 0.081 | 0 | UGEC1 | SPECC1 |
| 0 | 0.64073659 | 0.402 | 0.038 | 0 | UGEC1 | PROM2 |
| 0 | 0.63941005 | 0.416 | 0.058 | 0 | UGEC1 | MUC20 |
| 0 | 0.63926841 | 0.405 | 0.037 | 0 | UGEC1 | TMCO4 |
| 0 | 0.638899 | 0.889 | 0.569 | 0 | UGEC1 | DSTN |
| 0 | 0.63785156 | 0.425 | 0.087 | 0 | UGEC1 | SLC44A4 |
| 0 | 0.63445593 | 0.654 | 0.316 | 0 | UGEC1 | METRNL |
| 0 | 0.63162526 | 0.424 | 0.086 | 0 | UGEC1 | FUCA1 |
| 0 | 0.63008552 | 0.609 | 0.256 | 0 | UGEC1 | RAB11A |
| 0 | 0.62996873 | 0.441 | 0.067 | 0 | UGEC1 | LMTK2 |
| 0 | 0.62606748 | 0.392 | 0.138 | 0 | UGEC1 | SPTSSA |
| 0 | 0.62570998 | 0.237 | 0.004 | 0 | UGEC1 | B3GALT5-AS1 |
| 0 | 0.62240953 | 0.598 | 0.226 | 0 | UGEC1 | CHMP2B |
| 0 | 0.61682772 | 0.38 | 0.027 | 0 | UGEC1 | LNX1 |
| 0 | 0.61622926 | 0.518 | 0.187 | 0 | UGEC1 | CFDP1 |
| 0 | 0.61500641 | 0.981 | 0.947 | 0 | UGEC1 | H3F3B |
| 0 | 0.61249629 | 0.162 | 0.029 | 0 | UGEC1 | EDN2 |
| 0 | 0.61134271 | 0.331 | 0.044 | 0 | UGEC1 | SORD |
| 0 | 0.60828055 | 0.476 | 0.277 | 0 | UGEC1 | SMS |
| 0 | 0.60721882 | 0.424 | 0.104 | 0 | UGEC1 | SDC1 |
| 0 | 0.60531881 | 0.273 | 0.004 | 0 | UGEC1 | PAX9 |
| 0 | 0.60264202 | 0.344 | 0.072 | 0 | UGEC1 | ASS1 |
| 0 | 0.6006557 | 0.568 | 0.294 | 0 | UGEC1 | NDFIP1 |
| 0 | 0.599597 | 0.367 | 0.037 | 0 | UGEC1 | SCNN1B |
| 0 | 0.59829072 | 0.364 | 0.018 | 0 | UGEC1 | ACHE |
| 0 | 0.59780628 | 0.388 | 0.068 | 0 | UGEC1 | PTPN3 |
| 0 | 0.59315597 | 0.414 | 0.088 | 0 | UGEC1 | CCDC69 |
| 0 | 0.59305562 | 0.275 | 0.075 | 0 | UGEC1 | MTAP |
| 0 | 0.58814287 | 0.115 | 0.002 | 0 | UGEC1 | RPTN |
| 0 | 0.58683535 | 0.384 | 0.027 | 0 | UGEC1 | VGLL1 |
| 0 | 0.58511811 | 0.373 | 0.034 | 0 | UGEC1 | MLPH |
| 0 | 0.58373563 | 0.285 | 0.028 | 0 | UGEC1 | RIPK4 |
| 0 | 0.58258401 | 0.393 | 0.076 | 0 | UGEC1 | C1orf116 |
| 0 | 0.58140417 | 0.349 | 0.063 | 0 | UGEC1 | HIST1H2BG |
| 0 | 0.57981626 | 0.442 | 0.159 | 0 | UGEC1 | MACC1 |
| 0 | 0.57850581 | 0.742 | 0.454 | 0 | UGEC1 | DNAJB1 |
| 0 | 0.57751383 | 0.637 | 0.225 | 0 | UGEC1 | ADIRF |

| | | | | | | |
|---|---|---|---|---|---|---|
| 0 | 0.57714591 | 0.555 | 0.216 | 0 | UGEC1 | JUP |
| 0 | 0.57662464 | 0.83 | 0.486 | 0 | UGEC1 | KRT7 |
| 0 | 0.57612841 | 0.381 | 0.056 | 0 | UGEC1 | DENND2C |
| 0 | 0.57437891 | 0.527 | 0.206 | 0 | UGEC1 | CHP1 |
| 0 | 0.57380663 | 0.443 | 0.097 | 0 | UGEC1 | FBXO34 |
| 0 | 0.57309536 | 0.343 | 0.018 | 0 | UGEC1 | CRYBG2 |
| 0 | 0.56947818 | 0.503 | 0.236 | 0 | UGEC1 | TMEM30A |
| 0 | 0.56925195 | 0.685 | 0.458 | 0 | UGEC1 | VDAC2 |
| 0 | 0.56924936 | 0.46 | 0.149 | 0 | UGEC1 | MAP3K13 |
| 0 | 0.56695305 | 0.532 | 0.226 | 0 | UGEC1 | ADAM9 |
| 0 | 0.56665645 | 0.528 | 0.253 | 0 | UGEC1 | PICALM |
| 0 | 0.56429713 | 0.365 | 0.104 | 0 | UGEC1 | RALGDS |
| 0 | 0.5629081 | 0.295 | 0.094 | 0 | UGEC1 | GAN |
| 0 | 0.56210255 | 0.43 | 0.106 | 0 | UGEC1 | GALNT3 |
| 0 | 0.55903686 | 0.568 | 0.269 | 0 | UGEC1 | PERP |
| 0 | 0.55894442 | 0.478 | 0.227 | 0 | UGEC1 | QSOX1 |
| 0 | 0.55876354 | 0.216 | 0.011 | 0 | UGEC1 | TPRG1 |
| 0 | 0.55763989 | 0.518 | 0.203 | 0 | UGEC1 | LYPLA1 |
| 0 | 0.55580267 | 0.304 | 0.009 | 0 | UGEC1 | TTYH1 |
| 0 | 0.55402479 | 0.642 | 0.322 | 0 | UGEC1 | GCC2 |
| 0 | 0.55300009 | 0.644 | 0.406 | 0 | UGEC1 | DNAJB6 |
| 0 | 0.55177935 | 0.497 | 0.158 | 0 | UGEC1 | ADIPOR1 |
| 0 | 0.55077815 | 0.439 | 0.14 | 0 | UGEC1 | F11R |
| 0 | 0.55063484 | 0.378 | 0.045 | 0 | UGEC1 | ZNF185 |
| 0 | 0.5492775 | 0.563 | 0.282 | 0 | UGEC1 | RBM47 |
| 0 | 0.54880573 | 0.438 | 0.148 | 0 | UGEC1 | SECISBP2L |
| 0 | 0.54864357 | 0.31 | 0.008 | 0 | UGEC1 | XDH |
| 0 | 0.54779243 | 0.374 | 0.079 | 0 | UGEC1 | ERV3-1 |
| 0 | 0.54678484 | 0.489 | 0.204 | 0 | UGEC1 | PSMC6 |
| 0 | 0.54654719 | 0.396 | 0.097 | 0 | UGEC1 | NAPRT |
| 0 | 0.54602616 | 0.163 | 0.027 | 0 | UGEC1 | PPP1R3C |
| 0 | 0.54579652 | 0.381 | 0.08 | 0 | UGEC1 | SLCO4A1 |
| 0 | 0.54330819 | 0.312 | 0.075 | 0 | UGEC1 | OAS1 |
| 0 | 0.54170201 | 0.31 | 0.011 | 0 | UGEC1 | MUC3A |
| 0 | 0.54034397 | 0.43 | 0.1 | 0 | UGEC1 | CTTNBP2NL |
| 0 | 0.53801492 | 0.364 | 0.057 | 0 | UGEC1 | DTX2 |
| 0 | 0.53736457 | 0.379 | 0.065 | 0 | UGEC1 | GRB7 |
| 0 | 0.53685179 | 0.385 | 0.067 | 0 | UGEC1 | TC2N |
| 0 | 0.53626641 | 0.312 | 0.006 | 0 | UGEC1 | SH2D1B |
| 0 | 0.53236486 | 0.329 | 0.073 | 0 | UGEC1 | VASN |
| 0 | 0.52894082 | 0.383 | 0.065 | 0 | UGEC1 | ARNTL2 |
| 0 | 0.52886953 | 0.22 | 0.008 | 0 | UGEC1 | FAM83D |
| 0 | 0.52880845 | 0.399 | 0.061 | 0 | UGEC1 | CDS1 |
| 0 | 0.52733837 | 0.39 | 0.156 | 0 | UGEC1 | TSPAN3 |
| 0 | 0.52684207 | 0.347 | 0.072 | 0 | UGEC1 | TGFA |
| 0 | 0.52366914 | 0.594 | 0.296 | 0 | UGEC1 | IER5 |
| 0 | 0.52242139 | 0.559 | 0.294 | 0 | UGEC1 | YTHDC1 |
| 0 | 0.52091284 | 0.416 | 0.107 | 0 | UGEC1 | SSBP3 |
| 0 | 0.51990794 | 0.675 | 0.354 | 0 | UGEC1 | CD59 |
| 0 | 0.51871638 | 0.505 | 0.242 | 0 | UGEC1 | CLINT1 |

| | | | | | | |
|---|---|---|---|---|---|---|
| 0 | 0.5185882 | 0.434 | 0.185 | 0 | UGEC1 | TRIM56 |
| 0 | 0.51818396 | 0.391 | 0.099 | 0 | UGEC1 | ABHD3 |
| 0 | 0.51767289 | 0.192 | 0.005 | 0 | UGEC1 | TCN1 |
| 0 | 0.51742105 | 0.327 | 0.031 | 0 | UGEC1 | CNGA1 |
| 0 | 0.51529 | 0.341 | 0.067 | 0 | UGEC1 | TRIM16 |
| 0 | 0.51469766 | 0.45 | 0.183 | 0 | UGEC1 | PGD |
| 0 | 0.51401311 | 0.946 | 0.86 | 0 | UGEC1 | MYL12A |
| 0 | 0.51310668 | 0.374 | 0.182 | 0 | UGEC1 | ARL4C |
| 0 | 0.51250729 | 0.234 | 0.058 | 0 | UGEC1 | ZNF426 |
| 0 | 0.51089731 | 0.461 | 0.127 | 0 | UGEC1 | EPS8L2 |
| 0 | 0.50951285 | 0.323 | 0.012 | 0 | UGEC1 | FUT6 |
| 0 | 0.50573367 | 0.306 | 0.032 | 0 | UGEC1 | FOXA1 |
| 0 | 0.50470342 | 0.499 | 0.273 | 0 | UGEC1 | TMEM165 |
| 0 | 0.50343478 | 0.546 | 0.302 | 0 | UGEC1 | CPEB4 |
| 0 | 0.50286674 | 0.337 | 0.086 | 0 | UGEC1 | INSR |
| 0 | 0.50279153 | 0.328 | 0.079 | 0 | UGEC1 | HIST1H2BD |
| 0 | 0.49932416 | 0.325 | 0.068 | 0 | UGEC1 | MBOAT2 |
| 0 | 0.49759436 | 0.661 | 0.457 | 0 | UGEC1 | SELENOW |
| 0 | 0.49735516 | 0.302 | 0.105 | 0 | UGEC1 | FAM45A |
| 0 | 0.49692263 | 0.424 | 0.154 | 0 | UGEC1 | NT5C2 |
| 0 | 0.49669769 | 0.355 | 0.086 | 0 | UGEC1 | MPZL2 |
| 0 | 0.49596239 | 0.31 | 0.025 | 0 | UGEC1 | EPHX3 |
| 0 | 0.49400945 | 0.367 | 0.094 | 0 | UGEC1 | RABGAP1L |
| 0 | 0.49314444 | 0.265 | 0.018 | 0 | UGEC1 | NAT1 |
| 0 | 0.49141812 | 0.429 | 0.197 | 0 | UGEC1 | ARID5B |
| 0 | 0.49060872 | 0.155 | 0.001 | 0 | UGEC1 | SLURP1 |
| 0 | 0.48995374 | 0.415 | 0.141 | 0 | UGEC1 | MPRIP |
| 0 | 0.4875787 | 0.473 | 0.201 | 0 | UGEC1 | PKN2 |
| 0 | 0.48745458 | 0.504 | 0.21 | 0 | UGEC1 | UBE2R2 |
| 0 | 0.48659937 | 0.364 | 0.156 | 0 | UGEC1 | ITPRIPL2 |
| 0 | 0.48445695 | 0.431 | 0.122 | 0 | UGEC1 | TRAF4 |
| 0 | 0.48386037 | 0.316 | 0.052 | 0 | UGEC1 | ABHD17C |
| 0 | 0.48341813 | 0.465 | 0.223 | 0 | UGEC1 | TSPAN1 |
| 0 | 0.48183942 | 0.275 | 0.019 | 0 | UGEC1 | TRIM29 |
| 0 | 0.48166767 | 0.447 | 0.201 | 0 | UGEC1 | ITGB8 |
| 0 | 0.47886663 | 0.402 | 0.131 | 0 | UGEC1 | PIK3IP1 |
| 0 | 0.47875602 | 0.363 | 0.158 | 0 | UGEC1 | B4GALT5 |
| 0 | 0.47760741 | 0.315 | 0.018 | 0 | UGEC1 | PTK6 |
| 0 | 0.47718018 | 0.285 | 0.008 | 0 | UGEC1 | ATP10B |
| 0 | 0.47210787 | 0.444 | 0.147 | 0 | UGEC1 | TWF1 |
| 0 | 0.47059403 | 0.278 | 0.057 | 0 | UGEC1 | SECTM1 |
| 0 | 0.46912011 | 0.414 | 0.131 | 0 | UGEC1 | TEAD1 |
| 0 | 0.46780514 | 0.529 | 0.229 | 0 | UGEC1 | LYN |
| 0 | 0.46580349 | 0.339 | 0.113 | 0 | UGEC1 | HK2 |
| 0 | 0.46564275 | 0.424 | 0.177 | 0 | UGEC1 | KLHL24 |
| 0 | 0.46255283 | 0.255 | 0.101 | 0 | UGEC1 | SGTA |
| 0 | 0.45845611 | 0.27 | 0.019 | 0 | UGEC1 | ELF3-AS1 |
| 0 | 0.45774019 | 0.25 | 0.026 | 0 | UGEC1 | IRS1 |
| 0 | 0.45643961 | 0.351 | 0.073 | 0 | UGEC1 | CASP7 |
| 0 | 0.45514134 | 0.367 | 0.097 | 0 | UGEC1 | UBE2G1 |

| | | | | | | |
|---|---|---|---|---|---|---|
| 0 | 0.45421174 | 0.237 | 0.057 | 0 | UGEC1 | CLDN23 |
| 0 | 0.45336373 | 0.42 | 0.209 | 0 | UGEC1 | SNX9 |
| 0 | 0.45156219 | 0.397 | 0.13 | 0 | UGEC1 | TMEM54 |
| 0 | 0.45130588 | 0.49 | 0.221 | 0 | UGEC1 | TOR1AIP2 |
| 0 | 0.45109121 | 0.252 | 0.01 | 0 | UGEC1 | TMC1 |
| 0 | 0.44923909 | 0.318 | 0.096 | 0 | UGEC1 | GPAT3 |
| 0 | 0.4481613 | 0.224 | 0.042 | 0 | UGEC1 | TMEM80 |
| 0 | 0.44647215 | 0.198 | 0.049 | 0 | UGEC1 | GPX2 |
| 0 | 0.44566545 | 0.252 | 0.073 | 0 | UGEC1 | AVPI1 |
| 0 | 0.44542799 | 0.438 | 0.189 | 0 | UGEC1 | BCL10 |
| 0 | 0.44494733 | 0.319 | 0.091 | 0 | UGEC1 | FOXC1 |
| 0 | 0.44370682 | 0.409 | 0.138 | 0 | UGEC1 | GAK |
| 0 | 0.44353729 | 0.234 | 0.039 | 0 | UGEC1 | ABHD17B |
| 0 | 0.44264443 | 0.281 | 0.01 | 0 | UGEC1 | PTPRH |
| 0 | 0.44232234 | 0.242 | 0.05 | 0 | UGEC1 | RHOV |
| 0 | 0.44201386 | 0.492 | 0.197 | 0 | UGEC1 | CCDC186 |
| 0 | 0.44099655 | 0.233 | 0.011 | 0 | UGEC1 | AL121761.2 |
| 0 | 0.43694711 | 0.395 | 0.174 | 0 | UGEC1 | SREK1IP1 |
| 0 | 0.43694474 | 0.495 | 0.223 | 0 | UGEC1 | PDCD6IP |
| 0 | 0.43589543 | 0.551 | 0.326 | 0 | UGEC1 | WASF2 |
| 0 | 0.43535316 | 0.29 | 0.037 | 0 | UGEC1 | MYEOV |
| 0 | 0.43313799 | 0.191 | 0.015 | 0 | UGEC1 | MSLN |
| 0 | 0.43192558 | 0.427 | 0.183 | 0 | UGEC1 | KYNU |
| 0 | 0.43058198 | 0.191 | 0.013 | 0 | UGEC1 | CTSV |
| 0 | 0.42909404 | 0.349 | 0.07 | 0 | UGEC1 | GMDS |
| 0 | 0.42895615 | 0.454 | 0.16 | 0 | UGEC1 | MMADHC |
| 0 | 0.42850636 | 0.251 | 0.066 | 0 | UGEC1 | DUSP10 |
| 0 | 0.42834203 | 0.281 | 0.067 | 0 | UGEC1 | CEP135 |
| 0 | 0.42828408 | 0.832 | 0.602 | 0 | UGEC1 | IER3 |
| 0 | 0.42812772 | 0.394 | 0.135 | 0 | UGEC1 | FNBP1L |
| 0 | 0.42659767 | 0.479 | 0.18 | 0 | UGEC1 | TJP1 |
| 0 | 0.4229634 | 0.502 | 0.247 | 0 | UGEC1 | STK24 |
| 0 | 0.42277685 | 0.395 | 0.139 | 0 | UGEC1 | CCNDBP1 |
| 0 | 0.42271351 | 0.362 | 0.105 | 0 | UGEC1 | PLS3 |
| 0 | 0.42228698 | 0.407 | 0.175 | 0 | UGEC1 | CKB |
| 0 | 0.42118352 | 0.368 | 0.125 | 0 | UGEC1 | SLC12A6 |
| 0 | 0.41960929 | 0.42 | 0.165 | 0 | UGEC1 | NECTIN2 |
| 0 | 0.41950978 | 0.182 | 0.041 | 0 | UGEC1 | IL23A |
| 0 | 0.41928308 | 0.353 | 0.112 | 0 | UGEC1 | ZMYND8 |
| 0 | 0.41806747 | 0.391 | 0.124 | 0 | UGEC1 | GNG12 |
| 0 | 0.41737498 | 0.337 | 0.107 | 0 | UGEC1 | NR1D2 |
| 0 | 0.41652009 | 0.271 | 0.04 | 0 | UGEC1 | VWA1 |
| 0 | 0.41570174 | 0.283 | 0.058 | 0 | UGEC1 | RDH13 |
| 0 | 0.41488346 | 0.375 | 0.122 | 0 | UGEC1 | DDR1 |
| 0 | 0.414326 | 0.255 | 0.067 | 0 | UGEC1 | NET1 |
| 0 | 0.41325607 | 0.411 | 0.155 | 0 | UGEC1 | CAPN1 |
| 0 | 0.41294357 | 0.389 | 0.121 | 0 | UGEC1 | CXADR |
| 0 | 0.41249906 | 0.341 | 0.069 | 0 | UGEC1 | SPINT1-AS1 |
| 0 | 0.41190543 | 0.277 | 0.018 | 0 | UGEC1 | AGFG2 |
| 0 | 0.41067233 | 0.369 | 0.123 | 0 | UGEC1 | SMAD3 |

| | | | | | | |
|---|---|---|---|---|---|---|
| 0 | 0.41014134 | 0.497 | 0.242 | 0 | UGEC1 | ATP6V1D |
| 0 | 0.41007851 | 0.343 | 0.133 | 0 | UGEC1 | VPS37B |
| 0 | 0.40921989 | 0.16 | 0.006 | 0 | UGEC1 | NDUFA4L2 |
| 0 | 0.40869723 | 0.405 | 0.189 | 0 | UGEC1 | IER5L |
| 0 | 0.40389482 | 0.463 | 0.229 | 0 | UGEC1 | UBE2D1 |
| 0 | 0.40341147 | 0.259 | 0.031 | 0 | UGEC1 | SORBS2 |
| 0 | 0.40335089 | 0.309 | 0.069 | 0 | UGEC1 | SLC25A23 |
| 0 | 0.40224173 | 0.389 | 0.141 | 0 | UGEC1 | TSC22D4 |
| 0 | 0.4022188 | 0.386 | 0.184 | 0 | UGEC1 | SPATS2L |
| 0 | 0.39963488 | 0.294 | 0.068 | 0 | UGEC1 | RASAL2 |
| 0 | 0.39924191 | 0.174 | 0.015 | 0 | UGEC1 | SLC12A8 |
| 0 | 0.39775864 | 0.365 | 0.148 | 0 | UGEC1 | PHACTR2 |
| 0 | 0.39741781 | 0.443 | 0.21 | 0 | UGEC1 | MIEN1 |
| 0 | 0.39710005 | 0.332 | 0.074 | 0 | UGEC1 | C19orf33 |
| 0 | 0.39530371 | 0.254 | 0.011 | 0 | UGEC1 | MYZAP |
| 0 | 0.39383856 | 0.448 | 0.186 | 0 | UGEC1 | RNF10 |
| 0 | 0.39346584 | 0.307 | 0.085 | 0 | UGEC1 | SH3GL1 |
| 0 | 0.39223476 | 0.254 | 0.039 | 0 | UGEC1 | AL031777.3 |
| 0 | 0.39077465 | 0.405 | 0.166 | 0 | UGEC1 | EIF4EBP2 |
| 0 | 0.39055017 | 0.259 | 0.036 | 0 | UGEC1 | SH3BGRL2 |
| 0 | 0.38953033 | 0.355 | 0.113 | 0 | UGEC1 | DNAJC5 |
| 0 | 0.38913203 | 0.323 | 0.136 | 0 | UGEC1 | TP53BP2 |
| 0 | 0.38833165 | 0.29 | 0.071 | 0 | UGEC1 | STK39 |
| 0 | 0.38731317 | 0.44 | 0.233 | 0 | UGEC1 | AGFG1 |
| 0 | 0.38580638 | 0.285 | 0.079 | 0 | UGEC1 | RAI14 |
| 0 | 0.38513657 | 0.251 | 0.013 | 0 | UGEC1 | SLC15A2 |
| 0 | 0.38445602 | 0.409 | 0.204 | 0 | UGEC1 | KLF3 |
| 0 | 0.38313434 | 0.247 | 0.051 | 0 | UGEC1 | CYP3A5 |
| 0 | 0.38269135 | 0.283 | 0.088 | 0 | UGEC1 | FOXQ1 |
| 0 | 0.3820211 | 0.365 | 0.123 | 0 | UGEC1 | SLMAP |
| 0 | 0.38156785 | 0.229 | 0.012 | 0 | UGEC1 | CRYM |
| 0 | 0.38118027 | 0.294 | 0.08 | 0 | UGEC1 | ATMIN |
| 0 | 0.38010989 | 0.227 | 0.031 | 0 | UGEC1 | SFTA2 |
| 0 | 0.37975317 | 0.391 | 0.178 | 0 | UGEC1 | CNN3 |
| 0 | 0.37846835 | 0.396 | 0.145 | 0 | UGEC1 | NBR1 |
| 0 | 0.37842396 | 0.393 | 0.156 | 0 | UGEC1 | SWAP70 |
| 0 | 0.37671755 | 0.327 | 0.093 | 0 | UGEC1 | RASSF7 |
| 0 | 0.37644597 | 0.308 | 0.087 | 0 | UGEC1 | GALE |
| 0 | 0.3757248 | 0.478 | 0.238 | 0 | UGEC1 | TMF1 |
| 0 | 0.37547339 | 0.468 | 0.228 | 0 | UGEC1 | NPEPPS |
| 0 | 0.37447483 | 0.323 | 0.08 | 0 | UGEC1 | GSKIP |
| 0 | 0.37211504 | 0.243 | 0.017 | 0 | UGEC1 | SLC52A3 |
| 0 | 0.37172634 | 0.285 | 0.047 | 0 | UGEC1 | PCDH1 |
| 0 | 0.37162233 | 0.29 | 0.058 | 0 | UGEC1 | DIRC2 |
| 0 | 0.37127532 | 0.295 | 0.077 | 0 | UGEC1 | MYH14 |
| 0 | 0.37064022 | 0.251 | 0.061 | 0 | UGEC1 | TMEM173 |
| 0 | 0.36999098 | 0.264 | 0.094 | 0 | UGEC1 | ASCC2 |
| 0 | 0.36961584 | 0.482 | 0.255 | 0 | UGEC1 | SYAP1 |
| 0 | 0.36788907 | 0.306 | 0.114 | 0 | UGEC1 | YPEL2 |
| 0 | 0.36673345 | 0.118 | 0.007 | 0 | UGEC1 | CPA4 |

| | | | | | | |
|---|---|---|---|---|---|---|
| 0 | 0.36663705 | 0.328 | 0.096 | 0 | UGEC1 | SRGAP1 |
| 0 | 0.36653559 | 0.258 | 0.032 | 0 | UGEC1 | ACAD8 |
| 0 | 0.36609009 | 0.319 | 0.088 | 0 | UGEC1 | MYO5B |
| 0 | 0.36592865 | 0.364 | 0.122 | 0 | UGEC1 | MARCH5 |
| 0 | 0.36589203 | 0.404 | 0.188 | 0 | UGEC1 | EMC3 |
| 0 | 0.36581657 | 0.484 | 0.249 | 0 | UGEC1 | CLIP1 |
| 0 | 0.36570544 | 0.371 | 0.152 | 0 | UGEC1 | N4BP1 |
| 0 | 0.36482637 | 0.285 | 0.104 | 0 | UGEC1 | CBX4 |
| 0 | 0.36133858 | 0.301 | 0.077 | 0 | UGEC1 | RANBP9 |
| 0 | 0.36126003 | 0.353 | 0.131 | 0 | UGEC1 | CACUL1 |
| 0 | 0.35991057 | 0.185 | 0.001 | 0 | UGEC1 | KLK8 |
| 0 | 0.35822101 | 0.361 | 0.14 | 0 | UGEC1 | ANKRD13A |
| 0 | 0.35702062 | 0.352 | 0.154 | 0 | UGEC1 | PMEPA1 |
| 0 | 0.35700889 | 0.262 | 0.057 | 0 | UGEC1 | FSTL1 |
| 0 | 0.35638991 | 0.426 | 0.189 | 0 | UGEC1 | RAB18 |
| 0 | 0.35597556 | 0.53 | 0.296 | 0 | UGEC1 | CHMP5 |
| 0 | 0.3553752 | 0.296 | 0.066 | 0 | UGEC1 | DHX32 |
| 0 | 0.35492898 | 0.256 | 0.028 | 0 | UGEC1 | CFH |
| 0 | 0.35473556 | 0.289 | 0.065 | 0 | UGEC1 | TRIM11 |
| 0 | 0.35411129 | 0.368 | 0.151 | 0 | UGEC1 | UBA6 |
| 0 | 0.35402962 | 0.284 | 0.062 | 0 | UGEC1 | ABTB2 |
| 0 | 0.35401828 | 0.286 | 0.05 | 0 | UGEC1 | GOLPH3L |
| 0 | 0.35336666 | 0.409 | 0.151 | 0 | UGEC1 | RBPMS |
| 0 | 0.35015597 | 0.234 | 0.056 | 0 | UGEC1 | DEPP1 |
| 0 | 0.34856873 | 0.217 | 0.018 | 0 | UGEC1 | AL512274.1 |
| 0 | 0.34561697 | 0.445 | 0.199 | 0 | UGEC1 | OSTF1 |
| 0 | 0.3453505 | 0.219 | 0.014 | 0 | UGEC1 | MEIS2 |
| 0 | 0.34529907 | 0.214 | 0.028 | 0 | UGEC1 | HRH1 |
| 0 | 0.34511891 | 0.202 | 0.047 | 0 | UGEC1 | TP53I11 |
| 0 | 0.3440873 | 0.243 | 0.069 | 0 | UGEC1 | TRIM2 |
| 0 | 0.34309522 | 0.291 | 0.102 | 0 | UGEC1 | TM2D2 |
| 0 | 0.34170998 | 0.367 | 0.158 | 0 | UGEC1 | KIFC3 |
| 0 | 0.34092111 | 0.248 | 0.032 | 0 | UGEC1 | CAPN5 |
| 0 | 0.34023479 | 0.261 | 0.068 | 0 | UGEC1 | ENPP4 |
| 0 | 0.34005368 | 0.238 | 0.026 | 0 | UGEC1 | RARG |
| 0 | 0.3396091 | 0.293 | 0.097 | 0 | UGEC1 | NTN4 |
| 0 | 0.33904436 | 0.273 | 0.089 | 0 | UGEC1 | NT5C3A |
| 0 | 0.33572414 | 0.241 | 0.046 | 0 | UGEC1 | TSPAN13 |
| 0 | 0.33512731 | 0.298 | 0.102 | 0 | UGEC1 | THAP9-AS1 |
| 0 | 0.33477661 | 0.37 | 0.16 | 0 | UGEC1 | USP53 |
| 0 | 0.33418754 | 0.183 | 0.027 | 0 | UGEC1 | HIST1H2BH |
| 0 | 0.33336409 | 0.236 | 0.055 | 0 | UGEC1 | FRMD8 |
| 0 | 0.33306016 | 0.241 | 0.022 | 0 | UGEC1 | RHBDL2 |
| 0 | 0.33229715 | 0.251 | 0.072 | 0 | UGEC1 | TICAM1 |
| 0 | 0.33224831 | 0.274 | 0.067 | 0 | UGEC1 | BAIAP2L1 |
| 0 | 0.33207835 | 0.283 | 0.07 | 0 | UGEC1 | TST |
| 0 | 0.33166768 | 0.301 | 0.122 | 0 | UGEC1 | TMEM127 |
| 0 | 0.3316529 | 0.337 | 0.109 | 0 | UGEC1 | RALGPS2 |
| 0 | 0.33156041 | 0.142 | 0.001 | 0 | UGEC1 | KLK13 |
| 0 | 0.33043072 | 0.23 | 0.051 | 0 | UGEC1 | TRPM4 |

| | | | | | | |
|---|---|---|---|---|---|---|
| 0 | 0.32996248 | 0.929 | 0.63 | 0 | UGEC1 | NAMPT |
| 0 | 0.32853961 | 0.24 | 0.049 | 0 | UGEC1 | SH3D19 |
| 0 | 0.3281327 | 0.164 | 0.008 | 0 | UGEC1 | RAPGEF5 |
| 0 | 0.32802262 | 0.237 | 0.055 | 0 | UGEC1 | EXPH5 |
| 0 | 0.32788214 | 0.305 | 0.095 | 0 | UGEC1 | PALLD |
| 0 | 0.32762646 | 0.212 | 0.037 | 0 | UGEC1 | TRIB2 |
| 0 | 0.32758738 | 0.296 | 0.088 | 0 | UGEC1 | RNF141 |
| 0 | 0.3258611 | 0.184 | 0.008 | 0 | UGEC1 | LEMD1 |
| 0 | 0.32485054 | 0.206 | 0.014 | 0 | UGEC1 | LINC00342 |
| 0 | 0.32484395 | 0.243 | 0.043 | 0 | UGEC1 | MPZL3 |
| 0 | 0.32466311 | 0.233 | 0.05 | 0 | UGEC1 | LIMCH1 |
| 0 | 0.32407588 | 0.262 | 0.065 | 0 | UGEC1 | SMAGP |
| 0 | 0.32324622 | 0.325 | 0.111 | 0 | UGEC1 | STK38 |
| 0 | 0.32322558 | 0.291 | 0.082 | 0 | UGEC1 | SLC35A3 |
| 0 | 0.32316148 | 0.23 | 0.034 | 0 | UGEC1 | MAGI3 |
| 0 | 0.32295678 | 0.202 | 0.011 | 0 | UGEC1 | RBPMS-AS1 |
| 0 | 0.32274241 | 0.319 | 0.121 | 0 | UGEC1 | RNF169 |
| 0 | 0.32255284 | 0.27 | 0.099 | 0 | UGEC1 | ADIPOR2 |
| 0 | 0.32229365 | 0.309 | 0.112 | 0 | UGEC1 | FAM102A |
| 0 | 0.32214794 | 0.125 | 0.002 | 0 | UGEC1 | CALML3 |
| 0 | 0.32064325 | 0.183 | 0.022 | 0 | UGEC1 | HIST1H2BF |
| 0 | 0.32034037 | 0.324 | 0.123 | 0 | UGEC1 | GOLPH3 |
| 0 | 0.319299 | 0.269 | 0.078 | 0 | UGEC1 | TMEM45B |
| 0 | 0.31747534 | 0.268 | 0.098 | 0 | UGEC1 | PLEKHM1 |
| 0 | 0.3149628 | 0.254 | 0.073 | 0 | UGEC1 | LRRC8A |
| 0 | 0.31458521 | 0.153 | 0.001 | 0 | UGEC1 | SERPINB11 |
| 0 | 0.31420125 | 0.329 | 0.145 | 0 | UGEC1 | SPSB3 |
| 0 | 0.31389397 | 0.364 | 0.133 | 0 | UGEC1 | XIST |
| 0 | 0.31018007 | 0.292 | 0.092 | 0 | UGEC1 | TRIP6 |
| 0 | 0.30834159 | 0.232 | 0.038 | 0 | UGEC1 | A4GALT |
| 0 | 0.30791194 | 0.211 | 0.03 | 0 | UGEC1 | SH3PXD2A |
| 0 | 0.30791058 | 0.388 | 0.181 | 0 | UGEC1 | SMIM14 |
| 0 | 0.30786122 | 0.284 | 0.09 | 0 | UGEC1 | NCK2 |
| 0 | 0.3074317 | 0.247 | 0.05 | 0 | UGEC1 | SPAG1 |
| 0 | 0.30577172 | 0.195 | 0.018 | 0 | UGEC1 | GDPD3 |
| 0 | 0.3051037 | 0.198 | 0.011 | 0 | UGEC1 | GPR153 |
| 0 | 0.30397666 | 0.191 | 0.021 | 0 | UGEC1 | WNT7B |
| 0 | 0.30355035 | 0.201 | 0.011 | 0 | UGEC1 | AC004816.1 |
| 0 | 0.30236865 | 0.167 | 0.001 | 0 | UGEC1 | AC016717.2 |
| 0 | 0.30193018 | 0.206 | 0.037 | 0 | UGEC1 | AC006333.2 |
| 0 | 0.30187666 | 0.257 | 0.081 | 0 | UGEC1 | TUFT1 |
| 0 | 0.30113956 | 0.252 | 0.061 | 0 | UGEC1 | ZSWIM4 |
| 0 | 0.30111572 | 0.17 | 0.005 | 0 | UGEC1 | TRIM17 |
| 0 | 0.30014784 | 0.245 | 0.076 | 0 | UGEC1 | ARHGAP27 |
| 0 | 0.29957626 | 0.219 | 0.071 | 0 | UGEC1 | ALDH3A2 |
| 0 | 0.29755768 | 0.197 | 0.017 | 0 | UGEC1 | UPK3B |
| 0 | 0.29636863 | 0.668 | 0.404 | 0 | UGEC1 | MARCKS |
| 0 | 0.29618719 | 0.315 | 0.106 | 0 | UGEC1 | BACE2 |
| 0 | 0.29582725 | 0.273 | 0.085 | 0 | UGEC1 | DAAM1 |
| 0 | 0.29506155 | 0.284 | 0.11 | 0 | UGEC1 | RAB25 |

| | | | | | | |
|---|---|---|---|---|---|---|
| 0 | 0.29490221 | 0.172 | 0.013 | 0 | UGEC1 | ANXA9 |
| 0 | 0.29484745 | 0.29 | 0.103 | 0 | UGEC1 | HSDL2 |
| 0 | 0.2945922 | 0.286 | 0.107 | 0 | UGEC1 | LGALS9 |
| 0 | 0.29404158 | 0.235 | 0.057 | 0 | UGEC1 | TBX3 |
| 0 | 0.29377311 | 0.226 | 0.042 | 0 | UGEC1 | ARHGAP32 |
| 0 | 0.29357844 | 0.253 | 0.075 | 0 | UGEC1 | LMO2 |
| 0 | 0.29324187 | 0.205 | 0.042 | 0 | UGEC1 | CYP4B1 |
| 0 | 0.29299196 | 0.321 | 0.122 | 0 | UGEC1 | CYTH2 |
| 0 | 0.29075254 | 0.276 | 0.101 | 0 | UGEC1 | ODF2L |
| 0 | 0.28979295 | 0.336 | 0.151 | 0 | UGEC1 | DDA1 |
| 0 | 0.28843098 | 0.18 | 0.023 | 0 | UGEC1 | MROH6 |
| 0 | 0.28732081 | 0.185 | 0.013 | 0 | UGEC1 | TFAP2C |
| 0 | 0.28728325 | 0.25 | 0.092 | 0 | UGEC1 | USP38 |
| 0 | 0.28550461 | 0.309 | 0.132 | 0 | UGEC1 | RCOR1 |
| 0 | 0.28514713 | 0.218 | 0.045 | 0 | UGEC1 | PWWP2B |
| 0 | 0.28469354 | 0.256 | 0.083 | 0 | UGEC1 | KCNK6 |
| 0 | 0.28332917 | 0.149 | 0.003 | 0 | UGEC1 | LINC01559 |
| 0 | 0.28257389 | 0.249 | 0.09 | 0 | UGEC1 | MAPK8 |
| 0 | 0.28219897 | 0.171 | 0.018 | 0 | UGEC1 | MARCH3 |
| 0 | 0.28212538 | 0.231 | 0.047 | 0 | UGEC1 | MCU |
| 0 | 0.28107668 | 0.234 | 0.073 | 0 | UGEC1 | RNF103 |
| 0 | 0.27786533 | 0.245 | 0.076 | 0 | UGEC1 | OSBPL3 |
| 0 | 0.277607 | 0.292 | 0.11 | 0 | UGEC1 | SNCG |
| 0 | 0.27698006 | 0.315 | 0.131 | 0 | UGEC1 | ITCH |
| 0 | 0.27690259 | 0.237 | 0.075 | 0 | UGEC1 | MAP3K4 |
| 0 | 0.27636744 | 0.201 | 0.036 | 0 | UGEC1 | RTKN2 |
| 0 | 0.27391937 | 0.166 | 0.007 | 0 | UGEC1 | AL355312.4 |
| 0 | 0.27359661 | 0.173 | 0.016 | 0 | UGEC1 | MFSD4A |
| 0 | 0.27241093 | 0.208 | 0.032 | 0 | UGEC1 | ABAT |
| 0 | 0.27180074 | 0.191 | 0.02 | 0 | UGEC1 | SRPX2 |
| 0 | 0.2713512 | 0.162 | 0.001 | 0 | UGEC1 | LEXM |
| 0 | 0.27058694 | 0.252 | 0.067 | 0 | UGEC1 | STAP2 |
| 0 | 0.26928321 | 0.154 | 0.04 | 0 | UGEC1 | AC092683.1 |
| 0 | 0.26677893 | 0.237 | 0.081 | 0 | UGEC1 | ABCG1 |
| 0 | 0.26543697 | 0.195 | 0.028 | 0 | UGEC1 | CORO2A |
| 0 | 0.26543164 | 0.237 | 0.061 | 0 | UGEC1 | MED16 |
| 0 | 0.26520267 | 0.171 | 0.021 | 0 | UGEC1 | CENPM |
| 0 | 0.26473154 | 0.289 | 0.105 | 0 | UGEC1 | SCNN1A |
| 0 | 0.26438765 | 0.199 | 0.051 | 0 | UGEC1 | CGNL1 |
| 0 | 0.26341312 | 0.231 | 0.078 | 0 | UGEC1 | TNFRSF21 |
| 0 | 0.26301002 | 0.173 | 0.045 | 0 | UGEC1 | GNE |
| 0 | 0.26286706 | 0.174 | 0.009 | 0 | UGEC1 | FUT2 |
| 0 | 0.26151057 | 0.219 | 0.053 | 0 | UGEC1 | ZDHHC13 |
| 0 | 0.26100321 | 0.209 | 0.034 | 0 | UGEC1 | HOMEZ |
| 0 | 0.25934606 | 0.169 | 0.022 | 0 | UGEC1 | NXN |
| 0 | 0.25845087 | 0.134 | 0.012 | 0 | UGEC1 | PORCN |
| 0 | 0.25837667 | 0.196 | 0.045 | 0 | UGEC1 | HIST1H4H |
| 0 | 0.25745815 | 0.173 | 0.009 | 0 | UGEC1 | STYK1 |
| 0 | 0.2573561 | 0.192 | 0.025 | 0 | UGEC1 | LINC00511 |
| 0 | 0.25631351 | 0.172 | 0.037 | 0 | UGEC1 | HIST1H2BJ |

| | | | | | | |
|---|---|---|---|---|---|---|
| 0 | 0.25604399 | 0.201 | 0.042 | 0 | UGEC1 | B3GALT4 |
| 0 | 0.2553355 | 0.264 | 0.098 | 0 | UGEC1 | SELENOP |
| 0 | -0.283566 | 0.01 | 0.189 | 0 | UGEC1 | ESD |
| 0 | -0.3054886 | 0.006 | 0.18 | 0 | UGEC1 | SHTN1 |
| 0 | -0.3176252 | 0.006 | 0.193 | 0 | UGEC1 | RAB32 |
| 0 | -0.3289385 | 0.03 | 0.23 | 0 | UGEC1 | C1QBP |
| 0 | -0.3330427 | 0.004 | 0.177 | 0 | UGEC1 | DDX3Y |
| 0 | -0.3420821 | 0.033 | 0.236 | 0 | UGEC1 | C19orf70 |
| 0 | -0.3439305 | 0.038 | 0.235 | 0 | UGEC1 | PHB |
| 0 | -0.3453434 | 0.067 | 0.275 | 0 | UGEC1 | C8orf59 |
| 0 | -0.3460042 | 0.063 | 0.271 | 0 | UGEC1 | BANF1 |
| 0 | -0.3461435 | 0.02 | 0.221 | 0 | UGEC1 | GTF3A |
| 0 | -0.3493511 | 0.065 | 0.273 | 0 | UGEC1 | SNRPE |
| 0 | -0.3505757 | 0.024 | 0.227 | 0 | UGEC1 | RHOQ |
| 0 | -0.3549261 | 0.042 | 0.248 | 0 | UGEC1 | RPL17 |
| 0 | -0.3553521 | 0.043 | 0.246 | 0 | UGEC1 | CELF1 |
| 0 | -0.3554056 | 0.003 | 0.176 | 0 | UGEC1 | PLEKHO1 |
| 0 | -0.3560293 | 0.005 | 0.192 | 0 | UGEC1 | AP1S2 |
| 0 | -0.356751 | 0.01 | 0.224 | 0 | UGEC1 | AP2B1 |
| 0 | -0.3597465 | 0.016 | 0.209 | 0 | UGEC1 | PPP1R18 |
| 0 | -0.3608384 | 0.044 | 0.262 | 0 | UGEC1 | TMED5 |
| 0 | -0.3626674 | 0.007 | 0.19 | 0 | UGEC1 | MYADM |
| 0 | -0.3647813 | 0.068 | 0.294 | 0 | UGEC1 | EIF3L |
| 0 | -0.3699037 | 0.092 | 0.31 | 0 | UGEC1 | ATP5F1C |
| 0 | -0.371467 | 0.034 | 0.244 | 0 | UGEC1 | TCEAL4 |
| 0 | -0.3728097 | 0.054 | 0.286 | 0 | UGEC1 | CCT8 |
| 0 | -0.3731934 | 0.06 | 0.296 | 0 | UGEC1 | EIF3I |
| 0 | -0.3746197 | 0.031 | 0.259 | 0 | UGEC1 | SNRPF |
| 0 | -0.3750018 | 0.048 | 0.263 | 0 | UGEC1 | MRPL52 |
| 0 | -0.3761262 | 0.054 | 0.274 | 0 | UGEC1 | ATP5F1A |
| 0 | -0.3765217 | 0.049 | 0.262 | 0 | UGEC1 | COX20 |
| 0 | -0.3770635 | 0.009 | 0.187 | 0 | UGEC1 | OGFRL1 |
| 0 | -0.3781253 | 0.054 | 0.282 | 0 | UGEC1 | LSM7 |
| 0 | -0.3787647 | 0.001 | 0.193 | 0 | UGEC1 | WIPF1 |
| 0 | -0.3824792 | 0.033 | 0.255 | 0 | UGEC1 | MLEC |
| 0 | -0.3830386 | 0.064 | 0.291 | 0 | UGEC1 | EIF3M |
| 0 | -0.3845205 | 0.053 | 0.27 | 0 | UGEC1 | M6PR |
| 0 | -0.3864823 | 0.015 | 0.215 | 0 | UGEC1 | RILPL2 |
| 0 | -0.3865442 | 0.081 | 0.311 | 0 | UGEC1 | NDUFB8 |
| 0 | -0.3875123 | 0.125 | 0.354 | 0 | UGEC1 | EIF3F |
| 0 | -0.3881665 | 0.087 | 0.315 | 0 | UGEC1 | CCT5 |
| 0 | -0.3885866 | 0.002 | 0.182 | 0 | UGEC1 | CPVL |
| 0 | -0.3899015 | 0.055 | 0.267 | 0 | UGEC1 | FLOT1 |
| 0 | -0.3931027 | 0.023 | 0.22 | 0 | UGEC1 | GPSM3 |
| 0 | -0.3933178 | 0.033 | 0.233 | 0 | UGEC1 | CREG1 |
| 0 | -0.3934141 | 0.091 | 0.314 | 0 | UGEC1 | MRPL51 |
| 0 | -0.3934818 | 0.022 | 0.221 | 0 | UGEC1 | RNF130 |
| 0 | -0.3934883 | 0.004 | 0.201 | 0 | UGEC1 | GLIPR2 |
| 0 | -0.3953079 | 0.043 | 0.259 | 0 | UGEC1 | DDX18 |
| 0 | -0.3959209 | 0.055 | 0.262 | 0 | UGEC1 | TPP1 |

| | | | | | | |
|---|---|---|---|---|---|---|
| 0 | -0.397414 | 0.014 | 0.237 | 0 | UGEC1 | PRNP |
| 0 | -0.4001094 | 0.026 | 0.248 | 0 | UGEC1 | TRIM44 |
| 0 | -0.4002126 | 0.004 | 0.179 | 0 | UGEC1 | DMXL2 |
| 0 | -0.40167 | 0.07 | 0.291 | 0 | UGEC1 | DHRS7 |
| 0 | -0.4020313 | 0.029 | 0.265 | 0 | UGEC1 | RSL1D1 |
| 0 | -0.4023756 | 0.002 | 0.175 | 0 | UGEC1 | C1orf162 |
| 0 | -0.40568 | 0.003 | 0.195 | 0 | UGEC1 | FERMT3 |
| 0 | -0.4057623 | 0.105 | 0.341 | 0 | UGEC1 | SF3B5 |
| 0 | -0.4068789 | 0.053 | 0.274 | 0 | UGEC1 | PABPC4 |
| 0 | -0.407014 | 0.002 | 0.179 | 0 | UGEC1 | PLXDC2 |
| 0 | -0.4083681 | 0.1 | 0.319 | 0 | UGEC1 | LRPAP1 |
| 0 | -0.4092254 | 0.002 | 0.196 | 0 | UGEC1 | MYO1F |
| 0 | -0.4106966 | 0.092 | 0.336 | 0 | UGEC1 | EIF3G |
| 0 | -0.4111663 | 0.077 | 0.316 | 0 | UGEC1 | RSL24D1 |
| 0 | -0.4140691 | 0.088 | 0.303 | 0 | UGEC1 | ATOX1 |
| 0 | -0.4156689 | 0.004 | 0.198 | 0 | UGEC1 | CEP170 |
| 0 | -0.415844 | 0.023 | 0.272 | 0 | UGEC1 | DNAJC15 |
| 0 | -0.4162622 | 0.07 | 0.31 | 0 | UGEC1 | NOL7 |
| 0 | -0.4200519 | 0.082 | 0.306 | 0 | UGEC1 | EID1 |
| 0 | -0.420907 | 0.041 | 0.291 | 0 | UGEC1 | RPS17 |
| 0 | -0.4242172 | 0.001 | 0.173 | 0 | UGEC1 | TLR2 |
| 0 | -0.4242372 | 0.011 | 0.23 | 0 | UGEC1 | ACSL4 |
| 0 | -0.4279643 | 0.02 | 0.235 | 0 | UGEC1 | ARHGAP18 |
| 0 | -0.4290038 | 0.142 | 0.375 | 0 | UGEC1 | RBM25 |
| 0 | -0.4294509 | 0.003 | 0.215 | 0 | UGEC1 | FMNL1 |
| 0 | -0.429882 | 0.068 | 0.315 | 0 | UGEC1 | MINOS1 |
| 0 | -0.4304065 | 0.007 | 0.191 | 0 | UGEC1 | SELENOM |
| 0 | -0.4307221 | 0.021 | 0.223 | 0 | UGEC1 | PYCARD |
| 0 | -0.4331443 | 0.002 | 0.182 | 0 | UGEC1 | EVI2B |
| 0 | -0.4389055 | 0.105 | 0.34 | 0 | UGEC1 | EIF4A1 |
| 0 | -0.43966 | 0.075 | 0.332 | 0 | UGEC1 | CBX3 |
| 0 | -0.4422921 | 0.001 | 0.176 | 0 | UGEC1 | TREM1 |
| 0 | -0.442889 | 0.114 | 0.362 | 0 | UGEC1 | FKBP1A |
| 0 | -0.4445367 | 0.104 | 0.335 | 0 | UGEC1 | SELENOS |
| 0 | -0.4452207 | 0.671 | 0.809 | 0 | UGEC1 | BTF3 |
| 0 | -0.4457605 | 0.12 | 0.372 | 0 | UGEC1 | COPS9 |
| 0 | -0.4461124 | 0.588 | 0.75 | 0 | UGEC1 | YWHAB |
| 0 | -0.4475975 | 0.139 | 0.39 | 0 | UGEC1 | ENY2 |
| 0 | -0.4480963 | 0.16 | 0.388 | 0 | UGEC1 | FKBP2 |
| 0 | -0.4486547 | 0.088 | 0.304 | 0 | UGEC1 | HSPA9 |
| 0 | -0.4502077 | 0.018 | 0.211 | 0 | UGEC1 | LTA4H |
| 0 | -0.4502204 | 0.003 | 0.225 | 0 | UGEC1 | HCLS1 |
| 0 | -0.4507844 | 0.103 | 0.35 | 0 | UGEC1 | HNRNPM |
| 0 | -0.4508931 | 0.004 | 0.215 | 0 | UGEC1 | TMEM176B |
| 0 | -0.4541747 | 0.423 | 0.643 | 0 | UGEC1 | SUB1 |
| 0 | -0.4563851 | 0.057 | 0.296 | 0 | UGEC1 | RPL22L1 |
| 0 | -0.4590624 | 0.002 | 0.213 | 0 | UGEC1 | CCDC88A |
| 0 | -0.4607256 | 0.018 | 0.223 | 0 | UGEC1 | RAC2 |
| 0 | -0.4614562 | 0.139 | 0.393 | 0 | UGEC1 | TMEM258 |
| 0 | -0.4635856 | 0.07 | 0.298 | 0 | UGEC1 | MBNL1 |

| | | | | | | |
|---|---|---|---|---|---|---|
| 0 | -0.4646841 | 0.132 | 0.367 | 0 | UGEC1 | EIF1B |
| 0 | -0.4654278 | 0.002 | 0.174 | 0 | UGEC1 | SDC2 |
| 0 | -0.4676893 | 0.12 | 0.383 | 0 | UGEC1 | ARL6IP4 |
| 0 | -0.4677849 | 0.106 | 0.33 | 0 | UGEC1 | HNRNPAB |
| 0 | -0.4685155 | 0.031 | 0.26 | 0 | UGEC1 | ACTN1 |
| 0 | -0.4688814 | 0.004 | 0.209 | 0 | UGEC1 | KCTD12 |
| 0 | -0.4707732 | 0.026 | 0.289 | 0 | UGEC1 | PDCD5 |
| 0 | -0.4720552 | 0.118 | 0.38 | 0 | UGEC1 | EIF4B |
| 0 | -0.4731745 | 0.002 | 0.204 | 0 | UGEC1 | CD48 |
| 0 | -0.4732041 | 0.077 | 0.309 | 0 | UGEC1 | PHPT1 |
| 0 | -0.4754132 | 0.177 | 0.434 | 0 | UGEC1 | ATP5MPL |
| 0 | -0.4758745 | 0.085 | 0.317 | 0 | UGEC1 | MYDGF |
| 0 | -0.4768272 | 0.146 | 0.4 | 0 | UGEC1 | TRMT112 |
| 0 | -0.4772227 | 0.096 | 0.323 | 0 | UGEC1 | CXCL16 |
| 0 | -0.4779084 | 0.112 | 0.331 | 0 | UGEC1 | FXYD5 |
| 0 | -0.4780897 | 0.051 | 0.282 | 0 | UGEC1 | RBPJ |
| 0 | -0.4781826 | 0.148 | 0.413 | 0 | UGEC1 | SEC11A |
| 0 | -0.479633 | 0.04 | 0.323 | 0 | UGEC1 | SSB |
| 0 | -0.4809221 | 0.731 | 0.83 | 0 | UGEC1 | RPL36AL |
| 0 | -0.4813892 | 0.026 | 0.283 | 0 | UGEC1 | SP100 |
| 0 | -0.4824309 | 0.002 | 0.175 | 0 | UGEC1 | MS4A7 |
| 0 | -0.4824581 | 0.041 | 0.314 | 0 | UGEC1 | GADD45GIP1 |
| 0 | -0.4831535 | 0.002 | 0.215 | 0 | UGEC1 | CD37 |
| 0 | -0.4863653 | 0.129 | 0.389 | 0 | UGEC1 | LAMTOR4 |
| 0 | -0.4927665 | 0.002 | 0.233 | 0 | UGEC1 | SNX10 |
| 0 | -0.4938747 | 0.104 | 0.392 | 0 | UGEC1 | SSR2 |
| 0 | -0.4946829 | 0.089 | 0.35 | 0 | UGEC1 | NOP53 |
| 0 | -0.4974504 | 0.16 | 0.44 | 0 | UGEC1 | SERBP1 |
| 0 | -0.5003583 | 0.666 | 0.829 | 0 | UGEC1 | HMGB1 |
| 0 | -0.5012739 | 0.233 | 0.475 | 0 | UGEC1 | ATP5MF |
| 0 | -0.5023738 | 0 | 0.175 | 0 | UGEC1 | FPR1 |
| 0 | -0.5032172 | 0.104 | 0.342 | 0 | UGEC1 | DDX24 |
| 0 | -0.5042293 | 0.01 | 0.191 | 0 | UGEC1 | SERPINB9 |
| 0 | -0.5047372 | 0.002 | 0.198 | 0 | UGEC1 | LCP2 |
| 0 | -0.5049459 | 0.061 | 0.336 | 0 | UGEC1 | NDUFB9 |
| 0 | -0.505579 | 0.184 | 0.461 | 0 | UGEC1 | PCBP2 |
| 0 | -0.5060605 | 0.031 | 0.295 | 0 | UGEC1 | SH3BGRL |
| 0 | -0.5062774 | 0.052 | 0.307 | 0 | UGEC1 | TUBB |
| 0 | -0.5065138 | 0.106 | 0.344 | 0 | UGEC1 | SRSF2 |
| 0 | -0.5071361 | 0.023 | 0.264 | 0 | UGEC1 | GRINA |
| 0 | -0.5078055 | 0.156 | 0.392 | 0 | UGEC1 | TAF1D |
| 0 | -0.5083057 | 0.099 | 0.344 | 0 | UGEC1 | PSME2 |
| 0 | -0.5085509 | 0.34 | 0.597 | 0 | UGEC1 | HNRNPA3 |
| 0 | -0.5097518 | 0.165 | 0.43 | 0 | UGEC1 | HNRNPH1 |
| 0 | -0.509819 | 0.09 | 0.38 | 0 | UGEC1 | FAM96B |
| 0 | -0.5099371 | 0.106 | 0.387 | 0 | UGEC1 | SNU13 |
| 0 | -0.5103297 | 0.184 | 0.426 | 0 | UGEC1 | DEK |
| 0 | -0.5152544 | 0.711 | 0.817 | 0 | UGEC1 | ARPC2 |
| 0 | -0.5167339 | 0.083 | 0.327 | 0 | UGEC1 | YWHAH |
| 0 | -0.5168072 | 0.047 | 0.338 | 0 | UGEC1 | APRT |

| | | | | | | |
|---|---|---|---|---|---|---|
| 0 | -0.5180278 | 0.174 | 0.43 | 0 | UGEC1 | PSME1 |
| 0 | -0.5182677 | 0.082 | 0.354 | 0 | UGEC1 | PFDN2 |
| 0 | -0.519053 | 0.113 | 0.41 | 0 | UGEC1 | TOMM20 |
| 0 | -0.5213037 | 0.323 | 0.575 | 0 | UGEC1 | COX6C |
| 0 | -0.5240467 | 0.035 | 0.276 | 0 | UGEC1 | SNHG7 |
| 0 | -0.5245591 | 0.092 | 0.334 | 0 | UGEC1 | TUBA1C |
| 0 | -0.5256315 | 0.544 | 0.749 | 0 | UGEC1 | ATP5MG |
| 0 | -0.527082 | 0.141 | 0.418 | 0 | UGEC1 | ATP5MC3 |
| 0 | -0.5279223 | 0.736 | 0.898 | 0 | UGEC1 | HLA-C |
| 0 | -0.5292393 | 0.001 | 0.202 | 0 | UGEC1 | SLA |
| 0 | -0.5292659 | 0.143 | 0.386 | 0 | UGEC1 | NORAD |
| 0 | -0.5318791 | 0.058 | 0.294 | 0 | UGEC1 | ODC1 |
| 0 | -0.5326848 | 0.007 | 0.223 | 0 | UGEC1 | PTPRE |
| 0 | -0.5344939 | 0.176 | 0.423 | 0 | UGEC1 | N4BP2L2 |
| 0 | -0.5351287 | 0.328 | 0.587 | 0 | UGEC1 | POMP |
| 0 | -0.5369244 | 0.054 | 0.328 | 0 | UGEC1 | RBM3 |
| 0 | -0.5402934 | 0.01 | 0.302 | 0 | UGEC1 | C4orf48 |
| 0 | -0.541047 | 0.039 | 0.325 | 0 | UGEC1 | ROCK1 |
| 0 | -0.5424248 | 0.077 | 0.359 | 0 | UGEC1 | MZT2B |
| 0 | -0.5443898 | 0.109 | 0.406 | 0 | UGEC1 | CYCS |
| 0 | -0.5450675 | 0.045 | 0.323 | 0 | UGEC1 | DDX21 |
| 0 | -0.5471249 | 0.136 | 0.437 | 0 | UGEC1 | CNBP |
| 0 | -0.5500141 | 0.001 | 0.194 | 0 | UGEC1 | AQP9 |
| 0 | -0.5517985 | 0.144 | 0.429 | 0 | UGEC1 | EIF5A |
| 0 | -0.5523518 | 0.019 | 0.315 | 0 | UGEC1 | PLP2 |
| 0 | -0.5524164 | 0.013 | 0.259 | 0 | UGEC1 | DSE |
| 0 | -0.5526872 | 0.002 | 0.208 | 0 | UGEC1 | SMIM25 |
| 0 | -0.5527744 | 0.314 | 0.612 | 0 | UGEC1 | SET |
| 0 | -0.5530428 | 0.21 | 0.498 | 0 | UGEC1 | ATP5MD |
| 0 | -0.5559583 | 0.343 | 0.599 | 0 | UGEC1 | NDUFS5 |
| 0 | -0.5572259 | 0.088 | 0.407 | 0 | UGEC1 | EIF3E |
| 0 | -0.5600195 | 0.244 | 0.495 | 0 | UGEC1 | SEC61G |
| 0 | -0.5629874 | 0.107 | 0.405 | 0 | UGEC1 | SEPT7 |
| 0 | -0.5706181 | 0.078 | 0.338 | 0 | UGEC1 | CHCHD10 |
| 0 | -0.5736518 | 0.029 | 0.28 | 0 | UGEC1 | RNASET2 |
| 0 | -0.5741376 | 0.195 | 0.461 | 0 | UGEC1 | FUS |
| 0 | -0.5766288 | 0.015 | 0.252 | 0 | UGEC1 | NINJ1 |
| 0 | -0.5772302 | 0.433 | 0.694 | 0 | UGEC1 | UQCRH |
| 0 | -0.5791348 | 0.054 | 0.355 | 0 | UGEC1 | RPL36A |
| 0 | -0.5814924 | 0.001 | 0.174 | 0 | UGEC1 | CD93 |
| 0 | -0.5816845 | 0.019 | 0.245 | 0 | UGEC1 | SLC43A2 |
| 0 | -0.582935 | 0.135 | 0.402 | 0 | UGEC1 | TKT |
| 0 | -0.5832727 | 0.05 | 0.248 | 0 | UGEC1 | CREM |
| 0 | -0.5836401 | 0.009 | 0.244 | 0 | UGEC1 | CTSC |
| 0 | -0.5857432 | 0.396 | 0.655 | 0 | UGEC1 | SEC61B |
| 0 | -0.5869994 | 0.077 | 0.406 | 0 | UGEC1 | VDAC1 |
| 0 | -0.588662 | 0.001 | 0.182 | 0 | UGEC1 | CLEC5A |
| 0 | -0.5894538 | 0.107 | 0.413 | 0 | UGEC1 | MTDH |
| 0 | -0.59246 | 0.272 | 0.556 | 0 | UGEC1 | HNRNPDL |
| 0 | -0.5924992 | 0.122 | 0.388 | 0 | UGEC1 | WSB1 |

| | | | | | | |
|---|---|---|---|---|---|---|
| 0 | -0.5955119 | 0.144 | 0.416 | 0 | UGEC1 | PRDX6 |
| 0 | -0.596545 | 0.044 | 0.35 | 0 | UGEC1 | ARL6IP5 |
| 0 | -0.5991783 | 0.054 | 0.35 | 0 | UGEC1 | NUCKS1 |
| 0 | -0.6006766 | 0.178 | 0.479 | 0 | UGEC1 | SEC62 |
| 0 | -0.6042992 | 0.127 | 0.432 | 0 | UGEC1 | PRELID1 |
| 0 | -0.6055462 | 0.189 | 0.46 | 0 | UGEC1 | SFPQ |
| 0 | -0.6062642 | 0.326 | 0.6 | 0 | UGEC1 | SSR4 |
| 0 | -0.6093656 | 0.036 | 0.289 | 0 | UGEC1 | PTMS |
| 0 | -0.6136487 | 0.413 | 0.695 | 0 | UGEC1 | ATP5MC2 |
| 0 | -0.6168367 | 0.003 | 0.205 | 0 | UGEC1 | CXCR4 |
| 0 | -0.6200705 | 0.127 | 0.454 | 0 | UGEC1 | NDUFA1 |
| 0 | -0.6229132 | 0.051 | 0.305 | 0 | UGEC1 | RHOC |
| 0 | -0.6229262 | 0.001 | 0.219 | 0 | UGEC1 | SAMSN1 |
| 0 | -0.6235655 | 0.096 | 0.419 | 0 | UGEC1 | ATP5F1D |
| 0 | -0.6252893 | 0.032 | 0.264 | 0 | UGEC1 | OCIAD2 |
| 0 | -0.6260774 | 0.18 | 0.455 | 0 | UGEC1 | ATF4 |
| 0 | -0.627348 | 0.027 | 0.254 | 0 | UGEC1 | TUBA1A |
| 0 | -0.6298221 | 0.004 | 0.227 | 0 | UGEC1 | FCGR2A |
| 0 | -0.6304051 | 0.068 | 0.418 | 0 | UGEC1 | SNRPD2 |
| 0 | -0.6315211 | 0.138 | 0.471 | 0 | UGEC1 | EIF3K |
| 0 | -0.6338526 | 0.659 | 0.842 | 0 | UGEC1 | DDX5 |
| 0 | -0.6339348 | 0.038 | 0.268 | 0 | UGEC1 | IVNS1ABP |
| 0 | -0.6366571 | 0.003 | 0.242 | 0 | UGEC1 | NCF2 |
| 0 | -0.6420951 | 0.037 | 0.255 | 0 | UGEC1 | ZBTB20 |
| 0 | -0.6431722 | 0.223 | 0.541 | 0 | UGEC1 | CANX |
| 0 | -0.6440097 | 0.142 | 0.391 | 0 | UGEC1 | PEBP1 |
| 0 | -0.6454126 | 0.125 | 0.43 | 0 | UGEC1 | POLR2L |
| 0 | -0.6470097 | 0.002 | 0.205 | 0 | UGEC1 | TGFBI |
| 0 | -0.6470277 | 0.006 | 0.24 | 0 | UGEC1 | ITGAX |
| 0 | -0.6474036 | 0.442 | 0.682 | 0 | UGEC1 | TXNIP |
| 0 | -0.6476557 | 0.09 | 0.446 | 0 | UGEC1 | TBCA |
| 0 | -0.6504411 | 0.042 | 0.333 | 0 | UGEC1 | LDHB |
| 0 | -0.6516106 | 0.046 | 0.363 | 0 | UGEC1 | ARPC1B |
| 0 | -0.6517495 | 0.853 | 0.934 | 0 | UGEC1 | NACA |
| 0 | -0.6540089 | 0.001 | 0.19 | 0 | UGEC1 | CD300E |
| 0 | -0.6561891 | 0.001 | 0.227 | 0 | UGEC1 | FYB1 |
| 0 | -0.6573162 | 0.002 | 0.259 | 0 | UGEC1 | SPI1 |
| 0 | -0.6608954 | 0.044 | 0.339 | 0 | UGEC1 | ATP1B3 |
| 0 | -0.6615786 | 0.017 | 0.224 | 0 | UGEC1 | HBEGF |
| 0 | -0.6624925 | 0.07 | 0.379 | 0 | UGEC1 | CAPG |
| 0 | -0.6637637 | 0.102 | 0.411 | 0 | UGEC1 | ATP1A1 |
| 0 | -0.6666784 | 0.307 | 0.547 | 0 | UGEC1 | MT-ND4L |
| 0 | -0.6688407 | 0.003 | 0.212 | 0 | UGEC1 | ANPEP |
| 0 | -0.6715392 | 0.085 | 0.435 | 0 | UGEC1 | ATP5ME |
| 0 | -0.6763701 | 0.167 | 0.496 | 0 | UGEC1 | PDIA3 |
| 0 | -0.6786951 | 0.096 | 0.392 | 0 | UGEC1 | PDIA6 |
| 0 | -0.6790196 | 0.068 | 0.307 | 0 | UGEC1 | ID2 |
| 0 | -0.6819735 | 0.002 | 0.22 | 0 | UGEC1 | IGSF6 |
| 0 | -0.6838493 | 0.102 | 0.422 | 0 | UGEC1 | MYH9 |
| 0 | -0.6848749 | 0.654 | 0.871 | 0 | UGEC1 | PPIA |

| | | | | | | |
|---|---|---|---|---|---|---|
| 0 | -0.6848762 | 0.909 | 0.964 | 0 | UGEC1 | SERF2 |
| 0 | -0.6863046 | 0.703 | 0.936 | 0 | UGEC1 | HLA-A |
| 0 | -0.6891474 | 0.107 | 0.41 | 0 | UGEC1 | MIF |
| 0 | -0.6898436 | 0.538 | 0.819 | 0 | UGEC1 | TMA7 |
| 0 | -0.6904316 | 0.179 | 0.396 | 0 | UGEC1 | TUBA1B |
| 0 | -0.6905954 | 0.334 | 0.689 | 0 | UGEC1 | OST4 |
| 0 | -0.6911812 | 0.073 | 0.437 | 0 | UGEC1 | PARK7 |
| 0 | -0.6924735 | 0.031 | 0.335 | 0 | UGEC1 | CTSZ |
| 0 | -0.695475 | 0.662 | 0.872 | 0 | UGEC1 | COX4I1 |
| 0 | -0.7045783 | 0.948 | 0.929 | 0 | UGEC1 | MT-ND4 |
| 0 | -0.7075932 | 0.601 | 0.776 | 0 | UGEC1 | PRDX1 |
| 0 | -0.7090546 | 0.012 | 0.262 | 0 | UGEC1 | ITGB2 |
| 0 | -0.7112066 | 0.003 | 0.221 | 0 | UGEC1 | C5AR1 |
| 0 | -0.7148748 | 0.037 | 0.383 | 0 | UGEC1 | MGST3 |
| 0 | -0.7151873 | 0.523 | 0.721 | 0 | UGEC1 | BRI3 |
| 0 | -0.7168722 | 0.095 | 0.416 | 0 | UGEC1 | GNAI2 |
| 0 | -0.7234005 | 0.003 | 0.296 | 0 | UGEC1 | GMFG |
| 0 | -0.7247483 | 0.113 | 0.473 | 0 | UGEC1 | CDV3 |
| 0 | -0.7278845 | 0.427 | 0.742 | 0 | UGEC1 | PSMA7 |
| 0 | -0.72791 | 0.495 | 0.746 | 0 | UGEC1 | TPM3 |
| 0 | -0.7348899 | 0.095 | 0.376 | 0 | UGEC1 | SLC38A2 |
| 0 | -0.7362237 | 0.117 | 0.482 | 0 | UGEC1 | RAN |
| 0 | -0.7391667 | 0.216 | 0.469 | 0 | UGEC1 | SLC3A2 |
| 0 | -0.7436998 | 0.141 | 0.457 | 0 | UGEC1 | SOD1 |
| 0 | -0.7469524 | 0.429 | 0.703 | 0 | UGEC1 | ZFAS1 |
| 0 | -0.7491742 | 0.198 | 0.555 | 0 | UGEC1 | NCL |
| 0 | -0.7492067 | 0.219 | 0.548 | 0 | UGEC1 | TAGLN2 |
| 0 | -0.7517059 | 0.002 | 0.25 | 0 | UGEC1 | LST1 |
| 0 | -0.7524289 | 0.211 | 0.568 | 0 | UGEC1 | DBI |
| 0 | -0.7574142 | 0.216 | 0.58 | 0 | UGEC1 | NAP1L1 |
| 0 | -0.7590589 | 0.068 | 0.462 | 0 | UGEC1 | COMMD6 |
| 0 | -0.7613868 | 0.012 | 0.34 | 0 | UGEC1 | RPS4Y1 |
| 0 | -0.7690901 | 0.392 | 0.725 | 0 | UGEC1 | RTN4 |
| 0 | -0.7731849 | 0.134 | 0.362 | 0 | UGEC1 | VEGFA |
| 0 | -0.7785994 | 0.514 | 0.814 | 0 | UGEC1 | COX7C |
| 0 | -0.7787984 | 0.073 | 0.46 | 0 | UGEC1 | NDUFB2 |
| 0 | -0.7818282 | 0.231 | 0.632 | 0 | UGEC1 | SLC25A6 |
| 0 | -0.7864635 | 0.036 | 0.394 | 0 | UGEC1 | SNHG8 |
| 0 | -0.7887421 | 0.007 | 0.398 | 0 | UGEC1 | MSN |
| 0 | -0.7897106 | 0.272 | 0.679 | 0 | UGEC1 | HNRNPA1 |
| 0 | -0.7916743 | 0.019 | 0.398 | 0 | UGEC1 | TLN1 |
| 0 | -0.7980991 | 0.682 | 0.932 | 0 | UGEC1 | HLA-B |
| 0 | -0.7990316 | 0.239 | 0.61 | 0 | UGEC1 | RPS10 |
| 0 | -0.8014248 | 0.243 | 0.61 | 0 | UGEC1 | PPIB |
| 0 | -0.8074874 | 0.235 | 0.629 | 0 | UGEC1 | TOMM7 |
| 0 | -0.8097283 | 0.002 | 0.282 | 0 | UGEC1 | HCST |
| 0 | -0.8139477 | 0.068 | 0.376 | 0 | UGEC1 | LIMS1 |
| 0 | -0.8142112 | 0.182 | 0.489 | 0 | UGEC1 | CYSTM1 |
| 0 | -0.8159462 | 0.003 | 0.217 | 0 | UGEC1 | OLR1 |
| 0 | -0.8202296 | 0.18 | 0.438 | 0 | UGEC1 | HIF1A |

| | | | | | | |
|---|---|---|---|---|---|---|
| 0 | -0.8236939 | 0.389 | 0.713 | 0 | UGEC1 | CALM2 |
| 0 | -0.8298115 | 0.896 | 0.96 | 0 | UGEC1 | RPL18 |
| 0 | -0.832524 | 0.002 | 0.322 | 0 | UGEC1 | CD53 |
| 0 | -0.8358673 | 0.193 | 0.533 | 0 | UGEC1 | CD81 |
| 0 | -0.8394147 | 0.387 | 0.7 | 0 | UGEC1 | LDHA |
| 0 | -0.8409791 | 0.113 | 0.341 | 0 | UGEC1 | SERPINA1 |
| 0 | -0.8429223 | 0.002 | 0.242 | 0 | UGEC1 | CYBB |
| 0 | -0.8429587 | 0.045 | 0.344 | 0 | UGEC1 | ATP13A3 |
| 0 | -0.8433042 | 0.002 | 0.253 | 0 | UGEC1 | TNFRSF1B |
| 0 | -0.8451552 | 0.458 | 0.83 | 0 | UGEC1 | RPS26 |
| 0 | -0.8460247 | 0.385 | 0.714 | 0 | UGEC1 | NDUFA4 |
| 0 | -0.8513383 | 0.002 | 0.239 | 0 | UGEC1 | CD52 |
| 0 | -0.8666111 | 0.002 | 0.274 | 0 | UGEC1 | ZEB2 |
| 0 | -0.8725792 | 0.002 | 0.21 | 0 | UGEC1 | PLEK |
| 0 | -0.8744653 | 0.331 | 0.764 | 0 | UGEC1 | EEF2 |
| 0 | -0.8763538 | 0.31 | 0.662 | 0 | UGEC1 | HINT1 |
| 0 | -0.8884211 | 0.034 | 0.277 | 0 | UGEC1 | EPCAM |
| 0 | -0.8900421 | 0.005 | 0.225 | 0 | UGEC1 | AC020656.1 |
| 0 | -0.8949236 | 0.188 | 0.424 | 0 | UGEC1 | TYMP |
| 0 | -0.8965196 | 0.928 | 0.964 | 0 | UGEC1 | RPL15 |
| 0 | -0.9151553 | 0.001 | 0.204 | 0 | UGEC1 | FCN1 |
| 0 | -0.9223535 | 0.868 | 0.963 | 0 | UGEC1 | RPS16 |
| 0 | -0.9285038 | 0.341 | 0.779 | 0 | UGEC1 | RPL4 |
| 0 | -0.9368889 | 0.528 | 0.893 | 0 | UGEC1 | HNRNPA2B1 |
| 0 | -0.9453042 | 0.816 | 0.972 | 0 | UGEC1 | PTMA |
| 0 | -0.9525796 | 0.43 | 0.747 | 0 | UGEC1 | HSP90B1 |
| 0 | -0.9576193 | 0.04 | 0.273 | 0 | UGEC1 | PLIN2 |
| 0 | -0.9673257 | 0.39 | 0.774 | 0 | UGEC1 | GPX4 |
| 0 | -0.9709602 | 0.043 | 0.304 | 0 | UGEC1 | TPM1 |
| 0 | -0.9812309 | 0.298 | 0.572 | 0 | UGEC1 | CEBPB |
| 0 | -0.9815881 | 0.436 | 0.859 | 0 | UGEC1 | RPL31 |
| 0 | -0.9816238 | 0.028 | 0.268 | 0 | UGEC1 | IL32 |
| 0 | -0.9964915 | 0.339 | 0.757 | 0 | UGEC1 | NPM1 |
| 0 | -0.9970154 | 0.582 | 0.814 | 0 | UGEC1 | CTSD |
| 0 | -0.9975518 | 0.434 | 0.812 | 0 | UGEC1 | CLIC1 |
| 0 | -1.0066181 | 0.784 | 0.968 | 0 | UGEC1 | S100A6 |
| 0 | -1.015794 | 0.053 | 0.378 | 0 | UGEC1 | IFITM3 |
| 0 | -1.0223954 | 0.763 | 0.949 | 0 | UGEC1 | RPS20 |
| 0 | -1.0231821 | 0.734 | 0.897 | 0 | UGEC1 | RPLP0 |
| 0 | -1.0236432 | 0.033 | 0.416 | 0 | UGEC1 | GLS |
| 0 | -1.0291083 | 0.56 | 0.806 | 0 | UGEC1 | BTG1 |
| 0 | -1.0301068 | 0.841 | 0.967 | 0 | UGEC1 | RPL13A |
| 0 | -1.0315489 | 0.515 | 0.883 | 0 | UGEC1 | RPL27 |
| 0 | -1.0455315 | 0.69 | 0.945 | 0 | UGEC1 | RPL24 |
| 0 | -1.0467633 | 0.015 | 0.232 | 0 | UGEC1 | C12orf75 |
| 0 | -1.061044 | 0.682 | 0.944 | 0 | UGEC1 | RPL27A |
| 0 | -1.061121 | 0.049 | 0.363 | 0 | UGEC1 | HIST1H4C |
| 0 | -1.0633118 | 0.034 | 0.437 | 0 | UGEC1 | COTL1 |
| 0 | -1.0663751 | 0.436 | 0.856 | 0 | UGEC1 | EEF1D |
| 0 | -1.0753389 | 0.084 | 0.505 | 0 | UGEC1 | GSTO1 |

| | | | | | | |
|---|---|---|---|---|---|---|
| 0 | -1.0806137 | 0.385 | 0.821 | 0 | UGEC1 | EEF1B2 |
| 0 | -1.0826043 | 0.004 | 0.294 | 0 | UGEC1 | SLC11A1 |
| 0 | -1.0920351 | 0.9 | 0.969 | 0 | UGEC1 | RPS18 |
| 0 | -1.1000489 | 0.014 | 0.262 | 0 | UGEC1 | CD14 |
| 0 | -1.1208784 | 0.749 | 0.904 | 0 | UGEC1 | MT-ND3 |
| 0 | -1.1221896 | 0.356 | 0.803 | 0 | UGEC1 | ENO1 |
| 0 | -1.1234994 | 0.003 | 0.35 | 0 | UGEC1 | LCP1 |
| 0 | -1.1256851 | 0.536 | 0.927 | 0 | UGEC1 | RPL23 |
| 0 | -1.1309878 | 0.676 | 0.947 | 0 | UGEC1 | RPS11 |
| 0 | -1.1311393 | 0.779 | 0.962 | 0 | UGEC1 | FAU |
| 0 | -1.1318324 | 0.032 | 0.337 | 0 | UGEC1 | SLC25A37 |
| 0 | -1.1397747 | 0.374 | 0.801 | 0 | UGEC1 | PKM |
| 0 | -1.1454401 | 0.021 | 0.482 | 0 | UGEC1 | FLNA |
| 0 | -1.1474838 | 0.565 | 0.918 | 0 | UGEC1 | RPL7 |
| 0 | -1.1522331 | 0.46 | 0.889 | 0 | UGEC1 | PFDN5 |
| 0 | -1.1572136 | 0.459 | 0.888 | 0 | UGEC1 | YBX1 |
| 0 | -1.1668106 | 0.603 | 0.933 | 0 | UGEC1 | RPL23A |
| 0 | -1.1704399 | 0.033 | 0.234 | 0 | UGEC1 | THBS1 |
| 0 | -1.1828286 | 0.237 | 0.6 | 0 | UGEC1 | UPP1 |
| 0 | -1.1857735 | 0.95 | 0.986 | 0 | UGEC1 | RPL41 |
| 0 | -1.1865591 | 0.667 | 0.946 | 0 | UGEC1 | RPS25 |
| 0 | -1.186702 | 0.7 | 0.932 | 0 | UGEC1 | ATP5F1E |
| 0 | -1.1907582 | 0.73 | 0.957 | 0 | UGEC1 | RPL9 |
| 0 | -1.2085129 | 0.556 | 0.956 | 0 | UGEC1 | RPS6 |
| 0 | -1.210981 | 0.498 | 0.921 | 0 | UGEC1 | RPL38 |
| 0 | -1.2133501 | 0.857 | 0.955 | 0 | UGEC1 | TMSB4X |
| 0 | -1.2228572 | 0.002 | 0.264 | 0 | UGEC1 | ALOX5AP |
| 0 | -1.2245526 | 0.758 | 0.95 | 0 | UGEC1 | UBA52 |
| 0 | -1.2290727 | 0.01 | 0.338 | 0 | UGEC1 | IFITM2 |
| 0 | -1.2308117 | 0.657 | 0.948 | 0 | UGEC1 | RPL29 |
| 0 | -1.2421843 | 0.007 | 0.283 | 0 | UGEC1 | BCL2A1 |
| 0 | -1.2474648 | 0.002 | 0.281 | 0 | UGEC1 | SLC2A3 |
| 0 | -1.2475691 | 0.572 | 0.933 | 0 | UGEC1 | RPL14 |
| 0 | -1.2512608 | 0.796 | 0.962 | 0 | UGEC1 | RPL34 |
| 0 | -1.2607541 | 0.471 | 0.78 | 0 | UGEC1 | CYBA |
| 0 | -1.2663072 | 0.006 | 0.221 | 0 | UGEC1 | HLA-DPB1 |
| 0 | -1.2664941 | 0.652 | 0.947 | 0 | UGEC1 | RPL35A |
| 0 | -1.2763344 | 0.354 | 0.87 | 0 | UGEC1 | RPL10A |
| 0 | -1.2765414 | 0.002 | 0.339 | 0 | UGEC1 | PTPRC |
| 0 | -1.2818974 | 0.475 | 0.914 | 0 | UGEC1 | RACK1 |
| 0 | -1.2952986 | 0.765 | 0.957 | 0 | UGEC1 | RPL18A |
| 0 | -1.2979856 | 0.681 | 0.947 | 0 | UGEC1 | RPS7 |
| 0 | -1.3026749 | 0.433 | 0.876 | 0 | UGEC1 | RPS5 |
| 0 | -1.3216895 | 0.942 | 0.985 | 0 | UGEC1 | RPL28 |
| 0 | -1.3268181 | 0.035 | 0.261 | 0 | UGEC1 | G0S2 |
| 0 | -1.3315906 | 0.401 | 0.671 | 0 | UGEC1 | S100A4 |
| 0 | -1.3339919 | 0.716 | 0.966 | 0 | UGEC1 | RPL8 |
| 0 | -1.3373619 | 0.607 | 0.945 | 0 | UGEC1 | RPS3 |
| 0 | -1.3418609 | 0.657 | 0.942 | 0 | UGEC1 | RPS15A |
| 0 | -1.3432987 | 0.37 | 0.835 | 0 | UGEC1 | RPSA |

| | | | | | | |
|---|---|---|---|---|---|---|
| 0 | -1.3519226 | 0.676 | 0.952 | 0 | UGEC1 | RPL7A |
| 0 | -1.3570915 | 0.6 | 0.943 | 0 | UGEC1 | RPS13 |
| 0 | -1.3587578 | 0.545 | 0.959 | 0 | UGEC1 | RPL6 |
| 0 | -1.363891 | 0.426 | 0.915 | 0 | UGEC1 | RPL22 |
| 0 | -1.36629 | 0.78 | 0.965 | 0 | UGEC1 | RPS14 |
| 0 | -1.367654 | 0.821 | 0.955 | 0 | UGEC1 | GAPDH |
| 0 | -1.3704277 | 0.566 | 0.939 | 0 | UGEC1 | RPL26 |
| 0 | -1.3797756 | 0.532 | 0.914 | 0 | UGEC1 | RPS4X |
| 0 | -1.3863225 | 0.371 | 0.879 | 0 | UGEC1 | RPS21 |
| 0 | -1.3947398 | 0.481 | 0.922 | 0 | UGEC1 | RPL35 |
| 0 | -1.4078311 | 0.006 | 0.33 | 0 | UGEC1 | AIF1 |
| 0 | -1.4095258 | 0.057 | 0.322 | 0 | UGEC1 | HLA-DRB1 |
| 0 | -1.4150842 | 0.756 | 0.963 | 0 | UGEC1 | RPL30 |
| 0 | -1.420644 | 0.012 | 0.243 | 0 | UGEC1 | HLA-DPA1 |
| 0 | -1.4387569 | 0.011 | 0.402 | 0 | UGEC1 | LAPTM5 |
| 0 | -1.4409549 | 0.58 | 0.937 | 0 | UGEC1 | RPL3 |
| 0 | -1.4469928 | 0.396 | 0.811 | 0 | UGEC1 | PSAP |
| 0 | -1.4477089 | 0.084 | 0.495 | 0 | UGEC1 | EMP3 |
| 0 | -1.4555943 | 0.399 | 0.904 | 0 | UGEC1 | RPL5 |
| 0 | -1.461864 | 0.569 | 0.944 | 0 | UGEC1 | RPS29 |
| 0 | -1.4694529 | 0.618 | 0.955 | 0 | UGEC1 | RPS3A |
| 0 | -1.4784735 | 0.23 | 0.602 | 0 | UGEC1 | PLAUR |
| 0 | -1.4835398 | 0.724 | 0.969 | 0 | UGEC1 | RPL19 |
| 0 | -1.4930871 | 0.778 | 0.979 | 0 | UGEC1 | RPS19 |
| 0 | -1.4946959 | 0.686 | 0.971 | 0 | UGEC1 | RPL11 |
| 0 | -1.4973853 | 0.595 | 0.968 | 0 | UGEC1 | RPL37 |
| 0 | -1.5104566 | 0.091 | 0.397 | 0 | UGEC1 | CTSL |
| 0 | -1.518303 | 0.68 | 0.974 | 0 | UGEC1 | RPS24 |
| 0 | -1.5211964 | 0.668 | 0.972 | 0 | UGEC1 | RPS15 |
| 0 | -1.5237273 | 0.766 | 0.975 | 0 | UGEC1 | RPS28 |
| 0 | -1.5253603 | 0.559 | 0.965 | 0 | UGEC1 | RPL37A |
| 0 | -1.5349143 | 0.216 | 0.694 | 0 | UGEC1 | CD44 |
| 0 | -1.5349744 | 0.517 | 0.95 | 0 | UGEC1 | RPL36 |
| 0 | -1.5355263 | 0.598 | 0.961 | 0 | UGEC1 | RPL21 |
| 0 | -1.5392069 | 0.222 | 0.759 | 0 | UGEC1 | ANXA5 |
| 0 | -1.5440482 | 0.05 | 0.328 | 0 | UGEC1 | MT1E |
| 0 | -1.5584374 | 0.551 | 0.949 | 0 | UGEC1 | RPL12 |
| 0 | -1.5648786 | 0.773 | 0.978 | 0 | UGEC1 | RPS27 |
| 0 | -1.5723543 | 0.717 | 0.966 | 0 | UGEC1 | RPS27A |
| 0 | -1.5757029 | 0.639 | 0.953 | 0 | UGEC1 | RPL39 |
| 0 | -1.5764096 | 0.632 | 0.97 | 0 | UGEC1 | RPS8 |
| 0 | -1.5846215 | 0.373 | 0.877 | 0 | UGEC1 | PFN1 |
| 0 | -1.5854692 | 0.686 | 0.963 | 0 | UGEC1 | RPL32 |
| 0 | -1.5873083 | 0.836 | 0.971 | 0 | UGEC1 | RPL10 |
| 0 | -1.6171446 | 0.603 | 0.966 | 0 | UGEC1 | RPLP2 |
| 0 | -1.6172469 | 0.707 | 0.89 | 0 | UGEC1 | SH3BGRL3 |
| 0 | -1.6264881 | 0.855 | 0.987 | 0 | UGEC1 | RPL13 |
| 0 | -1.6346795 | 0.101 | 0.384 | 0 | UGEC1 | IGFBP7 |
| 0 | -1.654032 | 0.555 | 0.966 | 0 | UGEC1 | RPS9 |
| 0 | -1.6776209 | 0.979 | 0.988 | 0 | UGEC1 | FTL |

| | | | | | | |
|---|---|---|---|---|---|---|
| 0 | -1.6801261 | 0.912 | 0.995 | 0 | UGEC1 | EEF1A1 |
| 0 | -1.7989364 | 0.89 | 0.989 | 0 | UGEC1 | RPLP1 |
| 0 | -1.7997129 | 0.545 | 0.96 | 0 | UGEC1 | RPS23 |
| 0 | -1.8347164 | 0.104 | 0.443 | 0 | UGEC1 | FCER1G |
| 0 | -1.8619308 | 0.742 | 0.973 | 0 | UGEC1 | RPS12 |
| 0 | -1.877495 | 0.143 | 0.501 | 0 | UGEC1 | TYROBP |
| 0 | -1.8960765 | 0.001 | 0.268 | 0 | UGEC1 | VCAN |
| 0 | -1.9008789 | 0.206 | 0.55 | 0 | UGEC1 | CD74 |
| 0 | -1.9241681 | 0.471 | 0.97 | 0 | UGEC1 | RPS2 |
| 0 | -1.9332711 | 0.048 | 0.327 | 0 | UGEC1 | MT1X |
| 0 | -2.0013327 | 0.234 | 0.696 | 0 | UGEC1 | LGALS1 |
| 0 | -2.0895242 | 0.145 | 0.448 | 0 | UGEC1 | HLA-DRA |
| 0 | -2.1183376 | 0.07 | 0.292 | 0 | UGEC1 | MT1G |
| 0 | -2.4119163 | 0.005 | 0.211 | 0 | UGEC1 | IL1B |
| 0 | -2.4753624 | 0.216 | 0.515 | 0 | UGEC1 | LYZ |
| 0 | -2.7614107 | 0.198 | 0.505 | 0 | UGEC1 | SPP1 |
| 0 | -2.8119835 | 0.235 | 0.639 | 0 | UGEC1 | SRGN |
| 0 | -2.8759694 | 0.293 | 0.832 | 0 | UGEC1 | VIM |
| 0 | -2.9701033 | 0.326 | 0.728 | 0 | UGEC1 | MT2A |
| 0 | -3.954958 | 0.294 | 0.755 | 0 | UGEC1 | TIMP1 |
| 0 | -4.0366045 | 0.348 | 0.582 | 0 | UGEC1 | PLCG2 |
| 5.13E-308 | -0.4761432 | 0.1 | 0.312 | 1.212E-303 | UGEC1 | VASP |
| 6.841E-308 | -0.3419844 | 0.037 | 0.23 | 1.616E-303 | UGEC1 | ATP5MC1 |
| 1.245E-307 | -0.3570804 | 0.094 | 0.309 | 2.942E-303 | UGEC1 | UXT |
| 1.992E-307 | -0.8473107 | 0.475 | 0.68 | 4.707E-303 | UGEC1 | HSPA5 |
| 1.949E-306 | 0.2641943 | 0.256 | 0.099 | 4.605E-302 | UGEC1 | CMAS |
| 2.908E-306 | 0.46543835 | 0.886 | 0.782 | 6.87E-302 | UGEC1 | GNG5 |
| 1.531E-305 | -0.3288333 | 0.005 | 0.176 | 3.616E-301 | UGEC1 | ALDH2 |
| 2.303E-305 | 0.30773031 | 0.426 | 0.218 | 5.441E-301 | UGEC1 | SCP2 |
| 2.448E-305 | 0.25319882 | 0.276 | 0.109 | 5.782E-301 | UGEC1 | MGST2 |
| 4.994E-305 | -0.3546391 | 0.085 | 0.303 | 1.18E-300 | UGEC1 | TMEM123 |
| 5.42E-305 | -0.4790075 | 0.302 | 0.534 | 1.28E-300 | UGEC1 | SRSF3 |
| 6.698E-305 | 0.36839856 | 0.554 | 0.328 | 1.582E-300 | UGEC1 | PNRC2 |
| 8.368E-305 | -0.3116266 | 0.007 | 0.179 | 1.977E-300 | UGEC1 | ASAP1 |
| 1.184E-304 | -0.3742224 | 0.085 | 0.298 | 2.797E-300 | UGEC1 | EIF3J |
| 1.596E-303 | -0.3303386 | 0.038 | 0.229 | 3.771E-299 | UGEC1 | GNS |
| 4.29E-303 | -0.747257 | 0.012 | 0.186 | 1.013E-298 | UGEC1 | MT1F |
| 6.815E-303 | -0.5727972 | 0.15 | 0.368 | 1.61E-298 | UGEC1 | PPP1R14B |
| 3.662E-302 | -0.3578686 | 0.026 | 0.212 | 8.652E-298 | UGEC1 | WARS |
| 7.718E-302 | -0.3818227 | 0.019 | 0.199 | 1.823E-297 | UGEC1 | SLC38A1 |
| 1.145E-301 | 0.44494087 | 0.537 | 0.334 | 2.704E-297 | UGEC1 | TXNDC17 |
| 2.578E-301 | -0.3265581 | 0.001 | 0.165 | 6.091E-297 | UGEC1 | EVI2A |
| 2.965E-301 | -0.3246476 | 0.001 | 0.167 | 7.004E-297 | UGEC1 | EMILIN2 |
| 2.441E-300 | 0.42794821 | 0.543 | 0.33 | 5.766E-296 | UGEC1 | AES |
| 3.925E-300 | -0.8734912 | 0.107 | 0.318 | 9.272E-296 | UGEC1 | MAFB |
| 5.557E-300 | -0.3473872 | 0.078 | 0.287 | 1.313E-295 | UGEC1 | NDUFAB1 |
| 1.158E-299 | -0.4290272 | 0.144 | 0.361 | 2.735E-295 | UGEC1 | TGOLN2 |
| 2.549E-299 | -0.3153206 | 0.045 | 0.239 | 6.022E-295 | UGEC1 | SUGT1 |
| 4.131E-299 | -0.4064885 | 0.105 | 0.319 | 9.76E-295 | UGEC1 | DYNC1H1 |
| 1.49E-298 | -0.3726808 | 0.1 | 0.312 | 3.52E-294 | UGEC1 | ATP6AP2 |

| | | | | | | |
|---|---|---|---|---|---|---|
| 1.824E-298 | -0.340059 | 0.018 | 0.196 | 4.31E-294 | UGEC1 | EPB41L4A-AS1 |
| 2.888E-298 | -0.5077299 | 0.001 | 0.164 | 6.823E-294 | UGEC1 | MCEMP1 |
| 3.078E-298 | -0.4007816 | 0.111 | 0.324 | 7.271E-294 | UGEC1 | HM13 |
| 3.096E-298 | -0.3329909 | 0.013 | 0.185 | 7.313E-294 | UGEC1 | CSGALNACT2 |
| 1.053E-297 | -0.3888536 | 0.12 | 0.339 | 2.487E-293 | UGEC1 | PGAM1 |
| 2.314E-297 | -0.3303682 | 0.004 | 0.169 | 5.466E-293 | UGEC1 | LRP1 |
| 6.665E-297 | -0.3490796 | 0.001 | 0.164 | 1.574E-292 | UGEC1 | HCK |
| 6.537E-296 | -1.5017435 | 0.005 | 0.171 | 1.544E-291 | UGEC1 | CCL2 |
| 5.716E-295 | 0.29756485 | 0.387 | 0.182 | 1.35E-290 | UGEC1 | DSP |
| 3.289E-294 | -0.3544711 | 0.048 | 0.241 | 7.77E-290 | UGEC1 | MAP4 |
| 3.444E-294 | -0.2989876 | 0.031 | 0.214 | 8.136E-290 | UGEC1 | POLE4 |
| 5.565E-294 | -0.3360983 | 0.078 | 0.283 | 1.315E-289 | UGEC1 | TRAPPC1 |
| 1.083E-293 | -0.3294787 | 0.001 | 0.162 | 2.559E-289 | UGEC1 | FGR |
| 1.943E-293 | -0.5355388 | 0.342 | 0.549 | 4.591E-289 | UGEC1 | EIF5 |
| 5.529E-293 | -0.2976963 | 0.022 | 0.199 | 1.306E-288 | UGEC1 | HEXB |
| 5.667E-293 | -0.3392061 | 0.002 | 0.164 | 1.339E-288 | UGEC1 | ITGA5 |
| 6.625E-293 | -0.3019037 | 0.001 | 0.162 | 1.565E-288 | UGEC1 | NCKAP1L |
| 1.121E-292 | 0.3196545 | 0.387 | 0.191 | 2.649E-288 | UGEC1 | ST14 |
| 2.186E-292 | 0.35591334 | 0.545 | 0.32 | 5.165E-288 | UGEC1 | PLSCR1 |
| 4.313E-292 | 0.35930307 | 0.303 | 0.134 | 1.019E-287 | UGEC1 | CDKN1B |
| 1.114E-291 | -0.3010601 | 0.028 | 0.209 | 2.631E-287 | UGEC1 | REX1BD |
| 1.353E-291 | -0.5468885 | 0.416 | 0.605 | 3.196E-287 | UGEC1 | TSPO |
| 5.756E-291 | -0.3340111 | 0.001 | 0.161 | 1.36E-286 | UGEC1 | CYTIP |
| 7.897E-291 | -0.3830211 | 0.079 | 0.281 | 1.865E-286 | UGEC1 | CD47 |
| 2.062E-290 | -0.3331311 | 0.082 | 0.288 | 4.871E-286 | UGEC1 | HADHA |
| 1.051E-289 | -0.3833028 | 0.07 | 0.271 | 2.484E-285 | UGEC1 | PCBD1 |
| 5.368E-288 | -0.3560182 | 0.005 | 0.167 | 1.268E-283 | UGEC1 | IFI30 |
| 7.034E-288 | 0.75756549 | 0.594 | 0.423 | 1.662E-283 | UGEC1 | HSPH1 |
| 1.213E-287 | -0.4038721 | 0.058 | 0.25 | 2.866E-283 | UGEC1 | CDK2AP2 |
| 6.04E-286 | 0.34941182 | 0.661 | 0.446 | 1.427E-281 | UGEC1 | TMEM50A |
| 1.241E-285 | 0.34887366 | 0.293 | 0.13 | 2.931E-281 | UGEC1 | SUSD6 |
| 1.493E-285 | -0.3861559 | 0.055 | 0.247 | 3.527E-281 | UGEC1 | NKTR |
| 3.995E-285 | 0.25289172 | 0.368 | 0.172 | 9.437E-281 | UGEC1 | CTTN |
| 7.828E-285 | -0.3082123 | 0.001 | 0.157 | 1.849E-280 | UGEC1 | APBB1IP |
| 1.108E-284 | -0.3990421 | 0.013 | 0.18 | 2.618E-280 | UGEC1 | ITGA3 |
| 1.58E-284 | -0.3449427 | 0.044 | 0.23 | 3.732E-280 | UGEC1 | ITGAV |
| 2.203E-284 | -0.349708 | 0.048 | 0.236 | 5.205E-280 | UGEC1 | C6orf48 |
| 6.875E-284 | -0.5837863 | 0.08 | 0.276 | 1.624E-279 | UGEC1 | ATP2B1 |
| 8.985E-284 | -0.3010828 | 0.044 | 0.231 | 2.123E-279 | UGEC1 | SSR1 |
| 1.941E-283 | 0.78413387 | 0.992 | 0.93 | 4.586E-279 | UGEC1 | NEAT1 |
| 9.096E-283 | -0.4704958 | 0.009 | 0.17 | 2.149E-278 | UGEC1 | PAX8 |
| 6.453E-282 | -0.3463909 | 0.085 | 0.288 | 1.524E-277 | UGEC1 | PA2G4 |
| 2.859E-281 | -0.3789511 | 0.056 | 0.244 | 6.754E-277 | UGEC1 | VPS13C |
| 3.527E-281 | -0.4012387 | 0.089 | 0.287 | 8.332E-277 | UGEC1 | C1orf122 |
| 6.098E-281 | -0.3264813 | 0.052 | 0.24 | 1.44E-276 | UGEC1 | RBMX |
| 2.554E-280 | -0.3418731 | 0.097 | 0.302 | 6.033E-276 | UGEC1 | ERH |
| 3.11E-280 | -0.3507889 | 0.004 | 0.162 | 7.348E-276 | UGEC1 | COL18A1 |
| 2.255E-279 | -0.3085063 | 0.003 | 0.159 | 5.327E-275 | UGEC1 | PILRA |
| 4.515E-279 | -0.4326848 | 0.03 | 0.202 | 1.067E-274 | UGEC1 | CORO1A |
| 6.796E-279 | -0.6707142 | 0.159 | 0.365 | 1.605E-274 | UGEC1 | HERPUD1 |

| | | | | | | |
|---|---|---|---|---|---|---|
| 1.246E-278 | -0.4411364 | 0.23 | 0.447 | 2.944E-274 | UGEC1 | ATP5PF |
| 1.457E-278 | -0.3189983 | 0.002 | 0.156 | 3.442E-274 | UGEC1 | CELF2 |
| 1.508E-278 | -0.4388128 | 0.501 | 0.672 | 3.563E-274 | UGEC1 | EDF1 |
| 5.016E-278 | 0.35285029 | 0.345 | 0.169 | 1.185E-273 | UGEC1 | SMNDC1 |
| 1.049E-277 | -0.5812736 | 0.415 | 0.622 | 2.478E-273 | UGEC1 | KLF6 |
| 3.831E-277 | -0.3203134 | 0.023 | 0.192 | 9.051E-273 | UGEC1 | MTHFD2 |
| 6.451E-277 | 0.30703351 | 0.315 | 0.143 | 1.524E-272 | UGEC1 | GNL3 |
| 6.677E-277 | -0.341613 | 0.019 | 0.185 | 1.577E-272 | UGEC1 | SH3BP5 |
| 4.586E-276 | -0.3929848 | 0.218 | 0.448 | 1.083E-271 | UGEC1 | PTP4A2 |
| 8.855E-276 | -0.3339712 | 0.003 | 0.157 | 2.092E-271 | UGEC1 | LILRB3 |
| 5.56E-275 | 1.02960173 | 0.326 | 0.166 | 1.314E-270 | UGEC1 | HSPA6 |
| 9.518E-275 | -0.5954997 | 0.465 | 0.625 | 2.248E-270 | UGEC1 | NPC2 |
| 1.043E-274 | -0.3831597 | 0.147 | 0.361 | 2.464E-270 | UGEC1 | NSA2 |
| 1.056E-274 | -0.3153167 | 0.003 | 0.158 | 2.494E-270 | UGEC1 | TMEM176A |
| 3.658E-274 | -0.3216342 | 0.05 | 0.233 | 8.642E-270 | UGEC1 | NDUFS8 |
| 8.158E-274 | -0.3398874 | 0.108 | 0.313 | 1.927E-269 | UGEC1 | GDI2 |
| 8.846E-274 | -0.5583211 | 0.215 | 0.424 | 2.09E-269 | UGEC1 | AHNAK |
| 9.767E-274 | 0.33571879 | 0.321 | 0.15 | 2.307E-269 | UGEC1 | PHACTR4 |
| 1.594E-273 | -0.3106861 | 0.06 | 0.247 | 3.765E-269 | UGEC1 | RAD23A |
| 2.451E-273 | -0.7398792 | 0.95 | 0.974 | 5.789E-269 | UGEC1 | TMSB10 |
| 2.828E-273 | -0.3371549 | 0.005 | 0.16 | 6.682E-269 | UGEC1 | MAP3K20 |
| 3.656E-273 | -0.5609618 | 0.209 | 0.429 | 8.636E-269 | UGEC1 | SRSF7 |
| 1.372E-272 | -0.3318891 | 0.001 | 0.153 | 3.24E-268 | UGEC1 | IRAK3 |
| 1.914E-272 | -1.3466922 | 0.004 | 0.157 | 4.521E-268 | UGEC1 | APOE |
| 2.696E-272 | 1.14546274 | 0.837 | 0.773 | 6.37E-268 | UGEC1 | TXN |
| 3.443E-272 | -0.3143275 | 0.028 | 0.198 | 8.133E-268 | UGEC1 | GARS |
| 6.861E-272 | -0.3863076 | 0.058 | 0.244 | 1.621E-267 | UGEC1 | BDP1 |
| 7.448E-272 | -0.3367347 | 0.032 | 0.205 | 1.759E-267 | UGEC1 | ATXN1 |
| 3.282E-271 | 0.28714307 | 0.354 | 0.169 | 7.752E-267 | UGEC1 | ZNF91 |
| 4.729E-271 | -0.5413665 | 0.041 | 0.215 | 1.117E-266 | UGEC1 | ACSL1 |
| 8.141E-271 | -0.4592706 | 0.008 | 0.164 | 1.923E-266 | UGEC1 | DCDC2 |
| 8.231E-271 | -0.3252774 | 0.086 | 0.283 | 1.944E-266 | UGEC1 | MRPL20 |
| 3.865E-270 | -0.4191943 | 0.001 | 0.152 | 9.131E-266 | UGEC1 | MARCO |
| 7.557E-270 | -0.2922369 | 0.045 | 0.226 | 1.785E-265 | UGEC1 | SNRPD1 |
| 1.125E-269 | -0.2728008 | 0.02 | 0.186 | 2.657E-265 | UGEC1 | TMEM205 |
| 1.59E-269 | -0.2529115 | 0.009 | 0.165 | 3.755E-265 | UGEC1 | DEGS1 |
| 3.278E-269 | -0.3166746 | 0.028 | 0.198 | 7.743E-265 | UGEC1 | EPRS |
| 5.151E-269 | -0.3572837 | 0.135 | 0.344 | 1.217E-264 | UGEC1 | SSBP1 |
| 1.024E-268 | -0.3258765 | 0.08 | 0.272 | 2.42E-264 | UGEC1 | MRPS21 |
| 1.345E-268 | 0.34888184 | 0.552 | 0.339 | 3.177E-264 | UGEC1 | ARF4 |
| 4.16E-268 | -0.2857209 | 0.04 | 0.216 | 9.826E-264 | UGEC1 | PCMT1 |
| 4.465E-268 | -0.2969242 | 0.027 | 0.196 | 1.055E-263 | UGEC1 | HMGN3 |
| 5.167E-268 | -0.2893216 | 0.033 | 0.204 | 1.221E-263 | UGEC1 | IFI27L2 |
| 1.027E-267 | -0.3267293 | 0.104 | 0.304 | 2.425E-263 | UGEC1 | LMAN2 |
| 6.861E-267 | -0.3375986 | 0.084 | 0.277 | 1.621E-262 | UGEC1 | OSTC |
| 8.592E-267 | -0.37667 | 0.121 | 0.323 | 2.03E-262 | UGEC1 | NDUFB7 |
| 1.275E-266 | -0.3742967 | 0.062 | 0.246 | 3.012E-262 | UGEC1 | EML4 |
| 1.295E-266 | 0.34072662 | 0.29 | 0.13 | 3.06E-262 | UGEC1 | HOTAIRM1 |
| 1.923E-266 | -0.6754929 | 0.074 | 0.269 | 4.543E-262 | UGEC1 | FABP5 |
| 2.607E-266 | 0.30898515 | 0.362 | 0.182 | 6.158E-262 | UGEC1 | RNF145 |

| | | | | | | |
|---|---|---|---|---|---|---|
| 4.315E-266 | -0.4714493 | 0.016 | 0.175 | 1.019E-261 | UGEC1 | RGS2 |
| 7.332E-266 | -0.3452788 | 0.009 | 0.163 | 1.732E-261 | UGEC1 | CLEC7A |
| 8.676E-266 | -0.4201994 | 0.082 | 0.271 | 2.049E-261 | UGEC1 | C12orf57 |
| 1.356E-265 | -0.2673798 | 0.013 | 0.171 | 3.204E-261 | UGEC1 | IRAK1 |
| 1.898E-265 | -0.2915367 | 0.033 | 0.203 | 4.483E-261 | UGEC1 | VKORC1 |
| 2.663E-265 | -0.2644591 | 0.007 | 0.159 | 6.291E-261 | UGEC1 | PDLIM7 |
| 3.886E-265 | -1.3037894 | 0.001 | 0.148 | 9.179E-261 | UGEC1 | CCL3 |
| 1.127E-264 | -0.386503 | 0.188 | 0.403 | 2.662E-260 | UGEC1 | REEP5 |
| 2.014E-264 | -0.334385 | 0.006 | 0.157 | 4.758E-260 | UGEC1 | EPB41L3 |
| 5.879E-264 | -0.5158199 | 0.008 | 0.16 | 1.389E-259 | UGEC1 | UCHL1 |
| 1.308E-263 | 0.27429242 | 0.367 | 0.182 | 3.09E-259 | UGEC1 | OSER1 |
| 1.511E-263 | -0.3322895 | 0.074 | 0.261 | 3.569E-259 | UGEC1 | PJA2 |
| 2.807E-263 | -0.4249273 | 0.013 | 0.169 | 6.63E-259 | UGEC1 | NCF1 |
| 7.13E-263 | -0.6476266 | 0.092 | 0.273 | 1.684E-258 | UGEC1 | S100A13 |
| 9.847E-263 | -0.3249499 | 0.071 | 0.259 | 2.326E-258 | UGEC1 | SNRNP70 |
| 4.195E-262 | 0.31689128 | 0.279 | 0.122 | 9.909E-258 | UGEC1 | RASGEF1B |
| 6.485E-262 | -0.5531288 | 0.076 | 0.261 | 1.532E-257 | UGEC1 | PPIF |
| 1.227E-261 | -0.3278383 | 0.075 | 0.261 | 2.898E-257 | UGEC1 | SAT2 |
| 1.24E-261 | -0.3546386 | 0.093 | 0.285 | 2.929E-257 | UGEC1 | SRSF10 |
| 1.986E-261 | -0.2537849 | 0.019 | 0.178 | 4.691E-257 | UGEC1 | BCL7C |
| 5.743E-261 | 0.30462096 | 0.343 | 0.171 | 1.357E-256 | UGEC1 | NANS |
| 6.248E-261 | -0.294045 | 0.02 | 0.18 | 1.476E-256 | UGEC1 | EIF4EBP1 |
| 9.804E-261 | -0.3462369 | 0.034 | 0.202 | 2.316E-256 | UGEC1 | TNFAIP8 |
| 3.717E-260 | -0.3473404 | 0.018 | 0.178 | 8.781E-256 | UGEC1 | SORL1 |
| 4.895E-260 | -0.2605921 | 0.026 | 0.19 | 1.156E-255 | UGEC1 | FAAP20 |
| 5.57E-260 | 0.33619942 | 0.505 | 0.292 | 1.316E-255 | UGEC1 | ASPH |
| 6.986E-260 | -0.4277861 | 0.01 | 0.162 | 1.65E-255 | UGEC1 | ARHGAP29 |
| 8.285E-260 | -0.3515161 | 0.002 | 0.148 | 1.957E-255 | UGEC1 | PHACTR1 |
| 8.51E-260 | -0.2604511 | 0.001 | 0.145 | 2.01E-255 | UGEC1 | WAS |
| 1.253E-259 | -0.3320739 | 0.092 | 0.285 | 2.961E-255 | UGEC1 | FIS1 |
| 7.521E-259 | -0.2976892 | 0.048 | 0.225 | 1.777E-254 | UGEC1 | CCT2 |
| 1.627E-258 | -0.2816331 | 0.017 | 0.175 | 3.842E-254 | UGEC1 | NME1 |
| 2.075E-258 | -0.3563321 | 0.001 | 0.145 | 4.901E-254 | UGEC1 | LILRB2 |
| 2.203E-258 | -0.3938361 | 0.124 | 0.321 | 5.205E-254 | UGEC1 | ACADVL |
| 6.179E-258 | 0.41127549 | 0.637 | 0.456 | 1.46E-253 | UGEC1 | DYNLT1 |
| 8.142E-258 | -0.4369733 | 0.013 | 0.166 | 1.923E-253 | UGEC1 | GPNMB |
| 9.878E-258 | -0.323698 | 0.016 | 0.172 | 2.333E-253 | UGEC1 | CARD16 |
| 1.879E-257 | -0.291399 | 0.052 | 0.229 | 4.438E-253 | UGEC1 | TIMM13 |
| 2.594E-257 | 0.31447206 | 0.563 | 0.354 | 6.127E-253 | UGEC1 | CHMP2A |
| 3.045E-257 | 0.26168485 | 0.397 | 0.208 | 7.193E-253 | UGEC1 | CASP4 |
| 6.255E-257 | -0.3552629 | 0.127 | 0.326 | 1.478E-252 | UGEC1 | NDUFS6 |
| 6.679E-257 | -0.3274071 | 0.026 | 0.187 | 1.578E-252 | UGEC1 | TUT4 |
| 8.047E-257 | -0.3006608 | 0.076 | 0.264 | 1.901E-252 | UGEC1 | EIF2S3 |
| 2.191E-256 | -0.2833891 | 0.002 | 0.147 | 5.176E-252 | UGEC1 | DOCK10 |
| 2.544E-256 | -0.2798526 | 0.027 | 0.189 | 6.009E-252 | UGEC1 | BLOC1S2 |
| 2.75E-256 | -0.2652079 | 0.03 | 0.194 | 6.496E-252 | UGEC1 | MCRIP1 |
| 4.858E-255 | -0.4412732 | 0.203 | 0.405 | 1.148E-250 | UGEC1 | CIRBP |
| 6.24E-255 | -0.3281924 | 0.106 | 0.299 | 1.474E-250 | UGEC1 | RNH1 |
| 6.92E-255 | -0.257765 | 0.025 | 0.185 | 1.635E-250 | UGEC1 | FUNDC2 |
| 1.007E-254 | -0.3275286 | 0.098 | 0.291 | 2.378E-250 | UGEC1 | CCT3 |

| | | | | | | |
|---|---|---|---|---|---|---|
| 2.315E-254 | -0.2994013 | 0.058 | 0.235 | 5.469E-250 | UGEC1 | SLIRP |
| 3.113E-254 | -0.3140611 | 0.001 | 0.144 | 7.353E-250 | UGEC1 | CSF1R |
| 6.964E-254 | 0.26736439 | 0.341 | 0.169 | 1.645E-249 | UGEC1 | AKIRIN2 |
| 1.469E-253 | -0.4199442 | 0.052 | 0.224 | 3.471E-249 | UGEC1 | SDF2L1 |
| 1.553E-253 | -0.2929152 | 0.01 | 0.159 | 3.669E-249 | UGEC1 | PDE4DIP |
| 2.401E-253 | -0.2682014 | 0.021 | 0.177 | 5.671E-249 | UGEC1 | LPGAT1 |
| 2.824E-253 | -0.3215287 | 0.039 | 0.207 | 6.67E-249 | UGEC1 | ERICH1 |
| 4.421E-253 | -0.2863826 | 0.014 | 0.165 | 1.044E-248 | UGEC1 | ITPR2 |
| 1.553E-252 | -0.4412688 | 0.002 | 0.143 | 3.668E-248 | UGEC1 | FCGR3A |
| 4.599E-252 | -0.3089679 | 0.018 | 0.172 | 1.086E-247 | UGEC1 | UCP2 |
| 6.599E-252 | -0.2689935 | 0.002 | 0.143 | 1.559E-247 | UGEC1 | MPP1 |
| 1.893E-251 | -0.3950926 | 0.659 | 0.78 | 4.472E-247 | UGEC1 | UQCRB |
| 3.216E-251 | -0.4736229 | 0.13 | 0.32 | 7.597E-247 | UGEC1 | MACF1 |
| 3.941E-251 | -0.3143002 | 0.046 | 0.215 | 9.31E-247 | UGEC1 | GPATCH2L |
| 6.03E-251 | -0.2663487 | 0.042 | 0.21 | 1.424E-246 | UGEC1 | POLR2E |
| 9.296E-251 | -0.4605239 | 0.869 | 0.912 | 2.196E-246 | UGEC1 | CFL1 |
| 1.104E-250 | -0.3422175 | 0.066 | 0.242 | 2.607E-246 | UGEC1 | RNF13 |
| 1.177E-250 | -0.253915 | 0.003 | 0.145 | 2.78E-246 | UGEC1 | DENND5A |
| 1.229E-250 | 0.32364457 | 0.333 | 0.166 | 2.903E-246 | UGEC1 | GLTP |
| 1.887E-250 | -0.2705762 | 0.012 | 0.16 | 4.457E-246 | UGEC1 | SLC6A6 |
| 3.279E-250 | -0.2723488 | 0.001 | 0.141 | 7.746E-246 | UGEC1 | MYO1G |
| 4.017E-250 | -0.3756013 | 0.143 | 0.346 | 9.489E-246 | UGEC1 | CTNNB1 |
| 4.534E-250 | -0.4062069 | 0.191 | 0.399 | 1.071E-245 | UGEC1 | SLC25A5 |
| 5.319E-250 | -0.3319438 | 0.054 | 0.228 | 1.257E-245 | UGEC1 | MDM4 |
| 1.199E-249 | 0.29802275 | 0.269 | 0.12 | 2.834E-245 | UGEC1 | TUBB6 |
| 2.797E-249 | -0.2766573 | 0.008 | 0.153 | 6.607E-245 | UGEC1 | UTRN |
| 1.716E-248 | -0.3112919 | 0.022 | 0.178 | 4.054E-244 | UGEC1 | SLC44A1 |
| 1.72E-248 | -0.5914806 | 0.101 | 0.282 | 4.064E-244 | UGEC1 | SNHG12 |
| 5.545E-248 | -0.3753596 | 0.026 | 0.182 | 1.31E-243 | UGEC1 | ADGRE5 |
| 1.044E-247 | -0.2900609 | 0.003 | 0.144 | 2.467E-243 | UGEC1 | LIMD2 |
| 1.337E-247 | 0.33957052 | 0.769 | 0.584 | 3.159E-243 | UGEC1 | CAP1 |
| 1.386E-247 | -0.2904351 | 0.001 | 0.139 | 3.274E-243 | UGEC1 | C3AR1 |
| 6.165E-247 | -0.2702383 | 0.028 | 0.187 | 1.456E-242 | UGEC1 | DUT |
| 1.581E-246 | 0.28846096 | 0.268 | 0.119 | 3.734E-242 | UGEC1 | MXI1 |
| 2.012E-246 | -0.8494935 | 0.129 | 0.312 | 4.753E-242 | UGEC1 | CLU |
| 2.728E-246 | 0.57918625 | 0.704 | 0.505 | 6.445E-242 | UGEC1 | CLDN4 |
| 4.077E-246 | -0.2808467 | 0.015 | 0.164 | 9.63E-242 | UGEC1 | PTPN6 |
| 6.489E-246 | 0.28395863 | 0.417 | 0.226 | 1.533E-241 | UGEC1 | ERBIN |
| 8.588E-246 | -0.2595848 | 0.001 | 0.138 | 2.029E-241 | UGEC1 | ITGAM |
| 1.022E-245 | -0.2880126 | 0.052 | 0.224 | 2.415E-241 | UGEC1 | IPO7 |
| 1.408E-245 | 0.25289454 | 0.284 | 0.13 | 3.325E-241 | UGEC1 | R3HDM4 |
| 2.038E-245 | -0.3051597 | 0.043 | 0.208 | 4.814E-241 | UGEC1 | GNA13 |
| 3.078E-245 | -0.3859704 | 0.218 | 0.42 | 7.27E-241 | UGEC1 | NEDD8 |
| 8.318E-245 | -0.4119006 | 0.002 | 0.139 | 1.965E-240 | UGEC1 | CD163 |
| 1.256E-244 | -0.3484498 | 0.135 | 0.332 | 2.967E-240 | UGEC1 | ROMO1 |
| 1.645E-244 | -0.3813509 | 0.175 | 0.374 | 3.886E-240 | UGEC1 | ANP32B |
| 2.148E-244 | -0.2872603 | 0.061 | 0.235 | 5.075E-240 | UGEC1 | CHURC1 |
| 2.271E-244 | -1.9229219 | 0.023 | 0.178 | 5.364E-240 | UGEC1 | MT1H |
| 3.025E-244 | -0.2542438 | 0.036 | 0.199 | 7.146E-240 | UGEC1 | PHB2 |
| 3.736E-244 | -0.6662159 | 0.002 | 0.14 | 8.825E-240 | UGEC1 | MNDA |

| | | | | | | |
|---|---|---|---|---|---|---|
| 1.227E-243 | 0.26240237 | 0.181 | 0.063 | 2.899E-239 | UGEC1 | HS3ST1 |
| 1.955E-243 | -0.4513775 | 0.127 | 0.315 | 4.618E-239 | UGEC1 | CD151 |
| 5.51E-243 | -0.2838703 | 0.024 | 0.178 | 1.302E-238 | UGEC1 | LBR |
| 6.28E-243 | 0.34327043 | 0.224 | 0.092 | 1.483E-238 | UGEC1 | GCHFR |
| 6.599E-243 | 0.28884166 | 0.563 | 0.35 | 1.559E-238 | UGEC1 | RAB2A |
| 7.387E-242 | -0.2736127 | 0.008 | 0.149 | 1.745E-237 | UGEC1 | ATM |
| 2.246E-241 | -0.4245471 | 0.694 | 0.787 | 5.305E-237 | UGEC1 | RHOA |
| 2.284E-240 | 0.252364 | 0.299 | 0.141 | 5.394E-236 | UGEC1 | CHIC2 |
| 4.853E-240 | -0.4202163 | 0.18 | 0.376 | 1.146E-235 | UGEC1 | ATP5IF1 |
| 5.066E-240 | -0.3493997 | 0.186 | 0.386 | 1.197E-235 | UGEC1 | SNX3 |
| 5.469E-240 | -0.2531719 | 0.006 | 0.145 | 1.292E-235 | UGEC1 | LPCAT1 |
| 7.228E-240 | -0.3450915 | 0.142 | 0.335 | 1.708E-235 | UGEC1 | PRKAR1A |
| 8.005E-240 | -0.285045 | 0.043 | 0.207 | 1.891E-235 | UGEC1 | GPI |
| 1.859E-239 | -0.2720015 | 0.034 | 0.191 | 4.392E-235 | UGEC1 | CNPY3 |
| 2.546E-239 | -0.2828599 | 0.001 | 0.136 | 6.015E-235 | UGEC1 | FYN |
| 3.368E-239 | -0.3210469 | 0.017 | 0.164 | 7.956E-235 | UGEC1 | GCA |
| 4.375E-239 | -0.4926085 | 0.038 | 0.196 | 1.033E-234 | UGEC1 | AKR1B1 |
| 1.284E-238 | -0.3375556 | 0.07 | 0.244 | 3.033E-234 | UGEC1 | LMAN1 |
| 2.678E-238 | -0.3669692 | 0.169 | 0.364 | 6.325E-234 | UGEC1 | PSMB3 |
| 3.519E-238 | -0.3335969 | 0.025 | 0.175 | 8.312E-234 | UGEC1 | SMAP2 |
| 6.137E-238 | -0.4161026 | 0.229 | 0.421 | 1.45E-233 | UGEC1 | UQCR11 |
| 1.146E-237 | -0.3372038 | 0.106 | 0.292 | 2.708E-233 | UGEC1 | CUTA |
| 2.16E-237 | -0.3129457 | 0.002 | 0.136 | 5.102E-233 | UGEC1 | PECAM1 |
| 5.164E-237 | 0.32570987 | 0.49 | 0.291 | 1.22E-232 | UGEC1 | FOSL2 |
| 8.23E-237 | 0.27270883 | 0.384 | 0.201 | 1.944E-232 | UGEC1 | BAZ2B |
| 9.087E-237 | 0.26332556 | 0.321 | 0.156 | 2.147E-232 | UGEC1 | CDCP1 |
| 1.045E-236 | -0.2947645 | 0.108 | 0.295 | 2.469E-232 | UGEC1 | PSMA1 |
| 2.211E-236 | 0.25635307 | 0.32 | 0.156 | 5.222E-232 | UGEC1 | GTF2B |
| 3.601E-236 | -0.3504194 | 0.13 | 0.32 | 8.507E-232 | UGEC1 | NDUFC1 |
| 8.697E-236 | -0.3668386 | 0.198 | 0.397 | 2.054E-231 | UGEC1 | C19orf53 |
| 1.242E-235 | -0.8698267 | 0.002 | 0.136 | 2.934E-231 | UGEC1 | IL7R |
| 2.043E-235 | -0.3301901 | 0.152 | 0.348 | 4.826E-231 | UGEC1 | RBX1 |
| 2.137E-235 | -0.3771199 | 0.01 | 0.15 | 5.047E-231 | UGEC1 | SYNE1 |
| 3.184E-235 | -0.2957861 | 0.057 | 0.223 | 7.521E-231 | UGEC1 | PHC2 |
| 5.45E-235 | -0.2879583 | 0.068 | 0.241 | 1.287E-230 | UGEC1 | RNPS1 |
| 5.973E-235 | 0.29513584 | 0.446 | 0.252 | 1.411E-230 | UGEC1 | RB1CC1 |
| 1.542E-234 | 0.31294424 | 0.339 | 0.174 | 3.644E-230 | UGEC1 | MUC20-OT1 |
| 4.777E-234 | -0.2758811 | 0.022 | 0.169 | 1.128E-229 | UGEC1 | PCNX1 |
| 5.406E-234 | 0.27322053 | 0.331 | 0.167 | 1.277E-229 | UGEC1 | RNF11 |
| 6.029E-234 | -0.3818806 | 0.085 | 0.257 | 1.424E-229 | UGEC1 | FCGRT |
| 7.61E-234 | -0.3772957 | 0.063 | 0.23 | 1.798E-229 | UGEC1 | NFKB1 |
| 7.762E-234 | -0.3780912 | 0.195 | 0.393 | 1.834E-229 | UGEC1 | SRSF11 |
| 8.015E-234 | -0.780465 | 0.868 | 0.93 | 1.893E-229 | UGEC1 | S100A10 |
| 8.733E-234 | -0.2871905 | 0.063 | 0.233 | 2.063E-229 | UGEC1 | NENF |
| 2.509E-233 | -0.3924824 | 0.709 | 0.857 | 5.927E-229 | UGEC1 | UBB |
| 4.879E-233 | -0.3359774 | 0.124 | 0.311 | 1.153E-228 | UGEC1 | MRPL41 |
| 9.785E-233 | -0.3285423 | 0.049 | 0.211 | 2.312E-228 | UGEC1 | LMO4 |
| 1.102E-232 | 0.84279765 | 0.729 | 0.642 | 2.603E-228 | UGEC1 | HSPA1A |
| 2.671E-232 | 0.9564293 | 0.397 | 0.25 | 6.309E-228 | UGEC1 | CAMK2N1 |
| 4.299E-232 | -0.2566215 | 0.025 | 0.175 | 1.016E-227 | UGEC1 | LARS |

| | | | | | | |
|---|---|---|---|---|---|---|
| 4.436E-232 | -0.2538449 | 0.038 | 0.194 | 1.048E-227 | UGEC1 | TBCB |
| 2.444E-231 | -0.2637059 | 0.02 | 0.166 | 5.773E-227 | UGEC1 | NFIC |
| 4.408E-231 | -0.3512335 | 0.193 | 0.391 | 1.041E-226 | UGEC1 | ATP5PD |
| 9.61E-231 | -0.3092223 | 0.006 | 0.14 | 2.27E-226 | UGEC1 | CD86 |
| 1.014E-230 | -0.5066234 | 0.561 | 0.696 | 2.395E-226 | UGEC1 | CALR |
| 2.211E-230 | -0.2557448 | 0.038 | 0.195 | 5.223E-226 | UGEC1 | METTL26 |
| 2.27E-230 | -0.4022532 | 0.174 | 0.369 | 5.363E-226 | UGEC1 | LUC7L3 |
| 2.487E-230 | -0.2566815 | 0.041 | 0.2 | 5.875E-226 | UGEC1 | ATRAID |
| 2.495E-230 | -0.2521207 | 0.002 | 0.133 | 5.895E-226 | UGEC1 | LAIR1 |
| 4.955E-230 | -0.2702063 | 0.047 | 0.208 | 1.171E-225 | UGEC1 | RBCK1 |
| 1.553E-229 | 0.25985576 | 0.227 | 0.094 | 3.669E-225 | UGEC1 | TRNP1 |
| 2.187E-229 | 0.29554745 | 0.381 | 0.206 | 5.166E-225 | UGEC1 | PNPLA2 |
| 4.462E-229 | -0.2859836 | 0.076 | 0.249 | 1.054E-224 | UGEC1 | MDH2 |
| 1.193E-227 | -0.2958954 | 0.082 | 0.255 | 2.819E-223 | UGEC1 | ARPP19 |
| 2.57E-227 | 0.26726358 | 0.29 | 0.138 | 6.072E-223 | UGEC1 | CDYL |
| 4.271E-227 | -0.25768 | 0.031 | 0.184 | 1.009E-222 | UGEC1 | PRKDC |
| 4.445E-227 | -0.2697543 | 0.008 | 0.142 | 1.05E-222 | UGEC1 | FKBP5 |
| 5.14E-227 | -0.3457925 | 0.112 | 0.297 | 1.214E-222 | UGEC1 | XBP1 |
| 1.208E-226 | -0.2704057 | 0.048 | 0.207 | 2.853E-222 | UGEC1 | ZFR |
| 4.973E-226 | -0.3191841 | 0.112 | 0.293 | 1.175E-221 | UGEC1 | HNRNPR |
| 5.637E-226 | 0.28457582 | 0.947 | 0.85 | 1.332E-221 | UGEC1 | ARPC3 |
| 7.388E-226 | -0.6973955 | 0.014 | 0.152 | 1.745E-221 | UGEC1 | TFPI2 |
| 9.272E-226 | -0.3833861 | 0.455 | 0.617 | 2.19E-221 | UGEC1 | SLC25A3 |
| 2.333E-225 | -0.2555214 | 0.044 | 0.203 | 5.512E-221 | UGEC1 | TMEM147 |
| 2.431E-225 | -0.3877522 | 0.003 | 0.132 | 5.744E-221 | UGEC1 | TNF |
| 2.665E-225 | -0.404307 | 0.169 | 0.351 | 6.296E-221 | UGEC1 | AP2S1 |
| 3.591E-225 | -0.2650058 | 0.001 | 0.129 | 8.484E-221 | UGEC1 | PIK3AP1 |
| 7.084E-225 | -0.3117583 | 0.002 | 0.13 | 1.673E-220 | UGEC1 | PDE4B |
| 1.467E-224 | -0.324148 | 0.133 | 0.319 | 3.467E-220 | UGEC1 | MRFAP1 |
| 2.44E-224 | -0.2569763 | 0.043 | 0.2 | 5.764E-220 | UGEC1 | RPL7L1 |
| 3.508E-224 | -1.2132025 | 0.002 | 0.129 | 8.286E-220 | UGEC1 | CCL4 |
| 3.882E-224 | -0.3347856 | 0.157 | 0.346 | 9.17E-220 | UGEC1 | ARHGDIA |
| 2.969E-223 | -0.3711939 | 0.002 | 0.13 | 7.015E-219 | UGEC1 | FGL2 |
| 3.244E-223 | -0.2873228 | 0.071 | 0.238 | 7.664E-219 | UGEC1 | ILF3 |
| 5.317E-223 | -0.2509582 | 0.005 | 0.134 | 1.256E-218 | UGEC1 | LIPA |
| 1.259E-222 | -0.2569237 | 0.001 | 0.127 | 2.974E-218 | UGEC1 | GPRIN3 |
| 2.9E-222 | -0.3379857 | 0.002 | 0.128 | 6.851E-218 | UGEC1 | MS4A6A |
| 3.207E-222 | -0.2892191 | 0.102 | 0.28 | 7.576E-218 | UGEC1 | ATP5PB |
| 3.897E-222 | -0.2851667 | 0.103 | 0.282 | 9.207E-218 | UGEC1 | XRCC6 |
| 2.38E-221 | -0.2502289 | 0.002 | 0.128 | 5.623E-217 | UGEC1 | HAVCR2 |
| 4.068E-221 | -0.4102758 | 0.004 | 0.132 | 9.61E-217 | UGEC1 | FXYD2 |
| 5.342E-221 | -0.2656908 | 0.056 | 0.216 | 1.262E-216 | UGEC1 | PRDX3 |
| 1.124E-220 | -0.3093694 | 0.077 | 0.243 | 2.655E-216 | UGEC1 | ZYX |
| 5.954E-220 | 0.40544445 | 0.327 | 0.171 | 1.407E-215 | UGEC1 | PTGR1 |
| 7.955E-220 | -0.3972949 | 0.002 | 0.127 | 1.879E-215 | UGEC1 | CSF3R |
| 1.227E-219 | -0.294683 | 0.103 | 0.278 | 2.898E-215 | UGEC1 | EIF3D |
| 2.473E-219 | 0.37607783 | 0.257 | 0.119 | 5.841E-215 | UGEC1 | ARL4A |
| 3.833E-219 | -0.2564549 | 0.06 | 0.221 | 9.055E-215 | UGEC1 | MRPL54 |
| 5.892E-219 | -0.2687761 | 0.001 | 0.124 | 1.392E-214 | UGEC1 | CCR1 |
| 7.387E-219 | -0.366132 | 0.01 | 0.142 | 1.745E-214 | UGEC1 | AC007952.4 |

| | | | | | | |
|---|---|---|---|---|---|---|
| 1.736E-218 | -0.2850339 | 0.078 | 0.244 | 4.102E-214 | UGEC1 | RWDD1 |
| 3.139E-218 | -0.4640713 | 0.239 | 0.419 | 7.415E-214 | UGEC1 | HMGN2 |
| 8.488E-218 | -0.2587269 | 0.054 | 0.211 | 2.005E-213 | UGEC1 | DPP7 |
| 1.214E-217 | -0.2886436 | 0.056 | 0.213 | 2.867E-213 | UGEC1 | PRMT2 |
| 1.348E-217 | -0.2947761 | 0.028 | 0.173 | 3.185E-213 | UGEC1 | SCCPDH |
| 2.713E-217 | -0.2939258 | 0.004 | 0.13 | 6.408E-213 | UGEC1 | ACP5 |
| 6.62E-217 | -0.2591761 | 0.064 | 0.226 | 1.564E-212 | UGEC1 | TMEM230 |
| 1.039E-216 | -0.2570648 | 0.015 | 0.149 | 2.455E-212 | UGEC1 | ADAM8 |
| 1.379E-216 | -0.397664 | 0.184 | 0.365 | 3.257E-212 | UGEC1 | SSR3 |
| 3.241E-216 | -0.8273966 | 0.004 | 0.13 | 7.657E-212 | UGEC1 | APOC1 |
| 3.469E-216 | -0.2742848 | 0.041 | 0.191 | 8.195E-212 | UGEC1 | CDC42SE2 |
| 5.282E-216 | -0.2851564 | 0.045 | 0.196 | 1.248E-211 | UGEC1 | FNBP1 |
| 5.668E-216 | -0.3256679 | 0.164 | 0.352 | 1.339E-211 | UGEC1 | ZNF706 |
| 6.209E-216 | -0.2531497 | 0.015 | 0.15 | 1.467E-211 | UGEC1 | SLC4A7 |
| 7.445E-216 | 0.36798053 | 0.727 | 0.572 | 1.759E-211 | UGEC1 | VAMP8 |
| 1.356E-215 | -0.2857842 | 0.073 | 0.239 | 3.204E-211 | UGEC1 | G3BP1 |
| 1.385E-215 | -0.2775169 | 0.038 | 0.185 | 3.273E-211 | UGEC1 | CPD |
| 2.624E-215 | 0.25320699 | 0.261 | 0.12 | 6.198E-211 | UGEC1 | ZNF431 |
| 4.282E-215 | -0.2667757 | 0.001 | 0.122 | 1.012E-210 | UGEC1 | CLEC4E |
| 4.955E-215 | -0.2673902 | 0.003 | 0.126 | 1.17E-210 | UGEC1 | ENG |
| 9.242E-215 | -0.2838255 | 0.04 | 0.191 | 2.183E-210 | UGEC1 | FLNB |
| 1.799E-214 | -0.3099003 | 0.064 | 0.223 | 4.25E-210 | UGEC1 | TNRC6B |
| 3.128E-214 | -0.2611695 | 0.049 | 0.203 | 7.39E-210 | UGEC1 | SENP6 |
| 5.069E-214 | -0.2991206 | 0.09 | 0.259 | 1.197E-209 | UGEC1 | MRPL57 |
| 1.939E-213 | -0.360716 | 0.21 | 0.395 | 4.58E-209 | UGEC1 | PSMB1 |
| 2.081E-213 | 0.27339289 | 0.262 | 0.124 | 4.915E-209 | UGEC1 | BCAP29 |
| 2.154E-213 | -0.2912298 | 0.066 | 0.227 | 5.088E-209 | UGEC1 | TTC3 |
| 3.309E-213 | -0.2516968 | 0.03 | 0.173 | 7.818E-209 | UGEC1 | NOP56 |
| 5.545E-213 | -0.3255564 | 0.002 | 0.124 | 1.31E-208 | UGEC1 | LILRB1 |
| 5.579E-213 | -0.5102913 | 0.022 | 0.158 | 1.318E-208 | UGEC1 | HLA-DRB5 |
| 7.462E-213 | -0.2555556 | 0.048 | 0.201 | 1.763E-208 | UGEC1 | CYC1 |
| 8.074E-213 | -0.4455045 | 0.479 | 0.633 | 1.907E-208 | UGEC1 | SON |
| 1.348E-212 | -0.5639899 | 0 | 0.12 | 3.185E-208 | UGEC1 | RETN |
| 1.422E-212 | -0.2590567 | 0.035 | 0.181 | 3.359E-208 | UGEC1 | RIF1 |
| 3.499E-212 | -0.8179751 | 0.002 | 0.125 | 8.265E-208 | UGEC1 | CCL5 |
| 5.969E-212 | -0.2579822 | 0.001 | 0.122 | 1.41E-207 | UGEC1 | SLC8A1 |
| 6.204E-212 | -0.3483127 | 0.461 | 0.624 | 1.466E-207 | UGEC1 | HNRNPK |
| 6.344E-211 | -0.2990531 | 0.113 | 0.287 | 1.499E-206 | UGEC1 | RPS19BP1 |
| 1.546E-210 | -0.6202143 | 0.05 | 0.197 | 3.652E-206 | UGEC1 | HLA-DQB1 |
| 1.712E-210 | -0.2814832 | 0.098 | 0.268 | 4.044E-206 | UGEC1 | TUFM |
| 2.505E-210 | -0.3381052 | 0.058 | 0.212 | 5.919E-206 | UGEC1 | CKLF |
| 3.157E-210 | -0.4040558 | 0.005 | 0.127 | 7.458E-206 | UGEC1 | SLC16A10 |
| 3.171E-210 | -0.2617669 | 0.076 | 0.239 | 7.491E-206 | UGEC1 | ANXA7 |
| 1.266E-209 | -0.3112794 | 0.058 | 0.212 | 2.99E-205 | UGEC1 | HIPK2 |
| 2.164E-209 | -0.3197572 | 0.137 | 0.312 | 5.111E-205 | UGEC1 | COX7A2L |
| 6.108E-209 | -0.2790385 | 0.003 | 0.125 | 1.443E-204 | UGEC1 | DPP4 |
| 6.497E-209 | -0.3589542 | 0.17 | 0.352 | 1.535E-204 | UGEC1 | KRT10 |
| 5.032E-208 | -0.3290479 | 0.087 | 0.25 | 1.189E-203 | UGEC1 | MAP3K2 |
| 5.661E-208 | 0.32468074 | 0.723 | 0.545 | 1.337E-203 | UGEC1 | CD55 |
| 9.192E-208 | -0.3552946 | 0.076 | 0.235 | 2.171E-203 | UGEC1 | STK17B |

| | | | | | | |
|---|---|---|---|---|---|---|
| 1.224E-207 | -0.2683724 | 0.074 | 0.236 | 2.892E-203 | UGEC1 | CNIH4 |
| 5.48E-207 | -0.2726191 | 0 | 0.117 | 1.295E-202 | UGEC1 | IKZF1 |
| 5.511E-207 | 0.27031473 | 0.335 | 0.178 | 1.302E-202 | UGEC1 | CNOT1 |
| 2.349E-206 | -0.2508981 | 0.004 | 0.124 | 5.549E-202 | UGEC1 | TRIB3 |
| 2.925E-206 | -0.2615599 | 0.04 | 0.183 | 6.909E-202 | UGEC1 | PPT1 |
| 6.251E-206 | -0.5461498 | 0.131 | 0.298 | 1.477E-201 | UGEC1 | TSC22D3 |
| 1.374E-205 | -0.3587292 | 0.694 | 0.777 | 3.246E-201 | UGEC1 | UBL5 |
| 2.735E-205 | -0.3074842 | 0.019 | 0.151 | 6.461E-201 | UGEC1 | BEX2 |
| 3.288E-205 | -0.3988231 | 0.195 | 0.373 | 7.768E-201 | UGEC1 | BSG |
| 5.993E-205 | 0.2882874 | 0.846 | 0.684 | 1.416E-200 | UGEC1 | UBE2D3 |
| 1.099E-204 | -0.2581063 | 0.058 | 0.212 | 2.596E-200 | UGEC1 | NHP2 |
| 1.617E-204 | 0.25328523 | 0.337 | 0.18 | 3.821E-200 | UGEC1 | SNAP23 |
| 2.007E-204 | -0.2848804 | 0.123 | 0.295 | 4.74E-200 | UGEC1 | VPS29 |
| 2.094E-204 | -0.3395001 | 0.217 | 0.399 | 4.946E-200 | UGEC1 | GNB1 |
| 2.422E-204 | 0.28511785 | 0.882 | 0.749 | 5.721E-200 | UGEC1 | EIF4G2 |
| 2.584E-204 | -0.3352767 | 0.123 | 0.296 | 6.104E-200 | UGEC1 | SMG1 |
| 3.387E-204 | -0.2833632 | 0.024 | 0.16 | 8.001E-200 | UGEC1 | SLC30A1 |
| 8.316E-204 | -0.5225279 | 0.862 | 0.898 | 1.965E-199 | UGEC1 | CD63 |
| 2.748E-203 | -0.3975438 | 0.172 | 0.346 | 6.493E-199 | UGEC1 | BOD1L1 |
| 3.523E-203 | 0.30508868 | 0.557 | 0.376 | 8.322E-199 | UGEC1 | NDUFB4 |
| 3.345E-202 | -0.4429605 | 0.736 | 0.806 | 7.901E-198 | UGEC1 | CHCHD2 |
| 4.607E-202 | -0.2575977 | 0.074 | 0.232 | 1.088E-197 | UGEC1 | RPN1 |
| 8.323E-202 | -0.3646713 | 0.026 | 0.159 | 1.966E-197 | UGEC1 | HLA-DMA |
| 1.738E-201 | -0.265286 | 0.055 | 0.204 | 4.105E-197 | UGEC1 | RANBP2 |
| 2.528E-201 | -0.2591672 | 0.076 | 0.235 | 5.971E-197 | UGEC1 | COX14 |
| 2.723E-201 | 0.31372712 | 0.708 | 0.53 | 6.432E-197 | UGEC1 | CSNK1A1 |
| 5.658E-201 | -0.2824516 | 0.046 | 0.19 | 1.336E-196 | UGEC1 | MYCBP2 |
| 5.703E-201 | -0.3003184 | 0.117 | 0.285 | 1.347E-196 | UGEC1 | NDUFB10 |
| 7.001E-201 | -0.2517895 | 0.055 | 0.205 | 1.654E-196 | UGEC1 | RANBP1 |
| 1.02E-200 | -0.3235498 | 0.151 | 0.326 | 2.41E-196 | UGEC1 | RABAC1 |
| 1.306E-200 | -0.2733 | 0.047 | 0.193 | 3.086E-196 | UGEC1 | NRIP1 |
| 1.594E-200 | -0.2530685 | 0.036 | 0.174 | 3.766E-196 | UGEC1 | CLIC4 |
| 2.02E-200 | -0.3931071 | 0.077 | 0.232 | 4.771E-196 | UGEC1 | UBE2S |
| 2.297E-200 | -1.4450456 | 0.012 | 0.136 | 5.425E-196 | UGEC1 | MT1M |
| 2.898E-200 | -2.0954608 | 0.207 | 0.366 | 6.847E-196 | UGEC1 | MMP7 |
| 3.51E-200 | -0.25246 | 0.074 | 0.231 | 8.292E-196 | UGEC1 | TXNL4A |
| 4.507E-200 | -0.2999676 | 0.068 | 0.22 | 1.065E-195 | UGEC1 | SEC11C |
| 1.102E-199 | -0.3448256 | 0.087 | 0.244 | 2.604E-195 | UGEC1 | STK4 |
| 1.442E-199 | -0.2699354 | 0.04 | 0.18 | 3.407E-195 | UGEC1 | NFE2L1 |
| 2.148E-199 | 0.416997 | 0.699 | 0.55 | 5.073E-195 | UGEC1 | PRDX5 |
| 4.505E-199 | -0.2755387 | 0.102 | 0.271 | 1.064E-194 | UGEC1 | COMT |
| 5.161E-199 | -0.2942756 | 0.145 | 0.32 | 1.219E-194 | UGEC1 | HIGD2A |
| 5.793E-199 | -0.4997553 | 0.006 | 0.124 | 1.368E-194 | UGEC1 | LTB |
| 9.105E-199 | -0.3703643 | 0.075 | 0.229 | 2.151E-194 | UGEC1 | LSP1 |
| 1.114E-198 | -0.6695951 | 0.004 | 0.121 | 2.632E-194 | UGEC1 | HLA-DQA1 |
| 3.544E-198 | -0.2965184 | 0.115 | 0.283 | 8.372E-194 | UGEC1 | CCT6A |
| 3.942E-198 | -0.2632019 | 0.023 | 0.153 | 9.313E-194 | UGEC1 | TMX4 |
| 1.245E-197 | -0.4298316 | 0.013 | 0.136 | 2.94E-193 | UGEC1 | RRAD |
| 7.703E-197 | -0.3135784 | 0.163 | 0.337 | 1.82E-192 | UGEC1 | ANAPC16 |
| 6.668E-196 | 0.28668671 | 0.392 | 0.228 | 1.575E-191 | UGEC1 | TACC1 |

| | | | | | | |
|---|---|---|---|---|---|---|
| 2.981E-195 | -1.3556292 | 0.038 | 0.175 | 7.043E-191 | UGEC1 | CCL20 |
| 3.063E-195 | -0.5524829 | 0.017 | 0.141 | 7.235E-191 | UGEC1 | TCIM |
| 1.006E-194 | 0.29812169 | 0.457 | 0.289 | 2.377E-190 | UGEC1 | RER1 |
| 1.321E-194 | -0.3110564 | 0.082 | 0.237 | 3.121E-190 | UGEC1 | NFAT5 |
| 1.367E-194 | -0.36784 | 0.201 | 0.375 | 3.23E-190 | UGEC1 | SCAF11 |
| 1.368E-194 | 0.28221063 | 0.459 | 0.281 | 3.231E-190 | UGEC1 | SBDS |
| 1.724E-194 | -0.3375048 | 0.09 | 0.25 | 4.072E-190 | UGEC1 | MRPS6 |
| 2.019E-194 | 0.28067491 | 0.819 | 0.68 | 4.77E-190 | UGEC1 | SUMO2 |
| 2.908E-194 | -0.3889386 | 0.09 | 0.257 | 6.87E-190 | UGEC1 | CYB5A |
| 5.556E-194 | -0.2868294 | 0.149 | 0.323 | 1.313E-189 | UGEC1 | SEPT2 |
| 5.938E-194 | 0.48759849 | 0.23 | 0.109 | 1.403E-189 | UGEC1 | AIF1L |
| 2.404E-193 | -0.2514575 | 0.049 | 0.192 | 5.68E-189 | UGEC1 | NUP58 |
| 4.246E-193 | -0.2683855 | 0.094 | 0.253 | 1.003E-188 | UGEC1 | PSMB2 |
| 1.24E-192 | -0.2518484 | 0.062 | 0.209 | 2.929E-188 | UGEC1 | UBE2Z |
| 2.758E-192 | -0.2540635 | 0.079 | 0.234 | 6.514E-188 | UGEC1 | FAM50A |
| 6.222E-192 | -0.3569477 | 0.208 | 0.387 | 1.47E-187 | UGEC1 | UQCRQ |
| 9.436E-191 | -0.2984565 | 0.035 | 0.168 | 2.229E-186 | UGEC1 | CD40 |
| 1.476E-190 | -0.3315854 | 0.221 | 0.397 | 3.486E-186 | UGEC1 | WDR83OS |
| 2.079E-190 | -0.3034193 | 0.062 | 0.209 | 4.91E-186 | UGEC1 | DAB2 |
| 2.622E-190 | -0.4251934 | 0.191 | 0.368 | 6.195E-186 | UGEC1 | NDRG1 |
| 5.143E-190 | -0.2906596 | 0.105 | 0.265 | 1.215E-185 | UGEC1 | NCOR1 |
| 1.488E-189 | -0.8387721 | 0.358 | 0.519 | 3.516E-185 | UGEC1 | MTRNR2L8 |
| 3.308E-189 | 0.28407088 | 0.288 | 0.147 | 7.815E-185 | UGEC1 | B3GNT5 |
| 9.69E-189 | 0.30894268 | 0.54 | 0.359 | 2.289E-184 | UGEC1 | STAT3 |
| 1.528E-188 | -0.3860105 | 0.066 | 0.21 | 3.609E-184 | UGEC1 | SEPHS2 |
| 1.626E-188 | -0.4006097 | 0.057 | 0.195 | 3.841E-184 | UGEC1 | SAMHD1 |
| 2.93E-188 | -0.2669042 | 0.072 | 0.221 | 6.923E-184 | UGEC1 | ZC3H13 |
| 8.76E-188 | -0.5838819 | 0.025 | 0.151 | 2.069E-183 | UGEC1 | AKAP12 |
| 1.544E-187 | -0.2962255 | 0.149 | 0.316 | 3.648E-183 | UGEC1 | TMED10 |
| 2.156E-187 | -0.2754489 | 0.097 | 0.253 | 5.094E-183 | UGEC1 | XRN2 |
| 4.192E-187 | -0.3151432 | 0.203 | 0.377 | 9.903E-183 | UGEC1 | SNRPB |
| 4.268E-187 | -0.2501295 | 0.002 | 0.111 | 1.008E-182 | UGEC1 | TNFSF13B |
| 1.83E-186 | -0.3227855 | 0.004 | 0.114 | 4.322E-182 | UGEC1 | GPR183 |
| 6.07E-185 | 0.51777385 | 0.384 | 0.233 | 1.434E-180 | UGEC1 | PMAIP1 |
| 9.18E-185 | -0.3201733 | 0.217 | 0.389 | 2.169E-180 | UGEC1 | RBM8A |
| 2.422E-184 | -0.2687801 | 0.062 | 0.204 | 5.722E-180 | UGEC1 | SARS |
| 1.609E-183 | -0.3597211 | 0.027 | 0.152 | 3.802E-179 | UGEC1 | CD83 |
| 2.67E-183 | -0.2729668 | 0.123 | 0.284 | 6.307E-179 | UGEC1 | SF3B2 |
| 2.932E-183 | -0.283987 | 0.124 | 0.283 | 6.927E-179 | UGEC1 | DNAJC8 |
| 1.437E-182 | -0.3324256 | 0.2 | 0.372 | 3.395E-178 | UGEC1 | ATP5F1B |
| 5.188E-182 | -0.2913293 | 0.14 | 0.303 | 1.226E-177 | UGEC1 | SNX6 |
| 2.748E-181 | 0.58576389 | 0.248 | 0.13 | 6.492E-177 | UGEC1 | LPIN1 |
| 1.346E-179 | -0.3381829 | 0.106 | 0.256 | 3.181E-175 | UGEC1 | ATP6V1B2 |
| 3.793E-179 | -0.3091923 | 0.171 | 0.335 | 8.959E-175 | UGEC1 | SPCS3 |
| 4.102E-179 | -0.2524523 | 0.058 | 0.196 | 9.69E-175 | UGEC1 | CEBPZ |
| 5.358E-179 | -0.2556808 | 0.103 | 0.261 | 1.266E-174 | UGEC1 | SYPL1 |
| 1.693E-178 | -0.2517916 | 0.061 | 0.2 | 3.999E-174 | UGEC1 | MBOAT7 |
| 3.053E-178 | -0.310726 | 0.155 | 0.319 | 7.212E-174 | UGEC1 | HNRNPH3 |
| 3.104E-178 | -0.2640919 | 0.115 | 0.272 | 7.332E-174 | UGEC1 | VPS35 |
| 4.333E-178 | -0.3247578 | 0.189 | 0.353 | 1.024E-173 | UGEC1 | CDC37 |

| | | | | | | |
|---|---|---|---|---|---|---|
| 5.746E-178 | -0.2752398 | 0.162 | 0.33 | 1.357E-173 | UGEC1 | ATP5PO |
| 1.055E-177 | -0.2506873 | 0.068 | 0.21 | 2.492E-173 | UGEC1 | DYNLT3 |
| 3.59E-177 | -0.2612789 | 0.116 | 0.274 | 8.481E-173 | UGEC1 | CCT4 |
| 4.711E-176 | -0.3097789 | 0.135 | 0.293 | 1.113E-171 | UGEC1 | DDX46 |
| 1.286E-175 | 0.33822176 | 0.76 | 0.611 | 3.038E-171 | UGEC1 | GADD45B |
| 2.82E-175 | 0.25032931 | 0.162 | 0.063 | 6.661E-171 | UGEC1 | SOX9 |
| 3.074E-175 | 0.2897806 | 0.602 | 0.416 | 7.261E-171 | UGEC1 | ACTN4 |
| 4.004E-174 | -0.2626336 | 0.091 | 0.239 | 9.458E-170 | UGEC1 | METAP2 |
| 5.161E-174 | -0.2688897 | 0.055 | 0.189 | 1.219E-169 | UGEC1 | CCDC50 |
| 3.882E-173 | -0.318908 | 0.008 | 0.114 | 9.17E-169 | UGEC1 | IFIT2 |
| 4.661E-173 | -0.2703645 | 0.039 | 0.166 | 1.101E-168 | UGEC1 | GABPB1-AS1 |
| 7.654E-173 | -0.3386363 | 0.031 | 0.153 | 1.808E-168 | UGEC1 | RGCC |
| 9.115E-173 | -0.2948248 | 0.075 | 0.214 | 2.153E-168 | UGEC1 | CHD9 |
| 9.695E-173 | -0.2530779 | 0.05 | 0.183 | 2.29E-168 | UGEC1 | ZFHX3 |
| 2.714E-172 | -0.3046432 | 0.006 | 0.111 | 6.411E-168 | UGEC1 | ADAMTS9 |
| 7.354E-172 | 0.34515562 | 0.276 | 0.146 | 1.737E-167 | UGEC1 | TINAGL1 |
| 1.752E-171 | -0.2581398 | 0.142 | 0.303 | 4.139E-167 | UGEC1 | SNRPD3 |
| 2.543E-171 | -0.2867439 | 0.039 | 0.163 | 6.008E-167 | UGEC1 | PIK3R1 |
| 2.799E-171 | 0.33017174 | 0.386 | 0.234 | 6.612E-167 | UGEC1 | ARRDC3 |
| 5.998E-171 | -0.2758045 | 0.126 | 0.28 | 1.417E-166 | UGEC1 | RPN2 |
| 1.015E-170 | -0.3193013 | 0.186 | 0.348 | 2.397E-166 | UGEC1 | ERP29 |
| 1.025E-170 | -0.3761989 | 0.061 | 0.195 | 2.421E-166 | UGEC1 | STMN1 |
| 2.714E-170 | 0.7328021 | 0.93 | 0.874 | 6.412E-166 | UGEC1 | CSTB |
| 6.975E-170 | -0.2500067 | 0.021 | 0.135 | 1.648E-165 | UGEC1 | ADGRE2 |
| 9.765E-170 | -0.2770081 | 0.09 | 0.234 | 2.307E-165 | UGEC1 | PNPLA8 |
| 1.509E-169 | -1.0944879 | 0.037 | 0.157 | 3.564E-165 | UGEC1 | DEFB1 |
| 6.529E-169 | -1.0666543 | 0.653 | 0.714 | 1.542E-164 | UGEC1 | SOD2 |
| 7.701E-169 | -0.3773465 | 0.565 | 0.679 | 1.819E-164 | UGEC1 | RBM39 |
| 6.509E-168 | -0.5510302 | 0.754 | 0.837 | 1.538E-163 | UGEC1 | MT-ND5 |
| 7.624E-168 | -0.3562969 | 0.256 | 0.413 | 1.801E-163 | UGEC1 | DAD1 |
| 1.03E-167 | -0.3053492 | 0.165 | 0.323 | 2.433E-163 | UGEC1 | TRAM1 |
| 2.096E-167 | -0.3278654 | 0.106 | 0.25 | 4.951E-163 | UGEC1 | PNP |
| 3.533E-167 | -0.2506413 | 0.033 | 0.153 | 8.345E-163 | UGEC1 | SQLE |
| 9.181E-167 | -0.2837087 | 0.081 | 0.221 | 2.169E-162 | UGEC1 | CCPG1 |
| 1.522E-166 | -0.2958193 | 0.146 | 0.302 | 3.595E-162 | UGEC1 | PPIG |
| 2.177E-166 | -0.4354066 | 0.239 | 0.395 | 5.142E-162 | UGEC1 | ANKRD11 |
| 2.453E-166 | 0.27155505 | 0.418 | 0.262 | 5.796E-162 | UGEC1 | RALA |
| 5.281E-166 | -0.7397955 | 0.976 | 0.973 | 1.247E-161 | UGEC1 | ACTB |
| 5.613E-166 | -0.2838557 | 0.112 | 0.262 | 1.326E-161 | UGEC1 | PRPF38B |
| 5.787E-166 | 0.30269548 | 0.201 | 0.092 | 1.367E-161 | UGEC1 | GM2A |
| 6.091E-166 | -0.2902213 | 0.106 | 0.252 | 1.439E-161 | UGEC1 | SKAP2 |
| 1.415E-165 | -0.3866618 | 0.004 | 0.103 | 3.343E-161 | UGEC1 | ADAMTS1 |
| 8.292E-165 | 0.29740952 | 0.533 | 0.366 | 1.959E-160 | UGEC1 | NDUFB11 |
| 4E-164 | -0.5775462 | 0.146 | 0.291 | 9.45E-160 | UGEC1 | DDIT3 |
| 5.372E-164 | -0.2739313 | 0.037 | 0.156 | 1.269E-159 | UGEC1 | DDX60L |
| 9.838E-164 | -0.2993758 | 0.195 | 0.355 | 2.324E-159 | UGEC1 | LAMP1 |
| 8.894E-163 | -0.2521035 | 0.007 | 0.106 | 2.101E-158 | UGEC1 | DCBLD2 |
| 1.276E-162 | -0.3909798 | 0.306 | 0.453 | 3.015E-158 | UGEC1 | IQGAP1 |
| 2.933E-162 | -0.2668254 | 0.121 | 0.269 | 6.928E-158 | UGEC1 | NDUFA3 |
| 3.211E-162 | 0.31480608 | 0.33 | 0.194 | 7.585E-158 | UGEC1 | FOXO3 |

| | | | | | | |
|---|---|---|---|---|---|---|
| 1.754E-161 | -0.2712563 | 0.124 | 0.272 | 4.144E-157 | UGEC1 | SMARCA5 |
| 3.643E-161 | -0.3051733 | 0.004 | 0.101 | 8.607E-157 | UGEC1 | GBP5 |
| 1.306E-160 | -0.2972538 | 0.157 | 0.308 | 3.084E-156 | UGEC1 | RTN3 |
| 3.187E-160 | -0.5686559 | 0.025 | 0.135 | 7.529E-156 | UGEC1 | HSPA2 |
| 2.335E-159 | -1.5275104 | 0.045 | 0.165 | 5.516E-155 | UGEC1 | CXCL3 |
| 4.086E-159 | -0.2537084 | 0.138 | 0.291 | 9.653E-155 | UGEC1 | PSMA4 |
| 2.476E-158 | -0.2939132 | 0.08 | 0.214 | 5.849E-154 | UGEC1 | FNDC3A |
| 4.825E-158 | -0.2574697 | 0.12 | 0.267 | 1.14E-153 | UGEC1 | NUDC |
| 2.286E-157 | -0.2876857 | 0.149 | 0.3 | 5.401E-153 | UGEC1 | SF1 |
| 3.198E-157 | -0.2983431 | 0.219 | 0.374 | 7.555E-153 | UGEC1 | PSMD8 |
| 2.435E-155 | 0.30164672 | 0.808 | 0.663 | 5.752E-151 | UGEC1 | CAST |
| 3.26E-155 | -0.5006353 | 0.173 | 0.325 | 7.701E-151 | UGEC1 | BTG2 |
| 4.221E-155 | -0.2542553 | 0.137 | 0.285 | 9.972E-151 | UGEC1 | OCIAD1 |
| 4.36E-155 | -0.2618677 | 0.091 | 0.226 | 1.03E-150 | UGEC1 | RHOG |
| 1.125E-154 | -0.4870768 | 0.696 | 0.751 | 2.658E-150 | UGEC1 | APLP2 |
| 2.757E-154 | -0.2677457 | 0.158 | 0.31 | 6.514E-150 | UGEC1 | PRPF40A |
| 1.142E-153 | -0.4692412 | 0.079 | 0.207 | 2.698E-149 | UGEC1 | KLF2 |
| 1.445E-153 | 0.25132861 | 0.346 | 0.208 | 3.413E-149 | UGEC1 | LASP1 |
| 1.218E-152 | -0.6641253 | 0.013 | 0.112 | 2.878E-148 | UGEC1 | PTGS2 |
| 3.252E-152 | 0.38570209 | 0.784 | 0.668 | 7.681E-148 | UGEC1 | DNAJA1 |
| 1.85E-151 | 0.25759734 | 0.428 | 0.274 | 4.37E-147 | UGEC1 | SERTAD1 |
| 2.674E-151 | -0.2636456 | 0.014 | 0.114 | 6.318E-147 | UGEC1 | AC025164.1 |
| 4.799E-151 | 0.36508967 | 0.932 | 0.856 | 1.134E-146 | UGEC1 | PABPC1 |
| 2.53E-150 | -0.4190416 | 0.021 | 0.124 | 5.976E-146 | UGEC1 | THBD |
| 3.26E-150 | -0.2624253 | 0.064 | 0.188 | 7.7E-146 | UGEC1 | CHCHD7 |
| 6.902E-150 | -0.2599507 | 0.14 | 0.283 | 1.63E-145 | UGEC1 | PGLS |
| 6.998E-150 | -0.3545832 | 0.112 | 0.246 | 1.653E-145 | UGEC1 | STXBP2 |
| 1.067E-149 | -0.2896418 | 0.18 | 0.329 | 2.519E-145 | UGEC1 | TMED9 |
| 1.526E-149 | -0.3104364 | 0.628 | 0.716 | 3.604E-145 | UGEC1 | COX6B1 |
| 2.571E-149 | -0.2661156 | 0.161 | 0.309 | 6.073E-145 | UGEC1 | PSMB6 |
| 2.542E-147 | 0.79519419 | 0.168 | 0.077 | 6.005E-143 | UGEC1 | SAA1 |
| 4.278E-147 | -0.9240669 | 0.042 | 0.153 | 1.011E-142 | UGEC1 | EREG |
| 5.497E-147 | -0.5942931 | 0.97 | 0.938 | 1.299E-142 | UGEC1 | MT-CO3 |
| 6.075E-147 | -0.2642314 | 0.008 | 0.1 | 1.435E-142 | UGEC1 | CALD1 |
| 9.38E-146 | 0.31131438 | 0.21 | 0.106 | 2.216E-141 | UGEC1 | PER1 |
| 2.375E-144 | -0.2697607 | 0.109 | 0.243 | 5.61E-140 | UGEC1 | PCM1 |
| 7.442E-143 | -0.2695029 | 0.157 | 0.299 | 1.758E-138 | UGEC1 | PDXK |
| 2.235E-142 | -0.2767193 | 0.046 | 0.157 | 5.281E-138 | UGEC1 | ETS1 |
| 4.139E-142 | -0.3253054 | 0.096 | 0.224 | 9.779E-138 | UGEC1 | MANF |
| 4.824E-142 | 0.36004201 | 0.351 | 0.214 | 1.14E-137 | UGEC1 | CITED2 |
| 6.853E-141 | -0.3415711 | 0.493 | 0.602 | 1.619E-136 | UGEC1 | NOP10 |
| 8.045E-141 | -0.2550586 | 0.092 | 0.219 | 1.9E-136 | UGEC1 | MYO9B |
| 1.04E-140 | -0.2877048 | 0.076 | 0.199 | 2.458E-136 | UGEC1 | STAT1 |
| 7.451E-140 | -0.5191863 | 0.166 | 0.296 | 1.76E-135 | UGEC1 | TNFRSF12A |
| 7.297E-138 | -0.4242596 | 0.127 | 0.261 | 1.724E-133 | UGEC1 | TNFAIP3 |
| 4.62E-137 | -0.2706253 | 0.048 | 0.157 | 1.091E-132 | UGEC1 | MGLL |
| 5.085E-137 | 0.45760359 | 0.463 | 0.33 | 1.201E-132 | UGEC1 | ATF3 |
| 5.29E-137 | -0.2601294 | 0.094 | 0.219 | 1.25E-132 | UGEC1 | CHD1 |
| 5.38E-137 | -0.2567676 | 0.143 | 0.284 | 1.271E-132 | UGEC1 | TRA2B |
| 6.488E-136 | -0.2510279 | 0.034 | 0.137 | 1.533E-131 | UGEC1 | ATP2B1-AS1 |

| | | | | | | |
|---|---|---|---|---|---|---|
| 7.225E-136 | -0.5519438 | 0.932 | 0.911 | 1.707E-131 | UGEC1 | MT-ND1 |
| 3.496E-135 | -0.2717561 | 0.107 | 0.235 | 8.259E-131 | UGEC1 | DNTTIP2 |
| 9.877E-135 | -0.4057774 | 0.084 | 0.204 | 2.333E-130 | UGEC1 | SGK1 |
| 1.979E-134 | -0.2654223 | 0.057 | 0.169 | 4.675E-130 | UGEC1 | FAM129A |
| 6.666E-133 | -0.2863167 | 0.151 | 0.285 | 1.575E-128 | UGEC1 | MPHOSPH8 |
| 6.914E-131 | -0.2501455 | 0.121 | 0.249 | 1.633E-126 | UGEC1 | DHX36 |
| 8.338E-131 | 0.27529482 | 0.239 | 0.133 | 1.97E-126 | UGEC1 | FAM89B |
| 5.791E-130 | -0.2951665 | 0.182 | 0.32 | 1.368E-125 | UGEC1 | EIF5B |
| 6.631E-130 | -0.2598686 | 0.114 | 0.243 | 1.566E-125 | UGEC1 | ECHS1 |
| 1.487E-129 | -0.4436852 | 0.228 | 0.371 | 3.512E-125 | UGEC1 | ZFP36L2 |
| 4.451E-129 | -0.2560098 | 0.141 | 0.274 | 1.051E-124 | UGEC1 | BCLAF1 |
| 2.323E-128 | -0.3896514 | 0.223 | 0.353 | 5.487E-124 | UGEC1 | APP |
| 3.272E-128 | -0.4088977 | 0.216 | 0.35 | 7.73E-124 | UGEC1 | LMNA |
| 5.778E-128 | -0.3072684 | 0.101 | 0.222 | 1.365E-123 | UGEC1 | AMD1 |
| 4.133E-127 | -0.3052163 | 0.113 | 0.234 | 9.764E-123 | UGEC1 | ZFYVE16 |
| 5.006E-127 | -0.3440829 | 0.088 | 0.209 | 1.183E-122 | UGEC1 | SLC5A3 |
| 6.897E-126 | -0.2868955 | 0.125 | 0.248 | 1.629E-121 | UGEC1 | RGS10 |
| 9.568E-126 | -0.3625244 | 0.522 | 0.632 | 2.26E-121 | UGEC1 | SRRM2 |
| 1.084E-125 | -0.2775508 | 0.124 | 0.248 | 2.561E-121 | UGEC1 | IFNGR1 |
| 5.11E-125 | -0.254391 | 0.64 | 0.753 | 1.207E-120 | UGEC1 | GNAS |
| 1.354E-124 | -0.6972319 | 0.526 | 0.649 | 3.198E-120 | UGEC1 | JUN |
| 2.717E-124 | -0.2700273 | 0.066 | 0.175 | 6.418E-120 | UGEC1 | FNIP2 |
| 1.319E-123 | -0.2849982 | 0.059 | 0.165 | 3.116E-119 | UGEC1 | GSTM3 |
| 2.029E-123 | -0.306869 | 0.978 | 0.985 | 4.793E-119 | UGEC1 | EIF1 |
| 5.58E-123 | 0.37592515 | 0.996 | 0.977 | 1.318E-118 | UGEC1 | MALAT1 |
| 2.045E-122 | 0.28502596 | 0.303 | 0.191 | 4.832E-118 | UGEC1 | NBDY |
| 6.339E-121 | 0.30302177 | 0.267 | 0.156 | 1.498E-116 | UGEC1 | NR4A1 |
| 1.808E-120 | -0.3142904 | 0.544 | 0.631 | 4.27E-116 | UGEC1 | COX8A |
| 7.024E-120 | -0.3250104 | 0.162 | 0.286 | 1.659E-115 | UGEC1 | BACH1 |
| 8.934E-120 | 0.2880051 | 0.313 | 0.192 | 2.11E-115 | UGEC1 | LAMB3 |
| 1.428E-119 | -0.3045696 | 0.132 | 0.256 | 3.372E-115 | UGEC1 | MT-ATP8 |
| 4.173E-119 | -0.272406 | 0.236 | 0.368 | 9.858E-115 | UGEC1 | MTPN |
| 5.342E-119 | -0.2848524 | 0.067 | 0.172 | 1.262E-114 | UGEC1 | HIST1H1E |
| 8.089E-119 | -0.5389739 | 0.155 | 0.27 | 1.911E-114 | UGEC1 | SYNE2 |
| 1.154E-117 | -0.2956716 | 0.35 | 0.476 | 2.725E-113 | UGEC1 | ST13 |
| 1.272E-117 | -0.3485379 | 0.287 | 0.416 | 3.006E-113 | UGEC1 | PNISR |
| 5.511E-117 | -0.5015833 | 0.226 | 0.329 | 1.302E-112 | UGEC1 | CD68 |
| 1.447E-116 | -0.2684848 | 0.099 | 0.214 | 3.417E-112 | UGEC1 | IRS2 |
| 7.947E-116 | -0.2711039 | 0.148 | 0.271 | 1.877E-111 | UGEC1 | DNAJC3 |
| 7.115E-115 | 0.48935294 | 0.998 | 0.988 | 1.681E-110 | UGEC1 | B2M |
| 9.422E-115 | 0.55201904 | 0.342 | 0.226 | 2.226E-110 | UGEC1 | PLAU |
| 3.534E-113 | -0.3639588 | 0.211 | 0.337 | 8.348E-109 | UGEC1 | HMGB2 |
| 1.923E-112 | -0.2641814 | 0.517 | 0.618 | 4.543E-108 | UGEC1 | YWHAE |
| 1.453E-111 | -0.2802525 | 0.191 | 0.316 | 3.431E-107 | UGEC1 | CD99 |
| 5.117E-111 | -0.2717098 | 0.455 | 0.571 | 1.209E-106 | UGEC1 | UQCR10 |
| 3.703E-110 | -0.2845324 | 0.137 | 0.253 | 8.747E-106 | UGEC1 | KMT2A |
| 8.272E-110 | -0.251829 | 0.343 | 0.506 | 1.954E-105 | UGEC1 | HSPD1 |
| 3.758E-109 | -0.2995326 | 0.121 | 0.234 | 8.878E-105 | UGEC1 | KDM6B |
| 9.957E-109 | -0.4111194 | 0.066 | 0.165 | 2.352E-104 | UGEC1 | RASD1 |
| 3.37E-108 | 0.39224115 | 0.341 | 0.224 | 7.96E-104 | UGEC1 | KIAA1551 |

| | | | | | | |
|---|---|---|---|---|---|---|
| 1.197E-107 | -0.260971 | 0.212 | 0.339 | 2.829E-103 | UGEC1 | CMTM6 |
| 3.549E-107 | -0.2598086 | 0.088 | 0.193 | 8.383E-103 | UGEC1 | P4HA1 |
| 1.11E-105 | -0.3164695 | 0.092 | 0.201 | 2.622E-101 | UGEC1 | INSIG1 |
| 1.304E-104 | -0.2533239 | 0.113 | 0.223 | 3.081E-100 | UGEC1 | MAT2A |
| 3.641E-103 | -0.2801044 | 0.073 | 0.173 | 8.6E-99 | UGEC1 | CLDN3 |
| 4.691E-101 | -0.3182909 | 0.219 | 0.328 | 1.1083E-96 | UGEC1 | RAB13 |
| 5.409E-101 | -0.3262918 | 0.079 | 0.177 | 1.2777E-96 | UGEC1 | SOCS1 |
| 3.413E-100 | -0.3816287 | 0.725 | 0.786 | 8.0634E-96 | UGEC1 | TPI1 |
| 7.739E-100 | -0.2770441 | 0.679 | 0.734 | 1.8281E-95 | UGEC1 | COX6A1 |
| 1.432E-99 | -0.5287233 | 0.06 | 0.153 | 3.3828E-95 | UGEC1 | NUPR1 |
| 8.87E-99 | 0.28190485 | 0.356 | 0.238 | 2.0953E-94 | UGEC1 | TSC22D1 |
| 9.3774E-97 | 0.25277992 | 0.301 | 0.196 | 2.2152E-92 | UGEC1 | MKNK2 |
| 6.7708E-96 | -1.5957156 | 0.027 | 0.102 | 1.5995E-91 | UGEC1 | HBB |
| 4.4643E-95 | -0.3019015 | 0.122 | 0.227 | 1.0546E-90 | UGEC1 | SOCS3 |
| 1.5606E-94 | -0.2712282 | 0.156 | 0.266 | 3.6867E-90 | UGEC1 | CREBRF |
| 6.5323E-94 | -0.4148857 | 0.426 | 0.538 | 1.5431E-89 | UGEC1 | H1FX |
| 2.1482E-93 | -0.3403288 | 0.19 | 0.303 | 5.0747E-89 | UGEC1 | MARCKSL1 |
| 4.6211E-92 | -0.2800841 | 0.181 | 0.289 | 1.0916E-87 | UGEC1 | OSBPL8 |
| 9.7913E-92 | -0.9443165 | 0.049 | 0.132 | 2.313E-87 | UGEC1 | CXCL2 |
| 3.4254E-89 | -0.8846453 | 0.165 | 0.265 | 8.0918E-85 | UGEC1 | CRYAB |
| 1.8908E-86 | -0.3677877 | 0.874 | 0.887 | 4.4667E-82 | UGEC1 | MYL12B |
| 2.2292E-85 | -0.25615 | 0.225 | 0.338 | 5.2661E-81 | UGEC1 | C6orf62 |
| 2.5821E-85 | -0.2791755 | 0.44 | 0.526 | 6.0998E-81 | UGEC1 | COX7B |
| 8.7422E-85 | -0.3874782 | 0.973 | 0.93 | 2.0652E-80 | UGEC1 | MT-ATP6 |
| 2.1464E-84 | 0.25355377 | 0.325 | 0.227 | 5.0704E-80 | UGEC1 | CD82 |
| 1.0861E-83 | -0.3915473 | 0.669 | 0.695 | 2.5658E-79 | UGEC1 | PNRC1 |
| 1.9838E-83 | -0.3774116 | 0.455 | 0.532 | 4.6864E-79 | UGEC1 | H2AFZ |
| 1.0553E-82 | -0.4045608 | 0.16 | 0.266 | 2.493E-78 | UGEC1 | EGR1 |
| 3.3969E-82 | -0.3482453 | 0.193 | 0.292 | 8.0245E-78 | UGEC1 | ETS2 |
| 1.3879E-81 | -0.3088804 | 0.531 | 0.607 | 3.2788E-77 | UGEC1 | DDX17 |
| 3.9566E-79 | 0.48002987 | 0.424 | 0.328 | 9.3466E-75 | UGEC1 | GDF15 |
| 5.9349E-79 | -0.2874466 | 0.449 | 0.535 | 1.402E-74 | UGEC1 | ATP6V1F |
| 7.4736E-77 | 0.28099829 | 0.315 | 0.223 | 1.7655E-72 | UGEC1 | FKBP4 |
| 2.5543E-75 | -0.2725901 | 0.247 | 0.351 | 6.0341E-71 | UGEC1 | GOLGA4 |
| 7.4353E-72 | -0.2881198 | 0.449 | 0.517 | 1.7564E-67 | UGEC1 | JAK1 |
| 5.8662E-70 | -0.3374618 | 0.111 | 0.194 | 1.3858E-65 | UGEC1 | DST |
| 3.9881E-69 | 0.38874257 | 0.268 | 0.181 | 9.421E-65 | UGEC1 | SFN |
| 1.0117E-66 | -0.2831979 | 0.426 | 0.508 | 2.3899E-62 | UGEC1 | RSRP1 |
| 1.5568E-64 | -0.4071147 | 0.098 | 0.174 | 3.6775E-60 | UGEC1 | CSKMT |
| 5.8147E-64 | -0.6115678 | 0.991 | 0.994 | 1.3736E-59 | UGEC1 | FTH1 |
| 2.7658E-63 | -0.2758103 | 0.295 | 0.383 | 6.5336E-59 | UGEC1 | SPAG9 |
| 1.0223E-62 | -0.349512 | 0.205 | 0.297 | 2.4149E-58 | UGEC1 | RHOB |
| 1.5157E-62 | -0.2685596 | 0.463 | 0.542 | 3.5806E-58 | UGEC1 | EIF4A2 |
| 5.938E-62 | -0.2602742 | 0.274 | 0.366 | 1.4027E-57 | UGEC1 | PNN |
| 4.0409E-56 | -0.354861 | 0.146 | 0.219 | 9.5457E-52 | UGEC1 | GK |
| 1.1785E-52 | -0.3952936 | 0.577 | 0.608 | 2.784E-48 | UGEC1 | MCL1 |
| 1.5827E-51 | 0.28012969 | 0.368 | 0.29 | 3.7389E-47 | UGEC1 | BLVRB |
| 9.5237E-51 | -0.3746921 | 0.919 | 0.906 | 2.2498E-46 | UGEC1 | MT-ND2 |
| 2.846E-47 | -0.348563 | 0.637 | 0.64 | 6.7232E-43 | UGEC1 | ARPC5 |
| 1.377E-46 | -1.3567219 | 0.216 | 0.277 | 3.2529E-42 | UGEC1 | S100A2 |

| | | | | | | |
|---|---|---|---|---|---|---|
| 1.4804E-40 | 0.36374523 | 0.756 | 0.726 | 3.4971E-36 | UGEC1 | HSPA8 |
| 5.2548E-37 | -0.3639307 | 0.749 | 0.77 | 1.2413E-32 | UGEC1 | CST3 |
| 2.1433E-35 | -0.6281496 | 0.141 | 0.19 | 5.0631E-31 | UGEC1 | LAMC2 |
| 1.3107E-33 | -0.4644826 | 0.339 | 0.4 | 3.0962E-29 | UGEC1 | FOS |
| 2.4682E-33 | -0.4390046 | 0.178 | 0.23 | 5.8306E-29 | UGEC1 | CCND1 |
| 1.0429E-32 | -1.0176405 | 0.543 | 0.477 | 2.4637E-28 | UGEC1 | CXCL8 |
| 2.8131E-30 | 0.37416116 | 0.105 | 0.07 | 6.6454E-26 | UGEC1 | PAPSS2 |
| 1.3022E-22 | -0.3618647 | 0.247 | 0.279 | 3.0762E-18 | UGEC1 | MAL2 |
| 1.3929E-21 | -0.309339 | 0.367 | 0.412 | 3.2904E-17 | UGEC1 | RNF213 |
| 6.2768E-21 | -0.3104325 | 0.567 | 0.609 | 1.4828E-16 | UGEC1 | NFKBIA |
| 2.0917E-19 | -0.2948524 | 0.287 | 0.325 | 4.9412E-15 | UGEC1 | GADD45A |
| 1.2965E-16 | 0.49886944 | 0.31 | 0.292 | 3.0627E-12 | UGEC1 | GLRX |
| 3.3405E-11 | -0.5434658 | 0.56 | 0.524 | 7.8914E-07 | UGEC1 | CTSS |
| 4.5005E-10 | -0.3391338 | 0.641 | 0.541 | 1.0632E-05 | UGEC1 | KRT18 |
| 2.5544E-08 | -0.3739925 | 0.895 | 0.862 | 0.00060344 | UGEC1 | ANXA2 |
| 8.6656E-07 | -0.2707315 | 0.214 | 0.228 | 0.02047065 | UGEC1 | PPP1R10 |
| 0.00015861 | -0.2692408 | 0.256 | 0.221 | 1 | UGEC1 | HEXIM1 |
| 0.00413359 | -0.4305493 | 0.558 | 0.48 | 1 | UGEC1 | KRT8 |
| 0 | 2.853461 | 0.891 | 0.098 | 0 | UGEC2 | SPINK1 |
| 0 | 2.73702953 | 0.923 | 0.223 | 0 | UGEC2 | ADIRF |
| 0 | 2.1504022 | 0.771 | 0.116 | 0 | UGEC2 | AQP3 |
| 0 | 1.91841753 | 0.683 | 0.078 | 0 | UGEC2 | SNCG |
| 0 | 1.90311632 | 0.647 | 0.034 | 0 | UGEC2 | DHRS2 |
| 0 | 1.84039651 | 0.995 | 0.34 | 0 | UGEC2 | S100P |
| 0 | 1.62147158 | 0.658 | 0.111 | 0 | UGEC2 | FXYD3 |
| 0 | 1.5264854 | 0.961 | 0.496 | 0 | UGEC2 | KRT7 |
| 0 | 1.51706999 | 0.655 | 0.217 | 0 | UGEC2 | AREG |
| 0 | 1.5103456 | 0.917 | 0.275 | 0 | UGEC2 | PSCA |
| 0 | 1.49236623 | 0.997 | 0.875 | 0 | UGEC2 | RPS6 |
| 0 | 1.44405372 | 0.535 | 0.024 | 0 | UGEC2 | UPK1B |
| 0 | 1.42601383 | 0.647 | 0.22 | 0 | UGEC2 | MGST1 |
| 0 | 1.4214149 | 0.215 | 0.018 | 0 | UGEC2 | FABP4 |
| 0 | 1.41415939 | 0.485 | 0.016 | 0 | UGEC2 | KRT20 |
| 0 | 1.37856231 | 0.545 | 0.234 | 0 | UGEC2 | S100A2 |
| 0 | 1.31174247 | 0.96 | 0.491 | 0 | UGEC2 | CLDN4 |
| 0 | 1.28470513 | 0.918 | 0.569 | 0 | UGEC2 | EZR |
| 0 | 1.22287909 | 0.433 | 0.031 | 0 | UGEC2 | UPK1A |
| 0 | 1.20739696 | 0.99 | 0.531 | 0 | UGEC2 | KRT19 |
| 0 | 1.19718211 | 0.438 | 0.023 | 0 | UGEC2 | UPK2 |
| 0 | 1.08827833 | 0.546 | 0.192 | 0 | UGEC2 | CYB5A |
| 0 | 1.03642962 | 0.919 | 0.444 | 0 | UGEC2 | KRT8 |
| 0 | 1.02718071 | 0.966 | 0.523 | 0 | UGEC2 | TACSTD2 |
| 0 | 1.02349297 | 0.925 | 0.277 | 0 | UGEC2 | HPGD |
| 0 | 1.00740503 | 0.817 | 0.458 | 0 | UGEC2 | CD9 |
| 0 | 0.99666155 | 0.585 | 0.258 | 0 | UGEC2 | GPRC5A |
| 0 | 0.96070248 | 0.4 | 0.022 | 0 | UGEC2 | FBLN1 |
| 0 | 0.95308293 | 0.957 | 0.512 | 0 | UGEC2 | KRT18 |
| 0 | 0.94943714 | 0.472 | 0.123 | 0 | UGEC2 | SDC1 |
| 0 | 0.94561388 | 0.442 | 0.106 | 0 | UGEC2 | ID1 |
| 0 | 0.91915068 | 0.357 | 0.013 | 0 | UGEC2 | C10orf99 |

| | | | | | | |
|---|---|---|---|---|---|---|
| 0 | 0.91775251 | 1 | 0.93 | 0 | UGEC2 | S100A6 |
| 0 | 0.90421002 | 0.479 | 0.208 | 0 | UGEC2 | FABP5 |
| 0 | 0.89930249 | 0.547 | 0.158 | 0 | UGEC2 | KRT17 |
| 0 | 0.89781088 | 0.996 | 0.87 | 0 | UGEC2 | CSTB |
| 0 | 0.8808799 | 0.99 | 0.872 | 0 | UGEC2 | HSP90AB1 |
| 0 | 0.85559139 | 0.989 | 0.845 | 0 | UGEC2 | RPL23 |
| 0 | 0.84413946 | 0.348 | 0.029 | 0 | UGEC2 | TMEM97 |
| 0 | 0.80339918 | 0.994 | 0.863 | 0 | UGEC2 | RPL23A |
| 0 | 0.79941621 | 0.974 | 0.805 | 0 | UGEC2 | MT-ND5 |
| 0 | 0.78374551 | 0.997 | 0.953 | 0 | UGEC2 | RPS18 |
| 0 | 0.7751356 | 0.346 | 0.014 | 0 | UGEC2 | SLC14A1 |
| 0 | 0.77498461 | 0.996 | 0.939 | 0 | UGEC2 | RPL13A |
| 0 | 0.76979026 | 0.993 | 0.923 | 0 | UGEC2 | MT-CYB |
| 0 | 0.76159834 | 0.416 | 0.145 | 0 | UGEC2 | XIST |
| 0 | 0.75500553 | 0.303 | 0.048 | 0 | UGEC2 | GPX2 |
| 0 | 0.73638332 | 0.987 | 0.843 | 0 | UGEC2 | RPL7 |
| 0 | 0.71250841 | 0.817 | 0.592 | 0 | UGEC2 | KTN1 |
| 0 | 0.70665838 | 0.352 | 0.086 | 0 | UGEC2 | PPARG |
| 0 | 0.70319407 | 0.987 | 0.758 | 0 | UGEC2 | RPL10A |
| 0 | 0.69934993 | 0.353 | 0.11 | 0 | UGEC2 | IDH1 |
| 0 | 0.69235078 | 0.992 | 0.931 | 0 | UGEC2 | MT-ATP6 |
| 0 | 0.67412111 | 0.997 | 0.902 | 0 | UGEC2 | RPS8 |
| 0 | 0.6729082 | 0.991 | 0.832 | 0 | UGEC2 | RPS4X |
| 0 | 0.67094855 | 0.963 | 0.744 | 0 | UGEC2 | RPS26 |
| 0 | 0.66607422 | 0.992 | 0.888 | 0 | UGEC2 | RPL27A |
| 0 | 0.66034299 | 0.992 | 0.939 | 0 | UGEC2 | MT-CO2 |
| 0 | 0.65765108 | 0.79 | 0.55 | 0 | UGEC2 | CD55 |
| 0 | 0.65348991 | 0.995 | 0.861 | 0 | UGEC2 | RPL26 |
| 0 | 0.64456052 | 0.414 | 0.148 | 0 | UGEC2 | IGFBP3 |
| 0 | 0.63678757 | 0.992 | 0.887 | 0 | UGEC2 | RPL29 |
| 0 | 0.63640097 | 0.993 | 0.911 | 0 | UGEC2 | NACA |
| 0 | 0.63524774 | 0.361 | 0.094 | 0 | UGEC2 | AGR2 |
| 0 | 0.63515564 | 0.993 | 0.958 | 0 | UGEC2 | HSP90AA1 |
| 0 | 0.63183255 | 0.893 | 0.499 | 0 | UGEC2 | CD24 |
| 0 | 0.6316384 | 0.715 | 0.426 | 0 | UGEC2 | RAB11FIP1 |
| 0 | 0.62756681 | 0.848 | 0.626 | 0 | UGEC2 | UQCRH |
| 0 | 0.61919124 | 0.978 | 0.693 | 0 | UGEC2 | HSPB1 |
| 0 | 0.61089935 | 0.301 | 0.043 | 0 | UGEC2 | CYP4B1 |
| 0 | 0.60676681 | 0.269 | 0.016 | 0 | UGEC2 | SPTSSB |
| 0 | 0.60489048 | 0.303 | 0.086 | 0 | UGEC2 | GATA3 |
| 0 | 0.60097259 | 0.363 | 0.121 | 0 | UGEC2 | VSIG2 |
| 0 | 0.59618969 | 0.729 | 0.417 | 0 | UGEC2 | MARCKS |
| 0 | 0.59554469 | 0.986 | 0.856 | 0 | UGEC2 | RPLP0 |
| 0 | 0.59496536 | 0.997 | 0.895 | 0 | UGEC2 | RPL7A |
| 0 | 0.59073187 | 0.994 | 0.862 | 0 | UGEC2 | RPL3 |
| 0 | 0.58868186 | 0.996 | 0.875 | 0 | UGEC2 | RPL6 |
| 0 | 0.58612936 | 0.994 | 0.891 | 0 | UGEC2 | RPL24 |
| 0 | 0.57622839 | 0.99 | 0.891 | 0 | UGEC2 | RPS11 |
| 0 | 0.56824265 | 0.996 | 0.886 | 0 | UGEC2 | RPS3A |
| 0 | 0.56535453 | 0.213 | 0.021 | 0 | UGEC2 | AKR1C2 |

| | | | | | | |
|---|---|---|---|---|---|---|
| 0 | 0.55999159 | 0.317 | 0.104 | 0 | UGEC2 | MAST4 |
| 0 | 0.54380418 | 0.292 | 0.076 | 0 | UGEC2 | LIPH |
| 0 | 0.54215306 | 0.995 | 0.857 | 0 | UGEC2 | RPL14 |
| 0 | 0.54077729 | 0.342 | 0.116 | 0 | UGEC2 | EMP2 |
| 0 | 0.50322122 | 0.199 | 0.018 | 0 | UGEC2 | UPK3A |
| 0 | 0.50301913 | 0.279 | 0.097 | 0 | UGEC2 | OAS1 |
| 0 | 0.4999821 | 0.231 | 0.009 | 0 | UGEC2 | BTBD16 |
| 0 | 0.49210271 | 0.243 | 0.042 | 0 | UGEC2 | LYPD3 |
| 0 | 0.4849036 | 0.998 | 0.926 | 0 | UGEC2 | RPL34 |
| 0 | 0.47713054 | 0.219 | 0.009 | 0 | UGEC2 | SNX31 |
| 0 | 0.47583484 | 0.274 | 0.079 | 0 | UGEC2 | APOL1 |
| 0 | 0.46716528 | 0.232 | 0.044 | 0 | UGEC2 | AKR1C1 |
| 0 | 0.46637712 | 0.252 | 0.072 | 0 | UGEC2 | CMYA5 |
| 0 | 0.46307942 | 0.266 | 0.086 | 0 | UGEC2 | FAM3B |
| 0 | 0.45840197 | 0.213 | 0.017 | 0 | UGEC2 | BHMT |
| 0 | 0.44333527 | 0.994 | 0.945 | 0 | UGEC2 | MT-CO1 |
| 0 | 0.43945972 | 0.246 | 0.066 | 0 | UGEC2 | CYP3A5 |
| 0 | 0.42440829 | 0.219 | 0.052 | 0 | UGEC2 | TESC |
| 0 | 0.42350409 | 0.208 | 0.027 | 0 | UGEC2 | ARL14 |
| 0 | 0.42160219 | 0.189 | 0.011 | 0 | UGEC2 | CRTAC1 |
| 0 | 0.41909469 | 0.184 | 0.02 | 0 | UGEC2 | SPOCD1 |
| 0 | 0.39377979 | 0.933 | 0.511 | 0 | UGEC2 | ELF3 |
| 0 | 0.38100304 | 0.183 | 0.009 | 0 | UGEC2 | SYT8 |
| 0 | 0.36163713 | 0.137 | 0.005 | 0 | UGEC2 | ANXA10 |
| 0 | 0.35207698 | 0.153 | 0.007 | 0 | UGEC2 | SERPINB5 |
| 0 | 0.32472502 | 0.161 | 0.007 | 0 | UGEC2 | HSD17B2 |
| 0 | 0.31630777 | 0.154 | 0.017 | 0 | UGEC2 | ERP27 |
| 0 | 0.29241877 | 0.168 | 0.029 | 0 | UGEC2 | POF1B |
| 0 | -0.5456151 | 0.65 | 0.303 | 0 | UGEC2 | MMP7 |
| 0 | -0.6060699 | 0.074 | 0.379 | 0 | UGEC2 | TIMP2 |
| 0 | -0.6560626 | 0.038 | 0.343 | 0 | UGEC2 | CYTOR |
| 0 | -0.6744853 | 0.028 | 0.366 | 0 | UGEC2 | MSN |
| 0 | -0.7667818 | 0.001 | 0.261 | 0 | UGEC2 | HCST |
| 0 | -0.7727148 | 0.035 | 0.344 | 0 | UGEC2 | CD68 |
| 0 | -0.7793845 | 0.002 | 0.298 | 0 | UGEC2 | CD53 |
| 0 | -0.8660347 | 0.08 | 0.401 | 0 | UGEC2 | COTL1 |
| 0 | -0.887739 | 0.024 | 0.334 | 0 | UGEC2 | SERPINA1 |
| 0 | -0.8931886 | 0.081 | 0.44 | 0 | UGEC2 | FLNA |
| 0 | -0.9829805 | 0.051 | 0.339 | 0 | UGEC2 | HIST1H4C |
| 0 | -1.0144784 | 0.007 | 0.272 | 0 | UGEC2 | SLC11A1 |
| 0 | -1.0530344 | 0.93 | 0.918 | 0 | UGEC2 | S100A10 |
| 0 | -1.0537474 | 0.007 | 0.324 | 0 | UGEC2 | LCP1 |
| 0 | -1.0832241 | 0.997 | 0.989 | 0 | UGEC2 | B2M |
| 0 | -1.0840201 | 0.893 | 0.889 | 0 | UGEC2 | HLA-B |
| 0 | -1.1427385 | 0.019 | 0.312 | 0 | UGEC2 | IFITM2 |
| 0 | -1.1829465 | 0.002 | 0.26 | 0 | UGEC2 | SLC2A3 |
| 0 | -1.213007 | 0.001 | 0.313 | 0 | UGEC2 | PTPRC |
| 0 | -1.3481237 | 0.003 | 0.306 | 0 | UGEC2 | AIF1 |
| 0 | -1.3521557 | 0.022 | 0.372 | 0 | UGEC2 | LAPTM5 |
| 0 | -1.3737077 | 0.027 | 0.305 | 0 | UGEC2 | HLA-DRB1 |

| | | | | | | |
|---|---|---|---|---|---|---|
| 0 | -1.605132 | 0.829 | 0.352 | 0 | UGEC2 | SPRR3 |
| 0 | -2.0199589 | 0.998 | 0.993 | 0 | UGEC2 | FTH1 |
| 0 | -2.2549192 | 0.993 | 0.986 | 0 | UGEC2 | FTL |
| 0 | -2.2690222 | 0.713 | 0.743 | 0 | UGEC2 | VIM |
| 2.118E-307 | -1.1751444 | 0.007 | 0.262 | 5.003E-303 | UGEC2 | BCL2A1 |
| 2.547E-305 | -0.9787913 | 0.142 | 0.412 | 6.016E-301 | UGEC2 | TYMP |
| 1.306E-303 | -0.7132761 | 0.031 | 0.297 | 3.086E-299 | UGEC2 | TSC22D3 |
| 5.197E-303 | -1.2203499 | 0.206 | 0.45 | 1.228E-298 | UGEC2 | EMP3 |
| 3.141E-302 | 0.50517665 | 0.998 | 0.915 | 7.421E-298 | UGEC2 | RPS24 |
| 1.256E-300 | 0.47873722 | 0.301 | 0.112 | 2.967E-296 | UGEC2 | MPZL2 |
| 2.135E-300 | -0.6140765 | 0.064 | 0.344 | 5.044E-296 | UGEC2 | HMGB2 |
| 3.553E-300 | 0.47913081 | 0.992 | 0.875 | 8.392E-296 | UGEC2 | RPS3 |
| 3.773E-299 | -0.8080975 | 0.005 | 0.253 | 8.914E-295 | UGEC2 | ZEB2 |
| 5.04E-298 | -1.8132258 | 0.002 | 0.248 | 1.191E-293 | UGEC2 | VCAN |
| 3.091E-296 | -0.6448581 | 0.018 | 0.272 | 7.302E-292 | UGEC2 | GMFG |
| 2.751E-295 | 0.50869071 | 0.994 | 0.926 | 6.499E-291 | UGEC2 | RPS14 |
| 2.386E-294 | 0.41648557 | 0.246 | 0.08 | 5.636E-290 | UGEC2 | CAPS |
| 5.157E-294 | 0.45756537 | 0.4 | 0.173 | 1.218E-289 | UGEC2 | TM4SF1 |
| 1.383E-292 | 0.76905334 | 0.387 | 0.174 | 3.267E-288 | UGEC2 | SFN |
| 1.792E-292 | -1.0645696 | 0.002 | 0.245 | 4.233E-288 | UGEC2 | CD14 |
| 1.079E-291 | -1.1599376 | 0.002 | 0.244 | 2.55E-287 | UGEC2 | ALOX5AP |
| 2.983E-291 | -0.6812735 | 0.002 | 0.244 | 7.047E-287 | UGEC2 | ITGB2 |
| 6.986E-287 | 0.37488148 | 0.998 | 0.95 | 1.65E-282 | UGEC2 | ACTG1 |
| 1.025E-286 | -0.6186165 | 0.002 | 0.24 | 2.421E-282 | UGEC2 | SPI1 |
| 2.009E-285 | -1.8018002 | 0.048 | 0.306 | 4.745E-281 | UGEC2 | MT1X |
| 1.589E-283 | 0.32198966 | 0.184 | 0.049 | 3.755E-279 | UGEC2 | ST3GAL5 |
| 7.149E-281 | -0.5217195 | 0.067 | 0.336 | 1.689E-276 | UGEC2 | ARPC1B |
| 2.288E-279 | 0.49845598 | 0.992 | 0.954 | 5.405E-275 | UGEC2 | RPL15 |
| 8.286E-279 | 0.3611329 | 0.218 | 0.066 | 1.957E-274 | UGEC2 | AKR1C3 |
| 1.601E-278 | -0.5908972 | 0.052 | 0.31 | 3.781E-274 | UGEC2 | RPS4Y1 |
| 2.119E-278 | 0.31829948 | 0.198 | 0.056 | 5.006E-274 | UGEC2 | MAOA |
| 1.081E-275 | -0.7116017 | 0.001 | 0.232 | 2.553E-271 | UGEC2 | LST1 |
| 1.357E-273 | 0.58787127 | 0.944 | 0.767 | 3.206E-269 | UGEC2 | BTF3 |
| 1.004E-271 | -0.5544399 | 0.08 | 0.347 | 2.371E-267 | UGEC2 | SLC16A3 |
| 5.299E-271 | 0.5656598 | 0.357 | 0.169 | 1.252E-266 | UGEC2 | SH3YL1 |
| 1.196E-269 | -1.3510649 | 0.855 | 0.485 | 2.826E-265 | UGEC2 | LCN2 |
| 1.784E-267 | 0.42261305 | 0.303 | 0.115 | 4.215E-263 | UGEC2 | BCAS1 |
| 5.295E-267 | -1.3676246 | 0.056 | 0.306 | 1.251E-262 | UGEC2 | MT1E |
| 3.864E-265 | 0.5660931 | 0.589 | 0.366 | 9.128E-261 | UGEC2 | FOS |
| 4.497E-265 | 0.44907991 | 0.995 | 0.868 | 1.062E-260 | UGEC2 | RPL12 |
| 1.19E-264 | -0.7827461 | 0.007 | 0.233 | 2.81E-260 | UGEC2 | TNFRSF1B |
| 4.418E-264 | -0.6000306 | 0.001 | 0.224 | 1.044E-259 | UGEC2 | NCF2 |
| 1.99E-262 | -0.7928511 | 0.002 | 0.224 | 4.702E-258 | UGEC2 | CYBB |
| 3.713E-261 | 0.47216756 | 0.991 | 0.872 | 8.77E-257 | UGEC2 | RPS13 |
| 2.342E-260 | -0.6147044 | 0.002 | 0.223 | 5.532E-256 | UGEC2 | ITGAX |
| 9.609E-260 | -0.8079642 | 0.001 | 0.221 | 2.27E-255 | UGEC2 | CD52 |
| 3.775E-259 | -0.4716287 | 0.008 | 0.232 | 8.919E-255 | UGEC2 | ARRB2 |
| 4.429E-258 | 0.46339367 | 0.704 | 0.423 | 1.046E-253 | UGEC2 | CLDN7 |
| 1.039E-257 | -0.5260865 | 0.095 | 0.361 | 2.454E-253 | UGEC2 | TLN1 |
| 2.537E-256 | 0.36328026 | 0.991 | 0.926 | 5.994E-252 | UGEC2 | MT-ND4 |

| | | | | | | |
|---|---|---|---|---|---|---|
| 9.314E-256 | -0.6054592 | 0.013 | 0.238 | 2.2E-251 | UGEC2 | TUBA1A |
| 8.464E-255 | 0.49585127 | 0.913 | 0.659 | 1.999E-250 | UGEC2 | NPM1 |
| 5.843E-254 | -0.8137754 | 0.095 | 0.349 | 1.38E-249 | UGEC2 | IFITM3 |
| 9.992E-254 | 0.39902973 | 0.275 | 0.104 | 2.36E-249 | UGEC2 | FOXQ1 |
| 5.482E-252 | -0.9174859 | 0.996 | 0.967 | 1.295E-247 | UGEC2 | TMSB10 |
| 1.236E-251 | -0.4068374 | 0.153 | 0.443 | 2.92E-247 | UGEC2 | DAZAP2 |
| 8.311E-251 | -0.5158416 | 0.06 | 0.305 | 1.963E-246 | UGEC2 | LYN |
| 2.456E-249 | 0.46873851 | 0.563 | 0.319 | 5.801E-245 | UGEC2 | GDF15 |
| 2.496E-249 | -0.9196562 | 0.024 | 0.251 | 5.896E-245 | UGEC2 | IL32 |
| 8.945E-249 | -0.985814 | 0.066 | 0.307 | 2.113E-244 | UGEC2 | MAFB |
| 3.459E-248 | 0.62281604 | 0.655 | 0.473 | 8.171E-244 | UGEC2 | SELENOW |
| 7.014E-248 | -0.6290326 | 0.095 | 0.349 | 1.657E-243 | UGEC2 | LIMS1 |
| 7.008E-247 | -0.6713805 | 0.921 | 0.893 | 1.656E-242 | UGEC2 | HLA-A |
| 2.1E-245 | 0.52690428 | 0.82 | 0.608 | 4.961E-241 | UGEC2 | TAX1BP1 |
| 2.175E-245 | -0.6065068 | 0.019 | 0.241 | 5.138E-241 | UGEC2 | TGM2 |
| 4.708E-245 | -0.5947452 | 0.001 | 0.21 | 1.112E-240 | UGEC2 | FCGR2A |
| 8.361E-243 | -0.6144335 | 0.002 | 0.21 | 1.975E-238 | UGEC2 | FYB1 |
| 1.375E-242 | -0.4367473 | 0.029 | 0.254 | 3.249E-238 | UGEC2 | ATP6V1B2 |
| 2.37E-242 | 0.76580065 | 0.509 | 0.35 | 5.6E-238 | UGEC2 | RALBP1 |
| 4.618E-242 | -0.5484028 | 0.013 | 0.229 | 1.091E-237 | UGEC2 | SLC43A2 |
| 1.324E-241 | -0.6333532 | 0.884 | 0.869 | 3.127E-237 | UGEC2 | HLA-C |
| 5.683E-241 | 0.53644502 | 0.955 | 0.767 | 1.343E-236 | UGEC2 | RPL31 |
| 1.979E-240 | 0.38667926 | 0.231 | 0.082 | 4.675E-236 | UGEC2 | PDLIM1 |
| 3.069E-240 | -0.5283299 | 0.127 | 0.388 | 7.249E-236 | UGEC2 | GNAI2 |
| 3.463E-240 | -0.8426641 | 0.002 | 0.209 | 8.181E-236 | UGEC2 | AC020656.1 |
| 3.915E-240 | 0.39692273 | 0.998 | 0.96 | 9.249E-236 | UGEC2 | RPL13 |
| 1.16E-239 | 0.4461471 | 0.995 | 0.891 | 2.741E-235 | UGEC2 | RPS7 |
| 8.659E-239 | 0.45847291 | 0.984 | 0.798 | 2.046E-234 | UGEC2 | RPL5 |
| 9.82E-239 | 0.35667876 | 0.999 | 0.978 | 2.32E-234 | UGEC2 | RPL41 |
| 1.576E-238 | -0.4480946 | 0.007 | 0.215 | 3.722E-234 | UGEC2 | SNX10 |
| 1.596E-238 | -1.3462353 | 0.013 | 0.226 | 3.77E-234 | UGEC2 | HLA-DPA1 |
| 6.317E-238 | 0.54110689 | 0.972 | 0.857 | 1.492E-233 | UGEC2 | PABPC1 |
| 7.055E-237 | 0.41453122 | 0.994 | 0.872 | 1.667E-232 | UGEC2 | RPS2 |
| 2.296E-235 | -0.5850524 | 0 | 0.202 | 5.425E-231 | UGEC2 | SAMSN1 |
| 6.716E-235 | 0.55208512 | 0.779 | 0.578 | 1.587E-230 | UGEC2 | VAMP8 |
| 1.217E-233 | -0.6418716 | 0.002 | 0.203 | 2.876E-229 | UGEC2 | IGSF6 |
| 2.694E-232 | -0.4164123 | 0.005 | 0.208 | 6.364E-228 | UGEC2 | HCLS1 |
| 1.302E-231 | 0.42503102 | 0.913 | 0.71 | 3.076E-227 | UGEC2 | HSPA8 |
| 1.641E-231 | -0.666006 | 0.004 | 0.205 | 3.876E-227 | UGEC2 | C5AR1 |
| 6.054E-231 | -0.4527059 | 0.001 | 0.199 | 1.43E-226 | UGEC2 | CD37 |
| 3.789E-230 | -1.0093149 | 0.97 | 0.936 | 8.95E-226 | UGEC2 | SAT1 |
| 2.1E-229 | -0.4040154 | 0.002 | 0.2 | 4.961E-225 | UGEC2 | FMNL1 |
| 8.506E-229 | 0.33352036 | 0.2 | 0.065 | 2.009E-224 | UGEC2 | INPP4B |
| 1.041E-228 | 0.40562873 | 0.998 | 0.914 | 2.458E-224 | UGEC2 | RPL11 |
| 1.811E-228 | 0.33806531 | 0.215 | 0.073 | 4.277E-224 | UGEC2 | TBX3 |
| 1.038E-226 | -1.5626913 | 0.051 | 0.271 | 2.452E-222 | UGEC2 | CRYAB |
| 1.04E-226 | 0.43271382 | 0.994 | 0.91 | 2.456E-222 | UGEC2 | RPL9 |
| 7.016E-226 | 0.5403924 | 0.426 | 0.229 | 1.657E-221 | UGEC2 | KLF5 |
| 9.544E-226 | -1.2060392 | 0.006 | 0.205 | 2.255E-221 | UGEC2 | HLA-DPB1 |
| 1.429E-225 | -0.3872883 | 0.033 | 0.249 | 3.377E-221 | UGEC2 | TPP1 |

| | | | | | | |
|---|---|---|---|---|---|---|
| 5.684E-225 | 0.44148999 | 0.991 | 0.888 | 1.343E-220 | UGEC2 | RPS25 |
| 2.341E-224 | -0.4203096 | 0.003 | 0.199 | 5.531E-220 | UGEC2 | TMEM176B |
| 1.168E-223 | 0.28386334 | 0.148 | 0.039 | 2.758E-219 | UGEC2 | UCA1 |
| 2.062E-223 | -0.6288874 | 0.002 | 0.197 | 4.871E-219 | UGEC2 | ANPEP |
| 1.296E-222 | -0.8236142 | 0.001 | 0.195 | 3.063E-218 | UGEC2 | PLEK |
| 5.259E-222 | -0.508587 | 0.026 | 0.233 | 1.242E-217 | UGEC2 | NINJ1 |
| 7.005E-222 | -0.5185031 | 0.001 | 0.193 | 1.655E-217 | UGEC2 | SMIM25 |
| 2.767E-221 | -0.4020695 | 0.035 | 0.249 | 6.537E-217 | UGEC2 | RGS10 |
| 1.946E-219 | 0.49565498 | 0.278 | 0.118 | 4.597E-215 | UGEC2 | GCLC |
| 2.692E-217 | -0.4875201 | 0.05 | 0.27 | 6.36E-213 | UGEC2 | GPCPD1 |
| 7.884E-217 | 0.52348794 | 0.642 | 0.459 | 1.862E-212 | UGEC2 | HSPD1 |
| 2.967E-216 | -0.4535408 | 0.003 | 0.193 | 7.01E-212 | UGEC2 | CORO1A |
| 5.502E-216 | -0.8654357 | 0.001 | 0.189 | 1.3E-211 | UGEC2 | FCN1 |
| 1.467E-215 | 0.49337277 | 0.86 | 0.668 | 3.464E-211 | UGEC2 | CAST |
| 2.452E-214 | -0.444219 | 0.001 | 0.188 | 5.793E-210 | UGEC2 | CD48 |
| 2.344E-213 | -0.3731284 | 0.001 | 0.187 | 5.538E-209 | UGEC2 | GLIPR2 |
| 5.951E-213 | -0.4433208 | 0.133 | 0.384 | 1.406E-208 | UGEC2 | ATP1A1 |
| 1.747E-212 | 0.4185279 | 0.996 | 0.944 | 4.126E-208 | UGEC2 | RPL18 |
| 3.501E-212 | 0.3969736 | 0.995 | 0.914 | 8.271E-208 | UGEC2 | RPL8 |
| 1.046E-211 | -1.1133323 | 0.037 | 0.244 | 2.471E-207 | UGEC2 | G0S2 |
| 1.642E-211 | -0.4927574 | 0.002 | 0.187 | 3.878E-207 | UGEC2 | SLA |
| 5.061E-211 | 0.47347029 | 0.637 | 0.414 | 1.196E-206 | UGEC2 | ATP1B1 |
| 9.19E-211 | 0.44689418 | 0.314 | 0.144 | 2.171E-206 | UGEC2 | SYTL2 |
| 1.276E-209 | -0.4744513 | 0 | 0.183 | 3.015E-205 | UGEC2 | LCP2 |
| 3.752E-207 | -0.4228456 | 0.016 | 0.207 | 8.864E-203 | UGEC2 | RAC2 |
| 3.842E-207 | -0.4461129 | 0.063 | 0.283 | 9.076E-203 | UGEC2 | ARRDC3 |
| 3.642E-205 | 0.44453705 | 0.703 | 0.503 | 8.602E-201 | UGEC2 | HSPE1 |
| 1.671E-204 | -0.6697466 | 0.072 | 0.289 | 3.948E-200 | UGEC2 | DDIT3 |
| 3.378E-204 | 0.44700713 | 0.272 | 0.122 | 7.979E-200 | UGEC2 | MGST2 |
| 3.748E-204 | 0.33206129 | 0.968 | 0.731 | 8.855E-200 | UGEC2 | RPSA |
| 5.216E-204 | -0.515811 | 0 | 0.179 | 1.232E-199 | UGEC2 | AQP9 |
| 1.439E-203 | -0.4601373 | 0.007 | 0.19 | 3.398E-199 | UGEC2 | SAMHD1 |
| 3.542E-203 | -0.4063617 | 0.044 | 0.249 | 8.366E-199 | UGEC2 | FCGRT |
| 5.701E-203 | -0.5972703 | 0.007 | 0.189 | 1.347E-198 | UGEC2 | TGFBI |
| 1.432E-202 | -0.730738 | 0.013 | 0.199 | 3.383E-198 | UGEC2 | OLR1 |
| 1.927E-202 | -0.3794066 | 0.002 | 0.18 | 4.552E-198 | UGEC2 | FERMT3 |
| 1.972E-202 | -0.5893457 | 0.104 | 0.333 | 4.658E-198 | UGEC2 | IRF1 |
| 5.573E-202 | -2.3067578 | 0.011 | 0.194 | 1.317E-197 | UGEC2 | IL1B |
| 6.713E-202 | -0.6320203 | 0.092 | 0.318 | 1.586E-197 | UGEC2 | GLRX |
| 2.711E-201 | 0.40546292 | 0.336 | 0.164 | 6.403E-197 | UGEC2 | FBP1 |
| 5.335E-201 | 0.42030745 | 0.264 | 0.113 | 1.26E-196 | UGEC2 | GAN |
| 8.915E-201 | -0.3702321 | 0.027 | 0.221 | 2.106E-196 | UGEC2 | NR3C1 |
| 1.91E-200 | -0.6149435 | 0 | 0.176 | 4.512E-196 | UGEC2 | CD300E |
| 1.113E-199 | -0.4154887 | 0.085 | 0.308 | 2.629E-195 | UGEC2 | CXCL16 |
| 1.16E-199 | 0.42096504 | 0.277 | 0.124 | 2.74E-195 | UGEC2 | RAB25 |
| 1.213E-199 | 0.4032314 | 0.973 | 0.781 | 2.866E-195 | UGEC2 | RPS5 |
| 1.686E-199 | -0.349624 | 0.002 | 0.178 | 3.983E-195 | UGEC2 | WIPF1 |
| 1.913E-199 | 0.46633296 | 0.308 | 0.151 | 4.519E-195 | UGEC2 | CXADR |
| 1.938E-199 | -0.4308501 | 0.056 | 0.266 | 4.579E-195 | UGEC2 | RAB31 |
| 7.218E-198 | 0.29934964 | 0.466 | 0.239 | 1.705E-193 | UGEC2 | LMO7 |

| | | | | | | |
|---|---|---|---|---|---|---|
| 6.424E-197 | -0.3751074 | 0.16 | 0.416 | 1.518E-192 | UGEC2 | STAT3 |
| 1.597E-196 | -0.3603232 | 0.129 | 0.371 | 3.773E-192 | UGEC2 | CAPN2 |
| 6.252E-196 | -0.3728875 | 0.005 | 0.181 | 1.477E-191 | UGEC2 | MYO1F |
| 9.733E-196 | -0.4425841 | 0.04 | 0.238 | 2.299E-191 | UGEC2 | DSE |
| 2.067E-195 | -0.3560057 | 0.02 | 0.206 | 4.883E-191 | UGEC2 | RNF130 |
| 2.098E-195 | -0.3974696 | 0.096 | 0.322 | 4.955E-191 | UGEC2 | TNIP1 |
| 5.554E-195 | -0.3617831 | 0.018 | 0.202 | 1.312E-190 | UGEC2 | PSMB9 |
| 6.404E-195 | -0.8714597 | 0.095 | 0.307 | 1.513E-190 | UGEC2 | SLC25A37 |
| 2.994E-194 | -0.4337461 | 0.09 | 0.312 | 7.072E-190 | UGEC2 | MIR4435-2HG |
| 6.415E-194 | -0.3855142 | 0.126 | 0.363 | 1.515E-189 | UGEC2 | GRB2 |
| 3.513E-192 | -0.411297 | 0.089 | 0.308 | 8.3E-188 | UGEC2 | YWHAH |
| 5.287E-192 | -0.3490943 | 0.018 | 0.2 | 1.249E-187 | UGEC2 | RILPL2 |
| 6.898E-192 | 0.63791999 | 0.252 | 0.105 | 1.629E-187 | UGEC2 | HILPDA |
| 1.859E-191 | -0.3941769 | 0.016 | 0.196 | 4.392E-187 | UGEC2 | CCDC88A |
| 6.193E-191 | -1.0584423 | 0.044 | 0.235 | 1.463E-186 | UGEC2 | S100A12 |
| 7.567E-191 | -0.3146756 | 0.009 | 0.185 | 1.788E-186 | UGEC2 | SIRPA |
| 1.403E-190 | -0.4079103 | 0 | 0.169 | 3.313E-186 | UGEC2 | EVI2B |
| 2.148E-190 | -0.5528562 | 0.012 | 0.189 | 5.075E-186 | UGEC2 | CXCR4 |
| 2.88E-190 | -0.5522379 | 0 | 0.169 | 6.804E-186 | UGEC2 | CLEC5A |
| 4.662E-190 | -0.3587197 | 0.004 | 0.174 | 1.101E-185 | UGEC2 | OGFRL1 |
| 1.053E-189 | 0.38651578 | 0.255 | 0.109 | 2.487E-185 | UGEC2 | FAT1 |
| 1.451E-189 | -0.4077041 | 0.014 | 0.193 | 3.428E-185 | UGEC2 | KCTD12 |
| 2.358E-189 | 0.32112323 | 0.995 | 0.92 | 5.57E-185 | UGEC2 | RPL30 |
| 2.59E-189 | 0.51467868 | 0.71 | 0.544 | 6.118E-185 | UGEC2 | SET |
| 2.71E-189 | -0.3039505 | 0.098 | 0.325 | 6.403E-185 | UGEC2 | H2AFV |
| 7.189E-189 | -0.4004072 | 0.006 | 0.177 | 1.698E-184 | UGEC2 | SELENOM |
| 1.905E-188 | -0.4293341 | 0.02 | 0.202 | 4.499E-184 | UGEC2 | ICAM1 |
| 7.642E-187 | -0.352952 | 0.024 | 0.205 | 1.805E-182 | UGEC2 | GPSM3 |
| 1.219E-186 | -0.4533985 | 0.095 | 0.311 | 2.881E-182 | UGEC2 | ATP1B3 |
| 2.342E-186 | 0.63833243 | 0.627 | 0.478 | 5.531E-182 | UGEC2 | NCL |
| 9.951E-186 | -0.4159722 | 0.069 | 0.275 | 2.351E-181 | UGEC2 | ODC1 |
| 8.597E-185 | -0.3679879 | 0.011 | 0.183 | 2.031E-180 | UGEC2 | CEP170 |
| 1.267E-184 | -0.8865884 | 0.639 | 0.671 | 2.992E-180 | UGEC2 | ANXA5 |
| 3.006E-184 | 0.30966196 | 0.203 | 0.077 | 7.102E-180 | UGEC2 | ACSL5 |
| 3.567E-184 | -0.5816109 | 0.236 | 0.448 | 8.426E-180 | UGEC2 | SLC3A2 |
| 4.633E-184 | -0.6397765 | 0.016 | 0.19 | 1.094E-179 | UGEC2 | HLA-DQB1 |
| 5.748E-184 | -0.3102648 | 0.054 | 0.254 | 1.358E-179 | UGEC2 | CTSA |
| 2.309E-183 | -0.3569892 | 0.003 | 0.169 | 5.455E-179 | UGEC2 | CPVL |
| 1.916E-182 | -0.3561134 | 0.065 | 0.266 | 4.526E-178 | UGEC2 | PTPN12 |
| 2.485E-182 | -1.2593057 | 0.17 | 0.365 | 5.869E-178 | UGEC2 | CTSL |
| 2.671E-182 | 0.34127602 | 0.993 | 0.888 | 6.31E-178 | UGEC2 | RPL21 |
| 1.088E-181 | -0.4695799 | 0 | 0.162 | 2.571E-177 | UGEC2 | FPR1 |
| 1.31E-181 | -0.3714014 | 0.064 | 0.265 | 3.096E-177 | UGEC2 | CD82 |
| 5.654E-181 | -0.3051591 | 0.073 | 0.28 | 1.336E-176 | UGEC2 | ATP6V0D1 |
| 2.805E-180 | 0.30807181 | 0.206 | 0.08 | 6.626E-176 | UGEC2 | ATP8B1 |
| 3.492E-180 | -0.4655418 | 0.038 | 0.223 | 8.25E-176 | UGEC2 | CTSC |
| 8.395E-180 | -0.5636954 | 0.027 | 0.203 | 1.983E-175 | UGEC2 | KLF2 |
| 1.72E-179 | -0.3180546 | 0.02 | 0.194 | 4.063E-175 | UGEC2 | PPP1R18 |
| 8.882E-179 | -0.3765084 | 0.002 | 0.162 | 2.098E-174 | UGEC2 | C1orf162 |
| 9.196E-179 | -0.3978909 | 0.087 | 0.296 | 2.172E-174 | UGEC2 | PLIN3 |

| | | | | | | |
|---|---|---|---|---|---|---|
| 1.684E-178 | -0.3196957 | 0.01 | 0.177 | 3.978E-174 | UGEC2 | AP1S2 |
| 1.688E-178 | -0.4631496 | 0.011 | 0.178 | 3.988E-174 | UGEC2 | SERPINB9 |
| 2.787E-178 | -0.5433983 | 0.001 | 0.161 | 6.585E-174 | UGEC2 | CD93 |
| 5.864E-178 | 0.33898406 | 0.996 | 0.893 | 1.385E-173 | UGEC2 | RPL37 |
| 4.203E-177 | 0.44333425 | 0.968 | 0.803 | 9.928E-173 | UGEC2 | RPL27 |
| 1.449E-176 | -0.2805465 | 0.085 | 0.296 | 3.424E-172 | UGEC2 | LAMP2 |
| 2.405E-176 | -0.4079514 | 0.003 | 0.163 | 5.682E-172 | UGEC2 | TREM1 |
| 2.585E-176 | 0.34700254 | 0.997 | 0.919 | 6.106E-172 | UGEC2 | RPL19 |
| 4.481E-176 | -0.3930084 | 0.002 | 0.16 | 1.058E-171 | UGEC2 | TLR2 |
| 4.556E-176 | -0.5749631 | 0.096 | 0.302 | 1.076E-171 | UGEC2 | H1F0 |
| 1.355E-175 | -0.4479968 | 0.003 | 0.162 | 3.2E-171 | UGEC2 | MS4A7 |
| 1.635E-175 | -0.363901 | 0.051 | 0.24 | 3.862E-171 | UGEC2 | ACTN1 |
| 3.907E-175 | 0.37389979 | 0.996 | 0.884 | 9.23E-171 | UGEC2 | RPL37A |
| 4.556E-175 | -0.4320625 | 0.025 | 0.198 | 1.076E-170 | UGEC2 | WARS |
| 1.455E-174 | -0.530524 | 0.057 | 0.246 | 3.438E-170 | UGEC2 | IER5L |
| 5.333E-174 | -0.4427992 | 0.002 | 0.159 | 1.26E-169 | UGEC2 | NCF1 |
| 5.447E-174 | 0.30785628 | 0.996 | 0.906 | 1.287E-169 | UGEC2 | RPL32 |
| 1.534E-173 | -0.3808397 | 0.111 | 0.325 | 3.625E-169 | UGEC2 | BLVRB |
| 1.874E-172 | 0.33115212 | 0.993 | 0.916 | 4.428E-168 | UGEC2 | RPL18A |
| 3.359E-172 | -0.9791107 | 0.564 | 0.618 | 7.935E-168 | UGEC2 | CD44 |
| 3.719E-172 | -0.4417328 | 0.031 | 0.204 | 8.786E-168 | UGEC2 | PTPRE |
| 7.577E-171 | -0.8988381 | 0.038 | 0.213 | 1.79E-166 | UGEC2 | C12orf75 |
| 3.057E-170 | 0.36950496 | 0.941 | 0.724 | 7.221E-166 | UGEC2 | EEF1B2 |
| 3.465E-170 | -0.8837365 | 0.766 | 0.738 | 8.185E-166 | UGEC2 | PSAP |
| 2.371E-169 | -0.3031525 | 0.001 | 0.153 | 5.602E-165 | UGEC2 | EVI2A |
| 2.468E-169 | -0.6438809 | 0.209 | 0.416 | 5.831E-165 | UGEC2 | HIF1A |
| 2.522E-169 | -0.3273197 | 0 | 0.151 | 5.957E-165 | UGEC2 | HCK |
| 7.422E-169 | 0.31736913 | 0.998 | 0.975 | 1.753E-164 | UGEC2 | RPL28 |
| 7.656E-169 | -0.4750749 | 0 | 0.152 | 1.809E-164 | UGEC2 | MCEMP1 |
| 9.4E-169 | -0.5185847 | 0.053 | 0.233 | 2.221E-164 | UGEC2 | CREM |
| 2.788E-168 | 0.3414681 | 0.842 | 0.599 | 6.585E-164 | UGEC2 | DSTN |
| 4.079E-168 | 0.43557098 | 0.961 | 0.82 | 9.635E-164 | UGEC2 | PPIA |
| 8.007E-168 | -0.299921 | 0.037 | 0.213 | 1.892E-163 | UGEC2 | HLA-F |
| 1.056E-167 | -0.3360159 | 0.001 | 0.153 | 2.494E-163 | UGEC2 | CLEC7A |
| 2.968E-167 | -0.3349178 | 0.003 | 0.155 | 7.012E-163 | UGEC2 | IFI30 |
| 3.08E-167 | -0.5030089 | 0.031 | 0.201 | 7.277E-163 | UGEC2 | SGK1 |
| 8.041E-167 | -0.299021 | 0.007 | 0.163 | 1.899E-162 | UGEC2 | ALDH2 |
| 5.449E-166 | -0.2715226 | 0.025 | 0.192 | 1.287E-161 | UGEC2 | IFI27L2 |
| 5.764E-166 | -0.2818722 | 0 | 0.149 | 1.362E-161 | UGEC2 | NCKAP1L |
| 9.426E-166 | -0.3128035 | 0 | 0.149 | 2.227E-161 | UGEC2 | CYTIP |
| 1.171E-165 | -0.710424 | 0.014 | 0.173 | 2.766E-161 | UGEC2 | MT1F |
| 1.19E-165 | -0.361398 | 0.009 | 0.165 | 2.81E-161 | UGEC2 | PLXDC2 |
| 4.553E-165 | -0.2965375 | 0.003 | 0.154 | 1.076E-160 | UGEC2 | EMILIN2 |
| 6.46E-165 | -0.314638 | 0.002 | 0.152 | 1.526E-160 | UGEC2 | ITGA5 |
| 1.335E-163 | -0.3590653 | 0.109 | 0.315 | 3.154E-159 | UGEC2 | FXYD5 |
| 3.852E-163 | -0.5040455 | 0.065 | 0.247 | 9.1E-159 | UGEC2 | IVNS1ABP |
| 3.892E-163 | 0.26986055 | 0.184 | 0.07 | 9.193E-159 | UGEC2 | LAD1 |
| 4.161E-163 | -0.5320199 | 0.002 | 0.15 | 9.829E-159 | UGEC2 | HLA-DRB5 |
| 9.109E-163 | -0.3115778 | 0.093 | 0.294 | 2.152E-158 | UGEC2 | FAM49B |
| 1.329E-162 | 0.32406381 | 0.216 | 0.091 | 3.138E-158 | UGEC2 | TNFRSF21 |

| | | | | | | |
|---|---|---|---|---|---|---|
| 1.535E-162 | -0.3220428 | 0.079 | 0.272 | 3.625E-158 | UGEC2 | C4orf48 |
| 2.82E-162 | -0.2891563 | 0 | 0.146 | 6.663E-158 | UGEC2 | APBB1IP |
| 3.078E-162 | -0.3038326 | 0.002 | 0.149 | 7.272E-158 | UGEC2 | FGR |
| 4.12E-162 | -0.2907477 | 0.001 | 0.147 | 9.732E-158 | UGEC2 | PILRA |
| 4.848E-162 | 0.27649098 | 0.998 | 0.915 | 1.145E-157 | UGEC2 | RPS27A |
| 7.541E-162 | 0.31615668 | 0.199 | 0.08 | 1.781E-157 | UGEC2 | SEMA3C |
| 9.275E-162 | -0.3672721 | 0.06 | 0.242 | 2.191E-157 | UGEC2 | GRINA |
| 2.584E-161 | -0.7075423 | 0.072 | 0.253 | 6.105E-157 | UGEC2 | PDZK1IP1 |
| 5.621E-161 | -0.5705806 | 0.161 | 0.372 | 1.328E-156 | UGEC2 | GLS |
| 5.935E-161 | -0.4175307 | 0.009 | 0.161 | 1.402E-156 | UGEC2 | SDC2 |
| 1.221E-160 | 0.49559564 | 0.386 | 0.231 | 2.884E-156 | UGEC2 | SPTBN1 |
| 1.488E-160 | -0.3838728 | 0.161 | 0.383 | 3.514E-156 | UGEC2 | CFLAR |
| 2.758E-160 | -0.2997014 | 0.046 | 0.221 | 6.514E-156 | UGEC2 | RHOG |
| 4.703E-160 | -0.6152256 | 0.273 | 0.452 | 1.111E-155 | UGEC2 | GSTO1 |
| 1.501E-159 | -0.3149434 | 0.001 | 0.145 | 3.545E-155 | UGEC2 | LILRB3 |
| 3.414E-159 | -0.2876428 | 0.071 | 0.26 | 8.064E-155 | UGEC2 | CASP4 |
| 7.163E-159 | -0.2596172 | 0.018 | 0.175 | 1.692E-154 | UGEC2 | DDAH2 |
| 7.844E-159 | -0.4835846 | 0.077 | 0.264 | 1.853E-154 | UGEC2 | KIAA1551 |
| 9.26E-158 | -0.4082327 | 0.225 | 0.437 | 2.188E-153 | UGEC2 | GRN |
| 3.348E-157 | -0.3031432 | 0.035 | 0.203 | 7.908E-153 | UGEC2 | HIPK2 |
| 3.967E-157 | -0.2970818 | 0.002 | 0.145 | 9.371E-153 | UGEC2 | CELF2 |
| 6.508E-157 | -0.4177527 | 0.021 | 0.177 | 1.537E-152 | UGEC2 | SOCS1 |
| 1.658E-156 | 0.2524814 | 0.181 | 0.069 | 3.917E-152 | UGEC2 | TMEM154 |
| 6.812E-156 | -1.2786897 | 0.003 | 0.145 | 1.609E-151 | UGEC2 | APOE |
| 7.849E-156 | -1.4075409 | 0.01 | 0.158 | 1.854E-151 | UGEC2 | CCL2 |
| 1.436E-155 | -0.292709 | 0.046 | 0.217 | 3.393E-151 | UGEC2 | PRNP |
| 3.379E-155 | -0.3032223 | 0.108 | 0.309 | 7.981E-151 | UGEC2 | RALA |
| 5.126E-155 | -0.2685318 | 0.012 | 0.163 | 1.211E-150 | UGEC2 | SH3KBP1 |
| 9.029E-155 | -0.3382806 | 0.185 | 0.408 | 2.133E-150 | UGEC2 | DEK |
| 1.265E-154 | -1.5734155 | 0.012 | 0.16 | 2.988E-150 | UGEC2 | CXCL3 |
| 2.141E-154 | -1.5552088 | 0.268 | 0.399 | 5.057E-150 | UGEC2 | FCER1G |
| 5.737E-154 | -0.3577419 | 0.187 | 0.412 | 1.355E-149 | UGEC2 | HMGN2 |
| 7.385E-154 | -0.3924773 | 0.001 | 0.14 | 1.745E-149 | UGEC2 | MARCO |
| 9.387E-154 | -0.2826606 | 0.101 | 0.298 | 2.217E-149 | UGEC2 | PLEKHB2 |
| 2.527E-153 | -0.4410948 | 0.014 | 0.163 | 5.97E-149 | UGEC2 | RGS2 |
| 2.808E-153 | -2.0048992 | 0.017 | 0.167 | 6.633E-149 | UGEC2 | MT1H |
| 2.917E-153 | 0.27013114 | 0.99 | 0.882 | 6.89E-149 | UGEC2 | RPS15A |
| 5.439E-153 | -0.2898234 | 0.004 | 0.146 | 1.285E-148 | UGEC2 | TMEM176A |
| 4.22E-152 | -0.3187945 | 0.047 | 0.216 | 9.969E-148 | UGEC2 | ARHGAP18 |
| 4.327E-152 | -0.3537445 | 0.174 | 0.392 | 1.022E-147 | UGEC2 | PRDX6 |
| 8.068E-152 | -0.3847856 | 0.072 | 0.251 | 1.906E-147 | UGEC2 | PARP14 |
| 2.549E-151 | -0.2550015 | 0.041 | 0.208 | 6.021E-147 | UGEC2 | AGTRAP |
| 1.672E-150 | -0.3189393 | 0.007 | 0.149 | 3.951E-146 | UGEC2 | COL18A1 |
| 2.614E-150 | -0.6083282 | 0.077 | 0.253 | 6.176E-146 | UGEC2 | CDKN1C |
| 5.396E-150 | -0.3315563 | 0.016 | 0.164 | 1.275E-145 | UGEC2 | BCAT1 |
| 1.947E-149 | -0.2769694 | 0.046 | 0.212 | 4.6E-145 | UGEC2 | PHC2 |
| 2.578E-149 | -0.2585546 | 0.017 | 0.166 | 6.091E-145 | UGEC2 | CLIC4 |
| 3.017E-149 | -0.3364563 | 0.034 | 0.193 | 7.127E-145 | UGEC2 | GLIPR1 |
| 3.552E-149 | 0.3854324 | 0.978 | 0.823 | 8.391E-145 | UGEC2 | RACK1 |
| 4.395E-149 | -0.3047633 | 0.003 | 0.141 | 1.038E-144 | UGEC2 | IRAK3 |

| | | | | | | |
|---|---|---|---|---|---|---|
| 1.235E-148 | -0.3404368 | 0.017 | 0.165 | 2.918E-144 | UGEC2 | DMXL2 |
| 1.407E-148 | -0.2861845 | 0.067 | 0.243 | 3.323E-144 | UGEC2 | ARFGAP3 |
| 1.536E-148 | -0.507555 | 0.089 | 0.27 | 3.629E-144 | UGEC2 | SNHG12 |
| 5.623E-148 | -0.3341377 | 0 | 0.134 | 1.328E-143 | UGEC2 | LILRB2 |
| 8.841E-148 | -0.2651805 | 0.001 | 0.136 | 2.088E-143 | UGEC2 | DOCK10 |
| 1.182E-147 | -0.5328439 | 0.15 | 0.35 | 2.792E-143 | UGEC2 | HERPUD1 |
| 1.204E-147 | -0.3330605 | 0.081 | 0.261 | 2.845E-143 | UGEC2 | RBPJ |
| 1.257E-147 | -0.3840653 | 0.006 | 0.145 | 2.969E-143 | UGEC2 | CD83 |
| 1.751E-147 | -0.2810516 | 0.012 | 0.155 | 4.137E-143 | UGEC2 | IL2RG |
| 1.957E-147 | -0.3043727 | 0.016 | 0.161 | 4.624E-143 | UGEC2 | PLEKHO1 |
| 1.294E-146 | -1.2342596 | 0.002 | 0.136 | 3.058E-142 | UGEC2 | CCL3 |
| 4.676E-146 | -0.3518057 | 0.097 | 0.284 | 1.105E-141 | UGEC2 | BACH1 |
| 2.905E-145 | 0.54982294 | 0.443 | 0.306 | 6.863E-141 | UGEC2 | PERP |
| 3.344E-145 | -0.2904746 | 0.091 | 0.275 | 7.899E-141 | UGEC2 | VAMP2 |
| 4.017E-145 | -0.3003192 | 0.174 | 0.393 | 9.489E-141 | UGEC2 | CIRBP |
| 5.115E-145 | -0.2940808 | 0.001 | 0.133 | 1.208E-140 | UGEC2 | CSF1R |
| 6.238E-145 | -0.2640799 | 0.042 | 0.204 | 1.474E-140 | UGEC2 | RAPGEF1 |
| 7.815E-145 | -0.4141544 | 0 | 0.133 | 1.846E-140 | UGEC2 | FCGR3A |
| 1.378E-144 | -0.2793443 | 0.022 | 0.171 | 3.256E-140 | UGEC2 | CSGALNACT2 |
| 1.423E-144 | -0.3121364 | 0.019 | 0.165 | 3.361E-140 | UGEC2 | FAM129A |
| 1.02E-143 | -0.2507482 | 0.001 | 0.132 | 2.41E-139 | UGEC2 | MPP1 |
| 2.1E-143 | -0.643784 | 0.089 | 0.264 | 4.96E-139 | UGEC2 | CST6 |
| 3.003E-143 | -0.3485062 | 0.024 | 0.17 | 7.094E-139 | UGEC2 | ADGRE5 |
| 4.9E-143 | 0.43953798 | 0.941 | 0.818 | 1.157E-138 | UGEC2 | YWHAZ |
| 1.133E-142 | -0.2632849 | 0.059 | 0.228 | 2.677E-138 | UGEC2 | CYTH1 |
| 1.285E-142 | -0.2986684 | 0 | 0.131 | 3.036E-138 | UGEC2 | CD86 |
| 1.387E-142 | -0.2542393 | 0 | 0.131 | 3.276E-138 | UGEC2 | MYO1G |
| 1.593E-142 | -0.2779588 | 0.054 | 0.22 | 3.762E-138 | UGEC2 | NEU1 |
| 1.806E-142 | 0.35016579 | 0.743 | 0.553 | 4.266E-138 | UGEC2 | TSPO |
| 2.28E-142 | -2.6431563 | 0.108 | 0.271 | 5.387E-138 | UGEC2 | MT1G |
| 2.431E-142 | -0.2919491 | 0.076 | 0.249 | 5.743E-138 | UGEC2 | FLOT1 |
| 1.175E-141 | -0.2722555 | 0.139 | 0.345 | 2.776E-137 | UGEC2 | TGOLN2 |
| 1.617E-141 | 0.44831149 | 0.31 | 0.164 | 3.82E-137 | UGEC2 | DST |
| 2.167E-141 | -0.6295345 | 0 | 0.13 | 5.118E-137 | UGEC2 | MNDA |
| 1.537E-140 | -0.282902 | 0.016 | 0.156 | 3.63E-136 | UGEC2 | SERPING1 |
| 2.015E-140 | -0.2629145 | 0.054 | 0.217 | 4.759E-136 | UGEC2 | SEC14L1 |
| 2.227E-140 | -0.2705411 | 0 | 0.129 | 5.261E-136 | UGEC2 | C3AR1 |
| 4.106E-140 | -0.3201253 | 0.044 | 0.201 | 9.7E-136 | UGEC2 | KLF10 |
| 1.488E-139 | -0.2704853 | 0.042 | 0.198 | 3.516E-135 | UGEC2 | CYLD |
| 1.755E-139 | -0.3856677 | 0.001 | 0.129 | 4.145E-135 | UGEC2 | CD163 |
| 2.151E-139 | 0.36224227 | 0.847 | 0.67 | 5.081E-135 | UGEC2 | DNAJA1 |
| 2.263E-139 | -0.315727 | 0.104 | 0.29 | 5.347E-135 | UGEC2 | OSBPL8 |
| 3.692E-139 | -0.3671775 | 0.118 | 0.307 | 8.721E-135 | UGEC2 | MACF1 |
| 5.148E-139 | 0.4234331 | 0.384 | 0.233 | 1.216E-134 | UGEC2 | EGR1 |
| 6.817E-139 | 0.38195616 | 0.923 | 0.744 | 1.61E-134 | UGEC2 | COX7C |
| 1.811E-138 | -0.2803832 | 0.066 | 0.232 | 4.277E-134 | UGEC2 | ZYX |
| 3.52E-138 | 0.29571619 | 0.703 | 0.502 | 8.315E-134 | UGEC2 | HEBP2 |
| 7.48E-138 | -0.7882972 | 0.424 | 0.537 | 1.767E-133 | UGEC2 | CEBPB |
| 1.275E-137 | -0.2615408 | 0.057 | 0.221 | 3.012E-133 | UGEC2 | IGF2R |
| 1.544E-137 | -0.2666812 | 0.004 | 0.133 | 3.648E-133 | UGEC2 | LIMD2 |

| | | | | | | |
|---|---|---|---|---|---|---|
| 2.88E-137 | -0.3030911 | 0.084 | 0.257 | 6.803E-133 | UGEC2 | SP100 |
| 4.032E-137 | -0.3640187 | 0.074 | 0.241 | 9.525E-133 | UGEC2 | STXBP2 |
| 5.361E-137 | -0.5137832 | 0.049 | 0.204 | 1.266E-132 | UGEC2 | HBEGF |
| 6.96E-137 | -0.2937486 | 0 | 0.126 | 1.644E-132 | UGEC2 | PECAM1 |
| 7.224E-137 | -0.3164942 | 0.006 | 0.136 | 1.706E-132 | UGEC2 | PHACTR1 |
| 1.616E-136 | -0.2990717 | 0.213 | 0.434 | 3.818E-132 | UGEC2 | CDV3 |
| 1.914E-136 | 0.46453994 | 0.304 | 0.17 | 4.522E-132 | UGEC2 | ERRFI1 |
| 2.179E-136 | -0.2556584 | 0.016 | 0.153 | 5.147E-132 | UGEC2 | PTPN6 |
| 3.305E-136 | -0.3029572 | 0.041 | 0.194 | 7.808E-132 | UGEC2 | EHD1 |
| 4.338E-136 | 0.25226636 | 0.172 | 0.069 | 1.025E-131 | UGEC2 | TRAK1 |
| 4.979E-136 | -0.2851599 | 0.029 | 0.174 | 1.176E-131 | UGEC2 | MYADM |
| 1.742E-135 | -0.6782472 | 0.12 | 0.3 | 4.114E-131 | UGEC2 | CLU |
| 2.871E-135 | -0.3136167 | 0.028 | 0.171 | 6.781E-131 | UGEC2 | FNIP2 |
| 3.068E-135 | -0.3308801 | 0.091 | 0.265 | 7.247E-131 | UGEC2 | CREBRF |
| 5.61E-135 | 0.37475938 | 0.299 | 0.162 | 1.325E-130 | UGEC2 | SMIM22 |
| 5.968E-135 | 0.36430404 | 0.308 | 0.168 | 1.41E-130 | UGEC2 | CDH1 |
| 1.724E-134 | -1.3358266 | 0.201 | 0.352 | 4.072E-130 | UGEC2 | IGFBP7 |
| 4.032E-134 | -0.2511787 | 0.012 | 0.144 | 9.525E-130 | UGEC2 | IL6R |
| 1.952E-133 | -0.2605823 | 0.119 | 0.307 | 4.61E-129 | UGEC2 | HM13 |
| 3.153E-133 | -0.3975496 | 0.02 | 0.157 | 7.449E-129 | UGEC2 | PAX8 |
| 3.384E-133 | -0.291785 | 0.093 | 0.268 | 7.994E-129 | UGEC2 | SH3BGRL |
| 6.076E-133 | -0.4549686 | 0.014 | 0.148 | 1.435E-128 | UGEC2 | UCHL1 |
| 7.305E-133 | -0.2652116 | 0.13 | 0.321 | 1.726E-128 | UGEC2 | TAPBP |
| 4.742E-132 | -0.295255 | 0.025 | 0.164 | 1.12E-127 | UGEC2 | SMAP2 |
| 7.053E-132 | 0.380959 | 0.804 | 0.628 | 1.666E-127 | UGEC2 | ATP5MC2 |
| 1.155E-131 | -0.2597358 | 0.002 | 0.125 | 2.727E-127 | UGEC2 | FYN |
| 1.438E-131 | -0.3234382 | 0.013 | 0.143 | 3.398E-127 | UGEC2 | AIF1L |
| 1.554E-131 | -0.2613558 | 0.004 | 0.128 | 3.671E-127 | UGEC2 | ADGRE2 |
| 2.974E-131 | -0.2798405 | 0.022 | 0.159 | 7.024E-127 | UGEC2 | CARD16 |
| 2.986E-131 | -0.2949375 | 0 | 0.121 | 7.054E-127 | UGEC2 | PDE4B |
| 6.07E-131 | -0.3266302 | 0.029 | 0.169 | 1.434E-126 | UGEC2 | HIST1H1E |
| 6.67E-131 | -0.2761225 | 0.077 | 0.246 | 1.576E-126 | UGEC2 | PGD |
| 1.126E-130 | -0.2759688 | 0.097 | 0.272 | 2.659E-126 | UGEC2 | GABARAPL1 |
| 1.858E-130 | -0.2518688 | 0.008 | 0.134 | 4.388E-126 | UGEC2 | PTAFR |
| 1.963E-130 | -0.2704743 | 0.07 | 0.232 | 4.638E-126 | UGEC2 | TGFB1 |
| 2.248E-130 | -0.3463359 | 0.046 | 0.197 | 5.311E-126 | UGEC2 | PFKFB3 |
| 5.809E-130 | 0.40638946 | 0.575 | 0.41 | 1.372E-125 | UGEC2 | SPINT2 |
| 8.072E-130 | -0.2879361 | 0.054 | 0.209 | 1.907E-125 | UGEC2 | ACSL4 |
| 9.209E-130 | 0.34630621 | 0.981 | 0.91 | 2.176E-125 | UGEC2 | RPS20 |
| 2.077E-129 | -0.2603843 | 0.108 | 0.287 | 4.906E-125 | UGEC2 | AGFG1 |
| 5.487E-129 | -0.8157657 | 0.004 | 0.125 | 1.296E-124 | UGEC2 | IL7R |
| 6.636E-129 | 0.30903201 | 0.22 | 0.104 | 1.568E-124 | UGEC2 | DSG2 |
| 9.313E-129 | -0.6539796 | 0.019 | 0.151 | 2.2E-124 | UGEC2 | NUPR1 |
| 1.008E-128 | -0.2503296 | 0.02 | 0.153 | 2.382E-124 | UGEC2 | ITPR2 |
| 1.474E-128 | -0.2806569 | 0.023 | 0.16 | 3.482E-124 | UGEC2 | CD40 |
| 4.427E-128 | -1.1521045 | 0.001 | 0.12 | 1.046E-123 | UGEC2 | CCL4 |
| 6.759E-128 | -0.362757 | 0.002 | 0.122 | 1.597E-123 | UGEC2 | TNF |
| 1.317E-127 | -0.3176057 | 0.141 | 0.331 | 3.112E-123 | UGEC2 | SMS |
| 2.362E-127 | -0.3164818 | 0.001 | 0.119 | 5.581E-123 | UGEC2 | MS4A6A |
| 1.772E-126 | -0.3830122 | 0.001 | 0.119 | 4.186E-122 | UGEC2 | SLC16A10 |

| | | | | | | |
|---|---|---|---|---|---|---|
| 3.287E-126 | -0.2857237 | 0.016 | 0.144 | 7.765E-122 | UGEC2 | EPB41L3 |
| 1.464E-125 | -0.2604807 | 0.027 | 0.161 | 3.457E-121 | UGEC2 | DDX3Y |
| 3.373E-125 | -0.2630806 | 0.058 | 0.213 | 7.968E-121 | UGEC2 | CCPG1 |
| 3.657E-125 | -0.3786052 | 0.004 | 0.122 | 8.639E-121 | UGEC2 | FXYD2 |
| 9.576E-125 | -0.3711422 | 0.001 | 0.118 | 2.262E-120 | UGEC2 | CSF3R |
| 1.697E-124 | -0.3432343 | 0.003 | 0.12 | 4.008E-120 | UGEC2 | FGL2 |
| 2.269E-124 | -1.0261841 | 0.009 | 0.13 | 5.359E-120 | UGEC2 | CXCL2 |
| 2.345E-124 | -0.2507405 | 0 | 0.115 | 5.541E-120 | UGEC2 | CCR1 |
| 5.186E-124 | -0.3026237 | 0.038 | 0.178 | 1.225E-119 | UGEC2 | CLEC2B |
| 9.131E-124 | -0.305803 | 0 | 0.115 | 2.157E-119 | UGEC2 | LILRB1 |
| 1.342E-123 | -0.2840545 | 0.075 | 0.234 | 3.17E-119 | UGEC2 | STK4 |
| 1.451E-123 | -0.2537154 | 0.058 | 0.209 | 3.429E-119 | UGEC2 | KIFC3 |
| 2.818E-123 | -1.5221321 | 0.377 | 0.447 | 6.656E-119 | UGEC2 | TYROBP |
| 4.843E-123 | 0.32787542 | 0.932 | 0.767 | 1.144E-118 | UGEC2 | DYNLL1 |
| 2.586E-122 | -0.34609 | 0.018 | 0.145 | 6.11E-118 | UGEC2 | RGCC |
| 4.405E-122 | -0.9248414 | 0.02 | 0.147 | 1.041E-117 | UGEC2 | EREG |
| 4.882E-122 | -0.3391014 | 0.011 | 0.132 | 1.153E-117 | UGEC2 | AC007952.4 |
| 1.275E-121 | -0.265786 | 0.063 | 0.213 | 3.011E-117 | UGEC2 | MYO9B |
| 1.923E-121 | -0.3813346 | 0.076 | 0.231 | 4.543E-117 | UGEC2 | ARL4C |
| 4.131E-121 | -0.2840431 | 0.02 | 0.147 | 9.758E-117 | UGEC2 | MAP3K20 |
| 5.103E-120 | -0.2554565 | 0.025 | 0.154 | 1.206E-115 | UGEC2 | LRP1 |
| 1.099E-119 | 0.33615705 | 0.884 | 0.683 | 2.595E-115 | UGEC2 | RPL4 |
| 1.426E-119 | -0.6862059 | 0.117 | 0.276 | 3.369E-115 | UGEC2 | TPM1 |
| 1.938E-119 | -0.2767967 | 0.026 | 0.155 | 4.579E-115 | UGEC2 | OTUD1 |
| 2.393E-119 | -0.5279859 | 0 | 0.111 | 5.654E-115 | UGEC2 | RETN |
| 3.372E-119 | -0.2717925 | 0.111 | 0.282 | 7.965E-115 | UGEC2 | GSN |
| 7.072E-119 | 0.31002772 | 0.983 | 0.814 | 1.671E-114 | UGEC2 | RPL22 |
| 1.35E-118 | 0.3758266 | 0.28 | 0.16 | 3.19E-114 | UGEC2 | GNL3 |
| 3.505E-118 | -0.7599314 | 0.003 | 0.115 | 8.279E-114 | UGEC2 | CCL5 |
| 4.43E-118 | -0.3712225 | 0.101 | 0.262 | 1.047E-113 | UGEC2 | PTMS |
| 1.801E-117 | -1.2680549 | 0.648 | 0.613 | 4.254E-113 | UGEC2 | LGALS1 |
| 2.599E-117 | -0.3755531 | 0.024 | 0.15 | 6.14E-113 | UGEC2 | DCDC2 |
| 3.218E-117 | -0.2540478 | 0 | 0.109 | 7.602E-113 | UGEC2 | IKZF1 |
| 4.932E-117 | -0.6370076 | 0.002 | 0.113 | 1.165E-112 | UGEC2 | HLA-DQA1 |
| 9.357E-117 | -0.4745059 | 0.183 | 0.366 | 2.21E-112 | UGEC2 | IER5 |
| 1.548E-116 | 0.26343446 | 0.279 | 0.144 | 3.656E-112 | UGEC2 | MALL |
| 8.705E-116 | -0.2677868 | 0.024 | 0.148 | 2.056E-111 | UGEC2 | DDX60L |
| 9.419E-116 | 0.28530388 | 0.992 | 0.941 | 2.225E-111 | UGEC2 | RPS16 |
| 1.094E-115 | 0.42178388 | 0.419 | 0.285 | 2.585E-111 | UGEC2 | BTG2 |
| 1.109E-115 | -0.2812424 | 0.043 | 0.18 | 2.62E-111 | UGEC2 | CFD |
| 3.847E-115 | 0.27050462 | 0.22 | 0.106 | 9.087E-111 | UGEC2 | ASS1 |
| 4.312E-115 | -0.4136977 | 0.011 | 0.127 | 1.019E-110 | UGEC2 | RRAD |
| 5.02E-115 | -0.2634381 | 0.007 | 0.121 | 1.186E-110 | UGEC2 | ACP5 |
| 5.294E-115 | -0.2597262 | 0.157 | 0.339 | 1.251E-110 | UGEC2 | AP2S1 |
| 6.91E-115 | -0.2676988 | 0.124 | 0.294 | 1.632E-110 | UGEC2 | ROCK1 |
| 7.458E-115 | -0.326033 | 0.071 | 0.218 | 1.762E-110 | UGEC2 | LSP1 |
| 7.76E-115 | -0.2545706 | 0.102 | 0.263 | 1.833E-110 | UGEC2 | CD47 |
| 3.964E-114 | 0.33834329 | 0.224 | 0.115 | 9.364E-110 | UGEC2 | LRRC75A |
| 6.424E-114 | -0.8761661 | 0.063 | 0.202 | 1.518E-109 | UGEC2 | RARRES3 |
| 3.221E-113 | 0.34178939 | 0.62 | 0.466 | 7.609E-109 | UGEC2 | CD46 |

| | | | | | | |
|---|---|---|---|---|---|---|
| 6.445E-113 | -0.3919922 | 0.887 | 0.631 | 1.522E-108 | UGEC2 | HSPA1A |
| 7.211E-113 | -1.4095882 | 0.012 | 0.127 | 1.703E-108 | UGEC2 | MT1M |
| 7.463E-113 | -0.3511628 | 0.133 | 0.3 | 1.763E-108 | UGEC2 | CTSZ |
| 8.728E-113 | -0.3406713 | 0.097 | 0.253 | 2.062E-108 | UGEC2 | RNASET2 |
| 1.17E-112 | -0.2625897 | 0.015 | 0.132 | 2.763E-108 | UGEC2 | ATP2B1-AS1 |
| 3.106E-112 | -0.2842935 | 0.078 | 0.228 | 7.337E-108 | UGEC2 | ZFYVE16 |
| 4.139E-112 | -0.3050868 | 0.001 | 0.106 | 9.778E-108 | UGEC2 | GPR183 |
| 7.827E-112 | -0.8119746 | 0.768 | 0.734 | 1.849E-107 | UGEC2 | CTSB |
| 9.507E-112 | -0.2574219 | 0.109 | 0.271 | 2.246E-107 | UGEC2 | MGAT1 |
| 2.957E-111 | -0.4468141 | 0.006 | 0.116 | 6.986E-107 | UGEC2 | LTB |
| 8.788E-111 | -0.4514508 | 0.035 | 0.161 | 2.076E-106 | UGEC2 | RASD1 |
| 3.647E-110 | -0.2705796 | 0.091 | 0.243 | 8.616E-106 | UGEC2 | NUMB |
| 7.87E-110 | -0.2721432 | 0.039 | 0.168 | 1.859E-105 | UGEC2 | SESN2 |
| 1.374E-109 | 0.30141262 | 0.719 | 0.555 | 3.246E-105 | UGEC2 | VMP1 |
| 6.785E-109 | 0.28060299 | 0.187 | 0.088 | 1.603E-104 | UGEC2 | DGKH |
| 1.418E-108 | -0.2721177 | 0.115 | 0.276 | 3.35E-104 | UGEC2 | MBNL1 |
| 8.103E-108 | -0.3494272 | 0.162 | 0.337 | 1.914E-103 | UGEC2 | BRD2 |
| 2.593E-107 | -0.2607121 | 0.03 | 0.151 | 6.125E-103 | UGEC2 | GCA |
| 3.402E-107 | 0.32495822 | 0.22 | 0.115 | 8.036E-103 | UGEC2 | FAM213A |
| 4.081E-107 | 0.25205747 | 0.996 | 0.911 | 9.64E-103 | UGEC2 | RPS15 |
| 4.95E-107 | 0.37325628 | 0.732 | 0.598 | 1.169E-102 | UGEC2 | PTGES3 |
| 5.561E-107 | -0.3954554 | 0.031 | 0.152 | 1.314E-102 | UGEC2 | GADD45G |
| 6.924E-107 | -0.7138391 | 0.814 | 0.758 | 1.636E-102 | UGEC2 | BTG1 |
| 1.318E-106 | -0.278704 | 0.069 | 0.208 | 3.113E-102 | UGEC2 | IRS2 |
| 4.033E-106 | -0.298628 | 0.058 | 0.192 | 9.528E-102 | UGEC2 | STAT1 |
| 5.237E-106 | -0.267956 | 0.158 | 0.335 | 1.237E-101 | UGEC2 | BOD1L1 |
| 9.259E-106 | -0.2614892 | 0.005 | 0.107 | 2.187E-101 | UGEC2 | AC025164.1 |
| 2.025E-105 | -0.2810665 | 0.063 | 0.2 | 4.784E-101 | UGEC2 | CKLF |
| 3.088E-105 | 0.36869658 | 0.713 | 0.588 | 7.294E-101 | UGEC2 | YWHAE |
| 4.244E-105 | -0.2674022 | 0.092 | 0.241 | 1.003E-100 | UGEC2 | PNP |
| 5.974E-105 | -0.3547364 | 0.175 | 0.342 | 1.411E-100 | UGEC2 | ARHGDIB |
| 5.092E-104 | -0.4003022 | 0.062 | 0.196 | 1.203E-99 | UGEC2 | INSIG1 |
| 5.842E-104 | -0.2537415 | 0.105 | 0.258 | 1.38E-99 | UGEC2 | MAFG |
| 6.8E-103 | -0.3684077 | 0.956 | 0.926 | 1.6063E-98 | UGEC2 | OAZ1 |
| 7.694E-103 | -0.3204515 | 0.064 | 0.199 | 1.8176E-98 | UGEC2 | SEPHS2 |
| 8.482E-103 | -0.5988493 | 0.143 | 0.302 | 2.0038E-98 | UGEC2 | HMOX1 |
| 1.17E-102 | -1.1727687 | 0.743 | 0.699 | 2.7646E-98 | UGEC2 | SOD2 |
| 1.307E-102 | -0.5372763 | 0.004 | 0.103 | 3.0868E-98 | UGEC2 | SAA1 |
| 1.564E-102 | -0.267832 | 0.077 | 0.216 | 3.6941E-98 | UGEC2 | IFI16 |
| 2.283E-102 | 0.31842733 | 0.21 | 0.108 | 5.3933E-98 | UGEC2 | C19orf33 |
| 5.294E-102 | -0.2585302 | 0.026 | 0.14 | 1.2506E-97 | UGEC2 | BEX2 |
| 1.962E-101 | -1.0148979 | 0.034 | 0.149 | 4.6337E-97 | UGEC2 | DEFB1 |
| 1.06E-100 | 0.31365609 | 0.22 | 0.118 | 2.505E-96 | UGEC2 | HACD3 |
| 1.33E-100 | -0.2637134 | 0.089 | 0.232 | 3.1419E-96 | UGEC2 | CDK2AP2 |
| 1.519E-100 | -0.544798 | 0.02 | 0.128 | 3.588E-96 | UGEC2 | HSPA2 |
| 2.136E-100 | -0.3104281 | 0.37 | 0.521 | 5.0462E-96 | UGEC2 | JAK1 |
| 1.581E-99 | -0.2632052 | 0.129 | 0.286 | 3.7346E-95 | UGEC2 | PHPT1 |
| 8.055E-99 | -0.7737549 | 0.795 | 0.506 | 1.9028E-94 | UGEC2 | WFDC2 |
| 1.2989E-98 | -0.3454786 | 0.034 | 0.148 | 3.0685E-94 | UGEC2 | ARHGAP29 |
| 1.3034E-98 | 0.26158321 | 0.991 | 0.885 | 3.0791E-94 | UGEC2 | RPL35A |

| | | | | | | |
|---|---|---|---|---|---|---|
| 1.5769E-97 | 0.4251169 | 0.377 | 0.264 | 3.7251E-93 | UGEC2 | SLK |
| 7.4587E-97 | -0.5592895 | 0.984 | 0.949 | 1.762E-92 | UGEC2 | H3F3B |
| 9.4697E-97 | -0.2848407 | 0.043 | 0.162 | 2.237E-92 | UGEC2 | LPIN1 |
| 1.7245E-96 | -0.4319654 | 0.104 | 0.244 | 4.0737E-92 | UGEC2 | PPIF |
| 3.2163E-96 | -0.298255 | 0.098 | 0.239 | 7.5978E-92 | UGEC2 | KYNU |
| 6.8217E-96 | -0.385517 | 0.193 | 0.363 | 1.6115E-91 | UGEC2 | MIDN |
| 1.0283E-95 | -0.3810705 | 0.073 | 0.202 | 2.4291E-91 | UGEC2 | DUSP6 |
| 5.2418E-95 | -0.2975632 | 0.064 | 0.191 | 1.2383E-90 | UGEC2 | LTA4H |
| 5.915E-95 | -0.4049147 | 0.108 | 0.253 | 1.3973E-90 | UGEC2 | TNFAIP3 |
| 6.3034E-95 | -0.4156866 | 0.794 | 0.779 | 1.489E-90 | UGEC2 | MAP1LC3B |
| 7.2682E-95 | 0.31556821 | 0.249 | 0.143 | 1.717E-90 | UGEC2 | AHR |
| 8.8462E-95 | -2.4310391 | 0.736 | 0.65 | 2.0897E-90 | UGEC2 | MT2A |
| 9.6501E-95 | -0.4026656 | 0.1 | 0.24 | 2.2797E-90 | UGEC2 | PPP1R10 |
| 9.7431E-95 | -0.256381 | 0.036 | 0.15 | 2.3016E-90 | UGEC2 | MGLL |
| 1.7917E-94 | -0.411949 | 0.173 | 0.331 | 4.2326E-90 | UGEC2 | HIST1H2AC |
| 8.6446E-94 | -0.2815525 | 0.008 | 0.103 | 2.0421E-89 | UGEC2 | ADAMTS9 |
| 3.2672E-93 | -0.2837574 | 0.077 | 0.208 | 7.7181E-89 | UGEC2 | SDF2L1 |
| 9.4735E-93 | -0.3038724 | 0.036 | 0.148 | 2.2379E-88 | UGEC2 | HLA-DMA |
| 1.0615E-92 | -0.2806672 | 0.083 | 0.215 | 2.5075E-88 | UGEC2 | AMD1 |
| 2.3846E-91 | -0.2597654 | 0.074 | 0.201 | 5.6333E-87 | UGEC2 | PYCARD |
| 3.46E-91 | -0.2648971 | 0.068 | 0.193 | 8.1735E-87 | UGEC2 | ANXA3 |
| 1.3944E-90 | -0.343391 | 0.06 | 0.182 | 3.294E-86 | UGEC2 | AKR1B1 |
| 2.5809E-90 | -0.3291476 | 0.133 | 0.276 | 6.0969E-86 | UGEC2 | RHOC |
| 5.3918E-90 | -0.2675755 | 0.089 | 0.221 | 1.2737E-85 | UGEC2 | STK17B |
| 6.3974E-90 | 0.39288408 | 0.193 | 0.101 | 1.5113E-85 | UGEC2 | SLC2A1 |
| 1.0982E-89 | -0.4370095 | 0.67 | 0.693 | 2.5942E-85 | UGEC2 | PNRC1 |
| 1.9771E-89 | -0.2580049 | 0.058 | 0.177 | 4.6704E-85 | UGEC2 | JARID2 |
| 6.2122E-89 | -0.2510108 | 0.205 | 0.376 | 1.4675E-84 | UGEC2 | MIF |
| 1.4699E-88 | -0.287194 | 0.104 | 0.24 | 3.4723E-84 | UGEC2 | RAB9A |
| 2.0167E-88 | 0.31698471 | 0.468 | 0.338 | 4.764E-84 | UGEC2 | PHLDA2 |
| 3.368E-88 | -0.3234 | 0.041 | 0.152 | 7.9561E-84 | UGEC2 | GPNMB |
| 5.4131E-88 | -0.420721 | 0.188 | 0.342 | 1.2787E-83 | UGEC2 | CEBPD |
| 1.2957E-87 | -0.4392353 | 0.058 | 0.173 | 3.0609E-83 | UGEC2 | CSKMT |
| 3.1526E-87 | 0.32289619 | 0.78 | 0.647 | 7.4474E-83 | UGEC2 | HNRNPU |
| 3.3798E-87 | -0.2574755 | 0.144 | 0.291 | 7.9841E-83 | UGEC2 | VASP |
| 6.8994E-87 | -0.7342816 | 0.022 | 0.119 | 1.6298E-82 | UGEC2 | APOC1 |
| 7.3304E-87 | 0.34758978 | 0.349 | 0.236 | 1.7316E-82 | UGEC2 | SERINC2 |
| 2.5406E-86 | 0.41131183 | 0.439 | 0.338 | 6.0016E-82 | UGEC2 | PEBP1 |
| 3.1184E-86 | -1.8129558 | 0.33 | 0.404 | 7.3667E-82 | UGEC2 | HLA-DRA |
| 7.0462E-86 | 0.32280798 | 0.82 | 0.69 | 1.6645E-81 | UGEC2 | SUMO2 |
| 1.7983E-84 | -0.3996271 | 0.036 | 0.139 | 4.2481E-80 | UGEC2 | GBP1 |
| 1.8107E-84 | -0.3375744 | 0.859 | 0.833 | 4.2774E-80 | UGEC2 | HLA-E |
| 3.3982E-84 | -0.258666 | 0.037 | 0.141 | 8.0275E-80 | UGEC2 | HIST1H2BC |
| 3.8842E-84 | -0.3815524 | 0.077 | 0.199 | 9.1758E-80 | UGEC2 | TNFSF10 |
| 1.4821E-83 | -0.2813265 | 0.174 | 0.327 | 3.5011E-79 | UGEC2 | FNDC3B |
| 1.7501E-83 | -0.351745 | 0.036 | 0.139 | 4.1342E-79 | UGEC2 | AC023157.3 |
| 1.2253E-82 | -0.4443946 | 0.7 | 0.686 | 2.8946E-78 | UGEC2 | BRI3 |
| 2.381E-82 | -0.3610131 | 0.216 | 0.376 | 5.6246E-78 | UGEC2 | TUBA1B |
| 5.7847E-82 | 0.28716856 | 0.212 | 0.121 | 1.3665E-77 | UGEC2 | AIG1 |
| 3.4792E-81 | -0.470193 | 0.136 | 0.267 | 8.2189E-77 | UGEC2 | CARHSP1 |

| | | | | | | |
|---|---|---|---|---|---|---|
| 1.2486E-79 | 0.3078455 | 0.947 | 0.823 | 2.9496E-75 | UGEC2 | COX4I1 |
| 1.5491E-78 | -0.2659684 | 0.017 | 0.105 | 3.6594E-74 | UGEC2 | IFIT2 |
| 2.0246E-78 | -0.4532216 | 0.16 | 0.296 | 4.7826E-74 | UGEC2 | RHOB |
| 1.9071E-77 | -3.1919344 | 0.783 | 0.664 | 4.5052E-73 | UGEC2 | TIMP1 |
| 1.9801E-76 | -0.2549518 | 0.029 | 0.122 | 4.6777E-72 | UGEC2 | HIST1H2BG |
| 7.052E-75 | -1.2528943 | 0.06 | 0.163 | 1.6659E-70 | UGEC2 | CCL20 |
| 6.8482E-74 | 0.26823606 | 0.229 | 0.134 | 1.6178E-69 | UGEC2 | CLDN1 |
| 1.1002E-73 | -0.3785461 | 0.131 | 0.255 | 2.5989E-69 | UGEC2 | ATP2B1 |
| 1.4138E-73 | 0.27351044 | 0.103 | 0.042 | 3.3398E-69 | UGEC2 | C11orf96 |
| 7.8554E-73 | 0.27022235 | 0.224 | 0.133 | 1.8557E-68 | UGEC2 | OCLN |
| 1.2085E-72 | -0.7245377 | 0.052 | 0.149 | 2.8549E-68 | UGEC2 | CXCL1 |
| 1.3808E-72 | 0.45507962 | 0.386 | 0.309 | 3.2618E-68 | UGEC2 | RAB11A |
| 1.9499E-72 | -0.3624932 | 0.704 | 0.71 | 4.6064E-68 | UGEC2 | SDCBP |
| 2.4745E-72 | -0.3638258 | 0.063 | 0.165 | 5.8456E-68 | UGEC2 | CYR61 |
| 4.1493E-72 | -0.2906815 | 0.103 | 0.221 | 9.8018E-68 | UGEC2 | SOCS3 |
| 6.1559E-72 | -0.2604864 | 0.041 | 0.136 | 1.4542E-67 | UGEC2 | SYNE1 |
| 6.3357E-72 | -0.2932043 | 0.912 | 0.871 | 1.4967E-67 | UGEC2 | MYL12A |
| 8.8234E-72 | -0.2853159 | 0.075 | 0.183 | 2.0843E-67 | UGEC2 | STMN1 |
| 1.5967E-71 | -0.355411 | 0.132 | 0.255 | 3.7718E-67 | UGEC2 | PRSS8 |
| 2.7913E-70 | -0.3762796 | 0.088 | 0.197 | 6.594E-66 | UGEC2 | ACSL1 |
| 1.139E-68 | -0.2695514 | 0.021 | 0.101 | 2.6907E-64 | UGEC2 | CTGF |
| 1.8271E-68 | 0.26225914 | 0.835 | 0.7 | 4.3163E-64 | UGEC2 | SKP1 |
| 2.0491E-68 | 0.26661087 | 0.925 | 0.784 | 4.8407E-64 | UGEC2 | ARPC2 |
| 3.9771E-68 | -0.5825612 | 0.162 | 0.276 | 9.3952E-64 | UGEC2 | TSPAN1 |
| 7.1156E-68 | 0.26053079 | 0.185 | 0.103 | 1.6809E-63 | UGEC2 | DENND2C |
| 1.2887E-67 | 0.39064814 | 0.331 | 0.248 | 3.0442E-63 | UGEC2 | LYPLA1 |
| 1.5248E-67 | 0.31349357 | 0.275 | 0.181 | 3.6021E-63 | UGEC2 | LIMA1 |
| 1.4758E-66 | 0.34452925 | 0.335 | 0.243 | 3.4863E-62 | UGEC2 | CD2AP |
| 2.6769E-66 | -0.3432459 | 0.063 | 0.161 | 6.3237E-62 | UGEC2 | SLC7A11 |
| 5.0048E-66 | 0.26159204 | 0.932 | 0.797 | 1.1823E-61 | UGEC2 | DDX5 |
| 5.1785E-66 | -0.4214584 | 0.144 | 0.253 | 1.2233E-61 | UGEC2 | S100A13 |
| 3.5135E-65 | -0.3277003 | 0.107 | 0.218 | 8.2998E-61 | UGEC2 | GK |
| 8.8022E-65 | -0.3051072 | 0.098 | 0.208 | 2.0793E-60 | UGEC2 | BIRC3 |
| 8.9635E-65 | 0.27907503 | 0.229 | 0.144 | 2.1174E-60 | UGEC2 | RCN1 |
| 3.206E-64 | -0.7340064 | 0.514 | 0.541 | 7.5736E-60 | UGEC2 | UPP1 |
| 2.4756E-63 | -0.256016 | 0.069 | 0.166 | 5.8482E-59 | UGEC2 | CLDN3 |
| 3.1367E-63 | -0.3474045 | 0.114 | 0.224 | 7.4099E-59 | UGEC2 | BASP1 |
| 1.1865E-62 | -0.4239075 | 0.088 | 0.192 | 2.8029E-58 | UGEC2 | LAMC2 |
| 7.4336E-62 | -0.5677325 | 0.13 | 0.242 | 1.756E-57 | UGEC2 | ISG15 |
| 5.396E-61 | -0.4831444 | 0.536 | 0.583 | 1.2747E-56 | UGEC2 | LITAF |
| 1.7367E-60 | -2.1838933 | 0.655 | 0.56 | 4.1027E-56 | UGEC2 | SRGN |
| 4.1823E-60 | -0.3151086 | 0.121 | 0.056 | 9.8799E-56 | UGEC2 | FLG |
| 1.1979E-59 | 0.25375611 | 0.725 | 0.596 | 2.8299E-55 | UGEC2 | HNRNPA1 |
| 1.7414E-59 | 0.29001653 | 0.221 | 0.14 | 4.1138E-55 | UGEC2 | RALGPS2 |
| 2.7743E-59 | 0.26523904 | 0.839 | 0.709 | 6.5538E-55 | UGEC2 | YWHAB |
| 7.1971E-59 | -0.2552965 | 0.123 | 0.234 | 1.7002E-54 | UGEC2 | POLR2A |
| 1.0444E-58 | -0.2779697 | 0.135 | 0.248 | 2.4672E-54 | UGEC2 | NFKBIZ |
| 4.7776E-58 | 0.27043413 | 0.315 | 0.225 | 1.1286E-53 | UGEC2 | TOB1 |
| 5.2586E-58 | 0.29560308 | 0.361 | 0.274 | 1.2422E-53 | UGEC2 | LDHB |
| 1.2505E-57 | -0.4507637 | 0.147 | 0.257 | 2.9541E-53 | UGEC2 | PLAU |

| | | | | | | |
|---|---|---|---|---|---|---|
| 1.8116E-57 | -0.2515087 | 0.432 | 0.53 | 4.2796E-53 | UGEC2 | ATP6V1F |
| 1.6892E-56 | -0.4269024 | 0.285 | 0.418 | 3.9904E-52 | UGEC2 | RNF213 |
| 2.4553E-55 | -1.5921261 | 0.487 | 0.492 | 5.8002E-51 | UGEC2 | CD74 |
| 1.4772E-54 | 0.43324691 | 0.443 | 0.391 | 3.4896E-50 | UGEC2 | ARGLU1 |
| 9.3232E-54 | 0.31220531 | 0.626 | 0.545 | 2.2024E-49 | UGEC2 | HNRNPA3 |
| 1.2474E-53 | 0.30589761 | 0.242 | 0.166 | 2.9468E-49 | UGEC2 | FAM162A |
| 1.4945E-53 | -0.2830639 | 0.032 | 0.104 | 3.5304E-49 | UGEC2 | KLK10 |
| 2.9475E-52 | -0.3902634 | 0.094 | 0.184 | 6.9628E-48 | UGEC2 | SPNS2 |
| 1.1041E-51 | 0.29259671 | 0.949 | 0.868 | 2.6082E-47 | UGEC2 | TMBIM6 |
| 5.7316E-51 | -0.2718493 | 0.266 | 0.405 | 1.354E-46 | UGEC2 | SRSF7 |
| 6.2673E-51 | -0.451722 | 0.046 | 0.12 | 1.4805E-46 | UGEC2 | CYP1B1 |
| 1.2655E-50 | -0.7665236 | 0.954 | 0.848 | 2.9896E-46 | UGEC2 | SH3BGRL3 |
| 3.409E-50 | -0.2726198 | 0.043 | 0.116 | 8.053E-46 | UGEC2 | NR1D1 |
| 7.8658E-50 | 0.26199909 | 0.934 | 0.819 | 1.8581E-45 | UGEC2 | HNRNPA2B1 |
| 8.1165E-50 | 0.30090384 | 0.267 | 0.189 | 1.9174E-45 | UGEC2 | PTGR1 |
| 1.078E-49 | -0.3185898 | 0.168 | 0.278 | 2.5467E-45 | UGEC2 | ID2 |
| 1.3812E-49 | 0.30527938 | 0.567 | 0.493 | 3.2628E-45 | UGEC2 | ASAH1 |
| 1.9454E-49 | -0.3874944 | 0.848 | 0.599 | 4.5956E-45 | UGEC2 | S100A4 |
| 1.1046E-48 | -0.3939852 | 0.382 | 0.474 | 2.6095E-44 | UGEC2 | CDKN1A |
| 1.2838E-48 | 0.26673986 | 0.216 | 0.145 | 3.0328E-44 | UGEC2 | MT-ND6 |
| 2.9426E-48 | -2.1170182 | 0.491 | 0.461 | 6.9512E-44 | UGEC2 | LYZ |
| 5.5039E-48 | 0.43301803 | 0.393 | 0.352 | 1.3002E-43 | UGEC2 | UQCRQ |
| 1.5488E-46 | -0.3359109 | 0.069 | 0.147 | 3.6587E-42 | UGEC2 | PLAC8 |
| 1.7688E-46 | -0.3790813 | 0.075 | 0.156 | 4.1784E-42 | UGEC2 | SMC4 |
| 3.1098E-46 | -0.4611666 | 0.16 | 0.244 | 7.3463E-42 | UGEC2 | EPCAM |
| 5.2282E-46 | -0.2969791 | 0.148 | 0.248 | 1.2351E-41 | UGEC2 | CITED2 |
| 2.2509E-45 | -0.3099161 | 0.135 | 0.227 | 5.3173E-41 | UGEC2 | ZBTB20 |
| 1.28E-44 | -0.3176296 | 0.198 | 0.304 | 3.0236E-40 | UGEC2 | ATP13A3 |
| 6.4257E-44 | -1.4807982 | 0.874 | 0.71 | 1.5179E-39 | UGEC2 | S100A9 |
| 2.6485E-42 | -0.5284709 | 0.15 | 0.243 | 6.2566E-38 | UGEC2 | PLIN2 |
| 4.4886E-42 | -0.2546917 | 0.434 | 0.523 | 1.0603E-37 | UGEC2 | EIF5 |
| 2.7899E-41 | -0.3930158 | 0.059 | 0.127 | 6.5907E-37 | UGEC2 | TCIM |
| 3.6721E-41 | 0.29244785 | 0.295 | 0.23 | 8.6747E-37 | UGEC2 | PCBD1 |
| 4.4116E-41 | -0.9244382 | 0.721 | 0.632 | 1.0422E-36 | UGEC2 | IER3 |
| 9.8583E-40 | -0.3368963 | 0.164 | 0.255 | 2.3288E-35 | UGEC2 | CSTA |
| 1.3876E-38 | -0.3124353 | 0.051 | 0.113 | 3.2779E-34 | UGEC2 | TMC5 |
| 3.1046E-38 | -0.3127189 | 0.119 | 0.199 | 7.3341E-34 | UGEC2 | PLK2 |
| 1.837E-37 | 0.26674397 | 0.227 | 0.161 | 4.3396E-33 | UGEC2 | TINAGL1 |
| 2.3131E-37 | -0.6514515 | 0.995 | 0.931 | 5.4642E-33 | UGEC2 | TMSB4X |
| 5.546E-37 | -0.4346197 | 0.068 | 0.135 | 1.3101E-32 | UGEC2 | TFPI2 |
| 3.0263E-36 | 0.29433272 | 0.378 | 0.322 | 7.149E-32 | UGEC2 | LMNA |
| 3.1855E-36 | 0.33485179 | 0.288 | 0.237 | 7.525E-32 | UGEC2 | COMT |
| 1.633E-35 | -0.4942078 | 0.428 | 0.475 | 3.8576E-31 | UGEC2 | DUSP1 |
| 4.4357E-35 | 0.26165034 | 0.262 | 0.202 | 1.0479E-30 | UGEC2 | TCEAL4 |
| 4.0744E-34 | 0.26814628 | 0.271 | 0.212 | 9.6249E-30 | UGEC2 | SPATS2L |
| 9.9748E-34 | -1.517013 | 0.854 | 0.65 | 2.3564E-29 | UGEC2 | S100A8 |
| 3.502E-33 | 0.27031183 | 0.225 | 0.17 | 8.2728E-29 | UGEC2 | TMEM147 |
| 6.5841E-33 | -0.7014904 | 0.055 | 0.11 | 1.5554E-28 | UGEC2 | ZG16B |
| 1.0552E-32 | 0.2954782 | 0.288 | 0.237 | 2.4927E-28 | UGEC2 | SRI |
| 3.8517E-32 | -0.3112797 | 0.654 | 0.638 | 9.0989E-28 | UGEC2 | ARPC5 |

| | | | | | | |
|---|---|---|---|---|---|---|
| 9.8771E-32 | -0.2722581 | 0.167 | 0.252 | 2.3333E-27 | UGEC2 | ITGB8 |
| 1.7739E-31 | -0.8415785 | 0.575 | 0.534 | 4.1906E-27 | UGEC2 | PLAUR |
| 4.8173E-31 | -0.2906195 | 0.542 | 0.572 | 1.138E-26 | UGEC2 | RPS27L |
| 8.2026E-31 | -0.3378029 | 0.181 | 0.259 | 1.9377E-26 | UGEC2 | SYNE2 |
| 1.151E-30 | 0.27266717 | 0.279 | 0.229 | 2.719E-26 | UGEC2 | SYPL1 |
| 5.3E-30 | 0.33063591 | 0.381 | 0.346 | 1.252E-25 | UGEC2 | PNN |
| 6.213E-30 | -0.6232384 | 0.606 | 0.433 | 1.4677E-25 | UGEC2 | SOX4 |
| 1.4811E-29 | -0.4518057 | 0.909 | 0.777 | 3.4989E-25 | UGEC2 | PFN1 |
| 5.3096E-29 | 0.33752919 | 0.39 | 0.368 | 1.2543E-24 | UGEC2 | ATP5MC3 |
| 1.1569E-27 | -0.6154586 | 0.538 | 0.53 | 2.733E-23 | UGEC2 | CTSS |
| 1.5833E-27 | -0.283901 | 0.06 | 0.112 | 3.7401E-23 | UGEC2 | THBD |
| 1.6025E-27 | -0.3563284 | 0.926 | 0.827 | 3.7855E-23 | UGEC2 | SQSTM1 |
| 2.6929E-27 | 0.34923935 | 0.375 | 0.35 | 6.3614E-23 | UGEC2 | EIF3E |
| 2.9928E-27 | -0.4557526 | 0.986 | 0.926 | 7.07E-23 | UGEC2 | GAPDH |
| 5.4511E-27 | -0.3959838 | 0.163 | 0.234 | 1.2877E-22 | UGEC2 | HEXIM1 |
| 6.2963E-27 | -0.426484 | 0.093 | 0.148 | 1.4874E-22 | UGEC2 | CDKN2A |
| 8.3397E-27 | 0.28663064 | 0.365 | 0.324 | 1.9701E-22 | UGEC2 | ASPH |
| 9.0422E-27 | 0.26391017 | 0.303 | 0.256 | 2.136E-22 | UGEC2 | PRDX2 |
| 6.8668E-26 | -1.1354739 | 0.07 | 0.118 | 1.6222E-21 | UGEC2 | SPRR1B |
| 4.0321E-25 | 0.29925666 | 0.355 | 0.322 | 9.5251E-21 | UGEC2 | PPA1 |
| 1.184E-24 | -0.348845 | 0.139 | 0.198 | 2.7971E-20 | UGEC2 | FBXO32 |
| 1.7746E-24 | 0.25724468 | 0.277 | 0.233 | 4.1922E-20 | UGEC2 | TSTD1 |
| 2.9862E-24 | -0.7337212 | 0.085 | 0.139 | 7.0543E-20 | UGEC2 | SERPINB2 |
| 5.9067E-24 | 0.28430934 | 0.375 | 0.342 | 1.3953E-19 | UGEC2 | GOLGB1 |
| 1.2852E-23 | -0.6250491 | 0.47 | 0.357 | 3.036E-19 | UGEC2 | MAL |
| 3.597E-23 | 0.29745975 | 0.338 | 0.308 | 8.4972E-19 | UGEC2 | TMEM165 |
| 4.2563E-22 | -0.413289 | 0.262 | 0.175 | 1.0055E-17 | UGEC2 | DUOX2 |
| 8.3982E-22 | -0.4920715 | 0.426 | 0.322 | 1.9839E-17 | UGEC2 | IL1RN |
| 9.8641E-22 | -1.9369051 | 0.665 | 0.468 | 2.3302E-17 | UGEC2 | CXCL8 |
| 1.2398E-21 | -0.2606705 | 0.082 | 0.13 | 2.9287E-17 | UGEC2 | SRD5A3 |
| 1.8413E-21 | -0.3133706 | 0.225 | 0.152 | 4.3498E-17 | UGEC2 | CEACAM5 |
| 2.5114E-21 | -0.442297 | 0.09 | 0.136 | 5.9328E-17 | UGEC2 | GABRP |
| 1.4293E-20 | -0.4521163 | 0.074 | 0.116 | 3.3765E-16 | UGEC2 | KRT78 |
| 1.3442E-19 | 0.2625559 | 0.316 | 0.283 | 3.1754E-15 | UGEC2 | CLINT1 |
| 1.3135E-18 | -0.2904931 | 0.161 | 0.213 | 3.103E-14 | UGEC2 | CDKN2B |
| 1.3495E-18 | -0.2858642 | 0.075 | 0.116 | 3.1879E-14 | UGEC2 | TMPRSS11A |
| 1.7076E-18 | -0.4905241 | 0.866 | 0.66 | 4.0339E-14 | UGEC2 | NAMPT |
| 2.2515E-18 | -0.3332888 | 0.187 | 0.128 | 5.3187E-14 | UGEC2 | TMPRSS11B |
| 2.4771E-18 | 0.28057077 | 0.292 | 0.266 | 5.8516E-14 | UGEC2 | BAG1 |
| 1.8237E-16 | -2.6848425 | 0.712 | 0.522 | 4.3081E-12 | UGEC2 | PLCG2 |
| 6.7285E-16 | -0.301588 | 0.532 | 0.536 | 1.5895E-11 | UGEC2 | SERPINB1 |
| 6.9029E-15 | -0.3999412 | 0.992 | 0.979 | 1.6307E-10 | UGEC2 | MALAT1 |
| 1.1896E-14 | -0.2710027 | 0.621 | 0.595 | 2.8102E-10 | UGEC2 | NPC2 |
| 3.0055E-14 | -0.4008792 | 0.397 | 0.425 | 7.0998E-10 | UGEC2 | TXNRD1 |
| 3.4301E-14 | -0.4845744 | 0.689 | 0.624 | 8.1028E-10 | UGEC2 | ZFP36L1 |
| 5.0185E-13 | -0.5282098 | 0.995 | 0.971 | 1.1855E-08 | UGEC2 | ACTB |
| 2.8642E-12 | -1.3973669 | 0.115 | 0.147 | 6.7662E-08 | UGEC2 | SPRR2A |
| 6.611E-12 | -0.5516072 | 0.424 | 0.434 | 1.5617E-07 | UGEC2 | HIST1H1C |
| 2.7746E-11 | -0.2888846 | 0.991 | 0.962 | 6.5544E-07 | UGEC2 | UBC |
| 8.4334E-11 | -0.5214645 | 0.659 | 0.595 | 1.9922E-06 | UGEC2 | NFKBIA |

| | | | | | | |
|---|---|---|---|---|---|---|
| 1.1966E-10 | -0.2556819 | 0.642 | 0.514 | 2.8267E-06 | UGEC2 | ZFP36 |
| 1.6353E-10 | -0.2926957 | 0.778 | 0.62 | 3.8631E-06 | UGEC2 | GADD45B |
| 2.8666E-10 | -1.0561382 | 0.141 | 0.102 | 6.7718E-06 | UGEC2 | CRCT1 |
| 4.4793E-10 | -0.3330849 | 0.105 | 0.134 | 1.0581E-05 | UGEC2 | DHRS9 |
| 5.0356E-10 | -0.4650476 | 0.121 | 0.147 | 1.1896E-05 | UGEC2 | TMPRSS11E |
| 5.1192E-10 | 0.30468908 | 0.343 | 0.354 | 1.2093E-05 | UGEC2 | SPCS1 |
| 2.6966E-09 | -0.752664 | 0.529 | 0.391 | 6.3703E-05 | UGEC2 | SLPI |
| 2.8796E-09 | -2.1584138 | 0.524 | 0.445 | 6.8024E-05 | UGEC2 | SPP1 |
| 8.582E-09 | -0.4239492 | 0.128 | 0.151 | 0.00020273 | UGEC2 | TMPRSS11D |
| 1.6923E-08 | -0.524603 | 0.177 | 0.136 | 0.00039977 | UGEC2 | CXCL6 |
| 2.632E-08 | -0.4169214 | 0.865 | 0.711 | 0.00062175 | UGEC2 | CYBA |
| 3.6886E-08 | -0.2844368 | 0.232 | 0.182 | 0.00087137 | UGEC2 | TMPRSS2 |
| 3.9337E-08 | 0.31457685 | 0.351 | 0.371 | 0.00092926 | UGEC2 | TXNDC17 |
| 4.2493E-07 | -0.3259346 | 0.218 | 0.19 | 0.01003813 | UGEC2 | HSPA6 |
| 1.0037E-06 | -0.4590736 | 0.725 | 0.632 | 0.02371099 | UGEC2 | TXNIP |
| 1.5447E-06 | -0.4012414 | 0.137 | 0.155 | 0.03649001 | UGEC2 | SCEL |
| 2.1092E-06 | -0.3268849 | 0.192 | 0.151 | 0.04982514 | UGEC2 | KRT15 |
| 1.2073E-05 | -0.6219159 | 0.118 | 0.134 | 0.28520681 | UGEC2 | RHCG |
| 3.8644E-05 | -0.2753509 | 0.535 | 0.517 | 0.91288002 | UGEC2 | H1FX |
| 4.0554E-05 | -0.6439091 | 0.205 | 0.161 | 0.95800474 | UGEC2 | DUOXA2 |
| 6.9448E-05 | -0.3228077 | 0.176 | 0.143 | 1 | UGEC2 | CEACAM6 |
| 0.00012116 | -0.7659419 | 0.16 | 0.128 | 1 | UGEC2 | PLAT |
| 0.00014395 | -0.2814957 | 0.809 | 0.711 | 1 | UGEC2 | JUND |
| 0.0008622 | -0.584424 | 0.805 | 0.625 | 1 | UGEC2 | GLUL |
| 0.00087455 | 0.25386678 | 0.308 | 0.333 | 1 | UGEC2 | HNRNPF |
| 0.00207712 | -0.5789806 | 0.151 | 0.155 | 1 | UGEC2 | PITX1 |
| 0.00224989 | -1.2274018 | 0.098 | 0.108 | 1 | UGEC2 | CNFN |
| 0.00483609 | -0.6828771 | 0.163 | 0.16 | 1 | UGEC2 | MUC21 |
| 0.00506327 | -0.2644881 | 0.147 | 0.156 | 1 | UGEC2 | MUC4 |
| 0.00940823 | -1.2634923 | 0.13 | 0.133 | 1 | UGEC2 | LYPD2 |
