## Supplemental Table 3 for "Urinary single-cell sequencing captures intrarenal injury and repair processes in human acute kidney injury"

| p_val | avg_log2FC | pct.1 | pct.2 | p_val_adj | cluster | gene |
|---|---|---|---|---|---|---|
| 4.216E-163 | 2.69232753 | 1 | 0.59 | 9.001E-159 | ATL | SPP1 |
| 1.0046E-56 | 1.13743311 | 0.97 | 0.811 | 2.1449E-52 | ATL | IGFBP7 |
| 1.643E-19 | 1.05318646 | 0.301 | 0.131 | 3.5077E-15 | ATL | SERPINA1 |
| 1.4545E-88 | 0.98696889 | 0.408 | 0.083 | 3.1054E-84 | ATL | CLDN10 |
| 4.5018E-97 | 0.91492885 | 0.355 | 0.057 | 9.6114E-93 | ATL | SFRP1 |
| 1.5709E-26 | 0.84600899 | 0.535 | 0.279 | 3.3539E-22 | ATL | AKR1B1 |
| 3.8034E-12 | 0.76642809 | 0.726 | 0.643 | 8.1202E-08 | ATL | WFDC2 |
| 8.2019E-49 | 0.74831632 | 0.742 | 0.365 | 1.7511E-44 | ATL | LDHB |
| 3.3221E-41 | 0.73004374 | 0.572 | 0.226 | 7.0928E-37 | ATL | FXYD2 |
| 2.8959E-37 | 0.71257761 | 0.99 | 0.874 | 6.1828E-33 | ATL | ITM2B |
| 1.998E-31 | 0.7038238 | 0.609 | 0.311 | 4.2658E-27 | ATL | UGCG |
| 2.4776E-68 | 0.700562 | 0.341 | 0.07 | 5.2898E-64 | ATL | UMOD |
| 2.9209E-51 | 0.68443079 | 0.482 | 0.164 | 6.2361E-47 | ATL | SERPING1 |
| 1.3985E-87 | 0.65460714 | 0.237 | 0.029 | 2.9858E-83 | ATL | NNMT |
| 1.0697E-14 | 0.64805927 | 0.749 | 0.602 | 2.2838E-10 | ATL | SOD2 |
| 1.6477E-27 | 0.64781548 | 0.923 | 0.798 | 3.5179E-23 | ATL | CD24 |
| 1.0782E-26 | 0.64358294 | 0.538 | 0.277 | 2.3019E-22 | ATL | UCHL1 |
| 3.8896E-35 | 0.64123162 | 0.903 | 0.724 | 8.3044E-31 | ATL | ENO1 |
| 3.6245E-30 | 0.63338264 | 0.609 | 0.308 | 7.7384E-26 | ATL | MAL |
| 7.4243E-09 | 0.61398191 | 0.518 | 0.389 | 0.00015851 | ATL | TXNRD1 |
| 1.7058E-33 | 0.61195207 | 0.916 | 0.761 | 3.6418E-29 | ATL | PKM |
| 3.2917E-30 | 0.61148399 | 0.906 | 0.785 | 7.0278E-26 | ATL | NPM1 |
| 1.1884E-30 | 0.60518872 | 0.799 | 0.538 | 2.5373E-26 | ATL | CD59 |
| 2.6643E-65 | 0.59030174 | 0.502 | 0.146 | 5.6882E-61 | ATL | CPM |
| 1.6781E-18 | 0.58930897 | 0.686 | 0.482 | 3.5827E-14 | ATL | LDHA |
| 2.9205E-24 | 0.58775304 | 0.582 | 0.301 | 6.2353E-20 | ATL | PLAU |
| 1.7942E-30 | 0.57751248 | 0.361 | 0.132 | 3.8307E-26 | ATL | TM4SF1 |
| 5.3538E-16 | 0.56216324 | 0.291 | 0.135 | 1.143E-11 | ATL | TNFSF10 |
| 4.5579E-30 | 0.56211972 | 0.91 | 0.742 | 9.731E-26 | ATL | HINT1 |
| 4.6974E-24 | 0.56125099 | 0.9 | 0.865 | 1.0029E-19 | ATL | RPS4X |
| 2.5962E-22 | 0.5246114 | 0.555 | 0.314 | 5.5429E-18 | ATL | TKT |
| 2.1645E-22 | 0.52346279 | 0.575 | 0.324 | 4.6213E-18 | ATL | TIMP1 |
| 2.2182E-24 | 0.52335179 | 0.93 | 0.835 | 4.7358E-20 | ATL | PRDX1 |
| 1.443E-27 | 0.52070625 | 0.696 | 0.413 | 3.0808E-23 | ATL | PTTG1IP |
| 5.65E-27 | 0.5133683 | 0.97 | 0.87 | 1.2063E-22 | ATL | RACK1 |
| 3.7336E-14 | 0.50514511 | 0.381 | 0.212 | 7.9713E-10 | ATL | GCLM |
| 1.4262E-30 | 0.50380127 | 0.428 | 0.179 | 3.045E-26 | ATL | CBR1 |
| 3.4263E-48 | 0.50337313 | 0.462 | 0.152 | 7.3151E-44 | ATL | GOLM1 |
| 5.6438E-28 | 0.50285221 | 0.967 | 0.872 | 1.205E-23 | ATL | RPL7 |
| 7.2817E-28 | 0.50189036 | 0.963 | 0.837 | 1.5547E-23 | ATL | CD63 |
| 8.6477E-23 | 0.49513599 | 0.742 | 0.48 | 1.8463E-18 | ATL | ATP1A1 |
| 3.9967E-40 | 0.4860817 | 0.418 | 0.145 | 8.5329E-36 | ATL | TFPI |
| 9.859E-22 | 0.48355096 | 0.967 | 0.864 | 2.1049E-17 | ATL | RPL23 |
| 2.0613E-16 | 0.48180219 | 0.742 | 0.56 | 4.4009E-12 | ATL | CST3 |
| 0.00025055 | 0.47846126 | 0.338 | 0.251 | 1 | ATL | AKAP12 |
| 2.0702E-31 | 0.47335838 | 0.532 | 0.242 | 4.4199E-27 | ATL | MGLL |
| 2.0855E-24 | 0.46463115 | 0.95 | 0.84 | 4.4526E-20 | ATL | RPSA |
| 4.2272E-25 | 0.46276919 | 0.997 | 0.937 | 9.0251E-21 | ATL | RPL15 |
| 2.053E-14 | 0.45776853 | 0.739 | 0.579 | 4.3832E-10 | ATL | APP |

| | | | | | | |
|---|---|---|---|---|---|---|
| 3.0161E-23 | 0.45765506 | 0.997 | 0.987 | 6.4394E-19 | ATL | EEF1A1 |
| 1.9451E-18 | 0.44970399 | 0.585 | 0.354 | 4.1527E-14 | ATL | TALDO1 |
| 4.8823E-48 | 0.44262741 | 0.271 | 0.06 | 1.0424E-43 | ATL | SOD3 |
| 3.6474E-25 | 0.44106658 | 0.943 | 0.75 | 7.7872E-21 | ATL | RPL4 |
| 6.7885E-24 | 0.43874381 | 0.983 | 0.92 | 1.4494E-19 | ATL | RPS3A |
| 7.6109E-22 | 0.43865951 | 0.99 | 0.972 | 1.6249E-17 | ATL | TPT1 |
| 2.1453E-15 | 0.43831771 | 0.886 | 0.766 | 4.5802E-11 | ATL | COX7C |
| 7.9408E-18 | 0.43698719 | 0.485 | 0.266 | 1.6954E-13 | ATL | CKB |
| 2.9372E-74 | 0.43688792 | 0.204 | 0.025 | 6.2708E-70 | ATL | CLDN16 |
| 2.9219E-23 | 0.43283692 | 0.99 | 0.908 | 6.2383E-19 | ATL | RPL24 |
| 6.9838E-21 | 0.42770462 | 0.645 | 0.392 | 1.491E-16 | ATL | MGST3 |
| 2.1146E-18 | 0.41986056 | 0.876 | 0.72 | 4.5147E-14 | ATL | CTSD |
| 4.2452E-93 | 0.41949734 | 0.224 | 0.024 | 9.0635E-89 | ATL | SLC34A2 |
| 8.1034E-23 | 0.41431989 | 0.94 | 0.833 | 1.7301E-18 | ATL | PPIA |
| 8.0557E-08 | 0.4132714 | 0.926 | 0.91 | 0.0017199 | ATL | MT-CO3 |
| 2.319E-20 | 0.41304773 | 0.973 | 0.894 | 4.9511E-16 | ATL | RPS13 |
| 2.6753E-26 | 0.41054238 | 0.652 | 0.354 | 5.7117E-22 | ATL | ATP5MC3 |
| 3.0171E-12 | 0.40944922 | 0.395 | 0.217 | 6.4414E-08 | ATL | SFN |
| 9.7533E-23 | 0.40471763 | 0.542 | 0.29 | 2.0823E-18 | ATL | LEPROT |
| 1.9766E-16 | 0.40177113 | 0.452 | 0.245 | 4.2201E-12 | ATL | MRPS6 |
| 1.4174E-18 | 0.39951952 | 0.816 | 0.626 | 3.0262E-14 | ATL | ANXA5 |
| 4.8534E-18 | 0.39793494 | 0.983 | 0.928 | 1.0362E-13 | ATL | RPS14 |
| 4.4591E-22 | 0.38949402 | 0.987 | 0.931 | 9.5203E-18 | ATL | RPS6 |
| 6.614E-19 | 0.38809429 | 0.625 | 0.384 | 1.4121E-14 | ATL | POMP |
| 3.3668E-16 | 0.38458241 | 0.973 | 0.897 | 7.188E-12 | ATL | RPL5 |
| 2.1563E-25 | 0.38409684 | 0.515 | 0.251 | 4.6036E-21 | ATL | ATP5F1A |
| 1.1255E-54 | 0.38373909 | 0.328 | 0.076 | 2.403E-50 | ATL | PPP1R1A |
| 2.4941E-17 | 0.38364642 | 0.98 | 0.929 | 5.3249E-13 | ATL | RPL7A |
| 2.0308E-08 | 0.38235595 | 0.552 | 0.414 | 0.00043358 | ATL | CCND1 |
| 9.8975E-25 | 0.38219573 | 0.368 | 0.153 | 2.1131E-20 | ATL | FHL2 |
| 1.1201E-16 | 0.381259 | 0.656 | 0.432 | 2.3914E-12 | ATL | NAP1L1 |
| 4.3925E-33 | 0.37949193 | 0.264 | 0.075 | 9.378E-29 | ATL | CYS1 |
| 1.5813E-20 | 0.3784955 | 0.585 | 0.327 | 3.376E-16 | ATL | EIF3E |
| 3.6896E-21 | 0.37755428 | 0.528 | 0.279 | 7.8772E-17 | ATL | ATP5PO |
| 8.8832E-13 | 0.37615879 | 0.756 | 0.604 | 1.8966E-08 | ATL | APLP2 |
| 2.0941E-18 | 0.37521133 | 0.328 | 0.148 | 4.4709E-14 | ATL | NQO1 |
| 2.9677E-22 | 0.3747716 | 0.425 | 0.2 | 6.3361E-18 | ATL | MPC2 |
| 2.0414E-23 | 0.37399288 | 0.351 | 0.144 | 4.3583E-19 | ATL | ITM2C |
| 1.4649E-17 | 0.37380778 | 0.853 | 0.677 | 3.1275E-13 | ATL | CLIC1 |
| 2.2881E-19 | 0.37096431 | 0.348 | 0.16 | 4.885E-15 | ATL | CAVIN1 |
| 1.8292E-16 | 0.36750519 | 0.95 | 0.913 | 3.9054E-12 | ATL | HSP90AB1 |
| 7.8314E-17 | 0.36691606 | 0.836 | 0.659 | 1.672E-12 | ATL | RAC1 |
| 9.6052E-15 | 0.366529 | 0.833 | 0.639 | 2.0507E-10 | ATL | PEBP1 |
| 1.9077E-15 | 0.36602339 | 0.967 | 0.903 | 4.0729E-11 | ATL | RPS3 |
| 3.4375E-15 | 0.36582572 | 0.525 | 0.311 | 7.3391E-11 | ATL | PRDX2 |
| 5.4326E-12 | 0.36518648 | 0.806 | 0.713 | 1.1599E-07 | ATL | TXN |
| 3.3898E-15 | 0.36387286 | 0.498 | 0.297 | 7.2372E-11 | ATL | CIB1 |
| 3.892E-17 | 0.36337343 | 0.993 | 0.932 | 8.3095E-13 | ATL | RPS27A |
| 7.6632E-16 | 0.36184555 | 0.746 | 0.546 | 1.6361E-11 | ATL | CD9 |
| 3.2371E-20 | 0.36153722 | 0.94 | 0.746 | 6.9113E-16 | ATL | BTF3 |

| | | | | | | |
|---|---|---|---|---|---|---|
| 7.0595E-14 | 0.36062933 | 0.829 | 0.658 | 1.5072E-09 | ATL | UQCRB |
| 5.0313E-21 | 0.35983798 | 0.388 | 0.18 | 1.0742E-16 | ATL | ATP1B3 |
| 8.5311E-15 | 0.35884646 | 0.676 | 0.462 | 1.8214E-10 | ATL | IFITM3 |
| 4.178E-21 | 0.35878172 | 0.462 | 0.231 | 8.9201E-17 | ATL | ANXA4 |
| 5.6441E-16 | 0.35628957 | 0.983 | 0.932 | 1.205E-11 | ATL | RPL3 |
| 1.1762E-19 | 0.3554193 | 0.482 | 0.248 | 2.5113E-15 | ATL | LAPTM4B |
| 8.3282E-25 | 0.35486578 | 0.552 | 0.271 | 1.7781E-20 | ATL | OCIAD1 |
| 4.5042E-19 | 0.35442327 | 0.983 | 0.903 | 9.6164E-15 | ATL | RPS7 |
| 2.2623E-19 | 0.34971737 | 0.569 | 0.323 | 4.8301E-15 | ATL | EIF3H |
| 1.2346E-07 | 0.34837814 | 0.719 | 0.577 | 0.00263585 | ATL | ATP6V0E1 |
| 1.1682E-14 | 0.34766405 | 0.99 | 0.978 | 2.4942E-10 | ATL | FTH1 |
| 1.4585E-16 | 0.34763398 | 0.983 | 0.907 | 3.1139E-12 | ATL | RPL26 |
| 6.2959E-11 | 0.34721247 | 0.582 | 0.393 | 1.3442E-06 | ATL | DBI |
| 4.5285E-12 | 0.34505519 | 0.742 | 0.6 | 9.6683E-08 | ATL | CLTA |
| 3.5802E-18 | 0.34498588 | 0.629 | 0.382 | 7.6438E-14 | ATL | SUB1 |
| 3.7291E-14 | 0.34455661 | 0.438 | 0.241 | 7.9615E-10 | ATL | PDZK1IP1 |
| 2.0761E-18 | 0.34373601 | 0.983 | 0.918 | 4.4325E-14 | ATL | RPL9 |
| 1.4249E-14 | 0.34292229 | 0.953 | 0.903 | 3.0422E-10 | ATL | RPS20 |
| 2.5571E-12 | 0.34086489 | 0.508 | 0.315 | 5.4595E-08 | ATL | DCDC2 |
| 1.7967E-16 | 0.34018296 | 0.562 | 0.334 | 3.8359E-12 | ATL | SEC11A |
| 1.1179E-20 | 0.33929366 | 0.726 | 0.464 | 2.3867E-16 | ATL | PRDX6 |
| 2.5913E-15 | 0.33745043 | 0.398 | 0.215 | 5.5323E-11 | ATL | CTNNAL1 |
| 6.8741E-35 | 0.33687955 | 0.301 | 0.089 | 1.4676E-30 | ATL | KCNJ16 |
| 2.2403E-18 | 0.33685119 | 0.361 | 0.173 | 4.783E-14 | ATL | BCL2L1 |
| 1.1385E-15 | 0.33570195 | 0.806 | 0.604 | 2.4308E-11 | ATL | ATP5MG |
| 1.1869E-12 | 0.33475782 | 0.612 | 0.421 | 2.5341E-08 | ATL | TMEM59 |
| 2.7827E-13 | 0.3344814 | 0.271 | 0.123 | 5.9411E-09 | ATL | GPX3 |
| 6.4583E-17 | 0.33411875 | 0.485 | 0.272 | 1.3788E-12 | ATL | CCT5 |
| 1.1005E-13 | 0.33391426 | 0.87 | 0.756 | 2.3496E-09 | ATL | CALM2 |
| 3.1359E-16 | 0.33382699 | 0.98 | 0.927 | 6.6951E-12 | ATL | RPL21 |
| 9.0624E-14 | 0.33259812 | 0.649 | 0.443 | 1.9348E-09 | ATL | VDAC2 |
| 2.0051E-15 | 0.33197921 | 0.592 | 0.376 | 4.281E-11 | ATL | HSBP1 |
| 1.129E-14 | 0.33192105 | 0.87 | 0.724 | 2.4103E-10 | ATL | GNAS |
| 2.3185E-11 | 0.33063908 | 0.585 | 0.386 | 4.9499E-07 | ATL | CD151 |
| 2.9072E-15 | 0.33017979 | 0.569 | 0.361 | 6.2069E-11 | ATL | ATP5F1B |
| 8.6777E-16 | 0.33001306 | 0.609 | 0.378 | 1.8527E-11 | ATL | MRPL33 |
| 2.7499E-18 | 0.32829932 | 0.498 | 0.267 | 5.8711E-14 | ATL | NDUFV2 |
| 5.3317E-21 | 0.32532319 | 0.278 | 0.106 | 1.1383E-16 | ATL | ALDH1A1 |
| 6.3261E-14 | 0.3249607 | 0.913 | 0.823 | 1.3506E-09 | ATL | H3F3A |
| 1.4665E-16 | 0.32485806 | 0.582 | 0.351 | 3.131E-12 | ATL | EIF3K |
| 6.5963E-11 | 0.32378845 | 0.702 | 0.553 | 1.4083E-06 | ATL | LRRFIP1 |
| 3.1942E-17 | 0.32351228 | 0.431 | 0.222 | 6.8197E-13 | ATL | IFITM2 |
| 5.9065E-15 | 0.32338354 | 0.826 | 0.644 | 1.261E-10 | ATL | UQCRH |
| 1.248E-12 | 0.32292748 | 0.522 | 0.32 | 2.6645E-08 | ATL | CYBA |
| 3.3141E-12 | 0.32128071 | 0.365 | 0.203 | 7.0756E-08 | ATL | DNTTIP2 |
| 7.0985E-16 | 0.32072195 | 0.291 | 0.13 | 1.5155E-11 | ATL | CDH16 |
| 1.128E-11 | 0.31967088 | 0.696 | 0.521 | 2.4084E-07 | ATL | ATP5MC2 |
| 1.524E-16 | 0.31927165 | 0.93 | 0.824 | 3.2538E-12 | ATL | TMBIM6 |
| 4.969E-14 | 0.31903258 | 0.973 | 0.932 | 1.0609E-09 | ATL | RPS9 |
| 1.7641E-19 | 0.31852382 | 0.482 | 0.25 | 3.7663E-15 | ATL | TMEM123 |

| | | | | | | |
|---|---|---|---|---|---|---|
| 1.9415E-13 | 0.31850178 | 0.98 | 0.954 | 4.1452E-09 | ATL | S100A6 |
| 1.9302E-12 | 0.31802308 | 0.946 | 0.867 | 4.121E-08 | ATL | RPL10A |
| 7.69E-16 | 0.31752871 | 0.515 | 0.305 | 1.6418E-11 | ATL | NDUFC2 |
| 1.1495E-13 | 0.3168355 | 0.983 | 0.93 | 2.4542E-09 | ATL | RPL18 |
| 2.6504E-15 | 0.31598856 | 0.602 | 0.373 | 5.6585E-11 | ATL | REEP5 |
| 1.6343E-12 | 0.31448837 | 0.97 | 0.921 | 3.4892E-08 | ATL | RPL14 |
| 2.5606E-14 | 0.3133401 | 0.545 | 0.332 | 5.4669E-10 | ATL | EIF4B |
| 2.2131E-32 | 0.31328727 | 0.321 | 0.104 | 4.7249E-28 | ATL | CPVL |
| 6.7033E-29 | 0.31302728 | 0.311 | 0.106 | 1.4312E-24 | ATL | EFHD1 |
| 1.4812E-14 | 0.3129257 | 0.993 | 0.939 | 3.1625E-10 | ATL | RPL19 |
| 8.1327E-14 | 0.312375 | 0.983 | 0.945 | 1.7363E-09 | ATL | RPL11 |
| 5.0082E-13 | 0.31040549 | 0.759 | 0.583 | 1.0693E-08 | ATL | S100A13 |
| 1.1334E-14 | 0.30931449 | 0.88 | 0.728 | 2.4199E-10 | ATL | NDUFA4 |
| 5.5778E-14 | 0.3077429 | 0.535 | 0.332 | 1.1909E-09 | ATL | PSMB1 |
| 2.8477E-13 | 0.30694223 | 0.993 | 0.936 | 6.0798E-09 | ATL | RPS23 |
| 2.7216E-15 | 0.30672301 | 0.485 | 0.277 | 5.8107E-11 | ATL | LITAF |
| 2.0677E-16 | 0.30669956 | 0.95 | 0.85 | 4.4145E-12 | ATL | MYL12B |
| 8.5835E-15 | 0.3058363 | 0.639 | 0.405 | 1.8326E-10 | ATL | UQCR10 |
| 6.2403E-16 | 0.30352767 | 0.528 | 0.311 | 1.3323E-11 | ATL | SRP9 |
| 2.1698E-49 | 0.30307644 | 0.184 | 0.029 | 4.6326E-45 | ATL | TSPAN8 |
| 1.0562E-08 | 0.30288779 | 0.903 | 0.924 | 0.00022551 | ATL | RPL10 |
| 9.8061E-16 | 0.30219048 | 0.575 | 0.344 | 2.0936E-11 | ATL | COMMD6 |
| 3.025E-13 | 0.30058754 | 0.776 | 0.581 | 6.4583E-09 | ATL | EZR |
| 7.1223E-13 | 0.30031419 | 0.599 | 0.387 | 1.5206E-08 | ATL | CAPN2 |
| 2.0607E-13 | 0.30024422 | 0.973 | 0.888 | 4.3997E-09 | ATL | NACA |
| 7.2139E-15 | 0.29982658 | 0.923 | 0.778 | 1.5402E-10 | ATL | COX4I1 |
| 9.2611E-19 | 0.29869611 | 0.288 | 0.12 | 1.9773E-14 | ATL | PRSS23 |
| 9.1843E-10 | 0.2980163 | 0.438 | 0.283 | 1.9609E-05 | ATL | RABAC1 |
| 2.5074E-12 | 0.29764684 | 0.856 | 0.746 | 5.3534E-08 | ATL | RTN4 |
| 2.2009E-15 | 0.29762929 | 0.963 | 0.891 | 4.6989E-11 | ATL | GAPDH |
| 2.6934E-11 | 0.29747773 | 0.492 | 0.317 | 5.7505E-07 | ATL | NPC2 |
| 3.1527E-12 | 0.29560473 | 0.452 | 0.269 | 6.731E-08 | ATL | HIF1A |
| 2.9617E-11 | 0.29522925 | 0.518 | 0.335 | 6.3232E-07 | ATL | ERP29 |
| 1.611E-12 | 0.29494044 | 0.652 | 0.448 | 3.4395E-08 | ATL | ATP5PF |
| 7.3089E-11 | 0.29485591 | 0.595 | 0.412 | 1.5605E-06 | ATL | DYNLRB1 |
| 1.4816E-16 | 0.29483985 | 0.485 | 0.271 | 3.1632E-12 | ATL | CCT6A |
| 4.2744E-17 | 0.29408471 | 0.886 | 0.675 | 9.1259E-13 | ATL | EPCAM |
| 2.9788E-15 | 0.29318879 | 0.522 | 0.306 | 6.3597E-11 | ATL | RAB13 |
| 3.4195E-11 | 0.29308535 | 0.448 | 0.269 | 7.3007E-07 | ATL | MUC1 |
| 5.6439E-13 | 0.29253308 | 0.421 | 0.243 | 1.205E-08 | ATL | TES |
| 3.6987E-12 | 0.29219112 | 0.753 | 0.555 | 7.8966E-08 | ATL | RHOA |
| 8.8026E-17 | 0.29179614 | 0.622 | 0.37 | 1.8794E-12 | ATL | VAMP8 |
| 1.5391E-13 | 0.29174364 | 0.525 | 0.325 | 3.2859E-09 | ATL | SF3B6 |
| 1.064E-08 | 0.29122546 | 0.729 | 0.579 | 0.00022717 | ATL | COX5B |
| 1.3649E-17 | 0.29054469 | 0.291 | 0.124 | 2.9141E-13 | ATL | IMPA2 |
| 1.9888E-15 | 0.29049615 | 0.548 | 0.322 | 4.2461E-11 | ATL | NDUFC1 |
| 5.4482E-14 | 0.29016417 | 0.508 | 0.31 | 1.1632E-09 | ATL | MRFAP1 |
| 2.9384E-07 | 0.29008401 | 0.669 | 0.556 | 0.00627357 | ATL | HMGA1 |
| 1.8689E-12 | 0.28965559 | 0.749 | 0.583 | 3.99E-08 | ATL | YWHAB |
| 1.7452E-15 | 0.28880993 | 0.923 | 0.779 | 3.726E-11 | ATL | EEF1B2 |

| | | | | | | |
|---|---|---|---|---|---|---|
| 5.193E-08 | 0.28855325 | 0.522 | 0.357 | 0.00110871 | ATL | COX7B |
| 1.9247E-15 | 0.28827905 | 0.652 | 0.411 | 4.1092E-11 | ATL | PARK7 |
| 2.3798E-12 | 0.28810388 | 0.893 | 0.744 | 5.081E-08 | ATL | DSTN |
| 2.1887E-16 | 0.28757369 | 0.428 | 0.228 | 4.6729E-12 | ATL | MRPL20 |
| 1.5882E-14 | 0.28626182 | 0.425 | 0.238 | 3.3909E-10 | ATL | HIGD2A |
| 2.8225E-16 | 0.2856686 | 0.388 | 0.197 | 6.0261E-12 | ATL | NDUFB5 |
| 4.8838E-12 | 0.2842211 | 0.515 | 0.328 | 1.0427E-07 | ATL | SLC25A5 |
| 5.6771E-13 | 0.28351786 | 0.993 | 0.932 | 1.2121E-08 | ATL | RPL32 |
| 2.6611E-15 | 0.28329338 | 0.498 | 0.284 | 5.6815E-11 | ATL | SERINC1 |
| 3.9339E-40 | 0.28219334 | 0.227 | 0.051 | 8.3989E-36 | ATL | KLHL13 |
| 1.431E-08 | 0.28122419 | 0.936 | 0.926 | 0.00030552 | ATL | ACTG1 |
| 4.9093E-14 | 0.28081412 | 0.515 | 0.308 | 1.0481E-09 | ATL | PCBD1 |
| 1.8107E-11 | 0.28056427 | 0.957 | 0.914 | 3.8658E-07 | ATL | RPLP0 |
| 2.272E-16 | 0.28049179 | 0.271 | 0.117 | 4.8508E-12 | ATL | FBXO2 |
| 4.4798E-13 | 0.27998549 | 0.692 | 0.472 | 9.5643E-09 | ATL | MORF4L1 |
| 1.1066E-17 | 0.27958761 | 0.465 | 0.245 | 2.3626E-13 | ATL | AP2M1 |
| 1.4765E-11 | 0.27930159 | 0.896 | 0.817 | 3.1524E-07 | ATL | YBX1 |
| 1.5033E-09 | 0.27828643 | 0.331 | 0.192 | 3.2095E-05 | ATL | NCOA7 |
| 4.1438E-12 | 0.27691424 | 0.518 | 0.327 | 8.847E-08 | ATL | NDUFB4 |
| 1.5762E-09 | 0.2763806 | 0.465 | 0.307 | 3.3652E-05 | ATL | PGK1 |
| 2.4935E-11 | 0.27608072 | 0.987 | 0.955 | 5.3235E-07 | ATL | RPS2 |
| 3.6471E-10 | 0.27499201 | 0.977 | 0.919 | 7.7866E-06 | ATL | RPL34 |
| 9.6207E-16 | 0.27473367 | 0.395 | 0.203 | 2.054E-11 | ATL | PRDX3 |
| 1.864E-15 | 0.27312574 | 0.365 | 0.185 | 3.9796E-11 | ATL | TMEM205 |
| 9.6732E-11 | 0.273101 | 0.796 | 0.67 | 2.0652E-06 | ATL | HNRNPA1 |
| 5.9826E-14 | 0.27300622 | 0.395 | 0.216 | 1.2773E-09 | ATL | GDI2 |
| 1.9529E-12 | 0.27267277 | 0.535 | 0.336 | 4.1695E-08 | ATL | SPCS1 |
| 2.0064E-12 | 0.27246855 | 0.595 | 0.394 | 4.2837E-08 | ATL | YWHAQ |
| 7.9509E-09 | 0.27241299 | 0.579 | 0.41 | 0.00016975 | ATL | HNRNPK |
| 6.8333E-08 | 0.27112068 | 0.692 | 0.548 | 0.00145892 | ATL | CANX |
| 4.6962E-09 | 0.27094021 | 0.936 | 0.868 | 0.00010026 | ATL | RPS5 |
| 3.9294E-12 | 0.26989427 | 0.512 | 0.317 | 8.3892E-08 | ATL | ATP5MPL |
| 2.4596E-09 | 0.26964577 | 0.719 | 0.564 | 5.2511E-05 | ATL | HNRNPC |
| 1.6233E-12 | 0.26921543 | 0.876 | 0.771 | 3.4658E-08 | ATL | EEF2 |
| 4.7917E-13 | 0.26860195 | 0.478 | 0.289 | 1.023E-08 | ATL | MRPL51 |
| 1.0821E-06 | 0.26839431 | 0.619 | 0.494 | 0.02310241 | ATL | TNFRSF12A |
| 3.7104E-11 | 0.26806453 | 0.963 | 0.863 | 7.9217E-07 | ATL | UBA52 |
| 2.4415E-12 | 0.26757572 | 0.592 | 0.377 | 5.2127E-08 | ATL | DYNLT1 |
| 2.1589E-08 | 0.26748534 | 0.482 | 0.327 | 0.00046093 | ATL | TUBA1C |
| 1.3466E-13 | 0.26669391 | 0.408 | 0.224 | 2.875E-09 | ATL | MDH1 |
| 2.7857E-10 | 0.26667851 | 0.465 | 0.301 | 5.9475E-06 | ATL | EIF3F |
| 1.1298E-15 | 0.26640539 | 0.311 | 0.144 | 2.4122E-11 | ATL | SNX10 |
| 2.8068E-13 | 0.26625985 | 0.472 | 0.278 | 5.9925E-09 | ATL | TMEM14C |
| 4.0197E-11 | 0.26477487 | 0.445 | 0.272 | 8.5822E-07 | ATL | SMS |
| 3.0221E-13 | 0.26448923 | 0.418 | 0.238 | 6.4521E-09 | ATL | RAB11A |
| 8.5885E-17 | 0.26393136 | 0.405 | 0.205 | 1.8336E-12 | ATL | TMBIM4 |
| 2.2191E-09 | 0.26377282 | 0.609 | 0.424 | 4.7378E-05 | ATL | BSG |
| 6.9511E-16 | 0.26332445 | 0.344 | 0.165 | 1.4841E-11 | ATL | PON2 |
| 2.1228E-10 | 0.26320747 | 0.783 | 0.6 | 4.5321E-06 | ATL | PSMA7 |
| 3.2233E-13 | 0.26179581 | 0.502 | 0.306 | 6.8817E-09 | ATL | ATP5PD |

| | | | | | | |
|---|---|---|---|---|---|---|
| 2.0795E-11 | 0.25978016 | 0.448 | 0.27 | 4.4397E-07 | ATL | CAPNS1 |
| 1.2689E-10 | 0.25966139 | 0.977 | 0.947 | 2.7091E-06 | ATL | RPL13A |
| 3.7015E-12 | 0.25961432 | 0.378 | 0.211 | 7.9027E-08 | ATL | PSMC5 |
| 1.0487E-14 | 0.25944191 | 0.405 | 0.219 | 2.2389E-10 | ATL | ANXA7 |
| 5.3914E-10 | 0.25905823 | 0.344 | 0.199 | 1.1511E-05 | ATL | DPP4 |
| 1.2552E-13 | 0.25891826 | 0.676 | 0.448 | 2.6798E-09 | ATL | LAPTM4A |
| 1.3616E-14 | 0.25869832 | 0.355 | 0.18 | 2.9071E-10 | ATL | PLEKHA1 |
| 5.516E-12 | 0.25863193 | 0.488 | 0.306 | 1.1777E-07 | ATL | GABARAPL2 |
| 3.828E-12 | 0.2581887 | 0.997 | 0.971 | 8.1728E-08 | ATL | RPLP1 |
| 1.2887E-12 | 0.2577157 | 0.522 | 0.318 | 2.7515E-08 | ATL | COX5A |
| 4.2161E-09 | 0.25712908 | 0.997 | 0.966 | 9.0014E-05 | ATL | RPL41 |
| 1.7485E-12 | 0.25674318 | 0.492 | 0.295 | 3.733E-08 | ATL | ACTR3 |
| 3.3897E-12 | 0.25567122 | 0.515 | 0.315 | 7.237E-08 | ATL | DEK |
| 2.0165E-15 | 0.25447717 | 0.435 | 0.231 | 4.3053E-11 | ATL | PPP1CA |
| 2.6459E-46 | 0.25383117 | 0.224 | 0.044 | 5.6489E-42 | ATL | GYPC |
| 1.0998E-11 | 0.25205971 | 0.525 | 0.329 | 2.348E-07 | ATL | RAB2A |
| 9.3102E-11 | 0.25196876 | 0.462 | 0.281 | 1.9877E-06 | ATL | NDUFB11 |
| 2.1004E-13 | 0.2514611 | 0.431 | 0.242 | 4.4844E-09 | ATL | BCAP31 |
| 1.0846E-12 | 0.25127505 | 0.431 | 0.247 | 2.3156E-08 | ATL | LMAN2 |
| 2.1115E-13 | 0.25071753 | 0.472 | 0.274 | 4.5082E-09 | ATL | CNN3 |
| 3.8169E-15 | 0.2506482 | 0.378 | 0.194 | 8.149E-11 | ATL | CCDC47 |
| 9.6482E-20 | 0.25050197 | 0.301 | 0.124 | 2.0599E-15 | ATL | UXS1 |
| 3.8445E-09 | 0.25048173 | 0.515 | 0.357 | 8.2081E-05 | ATL | EIF3A |
| 6.3572E-07 | 0.25043699 | 0.572 | 0.431 | 0.01357257 | ATL | ST13 |
| 5.2144E-17 | 0.25030262 | 0.244 | 0.098 | 1.1133E-12 | ATL | SLC48A1 |
| 0.0063045 | -0.2505137 | 0.124 | 0.177 | 1 | ATL | FOXC1 |
| 0.0001535 | -0.2513193 | 0.094 | 0.174 | 1 | ATL | THSD7A |
| 0.00134148 | -0.2533295 | 0.077 | 0.139 | 1 | ATL | ISG15 |
| 0.0008037 | -0.2559701 | 0.204 | 0.274 | 1 | ATL | SMG1 |
| 3.0564E-06 | -0.2605817 | 0.02 | 0.1 | 0.06525381 | ATL | MAFA |
| 2.4981E-07 | -0.2613521 | 0.08 | 0.197 | 0.00533334 | ATL | EPS8 |
| 0.00106589 | -0.2618426 | 0.084 | 0.147 | 1 | ATL | ANKRD37 |
| 0.00777696 | -0.2627612 | 0.301 | 0.35 | 1 | ATL | DDX24 |
| 0.00048859 | -0.2650139 | 0.07 | 0.135 | 1 | ATL | CSRNP2 |
| 6.4566E-05 | -0.2682308 | 0.077 | 0.158 | 1 | ATL | CDKN2B |
| 0.00153339 | -0.2692837 | 0.211 | 0.275 | 1 | ATL | SPTAN1 |
| 0.00723321 | -0.277421 | 0.06 | 0.106 | 1 | ATL | ARC |
| 0.00019887 | -0.2778964 | 0.06 | 0.129 | 1 | ATL | HOMER1 |
| 8.7173E-06 | -0.2794685 | 0.03 | 0.109 | 0.18611361 | ATL | AL355075.4 |
| 0.00033897 | -0.2827361 | 0.214 | 0.291 | 1 | ATL | HP1BP3 |
| 0.00557396 | -0.2877464 | 0.251 | 0.303 | 1 | ATL | C6orf62 |
| 0.00611781 | -0.2895812 | 0.311 | 0.363 | 1 | ATL | AKAP9 |
| 0.00388871 | -0.2964749 | 0.234 | 0.286 | 1 | ATL | NUFIP2 |
| 5.8843E-05 | -0.2979901 | 0.074 | 0.154 | 1 | ATL | PFKFB3 |
| 0.00142375 | -0.2987821 | 0.047 | 0.101 | 1 | ATL | CXCL3 |
| 0.00547425 | -0.2995796 | 0.341 | 0.392 | 1 | ATL | NDRG1 |
| 0.00019476 | -0.3031468 | 0.134 | 0.216 | 1 | ATL | ZFP36 |
| 1.0989E-07 | -0.3042861 | 0.017 | 0.113 | 0.00234618 | ATL | HIST4H4 |
| 0.00307696 | -0.3096614 | 0.09 | 0.145 | 1 | ATL | ID1 |
| 0.00018788 | -0.3161357 | 0.154 | 0.235 | 1 | ATL | CDKN2AIP |

| | | | | | | |
|---|---|---|---|---|---|---|
| 2.3391E-05 | -0.3170158 | 0.08 | 0.167 | 0.49939869 | ATL | HIST1H1E |
| 0.00089328 | -0.3229019 | 0.184 | 0.254 | 1 | ATL | IRF1 |
| 2.1447E-05 | -0.3251635 | 0.181 | 0.275 | 0.45789569 | ATL | TCF25 |
| 8.9888E-07 | -0.3272865 | 0.023 | 0.113 | 0.01919104 | ATL | AC023157.3 |
| 0.00549901 | -0.3309229 | 0.214 | 0.266 | 1 | ATL | AMD1 |
| 2.6023E-06 | -0.3341409 | 0.057 | 0.152 | 0.05555985 | ATL | AC007952.4 |
| 0.0037594 | -0.3352434 | 0.572 | 0.669 | 1 | ATL | CRYAB |
| 0.00851 | -0.343652 | 0.258 | 0.303 | 1 | ATL | DNAJB1 |
| 6.2989E-06 | -0.3443235 | 0.157 | 0.264 | 0.13448163 | ATL | HIST1H2AC |
| 0.0071211 | -0.3486682 | 0.124 | 0.173 | 1 | ATL | FOSB |
| 0.00636582 | -0.3498262 | 0.435 | 0.482 | 1 | ATL | H1FX |
| 0.00012444 | -0.3582016 | 0.194 | 0.278 | 1 | ATL | VEGFA |
| 4.9791E-05 | -0.3597331 | 0.087 | 0.177 | 1 | ATL | IGFBP3 |
| 5.9262E-06 | -0.3629462 | 0.03 | 0.114 | 0.12652358 | ATL | C15orf48 |
| 2.0147E-05 | -0.3639835 | 0.104 | 0.194 | 0.43014542 | ATL | H2AFX |
| 0.00024515 | -0.3695312 | 0.157 | 0.235 | 1 | ATL | NFKBIZ |
| 1.4025E-07 | -0.3700016 | 0.01 | 0.102 | 0.00299436 | ATL | AP001160.1 |
| 4.6375E-05 | -0.3702977 | 0.268 | 0.372 | 0.99009867 | ATL | DDIT3 |
| 0.00087041 | -0.3733783 | 0.147 | 0.22 | 1 | ATL | TNFAIP2 |
| 3.1836E-09 | -0.3762853 | 0.054 | 0.185 | 6.797E-05 | ATL | ERN1 |
| 2.3019E-09 | -0.3786398 | 0.013 | 0.129 | 4.9146E-05 | ATL | HIST2H2BF |
| 0.00147732 | -0.3791478 | 0.207 | 0.272 | 1 | ATL | SEPHS2 |
| 0.00047525 | -0.3796416 | 0.047 | 0.11 | 1 | ATL | S100A9 |
| 0.00129213 | -0.3841011 | 0.278 | 0.344 | 1 | ATL | ZFP36L2 |
| 0.00601223 | -0.3855292 | 0.234 | 0.29 | 1 | ATL | RND3 |
| 7.0243E-05 | -0.3862424 | 0.274 | 0.36 | 1 | ATL | ARGLU1 |
| 4.7298E-05 | -0.3927153 | 0.191 | 0.288 | 1 | ATL | CEBPD |
| 0.0030061 | -0.3948828 | 0.077 | 0.129 | 1 | ATL | COL7A1 |
| 6.7897E-09 | -0.4043409 | 0.023 | 0.138 | 0.00014496 | ATL | AC025164.1 |
| 1.5975E-06 | -0.4050765 | 0.09 | 0.193 | 0.0341056 | ATL | TLE4 |
| 0.00179938 | -0.406027 | 0.174 | 0.234 | 1 | ATL | SYNE1 |
| 5.0246E-09 | -0.4084199 | 0.582 | 0.717 | 0.00010727 | ATL | HSPB1 |
| 4.4022E-05 | -0.4132552 | 0.298 | 0.38 | 0.93986749 | ATL | RSRP1 |
| 2.8146E-05 | -0.4181309 | 0.204 | 0.303 | 0.60091896 | ATL | DDIT4 |
| 5.1291E-05 | -0.4195107 | 0.171 | 0.266 | 1 | ATL | DUSP1 |
| 3.8727E-06 | -0.4204903 | 0.441 | 0.595 | 0.08268269 | ATL | KRT19 |
| 0.0015513 | -0.4351048 | 0.247 | 0.307 | 1 | ATL | HSPA1B |
| 0.0020867 | -0.4419014 | 0.351 | 0.414 | 1 | ATL | PHLDA2 |
| 3.014E-07 | -0.4498063 | 0.181 | 0.302 | 0.0064349 | ATL | MARCKSL1 |
| 0.00026269 | -0.4588197 | 0.181 | 0.255 | 1 | ATL | IER5 |
| 0.00017966 | -0.4588278 | 0.164 | 0.244 | 1 | ATL | ID4 |
| 0.00114139 | -0.4718758 | 0.167 | 0.244 | 1 | ATL | TFPI2 |
| 7.4594E-07 | -0.4851798 | 0.217 | 0.33 | 0.01592573 | ATL | CEBPB |
| 6.368E-08 | -0.4883146 | 0.452 | 0.564 | 0.00135958 | ATL | SRRM2 |
| 3.93E-05 | -0.4889558 | 0.187 | 0.279 | 0.83906297 | ATL | RASD1 |
| 0.00539695 | -0.4946655 | 0.278 | 0.328 | 1 | ATL | IL32 |
| 0.00013981 | -0.5100065 | 0.161 | 0.239 | 1 | ATL | INTS6 |
| 2.2869E-07 | -0.5274639 | 0.789 | 0.873 | 0.0048825 | ATL | SAT1 |
| 6.4717E-07 | -0.5289952 | 0.916 | 0.941 | 0.01381714 | ATL | MALAT1 |
| 2.5831E-06 | -0.5301944 | 0.187 | 0.286 | 0.05514929 | ATL | POLR2A |

| | | | | | | |
|---|---|---|---|---|---|---|
| 1.7629E-08 | -0.5316536 | 0.054 | 0.171 | 0.00037638 | ATL | EGLN3 |
| 1.0927E-08 | -0.5393077 | 0.462 | 0.603 | 0.00023328 | ATL | CLU |
| 0.00490816 | -0.5461741 | 0.217 | 0.273 | 1 | ATL | TCIM |
| 0.00016193 | -0.5494919 | 0.314 | 0.393 | 1 | ATL | NFKBIA |
| 3.3456E-07 | -0.5495008 | 0.107 | 0.216 | 0.00714282 | ATL | SOCS1 |
| 3.6536E-05 | -0.5532471 | 0.873 | 0.858 | 0.78004445 | ATL | H3F3B |
| 5.37E-05 | -0.5556814 | 0.311 | 0.388 | 1 | ATL | JUNB |
| 6.8076E-11 | -0.5585388 | 0.057 | 0.206 | 1.4534E-06 | ATL | SGK1 |
| 5.0441E-07 | -0.5585619 | 0.144 | 0.257 | 0.01076925 | ATL | IER5L |
| 1.0283E-08 | -0.570169 | 0.512 | 0.612 | 0.00021953 | ATL | BTG1 |
| 4.1381E-09 | -0.5802604 | 0.06 | 0.188 | 8.8349E-05 | ATL | PIK3R3 |
| 1.3915E-07 | -0.5937918 | 0.528 | 0.625 | 0.00297084 | ATL | PPP1R15A |
| 8.2445E-13 | -0.5948898 | 0.224 | 0.425 | 1.7602E-08 | ATL | TPM4 |
| 7.0928E-06 | -0.5972254 | 0.512 | 0.626 | 0.15143173 | ATL | VIM |
| 3.0806E-05 | -0.6014194 | 0.756 | 0.871 | 0.65771429 | ATL | TMSB4X |
| 4.0639E-10 | -0.6078359 | 0.13 | 0.284 | 8.6764E-06 | ATL | BTG2 |
| 3.3495E-12 | -0.6276019 | 0.134 | 0.316 | 7.1511E-08 | ATL | CDKN1C |
| 0.00052624 | -0.6280644 | 0.271 | 0.333 | 1 | ATL | RHOB |
| 2.2847E-07 | -0.6352619 | 0.258 | 0.377 | 0.00487783 | ATL | SLC38A2 |
| 4.2792E-07 | -0.6681307 | 0.204 | 0.314 | 0.00913606 | ATL | GLUL |
| 0.00096955 | -0.668232 | 0.234 | 0.291 | 1 | ATL | FOS |
| 7.0165E-05 | -0.6731251 | 0.548 | 0.606 | 1 | ATL | GADD45B |
| 1.2774E-11 | -0.6790943 | 0.181 | 0.345 | 2.7272E-07 | ATL | SNHG12 |
| 1.9178E-09 | -0.6827153 | 0.137 | 0.28 | 4.0945E-05 | ATL | PPP1R10 |
| 2.0342E-06 | -0.6973205 | 0.023 | 0.108 | 0.04342948 | ATL | FXYD4 |
| 1.3674E-08 | -0.7076332 | 0.147 | 0.28 | 0.00029193 | ATL | HES1 |
| 6.7203E-05 | -0.7424433 | 0.508 | 0.565 | 1 | ATL | IER2 |
| 0.00033778 | -0.757979 | 0.268 | 0.332 | 1 | ATL | LAMC2 |
| 2.0917E-07 | -0.7656285 | 0.291 | 0.409 | 0.00446571 | ATL | RNF213 |
| 2.771E-09 | -0.8248951 | 0.104 | 0.237 | 5.9161E-05 | ATL | CSKMT |
| 0.0037705 | -0.8359263 | 0.191 | 0.257 | 1 | ATL | LCN2 |
| 1.7328E-12 | -0.837325 | 0.518 | 0.642 | 3.6995E-08 | ATL | JUND |
| 5.7702E-05 | -0.840397 | 0.197 | 0.272 | 1 | ATL | HSPA2 |
| 3.5507E-08 | -0.9275891 | 0.137 | 0.264 | 0.00075807 | ATL | NUPR1 |
| 2.2846E-10 | -1.0154505 | 0.157 | 0.301 | 4.8776E-06 | ATL | HEXIM1 |
| 2.8303E-15 | -1.020936 | 0.431 | 0.59 | 6.0428E-11 | ATL | ELF3 |
| 8.0904E-08 | -1.0639678 | 0.274 | 0.38 | 0.00172729 | ATL | HIST1H4C |
| 1.3273E-15 | -1.2446222 | 0.087 | 0.287 | 2.8339E-11 | ATL | MAFB |
| 6.1914E-06 | -1.3033419 | 0.659 | 0.679 | 0.13218579 | ATL | MTRNR2L12 |
| 1.3852E-16 | -1.3167847 | 0.468 | 0.637 | 2.9574E-12 | ATL | JUN |
| 6.4376E-11 | -1.3361256 | 0.421 | 0.542 | 1.3744E-06 | ATL | SOX4 |
| 2.689E-16 | -1.3465625 | 0.783 | 0.816 | 5.7409E-12 | ATL | NEAT1 |
| 9.8602E-26 | -4.2860055 | 0.391 | 0.602 | 2.1052E-21 | ATL | PLCG2 |
| 0 | 3.16687782 | 0.683 | 0.022 | 0 | CD-IC | IGFBP5 |
| 6.831E-129 | 2.02986594 | 0.71 | 0.108 | 1.459E-124 | CD-IC | RARRES3 |
| 9.734E-212 | 1.79904851 | 0.648 | 0.052 | 2.078E-207 | CD-IC | TMEM213 |
| 2.3207E-08 | 1.77735453 | 0.179 | 0.067 | 0.00049548 | CD-IC | SPINK1 |
| 0 | 1.76101825 | 0.759 | 0.03 | 0 | CD-IC | ATP6V0D2 |
| 2.1442E-19 | 1.61640354 | 0.207 | 0.048 | 4.5779E-15 | CD-IC | TAC1 |
| 6.291E-274 | 1.61192155 | 0.731 | 0.052 | 1.343E-269 | CD-IC | IL18 |

| | | | | | | |
|---|---|---|---|---|---|---|
| 3.2558E-27 | 1.46083898 | 0.655 | 0.329 | 6.9512E-23 | CD-IC | CHCHD10 |
| 8.1386E-20 | 1.44161358 | 0.517 | 0.222 | 1.7376E-15 | CD-IC | DEFB1 |
| 1.8587E-70 | 1.34953091 | 0.676 | 0.162 | 3.9684E-66 | CD-IC | CA12 |
| 9.2341E-06 | 1.34664724 | 0.69 | 0.645 | 0.1971471 | CD-IC | WFDC2 |
| 0 | 1.30110136 | 0.524 | 0.007 | 0 | CD-IC | FOXI1 |
| 0 | 1.26493072 | 0.51 | 0.01 | 0 | CD-IC | LINC01187 |
| 2.9694E-36 | 1.22771952 | 0.807 | 0.415 | 6.3397E-32 | CD-IC | LGALS3 |
| 2.846E-102 | 1.2257096 | 0.69 | 0.117 | 6.0762E-98 | CD-IC | PTGER3 |
| 0 | 1.22090133 | 0.648 | 0.033 | 0 | CD-IC | ATP6V0A4 |
| 1.5348E-14 | 1.18118086 | 0.724 | 0.548 | 3.2768E-10 | CD-IC | CD9 |
| 0 | 1.14525744 | 0.276 | 0.004 | 0 | CD-IC | SPARCL1 |
| 7.7825E-07 | 1.11214546 | 0.71 | 0.617 | 0.01661556 | CD-IC | MTRNR2L8 |
| 1.1515E-43 | 1.08302723 | 0.648 | 0.222 | 2.4585E-39 | CD-IC | ATP6AP2 |
| 0 | 1.02467964 | 0.303 | 0.004 | 0 | CD-IC | ADGRF5 |
| 2.2586E-36 | 0.9856281 | 0.8 | 0.422 | 4.8222E-32 | CD-IC | ATP6V0B |
| 8.57E-113 | 0.97237853 | 0.303 | 0.02 | 1.83E-108 | CD-IC | SLC26A7 |
| 6.4525E-28 | 0.93739667 | 0.538 | 0.195 | 1.3776E-23 | CD-IC | MARCKS |
| 1.1546E-27 | 0.93626459 | 0.628 | 0.278 | 2.4651E-23 | CD-IC | LITAF |
| 8.9023E-29 | 0.90331821 | 0.552 | 0.219 | 1.9006E-24 | CD-IC | DYNLT3 |
| 3.1023E-32 | 0.89141428 | 0.759 | 0.356 | 6.6233E-28 | CD-IC | COX7B |
| 0 | 0.88721939 | 0.393 | 0.007 | 0 | CD-IC | DMRT2 |
| 0 | 0.88182288 | 0.338 | 0.004 | 0 | CD-IC | ATP6V1G3 |
| 0.00031189 | 0.87033059 | 0.738 | 0.678 | 1 | CD-IC | MTRNR2L12 |
| 4.1202E-42 | 0.82853814 | 0.517 | 0.142 | 8.7967E-38 | CD-IC | ATP6V1A |
| 9.8297E-21 | 0.82843883 | 0.648 | 0.329 | 2.0986E-16 | CD-IC | SLC25A5 |
| 6.8169E-50 | 0.8269268 | 0.662 | 0.189 | 1.4554E-45 | CD-IC | EPS8 |
| 2.4607E-22 | 0.81915662 | 0.441 | 0.161 | 5.2536E-18 | CD-IC | ADGRF1 |
| 2.2955E-22 | 0.80101923 | 0.517 | 0.217 | 4.9009E-18 | CD-IC | LIMS1 |
| 1.3185E-41 | 0.79351679 | 0.6 | 0.18 | 2.815E-37 | CD-IC | MUC20-OT1 |
| 2.0854E-18 | 0.78637216 | 0.49 | 0.214 | 4.4522E-14 | CD-IC | PTGR1 |
| 6.6752E-09 | 0.78191005 | 0.938 | 0.889 | 0.00014252 | CD-IC | MT-ND2 |
| 3.861E-22 | 0.77802243 | 0.938 | 0.78 | 8.2432E-18 | CD-IC | COX4I1 |
| 0 | 0.76855936 | 0.434 | 0.006 | 0 | CD-IC | STAC2 |
| 6.3425E-23 | 0.76752801 | 0.655 | 0.319 | 1.3541E-18 | CD-IC | COX5A |
| 1.1409E-94 | 0.76470719 | 0.262 | 0.018 | 2.4358E-90 | CD-IC | ERP27 |
| 2.7338E-32 | 0.76014626 | 0.51 | 0.165 | 5.8366E-28 | CD-IC | MAP1LC3A |
| 2.7959E-19 | 0.75528616 | 0.931 | 0.767 | 5.9692E-15 | CD-IC | COX7C |
| 1.8306E-33 | 0.74945027 | 0.641 | 0.244 | 3.9084E-29 | CD-IC | DHRS7 |
| 1.3183E-16 | 0.74419213 | 0.441 | 0.193 | 2.8146E-12 | CD-IC | NCOA7 |
| 5.3197E-18 | 0.74347369 | 0.648 | 0.358 | 1.1358E-13 | CD-IC | ATP5MC3 |
| 1.2609E-71 | 0.7367842 | 0.372 | 0.048 | 2.6919E-67 | CD-IC | CDA |
| 1.753E-151 | 0.73329259 | 0.428 | 0.03 | 3.742E-147 | CD-IC | RCAN2 |
| 1.1768E-07 | 0.73005364 | 0.924 | 0.891 | 0.0025124 | CD-IC | MT-ND1 |
| 7.4648E-18 | 0.7164421 | 0.8 | 0.526 | 1.5937E-13 | CD-IC | COX6C |
| 2.8141E-14 | 0.70858776 | 0.772 | 0.58 | 6.0081E-10 | CD-IC | COX5B |
| 1.1036E-30 | 0.70437808 | 0.393 | 0.098 | 2.3562E-26 | CD-IC | APOE |
| 3.788E-10 | 0.70350871 | 0.586 | 0.371 | 8.0875E-06 | CD-IC | ATF3 |
| 8.014E-50 | 0.69952188 | 0.248 | 0.031 | 1.711E-45 | CD-IC | TSPAN8 |
| 7.0199E-17 | 0.68105397 | 0.579 | 0.297 | 1.4987E-12 | CD-IC | DDT |
| 1.5518E-08 | 0.68088581 | 0.759 | 0.612 | 0.00033131 | CD-IC | MT-ND4L |

| | | | | | | |
|---|---|---|---|---|---|---|
| 6.4473E-16 | 0.68011162 | 0.814 | 0.624 | 1.3765E-11 | CD-IC | SLC25A6 |
| 9.5216E-18 | 0.67742652 | 0.986 | 0.957 | 2.0329E-13 | CD-IC | EIF1 |
| 1.063E-07 | 0.66162725 | 0.883 | 0.828 | 0.00226957 | CD-IC | MT-ND5 |
| 1.306E-212 | 0.66090292 | 0.379 | 0.016 | 2.787E-208 | CD-IC | ATP6V1C2 |
| 6.4745E-16 | 0.65485807 | 0.738 | 0.458 | 1.3823E-11 | CD-IC | COX8A |
| 0 | 0.652073 | 0.193 | 0 | 0 | CD-IC | TRGC1 |
| 4.0909E-20 | 0.64793033 | 0.566 | 0.253 | 8.7342E-16 | CD-IC | NEDD4L |
| 1.067E-14 | 0.63439488 | 0.545 | 0.268 | 2.2781E-10 | CD-IC | CKB |
| 1.0894E-14 | 0.63214241 | 0.924 | 0.826 | 2.3258E-10 | CD-IC | HLA-B |
| 0 | 0.6270198 | 0.372 | 0.006 | 0 | CD-IC | CLNK |
| 1.1913E-13 | 0.62148989 | 0.669 | 0.391 | 2.5435E-09 | CD-IC | ATP5F1D |
| 1.1968E-26 | 0.62106662 | 0.621 | 0.254 | 2.5552E-22 | CD-IC | NDUFA6 |
| 6.5186E-21 | 0.62010213 | 0.448 | 0.169 | 1.3917E-16 | CD-IC | MPC1 |
| 5.8147E-16 | 0.6136582 | 0.559 | 0.288 | 1.2414E-11 | CD-IC | YWHAH |
| 3.6731E-41 | 0.60731862 | 0.331 | 0.061 | 7.8421E-37 | CD-IC | RNF152 |
| 1.3471E-09 | 0.59975975 | 0.959 | 0.926 | 2.8762E-05 | CD-IC | MT-CO1 |
| 8.7255E-19 | 0.59494015 | 0.262 | 0.073 | 1.8629E-14 | CD-IC | JAG1 |
| 2.7708E-12 | 0.59315773 | 0.752 | 0.569 | 5.9156E-08 | CD-IC | COX7A2 |
| 2.7379E-11 | 0.59179118 | 0.841 | 0.678 | 5.8453E-07 | CD-IC | COX6A1 |
| 1.9338E-20 | 0.58769615 | 0.414 | 0.146 | 4.1288E-16 | CD-IC | TFCP2L1 |
| 9.5811E-15 | 0.58557222 | 0.559 | 0.285 | 2.0456E-10 | CD-IC | MT-ATP8 |
| 2.1253E-13 | 0.5827276 | 0.89 | 0.729 | 4.5375E-09 | CD-IC | NDUFA4 |
| 9.7582E-20 | 0.58021746 | 0.469 | 0.183 | 2.0834E-15 | CD-IC | ARHGAP18 |
| 2.5826E-36 | 0.57889298 | 0.29 | 0.053 | 5.5139E-32 | CD-IC | OXCT1 |
| 3.6926E-28 | 0.57679784 | 0.379 | 0.104 | 7.8838E-24 | CD-IC | MCRIP2 |
| 1.876E-230 | 0.57304975 | 0.138 | 0.001 | 4.004E-226 | CD-IC | HMX2 |
| 3.3333E-12 | 0.56611194 | 0.6 | 0.363 | 7.1166E-08 | CD-IC | ATP5F1B |
| 5.7119E-13 | 0.56552064 | 0.917 | 0.807 | 1.2195E-08 | CD-IC | ATP5F1E |
| 1.4755E-15 | 0.56213424 | 0.69 | 0.413 | 3.1502E-11 | CD-IC | DYNLRB1 |
| 1.0312E-09 | 0.56205923 | 0.972 | 0.949 | 2.2016E-05 | CD-IC | B2M |
| 4.7018E-11 | 0.55394839 | 0.6 | 0.373 | 1.0038E-06 | CD-IC | NDUFA1 |
| 1.9988E-26 | 0.55315263 | 0.421 | 0.129 | 4.2674E-22 | CD-IC | FOXP1 |
| 1.6024E-08 | 0.54058458 | 0.738 | 0.579 | 0.00034211 | CD-IC | ATP6V0E1 |
| 2.1466E-12 | 0.53258594 | 0.241 | 0.083 | 4.5829E-08 | CD-IC | ANKRD36C |
| 2.3254E-14 | 0.52998974 | 0.641 | 0.373 | 4.9648E-10 | CD-IC | VAMP8 |
| 1.729E-12 | 0.52737436 | 0.579 | 0.315 | 3.6915E-08 | CD-IC | FAM107B |
| 1.5779E-43 | 0.52655341 | 0.241 | 0.032 | 3.3688E-39 | CD-IC | RHCG |
| 1.5622E-20 | 0.52625414 | 0.386 | 0.13 | 3.3353E-16 | CD-IC | ACAT1 |
| 7.3581E-15 | 0.52489129 | 0.455 | 0.201 | 1.571E-10 | CD-IC | ERO1A |
| 8.4732E-27 | 0.52349872 | 0.366 | 0.099 | 1.809E-22 | CD-IC | MAP3K8 |
| 7.145E-09 | 0.52126372 | 0.697 | 0.488 | 0.00015255 | CD-IC | DNAJA1 |
| 5.7231E-13 | 0.51674602 | 0.779 | 0.602 | 1.2219E-08 | CD-IC | PSMA7 |
| 3.6575E-27 | 0.51595947 | 0.345 | 0.089 | 7.8088E-23 | CD-IC | SDCBP2 |
| 1.562E-10 | 0.51275955 | 0.621 | 0.392 | 3.3348E-06 | CD-IC | FKBP2 |
| 3.2199E-15 | 0.50872562 | 0.634 | 0.346 | 6.8744E-11 | CD-IC | VAPA |
| 4.5446E-06 | 0.50693617 | 0.952 | 0.912 | 0.09702817 | CD-IC | MT-ND4 |
| 3.3806E-11 | 0.50652158 | 0.676 | 0.45 | 7.2177E-07 | CD-IC | ATP5PF |
| 8.9232E-62 | 0.5035711 | 0.317 | 0.04 | 1.9051E-57 | CD-IC | ARSJ |
| 6.5392E-16 | 0.50330518 | 0.524 | 0.247 | 1.3961E-11 | CD-IC | MRPS6 |
| 5.0705E-16 | 0.50076669 | 0.717 | 0.407 | 1.0826E-11 | CD-IC | UQCR10 |

| | | | | | | |
|---|---|---|---|---|---|---|
| 2.4685E-18 | 0.50007458 | 0.269 | 0.078 | 5.2703E-14 | CD-IC | GNPNAT1 |
| 2.214E-22 | 0.49571853 | 0.366 | 0.109 | 4.7268E-18 | CD-IC | RARRES2 |
| 2.2174E-70 | 0.49386635 | 0.283 | 0.028 | 4.7342E-66 | CD-IC | ITIH5 |
| 1.3892E-69 | 0.49279005 | 0.207 | 0.015 | 2.966E-65 | CD-IC | CALCA |
| 1.1782E-52 | 0.48985599 | 0.352 | 0.055 | 2.5155E-48 | CD-IC | SMIM24 |
| 4.1082E-17 | 0.48979419 | 0.421 | 0.163 | 8.771E-13 | CD-IC | SELENOP |
| 1.0634E-07 | 0.48738374 | 0.855 | 0.722 | 0.00227046 | CD-IC | CTSD |
| 3.4824E-24 | 0.48731026 | 0.379 | 0.113 | 7.435E-20 | CD-IC | PRKAA2 |
| 2.2121E-20 | 0.48149416 | 0.51 | 0.203 | 4.7229E-16 | CD-IC | GABARAPL1 |
| 5.2537E-18 | 0.48039001 | 0.497 | 0.215 | 1.1217E-13 | CD-IC | SRI |
| 0.00439034 | 0.48019141 | 0.269 | 0.189 | 1 | CD-IC | THBS1 |
| 1.5692E-36 | 0.47782256 | 0.29 | 0.052 | 3.3502E-32 | CD-IC | IQGAP2 |
| 1.651E-09 | 0.4734232 | 0.855 | 0.69 | 3.5249E-05 | CD-IC | GPX4 |
| 2.7447E-16 | 0.47315384 | 0.4 | 0.16 | 5.86E-12 | CD-IC | PPP2R5A |
| 1.7744E-13 | 0.47308699 | 0.634 | 0.355 | 3.7883E-09 | CD-IC | SPAG9 |
| 4.8019E-09 | 0.47090933 | 0.593 | 0.379 | 0.00010252 | CD-IC | NDUFB2 |
| 1.47E-251 | 0.46845924 | 0.248 | 0.005 | 3.138E-247 | CD-IC | KIT |
| 5.535E-153 | 0.46703968 | 0.269 | 0.011 | 1.182E-148 | CD-IC | CCDC68 |
| 5.8022E-06 | 0.46675014 | 0.517 | 0.357 | 0.12387652 | CD-IC | ATP5ME |
| 5.2154E-71 | 0.46588868 | 0.331 | 0.038 | 1.1135E-66 | CD-IC | SNTB1 |
| 1.6191E-11 | 0.46549125 | 0.51 | 0.276 | 3.4568E-07 | CD-IC | MRPL41 |
| 1.5853E-13 | 0.46181106 | 0.407 | 0.182 | 3.3846E-09 | CD-IC | HADHB |
| 3.0642E-10 | 0.45909845 | 0.386 | 0.191 | 6.5422E-06 | CD-IC | HIPK2 |
| 2.1426E-13 | 0.45877381 | 0.641 | 0.353 | 4.5745E-09 | CD-IC | EIF3K |
| 1.476E-118 | 0.4566839 | 0.159 | 0.005 | 3.151E-114 | CD-IC | AQP6 |
| 2.1528E-13 | 0.45618061 | 0.49 | 0.232 | 4.5963E-09 | CD-IC | SCIN |
| 0.0001224 | 0.45552262 | 0.448 | 0.33 | 1 | CD-IC | TUBA1C |
| 3.0496E-09 | 0.45534054 | 0.759 | 0.607 | 6.5109E-05 | CD-IC | ATP5MG |
| 2.9299E-14 | 0.45345291 | 0.441 | 0.195 | 6.2554E-10 | CD-IC | BNIP3 |
| 1.7336E-23 | 0.45314312 | 0.379 | 0.115 | 3.7012E-19 | CD-IC | TMEM106C |
| 5.1547E-06 | 0.45278014 | 0.821 | 0.727 | 0.11005316 | CD-IC | GNAS |
| 2.8034E-15 | 0.45257026 | 0.614 | 0.324 | 5.9853E-11 | CD-IC | NDUFC1 |
| 4.6097E-11 | 0.45181224 | 0.648 | 0.423 | 9.8417E-07 | CD-IC | TMEM59 |
| 1.1166E-70 | 0.45167215 | 0.269 | 0.025 | 2.3839E-66 | CD-IC | CA8 |
| 4.5519E-05 | 0.4513744 | 0.752 | 0.661 | 0.97183498 | CD-IC | UQCRB |
| 4.0284E-20 | 0.44947215 | 0.483 | 0.189 | 8.6006E-16 | CD-IC | TBC1D1 |
| 7.4922E-09 | 0.44875383 | 0.441 | 0.262 | 0.00015996 | CD-IC | CAPZA2 |
| 2.4258E-33 | 0.44776124 | 0.359 | 0.081 | 5.179E-29 | CD-IC | RAB40B |
| 9.3556E-33 | 0.44717076 | 0.421 | 0.107 | 1.9974E-28 | CD-IC | EFHD1 |
| 7.2448E-10 | 0.44600291 | 0.628 | 0.394 | 1.5468E-05 | CD-IC | CYCS |
| 1.4344E-09 | 0.44584216 | 0.317 | 0.141 | 3.0625E-05 | CD-IC | ID3 |
| 2.6765E-10 | 0.44573432 | 0.662 | 0.426 | 5.7142E-06 | CD-IC | BSG |
| 5.3614E-07 | 0.44284568 | 0.31 | 0.162 | 0.01144666 | CD-IC | KATNBL1 |
| 1.2415E-08 | 0.44156081 | 0.91 | 0.797 | 0.00026506 | CD-IC | HLA-C |
| 8.6929E-49 | 0.43760603 | 0.269 | 0.036 | 1.8559E-44 | CD-IC | ABHD10 |
| 3.0308E-10 | 0.43442423 | 0.379 | 0.188 | 6.4707E-06 | CD-IC | NDUFS7 |
| 5.994E-15 | 0.43421859 | 0.448 | 0.194 | 1.2797E-10 | CD-IC | CYC1 |
| 2.8119E-13 | 0.43420228 | 0.772 | 0.489 | 6.0034E-09 | CD-IC | ATP1B1 |
| 1.1213E-12 | 0.4329237 | 0.531 | 0.282 | 2.3941E-08 | CD-IC | ATP5PO |
| 5.275E-08 | 0.43214842 | 0.579 | 0.386 | 0.00112621 | CD-IC | HSPE1 |

| | | | | | | |
|---|---|---|---|---|---|---|
| 5.337E-08 | 0.43173213 | 0.648 | 0.444 | 0.00113945 | CD-IC | ATP5MD |
| 4.2917E-12 | 0.42890015 | 0.407 | 0.188 | 9.1629E-08 | CD-IC | ECI1 |
| 1.7765E-14 | 0.42864839 | 0.428 | 0.186 | 3.7927E-10 | CD-IC | PNPLA8 |
| 6.2013E-10 | 0.42674023 | 0.6 | 0.376 | 1.324E-05 | CD-IC | REEP5 |
| 2.1891E-75 | 0.42618715 | 0.262 | 0.022 | 4.6738E-71 | CD-IC | CELF2 |
| 4.5697E-12 | 0.42614462 | 0.476 | 0.24 | 9.7563E-08 | CD-IC | HIGD2A |
| 3.2982E-19 | 0.42497862 | 0.428 | 0.159 | 7.0416E-15 | CD-IC | CTSA |
| 2.1373E-08 | 0.42384424 | 0.448 | 0.262 | 0.00045631 | CD-IC | SELENOM |
| 1.2406E-08 | 0.42384121 | 0.517 | 0.308 | 0.00026486 | CD-IC | ISCU |
| 1.0047E-14 | 0.42288414 | 0.469 | 0.21 | 2.1451E-10 | CD-IC | SLIRP |
| 4.2059E-05 | 0.4211349 | 0.29 | 0.17 | 0.89795784 | CD-IC | SERPING1 |
| 2.1426E-15 | 0.41305818 | 0.393 | 0.156 | 4.5744E-11 | CD-IC | MRPL34 |
| 1.222E-17 | 0.41300313 | 0.386 | 0.143 | 2.6091E-13 | CD-IC | CRB3 |
| 8.5318E-24 | 0.41107023 | 0.331 | 0.089 | 1.8215E-19 | CD-IC | SEMA3C |
| 3.1152E-12 | 0.4088498 | 0.586 | 0.322 | 6.6509E-08 | CD-IC | NDUFB7 |
| 1.0444E-09 | 0.40865559 | 0.945 | 0.892 | 2.2299E-05 | CD-IC | RPS11 |
| 7.5376E-10 | 0.4065189 | 0.455 | 0.234 | 1.6093E-05 | CD-IC | LPP |
| 4.2816E-06 | 0.40613648 | 0.552 | 0.368 | 0.09141266 | CD-IC | SPTBN1 |
| 6.7512E-14 | 0.40173822 | 0.517 | 0.251 | 1.4414E-09 | CD-IC | ATP6V1D |
| 5.1736E-18 | 0.39989067 | 0.393 | 0.145 | 1.1046E-13 | CD-IC | NDUFS2 |
| 2.7704E-12 | 0.39839401 | 0.393 | 0.178 | 5.9147E-08 | CD-IC | SLC25A4 |
| 2.045E-08 | 0.39806206 | 0.503 | 0.314 | 0.0004366 | CD-IC | SNX3 |
| 0 | 0.3971792 | 0.152 | 0 | 0 | CD-IC | GPRC6A |
| 1.0576E-36 | 0.39313432 | 0.324 | 0.063 | 2.258E-32 | CD-IC | TSPAN5 |
| 3.0547E-08 | 0.39051796 | 0.379 | 0.201 | 0.00065217 | CD-IC | NDUFB1 |
| 5.6904E-12 | 0.38940995 | 0.586 | 0.324 | 1.2149E-07 | CD-IC | VDAC1 |
| 1.2523E-06 | 0.38799771 | 0.717 | 0.559 | 0.02673628 | CD-IC | COX6B1 |
| 6.7468E-07 | 0.38637486 | 0.352 | 0.197 | 0.01440435 | CD-IC | TMEM141 |
| 5.1539E-12 | 0.38440243 | 0.476 | 0.24 | 1.1004E-07 | CD-IC | PLEKHB2 |
| 3.8689E-52 | 0.38199456 | 0.193 | 0.018 | 8.2602E-48 | CD-IC | LINC01564 |
| 8.1993E-12 | 0.38192215 | 0.428 | 0.209 | 1.7506E-07 | CD-IC | MIEN1 |
| 9.9388E-06 | 0.38164615 | 0.662 | 0.534 | 0.21219433 | CD-IC | HLA-E |
| 7.5176E-13 | 0.38079435 | 0.269 | 0.093 | 1.605E-08 | CD-IC | CRYBG3 |
| 5.5776E-09 | 0.37787387 | 0.607 | 0.385 | 0.00011908 | CD-IC | SUB1 |
| 4.5309E-19 | 0.37637679 | 0.352 | 0.115 | 9.6735E-15 | CD-IC | PBDC1 |
| 2.2433E-14 | 0.37631196 | 0.372 | 0.145 | 4.7894E-10 | CD-IC | RETREG1 |
| 3.9235E-10 | 0.37595369 | 0.566 | 0.339 | 8.3768E-06 | CD-IC | GHITM |
| 1.4567E-12 | 0.37542203 | 0.462 | 0.221 | 3.1101E-08 | CD-IC | C19orf70 |
| 9.3987E-05 | 0.37536331 | 0.138 | 0.062 | 1 | CD-IC | COX7A1 |
| 5.2305E-13 | 0.37533562 | 0.359 | 0.149 | 1.1167E-08 | CD-IC | TIMM17A |
| 1.2717E-12 | 0.37488604 | 0.552 | 0.282 | 2.7152E-08 | CD-IC | ROMO1 |
| 1.9144E-06 | 0.37410582 | 0.434 | 0.27 | 0.04087323 | CD-IC | NDUFV2 |
| 1.2985E-08 | 0.37228974 | 0.393 | 0.215 | 0.00027723 | CD-IC | MED10 |
| 6.476E-117 | 0.37068724 | 0.159 | 0.005 | 1.383E-112 | CD-IC | SLC4A9 |
| 1.953E-134 | 0.36912766 | 0.152 | 0.004 | 4.17E-130 | CD-IC | SLC26A4 |
| 4.8068E-12 | 0.36686022 | 0.469 | 0.228 | 1.0263E-07 | CD-IC | NDUFA12 |
| 1.7838E-19 | 0.36607744 | 0.352 | 0.112 | 3.8084E-15 | CD-IC | KIF21A |
| 8.9812E-24 | 0.36597593 | 0.345 | 0.096 | 1.9175E-19 | CD-IC | ABHD3 |
| 2.8058E-13 | 0.36506348 | 0.51 | 0.251 | 5.9903E-09 | CD-IC | SH3GLB1 |
| 4.7768E-12 | 0.36429639 | 0.49 | 0.244 | 1.0199E-07 | CD-IC | LAMTOR4 |

| | | | | | | |
|---|---|---|---|---|---|---|
| 8.0432E-08 | 0.36422134 | 0.517 | 0.314 | 0.00171721 | CD-IC | PRDX2 |
| 6.889E-17 | 0.36080582 | 0.469 | 0.191 | 1.4708E-12 | CD-IC | RALA |
| 3.9227E-18 | 0.35945026 | 0.317 | 0.098 | 8.3749E-14 | CD-IC | MUC20 |
| 9.0045E-05 | 0.35912216 | 0.966 | 0.955 | 1 | CD-IC | RPS27 |
| 1.3724E-06 | 0.35793527 | 0.552 | 0.369 | 0.02930026 | CD-IC | CAMK2N1 |
| 2.8135E-09 | 0.35727091 | 0.503 | 0.283 | 6.0069E-05 | CD-IC | NDUFB11 |
| 3.8274E-08 | 0.35721932 | 0.366 | 0.194 | 0.00081715 | CD-IC | SLC25A39 |
| 6.6691E-08 | 0.35437587 | 0.407 | 0.223 | 0.00142386 | CD-IC | CLIP1 |
| 6.2074E-07 | 0.35417637 | 0.379 | 0.214 | 0.01325281 | CD-IC | ATP5MC1 |
| 0.0002257 | 0.35276586 | 0.159 | 0.078 | 1 | CD-IC | LY6K |
| 2.1536E-19 | 0.35220739 | 0.428 | 0.149 | 4.598E-15 | CD-IC | CA2 |
| 4.7988E-08 | 0.35124306 | 0.414 | 0.232 | 0.00102455 | CD-IC | LRRFIP2 |
| 3.9255E-12 | 0.35059572 | 0.29 | 0.111 | 8.3808E-08 | CD-IC | BPGM |
| 1.7159E-40 | 0.34966969 | 0.166 | 0.017 | 3.6634E-36 | CD-IC | RGCC |
| 1.1918E-07 | 0.34955155 | 0.262 | 0.122 | 0.00254454 | CD-IC | PRSS23 |
| 5.7459E-05 | 0.34461819 | 0.959 | 0.909 | 1 | CD-IC | MT-ATP6 |
| 2.841E-05 | 0.34448659 | 0.641 | 0.502 | 0.60654445 | CD-IC | CLDN7 |
| 1.7311E-10 | 0.34329046 | 0.338 | 0.151 | 3.6959E-06 | CD-IC | UCHL3 |
| 2.5631E-09 | 0.34287484 | 0.566 | 0.316 | 5.4721E-05 | CD-IC | NDUFB9 |
| 2.429E-10 | 0.34210527 | 0.462 | 0.244 | 5.186E-06 | CD-IC | HADHA |
| 5.7752E-07 | 0.33987703 | 0.448 | 0.266 | 0.01233002 | CD-IC | BLVRB |
| 5.8898E-20 | 0.33958555 | 0.241 | 0.06 | 1.2575E-15 | CD-IC | GOLT1A |
| 1.1019E-06 | 0.33775137 | 0.476 | 0.304 | 0.02352505 | CD-IC | WDR1 |
| 5.9863E-10 | 0.33635946 | 0.359 | 0.164 | 1.2781E-05 | CD-IC | MIR22HG |
| 2.2846E-13 | 0.3352044 | 0.372 | 0.15 | 4.8776E-09 | CD-IC | ITGA6 |
| 3.7943E-18 | 0.3350843 | 0.359 | 0.12 | 8.1008E-14 | CD-IC | SDC2 |
| 1.4875E-06 | 0.33467553 | 0.469 | 0.287 | 0.03175839 | CD-IC | NFE2L2 |
| 6.063E-20 | 0.33149409 | 0.221 | 0.05 | 1.2945E-15 | CD-IC | CLTRN |
| 2.7821E-05 | 0.33123103 | 0.338 | 0.206 | 0.59398746 | CD-IC | NDUFA3 |
| 5.7237E-10 | 0.33109056 | 0.503 | 0.271 | 1.222E-05 | CD-IC | COX17 |
| 3.7404E-20 | 0.33100897 | 0.345 | 0.106 | 7.9857E-16 | CD-IC | NAXE |
| 2.4999E-31 | 0.33074616 | 0.214 | 0.034 | 5.3372E-27 | CD-IC | CLCNKB |
| 1.618E-17 | 0.32954388 | 0.331 | 0.108 | 3.4545E-13 | CD-IC | ESRRA |
| 5.1863E-05 | 0.32928924 | 0.462 | 0.312 | 1 | CD-IC | UQCR11 |
| 7.0955E-05 | 0.32808889 | 0.945 | 0.91 | 1 | CD-IC | MT-CO2 |
| 9.1502E-12 | 0.32789891 | 0.372 | 0.165 | 1.9536E-07 | CD-IC | CISD1 |
| 1.1983E-09 | 0.32775327 | 0.441 | 0.234 | 2.5584E-05 | CD-IC | ANXA4 |
| 3.4535E-06 | 0.32711268 | 0.517 | 0.338 | 0.07373172 | CD-IC | KRT10 |
| 1.489E-11 | 0.32308004 | 0.503 | 0.254 | 3.1791E-07 | CD-IC | SMDT1 |
| 1.4438E-15 | 0.322994 | 0.359 | 0.131 | 3.0825E-11 | CD-IC | RRAGD |
| 0.00034274 | 0.32255263 | 0.669 | 0.573 | 1 | CD-IC | SLC3A2 |
| 4.9356E-10 | 0.31986717 | 0.366 | 0.171 | 1.0538E-05 | CD-IC | DSG2 |
| 9.997E-12 | 0.31974461 | 0.407 | 0.184 | 2.1344E-07 | CD-IC | MRPS5 |
| 4.73E-08 | 0.31929661 | 0.462 | 0.275 | 0.00100985 | CD-IC | SSB |
| 0.00020386 | 0.31922321 | 0.607 | 0.495 | 1 | CD-IC | HSPD1 |
| 9.3373E-14 | 0.31919137 | 0.234 | 0.073 | 1.9935E-09 | CD-IC | MAGI1 |
| 2.7764E-11 | 0.31813523 | 0.317 | 0.131 | 5.9277E-07 | CD-IC | RALGPS2 |
| 2.0361E-07 | 0.31688612 | 0.448 | 0.261 | 0.004347 | CD-IC | NDUFB10 |
| 4.132E-259 | 0.31608205 | 0.193 | 0.002 | 8.823E-255 | CD-IC | HEPACAM2 |
| 1.342E-06 | 0.31582267 | 0.483 | 0.295 | 0.02865119 | CD-IC | SELENOS |

| | | | | | | |
|---|---|---|---|---|---|---|
| 7.8615E-11 | 0.31347509 | 0.366 | 0.166 | 1.6784E-06 | CD-IC | PDHA1 |
| 1.2504E-08 | 0.31234474 | 0.483 | 0.273 | 0.00026696 | CD-IC | NDUFS6 |
| 1.3522E-06 | 0.31207427 | 0.421 | 0.249 | 0.02886973 | CD-IC | PTBP3 |
| 2.0609E-13 | 0.31039842 | 0.407 | 0.17 | 4.4E-09 | CD-IC | SHTN1 |
| 1.6557E-29 | 0.31029278 | 0.159 | 0.02 | 3.535E-25 | CD-IC | PVALB |
| 2.2705E-08 | 0.30880671 | 0.241 | 0.103 | 0.00048475 | CD-IC | OGFRL1 |
| 4.7639E-08 | 0.30853185 | 0.517 | 0.307 | 0.0010171 | CD-IC | NDUFC2 |
| 6.64E-22 | 0.30832746 | 0.255 | 0.061 | 1.4176E-17 | CD-IC | SBF2-AS1 |
| 1.0941E-15 | 0.30642058 | 0.372 | 0.139 | 2.336E-11 | CD-IC | MRPS35 |
| 0.00047977 | 0.30629311 | 0.938 | 0.898 | 1 | CD-IC | MT-CYB |
| 1.95E-23 | 0.3060647 | 0.186 | 0.033 | 4.1633E-19 | CD-IC | FYB2 |
| 8.137E-07 | 0.30598936 | 0.441 | 0.267 | 0.01737257 | CD-IC | SNRPG |
| 3.2543E-09 | 0.30581735 | 0.366 | 0.18 | 6.9479E-05 | CD-IC | ALKBH7 |
| 4.8931E-88 | 0.3052453 | 0.221 | 0.013 | 1.0447E-83 | CD-IC | PDE1C |
| 9.0041E-05 | 0.30502893 | 0.559 | 0.395 | 1 | CD-IC | DBI |
| 3.5881E-10 | 0.30499986 | 0.366 | 0.17 | 7.6607E-06 | CD-IC | PLEKHJ1 |
| 1.4683E-35 | 0.30477671 | 0.234 | 0.036 | 3.1348E-31 | CD-IC | MINDY1 |
| 1.4824E-13 | 0.30453034 | 0.2 | 0.057 | 3.1649E-09 | CD-IC | NEXN |
| 1.2918E-05 | 0.30410581 | 0.145 | 0.059 | 0.27580126 | CD-IC | CD83 |
| 4.9274E-12 | 0.30212537 | 0.31 | 0.122 | 1.052E-07 | CD-IC | PRKAG2 |
| 1.4076E-08 | 0.3019697 | 0.462 | 0.256 | 0.00030051 | CD-IC | OPTN |
| 6.9551E-12 | 0.30152625 | 0.303 | 0.119 | 1.4849E-07 | CD-IC | HLA-F |
| 3.2307E-24 | 0.30066215 | 0.234 | 0.049 | 6.8976E-20 | CD-IC | OPN3 |
| 3.626E-15 | 0.30047948 | 0.262 | 0.082 | 7.7416E-11 | CD-IC | NDUFAF4 |
| 1.0068E-05 | 0.30025696 | 0.469 | 0.309 | 0.21496187 | CD-IC | ATP5PD |
| 0.00655672 | 0.29938705 | 0.124 | 0.07 | 1 | CD-IC | MALL |
| 9.3572E-23 | 0.29831179 | 0.145 | 0.022 | 1.9978E-18 | CD-IC | PCP4 |
| 3.9632E-08 | 0.29807334 | 0.545 | 0.332 | 0.00084615 | CD-IC | SPCS2 |
| 2.5998E-08 | 0.29691378 | 0.366 | 0.19 | 0.00055505 | CD-IC | FUNDC2 |
| 2.8149E-05 | 0.29625438 | 0.221 | 0.113 | 0.60097413 | CD-IC | CNKSR3 |
| 3.6634E-70 | 0.29620746 | 0.11 | 0.004 | 7.8214E-66 | CD-IC | CFTR |
| 1.7067E-08 | 0.29555277 | 0.29 | 0.134 | 0.00036438 | CD-IC | COMTD1 |
| 8.1472E-08 | 0.29472338 | 0.531 | 0.319 | 0.00173943 | CD-IC | ATP5MPL |
| 1.7291E-08 | 0.29380229 | 0.393 | 0.205 | 0.00036917 | CD-IC | SINHCAF |
| 1.3963E-07 | 0.29346719 | 0.241 | 0.109 | 0.00298108 | CD-IC | NUDT9 |
| 4.3773E-07 | 0.29325212 | 0.414 | 0.237 | 0.00934546 | CD-IC | COX20 |
| 4.9049E-06 | 0.29320782 | 0.69 | 0.54 | 0.1047191 | CD-IC | GNG5 |
| 0.0002492 | 0.29316876 | 0.29 | 0.174 | 1 | CD-IC | PHLDA1 |
| 2.2615E-09 | 0.29295191 | 0.331 | 0.154 | 4.8283E-05 | CD-IC | COPRS |
| 3.9126E-06 | 0.2929351 | 0.883 | 0.791 | 0.08353367 | CD-IC | PPDPF |
| 6.6238E-07 | 0.29158064 | 0.524 | 0.324 | 0.01414182 | CD-IC | EIF1B |
| 5.9246E-09 | 0.29118892 | 0.503 | 0.281 | 0.00012649 | CD-IC | FIS1 |
| 1.5938E-05 | 0.29102451 | 0.428 | 0.263 | 0.34028269 | CD-IC | LIMA1 |
| 1.3616E-13 | 0.29083427 | 0.234 | 0.074 | 2.907E-09 | CD-IC | AIFM2 |
| 4.6497E-12 | 0.29040186 | 0.386 | 0.166 | 9.9271E-08 | CD-IC | METRNL |
| 4.4258E-08 | 0.29009046 | 0.324 | 0.159 | 0.00094491 | CD-IC | FBXW5 |
| 1.1618E-09 | 0.28943496 | 0.283 | 0.119 | 2.4805E-05 | CD-IC | SH2D4A |
| 4.6304E-09 | 0.28903842 | 0.31 | 0.141 | 9.8858E-05 | CD-IC | KIF13B |
| 0.00053711 | 0.28792423 | 0.51 | 0.376 | 1 | CD-IC | SRSF5 |
| 8.4293E-10 | 0.28760974 | 0.366 | 0.175 | 1.7997E-05 | CD-IC | SDHD |

| | | | | | | |
|---|---|---|---|---|---|---|
| 2.796E-08 | 0.28573049 | 0.428 | 0.232 | 0.00059694 | CD-IC | NDUFS8 |
| 0.00016612 | 0.28537552 | 0.945 | 0.91 | 1 | CD-IC | MT-CO3 |
| 5.1654E-33 | 0.28524086 | 0.248 | 0.042 | 1.1028E-28 | CD-IC | MR1 |
| 4.3061E-08 | 0.28497439 | 0.379 | 0.197 | 0.00091936 | CD-IC | PHLDB2 |
| 6.1914E-12 | 0.28400067 | 0.345 | 0.144 | 1.3219E-07 | CD-IC | TRAPPC2L |
| 0.00069387 | 0.28350489 | 0.648 | 0.532 | 1 | CD-IC | SEC61B |
| 0.00039895 | 0.28336294 | 0.497 | 0.357 | 1 | CD-IC | KLF6 |
| 3.4347E-05 | 0.28304523 | 0.421 | 0.271 | 0.73330278 | CD-IC | ATP5F1C |
| 1.2174E-09 | 0.2817318 | 0.276 | 0.115 | 2.5992E-05 | CD-IC | CFL2 |
| 4.3989E-18 | 0.28120441 | 0.255 | 0.069 | 9.3917E-14 | CD-IC | CARMIL1 |
| 2.308E-171 | 0.28078791 | 0.172 | 0.003 | 4.928E-167 | CD-IC | DGKI |
| 8.3526E-12 | 0.28063605 | 0.4 | 0.178 | 1.7833E-07 | CD-IC | NDUFS4 |
| 1.4395E-15 | 0.28013039 | 0.324 | 0.11 | 3.0733E-11 | CD-IC | CREM |
| 2.2845E-19 | 0.28006463 | 0.207 | 0.047 | 4.8773E-15 | CD-IC | SMIM6 |
| 4.404E-06 | 0.27927327 | 0.807 | 0.699 | 0.09402465 | CD-IC | MAP1LC3B |
| 5.1802E-06 | 0.27921079 | 0.421 | 0.246 | 0.11059622 | CD-IC | RAP2B |
| 5.9546E-12 | 0.27915839 | 0.303 | 0.119 | 1.2713E-07 | CD-IC | EGLN1 |
| 8.4542E-06 | 0.27887811 | 0.91 | 0.839 | 0.18049669 | CD-IC | CD63 |
| 8.6193E-08 | 0.27784199 | 0.455 | 0.258 | 0.00184023 | CD-IC | LAMTOR1 |
| 5.4695E-07 | 0.27743935 | 0.497 | 0.301 | 0.01167738 | CD-IC | GADD45GIP1 |
| 4.989E-07 | 0.27563856 | 0.476 | 0.298 | 0.01065153 | CD-IC | NDFIP1 |
| 1.2008E-06 | 0.27272697 | 0.545 | 0.347 | 0.02563622 | CD-IC | COMMD6 |
| 4.5218E-08 | 0.27267981 | 0.386 | 0.203 | 0.0009654 | CD-IC | ADI1 |
| 4.7909E-06 | 0.27141274 | 0.434 | 0.267 | 0.10228577 | CD-IC | NDUFB8 |
| 2.6701E-21 | 0.27075972 | 0.138 | 0.021 | 5.7008E-17 | CD-IC | EEF1A2 |
| 7.1092E-06 | 0.27019301 | 0.283 | 0.155 | 0.15178224 | CD-IC | AKIRIN1 |
| 3.6631E-25 | 0.26980095 | 0.152 | 0.022 | 7.8207E-21 | CD-IC | TBXAS1 |
| 1.7157E-06 | 0.26943445 | 0.407 | 0.24 | 0.03663007 | CD-IC | GTF2A2 |
| 2.1651E-06 | 0.26916799 | 0.269 | 0.137 | 0.0462243 | CD-IC | SPIRE1 |
| 9.2873E-13 | 0.26910679 | 0.331 | 0.129 | 1.9828E-08 | CD-IC | PPP2R5E |
| 6.986E-12 | 0.26783072 | 0.324 | 0.13 | 1.4915E-07 | CD-IC | NIPSNAP2 |
| 3.5352E-05 | 0.26732285 | 0.145 | 0.063 | 0.75476587 | CD-IC | CTTNBP2 |
| 9.9059E-06 | 0.26664084 | 0.462 | 0.287 | 0.21149104 | CD-IC | FAM3C |
| 2.8622E-08 | 0.26591328 | 0.317 | 0.153 | 0.00061108 | CD-IC | SDHB |
| 2.6357E-05 | 0.26412076 | 0.421 | 0.265 | 0.56272092 | CD-IC | MINOS1 |
| 4.4577E-05 | 0.26402577 | 0.448 | 0.298 | 0.95172839 | CD-IC | ACTR3 |
| 2.6585E-14 | 0.2639216 | 0.248 | 0.077 | 5.6759E-10 | CD-IC | TBC1D4 |
| 4.025E-11 | 0.26381357 | 0.317 | 0.13 | 8.5933E-07 | CD-IC | SCRN1 |
| 1.5798E-07 | 0.26306639 | 0.345 | 0.18 | 0.00337291 | CD-IC | MAPRE1 |
| 2.241E-10 | 0.26233298 | 0.303 | 0.127 | 4.7845E-06 | CD-IC | CMC2 |
| 6.2384E-06 | 0.26185887 | 0.386 | 0.233 | 0.13319089 | CD-IC | CALM3 |
| 7.564E-08 | 0.26178079 | 0.628 | 0.391 | 0.00161492 | CD-IC | TAX1BP1 |
| 9.1476E-08 | 0.26176858 | 0.324 | 0.16 | 0.001953 | CD-IC | ETFB |
| 5.9595E-16 | 0.26112619 | 0.193 | 0.048 | 1.2724E-11 | CD-IC | CERS4 |
| 1.7683E-07 | 0.26089521 | 0.324 | 0.165 | 0.00377533 | CD-IC | UQCRC1 |
| 7.2206E-06 | 0.25985066 | 0.421 | 0.258 | 0.15416 | CD-IC | NDUFB3 |
| 0.00169557 | 0.25980698 | 0.959 | 0.947 | 1 | CD-IC | RPL13A |
| 4.733E-05 | 0.25951852 | 0.49 | 0.33 | 1 | CD-IC | ATP6V1F |
| 7.637E-08 | 0.25897654 | 0.29 | 0.138 | 0.00163051 | CD-IC | PITHD1 |
| 9.3496E-07 | 0.25860108 | 0.179 | 0.074 | 0.01996145 | CD-IC | TNFRSF21 |

| | | | | | | |
|---|---|---|---|---|---|---|
| 1.9805E-08 | 0.25808019 | 0.248 | 0.105 | 0.00042283 | CD-IC | FRK |
| 1.0611E-07 | 0.25799748 | 0.455 | 0.255 | 0.00226537 | CD-IC | ATP5F1A |
| 1.9018E-10 | 0.25786491 | 0.338 | 0.146 | 4.0604E-06 | CD-IC | SGMS2 |
| 2.0304E-09 | 0.25756427 | 0.386 | 0.188 | 4.335E-05 | CD-IC | FAM50A |
| 1.9964E-17 | 0.25657603 | 0.234 | 0.062 | 4.2624E-13 | CD-IC | PLEK2 |
| 2.0078E-10 | 0.2564246 | 0.71 | 0.468 | 4.2867E-06 | CD-IC | C12orf75 |
| 1.5138E-06 | 0.25640288 | 0.469 | 0.281 | 0.03231904 | CD-IC | PDCD5 |
| 6.8035E-70 | 0.25639381 | 0.159 | 0.009 | 1.4525E-65 | CD-IC | AVPR1A |
| 2.5518E-16 | 0.2562833 | 0.193 | 0.047 | 5.4481E-12 | CD-IC | HOXB-AS3 |
| 2.6083E-05 | 0.25621393 | 0.717 | 0.578 | 0.55686381 | CD-IC | PRDX5 |
| 0.00011683 | 0.25613455 | 0.331 | 0.205 | 1 | CD-IC | DNTTIP2 |
| 1.1288E-21 | 0.25605053 | 0.186 | 0.036 | 2.4099E-17 | CD-IC | COLCA1 |
| 1.11E-17 | 0.25533269 | 0.186 | 0.042 | 2.3698E-13 | CD-IC | MPP7 |
| 8.687E-06 | 0.25512904 | 0.345 | 0.204 | 0.18546796 | CD-IC | ZFAND6 |
| 1.7407E-11 | 0.25494791 | 0.324 | 0.133 | 3.7164E-07 | CD-IC | ACTR10 |
| 3.975E-11 | 0.25433095 | 0.366 | 0.157 | 8.4866E-07 | CD-IC | KIAA1217 |
| 1.2488E-07 | 0.25335149 | 0.324 | 0.164 | 0.00266614 | CD-IC | HTATIP2 |
| 1.353E-164 | 0.25324405 | 0.103 | 0.001 | 2.888E-160 | CD-IC | GLDN |
| 8.6038E-14 | 0.25302395 | 0.228 | 0.069 | 1.8369E-09 | CD-IC | PIGH |
| 1.2374E-07 | 0.2528915 | 0.338 | 0.174 | 0.0026419 | CD-IC | UQCRC2 |
| 0.00016713 | 0.25229851 | 0.648 | 0.532 | 1 | CD-IC | NDUFS5 |
| 1.4834E-06 | 0.25227984 | 0.276 | 0.137 | 0.03167134 | CD-IC | GCLC |
| 2.7616E-09 | 0.25217682 | 0.441 | 0.225 | 5.896E-05 | CD-IC | SEC11C |
| 5.1471E-11 | 0.25198768 | 0.262 | 0.098 | 1.0989E-06 | CD-IC | RELB |
| 0.00011321 | 0.25173598 | 0.483 | 0.327 | 1 | CD-IC | UQCRQ |
| 4.6502E-09 | 0.25155799 | 0.428 | 0.22 | 9.9283E-05 | CD-IC | RWDD1 |
| 7.8557E-07 | 0.25148514 | 0.324 | 0.171 | 0.01677197 | CD-IC | COA3 |
| 0.00137869 | 0.25098428 | 0.862 | 0.804 | 1 | CD-IC | RPL31 |
| 2.2292E-11 | 0.25084331 | 0.221 | 0.073 | 4.7592E-07 | CD-IC | GAS6 |
| 2.4434E-09 | 0.25076182 | 0.31 | 0.138 | 5.2166E-05 | CD-IC | GLUD1 |
| 7.7504E-35 | 0.25056292 | 0.138 | 0.014 | 1.6547E-30 | CD-IC | PACRG |
| 0.00077313 | -0.2501598 | 0.138 | 0.255 | 1 | CD-IC | LAPTM4B |
| 0.00024351 | -0.2504082 | 0.055 | 0.168 | 1 | CD-IC | DAB2 |
| 1.3675E-05 | -0.2521158 | 0.007 | 0.127 | 0.29196345 | CD-IC | SPNS2 |
| 9.1605E-05 | -0.2528324 | 0.055 | 0.178 | 1 | CD-IC | ACTN1 |
| 0.00078953 | -0.2540415 | 0.062 | 0.163 | 1 | CD-IC | PCSK1N |
| 5.0559E-05 | -0.2544545 | 0.034 | 0.158 | 1 | CD-IC | RAPGEF3 |
| 8.0849E-05 | -0.2547303 | 0.076 | 0.208 | 1 | CD-IC | ATP13A3 |
| 0.00038597 | -0.2547419 | 0.083 | 0.195 | 1 | CD-IC | FAM129B |
| 0.00121047 | -0.2578686 | 0.09 | 0.192 | 1 | CD-IC | ZNF83 |
| 0.00104367 | -0.2585052 | 0.034 | 0.123 | 1 | CD-IC | MT1F |
| 8.0409E-05 | -0.2605324 | 0.076 | 0.206 | 1 | CD-IC | AIF1L |
| 3.2347E-05 | -0.2609782 | 0.034 | 0.161 | 0.69061201 | CD-IC | ETS1 |
| 2.341E-05 | -0.2624217 | 0.034 | 0.166 | 0.49979409 | CD-IC | CAVIN1 |
| 3.1012E-06 | -0.262905 | 0.386 | 0.602 | 0.06621021 | CD-IC | SPP1 |
| 0.00290478 | -0.2655834 | 0.572 | 0.674 | 1 | CD-IC | HNRNPA1 |
| 2.7504E-05 | -0.2661723 | 0.014 | 0.132 | 0.58720823 | CD-IC | AL353751.1 |
| 0.00127813 | -0.2681958 | 0.055 | 0.153 | 1 | CD-IC | PFKFB3 |
| 0.00011205 | -0.272252 | 0.083 | 0.213 | 1 | CD-IC | PPFIBP1 |
| 9.8682E-06 | -0.2731922 | 0.021 | 0.153 | 0.2106869 | CD-IC | SLC7A5 |

| | | | | | | |
|---|---|---|---|---|---|---|
| 0.00010735 | -0.2733395 | 0.076 | 0.202 | 1 | CD-IC | FOXO3 |
| 0.00011357 | -0.2750735 | 0.076 | 0.201 | 1 | CD-IC | AGRN |
| 0.00031885 | -0.2751045 | 0.407 | 0.558 | 1 | CD-IC | LRRFIP1 |
| 4.6936E-05 | -0.2753648 | 0.034 | 0.158 | 1 | CD-IC | QPRT |
| 0.00295938 | -0.2810915 | 0.172 | 0.269 | 1 | CD-IC | MET |
| 0.00127205 | -0.2826692 | 0.124 | 0.232 | 1 | CD-IC | RPL22L1 |
| 7.9627E-06 | -0.2870029 | 0.007 | 0.133 | 0.1700043 | CD-IC | COL4A1 |
| 0.00498961 | -0.2880985 | 0.069 | 0.148 | 1 | CD-IC | AL118516.1 |
| 0.00072218 | -0.2910622 | 0.2 | 0.323 | 1 | CD-IC | RRBP1 |
| 5.6692E-06 | -0.2916321 | 0.028 | 0.167 | 0.12103636 | CD-IC | NTN4 |
| 0.00059043 | -0.2922384 | 0.041 | 0.139 | 1 | CD-IC | TNFSF10 |
| 0.00030228 | -0.2926204 | 0.048 | 0.154 | 1 | CD-IC | TSC22D3 |
| 0.00160294 | -0.2937049 | 0.145 | 0.25 | 1 | CD-IC | MGLL |
| 0.00493159 | -0.2944822 | 0.076 | 0.162 | 1 | CD-IC | G0S2 |
| 1.3354E-06 | -0.295254 | 0 | 0.14 | 0.02851102 | CD-IC | TMEM176A |
| 0.00233922 | -0.2952875 | 0.4 | 0.501 | 1 | CD-IC | HNRNPDL |
| 6.712E-07 | -0.2995958 | 0 | 0.147 | 0.01433017 | CD-IC | COL4A2 |
| 3.3782E-05 | -0.3028705 | 0.028 | 0.149 | 0.72125156 | CD-IC | ARL4C |
| 0.00075466 | -0.3107459 | 0.034 | 0.127 | 1 | CD-IC | PCDH9 |
| 0.00224879 | -0.3108694 | 0.069 | 0.16 | 1 | CD-IC | MGST1 |
| 0.00064276 | -0.3117638 | 0.034 | 0.131 | 1 | CD-IC | TNFRSF11B |
| 0.00706307 | -0.3119421 | 0.034 | 0.1 | 1 | CD-IC | AP001160.1 |
| 1.2919E-06 | -0.3150066 | 0.007 | 0.151 | 0.02758239 | CD-IC | FXYD5 |
| 0.00702086 | -0.3152702 | 0.634 | 0.701 | 1 | CD-IC | HMGB1 |
| 3.1459E-06 | -0.3168933 | 0.014 | 0.153 | 0.06716444 | CD-IC | KITLG |
| 0.00033679 | -0.3202235 | 0.097 | 0.213 | 1 | CD-IC | PLEC |
| 0.0009255 | -0.3221021 | 0.021 | 0.105 | 1 | CD-IC | SERPINE2 |
| 3.0955E-07 | -0.322836 | 0 | 0.154 | 0.00660893 | CD-IC | PROM1 |
| 0.00029637 | -0.3267261 | 0.069 | 0.181 | 1 | CD-IC | DCBLD2 |
| 0.00065344 | -0.3269102 | 0.241 | 0.375 | 1 | CD-IC | ODC1 |
| 0.00166182 | -0.329719 | 0.517 | 0.61 | 1 | CD-IC | EIF4A2 |
| 8.4594E-05 | -0.3309791 | 0.124 | 0.265 | 1 | CD-IC | PLXNB2 |
| 0.00146775 | -0.3310946 | 0.055 | 0.148 | 1 | CD-IC | MYL9 |
| 0.00207508 | -0.3313844 | 0.062 | 0.151 | 1 | CD-IC | AC007952.4 |
| 0.00086134 | -0.3373394 | 0.117 | 0.234 | 1 | CD-IC | SYNE1 |
| 1.37E-07 | -0.3389022 | 0 | 0.162 | 0.00292504 | CD-IC | TMEM176B |
| 0.00015059 | -0.3411623 | 0.317 | 0.483 | 1 | CD-IC | H1FX |
| 5.3989E-05 | -0.3428676 | 0.117 | 0.255 | 1 | CD-IC | PNPLA2 |
| 0.0001501 | -0.3441411 | 0.083 | 0.21 | 1 | CD-IC | ARID5B |
| 0.00433601 | -0.3443584 | 0.034 | 0.109 | 1 | CD-IC | S100A9 |
| 0.00060359 | -0.3445967 | 0.034 | 0.127 | 1 | CD-IC | HIST2H2BF |
| 0.00011968 | -0.3521227 | 0.4 | 0.566 | 1 | CD-IC | CST3 |
| 0.00016144 | -0.3537643 | 0.531 | 0.632 | 1 | CD-IC | PABPC1 |
| 6.2156E-07 | -0.3585839 | 0.028 | 0.191 | 0.01327039 | CD-IC | EMP1 |
| 0.0001057 | -0.3598733 | 0.062 | 0.183 | 1 | CD-IC | ERN1 |
| 0.0053403 | -0.3686279 | 0.172 | 0.263 | 1 | CD-IC | HIST1H2AC |
| 0.00929122 | -0.3699654 | 0.51 | 0.569 | 1 | CD-IC | RPS27L |
| 0.00033109 | -0.3781681 | 0.297 | 0.424 | 1 | CD-IC | SNHG8 |
| 0.00178607 | -0.3876267 | 0.186 | 0.289 | 1 | CD-IC | STMN1 |
| 0.00616441 | -0.3912554 | 0.214 | 0.331 | 1 | CD-IC | TIMP1 |

| | | | | | | |
|---|---|---|---|---|---|---|
| 5.038E-06 | -0.3925351 | 0.028 | 0.169 | 0.10756159 | CD-IC | FGF2 |
| 0.00112575 | -0.398829 | 0.028 | 0.113 | 1 | CD-IC | C15orf48 |
| 5.4375E-05 | -0.4001268 | 0.179 | 0.34 | 1 | CD-IC | PAX8 |
| 1.8841E-07 | -0.4028645 | 0.028 | 0.201 | 0.00402246 | CD-IC | FAT1 |
| 7.6658E-05 | -0.402931 | 0.566 | 0.668 | 1 | CD-IC | SPINT2 |
| 3.8425E-08 | -0.4048608 | 0.021 | 0.205 | 0.00082037 | CD-IC | DPP4 |
| 0.00239631 | -0.4081327 | 0.628 | 0.703 | 1 | CD-IC | CLDN4 |
| 6.8652E-07 | -0.4190186 | 0.007 | 0.157 | 0.01465712 | CD-IC | IGFBP2 |
| 2.548E-06 | -0.4234982 | 0.048 | 0.203 | 0.05440055 | CD-IC | CALD1 |
| 0.00139461 | -0.4258292 | 0.103 | 0.204 | 1 | CD-IC | SGK1 |
| 1.5239E-06 | -0.4286979 | 0.034 | 0.19 | 0.03253522 | CD-IC | SYTL2 |
| 0.00010963 | -0.4293425 | 0.028 | 0.139 | 1 | CD-IC | TM4SF1 |
| 1.599E-06 | -0.4310152 | 0.683 | 0.804 | 0.03413909 | CD-IC | CALM1 |
| 0.00449234 | -0.4326096 | 0.172 | 0.268 | 1 | CD-IC | TUBA1A |
| 0.00021606 | -0.4373478 | 0.159 | 0.282 | 1 | CD-IC | CLDN3 |
| 7.8651E-07 | -0.4373666 | 0.021 | 0.177 | 0.01679188 | CD-IC | FOXC1 |
| 8.221E-08 | -0.4387378 | 0.462 | 0.659 | 0.00175518 | CD-IC | PSAP |
| 3.1723E-06 | -0.4391934 | 0.103 | 0.268 | 0.06772805 | CD-IC | GSTM3 |
| 3.2124E-07 | -0.4454366 | 0.097 | 0.284 | 0.00685842 | CD-IC | ASPH |
| 0.00013968 | -0.4470247 | 0.262 | 0.394 | 1 | CD-IC | TAF1D |
| 0.00030722 | -0.4487902 | 0.117 | 0.235 | 1 | CD-IC | NFKBIZ |
| 1.5871E-05 | -0.4552242 | 0.345 | 0.497 | 0.33885005 | CD-IC | ACTN4 |
| 0.00115647 | -0.4609491 | 0.11 | 0.215 | 1 | CD-IC | SOCS1 |
| 0.00725006 | -0.4646567 | 0.062 | 0.136 | 1 | CD-IC | SERPINA1 |
| 0.00032415 | -0.4678813 | 0.028 | 0.124 | 1 | CD-IC | CXCL14 |
| 0.00010551 | -0.4741063 | 0.262 | 0.392 | 1 | CD-IC | NDRG1 |
| 9.462E-11 | -0.4757581 | 0.021 | 0.257 | 2.0201E-06 | CD-IC | COTL1 |
| 0.00059841 | -0.4767543 | 0.159 | 0.277 | 1 | CD-IC | CITED2 |
| 0.0001777 | -0.4771971 | 0.055 | 0.17 | 1 | CD-IC | EGLN3 |
| 1.6036E-07 | -0.4878341 | 0.428 | 0.617 | 0.00342365 | CD-IC | CD81 |
| 1.7635E-08 | -0.497066 | 0.083 | 0.291 | 0.00037651 | CD-IC | ITGA3 |
| 2.9198E-06 | -0.4988705 | 0.062 | 0.223 | 0.06233801 | CD-IC | SFN |
| 2.4124E-06 | -0.5005805 | 0.034 | 0.188 | 0.0515042 | CD-IC | ADIRF |
| 1.8487E-06 | -0.5032148 | 0.076 | 0.241 | 0.0394696 | CD-IC | ITGB6 |
| 9.0042E-07 | -0.511487 | 0.193 | 0.375 | 0.01922398 | CD-IC | FLNA |
| 3.6783E-06 | -0.5164565 | 0 | 0.13 | 0.07853157 | CD-IC | COL7A1 |
| 9.124E-07 | -0.5206912 | 0.172 | 0.361 | 0.01947981 | CD-IC | CD44 |
| 0.00029529 | -0.5282308 | 0.097 | 0.214 | 1 | CD-IC | GLRX |
| 0.00012878 | -0.5298052 | 0.097 | 0.22 | 1 | CD-IC | TNFAIP2 |
| 2.1747E-06 | -0.5309962 | 0.11 | 0.288 | 0.04643079 | CD-IC | PLAUR |
| 3.2504E-05 | -0.5313116 | 0.138 | 0.29 | 0.69395021 | CD-IC | CYR61 |
| 0.00012235 | -0.5378103 | 0.014 | 0.116 | 1 | CD-IC | TAGLN |
| 6.8617E-05 | -0.5450004 | 0.069 | 0.201 | 1 | CD-IC | ANXA1 |
| 1.2511E-06 | -0.5516735 | 0.09 | 0.261 | 0.02671086 | CD-IC | LAMB3 |
| 4.4964E-09 | -0.5646443 | 0.11 | 0.347 | 9.5998E-05 | CD-IC | S100A16 |
| 0.00060692 | -0.5655685 | 0.028 | 0.118 | 1 | CD-IC | NEFM |
| 9.1211E-05 | -0.5659272 | 0.207 | 0.343 | 1 | CD-IC | SNHG12 |
| 0.00099767 | -0.5725565 | 0.166 | 0.279 | 1 | CD-IC | HES1 |
| 0.00021836 | -0.5847332 | 0.393 | 0.512 | 1 | CD-IC | CDKN1A |
| 7.0278E-10 | -0.5854165 | 0 | 0.211 | 1.5004E-05 | CD-IC | ADAMTS9 |

| | | | | | | |
|---|---|---|---|---|---|---|
| 0.00015322 | -0.5888438 | 0.166 | 0.297 | 1 | CD-IC | ITGB8 |
| 2.9816E-05 | -0.58896 | 0.29 | 0.446 | 0.6365642 | CD-IC | ZFP36L1 |
| 1.4635E-07 | -0.5946837 | 0.09 | 0.285 | 0.00312462 | CD-IC | UCHL1 |
| 9.3488E-08 | -0.6037468 | 0.007 | 0.176 | 0.00199597 | CD-IC | IGFBP3 |
| 5.954E-05 | -0.6078872 | 0.159 | 0.303 | 1 | CD-IC | PLK2 |
| 4.8631E-05 | -0.6288944 | 0.166 | 0.321 | 1 | CD-IC | ITGA2 |
| 0.00198337 | -0.6309289 | 0.138 | 0.235 | 1 | CD-IC | CSKMT |
| 2.792E-05 | -0.6335414 | 0.117 | 0.256 | 0.59609737 | CD-IC | IER5L |
| 3.9495E-10 | -0.636302 | 0.014 | 0.235 | 8.4321E-06 | CD-IC | TGM2 |
| 0.00031106 | -0.6442056 | 0.124 | 0.248 | 1 | CD-IC | MT1X |
| 1.3601E-06 | -0.6449178 | 0.083 | 0.255 | 0.02903803 | CD-IC | AKAP12 |
| 5.4216E-06 | -0.651125 | 0.793 | 0.878 | 0.11575155 | CD-IC | ITM2B |
| 6.4514E-11 | -0.6532407 | 0.138 | 0.388 | 1.3774E-06 | CD-IC | OCIAD2 |
| 0.00010227 | -0.6553269 | 0.145 | 0.278 | 1 | CD-IC | PPP1R10 |
| 3.4331E-07 | -0.668437 | 0.49 | 0.664 | 0.00732972 | CD-IC | TACSTD2 |
| 4.8369E-08 | -0.6817201 | 0.09 | 0.287 | 0.00103269 | CD-IC | AKR1B1 |
| 2.6966E-07 | -0.6853012 | 0.117 | 0.303 | 0.00575714 | CD-IC | DDIT4 |
| 0.00072698 | -0.6882381 | 0.228 | 0.333 | 1 | CD-IC | RHOB |
| 5.2289E-05 | -0.7048861 | 0.379 | 0.541 | 1 | CD-IC | SOX4 |
| 1.5234E-12 | -0.7147915 | 0.034 | 0.307 | 3.2524E-08 | CD-IC | ARHGAP29 |
| 1.3349E-11 | -0.7174001 | 0.29 | 0.561 | 2.8501E-07 | CD-IC | HMGA1 |
| 1.2543E-10 | -0.7226681 | 0.897 | 0.935 | 2.6778E-06 | CD-IC | PTMA |
| 1.2202E-11 | -0.724321 | 0.566 | 0.767 | 2.6052E-07 | CD-IC | PKM |
| 2.0161E-07 | -0.732052 | 0.869 | 0.924 | 0.00430437 | CD-IC | TMSB10 |
| 4.667E-08 | -0.7563241 | 0.09 | 0.291 | 0.0009964 | CD-IC | RND3 |
| 8.5593E-11 | -0.7627932 | 0.014 | 0.248 | 1.8274E-06 | CD-IC | PDZK1IP1 |
| 7.1443E-07 | -0.7672691 | 0.31 | 0.487 | 0.01525309 | CD-IC | TSPAN1 |
| 1.2155E-13 | -0.7703971 | 0.034 | 0.323 | 2.5951E-09 | CD-IC | DCDC2 |
| 1.3125E-05 | -0.8158102 | 0.545 | 0.673 | 0.28022121 | CD-IC | SH3BGRL3 |
| 6.0686E-10 | -0.8213584 | 0.621 | 0.833 | 1.2956E-05 | CD-IC | ANXA2 |
| 0.00194312 | -0.8314509 | 0.145 | 0.257 | 1 | CD-IC | LCN2 |
| 2.9584E-08 | -0.8442176 | 0.041 | 0.238 | 0.00063162 | CD-IC | CST6 |
| 3.4573E-12 | -0.8638471 | 0.145 | 0.424 | 7.3814E-08 | CD-IC | TPM4 |
| 7.6349E-09 | -0.8694021 | 0.607 | 0.817 | 0.00016301 | CD-IC | IGFBP7 |
| 2.5702E-15 | -0.8759247 | 0.414 | 0.707 | 5.4873E-11 | CD-IC | ITGB1 |
| 1.7397E-11 | -0.8863579 | 0.152 | 0.42 | 3.7143E-07 | CD-IC | CCND1 |
| 2.0045E-15 | -0.8915365 | 0.379 | 0.671 | 4.2796E-11 | CD-IC | GLS |
| 9.6261E-08 | -0.8919377 | 0.503 | 0.632 | 0.00205517 | CD-IC | TPM1 |
| 1.1691E-17 | -0.8989963 | 0.234 | 0.569 | 2.496E-13 | CD-IC | MAL2 |
| 5.7257E-10 | -0.9235202 | 0.807 | 0.872 | 1.2224E-05 | CD-IC | SAT1 |
| 4.8184E-07 | -0.9383933 | 0.09 | 0.272 | 0.01028723 | CD-IC | HSPA2 |
| 2.2752E-11 | -0.9429626 | 0.048 | 0.31 | 4.8575E-07 | CD-IC | PLAU |
| 2.5607E-06 | -1.0091596 | 0.8 | 0.815 | 0.05467114 | CD-IC | NEAT1 |
| 1.7568E-10 | -1.0546123 | 0.014 | 0.245 | 3.7508E-06 | CD-IC | TFPI2 |
| 2.1657E-05 | -1.0798362 | 0.441 | 0.639 | 0.46236681 | CD-IC | MMP7 |
| 1.4601E-11 | -1.2199077 | 0.076 | 0.333 | 3.1172E-07 | CD-IC | LAMC2 |
| 5.2358E-05 | -1.2690233 | 0.069 | 0.199 | 1 | CD-IC | MT1G |
| 1.2099E-19 | -1.4525217 | 0.29 | 0.671 | 2.5831E-15 | CD-IC | CRYAB |
| 2.5799E-24 | -1.6349297 | 0.172 | 0.596 | 5.5081E-20 | CD-IC | KRT19 |
| 1.1218E-26 | -1.6854206 | 0.193 | 0.628 | 2.3951E-22 | CD-IC | VIM |

| | | | | | | |
|---|---|---|---|---|---|---|
| 6.6777E-15 | -1.8524784 | 0.083 | 0.395 | 1.4257E-10 | CD-IC | SLPI |
| 0 | 3.09353241 | 0.719 | 0.039 | 0 | CD-PC | AQP2 |
| 0 | 3.08630503 | 0.843 | 0.091 | 0 | CD-PC | FXYD4 |
| 1.905E-104 | 2.36550025 | 0.767 | 0.251 | 4.067E-100 | CD-PC | NUPR1 |
| 8.028E-108 | 2.29538752 | 0.771 | 0.245 | 1.714E-103 | CD-PC | LCN2 |
| 1.76E-192 | 2.15611476 | 0.562 | 0.07 | 3.757E-188 | CD-PC | PDK4 |
| 1.343E-168 | 1.95951327 | 0.622 | 0.096 | 2.868E-164 | CD-PC | AQP3 |
| 9.669E-116 | 1.95511436 | 0.843 | 0.323 | 2.064E-111 | CD-PC | RALBP1 |
| 3.3045E-63 | 1.93079452 | 0.847 | 0.56 | 7.055E-59 | CD-PC | GDF15 |
| 1.736E-113 | 1.83453342 | 0.639 | 0.159 | 3.706E-109 | CD-PC | FAM102A |
| 2.1002E-73 | 1.68111017 | 0.655 | 0.217 | 4.4839E-69 | CD-PC | DEFB1 |
| 1.9391E-66 | 1.66153922 | 0.759 | 0.351 | 4.1399E-62 | CD-PC | KLF6 |
| 3.982E-133 | 1.6043023 | 0.663 | 0.142 | 8.501E-129 | CD-PC | PFKFB3 |
| 1.3088E-62 | 1.49579988 | 0.912 | 0.594 | 2.7942E-58 | CD-PC | CLU |
| 7.936E-185 | 1.48593194 | 0.345 | 0.025 | 1.694E-180 | CD-PC | RARRES1 |
| 5.948E-240 | 1.43159837 | 0.554 | 0.054 | 1.27E-235 | CD-PC | PPARGC1A |
| 2.1195E-47 | 1.40955131 | 0.767 | 0.437 | 4.5251E-43 | CD-PC | ZFP36L1 |
| 5.104E-214 | 1.35196966 | 0.518 | 0.053 | 1.09E-209 | CD-PC | PPP1R3B |
| 2.5077E-84 | 1.3174935 | 0.474 | 0.108 | 5.3539E-80 | CD-PC | OTUD1 |
| 1.3197E-35 | 1.29767742 | 0.57 | 0.27 | 2.8176E-31 | CD-PC | CITED2 |
| 1.9577E-86 | 1.29400033 | 0.759 | 0.264 | 4.1797E-82 | CD-PC | MUC1 |
| 1.881E-92 | 1.25437156 | 0.586 | 0.157 | 4.0159E-88 | CD-PC | SPTSSA |
| 1.7763E-66 | 1.22380042 | 0.695 | 0.268 | 3.7923E-62 | CD-PC | CYB5A |
| 1.9097E-91 | 1.21230185 | 0.775 | 0.268 | 4.0773E-87 | CD-PC | COBLL1 |
| 8.0491E-79 | 1.09333594 | 0.45 | 0.099 | 1.7185E-74 | CD-PC | PIM1 |
| 0 | 1.06120674 | 0.478 | 0.025 | 0 | CD-PC | AVPR2 |
| 1.7072E-44 | 1.03777819 | 0.723 | 0.395 | 3.6449E-40 | CD-PC | CDV3 |
| 7.164E-219 | 1.03535965 | 0.494 | 0.046 | 1.53E-214 | CD-PC | FAM83F |
| 1.009E-116 | 1.01978314 | 0.791 | 0.222 | 2.155E-112 | CD-PC | ASAH1 |
| 6.297E-103 | 1.00869727 | 0.631 | 0.154 | 1.3445E-98 | CD-PC | AFF1 |
| 1.6056E-73 | 0.99113132 | 0.618 | 0.192 | 3.4279E-69 | CD-PC | MBP |
| 2.7763E-46 | 0.98999814 | 0.779 | 0.384 | 5.9274E-42 | CD-PC | SLPI |
| 2.852E-102 | 0.97840779 | 0.578 | 0.124 | 6.0884E-98 | CD-PC | RDH10 |
| 1.1472E-88 | 0.97513721 | 0.598 | 0.153 | 2.4493E-84 | CD-PC | FNIP2 |
| 8.249E-198 | 0.97191531 | 0.566 | 0.067 | 1.761E-193 | CD-PC | TBX2 |
| 8.8147E-63 | 0.96189366 | 0.506 | 0.147 | 1.8819E-58 | CD-PC | IVNS1ABP |
| 1.8496E-56 | 0.96146815 | 0.667 | 0.265 | 3.9489E-52 | CD-PC | HIF1A |
| 2.9684E-43 | 0.95371679 | 0.494 | 0.168 | 6.3374E-39 | CD-PC | IGFBP3 |
| 8.569E-150 | 0.939022 | 0.53 | 0.077 | 1.829E-145 | CD-PC | TOX3 |
| 5.602E-215 | 0.93624391 | 0.578 | 0.063 | 1.196E-210 | CD-PC | GATA3 |
| 1.0217E-61 | 0.86446042 | 0.594 | 0.196 | 2.1814E-57 | CD-PC | PIM3 |
| 9.5982E-35 | 0.85402328 | 0.888 | 0.586 | 2.0492E-30 | CD-PC | KRT19 |
| 5.1944E-80 | 0.83982854 | 0.679 | 0.204 | 1.109E-75 | CD-PC | CTSH |
| 1.1606E-77 | 0.83761983 | 0.514 | 0.129 | 2.4778E-73 | CD-PC | L1CAM |
| 1.0442E-52 | 0.83381547 | 0.727 | 0.31 | 2.2295E-48 | CD-PC | FAM107B |
| 8.16E-100 | 0.83319886 | 0.558 | 0.122 | 1.7421E-95 | CD-PC | KIAA1522 |
| 3.1885E-81 | 0.83174255 | 0.426 | 0.085 | 6.8075E-77 | CD-PC | BHLHE41 |
| 2.2653E-92 | 0.8308258 | 0.333 | 0.046 | 4.8365E-88 | CD-PC | AGR2 |
| 9.5602E-95 | 0.8073226 | 0.546 | 0.121 | 2.0411E-90 | CD-PC | MYH10 |
| 4.5979E-44 | 0.80361111 | 0.847 | 0.485 | 9.8165E-40 | CD-PC | ATP1B1 |

| | | | | | | |
|---|---|---|---|---|---|---|
| 5.2734E-45 | 0.80116405 | 0.639 | 0.259 | 1.1259E-40 | CD-PC | GSTM3 |
| 4.4014E-47 | 0.80108254 | 0.719 | 0.322 | 9.397E-43 | CD-PC | CD2AP |
| 1.6668E-72 | 0.79975981 | 0.534 | 0.144 | 3.5586E-68 | CD-PC | POR |
| 9.542E-109 | 0.78142163 | 0.494 | 0.086 | 2.037E-104 | CD-PC | ZNF503 |
| 9.5428E-87 | 0.77834041 | 0.49 | 0.107 | 2.0374E-82 | CD-PC | TWSG1 |
| 6.7048E-20 | 0.77464878 | 0.988 | 0.939 | 1.4315E-15 | CD-PC | MALAT1 |
| 2.5034E-30 | 0.77438388 | 0.526 | 0.24 | 5.3448E-26 | CD-PC | IRS2 |
| 2.582E-148 | 0.77181906 | 0.438 | 0.052 | 5.512E-144 | CD-PC | PAPPA |
| 3.8731E-47 | 0.75752303 | 0.582 | 0.223 | 8.2692E-43 | CD-PC | METAP2 |
| 5.3254E-76 | 0.75594089 | 0.133 | 0.009 | 1.137E-71 | CD-PC | TFF3 |
| 2.7989E-67 | 0.75487671 | 0.622 | 0.192 | 5.9757E-63 | CD-PC | EHF |
| 8.29E-54 | 0.75444299 | 0.606 | 0.216 | 1.7699E-49 | CD-PC | GPRC5C |
| 1.3347E-49 | 0.74614584 | 0.538 | 0.191 | 2.8496E-45 | CD-PC | ADGRG1 |
| 9.9648E-53 | 0.7437279 | 0.631 | 0.232 | 2.1275E-48 | CD-PC | KLF5 |
| 1.0206E-49 | 0.74262519 | 0.627 | 0.244 | 2.1789E-45 | CD-PC | PTBP3 |
| 3.8117E-30 | 0.74244541 | 0.562 | 0.258 | 8.138E-26 | CD-PC | DUSP1 |
| 1.5961E-37 | 0.74081503 | 0.586 | 0.266 | 3.4077E-33 | CD-PC | BAG1 |
| 2.4165E-40 | 0.72705087 | 0.474 | 0.167 | 5.1592E-36 | CD-PC | PLA2G16 |
| 6.0955E-71 | 0.72600192 | 0.538 | 0.139 | 1.3014E-66 | CD-PC | SLC25A29 |
| 3.914E-118 | 0.72539091 | 0.434 | 0.063 | 8.356E-114 | CD-PC | HS6ST1 |
| 1.4559E-53 | 0.72344915 | 0.442 | 0.125 | 3.1083E-49 | CD-PC | SLC2A1 |
| 2.2751E-35 | 0.7226396 | 0.679 | 0.331 | 4.8573E-31 | CD-PC | PAX8 |
| 7.9146E-93 | 0.71458099 | 0.542 | 0.118 | 1.6898E-88 | CD-PC | KAZN |
| 4.5461E-67 | 0.71253444 | 0.554 | 0.161 | 9.7059E-63 | CD-PC | METRNL |
| 8.2261E-16 | 0.70966608 | 0.478 | 0.266 | 1.7563E-11 | CD-PC | SEPHS2 |
| 3.5244E-22 | 0.69407059 | 0.554 | 0.281 | 7.5246E-18 | CD-PC | CEBPD |
| 5.0887E-52 | 0.69172809 | 0.245 | 0.042 | 1.0864E-47 | CD-PC | LTF |
| 2.3592E-38 | 0.68679907 | 0.699 | 0.345 | 5.0368E-34 | CD-PC | NUCKS1 |
| 8.613E-77 | 0.68554798 | 0.55 | 0.137 | 1.8389E-72 | CD-PC | LPIN2 |
| 1.234E-93 | 0.68224152 | 0.402 | 0.067 | 2.6345E-89 | CD-PC | KLF11 |
| 6.3359E-94 | 0.67322011 | 0.454 | 0.085 | 1.3527E-89 | CD-PC | SLC44A1 |
| 1.7243E-51 | 0.66936946 | 0.45 | 0.132 | 3.6814E-47 | CD-PC | GNA13 |
| 4.7158E-74 | 0.66519534 | 0.406 | 0.08 | 1.0068E-69 | CD-PC | HLA-DRA |
| 9.3038E-25 | 0.65617047 | 0.566 | 0.283 | 1.9864E-20 | CD-PC | RND3 |
| 6.5527E-93 | 0.65023976 | 0.498 | 0.099 | 1.399E-88 | CD-PC | CAPS |
| 2.046E-102 | 0.64632309 | 0.39 | 0.057 | 4.3678E-98 | CD-PC | ST6GAL1 |
| 7.017E-140 | 0.64482389 | 0.446 | 0.056 | 1.498E-135 | CD-PC | FAM20C |
| 1.374E-50 | 0.64346906 | 0.55 | 0.19 | 2.9336E-46 | CD-PC | SNX2 |
| 5.4987E-73 | 0.64003903 | 0.426 | 0.091 | 1.174E-68 | CD-PC | SASH1 |
| 2.8375E-52 | 0.63674969 | 0.498 | 0.158 | 6.0581E-48 | CD-PC | HSBP1L1 |
| 1.9933E-62 | 0.63123022 | 0.55 | 0.158 | 4.2558E-58 | CD-PC | NTN4 |
| 7.6756E-41 | 0.62809713 | 0.373 | 0.109 | 1.6387E-36 | CD-PC | NUAK2 |
| 7.3083E-30 | 0.62797246 | 0.96 | 0.797 | 1.5603E-25 | CD-PC | CD24 |
| 3.6811E-29 | 0.62338828 | 0.831 | 0.56 | 7.8592E-25 | CD-PC | MAL2 |
| 5.7378E-46 | 0.62211742 | 0.418 | 0.12 | 1.225E-41 | CD-PC | SPNS2 |
| 3.552E-207 | 0.62148829 | 0.41 | 0.032 | 7.583E-203 | CD-PC | MLPH |
| 5.1005E-35 | 0.61306221 | 0.418 | 0.146 | 1.0889E-30 | CD-PC | KITLG |
| 2.0731E-75 | 0.59921332 | 0.486 | 0.112 | 4.4261E-71 | CD-PC | RAB11FIP2 |
| 1.603E-22 | 0.59689085 | 0.699 | 0.426 | 3.4224E-18 | CD-PC | ZFAND5 |
| 5.8939E-54 | 0.59414073 | 0.558 | 0.181 | 1.2584E-49 | CD-PC | LLGL2 |

| | | | | | | |
|---|---|---|---|---|---|---|
| 8.1828E-79 | 0.59141224 | 0.394 | 0.073 | 1.747E-74 | CD-PC | TNFRSF14 |
| 4.8097E-40 | 0.58831534 | 0.57 | 0.224 | 1.0269E-35 | CD-PC | KMT2E |
| 2.003E-40 | 0.58799047 | 0.53 | 0.198 | 4.2765E-36 | CD-PC | AIF1L |
| 1.6008E-38 | 0.5869547 | 0.61 | 0.256 | 3.4178E-34 | CD-PC | PLXNB2 |
| 4.2626E-39 | 0.58288958 | 0.627 | 0.258 | 9.1007E-35 | CD-PC | LIMA1 |
| 7.4719E-67 | 0.5818194 | 0.422 | 0.094 | 1.5953E-62 | CD-PC | MUC20 |
| 2.979E-109 | 0.5758191 | 0.426 | 0.063 | 6.361E-105 | CD-PC | GCNT3 |
| 1.1507E-37 | 0.5755237 | 0.622 | 0.267 | 2.4568E-33 | CD-PC | CREBRF |
| 3.8039E-43 | 0.57403195 | 0.426 | 0.133 | 8.1213E-39 | CD-PC | SPIRE1 |
| 7.1774E-52 | 0.57164756 | 0.365 | 0.089 | 1.5324E-47 | CD-PC | MTMR10 |
| 1.583E-117 | 0.57046694 | 0.418 | 0.056 | 3.381E-113 | CD-PC | TBX3 |
| 3.2394E-25 | 0.56931966 | 0.631 | 0.328 | 6.9162E-21 | CD-PC | ZNF292 |
| 1.1561E-63 | 0.56522371 | 0.49 | 0.127 | 2.4682E-59 | CD-PC | SMIM22 |
| 8.6705E-45 | 0.56412736 | 0.349 | 0.089 | 1.8512E-40 | CD-PC | AKR1C3 |
| 2.3875E-40 | 0.56316663 | 0.454 | 0.151 | 5.0973E-36 | CD-PC | RAPGEF3 |
| 2.8205E-93 | 0.55928005 | 0.41 | 0.069 | 6.0218E-89 | CD-PC | SLC16A7 |
| 5.1611E-29 | 0.55712439 | 0.719 | 0.405 | 1.1019E-24 | CD-PC | UQCR10 |
| 2.8172E-36 | 0.55311041 | 0.518 | 0.199 | 6.0148E-32 | CD-PC | FOSL2 |
| 4.7532E-37 | 0.55205131 | 0.498 | 0.186 | 1.0148E-32 | CD-PC | PHLDA3 |
| 6.5075E-38 | 0.54717244 | 0.647 | 0.282 | 1.3893E-33 | CD-PC | NFE2L2 |
| 1.9557E-62 | 0.54618425 | 0.361 | 0.076 | 4.1754E-58 | CD-PC | SH3BP5 |
| 1.19E-111 | 0.54604357 | 0.398 | 0.055 | 2.541E-107 | CD-PC | BLNK |
| 3.3262E-88 | 0.54468656 | 0.43 | 0.078 | 7.1015E-84 | CD-PC | SLC16A4 |
| 1.9298E-18 | 0.54421064 | 0.454 | 0.221 | 4.1202E-14 | CD-PC | CD74 |
| 9.7885E-25 | 0.53929946 | 0.876 | 0.641 | 2.0898E-20 | CD-PC | WFDC2 |
| 1.2533E-42 | 0.53367237 | 0.562 | 0.204 | 2.6758E-38 | CD-PC | SCCPDH |
| 6.3629E-41 | 0.53332722 | 0.586 | 0.231 | 1.3585E-36 | CD-PC | CLINT1 |
| 4.1133E-41 | 0.53071093 | 0.546 | 0.204 | 8.7819E-37 | CD-PC | SERINC2 |
| 8.0833E-41 | 0.53043241 | 0.402 | 0.12 | 1.7258E-36 | CD-PC | GPNMB |
| 1.5205E-56 | 0.52894604 | 0.398 | 0.096 | 3.2462E-52 | CD-PC | PLEKHG3 |
| 6.939E-158 | 0.5271797 | 0.329 | 0.027 | 1.481E-153 | CD-PC | PQLC2L |
| 1.0388E-91 | 0.52694706 | 0.418 | 0.07 | 2.2179E-87 | CD-PC | NOV |
| 6.2576E-30 | 0.52413429 | 0.635 | 0.316 | 1.336E-25 | CD-PC | AES |
| 9.9089E-63 | 0.52041051 | 0.478 | 0.122 | 2.1156E-58 | CD-PC | ADSS |
| 6.1375E-40 | 0.51770224 | 0.418 | 0.128 | 1.3104E-35 | CD-PC | CDH16 |
| 7.3357E-23 | 0.51664377 | 0.474 | 0.223 | 1.5662E-18 | CD-PC | PRSS8 |
| 1.0387E-60 | 0.51658188 | 0.365 | 0.079 | 2.2176E-56 | CD-PC | RAB40B |
| 2.5794E-21 | 0.51318353 | 0.478 | 0.233 | 5.507E-17 | CD-PC | HECTD1 |
| 4.5357E-22 | 0.51070878 | 0.843 | 0.582 | 9.6837E-18 | CD-PC | ELF3 |
| 2.418E-26 | 0.50866249 | 0.659 | 0.354 | 5.1624E-22 | CD-PC | SEC62 |
| 1.803E-133 | 0.50748178 | 0.365 | 0.039 | 3.85E-129 | CD-PC | ARHGAP6 |
| 6.9616E-68 | 0.50532126 | 0.382 | 0.075 | 1.4863E-63 | CD-PC | AC019197.1 |
| 3.5085E-16 | 0.50518445 | 0.414 | 0.198 | 7.4907E-12 | CD-PC | SGK1 |
| 6.6214E-76 | 0.50394719 | 0.41 | 0.08 | 1.4137E-71 | CD-PC | MTUS1 |
| 5.756E-154 | 0.50298932 | 0.333 | 0.028 | 1.229E-149 | CD-PC | CLDN8 |
| 1.1446E-52 | 0.50236598 | 0.378 | 0.092 | 2.4437E-48 | CD-PC | STAM |
| 3.9539E-80 | 0.50201517 | 0.321 | 0.049 | 8.4417E-76 | CD-PC | SMIM5 |
| 8.555E-190 | 0.49979373 | 0.309 | 0.019 | 1.826E-185 | CD-PC | MXRA5 |
| 2.8773E-66 | 0.49878016 | 0.301 | 0.05 | 6.143E-62 | CD-PC | HSD11B2 |
| 4.7591E-33 | 0.49787468 | 0.422 | 0.154 | 1.0161E-28 | CD-PC | RRAGC |

| | | | | | | |
|---|---|---|---|---|---|---|
| 4.0312E-39 | 0.49702034 | 0.554 | 0.214 | 8.6067E-35 | CD-PC | PCMTD1 |
| 5.5859E-51 | 0.49608106 | 0.365 | 0.089 | 1.1926E-46 | CD-PC | MPZL2 |
| 1.2912E-48 | 0.49605087 | 0.434 | 0.122 | 2.7568E-44 | CD-PC | CRIP2 |
| 8.3145E-22 | 0.49304217 | 0.606 | 0.334 | 1.7751E-17 | CD-PC | KRT10 |
| 4.3676E-66 | 0.49249576 | 0.39 | 0.082 | 9.3248E-62 | CD-PC | PRRG1 |
| 2.2771E-37 | 0.49081671 | 0.45 | 0.158 | 4.8617E-33 | CD-PC | XIAP |
| 7.8089E-46 | 0.48889054 | 0.502 | 0.157 | 1.6672E-41 | CD-PC | ADGRF1 |
| 5.5812E-25 | 0.4887042 | 0.627 | 0.327 | 1.1916E-20 | CD-PC | ANKRD12 |
| 2.8646E-34 | 0.48723145 | 0.442 | 0.158 | 6.116E-30 | CD-PC | ESF1 |
| 9.0401E-38 | 0.4872212 | 0.369 | 0.11 | 1.9301E-33 | CD-PC | NEDD9 |
| 1.97E-66 | 0.4872097 | 0.406 | 0.088 | 4.206E-62 | CD-PC | RNF24 |
| 3.387E-111 | 0.48452857 | 0.325 | 0.037 | 7.231E-107 | CD-PC | PALM3 |
| 1.3165E-13 | 0.48423139 | 0.823 | 0.722 | 2.8108E-09 | CD-PC | CTSD |
| 4.2073E-59 | 0.48359172 | 0.373 | 0.082 | 8.9827E-55 | CD-PC | TRIM24 |
| 2.8678E-47 | 0.48339606 | 0.414 | 0.117 | 6.1227E-43 | CD-PC | TPD52L1 |
| 3.1624E-26 | 0.48289275 | 0.502 | 0.225 | 6.7517E-22 | CD-PC | NFE2L1 |
| 2.8817E-35 | 0.4795459 | 0.446 | 0.157 | 6.1524E-31 | CD-PC | KIAA1109 |
| 1.0886E-42 | 0.47949935 | 0.345 | 0.093 | 2.3241E-38 | CD-PC | BCAR3 |
| 5.9472E-53 | 0.47930409 | 0.434 | 0.117 | 1.2697E-48 | CD-PC | CHPT1 |
| 9.6711E-63 | 0.47923 | 0.406 | 0.092 | 2.0648E-58 | CD-PC | ACBD5 |
| 4.0703E-14 | 0.4772752 | 0.703 | 0.543 | 8.6901E-10 | CD-PC | BRI3 |
| 2.7314E-36 | 0.47389103 | 0.55 | 0.212 | 5.8315E-32 | CD-PC | EPS8L2 |
| 2.37E-252 | 0.4735653 | 0.317 | 0.014 | 5.061E-248 | CD-PC | FAM20A |
| 4.0081E-65 | 0.47249969 | 0.382 | 0.079 | 8.5572E-61 | CD-PC | ATP8B1 |
| 9.7864E-43 | 0.47190973 | 0.47 | 0.151 | 2.0894E-38 | CD-PC | CXXC5 |
| 8.3539E-08 | 0.47068692 | 0.924 | 0.813 | 0.00178355 | CD-PC | NEAT1 |
| 2.0529E-31 | 0.47047439 | 0.482 | 0.193 | 4.3829E-27 | CD-PC | NFIC |
| 1.065E-244 | 0.46662364 | 0.301 | 0.013 | 2.274E-240 | CD-PC | KCNK3 |
| 1.2205E-65 | 0.46586134 | 0.353 | 0.069 | 2.6058E-61 | CD-PC | CRYBG1 |
| 3.4293E-30 | 0.465138 | 0.506 | 0.209 | 7.3216E-26 | CD-PC | PARD6B |
| 4.7966E-33 | 0.4636069 | 0.53 | 0.215 | 1.0241E-28 | CD-PC | RBM47 |
| 2.5379E-41 | 0.46272068 | 0.466 | 0.148 | 5.4184E-37 | CD-PC | CHD3 |
| 1.693E-179 | 0.46009305 | 0.253 | 0.013 | 3.615E-175 | CD-PC | SLC14A2 |
| 2.612E-25 | 0.45791042 | 0.434 | 0.18 | 5.5766E-21 | CD-PC | MUC20-OT1 |
| 1.6963E-26 | 0.45705947 | 0.285 | 0.092 | 3.6216E-22 | CD-PC | PXDC1 |
| 3.684E-32 | 0.45204341 | 0.414 | 0.148 | 7.8654E-28 | CD-PC | ROCK2 |
| 4.5338E-28 | 0.45127252 | 0.546 | 0.247 | 9.6796E-24 | CD-PC | PTP4A1 |
| 2.28E-87 | 0.45042112 | 0.365 | 0.057 | 4.8678E-83 | CD-PC | SLC16A5 |
| 3.2384E-28 | 0.44945216 | 0.554 | 0.253 | 6.9139E-24 | CD-PC | NDUFA5 |
| 1.6828E-44 | 0.44575424 | 0.414 | 0.121 | 3.5928E-40 | CD-PC | PKN1 |
| 3.8665E-23 | 0.44553125 | 0.618 | 0.329 | 8.255E-19 | CD-PC | PTP4A2 |
| 2.4351E-24 | 0.44525541 | 0.578 | 0.293 | 5.199E-20 | CD-PC | TM9SF3 |
| 8.646E-114 | 0.44442731 | 0.309 | 0.033 | 1.846E-109 | CD-PC | PCDH7 |
| 4.0216E-21 | 0.44432752 | 0.422 | 0.187 | 8.5862E-17 | CD-PC | BHLHE40 |
| 1.7597E-46 | 0.44395852 | 0.394 | 0.107 | 3.7569E-42 | CD-PC | GSR |
| 2.1163E-27 | 0.44329686 | 0.582 | 0.273 | 4.5182E-23 | CD-PC | YY1 |
| 2.4885E-18 | 0.44160807 | 0.618 | 0.361 | 5.313E-14 | CD-PC | PPP1CB |
| 2.4727E-43 | 0.44146305 | 0.43 | 0.133 | 5.2792E-39 | CD-PC | FDX1 |
| 4.1146E-49 | 0.44075014 | 0.373 | 0.09 | 8.7846E-45 | CD-PC | ALDH1A3 |
| 1.0002E-64 | 0.43465642 | 0.378 | 0.078 | 2.1354E-60 | CD-PC | ARRDC1 |

| | | | | | | |
|---|---|---|---|---|---|---|
| 8.4047E-18 | 0.43271717 | 0.618 | 0.337 | 1.7944E-13 | CD-PC | ZFP36L2 |
| 3.098E-120 | 0.43232872 | 0.289 | 0.027 | 6.614E-116 | CD-PC | C4orf19 |
| 2.8022E-26 | 0.43175649 | 0.414 | 0.165 | 5.9828E-22 | CD-PC | NRIP1 |
| 1.5975E-41 | 0.43068296 | 0.345 | 0.093 | 3.4106E-37 | CD-PC | SLC12A7 |
| 2.201E-42 | 0.43033282 | 0.369 | 0.103 | 4.6991E-38 | CD-PC | KMT5A |
| 2.1801E-24 | 0.43032563 | 0.43 | 0.183 | 4.6544E-20 | CD-PC | KLHL24 |
| 1.1367E-16 | 0.42865962 | 0.602 | 0.365 | 2.4269E-12 | CD-PC | SPTBN1 |
| 9.5835E-18 | 0.42838844 | 0.936 | 0.867 | 2.0461E-13 | CD-PC | TMSB4X |
| 7.5731E-19 | 0.42835422 | 0.414 | 0.19 | 1.6169E-14 | CD-PC | TOB1 |
| 1.0133E-39 | 0.42796005 | 0.39 | 0.118 | 2.1634E-35 | CD-PC | TRIM2 |
| 1.0798E-29 | 0.42752827 | 0.426 | 0.159 | 2.3053E-25 | CD-PC | CD82 |
| 3.9507E-50 | 0.42624091 | 0.369 | 0.089 | 8.4347E-46 | CD-PC | KCNJ16 |
| 2.2306E-49 | 0.42583931 | 0.273 | 0.054 | 4.7623E-45 | CD-PC | EMX1 |
| 2.0335E-65 | 0.4254513 | 0.333 | 0.062 | 4.3416E-61 | CD-PC | VAV2 |
| 3.0897E-35 | 0.42531761 | 0.378 | 0.12 | 6.5965E-31 | CD-PC | VEGFB |
| 2.7804E-38 | 0.42528265 | 0.446 | 0.149 | 5.9362E-34 | CD-PC | TMEM87A |
| 1.5723E-26 | 0.42424474 | 0.526 | 0.233 | 3.3568E-22 | CD-PC | NSD3 |
| 9.4615E-24 | 0.42338506 | 0.486 | 0.226 | 2.02E-19 | CD-PC | SCARB2 |
| 1.08E-131 | 0.42338124 | 0.309 | 0.028 | 2.306E-127 | CD-PC | VEPH1 |
| 1.0266E-85 | 0.42272205 | 0.293 | 0.038 | 2.1917E-81 | CD-PC | RORC |
| 8.8723E-23 | 0.42236619 | 0.486 | 0.232 | 1.8942E-18 | CD-PC | SMIM26 |
| 1.0519E-55 | 0.42196431 | 0.353 | 0.077 | 2.2459E-51 | CD-PC | SH3BP2 |
| 3.8713E-40 | 0.42086255 | 0.41 | 0.126 | 8.2651E-36 | CD-PC | AFG3L2 |
| 3.1284E-71 | 0.42048378 | 0.225 | 0.027 | 6.6792E-67 | CD-PC | AC129507.4 |
| 1.9712E-54 | 0.41930788 | 0.349 | 0.078 | 4.2084E-50 | CD-PC | DHRS3 |
| 1.4601E-56 | 0.418666 | 0.337 | 0.07 | 3.1173E-52 | CD-PC | SPTBN2 |
| 5.3717E-27 | 0.41755739 | 0.554 | 0.255 | 1.1469E-22 | CD-PC | PAK2 |
| 1.7472E-72 | 0.41431188 | 0.301 | 0.047 | 3.7303E-68 | CD-PC | AOC1 |
| 2.2141E-29 | 0.4131249 | 0.474 | 0.189 | 4.7271E-25 | CD-PC | TBL1XR1 |
| 7.287E-183 | 0.41219643 | 0.253 | 0.013 | 1.556E-178 | CD-PC | CREB3L1 |
| 6.4215E-26 | 0.41204227 | 0.442 | 0.183 | 1.371E-21 | CD-PC | SREK1 |
| 1.403E-28 | 0.41180299 | 0.51 | 0.218 | 2.9953E-24 | CD-PC | TMED4 |
| 2.3782E-19 | 0.4109348 | 0.631 | 0.372 | 5.0774E-15 | CD-PC | CD164 |
| 1.7923E-22 | 0.41038438 | 0.538 | 0.26 | 3.8265E-18 | CD-PC | SLC38A1 |
| 1.158E-33 | 0.4083879 | 0.426 | 0.148 | 2.4724E-29 | CD-PC | ZNF148 |
| 6.5428E-60 | 0.40838656 | 0.349 | 0.072 | 1.3969E-55 | CD-PC | SAMD12 |
| 2.1548E-30 | 0.40599454 | 0.394 | 0.139 | 4.6005E-26 | CD-PC | PTPRF |
| 2.2461E-78 | 0.40513115 | 0.285 | 0.039 | 4.7954E-74 | CD-PC | PPFIBP2 |
| 1.0733E-40 | 0.40382512 | 0.369 | 0.105 | 2.2915E-36 | CD-PC | ESRRA |
| 1.0218E-28 | 0.40362518 | 0.458 | 0.182 | 2.1816E-24 | CD-PC | TMED5 |
| 8.6145E-22 | 0.40320668 | 0.53 | 0.267 | 1.8392E-17 | CD-PC | NDUFV2 |
| 1.78E-151 | 0.4021851 | 0.273 | 0.019 | 3.8E-147 | CD-PC | EYA4 |
| 3.5878E-20 | 0.40164523 | 0.578 | 0.305 | 7.6599E-16 | CD-PC | ARHGAP5 |
| 1.356E-57 | 0.40081581 | 0.325 | 0.066 | 2.895E-53 | CD-PC | C16orf87 |
| 5.5258E-63 | 0.40078424 | 0.341 | 0.065 | 1.1798E-58 | CD-PC | VAV3 |
| 1.4899E-13 | 0.40078424 | 0.446 | 0.233 | 3.181E-09 | CD-PC | SLC5A3 |
| 2.4309E-34 | 0.40048566 | 0.41 | 0.138 | 5.1899E-30 | CD-PC | KIF13B |
| 6.0567E-43 | 0.40042681 | 0.289 | 0.066 | 1.2931E-38 | CD-PC | NUPR2 |
| 8.31E-27 | 0.39973952 | 0.382 | 0.144 | 1.7742E-22 | CD-PC | YAP1 |
| 2.0397E-18 | 0.39948707 | 0.884 | 0.766 | 4.3547E-14 | CD-PC | COX7C |

| | | | | | | |
|---|---|---|---|---|---|---|
| 4.3071E-45 | 0.39940023 | 0.149 | 0.019 | 9.1957E-41 | CD-PC | NR0B2 |
| 2.9728E-26 | 0.39867098 | 0.406 | 0.159 | 6.3469E-22 | CD-PC | LARP1 |
| 6.3279E-40 | 0.3963389 | 0.406 | 0.124 | 1.351E-35 | CD-PC | TRIP6 |
| 3.8405E-62 | 0.39539963 | 0.293 | 0.051 | 8.1995E-58 | CD-PC | FGFR2 |
| 4.4564E-47 | 0.39406477 | 0.305 | 0.068 | 9.5145E-43 | CD-PC | PCDH1 |
| 2.636E-29 | 0.39390889 | 0.345 | 0.115 | 5.6279E-25 | CD-PC | MAN1A1 |
| 3.8932E-32 | 0.39384505 | 0.426 | 0.153 | 8.312E-28 | CD-PC | ZFAND3 |
| 2.2597E-34 | 0.3929679 | 0.39 | 0.129 | 4.8244E-30 | CD-PC | NDUFA8 |
| 1.0206E-18 | 0.39282803 | 0.454 | 0.22 | 2.1791E-14 | CD-PC | NFAT5 |
| 8.9688E-23 | 0.39163417 | 0.446 | 0.198 | 1.9148E-18 | CD-PC | PLSCR1 |
| 2.5885E-29 | 0.39104462 | 0.414 | 0.151 | 5.5264E-25 | CD-PC | NUDT4 |
| 2.1545E-36 | 0.39090328 | 0.369 | 0.112 | 4.5998E-32 | CD-PC | GPR160 |
| 1.0259E-21 | 0.39056871 | 0.221 | 0.065 | 2.1903E-17 | CD-PC | SPINK1 |
| 1.1211E-35 | 0.39017823 | 0.39 | 0.123 | 2.3935E-31 | CD-PC | RASSF7 |
| 4.2804E-17 | 0.38985737 | 0.237 | 0.087 | 9.1385E-13 | CD-PC | ASS1 |
| 1.3805E-21 | 0.38970469 | 0.466 | 0.213 | 2.9473E-17 | CD-PC | PTGR1 |
| 6.0474E-22 | 0.38863399 | 0.418 | 0.181 | 1.2911E-17 | CD-PC | MSI2 |
| 3.4779E-37 | 0.38844339 | 0.418 | 0.134 | 7.4253E-33 | CD-PC | PACSIN2 |
| 2.7049E-46 | 0.3883714 | 0.378 | 0.098 | 5.7749E-42 | CD-PC | ERBB2 |
| 9.2478E-18 | 0.38675962 | 0.647 | 0.398 | 1.9744E-13 | CD-PC | LAMTOR5 |
| 1.1065E-40 | 0.38644509 | 0.329 | 0.086 | 2.3623E-36 | CD-PC | EMX2OS |
| 1.8323E-87 | 0.3864347 | 0.261 | 0.03 | 3.912E-83 | CD-PC | CNGA1 |
| 8.1002E-63 | 0.38582052 | 0.289 | 0.049 | 1.7294E-58 | CD-PC | FAM213B |
| 7.2712E-21 | 0.38539141 | 0.558 | 0.28 | 1.5524E-16 | CD-PC | SRRM1 |
| 3.1461E-19 | 0.38509872 | 0.502 | 0.262 | 6.717E-15 | CD-PC | MINOS1 |
| 8.9118E-20 | 0.38441396 | 0.418 | 0.19 | 1.9027E-15 | CD-PC | CCNG2 |
| 6.7572E-68 | 0.38403853 | 0.317 | 0.054 | 1.4427E-63 | CD-PC | KLHL8 |
| 8.3206E-37 | 0.38350951 | 0.378 | 0.111 | 1.7765E-32 | CD-PC | ANK3 |
| 1.5954E-72 | 0.38193401 | 0.281 | 0.041 | 3.4061E-68 | CD-PC | FXYD3 |
| 5.6015E-16 | 0.37987082 | 0.442 | 0.219 | 1.1959E-11 | CD-PC | GPRC5A |
| 4.2942E-26 | 0.37979317 | 0.486 | 0.202 | 9.1682E-22 | CD-PC | VPS13C |
| 1.9645E-19 | 0.3788665 | 0.418 | 0.194 | 4.1943E-15 | CD-PC | SLK |
| 3.8387E-18 | 0.37752356 | 0.502 | 0.264 | 8.1956E-14 | CD-PC | RTN3 |
| 1.8299E-25 | 0.37737888 | 0.546 | 0.243 | 3.9068E-21 | CD-PC | MACC1 |
| 2.1057E-12 | 0.37645264 | 0.771 | 0.553 | 4.4956E-08 | CD-PC | LRRFIP1 |
| 7.4188E-26 | 0.37566367 | 0.627 | 0.293 | 1.5839E-21 | CD-PC | CDH1 |
| 4.6268E-23 | 0.37535572 | 0.402 | 0.168 | 9.8781E-19 | CD-PC | COA3 |
| 4.6274E-19 | 0.3751586 | 0.622 | 0.344 | 9.8796E-15 | CD-PC | VAPA |
| 6.9102E-23 | 0.37344556 | 0.353 | 0.134 | 1.4753E-18 | CD-PC | EFNB2 |
| 1.8161E-15 | 0.37317885 | 0.598 | 0.354 | 3.8773E-11 | CD-PC | ATP5ME |
| 3.3878E-32 | 0.37301677 | 0.309 | 0.09 | 7.2329E-28 | CD-PC | PDP1 |
| 5.8154E-32 | 0.3727322 | 0.373 | 0.124 | 1.2416E-27 | CD-PC | HOXB2 |
| 7.4144E-24 | 0.37111796 | 0.478 | 0.213 | 1.583E-19 | CD-PC | SRI |
| 1.7509E-34 | 0.37047147 | 0.353 | 0.107 | 3.7382E-30 | CD-PC | SHROOM3 |
| 2.1738E-51 | 0.37040754 | 0.189 | 0.026 | 4.641E-47 | CD-PC | AJAP1 |
| 9.0377E-17 | 0.37012929 | 0.357 | 0.164 | 1.9295E-12 | CD-PC | AMOTL2 |
| 2.182E-33 | 0.36990705 | 0.349 | 0.106 | 4.6586E-29 | CD-PC | DDX3Y |
| 2.0137E-25 | 0.36955051 | 0.361 | 0.133 | 4.2993E-21 | CD-PC | RBBP8 |
| 7.9448E-28 | 0.36942238 | 0.474 | 0.193 | 1.6962E-23 | CD-PC | BBX |
| 6.7938E-52 | 0.36895885 | 0.317 | 0.067 | 1.4505E-47 | CD-PC | SNX18 |

| | | | | | | |
|---|---|---|---|---|---|---|
| 2.7605E-44 | 0.36799274 | 0.241 | 0.047 | 5.8937E-40 | CD-PC | HLA-DPA1 |
| 5.755E-23 | 0.36788821 | 0.378 | 0.151 | 1.2287E-18 | CD-PC | SMAD3 |
| 7.1405E-20 | 0.36788621 | 0.458 | 0.222 | 1.5245E-15 | CD-PC | PDXK |
| 7.2145E-24 | 0.36664153 | 0.337 | 0.121 | 1.5403E-19 | CD-PC | CMYA5 |
| 2.3924E-18 | 0.36631656 | 0.542 | 0.283 | 5.1078E-14 | CD-PC | TGOLN2 |
| 2.2139E-24 | 0.36626467 | 0.394 | 0.155 | 4.7267E-20 | CD-PC | TRAF4 |
| 2.9617E-26 | 0.36546493 | 0.373 | 0.137 | 6.3232E-22 | CD-PC | FNDC3A |
| 3.9063E-23 | 0.36497932 | 0.337 | 0.128 | 8.3399E-19 | CD-PC | VASN |
| 1.529E-29 | 0.36456006 | 0.402 | 0.143 | 3.2645E-25 | CD-PC | RETREG1 |
| 1.4871E-18 | 0.36441441 | 0.59 | 0.317 | 3.175E-14 | CD-PC | RSRC2 |
| 1.0545E-19 | 0.36429219 | 0.635 | 0.342 | 2.2514E-15 | CD-PC | UBE2H |
| 4.6605E-29 | 0.36357663 | 0.325 | 0.104 | 9.9502E-25 | CD-PC | POGZ |
| 8.3609E-37 | 0.3629583 | 0.329 | 0.091 | 1.7851E-32 | CD-PC | PPL |
| 3.2594E-26 | 0.36265491 | 0.402 | 0.154 | 6.9588E-22 | CD-PC | MRPL34 |
| 2.451E-19 | 0.36249294 | 0.859 | 0.635 | 5.233E-15 | CD-PC | JUND |
| 1.2613E-20 | 0.36155256 | 0.494 | 0.237 | 2.6929E-16 | CD-PC | TOR1AIP2 |
| 6.6301E-15 | 0.3604912 | 0.49 | 0.276 | 1.4155E-10 | CD-PC | GRN |
| 3.0429E-36 | 0.36036417 | 0.325 | 0.09 | 6.4967E-32 | CD-PC | MAP3K1 |
| 1.159E-148 | 0.3601029 | 0.265 | 0.018 | 2.475E-144 | CD-PC | ELF5 |
| 1.2683E-36 | 0.35996747 | 0.337 | 0.096 | 2.7078E-32 | CD-PC | TMEM45B |
| 5.5102E-20 | 0.35974421 | 0.434 | 0.199 | 1.1764E-15 | CD-PC | RTF1 |
| 2.974E-93 | 0.35974368 | 0.281 | 0.033 | 6.3495E-89 | CD-PC | TJP3 |
| 1.7713E-17 | 0.35954529 | 0.406 | 0.195 | 3.7818E-13 | CD-PC | CDC42BPB |
| 2.1178E-20 | 0.35929047 | 0.458 | 0.214 | 4.5214E-16 | CD-PC | FNBP1L |
| 1.2997E-21 | 0.35900733 | 0.502 | 0.235 | 2.7748E-17 | CD-PC | AAK1 |
| 1.4785E-34 | 0.35894035 | 0.329 | 0.095 | 3.1566E-30 | CD-PC | AC118549.1 |
| 1.71E-260 | 0.35866497 | 0.205 | 0.004 | 3.651E-256 | CD-PC | ADH1C |
| 1.3302E-51 | 0.35807452 | 0.325 | 0.07 | 2.84E-47 | CD-PC | SORL1 |
| 4.4544E-20 | 0.35711803 | 0.394 | 0.173 | 9.5102E-16 | CD-PC | BACE2 |
| 8.9562E-16 | 0.35697952 | 0.414 | 0.21 | 1.9122E-11 | CD-PC | BTG3 |
| 9.8458E-33 | 0.35683976 | 0.321 | 0.095 | 2.1021E-28 | CD-PC | TM7SF3 |
| 5.627E-21 | 0.35649233 | 0.514 | 0.252 | 1.2014E-16 | CD-PC | PRKAR1A |
| 1.2549E-22 | 0.35569925 | 0.51 | 0.239 | 2.6792E-18 | CD-PC | RBPMS |
| 3.063E-44 | 0.35473559 | 0.305 | 0.071 | 6.5395E-40 | CD-PC | KCNQ1 |
| 6.8988E-33 | 0.35462788 | 0.353 | 0.112 | 1.4729E-28 | CD-PC | PLEKHM2 |
| 5.7347E-25 | 0.35450392 | 0.289 | 0.093 | 1.2244E-20 | CD-PC | HLA-DRB1 |
| 9.9777E-28 | 0.35349629 | 0.43 | 0.165 | 2.1302E-23 | CD-PC | CBX5 |
| 5.8137E-30 | 0.35325308 | 0.394 | 0.138 | 1.2412E-25 | CD-PC | SH3GLB2 |
| 1.6221E-69 | 0.35325308 | 0.229 | 0.029 | 3.4633E-65 | CD-PC | DOK7 |
| 1.0395E-12 | 0.35323577 | 0.43 | 0.236 | 2.2194E-08 | CD-PC | MDM2 |
| 3.1205E-29 | 0.35301876 | 0.325 | 0.102 | 6.6623E-25 | CD-PC | RNF144B |
| 2.306E-55 | 0.35263383 | 0.285 | 0.053 | 4.9232E-51 | CD-PC | EMP2 |
| 2.5565E-19 | 0.35197933 | 0.486 | 0.23 | 5.4582E-15 | CD-PC | SCIN |
| 2.5036E-30 | 0.35163582 | 0.329 | 0.103 | 5.3453E-26 | CD-PC | MCRIP2 |
| 6.7418E-31 | 0.35029349 | 0.333 | 0.103 | 1.4394E-26 | CD-PC | MTR |
| 9.7575E-69 | 0.35029165 | 0.217 | 0.026 | 2.0832E-64 | CD-PC | DEGS2 |
| 2.8454E-19 | 0.35023719 | 0.434 | 0.207 | 6.075E-15 | CD-PC | RAB14 |
| 7.0085E-28 | 0.34887078 | 0.398 | 0.142 | 1.4963E-23 | CD-PC | EMX2 |
| 5.8104E-28 | 0.34827714 | 0.373 | 0.13 | 1.2405E-23 | CD-PC | PTPN13 |
| 1.031E-35 | 0.3481984 | 0.394 | 0.122 | 2.2011E-31 | CD-PC | TULP4 |

| | | | | | | |
|---|---|---|---|---|---|---|
| 1.8755E-39 | 0.34773637 | 0.301 | 0.075 | 4.0042E-35 | CD-PC | DENND4C |
| 2.928E-31 | 0.3476568 | 0.325 | 0.099 | 6.2512E-27 | CD-PC | MECOM |
| 1.2023E-27 | 0.34718952 | 0.373 | 0.133 | 2.567E-23 | CD-PC | SEPT9 |
| 9.8132E-37 | 0.34691606 | 0.321 | 0.087 | 2.0951E-32 | CD-PC | SLC25A25 |
| 2.8093E-22 | 0.34668185 | 0.434 | 0.186 | 5.9978E-18 | CD-PC | ARID1B |
| 4.4839E-44 | 0.3457937 | 0.297 | 0.068 | 9.5732E-40 | CD-PC | TRAK1 |
| 6.3972E-34 | 0.3453809 | 0.325 | 0.094 | 1.3658E-29 | CD-PC | CAT |
| 7.1102E-37 | 0.3446369 | 0.349 | 0.1 | 1.518E-32 | CD-PC | LTBR |
| 4.6409E-19 | 0.34376397 | 0.598 | 0.315 | 9.9082E-15 | CD-PC | CCNL1 |
| 4.8496E-41 | 0.34318203 | 0.337 | 0.088 | 1.0354E-36 | CD-PC | RCOR3 |
| 1.8548E-15 | 0.34317678 | 0.783 | 0.565 | 3.9599E-11 | CD-PC | RPS27L |
| 5.4405E-26 | 0.34299113 | 0.357 | 0.127 | 1.1616E-21 | CD-PC | PBRM1 |
| 2.8185E-13 | 0.3424125 | 0.538 | 0.328 | 6.0174E-09 | CD-PC | ATP6V1F |
| 9.0137E-74 | 0.34232173 | 0.233 | 0.028 | 1.9244E-69 | CD-PC | AMIGO2 |
| 1.2004E-35 | 0.34184214 | 0.309 | 0.084 | 2.5629E-31 | CD-PC | BRD3 |
| 1.1036E-24 | 0.34108097 | 0.373 | 0.142 | 2.3563E-20 | CD-PC | RRM2B |
| 1.9436E-29 | 0.34089761 | 0.289 | 0.085 | 4.1495E-25 | CD-PC | FAM241A |
| 1.6864E-39 | 0.34019765 | 0.265 | 0.06 | 3.6005E-35 | CD-PC | SLC45A4 |
| 3.0525E-54 | 0.33977136 | 0.277 | 0.051 | 6.5171E-50 | CD-PC | ARHGDIB |
| 6.8499E-38 | 0.33966401 | 0.337 | 0.093 | 1.4625E-33 | CD-PC | TBC1D9 |
| 1.3062E-19 | 0.33936285 | 0.598 | 0.322 | 2.7888E-15 | CD-PC | NDUFC1 |
| 3.2228E-21 | 0.33871767 | 0.394 | 0.168 | 6.8806E-17 | CD-PC | TWF1 |
| 1.6742E-21 | 0.33794323 | 0.442 | 0.199 | 3.5744E-17 | CD-PC | SVIP |
| 1.6922E-17 | 0.33782416 | 0.494 | 0.258 | 3.6128E-13 | CD-PC | PPP2CB |
| 7.1503E-79 | 0.33778442 | 0.249 | 0.03 | 1.5266E-74 | CD-PC | RASAL1 |
| 4.7414E-68 | 0.33725423 | 0.277 | 0.042 | 1.0123E-63 | CD-PC | ADAP1 |
| 8.25E-13 | 0.33685338 | 0.635 | 0.386 | 1.7614E-08 | CD-PC | RAB11FIP1 |
| 8.3074E-29 | 0.33444233 | 0.434 | 0.161 | 1.7736E-24 | CD-PC | SELENOP |
| 4.3574E-14 | 0.33395927 | 0.486 | 0.275 | 9.3031E-10 | CD-PC | MRPL41 |
| 4.3735E-21 | 0.3331063 | 0.526 | 0.251 | 9.3374E-17 | CD-PC | SF1 |
| 3.7591E-31 | 0.33289305 | 0.386 | 0.129 | 8.0256E-27 | CD-PC | SUDS3 |
| 1.9285E-52 | 0.33225427 | 0.301 | 0.061 | 4.1174E-48 | CD-PC | ZNRF1 |
| 1.88E-116 | 0.33210855 | 0.217 | 0.015 | 4.015E-112 | CD-PC | EPN3 |
| 1.0786E-16 | 0.3310023 | 0.446 | 0.227 | 2.3028E-12 | CD-PC | DNAJC15 |
| 6.0661E-17 | 0.33069555 | 0.466 | 0.231 | 1.2951E-12 | CD-PC | MIDN |
| 1.2868E-20 | 0.32977038 | 0.442 | 0.198 | 2.7472E-16 | CD-PC | GIPC1 |
| 1.5634E-17 | 0.32897733 | 0.631 | 0.353 | 3.3378E-13 | CD-PC | GCC2 |
| 1.3992E-83 | 0.3286853 | 0.253 | 0.029 | 2.9873E-79 | CD-PC | TNFRSF14-AS1 |
| 5.2337E-12 | 0.32865993 | 0.414 | 0.233 | 1.1174E-07 | CD-PC | AKAP13 |
| 1.4338E-33 | 0.32796868 | 0.309 | 0.087 | 3.0612E-29 | CD-PC | TMEM127 |
| 9.0421E-18 | 0.3277221 | 0.382 | 0.173 | 1.9305E-13 | CD-PC | MYO10 |
| 4.5052E-43 | 0.32769137 | 0.265 | 0.056 | 9.6186E-39 | CD-PC | NIPAL1 |
| 2.07E-16 | 0.32769137 | 0.434 | 0.213 | 4.4195E-12 | CD-PC | USP53 |
| 2.7747E-21 | 0.32727097 | 0.442 | 0.197 | 5.9239E-17 | CD-PC | COX14 |
| 5.8252E-24 | 0.32720677 | 0.329 | 0.117 | 1.2437E-19 | CD-PC | COL4A4 |
| 5.7541E-26 | 0.3265531 | 0.297 | 0.094 | 1.2285E-21 | CD-PC | EXPH5 |
| 1.2016E-18 | 0.32620175 | 0.402 | 0.184 | 2.5655E-14 | CD-PC | PPP4R2 |
| 1.6888E-24 | 0.32538351 | 0.422 | 0.171 | 3.6055E-20 | CD-PC | FAM213A |
| 2.4848E-18 | 0.32537485 | 0.506 | 0.253 | 5.305E-14 | CD-PC | BRD4 |
| 3.8528E-21 | 0.32514722 | 0.446 | 0.195 | 8.2256E-17 | CD-PC | DDR1 |

| | | | | | | |
|---|---|---|---|---|---|---|
| 2.654E-35 | 0.32474826 | 0.313 | 0.086 | 5.6664E-31 | CD-PC | PTPRJ |
| 2.2837E-25 | 0.32434496 | 0.337 | 0.117 | 4.8758E-21 | CD-PC | HOXB6 |
| 3.6923E-32 | 0.32425817 | 0.241 | 0.06 | 7.8831E-28 | CD-PC | GLIS2 |
| 4.8694E-48 | 0.32405778 | 0.269 | 0.053 | 1.0396E-43 | CD-PC | RAI1 |
| 1.5103E-26 | 0.32401442 | 0.321 | 0.108 | 3.2245E-22 | CD-PC | NEK7 |
| 6.203E-27 | 0.32371152 | 0.357 | 0.126 | 1.3243E-22 | CD-PC | ADNP |
| 3.6158E-12 | 0.32264804 | 0.514 | 0.319 | 7.7197E-08 | CD-PC | COX5A |
| 9.5809E-28 | 0.32240192 | 0.317 | 0.102 | 2.0455E-23 | CD-PC | FRK |
| 2.5107E-50 | 0.32222942 | 0.265 | 0.05 | 5.3604E-46 | CD-PC | C15orf39 |
| 4.222E-27 | 0.32199877 | 0.297 | 0.094 | 9.014E-23 | CD-PC | NAPRT |
| 2.9595E-21 | 0.32197834 | 0.386 | 0.161 | 6.3186E-17 | CD-PC | IFNAR1 |
| 7.3144E-50 | 0.32179624 | 0.257 | 0.048 | 1.5616E-45 | CD-PC | PPARG |
| 1.004E-18 | 0.32179228 | 0.289 | 0.113 | 2.1435E-14 | CD-PC | PDCD4 |
| 2.1769E-22 | 0.3217421 | 0.434 | 0.185 | 4.6477E-18 | CD-PC | ARF6 |
| 3.0709E-18 | 0.32108259 | 0.438 | 0.21 | 6.5564E-14 | CD-PC | RSF1 |
| 4.393E-119 | 0.32039 | 0.205 | 0.013 | 9.378E-115 | CD-PC | ALDH8A1 |
| 4.7532E-20 | 0.31952902 | 0.992 | 0.917 | 1.0148E-15 | CD-PC | S100A11 |
| 4.5635E-16 | 0.31940441 | 0.574 | 0.321 | 9.7431E-12 | CD-PC | ENY2 |
| 4.7885E-17 | 0.31924588 | 0.518 | 0.268 | 1.0224E-12 | CD-PC | COX17 |
| 6.0773E-24 | 0.31882367 | 0.422 | 0.173 | 1.2975E-19 | CD-PC | DPY30 |
| 1.4413E-17 | 0.31841761 | 0.414 | 0.198 | 3.0772E-13 | CD-PC | SMIM14 |
| 6.6036E-59 | 0.31757024 | 0.253 | 0.04 | 1.4099E-54 | CD-PC | REPS2 |
| 1.3404E-33 | 0.31727024 | 0.293 | 0.079 | 2.8618E-29 | CD-PC | TMPRSS2 |
| 5.7276E-53 | 0.31643596 | 0.253 | 0.044 | 1.2228E-48 | CD-PC | GPRC5B |
| 5.8672E-17 | 0.31551717 | 0.386 | 0.178 | 1.2526E-12 | CD-PC | ZFHX3 |
| 2.5865E-11 | 0.31544545 | 0.514 | 0.318 | 5.5221E-07 | CD-PC | PAWR |
| 1.2957E-14 | 0.3153523 | 0.643 | 0.386 | 2.7664E-10 | CD-PC | EIF2S2 |
| 9.1243E-20 | 0.31532971 | 0.414 | 0.184 | 1.948E-15 | CD-PC | ACBD3 |
| 3.2501E-29 | 0.31520574 | 0.317 | 0.099 | 6.939E-25 | CD-PC | UBQLN2 |
| 1.9496E-18 | 0.3144741 | 0.422 | 0.196 | 4.1625E-14 | CD-PC | ARHGEF12 |
| 4.5903E-21 | 0.31432116 | 0.402 | 0.168 | 9.8002E-17 | CD-PC | TACC1 |
| 4.5188E-06 | 0.31398909 | 0.418 | 0.282 | 0.09647697 | CD-PC | AKR1B1 |
| 3.9255E-18 | 0.31397336 | 0.474 | 0.229 | 8.381E-14 | CD-PC | PHF3 |
| 2.4929E-17 | 0.31377985 | 0.353 | 0.157 | 5.3224E-13 | CD-PC | RNF187 |
| 8.9407E-36 | 0.31328549 | 0.289 | 0.075 | 1.9088E-31 | CD-PC | RDH14 |
| 2.9389E-27 | 0.31149903 | 0.325 | 0.107 | 6.2745E-23 | CD-PC | TRIM33 |
| 2.1303E-33 | 0.31139401 | 0.253 | 0.063 | 4.5482E-29 | CD-PC | RPS6KA1 |
| 8.237E-53 | 0.31129011 | 0.265 | 0.048 | 1.7586E-48 | CD-PC | NPAS2 |
| 2.1699E-31 | 0.31101263 | 0.301 | 0.086 | 4.6328E-27 | CD-PC | IRF2BPL |
| 4.82E-18 | 0.31038205 | 0.349 | 0.152 | 1.0291E-13 | CD-PC | GNS |
| 5.1901E-17 | 0.31030929 | 0.43 | 0.211 | 1.1081E-12 | CD-PC | SAT2 |
| 3.8276E-19 | 0.31023624 | 0.386 | 0.166 | 8.172E-15 | CD-PC | BICC1 |
| 5.6624E-11 | 0.30931736 | 0.566 | 0.37 | 1.2089E-06 | CD-PC | HEBP2 |
| 7.2249E-17 | 0.30930973 | 0.39 | 0.182 | 1.5425E-12 | CD-PC | MRPS5 |
| 3.2747E-90 | 0.30826436 | 0.249 | 0.026 | 6.9916E-86 | CD-PC | PRR15 |
| 4.6371E-18 | 0.30758288 | 0.309 | 0.126 | 9.9003E-14 | CD-PC | ITPR3 |
| 1.1451E-27 | 0.30740898 | 0.39 | 0.139 | 2.4448E-23 | CD-PC | GALNT3 |
| 8.0326E-15 | 0.30738635 | 0.361 | 0.172 | 1.715E-10 | CD-PC | SP100 |
| 2.8722E-19 | 0.30725139 | 0.297 | 0.112 | 6.1321E-15 | CD-PC | PAX2 |
| 2.1374E-15 | 0.30691543 | 0.514 | 0.277 | 4.5634E-11 | CD-PC | LITAF |

| | | | | | | |
|---|---|---|---|---|---|---|
| 1.6616E-29 | 0.30624729 | 0.333 | 0.105 | 3.5474E-25 | CD-PC | KAT6B |
| 1.8603E-20 | 0.30619206 | 0.341 | 0.137 | 3.9717E-16 | CD-PC | PCGF5 |
| 1.4285E-12 | 0.30554463 | 0.534 | 0.323 | 3.0499E-08 | CD-PC | TRAM1 |
| 2.5274E-43 | 0.30414771 | 0.269 | 0.058 | 5.396E-39 | CD-PC | PLEKHA6 |
| 4.6351E-31 | 0.30382224 | 0.169 | 0.032 | 9.8959E-27 | CD-PC | PIK3C2G |
| 1.2221E-14 | 0.30363477 | 0.47 | 0.245 | 2.6092E-10 | CD-PC | KANSL1 |
| 3.3734E-18 | 0.3035555 | 0.325 | 0.135 | 7.2022E-14 | CD-PC | ZBTB7A |
| 2.5855E-34 | 0.30350866 | 0.269 | 0.068 | 5.52E-30 | CD-PC | ZNF639 |
| 3.357E-29 | 0.30310896 | 0.293 | 0.087 | 7.1671E-25 | CD-PC | LACTB |
| 1.4135E-24 | 0.30285115 | 0.337 | 0.12 | 3.0179E-20 | CD-PC | KLF13 |
| 4.7374E-15 | 0.30232928 | 0.333 | 0.157 | 1.0114E-10 | CD-PC | PNKD |
| 6.1447E-55 | 0.30208051 | 0.269 | 0.048 | 1.3119E-50 | CD-PC | PWWP2B |
| 1.9924E-16 | 0.30201702 | 0.333 | 0.148 | 4.2537E-12 | CD-PC | VEZF1 |
| 1.063E-39 | 0.30191552 | 0.277 | 0.064 | 2.2695E-35 | CD-PC | THRA |
| 5.6891E-62 | 0.30156896 | 0.157 | 0.015 | 1.2146E-57 | CD-PC | PTGER1 |
| 6.8378E-14 | 0.30134165 | 0.522 | 0.294 | 1.4599E-09 | CD-PC | CAPG |
| 6.3846E-26 | 0.30127288 | 0.301 | 0.098 | 1.3631E-21 | CD-PC | NR2F6 |
| 7.7525E-26 | 0.30103308 | 0.277 | 0.086 | 1.6552E-21 | CD-PC | ARHGAP35 |
| 4.2834E-09 | 0.29966952 | 0.843 | 0.724 | 9.145E-05 | CD-PC | CSTB |
| 4.3972E-13 | 0.29965972 | 0.329 | 0.157 | 9.3879E-09 | CD-PC | TMEM176B |
| 6.6046E-33 | 0.29951705 | 0.289 | 0.079 | 1.4101E-28 | CD-PC | AP1AR |
| 9.0749E-15 | 0.29936113 | 0.55 | 0.313 | 1.9375E-10 | CD-PC | PCBP1 |
| 1.3702E-29 | 0.29926419 | 0.261 | 0.072 | 2.9254E-25 | CD-PC | AIFM2 |
| 1.7428E-16 | 0.29869537 | 0.333 | 0.145 | 3.721E-12 | CD-PC | RHOF |
| 1.5691E-31 | 0.29861316 | 0.317 | 0.093 | 3.3501E-27 | CD-PC | TGFBR2 |
| 6.5036E-17 | 0.29842103 | 0.285 | 0.116 | 1.3885E-12 | CD-PC | TTC37 |
| 2.9454E-24 | 0.29789041 | 0.341 | 0.123 | 6.2883E-20 | CD-PC | TSC22D4 |
| 4.6429E-28 | 0.29754944 | 0.361 | 0.123 | 9.9126E-24 | CD-PC | DNM2 |
| 2.8829E-22 | 0.29726378 | 0.386 | 0.154 | 6.1551E-18 | CD-PC | ECH1 |
| 1.4989E-34 | 0.29716734 | 0.273 | 0.069 | 3.2002E-30 | CD-PC | TLE2 |
| 8.229E-25 | 0.297157 | 0.341 | 0.12 | 1.7569E-20 | CD-PC | NFIB |
| 8.4065E-41 | 0.29704116 | 0.265 | 0.058 | 1.7948E-36 | CD-PC | DTNBP1 |
| 2.8118E-09 | 0.29687224 | 0.474 | 0.291 | 6.0032E-05 | CD-PC | CD55 |
| 3.8043E-12 | 0.2965665 | 0.546 | 0.332 | 8.1222E-08 | CD-PC | SYF2 |
| 7.9148E-13 | 0.29649536 | 0.454 | 0.26 | 1.6898E-08 | CD-PC | CAPZA2 |
| 3.7457E-30 | 0.29616024 | 0.257 | 0.068 | 7.997E-26 | CD-PC | PTGES |
| 3.8581E-26 | 0.2961163 | 0.297 | 0.095 | 8.2371E-22 | CD-PC | MOB1B |
| 9.8041E-26 | 0.29521935 | 0.309 | 0.101 | 2.0932E-21 | CD-PC | SIAH1 |
| 7.5598E-20 | 0.29488134 | 0.345 | 0.14 | 1.614E-15 | CD-PC | MAPK6 |
| 4.0708E-10 | 0.29473723 | 0.663 | 0.469 | 8.6912E-06 | CD-PC | SUMO2 |
| 2.21E-212 | 0.29421649 | 0.173 | 0.004 | 4.719E-208 | CD-PC | AGR3 |
| 1.3332E-18 | 0.2941062 | 0.301 | 0.119 | 2.8464E-14 | CD-PC | GSE1 |
| 8.3019E-20 | 0.29295107 | 0.39 | 0.168 | 1.7725E-15 | CD-PC | PPP1R12A |
| 2.1152E-43 | 0.29246806 | 0.189 | 0.03 | 4.5159E-39 | CD-PC | MSLN |
| 1.6316E-14 | 0.292389 | 0.361 | 0.173 | 3.4835E-10 | CD-PC | SECISBP2L |
| 3.1128E-15 | 0.29191088 | 0.321 | 0.142 | 6.6458E-11 | CD-PC | THOC2 |
| 1.0137E-34 | 0.29123394 | 0.257 | 0.062 | 2.1643E-30 | CD-PC | GATA2 |
| 8.3829E-16 | 0.2912259 | 0.394 | 0.19 | 1.7897E-11 | CD-PC | TSPAN3 |
| 1.2887E-33 | 0.29057752 | 0.301 | 0.083 | 2.7513E-29 | CD-PC | HINT3 |
| 9.8476E-23 | 0.29040422 | 0.349 | 0.132 | 2.1025E-18 | CD-PC | COMTD1 |

| | | | | | | |
|---|---|---|---|---|---|---|
| 3.0624E-15 | 0.28954252 | 0.305 | 0.136 | 6.5382E-11 | CD-PC | RAB25 |
| 2.2757E-17 | 0.28953682 | 0.402 | 0.188 | 4.8586E-13 | CD-PC | RAD23B |
| 1.9902E-20 | 0.28950917 | 0.349 | 0.141 | 4.2491E-16 | CD-PC | STX7 |
| 1.6692E-44 | 0.28906855 | 0.281 | 0.06 | 3.5638E-40 | CD-PC | TC2N |
| 1.0751E-24 | 0.28823577 | 0.293 | 0.096 | 2.2953E-20 | CD-PC | DMXL1 |
| 2.5748E-15 | 0.28822263 | 0.502 | 0.271 | 5.4972E-11 | CD-PC | ZNF706 |
| 2.7537E-18 | 0.28814985 | 0.454 | 0.216 | 5.8793E-14 | CD-PC | SSU72 |
| 2.1197E-29 | 0.28810546 | 0.297 | 0.088 | 4.5255E-25 | CD-PC | MARK2 |
| 9.3142E-13 | 0.28805766 | 0.446 | 0.249 | 1.9886E-08 | CD-PC | C4orf3 |
| 1.5684E-21 | 0.28786702 | 0.289 | 0.102 | 3.3485E-17 | CD-PC | MORC3 |
| 1.5785E-15 | 0.28750282 | 0.309 | 0.134 | 3.3701E-11 | CD-PC | LRBA |
| 8.8031E-26 | 0.28676086 | 0.269 | 0.082 | 1.8795E-21 | CD-PC | GNPTAB |
| 1.0008E-29 | 0.28655268 | 0.281 | 0.08 | 2.1368E-25 | CD-PC | FOXK1 |
| 3.2963E-21 | 0.28655268 | 0.365 | 0.145 | 7.0375E-17 | CD-PC | ZBTB44 |
| 1.1815E-12 | 0.28628252 | 0.325 | 0.153 | 2.5224E-08 | CD-PC | SLC20A1 |
| 1.4864E-18 | 0.28622475 | 0.373 | 0.161 | 3.1736E-14 | CD-PC | SREK1IP1 |
| 2.242E-16 | 0.28614214 | 0.39 | 0.183 | 4.7867E-12 | CD-PC | PRRC2B |
| 1.1938E-37 | 0.2852657 | 0.261 | 0.06 | 2.5488E-33 | CD-PC | ETV3 |
| 1.0643E-17 | 0.28521623 | 0.281 | 0.108 | 2.2723E-13 | CD-PC | ZMAT3 |
| 2.3088E-31 | 0.28474857 | 0.309 | 0.09 | 4.9293E-27 | CD-PC | PLEKHF2 |
| 1.6964E-32 | 0.28414796 | 0.241 | 0.058 | 3.6217E-28 | CD-PC | DCPS |
| 8.3838E-12 | 0.28411114 | 0.321 | 0.162 | 1.7899E-07 | CD-PC | SEMA3B |
| 3.7526E-22 | 0.28302 | 0.317 | 0.117 | 8.0119E-18 | CD-PC | RAB4A |
| 1.9449E-40 | 0.28290655 | 0.233 | 0.047 | 4.1524E-36 | CD-PC | CDK18 |
| 5.3553E-15 | 0.28262745 | 0.414 | 0.208 | 1.1434E-10 | CD-PC | PUM1 |
| 4.1869E-34 | 0.28247475 | 0.289 | 0.076 | 8.9391E-30 | CD-PC | TFAP2A |
| 1.9116E-16 | 0.2823436 | 0.353 | 0.158 | 4.0813E-12 | CD-PC | CTSA |
| 1.557E-17 | 0.28217085 | 0.474 | 0.233 | 3.3242E-13 | CD-PC | WASL |
| 9.1352E-18 | 0.2818652 | 0.297 | 0.119 | 1.9504E-13 | CD-PC | PUM2 |
| 1.0278E-18 | 0.28182896 | 0.309 | 0.124 | 2.1943E-14 | CD-PC | CASP4 |
| 4.8476E-17 | 0.28151513 | 0.373 | 0.168 | 1.035E-12 | CD-PC | ERBIN |
| 2.0036E-76 | 0.28114148 | 0.225 | 0.025 | 4.2777E-72 | CD-PC | LYPD6B |
| 8.2613E-17 | 0.28095649 | 0.382 | 0.175 | 1.7638E-12 | CD-PC | MIER1 |
| 9.7138E-15 | 0.28034774 | 0.313 | 0.142 | 2.0739E-10 | CD-PC | EFCAB14 |
| 1.4153E-22 | 0.28029235 | 0.281 | 0.095 | 3.0217E-18 | CD-PC | AKT2 |
| 2.623E-43 | 0.28027476 | 0.273 | 0.058 | 5.6001E-39 | CD-PC | SHMT1 |
| 2.7133E-20 | 0.27986029 | 0.345 | 0.139 | 5.7929E-16 | CD-PC | PAFAH1B2 |
| 1.1346E-18 | 0.27965383 | 0.382 | 0.163 | 2.4225E-14 | CD-PC | ANKRD13A |
| 5.1987E-15 | 0.27943202 | 0.458 | 0.231 | 1.1099E-10 | CD-PC | CHD9 |
| 1.0814E-27 | 0.27890451 | 0.301 | 0.093 | 2.3088E-23 | CD-PC | FBXO34 |
| 1.1732E-29 | 0.27870204 | 0.265 | 0.073 | 2.5049E-25 | CD-PC | KIZ |
| 3.8046E-20 | 0.27689601 | 0.349 | 0.14 | 8.1228E-16 | CD-PC | TLK1 |
| 9.7346E-21 | 0.27662537 | 0.289 | 0.104 | 2.0783E-16 | CD-PC | TNKS1BP1 |
| 6.2467E-29 | 0.27594537 | 0.217 | 0.053 | 1.3337E-24 | CD-PC | OSBPL11 |
| 1.1989E-16 | 0.27564631 | 0.325 | 0.139 | 2.5597E-12 | CD-PC | ARG2 |
| 1.0398E-26 | 0.27538522 | 0.305 | 0.097 | 2.2199E-22 | CD-PC | SLC2A4RG |
| 8.3929E-26 | 0.27522008 | 0.265 | 0.08 | 1.7919E-21 | CD-PC | NCOA2 |
| 3.5274E-16 | 0.27480059 | 0.321 | 0.139 | 7.531E-12 | CD-PC | AGPAT2 |
| 5.9216E-16 | 0.27469919 | 0.406 | 0.194 | 1.2643E-11 | CD-PC | MINDY2 |
| 1.7059E-14 | 0.27466044 | 0.369 | 0.179 | 3.6422E-10 | CD-PC | CELF1 |

| | | | | | | |
|---|---|---|---|---|---|---|
| 4.4881E-23 | 0.274577 | 0.269 | 0.088 | 9.5821E-19 | CD-PC | ISCA2 |
| 8.0458E-18 | 0.27429276 | 0.434 | 0.202 | 1.7178E-13 | CD-PC | SINHCAF |
| 2.2765E-14 | 0.27416759 | 0.526 | 0.289 | 4.8603E-10 | CD-PC | NOL7 |
| 2.3184E-13 | 0.27404235 | 0.47 | 0.252 | 4.9499E-09 | CD-PC | MPHOSPH8 |
| 5.6753E-34 | 0.27393264 | 0.261 | 0.064 | 1.2117E-29 | CD-PC | FLCN |
| 5.2561E-14 | 0.27381146 | 0.317 | 0.145 | 1.1222E-09 | CD-PC | HCFC1R1 |
| 7.429E-36 | 0.27323946 | 0.245 | 0.056 | 1.5861E-31 | CD-PC | CELSR2 |
| 2.764E-20 | 0.27300893 | 0.293 | 0.107 | 5.9011E-16 | CD-PC | ALDH1A1 |
| 1.4882E-12 | 0.27277075 | 0.478 | 0.275 | 3.1773E-08 | CD-PC | SEPT7 |
| 2.4273E-34 | 0.2724295 | 0.277 | 0.07 | 5.1824E-30 | CD-PC | MAP3K21 |
| 1.2656E-14 | 0.27208867 | 0.406 | 0.201 | 2.7021E-10 | CD-PC | REST |
| 1.1612E-19 | 0.27164232 | 0.337 | 0.134 | 2.4792E-15 | CD-PC | ALDH2 |
| 1.7581E-11 | 0.27148656 | 0.249 | 0.112 | 3.7536E-07 | CD-PC | RARRES3 |
| 2.6307E-17 | 0.27144163 | 0.301 | 0.121 | 5.6166E-13 | CD-PC | PKP4 |
| 1.2571E-30 | 0.27138358 | 0.277 | 0.077 | 2.684E-26 | CD-PC | UBAC2 |
| 3.9294E-17 | 0.2706717 | 0.357 | 0.155 | 8.3893E-13 | CD-PC | NUMA1 |
| 1.2302E-12 | 0.27032019 | 0.606 | 0.374 | 2.6264E-08 | CD-PC | REEP5 |
| 6.7738E-11 | 0.27014018 | 0.382 | 0.208 | 1.4462E-06 | CD-PC | GFPT1 |
| 2.0638E-50 | 0.2699333 | 0.225 | 0.037 | 4.4062E-46 | CD-PC | ESRP2 |
| 1.0812E-20 | 0.26977844 | 0.289 | 0.103 | 2.3083E-16 | CD-PC | FBXL5 |
| 8.7138E-23 | 0.2695778 | 0.305 | 0.107 | 1.8604E-18 | CD-PC | ELF2 |
| 6.7557E-09 | 0.2695037 | 0.691 | 0.515 | 0.00014423 | CD-PC | OST4 |
| 4.2718E-13 | 0.26890538 | 0.357 | 0.179 | 9.1202E-09 | CD-PC | HACD3 |
| 3.9771E-15 | 0.26884322 | 0.45 | 0.231 | 8.491E-11 | CD-PC | ARPC5L |
| 7.851E-15 | 0.26879502 | 0.333 | 0.153 | 1.6762E-10 | CD-PC | DENR |
| 3.6728E-12 | 0.26869749 | 0.57 | 0.341 | 7.8414E-08 | CD-PC | MTDH |
| 4.774E-18 | 0.26864039 | 0.418 | 0.191 | 1.0193E-13 | CD-PC | SLC25A36 |
| 3.0093E-33 | 0.26822139 | 0.281 | 0.074 | 6.4249E-29 | CD-PC | CDC42EP4 |
| 6.7546E-34 | 0.26788855 | 0.257 | 0.063 | 1.4421E-29 | CD-PC | RFX1 |
| 7.9732E-14 | 0.26785205 | 0.329 | 0.155 | 1.7023E-09 | CD-PC | ASAP2 |
| 1.0333E-15 | 0.26783666 | 0.398 | 0.19 | 2.206E-11 | CD-PC | ARFGEF1 |
| 7.232E-17 | 0.26776143 | 0.365 | 0.165 | 1.544E-12 | CD-PC | VGLL4 |
| 5.1223E-31 | 0.26736172 | 0.253 | 0.065 | 1.0936E-26 | CD-PC | FOXN3 |
| 1.4497E-18 | 0.26734621 | 0.253 | 0.09 | 3.0951E-14 | CD-PC | NR1D1 |
| 7.0667E-20 | 0.2672254 | 0.269 | 0.096 | 1.5087E-15 | CD-PC | PIAS1 |
| 1.0543E-19 | 0.26721516 | 0.265 | 0.091 | 2.2509E-15 | CD-PC | FOXQ1 |
| 2.8068E-25 | 0.26696553 | 0.257 | 0.077 | 5.9926E-21 | CD-PC | SCAMP1 |
| 4.8533E-43 | 0.26680017 | 0.245 | 0.048 | 1.0362E-38 | CD-PC | NFIA |
| 2.699E-16 | 0.26674497 | 0.502 | 0.261 | 5.7624E-12 | CD-PC | SYPL1 |
| 4.8538E-14 | 0.26652243 | 0.446 | 0.235 | 1.0363E-09 | CD-PC | MBD2 |
| 2.3217E-18 | 0.26579412 | 0.301 | 0.118 | 4.9568E-14 | CD-PC | MAP4K5 |
| 2.7211E-17 | 0.2657659 | 0.325 | 0.136 | 5.8095E-13 | CD-PC | VEZT |
| 1.7686E-20 | 0.26555238 | 0.297 | 0.109 | 3.776E-16 | CD-PC | LAD1 |
| 3.6621E-28 | 0.26512028 | 0.249 | 0.068 | 7.8186E-24 | CD-PC | RANBP9 |
| 7.2562E-16 | 0.26470842 | 0.305 | 0.13 | 1.5492E-11 | CD-PC | ZMYM2 |
| 1.2638E-19 | 0.26467533 | 0.301 | 0.113 | 2.6982E-15 | CD-PC | TMEM51 |
| 1.0331E-23 | 0.26450886 | 0.273 | 0.088 | 2.2057E-19 | CD-PC | RREB1 |
| 3.0782E-18 | 0.26444615 | 0.309 | 0.122 | 6.5719E-14 | CD-PC | GNPDA1 |
| 2.0998E-08 | 0.26357826 | 0.562 | 0.365 | 0.00044831 | CD-PC | DDIT3 |
| 6.0933E-33 | 0.26321337 | 0.281 | 0.073 | 1.3009E-28 | CD-PC | CSAD |

| | | | | | | |
|---|---|---|---|---|---|---|
| 4.4696E-69 | 0.26298157 | 0.181 | 0.018 | 9.5426E-65 | CD-PC | KSR2 |
| 1.6718E-14 | 0.26291354 | 0.341 | 0.16 | 3.5692E-10 | CD-PC | CXADR |
| 7.426E-24 | 0.26275156 | 0.273 | 0.088 | 1.5854E-19 | CD-PC | OSGIN2 |
| 1.9868E-15 | 0.26250648 | 0.43 | 0.214 | 4.2418E-11 | CD-PC | CCPG1 |
| 2.3846E-16 | 0.26146812 | 0.378 | 0.174 | 5.0911E-12 | CD-PC | RYBP |
| 2.4648E-28 | 0.2614177 | 0.269 | 0.076 | 5.2624E-24 | CD-PC | RAB17 |
| 1.3302E-15 | 0.26127508 | 0.253 | 0.1 | 2.8399E-11 | CD-PC | FUCA1 |
| 1.7966E-12 | 0.26113855 | 0.51 | 0.296 | 3.8357E-08 | CD-PC | NDFIP1 |
| 3.0009E-11 | 0.26099259 | 0.522 | 0.308 | 6.407E-07 | CD-PC | WSB1 |
| 6.4696E-19 | 0.26083785 | 0.297 | 0.113 | 1.3813E-14 | CD-PC | PISD |
| 1.1162E-14 | 0.26059637 | 0.369 | 0.178 | 2.3831E-10 | CD-PC | CAPN1 |
| 2.8423E-16 | 0.26023849 | 0.361 | 0.162 | 6.0684E-12 | CD-PC | SNRPN |
| 1.8952E-16 | 0.25969676 | 0.241 | 0.089 | 4.0463E-12 | CD-PC | NBEAL2 |
| 6.6879E-15 | 0.25920699 | 0.281 | 0.12 | 1.4279E-10 | CD-PC | BCAP29 |
| 2.2179E-22 | 0.25914762 | 0.309 | 0.11 | 4.7353E-18 | CD-PC | IFNGR1 |
| 3.9673E-27 | 0.2587932 | 0.241 | 0.066 | 8.4702E-23 | CD-PC | ZNF768 |
| 1.684E-13 | 0.25875155 | 0.285 | 0.128 | 3.5954E-09 | CD-PC | C7orf50 |
| 2.2211E-54 | 0.25871847 | 0.213 | 0.031 | 4.7421E-50 | CD-PC | LIN7A |
| 1.4689E-27 | 0.25783442 | 0.241 | 0.065 | 3.1362E-23 | CD-PC | BAG4 |
| 4.5768E-12 | 0.25768671 | 0.349 | 0.18 | 9.7716E-08 | CD-PC | KLF3 |
| 1.3157E-10 | 0.25704311 | 0.566 | 0.358 | 2.8089E-06 | CD-PC | CSDE1 |
| 5.0169E-26 | 0.25669155 | 0.217 | 0.057 | 1.0711E-21 | CD-PC | MATR3.1 |
| 7.5025E-16 | 0.2566158 | 0.253 | 0.097 | 1.6018E-11 | CD-PC | DCAF5 |
| 7.7448E-16 | 0.25647496 | 0.305 | 0.128 | 1.6535E-11 | CD-PC | HDAC7 |
| 3.0555E-11 | 0.25639538 | 0.534 | 0.311 | 6.5236E-07 | CD-PC | MAL |
| 8.1442E-11 | 0.25637367 | 0.365 | 0.197 | 1.7388E-06 | CD-PC | FOXO3 |
| 2.2124E-21 | 0.25626789 | 0.253 | 0.083 | 4.7236E-17 | CD-PC | SLC35A3 |
| 1.4552E-32 | 0.25594219 | 0.245 | 0.059 | 3.1069E-28 | CD-PC | PLLP |
| 2.9798E-21 | 0.25588248 | 0.305 | 0.111 | 6.362E-17 | CD-PC | COMMD2 |
| 1.1262E-25 | 0.25577314 | 0.205 | 0.053 | 2.4044E-21 | CD-PC | BCAS4 |
| 4.7431E-11 | 0.25554819 | 0.574 | 0.358 | 1.0126E-06 | CD-PC | AKAP9 |
| 1.0863E-12 | 0.25552868 | 0.394 | 0.204 | 2.3193E-08 | CD-PC | TMEM30A |
| 7.1193E-07 | 0.25544282 | 0.43 | 0.278 | 0.01519967 | CD-PC | CLDN3 |
| 2.5133E-24 | 0.25512696 | 0.289 | 0.094 | 5.366E-20 | CD-PC | PPP4R1 |
| 8.3609E-22 | 0.2550242 | 0.281 | 0.097 | 1.7851E-17 | CD-PC | ZDHHC3 |
| 6.5471E-83 | 0.25501399 | 0.181 | 0.015 | 1.3978E-78 | CD-PC | OTULINL |
| 3.6878E-19 | 0.25495085 | 0.321 | 0.126 | 7.8734E-15 | CD-PC | PHAX |
| 2.568E-34 | 0.25487233 | 0.241 | 0.055 | 5.4828E-30 | CD-PC | GALM |
| 1.3289E-31 | 0.25459677 | 0.201 | 0.044 | 2.8373E-27 | CD-PC | DOC2B |
| 8.6319E-25 | 0.25420238 | 0.277 | 0.087 | 1.8429E-20 | CD-PC | SCPEP1 |
| 1.5383E-15 | 0.25411005 | 0.305 | 0.131 | 3.2842E-11 | CD-PC | FCHO2 |
| 3.0703E-31 | 0.25405298 | 0.265 | 0.07 | 6.555E-27 | CD-PC | NBN |
| 1.9665E-15 | 0.2540176 | 0.386 | 0.184 | 4.1985E-11 | CD-PC | CNDP2 |
| 9.6833E-20 | 0.25389432 | 0.265 | 0.093 | 2.0674E-15 | CD-PC | SIGIRR |
| 2.4063E-19 | 0.25377359 | 0.233 | 0.077 | 5.1374E-15 | CD-PC | AKAP11 |
| 8.6667E-15 | 0.25343437 | 0.333 | 0.151 | 1.8503E-10 | CD-PC | CFAP97 |
| 2.3591E-16 | 0.25310534 | 0.269 | 0.106 | 5.0367E-12 | CD-PC | RORA |
| 3.9842E-29 | 0.25308926 | 0.273 | 0.076 | 8.5063E-25 | CD-PC | ERMP1 |
| 2.8288E-16 | 0.2528446 | 0.309 | 0.131 | 6.0395E-12 | CD-PC | UBE2V2 |
| 8.659E-13 | 0.25278041 | 0.373 | 0.19 | 1.8487E-08 | CD-PC | RHOBTB3 |

| | | | | | | |
|---|---|---|---|---|---|---|
| 9.0291E-32 | 0.25273932 | 0.277 | 0.072 | 1.9277E-27 | CD-PC | PGGHG |
| 2.8912E-26 | 0.25256679 | 0.241 | 0.066 | 6.1727E-22 | CD-PC | SRGAP3 |
| 8.9163E-19 | 0.25255587 | 0.281 | 0.105 | 1.9036E-14 | CD-PC | OGDH |
| 1.3995E-17 | 0.25248888 | 0.325 | 0.134 | 2.9878E-13 | CD-PC | ATF7IP |
| 4.2076E-20 | 0.25230101 | 0.309 | 0.116 | 8.9833E-16 | CD-PC | WWP1 |
| 4.9579E-11 | 0.25228012 | 0.462 | 0.264 | 1.0585E-06 | CD-PC | MET |
| 9.1845E-22 | 0.25220117 | 0.309 | 0.109 | 1.9609E-17 | CD-PC | NECTIN4 |
| 2.0413E-13 | 0.25215389 | 0.337 | 0.162 | 4.3582E-09 | CD-PC | IST1 |
| 1.7553E-09 | 0.25194569 | 0.498 | 0.306 | 3.7476E-05 | CD-PC | ISCU |
| 6.6387E-15 | 0.25177809 | 0.345 | 0.16 | 1.4174E-10 | CD-PC | RANBP2 |
| 6.8535E-09 | 0.25171882 | 0.349 | 0.201 | 0.00014632 | CD-PC | ERO1A |
| 2.9063E-15 | 0.25162216 | 0.414 | 0.204 | 6.205E-11 | CD-PC | SCP2 |
| 1.3715E-17 | 0.25155044 | 0.285 | 0.111 | 2.9281E-13 | CD-PC | INSR |
| 4.6826E-19 | 0.2514769 | 0.257 | 0.091 | 9.9974E-15 | CD-PC | REPIN1 |
| 1.0491E-17 | 0.25147246 | 0.229 | 0.079 | 2.2397E-13 | CD-PC | YOD1 |
| 2.7309E-41 | 0.25132542 | 0.193 | 0.033 | 5.8305E-37 | CD-PC | GATA3-AS1 |
| 1.2618E-26 | 0.25124676 | 0.245 | 0.069 | 2.694E-22 | CD-PC | SLC29A1 |
| 3.8058E-14 | 0.25111241 | 0.369 | 0.18 | 8.1254E-10 | CD-PC | FNIP1 |
| 6.3431E-14 | 0.25108251 | 0.337 | 0.158 | 1.3542E-09 | CD-PC | PLS3 |
| 1.8703E-14 | 0.2510175 | 0.285 | 0.123 | 3.9931E-10 | CD-PC | INO80D |
| 5.1275E-09 | 0.25100407 | 0.438 | 0.271 | 0.00010947 | CD-PC | VPS28 |
| 2.0052E-22 | 0.25095979 | 0.261 | 0.084 | 4.281E-18 | CD-PC | HMGXB4 |
| 1.1547E-15 | 0.25086296 | 0.438 | 0.214 | 2.4654E-11 | CD-PC | CCDC186 |
| 1.4354E-15 | 0.25079151 | 0.317 | 0.139 | 3.0646E-11 | CD-PC | EEF1E1 |
| 1.7314E-14 | 0.2502846 | 0.333 | 0.154 | 3.6964E-10 | CD-PC | KRAS |
| 6.5609E-20 | 0.2502613 | 0.309 | 0.116 | 1.4008E-15 | CD-PC | GAK |
| 2.0577E-40 | 0.25001278 | 0.217 | 0.041 | 4.3931E-36 | CD-PC | GRHL1 |
| 0.00244573 | -0.2515441 | 0.382 | 0.452 | 1 | CD-PC | RPS10 |
| 0.00173997 | -0.2523112 | 0.189 | 0.262 | 1 | CD-PC | GSTO1 |
| 0.00058528 | -0.2524699 | 0.333 | 0.427 | 1 | CD-PC | SSR4 |
| 0.00261324 | -0.2528684 | 0.221 | 0.299 | 1 | CD-PC | HSPH1 |
| 1.8718E-06 | -0.2537292 | 0.08 | 0.2 | 0.03996344 | CD-PC | MAP2K3 |
| 0.00200091 | -0.2540589 | 0.679 | 0.776 | 1 | CD-PC | MYL12A |
| 0.00115436 | -0.254758 | 0.892 | 0.942 | 1 | CD-PC | RPL19 |
| 0.00071523 | -0.2552621 | 0.108 | 0.191 | 1 | CD-PC | THBS1 |
| 6.4688E-08 | -0.2585746 | 0.016 | 0.134 | 0.00138108 | CD-PC | COL4A1 |
| 0.00961015 | -0.2590953 | 0.157 | 0.216 | 1 | CD-PC | ZFP36 |
| 0.00102131 | -0.2635314 | 0.51 | 0.57 | 1 | CD-PC | RAN |
| 7.8047E-08 | -0.2645164 | 0.04 | 0.167 | 0.0016663 | CD-PC | CAVIN1 |
| 9.8045E-05 | -0.2648803 | 0.189 | 0.291 | 1 | CD-PC | TJP1 |
| 5.5813E-05 | -0.2671381 | 0.108 | 0.208 | 1 | CD-PC | THUMPD3-AS1 |
| 5.173E-08 | -0.2675552 | 0.016 | 0.133 | 0.00110444 | CD-PC | SLC6A8 |
| 4.6272E-08 | -0.2702491 | 0.032 | 0.157 | 0.0009879 | CD-PC | CPM |
| 0.00049895 | -0.2713466 | 0.281 | 0.372 | 1 | CD-PC | TAGLN2 |
| 2.4085E-09 | -0.2716182 | 0.032 | 0.176 | 5.1422E-05 | CD-PC | DSG2 |
| 5.579E-09 | -0.2726792 | 0.016 | 0.147 | 0.00011911 | CD-PC | COL4A2 |
| 0.00527213 | -0.272719 | 0.072 | 0.127 | 1 | CD-PC | HIST2H2BF |
| 0.00077148 | -0.2776256 | 0.221 | 0.302 | 1 | CD-PC | MYDGF |
| 6.6485E-06 | -0.2776896 | 0.916 | 0.968 | 0.14194457 | CD-PC | RPL41 |
| 8.8044E-05 | -0.277809 | 0.052 | 0.135 | 1 | CD-PC | CSRNP2 |

| | | | | | | |
|---|---|---|---|---|---|---|
| 8.2717E-08 | -0.2778472 | 0.008 | 0.118 | 0.00176601 | CD-PC | FSCN1 |
| 0.00012948 | -0.279772 | 0.173 | 0.272 | 1 | CD-PC | RBM3 |
| 1.5226E-05 | -0.280329 | 0.024 | 0.109 | 0.32506696 | CD-PC | AL355075.4 |
| 2.4021E-14 | -0.2809418 | 0.333 | 0.619 | 5.1284E-10 | CD-PC | MT-ND4L |
| 8.9568E-06 | -0.2811388 | 0.064 | 0.169 | 0.19122851 | CD-PC | TSPYL2 |
| 0.00021929 | -0.2814077 | 0.129 | 0.22 | 1 | CD-PC | SERTAD1 |
| 0.00148891 | -0.2817275 | 0.197 | 0.28 | 1 | CD-PC | SERPINB1 |
| 0.00186185 | -0.2832062 | 0.494 | 0.566 | 1 | CD-PC | CST3 |
| 0.0017309 | -0.2840742 | 0.261 | 0.332 | 1 | CD-PC | TUBA1C |
| 7.3927E-09 | -0.2842434 | 0.008 | 0.134 | 0.00015783 | CD-PC | CSRP2 |
| 8.6793E-07 | -0.2869444 | 0.08 | 0.205 | 0.01853033 | CD-PC | DPP4 |
| 5.9547E-07 | -0.2870584 | 0.47 | 0.606 | 0.01271318 | CD-PC | CLTA |
| 1.194E-05 | -0.2884713 | 0.048 | 0.146 | 0.25491219 | CD-PC | CKS2 |
| 1.4521E-07 | -0.2887676 | 0 | 0.1 | 0.00310028 | CD-PC | MAFA |
| 0.00901249 | -0.2901414 | 0.671 | 0.728 | 1 | CD-PC | KRT8 |
| 1.0535E-06 | -0.2909769 | 0.815 | 0.912 | 0.02249135 | CD-PC | FAU |
| 0.00380713 | -0.2911955 | 0.189 | 0.257 | 1 | CD-PC | S100A14 |
| 0.00565742 | -0.2913344 | 0.08 | 0.136 | 1 | CD-PC | AC025164.1 |
| 2.2579E-06 | -0.2916697 | 0.088 | 0.205 | 0.04820535 | CD-PC | ZFC3H1 |
| 9.8274E-05 | -0.2927077 | 0.835 | 0.872 | 1 | CD-PC | SAT1 |
| 0.00033146 | -0.2939927 | 0.032 | 0.101 | 1 | CD-PC | CXCL3 |
| 0.00493572 | -0.2944379 | 0.056 | 0.113 | 1 | CD-PC | C15orf48 |
| 8.4564E-07 | -0.2965734 | 0.651 | 0.784 | 0.01805441 | CD-PC | COX4I1 |
| 1.4958E-05 | -0.2977046 | 0.349 | 0.473 | 0.31934693 | CD-PC | PRDX6 |
| 1.5702E-06 | -0.299015 | 0.04 | 0.149 | 0.03352369 | CD-PC | AL118516.1 |
| 1.4919E-07 | -0.3007568 | 0.032 | 0.151 | 0.00318526 | CD-PC | HBEGF |
| 7.8513E-08 | -0.3009707 | 0.052 | 0.185 | 0.00167625 | CD-PC | CD40 |
| 0.00319019 | -0.3050189 | 0.133 | 0.207 | 1 | CD-PC | RRAD |
| 8.1446E-06 | -0.3068206 | 0.076 | 0.182 | 0.17388685 | CD-PC | DCBLD2 |
| 9.0645E-05 | -0.3091663 | 0.578 | 0.631 | 1 | CD-PC | PABPC1 |
| 3.7881E-06 | -0.3098677 | 0.265 | 0.394 | 0.08087593 | CD-PC | CAPN2 |
| 3.8835E-05 | -0.3142868 | 0.104 | 0.203 | 0.82911891 | CD-PC | CHCHD7 |
| 5.1633E-05 | -0.3149044 | 0.631 | 0.717 | 1 | CD-PC | TXN |
| 2.3939E-05 | -0.3151381 | 0.317 | 0.43 | 0.51110358 | CD-PC | PDIA3 |
| 0.00015646 | -0.3155113 | 0.9 | 0.934 | 1 | CD-PC | RPS27A |
| 6.2531E-06 | -0.3155438 | 0.1 | 0.224 | 0.13350268 | CD-PC | SFN |
| 4.8924E-07 | -0.3162851 | 0.494 | 0.638 | 0.01044524 | CD-PC | CCNI |
| 0.00053465 | -0.319474 | 0.9 | 0.938 | 1 | CD-PC | RPS23 |
| 6.2523E-05 | -0.3199853 | 0.1 | 0.199 | 1 | CD-PC | MT-ND6 |
| 3.704E-05 | -0.3205925 | 0.084 | 0.18 | 0.79081018 | CD-PC | KLF4 |
| 0.00024004 | -0.3256445 | 0.032 | 0.101 | 1 | CD-PC | AP001160.1 |
| 3.713E-16 | -0.3262143 | 0.699 | 0.878 | 7.9274E-12 | CD-PC | HLA-A |
| 0.00430566 | -0.3277424 | 0.333 | 0.385 | 1 | CD-PC | MYH9 |
| 3.406E-05 | -0.3285198 | 0.606 | 0.702 | 0.72718933 | CD-PC | MAP1LC3B |
| 8.5065E-06 | -0.3289551 | 0.024 | 0.112 | 0.18161426 | CD-PC | AC023157.3 |
| 1.0504E-05 | -0.328958 | 0.446 | 0.564 | 0.22425234 | CD-PC | GUK1 |
| 2.9881E-06 | -0.3299584 | 0.406 | 0.518 | 0.06379674 | CD-PC | TPM3 |
| 0.00027726 | -0.3299938 | 0.803 | 0.838 | 1 | CD-PC | PRDX1 |
| 0.0003538 | -0.3329419 | 0.325 | 0.408 | 1 | CD-PC | SDC4 |
| 1.5359E-06 | -0.3359997 | 0.145 | 0.289 | 0.0327914 | CD-PC | PLAUR |

| | | | | | | |
|---|---|---|---|---|---|---|
| 0.00064334 | -0.3374534 | 0.217 | 0.302 | 1 | CD-PC | DDIT4 |
| 0.00068664 | -0.3382036 | 0.205 | 0.29 | 1 | CD-PC | STMN1 |
| 1.6267E-08 | -0.3384079 | 0.659 | 0.776 | 0.0003473 | CD-PC | EEF2 |
| 0.00015349 | -0.3445517 | 0.916 | 0.932 | 1 | CD-PC | RPS6 |
| 0.00072716 | -0.3451809 | 0.871 | 0.9 | 1 | CD-PC | SERF2 |
| 2.2521E-06 | -0.3465344 | 0.847 | 0.911 | 0.04808189 | CD-PC | RPL24 |
| 7.2932E-07 | -0.3491552 | 0.169 | 0.305 | 0.01557109 | CD-PC | EIF4A1 |
| 2.0852E-09 | -0.3502157 | 0.506 | 0.655 | 4.4519E-05 | CD-PC | SLC25A3 |
| 1.8806E-05 | -0.3512777 | 0.108 | 0.211 | 0.40151753 | CD-PC | MAT2A |
| 1.4805E-07 | -0.3513605 | 0.145 | 0.291 | 0.00316096 | CD-PC | CDK2AP2 |
| 1.0944E-06 | -0.3561319 | 0.831 | 0.898 | 0.02336459 | CD-PC | RPL35A |
| 1.2832E-07 | -0.3563459 | 0.867 | 0.933 | 0.00273956 | CD-PC | RPL18 |
| 1.7024E-08 | -0.3570452 | 0.1 | 0.252 | 0.00036345 | CD-PC | MGLL |
| 8.6988E-06 | -0.3587835 | 0.892 | 0.922 | 0.18572042 | CD-PC | RPL14 |
| 8.8348E-06 | -0.3592081 | 0.02 | 0.106 | 0.18862335 | CD-PC | ARC |
| 3.8586E-07 | -0.3601009 | 0.205 | 0.353 | 0.00823801 | CD-PC | SELENOK |
| 0.00047204 | -0.3602238 | 0.213 | 0.304 | 1 | CD-PC | DNAJB1 |
| 2.1924E-05 | -0.3605225 | 0.056 | 0.149 | 0.46807889 | CD-PC | MYL9 |
| 5.6772E-07 | -0.3615209 | 0.542 | 0.696 | 0.0121208 | CD-PC | ZFAS1 |
| 1.6214E-05 | -0.3618322 | 0.249 | 0.361 | 0.34616901 | CD-PC | SPAG9 |
| 3.515E-06 | -0.361937 | 0.249 | 0.386 | 0.07504468 | CD-PC | PTMS |
| 1.3281E-08 | -0.3653836 | 0.908 | 0.966 | 0.00028355 | CD-PC | RPS19 |
| 0.00021391 | -0.3655146 | 0.293 | 0.394 | 1 | CD-PC | TXNRD1 |
| 0.00501881 | -0.3656051 | 0.157 | 0.219 | 1 | CD-PC | TNFAIP2 |
| 5.9177E-11 | -0.3656609 | 0.016 | 0.174 | 1.2634E-06 | CD-PC | EMP3 |
| 3.2327E-09 | -0.3677667 | 0.807 | 0.898 | 6.9017E-05 | CD-PC | RPS13 |
| 1.3995E-05 | -0.3710035 | 0.779 | 0.895 | 0.29879981 | CD-PC | GAPDH |
| 9.4832E-08 | -0.3715696 | 0.747 | 0.874 | 0.00202467 | CD-PC | RACK1 |
| 1.4673E-07 | -0.3726307 | 0.94 | 0.935 | 0.0031326 | CD-PC | PTMA |
| 7.0953E-08 | -0.3757858 | 0.671 | 0.807 | 0.00151484 | CD-PC | RPL31 |
| 1.5327E-09 | -0.3770453 | 0.514 | 0.684 | 3.2724E-05 | CD-PC | CLIC1 |
| 8.2865E-08 | -0.377743 | 0.1 | 0.244 | 0.00176916 | CD-PC | HES4 |
| 0.00195888 | -0.378313 | 0.699 | 0.732 | 1 | CD-PC | PFN1 |
| 3.1636E-07 | -0.3803346 | 0.345 | 0.494 | 0.00675429 | CD-PC | DNAJA1 |
| 1.837E-07 | -0.380907 | 0.863 | 0.931 | 0.00392191 | CD-PC | RPS14 |
| 1.1714E-07 | -0.3815587 | 0.112 | 0.256 | 0.00250095 | CD-PC | IRF1 |
| 2.3123E-07 | -0.3841609 | 0.871 | 0.931 | 0.00493672 | CD-PC | RPL7A |
| 9.2349E-06 | -0.3848665 | 0.124 | 0.248 | 0.19716596 | CD-PC | PDZK1IP1 |
| 4.9437E-09 | -0.3878971 | 0.683 | 0.802 | 0.00010555 | CD-PC | RPL27 |
| 1.5125E-09 | -0.3917286 | 0 | 0.129 | 3.2291E-05 | CD-PC | PCDH9 |
| 0.00407395 | -0.3924419 | 0.55 | 0.625 | 1 | CD-PC | PPP1R15A |
| 4.7451E-08 | -0.3928807 | 0.98 | 0.988 | 0.00101308 | CD-PC | EEF1A1 |
| 1.1445E-06 | -0.3944422 | 0.888 | 0.947 | 0.02443604 | CD-PC | RPL11 |
| 0.0058904 | -0.3950456 | 0.365 | 0.444 | 1 | CD-PC | UPP1 |
| 7.1348E-07 | -0.3970498 | 0.643 | 0.704 | 0.01523287 | CD-PC | EIF4G2 |
| 6.5056E-07 | -0.3988069 | 0.072 | 0.194 | 0.01388942 | CD-PC | H2AFX |
| 4.2399E-06 | -0.3998982 | 0.06 | 0.168 | 0.09052188 | CD-PC | CTGF |
| 1.814E-05 | -0.4003026 | 0.261 | 0.376 | 0.38728926 | CD-PC | ATF3 |
| 4.8902E-07 | -0.4009755 | 0.008 | 0.106 | 0.01044053 | CD-PC | SERPINE2 |
| 1.3898E-05 | -0.4012186 | 0.916 | 0.935 | 0.29671298 | CD-PC | RPL37 |

| | | | | | | |
|---|---|---|---|---|---|---|
| 1.5817E-06 | -0.4044971 | 0.265 | 0.387 | 0.03376856 | CD-PC | OCIAD2 |
| 1.1742E-09 | -0.4110047 | 0.783 | 0.925 | 2.507E-05 | CD-PC | RPL30 |
| 2.5854E-08 | -0.4112993 | 0.799 | 0.9 | 0.00055198 | CD-PC | RPL23A |
| 1.9982E-07 | -0.413505 | 0.157 | 0.302 | 0.00426613 | CD-PC | MARCKSL1 |
| 0.0002658 | -0.415852 | 0.233 | 0.314 | 1 | CD-PC | SRSF2 |
| 1.3258E-09 | -0.4166178 | 0.779 | 0.901 | 2.8306E-05 | CD-PC | RPL5 |
| 1.4347E-09 | -0.4170309 | 0.032 | 0.178 | 3.063E-05 | CD-PC | FOXC1 |
| 2.2841E-10 | -0.4189358 | 0.293 | 0.483 | 4.8766E-06 | CD-PC | P4HB |
| 1.9488E-08 | -0.4192856 | 0.173 | 0.329 | 0.00041606 | CD-PC | CTSB |
| 0.00011975 | -0.4221142 | 0.233 | 0.322 | 1 | CD-PC | DCDC2 |
| 2.896E-10 | -0.4250638 | 0.044 | 0.204 | 6.1829E-06 | CD-PC | CALD1 |
| 0.00107655 | -0.4257448 | 0.261 | 0.328 | 1 | CD-PC | CEBPB |
| 2.4102E-05 | -0.4284957 | 0.241 | 0.339 | 0.5145842 | CD-PC | HNRNPAB |
| 1.3986E-10 | -0.428716 | 0.751 | 0.9 | 2.9859E-06 | CD-PC | RPS15A |
| 2.1297E-09 | -0.4305558 | 0.779 | 0.856 | 4.5469E-05 | CD-PC | RPL38 |
| 7.205E-07 | -0.4311023 | 0.446 | 0.568 | 0.01538258 | CD-PC | PPIB |
| 6.7814E-09 | -0.434323 | 0.55 | 0.666 | 0.00014478 | CD-PC | SOD1 |
| 8.8943E-15 | -0.4351079 | 0.462 | 0.677 | 1.8989E-10 | CD-PC | HNRNPA1 |
| 1.7064E-09 | -0.4362318 | 0.016 | 0.153 | 3.6431E-05 | CD-PC | AC007952.4 |
| 6.8337E-06 | -0.439264 | 0.257 | 0.368 | 0.14589899 | CD-PC | PNN |
| 0.00036871 | -0.4399459 | 0.313 | 0.379 | 1 | CD-PC | RSRP1 |
| 1.1831E-05 | -0.4402545 | 0.161 | 0.267 | 0.25260202 | CD-PC | AMD1 |
| 1.024E-06 | -0.441406 | 0.189 | 0.32 | 0.02186173 | CD-PC | UGCG |
| 7.9973E-09 | -0.4467806 | 0.671 | 0.761 | 0.00017074 | CD-PC | CALM2 |
| 2.4575E-06 | -0.4477943 | 0.916 | 0.939 | 0.05246734 | CD-PC | RPS28 |
| 0.00012832 | -0.4503099 | 0.723 | 0.635 | 1 | CD-PC | MMP7 |
| 7.0913E-13 | -0.451124 | 0.602 | 0.78 | 1.514E-08 | CD-PC | OAZ1 |
| 4.696E-09 | -0.4514058 | 0.048 | 0.193 | 0.00010026 | CD-PC | TLE4 |
| 2.0462E-09 | -0.4515162 | 0.779 | 0.915 | 4.3686E-05 | CD-PC | RPL29 |
| 2.3746E-08 | -0.4533762 | 0.016 | 0.14 | 0.00050699 | CD-PC | TM4SF1 |
| 3.747E-05 | -0.4566929 | 0.024 | 0.103 | 0.79998999 | CD-PC | APOE |
| 3.3909E-15 | -0.4591879 | 0.779 | 0.942 | 7.2395E-11 | CD-PC | RPL15 |
| 7.8689E-06 | -0.4604239 | 0.502 | 0.608 | 0.16800086 | CD-PC | GADD45A |
| 7.3646E-09 | -0.4609999 | 0.795 | 0.907 | 0.00015723 | CD-PC | RPS3 |
| 2.5885E-10 | -0.4620231 | 0.386 | 0.555 | 5.5264E-06 | CD-PC | EIF5 |
| 5.0096E-08 | -0.463578 | 0.08 | 0.219 | 0.00106956 | CD-PC | UBE2S |
| 2.4019E-05 | -0.4641751 | 0.767 | 0.796 | 0.51279802 | CD-PC | HNRNPA2B1 |
| 2.7714E-06 | -0.4642424 | 0.309 | 0.425 | 0.05917017 | CD-PC | SNHG8 |
| 1.1286E-06 | -0.4678619 | 0.787 | 0.841 | 0.02409632 | CD-PC | CD63 |
| 3.3684E-09 | -0.4716463 | 0.177 | 0.339 | 7.1915E-05 | CD-PC | LUC7L3 |
| 1.3653E-06 | -0.4722364 | 0.237 | 0.36 | 0.02914981 | CD-PC | ARGLU1 |
| 0.00120318 | -0.4727201 | 0.201 | 0.279 | 1 | CD-PC | RASD1 |
| 6.5322E-11 | -0.4799939 | 0.759 | 0.876 | 1.3946E-06 | CD-PC | RPL7 |
| 4.8022E-08 | -0.480267 | 0.795 | 0.888 | 0.00102527 | CD-PC | RPS29 |
| 3.5741E-09 | -0.4803814 | 0.823 | 0.911 | 7.6307E-05 | CD-PC | RPL26 |
| 2.5774E-05 | -0.48141 | 0.141 | 0.245 | 0.55026676 | CD-PC | ID4 |
| 1.4808E-11 | -0.4841718 | 0.783 | 0.924 | 3.1615E-07 | CD-PC | RPS3A |
| 1.144E-09 | -0.4894049 | 0.028 | 0.174 | 2.4425E-05 | CD-PC | FOSB |
| 1.2022E-13 | -0.4908446 | 0.799 | 0.936 | 2.5667E-09 | CD-PC | RPL3 |
| 0.00022906 | -0.4914642 | 0.317 | 0.387 | 1 | CD-PC | BRD2 |

| | | | | | | |
|---|---|---|---|---|---|---|
| 3.4786E-10 | -0.4946372 | 0.735 | 0.868 | 7.4267E-06 | CD-PC | UBA52 |
| 6.2986E-17 | -0.4953363 | 0.562 | 0.758 | 1.3447E-12 | CD-PC | RPL4 |
| 4.2852E-09 | -0.4955458 | 0.044 | 0.189 | 9.1489E-05 | CD-PC | ADIRF |
| 6.5645E-12 | -0.4959401 | 0.795 | 0.918 | 1.4015E-07 | CD-PC | RPLP0 |
| 2.9033E-12 | -0.4999125 | 0.863 | 0.929 | 6.1986E-08 | CD-PC | RPL36 |
| 1.673E-07 | -0.5016794 | 0.161 | 0.308 | 0.00357179 | CD-PC | PMAIP1 |
| 1.4464E-11 | -0.5033801 | 0.185 | 0.377 | 3.0881E-07 | CD-PC | ODC1 |
| 7.4198E-09 | -0.5051233 | 0.004 | 0.126 | 0.00015841 | CD-PC | CXCL14 |
| 9.3607E-09 | -0.5083329 | 0.008 | 0.131 | 0.00019985 | CD-PC | COL7A1 |
| 8.6417E-10 | -0.5106624 | 0.173 | 0.348 | 1.845E-05 | CD-PC | S100A16 |
| 7.9237E-06 | -0.5108288 | 0.321 | 0.424 | 0.16917016 | CD-PC | TUBB4B |
| 2.9081E-11 | -0.5115734 | 0.185 | 0.377 | 6.2089E-07 | CD-PC | FLNA |
| 0.00329468 | -0.5122606 | 0.333 | 0.382 | 1 | CD-PC | HERPUD1 |
| 3.0407E-10 | -0.514855 | 0.863 | 0.925 | 6.4919E-06 | CD-PC | RPLP2 |
| 1.7363E-10 | -0.5150365 | 0.851 | 0.93 | 3.707E-06 | CD-PC | RPL21 |
| 5.7777E-11 | -0.5175911 | 0.896 | 0.946 | 1.2335E-06 | CD-PC | RPS15 |
| 9.3964E-14 | -0.5214982 | 0.454 | 0.664 | 2.0061E-09 | CD-PC | CALR |
| 5.1737E-17 | -0.5218258 | 0.325 | 0.593 | 1.1046E-12 | CD-PC | S100A13 |
| 1.3997E-07 | -0.5222229 | 0.04 | 0.163 | 0.00298826 | CD-PC | G0S2 |
| 2.0495E-05 | -0.5251549 | 0.165 | 0.269 | 0.43756816 | CD-PC | TUBA1A |
| 1.7978E-12 | -0.5300213 | 0.747 | 0.896 | 3.8383E-08 | CD-PC | RPS11 |
| 1.1021E-07 | -0.5307639 | 0.064 | 0.195 | 0.00235297 | CD-PC | GADD45G |
| 8.5726E-14 | -0.5315146 | 0.815 | 0.946 | 1.8302E-09 | CD-PC | RPL8 |
| 1.0015E-08 | -0.5316981 | 0.036 | 0.171 | 0.00021382 | CD-PC | EGLN3 |
| 4.646E-06 | -0.5323946 | 0.08 | 0.187 | 0.09919128 | CD-PC | PIK3R3 |
| 0.00307136 | -0.533735 | 0.337 | 0.387 | 1 | CD-PC | JUNB |
| 1.8053E-06 | -0.5350602 | 0.02 | 0.117 | 0.03854287 | CD-PC | TAGLN |
| 4.2223E-09 | -0.53859 | 0.181 | 0.362 | 9.0145E-05 | CD-PC | HSPA1A |
| 2.6679E-13 | -0.5393385 | 0.827 | 0.915 | 5.6959E-09 | CD-PC | HSP90AB1 |
| 3.2532E-13 | -0.5427105 | 0.904 | 0.973 | 6.9455E-09 | CD-PC | TPT1 |
| 1.4807E-06 | -0.544336 | 0.149 | 0.269 | 0.03161239 | CD-PC | KCNQ1OT1 |
| 0.00010646 | -0.5451697 | 0.161 | 0.255 | 1 | CD-PC | AKAP12 |
| 3.9123E-16 | -0.5460658 | 0.534 | 0.762 | 8.3527E-12 | CD-PC | DYNLL1 |
| 9.1687E-14 | -0.5477345 | 0.719 | 0.873 | 1.9575E-09 | CD-PC | RPS5 |
| 1.8973E-12 | -0.5491125 | 0.815 | 0.914 | 4.0506E-08 | CD-PC | RPL35 |
| 7.3548E-14 | -0.5503025 | 0.341 | 0.548 | 1.5703E-09 | CD-PC | CD59 |
| 5.9656E-11 | -0.5553305 | 0.747 | 0.868 | 1.2737E-06 | CD-PC | RPS4X |
| 4.3759E-11 | -0.5562137 | 0.871 | 0.941 | 9.3426E-07 | CD-PC | RPL37A |
| 1.8041E-11 | -0.5576792 | 0.49 | 0.672 | 3.8517E-07 | CD-PC | TPI1 |
| 2.1786E-16 | -0.5620613 | 0.631 | 0.821 | 4.6514E-12 | CD-PC | UBB |
| 4.0113E-07 | -0.5640571 | 0.277 | 0.416 | 0.00856414 | CD-PC | PHLDA2 |
| 0.00305396 | -0.5662501 | 0.221 | 0.278 | 1 | CD-PC | PPP1R10 |
| 3.1772E-10 | -0.5673365 | 0.735 | 0.845 | 6.7834E-06 | CD-PC | RPSA |
| 1.5704E-14 | -0.5674018 | 0.936 | 0.974 | 3.3529E-10 | CD-PC | RPL13 |
| 6.191E-12 | -0.5674063 | 0.863 | 0.943 | 1.3218E-07 | CD-PC | RPS24 |
| 1.5257E-14 | -0.5675087 | 0.763 | 0.904 | 3.2574E-10 | CD-PC | RPL27A |
| 2.7528E-14 | -0.5714823 | 0.751 | 0.869 | 5.8772E-10 | CD-PC | RPL23 |
| 1.0077E-09 | -0.5716363 | 0.353 | 0.513 | 2.1514E-05 | CD-PC | H2AFZ |
| 6.7622E-11 | -0.5723716 | 0.952 | 0.961 | 1.4437E-06 | CD-PC | RPL28 |
| 1.7786E-10 | -0.576345 | 0.88 | 0.953 | 3.7973E-06 | CD-PC | RPS12 |

| | | | | | | |
|---|---|---|---|---|---|---|
| 1.4458E-14 | -0.5794951 | 0.867 | 0.952 | 3.0867E-10 | CD-PC | RPS8 |
| 1.6298E-07 | -0.586141 | 0.157 | 0.286 | 0.00347955 | CD-PC | POLR2A |
| 2.3719E-16 | -0.5888288 | 0.823 | 0.936 | 5.064E-12 | CD-PC | RPL6 |
| 4.2778E-14 | -0.5891756 | 0.888 | 0.949 | 9.1332E-10 | CD-PC | RPL13A |
| 1.942E-12 | -0.5902182 | 0.815 | 0.925 | 4.1461E-08 | CD-PC | RPL10 |
| 2.9054E-15 | -0.5927493 | 0.799 | 0.907 | 6.203E-11 | CD-PC | RPS20 |
| 1.2099E-09 | -0.5962817 | 0.056 | 0.214 | 2.5831E-05 | CD-PC | ADAMTS1 |
| 1.4519E-17 | -0.5976073 | 0.606 | 0.827 | 3.0999E-13 | CD-PC | RPS21 |
| 6.929E-10 | -0.6075945 | 0.309 | 0.484 | 1.4793E-05 | CD-PC | H1FX |
| 8.276E-06 | -0.6081943 | 0.137 | 0.24 | 0.17669285 | CD-PC | INTS6 |
| 2.1224E-08 | -0.6131168 | 0.004 | 0.119 | 0.00045312 | CD-PC | NEFM |
| 2.4845E-18 | -0.6179516 | 0.707 | 0.872 | 5.3044E-14 | CD-PC | RPL10A |
| 6.1958E-06 | -0.6202332 | 0.149 | 0.262 | 0.13228121 | CD-PC | EGR1 |
| 2.2291E-17 | -0.6215244 | 0.867 | 0.963 | 4.7591E-13 | CD-PC | RPS18 |
| 6.1712E-13 | -0.6218489 | 0.161 | 0.363 | 1.3175E-08 | CD-PC | CD44 |
| 3.0239E-16 | -0.6222331 | 0.313 | 0.563 | 6.456E-12 | CD-PC | HMGA1 |
| 3.8491E-14 | -0.6236414 | 0.205 | 0.425 | 8.2179E-10 | CD-PC | SFPQ |
| 2.9693E-17 | -0.6265286 | 0.594 | 0.808 | 6.3394E-13 | CD-PC | CFL1 |
| 7.7905E-18 | -0.6283617 | 0.703 | 0.911 | 1.6633E-13 | CD-PC | MYL6 |
| 0.00323469 | -0.6385316 | 0.289 | 0.333 | 1 | CD-PC | RHOB |
| 3.536E-06 | -0.6409045 | 0.116 | 0.238 | 0.07549271 | CD-PC | CST6 |
| 3.8965E-19 | -0.6428315 | 0.763 | 0.908 | 8.3191E-15 | CD-PC | RPS7 |
| 1.9372E-24 | -0.6437916 | 0.526 | 0.81 | 4.1359E-20 | CD-PC | EEF1D |
| 6.527E-12 | -0.6441635 | 0.57 | 0.707 | 1.3935E-07 | CD-PC | ITGB1 |
| 7.4184E-22 | -0.6462749 | 0.55 | 0.824 | 1.5838E-17 | CD-PC | YBX1 |
| 1.8125E-15 | -0.6468566 | 0.823 | 0.936 | 3.8697E-11 | CD-PC | RPL32 |
| 1.6213E-06 | -0.6483591 | 0.129 | 0.249 | 0.03461473 | CD-PC | MT1X |
| 1.2481E-19 | -0.6499691 | 0.747 | 0.91 | 2.6647E-15 | CD-PC | RPS16 |
| 4.2624E-09 | -0.6523772 | 0.237 | 0.393 | 9.1002E-05 | CD-PC | TUBA1B |
| 2.532E-11 | -0.6533933 | 0.076 | 0.257 | 5.4059E-07 | CD-PC | IER5 |
| 1.6446E-06 | -0.6543489 | 0.233 | 0.343 | 0.03511153 | CD-PC | SNHG12 |
| 1.944E-19 | -0.655013 | 0.759 | 0.923 | 4.1505E-15 | CD-PC | RPL9 |
| 2.8361E-05 | -0.6629482 | 0.048 | 0.137 | 0.60551617 | CD-PC | SERPINA1 |
| 2.3056E-11 | -0.6665999 | 0.116 | 0.309 | 4.9225E-07 | CD-PC | HSPA1B |
| 2.6696E-08 | -0.6678641 | 0.133 | 0.293 | 0.00056997 | CD-PC | FOS |
| 3.0341E-17 | -0.6711073 | 0.695 | 0.886 | 6.4778E-13 | CD-PC | RPL39 |
| 9.2515E-17 | -0.6848603 | 0.012 | 0.239 | 1.9752E-12 | CD-PC | FXYD2 |
| 1.1555E-21 | -0.689739 | 0.53 | 0.788 | 2.4669E-17 | CD-PC | EEF1B2 |
| 2.0566E-20 | -0.697064 | 0.759 | 0.931 | 4.3908E-16 | CD-PC | RPL18A |
| 4.9762E-13 | -0.6985084 | 0.06 | 0.256 | 1.0624E-08 | CD-PC | ID2 |
| 9.2703E-29 | -0.7047648 | 0.51 | 0.875 | 1.9792E-24 | CD-PC | MT-ND3 |
| 1.1979E-14 | -0.7213441 | 0.072 | 0.287 | 2.5575E-10 | CD-PC | UCHL1 |
| 1.9056E-17 | -0.7381516 | 0.9 | 0.973 | 4.0685E-13 | CD-PC | RPLP1 |
| 2.4264E-25 | -0.7405019 | 0.466 | 0.836 | 5.1804E-21 | CD-PC | MT-ND5 |
| 2.0083E-14 | -0.7426576 | 0.442 | 0.676 | 4.2878E-10 | CD-PC | SH3BGRL3 |
| 1.2478E-11 | -0.7547641 | 0.116 | 0.305 | 2.6641E-07 | CD-PC | PLK2 |
| 2.4938E-11 | -0.7557094 | 0.137 | 0.316 | 5.3242E-07 | CD-PC | CDKN1C |
| 0.00010059 | -0.7694577 | 0.197 | 0.279 | 1 | CD-PC | HES1 |
| 2.0353E-12 | -0.7821243 | 0.502 | 0.687 | 4.3455E-08 | CD-PC | HSPA5 |
| 1.1461E-24 | -0.7839715 | 0.819 | 0.936 | 2.4469E-20 | CD-PC | HSP90AA1 |

| | | | | | | |
|---|---|---|---|---|---|---|
| 0.0099883 | -0.7940977 | 0.076 | 0.128 | 1 | CD-PC | GPX3 |
| 7.1926E-27 | -0.8036187 | 0.321 | 0.663 | 1.5356E-22 | CD-PC | HSPA8 |
| 2.6261E-09 | -0.8049196 | 0.161 | 0.333 | 5.6068E-05 | CD-PC | TIMP1 |
| 5.1965E-13 | -0.8236398 | 0.12 | 0.331 | 1.1095E-08 | CD-PC | MT1E |
| 1.856E-33 | -0.8258291 | 0.566 | 0.92 | 3.9626E-29 | CD-PC | MT-ND4 |
| 7.7356E-26 | -0.8299762 | 0.277 | 0.624 | 1.6516E-21 | CD-PC | MTRNR2L8 |
| 3.9537E-31 | -0.8327105 | 0.293 | 0.637 | 8.4412E-27 | CD-PC | ANXA5 |
| 8.6335E-29 | -0.833171 | 0.751 | 0.937 | 1.8433E-24 | CD-PC | RPS9 |
| 3.2463E-33 | -0.8577265 | 0.526 | 0.906 | 6.9309E-29 | CD-PC | MT-CYB |
| 8.7898E-16 | -0.8635115 | 0.092 | 0.331 | 1.8766E-11 | CD-PC | IL32 |
| 2.7901E-30 | -0.8839657 | 0.731 | 0.934 | 5.957E-26 | CD-PC | RPL12 |
| 1.0944E-22 | -0.8842204 | 0.518 | 0.778 | 2.3365E-18 | CD-PC | HSP90B1 |
| 9.0437E-14 | -0.9014163 | 0.036 | 0.238 | 1.9308E-09 | CD-PC | CSKMT |
| 3.9863E-33 | -0.9025581 | 0.578 | 0.933 | 8.5108E-29 | CD-PC | MT-CO1 |
| 1.7218E-33 | -0.9157463 | 0.526 | 0.918 | 3.676E-29 | CD-PC | MT-CO2 |
| 6.8171E-08 | -0.9158814 | 0.197 | 0.333 | 0.00145546 | CD-PC | LAMC2 |
| 3.9517E-34 | -0.929441 | 0.562 | 0.916 | 8.4368E-30 | CD-PC | MT-ATP6 |
| 4.731E-18 | -0.9466326 | 0.426 | 0.634 | 1.0101E-13 | CD-PC | TPM1 |
| 3.277E-19 | -0.964077 | 0.651 | 0.834 | 6.9964E-15 | CD-PC | ANXA2 |
| 2.2851E-33 | -0.970915 | 0.815 | 0.959 | 4.8787E-29 | CD-PC | RPS2 |
| 8.9449E-13 | -0.9738314 | 0.08 | 0.282 | 1.9097E-08 | CD-PC | LGALS1 |
| 6.2571E-33 | -0.9849793 | 0.546 | 0.896 | 1.3359E-28 | CD-PC | MT-ND2 |
| 7.9317E-07 | -0.9969704 | 0.542 | 0.635 | 0.01693422 | CD-PC | JUN |
| 9.6738E-36 | -1.0022172 | 0.55 | 0.917 | 2.0654E-31 | CD-PC | MT-CO3 |
| 6.2969E-35 | -1.009632 | 0.526 | 0.899 | 1.3444E-30 | CD-PC | MT-ND1 |
| 2.145E-10 | -1.0254752 | 0.273 | 0.443 | 4.5796E-06 | CD-PC | IER3 |
| 2.9855E-10 | -1.0289117 | 0.108 | 0.273 | 6.3741E-06 | CD-PC | HSPA2 |
| 1.3943E-20 | -1.0452045 | 0.036 | 0.313 | 2.9768E-16 | CD-PC | PLAU |
| 5.5673E-25 | -1.0480053 | 0.329 | 0.685 | 1.1886E-20 | CD-PC | MTRNR2L12 |
| 2.7791E-26 | -1.0539247 | 0.779 | 0.927 | 5.9335E-22 | CD-PC | TMSB10 |
| 2.0353E-11 | -1.0641121 | 0.124 | 0.301 | 4.3453E-07 | CD-PC | HEXIM1 |
| 2.355E-18 | -1.0912606 | 0.386 | 0.628 | 5.0279E-14 | CD-PC | VIM |
| 1.575E-20 | -1.1077557 | 0.301 | 0.569 | 3.3625E-16 | CD-PC | IER2 |
| 8.7007E-25 | -1.13915 | 0.679 | 0.874 | 1.8576E-20 | CD-PC | S100A10 |
| 1.452E-24 | -1.1790178 | 0.309 | 0.611 | 3.1E-20 | CD-PC | GADD45B |
| 1.1221E-39 | -1.1999407 | 0.402 | 0.823 | 2.3957E-35 | CD-PC | IGFBP7 |
| 2.8937E-10 | -1.2186452 | 0.928 | 0.962 | 6.178E-06 | CD-PC | FTL |
| 1.0423E-08 | -1.243545 | 0.056 | 0.2 | 0.00022253 | CD-PC | MT1G |
| 4.5437E-14 | -1.2664858 | 0.177 | 0.381 | 9.7007E-10 | CD-PC | HIST1H4C |
| 6.6637E-07 | -1.3209611 | 0.466 | 0.541 | 0.01422692 | CD-PC | SOX4 |
| 9.1505E-18 | -1.3839152 | 0.044 | 0.287 | 1.9536E-13 | CD-PC | MAFB |
| 4.9936E-32 | -1.508686 | 0.357 | 0.718 | 1.0661E-27 | CD-PC | MT2A |
| 1.2821E-27 | -1.8215443 | 0.253 | 0.606 | 2.7373E-23 | CD-PC | SPP1 |
| 3.373E-15 | -1.8322536 | 0.269 | 0.496 | 7.2013E-11 | CD-PC | S100A2 |
| 0.00311629 | -2.1103036 | 0.807 | 0.593 | 1 | CD-PC | PLCG2 |
| 0 | 4.16086744 | 0.821 | 0.027 | 0 | CNT/CD | CALB1 |
| 0 | 3.68839447 | 0.929 | 0.04 | 0 | CNT/CD | AQP2 |
| 1.554E-153 | 3.19905259 | 0.924 | 0.215 | 3.317E-149 | CNT/CD | DEFB1 |
| 0 | 2.79931387 | 0.777 | 0.056 | 0 | CNT/CD | STC1 |
| 0 | 2.70943994 | 0.832 | 0.043 | 0 | CNT/CD | HSD11B2 |

| | | | | | | |
|---|---|---|---|---|---|---|
| 3.895E-277 | 2.49763114 | 0.859 | 0.096 | 8.317E-273 | CNT/CD | AQP3 |
| 3.169E-281 | 2.44656677 | 0.886 | 0.095 | 6.766E-277 | CNT/CD | FXYD4 |
| 9.198E-226 | 2.2445051 | 0.908 | 0.141 | 1.964E-221 | CNT/CD | CA2 |
| 6.0544E-74 | 1.97472217 | 0.929 | 0.486 | 1.2926E-69 | CNT/CD | ATP1B1 |
| 1.163E-150 | 1.91225197 | 0.793 | 0.143 | 2.482E-146 | CNT/CD | RCAN1 |
| 6.664E-196 | 1.71768495 | 0.821 | 0.124 | 1.423E-191 | CNT/CD | CDH16 |
| 1.187E-276 | 1.64025266 | 0.821 | 0.083 | 2.535E-272 | CNT/CD | PIGR |
| 0 | 1.61239275 | 0.614 | 0.025 | 0 | CNT/CD | SLC8A1 |
| 0 | 1.60787661 | 0.679 | 0.022 | 0 | CNT/CD | TMEM52B |
| 1.6514E-55 | 1.58434917 | 0.984 | 0.812 | 3.5258E-51 | CNT/CD | IGFBP7 |
| 7.2446E-96 | 1.53907665 | 0.837 | 0.258 | 1.5467E-91 | CNT/CD | GSTM3 |
| 2.81E-149 | 1.53903915 | 0.832 | 0.158 | 6E-145 | CNT/CD | CA12 |
| 3.1075E-71 | 1.53616413 | 0.875 | 0.366 | 6.6345E-67 | CNT/CD | ATF3 |
| 7.8164E-76 | 1.52594913 | 0.5 | 0.098 | 1.6688E-71 | CNT/CD | RSAD2 |
| 0 | 1.51857457 | 0.609 | 0.007 | 0 | CNT/CD | HMGCS2 |
| 7.952E-100 | 1.49578771 | 0.696 | 0.161 | 1.6978E-95 | CNT/CD | WNK1 |
| 5.0803E-58 | 1.48098686 | 0.739 | 0.284 | 1.0846E-53 | CNT/CD | MAFF |
| 0 | 1.47798881 | 0.609 | 0.013 | 0 | CNT/CD | PVALB |
| 0 | 1.45400345 | 0.549 | 0.027 | 0 | CNT/CD | RHCG |
| 4.221E-111 | 1.40633511 | 0.63 | 0.113 | 9.012E-107 | CNT/CD | NEURL3 |
| 3.5416E-26 | 1.38292257 | 0.815 | 0.487 | 7.5613E-22 | CNT/CD | S100A2 |
| 7.8147E-70 | 1.34877277 | 0.832 | 0.308 | 1.6684E-65 | CNT/CD | MAL |
| 8.2438E-89 | 1.34059288 | 0.641 | 0.142 | 1.76E-84 | CNT/CD | TIMP3 |
| 6.308E-224 | 1.34023037 | 0.728 | 0.081 | 1.347E-219 | CNT/CD | SCPEP1 |
| 4.5627E-25 | 1.28863524 | 0.696 | 0.409 | 9.7413E-21 | CNT/CD | PHLDA2 |
| 1.2927E-82 | 1.27445186 | 0.707 | 0.188 | 2.7599E-78 | CNT/CD | NCOA7 |
| 3.3479E-23 | 1.26460164 | 0.538 | 0.242 | 7.1478E-19 | CNT/CD | MT1X |
| 4.1983E-65 | 1.24511148 | 0.837 | 0.354 | 8.9633E-61 | CNT/CD | COX7B |
| 2.1107E-65 | 1.22485505 | 0.793 | 0.311 | 4.5063E-61 | CNT/CD | FAM107B |
| 5.9975E-88 | 1.22405682 | 0.668 | 0.153 | 1.2805E-83 | CNT/CD | TNFAIP3 |
| 2.6089E-75 | 1.19431955 | 0.848 | 0.325 | 5.57E-71 | CNT/CD | CHCHD10 |
| 1.8391E-50 | 1.18461062 | 0.946 | 0.728 | 3.9264E-46 | CNT/CD | NDUFA4 |
| 5.4666E-69 | 1.12835636 | 0.821 | 0.27 | 1.1671E-64 | CNT/CD | COBLL1 |
| 1.9159E-64 | 1.11298675 | 0.875 | 0.367 | 4.0904E-60 | CNT/CD | LDHB |
| 5.012E-142 | 1.1017344 | 0.663 | 0.101 | 1.07E-137 | CNT/CD | CPVL |
| 4.3533E-81 | 1.07503673 | 0.728 | 0.208 | 9.2942E-77 | CNT/CD | ATP5MC1 |
| 2.7252E-47 | 1.06637008 | 0.62 | 0.2 | 5.8184E-43 | CNT/CD | RRAD |
| 3.185E-78 | 1.04931453 | 0.832 | 0.268 | 6.8E-74 | CNT/CD | CYB5A |
| 3.2307E-60 | 1.0485918 | 0.832 | 0.359 | 6.8975E-56 | CNT/CD | ATP5F1B |
| 3.6135E-40 | 1.04820912 | 0.75 | 0.356 | 7.7149E-36 | CNT/CD | ATP5MC3 |
| 1.0627E-79 | 1.043245 | 0.728 | 0.203 | 2.2688E-75 | CNT/CD | BEX3 |
| 4.6388E-59 | 1.04247431 | 0.783 | 0.309 | 9.9038E-55 | CNT/CD | PRDX2 |
| 3.8508E-16 | 1.03830181 | 0.527 | 0.294 | 8.2215E-12 | CNT/CD | CYTOR |
| 8.0792E-50 | 1.038054 | 0.962 | 0.641 | 1.7249E-45 | CNT/CD | WFDC2 |
| 2.1928E-18 | 1.03745261 | 0.864 | 0.709 | 4.6816E-14 | CNT/CD | MT2A |
| 1.7282E-63 | 1.03727737 | 0.886 | 0.39 | 3.6898E-59 | CNT/CD | CYCS |
| 4.9347E-50 | 1.03196491 | 0.848 | 0.351 | 1.0536E-45 | CNT/CD | KLF6 |
| 1.4382E-46 | 1.02923305 | 0.75 | 0.299 | 3.0706E-42 | CNT/CD | PMAIP1 |
| 1.715E-122 | 1.0268219 | 0.478 | 0.059 | 3.661E-118 | CNT/CD | PRPS2 |
| 3.0137E-92 | 1.01957571 | 0.761 | 0.196 | 6.4343E-88 | CNT/CD | AIF1L |

| | | | | | | |
|---|---|---|---|---|---|---|
| 5.6761E-24 | 1.01145512 | 0.587 | 0.291 | 1.2118E-19 | CNT/CD | MIR4435-2HG |
| 4.7912E-85 | 1.01078035 | 0.793 | 0.231 | 1.0229E-80 | CNT/CD | COX20 |
| 1.3879E-10 | 1.0068274 | 0.38 | 0.195 | 2.9631E-06 | CNT/CD | MT1G |
| 2.14E-41 | 1.00454329 | 0.951 | 0.72 | 4.569E-37 | CNT/CD | CTSD |
| 2.33E-56 | 1.00362139 | 0.799 | 0.327 | 4.9746E-52 | CNT/CD | RALBP1 |
| 1.157E-115 | 0.99444164 | 0.565 | 0.085 | 2.471E-111 | CNT/CD | NR4A1 |
| 2.0322E-25 | 0.98773915 | 0.685 | 0.321 | 4.3387E-21 | CNT/CD | MT1E |
| 1.2523E-55 | 0.9800477 | 0.957 | 0.676 | 2.6736E-51 | CNT/CD | EPCAM |
| 1.259E-248 | 0.97562057 | 0.446 | 0.024 | 2.688E-244 | CNT/CD | KCNJ1 |
| 8.088E-192 | 0.96726484 | 0.538 | 0.047 | 1.727E-187 | CNT/CD | AC092803.2 |
| 7.196E-84 | 0.96464938 | 0.337 | 0.041 | 1.5364E-79 | CNT/CD | ARL14 |
| 8.089E-145 | 0.96426243 | 0.565 | 0.07 | 1.727E-140 | CNT/CD | FOLR1 |
| 1.4961E-72 | 0.96056718 | 0.674 | 0.164 | 3.1942E-68 | CNT/CD | FOSB |
| 3.7464E-44 | 0.96041881 | 0.989 | 0.746 | 7.9985E-40 | CNT/CD | RTN4 |
| 3.8322E-72 | 0.95836625 | 0.707 | 0.196 | 8.1818E-68 | CNT/CD | PNP |
| 1.1649E-29 | 0.94545719 | 0.913 | 0.601 | 2.4871E-25 | CNT/CD | GADD45B |
| 3.5565E-07 | 0.93772354 | 0.69 | 0.667 | 0.00759317 | CNT/CD | CRYAB |
| 7.6919E-29 | 0.92721813 | 0.652 | 0.284 | 1.6422E-24 | CNT/CD | FOS |
| 1.0547E-72 | 0.91672444 | 0.495 | 0.094 | 2.2517E-68 | CNT/CD | CXCL3 |
| 2.229E-205 | 0.90737163 | 0.435 | 0.028 | 4.76E-201 | CNT/CD | SERPINA5 |
| 6.1333E-75 | 0.90480848 | 0.712 | 0.199 | 1.3095E-70 | CNT/CD | GABARAPL1 |
| 1.4922E-94 | 0.90118839 | 0.625 | 0.128 | 3.1858E-90 | CNT/CD | DECR1 |
| 7.9345E-83 | 0.89883483 | 0.668 | 0.161 | 1.694E-78 | CNT/CD | PDHA1 |
| 4.8494E-10 | 0.88931486 | 0.576 | 0.355 | 1.0353E-05 | CNT/CD | HSPA1A |
| 1.1511E-81 | 0.88916228 | 0.674 | 0.165 | 2.4576E-77 | CNT/CD | MPC1 |
| 5.7376E-50 | 0.88914047 | 0.473 | 0.118 | 1.225E-45 | CNT/CD | IFIT3 |
| 1.2417E-30 | 0.87840132 | 0.989 | 0.926 | 2.651E-26 | CNT/CD | MT-CO1 |
| 1.6747E-56 | 0.8686557 | 0.636 | 0.185 | 3.5755E-52 | CNT/CD | BHLHE40 |
| 9.9926E-29 | 0.86818506 | 0.658 | 0.292 | 2.1334E-24 | CNT/CD | HSPH1 |
| 1.8285E-38 | 0.86668372 | 0.929 | 0.658 | 3.9038E-34 | CNT/CD | UQCRB |
| 0 | 0.86103097 | 0.549 | 0.028 | 0 | CNT/CD | CLCNKB |
| 2.7047E-61 | 0.85972123 | 0.543 | 0.136 | 5.7745E-57 | CNT/CD | ARG2 |
| 4.967E-28 | 0.85775357 | 0.989 | 0.897 | 1.0604E-23 | CNT/CD | MT-CYB |
| 4.1841E-68 | 0.85272568 | 0.717 | 0.206 | 8.9331E-64 | CNT/CD | CTSH |
| 1.7043E-53 | 0.84495643 | 0.565 | 0.167 | 3.6387E-49 | CNT/CD | COA3 |
| 3.7455E-33 | 0.8358482 | 0.864 | 0.578 | 7.9967E-29 | CNT/CD | COX5B |
| 1.608E-105 | 0.83519273 | 0.19 | 0.01 | 3.433E-101 | CNT/CD | KLK1 |
| 1.0327E-47 | 0.82038192 | 0.63 | 0.211 | 2.2048E-43 | CNT/CD | PTGR1 |
| 6.6147E-37 | 0.81785807 | 0.826 | 0.447 | 1.4122E-32 | CNT/CD | ATP5PF |
| 1.605E-110 | 0.81448597 | 0.418 | 0.049 | 3.426E-106 | CNT/CD | SMIM5 |
| 1.9822E-49 | 0.81413364 | 0.614 | 0.2 | 4.2321E-45 | CNT/CD | CSRP1 |
| 3.342E-178 | 0.8038569 | 0.549 | 0.052 | 7.135E-174 | CNT/CD | OGDHL |
| 9.1106E-38 | 0.80067202 | 0.815 | 0.391 | 1.9451E-33 | CNT/CD | DBI |
| 1.5281E-43 | 0.79575704 | 0.821 | 0.377 | 3.2624E-39 | CNT/CD | MRPL33 |
| 2.3831E-72 | 0.79523249 | 0.451 | 0.081 | 5.0879E-68 | CNT/CD | HLA-DRA |
| 9.454E-153 | 0.78476565 | 0.516 | 0.053 | 2.018E-148 | CNT/CD | TMEM213 |
| 1.064E-144 | 0.78363362 | 0.446 | 0.043 | 2.272E-140 | CNT/CD | SMIM6 |
| 2.7222E-81 | 0.77849209 | 0.614 | 0.138 | 5.812E-77 | CNT/CD | SH3BGRL |
| 2.2496E-70 | 0.77790981 | 0.473 | 0.091 | 4.8028E-66 | CNT/CD | HLA-DRB1 |
| 1.1739E-48 | 0.77581091 | 0.679 | 0.251 | 2.5062E-44 | CNT/CD | ATP5F1A |

| | | | | | | |
|---|---|---|---|---|---|---|
| 6.5344E-53 | 0.77378268 | 0.701 | 0.252 | 1.3951E-48 | CNT/CD | NDUFA5 |
| 1.878E-126 | 0.76891689 | 0.473 | 0.054 | 4.01E-122 | CNT/CD | CD83 |
| 1.335E-182 | 0.76830424 | 0.516 | 0.046 | 2.849E-178 | CNT/CD | SCNN1B |
| 5.394E-66 | 0.76781275 | 0.56 | 0.132 | 1.1516E-61 | CNT/CD | FDX1 |
| 0 | 0.7641991 | 0.495 | 0.011 | 0 | CNT/CD | PTGER1 |
| 9.0949E-78 | 0.76340585 | 0.255 | 0.026 | 1.9418E-73 | CNT/CD | LINC02154 |
| 1.6469E-47 | 0.76104949 | 0.495 | 0.139 | 3.5161E-43 | CNT/CD | IDH2 |
| 2.3649E-36 | 0.7582383 | 0.837 | 0.455 | 5.049E-32 | CNT/CD | COX8A |
| 9.4773E-35 | 0.75548147 | 0.864 | 0.524 | 2.0234E-30 | CNT/CD | COX6C |
| 4.6614E-17 | 0.75461911 | 0.337 | 0.131 | 9.9521E-13 | CNT/CD | IFIT2 |
| 1.2655E-60 | 0.753715 | 0.609 | 0.169 | 2.7019E-56 | CNT/CD | UQCRC2 |
| 3.4401E-26 | 0.75079527 | 0.723 | 0.378 | 7.3446E-22 | CNT/CD | PTMS |
| 2.1951E-42 | 0.75038822 | 0.957 | 0.595 | 4.6866E-38 | CNT/CD | CLU |
| 8.0839E-96 | 0.75026195 | 0.582 | 0.104 | 1.7259E-91 | CNT/CD | ALDH1A1 |
| 2.1981E-27 | 0.74588792 | 0.94 | 0.816 | 4.693E-23 | CNT/CD | UBB |
| 2.1164E-34 | 0.74430059 | 0.576 | 0.209 | 4.5185E-30 | CNT/CD | ZFP36 |
| 1.6538E-65 | 0.73515366 | 0.603 | 0.156 | 3.5309E-61 | CNT/CD | SUCLG1 |
| 8.0236E-23 | 0.73308805 | 0.75 | 0.415 | 1.713E-18 | CNT/CD | LGALS3 |
| 7.0665E-77 | 0.72532257 | 0.408 | 0.064 | 1.5087E-72 | CNT/CD | GBP1 |
| 1.4047E-24 | 0.72370662 | 0.967 | 0.866 | 2.9991E-20 | CNT/CD | MT-ND3 |
| 3.2155E-31 | 0.71729439 | 0.815 | 0.415 | 6.865E-27 | CNT/CD | TPM4 |
| 1.6061E-54 | 0.71566227 | 0.576 | 0.156 | 3.429E-50 | CNT/CD | PCSK1N |
| 4.0261E-98 | 0.71471704 | 0.435 | 0.058 | 8.5957E-94 | CNT/CD | ST6GAL1 |
| 5.9865E-52 | 0.71346838 | 0.63 | 0.188 | 1.2781E-47 | CNT/CD | BCAM |
| 2.4045E-55 | 0.71219343 | 0.255 | 0.034 | 5.1336E-51 | CNT/CD | KNG1 |
| 1.5461E-33 | 0.70739999 | 0.924 | 0.686 | 3.301E-29 | CNT/CD | CYSTM1 |
| 3.3446E-27 | 0.70728484 | 0.891 | 0.61 | 7.1406E-23 | CNT/CD | MT-ND4L |
| 1.7775E-55 | 0.70328451 | 0.755 | 0.26 | 3.795E-51 | CNT/CD | MINOS1 |
| 1.044E-101 | 0.70215169 | 0.467 | 0.065 | 2.2296E-97 | CNT/CD | ABCB1 |
| 7.9664E-24 | 0.70061354 | 0.467 | 0.182 | 1.7008E-19 | CNT/CD | BIRC3 |
| 6.7842E-35 | 0.69511576 | 0.772 | 0.37 | 1.4484E-30 | CNT/CD | NDUFA1 |
| 2.2549E-21 | 0.69145813 | 0.989 | 0.909 | 4.8143E-17 | CNT/CD | MT-CO3 |
| 1.6964E-22 | 0.69066794 | 0.435 | 0.171 | 3.6218E-18 | CNT/CD | PHLDA1 |
| 1.5932E-27 | 0.68492795 | 0.473 | 0.184 | 3.4015E-23 | CNT/CD | B4GALT1 |
| 6.0939E-37 | 0.68385308 | 0.326 | 0.076 | 1.301E-32 | CNT/CD | PDK4 |
| 1.1907E-68 | 0.68268746 | 0.484 | 0.097 | 2.5422E-64 | CNT/CD | MAP3K8 |
| 1.1649E-57 | 0.67223103 | 0.576 | 0.153 | 2.487E-53 | CNT/CD | ECH1 |
| 2.779E-116 | 0.67050414 | 0.446 | 0.052 | 5.934E-112 | CNT/CD | EMX1 |
| 0 | 0.66755546 | 0.435 | 0.002 | 0 | CNT/CD | TEX41 |
| 8.7244E-42 | 0.66269508 | 0.777 | 0.326 | 1.8627E-37 | CNT/CD | ATP6V1F |
| 4.2029E-60 | 0.6601944 | 0.658 | 0.19 | 8.9731E-56 | CNT/CD | SNX2 |
| 4.4524E-52 | 0.65844957 | 0.625 | 0.195 | 9.5058E-48 | CNT/CD | NDUFB5 |
| 1.1263E-20 | 0.65702791 | 0.989 | 0.91 | 2.4047E-16 | CNT/CD | MT-CO2 |
| 6.083E-148 | 0.65669496 | 0.293 | 0.017 | 1.299E-143 | CNT/CD | MEST |
| 7.5157E-30 | 0.65660122 | 0.886 | 0.604 | 1.6046E-25 | CNT/CD | ATP5MG |
| 1.0073E-32 | 0.65257768 | 0.614 | 0.268 | 2.1505E-28 | CNT/CD | ATP5F1C |
| 2.8042E-22 | 0.64994616 | 0.989 | 0.908 | 5.987E-18 | CNT/CD | MT-ATP6 |
| 1.0035E-32 | 0.64731067 | 0.717 | 0.323 | 2.1426E-28 | CNT/CD | UQCRQ |
| 1.4467E-23 | 0.64695673 | 0.984 | 0.912 | 3.0886E-19 | CNT/CD | MT-ND4 |
| 1.9668E-28 | 0.64482065 | 0.685 | 0.327 | 4.1992E-24 | CNT/CD | SLC25A5 |

| | | | | | | |
|---|---|---|---|---|---|---|
| 4.5401E-84 | 0.64396721 | 0.462 | 0.074 | 9.6931E-80 | CNT/CD | CYS1 |
| 3.885E-45 | 0.6417548 | 0.62 | 0.204 | 8.2945E-41 | CNT/CD | RAB3IP |
| 1.6765E-19 | 0.64151705 | 0.533 | 0.257 | 3.5794E-15 | CNT/CD | VCL |
| 1.3958E-92 | 0.63986647 | 0.348 | 0.04 | 2.98E-88 | CNT/CD | AC026250.1 |
| 2.0263E-59 | 0.6398571 | 0.592 | 0.157 | 4.3261E-55 | CNT/CD | NDUFV1 |
| 2.111E-11 | 0.63887315 | 0.685 | 0.495 | 4.5069E-07 | CNT/CD | TNFRSF12A |
| 3.593E-31 | 0.63755715 | 0.668 | 0.305 | 7.6712E-27 | CNT/CD | ATP5PD |
| 5.828E-161 | 0.63691787 | 0.402 | 0.03 | 1.244E-156 | CNT/CD | GATA3-AS1 |
| 6.5763E-74 | 0.63651789 | 0.533 | 0.106 | 1.404E-69 | CNT/CD | SULT1C2 |
| 1.6345E-16 | 0.6303373 | 0.516 | 0.261 | 3.4896E-12 | CNT/CD | LIMA1 |
| 3.2741E-47 | 0.62846035 | 0.674 | 0.226 | 6.9902E-43 | CNT/CD | ASAH1 |
| 1.4513E-25 | 0.62437682 | 0.543 | 0.221 | 3.0985E-21 | CNT/CD | CD74 |
| 2.1962E-46 | 0.61995348 | 0.609 | 0.2 | 4.6889E-42 | CNT/CD | METTL9 |
| 2.676E-26 | 0.61983158 | 0.587 | 0.268 | 5.7132E-22 | CNT/CD | NDUFV2 |
| 6.0858E-48 | 0.61920269 | 0.364 | 0.073 | 1.2993E-43 | CNT/CD | CXCL2 |
| 1.9815E-52 | 0.61861165 | 0.413 | 0.089 | 4.2305E-48 | CNT/CD | ELMSAN1 |
| 4.2185E-54 | 0.61539797 | 0.582 | 0.16 | 9.0065E-50 | CNT/CD | SNRPN |
| 1.6023E-12 | 0.61478568 | 0.69 | 0.465 | 3.4209E-08 | CNT/CD | LMNA |
| 7.87E-35 | 0.61448633 | 0.609 | 0.23 | 1.6802E-30 | CNT/CD | SCIN |
| 4.8215E-17 | 0.61072247 | 0.891 | 0.712 | 1.0294E-12 | CNT/CD | HSPB1 |
| 3.3232E-30 | 0.60951138 | 0.658 | 0.288 | 7.095E-26 | CNT/CD | TOP1 |
| 8.263E-204 | 0.60915408 | 0.391 | 0.022 | 1.764E-199 | CNT/CD | SCNN1G |
| 5.5738E-41 | 0.60888499 | 0.565 | 0.184 | 1.19E-36 | CNT/CD | RAB9A |
| 2.5223E-15 | 0.60818464 | 0.658 | 0.384 | 5.3851E-11 | CNT/CD | HSPE1 |
| 2.7832E-17 | 0.6081036 | 0.989 | 0.925 | 5.942E-13 | CNT/CD | ACTG1 |
| 1.288E-25 | 0.60569858 | 0.929 | 0.766 | 2.7499E-21 | CNT/CD | COX7C |
| 1.1774E-27 | 0.60362329 | 0.766 | 0.386 | 2.5138E-23 | CNT/CD | NDRG1 |
| 5.8012E-34 | 0.60349093 | 0.679 | 0.283 | 1.2386E-29 | CNT/CD | NFE2L2 |
| 1.2615E-60 | 0.6014381 | 0.549 | 0.132 | 2.6932E-56 | CNT/CD | ALDH2 |
| 2.5291E-38 | 0.59873789 | 0.63 | 0.233 | 5.3995E-34 | CNT/CD | TCEAL9 |
| 8.1838E-20 | 0.59396564 | 0.234 | 0.066 | 1.7472E-15 | CNT/CD | LIF |
| 4.147E-157 | 0.59382525 | 0.337 | 0.022 | 8.853E-153 | CNT/CD | FGL2 |
| 1.5713E-26 | 0.59195592 | 0.609 | 0.28 | 3.3547E-22 | CNT/CD | ATP5PO |
| 7.5955E-08 | 0.59181474 | 0.723 | 0.605 | 0.00162164 | CNT/CD | GADD45A |
| 3.4257E-25 | 0.59067548 | 0.658 | 0.318 | 7.3139E-21 | CNT/CD | COX5A |
| 1.0544E-26 | 0.59038575 | 0.484 | 0.176 | 2.251E-22 | CNT/CD | HMGCS1 |
| 8.3367E-17 | 0.59009844 | 0.772 | 0.487 | 1.7799E-12 | CNT/CD | DNAJA1 |
| 1.7692E-27 | 0.58978542 | 0.783 | 0.42 | 3.7773E-23 | CNT/CD | TMEM59 |
| 1.6289E-24 | 0.58537187 | 0.592 | 0.266 | 3.4777E-20 | CNT/CD | CKB |
| 3.0295E-29 | 0.58243464 | 0.658 | 0.295 | 6.468E-25 | CNT/CD | DDT |
| 4.4601E-15 | 0.57912818 | 0.761 | 0.563 | 9.5223E-11 | CNT/CD | GDF15 |
| 1.189E-239 | 0.57796206 | 0.261 | 0.007 | 2.539E-235 | CNT/CD | CXCL11 |
| 2.8958E-55 | 0.57602872 | 0.457 | 0.102 | 6.1825E-51 | CNT/CD | CAPS |
| 2.6638E-90 | 0.57535879 | 0.353 | 0.041 | 5.6872E-86 | CNT/CD | THY1 |
| 3.7472E-21 | 0.57423795 | 0.989 | 0.921 | 8.0002E-17 | CNT/CD | ACTB |
| 1.451E-215 | 0.57136409 | 0.353 | 0.016 | 3.097E-211 | CNT/CD | NR0B2 |
| 9.4677E-67 | 0.56895022 | 0.467 | 0.092 | 2.0213E-62 | CNT/CD | ACADM |
| 7.8996E-94 | 0.56851209 | 0.37 | 0.044 | 1.6866E-89 | CNT/CD | AC004130.1 |
| 2.2136E-67 | 0.56397448 | 0.37 | 0.059 | 4.7259E-63 | CNT/CD | COX7A1 |
| 3.98E-113 | 0.56011335 | 0.353 | 0.034 | 8.496E-109 | CNT/CD | AC025263.1 |

| | | | | | | |
|---|---|---|---|---|---|---|
| 8.154E-19 | 0.55926472 | 0.973 | 0.89 | 1.7409E-14 | CNT/CD | MT-ND1 |
| 1.3359E-22 | 0.5582345 | 0.701 | 0.383 | 2.8521E-18 | CNT/CD | ATP5IF1 |
| 6.4005E-14 | 0.55744512 | 0.397 | 0.187 | 1.3665E-09 | CNT/CD | THBS1 |
| 7.7704E-45 | 0.55712889 | 0.56 | 0.172 | 1.659E-40 | CNT/CD | SDHD |
| 1.2983E-78 | 0.55507081 | 0.353 | 0.048 | 2.7719E-74 | CNT/CD | PINX1.1 |
| 1.0098E-22 | 0.55445463 | 0.614 | 0.284 | 2.156E-18 | CNT/CD | MT-ATP8 |
| 5.5339E-56 | 0.5540778 | 0.429 | 0.089 | 1.1815E-51 | CNT/CD | KCNJ16 |
| 3.259E-132 | 0.55239436 | 0.391 | 0.035 | 6.958E-128 | CNT/CD | EDNRB |
| 1.2333E-20 | 0.5521205 | 0.652 | 0.307 | 2.6331E-16 | CNT/CD | WSB1 |
| 8.9774E-22 | 0.55105913 | 0.897 | 0.645 | 1.9167E-17 | CNT/CD | UQCRH |
| 2.325E-31 | 0.54978126 | 0.527 | 0.194 | 4.9639E-27 | CNT/CD | TMEM141 |
| 9.0777E-32 | 0.54910119 | 0.538 | 0.201 | 1.9381E-27 | CNT/CD | MPC2 |
| 1.9407E-68 | 0.54817588 | 0.435 | 0.076 | 4.1434E-64 | CNT/CD | AC019197.1 |
| 1.1394E-81 | 0.54712136 | 0.424 | 0.062 | 2.4327E-77 | CNT/CD | MGP |
| 2.7101E-54 | 0.54497326 | 0.413 | 0.084 | 5.7861E-50 | CNT/CD | AC015912.3 |
| 2.3382E-48 | 0.54374831 | 0.272 | 0.044 | 4.9921E-44 | CNT/CD | RAET1L |
| 2.5834E-33 | 0.54229495 | 0.571 | 0.209 | 5.5157E-29 | CNT/CD | BTG3 |
| 2.571E-162 | 0.54215161 | 0.402 | 0.03 | 5.489E-158 | CNT/CD | PIK3C2G |
| 4.1821E-11 | 0.54119836 | 0.293 | 0.133 | 8.9287E-07 | CNT/CD | KIAA1551 |
| 6.6856E-20 | 0.54037506 | 0.832 | 0.557 | 1.4274E-15 | CNT/CD | COX6B1 |
| 3.6054E-26 | 0.5402833 | 0.707 | 0.352 | 7.6974E-22 | CNT/CD | EIF3K |
| 1.8649E-34 | 0.53794522 | 0.658 | 0.262 | 3.9815E-30 | CNT/CD | TSTD1 |
| 1.8145E-11 | 0.53532026 | 0.5 | 0.277 | 3.874E-07 | CNT/CD | BTG2 |
| 4.3293E-30 | 0.53480306 | 0.533 | 0.198 | 9.243E-26 | CNT/CD | NDUFB1 |
| 1.38E-53 | 0.53471057 | 0.38 | 0.076 | 2.9462E-49 | CNT/CD | HOXD8 |
| 4.268E-28 | 0.53274437 | 0.489 | 0.178 | 9.1122E-24 | CNT/CD | BCL10 |
| 4.4879E-44 | 0.53171543 | 0.337 | 0.07 | 9.5818E-40 | CNT/CD | GATA3 |
| 7.3383E-46 | 0.53158723 | 0.582 | 0.177 | 1.5667E-41 | CNT/CD | HACD3 |
| 3.7109E-38 | 0.53108968 | 0.554 | 0.184 | 7.9227E-34 | CNT/CD | NDUFS7 |
| 3.1082E-15 | 0.52995006 | 0.989 | 0.913 | 6.636E-11 | CNT/CD | HSP90AB1 |
| 2.3656E-08 | 0.52937298 | 0.978 | 0.933 | 0.00050505 | CNT/CD | HSP90AA1 |
| 5.845E-35 | 0.52576482 | 0.418 | 0.12 | 1.2479E-30 | CNT/CD | WEE1 |
| 2.7254E-43 | 0.52494685 | 0.543 | 0.161 | 5.8188E-39 | CNT/CD | SELENOP |
| 6.172E-125 | 0.52443057 | 0.348 | 0.029 | 1.318E-120 | CNT/CD | CLDN8 |
| 5.4406E-23 | 0.5221685 | 0.734 | 0.393 | 1.1616E-18 | CNT/CD | MGST3 |
| 2.4685E-29 | 0.52215209 | 0.598 | 0.239 | 5.2702E-25 | CNT/CD | MMP24OS |
| 1.429E-08 | 0.51918815 | 0.837 | 0.7 | 0.00030509 | CNT/CD | CLDN4 |
| 5.2106E-45 | 0.51910567 | 0.31 | 0.059 | 1.1125E-40 | CNT/CD | LGALSL |
| 2.1842E-32 | 0.51774612 | 0.625 | 0.247 | 4.6633E-28 | CNT/CD | ECHS1 |
| 8.9248E-44 | 0.51758484 | 0.489 | 0.135 | 1.9054E-39 | CNT/CD | CLN5 |
| 2.431E-28 | 0.51489774 | 0.924 | 0.64 | 5.1903E-24 | CNT/CD | PEBP1 |
| 7.3876E-34 | 0.50942327 | 0.38 | 0.104 | 1.5772E-29 | CNT/CD | AL121944.1 |
| 3.229E-70 | 0.50548731 | 0.348 | 0.05 | 6.8938E-66 | CNT/CD | AL139274.2 |
| 5.8689E-68 | 0.5037092 | 0.332 | 0.048 | 1.253E-63 | CNT/CD | NR1H4 |
| 3.0752E-19 | 0.50366515 | 0.386 | 0.153 | 6.5656E-15 | CNT/CD | PALLD |
| 3.4902E-10 | 0.50251046 | 0.489 | 0.257 | 7.4515E-06 | CNT/CD | EGR1 |
| 2.4946E-49 | 0.50052425 | 0.451 | 0.109 | 5.326E-45 | CNT/CD | ECI2 |
| 1.8924E-49 | 0.4978641 | 0.397 | 0.086 | 4.0402E-45 | CNT/CD | COQ9 |
| 2.6676E-37 | 0.49672113 | 0.473 | 0.143 | 5.6954E-33 | CNT/CD | NDUFS2 |
| 5.941E-55 | 0.49608695 | 0.321 | 0.054 | 1.2684E-50 | CNT/CD | GMNN |

| | | | | | | |
|---|---|---|---|---|---|---|
| 1.0376E-21 | 0.49570288 | 0.728 | 0.377 | 2.2153E-17 | CNT/CD | NDUFB2 |
| 7.9062E-54 | 0.49567744 | 0.136 | 0.01 | 1.688E-49 | CNT/CD | CAVIN2 |
| 8.3982E-53 | 0.49466052 | 0.391 | 0.079 | 1.793E-48 | CNT/CD | PLPP3 |
| 2.051E-101 | 0.49440211 | 0.413 | 0.049 | 4.3793E-97 | CNT/CD | IQGAP2 |
| 2.3148E-16 | 0.49395439 | 0.755 | 0.439 | 4.942E-12 | CNT/CD | ZFP36L1 |
| 5.818E-36 | 0.49374052 | 0.353 | 0.088 | 1.2421E-31 | CNT/CD | BHLHE41 |
| 7.2533E-17 | 0.49296704 | 0.315 | 0.119 | 1.5486E-12 | CNT/CD | MT1F |
| 2.1022E-53 | 0.49286159 | 0.516 | 0.127 | 4.4882E-49 | CNT/CD | ACAT1 |
| 4.1718E-37 | 0.49273698 | 0.168 | 0.022 | 8.9068E-33 | CNT/CD | AL358472.4 |
| 9.9177E-67 | 0.49171065 | 0.332 | 0.048 | 2.1174E-62 | CNT/CD | SOCS2 |
| 1.3017E-32 | 0.48983479 | 0.538 | 0.194 | 2.7792E-28 | CNT/CD | GLRX5 |
| 2.1762E-17 | 0.48682286 | 0.674 | 0.365 | 4.6463E-13 | CNT/CD | DDIT3 |
| 1.3014E-22 | 0.48678604 | 0.929 | 0.745 | 2.7784E-18 | CNT/CD | DSTN |
| 1.029E-146 | 0.48575216 | 0.359 | 0.027 | 2.197E-142 | CNT/CD | CKMT2 |
| 1.8214E-22 | 0.48138841 | 0.87 | 0.576 | 3.8887E-18 | CNT/CD | ATP6V0E1 |
| 9.4458E-28 | 0.47912244 | 0.5 | 0.185 | 2.0167E-23 | CNT/CD | FERMT2 |
| 1.3748E-25 | 0.47885057 | 0.277 | 0.074 | 2.9352E-21 | CNT/CD | NOV |
| 4.4346E-31 | 0.47582991 | 0.527 | 0.188 | 9.4679E-27 | CNT/CD | RHOBTB3 |
| 6.7817E-38 | 0.47494512 | 0.408 | 0.11 | 1.4479E-33 | CNT/CD | TWSG1 |
| 1.1551E-56 | 0.47483893 | 0.38 | 0.071 | 2.4661E-52 | CNT/CD | SORL1 |
| 4.4339E-19 | 0.47450469 | 0.543 | 0.252 | 9.4663E-15 | CNT/CD | NEDD4L |
| 2.7111E-27 | 0.4717899 | 0.81 | 0.386 | 5.7881E-23 | CNT/CD | CIRBP |
| 3.4725E-34 | 0.47124706 | 0.446 | 0.134 | 7.4137E-30 | CNT/CD | PHYH |
| 6.4185E-39 | 0.4709317 | 0.511 | 0.155 | 1.3704E-34 | CNT/CD | PNKD |
| 6.8931E-26 | 0.46950405 | 0.495 | 0.193 | 1.4717E-21 | CNT/CD | CYC1 |
| 2.0508E-10 | 0.46867024 | 0.978 | 0.911 | 4.3784E-06 | CNT/CD | UBC |
| 6.2349E-36 | 0.4669463 | 0.348 | 0.086 | 1.3311E-31 | CNT/CD | ZNRD1 |
| 1.3632E-20 | 0.46513631 | 0.288 | 0.09 | 2.9105E-16 | CNT/CD | SNAPC1 |
| 7.7711E-35 | 0.46444947 | 0.549 | 0.183 | 1.6591E-30 | CNT/CD | HMGN3 |
| 1.3968E-34 | 0.4620669 | 0.5 | 0.163 | 2.9821E-30 | CNT/CD | CISD1 |
| 1.3316E-23 | 0.46062187 | 0.598 | 0.267 | 2.8429E-19 | CNT/CD | BAG1 |
| 4.3826E-43 | 0.46057767 | 0.38 | 0.087 | 9.3569E-39 | CNT/CD | SLC25A25 |
| 2.2994E-38 | 0.45940963 | 0.429 | 0.118 | 4.9093E-34 | CNT/CD | CHPT1 |
| 3.9057E-35 | 0.4560623 | 0.505 | 0.162 | 8.3386E-31 | CNT/CD | GSTK1 |
| 1.771E-12 | 0.45536922 | 0.353 | 0.164 | 3.7811E-08 | CNT/CD | STAT1 |
| 4.1928E-31 | 0.45470592 | 0.582 | 0.223 | 8.9517E-27 | CNT/CD | MDH1 |
| 2.383E-125 | 0.45434575 | 0.288 | 0.02 | 5.088E-121 | CNT/CD | FOLR3 |
| 2.2549E-29 | 0.45414254 | 0.413 | 0.127 | 4.8142E-25 | CNT/CD | LDLR |
| 2.55E-24 | 0.45092613 | 0.609 | 0.263 | 5.4443E-20 | CNT/CD | BLVRB |
| 1.2392E-31 | 0.45036833 | 0.641 | 0.253 | 2.6458E-27 | CNT/CD | NDUFA6 |
| 4.2425E-11 | 0.45021382 | 0.397 | 0.206 | 9.0578E-07 | CNT/CD | ARID5B |
| 3.5816E-21 | 0.44986575 | 0.402 | 0.15 | 7.6467E-17 | CNT/CD | INSIG1 |
| 4.9475E-19 | 0.44882442 | 0.908 | 0.677 | 1.0563E-14 | CNT/CD | COX6A1 |
| 2.732E-25 | 0.4486903 | 0.641 | 0.285 | 5.8329E-21 | CNT/CD | LRPAP1 |
| 8.2712E-31 | 0.44794426 | 0.995 | 0.875 | 1.7659E-26 | CNT/CD | ITM2B |
| 3.1504E-26 | 0.44619234 | 0.478 | 0.176 | 6.726E-22 | CNT/CD | SLC25A4 |
| 7.5218E-94 | 0.44581389 | 0.299 | 0.029 | 1.6059E-89 | CNT/CD | LINC00982 |
| 5.9271E-37 | 0.44524627 | 0.375 | 0.097 | 1.2654E-32 | CNT/CD | TEX264 |
| 2.1617E-30 | 0.44251772 | 0.38 | 0.107 | 4.6152E-26 | CNT/CD | AC023157.3 |
| 3.2157E-21 | 0.44216509 | 0.641 | 0.309 | 6.8656E-17 | CNT/CD | UQCR11 |

| | | | | | | |
|---|---|---|---|---|---|---|
| 7.4265E-38 | 0.44164254 | 0.489 | 0.144 | 1.5856E-33 | CNT/CD | ITM2C |
| 2.3008E-15 | 0.44066894 | 0.685 | 0.407 | 4.9123E-11 | CNT/CD | UQCR10 |
| 4.2132E-27 | 0.44051204 | 0.353 | 0.106 | 8.9952E-23 | CNT/CD | FOSL1 |
| 1.8236E-17 | 0.43869648 | 0.353 | 0.139 | 3.8934E-13 | CNT/CD | EFNA1 |
| 2.225E-105 | 0.43841032 | 0.321 | 0.029 | 4.751E-101 | CNT/CD | VEPH1 |
| 1.661E-111 | 0.43802842 | 0.261 | 0.018 | 3.546E-107 | CNT/CD | NR4A3 |
| 3.0263E-15 | 0.43787662 | 0.576 | 0.309 | 6.4611E-11 | CNT/CD | DNAJB6 |
| 1.8693E-13 | 0.43702689 | 0.734 | 0.482 | 3.9911E-09 | CNT/CD | ATP1A1 |
| 0.00033826 | 0.43690572 | 0.261 | 0.171 | 1 | CNT/CD | SETX |
| 1.013E-110 | 0.43666464 | 0.332 | 0.03 | 2.163E-106 | CNT/CD | MPP1 |
| 4.9818E-28 | 0.4361989 | 0.641 | 0.249 | 1.0636E-23 | CNT/CD | RPS4Y1 |
| 3.6783E-88 | 0.43537767 | 0.158 | 0.008 | 7.8531E-84 | CNT/CD | AL031429.1 |
| 1.3209E-29 | 0.43522417 | 0.576 | 0.217 | 2.8201E-25 | CNT/CD | BEX2 |
| 1.6548E-26 | 0.43484328 | 0.62 | 0.264 | 3.5331E-22 | CNT/CD | NDUFB8 |
| 2.8832E-32 | 0.4344083 | 0.424 | 0.129 | 6.1557E-28 | CNT/CD | NDUFA8 |
| 1.3877E-16 | 0.43323373 | 0.625 | 0.327 | 2.9627E-12 | CNT/CD | TUBA1C |
| 7.4693E-26 | 0.43311118 | 0.527 | 0.199 | 1.5947E-21 | CNT/CD | PIM3 |
| 3.2208E-71 | 0.43205633 | 0.332 | 0.045 | 6.8765E-67 | CNT/CD | RASSF9 |
| 2.9142E-63 | 0.43077929 | 0.359 | 0.058 | 6.2217E-59 | CNT/CD | CTSF |
| 1.2301E-17 | 0.43044371 | 0.929 | 0.726 | 2.6263E-13 | CNT/CD | ENO1 |
| 3.3792E-44 | 0.42962866 | 0.31 | 0.06 | 7.2145E-40 | CNT/CD | STIM2 |
| 5.184E-34 | 0.42932258 | 0.533 | 0.179 | 1.1068E-29 | CNT/CD | HADHB |
| 9.654E-18 | 0.4279851 | 0.832 | 0.568 | 2.0611E-13 | CNT/CD | COX7A2 |
| 3.0551E-16 | 0.42792167 | 0.935 | 0.779 | 6.5226E-12 | CNT/CD | COX4I1 |
| 1.0275E-27 | 0.42652146 | 0.505 | 0.191 | 2.1937E-23 | CNT/CD | NDUFA10 |
| 1.7324E-22 | 0.42628812 | 0.245 | 0.066 | 3.6986E-18 | CNT/CD | GLA |
| 2.1548E-84 | 0.42586302 | 0.283 | 0.028 | 4.6006E-80 | CNT/CD | AC007032.1 |
| 8.5736E-40 | 0.42566253 | 0.326 | 0.07 | 1.8305E-35 | CNT/CD | TCEAL2 |
| 1.6819E-21 | 0.42314935 | 0.658 | 0.322 | 3.5909E-17 | CNT/CD | VDAC1 |
| 8.7243E-16 | 0.42296004 | 0.674 | 0.39 | 1.8626E-11 | CNT/CD | ATP5F1D |
| 3.8003E-32 | 0.42118586 | 0.435 | 0.132 | 8.1137E-28 | CNT/CD | TUBB2A |
| 2.7735E-21 | 0.42099566 | 0.38 | 0.138 | 5.9213E-17 | CNT/CD | NFKB1 |
| 4.3769E-42 | 0.42084122 | 0.451 | 0.118 | 9.3446E-38 | CNT/CD | SUCLG2 |
| 7.3837E-49 | 0.41982824 | 0.217 | 0.029 | 1.5764E-44 | CNT/CD | AC129507.4 |
| 7.5827E-16 | 0.41967608 | 0.636 | 0.34 | 1.6189E-11 | CNT/CD | TSC22D1 |
| 2.9623E-40 | 0.41914313 | 0.522 | 0.154 | 6.3245E-36 | CNT/CD | TXN2 |
| 1.9337E-19 | 0.41865456 | 0.511 | 0.23 | 4.1285E-15 | CNT/CD | NDUFS8 |
| 2.9543E-64 | 0.41566388 | 0.201 | 0.019 | 6.3074E-60 | CNT/CD | FIBIN |
| 1.3413E-13 | 0.41555685 | 0.62 | 0.345 | 2.8637E-09 | CNT/CD | DDX24 |
| 8.8489E-27 | 0.41476553 | 0.527 | 0.203 | 1.8892E-22 | CNT/CD | PRDX3 |
| 2.5962E-71 | 0.41403669 | 0.283 | 0.033 | 5.5428E-67 | CNT/CD | KCNJ15 |
| 7.5594E-24 | 0.41399727 | 0.364 | 0.123 | 1.6139E-19 | CNT/CD | GNPDA1 |
| 6.557E-26 | 0.41378225 | 0.696 | 0.304 | 1.3999E-21 | CNT/CD | ISCU |
| 2.3683E-19 | 0.41329749 | 0.435 | 0.181 | 5.0563E-15 | CNT/CD | CBR1 |
| 1.1791E-09 | 0.41329192 | 0.766 | 0.547 | 2.5173E-05 | CNT/CD | CD9 |
| 2.063E-55 | 0.41154378 | 0.364 | 0.066 | 4.4046E-51 | CNT/CD | NUPR2 |
| 1.1885E-05 | 0.41100759 | 0.467 | 0.303 | 0.25373676 | CNT/CD | HSPA1B |
| 1.6828E-32 | 0.41085043 | 0.408 | 0.119 | 3.5929E-28 | CNT/CD | ACO2 |
| 8.603E-62 | 0.40966245 | 0.332 | 0.05 | 1.8367E-57 | CNT/CD | MNS1 |
| 7.5846E-44 | 0.40823819 | 0.359 | 0.078 | 1.6193E-39 | CNT/CD | HIBCH |

| | | | | | | |
|---|---|---|---|---|---|---|
| 1.311E-158 | 0.40801284 | 0.315 | 0.018 | 2.799E-154 | CNT/CD | ELF5 |
| 8.1483E-19 | 0.40765069 | 0.592 | 0.287 | 1.7397E-14 | CNT/CD | YWHAH |
| 2.773E-121 | 0.4059929 | 0.337 | 0.028 | 5.92E-117 | CNT/CD | CLCNKA |
| 7.2614E-35 | 0.4054322 | 0.321 | 0.075 | 1.5503E-30 | CNT/CD | NNT |
| 2.1189E-20 | 0.40510116 | 0.484 | 0.204 | 4.5239E-16 | CNT/CD | NDUFA3 |
| 2.2879E-96 | 0.40425788 | 0.359 | 0.04 | 4.8846E-92 | CNT/CD | ALDH4A1 |
| 3.0419E-12 | 0.40309178 | 0.772 | 0.564 | 6.4944E-08 | CNT/CD | HNRNPC |
| 1.0351E-26 | 0.40184919 | 0.168 | 0.03 | 2.21E-22 | CNT/CD | AC007906.2 |
| 3.0804E-35 | 0.40151203 | 0.402 | 0.11 | 6.5767E-31 | CNT/CD | FCGRT |
| 3.407E-20 | 0.40146594 | 0.723 | 0.366 | 7.274E-16 | CNT/CD | CAMK2N1 |
| 1.9853E-34 | 0.40106977 | 0.37 | 0.097 | 4.2387E-30 | CNT/CD | SLC19A2 |
| 6.7038E-25 | 0.4001802 | 0.44 | 0.157 | 1.4313E-20 | CNT/CD | PLS3 |
| 2.4358E-15 | 0.3997751 | 0.967 | 0.888 | 5.2005E-11 | CNT/CD | MT-ND2 |
| 2.5057E-18 | 0.39974176 | 0.668 | 0.345 | 5.3496E-14 | CNT/CD | ACADVL |
| 8.639E-30 | 0.39894101 | 0.332 | 0.09 | 1.8444E-25 | CNT/CD | MLF1 |
| 7.9906E-26 | 0.39883831 | 0.293 | 0.08 | 1.706E-21 | CNT/CD | ZNF165 |
| 1.5674E-14 | 0.39841062 | 0.489 | 0.249 | 3.3464E-10 | CNT/CD | NDUFAB1 |
| 2.7663E-33 | 0.39713553 | 0.478 | 0.151 | 5.9061E-29 | CNT/CD | AIG1 |
| 9.036E-118 | 0.39690238 | 0.31 | 0.024 | 1.929E-113 | CNT/CD | LCN12 |
| 9.6504E-23 | 0.39634499 | 0.375 | 0.129 | 2.0604E-18 | CNT/CD | ABHD11 |
| 1.2272E-16 | 0.39596519 | 0.848 | 0.588 | 2.6202E-12 | CNT/CD | KRT19 |
| 1.4637E-21 | 0.39510147 | 0.533 | 0.232 | 3.1249E-17 | CNT/CD | MDH2 |
| 4.9427E-27 | 0.39508577 | 0.473 | 0.168 | 1.0553E-22 | CNT/CD | GPI |
| 9.7496E-25 | 0.39414806 | 0.386 | 0.128 | 2.0815E-20 | CNT/CD | KLF10 |
| 1.1229E-35 | 0.39381489 | 0.353 | 0.088 | 2.3974E-31 | CNT/CD | IFIT5 |
| 8.9824E-15 | 0.39373343 | 0.745 | 0.442 | 1.9177E-10 | CNT/CD | ATP5MD |
| 1.0419E-65 | 0.3936535 | 0.261 | 0.031 | 2.2244E-61 | CNT/CD | AVPR2 |
| 1.0508E-28 | 0.39220481 | 0.582 | 0.222 | 2.2436E-24 | CNT/CD | ATP6AP2 |
| 8.3331E-16 | 0.39217737 | 0.69 | 0.39 | 1.7791E-11 | CNT/CD | FKBP2 |
| 0.00448026 | 0.39178573 | 0.245 | 0.168 | 1 | CNT/CD | ERRFI1 |
| 4.4505E-30 | 0.3916573 | 0.375 | 0.109 | 9.5018E-26 | CNT/CD | CREM |
| 1.3704E-26 | 0.39161041 | 0.5 | 0.184 | 2.9257E-22 | CNT/CD | BEX4 |
| 7.2659E-17 | 0.39105903 | 0.625 | 0.317 | 1.5513E-12 | CNT/CD | NPC2 |
| 1.2514E-23 | 0.39010687 | 0.353 | 0.113 | 2.6718E-19 | CNT/CD | CRABP2 |
| 6.3918E-85 | 0.38944992 | 0.31 | 0.034 | 1.3647E-80 | CNT/CD | CLYBL |
| 4.6349E-26 | 0.38887161 | 0.625 | 0.262 | 9.8956E-22 | CNT/CD | TCEAL4 |
| 2.6E-16 | 0.38877468 | 0.446 | 0.198 | 5.551E-12 | CNT/CD | IRF2BP2 |
| 3.1485E-17 | 0.3885807 | 0.473 | 0.211 | 6.7221E-13 | CNT/CD | PARD6B |
| 9.4365E-85 | 0.3885719 | 0.304 | 0.033 | 2.0147E-80 | CNT/CD | UGT3A1 |
| 8.5232E-48 | 0.38803201 | 0.25 | 0.038 | 1.8197E-43 | CNT/CD | ZNF556 |
| 1.0563E-28 | 0.38794477 | 0.31 | 0.082 | 2.2553E-24 | CNT/CD | SLC16A4 |
| 5.7277E-35 | 0.38663483 | 0.342 | 0.085 | 1.2229E-30 | CNT/CD | TMEM251 |
| 1.2781E-17 | 0.3864792 | 0.859 | 0.575 | 2.7287E-13 | CNT/CD | PRDX5 |
| 1.4519E-27 | 0.38588964 | 0.44 | 0.145 | 3.0998E-23 | CNT/CD | NUCB2 |
| 0.00944886 | 0.38502188 | 0.364 | 0.285 | 1 | CNT/CD | PLAUR |
| 6.1845E-35 | 0.38495654 | 0.288 | 0.062 | 1.3204E-30 | CNT/CD | GATA2 |
| 2.3896E-25 | 0.38463351 | 0.304 | 0.086 | 5.1019E-21 | CNT/CD | CADPS2 |
| 1.0953E-30 | 0.38401923 | 0.44 | 0.139 | 2.3386E-26 | CNT/CD | ACAA2 |
| 6.3548E-11 | 0.38390592 | 0.489 | 0.273 | 1.3567E-06 | CNT/CD | SMS |
| 6.127E-25 | 0.381807 | 0.429 | 0.152 | 1.3081E-20 | CNT/CD | FAM162A |

| | | | | | | |
|---|---|---|---|---|---|---|
| 2.9465E-14 | 0.38075266 | 0.745 | 0.458 | 6.2908E-10 | CNT/CD | SERP1 |
| 9.5305E-15 | 0.37956545 | 0.25 | 0.088 | 2.0348E-10 | CNT/CD | TRIB1 |
| 1.4018E-16 | 0.37856997 | 0.679 | 0.366 | 2.9929E-12 | CNT/CD | TAGLN2 |
| 6.0788E-20 | 0.3784214 | 0.592 | 0.275 | 1.2978E-15 | CNT/CD | ANAPC16 |
| 7.6859E-14 | 0.37839828 | 0.533 | 0.282 | 1.6409E-09 | CNT/CD | NDUFB11 |
| 1.1702E-51 | 0.37838686 | 0.299 | 0.048 | 2.4984E-47 | CNT/CD | ZNF555 |
| 0.00761241 | 0.3774451 | 0.13 | 0.077 | 1 | CNT/CD | HSPA6 |
| 4.0812E-05 | 0.37692705 | 0.701 | 0.604 | 0.87133248 | CNT/CD | SOD2 |
| 8.4116E-17 | 0.3768656 | 0.701 | 0.398 | 1.7959E-12 | CNT/CD | LAMTOR5 |
| 8.5463E-13 | 0.37514158 | 0.717 | 0.427 | 1.8246E-08 | CNT/CD | ZFAND5 |
| 8.8098E-15 | 0.37457163 | 0.701 | 0.402 | 1.8809E-10 | CNT/CD | SDC4 |
| 7.7471E-21 | 0.37417813 | 0.44 | 0.171 | 1.654E-16 | CNT/CD | ZNF217 |
| 0 | 0.37400622 | 0.288 | 0.002 | 0 | CNT/CD | PRR35 |
| 1.65E-286 | 0.37345529 | 0.272 | 0.006 | 3.522E-282 | CNT/CD | AL031710.1 |
| 1.773E-14 | 0.37219015 | 0.565 | 0.3 | 3.7854E-10 | CNT/CD | PNRC2 |
| 1.6944E-12 | 0.371855 | 0.592 | 0.331 | 3.6175E-08 | CNT/CD | ZNF292 |
| 1.9671E-23 | 0.37185129 | 0.277 | 0.077 | 4.1997E-19 | CNT/CD | AGPAT5 |
| 1.4813E-15 | 0.37143477 | 0.332 | 0.132 | 3.1626E-11 | CNT/CD | ITPKC |
| 1.5851E-29 | 0.37109153 | 0.234 | 0.049 | 3.3842E-25 | CNT/CD | AOC1 |
| 1.5824E-33 | 0.3709294 | 0.272 | 0.058 | 3.3785E-29 | CNT/CD | SPHK1 |
| 3.8689E-15 | 0.37008156 | 0.174 | 0.048 | 8.2601E-11 | CNT/CD | TAC1 |
| 3.2571E-19 | 0.37000078 | 0.505 | 0.225 | 6.954E-15 | CNT/CD | ATP5PB |
| 2.4272E-40 | 0.36824191 | 0.332 | 0.072 | 5.1822E-36 | CNT/CD | IVD |
| 2.3334E-22 | 0.36811949 | 0.467 | 0.185 | 4.9817E-18 | CNT/CD | TMEM205 |
| 4.3258E-63 | 0.36768782 | 0.288 | 0.039 | 9.2355E-59 | CNT/CD | SGK2 |
| 8.5068E-46 | 0.36675034 | 0.125 | 0.01 | 1.8162E-41 | CNT/CD | CXCL10 |
| 1.0568E-59 | 0.36605353 | 0.288 | 0.04 | 2.2563E-55 | CNT/CD | ARHGAP5-AS1 |
| 1.2102E-61 | 0.36585082 | 0.266 | 0.034 | 2.5837E-57 | CNT/CD | COLCA1 |
| 3.5239E-17 | 0.36518234 | 0.533 | 0.256 | 7.5236E-13 | CNT/CD | NDUFB3 |
| 4.391E-57 | 0.36436825 | 0.261 | 0.035 | 9.3749E-53 | CNT/CD | ZNF554 |
| 1.2242E-16 | 0.36420714 | 0.641 | 0.337 | 2.6137E-12 | CNT/CD | GHITM |
| 9.8088E-44 | 0.36331467 | 0.315 | 0.06 | 2.0942E-39 | CNT/CD | PPARGC1A |
| 3.5622E-90 | 0.36253822 | 0.223 | 0.016 | 7.6053E-86 | CNT/CD | LRRTM1 |
| 3.9678E-17 | 0.36226427 | 0.63 | 0.323 | 8.4713E-13 | CNT/CD | NDUFC1 |
| 2.2304E-25 | 0.36219911 | 0.408 | 0.137 | 4.7618E-21 | CNT/CD | TCEAL3 |
| 7.2355E-39 | 0.36179283 | 0.348 | 0.08 | 1.5448E-34 | CNT/CD | TSPAN6 |
| 2.6805E-56 | 0.36107667 | 0.31 | 0.048 | 5.7228E-52 | CNT/CD | DZIP3 |
| 4.0874E-26 | 0.3608383 | 0.299 | 0.082 | 8.7267E-22 | CNT/CD | IFIH1 |
| 1.0825E-27 | 0.36024563 | 0.397 | 0.12 | 2.311E-23 | CNT/CD | PTGER3 |
| 9.9337E-66 | 0.3591709 | 0.288 | 0.037 | 2.1208E-61 | CNT/CD | HLA-DMA |
| 2.3598E-46 | 0.35907707 | 0.332 | 0.064 | 5.0382E-42 | CNT/CD | TRIM69 |
| 1.5555E-27 | 0.359049 | 0.293 | 0.076 | 3.3211E-23 | CNT/CD | HNF1B |
| 1.5863E-27 | 0.35819041 | 0.375 | 0.115 | 3.3867E-23 | CNT/CD | NDUFA9 |
| 5.7759E-42 | 0.35806986 | 0.359 | 0.079 | 1.2332E-37 | CNT/CD | PRCP |
| 2.9816E-18 | 0.35728297 | 0.435 | 0.181 | 6.3657E-14 | CNT/CD | ATP1B3 |
| 1.1881E-79 | 0.35717739 | 0.272 | 0.028 | 2.5367E-75 | CNT/CD | FAM107A |
| 8.4834E-33 | 0.3568536 | 0.359 | 0.095 | 1.8112E-28 | CNT/CD | CAT |
| 2.575E-100 | 0.35541127 | 0.283 | 0.024 | 5.4981E-96 | CNT/CD | LINC00346 |
| 3.5961E-24 | 0.35536145 | 0.424 | 0.151 | 7.6776E-20 | CNT/CD | SDHB |
| 6.8034E-17 | 0.35515976 | 0.582 | 0.296 | 1.4525E-12 | CNT/CD | NDFIP1 |

| | | | | | | |
|---|---|---|---|---|---|---|
| 1.2045E-27 | 0.35344562 | 0.321 | 0.089 | 2.5716E-23 | CNT/CD | FAM3B |
| 2.6417E-13 | 0.3534124 | 0.418 | 0.197 | 5.64E-09 | CNT/CD | EHF |
| 1.2821E-48 | 0.35267078 | 0.31 | 0.054 | 2.7372E-44 | CNT/CD | SMIM24 |
| 8.6672E-19 | 0.35212718 | 0.446 | 0.185 | 1.8504E-14 | CNT/CD | ATRAID |
| 5.227E-17 | 0.35120863 | 0.571 | 0.269 | 1.116E-12 | CNT/CD | MUC1 |
| 1.0656E-31 | 0.35056557 | 0.353 | 0.094 | 2.275E-27 | CNT/CD | SPINT1-AS1 |
| 1.1113E-07 | 0.35047501 | 0.332 | 0.183 | 0.0023726 | CNT/CD | MRPL18 |
| 6.4754E-63 | 0.35012586 | 0.255 | 0.031 | 1.3825E-58 | CNT/CD | MUC15 |
| 5.3084E-35 | 0.34971768 | 0.272 | 0.056 | 1.1334E-30 | CNT/CD | RHOV |
| 7.9976E-49 | 0.34914866 | 0.266 | 0.042 | 1.7075E-44 | CNT/CD | MAP3K14 |
| 4.3887E-07 | 0.34871415 | 0.234 | 0.113 | 0.00936989 | CNT/CD | ZC3HAV1 |
| 2.724E-21 | 0.34848409 | 0.473 | 0.191 | 5.8156E-17 | CNT/CD | NUCB1 |
| 3.6083E-19 | 0.34834829 | 0.478 | 0.204 | 7.7037E-15 | CNT/CD | SCP2 |
| 6.9949E-30 | 0.34758831 | 0.255 | 0.057 | 1.4934E-25 | CNT/CD | PDE4D |
| 7.3478E-19 | 0.34693966 | 0.413 | 0.165 | 1.5688E-14 | CNT/CD | MAP1LC3A |
| 2.254E-47 | 0.34675396 | 0.272 | 0.044 | 4.8122E-43 | CNT/CD | PFKM |
| 6.3682E-14 | 0.3457347 | 0.527 | 0.267 | 1.3596E-09 | CNT/CD | SEPHS2 |
| 3.7661E-18 | 0.34567264 | 0.348 | 0.127 | 8.0406E-14 | CNT/CD | KDM6B |
| 1.5689E-14 | 0.34510386 | 0.701 | 0.397 | 3.3497E-10 | CNT/CD | CDV3 |
| 6.1121E-43 | 0.3447365 | 0.266 | 0.046 | 1.3049E-38 | CNT/CD | PHACTR1 |
| 4.325E-36 | 0.3442742 | 0.288 | 0.061 | 9.2339E-32 | CNT/CD | SYNGR1 |
| 8.8036E-14 | 0.34383087 | 0.348 | 0.152 | 1.8796E-09 | CNT/CD | LGALS3BP |
| 5.981E-31 | 0.34359503 | 0.348 | 0.093 | 1.2769E-26 | CNT/CD | MOCS2 |
| 1.1111E-13 | 0.34317795 | 0.413 | 0.194 | 2.3721E-09 | CNT/CD | MAP2K3 |
| 7.3516E-12 | 0.34272908 | 0.495 | 0.268 | 1.5696E-07 | CNT/CD | NDUFA2 |
| 3.1949E-11 | 0.34254422 | 0.288 | 0.127 | 6.8212E-07 | CNT/CD | SLC30A1 |
| 4.8055E-36 | 0.34235227 | 0.326 | 0.076 | 1.026E-31 | CNT/CD | ABHD14B |
| 1.7447E-20 | 0.34126908 | 0.386 | 0.142 | 3.7248E-16 | CNT/CD | MAGED2 |
| 7.7449E-26 | 0.34119773 | 0.277 | 0.073 | 1.6535E-21 | CNT/CD | BABAM2 |
| 2.1713E-18 | 0.33967297 | 0.402 | 0.163 | 4.6357E-14 | CNT/CD | UQCRC1 |
| 1.1085E-29 | 0.3386032 | 0.332 | 0.089 | 2.3667E-25 | CNT/CD | KIF9 |
| 1.1428E-27 | 0.33758644 | 0.261 | 0.062 | 2.4399E-23 | CNT/CD | SERTAD4-AS1 |
| 1.5456E-58 | 0.33537917 | 0.288 | 0.041 | 3.2998E-54 | CNT/CD | ACSM3 |
| 5.0403E-24 | 0.33411144 | 0.386 | 0.129 | 1.0761E-19 | CNT/CD | NIPSNAP2 |
| 2.5076E-20 | 0.333347 | 0.342 | 0.117 | 5.3538E-16 | CNT/CD | GNG11 |
| 4.898E-15 | 0.33317336 | 0.37 | 0.157 | 1.0457E-10 | CNT/CD | TP53BP2 |
| 5.9397E-89 | 0.33260486 | 0.266 | 0.024 | 1.2681E-84 | CNT/CD | TRPM3 |
| 3.4837E-21 | 0.33195383 | 0.315 | 0.1 | 7.4376E-17 | CNT/CD | MYO1B |
| 1.7664E-16 | 0.3315421 | 0.603 | 0.302 | 3.7713E-12 | CNT/CD | EID1 |
| 2.484E-18 | 0.33078939 | 0.348 | 0.13 | 5.3034E-14 | CNT/CD | NDUFS3 |
| 1.7512E-18 | 0.33056621 | 0.326 | 0.117 | 3.7388E-14 | CNT/CD | SYNJ2BP |
| 1.1581E-24 | 0.33035656 | 0.5 | 0.183 | 2.4726E-20 | CNT/CD | SCNN1A |
| 1.2721E-21 | 0.33032448 | 0.375 | 0.132 | 2.716E-17 | CNT/CD | HEXA |
| 5.5577E-22 | 0.33013072 | 0.424 | 0.158 | 1.1866E-17 | CNT/CD | ETFB |
| 1.768E-11 | 0.32978341 | 0.598 | 0.355 | 3.7747E-07 | CNT/CD | ATP5ME |
| 2.374E-28 | 0.3284718 | 0.288 | 0.072 | 5.0685E-24 | CNT/CD | SLC16A7 |
| 2.4116E-53 | 0.32845361 | 0.261 | 0.037 | 5.1487E-49 | CNT/CD | FRAS1 |
| 1.1409E-13 | 0.32730069 | 0.261 | 0.098 | 2.4359E-09 | CNT/CD | CD68 |
| 6.2506E-89 | 0.32654068 | 0.25 | 0.021 | 1.3345E-84 | CNT/CD | SLC9A4 |
| 1.3375E-14 | 0.32643034 | 0.582 | 0.306 | 2.8556E-10 | CNT/CD | GABARAPL2 |

| | | | | | | |
|---|---|---|---|---|---|---|
| 1.7855E-14 | 0.32637853 | 0.668 | 0.338 | 3.812E-10 | CNT/CD | ZFP36L2 |
| 6.899E-213 | 0.32557031 | 0.125 | 0.001 | 1.473E-208 | CNT/CD | AC087379.2 |
| 5.5807E-13 | 0.32443517 | 0.473 | 0.243 | 1.1915E-08 | CNT/CD | TUFM |
| 1.2153E-62 | 0.32420312 | 0.266 | 0.033 | 2.5947E-58 | CNT/CD | TBX2-AS1 |
| 2.1299E-24 | 0.32388845 | 0.321 | 0.095 | 4.5474E-20 | CNT/CD | SLC25A11 |
| 2.5515E-16 | 0.32374598 | 0.446 | 0.201 | 5.4474E-12 | CNT/CD | ADI1 |
| 4.2522E-34 | 0.32304451 | 0.304 | 0.069 | 9.0784E-30 | CNT/CD | LIMCH1 |
| 1.5024E-22 | 0.32293658 | 0.413 | 0.149 | 3.2076E-18 | CNT/CD | ATP6AP1 |
| 3.8446E-12 | 0.32111988 | 0.293 | 0.123 | 8.2083E-08 | CNT/CD | GPNMB |
| 6.4548E-10 | 0.320371 | 0.571 | 0.338 | 1.3781E-05 | CNT/CD | ARF4 |
| 4.687E-224 | 0.32035096 | 0.223 | 0.005 | 1.001E-219 | CNT/CD | AC148477.4 |
| 1.7492E-11 | 0.32001571 | 0.614 | 0.366 | 3.7346E-07 | CNT/CD | SDCBP |
| 7.8619E-20 | 0.31984331 | 0.342 | 0.119 | 1.6785E-15 | CNT/CD | TUFT1 |
| 3.7708E-13 | 0.31943199 | 0.429 | 0.202 | 8.0506E-09 | CNT/CD | FOSL2 |
| 1.0662E-33 | 0.31879192 | 0.223 | 0.041 | 2.2763E-29 | CNT/CD | RASL11A |
| 2.1443E-40 | 0.31870918 | 0.326 | 0.068 | 4.5781E-36 | CNT/CD | TMC4 |
| 3.5768E-20 | 0.31794621 | 0.348 | 0.123 | 7.6366E-16 | CNT/CD | CRYZ |
| 2.647E-22 | 0.31747311 | 0.283 | 0.082 | 5.6513E-18 | CNT/CD | ALAS1 |
| 1.6563E-19 | 0.31612781 | 0.364 | 0.134 | 3.5361E-15 | CNT/CD | HDDC2 |
| 6.1085E-21 | 0.31554138 | 0.359 | 0.125 | 1.3042E-16 | CNT/CD | CMC2 |
| 4.279E-62 | 0.31548616 | 0.25 | 0.03 | 9.1356E-58 | CNT/CD | NFATC1 |
| 2.1025E-18 | 0.31544005 | 0.391 | 0.154 | 4.4888E-14 | CNT/CD | MFF |
| 3.3685E-18 | 0.3152226 | 0.446 | 0.186 | 7.1918E-14 | CNT/CD | SDHC |
| 6.999E-15 | 0.31518926 | 0.353 | 0.146 | 1.4943E-10 | CNT/CD | TFCP2L1 |
| 7.6295E-46 | 0.31515889 | 0.261 | 0.042 | 1.6289E-41 | CNT/CD | AL162377.1 |
| 2.5145E-14 | 0.3143285 | 0.587 | 0.309 | 5.3684E-10 | CNT/CD | PCBD1 |
| 3.1221E-20 | 0.31419638 | 0.114 | 0.018 | 6.6657E-16 | CNT/CD | HPD |
| 1.058E-11 | 0.31407921 | 0.484 | 0.249 | 2.2589E-07 | CNT/CD | IRF1 |
| 4.0379E-21 | 0.31384327 | 0.288 | 0.088 | 8.6209E-17 | CNT/CD | EMX2OS |
| 2.7711E-40 | 0.31286443 | 0.277 | 0.052 | 5.9164E-36 | CNT/CD | SCML1 |
| 1.2941E-23 | 0.31282019 | 0.255 | 0.067 | 2.7629E-19 | CNT/CD | OVOL1 |
| 9.9532E-51 | 0.31233752 | 0.266 | 0.04 | 2.125E-46 | CNT/CD | IQUB |
| 4.4052E-55 | 0.31153934 | 0.179 | 0.018 | 9.4052E-51 | CNT/CD | LAMA2 |
| 2.1126E-14 | 0.31119195 | 0.571 | 0.297 | 4.5104E-10 | CNT/CD | CUTA |
| 1.3595E-46 | 0.31114288 | 0.31 | 0.056 | 2.9024E-42 | CNT/CD | NDRG2 |
| 4.3573E-63 | 0.30903902 | 0.223 | 0.024 | 9.3029E-59 | CNT/CD | GABRA2 |
| 1.2224E-09 | 0.30893356 | 0.342 | 0.169 | 2.6097E-05 | CNT/CD | SH3BP4 |
| 5.0152E-18 | 0.30882456 | 0.359 | 0.133 | 1.0707E-13 | CNT/CD | L1CAM |
| 2.0424E-75 | 0.30819512 | 0.228 | 0.021 | 4.3606E-71 | CNT/CD | BARX2 |
| 2.1408E-23 | 0.3078465 | 0.163 | 0.031 | 4.5706E-19 | CNT/CD | KL |
| 2.0412E-11 | 0.30776888 | 0.332 | 0.149 | 4.358E-07 | CNT/CD | KITLG |
| 1.5692E-42 | 0.30759307 | 0.304 | 0.058 | 3.3502E-38 | CNT/CD | BLNK |
| 0 | 0.30705233 | 0.185 | 0.001 | 0 | CNT/CD | RBP2 |
| 4.989E-113 | 0.30645065 | 0.234 | 0.014 | 1.065E-108 | CNT/CD | RHBG |
| 4.5096E-13 | 0.30567419 | 0.766 | 0.487 | 9.628E-09 | CNT/CD | TOMM7 |
| 1.438E-205 | 0.30486397 | 0.212 | 0.005 | 3.07E-201 | CNT/CD | ADH1C |
| 2.3889E-09 | 0.30484036 | 0.603 | 0.377 | 5.1004E-05 | CNT/CD | RHEB |
| 2.7782E-27 | 0.30391697 | 0.255 | 0.061 | 5.9316E-23 | CNT/CD | SBF2-AS1 |
| 7.3667E-47 | 0.3037986 | 0.266 | 0.042 | 1.5728E-42 | CNT/CD | FXYD3 |
| 4.9119E-51 | 0.3031284 | 0.255 | 0.037 | 1.0487E-46 | CNT/CD | ARL15 |

| | | | | | | |
|---|---|---|---|---|---|---|
| 1.829E-18 | 0.30308019 | 0.315 | 0.109 | 3.9049E-14 | CNT/CD | BDH2 |
| 8.874E-40 | 0.30274069 | 0.255 | 0.045 | 1.8946E-35 | CNT/CD | NOCT |
| 6.0011E-14 | 0.30204678 | 0.457 | 0.22 | 1.2812E-09 | CNT/CD | TMED4 |
| 6.2207E-53 | 0.30194022 | 0.245 | 0.033 | 1.3281E-48 | CNT/CD | DYRK3 |
| 0.0029857 | 0.30168886 | 0.147 | 0.085 | 1 | CNT/CD | MT1H |
| 1.1377E-06 | 0.30104785 | 0.299 | 0.166 | 0.02428913 | CNT/CD | FGF2 |
| 2.0979E-23 | 0.30065622 | 0.31 | 0.091 | 4.479E-19 | CNT/CD | NFKBIE |
| 9.9493E-28 | 0.30058995 | 0.185 | 0.034 | 2.1242E-23 | CNT/CD | KLHDC7B |
| 8.395E-32 | 0.30026382 | 0.255 | 0.053 | 1.7923E-27 | CNT/CD | HRK |
| 9.9345E-37 | 0.29938685 | 0.266 | 0.052 | 2.121E-32 | CNT/CD | PROM2 |
| 1.8705E-10 | 0.29877373 | 0.272 | 0.118 | 3.9935E-06 | CNT/CD | FBXO32 |
| 1.164E-06 | 0.29839528 | 0.62 | 0.44 | 0.02485154 | CNT/CD | UPP1 |
| 4.1305E-12 | 0.29767588 | 0.478 | 0.254 | 8.8187E-08 | CNT/CD | PRKAR1A |
| 1.258E-126 | 0.29719844 | 0.217 | 0.011 | 2.685E-122 | CNT/CD | TMEM72 |
| 1.8092E-09 | 0.2963579 | 0.88 | 0.733 | 3.8626E-05 | CNT/CD | SKP1 |
| 1.8502E-14 | 0.2960268 | 0.293 | 0.111 | 3.9503E-10 | CNT/CD | MYADM |
| 6.9369E-17 | 0.29580148 | 0.364 | 0.143 | 1.481E-12 | CNT/CD | ATP6V1A |
| 8.1225E-60 | 0.29568837 | 0.261 | 0.033 | 1.7341E-55 | CNT/CD | SELENBP1 |
| 1.8624E-26 | 0.29533294 | 0.261 | 0.064 | 3.9762E-22 | CNT/CD | AC093673.1 |
| 4.2286E-43 | 0.29507046 | 0.266 | 0.046 | 9.028E-39 | CNT/CD | EPHX2 |
| 1.2052E-16 | 0.29485614 | 0.288 | 0.102 | 2.5731E-12 | CNT/CD | TCEAL8 |
| 1.5159E-14 | 0.29454264 | 0.364 | 0.153 | 3.2365E-10 | CNT/CD | TGIF1 |
| 9.5616E-19 | 0.29369141 | 0.288 | 0.095 | 2.0414E-14 | CNT/CD | TBC1D9 |
| 1.888E-09 | 0.29312048 | 0.495 | 0.282 | 4.0308E-05 | CNT/CD | HIST1H1C |
| 6.9168E-05 | 0.29312048 | 0.255 | 0.15 | 1 | CNT/CD | BAG3 |
| 2.3085E-13 | 0.2923845 | 0.31 | 0.129 | 4.9287E-09 | CNT/CD | TANK |
| 1.4536E-08 | 0.2921794 | 0.31 | 0.156 | 0.00031034 | CNT/CD | FHL2 |
| 9.8121E-16 | 0.29188967 | 0.353 | 0.142 | 2.0949E-11 | CNT/CD | CRB3 |
| 2.2645E-07 | 0.29150316 | 0.375 | 0.218 | 0.00483473 | CNT/CD | WTAP |
| 1.988E-18 | 0.28950441 | 0.397 | 0.155 | 4.2444E-14 | CNT/CD | MAP3K13 |
| 5.6097E-27 | 0.28932004 | 0.217 | 0.046 | 1.1977E-22 | CNT/CD | TIFA |
| 3.3193E-18 | 0.28903955 | 0.261 | 0.079 | 7.0867E-14 | CNT/CD | CXCL1 |
| 0.000344 | 0.28880513 | 0.63 | 0.494 | 1 | CNT/CD | HSPD1 |
| 2.4421E-14 | 0.28848109 | 0.353 | 0.148 | 5.2138E-10 | CNT/CD | ETS2 |
| 3.1528E-15 | 0.28832147 | 0.408 | 0.175 | 6.7311E-11 | CNT/CD | RYBP |
| 3.4376E-13 | 0.28808176 | 0.332 | 0.143 | 7.3394E-09 | CNT/CD | EFCAB14 |
| 3.7738E-10 | 0.28764946 | 0.462 | 0.26 | 8.057E-06 | CNT/CD | NDUFB10 |
| 4.7659E-26 | 0.28747342 | 0.288 | 0.076 | 1.0175E-21 | CNT/CD | ATPAF1 |
| 5.4237E-60 | 0.28703618 | 0.25 | 0.031 | 1.158E-55 | CNT/CD | C1orf54 |
| 7.8434E-23 | 0.28699481 | 0.255 | 0.068 | 1.6746E-18 | CNT/CD | SIM1 |
| 2.4665E-17 | 0.2866497 | 0.332 | 0.122 | 5.2661E-13 | CNT/CD | VEGFB |
| 7.4147E-09 | 0.28655944 | 0.424 | 0.237 | 0.0001583 | CNT/CD | KLF5 |
| 1.6415E-18 | 0.28567005 | 0.342 | 0.124 | 3.5047E-14 | CNT/CD | MRPL19 |
| 2.1393E-17 | 0.28527411 | 0.348 | 0.131 | 4.5674E-13 | CNT/CD | RRAGD |
| 5.9667E-17 | 0.28518484 | 0.473 | 0.208 | 1.2739E-12 | CNT/CD | RAB14 |
| 1.8846E-17 | 0.28507771 | 0.342 | 0.129 | 4.0236E-13 | CNT/CD | NDUFS1 |
| 3.9791E-55 | 0.28505401 | 0.239 | 0.03 | 8.4954E-51 | CNT/CD | AL353708.3 |
| 9.8729E-54 | 0.2850025 | 0.245 | 0.032 | 2.1079E-49 | CNT/CD | FYB2 |
| 1.1297E-18 | 0.28488133 | 0.353 | 0.129 | 2.4118E-14 | CNT/CD | ECHDC2 |
| 5.838E-07 | 0.28422225 | 0.446 | 0.272 | 0.01246422 | CNT/CD | VMP1 |

| | | | | | | |
|---|---|---|---|---|---|---|
| 1.384E-24 | 0.28401887 | 0.283 | 0.077 | 2.9548E-20 | CNT/CD | GAA |
| 4.6057E-10 | 0.28326456 | 0.821 | 0.662 | 9.8332E-06 | CNT/CD | SOD1 |
| 5.8904E-18 | 0.28319014 | 0.288 | 0.096 | 1.2576E-13 | CNT/CD | DLGAP1-AS1 |
| 2.9174E-11 | 0.28289414 | 0.326 | 0.15 | 6.2288E-07 | CNT/CD | USP12 |
| 1.7711E-17 | 0.28262097 | 0.283 | 0.096 | 3.7814E-13 | CNT/CD | METTL23 |
| 1.3803E-23 | 0.28261889 | 0.288 | 0.081 | 2.9469E-19 | CNT/CD | HIBADH |
| 1.1841E-36 | 0.28156875 | 0.228 | 0.039 | 2.528E-32 | CNT/CD | AL355353.1 |
| 7.8663E-56 | 0.28117224 | 0.223 | 0.026 | 1.6794E-51 | CNT/CD | CYBRD1 |
| 1.1219E-17 | 0.28041631 | 0.38 | 0.149 | 2.3952E-13 | CNT/CD | RDH11 |
| 3.4849E-13 | 0.28019538 | 0.348 | 0.153 | 7.4402E-09 | CNT/CD | GNS |
| 6.0482E-13 | 0.2796328 | 0.549 | 0.284 | 1.2913E-08 | CNT/CD | TGOLN2 |
| 1.0016E-12 | 0.27948248 | 0.576 | 0.313 | 2.1383E-08 | CNT/CD | SARAF |
| 4.9272E-13 | 0.27845635 | 0.174 | 0.053 | 1.052E-08 | CNT/CD | RASL11B |
| 3.541E-13 | 0.27835491 | 0.397 | 0.183 | 7.5601E-09 | CNT/CD | TNFRSF10B |
| 4.7301E-35 | 0.27795834 | 0.239 | 0.044 | 1.0099E-30 | CNT/CD | ACADSB |
| 7.649E-09 | 0.27784982 | 0.261 | 0.122 | 0.00016331 | CNT/CD | ARL5B |
| 1.0901E-16 | 0.27714568 | 0.337 | 0.127 | 2.3274E-12 | CNT/CD | HINT2 |
| 2.758E-170 | 0.27695689 | 0.217 | 0.007 | 5.887E-166 | CNT/CD | C9orf24 |
| 3.7035E-15 | 0.27678765 | 0.315 | 0.121 | 7.907E-11 | CNT/CD | PRKAG2 |
| 6.4197E-38 | 0.27646224 | 0.25 | 0.045 | 1.3706E-33 | CNT/CD | GPRC5B |
| 3.3989E-13 | 0.27636463 | 0.418 | 0.199 | 7.2567E-09 | CNT/CD | TMEM230 |
| 3.0237E-12 | 0.27533526 | 0.565 | 0.307 | 6.4557E-08 | CNT/CD | PGK1 |
| 7.9118E-29 | 0.27479478 | 0.13 | 0.017 | 1.6892E-24 | CNT/CD | KIF5C |
| 2.1441E-31 | 0.27433256 | 0.163 | 0.025 | 4.5777E-27 | CNT/CD | HPSE |
| 4.9764E-41 | 0.27431723 | 0.223 | 0.034 | 1.0625E-36 | CNT/CD | LHX1-DT |
| 1.1895E-09 | 0.27381469 | 0.272 | 0.122 | 2.5396E-05 | CNT/CD | HOTAIRM1 |
| 4.3838E-15 | 0.27322678 | 0.353 | 0.144 | 9.3595E-11 | CNT/CD | LGMN |
| 4.4472E-12 | 0.27317308 | 0.228 | 0.085 | 9.4947E-08 | CNT/CD | BAMBI |
| 8.3451E-13 | 0.27210791 | 0.402 | 0.188 | 1.7817E-08 | CNT/CD | ECI1 |
| 2.2858E-09 | 0.27202031 | 0.598 | 0.341 | 4.8801E-05 | CNT/CD | MIF |
| 3.5889E-15 | 0.27156025 | 0.543 | 0.26 | 7.6624E-11 | CNT/CD | SELENOM |
| 4.2075E-12 | 0.27146831 | 0.674 | 0.39 | 8.9831E-08 | CNT/CD | SRSF3 |
| 2.5958E-13 | 0.27146303 | 0.293 | 0.119 | 5.542E-09 | CNT/CD | SLC39A6 |
| 1.6587E-22 | 0.27061742 | 0.315 | 0.096 | 3.5413E-18 | CNT/CD | HSD17B4 |
| 4.4868E-20 | 0.2704554 | 0.239 | 0.066 | 9.5793E-16 | CNT/CD | FAM110B |
| 9.2133E-32 | 0.27044659 | 0.261 | 0.055 | 1.967E-27 | CNT/CD | METTL7A |
| 2.0373E-21 | 0.27034632 | 0.337 | 0.11 | 4.3496E-17 | CNT/CD | MORN2 |
| 2.8196E-41 | 0.27006677 | 0.255 | 0.044 | 6.02E-37 | CNT/CD | RASGRP1 |
| 1.3545E-76 | 0.27000085 | 0.223 | 0.019 | 2.8919E-72 | CNT/CD | INPP5J |
| 1.0901E-08 | 0.26977029 | 0.533 | 0.328 | 0.00023274 | CNT/CD | NDUFB4 |
| 1.9461E-64 | 0.26955571 | 0.217 | 0.022 | 4.1548E-60 | CNT/CD | PLEKHB1 |
| 1.0868E-13 | 0.26930089 | 0.418 | 0.194 | 2.3203E-09 | CNT/CD | HSD17B12 |
| 3.3265E-24 | 0.26883007 | 0.245 | 0.061 | 7.102E-20 | CNT/CD | SGPP1 |
| 6.5252E-25 | 0.26785187 | 0.245 | 0.06 | 1.3931E-20 | CNT/CD | FAM229B |
| 3.5136E-18 | 0.26755152 | 0.234 | 0.068 | 7.5015E-14 | CNT/CD | VAV3 |
| 2.7729E-10 | 0.26740689 | 0.859 | 0.632 | 5.9201E-06 | CNT/CD | UBL5 |
| 2.871E-12 | 0.26723534 | 0.473 | 0.243 | 6.1296E-08 | CNT/CD | HADHA |
| 5.5578E-21 | 0.26715698 | 0.272 | 0.079 | 1.1866E-16 | CNT/CD | RGP1 |
| 1.2592E-13 | 0.26709284 | 0.304 | 0.124 | 2.6883E-09 | CNT/CD | HPRT1 |
| 3.544E-18 | 0.2666865 | 0.255 | 0.078 | 7.5665E-14 | CNT/CD | BCL2 |

| | | | | | | |
|---|---|---|---|---|---|---|
| 1.0799E-53 | 0.26655461 | 0.207 | 0.024 | 2.3056E-49 | CNT/CD | AL031058.1 |
| 6.6464E-14 | 0.26624929 | 0.239 | 0.084 | 1.419E-09 | CNT/CD | RIPK2 |
| 0.0026231 | 0.26605243 | 0.375 | 0.265 | 1 | CNT/CD | KCNQ1OT1 |
| 1.4057E-10 | 0.26575484 | 0.451 | 0.243 | 3.0011E-06 | CNT/CD | CAMTA1 |
| 8.0634E-11 | 0.26571862 | 0.353 | 0.169 | 1.7215E-06 | CNT/CD | NAA38 |
| 7.967E-107 | 0.26512473 | 0.207 | 0.012 | 1.701E-102 | CNT/CD | CCDC151 |
| 1.6104E-34 | 0.26491355 | 0.217 | 0.038 | 3.4383E-30 | CNT/CD | LINC01503 |
| 1.368E-11 | 0.26458261 | 0.37 | 0.174 | 2.9206E-07 | CNT/CD | SLC39A1 |
| 8.5905E-31 | 0.26436591 | 0.245 | 0.051 | 1.8341E-26 | CNT/CD | CFI |
| 5.2391E-10 | 0.26418247 | 0.315 | 0.149 | 1.1185E-05 | CNT/CD | ACSL4 |
| 6.4944E-30 | 0.26397006 | 0.228 | 0.046 | 1.3866E-25 | CNT/CD | NME5 |
| 4.515E-06 | 0.26362117 | 0.342 | 0.209 | 0.09639553 | CNT/CD | SNX9 |
| 3.3917E-13 | 0.26354533 | 0.353 | 0.151 | 7.2413E-09 | CNT/CD | IVNS1ABP |
| 3.9634E-11 | 0.26309105 | 0.337 | 0.156 | 8.4619E-07 | CNT/CD | MRPL34 |
| 8.5165E-13 | 0.26175631 | 0.293 | 0.12 | 1.8183E-08 | CNT/CD | MSRB2 |
| 3.2474E-20 | 0.26162089 | 0.364 | 0.125 | 6.9333E-16 | CNT/CD | CCDC198 |
| 5.0888E-11 | 0.26084939 | 0.364 | 0.177 | 1.0865E-06 | CNT/CD | NDUFS4 |
| 3.4289E-20 | 0.26060651 | 0.212 | 0.055 | 7.3206E-16 | CNT/CD | LINC01116 |
| 0.00279804 | 0.26021907 | 0.141 | 0.083 | 1 | CNT/CD | NMB |
| 5.197E-31 | 0.25996303 | 0.25 | 0.052 | 1.1096E-26 | CNT/CD | DUSP15 |
| 3.5357E-14 | 0.25926365 | 0.391 | 0.17 | 7.5486E-10 | CNT/CD | ARL2 |
| 6.8399E-35 | 0.25892253 | 0.212 | 0.036 | 1.4603E-30 | CNT/CD | LETM2 |
| 9.5876E-16 | 0.25793144 | 0.321 | 0.123 | 2.0469E-11 | CNT/CD | CETN2 |
| 5.8311E-30 | 0.2571779 | 0.25 | 0.054 | 1.2449E-25 | CNT/CD | BCL3 |
| 8.8578E-20 | 0.25702064 | 0.152 | 0.031 | 1.8911E-15 | CNT/CD | EGR3 |
| 2.6603E-30 | 0.25693805 | 0.239 | 0.05 | 5.6798E-26 | CNT/CD | AL035661.1 |
| 4.0315E-30 | 0.25692336 | 0.272 | 0.062 | 8.6073E-26 | CNT/CD | PCCB |
| 7.7736E-12 | 0.25678727 | 0.44 | 0.218 | 1.6597E-07 | CNT/CD | RBM47 |
| 3.6851E-11 | 0.25665242 | 0.962 | 0.827 | 7.8678E-07 | CNT/CD | MT-ND5 |
| 1.4078E-08 | 0.25639397 | 0.837 | 0.649 | 0.00030057 | CNT/CD | SLC25A3 |
| 2.9574E-08 | 0.256268 | 0.397 | 0.221 | 0.00063141 | CNT/CD | GPRC5C |
| 1.8237E-33 | 0.2557514 | 0.239 | 0.046 | 3.8937E-29 | CNT/CD | HOXB-AS3 |
| 6.3403E-22 | 0.25565551 | 0.299 | 0.089 | 1.3537E-17 | CNT/CD | GEM |
| 2.5489E-42 | 0.25512036 | 0.179 | 0.022 | 5.4418E-38 | CNT/CD | LPP-AS2 |
| 1.2589E-14 | 0.2550952 | 0.386 | 0.165 | 2.6878E-10 | CNT/CD | SPAG7 |
| 2.0959E-24 | 0.25486891 | 0.217 | 0.049 | 4.4747E-20 | CNT/CD | BEX1 |
| 4.2834E-51 | 0.25469049 | 0.239 | 0.032 | 9.1451E-47 | CNT/CD | RGN |
| 3.0599E-11 | 0.25466842 | 0.467 | 0.243 | 6.5328E-07 | CNT/CD | PGAM1 |
| 4.9331E-07 | 0.25438045 | 0.44 | 0.277 | 0.01053208 | CNT/CD | MRPL41 |
| 1.4472E-30 | 0.25416373 | 0.223 | 0.043 | 3.0897E-26 | CNT/CD | ESRRG |
| 7.4918E-51 | 0.25409097 | 0.212 | 0.026 | 1.5995E-46 | CNT/CD | AC027682.6 |
| 3.1595E-38 | 0.25409097 | 0.185 | 0.026 | 6.7455E-34 | CNT/CD | AC010329.1 |
| 9.9711E-08 | 0.2538315 | 0.413 | 0.245 | 0.00212883 | CNT/CD | LAMTOR4 |
| 3.4673E-09 | 0.25355955 | 0.663 | 0.423 | 7.4027E-05 | CNT/CD | ATP6V0B |
| 7.4263E-20 | 0.25337268 | 0.304 | 0.098 | 1.5855E-15 | CNT/CD | TMEM45B |
| 4.181E-29 | 0.25304538 | 0.25 | 0.055 | 8.9265E-25 | CNT/CD | TMEM116 |
| 1.6749E-27 | 0.25270721 | 0.283 | 0.07 | 3.5759E-23 | CNT/CD | ZSCAN16-AS1 |
| 5.1906E-19 | 0.25262052 | 0.272 | 0.084 | 1.1082E-14 | CNT/CD | TACC2 |
| 6.8877E-21 | 0.25234928 | 0.239 | 0.065 | 1.4705E-16 | CNT/CD | TMEM136 |
| 8.4709E-12 | 0.25214352 | 0.717 | 0.424 | 1.8085E-07 | CNT/CD | BSG |

| | | | | | | |
|---|---|---|---|---|---|---|
| 6.3384E-90 | 0.25139901 | 0.196 | 0.013 | 1.3532E-85 | CNT/CD | TSPAN7 |
| 2.0403E-17 | 0.25124891 | 0.293 | 0.101 | 4.3561E-13 | CNT/CD | DLD |
| 6.4151E-18 | 0.25117297 | 0.293 | 0.097 | 1.3696E-13 | CNT/CD | COL6A1 |
| 2.3623E-15 | 0.25030367 | 0.342 | 0.136 | 5.0436E-11 | CNT/CD | DMAC1 |
| 5.5217E-10 | 0.25024996 | 0.212 | 0.085 | 1.1789E-05 | CNT/CD | FAM210A |
| 0.00027857 | -0.2502003 | 0.098 | 0.199 | 1 | CNT/CD | CDK12 |
| 6.7017E-05 | -0.2520989 | 0.12 | 0.241 | 1 | CNT/CD | EPRS |
| 0.00102504 | -0.2521839 | 0.109 | 0.198 | 1 | CNT/CD | UPF2 |
| 7.2384E-05 | -0.2523502 | 0.125 | 0.245 | 1 | CNT/CD | NCKAP1 |
| 0.00051788 | -0.2525024 | 0.141 | 0.245 | 1 | CNT/CD | DNAJC3 |
| 0.00963868 | -0.2527887 | 0.103 | 0.167 | 1 | CNT/CD | TSPYL2 |
| 3.7506E-05 | -0.254401 | 0.147 | 0.28 | 0.80074891 | CNT/CD | SERPINB1 |
| 0.00058954 | -0.254901 | 0.098 | 0.193 | 1 | CNT/CD | SESN2 |
| 0.00049867 | -0.2555478 | 0.098 | 0.196 | 1 | CNT/CD | SDF2L1 |
| 1.4003E-06 | -0.2558834 | 0.054 | 0.194 | 0.02989689 | CNT/CD | BNIP3L |
| 7.4385E-05 | -0.256223 | 0.136 | 0.259 | 1 | CNT/CD | TLN1 |
| 3.3033E-07 | -0.2564218 | 0.022 | 0.162 | 0.00705252 | CNT/CD | MTHFD2 |
| 0.00114584 | -0.2574765 | 0.13 | 0.221 | 1 | CNT/CD | TIMP2 |
| 0.00025868 | -0.2580941 | 0.092 | 0.196 | 1 | CNT/CD | SLC25A37 |
| 0.00014033 | -0.2581162 | 0.13 | 0.247 | 1 | CNT/CD | SLTM |
| 1.077E-05 | -0.2591744 | 0.038 | 0.154 | 0.22993759 | CNT/CD | SLC7A5 |
| 7.7688E-05 | -0.2595048 | 0.13 | 0.256 | 1 | CNT/CD | COTL1 |
| 4.1938E-06 | -0.2602445 | 0.049 | 0.179 | 0.08953848 | CNT/CD | SLC1A5 |
| 0.00060862 | -0.2616215 | 0.87 | 0.89 | 1 | CNT/CD | NACA |
| 0.0042311 | -0.2648174 | 0.196 | 0.277 | 1 | CNT/CD | ATRX |
| 1.147E-06 | -0.2673801 | 0.087 | 0.24 | 0.02448876 | CNT/CD | RNPS1 |
| 0.00372606 | -0.2678635 | 0.141 | 0.219 | 1 | CNT/CD | EPS8L2 |
| 0.0013592 | -0.2685993 | 0.223 | 0.314 | 1 | CNT/CD | NOP53 |
| 1.1065E-05 | -0.2694151 | 0.092 | 0.225 | 0.23623409 | CNT/CD | ILF3 |
| 0.00022513 | -0.2698395 | 0.114 | 0.219 | 1 | CNT/CD | ASH1L |
| 0.00016061 | -0.2701393 | 0.185 | 0.299 | 1 | CNT/CD | UBXN1 |
| 4.6602E-05 | -0.2702612 | 0.092 | 0.214 | 0.99495954 | CNT/CD | KMT2C |
| 2.3221E-05 | -0.2702993 | 0.06 | 0.175 | 0.49576567 | CNT/CD | CCNL2 |
| 0.00142042 | -0.2703015 | 0.12 | 0.204 | 1 | CNT/CD | SGK1 |
| 6.5263E-06 | -0.2714053 | 0.033 | 0.151 | 0.13933705 | CNT/CD | FXYD5 |
| 0.00042543 | -0.2738015 | 0.109 | 0.21 | 1 | CNT/CD | SLFN5 |
| 1.2895E-05 | -0.2748749 | 0.011 | 0.113 | 0.2753121 | CNT/CD | SLC7A11 |
| 0.00521963 | -0.2751322 | 0.266 | 0.34 | 1 | CNT/CD | SSR3 |
| 0.00381038 | -0.276662 | 0.19 | 0.267 | 1 | CNT/CD | ZRANB2 |
| 0.00094269 | -0.2769246 | 0.158 | 0.25 | 1 | CNT/CD | TRIM44 |
| 0.00060032 | -0.2792351 | 0.201 | 0.305 | 1 | CNT/CD | YTHDC1 |
| 0.00010088 | -0.2810771 | 0.201 | 0.323 | 1 | CNT/CD | GNB1 |
| 2.7073E-05 | -0.2817447 | 0.103 | 0.228 | 0.5780122 | CNT/CD | SNRNP70 |
| 0.0023122 | -0.2821004 | 0.467 | 0.547 | 1 | CNT/CD | BRI3 |
| 0.00093465 | -0.2828175 | 0.174 | 0.271 | 1 | CNT/CD | TPR |
| 0.00014336 | -0.2838114 | 0.12 | 0.235 | 1 | CNT/CD | PKN2 |
| 0.00180043 | -0.284081 | 0.136 | 0.219 | 1 | CNT/CD | RERE |
| 0.00072623 | -0.2841097 | 0.533 | 0.613 | 1 | CNT/CD | SAP18 |
| 1.1789E-06 | -0.2842323 | 0.109 | 0.265 | 0.02516955 | CNT/CD | RPS19BP1 |
| 3.0797E-06 | -0.2849644 | 0.076 | 0.214 | 0.06575148 | CNT/CD | BAX |

| | | | | | | |
|---|---|---|---|---|---|---|
| 0.00160539 | -0.285451 | 0.19 | 0.284 | 1 | CNT/CD | ASPH |
| 1.251E-06 | -0.288743 | 0.076 | 0.223 | 0.02670943 | CNT/CD | GARS |
| 1.9901E-05 | -0.2893158 | 0.114 | 0.246 | 0.42489002 | CNT/CD | TNIP1 |
| 0.00055982 | -0.2905295 | 0.239 | 0.338 | 1 | CNT/CD | PSMB1 |
| 0.00187103 | -0.2920388 | 0.364 | 0.439 | 1 | CNT/CD | HMGN1 |
| 0.00365364 | -0.293642 | 0.201 | 0.277 | 1 | CNT/CD | BPTF |
| 9.8439E-05 | -0.2939971 | 0.266 | 0.386 | 1 | CNT/CD | CTNNA1 |
| 8.7549E-07 | -0.2942818 | 0.054 | 0.196 | 0.01869162 | CNT/CD | FAM129B |
| 8.0304E-05 | -0.2946088 | 0.022 | 0.112 | 1 | CNT/CD | HIST4H4 |
| 4.2467E-08 | -0.2947073 | 0.027 | 0.184 | 0.00090668 | CNT/CD | CLIC4 |
| 0.00377484 | -0.2951431 | 0.125 | 0.203 | 1 | CNT/CD | MANF |
| 4.3865E-07 | -0.2960688 | 0.011 | 0.14 | 0.00936524 | CNT/CD | TRIB3 |
| 1.5848E-07 | -0.2965583 | 0.065 | 0.224 | 0.0033836 | CNT/CD | GNAI2 |
| 9.6942E-08 | -0.2974019 | 0.033 | 0.184 | 0.00206971 | CNT/CD | RBM33 |
| 0.00011072 | -0.2979026 | 0.168 | 0.284 | 1 | CNT/CD | CDC37 |
| 1.5825E-05 | -0.2983539 | 0.467 | 0.603 | 0.33786653 | CNT/CD | CAST |
| 6.7205E-06 | -0.2986346 | 0.087 | 0.22 | 0.14348357 | CNT/CD | USP34 |
| 6.0823E-07 | -0.2993409 | 0.049 | 0.194 | 0.0129857 | CNT/CD | PHLDA3 |
| 1.9392E-05 | -0.3009794 | 0.082 | 0.205 | 0.41400981 | CNT/CD | ZFC3H1 |
| 5.0673E-05 | -0.3009865 | 0.13 | 0.255 | 1 | CNT/CD | PNPLA2 |
| 5.5812E-05 | -0.3010798 | 0.087 | 0.207 | 1 | CNT/CD | MDK |
| 5.2862E-06 | -0.3019397 | 0.141 | 0.291 | 0.11285979 | CNT/CD | FAM3C |
| 4.6616E-05 | -0.303632 | 0.136 | 0.266 | 0.99524553 | CNT/CD | RB1CC1 |
| 2.0574E-05 | -0.3036942 | 0.103 | 0.232 | 0.43926057 | CNT/CD | RPL22L1 |
| 9.4624E-08 | -0.304203 | 0.038 | 0.193 | 0.00202021 | CNT/CD | CARHSP1 |
| 0.00329511 | -0.3044906 | 0.446 | 0.496 | 1 | CNT/CD | ACTN4 |
| 1.538E-05 | -0.3059834 | 0.071 | 0.193 | 0.32836859 | CNT/CD | TUT4 |
| 7.3937E-07 | -0.3067639 | 0.065 | 0.213 | 0.01578552 | CNT/CD | GFPT1 |
| 0.00012533 | -0.3069603 | 0.217 | 0.333 | 1 | CNT/CD | KIF5B |
| 7.9503E-08 | -0.3084933 | 0.011 | 0.154 | 0.0016974 | CNT/CD | TFPI |
| 8.0757E-05 | -0.3090769 | 0.038 | 0.139 | 1 | CNT/CD | TM4SF1 |
| 6.3076E-07 | -0.3095289 | 0.076 | 0.231 | 0.01346681 | CNT/CD | ST14 |
| 7.124E-06 | -0.3102887 | 0.071 | 0.206 | 0.1520965 | CNT/CD | CLDN1 |
| 1.3639E-08 | -0.3114517 | 0.016 | 0.176 | 0.0002912 | CNT/CD | EIF4EBP1 |
| 2.2007E-05 | -0.3115433 | 0.114 | 0.241 | 0.46985545 | CNT/CD | ANKLE2 |
| 4.7637E-05 | -0.3126596 | 0.201 | 0.329 | 1 | CNT/CD | TMED9 |
| 3.6942E-05 | -0.3150612 | 0.033 | 0.135 | 0.78871079 | CNT/CD | CSRNP2 |
| 1.1778E-07 | -0.3155756 | 0.076 | 0.241 | 0.00251451 | CNT/CD | RPS17 |
| 1.0268E-05 | -0.316942 | 0.027 | 0.14 | 0.21922217 | CNT/CD | TNFSF10 |
| 8.0907E-06 | -0.3171347 | 0.071 | 0.201 | 0.17273703 | CNT/CD | MARCKS |
| 0.00101294 | -0.3176363 | 0.261 | 0.346 | 1 | CNT/CD | CAP1 |
| 1.0933E-07 | -0.3177658 | 0.087 | 0.257 | 0.00233424 | CNT/CD | SARS |
| 0.00010852 | -0.3187814 | 0.549 | 0.637 | 1 | CNT/CD | CCNI |
| 4.5122E-05 | -0.318827 | 0.092 | 0.209 | 0.96334686 | CNT/CD | VPS13C |
| 1.2167E-06 | -0.3196543 | 0.087 | 0.236 | 0.0259756 | CNT/CD | PHF3 |
| 3.0679E-05 | -0.320238 | 0.092 | 0.223 | 0.6550039 | CNT/CD | SFN |
| 3.4773E-08 | -0.3227226 | 0.043 | 0.208 | 0.00074241 | CNT/CD | MAP4K4 |
| 8.354E-06 | -0.3232766 | 0.082 | 0.214 | 0.17835853 | CNT/CD | PLEC |
| 0.00073703 | -0.3255244 | 0.196 | 0.292 | 1 | CNT/CD | KMT2A |
| 0.00408995 | -0.3262547 | 0.554 | 0.583 | 1 | CNT/CD | ARPC3 |

| | | | | | | |
|---|---|---|---|---|---|---|
| 0.00595866 | -0.3327635 | 0.293 | 0.359 | 1 | CNT/CD | SPAG9 |
| 4.9759E-06 | -0.3359745 | 0.06 | 0.19 | 0.10623552 | CNT/CD | PDIA4 |
| 3.8238E-05 | -0.3375675 | 0.152 | 0.274 | 0.816386 | CNT/CD | SMG1 |
| 1.0023E-08 | -0.3379156 | 0.06 | 0.237 | 0.00021398 | CNT/CD | MGAT4B |
| 2.7136E-05 | -0.3382458 | 0.674 | 0.778 | 0.57935975 | CNT/CD | OAZ1 |
| 0.00018698 | -0.3390322 | 0.668 | 0.719 | 1 | CNT/CD | RPL36AL |
| 0.00205479 | -0.3393388 | 0.31 | 0.386 | 1 | CNT/CD | MYO6 |
| 5.5291E-08 | -0.3398101 | 0.071 | 0.24 | 0.00118046 | CNT/CD | CRIM1 |
| 7.7003E-07 | -0.3423442 | 0.038 | 0.175 | 0.01644016 | CNT/CD | PLA2G16 |
| 0.00364428 | -0.3458947 | 0.967 | 0.937 | 1 | CNT/CD | RPS23 |
| 8.8155E-07 | -0.3477683 | 0.12 | 0.274 | 0.0188212 | CNT/CD | OGA |
| 4.8075E-07 | -0.3496001 | 0.038 | 0.182 | 0.01026404 | CNT/CD | DCBLD2 |
| 0.00020446 | -0.3498476 | 0.478 | 0.563 | 1 | CNT/CD | GUK1 |
| 7.7103E-06 | -0.3549741 | 0.13 | 0.27 | 0.16461537 | CNT/CD | TUBB |
| 1.4813E-05 | -0.3558856 | 0.022 | 0.127 | 0.3162593 | CNT/CD | HIST2H2BF |
| 4.9727E-05 | -0.3573 | 0.147 | 0.265 | 1 | CNT/CD | PLXNB2 |
| 0.0004737 | -0.3581068 | 0.223 | 0.315 | 1 | CNT/CD | AFDN |
| 2.5482E-08 | -0.3586189 | 0.038 | 0.205 | 0.00054404 | CNT/CD | DPP4 |
| 0.00010445 | -0.3586762 | 0.603 | 0.646 | 1 | CNT/CD | EDF1 |
| 2.1551E-05 | -0.3598424 | 0.19 | 0.321 | 0.46011329 | CNT/CD | TKT |
| 1.3028E-06 | -0.3608066 | 0.109 | 0.257 | 0.02781512 | CNT/CD | DDX46 |
| 6.7601E-07 | -0.3613039 | 0.136 | 0.303 | 0.01443276 | CNT/CD | MYDGF |
| 3.9335E-07 | -0.362936 | 0.109 | 0.271 | 0.00839808 | CNT/CD | PABPC4 |
| 3.1433E-05 | -0.3638843 | 0.304 | 0.421 | 0.67109836 | CNT/CD | PTTG1IP |
| 2.6189E-07 | -0.3652427 | 0.467 | 0.606 | 0.00559132 | CNT/CD | CLTA |
| 1.7723E-06 | -0.3657069 | 0.12 | 0.272 | 0.03783867 | CNT/CD | RBM3 |
| 0.00026871 | -0.3665889 | 0.348 | 0.436 | 1 | CNT/CD | PRRC2C |
| 0.00445096 | -0.3723395 | 0.207 | 0.277 | 1 | CNT/CD | VEGFA |
| 1.1295E-05 | -0.3731499 | 0.908 | 0.91 | 0.24115102 | CNT/CD | FAU |
| 0.00384765 | -0.3735044 | 0.158 | 0.235 | 1 | CNT/CD | NFKBIZ |
| 3.4996E-06 | -0.3745808 | 0.913 | 0.909 | 0.07471711 | CNT/CD | RPL24 |
| 0.0023213 | -0.3750644 | 0.19 | 0.29 | 1 | CNT/CD | RND3 |
| 4.3506E-06 | -0.3763464 | 0.239 | 0.376 | 0.09288464 | CNT/CD | HEBP2 |
| 5.1679E-05 | -0.3785767 | 0.207 | 0.323 | 1 | CNT/CD | RRBP1 |
| 8.8124E-06 | -0.3813417 | 0.076 | 0.203 | 0.18814463 | CNT/CD | CALD1 |
| 0.00014018 | -0.3825058 | 0.549 | 0.61 | 1 | CNT/CD | EIF4A2 |
| 0.00013081 | -0.3844301 | 0.212 | 0.319 | 1 | CNT/CD | BOD1L1 |
| 1.7484E-07 | -0.3854474 | 0.082 | 0.242 | 0.00373292 | CNT/CD | LMO4 |
| 1.3988E-05 | -0.386719 | 0.087 | 0.211 | 0.29864674 | CNT/CD | MAT2A |
| 5.7007E-05 | -0.3940228 | 0.25 | 0.361 | 1 | CNT/CD | TALDO1 |
| 2.7485E-05 | -0.3950783 | 0.011 | 0.106 | 0.58680246 | CNT/CD | SERPINE2 |
| 2.0078E-05 | -0.3974105 | 0.158 | 0.287 | 0.42866645 | CNT/CD | NUFIP2 |
| 0.00028144 | -0.3986565 | 0.163 | 0.266 | 1 | CNT/CD | AMD1 |
| 0.00015396 | -0.4051041 | 0.293 | 0.398 | 1 | CNT/CD | RHOC |
| 1.0312E-05 | -0.4053463 | 0.266 | 0.387 | 0.22015131 | CNT/CD | OCIAD2 |
| 0.0008998 | -0.4109799 | 0.266 | 0.357 | 1 | CNT/CD | DDX3X |
| 0.00014506 | -0.4115487 | 0.223 | 0.327 | 1 | CNT/CD | CTSB |
| 0.00057544 | -0.4132219 | 0.641 | 0.66 | 1 | CNT/CD | CALR |
| 2.7623E-07 | -0.4135907 | 0.734 | 0.805 | 0.00589754 | CNT/CD | RPL31 |
| 0.00060663 | -0.4160781 | 0.299 | 0.385 | 1 | CNT/CD | MYH9 |

| | | | | | | |
|---|---|---|---|---|---|---|
| 1.4661E-06 | -0.4169593 | 0.212 | 0.363 | 0.03130028 | CNT/CD | EIF3A |
| 7.6537E-05 | -0.4176522 | 0.935 | 0.93 | 1 | CNT/CD | RPL7A |
| 1.1334E-06 | -0.4182534 | 0.13 | 0.284 | 0.02419788 | CNT/CD | PRPF4B |
| 0.00651316 | -0.4189268 | 0.141 | 0.214 | 1 | CNT/CD | GLRX |
| 6.1512E-06 | -0.419078 | 0.413 | 0.517 | 0.13132913 | CNT/CD | TPM3 |
| 0.00016164 | -0.4195717 | 0.201 | 0.305 | 1 | CNT/CD | SNHG7 |
| 1.0773E-08 | -0.4211605 | 0.212 | 0.401 | 0.00023 | CNT/CD | YWHAQ |
| 0.00831906 | -0.4222656 | 0.212 | 0.278 | 1 | CNT/CD | RASD1 |
| 2.7771E-11 | -0.4233823 | 0.016 | 0.22 | 5.929E-07 | CNT/CD | SVIL |
| 0.00345168 | -0.4238257 | 0.549 | 0.611 | 1 | CNT/CD | BTG1 |
| 0.00034799 | -0.4271948 | 0.962 | 0.933 | 1 | CNT/CD | RPS9 |
| 5.3506E-08 | -0.4289301 | 0.163 | 0.34 | 0.00114235 | CNT/CD | PRELID1 |
| 0.00016402 | -0.4296006 | 0.929 | 0.92 | 1 | CNT/CD | RPL34 |
| 5.4473E-05 | -0.4307242 | 0.277 | 0.392 | 1 | CNT/CD | CD151 |
| 7.6238E-06 | -0.4310064 | 0.913 | 0.898 | 0.1627675 | CNT/CD | RPL5 |
| 1.1926E-09 | -0.4335601 | 0.114 | 0.321 | 2.5461E-05 | CNT/CD | DYNC1H1 |
| 0.00087139 | -0.4361767 | 0.359 | 0.422 | 1 | CNT/CD | SFPQ |
| 9.9453E-06 | -0.4367901 | 0.652 | 0.756 | 0.21233111 | CNT/CD | RPS26 |
| 2.8752E-06 | -0.4387894 | 0.06 | 0.192 | 0.06138657 | CNT/CD | TLE4 |
| 2.3457E-07 | -0.438898 | 0.538 | 0.669 | 0.00500806 | CNT/CD | SPINT2 |
| 0.0005895 | -0.4419244 | 0.946 | 0.934 | 1 | CNT/CD | RPS27A |
| 0.00042512 | -0.4432649 | 0.179 | 0.285 | 1 | CNT/CD | UCHL1 |
| 0.00058272 | -0.4562708 | 0.967 | 0.938 | 1 | CNT/CD | RPS28 |
| 0.00015879 | -0.4571226 | 0.935 | 0.943 | 1 | CNT/CD | RPL8 |
| 0.00106442 | -0.4582048 | 0.147 | 0.238 | 1 | CNT/CD | SLC5A3 |
| 0.00109233 | -0.4603572 | 0.087 | 0.17 | 1 | CNT/CD | EGLN3 |
| 0.00032741 | -0.4611718 | 0.196 | 0.301 | 1 | CNT/CD | MARCKSL1 |
| 1.4116E-05 | -0.461832 | 0.315 | 0.44 | 0.30136886 | CNT/CD | PPP1R14B |
| 1.2923E-09 | -0.4624417 | 0.005 | 0.178 | 2.759E-05 | CNT/CD | FOXC1 |
| 7.9456E-07 | -0.4629991 | 0.098 | 0.255 | 0.0169639 | CNT/CD | AKAP12 |
| 0.00674076 | -0.4659968 | 0.185 | 0.256 | 1 | CNT/CD | IER5L |
| 2.6923E-07 | -0.4687711 | 0.87 | 0.888 | 0.00574803 | CNT/CD | RPL22 |
| 0.00051005 | -0.4694021 | 0.364 | 0.434 | 1 | CNT/CD | DDX17 |
| 8.4888E-06 | -0.4696959 | 0.016 | 0.125 | 0.18123608 | CNT/CD | CXCL14 |
| 3.1581E-07 | -0.4708324 | 0.935 | 0.902 | 0.00674255 | CNT/CD | RPS25 |
| 8.9269E-06 | -0.4710723 | 0.913 | 0.93 | 0.19058847 | CNT/CD | RPS14 |
| 0.00094014 | -0.4719758 | 0.038 | 0.116 | 1 | CNT/CD | TAGLN |
| 3.8195E-08 | -0.4754013 | 0.902 | 0.934 | 0.00081546 | CNT/CD | RPL6 |
| 1.4398E-05 | -0.4764874 | 0.946 | 0.932 | 0.30740579 | CNT/CD | RPS6 |
| 1.5141E-05 | -0.4768233 | 0.973 | 0.941 | 0.32325717 | CNT/CD | RPS24 |
| 8.0951E-08 | -0.4769815 | 0.201 | 0.376 | 0.00172831 | CNT/CD | FLNA |
| 8.8199E-05 | -0.4770114 | 0.326 | 0.434 | 1 | CNT/CD | ZBTB20 |
| 4.0038E-05 | -0.4776867 | 0.467 | 0.545 | 0.85480722 | CNT/CD | CD59 |
| 1.1867E-09 | -0.4848613 | 0.076 | 0.27 | 2.5335E-05 | CNT/CD | BDP1 |
| 5.5346E-07 | -0.4854258 | 0.103 | 0.262 | 0.01181628 | CNT/CD | LAMB3 |
| 0.00175096 | -0.4865018 | 0.342 | 0.4 | 1 | CNT/CD | JPT1 |
| 1.6674E-06 | -0.4869713 | 0.19 | 0.339 | 0.03559846 | CNT/CD | HNRNPAB |
| 5.362E-05 | -0.4892984 | 0.408 | 0.512 | 1 | CNT/CD | H2AFZ |
| 1.8217E-05 | -0.4935101 | 0.011 | 0.11 | 0.38892446 | CNT/CD | S100A9 |
| 6.9593E-08 | -0.4959655 | 0.842 | 0.869 | 0.00148581 | CNT/CD | RPL10A |

| | | | | | | |
|---|---|---|---|---|---|---|
| 0.00976832 | -0.4960454 | 0.185 | 0.243 | 1 | CNT/CD | ID4 |
| 0.00204773 | -0.4966546 | 0.185 | 0.268 | 1 | CNT/CD | TUBA1A |
| 5.8745E-07 | -0.497782 | 0.212 | 0.37 | 0.01254214 | CNT/CD | GOLGA4 |
| 4.3419E-06 | -0.4980834 | 0.973 | 0.972 | 0.0927 | CNT/CD | TPT1 |
| 8.1946E-09 | -0.5007099 | 0.842 | 0.887 | 0.00017496 | CNT/CD | RPS29 |
| 9.476E-09 | -0.5015598 | 0.435 | 0.59 | 0.00020231 | CNT/CD | S100A13 |
| 3.892E-05 | -0.5016998 | 0.908 | 0.909 | 0.83094503 | CNT/CD | RPL26 |
| 0.00074621 | -0.501973 | 0.913 | 0.882 | 1 | CNT/CD | RPL39 |
| 1.8883E-11 | -0.5039003 | 0.065 | 0.292 | 4.0316E-07 | CNT/CD | ITGA3 |
| 0.00458607 | -0.5044182 | 0.745 | 0.773 | 1 | CNT/CD | HSP90B1 |
| 5.1692E-07 | -0.5098932 | 0.005 | 0.13 | 0.01103628 | CNT/CD | COL7A1 |
| 6.9191E-07 | -0.5153081 | 0.929 | 0.919 | 0.01477235 | CNT/CD | RPL9 |
| 1.2176E-07 | -0.5192839 | 0.891 | 0.897 | 0.00259959 | CNT/CD | RPL35A |
| 2.9069E-12 | -0.5210109 | 0.625 | 0.776 | 6.2063E-08 | CNT/CD | EEF2 |
| 3.4106E-07 | -0.5226138 | 0.266 | 0.426 | 0.00728153 | CNT/CD | AHNAK |
| 0.00478859 | -0.523647 | 0.255 | 0.328 | 1 | CNT/CD | IL32 |
| 8.6804E-06 | -0.5275116 | 0.234 | 0.359 | 0.18532583 | CNT/CD | ARGLU1 |
| 3.0631E-08 | -0.5296604 | 0.223 | 0.395 | 0.00065398 | CNT/CD | TAF1D |
| 6.4032E-08 | -0.5352327 | 0.054 | 0.221 | 0.00136709 | CNT/CD | TNFAIP2 |
| 0.0011977 | -0.5392952 | 0.043 | 0.118 | 1 | CNT/CD | NEFM |
| 0.00770553 | -0.5406365 | 0.266 | 0.333 | 1 | CNT/CD | RHOB |
| 4.4418E-13 | -0.5414461 | 0.587 | 0.757 | 9.4832E-09 | CNT/CD | RPL4 |
| 2.061E-05 | -0.5446532 | 0.946 | 0.928 | 0.4400314 | CNT/CD | RPL21 |
| 3.7311E-09 | -0.5531141 | 0.842 | 0.872 | 7.966E-05 | CNT/CD | RACK1 |
| 2.9755E-09 | -0.5587346 | 0.913 | 0.898 | 6.3526E-05 | CNT/CD | RPL23A |
| 1.4277E-06 | -0.560374 | 0.109 | 0.255 | 0.03048135 | CNT/CD | IER5 |
| 3.7751E-09 | -0.5606059 | 0.951 | 0.94 | 8.0598E-05 | CNT/CD | RPL19 |
| 1.6028E-06 | -0.5620867 | 0.179 | 0.327 | 0.03421996 | CNT/CD | CYBA |
| 3.0761E-14 | -0.5622021 | 0.37 | 0.629 | 6.5675E-10 | CNT/CD | ATF4 |
| 3.1073E-12 | -0.5661292 | 0.668 | 0.802 | 6.6341E-08 | CNT/CD | RPL27 |
| 3.4278E-07 | -0.5665159 | 0.272 | 0.415 | 0.00731826 | CNT/CD | GOLGB1 |
| 4.5811E-08 | -0.5717419 | 0.152 | 0.33 | 0.00097806 | CNT/CD | CEBPB |
| 1.5224E-08 | -0.575032 | 0.88 | 0.905 | 0.00032504 | CNT/CD | RPS3 |
| 0.00318072 | -0.5785892 | 0.842 | 0.871 | 1 | CNT/CD | S100A10 |
| 6.1082E-10 | -0.5807586 | 0.043 | 0.235 | 1.3041E-05 | CNT/CD | TGM2 |
| 1.6469E-07 | -0.5852235 | 0.082 | 0.248 | 0.00351618 | CNT/CD | PDZK1IP1 |
| 1.2794E-16 | -0.5870523 | 0.353 | 0.634 | 2.7315E-12 | CNT/CD | PABPC1 |
| 7.5907E-09 | -0.588924 | 0.185 | 0.371 | 0.00016206 | CNT/CD | MACF1 |
| 1.7361E-12 | -0.5930859 | 0.245 | 0.483 | 3.7066E-08 | CNT/CD | P4HB |
| 4.5816E-08 | -0.5959015 | 0.962 | 0.938 | 0.00097818 | CNT/CD | RPL15 |
| 1.1358E-12 | -0.6009941 | 0.924 | 0.932 | 2.425E-08 | CNT/CD | RPL18 |
| 0.00068843 | -0.6021853 | 0.571 | 0.685 | 1 | CNT/CD | HSPA5 |
| 3.7243E-15 | -0.6108334 | 0.467 | 0.719 | 7.9515E-11 | CNT/CD | TXN |
| 3.8826E-09 | -0.6110578 | 0.082 | 0.264 | 8.2893E-05 | CNT/CD | HIST1H2AC |
| 2.3102E-09 | -0.6148577 | 0.908 | 0.912 | 4.9324E-05 | CNT/CD | RPL29 |
| 2.1461E-08 | -0.6207946 | 0.94 | 0.922 | 0.00045818 | CNT/CD | RPL30 |
| 3.0599E-09 | -0.6224674 | 0.967 | 0.946 | 6.533E-05 | CNT/CD | RPL11 |
| 2.8504E-06 | -0.6352396 | 0.076 | 0.213 | 0.0608551 | CNT/CD | ADAMTS1 |
| 9.1456E-14 | -0.6455579 | 0.864 | 0.898 | 1.9526E-09 | CNT/CD | RPS15A |
| 1.8677E-05 | -0.6457058 | 0.174 | 0.297 | 0.39874353 | CNT/CD | ITGB8 |

| | | | | | | |
|---|---|---|---|---|---|---|
| 5.7082E-12 | -0.646687 | 0.125 | 0.362 | 1.2187E-07 | CNT/CD | CD44 |
| 1.587E-15 | -0.6544009 | 0.734 | 0.867 | 3.3883E-11 | CNT/CD | UBA52 |
| 3.8864E-16 | -0.6555413 | 0.696 | 0.857 | 8.2975E-12 | CNT/CD | RPL38 |
| 4.0906E-13 | -0.6568743 | 0.799 | 0.867 | 8.7335E-09 | CNT/CD | RPL23 |
| 2.0465E-14 | -0.6616621 | 0.462 | 0.696 | 4.3692E-10 | CNT/CD | ZFAS1 |
| 1.6456E-12 | -0.6639138 | 0.902 | 0.941 | 3.5134E-08 | CNT/CD | RPL37A |
| 2.2154E-07 | -0.6652554 | 0.234 | 0.388 | 0.00472983 | CNT/CD | BRD2 |
| 2.3825E-11 | -0.6747586 | 0.929 | 0.935 | 5.0866E-07 | CNT/CD | RPL37 |
| 5.0569E-08 | -0.6771801 | 0.967 | 0.972 | 0.00107965 | CNT/CD | RPLP1 |
| 1.4543E-11 | -0.687806 | 0.37 | 0.564 | 3.1049E-07 | CNT/CD | SRRM2 |
| 6.4897E-14 | -0.6887847 | 0.967 | 0.955 | 1.3856E-09 | CNT/CD | RPS27 |
| 1.3631E-15 | -0.6957325 | 0.668 | 0.821 | 2.9103E-11 | CNT/CD | YBX1 |
| 8.3837E-06 | -0.7072465 | 0.272 | 0.392 | 0.17899186 | CNT/CD | TUBA1B |
| 1.0013E-11 | -0.7102883 | 0.902 | 0.896 | 2.1379E-07 | CNT/CD | RPS13 |
| 7.932E-10 | -0.7109817 | 0.728 | 0.839 | 1.6935E-05 | CNT/CD | PRDX1 |
| 1.9247E-13 | -0.7137652 | 0.147 | 0.401 | 4.1092E-09 | CNT/CD | ANKRD11 |
| 1.1655E-07 | -0.7159117 | 0.957 | 0.951 | 0.00248837 | CNT/CD | RPS12 |
| 7.5585E-11 | -0.7260667 | 0.12 | 0.344 | 1.6137E-06 | CNT/CD | SNHG12 |
| 3.9738E-13 | -0.7312841 | 0.929 | 0.93 | 8.484E-09 | CNT/CD | RPL12 |
| 8.9515E-12 | -0.7362438 | 0.326 | 0.52 | 1.9111E-07 | CNT/CD | SON |
| 1.442E-11 | -0.7388317 | 0.918 | 0.934 | 3.0788E-07 | CNT/CD | RPL32 |
| 3.0064E-17 | -0.7411289 | 0.652 | 0.825 | 6.4187E-13 | CNT/CD | RPS21 |
| 6.99E-07 | -0.741371 | 0.011 | 0.137 | 0.01492373 | CNT/CD | SERPINA1 |
| 5.246E-13 | -0.7435247 | 0.049 | 0.287 | 1.12E-08 | CNT/CD | POLR2A |
| 2.0494E-13 | -0.7469044 | 0.054 | 0.299 | 4.3755E-09 | CNT/CD | DST |
| 4.6227E-09 | -0.7580011 | 0.114 | 0.304 | 9.8695E-05 | CNT/CD | PLK2 |
| 2.8416E-19 | -0.7792144 | 0.56 | 0.786 | 6.0668E-15 | CNT/CD | EEF1B2 |
| 5.482E-06 | -0.7796211 | 0.859 | 0.873 | 0.11704047 | CNT/CD | GSTP1 |
| 1.9144E-16 | -0.7848912 | 0.848 | 0.906 | 4.0873E-12 | CNT/CD | RPS7 |
| 3.2156E-09 | -0.793194 | 0.125 | 0.314 | 6.8654E-05 | CNT/CD | GLUL |
| 2.2869E-12 | -0.8006105 | 0.984 | 0.961 | 4.8826E-08 | CNT/CD | RPS18 |
| 8.8133E-18 | -0.81666 | 0.87 | 0.929 | 1.8816E-13 | CNT/CD | RPL36 |
| 1.1876E-21 | -0.8177623 | 0.755 | 0.907 | 2.5355E-17 | CNT/CD | RPS20 |
| 6.5789E-19 | -0.8208094 | 0.777 | 0.895 | 1.4046E-14 | CNT/CD | RPS11 |
| 8.3105E-05 | -0.8280415 | 0.989 | 0.978 | 1 | CNT/CD | FTH1 |
| 2.1156E-06 | -0.8339155 | 0.266 | 0.408 | 0.04516773 | CNT/CD | RNF213 |
| 1.187E-27 | -0.8354731 | 0.342 | 0.708 | 2.5343E-23 | CNT/CD | EIF4G2 |
| 2.7654E-22 | -0.8383719 | 0.772 | 0.903 | 5.9042E-18 | CNT/CD | RPL27A |
| 1.1965E-05 | -0.8442969 | 0.147 | 0.273 | 0.25545055 | CNT/CD | TCIM |
| 2.6268E-21 | -0.8450423 | 0.891 | 0.922 | 5.6082E-17 | CNT/CD | RPL14 |
| 6.5025E-22 | -0.8488354 | 0.821 | 0.914 | 1.3883E-17 | CNT/CD | RPL35 |
| 5.8459E-19 | -0.8504044 | 0.266 | 0.573 | 1.2481E-14 | CNT/CD | RPS27L |
| 2.5551E-09 | -0.8590923 | 0.19 | 0.377 | 5.4551E-05 | CNT/CD | SLC38A2 |
| 2.4481E-11 | -0.8644058 | 0.027 | 0.239 | 5.2266E-07 | CNT/CD | CST6 |
| 2.5228E-10 | -0.8790295 | 0.842 | 0.872 | 5.3861E-06 | CNT/CD | SAT1 |
| 3.3443E-05 | -0.8804641 | 0.027 | 0.128 | 0.71400543 | CNT/CD | GPX3 |
| 6.3175E-09 | -0.881818 | 0.06 | 0.237 | 0.00013488 | CNT/CD | CSKMT |
| 2.886E-21 | -0.8837129 | 0.897 | 0.946 | 6.1616E-17 | CNT/CD | RPS15 |
| 4.53E-20 | -0.9081037 | 0.424 | 0.708 | 9.6716E-16 | CNT/CD | ITGB1 |
| 2.2458E-21 | -0.9143709 | 0.712 | 0.872 | 4.7948E-17 | CNT/CD | RPS5 |

| | | | | | | |
|---|---|---|---|---|---|---|
| 9.2343E-36 | -0.9182922 | 0.424 | 0.81 | 1.9715E-31 | CNT/CD | EEF1D |
| 2.6123E-09 | -0.9233254 | 0.071 | 0.264 | 5.5773E-05 | CNT/CD | NUPR1 |
| 4.7888E-20 | -0.9463365 | 0.652 | 0.846 | 1.0224E-15 | CNT/CD | RPSA |
| 2.6402E-24 | -0.9568535 | 0.908 | 0.951 | 5.6369E-20 | CNT/CD | RPS8 |
| 5.8034E-06 | -0.9583742 | 0.114 | 0.258 | 0.12390199 | CNT/CD | LCN2 |
| 7.2358E-17 | -0.9588116 | 0.087 | 0.397 | 1.5448E-12 | CNT/CD | TXNRD1 |
| 1.0423E-05 | -0.9607803 | 0.147 | 0.272 | 0.22253222 | CNT/CD | HSPA2 |
| 1.2515E-24 | -0.9678694 | 0.864 | 0.949 | 2.672E-20 | CNT/CD | RPL13A |
| 1.0824E-24 | -0.9800607 | 0.973 | 0.973 | 2.3109E-20 | CNT/CD | RPL13 |
| 1.1733E-24 | -0.9843524 | 0.821 | 0.925 | 2.505E-20 | CNT/CD | RPLP2 |
| 4.5107E-14 | -0.9848702 | 0.185 | 0.432 | 9.6304E-10 | CNT/CD | SYNE2 |
| 9.8146E-28 | -0.9935143 | 0.783 | 0.909 | 2.0954E-23 | CNT/CD | RPS16 |
| 1.6538E-09 | -0.9999011 | 0.272 | 0.473 | 3.5309E-05 | CNT/CD | C12orf75 |
| 2.719E-07 | -1.0071758 | 0.179 | 0.333 | 0.005805 | CNT/CD | LAMC2 |
| 3.5158E-14 | -1.0666526 | 0.076 | 0.334 | 7.5063E-10 | CNT/CD | TIMP1 |
| 2.8507E-27 | -1.0824886 | 0.935 | 0.961 | 6.0863E-23 | CNT/CD | RPL28 |
| 2.7729E-27 | -1.0844499 | 0.87 | 0.929 | 5.9202E-23 | CNT/CD | RPL18A |
| 2.3661E-27 | -1.0934384 | 0.913 | 0.957 | 5.0516E-23 | CNT/CD | RPS2 |
| 5.0203E-09 | -1.1144991 | 0.125 | 0.3 | 0.00010718 | CNT/CD | HEXIM1 |
| 5.0638E-17 | -1.1239196 | 0.582 | 0.835 | 1.0811E-12 | CNT/CD | ANXA2 |
| 1.0366E-25 | -1.1576716 | 0.212 | 0.629 | 2.213E-21 | CNT/CD | VIM |
| 8.1014E-38 | -1.1819505 | 0.63 | 0.919 | 1.7297E-33 | CNT/CD | RPLP0 |
| 1.0416E-07 | -1.1821078 | 0.951 | 0.961 | 0.00222392 | CNT/CD | FTL |
| 3.9583E-23 | -1.1958633 | 0.321 | 0.635 | 8.451E-19 | CNT/CD | TPM1 |
| 1.7724E-05 | -1.2463124 | 0.462 | 0.639 | 0.37841056 | CNT/CD | MMP7 |
| 1.3654E-21 | -1.2477246 | 0.364 | 0.676 | 2.915E-17 | CNT/CD | SH3BGRL3 |
| 2.733E-27 | -1.3328635 | 0.25 | 0.655 | 5.835E-23 | CNT/CD | KRT7 |
| 2.7958E-11 | -1.3487228 | 0.076 | 0.286 | 5.969E-07 | CNT/CD | MAFB |
| 2.5432E-36 | -1.3494082 | 0.924 | 0.965 | 5.4298E-32 | CNT/CD | RPS19 |
| 2.3563E-13 | -1.4789209 | 0.332 | 0.603 | 5.0307E-09 | CNT/CD | SPP1 |
| 8.1487E-32 | -1.5629343 | 0.826 | 0.925 | 1.7398E-27 | CNT/CD | TMSB10 |
| 4.2805E-10 | -1.5637142 | 0.375 | 0.542 | 9.1389E-06 | CNT/CD | SOX4 |
| 8.7447E-18 | -1.6722365 | 0.087 | 0.396 | 1.867E-13 | CNT/CD | SLPI |
| 2.7337E-28 | -1.8682238 | 0.636 | 0.818 | 5.8364E-24 | CNT/CD | NEAT1 |
| 9.1178E-14 | -4.6975142 | 0.462 | 0.599 | 1.9466E-09 | CNT/CD | PLCG2 |
| 6.945E-227 | 3.14373992 | 0.907 | 0.474 | 1.483E-222 | TAL/DCT | S100A2 |
| 6.446E-184 | 2.92477688 | 0.387 | 0.063 | 1.376E-179 | TAL/DCT | UMOD |
| 1.114E-138 | 2.37601158 | 0.624 | 0.208 | 2.378E-134 | TAL/DCT | DEFB1 |
| 7.704E-122 | 1.63548048 | 0.806 | 0.427 | 1.645E-117 | TAL/DCT | UPP1 |
| 3.3085E-50 | 1.48295715 | 0.425 | 0.19 | 7.0636E-46 | TAL/DCT | ANXA1 |
| 4.921E-151 | 1.34759762 | 0.43 | 0.09 | 1.051E-146 | TAL/DCT | RSAD2 |
| 4.9984E-75 | 1.32829123 | 0.387 | 0.127 | 1.0672E-70 | TAL/DCT | TM4SF1 |
| 1.815E-133 | 1.1367395 | 0.717 | 0.277 | 3.874E-129 | TAL/DCT | MIR4435-2HG |
| 2.2081E-18 | 1.12891771 | 0.296 | 0.166 | 4.7142E-14 | TAL/DCT | SETX |
| 3.497E-135 | 1.07325666 | 0.721 | 0.279 | 7.467E-131 | TAL/DCT | CYTOR |
| 5.2891E-71 | 1.05682744 | 0.622 | 0.294 | 1.1292E-66 | TAL/DCT | PLAU |
| 1.0472E-58 | 1.05677169 | 0.866 | 0.704 | 2.2358E-54 | TAL/DCT | MT2A |
| 1.109E-115 | 1.00538392 | 0.508 | 0.153 | 2.368E-111 | TAL/DCT | FGF2 |
| 1.196E-143 | 0.98068227 | 0.361 | 0.066 | 2.553E-139 | TAL/DCT | LY6K |
| 1.002E-167 | 0.91796641 | 0.231 | 0.022 | 2.139E-163 | TAL/DCT | VGF |

| | | | | | | |
|---|---|---|---|---|---|---|
| 1.0242E-35 | 0.87321654 | 0.775 | 0.64 | 2.1868E-31 | TAL/DCT | WFDC2 |
| 1.3114E-70 | 0.86563361 | 0.57 | 0.264 | 2.7999E-66 | TAL/DCT | SMS |
| 2.1902E-33 | 0.86497379 | 0.246 | 0.091 | 4.6762E-29 | TAL/DCT | SLC12A1 |
| 4.8392E-45 | 0.84589099 | 0.669 | 0.402 | 1.0332E-40 | TAL/DCT | PHLDA2 |
| 1.2E-195 | 0.83275535 | 0.251 | 0.022 | 2.562E-191 | TAL/DCT | KCNMA1 |
| 4.4013E-20 | 0.82892429 | 0.209 | 0.094 | 9.3967E-16 | TAL/DCT | MAEA |
| 3.1047E-37 | 0.82222662 | 0.352 | 0.155 | 6.6286E-33 | TAL/DCT | KATNBL1 |
| 2.2958E-71 | 0.81852677 | 0.868 | 0.62 | 4.9015E-67 | TAL/DCT | ANXA5 |
| 8.3797E-46 | 0.81690335 | 0.261 | 0.085 | 1.7891E-41 | TAL/DCT | SNAPC1 |
| 1.3404E-49 | 0.81444909 | 0.587 | 0.323 | 2.8617E-45 | TAL/DCT | ZNF292 |
| 3.5567E-79 | 0.79272448 | 0.976 | 0.866 | 7.5936E-75 | TAL/DCT | S100A10 |
| 4.4776E-20 | 0.78243499 | 0.445 | 0.278 | 9.5596E-16 | TAL/DCT | HIST1H1C |
| 4.534E-100 | 0.77684565 | 0.523 | 0.174 | 9.6792E-96 | TAL/DCT | EMP1 |
| 2.7235E-37 | 0.77321019 | 0.723 | 0.501 | 5.8146E-33 | TAL/DCT | CDKN1A |
| 1.89E-123 | 0.75776014 | 0.281 | 0.046 | 4.036E-119 | TAL/DCT | HRK |
| 2.1961E-58 | 0.75327763 | 0.514 | 0.237 | 4.6887E-54 | TAL/DCT | MGLL |
| 1.2618E-07 | 0.74135739 | 0.669 | 0.667 | 0.00269398 | TAL/DCT | CRYAB |
| 3.1447E-31 | 0.73994934 | 0.931 | 0.921 | 6.714E-27 | TAL/DCT | ACTB |
| 6.0115E-61 | 0.70613484 | 0.704 | 0.393 | 1.2835E-56 | TAL/DCT | SDC4 |
| 1.1446E-47 | 0.69479871 | 0.68 | 0.408 | 2.4436E-43 | TAL/DCT | LGALS3 |
| 7.5642E-95 | 0.69449965 | 0.378 | 0.101 | 1.615E-90 | TAL/DCT | SULT1C2 |
| 7.7644E-47 | 0.67837113 | 0.756 | 0.537 | 1.6577E-42 | TAL/DCT | BRI3 |
| 7.7434E-50 | 0.66271031 | 0.341 | 0.127 | 1.6532E-45 | TAL/DCT | KIAA1551 |
| 6.9373E-27 | 0.64394382 | 0.471 | 0.266 | 1.4811E-22 | TAL/DCT | VMP1 |
| 6.3443E-31 | 0.63990844 | 0.95 | 0.925 | 1.3545E-26 | TAL/DCT | ACTG1 |
| 1.6862E-67 | 0.63403436 | 0.371 | 0.12 | 3.6E-63 | TAL/DCT | AL353751.1 |
| 2.277E-51 | 0.6243139 | 0.81 | 0.547 | 4.8614E-47 | TAL/DCT | HMGA1 |
| 9.6465E-49 | 0.61872105 | 0.626 | 0.332 | 2.0595E-44 | TAL/DCT | S100A16 |
| 2.738E-112 | 0.61761507 | 0.378 | 0.088 | 5.846E-108 | TAL/DCT | CD68 |
| 6.1622E-68 | 0.61597872 | 0.611 | 0.288 | 1.3156E-63 | TAL/DCT | CUTA |
| 1.3251E-41 | 0.60187023 | 0.59 | 0.315 | 2.8292E-37 | TAL/DCT | MT1E |
| 3.0584E-58 | 0.59950514 | 0.685 | 0.379 | 6.5296E-54 | TAL/DCT | CAPN2 |
| 4.2414E-37 | 0.59858671 | 0.741 | 0.542 | 9.0553E-33 | TAL/DCT | CD9 |
| 7.2318E-57 | 0.59811214 | 0.583 | 0.278 | 1.544E-52 | TAL/DCT | MAFF |
| 1.3096E-45 | 0.5937018 | 0.503 | 0.25 | 2.796E-41 | TAL/DCT | GSTO1 |
| 3.2709E-43 | 0.57320916 | 0.749 | 0.486 | 6.9833E-39 | TAL/DCT | TNFRSF12A |
| 6.0863E-40 | 0.57213483 | 0.613 | 0.36 | 1.2994E-35 | TAL/DCT | CAMK2N1 |
| 4.1842E-52 | 0.55947932 | 0.216 | 0.056 | 8.9332E-48 | TAL/DCT | FABP3 |
| 6.0662E-51 | 0.55118288 | 0.657 | 0.369 | 1.2951E-46 | TAL/DCT | RHEB |
| 2.4531E-29 | 0.55071347 | 0.326 | 0.154 | 5.2375E-25 | TAL/DCT | FNIP2 |
| 1.0165E-78 | 0.55024022 | 0.188 | 0.031 | 2.1703E-74 | TAL/DCT | KNG1 |
| 4.2055E-44 | 0.54503708 | 0.896 | 0.743 | 8.9787E-40 | TAL/DCT | RTN4 |
| 1.1638E-27 | 0.54196707 | 0.432 | 0.226 | 2.4847E-23 | TAL/DCT | FXYD2 |
| 2.4043E-50 | 0.52493747 | 0.449 | 0.195 | 5.1331E-46 | TAL/DCT | CSRP1 |
| 1.1209E-36 | 0.5239579 | 0.432 | 0.213 | 2.3931E-32 | TAL/DCT | BEX2 |
| 9.7759E-28 | 0.52160768 | 0.894 | 0.811 | 2.0871E-23 | TAL/DCT | IGFBP7 |
| 1.856E-121 | 0.52145826 | 0.268 | 0.042 | 3.963E-117 | TAL/DCT | BEX1 |
| 3.5428E-39 | 0.51876966 | 0.488 | 0.245 | 7.5639E-35 | TAL/DCT | S100A14 |
| 3.727E-47 | 0.51608206 | 0.315 | 0.112 | 7.9571E-43 | TAL/DCT | NEURL3 |
| 3.3569E-72 | 0.51287997 | 0.203 | 0.039 | 7.167E-68 | TAL/DCT | ARL14 |

| | | | | | | |
|---|---|---|---|---|---|---|
| 4.748E-38 | 0.50723816 | 0.892 | 0.769 | 1.0137E-33 | TAL/DCT | MYL12A |
| 3.1488E-42 | 0.50526297 | 0.98 | 0.956 | 6.7226E-38 | TAL/DCT | EIF1 |
| 4.4979E-44 | 0.49923265 | 0.661 | 0.373 | 9.6029E-40 | TAL/DCT | OCIAD2 |
| 5.5624E-45 | 0.49654347 | 0.318 | 0.117 | 1.1876E-40 | TAL/DCT | GPNMB |
| 1.0513E-24 | 0.49357801 | 0.4 | 0.213 | 2.2445E-20 | TAL/DCT | SFN |
| 4.1421E-31 | 0.49254657 | 0.544 | 0.308 | 8.8433E-27 | TAL/DCT | UGCG |
| 1.0304E-50 | 0.49157992 | 0.384 | 0.149 | 2.1999E-46 | TAL/DCT | FHL2 |
| 9.7715E-31 | 0.49102508 | 0.842 | 0.694 | 2.0862E-26 | TAL/DCT | MAP1LC3B |
| 1.4101E-14 | 0.49007767 | 0.521 | 0.362 | 3.0105E-10 | TAL/DCT | DDIT3 |
| 4.0707E-22 | 0.4846356 | 0.594 | 0.41 | 8.6909E-18 | TAL/DCT | CCND1 |
| 1.0903E-35 | 0.48406254 | 0.615 | 0.363 | 2.3278E-31 | TAL/DCT | ODC1 |
| 2.2521E-38 | 0.48361248 | 0.561 | 0.308 | 4.8083E-34 | TAL/DCT | NDUFB9 |
| 7.2765E-40 | 0.48262126 | 0.773 | 0.558 | 1.5535E-35 | TAL/DCT | HNRNPC |
| 6.1082E-65 | 0.47993951 | 0.218 | 0.048 | 1.3041E-60 | TAL/DCT | CLDN14 |
| 6.3111E-40 | 0.46981698 | 0.633 | 0.358 | 1.3474E-35 | TAL/DCT | SDCBP |
| 3.605E-53 | 0.46810329 | 0.425 | 0.172 | 7.6966E-49 | TAL/DCT | BCL10 |
| 8.2705E-05 | 0.46631948 | 0.628 | 0.604 | 1 | TAL/DCT | SOD2 |
| 2.4873E-23 | 0.46328818 | 0.728 | 0.581 | 5.3104E-19 | TAL/DCT | S100A13 |
| 2.986E-24 | 0.46259227 | 0.814 | 0.72 | 6.375E-20 | TAL/DCT | CTSD |
| 2.4154E-55 | 0.46253334 | 0.421 | 0.166 | 5.1569E-51 | TAL/DCT | SLC39A1 |
| 1.3416E-57 | 0.4624472 | 0.253 | 0.069 | 2.8644E-53 | TAL/DCT | NOV |
| 3.8642E-38 | 0.46076841 | 0.492 | 0.251 | 8.2501E-34 | TAL/DCT | VCL |
| 1.3014E-40 | 0.45901295 | 0.983 | 0.911 | 2.7784E-36 | TAL/DCT | HSP90AB1 |
| 2.3027E-42 | 0.45667254 | 0.231 | 0.073 | 4.9164E-38 | TAL/DCT | SYNJ2 |
| 4.0731E-38 | 0.45526142 | 0.583 | 0.329 | 8.6961E-34 | TAL/DCT | EIF1AX |
| 5.7505E-40 | 0.45428684 | 0.389 | 0.172 | 1.2277E-35 | TAL/DCT | LINC00958 |
| 2.4414E-47 | 0.45284238 | 0.613 | 0.326 | 5.2124E-43 | TAL/DCT | NOP10 |
| 1.7112E-51 | 0.45083518 | 0.188 | 0.044 | 3.6533E-47 | TAL/DCT | TAC1 |
| 4.6446E-43 | 0.44752633 | 0.419 | 0.188 | 9.9161E-39 | TAL/DCT | MAP2K3 |
| 8.8293E-74 | 0.44671892 | 0.291 | 0.074 | 1.8851E-69 | TAL/DCT | PLPP3 |
| 7.304E-27 | 0.4465161 | 0.864 | 0.754 | 1.5594E-22 | TAL/DCT | CALM2 |
| 1.3632E-50 | 0.44593132 | 0.382 | 0.145 | 2.9105E-46 | TAL/DCT | CPM |
| 1.0391E-30 | 0.44581128 | 0.829 | 0.674 | 2.2184E-26 | TAL/DCT | CLIC1 |
| 5.3338E-32 | 0.44419003 | 0.475 | 0.25 | 1.1388E-27 | TAL/DCT | LAMB3 |
| 2.2906E-57 | 0.44385128 | 0.367 | 0.129 | 4.8905E-53 | TAL/DCT | C12orf49 |
| 1.1602E-12 | 0.44375511 | 0.769 | 0.699 | 2.477E-08 | TAL/DCT | CLDN4 |
| 1.0002E-31 | 0.44258964 | 0.298 | 0.127 | 2.1355E-27 | TAL/DCT | PPIF |
| 7.0661E-38 | 0.44045198 | 0.533 | 0.291 | 1.5086E-33 | TAL/DCT | RAP1B |
| 2.8574E-40 | 0.43827915 | 0.412 | 0.182 | 6.1005E-36 | TAL/DCT | BHLHE40 |
| 5.2624E-37 | 0.43504998 | 0.238 | 0.082 | 1.1235E-32 | TAL/DCT | AC015912.3 |
| 2.6426E-55 | 0.43425892 | 0.311 | 0.1 | 5.642E-51 | TAL/DCT | FOSL1 |
| 3.5446E-40 | 0.42905542 | 0.378 | 0.163 | 7.5676E-36 | TAL/DCT | SH3BP4 |
| 2.0511E-41 | 0.42878709 | 0.432 | 0.201 | 4.3792E-37 | TAL/DCT | SNX9 |
| 4.7665E-23 | 0.42762358 | 0.432 | 0.249 | 1.0177E-18 | TAL/DCT | NEDD4L |
| 7.1589E-12 | 0.42663479 | 0.358 | 0.234 | 1.5284E-07 | TAL/DCT | MDM2 |
| 1.0882E-42 | 0.42373919 | 0.205 | 0.058 | 2.3234E-38 | TAL/DCT | SFRP1 |
| 1.0259E-60 | 0.42221307 | 0.231 | 0.056 | 2.1902E-56 | TAL/DCT | SERPINE1 |
| 4.5259E-43 | 0.41938058 | 0.439 | 0.199 | 9.6628E-39 | TAL/DCT | GULP1 |
| 1.579E-35 | 0.41615604 | 0.622 | 0.348 | 3.3712E-31 | TAL/DCT | CD44 |
| 8.071E-20 | 0.41527312 | 0.402 | 0.234 | 1.7232E-15 | TAL/DCT | HES4 |

| | | | | | | |
|---|---|---|---|---|---|---|
| 1.0297E-38 | 0.41306033 | 0.918 | 0.836 | 2.1984E-34 | TAL/DCT | CD63 |
| 1.2527E-44 | 0.41280482 | 0.304 | 0.108 | 2.6746E-40 | TAL/DCT | CRABP2 |
| 3.893E-101 | 0.41043354 | 0.203 | 0.029 | 8.3116E-97 | TAL/DCT | FXYD6 |
| 4.1876E-31 | 0.40982415 | 0.415 | 0.214 | 8.9406E-27 | TAL/DCT | TMEM167A |
| 1.5442E-20 | 0.40912844 | 0.3 | 0.15 | 3.2969E-16 | TAL/DCT | SLC20A1 |
| 7.5613E-26 | 0.40761957 | 0.462 | 0.263 | 1.6143E-21 | TAL/DCT | CKB |
| 1.0658E-17 | 0.40473617 | 0.931 | 0.906 | 2.2754E-13 | TAL/DCT | MYL6 |
| 4.0225E-26 | 0.40160342 | 0.587 | 0.356 | 8.5881E-22 | TAL/DCT | PPP1CB |
| 3.4294E-53 | 0.39645505 | 0.471 | 0.199 | 7.3217E-49 | TAL/DCT | PEA15 |
| 5.478E-25 | 0.39522815 | 0.585 | 0.349 | 1.1695E-20 | TAL/DCT | KLF6 |
| 5.9385E-66 | 0.39332016 | 0.378 | 0.123 | 1.2679E-61 | TAL/DCT | RRAGD |
| 1.0094E-23 | 0.39292226 | 0.475 | 0.278 | 2.155E-19 | TAL/DCT | PLAUR |
| 6.4029E-70 | 0.39139308 | 0.317 | 0.088 | 1.367E-65 | TAL/DCT | C1orf116 |
| 8.4655E-31 | 0.39071568 | 0.348 | 0.16 | 1.8074E-26 | TAL/DCT | CA12 |
| 5.3022E-10 | 0.39039242 | 0.304 | 0.201 | 1.132E-05 | TAL/DCT | MDK |
| 6.9284E-19 | 0.38988084 | 0.263 | 0.128 | 1.4792E-14 | TAL/DCT | IFIT2 |
| 3.1485E-30 | 0.38956693 | 0.343 | 0.162 | 6.7221E-26 | TAL/DCT | CDYL |
| 3.5933E-45 | 0.38917739 | 0.317 | 0.115 | 7.6718E-41 | TAL/DCT | PRSS23 |
| 5.7108E-30 | 0.38851076 | 0.663 | 0.422 | 1.2193E-25 | TAL/DCT | BZW1 |
| 9.3526E-11 | 0.38798005 | 0.374 | 0.247 | 1.9968E-06 | TAL/DCT | AKAP12 |
| 1.6589E-35 | 0.38561213 | 0.497 | 0.258 | 3.5418E-31 | TAL/DCT | MET |
| 3.6162E-19 | 0.38553361 | 0.788 | 0.66 | 7.7207E-15 | TAL/DCT | TMA7 |
| 1.8386E-32 | 0.38482075 | 0.78 | 0.572 | 3.9254E-28 | TAL/DCT | ATP6V0E1 |
| 1.1286E-34 | 0.38410781 | 0.283 | 0.113 | 2.4097E-30 | TAL/DCT | FBXO32 |
| 1.471E-20 | 0.38399334 | 0.689 | 0.529 | 3.1405E-16 | TAL/DCT | HLA-E |
| 6.2127E-25 | 0.37843015 | 0.536 | 0.322 | 1.3264E-20 | TAL/DCT | TUBA1C |
| 1.4482E-29 | 0.37769789 | 0.385 | 0.192 | 3.0919E-25 | TAL/DCT | TMEM106B |
| 6.2796E-93 | 0.37757481 | 0.162 | 0.02 | 1.3407E-88 | TAL/DCT | C2orf54 |
| 7.0277E-53 | 0.37693866 | 0.231 | 0.061 | 1.5004E-48 | TAL/DCT | TRIM55 |
| 7.2084E-34 | 0.37610256 | 0.987 | 0.953 | 1.539E-29 | TAL/DCT | S100A6 |
| 2.5363E-43 | 0.37493846 | 0.35 | 0.139 | 5.415E-39 | TAL/DCT | SGMS2 |
| 9.6202E-11 | 0.37452552 | 0.415 | 0.277 | 2.0539E-06 | TAL/DCT | UCHL1 |
| 1.4068E-39 | 0.37137777 | 0.251 | 0.086 | 3.0035E-35 | TAL/DCT | SDC1 |
| 3.5636E-11 | 0.36996115 | 0.877 | 0.868 | 7.6082E-07 | TAL/DCT | TMSB4X |
| 1.3737E-27 | 0.36886243 | 0.367 | 0.187 | 2.9328E-23 | TAL/DCT | EMC4 |
| 5.1475E-14 | 0.3686549 | 0.477 | 0.325 | 1.099E-09 | TAL/DCT | H1F0 |
| 1.0598E-17 | 0.36859728 | 0.294 | 0.155 | 2.2626E-13 | TAL/DCT | G0S2 |
| 3.9583E-35 | 0.36850125 | 0.503 | 0.265 | 8.451E-31 | TAL/DCT | ZNF706 |
| 1.7835E-42 | 0.36764303 | 0.425 | 0.188 | 3.8079E-38 | TAL/DCT | WBP2 |
| 2.9522E-69 | 0.36726662 | 0.186 | 0.034 | 6.303E-65 | TAL/DCT | ZNF556 |
| 5.2628E-28 | 0.3665002 | 0.291 | 0.13 | 1.1236E-23 | TAL/DCT | RMND5A |
| 2.104E-21 | 0.36606785 | 0.762 | 0.577 | 4.4919E-17 | TAL/DCT | EZR |
| 2.4236E-20 | 0.36527941 | 0.363 | 0.202 | 5.1745E-16 | TAL/DCT | ARID5B |
| 3.1064E-23 | 0.36344658 | 0.315 | 0.154 | 6.6321E-19 | TAL/DCT | TNFAIP3 |
| 1.2352E-30 | 0.36317361 | 0.38 | 0.184 | 2.6371E-26 | TAL/DCT | KLHL21 |
| 6.5859E-14 | 0.3600594 | 0.225 | 0.118 | 1.4061E-09 | TAL/DCT | IFIT3 |
| 2.2749E-27 | 0.35930042 | 0.508 | 0.295 | 4.8568E-23 | TAL/DCT | PNRC2 |
| 8.3935E-21 | 0.35838712 | 0.46 | 0.27 | 1.792E-16 | TAL/DCT | SERPINB1 |
| 5.1094E-91 | 0.35787014 | 0.102 | 0.008 | 1.0909E-86 | TAL/DCT | TMEM233 |
| 7.5794E-30 | 0.35738527 | 0.603 | 0.36 | 1.6182E-25 | TAL/DCT | TAGLN2 |

| | | | | | | |
|---|---|---|---|---|---|---|
| 1.9649E-21 | 0.3571025 | 0.164 | 0.061 | 4.1951E-17 | TAL/DCT | SOD3 |
| 5.4391E-25 | 0.35678176 | 0.998 | 0.987 | 1.1613E-20 | TAL/DCT | EEF1A1 |
| 1.1214E-96 | 0.35386948 | 0.181 | 0.024 | 2.3942E-92 | TAL/DCT | EEPD1 |
| 8.4021E-21 | 0.35195904 | 0.929 | 0.863 | 1.7939E-16 | TAL/DCT | RPS4X |
| 2.8539E-15 | 0.35071627 | 0.151 | 0.065 | 6.0932E-11 | TAL/DCT | SPINK1 |
| 4.2495E-39 | 0.35068425 | 0.385 | 0.169 | 9.0728E-35 | TAL/DCT | BACE2 |
| 6.538E-26 | 0.34969598 | 0.89 | 0.782 | 1.3959E-21 | TAL/DCT | CHCHD2 |
| 8.519E-81 | 0.34965404 | 0.216 | 0.04 | 1.8188E-76 | TAL/DCT | RAET1L |
| 3.9867E-24 | 0.34948519 | 0.935 | 0.849 | 8.5116E-20 | TAL/DCT | MYL12B |
| 3.0157E-33 | 0.34808928 | 0.533 | 0.289 | 6.4386E-29 | TAL/DCT | ACTR3 |
| 1.3453E-21 | 0.34735496 | 0.749 | 0.574 | 2.8723E-17 | TAL/DCT | ATP6V1G1 |
| 4.7591E-33 | 0.34613005 | 0.302 | 0.124 | 1.0161E-28 | TAL/DCT | LDLR |
| 2.051E-37 | 0.34499924 | 0.417 | 0.194 | 4.3788E-33 | TAL/DCT | SUN1 |
| 5.45E-30 | 0.3449748 | 0.512 | 0.285 | 1.1636E-25 | TAL/DCT | EIF6 |
| 1.9886E-21 | 0.34486835 | 0.965 | 0.932 | 4.2458E-17 | TAL/DCT | RPL3 |
| 2.4824E-22 | 0.34420308 | 0.313 | 0.158 | 5.3E-18 | TAL/DCT | SPTSSA |
| 1.5301E-30 | 0.34298098 | 0.458 | 0.246 | 3.2667E-26 | TAL/DCT | ATP6V1D |
| 3.635E-30 | 0.34113559 | 0.359 | 0.17 | 7.7607E-26 | TAL/DCT | BCL2L1 |
| 0.00017944 | 0.34099224 | 0.164 | 0.113 | 1 | TAL/DCT | TAGLN |
| 4.1961E-22 | 0.33894218 | 0.831 | 0.722 | 8.9586E-18 | TAL/DCT | CSTB |
| 5.9103E-27 | 0.33882709 | 0.367 | 0.178 | 1.2619E-22 | TAL/DCT | ADIRF |
| 2.0429E-32 | 0.33788681 | 0.454 | 0.234 | 4.3615E-28 | TAL/DCT | PLEKHB2 |
| 2.5486E-22 | 0.33767366 | 0.281 | 0.137 | 5.4413E-18 | TAL/DCT | ANKRD10 |
| 2.8851E-44 | 0.33677558 | 0.263 | 0.086 | 6.1597E-40 | TAL/DCT | NR1D1 |
| 1.385E-100 | 0.33658547 | 0.194 | 0.026 | 2.9576E-96 | TAL/DCT | CPXM2 |
| 3.6745E-33 | 0.33397885 | 0.369 | 0.17 | 7.845E-29 | TAL/DCT | RYBP |
| 1.5038E-16 | 0.33324804 | 0.471 | 0.304 | 3.2107E-12 | TAL/DCT | RAB13 |
| 2.4358E-24 | 0.32951471 | 0.425 | 0.231 | 5.2005E-20 | TAL/DCT | TCEAL9 |
| 3.9122E-19 | 0.327385 | 0.568 | 0.377 | 8.3526E-15 | TAL/DCT | PERP |
| 8.6776E-38 | 0.32593935 | 0.304 | 0.118 | 1.8527E-33 | TAL/DCT | GNPDA1 |
| 9.4069E-25 | 0.323007 | 0.683 | 0.461 | 2.0084E-20 | TAL/DCT | PRDX6 |
| 1.7515E-30 | 0.32250054 | 0.525 | 0.294 | 3.7394E-26 | TAL/DCT | WASF2 |
| 3.045E-18 | 0.32124149 | 0.952 | 0.922 | 6.5011E-14 | TAL/DCT | RPL10 |
| 2.0116E-60 | 0.31980926 | 0.231 | 0.056 | 4.2947E-56 | TAL/DCT | SBF2-AS1 |
| 4.5549E-24 | 0.31973605 | 0.749 | 0.535 | 9.7247E-20 | TAL/DCT | CD59 |
| 9.2701E-43 | 0.31832275 | 0.162 | 0.039 | 1.9792E-38 | TAL/DCT | AC026250.1 |
| 9.1642E-19 | 0.31688871 | 0.799 | 0.673 | 1.9566E-14 | TAL/DCT | YWHAZ |
| 1.524E-33 | 0.31612432 | 0.363 | 0.165 | 3.2538E-29 | TAL/DCT | DSG2 |
| 4.7806E-19 | 0.31597651 | 0.179 | 0.075 | 1.0206E-14 | TAL/DCT | AGPAT5 |
| 1.3605E-16 | 0.31584481 | 0.451 | 0.288 | 2.9047E-12 | TAL/DCT | CD55 |
| 9.398E-25 | 0.3139779 | 0.523 | 0.315 | 2.0065E-20 | TAL/DCT | FAM96B |
| 3.8249E-22 | 0.31190106 | 0.406 | 0.23 | 8.1662E-18 | TAL/DCT | HECTD1 |
| 1.0465E-20 | 0.31170179 | 0.914 | 0.827 | 2.2343E-16 | TAL/DCT | ANXA2 |
| 3.6615E-16 | 0.31099269 | 0.629 | 0.461 | 7.8174E-12 | TAL/DCT | LMNA |
| 1.1119E-20 | 0.31072514 | 0.441 | 0.265 | 2.374E-16 | TAL/DCT | NDUFV2 |
| 3.8354E-32 | 0.31026416 | 0.298 | 0.124 | 8.1885E-28 | TAL/DCT | SLC2A1 |
| 7.2407E-19 | 0.31023284 | 0.372 | 0.213 | 1.5459E-14 | TAL/DCT | TIMP2 |
| 2.8416E-46 | 0.30968317 | 0.201 | 0.054 | 6.0668E-42 | TAL/DCT | DOK5 |
| 1.0047E-25 | 0.30965015 | 0.508 | 0.297 | 2.1451E-21 | TAL/DCT | WDR1 |
| 4.2474E-05 | 0.3095163 | 0.676 | 0.648 | 0.9068294 | TAL/DCT | KRT7 |

| | | | | | | |
|---|---|---|---|---|---|---|
| 5.8039E-23 | 0.30932046 | 0.752 | 0.564 | 1.2391E-18 | TAL/DCT | COX7A2 |
| 1.5987E-15 | 0.30844688 | 0.67 | 0.497 | 3.4132E-11 | TAL/DCT | CLDN7 |
| 7.3732E-33 | 0.30830629 | 0.225 | 0.08 | 1.5742E-28 | TAL/DCT | FAM210A |
| 1.0852E-39 | 0.30594499 | 0.317 | 0.121 | 2.3168E-35 | TAL/DCT | EPHA2 |
| 1.358E-20 | 0.30465571 | 0.832 | 0.684 | 2.8993E-16 | TAL/DCT | CYSTM1 |
| 3.2949E-97 | 0.30457059 | 0.112 | 0.008 | 7.0346E-93 | TAL/DCT | TCEAL5 |
| 1.773E-14 | 0.30395759 | 0.831 | 0.727 | 3.7855E-10 | TAL/DCT | PFN1 |
| 1.6366E-25 | 0.30359616 | 0.251 | 0.106 | 3.4942E-21 | TAL/DCT | ISG20 |
| 6.7313E-20 | 0.30341829 | 0.507 | 0.318 | 1.4371E-15 | TAL/DCT | EIF1B |
| 1.4088E-23 | 0.303382 | 0.691 | 0.479 | 3.0078E-19 | TAL/DCT | UBE2D3 |
| 8.9052E-23 | 0.30179619 | 0.309 | 0.153 | 1.9013E-18 | TAL/DCT | GOLM1 |
| 9.3654E-12 | 0.30129896 | 0.201 | 0.107 | 1.9995E-07 | TAL/DCT | AC023157.3 |
| 1.0976E-20 | 0.29921945 | 0.762 | 0.571 | 2.3434E-16 | TAL/DCT | PRDX5 |
| 1.631E-35 | 0.29916935 | 0.335 | 0.14 | 3.4821E-31 | TAL/DCT | SNX10 |
| 1.0107E-06 | 0.29884236 | 0.907 | 0.889 | 0.02157738 | TAL/DCT | MT-ND2 |
| 2.2832E-24 | 0.29811956 | 0.54 | 0.323 | 4.8746E-20 | TAL/DCT | ATP6V1F |
| 1.2968E-12 | 0.29727947 | 0.311 | 0.191 | 2.7688E-08 | TAL/DCT | NCOA7 |
| 1.9027E-15 | 0.29723749 | 0.734 | 0.598 | 4.0622E-11 | TAL/DCT | PSMA7 |
| 1.2031E-37 | 0.29618981 | 0.248 | 0.085 | 2.5686E-33 | TAL/DCT | KIF9 |
| 1.0249E-29 | 0.29593091 | 0.343 | 0.156 | 2.1882E-25 | TAL/DCT | KHDRBS3 |
| 6.603E-113 | 0.29415611 | 0.151 | 0.014 | 1.41E-108 | TAL/DCT | LRRTM1 |
| 8.1264E-27 | 0.29374879 | 0.289 | 0.128 | 1.735E-22 | TAL/DCT | CREB5 |
| 5.024E-25 | 0.29344831 | 0.413 | 0.219 | 1.0726E-20 | TAL/DCT | PRSS8 |
| 1.1326E-18 | 0.29245651 | 0.337 | 0.184 | 2.4181E-14 | TAL/DCT | ZBTB43 |
| 7.3206E-20 | 0.29126349 | 0.287 | 0.145 | 1.5629E-15 | TAL/DCT | ACSL4 |
| 1.2905E-39 | 0.29116637 | 0.253 | 0.085 | 2.7551E-35 | TAL/DCT | CAV1 |
| 4.0413E-67 | 0.28971735 | 0.177 | 0.032 | 8.6281E-63 | TAL/DCT | ZNF554 |
| 7.1565E-23 | 0.28944584 | 0.21 | 0.086 | 1.5279E-18 | TAL/DCT | GEM |
| 3.903E-26 | 0.28736257 | 0.467 | 0.258 | 8.3329E-22 | TAL/DCT | MINOS1 |
| 4.5571E-45 | 0.28570806 | 0.158 | 0.036 | 9.7293E-41 | TAL/DCT | CAPN10-DT |
| 3.9248E-47 | 0.285627 | 0.259 | 0.081 | 8.3794E-43 | TAL/DCT | TMEM251 |
| 1.3191E-16 | 0.28431826 | 0.447 | 0.284 | 2.8163E-12 | TAL/DCT | YWHAH |
| 1.4923E-26 | 0.28350774 | 0.408 | 0.214 | 3.186E-22 | TAL/DCT | ARPC4 |
| 1.8621E-46 | 0.28342261 | 0.196 | 0.051 | 3.9756E-42 | TAL/DCT | RASGRP3 |
| 4.2824E-41 | 0.28319395 | 0.21 | 0.063 | 9.1429E-37 | TAL/DCT | TEX30 |
| 3.882E-05 | 0.28290167 | 0.659 | 0.604 | 0.82881762 | TAL/DCT | GADD45A |
| 1.9019E-18 | 0.28268355 | 0.989 | 0.971 | 4.0606E-14 | TAL/DCT | TPT1 |
| 2.3663E-45 | 0.28245999 | 0.201 | 0.054 | 5.0521E-41 | TAL/DCT | TMPRSS4 |
| 3.4479E-29 | 0.28159136 | 0.354 | 0.166 | 7.3613E-25 | TAL/DCT | PLIN3 |
| 1.8832E-22 | 0.28138704 | 0.475 | 0.275 | 4.0207E-18 | TAL/DCT | MSN |
| 9.9286E-21 | 0.2811887 | 0.339 | 0.18 | 2.1197E-16 | TAL/DCT | ARHGAP18 |
| 4.2157E-61 | 0.28107704 | 0.205 | 0.045 | 9.0005E-57 | TAL/DCT | BACH2 |
| 2.6183E-19 | 0.2793707 | 0.423 | 0.247 | 5.5901E-15 | TAL/DCT | COTL1 |
| 5.5931E-15 | 0.27912161 | 0.568 | 0.389 | 1.1941E-10 | TAL/DCT | RHOC |
| 3.1575E-17 | 0.2790894 | 0.572 | 0.376 | 6.7412E-13 | TAL/DCT | MYH9 |
| 1.055E-18 | 0.27753527 | 0.644 | 0.439 | 2.2524E-14 | TAL/DCT | VDAC2 |
| 4.5122E-26 | 0.27751213 | 0.203 | 0.076 | 9.6336E-22 | TAL/DCT | PTH1R |
| 1.5905E-36 | 0.27577114 | 0.153 | 0.039 | 3.3958E-32 | TAL/DCT | IL1R2 |
| 1.0997E-11 | 0.27527241 | 0.292 | 0.176 | 2.3479E-07 | TAL/DCT | HMGCS1 |
| 8.054E-24 | 0.27521362 | 0.454 | 0.259 | 1.7195E-19 | TAL/DCT | PSMB7 |

| | | | | | | |
|---|---|---|---|---|---|---|
| 1.4254E-23 | 0.27502566 | 0.145 | 0.048 | 3.0432E-19 | TAL/DCT | IGFBP6 |
| 1.9555E-19 | 0.27441041 | 0.248 | 0.119 | 4.175E-15 | TAL/DCT | WEE1 |
| 7.8491E-19 | 0.27417062 | 0.57 | 0.373 | 1.6758E-14 | TAL/DCT | DYNLT1 |
| 1.0869E-24 | 0.27227418 | 0.372 | 0.192 | 2.3205E-20 | TAL/DCT | CHMP1B |
| 1.7988E-19 | 0.27129365 | 0.659 | 0.454 | 3.8404E-15 | TAL/DCT | SERP1 |
| 3.1357E-23 | 0.271109 | 0.406 | 0.219 | 6.6948E-19 | TAL/DCT | RNMT |
| 1.8038E-17 | 0.27099938 | 0.35 | 0.199 | 3.8512E-13 | TAL/DCT | OSER1 |
| 2.5729E-23 | 0.2703685 | 0.41 | 0.229 | 5.4932E-19 | TAL/DCT | TMEM14B |
| 9.7028E-40 | 0.26806918 | 0.264 | 0.092 | 2.0715E-35 | TAL/DCT | DLGAP1-AS1 |
| 2.712E-44 | 0.26748615 | 0.205 | 0.057 | 5.7901E-40 | TAL/DCT | IL6R |
| 3.7795E-16 | 0.26726218 | 0.689 | 0.508 | 8.0692E-12 | TAL/DCT | TPM3 |
| 2.3933E-18 | 0.26667862 | 0.196 | 0.086 | 5.1098E-14 | TAL/DCT | CLDN10 |
| 2.2634E-15 | 0.26656872 | 0.588 | 0.409 | 4.8324E-11 | TAL/DCT | DYNLRB1 |
| 2.5702E-20 | 0.2664178 | 0.408 | 0.236 | 5.4873E-16 | TAL/DCT | PFKP |
| 1.172E-44 | 0.26635328 | 0.276 | 0.091 | 2.5022E-40 | TAL/DCT | SMIM29 |
| 1.4379E-31 | 0.26609812 | 0.302 | 0.126 | 3.0699E-27 | TAL/DCT | NPC1 |
| 2.7382E-15 | 0.26565934 | 0.704 | 0.558 | 5.846E-11 | TAL/DCT | CST3 |
| 1.9533E-20 | 0.26491767 | 0.493 | 0.302 | 4.1704E-16 | TAL/DCT | GABARAPL2 |
| 3.4611E-18 | 0.26417697 | 0.298 | 0.158 | 7.3893E-14 | TAL/DCT | CYLD |
| 1.4496E-23 | 0.26390782 | 0.292 | 0.14 | 3.095E-19 | TAL/DCT | ATP6V1A |
| 7.1599E-27 | 0.26276643 | 0.277 | 0.121 | 1.5286E-22 | TAL/DCT | TOR1AIP1 |
| 2.1025E-15 | 0.26270078 | 0.939 | 0.871 | 4.4888E-11 | TAL/DCT | RPL7 |
| 4.9679E-14 | 0.26262701 | 0.529 | 0.363 | 1.0607E-09 | TAL/DCT | SPTBN1 |
| 7.3902E-25 | 0.26159275 | 0.479 | 0.272 | 1.5778E-20 | TAL/DCT | MTCH1 |
| 3.4441E-10 | 0.26016511 | 0.277 | 0.171 | 7.3531E-06 | TAL/DCT | PHLDA1 |
| 2.3702E-26 | 0.2589518 | 0.637 | 0.392 | 5.0604E-22 | TAL/DCT | LAMTOR5 |
| 4.6617E-22 | 0.2579921 | 0.326 | 0.164 | 9.9528E-18 | TAL/DCT | SERPING1 |
| 1.3267E-17 | 0.25798844 | 0.549 | 0.362 | 2.8326E-13 | TAL/DCT | UBE2B |
| 2.3178E-16 | 0.2576627 | 0.274 | 0.146 | 4.9486E-12 | TAL/DCT | TFPI |
| 5.7262E-16 | 0.25744815 | 0.98 | 0.902 | 1.2225E-11 | TAL/DCT | RPS7 |
| 1.6134E-17 | 0.25726128 | 0.296 | 0.159 | 3.4445E-13 | TAL/DCT | CAVIN1 |
| 1.2874E-23 | 0.25703394 | 0.274 | 0.125 | 2.7486E-19 | TAL/DCT | SLC6A8 |
| 8.0723E-35 | 0.25679487 | 0.229 | 0.078 | 1.7234E-30 | TAL/DCT | HDAC9 |
| 2.9726E-18 | 0.25570941 | 0.343 | 0.191 | 6.3465E-14 | TAL/DCT | ADGRG1 |
| 7.7036E-18 | 0.25562901 | 0.972 | 0.906 | 1.6447E-13 | TAL/DCT | RPL26 |
| 3.1879E-31 | 0.25464019 | 0.283 | 0.115 | 6.8061E-27 | TAL/DCT | LMTK3 |
| 1.366E-22 | 0.25453358 | 0.369 | 0.193 | 2.9164E-18 | TAL/DCT | FOXO3 |
| 2.5786E-34 | 0.25449223 | 0.238 | 0.085 | 5.5053E-30 | TAL/DCT | SDSL |
| 2.996E-16 | 0.25445744 | 0.711 | 0.595 | 6.3964E-12 | TAL/DCT | SPP1 |
| 2.97E-20 | 0.25301186 | 0.25 | 0.119 | 6.3409E-16 | TAL/DCT | PRKAG2 |
| 1.4661E-13 | 0.25296961 | 0.672 | 0.509 | 3.1302E-09 | TAL/DCT | ARPC2 |
| 2.4958E-33 | 0.25250624 | 0.326 | 0.137 | 5.3285E-29 | TAL/DCT | CRB3 |
| 1.4172E-18 | 0.25172134 | 0.453 | 0.272 | 3.0258E-14 | TAL/DCT | BUD31 |
| 8.8259E-13 | 0.25112801 | 0.261 | 0.148 | 1.8843E-08 | TAL/DCT | RCAN1 |
| 4.5671E-22 | 0.25106305 | 0.35 | 0.181 | 9.7508E-18 | TAL/DCT | BEX4 |
| 4.9406E-29 | 0.25054453 | 0.277 | 0.115 | 1.0548E-24 | TAL/DCT | IRF6 |
| 1.2905E-12 | 0.25007373 | 0.581 | 0.432 | 2.7552E-08 | TAL/DCT | PPP1R14B |
| 2.6541E-06 | -0.2502792 | 0.11 | 0.182 | 0.05666526 | TAL/DCT | MYO9A |
| 0.00069522 | -0.2519516 | 0.201 | 0.255 | 1 | TAL/DCT | IRF1 |
| 0.00010219 | -0.2541716 | 0.093 | 0.148 | 1 | TAL/DCT | ANKRD37 |

| | | | | | | |
|---|---|---|---|---|---|---|
| 0.00055834 | -0.2549415 | 0.134 | 0.184 | 1 | TAL/DCT | ERN1 |
| 1.9924E-08 | -0.2549724 | 0.125 | 0.222 | 0.00042538 | TAL/DCT | EPS8L2 |
| 0.00393245 | -0.255453 | 0.356 | 0.384 | 1 | TAL/DCT | N4BP2L2 |
| 0.00080906 | -0.2567261 | 0.214 | 0.269 | 1 | TAL/DCT | TUBA1A |
| 6.0694E-06 | -0.2585809 | 0.073 | 0.136 | 0.1295827 | TAL/DCT | CSRNP2 |
| 2.1071E-09 | -0.2603431 | 0.138 | 0.244 | 4.4987E-05 | TAL/DCT | NSD3 |
| 6.613E-09 | -0.2610544 | 0.035 | 0.114 | 0.00014119 | TAL/DCT | HIST4H4 |
| 1.1208E-07 | -0.2668285 | 0.615 | 0.704 | 0.00239284 | TAL/DCT | HMGB1 |
| 1.5777E-07 | -0.2676345 | 0.177 | 0.267 | 0.0033683 | TAL/DCT | PLXNB2 |
| 2.0929E-08 | -0.2681903 | 0.035 | 0.11 | 0.00044683 | TAL/DCT | AL355075.4 |
| 2.731E-12 | -0.2688812 | 0.047 | 0.155 | 5.8308E-08 | TAL/DCT | AC007952.4 |
| 5.1301E-09 | -0.2699507 | 0.071 | 0.161 | 0.00010953 | TAL/DCT | PLEKHH2 |
| 0.00048405 | -0.2702061 | 0.089 | 0.137 | 1 | TAL/DCT | AC025164.1 |
| 1.7508E-09 | -0.2705793 | 0.08 | 0.175 | 3.7379E-05 | TAL/DCT | AHI1 |
| 0.00085823 | -0.2729204 | 0.233 | 0.287 | 1 | TAL/DCT | AKR1B1 |
| 0.00057118 | -0.2740562 | 0.304 | 0.358 | 1 | TAL/DCT | DDX3X |
| 7.933E-06 | -0.2766044 | 0.089 | 0.155 | 0.16936994 | TAL/DCT | PFKFB3 |
| 1.4624E-06 | -0.2803232 | 0.631 | 0.71 | 0.03122185 | TAL/DCT | DDX5 |
| 0.00402091 | -0.2821375 | 0.274 | 0.314 | 1 | TAL/DCT | CDKN1C |
| 4.2595E-12 | -0.2954034 | 0.08 | 0.196 | 9.0941E-08 | TAL/DCT | TUT4 |
| 0.00082478 | -0.2964256 | 0.22 | 0.263 | 1 | TAL/DCT | HIST1H2AC |
| 6.7784E-05 | -0.2995703 | 0.052 | 0.102 | 1 | TAL/DCT | AP001160.1 |
| 5.6661E-08 | -0.3003134 | 0.345 | 0.438 | 0.0012097 | TAL/DCT | PNRC1 |
| 5.1405E-08 | -0.3029133 | 0.199 | 0.293 | 0.00109749 | TAL/DCT | HP1BP3 |
| 0.00024463 | -0.3071435 | 0.32 | 0.37 | 1 | TAL/DCT | MACF1 |
| 1.2741E-12 | -0.3080971 | 0.089 | 0.212 | 2.7202E-08 | TAL/DCT | VPS13C |
| 7.4821E-07 | -0.3101823 | 0.218 | 0.307 | 0.01597425 | TAL/DCT | ARHGAP29 |
| 1.8699E-12 | -0.311683 | 0.43 | 0.565 | 3.9922E-08 | TAL/DCT | KTN1 |
| 0.00481119 | -0.3139747 | 0.08 | 0.119 | 1 | TAL/DCT | CXCL8 |
| 9.5658E-07 | -0.3142159 | 0.302 | 0.38 | 0.0204229 | TAL/DCT | PNISR |
| 0.00014919 | -0.3192073 | 0.775 | 0.797 | 1 | TAL/DCT | HNRNPA2B1 |
| 0.00010483 | -0.3233562 | 0.076 | 0.13 | 1 | TAL/DCT | COL7A1 |
| 2.2298E-08 | -0.3235198 | 0.642 | 0.778 | 0.00047606 | TAL/DCT | HSP90B1 |
| 2.2111E-09 | -0.3237858 | 0.045 | 0.129 | 4.7207E-05 | TAL/DCT | HIST2H2BF |
| 6.832E-07 | -0.3252469 | 0.35 | 0.424 | 0.0145863 | TAL/DCT | SFPQ |
| 1.386E-07 | -0.326947 | 0.199 | 0.286 | 0.00295911 | TAL/DCT | PRPF4B |
| 0.00021296 | -0.3279185 | 0.499 | 0.52 | 1 | TAL/DCT | HNRNPU |
| 6.2569E-09 | -0.3344463 | 0.57 | 0.672 | 0.00013359 | TAL/DCT | GLS |
| 1.6249E-11 | -0.3426785 | 0.153 | 0.272 | 3.4692E-07 | TAL/DCT | BDP1 |
| 3.438E-13 | -0.3431478 | 0.143 | 0.278 | 7.3401E-09 | TAL/DCT | TCF25 |
| 3.022E-06 | -0.344134 | 0.348 | 0.416 | 0.06451907 | TAL/DCT | GOLGB1 |
| 8.0055E-07 | -0.3506472 | 0.43 | 0.514 | 0.01709171 | TAL/DCT | H2AFZ |
| 2.8173E-08 | -0.3524768 | 0.065 | 0.147 | 0.00060149 | TAL/DCT | ID1 |
| 4.9754E-07 | -0.360583 | 0.233 | 0.316 | 0.01062238 | TAL/DCT | SRSF2 |
| 0.00020388 | -0.3724935 | 0.235 | 0.286 | 1 | TAL/DCT | POLR2A |
| 1.2136E-08 | -0.3734643 | 0.242 | 0.341 | 0.0002591 | TAL/DCT | HNRNPAB |
| 0.00140124 | -0.378974 | 0.223 | 0.266 | 1 | TAL/DCT | DUSP1 |
| 1.7093E-06 | -0.3795564 | 0.251 | 0.33 | 0.03649409 | TAL/DCT | CEBPB |
| 7.5246E-14 | -0.3796684 | 0.518 | 0.666 | 1.6065E-09 | TAL/DCT | CALR |
| 0.00177503 | -0.3827269 | 0.177 | 0.215 | 1 | TAL/DCT | SOCS1 |

| | | | | | | |
|---|---|---|---|---|---|---|
| 2.6146E-06 | -0.386968 | 0.46 | 0.52 | 0.05582123 | TAL/DCT | SON |
| 0.00023848 | -0.3884229 | 0.201 | 0.255 | 1 | TAL/DCT | ID2 |
| 3.2812E-08 | -0.3921439 | 0.177 | 0.268 | 0.00070053 | TAL/DCT | AMD1 |
| 6.0405E-11 | -0.3937238 | 0.089 | 0.197 | 1.2896E-06 | TAL/DCT | H2AFX |
| 5.8529E-13 | -0.4010521 | 0.074 | 0.195 | 1.2496E-08 | TAL/DCT | TLE4 |
| 0.00255685 | -0.4031413 | 0.257 | 0.195 | 1 | TAL/DCT | MT1G |
| 5.2782E-09 | -0.408292 | 0.304 | 0.402 | 0.00011269 | TAL/DCT | ANKRD11 |
| 0.00037734 | -0.4154732 | 0.086 | 0.138 | 1 | TAL/DCT | SERPINA1 |
| 1.4307E-05 | -0.4169492 | 0.128 | 0.195 | 0.30545699 | TAL/DCT | GADD45G |
| 3.3078E-16 | -0.4258044 | 0.492 | 0.663 | 7.0621E-12 | TAL/DCT | HSPA8 |
| 0.00458334 | -0.431108 | 0.51 | 0.642 | 1 | TAL/DCT | MMP7 |
| 3.7969E-10 | -0.4317906 | 0.173 | 0.281 | 8.1063E-06 | TAL/DCT | VEGFA |
| 9.8464E-08 | -0.4371994 | 0.257 | 0.345 | 0.0021022 | TAL/DCT | SNHG12 |
| 7.9954E-08 | -0.4428333 | 0.309 | 0.381 | 0.00170701 | TAL/DCT | RSRP1 |
| 3.6792E-05 | -0.4437177 | 0.063 | 0.119 | 0.78551878 | TAL/DCT | NEFM |
| 1.1974E-06 | -0.4606065 | 0.952 | 0.94 | 0.02556523 | TAL/DCT | MALAT1 |
| 3.4361E-10 | -0.4700954 | 0.155 | 0.259 | 7.3361E-06 | TAL/DCT | IER5L |
| 1.4286E-16 | -0.4707599 | 0.21 | 0.368 | 3.05E-12 | TAL/DCT | AKAP9 |
| 4.4448E-10 | -0.4716842 | 0.134 | 0.241 | 9.4898E-06 | TAL/DCT | SLC5A3 |
| 2.1483E-06 | -0.4756491 | 0.858 | 0.872 | 0.04586712 | TAL/DCT | SAT1 |
| 9.0561E-13 | -0.4779462 | 0.151 | 0.281 | 1.9335E-08 | TAL/DCT | CITED2 |
| 2.044E-10 | -0.4799716 | 0.196 | 0.307 | 4.364E-06 | TAL/DCT | DNAJB1 |
| 7.1427E-11 | -0.4839577 | 0.233 | 0.343 | 1.525E-06 | TAL/DCT | PAX8 |
| 0.00061666 | -0.4857248 | 0.225 | 0.273 | 1 | TAL/DCT | TCIM |
| 4.0892E-07 | -0.4862145 | 0.227 | 0.309 | 0.00873039 | TAL/DCT | HSPA1B |
| 0.00037628 | -0.4920016 | 0.562 | 0.601 | 1 | TAL/DCT | CLU |
| 1.37E-05 | -0.4939425 | 0.289 | 0.361 | 0.29248651 | TAL/DCT | HSPA1A |
| 1.8744E-08 | -0.4988284 | 0.166 | 0.257 | 0.00040017 | TAL/DCT | IER5 |
| 1.0955E-10 | -0.4993466 | 0.231 | 0.348 | 2.339E-06 | TAL/DCT | ZFP36L2 |
| 1.4831E-07 | -0.5026512 | 0.201 | 0.284 | 0.00316637 | TAL/DCT | BTG2 |
| 1.0297E-08 | -0.5081632 | 0.557 | 0.689 | 0.00021983 | TAL/DCT | HSPA5 |
| 4.9563E-12 | -0.5083425 | 0.065 | 0.173 | 1.0582E-07 | TAL/DCT | EGLN3 |
| 9.9008E-10 | -0.5089075 | 0.201 | 0.305 | 2.1138E-05 | TAL/DCT | DDIT4 |
| 1.6065E-16 | -0.50904 | 0.095 | 0.24 | 3.4298E-12 | TAL/DCT | NFKBIZ |
| 0.00019477 | -0.5138802 | 0.393 | 0.416 | 1 | TAL/DCT | SRSF7 |
| 7.4306E-15 | -0.5177956 | 0.553 | 0.673 | 1.5864E-10 | TAL/DCT | TPI1 |
| 1.3478E-07 | -0.5183192 | 0.209 | 0.289 | 0.00287749 | TAL/DCT | CEBPD |
| 1.5804E-09 | -0.5193392 | 0.296 | 0.39 | 3.3741E-05 | TAL/DCT | BRD2 |
| 2.0158E-15 | -0.5211553 | 0.3 | 0.439 | 4.3037E-11 | TAL/DCT | DDX17 |
| 8.3796E-12 | -0.5231765 | 0.128 | 0.248 | 1.789E-07 | TAL/DCT | ID4 |
| 1.9064E-05 | -0.535007 | 0.199 | 0.263 | 0.40702303 | TAL/DCT | EGR1 |
| 1.111E-17 | -0.5371231 | 0.205 | 0.364 | 2.372E-13 | TAL/DCT | ARGLU1 |
| 1.3515E-13 | -0.5479688 | 0.069 | 0.19 | 2.8855E-09 | TAL/DCT | PIK3R3 |
| 1.3941E-15 | -0.5631756 | 0.238 | 0.397 | 2.9765E-11 | TAL/DCT | TUBA1B |
| 7.0238E-18 | -0.5692317 | 0.061 | 0.209 | 1.4996E-13 | TAL/DCT | SGK1 |
| 2.74E-14 | -0.5724372 | 0.11 | 0.243 | 5.8499E-10 | TAL/DCT | INTS6 |
| 5.7427E-12 | -0.5739312 | 0.099 | 0.216 | 1.2261E-07 | TAL/DCT | ADAMTS1 |
| 3.5449E-07 | -0.6128408 | 0.328 | 0.389 | 0.00756831 | TAL/DCT | JUNB |
| 6.0557E-09 | -0.6156001 | 0.188 | 0.281 | 0.00012929 | TAL/DCT | PPP1R10 |
| 1.1597E-22 | -0.6416183 | 0.406 | 0.568 | 2.476E-18 | TAL/DCT | SRRM2 |

| | | | | | | |
|---|---|---|---|---|---|---|
| 1.9926E-13 | -0.6549922 | 0.186 | 0.317 | 4.2543E-09 | TAL/DCT | GLUL |
| 2.6831E-07 | -0.6568651 | 0.21 | 0.293 | 0.00572834 | TAL/DCT | FOS |
| 3.148E-05 | -0.6683283 | 0.544 | 0.565 | 0.67210058 | TAL/DCT | IER2 |
| 9.1318E-17 | -0.725403 | 0.134 | 0.283 | 1.9496E-12 | TAL/DCT | RASD1 |
| 3.7259E-10 | -0.7403826 | 0.311 | 0.41 | 7.9549E-06 | TAL/DCT | RNF213 |
| 1.8216E-14 | -0.7438448 | 0.201 | 0.337 | 3.8891E-10 | TAL/DCT | RHOB |
| 2.2514E-14 | -0.7568013 | 0.145 | 0.283 | 4.8067E-10 | TAL/DCT | HES1 |
| 9.7712E-15 | -0.7626565 | 0.106 | 0.24 | 2.0862E-10 | TAL/DCT | CSKMT |
| 0.00853586 | -0.7650767 | 0.093 | 0.128 | 1 | TAL/DCT | GPX3 |
| 1.4361E-20 | -0.7680548 | 0.136 | 0.308 | 3.0661E-16 | TAL/DCT | PLK2 |
| 7.4615E-22 | -0.7804556 | 0.257 | 0.436 | 1.593E-17 | TAL/DCT | SYNE2 |
| 9.0494E-08 | -0.8160917 | 0.372 | 0.443 | 0.00193204 | TAL/DCT | IER3 |
| 1.0137E-19 | -0.8200042 | 0.222 | 0.381 | 2.1642E-15 | TAL/DCT | SLC38A2 |
| 8.1875E-32 | -0.8744891 | 0.439 | 0.648 | 1.748E-27 | TAL/DCT | JUND |
| 9.8993E-22 | -0.9326043 | 0.088 | 0.268 | 2.1135E-17 | TAL/DCT | NUPR1 |
| 4.7285E-28 | -0.9610745 | 0.728 | 0.819 | 1.0095E-23 | TAL/DCT | NEAT1 |
| 3.1055E-23 | -0.977064 | 0.441 | 0.593 | 6.6302E-19 | TAL/DCT | ELF3 |
| 2.5027E-06 | -0.980596 | 0.503 | 0.541 | 0.05343166 | TAL/DCT | SOX4 |
| 3.5545E-15 | -0.9983051 | 0.143 | 0.276 | 7.5888E-11 | TAL/DCT | HSPA2 |
| 4.1938E-26 | -1.0913701 | 0.106 | 0.306 | 8.9538E-22 | TAL/DCT | HEXIM1 |
| 8.584E-18 | -1.1776546 | 0.533 | 0.638 | 1.8327E-13 | TAL/DCT | JUN |
| 4.7504E-21 | -1.2066201 | 0.121 | 0.29 | 1.0142E-16 | TAL/DCT | MAFB |
| 2.5469E-38 | -3.9694449 | 0.426 | 0.605 | 5.4376E-34 | TAL/DCT | PLCG2 |
| 0 | 1.46052699 | 0.383 | 0.015 | 0 | DTL | VCAM1 |
| 9.2292E-83 | 1.2259078 | 0.513 | 0.13 | 1.9704E-78 | DTL | TNFSF10 |
| 1.5534E-36 | 1.20811057 | 0.538 | 0.23 | 3.3166E-32 | DTL | SLC5A3 |
| 3.9979E-55 | 1.09926615 | 0.318 | 0.072 | 8.5354E-51 | DTL | AKR1C1 |
| 1.6611E-64 | 1.07967276 | 0.841 | 0.424 | 3.5464E-60 | DTL | ZBTB20 |
| 7.576E-300 | 1.06464225 | 0.397 | 0.021 | 1.618E-295 | DTL | SPON2 |
| 2.652E-160 | 1.04930918 | 0.245 | 0.015 | 5.663E-156 | DTL | AQP1 |
| 1.393E-224 | 1.04853871 | 0.245 | 0.01 | 2.975E-220 | DTL | RBP4 |
| 2.607E-171 | 1.04818555 | 0.318 | 0.025 | 5.567E-167 | DTL | CPE |
| 2.4004E-48 | 1.03203495 | 0.632 | 0.253 | 5.1249E-44 | DTL | NUPR1 |
| 6.966E-181 | 1.01694382 | 0.231 | 0.012 | 1.487E-176 | DTL | CFH |
| 4.5534E-21 | 1.01615454 | 0.202 | 0.064 | 9.7215E-17 | DTL | MGP |
| 5.8023E-50 | 1.0034858 | 0.801 | 0.42 | 1.2388E-45 | DTL | SYNE2 |
| 3.1666E-44 | 0.99957436 | 0.971 | 0.875 | 6.7608E-40 | DTL | ITM2B |
| 5.6867E-47 | 0.98150527 | 0.509 | 0.188 | 1.2141E-42 | DTL | BCAM |
| 2.7902E-77 | 0.96240313 | 0.563 | 0.164 | 5.9571E-73 | DTL | PLA2G16 |
| 4.1247E-14 | 0.95328293 | 0.422 | 0.246 | 8.8063E-10 | DTL | MRPS6 |
| 1.6882E-13 | 0.91938892 | 0.715 | 0.622 | 3.6044E-09 | DTL | VIM |
| 4.6062E-06 | 0.91751034 | 0.549 | 0.465 | 0.09834131 | DTL | IFITM3 |
| 1.1655E-42 | 0.86209105 | 0.614 | 0.274 | 2.4884E-38 | DTL | CLDN3 |
| 8.8603E-54 | 0.85854054 | 0.462 | 0.146 | 1.8917E-49 | DTL | PROM1 |
| 5.4849E-39 | 0.79358645 | 0.819 | 0.577 | 1.171E-34 | DTL | APP |
| 3.3504E-23 | 0.79260056 | 0.621 | 0.38 | 7.1532E-19 | DTL | PERP |
| 1.2081E-45 | 0.78599206 | 0.711 | 0.33 | 2.5792E-41 | DTL | PAX8 |
| 2.7E-67 | 0.7651571 | 0.509 | 0.144 | 5.7645E-63 | DTL | TFPI |
| 6.0018E-33 | 0.76073803 | 0.614 | 0.288 | 1.2814E-28 | DTL | ITGB8 |
| 9.2563E-22 | 0.75903403 | 0.928 | 0.812 | 1.9762E-17 | DTL | IGFBP7 |

| | | | | | | |
|---|---|---|---|---|---|---|
| 1.874E-40 | 0.75452786 | 0.415 | 0.142 | 4.0011E-36 | DTL | RETREG1 |
| 6.3699E-45 | 0.74911551 | 0.451 | 0.154 | 1.36E-40 | DTL | NR2F2 |
| 2.2668E-58 | 0.74571833 | 0.679 | 0.269 | 4.8396E-54 | DTL | GTF2I |
| 1.8544E-97 | 0.74360361 | 0.314 | 0.043 | 3.9591E-93 | DTL | CYP1B1 |
| 1.4414E-41 | 0.72444535 | 0.65 | 0.296 | 3.0773E-37 | DTL | ARHGAP29 |
| 9.4656E-31 | 0.717575 | 0.733 | 0.417 | 2.0209E-26 | DTL | AHNAK |
| 1.2106E-37 | 0.70628192 | 0.166 | 0.029 | 2.5846E-33 | DTL | AKR1C2 |
| 5.8427E-39 | 0.70241238 | 0.181 | 0.033 | 1.2474E-34 | DTL | PTGDS |
| 0.00885591 | 0.70012223 | 0.217 | 0.165 | 1 | DTL | CTGF |
| 1.2254E-34 | 0.68181937 | 0.57 | 0.246 | 2.6163E-30 | DTL | AKAP12 |
| 1.0137E-39 | 0.67705486 | 0.509 | 0.194 | 2.1642E-35 | DTL | CALD1 |
| 1.611E-220 | 0.67025873 | 0.375 | 0.027 | 3.439E-216 | DTL | PRUNE2 |
| 1.4205E-70 | 0.66814207 | 0.31 | 0.055 | 3.0327E-66 | DTL | IGFBP4 |
| 0.00394842 | 0.65435148 | 0.35 | 0.283 | 1 | DTL | AKR1B1 |
| 1.4537E-19 | 0.63519715 | 0.556 | 0.312 | 3.1036E-15 | DTL | UGCG |
| 2.6197E-50 | 0.62716272 | 0.336 | 0.083 | 5.593E-46 | DTL | PXDN |
| 3.858E-18 | 0.62677941 | 0.816 | 0.626 | 8.2368E-14 | DTL | TPM1 |
| 5.2308E-48 | 0.62481072 | 0.332 | 0.086 | 1.1168E-43 | DTL | LY6E |
| 1.7065E-13 | 0.60641454 | 0.22 | 0.092 | 3.6435E-09 | DTL | CCL2 |
| 4.5625E-44 | 0.60154905 | 0.368 | 0.109 | 9.741E-40 | DTL | CTSC |
| 1.675E-187 | 0.59238207 | 0.318 | 0.022 | 3.576E-183 | DTL | AEBP1 |
| 1.5379E-52 | 0.5869729 | 0.523 | 0.174 | 3.2835E-48 | DTL | CLIC4 |
| 1.4736E-71 | 0.58116862 | 0.285 | 0.047 | 3.1461E-67 | DTL | CLTRN |
| 7.2933E-45 | 0.58047885 | 0.527 | 0.192 | 1.5571E-40 | DTL | FAT1 |
| 5.6903E-52 | 0.57726233 | 0.502 | 0.163 | 1.2149E-47 | DTL | BICC1 |
| 8.0314E-11 | 0.57714172 | 0.661 | 0.489 | 1.7147E-06 | DTL | ATP1B1 |
| 8.6354E-14 | 0.57331286 | 0.953 | 0.866 | 1.8437E-09 | DTL | MT-ND3 |
| 1.4244E-22 | 0.57227808 | 0.628 | 0.364 | 3.041E-18 | DTL | SPTBN1 |
| 6.6073E-23 | 0.55637572 | 0.477 | 0.227 | 1.4107E-18 | DTL | SYNE1 |
| 7.5023E-22 | 0.55360349 | 0.682 | 0.448 | 1.6017E-17 | DTL | LAPTM4A |
| 2.0883E-24 | 0.55315377 | 0.859 | 0.663 | 4.4585E-20 | DTL | CRYAB |
| 3.1568E-86 | 0.54311748 | 0.339 | 0.055 | 6.7398E-82 | DTL | LINC01320 |
| 4.6917E-27 | 0.54130181 | 0.455 | 0.198 | 1.0017E-22 | DTL | CLDN1 |
| 4.4637E-21 | 0.52210724 | 0.422 | 0.204 | 9.53E-17 | DTL | ARID5B |
| 8.7498E-38 | 0.51965412 | 0.379 | 0.122 | 1.8681E-33 | DTL | CCDC198 |
| 8.1153E-23 | 0.51607752 | 0.43 | 0.195 | 1.7326E-18 | DTL | ANXA1 |
| 1.0686E-24 | 0.51572828 | 0.625 | 0.354 | 2.2814E-20 | DTL | SEC62 |
| 5.4076E-90 | 0.50691555 | 0.314 | 0.046 | 1.1545E-85 | DTL | NFIA |
| 8.5066E-33 | 0.50400849 | 0.235 | 0.061 | 1.8162E-28 | DTL | PDE1A |
| 6.0156E-51 | 0.5005606 | 0.184 | 0.028 | 1.2843E-46 | DTL | CYP3A5 |
| 5.5622E-92 | 0.49718526 | 0.256 | 0.03 | 1.1875E-87 | DTL | IRX3 |
| 1.162E-256 | 0.49651036 | 0.307 | 0.014 | 2.481E-252 | DTL | CDH2 |
| 2.2347E-74 | 0.49309658 | 0.329 | 0.059 | 4.7711E-70 | DTL | CCL28 |
| 6.1124E-40 | 0.49252376 | 0.253 | 0.059 | 1.305E-35 | DTL | CTTNBP2 |
| 1.6194E-30 | 0.49135531 | 0.592 | 0.289 | 3.4573E-26 | DTL | LEPROT |
| 5.2859E-16 | 0.48527385 | 0.581 | 0.354 | 1.1285E-11 | DTL | KLF6 |
| 8.5351E-21 | 0.48406337 | 0.505 | 0.262 | 1.8222E-16 | DTL | MET |
| 2.6098E-32 | 0.48140538 | 0.495 | 0.205 | 5.572E-28 | DTL | PPFIBP1 |
| 9.4383E-33 | 0.48128626 | 0.332 | 0.107 | 2.0151E-28 | DTL | RARRES2 |
| 1.084E-44 | 0.48060042 | 0.339 | 0.089 | 2.3144E-40 | DTL | KCNJ16 |

| | | | | | | |
|---|---|---|---|---|---|---|
| 4.247E-28 | 0.4792056 | 0.495 | 0.224 | 9.0673E-24 | DTL | NFE2L1 |
| 3.8263E-17 | 0.4746165 | 0.809 | 0.64 | 8.1691E-13 | DTL | PEBP1 |
| 1.0371E-09 | 0.4737318 | 0.477 | 0.323 | 2.2142E-05 | DTL | CTSB |
| 1.1093E-29 | 0.47333885 | 0.379 | 0.143 | 2.3683E-25 | DTL | HCFC1R1 |
| 1.0267E-24 | 0.4662118 | 0.52 | 0.257 | 2.1921E-20 | DTL | IL6ST |
| 1.7332E-24 | 0.46361607 | 0.314 | 0.117 | 3.7004E-20 | DTL | MT1F |
| 5.36E-129 | 0.46069539 | 0.256 | 0.021 | 1.144E-124 | DTL | EDIL3 |
| 1.3307E-37 | 0.45552557 | 0.39 | 0.126 | 2.841E-33 | DTL | COL4A1 |
| 6.3598E-88 | 0.45530226 | 0.206 | 0.02 | 1.3578E-83 | DTL | SPARC |
| 3.5713E-37 | 0.45288707 | 0.412 | 0.139 | 7.6247E-33 | DTL | COL4A2 |
| 2.7254E-29 | 0.44940518 | 0.484 | 0.206 | 5.8188E-25 | DTL | ADAM10 |
| 2.2348E-45 | 0.44928286 | 0.321 | 0.082 | 4.7713E-41 | DTL | DSC2 |
| 3.846E-48 | 0.44795545 | 0.383 | 0.105 | 8.2111E-44 | DTL | PLOD2 |
| 4.9732E-81 | 0.44739001 | 0.264 | 0.036 | 1.0618E-76 | DTL | SLC4A11 |
| 4.519E-112 | 0.44615454 | 0.278 | 0.029 | 9.648E-108 | DTL | SULT1C4 |
| 2.2303E-58 | 0.44341109 | 0.347 | 0.078 | 4.7616E-54 | DTL | PKD2 |
| 8.0833E-21 | 0.44316137 | 0.617 | 0.353 | 1.7258E-16 | DTL | GCC2 |
| 1.1234E-17 | 0.44158537 | 0.44 | 0.234 | 2.3986E-13 | DTL | CRIM1 |
| 1.5337E-30 | 0.44012643 | 0.545 | 0.249 | 3.2745E-26 | DTL | TMEM123 |
| 2.3199E-18 | 0.43979121 | 0.415 | 0.207 | 4.9529E-14 | DTL | SCCPDH |
| 3.379E-23 | 0.43975091 | 0.379 | 0.161 | 7.2141E-19 | DTL | NTN4 |
| 2.4587E-80 | 0.4394765 | 0.307 | 0.048 | 5.2494E-76 | DTL | GPC4 |
| 2.0955E-15 | 0.43480331 | 0.505 | 0.281 | 4.4739E-11 | DTL | CEBPD |
| 3.2575E-24 | 0.42797244 | 0.426 | 0.183 | 6.9548E-20 | DTL | SYTL2 |
| 5.2608E-52 | 0.4257961 | 0.325 | 0.075 | 1.1232E-47 | DTL | BCL2 |
| 3.5832E-26 | 0.42460051 | 0.523 | 0.245 | 7.6502E-22 | DTL | PTBP3 |
| 7.4328E-22 | 0.42324567 | 0.56 | 0.29 | 1.5869E-17 | DTL | DST |
| 8.7047E-16 | 0.42184442 | 0.545 | 0.315 | 1.8585E-11 | DTL | DCDC2 |
| 3.3617E-82 | 0.41850711 | 0.227 | 0.026 | 7.1773E-78 | DTL | RERG |
| 1.5639E-18 | 0.41194222 | 0.56 | 0.317 | 3.3389E-14 | DTL | PAWR |
| 0.00374823 | 0.41129387 | 0.105 | 0.062 | 1 | DTL | LTB |
| 2.1581E-24 | 0.4108052 | 0.534 | 0.264 | 4.6076E-20 | DTL | FARP1 |
| 1.7577E-30 | 0.40975539 | 0.249 | 0.07 | 3.7527E-26 | DTL | IL1R1 |
| 5.4546E-29 | 0.40576009 | 0.455 | 0.187 | 1.1646E-24 | DTL | HIPK2 |
| 6.4634E-27 | 0.39863369 | 0.44 | 0.185 | 1.3799E-22 | DTL | CARHSP1 |
| 9.1497E-21 | 0.39692717 | 0.249 | 0.089 | 1.9535E-16 | DTL | CAV1 |
| 4.6946E-46 | 0.39682141 | 0.249 | 0.052 | 1.0023E-41 | DTL | ABCC4 |
| 2.3985E-16 | 0.39616651 | 0.321 | 0.149 | 5.1209E-12 | DTL | MKLN1 |
| 1.0017E-22 | 0.39616608 | 0.354 | 0.144 | 2.1387E-18 | DTL | LAMB1 |
| 1.8709E-51 | 0.39339897 | 0.238 | 0.044 | 3.9943E-47 | DTL | SLC6A12 |
| 8.1939E-31 | 0.39251479 | 0.188 | 0.042 | 1.7494E-26 | DTL | THY1 |
| 0.0002562 | 0.38722982 | 0.798 | 0.8 | 1 | DTL | CD24 |
| 8.5144E-29 | 0.38448171 | 0.292 | 0.093 | 1.8178E-24 | DTL | SASH1 |
| 6.187E-18 | 0.38258537 | 0.801 | 0.603 | 1.3209E-13 | DTL | APLP2 |
| 4.985E-156 | 0.38228402 | 0.227 | 0.013 | 1.064E-151 | DTL | LRP2 |
| 3.5507E-14 | 0.38132662 | 0.603 | 0.387 | 7.5807E-10 | DTL | CIRBP |
| 1.0042E-20 | 0.37891473 | 0.282 | 0.107 | 2.1439E-16 | DTL | ALDH1A1 |
| 1.4364E-16 | 0.37511805 | 0.975 | 0.934 | 3.0667E-12 | DTL | PTMA |
| 9.4331E-33 | 0.37457352 | 0.372 | 0.128 | 2.014E-28 | DTL | IGF1R |
| 8.1265E-41 | 0.37389243 | 0.375 | 0.11 | 1.735E-36 | DTL | PAX2 |

| | | | | | | |
|---|---|---|---|---|---|---|
| 8.4992E-24 | 0.37387233 | 0.404 | 0.175 | 1.8146E-19 | DTL | CTSZ |
| 1.088E-30 | 0.37362781 | 0.397 | 0.148 | 2.3228E-26 | DTL | GLG1 |
| 1.0127E-43 | 0.37299802 | 0.343 | 0.092 | 2.1621E-39 | DTL | TGFBR2 |
| 1.2864E-39 | 0.37061975 | 0.329 | 0.092 | 2.7465E-35 | DTL | C11orf54 |
| 5.2131E-20 | 0.37031691 | 0.527 | 0.284 | 1.113E-15 | DTL | SERINC1 |
| 7.2167E-21 | 0.36953124 | 0.473 | 0.231 | 1.5408E-16 | DTL | LPP |
| 2.5062E-19 | 0.36839696 | 0.614 | 0.352 | 5.3508E-15 | DTL | CD46 |
| 1.4846E-37 | 0.36794453 | 0.206 | 0.043 | 3.1696E-33 | DTL | FHL1 |
| 5.3631E-47 | 0.36790451 | 0.264 | 0.056 | 1.145E-42 | DTL | LRRK2 |
| 8.8506E-14 | 0.36647533 | 0.231 | 0.098 | 1.8896E-09 | DTL | PIK3R1 |
| 1.4099E-31 | 0.36569995 | 0.314 | 0.099 | 3.0101E-27 | DTL | ERBB2 |
| 3.4763E-23 | 0.36459625 | 0.426 | 0.191 | 7.4218E-19 | DTL | IGF2R |
| 5.6463E-24 | 0.36442964 | 0.314 | 0.116 | 1.2055E-19 | DTL | POU3F3 |
| 4.1549E-18 | 0.36389415 | 0.339 | 0.153 | 8.8708E-14 | DTL | PALLD |
| 8.4093E-30 | 0.36275516 | 0.361 | 0.127 | 1.7954E-25 | DTL | UACA |
| 8.3904E-67 | 0.36139113 | 0.227 | 0.032 | 1.7913E-62 | DTL | PAPLN |
| 4.4506E-13 | 0.36108798 | 0.549 | 0.34 | 9.502E-09 | DTL | TSC22D1 |
| 1.0016E-14 | 0.35694828 | 0.534 | 0.332 | 2.1385E-10 | DTL | EIF4B |
| 1.8509E-29 | 0.35686708 | 0.39 | 0.147 | 3.9517E-25 | DTL | SLC35F5 |
| 6.0207E-25 | 0.35631244 | 0.433 | 0.187 | 1.2854E-20 | DTL | BNIP3L |
| 5.4533E-21 | 0.35566481 | 0.448 | 0.21 | 1.1643E-16 | DTL | PARD6B |
| 3.3132E-28 | 0.35541467 | 0.267 | 0.082 | 7.0736E-24 | DTL | AHR |
| 2.4487E-17 | 0.35496931 | 0.592 | 0.351 | 5.2281E-13 | DTL | UBXN4 |
| 1.3926E-28 | 0.35453978 | 0.404 | 0.151 | 2.9731E-24 | DTL | PLEKHH2 |
| 2.7735E-20 | 0.35381195 | 0.581 | 0.308 | 5.9214E-16 | DTL | AFDN |
| 1.2407E-23 | 0.35282205 | 0.148 | 0.035 | 2.6489E-19 | DTL | C1R |
| 2.1016E-16 | 0.35155986 | 0.35 | 0.17 | 4.487E-12 | DTL | DSG2 |
| 3.881E-26 | 0.35114977 | 0.231 | 0.068 | 8.286E-22 | DTL | SLC3A1 |
| 3.3033E-34 | 0.34889576 | 0.303 | 0.089 | 7.0526E-30 | DTL | FNBP1 |
| 1.3311E-18 | 0.34836036 | 0.484 | 0.256 | 2.842E-14 | DTL | PAK2 |
| 1.4002E-21 | 0.348016 | 0.401 | 0.179 | 2.9895E-17 | DTL | PLEKHA1 |
| 7.6883E-43 | 0.34788984 | 0.26 | 0.059 | 1.6414E-38 | DTL | TSPAN12 |
| 9.9354E-48 | 0.34782039 | 0.289 | 0.064 | 2.1212E-43 | DTL | TNS1 |
| 4.3568E-31 | 0.34668195 | 0.31 | 0.098 | 9.3018E-27 | DTL | UGT2B7 |
| 2.2941E-18 | 0.34657644 | 0.318 | 0.138 | 4.8979E-14 | DTL | EEF1E1 |
| 5.5888E-42 | 0.34637165 | 0.17 | 0.028 | 1.1932E-37 | DTL | TGFB2 |
| 2.6711E-53 | 0.34611963 | 0.202 | 0.031 | 5.7028E-49 | DTL | ENPP1 |
| 5.6781E-16 | 0.34580712 | 0.603 | 0.356 | 1.2123E-11 | DTL | AKAP9 |
| 8.239E-107 | 0.34578123 | 0.217 | 0.018 | 1.759E-102 | DTL | NPR1 |
| 6.4114E-14 | 0.34551246 | 0.43 | 0.241 | 1.3688E-09 | DTL | RBPMS |
| 8.7465E-32 | 0.34542388 | 0.285 | 0.085 | 1.8674E-27 | DTL | TMEM43 |
| 3.935E-13 | 0.34457629 | 0.419 | 0.23 | 8.4013E-09 | DTL | FXYD2 |
| 1.9042E-32 | 0.34420198 | 0.339 | 0.109 | 4.0655E-28 | DTL | SLCO3A1 |
| 1.5483E-16 | 0.34254403 | 0.354 | 0.167 | 3.3056E-12 | DTL | EMP3 |
| 5.2528E-17 | 0.34190477 | 0.394 | 0.19 | 1.1215E-12 | DTL | SLC25A37 |
| 1.8168E-18 | 0.34161826 | 0.325 | 0.142 | 3.8789E-14 | DTL | LPIN2 |
| 6.3873E-07 | 0.34061369 | 0.318 | 0.2 | 0.01363696 | DTL | DPP4 |
| 4.2499E-17 | 0.34037064 | 0.531 | 0.285 | 9.0734E-13 | DTL | KMT2A |
| 6.3424E-47 | 0.34016429 | 0.217 | 0.039 | 1.3541E-42 | DTL | SPOCK2 |
| 1.0312E-16 | 0.33978521 | 0.235 | 0.091 | 2.2017E-12 | DTL | PIGR |

| | | | | | | |
|---|---|---|---|---|---|---|
| 3.7416E-45 | 0.3390108 | 0.267 | 0.059 | 7.9883E-41 | DTL | BLOC1S5 |
| 5.093E-14 | 0.33827775 | 0.783 | 0.554 | 1.0874E-09 | DTL | KTN1 |
| 1.274E-09 | 0.33797917 | 0.996 | 0.939 | 2.72E-05 | DTL | MALAT1 |
| 1.0642E-06 | 0.33568238 | 0.339 | 0.224 | 0.02272019 | DTL | IFITM2 |
| 3.1132E-20 | 0.33564473 | 0.458 | 0.22 | 6.6466E-16 | DTL | NFAT5 |
| 7.7873E-11 | 0.334972 | 0.502 | 0.311 | 1.6626E-06 | DTL | MAL |
| 5.9803E-27 | 0.33481656 | 0.339 | 0.122 | 1.2768E-22 | DTL | KAZN |
| 1.0594E-29 | 0.33461688 | 0.307 | 0.098 | 2.2619E-25 | DTL | LGR4 |
| 7.9035E-14 | 0.33384551 | 0.397 | 0.214 | 1.6874E-09 | DTL | FNBP1L |
| 8.3133E-15 | 0.33340879 | 0.48 | 0.272 | 1.7749E-10 | DTL | ATRX |
| 4.1437E-29 | 0.33141016 | 0.152 | 0.031 | 8.8467E-25 | DTL | MSLN |
| 1.2063E-23 | 0.3307164 | 0.321 | 0.123 | 2.5754E-19 | DTL | VAT1 |
| 5.1898E-12 | 0.33068056 | 0.321 | 0.168 | 1.108E-07 | DTL | SERPING1 |
| 1.3554E-30 | 0.32847975 | 0.224 | 0.057 | 2.8938E-26 | DTL | MT1M |
| 2.9619E-29 | 0.32818246 | 0.397 | 0.148 | 6.3237E-25 | DTL | ZNF148 |
| 3.047E-23 | 0.32687995 | 0.321 | 0.121 | 6.5054E-19 | DTL | MICAL3 |
| 1.6473E-12 | 0.32683524 | 0.31 | 0.157 | 3.5169E-08 | DTL | TMEM176B |
| 3.494E-08 | 0.32672554 | 0.477 | 0.323 | 0.00074597 | DTL | MT1E |
| 4.2667E-35 | 0.3253763 | 0.238 | 0.058 | 9.1094E-31 | DTL | NAPEPLD |
| 1.0383E-19 | 0.32409088 | 0.285 | 0.11 | 2.2167E-15 | DTL | CNKSR3 |
| 1.6905E-26 | 0.32344137 | 0.336 | 0.123 | 3.6092E-22 | DTL | CDC42SE2 |
| 5.6987E-16 | 0.3234141 | 0.401 | 0.2 | 1.2167E-11 | DTL | AIF1L |
| 1.195E-21 | 0.3229109 | 0.343 | 0.142 | 2.5514E-17 | DTL | CLSTN1 |
| 8.6755E-23 | 0.32269125 | 0.162 | 0.042 | 1.8522E-18 | DTL | ACSM3 |
| 5.0368E-16 | 0.32114579 | 0.343 | 0.161 | 1.0754E-11 | DTL | CAVIN1 |
| 8.7743E-20 | 0.32098904 | 0.469 | 0.231 | 1.8733E-15 | DTL | ANXA4 |
| 3.3078E-16 | 0.32031896 | 0.267 | 0.114 | 7.0622E-12 | DTL | TMEM51 |
| 1.017E-21 | 0.32010415 | 0.448 | 0.202 | 2.1713E-17 | DTL | VPS13C |
| 4.6633E-27 | 0.31984055 | 0.372 | 0.14 | 9.9561E-23 | DTL | SKAP2 |
| 3.0602E-29 | 0.31958347 | 0.347 | 0.12 | 6.5336E-25 | DTL | NFIB |
| 8.518E-20 | 0.31926391 | 0.469 | 0.229 | 1.8186E-15 | DTL | PHF3 |
| 1.5375E-18 | 0.31925012 | 0.282 | 0.112 | 3.2826E-14 | DTL | NEDD9 |
| 4.5072E-15 | 0.31886296 | 0.419 | 0.221 | 9.6229E-11 | DTL | TTC3 |
| 1.5675E-11 | 0.31869063 | 0.365 | 0.207 | 3.3467E-07 | DTL | BEX3 |
| 5.4599E-16 | 0.31712797 | 0.574 | 0.347 | 1.1657E-11 | DTL | NUCKS1 |
| 6.5622E-27 | 0.31657991 | 0.274 | 0.087 | 1.401E-22 | DTL | BHLHE41 |
| 3.3522E-07 | 0.31609579 | 0.318 | 0.196 | 0.00715693 | DTL | TINAGL1 |
| 1.6507E-25 | 0.31506522 | 0.357 | 0.139 | 3.5243E-21 | DTL | CERS2 |
| 1.4722E-34 | 0.31497859 | 0.249 | 0.063 | 3.143E-30 | DTL | SEMA4D |
| 2.8795E-33 | 0.31492118 | 0.303 | 0.09 | 6.1477E-29 | DTL | MAP3K1 |
| 8.2118E-15 | 0.31443227 | 0.682 | 0.439 | 1.7532E-10 | DTL | ZFP36L1 |
| 7.299E-61 | 0.31409525 | 0.184 | 0.023 | 1.5583E-56 | DTL | COL6A2 |
| 3.2121E-40 | 0.31391833 | 0.285 | 0.072 | 6.8578E-36 | DTL | TBL1X |
| 2.6947E-70 | 0.31279898 | 0.155 | 0.014 | 5.7532E-66 | DTL | BGN |
| 8.914E-107 | 0.31259929 | 0.144 | 0.008 | 1.903E-102 | DTL | F5 |
| 8.4865E-39 | 0.31232456 | 0.296 | 0.077 | 1.8119E-34 | DTL | SOX6 |
| 5.0665E-10 | 0.31178818 | 0.368 | 0.219 | 1.0817E-05 | DTL | ARL6IP5 |
| 2.1093E-43 | 0.31170732 | 0.264 | 0.059 | 4.5033E-39 | DTL | BIN1 |
| 2.2686E-51 | 0.31162051 | 0.202 | 0.032 | 4.8434E-47 | DTL | PHYHIPL |
| 9.8313E-46 | 0.31160015 | 0.26 | 0.056 | 2.099E-41 | DTL | TMEM98 |

| | | | | | | |
|---|---|---|---|---|---|---|
| 1.8317E-36 | 0.31149411 | 0.292 | 0.079 | 3.9106E-32 | DTL | ARHGAP24 |
| 1.1266E-24 | 0.31140232 | 0.383 | 0.152 | 2.4052E-20 | DTL | CXXC5 |
| 6.819E-16 | 0.31088923 | 0.394 | 0.195 | 1.4559E-11 | DTL | AGRN |
| 8.2862E-12 | 0.30935757 | 0.451 | 0.275 | 1.7691E-07 | DTL | CNN3 |
| 2.879E-38 | 0.30899884 | 0.206 | 0.042 | 6.1467E-34 | DTL | MAF |
| 3.1293E-17 | 0.30833784 | 0.336 | 0.147 | 6.6811E-13 | DTL | DAPK1 |
| 1.7265E-09 | 0.30782311 | 0.527 | 0.355 | 3.686E-05 | DTL | SPAG9 |
| 5.6637E-15 | 0.30765009 | 0.383 | 0.194 | 1.2092E-10 | DTL | MINDY2 |
| 1.1608E-22 | 0.30754082 | 0.354 | 0.142 | 2.4783E-18 | DTL | LNPEP |
| 4.7297E-19 | 0.30751929 | 0.386 | 0.174 | 1.0098E-14 | DTL | MYO9A |
| 2.5507E-45 | 0.30744037 | 0.199 | 0.035 | 5.4458E-41 | DTL | RHEX |
| 3.265E-48 | 0.30583404 | 0.256 | 0.052 | 6.9707E-44 | DTL | PBX1 |
| 2.239E-140 | 0.30525467 | 0.173 | 0.008 | 4.781E-136 | DTL | CDH6 |
| 3.712E-124 | 0.30446051 | 0.159 | 0.008 | 7.926E-120 | DTL | CRB2 |
| 4.2056E-12 | 0.3039841 | 0.628 | 0.408 | 8.979E-08 | DTL | GOLGB1 |
| 2.3798E-41 | 0.30352371 | 0.242 | 0.053 | 5.0809E-37 | DTL | CRNDE |
| 3.7166E-09 | 0.30307401 | 0.412 | 0.258 | 7.9349E-05 | DTL | CFLAR |
| 1.9023E-23 | 0.30302888 | 0.329 | 0.123 | 4.0615E-19 | DTL | FTX |
| 5.8963E-18 | 0.30250745 | 0.339 | 0.153 | 1.2589E-13 | DTL | CASC4 |
| 2.5769E-23 | 0.30238797 | 0.289 | 0.103 | 5.5016E-19 | DTL | ZMYND8 |
| 1.6461E-36 | 0.30222147 | 0.278 | 0.072 | 3.5144E-32 | DTL | EVC |
| 3.79E-12 | 0.30216073 | 0.606 | 0.378 | 8.0916E-08 | DTL | N4BP2L2 |
| 1.6722E-10 | 0.30200502 | 0.426 | 0.262 | 3.5701E-06 | DTL | TAPBP |
| 1.9423E-16 | 0.30168861 | 0.401 | 0.199 | 4.1468E-12 | DTL | FLNB |
| 4.9069E-27 | 0.30115016 | 0.141 | 0.029 | 1.0476E-22 | DTL | DPYSL3 |
| 9.3828E-25 | 0.30037058 | 0.264 | 0.087 | 2.0032E-20 | DTL | MEGF9 |
| 1.2752E-19 | 0.300172 | 0.289 | 0.114 | 2.7226E-15 | DTL | C9orf3 |
| 3.9911E-16 | 0.30006 | 0.3 | 0.134 | 8.521E-12 | DTL | RBBP8 |
| 4.3663E-14 | 0.29976868 | 0.375 | 0.196 | 9.3219E-10 | DTL | DDR1 |
| 5.2457E-21 | 0.29976657 | 0.285 | 0.108 | 1.12E-16 | DTL | CRTAP |
| 4.3587E-15 | 0.29922152 | 0.542 | 0.315 | 9.3059E-11 | DTL | DEK |
| 3.6157E-25 | 0.29807745 | 0.314 | 0.112 | 7.7195E-21 | DTL | ARHGEF28 |
| 1.16E-64 | 0.29727836 | 0.209 | 0.028 | 2.4766E-60 | DTL | IRX5 |
| 4.031E-101 | 0.29620751 | 0.148 | 0.009 | 8.6066E-97 | DTL | ALPK2 |
| 1.8949E-15 | 0.29581564 | 0.206 | 0.076 | 4.0456E-11 | DTL | TBC1D4 |
| 4.5827E-26 | 0.29560155 | 0.213 | 0.058 | 9.7842E-22 | DTL | S100A1 |
| 3.1028E-18 | 0.29528697 | 0.325 | 0.143 | 6.6244E-14 | DTL | STAG2 |
| 2.4478E-08 | 0.29484798 | 0.3 | 0.169 | 0.0005226 | DTL | THSD7A |
| 6.1574E-14 | 0.29477077 | 0.282 | 0.126 | 1.3146E-09 | DTL | TNFRSF11B |
| 3.0354E-15 | 0.29314899 | 0.336 | 0.159 | 6.4806E-11 | DTL | AFF1 |
| 1.5716E-18 | 0.29193635 | 0.336 | 0.144 | 3.3553E-14 | DTL | ZBTB38 |
| 5.1632E-11 | 0.29088791 | 0.888 | 0.823 | 1.1023E-06 | DTL | H3F3A |
| 1.8427E-14 | 0.29060425 | 0.282 | 0.127 | 3.9342E-10 | DTL | NCEH1 |
| 9.8879E-40 | 0.29045841 | 0.253 | 0.059 | 2.1111E-35 | DTL | DPYSL2 |
| 2.4544E-12 | 0.29036074 | 0.509 | 0.305 | 5.2401E-08 | DTL | DSP |
| 6.124E-16 | 0.29030075 | 0.354 | 0.168 | 1.3075E-11 | DTL | PPP1R12A |
| 4.0711E-14 | 0.28956879 | 0.422 | 0.23 | 8.6919E-10 | DTL | MGAT4B |
| 7.2392E-33 | 0.2889733 | 0.235 | 0.059 | 1.5456E-28 | DTL | KANK2 |
| 6.7125E-23 | 0.28872274 | 0.292 | 0.105 | 1.4331E-18 | DTL | RORA |
| 1.1376E-33 | 0.28832543 | 0.202 | 0.045 | 2.4289E-29 | DTL | ALDH1A2 |

| | | | | | | |
|---|---|---|---|---|---|---|
| 1.1739E-18 | 0.28813521 | 0.357 | 0.16 | 2.5063E-14 | DTL | SUPT16H |
| 1.1259E-06 | 0.28795096 | 0.978 | 0.908 | 0.02403876 | DTL | MT-ATP6 |
| 1.3733E-10 | 0.28669655 | 0.361 | 0.206 | 2.932E-06 | DTL | RAB3IP |
| 6.4856E-30 | 0.28600114 | 0.264 | 0.076 | 1.3847E-25 | DTL | PBXIP1 |
| 3.8415E-09 | 0.28543253 | 0.39 | 0.24 | 8.2017E-05 | DTL | PFKP |
| 2.1341E-44 | 0.28525725 | 0.235 | 0.047 | 4.5564E-40 | DTL | ADGRG6 |
| 4.5764E-51 | 0.28522936 | 0.217 | 0.036 | 9.7705E-47 | DTL | SLC22A2 |
| 1.0851E-12 | 0.28448114 | 0.516 | 0.312 | 2.3167E-08 | DTL | EIF5B |
| 3.1121E-13 | 0.28385856 | 0.458 | 0.26 | 6.6444E-09 | DTL | SCAF11 |
| 7.129E-19 | 0.283846 | 0.307 | 0.124 | 1.522E-14 | DTL | UTRN |
| 1.4904E-16 | 0.28357493 | 0.422 | 0.213 | 3.182E-12 | DTL | THRAP3 |
| 1.0459E-12 | 0.28332755 | 0.17 | 0.063 | 2.233E-08 | DTL | SOD3 |
| 7.5939E-15 | 0.28326404 | 0.329 | 0.158 | 1.6213E-10 | DTL | DDAH2 |
| 6.7973E-16 | 0.28278634 | 0.264 | 0.111 | 1.4512E-11 | DTL | INSR |
| 1.2749E-21 | 0.28201176 | 0.256 | 0.089 | 2.7219E-17 | DTL | NRP1 |
| 3.2983E-33 | 0.28130097 | 0.274 | 0.076 | 7.0418E-29 | DTL | MTMR12 |
| 4.2537E-15 | 0.28096337 | 0.43 | 0.222 | 9.0817E-11 | DTL | TRIM8 |
| 7.0756E-20 | 0.28058028 | 0.3 | 0.118 | 1.5106E-15 | DTL | SDC2 |
| 2.3691E-15 | 0.28030134 | 0.495 | 0.275 | 5.0581E-11 | DTL | LMAN1 |
| 8.5248E-40 | 0.28027882 | 0.242 | 0.054 | 1.8201E-35 | DTL | ZNF462 |
| 1.8645E-14 | 0.28007905 | 0.3 | 0.139 | 3.9808E-10 | DTL | PHACTR2 |
| 5.5783E-17 | 0.28003717 | 0.354 | 0.165 | 1.191E-12 | DTL | VGLL4 |
| 9.0735E-21 | 0.27977341 | 0.332 | 0.134 | 1.9372E-16 | DTL | YPEL2 |
| 1.4588E-21 | 0.27830564 | 0.292 | 0.108 | 3.1144E-17 | DTL | GNAI1 |
| 1.0559E-13 | 0.27665276 | 0.274 | 0.125 | 2.2542E-09 | DTL | CRIP2 |
| 8.5558E-17 | 0.27652346 | 0.357 | 0.164 | 1.8267E-12 | DTL | PRMT2 |
| 5.1715E-14 | 0.27616666 | 0.401 | 0.21 | 1.1041E-09 | DTL | TRIP11 |
| 5.1497E-17 | 0.27616047 | 0.336 | 0.152 | 1.0995E-12 | DTL | MARCH6 |
| 2.2535E-23 | 0.27520758 | 0.264 | 0.089 | 4.8111E-19 | DTL | MITF |
| 6.0865E-08 | 0.2750624 | 0.996 | 0.987 | 0.00129948 | DTL | EEF1A1 |
| 9.3065E-20 | 0.27476174 | 0.336 | 0.14 | 1.9869E-15 | DTL | MIA2 |
| 6.0776E-09 | 0.27475493 | 0.505 | 0.323 | 0.00012976 | DTL | IL32 |
| 1.0455E-08 | 0.27444233 | 0.621 | 0.469 | 0.00022321 | DTL | SUMO2 |
| 1.4828E-40 | 0.27413636 | 0.217 | 0.044 | 3.1658E-36 | DTL | SORBS2 |
| 6.848E-12 | 0.27347188 | 0.552 | 0.341 | 1.4621E-07 | DTL | MTDH |
| 1.1187E-24 | 0.27327254 | 0.267 | 0.088 | 2.3884E-20 | DTL | CDS1 |
| 2.2189E-15 | 0.27322991 | 0.401 | 0.201 | 4.7373E-11 | DTL | TNRC6B |
| 3.3222E-11 | 0.27299422 | 0.466 | 0.28 | 7.0928E-07 | DTL | COL18A1 |
| 4.7617E-13 | 0.27235245 | 0.253 | 0.111 | 1.0166E-08 | DTL | RARRES3 |
| 4.4466E-05 | 0.27141658 | 0.671 | 0.612 | 0.94934599 | DTL | MT-ND4L |
| 3.5078E-97 | 0.27130563 | 0.191 | 0.016 | 7.4892E-93 | DTL | PROS1 |
| 3.6377E-10 | 0.2704018 | 0.549 | 0.358 | 7.7666E-06 | DTL | CSDE1 |
| 2.3328E-12 | 0.27012575 | 0.69 | 0.472 | 4.9804E-08 | DTL | MORF4L1 |
| 1.7371E-14 | 0.26960178 | 0.39 | 0.198 | 3.7088E-10 | DTL | IRF2BP2 |
| 1.1417E-23 | 0.26924132 | 0.126 | 0.026 | 2.4375E-19 | DTL | IFITM1 |
| 1.1662E-14 | 0.26919191 | 0.477 | 0.258 | 2.4899E-10 | DTL | PLXNB2 |
| 2.2558E-14 | 0.26880204 | 0.412 | 0.219 | 4.8162E-10 | DTL | ANXA7 |
| 9.0118E-13 | 0.26850162 | 0.469 | 0.275 | 1.924E-08 | DTL | SEPT7 |
| 1.1215E-15 | 0.2676298 | 0.375 | 0.184 | 2.3944E-11 | DTL | HMGN3 |
| 5.6097E-22 | 0.26748534 | 0.253 | 0.086 | 1.1977E-17 | DTL | MGAT4A |

| | | | | | | |
|---|---|---|---|---|---|---|
| 1.4665E-25 | 0.26707074 | 0.314 | 0.11 | 3.1311E-21 | DTL | HOOK1 |
| 4.4214E-16 | 0.26641392 | 0.422 | 0.213 | 9.4396E-12 | DTL | PJA2 |
| 8.4264E-19 | 0.26628759 | 0.318 | 0.133 | 1.799E-14 | DTL | C5orf24 |
| 2.3452E-25 | 0.26610014 | 0.116 | 0.021 | 5.0069E-21 | DTL | PCP4 |
| 1.306E-44 | 0.26607848 | 0.235 | 0.047 | 2.7884E-40 | DTL | PDZK1 |
| 4.726E-24 | 0.26587713 | 0.271 | 0.091 | 1.009E-19 | DTL | ZNF22 |
| 1.7756E-17 | 0.26583637 | 0.278 | 0.113 | 3.7909E-13 | DTL | MXI1 |
| 9.0337E-15 | 0.2643115 | 0.404 | 0.21 | 1.9287E-10 | DTL | RAD21 |
| 1.5671E-14 | 0.26368538 | 0.357 | 0.176 | 3.3458E-10 | DTL | ZC3H13 |
| 6.4772E-18 | 0.26320749 | 0.289 | 0.119 | 1.3829E-13 | DTL | TRIOBP |
| 2.0388E-09 | 0.26302765 | 0.357 | 0.217 | 4.3527E-05 | DTL | LAMP2 |
| 6.504E-14 | 0.26197689 | 0.375 | 0.187 | 1.3886E-09 | DTL | TUT4 |
| 2.4612E-10 | 0.26185302 | 0.357 | 0.206 | 5.2546E-06 | DTL | TMBIM4 |
| 2.1476E-13 | 0.26165249 | 0.329 | 0.163 | 4.585E-09 | DTL | SELENOP |
| 5.272E-14 | 0.26164064 | 0.332 | 0.163 | 1.1256E-09 | DTL | DAB2 |
| 1.9045E-18 | 0.26122394 | 0.296 | 0.12 | 4.0661E-14 | DTL | SSFA2 |
| 4.6793E-15 | 0.26085227 | 0.368 | 0.18 | 9.9903E-11 | DTL | SPEN |
| 1.4089E-26 | 0.26081185 | 0.253 | 0.076 | 3.0079E-22 | DTL | DYNC2LI1 |
| 1.2798E-10 | 0.26072588 | 0.469 | 0.285 | 2.7325E-06 | DTL | NFE2L2 |
| 1.0611E-15 | 0.2605074 | 0.256 | 0.106 | 2.2655E-11 | DTL | CPVL |
| 2.6622E-27 | 0.25998539 | 0.292 | 0.094 | 5.6839E-23 | DTL | TNS3 |
| 3.063E-06 | 0.2594832 | 0.675 | 0.548 | 0.06539606 | DTL | CD9 |
| 2.1852E-10 | 0.25946958 | 0.755 | 0.569 | 4.6653E-06 | DTL | SET |
| 4.1658E-13 | 0.25917613 | 0.343 | 0.173 | 8.894E-09 | DTL | ACTN1 |
| 5.4834E-08 | 0.25869175 | 0.296 | 0.17 | 0.00117071 | DTL | TACC1 |
| 1.1915E-16 | 0.25832373 | 0.26 | 0.104 | 2.5438E-12 | DTL | VPS13A |
| 5.8509E-33 | 0.25797839 | 0.238 | 0.06 | 1.2492E-28 | DTL | VAMP5 |
| 4.1389E-19 | 0.25772056 | 0.285 | 0.112 | 8.8365E-15 | DTL | PDCD4 |
| 1.0159E-22 | 0.25725368 | 0.177 | 0.047 | 2.1689E-18 | DTL | RASSF4 |
| 3.4776E-17 | 0.25681098 | 0.339 | 0.151 | 7.4247E-13 | DTL | CFAP97 |
| 1.1933E-13 | 0.25633716 | 0.329 | 0.162 | 2.5477E-09 | DTL | IFNAR1 |
| 2.8463E-33 | 0.2558408 | 0.184 | 0.038 | 6.0769E-29 | DTL | MCAM |
| 3.1406E-06 | 0.25571795 | 0.729 | 0.597 | 0.06705235 | DTL | CLU |
| 1.025E-20 | 0.2556315 | 0.303 | 0.116 | 2.1885E-16 | DTL | LBR |
| 3.7866E-09 | 0.25556829 | 0.422 | 0.269 | 8.0844E-05 | DTL | ARPC5 |
| 7.2986E-15 | 0.25551254 | 0.354 | 0.171 | 1.5583E-10 | DTL | ZKSCAN1 |
| 8.4888E-16 | 0.25549411 | 0.52 | 0.281 | 1.8124E-11 | DTL | SRRM1 |
| 9.733E-32 | 0.25528594 | 0.177 | 0.037 | 2.078E-27 | DTL | ABCG1 |
| 2.314E-36 | 0.25497633 | 0.213 | 0.046 | 4.9405E-32 | DTL | DEPTOR |
| 1.5309E-09 | 0.25464474 | 0.26 | 0.135 | 3.2686E-05 | DTL | TMEM176A |
| 9.837E-27 | 0.25456029 | 0.184 | 0.045 | 2.1002E-22 | DTL | TNC |
| 2.8676E-06 | 0.25440147 | 0.718 | 0.634 | 0.06122335 | DTL | CCNI |
| 3.2119E-10 | 0.25439978 | 0.469 | 0.295 | 6.8574E-06 | DTL | TM9SF3 |
| 3.432E-15 | 0.25400654 | 0.408 | 0.206 | 7.3272E-11 | DTL | PURA |
| 5.6492E-21 | 0.25388383 | 0.289 | 0.108 | 1.2061E-16 | DTL | PARP4 |
| 2.041E-11 | 0.25366208 | 0.209 | 0.091 | 4.3576E-07 | DTL | AKR1C3 |
| 0.00933377 | 0.25333465 | 0.415 | 0.348 | 1 | DTL | TXNIP |
| 4.9176E-09 | 0.25326852 | 0.379 | 0.231 | 0.00010499 | DTL | ITGAV |
| 1.7242E-12 | 0.2530835 | 0.375 | 0.201 | 3.6811E-08 | DTL | OCLN |
| 1.0892E-26 | 0.25301918 | 0.235 | 0.067 | 2.3254E-22 | DTL | ABCB1 |

| | | | | | | |
|---|---|---|---|---|---|---|
| 2.9582E-14 | 0.25274813 | 0.401 | 0.21 | 6.3157E-10 | DTL | RSF1 |
| 3.305E-19 | 0.25271328 | 0.112 | 0.025 | 7.0562E-15 | DTL | NAPSA |
| 4.113E-09 | 0.2525252 | 0.26 | 0.14 | 8.7813E-05 | DTL | NBL1 |
| 6.9807E-15 | 0.25230127 | 0.318 | 0.15 | 1.4904E-10 | DTL | SMC3 |
| 5.7741E-10 | 0.2522877 | 0.603 | 0.41 | 1.2328E-05 | DTL | HNRNPK |
| 4.8793E-13 | 0.25131632 | 0.289 | 0.139 | 1.0417E-08 | DTL | GNAQ |
| 1.1865E-14 | 0.25120409 | 0.285 | 0.128 | 2.5331E-10 | DTL | ELK4 |
| 4.617E-19 | 0.25097209 | 0.343 | 0.143 | 9.8573E-15 | DTL | EMX2 |
| 5.6205E-10 | 0.25092204 | 0.307 | 0.167 | 1.2E-05 | DTL | PHIP |
| 4.7554E-20 | 0.25045166 | 0.307 | 0.12 | 1.0153E-15 | DTL | KLF13 |
| 1.8942E-14 | 0.25011571 | 0.321 | 0.151 | 4.0442E-10 | DTL | CUX1 |
| 6.1211E-22 | 0.25003763 | 0.235 | 0.076 | 1.3069E-17 | DTL | FRY |
| 0.00065245 | -0.2510711 | 0.126 | 0.202 | 1 | DTL | CHCHD7 |
| 0.00081081 | -0.2541968 | 0.065 | 0.131 | 1 | DTL | PRSS22 |
| 0.00070077 | -0.2552203 | 0.686 | 0.766 | 1 | DTL | PKM |
| 0.00035027 | -0.2590251 | 0.592 | 0.667 | 1 | DTL | TMA7 |
| 0.00456781 | -0.2595622 | 0.819 | 0.888 | 1 | DTL | RPS29 |
| 7.9917E-06 | -0.2596329 | 0.101 | 0.206 | 0.17062278 | DTL | PNP |
| 7.5721E-05 | -0.2599766 | 0.036 | 0.109 | 1 | DTL | AL355075.4 |
| 3.4223E-05 | -0.2602833 | 0.682 | 0.778 | 0.73066036 | DTL | OAZ1 |
| 0.00452503 | -0.260742 | 0.628 | 0.716 | 1 | DTL | HSPB1 |
| 4.208E-05 | -0.2608 | 0.036 | 0.113 | 0.8984145 | DTL | HIST4H4 |
| 0.00076316 | -0.2615119 | 0.913 | 0.944 | 1 | DTL | RPL8 |
| 3.4069E-07 | -0.2615142 | 0.014 | 0.11 | 0.00727365 | DTL | RGS2 |
| 2.9049E-05 | -0.2617513 | 0.065 | 0.153 | 0.62019032 | DTL | SLC16A3 |
| 0.00019212 | -0.2618911 | 0.061 | 0.135 | 1 | DTL | CSRNP2 |
| 0.00025163 | -0.2625923 | 0.112 | 0.196 | 1 | DTL | SDF2L1 |
| 3.3691E-05 | -0.26267 | 0.069 | 0.158 | 0.71931187 | DTL | CDKN2B |
| 7.0072E-05 | -0.2627452 | 0.191 | 0.293 | 1 | DTL | MAFF |
| 0.00012305 | -0.2636878 | 0.942 | 0.965 | 1 | DTL | RPS19 |
| 0.00032567 | -0.2655812 | 0.242 | 0.334 | 1 | DTL | SLC25A5 |
| 0.0007902 | -0.2668333 | 0.56 | 0.613 | 1 | DTL | SAP18 |
| 0.00044144 | -0.2676098 | 0.43 | 0.506 | 1 | DTL | CLDN7 |
| 0.00385745 | -0.2685463 | 0.339 | 0.398 | 1 | DTL | DBI |
| 0.00045668 | -0.270292 | 0.087 | 0.162 | 1 | DTL | TNFAIP3 |
| 0.00038343 | -0.2706504 | 0.166 | 0.252 | 1 | DTL | IDI1 |
| 0.00339095 | -0.2707679 | 0.78 | 0.872 | 1 | DTL | RPS5 |
| 0.0008973 | -0.2762495 | 0.899 | 0.934 | 1 | DTL | RPS9 |
| 7.85E-05 | -0.2773263 | 0.13 | 0.225 | 1 | DTL | GPRC5A |
| 0.00049525 | -0.2782879 | 0.264 | 0.34 | 1 | DTL | SSR3 |
| 0.00676691 | -0.2797679 | 0.274 | 0.334 | 1 | DTL | CHCHD10 |
| 1.7264E-06 | -0.2865095 | 0.119 | 0.24 | 0.03685928 | DTL | HIGD1A |
| 0.00623703 | -0.2873018 | 0.419 | 0.462 | 1 | DTL | COX8A |
| 0.00150759 | -0.2899725 | 0.787 | 0.818 | 1 | DTL | UBB |
| 0.00060661 | -0.2915554 | 0.069 | 0.136 | 1 | DTL | AC025164.1 |
| 5.993E-05 | -0.2919243 | 0.137 | 0.235 | 1 | DTL | CDKN2AIP |
| 0.00019339 | -0.2944574 | 0.292 | 0.377 | 1 | DTL | SEC61G |
| 0.00157269 | -0.294707 | 0.906 | 0.929 | 1 | DTL | RPL36 |
| 0.00196807 | -0.2978055 | 0.44 | 0.512 | 1 | DTL | H2AFZ |
| 0.00021452 | -0.3002781 | 0.516 | 0.581 | 1 | DTL | PRDX5 |

| | | | | | | |
|---|---|---|---|---|---|---|
| 6.0677E-06 | -0.3008809 | 0.614 | 0.757 | 0.12954454 | DTL | RPS26 |
| 0.00010617 | -0.3023152 | 0.274 | 0.37 | 1 | DTL | SELENOW |
| 0.00020905 | -0.3065417 | 0.101 | 0.179 | 1 | DTL | KLF4 |
| 1.586E-05 | -0.3079199 | 0.469 | 0.571 | 0.33860869 | DTL | RAN |
| 3.2676E-05 | -0.3107554 | 0.722 | 0.81 | 0.69762916 | DTL | ATP5F1E |
| 9.2931E-05 | -0.3129441 | 0.372 | 0.464 | 1 | DTL | SERP1 |
| 3.5539E-07 | -0.3138946 | 0.264 | 0.4 | 0.00758765 | DTL | CYCS |
| 4.9215E-06 | -0.3151461 | 0.054 | 0.149 | 0.10507442 | DTL | AL118516.1 |
| 0.00382896 | -0.3155608 | 0.531 | 0.566 | 1 | DTL | PPIB |
| 3.6174E-05 | -0.3166654 | 0.087 | 0.177 | 0.77232444 | DTL | PHLDA1 |
| 9.8567E-06 | -0.3171697 | 0.224 | 0.338 | 0.2104413 | DTL | H2AFJ |
| 8.8988E-05 | -0.3172112 | 0.069 | 0.148 | 1 | DTL | ANKRD37 |
| 2.1323E-05 | -0.3175452 | 0.57 | 0.666 | 0.45525069 | DTL | SOD1 |
| 3.5175E-06 | -0.3206914 | 0.606 | 0.704 | 0.0750996 | DTL | EIF4G2 |
| 6.3736E-06 | -0.3211859 | 0.83 | 0.906 | 0.13607672 | DTL | RPS20 |
| 1.1297E-05 | -0.3214674 | 0.245 | 0.351 | 0.24118142 | DTL | ACADVL |
| 0.00093581 | -0.3241924 | 0.827 | 0.866 | 1 | DTL | UBA52 |
| 0.00057996 | -0.3266962 | 0.184 | 0.263 | 1 | DTL | HMGB2 |
| 4.1506E-07 | -0.3278077 | 0.704 | 0.807 | 0.00886162 | DTL | EEF1D |
| 2.1269E-06 | -0.3285427 | 0.184 | 0.305 | 0.04540942 | DTL | EIF4A1 |
| 1.8267E-06 | -0.3337779 | 0.076 | 0.184 | 0.03900008 | DTL | ERN1 |
| 0.00019855 | -0.3349518 | 0.776 | 0.885 | 1 | DTL | RPL39 |
| 6.7456E-07 | -0.3353232 | 0.029 | 0.128 | 0.01440193 | DTL | HIST2H2BF |
| 1.9473E-06 | -0.3374529 | 0.368 | 0.482 | 0.04157477 | DTL | P4HB |
| 0.00161937 | -0.3404373 | 0.419 | 0.482 | 1 | DTL | H1FX |
| 1.5529E-05 | -0.3408744 | 0.451 | 0.535 | 0.33153699 | DTL | SEC61B |
| 1.4672E-06 | -0.3410721 | 0.17 | 0.29 | 0.03132404 | DTL | CDK2AP2 |
| 0.00243972 | -0.3438042 | 0.682 | 0.733 | 1 | DTL | PFN1 |
| 0.00012344 | -0.3452491 | 0.884 | 0.947 | 1 | DTL | RPS15 |
| 0.0001139 | -0.3453698 | 0.032 | 0.101 | 1 | DTL | CXCL3 |
| 5.1635E-08 | -0.3469986 | 0.126 | 0.276 | 0.00110242 | DTL | MUC1 |
| 1.133E-07 | -0.3470937 | 0.505 | 0.628 | 0.00241892 | DTL | ATF4 |
| 1.8178E-06 | -0.3479353 | 0.018 | 0.106 | 0.03881042 | DTL | ARC |
| 1.0837E-05 | -0.3524291 | 0.191 | 0.3 | 0.23136102 | DTL | CYTOR |
| 2.5053E-05 | -0.3558466 | 0.755 | 0.82 | 0.53489153 | DTL | YBX1 |
| 0.00060042 | -0.3562577 | 0.412 | 0.47 | 1 | DTL | PDIA6 |
| 0.00711214 | -0.3573827 | 0.278 | 0.328 | 1 | DTL | CEBPB |
| 2.6807E-07 | -0.3575664 | 0.538 | 0.651 | 0.00572326 | DTL | UQCRH |
| 1.3649E-07 | -0.3581021 | 0.17 | 0.306 | 0.00291405 | DTL | C9orf16 |
| 0.00121566 | -0.3596577 | 0.874 | 0.929 | 1 | DTL | RPL18A |
| 0.00017599 | -0.3631712 | 0.643 | 0.725 | 1 | DTL | CTSD |
| 3.2278E-10 | -0.3656656 | 0.408 | 0.575 | 6.8914E-06 | DTL | COX7A2 |
| 0.00015472 | -0.3727182 | 0.368 | 0.44 | 1 | DTL | PPP1R14B |
| 6.9257E-06 | -0.3733082 | 0.148 | 0.262 | 0.14786402 | DTL | LAMB3 |
| 1.8849E-07 | -0.373523 | 0.007 | 0.102 | 0.00402416 | DTL | AP001160.1 |
| 8.6715E-07 | -0.3741751 | 0.671 | 0.785 | 0.01851374 | DTL | EEF1B2 |
| 1.1126E-06 | -0.3754111 | 0.274 | 0.394 | 0.02375312 | DTL | NDRG1 |
| 0.00014646 | -0.3773413 | 0.903 | 0.924 | 1 | DTL | RPLP2 |
| 5.7101E-08 | -0.380833 | 0.697 | 0.806 | 0.0012191 | DTL | CFL1 |
| 0.0025463 | -0.3824098 | 0.238 | 0.301 | 1 | DTL | MARCKSL1 |

| | | | | | | |
|---|---|---|---|---|---|---|
| 1.2444E-05 | -0.3876313 | 0.13 | 0.235 | 0.26567093 | DTL | TGM2 |
| 0.00018141 | -0.3896443 | 0.231 | 0.314 | 1 | DTL | SRSF2 |
| 4.8206E-09 | -0.3904354 | 0.426 | 0.577 | 0.00010292 | DTL | SLC3A2 |
| 8.1137E-06 | -0.3914884 | 0.296 | 0.42 | 0.17322794 | DTL | CCND1 |
| 0.00033316 | -0.3941351 | 0.928 | 0.934 | 1 | DTL | HSP90AA1 |
| 3.8687E-05 | -0.3974302 | 0.264 | 0.371 | 0.82597412 | DTL | DDIT3 |
| 1.2826E-08 | -0.3995474 | 0.083 | 0.225 | 0.00027383 | DTL | ADM |
| 1.4551E-05 | -0.402952 | 0.079 | 0.174 | 0.31066627 | DTL | FOSB |
| 1.4918E-06 | -0.4037274 | 0.628 | 0.734 | 0.03185068 | DTL | NDUFA4 |
| 6.1737E-08 | -0.4055563 | 0.448 | 0.564 | 0.00131808 | DTL | GUK1 |
| 3.1748E-06 | -0.4070836 | 0.798 | 0.83 | 0.0677816 | DTL | SQSTM1 |
| 1.5728E-05 | -0.4079763 | 0.26 | 0.376 | 0.33578826 | DTL | ATF3 |
| 1.5967E-08 | -0.4093801 | 0.679 | 0.789 | 0.0003409 | DTL | CHCHD2 |
| 9.475E-06 | -0.4094042 | 0.177 | 0.286 | 0.20229227 | DTL | UCHL1 |
| 2.7818E-06 | -0.4126456 | 0.166 | 0.281 | 0.05939199 | DTL | SERPINB1 |
| 4.5812E-06 | -0.4138716 | 0.928 | 0.962 | 0.0978081 | DTL | RPS18 |
| 2.158E-07 | -0.415759 | 0.556 | 0.662 | 0.00460737 | DTL | CALR |
| 0.00136304 | -0.4182731 | 0.209 | 0.291 | 1 | DTL | FOS |
| 0.00903712 | -0.4197839 | 0.336 | 0.382 | 1 | DTL | HERPUD1 |
| 0.00092352 | -0.4220167 | 0.556 | 0.651 | 1 | DTL | KRT7 |
| 4.4465E-08 | -0.4224508 | 0.069 | 0.195 | 0.00094934 | DTL | H2AFX |
| 1.0304E-08 | -0.4258141 | 0.144 | 0.299 | 0.00021999 | DTL | MIR4435-2HG |
| 1.9165E-05 | -0.4308402 | 0.841 | 0.901 | 0.40917004 | DTL | SERF2 |
| 2.3428E-05 | -0.4337161 | 0.047 | 0.13 | 0.50017743 | DTL | COL7A1 |
| 0.00919328 | -0.437743 | 0.209 | 0.256 | 1 | DTL | IER5L |
| 7.7877E-07 | -0.4402265 | 0.303 | 0.425 | 0.01662682 | DTL | SNHG8 |
| 0.00034035 | -0.4463926 | 0.35 | 0.422 | 1 | DTL | TPM4 |
| 6.0581E-05 | -0.4470658 | 0.087 | 0.17 | 1 | DTL | EGLN3 |
| 0.00164745 | -0.4484868 | 0.22 | 0.29 | 1 | DTL | RND3 |
| 0.00104788 | -0.4498144 | 0.17 | 0.239 | 1 | DTL | INTS6 |
| 1.4752E-05 | -0.4505675 | 0.097 | 0.194 | 0.31494729 | DTL | GADD45G |
| 3.8083E-05 | -0.4539134 | 0.191 | 0.286 | 0.8130665 | DTL | POLR2A |
| 1.5359E-06 | -0.4548024 | 0.094 | 0.205 | 0.03279048 | DTL | SGK1 |
| 5.8521E-06 | -0.4559507 | 0.191 | 0.31 | 0.12494229 | DTL | PLAU |
| 0.00205845 | -0.456286 | 0.895 | 0.922 | 1 | DTL | ACTB |
| 0.00885918 | -0.4624067 | 0.069 | 0.118 | 1 | DTL | CXCL8 |
| 0.00048336 | -0.4653694 | 0.325 | 0.387 | 1 | DTL | BRD2 |
| 5.506E-08 | -0.4702473 | 0.856 | 0.914 | 0.00117554 | DTL | RPL35 |
| 0.00863331 | -0.4749248 | 0.339 | 0.392 | 1 | DTL | NFKBIA |
| 2.3215E-05 | -0.4820228 | 0.343 | 0.424 | 0.49564443 | DTL | TUBB4B |
| 8.6811E-05 | -0.4830261 | 0.166 | 0.254 | 1 | DTL | ID2 |
| 1.1033E-08 | -0.4838539 | 0.083 | 0.219 | 0.00023555 | DTL | UBE2S |
| 1.9657E-09 | -0.48501 | 0.531 | 0.671 | 4.1968E-05 | DTL | TPI1 |
| 4.2659E-06 | -0.4872257 | 0.964 | 0.972 | 0.09107724 | DTL | RPLP1 |
| 1.74E-06 | -0.4954514 | 0.895 | 0.953 | 0.03714897 | DTL | RPS12 |
| 1.7214E-09 | -0.5004053 | 0.076 | 0.224 | 3.6753E-05 | DTL | SFN |
| 2.8676E-12 | -0.5074008 | 0.968 | 0.957 | 6.1223E-08 | DTL | EIF1 |
| 6.8781E-10 | -0.5085538 | 0.899 | 0.916 | 1.4685E-05 | DTL | RPLP0 |
| 1.3399E-08 | -0.5137073 | 0.852 | 0.932 | 0.00028606 | DTL | RPL12 |
| 2.7104E-09 | -0.5139111 | 0.148 | 0.304 | 5.7868E-05 | DTL | DDIT4 |

| | | | | | | |
|---|---|---|---|---|---|---|
| 5.5023E-07 | -0.5179339 | 0.195 | 0.322 | 0.01174733 | DTL | ITGA2 |
| 0.00027008 | -0.5196257 | 0.199 | 0.278 | 1 | DTL | PPP1R10 |
| 3.6502E-07 | -0.5255329 | 0.155 | 0.273 | 0.00779308 | DTL | SEPHS2 |
| 0.00629919 | -0.5262558 | 0.213 | 0.273 | 1 | DTL | TCIM |
| 1.2174E-06 | -0.5282415 | 0.079 | 0.187 | 0.02599069 | DTL | PIK3R3 |
| 3.0728E-09 | -0.5380208 | 0.152 | 0.308 | 6.5605E-05 | DTL | PMAIP1 |
| 0.00034656 | -0.5419806 | 0.325 | 0.392 | 1 | DTL | TUBA1B |
| 5.372E-13 | -0.5436662 | 0.596 | 0.761 | 1.1469E-08 | DTL | DYNLL1 |
| 3.4613E-16 | -0.5443543 | 0.498 | 0.705 | 7.39E-12 | DTL | MAP1LC3B |
| 7.5033E-08 | -0.5474701 | 0.805 | 0.875 | 0.00160196 | DTL | GSTP1 |
| 6.6386E-06 | -0.5479154 | 0.032 | 0.119 | 0.14173475 | DTL | NEFM |
| 1.604E-09 | -0.5493983 | 0.448 | 0.61 | 3.4246E-05 | DTL | GADD45A |
| 3.1849E-09 | -0.5537085 | 0.336 | 0.514 | 6.7998E-05 | DTL | CDKN1A |
| 2.6758E-08 | -0.5665432 | 0.166 | 0.305 | 0.00057128 | DTL | DNAJB1 |
| 1.9301E-09 | -0.5776325 | 0.134 | 0.289 | 4.1209E-05 | DTL | PLAUR |
| 1.5215E-07 | -0.5844734 | 0.339 | 0.488 | 0.00324839 | DTL | TSPAN1 |
| 6.1955E-11 | -0.5886387 | 0.058 | 0.217 | 1.3227E-06 | DTL | SOCS1 |
| 2.2082E-05 | -0.5968378 | 0.217 | 0.307 | 0.47144873 | DTL | HSPA1B |
| 1.3327E-07 | -0.5972327 | 0.675 | 0.775 | 0.00284536 | DTL | HSP90B1 |
| 2.224E-09 | -0.598628 | 0.108 | 0.256 | 4.7483E-05 | DTL | IER5 |
| 4.0222E-12 | -0.5988777 | 0.928 | 0.961 | 8.5874E-08 | DTL | RPL28 |
| 7.7432E-11 | -0.600718 | 0.527 | 0.659 | 1.6532E-06 | DTL | HSPA8 |
| 3.0728E-07 | -0.621562 | 0.155 | 0.28 | 0.00656047 | DTL | RASD1 |
| 0.00031344 | -0.6250065 | 0.791 | 0.87 | 1 | DTL | TMSB4X |
| 1.0402E-16 | -0.6295274 | 0.321 | 0.564 | 2.2208E-12 | DTL | HMGA1 |
| 1.99E-06 | -0.6404114 | 0.195 | 0.304 | 0.04248582 | DTL | PLK2 |
| 2.0864E-07 | -0.6492168 | 0.051 | 0.163 | 0.00445453 | DTL | G0S2 |
| 5.1246E-12 | -0.6540994 | 0.921 | 0.911 | 1.0941E-07 | DTL | UBC |
| 7.7303E-15 | -0.660582 | 0.903 | 0.957 | 1.6504E-10 | DTL | RPS2 |
| 9.9498E-15 | -0.6727482 | 0.195 | 0.403 | 2.1243E-10 | DTL | JPT1 |
| 3.9833E-19 | -0.6805236 | 0.108 | 0.365 | 8.5043E-15 | DTL | CD44 |
| 2.1421E-07 | -0.6960439 | 0.148 | 0.28 | 0.00457337 | DTL | HES1 |
| 3.6062E-06 | -0.7178826 | 0.477 | 0.566 | 0.0769934 | DTL | IER2 |
| 1.6705E-07 | -0.7287024 | 0.011 | 0.108 | 0.00356655 | DTL | FXYD4 |
| 6.9409E-13 | -0.741908 | 0.155 | 0.345 | 1.4819E-08 | DTL | SNHG12 |
| 7.6877E-05 | -0.7458314 | 0.563 | 0.587 | 1 | DTL | ELF3 |
| 0.00090349 | -0.7490077 | 0.823 | 0.798 | 1 | DTL | KRT18 |
| 1.3122E-05 | -0.7571038 | 0.141 | 0.245 | 0.28016193 | DTL | TFPI2 |
| 9.0621E-11 | -0.7655352 | 0.076 | 0.239 | 1.9348E-06 | DTL | CST6 |
| 1.1187E-15 | -0.7741432 | 0.722 | 0.875 | 2.3885E-11 | DTL | SAT1 |
| 2.2418E-07 | -0.7861602 | 0.495 | 0.62 | 0.00478617 | DTL | MTRNR2L8 |
| 0.00014287 | -0.7879159 | 0.227 | 0.299 | 1 | DTL | HEXIM1 |
| 1.3355E-15 | -0.7909171 | 0.444 | 0.627 | 2.8513E-11 | DTL | PPP1R15A |
| 1.5421E-08 | -0.8095759 | 0.105 | 0.237 | 0.00032923 | DTL | CSKMT |
| 1.1864E-15 | -0.8404826 | 0.791 | 0.895 | 2.533E-11 | DTL | GAPDH |
| 8.2435E-06 | -0.8471749 | 0.964 | 0.961 | 0.17599795 | DTL | FTL |
| 8.2362E-07 | -0.8786748 | 0.159 | 0.273 | 0.01758435 | DTL | HSPA2 |
| 1.5669E-09 | -0.8825604 | 0.61 | 0.73 | 3.3453E-05 | DTL | KRT8 |
| 3.7486E-10 | -0.8973301 | 0.74 | 0.833 | 8.0032E-06 | DTL | ANXA2 |
| 3.8154E-11 | -0.9111851 | 0.534 | 0.687 | 8.1459E-07 | DTL | HSPA5 |

| | | | | | | |
|---|---|---|---|---|---|---|
| 1.1208E-10 | -0.9274857 | 0.495 | 0.649 | 2.3929E-06 | DTL | WFDC2 |
| 5.2566E-10 | -0.9408973 | 0.181 | 0.334 | 1.1223E-05 | DTL | LAMC2 |
| 9.3344E-10 | -0.9547215 | 0.285 | 0.443 | 1.9929E-05 | DTL | IER3 |
| 0.00395534 | -0.970633 | 0.646 | 0.679 | 1 | DTL | MTRNR2L12 |
| 1.5102E-23 | -0.9836165 | 0.126 | 0.419 | 3.2242E-19 | DTL | PHLDA2 |
| 1.6297E-09 | -0.9893609 | 0.339 | 0.473 | 3.4794E-05 | DTL | C12orf75 |
| 0.0030521 | -1.0209073 | 0.162 | 0.227 | 1 | DTL | DEFB1 |
| 4.8739E-19 | -1.0625175 | 0.195 | 0.448 | 1.0406E-14 | DTL | UPP1 |
| 9.5728E-21 | -1.0654602 | 0.437 | 0.677 | 2.0438E-16 | DTL | SH3BGRL3 |
| 2.5455E-10 | -1.0926716 | 0.094 | 0.259 | 5.4346E-06 | DTL | LCN2 |
| 3.0876E-29 | -1.2270732 | 0.296 | 0.612 | 6.5921E-25 | DTL | GADD45B |
| 2.2831E-16 | -1.2756436 | 0.069 | 0.287 | 4.8745E-12 | DTL | MAFB |
| 5.6294E-09 | -1.4925179 | 0.473 | 0.64 | 0.00012019 | DTL | MMP7 |
| 1.8199E-24 | -1.886737 | 0.195 | 0.498 | 3.8855E-20 | DTL | S100A2 |
| 4.7514E-08 | -2.8430728 | 0.538 | 0.599 | 0.00101442 | DTL | PLCG2 |
| 6.863E-305 | 3.79074946 | 0.627 | 0.107 | 1.465E-300 | PCT | GPX3 |
| 0 | 3.5438234 | 0.379 | 0.005 | 0 | PCT | ALDOB |
| 0 | 3.52842714 | 0.792 | 0.173 | 0 | PCT | MT1G |
| 0 | 2.67166947 | 0.617 | 0.081 | 0 | PCT | APOE |
| 1.006E-292 | 2.37489749 | 0.523 | 0.068 | 2.149E-288 | PCT | MT1H |
| 1.369E-140 | 2.2454109 | 0.988 | 0.96 | 2.922E-136 | PCT | FTL |
| 0 | 2.02937635 | 0.317 | 0.01 | 0 | PCT | GSTA1 |
| 3.303E-210 | 1.96208942 | 0.371 | 0.047 | 7.052E-206 | PCT | GATM |
| 0 | 1.89925356 | 0.238 | 0.002 | 0 | PCT | FABP1 |
| 0 | 1.87144159 | 0.345 | 0.007 | 0 | PCT | MIOX |
| 1.598E-140 | 1.72286687 | 0.643 | 0.229 | 3.411E-136 | PCT | PDZK1IP1 |
| 0 | 1.54227352 | 0.214 | 0.001 | 0 | PCT | GSTA2 |
| 5.838E-162 | 1.54162326 | 0.495 | 0.108 | 1.246E-157 | PCT | CXCL14 |
| 5.2756E-65 | 1.47802696 | 0.838 | 0.66 | 1.1263E-60 | PCT | CRYAB |
| 2.57E-102 | 1.47591269 | 0.695 | 0.312 | 5.4859E-98 | PCT | MT1E |
| 1.1876E-95 | 1.43526624 | 0.595 | 0.232 | 2.5356E-91 | PCT | MT1X |
| 4.992E-67 | 1.4118429 | 0.561 | 0.265 | 1.0658E-62 | PCT | CYB5A |
| 0 | 1.4049434 | 0.365 | 0.011 | 0 | PCT | RBP5 |
| 3.049E-166 | 1.38094701 | 0.567 | 0.142 | 6.509E-162 | PCT | MGST1 |
| 4.1392E-69 | 1.36310484 | 0.872 | 0.704 | 8.8372E-65 | PCT | MT2A |
| 9.459E-112 | 1.35099076 | 0.996 | 0.977 | 2.02E-107 | PCT | FTH1 |
| 2.654E-176 | 1.32641381 | 0.415 | 0.072 | 5.665E-172 | PCT | CMBL |
| 1.5453E-85 | 1.2892343 | 0.443 | 0.151 | 3.2992E-81 | PCT | SUCLG1 |
| 1.7528E-51 | 1.25456935 | 0.808 | 0.637 | 3.7423E-47 | PCT | PEBP1 |
| 6.3903E-82 | 1.24492484 | 0.206 | 0.035 | 1.3643E-77 | PCT | PCK1 |
| 0 | 1.23441812 | 0.315 | 0.008 | 0 | PCT | HPD |
| 3.1576E-76 | 1.2274985 | 0.541 | 0.241 | 6.7416E-72 | PCT | ECHS1 |
| 3.053E-102 | 1.1309825 | 0.451 | 0.123 | 6.5174E-98 | PCT | SERPINA1 |
| 7.6181E-73 | 1.10403188 | 0.381 | 0.117 | 1.6265E-68 | PCT | HMOX1 |
| 6.101E-53 | 1.10034816 | 0.425 | 0.181 | 1.3026E-48 | PCT | DCXR |
| 3.84E-64 | 1.09542963 | 0.631 | 0.314 | 8.1984E-60 | PCT | IL32 |
| 2.833E-156 | 1.08565911 | 0.311 | 0.044 | 6.048E-152 | PCT | AK4 |
| 8.1596E-76 | 1.08308493 | 0.675 | 0.362 | 1.7421E-71 | PCT | LDHB |
| 9.18E-133 | 1.07746652 | 0.401 | 0.087 | 1.96E-128 | PCT | PEPD |
| 5.505E-129 | 1.07624081 | 0.455 | 0.108 | 1.175E-124 | PCT | MT1F |

| | | | | | | |
|---|---|---|---|---|---|---|
| 8.18E-209 | 1.05851337 | 0.343 | 0.04 | 1.747E-204 | PCT | RIDA |
| 2.76E-271 | 1.03681648 | 0.279 | 0.017 | 5.893E-267 | PCT | BHMT |
| 2.6997E-90 | 1.03517535 | 0.409 | 0.12 | 5.7639E-86 | PCT | PLIN2 |
| 1.046E-92 | 1.01833675 | 0.333 | 0.08 | 2.2332E-88 | PCT | ASS1 |
| 1.8265E-69 | 0.98837754 | 0.473 | 0.194 | 3.8997E-65 | PCT | ADI1 |
| 9.4803E-52 | 0.96969902 | 0.637 | 0.387 | 2.024E-47 | PCT | DBI |
| 1.6073E-98 | 0.94961809 | 0.369 | 0.092 | 3.4315E-94 | PCT | UGT2B7 |
| 4.197E-286 | 0.9493712 | 0.238 | 0.011 | 8.96E-282 | PCT | BBOX1 |
| 1.0581E-90 | 0.94579022 | 0.305 | 0.069 | 2.259E-86 | PCT | ALDH6A1 |
| 0 | 0.94173862 | 0.349 | 0.022 | 0 | PCT | ANPEP |
| 1.3489E-60 | 0.93610083 | 0.629 | 0.35 | 2.8799E-56 | PCT | ATP5MC3 |
| 1.4438E-68 | 0.9359846 | 0.309 | 0.089 | 3.0825E-64 | PCT | C11orf54 |
| 9.387E-52 | 0.93582723 | 0.651 | 0.393 | 2.0041E-47 | PCT | LAMTOR5 |
| 1.0895E-52 | 0.92313078 | 0.643 | 0.378 | 2.3261E-48 | PCT | HSPE1 |
| 1.5388E-61 | 0.92219022 | 0.601 | 0.322 | 3.2853E-57 | PCT | CHCHD10 |
| 6.189E-74 | 0.88890149 | 0.439 | 0.155 | 1.3214E-69 | PCT | DAB2 |
| 1.889E-125 | 0.8695126 | 0.279 | 0.043 | 4.034E-121 | PCT | FBP1 |
| 7.5512E-80 | 0.86947922 | 0.403 | 0.127 | 1.6122E-75 | PCT | ALDH2 |
| 2.6565E-46 | 0.86553358 | 0.826 | 0.686 | 5.6716E-42 | PCT | GPX4 |
| 0 | 0.86234499 | 0.265 | 0.012 | 0 | PCT | KHK |
| 0 | 0.86029166 | 0.281 | 0.005 | 0 | PCT | LGALS2 |
| 1.567E-129 | 0.8574109 | 0.393 | 0.082 | 3.347E-125 | PCT | AKR1C3 |
| 8.8275E-88 | 0.85288387 | 0.435 | 0.139 | 1.8847E-83 | PCT | AKR1A1 |
| 0 | 0.85087433 | 0.253 | 0.002 | 0 | PCT | NAT8 |
| 6.2366E-46 | 0.84394659 | 0.856 | 0.71 | 1.3315E-41 | PCT | TXN |
| 1.3484E-45 | 0.83052707 | 0.563 | 0.319 | 2.8788E-41 | PCT | UQCRQ |
| 1.0805E-91 | 0.82014688 | 0.365 | 0.095 | 2.3068E-87 | PCT | SLC9A3R1 |
| 0 | 0.81566372 | 0.208 | 0.001 | 0 | PCT | ALB |
| 6.732E-155 | 0.80406687 | 0.301 | 0.041 | 1.437E-150 | PCT | PDZK1 |
| 4.4526E-43 | 0.77891377 | 0.872 | 0.764 | 9.5063E-39 | PCT | COX7C |
| 4.1281E-65 | 0.77142272 | 0.409 | 0.151 | 8.8135E-61 | PCT | ETFB |
| 1.9545E-66 | 0.77075339 | 0.369 | 0.123 | 4.1729E-62 | PCT | ACAT1 |
| 1.3172E-36 | 0.76827748 | 0.271 | 0.098 | 2.8121E-32 | PCT | SERPINE2 |
| 1.2619E-47 | 0.75030234 | 0.623 | 0.372 | 2.6942E-43 | PCT | NDUFB2 |
| 3.9541E-53 | 0.74382331 | 0.325 | 0.114 | 8.442E-49 | PCT | ACAA1 |
| 1.121E-114 | 0.74091227 | 0.369 | 0.081 | 2.393E-110 | PCT | BHMT2 |
| 9.478E-84 | 0.73677849 | 0.377 | 0.104 | 2.0236E-79 | PCT | RARRES3 |
| 1.8667E-33 | 0.7350203 | 0.557 | 0.353 | 3.9854E-29 | PCT | COX7B |
| 2.5673E-60 | 0.73231572 | 0.425 | 0.169 | 5.4811E-56 | PCT | DNPH1 |
| 3.247E-51 | 0.7309145 | 0.994 | 0.971 | 6.9323E-47 | PCT | TPT1 |
| 1.2401E-48 | 0.72184856 | 0.974 | 0.89 | 2.6476E-44 | PCT | GAPDH |
| 5.5074E-41 | 0.71815672 | 0.904 | 0.804 | 1.1758E-36 | PCT | ATP5F1E |
| 4.6839E-46 | 0.7166195 | 0.964 | 0.897 | 1E-41 | PCT | SERF2 |
| 1.6335E-42 | 0.70687638 | 0.471 | 0.225 | 3.4875E-38 | PCT | FXYD2 |
| 2.7235E-75 | 0.70451797 | 0.401 | 0.127 | 5.8147E-71 | PCT | TMEM176A |
| 4.3223E-65 | 0.69865827 | 0.299 | 0.085 | 9.2281E-61 | PCT | CRYL1 |
| 5.8887E-25 | 0.69670903 | 0.776 | 0.598 | 1.2572E-20 | PCT | SOD2 |
| 1.7028E-43 | 0.6933475 | 0.597 | 0.349 | 3.6356E-39 | PCT | ATP5ME |
| 9.9224E-48 | 0.6930776 | 0.9 | 0.74 | 2.1184E-43 | PCT | HINT1 |
| 0 | 0.69251549 | 0.208 | 0.002 | 0 | PCT | GLYAT |

| | | | | | | |
|---|---|---|---|---|---|---|
| 2.3214E-43 | 0.68963863 | 0.445 | 0.203 | 4.9562E-39 | PCT | GLRX |
| 4.6916E-70 | 0.68681315 | 0.244 | 0.054 | 1.0017E-65 | PCT | S100A1 |
| 7.1838E-43 | 0.6839261 | 0.699 | 0.451 | 1.5337E-38 | PCT | COX8A |
| 4.3151E-37 | 0.68369631 | 0.657 | 0.438 | 9.2128E-33 | PCT | ATP5MD |
| 1.6774E-39 | 0.67988003 | 0.601 | 0.366 | 3.5813E-35 | PCT | NDUFA1 |
| 5.1653E-56 | 0.67926741 | 0.453 | 0.195 | 1.1028E-51 | PCT | MPC2 |
| 1.9689E-51 | 0.67805474 | 0.984 | 0.937 | 4.2037E-47 | PCT | RPS28 |
| 8.9303E-51 | 0.67698874 | 0.601 | 0.326 | 1.9066E-46 | PCT | PRELID1 |
| 1.1E-44 | 0.67684272 | 0.483 | 0.243 | 2.3484E-40 | PCT | ATOX1 |
| 6.6587E-39 | 0.67105858 | 0.615 | 0.377 | 1.4216E-34 | PCT | POLR2L |
| 0 | 0.66655316 | 0.263 | 0.002 | 0 | PCT | DPYS |
| 4.893E-36 | 0.65990087 | 0.705 | 0.488 | 1.0447E-31 | PCT | HSPD1 |
| 1.9758E-40 | 0.65672139 | 0.425 | 0.207 | 4.2184E-36 | PCT | SAT2 |
| 2.3068E-33 | 0.652466 | 0.645 | 0.445 | 4.9251E-29 | PCT | ATP5PF |
| 2.3817E-43 | 0.64861201 | 0.866 | 0.673 | 5.085E-39 | PCT | COX6A1 |
| 6.622E-193 | 0.64690863 | 0.222 | 0.016 | 1.414E-188 | PCT | AKR7A3 |
| 1.9693E-49 | 0.64033964 | 0.577 | 0.311 | 4.2044E-45 | PCT | ATP5MPL |
| 1.2211E-91 | 0.64024035 | 0.377 | 0.099 | 2.607E-87 | PCT | ISOC2 |
| 7.3334E-53 | 0.63721836 | 0.385 | 0.147 | 1.5657E-48 | PCT | QPRT |
| 1.0038E-59 | 0.63514279 | 0.401 | 0.149 | 2.1431E-55 | PCT | ECH1 |
| 1.5567E-26 | 0.63451734 | 0.772 | 0.659 | 3.3236E-22 | PCT | SOD1 |
| 0 | 0.62654477 | 0.273 | 0.005 | 0 | PCT | DPEP1 |
| 5.7363E-51 | 0.62107184 | 0.994 | 0.987 | 1.2247E-46 | PCT | EEF1A1 |
| 9.8461E-39 | 0.62052531 | 0.772 | 0.575 | 2.1021E-34 | PCT | COX5B |
| 9.4803E-46 | 0.62042858 | 0.407 | 0.179 | 2.0241E-41 | PCT | CNDP2 |
| 4.0549E-42 | 0.6185186 | 0.571 | 0.317 | 8.6573E-38 | PCT | NDUFC1 |
| 4.8531E-32 | 0.61572374 | 0.533 | 0.324 | 1.0361E-27 | PCT | SLC25A5 |
| 2.3002E-54 | 0.61548778 | 0.597 | 0.298 | 4.9109E-50 | PCT | ISCU |
| 5.688E-26 | 0.61448484 | 0.565 | 0.38 | 1.2144E-21 | PCT | ATP5IF1 |
| 8.1563E-35 | 0.61059142 | 0.603 | 0.386 | 1.7414E-30 | PCT | ATP5F1D |
| 4.1191E-80 | 0.60989505 | 0.246 | 0.05 | 8.7942E-76 | PCT | SMIM24 |
| 4.582E-49 | 0.60842096 | 0.391 | 0.157 | 9.7825E-45 | PCT | SELENOP |
| 1.382E-103 | 0.59650962 | 0.285 | 0.053 | 2.951E-99 | PCT | MSRA |
| 2.7774E-54 | 0.59569895 | 0.395 | 0.151 | 5.9297E-50 | PCT | TMEM176B |
| 4.3717E-33 | 0.59296062 | 0.507 | 0.292 | 9.3336E-29 | PCT | DDT |
| 1.0419E-49 | 0.59126128 | 0.355 | 0.135 | 2.2244E-45 | PCT | ACAA2 |
| 6.9627E-22 | 0.58816465 | 0.497 | 0.319 | 1.4865E-17 | PCT | CTSB |
| 3.8882E-33 | 0.58737837 | 0.613 | 0.389 | 8.3014E-29 | PCT | MGST3 |
| 4.3172E-27 | 0.57932323 | 0.543 | 0.359 | 9.2173E-23 | PCT | ATP5F1B |
| 8.225E-50 | 0.57638533 | 0.335 | 0.12 | 1.756E-45 | PCT | IMPA2 |
| 1.5079E-34 | 0.57265188 | 0.888 | 0.749 | 3.2193E-30 | PCT | RPS26 |
| 1.4292E-33 | 0.57130605 | 0.343 | 0.154 | 3.0514E-29 | PCT | PCSK1N |
| 1.5101E-78 | 0.57115726 | 0.263 | 0.057 | 3.2241E-74 | PCT | DHRS4L2 |
| 2.8981E-32 | 0.57045126 | 0.577 | 0.359 | 6.1875E-28 | PCT | ATP5MF |
| 1.0698E-37 | 0.55846517 | 0.978 | 0.939 | 2.2839E-33 | PCT | RPL19 |
| 4.6742E-41 | 0.55665922 | 0.497 | 0.256 | 9.9795E-37 | PCT | TXNDC17 |
| 3.1838E-92 | 0.55414555 | 0.273 | 0.054 | 6.7974E-88 | PCT | SHMT1 |
| 1.6229E-27 | 0.55192995 | 0.703 | 0.522 | 3.4649E-23 | PCT | COX6C |
| 2.6428E-26 | 0.54827912 | 0.489 | 0.307 | 5.6423E-22 | PCT | UQCR11 |
| 1.3683E-90 | 0.54815 | 0.222 | 0.037 | 2.9213E-86 | PCT | ALDH4A1 |

| | | | | | | |
|---|---|---|---|---|---|---|
| 1.2429E-42 | 0.54669194 | 0.992 | 0.966 | 2.6535E-38 | PCT | RPL41 |
| 1.2566E-50 | 0.54418846 | 0.375 | 0.145 | 2.6829E-46 | PCT | DUSP23 |
| 2.4396E-35 | 0.54299641 | 0.335 | 0.145 | 5.2086E-31 | PCT | NQO1 |
| 5.6507E-32 | 0.54004243 | 0.391 | 0.195 | 1.2064E-27 | PCT | NDUFB1 |
| 1.7405E-31 | 0.537907 | 0.786 | 0.601 | 3.7159E-27 | PCT | ATP5MG |
| 1.952E-224 | 0.5369322 | 0.218 | 0.012 | 4.167E-220 | PCT | AQP1 |
| 1.7872E-39 | 0.53638582 | 0.922 | 0.776 | 3.8158E-35 | PCT | COX4I1 |
| 6.5389E-37 | 0.53570389 | 0.731 | 0.517 | 1.3961E-32 | PCT | ATP5MC2 |
| 2.2621E-92 | 0.53476847 | 0.246 | 0.044 | 4.8296E-88 | PCT | RAB11FIP3 |
| 2.9347E-27 | 0.53464193 | 0.8 | 0.656 | 6.2655E-23 | PCT | UQCRB |
| 3.088E-45 | 0.53270165 | 0.393 | 0.164 | 6.593E-41 | PCT | PLA2G16 |
| 2.491E-32 | 0.53135674 | 0.415 | 0.209 | 5.3182E-28 | PCT | PTGR1 |
| 1.2216E-85 | 0.53021059 | 0.196 | 0.031 | 2.608E-81 | PCT | PBLD |
| 0 | 0.52934339 | 0.182 | 0.002 | 0 | PCT | PRODH2 |
| 0 | 0.52833397 | 0.208 | 0.005 | 0 | PCT | AGMAT |
| 3.0343E-24 | 0.52828864 | 0.511 | 0.318 | 6.4782E-20 | PCT | CYBA |
| 1.1973E-34 | 0.5244592 | 0.756 | 0.564 | 2.5563E-30 | PCT | COX7A2 |
| 3.4771E-32 | 0.52381641 | 0.519 | 0.305 | 7.4237E-28 | PCT | PCBD1 |
| 1.9483E-33 | 0.52379853 | 0.954 | 0.887 | 4.1597E-29 | PCT | NACA |
| 6.3799E-34 | 0.51784538 | 0.735 | 0.519 | 1.3621E-29 | PCT | ELOB |
| 1.5146E-53 | 0.51255199 | 0.317 | 0.105 | 3.2336E-49 | PCT | ECI2 |
| 5.9227E-31 | 0.5121501 | 0.417 | 0.221 | 1.2645E-26 | PCT | MDH1 |
| 5.3333E-22 | 0.51198104 | 0.575 | 0.404 | 1.1387E-17 | PCT | UQCR10 |
| 1.5862E-43 | 0.51072789 | 0.339 | 0.133 | 3.3866E-39 | PCT | GCSH |
| 2.0702E-33 | 0.5089 | 0.461 | 0.249 | 4.4199E-29 | PCT | ATP5F1A |
| 3.2135E-47 | 0.50524342 | 0.435 | 0.192 | 6.8608E-43 | PCT | COX14 |
| 0 | 0.49836813 | 0.214 | 0.006 | 0 | PCT | ACSM2B |
| 2.611E-31 | 0.49807264 | 0.749 | 0.538 | 5.5744E-27 | PCT | BRI3 |
| 1.0457E-14 | 0.49804408 | 0.409 | 0.255 | 2.2326E-10 | PCT | NUPR1 |
| 7.0282E-25 | 0.49769029 | 0.711 | 0.555 | 1.5005E-20 | PCT | COX6B1 |
| 7.3031E-29 | 0.49569718 | 0.952 | 0.883 | 1.5592E-24 | PCT | RPS29 |
| 3.02E-33 | 0.49483584 | 0.489 | 0.271 | 6.4478E-29 | PCT | MRPL41 |
| 1.5542E-31 | 0.49274126 | 0.461 | 0.254 | 3.3183E-27 | PCT | PRR13 |
| 1.9133E-70 | 0.49172134 | 0.222 | 0.045 | 4.0848E-66 | PCT | CLTRN |
| 4.4888E-23 | 0.48937516 | 0.92 | 0.864 | 9.5837E-19 | PCT | RPL23 |
| 1.1009E-22 | 0.48654626 | 0.349 | 0.18 | 2.3504E-18 | PCT | ADIRF |
| 3.3942E-26 | 0.48332301 | 0.427 | 0.246 | 7.2467E-22 | PCT | C4orf3 |
| 1.8244E-41 | 0.48258321 | 0.289 | 0.105 | 3.8952E-37 | PCT | BDH2 |
| 2.2229E-27 | 0.48205979 | 0.914 | 0.795 | 4.7458E-23 | PCT | RPL27 |
| 4.2459E-20 | 0.48133743 | 0.549 | 0.378 | 9.0649E-16 | PCT | OCIAD2 |
| 1.8695E-33 | 0.48097864 | 0.395 | 0.19 | 3.9915E-29 | PCT | BNIP3 |
| 9.9651E-24 | 0.48074867 | 0.974 | 0.933 | 2.1276E-19 | PCT | RPL37 |
| 1.9639E-60 | 0.4798072 | 0.253 | 0.064 | 4.193E-56 | PCT | SLC3A1 |
| 1.7462E-27 | 0.47597177 | 0.433 | 0.245 | 3.7281E-23 | PCT | NDUFAB1 |
| 8.3094E-26 | 0.47329012 | 0.715 | 0.526 | 1.7741E-21 | PCT | NDUFS5 |
| 3.6031E-41 | 0.47229603 | 0.796 | 0.592 | 7.6927E-37 | PCT | SPP1 |
| 6.8521E-81 | 0.46856909 | 0.259 | 0.052 | 1.4629E-76 | PCT | IGFBP4 |
| 1.377E-45 | 0.46854987 | 0.325 | 0.117 | 2.9398E-41 | PCT | NQO2 |
| 0 | 0.46845365 | 0.16 | 0.002 | 0 | PCT | PAH |
| 1.4639E-30 | 0.46666664 | 0.964 | 0.903 | 3.1254E-26 | PCT | RPS7 |

| | | | | | | |
|---|---|---|---|---|---|---|
| 0 | 0.46622638 | 0.224 | 0.007 | 0 | PCT | SLC7A7 |
| 3.7994E-26 | 0.46617116 | 0.497 | 0.303 | 8.1118E-22 | PCT | ATP5PD |
| 4.128E-27 | 0.46611868 | 0.537 | 0.333 | 8.8132E-23 | PCT | GHITM |
| 1.315E-86 | 0.46540779 | 0.244 | 0.046 | 2.8076E-82 | PCT | GAMT |
| 6.3606E-35 | 0.4631072 | 0.467 | 0.243 | 1.358E-30 | PCT | PHB |
| 1.9419E-30 | 0.46241395 | 0.982 | 0.918 | 4.1459E-26 | PCT | RPL34 |
| 4.8363E-32 | 0.46203043 | 0.457 | 0.249 | 1.0326E-27 | PCT | NDUFA11 |
| 2.2737E-29 | 0.46086955 | 0.389 | 0.201 | 4.8543E-25 | PCT | NDUFA3 |
| 1.4579E-40 | 0.45661687 | 0.198 | 0.055 | 3.1126E-36 | PCT | MT1M |
| 1.7358E-22 | 0.45472415 | 0.796 | 0.643 | 3.706E-18 | PCT | UQCRH |
| 9.4134E-49 | 0.4538734 | 0.307 | 0.102 | 2.0098E-44 | PCT | ALDH1A1 |
| 2.0577E-64 | 0.4527603 | 0.236 | 0.054 | 4.3931E-60 | PCT | MAOA |
| 0 | 0.45110792 | 0.194 | 0.002 | 0 | PCT | DDC |
| 4.394E-29 | 0.44680755 | 0.18 | 0.059 | 9.3813E-25 | PCT | RNF152 |
| 4.5159E-22 | 0.44354445 | 0.928 | 0.852 | 9.6415E-18 | PCT | RPL38 |
| 1.4781E-25 | 0.44293383 | 0.505 | 0.308 | 3.1557E-21 | PCT | PRDX2 |
| 7.9067E-32 | 0.44108577 | 0.411 | 0.208 | 1.6881E-27 | PCT | ATP5MC1 |
| 4.9924E-26 | 0.44085414 | 0.998 | 0.972 | 1.0659E-21 | PCT | RPL13 |
| 1.25E-127 | 0.44071064 | 0.18 | 0.017 | 2.669E-123 | PCT | GPD1 |
| 2.1329E-26 | 0.43959004 | 0.974 | 0.936 | 4.5538E-22 | PCT | RPS23 |
| 3.0792E-20 | 0.43867741 | 0.952 | 0.905 | 6.5741E-16 | PCT | RPS16 |
| 1.4138E-15 | 0.43804434 | 0.321 | 0.186 | 3.0184E-11 | PCT | CTSL |
| 0 | 0.43669943 | 0.158 | 0.001 | 0 | PCT | AGXT2 |
| 1.6879E-22 | 0.43621647 | 0.974 | 0.91 | 3.6036E-18 | PCT | RPL29 |
| 1.2125E-22 | 0.43533778 | 0.643 | 0.443 | 2.5886E-18 | PCT | RPS10 |
| 1.0632E-26 | 0.43447789 | 0.968 | 0.939 | 2.2699E-22 | PCT | RPL37A |
| 3.0317E-22 | 0.43414824 | 0.86 | 0.723 | 6.4727E-18 | PCT | ENO1 |
| 2.2792E-18 | 0.43255601 | 0.519 | 0.343 | 4.8662E-14 | PCT | SELENOK |
| 2.0392E-20 | 0.43250633 | 0.415 | 0.248 | 4.3537E-16 | PCT | RPS4Y1 |
| 1.1872E-22 | 0.43192922 | 0.776 | 0.647 | 2.5347E-18 | PCT | SLC25A3 |
| 2.1703E-23 | 0.43068478 | 0.972 | 0.926 | 4.6337E-19 | PCT | RPL21 |
| 1.0444E-24 | 0.43050317 | 0.511 | 0.324 | 2.2298E-20 | PCT | NDUFB4 |
| 1.3265E-20 | 0.42965141 | 0.489 | 0.313 | 2.8321E-16 | PCT | TKT |
| 1.5611E-27 | 0.42737595 | 0.355 | 0.178 | 3.3329E-23 | PCT | CBR1 |
| 1.8568E-22 | 0.4262235 | 0.92 | 0.818 | 3.9642E-18 | PCT | RPS21 |
| 9.807E-66 | 0.42461915 | 0.244 | 0.058 | 2.0938E-61 | PCT | QDPR |
| 1.4642E-24 | 0.42455924 | 0.968 | 0.92 | 3.126E-20 | PCT | RPS3A |
| 8.5176E-21 | 0.42337001 | 0.415 | 0.254 | 1.8185E-16 | PCT | NDUFB3 |
| 1.5851E-16 | 0.42323612 | 0.567 | 0.429 | 3.3841E-12 | PCT | ST13 |
| 7.6007E-26 | 0.42086239 | 0.431 | 0.244 | 1.6227E-21 | PCT | UQCRFS1 |
| 2.2168E-18 | 0.41908234 | 0.948 | 0.895 | 4.7329E-14 | PCT | RPL35A |
| 3.06E-151 | 0.41844745 | 0.198 | 0.017 | 6.533E-147 | PCT | APOM |
| 2.5518E-20 | 0.4174332 | 0.497 | 0.325 | 5.448E-16 | PCT | ATP6V1F |
| 8.1821E-35 | 0.41728934 | 0.285 | 0.113 | 1.7469E-30 | PCT | GRHPR |
| 2.8161E-21 | 0.4171494 | 0.866 | 0.726 | 6.0123E-17 | PCT | NDUFA4 |
| 4.3024E-26 | 0.4169929 | 0.417 | 0.227 | 9.1857E-22 | PCT | NDUFS8 |
| 3.1422E-65 | 0.41503396 | 0.232 | 0.053 | 6.7086E-61 | PCT | AIFM1 |
| 1.8946E-23 | 0.4111471 | 0.982 | 0.942 | 4.045E-19 | PCT | RPL8 |
| 3.725E-283 | 0.41039279 | 0.13 | 0.001 | 7.953E-279 | PCT | GLYATL1 |
| 1.4207E-62 | 0.41037784 | 0.196 | 0.039 | 3.0332E-58 | PCT | MAF |

| | | | | | | |
|---|---|---|---|---|---|---|
| 7.8182E-18 | 0.41011657 | 0.475 | 0.319 | 1.6692E-13 | PCT | NDUFB7 |
| 7.767E-173 | 0.40935 | 0.18 | 0.011 | 1.658E-168 | PCT | LRP2 |
| 1.7036E-54 | 0.40900187 | 0.23 | 0.059 | 3.6372E-50 | PCT | GCHFR |
| 1.0734E-55 | 0.40747548 | 0.224 | 0.056 | 2.2916E-51 | PCT | MSRB1 |
| 7.1329E-21 | 0.40740219 | 0.958 | 0.908 | 1.5229E-16 | PCT | RPL24 |
| 6.1776E-29 | 0.40649853 | 0.405 | 0.21 | 1.3189E-24 | PCT | BLOC1S1 |
| 7.2645E-75 | 0.40598818 | 0.212 | 0.04 | 1.551E-70 | PCT | ALDH1L1 |
| 2.0699E-22 | 0.40578506 | 0.974 | 0.932 | 4.4191E-18 | PCT | RPL32 |
| 1.2944E-22 | 0.40468355 | 0.529 | 0.331 | 2.7636E-18 | PCT | NOP10 |
| 3.3542E-29 | 0.40400024 | 0.553 | 0.329 | 7.1613E-25 | PCT | AURKAIP1 |
| 3.0557E-69 | 0.40393195 | 0.23 | 0.049 | 6.524E-65 | PCT | AC245595.1 |
| 8.9506E-17 | 0.40197778 | 0.551 | 0.399 | 1.9109E-12 | PCT | TBCA |
| 6.3238E-20 | 0.39884018 | 0.974 | 0.928 | 1.3501E-15 | PCT | RPS14 |
| 6.115E-21 | 0.39882279 | 0.497 | 0.32 | 1.3056E-16 | PCT | VDAC1 |
| 9.4994E-22 | 0.39802634 | 0.96 | 0.893 | 2.0281E-17 | PCT | RPS13 |
| 4.5013E-25 | 0.39731559 | 0.471 | 0.277 | 9.6102E-21 | PCT | ATP5PO |
| 4.2005E-57 | 0.3962508 | 0.22 | 0.053 | 8.968E-53 | PCT | RAB29 |
| 1.0348E-24 | 0.39558106 | 0.956 | 0.869 | 2.2093E-20 | PCT | RACK1 |
| 1.643E-298 | 0.39556058 | 0.198 | 0.006 | 3.509E-294 | PCT | ACSM2A |
| 4.02E-25 | 0.3929993 | 0.511 | 0.315 | 8.5827E-21 | PCT | COX5A |
| 4.3896E-16 | 0.39296361 | 0.739 | 0.622 | 9.3719E-12 | PCT | SLC25A6 |
| 2.4063E-40 | 0.39274056 | 0.289 | 0.105 | 5.1374E-36 | PCT | AKR7A2 |
| 5.1382E-24 | 0.39144208 | 0.453 | 0.265 | 1.097E-19 | PCT | NDUFV2 |
| 8.9168E-22 | 0.390296 | 0.547 | 0.361 | 1.9037E-17 | PCT | ARL6IP4 |
| 2.1105E-16 | 0.39012677 | 0.926 | 0.872 | 4.5058E-12 | PCT | RPL7 |
| 1.1842E-30 | 0.38880567 | 0.24 | 0.092 | 2.5282E-26 | PCT | ACADM |
| 1.7927E-28 | 0.38811652 | 0.371 | 0.181 | 3.8273E-24 | PCT | HMGN3 |
| 1.7996E-20 | 0.3856304 | 0.683 | 0.512 | 3.8421E-16 | PCT | OST4 |
| 7.1118E-84 | 0.38423837 | 0.194 | 0.03 | 1.5184E-79 | PCT | EHHADH |
| 4.713E-226 | 0.38394891 | 0.186 | 0.008 | 1.006E-221 | PCT | ACMSD |
| 9.7041E-20 | 0.3836404 | 0.956 | 0.929 | 2.0718E-15 | PCT | RPL7A |
| 5.232E-20 | 0.38344295 | 0.439 | 0.277 | 1.117E-15 | PCT | FIS1 |
| 1.4405E-20 | 0.38331505 | 0.317 | 0.171 | 3.0754E-16 | PCT | SDHD |
| 4.1759E-21 | 0.38209061 | 0.922 | 0.833 | 8.9156E-17 | PCT | PRDX1 |
| 2.5462E-21 | 0.38044617 | 0.435 | 0.257 | 5.4361E-17 | PCT | APRT |
| 5.4629E-13 | 0.379571 | 0.16 | 0.074 | 1.1663E-08 | PCT | AKR1C1 |
| 1.1546E-17 | 0.37936515 | 0.451 | 0.295 | 2.4652E-13 | PCT | CIB1 |
| 1.8227E-22 | 0.37884476 | 0.96 | 0.884 | 3.8915E-18 | PCT | RPL22 |
| 7.4437E-17 | 0.37864514 | 0.946 | 0.907 | 1.5892E-12 | PCT | RPL26 |
| 1.013E-301 | 0.3781395 | 0.162 | 0.003 | 2.163E-297 | PCT | SLC17A3 |
| 1.4993E-17 | 0.37636202 | 0.341 | 0.202 | 3.201E-13 | PCT | SCP2 |
| 2.3759E-19 | 0.37613411 | 0.948 | 0.895 | 5.0724E-15 | PCT | RPS15A |
| 5.048E-270 | 0.37560156 | 0.184 | 0.006 | 1.078E-265 | PCT | AZGP1 |
| 1.9264E-19 | 0.37521672 | 0.234 | 0.107 | 4.1128E-15 | PCT | RARRES2 |
| 2.2007E-39 | 0.37451088 | 0.281 | 0.101 | 4.6986E-35 | PCT | LACTB2 |
| 1.584E-20 | 0.37411088 | 0.437 | 0.268 | 3.3818E-16 | PCT | NDUFS6 |
| 1.9265E-23 | 0.37249696 | 0.986 | 0.921 | 4.1131E-19 | PCT | RPLP2 |
| 3.8849E-22 | 0.37246534 | 0.345 | 0.184 | 8.2942E-18 | PCT | NDUFS7 |
| 2.1833E-30 | 0.37240677 | 0.325 | 0.148 | 4.6613E-26 | PCT | SDHB |
| 2.2213E-46 | 0.37224326 | 0.226 | 0.064 | 4.7425E-42 | PCT | ECHDC3 |

| | | | | | | |
|---|---|---|---|---|---|---|
| 1.5026E-23 | 0.37105486 | 0.994 | 0.971 | 3.2081E-19 | PCT | RPLP1 |
| 2.4918E-13 | 0.37099018 | 0.966 | 0.947 | 5.32E-09 | PCT | RPL13A |
| 5.1436E-21 | 0.37012392 | 0.453 | 0.28 | 1.0982E-16 | PCT | COPS9 |
| 4.2666E-22 | 0.36968851 | 0.98 | 0.932 | 9.1092E-18 | PCT | RPS27A |
| 1.8577E-15 | 0.36809336 | 0.473 | 0.317 | 3.9662E-11 | PCT | PHPT1 |
| 5.8399E-44 | 0.36710926 | 0.307 | 0.107 | 1.2468E-39 | PCT | FAHD1 |
| 7.8537E-31 | 0.36680127 | 0.279 | 0.115 | 1.6768E-26 | PCT | TMEM256 |
| 4.2241E-18 | 0.36623358 | 0.413 | 0.255 | 9.0184E-14 | PCT | GSTO1 |
| 1.496E-23 | 0.36531866 | 0.407 | 0.226 | 3.194E-19 | PCT | RNF181 |
| 1.112E-18 | 0.36475932 | 0.962 | 0.92 | 2.3742E-14 | PCT | RPL14 |
| 6.0556E-39 | 0.36418626 | 0.236 | 0.077 | 1.2929E-34 | PCT | TST |
| 3.1503E-16 | 0.36380846 | 0.974 | 0.955 | 6.7259E-12 | PCT | RPS27 |
| 1.1579E-20 | 0.36355694 | 0.491 | 0.317 | 2.4721E-16 | PCT | FAM96B |
| 3.8085E-15 | 0.36323175 | 0.539 | 0.39 | 8.1312E-11 | PCT | DAD1 |
| 4.5358E-18 | 0.36189724 | 0.906 | 0.814 | 9.6839E-14 | PCT | UBB |
| 5.5565E-31 | 0.36159173 | 0.253 | 0.099 | 1.1863E-26 | PCT | NIT2 |
| 2.6584E-22 | 0.36127923 | 0.311 | 0.157 | 5.6756E-18 | PCT | NDUFV1 |
| 7.6711E-22 | 0.36122733 | 0.766 | 0.598 | 1.6378E-17 | PCT | PSMA7 |
| 3.9703E-30 | 0.36108456 | 0.337 | 0.155 | 8.4767E-26 | PCT | ADH5 |
| 3.3146E-19 | 0.36102161 | 0.433 | 0.267 | 7.0767E-15 | PCT | ATP5F1C |
| 2.0109E-20 | 0.36072949 | 0.952 | 0.861 | 4.2933E-16 | PCT | UBA52 |
| 2.5174E-21 | 0.36061214 | 0.491 | 0.309 | 5.3746E-17 | PCT | SNX3 |
| 1.169E-21 | 0.36061013 | 0.435 | 0.26 | 2.4957E-17 | PCT | MINOS1 |
| 5.8999E-12 | 0.36033737 | 0.946 | 0.911 | 1.2596E-07 | PCT | RPL35 |
| 9.6986E-19 | 0.35950915 | 0.375 | 0.223 | 2.0706E-14 | PCT | ATP5PB |
| 2.2876E-18 | 0.359018 | 0.625 | 0.464 | 4.884E-14 | PCT | PRDX6 |
| 1.7407E-49 | 0.35792746 | 0.186 | 0.044 | 3.7164E-45 | PCT | PXMP2 |
| 2.9047E-21 | 0.35788384 | 0.982 | 0.944 | 6.2014E-17 | PCT | RPL11 |
| 1.1138E-21 | 0.35748351 | 0.986 | 0.917 | 2.378E-17 | PCT | RPL9 |
| 9.3193E-18 | 0.35733027 | 0.379 | 0.229 | 1.9897E-13 | PCT | CALM3 |
| 1.7117E-21 | 0.35700997 | 0.407 | 0.237 | 3.6545E-17 | PCT | LSM3 |
| 1.9603E-14 | 0.35625323 | 0.259 | 0.143 | 4.1852E-10 | PCT | LGMN |
| 3.0284E-26 | 0.35618768 | 0.373 | 0.19 | 6.4657E-22 | PCT | CYC1 |
| 3.4623E-21 | 0.35586709 | 0.413 | 0.241 | 7.392E-17 | PCT | ZNHIT1 |
| 5.3899E-24 | 0.35582729 | 0.307 | 0.149 | 1.1507E-19 | PCT | COA4 |
| 0 | 0.35572548 | 0.154 | 0 | 0 | PCT | SLC22A12 |
| 3.6207E-20 | 0.3552788 | 0.429 | 0.256 | 7.7302E-16 | PCT | NDUFB10 |
| 2.874E-128 | 0.35273252 | 0.144 | 0.01 | 6.137E-124 | PCT | CUBN |
| 7.7657E-18 | 0.35192855 | 0.976 | 0.932 | 1.658E-13 | PCT | RPL6 |
| 1.3015E-36 | 0.35161389 | 0.206 | 0.064 | 2.7788E-32 | PCT | BPHL |
| 1.1728E-18 | 0.34982298 | 0.972 | 0.896 | 2.504E-14 | PCT | RPL5 |
| 1.9376E-37 | 0.34970189 | 0.232 | 0.076 | 4.1367E-33 | PCT | PSAT1 |
| 6.5816E-14 | 0.34939228 | 0.647 | 0.528 | 1.4052E-09 | PCT | SEC61B |
| 1.386E-238 | 0.34889378 | 0.172 | 0.006 | 2.958E-234 | PCT | FTCD |
| 5.8885E-15 | 0.34799535 | 0.701 | 0.563 | 1.2572E-10 | PCT | RPS27L |
| 1.8535E-19 | 0.34792179 | 0.427 | 0.262 | 3.9573E-15 | PCT | NDUFB8 |
| 8.3693E-22 | 0.34736065 | 0.313 | 0.161 | 1.7869E-17 | PCT | UQCRC1 |
| 5.0886E-18 | 0.34416019 | 0.962 | 0.931 | 1.0864E-13 | PCT | RPS6 |
| 3.092E-16 | 0.34333804 | 0.948 | 0.891 | 6.6015E-12 | PCT | RPS11 |
| 5.95E-176 | 0.34150527 | 0.172 | 0.01 | 1.27E-171 | PCT | SLC6A13 |

| | | | | | | |
|---|---|---|---|---|---|---|
| 4.4735E-16 | 0.34063148 | 0.948 | 0.896 | 9.5509E-12 | PCT | RPL23A |
| 2.3009E-22 | 0.34059082 | 0.311 | 0.158 | 4.9123E-18 | PCT | AK2 |
| 1.5434E-21 | 0.3404644 | 0.339 | 0.176 | 3.2951E-17 | PCT | CLIC4 |
| 3.5335E-14 | 0.34013478 | 0.471 | 0.315 | 7.5439E-10 | PCT | NPC2 |
| 1.7104E-24 | 0.33976942 | 0.281 | 0.131 | 3.6517E-20 | PCT | ETFA |
| 7.469E-35 | 0.33964702 | 0.293 | 0.116 | 1.5946E-30 | PCT | MSRB2 |
| 1.4032E-15 | 0.33962908 | 0.633 | 0.485 | 2.9958E-11 | PCT | TOMM7 |
| 8.682E-21 | 0.33937749 | 0.369 | 0.207 | 1.8536E-16 | PCT | SLIRP |
| 1.2412E-12 | 0.33905951 | 0.174 | 0.085 | 2.6499E-08 | PCT | SLC7A8 |
| 2.3185E-19 | 0.33891299 | 0.244 | 0.119 | 4.95E-15 | PCT | ACO2 |
| 8.4737E-15 | 0.33887108 | 0.487 | 0.344 | 1.8091E-10 | PCT | COMMD6 |
| 9.6154E-17 | 0.33811946 | 0.97 | 0.899 | 2.0529E-12 | PCT | RPS25 |
| 6.6499E-37 | 0.33799601 | 0.263 | 0.093 | 1.4197E-32 | PCT | NINJ1 |
| 1.6103E-28 | 0.33759683 | 0.255 | 0.103 | 3.438E-24 | PCT | MPST |
| 1.1801E-13 | 0.33665423 | 0.349 | 0.211 | 2.5195E-09 | PCT | GCLM |
| 7.8772E-72 | 0.33638751 | 0.194 | 0.034 | 1.6818E-67 | PCT | SLC22A2 |
| 0.00232526 | 0.33557439 | 0.132 | 0.093 | 1 | PCT | CCL2 |
| 2.8471E-20 | 0.33553946 | 0.267 | 0.131 | 6.0785E-16 | PCT | HAGH |
| 4.9892E-16 | 0.33527998 | 0.409 | 0.266 | 1.0652E-11 | PCT | NDUFA2 |
| 2.4195E-49 | 0.3344305 | 0.174 | 0.039 | 5.1656E-45 | PCT | ETNK2 |
| 6.9475E-20 | 0.33432244 | 0.713 | 0.535 | 1.4833E-15 | PCT | GNG5 |
| 1.0963E-35 | 0.33418965 | 0.2 | 0.062 | 2.3406E-31 | PCT | SORD |
| 0 | 0.33409491 | 0.142 | 0.001 | 0 | PCT | PRAP1 |
| 5.376E-256 | 0.33343014 | 0.166 | 0.005 | 1.148E-251 | PCT | SLC13A3 |
| 1.243E-16 | 0.33335206 | 0.511 | 0.35 | 2.6538E-12 | PCT | EIF3K |
| 3.1374E-22 | 0.3332103 | 0.347 | 0.184 | 6.6983E-18 | PCT | ECI1 |
| 6.069E-20 | 0.33274683 | 0.453 | 0.278 | 1.2957E-15 | PCT | PSMB6 |
| 1.9721E-17 | 0.33270198 | 0.391 | 0.24 | 4.2105E-13 | PCT | TUFM |
| 4.3924E-15 | 0.33264478 | 0.99 | 0.96 | 9.3778E-11 | PCT | RPS18 |
| 8.3415E-14 | 0.33259347 | 0.956 | 0.927 | 1.7809E-09 | PCT | RPL36 |
| 2.0332E-17 | 0.33118737 | 0.509 | 0.344 | 4.3409E-13 | PCT | C19orf53 |
| 7.9539E-18 | 0.33116292 | 0.323 | 0.185 | 1.6982E-13 | PCT | SDHC |
| 2.2043E-14 | 0.33080593 | 0.852 | 0.773 | 4.7061E-10 | PCT | OAZ1 |
| 1.4602E-14 | 0.33044592 | 0.389 | 0.248 | 3.1175E-10 | PCT | COTL1 |
| 2.9813E-19 | 0.33034013 | 0.353 | 0.201 | 6.3651E-15 | PCT | PRDX3 |
| 1.1604E-25 | 0.3302774 | 0.369 | 0.188 | 2.4774E-21 | PCT | C1QBP |
| 6.2656E-18 | 0.33026063 | 0.413 | 0.257 | 1.3377E-13 | PCT | RPS19BP1 |
| 1.2218E-23 | 0.33012867 | 0.385 | 0.209 | 2.6085E-19 | PCT | TIMM13 |
| 6.5026E-15 | 0.32960829 | 0.349 | 0.215 | 1.3883E-10 | PCT | NDUFA13 |
| 4.3833E-52 | 0.32859368 | 0.22 | 0.056 | 9.3584E-48 | PCT | FUOM |
| 1.1567E-63 | 0.32769539 | 0.194 | 0.038 | 2.4695E-59 | PCT | TTC38 |
| 3.6696E-12 | 0.32755283 | 0.86 | 0.802 | 7.8346E-08 | PCT | RPL31 |
| 1.4362E-23 | 0.32682046 | 0.331 | 0.166 | 3.0663E-19 | PCT | COA3 |
| 1.9713E-37 | 0.32677607 | 0.269 | 0.096 | 4.2088E-33 | PCT | COMMD8 |
| 6.1358E-14 | 0.32607087 | 0.469 | 0.331 | 1.31E-09 | PCT | PSMB1 |
| 2.631E-108 | 0.32561568 | 0.196 | 0.024 | 5.617E-104 | PCT | AMN |
| 2.1282E-18 | 0.3244687 | 0.291 | 0.154 | 4.5438E-14 | PCT | MRPL34 |
| 3.207E-18 | 0.32395734 | 0.415 | 0.254 | 6.8469E-14 | PCT | LAMTOR1 |
| 1.222E-234 | 0.32315779 | 0.13 | 0.002 | 2.61E-230 | PCT | ACY3 |
| 1.8845E-14 | 0.32302464 | 0.86 | 0.75 | 4.0235E-10 | PCT | RPL4 |

| | | | | | | |
|---|---|---|---|---|---|---|
| 1.4747E-15 | 0.32250657 | 0.864 | 0.78 | 3.1484E-11 | PCT | EEF1B2 |
| 1.4438E-85 | 0.32209372 | 0.186 | 0.027 | 3.0826E-81 | PCT | HGD |
| 5.0881E-15 | 0.32144276 | 0.752 | 0.641 | 1.0863E-10 | PCT | EDF1 |
| 3.8615E-19 | 0.32135627 | 0.351 | 0.2 | 8.2444E-15 | PCT | METTL9 |
| 1.1711E-52 | 0.32111875 | 0.224 | 0.056 | 2.5003E-48 | PCT | FABP3 |
| 6.337E-11 | 0.32111568 | 0.397 | 0.281 | 1.353E-06 | PCT | NDUFB11 |
| 5.8914E-17 | 0.32061893 | 0.563 | 0.39 | 1.2578E-12 | PCT | RHOC |
| 5.407E-243 | 0.32024925 | 0.152 | 0.004 | 1.154E-238 | PCT | PIPOX |
| 4.9506E-12 | 0.32020775 | 0.471 | 0.34 | 1.057E-07 | PCT | MIF |
| 3.1972E-15 | 0.31787499 | 0.499 | 0.343 | 6.826E-11 | PCT | EIF5A |
| 2.4796E-63 | 0.31737452 | 0.184 | 0.035 | 5.2939E-59 | PCT | CHDH |
| 5.2647E-16 | 0.31732305 | 0.291 | 0.163 | 1.124E-11 | PCT | CISD1 |
| 3.6242E-16 | 0.31671728 | 0.79 | 0.66 | 7.7376E-12 | PCT | TMA7 |
| 2.403E-16 | 0.31623049 | 0.381 | 0.234 | 5.1303E-12 | PCT | MRPL57 |
| 1.893E-20 | 0.31560104 | 0.212 | 0.094 | 4.0416E-16 | PCT | CAT |
| 9.8237E-21 | 0.31532525 | 0.341 | 0.185 | 2.0973E-16 | PCT | MRPS36 |
| 2.1384E-17 | 0.31517294 | 0.729 | 0.573 | 4.5655E-13 | PCT | PRDX5 |
| 2.2557E-15 | 0.31436751 | 0.93 | 0.867 | 4.816E-11 | PCT | RPS5 |
| 0.00125428 | 0.31356367 | 0.126 | 0.085 | 1 | PCT | SAA1 |
| 0.00064234 | 0.31321355 | 0.251 | 0.195 | 1 | PCT | MT-ND6 |
| 2.166E-123 | 0.31153516 | 0.17 | 0.015 | 4.624E-119 | PCT | GIPC2 |
| 1.0802E-24 | 0.30975038 | 0.297 | 0.139 | 2.3061E-20 | PCT | GRPEL1 |
| 1.5261E-13 | 0.30947884 | 0.295 | 0.175 | 3.2582E-09 | PCT | CTSZ |
| 1.0711E-13 | 0.30915813 | 0.96 | 0.927 | 2.2868E-09 | PCT | RPL18A |
| 6.0961E-18 | 0.3090328 | 0.523 | 0.342 | 1.3015E-13 | PCT | MZT2B |
| 2.0819E-41 | 0.30862262 | 0.168 | 0.04 | 4.445E-37 | PCT | THY1 |
| 5.2868E-16 | 0.30746656 | 0.822 | 0.715 | 1.1287E-11 | PCT | RPL36AL |
| 2.2261E-89 | 0.30729774 | 0.178 | 0.024 | 4.7528E-85 | PCT | ABAT |
| 1.207E-26 | 0.30677563 | 0.15 | 0.045 | 2.5769E-22 | PCT | ANK2 |
| 5.7691E-58 | 0.30621552 | 0.128 | 0.019 | 1.2317E-53 | PCT | VCAM1 |
| 3.179E-25 | 0.30606692 | 0.216 | 0.085 | 6.7871E-21 | PCT | ACSL1 |
| 3.9725E-19 | 0.30595059 | 0.481 | 0.296 | 8.4813E-15 | PCT | GADD45GIP1 |
| 3.6445E-84 | 0.30488313 | 0.146 | 0.017 | 7.781E-80 | PCT | CRYM |
| 4.779E-31 | 0.30459138 | 0.196 | 0.065 | 1.0203E-26 | PCT | ACO1 |
| 2.7452E-22 | 0.30302525 | 0.281 | 0.135 | 5.8611E-18 | PCT | MRPS23 |
| 0.00011706 | 0.30264202 | 0.347 | 0.282 | 1 | PCT | AKR1B1 |
| 3.2067E-24 | 0.30247668 | 0.263 | 0.116 | 6.8464E-20 | PCT | LAP3 |
| 7.5403E-18 | 0.3022627 | 0.317 | 0.175 | 1.6098E-13 | PCT | TIMM8B |
| 1.2276E-33 | 0.30193698 | 0.281 | 0.109 | 2.621E-29 | PCT | MTCH2 |
| 1.8889E-17 | 0.30125514 | 0.309 | 0.167 | 4.0329E-13 | PCT | MPC1 |
| 4.4207E-44 | 0.3006927 | 0.228 | 0.066 | 9.4382E-40 | PCT | ALDH7A1 |
| 2.6401E-10 | 0.29990322 | 0.924 | 0.881 | 5.6367E-06 | PCT | RPL39 |
| 9.2256E-36 | 0.29984939 | 0.194 | 0.059 | 1.9697E-31 | PCT | HADH |
| 2.1576E-56 | 0.2997488 | 0.164 | 0.031 | 4.6065E-52 | PCT | SLC4A4 |
| 5.9839E-28 | 0.29925405 | 0.146 | 0.043 | 1.2776E-23 | PCT | ACADSB |
| 2.6671E-07 | 0.29920277 | 0.275 | 0.197 | 0.00569421 | PCT | SH3YL1 |
| 6.1335E-30 | 0.29909631 | 0.216 | 0.076 | 1.3095E-25 | PCT | PTH1R |
| 2.6804E-20 | 0.29903135 | 0.251 | 0.118 | 5.7226E-16 | PCT | SUCLG2 |
| 1.375E-277 | 0.29897973 | 0.128 | 0.001 | 2.936E-273 | PCT | SLC5A12 |
| 7.9648E-19 | 0.2989351 | 0.351 | 0.198 | 1.7005E-14 | PCT | MRPL52 |

| | | | | | | |
|---|---|---|---|---|---|---|
| 9.3725E-10 | 0.29784109 | 0.259 | 0.157 | 2.001E-05 | PCT | G0S2 |
| 9.583E-19 | 0.29777235 | 0.234 | 0.11 | 2.046E-14 | PCT | CTSC |
| 1.1848E-16 | 0.29700025 | 0.353 | 0.21 | 2.5295E-12 | PCT | PSMA5 |
| 1.8342E-20 | 0.29679695 | 0.363 | 0.203 | 3.916E-16 | PCT | RAB5C |
| 1.6017E-14 | 0.2964644 | 0.397 | 0.256 | 3.4195E-10 | PCT | SCAND1 |
| 1.6029E-13 | 0.29622165 | 0.557 | 0.411 | 3.4222E-09 | PCT | PARK7 |
| 6.8246E-13 | 0.2944299 | 0.425 | 0.288 | 1.457E-08 | PCT | SF3B5 |
| 1.7623E-20 | 0.2936715 | 0.279 | 0.138 | 3.7625E-16 | PCT | AP1S1 |
| 1.4189E-14 | 0.29311131 | 0.98 | 0.931 | 3.0294E-10 | PCT | RPS9 |
| 4.633E-180 | 0.29288729 | 0.174 | 0.01 | 9.891E-176 | PCT | USH1C |
| 8.3248E-14 | 0.29191026 | 0.956 | 0.921 | 1.7773E-09 | PCT | RPL30 |
| 8.61E-11 | 0.29103458 | 0.333 | 0.221 | 1.8382E-06 | PCT | SPATS2L |
| 1.1861E-13 | 0.29076104 | 0.828 | 0.749 | 2.5323E-09 | PCT | PFDN5 |
| 8.1441E-21 | 0.29072955 | 0.323 | 0.166 | 1.7388E-16 | PCT | NAA38 |
| 8.313E-307 | 0.29061523 | 0.128 | 0.001 | 1.775E-302 | PCT | ASPDH |
| 7.1355E-09 | 0.29005204 | 0.834 | 0.771 | 0.00015234 | PCT | EEF2 |
| 1.5836E-13 | 0.28959992 | 0.485 | 0.336 | 3.381E-09 | PCT | TRIR |
| 2.034E-149 | 0.28898921 | 0.14 | 0.008 | 4.342E-145 | PCT | AOX1 |
| 4.5644E-24 | 0.28869659 | 0.267 | 0.119 | 9.745E-20 | PCT | ABRACL |
| 1.8557E-16 | 0.2886309 | 0.363 | 0.214 | 3.9619E-12 | PCT | GTF3C6 |
| 1.0107E-34 | 0.28810704 | 0.184 | 0.054 | 2.1579E-30 | PCT | GALM |
| 3.0325E-08 | 0.28773083 | 0.503 | 0.399 | 0.00064743 | PCT | PCBP2 |
| 8.326E-07 | 0.28755523 | 0.876 | 0.827 | 0.01777606 | PCT | SQSTM1 |
| 1.9922E-37 | 0.28726893 | 0.253 | 0.085 | 4.2533E-33 | PCT | LY6E |
| 2.187E-289 | 0.28708599 | 0.128 | 0.001 | 4.669E-285 | PCT | SLC22A8 |
| 2.9052E-27 | 0.28663838 | 0.244 | 0.099 | 6.2025E-23 | PCT | TPMT |
| 1.0125E-25 | 0.28591307 | 0.228 | 0.091 | 2.1617E-21 | PCT | SIGIRR |
| 3.549E-23 | 0.28569513 | 0.218 | 0.09 | 7.577E-19 | PCT | PPP1R16A |
| 1.4642E-14 | 0.28479738 | 0.353 | 0.219 | 3.1261E-10 | PCT | C19orf70 |
| 1.9156E-19 | 0.28382104 | 0.283 | 0.144 | 4.0899E-15 | PCT | RAB5IF |
| 4.2881E-44 | 0.28379978 | 0.182 | 0.045 | 9.1551E-40 | PCT | RASSF4 |
| 1.1827E-27 | 0.28339201 | 0.251 | 0.102 | 2.525E-23 | PCT | HMGCL |
| 7.7512E-14 | 0.28244948 | 0.974 | 0.944 | 1.6549E-09 | PCT | RPS15 |
| 2.9864E-12 | 0.28167044 | 0.521 | 0.376 | 6.3759E-08 | PCT | DYNLT1 |
| 3.2152E-18 | 0.28137946 | 0.261 | 0.13 | 6.8645E-14 | PCT | DECR1 |
| 5.5719E-12 | 0.28110135 | 0.972 | 0.941 | 1.1896E-07 | PCT | RPS24 |
| 2.2458E-28 | 0.28046765 | 0.224 | 0.084 | 4.7949E-24 | PCT | CISD3 |
| 2.3851E-26 | 0.2804245 | 0.283 | 0.124 | 5.0922E-22 | PCT | HINT2 |
| 3.6338E-19 | 0.27962847 | 0.357 | 0.198 | 7.7582E-15 | PCT | AP2S1 |
| 4.1558E-13 | 0.27906925 | 0.365 | 0.233 | 8.8726E-09 | PCT | RPS17 |
| 4.1286E-16 | 0.27873526 | 0.533 | 0.354 | 8.8146E-12 | PCT | NEDD8 |
| 5.6242E-14 | 0.27849057 | 0.758 | 0.63 | 1.2008E-09 | PCT | UBL5 |
| 3.1979E-14 | 0.27844373 | 0.964 | 0.932 | 6.8275E-10 | PCT | RPL3 |
| 4.4081E-12 | 0.27764763 | 0.413 | 0.28 | 9.4112E-08 | PCT | ROMO1 |
| 5.0967E-14 | 0.27718531 | 0.978 | 0.937 | 1.0882E-09 | PCT | RPL15 |
| 4.3026E-08 | 0.27590677 | 0.124 | 0.063 | 0.0009186 | PCT | CCL20 |
| 7.65E-13 | 0.27530334 | 0.958 | 0.922 | 1.6333E-08 | PCT | RPL10 |
| 6.223E-24 | 0.27451457 | 0.24 | 0.102 | 1.3286E-19 | PCT | MCRIP2 |
| 3.2196E-41 | 0.27410154 | 0.206 | 0.058 | 6.8739E-37 | PCT | BIN1 |
| 3.882E-14 | 0.27368223 | 0.381 | 0.245 | 8.288E-10 | PCT | RAB1A |

| | | | | | | |
|---|---|---|---|---|---|---|
| 9.6334E-64 | 0.27353396 | 0.192 | 0.037 | 2.0567E-59 | PCT | FAH |
| 2.685E-16 | 0.27268584 | 0.385 | 0.233 | 5.7326E-12 | PCT | VPS29 |
| 5.6212E-95 | 0.27167517 | 0.172 | 0.021 | 1.2001E-90 | PCT | NAPSA |
| 5.7041E-15 | 0.27167163 | 0.383 | 0.241 | 1.2178E-10 | PCT | MRPS21 |
| 6.5072E-10 | 0.27149878 | 0.816 | 0.748 | 1.3893E-05 | PCT | BTF3 |
| 6.333E-258 | 0.2714842 | 0.116 | 0.001 | 1.352E-253 | PCT | SMIM32 |
| 1.6822E-36 | 0.27092367 | 0.19 | 0.055 | 3.5914E-32 | PCT | MACROD1 |
| 1.7809E-18 | 0.27065308 | 0.279 | 0.142 | 3.8022E-14 | PCT | NDUFS2 |
| 7.4224E-12 | 0.27029542 | 0.467 | 0.334 | 1.5847E-07 | PCT | ERP29 |
| 5.0455E-13 | 0.27021058 | 0.363 | 0.231 | 1.0772E-08 | PCT | SMIM26 |
| 3.782E-304 | 0.26850656 | 0.132 | 0.001 | 8.075E-300 | PCT | SLC22A6 |
| 2.7452E-11 | 0.26848486 | 0.521 | 0.384 | 5.8611E-07 | PCT | POMP |
| 2.761E-187 | 0.26811904 | 0.144 | 0.006 | 5.895E-183 | PCT | AGT |
| 1.9972E-26 | 0.26809338 | 0.24 | 0.097 | 4.264E-22 | PCT | PMVK |
| 3.0882E-11 | 0.26800289 | 0.952 | 0.899 | 6.5934E-07 | PCT | RPL27A |
| 7.5754E-14 | 0.26800129 | 0.363 | 0.231 | 1.6174E-09 | PCT | MDH2 |
| 1.4575E-18 | 0.26760992 | 0.313 | 0.168 | 3.1117E-14 | PCT | LAMTOR2 |
| 5.0411E-12 | 0.26734051 | 0.782 | 0.686 | 1.0763E-07 | PCT | CYSTM1 |
| 2.501E-12 | 0.26678518 | 0.367 | 0.237 | 5.3397E-08 | PCT | HIGD2A |
| 8.6441E-14 | 0.26630433 | 0.321 | 0.194 | 1.8455E-09 | PCT | GLRX5 |
| 1.267E-29 | 0.2661775 | 0.23 | 0.085 | 2.7051E-25 | PCT | MRPL12 |
| 1.4173E-70 | 0.26611889 | 0.198 | 0.036 | 3.0259E-66 | PCT | HPN |
| 8.8713E-16 | 0.26600054 | 0.289 | 0.159 | 1.894E-11 | PCT | SSNA1 |
| 1.3767E-25 | 0.26487368 | 0.182 | 0.064 | 2.9392E-21 | PCT | TSPAN33 |
| 4.8514E-12 | 0.26469787 | 0.399 | 0.268 | 1.0358E-07 | PCT | ANAPC11 |
| 1.3265E-12 | 0.26443789 | 0.832 | 0.728 | 2.832E-08 | PCT | PFN1 |
| 1.3993E-77 | 0.26415388 | 0.168 | 0.024 | 2.9874E-73 | PCT | HSD17B14 |
| 2.9716E-09 | 0.26397811 | 0.232 | 0.143 | 6.3444E-05 | PCT | DDAH1 |
| 3.4342E-12 | 0.26363284 | 0.445 | 0.31 | 7.3321E-08 | PCT | COX7A2L |
| 4.4027E-13 | 0.2632776 | 0.321 | 0.198 | 9.3998E-09 | PCT | NDUFAF3 |
| 1.9895E-08 | 0.26274568 | 0.373 | 0.275 | 0.00042477 | PCT | CNN3 |
| 3.124E-77 | 0.262396 | 0.134 | 0.016 | 6.6697E-73 | PCT | ACSF2 |
| 0.00209159 | 0.26233995 | 0.333 | 0.286 | 1 | PCT | CDK2AP2 |
| 2.0498E-10 | 0.26174083 | 0.415 | 0.302 | 4.3764E-06 | PCT | TMEM258 |
| 1.105E-15 | 0.26164134 | 0.343 | 0.201 | 2.3592E-11 | PCT | TXNL4A |
| 6.4358E-11 | 0.26138397 | 0.381 | 0.263 | 1.374E-06 | PCT | PSMB7 |
| 9.9711E-16 | 0.26059644 | 0.176 | 0.079 | 2.1288E-11 | PCT | STK19 |
| 1.0885E-15 | 0.26031906 | 0.361 | 0.216 | 2.3239E-11 | PCT | PSMA4 |
| 2.048E-08 | 0.26026383 | 0.962 | 0.956 | 0.00043725 | PCT | RPS2 |
| 2.5623E-14 | 0.25996681 | 0.253 | 0.139 | 5.4705E-10 | PCT | BOLA3 |
| 3.4441E-10 | 0.25995913 | 0.228 | 0.138 | 7.3532E-06 | PCT | AP1M2 |
| 2.3098E-11 | 0.25858895 | 0.493 | 0.356 | 4.9313E-07 | PCT | TRMT112 |
| 5.892E-29 | 0.25826796 | 0.21 | 0.075 | 1.2579E-24 | PCT | ALDH3A2 |
| 2.7682E-29 | 0.25810384 | 0.176 | 0.056 | 5.91E-25 | PCT | ACOT7 |
| 2.8072E-13 | 0.25783929 | 0.463 | 0.313 | 5.9934E-09 | PCT | NDUFB9 |
| 1.0649E-11 | 0.25715889 | 0.389 | 0.263 | 2.2735E-07 | PCT | BLVRB |
| 2.1304E-26 | 0.25687648 | 0.226 | 0.089 | 4.5484E-22 | PCT | POLDIP2 |
| 3.0905E-13 | 0.25656045 | 0.938 | 0.863 | 6.5982E-09 | PCT | RPS4X |
| 1.585E-20 | 0.25639116 | 0.259 | 0.121 | 3.3839E-16 | PCT | GNPDA1 |
| 6.6149E-09 | 0.25559063 | 0.495 | 0.379 | 0.00014123 | PCT | PTMS |

| | | | | | | |
|---|---|---|---|---|---|---|
| 3.341E-17 | 0.25547532 | 0.226 | 0.108 | 7.1329E-13 | PCT | SMIM37 |
| 4.5095E-23 | 0.25491085 | 0.23 | 0.097 | 9.6278E-19 | PCT | ACOT13 |
| 2.031E-09 | 0.25486499 | 0.944 | 0.909 | 4.3361E-05 | PCT | FAU |
| 5.0073E-15 | 0.2543926 | 0.283 | 0.157 | 1.0691E-10 | PCT | CTSA |
| 6.7429E-05 | 0.25404466 | 0.882 | 0.877 | 1 | PCT | ITM2B |
| 0.00407536 | 0.25318791 | 0.267 | 0.221 | 1 | PCT | ARL6IP5 |
| 4.125E-14 | 0.25159388 | 0.283 | 0.161 | 8.8069E-10 | PCT | NDUFB6 |
| 4.3221E-09 | 0.25131051 | 0.509 | 0.39 | 9.2277E-05 | PCT | FKBP2 |
| 1.4902E-16 | 0.25114267 | 0.261 | 0.135 | 3.1816E-12 | PCT | PPCS |
| 3.835E-05 | -0.2503974 | 0.727 | 0.776 | 0.81876589 | PCT | MYL12A |
| 4.2649E-08 | -0.2512448 | 0.154 | 0.255 | 0.00091056 | PCT | ARID4B |
| 4.132E-08 | -0.2513363 | 0.184 | 0.293 | 0.00088218 | PCT | TJP1 |
| 2.7936E-07 | -0.2547082 | 0.074 | 0.159 | 0.00596442 | PCT | IGFBP2 |
| 1.6429E-07 | -0.2550765 | 0.158 | 0.259 | 0.00350767 | PCT | S100A14 |
| 1.6996E-05 | -0.2568536 | 0.094 | 0.163 | 0.36286436 | PCT | TNFAIP3 |
| 2.0468E-05 | -0.2570451 | 0.176 | 0.253 | 0.43699303 | PCT | IDI1 |
| 2.4325E-07 | -0.2572911 | 0.13 | 0.225 | 0.00519349 | PCT | ADM |
| 7.5732E-10 | -0.2598675 | 0.168 | 0.293 | 1.6169E-05 | PCT | STMN1 |
| 2.0429E-05 | -0.2604194 | 0.311 | 0.396 | 0.43615509 | PCT | TAF1D |
| 3.2679E-07 | -0.2608923 | 0.108 | 0.198 | 0.006977 | PCT | BCAM |
| 5.8891E-05 | -0.2609177 | 0.09 | 0.15 | 1 | PCT | AL118516.1 |
| 2.0502E-07 | -0.2619265 | 0.146 | 0.245 | 0.00437709 | PCT | HES4 |
| 8.1266E-08 | -0.2621355 | 0.198 | 0.303 | 0.00173502 | PCT | CAPG |
| 6.1808E-11 | -0.2622167 | 0.156 | 0.292 | 1.3196E-06 | PCT | ANXA3 |
| 1.8719E-09 | -0.2623451 | 0.13 | 0.241 | 3.9964E-05 | PCT | AKAP13 |
| 9.9972E-10 | -0.2636192 | 0.048 | 0.142 | 2.1344E-05 | PCT | EFNB2 |
| 1.2629E-09 | -0.2647452 | 0.098 | 0.207 | 2.6963E-05 | PCT | CLK1 |
| 4.393E-08 | -0.265104 | 0.128 | 0.231 | 0.00093791 | PCT | IFITM2 |
| 3.6068E-05 | -0.2660777 | 0.248 | 0.325 | 0.7700558 | PCT | RSRC2 |
| 9.704E-06 | -0.2662648 | 0.299 | 0.38 | 0.20717984 | PCT | SRSF5 |
| 9.0644E-06 | -0.2673545 | 0.044 | 0.104 | 0.19352589 | PCT | TNF |
| 5.9643E-09 | -0.2682183 | 0.164 | 0.277 | 0.00012734 | PCT | SMG1 |
| 4.9061E-08 | -0.2684803 | 0.218 | 0.33 | 0.00104745 | PCT | HNRNPH1 |
| 2.0052E-11 | -0.2701839 | 0.056 | 0.168 | 4.281E-07 | PCT | ADGRF1 |
| 2.0814E-11 | -0.2705605 | 0.052 | 0.161 | 4.4438E-07 | PCT | RNPC3 |
| 8.5246E-10 | -0.2708059 | 0.05 | 0.147 | 1.82E-05 | PCT | ID3 |
| 0.00126287 | -0.2722958 | 0.084 | 0.131 | 1 | PCT | TNFRSF11B |
| 7.9211E-07 | -0.2727668 | 0.058 | 0.13 | 0.01691165 | PCT | HOMER1 |
| 1.772E-11 | -0.2730619 | 0.042 | 0.148 | 3.7831E-07 | PCT | ID1 |
| 1.2469E-10 | -0.2752676 | 0.126 | 0.247 | 2.6621E-06 | PCT | FNBP4 |
| 2.7256E-10 | -0.2753462 | 0.15 | 0.274 | 5.8192E-06 | PCT | TPR |
| 7.3356E-05 | -0.2754943 | 0.269 | 0.34 | 1 | PCT | HNRNPAB |
| 2.3086E-11 | -0.2783455 | 0.224 | 0.363 | 4.9288E-07 | PCT | CD46 |
| 4.5453E-10 | -0.2786386 | 0.146 | 0.266 | 9.7042E-06 | PCT | VCL |
| 1.5172E-08 | -0.27896 | 0.136 | 0.24 | 0.00032392 | PCT | CHD9 |
| 2.9758E-10 | -0.279894 | 0.263 | 0.397 | 6.3532E-06 | PCT | CAPN2 |
| 4.8693E-09 | -0.2799449 | 0.251 | 0.378 | 0.00010396 | PCT | FLNA |
| 1.9114E-05 | -0.2800592 | 0.844 | 0.9 | 0.40808111 | PCT | MT-CYB |
| 6.4317E-10 | -0.2805206 | 0.078 | 0.183 | 1.3732E-05 | PCT | MYO9A |
| 2.0822E-05 | -0.2809108 | 0.253 | 0.337 | 0.44455257 | PCT | RALBP1 |

| | | | | | | |
|---|---|---|---|---|---|---|
| 1.8203E-08 | -0.2821343 | 0.391 | 0.504 | 0.00038864 | PCT | HNRNPDL |
| 1.9472E-07 | -0.2853541 | 0.236 | 0.337 | 0.00415731 | PCT | ANKRD12 |
| 6.0545E-10 | -0.2859037 | 0.042 | 0.137 | 1.2926E-05 | PCT | CSRNP2 |
| 1.8852E-12 | -0.286498 | 0.098 | 0.23 | 4.0249E-08 | PCT | ZNF91 |
| 3.8583E-09 | -0.2867122 | 0.19 | 0.305 | 8.2375E-05 | PCT | GPBP1 |
| 7.7043E-12 | -0.2881444 | 0.076 | 0.195 | 1.6449E-07 | PCT | ZNF83 |
| 0.00299998 | -0.2882832 | 0.557 | 0.62 | 1 | PCT | MTRNR2L8 |
| 8.7627E-08 | -0.288298 | 0.188 | 0.295 | 0.00187084 | PCT | MAFF |
| 2.7328E-06 | -0.2884297 | 0.212 | 0.301 | 0.05834484 | PCT | CYTOR |
| 6.1393E-09 | -0.2884654 | 0.03 | 0.11 | 0.00013107 | PCT | AL355075.4 |
| 1.1363E-06 | -0.2890243 | 0.04 | 0.107 | 0.02426025 | PCT | ARC |
| 2.5635E-12 | -0.2897434 | 0.122 | 0.257 | 5.4731E-08 | PCT | CD47 |
| 4.4057E-12 | -0.28982 | 0.122 | 0.256 | 9.4061E-08 | PCT | JMJD1C |
| 1.7597E-13 | -0.2918574 | 0.074 | 0.208 | 3.7569E-09 | PCT | ZFC3H1 |
| 1.5761E-10 | -0.2922029 | 0.024 | 0.115 | 3.365E-06 | PCT | HIST4H4 |
| 7.1256E-09 | -0.2940385 | 0.092 | 0.194 | 0.00015213 | PCT | THBS1 |
| 4.5183E-09 | -0.296136 | 0.078 | 0.176 | 9.6466E-05 | PCT | THSD7A |
| 0.00091307 | -0.2974754 | 0.068 | 0.113 | 1 | PCT | C15orf48 |
| 7.637E-09 | -0.3005653 | 0.265 | 0.389 | 0.00016305 | PCT | MYH9 |
| 8.4435E-11 | -0.3017824 | 0.054 | 0.16 | 1.8027E-06 | PCT | CDKN2B |
| 0.00023348 | -0.3019357 | 0.447 | 0.513 | 1 | PCT | CDKN1A |
| 3.2883E-11 | -0.3026178 | 0.453 | 0.588 | 7.0205E-07 | PCT | APP |
| 7.2359E-08 | -0.3052096 | 0.092 | 0.181 | 0.00154487 | PCT | KLF4 |
| 1.5334E-12 | -0.3108644 | 0.046 | 0.161 | 3.2739E-08 | PCT | PLEKHH2 |
| 0.00010628 | -0.3112505 | 0.134 | 0.194 | 1 | PCT | H2AFX |
| 3.0614E-08 | -0.3119317 | 0.036 | 0.114 | 0.0006536 | PCT | AC023157.3 |
| 0.0001295 | -0.3136219 | 0.132 | 0.195 | 1 | PCT | GADD45G |
| 1.0745E-10 | -0.3148543 | 0.16 | 0.289 | 2.294E-06 | PCT | COL18A1 |
| 4.1967E-12 | -0.316737 | 0.088 | 0.212 | 8.9599E-08 | PCT | VPS13C |
| 5.5887E-10 | -0.3172634 | 0.066 | 0.169 | 1.1932E-05 | PCT | HIST1H1E |
| 7.9878E-11 | -0.3180103 | 0.144 | 0.271 | 1.7054E-06 | PCT | GSTM3 |
| 1.4191E-09 | -0.318648 | 0.24 | 0.363 | 3.0297E-05 | PCT | GCC2 |
| 8.3119E-06 | -0.3210553 | 0.214 | 0.3 | 0.17745802 | PCT | HSPH1 |
| 1.6932E-09 | -0.3223987 | 0.078 | 0.179 | 3.615E-05 | PCT | FOXC1 |
| 8.9987E-12 | -0.3244726 | 0.206 | 0.346 | 1.9212E-07 | PCT | JUP |
| 1.5809E-07 | -0.3248864 | 0.082 | 0.169 | 0.00337513 | PCT | CTGF |
| 1.7885E-07 | -0.3255244 | 0.162 | 0.263 | 0.0038184 | PCT | LAMB3 |
| 1.4737E-11 | -0.3261728 | 0.146 | 0.279 | 3.1464E-07 | PCT | SPTAN1 |
| 7.7466E-07 | -0.3275851 | 0.419 | 0.522 | 0.01653896 | PCT | SON |
| 4.3509E-11 | -0.3295356 | 0.218 | 0.365 | 9.2891E-07 | PCT | CD44 |
| 1.5429E-08 | -0.3316467 | 0.026 | 0.103 | 0.00032942 | PCT | AP001160.1 |
| 6.254E-11 | -0.3319395 | 0.13 | 0.251 | 1.3352E-06 | PCT | IRS2 |
| 1.9352E-11 | -0.3324727 | 0.188 | 0.322 | 4.1316E-07 | PCT | BOD1L1 |
| 0.00104449 | -0.3330936 | 0.251 | 0.304 | 1 | PCT | DNAJB1 |
| 1.3736E-13 | -0.3337878 | 0.134 | 0.282 | 2.9326E-09 | PCT | ATRX |
| 2.183E-11 | -0.3344603 | 0.03 | 0.13 | 4.6608E-07 | PCT | PCDH9 |
| 3.6607E-08 | -0.3349582 | 0.156 | 0.256 | 0.00078156 | PCT | ID2 |
| 5.3232E-06 | -0.3353255 | 0.279 | 0.362 | 0.11365039 | PCT | MCL1 |
| 1.654E-10 | -0.3357293 | 0.054 | 0.156 | 3.5312E-06 | PCT | PFKFB3 |
| 3.708E-08 | -0.3363641 | 0.202 | 0.308 | 0.00079167 | PCT | ARHGAP29 |

| | | | | | | |
|---|---|---|---|---|---|---|
| 1.9216E-07 | -0.3365279 | 0.439 | 0.522 | 0.00410262 | PCT | HNRNPU |
| 4.8563E-10 | -0.3374985 | 0.499 | 0.62 | 1.0368E-05 | PCT | CD81 |
| 1.2088E-09 | -0.3399059 | 0.174 | 0.29 | 2.5807E-05 | PCT | NUFIP2 |
| 2.4767E-08 | -0.3420624 | 0.082 | 0.178 | 0.00052877 | PCT | IGFBP3 |
| 2.3564E-12 | -0.34296 | 0.146 | 0.281 | 5.0308E-08 | PCT | BPTF |
| 2.4602E-14 | -0.3435543 | 0.096 | 0.239 | 5.2525E-10 | PCT | PKN2 |
| 3.7049E-11 | -0.3438681 | 0.301 | 0.441 | 7.91E-07 | PCT | PRRC2C |
| 1.126E-15 | -0.345165 | 0.092 | 0.246 | 2.404E-11 | PCT | KLF5 |
| 5.5526E-13 | -0.3452972 | 0.13 | 0.272 | 1.1855E-08 | PCT | BDP1 |
| 4.0918E-09 | -0.3469727 | 0.866 | 0.913 | 8.7359E-05 | PCT | MT-CO2 |
| 4.8858E-05 | -0.3470802 | 0.18 | 0.256 | 1 | PCT | AKAP12 |
| 2.4729E-11 | -0.3472579 | 0.259 | 0.39 | 5.2796E-07 | PCT | MYO6 |
| 1.1326E-14 | -0.3519334 | 0.12 | 0.271 | 2.4181E-10 | PCT | LIMA1 |
| 1.2544E-06 | -0.3535929 | 0.238 | 0.335 | 0.0267812 | PCT | H1F0 |
| 5.695E-06 | -0.3549531 | 0.279 | 0.359 | 0.12158893 | PCT | DDX3X |
| 0.00016254 | -0.3550751 | 0.206 | 0.264 | 1 | PCT | HIST1H2AC |
| 3.0555E-07 | -0.3570139 | 0.888 | 0.928 | 0.00652348 | PCT | ACTG1 |
| 9.7898E-09 | -0.3618554 | 0.291 | 0.403 | 0.00020901 | PCT | JPT1 |
| 2.0872E-07 | -0.362252 | 0.186 | 0.28 | 0.00445621 | PCT | VEGFA |
| 1.7423E-06 | -0.3634121 | 0.327 | 0.421 | 0.03719752 | PCT | CCND1 |
| 1.0944E-11 | -0.3649328 | 0.176 | 0.321 | 2.3366E-07 | PCT | MAL |
| 8.272E-15 | -0.3671271 | 0.383 | 0.557 | 1.7661E-10 | PCT | CD9 |
| 5.4446E-11 | -0.3671316 | 0.154 | 0.279 | 1.1624E-06 | PCT | IFRD1 |
| 5.0457E-10 | -0.3673849 | 0.315 | 0.438 | 1.0773E-05 | PCT | ZBTB20 |
| 2.1583E-13 | -0.3684398 | 0.02 | 0.13 | 4.6081E-09 | PCT | HIST2H2BF |
| 1.7844E-09 | -0.3691316 | 0.365 | 0.485 | 3.8097E-05 | PCT | H1FX |
| 2.9987E-06 | -0.3695212 | 0.259 | 0.341 | 0.06402152 | PCT | PAX8 |
| 1.2764E-09 | -0.3695703 | 0.074 | 0.175 | 2.7251E-05 | PCT | FOSB |
| 2.011E-13 | -0.3697879 | 0.06 | 0.187 | 4.2935E-09 | PCT | ERN1 |
| 0.00208916 | -0.3703142 | 0.697 | 0.728 | 1 | PCT | KRT8 |
| 2.3059E-16 | -0.3714256 | 0.086 | 0.241 | 4.9231E-12 | PCT | SCIN |
| 4.4142E-07 | -0.3718867 | 0.226 | 0.316 | 0.00942422 | PCT | SRSF2 |
| 4.5239E-16 | -0.3723197 | 0.399 | 0.566 | 9.6585E-12 | PCT | KTN1 |
| 1.1576E-13 | -0.3727551 | 0.09 | 0.229 | 2.4714E-09 | PCT | GPRC5A |
| 1.4527E-11 | -0.3730484 | 0.124 | 0.254 | 3.1015E-07 | PCT | MACC1 |
| 4.0599E-12 | -0.3741172 | 0.154 | 0.287 | 8.6679E-08 | PCT | PRPF4B |
| 4.5289E-06 | -0.3752559 | 0.301 | 0.389 | 0.09669284 | PCT | BRD2 |
| 1.4722E-06 | -0.3777746 | 0.244 | 0.33 | 0.03143074 | PCT | CEBPB |
| 1.1249E-11 | -0.3779256 | 0.046 | 0.154 | 2.4016E-07 | PCT | AC007952.4 |
| 1.4398E-12 | -0.3785212 | 0.467 | 0.597 | 3.074E-08 | PCT | RBM39 |
| 3.8085E-12 | -0.3818496 | 0.122 | 0.253 | 8.1311E-08 | PCT | RAP2B |
| 1.1999E-12 | -0.3831522 | 0.03 | 0.139 | 2.5617E-08 | PCT | AC025164.1 |
| 0.00020073 | -0.3832347 | 0.83 | 0.875 | 1 | PCT | GSTP1 |
| 6.0445E-13 | -0.383448 | 0.144 | 0.284 | 1.2905E-08 | PCT | COBLL1 |
| 7.1251E-08 | -0.3840201 | 0.301 | 0.402 | 0.00152121 | PCT | ANKRD11 |
| 0.00040035 | -0.3857007 | 0.066 | 0.117 | 1 | PCT | TAGLN |
| 0.00021825 | -0.386803 | 0.772 | 0.816 | 1 | PCT | IGFBP7 |
| 2.3709E-09 | -0.3869237 | 0.18 | 0.3 | 5.0618E-05 | PCT | DST |
| 1.7449E-12 | -0.3876579 | 0.273 | 0.43 | 3.7253E-08 | PCT | AHNAK |
| 2.415E-06 | -0.38843 | 0.261 | 0.353 | 0.05155955 | PCT | TXNIP |

| | | | | | | |
|---|---|---|---|---|---|---|
| 0.00041669 | -0.3932181 | 0.07 | 0.119 | 1 | PCT | NEFM |
| 7.2632E-11 | -0.3978754 | 0.303 | 0.426 | 1.5507E-06 | PCT | SFPQ |
| 3.1997E-13 | -0.4010049 | 0.567 | 0.684 | 6.8314E-09 | PCT | EPCAM |
| 6.811E-14 | -0.4049725 | 0.19 | 0.342 | 1.4541E-09 | PCT | LUC7L3 |
| 1.2824E-16 | -0.4071706 | 0.048 | 0.194 | 2.738E-12 | PCT | SYTL2 |
| 4.0116E-17 | -0.4095945 | 0.128 | 0.297 | 8.5648E-13 | PCT | KMT2A |
| 9.0756E-09 | -0.4107048 | 0.595 | 0.671 | 0.00019376 | PCT | GLS |
| 3.4781E-10 | -0.4110664 | 0.202 | 0.325 | 7.4257E-06 | PCT | CCNL1 |
| 8.5495E-14 | -0.4122003 | 0.228 | 0.383 | 1.8253E-09 | PCT | PNISR |
| 2.5504E-11 | -0.4148316 | 0.094 | 0.213 | 5.4451E-07 | PCT | ADAMTS9 |
| 5.666E-14 | -0.4201165 | 0.236 | 0.389 | 1.2097E-09 | PCT | N4BP2L2 |
| 2.1973E-12 | -0.4208997 | 0.074 | 0.195 | 4.6913E-08 | PCT | TLE4 |
| 2.9758E-18 | -0.423896 | 0.144 | 0.321 | 6.3533E-14 | PCT | AFDN |
| 3.1273E-22 | -0.4249784 | 0.106 | 0.307 | 6.6768E-18 | PCT | CDH1 |
| 3.4377E-08 | -0.4266327 | 0.146 | 0.246 | 0.00073394 | PCT | ID4 |
| 7.0163E-14 | -0.4298296 | 0.216 | 0.374 | 1.498E-09 | PCT | MACF1 |
| 5.4086E-07 | -0.4298574 | 0.289 | 0.395 | 0.01154744 | PCT | NFKBIA |
| 1.4486E-08 | -0.4325501 | 0.188 | 0.299 | 0.00030927 | PCT | ITGB8 |
| 5.8009E-11 | -0.4335146 | 0.118 | 0.237 | 1.2385E-06 | PCT | SYNE1 |
| 1.4573E-13 | -0.4351486 | 0.607 | 0.71 | 3.1114E-09 | PCT | DDX5 |
| 2.5816E-09 | -0.4381933 | 0.178 | 0.293 | 5.5117E-05 | PCT | CYR61 |
| 6.5181E-14 | -0.4426789 | 0.265 | 0.419 | 1.3916E-09 | PCT | GOLGB1 |
| 4.366E-15 | -0.4438493 | 0.164 | 0.326 | 9.3214E-11 | PCT | DCDC2 |
| 1.1777E-16 | -0.4451919 | 0.15 | 0.318 | 2.5143E-12 | PCT | WSB1 |
| 1.6622E-16 | -0.4452938 | 0.086 | 0.239 | 3.5489E-12 | PCT | CDKN2AIP |
| 4.0405E-20 | -0.4488346 | 0.315 | 0.512 | 8.6265E-16 | PCT | CLDN7 |
| 8.4556E-22 | -0.4521143 | 0.184 | 0.393 | 1.8053E-17 | PCT | PERP |
| 1.505E-14 | -0.4564733 | 0.218 | 0.372 | 3.2132E-10 | PCT | PNN |
| 7.9218E-10 | -0.4588649 | 0.15 | 0.268 | 1.6913E-05 | PCT | DUSP1 |
| 1.088E-12 | -0.4624778 | 0.2 | 0.34 | 2.3229E-08 | PCT | ZNF292 |
| 3.9156E-14 | -0.4652 | 0.257 | 0.407 | 8.3598E-10 | PCT | FUS |
| 4.7319E-09 | -0.4704276 | 0.184 | 0.288 | 0.00010103 | PCT | POLR2A |
| 8.3488E-08 | -0.4707811 | 0.206 | 0.309 | 0.00178246 | PCT | HSPA1B |
| 6.6021E-09 | -0.4724049 | 0.345 | 0.448 | 0.00014096 | PCT | ZFP36L1 |
| 1.808E-12 | -0.4733919 | 0.026 | 0.132 | 3.8601E-08 | PCT | COL7A1 |
| 1.1773E-10 | -0.4742658 | 0.096 | 0.207 | 2.5135E-06 | PCT | SGK1 |
| 9.9343E-19 | -0.475163 | 0.687 | 0.8 | 2.121E-14 | PCT | HNRNPA2B1 |
| 5.3605E-12 | -0.4759051 | 0.154 | 0.291 | 1.1445E-07 | PCT | PLAUR |
| 6.6882E-22 | -0.4766375 | 0.084 | 0.281 | 1.4279E-17 | PCT | MUC1 |
| 2.9831E-18 | -0.4852025 | 0.395 | 0.572 | 6.369E-14 | PCT | MAL2 |
| 1.5662E-14 | -0.487091 | 0.098 | 0.239 | 3.3439E-10 | PCT | NFKBIZ |
| 2.0481E-10 | -0.4918922 | 0.166 | 0.28 | 4.3728E-06 | PCT | CITED2 |
| 2.8305E-19 | -0.4961249 | 0.184 | 0.369 | 6.0431E-15 | PCT | AKAP9 |
| 7.2296E-12 | -0.4962848 | 0.06 | 0.173 | 1.5435E-07 | PCT | EGLN3 |
| 1.2898E-11 | -0.5040657 | 0.511 | 0.614 | 2.7537E-07 | PCT | BTG1 |
| 3.3466E-15 | -0.5041314 | 0.094 | 0.243 | 7.1451E-11 | PCT | SLC5A3 |
| 1.0101E-20 | -0.5077089 | 0.066 | 0.246 | 2.1565E-16 | PCT | ITGB6 |
| 4.0095E-15 | -0.5079079 | 0.21 | 0.364 | 8.5602E-11 | PCT | ARGLU1 |
| 2.0773E-11 | -0.5083375 | 0.176 | 0.306 | 4.435E-07 | PCT | DDIT4 |
| 2.3265E-18 | -0.5093949 | 0.537 | 0.711 | 4.9671E-14 | PCT | ITGB1 |

| | | | | | | |
|---|---|---|---|---|---|---|
| 1.5168E-07 | -0.5105897 | 0.086 | 0.175 | 0.00323846 | PCT | S100A4 |
| 1.4782E-10 | -0.5113432 | 0.293 | 0.418 | 3.156E-06 | PCT | PHLDA2 |
| 4.3873E-10 | -0.5182954 | 0.154 | 0.271 | 9.3669E-06 | PCT | TUBA1A |
| 1.4242E-13 | -0.5197975 | 0.86 | 0.929 | 3.0406E-09 | PCT | MT-CO1 |
| 1.6893E-18 | -0.5219218 | 0.128 | 0.306 | 3.6066E-14 | PCT | MARCKSL1 |
| 2.1271E-18 | -0.5244367 | 0.253 | 0.441 | 4.5414E-14 | PCT | DDX17 |
| 1.2706E-06 | -0.5283404 | 0.764 | 0.834 | 0.02712821 | PCT | ANXA2 |
| 4.8967E-12 | -0.5293559 | 0.008 | 0.101 | 1.0455E-07 | PCT | SLC12A1 |
| 3.8515E-13 | -0.5308035 | 0.09 | 0.219 | 8.2229E-09 | PCT | SOCS1 |
| 5.0184E-11 | -0.547148 | 0.18 | 0.306 | 1.0714E-06 | PCT | PLK2 |
| 2.3292E-17 | -0.549038 | 0.106 | 0.273 | 4.9729E-13 | PCT | KCNQ1OT1 |
| 8.5237E-21 | -0.5538445 | 0.188 | 0.385 | 1.8198E-16 | PCT | RSRP1 |
| 4.1243E-06 | -0.5564528 | 0.864 | 0.921 | 0.08805442 | PCT | S100A11 |
| 4.6328E-14 | -0.5615666 | 0.192 | 0.347 | 9.8911E-10 | PCT | SNHG12 |
| 6.862E-11 | -0.56418 | 0.533 | 0.634 | 1.465E-06 | PCT | TPM1 |
| 2.4554E-12 | -0.5644249 | 0.182 | 0.317 | 5.2422E-08 | PCT | CDKN1C |
| 1.1394E-15 | -0.5708938 | 0.106 | 0.259 | 2.4326E-11 | PCT | IER5 |
| 6.5802E-10 | -0.5715076 | 0.124 | 0.24 | 1.4049E-05 | PCT | CST6 |
| 1.694E-10 | -0.5788708 | 0.104 | 0.215 | 3.6167E-06 | PCT | ADAMTS1 |
| 4.7803E-18 | -0.589633 | 0.427 | 0.567 | 1.0206E-13 | PCT | SRRM2 |
| 1.7595E-21 | -0.5923728 | 0.196 | 0.398 | 3.7566E-17 | PCT | RAB11FIP1 |
| 1.2146E-17 | -0.6042784 | 0.096 | 0.261 | 2.5932E-13 | PCT | IER5L |
| 8.4044E-08 | -0.6053531 | 0.24 | 0.335 | 0.00179434 | PCT | RHOB |
| 4.1599E-19 | -0.6076489 | 0.034 | 0.191 | 8.8815E-15 | PCT | PIK3R3 |
| 8.9595E-18 | -0.6138051 | 0.116 | 0.287 | 1.9128E-13 | PCT | BTG2 |
| 9.9696E-08 | -0.6141642 | 0.305 | 0.41 | 0.0021285 | PCT | RNF213 |
| 1.3803E-10 | -0.614471 | 0.164 | 0.281 | 2.9469E-06 | PCT | PPP1R10 |
| 5.454E-16 | -0.636728 | 0.094 | 0.243 | 1.1644E-11 | PCT | INTS6 |
| 4.1815E-16 | -0.6407905 | 0.19 | 0.349 | 8.9275E-12 | PCT | ZFP36L2 |
| 4.5385E-14 | -0.644941 | 0.12 | 0.266 | 9.6897E-10 | PCT | EGR1 |
| 4.6809E-20 | -0.6628085 | 0.465 | 0.63 | 9.9938E-16 | PCT | PPP1R15A |
| 1.1063E-13 | -0.6687059 | 0.176 | 0.317 | 2.362E-09 | PCT | GLUL |
| 1.493E-22 | -0.6763607 | 0.116 | 0.313 | 3.1875E-18 | PCT | PMAIP1 |
| 3.7205E-22 | -0.6855856 | 0.389 | 0.6 | 7.9433E-18 | PCT | KRT19 |
| 1.2515E-22 | -0.6957409 | 0.226 | 0.429 | 2.672E-18 | PCT | TPM4 |
| 7.1421E-29 | -0.701253 | 0.499 | 0.71 | 1.5248E-24 | PCT | CLDN4 |
| 8.9778E-09 | -0.7024909 | 0.03 | 0.109 | 0.00019168 | PCT | FXYD4 |
| 1.584E-10 | -0.7031657 | 0.505 | 0.604 | 3.3819E-06 | PCT | CLU |
| 3.007E-13 | -0.7060112 | 0.17 | 0.313 | 6.4199E-09 | PCT | PLAU |
| 9.6563E-32 | -0.7061868 | 0.431 | 0.672 | 2.0616E-27 | PCT | TACSTD2 |
| 4.968E-17 | -0.7075879 | 0.212 | 0.381 | 1.0607E-12 | PCT | SLC38A2 |
| 1.2972E-15 | -0.7173408 | 0.136 | 0.296 | 2.7696E-11 | PCT | FOS |
| 5.9567E-17 | -0.7180988 | 0.22 | 0.393 | 1.2718E-12 | PCT | JUNB |
| 8.8114E-14 | -0.7201807 | 0.543 | 0.643 | 1.8812E-09 | PCT | JUND |
| 9.2648E-22 | -0.727043 | 0.238 | 0.436 | 1.978E-17 | PCT | SYNE2 |
| 2.8005E-19 | -0.7286025 | 0.14 | 0.326 | 5.9791E-15 | PCT | ITGA2 |
| 4.2967E-15 | -0.7390762 | 0.846 | 0.859 | 9.1735E-11 | PCT | H3F3B |
| 4.4705E-22 | -0.7456032 | 0.104 | 0.296 | 9.5445E-18 | PCT | RND3 |
| 2.9163E-10 | -0.7487378 | 0.128 | 0.247 | 6.2262E-06 | PCT | TFPI2 |
| 3.7618E-21 | -0.769436 | 0.232 | 0.422 | 8.0313E-17 | PCT | SRSF7 |

| | | | | | | |
|---|---|---|---|---|---|---|
| 1.2858E-20 | -0.7987742 | 0.104 | 0.284 | 2.7452E-16 | PCT | RASD1 |
| 6.2691E-06 | -0.8049799 | 0.174 | 0.259 | 0.13384619 | PCT | LCN2 |
| 2.4386E-30 | -0.8072683 | 0.425 | 0.658 | 5.2064E-26 | PCT | KRT7 |
| 4.495E-16 | -0.820588 | 0.487 | 0.61 | 9.5968E-12 | PCT | GADD45B |
| 1.2185E-16 | -0.8293204 | 0.168 | 0.337 | 2.6016E-12 | PCT | LAMC2 |
| 1.6203E-18 | -0.843072 | 0.078 | 0.24 | 3.4593E-14 | PCT | CSKMT |
| 3.9794E-11 | -0.8768414 | 0.261 | 0.382 | 8.4961E-07 | PCT | HIST1H4C |
| 1.146E-28 | -0.8853924 | 0.77 | 0.875 | 2.4466E-24 | PCT | SAT1 |
| 2.6685E-09 | -0.9021125 | 0.271 | 0.396 | 5.6972E-05 | PCT | SLPI |
| 6.4962E-24 | -0.9261063 | 0.387 | 0.571 | 1.3869E-19 | PCT | IER2 |
| 6.279E-14 | -0.9306492 | 0.168 | 0.302 | 1.3406E-09 | PCT | HEXIM1 |
| 3.8811E-21 | -0.936266 | 0.104 | 0.284 | 8.2862E-17 | PCT | HES1 |
| 7.5237E-33 | -0.9387031 | 0.677 | 0.82 | 1.6063E-28 | PCT | NEAT1 |
| 1.3739E-06 | -0.9607172 | 0.142 | 0.229 | 0.02933249 | PCT | DEFB1 |
| 7.0369E-35 | -0.9866723 | 0.661 | 0.877 | 1.5024E-30 | PCT | TMSB4X |
| 2.1431E-26 | -0.9907814 | 0.068 | 0.279 | 4.5755E-22 | PCT | TCIM |
| 1.283E-39 | -1.0198434 | 0.764 | 0.962 | 2.7393E-35 | PCT | S100A6 |
| 1.6583E-18 | -1.0432031 | 0.279 | 0.446 | 3.5405E-14 | PCT | IER3 |
| 5.1374E-21 | -1.0720184 | 0.497 | 0.639 | 1.0968E-16 | PCT | JUN |
| 6.6713E-24 | -1.0826048 | 0.082 | 0.278 | 1.4243E-19 | PCT | HSPA2 |
| 8.2363E-14 | -1.1087583 | 0.575 | 0.683 | 1.7584E-09 | PCT | MTRNR2L12 |
| 4.159E-31 | -1.1127827 | 0.397 | 0.594 | 8.8794E-27 | PCT | ELF3 |
| 1.2152E-60 | -1.1517138 | 0.794 | 0.946 | 2.5945E-56 | PCT | MALAT1 |
| 5.608E-37 | -1.2957828 | 0.403 | 0.655 | 1.1973E-32 | PCT | WFDC2 |
| 2.7903E-28 | -1.3266893 | 0.074 | 0.291 | 5.9573E-24 | PCT | MAFB |
| 1.9415E-26 | -1.4074144 | 0.361 | 0.547 | 4.1451E-22 | PCT | SOX4 |
| 7.266E-26 | -1.7266574 | 0.287 | 0.5 | 1.5513E-21 | PCT | S100A2 |
| 2.7637E-29 | -1.8162861 | 0.421 | 0.645 | 5.9005E-25 | PCT | MMP7 |
| 7.859E-16 | -2.8985042 | 0.543 | 0.6 | 1.6779E-11 | PCT | PLCG2 |
| 0 | 4.41825874 | 0.895 | 0.02 | 0 | PDC | PTGDS |
| 0 | 4.1591045 | 0.949 | 0.028 | 0 | PDC | PODXL |
| 0 | 3.34249266 | 0.899 | 0.032 | 0 | PDC | MME |
| 0 | 3.20019135 | 0.768 | 0.018 | 0 | PDC | CHI3L1 |
| 0 | 3.12911428 | 0.895 | 0.028 | 0 | PDC | HTRA1 |
| 0 | 3.07156063 | 0.802 | 0.008 | 0 | PDC | NPHS2 |
| 0 | 2.88117309 | 0.785 | 0.013 | 0 | PDC | DCN |
| 0 | 2.62089076 | 0.844 | 0.02 | 0 | PDC | CLIC5 |
| 0 | 2.5155509 | 0.848 | 0.014 | 0 | PDC | PTPRO |
| 0 | 2.48652118 | 0.624 | 0.012 | 0 | PDC | AIF1 |
| 0 | 2.48513359 | 0.831 | 0.025 | 0 | PDC | PLA2R1 |
| 0 | 2.46993188 | 0.654 | 0.011 | 0 | PDC | TPPP3 |
| 0 | 2.36284699 | 0.819 | 0.094 | 0 | PDC | BST2 |
| 0 | 2.35905862 | 0.797 | 0.015 | 0 | PDC | ENPEP |
| 1.847E-251 | 2.32915899 | 0.848 | 0.134 | 3.944E-247 | PDC | MYL9 |
| 0 | 2.19580913 | 0.7 | 0.013 | 0 | PDC | TCF21 |
| 2.5822E-62 | 2.04023332 | 0.789 | 0.321 | 5.513E-58 | PDC | TIMP1 |
| 0 | 1.96674837 | 0.793 | 0.084 | 0 | PDC | NPNT |
| 0 | 1.8791983 | 0.641 | 0.02 | 0 | PDC | NES |
| 0 | 1.86757752 | 0.785 | 0.053 | 0 | PDC | TGFBR3 |
| 1.2805E-97 | 1.85782318 | 0.949 | 0.609 | 2.7338E-93 | PDC | CD81 |

| | | | | | | |
|---|---|---|---|---|---|---|
| 3.006E-153 | 1.84242605 | 0.722 | 0.145 | 6.417E-149 | PDC | IGFBP2 |
| 0 | 1.8361236 | 0.726 | 0.019 | 0 | PDC | DACH1 |
| 3.28E-300 | 1.81869972 | 0.772 | 0.088 | 7.003E-296 | PDC | GMDS |
| 8.8546E-52 | 1.73656227 | 0.519 | 0.165 | 1.8905E-47 | PDC | S100A4 |
| 0 | 1.67991258 | 0.7 | 0.025 | 0 | PDC | FGF1 |
| 0 | 1.67518224 | 0.734 | 0.026 | 0 | PDC | SPOCK1 |
| 0 | 1.66864324 | 0.675 | 0.008 | 0 | PDC | NDNF |
| 0 | 1.64091618 | 0.675 | 0.035 | 0 | PDC | SBSPON |
| 2.338E-189 | 1.63905751 | 0.591 | 0.075 | 4.992E-185 | PDC | HPGD |
| 0 | 1.61221322 | 0.709 | 0.009 | 0 | PDC | WT1 |
| 1.8417E-85 | 1.58662462 | 0.759 | 0.267 | 3.932E-81 | PDC | VEGFA |
| 0 | 1.57568442 | 0.743 | 0.03 | 0 | PDC | SPOCK2 |
| 0 | 1.55343498 | 0.705 | 0.042 | 0 | PDC | IQGAP2 |
| 2.8424E-72 | 1.53667362 | 0.945 | 0.617 | 6.0686E-68 | PDC | VIM |
| 2.886E-117 | 1.50935219 | 0.835 | 0.279 | 6.161E-113 | PDC | TJP1 |
| 0 | 1.49965405 | 0.633 | 0.02 | 0 | PDC | GAS1 |
| 1.364E-171 | 1.48697734 | 0.81 | 0.164 | 2.911E-167 | PDC | FOXC1 |
| 1.414E-150 | 1.48337496 | 0.705 | 0.144 | 3.019E-146 | PDC | VKORC1 |
| 9.7236E-46 | 1.47577519 | 0.937 | 0.812 | 2.076E-41 | PDC | IGFBP7 |
| 0 | 1.47536497 | 0.696 | 0.041 | 0 | PDC | NTNG1 |
| 7.055E-176 | 1.47309579 | 0.722 | 0.128 | 1.506E-171 | PDC | CDC42EP3 |
| 1.507E-127 | 1.45754646 | 0.667 | 0.143 | 3.217E-123 | PDC | TSC22D3 |
| 0 | 1.45104494 | 0.713 | 0.022 | 0 | PDC | F2R |
| 0 | 1.44966118 | 0.633 | 0.013 | 0 | PDC | SPARC |
| 5.584E-73 | 1.44535603 | 0.895 | 0.459 | 1.1922E-68 | PDC | IFITM3 |
| 0 | 1.4410399 | 0.489 | 0.017 | 0 | PDC | PLTP |
| 4.7523E-87 | 1.4363439 | 0.65 | 0.186 | 1.0146E-82 | PDC | BCAM |
| 0 | 1.42499028 | 0.553 | 0.021 | 0 | PDC | APOD |
| 0 | 1.42491026 | 0.713 | 0.017 | 0 | PDC | ARHGAP28 |
| 1.777E-130 | 1.41767736 | 0.814 | 0.244 | 3.793E-126 | PDC | EIF3M |
| 8.8E-201 | 1.41442447 | 0.768 | 0.127 | 1.879E-196 | PDC | PDE4DIP |
| 1.6874E-49 | 1.40099472 | 0.979 | 0.866 | 3.6026E-45 | PDC | TMSB4X |
| 1.006E-174 | 1.39491352 | 0.624 | 0.094 | 2.147E-170 | PDC | ZDHHC6 |
| 0 | 1.37830011 | 0.654 | 0.014 | 0 | PDC | MXRA8 |
| 3.419E-163 | 1.37251262 | 0.743 | 0.15 | 7.299E-159 | PDC | PLS3 |
| 2.427E-286 | 1.37004971 | 0.7 | 0.072 | 5.181E-282 | PDC | GOLIM4 |
| 0.0005751 | 1.36185586 | 0.363 | 0.288 | 1 | PDC | FOS |
| 0 | 1.33857205 | 0.692 | 0.052 | 0 | PDC | VAMP5 |
| 0 | 1.33435288 | 0.662 | 0.051 | 0 | PDC | SULF2 |
| 0 | 1.33377258 | 0.405 | 0.001 | 0 | PDC | DDN |
| 7.623E-157 | 1.32590803 | 0.565 | 0.082 | 1.627E-152 | PDC | F3 |
| 1.3355E-75 | 1.32458017 | 0.996 | 0.875 | 2.8514E-71 | PDC | ITM2B |
| 3.41E-280 | 1.32027134 | 0.764 | 0.088 | 7.281E-276 | PDC | NEBL |
| 0 | 1.29976338 | 0.662 | 0.018 | 0 | PDC | PTPRD |
| 0 | 1.28126738 | 0.679 | 0.049 | 0 | PDC | ARHGEF3 |
| 1.48E-243 | 1.26756982 | 0.713 | 0.088 | 3.159E-239 | PDC | MTSS1 |
| 4.8133E-83 | 1.23439438 | 0.806 | 0.286 | 1.0276E-78 | PDC | DST |
| 1.502E-161 | 1.22813452 | 0.751 | 0.145 | 3.207E-157 | PDC | PALLD |
| 0 | 1.2059704 | 0.646 | 0.044 | 0 | PDC | SLC9A3R2 |
| 0 | 1.19187459 | 0.329 | 0.004 | 0 | PDC | C1QL1 |

| | | | | | | |
|---|---|---|---|---|---|---|
| 1.42E-56 | 1.1795779 | 0.848 | 0.489 | 3.0316E-52 | PDC | ACTN4 |
| 4.378E-137 | 1.16476178 | 0.806 | 0.19 | 9.348E-133 | PDC | CALD1 |
| 2.0833E-26 | 1.15593611 | 0.523 | 0.241 | 4.4479E-22 | PDC | MT1X |
| 1.928E-120 | 1.15298518 | 0.713 | 0.162 | 4.116E-116 | PDC | THSD7A |
| 7.278E-134 | 1.14740133 | 0.717 | 0.156 | 1.554E-129 | PDC | SELENOP |
| 3.628E-288 | 1.13893775 | 0.633 | 0.055 | 7.746E-284 | PDC | PDLIM2 |
| 0 | 1.13291806 | 0.557 | 0.006 | 0 | PDC | NPHS1 |
| 3.585E-52 | 1.1297903 | 0.692 | 0.27 | 7.654E-48 | PDC | LGALS1 |
| 4.3105E-17 | 1.12909278 | 0.477 | 0.26 | 9.2029E-13 | PDC | DUSP1 |
| 1.521E-110 | 1.12805418 | 0.785 | 0.232 | 3.248E-106 | PDC | VAMP2 |
| 0 | 1.12637477 | 0.624 | 0.039 | 0 | PDC | SEMA5A |
| 1.737E-279 | 1.1222126 | 0.591 | 0.049 | 3.709E-275 | PDC | NDRG2 |
| 2.5249E-60 | 1.11580407 | 0.57 | 0.179 | 5.3906E-56 | PDC | ADIRF |
| 0 | 1.11575282 | 0.557 | 0.004 | 0 | PDC | MRGPRF |
| 0 | 1.10270556 | 0.637 | 0.005 | 0 | PDC | SULF1 |
| 0 | 1.09873098 | 0.578 | 0.019 | 0 | PDC | PCOLCE2 |
| 0 | 1.08687088 | 0.62 | 0.013 | 0 | PDC | NECAB1 |
| 0 | 1.07819183 | 0.468 | 0.007 | 0 | PDC | RAMP1 |
| 6.4613E-76 | 1.07623574 | 0.692 | 0.218 | 1.3795E-71 | PDC | IFITM2 |
| 0 | 1.074806 | 0.667 | 0.017 | 0 | PDC | SNCA |
| 0 | 1.06626007 | 0.557 | 0.022 | 0 | PDC | TYRO3 |
| 6.1055E-44 | 1.06016727 | 0.287 | 0.063 | 1.3035E-39 | PDC | MGP |
| 1.085E-197 | 1.0591439 | 0.629 | 0.082 | 2.316E-193 | PDC | MGAT5 |
| 1.822E-209 | 1.05587743 | 0.608 | 0.072 | 3.891E-205 | PDC | KLHL42 |
| 4.654E-108 | 1.03666469 | 0.717 | 0.189 | 9.936E-104 | PDC | NFIC |
| 6.349E-108 | 1.03588112 | 0.705 | 0.179 | 1.356E-103 | PDC | BEX4 |
| 2.198E-117 | 1.03438182 | 0.591 | 0.119 | 4.693E-113 | PDC | MPP5 |
| 0 | 1.03422182 | 0.582 | 0.039 | 0 | PDC | ARMH4 |
| 2.8759E-69 | 1.02820454 | 0.333 | 0.058 | 6.14E-65 | PDC | KLF2 |
| 4.071E-146 | 1.02756392 | 0.523 | 0.073 | 8.691E-142 | PDC | PTH1R |
| 6.456E-162 | 1.01810696 | 0.515 | 0.066 | 1.378E-157 | PDC | TCEAL2 |
| 2.0925E-86 | 1.0169065 | 0.802 | 0.297 | 4.4675E-82 | PDC | EID1 |
| 6.1585E-96 | 1.01681799 | 0.679 | 0.186 | 1.3148E-91 | PDC | NUCB1 |
| 0 | 1.01647143 | 0.629 | 0.03 | 0 | PDC | PLCE1 |
| 1.63E-303 | 1.01103069 | 0.485 | 0.029 | 3.48E-299 | PDC | C1R |
| 4.6533E-97 | 0.99151091 | 0.692 | 0.19 | 9.9347E-93 | PDC | SEPT11 |
| 1.1161E-38 | 0.98911014 | 0.717 | 0.319 | 2.3829E-34 | PDC | MT1E |
| 3.5842E-74 | 0.988053 | 0.658 | 0.191 | 7.6524E-70 | PDC | ANXA1 |
| 1.251E-203 | 0.98746407 | 0.62 | 0.073 | 2.67E-199 | PDC | MAP1B |
| 0 | 0.97955747 | 0.565 | 0.03 | 0 | PDC | SYNPO |
| 2.694E-130 | 0.97624058 | 0.684 | 0.138 | 5.752E-126 | PDC | RETREG1 |
| 3.9815E-60 | 0.96655872 | 0.751 | 0.274 | 8.5006E-56 | PDC | MAFB |
| 0 | 0.95207275 | 0.586 | 0.009 | 0 | PDC | CCBE1 |
| 7.276E-256 | 0.95199711 | 0.549 | 0.046 | 1.553E-251 | PDC | LTBP4 |
| 4.462E-160 | 0.94624249 | 0.515 | 0.064 | 9.527E-156 | PDC | COL4A3 |
| 0 | 0.94422137 | 0.591 | 0.02 | 0 | PDC | WIPF3 |
| 4.0844E-54 | 0.94198605 | 0.696 | 0.253 | 8.7201E-50 | PDC | NUPR1 |
| 8.9035E-25 | 0.94147614 | 0.819 | 0.602 | 1.9009E-20 | PDC | GADD45A |
| 4.1844E-90 | 0.94127783 | 0.679 | 0.191 | 8.9336E-86 | PDC | ARHGEF12 |
| 0 | 0.93931015 | 0.591 | 0.028 | 0 | PDC | ST3GAL6 |

| | | | | | | |
|---|---|---|---|---|---|---|
| 2.2931E-75 | 0.931299 | 0.734 | 0.271 | 4.8959E-71 | PDC | ANAPC16 |
| 1.2905E-56 | 0.93022532 | 0.591 | 0.207 | 2.7552E-52 | PDC | BTG3 |
| 9.108E-180 | 0.92887215 | 0.527 | 0.06 | 1.945E-175 | PDC | MAGI2 |
| 2.949E-113 | 0.92504349 | 0.658 | 0.147 | 6.296E-109 | PDC | RRAS |
| 1.1855E-92 | 0.92405376 | 0.447 | 0.08 | 2.531E-88 | PDC | MAP3K20 |
| 4.46E-107 | 0.92186943 | 0.641 | 0.149 | 9.522E-103 | PDC | PDLIM5 |
| 2.4144E-64 | 0.91637386 | 0.637 | 0.213 | 5.1548E-60 | PDC | PCMTD1 |
| 1.4476E-80 | 0.91091007 | 0.696 | 0.211 | 3.0907E-76 | PDC | TIMP2 |
| 0 | 0.91053662 | 0.544 | 0.021 | 0 | PDC | RERG |
| 3.7625E-52 | 0.89795849 | 0.793 | 0.384 | 8.033E-48 | PDC | CIRBP |
| 8.947E-112 | 0.89413464 | 0.549 | 0.105 | 1.91E-107 | PDC | RNF145 |
| 0 | 0.891042 | 0.485 | 0.012 | 0 | PDC | FXYD1 |
| 2.7791E-91 | 0.88614567 | 0.502 | 0.103 | 5.9335E-87 | PDC | RIPOR1 |
| 2.0094E-81 | 0.86972311 | 0.675 | 0.201 | 4.2901E-77 | PDC | BEX3 |
| 4.415E-197 | 0.86966071 | 0.519 | 0.054 | 9.426E-193 | PDC | VWA1 |
| 5.756E-230 | 0.86808501 | 0.468 | 0.036 | 1.229E-225 | PDC | FAM89A |
| 1.225E-199 | 0.86537128 | 0.544 | 0.058 | 2.615E-195 | PDC | MYLK |
| 2.2714E-49 | 0.859839 | 0.903 | 0.63 | 4.8495E-45 | PDC | CCNI |
| 1.392E-108 | 0.85759092 | 0.397 | 0.054 | 2.972E-104 | PDC | IGFBP4 |
| 9.9229E-97 | 0.85558752 | 0.54 | 0.111 | 2.1185E-92 | PDC | SRGAP1 |
| 1.5359E-56 | 0.85512253 | 0.814 | 0.375 | 3.2791E-52 | PDC | PTMS |
| 5.297E-188 | 0.84682783 | 0.498 | 0.052 | 1.131E-183 | PDC | TMEM98 |
| 3.678E-143 | 0.84403672 | 0.257 | 0.017 | 7.852E-139 | PDC | IGFBP1 |
| 5.622E-182 | 0.83210761 | 0.549 | 0.065 | 1.2E-177 | PDC | PAK1 |
| 6.6625E-60 | 0.83190754 | 0.789 | 0.344 | 1.4224E-55 | PDC | NUCKS1 |
| 3.5E-68 | 0.83146613 | 0.616 | 0.191 | 7.4726E-64 | PDC | SLK |
| 9.3743E-74 | 0.83016469 | 0.57 | 0.15 | 2.0014E-69 | PDC | PMEPA1 |
| 0 | 0.82922239 | 0.502 | 0.022 | 0 | PDC | ITIH5 |
| 3.0866E-63 | 0.82805238 | 0.591 | 0.185 | 6.5898E-59 | PDC | HIPK2 |
| 0 | 0.82341501 | 0.506 | 0.019 | 0 | PDC | LRRC2 |
| 2.6185E-45 | 0.82138862 | 0.932 | 0.652 | 5.5905E-41 | PDC | PSAP |
| 1.655E-229 | 0.81720888 | 0.321 | 0.016 | 3.533E-225 | PDC | ECM1 |
| 2.8254E-50 | 0.81189335 | 0.941 | 0.744 | 6.0321E-46 | PDC | DSTN |
| 0 | 0.81078629 | 0.519 | 0.007 | 0 | PDC | ST6GALNAC3 |
| 1.557E-95 | 0.80723593 | 0.544 | 0.114 | 3.3243E-91 | PDC | SDC2 |
| 2.404E-273 | 0.80710045 | 0.549 | 0.041 | 5.132E-269 | PDC | PTGER4 |
| 1.7253E-74 | 0.80190448 | 0.62 | 0.18 | 3.6836E-70 | PDC | HMGN3 |
| 0 | 0.79772664 | 0.388 | 0.01 | 0 | PDC | BGN |
| 7.488E-139 | 0.79081508 | 0.553 | 0.083 | 1.599E-134 | PDC | LY6E |
| 1.111E-191 | 0.78847534 | 0.561 | 0.064 | 2.373E-187 | PDC | FYN |
| 4.6419E-28 | 0.784923 | 0.983 | 0.948 | 9.9105E-24 | PDC | B2M |
| 1.802E-122 | 0.78445173 | 0.536 | 0.09 | 3.846E-118 | PDC | SUN2 |
| 0 | 0.78059228 | 0.473 | 0.009 | 0 | PDC | PHYHIP |
| 5.0056E-75 | 0.77880072 | 0.477 | 0.105 | 1.0687E-70 | PDC | RARRES2 |
| 1.0542E-34 | 0.77839016 | 0.658 | 0.328 | 2.2507E-30 | PDC | PPA1 |
| 1.1194E-87 | 0.77803425 | 0.54 | 0.118 | 2.3899E-83 | PDC | CMYA5 |
| 0 | 0.77743798 | 0.43 | 0.016 | 0 | PDC | LINC00839 |
| 2.97E-248 | 0.77719644 | 0.532 | 0.044 | 6.34E-244 | PDC | SRGAP2C |
| 2.6504E-86 | 0.7761465 | 0.608 | 0.152 | 5.6586E-82 | PDC | RFC1 |
| 8.296E-106 | 0.77263774 | 0.494 | 0.088 | 1.771E-101 | PDC | DAG1 |

| | | | | | | |
|---|---|---|---|---|---|---|
| 0 | 0.76860832 | 0.477 | 0.013 | 0 | PDC | SAMD11 |
| 1.3081E-37 | 0.76352861 | 1 | 0.933 | 2.7928E-33 | PDC | PTMA |
| 0 | 0.7564299 | 0.384 | 0.001 | 0 | PDC | AC092813.2 |
| 0 | 0.74579527 | 0.392 | 0.004 | 0 | PDC | ROBO2 |
| 7.1189E-28 | 0.74534863 | 0.679 | 0.385 | 1.5199E-23 | PDC | CD151 |
| 1.5856E-74 | 0.74242758 | 0.629 | 0.182 | 3.3852E-70 | PDC | FERMT2 |
| 1.008E-165 | 0.74024043 | 0.527 | 0.065 | 2.151E-161 | PDC | RGS10 |
| 2.9899E-52 | 0.73923062 | 0.363 | 0.084 | 6.3835E-48 | PDC | IFI6 |
| 2.007E-186 | 0.73790432 | 0.481 | 0.048 | 4.286E-182 | PDC | CBLB |
| 1.848E-89 | 0.7278622 | 0.608 | 0.141 | 3.9455E-85 | PDC | TIMP3 |
| 8.8025E-50 | 0.72433764 | 0.722 | 0.302 | 1.8793E-45 | PDC | ISCU |
| 8.313E-308 | 0.72391709 | 0.447 | 0.023 | 1.775E-303 | PDC | TMEM150C |
| 0 | 0.720488 | 0.3 | 0.003 | 0 | PDC | MFAP4 |
| 0 | 0.71579109 | 0.506 | 0.022 | 0 | PDC | DPYSL3 |
| 8.9864E-46 | 0.71071225 | 0.755 | 0.355 | 1.9186E-41 | PDC | HMGN2 |
| 1.5747E-52 | 0.71001087 | 0.675 | 0.271 | 3.3621E-48 | PDC | SEPT7 |
| 9.902E-52 | 0.70957351 | 0.599 | 0.211 | 2.1141E-47 | PDC | CCPG1 |
| 0 | 0.70929058 | 0.376 | 0.003 | 0 | PDC | SEMA3G |
| 0 | 0.70905616 | 0.439 | 0.019 | 0 | PDC | NLRP1 |
| 2.4727E-41 | 0.70067626 | 0.827 | 0.464 | 5.2793E-37 | PDC | PRDX6 |
| 5.4148E-58 | 0.70030118 | 0.65 | 0.226 | 1.1561E-53 | PDC | GSN |
| 2.3746E-38 | 0.69394702 | 0.907 | 0.602 | 5.0698E-34 | PDC | APLP2 |
| 3.7026E-59 | 0.69170711 | 0.468 | 0.119 | 7.9051E-55 | PDC | GPNMB |
| 6.6171E-28 | 0.69075431 | 0.928 | 0.756 | 1.4127E-23 | PDC | CALM2 |
| 0 | 0.68801387 | 0.477 | 0.008 | 0 | PDC | TRIM54 |
| 5.6887E-61 | 0.68728893 | 0.494 | 0.134 | 1.2145E-56 | PDC | TCEAL3 |
| 0 | 0.6822959 | 0.321 | 0.004 | 0 | PDC | PHACTR3 |
| 9.8463E-41 | 0.68107441 | 0.435 | 0.144 | 2.1022E-36 | PDC | ITM2C |
| 1.873E-158 | 0.6800264 | 0.435 | 0.046 | 3.999E-154 | PDC | CDC14A |
| 1.5299E-40 | 0.67768049 | 0.819 | 0.466 | 3.2662E-36 | PDC | SUMO2 |
| 2.391E-105 | 0.67733009 | 0.316 | 0.036 | 5.104E-101 | PDC | TPM2 |
| 2.197E-114 | 0.67320995 | 0.443 | 0.064 | 4.691E-110 | PDC | ALS2CL |
| 5.45E-10 | 0.67277439 | 0.827 | 0.709 | 1.1636E-05 | PDC | MT2A |
| 7.4091E-48 | 0.66821962 | 0.608 | 0.223 | 1.5818E-43 | PDC | NFE2L1 |
| 5.1655E-43 | 0.66765991 | 0.586 | 0.231 | 1.1028E-38 | PDC | WASL |
| 1.5354E-63 | 0.66585971 | 0.523 | 0.14 | 3.278E-59 | PDC | RASSF8 |
| 1.703E-45 | 0.66501661 | 0.684 | 0.303 | 3.6359E-41 | PDC | GABARAPL2 |
| 4.044E-112 | 0.66489494 | 0.414 | 0.057 | 8.634E-108 | PDC | ST6GAL1 |
| 8.107E-44 | 0.66453039 | 0.675 | 0.289 | 1.7309E-39 | PDC | LEPROT |
| 1.9773E-12 | 0.66350278 | 0.236 | 0.102 | 4.2216E-08 | PDC | SERPINE2 |
| 6.2778E-74 | 0.66324898 | 0.523 | 0.127 | 1.3403E-69 | PDC | PSIP1 |
| 1.9731E-71 | 0.6631453 | 0.62 | 0.18 | 4.2126E-67 | PDC | ARL3 |
| 1.3276E-83 | 0.66295998 | 0.443 | 0.081 | 2.8344E-79 | PDC | SLC7A8 |
| 8.158E-262 | 0.66060375 | 0.405 | 0.023 | 1.742E-257 | PDC | C1orf21 |
| 7.6633E-23 | 0.65916641 | 0.65 | 0.344 | 1.6361E-18 | PDC | TXNIP |
| 4.266E-154 | 0.65882832 | 0.473 | 0.055 | 9.109E-150 | PDC | KANK2 |
| 9.83E-164 | 0.65165264 | 0.414 | 0.04 | 2.099E-159 | PDC | FHL1 |
| 1.0835E-49 | 0.65129958 | 0.646 | 0.244 | 2.3132E-45 | PDC | PTBP3 |
| 9.1412E-50 | 0.65068703 | 0.557 | 0.185 | 1.9516E-45 | PDC | CTSL |
| 4.196E-161 | 0.6486455 | 0.312 | 0.022 | 8.958E-157 | PDC | BASP1 |

| | | | | | | |
|---|---|---|---|---|---|---|
| 2.862E-138 | 0.6479224 | 0.485 | 0.064 | 6.111E-134 | PDC | PARD3B |
| 7.0286E-76 | 0.64705285 | 0.506 | 0.114 | 1.5006E-71 | PDC | COL4A4 |
| 1.934E-168 | 0.64686883 | 0.447 | 0.045 | 4.129E-164 | PDC | FKBP5 |
| 7.283E-255 | 0.64330343 | 0.414 | 0.024 | 1.555E-250 | PDC | FRMD3 |
| 2.478E-263 | 0.64234288 | 0.241 | 0.006 | 5.292E-259 | PDC | ADCYAP1 |
| 1.532E-148 | 0.6421256 | 0.447 | 0.051 | 3.271E-144 | PDC | CERS6 |
| 0 | 0.64026474 | 0.485 | 0.021 | 0 | PDC | MSRB3 |
| 3.516E-290 | 0.63908693 | 0.346 | 0.014 | 7.507E-286 | PDC | MIR503HG |
| 7.9988E-80 | 0.63789274 | 0.464 | 0.095 | 1.7077E-75 | PDC | P3H2 |
| 3.7832E-48 | 0.63538534 | 0.527 | 0.175 | 8.0772E-44 | PDC | RBMS1 |
| 1.1842E-42 | 0.63397044 | 0.544 | 0.202 | 2.5284E-38 | PDC | REXO2 |
| 5.737E-103 | 0.63389068 | 0.506 | 0.092 | 1.2248E-98 | PDC | PCYOX1 |
| 1.646E-143 | 0.63383836 | 0.401 | 0.042 | 3.514E-139 | PDC | FOXD1 |
| 1.108E-74 | 0.62836279 | 0.586 | 0.154 | 2.3657E-70 | PDC | ADH5 |
| 5.8807E-70 | 0.62629961 | 0.485 | 0.116 | 1.2555E-65 | PDC | PLD3 |
| 7.2993E-37 | 0.62461597 | 0.819 | 0.471 | 1.5584E-32 | PDC | MORF4L1 |
| 0 | 0.62375821 | 0.186 | 0.002 | 0 | PDC | SOST |
| 4.128E-170 | 0.62306791 | 0.451 | 0.045 | 8.814E-166 | PDC | FLRT3 |
| 1.6726E-54 | 0.61983489 | 0.523 | 0.153 | 3.571E-50 | PDC | NR2F2 |
| 2.5538E-47 | 0.61815809 | 0.637 | 0.245 | 5.4523E-43 | PDC | C4orf3 |
| 2.4814E-35 | 0.61813839 | 0.713 | 0.361 | 5.2979E-31 | PDC | SBDS |
| 4.174E-258 | 0.61592276 | 0.418 | 0.025 | 8.911E-254 | PDC | FGFR4 |
| 1.1649E-57 | 0.6149626 | 0.591 | 0.188 | 2.487E-53 | PDC | RDX |
| 1.254E-167 | 0.61483628 | 0.304 | 0.02 | 2.677E-163 | PDC | CHST2 |
| 5.4636E-46 | 0.614326 | 0.435 | 0.129 | 1.1665E-41 | PDC | ALCAM |
| 1.097E-105 | 0.61301648 | 0.422 | 0.063 | 2.342E-101 | PDC | KANK1 |
| 1.6126E-40 | 0.61141413 | 0.73 | 0.304 | 3.4428E-36 | PDC | CDKN1C |
| 1.193E-42 | 0.61084128 | 0.743 | 0.338 | 2.5472E-38 | PDC | CNBP |
| 2.8486E-43 | 0.60423473 | 0.502 | 0.164 | 6.0819E-39 | PDC | EMP3 |
| 4.3493E-26 | 0.60385226 | 0.557 | 0.271 | 9.2858E-22 | PDC | TCEA1 |
| 8.0303E-52 | 0.60131941 | 0.506 | 0.157 | 1.7145E-47 | PDC | CXADR |
| 1.187E-103 | 0.59657229 | 0.439 | 0.069 | 2.534E-99 | PDC | PLPP1 |
| 9.86E-229 | 0.59395353 | 0.401 | 0.026 | 2.105E-224 | PDC | NR2F1 |
| 1.068E-265 | 0.59391566 | 0.367 | 0.018 | 2.279E-261 | PDC | CSDC2 |
| 2.6694E-35 | 0.59199181 | 0.485 | 0.18 | 5.6991E-31 | PDC | MSI2 |
| 5.117E-160 | 0.59148707 | 0.236 | 0.012 | 1.093E-155 | PDC | CFH |
| 3.5975E-74 | 0.59059544 | 0.498 | 0.116 | 7.6807E-70 | PDC | SORT1 |
| 1.047E-99 | 0.58883737 | 0.451 | 0.074 | 2.2364E-95 | PDC | SH3BP5 |
| 1.0532E-81 | 0.58709747 | 0.414 | 0.075 | 2.2486E-77 | PDC | UBE2L6 |
| 0 | 0.58674872 | 0.295 | 0.005 | 0 | PDC | XKR4 |
| 1.094E-231 | 0.58652661 | 0.426 | 0.029 | 2.335E-227 | PDC | CORO2B |
| 4.4005E-29 | 0.58579395 | 0.857 | 0.597 | 9.3951E-25 | PDC | CAST |
| 0 | 0.58408479 | 0.371 | 0.014 | 0 | PDC | GJA1 |
| 2.9032E-16 | 0.58047862 | 0.886 | 0.721 | 6.1983E-12 | PDC | CTSD |
| 1.434E-38 | 0.57818422 | 0.629 | 0.265 | 3.0615E-34 | PDC | ARPC5 |
| 2.0295E-14 | 0.57786916 | 0.92 | 0.796 | 4.3329E-10 | PDC | HLA-C |
| 8.46E-45 | 0.57784263 | 0.519 | 0.18 | 1.8062E-40 | PDC | RHOQ |
| 3.4212E-73 | 0.57280303 | 0.46 | 0.099 | 7.3043E-69 | PDC | ADD3 |
| 2.9812E-23 | 0.57112979 | 0.519 | 0.249 | 6.3649E-19 | PDC | RPS4Y1 |
| 0 | 0.57021418 | 0.329 | 0.003 | 0 | PDC | TRPC6 |

| | | | | | | |
|---|---|---|---|---|---|---|
| 1.6603E-32 | 0.56862517 | 0.418 | 0.147 | 3.5447E-28 | PDC | NQO1 |
| 0 | 0.56792584 | 0.325 | 0.005 | 0 | PDC | CRB2 |
| 1.7755E-39 | 0.56731216 | 0.46 | 0.155 | 3.7907E-35 | PDC | RNF187 |
| 6.0237E-53 | 0.56673774 | 0.599 | 0.202 | 1.2861E-48 | PDC | TMBIM4 |
| 2.7068E-41 | 0.56650229 | 0.532 | 0.19 | 5.779E-37 | PDC | EPB41L5 |
| 1.5181E-27 | 0.5657567 | 0.489 | 0.203 | 3.2412E-23 | PDC | ARID5B |
| 0 | 0.56373596 | 0.376 | 0.015 | 0 | PDC | PARD6G |
| 0 | 0.56202995 | 0.287 | 0.006 | 0 | PDC | FAM180A |
| 7.329E-103 | 0.56042231 | 0.451 | 0.072 | 1.5647E-98 | PDC | SRGAP2 |
| 4.7133E-80 | 0.55969743 | 0.511 | 0.111 | 1.0063E-75 | PDC | C9orf3 |
| 0 | 0.55616888 | 0.228 | 0.002 | 0 | PDC | CLDN5 |
| 2.1044E-35 | 0.55555884 | 0.295 | 0.075 | 4.4929E-31 | PDC | PDK4 |
| 7.276E-227 | 0.55501153 | 0.371 | 0.022 | 1.553E-222 | PDC | PTP4A3 |
| 1.8278E-10 | 0.55484071 | 0.873 | 0.826 | 3.9024E-06 | PDC | HLA-B |
| 2.0577E-27 | 0.5539372 | 0.772 | 0.486 | 4.3932E-23 | PDC | TOMM7 |
| 0 | 0.54781057 | 0.278 | 0.004 | 0 | PDC | TCEAL6 |
| 1.4235E-47 | 0.54610914 | 0.603 | 0.21 | 3.0392E-43 | PDC | PJA2 |
| 4.3261E-20 | 0.54309051 | 0.549 | 0.281 | 9.2362E-16 | PDC | CEBPD |
| 0 | 0.54127008 | 0.388 | 0.006 | 0 | PDC | LMX1B |
| 1.7361E-31 | 0.5407851 | 0.624 | 0.297 | 3.7066E-27 | PDC | WASF2 |
| 7.241E-158 | 0.54069488 | 0.371 | 0.033 | 1.546E-153 | PDC | TSHZ1 |
| 1.0969E-25 | 0.53925934 | 0.608 | 0.322 | 2.3418E-21 | PDC | NDUFC1 |
| 3.7084E-94 | 0.53692598 | 0.35 | 0.048 | 7.9173E-90 | PDC | ST3GAL1 |
| 1.0403E-35 | 0.53621889 | 0.481 | 0.184 | 2.2211E-31 | PDC | ATRAID |
| 3.25E-192 | 0.53604922 | 0.363 | 0.025 | 6.939E-188 | PDC | TNS2 |
| 3.0621E-50 | 0.5360087 | 0.481 | 0.141 | 6.5375E-46 | PDC | ZBTB38 |
| 6.3589E-39 | 0.53591689 | 0.527 | 0.188 | 1.3576E-34 | PDC | SLC25A37 |
| 2.3268E-35 | 0.53562063 | 0.671 | 0.311 | 4.9677E-31 | PDC | SARAF |
| 1.606E-216 | 0.53530221 | 0.354 | 0.021 | 3.43E-212 | PDC | ZEB2 |
| 8.2444E-40 | 0.53407981 | 0.532 | 0.196 | 1.7602E-35 | PDC | NDUFAF3 |
| 2.2729E-33 | 0.5322583 | 0.38 | 0.124 | 4.8526E-29 | PDC | CRIP2 |
| 0 | 0.53183568 | 0.304 | 0.001 | 0 | PDC | PTPRQ |
| 9.737E-147 | 0.53155795 | 0.388 | 0.039 | 2.079E-142 | PDC | MCC |
| 2.0501E-29 | 0.53131165 | 0.523 | 0.229 | 4.3769E-25 | PDC | ITGAV |
| 0 | 0.53009205 | 0.367 | 0.007 | 0 | PDC | FOXD2 |
| 4.6367E-44 | 0.52978967 | 0.498 | 0.16 | 9.8993E-40 | PDC | DAB2 |
| 3.926E-140 | 0.52662828 | 0.397 | 0.043 | 8.381E-136 | PDC | NDN |
| 1.125E-127 | 0.52638692 | 0.422 | 0.052 | 2.402E-123 | PDC | SSBP2 |
| 1.9111E-35 | 0.52421455 | 0.641 | 0.28 | 4.0801E-31 | PDC | RABAC1 |
| 4.8135E-83 | 0.52360529 | 0.35 | 0.053 | 1.0277E-78 | PDC | BMP2 |
| 1.4505E-51 | 0.52224789 | 0.549 | 0.174 | 3.0967E-47 | PDC | STUB1 |
| 1.156E-223 | 0.5219476 | 0.287 | 0.013 | 2.469E-219 | PDC | POLA2 |
| 9.6941E-52 | 0.51945443 | 0.511 | 0.151 | 2.0697E-47 | PDC | CXXC5 |
| 6.163E-184 | 0.51932179 | 0.376 | 0.029 | 1.316E-179 | PDC | TCF7L1 |
| 5.848E-100 | 0.51832204 | 0.397 | 0.058 | 1.2486E-95 | PDC | ZNRF3 |
| 2.079E-135 | 0.51815801 | 0.359 | 0.036 | 4.439E-131 | PDC | KIRREL1 |
| 1.8347E-74 | 0.51600269 | 0.392 | 0.073 | 3.917E-70 | PDC | VTI1A |
| 1.107E-124 | 0.51574107 | 0.35 | 0.037 | 2.363E-120 | PDC | SRGAP2B |
| 2.2873E-32 | 0.51549631 | 0.553 | 0.232 | 4.8834E-28 | PDC | CRIM1 |
| 1.9998E-23 | 0.51450401 | 0.789 | 0.523 | 4.2697E-19 | PDC | ELOB |

| | | | | | | |
|---|---|---|---|---|---|---|
| 1.0129E-25 | 0.51356979 | 0.903 | 0.732 | 2.1626E-21 | PDC | SKP1 |
| 1.3671E-06 | 0.51334017 | 0.316 | 0.195 | 0.02918733 | PDC | MT1G |
| 7.6409E-28 | 0.51331238 | 0.734 | 0.41 | 1.6313E-23 | PDC | PARK7 |
| 2.3484E-30 | 0.51258366 | 0.709 | 0.353 | 5.0139E-26 | PDC | SEC62 |
| 7.866E-207 | 0.51227939 | 0.316 | 0.017 | 1.679E-202 | PDC | NPR1 |
| 8.0151E-50 | 0.51154923 | 0.519 | 0.161 | 1.7112E-45 | PDC | ARMCX3 |
| 9.975E-149 | 0.51150382 | 0.384 | 0.037 | 2.13E-144 | PDC | DBP |
| 2.504E-164 | 0.51080115 | 0.376 | 0.032 | 5.347E-160 | PDC | CA11 |
| 8.294E-31 | 0.51074051 | 0.62 | 0.283 | 1.7708E-26 | PDC | SERINC1 |
| 6.6608E-39 | 0.50957734 | 0.519 | 0.185 | 1.4221E-34 | PDC | SEL1L |
| 4.3388E-66 | 0.50880358 | 0.414 | 0.087 | 9.2634E-62 | PDC | FNBP1 |
| 5.347E-299 | 0.50784703 | 0.245 | 0.005 | 1.142E-294 | PDC | CFAP45 |
| 6.4941E-49 | 0.50706833 | 0.515 | 0.16 | 1.3865E-44 | PDC | SNRPN |
| 6.3843E-49 | 0.5055549 | 0.422 | 0.115 | 1.363E-44 | PDC | MIB1 |
| 2.1337E-42 | 0.50505711 | 0.397 | 0.115 | 4.5553E-38 | PDC | FBXO2 |
| 4.9622E-78 | 0.50434264 | 0.38 | 0.066 | 1.0594E-73 | PDC | TGFB1I1 |
| 1.7878E-91 | 0.50395092 | 0.325 | 0.042 | 3.817E-87 | PDC | PPM1L |
| 9.0716E-66 | 0.5023123 | 0.43 | 0.094 | 1.9368E-61 | PDC | CDKN1B |
| 2.9816E-73 | 0.50197227 | 0.388 | 0.07 | 6.3658E-69 | PDC | FAM43A |
| 6.0176E-54 | 0.49989139 | 0.464 | 0.125 | 1.2848E-49 | PDC | C7orf50 |
| 2.1716E-28 | 0.49850722 | 0.481 | 0.201 | 4.6365E-24 | PDC | CSRP1 |
| 3.607E-39 | 0.49786615 | 0.515 | 0.185 | 7.7009E-35 | PDC | PHACTR4 |
| 4.6407E-36 | 0.4966629 | 0.502 | 0.187 | 9.9079E-32 | PDC | BNIP3L |
| 0 | 0.49632785 | 0.342 | 0.009 | 0 | PDC | DPP6 |
| 4.2724E-52 | 0.49553236 | 0.494 | 0.141 | 9.1215E-48 | PDC | NME3 |
| 5.2688E-91 | 0.49536406 | 0.397 | 0.063 | 1.1249E-86 | PDC | FAM98A |
| 4.5514E-59 | 0.49425731 | 0.409 | 0.093 | 9.7173E-55 | PDC | LAMB2 |
| 2.5661E-68 | 0.49403349 | 0.43 | 0.093 | 5.4785E-64 | PDC | PARVA |
| 4.6584E-48 | 0.49348642 | 0.456 | 0.128 | 9.9456E-44 | PDC | PTPN13 |
| 3.0752E-05 | 0.49305087 | 0.186 | 0.104 | 0.65656067 | PDC | AREG |
| 1.8007E-28 | 0.49178786 | 0.591 | 0.278 | 3.8444E-24 | PDC | FIS1 |
| 1.7447E-17 | 0.49154605 | 0.916 | 0.823 | 3.7248E-13 | PDC | H3F3A |
| 6.3224E-60 | 0.49098723 | 0.464 | 0.117 | 1.3498E-55 | PDC | CHPT1 |
| 5.7813E-79 | 0.4907758 | 0.426 | 0.081 | 1.2343E-74 | PDC | CTDSPL |
| 4.1936E-25 | 0.49058793 | 0.696 | 0.39 | 8.9534E-21 | PDC | DAD1 |
| 1.7738E-74 | 0.49023637 | 0.354 | 0.061 | 3.7871E-70 | PDC | TSPAN5 |
| 5.5918E-32 | 0.48920094 | 0.616 | 0.277 | 1.1939E-27 | PDC | LAMP1 |
| 6.494E-204 | 0.48918535 | 0.367 | 0.024 | 1.386E-199 | PDC | GPC6 |
| 1.2809E-87 | 0.48896651 | 0.329 | 0.044 | 2.7348E-83 | PDC | KLK6 |
| 4.429E-268 | 0.48753964 | 0.27 | 0.008 | 9.456E-264 | PDC | TRAC |
| 9.0097E-86 | 0.4860841 | 0.401 | 0.067 | 1.9236E-81 | PDC | CPQ |
| 1.769E-104 | 0.48462373 | 0.409 | 0.057 | 3.776E-100 | PDC | CTTNBP2 |
| 0 | 0.4843593 | 0.283 | 0.004 | 0 | PDC | OLFML2A |
| 7.566E-99 | 0.48352607 | 0.418 | 0.064 | 1.6154E-94 | PDC | MDFIC |
| 1.166E-163 | 0.4829403 | 0.304 | 0.021 | 2.49E-159 | PDC | TMOD1 |
| 1.4578E-48 | 0.48234684 | 0.456 | 0.133 | 3.1123E-44 | PDC | DMAC1 |
| 2.1723E-36 | 0.48167619 | 0.304 | 0.08 | 4.6378E-32 | PDC | RNF220 |
| 5.3197E-24 | 0.47831134 | 0.414 | 0.167 | 1.1358E-19 | PDC | SERPING1 |
| 1.5355E-20 | 0.47791464 | 0.903 | 0.79 | 3.2782E-16 | PDC | PPDPF |
| 4.6054E-40 | 0.4765042 | 0.481 | 0.163 | 9.8326E-36 | PDC | VGLL4 |

| | | | | | | |
|---|---|---|---|---|---|---|
| 6.3111E-30 | 0.47633149 | 0.65 | 0.308 | 1.3474E-25 | PDC | MRFAP1 |
| 5.7333E-29 | 0.47568839 | 0.519 | 0.216 | 1.2241E-24 | PDC | BEX2 |
| 3.1217E-49 | 0.47541841 | 0.443 | 0.125 | 6.6648E-45 | PDC | FEZ2 |
| 4.3508E-43 | 0.47531579 | 0.194 | 0.032 | 9.289E-39 | PDC | RFLNB |
| 3.6896E-44 | 0.47492289 | 0.511 | 0.17 | 7.8773E-40 | PDC | ESD |
| 8.76E-101 | 0.47337174 | 0.363 | 0.048 | 1.8702E-96 | PDC | HOXA7 |
| 1.387E-75 | 0.47315522 | 0.451 | 0.091 | 2.9612E-71 | PDC | SASH1 |
| 0 | 0.47279227 | 0.333 | 0.01 | 0 | PDC | ABLIM2 |
| 1.119E-29 | 0.47221588 | 0.519 | 0.22 | 2.3892E-25 | PDC | SPATS2L |
| 1.032E-157 | 0.47163136 | 0.367 | 0.032 | 2.203E-153 | PDC | GSTA4 |
| 3.108E-209 | 0.47000453 | 0.359 | 0.022 | 6.635E-205 | PDC | B4GAT1 |
| 5.9535E-86 | 0.46922009 | 0.329 | 0.046 | 1.2711E-81 | PDC | PLXDC2 |
| 1.0232E-22 | 0.46902697 | 0.709 | 0.411 | 2.1846E-18 | PDC | DYNLRB1 |
| 1.3604E-31 | 0.46885553 | 0.675 | 0.311 | 2.9044E-27 | PDC | PCBP1 |
| 9.8631E-65 | 0.46601346 | 0.228 | 0.029 | 2.1058E-60 | PDC | TSPAN8 |
| 2.4384E-64 | 0.46540317 | 0.392 | 0.082 | 5.206E-60 | PDC | AC245297.3 |
| 5.0437E-29 | 0.46465535 | 0.43 | 0.167 | 1.0768E-24 | PDC | NECTIN2 |
| 3.3275E-72 | 0.46443592 | 0.27 | 0.037 | 7.1042E-68 | PDC | SHISA2 |
| 1.464E-120 | 0.46384911 | 0.308 | 0.029 | 3.125E-116 | PDC | NPR3 |
| 1.0647E-64 | 0.46321593 | 0.376 | 0.074 | 2.2732E-60 | PDC | FRY |
| 2.2913E-50 | 0.463206 | 0.392 | 0.098 | 4.8919E-46 | PDC | MYO1B |
| 6.674E-37 | 0.46258776 | 0.494 | 0.177 | 1.4249E-32 | PDC | DUT |
| 1.3052E-46 | 0.45835643 | 0.363 | 0.091 | 2.7867E-42 | PDC | EIF1AY |
| 2.301E-26 | 0.4580339 | 0.502 | 0.222 | 4.9126E-22 | PDC | COMT |
| 0 | 0.45790006 | 0.295 | 0.004 | 0 | PDC | CNTN1 |
| 3.8327E-83 | 0.45762069 | 0.367 | 0.058 | 8.1829E-79 | PDC | FAM20C |
| 2.2293E-61 | 0.45615248 | 0.295 | 0.051 | 4.7595E-57 | PDC | STON2 |
| 1.707E-183 | 0.45525961 | 0.35 | 0.025 | 3.644E-179 | PDC | PRKAR2B |
| 0 | 0.45448889 | 0.274 | 0.007 | 0 | PDC | AC008264.2 |
| 4.214E-103 | 0.45273211 | 0.354 | 0.045 | 8.996E-99 | PDC | A1BG |
| 9.936E-114 | 0.45244841 | 0.304 | 0.03 | 2.121E-109 | PDC | ADAMTS9-AS1 |
| 1.1714E-19 | 0.45120101 | 0.599 | 0.331 | 2.501E-15 | PDC | H2AFJ |
| 0 | 0.45086544 | 0.295 | 0.006 | 0 | PDC | FRMD1 |
| 4.4793E-36 | 0.45066741 | 0.426 | 0.141 | 9.5634E-32 | PDC | MAGED2 |
| 1.8027E-35 | 0.44842269 | 0.384 | 0.12 | 3.8487E-31 | PDC | FBXO21 |
| 1.6035E-31 | 0.44755888 | 0.494 | 0.196 | 3.4236E-27 | PDC | COX14 |
| 6.6916E-31 | 0.4471448 | 0.586 | 0.254 | 1.4287E-26 | PDC | ERH |
| 3.1088E-34 | 0.44664153 | 0.43 | 0.149 | 6.6373E-30 | PDC | DUSP23 |
| 0 | 0.44541085 | 0.27 | 0.006 | 0 | PDC | CLDN11 |
| 2.4011E-40 | 0.44500305 | 0.451 | 0.142 | 5.1264E-36 | PDC | EEA1 |
| 9.6055E-30 | 0.44472635 | 0.578 | 0.252 | 2.0508E-25 | PDC | TLN1 |
| 5.2155E-26 | 0.44449909 | 0.62 | 0.309 | 1.1135E-21 | PDC | COX7A2L |
| 2.028E-39 | 0.44415829 | 0.506 | 0.176 | 4.3297E-35 | PDC | ALKBH7 |
| 2.4588E-36 | 0.44378589 | 0.494 | 0.177 | 5.2496E-32 | PDC | TBCB |
| 1.7744E-29 | 0.44309422 | 0.511 | 0.212 | 3.7884E-25 | PDC | BLOC1S1 |
| 3.5367E-48 | 0.44280493 | 0.405 | 0.107 | 7.5508E-44 | PDC | PAM |
| 2.778E-200 | 0.44263182 | 0.316 | 0.018 | 5.931E-196 | PDC | MAP6 |
| 1.069E-27 | 0.44180664 | 0.916 | 0.696 | 2.2824E-23 | PDC | HMGB1 |
| 7.9698E-30 | 0.44152465 | 0.57 | 0.246 | 1.7016E-25 | PDC | ATOX1 |
| 2.4578E-22 | 0.44025946 | 0.549 | 0.272 | 5.2474E-18 | PDC | SSB |

| | | | | | | |
|---|---|---|---|---|---|---|
| 5.518E-124 | 0.43974093 | 0.304 | 0.028 | 1.178E-119 | PDC | C1S |
| 3.8492E-28 | 0.43856822 | 0.536 | 0.239 | 8.2179E-24 | PDC | RBPMS |
| 1.864E-69 | 0.437703 | 0.38 | 0.072 | 3.9796E-65 | PDC | LZTS2 |
| 1.5168E-33 | 0.43761091 | 0.388 | 0.127 | 3.2383E-29 | PDC | VASN |
| 1.442E-280 | 0.43737101 | 0.316 | 0.012 | 3.079E-276 | PDC | ANKRD65 |
| 2.8292E-55 | 0.43623496 | 0.384 | 0.087 | 6.0404E-51 | PDC | WASHC1 |
| 1.2256E-23 | 0.43504863 | 0.477 | 0.216 | 2.6166E-19 | PDC | TMOD3 |
| 7.082E-278 | 0.43461993 | 0.308 | 0.011 | 1.512E-273 | PDC | CLIP3 |
| 3.024E-224 | 0.43331152 | 0.304 | 0.014 | 6.456E-220 | PDC | SMCO3 |
| 1.1769E-51 | 0.43278613 | 0.354 | 0.08 | 2.5127E-47 | PDC | GNPTAB |
| 4.2235E-66 | 0.4327006 | 0.321 | 0.055 | 9.0172E-62 | PDC | DOCK5 |
| 6.8117E-14 | 0.4320247 | 0.937 | 0.873 | 1.4543E-09 | PDC | HLA-A |
| 3.5457E-23 | 0.43058905 | 0.658 | 0.357 | 7.57E-19 | PDC | CSDE1 |
| 8.9385E-35 | 0.43053856 | 0.392 | 0.123 | 1.9084E-30 | PDC | UTRN |
| 1.9325E-30 | 0.42917596 | 0.46 | 0.178 | 4.1259E-26 | PDC | CPNE3 |
| 6.3633E-52 | 0.42903886 | 0.557 | 0.166 | 1.3586E-47 | PDC | PLA2G16 |
| 8.7985E-28 | 0.42894833 | 0.426 | 0.162 | 1.8785E-23 | PDC | STAT1 |
| 3.7878E-39 | 0.42884531 | 0.456 | 0.149 | 8.087E-35 | PDC | TPD52 |
| 3.135E-129 | 0.42872324 | 0.316 | 0.029 | 6.694E-125 | PDC | LOXL1 |
| 6.366E-11 | 0.42748734 | 0.16 | 0.058 | 1.3591E-06 | PDC | MT1M |
| 2.164E-149 | 0.42748609 | 0.304 | 0.023 | 4.62E-145 | PDC | CARD16 |
| 2.187E-281 | 0.4270441 | 0.274 | 0.008 | 4.67E-277 | PDC | FOXC2 |
| 4.8893E-19 | 0.42593484 | 0.764 | 0.514 | 1.0439E-14 | PDC | OST4 |
| 4.9673E-33 | 0.42436843 | 0.439 | 0.154 | 1.0605E-28 | PDC | ECH1 |
| 4.7547E-76 | 0.42411885 | 0.359 | 0.06 | 1.0151E-71 | PDC | HSPA12A |
| 1.0287E-18 | 0.42373504 | 0.46 | 0.228 | 2.1963E-14 | PDC | C1orf122 |
| 2.3291E-47 | 0.42330113 | 0.354 | 0.085 | 4.9727E-43 | PDC | EFEMP1 |
| 9.6376E-27 | 0.42255582 | 0.397 | 0.15 | 2.0576E-22 | PDC | CUX1 |
| 2.7638E-31 | 0.42060883 | 0.502 | 0.195 | 5.9007E-27 | PDC | TMEM106B |
| 6.6084E-55 | 0.42055423 | 0.359 | 0.078 | 1.4109E-50 | PDC | ZBTB4 |
| 1.6528E-30 | 0.42051611 | 0.38 | 0.128 | 3.5286E-26 | PDC | UACA |
| 1.263E-191 | 0.42027035 | 0.27 | 0.013 | 2.697E-187 | PDC | TMEM178A |
| 2.5149E-72 | 0.42022687 | 0.371 | 0.067 | 5.3693E-68 | PDC | TCEA2 |
| 0 | 0.41920904 | 0.304 | 0.006 | 0 | PDC | PDPN |
| 2.4078E-24 | 0.41911008 | 0.692 | 0.369 | 5.1406E-20 | PDC | C11orf58 |
| 2.8652E-22 | 0.4189772 | 0.65 | 0.356 | 6.1171E-18 | PDC | TRMT112 |
| 4.3426E-41 | 0.41869247 | 0.371 | 0.104 | 9.2715E-37 | PDC | ITGB5 |
| 1.0237E-73 | 0.41866179 | 0.325 | 0.052 | 2.1857E-69 | PDC | ZADH2 |
| 2.8947E-25 | 0.41831939 | 0.595 | 0.288 | 6.1802E-21 | PDC | SEPT2 |
| 0 | 0.41810994 | 0.257 | 0.001 | 0 | PDC | NRIP2 |
| 2.8625E-81 | 0.41787166 | 0.363 | 0.058 | 6.1114E-77 | PDC | C1orf115 |
| 1.2688E-23 | 0.41697289 | 0.743 | 0.422 | 2.7089E-19 | PDC | PDIA3 |
| 1.2184E-21 | 0.41557156 | 0.641 | 0.352 | 2.6013E-17 | PDC | CD46 |
| 5.717E-200 | 0.41522906 | 0.249 | 0.01 | 1.221E-195 | PDC | BMP7 |
| 6.1151E-39 | 0.41452621 | 0.35 | 0.095 | 1.3056E-34 | PDC | PSMB9 |
| 1.9571E-40 | 0.41389504 | 0.401 | 0.117 | 4.1784E-36 | PDC | RTL8A |
| 2.4645E-30 | 0.41377739 | 0.515 | 0.205 | 5.2618E-26 | PDC | PURA |
| 6.1852E-41 | 0.41369501 | 0.405 | 0.119 | 1.3205E-36 | PDC | BORCS7 |
| 3.2539E-63 | 0.41345953 | 0.363 | 0.071 | 6.9471E-59 | PDC | UFSP2 |
| 8.9E-59 | 0.41272504 | 0.287 | 0.049 | 1.9002E-54 | PDC | IFI16 |

| | | | | | | |
|---|---|---|---|---|---|---|
| 1.0919E-80 | 0.41197456 | 0.325 | 0.047 | 2.3311E-76 | PDC | SERPINI1 |
| 9.8282E-24 | 0.41128545 | 0.549 | 0.258 | 2.0983E-19 | PDC | SELENOM |
| 3.33E-152 | 0.4111794 | 0.3 | 0.022 | 7.11E-148 | PDC | SFMBT2 |
| 2.699E-295 | 0.4108219 | 0.27 | 0.007 | 5.763E-291 | PDC | PALMD |
| 7.0787E-45 | 0.41024777 | 0.384 | 0.102 | 1.5113E-40 | PDC | TMEM9B |
| 9.2946E-44 | 0.40949193 | 0.35 | 0.088 | 1.9844E-39 | PDC | FAM114A1 |
| 2.4951E-26 | 0.40908706 | 0.456 | 0.186 | 5.327E-22 | PDC | PSMB8 |
| 7.7305E-24 | 0.40841955 | 0.388 | 0.154 | 1.6505E-19 | PDC | IP6K2 |
| 3.8048E-31 | 0.40700727 | 0.329 | 0.1 | 8.1232E-27 | PDC | OGFRL1 |
| 1.5203E-83 | 0.40612903 | 0.194 | 0.017 | 3.2458E-79 | PDC | PMP22 |
| 1.1607E-24 | 0.40603661 | 0.443 | 0.189 | 2.478E-20 | PDC | TSPAN3 |
| 5.1886E-24 | 0.40439793 | 0.527 | 0.242 | 1.1078E-19 | PDC | LAMTOR4 |
| 4.1788E-21 | 0.40410103 | 0.616 | 0.328 | 8.9218E-17 | PDC | RAB2A |
| 1.663E-41 | 0.40403637 | 0.325 | 0.081 | 3.5505E-37 | PDC | TNFRSF1A |
| 1.2676E-36 | 0.40400367 | 0.456 | 0.154 | 2.7064E-32 | PDC | PYURF |
| 1.6938E-14 | 0.4035383 | 0.582 | 0.34 | 3.6163E-10 | PDC | TSC22D1 |
| 2.7026E-23 | 0.4033279 | 0.46 | 0.194 | 5.77E-19 | PDC | FAT1 |
| 1.055E-301 | 0.40239968 | 0.287 | 0.008 | 2.252E-297 | PDC | GRK5 |
| 9.0177E-66 | 0.40199672 | 0.342 | 0.062 | 1.9253E-61 | PDC | NOL3 |
| 1.9939E-30 | 0.40160997 | 0.443 | 0.162 | 4.2569E-26 | PDC | TROVE2 |
| 8.3157E-33 | 0.40103358 | 0.489 | 0.182 | 1.7754E-28 | PDC | PCMT1 |
| 2.43E-23 | 0.40062107 | 0.468 | 0.207 | 5.188E-19 | PDC | RAD23A |
| 3.92E-45 | 0.39987634 | 0.367 | 0.093 | 8.3693E-41 | PDC | NOTCH2 |
| 1.2397E-71 | 0.39869464 | 0.211 | 0.023 | 2.6467E-67 | PDC | EDIL3 |
| 1.765E-185 | 0.39667917 | 0.274 | 0.014 | 3.768E-181 | PDC | CA10 |
| 1.6954E-43 | 0.39613516 | 0.342 | 0.085 | 3.6196E-39 | PDC | RASSF8-AS1 |
| 2.0973E-33 | 0.39590664 | 0.397 | 0.128 | 4.4778E-29 | PDC | EPHX1 |
| 3.1883E-29 | 0.39589646 | 0.595 | 0.261 | 6.8071E-25 | PDC | TCEAL4 |
| 3.149E-65 | 0.39319393 | 0.338 | 0.061 | 6.7231E-61 | PDC | TXNDC15 |
| 5.6709E-62 | 0.39314402 | 0.338 | 0.064 | 1.2107E-57 | PDC | ACTR1B |
| 1.0563E-38 | 0.39260409 | 0.325 | 0.085 | 2.2552E-34 | PDC | SGCB |
| 1.204E-24 | 0.39225785 | 0.57 | 0.272 | 2.5704E-20 | PDC | OCIAD1 |
| 7.3371E-22 | 0.39095255 | 0.616 | 0.326 | 1.5665E-17 | PDC | NDUFB4 |
| 1.4817E-33 | 0.39077595 | 0.354 | 0.107 | 3.1635E-29 | PDC | PLOD2 |
| 4.9047E-22 | 0.39008087 | 0.616 | 0.31 | 1.0471E-17 | PDC | PRDX2 |
| 3.2203E-29 | 0.38996754 | 0.228 | 0.056 | 6.8753E-25 | PDC | PAPPA |
| 0 | 0.38848688 | 0.181 | 0.002 | 0 | PDC | COL3A1 |
| 8.5317E-51 | 0.38834421 | 0.359 | 0.083 | 1.8215E-46 | PDC | ZFYVE21 |
| 4.4115E-22 | 0.38755764 | 0.603 | 0.304 | 9.4185E-18 | PDC | NDUFC2 |
| 1.4464E-18 | 0.38692179 | 0.717 | 0.448 | 3.0882E-14 | PDC | LAPTM4A |
| 2.9204E-82 | 0.38685572 | 0.245 | 0.027 | 6.2351E-78 | PDC | GPM6B |
| 3.6204E-33 | 0.38678895 | 0.346 | 0.102 | 7.7296E-29 | PDC | CGNL1 |
| 4.7064E-25 | 0.38672847 | 0.675 | 0.335 | 1.0048E-20 | PDC | SPCS1 |
| 1.1957E-27 | 0.38581007 | 0.498 | 0.206 | 2.5527E-23 | PDC | RAB14 |
| 5.052E-18 | 0.38546671 | 0.776 | 0.521 | 1.0786E-13 | PDC | ATP5MC2 |
| 2.098E-36 | 0.38467244 | 0.333 | 0.092 | 4.4792E-32 | PDC | P4HTM |
| 2.8575E-05 | 0.38425806 | 0.16 | 0.085 | 0.61006849 | PDC | MT1H |
| 3.445E-23 | 0.38401007 | 0.489 | 0.223 | 7.355E-19 | PDC | TMEM219 |
| 4.4964E-18 | 0.38368773 | 0.688 | 0.42 | 9.5999E-14 | PDC | TMEM59 |
| 3.9858E-85 | 0.38361765 | 0.329 | 0.046 | 8.5098E-81 | PDC | UNG |

| | | | | | | |
|---|---|---|---|---|---|---|
| 1.2425E-23 | 0.38334423 | 0.371 | 0.141 | 2.6527E-19 | PDC | PTPN14 |
| 1.6076E-55 | 0.3832453 | 0.363 | 0.078 | 3.4322E-51 | PDC | GTF3C1 |
| 2.0289E-42 | 0.38319512 | 0.38 | 0.103 | 4.3318E-38 | PDC | UNC50 |
| 3.3191E-42 | 0.38308368 | 0.346 | 0.089 | 7.0864E-38 | PDC | SQOR |
| 2.8249E-28 | 0.38275133 | 0.316 | 0.099 | 6.0311E-24 | PDC | INF2 |
| 1.1032E-38 | 0.38237808 | 0.346 | 0.093 | 2.3554E-34 | PDC | TNS3 |
| 7.2309E-60 | 0.38109004 | 0.338 | 0.064 | 1.5438E-55 | PDC | PHLDB1 |
| 3.9065E-65 | 0.38107687 | 0.316 | 0.054 | 8.3404E-61 | PDC | TMEM47 |
| 2.6945E-44 | 0.38088394 | 0.367 | 0.095 | 5.7528E-40 | PDC | LGALS8 |
| 3.439E-223 | 0.38073753 | 0.262 | 0.01 | 7.343E-219 | PDC | NPAS3 |
| 2.0289E-25 | 0.38046327 | 0.903 | 0.639 | 4.3317E-21 | PDC | PEBP1 |
| 3.2742E-30 | 0.38013847 | 0.468 | 0.178 | 6.9905E-26 | PDC | SERINC3 |
| 9.7044E-21 | 0.37972189 | 0.333 | 0.131 | 2.0719E-16 | PDC | EVA1B |
| 3.7934E-73 | 0.37936439 | 0.329 | 0.053 | 8.0989E-69 | PDC | PRKACB |
| 3.4294E-56 | 0.37897553 | 0.215 | 0.03 | 7.3218E-52 | PDC | MSLN |
| 6.8907E-31 | 0.37800595 | 0.312 | 0.093 | 1.4712E-26 | PDC | SERINC5 |
| 1.793E-35 | 0.37635941 | 0.274 | 0.066 | 3.8281E-31 | PDC | SLC40A1 |
| 5.5643E-32 | 0.37585361 | 0.321 | 0.093 | 1.188E-27 | PDC | KLF9 |
| 1.1204E-40 | 0.37446159 | 0.401 | 0.115 | 2.392E-36 | PDC | MED4 |
| 7.7058E-20 | 0.37437005 | 0.549 | 0.276 | 1.6452E-15 | PDC | MAP4 |
| 2.5682E-98 | 0.37425456 | 0.321 | 0.039 | 5.483E-94 | PDC | SNTA1 |
| 5.4441E-59 | 0.37378683 | 0.316 | 0.058 | 1.1623E-54 | PDC | TMCC3 |
| 1.5947E-37 | 0.37358275 | 0.228 | 0.046 | 3.4047E-33 | PDC | CYP1B1 |
| 8.503E-41 | 0.37311542 | 0.321 | 0.079 | 1.8154E-36 | PDC | CLIP4 |
| 5.2032E-17 | 0.37262229 | 0.684 | 0.406 | 1.1109E-12 | PDC | UQCR10 |
| 7.7899E-64 | 0.37244234 | 0.3 | 0.05 | 1.6631E-59 | PDC | CCDC85C |
| 3.2784E-23 | 0.37171784 | 0.456 | 0.193 | 6.9993E-19 | PDC | BBX |
| 5.814E-20 | 0.37163004 | 0.515 | 0.258 | 1.2413E-15 | PDC | NDUFB10 |
| 3.8472E-22 | 0.36990209 | 0.363 | 0.143 | 8.2137E-18 | PDC | MXRA7 |
| 8.0688E-22 | 0.36899776 | 0.397 | 0.164 | 1.7227E-17 | PDC | PON2 |
| 4.0445E-69 | 0.36888547 | 0.283 | 0.042 | 8.635E-65 | PDC | RBM43 |
| 4.3172E-46 | 0.36834792 | 0.333 | 0.078 | 9.2172E-42 | PDC | EXOSC7 |
| 1.4683E-19 | 0.36780175 | 0.667 | 0.376 | 3.1348E-15 | PDC | NDUFB2 |
| 7.6471E-37 | 0.36647881 | 0.367 | 0.107 | 1.6326E-32 | PDC | RTL8C |
| 2.5213E-17 | 0.36632692 | 0.536 | 0.285 | 5.3829E-13 | PDC | LRPAP1 |
| 2.0491E-44 | 0.36620596 | 0.342 | 0.083 | 4.3748E-40 | PDC | SNX17 |
| 1.1576E-30 | 0.36548296 | 0.388 | 0.131 | 2.4716E-26 | PDC | TUSC3 |
| 9.0857E-34 | 0.36531766 | 0.397 | 0.127 | 1.9398E-29 | PDC | IFI27L2 |
| 9.0002E-22 | 0.36512099 | 0.35 | 0.137 | 1.9215E-17 | PDC | RAB34 |
| 2.215E-17 | 0.36488339 | 0.519 | 0.275 | 4.7289E-13 | PDC | LMAN1 |
| 1.454E-24 | 0.36468131 | 0.418 | 0.165 | 3.1042E-20 | PDC | CBX5 |
| 4.4546E-36 | 0.36468104 | 0.342 | 0.096 | 9.5106E-32 | PDC | AP3S1 |
| 2.3499E-36 | 0.36457442 | 0.312 | 0.082 | 5.0171E-32 | PDC | TACC2 |
| 1.3828E-61 | 0.36384478 | 0.316 | 0.056 | 2.9523E-57 | PDC | LCA5 |
| 6.8292E-34 | 0.36353629 | 0.418 | 0.137 | 1.458E-29 | PDC | LYRM2 |
| 3.3314E-39 | 0.36256081 | 0.333 | 0.088 | 7.1124E-35 | PDC | NSMCE1 |
| 1.297E-48 | 0.36253696 | 0.304 | 0.064 | 2.7691E-44 | PDC | NLK |
| 3.7094E-26 | 0.36135261 | 0.405 | 0.153 | 7.9195E-22 | PDC | PIN1 |
| 7.2562E-32 | 0.36097322 | 0.38 | 0.124 | 1.5492E-27 | PDC | TRAPPC4 |
| 0 | 0.36083791 | 0.19 | 0.002 | 0 | PDC | RAMP3 |

| | | | | | | |
|---|---|---|---|---|---|---|
| 5.7062E-41 | 0.35995815 | 0.346 | 0.09 | 1.2183E-36 | PDC | CREB3L2 |
| 1.4578E-36 | 0.35888112 | 0.35 | 0.098 | 3.1123E-32 | PDC | NR3C1 |
| 3.9517E-46 | 0.35849341 | 0.3 | 0.064 | 8.4369E-42 | PDC | TOM1L2 |
| 1.8787E-12 | 0.35835679 | 0.101 | 0.026 | 4.011E-08 | PDC | MT1A |
| 5.609E-108 | 0.35812952 | 0.283 | 0.028 | 1.198E-103 | PDC | SMIM1 |
| 2.8238E-20 | 0.3581095 | 0.443 | 0.199 | 6.0289E-16 | PDC | ROCK1 |
| 2.8725E-21 | 0.35752402 | 0.549 | 0.27 | 6.1329E-17 | PDC | NDUFS6 |
| 1.3238E-23 | 0.35722919 | 0.578 | 0.279 | 2.8263E-19 | PDC | C12orf57 |
| 2.5817E-19 | 0.35722587 | 0.473 | 0.228 | 5.5119E-15 | PDC | CHD4 |
| 2.3879E-36 | 0.35615086 | 0.295 | 0.074 | 5.0981E-32 | PDC | APOL1 |
| 1.628E-59 | 0.35551822 | 0.3 | 0.052 | 3.4759E-55 | PDC | PLEKHO1 |
| 1.7786E-05 | 0.35537377 | 0.215 | 0.126 | 0.37973076 | PDC | SOCS3 |
| 3.1593E-54 | 0.35508753 | 0.3 | 0.057 | 6.7452E-50 | PDC | POLR3GL |
| 6.0065E-25 | 0.35500135 | 0.392 | 0.149 | 1.2824E-20 | PDC | FAM200B |
| 1.2829E-14 | 0.35486053 | 0.882 | 0.743 | 2.7389E-10 | PDC | HINT1 |
| 5.016E-23 | 0.3546673 | 0.384 | 0.15 | 1.0709E-18 | PDC | CFAP97 |
| 4.6703E-27 | 0.35457611 | 0.367 | 0.129 | 9.9711E-23 | PDC | IGF1R |
| 2.2816E-18 | 0.35453759 | 0.346 | 0.145 | 4.8712E-14 | PDC | HCFC1R1 |
| 5.2986E-21 | 0.35449171 | 0.473 | 0.219 | 1.1313E-16 | PDC | TMED4 |
| 9.4191E-47 | 0.35436403 | 0.287 | 0.059 | 2.011E-42 | PDC | CPNE8 |
| 2.0079E-21 | 0.35420587 | 0.371 | 0.153 | 4.2868E-17 | PDC | NAPA |
| 1.6144E-16 | 0.35394762 | 0.688 | 0.434 | 3.4468E-12 | PDC | HMGN1 |
| 4.0895E-28 | 0.3534643 | 0.405 | 0.145 | 8.731E-24 | PDC | NUCB2 |
| 1.8836E-19 | 0.3524536 | 0.586 | 0.311 | 4.0216E-15 | PDC | SNX3 |
| 1.8808E-77 | 0.3512349 | 0.274 | 0.036 | 4.0155E-73 | PDC | ANKH |
| 8.7858E-25 | 0.35114404 | 0.262 | 0.08 | 1.8758E-20 | PDC | SLC6A6 |
| 1.2933E-17 | 0.35091109 | 0.371 | 0.166 | 2.7612E-13 | PDC | NRIP1 |
| 4.265E-50 | 0.35025443 | 0.325 | 0.07 | 9.1058E-46 | PDC | TMEM109 |
| 4.5887E-22 | 0.34966082 | 0.3 | 0.105 | 9.7969E-18 | PDC | CCDC85B |
| 2.6401E-29 | 0.34932832 | 0.38 | 0.13 | 5.6367E-25 | PDC | ACTR10 |
| 0 | 0.34874695 | 0.194 | 0.001 | 0 | PDC | AL118508.1 |
| 1.2791E-86 | 0.34850599 | 0.283 | 0.034 | 2.7309E-82 | PDC | DLC1 |
| 5.0567E-64 | 0.34848974 | 0.304 | 0.05 | 1.0796E-59 | PDC | SIRPA |
| 3.4567E-14 | 0.34840718 | 0.367 | 0.18 | 7.3801E-10 | PDC | SLC44A2 |
| 0 | 0.34794719 | 0.236 | 0.001 | 0 | PDC | CR1 |
| 3.7333E-25 | 0.34697521 | 0.371 | 0.137 | 7.9705E-21 | PDC | CLCN3 |
| 1.1405E-35 | 0.34570514 | 0.333 | 0.093 | 2.4349E-31 | PDC | MOCS2 |
| 6.5053E-15 | 0.34538043 | 0.97 | 0.925 | 1.3889E-10 | PDC | ACTG1 |
| 3.0682E-81 | 0.34532329 | 0.27 | 0.033 | 6.5507E-77 | PDC | CHST12 |
| 4.6263E-30 | 0.34521486 | 0.346 | 0.109 | 9.8771E-26 | PDC | FCGRT |
| 0 | 0.34509455 | 0.241 | 0.003 | 0 | PDC | TENM2 |
| 3.0044E-55 | 0.34408196 | 0.266 | 0.045 | 6.4145E-51 | PDC | PAQR7 |
| 2.7524E-36 | 0.34325599 | 0.325 | 0.088 | 5.8764E-32 | PDC | PKIG |
| 2.6726E-52 | 0.34321129 | 0.262 | 0.046 | 5.7059E-48 | PDC | PLBD1 |
| 4.2482E-69 | 0.34278169 | 0.278 | 0.041 | 9.0699E-65 | PDC | B9D1 |
| 1.145E-24 | 0.34266559 | 0.401 | 0.156 | 2.4446E-20 | PDC | PNKD |
| 4.3294E-20 | 0.34223267 | 0.549 | 0.278 | 9.2432E-16 | PDC | TMEM14C |
| 5.2385E-25 | 0.34181112 | 0.418 | 0.164 | 1.1184E-20 | PDC | COPS6 |
| 5.1379E-21 | 0.34172784 | 0.464 | 0.21 | 1.0969E-16 | PDC | PSMC5 |
| 1.1492E-19 | 0.34135741 | 0.591 | 0.311 | 2.4535E-15 | PDC | SRP9 |

| | | | | | | |
|---|---|---|---|---|---|---|
| 1.985E-147 | 0.34088634 | 0.186 | 0.008 | 4.237E-143 | PDC | VSIG8 |
| 1.7779E-17 | 0.34072519 | 0.498 | 0.257 | 3.7958E-13 | PDC | JTB |
| 2.175E-272 | 0.33973964 | 0.198 | 0.004 | 4.643E-268 | PDC | NFASC |
| 2.33E-23 | 0.33973844 | 0.464 | 0.2 | 4.9746E-19 | PDC | POLR2I |
| 7.4942E-20 | 0.33932517 | 0.338 | 0.133 | 1.6E-15 | PDC | TUBB2A |
| 3.702E-177 | 0.33930121 | 0.228 | 0.01 | 7.904E-173 | PDC | HS3ST3B1 |
| 1.8863E-23 | 0.33857328 | 0.359 | 0.136 | 4.0272E-19 | PDC | TMEM245 |
| 5.1855E-46 | 0.33835721 | 0.262 | 0.05 | 1.1071E-41 | PDC | TRIB2 |
| 8.8228E-14 | 0.33703624 | 0.928 | 0.838 | 1.8837E-09 | PDC | CD63 |
| 8.848E-20 | 0.33702062 | 0.384 | 0.161 | 1.889E-15 | PDC | SEMA3B |
| 4.403E-159 | 0.33645699 | 0.203 | 0.009 | 9.401E-155 | PDC | TMIE |
| 7.3083E-45 | 0.33639433 | 0.203 | 0.033 | 1.5603E-40 | PDC | UNC5B |
| 3.6626E-47 | 0.33631322 | 0.359 | 0.085 | 7.8197E-43 | PDC | CCDC115 |
| 5.8135E-19 | 0.3358068 | 0.494 | 0.242 | 1.2412E-14 | PDC | MRPS21 |
| 1.4361E-87 | 0.33580514 | 0.257 | 0.028 | 3.066E-83 | PDC | AMIGO2 |
| 1.691E-32 | 0.335702 | 0.35 | 0.107 | 3.6102E-28 | PDC | EMC2 |
| 3.3711E-16 | 0.33563368 | 0.586 | 0.326 | 7.1972E-12 | PDC | KIF5B |
| 1.083E-145 | 0.33548 | 0.241 | 0.014 | 2.311E-141 | PDC | SCN9A |
| 1.8171E-35 | 0.33540811 | 0.295 | 0.075 | 3.8796E-31 | PDC | KLF7 |
| 5.3861E-12 | 0.33466079 | 0.616 | 0.387 | 1.1499E-07 | PDC | CAPN2 |
| 8.4959E-60 | 0.3345721 | 0.266 | 0.042 | 1.8139E-55 | PDC | CMTM3 |
| 0 | 0.33448293 | 0.232 | 0.004 | 0 | PDC | FMN2 |
| 0 | 0.33448293 | 0.245 | 0.004 | 0 | PDC | KCNH2 |
| 2.012E-21 | 0.33325045 | 0.401 | 0.169 | 4.2956E-17 | PDC | ARL2 |
| 1.8565E-14 | 0.33304699 | 0.468 | 0.253 | 3.9636E-10 | PDC | PRKAR1A |
| 6.072E-27 | 0.33193891 | 0.346 | 0.117 | 1.2964E-22 | PDC | TM2D1 |
| 2.7559E-19 | 0.33160561 | 0.561 | 0.283 | 5.8839E-15 | PDC | RBX1 |
| 4.037E-161 | 0.33136692 | 0.232 | 0.012 | 8.619E-157 | PDC | SLC2A12 |
| 4.225E-134 | 0.33117651 | 0.236 | 0.015 | 9.019E-130 | PDC | TSPAN2 |
| 6.7734E-27 | 0.33110963 | 0.405 | 0.15 | 1.4461E-22 | PDC | CFAP36 |
| 5.108E-69 | 0.33094706 | 0.215 | 0.025 | 1.0906E-64 | PDC | GRK3 |
| 3.6952E-67 | 0.33090727 | 0.241 | 0.032 | 7.8893E-63 | PDC | TMOD2 |
| 1.3589E-24 | 0.32979371 | 0.376 | 0.14 | 2.9013E-20 | PDC | SH3BGRL |
| 4.6747E-43 | 0.32922808 | 0.291 | 0.064 | 9.9804E-39 | PDC | THRA |
| 2.5698E-24 | 0.32900017 | 0.43 | 0.174 | 5.4864E-20 | PDC | UBE2M |
| 5.991E-283 | 0.32875491 | 0.211 | 0.004 | 1.279E-278 | PDC | ATP10A |
| 7.088E-146 | 0.32854684 | 0.207 | 0.01 | 1.513E-141 | PDC | KLK7 |
| 5.5615E-45 | 0.32768264 | 0.156 | 0.02 | 1.1874E-40 | PDC | IFI27 |
| 4.4779E-15 | 0.32753628 | 0.684 | 0.409 | 9.5603E-11 | PDC | HNRNPK |
| 5.0151E-18 | 0.32747365 | 0.553 | 0.289 | 1.0707E-13 | PDC | BRK1 |
| 6.378E-18 | 0.32728758 | 0.308 | 0.123 | 1.3617E-13 | PDC | HSPA13 |
| 6.8809E-84 | 0.32686654 | 0.287 | 0.036 | 1.4691E-79 | PDC | ARHGAP23 |
| 3.4422E-18 | 0.32683095 | 0.595 | 0.315 | 7.349E-14 | PDC | XRCC5 |
| 9.9422E-27 | 0.32659981 | 0.363 | 0.126 | 2.1227E-22 | PDC | HINT2 |
| 1.42E-22 | 0.32632238 | 0.329 | 0.119 | 3.0317E-18 | PDC | TPP1 |
| 3.2106E-22 | 0.32623948 | 0.304 | 0.107 | 6.8546E-18 | PDC | TERF1 |
| 3.7042E-17 | 0.32602958 | 0.612 | 0.329 | 7.9085E-13 | PDC | SPCS2 |
| 2.1628E-17 | 0.32487617 | 0.464 | 0.229 | 4.6175E-13 | PDC | FXR1 |
| 5.9825E-11 | 0.32480018 | 0.591 | 0.366 | 1.2773E-06 | PDC | TAGLN2 |
| 6.1862E-20 | 0.32465027 | 0.43 | 0.192 | 1.3208E-15 | PDC | ZCRB1 |

| | | | | | | |
|---|---|---|---|---|---|---|
| 2.418E-125 | 0.32427085 | 0.241 | 0.017 | 5.162E-121 | PDC | LHFPL6 |
| 2.9609E-28 | 0.32419334 | 0.38 | 0.131 | 6.3216E-24 | PDC | ANP32A |
| 3.6502E-14 | 0.32370999 | 0.637 | 0.379 | 7.7932E-10 | PDC | MYH9 |
| 1.7229E-23 | 0.32339195 | 0.414 | 0.164 | 3.6784E-19 | PDC | PRMT2 |
| 8.8259E-38 | 0.32284349 | 0.3 | 0.073 | 1.8843E-33 | PDC | MUM1 |
| 4.8147E-60 | 0.32278862 | 0.194 | 0.023 | 1.0279E-55 | PDC | CLGN |
| 2.457E-100 | 0.32267005 | 0.101 | 0.003 | 5.2459E-96 | PDC | COL1A2 |
| 1.4098E-69 | 0.32225343 | 0.278 | 0.04 | 3.0099E-65 | PDC | MANEA |
| 2.7794E-43 | 0.32199781 | 0.283 | 0.061 | 5.934E-39 | PDC | LIMK2 |
| 5.173E-126 | 0.32141982 | 0.262 | 0.02 | 1.104E-121 | PDC | FIGN |
| 1.7375E-15 | 0.32102366 | 0.338 | 0.152 | 3.7097E-11 | PDC | LGALS3BP |
| 9.7857E-61 | 0.32089605 | 0.27 | 0.043 | 2.0892E-56 | PDC | NTHL1 |
| 1.7713E-19 | 0.32083656 | 0.426 | 0.19 | 3.7817E-15 | PDC | RAB18 |
| 4.8201E-35 | 0.32077747 | 0.316 | 0.085 | 1.0291E-30 | PDC | CISD3 |
| 1.95E-36 | 0.32066167 | 0.295 | 0.075 | 4.1632E-32 | PDC | BLVRA |
| 1.5287E-18 | 0.32059577 | 0.485 | 0.238 | 3.2638E-14 | PDC | RNH1 |
| 6.4113E-26 | 0.32058357 | 0.333 | 0.11 | 1.3688E-21 | PDC | HSPB8 |
| 4.188E-154 | 0.32049738 | 0.253 | 0.015 | 8.941E-150 | PDC | STON1 |
| 5.3624E-16 | 0.32042921 | 0.418 | 0.204 | 1.1449E-11 | PDC | TMEM30A |
| 1.9099E-43 | 0.32014349 | 0.278 | 0.059 | 4.0777E-39 | PDC | IL6R |
| 1.505E-38 | 0.3200452 | 0.257 | 0.056 | 3.2131E-34 | PDC | ZBED3 |
| 2.1736E-31 | 0.31991691 | 0.3 | 0.084 | 4.6406E-27 | PDC | MVB12A |
| 9.2802E-27 | 0.31951105 | 0.388 | 0.139 | 1.9813E-22 | PDC | ERLEC1 |
| 1.5716E-20 | 0.31945247 | 0.477 | 0.22 | 3.3554E-16 | PDC | C1orf43 |
| 1.699E-200 | 0.31911081 | 0.245 | 0.01 | 3.626E-196 | PDC | LIMS2 |
| 3.4573E-16 | 0.31905 | 0.257 | 0.099 | 7.3812E-12 | PDC | KIAA0040 |
| 5.029E-166 | 0.31846775 | 0.194 | 0.007 | 1.074E-161 | PDC | ADORA1 |
| 1.4309E-31 | 0.31833516 | 0.346 | 0.106 | 3.0549E-27 | PDC | ALDH9A1 |
| 1.8014E-28 | 0.31789671 | 0.321 | 0.1 | 3.846E-24 | PDC | TCEAL8 |
| 1.011E-30 | 0.31778228 | 0.304 | 0.086 | 2.1585E-26 | PDC | IRF2BPL |
| 5.1971E-18 | 0.31717267 | 0.376 | 0.166 | 1.1096E-13 | PDC | FAM204A |
| 0 | 0.31694135 | 0.194 | 0.002 | 0 | PDC | CDH11 |
| 5.0341E-17 | 0.31682438 | 0.477 | 0.238 | 1.0748E-12 | PDC | GTF2A2 |
| 2.074E-102 | 0.3165674 | 0.219 | 0.017 | 4.429E-98 | PDC | PRICKLE1 |
| 1.331E-29 | 0.31560918 | 0.266 | 0.072 | 2.8417E-25 | PDC | SORL1 |
| 3.0025E-82 | 0.31559432 | 0.249 | 0.028 | 6.4103E-78 | PDC | GPSM1 |
| 1.0701E-77 | 0.31556922 | 0.257 | 0.031 | 2.2847E-73 | PDC | LIX1L |
| 2.2913E-45 | 0.31539803 | 0.249 | 0.046 | 4.8919E-41 | PDC | ADAMTS9-AS2 |
| 2.9138E-13 | 0.31537174 | 0.629 | 0.384 | 6.2209E-09 | PDC | TOMM20 |
| 3.7616E-26 | 0.31536715 | 0.333 | 0.112 | 8.031E-22 | PDC | WDR13 |
| 4.3007E-32 | 0.31349442 | 0.367 | 0.114 | 9.1819E-28 | PDC | CCNDBP1 |
| 4.5524E-44 | 0.31347311 | 0.308 | 0.069 | 9.7193E-40 | PDC | TMEM14A |
| 1.1819E-22 | 0.31327744 | 0.19 | 0.049 | 2.5234E-18 | PDC | BEX1 |
| 3.2787E-57 | 0.3125265 | 0.274 | 0.046 | 7E-53 | PDC | STN1 |
| 3.578E-15 | 0.31216786 | 0.62 | 0.364 | 7.6391E-11 | PDC | ARL6IP4 |
| 1.3865E-21 | 0.31208683 | 0.367 | 0.143 | 2.9602E-17 | PDC | PTPRA |
| 3.8373E-37 | 0.31207957 | 0.266 | 0.06 | 8.1926E-33 | PDC | HOXA9 |
| 4.0919E-39 | 0.31166808 | 0.295 | 0.071 | 8.7361E-35 | PDC | WRB |
| 1.5858E-15 | 0.31107057 | 0.376 | 0.177 | 3.3856E-11 | PDC | CD99 |
| 1.8029E-19 | 0.31105108 | 0.198 | 0.059 | 3.8493E-15 | PDC | SESN3 |

| | | | | | | |
|---|---|---|---|---|---|---|
| 2.259E-120 | 0.31099182 | 0.219 | 0.015 | 4.823E-116 | PDC | ANTXR2 |
| 4.554E-31 | 0.31098379 | 0.274 | 0.072 | 9.7228E-27 | PDC | ANKRD33B |
| 4.6159E-14 | 0.31062183 | 0.574 | 0.332 | 9.855E-10 | PDC | PSMB1 |
| 4.104E-36 | 0.31038763 | 0.291 | 0.072 | 8.762E-32 | PDC | SGCE |
| 1.062E-80 | 0.30973758 | 0.241 | 0.027 | 2.2673E-76 | PDC | VSIR |
| 3.8358E-31 | 0.30963747 | 0.38 | 0.122 | 8.1894E-27 | PDC | MRPS26 |
| 2.1053E-16 | 0.3093422 | 0.401 | 0.191 | 4.4948E-12 | PDC | EMC7 |
| 1.7549E-18 | 0.30777857 | 0.435 | 0.202 | 3.7466E-14 | PDC | RCN2 |
| 3.9537E-24 | 0.30776613 | 0.392 | 0.151 | 8.4411E-20 | PDC | TAL/DCTN3 |
| 1.713E-121 | 0.30773454 | 0.245 | 0.018 | 3.658E-117 | PDC | RAB3IL1 |
| 1.0352E-11 | 0.30748199 | 0.764 | 0.579 | 2.2102E-07 | PDC | COX5B |
| 2.298E-15 | 0.30737192 | 0.481 | 0.253 | 4.9062E-11 | PDC | NDUFA11 |
| 4.693E-196 | 0.30693917 | 0.228 | 0.009 | 1.002E-191 | PDC | ZBTB7C |
| 3.7891E-49 | 0.30634163 | 0.287 | 0.056 | 8.0897E-45 | PDC | SLC17A5 |
| 3.3372E-18 | 0.30628236 | 0.397 | 0.178 | 7.1249E-14 | PDC | SIVA1 |
| 5.3345E-13 | 0.30620157 | 0.612 | 0.384 | 1.1389E-08 | PDC | SUB1 |
| 1.0914E-27 | 0.30617206 | 0.27 | 0.077 | 2.3301E-23 | PDC | ENDOD1 |
| 7.7774E-22 | 0.30615193 | 0.321 | 0.118 | 1.6605E-17 | PDC | APPL1 |
| 8.3237E-80 | 0.30541441 | 0.262 | 0.031 | 1.7771E-75 | PDC | AC068888.1 |
| 0 | 0.30521335 | 0.169 | 0.001 | 0 | PDC | RAET1E |
| 1.4029E-50 | 0.30511101 | 0.241 | 0.04 | 2.9953E-46 | PDC | PLCD3 |
| 2.274E-67 | 0.30496246 | 0.219 | 0.026 | 4.855E-63 | PDC | HSD17B14 |
| 6.3729E-48 | 0.30496069 | 0.291 | 0.059 | 1.3606E-43 | PDC | DPYSL2 |
| 7.2758E-24 | 0.3040112 | 0.321 | 0.112 | 1.5534E-19 | PDC | ARL2BP |
| 5.7663E-30 | 0.30383393 | 0.363 | 0.116 | 1.2311E-25 | PDC | ZNF667-AS1 |
| 2.1153E-38 | 0.30367763 | 0.274 | 0.063 | 4.5162E-34 | PDC | EFEMP2 |
| 4.9076E-29 | 0.30321895 | 0.266 | 0.072 | 1.0478E-24 | PDC | RABGAP1L |
| 1.1168E-20 | 0.30318805 | 0.333 | 0.128 | 2.3843E-16 | PDC | TMEM179B |
| 2.8772E-26 | 0.30294015 | 0.316 | 0.103 | 6.1427E-22 | PDC | WASHC3 |
| 3.2501E-26 | 0.30280645 | 0.291 | 0.089 | 6.9389E-22 | PDC | MAP9 |
| 0 | 0.30247297 | 0.203 | 0.002 | 0 | PDC | ADAMTS19 |
| 1.1286E-16 | 0.30242177 | 0.456 | 0.227 | 2.4095E-12 | PDC | NDUFA12 |
| 3.4409E-15 | 0.30169742 | 0.489 | 0.259 | 7.3464E-11 | PDC | MLEC |
| 7.0026E-23 | 0.30169401 | 0.359 | 0.133 | 1.4951E-18 | PDC | RNASEH2C |
| 2.1691E-16 | 0.3008506 | 0.312 | 0.131 | 4.631E-12 | PDC | EHD4 |
| 3.6114E-22 | 0.30074445 | 0.278 | 0.091 | 7.7102E-18 | PDC | ZNF503 |
| 9.1924E-26 | 0.30073655 | 0.325 | 0.108 | 1.9626E-21 | PDC | CRTAP |
| 3.0869E-36 | 0.30041639 | 0.283 | 0.069 | 6.5905E-32 | PDC | LRP11 |
| 1.3849E-40 | 0.30036832 | 0.27 | 0.058 | 2.9567E-36 | PDC | AHDC1 |
| 4.1021E-16 | 0.30034963 | 0.388 | 0.181 | 8.7581E-12 | PDC | TSPYL1 |
| 7.3699E-22 | 0.30031044 | 0.397 | 0.161 | 1.5735E-17 | PDC | CHURC1 |
| 1.1165E-16 | 0.30000923 | 0.46 | 0.226 | 2.3836E-12 | PDC | ENSA |
| 1.3711E-50 | 0.29978563 | 0.262 | 0.047 | 2.9272E-46 | PDC | LRFN4 |
| 5.8965E-34 | 0.29956965 | 0.304 | 0.08 | 1.2589E-29 | PDC | EFNB1 |
| 9.3607E-58 | 0.29931936 | 0.232 | 0.034 | 1.9985E-53 | PDC | TRAM2 |
| 7.8106E-49 | 0.29908279 | 0.278 | 0.054 | 1.6676E-44 | PDC | C6orf203 |
| 5.4337E-28 | 0.29894207 | 0.283 | 0.082 | 1.1601E-23 | PDC | MFSD6 |
| 1.3726E-19 | 0.29872963 | 0.316 | 0.121 | 2.9306E-15 | PDC | PLEKHA5 |
| 5.0711E-36 | 0.29872209 | 0.287 | 0.071 | 1.0827E-31 | PDC | PNMA1 |
| 1.2633E-49 | 0.29856024 | 0.257 | 0.046 | 2.6971E-45 | PDC | WFS1 |

| | | | | | | |
|---|---|---|---|---|---|---|
| 2.582E-16 | 0.29845962 | 0.549 | 0.284 | 5.5126E-12 | PDC | NFE2L2 |
| 9.8974E-39 | 0.29840463 | 0.236 | 0.048 | 2.1131E-34 | PDC | TMEM80 |
| 6.7655E-55 | 0.29816972 | 0.241 | 0.037 | 1.4444E-50 | PDC | SEPT1 |
| 7.9548E-35 | 0.29797049 | 0.308 | 0.082 | 1.6983E-30 | PDC | PPT1 |
| 2.8797E-38 | 0.2978211 | 0.262 | 0.058 | 6.1481E-34 | PDC | CBY1 |
| 6.9412E-29 | 0.2977794 | 0.329 | 0.102 | 1.4819E-24 | PDC | SLC9A3R1 |
| 2.3264E-20 | 0.29739721 | 0.278 | 0.098 | 4.9668E-16 | PDC | SLC48A1 |
| 1.9903E-20 | 0.29700974 | 0.371 | 0.15 | 4.2492E-16 | PDC | GLG1 |
| 3.7513E-14 | 0.29654592 | 0.582 | 0.332 | 8.009E-10 | PDC | EIF4B |
| 1.1559E-29 | 0.29640358 | 0.304 | 0.089 | 2.4678E-25 | PDC | TMEM18 |
| 5.4804E-12 | 0.29635119 | 0.38 | 0.201 | 1.1701E-07 | PDC | AIF1L |
| 6.4831E-44 | 0.2961891 | 0.308 | 0.068 | 1.3841E-39 | PDC | IFIT1 |
| 1.5192E-16 | 0.29617603 | 0.46 | 0.226 | 3.2435E-12 | PDC | SCARB2 |
| 7.5316E-50 | 0.29514561 | 0.257 | 0.046 | 1.608E-45 | PDC | TNFSF12 |
| 1.2473E-36 | 0.29502274 | 0.3 | 0.075 | 2.6629E-32 | PDC | STARD3NL |
| 1.1873E-21 | 0.29450133 | 0.329 | 0.121 | 2.5348E-17 | PDC | SMAD5 |
| 7.4439E-40 | 0.29425966 | 0.287 | 0.066 | 1.5893E-35 | PDC | SNAPIN |
| 3.3587E-30 | 0.29407301 | 0.266 | 0.07 | 7.1708E-26 | PDC | TANC1 |
| 2.4053E-18 | 0.29372891 | 0.646 | 0.343 | 5.1354E-14 | PDC | DDX24 |
| 2.388E-228 | 0.29343984 | 0.194 | 0.005 | 5.098E-224 | PDC | FLRT2 |
| 3.9377E-19 | 0.2931239 | 0.312 | 0.121 | 8.4069E-15 | PDC | OSBPL9 |
| 8.8559E-22 | 0.29297331 | 0.38 | 0.151 | 1.8907E-17 | PDC | METTL5 |
| 2.0305E-43 | 0.29219875 | 0.249 | 0.048 | 4.335E-39 | PDC | CDC25B |
| 1.1149E-19 | 0.29196722 | 0.342 | 0.135 | 2.3804E-15 | PDC | MAGT1 |
| 5.2862E-65 | 0.29188634 | 0.241 | 0.032 | 1.1286E-60 | PDC | SULT1A1 |
| 1.6772E-46 | 0.29171098 | 0.278 | 0.056 | 3.5808E-42 | PDC | METTL21A |
| 4.1682E-16 | 0.29123822 | 0.481 | 0.242 | 8.899E-12 | PDC | PPIG |
| 1.1512E-21 | 0.29099267 | 0.397 | 0.161 | 2.4578E-17 | PDC | RTF2 |
| 7.149E-171 | 0.29048536 | 0.186 | 0.006 | 1.526E-166 | PDC | ARHGAP20 |
| 6.9253E-54 | 0.29024074 | 0.257 | 0.043 | 1.4785E-49 | PDC | EHD2 |
| 1.1299E-37 | 0.28988104 | 0.253 | 0.055 | 2.4122E-33 | PDC | C2orf74 |
| 7.8545E-12 | 0.28969564 | 0.435 | 0.244 | 1.6769E-07 | PDC | BANF1 |
| 4.3275E-22 | 0.28961308 | 0.359 | 0.138 | 9.2392E-18 | PDC | ZNF428 |
| 4.7567E-28 | 0.28931235 | 0.304 | 0.092 | 1.0155E-23 | PDC | TADA3 |
| 2.4904E-25 | 0.28920252 | 0.316 | 0.104 | 5.317E-21 | PDC | USP1 |
| 6.7485E-40 | 0.28917514 | 0.249 | 0.052 | 1.4408E-35 | PDC | ITGA1 |
| 5.6691E-13 | 0.2890654 | 0.473 | 0.261 | 1.2103E-08 | PDC | TAPBP |
| 1.1448E-36 | 0.28898923 | 0.278 | 0.067 | 2.4442E-32 | PDC | SORBS3 |
| 3.4397E-18 | 0.28885786 | 0.354 | 0.149 | 7.3438E-14 | PDC | ATP6AP1 |
| 3.5638E-36 | 0.28868133 | 0.236 | 0.051 | 7.6088E-32 | PDC | GPX8 |
| 1.3997E-16 | 0.28839076 | 0.857 | 0.605 | 2.9883E-12 | PDC | EIF4A2 |
| 1.8962E-13 | 0.28806876 | 0.464 | 0.253 | 4.0485E-09 | PDC | MTPN |
| 5.7868E-33 | 0.2880303 | 0.3 | 0.081 | 1.2355E-28 | PDC | FAM210B |
| 2.2576E-54 | 0.28783423 | 0.186 | 0.023 | 4.82E-50 | PDC | UNC5B-AS1 |
| 9.748E-160 | 0.28783195 | 0.228 | 0.011 | 2.081E-155 | PDC | SMIM10 |
| 4.242E-12 | 0.28698737 | 0.544 | 0.319 | 9.0566E-08 | PDC | PHPT1 |
| 1.7587E-12 | 0.28662384 | 0.675 | 0.422 | 3.7548E-08 | PDC | ATP6V0B |
| 8.4093E-22 | 0.28639934 | 0.316 | 0.115 | 1.7954E-17 | PDC | CCDC91 |
| 7.956E-100 | 0.28621277 | 0.203 | 0.015 | 1.6986E-95 | PDC | PC |
| 9.4441E-25 | 0.28620651 | 0.304 | 0.099 | 2.0163E-20 | PDC | PMVK |

| | | | | | | |
|---|---|---|---|---|---|---|
| 2.5233E-14 | 0.28602753 | 0.409 | 0.204 | 5.3871E-10 | PDC | MYO1E |
| 3.5419E-13 | 0.28584974 | 0.62 | 0.383 | 7.5619E-09 | PDC | ATP5IF1 |
| 8.8009E-12 | 0.28584403 | 0.738 | 0.517 | 1.879E-07 | PDC | ARF1 |
| 1.3985E-31 | 0.28565831 | 0.283 | 0.076 | 2.9858E-27 | PDC | NME4 |
| 2.2535E-28 | 0.2854879 | 0.304 | 0.091 | 4.8112E-24 | PDC | FRA10AC1 |
| 3.0165E-24 | 0.285418 | 0.278 | 0.088 | 6.4402E-20 | PDC | TM2D3 |
| 1.6774E-23 | 0.28535537 | 0.287 | 0.092 | 3.5813E-19 | PDC | SLF2 |
| 1.027E-265 | 0.28534552 | 0.203 | 0.004 | 2.193E-261 | PDC | SLC45A1 |
| 7.6615E-15 | 0.28518368 | 0.359 | 0.166 | 1.6357E-10 | PDC | PHIP |
| 1.7435E-25 | 0.28502471 | 0.316 | 0.103 | 3.7223E-21 | PDC | FBXL5 |
| 1.9115E-16 | 0.28469601 | 0.473 | 0.235 | 4.0811E-12 | PDC | MRPL57 |
| 5.2981E-21 | 0.28465333 | 0.418 | 0.172 | 1.1311E-16 | PDC | DNAJB9 |
| 2.5366E-35 | 0.28399371 | 0.283 | 0.07 | 5.4156E-31 | PDC | ZNF32 |
| 6.3434E-29 | 0.2839622 | 0.127 | 0.02 | 1.3543E-24 | PDC | GSTA1 |
| 1.8183E-47 | 0.28388063 | 0.257 | 0.048 | 3.882E-43 | PDC | CBX7 |
| 1.1372E-69 | 0.28370204 | 0.249 | 0.032 | 2.4279E-65 | PDC | ROBO1 |
| 1.5699E-13 | 0.28356229 | 0.414 | 0.213 | 3.3518E-09 | PDC | THRAP3 |
| 1.1288E-33 | 0.28341283 | 0.262 | 0.063 | 2.4099E-29 | PDC | MAPK3 |
| 0 | 0.2833264 | 0.177 | 0.002 | 0 | PDC | ATP1A2 |
| 9.3606E-23 | 0.28313563 | 0.35 | 0.129 | 1.9985E-18 | PDC | MRPL40 |
| 2.1131E-19 | 0.28283612 | 0.3 | 0.112 | 4.5115E-15 | PDC | MYO1D |
| 4.9253E-14 | 0.28281153 | 0.384 | 0.187 | 1.0515E-09 | PDC | CARHSP1 |
| 6.1652E-17 | 0.28277155 | 0.342 | 0.146 | 1.3163E-12 | PDC | MXD4 |
| 4.0171E-23 | 0.28225414 | 0.316 | 0.109 | 8.5764E-19 | PDC | ESYT1 |
| 4.9771E-25 | 0.28199226 | 0.257 | 0.076 | 1.0626E-20 | PDC | ATP2B4 |
| 1.4971E-26 | 0.28169518 | 0.295 | 0.091 | 3.1963E-22 | PDC | ZNF580 |
| 1.4576E-13 | 0.28108142 | 0.485 | 0.272 | 3.112E-09 | PDC | CYB5A |
| 5.3747E-22 | 0.28099766 | 0.287 | 0.097 | 1.1475E-17 | PDC | SLC2A4RG |
| 4.7265E-52 | 0.28079581 | 0.253 | 0.043 | 1.0091E-47 | PDC | RELL1 |
| 1.2057E-18 | 0.28069657 | 0.363 | 0.152 | 2.5741E-14 | PDC | COA4 |
| 3.1661E-18 | 0.28057892 | 0.354 | 0.147 | 6.7596E-14 | PDC | ETS2 |
| 8.7456E-21 | 0.28053646 | 0.287 | 0.1 | 1.8672E-16 | PDC | JADE1 |
| 0 | 0.28049579 | 0.169 | 0 | 0 | PDC | AC109466.1 |
| 1.86E-271 | 0.28036212 | 0.203 | 0.004 | 3.971E-267 | PDC | APLP1 |
| 4.394E-19 | 0.28023586 | 0.397 | 0.171 | 9.3813E-15 | PDC | RNF11 |
| 5.5193E-40 | 0.27999567 | 0.215 | 0.039 | 1.1784E-35 | PDC | PPP1R3C |
| 1.1147E-30 | 0.2798453 | 0.274 | 0.073 | 2.3799E-26 | PDC | APPBP2 |
| 2.6334E-19 | 0.2796249 | 0.316 | 0.121 | 5.6224E-15 | PDC | VEGFB |
| 1.308E-12 | 0.27952068 | 0.414 | 0.219 | 2.7927E-08 | PDC | ARL6IP5 |
| 5.2082E-28 | 0.27941192 | 0.338 | 0.107 | 1.112E-23 | PDC | IMP3 |
| 2.8678E-74 | 0.27935634 | 0.215 | 0.023 | 6.1227E-70 | PDC | IL1RAP |
| 0 | 0.27901065 | 0.173 | 0.002 | 0 | PDC | SCN7A |
| 1.8708E-51 | 0.27888264 | 0.203 | 0.029 | 3.9941E-47 | PDC | PGAP1 |
| 2.4641E-32 | 0.27876558 | 0.287 | 0.075 | 5.2609E-28 | PDC | SPATA13 |
| 0 | 0.27775519 | 0.19 | 0.002 | 0 | PDC | GJA3 |
| 8.8312E-16 | 0.27737541 | 0.333 | 0.147 | 1.8855E-11 | PDC | MAP3K2 |
| 2.6023E-12 | 0.27694836 | 0.346 | 0.177 | 5.5559E-08 | PDC | NDUFS4 |
| 2.1165E-12 | 0.27626745 | 0.405 | 0.215 | 4.5187E-08 | PDC | FNBP1L |
| 5.4143E-60 | 0.27611747 | 0.181 | 0.02 | 1.156E-55 | PDC | GPD1 |
| 7.5579E-34 | 0.2758605 | 0.257 | 0.062 | 1.6136E-29 | PDC | AP1S2 |

| | | | | | | |
|---|---|---|---|---|---|---|
| 3.7391E-26 | 0.27540892 | 0.304 | 0.095 | 7.983E-22 | PDC | MRPL17 |
| 7.1523E-20 | 0.27499263 | 0.308 | 0.116 | 1.527E-15 | PDC | GRHPR |
| 1.1298E-17 | 0.27497999 | 0.363 | 0.155 | 2.412E-13 | PDC | MRPL34 |
| 7.3213E-19 | 0.27488367 | 0.397 | 0.168 | 1.5631E-14 | PDC | OTUD6B-AS1 |
| 1.7497E-21 | 0.27469025 | 0.3 | 0.105 | 3.7356E-17 | PDC | CCDC107 |
| 1.1281E-21 | 0.27429234 | 0.316 | 0.113 | 2.4084E-17 | PDC | CFL2 |
| 1.4156E-12 | 0.27423853 | 0.73 | 0.475 | 3.0224E-08 | PDC | PTGES3 |
| 3.3983E-16 | 0.27383924 | 0.354 | 0.157 | 7.2554E-12 | PDC | EAPP |
| 1.5245E-20 | 0.27370612 | 0.342 | 0.132 | 3.2548E-16 | PDC | ATXN10 |
| 2.6789E-24 | 0.27365475 | 0.278 | 0.087 | 5.7195E-20 | PDC | MEGF9 |
| 5.8717E-18 | 0.27328078 | 0.523 | 0.251 | 1.2536E-13 | PDC | MPHOSPH8 |
| 3.04E-22 | 0.27319838 | 0.333 | 0.122 | 6.4903E-18 | PDC | CETN2 |
| 5.6383E-18 | 0.27279184 | 0.371 | 0.159 | 1.2038E-13 | PDC | HAX1 |
| 3.9649E-29 | 0.27268503 | 0.274 | 0.076 | 8.465E-25 | PDC | ETFRF1 |
| 5.9506E-15 | 0.27228748 | 0.253 | 0.098 | 1.2705E-10 | PDC | PIK3R1 |
| 1.1198E-17 | 0.27204535 | 0.278 | 0.106 | 2.3907E-13 | PDC | TIMM17B |
| 1.7605E-13 | 0.27175305 | 0.464 | 0.245 | 3.7587E-09 | PDC | HNRNPR |
| 3.085E-14 | 0.27169506 | 0.35 | 0.164 | 6.5864E-10 | PDC | PRKDC |
| 0 | 0.27112809 | 0.114 | 0 | 0 | PDC | MYBPH |
| 7.6858E-61 | 0.271018 | 0.224 | 0.03 | 1.6409E-56 | PDC | EIF5A2 |
| 3.574E-140 | 0.27045726 | 0.127 | 0.003 | 7.63E-136 | PDC | CPXM1 |
| 4.8007E-28 | 0.27044804 | 0.257 | 0.07 | 1.025E-23 | PDC | NBN |
| 7.0333E-12 | 0.26924364 | 0.759 | 0.54 | 1.5016E-07 | PDC | CD59 |
| 2.4189E-23 | 0.26920725 | 0.367 | 0.135 | 5.1643E-19 | PDC | PPCS |
| 2.6178E-64 | 0.26909612 | 0.207 | 0.024 | 5.5889E-60 | PDC | IQSEC2 |
| 2.7452E-72 | 0.26907665 | 0.198 | 0.02 | 5.861E-68 | PDC | TBXAS1 |
| 5.3892E-14 | 0.26888672 | 0.291 | 0.124 | 1.1506E-09 | PDC | FTX |
| 1.3856E-13 | 0.26864043 | 0.544 | 0.299 | 2.9583E-09 | PDC | ARHGAP29 |
| 9.2545E-17 | 0.26853309 | 0.363 | 0.158 | 1.9758E-12 | PDC | DDAH2 |
| 1.9887E-15 | 0.26822997 | 0.325 | 0.141 | 4.2458E-11 | PDC | SKAP2 |
| 1.3289E-19 | 0.26773932 | 0.232 | 0.073 | 2.8372E-15 | PDC | DDIT4L |
| 6.5033E-14 | 0.26748253 | 0.338 | 0.158 | 1.3884E-09 | PDC | ETFB |
| 1.3229E-26 | 0.26687775 | 0.291 | 0.088 | 2.8243E-22 | PDC | TMED1 |
| 1.199E-32 | 0.26641181 | 0.287 | 0.074 | 2.5599E-28 | PDC | TMEM107 |
| 2.1722E-23 | 0.26588685 | 0.266 | 0.083 | 4.6377E-19 | PDC | IL13RA1 |
| 6.7181E-21 | 0.26582957 | 0.232 | 0.071 | 1.4343E-16 | PDC | GAS6 |
| 7.8018E-94 | 0.2657752 | 0.203 | 0.016 | 1.6657E-89 | PDC | PROS1 |
| 1.072E-15 | 0.26560568 | 0.342 | 0.152 | 2.2886E-11 | PDC | COPRS |
| 2.6139E-38 | 0.26548503 | 0.253 | 0.054 | 5.5807E-34 | PDC | LANCL1 |
| 6.6732E-18 | 0.26544048 | 0.363 | 0.153 | 1.4247E-13 | PDC | CASC4 |
| 7.17E-12 | 0.26535277 | 0.401 | 0.216 | 1.5308E-07 | PDC | LAMP2 |
| 1.9343E-18 | 0.26496091 | 0.397 | 0.171 | 4.1297E-14 | PDC | PSMD1 |
| 8.7562E-18 | 0.2648386 | 0.304 | 0.119 | 1.8694E-13 | PDC | MSRB2 |
| 9.7666E-18 | 0.2647743 | 0.388 | 0.17 | 2.0852E-13 | PDC | CIR1 |
| 5.1966E-21 | 0.2646487 | 0.354 | 0.137 | 1.1095E-16 | PDC | GCSH |
| 1.2036E-17 | 0.26424384 | 0.274 | 0.103 | 2.5698E-13 | PDC | LMBRD1 |
| 6.0296E-19 | 0.26423451 | 0.329 | 0.13 | 1.2873E-14 | PDC | IER3IP1 |
| 6.5923E-17 | 0.26423451 | 0.304 | 0.122 | 1.4075E-12 | PDC | ABRACL |
| 2.7779E-14 | 0.26406938 | 0.489 | 0.264 | 5.9309E-10 | PDC | NUDC |
| 3.9043E-24 | 0.26400707 | 0.278 | 0.087 | 8.3357E-20 | PDC | SIK3 |

| | | | | | | |
|---|---|---|---|---|---|---|
| 1.322E-11 | 0.26377937 | 0.553 | 0.338 | 2.8226E-07 | PDC | TRIR |
| 1.7227E-26 | 0.26369108 | 0.245 | 0.067 | 3.678E-22 | PDC | C15orf61 |
| 2.1789E-13 | 0.26368353 | 0.498 | 0.272 | 4.6519E-09 | PDC | CHMP2A |
| 4.1505E-11 | 0.26354815 | 0.544 | 0.331 | 8.8613E-07 | PDC | KDELR1 |
| 2.932E-16 | 0.26351641 | 0.346 | 0.152 | 6.2598E-12 | PDC | ARF5 |
| 9.0755E-08 | 0.26326205 | 0.439 | 0.272 | 0.00193762 | PDC | CITED2 |
| 3.8932E-43 | 0.26288831 | 0.241 | 0.045 | 8.3119E-39 | PDC | PLEKHG1 |
| 6.3806E-15 | 0.26281944 | 0.494 | 0.258 | 1.3623E-10 | PDC | PPP2CB |
| 2.3749E-10 | 0.26240589 | 0.578 | 0.363 | 5.0704E-06 | PDC | WDR83OS |
| 2.6533E-14 | 0.26145195 | 0.485 | 0.255 | 5.6647E-10 | PDC | NDUFA5 |
| 1.7427E-23 | 0.2614265 | 0.287 | 0.093 | 3.7206E-19 | PDC | EBAG9 |
| 1.2079E-26 | 0.26129146 | 0.253 | 0.07 | 2.5788E-22 | PDC | HPF1 |
| 1.9666E-38 | 0.26116713 | 0.215 | 0.041 | 4.1987E-34 | PDC | SEZ6L2 |
| 1.1634E-36 | 0.26102363 | 0.241 | 0.051 | 2.4838E-32 | PDC | EVL |
| 3.9896E-22 | 0.26092382 | 0.304 | 0.105 | 8.5178E-18 | PDC | CDC40 |
| 1.8651E-35 | 0.26074641 | 0.253 | 0.058 | 3.9821E-31 | PDC | THYN1 |
| 5.6212E-34 | 0.26063889 | 0.245 | 0.056 | 1.2001E-29 | PDC | ARL13B |
| 1.8827E-18 | 0.26048748 | 0.338 | 0.135 | 4.0195E-14 | PDC | SEC14L1 |
| 8.5566E-09 | 0.26036844 | 0.738 | 0.556 | 0.00018268 | PDC | RHOA |
| 7.9169E-11 | 0.26031091 | 0.46 | 0.265 | 1.6903E-06 | PDC | NDUFB8 |
| 2.2854E-37 | 0.26010143 | 0.257 | 0.057 | 4.8793E-33 | PDC | TCTN1 |
| 6.7488E-14 | 0.25991451 | 0.329 | 0.148 | 1.4409E-09 | PDC | TFPI |
| 3.4467E-14 | 0.25991118 | 0.308 | 0.137 | 7.3587E-10 | PDC | USO1 |
| 7.1163E-55 | 0.25937271 | 0.228 | 0.034 | 1.5193E-50 | PDC | GALC |
| 2.3559E-19 | 0.259301 | 0.287 | 0.104 | 5.0299E-15 | PDC | SPAG16 |
| 3.598E-35 | 0.25913881 | 0.241 | 0.053 | 7.6817E-31 | PDC | AKIP1 |
| 2.5704E-12 | 0.25913239 | 0.481 | 0.269 | 5.4879E-08 | PDC | COX17 |
| 2.754E-26 | 0.25911789 | 0.27 | 0.078 | 5.8797E-22 | PDC | FMC1 |
| 5.8632E-21 | 0.25884161 | 0.3 | 0.105 | 1.2518E-16 | PDC | FGFR1 |
| 8.6503E-31 | 0.25849026 | 0.245 | 0.06 | 1.8468E-26 | PDC | TRIQK |
| 7.4292E-16 | 0.25799913 | 0.321 | 0.137 | 1.5861E-11 | PDC | GANAB |
| 3.5745E-39 | 0.25789403 | 0.245 | 0.05 | 7.6316E-35 | PDC | NIPSNAP3A |
| 2.9361E-54 | 0.25788595 | 0.228 | 0.034 | 6.2685E-50 | PDC | TTLL7 |
| 1.5004E-27 | 0.25775363 | 0.27 | 0.076 | 3.2033E-23 | PDC | USP46 |
| 9.6389E-18 | 0.2576157 | 0.262 | 0.095 | 2.0579E-13 | PDC | PIGP |
| 3.4037E-16 | 0.25758735 | 0.354 | 0.156 | 7.2669E-12 | PDC | HRAS |
| 6.2702E-49 | 0.25667808 | 0.236 | 0.04 | 1.3387E-44 | PDC | MANSC1 |
| 5.2485E-21 | 0.25666713 | 0.253 | 0.082 | 1.1206E-16 | PDC | ATP8B1 |
| 4.195E-133 | 0.25633809 | 0.173 | 0.008 | 8.957E-129 | PDC | F5 |
| 9.7055E-42 | 0.25628394 | 0.266 | 0.055 | 2.0721E-37 | PDC | THAP7 |
| 3.4471E-13 | 0.25610202 | 0.418 | 0.214 | 7.3596E-09 | PDC | DHX36 |
| 2.462E-19 | 0.25603325 | 0.304 | 0.113 | 5.2563E-15 | PDC | FAM120AOS |
| 4.5073E-28 | 0.25601073 | 0.257 | 0.069 | 9.623E-24 | PDC | ZSCAN16-AS1 |
| 2.9388E-70 | 0.25596841 | 0.19 | 0.019 | 6.2742E-66 | PDC | CD200 |
| 1.2215E-11 | 0.25587811 | 0.414 | 0.229 | 2.608E-07 | PDC | MRPL20 |
| 9.4007E-42 | 0.25581835 | 0.236 | 0.045 | 2.0071E-37 | PDC | RAB28 |
| 1.9608E-12 | 0.25555407 | 0.414 | 0.219 | 4.1863E-08 | PDC | ANXA7 |
| 2.2754E-14 | 0.25552528 | 0.388 | 0.186 | 4.858E-10 | PDC | ARF6 |
| 2.1332E-15 | 0.25551069 | 0.439 | 0.213 | 4.5544E-11 | PDC | QKI |
| 1.5144E-25 | 0.25520961 | 0.224 | 0.058 | 3.2333E-21 | PDC | MAOA |

| | | | | | | |
|---|---|---|---|---|---|---|
| 1.0028E-28 | 0.25499399 | 0.236 | 0.06 | 2.1409E-24 | PDC | C1GALT1C1 |
| 1.9017E-13 | 0.25496699 | 0.477 | 0.259 | 4.0602E-09 | PDC | TMCO1 |
| 3.3244E-76 | 0.25480579 | 0.219 | 0.023 | 7.0976E-72 | PDC | PRMT6 |
| 1.3893E-21 | 0.25470587 | 0.253 | 0.081 | 2.9662E-17 | PDC | OSTF1 |
| 7.6939E-17 | 0.25458951 | 0.325 | 0.134 | 1.6426E-12 | PDC | PREPL |
| 2.6634E-10 | 0.25429737 | 0.266 | 0.129 | 5.6863E-06 | PDC | XPOT |
| 1.6428E-20 | 0.25426326 | 0.278 | 0.096 | 3.5075E-16 | PDC | MALSU1 |
| 4.2879E-15 | 0.25424546 | 0.371 | 0.173 | 9.1546E-11 | PDC | METTL26 |
| 6.8014E-13 | 0.25419262 | 0.806 | 0.577 | 1.4521E-08 | PDC | ATP6V1G1 |
| 9.6057E-11 | 0.2537838 | 0.464 | 0.27 | 2.0508E-06 | PDC | VPS28 |
| 2.9645E-17 | 0.25373046 | 0.295 | 0.116 | 6.3293E-13 | PDC | DPM3 |
| 3.6176E-37 | 0.2532795 | 0.207 | 0.039 | 7.7236E-33 | PDC | CERK |
| 9.6434E-12 | 0.25312886 | 0.312 | 0.153 | 2.0589E-07 | PDC | MARCH6 |
| 2.7003E-14 | 0.25304807 | 0.612 | 0.345 | 5.7652E-10 | PDC | VAPA |
| 2.8325E-06 | 0.25289947 | 0.177 | 0.09 | 0.06047356 | PDC | NR4A1 |
| 9.0936E-14 | 0.25264256 | 0.473 | 0.254 | 1.9415E-09 | PDC | ATP6V1E1 |
| 6.3792E-10 | 0.25261212 | 0.249 | 0.118 | 1.362E-05 | PDC | FBXO32 |
| 4.0877E-10 | 0.25255671 | 0.646 | 0.433 | 8.7273E-06 | PDC | NAP1L1 |
| 1.8282E-19 | 0.25236839 | 0.325 | 0.125 | 3.9032E-15 | PDC | FBXO7 |
| 8.0241E-20 | 0.25228468 | 0.3 | 0.11 | 1.7132E-15 | PDC | TAL/DCTD |
| 1.9495E-20 | 0.25225349 | 0.295 | 0.105 | 4.1622E-16 | PDC | PARP1 |
| 4.0347E-28 | 0.25172547 | 0.236 | 0.06 | 8.614E-24 | PDC | COX7A1 |
| 5.2479E-22 | 0.2516949 | 0.249 | 0.077 | 1.1204E-17 | PDC | JMJD8 |
| 1.8795E-18 | 0.25148798 | 0.304 | 0.116 | 4.0128E-14 | PDC | TSR3 |
| 9.191E-12 | 0.25141574 | 0.439 | 0.245 | 1.9623E-07 | PDC | PGRMC1 |
| 1.8739E-28 | 0.25128598 | 0.253 | 0.067 | 4.0008E-24 | PDC | FAM20B |
| 1.6273E-21 | 0.25125252 | 0.283 | 0.095 | 3.4742E-17 | PDC | THAP5 |
| 3.0906E-09 | 0.25088723 | 0.726 | 0.539 | 6.5984E-05 | PDC | GNG5 |
| 2.5316E-39 | 0.25076314 | 0.219 | 0.042 | 5.4051E-35 | PDC | CDC42EP2 |
| 4.8918E-36 | 0.25051684 | 0.215 | 0.043 | 1.0444E-31 | PDC | MAP7D3 |
| 1.2787E-17 | 0.25035723 | 0.287 | 0.11 | 2.73E-13 | PDC | TMEM126B |
| 2.0789E-58 | 0.25026848 | 0.19 | 0.023 | 4.4384E-54 | PDC | ANKRD44 |
| 5.9453E-05 | -0.250123 | 0.152 | 0.26 | 1 | PDC | HM13 |
| 1.1036E-05 | -0.2511713 | 0.021 | 0.109 | 0.23562372 | PDC | RGS2 |
| 0.00558635 | -0.2528995 | 0.819 | 0.826 | 1 | PDC | TMBIM6 |
| 7.3292E-05 | -0.2539858 | 0.059 | 0.149 | 1 | PDC | AL118516.1 |
| 0.00190235 | -0.2543333 | 0.122 | 0.194 | 1 | PDC | ERICH1 |
| 2.4607E-05 | -0.2549254 | 0.127 | 0.236 | 0.52536578 | PDC | LRRFIP2 |
| 0.00010371 | -0.2550773 | 0.063 | 0.154 | 1 | PDC | CA2 |
| 2.5205E-05 | -0.2554146 | 0.089 | 0.194 | 0.53812213 | PDC | KLHL21 |
| 7.8721E-07 | -0.2555092 | 0.084 | 0.214 | 0.0168069 | PDC | ADAM10 |
| 0.00867655 | -0.2562774 | 0.207 | 0.268 | 1 | PDC | BDP1 |
| 5.1239E-06 | -0.2566954 | 0.051 | 0.158 | 0.10939449 | PDC | SLC20A1 |
| 0.00036025 | -0.2567967 | 0.152 | 0.241 | 1 | PDC | ANKLE2 |
| 0.00061282 | -0.2572033 | 0.114 | 0.197 | 1 | PDC | NCOA7 |
| 1.2001E-08 | -0.257703 | 0.004 | 0.128 | 0.00025622 | PDC | SPNS2 |
| 3.2238E-09 | -0.2578935 | 0.004 | 0.137 | 6.8829E-05 | PDC | SMIM22 |
| 4.2334E-05 | -0.2579305 | 0.021 | 0.1 | 0.90383488 | PDC | MAFA |
| 7.5962E-08 | -0.2582718 | 0.059 | 0.2 | 0.00162178 | PDC | MAP2K3 |
| 0.00654849 | -0.2582892 | 0.945 | 0.946 | 1 | PDC | RPL11 |

| | | | | | | |
|---|---|---|---|---|---|---|
| 1.3184E-10 | -0.2583382 | 0 | 0.149 | 2.8147E-06 | PDC | DDAH1 |
| 0.0006458 | -0.2583709 | 0.734 | 0.782 | 1 | PDC | COX4I1 |
| 2.1516E-10 | -0.259294 | 0 | 0.146 | 4.5936E-06 | PDC | GALNT3 |
| 2.1664E-05 | -0.2592962 | 0.084 | 0.188 | 0.46252099 | PDC | TNFRSF10B |
| 0.00057227 | -0.2596219 | 0.228 | 0.321 | 1 | PDC | TKT |
| 6.8297E-07 | -0.2596659 | 0.03 | 0.141 | 0.01458133 | PDC | SNX29 |
| 1.5132E-08 | -0.2603664 | 0.076 | 0.234 | 0.00032308 | PDC | DMKN |
| 0.00068584 | -0.2606851 | 0.143 | 0.227 | 1 | PDC | ZNF91 |
| 0.00037774 | -0.2612315 | 0.249 | 0.34 | 1 | PDC | NOP10 |
| 4.929E-10 | -0.2620777 | 0.004 | 0.148 | 1.0524E-05 | PDC | CDCP1 |
| 1.3111E-06 | -0.2624911 | 0.152 | 0.291 | 0.0279927 | PDC | MT-ATP8 |
| 0.00036556 | -0.263297 | 0.063 | 0.146 | 1 | PDC | ID1 |
| 5.2827E-10 | -0.2634457 | 0.013 | 0.163 | 1.1279E-05 | PDC | WLS |
| 2.0872E-09 | -0.2640199 | 0.021 | 0.167 | 4.4563E-05 | PDC | CD82 |
| 4.2219E-07 | -0.2644876 | 0.021 | 0.134 | 0.00901365 | PDC | CSRP2 |
| 8.2784E-08 | -0.2645134 | 0.046 | 0.179 | 0.00176744 | PDC | ACTN1 |
| 1.1E-07 | -0.2652286 | 0.034 | 0.16 | 0.0023485 | PDC | HOOK2 |
| 3.2176E-09 | -0.2654037 | 0.013 | 0.151 | 6.8697E-05 | PDC | LAMB1 |
| 1.0235E-07 | -0.2658679 | 0.025 | 0.149 | 0.00218528 | PDC | SLC25A29 |
| 0.001587 | -0.2665462 | 0.27 | 0.342 | 1 | PDC | ARF4 |
| 9.6692E-06 | -0.2667446 | 0.063 | 0.17 | 0.20643638 | PDC | FAM102A |
| 0.00293464 | -0.2670098 | 0.063 | 0.128 | 1 | PDC | HMOX1 |
| 0.00041401 | -0.2675572 | 0.059 | 0.135 | 1 | PDC | CSRNP2 |
| 1.6219E-06 | -0.2690055 | 0.072 | 0.193 | 0.0346266 | PDC | ZBTB43 |
| 5.8113E-06 | -0.2701718 | 0.114 | 0.243 | 0.12407019 | PDC | HES4 |
| 4.3214E-09 | -0.2718859 | 0.021 | 0.162 | 9.2263E-05 | PDC | GOLM1 |
| 2.8746E-09 | -0.2733655 | 0 | 0.13 | 6.1373E-05 | PDC | CCDC198 |
| 6.5433E-08 | -0.2734844 | 0.034 | 0.162 | 0.001397 | PDC | ETS1 |
| 0.00014542 | -0.2737105 | 0.785 | 0.856 | 1 | PDC | RPL38 |
| 8.3979E-09 | -0.275354 | 0.051 | 0.199 | 0.0001793 | PDC | STX3 |
| 5.671E-08 | -0.2775692 | 0.063 | 0.206 | 0.00121075 | PDC | ERO1A |
| 5.8065E-08 | -0.2784939 | 0.008 | 0.126 | 0.00123968 | PDC | PTGER3 |
| 0.00207519 | -0.2788663 | 0.122 | 0.198 | 1 | PDC | MT-ND6 |
| 5.1037E-07 | -0.2793488 | 0.025 | 0.137 | 0.01089637 | PDC | ITPKC |
| 1.7857E-07 | -0.2798425 | 0.03 | 0.149 | 0.00381244 | PDC | EMX2 |
| 1.5239E-08 | -0.2819346 | 0.013 | 0.14 | 0.00032535 | PDC | NABP1 |
| 1.7802E-05 | -0.2826557 | 0.591 | 0.68 | 0.38008259 | PDC | YWHAZ |
| 3.5425E-10 | -0.2830273 | 0.004 | 0.151 | 7.5632E-06 | PDC | TFCP2L1 |
| 1.3804E-10 | -0.283193 | 0.004 | 0.156 | 2.9472E-06 | PDC | TRNP1 |
| 3.6811E-05 | -0.2845921 | 0.03 | 0.113 | 0.78590763 | PDC | HIST4H4 |
| 2.5538E-06 | -0.2854288 | 0.097 | 0.221 | 0.05452389 | PDC | EPS8L2 |
| 1.2352E-05 | -0.2857296 | 0.139 | 0.256 | 0.26371428 | PDC | COTL1 |
| 3.4578E-06 | -0.2861027 | 0.118 | 0.242 | 0.07382393 | PDC | AAK1 |
| 2.3425E-07 | -0.2861896 | 0.046 | 0.172 | 0.0050013 | PDC | TYMP |
| 3.8682E-07 | -0.2872525 | 0.042 | 0.164 | 0.0082586 | PDC | FNIP2 |
| 6.7495E-10 | -0.2874041 | 0.013 | 0.16 | 1.441E-05 | PDC | KIFC3 |
| 1.1551E-06 | -0.2888165 | 0.08 | 0.205 | 0.02466212 | PDC | CLK1 |
| 4.2629E-05 | -0.2890005 | 0.139 | 0.255 | 0.91012101 | PDC | PTP4A1 |
| 1.6149E-09 | -0.2896799 | 0.059 | 0.226 | 3.4477E-05 | PDC | CXCL16 |
| 5.5142E-10 | -0.2909728 | 0.021 | 0.175 | 1.1773E-05 | PDC | SH3BP4 |

| | | | | | | |
|---|---|---|---|---|---|---|
| 3.4735E-13 | -0.2915454 | 0.287 | 0.124 | 7.4159E-09 | PDC | GPX3 |
| 1.4233E-06 | -0.291865 | 0.135 | 0.268 | 0.03038817 | PDC | SLC38A1 |
| 8.3371E-10 | -0.2924662 | 0.017 | 0.165 | 1.78E-05 | PDC | HSPG2 |
| 0.00038373 | -0.2928253 | 0.793 | 0.871 | 1 | PDC | RPS5 |
| 2.1231E-07 | -0.2930394 | 0.089 | 0.224 | 0.00453271 | PDC | LSR |
| 0.00185577 | -0.2941383 | 0.211 | 0.287 | 1 | PDC | NUFIP2 |
| 0.00473412 | -0.2944832 | 0.148 | 0.21 | 1 | PDC | MAT2A |
| 1.7827E-09 | -0.295584 | 0.025 | 0.173 | 3.806E-05 | PDC | BICC1 |
| 0.00368191 | -0.2960996 | 0.312 | 0.371 | 1 | PDC | SPTBN1 |
| 6.0335E-11 | -0.297233 | 0.017 | 0.181 | 1.2882E-06 | PDC | BACE2 |
| 1.1698E-08 | -0.2977272 | 0.042 | 0.185 | 0.00024976 | PDC | TMEM238 |
| 1.7855E-09 | -0.2979441 | 0 | 0.133 | 3.812E-05 | PDC | AL353751.1 |
| 6.6499E-09 | -0.3001309 | 0.072 | 0.227 | 0.00014197 | PDC | GPRC5C |
| 2.5933E-06 | -0.3012243 | 0.038 | 0.146 | 0.05536733 | PDC | CKS2 |
| 0.00287728 | -0.3014216 | 0.911 | 0.914 | 1 | PDC | HSP90AB1 |
| 0.0031365 | -0.3018374 | 0.911 | 0.93 | 1 | PDC | RPS14 |
| 1.0514E-08 | -0.3020074 | 0.008 | 0.136 | 0.00022448 | PDC | CDH16 |
| 2.972E-09 | -0.3023093 | 0 | 0.13 | 6.3453E-05 | PDC | PKHD1 |
| 3.6357E-05 | -0.3036305 | 0.114 | 0.219 | 0.7762308 | PDC | PTGR1 |
| 3.0072E-05 | -0.3049116 | 0.16 | 0.27 | 0.64203244 | PDC | PABPC4 |
| 0.00052517 | -0.3058215 | 0.219 | 0.304 | 1 | PDC | EIF4A1 |
| 6.3773E-09 | -0.3059474 | 0.059 | 0.212 | 0.00013616 | PDC | RAB3IP |
| 1.8721E-09 | -0.3064637 | 0.042 | 0.197 | 3.9968E-05 | PDC | FAM129B |
| 0.00172799 | -0.3067427 | 0.287 | 0.36 | 1 | PDC | SPAG9 |
| 0.00013182 | -0.3077672 | 0.089 | 0.179 | 1 | PDC | KLF4 |
| 1.7601E-06 | -0.3079829 | 0.013 | 0.109 | 0.03757907 | PDC | AL355075.4 |
| 1.1077E-08 | -0.3081 | 0.017 | 0.15 | 0.00023648 | PDC | ARL4C |
| 0.00363992 | -0.3084065 | 0.236 | 0.303 | 1 | PDC | C6orf62 |
| 1.011E-08 | -0.3086016 | 0 | 0.122 | 0.00021584 | PDC | NEURL3 |
| 0.00016201 | -0.3090509 | 0.186 | 0.284 | 1 | PDC | ASPH |
| 1.1648E-07 | -0.3095457 | 0.03 | 0.153 | 0.0024869 | PDC | SLC16A3 |
| 7.2127E-07 | -0.309656 | 0.072 | 0.196 | 0.01539903 | PDC | POLR2J3.1 |
| 1.0805E-10 | -0.3101704 | 0.004 | 0.158 | 2.3068E-06 | PDC | CPM |
| 1.2644E-09 | -0.3107974 | 0.017 | 0.164 | 2.6995E-05 | PDC | PCSK1N |
| 1.2938E-07 | -0.3109968 | 0.004 | 0.113 | 0.00276225 | PDC | EGOT |
| 2.5902E-05 | -0.3144578 | 0.135 | 0.248 | 0.55301495 | PDC | IRS2 |
| 0.00854444 | -0.3147896 | 0.278 | 0.334 | 1 | PDC | RALBP1 |
| 5.34E-07 | -0.3148048 | 0.084 | 0.214 | 0.01140098 | PDC | SCCPDH |
| 3.4449E-06 | -0.3149547 | 0.068 | 0.183 | 0.07354758 | PDC | HMGCS1 |
| 0.00014557 | -0.316505 | 0.886 | 0.898 | 1 | PDC | MT-CYB |
| 7.0262E-06 | -0.3168213 | 0.173 | 0.291 | 0.15000999 | PDC | FAM3C |
| 2.1023E-05 | -0.317356 | 0.097 | 0.203 | 0.44884113 | PDC | CHCHD7 |
| 1.4028E-11 | -0.3177417 | 0.004 | 0.171 | 2.9949E-07 | PDC | ELK3 |
| 8.1416E-06 | -0.3182667 | 0.034 | 0.129 | 0.17382366 | PDC | HOMER1 |
| 1.6013E-10 | -0.3187781 | 0.004 | 0.155 | 3.4188E-06 | PDC | PROM1 |
| 0.00047122 | -0.3211844 | 0.089 | 0.166 | 1 | PDC | HIST1H1E |
| 6.0619E-09 | -0.324613 | 0.042 | 0.192 | 0.00012942 | PDC | EMP1 |
| 6.7521E-13 | -0.32467 | 0 | 0.181 | 1.4416E-08 | PDC | MYO10 |
| 1.5342E-07 | -0.3259809 | 0.105 | 0.246 | 0.00327557 | PDC | TNIP1 |
| 4.9259E-05 | -0.3288495 | 0.025 | 0.105 | 1 | PDC | RSAD2 |

| | | | | | | |
|---|---|---|---|---|---|---|
| 1.793E-05 | -0.3325034 | 0.097 | 0.207 | 0.38281342 | PDC | MDK |
| 1.9213E-10 | -0.3328604 | 0.004 | 0.154 | 4.102E-06 | PDC | KITLG |
| 3.229E-09 | -0.3337744 | 0.046 | 0.197 | 6.8939E-05 | PDC | EPS8 |
| 9.8082E-06 | -0.3350307 | 0.207 | 0.327 | 0.20940506 | PDC | ARL6IP1 |
| 2.3275E-06 | -0.3351326 | 0.489 | 0.616 | 0.04969319 | PDC | MT-ND4L |
| 1.7545E-06 | -0.3364769 | 0.278 | 0.41 | 0.03745872 | PDC | JAK1 |
| 1.9073E-12 | -0.3373043 | 0.025 | 0.214 | 4.0721E-08 | PDC | SERINC2 |
| 1.5185E-06 | -0.3382481 | 0.008 | 0.103 | 0.0324199 | PDC | TNF |
| 1.8964E-09 | -0.3387325 | 0.03 | 0.178 | 4.0488E-05 | PDC | ZNF217 |
| 0.00188664 | -0.3400155 | 0.101 | 0.173 | 1 | PDC | FOSB |
| 9.1325E-13 | -0.3422789 | 0.008 | 0.192 | 1.9498E-08 | PDC | LLGL2 |
| 2.0679E-07 | -0.3427848 | 0.131 | 0.276 | 0.00441489 | PDC | HIF1A |
| 1.9457E-06 | -0.3433301 | 0.139 | 0.265 | 0.04154123 | PDC | CHD2 |
| 3.6598E-07 | -0.3438437 | 0.051 | 0.177 | 0.00781371 | PDC | PHLDA1 |
| 0.00447415 | -0.3439134 | 0.785 | 0.844 | 1 | PDC | RPSA |
| 0.00081266 | -0.3444522 | 0.831 | 0.866 | 1 | PDC | UBA52 |
| 0.00013448 | -0.3450642 | 0.949 | 0.933 | 1 | PDC | RPS9 |
| 0.00208301 | -0.3460579 | 0.802 | 0.804 | 1 | PDC | CFL1 |
| 0.00293972 | -0.3461927 | 0.865 | 0.886 | 1 | PDC | RPS29 |
| 1.9474E-06 | -0.3462938 | 0.515 | 0.628 | 0.04157653 | PDC | ATF4 |
| 0.00013416 | -0.3464494 | 0.312 | 0.398 | 1 | PDC | CYCS |
| 0.00120817 | -0.3475303 | 0.139 | 0.217 | 1 | PDC | GCLM |
| 6.5207E-12 | -0.3478381 | 0.021 | 0.201 | 1.3922E-07 | PDC | ADGRG1 |
| 0.00018197 | -0.3485894 | 0.873 | 0.913 | 1 | PDC | RPL29 |
| 6.2233E-13 | -0.3502676 | 0.017 | 0.208 | 1.3287E-08 | PDC | OCLN |
| 4.1175E-07 | -0.3508504 | 0.008 | 0.113 | 0.00879076 | PDC | AC023157.3 |
| 9.9691E-06 | -0.3513139 | 0.924 | 0.913 | 0.21284104 | PDC | MT-ND4 |
| 5.1179E-05 | -0.3516201 | 0.911 | 0.942 | 1 | PDC | RPS24 |
| 7.9384E-06 | -0.3519992 | 0.743 | 0.806 | 0.16948409 | PDC | EEF1D |
| 0.00830701 | -0.3520506 | 0.38 | 0.423 | 1 | PDC | SNHG8 |
| 1.1907E-07 | -0.3526109 | 0.228 | 0.376 | 0.00254215 | PDC | HEBP2 |
| 3.205E-08 | -0.3529189 | 0.105 | 0.256 | 0.00068427 | PDC | PNPLA2 |
| 5.6156E-05 | -0.3548646 | 0.173 | 0.276 | 1 | PDC | IFRD1 |
| 2.3904E-09 | -0.3555648 | 0 | 0.132 | 5.1034E-05 | PDC | PRSS22 |
| 0.00396406 | -0.3578191 | 0.215 | 0.287 | 1 | PDC | HIST1H1C |
| 1.3928E-05 | -0.3585115 | 0.017 | 0.101 | 0.29737144 | PDC | AP001160.1 |
| 0.00162157 | -0.3589155 | 0.709 | 0.716 | 1 | PDC | TXN |
| 8.1802E-07 | -0.3593809 | 0.152 | 0.293 | 0.01746481 | PDC | MAFF |
| 1.4066E-07 | -0.3601334 | 0.219 | 0.373 | 0.00300299 | PDC | SDCBP |
| 1.861E-09 | -0.3605 | 0.004 | 0.141 | 3.9732E-05 | PDC | TNFSF10 |
| 0.00018081 | -0.3616142 | 0.785 | 0.823 | 1 | PDC | RPS21 |
| 3.2691E-09 | -0.3617353 | 0.03 | 0.174 | 6.9795E-05 | PDC | DUSP5 |
| 3.4945E-05 | -0.362068 | 0.19 | 0.299 | 0.74607735 | PDC | HSPH1 |
| 5.2091E-05 | -0.3624894 | 0.435 | 0.567 | 1 | PDC | IER2 |
| 3.1422E-09 | -0.3639821 | 0.173 | 0.353 | 6.7085E-05 | PDC | ANXA11 |
| 1.979E-06 | -0.3648681 | 0.228 | 0.35 | 0.04225167 | PDC | UBE2H |
| 6.5147E-07 | -0.3658956 | 0.021 | 0.128 | 0.01390886 | PDC | HIST2H2BF |
| 1.7966E-08 | -0.3663014 | 0.025 | 0.161 | 0.00038357 | PDC | MGST1 |
| 2.2746E-11 | -0.3663724 | 0.004 | 0.167 | 4.8563E-07 | PDC | ADGRF1 |
| 0.00010559 | -0.3669411 | 0.266 | 0.36 | 1 | PDC | GCC2 |

| | | | | | | |
|---|---|---|---|---|---|---|
| 6.0702E-09 | -0.3675458 | 0.021 | 0.158 | 0.0001296 | PDC | CDKN2B |
| 2.0406E-05 | -0.3676248 | 0.852 | 0.894 | 0.43567258 | PDC | RPS11 |
| 0.00038316 | -0.3676478 | 0.059 | 0.135 | 1 | PDC | IFIT2 |
| 9.1403E-14 | -0.3681895 | 0.004 | 0.2 | 1.9515E-09 | PDC | WWC1 |
| 2.8116E-06 | -0.36926 | 0.013 | 0.106 | 0.06002833 | PDC | ARC |
| 2.113E-07 | -0.3697706 | 0.03 | 0.148 | 0.00451115 | PDC | ANKRD37 |
| 1.2065E-11 | -0.3704553 | 0.03 | 0.209 | 2.5759E-07 | PDC | ABHD5 |
| 8.5427E-05 | -0.3708415 | 0.713 | 0.784 | 1 | PDC | EEF1B2 |
| 0.00011819 | -0.3724327 | 0.156 | 0.251 | 1 | PDC | IDI1 |
| 0.00020725 | -0.3751249 | 0.127 | 0.218 | 1 | PDC | UBE2S |
| 0.001475 | -0.3754866 | 0.527 | 0.552 | 1 | PDC | EIF5 |
| 1.2641E-06 | -0.3757622 | 0.177 | 0.306 | 0.02698819 | PDC | YTHDC1 |
| 1.2119E-13 | -0.3766808 | 0 | 0.191 | 2.5874E-09 | PDC | SCNN1A |
| 1.3701E-05 | -0.3769011 | 0.017 | 0.102 | 0.2925185 | PDC | CXCL3 |
| 1.8172E-13 | -0.3774337 | 0.013 | 0.207 | 3.8797E-09 | PDC | FLNB |
| 1.3806E-07 | -0.3785271 | 0.038 | 0.163 | 0.00294753 | PDC | TNFAIP3 |
| 0.00015153 | -0.3803284 | 0.97 | 0.957 | 1 | PDC | EIF1 |
| 1.1472E-09 | -0.3830123 | 0.105 | 0.28 | 2.4493E-05 | PDC | SMS |
| 3.2229E-15 | -0.3836492 | 0.008 | 0.225 | 6.8809E-11 | PDC | RBM47 |
| 2.975E-08 | -0.3839188 | 0.405 | 0.609 | 0.00063516 | PDC | GADD45B |
| 0.00014246 | -0.3876035 | 0.173 | 0.266 | 1 | PDC | AMD1 |
| 0.00015464 | -0.3881857 | 0.675 | 0.707 | 1 | PDC | DDX5 |
| 4.3461E-14 | -0.3885499 | 0.004 | 0.203 | 9.2789E-10 | PDC | TINAGL1 |
| 6.1568E-06 | -0.38941 | 0.814 | 0.873 | 0.13144845 | PDC | RACK1 |
| 1.6153E-07 | -0.3909454 | 0.207 | 0.352 | 0.00344863 | PDC | ACADVL |
| 2.867E-12 | -0.3931094 | 0.034 | 0.229 | 6.121E-08 | PDC | CD74 |
| 7.5367E-13 | -0.3937127 | 0.034 | 0.229 | 1.6091E-08 | PDC | C4orf48 |
| 4.3899E-07 | -0.3938763 | 0.025 | 0.137 | 0.00937251 | PDC | AC025164.1 |
| 2.0446E-08 | -0.3950593 | 0.73 | 0.804 | 0.00043652 | PDC | CALM1 |
| 7.3286E-06 | -0.3957032 | 0.886 | 0.905 | 0.15646484 | PDC | RPS7 |
| 0.00085568 | -0.4014831 | 0.641 | 0.656 | 1 | PDC | HSPA8 |
| 2.9932E-12 | -0.4015124 | 0.008 | 0.185 | 6.3906E-08 | PDC | CD40 |
| 1.6935E-08 | -0.4024073 | 0.848 | 0.923 | 0.00036155 | PDC | RPL14 |
| 6.3123E-12 | -0.4025272 | 0.051 | 0.243 | 1.3477E-07 | PDC | KLF5 |
| 1.6861E-12 | -0.4025344 | 0.021 | 0.207 | 3.5999E-08 | PDC | PNP |
| 6.578E-06 | -0.40254 | 0.916 | 0.934 | 0.14043959 | PDC | RPL6 |
| 2.9365E-10 | -0.4041025 | 0.084 | 0.259 | 6.2693E-06 | PDC | NEDD4L |
| 5.2875E-07 | -0.404662 | 0.038 | 0.152 | 0.01128875 | PDC | AC007952.4 |
| 5.6996E-10 | -0.404816 | 0.008 | 0.155 | 1.2169E-05 | PDC | PFKFB3 |
| 3.603E-14 | -0.4053277 | 0.004 | 0.204 | 7.6924E-10 | PDC | EHF |
| 3.1514E-11 | -0.4090064 | 0.038 | 0.217 | 6.7283E-07 | PDC | CTSH |
| 0.00036943 | -0.41045 | 0.291 | 0.368 | 1 | PDC | PNN |
| 1.0715E-10 | -0.4117933 | 0.013 | 0.17 | 2.2876E-06 | PDC | CA12 |
| 0.00255045 | -0.4150301 | 0.819 | 0.795 | 1 | PDC | HNRNPA2B1 |
| 5.9367E-13 | -0.4171179 | 0.013 | 0.202 | 1.2675E-08 | PDC | MARCKS |
| 0.00013425 | -0.4194054 | 0.321 | 0.402 | 1 | PDC | FUS |
| 0.00037531 | -0.4196611 | 0.038 | 0.11 | 1 | PDC | S100A9 |
| 0.00318053 | -0.419741 | 0.241 | 0.307 | 1 | PDC | HSPA1B |
| 1.1262E-05 | -0.4201757 | 0.101 | 0.211 | 0.24043575 | PDC | ADAMTS9 |
| 4.7079E-06 | -0.4214347 | 0.063 | 0.177 | 0.10051356 | PDC | IGFBP3 |

| | | | | | | |
|---|---|---|---|---|---|---|
| 1.477E-16 | -0.4234782 | 0.013 | 0.247 | 3.1533E-12 | PDC | PFKP |
| 6.6965E-15 | -0.4239791 | 0.017 | 0.232 | 1.4297E-10 | PDC | ST14 |
| 1.4814E-11 | -0.426236 | 0.004 | 0.171 | 3.1629E-07 | PDC | FGF2 |
| 1.3188E-09 | -0.4299649 | 0.034 | 0.185 | 2.8156E-05 | PDC | ERN1 |
| 0.00067427 | -0.4319394 | 0.245 | 0.314 | 1 | PDC | SRSF2 |
| 3.6751E-05 | -0.4323294 | 0.852 | 0.883 | 0.78464012 | PDC | RPL39 |
| 1.2653E-06 | -0.4331407 | 0.907 | 0.944 | 0.02701494 | PDC | RPL8 |
| 0.00214592 | -0.4333536 | 0.485 | 0.511 | 1 | PDC | H2AFZ |
| 7.6272E-05 | -0.4356508 | 0.405 | 0.494 | 1 | PDC | ATP1B1 |
| 2.3787E-06 | -0.4362001 | 0.017 | 0.113 | 0.05078542 | PDC | C15orf48 |
| 2.7569E-08 | -0.4363787 | 0.544 | 0.65 | 0.00058861 | PDC | UQCRH |
| 3.4464E-11 | -0.4367705 | 0.118 | 0.318 | 7.358E-07 | PDC | AFDN |
| 7.4407E-11 | -0.4481299 | 0.025 | 0.191 | 1.5886E-06 | PDC | SYTL2 |
| 1.9499E-07 | -0.4489209 | 0.869 | 0.913 | 0.00416301 | PDC | RPL35 |
| 6.4047E-08 | -0.4491669 | 0.747 | 0.82 | 0.00136741 | PDC | YBX1 |
| 1.5595E-06 | -0.4498914 | 0.907 | 0.911 | 0.0332947 | PDC | MT-CO2 |
| 7.3846E-07 | -0.4558103 | 0.139 | 0.277 | 0.01576618 | PDC | VMP1 |
| 5.0689E-12 | -0.4563667 | 0.072 | 0.269 | 1.0822E-07 | PDC | LIMA1 |
| 6.36E-16 | -0.4590795 | 0.021 | 0.252 | 1.3579E-11 | PDC | TES |
| 7.9565E-07 | -0.4613132 | 0.224 | 0.377 | 0.01698722 | PDC | ATF3 |
| 3.2713E-06 | -0.4640459 | 0.928 | 0.946 | 0.0698421 | PDC | RPS15 |
| 1.4822E-05 | -0.4651295 | 0.038 | 0.13 | 0.31644047 | PDC | COL7A1 |
| 1.7607E-09 | -0.4667144 | 0.042 | 0.195 | 3.759E-05 | PDC | H2AFX |
| 7.8912E-11 | -0.4683323 | 0.684 | 0.807 | 1.6848E-06 | PDC | RPL31 |
| 8.3947E-09 | -0.4701046 | 0.143 | 0.31 | 0.00017923 | PDC | YME1L1 |
| 5.6581E-10 | -0.4748177 | 0.152 | 0.329 | 1.208E-05 | PDC | CTSB |
| 5.8777E-14 | -0.4785703 | 0.008 | 0.207 | 1.2549E-09 | PDC | CLDN1 |
| 2.1017E-16 | -0.4793196 | 0.004 | 0.232 | 4.4872E-12 | PDC | PRSS8 |
| 1.2432E-05 | -0.4814892 | 0.253 | 0.358 | 0.26541759 | PDC | DDX3X |
| 4.2675E-12 | -0.4852234 | 0.139 | 0.344 | 9.1112E-08 | PDC | JUP |
| 4.0422E-05 | -0.4867977 | 0.295 | 0.379 | 0.86301417 | PDC | RSRP1 |
| 4.7813E-13 | -0.4905527 | 0.481 | 0.697 | 1.0208E-08 | PDC | ZFAS1 |
| 2.222E-11 | -0.4925325 | 0.055 | 0.237 | 4.744E-07 | PDC | CDKN2AIP |
| 2.5775E-08 | -0.4927928 | 0.928 | 0.948 | 0.0005503 | PDC | RPL13A |
| 5.2664E-07 | -0.4953371 | 0.92 | 0.928 | 0.01124382 | PDC | RPL18A |
| 1.7248E-18 | -0.4953736 | 0.025 | 0.288 | 3.6826E-14 | PDC | SPINT1 |
| 0.00033945 | -0.4968485 | 0.16 | 0.244 | 1 | PDC | ID4 |
| 2.9551E-16 | -0.4978503 | 0.021 | 0.253 | 6.3092E-12 | PDC | MGLL |
| 1.4841E-11 | -0.5015672 | 0.186 | 0.389 | 3.1685E-07 | PDC | MYO6 |
| 6.69E-07 | -0.5023462 | 0.19 | 0.323 | 0.01428315 | PDC | CCNL1 |
| 2.7698E-07 | -0.5038745 | 0.954 | 0.972 | 0.00591345 | PDC | RPLP1 |
| 1.0576E-11 | -0.5065344 | 0.557 | 0.705 | 2.258E-07 | PDC | EIF4G2 |
| 2.9773E-08 | -0.506974 | 0.224 | 0.377 | 0.00063565 | PDC | LDHB |
| 1.1667E-11 | -0.5070892 | 0.063 | 0.251 | 2.491E-07 | PDC | RAP2B |
| 1.8747E-07 | -0.5081058 | 0.131 | 0.273 | 0.00400243 | PDC | SEPHS2 |
| 7.3423E-10 | -0.5143981 | 0.034 | 0.189 | 1.5676E-05 | PDC | BIRC3 |
| 5.6239E-16 | -0.5159396 | 0.068 | 0.314 | 1.2007E-11 | PDC | DSP |
| 1.4852E-17 | -0.5175433 | 0 | 0.239 | 3.1708E-13 | PDC | SCIN |
| 0.0009341 | -0.5175815 | 0.827 | 0.837 | 1 | PDC | PRDX1 |
| 4.8549E-19 | -0.5191065 | 0.021 | 0.291 | 1.0365E-14 | PDC | ANXA3 |

| | | | | | | |
|---|---|---|---|---|---|---|
| 1.7869E-12 | -0.5194462 | 0.405 | 0.591 | 3.8151E-08 | PDC | S100A13 |
| 1.0518E-10 | -0.5219654 | 0.928 | 0.973 | 2.2457E-06 | PDC | TPT1 |
| 1.184E-13 | -0.5233138 | 0.059 | 0.27 | 2.5278E-09 | PDC | GSTM3 |
| 1.0575E-09 | -0.5240602 | 0.046 | 0.209 | 2.2578E-05 | PDC | RRAD |
| 3.2468E-08 | -0.5251639 | 0.92 | 0.934 | 0.0006932 | PDC | HSP90AA1 |
| 1.7877E-07 | -0.5254828 | 0.654 | 0.733 | 0.00381676 | PDC | PFN1 |
| 3.985E-11 | -0.5277551 | 0.962 | 0.974 | 8.508E-07 | PDC | RPL13 |
| 4.1749E-10 | -0.528706 | 0.16 | 0.34 | 8.9134E-06 | PDC | HNRNPAB |
| 1.1903E-05 | -0.532717 | 0.017 | 0.103 | 0.25412572 | PDC | APOE |
| 3.5519E-12 | -0.5328919 | 0.844 | 0.902 | 7.5834E-08 | PDC | RPL27A |
| 3.3109E-07 | -0.5346958 | 0 | 0.1 | 0.00706886 | PDC | SLC12A1 |
| 3.2425E-08 | -0.5377455 | 0.287 | 0.424 | 0.00069228 | PDC | SFPQ |
| 1.068E-07 | -0.5437481 | 0.143 | 0.286 | 0.00228017 | PDC | POLR2A |
| 0.00019824 | -0.5443567 | 0.806 | 0.815 | 1 | PDC | NEAT1 |
| 1.8431E-20 | -0.549695 | 0 | 0.273 | 3.935E-16 | PDC | MET |
| 9.5187E-11 | -0.5547461 | 0.38 | 0.554 | 2.0323E-06 | PDC | CD9 |
| 9.3273E-13 | -0.5624172 | 0.089 | 0.306 | 1.9914E-08 | PDC | DNAJB1 |
| 2.8346E-13 | -0.5661506 | 0.089 | 0.302 | 6.0519E-09 | PDC | CYTOR |
| 6.7998E-12 | -0.5713327 | 0.852 | 0.936 | 1.4518E-07 | PDC | RPL37 |
| 2.3589E-13 | -0.5720219 | 0.852 | 0.932 | 5.0362E-09 | PDC | RPL12 |
| 3.4631E-08 | -0.5766454 | 0.08 | 0.221 | 0.00073938 | PDC | TNFAIP2 |
| 2.2535E-11 | -0.5823729 | 0.852 | 0.89 | 4.8113E-07 | PDC | MT-ND2 |
| 2.9248E-17 | -0.584707 | 0.076 | 0.336 | 6.2445E-13 | PDC | TUBA1C |
| 2.9077E-08 | -0.5867964 | 0.19 | 0.337 | 0.0006208 | PDC | ZNF292 |
| 3.2674E-11 | -0.5872535 | 0.008 | 0.172 | 6.9758E-07 | PDC | EGLN3 |
| 1.3633E-18 | -0.5879076 | 0.198 | 0.503 | 2.9107E-14 | PDC | TNFRSF12A |
| 4.4742E-08 | -0.5964666 | 0.802 | 0.83 | 0.00095525 | PDC | SQSTM1 |
| 3.3383E-21 | -0.5975076 | 0.017 | 0.304 | 7.1273E-17 | PDC | CAPG |
| 3.409E-16 | -0.6028406 | 0.055 | 0.3 | 7.2781E-12 | PDC | MIR4435-2HG |
| 4.4355E-13 | -0.6046788 | 0.932 | 0.962 | 9.4697E-09 | PDC | RPS18 |
| 6.7844E-16 | -0.6057569 | 0.498 | 0.693 | 1.4485E-11 | PDC | CYSTM1 |
| 1.3896E-07 | -0.6062224 | 0.008 | 0.119 | 0.00296672 | PDC | NEFM |
| 4.0724E-05 | -0.6070678 | 0.034 | 0.119 | 0.86946331 | PDC | CXCL8 |
| 3.1442E-11 | -0.6096661 | 0.257 | 0.442 | 6.7128E-07 | PDC | PPP1R14B |
| 2.0037E-22 | -0.6122437 | 0.008 | 0.305 | 4.2779E-18 | PDC | CDH1 |
| 2.2289E-09 | -0.6128645 | 0.093 | 0.256 | 4.7587E-05 | PDC | IER5 |
| 7.6856E-19 | -0.6141698 | 0.738 | 0.908 | 1.6409E-14 | PDC | RPS20 |
| 3.1035E-16 | -0.6142576 | 0.046 | 0.285 | 6.626E-12 | PDC | CLDN3 |
| 6.1018E-18 | -0.6149077 | 0.498 | 0.704 | 1.3027E-13 | PDC | MAP1LC3B |
| 1.4314E-13 | -0.6150419 | 0.034 | 0.237 | 3.056E-09 | PDC | NFKBIZ |
| 1.602E-07 | -0.6213085 | 0.819 | 0.868 | 0.00342018 | PDC | MT-ND3 |
| 6.6257E-08 | -0.6224959 | 0.156 | 0.304 | 0.00141458 | PDC | DDIT4 |
| 2.1164E-16 | -0.622595 | 0.013 | 0.243 | 4.5186E-12 | PDC | ITGB6 |
| 8.3869E-19 | -0.6239935 | 0.004 | 0.26 | 1.7906E-14 | PDC | S100A14 |
| 8.9187E-15 | -0.6248442 | 0.165 | 0.397 | 1.9041E-10 | PDC | TAF1D |
| 5.4968E-18 | -0.6275044 | 0.03 | 0.283 | 1.1736E-13 | PDC | COBLL1 |
| 1.0924E-15 | -0.6295672 | 0.013 | 0.237 | 2.3322E-11 | PDC | TGM2 |
| 1.0563E-18 | -0.6405223 | 0 | 0.253 | 2.2553E-14 | PDC | MACC1 |
| 3.3994E-13 | -0.6415263 | 0.945 | 0.956 | 7.2577E-09 | PDC | RPS2 |
| 4.4165E-16 | -0.6468808 | 0.852 | 0.925 | 9.4293E-12 | PDC | RPLP2 |

| | | | | | | |
|---|---|---|---|---|---|---|
| 7.7109E-15 | -0.6484407 | 0.013 | 0.225 | 1.6463E-10 | PDC | SFN |
| 1.3027E-06 | -0.6496065 | 0.236 | 0.361 | 0.02781303 | PDC | KLF6 |
| 2.6961E-08 | -0.6554025 | 0.253 | 0.388 | 0.00057561 | PDC | BRD2 |
| 4.9383E-12 | -0.6586661 | 0.013 | 0.188 | 1.0543E-07 | PDC | PIK3R3 |
| 9.556E-20 | -0.6634997 | 0.008 | 0.278 | 2.0402E-15 | PDC | MUC1 |
| 6.1685E-14 | -0.6636838 | 0.878 | 0.929 | 1.317E-09 | PDC | RPL36 |
| 4.1832E-23 | -0.6680833 | 0.059 | 0.377 | 8.9312E-19 | PDC | CAMK2N1 |
| 2.688E-18 | -0.6816902 | 0.127 | 0.39 | 5.7388E-14 | PDC | OCIAD2 |
| 5.3046E-13 | -0.6861201 | 0.021 | 0.216 | 1.1325E-08 | PDC | GLRX |
| 2.8443E-14 | -0.6888355 | 0.013 | 0.218 | 6.0725E-10 | PDC | SOCS1 |
| 4.826E-23 | -0.6918339 | 0.017 | 0.324 | 1.0303E-18 | PDC | FAM107B |
| 1.6903E-16 | -0.6953362 | 0.008 | 0.239 | 3.6089E-12 | PDC | FXYD2 |
| 2.4561E-05 | -0.7043759 | 0.523 | 0.636 | 0.52438303 | PDC | JUN |
| 8.4621E-07 | -0.7121419 | 0.008 | 0.108 | 0.0180666 | PDC | FXYD4 |
| 0.00192659 | -0.7127199 | 0.338 | 0.392 | 1 | PDC | NFKBIA |
| 3.3497E-18 | -0.7164377 | 0.013 | 0.264 | 7.1516E-14 | PDC | LAMB3 |
| 1.6303E-14 | -0.7225052 | 0.068 | 0.289 | 3.4807E-10 | PDC | AKR1B1 |
| 9.6261E-12 | -0.7252079 | 0.072 | 0.258 | 2.0552E-07 | PDC | IER5L |
| 3.568E-17 | -0.7271421 | 0.165 | 0.424 | 7.6178E-13 | PDC | LGALS3 |
| 1.4703E-15 | -0.7303725 | 0.055 | 0.293 | 3.1392E-11 | PDC | CYR61 |
| 5.6707E-12 | -0.7312695 | 0.055 | 0.241 | 1.2107E-07 | PDC | INTS6 |
| 6.3031E-07 | -0.7322383 | 0.253 | 0.377 | 0.0134572 | PDC | SLC38A2 |
| 2.1735E-19 | -0.7346751 | 0.051 | 0.325 | 4.6404E-15 | PDC | DCDC2 |
| 8.8645E-15 | -0.7445133 | 0.038 | 0.256 | 1.8926E-10 | PDC | ID2 |
| 3.7107E-18 | -0.7533873 | 0.92 | 0.966 | 7.9223E-14 | PDC | RPS19 |
| 7.499E-13 | -0.753663 | 0.869 | 0.91 | 1.601E-08 | PDC | MT-ATP6 |
| 4.5264E-17 | -0.757132 | 0.473 | 0.671 | 9.6638E-13 | PDC | GLS |
| 2.8577E-09 | -0.7629183 | 0.004 | 0.138 | 6.1012E-05 | PDC | SERPINA1 |
| 1.9172E-13 | -0.763006 | 0.017 | 0.215 | 4.0933E-09 | PDC | ADAMTS1 |
| 5.4238E-17 | -0.7684029 | 0.042 | 0.29 | 1.158E-12 | PDC | PLAUR |
| 2.7345E-08 | -0.7712224 | 0.139 | 0.279 | 0.00058381 | PDC | PPP1R10 |
| 6.3629E-21 | -0.7788256 | 0.046 | 0.344 | 1.3585E-16 | PDC | PAX8 |
| 3.0843E-18 | -0.7832407 | 0.477 | 0.672 | 6.5849E-14 | PDC | TPI1 |
| 4.0273E-18 | -0.7951653 | 0.131 | 0.396 | 8.5984E-14 | PDC | RAB11FIP1 |
| 2.6768E-23 | -0.7952586 | 0.776 | 0.918 | 5.715E-19 | PDC | RPLP0 |
| 1.3252E-09 | -0.8033683 | 0.27 | 0.418 | 2.8293E-05 | PDC | SRSF7 |
| 2.1841E-20 | -0.8083008 | 0.025 | 0.305 | 4.6632E-16 | PDC | MARCKSL1 |
| 1.0209E-23 | -0.8134053 | 0.046 | 0.365 | 2.1796E-19 | PDC | CD44 |
| 8.8074E-11 | -0.8149834 | 0.143 | 0.33 | 1.8804E-06 | PDC | IL32 |
| 1.6873E-22 | -0.8209889 | 0.017 | 0.321 | 3.6023E-18 | PDC | MAL |
| 1.186E-27 | -0.8331518 | 0.494 | 0.77 | 2.5321E-23 | PDC | PKM |
| 2.3644E-20 | -0.8338786 | 0.342 | 0.636 | 5.048E-16 | PDC | TPM1 |
| 1.4259E-20 | -0.8412399 | 0.941 | 0.961 | 3.0444E-16 | PDC | RPL28 |
| 7.4772E-13 | -0.8436349 | 0.709 | 0.831 | 1.5964E-08 | PDC | MT-ND5 |
| 1.3436E-15 | -0.8459853 | 0.03 | 0.257 | 2.8687E-11 | PDC | AKAP12 |
| 6.6129E-17 | -0.8460442 | 0.844 | 0.892 | 1.4119E-12 | PDC | MT-ND1 |
| 5.9511E-12 | -0.8465195 | 0.743 | 0.873 | 1.2706E-07 | PDC | S100A10 |
| 2.5605E-17 | -0.85682 | 0.042 | 0.293 | 5.4667E-13 | PDC | RND3 |
| 3.2706E-14 | -0.878597 | 0.781 | 0.875 | 6.9828E-10 | PDC | GSTP1 |
| 1.9852E-23 | -0.8883819 | 0.004 | 0.311 | 4.2384E-19 | PDC | PMAIP1 |

| | | | | | | |
|---|---|---|---|---|---|---|
| 8.9562E-19 | -0.8918421 | 0.076 | 0.346 | 1.9122E-14 | PDC | SNHG12 |
| 6.3554E-15 | -0.8931202 | 0.899 | 0.91 | 1.3569E-10 | PDC | MT-CO3 |
| 6.4617E-27 | -0.896305 | 0.051 | 0.405 | 1.3796E-22 | PDC | JPT1 |
| 4.3721E-19 | -0.9007643 | 0.401 | 0.614 | 9.3343E-15 | PDC | BTG1 |
| 6.5969E-15 | -0.9020224 | 0.021 | 0.24 | 1.4084E-10 | PDC | CST6 |
| 2.9368E-36 | -0.9097524 | 0.367 | 0.735 | 6.2701E-32 | PDC | ENO1 |
| 7.8536E-13 | -0.9239311 | 0.042 | 0.238 | 1.6767E-08 | PDC | CSKMT |
| 1.0506E-20 | -0.931507 | 0.021 | 0.307 | 2.243E-16 | PDC | PLK2 |
| 2.8054E-24 | -0.9380105 | 0.321 | 0.629 | 5.9895E-20 | PDC | PPP1R15A |
| 3.8734E-12 | -0.9428741 | 0.084 | 0.281 | 8.2697E-08 | PDC | HES1 |
| 1.0054E-13 | -0.9945696 | 0.034 | 0.246 | 2.1466E-09 | PDC | TFPI2 |
| 3.8882E-38 | -0.9965426 | 0.076 | 0.512 | 8.3013E-34 | PDC | CLDN7 |
| 9.2856E-21 | -1.0040673 | 0.004 | 0.282 | 1.9825E-16 | PDC | RASD1 |
| 1.2135E-10 | -1.0073877 | 0.097 | 0.274 | 2.5908E-06 | PDC | HSPA2 |
| 2.2591E-15 | -1.0331427 | 0.084 | 0.312 | 4.8233E-11 | PDC | PLAU |
| 1.0582E-16 | -1.0343603 | 0.16 | 0.411 | 2.2592E-12 | PDC | RNF213 |
| 1.3866E-23 | -1.0347333 | 0.101 | 0.423 | 2.9603E-19 | PDC | CCND1 |
| 2.3775E-09 | -1.0468127 | 0.148 | 0.3 | 5.076E-05 | PDC | HEXIM1 |
| 3.0369E-19 | -1.0724652 | 0.865 | 0.925 | 6.4837E-15 | PDC | TMSB10 |
| 2.1065E-21 | -1.0771412 | 0.034 | 0.325 | 4.4973E-17 | PDC | ITGA2 |
| 3.0407E-20 | -1.1033048 | 0.321 | 0.623 | 6.4918E-16 | PDC | MTRNR2L8 |
| 3.4574E-27 | -1.1063735 | 0.274 | 0.611 | 7.3815E-23 | PDC | SOD2 |
| 7.4728E-20 | -1.1097824 | 0.439 | 0.676 | 1.5954E-15 | PDC | SH3BGRL3 |
| 3.2367E-18 | -1.1109079 | 0.021 | 0.276 | 6.9103E-14 | PDC | TCIM |
| 1.7081E-27 | -1.1176928 | 0.063 | 0.42 | 3.6469E-23 | PDC | PHLDA2 |
| 1.3306E-41 | -1.1244044 | 0.114 | 0.567 | 2.8409E-37 | PDC | HMGA1 |
| 8.2888E-18 | -1.1862052 | 0.291 | 0.571 | 1.7697E-13 | PDC | GDF15 |
| 2.2999E-20 | -1.2421203 | 0.051 | 0.336 | 4.9102E-16 | PDC | LAMC2 |
| 1.6761E-32 | -1.2482016 | 0.097 | 0.492 | 3.5785E-28 | PDC | TSPAN1 |
| 1.3027E-27 | -1.2756198 | 0.903 | 0.927 | 2.7812E-23 | PDC | MT-CO1 |
| 3.1654E-65 | -1.2803041 | 0.101 | 0.69 | 6.7581E-61 | PDC | EPCAM |
| 5.0339E-26 | -1.2829839 | 0.118 | 0.449 | 1.0747E-21 | PDC | UPP1 |
| 7.607E-14 | -1.3407411 | 0.046 | 0.259 | 1.6241E-09 | PDC | LCN2 |
| 1.4451E-30 | -1.3553057 | 0.093 | 0.477 | 3.0853E-26 | PDC | C12orf75 |
| 6.5536E-24 | -1.4182523 | 0.122 | 0.446 | 1.3992E-19 | PDC | IER3 |
| 1.2994E-15 | -1.5005443 | 0.008 | 0.23 | 2.7743E-11 | PDC | DEFB1 |
| 6.3739E-42 | -1.5039528 | 0.152 | 0.608 | 1.3608E-37 | PDC | CLU |
| 3.0429E-45 | -1.6105027 | 0.118 | 0.6 | 6.4966E-41 | PDC | KRT19 |
| 8.5475E-60 | -1.6123787 | 0.11 | 0.673 | 1.8249E-55 | PDC | TACSTD2 |
| 3.5051E-41 | -1.6136293 | 0.7 | 0.923 | 7.4834E-37 | PDC | S100A11 |
| 1.4105E-19 | -1.6414071 | 0.426 | 0.683 | 3.0115E-15 | PDC | MTRNR2L12 |
| 9.6089E-62 | -1.6570862 | 0.148 | 0.738 | 2.0515E-57 | PDC | KRT8 |
| 2.357E-55 | -1.6784093 | 0.118 | 0.659 | 5.0321E-51 | PDC | KRT7 |
| 6.8966E-59 | -1.7155981 | 0.494 | 0.878 | 1.4724E-54 | PDC | SAT1 |
| 5.1241E-48 | -1.7269134 | 0.101 | 0.596 | 1.094E-43 | PDC | ELF3 |
| 2.1805E-66 | -1.7693029 | 0.118 | 0.713 | 4.6553E-62 | PDC | CLDN4 |
| 7.8801E-22 | -1.7727953 | 0.093 | 0.397 | 1.6824E-17 | PDC | SLPI |
| 9.2275E-68 | -1.7887012 | 0.236 | 0.81 | 1.9701E-63 | PDC | KRT18 |
| 1.2918E-86 | -1.7948849 | 0.131 | 0.813 | 2.7579E-82 | PDC | CD24 |
| 2.4123E-51 | -1.8847833 | 0.131 | 0.655 | 5.1502E-47 | PDC | WFDC2 |

| | | | | | | |
|---|---|---|---|---|---|---|
| 6.8463E-39 | -2.2224188 | 0.118 | 0.547 | 1.4617E-34 | PDC | SOX4 |
| 5.2723E-31 | -2.349017 | 0.122 | 0.499 | 1.1256E-26 | PDC | S100A2 |
| 1.6763E-44 | -2.608642 | 0.135 | 0.608 | 3.5789E-40 | PDC | SPP1 |
| 4.6802E-50 | -3.1161391 | 0.139 | 0.646 | 9.9923E-46 | PDC | MMP7 |
| 2.007E-23 | -4.9521379 | 0.38 | 0.602 | 4.2848E-19 | PDC | PLCG2 |
| 0 | 2.57898246 | 0.854 | 0.088 | 0 | PDC_inj | SERPINE2 |
| 3.382E-285 | 2.38059808 | 0.683 | 0.073 | 7.221E-281 | PDC_inj | MT1H |
| 0 | 2.34360163 | 0.787 | 0.014 | 0 | PDC_inj | APOD |
| 1.298E-201 | 2.27662694 | 0.933 | 0.246 | 2.771E-197 | PDC_inj | NUPR1 |
| 0 | 2.25968936 | 0.53 | 0.01 | 0 | PDC_inj | IGFBP1 |
| 1.355E-143 | 2.16956591 | 0.761 | 0.185 | 2.892E-139 | PDC_inj | MT1G |
| 0 | 1.80659095 | 0.787 | 0.093 | 0 | PDC_inj | BST2 |
| 2.081E-267 | 1.76060495 | 0.84 | 0.132 | 4.443E-263 | PDC_inj | MYL9 |
| 0 | 1.72758431 | 0.791 | 0.072 | 0 | PDC_inj | MAP3K20 |
| 6.681E-144 | 1.71808388 | 0.862 | 0.271 | 1.426E-139 | PDC_inj | UCHL1 |
| 3.0447E-68 | 1.62919339 | 0.664 | 0.238 | 6.5004E-64 | PDC_inj | MT1X |
| 4.167E-269 | 1.61338286 | 0.433 | 0.027 | 8.897E-265 | PDC_inj | PTGDS |
| 3.338E-100 | 1.59369153 | 0.996 | 0.978 | 7.1274E-96 | PDC_inj | FTH1 |
| 3.691E-178 | 1.58066972 | 0.836 | 0.209 | 7.88E-174 | PDC_inj | BEX2 |
| 0 | 1.5328337 | 0.507 | 0.022 | 0 | PDC_inj | CHI3L1 |
| 2.005E-163 | 1.50510446 | 0.78 | 0.179 | 4.28E-159 | PDC_inj | CTSL |
| 0 | 1.49825961 | 0.784 | 0.032 | 0 | PDC_inj | MME |
| 6.49E-172 | 1.49755527 | 0.892 | 0.263 | 1.386E-167 | PDC_inj | TCEA1 |
| 3.5352E-89 | 1.43420083 | 0.996 | 0.961 | 7.5477E-85 | PDC_inj | FTL |
| 7.291E-98 | 1.42623986 | 0.955 | 0.651 | 1.5566E-93 | PDC_inj | PSAP |
| 1.689E-171 | 1.42077627 | 0.821 | 0.205 | 3.606E-167 | PDC_inj | CCPG1 |
| 0 | 1.41576175 | 0.728 | 0.046 | 0 | PDC_inj | IGFBP4 |
| 3.473E-104 | 1.3568602 | 0.896 | 0.358 | 7.415E-100 | PDC_inj | DDIT3 |
| 0 | 1.27540487 | 0.739 | 0.069 | 0 | PDC_inj | MAP1B |
| 6.74E-261 | 1.2507589 | 0.743 | 0.1 | 1.439E-256 | PDC_inj | GBE1 |
| 1.9463E-85 | 1.24167629 | 0.985 | 0.718 | 4.1554E-81 | PDC_inj | CTSD |
| 1.318E-117 | 1.1600395 | 0.847 | 0.299 | 2.813E-113 | PDC_inj | ISCU |
| 0 | 1.14606747 | 0.634 | 0.04 | 0 | PDC_inj | SERPINI1 |
| 0 | 1.13555377 | 0.631 | 0.055 | 0 | PDC_inj | TGFBR3 |
| 0 | 1.12837998 | 0.638 | 0.035 | 0 | PDC_inj | PPM1L |
| 8.3336E-81 | 1.12270894 | 0.985 | 0.931 | 1.7792E-76 | PDC_inj | RPS6 |
| 9.867E-71 | 1.11178847 | 0.799 | 0.316 | 2.1066E-66 | PDC_inj | MT1E |
| 2.4817E-81 | 1.10763829 | 0.925 | 0.744 | 5.2984E-77 | PDC_inj | DSTN |
| 1.1745E-63 | 1.10739984 | 0.993 | 0.95 | 2.5076E-59 | PDC_inj | RPS12 |
| 1.953E-174 | 1.09608752 | 0.672 | 0.114 | 4.17E-170 | PDC_inj | GPNMB |
| 1.5855E-88 | 1.06766051 | 0.687 | 0.213 | 3.3851E-84 | PDC_inj | NAMPT |
| 2.5768E-68 | 1.03226563 | 0.966 | 0.921 | 5.5015E-64 | PDC_inj | RPL30 |
| 1.9743E-88 | 1.02956198 | 0.955 | 0.629 | 4.2152E-84 | PDC_inj | CCNI |
| 0 | 1.02649848 | 0.601 | 0.014 | 0 | PDC_inj | CLGN |
| 7.973E-157 | 1.02575604 | 0.698 | 0.138 | 1.702E-152 | PDC_inj | NUCB2 |
| 2.6227E-61 | 1.01223652 | 0.847 | 0.459 | 5.5994E-57 | PDC_inj | IFITM3 |
| 1.578E-54 | 1.01152067 | 0.959 | 0.706 | 3.3691E-50 | PDC_inj | MT2A |
| 1.099E-180 | 1.0024972 | 0.761 | 0.149 | 2.347E-176 | PDC_inj | ADH5 |
| 6.5963E-64 | 0.993838 | 0.993 | 0.922 | 1.4083E-59 | PDC_inj | RPL10 |
| 1.0349E-48 | 0.99375389 | 0.918 | 0.751 | 2.2094E-44 | PDC_inj | RPS26 |

| | | | | | | |
|---|---|---|---|---|---|---|
| 2.455E-244 | 0.98502648 | 0.619 | 0.07 | 5.241E-240 | PDC_inj | SH3BP5 |
| 0 | 0.98433686 | 0.496 | 0.024 | 0 | PDC_inj | TMEM91 |
| 5.804E-196 | 0.98364736 | 0.713 | 0.119 | 1.239E-191 | PDC_inj | C7orf50 |
| 3.8392E-56 | 0.95890954 | 0.612 | 0.247 | 8.1967E-52 | PDC_inj | RPS4Y1 |
| 3.9563E-62 | 0.9567847 | 0.993 | 0.95 | 8.4468E-58 | PDC_inj | RPS8 |
| 7.1172E-50 | 0.95139818 | 0.907 | 0.679 | 1.5195E-45 | PDC_inj | HSPA5 |
| 8.353E-195 | 0.94890484 | 0.631 | 0.089 | 1.783E-190 | PDC_inj | NEBL |
| 2.2986E-91 | 0.94394624 | 0.69 | 0.218 | 4.9074E-87 | PDC_inj | SEC11C |
| 8.716E-133 | 0.93173178 | 0.787 | 0.189 | 1.861E-128 | PDC_inj | CALD1 |
| 9.78E-112 | 0.92713906 | 0.619 | 0.144 | 2.088E-107 | PDC_inj | DUSP23 |
| 9.9581E-75 | 0.91344233 | 1 | 0.937 | 2.126E-70 | PDC_inj | RPS28 |
| 2.999E-63 | 0.90824817 | 1 | 0.955 | 6.4029E-59 | PDC_inj | RPS27 |
| 4.327E-119 | 0.89883432 | 0.713 | 0.164 | 9.238E-115 | PDC_inj | FOXC1 |
| 4.845E-111 | 0.8984421 | 0.634 | 0.146 | 1.034E-106 | PDC_inj | PALLD |
| 1.7879E-60 | 0.89101031 | 0.996 | 0.939 | 3.8171E-56 | PDC_inj | RPL19 |
| 7.82E-139 | 0.88950513 | 0.698 | 0.149 | 1.67E-134 | PDC_inj | RNF187 |
| 1.136E-170 | 0.88893707 | 0.672 | 0.117 | 2.426E-166 | PDC_inj | UQCC2 |
| 3.686E-213 | 0.87570015 | 0.687 | 0.096 | 7.87E-209 | PDC_inj | FAM129A |
| 1.1275E-81 | 0.87197783 | 0.787 | 0.284 | 2.4072E-77 | PDC_inj | CD55 |
| 4.4961E-61 | 0.86850967 | 1 | 0.939 | 9.5992E-57 | PDC_inj | RPL37A |
| 1.058E-111 | 0.86502783 | 0.806 | 0.243 | 2.258E-107 | PDC_inj | EIF3M |
| 1.9837E-98 | 0.84934246 | 0.799 | 0.268 | 4.2352E-94 | PDC_inj | ANAPC16 |
| 9.886E-155 | 0.84515967 | 0.567 | 0.09 | 2.111E-150 | PDC_inj | PSMB9 |
| 1.1695E-58 | 0.8361719 | 0.985 | 0.918 | 2.4968E-54 | PDC_inj | RPL9 |
| 0 | 0.83609903 | 0.407 | 0.008 | 0 | PDC_inj | CES1 |
| 2.9276E-71 | 0.82957713 | 0.806 | 0.352 | 6.2505E-67 | PDC_inj | TRMT112 |
| 8.9439E-56 | 0.82579991 | 0.978 | 0.905 | 1.9095E-51 | PDC_inj | RPS16 |
| 1.072E-91 | 0.82367868 | 0.731 | 0.229 | 2.2887E-87 | PDC_inj | C6orf48 |
| 6.734E-137 | 0.82031 | 0.627 | 0.121 | 1.438E-132 | PDC_inj | XPOT |
| 4.0534E-59 | 0.81564777 | 0.974 | 0.901 | 8.6541E-55 | PDC_inj | RPS25 |
| 5.9269E-85 | 0.81533268 | 0.41 | 0.077 | 1.2654E-80 | PDC_inj | HPGD |
| 0 | 0.81501931 | 0.41 | 0.018 | 0 | PDC_inj | ADAMTS5 |
| 3.678E-56 | 0.81384111 | 0.847 | 0.484 | 7.8526E-52 | PDC_inj | TOMM7 |
| 7.3493E-71 | 0.81101456 | 0.821 | 0.329 | 1.5691E-66 | PDC_inj | HSPA9 |
| 4.651E-205 | 0.8107427 | 0.522 | 0.057 | 9.929E-201 | PDC_inj | MYLK |
| 2.483E-174 | 0.80916731 | 0.459 | 0.052 | 5.302E-170 | PDC_inj | CAMTA2 |
| 1.602E-116 | 0.79643855 | 0.582 | 0.116 | 3.42E-112 | PDC_inj | CMYA5 |
| 7.27E-106 | 0.79479197 | 0.683 | 0.178 | 1.552E-101 | PDC_inj | BEX4 |
| 9.854E-51 | 0.79386119 | 0.959 | 0.794 | 2.1038E-46 | PDC_inj | HLA-C |
| 3.3605E-74 | 0.77952882 | 0.724 | 0.258 | 7.1747E-70 | PDC_inj | BLVRB |
| 3.144E-125 | 0.77895835 | 0.649 | 0.137 | 6.713E-121 | PDC_inj | RASSF8 |
| 0 | 0.77704827 | 0.593 | 0.032 | 0 | PDC_inj | HTRA1 |
| 8.3088E-86 | 0.77035263 | 0.638 | 0.181 | 1.7739E-81 | PDC_inj | SEL1L |
| 7.9746E-52 | 0.76931739 | 0.993 | 0.927 | 1.7026E-47 | PDC_inj | RPL21 |
| 2.1058E-77 | 0.76575846 | 0.757 | 0.274 | 4.496E-73 | PDC_inj | C12orf57 |
| 1.798E-119 | 0.76563725 | 0.582 | 0.117 | 3.84E-115 | PDC_inj | HSPA13 |
| 5.0846E-80 | 0.75855918 | 0.761 | 0.253 | 1.0856E-75 | PDC_inj | SELENOM |
| 3.9253E-54 | 0.7570235 | 0.918 | 0.595 | 8.3804E-50 | PDC_inj | CAST |
| 6.8993E-63 | 0.75604142 | 0.642 | 0.221 | 1.473E-58 | PDC_inj | NFE2L1 |
| 1.2966E-49 | 0.75382002 | 0.989 | 0.932 | 2.7682E-45 | PDC_inj | RPL3 |

| | | | | | | |
|---|---|---|---|---|---|---|
| 2.7278E-67 | 0.75307466 | 0.795 | 0.332 | 5.8238E-63 | PDC_inj | GHITM |
| 6.17E-79 | 0.74264183 | 0.664 | 0.202 | 1.3173E-74 | PDC_inj | GFPT1 |
| 1.4072E-47 | 0.74066637 | 0.933 | 0.789 | 3.0044E-43 | PDC_inj | PPDPF |
| 3.497E-107 | 0.73964437 | 0.616 | 0.145 | 7.467E-103 | PDC_inj | CFAP36 |
| 0 | 0.73359336 | 0.47 | 0.02 | 0 | PDC_inj | PTPRO |
| 1.0121E-43 | 0.73252952 | 0.951 | 0.824 | 2.1609E-39 | PDC_inj | HLA-B |
| 7.2239E-53 | 0.72456836 | 0.66 | 0.261 | 1.5423E-48 | PDC_inj | RPL36A |
| 0 | 0.72398621 | 0.422 | 0.019 | 0 | PDC_inj | DCN |
| 2.0906E-76 | 0.72363232 | 0.716 | 0.252 | 4.4634E-72 | PDC_inj | JTB |
| 1.4877E-50 | 0.71955545 | 0.978 | 0.895 | 3.1761E-46 | PDC_inj | RPL35A |
| 0 | 0.71381665 | 0.504 | 0.017 | 0 | PDC_inj | LRRC2 |
| 2.5332E-42 | 0.71279311 | 1 | 0.971 | 5.4083E-38 | PDC_inj | RPLP1 |
| 0 | 0.70969047 | 0.481 | 0.026 | 0 | PDC_inj | CLIC5 |
| 1.156E-142 | 0.70919164 | 0.463 | 0.064 | 2.468E-138 | PDC_inj | PARD3B |
| 1.995E-156 | 0.70408498 | 0.515 | 0.072 | 4.259E-152 | PDC_inj | UBE2L6 |
| 1.1212E-72 | 0.70393346 | 0.731 | 0.263 | 2.3938E-68 | PDC_inj | COX17 |
| 8.1885E-42 | 0.70327784 | 0.978 | 0.908 | 1.7482E-37 | PDC_inj | RPL26 |
| 3.204E-43 | 0.70206136 | 0.978 | 0.867 | 6.8405E-39 | PDC_inj | RPS5 |
| 4.1298E-83 | 0.69561781 | 0.664 | 0.199 | 8.817E-79 | PDC_inj | REXO2 |
| 4.449E-116 | 0.69504955 | 0.586 | 0.12 | 9.498E-112 | PDC_inj | HINT2 |
| 4.5393E-75 | 0.69244768 | 0.769 | 0.275 | 9.6914E-71 | PDC_inj | ANP32B |
| 4.0722E-63 | 0.69117137 | 0.84 | 0.374 | 8.6941E-59 | PDC_inj | PTMS |
| 5.9017E-56 | 0.69013289 | 0.69 | 0.273 | 1.26E-51 | PDC_inj | MAP4 |
| 2.6247E-43 | 0.6897476 | 0.981 | 0.937 | 5.6038E-39 | PDC_inj | RPS23 |
| 1.115E-234 | 0.68950029 | 0.504 | 0.045 | 2.379E-230 | PDC_inj | IQGAP2 |
| 1.3328E-40 | 0.68672141 | 0.914 | 0.688 | 2.8456E-36 | PDC_inj | ZFAS1 |
| 5.5408E-62 | 0.68490643 | 0.321 | 0.065 | 1.183E-57 | PDC_inj | SLC40A1 |
| 2.1108E-42 | 0.68358557 | 0.974 | 0.933 | 4.5066E-38 | PDC_inj | RPL32 |
| 5.4105E-57 | 0.68149121 | 0.683 | 0.271 | 1.1551E-52 | PDC_inj | LMAN1 |
| 3.0339E-83 | 0.68113721 | 0.623 | 0.176 | 6.4773E-79 | PDC_inj | DNAJC1 |
| 2.7692E-80 | 0.67330442 | 0.653 | 0.193 | 5.9123E-76 | PDC_inj | NDUFAF3 |
| 4.891E-104 | 0.67264288 | 0.575 | 0.125 | 1.044E-99 | PDC_inj | PSIP1 |
| 1.368E-119 | 0.67172626 | 0.485 | 0.082 | 2.921E-115 | PDC_inj | EFEMP1 |
| 6.4892E-82 | 0.67077559 | 0.668 | 0.202 | 1.3854E-77 | PDC_inj | RAD23A |
| 2.4703E-83 | 0.6706375 | 0.56 | 0.144 | 5.2742E-79 | PDC_inj | WARS |
| 1.7821E-74 | 0.67008908 | 0.425 | 0.093 | 3.8048E-70 | PDC_inj | GMDS |
| 3.1053E-62 | 0.66728916 | 0.821 | 0.359 | 6.6299E-58 | PDC_inj | ARL6IP4 |
| 1.2642E-98 | 0.66722344 | 0.646 | 0.163 | 2.699E-94 | PDC_inj | ARL2 |
| 1.8093E-71 | 0.66376392 | 0.709 | 0.245 | 3.8629E-67 | PDC_inj | SARS |
| 9.0199E-39 | 0.66302501 | 0.959 | 0.894 | 1.9257E-34 | PDC_inj | RPS13 |
| 1.582E-162 | 0.66117701 | 0.41 | 0.044 | 3.377E-158 | PDC_inj | PLXDC2 |
| 1.4108E-40 | 0.66060115 | 0.97 | 0.896 | 3.012E-36 | PDC_inj | RPS15A |
| 7.9098E-40 | 0.65888102 | 0.981 | 0.933 | 1.6887E-35 | PDC_inj | RPS27A |
| 5.5993E-59 | 0.65718355 | 0.608 | 0.213 | 1.1955E-54 | PDC_inj | PCMTD1 |
| 1.0117E-79 | 0.65242697 | 0.627 | 0.182 | 2.1599E-75 | PDC_inj | PSMB8 |
| 3.5992E-48 | 0.64975087 | 0.731 | 0.327 | 7.6843E-44 | PDC_inj | H2AFJ |
| 4.4431E-46 | 0.64709763 | 0.795 | 0.383 | 9.4859E-42 | PDC_inj | CIRBP |
| 1.5012E-90 | 0.64634163 | 0.604 | 0.153 | 3.205E-86 | PDC_inj | HAX1 |
| 1.3247E-67 | 0.64488178 | 0.537 | 0.147 | 2.8281E-63 | PDC_inj | IGFBP2 |
| 1.7557E-57 | 0.64343038 | 0.739 | 0.306 | 3.7484E-53 | PDC_inj | COX7A2L |

| | | | | | | |
|---|---|---|---|---|---|---|
| 3.464E-43 | 0.63967252 | 0.854 | 0.526 | 7.3956E-39 | PDC_inj | SEC61B |
| 7.5437E-45 | 0.63834327 | 0.772 | 0.387 | 1.6106E-40 | PDC_inj | FKBP2 |
| 6.4554E-39 | 0.63800144 | 1 | 0.973 | 1.3782E-34 | PDC_inj | RPL13 |
| 1.197E-83 | 0.63734433 | 0.541 | 0.132 | 2.5557E-79 | PDC_inj | CLCN3 |
| 3.9079E-43 | 0.63721148 | 0.993 | 0.897 | 8.3434E-39 | PDC_inj | RPL5 |
| 6.2038E-36 | 0.62882369 | 0.996 | 0.961 | 1.3245E-31 | PDC_inj | RPS18 |
| 6.7392E-39 | 0.62866957 | 0.866 | 0.545 | 1.4388E-34 | PDC_inj | CANX |
| 3.1793E-74 | 0.62603859 | 0.59 | 0.168 | 6.7878E-70 | PDC_inj | DNAJB9 |
| 5.2663E-41 | 0.62542505 | 0.993 | 0.92 | 1.1244E-36 | PDC_inj | RPL14 |
| 2.422E-104 | 0.62523241 | 0.556 | 0.116 | 5.171E-100 | PDC_inj | VEGFB |
| 6.0362E-71 | 0.62411227 | 0.701 | 0.232 | 1.2887E-66 | PDC_inj | VAMP2 |
| 3.7957E-74 | 0.62389524 | 0.604 | 0.18 | 8.1037E-70 | PDC_inj | ATRAID |
| 2.9396E-65 | 0.62361601 | 0.683 | 0.243 | 6.276E-61 | PDC_inj | PSMA3 |
| 7.8579E-93 | 0.62347091 | 0.575 | 0.137 | 1.6777E-88 | PDC_inj | NDUFAF2 |
| 1.6016E-40 | 0.62194898 | 0.948 | 0.889 | 3.4195E-36 | PDC_inj | NACA |
| 4.1473E-46 | 0.62134355 | 0.84 | 0.462 | 8.8546E-42 | PDC_inj | PRDX6 |
| 3.7592E-44 | 0.61743853 | 0.854 | 0.518 | 8.0259E-40 | PDC_inj | ATP5MC2 |
| 3.8455E-35 | 0.61475151 | 0.974 | 0.92 | 8.2102E-31 | PDC_inj | RPS3A |
| 2.5039E-56 | 0.61448163 | 0.776 | 0.337 | 5.3458E-52 | PDC_inj | CNBP |
| 5.8162E-41 | 0.61388175 | 0.981 | 0.903 | 1.2418E-36 | PDC_inj | RPS3 |
| 3.3751E-39 | 0.61327562 | 0.97 | 0.852 | 7.2058E-35 | PDC_inj | RPL38 |
| 3.0744E-72 | 0.60754188 | 0.731 | 0.25 | 6.5639E-68 | PDC_inj | ERH |
| 2.0849E-68 | 0.60493772 | 0.549 | 0.152 | 4.4512E-64 | PDC_inj | HYOU1 |
| 8.422E-127 | 0.60388876 | 0.448 | 0.065 | 1.798E-122 | PDC_inj | RGS10 |
| 2.042E-109 | 0.60315319 | 0.451 | 0.075 | 4.36E-105 | PDC_inj | GOLIM4 |
| 2.8619E-38 | 0.60217489 | 0.981 | 0.919 | 6.1101E-34 | PDC_inj | RPL34 |
| 5.664E-99 | 0.60093471 | 0.537 | 0.114 | 1.2093E-94 | PDC_inj | TPP1 |
| 8.851E-101 | 0.59953421 | 0.541 | 0.114 | 1.8897E-96 | PDC_inj | CHPT1 |
| 7.8385E-36 | 0.59777016 | 1 | 0.942 | 1.6735E-31 | PDC_inj | RPL8 |
| 1.4686E-72 | 0.5975877 | 0.642 | 0.2 | 3.1355E-68 | PDC_inj | TMBIM4 |
| 2.3624E-80 | 0.59557104 | 0.537 | 0.13 | 5.0438E-76 | PDC_inj | TRIB3 |
| 2.4284E-48 | 0.59500779 | 0.761 | 0.354 | 5.1846E-44 | PDC_inj | CSDE1 |
| 2.0973E-38 | 0.59403254 | 0.974 | 0.934 | 4.4778E-34 | PDC_inj | RPL37 |
| 0 | 0.59313958 | 0.437 | 0.016 | 0 | PDC_inj | TCF21 |
| 3.508E-145 | 0.59280973 | 0.396 | 0.045 | 7.49E-141 | PDC_inj | SRGAP2C |
| 2.925E-270 | 0.58954308 | 0.399 | 0.023 | 6.244E-266 | PDC_inj | DPYSL3 |
| 6.7213E-66 | 0.58927185 | 0.612 | 0.197 | 1.435E-61 | PDC_inj | POLR2I |
| 1.3662E-42 | 0.58920115 | 0.925 | 0.62 | 2.9168E-38 | PDC_inj | SLC25A6 |
| 2.325E-146 | 0.58724543 | 0.418 | 0.05 | 4.964E-142 | PDC_inj | PRKACB |
| 1.2116E-81 | 0.58498696 | 0.582 | 0.154 | 2.5868E-77 | PDC_inj | BTF3L4 |
| 1.7301E-53 | 0.5848891 | 0.728 | 0.303 | 3.6937E-49 | PDC_inj | NOP53 |
| 7.0775E-33 | 0.58375975 | 0.94 | 0.865 | 1.511E-28 | PDC_inj | RPL23 |
| 7.072E-215 | 0.58144357 | 0.384 | 0.028 | 1.51E-210 | PDC_inj | F2R |
| 5.559E-44 | 0.58003776 | 1 | 0.987 | 1.1868E-39 | PDC_inj | EEF1A1 |
| 8.8986E-41 | 0.57970472 | 0.802 | 0.42 | 1.8999E-36 | PDC_inj | PDIA3 |
| 1.257E-166 | 0.5788597 | 0.422 | 0.044 | 2.683E-162 | PDC_inj | FKBP5 |
| 1.071E-242 | 0.57852628 | 0.437 | 0.031 | 2.286E-238 | PDC_inj | PLA2R1 |
| 4.938E-262 | 0.57812409 | 0.481 | 0.035 | 1.054E-257 | PDC_inj | PODXL |
| 1.7163E-37 | 0.57773328 | 0.955 | 0.796 | 3.6642E-33 | PDC_inj | RPL27 |
| 1.7227E-43 | 0.57683599 | 0.765 | 0.372 | 3.6779E-39 | PDC_inj | RHEB |

| | | | | | | |
|---|---|---|---|---|---|---|
| 3.952E-104 | 0.57515382 | 0.44 | 0.076 | 8.437E-100 | PDC_inj | CLIP4 |
| 4.33E-146 | 0.57410546 | 0.422 | 0.051 | 9.245E-142 | PDC_inj | CERS6 |
| 7.3249E-75 | 0.57177577 | 0.601 | 0.172 | 1.5639E-70 | PDC_inj | STUB1 |
| 6.247E-211 | 0.57083136 | 0.425 | 0.035 | 1.334E-206 | PDC_inj | SPOCK2 |
| 3.0851E-35 | 0.5688923 | 0.53 | 0.22 | 6.5867E-31 | PDC_inj | IFITM2 |
| 6.8287E-31 | 0.56689951 | 0.981 | 0.928 | 1.4579E-26 | PDC_inj | RPS14 |
| 5.1763E-68 | 0.56146825 | 0.616 | 0.189 | 1.1051E-63 | PDC_inj | BNIP3 |
| 2.1054E-53 | 0.56071451 | 0.713 | 0.288 | 4.4951E-49 | PDC_inj | UBXN1 |
| 7.8551E-54 | 0.55882579 | 0.507 | 0.159 | 1.6771E-49 | PDC_inj | DAB2 |
| 4.5492E-34 | 0.55775668 | 0.97 | 0.891 | 9.7125E-30 | PDC_inj | RPS11 |
| 2.8374E-52 | 0.55672488 | 0.563 | 0.191 | 6.0578E-48 | PDC_inj | NFIC |
| 9.1981E-36 | 0.55445442 | 0.996 | 0.93 | 1.9638E-31 | PDC_inj | RPL18 |
| 2.9971E-44 | 0.55271305 | 0.705 | 0.325 | 6.3989E-40 | PDC_inj | EIF3E |
| 6.5879E-35 | 0.55035385 | 0.996 | 0.955 | 1.4065E-30 | PDC_inj | RPS2 |
| 1.9951E-40 | 0.54690214 | 0.66 | 0.299 | 4.2596E-36 | PDC_inj | TMEM258 |
| 5.1225E-34 | 0.54587886 | 0.989 | 0.948 | 1.0936E-29 | PDC_inj | B2M |
| 6.7684E-60 | 0.54125961 | 0.493 | 0.138 | 1.4451E-55 | PDC_inj | PTPN14 |
| 4.6992E-39 | 0.54066058 | 0.72 | 0.349 | 1.0033E-34 | PDC_inj | EIF3K |
| 4.01E-262 | 0.54023376 | 0.384 | 0.022 | 8.56E-258 | PDC_inj | C1orf21 |
| 1.061E-75 | 0.53922505 | 0.187 | 0.019 | 2.2653E-71 | PDC_inj | IFI27 |
| 0 | 0.5391371 | 0.392 | 0.006 | 0 | PDC_inj | LRRN4CL |
| 1.4347E-35 | 0.53823205 | 0.799 | 0.443 | 3.0631E-31 | PDC_inj | RPS10 |
| 1.782E-95 | 0.53653269 | 0.534 | 0.114 | 3.8045E-91 | PDC_inj | RTL8A |
| 2.5007E-76 | 0.53568236 | 0.526 | 0.133 | 5.3389E-72 | PDC_inj | DNAJC19 |
| 2.7001E-62 | 0.53542513 | 0.504 | 0.14 | 5.7647E-58 | PDC_inj | RETREG1 |
| 7.5061E-50 | 0.53429145 | 0.537 | 0.182 | 1.6026E-45 | PDC_inj | FERMT2 |
| 7.1461E-53 | 0.53418336 | 0.66 | 0.252 | 1.5257E-48 | PDC_inj | LAMTOR1 |
| 2.033E-307 | 0.53407129 | 0.41 | 0.021 | 4.341E-303 | PDC_inj | SNCA |
| 2.5907E-48 | 0.5331898 | 0.683 | 0.278 | 5.5311E-44 | PDC_inj | RABAC1 |
| 4.0676E-47 | 0.53234847 | 0.705 | 0.298 | 8.6844E-43 | PDC_inj | EID1 |
| 6.3376E-45 | 0.53174725 | 0.575 | 0.218 | 1.3531E-40 | PDC_inj | SPATS2L |
| 2.2622E-36 | 0.52957811 | 1 | 0.966 | 4.8299E-32 | PDC_inj | RPL41 |
| 2.5656E-42 | 0.52754996 | 0.646 | 0.275 | 5.4776E-38 | PDC_inj | LAMP1 |
| 5.5631E-54 | 0.52665007 | 0.638 | 0.235 | 1.1877E-49 | PDC_inj | RNH1 |
| 5.2855E-90 | 0.52287055 | 0.463 | 0.091 | 1.1285E-85 | PDC_inj | MTSS1 |
| 8.739E-195 | 0.52217619 | 0.388 | 0.032 | 1.866E-190 | PDC_inj | BAG2 |
| 6.1218E-60 | 0.52129426 | 0.563 | 0.175 | 1.307E-55 | PDC_inj | DUT |
| 5.9693E-61 | 0.52045977 | 0.515 | 0.151 | 1.2744E-56 | PDC_inj | KRTCAP2 |
| 4.8223E-30 | 0.51822559 | 0.28 | 0.085 | 1.0296E-25 | PDC_inj | IFI6 |
| 7.7317E-48 | 0.51763537 | 0.571 | 0.202 | 1.6507E-43 | PDC_inj | BEX3 |
| 1.7652E-28 | 0.51509863 | 0.97 | 0.929 | 3.7686E-24 | PDC_inj | RPL7A |
| 9.3568E-47 | 0.5141523 | 0.604 | 0.231 | 1.9977E-42 | PDC_inj | EPRS |
| 3.2093E-47 | 0.51179394 | 0.604 | 0.229 | 6.8519E-43 | PDC_inj | SMIM26 |
| 3.8653E-52 | 0.5113445 | 0.541 | 0.179 | 8.2524E-48 | PDC_inj | RHOQ |
| 1.0331E-36 | 0.50930171 | 0.698 | 0.342 | 2.2056E-32 | PDC_inj | COMMD6 |
| 7.696E-242 | 0.50915793 | 0.362 | 0.021 | 1.643E-237 | PDC_inj | PTP4A3 |
| 9.8525E-61 | 0.50724617 | 0.47 | 0.13 | 2.1035E-56 | PDC_inj | HDDC2 |
| 7.438E-45 | 0.50682893 | 0.586 | 0.225 | 1.588E-40 | PDC_inj | MRPL20 |
| 1.2286E-65 | 0.50646924 | 0.493 | 0.132 | 2.6232E-61 | PDC_inj | PPCS |
| 2.573E-102 | 0.50629034 | 0.381 | 0.057 | 5.4927E-98 | PDC_inj | VAMP5 |

| | | | | | | |
|---|---|---|---|---|---|---|
| 5.0086E-28 | 0.50443516 | 0.933 | 0.867 | 1.0693E-23 | PDC_inj | RPL10A |
| 5.1713E-38 | 0.50275415 | 0.687 | 0.315 | 1.1041E-33 | PDC_inj | PHPT1 |
| 5.3352E-46 | 0.50238017 | 0.754 | 0.332 | 1.1391E-41 | PDC_inj | SPCS1 |
| 1.86E-175 | 0.50213293 | 0.369 | 0.032 | 3.972E-171 | PDC_inj | SPOCK1 |
| 2.7299E-60 | 0.50128316 | 0.556 | 0.165 | 5.8283E-56 | PDC_inj | EIF4EBP1 |
| 1.0967E-35 | 0.49978413 | 0.631 | 0.282 | 2.3414E-31 | PDC_inj | TJP1 |
| 4.5964E-79 | 0.4992862 | 0.437 | 0.093 | 9.8132E-75 | PDC_inj | CDK4 |
| 1.4886E-46 | 0.4971921 | 0.623 | 0.245 | 3.1783E-42 | PDC_inj | NDUFAB1 |
| 2.057E-179 | 0.49684121 | 0.399 | 0.037 | 4.391E-175 | PDC_inj | FAM89A |
| 1.4184E-73 | 0.49543002 | 0.507 | 0.127 | 3.0283E-69 | PDC_inj | SURF1 |
| 4.9725E-76 | 0.49541948 | 0.414 | 0.085 | 1.0616E-71 | PDC_inj | LY6E |
| 9.9895E-25 | 0.4944783 | 0.869 | 0.716 | 2.1328E-20 | PDC_inj | RPL36AL |
| 1.1203E-30 | 0.49435069 | 0.966 | 0.93 | 2.3919E-26 | PDC_inj | RPL12 |
| 6.4885E-43 | 0.49392321 | 0.257 | 0.056 | 1.3853E-38 | PDC_inj | MT1M |
| 5.6076E-58 | 0.49339742 | 0.437 | 0.115 | 1.1972E-53 | PDC_inj | NKAP |
| 2.6645E-48 | 0.49249766 | 0.653 | 0.255 | 5.6887E-44 | PDC_inj | NDUFB10 |
| 6.083E-166 | 0.49170302 | 0.328 | 0.026 | 1.299E-161 | PDC_inj | DACH1 |
| 7.9763E-54 | 0.49136292 | 0.653 | 0.239 | 1.7029E-49 | PDC_inj | LAMTOR4 |
| 5.0855E-64 | 0.49061223 | 0.504 | 0.138 | 1.0857E-59 | PDC_inj | TRAPPC2L |
| 2.0042E-30 | 0.48987666 | 0.854 | 0.576 | 4.279E-26 | PDC_inj | COX5B |
| 3.7604E-48 | 0.48971378 | 0.56 | 0.196 | 8.0284E-44 | PDC_inj | MRPL52 |
| 1.2352E-28 | 0.48585868 | 0.981 | 0.927 | 2.6371E-24 | PDC_inj | RPL36 |
| 4.773E-293 | 0.48508754 | 0.239 | 0.006 | 1.019E-288 | PDC_inj | ADCYAP1 |
| 3.2978E-82 | 0.4847352 | 0.481 | 0.106 | 7.0408E-78 | PDC_inj | TMEM126B |
| 2.116E-229 | 0.48360839 | 0.362 | 0.023 | 4.517E-225 | PDC_inj | HSD17B14 |
| 1.9853E-47 | 0.48324388 | 0.552 | 0.191 | 4.2386E-43 | PDC_inj | SLK |
| 3.8543E-44 | 0.48034397 | 0.575 | 0.214 | 8.2289E-40 | PDC_inj | SPCS3 |
| 1.7536E-66 | 0.4787325 | 0.466 | 0.118 | 3.7439E-62 | PDC_inj | DTD1 |
| 1.1636E-25 | 0.47754702 | 0.966 | 0.863 | 2.4842E-21 | PDC_inj | RPS4X |
| 5.5093E-63 | 0.47739766 | 0.549 | 0.161 | 1.1762E-58 | PDC_inj | COPS6 |
| 5.069E-202 | 0.47734361 | 0.347 | 0.024 | 1.082E-197 | PDC_inj | ITIH5 |
| 1.888E-56 | 0.47651281 | 0.515 | 0.154 | 4.0309E-52 | PDC_inj | PLS3 |
| 1.161E-286 | 0.47552625 | 0.306 | 0.012 | 2.478E-282 | PDC_inj | PHYHIP |
| 2.3651E-57 | 0.47483651 | 0.522 | 0.157 | 5.0495E-53 | PDC_inj | ORMDL1 |
| 3.1589E-28 | 0.47221895 | 0.981 | 0.947 | 6.7443E-24 | PDC_inj | RPL13A |
| 8.2985E-64 | 0.47179852 | 0.478 | 0.126 | 1.7717E-59 | PDC_inj | MRPL40 |
| 3.3143E-34 | 0.46882228 | 0.694 | 0.331 | 7.0761E-30 | PDC_inj | SSR3 |
| 1.0248E-26 | 0.46736779 | 0.601 | 0.289 | 2.188E-22 | PDC_inj | DST |
| 1.2789E-28 | 0.46701803 | 0.978 | 0.908 | 2.7305E-24 | PDC_inj | RPL24 |
| 7.8723E-44 | 0.46618344 | 0.683 | 0.285 | 1.6807E-39 | PDC_inj | NOL7 |
| 6.911E-75 | 0.46534309 | 0.362 | 0.068 | 1.4755E-70 | PDC_inj | TCEAL2 |
| 1.2614E-43 | 0.46506181 | 0.567 | 0.218 | 2.6931E-39 | PDC_inj | C1orf43 |
| 9.4153E-43 | 0.46492641 | 0.698 | 0.293 | 2.0102E-38 | PDC_inj | CUTA |
| 8.1315E-54 | 0.46353707 | 0.485 | 0.145 | 1.7361E-49 | PDC_inj | MRPS18C |
| 8.7382E-27 | 0.46353629 | 0.951 | 0.885 | 1.8656E-22 | PDC_inj | RPS29 |
| 1.074E-212 | 0.46329282 | 0.347 | 0.023 | 2.294E-208 | PDC_inj | MSRB3 |
| 3.9394E-46 | 0.46234159 | 0.526 | 0.183 | 8.4105E-42 | PDC_inj | MRPL54 |
| 5.2053E-43 | 0.46227888 | 0.466 | 0.154 | 1.1113E-38 | PDC_inj | RFC1 |
| 9.6139E-49 | 0.46227089 | 0.5 | 0.161 | 2.0526E-44 | PDC_inj | DNAJC10 |
| 2.71E-63 | 0.46112637 | 0.534 | 0.151 | 5.7858E-59 | PDC_inj | PYURF |

| | | | | | | |
|---|---|---|---|---|---|---|
| 5.7776E-39 | 0.46102672 | 0.575 | 0.232 | 1.2335E-34 | PDC_inj | MRPL57 |
| 4.317E-189 | 0.46043293 | 0.328 | 0.023 | 9.216E-185 | PDC_inj | PCOLCE2 |
| 2.1113E-27 | 0.46036228 | 0.981 | 0.899 | 4.5076E-23 | PDC_inj | RPL27A |
| 1.3467E-36 | 0.46029059 | 0.366 | 0.115 | 2.8751E-32 | PDC_inj | FBXO32 |
| 2.7205E-34 | 0.45978441 | 0.698 | 0.323 | 5.8082E-30 | PDC_inj | KIF5B |
| 8.2622E-61 | 0.45936018 | 0.392 | 0.092 | 1.764E-56 | PDC_inj | CHPF |
| 1.2648E-31 | 0.45933821 | 0.769 | 0.418 | 2.7003E-27 | PDC_inj | SSR4 |
| 3.3316E-58 | 0.45602189 | 0.44 | 0.116 | 7.113E-54 | PDC_inj | PLD3 |
| 1.6025E-38 | 0.45590012 | 0.739 | 0.344 | 3.4213E-34 | PDC_inj | NUCKS1 |
| 1.8731E-70 | 0.45565949 | 0.418 | 0.093 | 3.9991E-66 | PDC_inj | STX8 |
| 1.7886E-75 | 0.45516568 | 0.384 | 0.075 | 3.8188E-71 | PDC_inj | KLHL42 |
| 0 | 0.45492737 | 0.28 | 0.006 | 0 | PDC_inj | FAM180A |
| 2.4489E-35 | 0.45432649 | 0.757 | 0.381 | 5.2283E-31 | PDC_inj | TOMM20 |
| 2.3627E-37 | 0.45412673 | 0.683 | 0.309 | 5.0443E-33 | PDC_inj | SARAF |
| 1.648E-27 | 0.45235685 | 0.963 | 0.909 | 3.5184E-23 | PDC_inj | FAU |
| 1.8984E-48 | 0.45185319 | 0.5 | 0.16 | 4.053E-44 | PDC_inj | ARMCX3 |
| 3.0721E-79 | 0.45175381 | 0.269 | 0.037 | 6.559E-75 | PDC_inj | SH3BGR |
| 2.805E-200 | 0.45145426 | 0.321 | 0.021 | 5.989E-196 | PDC_inj | SFMBT2 |
| 2.8533E-52 | 0.45076907 | 0.455 | 0.132 | 6.0918E-48 | PDC_inj | ATF6 |
| 6.864E-42 | 0.45073012 | 0.392 | 0.116 | 1.4655E-37 | PDC_inj | MT1F |
| 1.7024E-21 | 0.44997857 | 0.944 | 0.873 | 3.6347E-17 | PDC_inj | RPL7 |
| 6.124E-38 | 0.44996329 | 0.541 | 0.214 | 1.3075E-33 | PDC_inj | RSL1D1 |
| 7.8727E-35 | 0.44984984 | 0.425 | 0.152 | 1.6808E-30 | PDC_inj | PDLIM5 |
| 8.483E-228 | 0.44973849 | 0.366 | 0.023 | 1.811E-223 | PDC_inj | ARHGAP28 |
| 8.679E-147 | 0.44912489 | 0.325 | 0.03 | 1.853E-142 | PDC_inj | LAMP3 |
| 1.5203E-73 | 0.44772081 | 0.451 | 0.102 | 3.2458E-69 | PDC_inj | CCDC107 |
| 6.7238E-62 | 0.44748439 | 0.388 | 0.089 | 1.4355E-57 | PDC_inj | EIF1AY |
| 2.3198E-26 | 0.44740278 | 0.985 | 0.941 | 4.9528E-22 | PDC_inj | RPS24 |
| 8.5568E-53 | 0.44739496 | 0.433 | 0.12 | 1.8269E-48 | PDC_inj | PKN1 |
| 2.6488E-53 | 0.44702395 | 0.504 | 0.152 | 5.6551E-49 | PDC_inj | MRPL34 |
| 2.997E-49 | 0.44631166 | 0.519 | 0.171 | 6.3986E-45 | PDC_inj | UBE2M |
| 2.6503E-38 | 0.44623215 | 0.608 | 0.251 | 5.6584E-34 | PDC_inj | OPTN |
| 1.878E-47 | 0.44604563 | 0.478 | 0.151 | 4.0095E-43 | PDC_inj | REX1BD |
| 5.3795E-35 | 0.44571709 | 0.549 | 0.227 | 1.1485E-30 | PDC_inj | RNF181 |
| 1.0478E-57 | 0.44224389 | 0.414 | 0.105 | 2.237E-53 | PDC_inj | ARMC10 |
| 2.7857E-13 | 0.44209611 | 0.582 | 0.369 | 5.9474E-09 | PDC_inj | FLNA |
| 3.2245E-64 | 0.44152204 | 0.47 | 0.119 | 6.8844E-60 | PDC_inj | MRPS26 |
| 1.064E-126 | 0.44010443 | 0.369 | 0.044 | 2.273E-122 | PDC_inj | A1BG |
| 5.8415E-54 | 0.43898075 | 0.485 | 0.141 | 1.2472E-49 | PDC_inj | DPP7 |
| 3.118E-124 | 0.43750181 | 0.276 | 0.025 | 6.657E-120 | PDC_inj | ENPEP |
| 1.3726E-30 | 0.43594504 | 0.776 | 0.419 | 2.9306E-26 | PDC_inj | ATP6V0B |
| 1.526E-101 | 0.43465349 | 0.343 | 0.047 | 3.2571E-97 | PDC_inj | NTNG1 |
| 7.387E-209 | 0.43450015 | 0.28 | 0.014 | 1.577E-204 | PDC_inj | MSC-AS1 |
| 3.3264E-25 | 0.43376389 | 0.784 | 0.49 | 7.1019E-21 | PDC_inj | HSPD1 |
| 1.1445E-58 | 0.43350977 | 0.396 | 0.097 | 2.4435E-54 | PDC_inj | HIKESHI |
| 3.852E-127 | 0.43237899 | 0.336 | 0.037 | 8.224E-123 | PDC_inj | SRGAP2B |
| 8.3361E-22 | 0.43110015 | 0.481 | 0.233 | 1.7798E-17 | PDC_inj | CRIM1 |
| 7.0258E-25 | 0.43088157 | 0.944 | 0.785 | 1.5E-20 | PDC_inj | NPM1 |
| 7.0697E-41 | 0.43072079 | 0.53 | 0.194 | 1.5094E-36 | PDC_inj | COX14 |
| 7.368E-89 | 0.42959463 | 0.377 | 0.063 | 1.5731E-84 | PDC_inj | PSMG1 |

| | | | | | | |
|---|---|---|---|---|---|---|
| 0 | 0.42936294 | 0.306 | 0.009 | 0 | PDC_inj | DPP6 |
| 1.1514E-37 | 0.42925129 | 0.549 | 0.215 | 2.4582E-33 | PDC_inj | RWDD1 |
| 2.9685E-48 | 0.42730326 | 0.448 | 0.136 | 6.3378E-44 | PDC_inj | EMD |
| 1.3012E-38 | 0.42589618 | 0.56 | 0.224 | 2.778E-34 | PDC_inj | NDUFA12 |
| 1.6845E-52 | 0.42504865 | 0.422 | 0.115 | 3.5963E-48 | PDC_inj | TMEM256 |
| 2.835E-22 | 0.42444353 | 0.903 | 0.78 | 6.0527E-18 | PDC_inj | EEF1B2 |
| 1.112E-135 | 0.42371742 | 0.34 | 0.035 | 2.374E-131 | PDC_inj | PLCE1 |
| 2.9438E-24 | 0.42363733 | 0.675 | 0.369 | 6.2851E-20 | PDC_inj | SEC61G |
| 8.0897E-53 | 0.42356053 | 0.493 | 0.149 | 1.7271E-48 | PDC_inj | COPRS |
| 1.0456E-44 | 0.42184378 | 0.515 | 0.175 | 2.2324E-40 | PDC_inj | NDUFAF8 |
| 1.4407E-32 | 0.42170729 | 0.5 | 0.208 | 3.0758E-28 | PDC_inj | MCFD2 |
| 1.2387E-65 | 0.42123768 | 0.381 | 0.081 | 2.6446E-61 | PDC_inj | PCK2 |
| 2.1342E-35 | 0.41926811 | 0.597 | 0.255 | 4.5566E-31 | PDC_inj | RPS19BP1 |
| 2.0365E-68 | 0.41921646 | 0.384 | 0.081 | 4.3479E-64 | PDC_inj | CMSS1 |
| 2.9799E-21 | 0.41895526 | 0.951 | 0.897 | 6.3621E-17 | PDC_inj | RPL23A |
| 1.9221E-28 | 0.41895502 | 0.877 | 0.562 | 4.1037E-24 | PDC_inj | RPS27L |
| 7.5315E-52 | 0.41843542 | 0.507 | 0.152 | 1.608E-47 | PDC_inj | MTHFD2 |
| 7.478E-34 | 0.41727216 | 0.623 | 0.276 | 1.5965E-29 | PDC_inj | PDCD5 |
| 1.181E-104 | 0.41664917 | 0.351 | 0.048 | 2.521E-100 | PDC_inj | TMEM140 |
| 2.9742E-82 | 0.41658008 | 0.332 | 0.054 | 6.35E-78 | PDC_inj | AC084033.3 |
| 7.7358E-40 | 0.41600505 | 0.448 | 0.147 | 1.6516E-35 | PDC_inj | TSC22D3 |
| 4.4236E-50 | 0.41594645 | 0.571 | 0.186 | 9.4445E-46 | PDC_inj | SLC25A37 |
| 2.6663E-42 | 0.41405613 | 0.511 | 0.181 | 5.6926E-38 | PDC_inj | PCMT1 |
| 8.8001E-35 | 0.41212481 | 0.526 | 0.213 | 1.8788E-30 | PDC_inj | LAMP2 |
| 1.6184E-39 | 0.41178889 | 0.478 | 0.169 | 3.4552E-35 | PDC_inj | ESD |
| 1.9636E-24 | 0.41034909 | 0.996 | 0.927 | 4.1922E-20 | PDC_inj | RPL18A |
| 2.393E-156 | 0.41000364 | 0.306 | 0.024 | 5.108E-152 | PDC_inj | PTPRD |
| 5.1E-175 | 0.40973494 | 0.257 | 0.015 | 1.089E-170 | PDC_inj | CCBE1 |
| 3.879E-27 | 0.40835275 | 0.657 | 0.33 | 8.2816E-23 | PDC_inj | EIF4B |
| 4.2415E-98 | 0.40801952 | 0.325 | 0.043 | 9.0556E-94 | PDC_inj | FGF13 |
| 5.1417E-52 | 0.40751146 | 0.448 | 0.128 | 1.0978E-47 | PDC_inj | ACTR10 |
| 1.0644E-43 | 0.4074678 | 0.53 | 0.183 | 2.2725E-39 | PDC_inj | FAM50A |
| 6.384E-174 | 0.40616565 | 0.28 | 0.018 | 1.363E-169 | PDC_inj | DIRAS1 |
| 4.5563E-40 | 0.40573438 | 0.463 | 0.159 | 9.7278E-36 | PDC_inj | SEC63 |
| 1.9844E-28 | 0.40562357 | 0.504 | 0.223 | 4.2367E-24 | PDC_inj | UXT |
| 9.623E-51 | 0.40550973 | 0.593 | 0.191 | 2.0545E-46 | PDC_inj | ANXA1 |
| 1.6282E-49 | 0.40544807 | 0.396 | 0.107 | 3.4762E-45 | PDC_inj | SRM |
| 1.298E-33 | 0.4041596 | 0.601 | 0.26 | 2.7713E-29 | PDC_inj | TCEAL4 |
| 1.5031E-29 | 0.40302008 | 0.668 | 0.332 | 3.2091E-25 | PDC_inj | ERP29 |
| 1.6619E-72 | 0.4023679 | 0.358 | 0.067 | 3.5482E-68 | PDC_inj | FYN |
| 1.2187E-39 | 0.40175271 | 0.407 | 0.127 | 2.6018E-35 | PDC_inj | EPHX1 |
| 2.6875E-27 | 0.40165836 | 0.631 | 0.311 | 5.7378E-23 | PDC_inj | PCBP1 |
| 4.6548E-34 | 0.40120497 | 0.504 | 0.188 | 9.938E-30 | PDC_inj | SDF2L1 |
| 2.26E-37 | 0.40081521 | 0.47 | 0.17 | 4.825E-33 | PDC_inj | METTL26 |
| 5.191E-32 | 0.40059619 | 0.575 | 0.25 | 1.1083E-27 | PDC_inj | SMDT1 |
| 7.6131E-51 | 0.40017399 | 0.459 | 0.134 | 1.6254E-46 | PDC_inj | NELFE |
| 3.562E-182 | 0.40012439 | 0.291 | 0.018 | 7.604E-178 | PDC_inj | NECAB1 |
| 6.2898E-51 | 0.4000786 | 0.354 | 0.086 | 1.3429E-46 | PDC_inj | MGAT5 |
| 1.141E-36 | 0.39859911 | 0.549 | 0.22 | 2.4361E-32 | PDC_inj | PCNP |
| 3.7051E-62 | 0.39751909 | 0.377 | 0.084 | 7.9103E-58 | PDC_inj | CISD3 |

| | | | | | | |
|---|---|---|---|---|---|---|
| 2.9572E-32 | 0.39722766 | 0.623 | 0.278 | 6.3136E-28 | PDC_inj | PSMB6 |
| 4.3782E-27 | 0.39691369 | 0.843 | 0.54 | 9.3475E-23 | PDC_inj | BRI3 |
| 5.4226E-30 | 0.39682392 | 0.638 | 0.309 | 1.1577E-25 | PDC_inj | SNX3 |
| 2.2391E-75 | 0.39587932 | 0.28 | 0.042 | 4.7804E-71 | PDC_inj | TSLP |
| 6.0063E-59 | 0.39489904 | 0.396 | 0.094 | 1.2823E-54 | PDC_inj | SHMT2 |
| 1.4213E-38 | 0.39441145 | 0.649 | 0.267 | 3.0344E-34 | PDC_inj | NDUFS6 |
| 2.8907E-38 | 0.39396768 | 0.437 | 0.149 | 6.1716E-34 | PDC_inj | METTL5 |
| 2.6227E-27 | 0.39368884 | 0.817 | 0.473 | 5.5995E-23 | PDC_inj | PTGES3 |
| 4.2826E-41 | 0.39350272 | 0.522 | 0.188 | 9.1433E-37 | PDC_inj | C1QBP |
| 7.2343E-25 | 0.39325294 | 0.944 | 0.75 | 1.5445E-20 | PDC_inj | RPL4 |
| 9.7028E-21 | 0.39225918 | 0.97 | 0.87 | 2.0715E-16 | PDC_inj | RACK1 |
| 2.7964E-21 | 0.3919732 | 0.925 | 0.82 | 5.9702E-17 | PDC_inj | RPS21 |
| 1.6385E-35 | 0.39121686 | 0.478 | 0.178 | 3.4982E-31 | PDC_inj | HADHB |
| 3.0256E-44 | 0.39088771 | 0.433 | 0.134 | 6.4596E-40 | PDC_inj | GANAB |
| 1.8899E-84 | 0.39069828 | 0.31 | 0.046 | 4.0348E-80 | PDC_inj | HOXD-AS2 |
| 1.5129E-62 | 0.39038062 | 0.347 | 0.072 | 3.23E-58 | PDC_inj | SLC35F6 |
| 1.9016E-34 | 0.38866872 | 0.455 | 0.169 | 4.0598E-30 | PDC_inj | SRP72 |
| 2.3274E-24 | 0.38859247 | 0.679 | 0.352 | 4.9689E-20 | PDC_inj | TALDO1 |
| 1.2469E-50 | 0.3884478 | 0.313 | 0.071 | 2.662E-46 | PDC_inj | CDK5RAP2 |
| 1.4694E-61 | 0.38553096 | 0.336 | 0.069 | 3.1372E-57 | PDC_inj | WDR41 |
| 3.3087E-40 | 0.38501535 | 0.951 | 0.617 | 7.0641E-36 | PDC_inj | VIM |
| 1.423E-216 | 0.38498024 | 0.302 | 0.016 | 3.037E-212 | PDC_inj | RAB3IL1 |
| 3.0768E-44 | 0.38429204 | 0.347 | 0.092 | 6.5689E-40 | PDC_inj | NPNT |
| 1.2096E-43 | 0.38350163 | 0.433 | 0.134 | 2.5826E-39 | PDC_inj | LARP6 |
| 3.9113E-19 | 0.38320271 | 0.616 | 0.327 | 8.3507E-15 | PDC_inj | CHCHD10 |
| 5.5725E-31 | 0.38315706 | 0.612 | 0.279 | 1.1897E-26 | PDC_inj | EIF3G |
| 3.426E-19 | 0.38302198 | 0.974 | 0.911 | 7.3144E-15 | PDC_inj | RPL29 |
| 1.338E-170 | 0.38249804 | 0.228 | 0.012 | 2.857E-166 | PDC_inj | ABLIM2 |
| 2.7261E-32 | 0.38213309 | 0.575 | 0.245 | 5.8203E-28 | PDC_inj | TAF10 |
| 3.907E-197 | 0.38164777 | 0.28 | 0.015 | 8.341E-193 | PDC_inj | NDNF |
| 5.6016E-35 | 0.38073994 | 0.575 | 0.235 | 1.1959E-30 | PDC_inj | NARS |
| 1.2956E-29 | 0.3803626 | 0.959 | 0.84 | 2.7662E-25 | PDC_inj | RPSA |
| 7.139E-44 | 0.3803392 | 0.429 | 0.132 | 1.5242E-39 | PDC_inj | DMAC1 |
| 1.1224E-41 | 0.37997514 | 0.504 | 0.173 | 2.3963E-37 | PDC_inj | CEBPG |
| 2.0923E-47 | 0.37937127 | 0.414 | 0.118 | 4.4671E-43 | PDC_inj | APMAP |
| 3.4604E-34 | 0.37893685 | 0.478 | 0.182 | 7.3879E-30 | PDC_inj | ARL3 |
| 6.369E-29 | 0.37848324 | 0.608 | 0.287 | 1.3598E-24 | PDC_inj | TMED10 |
| 2.2784E-18 | 0.37819699 | 0.444 | 0.229 | 4.8644E-14 | PDC_inj | GSN |
| 1.9995E-25 | 0.37811021 | 0.631 | 0.321 | 4.2689E-21 | PDC_inj | NDUFC1 |
| 1.6131E-33 | 0.37584387 | 0.537 | 0.214 | 3.4441E-29 | PDC_inj | GARS |
| 2.543E-50 | 0.37577646 | 0.433 | 0.119 | 5.4293E-46 | PDC_inj | NQO2 |
| 9.0972E-27 | 0.37552402 | 0.582 | 0.277 | 1.9423E-22 | PDC_inj | FIS1 |
| 9.2142E-27 | 0.37549821 | 0.896 | 0.657 | 1.9672E-22 | PDC_inj | UQCRB |
| 2.0562E-39 | 0.37521136 | 0.396 | 0.124 | 4.39E-35 | PDC_inj | YARS |
| 9.1944E-30 | 0.3747699 | 0.336 | 0.112 | 1.963E-25 | PDC_inj | EXOSC6 |
| 1.7142E-45 | 0.37471768 | 0.381 | 0.105 | 3.6599E-41 | PDC_inj | EMC2 |
| 1.4293E-30 | 0.37457464 | 0.522 | 0.221 | 3.0515E-26 | PDC_inj | TMEM219 |
| 1.9929E-28 | 0.37348216 | 0.563 | 0.256 | 4.2549E-24 | PDC_inj | PPP2CB |
| 1.9227E-42 | 0.37320562 | 0.455 | 0.147 | 4.1049E-38 | PDC_inj | VKORC1 |
| 2.3708E-45 | 0.37215723 | 0.396 | 0.112 | 5.0617E-41 | PDC_inj | CCDC91 |

| | | | | | | |
|---|---|---|---|---|---|---|
| 8.892E-118 | 0.37143076 | 0.272 | 0.026 | 1.898E-113 | PDC_inj | RERG |
| 6.2652E-40 | 0.37020113 | 0.481 | 0.166 | 1.3376E-35 | PDC_inj | CNPY2 |
| 6.8967E-47 | 0.36996389 | 0.272 | 0.059 | 1.4725E-42 | PDC_inj | KANK2 |
| 1.8122E-52 | 0.36989979 | 0.373 | 0.093 | 3.8691E-48 | PDC_inj | SLC25A11 |
| 1.0173E-43 | 0.36937514 | 0.403 | 0.119 | 2.172E-39 | PDC_inj | ABRACL |
| 6.9682E-32 | 0.36919514 | 0.451 | 0.174 | 1.4877E-27 | PDC_inj | NDUFS4 |
| 1.6258E-69 | 0.36865682 | 0.272 | 0.042 | 3.4712E-65 | PDC_inj | FHL1 |
| 1.8423E-33 | 0.36839582 | 0.489 | 0.19 | 3.9334E-29 | PDC_inj | ZCRB1 |
| 1.0701E-62 | 0.36802202 | 0.269 | 0.045 | 2.2846E-58 | PDC_inj | SEMA5A |
| 3.3447E-22 | 0.36793673 | 0.224 | 0.069 | 7.141E-18 | PDC_inj | FABP5 |
| 3.596E-109 | 0.36785936 | 0.28 | 0.029 | 7.678E-105 | PDC_inj | NPR3 |
| 3.5112E-50 | 0.36785936 | 0.201 | 0.032 | 7.4965E-46 | PDC_inj | UNC5B |
| 1.7068E-34 | 0.36674323 | 0.362 | 0.116 | 3.644E-30 | PDC_inj | HLA-F |
| 2.3944E-25 | 0.36651439 | 0.989 | 0.922 | 5.1121E-21 | PDC_inj | RPLP2 |
| 2.8688E-35 | 0.36646208 | 0.459 | 0.165 | 6.1249E-31 | PDC_inj | CACYBP |
| 1.0578E-39 | 0.36638883 | 0.336 | 0.094 | 2.2584E-35 | PDC_inj | OSTM1 |
| 1.3969E-58 | 0.36350385 | 0.239 | 0.039 | 2.9825E-54 | PDC_inj | SLC2A13 |
| 9.967E-169 | 0.36208886 | 0.228 | 0.012 | 2.128E-164 | PDC_inj | TENM4 |
| 2.7905E-34 | 0.36203387 | 0.47 | 0.173 | 5.9577E-30 | PDC_inj | DNPH1 |
| 3.9918E-36 | 0.36190488 | 0.466 | 0.168 | 8.5224E-32 | PDC_inj | GTF3A |
| 1.5671E-31 | 0.36132041 | 0.407 | 0.15 | 3.3458E-27 | PDC_inj | TPD52 |
| 7.3495E-29 | 0.3602752 | 0.399 | 0.147 | 1.5691E-24 | PDC_inj | NQO1 |
| 9.646E-127 | 0.35955666 | 0.299 | 0.029 | 2.059E-122 | PDC_inj | KIAA1841 |
| 1.9917E-37 | 0.35893494 | 0.373 | 0.116 | 4.2522E-33 | PDC_inj | RPA3 |
| 3.722E-23 | 0.35875928 | 0.784 | 0.462 | 7.9464E-19 | PDC_inj | PDIA6 |
| 2.3232E-21 | 0.35875588 | 0.862 | 0.599 | 4.96E-17 | PDC_inj | PSMA7 |
| 3.0755E-48 | 0.35713298 | 0.399 | 0.109 | 6.5663E-44 | PDC_inj | RPF2 |
| 9.6263E-64 | 0.35597958 | 0.321 | 0.061 | 2.0552E-59 | PDC_inj | PDLIM2 |
| 5.2103E-72 | 0.35592267 | 0.321 | 0.056 | 1.1124E-67 | PDC_inj | THYN1 |
| 1.029E-36 | 0.35576954 | 0.388 | 0.125 | 2.1968E-32 | PDC_inj | FEZ2 |
| 9.2963E-35 | 0.35525576 | 0.399 | 0.135 | 1.9848E-30 | PDC_inj | USO1 |
| 1.329E-21 | 0.35525362 | 0.485 | 0.236 | 2.8373E-17 | PDC_inj | OSTC |
| 3.127E-26 | 0.35516972 | 0.631 | 0.308 | 6.6761E-22 | PDC_inj | MRFAP1 |
| 2.4311E-52 | 0.35480499 | 0.343 | 0.08 | 5.1904E-48 | PDC_inj | TNFRSF1A |
| 1.0432E-35 | 0.35475759 | 0.433 | 0.148 | 2.2272E-31 | PDC_inj | IARS |
| 1.7971E-48 | 0.35406935 | 0.362 | 0.092 | 3.8369E-44 | PDC_inj | PIGP |
| 2.623E-42 | 0.35393944 | 0.328 | 0.086 | 5.6E-38 | PDC_inj | EMX2OS |
| 2.4347E-33 | 0.35322526 | 0.433 | 0.157 | 5.198E-29 | PDC_inj | CHCHD5 |
| 2.7143E-25 | 0.35208582 | 0.552 | 0.257 | 5.795E-21 | PDC_inj | APRT |
| 7.4117E-28 | 0.3517654 | 0.843 | 0.56 | 1.5824E-23 | PDC_inj | GDF15 |
| 1.5285E-38 | 0.3517142 | 0.384 | 0.117 | 3.2634E-34 | PDC_inj | SDC2 |
| 1.3375E-31 | 0.35157445 | 0.549 | 0.227 | 2.8555E-27 | PDC_inj | NDUFS8 |
| 2.5173E-30 | 0.35088645 | 0.619 | 0.279 | 5.3744E-26 | PDC_inj | RSL24D1 |
| 1.6922E-19 | 0.3498932 | 0.414 | 0.192 | 3.6127E-15 | PDC_inj | IGF2R |
| 5.787E-32 | 0.34965967 | 0.388 | 0.134 | 1.2355E-27 | PDC_inj | CDC42EP3 |
| 1.3865E-34 | 0.34902493 | 0.276 | 0.073 | 2.9602E-30 | PDC_inj | AKR1C1 |
| 3.2945E-22 | 0.34874787 | 0.623 | 0.328 | 7.0337E-18 | PDC_inj | SPCS2 |
| 1.5834E-39 | 0.34872156 | 0.478 | 0.164 | 3.3806E-35 | PDC_inj | THOC7 |
| 2.1204E-26 | 0.34840416 | 0.549 | 0.252 | 4.527E-22 | PDC_inj | ATP6V1E1 |
| 1.1562E-49 | 0.34833206 | 0.306 | 0.069 | 2.4684E-45 | PDC_inj | HPF1 |

| | | | | | | |
|---|---|---|---|---|---|---|
| 1.4824E-29 | 0.34808168 | 0.466 | 0.186 | 3.1649E-25 | PDC_inj | LARS |
| 5.8043E-41 | 0.34775955 | 0.403 | 0.123 | 1.2392E-36 | PDC_inj | TRAPPC4 |
| 9.7285E-27 | 0.34773557 | 0.5 | 0.218 | 2.077E-22 | PDC_inj | C19orf70 |
| 1.734E-55 | 0.34719048 | 0.306 | 0.063 | 3.7021E-51 | PDC_inj | LPCAT2 |
| 7.1481E-20 | 0.34715538 | 0.91 | 0.802 | 1.5261E-15 | PDC_inj | RPL31 |
| 2.5167E-34 | 0.34711258 | 0.493 | 0.188 | 5.3731E-30 | PDC_inj | EMC7 |
| 3.5327E-17 | 0.34698 | 0.582 | 0.329 | 7.5423E-13 | PDC_inj | PPA1 |
| 5.8237E-26 | 0.34694315 | 0.541 | 0.253 | 1.2434E-21 | PDC_inj | RTRAF |
| 4.553E-22 | 0.34689119 | 0.463 | 0.216 | 9.7206E-18 | PDC_inj | PSMA4 |
| 4.9807E-57 | 0.34634771 | 0.291 | 0.056 | 1.0634E-52 | PDC_inj | LAGE3 |
| 1.9721E-27 | 0.34604704 | 0.66 | 0.318 | 4.2104E-23 | PDC_inj | NDUFB7 |
| 1.8757E-42 | 0.34588633 | 0.396 | 0.117 | 4.0046E-38 | PDC_inj | MSRB2 |
| 2.0705E-80 | 0.34552818 | 0.269 | 0.036 | 4.4206E-76 | PDC_inj | AC007405.3 |
| 4.8901E-45 | 0.34542316 | 0.354 | 0.094 | 1.044E-40 | PDC_inj | MRPL17 |
| 4.4579E-42 | 0.34538324 | 0.384 | 0.111 | 9.5175E-38 | PDC_inj | MXI1 |
| 1.5228E-41 | 0.34501117 | 0.433 | 0.132 | 3.2512E-37 | PDC_inj | PDE4DIP |
| 1.6174E-43 | 0.34468549 | 0.354 | 0.095 | 3.4531E-39 | PDC_inj | CDKN1B |
| 2.4265E-24 | 0.34466742 | 0.474 | 0.212 | 5.1807E-20 | PDC_inj | BLOC1S1 |
| 7.5737E-29 | 0.34414092 | 0.504 | 0.213 | 1.617E-24 | PDC_inj | GTF3C6 |
| 1.3392E-21 | 0.34393773 | 0.75 | 0.446 | 2.8592E-17 | PDC_inj | ATP5PF |
| 4.0573E-21 | 0.34354323 | 0.53 | 0.27 | 8.6624E-17 | PDC_inj | CHMP2A |
| 1.4984E-35 | 0.34346893 | 0.351 | 0.108 | 3.1991E-31 | PDC_inj | ESYT1 |
| 4.8855E-30 | 0.34275341 | 0.451 | 0.177 | 1.0431E-25 | PDC_inj | SIVA1 |
| 2.9272E-61 | 0.34234676 | 0.269 | 0.046 | 6.2496E-57 | PDC_inj | SGTB |
| 2.1426E-37 | 0.34230365 | 0.295 | 0.079 | 4.5744E-33 | PDC_inj | P4HA2 |
| 3.377E-34 | 0.34220212 | 0.388 | 0.128 | 7.2099E-30 | PDC_inj | RRAGD |
| 3.9245E-28 | 0.34182942 | 0.381 | 0.141 | 8.3788E-24 | PDC_inj | MAGED2 |
| 3.6476E-32 | 0.34177064 | 0.713 | 0.319 | 7.7876E-28 | PDC_inj | CEBPB |
| 2.1856E-26 | 0.34175939 | 0.511 | 0.225 | 4.6663E-22 | PDC_inj | VPS35 |
| 4.8484E-64 | 0.34169228 | 0.276 | 0.047 | 1.0351E-59 | PDC_inj | UNG |
| 2.4125E-32 | 0.34166784 | 0.414 | 0.148 | 5.1507E-28 | PDC_inj | SMC3 |
| 4.2583E-23 | 0.34155686 | 0.646 | 0.331 | 9.0914E-19 | PDC_inj | AURKAIP1 |
| 3.2491E-24 | 0.34140824 | 0.515 | 0.242 | 6.9368E-20 | PDC_inj | RPN2 |
| 3.5333E-15 | 0.34111564 | 0.918 | 0.769 | 7.5437E-11 | PDC_inj | HSP90B1 |
| 4.6338E-33 | 0.34065237 | 0.455 | 0.166 | 9.8931E-29 | PDC_inj | OTUD6B-AS1 |
| 2.5092E-26 | 0.34042774 | 0.437 | 0.182 | 5.3571E-22 | PDC_inj | TMEM147 |
| 4.772E-182 | 0.34002634 | 0.243 | 0.012 | 1.019E-177 | PDC_inj | LPL |
| 4.5865E-19 | 0.33954911 | 0.881 | 0.732 | 9.7922E-15 | PDC_inj | SKP1 |
| 8.523E-116 | 0.33941298 | 0.269 | 0.026 | 1.82E-111 | PDC_inj | PRKAR2B |
| 6.2615E-50 | 0.33936284 | 0.343 | 0.083 | 1.3368E-45 | PDC_inj | MVB12A |
| 3.6552E-22 | 0.3392697 | 0.668 | 0.365 | 7.8038E-18 | PDC_inj | SNU13 |
| 2.4573E-52 | 0.33898274 | 0.284 | 0.057 | 5.2463E-48 | PDC_inj | MAOA |
| 6.3859E-21 | 0.3383988 | 0.604 | 0.32 | 1.3634E-16 | PDC_inj | EIF1B |
| 3.6308E-21 | 0.338303 | 0.903 | 0.749 | 7.7518E-17 | PDC_inj | PFDN5 |
| 4.014E-23 | 0.33804973 | 0.627 | 0.322 | 8.57E-19 | PDC_inj | EIF3H |
| 2.323E-23 | 0.33754742 | 0.478 | 0.212 | 4.9597E-19 | PDC_inj | EPB41L4A-AS1 |
| 5.184E-24 | 0.33749054 | 0.75 | 0.409 | 1.1068E-19 | PDC_inj | PARK7 |
| 1.0312E-17 | 0.33699095 | 0.933 | 0.802 | 2.2016E-13 | PDC_inj | EEF1D |
| 5.7506E-35 | 0.33695329 | 0.41 | 0.139 | 1.2277E-30 | PDC_inj | MRPS18B |
| 3.1194E-26 | 0.33691244 | 0.444 | 0.186 | 6.6598E-22 | PDC_inj | VTI1B |

| | | | | | | |
|---|---|---|---|---|---|---|
| 2.7188E-15 | 0.33678705 | 0.701 | 0.428 | 5.8047E-11 | PDC_inj | PNRC1 |
| 1.1807E-24 | 0.33659088 | 0.713 | 0.375 | 2.5207E-20 | PDC_inj | NDUFB2 |
| 8.762E-24 | 0.33628922 | 0.381 | 0.154 | 1.8707E-19 | PDC_inj | CCDC6 |
| 7.8729E-95 | 0.33616361 | 0.239 | 0.024 | 1.6809E-90 | PDC_inj | TCEAL7 |
| 1.1086E-14 | 0.33564323 | 0.974 | 0.925 | 2.3668E-10 | PDC_inj | ACTG1 |
| 2.0244E-33 | 0.33557812 | 0.44 | 0.16 | 4.3221E-29 | PDC_inj | RTF2 |
| 5.2924E-28 | 0.3353433 | 0.5 | 0.212 | 1.1299E-23 | PDC_inj | PJA2 |
| 1.0315E-29 | 0.33503157 | 0.396 | 0.144 | 2.2021E-25 | PDC_inj | SPART |
| 3.6134E-41 | 0.33501677 | 0.306 | 0.078 | 7.7145E-37 | PDC_inj | PSAT1 |
| 2.06E-45 | 0.33499303 | 0.313 | 0.075 | 4.3982E-41 | PDC_inj | FRY |
| 1.3751E-23 | 0.3344938 | 0.545 | 0.263 | 2.9359E-19 | PDC_inj | RTN3 |
| 1.0469E-32 | 0.33444393 | 0.31 | 0.093 | 2.2352E-28 | PDC_inj | PYGB |
| 1.826E-86 | 0.33372962 | 0.265 | 0.033 | 3.8986E-82 | PDC_inj | FGF1 |
| 1.1302E-17 | 0.33344783 | 0.396 | 0.182 | 2.4129E-13 | PDC_inj | ADIRF |
| 4.012E-109 | 0.33335091 | 0.261 | 0.026 | 8.565E-105 | PDC_inj | PIR |
| 1.867E-132 | 0.33318919 | 0.287 | 0.025 | 3.986E-128 | PDC_inj | WIPF3 |
| 1.8687E-48 | 0.33224593 | 0.302 | 0.068 | 3.9896E-44 | PDC_inj | ME2 |
| 4.512E-153 | 0.3322134 | 0.246 | 0.016 | 9.632E-149 | PDC_inj | ERICH2 |
| 1.1977E-25 | 0.33185042 | 0.56 | 0.262 | 2.5571E-21 | PDC_inj | NDUFB8 |
| 1.4533E-41 | 0.33160879 | 0.388 | 0.114 | 3.1027E-37 | PDC_inj | TSR3 |
| 1.9359E-34 | 0.33159841 | 0.373 | 0.121 | 4.1332E-30 | PDC_inj | MRPL36 |
| 7.4513E-20 | 0.33141784 | 0.601 | 0.325 | 1.5908E-15 | PDC_inj | NDUFB4 |
| 2.1252E-21 | 0.33112613 | 0.642 | 0.343 | 4.5374E-17 | PDC_inj | MZT2B |
| 1.1007E-26 | 0.33110029 | 0.351 | 0.128 | 2.3499E-22 | PDC_inj | IFI27L2 |
| 9.6467E-30 | 0.33109782 | 0.384 | 0.137 | 2.0596E-25 | PDC_inj | LYRM2 |
| 3.6783E-39 | 0.33052384 | 0.34 | 0.096 | 7.8531E-35 | PDC_inj | TMEM38B |
| 3.6095E-17 | 0.33050215 | 0.985 | 0.945 | 7.7064E-13 | PDC_inj | RPL11 |
| 1.2301E-19 | 0.32982456 | 0.933 | 0.742 | 2.6263E-15 | PDC_inj | HINT1 |
| 3.7013E-46 | 0.32982411 | 0.246 | 0.049 | 7.9023E-42 | PDC_inj | IFI16 |
| 0 | 0.3290698 | 0.246 | 0.004 | 0 | PDC_inj | SCUBE1 |
| 2.9794E-24 | 0.32883275 | 0.511 | 0.231 | 6.3609E-20 | PDC_inj | MKNK2 |
| 3.096E-129 | 0.32865362 | 0.235 | 0.017 | 6.61E-125 | PDC_inj | FIBIN |
| 3.8877E-85 | 0.32845098 | 0.216 | 0.022 | 8.3003E-81 | PDC_inj | EDIL3 |
| 9.3015E-42 | 0.32813393 | 0.325 | 0.085 | 1.9859E-37 | PDC_inj | CCDC115 |
| 9.9454E-20 | 0.32792438 | 0.41 | 0.19 | 2.1233E-15 | PDC_inj | NUCB1 |
| 6.9235E-33 | 0.32760027 | 0.422 | 0.149 | 1.4782E-28 | PDC_inj | CFAP97 |
| 1.54E-40 | 0.32729092 | 0.231 | 0.049 | 3.2878E-36 | PDC_inj | FLRT3 |
| 5.3919E-46 | 0.32721392 | 0.25 | 0.051 | 1.1512E-41 | PDC_inj | ITGA1 |
| 3.4449E-29 | 0.32675982 | 0.34 | 0.115 | 7.3548E-25 | PDC_inj | DRAM1 |
| 8.7736E-33 | 0.32585686 | 0.433 | 0.157 | 1.8732E-28 | PDC_inj | MRPS16 |
| 5.2452E-40 | 0.32561797 | 0.369 | 0.107 | 1.1199E-35 | PDC_inj | SMIM37 |
| 8.4999E-28 | 0.3254656 | 0.578 | 0.261 | 1.8147E-23 | PDC_inj | NUDC |
| 7.8176E-19 | 0.32529507 | 0.321 | 0.136 | 1.6691E-14 | PDC_inj | TMEM245 |
| 7.241E-30 | 0.32482217 | 0.377 | 0.133 | 1.5459E-25 | PDC_inj | ISG15 |
| 3.0236E-37 | 0.32442188 | 0.478 | 0.166 | 6.4553E-33 | PDC_inj | PLA2G16 |
| 2.8335E-34 | 0.32400092 | 0.328 | 0.099 | 6.0496E-30 | PDC_inj | CKS1B |
| 7.3609E-18 | 0.32397933 | 0.989 | 0.938 | 1.5715E-13 | PDC_inj | RPL15 |
| 1.1721E-32 | 0.32370425 | 0.354 | 0.115 | 2.5025E-28 | PDC_inj | UFL1 |
| 5.2621E-28 | 0.32341772 | 0.425 | 0.166 | 1.1235E-23 | PDC_inj | PHB2 |
| 9.4064E-41 | 0.32321408 | 0.332 | 0.09 | 2.0083E-36 | PDC_inj | FRA10AC1 |

| | | | | | | |
|---|---|---|---|---|---|---|
| 1.0352E-29 | 0.32176434 | 0.47 | 0.184 | 2.2101E-25 | PDC_inj | NDUFS7 |
| 1.1924E-26 | 0.31981524 | 0.403 | 0.157 | 2.5457E-22 | PDC_inj | CTSA |
| 6.9503E-34 | 0.31976971 | 0.399 | 0.135 | 1.4839E-29 | PDC_inj | GCSH |
| 3.5097E-18 | 0.319522 | 0.552 | 0.28 | 7.4932E-14 | PDC_inj | CEBPD |
| 1.3537E-19 | 0.31900295 | 0.616 | 0.331 | 2.8901E-15 | PDC_inj | PSMB1 |
| 7.6502E-30 | 0.31871251 | 0.437 | 0.168 | 1.6333E-25 | PDC_inj | VPS26A |
| 6.5689E-22 | 0.31838843 | 0.623 | 0.32 | 1.4025E-17 | PDC_inj | TMED9 |
| 1.3287E-21 | 0.31752533 | 0.978 | 0.911 | 2.8369E-17 | PDC_inj | RPL35 |
| 8.6831E-51 | 0.31692227 | 0.19 | 0.028 | 1.8538E-46 | PDC_inj | GAS1 |
| 8.6271E-33 | 0.3166476 | 0.287 | 0.082 | 1.8419E-28 | PDC_inj | WASHC2A |
| 1.2989E-16 | 0.31613082 | 0.519 | 0.284 | 2.7731E-12 | PDC_inj | PSME2 |
| 1.8481E-22 | 0.31603913 | 0.485 | 0.226 | 3.9456E-18 | PDC_inj | PA2G4 |
| 1.971E-120 | 0.31602228 | 0.228 | 0.017 | 4.208E-116 | PDC_inj | IL6 |
| 4.3012E-47 | 0.31589785 | 0.31 | 0.071 | 9.183E-43 | PDC_inj | KIZ |
| 4.623E-185 | 0.31574122 | 0.254 | 0.013 | 9.869E-181 | PDC_inj | ANTXR2 |
| 8.775E-61 | 0.31559666 | 0.209 | 0.029 | 1.8735E-56 | PDC_inj | AKR1C2 |
| 8.8354E-17 | 0.31538206 | 0.619 | 0.354 | 1.8864E-12 | PDC_inj | SEC62 |
| 6.736E-202 | 0.31517734 | 0.25 | 0.011 | 1.438E-197 | PDC_inj | ST6GALNAC3 |
| 1.7958E-19 | 0.31496266 | 0.567 | 0.297 | 3.8341E-15 | PDC_inj | GADD45GIP1 |
| 4.0365E-22 | 0.31490688 | 0.72 | 0.389 | 8.6179E-18 | PDC_inj | DAD1 |
| 1.348E-20 | 0.31451819 | 0.75 | 0.427 | 2.8779E-16 | PDC_inj | ST13 |
| 1.0748E-21 | 0.31412101 | 0.571 | 0.288 | 2.2947E-17 | PDC_inj | SSBP1 |
| 2.2739E-23 | 0.31396185 | 0.466 | 0.209 | 4.8548E-19 | PDC_inj | PSMC5 |
| 2.6614E-29 | 0.31347984 | 0.31 | 0.098 | 5.682E-25 | PDC_inj | NR3C1 |
| 9.7382E-32 | 0.31331944 | 0.336 | 0.107 | 2.0791E-27 | PDC_inj | CRTAP |
| 1.3787E-18 | 0.31043824 | 0.493 | 0.252 | 2.9435E-14 | PDC_inj | NDUFA11 |
| 1.3662E-41 | 0.31036145 | 0.34 | 0.091 | 2.9168E-37 | PDC_inj | EBAG9 |
| 4.3513E-33 | 0.30995182 | 0.343 | 0.108 | 9.2899E-29 | PDC_inj | HYI |
| 1.0038E-19 | 0.30988934 | 0.463 | 0.22 | 2.1432E-15 | PDC_inj | CLIP1 |
| 4.0846E-25 | 0.30984296 | 0.489 | 0.215 | 8.7207E-21 | PDC_inj | SSU72 |
| 1.4711E-22 | 0.30974636 | 0.608 | 0.304 | 3.1407E-18 | PDC_inj | ATP5PD |
| 1.1693E-29 | 0.30887505 | 0.306 | 0.096 | 2.4964E-25 | PDC_inj | SLC2A4RG |
| 5.7132E-22 | 0.30872672 | 0.586 | 0.296 | 1.2198E-17 | PDC_inj | SSR2 |
| 3.926E-21 | 0.30869301 | 0.582 | 0.295 | 8.3821E-17 | PDC_inj | MYDGF |
| 1.0957E-24 | 0.30855224 | 0.388 | 0.154 | 2.3394E-20 | PDC_inj | RABGGTB |
| 7.7535E-22 | 0.30824224 | 0.455 | 0.208 | 1.6554E-17 | PDC_inj | SLIRP |
| 2.9912E-24 | 0.30810941 | 0.571 | 0.271 | 6.3862E-20 | PDC_inj | SSB |
| 1.7913E-17 | 0.30792934 | 0.593 | 0.334 | 3.8245E-13 | PDC_inj | KRT10 |
| 1.968E-33 | 0.30765398 | 0.321 | 0.096 | 4.2016E-29 | PDC_inj | SLC48A1 |
| 7.0618E-31 | 0.30759656 | 0.392 | 0.137 | 1.5077E-26 | PDC_inj | ARG2 |
| 8.6863E-25 | 0.30744951 | 0.425 | 0.177 | 1.8545E-20 | PDC_inj | ALKBH7 |
| 7.2293E-23 | 0.30709342 | 0.392 | 0.162 | 1.5435E-18 | PDC_inj | PRKDC |
| 6.6155E-26 | 0.30703915 | 0.287 | 0.095 | 1.4124E-21 | PDC_inj | DLGAP1-AS1 |
| 7.662E-53 | 0.30699697 | 0.317 | 0.069 | 1.6358E-48 | PDC_inj | ZNF32 |
| 3.544E-59 | 0.3065811 | 0.269 | 0.046 | 7.5665E-55 | PDC_inj | PTGER4 |
| 2.7338E-33 | 0.30639771 | 0.377 | 0.124 | 5.8366E-29 | PDC_inj | MRPL55 |
| 6.8073E-45 | 0.30637193 | 0.254 | 0.053 | 1.4534E-40 | PDC_inj | PLEKHO1 |
| 9.6656E-18 | 0.30587662 | 0.963 | 0.881 | 2.0636E-13 | PDC_inj | RPL39 |
| 2.4154E-19 | 0.30504612 | 0.433 | 0.202 | 5.1569E-15 | PDC_inj | RPL17 |
| 4.0764E-40 | 0.3046828 | 0.358 | 0.102 | 8.7032E-36 | PDC_inj | WASHC3 |

| | | | | | | |
|---|---|---|---|---|---|---|
| 2.3802E-30 | 0.30382804 | 0.381 | 0.134 | 5.0817E-26 | PDC_inj | PSENEN |
| 3.0235E-40 | 0.30296013 | 0.351 | 0.097 | 6.4551E-36 | PDC_inj | ACOT13 |
| 2.1278E-24 | 0.30282924 | 0.444 | 0.189 | 4.5428E-20 | PDC_inj | RALA |
| 3.9655E-40 | 0.3012938 | 0.306 | 0.079 | 8.4662E-36 | PDC_inj | LSM10 |
| 1.8452E-43 | 0.30128237 | 0.265 | 0.058 | 3.9395E-39 | PDC_inj | SULF2 |
| 4.3965E-20 | 0.3010134 | 0.451 | 0.209 | 9.3865E-16 | PDC_inj | BTG3 |
| 2.1932E-24 | 0.30037443 | 0.31 | 0.11 | 4.6825E-20 | PDC_inj | INSR |
| 6.9137E-39 | 0.30024764 | 0.254 | 0.059 | 1.4761E-34 | PDC_inj | CPNE8 |
| 2.9076E-29 | 0.30021081 | 0.377 | 0.133 | 6.2077E-25 | PDC_inj | PREPL |
| 8.7127E-26 | 0.29980109 | 0.437 | 0.178 | 1.8602E-21 | PDC_inj | TBCB |
| 5.263E-119 | 0.29901256 | 0.235 | 0.019 | 1.124E-114 | PDC_inj | LINC00839 |
| 1.8724E-10 | 0.29873436 | 0.213 | 0.098 | 3.9976E-06 | PDC_inj | ATF5 |
| 1.2773E-19 | 0.29868579 | 0.213 | 0.069 | 2.727E-15 | PDC_inj | CSF1 |
| 2.7176E-35 | 0.29831303 | 0.284 | 0.076 | 5.802E-31 | PDC_inj | NUDT14 |
| 2.1262E-57 | 0.29825936 | 0.287 | 0.054 | 4.5394E-53 | PDC_inj | THAP7 |
| 5.424E-20 | 0.29813289 | 0.351 | 0.148 | 1.158E-15 | PDC_inj | POR |
| 3.0275E-42 | 0.29804046 | 0.287 | 0.068 | 6.4637E-38 | PDC_inj | WASHC2C |
| 4.336E-134 | 0.29800646 | 0.231 | 0.016 | 9.257E-130 | PDC_inj | FXYD1 |
| 4.1071E-17 | 0.29728269 | 0.556 | 0.305 | 8.7686E-13 | PDC_inj | NSA2 |
| 5.1118E-32 | 0.29726596 | 0.299 | 0.088 | 1.0914E-27 | PDC_inj | TMEM18 |
| 5.078E-283 | 0.29682395 | 0.224 | 0.005 | 1.084E-278 | PDC_inj | PTPRD-AS1 |
| 1.2166E-16 | 0.29639234 | 0.791 | 0.522 | 2.5974E-12 | PDC_inj | ELOB |
| 1.4713E-26 | 0.29608714 | 0.373 | 0.139 | 3.1412E-22 | PDC_inj | ERLEC1 |
| 8.91E-192 | 0.29605417 | 0.198 | 0.007 | 1.902E-187 | PDC_inj | CLDN11 |
| 6.845E-38 | 0.2953332 | 0.291 | 0.075 | 1.4614E-33 | PDC_inj | C11orf1 |
| 7.1418E-20 | 0.29511023 | 0.343 | 0.143 | 1.5248E-15 | PDC_inj | MRPL21 |
| 7.3889E-30 | 0.29485394 | 0.328 | 0.106 | 1.5775E-25 | PDC_inj | AC027644.3 |
| 6.85E-22 | 0.29419897 | 0.388 | 0.165 | 1.4625E-17 | PDC_inj | TAL/DCTN2 |
| 4.08E-181 | 0.29367021 | 0.201 | 0.008 | 8.712E-177 | PDC_inj | PALMD |
| 9.2476E-17 | 0.29351682 | 0.899 | 0.712 | 1.9744E-12 | PDC_inj | TXN |
| 2.0227E-29 | 0.29310197 | 0.354 | 0.122 | 4.3186E-25 | PDC_inj | PPP1R11 |
| 1.1787E-42 | 0.2928365 | 0.269 | 0.061 | 2.5165E-38 | PDC_inj | QDPR |
| 1.4658E-21 | 0.29264972 | 0.444 | 0.201 | 3.1294E-17 | PDC_inj | TXNL4A |
| 5.372E-18 | 0.29253012 | 0.343 | 0.145 | 1.1469E-13 | PDC_inj | TIMP3 |
| 9.5014E-27 | 0.29232211 | 0.396 | 0.15 | 2.0286E-22 | PDC_inj | COA4 |
| 1.7182E-18 | 0.29226598 | 0.549 | 0.288 | 3.6683E-14 | PDC_inj | SF3B5 |
| 1.1047E-33 | 0.29216418 | 0.403 | 0.133 | 2.3586E-29 | PDC_inj | PHYH |
| 5.6943E-21 | 0.2919539 | 0.328 | 0.129 | 1.2157E-16 | PDC_inj | TRIM38 |
| 4.3078E-23 | 0.29192588 | 0.362 | 0.144 | 9.1971E-19 | PDC_inj | URI1 |
| 6.3605E-21 | 0.29105512 | 0.381 | 0.163 | 1.358E-16 | PDC_inj | SSR1 |
| 4.3843E-36 | 0.29090337 | 0.347 | 0.103 | 9.3605E-32 | PDC_inj | USP1 |
| 6.1773E-15 | 0.290842 | 0.806 | 0.576 | 1.3189E-10 | PDC_inj | ATP6V1G1 |
| 2.0387E-29 | 0.29078843 | 0.381 | 0.136 | 4.3525E-25 | PDC_inj | MRPL3 |
| 6.5366E-24 | 0.29023915 | 0.451 | 0.195 | 1.3956E-19 | PDC_inj | PSMC3 |
| 1.6831E-22 | 0.28987357 | 0.444 | 0.197 | 3.5935E-18 | PDC_inj | TMEM230 |
| 1.3299E-23 | 0.28981241 | 0.384 | 0.156 | 2.8392E-19 | PDC_inj | DDOST |
| 3.138E-30 | 0.28978266 | 0.377 | 0.13 | 6.6995E-26 | PDC_inj | HEXA |
| 2.5799E-15 | 0.28962242 | 0.552 | 0.304 | 5.5081E-11 | PDC_inj | HDLBP |
| 1.1328E-16 | 0.28951261 | 0.407 | 0.196 | 2.4185E-12 | PDC_inj | TMEM106B |
| 2.8323E-25 | 0.28950786 | 0.448 | 0.185 | 6.0469E-21 | PDC_inj | DCXR |

| | | | | | | |
|---|---|---|---|---|---|---|
| 1.7006E-19 | 0.2892338 | 0.433 | 0.201 | 3.6308E-15 | PDC_inj | CALU |
| 1.3521E-58 | 0.2892051 | 0.19 | 0.025 | 2.8868E-54 | PDC_inj | XAF1 |
| 1.0381E-30 | 0.28867138 | 0.291 | 0.087 | 2.2163E-26 | PDC_inj | SDSL |
| 7.0943E-31 | 0.28852348 | 0.336 | 0.108 | 1.5146E-26 | PDC_inj | IMP4 |
| 3.9312E-24 | 0.2879378 | 0.332 | 0.123 | 8.3931E-20 | PDC_inj | ANAPC13 |
| 1.8733E-47 | 0.2878346 | 0.239 | 0.045 | 3.9995E-43 | PDC_inj | FOXD1 |
| 2.3474E-24 | 0.28719713 | 0.481 | 0.21 | 5.0116E-20 | PDC_inj | SAT2 |
| 9.4874E-19 | 0.28674868 | 0.522 | 0.268 | 2.0256E-14 | PDC_inj | ANAPC11 |
| 4.0076E-20 | 0.28663462 | 0.414 | 0.187 | 8.5562E-16 | PDC_inj | MRPS15 |
| 5.2815E-18 | 0.28531105 | 0.679 | 0.385 | 1.1276E-13 | PDC_inj | EIF2S2 |
| 1.8063E-21 | 0.28364456 | 0.422 | 0.184 | 3.8563E-17 | PDC_inj | TARS |
| 5.8998E-25 | 0.28360781 | 0.317 | 0.112 | 1.2596E-20 | PDC_inj | MCUR1 |
| 1.6786E-32 | 0.28342252 | 0.362 | 0.117 | 3.5839E-28 | PDC_inj | UBA2 |
| 1.0506E-19 | 0.28298179 | 0.418 | 0.191 | 2.243E-15 | PDC_inj | RDX |
| 2.0387E-18 | 0.28285891 | 0.619 | 0.341 | 4.3525E-14 | PDC_inj | PSMD8 |
| 1.0339E-24 | 0.28279946 | 0.407 | 0.163 | 2.2074E-20 | PDC_inj | ZNF593 |
| 1.3564E-19 | 0.28245361 | 0.347 | 0.146 | 2.8959E-15 | PDC_inj | MAP3K2 |
| 1.9417E-29 | 0.2819273 | 0.381 | 0.135 | 4.1455E-25 | PDC_inj | PSMF1 |
| 6.4719E-24 | 0.28129447 | 0.485 | 0.213 | 1.3817E-19 | PDC_inj | PSMD4 |
| 8.1856E-27 | 0.28123265 | 0.384 | 0.143 | 1.7476E-22 | PDC_inj | ZBTB38 |
| 2.5138E-20 | 0.28096308 | 0.466 | 0.22 | 5.367E-16 | PDC_inj | PSMA1 |
| 3.0852E-31 | 0.27989883 | 0.31 | 0.094 | 6.5868E-27 | PDC_inj | THAP5 |
| 9.1254E-28 | 0.27940751 | 0.332 | 0.113 | 1.9483E-23 | PDC_inj | CFL2 |
| 2.4835E-46 | 0.27919604 | 0.276 | 0.06 | 5.3023E-42 | PDC_inj | NUDT2 |
| 1.4409E-34 | 0.2789717 | 0.28 | 0.075 | 3.0762E-30 | PDC_inj | ETFRF1 |
| 2.9672E-22 | 0.27884747 | 0.328 | 0.127 | 6.335E-18 | PDC_inj | MRPS10 |
| 7.2882E-32 | 0.2787163 | 0.265 | 0.073 | 1.556E-27 | PDC_inj | UROS |
| 3.6671E-48 | 0.27870257 | 0.257 | 0.051 | 7.8292E-44 | PDC_inj | SLC9A3R2 |
| 8.578E-29 | 0.27854826 | 0.384 | 0.138 | 1.8314E-24 | PDC_inj | SAR1B |
| 2.4221E-19 | 0.27853429 | 0.313 | 0.126 | 5.1713E-15 | PDC_inj | LONP1 |
| 1.6575E-21 | 0.27852009 | 0.325 | 0.127 | 3.5388E-17 | PDC_inj | PPP2R5E |
| 5.1002E-19 | 0.27806891 | 0.246 | 0.088 | 1.0889E-14 | PDC_inj | MAP2 |
| 1.3509E-22 | 0.27804144 | 0.34 | 0.132 | 2.8841E-18 | PDC_inj | AGTRAP |
| 6.4153E-20 | 0.27785457 | 0.455 | 0.214 | 1.3697E-15 | PDC_inj | COPZ1 |
| 2.0577E-84 | 0.2774827 | 0.231 | 0.026 | 4.3931E-80 | PDC_inj | DTWD2 |
| 1.1673E-19 | 0.27713606 | 0.377 | 0.163 | 2.4922E-15 | PDC_inj | STAT1 |
| 1.4466E-13 | 0.27703478 | 0.631 | 0.386 | 3.0885E-09 | PDC_inj | CAPN2 |
| 5.5453E-31 | 0.27655009 | 0.354 | 0.116 | 1.1839E-26 | PDC_inj | APPL1 |
| 4.2869E-17 | 0.27637886 | 0.832 | 0.56 | 9.1525E-13 | PDC_inj | PPIB |
| 1.8165E-15 | 0.27552286 | 0.399 | 0.196 | 3.8783E-11 | PDC_inj | ARHGEF12 |
| 2.7186E-15 | 0.27545691 | 0.34 | 0.16 | 5.8041E-11 | PDC_inj | LARP1 |
| 4.3047E-32 | 0.27521833 | 0.291 | 0.084 | 9.1905E-28 | PDC_inj | SNX17 |
| 3.49E-19 | 0.27521517 | 0.552 | 0.288 | 7.4511E-15 | PDC_inj | MRPL51 |
| 1.1028E-32 | 0.27491262 | 0.228 | 0.055 | 2.3544E-28 | PDC_inj | TMEM47 |
| 5.0651E-34 | 0.27470638 | 0.272 | 0.073 | 1.0814E-29 | PDC_inj | ZDHHC4 |
| 2.4437E-28 | 0.27396324 | 0.246 | 0.07 | 5.2174E-24 | PDC_inj | EDEM1 |
| 7.2566E-23 | 0.27338604 | 0.888 | 0.639 | 1.5493E-18 | PDC_inj | PEBP1 |
| 3.0032E-18 | 0.27311304 | 0.653 | 0.353 | 6.4118E-14 | PDC_inj | ATP5ME |
| 1.8903E-20 | 0.27311056 | 0.358 | 0.149 | 4.0358E-16 | PDC_inj | FAM200B |
| 5.9704E-20 | 0.27277624 | 0.336 | 0.137 | 1.2747E-15 | PDC_inj | FNDC3A |

| | | | | | | |
|---|---|---|---|---|---|---|
| 1.5002E-19 | 0.2725336 | 0.351 | 0.148 | 3.2029E-15 | PDC_inj | CCDC90B |
| 4.4691E-20 | 0.27241707 | 0.261 | 0.094 | 9.5415E-16 | PDC_inj | TNS3 |
| 5.2139E-11 | 0.27206049 | 0.951 | 0.903 | 1.1132E-06 | PDC_inj | RPS20 |
| 4.4517E-30 | 0.27185808 | 0.284 | 0.083 | 9.5043E-26 | PDC_inj | UGGT2 |
| 8.9569E-34 | 0.27161959 | 0.276 | 0.074 | 1.9123E-29 | PDC_inj | SNAPC5 |
| 3.8808E-17 | 0.27159192 | 0.593 | 0.318 | 8.2855E-13 | PDC_inj | FAM96B |
| 3.7487E-31 | 0.27157935 | 0.34 | 0.109 | 8.0035E-27 | PDC_inj | LSM2 |
| 1.2007E-27 | 0.27131116 | 0.336 | 0.113 | 2.5636E-23 | PDC_inj | C9orf3 |
| 9.5777E-34 | 0.27122731 | 0.321 | 0.095 | 2.0448E-29 | PDC_inj | MALSU1 |
| 3.3187E-18 | 0.27121278 | 0.534 | 0.274 | 7.0854E-14 | PDC_inj | MRPL41 |
| 3.6843E-88 | 0.27115133 | 0.22 | 0.022 | 7.8661E-84 | PDC_inj | NLRP1 |
| 6.4522E-16 | 0.27113913 | 0.478 | 0.252 | 1.3775E-11 | PDC_inj | MTPN |
| 3.4277E-28 | 0.27111515 | 0.299 | 0.095 | 7.3181E-24 | PDC_inj | SMIM29 |
| 8.3079E-28 | 0.27095125 | 0.257 | 0.075 | 1.7737E-23 | PDC_inj | SRGAP2 |
| 1.1849E-22 | 0.27027969 | 0.444 | 0.192 | 2.5298E-18 | PDC_inj | CYC1 |
| 6.2775E-24 | 0.27016371 | 0.299 | 0.105 | 1.3402E-19 | PDC_inj | MPST |
| 1.0001E-26 | 0.26986372 | 0.31 | 0.103 | 2.1352E-22 | PDC_inj | SMIM30 |
| 1.6659E-32 | 0.26982447 | 0.272 | 0.075 | 3.5567E-28 | PDC_inj | AMZ2 |
| 1.1179E-24 | 0.26951913 | 0.399 | 0.156 | 2.3867E-20 | PDC_inj | ETFB |
| 3.4247E-29 | 0.26906966 | 0.291 | 0.089 | 7.3118E-25 | PDC_inj | SWI5 |
| 1.2764E-20 | 0.26905616 | 0.392 | 0.169 | 2.725E-16 | PDC_inj | CHMP4A |
| 5.2151E-25 | 0.26892363 | 0.34 | 0.124 | 1.1134E-20 | PDC_inj | TIMMDC1 |
| 3.6555E-88 | 0.2685969 | 0.22 | 0.022 | 7.8046E-84 | PDC_inj | IL1RAP |
| 4.2112E-21 | 0.26845224 | 0.474 | 0.219 | 8.9909E-17 | PDC_inj | DNAJC8 |
| 1.7262E-28 | 0.26834424 | 0.28 | 0.085 | 3.6855E-24 | PDC_inj | NFU1 |
| 4.7388E-18 | 0.26834072 | 0.239 | 0.087 | 1.0117E-13 | PDC_inj | MEGF9 |
| 7.9951E-15 | 0.26810978 | 0.362 | 0.177 | 1.7069E-10 | PDC_inj | RBMS1 |
| 9.8089E-86 | 0.26803728 | 0.22 | 0.023 | 2.0942E-81 | PDC_inj | ZEB2 |
| 1.0382E-22 | 0.26801663 | 0.414 | 0.172 | 2.2167E-18 | PDC_inj | SLC1A5 |
| 6.6924E-68 | 0.26792619 | 0.224 | 0.03 | 1.4288E-63 | PDC_inj | TSKU |
| 4.5709E-28 | 0.26751862 | 0.313 | 0.102 | 9.7588E-24 | PDC_inj | ZNF770 |
| 1.6845E-12 | 0.26744154 | 0.989 | 0.826 | 3.5965E-08 | PDC_inj | SQSTM1 |
| 1.4385E-19 | 0.26666122 | 0.347 | 0.147 | 3.0712E-15 | PDC_inj | SMIM7 |
| 3.261E-42 | 0.26651247 | 0.243 | 0.051 | 6.9623E-38 | PDC_inj | PIP4P2 |
| 7.849E-104 | 0.26617095 | 0.213 | 0.017 | 1.676E-99 | PDC_inj | WT1 |
| 7.6818E-14 | 0.26523329 | 0.354 | 0.177 | 1.6401E-09 | PDC_inj | AP3D1 |
| 9.3332E-15 | 0.2651585 | 0.515 | 0.289 | 1.9926E-10 | PDC_inj | RNF7 |
| 4.5672E-21 | 0.26499328 | 0.354 | 0.145 | 9.7509E-17 | PDC_inj | TMED7 |
| 5.375E-19 | 0.26495588 | 0.455 | 0.216 | 1.1476E-14 | PDC_inj | NDUFA13 |
| 1.4485E-29 | 0.26489722 | 0.34 | 0.112 | 3.0926E-25 | PDC_inj | AK6 |
| 2.7814E-19 | 0.26475067 | 0.358 | 0.154 | 5.9382E-15 | PDC_inj | EBNA1BP2 |
| 7.6696E-51 | 0.26474639 | 0.239 | 0.043 | 1.6375E-46 | PDC_inj | BTN3A2 |
| 1.3119E-24 | 0.26470882 | 0.377 | 0.146 | 2.8009E-20 | PDC_inj | STOML2 |
| 2.2245E-24 | 0.26379945 | 0.317 | 0.113 | 4.7494E-20 | PDC_inj | FAM120AOS |
| 3.9368E-30 | 0.26375232 | 0.284 | 0.083 | 8.4051E-26 | PDC_inj | AC245297.3 |
| 3.473E-34 | 0.26223008 | 0.287 | 0.078 | 7.4149E-30 | PDC_inj | COMMD3 |
| 2.0221E-35 | 0.2621413 | 0.295 | 0.08 | 4.3172E-31 | PDC_inj | USE1 |
| 3.3962E-46 | 0.26199323 | 0.224 | 0.041 | 7.2509E-42 | PDC_inj | B9D1 |
| 7.5881E-17 | 0.26172562 | 0.325 | 0.142 | 1.6201E-12 | PDC_inj | DLGAP4 |
| 1.8843E-22 | 0.2611894 | 0.362 | 0.145 | 4.023E-18 | PDC_inj | EIF2S1 |

| | | | | | | |
|---|---|---|---|---|---|---|
| 2.214E-138 | 0.26077316 | 0.19 | 0.01 | 4.727E-134 | PDC_inj | FOXD2 |
| 1.5285E-33 | 0.26061199 | 0.19 | 0.04 | 3.2634E-29 | PDC_inj | PPP1R3C |
| 3.2958E-22 | 0.26050771 | 0.351 | 0.138 | 7.0366E-18 | PDC_inj | EEF1E1 |
| 1.5898E-26 | 0.25998519 | 0.336 | 0.117 | 3.3941E-22 | PDC_inj | FAM173A |
| 6.1911E-21 | 0.25976253 | 0.306 | 0.117 | 1.3218E-16 | PDC_inj | CCDC25 |
| 4.6899E-69 | 0.25961858 | 0.201 | 0.024 | 1.0013E-64 | PDC_inj | MT1A |
| 3.0962E-20 | 0.2591453 | 0.478 | 0.224 | 6.6103E-16 | PDC_inj | POLR2K |
| 3.448E-16 | 0.25870766 | 0.459 | 0.234 | 7.3615E-12 | PDC_inj | VPS29 |
| 8.6452E-22 | 0.25868416 | 0.377 | 0.154 | 1.8458E-17 | PDC_inj | ECH1 |
| 8.4046E-26 | 0.25862867 | 0.257 | 0.079 | 1.7944E-21 | PDC_inj | LZIC |
| 1.4899E-39 | 0.25734896 | 0.235 | 0.051 | 3.181E-35 | PDC_inj | WDSUB1 |
| 8.9271E-51 | 0.25697022 | 0.194 | 0.029 | 1.9059E-46 | PDC_inj | NR2F1 |
| 1.0436E-26 | 0.25678804 | 0.276 | 0.086 | 2.2282E-22 | PDC_inj | ARHGAP35 |
| 3.3367E-19 | 0.25630634 | 0.381 | 0.166 | 7.1238E-15 | PDC_inj | MRPS34 |
| 5.6303E-20 | 0.25591367 | 0.313 | 0.124 | 1.2021E-15 | PDC_inj | VAT1 |
| 1.9524E-15 | 0.25571978 | 0.634 | 0.37 | 4.1684E-11 | PDC_inj | C11orf58 |
| 5.1335E-18 | 0.25564786 | 0.545 | 0.279 | 1.096E-13 | PDC_inj | ATP5PO |
| 2.5058E-20 | 0.25562878 | 0.746 | 0.415 | 5.3499E-16 | PDC_inj | SNHG8 |
| 3.1102E-62 | 0.2556107 | 0.194 | 0.024 | 6.6402E-58 | PDC_inj | CARD16 |
| 4.9195E-64 | 0.25542558 | 0.213 | 0.028 | 1.0503E-59 | PDC_inj | SMIM1 |
| 7.1278E-43 | 0.25510697 | 0.243 | 0.05 | 1.5218E-38 | PDC_inj | HOXA10 |
| 3.9609E-21 | 0.25465861 | 0.373 | 0.154 | 8.4566E-17 | PDC_inj | POLE4 |
| 2.994E-126 | 0.25458839 | 0.175 | 0.009 | 6.393E-122 | PDC_inj | ALDH1L2 |
| 2.3205E-22 | 0.25445555 | 0.332 | 0.127 | 4.9542E-18 | PDC_inj | BCAS2 |
| 2.8821E-15 | 0.25445018 | 0.597 | 0.345 | 6.1532E-11 | PDC_inj | C19orf53 |
| 7.3619E-27 | 0.25426642 | 0.235 | 0.066 | 1.5718E-22 | PDC_inj | KANK1 |
| 3.1877E-29 | 0.25423918 | 0.265 | 0.076 | 6.8057E-25 | PDC_inj | DGCR6L |
| 3.3428E-25 | 0.25385719 | 0.25 | 0.077 | 7.1369E-21 | PDC_inj | ENDOD1 |
| 8.7867E-19 | 0.25372787 | 0.466 | 0.224 | 1.876E-14 | PDC_inj | TXNL1 |
| 2.4501E-17 | 0.25370803 | 0.437 | 0.211 | 5.231E-13 | PDC_inj | ATP5MC1 |
| 2.9802E-28 | 0.25364041 | 0.287 | 0.088 | 6.3627E-24 | PDC_inj | C1orf123 |
| 1.2907E-14 | 0.25361656 | 0.392 | 0.196 | 2.7556E-10 | PDC_inj | ACSL3 |
| 6.8206E-18 | 0.25332409 | 0.291 | 0.117 | 1.4562E-13 | PDC_inj | SERTAD2 |
| 2.0551E-95 | 0.25331735 | 0.187 | 0.015 | 4.3877E-91 | PDC_inj | SCN9A |
| 5.5117E-18 | 0.25325589 | 0.496 | 0.248 | 1.1768E-13 | PDC_inj | SH3GLB1 |
| 5.0018E-65 | 0.25318229 | 0.183 | 0.021 | 1.0679E-60 | PDC_inj | FIGN |
| 8.5233E-43 | 0.25309538 | 0.216 | 0.041 | 1.8197E-38 | PDC_inj | CEP126 |
| 7.7777E-23 | 0.25296654 | 0.34 | 0.129 | 1.6605E-18 | PDC_inj | NDUFA8 |
| 1.327E-122 | 0.25232071 | 0.183 | 0.011 | 2.833E-118 | PDC_inj | LIMS2 |
| 2.1562E-22 | 0.25224978 | 0.201 | 0.058 | 4.6034E-18 | PDC_inj | ARHGEF3 |
| 1.1362E-21 | 0.25221647 | 0.302 | 0.112 | 2.4257E-17 | PDC_inj | PIN4 |
| 9.1595E-30 | 0.25188538 | 0.254 | 0.07 | 1.9556E-25 | PDC_inj | TSEN15 |
| 3.5884E-24 | 0.25185734 | 0.325 | 0.117 | 7.6612E-20 | PDC_inj | TM2D1 |
| 2.5559E-61 | 0.25179788 | 0.187 | 0.023 | 5.4567E-57 | PDC_inj | MXRA8 |
| 3.2833E-25 | 0.25075861 | 0.254 | 0.079 | 7.0099E-21 | PDC_inj | LETMD1 |
| 3.2456E-23 | 0.25069887 | 0.321 | 0.118 | 6.9294E-19 | PDC_inj | ATG3 |
| 9.8154E-50 | 0.25065487 | 0.183 | 0.027 | 2.0956E-45 | PDC_inj | IFI44L |
| 7.659E-152 | 0.25045537 | 0.19 | 0.009 | 1.635E-147 | PDC_inj | ZBTB7C |
| 2.8903E-20 | 0.25011046 | 0.254 | 0.089 | 6.1707E-16 | PDC_inj | SEMA3C |
| 4.4872E-05 | -0.2503174 | 0.123 | 0.22 | 0.95802403 | PDC_inj | FAM133B |

| | | | | | | |
|---|---|---|---|---|---|---|
| 1.4182E-07 | -0.2507696 | 0.078 | 0.207 | 0.00302786 | PDC_inj | WWTR1 |
| 5.8305E-10 | -0.250909 | 0.015 | 0.151 | 1.2448E-05 | PDC_inj | SGMS2 |
| 2.7158E-07 | -0.2512005 | 0.086 | 0.211 | 0.00579823 | PDC_inj | ADAM9 |
| 6.0202E-08 | -0.251534 | 0.097 | 0.237 | 0.00128532 | PDC_inj | TMBIM1 |
| 5.5134E-09 | -0.2524388 | 0.026 | 0.155 | 0.00011771 | PDC_inj | ITGA6 |
| 0.00407498 | -0.2524737 | 0.149 | 0.216 | 1 | PDC_inj | ZFP36 |
| 0.00338165 | -0.2534881 | 0.201 | 0.265 | 1 | PDC_inj | RB1CC1 |
| 9.7409E-09 | -0.2540265 | 0.034 | 0.162 | 0.00020797 | PDC_inj | KIAA1217 |
| 8.9422E-05 | -0.2544865 | 0.037 | 0.112 | 1 | PDC_inj | ICAM1 |
| 2.3616E-09 | -0.2546727 | 0.026 | 0.161 | 5.0421E-05 | PDC_inj | FHL2 |
| 1.7832E-10 | -0.254784 | 0.011 | 0.152 | 3.8071E-06 | PDC_inj | RHOF |
| 2.1868E-06 | -0.2548725 | 0.019 | 0.108 | 0.0466877 | PDC_inj | AC012306.2 |
| 2.0309E-05 | -0.2555945 | 0.134 | 0.239 | 0.4336022 | PDC_inj | AKAP13 |
| 6.9344E-05 | -0.2564314 | 0.078 | 0.165 | 1 | PDC_inj | KATNBL1 |
| 1.2603E-09 | -0.2568342 | 0.026 | 0.163 | 2.6908E-05 | PDC_inj | WLS |
| 6.1918E-09 | -0.2571835 | 0.004 | 0.119 | 0.00013219 | PDC_inj | CRABP2 |
| 5.2857E-07 | -0.2572984 | 0.078 | 0.198 | 0.01128492 | PDC_inj | BACH1 |
| 1.8587E-05 | -0.2577856 | 0.123 | 0.227 | 0.39684275 | PDC_inj | NFAT5 |
| 2.3595E-05 | -0.2584183 | 0.067 | 0.158 | 0.50376184 | PDC_inj | SLC20A1 |
| 9.2974E-06 | -0.2588055 | 0.09 | 0.193 | 0.19849888 | PDC_inj | ZNF83 |
| 2.6975E-10 | -0.2589075 | 0.019 | 0.161 | 5.7592E-06 | PDC_inj | CAV2 |
| 3.0891E-07 | -0.2597298 | 0.022 | 0.126 | 0.00659521 | PDC_inj | PTGER3 |
| 1.9363E-07 | -0.2598778 | 0.071 | 0.195 | 0.00413404 | PDC_inj | FDFT1 |
| 1.9339E-07 | -0.2599406 | 0.127 | 0.266 | 0.00412897 | PDC_inj | HDGF |
| 0.00039966 | -0.2603504 | 0.153 | 0.234 | 1 | PDC_inj | RBBP6 |
| 0.00020787 | -0.2605099 | 0.22 | 0.305 | 1 | PDC_inj | PNRC2 |
| 4.9784E-07 | -0.2611069 | 0.097 | 0.219 | 0.01062882 | PDC_inj | CHMP4B |
| 3.2167E-08 | -0.2617812 | 0.019 | 0.134 | 0.00068676 | PDC_inj | COL4A1 |
| 4.7083E-10 | -0.2621834 | 0.022 | 0.163 | 1.0052E-05 | PDC_inj | TRAF4 |
| 3.6286E-08 | -0.2621921 | 0.037 | 0.161 | 0.00077471 | PDC_inj | HOOK2 |
| 6.1054E-08 | -0.2622328 | 0.067 | 0.197 | 0.0013035 | PDC_inj | CCNG2 |
| 2.542E-11 | -0.2634144 | 0.022 | 0.181 | 5.4271E-07 | PDC_inj | BACE2 |
| 7.3494E-09 | -0.2640208 | 0.041 | 0.175 | 0.00015691 | PDC_inj | SH3BP4 |
| 2.1317E-05 | -0.2640537 | 0.086 | 0.182 | 0.45512099 | PDC_inj | SQLE |
| 0.00177595 | -0.2644107 | 0.134 | 0.203 | 1 | PDC_inj | ARRDC3 |
| 4.7869E-08 | -0.2644525 | 0.138 | 0.291 | 0.00102201 | PDC_inj | MT-ATP8 |
| 9.2108E-06 | -0.2645851 | 0.049 | 0.141 | 0.19664982 | PDC_inj | EFNB2 |
| 1.3175E-09 | -0.2658052 | 0.03 | 0.17 | 2.8129E-05 | PDC_inj | AMOTL2 |
| 1.1715E-05 | -0.266346 | 0.153 | 0.265 | 0.25010582 | PDC_inj | PLXNB2 |
| 4.0377E-07 | -0.2665882 | 0.082 | 0.203 | 0.0086204 | PDC_inj | FOXO3 |
| 6.3179E-06 | -0.2671129 | 0.049 | 0.145 | 0.13488623 | PDC_inj | ID3 |
| 2.5511E-05 | -0.2672374 | 0.183 | 0.291 | 0.54465231 | PDC_inj | FAM3C |
| 0.00011326 | -0.2672774 | 0.187 | 0.28 | 1 | PDC_inj | GTF2I |
| 3.5522E-07 | -0.2672848 | 0.078 | 0.2 | 0.00758387 | PDC_inj | CDK12 |
| 9.1503E-10 | -0.2682161 | 0.007 | 0.137 | 1.9536E-05 | PDC_inj | PDLIM4 |
| 1.9441E-10 | -0.268258 | 0.015 | 0.156 | 4.1508E-06 | PDC_inj | TRNP1 |
| 0.00015337 | -0.2686935 | 0.261 | 0.35 | 1 | PDC_inj | UBE2H |
| 5.305E-10 | -0.2688601 | 0.015 | 0.151 | 1.1326E-05 | PDC_inj | TFCP2L1 |
| 2.8788E-09 | -0.2697366 | 0.019 | 0.148 | 6.1462E-05 | PDC_inj | COL4A2 |
| 7.8595E-05 | -0.2697473 | 0.093 | 0.181 | 1 | PDC_inj | MYO9A |

| | | | | | | |
|---|---|---|---|---|---|---|
| 1.5186E-08 | -0.2697514 | 0.034 | 0.159 | 0.00032422 | PDC_inj | NUDT4 |
| 2.1385E-06 | -0.270925 | 0.082 | 0.193 | 0.04565612 | PDC_inj | PRPF38B |
| 9.6567E-09 | -0.271131 | 0.015 | 0.136 | 0.00020617 | PDC_inj | RDH10 |
| 8.1355E-08 | -0.2714772 | 0.007 | 0.11 | 0.00173693 | PDC_inj | RGS2 |
| 0.00012597 | -0.2719408 | 0.187 | 0.277 | 1 | PDC_inj | BPTF |
| 2.8158E-06 | -0.2719838 | 0.063 | 0.169 | 0.06011662 | PDC_inj | MIR22HG |
| 1.0599E-06 | -0.2721455 | 0.052 | 0.159 | 0.02262787 | PDC_inj | RNPC3 |
| 2.5681E-10 | -0.2739787 | 0 | 0.131 | 5.4828E-06 | PDC_inj | CCDC198 |
| 5.7345E-08 | -0.2740886 | 0.034 | 0.152 | 0.00122431 | PDC_inj | FXYD5 |
| 8.3867E-06 | -0.2741155 | 0.09 | 0.194 | 0.17905562 | PDC_inj | ERICH1 |
| 0.00025962 | -0.2748111 | 0.616 | 0.649 | 1 | PDC_inj | UQCRH |
| 2.8773E-05 | -0.2751255 | 0.045 | 0.128 | 0.61430023 | PDC_inj | AL135925.1 |
| 0.00015405 | -0.2759652 | 0.116 | 0.201 | 1 | PDC_inj | FAT1 |
| 7.7054E-08 | -0.2759677 | 0.022 | 0.134 | 0.00164511 | PDC_inj | CSRP2 |
| 1.2527E-07 | -0.2769656 | 0.049 | 0.168 | 0.00267447 | PDC_inj | PAXBP1 |
| 1.7948E-08 | -0.2773827 | 0.03 | 0.154 | 0.00038318 | PDC_inj | DAPK1 |
| 1.3504E-08 | -0.277702 | 0.06 | 0.196 | 0.00028832 | PDC_inj | RHOBTB3 |
| 2.7398E-10 | -0.2793762 | 0.019 | 0.16 | 5.8495E-06 | PDC_inj | KIFC3 |
| 7.905E-09 | -0.2794682 | 0.004 | 0.118 | 0.00016877 | PDC_inj | FSCN1 |
| 2.3414E-10 | -0.2796533 | 0.019 | 0.163 | 4.9989E-06 | PDC_inj | ETS1 |
| 0.00410895 | -0.27985 | 0.246 | 0.304 | 1 | PDC_inj | C9orf16 |
| 4.0575E-12 | -0.2802104 | 0.007 | 0.168 | 8.6629E-08 | PDC_inj | CD82 |
| 4.2049E-08 | -0.2804685 | 0.041 | 0.163 | 0.00089775 | PDC_inj | TUBA4A |
| 1.1604E-06 | -0.2809999 | 0.09 | 0.208 | 0.02477361 | PDC_inj | FOSL2 |
| 3.4505E-12 | -0.2811426 | 0.037 | 0.214 | 7.3669E-08 | PDC_inj | SERINC2 |
| 0.00289224 | -0.2813735 | 0.257 | 0.317 | 1 | PDC_inj | STAT3 |
| 1.2048E-09 | -0.282108 | 0.056 | 0.208 | 2.5721E-05 | PDC_inj | OCLN |
| 2.4782E-10 | -0.2834639 | 0.041 | 0.197 | 5.291E-06 | PDC_inj | FAM129B |
| 1.4021E-07 | -0.2836957 | 0.004 | 0.101 | 0.00299342 | PDC_inj | MAFA |
| 1.7772E-08 | -0.2841729 | 0.019 | 0.137 | 0.00037942 | PDC_inj | ITPKC |
| 4.2578E-06 | -0.2861087 | 0.075 | 0.182 | 0.09090298 | PDC_inj | DCBLD2 |
| 1.065E-06 | -0.2864762 | 0.175 | 0.304 | 0.02273779 | PDC_inj | RAP1B |
| 1.777E-06 | -0.2866785 | 0.101 | 0.215 | 0.03793989 | PDC_inj | ACIN1 |
| 4.8668E-06 | -0.2872478 | 0.123 | 0.238 | 0.10390598 | PDC_inj | MIDN |
| 1.393E-07 | -0.2874127 | 0.093 | 0.226 | 0.002974 | PDC_inj | TEAD1 |
| 7.055E-08 | -0.2882601 | 0.082 | 0.214 | 0.00150625 | PDC_inj | PPFIBP1 |
| 4.6379E-10 | -0.2889569 | 0.063 | 0.222 | 9.9019E-06 | PDC_inj | YPEL3 |
| 4.954E-05 | -0.2899709 | 0.078 | 0.17 | 1 | PDC_inj | FGF2 |
| 1.0116E-05 | -0.28998 | 0.116 | 0.22 | 0.21597888 | PDC_inj | RERE |
| 0.00129661 | -0.2904533 | 0.235 | 0.302 | 1 | PDC_inj | GPBP1 |
| 5.8916E-08 | -0.2905362 | 0.037 | 0.156 | 0.00125786 | PDC_inj | INSIG1 |
| 5.5828E-12 | -0.290677 | 0.034 | 0.204 | 1.1919E-07 | PDC_inj | SH3YL1 |
| 0.00638093 | -0.2909066 | 0.205 | 0.263 | 1 | PDC_inj | HMGB2 |
| 1.689E-11 | -0.2911819 | 0.004 | 0.152 | 3.606E-07 | PDC_inj | LAMB1 |
| 1.1929E-06 | -0.2919038 | 0.082 | 0.196 | 0.02546765 | PDC_inj | POLR2J3.1 |
| 1.6503E-11 | -0.292894 | 0.022 | 0.181 | 3.5233E-07 | PDC_inj | MYO10 |
| 4.5937E-10 | -0.2931407 | 0.004 | 0.134 | 9.8076E-06 | PDC_inj | AL353751.1 |
| 1.885E-05 | -0.2945259 | 0.049 | 0.135 | 0.40245466 | PDC_inj | CSRNP2 |
| 1.4268E-10 | -0.2952945 | 0.015 | 0.158 | 3.0461E-06 | PDC_inj | CPM |
| 2.4599E-06 | -0.2956422 | 0.16 | 0.283 | 0.0525187 | PDC_inj | ZNF207 |

| | | | | | | |
|---|---|---|---|---|---|---|
| 5.0498E-05 | -0.2982791 | 0.142 | 0.235 | 1 | PDC_inj | PKN2 |
| 4.5945E-06 | -0.2988622 | 0.09 | 0.196 | 0.09809259 | PDC_inj | TOB1 |
| 1.0432E-05 | -0.2993891 | 0.123 | 0.227 | 0.22272161 | PDC_inj | ZNF91 |
| 8.6525E-07 | -0.2996357 | 0.09 | 0.208 | 0.01847299 | PDC_inj | THUMPD3-AS1 |
| 3.7061E-11 | -0.3000091 | 0.026 | 0.185 | 7.9126E-07 | PDC_inj | LINC00958 |
| 4.2069E-07 | -0.3000717 | 0.123 | 0.252 | 0.00898171 | PDC_inj | KANSL1 |
| 2.418E-06 | -0.3003393 | 0.052 | 0.154 | 0.05162347 | PDC_inj | CA2 |
| 0.00045235 | -0.300743 | 0.373 | 0.434 | 1 | PDC_inj | BZW1 |
| 4.8747E-12 | -0.3013445 | 0.007 | 0.165 | 1.0408E-07 | PDC_inj | HSPG2 |
| 5.2023E-08 | -0.3015285 | 0.007 | 0.113 | 0.00111069 | PDC_inj | EGOT |
| 1.3956E-10 | -0.3016537 | 0.022 | 0.169 | 2.9796E-06 | PDC_inj | NTN4 |
| 2.2867E-09 | -0.3019365 | 0.022 | 0.154 | 4.8822E-05 | PDC_inj | KITLG |
| 6.238E-05 | -0.3020319 | 0.049 | 0.128 | 1 | PDC_inj | PCDH9 |
| 2.6666E-10 | -0.302981 | 0 | 0.131 | 5.6933E-06 | PDC_inj | PKHD1 |
| 5.8462E-06 | -0.3036573 | 0.16 | 0.276 | 0.12481665 | PDC_inj | CREBRF |
| 2.9387E-08 | -0.3041214 | 0.022 | 0.141 | 0.0006274 | PDC_inj | TNFSF10 |
| 2.7667E-08 | -0.3051893 | 0.03 | 0.151 | 0.00059068 | PDC_inj | HBEGF |
| 9.4439E-09 | -0.3056959 | 0.06 | 0.197 | 0.00020163 | PDC_inj | EPS8 |
| 2.1001E-06 | -0.3058674 | 0.138 | 0.253 | 0.04483734 | PDC_inj | ARID4B |
| 5.3462E-12 | -0.3062565 | 0.034 | 0.204 | 1.1414E-07 | PDC_inj | DDR1 |
| 0.00047511 | -0.3064398 | 0.537 | 0.569 | 1 | PDC_inj | RAN |
| 3.0088E-12 | -0.3064671 | 0.011 | 0.174 | 6.4238E-08 | PDC_inj | BICC1 |
| 5.3899E-08 | -0.3078817 | 0.019 | 0.132 | 0.00115075 | PDC_inj | PRSS22 |
| 2.6989E-07 | -0.3089618 | 0.153 | 0.286 | 0.00576209 | PDC_inj | MSN |
| 3.9477E-08 | -0.3090168 | 0.127 | 0.275 | 0.00084284 | PDC_inj | OGA |
| 1.2257E-11 | -0.3090355 | 0.015 | 0.171 | 2.617E-07 | PDC_inj | ELK3 |
| 1.0637E-09 | -0.3092858 | 0 | 0.123 | 2.2711E-05 | PDC_inj | NEURL3 |
| 2.8318E-10 | -0.3095451 | 0.004 | 0.136 | 6.0458E-06 | PDC_inj | CDH16 |
| 2.0051E-05 | -0.3098719 | 0.201 | 0.305 | 0.42807961 | PDC_inj | AFF4 |
| 0.00021144 | -0.3114442 | 0.519 | 0.558 | 1 | PDC_inj | LRRFIP1 |
| 1.2159E-07 | -0.3116115 | 0.011 | 0.113 | 0.002596 | PDC_inj | HIST4H4 |
| 1.8174E-06 | -0.3123053 | 0.082 | 0.194 | 0.03880221 | PDC_inj | BHLHE40 |
| 1.5234E-06 | -0.3126407 | 0.086 | 0.198 | 0.03252431 | PDC_inj | MSMO1 |
| 0.00160825 | -0.3126974 | 0.071 | 0.135 | 1 | PDC_inj | IFIT2 |
| 6.8358E-09 | -0.3129202 | 0.056 | 0.195 | 0.00014595 | PDC_inj | KLHL21 |
| 7.301E-09 | -0.3143793 | 0.067 | 0.208 | 0.00015588 | PDC_inj | ABHD5 |
| 1.8037E-05 | -0.3157573 | 0.104 | 0.203 | 0.38509515 | PDC_inj | CHCHD7 |
| 7.9308E-05 | -0.315998 | 0.093 | 0.179 | 1 | PDC_inj | KLF4 |
| 1.5684E-05 | -0.3162812 | 0.071 | 0.167 | 0.33485792 | PDC_inj | HIST1H1E |
| 6.2118E-06 | -0.3169174 | 0.149 | 0.258 | 0.13262296 | PDC_inj | NEDD4L |
| 5.2843E-11 | -0.3172986 | 0.052 | 0.215 | 1.1282E-06 | PDC_inj | ADAM10 |
| 5.0593E-05 | -0.318045 | 0.951 | 0.867 | 1 | PDC_inj | TMSB4X |
| 6.0718E-09 | -0.319905 | 0.071 | 0.214 | 0.00012963 | PDC_inj | SCCPDH |
| 5.9659E-11 | -0.320146 | 0.082 | 0.258 | 1.2737E-06 | PDC_inj | COTL1 |
| 1.788E-10 | -0.3206837 | 0.063 | 0.224 | 3.8173E-06 | PDC_inj | RBM47 |
| 0.0002561 | -0.3211053 | 0.56 | 0.581 | 1 | PDC_inj | ATP6V0E1 |
| 1.0721E-09 | -0.3215992 | 0.067 | 0.217 | 2.2889E-05 | PDC_inj | PATJ |
| 3.0185E-05 | -0.3216141 | 0.034 | 0.112 | 0.64444125 | PDC_inj | AC023157.3 |
| 3.1392E-08 | -0.3216587 | 0.004 | 0.109 | 0.00067022 | PDC_inj | AL355075.4 |
| 2.0571E-08 | -0.3217646 | 0.071 | 0.206 | 0.00043919 | PDC_inj | ZFC3H1 |

| | | | | | | |
|---|---|---|---|---|---|---|
| 3.0636E-07 | -0.3227012 | 0.116 | 0.249 | 0.00654083 | PDC_inj | IRS2 |
| 2.6617E-12 | -0.322888 | 0.045 | 0.226 | 5.6827E-08 | PDC_inj | LSR |
| 6.8759E-06 | -0.324325 | 0.157 | 0.269 | 0.1467994 | PDC_inj | BDP1 |
| 3.6953E-07 | -0.3245225 | 0.026 | 0.13 | 0.00788944 | PDC_inj | HOMER1 |
| 1.8849E-08 | -0.325574 | 0.041 | 0.17 | 0.00040243 | PDC_inj | CA12 |
| 2.7238E-12 | -0.3256604 | 0 | 0.156 | 5.8153E-08 | PDC_inj | PROM1 |
| 5.1037E-12 | -0.3257318 | 0.049 | 0.227 | 1.0896E-07 | PDC_inj | CXCL16 |
| 2.2902E-09 | -0.3264561 | 0.022 | 0.153 | 4.8896E-05 | PDC_inj | SLC16A3 |
| 3.2211E-07 | -0.3264972 | 0.037 | 0.146 | 0.00687702 | PDC_inj | CKS2 |
| 1.2686E-06 | -0.3271395 | 0.157 | 0.279 | 0.02708461 | PDC_inj | SMS |
| 5.9703E-13 | -0.3282394 | 0.019 | 0.192 | 1.2747E-08 | PDC_inj | LLGL2 |
| 0.00012536 | -0.3289029 | 0.101 | 0.184 | 1 | PDC_inj | ERN1 |
| 3.4856E-11 | -0.3299258 | 0.034 | 0.194 | 7.4417E-07 | PDC_inj | ZBTB43 |
| 4.8441E-07 | -0.3314729 | 0.16 | 0.288 | 0.01034208 | PDC_inj | RBM25 |
| 0.00064247 | -0.3315415 | 0.295 | 0.37 | 1 | PDC_inj | MACF1 |
| 0.00045019 | -0.3321749 | 0.146 | 0.223 | 1 | PDC_inj | ADM |
| 3.3316E-06 | -0.3337699 | 0.019 | 0.106 | 0.07112957 | PDC_inj | RSAD2 |
| 1.362E-11 | -0.3351748 | 0.037 | 0.203 | 2.9079E-07 | PDC_inj | AGRN |
| 4.1151E-13 | -0.335371 | 0.022 | 0.201 | 8.7857E-09 | PDC_inj | MAP2K3 |
| 7.6701E-05 | -0.3370702 | 0.034 | 0.108 | 1 | PDC_inj | AQP3 |
| 0.00016546 | -0.3378991 | 0.269 | 0.351 | 1 | PDC_inj | EIF5A |
| 2.3393E-07 | -0.3404113 | 0.007 | 0.104 | 0.00499438 | PDC_inj | TNF |
| 7.3618E-12 | -0.3413198 | 0.052 | 0.228 | 1.5718E-07 | PDC_inj | GPRC5C |
| 3.7811E-07 | -0.3430473 | 0.041 | 0.15 | 0.00807265 | PDC_inj | AL118516.1 |
| 4.4743E-12 | -0.3432843 | 0.007 | 0.165 | 9.5526E-08 | PDC_inj | PCSK1N |
| 9.2305E-07 | -0.3438459 | 0.082 | 0.197 | 0.01970719 | PDC_inj | BCAM |
| 2.1506E-13 | -0.3439294 | 0.03 | 0.215 | 4.5916E-09 | PDC_inj | UBALD2 |
| 2.0401E-10 | -0.3441582 | 0.082 | 0.247 | 4.3555E-06 | PDC_inj | TNIP1 |
| 1.1795E-12 | -0.3442979 | 0.026 | 0.2 | 2.5182E-08 | PDC_inj | WWC1 |
| 3.9269E-08 | -0.344567 | 0.127 | 0.265 | 0.00083839 | PDC_inj | CFLAR |
| 0.00380434 | -0.3448705 | 0.134 | 0.191 | 1 | PDC_inj | TLE4 |
| 1.1941E-13 | -0.3450247 | 0.011 | 0.191 | 2.5495E-09 | PDC_inj | SCNN1A |
| 5.7558E-06 | -0.3456727 | 0.194 | 0.305 | 0.12288533 | PDC_inj | EIF4A1 |
| 0.00507495 | -0.3465991 | 0.299 | 0.337 | 1 | PDC_inj | LUC7L3 |
| 0.00041529 | -0.3472426 | 0.231 | 0.309 | 1 | PDC_inj | YME1L1 |
| 4.0112E-09 | -0.347265 | 0.063 | 0.206 | 8.5638E-05 | PDC_inj | CLK1 |
| 2.5006E-07 | -0.3474159 | 0.216 | 0.352 | 0.00533874 | PDC_inj | ACADVL |
| 9.0624E-10 | -0.3504674 | 0.041 | 0.185 | 1.9348E-05 | PDC_inj | CD40 |
| 5.9452E-10 | -0.3521042 | 0.09 | 0.251 | 1.2693E-05 | PDC_inj | TES |
| 0.00272912 | -0.3529549 | 0.351 | 0.395 | 1 | PDC_inj | SRSF3 |
| 2.5648E-11 | -0.3530559 | 0.06 | 0.229 | 5.4758E-07 | PDC_inj | C4orf48 |
| 0.00808692 | -0.3532002 | 0.522 | 0.519 | 1 | PDC_inj | HNRNPU |
| 2.8727E-13 | -0.3535622 | 0.026 | 0.208 | 6.1333E-09 | PDC_inj | FLNB |
| 1.7147E-12 | -0.3549566 | 0.067 | 0.257 | 3.6608E-08 | PDC_inj | LAPTM4B |
| 3.7348E-10 | -0.3567704 | 0.03 | 0.175 | 7.9738E-06 | PDC_inj | DUSP5 |
| 8.3868E-12 | -0.3570184 | 0.011 | 0.167 | 1.7906E-07 | PDC_inj | ADGRF1 |
| 1.7414E-07 | -0.3600987 | 0.235 | 0.366 | 0.00371794 | PDC_inj | SRSF11 |
| 2.0609E-07 | -0.3602688 | 0.037 | 0.148 | 0.00439999 | PDC_inj | ANKRD37 |
| 2.8775E-12 | -0.360441 | 0.015 | 0.178 | 6.1435E-08 | PDC_inj | ZNF217 |
| 0.00743894 | -0.3618645 | 0.052 | 0.102 | 1 | PDC_inj | APOE |

| | | | | | | |
|---|---|---|---|---|---|---|
| 2.2832E-06 | -0.3621116 | 0.015 | 0.101 | 0.04874586 | PDC_inj | AP001160.1 |
| 1.4391E-11 | -0.3623358 | 0.134 | 0.33 | 3.0724E-07 | PDC_inj | CTSB |
| 1.4866E-05 | -0.3644505 | 0.022 | 0.102 | 0.31737942 | PDC_inj | CXCL3 |
| 1.6211E-07 | -0.3648384 | 0.022 | 0.128 | 0.00346109 | PDC_inj | HIST2H2BF |
| 6.8616E-06 | -0.3651456 | 0.213 | 0.319 | 0.14649484 | PDC_inj | BOD1L1 |
| 5.3721E-10 | -0.3654119 | 0.086 | 0.243 | 1.1469E-05 | PDC_inj | ANKLE2 |
| 4.2347E-07 | -0.3654817 | 0.81 | 0.825 | 0.00904106 | PDC_inj | H3F3A |
| 6.0135E-10 | -0.3695846 | 0.063 | 0.221 | 1.2839E-05 | PDC_inj | SERTAD1 |
| 5.967E-08 | -0.3710836 | 0.183 | 0.328 | 0.00127395 | PDC_inj | HNRNPH1 |
| 4.3509E-06 | -0.3711716 | 0.056 | 0.155 | 0.09289155 | PDC_inj | RCAN1 |
| 2.5335E-12 | -0.3727053 | 0.022 | 0.193 | 5.409E-08 | PDC_inj | EMP1 |
| 8.1456E-09 | -0.3730111 | 0.127 | 0.276 | 0.00017391 | PDC_inj | HIF1A |
| 0.00207371 | -0.3735001 | 0.056 | 0.116 | 1 | PDC_inj | TAGLN |
| 4.1935E-15 | -0.3749632 | 0.007 | 0.202 | 8.953E-11 | PDC_inj | ADGRG1 |
| 2.7725E-08 | -0.3763738 | 0.146 | 0.292 | 0.00059193 | PDC_inj | ITGA3 |
| 4.504E-06 | -0.3778238 | 0.037 | 0.131 | 0.09616095 | PDC_inj | TNFRSF11B |
| 2.0597E-06 | -0.3788485 | 0.127 | 0.243 | 0.04397454 | PDC_inj | HES4 |
| 3.0838E-15 | -0.3792405 | 0.007 | 0.203 | 6.5839E-11 | PDC_inj | TINAGL1 |
| 2.6777E-05 | -0.3798084 | 0.19 | 0.284 | 0.57169882 | PDC_inj | PRPF4B |
| 5.7103E-06 | -0.3813707 | 0.116 | 0.226 | 0.12191578 | PDC_inj | GPRC5A |
| 6.8314E-11 | -0.3830771 | 0.101 | 0.276 | 1.4585E-06 | PDC_inj | SMG1 |
| 2.3057E-11 | -0.3832277 | 0.075 | 0.246 | 4.9227E-07 | PDC_inj | FNBP4 |
| 1.8293E-12 | -0.3855824 | 0.034 | 0.207 | 3.9055E-08 | PDC_inj | PNP |
| 2.0165E-05 | -0.386389 | 0.799 | 0.803 | 0.43053081 | PDC_inj | CALM1 |
| 2.4314E-09 | -0.3872579 | 0.045 | 0.183 | 5.191E-05 | PDC_inj | HMGCS1 |
| 3.2521E-07 | -0.3873329 | 0.164 | 0.293 | 0.00694323 | PDC_inj | KMT2A |
| 4.9729E-08 | -0.3891425 | 0.004 | 0.107 | 0.00106172 | PDC_inj | ARC |
| 4.8605E-10 | -0.3915131 | 0.022 | 0.163 | 1.0377E-05 | PDC_inj | TNFAIP3 |
| 6.7574E-10 | -0.3916668 | 0.019 | 0.155 | 1.4427E-05 | PDC_inj | PFKFB3 |
| 2.5205E-10 | -0.3942042 | 0.022 | 0.169 | 5.3813E-06 | PDC_inj | CTGF |
| 3.0462E-06 | -0.394978 | 0.284 | 0.395 | 0.06503566 | PDC_inj | TAF1D |
| 1.0755E-16 | -0.4012864 | 0.026 | 0.247 | 2.2962E-12 | PDC_inj | PFKP |
| 6.4743E-12 | -0.4041234 | 0.131 | 0.317 | 1.3823E-07 | PDC_inj | DNAJB6 |
| 5.7079E-12 | -0.4068139 | 0.078 | 0.257 | 1.2186E-07 | PDC_inj | PNPLA2 |
| 6.3139E-16 | -0.4069104 | 0.004 | 0.204 | 1.348E-11 | PDC_inj | EHF |
| 8.73E-16 | -0.4070709 | 0.022 | 0.233 | 1.8638E-11 | PDC_inj | ST14 |
| 7.0404E-19 | -0.408202 | 0.004 | 0.236 | 1.5031E-14 | PDC_inj | DMKN |
| 1.4534E-12 | -0.4083309 | 0.037 | 0.217 | 3.1031E-08 | PDC_inj | CTSH |
| 6.7358E-07 | -0.4084449 | 0.291 | 0.416 | 0.01438104 | PDC_inj | NORAD |
| 2.1049E-09 | -0.4089091 | 0.377 | 0.519 | 4.494E-05 | PDC_inj | TPM3 |
| 0.00028445 | -0.4100117 | 0.231 | 0.305 | 1 | PDC_inj | ARHGAP29 |
| 1.8198E-05 | -0.4111654 | 0.224 | 0.324 | 0.38852991 | PDC_inj | RSRC2 |
| 4.9856E-07 | -0.4112892 | 0.022 | 0.125 | 0.0106443 | PDC_inj | CXCL14 |
| 3.3273E-08 | -0.4117474 | 0.034 | 0.153 | 0.00071038 | PDC_inj | AC007952.4 |
| 7.6046E-13 | -0.4130328 | 0.041 | 0.222 | 1.6236E-08 | PDC_inj | EPS8L2 |
| 5.9204E-12 | -0.4154253 | 0.795 | 0.915 | 1.264E-07 | PDC_inj | MT-ND4 |
| 7.7806E-12 | -0.4160202 | 0.093 | 0.278 | 1.6612E-07 | PDC_inj | SPTAN1 |
| 2.0904E-13 | -0.4161579 | 0.022 | 0.202 | 4.463E-09 | PDC_inj | MARCKS |
| 0.00440446 | -0.4166153 | 0.948 | 0.899 | 1 | PDC_inj | SERF2 |
| 1.2688E-08 | -0.4175518 | 0.127 | 0.274 | 0.00027088 | PDC_inj | CKB |

| | | | | | | |
|---|---|---|---|---|---|---|
| 8.7994E-07 | -0.4176616 | 0.646 | 0.701 | 0.01878677 | PDC_inj | MAP1LC3B |
| 0.00549689 | -0.4177794 | 0.422 | 0.439 | 1 | PDC_inj | PPP1R14B |
| 2.3849E-09 | -0.4198617 | 0.153 | 0.307 | 5.0917E-05 | PDC_inj | YTHDC1 |
| 2.5265E-11 | -0.4198654 | 0.022 | 0.178 | 5.3941E-07 | PDC_inj | PHLDA1 |
| 1.7561E-09 | -0.4210756 | 0.011 | 0.138 | 3.7493E-05 | PDC_inj | AC025164.1 |
| 3.6688E-07 | -0.4246113 | 0.235 | 0.361 | 0.00783287 | PDC_inj | GCC2 |
| 4.3799E-05 | -0.4250334 | 0.313 | 0.4 | 0.93510191 | PDC_inj | ANKRD11 |
| 5.273E-10 | -0.4287086 | 0.168 | 0.333 | 1.1258E-05 | PDC_inj | CD2AP |
| 1.1684E-11 | -0.4305264 | 0.03 | 0.193 | 2.4944E-07 | PDC_inj | THBS1 |
| 8.582E-13 | -0.432952 | 0.03 | 0.207 | 1.8323E-08 | PDC_inj | CLDN1 |
| 7.9569E-07 | -0.4345655 | 0.019 | 0.114 | 0.01698805 | PDC_inj | C15orf48 |
| 2.0086E-07 | -0.4370472 | 0.03 | 0.14 | 0.00428839 | PDC_inj | TM4SF1 |
| 4.6155E-07 | -0.4379052 | 0.511 | 0.593 | 0.00985415 | PDC_inj | RBM39 |
| 1.9769E-08 | -0.4382207 | 0.619 | 0.691 | 0.00042206 | PDC_inj | CYSTM1 |
| 2.715E-09 | -0.4388347 | 0.019 | 0.147 | 5.7966E-05 | PDC_inj | ID1 |
| 9.23E-16 | -0.4393764 | 0.037 | 0.253 | 1.9706E-11 | PDC_inj | MGLL |
| 3.5146E-08 | -0.4514271 | 0.194 | 0.336 | 0.00075037 | PDC_inj | RALBP1 |
| 1.6927E-13 | -0.4514995 | 0.071 | 0.267 | 3.614E-09 | PDC_inj | CHD2 |
| 1.5946E-05 | -0.4522787 | 0.175 | 0.276 | 0.3404533 | PDC_inj | VMP1 |
| 8.0348E-17 | -0.454057 | 0.015 | 0.232 | 1.7154E-12 | PDC_inj | PRSS8 |
| 0.00011999 | -0.4551759 | 0.522 | 0.562 | 1 | PDC_inj | SRRM2 |
| 8.0562E-07 | -0.4617882 | 0.149 | 0.267 | 0.01720005 | PDC_inj | AMD1 |
| 2.5896E-07 | -0.4630015 | 0.261 | 0.38 | 0.00552881 | PDC_inj | SRSF5 |
| 2.204E-05 | -0.4648147 | 0.384 | 0.483 | 0.47056462 | PDC_inj | H1FX |
| 1.9145E-11 | -0.4656839 | 0.075 | 0.251 | 4.0874E-07 | PDC_inj | RAP2B |
| 5.6815E-16 | -0.4670157 | 0.75 | 0.894 | 1.213E-11 | PDC_inj | MT-ND1 |
| 3.3145E-08 | -0.4674311 | 0.272 | 0.4 | 0.00070765 | PDC_inj | DBI |
| 3.4552E-09 | -0.468156 | 0.287 | 0.438 | 7.3769E-05 | PDC_inj | PRRC2C |
| 2.2004E-06 | -0.4697156 | 0.269 | 0.376 | 0.04697794 | PDC_inj | LDHB |
| 6.6155E-09 | -0.4707841 | 0.231 | 0.38 | 0.00014124 | PDC_inj | PNISR |
| 1.157E-06 | -0.4722067 | 0.396 | 0.489 | 0.02470251 | PDC_inj | LDHA |
| 3.5691E-15 | -0.4772579 | 0.104 | 0.33 | 7.62E-11 | PDC_inj | ARL6IP1 |
| 4.6732E-08 | -0.4805609 | 0.075 | 0.205 | 0.00099772 | PDC_inj | SGK1 |
| 1.7394E-14 | -0.485478 | 0.205 | 0.424 | 3.7136E-10 | PDC_inj | PTTG1IP |
| 1.7585E-17 | -0.4861911 | 0.015 | 0.24 | 3.7544E-13 | PDC_inj | SCIN |
| 1.1157E-07 | -0.4877241 | 0.213 | 0.34 | 0.00238202 | PDC_inj | HNRNPAB |
| 2.375E-15 | -0.4917181 | 0 | 0.192 | 5.0706E-11 | PDC_inj | SYTL2 |
| 3.7366E-18 | -0.4946776 | 0.015 | 0.245 | 7.9777E-14 | PDC_inj | KLF5 |
| 3.9376E-06 | -0.4961403 | 0.09 | 0.194 | 0.0840669 | PDC_inj | GADD45G |
| 8.493E-13 | -0.4963816 | 0.022 | 0.196 | 1.8133E-08 | PDC_inj | H2AFX |
| 3.0546E-11 | -0.4975587 | 0.31 | 0.49 | 6.5216E-07 | PDC_inj | ATP1A1 |
| 2.4871E-13 | -0.4985124 | 0.049 | 0.237 | 5.31E-09 | PDC_inj | CDKN2AIP |
| 6.4381E-09 | -0.4989999 | 0.011 | 0.131 | 0.00013745 | PDC_inj | COL7A1 |
| 1.7503E-10 | -0.5006796 | 0.037 | 0.189 | 3.7368E-06 | PDC_inj | BIRC3 |
| 2.0069E-13 | -0.501954 | 0.06 | 0.254 | 4.2847E-09 | PDC_inj | IDI1 |
| 4.1516E-21 | -0.5032801 | 0.022 | 0.289 | 8.8637E-17 | PDC_inj | SPINT1 |
| 4.6042E-07 | -0.5037158 | 0.787 | 0.836 | 0.00982997 | PDC_inj | PPIA |
| 5.9396E-13 | -0.5043261 | 0.097 | 0.289 | 1.2681E-08 | PDC_inj | NUFIP2 |
| 8.3586E-15 | -0.5063012 | 0.075 | 0.289 | 1.7846E-10 | PDC_inj | COL18A1 |
| 1.8872E-16 | -0.5088677 | 0.045 | 0.271 | 4.0292E-12 | PDC_inj | GSTM3 |

| | | | | | | |
|---|---|---|---|---|---|---|
| 0.0001106 | -0.509072 | 0.194 | 0.277 | 1 | PDC_inj | CITED2 |
| 1.8708E-09 | -0.5091675 | 0.22 | 0.369 | 3.9943E-05 | PDC_inj | PNN |
| 0.0001723 | -0.5125074 | 0.22 | 0.299 | 1 | PDC_inj | HSPH1 |
| 1.6961E-11 | -0.5155769 | 0.019 | 0.175 | 3.6211E-07 | PDC_inj | FOSB |
| 5.6104E-15 | -0.5159642 | 0.168 | 0.39 | 1.1978E-10 | PDC_inj | PERP |
| 8.0408E-05 | -0.517255 | 0.213 | 0.125 | 1 | PDC_inj | GPX3 |
| 5.4612E-18 | -0.5175209 | 0.034 | 0.27 | 1.166E-13 | PDC_inj | LIMA1 |
| 2.6394E-20 | -0.518408 | 0.019 | 0.273 | 5.6352E-16 | PDC_inj | MET |
| 2.6607E-16 | -0.5200698 | 0.131 | 0.375 | 5.6806E-12 | PDC_inj | SPTBN1 |
| 7.981E-10 | -0.52051 | 0.123 | 0.293 | 1.704E-05 | PDC_inj | FOS |
| 5.3463E-14 | -0.5217238 | 0.034 | 0.229 | 1.1414E-09 | PDC_inj | CD74 |
| 1.5368E-20 | -0.5290247 | 0.037 | 0.304 | 3.281E-16 | PDC_inj | CAPG |
| 2.7756E-17 | -0.5333827 | 0.299 | 0.529 | 5.9258E-13 | PDC_inj | CDC42 |
| 4.7753E-12 | -0.5354921 | 0.101 | 0.288 | 1.0195E-07 | PDC_inj | AKR1B1 |
| 5.4761E-08 | -0.5357959 | 0 | 0.1 | 0.00116914 | PDC_inj | SLC12A1 |
| 8.9133E-10 | -0.5362609 | 0.16 | 0.321 | 1.903E-05 | PDC_inj | UGCG |
| 7.1468E-14 | -0.5419709 | 0.19 | 0.411 | 1.5258E-09 | PDC_inj | SDC4 |
| 1.0285E-05 | -0.545039 | 0.045 | 0.137 | 0.21958303 | PDC_inj | SERPINA1 |
| 4.3293E-19 | -0.5517194 | 0.015 | 0.26 | 9.243E-15 | PDC_inj | S100A14 |
| 1.0857E-16 | -0.55648 | 0.09 | 0.319 | 2.3181E-12 | PDC_inj | AFDN |
| 1.2481E-20 | -0.5589533 | 0.045 | 0.315 | 2.6648E-16 | PDC_inj | DSP |
| 3.2597E-05 | -0.5634912 | 0.257 | 0.343 | 0.69595126 | PDC_inj | SNHG12 |
| 2.3666E-17 | -0.5654685 | 0.06 | 0.296 | 5.0527E-13 | PDC_inj | MAFF |
| 1.3081E-17 | -0.5667013 | 0.522 | 0.706 | 2.7927E-13 | PDC_inj | EIF4G2 |
| 5.6918E-05 | -0.5675974 | 0.172 | 0.257 | 1 | PDC_inj | IER5L |
| 3.0408E-15 | -0.5684336 | 0.015 | 0.213 | 6.4922E-11 | PDC_inj | ADAMTS9 |
| 1.0206E-12 | -0.5691731 | 0.153 | 0.348 | 2.179E-08 | PDC_inj | S100A16 |
| 7.7699E-12 | -0.5713444 | 0.086 | 0.265 | 1.6589E-07 | PDC_inj | HIST1H2AC |
| 2.7936E-10 | -0.5761386 | 0.201 | 0.361 | 5.9644E-06 | PDC_inj | ARGLU1 |
| 1.004E-23 | -0.5768545 | 0.418 | 0.684 | 2.1434E-19 | PDC_inj | YWHAZ |
| 1.5864E-22 | -0.5774419 | 0.026 | 0.305 | 3.3869E-18 | PDC_inj | CDH1 |
| 2.7759E-18 | -0.5793741 | 0.086 | 0.336 | 5.9265E-14 | PDC_inj | TUBA1C |
| 7.0864E-09 | -0.5805642 | 0.672 | 0.733 | 0.0001513 | PDC_inj | PFN1 |
| 7.4508E-14 | -0.5817999 | 0.69 | 0.776 | 1.5908E-09 | PDC_inj | MYL12A |
| 1.3267E-11 | -0.5823109 | 0.183 | 0.363 | 2.8326E-07 | PDC_inj | MCL1 |
| 1.0262E-20 | -0.5842727 | 0.075 | 0.346 | 2.191E-16 | PDC_inj | JUP |
| 2.5342E-18 | -0.5865111 | 0.041 | 0.283 | 5.4105E-14 | PDC_inj | COBLL1 |
| 1.9372E-24 | -0.5870215 | 0.004 | 0.292 | 4.136E-20 | PDC_inj | ANXA3 |
| 1.5432E-14 | -0.5872856 | 0.735 | 0.82 | 3.2948E-10 | PDC_inj | UBB |
| 1.3675E-07 | -0.5879631 | 0.586 | 0.657 | 0.00291959 | PDC_inj | HSPA8 |
| 2.9815E-17 | -0.5886501 | 0.019 | 0.244 | 6.3656E-13 | PDC_inj | ITGB6 |
| 1.262E-11 | -0.5903491 | 0.09 | 0.27 | 2.6944E-07 | PDC_inj | KCNQ1OT1 |
| 3.7084E-10 | -0.5909765 | 0.343 | 0.496 | 7.9174E-06 | PDC_inj | ATP1B1 |
| 1.4646E-17 | -0.5913997 | 0.06 | 0.3 | 3.127E-13 | PDC_inj | MIR4435-2HG |
| 2.8303E-18 | -0.5934815 | 0.019 | 0.253 | 6.0427E-14 | PDC_inj | MACC1 |
| 1.1862E-21 | -0.5939894 | 0.075 | 0.355 | 2.5325E-17 | PDC_inj | ANXA11 |
| 6.0425E-14 | -0.5954362 | 0.119 | 0.329 | 1.2901E-09 | PDC_inj | CYBA |
| 1.0824E-12 | -0.5959671 | 0.007 | 0.172 | 2.3109E-08 | PDC_inj | EGLN3 |
| 2.0282E-13 | -0.6029609 | 0.026 | 0.21 | 4.3301E-09 | PDC_inj | RRAD |
| 2.3034E-14 | -0.6049264 | 0.03 | 0.225 | 4.9178E-10 | PDC_inj | SFN |

| | | | | | | |
|---|---|---|---|---|---|---|
| 2.2796E-11 | -0.6080447 | 0.19 | 0.359 | 4.867E-07 | PDC_inj | DDX3X |
| 8.4872E-15 | -0.6126553 | 0.037 | 0.238 | 1.812E-10 | PDC_inj | NFKBIZ |
| 2.3595E-16 | -0.6130477 | 0.022 | 0.237 | 5.0375E-12 | PDC_inj | TGM2 |
| 8.4784E-06 | -0.6137466 | 0.854 | 0.837 | 0.18101479 | PDC_inj | PRDX1 |
| 1.1111E-05 | -0.6151543 | 0.034 | 0.119 | 0.23721194 | PDC_inj | CXCL8 |
| 8.0233E-11 | -0.6171421 | 0.381 | 0.521 | 1.713E-06 | PDC_inj | SON |
| 6.6322E-14 | -0.6176543 | 0.116 | 0.317 | 1.416E-09 | PDC_inj | SRSF2 |
| 1.9105E-09 | -0.6197334 | 0 | 0.119 | 4.0788E-05 | PDC_inj | NEFM |
| 5.4035E-23 | -0.6229279 | 0.082 | 0.377 | 1.1536E-18 | PDC_inj | CAMK2N1 |
| 8.7552E-19 | -0.6244918 | 0.056 | 0.303 | 1.8692E-14 | PDC_inj | CYTOR |
| 1.5417E-08 | -0.6265968 | 0.963 | 0.911 | 0.00032914 | PDC_inj | UBC |
| 1.0552E-07 | -0.6305155 | 0.179 | 0.314 | 0.00225286 | PDC_inj | GLUL |
| 8.7249E-11 | -0.631016 | 0.877 | 0.908 | 1.8628E-06 | PDC_inj | MYL6 |
| 2.9426E-06 | -0.6337582 | 0.019 | 0.108 | 0.06282471 | PDC_inj | FXYD4 |
| 4.6196E-15 | -0.6350013 | 0.082 | 0.292 | 9.8628E-11 | PDC_inj | STMN1 |
| 7.5969E-12 | -0.6359455 | 0.209 | 0.389 | 1.6219E-07 | PDC_inj | BRD2 |
| 8.4518E-16 | -0.6433233 | 0.183 | 0.406 | 1.8045E-11 | PDC_inj | FUS |
| 2.5165E-11 | -0.6434284 | 0.112 | 0.289 | 5.3727E-07 | PDC_inj | HIST1H1C |
| 2.9624E-20 | -0.6458496 | 0.03 | 0.286 | 6.3247E-16 | PDC_inj | CLDN3 |
| 3.9669E-23 | -0.6466888 | 0.093 | 0.391 | 8.4693E-19 | PDC_inj | OCIAD2 |
| 7.5644E-08 | -0.6483721 | 0.22 | 0.345 | 0.00161501 | PDC_inj | ZFP36L2 |
| 2.6543E-15 | -0.6491821 | 0.108 | 0.325 | 5.6669E-11 | PDC_inj | CCNL1 |
| 4.768E-16 | -0.6501731 | 0.026 | 0.239 | 1.018E-11 | PDC_inj | FXYD2 |
| 8.8762E-16 | -0.6534256 | 0.201 | 0.428 | 1.8951E-11 | PDC_inj | AHNAK |
| 3.0609E-09 | -0.6555137 | 0.134 | 0.284 | 6.535E-05 | PDC_inj | BTG2 |
| 2.7153E-21 | -0.6569074 | 0.015 | 0.279 | 5.7973E-17 | PDC_inj | MUC1 |
| 6.7431E-13 | -0.6590373 | 0.146 | 0.338 | 1.4397E-08 | PDC_inj | ZNF292 |
| 2.1046E-17 | -0.6617829 | 0.034 | 0.264 | 4.4932E-13 | PDC_inj | LAMB3 |
| 6.0478E-24 | -0.6657018 | 0.03 | 0.324 | 1.2912E-19 | PDC_inj | FAM107B |
| 1.4143E-07 | -0.6660536 | 0.888 | 0.893 | 0.00301949 | PDC_inj | GAPDH |
| 4.5011E-14 | -0.6674888 | 0.007 | 0.188 | 9.6099E-10 | PDC_inj | PIK3R3 |
| 3.7714E-13 | -0.6786358 | 0.034 | 0.217 | 8.0519E-09 | PDC_inj | GLRX |
| 1.3066E-16 | -0.680026 | 0.011 | 0.223 | 2.7895E-12 | PDC_inj | TNFAIP2 |
| 2.9115E-15 | -0.6822841 | 0.019 | 0.218 | 6.2161E-11 | PDC_inj | SOCS1 |
| 5.3105E-17 | -0.685576 | 0.142 | 0.383 | 1.1338E-12 | PDC_inj | RSRP1 |
| 7.7403E-24 | -0.6922403 | 0.093 | 0.391 | 1.6526E-19 | PDC_inj | MYO6 |
| 1.3415E-11 | -0.696229 | 0.071 | 0.241 | 2.864E-07 | PDC_inj | INTS6 |
| 2.2069E-16 | -0.7020024 | 0.037 | 0.257 | 4.7118E-12 | PDC_inj | ID2 |
| 2.1512E-09 | -0.7026955 | 0.157 | 0.309 | 4.5929E-05 | PDC_inj | HSPA1B |
| 4.1262E-10 | -0.7072667 | 0.53 | 0.612 | 8.8095E-06 | PDC_inj | BTG1 |
| 3.2466E-16 | -0.7138755 | 0.269 | 0.515 | 6.9314E-12 | PDC_inj | CDKN1A |
| 1.6286E-16 | -0.7144951 | 0.063 | 0.288 | 3.4771E-12 | PDC_inj | POLR2A |
| 6.1947E-14 | -0.7147724 | 0.063 | 0.257 | 1.3226E-09 | PDC_inj | IER5 |
| 7.3143E-14 | -0.7194205 | 0.06 | 0.264 | 1.5616E-09 | PDC_inj | EGR1 |
| 1.3763E-10 | -0.7222766 | 0.933 | 0.934 | 2.9385E-06 | PDC_inj | HSP90AA1 |
| 3.7702E-15 | -0.723959 | 0.09 | 0.307 | 8.0494E-11 | PDC_inj | DNAJB1 |
| 1.9714E-21 | -0.7293603 | 0.664 | 0.807 | 4.2089E-17 | PDC_inj | CFL1 |
| 4.4203E-20 | -0.7361252 | 0.597 | 0.834 | 9.4373E-16 | PDC_inj | MT-ND5 |
| 4.281E-14 | -0.7463554 | 0.093 | 0.299 | 9.1399E-10 | PDC_inj | ITGB8 |
| 1.0756E-11 | -0.7504523 | 0.049 | 0.214 | 2.2964E-07 | PDC_inj | ADAMTS1 |

| | | | | | | |
|---|---|---|---|---|---|---|
| 3.604E-24 | -0.7565484 | 0.03 | 0.326 | 7.6946E-20 | PDC_inj | DCDC2 |
| 6.3593E-19 | -0.7659507 | 0.06 | 0.31 | 1.3577E-14 | PDC_inj | PMAIP1 |
| 1.7009E-10 | -0.7705148 | 0.272 | 0.418 | 3.6314E-06 | PDC_inj | SRSF7 |
| 2.2499E-10 | -0.7717496 | 0.966 | 0.94 | 4.8036E-06 | PDC_inj | MALAT1 |
| 3.5442E-19 | -0.7818545 | 0.776 | 0.854 | 7.5669E-15 | PDC_inj | MYL12B |
| 1.2576E-30 | -0.7854806 | 0.448 | 0.735 | 2.6849E-26 | PDC_inj | ENO1 |
| 2.0231E-10 | -0.7935588 | 0.22 | 0.378 | 4.3194E-06 | PDC_inj | SLC38A2 |
| 3.0335E-20 | -0.7955262 | 0.034 | 0.291 | 6.4766E-16 | PDC_inj | PLAUR |
| 6.5806E-23 | -0.8000713 | 0.146 | 0.427 | 1.405E-18 | PDC_inj | SFPQ |
| 1.3911E-17 | -0.8003365 | 0.041 | 0.272 | 2.97E-13 | PDC_inj | TUBA1A |
| 6.1471E-30 | -0.8070176 | 0.205 | 0.551 | 1.3124E-25 | PDC_inj | CD59 |
| 2.4859E-15 | -0.8095452 | 0.078 | 0.293 | 5.3075E-11 | PDC_inj | RND3 |
| 6.1363E-28 | -0.8147415 | 0.034 | 0.366 | 1.3101E-23 | PDC_inj | CD44 |
| 3.6488E-31 | -0.8176382 | 0.25 | 0.592 | 7.7902E-27 | PDC_inj | EZR |
| 3.1887E-26 | -0.8201244 | 0.03 | 0.345 | 6.8079E-22 | PDC_inj | PAX8 |
| 5.8687E-22 | -0.8355748 | 0.022 | 0.294 | 1.253E-17 | PDC_inj | CYR61 |
| 8.4588E-18 | -0.8387445 | 0.026 | 0.258 | 1.806E-13 | PDC_inj | AKAP12 |
| 9.2029E-26 | -0.8393013 | 0.015 | 0.321 | 1.9648E-21 | PDC_inj | MAL |
| 2.5873E-25 | -0.8414677 | 0.007 | 0.306 | 5.5239E-21 | PDC_inj | MARCKSL1 |
| 8.8372E-21 | -0.8449851 | 0.164 | 0.427 | 1.8867E-16 | PDC_inj | TUBB4B |
| 1.8091E-19 | -0.847004 | 0.19 | 0.449 | 3.8625E-15 | PDC_inj | ZFP36L1 |
| 1.7536E-19 | -0.8502323 | 0.328 | 0.624 | 3.7439E-15 | PDC_inj | MTRNR2L8 |
| 2.9732E-19 | -0.8530394 | 0.41 | 0.628 | 6.3479E-15 | PDC_inj | PPP1R15A |
| 1.8334E-30 | -0.8545743 | 0.44 | 0.712 | 3.9143E-26 | PDC_inj | DDX5 |
| 6.8349E-07 | -0.8573773 | 0.243 | 0.361 | 0.01459258 | PDC_inj | HSPA1A |
| 6.718E-36 | -0.8666064 | 0.101 | 0.506 | 1.4343E-31 | PDC_inj | TNFRSF12A |
| 7.4843E-23 | -0.8688601 | 0.489 | 0.719 | 1.5979E-18 | PDC_inj | HSPB1 |
| 2.8093E-15 | -0.8701855 | 0.071 | 0.281 | 5.9978E-11 | PDC_inj | PPP1R10 |
| 4.5984E-43 | -0.8785939 | 0.254 | 0.676 | 9.8175E-39 | PDC_inj | SPINT2 |
| 4.2377E-19 | -0.8803454 | 0.358 | 0.61 | 9.0476E-15 | PDC_inj | SOD2 |
| 1.2782E-20 | -0.8836144 | 0.06 | 0.332 | 2.7289E-16 | PDC_inj | IL32 |
| 1.0053E-15 | -0.8836228 | 0.183 | 0.395 | 2.1463E-11 | PDC_inj | TUBA1B |
| 1.9267E-30 | -0.8909608 | 0.563 | 0.801 | 4.1134E-26 | PDC_inj | HNRNPA2B1 |
| 2.1638E-10 | -0.8978827 | 0.937 | 0.923 | 4.6198E-06 | PDC_inj | TMSB10 |
| 2.3465E-27 | -0.9004618 | 0.067 | 0.398 | 5.0097E-23 | PDC_inj | RAB11FIP1 |
| 3.0224E-15 | -0.9096628 | 0.119 | 0.336 | 6.4527E-11 | PDC_inj | RHOB |
| 3.9659E-31 | -0.9135512 | 0.045 | 0.406 | 8.4671E-27 | PDC_inj | JPT1 |
| 2.2183E-13 | -0.9138343 | 0.851 | 0.874 | 4.7361E-09 | PDC_inj | GSTP1 |
| 9.9574E-18 | -0.922707 | 0.119 | 0.364 | 2.1259E-13 | PDC_inj | KLF6 |
| 3.5955E-27 | -0.9284288 | 0.713 | 0.915 | 7.6763E-23 | PDC_inj | MT-CO2 |
| 1.148E-33 | -0.9303438 | 0.347 | 0.675 | 2.4511E-29 | PDC_inj | TPI1 |
| 4.2353E-18 | -0.9312537 | 0.142 | 0.391 | 9.0423E-14 | PDC_inj | JUNB |
| 3.831E-23 | -0.9536197 | 0.022 | 0.307 | 8.1792E-19 | PDC_inj | PLK2 |
| 1.2133E-18 | -0.9541165 | 0.007 | 0.241 | 2.5904E-14 | PDC_inj | CST6 |
| 1.4902E-15 | -0.9669463 | 0.034 | 0.238 | 3.1815E-11 | PDC_inj | CSKMT |
| 0.00786952 | -0.9712676 | 0.257 | 0.283 | 1 | PDC_inj | MAFB |
| 1.8142E-43 | -0.9722913 | 0.063 | 0.513 | 3.8733E-39 | PDC_inj | CLDN7 |
| 5.5759E-22 | -0.9903858 | 0.015 | 0.283 | 1.1905E-17 | PDC_inj | RASD1 |
| 3.5287E-29 | -0.9972118 | 0.612 | 0.819 | 7.5338E-25 | PDC_inj | IGFBP7 |
| 8.359E-24 | -1.0042851 | 0.03 | 0.325 | 1.7847E-19 | PDC_inj | ITGA2 |

| | | | | | | |
|---|---|---|---|---|---|---|
| 4.6843E-49 | -1.006351 | 0.127 | 0.597 | 1.0001E-44 | PDC_inj | S100A13 |
| 7.3869E-41 | -1.0120428 | 0.138 | 0.567 | 1.5771E-36 | PDC_inj | HMGA1 |
| 2.2388E-45 | -1.0297848 | 0.097 | 0.56 | 4.7799E-41 | PDC_inj | CD9 |
| 6.6676E-19 | -1.0303899 | 0.037 | 0.282 | 1.4235E-14 | PDC_inj | HES1 |
| 1.1885E-51 | -1.0414882 | 0.332 | 0.774 | 2.5374E-47 | PDC_inj | PKM |
| 0.00020096 | -1.0461712 | 0.34 | 0.378 | 1 | PDC_inj | HIST1H4C |
| 1.0552E-17 | -1.0578742 | 0.175 | 0.411 | 2.2528E-13 | PDC_inj | RNF213 |
| 1.4364E-17 | -1.0848892 | 0.019 | 0.247 | 3.0668E-13 | PDC_inj | TFPI2 |
| 1.8921E-37 | -1.0929187 | 0.522 | 0.875 | 4.0397E-33 | PDC_inj | MT-ND3 |
| 2.3658E-24 | -1.105702 | 0.019 | 0.314 | 5.0511E-20 | PDC_inj | PLAU |
| 2.2744E-30 | -1.1136757 | 0.313 | 0.637 | 4.8559E-26 | PDC_inj | TPM1 |
| 4.2737E-26 | -1.1271206 | 0.123 | 0.45 | 9.1244E-22 | PDC_inj | UPP1 |
| 1.091E-22 | -1.1316721 | 0.104 | 0.397 | 2.3293E-18 | PDC_inj | NFKBIA |
| 2.8315E-32 | -1.1329789 | 0.06 | 0.425 | 6.0453E-28 | PDC_inj | CCND1 |
| 1.0591E-22 | -1.1503445 | 0.004 | 0.277 | 2.2611E-18 | PDC_inj | TCIM |
| 5.4026E-35 | -1.1589588 | 0.664 | 0.903 | 1.1535E-30 | PDC_inj | MT-CYB |
| 9.0581E-33 | -1.1603425 | 0.052 | 0.421 | 1.9339E-28 | PDC_inj | PHLDA2 |
| 5.4392E-18 | -1.1727634 | 0.041 | 0.275 | 1.1613E-13 | PDC_inj | HSPA2 |
| 3.5455E-71 | -1.24168 | 0.104 | 0.692 | 7.5696E-67 | PDC_inj | EPCAM |
| 1.9652E-22 | -1.2431674 | 0.026 | 0.303 | 4.1956E-18 | PDC_inj | HEXIM1 |
| 7.0429E-57 | -1.2553637 | 0.172 | 0.678 | 1.5037E-52 | PDC_inj | GLS |
| 1.427E-26 | -1.3035377 | 0.022 | 0.337 | 3.0467E-22 | PDC_inj | LAMC2 |
| 5.4527E-16 | -1.3038699 | 0.041 | 0.26 | 1.1642E-11 | PDC_inj | LCN2 |
| 4.1852E-37 | -1.3050857 | 0.31 | 0.68 | 8.9354E-33 | PDC_inj | SH3BGRL3 |
| 1.5994E-41 | -1.3155626 | 0.06 | 0.494 | 3.4147E-37 | PDC_inj | TSPAN1 |
| 8.4619E-49 | -1.3213997 | 0.612 | 0.916 | 1.8066E-44 | PDC_inj | MT-ATP6 |
| 5.9442E-14 | -1.3451417 | 0.034 | 0.23 | 1.2691E-09 | PDC_inj | DEFB1 |
| 4.3917E-41 | -1.3579847 | 0.623 | 0.875 | 9.3763E-37 | PDC_inj | S100A10 |
| 7.8309E-39 | -1.4197427 | 0.164 | 0.573 | 1.6719E-34 | PDC_inj | IER2 |
| 2.3795E-23 | -1.4408364 | 0.966 | 0.954 | 5.0801E-19 | PDC_inj | S100A6 |
| 8.1699E-44 | -1.4634646 | 0.172 | 0.614 | 1.7443E-39 | PDC_inj | GADD45B |
| 4.0401E-39 | -1.4645323 | 0.063 | 0.479 | 8.6256E-35 | PDC_inj | C12orf75 |
| 1.8286E-49 | -1.5044501 | 0.616 | 0.916 | 3.904E-45 | PDC_inj | MT-CO3 |
| 5.4018E-38 | -1.5497545 | 0.832 | 0.921 | 1.1533E-33 | PDC_inj | S100A11 |
| 3.1759E-53 | -1.5704829 | 0.108 | 0.61 | 6.7805E-49 | PDC_inj | CLU |
| 4.3842E-34 | -1.5755224 | 0.063 | 0.448 | 9.3603E-30 | PDC_inj | IER3 |
| 7.521E-53 | -1.6028392 | 0.731 | 0.931 | 1.6057E-48 | PDC_inj | MT-CO1 |
| 1.8921E-69 | -1.6146302 | 0.086 | 0.675 | 4.0397E-65 | PDC_inj | TACSTD2 |
| 1.2919E-35 | -1.6327175 | 0.272 | 0.641 | 2.7583E-31 | PDC_inj | JUN |
| 3.3753E-92 | -1.6652091 | 0.131 | 0.815 | 7.2063E-88 | PDC_inj | CD24 |
| 3.4629E-54 | -1.6928481 | 0.093 | 0.602 | 7.3933E-50 | PDC_inj | KRT19 |
| 7.6568E-75 | -1.7242514 | 0.104 | 0.715 | 1.6347E-70 | PDC_inj | CLDN4 |
| 1.5364E-34 | -1.7334374 | 0.593 | 0.82 | 3.2801E-30 | PDC_inj | NEAT1 |
| 1.3281E-56 | -1.7640994 | 0.082 | 0.597 | 2.8354E-52 | PDC_inj | ELF3 |
| 1.2346E-73 | -1.7680161 | 0.142 | 0.74 | 2.6358E-69 | PDC_inj | KRT8 |
| 3.7958E-28 | -1.7757779 | 0.06 | 0.398 | 8.104E-24 | PDC_inj | SLPI |
| 2.2292E-82 | -1.7829638 | 0.168 | 0.812 | 4.7594E-78 | PDC_inj | KRT18 |
| 8.4102E-68 | -1.789705 | 0.09 | 0.661 | 1.7956E-63 | PDC_inj | KRT7 |
| 1.8651E-37 | -1.8764877 | 0.306 | 0.687 | 3.982E-33 | PDC_inj | MTRNR2L12 |
| 1.1276E-66 | -2.1127579 | 0.086 | 0.657 | 2.4075E-62 | PDC_inj | WFDC2 |

| | | | | | | |
|---|---|---|---|---|---|---|
| 5.3231E-93 | -2.2111048 | 0.358 | 0.882 | 1.1365E-88 | PDC_inj | SAT1 |
| 1.5642E-45 | -2.2676653 | 0.138 | 0.609 | 3.3395E-41 | PDC_inj | SPP1 |
| 6.5668E-48 | -2.288155 | 0.086 | 0.549 | 1.402E-43 | PDC_inj | SOX4 |
| 5.0901E-40 | -2.3089093 | 0.075 | 0.501 | 1.0867E-35 | PDC_inj | S100A2 |
| 2.8125E-59 | -2.964776 | 0.108 | 0.648 | 6.0047E-55 | PDC_inj | MMP7 |
| 6.1554E-37 | -5.1077951 | 0.272 | 0.604 | 1.3142E-32 | PDC_inj | PLCG2 |
| 1.711E-157 | 3.22249167 | 0.624 | 0.07 | 3.653E-153 | TAL | UMOD |
| 8.098E-171 | 2.83386251 | 0.72 | 0.09 | 1.729E-166 | TAL | SLC12A1 |
| 5.79E-231 | 1.50017824 | 0.522 | 0.032 | 1.236E-226 | TAL | KNG1 |
| 5.7413E-41 | 1.48062315 | 0.643 | 0.22 | 1.2258E-36 | TAL | DEFB1 |
| 3.5422E-57 | 1.36939989 | 0.879 | 0.368 | 7.5626E-53 | TAL | LDHB |
| 1.216E-101 | 1.349363 | 0.535 | 0.076 | 2.5958E-97 | TAL | PPP1R1A |
| 4.465E-114 | 1.30668334 | 0.592 | 0.084 | 9.532E-110 | TAL | CLDN10 |
| 2.1133E-29 | 1.19893786 | 0.822 | 0.488 | 4.5118E-25 | TAL | ATP1B1 |
| 7.668E-116 | 1.19470788 | 0.503 | 0.059 | 1.637E-111 | TAL | SFRP1 |
| 8.2472E-48 | 1.15101575 | 0.707 | 0.229 | 1.7608E-43 | TAL | FXYD2 |
| 9.5687E-58 | 1.04552489 | 0.618 | 0.162 | 2.0429E-53 | TAL | CA12 |
| 9.7145E-21 | 0.99434036 | 0.554 | 0.268 | 2.074E-16 | TAL | CKB |
| 3.2318E-26 | 0.94016966 | 0.828 | 0.482 | 6.9E-22 | TAL | ATP1A1 |
| 1.5574E-20 | 0.89418922 | 0.618 | 0.329 | 3.3251E-16 | TAL | CHCHD10 |
| 2.5426E-38 | 0.86076822 | 0.529 | 0.168 | 5.4284E-34 | TAL | MPC1 |
| 3.9771E-15 | 0.85829049 | 0.822 | 0.66 | 8.4911E-11 | TAL | UQCRB |
| 4.0631E-24 | 0.84994901 | 0.873 | 0.641 | 8.6748E-20 | TAL | PEBP1 |
| 2.1972E-44 | 0.84757254 | 0.306 | 0.052 | 4.691E-40 | TAL | CLDN14 |
| 2.5341E-30 | 0.80663571 | 0.713 | 0.31 | 5.4103E-26 | TAL | MAL |
| 2.2361E-40 | 0.80372148 | 0.325 | 0.062 | 4.774E-36 | TAL | SOD3 |
| 3.1748E-31 | 0.78545789 | 0.624 | 0.253 | 6.7782E-27 | TAL | ATP5F1A |
| 6.8177E-68 | 0.77157495 | 0.49 | 0.089 | 1.4556E-63 | TAL | KCNJ16 |
| 4.1904E-14 | 0.76695253 | 0.694 | 0.458 | 8.9464E-10 | TAL | COX8A |
| 4.1812E-12 | 0.75356001 | 0.752 | 0.58 | 8.9268E-08 | TAL | COX5B |
| 1.4893E-50 | 0.73053532 | 0.401 | 0.077 | 3.1796E-46 | TAL | PTH1R |
| 6.4146E-09 | 0.7174505 | 0.42 | 0.235 | 0.00013695 | TAL | SLC5A3 |
| 1.5981E-63 | 0.68344229 | 0.268 | 0.029 | 3.4119E-59 | TAL | CLCNKA |
| 1.6457E-15 | 0.67410716 | 0.631 | 0.363 | 3.5135E-11 | TAL | ATP5F1B |
| 7.3304E-38 | 0.67200507 | 0.414 | 0.104 | 1.565E-33 | TAL | CYFIP2 |
| 2.7933E-06 | 0.66553299 | 0.471 | 0.331 | 0.05963692 | TAL | SLC25A5 |
| 1.5948E-06 | 0.66166573 | 0.554 | 0.396 | 0.03404938 | TAL | ANKRD11 |
| 1.8245E-12 | 0.65616343 | 0.573 | 0.359 | 3.8953E-08 | TAL | ATP5MC3 |
| 1.231E-50 | 0.65198789 | 0.395 | 0.075 | 2.6281E-46 | TAL | CYS1 |
| 1.8162E-64 | 0.64345043 | 0.452 | 0.078 | 3.8777E-60 | TAL | IRX2 |
| 4.7801E-48 | 0.64031125 | 0.465 | 0.106 | 1.0206E-43 | TAL | EFHD1 |
| 3.1298E-41 | 0.6338723 | 0.478 | 0.124 | 6.6821E-37 | TAL | IMPA2 |
| 1.5326E-23 | 0.61978924 | 0.414 | 0.141 | 3.2721E-19 | TAL | IDH2 |
| 7.015E-20 | 0.6187069 | 0.433 | 0.169 | 1.4977E-15 | TAL | TACC1 |
| 3.621E-08 | 0.61522028 | 0.376 | 0.212 | 0.00077307 | TAL | TRIP11 |
| 5.3912E-12 | 0.59773517 | 0.363 | 0.167 | 1.151E-07 | TAL | WNK1 |
| 7.1965E-25 | 0.59687328 | 0.452 | 0.153 | 1.5364E-20 | TAL | NUDT4 |
| 9.4858E-12 | 0.59350696 | 0.873 | 0.767 | 2.0252E-07 | TAL | COX7C |
| 1.8376E-10 | 0.59262896 | 0.936 | 0.909 | 3.9233E-06 | TAL | MT-ATP6 |
| 2.5619E-85 | 0.59125736 | 0.293 | 0.027 | 5.4696E-81 | TAL | KCNJ1 |

| | | | | | | |
|---|---|---|---|---|---|---|
| 1.8001E-08 | 0.57686451 | 0.548 | 0.357 | 0.00038432 | TAL | ATP5ME |
| 1.4401E-05 | 0.57473053 | 0.51 | 0.366 | 0.30745376 | TAL | GOLGA4 |
| 4.3808E-11 | 0.56173088 | 0.79 | 0.647 | 9.3531E-07 | TAL | UQCRH |
| 2.2187E-43 | 0.55768699 | 0.318 | 0.057 | 4.7368E-39 | TAL | ERBB4 |
| 1.9555E-14 | 0.55746527 | 0.949 | 0.799 | 4.175E-10 | TAL | CD24 |
| 1.0787E-37 | 0.5557145 | 0.427 | 0.106 | 2.3031E-33 | TAL | ALDH1A1 |
| 4.0744E-19 | 0.55557073 | 0.847 | 0.596 | 8.6989E-15 | TAL | SPP1 |
| 6.3523E-12 | 0.55243457 | 0.688 | 0.45 | 1.3562E-07 | TAL | ATP5PF |
| 2.8275E-13 | 0.55091711 | 0.484 | 0.246 | 6.0368E-09 | TAL | MGLL |
| 9.4551E-31 | 0.5507236 | 0.529 | 0.176 | 2.0187E-26 | TAL | SLC25A4 |
| 2.9727E-08 | 0.54577374 | 0.936 | 0.867 | 0.00063467 | TAL | MT-ND3 |
| 2.821E-15 | 0.54276815 | 0.535 | 0.272 | 6.0228E-11 | TAL | NDUFS6 |
| 6.2189E-21 | 0.54250456 | 0.414 | 0.151 | 1.3277E-16 | TAL | IVNS1ABP |
| 1.1564E-13 | 0.54035918 | 0.497 | 0.247 | 2.4689E-09 | TAL | MRPS6 |
| 5.9032E-34 | 0.53647201 | 0.49 | 0.145 | 1.2603E-29 | TAL | ITM2C |
| 1.3676E-37 | 0.53277828 | 0.42 | 0.103 | 2.9199E-33 | TAL | CGNL1 |
| 2.5704E-10 | 0.53240862 | 0.834 | 0.73 | 5.4878E-06 | TAL | NDUFA4 |
| 9.2941E-22 | 0.53139944 | 0.427 | 0.151 | 1.9843E-17 | TAL | CPM |
| 1.3122E-48 | 0.53106522 | 0.325 | 0.054 | 2.8016E-44 | TAL | SLC9A3 |
| 2.5412E-32 | 0.52872135 | 0.503 | 0.155 | 5.4255E-28 | TAL | GOLM1 |
| 5.528E-21 | 0.5283619 | 0.357 | 0.118 | 1.1802E-16 | TAL | POU3F3 |
| 3.1052E-64 | 0.52716635 | 0.255 | 0.026 | 6.6297E-60 | TAL | CLDN16 |
| 3.1419E-13 | 0.51390051 | 0.599 | 0.354 | 6.708E-09 | TAL | EIF3K |
| 6.2834E-14 | 0.51268731 | 0.325 | 0.124 | 1.3415E-09 | TAL | PCDH9 |
| 3.4426E-12 | 0.50554269 | 0.408 | 0.2 | 7.35E-08 | TAL | NDUFB1 |
| 6.3431E-11 | 0.50510636 | 0.783 | 0.606 | 1.3542E-06 | TAL | ATP5MG |
| 1.7296E-27 | 0.50251697 | 0.408 | 0.121 | 3.6928E-23 | TAL | PKP4 |
| 3.57E-14 | 0.49065974 | 0.943 | 0.813 | 7.6219E-10 | TAL | IGFBP7 |
| 1.2307E-17 | 0.48744885 | 0.465 | 0.202 | 2.6275E-13 | TAL | MPC2 |
| 1.599E-09 | 0.48486522 | 0.72 | 0.526 | 3.4138E-05 | TAL | COX6C |
| 5.1803E-09 | 0.48041417 | 0.51 | 0.314 | 0.0001106 | TAL | PRDX2 |
| 1.3027E-32 | 0.48020435 | 0.312 | 0.068 | 2.7813E-28 | TAL | SLC3A1 |
| 0.00024182 | 0.47914397 | 0.446 | 0.332 | 1 | TAL | ANKRD12 |
| 2.3351E-25 | 0.47906491 | 0.261 | 0.061 | 4.9855E-21 | TAL | COX7A1 |
| 3.5589E-18 | 0.47641905 | 0.35 | 0.121 | 7.5983E-14 | TAL | CXCL14 |
| 2.1654E-07 | 0.47563533 | 0.949 | 0.91 | 0.00462305 | TAL | MT-CO3 |
| 2.0434E-17 | 0.47493162 | 0.465 | 0.191 | 4.3627E-13 | TAL | BCAM |
| 2.8186E-14 | 0.47107733 | 0.49 | 0.249 | 6.0177E-10 | TAL | UQCRFS1 |
| 4.0496E-48 | 0.47100836 | 0.35 | 0.061 | 8.646E-44 | TAL | PDE1A |
| 2.8213E-19 | 0.46983837 | 0.338 | 0.109 | 6.0235E-15 | TAL | SULT1C2 |
| 1.8556E-30 | 0.46577514 | 0.42 | 0.119 | 3.9616E-26 | TAL | TRIM2 |
| 3.0843E-09 | 0.46383323 | 0.745 | 0.569 | 6.585E-05 | TAL | COX7A2 |
| 6.349E-10 | 0.46030746 | 0.739 | 0.581 | 1.3555E-05 | TAL | APP |
| 8.4194E-12 | 0.45978598 | 0.643 | 0.391 | 1.7975E-07 | TAL | ATP5F1D |
| 1.2433E-07 | 0.45712149 | 0.51 | 0.323 | 0.00265441 | TAL | NDUFB7 |
| 1.3441E-31 | 0.45597748 | 0.287 | 0.06 | 2.8696E-27 | TAL | RAP1GAP |
| 5.2777E-33 | 0.44921467 | 0.312 | 0.067 | 1.1268E-28 | TAL | BICDL1 |
| 4.499E-09 | 0.44693851 | 0.312 | 0.15 | 9.6053E-05 | TAL | TFPI |
| 1.5182E-06 | 0.44610386 | 0.338 | 0.194 | 0.03241268 | TAL | NCOA7 |
| 1.1418E-07 | 0.44567032 | 0.268 | 0.13 | 0.00243773 | TAL | COL4A1 |

| | | | | | | |
|---|---|---|---|---|---|---|
| 1.9267E-66 | 0.44456341 | 0.363 | 0.05 | 4.1134E-62 | TAL | PLCB1 |
| 6.3576E-45 | 0.44433421 | 0.185 | 0.02 | 1.3573E-40 | TAL | PVALB |
| 1.1583E-13 | 0.4436741 | 0.35 | 0.146 | 2.473E-09 | TAL | TFCP2L1 |
| 1.1123E-20 | 0.44208846 | 0.369 | 0.121 | 2.3749E-16 | TAL | PTGER3 |
| 3.2364E-13 | 0.44006769 | 0.592 | 0.32 | 6.9097E-09 | TAL | PHPT1 |
| 7.4063E-40 | 0.43995043 | 0.318 | 0.061 | 1.5812E-35 | TAL | ST3GAL4 |
| 2.6956E-05 | 0.43893249 | 0.306 | 0.182 | 0.57550037 | TAL | SPEN |
| 8.6327E-74 | 0.43511083 | 0.338 | 0.04 | 1.8431E-69 | TAL | C5orf38 |
| 3.4598E-56 | 0.43502264 | 0.268 | 0.033 | 7.3867E-52 | TAL | CLCNKB |
| 1.4924E-54 | 0.43363858 | 0.248 | 0.029 | 3.1864E-50 | TAL | IRX1 |
| 7.7093E-22 | 0.43029872 | 0.395 | 0.13 | 1.6459E-17 | TAL | CDH16 |
| 5.8385E-16 | 0.42894275 | 0.389 | 0.159 | 1.2465E-11 | TAL | SUCLG1 |
| 6.2754E-15 | 0.42634864 | 0.433 | 0.194 | 1.3398E-10 | TAL | CYC1 |
| 6.4827E-10 | 0.42510901 | 0.395 | 0.208 | 1.384E-05 | TAL | BEX3 |
| 2.4524E-41 | 0.42508646 | 0.274 | 0.045 | 5.2359E-37 | TAL | PLCL1 |
| 0.00010111 | 0.42507384 | 0.363 | 0.238 | 1 | TAL | MDM2 |
| 4.0459E-13 | 0.42501982 | 0.58 | 0.316 | 8.638E-09 | TAL | NDUFB9 |
| 1.5559E-14 | 0.42339642 | 0.51 | 0.253 | 3.3218E-10 | TAL | SMDT1 |
| 3.689E-13 | 0.42323457 | 0.567 | 0.307 | 7.8761E-09 | TAL | ISCU |
| 9.0717E-14 | 0.42322007 | 0.535 | 0.282 | 1.9368E-09 | TAL | ATP5PO |
| 2.5629E-06 | 0.42320125 | 0.471 | 0.312 | 0.0547179 | TAL | UQCR11 |
| 0.00140481 | 0.42148561 | 0.471 | 0.374 | 1 | TAL | NDUFA1 |
| 1.7987E-12 | 0.41860243 | 0.452 | 0.224 | 3.8403E-08 | TAL | COMT |
| 7.1203E-08 | 0.41809353 | 0.93 | 0.913 | 0.00152018 | TAL | MT-ND4 |
| 1.167E-07 | 0.41691862 | 0.618 | 0.408 | 0.00249164 | TAL | UQCR10 |
| 2.0384E-30 | 0.41564931 | 0.401 | 0.108 | 4.3521E-26 | TAL | GNAI1 |
| 3.6676E-17 | 0.41513294 | 0.408 | 0.161 | 7.8303E-13 | TAL | MTURN |
| 2.9953E-12 | 0.41331745 | 0.51 | 0.267 | 6.3949E-08 | TAL | FARP1 |
| 3.8566E-08 | 0.41265316 | 0.382 | 0.214 | 0.00082338 | TAL | ATP5MC1 |
| 5.9052E-57 | 0.41173363 | 0.178 | 0.015 | 1.2608E-52 | TAL | ACPP |
| 1.8514E-09 | 0.41143198 | 0.376 | 0.187 | 3.9528E-05 | TAL | EMP1 |
| 1.4873E-38 | 0.40945564 | 0.217 | 0.031 | 3.1753E-34 | TAL | TMEM37 |
| 2.1522E-09 | 0.40766751 | 0.822 | 0.726 | 4.595E-05 | TAL | GNAS |
| 1.8226E-18 | 0.40457986 | 0.229 | 0.06 | 3.8914E-14 | TAL | FABP3 |
| 1.9483E-05 | 0.40451753 | 0.318 | 0.185 | 0.41596536 | TAL | ADIRF |
| 0.00028045 | 0.40188673 | 0.471 | 0.359 | 1 | TAL | COX7B |
| 1.3932E-27 | 0.4006714 | 0.299 | 0.072 | 2.9745E-23 | TAL | SLC16A7 |
| 1.1098E-23 | 0.39900686 | 0.255 | 0.061 | 2.3694E-19 | TAL | TSPAN12 |
| 0.00026398 | 0.39608412 | 0.975 | 0.94 | 1 | TAL | MALAT1 |
| 3.3601E-09 | 0.39599506 | 0.433 | 0.237 | 7.1738E-05 | TAL | COX20 |
| 3.2676E-11 | 0.39521378 | 0.42 | 0.205 | 6.9764E-07 | TAL | NDUFA3 |
| 4.7839E-15 | 0.39508702 | 0.401 | 0.168 | 1.0214E-10 | TAL | BICC1 |
| 6.0875E-16 | 0.39481208 | 0.414 | 0.173 | 1.2997E-11 | TAL | FAM213A |
| 3.517E-24 | 0.39352865 | 0.401 | 0.128 | 7.5087E-20 | TAL | NDUFS1 |
| 1.0288E-16 | 0.39200015 | 0.503 | 0.225 | 2.1965E-12 | TAL | MDH1 |
| 2.1613E-06 | 0.39140588 | 0.847 | 0.781 | 0.04614356 | TAL | COX4I1 |
| 1.3958E-28 | 0.39012351 | 0.268 | 0.057 | 2.98E-24 | TAL | TMEM213 |
| 6.5645E-21 | 0.38927699 | 0.312 | 0.094 | 1.4015E-16 | TAL | MOCS2 |
| 9.6677E-08 | 0.38720998 | 0.72 | 0.523 | 0.00206405 | TAL | ATP5MC2 |
| 4.8465E-06 | 0.38633147 | 0.694 | 0.559 | 0.10347336 | TAL | COX6B1 |

| | | | | | | |
|---|---|---|---|---|---|---|
| 1.375E-11 | 0.38550852 | 0.357 | 0.159 | 2.9356E-07 | TAL | PCSK1N |
| 2.9823E-98 | 0.38092196 | 0.229 | 0.014 | 6.3672E-94 | TAL | CXCL12 |
| 1.8037E-15 | 0.3804936 | 0.459 | 0.202 | 3.8508E-11 | TAL | ADI1 |
| 1.2462E-07 | 0.37845786 | 0.93 | 0.891 | 0.00266071 | TAL | MT-ND1 |
| 1.718E-06 | 0.37829749 | 0.93 | 0.911 | 0.03667954 | TAL | MT-CO2 |
| 2.4328E-49 | 0.37560547 | 0.299 | 0.045 | 5.1941E-45 | TAL | GYPC |
| 7.2089E-20 | 0.37498274 | 0.287 | 0.085 | 1.5391E-15 | TAL | CTDSPL |
| 2.3283E-14 | 0.37478342 | 0.414 | 0.182 | 4.9709E-10 | TAL | CBR1 |
| 6.2344E-12 | 0.37471768 | 0.522 | 0.286 | 1.331E-07 | TAL | SERINC1 |
| 6.9123E-24 | 0.37453076 | 0.299 | 0.079 | 1.4758E-19 | TAL | PCCA |
| 6.9953E-05 | 0.37116469 | 0.58 | 0.411 | 1 | TAL | GOLGB1 |
| 7.6947E-07 | 0.37039955 | 0.503 | 0.327 | 0.01642814 | TAL | UQCRQ |
| 9.9978E-10 | 0.36998206 | 0.79 | 0.605 | 2.1345E-05 | TAL | APLP2 |
| 8.0388E-05 | 0.36969629 | 0.701 | 0.626 | 1 | TAL | SLC25A6 |
| 1.0662E-39 | 0.36921031 | 0.344 | 0.068 | 2.2763E-35 | TAL | SIM2 |
| 1.5805E-06 | 0.36801779 | 0.299 | 0.16 | 0.03374343 | TAL | FNIP2 |
| 6.6831E-07 | 0.36452643 | 0.58 | 0.39 | 0.01426843 | TAL | CIRBP |
| 1.7358E-09 | 0.36421321 | 0.465 | 0.257 | 3.7059E-05 | TAL | NDUFB3 |
| 7.5498E-14 | 0.36400745 | 0.376 | 0.156 | 1.6119E-09 | TAL | MRPL34 |
| 9.4127E-12 | 0.36376527 | 0.318 | 0.137 | 2.0096E-07 | TAL | SEC14L1 |
| 5.9196E-10 | 0.36222685 | 0.312 | 0.143 | 1.2638E-05 | TAL | COL4A2 |
| 6.1544E-09 | 0.36156962 | 0.516 | 0.308 | 0.0001314 | TAL | ATP5PD |
| 5.2147E-12 | 0.36103808 | 0.127 | 0.031 | 1.1133E-07 | TAL | KCNMA1 |
| 5.5241E-07 | 0.360922 | 0.427 | 0.261 | 0.01179399 | TAL | NDUFB10 |
| 3.8512E-12 | 0.35964276 | 0.414 | 0.2 | 8.2224E-08 | TAL | TMEM230 |
| 6.6544E-11 | 0.35928504 | 0.497 | 0.264 | 1.4207E-06 | TAL | GSTM3 |
| 1.2963E-14 | 0.35882791 | 0.344 | 0.135 | 2.7676E-10 | TAL | AK3 |
| 1.4643E-05 | 0.3576767 | 0.637 | 0.489 | 0.31263707 | TAL | TOMM7 |
| 5.8855E-12 | 0.357154 | 0.376 | 0.17 | 1.2566E-07 | TAL | COA3 |
| 5.3708E-25 | 0.35708581 | 0.242 | 0.053 | 1.1467E-20 | TAL | OXCT1 |
| 2.652E-05 | 0.35705625 | 0.764 | 0.723 | 0.56621046 | TAL | CTSD |
| 2.117E-10 | 0.35508697 | 0.446 | 0.231 | 4.5198E-06 | TAL | NDUFS8 |
| 8.9613E-15 | 0.35477469 | 0.382 | 0.156 | 1.9132E-10 | TAL | ECH1 |
| 8.149E-05 | 0.35398298 | 0.675 | 0.564 | 1 | TAL | GDF15 |
| 4.4849E-13 | 0.35383863 | 0.427 | 0.198 | 9.5752E-09 | TAL | NDUFB5 |
| 3.3373E-32 | 0.35286323 | 0.185 | 0.027 | 7.1251E-28 | TAL | SLC34A2 |
| 4.1634E-21 | 0.35250258 | 0.248 | 0.063 | 8.8889E-17 | TAL | DNASE1 |
| 9.8218E-26 | 0.35226886 | 0.338 | 0.09 | 2.097E-21 | TAL | SDC1 |
| 1.6154E-24 | 0.35223362 | 0.338 | 0.095 | 3.449E-20 | TAL | ACADM |
| 3.1939E-23 | 0.35133544 | 0.344 | 0.1 | 6.8189E-19 | TAL | LGR4 |
| 9.831E-07 | 0.3511745 | 0.49 | 0.32 | 0.02098913 | TAL | COX5A |
| 1.1695E-17 | 0.34862678 | 0.248 | 0.071 | 2.4969E-13 | TAL | COL4A3 |
| 5.6543E-08 | 0.34860514 | 0.459 | 0.274 | 0.00120719 | TAL | CYB5A |
| 4.1657E-08 | 0.34585038 | 0.631 | 0.426 | 0.00088937 | TAL | BSG |
| 2.854E-13 | 0.34538063 | 0.573 | 0.285 | 6.0934E-09 | TAL | MT-ATP8 |
| 2.3592E-18 | 0.3425391 | 0.293 | 0.089 | 5.0368E-14 | TAL | TUBB2B |
| 6.4133E-14 | 0.34162166 | 0.363 | 0.147 | 1.3692E-09 | TAL | ABLIM1 |
| 4.3515E-24 | 0.34023291 | 0.229 | 0.05 | 9.2905E-20 | TAL | FKBP5 |
| 8.8546E-12 | 0.33976771 | 0.389 | 0.182 | 1.8904E-07 | TAL | HADHB |
| 1.376E-40 | 0.3394084 | 0.268 | 0.043 | 2.9377E-36 | TAL | ESRRG |

| | | | | | | |
|---|---|---|---|---|---|---|
| 2.6775E-14 | 0.33892386 | 0.338 | 0.131 | 5.7164E-10 | TAL | RRAGD |
| 1.9928E-05 | 0.33779526 | 0.522 | 0.38 | 0.42547128 | TAL | NDUFB2 |
| 1.6589E-05 | 0.33760482 | 0.783 | 0.679 | 0.35418048 | TAL | COX6A1 |
| 0.00010591 | 0.33579913 | 0.516 | 0.381 | 1 | TAL | N4BP2L2 |
| 1.3713E-19 | 0.33570077 | 0.274 | 0.078 | 2.9277E-15 | TAL | HOXD8 |
| 7.4515E-14 | 0.33477796 | 0.325 | 0.123 | 1.5909E-09 | TAL | GPNMB |
| 5.783E-19 | 0.33374791 | 0.268 | 0.076 | 1.2347E-14 | TAL | TBC1D4 |
| 2.8293E-06 | 0.33288354 | 0.503 | 0.331 | 0.06040482 | TAL | CPEB4 |
| 1.22E-08 | 0.33185797 | 0.637 | 0.401 | 0.00026048 | TAL | PCBP2 |
| 5.5758E-08 | 0.33156926 | 0.631 | 0.423 | 0.00119044 | TAL | TMEM59 |
| 9.0269E-13 | 0.33142616 | 0.439 | 0.195 | 1.9272E-08 | TAL | ADGRG1 |
| 2.0317E-12 | 0.33139862 | 0.382 | 0.173 | 4.3378E-08 | TAL | UQCRC2 |
| 4.746E-08 | 0.33105227 | 0.338 | 0.177 | 0.00101327 | TAL | SETD2 |
| 2.8469E-11 | 0.3307283 | 0.331 | 0.146 | 6.0781E-07 | TAL | SNX10 |
| 1.5914E-10 | 0.33045661 | 0.312 | 0.138 | 3.3977E-06 | TAL | PPCS |
| 2.5052E-12 | 0.33037949 | 0.357 | 0.154 | 5.3485E-08 | TAL | AKIRIN1 |
| 4.2833E-14 | 0.33030192 | 0.287 | 0.101 | 9.1449E-10 | TAL | MECOM |
| 2.2561E-17 | 0.32977181 | 0.325 | 0.111 | 4.8169E-13 | TAL | ECI2 |
| 4.4426E-12 | 0.32832172 | 0.318 | 0.133 | 9.4849E-08 | TAL | DECR1 |
| 8.0037E-09 | 0.32797914 | 0.446 | 0.256 | 0.00017088 | TAL | NDUFA5 |
| 2.5261E-19 | 0.32593063 | 0.28 | 0.081 | 5.3933E-15 | TAL | ARHGAP24 |
| 1.1692E-05 | 0.32536833 | 0.936 | 0.898 | 0.24962033 | TAL | MT-CYB |
| 0.00019607 | 0.32304409 | 0.873 | 0.807 | 1 | TAL | ATP5F1E |
| 3.3905E-39 | 0.32295937 | 0.153 | 0.016 | 7.2388E-35 | TAL | RANBP3L |
| 4.5771E-07 | 0.32260386 | 0.318 | 0.169 | 0.00977204 | TAL | SERPING1 |
| 1.4699E-09 | 0.32070124 | 0.42 | 0.224 | 3.1383E-05 | TAL | PCNP |
| 2.2813E-11 | 0.32032975 | 0.363 | 0.165 | 4.8705E-07 | TAL | CISD1 |
| 2.0829E-19 | 0.3193565 | 0.261 | 0.072 | 4.4469E-15 | TAL | AC027031.2 |
| 8.1634E-08 | 0.31885518 | 0.465 | 0.281 | 0.00174288 | TAL | FIS1 |
| 3.704E-15 | 0.31878424 | 0.185 | 0.048 | 7.9081E-11 | TAL | ANK2 |
| 5.2528E-08 | 0.31610293 | 0.331 | 0.168 | 0.00112148 | TAL | NRIP1 |
| 5.2467E-08 | 0.31517159 | 0.529 | 0.334 | 0.00112017 | TAL | EIF4B |
| 6.6116E-10 | 0.31509785 | 0.344 | 0.161 | 1.4116E-05 | TAL | RANBP2 |
| 0.00544444 | 0.31350703 | 0.306 | 0.207 | 1 | TAL | G3BP2 |
| 4.8874E-09 | 0.3134681 | 0.357 | 0.18 | 0.00010434 | TAL | NDUFAF8 |
| 1.5685E-06 | 0.31313764 | 1 | 0.987 | 0.03348767 | TAL | EEF1A1 |
| 8.3677E-32 | 0.3113483 | 0.191 | 0.029 | 1.7865E-27 | TAL | MFSD4A |
| 2.0337E-08 | 0.31080023 | 0.439 | 0.25 | 0.00043419 | TAL | NDUFAB1 |
| 6.4869E-06 | 0.30988028 | 0.764 | 0.612 | 0.1384953 | TAL | MT-ND4L |
| 1.2258E-23 | 0.3098786 | 0.255 | 0.061 | 2.617E-19 | TAL | LINC02381 |
| 3.9971E-10 | 0.30909728 | 0.427 | 0.222 | 8.5338E-06 | TAL | C1orf43 |
| 0.00026639 | 0.3074972 | 0.248 | 0.15 | 1 | TAL | ACSL4 |
| 4.8181E-20 | 0.3071254 | 0.261 | 0.07 | 1.0287E-15 | TAL | UGT8 |
| 1.6108E-11 | 0.30706482 | 0.478 | 0.243 | 3.439E-07 | TAL | TUFM |
| 5.2357E-07 | 0.30704195 | 0.439 | 0.265 | 0.0111783 | TAL | MINOS1 |
| 2.009E-19 | 0.30472784 | 0.248 | 0.067 | 4.2893E-15 | TAL | TSPAN33 |
| 7.0299E-09 | 0.3047003 | 0.465 | 0.266 | 0.00015009 | TAL | NDUFB8 |
| 5.1322E-39 | 0.30365918 | 0.178 | 0.021 | 1.0957E-34 | TAL | PCP4 |
| 5.8234E-06 | 0.30322712 | 0.42 | 0.271 | 0.12433013 | TAL | ATP5F1C |
| 1.1341E-11 | 0.30294179 | 0.274 | 0.105 | 2.4213E-07 | TAL | MCRIP2 |

| | | | | | | |
|---|---|---|---|---|---|---|
| 7.2238E-10 | 0.30294179 | 0.255 | 0.104 | 1.5423E-05 | TAL | PRR14L |
| 1.9298E-12 | 0.30134202 | 0.274 | 0.102 | 4.1202E-08 | TAL | PILRB |
| 2.8457E-17 | 0.30125406 | 0.312 | 0.104 | 6.0755E-13 | TAL | LMBRD1 |
| 3.3251E-08 | 0.30095735 | 0.599 | 0.374 | 0.0007099 | TAL | PNISR |
| 2.7151E-07 | 0.29919895 | 0.497 | 0.312 | 0.00579674 | TAL | MRFAP1 |
| 5.0992E-16 | 0.29857267 | 0.236 | 0.069 | 1.0887E-11 | TAL | VAV3 |
| 7.4898E-06 | 0.29792188 | 0.484 | 0.31 | 0.15990762 | TAL | WSB1 |
| 2.7157E-13 | 0.29772165 | 0.344 | 0.138 | 5.798E-09 | TAL | DNAJC19 |
| 2.7498E-06 | 0.29704991 | 0.924 | 0.889 | 0.05870782 | TAL | MT-ND2 |
| 2.5612E-55 | 0.29573703 | 0.197 | 0.019 | 5.4682E-51 | TAL | CASR |
| 4.3658E-16 | 0.29556002 | 0.261 | 0.082 | 9.3209E-12 | TAL | HIBADH |
| 1.634E-06 | 0.29423561 | 0.42 | 0.255 | 0.03488653 | TAL | NDUFA11 |
| 3.1246E-11 | 0.29360916 | 0.395 | 0.183 | 6.671E-07 | TAL | MSI2 |
| 3.7791E-11 | 0.29297264 | 0.299 | 0.126 | 8.0685E-07 | TAL | UXS1 |
| 0.00088485 | 0.29217657 | 0.433 | 0.326 | 1 | TAL | NDUFC1 |
| 9.0653E-10 | 0.29200328 | 0.28 | 0.122 | 1.9354E-05 | TAL | SSFA2 |
| 3.9905E-26 | 0.29169941 | 0.255 | 0.056 | 8.5197E-22 | TAL | METTL7A |
| 2.2055E-09 | 0.29083515 | 0.51 | 0.282 | 4.7088E-05 | TAL | C12orf57 |
| 0.00064424 | 0.29083133 | 0.573 | 0.445 | 1 | TAL | ATP5MD |
| 0.00020725 | 0.29036647 | 0.49 | 0.357 | 1 | TAL | GCC2 |
| 7.2469E-08 | 0.29022717 | 0.331 | 0.168 | 0.00154721 | TAL | ZNF644 |
| 4.4456E-07 | 0.28972859 | 0.178 | 0.073 | 0.00949144 | TAL | GAS6 |
| 2.3714E-18 | 0.28948564 | 0.223 | 0.057 | 5.063E-14 | TAL | GRB14 |
| 5.2223E-10 | 0.28827237 | 0.382 | 0.187 | 1.115E-05 | TAL | NDUFS7 |
| 8.4331E-07 | 0.28746998 | 0.229 | 0.11 | 0.01800465 | TAL | ZMAT3 |
| 3.7573E-07 | 0.28740934 | 0.408 | 0.24 | 0.00802188 | TAL | VAMP2 |
| 0.00018392 | 0.28640372 | 0.955 | 0.926 | 1 | TAL | MT-CO1 |
| 4.9143E-36 | 0.28627722 | 0.223 | 0.034 | 1.0492E-31 | TAL | NTRK2 |
| 2.3547E-11 | 0.28620984 | 0.369 | 0.169 | 5.0272E-07 | TAL | EIF2A |
| 3.3002E-20 | 0.2861282 | 0.204 | 0.046 | 7.0459E-16 | TAL | FECH |
| 2.9125E-25 | 0.28540286 | 0.178 | 0.031 | 6.2182E-21 | TAL | LINC00982 |
| 1.2358E-08 | 0.2845753 | 0.357 | 0.181 | 0.00026384 | TAL | ITGB1BP1 |
| 4.2119E-13 | 0.28408154 | 0.325 | 0.128 | 8.9923E-09 | TAL | HINT2 |
| 5.5043E-11 | 0.28118959 | 0.446 | 0.218 | 1.1752E-06 | TAL | NDUFA13 |
| 2.6E-06 | 0.2803683 | 0.242 | 0.123 | 0.05551081 | TAL | VEGFB |
| 2.0163E-48 | 0.27979879 | 0.242 | 0.03 | 4.3049E-44 | TAL | TSPAN8 |
| 2.208E-87 | 0.27902289 | 0.146 | 0.006 | 4.7141E-83 | TAL | SUCNR1 |
| 4.2312E-36 | 0.27900922 | 0.21 | 0.031 | 9.0335E-32 | TAL | PROX1 |
| 0.00054403 | 0.27889875 | 0.439 | 0.32 | 1 | TAL | ATP5MPL |
| 5.9985E-17 | 0.27864235 | 0.217 | 0.058 | 1.2807E-12 | TAL | OGDHL |
| 1.508E-14 | 0.2783629 | 0.242 | 0.077 | 3.2195E-10 | TAL | ATPAF1 |
| 2.8926E-14 | 0.27834857 | 0.376 | 0.153 | 6.1756E-10 | TAL | SDHB |
| 1.8148E-06 | 0.27788875 | 0.471 | 0.302 | 0.03874676 | TAL | PNRC2 |
| 0.00242743 | 0.27726859 | 0.357 | 0.261 | 1 | TAL | PRR13 |
| 1.2458E-08 | 0.27672651 | 0.401 | 0.213 | 0.00026597 | TAL | SAT2 |
| 5.2286E-06 | 0.27592088 | 0.459 | 0.299 | 0.11163098 | TAL | DDT |
| 2.2419E-10 | 0.2754975 | 0.287 | 0.119 | 4.7865E-06 | TAL | COL4A4 |
| 1.1714E-08 | 0.27516544 | 0.376 | 0.197 | 0.00025009 | TAL | GLRX5 |
| 4.5642E-15 | 0.27481773 | 0.274 | 0.091 | 9.7446E-11 | TAL | CRYL1 |
| 5.6123E-10 | 0.27427972 | 0.401 | 0.196 | 1.1982E-05 | TAL | NFIC |

| | | | | | | |
|---|---|---|---|---|---|---|
| 4.021E-06 | 0.2741523 | 0.382 | 0.23 | 0.08584849 | TAL | RBBP6 |
| 7.3882E-07 | 0.27411036 | 0.631 | 0.445 | 0.0157738 | TAL | VDAC2 |
| 1.5671E-14 | 0.27377099 | 0.299 | 0.106 | 3.3457E-10 | TAL | ESCO1 |
| 4.1411E-11 | 0.271751 | 0.35 | 0.155 | 8.8413E-07 | TAL | CXXC5 |
| 2.1769E-09 | 0.2717446 | 0.35 | 0.17 | 4.6477E-05 | TAL | NAA38 |
| 2.3082E-11 | 0.27167283 | 0.274 | 0.107 | 4.9281E-07 | TAL | CPVL |
| 6.2367E-10 | 0.27166108 | 0.338 | 0.157 | 1.3315E-05 | TAL | TXN2 |
| 2.2838E-10 | 0.27128035 | 0.318 | 0.143 | 4.8759E-06 | TAL | MRPS18B |
| 1.729E-06 | 0.2709493 | 0.395 | 0.234 | 0.03691449 | TAL | ANXA4 |
| 1.3627E-52 | 0.27092794 | 0.153 | 0.012 | 2.9094E-48 | TAL | DUSP9 |
| 3.228E-35 | 0.2704888 | 0.166 | 0.02 | 6.8918E-31 | TAL | TDGF1 |
| 4.3088E-07 | 0.2704863 | 0.471 | 0.28 | 0.00919932 | TAL | TMEM14C |
| 1.5331E-08 | 0.27012994 | 0.459 | 0.256 | 0.00032731 | TAL | NDUFA6 |
| 2.7633E-05 | 0.26994668 | 0.49 | 0.329 | 0.58995409 | TAL | NDUFB4 |
| 1.2864E-09 | 0.26960647 | 0.363 | 0.178 | 2.7465E-05 | TAL | NDUFS4 |
| 9.1029E-20 | 0.2682576 | 0.204 | 0.047 | 1.9435E-15 | TAL | ALDH1A2 |
| 1.3416E-18 | 0.26746673 | 0.159 | 0.032 | 2.8644E-14 | TAL | SERPINA5 |
| 0.00786153 | 0.26702809 | 0.21 | 0.137 | 1 | TAL | TM4SF1 |
| 6.1644E-09 | 0.26696133 | 0.287 | 0.131 | 0.00013161 | TAL | ACAT1 |
| 3.2133E-12 | 0.26593179 | 0.344 | 0.144 | 6.8605E-08 | TAL | ATP6V1A |
| 1.1675E-06 | 0.26499806 | 0.401 | 0.236 | 0.02492716 | TAL | CLINT1 |
| 2.4499E-07 | 0.26468663 | 0.312 | 0.164 | 0.00523061 | TAL | SNRPN |
| 1.0335E-09 | 0.26384083 | 0.306 | 0.136 | 2.2065E-05 | TAL | ALDH2 |
| 7.4958E-06 | 0.26282378 | 0.369 | 0.227 | 0.16003632 | TAL | ATP5PB |
| 6.0668E-10 | 0.26248047 | 0.306 | 0.135 | 1.2953E-05 | TAL | RBBP8 |
| 2.0108E-10 | 0.26245812 | 0.21 | 0.076 | 4.2931E-06 | TAL | CSAD |
| 3.2398E-16 | 0.26158202 | 0.115 | 0.02 | 6.9169E-12 | TAL | NR0B2 |
| 3.5573E-10 | 0.2614953 | 0.28 | 0.118 | 7.5948E-06 | TAL | SYNJ2BP |
| 4.674E-14 | 0.26126086 | 0.28 | 0.099 | 9.979E-10 | TAL | TMEM38B |
| 6.028E-14 | 0.26049668 | 0.261 | 0.088 | 1.287E-09 | TAL | ACSL1 |
| 5.9161E-23 | 0.2602665 | 0.172 | 0.031 | 1.2631E-18 | TAL | SLC16A12 |
| 2.7895E-08 | 0.25994947 | 0.389 | 0.208 | 0.00059557 | TAL | SERINC2 |
| 5.9184E-09 | 0.25984135 | 0.427 | 0.223 | 0.00012636 | TAL | TTC3 |
| 6.0472E-07 | 0.25935675 | 0.274 | 0.137 | 0.01291079 | TAL | C12orf49 |
| 4.5367E-06 | 0.25851432 | 0.834 | 0.747 | 0.09685787 | TAL | DSTN |
| 1.966E-24 | 0.25815336 | 0.261 | 0.061 | 4.1973E-20 | TAL | KCTD1 |
| 2.0562E-06 | 0.25814231 | 0.739 | 0.577 | 0.04390074 | TAL | PRDX5 |
| 1.5371E-40 | 0.25806169 | 0.248 | 0.037 | 3.2818E-36 | TAL | ATP6V0A4 |
| 0.00024973 | 0.25804811 | 0.306 | 0.197 | 1 | TAL | TMEM141 |
| 4.8558E-05 | 0.25735795 | 0.166 | 0.078 | 1 | TAL | LY6K |
| 1.0157E-22 | 0.25730762 | 0.217 | 0.047 | 2.1685E-18 | TAL | EPHX2 |
| 3.9035E-06 | 0.25717787 | 0.369 | 0.215 | 0.08333944 | TAL | QKI |
| 2.9475E-05 | 0.25704783 | 0.554 | 0.393 | 0.62928254 | TAL | FKBP2 |
| 0.00144024 | 0.25507579 | 0.898 | 0.817 | 1 | TAL | UBB |
| 2.5146E-09 | 0.25436142 | 0.369 | 0.182 | 5.3686E-05 | TAL | PLEKHA1 |
| 1.2694E-19 | 0.25421016 | 0.229 | 0.057 | 2.7101E-15 | TAL | NDRG2 |
| 5.991E-10 | 0.25367952 | 0.236 | 0.092 | 1.2791E-05 | TAL | ZNF44 |
| 1.6785E-05 | 0.25344738 | 0.42 | 0.275 | 0.35835265 | TAL | OCIAD1 |
| 2.4449E-06 | 0.25248016 | 0.357 | 0.205 | 0.05219831 | TAL | PRDX3 |
| 2.9179E-17 | 0.25129575 | 0.172 | 0.039 | 6.2298E-13 | TAL | SLC22A2 |

| | | | | | | |
|---|---|---|---|---|---|---|
| 6.7792E-08 | 0.25123582 | 0.401 | 0.224 | 0.00144736 | TAL | EIF3L |
| 5.1517E-08 | 0.25105921 | 0.299 | 0.146 | 0.00109988 | TAL | NDUFS2 |
| 1.3514E-09 | 0.25104974 | 0.312 | 0.141 | 2.8851E-05 | TAL | ACAA2 |
| 7.9289E-07 | 0.25065434 | 0.325 | 0.175 | 0.0169282 | TAL | SDHD |
| 8.731E-11 | 0.25052292 | 0.217 | 0.078 | 1.8641E-06 | TAL | NME4 |
| 2.9311E-09 | 0.25019311 | 0.318 | 0.147 | 6.2579E-05 | TAL | ZBTB44 |
| 1.2374E-08 | 0.25019311 | 0.318 | 0.155 | 0.00026418 | TAL | PLPP5 |
| 0.00256728 | -0.2501144 | 0.643 | 0.775 | 1 | TAL | MYL12A |
| 0.0068271 | -0.2502771 | 0.121 | 0.206 | 1 | TAL | MDK |
| 0.0006432 | -0.2569123 | 0.089 | 0.195 | 1 | TAL | SLC25A37 |
| 1.5463E-05 | -0.2586792 | 0.051 | 0.184 | 0.33012688 | TAL | QSOX1 |
| 0.00201155 | -0.2600924 | 0.096 | 0.189 | 1 | TAL | PDIA4 |
| 0.00574416 | -0.2666551 | 0.127 | 0.209 | 1 | TAL | SLFN5 |
| 0.00376102 | -0.2714646 | 0.076 | 0.157 | 1 | TAL | CDKN2B |
| 0.00533388 | -0.2735591 | 0.064 | 0.139 | 1 | TAL | ISG15 |
| 0.00014871 | -0.2837426 | 0.045 | 0.152 | 1 | TAL | SLC16A3 |
| 0.00991735 | -0.2853069 | 0.127 | 0.21 | 1 | TAL | ADAMTS9 |
| 0.00853742 | -0.2881278 | 0.108 | 0.187 | 1 | TAL | BIRC3 |
| 0.00344586 | -0.2886833 | 0.611 | 0.716 | 1 | TAL | HSPB1 |
| 0.00778321 | -0.2989046 | 0.331 | 0.402 | 1 | TAL | CDV3 |
| 5.0207E-05 | -0.3009598 | 0.083 | 0.213 | 1 | TAL | SCCPDH |
| 0.00391788 | -0.301152 | 0.274 | 0.374 | 1 | TAL | FLNA |
| 0.00016336 | -0.30294 | 0.083 | 0.203 | 1 | TAL | CALD1 |
| 0.00122225 | -0.3119356 | 0.025 | 0.103 | 1 | TAL | TNF |
| 4.0356E-06 | -0.3182282 | 0.013 | 0.14 | 0.08615933 | TAL | EFNB2 |
| 0.00319904 | -0.3182426 | 0.955 | 0.961 | 1 | TAL | RPL28 |
| 0.00522812 | -0.3232643 | 0.045 | 0.116 | 1 | TAL | TAGLN |
| 1.371E-07 | -0.3252877 | 0 | 0.151 | 0.00292718 | TAL | FXYD5 |
| 0.00150443 | -0.325352 | 0.599 | 0.704 | 1 | TAL | EIF4G2 |
| 0.00872562 | -0.3258713 | 0.911 | 0.924 | 1 | TAL | RPLP2 |
| 0.00035176 | -0.3262293 | 0.076 | 0.181 | 1 | TAL | DCBLD2 |
| 0.00492347 | -0.3276164 | 0.924 | 0.912 | 1 | TAL | RPL35 |
| 0.00236517 | -0.3280436 | 0.032 | 0.105 | 1 | TAL | ARC |
| 0.00225422 | -0.3320685 | 0.777 | 0.884 | 1 | TAL | RPL39 |
| 6.0345E-06 | -0.3325428 | 0.038 | 0.175 | 0.12883756 | TAL | PLA2G16 |
| 0.00023254 | -0.3333964 | 0.561 | 0.695 | 1 | TAL | ZFAS1 |
| 0.00289826 | -0.334474 | 0.338 | 0.429 | 1 | TAL | PDIA3 |
| 0.00020877 | -0.3418216 | 0.924 | 0.945 | 1 | TAL | RPS15 |
| 0.00480351 | -0.3426874 | 0.516 | 0.606 | 1 | TAL | GADD45B |
| 0.00108665 | -0.3430779 | 0.089 | 0.183 | 1 | TAL | ERN1 |
| 0.00332801 | -0.3462183 | 0.248 | 0.345 | 1 | TAL | S100A16 |
| 0.00918175 | -0.3493514 | 0.045 | 0.109 | 1 | TAL | S100A9 |
| 0.00933827 | -0.3527359 | 0.248 | 0.328 | 1 | TAL | CEBPB |
| 0.00260537 | -0.3531854 | 0.389 | 0.47 | 1 | TAL | PDIA6 |
| 6.2816E-05 | -0.3543827 | 0.134 | 0.267 | 1 | TAL | LIMA1 |
| 0.00197271 | -0.3591881 | 0.764 | 0.83 | 1 | TAL | SQSTM1 |
| 0.00050694 | -0.3663755 | 0.446 | 0.567 | 1 | TAL | PPIB |
| 0.00018584 | -0.3698935 | 0.127 | 0.249 | 1 | TAL | RAP2B |
| 0.00094015 | -0.3722188 | 0.096 | 0.193 | 1 | TAL | H2AFX |
| 1.4107E-05 | -0.3810013 | 0.07 | 0.205 | 0.30117433 | TAL | CLDN1 |

| | | | | | | |
|---|---|---|---|---|---|---|
| 0.0001614 | -0.3855639 | 0.261 | 0.398 | 1 | TAL | RHOC |
| 0.00030808 | -0.3861953 | 0.019 | 0.108 | 1 | TAL | AQP3 |
| 1.8102E-06 | -0.3907037 | 0.032 | 0.177 | 0.03864814 | TAL | PHLDA1 |
| 0.00313865 | -0.3943893 | 0.146 | 0.235 | 1 | TAL | NFKBIZ |
| 9.8188E-05 | -0.3960557 | 0.242 | 0.372 | 1 | TAL | TAGLN2 |
| 0.00017157 | -0.4041469 | 0.701 | 0.828 | 1 | TAL | HLA-B |
| 3.3354E-09 | -0.411539 | 0 | 0.184 | 7.121E-05 | TAL | CD40 |
| 0.0002393 | -0.4188822 | 0.159 | 0.28 | 1 | TAL | SERPINB1 |
| 2.2822E-05 | -0.4240856 | 0.089 | 0.225 | 0.48725442 | TAL | GPRC5A |
| 2.5133E-05 | -0.4275769 | 0.185 | 0.328 | 0.53659855 | TAL | CTSB |
| 1.5618E-06 | -0.4374276 | 0.662 | 0.805 | 0.03334443 | TAL | CALM1 |
| 0.00026523 | -0.4398636 | 0.102 | 0.215 | 1 | TAL | SOCS1 |
| 6.602E-07 | -0.4403089 | 0.561 | 0.729 | 0.01409526 | TAL | CSTB |
| 0.00618146 | -0.4448809 | 0.14 | 0.219 | 1 | TAL | TNFAIP2 |
| 0.00078654 | -0.4460897 | 0.159 | 0.263 | 1 | TAL | HIST1H2AC |
| 4.0912E-09 | -0.4517245 | 0.439 | 0.662 | 8.7348E-05 | TAL | CALR |
| 0.00263812 | -0.4529009 | 0.331 | 0.418 | 1 | TAL | CCND1 |
| 7.8812E-06 | -0.454965 | 0.669 | 0.805 | 0.16826306 | TAL | CFL1 |
| 1.3185E-06 | -0.4576438 | 0.376 | 0.561 | 0.02814992 | TAL | HMGA1 |
| 0.00194059 | -0.4596614 | 0.115 | 0.204 | 1 | TAL | SGK1 |
| 0.00242167 | -0.4605185 | 0.21 | 0.306 | 1 | TAL | PMAIP1 |
| 0.00391405 | -0.4617996 | 0.166 | 0.254 | 1 | TAL | IER5 |
| 0.00031733 | -0.4679797 | 0.369 | 0.482 | 1 | TAL | H1FX |
| 0.00655014 | -0.4742923 | 0.236 | 0.313 | 1 | TAL | CDKN1C |
| 0.0005738 | -0.4762603 | 0.28 | 0.392 | 1 | TAL | RAB11FIP1 |
| 0.0020893 | -0.4853238 | 0.21 | 0.302 | 1 | TAL | DDIT4 |
| 7.3724E-06 | -0.4988192 | 0.089 | 0.234 | 0.1574007 | TAL | TGM2 |
| 1.2933E-06 | -0.5066328 | 0.656 | 0.839 | 0.02761271 | TAL | PRDX1 |
| 5.7808E-06 | -0.5148876 | 0.586 | 0.733 | 0.12342011 | TAL | PFN1 |
| 0.00047191 | -0.5161911 | 0.497 | 0.566 | 1 | TAL | MAL2 |
| 7.7942E-05 | -0.5280576 | 0.49 | 0.625 | 1 | TAL | PPP1R15A |
| 1.5872E-05 | -0.5368051 | 0.102 | 0.247 | 0.33886939 | TAL | PDZK1IP1 |
| 0.00204203 | -0.539805 | 0.038 | 0.118 | 1 | TAL | CXCL8 |
| 0.00018474 | -0.5416127 | 0.822 | 0.923 | 1 | TAL | ACTB |
| 8.3635E-07 | -0.5453626 | 0.535 | 0.718 | 0.01785607 | TAL | TXN |
| 2.6159E-05 | -0.5537042 | 0.146 | 0.287 | 0.55850385 | TAL | PLAUR |
| 0.00059572 | -0.5539005 | 0.65 | 0.774 | 1 | TAL | HSP90B1 |
| 2.0855E-08 | -0.562237 | 0.484 | 0.707 | 0.00044525 | TAL | ITGB1 |
| 3.1004E-05 | -0.5701358 | 0.535 | 0.664 | 0.66192799 | TAL | TACSTD2 |
| 0.0014474 | -0.5756249 | 0.312 | 0.391 | 1 | TAL | TUBA1B |
| 3.6699E-06 | -0.5821217 | 0.248 | 0.401 | 0.07835214 | TAL | JPT1 |
| 4.7388E-05 | -0.5960145 | 0.936 | 0.949 | 1 | TAL | B2M |
| 0.0039216 | -0.5962002 | 0.299 | 0.375 | 1 | TAL | SLC38A2 |
| 0.00284465 | -0.6023625 | 0.471 | 0.565 | 1 | TAL | IER2 |
| 0.00080411 | -0.6031793 | 0.217 | 0.32 | 1 | TAL | ITGA2 |
| 1.2329E-07 | -0.6081522 | 0.115 | 0.302 | 0.00263221 | TAL | MARCKSL1 |
| 1.9617E-08 | -0.6195322 | 0.439 | 0.651 | 0.00041882 | TAL | KRT7 |
| 0.0001247 | -0.6283778 | 0.178 | 0.297 | 1 | TAL | ITGB8 |
| 0.0017416 | -0.6411721 | 0.159 | 0.262 | 1 | TAL | NUPR1 |
| 0.00600223 | -0.6481136 | 0.541 | 0.635 | 1 | TAL | JUN |

| | | | | | | |
|---|---|---|---|---|---|---|
| 0.00196473 | -0.6684453 | 0.153 | 0.248 | 1 | TAL | MT1X |
| 0.00086327 | -0.6755335 | 0.025 | 0.107 | 1 | TAL | FXYD4 |
| 3.141E-06 | -0.7071047 | 0.14 | 0.29 | 0.06706008 | TAL | RND3 |
| 3.2943E-12 | -0.7201066 | 0.433 | 0.671 | 7.0333E-08 | TAL | GLS |
| 5.0342E-05 | -0.7232884 | 0.108 | 0.237 | 1 | TAL | CST6 |
| 0.00031591 | -0.7316237 | 0.127 | 0.235 | 1 | TAL | CSKMT |
| 1.2545E-09 | -0.7563549 | 0.401 | 0.633 | 2.6784E-05 | TAL | TPM1 |
| 8.3939E-07 | -0.7563671 | 0.242 | 0.415 | 0.01792101 | TAL | PHLDA2 |
| 8.77E-06 | -0.7633406 | 0.478 | 0.67 | 0.18723999 | TAL | CRYAB |
| 6.1969E-08 | -0.7781941 | 0.159 | 0.343 | 0.00132303 | TAL | SNHG12 |
| 4.7821E-09 | -0.791565 | 0.268 | 0.488 | 0.0001021 | TAL | TSPAN1 |
| 2.8402E-06 | -0.8197818 | 0.529 | 0.685 | 0.0606391 | TAL | HSPA5 |
| 2.1303E-14 | -0.864508 | 0.662 | 0.874 | 4.5481E-10 | TAL | SAT1 |
| 1.3984E-10 | -0.8662925 | 0.516 | 0.704 | 2.9857E-06 | TAL | CLDN4 |
| 1.6898E-08 | -0.92618 | 0.21 | 0.409 | 0.00036076 | TAL | RNF213 |
| 5.0725E-17 | -0.9392196 | 0.752 | 0.926 | 1.083E-12 | TAL | TMSB10 |
| 0.00090268 | -0.9783838 | 0.191 | 0.284 | 1 | TAL | MAFB |
| 8.1502E-08 | -0.9996463 | 0.446 | 0.588 | 0.00174008 | TAL | ELF3 |
| 2.2575E-17 | -1.0127285 | 0.096 | 0.425 | 4.8197E-13 | TAL | TPM4 |
| 1.1913E-08 | -1.0281405 | 0.051 | 0.245 | 0.00025434 | TAL | TFPI2 |
| 3.1332E-11 | -1.0350257 | 0.586 | 0.802 | 6.6894E-07 | TAL | KRT18 |
| 1.2573E-12 | -1.0378294 | 0.07 | 0.33 | 2.6842E-08 | TAL | IL32 |
| 6.8277E-10 | -1.0776312 | 0.108 | 0.333 | 1.4577E-05 | TAL | LAMC2 |
| 3.1162E-12 | -1.0927937 | 0.758 | 0.921 | 6.6531E-08 | TAL | S100A11 |
| 3.0863E-10 | -1.0932351 | 0.242 | 0.473 | 6.5892E-06 | TAL | C12orf75 |
| 2.801E-09 | -1.1069051 | 0.07 | 0.28 | 5.9801E-05 | TAL | LGALS1 |
| 3.1458E-16 | -1.1132682 | 0.433 | 0.731 | 6.7163E-12 | TAL | KRT8 |
| 6.8837E-17 | -1.1263825 | 0.344 | 0.676 | 1.4697E-12 | TAL | SH3BGRL3 |
| 1.1355E-15 | -1.2024425 | 0.318 | 0.603 | 2.4243E-11 | TAL | CLU |
| 0.00175118 | -1.2220131 | 0.108 | 0.199 | 1 | TAL | MT1G |
| 1.6472E-19 | -1.2462017 | 0.248 | 0.628 | 3.5167E-15 | TAL | VIM |
| 4.9031E-08 | -1.2473326 | 0.07 | 0.258 | 0.00104682 | TAL | LCN2 |
| 4.0467E-20 | -1.3073586 | 0.49 | 0.835 | 8.6396E-16 | TAL | ANXA2 |
| 2.4201E-07 | -1.3843516 | 0.204 | 0.394 | 0.00516685 | TAL | SLPI |
| 6.1782E-25 | -1.5969452 | 0.592 | 0.872 | 1.319E-20 | TAL | TMSB4X |
| 3.7169E-30 | -1.6799647 | 0.121 | 0.597 | 7.9355E-26 | TAL | KRT19 |
| 2.8614E-26 | -2.7293687 | 0.204 | 0.642 | 6.1091E-22 | TAL | MMP7 |
| 1.067E-194 | 3.2027813 | 0.988 | 0.346 | 2.279E-190 | TEC_dmg | HSPA1A |
| 1.1036E-63 | 2.34753718 | 0.893 | 0.663 | 2.3562E-59 | TEC_dmg | CRYAB |
| 9.002E-290 | 2.3398115 | 0.663 | 0.066 | 1.922E-285 | TEC_dmg | HSPA6 |
| 8.76E-156 | 2.20994399 | 0.913 | 0.293 | 1.87E-151 | TEC_dmg | HSPA1B |
| 3.722E-140 | 2.06413897 | 0.877 | 0.285 | 7.946E-136 | TEC_dmg | HSPH1 |
| 1.6348E-84 | 1.76862477 | 0.984 | 0.933 | 3.4902E-80 | TEC_dmg | HSP90AA1 |
| 2.01E-110 | 1.66685136 | 0.841 | 0.292 | 4.291E-106 | TEC_dmg | DNAJB1 |
| 1.7115E-76 | 1.3291593 | 0.992 | 0.912 | 3.6541E-72 | TEC_dmg | HSP90AB1 |
| 2.816E-123 | 1.21514922 | 0.639 | 0.142 | 6.012E-119 | TEC_dmg | BAG3 |
| 7.3811E-23 | 1.2023336 | 0.575 | 0.326 | 1.5759E-18 | TEC_dmg | LAMC2 |
| 1.305E-45 | 1.19080593 | 0.909 | 0.679 | 2.7862E-41 | TEC_dmg | HSPA5 |
| 1.8615E-48 | 1.00983003 | 0.829 | 0.49 | 3.9744E-44 | TEC_dmg | HSPD1 |
| 3.9669E-46 | 1.00921238 | 0.774 | 0.381 | 8.4694E-42 | TEC_dmg | HSPE1 |

| | | | | | | |
|---|---|---|---|---|---|---|
| 2.324E-37 | 1.0065231 | 0.913 | 0.71 | 4.9618E-33 | TEC_dmg | HSPB1 |
| 3.5404E-43 | 0.91994415 | 0.821 | 0.484 | 7.5588E-39 | TEC_dmg | DNAJA1 |
| 3.154E-107 | 0.87887195 | 0.44 | 0.072 | 6.734E-103 | TEC_dmg | DNAJA4 |
| 2.744E-37 | 0.87176586 | 0.988 | 0.91 | 5.8584E-33 | TEC_dmg | UBC |
| 1.7134E-57 | 0.84298614 | 0.429 | 0.111 | 3.658E-53 | TEC_dmg | SERPINH1 |
| 5.1475E-16 | 0.7822801 | 0.496 | 0.291 | 1.099E-11 | TEC_dmg | ITGB8 |
| 3.8233E-27 | 0.75705877 | 0.877 | 0.652 | 8.1627E-23 | TEC_dmg | HSPA8 |
| 1.0393E-47 | 0.75512269 | 0.492 | 0.164 | 2.2189E-43 | TEC_dmg | CACYBP |
| 1.9168E-20 | 0.73511284 | 0.496 | 0.255 | 4.0923E-16 | TEC_dmg | LAMB3 |
| 2.8081E-18 | 0.7193568 | 0.913 | 0.816 | 5.9953E-14 | TEC_dmg | UBB |
| 5.9452E-23 | 0.69381301 | 0.591 | 0.314 | 1.2693E-18 | TEC_dmg | ITGA2 |
| 3.1524E-07 | 0.66951479 | 0.218 | 0.115 | 0.00673033 | TEC_dmg | CXCL8 |
| 8.7705E-11 | 0.63972075 | 0.492 | 0.323 | 1.8725E-06 | TEC_dmg | IL32 |
| 3.0382E-57 | 0.63764157 | 0.437 | 0.114 | 6.4866E-53 | TEC_dmg | DNAJB4 |
| 4.4058E-23 | 0.63531668 | 0.548 | 0.281 | 9.4064E-19 | TEC_dmg | PLAUR |
| 2.4497E-11 | 0.63472832 | 0.238 | 0.109 | 5.2301E-07 | TEC_dmg | C15orf48 |
| 7.7561E-34 | 0.63280098 | 0.635 | 0.307 | 1.6559E-29 | TEC_dmg | DNAJB6 |
| 2.2326E-31 | 0.61434552 | 0.444 | 0.18 | 4.7665E-27 | TEC_dmg | MRPL18 |
| 7.8654E-52 | 0.60898875 | 0.488 | 0.149 | 1.6793E-47 | TEC_dmg | CHORDC1 |
| 9.0522E-06 | 0.57263756 | 0.421 | 0.297 | 0.19326459 | TEC_dmg | MARCKSL1 |
| 1.8538E-19 | 0.56883635 | 0.627 | 0.376 | 3.9578E-15 | TEC_dmg | HERPUD1 |
| 3.3364E-13 | 0.56618172 | 0.706 | 0.562 | 7.1232E-09 | TEC_dmg | MAL2 |
| 1.4785E-06 | 0.54781873 | 0.802 | 0.7 | 0.03156585 | TEC_dmg | CLDN4 |
| 1.3207E-14 | 0.5420003 | 0.603 | 0.386 | 2.8198E-10 | TEC_dmg | RAB11FIP1 |
| 3.6246E-11 | 0.5371397 | 0.488 | 0.304 | 7.7385E-07 | TEC_dmg | PLAU |
| 4.3178E-15 | 0.52832069 | 0.5 | 0.29 | 9.2184E-11 | TEC_dmg | CD55 |
| 5.8279E-13 | 0.50583449 | 0.417 | 0.229 | 1.2443E-08 | TEC_dmg | TGM2 |
| 2.3865E-07 | 0.50553031 | 0.476 | 0.329 | 0.00509525 | TEC_dmg | H1F0 |
| 3.5872E-25 | 0.48689895 | 0.306 | 0.106 | 7.6586E-21 | TEC_dmg | ZFAND2A |
| 1.4859E-14 | 0.4757593 | 0.655 | 0.439 | 3.1723E-10 | TEC_dmg | UPP1 |
| 8.9232E-26 | 0.47336947 | 0.52 | 0.251 | 1.9051E-21 | TEC_dmg | CCT4 |
| 2.5651E-21 | 0.47196655 | 0.421 | 0.192 | 5.4766E-17 | TEC_dmg | FKBP4 |
| 2.7477E-34 | 0.47144456 | 0.464 | 0.173 | 5.8664E-30 | TEC_dmg | AHSA1 |
| 4.2286E-10 | 0.47035347 | 0.591 | 0.414 | 9.0281E-06 | TEC_dmg | CCND1 |
| 4.8678E-14 | 0.46940195 | 0.841 | 0.701 | 1.0393E-09 | TEC_dmg | ITGB1 |
| 3.5307E-34 | 0.46790071 | 0.448 | 0.165 | 7.5381E-30 | TEC_dmg | HSPA4 |
| 1.5901E-13 | 0.45885624 | 0.548 | 0.34 | 3.3949E-09 | TEC_dmg | S100A16 |
| 4.1742E-19 | 0.45811874 | 0.575 | 0.335 | 8.912E-15 | TEC_dmg | PFDN2 |
| 6.2539E-06 | 0.45692042 | 0.433 | 0.298 | 0.13352128 | TEC_dmg | DDIT4 |
| 1.2493E-17 | 0.45630496 | 0.611 | 0.365 | 2.6672E-13 | TEC_dmg | UBE2B |
| 1.3384E-10 | 0.4500648 | 0.587 | 0.417 | 2.8574E-06 | TEC_dmg | TPM4 |
| 6.3327E-07 | 0.44401865 | 0.611 | 0.478 | 0.0135203 | TEC_dmg | H1FX |
| 5.1846E-17 | 0.43120088 | 0.401 | 0.195 | 1.1069E-12 | TEC_dmg | MARCKS |
| 4.376E-13 | 0.43103737 | 0.575 | 0.354 | 9.3427E-09 | TEC_dmg | KLF6 |
| 5.8023E-11 | 0.42840469 | 0.885 | 0.77 | 1.2388E-06 | TEC_dmg | HSP90B1 |
| 1.8056E-14 | 0.42139794 | 0.476 | 0.269 | 3.855E-10 | TEC_dmg | TAF7 |
| 0.00014845 | 0.41684034 | 0.44 | 0.348 | 1 | TEC_dmg | TXNIP |
| 7.8605E-18 | 0.41663244 | 0.405 | 0.191 | 1.6782E-13 | TEC_dmg | MSMO1 |
| 6.2988E-08 | 0.41268795 | 0.536 | 0.366 | 0.00134479 | TEC_dmg | DDIT3 |
| 5.7155E-26 | 0.41262827 | 0.306 | 0.101 | 1.2203E-21 | TEC_dmg | VGLL3 |

| | | | | | | |
|---|---|---|---|---|---|---|
| 4.1308E-15 | 0.40870659 | 0.571 | 0.343 | 8.8193E-11 | TEC_dmg | UBE2H |
| 3.5977E-31 | 0.40768493 | 0.286 | 0.083 | 7.6811E-27 | TEC_dmg | HSPA4L |
| 6.2836E-12 | 0.40368612 | 0.44 | 0.245 | 1.3416E-07 | TEC_dmg | MACC1 |
| 1.037E-11 | 0.40122962 | 0.306 | 0.151 | 2.214E-07 | TEC_dmg | INSIG1 |
| 1.135E-25 | 0.39706964 | 0.333 | 0.116 | 2.4232E-21 | TEC_dmg | NEURL3 |
| 2.1053E-21 | 0.39705472 | 0.385 | 0.163 | 4.4947E-17 | TEC_dmg | ELK3 |
| 1.8976E-10 | 0.39307822 | 0.647 | 0.477 | 4.0515E-06 | TEC_dmg | PTGES3 |
| 9.6159E-15 | 0.38934281 | 0.571 | 0.346 | 2.053E-10 | TEC_dmg | SELENOK |
| 1.6439E-19 | 0.38316744 | 0.468 | 0.223 | 3.5097E-15 | TEC_dmg | PRSS8 |
| 1.7095E-23 | 0.38267722 | 0.274 | 0.088 | 3.6497E-19 | TEC_dmg | PARD6G-AS1 |
| 3.1797E-14 | 0.38099639 | 0.683 | 0.458 | 6.7887E-10 | TEC_dmg | SERP1 |
| 2.3436E-16 | 0.37922029 | 0.437 | 0.219 | 5.0035E-12 | TEC_dmg | TEAD1 |
| 1.4012E-10 | 0.3776215 | 0.548 | 0.367 | 2.9915E-06 | TEC_dmg | CAMK2N1 |
| 5.1352E-10 | 0.36707463 | 0.294 | 0.155 | 1.0964E-05 | TEC_dmg | PMEPA1 |
| 1.4875E-10 | 0.36315802 | 0.802 | 0.676 | 3.1758E-06 | TEC_dmg | YWHAZ |
| 0.00074812 | 0.36265349 | 0.802 | 0.757 | 1 | TEC_dmg | DYNLL1 |
| 6.5518E-26 | 0.36067183 | 0.194 | 0.048 | 1.3988E-21 | TEC_dmg | VNN1 |
| 4.8066E-13 | 0.36043724 | 0.548 | 0.334 | 1.0262E-08 | TEC_dmg | SSR3 |
| 6.1642E-09 | 0.35775568 | 0.258 | 0.135 | 0.0001316 | TEC_dmg | TM4SF1 |
| 1.7758E-09 | 0.35743785 | 0.56 | 0.388 | 3.7914E-05 | TEC_dmg | EIF2S2 |
| 2.868E-09 | 0.35660519 | 0.726 | 0.571 | 6.1232E-05 | TEC_dmg | SLC3A2 |
| 1.663E-21 | 0.35345554 | 0.385 | 0.164 | 3.5504E-17 | TEC_dmg | STIP1 |
| 1.7058E-09 | 0.35136753 | 0.786 | 0.62 | 3.6419E-05 | TEC_dmg | PPP1R15A |
| 2.4344E-13 | 0.35072674 | 0.302 | 0.14 | 5.1975E-09 | TEC_dmg | CKS2 |
| 5.8251E-12 | 0.34733998 | 0.472 | 0.28 | 1.2437E-07 | TEC_dmg | SPINT1 |
| 1.2143E-21 | 0.34688185 | 0.325 | 0.125 | 2.5924E-17 | TEC_dmg | KCNK1 |
| 2.1896E-14 | 0.34601229 | 0.44 | 0.235 | 4.6748E-10 | TEC_dmg | TCEAL9 |
| 6.6202E-10 | 0.34422521 | 0.825 | 0.698 | 1.4134E-05 | TEC_dmg | MAP1LC3B |
| 2.7584E-18 | 0.34326121 | 0.29 | 0.116 | 5.8891E-14 | TEC_dmg | DAPP1 |
| 1.9187E-06 | 0.34283779 | 0.29 | 0.178 | 0.04096412 | TEC_dmg | DCBLD2 |
| 1.988E-18 | 0.34206712 | 0.567 | 0.298 | 4.2443E-14 | TEC_dmg | PNRC2 |
| 1.016E-34 | 0.34049604 | 0.337 | 0.099 | 2.1692E-30 | TEC_dmg | DEDD2 |
| 2.0151E-11 | 0.34044448 | 0.627 | 0.428 | 4.3023E-07 | TEC_dmg | BZW1 |
| 6.7248E-18 | 0.34004307 | 0.294 | 0.119 | 1.4358E-13 | TEC_dmg | PIK3IP1 |
| 1.9861E-10 | 0.33976031 | 0.516 | 0.341 | 4.2404E-06 | TEC_dmg | CAP1 |
| 5.6813E-15 | 0.33967318 | 0.425 | 0.219 | 1.213E-10 | TEC_dmg | CXCL16 |
| 3.3142E-21 | 0.33938841 | 0.452 | 0.207 | 7.0758E-17 | TEC_dmg | YWHAG |
| 1.0985E-17 | 0.33810436 | 0.5 | 0.257 | 2.3453E-13 | TEC_dmg | HMGB2 |
| 4.805E-15 | 0.32950293 | 0.381 | 0.187 | 1.0259E-10 | TEC_dmg | ZBTB43 |
| 2.1332E-09 | 0.32770973 | 0.377 | 0.228 | 4.5544E-05 | TEC_dmg | RPL22L1 |
| 1.6119E-06 | 0.32694694 | 0.948 | 0.921 | 0.03441442 | TEC_dmg | ACTB |
| 1.193E-10 | 0.32643128 | 0.341 | 0.188 | 2.5471E-06 | TEC_dmg | BHLHE40 |
| 9.4315E-12 | 0.3240673 | 0.361 | 0.197 | 2.0136E-07 | TEC_dmg | FOXO3 |
| 1.4122E-08 | 0.32115505 | 0.69 | 0.549 | 0.00030151 | TEC_dmg | CANX |
| 6.0068E-09 | 0.32008743 | 0.302 | 0.165 | 0.00012825 | TEC_dmg | FGF2 |
| 8.26E-13 | 0.31369254 | 0.46 | 0.265 | 1.7635E-08 | TEC_dmg | TCP1 |
| 1.5085E-10 | 0.31032749 | 0.496 | 0.318 | 3.2207E-06 | TEC_dmg | GNB1 |
| 6.7556E-19 | 0.30675786 | 0.194 | 0.06 | 1.4423E-14 | TEC_dmg | SEMA7A |
| 5.4026E-20 | 0.30625562 | 0.294 | 0.11 | 1.1535E-15 | TEC_dmg | NECTIN4 |
| 1.2284E-10 | 0.3062222 | 0.468 | 0.286 | 2.6226E-06 | TEC_dmg | FAM3C |

| | | | | | | |
|---|---|---|---|---|---|---|
| 1.1852E-09 | 0.30610105 | 0.353 | 0.203 | 2.5305E-05 | TEC_dmg | ATP13A3 |
| 1.4183E-14 | 0.30590747 | 0.361 | 0.174 | 3.0281E-10 | TEC_dmg | RYBP |
| 4.3109E-13 | 0.30520449 | 0.357 | 0.178 | 9.2038E-09 | TEC_dmg | CD40 |
| 2.1031E-05 | 0.30472039 | 0.389 | 0.276 | 0.44900366 | TEC_dmg | SERPINB1 |
| 2.2126E-15 | 0.30093696 | 0.357 | 0.168 | 4.724E-11 | TEC_dmg | AZIN1 |
| 3.1134E-10 | 0.29916267 | 0.369 | 0.208 | 6.6471E-06 | TEC_dmg | MAFG |
| 3.91E-11 | 0.29904076 | 0.524 | 0.318 | 8.3478E-07 | TEC_dmg | RSRC2 |
| 1.0145E-10 | 0.2979617 | 0.385 | 0.218 | 2.1659E-06 | TEC_dmg | TMOD3 |
| 6.9843E-08 | 0.29789679 | 0.492 | 0.328 | 0.00149115 | TEC_dmg | TUBA1C |
| 0.00048891 | 0.29510331 | 0.702 | 0.604 | 1 | TEC_dmg | GADD45A |
| 1.6209E-07 | 0.28888469 | 0.417 | 0.271 | 0.00346065 | TEC_dmg | CREBRF |
| 1.4961E-05 | 0.28765005 | 0.321 | 0.204 | 0.31942269 | TEC_dmg | RRAD |
| 4.1091E-08 | 0.28724421 | 0.456 | 0.296 | 0.00087729 | TEC_dmg | CDH1 |
| 5.9024E-11 | 0.28606772 | 0.369 | 0.202 | 1.2602E-06 | TEC_dmg | OSER1 |
| 5.674E-12 | 0.28307136 | 0.246 | 0.109 | 1.2114E-07 | TEC_dmg | ARRDC4 |
| 1.7551E-10 | 0.28293895 | 0.397 | 0.23 | 3.7471E-06 | TEC_dmg | FXR1 |
| 3.4241E-10 | 0.28275403 | 0.262 | 0.128 | 7.3104E-06 | TEC_dmg | AL353751.1 |
| 0.00095571 | 0.28026229 | 0.758 | 0.726 | 1 | TEC_dmg | CSTB |
| 3.7799E-15 | 0.27826065 | 0.429 | 0.219 | 8.0701E-11 | TEC_dmg | RAB21 |
| 1.5332E-06 | 0.27750275 | 0.437 | 0.306 | 0.03273279 | TEC_dmg | DSP |
| 2.4126E-08 | 0.27678998 | 0.46 | 0.298 | 0.00051509 | TEC_dmg | RAP1B |
| 3.0355E-10 | 0.27445313 | 0.381 | 0.214 | 6.4807E-06 | TEC_dmg | USP53 |
| 4.8296E-15 | 0.27432593 | 0.337 | 0.157 | 1.0311E-10 | TEC_dmg | F11R |
| 1.1867E-09 | 0.27377938 | 0.421 | 0.256 | 2.5335E-05 | TEC_dmg | RAB7A |
| 5.6851E-09 | 0.27327159 | 0.29 | 0.157 | 0.00012138 | TEC_dmg | TP53BP2 |
| 1.4877E-14 | 0.27285065 | 0.425 | 0.219 | 3.1763E-10 | TEC_dmg | NAMPT |
| 2.1302E-12 | 0.27083136 | 0.282 | 0.133 | 4.548E-08 | TEC_dmg | DESI2 |
| 3.4936E-13 | 0.27041342 | 0.262 | 0.114 | 7.4589E-09 | TEC_dmg | LRIF1 |
| 8.2135E-10 | 0.26961157 | 0.488 | 0.306 | 1.7536E-05 | TEC_dmg | ARHGAP5 |
| 1.0141E-09 | 0.26920573 | 0.341 | 0.195 | 2.165E-05 | TEC_dmg | WBP2 |
| 5.1197E-07 | 0.26844773 | 0.956 | 0.961 | 0.01093047 | TEC_dmg | FTL |
| 5.3505E-12 | 0.26818863 | 0.258 | 0.119 | 1.1423E-07 | TEC_dmg | IRF6 |
| 5.2424E-09 | 0.26801313 | 0.56 | 0.377 | 0.00011192 | TEC_dmg | RHEB |
| 2.0946E-13 | 0.26758031 | 0.833 | 0.669 | 4.4719E-09 | TEC_dmg | SH3BGRL3 |
| 7.0148E-07 | 0.26647654 | 0.571 | 0.389 | 0.0149765 | TEC_dmg | TAF1D |
| 1.9753E-08 | 0.26515006 | 0.329 | 0.189 | 0.00042173 | TEC_dmg | SESN2 |
| 1.9168E-09 | 0.26496098 | 0.865 | 0.757 | 4.0923E-05 | TEC_dmg | CALM2 |
| 1.2655E-07 | 0.26220059 | 0.46 | 0.313 | 0.00270185 | TEC_dmg | SNX3 |
| 7.8118E-12 | 0.25958652 | 0.25 | 0.114 | 1.6678E-07 | TEC_dmg | SNAP23 |
| 1.0891E-06 | 0.25895806 | 0.437 | 0.298 | 0.02325261 | TEC_dmg | MYDGF |
| 0.00017799 | 0.25862625 | 0.357 | 0.251 | 1 | TEC_dmg | AKAP12 |
| 7.8549E-06 | 0.25774819 | 0.504 | 0.367 | 0.16770274 | TEC_dmg | SDCBP |
| 3.3934E-07 | 0.25654462 | 0.385 | 0.253 | 0.00724485 | TEC_dmg | PPP2R1A |
| 7.0721E-11 | 0.25634932 | 0.274 | 0.135 | 1.5099E-06 | TEC_dmg | CTSS |
| 8.5162E-09 | 0.25634496 | 0.492 | 0.327 | 0.00018182 | TEC_dmg | SF3B6 |
| 1.4379E-08 | 0.25612633 | 0.575 | 0.381 | 0.00030699 | TEC_dmg | PERP |
| 1.2033E-08 | 0.25519228 | 0.274 | 0.145 | 0.0002569 | TEC_dmg | ARL4C |
| 2.9052E-09 | 0.25488636 | 0.52 | 0.342 | 6.2025E-05 | TEC_dmg | CNBP |
| 6.1715E-06 | 0.25404566 | 0.79 | 0.66 | 0.1317609 | TEC_dmg | TACSTD2 |
| 1.1703E-09 | 0.25342091 | 0.222 | 0.103 | 2.4986E-05 | TEC_dmg | PIM1 |

| | | | | | | |
|---|---|---|---|---|---|---|
| 1.3538E-06 | 0.25330963 | 0.23 | 0.125 | 0.02890278 | TEC_dmg | HMOX1 |
| 7.2417E-07 | 0.25296724 | 0.115 | 0.047 | 0.01546102 | TEC_dmg | PI3 |
| 1.054E-09 | 0.25116062 | 0.476 | 0.294 | 2.2503E-05 | TEC_dmg | CYTOR |
| 1.1055E-07 | 0.25083449 | 0.746 | 0.606 | 0.00236029 | TEC_dmg | EIF4A2 |
| 2.631E-09 | 0.25021214 | 0.353 | 0.2 | 5.6172E-05 | TEC_dmg | CLK1 |
| 0.00036179 | -0.2530527 | 0.175 | 0.272 | 1 | TEC_dmg | SEPHS2 |
| 2.6213E-05 | -0.2540877 | 0.091 | 0.194 | 0.55964323 | TEC_dmg | ERICH1 |
| 2.2497E-05 | -0.2556883 | 0.079 | 0.181 | 0.48031508 | TEC_dmg | MYO9A |
| 1.0064E-06 | -0.2567436 | 0.083 | 0.206 | 0.02148685 | TEC_dmg | RTF1 |
| 2.6424E-06 | -0.2568033 | 0.032 | 0.13 | 0.05641606 | TEC_dmg | PKHD1 |
| 0.00276555 | -0.2576379 | 0.139 | 0.21 | 1 | TEC_dmg | MAT2A |
| 0.00393142 | -0.2576445 | 0.325 | 0.388 | 1 | TEC_dmg | POLR2L |
| 0.00011702 | -0.2584415 | 0.083 | 0.174 | 1 | TEC_dmg | THSD7A |
| 0.00167683 | -0.2635296 | 0.063 | 0.127 | 1 | TEC_dmg | HIST2H2BF |
| 0.00088535 | -0.2671485 | 0.206 | 0.29 | 1 | TEC_dmg | MT-ATP8 |
| 0.00038264 | -0.2686948 | 0.444 | 0.529 | 1 | TEC_dmg | ELOB |
| 3.04E-05 | -0.2696953 | 0.028 | 0.109 | 0.64903513 | TEC_dmg | AL355075.4 |
| 3.075E-05 | -0.2778513 | 0.163 | 0.275 | 0.65651109 | TEC_dmg | TCF25 |
| 0.00477501 | -0.2787317 | 0.687 | 0.732 | 1 | TEC_dmg | NDUFA4 |
| 0.00039474 | -0.2790094 | 0.853 | 0.899 | 1 | TEC_dmg | RPL23A |
| 2.5839E-06 | -0.293078 | 0.02 | 0.113 | 0.05516687 | TEC_dmg | HIST4H4 |
| 7.0549E-07 | -0.2932107 | 0.024 | 0.128 | 0.01506219 | TEC_dmg | AL135925.1 |
| 3.0212E-05 | -0.2964434 | 0.119 | 0.22 | 0.64502794 | TEC_dmg | EPS8L2 |
| 0.00357835 | -0.2977198 | 0.127 | 0.191 | 1 | TEC_dmg | TLE4 |
| 0.00881716 | -0.2988873 | 0.444 | 0.512 | 1 | TEC_dmg | H2AFZ |
| 8.7742E-05 | -0.2992063 | 0.202 | 0.315 | 1 | TEC_dmg | SRSF2 |
| 0.00499295 | -0.2992632 | 0.171 | 0.235 | 1 | TEC_dmg | NFKBIZ |
| 0.00257613 | -0.2993619 | 0.21 | 0.289 | 1 | TEC_dmg | STMN1 |
| 2.6066E-05 | -0.3019474 | 0.21 | 0.325 | 0.55650947 | TEC_dmg | PHPT1 |
| 0.00062777 | -0.3184136 | 0.877 | 0.9 | 1 | TEC_dmg | SERF2 |
| 0.00020141 | -0.3220874 | 0.087 | 0.173 | 1 | TEC_dmg | FOSB |
| 4.1018E-06 | -0.3264527 | 0.948 | 0.956 | 0.08757375 | TEC_dmg | RPS27 |
| 1.7093E-06 | -0.329211 | 0.028 | 0.128 | 0.0364938 | TEC_dmg | PCDH9 |
| 9.93E-05 | -0.3342099 | 0.175 | 0.268 | 1 | TEC_dmg | GSTM3 |
| 5.7409E-05 | -0.3358514 | 0.052 | 0.137 | 1 | TEC_dmg | AC025164.1 |
| 4.0453E-06 | -0.3379831 | 0.48 | 0.616 | 0.08636694 | TEC_dmg | MT-ND4L |
| 0.00675513 | -0.3398645 | 0.825 | 0.835 | 1 | TEC_dmg | PPIA |
| 0.00033517 | -0.3411412 | 0.246 | 0.339 | 1 | TEC_dmg | HNRNPAB |
| 2.4875E-08 | -0.3503247 | 0.536 | 0.683 | 0.00053108 | TEC_dmg | YBX3 |
| 6.1925E-06 | -0.3546252 | 0.016 | 0.101 | 0.13220962 | TEC_dmg | AP001160.1 |
| 0.00075477 | -0.359561 | 0.179 | 0.269 | 1 | TEC_dmg | TUBA1A |
| 0.00843515 | -0.3596487 | 0.286 | 0.339 | 1 | TEC_dmg | PAX8 |
| 0.00024601 | -0.3608721 | 0.893 | 0.89 | 1 | TEC_dmg | MT-ND2 |
| 6.579E-05 | -0.3678258 | 0.492 | 0.612 | 1 | TEC_dmg | BTG1 |
| 5.0727E-07 | -0.3685717 | 0.06 | 0.178 | 0.0108302 | TEC_dmg | FOXC1 |
| 2.0143E-05 | -0.3686742 | 0.46 | 0.571 | 0.43005419 | TEC_dmg | RPS27L |
| 2.3671E-06 | -0.3695082 | 0.048 | 0.152 | 0.05053681 | TEC_dmg | AC007952.4 |
| 0.0003038 | -0.3816786 | 0.056 | 0.13 | 1 | TEC_dmg | COL7A1 |
| 0.00795936 | -0.3928725 | 0.23 | 0.287 | 1 | TEC_dmg | CEBPD |
| 0.00206786 | -0.4064771 | 0.298 | 0.376 | 1 | TEC_dmg | SLC38A2 |

| | | | | | | |
|---|---|---|---|---|---|---|
| 0.0035438 | -0.4193966 | 0.278 | 0.344 | 1 | TEC_dmg | ZFP36L2 |
| 3.2292E-05 | -0.4209932 | 0.04 | 0.125 | 0.68944229 | TEC_dmg | CXCL14 |
| 4.727E-05 | -0.4305002 | 0.591 | 0.67 | 1 | TEC_dmg | TPI1 |
| 4.2162E-06 | -0.4388274 | 0.099 | 0.213 | 0.09001667 | TEC_dmg | ADAMTS1 |
| 5.7277E-05 | -0.4401072 | 0.925 | 0.911 | 1 | TEC_dmg | MT-CO2 |
| 8.9242E-05 | -0.4416854 | 0.083 | 0.17 | 1 | TEC_dmg | EGLN3 |
| 0.00021476 | -0.4639243 | 0.131 | 0.216 | 1 | TEC_dmg | SOCS1 |
| 1.3182E-05 | -0.4765157 | 0.952 | 0.909 | 0.28142703 | TEC_dmg | MT-ATP6 |
| 0.00013712 | -0.489561 | 0.155 | 0.244 | 1 | TEC_dmg | ID4 |
| 0.00155142 | -0.4901268 | 0.325 | 0.392 | 1 | TEC_dmg | TUBA1B |
| 1.4701E-07 | -0.4949482 | 0.131 | 0.28 | 0.0031387 | TEC_dmg | PPP1R10 |
| 0.00089309 | -0.5072517 | 0.345 | 0.416 | 1 | TEC_dmg | SRSF7 |
| 1.0638E-06 | -0.5130991 | 0.115 | 0.24 | 0.022713 | TEC_dmg | INTS6 |
| 1.4393E-07 | -0.5184475 | 0.06 | 0.187 | 0.00307301 | TEC_dmg | PIK3R3 |
| 0.00253152 | -0.5267782 | 0.206 | 0.273 | 1 | TEC_dmg | TCIM |
| 4.2428E-07 | -0.5271281 | 0.905 | 0.891 | 0.00905833 | TEC_dmg | MT-ND1 |
| 2.8231E-05 | -0.5274899 | 0.155 | 0.257 | 0.60273866 | TEC_dmg | IER5L |
| 7.1019E-08 | -0.5554411 | 0.849 | 0.868 | 0.00151626 | TEC_dmg | MT-ND3 |
| 8.4029E-10 | -0.5616928 | 0.052 | 0.206 | 1.794E-05 | TEC_dmg | SGK1 |
| 1.1781E-05 | -0.5642106 | 0.329 | 0.43 | 0.25151907 | TEC_dmg | SYNE2 |
| 3.1121E-05 | -0.5675781 | 0.202 | 0.303 | 0.664443 | TEC_dmg | PLK2 |
| 2.9241E-09 | -0.5778708 | 0.115 | 0.279 | 6.2429E-05 | TEC_dmg | CITED2 |
| 5.9742E-08 | -0.5879689 | 0.139 | 0.283 | 0.0012755 | TEC_dmg | BTG2 |
| 2.7665E-07 | -0.6251301 | 0.921 | 0.91 | 0.00590641 | TEC_dmg | MT-CO3 |
| 5.6493E-06 | -0.6398941 | 0.171 | 0.279 | 0.12061197 | TEC_dmg | RASD1 |
| 1.2582E-07 | -0.6568761 | 0.175 | 0.314 | 0.00268618 | TEC_dmg | GLUL |
| 1.0211E-05 | -0.6706663 | 0.575 | 0.64 | 0.21801486 | TEC_dmg | JUND |
| 4.0324E-07 | -0.6967796 | 0.444 | 0.567 | 0.00860915 | TEC_dmg | IER2 |
| 2.9345E-07 | -0.702863 | 0.726 | 0.816 | 0.00626511 | TEC_dmg | IGFBP7 |
| 4.4138E-09 | -0.7077048 | 0.79 | 0.829 | 9.4236E-05 | TEC_dmg | MT-ND5 |
| 5.0847E-12 | -0.7308439 | 0.881 | 0.899 | 1.0856E-07 | TEC_dmg | MT-CYB |
| 0.00394894 | -0.7384644 | 0.067 | 0.128 | 1 | TEC_dmg | GPX3 |
| 9.6609E-08 | -0.7714649 | 0.48 | 0.589 | 0.0020626 | TEC_dmg | ELF3 |
| 0.00020459 | -0.7787765 | 0.607 | 0.68 | 1 | TEC_dmg | MTRNR2L12 |
| 6.5288E-11 | -0.7993703 | 0.063 | 0.237 | 1.3939E-06 | TEC_dmg | CSKMT |
| 1.8713E-09 | -0.8698779 | 0.27 | 0.443 | 3.9951E-05 | TEC_dmg | IER3 |
| 1.0156E-09 | -0.8920915 | 0.135 | 0.3 | 2.1683E-05 | TEC_dmg | HEXIM1 |
| 7.9992E-10 | -0.9587884 | 0.115 | 0.273 | 1.7078E-05 | TEC_dmg | HSPA2 |
| 4.9437E-08 | -1.0431908 | 0.147 | 0.285 | 0.00105548 | TEC_dmg | MAFB |
| 1.2694E-09 | -1.0584226 | 0.222 | 0.38 | 2.7101E-05 | TEC_dmg | HIST1H4C |
| 4.4526E-05 | -1.0930299 | 0.48 | 0.541 | 0.95062532 | TEC_dmg | SOX4 |
| 3.2524E-13 | -3.2244916 | 0.464 | 0.6 | 6.9439E-09 | TEC_dmg | PLCG2 |
| 1.797E-108 | 1.71370653 | 0.955 | 0.802 | 3.836E-104 | TEC_GCLM | NEAT1 |
| 1.179E-121 | 1.00304507 | 0.997 | 0.935 | 2.518E-117 | TEC_GCLM | MALAT1 |
| 4.0635E-35 | 0.97987496 | 0.772 | 0.676 | 8.6756E-31 | TEC_GCLM | HSPA5 |
| 9.1301E-93 | 0.9566816 | 0.645 | 0.37 | 1.9493E-88 | TEC_GCLM | TXNRD1 |
| 9.3043E-88 | 0.80316354 | 0.337 | 0.127 | 1.9865E-83 | TEC_GCLM | XIST |
| 5.059E-114 | 0.71021666 | 0.663 | 0.341 | 1.08E-109 | TEC_GCLM | GOLGA4 |
| 5.2419E-89 | 0.66100856 | 0.448 | 0.195 | 1.1191E-84 | TEC_GCLM | GCLM |
| 5.272E-126 | 0.63489635 | 0.558 | 0.238 | 1.126E-121 | TEC_GCLM | RB1CC1 |

| | | | | | | |
|---|---|---|---|---|---|---|
| 3.3269E-74 | 0.58943413 | 0.582 | 0.312 | 7.103E-70 | TEC_GCLM | ZNF292 |
| 6.7525E-29 | 0.58610751 | 0.522 | 0.369 | 1.4417E-24 | TEC_GCLM | HERPUD1 |
| 6.423E-110 | 0.57665877 | 0.316 | 0.093 | 1.371E-105 | TEC_GCLM | SLC7A11 |
| 9.8457E-47 | 0.54238599 | 0.622 | 0.416 | 2.1021E-42 | TEC_GCLM | ZBTB20 |
| 1.0045E-84 | 0.53985341 | 0.676 | 0.373 | 2.1447E-80 | TEC_GCLM | ANKRD11 |
| 7.732E-99 | 0.5296704 | 0.454 | 0.19 | 1.6509E-94 | TEC_GCLM | MAFG |
| 7.5001E-12 | 0.52935047 | 0.785 | 0.771 | 1.6013E-07 | TEC_GCLM | HSP90B1 |
| 1.976E-31 | 0.52756697 | 0.421 | 0.261 | 4.2188E-27 | TEC_GCLM | VMP1 |
| 8.3026E-95 | 0.52086937 | 0.522 | 0.243 | 1.7726E-90 | TEC_GCLM | SLC38A1 |
| 1.1642E-95 | 0.50548234 | 0.424 | 0.17 | 2.4856E-91 | TEC_GCLM | ZBTB43 |
| 1.9459E-64 | 0.48979876 | 0.653 | 0.392 | 4.1544E-60 | TEC_GCLM | GOLGB1 |
| 8.5342E-78 | 0.47691421 | 0.513 | 0.252 | 1.8221E-73 | TEC_GCLM | CREBRF |
| 1.4492E-59 | 0.47555664 | 0.436 | 0.226 | 3.0941E-55 | TEC_GCLM | DNAJC3 |
| 1.2012E-77 | 0.46788072 | 0.508 | 0.251 | 2.5645E-73 | TEC_GCLM | OGA |
| 4.0068E-61 | 0.46493462 | 0.378 | 0.175 | 8.5544E-57 | TEC_GCLM | SESN2 |
| 2.3691E-42 | 0.46291135 | 0.415 | 0.234 | 5.058E-38 | TEC_GCLM | MACC1 |
| 8.0348E-56 | 0.45982347 | 0.738 | 0.541 | 1.7154E-51 | TEC_GCLM | LRRFIP1 |
| 3.4568E-54 | 0.45224214 | 0.6 | 0.363 | 7.3802E-50 | TEC_GCLM | N4BP2L2 |
| 1.0435E-25 | 0.44665745 | 0.457 | 0.307 | 2.228E-21 | TEC_GCLM | ITGA2 |
| 3.7839E-72 | 0.43598868 | 0.547 | 0.289 | 8.0786E-68 | TEC_GCLM | ARHGAP5 |
| 1.3417E-46 | 0.43530218 | 0.601 | 0.374 | 2.8645E-42 | TEC_GCLM | TAF1D |
| 7.3392E-74 | 0.43414511 | 0.498 | 0.242 | 1.5669E-69 | TEC_GCLM | CHD2 |
| 6.5923E-66 | 0.42753266 | 0.506 | 0.26 | 1.4075E-61 | TEC_GCLM | ZNF207 |
| 1.5542E-61 | 0.42608366 | 0.582 | 0.341 | 3.3182E-57 | TEC_GCLM | EIF3A |
| 5.0236E-64 | 0.41997974 | 0.407 | 0.194 | 1.0725E-59 | TEC_GCLM | GFPT1 |
| 6.1896E-60 | 0.41844441 | 0.395 | 0.19 | 1.3215E-55 | TEC_GCLM | DNTTIP2 |
| 4.7129E-84 | 0.40950019 | 0.399 | 0.164 | 1.0062E-79 | TEC_GCLM | CELF1 |
| 1.3116E-65 | 0.40738129 | 0.474 | 0.245 | 2.8002E-61 | TEC_GCLM | UFM1 |
| 1.8223E-50 | 0.40424014 | 0.541 | 0.315 | 3.8906E-46 | TEC_GCLM | ANKRD12 |
| 5.6653E-44 | 0.39750699 | 0.51 | 0.301 | 1.2095E-39 | TEC_GCLM | UGCG |
| 4.6417E-51 | 0.39665557 | 0.359 | 0.176 | 9.9101E-47 | TEC_GCLM | SEL1L |
| 4.8952E-42 | 0.39562796 | 0.398 | 0.222 | 1.0451E-37 | TEC_GCLM | LPP |
| 9.7923E-52 | 0.38943361 | 0.404 | 0.207 | 2.0907E-47 | TEC_GCLM | TEAD1 |
| 1.0465E-16 | 0.38495129 | 0.394 | 0.286 | 2.2343E-12 | TEC_GCLM | ITGB8 |
| 1.1916E-36 | 0.38432216 | 0.503 | 0.325 | 2.5441E-32 | TEC_GCLM | HSPA9 |
| 3.954E-68 | 0.38388388 | 0.411 | 0.188 | 8.4418E-64 | TEC_GCLM | ATP13A3 |
| 6.2168E-70 | 0.38300816 | 0.433 | 0.2 | 1.3273E-65 | TEC_GCLM | CCDC186 |
| 5.0239E-20 | 0.37411904 | 0.376 | 0.249 | 1.0726E-15 | TEC_GCLM | LAMB3 |
| 4.6359E-43 | 0.36989803 | 0.583 | 0.358 | 9.8977E-39 | TEC_GCLM | PNISR |
| 1.3135E-47 | 0.3693261 | 0.45 | 0.249 | 2.8043E-43 | TEC_GCLM | RIOK3 |
| 7.0061E-82 | 0.3650449 | 0.304 | 0.106 | 1.4958E-77 | TEC_GCLM | PGM3 |
| 1.2865E-16 | 0.36343068 | 0.447 | 0.341 | 2.7467E-12 | TEC_GCLM | SELENOK |
| 8.027E-48 | 0.36339317 | 0.393 | 0.206 | 1.7138E-43 | TEC_GCLM | GARS |
| 7.4667E-31 | 0.35879671 | 0.425 | 0.252 | 1.5941E-26 | TEC_GCLM | KCNQ1OT1 |
| 2.4471E-55 | 0.34610114 | 0.325 | 0.145 | 5.2245E-51 | TEC_GCLM | TRIM56 |
| 4.2398E-38 | 0.34390052 | 0.44 | 0.265 | 9.0519E-34 | TEC_GCLM | VCP |
| 2.6987E-28 | 0.33930462 | 0.336 | 0.197 | 5.7616E-24 | TEC_GCLM | SLFN5 |
| 4.2143E-15 | 0.33887597 | 0.549 | 0.455 | 8.9975E-11 | TEC_GCLM | SERP1 |
| 1.7184E-28 | 0.33793442 | 0.433 | 0.285 | 3.6687E-24 | TEC_GCLM | SELENOS |
| 2.7399E-25 | 0.33704786 | 0.561 | 0.426 | 5.8496E-21 | TEC_GCLM | NAP1L1 |

| | | | | | | |
|---|---|---|---|---|---|---|
| 2.5834E-43 | 0.33548423 | 0.513 | 0.3 | 5.5155E-39 | TEC_GCLM | BOD1L1 |
| 3.2496E-47 | 0.33386347 | 0.48 | 0.268 | 6.9379E-43 | TEC_GCLM | RBM25 |
| 3.0087E-38 | 0.33110198 | 0.407 | 0.233 | 6.4237E-34 | TEC_GCLM | TES |
| 3.3058E-89 | 0.3295068 | 0.3 | 0.098 | 7.0579E-85 | TEC_GCLM | AC005261.1 |
| 1.6541E-28 | 0.32934466 | 0.258 | 0.133 | 3.5314E-24 | TEC_GCLM | GABPB1-AS1 |
| 2.3672E-58 | 0.32731827 | 0.383 | 0.18 | 5.0539E-54 | TEC_GCLM | STX3 |
| 3.9886E-58 | 0.3259337 | 0.412 | 0.198 | 8.5156E-54 | TEC_GCLM | TMF1 |
| 7.0185E-52 | 0.32159688 | 0.362 | 0.172 | 1.4985E-47 | TEC_GCLM | KLHL24 |
| 3.427E-47 | 0.32041085 | 0.392 | 0.205 | 7.3166E-43 | TEC_GCLM | LIMS1 |
| 6.6478E-51 | 0.32038891 | 0.278 | 0.117 | 1.4193E-46 | TEC_GCLM | OGT |
| 9.7772E-24 | 0.31805487 | 0.427 | 0.285 | 2.0874E-19 | TEC_GCLM | HSPH1 |
| 4.7658E-42 | 0.31715804 | 0.425 | 0.236 | 1.0175E-37 | TEC_GCLM | BCLAF1 |
| 4.9925E-28 | 0.31702004 | 0.698 | 0.539 | 1.0659E-23 | TEC_GCLM | EIF5 |
| 1.0218E-20 | 0.31540295 | 0.461 | 0.33 | 2.1816E-16 | TEC_GCLM | ARF4 |
| 2.4114E-43 | 0.315381 | 0.474 | 0.268 | 5.1484E-39 | TEC_GCLM | NUFIP2 |
| 5.1991E-58 | 0.31514926 | 0.496 | 0.253 | 1.11E-53 | TEC_GCLM | SMG1 |
| 2.0261E-20 | 0.31409299 | 0.679 | 0.564 | 4.3258E-16 | TEC_GCLM | SLC3A2 |
| 1.5314E-47 | 0.31269303 | 0.375 | 0.19 | 3.2695E-43 | TEC_GCLM | OCLN |
| 7.1781E-40 | 0.31232633 | 0.354 | 0.187 | 1.5325E-35 | TEC_GCLM | FOXO3 |
| 2.169E-46 | 0.31228186 | 0.45 | 0.248 | 4.6309E-42 | TEC_GCLM | SCAF11 |
| 7.0241E-66 | 0.31101305 | 0.373 | 0.162 | 1.4996E-61 | TEC_GCLM | UHMK1 |
| 1.0762E-55 | 0.31004343 | 0.359 | 0.165 | 2.2977E-51 | TEC_GCLM | LINC00958 |
| 6.8979E-38 | 0.30835873 | 0.471 | 0.285 | 1.4727E-33 | TEC_GCLM | PAPOLA |
| 4.6099E-37 | 0.30802876 | 0.546 | 0.354 | 9.8421E-33 | TEC_GCLM | UBE2B |
| 2.1502E-39 | 0.30677939 | 0.461 | 0.274 | 4.5908E-35 | TEC_GCLM | SERINC1 |
| 8.5627E-34 | 0.30661744 | 0.565 | 0.369 | 1.8281E-29 | TEC_GCLM | MYO6 |
| 3.4658E-50 | 0.30531975 | 0.257 | 0.105 | 7.3996E-46 | TEC_GCLM | ABL2 |
| 9.871E-54 | 0.30503817 | 0.34 | 0.155 | 2.1075E-49 | TEC_GCLM | PRDM2 |
| 2.2147E-52 | 0.30432243 | 0.343 | 0.159 | 4.7285E-48 | TEC_GCLM | MED13 |
| 9.1521E-51 | 0.30427264 | 0.37 | 0.178 | 1.954E-46 | TEC_GCLM | GPATCH2L |
| 1.3523E-32 | 0.30389479 | 0.487 | 0.317 | 2.8872E-28 | TEC_GCLM | KIF5B |
| 1.0358E-53 | 0.3027812 | 0.293 | 0.124 | 2.2114E-49 | TEC_GCLM | AGO2 |
| 8.6057E-33 | 0.30248585 | 0.586 | 0.398 | 1.8373E-28 | TEC_GCLM | NORAD |
| 2.5706E-40 | 0.30143187 | 0.41 | 0.232 | 5.4881E-36 | TEC_GCLM | SEC31A |
| 2.4624E-33 | 0.29999731 | 0.508 | 0.331 | 5.2572E-29 | TEC_GCLM | MTDH |
| 1.6191E-29 | 0.29902425 | 0.318 | 0.179 | 3.4568E-25 | TEC_GCLM | ZNF83 |
| 7.1843E-42 | 0.29762102 | 0.438 | 0.247 | 1.5339E-37 | TEC_GCLM | IL6ST |
| 1.1313E-49 | 0.29747617 | 0.36 | 0.173 | 2.4154E-45 | TEC_GCLM | SREK1 |
| 7.5385E-48 | 0.29710452 | 0.365 | 0.178 | 1.6095E-43 | TEC_GCLM | POLR2J3.1 |
| 6.7676E-12 | 0.29270063 | 0.34 | 0.245 | 1.4449E-07 | TEC_GCLM | AKAP12 |
| 1.0261E-43 | 0.29212675 | 0.369 | 0.19 | 2.1907E-39 | TEC_GCLM | C16orf72 |
| 1.0394E-41 | 0.29156533 | 0.413 | 0.229 | 2.2192E-37 | TEC_GCLM | NCKAP1 |
| 9.1786E-53 | 0.29133927 | 0.33 | 0.148 | 1.9596E-48 | TEC_GCLM | XIAP |
| 2.6329E-31 | 0.29114903 | 0.646 | 0.473 | 5.6211E-27 | TEC_GCLM | UBE2D3 |
| 3.6097E-48 | 0.29099913 | 0.318 | 0.148 | 7.7067E-44 | TEC_GCLM | TAOK1 |
| 1.7539E-40 | 0.28865891 | 0.404 | 0.221 | 3.7445E-36 | TEC_GCLM | NIPBL |
| 6.8728E-30 | 0.28809766 | 0.527 | 0.344 | 1.4673E-25 | TEC_GCLM | GCC2 |
| 3.7826E-44 | 0.28772873 | 0.442 | 0.245 | 8.0758E-40 | TEC_GCLM | WAC |
| 1.7614E-23 | 0.28648848 | 0.667 | 0.541 | 3.7606E-19 | TEC_GCLM | CANX |
| 3.4127E-40 | 0.2862697 | 0.273 | 0.126 | 7.2861E-36 | TEC_GCLM | TRIB3 |

| | | | | | | |
|---|---|---|---|---|---|---|
| 1.0565E-42 | 0.28621427 | 0.416 | 0.226 | 2.2556E-38 | TEC_GCLM | TOR1AIP2 |
| 2.4391E-41 | 0.28576453 | 0.433 | 0.242 | 5.2075E-37 | TEC_GCLM | BRD4 |
| 1.5056E-44 | 0.28493207 | 0.395 | 0.208 | 3.2145E-40 | TEC_GCLM | RAB21 |
| 2.863E-35 | 0.28424748 | 0.361 | 0.204 | 6.1126E-31 | TEC_GCLM | MED10 |
| 2.8143E-40 | 0.28383017 | 0.325 | 0.165 | 6.0086E-36 | TEC_GCLM | SRP19 |
| 1.069E-44 | 0.2835492 | 0.347 | 0.174 | 2.2824E-40 | TEC_GCLM | PNPLA8 |
| 1.064E-37 | 0.28331473 | 0.344 | 0.182 | 2.2717E-33 | TEC_GCLM | BACH1 |
| 7.7542E-20 | 0.28309095 | 0.51 | 0.381 | 1.6555E-15 | TEC_GCLM | EIF2S2 |
| 8.6807E-40 | 0.28284925 | 0.39 | 0.216 | 1.8533E-35 | TEC_GCLM | GNAI3 |
| 1.8094E-29 | 0.28275053 | 0.3 | 0.165 | 3.863E-25 | TEC_GCLM | DNAJB9 |
| 2.9821E-42 | 0.28020213 | 0.473 | 0.273 | 6.3668E-38 | TEC_GCLM | TJP1 |
| 5.5536E-41 | 0.27977582 | 0.391 | 0.212 | 1.1857E-36 | TEC_GCLM | KPNA4 |
| 1.5416E-48 | 0.27973436 | 0.351 | 0.168 | 3.2914E-44 | TEC_GCLM | BTBD7 |
| 3.2646E-49 | 0.27913559 | 0.359 | 0.173 | 6.97E-45 | TEC_GCLM | DDX6 |
| 1.6719E-18 | 0.27819239 | 0.353 | 0.227 | 3.5695E-14 | TEC_GCLM | SLC5A3 |
| 9.4908E-54 | 0.27803725 | 0.284 | 0.117 | 2.0263E-49 | TEC_GCLM | ZNF791 |
| 2.6879E-50 | 0.27750296 | 0.311 | 0.14 | 5.7387E-46 | TEC_GCLM | KIF1B |
| 2.9627E-32 | 0.27640872 | 0.365 | 0.206 | 6.3255E-28 | TEC_GCLM | NKTR |
| 9.9594E-24 | 0.27615237 | 0.528 | 0.382 | 2.1263E-19 | TEC_GCLM | TAX1BP1 |
| 1.1133E-46 | 0.27536544 | 0.374 | 0.185 | 2.3769E-42 | TEC_GCLM | MDM4 |
| 8.107E-40 | 0.27441399 | 0.489 | 0.286 | 1.7308E-35 | TEC_GCLM | AFF4 |
| 5.4317E-41 | 0.2732267 | 0.368 | 0.194 | 1.1597E-36 | TEC_GCLM | CSNK2A1 |
| 2.4072E-15 | 0.27234684 | 0.528 | 0.414 | 5.1393E-11 | TEC_GCLM | AHNAK |
| 1.2359E-36 | 0.27186167 | 0.4 | 0.226 | 2.6386E-32 | TEC_GCLM | AAK1 |
| 4.1336E-34 | 0.2718048 | 0.362 | 0.205 | 8.8253E-30 | TEC_GCLM | SEC61A1 |
| 2.1298E-48 | 0.27174738 | 0.268 | 0.114 | 4.5472E-44 | TEC_GCLM | HSPA13 |
| 9.231E-11 | 0.27032119 | 0.273 | 0.196 | 1.9708E-06 | TEC_GCLM | MANF |
| 1.1946E-31 | 0.27025046 | 0.376 | 0.222 | 2.5504E-27 | TEC_GCLM | CHMP2B |
| 4.5134E-45 | 0.27009937 | 0.335 | 0.163 | 9.6362E-41 | TEC_GCLM | TNPO1 |
| 8.6383E-38 | 0.26975415 | 0.323 | 0.166 | 1.8443E-33 | TEC_GCLM | USP8 |
| 3.0714E-22 | 0.2692635 | 0.26 | 0.151 | 6.5573E-18 | TEC_GCLM | HYOU1 |
| 2.4406E-11 | 0.26896107 | 0.51 | 0.409 | 5.2106E-07 | TEC_GCLM | CCND1 |
| 2.5118E-13 | 0.26866833 | 0.468 | 0.349 | 5.3627E-09 | TEC_GCLM | KLF6 |
| 6.8886E-30 | 0.26860679 | 0.32 | 0.18 | 1.4707E-25 | TEC_GCLM | FNDC3B |
| 1.1262E-26 | 0.26768599 | 0.271 | 0.15 | 2.4044E-22 | TEC_GCLM | MTHFD2 |
| 6.2903E-46 | 0.26663542 | 0.295 | 0.135 | 1.343E-41 | TEC_GCLM | SAMD4B |
| 2.0375E-51 | 0.26656384 | 0.264 | 0.108 | 4.35E-47 | TEC_GCLM | SLC7A1 |
| 3.692E-47 | 0.26635559 | 0.293 | 0.131 | 7.8824E-43 | TEC_GCLM | TBC1D15 |
| 2.5391E-35 | 0.26628442 | 0.347 | 0.191 | 5.421E-31 | TEC_GCLM | IPO7 |
| 4.287E-48 | 0.26605505 | 0.317 | 0.145 | 9.1527E-44 | TEC_GCLM | RIF1 |
| 1.0931E-58 | 0.26601032 | 0.298 | 0.12 | 2.3339E-54 | TEC_GCLM | KHDC4 |
| 5.4523E-39 | 0.2657801 | 0.318 | 0.162 | 1.1641E-34 | TEC_GCLM | PRNP |
| 7.5925E-27 | 0.26400524 | 0.483 | 0.321 | 1.621E-22 | TEC_GCLM | PTP4A2 |
| 4.8003E-26 | 0.2638657 | 0.453 | 0.296 | 1.0249E-21 | TEC_GCLM | DSP |
| 1.5132E-26 | 0.26375606 | 0.47 | 0.309 | 3.2307E-22 | TEC_GCLM | PAWR |
| 2.0269E-43 | 0.26351697 | 0.317 | 0.153 | 4.3273E-39 | TEC_GCLM | UBE2J1 |
| 3.2601E-44 | 0.26307161 | 0.317 | 0.15 | 6.9602E-40 | TEC_GCLM | RANBP2 |
| 7.1047E-16 | 0.26287586 | 0.434 | 0.33 | 1.5168E-11 | TEC_GCLM | SSR3 |
| 6.5243E-32 | 0.26089435 | 0.471 | 0.288 | 1.3929E-27 | TEC_GCLM | YTHDC1 |
| 1.5551E-15 | 0.26069882 | 0.284 | 0.187 | 3.3201E-11 | TEC_GCLM | SDF2L1 |

| | | | | | | |
|---|---|---|---|---|---|---|
| 1.1826E-33 | 0.25896224 | 0.379 | 0.212 | 2.5248E-29 | TEC_GCLM | ZNF91 |
| 1.7027E-41 | 0.2585228 | 0.337 | 0.169 | 3.6353E-37 | TEC_GCLM | SRSF4 |
| 1.3927E-36 | 0.25836402 | 0.291 | 0.144 | 2.9735E-32 | TEC_GCLM | CHORDC1 |
| 2.1949E-32 | 0.2583101 | 0.31 | 0.167 | 4.6861E-28 | TEC_GCLM | SRP54 |
| 4.4026E-51 | 0.25790149 | 0.263 | 0.108 | 9.3996E-47 | TEC_GCLM | NUP58 |
| 2.7225E-52 | 0.2577954 | 0.306 | 0.131 | 5.8125E-48 | TEC_GCLM | DYRK1A |
| 6.5808E-37 | 0.25737495 | 0.351 | 0.188 | 1.405E-32 | TEC_GCLM | UBE2Z |
| 1.3073E-35 | 0.25697454 | 0.361 | 0.198 | 2.7911E-31 | TEC_GCLM | G3BP1 |
| 3.1735E-27 | 0.25673524 | 0.504 | 0.334 | 6.7754E-23 | TEC_GCLM | UBE2H |
| 8.7208E-50 | 0.25635319 | 0.226 | 0.085 | 1.8619E-45 | TEC_GCLM | MIR29B2CHG |
| 2.4062E-44 | 0.25629518 | 0.285 | 0.13 | 5.1373E-40 | TEC_GCLM | UBE2G2 |
| 9.3691E-30 | 0.25515701 | 0.291 | 0.155 | 2.0003E-25 | TEC_GCLM | MIR22HG |
| 1.0136E-25 | 0.25458093 | 0.34 | 0.203 | 2.164E-21 | TEC_GCLM | TRIP11 |
| 3.1557E-47 | 0.25454027 | 0.322 | 0.15 | 6.7373E-43 | TEC_GCLM | CYLD |
| 3.2568E-34 | 0.25439805 | 0.413 | 0.237 | 6.9533E-30 | TEC_GCLM | JMJD1C |
| 1.198E-61 | 0.25299682 | 0.2 | 0.062 | 2.5577E-57 | TEC_GCLM | EID3 |
| 3.4246E-33 | 0.25290734 | 0.368 | 0.204 | 7.3115E-29 | TEC_GCLM | USP53 |
| 4.3008E-34 | 0.25230624 | 0.403 | 0.232 | 9.1822E-30 | TEC_GCLM | SLTM |
| 5.7567E-29 | 0.25141302 | 0.53 | 0.343 | 1.229E-24 | TEC_GCLM | SPAG9 |
| 1.3648E-40 | 0.25135068 | 0.295 | 0.14 | 2.9139E-36 | TEC_GCLM | MKLN1 |
| 1.2786E-25 | 0.25108106 | 0.529 | 0.347 | 2.7297E-21 | TEC_GCLM | AKAP9 |
| 6.0388E-41 | 0.25107158 | 0.387 | 0.203 | 1.2893E-36 | TEC_GCLM | ASH1L |
| 2.7022E-41 | 0.25039294 | 0.333 | 0.167 | 5.7693E-37 | TEC_GCLM | CEBPG |
| 1.6204E-29 | 0.25022594 | 0.379 | 0.225 | 3.4596E-25 | TEC_GCLM | HECTD1 |
| 1.6552E-33 | 0.25010768 | 0.309 | 0.163 | 3.5338E-29 | TEC_GCLM | ZKSCAN1 |
| 1.357E-20 | -0.2513685 | 0.164 | 0.296 | 2.8973E-16 | TEC_GCLM | NDUFB11 |
| 6.3009E-19 | -0.2526773 | 0.067 | 0.169 | 1.3453E-14 | TEC_GCLM | TMEM176B |
| 6.4091E-12 | -0.2542194 | 0.062 | 0.132 | 1.3683E-07 | TEC_GCLM | AL135925.1 |
| 6.8412E-09 | -0.2542293 | 0.089 | 0.151 | 0.00014606 | TEC_GCLM | ANKRD37 |
| 8.8599E-09 | -0.2542293 | 0.189 | 0.258 | 0.00018916 | TEC_GCLM | IRF1 |
| 7.29E-09 | -0.2545159 | 0.266 | 0.338 | 0.00015564 | TEC_GCLM | SLC25A5 |
| 9.2398E-13 | -0.2546988 | 0.26 | 0.357 | 1.9727E-08 | TEC_GCLM | ACADVL |
| 8.8649E-19 | -0.2561883 | 0.107 | 0.219 | 1.8926E-14 | TEC_GCLM | BEX3 |
| 2.9191E-11 | -0.2566986 | 0.077 | 0.149 | 6.2322E-07 | TEC_GCLM | ID3 |
| 5.5986E-07 | -0.2574629 | 0.763 | 0.843 | 0.01195306 | TEC_GCLM | PRDX1 |
| 9.2616E-15 | -0.2587766 | 0.437 | 0.542 | 1.9774E-10 | TEC_GCLM | NDUFS5 |
| 3.6294E-11 | -0.2595046 | 0.065 | 0.133 | 7.7488E-07 | TEC_GCLM | HOMER1 |
| 3.2679E-05 | -0.2612634 | 0.15 | 0.194 | 0.6976978 | TEC_GCLM | TLE4 |
| 1.9996E-11 | -0.2628104 | 0.351 | 0.436 | 4.2692E-07 | TEC_GCLM | BSG |
| 1.2164E-05 | -0.2628236 | 0.305 | 0.348 | 0.25969582 | TEC_GCLM | TSC22D1 |
| 1.2538E-07 | -0.2637771 | 0.21 | 0.272 | 0.00267678 | TEC_GCLM | GSTM3 |
| 7.5592E-08 | -0.2638751 | 0.237 | 0.306 | 0.00161389 | TEC_GCLM | DDIT4 |
| 1.2731E-14 | -0.2662202 | 0.038 | 0.113 | 2.7182E-10 | TEC_GCLM | AL355075.4 |
| 3.7086E-20 | -0.2670378 | 0.027 | 0.119 | 7.918E-16 | TEC_GCLM | RARRES2 |
| 2.067E-23 | -0.267477 | 0.034 | 0.142 | 4.4131E-19 | TEC_GCLM | CDH16 |
| 8.711E-25 | -0.2676484 | 0.806 | 0.919 | 1.8598E-20 | TEC_GCLM | FAU |
| 1.8374E-13 | -0.2699567 | 0.277 | 0.376 | 3.9228E-09 | TEC_GCLM | ATP5MF |
| 0.00053199 | -0.2701909 | 0.075 | 0.107 | 1 | TEC_GCLM | SERPINE2 |
| 2.8794E-12 | -0.2702519 | 0.372 | 0.457 | 6.1474E-08 | TEC_GCLM | RPS10 |
| 1.0621E-13 | -0.2713168 | 0.286 | 0.384 | 2.2675E-09 | TEC_GCLM | VAMP8 |

| | | | | | | |
|---|---|---|---|---|---|---|
| 1.4956E-13 | -0.2716325 | 0.689 | 0.791 | 3.1931E-09 | TEC_GCLM | EEF1B2 |
| 1.7124E-18 | -0.2720784 | 0.032 | 0.122 | 3.6561E-14 | TEC_GCLM | RARRES3 |
| 5.9345E-09 | -0.2724698 | 0.216 | 0.287 | 0.0001267 | TEC_GCLM | CLDN3 |
| 1.9262E-11 | -0.2743486 | 0.162 | 0.244 | 4.1124E-07 | TEC_GCLM | HIGD1A |
| 0.00199862 | -0.2746268 | 0.189 | 0.221 | 1 | TEC_GCLM | TNFAIP2 |
| 1.0741E-15 | -0.2754553 | 0.945 | 0.991 | 2.2933E-11 | TEC_GCLM | EEF1A1 |
| 3.6182E-10 | -0.2763039 | 0.669 | 0.734 | 7.7249E-06 | TEC_GCLM | ENO1 |
| 3.6532E-14 | -0.2797169 | 0.208 | 0.307 | 7.7997E-10 | TEC_GCLM | CAPG |
| 4.1357E-17 | -0.2800397 | 0.311 | 0.426 | 8.8298E-13 | TEC_GCLM | PARK7 |
| 4.7876E-10 | -0.280359 | 0.481 | 0.556 | 1.0222E-05 | TEC_GCLM | CD9 |
| 1.796E-19 | -0.2804913 | 0.24 | 0.367 | 3.8345E-15 | TEC_GCLM | EIF3K |
| 1.1109E-09 | -0.2806241 | 0.049 | 0.106 | 2.3717E-05 | TEC_GCLM | TNF |
| 5.4667E-18 | -0.2809736 | 0.652 | 0.756 | 1.1671E-13 | TEC_GCLM | DSTN |
| 1.7287E-05 | -0.2812729 | 0.307 | 0.347 | 0.36908444 | TEC_GCLM | S100A16 |
| 4.6791E-16 | -0.2825713 | 0.155 | 0.261 | 9.9899E-12 | TEC_GCLM | ECHS1 |
| 1.1189E-23 | -0.2829338 | 0.513 | 0.646 | 2.3888E-19 | TEC_GCLM | UBL5 |
| 2.0746E-15 | -0.2830568 | 0.106 | 0.203 | 4.4293E-11 | TEC_GCLM | BCAM |
| 5.2075E-10 | -0.2837454 | 0.288 | 0.367 | 1.1118E-05 | TEC_GCLM | COX7B |
| 1.9212E-17 | -0.2848516 | 0.524 | 0.636 | 4.1017E-13 | TEC_GCLM | SLC25A6 |
| 4.0628E-15 | -0.2856143 | 0.815 | 0.912 | 8.674E-11 | TEC_GCLM | RPS3 |
| 1.6373E-09 | -0.2863234 | 0.235 | 0.319 | 3.4956E-05 | TEC_GCLM | CDKN1C |
| 2.2997E-18 | -0.2867834 | 0.031 | 0.118 | 4.91E-14 | TEC_GCLM | HIST4H4 |
| 1.0818E-08 | -0.287368 | 0.055 | 0.111 | 0.00023095 | TEC_GCLM | AQP3 |
| 3.4422E-14 | -0.2874785 | 0.242 | 0.344 | 7.3491E-10 | TEC_GCLM | H2AFJ |
| 4.4639E-21 | -0.2875291 | 0.777 | 0.897 | 9.5305E-17 | TEC_GCLM | RPL22 |
| 1.2763E-16 | -0.2875518 | 0.548 | 0.657 | 2.7249E-12 | TEC_GCLM | UQCRH |
| 8.5016E-18 | -0.2886982 | 0.513 | 0.617 | 1.8151E-13 | TEC_GCLM | ATP5MG |
| 9.7517E-11 | -0.28924 | 0.292 | 0.367 | 2.082E-06 | TEC_GCLM | ATP5MC3 |
| 3.8304E-13 | -0.2896903 | 0.792 | 0.902 | 8.178E-09 | TEC_GCLM | RPS11 |
| 0.00020074 | -0.2902766 | 0.156 | 0.195 | 1 | TEC_GCLM | GADD45G |
| 1.3661E-08 | -0.2910399 | 0.34 | 0.402 | 0.00029166 | TEC_GCLM | RHOC |
| 6.4866E-05 | -0.2923234 | 0.307 | 0.34 | 1 | TEC_GCLM | HNRNPAB |
| 9.2984E-22 | -0.2923356 | 0.194 | 0.335 | 1.9852E-17 | TEC_GCLM | PHPT1 |
| 2.2145E-05 | -0.2939529 | 0.09 | 0.132 | 0.47279211 | TEC_GCLM | COL7A1 |
| 1.4446E-13 | -0.296063 | 0.832 | 0.929 | 3.0842E-09 | TEC_GCLM | RPS3A |
| 1.8552E-12 | -0.2970791 | 0.742 | 0.801 | 3.9608E-08 | TEC_GCLM | HNRNPA2B1 |
| 6.3298E-20 | -0.2977689 | 0.794 | 0.906 | 1.3514E-15 | TEC_GCLM | RPS15A |
| 9.7037E-08 | -0.2983179 | 0.221 | 0.284 | 0.00207175 | TEC_GCLM | SERPINB1 |
| 7.832E-07 | -0.3005583 | 0.262 | 0.317 | 0.01672132 | TEC_GCLM | SRSF2 |
| 1.0094E-10 | -0.3010186 | 0.549 | 0.621 | 2.1551E-06 | TEC_GCLM | CD81 |
| 1.6273E-13 | -0.301589 | 0.719 | 0.828 | 3.4742E-09 | TEC_GCLM | YBX1 |
| 4.4278E-13 | -0.3015995 | 0.24 | 0.335 | 9.4534E-09 | TEC_GCLM | TSPO |
| 1.7448E-19 | -0.3022875 | 0.815 | 0.932 | 3.7251E-15 | TEC_GCLM | RPL30 |
| 4.0085E-13 | -0.303017 | 0.102 | 0.184 | 8.5581E-09 | TEC_GCLM | KLF4 |
| 1.4207E-18 | -0.3039279 | 0.834 | 0.93 | 3.0332E-14 | TEC_GCLM | RPL14 |
| 3.0527E-13 | -0.3040133 | 0.332 | 0.434 | 6.5176E-09 | TEC_GCLM | SSR4 |
| 1.4336E-15 | -0.3041379 | 0.845 | 0.937 | 3.0607E-11 | TEC_GCLM | RPS14 |
| 1.3129E-10 | -0.3043827 | 0.822 | 0.879 | 2.803E-06 | TEC_GCLM | HLA-A |
| 1.1044E-18 | -0.3056343 | 0.259 | 0.386 | 2.3579E-14 | TEC_GCLM | NDUFA1 |
| 1.0188E-13 | -0.3057577 | 0.855 | 0.939 | 2.1752E-09 | TEC_GCLM | RPS6 |

| | | | | | | |
|---|---|---|---|---|---|---|
| 1.3725E-12 | -0.3063599 | 0.649 | 0.73 | 2.9303E-08 | TEC_GCLM | CTSD |
| 3.3903E-18 | -0.3069495 | 0.731 | 0.811 | 7.2384E-14 | TEC_GCLM | EEF1D |
| 2.46E-17 | -0.3072015 | 0.808 | 0.91 | 5.2521E-13 | TEC_GCLM | RPS25 |
| 1.0296E-13 | -0.308381 | 0.866 | 0.94 | 2.1983E-09 | TEC_GCLM | RPS27A |
| 4.9727E-13 | -0.3099385 | 0.043 | 0.113 | 1.0617E-08 | TEC_GCLM | BST2 |
| 2.8157E-24 | -0.310312 | 0.761 | 0.862 | 6.0114E-20 | TEC_GCLM | SRP14 |
| 3.4286E-21 | -0.3126981 | 0.193 | 0.325 | 7.32E-17 | TEC_GCLM | UQCR11 |
| 2.1328E-06 | -0.3131451 | 0.148 | 0.196 | 0.04553613 | TEC_GCLM | H2AFX |
| 2.3114E-17 | -0.3141136 | 0.851 | 0.945 | 4.9348E-13 | TEC_GCLM | RPS23 |
| 6.8377E-19 | -0.3152033 | 0.479 | 0.597 | 1.4598E-14 | TEC_GCLM | S100A13 |
| 9.0004E-17 | -0.316132 | 0.294 | 0.407 | 1.9216E-12 | TEC_GCLM | MGST3 |
| 3.6316E-15 | -0.3167182 | 0.051 | 0.133 | 7.7534E-11 | TEC_GCLM | HIST2H2BF |
| 1.8454E-21 | -0.3179281 | 0.103 | 0.223 | 3.9398E-17 | TEC_GCLM | CTSH |
| 4.5683E-12 | -0.3197961 | 0.204 | 0.295 | 9.7533E-08 | TEC_GCLM | STMN1 |
| 0.00979485 | -0.3206842 | 0.287 | 0.309 | 1 | TEC_GCLM | PLAU |
| 1.7839E-06 | -0.321322 | 0.069 | 0.115 | 0.03808576 | TEC_GCLM | C15orf48 |
| 4.9256E-16 | -0.3217737 | 0.596 | 0.698 | 1.0516E-11 | TEC_GCLM | CYSTM1 |
| 9.054E-12 | -0.3223279 | 0.855 | 0.93 | 1.933E-07 | TEC_GCLM | RPLP2 |
| 8.2294E-16 | -0.3224601 | 0.394 | 0.494 | 1.757E-11 | TEC_GCLM | ATP1A1 |
| 3.6584E-20 | -0.3234125 | 0.246 | 0.379 | 7.8107E-16 | TEC_GCLM | SELENOW |
| 8.2642E-32 | -0.3255107 | 0.626 | 0.763 | 1.7644E-27 | TEC_GCLM | PFDN5 |
| 3.6227E-20 | -0.3264913 | 0.624 | 0.736 | 7.7346E-16 | TEC_GCLM | CSTB |
| 2.2688E-10 | -0.3268479 | 0.106 | 0.181 | 4.8439E-06 | TEC_GCLM | IGFBP3 |
| 1.2872E-13 | -0.3276707 | 0.039 | 0.11 | 2.7482E-09 | TEC_GCLM | ARC |
| 1.6896E-12 | -0.3285891 | 0.137 | 0.221 | 3.6074E-08 | TEC_GCLM | ZFP36 |
| 2.5664E-20 | -0.3296465 | 0.89 | 0.979 | 5.4792E-16 | TEC_GCLM | TPT1 |
| 5.3828E-12 | -0.3309319 | 0.707 | 0.77 | 1.1492E-07 | TEC_GCLM | PKM |
| 5.8481E-17 | -0.3328381 | 0.773 | 0.878 | 1.2486E-12 | TEC_GCLM | RPS5 |
| 1.0237E-13 | -0.3330663 | 0.072 | 0.154 | 2.1856E-09 | TEC_GCLM | MYL9 |
| 3.0108E-20 | -0.3336337 | 0.874 | 0.952 | 6.428E-16 | TEC_GCLM | RPL11 |
| 9.2407E-15 | -0.3344446 | 0.032 | 0.106 | 1.9729E-10 | TEC_GCLM | AP001160.1 |
| 2.5695E-14 | -0.334554 | 0.851 | 0.93 | 5.4859E-10 | TEC_GCLM | RPL10 |
| 1.3966E-21 | -0.3346459 | 0.234 | 0.37 | 2.9818E-17 | TEC_GCLM | ATP5ME |
| 4.3866E-20 | -0.3361604 | 0.841 | 0.927 | 9.3655E-16 | TEC_GCLM | RPL34 |
| 1.2761E-25 | -0.3365837 | 0.559 | 0.704 | 2.7246E-21 | TEC_GCLM | GPX4 |
| 8.6778E-17 | -0.3366693 | 0.289 | 0.396 | 1.8527E-12 | TEC_GCLM | ATP5IF1 |
| 1.1116E-21 | -0.3369268 | 0.413 | 0.535 | 2.3733E-17 | TEC_GCLM | ATP5MC2 |
| 1.8029E-18 | -0.3373582 | 0.195 | 0.312 | 3.8493E-14 | TEC_GCLM | C9orf16 |
| 6.9205E-14 | -0.3381441 | 0.103 | 0.194 | 1.4775E-09 | TEC_GCLM | ADIRF |
| 9.5285E-07 | -0.3396733 | 0.252 | 0.31 | 0.02034344 | TEC_GCLM | PMAIP1 |
| 1.0521E-22 | -0.3407217 | 0.783 | 0.907 | 2.2463E-18 | TEC_GCLM | RPL35A |
| 3.1625E-25 | -0.3431698 | 0.561 | 0.692 | 6.7519E-21 | TEC_GCLM | CLIC1 |
| 3.3323E-17 | -0.3455527 | 0.615 | 0.708 | 7.1146E-13 | TEC_GCLM | HMGB1 |
| 1.2211E-25 | -0.3456628 | 0.694 | 0.814 | 2.6072E-21 | TEC_GCLM | RPL31 |
| 5.462E-21 | -0.3465723 | 0.869 | 0.948 | 1.1661E-16 | TEC_GCLM | RPS24 |
| 3.2788E-19 | -0.3469201 | 0.079 | 0.184 | 7.0003E-15 | TEC_GCLM | FOXC1 |
| 2.0356E-05 | -0.3478935 | 0.471 | 0.494 | 0.43460406 | TEC_GCLM | ATP1B1 |
| 9.5685E-17 | -0.3492704 | 0.665 | 0.762 | 2.0429E-12 | TEC_GCLM | RPS26 |
| 1.7127E-24 | -0.3498051 | 0.401 | 0.54 | 3.6566E-20 | TEC_GCLM | COX6C |
| 2.0756E-23 | -0.3498471 | 0.05 | 0.165 | 4.4314E-19 | TEC_GCLM | IGFBP2 |

| | | | | | | |
|---|---|---|---|---|---|---|
| 5.9516E-23 | -0.351253 | 0.203 | 0.34 | 1.2707E-18 | TEC_GCLM | UQCRQ |
| 1.1542E-11 | -0.3516366 | 0.342 | 0.426 | 2.4643E-07 | TEC_GCLM | LGALS3 |
| 1.5064E-15 | -0.3518834 | 0.051 | 0.136 | 3.2161E-11 | TEC_GCLM | TNFRSF11B |
| 2.1625E-23 | -0.3519906 | 0.851 | 0.941 | 4.617E-19 | TEC_GCLM | RPL6 |
| 1.8866E-20 | -0.3521144 | 0.043 | 0.143 | 4.0279E-16 | TEC_GCLM | AC025164.1 |
| 0.00106744 | -0.3553601 | 0.391 | 0.415 | 1 | TEC_GCLM | PHLDA2 |
| 1.1159E-06 | -0.3556578 | 0.217 | 0.271 | 0.02382514 | TEC_GCLM | TUBA1A |
| 4.3388E-25 | -0.3556812 | 0.218 | 0.361 | 9.2632E-21 | TEC_GCLM | MZT2B |
| 2.1169E-14 | -0.3561024 | 0.307 | 0.398 | 4.5197E-10 | TEC_GCLM | CD151 |
| 4.9919E-07 | -0.3589522 | 0.108 | 0.165 | 0.01065779 | TEC_GCLM | G0S2 |
| 1.9798E-22 | -0.3597077 | 0.798 | 0.894 | 4.2268E-18 | TEC_GCLM | RPS29 |
| 2.4649E-10 | -0.3599595 | 0.041 | 0.1 | 5.2625E-06 | TEC_GCLM | CCL2 |
| 1.1966E-24 | -0.3628312 | 0.856 | 0.942 | 2.5547E-20 | TEC_GCLM | RPL37 |
| 1.0437E-18 | -0.3632947 | 0.495 | 0.59 | 2.2284E-14 | TEC_GCLM | ARPC3 |
| 1.9957E-24 | -0.3642642 | 0.25 | 0.399 | 4.2609E-20 | TEC_GCLM | POLR2L |
| 4.6579E-21 | -0.3647965 | 0.245 | 0.371 | 9.9446E-17 | TEC_GCLM | PSME1 |
| 2.6596E-14 | -0.3649139 | 0.842 | 0.922 | 5.6782E-10 | TEC_GCLM | RPLP0 |
| 1.2384E-21 | -0.3660579 | 0.854 | 0.937 | 2.6439E-17 | TEC_GCLM | RPL7A |
| 1.1506E-16 | -0.3678025 | 0.809 | 0.889 | 2.4566E-12 | TEC_GCLM | RPL39 |
| 2.1349E-16 | -0.3678618 | 0.724 | 0.807 | 4.5579E-12 | TEC_GCLM | CD24 |
| 1.2422E-16 | -0.3679076 | 0.028 | 0.106 | 2.6521E-12 | TEC_GCLM | CXCL3 |
| 2.1403E-29 | -0.3683054 | 0.538 | 0.673 | 4.5696E-25 | TEC_GCLM | UQCRB |
| 1.2016E-22 | -0.3690646 | 0.262 | 0.392 | 2.5653E-18 | TEC_GCLM | NDUFB2 |
| 4.0484E-27 | -0.3718785 | 0.671 | 0.791 | 8.6434E-23 | TEC_GCLM | COX4I1 |
| 2.8339E-23 | -0.373374 | 0.111 | 0.237 | 6.0504E-19 | TEC_GCLM | IFITM2 |
| 4.6161E-17 | -0.3741744 | 0.281 | 0.393 | 9.8554E-13 | TEC_GCLM | PTMS |
| 3.6767E-25 | -0.3742289 | 0.398 | 0.529 | 7.8497E-21 | TEC_GCLM | OST4 |
| 1.0128E-22 | -0.3745879 | 0.668 | 0.783 | 2.1623E-18 | TEC_GCLM | MYL12A |
| 9.3842E-21 | -0.3747074 | 0.256 | 0.38 | 2.0035E-16 | TEC_GCLM | TAGLN2 |
| 2.854E-24 | -0.3754813 | 0.843 | 0.941 | 6.0933E-20 | TEC_GCLM | RPS9 |
| 1.7904E-31 | -0.3756619 | 0.603 | 0.772 | 3.8225E-27 | TEC_GCLM | DYNLL1 |
| 1.08E-17 | -0.3773838 | 0.771 | 0.874 | 2.3059E-13 | TEC_GCLM | RPS4X |
| 9.3541E-12 | -0.3783936 | 0.121 | 0.207 | 1.9971E-07 | TEC_GCLM | ANXA1 |
| 5.8727E-25 | -0.378588 | 0.86 | 0.959 | 1.2538E-20 | TEC_GCLM | RPS8 |
| 5.2141E-14 | -0.3790908 | 0.762 | 0.833 | 1.1132E-09 | TEC_GCLM | HLA-B |
| 1.3029E-25 | -0.3818908 | 0.261 | 0.406 | 2.7817E-21 | TEC_GCLM | ATP5F1D |
| 7.0563E-23 | -0.3834229 | 0.532 | 0.654 | 1.5065E-18 | TEC_GCLM | PEBP1 |
| 3.4654E-30 | -0.3836049 | 0.423 | 0.573 | 7.3986E-26 | TEC_GCLM | COX6B1 |
| 1.1226E-13 | -0.3850537 | 0.226 | 0.323 | 2.3968E-09 | TEC_GCLM | MAL |
| 3.7336E-34 | -0.3853849 | 0.533 | 0.693 | 7.9713E-30 | TEC_GCLM | COX6A1 |
| 1.7153E-19 | -0.3871502 | 0.704 | 0.806 | 3.6621E-15 | TEC_GCLM | HLA-C |
| 8.5827E-21 | -0.3875308 | 0.115 | 0.235 | 1.8324E-16 | TEC_GCLM | CD74 |
| 4.7583E-11 | -0.3881285 | 0.765 | 0.85 | 1.0159E-06 | TEC_GCLM | RPSA |
| 1.6613E-29 | -0.3904641 | 0.749 | 0.879 | 3.547E-25 | TEC_GCLM | RPL10A |
| 2.8186E-14 | -0.3905421 | 0.501 | 0.614 | 6.0178E-10 | TEC_GCLM | SOD2 |
| 4.7066E-21 | -0.3932457 | 0.048 | 0.153 | 1.0049E-16 | TEC_GCLM | ID1 |
| 2.6763E-24 | -0.393372 | 0.801 | 0.922 | 5.7139E-20 | TEC_GCLM | RPL29 |
| 5.2001E-28 | -0.4067336 | 0.906 | 0.979 | 1.1102E-23 | TEC_GCLM | RPL13 |
| 2.2448E-19 | -0.4087082 | 0.075 | 0.18 | 4.7927E-15 | TEC_GCLM | FOSB |
| 1.5794E-28 | -0.4097878 | 0.323 | 0.473 | 3.372E-24 | TEC_GCLM | COX8A |

| | | | | | | |
|---|---|---|---|---|---|---|
| 2.0047E-11 | -0.412459 | 0.179 | 0.259 | 4.2801E-07 | TEC_GCLM | ID2 |
| 1.3899E-12 | -0.4126343 | 0.036 | 0.103 | 2.9675E-08 | TEC_GCLM | SLC12A1 |
| 1.4436E-16 | -0.4131312 | 0.302 | 0.407 | 3.0821E-12 | TEC_GCLM | JPT1 |
| 4.2292E-34 | -0.4150507 | 0.432 | 0.596 | 9.0294E-30 | TEC_GCLM | COX5B |
| 1.0536E-15 | -0.4184895 | 0.588 | 0.662 | 2.2495E-11 | TEC_GCLM | HSPA8 |
| 1.5187E-05 | -0.4191881 | 0.2 | 0.241 | 0.32423348 | TEC_GCLM | INTS6 |
| 4.6385E-24 | -0.4230818 | 0.837 | 0.942 | 9.9033E-20 | TEC_GCLM | RPL32 |
| 1.0955E-26 | -0.4234104 | 0.865 | 0.947 | 2.3389E-22 | TEC_GCLM | RPL37A |
| 1.4303E-22 | -0.4239454 | 0.754 | 0.862 | 3.0537E-18 | TEC_GCLM | MYL12B |
| 2.9039E-07 | -0.424559 | 0.945 | 0.981 | 0.00619983 | TEC_GCLM | FTH1 |
| 1.5287E-25 | -0.4257029 | 0.266 | 0.409 | 3.2637E-21 | TEC_GCLM | DBI |
| 1.0067E-30 | -0.425844 | 0.781 | 0.908 | 2.1493E-26 | TEC_GCLM | RPL23A |
| 7.365E-09 | -0.4266403 | 0.584 | 0.655 | 0.00015724 | TEC_GCLM | KRT7 |
| 3.213E-18 | -0.4342994 | 0.04 | 0.131 | 6.8598E-14 | TEC_GCLM | CXCL14 |
| 9.4535E-39 | -0.4373117 | 0.36 | 0.543 | 2.0183E-34 | TEC_GCLM | ELOB |
| 3.6338E-21 | -0.4374786 | 0.217 | 0.343 | 7.7581E-17 | TEC_GCLM | CHCHD10 |
| 7.4518E-13 | -0.437679 | 0.044 | 0.114 | 1.5909E-08 | TEC_GCLM | S100A9 |
| 1.4881E-37 | -0.4395788 | 0.516 | 0.677 | 3.177E-33 | TEC_GCLM | SOD1 |
| 7.1079E-38 | -0.4397911 | 0.882 | 0.939 | 1.5175E-33 | TEC_GCLM | PTMA |
| 6.5287E-37 | -0.4437811 | 0.852 | 0.954 | 1.3939E-32 | TEC_GCLM | RPS15 |
| 2.8416E-26 | -0.446082 | 0.884 | 0.968 | 6.0668E-22 | TEC_GCLM | RPS18 |
| 4.5245E-30 | -0.4468351 | 0.759 | 0.874 | 9.6599E-26 | TEC_GCLM | UBA52 |
| 8.3511E-09 | -0.4477918 | 0.197 | 0.258 | 0.0001783 | TEC_GCLM | IER5 |
| 2.8008E-33 | -0.4481322 | 0.815 | 0.921 | 5.9797E-29 | TEC_GCLM | RPL35 |
| 1.249E-20 | -0.4502739 | 0.267 | 0.384 | 2.6667E-16 | TEC_GCLM | LDHB |
| 1.9884E-31 | -0.4509206 | 0.198 | 0.358 | 4.2452E-27 | TEC_GCLM | MIF |
| 6.3234E-25 | -0.452129 | 0.859 | 0.932 | 1.3501E-20 | TEC_GCLM | ACTG1 |
| 1.3545E-30 | -0.4531033 | 0.906 | 0.97 | 2.8919E-26 | TEC_GCLM | RPS19 |
| 7.3088E-06 | -0.4549446 | 0.076 | 0.121 | 0.15604313 | TEC_GCLM | CXCL8 |
| 6.1436E-41 | -0.4553678 | 0.412 | 0.594 | 1.3117E-36 | TEC_GCLM | PRDX5 |
| 7.4118E-32 | -0.4593369 | 0.826 | 0.94 | 1.5824E-27 | TEC_GCLM | RPL12 |
| 8.4056E-24 | -0.4594036 | 0.838 | 0.936 | 1.7946E-19 | TEC_GCLM | RPL18A |
| 8.0056E-17 | -0.4613136 | 0.167 | 0.273 | 1.7092E-12 | TEC_GCLM | DUSP1 |
| 1.9298E-15 | -0.4634903 | 0.211 | 0.307 | 4.1201E-11 | TEC_GCLM | MARCKSL1 |
| 1.1274E-11 | -0.4658517 | 0.207 | 0.287 | 2.4069E-07 | TEC_GCLM | BTG2 |
| 1.5721E-41 | -0.46597 | 0.139 | 0.332 | 3.3565E-37 | TEC_GCLM | PRDX2 |
| 7.8068E-28 | -0.4667189 | 0.902 | 0.978 | 1.6668E-23 | TEC_GCLM | RPLP1 |
| 1.9794E-45 | -0.4722571 | 0.616 | 0.782 | 4.226E-41 | TEC_GCLM | COX7C |
| 1.5443E-37 | -0.4725733 | 0.627 | 0.756 | 3.2971E-33 | TEC_GCLM | HINT1 |
| 5.4389E-31 | -0.4755054 | 0.893 | 0.967 | 1.1612E-26 | TEC_GCLM | RPL28 |
| 1.7398E-28 | -0.4759884 | 0.711 | 0.812 | 3.7144E-24 | TEC_GCLM | CFL1 |
| 4.9921E-35 | -0.4767887 | 0.853 | 0.948 | 1.0658E-30 | TEC_GCLM | RPL19 |
| 6.2246E-27 | -0.4772164 | 0.143 | 0.288 | 1.329E-22 | TEC_GCLM | CYB5A |
| 7.5847E-07 | -0.4780378 | 0.087 | 0.14 | 0.01619324 | TEC_GCLM | SERPINA1 |
| 6.1591E-11 | -0.4797123 | 0.045 | 0.106 | 1.315E-06 | TEC_GCLM | APOE |
| 2.8453E-40 | -0.4805084 | 0.675 | 0.831 | 6.0747E-36 | TEC_GCLM | UBB |
| 1.0602E-37 | -0.4821094 | 0.836 | 0.942 | 2.2635E-33 | TEC_GCLM | RPL3 |
| 3.0055E-23 | -0.4835852 | 0.629 | 0.741 | 6.4168E-19 | TEC_GCLM | PFN1 |
| 2.9778E-24 | -0.4836432 | 0.112 | 0.245 | 6.3577E-20 | TEC_GCLM | FXYD2 |
| 1.1932E-35 | -0.484982 | 0.586 | 0.744 | 2.5475E-31 | TEC_GCLM | NDUFA4 |

| | | | | | | |
|---|---|---|---|---|---|---|
| 2.4434E-26 | -0.4904777 | 0.857 | 0.96 | 5.2166E-22 | TEC_GCLM | RPS12 |
| 4.0923E-43 | -0.4935394 | 0.832 | 0.937 | 8.7371E-39 | TEC_GCLM | RPL36 |
| 2.7055E-05 | -0.4974154 | 0.197 | 0.239 | 0.57761606 | TEC_GCLM | CST6 |
| 1.4303E-18 | -0.4977707 | 0.853 | 0.873 | 3.0536E-14 | TEC_GCLM | SAT1 |
| 7.912E-27 | -0.4996442 | 0.311 | 0.45 | 1.6892E-22 | TEC_GCLM | PPP1R14B |
| 1.3231E-14 | -0.5010864 | 0.159 | 0.25 | 2.8247E-10 | TEC_GCLM | ID4 |
| 2.9059E-51 | -0.5068844 | 0.843 | 0.966 | 6.204E-47 | TEC_GCLM | RPS27 |
| 2.3703E-20 | -0.5079991 | 0.526 | 0.64 | 5.0606E-16 | TEC_GCLM | TPM1 |
| 1.0893E-31 | -0.5137911 | 0.741 | 0.848 | 2.3256E-27 | TEC_GCLM | CD63 |
| 2.9463E-12 | -0.5201439 | 0.346 | 0.421 | 6.2903E-08 | TEC_GCLM | SRSF7 |
| 5.0187E-24 | -0.5246486 | 0.062 | 0.178 | 1.0715E-19 | TEC_GCLM | EGLN3 |
| 9.1236E-49 | -0.5250373 | 0.668 | 0.821 | 1.9479E-44 | TEC_GCLM | ATP5F1E |
| 2.3534E-47 | -0.5316557 | 0.379 | 0.581 | 5.0245E-43 | TEC_GCLM | CST3 |
| 3.0565E-21 | -0.5347591 | 0.083 | 0.194 | 6.5256E-17 | TEC_GCLM | PIK3R3 |
| 1.2308E-39 | -0.5392929 | 0.858 | 0.946 | 2.6278E-35 | TEC_GCLM | RPS28 |
| 4.1873E-39 | -0.5401577 | 0.849 | 0.952 | 8.9398E-35 | TEC_GCLM | RPL8 |
| 6.1801E-11 | -0.540779 | 0.248 | 0.333 | 1.3194E-06 | TEC_GCLM | MT1E |
| 1.4924E-24 | -0.5438658 | 0.152 | 0.286 | 3.1863E-20 | TEC_GCLM | CITED2 |
| 1.4513E-37 | -0.5467694 | 0.881 | 0.963 | 3.0985E-33 | TEC_GCLM | RPS2 |
| 1.2645E-06 | -0.5471563 | 0.209 | 0.265 | 0.02699688 | TEC_GCLM | NUPR1 |
| 1.2318E-27 | -0.5487443 | 0.189 | 0.337 | 2.6298E-23 | TEC_GCLM | CYBA |
| 9.1394E-13 | -0.5489591 | 0.322 | 0.396 | 1.9513E-08 | TEC_GCLM | TUBA1B |
| 2.1137E-41 | -0.5506963 | 0.812 | 0.916 | 4.5128E-37 | TEC_GCLM | MYL6 |
| 1.0717E-43 | -0.5512476 | 0.702 | 0.847 | 2.2881E-39 | TEC_GCLM | PPIA |
| 2.0757E-24 | -0.5585943 | 0.1 | 0.224 | 4.4316E-20 | TEC_GCLM | SOCS1 |
| 7.4559E-12 | -0.5625198 | 0.739 | 0.804 | 1.5918E-07 | TEC_GCLM | KRT18 |
| 1.4225E-20 | -0.5636321 | 0.146 | 0.27 | 3.037E-16 | TEC_GCLM | EGR1 |
| 8.2947E-11 | -0.5638474 | 0.259 | 0.333 | 1.7709E-06 | TEC_GCLM | IL32 |
| 3.7851E-19 | -0.5687753 | 0.111 | 0.22 | 8.0811E-15 | TEC_GCLM | ADAMTS1 |
| 1.5396E-23 | -0.5707975 | 0.128 | 0.256 | 3.2871E-19 | TEC_GCLM | PDZK1IP1 |
| 1.1876E-35 | -0.5724647 | 0.058 | 0.215 | 2.5355E-31 | TEC_GCLM | SGK1 |
| 3.3609E-11 | -0.5727308 | 0.24 | 0.307 | 7.1754E-07 | TEC_GCLM | PLK2 |
| 3.2725E-31 | -0.585266 | 0.191 | 0.356 | 6.9868E-27 | TEC_GCLM | ZFP36L2 |
| 4.7611E-13 | -0.5865076 | 0.094 | 0.179 | 1.0165E-08 | TEC_GCLM | S100A4 |
| 6.4473E-46 | -0.5873249 | 0.906 | 0.973 | 1.3765E-41 | TEC_GCLM | RPL41 |
| 2.815E-26 | -0.5919076 | 0.856 | 0.928 | 6.0101E-22 | TEC_GCLM | ACTB |
| 4.8429E-31 | -0.5951449 | 0.14 | 0.299 | 1.034E-26 | TEC_GCLM | CEBPD |
| 3.8175E-12 | -0.6020175 | 0.423 | 0.491 | 8.1504E-08 | TEC_GCLM | TSPAN1 |
| 1.0737E-13 | -0.6034349 | 0.162 | 0.254 | 2.2924E-09 | TEC_GCLM | MT1X |
| 5.1865E-18 | -0.6046308 | 0.18 | 0.285 | 1.1073E-13 | TEC_GCLM | PPP1R10 |
| 2.1377E-26 | -0.60907 | 0.13 | 0.266 | 4.564E-22 | TEC_GCLM | IER5L |
| 3.8548E-12 | -0.6342421 | 0.625 | 0.719 | 8.2301E-08 | TEC_GCLM | MT2A |
| 8.089E-53 | -0.6521865 | 0.501 | 0.683 | 1.727E-48 | TEC_GCLM | TPI1 |
| 1.0609E-08 | -0.6598716 | 0.808 | 0.833 | 0.00022651 | TEC_GCLM | ANXA2 |
| 1.96E-17 | -0.6627962 | 0.291 | 0.4 | 4.1847E-13 | TEC_GCLM | NFKBIA |
| 1.2954E-35 | -0.6725907 | 0.312 | 0.481 | 2.7657E-31 | TEC_GCLM | IFITM3 |
| 1.9217E-16 | -0.6863814 | 0.178 | 0.287 | 4.1028E-12 | TEC_GCLM | LGALS1 |
| 2.9161E-37 | -0.6866006 | 0.812 | 0.862 | 6.2258E-33 | TEC_GCLM | H3F3B |
| 7.2983E-26 | -0.6872342 | 0.187 | 0.323 | 1.5582E-21 | TEC_GCLM | GLUL |
| 8.9533E-09 | -0.7049606 | 0.072 | 0.132 | 0.00019115 | TEC_GCLM | GPX3 |

| | | | | | | |
|---|---|---|---|---|---|---|
| 3.302E-24 | -0.7089713 | 0.206 | 0.341 | 7.0498E-20 | TEC_GCLM | TIMP1 |
| 6.1159E-87 | -0.7185792 | 0.577 | 0.811 | 1.3058E-82 | TEC_GCLM | PPDPF |
| 1.2648E-26 | -0.7220171 | 0.261 | 0.397 | 2.7004E-22 | TEC_GCLM | JUNB |
| 4.8482E-17 | -0.7226772 | 0.031 | 0.113 | 1.0351E-12 | TEC_GCLM | FXYD4 |
| 3.3308E-20 | -0.7230218 | 0.23 | 0.341 | 7.1112E-16 | TEC_GCLM | RHOB |
| 5.1376E-47 | -0.7354169 | 0.892 | 0.954 | 1.0969E-42 | TEC_GCLM | B2M |
| 2.0189E-42 | -0.7403862 | 0.474 | 0.684 | 4.3103E-38 | TEC_GCLM | CRYAB |
| 2.714E-25 | -0.7433628 | 0.568 | 0.645 | 5.7943E-21 | TEC_GCLM | JUND |
| 8.3074E-26 | -0.7499981 | 0.79 | 0.877 | 1.7736E-21 | TEC_GCLM | S100A10 |
| 4.0805E-20 | -0.7540569 | 0.496 | 0.6 | 8.7119E-16 | TEC_GCLM | KRT19 |
| 0.00355712 | -0.7617308 | 0.23 | 0.258 | 1 | TEC_GCLM | LCN2 |
| 3.0591E-12 | -0.7640835 | 0.166 | 0.249 | 6.5311E-08 | TEC_GCLM | TFPI2 |
| 4.9579E-93 | -0.7705916 | 0.755 | 0.912 | 1.0585E-88 | TEC_GCLM | SERF2 |
| 1.9585E-17 | -0.7721294 | 0.553 | 0.61 | 4.1815E-13 | TEC_GCLM | GADD45B |
| 2.1031E-27 | -0.7889693 | 0.14 | 0.283 | 4.49E-23 | TEC_GCLM | TCIM |
| 1.3328E-36 | -0.7893834 | 0.124 | 0.291 | 2.8456E-32 | TEC_GCLM | RASD1 |
| 2.8263E-24 | -0.7977967 | 0.642 | 0.735 | 6.0341E-20 | TEC_GCLM | KRT8 |
| 4.5111E-26 | -0.798559 | 0.489 | 0.595 | 9.6312E-22 | TEC_GCLM | ELF3 |
| 2.4305E-18 | -0.7993448 | 0.504 | 0.57 | 5.1891E-14 | TEC_GCLM | IER2 |
| 5.4031E-24 | -0.8050019 | 0.571 | 0.681 | 1.1536E-19 | TEC_GCLM | SH3BGRL3 |
| 4.719E-62 | -0.8093344 | 0.756 | 0.888 | 1.0075E-57 | TEC_GCLM | ITM2B |
| 2.4541E-28 | -0.8096045 | 0.107 | 0.245 | 5.2396E-24 | TEC_GCLM | CSKMT |
| 7.6441E-19 | -0.8156396 | 0.372 | 0.479 | 1.632E-14 | TEC_GCLM | C12orf75 |
| 1.7023E-28 | -0.818885 | 0.145 | 0.289 | 3.6345E-24 | TEC_GCLM | HES1 |
| 4.7668E-54 | -0.8194096 | 0.819 | 0.933 | 1.0177E-49 | TEC_GCLM | TMSB10 |
| 9.7948E-80 | -0.8257745 | 0.476 | 0.735 | 2.0912E-75 | TEC_GCLM | HSPB1 |
| 3.7671E-57 | -0.8599322 | 0.748 | 0.884 | 8.0427E-53 | TEC_GCLM | GSTP1 |
| 1.7995E-35 | -0.9074854 | 0.136 | 0.303 | 3.842E-31 | TEC_GCLM | FOS |
| 1.3821E-59 | -0.9079576 | 0.778 | 0.903 | 2.9507E-55 | TEC_GCLM | GAPDH |
| 2.6366E-43 | -0.9144798 | 0.455 | 0.639 | 5.6291E-39 | TEC_GCLM | VIM |
| 4.4607E-27 | -0.9458605 | 0.174 | 0.308 | 9.5236E-23 | TEC_GCLM | HEXIM1 |
| 3.7741E-67 | -0.9604797 | 0.648 | 0.829 | 8.0577E-63 | TEC_GCLM | IGFBP7 |
| 6.5784E-38 | -0.9625887 | 0.859 | 0.924 | 1.4045E-33 | TEC_GCLM | S100A11 |
| 2.8048E-05 | -0.9737611 | 0.156 | 0.201 | 0.5988338 | TEC_GCLM | MT1G |
| 3.0115E-58 | -0.9780078 | 0.408 | 0.617 | 6.4296E-54 | TEC_GCLM | CLU |
| 1.9894E-35 | -1.0118245 | 0.293 | 0.453 | 4.2474E-31 | TEC_GCLM | IER3 |
| 1.1446E-56 | -1.0179508 | 0.913 | 0.958 | 2.4437E-52 | TEC_GCLM | S100A6 |
| 3.3755E-34 | -1.0252924 | 0.125 | 0.283 | 7.2066E-30 | TEC_GCLM | HSPA2 |
| 3.6839E-16 | -1.0328252 | 0.6 | 0.636 | 7.8651E-12 | TEC_GCLM | JUN |
| 1.3637E-50 | -1.0331551 | 0.749 | 0.879 | 2.9115E-46 | TEC_GCLM | TMSB4X |
| 4.3642E-13 | -1.0627909 | 0.636 | 0.682 | 9.3176E-09 | TEC_GCLM | MTRNR2L12 |
| 9.8109E-12 | -1.0870525 | 0.318 | 0.398 | 2.0946E-07 | TEC_GCLM | SLPI |
| 8.0983E-18 | -1.1020593 | 0.126 | 0.234 | 1.729E-13 | TEC_GCLM | DEFB1 |
| 1.9559E-59 | -1.1377272 | 0.446 | 0.663 | 4.1758E-55 | TEC_GCLM | WFDC2 |
| 1.3261E-31 | -1.1447104 | 0.147 | 0.295 | 2.8312E-27 | TEC_GCLM | MAFB |
| 1.6453E-44 | -1.1816922 | 0.204 | 0.393 | 3.5126E-40 | TEC_GCLM | HIST1H4C |
| 7.1675E-39 | -1.2133855 | 0.383 | 0.553 | 1.5303E-34 | TEC_GCLM | SOX4 |
| 6.2111E-26 | -1.4154376 | 0.47 | 0.611 | 1.3261E-21 | TEC_GCLM | SPP1 |
| 2.0256E-33 | -1.6203226 | 0.341 | 0.505 | 4.3247E-29 | TEC_GCLM | S100A2 |
| 1.2346E-25 | -1.7133522 | 0.514 | 0.648 | 2.6359E-21 | TEC_GCLM | MMP7 |

| | | | | | | |
|---|---|---|---|---|---|---|
| 2.2713E-40 | -3.4412669 | 0.506 | 0.606 | 4.8492E-36 | TEC_GCLM | PLCG2 |
| 1.0452E-57 | 2.56634007 | 0.708 | 0.131 | 2.2314E-53 | TEC_IFIT | IFIT2 |
| 3.0712E-53 | 2.42757316 | 0.694 | 0.135 | 6.557E-49 | TEC_IFIT | ISG15 |
| 2.0948E-74 | 2.28946694 | 0.778 | 0.119 | 4.4725E-70 | TEC_IFIT | IFIT3 |
| 5.406E-111 | 1.79042594 | 0.736 | 0.069 | 1.154E-106 | TEC_IFIT | IFIT1 |
| 2.3317E-28 | 1.3442326 | 0.153 | 0.011 | 4.9782E-24 | TEC_IFIT | CXCL10 |
| 3.7165E-50 | 1.30612261 | 0.639 | 0.109 | 7.9347E-46 | TEC_IFIT | ISG20 |
| 2.4557E-08 | 1.13075461 | 0.583 | 0.325 | 0.00052429 | TEC_IFIT | IL32 |
| 1.5605E-70 | 1.01968602 | 0.181 | 0.006 | 3.3316E-66 | TEC_IFIT | APOBEC3A |
| 1.0677E-09 | 1.00983208 | 0.833 | 0.701 | 2.2795E-05 | TEC_IFIT | CLDN4 |
| 3.6673E-33 | 0.9862905 | 0.528 | 0.102 | 7.8297E-29 | TEC_IFIT | RSAD2 |
| 5.5151E-26 | 0.93774124 | 0.375 | 0.068 | 1.1775E-21 | TEC_IFIT | HERC5 |
| 4.1182E-14 | 0.91711804 | 0.556 | 0.216 | 8.7924E-10 | TEC_IFIT | PARP14 |
| 1.0773E-09 | 0.90390919 | 0.986 | 0.949 | 2.3E-05 | TEC_IFIT | B2M |
| 1.8163E-11 | 0.88438767 | 0.694 | 0.389 | 3.8777E-07 | TEC_IFIT | RAB11FIP1 |
| 2.1741E-09 | 0.87336334 | 0.861 | 0.661 | 4.6417E-05 | TEC_IFIT | TACSTD2 |
| 5.9146E-10 | 0.86230541 | 0.389 | 0.151 | 1.2628E-05 | TEC_IFIT | WARS |
| 3.2024E-09 | 0.84908925 | 0.472 | 0.211 | 6.8371E-05 | TEC_IFIT | GLRX |
| 1.4483E-29 | 0.84213 | 0.472 | 0.093 | 3.0922E-25 | TEC_IFIT | MX1 |
| 1.7519E-06 | 0.82205966 | 0.792 | 0.604 | 0.03740331 | TEC_IFIT | SOD2 |
| 1.2348E-79 | 0.78682037 | 0.347 | 0.02 | 2.6362E-75 | TEC_IFIT | OASL |
| 1.0054E-16 | 0.77588887 | 0.417 | 0.113 | 2.1466E-12 | TEC_IFIT | RARRES3 |
| 1.9147E-09 | 0.76418763 | 0.333 | 0.113 | 4.0878E-05 | TEC_IFIT | ZC3HAV1 |
| 1.6357E-10 | 0.76129471 | 0.917 | 0.826 | 3.4921E-06 | TEC_IFIT | HLA-B |
| 1.3426E-16 | 0.75709942 | 0.111 | 0.01 | 2.8665E-12 | TEC_IFIT | CXCL11 |
| 0.00013521 | 0.73776874 | 0.5 | 0.306 | 1 | TEC_IFIT | PLAU |
| 1.7674E-11 | 0.7309513 | 0.611 | 0.284 | 3.7733E-07 | TEC_IFIT | PLAUR |
| 4.4039E-08 | 0.72261793 | 0.681 | 0.405 | 0.00094022 | TEC_IFIT | RNF213 |
| 3.7065E-07 | 0.69668436 | 0.542 | 0.298 | 0.00791344 | TEC_IFIT | MARCKSL1 |
| 1.6024E-06 | 0.69571272 | 0.444 | 0.22 | 0.03421136 | TEC_IFIT | SFN |
| 1.9227E-06 | 0.67550749 | 0.514 | 0.283 | 0.04105043 | TEC_IFIT | AKR1B1 |
| 1.8165E-08 | 0.66696563 | 0.431 | 0.185 | 0.00038782 | TEC_IFIT | BIRC3 |
| 6.5303E-16 | 0.66291257 | 0.361 | 0.09 | 1.3942E-11 | TEC_IFIT | IFIT5 |
| 0.00131308 | 0.66062881 | 0.75 | 0.606 | 1 | TEC_IFIT | GADD45A |
| 0.00011765 | 0.65974296 | 0.389 | 0.222 | 1 | TEC_IFIT | GPRC5A |
| 1.566E-09 | 0.65780008 | 0.403 | 0.151 | 3.3433E-05 | TEC_IFIT | RCAN1 |
| 1.7184E-16 | 0.65296387 | 0.431 | 0.124 | 3.6687E-12 | TEC_IFIT | APOL6 |
| 0.00022222 | 0.63424315 | 0.556 | 0.348 | 1 | TEC_IFIT | TXNIP |
| 1.5397E-11 | 0.62772053 | 0.5 | 0.194 | 3.2872E-07 | TEC_IFIT | NCOA7 |
| 2.5564E-08 | 0.62435135 | 0.556 | 0.287 | 0.0005458 | TEC_IFIT | PSME2 |
| 5.9549E-08 | 0.61897839 | 0.972 | 0.874 | 0.00127137 | TEC_IFIT | HLA-A |
| 1.8824E-06 | 0.60608018 | 0.792 | 0.6 | 0.04019004 | TEC_IFIT | CAST |
| 8.107E-07 | 0.60308317 | 0.722 | 0.534 | 0.01730843 | TEC_IFIT | HLA-E |
| 2.4112E-19 | 0.59281759 | 0.333 | 0.068 | 5.1479E-15 | TEC_IFIT | GBP1 |
| 1.4535E-06 | 0.57365402 | 0.625 | 0.357 | 0.03103175 | TEC_IFIT | KLF6 |
| 4.9445E-09 | 0.57229216 | 0.528 | 0.246 | 0.00010556 | TEC_IFIT | RAP2B |
| 4.1126E-05 | 0.56815231 | 0.889 | 0.797 | 0.87804779 | TEC_IFIT | HLA-C |
| 1.9497E-05 | 0.56697482 | 0.208 | 0.077 | 0.41625232 | TEC_IFIT | HSPA6 |
| 1.8254E-05 | 0.56297311 | 0.528 | 0.318 | 0.38972359 | TEC_IFIT | ITGA2 |
| 1.1579E-12 | 0.5607981 | 0.389 | 0.12 | 2.4721E-08 | TEC_IFIT | IRF6 |

| | | | | | | |
|---|---|---|---|---|---|---|
| 5.4928E-06 | 0.55582338 | 0.833 | 0.718 | 0.1172722 | TEC_IFIT | RPL36AL |
| 3.0068E-06 | 0.54711156 | 0.361 | 0.168 | 0.0641956 | TEC_IFIT | CDYL |
| 7.8295E-09 | 0.52088165 | 0.417 | 0.166 | 0.00016716 | TEC_IFIT | STAT1 |
| 0.00010853 | 0.51666576 | 0.861 | 0.715 | 1 | TEC_IFIT | TXN |
| 1.624E-06 | 0.51622642 | 0.847 | 0.677 | 0.03467337 | TEC_IFIT | YWHAZ |
| 1.1026E-25 | 0.51393981 | 0.181 | 0.017 | 2.3541E-21 | TEC_IFIT | GBP4 |
| 0.00525968 | 0.50947044 | 0.389 | 0.238 | 1 | TEC_IFIT | MDM2 |
| 3.39E-05 | 0.507311 | 0.458 | 0.252 | 0.72376607 | TEC_IFIT | IRF1 |
| 1.7131E-05 | 0.50629603 | 0.444 | 0.247 | 0.36575139 | TEC_IFIT | MACC1 |
| 2.2019E-11 | 0.50420157 | 0.486 | 0.19 | 4.701E-07 | TEC_IFIT | PSMB8 |
| 0.00711293 | 0.49315034 | 0.389 | 0.26 | 1 | TEC_IFIT | NUPR1 |
| 8.9481E-08 | 0.49162671 | 0.403 | 0.174 | 0.00191042 | TEC_IFIT | ZNF217 |
| 1.9835E-07 | 0.48646363 | 0.472 | 0.235 | 0.00423484 | TEC_IFIT | NAA20 |
| 6.2815E-09 | 0.47669043 | 0.639 | 0.332 | 0.00013411 | TEC_IFIT | RAB2A |
| 1.0367E-06 | 0.47376555 | 0.583 | 0.318 | 0.02213335 | TEC_IFIT | NDUFB9 |
| 0.00036059 | 0.47323383 | 0.583 | 0.416 | 1 | TEC_IFIT | CCND1 |
| 1.2539E-09 | 0.4720028 | 0.167 | 0.035 | 2.6771E-05 | TEC_IFIT | KLHDC7B |
| 3.5488E-16 | 0.47168564 | 0.347 | 0.084 | 7.5768E-12 | TEC_IFIT | APOL2 |
| 0.00181996 | 0.4672867 | 0.431 | 0.28 | 1 | TEC_IFIT | CLDN3 |
| 0.00065342 | 0.46519362 | 0.486 | 0.34 | 1 | TEC_IFIT | JUP |
| 6.41E-09 | 0.46498222 | 0.292 | 0.096 | 0.00013685 | TEC_IFIT | HLA-DRB1 |
| 0.00320166 | 0.46377285 | 0.306 | 0.182 | 1 | TEC_IFIT | SAR1A |
| 7.9567E-05 | 0.46221864 | 0.875 | 0.758 | 1 | TEC_IFIT | CALM2 |
| 0.00012174 | 0.45739165 | 0.431 | 0.261 | 1 | TEC_IFIT | CFLAR |
| 0.00062908 | 0.45722813 | 0.625 | 0.472 | 1 | TEC_IFIT | SUMO2 |
| 9.7502E-14 | 0.4539292 | 0.264 | 0.059 | 2.0817E-09 | TEC_IFIT | SAMD9 |
| 1.9669E-15 | 0.45289248 | 0.403 | 0.109 | 4.1992E-11 | TEC_IFIT | ZNFX1 |
| 0.00084424 | 0.45238691 | 0.611 | 0.447 | 1 | TEC_IFIT | VDAC2 |
| 1.3687E-05 | 0.45237233 | 0.5 | 0.293 | 0.29221671 | TEC_IFIT | EIF6 |
| 3.655E-85 | 0.44841176 | 0.153 | 0.003 | 7.8034E-81 | TEC_IFIT | IDO1 |
| 3.7425E-05 | 0.44353119 | 0.306 | 0.137 | 0.79903203 | TEC_IFIT | TNFSF10 |
| 0.00038179 | 0.44170059 | 0.167 | 0.063 | 1 | TEC_IFIT | LTB |
| 6.1222E-16 | 0.43846707 | 0.347 | 0.083 | 1.3071E-11 | TEC_IFIT | IFIH1 |
| 0.00021537 | 0.43478394 | 0.597 | 0.419 | 1 | TEC_IFIT | PTTG1IP |
| 6.2901E-05 | 0.43474894 | 0.681 | 0.491 | 1 | TEC_IFIT | ATP1B1 |
| 0.00091021 | 0.4346116 | 0.347 | 0.2 | 1 | TEC_IFIT | TMEM106B |
| 6.8668E-06 | 0.43457614 | 0.75 | 0.503 | 0.146607 | TEC_IFIT | CLDN7 |
| 5.9317E-09 | 0.43355496 | 0.278 | 0.088 | 0.00012664 | TEC_IFIT | IFI6 |
| 4.3501E-05 | 0.43080802 | 0.389 | 0.201 | 0.92875648 | TEC_IFIT | PLSCR1 |
| 5.3007E-23 | 0.42825994 | 0.264 | 0.038 | 1.1317E-18 | TEC_IFIT | IFI44 |
| 0.00893173 | 0.42422438 | 0.347 | 0.217 | 1 | TEC_IFIT | TNFAIP2 |
| 0.00027356 | 0.42245978 | 0.472 | 0.294 | 1 | TEC_IFIT | MIR4435-2HG |
| 2.4944E-06 | 0.42084137 | 0.708 | 0.465 | 0.05325497 | TEC_IFIT | IFITM3 |
| 3.1012E-06 | 0.42027098 | 0.806 | 0.584 | 0.06621163 | TEC_IFIT | EZR |
| 5.1758E-05 | 0.41864384 | 0.458 | 0.257 | 1 | TEC_IFIT | OPTN |
| 4.6823E-08 | 0.41525459 | 0.486 | 0.221 | 0.00099967 | TEC_IFIT | TEAD1 |
| 9.1421E-05 | 0.41234706 | 0.264 | 0.12 | 1 | TEC_IFIT | SH2D4A |
| 0.00116831 | 0.41214371 | 0.847 | 0.723 | 1 | TEC_IFIT | CTSD |
| 0.00207866 | 0.41043147 | 0.403 | 0.267 | 1 | TEC_IFIT | MET |
| 4.4309E-08 | 0.40664666 | 0.472 | 0.214 | 0.00094599 | TEC_IFIT | TMF1 |

| | | | | | | |
|---|---|---|---|---|---|---|
| 8.9235E-06 | 0.40529628 | 0.958 | 0.913 | 0.19051646 | TEC_IFIT | HSP90AB1 |
| 0.00013711 | 0.39888252 | 0.569 | 0.366 | 1 | TEC_IFIT | SBDS |
| 6.9434E-12 | 0.39781854 | 0.375 | 0.115 | 1.4824E-07 | TEC_IFIT | TMEM51 |
| 8.6623E-05 | 0.3957585 | 0.486 | 0.281 | 1 | TEC_IFIT | LITAF |
| 0.00331476 | 0.39505599 | 0.292 | 0.167 | 1 | TEC_IFIT | FGF2 |
| 0.0003449 | 0.39368207 | 0.486 | 0.308 | 1 | TEC_IFIT | DSP |
| 4.5336E-19 | 0.3914872 | 0.222 | 0.033 | 9.6793E-15 | TEC_IFIT | OAS1 |
| 9.4638E-05 | 0.39113944 | 0.431 | 0.24 | 1 | TEC_IFIT | HES4 |
| 2.39E-05 | 0.39025009 | 0.403 | 0.215 | 0.51026472 | TEC_IFIT | PSMA5 |
| 0.00623046 | 0.39019161 | 0.722 | 0.63 | 1 | TEC_IFIT | PABPC1 |
| 4.5727E-05 | 0.38985548 | 0.375 | 0.2 | 0.97626216 | TEC_IFIT | FOXO3 |
| 4.2774E-06 | 0.38812501 | 0.389 | 0.19 | 0.0913217 | TEC_IFIT | ZBTB43 |
| 0.00021696 | 0.38579481 | 0.708 | 0.489 | 1 | TEC_IFIT | DNAJA1 |
| 0.00080659 | 0.38483453 | 0.333 | 0.188 | 1 | TEC_IFIT | EMP1 |
| 3.9061E-13 | 0.38180673 | 0.264 | 0.06 | 8.3395E-09 | TEC_IFIT | DDX58 |
| 1.1139E-06 | 0.3806653 | 0.306 | 0.121 | 0.02378169 | TEC_IFIT | PIK3IP1 |
| 4.364E-05 | 0.37862451 | 0.431 | 0.243 | 0.931715 | TEC_IFIT | TNIP1 |
| 2.9443E-05 | 0.37834442 | 0.653 | 0.405 | 0.62861851 | TEC_IFIT | SDC4 |
| 2.61E-10 | 0.37776321 | 0.222 | 0.055 | 5.5723E-06 | TEC_IFIT | SAMD9L |
| 0.00024058 | 0.37452911 | 0.819 | 0.662 | 1 | TEC_IFIT | RAC1 |
| 0.00020773 | 0.37277675 | 0.444 | 0.267 | 1 | TEC_IFIT | NUDC |
| 0.00044923 | 0.37165143 | 0.514 | 0.33 | 1 | TEC_IFIT | TUBA1C |
| 5.276E-05 | 0.36880892 | 0.347 | 0.168 | 1 | TEC_IFIT | TYMP |
| 5.0977E-05 | 0.36766945 | 0.806 | 0.642 | 1 | TEC_IFIT | YWHAE |
| 9.9377E-06 | 0.36738452 | 0.833 | 0.71 | 0.21216963 | TEC_IFIT | MT2A |
| 2.1924E-07 | 0.3661756 | 0.375 | 0.158 | 0.00468077 | TEC_IFIT | BAZ1A |
| 5.4293E-05 | 0.36456021 | 0.403 | 0.22 | 1 | TEC_IFIT | RBCK1 |
| 1.7672E-05 | 0.36278416 | 0.278 | 0.118 | 0.3772987 | TEC_IFIT | DAPP1 |
| 0.0012108 | 0.36257741 | 0.708 | 0.545 | 1 | TEC_IFIT | BRI3 |
| 0.00235185 | 0.36257451 | 0.694 | 0.523 | 1 | TEC_IFIT | CDC42 |
| 8.6649E-05 | 0.36194777 | 0.472 | 0.282 | 1 | TEC_IFIT | SPINT1 |
| 0.00357499 | 0.36013896 | 0.333 | 0.207 | 1 | TEC_IFIT | SLFN5 |
| 0.00011547 | 0.35858769 | 0.431 | 0.252 | 1 | TEC_IFIT | SF3B2 |
| 4.8507E-09 | 0.35823534 | 0.25 | 0.072 | 0.00010356 | TEC_IFIT | SAA2 |
| 0.00171576 | 0.35792094 | 0.389 | 0.252 | 1 | TEC_IFIT | PSMA3 |
| 0.0013845 | 0.3575786 | 0.569 | 0.388 | 1 | TEC_IFIT | POMP |
| 8.0218E-07 | 0.35741748 | 0.375 | 0.16 | 0.01712663 | TEC_IFIT | BBC3 |
| 0.00022728 | 0.35717814 | 0.778 | 0.566 | 1 | TEC_IFIT | HNRNPC |
| 3.6009E-05 | 0.35713249 | 0.347 | 0.168 | 0.76879201 | TEC_IFIT | METRNL |
| 4.6025E-14 | 0.35679648 | 0.306 | 0.074 | 9.8263E-10 | TEC_IFIT | NKIRAS2 |
| 1.2571E-17 | 0.35673131 | 0.236 | 0.039 | 2.684E-13 | TEC_IFIT | CDKN2A |
| 4.6774E-06 | 0.35661782 | 0.347 | 0.159 | 0.09986302 | TEC_IFIT | TRIM56 |
| 1.2502E-12 | 0.35624011 | 0.25 | 0.057 | 2.6692E-08 | TEC_IFIT | IFI35 |
| 0.00792797 | 0.3548559 | 0.556 | 0.39 | 1 | TEC_IFIT | NFKBIA |
| 1.6726E-07 | 0.35397973 | 0.347 | 0.136 | 0.00357096 | TEC_IFIT | CTSS |
| 1.297E-06 | 0.35380274 | 0.653 | 0.391 | 0.02768992 | TEC_IFIT | TXNRD1 |
| 5.2235E-07 | 0.35336178 | 0.403 | 0.17 | 0.01115207 | TEC_IFIT | DUSP5 |
| 0.00304648 | 0.35300194 | 0.417 | 0.282 | 1 | TEC_IFIT | MSN |
| 0.00017709 | 0.35216177 | 0.208 | 0.086 | 1 | TEC_IFIT | HLA-DRA |
| 0.00057336 | 0.35201914 | 0.528 | 0.342 | 1 | TEC_IFIT | ZFP36L2 |

| | | | | | | |
|---|---|---|---|---|---|---|
| 0.00010101 | 0.35036527 | 0.375 | 0.2 | 1 | TEC_IFIT | UBE2Z |
| 2.3037E-05 | 0.35030345 | 0.278 | 0.121 | 0.49183763 | TEC_IFIT | TAP1 |
| 4.0968E-10 | 0.34922867 | 0.347 | 0.112 | 8.7467E-06 | TEC_IFIT | NECTIN4 |
| 1.9106E-08 | 0.34825229 | 0.347 | 0.125 | 0.00040792 | TEC_IFIT | FMR1 |
| 1.0104E-11 | 0.34787786 | 0.25 | 0.061 | 2.1572E-07 | TEC_IFIT | SP110 |
| 0.0001689 | 0.34677552 | 0.361 | 0.201 | 1 | TEC_IFIT | STK24 |
| 3.2862E-06 | 0.34541213 | 0.333 | 0.146 | 0.07016041 | TEC_IFIT | LGMN |
| 0.00021781 | 0.34408418 | 0.278 | 0.13 | 1 | TEC_IFIT | LDLR |
| 0.00214279 | 0.34178211 | 0.528 | 0.333 | 1 | TEC_IFIT | ZNF292 |
| 1.2027E-05 | 0.34039443 | 0.569 | 0.316 | 0.25676905 | TEC_IFIT | UGCG |
| 3.9988E-06 | 0.33858471 | 0.292 | 0.12 | 0.08537486 | TEC_IFIT | SGPP2 |
| 1.0173E-05 | 0.33766898 | 0.403 | 0.199 | 0.21719842 | TEC_IFIT | CHMP1B |
| 0.00043222 | 0.33731863 | 0.375 | 0.216 | 1 | TEC_IFIT | CHMP4B |
| 0.00292839 | 0.3368762 | 0.431 | 0.278 | 1 | TEC_IFIT | SERPINB1 |
| 4.9549E-08 | 0.33607881 | 0.333 | 0.119 | 0.00105788 | TEC_IFIT | NEURL3 |
| 0.00107568 | 0.33595702 | 0.625 | 0.433 | 1 | TEC_IFIT | PNRC1 |
| 1.9497E-10 | 0.33385855 | 0.306 | 0.09 | 4.1627E-06 | TEC_IFIT | DAB2IP |
| 2.4742E-05 | 0.32988533 | 0.222 | 0.085 | 0.52823575 | TEC_IFIT | TMEM30B |
| 3.3642E-06 | 0.32948463 | 0.292 | 0.12 | 0.07182618 | TEC_IFIT | GBP3 |
| 0.00076482 | 0.32865107 | 0.694 | 0.52 | 1 | TEC_IFIT | ARF1 |
| 0.00283142 | 0.32844277 | 0.667 | 0.496 | 1 | TEC_IFIT | TNFRSF12A |
| 0.00043917 | 0.32616065 | 0.611 | 0.431 | 1 | TEC_IFIT | BZW1 |
| 0.00088171 | 0.32479721 | 0.222 | 0.105 | 1 | TEC_IFIT | VGLL3 |
| 2.0686E-05 | 0.32401785 | 0.333 | 0.157 | 0.44165662 | TEC_IFIT | MAP3K13 |
| 4.5241E-08 | 0.32323675 | 0.292 | 0.099 | 0.00096589 | TEC_IFIT | PSMB9 |
| 0.00104273 | 0.32300584 | 0.597 | 0.406 | 1 | TEC_IFIT | JAK1 |
| 6.0018E-06 | 0.32251581 | 0.333 | 0.149 | 0.12813868 | TEC_IFIT | RHOD |
| 9.4064E-06 | 0.32173314 | 0.222 | 0.081 | 0.2008258 | TEC_IFIT | RIN2 |
| 9.7647E-06 | 0.32110996 | 0.472 | 0.248 | 0.20847714 | TEC_IFIT | EIF3D |
| 1.5161E-06 | 0.32091255 | 0.208 | 0.068 | 0.03236968 | TEC_IFIT | CX3CL1 |
| 0.00083493 | 0.32065227 | 0.347 | 0.207 | 1 | TEC_IFIT | MYO1E |
| 0.0004751 | 0.31973765 | 0.333 | 0.181 | 1 | TEC_IFIT | CD40 |
| 2.8541E-09 | 0.31880212 | 0.181 | 0.042 | 6.0935E-05 | TEC_IFIT | EPSTI1 |
| 4.0718E-16 | 0.31744963 | 0.264 | 0.051 | 8.6934E-12 | TEC_IFIT | NLRC5 |
| 6.4432E-08 | 0.31739189 | 0.194 | 0.052 | 0.00137563 | TEC_IFIT | IFI16 |
| 8.4245E-05 | 0.31719667 | 0.5 | 0.288 | 1 | TEC_IFIT | TJP1 |
| 0.00123133 | 0.31684449 | 0.361 | 0.214 | 1 | TEC_IFIT | PARD6B |
| 0.00022855 | 0.3162061 | 0.292 | 0.144 | 1 | TEC_IFIT | EIF2AK2 |
| 2.2427E-09 | 0.31599684 | 0.361 | 0.125 | 4.7881E-05 | TEC_IFIT | B4GALT5 |
| 2.1481E-09 | 0.31585676 | 0.306 | 0.096 | 4.5862E-05 | TEC_IFIT | CKAP4 |
| 2.5672E-08 | 0.31521686 | 0.292 | 0.096 | 0.0005481 | TEC_IFIT | C1orf116 |
| 0.0001241 | 0.3144871 | 0.333 | 0.173 | 1 | TEC_IFIT | CIR1 |
| 0.00027119 | 0.31341835 | 0.375 | 0.206 | 1 | TEC_IFIT | DNTTIP2 |
| 6.3539E-13 | 0.31265259 | 0.264 | 0.06 | 1.3566E-08 | TEC_IFIT | DDX60L |
| 5.1806E-05 | 0.31108398 | 0.333 | 0.159 | 1 | TEC_IFIT | TRAF4 |
| 0.00082059 | 0.31103549 | 0.25 | 0.129 | 1 | TEC_IFIT | NTMT1 |
| 0.00017764 | 0.30881586 | 0.819 | 0.603 | 1 | TEC_IFIT | PSMA7 |
| 0.00043431 | 0.30834131 | 0.403 | 0.226 | 1 | TEC_IFIT | PRSS8 |
| 0.0077333 | 0.30800831 | 0.306 | 0.197 | 1 | TEC_IFIT | MAP2K3 |
| 0.0007885 | 0.30758824 | 0.472 | 0.296 | 1 | TEC_IFIT | UBXN1 |

| | | | | | | |
|---|---|---|---|---|---|---|
| 5.5902E-05 | 0.30695402 | 0.569 | 0.335 | 1 | TEC_IFIT | PSMB1 |
| 8.4122E-15 | 0.30661094 | 0.222 | 0.04 | 1.796E-10 | TEC_IFIT | RTP4 |
| 5.5326E-05 | 0.30619769 | 0.472 | 0.271 | 1 | TEC_IFIT | ARPC5 |
| 4.3056E-05 | 0.30531115 | 0.486 | 0.264 | 0.91924779 | TEC_IFIT | RIOK3 |
| 1.6695E-05 | 0.30530499 | 0.292 | 0.125 | 0.35643366 | TEC_IFIT | ZNHIT3 |
| 0.0033153 | 0.30495488 | 0.431 | 0.286 | 1 | TEC_IFIT | ANXA3 |
| 0.0028345 | 0.30484047 | 0.431 | 0.271 | 1 | TEC_IFIT | OGA |
| 2.2636E-05 | 0.30421173 | 0.361 | 0.171 | 0.48328693 | TEC_IFIT | SH3BP4 |
| 1.2543E-05 | 0.30311598 | 0.319 | 0.143 | 0.26779374 | TEC_IFIT | DERL2 |
| 0.00012506 | 0.3017028 | 0.472 | 0.274 | 1 | TEC_IFIT | ZNF706 |
| 0.00137023 | 0.30140517 | 0.722 | 0.541 | 1 | TEC_IFIT | GNG5 |
| 1.6641E-05 | 0.30109916 | 0.431 | 0.22 | 0.35528744 | TEC_IFIT | RBM47 |
| 2.4303E-05 | 0.29996891 | 0.819 | 0.572 | 0.51885922 | TEC_IFIT | SET |
| 6.9443E-05 | 0.29918341 | 0.472 | 0.26 | 1 | TEC_IFIT | GSTO1 |
| 5.5555E-05 | 0.29883309 | 0.333 | 0.164 | 1 | TEC_IFIT | GRAMD2B |
| 4.4453E-05 | 0.29739737 | 0.375 | 0.189 | 0.94906888 | TEC_IFIT | RAB9A |
| 0.00275538 | 0.29718568 | 0.5 | 0.336 | 1 | TEC_IFIT | SYF2 |
| 4.0612E-07 | 0.29695486 | 0.375 | 0.154 | 0.00867067 | TEC_IFIT | TGIF1 |
| 0.00289718 | 0.29663344 | 0.306 | 0.176 | 1 | TEC_IFIT | SLC39A1 |
| 0.00017639 | 0.29629891 | 0.472 | 0.279 | 1 | TEC_IFIT | RAB10 |
| 0.00120543 | 0.29573852 | 0.319 | 0.177 | 1 | TEC_IFIT | BCL2L1 |
| 8.658E-12 | 0.29484577 | 0.236 | 0.054 | 1.8485E-07 | TEC_IFIT | DDX60 |
| 0.00155699 | 0.29470929 | 0.958 | 0.836 | 1 | TEC_IFIT | PRDX1 |
| 0.00010722 | 0.29327263 | 0.222 | 0.092 | 1 | TEC_IFIT | TRIM22 |
| 0.00067409 | 0.29025181 | 0.333 | 0.183 | 1 | TEC_IFIT | ELF1 |
| 0.00559376 | 0.28932477 | 0.486 | 0.348 | 1 | TEC_IFIT | ANXA11 |
| 0.00144879 | 0.28923204 | 0.944 | 0.926 | 1 | TEC_IFIT | ACTG1 |
| 5.1205E-06 | 0.28830158 | 0.264 | 0.102 | 0.10932279 | TEC_IFIT | GBP2 |
| 0.00020072 | 0.28828105 | 0.403 | 0.226 | 1 | TEC_IFIT | PSMD7 |
| 1.7534E-06 | 0.28705441 | 0.361 | 0.155 | 0.03743574 | TEC_IFIT | COPA |
| 0.00177889 | 0.28561489 | 0.583 | 0.413 | 1 | TEC_IFIT | NORAD |
| 5.1291E-06 | 0.28535036 | 0.389 | 0.181 | 0.10950532 | TEC_IFIT | BCL10 |
| 0.00239828 | 0.28510176 | 0.528 | 0.369 | 1 | TEC_IFIT | UBE2B |
| 3.5973E-14 | 0.28435445 | 0.181 | 0.029 | 7.6802E-10 | TEC_IFIT | IFI44L |
| 0.0056942 | 0.28405013 | 0.306 | 0.186 | 1 | TEC_IFIT | PPP2CA |
| 3.7372E-05 | 0.28381173 | 0.361 | 0.181 | 0.79790224 | TEC_IFIT | TSG101 |
| 2.6963E-06 | 0.28367869 | 0.25 | 0.091 | 0.05756555 | TEC_IFIT | SDCBP2 |
| 0.00104193 | 0.28355251 | 0.347 | 0.19 | 1 | TEC_IFIT | PHACTR4 |
| 0.00165438 | 0.28218597 | 0.972 | 0.934 | 1 | TEC_IFIT | HSP90AA1 |
| 1.8295E-05 | 0.28206756 | 0.208 | 0.076 | 0.39060724 | TEC_IFIT | VPS26C |
| 0.00437502 | 0.28203362 | 0.5 | 0.347 | 1 | TEC_IFIT | UBE2H |
| 1E-05 | 0.28147765 | 0.514 | 0.276 | 0.21350998 | TEC_IFIT | ARPP19 |
| 0.00155499 | 0.28043342 | 0.556 | 0.365 | 1 | TEC_IFIT | PPP1CB |
| 0.00161668 | 0.28042872 | 0.542 | 0.375 | 1 | TEC_IFIT | VAMP8 |
| 0.00033789 | 0.2803492 | 0.306 | 0.157 | 1 | TEC_IFIT | ERP44 |
| 0.00030272 | 0.27903598 | 0.986 | 0.957 | 1 | TEC_IFIT | EIF1 |
| 0.00531046 | 0.27847328 | 0.875 | 0.871 | 1 | TEC_IFIT | SAT1 |
| 3.3565E-07 | 0.27813703 | 0.278 | 0.099 | 0.00716605 | TEC_IFIT | PROSER2 |
| 5.7585E-06 | 0.27712972 | 0.264 | 0.101 | 0.12294311 | TEC_IFIT | MAP3K8 |
| 0.00456988 | 0.27688921 | 0.597 | 0.443 | 1 | TEC_IFIT | ZFP36L1 |

| | | | | | | |
|---|---|---|---|---|---|---|
| 0.00601661 | 0.27574274 | 0.778 | 0.679 | 1 | TEC_IFIT | EPCAM |
| 8.7653E-13 | 0.27542652 | 0.236 | 0.05 | 1.8714E-08 | TEC_IFIT | BTN2A2 |
| 0.00153546 | 0.27542069 | 0.333 | 0.195 | 1 | TEC_IFIT | SYNGR2 |
| 0.00186153 | 0.27526574 | 0.514 | 0.332 | 1 | TEC_IFIT | EIF3E |
| 0.00852346 | 0.27523956 | 0.653 | 0.56 | 1 | TEC_IFIT | NCL |
| 1.6E-05 | 0.27523767 | 0.375 | 0.179 | 0.34160409 | TEC_IFIT | FAM120A |
| 0.00100176 | 0.27469228 | 0.444 | 0.276 | 1 | TEC_IFIT | SSB |
| 0.00426984 | 0.27412637 | 0.444 | 0.3 | 1 | TEC_IFIT | PAPOLA |
| 0.00190021 | 0.27204464 | 0.5 | 0.336 | 1 | TEC_IFIT | PRELID1 |
| 4.9578E-07 | 0.27160468 | 0.25 | 0.085 | 0.0105849 | TEC_IFIT | RIPK2 |
| 6.2311E-06 | 0.27034193 | 0.208 | 0.071 | 0.13303361 | TEC_IFIT | SLC29A1 |
| 0.00681055 | 0.27030649 | 0.208 | 0.112 | 1 | TEC_IFIT | SULT1C2 |
| 0.00115431 | 0.27019397 | 0.569 | 0.393 | 1 | TEC_IFIT | TAX1BP1 |
| 0.00340081 | 0.2701719 | 0.333 | 0.205 | 1 | TEC_IFIT | IK |
| 0.0091497 | 0.26912468 | 0.306 | 0.195 | 1 | TEC_IFIT | MSMO1 |
| 0.00026166 | 0.26893626 | 0.375 | 0.204 | 1 | TEC_IFIT | CTBP2 |
| 6.7839E-05 | 0.26816104 | 0.431 | 0.233 | 1 | TEC_IFIT | PSMB2 |
| 0.00297208 | 0.26803226 | 0.306 | 0.174 | 1 | TEC_IFIT | IGFBP3 |
| 2.0633E-06 | 0.26762455 | 0.25 | 0.09 | 0.04405186 | TEC_IFIT | TTC9 |
| 0.00036118 | 0.26725372 | 0.5 | 0.3 | 1 | TEC_IFIT | GPBP1 |
| 0.00036083 | 0.26679999 | 0.458 | 0.273 | 1 | TEC_IFIT | CREBRF |
| 0.00340256 | 0.26671644 | 0.458 | 0.299 | 1 | TEC_IFIT | ACTR3 |
| 0.00587452 | 0.26517482 | 0.917 | 0.866 | 1 | TEC_IFIT | RPL23 |
| 1.2622E-06 | 0.26430441 | 0.361 | 0.152 | 0.02694849 | TEC_IFIT | AP1G1 |
| 0.00018834 | 0.26261868 | 0.306 | 0.153 | 1 | TEC_IFIT | MRPL14 |
| 5.2557E-06 | 0.26252402 | 0.347 | 0.154 | 0.1122093 | TEC_IFIT | RAMMET |
| 0.00191419 | 0.26224363 | 0.319 | 0.186 | 1 | TEC_IFIT | GRB2 |
| 0.0038447 | 0.26163862 | 0.347 | 0.216 | 1 | TEC_IFIT | MED10 |
| 0.00025974 | 0.26158508 | 0.319 | 0.157 | 1 | TEC_IFIT | MAN1A2 |
| 0.00463779 | 0.26145549 | 0.5 | 0.337 | 1 | TEC_IFIT | AURKAIP1 |
| 0.00129543 | 0.26086897 | 0.389 | 0.23 | 1 | TEC_IFIT | SCARB2 |
| 0.00934465 | 0.26036938 | 0.347 | 0.225 | 1 | TEC_IFIT | SPATS2L |
| 0.00040062 | 0.25977694 | 0.25 | 0.119 | 1 | TEC_IFIT | XRN1 |
| 3.8382E-06 | 0.25977176 | 0.236 | 0.085 | 0.08194471 | TEC_IFIT | TSPAN15 |
| 0.00226941 | 0.25967865 | 0.319 | 0.181 | 1 | TEC_IFIT | ZFHX3 |
| 0.00860221 | 0.25937171 | 0.972 | 0.938 | 1 | TEC_IFIT | RPL15 |
| 0.00163176 | 0.25904209 | 0.75 | 0.564 | 1 | TEC_IFIT | MAL2 |
| 0.00899586 | 0.25895112 | 0.514 | 0.388 | 1 | TEC_IFIT | TOMM20 |
| 3.3893E-07 | 0.25877536 | 0.222 | 0.07 | 0.00723606 | TEC_IFIT | THEMIS2 |
| 0.00023701 | 0.25777261 | 0.222 | 0.096 | 1 | TEC_IFIT | MAT2B |
| 0.00480727 | 0.25758556 | 0.444 | 0.292 | 1 | TEC_IFIT | NOL7 |
| 1.5502E-07 | 0.2571111 | 0.264 | 0.088 | 0.00330971 | TEC_IFIT | OGFR |
| 0.00014551 | 0.25594584 | 0.292 | 0.138 | 1 | TEC_IFIT | GNA13 |
| 0.00131326 | 0.2558507 | 0.375 | 0.224 | 1 | TEC_IFIT | PSMA1 |
| 2.068E-06 | 0.25463818 | 0.25 | 0.09 | 0.04415283 | TEC_IFIT | IRF2BPL |
| 0.00096366 | 0.25443619 | 0.347 | 0.189 | 1 | TEC_IFIT | PRPF38B |
| 3.2607E-05 | 0.25371801 | 0.264 | 0.112 | 0.69616073 | TEC_IFIT | PSMC2 |
| 0.00281707 | 0.25370183 | 0.736 | 0.586 | 1 | TEC_IFIT | YWHAB |
| 0.00038427 | 0.25352477 | 0.347 | 0.187 | 1 | TEC_IFIT | PSMC6 |
| 0.00061108 | 0.2531956 | 0.931 | 0.826 | 1 | TEC_IFIT | TMBIM6 |

| | | | | | | |
|---|---|---|---|---|---|---|
| 0.00871436 | 0.25315168 | 0.111 | 0.047 | 1 | TEC_IFIT | HLA-DRB5 |
| 0.00841801 | 0.25241358 | 0.167 | 0.082 | 1 | TEC_IFIT | CRNKL1 |
| 0.00025014 | 0.25079926 | 0.292 | 0.144 | 1 | TEC_IFIT | CRB3 |
| 0.00035845 | 0.25074676 | 0.167 | 0.064 | 1 | TEC_IFIT | CCL28 |
| 0.00084087 | 0.25048082 | 0.194 | 0.086 | 1 | TEC_IFIT | CES2 |
| 0.00427081 | -0.2583378 | 0.028 | 0.146 | 1 | TEC_IFIT | COL4A2 |
| 0.00972268 | -0.302638 | 0.014 | 0.108 | 1 | TEC_IFIT | AL355075.4 |
| 0.00999585 | -0.3299888 | 0.361 | 0.487 | 1 | TEC_IFIT | ATP1A1 |
| 0.00901477 | -0.3568224 | 0.042 | 0.151 | 1 | TEC_IFIT | AC007952.4 |
| 0.0072381 | -0.3593291 | 0.042 | 0.153 | 1 | TEC_IFIT | PFKFB3 |
| 0.00605432 | -0.3773497 | 0.083 | 0.21 | 1 | TEC_IFIT | MAT2A |
| 0.00014771 | -0.4632496 | 0.083 | 0.285 | 1 | TEC_IFIT | COL18A1 |
| 0.00159427 | -0.5462888 | 0.083 | 0.234 | 1 | TEC_IFIT | SYNE1 |
| 0.00518117 | -0.5555749 | 0.222 | 0.375 | 1 | TEC_IFIT | SLC38A2 |
| 0.00045527 | -0.6119217 | 0.042 | 0.204 | 1 | TEC_IFIT | SGK1 |
| 0.00118468 | -0.6186501 | 0.569 | 0.669 | 1 | TEC_IFIT | TPI1 |
| 0.00183324 | -0.6311062 | 0.097 | 0.243 | 1 | TEC_IFIT | ID4 |
| 0.00462179 | -0.6456053 | 0.139 | 0.291 | 1 | TEC_IFIT | FOS |
| 0.00052962 | -0.672612 | 0.069 | 0.239 | 1 | TEC_IFIT | INTS6 |
| 0.00457175 | -0.6841004 | 0.292 | 0.416 | 1 | TEC_IFIT | SRSF7 |
| 0.00377709 | -0.6940222 | 0.139 | 0.278 | 1 | TEC_IFIT | RASD1 |
| 0.00037086 | -0.7179723 | 0.042 | 0.212 | 1 | TEC_IFIT | ADAMTS1 |
| 0.00787031 | -0.7860549 | 0.528 | 0.6 | 1 | TEC_IFIT | CLU |
| 0.00018702 | -0.7926545 | 0.069 | 0.261 | 1 | TEC_IFIT | EGR1 |
| 0.00135251 | -0.8593863 | 0.125 | 0.278 | 1 | TEC_IFIT | HES1 |
| 0.00337153 | -0.9760757 | 0.167 | 0.298 | 1 | TEC_IFIT | HEXIM1 |
| 0.00367943 | -1.0400018 | 0.25 | 0.378 | 1 | TEC_IFIT | HIST1H4C |
| 0.00960834 | -1.1464161 | 0.611 | 0.679 | 1 | TEC_IFIT | MTRNR2L12 |
| 3.2714E-06 | -1.4025581 | 0.042 | 0.284 | 0.0698444 | TEC_IFIT | MAFB |
| 0.00032797 | -4.0538253 | 0.514 | 0.598 | 1 | TEC_IFIT | PLCG2 |
| 1.532E-106 | 1.59772228 | 0.248 | 0.054 | 3.27E-102 | TEC_IL32 | CCL20 |
| 4.982E-259 | 1.53908951 | 0.826 | 0.298 | 1.064E-254 | TEC_IL32 | LAMC2 |
| 2.586E-211 | 1.4969853 | 0.923 | 0.57 | 5.522E-207 | TEC_IL32 | KRT19 |
| 3.114E-240 | 1.45388808 | 0.783 | 0.289 | 6.649E-236 | TEC_IL32 | ITGA2 |
| 1.299E-158 | 1.35250815 | 0.955 | 0.789 | 2.774E-154 | TEC_IL32 | KRT18 |
| 5.9334E-48 | 1.27366315 | 0.27 | 0.107 | 1.2668E-43 | TEC_IL32 | CXCL8 |
| 1.392E-166 | 1.22477566 | 0.707 | 0.281 | 2.972E-162 | TEC_IL32 | PLAU |
| 1.0241E-56 | 1.21787242 | 0.822 | 0.674 | 2.1864E-52 | TEC_IL32 | HSPA5 |
| 2.469E-216 | 1.18518012 | 0.747 | 0.265 | 5.272E-212 | TEC_IL32 | ITGB8 |
| 1.036E-168 | 1.14366574 | 0.907 | 0.619 | 2.211E-164 | TEC_IL32 | MMP7 |
| 8.5372E-31 | 1.1403732 | 0.117 | 0.035 | 1.8227E-26 | TEC_IL32 | CCL5 |
| 1.141E-186 | 1.12980226 | 0.71 | 0.258 | 2.436E-182 | TEC_IL32 | PLAUR |
| 3.7714E-41 | 1.11031957 | 0.115 | 0.028 | 8.052E-37 | TEC_IL32 | MMP1 |
| 3.727E-158 | 1.10665211 | 0.952 | 0.712 | 7.957E-154 | TEC_IL32 | KRT8 |
| 1.237E-185 | 1.09718659 | 0.827 | 0.394 | 2.64E-181 | TEC_IL32 | TPM4 |
| 1.638E-60 | 1.05095333 | 0.256 | 0.084 | 3.4972E-56 | TEC_IL32 | CCL2 |
| 1.6094E-42 | 1.04366201 | 0.562 | 0.369 | 3.4361E-38 | TEC_IL32 | HERPUD1 |
| 3.291E-160 | 1.04334106 | 0.811 | 0.392 | 7.027E-156 | TEC_IL32 | CCND1 |
| 2.235E-101 | 1.04227109 | 0.335 | 0.097 | 4.7709E-97 | TEC_IL32 | C15orf48 |
| 2.339E-185 | 1.03268416 | 0.412 | 0.09 | 4.993E-181 | TEC_IL32 | ICAM1 |

| | | | | | | |
|---|---|---|---|---|---|---|
| 6.238E-103 | 1.02347895 | 0.369 | 0.114 | 1.3317E-98 | TEC_IL32 | TNFRSF11B |
| 2.372E-168 | 1.02279922 | 0.665 | 0.253 | 5.064E-164 | TEC_IL32 | SERPINB1 |
| 1.146E-141 | 1.00138664 | 0.705 | 0.302 | 2.448E-137 | TEC_IL32 | IL32 |
| 2.5734E-18 | 0.9967436 | 0.262 | 0.154 | 5.4943E-14 | TEC_IL32 | G0S2 |
| 3.387E-130 | 0.9942772 | 0.898 | 0.632 | 7.231E-126 | TEC_IL32 | KRT7 |
| 2.606E-114 | 0.95106951 | 0.359 | 0.099 | 5.563E-110 | TEC_IL32 | TAGLN |
| 3.85E-141 | 0.9378332 | 0.997 | 0.917 | 8.221E-137 | TEC_IL32 | ACTB |
| 1.741E-183 | 0.93401077 | 0.773 | 0.316 | 3.716E-179 | TEC_IL32 | S100A16 |
| 2.094E-138 | 0.9045405 | 0.617 | 0.236 | 4.47E-134 | TEC_IL32 | LAMB3 |
| 3.307E-186 | 0.85866345 | 0.649 | 0.205 | 7.061E-182 | TEC_IL32 | TGM2 |
| 1.854E-106 | 0.85806205 | 0.27 | 0.061 | 3.958E-102 | TEC_IL32 | SPRR2A |
| 4.908E-216 | 0.85798547 | 0.584 | 0.156 | 1.048E-211 | TEC_IL32 | CD40 |
| 1.846E-132 | 0.84832077 | 0.942 | 0.644 | 3.941E-128 | TEC_IL32 | TACSTD2 |
| 5.1745E-61 | 0.84206109 | 0.895 | 0.764 | 1.1047E-56 | TEC_IL32 | HSP90B1 |
| 1.051E-178 | 0.83913766 | 0.985 | 0.821 | 2.244E-174 | TEC_IL32 | ANXA2 |
| 6.339E-104 | 0.8330123 | 0.564 | 0.232 | 1.353E-99 | TEC_IL32 | AKAP12 |
| 8.46E-106 | 0.81191751 | 0.24 | 0.05 | 1.806E-101 | TEC_IL32 | KRTAP2-3 |
| 4.213E-125 | 0.80065646 | 0.924 | 0.689 | 8.995E-121 | TEC_IL32 | ITGB1 |
| 2.112E-118 | 0.79949583 | 0.741 | 0.368 | 4.509E-114 | TEC_IL32 | RAB11FIP1 |
| 2.94E-148 | 0.78557177 | 0.61 | 0.211 | 6.276E-144 | TEC_IL32 | CST6 |
| 7.19E-155 | 0.78160988 | 0.962 | 0.653 | 1.535E-150 | TEC_IL32 | SH3BGRL3 |
| 3.3507E-60 | 0.7802739 | 0.311 | 0.117 | 7.1536E-56 | TEC_IL32 | PRSS22 |
| 2.685E-135 | 0.77643767 | 0.868 | 0.48 | 5.732E-131 | TEC_IL32 | CLDN7 |
| 1.49E-125 | 0.76967464 | 0.887 | 0.537 | 3.181E-121 | TEC_IL32 | HMGA1 |
| 2.125E-108 | 0.76743542 | 0.987 | 0.922 | 4.537E-104 | TEC_IL32 | ACTG1 |
| 1.79E-141 | 0.76672585 | 0.498 | 0.159 | 3.821E-137 | TEC_IL32 | DCBLD2 |
| 6.5043E-81 | 0.75882185 | 0.562 | 0.259 | 1.3887E-76 | TEC_IL32 | LGALS1 |
| 6.652E-63 | 0.72859267 | 0.823 | 0.618 | 1.4202E-58 | TEC_IL32 | TPM1 |
| 3.5016E-69 | 0.71778286 | 0.905 | 0.688 | 7.4759E-65 | TEC_IL32 | CLDN4 |
| 1.0044E-90 | 0.7013907 | 0.836 | 0.547 | 2.1444E-86 | TEC_IL32 | MAL2 |
| 1.473E-150 | 0.70079292 | 0.883 | 0.459 | 3.146E-146 | TEC_IL32 | TSPAN1 |
| 6.7785E-56 | 0.6973558 | 0.648 | 0.398 | 1.4472E-51 | TEC_IL32 | PHLDA2 |
| 5.287E-127 | 0.69558986 | 0.96 | 0.717 | 1.129E-122 | TEC_IL32 | PFN1 |
| 1.6195E-67 | 0.69472383 | 0.426 | 0.187 | 3.4577E-63 | TEC_IL32 | MANF |
| 4.523E-105 | 0.6918755 | 0.527 | 0.201 | 9.658E-101 | TEC_IL32 | SFN |
| 6.941E-94 | 0.67894316 | 0.961 | 0.79 | 1.4819E-89 | TEC_IL32 | CD24 |
| 3.141E-99 | 0.66625533 | 0.646 | 0.298 | 6.7068E-95 | TEC_IL32 | DCDC2 |
| 6.135E-103 | 0.6662035 | 0.992 | 0.919 | 1.3098E-98 | TEC_IL32 | TMSB10 |
| 2.246E-162 | 0.66219816 | 0.444 | 0.114 | 4.796E-158 | TEC_IL32 | PDLIM4 |
| 2.9799E-49 | 0.65891269 | 0.614 | 0.378 | 6.3622E-45 | TEC_IL32 | TXNRD1 |
| 6.7711E-55 | 0.65888594 | 0.193 | 0.055 | 1.4456E-50 | TEC_IL32 | LTB |
| 8.817E-127 | 0.65869674 | 0.704 | 0.321 | 1.882E-122 | TEC_IL32 | CAP1 |
| 3.98E-118 | 0.64933201 | 0.707 | 0.326 | 8.497E-114 | TEC_IL32 | ANXA11 |
| 1.1737E-98 | 0.64347919 | 0.975 | 0.861 | 2.5058E-94 | TEC_IL32 | TMSB4X |
| 3.0268E-60 | 0.63637512 | 0.406 | 0.181 | 6.4622E-56 | TEC_IL32 | SDF2L1 |
| 3.1583E-62 | 0.6357669 | 0.564 | 0.282 | 6.7429E-58 | TEC_IL32 | MARCKSL1 |
| 3.5149E-68 | 0.62782173 | 0.45 | 0.197 | 7.5043E-64 | TEC_IL32 | GLRX |
| 8.1513E-43 | 0.6200483 | 0.658 | 0.429 | 1.7403E-38 | TEC_IL32 | UPP1 |
| 9.021E-108 | 0.5948273 | 0.508 | 0.184 | 1.926E-103 | TEC_IL32 | CLDN1 |
| 8.348E-141 | 0.58335306 | 0.999 | 0.914 | 1.782E-136 | TEC_IL32 | S100A11 |

| | | | | | | |
|---|---|---|---|---|---|---|
| 2.917E-115 | 0.57782662 | 0.571 | 0.222 | 6.227E-111 | TEC_IL32 | TNIP1 |
| 5.976E-100 | 0.57341327 | 0.682 | 0.318 | 1.2759E-95 | TEC_IL32 | JUP |
| 1.5206E-38 | 0.56882997 | 0.398 | 0.206 | 3.2466E-34 | TEC_IL32 | TNFAIP2 |
| 8.0313E-67 | 0.56672615 | 0.487 | 0.223 | 1.7147E-62 | TEC_IL32 | ITGB6 |
| 2.3733E-68 | 0.56278708 | 0.777 | 0.479 | 5.067E-64 | TEC_IL32 | TNFRSF12A |
| 1.114E-112 | 0.5582614 | 0.632 | 0.261 | 2.378E-108 | TEC_IL32 | SPINT1 |
| 2.65E-69 | 0.55373735 | 0.596 | 0.297 | 5.6577E-65 | TEC_IL32 | MAL |
| 7.8978E-75 | 0.55354222 | 0.657 | 0.339 | 1.6862E-70 | TEC_IL32 | CD44 |
| 2.1688E-93 | 0.55351243 | 0.733 | 0.375 | 4.6305E-89 | TEC_IL32 | RHOC |
| 1.1394E-56 | 0.55112904 | 0.541 | 0.288 | 2.4327E-52 | TEC_IL32 | ARHGAP29 |
| 8.2753E-91 | 0.54992676 | 0.852 | 0.569 | 1.7668E-86 | TEC_IL32 | YWHAB |
| 7.0863E-88 | 0.54945488 | 0.801 | 0.497 | 1.5129E-83 | TEC_IL32 | ARPC2 |
| 3.998E-105 | 0.54912646 | 0.926 | 0.646 | 8.536E-101 | TEC_IL32 | RAC1 |
| 5.1138E-84 | 0.54636568 | 0.613 | 0.289 | 1.0918E-79 | TEC_IL32 | DSP |
| 8.7526E-96 | 0.53545078 | 0.622 | 0.278 | 1.8687E-91 | TEC_IL32 | CDH1 |
| 1.0471E-78 | 0.53196222 | 0.628 | 0.31 | 2.2355E-74 | TEC_IL32 | CD2AP |
| 1.02E-151 | 0.53095176 | 0.409 | 0.1 | 2.178E-147 | TEC_IL32 | DAPP1 |
| 6.595E-128 | 0.53087466 | 0.357 | 0.089 | 1.408E-123 | TEC_IL32 | VGLL3 |
| 1.5831E-18 | 0.5299051 | 0.431 | 0.292 | 3.3799E-14 | TEC_IL32 | DDIT4 |
| 7.8358E-89 | 0.52620429 | 0.491 | 0.19 | 1.6729E-84 | TEC_IL32 | ADAMTS9 |
| 1.192E-113 | 0.52517889 | 0.542 | 0.202 | 2.545E-109 | TEC_IL32 | CXCL16 |
| 1.4446E-98 | 0.52251046 | 0.967 | 0.793 | 3.0842E-94 | TEC_IL32 | CFL1 |
| 3.5543E-73 | 0.52215236 | 0.512 | 0.225 | 7.5884E-69 | TEC_IL32 | TFPI2 |
| 1.0483E-66 | 0.51288272 | 0.343 | 0.124 | 2.2382E-62 | TEC_IL32 | TM4SF1 |
| 5.4316E-55 | 0.5095225 | 0.807 | 0.57 | 1.1597E-50 | TEC_IL32 | EZR |
| 1.1535E-81 | 0.50949488 | 0.458 | 0.182 | 2.4627E-77 | TEC_IL32 | MARCKS |
| 3.9639E-46 | 0.50714211 | 0.639 | 0.391 | 8.4629E-42 | TEC_IL32 | SDC4 |
| 2.847E-83 | 0.50286516 | 0.719 | 0.363 | 6.0783E-79 | TEC_IL32 | OCIAD2 |
| 1.578E-72 | 0.5019485 | 0.828 | 0.532 | 3.369E-68 | TEC_IL32 | CD9 |
| 1.9877E-55 | 0.4996026 | 0.485 | 0.24 | 4.2437E-51 | TEC_IL32 | S100A14 |
| 1.6783E-71 | 0.49320893 | 0.72 | 0.413 | 3.5832E-67 | TEC_IL32 | BZW1 |
| 2.7383E-55 | 0.4927083 | 0.527 | 0.273 | 5.8463E-51 | TEC_IL32 | ITGA3 |
| 8.233E-110 | 0.48913574 | 0.218 | 0.04 | 1.758E-105 | TEC_IL32 | VNN1 |
| 2.5336E-78 | 0.48601265 | 0.375 | 0.133 | 5.4093E-74 | TEC_IL32 | ARL4C |
| 1.5931E-39 | 0.48096548 | 0.31 | 0.142 | 3.4012E-35 | TEC_IL32 | RCAN1 |
| 1.3292E-93 | 0.48044242 | 0.668 | 0.305 | 2.8378E-89 | TEC_IL32 | TSPO |
| 2.6312E-63 | 0.47566557 | 0.875 | 0.665 | 5.6176E-59 | TEC_IL32 | YWHAZ |
| 1.4409E-66 | 0.47118885 | 0.54 | 0.266 | 3.0763E-62 | TEC_IL32 | MSN |
| 4.1278E-72 | 0.46761136 | 0.673 | 0.35 | 8.8129E-68 | TEC_IL32 | TAGLN2 |
| 2.744E-44 | 0.46496913 | 0.371 | 0.176 | 5.8585E-40 | TEC_IL32 | PDIA4 |
| 8.0753E-86 | 0.46471455 | 0.598 | 0.266 | 1.7241E-81 | TEC_IL32 | ANXA3 |
| 5.2723E-39 | 0.46468897 | 0.821 | 0.658 | 1.1256E-34 | TEC_IL32 | GLS |
| 2.5458E-67 | 0.46181821 | 0.69 | 0.389 | 5.4353E-63 | TEC_IL32 | JAK1 |
| 1.2723E-67 | 0.46155655 | 0.527 | 0.248 | 2.7164E-63 | TEC_IL32 | LIMA1 |
| 1.4179E-79 | 0.45498625 | 0.503 | 0.21 | 3.0272E-75 | TEC_IL32 | ST14 |
| 5.6679E-59 | 0.4519145 | 0.879 | 0.668 | 1.2101E-54 | TEC_IL32 | CLIC1 |
| 4.0075E-49 | 0.45033768 | 0.513 | 0.273 | 8.5559E-45 | TEC_IL32 | CDK2AP2 |
| 1.737E-54 | 0.44924774 | 0.929 | 0.754 | 3.7084E-50 | TEC_IL32 | PKM |
| 6.7344E-68 | 0.4472206 | 0.946 | 0.747 | 1.4378E-63 | TEC_IL32 | CALM2 |
| 1.9914E-42 | 0.4454119 | 0.329 | 0.149 | 4.2516E-38 | TEC_IL32 | TNFAIP3 |

| | | | | | | |
|---|---|---|---|---|---|---|
| 2.2745E-97 | 0.43746433 | 0.43 | 0.147 | 4.856E-93 | TEC_IL32 | GRAMD2B |
| 2.2055E-39 | 0.43713016 | 0.363 | 0.178 | 4.7088E-35 | TEC_IL32 | THBS1 |
| 1.421E-47 | 0.43674233 | 0.55 | 0.302 | 3.0339E-43 | TEC_IL32 | UGCG |
| 8.217E-112 | 0.43426095 | 0.828 | 0.447 | 1.754E-107 | TEC_IL32 | C12orf75 |
| 5.9961E-46 | 0.4335314 | 0.714 | 0.472 | 1.2802E-41 | TEC_IL32 | LDHA |
| 2.042E-73 | 0.43117821 | 0.462 | 0.195 | 4.3597E-69 | TEC_IL32 | UBALD2 |
| 5.3378E-52 | 0.43039328 | 0.784 | 0.551 | 1.1396E-47 | TEC_IL32 | PPIB |
| 6.9249E-79 | 0.42963988 | 0.383 | 0.137 | 1.4785E-74 | TEC_IL32 | PROM1 |
| 1.6775E-41 | 0.42836315 | 0.169 | 0.053 | 3.5816E-37 | TEC_IL32 | EDN1 |
| 2.3887E-77 | 0.42440844 | 0.535 | 0.233 | 5.0998E-73 | TEC_IL32 | CD47 |
| 1.5053E-90 | 0.42101656 | 0.508 | 0.203 | 3.2137E-86 | TEC_IL32 | GNAI2 |
| 1.6978E-80 | 0.42083842 | 0.351 | 0.117 | 3.6249E-76 | TEC_IL32 | AL353751.1 |
| 2.306E-99 | 0.4193484 | 0.441 | 0.15 | 4.9223E-95 | TEC_IL32 | ELK3 |
| 6.1239E-60 | 0.41903416 | 0.288 | 0.1 | 1.3074E-55 | TEC_IL32 | ISG20 |
| 9.7242E-64 | 0.41828146 | 0.46 | 0.206 | 2.0761E-59 | TEC_IL32 | LSR |
| 5.5779E-70 | 0.41812433 | 0.581 | 0.288 | 1.1909E-65 | TEC_IL32 | WDR1 |
| 1.1666E-57 | 0.41777976 | 0.61 | 0.332 | 2.4907E-53 | TEC_IL32 | EIF5A |
| 3.4315E-65 | 0.41671682 | 0.967 | 0.867 | 7.3263E-61 | TEC_IL32 | GSTP1 |
| 9.3486E-36 | 0.41623637 | 0.279 | 0.125 | 1.9959E-31 | TEC_IL32 | ITPKC |
| 3.0879E-12 | 0.41227894 | 0.689 | 0.6 | 6.5927E-08 | TEC_IL32 | SOD2 |
| 1.4628E-51 | 0.41221955 | 0.343 | 0.145 | 3.1231E-47 | TEC_IL32 | PMEPA1 |
| 3.6035E-62 | 0.41120571 | 0.559 | 0.283 | 7.6935E-58 | TEC_IL32 | ACTR3 |
| 7.9031E-20 | 0.40803643 | 0.433 | 0.279 | 1.6873E-15 | TEC_IL32 | CYR61 |
| 6.5002E-44 | 0.40534629 | 0.358 | 0.163 | 1.3878E-39 | TEC_IL32 | PHLDA1 |
| 4.469E-55 | 0.40179683 | 0.989 | 0.909 | 9.5414E-51 | TEC_IL32 | HSP90AB1 |
| 1.596E-66 | 0.39847416 | 0.73 | 0.419 | 3.4075E-62 | TEC_IL32 | PPP1R14B |
| 5.5162E-31 | 0.39609885 | 0.201 | 0.082 | 1.1777E-26 | TEC_IL32 | LAMA3 |
| 1.2974E-54 | 0.39372591 | 0.594 | 0.322 | 2.7699E-50 | TEC_IL32 | SSR3 |
| 1.3791E-55 | 0.39321663 | 0.567 | 0.305 | 2.9443E-51 | TEC_IL32 | GNB1 |
| 5.1178E-49 | 0.39248218 | 0.758 | 0.509 | 1.0926E-44 | TEC_IL32 | CDC42 |
| 2.9577E-47 | 0.39127415 | 0.701 | 0.453 | 6.3147E-43 | TEC_IL32 | PDIA6 |
| 5.9076E-42 | 0.38858076 | 0.343 | 0.158 | 1.2613E-37 | TEC_IL32 | TYMP |
| 1.8121E-45 | 0.38795885 | 0.952 | 0.819 | 3.8689E-41 | TEC_IL32 | HLA-B |
| 1.011E-56 | 0.38604887 | 0.956 | 0.816 | 2.1585E-52 | TEC_IL32 | H3F3A |
| 2.3918E-59 | 0.38406257 | 0.654 | 0.355 | 5.1066E-55 | TEC_IL32 | HEBP2 |
| 9.2862E-54 | 0.38341031 | 0.585 | 0.318 | 1.9826E-49 | TEC_IL32 | PTP4A2 |
| 5.2237E-69 | 0.38326935 | 0.392 | 0.155 | 1.1153E-64 | TEC_IL32 | PHC2 |
| 7.3495E-58 | 0.38282892 | 0.841 | 0.588 | 1.5691E-53 | TEC_IL32 | CLTA |
| 1.7345E-58 | 0.38226969 | 0.464 | 0.212 | 3.7031E-54 | TEC_IL32 | PRSS8 |
| 3.4773E-53 | 0.37958704 | 0.516 | 0.265 | 7.424E-49 | TEC_IL32 | CLTB |
| 7.6756E-52 | 0.37863259 | 0.503 | 0.253 | 1.6387E-47 | TEC_IL32 | TUBB |
| 3.1827E-69 | 0.37835753 | 0.435 | 0.18 | 6.7952E-65 | TEC_IL32 | SYNGR2 |
| 7.781E-61 | 0.37819882 | 0.498 | 0.231 | 1.6612E-56 | TEC_IL32 | TES |
| 3.9013E-46 | 0.37664311 | 0.813 | 0.573 | 8.3293E-42 | TEC_IL32 | S100A13 |
| 1.9247E-27 | 0.37637402 | 0.304 | 0.157 | 4.1093E-23 | TEC_IL32 | CTGF |
| 2.8518E-65 | 0.37616743 | 0.433 | 0.182 | 6.0885E-61 | TEC_IL32 | MAP2K3 |
| 8.4226E-69 | 0.37413169 | 0.858 | 0.565 | 1.7982E-64 | TEC_IL32 | ARPC3 |
| 5.7693E-47 | 0.37308063 | 0.465 | 0.234 | 1.2317E-42 | TEC_IL32 | MACC1 |
| 1.0774E-32 | 0.37290548 | 0.675 | 0.481 | 2.3002E-28 | TEC_IL32 | ATP1B1 |
| 3.8328E-63 | 0.37187787 | 0.238 | 0.072 | 8.183E-59 | TEC_IL32 | FSD1L |

| | | | | | | |
|---|---|---|---|---|---|---|
| 8.3331E-64 | 0.37099782 | 0.599 | 0.301 | 1.7791E-59 | TEC_IL32 | SNRPB |
| 8.2687E-58 | 0.36963273 | 0.556 | 0.284 | 1.7654E-53 | TEC_IL32 | RAP1B |
| 5.3323E-45 | 0.36934601 | 0.916 | 0.764 | 1.1385E-40 | TEC_IL32 | MYL12A |
| 3.249E-118 | 0.369043 | 0.253 | 0.05 | 6.937E-114 | TEC_IL32 | SEMA7A |
| 2.6251E-47 | 0.36857874 | 0.671 | 0.403 | 5.6046E-43 | TEC_IL32 | PTTG1IP |
| 3.0429E-41 | 0.36849243 | 0.451 | 0.241 | 6.4965E-37 | TEC_IL32 | COTL1 |
| 3.231E-59 | 0.36805487 | 0.97 | 0.831 | 6.8982E-55 | TEC_IL32 | CD63 |
| 1.6471E-51 | 0.3679311 | 0.453 | 0.22 | 3.5165E-47 | TEC_IL32 | ITGAV |
| 2.2139E-45 | 0.36769614 | 0.526 | 0.286 | 4.7267E-41 | TEC_IL32 | MYDGF |
| 2.9637E-47 | 0.36540052 | 0.435 | 0.209 | 6.3275E-43 | TEC_IL32 | GPRC5A |
| 3.3642E-37 | 0.36432349 | 0.823 | 0.649 | 7.1825E-33 | TEC_IL32 | CALR |
| 6.5671E-74 | 0.36365497 | 0.454 | 0.186 | 1.4021E-69 | TEC_IL32 | GIPC1 |
| 4.3071E-69 | 0.36137289 | 0.537 | 0.244 | 9.1956E-65 | TEC_IL32 | FKBP1A |
| 1.4025E-47 | 0.35968093 | 0.943 | 0.83 | 2.9942E-43 | TEC_IL32 | PRDX1 |
| 5.5367E-32 | 0.35962489 | 0.52 | 0.312 | 1.1821E-27 | TEC_IL32 | CYBA |
| 5.2908E-70 | 0.35894915 | 0.995 | 0.952 | 1.1296E-65 | TEC_IL32 | S100A6 |
| 6.5182E-76 | 0.35822907 | 0.333 | 0.112 | 1.3916E-71 | TEC_IL32 | IFNGR2 |
| 1.6294E-55 | 0.3579045 | 0.493 | 0.24 | 3.4787E-51 | TEC_IL32 | PPP2R1A |
| 1.6081E-49 | 0.35778503 | 0.523 | 0.274 | 3.4333E-45 | TEC_IL32 | FAM3C |
| 1.0552E-35 | 0.35743794 | 0.607 | 0.37 | 2.2529E-31 | TEC_IL32 | MYH9 |
| 5.6438E-33 | 0.35705545 | 0.74 | 0.545 | 1.205E-28 | TEC_IL32 | LRRFIP1 |
| 1.2587E-55 | 0.35606886 | 0.952 | 0.846 | 2.6872E-51 | TEC_IL32 | MYL12B |
| 6.786E-32 | 0.3559292 | 0.342 | 0.176 | 1.4488E-27 | TEC_IL32 | BIRC3 |
| 3.515E-37 | 0.35585595 | 0.494 | 0.281 | 7.5046E-33 | TEC_IL32 | CD55 |
| 1.2856E-50 | 0.35534839 | 0.411 | 0.191 | 2.7448E-46 | TEC_IL32 | WWTR1 |
| 1.2117E-42 | 0.35513987 | 0.185 | 0.061 | 2.587E-38 | TEC_IL32 | LIF |
| 5.9574E-49 | 0.35473751 | 0.67 | 0.381 | 1.2719E-44 | TEC_IL32 | JPT1 |
| 7.8632E-46 | 0.35454519 | 0.711 | 0.457 | 1.6788E-41 | TEC_IL32 | SUMO2 |
| 1.2476E-41 | 0.35287592 | 0.136 | 0.037 | 2.6636E-37 | TEC_IL32 | EDN2 |
| 1.6741E-41 | 0.35184567 | 0.89 | 0.718 | 3.5743E-37 | TEC_IL32 | ENO1 |
| 9.5344E-69 | 0.35159938 | 0.416 | 0.167 | 2.0356E-64 | TEC_IL32 | QSOX1 |
| 3.1402E-45 | 0.35045842 | 0.474 | 0.247 | 6.7042E-41 | TEC_IL32 | VCL |
| 4.1315E-58 | 0.34979742 | 0.908 | 0.651 | 8.8207E-54 | TEC_IL32 | SPINT2 |
| 4.0641E-43 | 0.34901866 | 0.66 | 0.413 | 8.6768E-39 | TEC_IL32 | PDIA3 |
| 3.6036E-53 | 0.34796912 | 0.552 | 0.282 | 7.6936E-49 | TEC_IL32 | CAPG |
| 1.0001E-48 | 0.34756023 | 0.565 | 0.302 | 2.1353E-44 | TEC_IL32 | FAM107B |
| 4.2473E-77 | 0.34727786 | 0.364 | 0.126 | 9.068E-73 | TEC_IL32 | SLC44A4 |
| 1.8024E-55 | 0.34642676 | 0.406 | 0.182 | 3.8481E-51 | TEC_IL32 | CMTM6 |
| 7.9596E-65 | 0.34547157 | 0.383 | 0.153 | 1.6994E-60 | TEC_IL32 | UBE2J1 |
| 3.3758E-77 | 0.34490335 | 0.989 | 0.862 | 7.2073E-73 | TEC_IL32 | S100A10 |
| 1.3856E-55 | 0.34405251 | 0.465 | 0.219 | 2.9584E-51 | TEC_IL32 | LRRFIP2 |
| 1.7899E-62 | 0.34300438 | 0.177 | 0.042 | 3.8215E-58 | TEC_IL32 | SPRR2D |
| 6.3623E-28 | 0.34278485 | 0.34 | 0.186 | 1.3584E-23 | TEC_IL32 | MSMO1 |
| 2.9616E-26 | 0.3427342 | 0.193 | 0.084 | 6.3231E-22 | TEC_IL32 | F3 |
| 3.5978E-46 | 0.34202278 | 0.918 | 0.738 | 7.6814E-42 | TEC_IL32 | RTN4 |
| 1.0675E-52 | 0.34121196 | 0.889 | 0.666 | 2.279E-48 | TEC_IL32 | EPCAM |
| 4.4154E-54 | 0.34093615 | 0.676 | 0.372 | 9.4269E-50 | TEC_IL32 | CD151 |
| 2.3015E-50 | 0.33383968 | 0.472 | 0.234 | 4.9138E-46 | TEC_IL32 | YPEL5 |
| 2.8512E-39 | 0.33368057 | 0.494 | 0.279 | 6.0874E-35 | TEC_IL32 | ACTR2 |
| 1.2434E-46 | 0.33367226 | 0.574 | 0.321 | 2.6546E-42 | TEC_IL32 | PRELID1 |

| | | | | | | |
|---|---|---|---|---|---|---|
| 2.5269E-47 | 0.33271901 | 0.363 | 0.165 | 5.3948E-43 | TEC_IL32 | SECISBP2L |
| 4.573E-62 | 0.33235067 | 0.293 | 0.101 | 9.7634E-58 | TEC_IL32 | BPGM |
| 3.472E-45 | 0.33190916 | 0.516 | 0.28 | 7.4128E-41 | TEC_IL32 | EIF6 |
| 4.1898E-52 | 0.33082873 | 0.404 | 0.18 | 8.9453E-48 | TEC_IL32 | FAM129B |
| 6.5098E-37 | 0.33074214 | 0.605 | 0.377 | 1.3898E-32 | TEC_IL32 | EIF2S2 |
| 2.7271E-27 | 0.32968871 | 0.265 | 0.13 | 5.8223E-23 | TEC_IL32 | TNFSF10 |
| 1.0201E-52 | 0.32943641 | 0.338 | 0.139 | 2.178E-48 | TEC_IL32 | CDC34 |
| 2.3826E-50 | 0.32908197 | 0.338 | 0.141 | 5.0869E-46 | TEC_IL32 | EGFR |
| 1.5838E-42 | 0.32856223 | 0.491 | 0.268 | 3.3815E-38 | TEC_IL32 | LITAF |
| 1.5816E-54 | 0.3280112 | 0.425 | 0.189 | 3.3768E-50 | TEC_IL32 | ERO1A |
| 5.5333E-29 | 0.32780892 | 0.318 | 0.162 | 1.1813E-24 | TEC_IL32 | DUSP5 |
| 1.4748E-45 | 0.32748033 | 0.412 | 0.198 | 3.1487E-41 | TEC_IL32 | PPFIBP1 |
| 4.5001E-36 | 0.32577417 | 0.957 | 0.904 | 9.6077E-32 | TEC_IL32 | MYL6 |
| 1.0673E-54 | 0.32442403 | 0.427 | 0.191 | 2.2787E-50 | TEC_IL32 | MAP4K4 |
| 6.8552E-38 | 0.32375269 | 0.734 | 0.502 | 1.4636E-33 | TEC_IL32 | TPM3 |
| 5.4002E-50 | 0.322873 | 0.44 | 0.209 | 1.1529E-45 | TEC_IL32 | TEAD1 |
| 5.814E-32 | 0.3198668 | 0.968 | 0.868 | 1.2413E-27 | TEC_IL32 | HLA-A |
| 6.3593E-41 | 0.31844552 | 0.434 | 0.223 | 1.3577E-36 | TEC_IL32 | LPP |
| 1.8133E-58 | 0.31799687 | 0.339 | 0.132 | 3.8714E-54 | TEC_IL32 | VASP |
| 5.0121E-68 | 0.31721193 | 0.322 | 0.111 | 1.0701E-63 | TEC_IL32 | CDK6 |
| 2.4399E-62 | 0.31709354 | 0.306 | 0.109 | 5.2092E-58 | TEC_IL32 | SGPP2 |
| 5.3592E-72 | 0.31623734 | 0.309 | 0.101 | 1.1442E-67 | TEC_IL32 | TMEM132A |
| 1.6481E-41 | 0.31533764 | 0.759 | 0.505 | 3.5186E-37 | TEC_IL32 | ARF1 |
| 1.2435E-42 | 0.31465298 | 0.406 | 0.199 | 2.6548E-38 | TEC_IL32 | SCCPDH |
| 1.5513E-64 | 0.31457829 | 0.332 | 0.121 | 3.312E-60 | TEC_IL32 | EHD4 |
| 3.2516E-44 | 0.31374091 | 0.615 | 0.354 | 6.9421E-40 | TEC_IL32 | SDCBP |
| 9.949E-37 | 0.31293484 | 0.99 | 0.931 | 2.1241E-32 | TEC_IL32 | PTMA |
| 8.0011E-35 | 0.31290452 | 0.392 | 0.206 | 1.7082E-30 | TEC_IL32 | SVIL |
| 2.6399E-33 | 0.31246627 | 0.98 | 0.926 | 5.6362E-29 | TEC_IL32 | RPS14 |
| 9.8728E-12 | 0.31231548 | 0.45 | 0.343 | 2.1078E-07 | TEC_IL32 | TXNIP |
| 4.4775E-51 | 0.31107001 | 0.522 | 0.262 | 9.5594E-47 | TEC_IL32 | CNN3 |
| 2.3265E-25 | 0.31095575 | 0.39 | 0.234 | 4.967E-21 | TEC_IL32 | DNAJC3 |
| 4.8583E-24 | 0.30942421 | 0.667 | 0.484 | 1.0372E-19 | TEC_IL32 | ACTN4 |
| 2.8283E-50 | 0.30744193 | 0.39 | 0.175 | 6.0384E-46 | TEC_IL32 | B4GALT1 |
| 2.1512E-48 | 0.30743343 | 0.451 | 0.222 | 4.5929E-44 | TEC_IL32 | PPP1CA |
| 7.6849E-46 | 0.30681464 | 0.293 | 0.119 | 1.6407E-41 | TEC_IL32 | NCEH1 |
| 8.351E-41 | 0.30645605 | 0.472 | 0.252 | 1.7829E-36 | TEC_IL32 | TAPBP |
| 3.6683E-34 | 0.30535456 | 0.984 | 0.919 | 7.8319E-30 | TEC_IL32 | RPL10 |
| 4.3892E-15 | 0.30483179 | 0.612 | 0.472 | 9.3708E-11 | TEC_IL32 | H1FX |
| 1.8031E-34 | 0.30473205 | 0.588 | 0.371 | 3.8496E-30 | TEC_IL32 | CTNNA1 |
| 8.707E-70 | 0.30465514 | 0.349 | 0.125 | 1.8589E-65 | TEC_IL32 | RAB25 |
| 5.1891E-52 | 0.30436752 | 0.426 | 0.196 | 1.1079E-47 | TEC_IL32 | SNX9 |
| 1.3428E-21 | 0.303972 | 0.32 | 0.186 | 2.867E-17 | TEC_IL32 | FDPS |
| 1.7608E-40 | 0.30300225 | 0.905 | 0.703 | 3.7594E-36 | TEC_IL32 | TXN |
| 1.157E-54 | 0.30260115 | 0.223 | 0.07 | 2.4702E-50 | TEC_IL32 | RBP1 |
| 4.3504E-31 | 0.30244332 | 0.256 | 0.117 | 9.2881E-27 | TEC_IL32 | SPNS2 |
| 9.8984E-31 | 0.30118663 | 0.356 | 0.188 | 2.1133E-26 | TEC_IL32 | PHLDB2 |
| 8.9874E-35 | 0.300729 | 0.979 | 0.887 | 1.9188E-30 | TEC_IL32 | GAPDH |
| 7.7521E-48 | 0.30061468 | 0.338 | 0.146 | 1.6551E-43 | TEC_IL32 | CAV2 |
| 1.293E-25 | 0.30010187 | 0.126 | 0.044 | 2.7606E-21 | TEC_IL32 | PI3 |

| | | | | | | |
|---|---|---|---|---|---|---|
| 3.2001E-41 | 0.29965037 | 0.303 | 0.131 | 6.8322E-37 | TEC_IL32 | NFKB1 |
| 1.12E-07 | 0.29844636 | 0.132 | 0.079 | 0.00239126 | TEC_IL32 | CXCL1 |
| 2.9286E-33 | 0.29804416 | 0.453 | 0.255 | 6.2526E-29 | TEC_IL32 | MET |
| 1.3137E-32 | 0.29763145 | 0.608 | 0.377 | 2.8048E-28 | TEC_IL32 | CAPN2 |
| 8.3181E-53 | 0.29728593 | 0.204 | 0.061 | 1.7759E-48 | TEC_IL32 | MALL |
| 3.0557E-37 | 0.29612836 | 0.598 | 0.361 | 6.5239E-33 | TEC_IL32 | VAMP8 |
| 2.6782E-37 | 0.29504405 | 0.506 | 0.287 | 5.718E-33 | TEC_IL32 | IQGAP1 |
| 4.6785E-50 | 0.29381279 | 0.356 | 0.149 | 9.9886E-46 | TEC_IL32 | HSPG2 |
| 9.2057E-41 | 0.29365381 | 0.532 | 0.297 | 1.9654E-36 | TEC_IL32 | TMEM50A |
| 1.8761E-58 | 0.29324972 | 0.499 | 0.228 | 4.0055E-54 | TEC_IL32 | PDZK1IP1 |
| 5.1001E-43 | 0.29268606 | 0.463 | 0.235 | 1.0889E-38 | TEC_IL32 | MGLL |
| 9.1861E-51 | 0.29261212 | 0.266 | 0.097 | 1.9612E-46 | TEC_IL32 | F2RL1 |
| 1.389E-36 | 0.29237626 | 0.97 | 0.866 | 2.9656E-32 | TEC_IL32 | RACK1 |
| 1.0976E-45 | 0.29230382 | 0.405 | 0.193 | 2.3434E-41 | TEC_IL32 | CSRP1 |
| 2.6215E-58 | 0.29212685 | 0.148 | 0.033 | 5.5968E-54 | TEC_IL32 | CDKN2A |
| 1.9227E-26 | 0.2915533 | 0.889 | 0.716 | 4.105E-22 | TEC_IL32 | CSTB |
| 2.1166E-44 | 0.2906552 | 0.419 | 0.205 | 4.5189E-40 | TEC_IL32 | SRI |
| 2.1959E-44 | 0.29059066 | 0.335 | 0.149 | 4.6883E-40 | TEC_IL32 | F11R |
| 1.5073E-38 | 0.28975379 | 0.42 | 0.212 | 3.218E-34 | TEC_IL32 | C4orf48 |
| 9.6373E-47 | 0.28964977 | 0.45 | 0.22 | 2.0576E-42 | TEC_IL32 | TMBIM1 |
| 8.9407E-40 | 0.28941039 | 0.338 | 0.156 | 1.9088E-35 | TEC_IL32 | AMOTL2 |
| 9.5692E-42 | 0.28861366 | 0.441 | 0.229 | 2.043E-37 | TEC_IL32 | RAB11A |
| 7.7535E-61 | 0.28729858 | 0.299 | 0.104 | 1.6554E-56 | TEC_IL32 | MYH14 |
| 4.7957E-33 | 0.28699517 | 0.705 | 0.465 | 1.0239E-28 | TEC_IL32 | PTGES3 |
| 4.1241E-40 | 0.28650343 | 0.391 | 0.19 | 8.8051E-36 | TEC_IL32 | DPP4 |
| 5.7374E-89 | 0.28649227 | 0.24 | 0.056 | 1.2249E-84 | TEC_IL32 | MYEOV |
| 4.8732E-39 | 0.28634179 | 0.357 | 0.172 | 1.0404E-34 | TEC_IL32 | SLC44A2 |
| 3.1298E-35 | 0.28565161 | 0.454 | 0.248 | 6.6821E-31 | TEC_IL32 | GSTO1 |
| 1.4372E-53 | 0.28528255 | 0.284 | 0.104 | 3.0684E-49 | TEC_IL32 | RAI14 |
| 7.4578E-43 | 0.28524161 | 0.421 | 0.209 | 1.5922E-38 | TEC_IL32 | ARPC4 |
| 7.7479E-68 | 0.28454587 | 0.182 | 0.042 | 1.6542E-63 | TEC_IL32 | RTKN2 |
| 2.0934E-34 | 0.28338319 | 0.46 | 0.26 | 4.4693E-30 | TEC_IL32 | ARPC5 |
| 1.1063E-27 | 0.28290904 | 0.657 | 0.45 | 2.362E-23 | TEC_IL32 | SERP1 |
| 5.8771E-46 | 0.28175342 | 0.396 | 0.187 | 1.2548E-41 | TEC_IL32 | PPP4C |
| 2.9523E-64 | 0.28169618 | 0.363 | 0.135 | 6.3031E-60 | TEC_IL32 | FXYD5 |
| 1.989E-30 | 0.28125709 | 0.544 | 0.337 | 4.2466E-26 | TEC_IL32 | SELENOK |
| 4.3362E-38 | 0.28084601 | 0.346 | 0.166 | 9.2577E-34 | TEC_IL32 | CAB39 |
| 6.3939E-35 | 0.28072329 | 0.48 | 0.269 | 1.3651E-30 | TEC_IL32 | ASPH |
| 6.0106E-30 | 0.28011833 | 0.612 | 0.387 | 1.2833E-25 | TEC_IL32 | CDV3 |
| 1.1004E-40 | 0.27951334 | 0.472 | 0.25 | 2.3493E-36 | TEC_IL32 | HDGF |
| 5.1047E-29 | 0.27947561 | 0.614 | 0.407 | 1.0899E-24 | TEC_IL32 | LGALS3 |
| 5.636E-32 | 0.27897531 | 0.252 | 0.111 | 1.2033E-27 | TEC_IL32 | NEURL3 |
| 8.0009E-52 | 0.27780215 | 0.317 | 0.125 | 1.7082E-47 | TEC_IL32 | L1CAM |
| 1.1371E-42 | 0.27739706 | 0.311 | 0.135 | 2.4277E-38 | TEC_IL32 | CDCP1 |
| 1.9987E-55 | 0.27719772 | 0.277 | 0.099 | 4.2672E-51 | TEC_IL32 | TGFA |
| 6.6892E-36 | 0.27639227 | 0.513 | 0.289 | 1.4281E-31 | TEC_IL32 | C9orf16 |
| 8.2969E-47 | 0.27626382 | 0.308 | 0.126 | 1.7714E-42 | TEC_IL32 | CTSS |
| 2.6516E-32 | 0.27571354 | 0.304 | 0.147 | 5.6611E-28 | TEC_IL32 | QPRT |
| 5.4439E-47 | 0.27552951 | 0.275 | 0.107 | 1.1623E-42 | TEC_IL32 | GPR160 |
| 5.6463E-19 | 0.27552335 | 0.344 | 0.207 | 1.2055E-14 | TEC_IL32 | GCLM |

| | | | | | | |
|---|---|---|---|---|---|---|
| 1.7546E-35 | 0.2752867 | 0.404 | 0.209 | 3.746E-31 | TEC_IL32 | GARS |
| 4.3306E-36 | 0.27461145 | 0.445 | 0.243 | 9.2457E-32 | TEC_IL32 | CAPZB |
| 1.536E-43 | 0.27422842 | 0.318 | 0.137 | 3.2794E-39 | TEC_IL32 | SNX10 |
| 8.7209E-32 | 0.27422665 | 0.576 | 0.345 | 1.8619E-27 | TEC_IL32 | TALDO1 |
| 1.4272E-43 | 0.27376635 | 0.811 | 0.545 | 3.047E-39 | TEC_IL32 | GUK1 |
| 2.8831E-28 | 0.27349863 | 0.834 | 0.633 | 6.1553E-24 | TEC_IL32 | EDF1 |
| 2.5235E-33 | 0.27335387 | 0.604 | 0.374 | 5.3876E-29 | TEC_IL32 | SUB1 |
| 1.8809E-33 | 0.27331003 | 0.54 | 0.32 | 4.0156E-29 | TEC_IL32 | PPA1 |
| 1.4289E-25 | 0.27330818 | 0.729 | 0.523 | 3.0506E-21 | TEC_IL32 | HLA-E |
| 2.031E-34 | 0.27325096 | 0.515 | 0.298 | 4.3362E-30 | TEC_IL32 | RAB13 |
| 4.2422E-17 | 0.27301824 | 0.286 | 0.167 | 9.0571E-13 | TEC_IL32 | IGFBP3 |
| 1.0134E-48 | 0.27186369 | 0.319 | 0.131 | 2.1636E-44 | TEC_IL32 | NBL1 |
| 9.7527E-38 | 0.27160372 | 0.544 | 0.313 | 2.0822E-33 | TEC_IL32 | TRAM1 |
| 2.2511E-37 | 0.27134071 | 0.561 | 0.316 | 4.806E-33 | TEC_IL32 | TUBA1C |
| 2.2534E-34 | 0.27133013 | 0.484 | 0.276 | 4.8109E-30 | TEC_IL32 | PSME2 |
| 6.5437E-40 | 0.27121935 | 0.362 | 0.174 | 1.3971E-35 | TEC_IL32 | PICALM |
| 1.7561E-41 | 0.27091605 | 0.47 | 0.248 | 3.7492E-37 | TEC_IL32 | GNB2 |
| 9.8895E-34 | 0.27048861 | 0.52 | 0.307 | 2.1114E-29 | TEC_IL32 | XRCC5 |
| 2.9922E-33 | 0.27048533 | 0.412 | 0.224 | 6.3883E-29 | TEC_IL32 | ARPC5L |
| 3.6308E-25 | 0.26985914 | 0.448 | 0.274 | 7.7517E-21 | TEC_IL32 | AKR1B1 |
| 1.5784E-34 | 0.26979599 | 0.617 | 0.375 | 3.3699E-30 | TEC_IL32 | POMP |
| 2.5996E-39 | 0.26915867 | 0.478 | 0.257 | 5.55E-35 | TEC_IL32 | UBE2I |
| 6.0114E-33 | 0.26859022 | 0.31 | 0.15 | 1.2834E-28 | TEC_IL32 | ETS1 |
| 7.2831E-32 | 0.26857675 | 0.749 | 0.528 | 1.5549E-27 | TEC_IL32 | GNG5 |
| 8.3594E-38 | 0.26827369 | 0.477 | 0.26 | 1.7847E-33 | TEC_IL32 | CAPNS1 |
| 1.2395E-39 | 0.26811803 | 0.369 | 0.178 | 2.6464E-35 | TEC_IL32 | RALY |
| 2.6089E-27 | 0.26789234 | 0.554 | 0.353 | 5.57E-23 | TEC_IL32 | PPP1CB |
| 1.2912E-58 | 0.26728432 | 0.291 | 0.102 | 2.7568E-54 | TEC_IL32 | NECTIN4 |
| 1.3509E-21 | 0.26714173 | 0.195 | 0.093 | 2.8842E-17 | TEC_IL32 | CRELD2 |
| 1.9197E-21 | 0.26608814 | 0.412 | 0.254 | 4.0985E-17 | TEC_IL32 | SELENOM |
| 1.3753E-28 | 0.26523615 | 0.473 | 0.281 | 2.9362E-24 | TEC_IL32 | TOP1 |
| 4.7363E-18 | 0.26433772 | 0.255 | 0.141 | 1.0112E-13 | TEC_IL32 | HBEGF |
| 3.9856E-44 | 0.2642214 | 0.368 | 0.169 | 8.5093E-40 | TEC_IL32 | MYO1C |
| 5.3293E-61 | 0.26412383 | 0.272 | 0.09 | 1.1378E-56 | TEC_IL32 | STAP2 |
| 3.3111E-37 | 0.2639483 | 0.438 | 0.23 | 7.0692E-33 | TEC_IL32 | PFKP |
| 1.2794E-38 | 0.26353773 | 0.343 | 0.163 | 2.7316E-34 | TEC_IL32 | PLIN3 |
| 1.854E-37 | 0.26347353 | 0.334 | 0.157 | 3.9583E-33 | TEC_IL32 | ANKRD13A |
| 3.6942E-21 | 0.26264164 | 0.338 | 0.196 | 7.8871E-17 | TEC_IL32 | MDK |
| 1.2259E-64 | 0.26248518 | 0.194 | 0.049 | 2.6172E-60 | TEC_IL32 | SERPINB9 |
| 7.1691E-40 | 0.2619761 | 0.598 | 0.345 | 1.5306E-35 | TEC_IL32 | PSME1 |
| 1.8082E-40 | 0.26126911 | 0.288 | 0.122 | 3.8604E-36 | TEC_IL32 | HDAC7 |
| 1.7828E-27 | 0.26124952 | 0.448 | 0.262 | 3.8064E-23 | TEC_IL32 | MUC1 |
| 3.4241E-33 | 0.26077823 | 0.586 | 0.358 | 7.3104E-29 | TEC_IL32 | SNU13 |
| 7.0221E-32 | 0.26035584 | 0.353 | 0.183 | 1.4992E-27 | TEC_IL32 | RALA |
| 7.4511E-50 | 0.25894989 | 0.301 | 0.118 | 1.5908E-45 | TEC_IL32 | KCNK1 |
| 2.5912E-33 | 0.25889369 | 0.607 | 0.367 | 5.5322E-29 | TEC_IL32 | DYNLT1 |
| 2.3077E-56 | 0.25875658 | 0.255 | 0.084 | 4.927E-52 | TEC_IL32 | MPZL2 |
| 3.4453E-47 | 0.25864026 | 0.285 | 0.112 | 7.3556E-43 | TEC_IL32 | LMTK3 |
| 1.025E-52 | 0.25798451 | 0.295 | 0.11 | 2.1885E-48 | TEC_IL32 | IRF6 |
| 3.0056E-32 | 0.25760998 | 0.955 | 0.86 | 6.417E-28 | TEC_IL32 | RPS4X |

| | | | | | | |
|---|---|---|---|---|---|---|
| 4.0214E-35 | 0.25759427 | 0.4 | 0.209 | 8.5857E-31 | TEC_IL32 | TMOD3 |
| 2.3371E-31 | 0.25755021 | 0.779 | 0.554 | 4.9897E-27 | TEC_IL32 | RAN |
| 3.135E-43 | 0.25720961 | 0.319 | 0.138 | 6.6931E-39 | TEC_IL32 | ZYX |
| 2.586E-41 | 0.25675407 | 0.185 | 0.061 | 5.5211E-37 | TEC_IL32 | CX3CL1 |
| 2.4117E-19 | 0.25663303 | 0.33 | 0.193 | 5.149E-15 | TEC_IL32 | CALD1 |
| 2.1494E-12 | 0.25628305 | 0.1 | 0.045 | 4.5889E-08 | TEC_IL32 | KRT17 |
| 3.3993E-41 | 0.25602233 | 0.356 | 0.167 | 7.2575E-37 | TEC_IL32 | RPN1 |
| 1.4199E-22 | 0.25443027 | 0.324 | 0.183 | 3.0316E-18 | TEC_IL32 | SESN2 |
| 8.3577E-29 | 0.25422212 | 0.342 | 0.181 | 1.7844E-24 | TEC_IL32 | CARHSP1 |
| 1.2431E-31 | 0.25407879 | 0.439 | 0.246 | 2.654E-27 | TEC_IL32 | HM13 |
| 1.3718E-34 | 0.25399298 | 0.411 | 0.22 | 2.9289E-30 | TEC_IL32 | VPS35 |
| 1.9528E-25 | 0.25394039 | 0.496 | 0.311 | 4.1693E-21 | TEC_IL32 | AES |
| 4.3859E-29 | 0.25373903 | 0.997 | 0.97 | 9.3638E-25 | TEC_IL32 | TPT1 |
| 1.3914E-24 | 0.25372083 | 0.55 | 0.359 | 2.9705E-20 | TEC_IL32 | CAMK2N1 |
| 1.2689E-30 | 0.25317493 | 0.443 | 0.25 | 2.7091E-26 | TEC_IL32 | CFLAR |
| 5.878E-34 | 0.25314511 | 0.376 | 0.193 | 1.2549E-29 | TEC_IL32 | MYOF |
| 3.1391E-55 | 0.25304382 | 0.243 | 0.078 | 6.702E-51 | TEC_IL32 | PXDN |
| 1.9926E-29 | 0.25228926 | 0.535 | 0.327 | 4.2541E-25 | TEC_IL32 | PFDN2 |
| 3.8243E-39 | 0.25225371 | 0.271 | 0.114 | 8.1649E-35 | TEC_IL32 | SCD5 |
| 1.3889E-41 | 0.25201636 | 0.305 | 0.131 | 2.9653E-37 | TEC_IL32 | ETHE1 |
| 2.1085E-05 | 0.25189686 | 0.184 | 0.132 | 0.45016418 | TEC_IL32 | SERPINA1 |
| 3.8102E-44 | 0.25167366 | 0.325 | 0.141 | 8.1347E-40 | TEC_IL32 | RAB32 |
| 1.7524E-14 | 0.25124785 | 0.327 | 0.211 | 3.7415E-10 | TEC_IL32 | SERTAD1 |
| 2.7684E-28 | 0.25114599 | 0.48 | 0.285 | 5.9104E-24 | TEC_IL32 | SELENOS |
| 5.5019E-27 | 0.25068213 | 0.419 | 0.244 | 1.1747E-22 | TEC_IL32 | RER1 |
| 1.1433E-24 | 0.25000773 | 0.971 | 0.9 | 2.441E-20 | TEC_IL32 | RPS3 |
| 2.9852E-08 | -0.2501688 | 0.139 | 0.212 | 0.00063734 | TEC_IL32 | VPS13C |
| 0.0001424 | -0.251075 | 0.861 | 0.914 | 1 | TEC_IL32 | MT-CO2 |
| 7.7051E-18 | -0.2514846 | 0.025 | 0.127 | 1.645E-13 | TEC_IL32 | LINC00472 |
| 5.656E-07 | -0.252479 | 0.168 | 0.232 | 0.01207563 | TEC_IL32 | TRA2B |
| 3.8783E-06 | -0.2538285 | 0.158 | 0.218 | 0.08280211 | TEC_IL32 | ZFP36 |
| 0.00938191 | -0.2545645 | 0.499 | 0.511 | 1 | TEC_IL32 | H2AFZ |
| 6.2503E-11 | -0.2551107 | 0.12 | 0.211 | 1.3344E-06 | TEC_IL32 | THUMPD3-AS1 |
| 1.4703E-11 | -0.2555267 | 0.111 | 0.203 | 3.139E-07 | TEC_IL32 | UPF2 |
| 2.1153E-16 | -0.2557193 | 0.033 | 0.131 | 4.5161E-12 | TEC_IL32 | GPNMB |
| 2.3505E-15 | -0.2565959 | 0.061 | 0.164 | 5.0184E-11 | TEC_IL32 | RNPC3 |
| 0.00486133 | -0.2573114 | 0.284 | 0.308 | 1 | TEC_IL32 | YME1L1 |
| 1.0557E-17 | -0.2583691 | 0.03 | 0.134 | 2.254E-13 | TEC_IL32 | PKHD1 |
| 2.0816E-15 | -0.2599012 | 0.055 | 0.158 | 4.4442E-11 | TEC_IL32 | SNHG25 |
| 6.2289E-17 | -0.2606611 | 0.045 | 0.151 | 1.3299E-12 | TEC_IL32 | ILF3-DT |
| 1.4692E-07 | -0.2608711 | 0.156 | 0.226 | 0.00313669 | TEC_IL32 | BEX2 |
| 0.0044976 | -0.2623456 | 0.352 | 0.362 | 1 | TEC_IL32 | AKAP9 |
| 1.4809E-13 | -0.262646 | 0.024 | 0.103 | 3.1617E-09 | TEC_IL32 | MAFA |
| 9.2984E-12 | -0.2635478 | 0.105 | 0.197 | 1.9852E-07 | TEC_IL32 | TUT4 |
| 2.1012E-09 | -0.2662456 | 0.313 | 0.397 | 4.4861E-05 | TEC_IL32 | CIRBP |
| 1.6354E-07 | -0.2683908 | 0.092 | 0.156 | 0.00349149 | TEC_IL32 | PFKFB3 |
| 8.0623E-11 | -0.2729724 | 0.117 | 0.206 | 1.7213E-06 | TEC_IL32 | MDM4 |
| 3.8444E-12 | -0.2751452 | 0.129 | 0.225 | 8.2078E-08 | TEC_IL32 | NKTR |
| 1.0277E-07 | -0.2769238 | 0.137 | 0.205 | 0.00219409 | TEC_IL32 | CHCHD7 |
| 0.00062763 | -0.2778654 | 0.32 | 0.346 | 1 | TEC_IL32 | TSC22D1 |

| | | | | | | |
|---|---|---|---|---|---|---|
| 0.00187925 | -0.2803048 | 0.846 | 0.828 | 1 | TEC_IL32 | SQSTM1 |
| 1.9412E-12 | -0.2806819 | 0.159 | 0.262 | 4.1446E-08 | TEC_IL32 | MPHOSPH8 |
| 9.8407E-05 | -0.2824682 | 0.266 | 0.304 | 1 | TEC_IL32 | C6orf62 |
| 0.00011773 | -0.2847023 | 0.148 | 0.203 | 1 | TEC_IL32 | ANXA1 |
| 1.4692E-16 | -0.28537 | 0.067 | 0.178 | 3.1368E-12 | TEC_IL32 | AHI1 |
| 8.9886E-06 | -0.2855578 | 0.569 | 0.593 | 0.19190694 | TEC_IL32 | RBM39 |
| 6.9944E-09 | -0.2863889 | 0.122 | 0.202 | 0.00014933 | TEC_IL32 | MT-ND6 |
| 0.00070336 | -0.2864883 | 0.071 | 0.106 | 1 | TEC_IL32 | SERPINE2 |
| 1.9058E-07 | -0.2866818 | 0.212 | 0.278 | 0.00406896 | TEC_IL32 | IFRD1 |
| 1.1719E-12 | -0.2870899 | 0.125 | 0.224 | 2.5019E-08 | TEC_IL32 | RERE |
| 5.8887E-14 | -0.2870947 | 0.076 | 0.174 | 1.2572E-09 | TEC_IL32 | TMX4 |
| 5.7361E-14 | -0.2872129 | 0.084 | 0.185 | 1.2246E-09 | TEC_IL32 | MYO9A |
| 2.1185E-07 | -0.2877651 | 0.212 | 0.277 | 0.00452291 | TEC_IL32 | TCF25 |
| 9.9237E-08 | -0.2879498 | 0.226 | 0.293 | 0.0021187 | TEC_IL32 | HP1BP3 |
| 0.00042829 | -0.2891824 | 0.352 | 0.379 | 1 | TEC_IL32 | RSRP1 |
| 2.402E-08 | -0.293292 | 0.644 | 0.665 | 0.00051283 | TEC_IL32 | SOD1 |
| 7.8616E-10 | -0.2971773 | 0.103 | 0.183 | 1.6785E-05 | TEC_IL32 | KLF4 |
| 1.9377E-07 | -0.3002227 | 0.303 | 0.362 | 0.00413694 | TEC_IL32 | SPAG9 |
| 0.00048109 | -0.3013292 | 0.298 | 0.337 | 1 | TEC_IL32 | ZNF292 |
| 4.7767E-08 | -0.301668 | 0.286 | 0.353 | 0.00101983 | TEC_IL32 | DDX24 |
| 8.8813E-08 | -0.3031133 | 0.221 | 0.289 | 0.00189616 | TEC_IL32 | NUFIP2 |
| 1.159E-14 | -0.3032768 | 0.045 | 0.139 | 2.4745E-10 | TEC_IL32 | CSRNP2 |
| 2.7358E-17 | -0.3038527 | 0.018 | 0.113 | 5.841E-13 | TEC_IL32 | AL355075.4 |
| 5.718E-19 | -0.3053888 | 0.025 | 0.133 | 1.2208E-14 | TEC_IL32 | AL135925.1 |
| 1.714E-08 | -0.3067773 | 0.349 | 0.427 | 0.00036593 | TEC_IL32 | SNHG8 |
| 1.8052E-05 | -0.3075693 | 0.28 | 0.33 | 0.38541114 | TEC_IL32 | CEBPB |
| 3.0879E-08 | -0.3139328 | 0.119 | 0.186 | 0.00065927 | TEC_IL32 | ERN1 |
| 1.2764E-05 | -0.3176326 | 0.216 | 0.267 | 0.27251929 | TEC_IL32 | DUSP1 |
| 5.0731E-11 | -0.3228202 | 0.203 | 0.296 | 1.0831E-06 | TEC_IL32 | KMT2A |
| 1.9752E-13 | -0.3241963 | 0.033 | 0.116 | 4.2171E-09 | TEC_IL32 | AC023157.3 |
| 1.6068E-07 | -0.3254384 | 0.13 | 0.196 | 0.00343054 | TEC_IL32 | H2AFX |
| 9.6904E-14 | -0.3302211 | 0.074 | 0.171 | 2.0689E-09 | TEC_IL32 | HIST1H1E |
| 2.4869E-07 | -0.3320385 | 0.284 | 0.34 | 0.00530946 | TEC_IL32 | LUC7L3 |
| 6.401E-13 | -0.3326147 | 0.188 | 0.295 | 1.3666E-08 | TEC_IL32 | MT-ATP8 |
| 3.2258E-23 | -0.3330223 | 0.004 | 0.118 | 6.887E-19 | TEC_IL32 | HIST4H4 |
| 6.4551E-16 | -0.3335684 | 0.052 | 0.154 | 1.3782E-11 | TEC_IL32 | AL118516.1 |
| 1.0151E-08 | -0.3348843 | 0.58 | 0.611 | 0.00021673 | TEC_IL32 | EIF4A2 |
| 4.1858E-17 | -0.341635 | 0.187 | 0.318 | 8.9367E-13 | TEC_IL32 | ISCU |
| 2.9688E-20 | -0.3439568 | 0.023 | 0.134 | 6.3384E-16 | TEC_IL32 | HOMER1 |
| 1.8251E-06 | -0.345386 | 0.643 | 0.67 | 0.03896531 | TEC_IL32 | TPI1 |
| 4.9745E-19 | -0.3487356 | 0.059 | 0.179 | 1.0621E-14 | TEC_IL32 | THSD7A |
| 2.6776E-08 | -0.3500849 | 0.317 | 0.381 | 0.00057167 | TEC_IL32 | PNISR |
| 4.5044E-10 | -0.3501409 | 0.073 | 0.149 | 9.6169E-06 | TEC_IL32 | ID1 |
| 3.4553E-06 | -0.3535602 | 0.362 | 0.403 | 0.07377063 | TEC_IL32 | FUS |
| 7.5492E-13 | -0.3591761 | 0.216 | 0.318 | 1.6118E-08 | TEC_IL32 | WSB1 |
| 1.3783E-19 | -0.3619385 | 0.024 | 0.133 | 2.9426E-15 | TEC_IL32 | PCDH9 |
| 3.4249E-16 | -0.3700374 | 0.016 | 0.105 | 7.3121E-12 | TEC_IL32 | AP001160.1 |
| 6.7158E-17 | -0.3713297 | 0.1 | 0.216 | 1.4338E-12 | TEC_IL32 | MAT2A |
| 2.8199E-06 | -0.3716214 | 0.874 | 0.93 | 0.06020499 | TEC_IL32 | MT-CO1 |
| 0.00685075 | -0.3726481 | 0.217 | 0.248 | 1 | TEC_IL32 | MT1X |

| | | | | | | |
|---|---|---|---|---|---|---|
| 1.6934E-10 | -0.373305 | 0.456 | 0.523 | 3.6154E-06 | TEC_IL32 | HNRNPU |
| 1.3841E-24 | -0.3804986 | 0.091 | 0.245 | 2.955E-20 | TEC_IL32 | CHD9 |
| 2.863E-10 | -0.3811271 | 0.285 | 0.364 | 6.1126E-06 | TEC_IL32 | MCL1 |
| 4.5621E-22 | -0.3820856 | 0.016 | 0.133 | 9.74E-18 | TEC_IL32 | HIST2H2BF |
| 9.9021E-14 | -0.3864918 | 0.235 | 0.339 | 2.1141E-09 | TEC_IL32 | CPEB4 |
| 1.0941E-05 | -0.3908696 | 0.083 | 0.131 | 0.23359724 | TEC_IL32 | COL7A1 |
| 8.5451E-05 | -0.3929958 | 0.275 | 0.33 | 1 | TEC_IL32 | MT1E |
| 2.607E-11 | -0.3938878 | 0.305 | 0.388 | 5.566E-07 | TEC_IL32 | N4BP2L2 |
| 1.7451E-06 | -0.4049138 | 0.291 | 0.346 | 0.03725729 | TEC_IL32 | ZFP36L2 |
| 1.1709E-13 | -0.4057135 | 0.962 | 0.955 | 2.4999E-09 | TEC_IL32 | RPS27 |
| 2.2803E-10 | -0.4081912 | 0.802 | 0.795 | 4.8685E-06 | TEC_IL32 | HNRNPA2B1 |
| 2.0186E-10 | -0.4125719 | 0.237 | 0.318 | 4.3098E-06 | TEC_IL32 | SRSF2 |
| 3.012E-26 | -0.4216351 | 0.01 | 0.143 | 6.4305E-22 | TEC_IL32 | AC025164.1 |
| 5.1012E-25 | -0.4224086 | 0.043 | 0.184 | 1.0891E-20 | TEC_IL32 | FOXC1 |
| 8.5547E-22 | -0.423942 | 0.145 | 0.291 | 1.8264E-17 | TEC_IL32 | PRPF4B |
| 5.9302E-07 | -0.427315 | 0.34 | 0.389 | 0.01266093 | TEC_IL32 | BRD2 |
| 1.1383E-11 | -0.4293452 | 0.29 | 0.371 | 2.4303E-07 | TEC_IL32 | PNN |
| 4.3809E-10 | -0.4308866 | 0.216 | 0.301 | 9.3532E-06 | TEC_IL32 | DST |
| 7.9551E-14 | -0.4357287 | 0.219 | 0.327 | 1.6984E-09 | TEC_IL32 | CCNL1 |
| 4.3752E-17 | -0.4358048 | 0.12 | 0.241 | 9.341E-13 | TEC_IL32 | NFKBIZ |
| 1.2917E-19 | -0.4435356 | 0.148 | 0.285 | 2.7579E-15 | TEC_IL32 | VEGFA |
| 1.623E-15 | -0.4446787 | 0.154 | 0.274 | 3.4652E-11 | TEC_IL32 | KCNQ1OT1 |
| 8.5442E-28 | -0.4451378 | 0.016 | 0.159 | 1.8242E-23 | TEC_IL32 | AC007952.4 |
| 1.2413E-16 | -0.4469046 | 0.03 | 0.129 | 2.6501E-12 | TEC_IL32 | CXCL14 |
| 3.4077E-22 | -0.449021 | 0.048 | 0.18 | 7.2754E-18 | TEC_IL32 | FOSB |
| 5.8819E-07 | -0.450983 | 0.596 | 0.611 | 0.01255782 | TEC_IL32 | BTG1 |
| 1.1081E-08 | -0.4601246 | 0.188 | 0.265 | 0.00023657 | TEC_IL32 | EGR1 |
| 1.178E-14 | -0.4620421 | 0.146 | 0.259 | 2.515E-10 | TEC_IL32 | ID2 |
| 3.842E-13 | -0.4626026 | 0.84 | 0.902 | 8.2027E-09 | TEC_IL32 | MT-CYB |
| 3.677E-18 | -0.4629075 | 0.146 | 0.272 | 7.8504E-14 | TEC_IL32 | AMD1 |
| 2.3134E-13 | -0.4644768 | 0.449 | 0.522 | 4.9391E-09 | TEC_IL32 | SON |
| 4.2729E-09 | -0.4737057 | 0.042 | 0.105 | 9.1225E-05 | TEC_IL32 | APOE |
| 1.9228E-17 | -0.4770404 | 0.119 | 0.245 | 4.1051E-13 | TEC_IL32 | SLC5A3 |
| 1.2332E-18 | -0.4861983 | 0.464 | 0.576 | 2.6328E-14 | TEC_IL32 | RPS27L |
| 1.1656E-07 | -0.4910084 | 0.276 | 0.335 | 0.00248846 | TEC_IL32 | RHOB |
| 3.9671E-15 | -0.4927313 | 0.132 | 0.246 | 8.4697E-11 | TEC_IL32 | MDM2 |
| 1.3791E-09 | -0.4962796 | 0.182 | 0.258 | 2.9444E-05 | TEC_IL32 | IER5 |
| 4.0794E-06 | -0.4999086 | 0.257 | 0.304 | 0.08709491 | TEC_IL32 | PLK2 |
| 2.0722E-14 | -0.5023912 | 0.172 | 0.282 | 4.4241E-10 | TEC_IL32 | CITED2 |
| 6.107E-22 | -0.503126 | 0.1 | 0.243 | 1.3038E-17 | TEC_IL32 | FXYD2 |
| 6.1355E-23 | -0.5038073 | 0.121 | 0.271 | 1.3099E-18 | TEC_IL32 | HIST1H2AC |
| 1.994E-17 | -0.5053264 | 0.859 | 0.916 | 4.2572E-13 | TEC_IL32 | MT-ND4 |
| 1.0765E-10 | -0.5080554 | 0.129 | 0.216 | 2.2984E-06 | TEC_IL32 | ADAMTS1 |
| 1.8571E-32 | -0.5087826 | 0.032 | 0.201 | 3.9649E-28 | TEC_IL32 | TLE4 |
| 3.5894E-19 | -0.5201433 | 0.058 | 0.176 | 7.6633E-15 | TEC_IL32 | EGLN3 |
| 0.00697336 | -0.5210249 | 0.628 | 0.604 | 1 | TEC_IL32 | GADD45B |
| 2.9E-18 | -0.5230238 | 0.246 | 0.365 | 6.1915E-14 | TEC_IL32 | ARGLU1 |
| 1.9344E-23 | -0.5414766 | 0.129 | 0.28 | 4.13E-19 | TEC_IL32 | SEPHS2 |
| 9.0613E-20 | -0.544315 | 0.516 | 0.62 | 1.9346E-15 | TEC_IL32 | MT-ND4L |
| 1.3608E-07 | -0.5464281 | 0.221 | 0.281 | 0.00290541 | TEC_IL32 | RASD1 |

| | | | | | | |
|---|---|---|---|---|---|---|
| 2.5886E-24 | -0.5464684 | 0.136 | 0.295 | 5.5267E-20 | TEC_IL32 | HIST1H1C |
| 3.3275E-20 | -0.5494569 | 0.004 | 0.104 | 7.1042E-16 | TEC_IL32 | SLC12A1 |
| 9.7639E-18 | -0.5528145 | 0.126 | 0.25 | 2.0846E-13 | TEC_IL32 | ID4 |
| 2.5698E-19 | -0.5641538 | 0.488 | 0.566 | 5.4864E-15 | TEC_IL32 | SRRM2 |
| 1.2771E-29 | -0.5647581 | 0.073 | 0.243 | 2.7267E-25 | TEC_IL32 | SYNE1 |
| 5.1078E-22 | -0.567214 | 0.086 | 0.222 | 1.0905E-17 | TEC_IL32 | SOCS1 |
| 5.8986E-23 | -0.5734368 | 0.145 | 0.295 | 1.2593E-18 | TEC_IL32 | CEBPD |
| 4.4973E-18 | -0.5758919 | 0.85 | 0.913 | 9.6017E-14 | TEC_IL32 | MT-ATP6 |
| 6.7124E-22 | -0.5775614 | 0.15 | 0.292 | 1.4331E-17 | TEC_IL32 | POLR2A |
| 1.4249E-08 | -0.5798315 | 0.333 | 0.39 | 0.00030422 | TEC_IL32 | JUNB |
| 1.5941E-17 | -0.5845172 | 0.946 | 0.94 | 3.4035E-13 | TEC_IL32 | MALAT1 |
| 1.055E-26 | -0.5886713 | 0.289 | 0.443 | 2.2525E-22 | TEC_IL32 | DDX17 |
| 5.1107E-13 | -0.5905493 | 0.348 | 0.433 | 1.0911E-08 | TEC_IL32 | SYNE2 |
| 1.1935E-12 | -0.6006946 | 0.352 | 0.445 | 2.5481E-08 | TEC_IL32 | IER3 |
| 3.8212E-20 | -0.6126767 | 0.153 | 0.289 | 8.1583E-16 | TEC_IL32 | BTG2 |
| 1.2728E-07 | -0.6214 | 0.561 | 0.588 | 0.00271746 | TEC_IL32 | ELF3 |
| 9.4047E-09 | -0.6300485 | 0.204 | 0.276 | 0.00020079 | TEC_IL32 | TCIM |
| 5.5441E-29 | -0.636841 | 0.039 | 0.194 | 1.1837E-24 | TEC_IL32 | PIK3R3 |
| 3.1869E-35 | -0.6415084 | 0.034 | 0.214 | 6.804E-31 | TEC_IL32 | SGK1 |
| 6.571E-12 | -0.6469665 | 0.166 | 0.267 | 1.4029E-07 | TEC_IL32 | NUPR1 |
| 1.061E-26 | -0.6646493 | 0.105 | 0.265 | 2.2651E-22 | TEC_IL32 | IER5L |
| 8.3658E-15 | -0.6670843 | 0.448 | 0.573 | 1.7861E-10 | TEC_IL32 | GDF15 |
| 3.7271E-20 | -0.6753123 | 0.185 | 0.32 | 7.9574E-16 | TEC_IL32 | GLUL |
| 2.4337E-17 | -0.6818392 | 0.27 | 0.381 | 5.196E-13 | TEC_IL32 | SLC38A2 |
| 4.9523E-33 | -0.7114825 | 0.068 | 0.249 | 1.0573E-28 | TEC_IL32 | INTS6 |
| 6.6631E-22 | -0.7313707 | 0.298 | 0.423 | 1.4226E-17 | TEC_IL32 | SRSF7 |
| 5.5321E-31 | -0.7353355 | 0.803 | 0.897 | 1.1811E-26 | TEC_IL32 | MT-ND1 |
| 9.4596E-31 | -0.7542873 | 0.765 | 0.874 | 2.0196E-26 | TEC_IL32 | MT-ND3 |
| 1.2924E-13 | -0.7651835 | 0.508 | 0.568 | 2.7593E-09 | TEC_IL32 | IER2 |
| 4.5128E-34 | -0.768206 | 0.788 | 0.896 | 9.6349E-30 | TEC_IL32 | MT-ND2 |
| 4.1057E-20 | -0.8097013 | 0.168 | 0.298 | 8.7656E-16 | TEC_IL32 | FOS |
| 2.968E-26 | -0.8106486 | 0.468 | 0.609 | 6.3367E-22 | TEC_IL32 | CLU |
| 1.0976E-16 | -0.8907096 | 0.033 | 0.133 | 2.3434E-12 | TEC_IL32 | GPX3 |
| 1.2502E-32 | -0.8914196 | 0.533 | 0.646 | 2.6692E-28 | TEC_IL32 | JUND |
| 3.1228E-32 | -0.898723 | 0.468 | 0.627 | 6.6671E-28 | TEC_IL32 | MTRNR2L8 |
| 1.0098E-41 | -0.9064943 | 0.706 | 0.837 | 2.1559E-37 | TEC_IL32 | MT-ND5 |
| 2.5996E-49 | -0.9104153 | 0.053 | 0.291 | 5.5503E-45 | TEC_IL32 | PPP1R10 |
| 1.0402E-32 | -0.9341923 | 0.106 | 0.289 | 2.2208E-28 | TEC_IL32 | HES1 |
| 2.6996E-42 | -0.9713921 | 0.039 | 0.247 | 5.7636E-38 | TEC_IL32 | CSKMT |
| 5.2719E-26 | -1.0258022 | 0.76 | 0.818 | 1.1255E-21 | TEC_IL32 | NEAT1 |
| 4.3402E-06 | -1.0383302 | 0.142 | 0.201 | 0.09266301 | TEC_IL32 | MT1G |
| 1.0096E-10 | -1.0932739 | 0.141 | 0.231 | 2.1556E-06 | TEC_IL32 | DEFB1 |
| 1.2208E-45 | -1.1746582 | 0.061 | 0.284 | 2.6063E-41 | TEC_IL32 | HSPA2 |
| 1.8663E-15 | -1.2072118 | 0.47 | 0.544 | 3.9846E-11 | TEC_IL32 | SOX4 |
| 9.2345E-58 | -1.2445718 | 0.049 | 0.314 | 1.9716E-53 | TEC_IL32 | HEXIM1 |
| 1.3407E-45 | -1.3031037 | 0.166 | 0.391 | 2.8624E-41 | TEC_IL32 | HIST1H4C |
| 1.3372E-42 | -1.3152966 | 0.081 | 0.296 | 2.855E-38 | TEC_IL32 | MAFB |
| 9.3334E-43 | -1.3932938 | 0.492 | 0.643 | 1.9927E-38 | TEC_IL32 | JUN |
| 1.1733E-89 | -1.7140044 | 0.396 | 0.697 | 2.505E-85 | TEC_IL32 | MTRNR2L12 |
| 3.592E-108 | -5.2813578 | 0.277 | 0.619 | 7.67E-104 | TEC_IL32 | PLCG2 |

| | | | | | | |
|---|---|---|---|---|---|---|
| 1.28E-128 | 1.99733702 | 0.894 | 0.308 | 2.734E-124 | TEC_LIF | ITGA2 |
| 2.728E-124 | 1.87419844 | 0.846 | 0.284 | 5.824E-120 | TEC_LIF | ITGB8 |
| 8.151E-100 | 1.51448194 | 0.898 | 0.38 | 1.7402E-95 | TEC_LIF | RAB11FIP1 |
| 6.1081E-84 | 1.50553015 | 0.835 | 0.32 | 1.3041E-79 | TEC_LIF | LAMC2 |
| 8.611E-118 | 1.49610448 | 0.799 | 0.237 | 1.838E-113 | TEC_LIF | MACC1 |
| 7.822E-130 | 1.46510455 | 0.768 | 0.197 | 1.67E-125 | TEC_LIF | ADAMTS9 |
| 5.0079E-83 | 1.43362862 | 0.846 | 0.349 | 1.0692E-78 | TEC_LIF | KLF6 |
| 4.3087E-68 | 1.35349949 | 0.744 | 0.299 | 9.1991E-64 | TEC_LIF | PLAU |
| 3.9808E-97 | 1.32402738 | 0.724 | 0.213 | 8.499E-93 | TEC_LIF | GPRC5A |
| 8.8162E-84 | 1.31459662 | 0.583 | 0.152 | 1.8823E-79 | TEC_LIF | TNFAIP3 |
| 8.603E-105 | 1.27694245 | 0.78 | 0.237 | 1.837E-100 | TEC_LIF | RAP2B |
| 2.0244E-76 | 1.23267519 | 0.76 | 0.276 | 4.3221E-72 | TEC_LIF | PLAUR |
| 8.742E-116 | 1.19792851 | 0.417 | 0.061 | 1.866E-111 | TEC_LIF | LIF |
| 4.8994E-49 | 1.1753921 | 0.957 | 0.697 | 1.046E-44 | TEC_LIF | CLDN4 |
| 1.6913E-22 | 1.15660349 | 0.303 | 0.113 | 3.611E-18 | TEC_LIF | CXCL8 |
| 1.2362E-44 | 1.12975248 | 0.602 | 0.246 | 2.6394E-40 | TEC_LIF | AKAP12 |
| 4.8301E-54 | 1.09241797 | 0.925 | 0.662 | 1.0312E-49 | TEC_LIF | GLS |
| 4.6559E-57 | 1.09043022 | 0.827 | 0.425 | 9.9404E-53 | TEC_LIF | ZBTB20 |
| 3.1431E-96 | 1.07917064 | 0.697 | 0.194 | 6.7104E-92 | TEC_LIF | CLDN1 |
| 2.6927E-47 | 1.06565638 | 0.772 | 0.41 | 5.749E-43 | TEC_LIF | CCND1 |
| 7.1153E-52 | 1.06202221 | 0.917 | 0.657 | 1.5191E-47 | TEC_LIF | TACSTD2 |
| 2.9575E-57 | 1.05271263 | 0.917 | 0.585 | 6.3142E-53 | TEC_LIF | KRT19 |
| 3.9195E-85 | 1.04071917 | 0.795 | 0.289 | 8.368E-81 | TEC_LIF | CDH1 |
| 1.1176E-51 | 1.00529907 | 0.516 | 0.173 | 2.386E-47 | TEC_LIF | DCBLD2 |
| 1.544E-109 | 1.00275915 | 0.516 | 0.097 | 3.297E-105 | TEC_LIF | VGLL3 |
| 1.6994E-41 | 0.9647711 | 0.642 | 0.297 | 3.6283E-37 | TEC_LIF | ARHGAP29 |
| 6.0524E-57 | 0.96332971 | 0.295 | 0.057 | 1.2922E-52 | TEC_LIF | KRTAP2-3 |
| 2.5413E-29 | 0.95676255 | 0.449 | 0.184 | 5.4257E-25 | TEC_LIF | THBS1 |
| 1.9456E-63 | 0.95485835 | 0.764 | 0.309 | 4.1538E-59 | TEC_LIF | UGCG |
| 9.1362E-48 | 0.95421284 | 0.799 | 0.413 | 1.9506E-43 | TEC_LIF | TPM4 |
| 2.4114E-23 | 0.95072557 | 0.547 | 0.283 | 5.1484E-19 | TEC_LIF | CYR61 |
| 2.8467E-34 | 0.93239783 | 0.579 | 0.268 | 6.0778E-30 | TEC_LIF | VMP1 |
| 1.1262E-64 | 0.92426998 | 0.559 | 0.164 | 2.4044E-60 | TEC_LIF | DUSP5 |
| 2.2582E-44 | 0.92051814 | 0.571 | 0.214 | 4.8212E-40 | TEC_LIF | SFN |
| 1.3469E-36 | 0.89978807 | 0.48 | 0.18 | 2.8757E-32 | TEC_LIF | BIRC3 |
| 4.3676E-63 | 0.89579907 | 0.776 | 0.332 | 9.3248E-59 | TEC_LIF | JUP |
| 4.1599E-62 | 0.89110974 | 0.732 | 0.301 | 8.8815E-58 | TEC_LIF | DSP |
| 2.2176E-43 | 0.88476063 | 0.614 | 0.252 | 4.7345E-39 | TEC_LIF | LAMB3 |
| 1.5496E-46 | 0.87889428 | 0.461 | 0.146 | 3.3084E-42 | TEC_LIF | RCAN1 |
| 8.8482E-85 | 0.87822153 | 0.701 | 0.213 | 1.8891E-80 | TEC_LIF | TEAD1 |
| 1.5355E-62 | 0.87662602 | 0.303 | 0.055 | 3.2783E-58 | TEC_LIF | EDN1 |
| 7.3693E-22 | 0.87448396 | 0.323 | 0.126 | 1.5734E-17 | TEC_LIF | TNFRSF11B |
| 6.706E-99 | 0.86213027 | 0.634 | 0.158 | 1.4317E-94 | TEC_LIF | ELK3 |
| 4.3768E-23 | 0.84376658 | 1 | 0.939 | 9.3444E-19 | TEC_LIF | MALAT1 |
| 1.8289E-52 | 0.82573548 | 0.681 | 0.281 | 3.9048E-48 | TEC_LIF | ITGA3 |
| 3.2448E-59 | 0.82573165 | 0.469 | 0.128 | 6.9277E-55 | TEC_LIF | ITPKC |
| 1.0621E-71 | 0.81343525 | 0.74 | 0.258 | 2.2675E-67 | TEC_LIF | MET |
| 4.3213E-40 | 0.80964242 | 0.626 | 0.271 | 9.2259E-36 | TEC_LIF | SERPINB1 |
| 2.443E-53 | 0.80594028 | 0.563 | 0.191 | 5.2158E-49 | TEC_LIF | PHLDB2 |
| 1.2943E-68 | 0.80178843 | 0.657 | 0.22 | 2.7634E-64 | TEC_LIF | ST14 |

| | | | | | | |
|---|---|---|---|---|---|---|
| 1.6421E-50 | 0.78047958 | 0.732 | 0.322 | 3.5058E-46 | TEC_LIF | CD2AP |
| 9.2642E-43 | 0.78011608 | 0.579 | 0.225 | 1.9779E-38 | TEC_LIF | TGM2 |
| 1.536E-25 | 0.77761142 | 0.748 | 0.438 | 3.2795E-21 | TEC_LIF | ZFP36L1 |
| 2.6721E-65 | 0.77525095 | 0.571 | 0.167 | 5.7049E-61 | TEC_LIF | ZNF217 |
| 1.1523E-41 | 0.77209642 | 0.472 | 0.163 | 2.4601E-37 | TEC_LIF | ERRFI1 |
| 2.4503E-72 | 0.76960696 | 0.48 | 0.116 | 5.2314E-68 | TEC_LIF | CDK6 |
| 1.6077E-97 | 0.76816344 | 0.5 | 0.1 | 3.4324E-93 | TEC_LIF | F2RL1 |
| 4.5318E-32 | 0.76355208 | 0.835 | 0.56 | 9.6754E-28 | TEC_LIF | MAL2 |
| 1.9246E-38 | 0.74942955 | 0.669 | 0.313 | 4.1089E-34 | TEC_LIF | DCDC2 |
| 2.1666E-40 | 0.74457447 | 0.831 | 0.489 | 4.6256E-36 | TEC_LIF | ACTN4 |
| 5.1066E-65 | 0.74326339 | 0.535 | 0.152 | 1.0903E-60 | TEC_LIF | TP53BP2 |
| 2.836E-31 | 0.74216098 | 0.839 | 0.58 | 6.0549E-27 | TEC_LIF | EZR |
| 2.6749E-72 | 0.73936797 | 0.398 | 0.082 | 5.7109E-68 | TEC_LIF | PXDN |
| 3.703E-23 | 0.73907501 | 0.917 | 0.798 | 7.9059E-19 | TEC_LIF | CD24 |
| 8.3345E-88 | 0.73445616 | 0.606 | 0.156 | 1.7794E-83 | TEC_LIF | GRAMD2B |
| 1.0083E-27 | 0.73179771 | 0.52 | 0.233 | 2.1527E-23 | TEC_LIF | ITGB6 |
| 2.8481E-70 | 0.7293872 | 0.5 | 0.126 | 6.0808E-66 | TEC_LIF | FGD6 |
| 2.7823E-15 | 0.7290849 | 0.508 | 0.297 | 5.9401E-11 | TEC_LIF | DDIT4 |
| 9.1402E-23 | 0.7279124 | 0.346 | 0.14 | 1.9514E-18 | TEC_LIF | XIST |
| 1.0662E-38 | 0.72254888 | 0.358 | 0.105 | 2.2764E-34 | TEC_LIF | ICAM1 |
| 7.7823E-43 | 0.71357983 | 0.705 | 0.342 | 1.6615E-38 | TEC_LIF | ANXA11 |
| 2.925E-104 | 0.71327331 | 0.531 | 0.105 | 6.244E-100 | TEC_LIF | NECTIN4 |
| 2.295E-41 | 0.71274014 | 0.433 | 0.142 | 4.8998E-37 | TEC_LIF | ARL4C |
| 4.5614E-64 | 0.71000552 | 0.669 | 0.228 | 9.7386E-60 | TEC_LIF | LPP |
| 2.4214E-45 | 0.70580935 | 0.528 | 0.185 | 5.1697E-41 | TEC_LIF | BHLHE40 |
| 6.9149E-55 | 0.70290956 | 0.638 | 0.227 | 1.4763E-50 | TEC_LIF | MIDN |
| 7.5723E-69 | 0.70260074 | 0.327 | 0.058 | 1.6167E-64 | TEC_LIF | ENC1 |
| 3.0801E-52 | 0.70134346 | 0.587 | 0.21 | 6.576E-48 | TEC_LIF | SVIL |
| 2.3079E-36 | 0.69687717 | 0.925 | 0.699 | 4.9273E-32 | TEC_LIF | ITGB1 |
| 9.832E-44 | 0.69504485 | 0.516 | 0.188 | 2.0991E-39 | TEC_LIF | SLC25A37 |
| 2.2327E-42 | 0.69316526 | 0.551 | 0.21 | 4.7668E-38 | TEC_LIF | USP53 |
| 6.0824E-47 | 0.69001542 | 0.571 | 0.207 | 1.2986E-42 | TEC_LIF | PARD6B |
| 1.9607E-55 | 0.68974896 | 0.449 | 0.124 | 4.1861E-51 | TEC_LIF | NCEH1 |
| 4.2557E-39 | 0.68769912 | 0.476 | 0.169 | 9.0859E-35 | TEC_LIF | PHLDA1 |
| 2.1036E-32 | 0.6844409 | 0.724 | 0.4 | 4.4912E-28 | TEC_LIF | RNF213 |
| 6.071E-47 | 0.68441906 | 0.488 | 0.157 | 1.2962E-42 | TEC_LIF | ADGRF1 |
| 2.6412E-49 | 0.67132995 | 0.626 | 0.244 | 5.6389E-45 | TEC_LIF | PTBP3 |
| 9.2115E-45 | 0.66775144 | 0.555 | 0.201 | 1.9667E-40 | TEC_LIF | SLFN5 |
| 2.5457E-14 | 0.66240907 | 0.197 | 0.072 | 5.4352E-10 | TEC_LIF | SPRR2A |
| 5.415E-30 | 0.66091833 | 0.689 | 0.352 | 1.1561E-25 | TEC_LIF | CD44 |
| 3.088E-49 | 0.65930725 | 0.555 | 0.194 | 6.5929E-45 | TEC_LIF | FOXO3 |
| 7.0832E-16 | 0.65801731 | 0.358 | 0.171 | 1.5123E-11 | TEC_LIF | IGFBP3 |
| 3.6775E-79 | 0.6571671 | 0.492 | 0.112 | 7.8515E-75 | TEC_LIF | DAPP1 |
| 5.0936E-58 | 0.65514681 | 0.575 | 0.186 | 1.0875E-53 | TEC_LIF | FAM129B |
| 1.8048E-40 | 0.65031361 | 0.516 | 0.192 | 3.8532E-36 | TEC_LIF | MARCKS |
| 2.0954E-31 | 0.649579 | 0.701 | 0.378 | 4.4736E-27 | TEC_LIF | MYH9 |
| 3.43E-73 | 0.64734831 | 0.563 | 0.152 | 7.3231E-69 | TEC_LIF | F11R |
| 2.2779E-43 | 0.64696139 | 0.689 | 0.302 | 4.8633E-39 | TEC_LIF | ARHGAP5 |
| 1.2587E-42 | 0.6468157 | 0.776 | 0.377 | 2.6873E-38 | TEC_LIF | MYO6 |
| 4.365E-48 | 0.64114037 | 0.484 | 0.153 | 9.3192E-44 | TEC_LIF | ETS1 |

| | | | | | | |
|---|---|---|---|---|---|---|
| 2.0014E-27 | 0.63765409 | 0.728 | 0.417 | 4.273E-23 | TEC_LIF | AHNAK |
| 1.1774E-51 | 0.63659476 | 0.555 | 0.19 | 2.5138E-47 | TEC_LIF | MAP2K3 |
| 4.2418E-26 | 0.63433398 | 0.346 | 0.125 | 9.0563E-22 | TEC_LIF | PRSS22 |
| 1.0963E-54 | 0.6299418 | 0.535 | 0.17 | 2.3405E-50 | TEC_LIF | BCL2L1 |
| 4.8212E-52 | 0.62920101 | 0.602 | 0.216 | 1.0293E-47 | TEC_LIF | NPEPPS |
| 2.0986E-51 | 0.62619733 | 0.291 | 0.06 | 4.4805E-47 | TEC_LIF | CLCF1 |
| 1.4024E-38 | 0.61767263 | 0.402 | 0.128 | 2.9942E-34 | TEC_LIF | MAST4 |
| 1.8617E-51 | 0.61762746 | 0.559 | 0.188 | 3.9747E-47 | TEC_LIF | TBL1XR1 |
| 2.9279E-76 | 0.61717412 | 0.335 | 0.057 | 6.2511E-72 | TEC_LIF | SEMA7A |
| 2.4757E-26 | 0.61678211 | 0.724 | 0.425 | 5.2857E-22 | TEC_LIF | ZFAND5 |
| 3.1706E-43 | 0.61658545 | 0.433 | 0.138 | 6.7691E-39 | TEC_LIF | LPIN1 |
| 1.9211E-53 | 0.61330225 | 0.531 | 0.17 | 4.1016E-49 | TEC_LIF | SECISBP2L |
| 3.2293E-62 | 0.60950093 | 0.543 | 0.16 | 6.8947E-58 | TEC_LIF | ANKRD13A |
| 1.1605E-39 | 0.60920977 | 0.571 | 0.228 | 2.4776E-35 | TEC_LIF | ITGAV |
| 1.7762E-23 | 0.60785468 | 0.394 | 0.163 | 3.7922E-19 | TEC_LIF | LMO7 |
| 1.2006E-31 | 0.60673918 | 0.717 | 0.361 | 2.5632E-27 | TEC_LIF | MACF1 |
| 1.8496E-32 | 0.60458547 | 0.37 | 0.126 | 3.9488E-28 | TEC_LIF | LDLR |
| 1.49E-46 | 0.6029062 | 0.669 | 0.276 | 3.1812E-42 | TEC_LIF | SPINT1 |
| 1.1578E-43 | 0.60157735 | 0.638 | 0.257 | 2.4719E-39 | TEC_LIF | LIMA1 |
| 6.5346E-48 | 0.5994314 | 0.421 | 0.123 | 1.3951E-43 | TEC_LIF | ELL2 |
| 2.43E-20 | 0.59857435 | 0.567 | 0.294 | 5.1882E-16 | TEC_LIF | MARCKSL1 |
| 3.2589E-45 | 0.59372516 | 0.543 | 0.195 | 6.9578E-41 | TEC_LIF | IRF2BP2 |
| 1.1722E-29 | 0.59316051 | 0.862 | 0.551 | 2.5026E-25 | TEC_LIF | LRRFIP1 |
| 9.8523E-39 | 0.58988354 | 0.61 | 0.255 | 2.1035E-34 | TEC_LIF | CFLAR |
| 4.7437E-36 | 0.58350815 | 0.484 | 0.183 | 1.0128E-31 | TEC_LIF | EMP1 |
| 7.435E-25 | 0.58219628 | 0.587 | 0.292 | 1.5874E-20 | TEC_LIF | CYTOR |
| 1.0994E-21 | 0.58003475 | 0.185 | 0.051 | 2.3472E-17 | TEC_LIF | IL11 |
| 3.2905E-35 | 0.5787725 | 0.587 | 0.254 | 7.0251E-31 | TEC_LIF | VCL |
| 1.6967E-45 | 0.5740243 | 0.421 | 0.125 | 3.6224E-41 | TEC_LIF | KDM6B |
| 4.7214E-48 | 0.57191223 | 0.622 | 0.229 | 1.008E-43 | TEC_LIF | AKAP13 |
| 6.0449E-45 | 0.5692637 | 0.547 | 0.198 | 1.2906E-40 | TEC_LIF | OCLN |
| 1.3193E-52 | 0.56889281 | 0.441 | 0.123 | 2.8167E-48 | TEC_LIF | EPHA2 |
| 3.9598E-35 | 0.56565648 | 0.748 | 0.379 | 8.4541E-31 | TEC_LIF | CSNK1A1 |
| 1.6238E-31 | 0.56459232 | 0.409 | 0.147 | 3.4668E-27 | TEC_LIF | ITGA6 |
| 8.6696E-39 | 0.56366261 | 0.539 | 0.203 | 1.851E-34 | TEC_LIF | RAB3IP |
| 1.6495E-48 | 0.56189437 | 0.567 | 0.2 | 3.5217E-44 | TEC_LIF | MYO1E |
| 2.9081E-19 | 0.55668924 | 0.453 | 0.213 | 6.2088E-15 | TEC_LIF | TNFAIP2 |
| 2.5813E-36 | 0.55637433 | 0.685 | 0.314 | 5.5111E-32 | TEC_LIF | RRBP1 |
| 6.3903E-43 | 0.5556951 | 0.555 | 0.204 | 1.3643E-38 | TEC_LIF | PPFIBP1 |
| 8.3694E-61 | 0.54944178 | 0.457 | 0.119 | 1.7869E-56 | TEC_LIF | B4GALT5 |
| 1.5589E-62 | 0.54917247 | 0.48 | 0.125 | 3.3282E-58 | TEC_LIF | HDAC7 |
| 1.1614E-24 | 0.5491495 | 0.669 | 0.364 | 2.4795E-20 | TEC_LIF | SPTBN1 |
| 1.3937E-56 | 0.54822237 | 0.323 | 0.067 | 2.9756E-52 | TEC_LIF | HS3ST1 |
| 4.5272E-10 | 0.54733233 | 0.205 | 0.093 | 9.6656E-06 | TEC_LIF | CCL2 |
| 3.2732E-31 | 0.54731578 | 0.476 | 0.195 | 6.9882E-27 | TEC_LIF | EHF |
| 4.0732E-51 | 0.54665141 | 0.39 | 0.1 | 8.6964E-47 | TEC_LIF | PIM1 |
| 5.4844E-39 | 0.54547915 | 0.551 | 0.217 | 1.1709E-34 | TEC_LIF | CXCL16 |
| 1.4761E-49 | 0.54326123 | 0.48 | 0.147 | 3.1515E-45 | TEC_LIF | EGFR |
| 1.2812E-34 | 0.54193339 | 0.709 | 0.341 | 2.7354E-30 | TEC_LIF | UBE2H |
| 5.8563E-42 | 0.54026539 | 0.551 | 0.205 | 1.2503E-37 | TEC_LIF | ADAM10 |

| | | | | | | |
|---|---|---|---|---|---|---|
| 1.7949E-26 | 0.53969154 | 0.642 | 0.338 | 3.832E-22 | TEC_LIF | S100A16 |
| 2.2525E-62 | 0.53602253 | 0.394 | 0.088 | 4.8091E-58 | TEC_LIF | PDP1 |
| 3.2414E-13 | 0.5318505 | 0.331 | 0.164 | 6.9203E-09 | TEC_LIF | FGF2 |
| 2.1128E-47 | 0.52836499 | 0.453 | 0.135 | 4.5109E-43 | TEC_LIF | NFKB1 |
| 5.2962E-48 | 0.52752376 | 0.484 | 0.152 | 1.1307E-43 | TEC_LIF | ASAP2 |
| 9.9918E-35 | 0.52599589 | 0.433 | 0.152 | 2.1332E-30 | TEC_LIF | PMEPA1 |
| 6.8282E-42 | 0.5250481 | 0.539 | 0.199 | 1.4578E-37 | TEC_LIF | MAP4K4 |
| 1.8649E-43 | 0.52344245 | 0.516 | 0.179 | 3.9815E-39 | TEC_LIF | PICALM |
| 1.8336E-36 | 0.52310032 | 0.571 | 0.231 | 3.9146E-32 | TEC_LIF | HECTD1 |
| 1.1566E-35 | 0.52124547 | 0.512 | 0.2 | 2.4693E-31 | TEC_LIF | ATP13A3 |
| 2.5121E-54 | 0.51761265 | 0.26 | 0.047 | 5.3633E-50 | TEC_LIF | RTKN2 |
| 8.1914E-35 | 0.51554697 | 0.575 | 0.241 | 1.7489E-30 | TEC_LIF | TES |
| 5.3851E-48 | 0.51523447 | 0.413 | 0.116 | 1.1497E-43 | TEC_LIF | IRF6 |
| 5.0545E-25 | 0.51395917 | 0.799 | 0.498 | 1.0791E-20 | TEC_LIF | CLDN7 |
| 1.1115E-48 | 0.51347124 | 0.402 | 0.109 | 2.3731E-44 | TEC_LIF | C6orf132 |
| 5.8337E-32 | 0.51194929 | 0.48 | 0.182 | 1.2455E-27 | TEC_LIF | SYTL2 |
| 8.1316E-25 | 0.51143018 | 0.913 | 0.746 | 1.7361E-20 | TEC_LIF | RTN4 |
| 1.5254E-32 | 0.51136872 | 0.626 | 0.293 | 3.2567E-28 | TEC_LIF | ACTR3 |
| 1.288E-55 | 0.50629842 | 0.409 | 0.104 | 2.7499E-51 | TEC_LIF | TGFA |
| 3.1641E-19 | 0.50603741 | 0.614 | 0.337 | 6.7553E-15 | TEC_LIF | ZFP36L2 |
| 8.3417E-27 | 0.50537335 | 0.5 | 0.222 | 1.781E-22 | TEC_LIF | PRSS8 |
| 4.4216E-39 | 0.50470732 | 0.469 | 0.161 | 9.44E-35 | TEC_LIF | AMOTL2 |
| 1.8436E-46 | 0.50376785 | 0.547 | 0.188 | 3.936E-42 | TEC_LIF | BACH1 |
| 4.6319E-34 | 0.49986995 | 0.677 | 0.307 | 9.8891E-30 | TEC_LIF | AFDN |
| 4.0814E-38 | 0.49926233 | 0.504 | 0.185 | 8.7139E-34 | TEC_LIF | FNDC3B |
| 3.8822E-60 | 0.49801329 | 0.394 | 0.091 | 8.2885E-56 | TEC_LIF | RASA2 |
| 8.828E-36 | 0.49753162 | 0.48 | 0.177 | 1.8848E-31 | TEC_LIF | CELF1 |
| 1.7321E-33 | 0.49721893 | 0.398 | 0.14 | 3.698E-29 | TEC_LIF | CBX6 |
| 2.6073E-41 | 0.49662469 | 0.457 | 0.149 | 5.5666E-37 | TEC_LIF | SMAD3 |
| 1.0105E-19 | 0.49566188 | 0.673 | 0.401 | 2.1573E-15 | TEC_LIF | SDC4 |
| 4.0347E-30 | 0.49542589 | 0.583 | 0.259 | 8.6142E-26 | TEC_LIF | SLC38A1 |
| 1.1797E-32 | 0.49205935 | 0.563 | 0.237 | 2.5186E-28 | TEC_LIF | TNIP1 |
| 1.2117E-39 | 0.49201356 | 0.429 | 0.14 | 2.587E-35 | TEC_LIF | MYO5B |
| 3.1572E-29 | 0.48934717 | 0.366 | 0.13 | 6.7407E-25 | TEC_LIF | CREB5 |
| 1.2461E-31 | 0.48780584 | 0.409 | 0.147 | 2.6605E-27 | TEC_LIF | PROM1 |
| 1.1857E-63 | 0.48641435 | 0.421 | 0.098 | 2.5314E-59 | TEC_LIF | TSPAN14 |
| 8.6183E-33 | 0.48638973 | 0.382 | 0.129 | 1.84E-28 | TEC_LIF | FAM84B |
| 8.7691E-36 | 0.48245583 | 0.508 | 0.198 | 1.8722E-31 | TEC_LIF | MYOF |
| 2.003E-19 | 0.48187411 | 0.102 | 0.02 | 4.2764E-15 | TEC_LIF | IL6 |
| 9.9944E-45 | 0.47796623 | 0.465 | 0.146 | 2.1338E-40 | TEC_LIF | TRIO |
| 3.6889E-16 | 0.47749727 | 0.382 | 0.188 | 7.8758E-12 | TEC_LIF | SESN2 |
| 1.1306E-14 | 0.47694753 | 0.591 | 0.369 | 2.4139E-10 | TEC_LIF | FLNA |
| 1.7731E-25 | 0.47627731 | 0.476 | 0.207 | 3.7855E-21 | TEC_LIF | PLEC |
| 7.4483E-36 | 0.47592354 | 0.48 | 0.176 | 1.5902E-31 | TEC_LIF | BCL10 |
| 1.1549E-37 | 0.47284997 | 0.421 | 0.14 | 2.4657E-33 | TEC_LIF | CDCP1 |
| 3.7653E-06 | 0.47152982 | 0.327 | 0.21 | 0.08038941 | TEC_LIF | GLRX |
| 2.7375E-36 | 0.47048604 | 0.476 | 0.175 | 5.8446E-32 | TEC_LIF | RBMS1 |
| 2.7793E-33 | 0.46992057 | 0.63 | 0.286 | 5.9337E-29 | TEC_LIF | ACTR2 |
| 2.1532E-14 | 0.46852947 | 0.484 | 0.269 | 4.597E-10 | TEC_LIF | MUC1 |
| 6.2939E-30 | 0.46754116 | 0.339 | 0.114 | 1.3438E-25 | TEC_LIF | LIPG |

| | | | | | | |
|---|---|---|---|---|---|---|
| 2.765E-29 | 0.46737086 | 0.622 | 0.294 | 5.9034E-25 | TEC_LIF | IQGAP1 |
| 5.6316E-36 | 0.4662012 | 0.406 | 0.133 | 1.2024E-31 | TEC_LIF | AGO2 |
| 1.7046E-08 | 0.46534575 | 0.453 | 0.285 | 0.00036393 | TEC_LIF | RND3 |
| 2.3725E-35 | 0.46396818 | 0.445 | 0.156 | 5.0653E-31 | TEC_LIF | BBC3 |
| 2.6468E-13 | 0.46070761 | 0.134 | 0.041 | 5.651E-09 | TEC_LIF | EDN2 |
| 2.9908E-11 | 0.4605565 | 0.78 | 0.627 | 6.3853E-07 | TEC_LIF | TPM1 |
| 2.9262E-18 | 0.46004221 | 0.843 | 0.645 | 6.2475E-14 | TEC_LIF | KRT7 |
| 1.1741E-35 | 0.45752419 | 0.508 | 0.195 | 2.5066E-31 | TEC_LIF | UBE2Z |
| 1.9807E-23 | 0.45643773 | 0.602 | 0.313 | 4.2287E-19 | TEC_LIF | FAM107B |
| 1.6145E-26 | 0.45542028 | 0.457 | 0.193 | 3.4471E-22 | TEC_LIF | MINDY2 |
| 4.9773E-38 | 0.45427 | 0.591 | 0.232 | 1.0627E-33 | TEC_LIF | ANKLE2 |
| 1.1996E-17 | 0.45033694 | 0.398 | 0.192 | 2.5612E-13 | TEC_LIF | NCOA7 |
| 6.6493E-38 | 0.4491151 | 0.453 | 0.156 | 1.4196E-33 | TEC_LIF | TAOK1 |
| 1.1623E-50 | 0.44636875 | 0.394 | 0.102 | 2.4814E-46 | TEC_LIF | IGF2BP2 |
| 5.8003E-34 | 0.44232938 | 0.492 | 0.19 | 1.2384E-29 | TEC_LIF | CMTM6 |
| 1.5253E-20 | 0.44034947 | 0.335 | 0.134 | 3.2565E-16 | TEC_LIF | NABP1 |
| 3.3079E-17 | 0.43959097 | 0.78 | 0.554 | 7.0624E-13 | TEC_LIF | HMGA1 |
| 1.1578E-37 | 0.43862645 | 0.339 | 0.096 | 2.472E-33 | TEC_LIF | ATXN1 |
| 1.3733E-11 | 0.4382817 | 0.575 | 0.37 | 2.9321E-07 | TEC_LIF | ATF3 |
| 2.4525E-36 | 0.43810504 | 0.358 | 0.108 | 5.236E-32 | TEC_LIF | SAMD4A |
| 7.4097E-24 | 0.4365779 | 0.665 | 0.36 | 1.582E-19 | TEC_LIF | PPP1CB |
| 2.2843E-13 | 0.43596798 | 0.354 | 0.177 | 4.877E-09 | TEC_LIF | HMGCS1 |
| 1.2875E-36 | 0.43557998 | 0.378 | 0.117 | 2.7488E-32 | TEC_LIF | PIK3IP1 |
| 1.1287E-14 | 0.43538016 | 0.504 | 0.274 | 2.4098E-10 | TEC_LIF | COBLL1 |
| 2.2232E-14 | 0.43532352 | 0.508 | 0.29 | 4.7466E-10 | TEC_LIF | CD55 |
| 4.0476E-28 | 0.43372435 | 0.583 | 0.266 | 8.6417E-24 | TEC_LIF | OGA |
| 2.1756E-18 | 0.43370471 | 0.24 | 0.083 | 4.6449E-14 | TEC_LIF | CCDC71L |
| 1.8437E-20 | 0.43273386 | 0.366 | 0.152 | 3.9363E-16 | TEC_LIF | RAPGEF3 |
| 1.3815E-32 | 0.43126322 | 0.457 | 0.171 | 2.9495E-28 | TEC_LIF | CAB39 |
| 2.4664E-07 | 0.43097346 | 0.484 | 0.347 | 0.00526578 | TEC_LIF | TXNIP |
| 1.3725E-56 | 0.43051439 | 0.358 | 0.08 | 2.9304E-52 | TEC_LIF | TICAM1 |
| 7.8597E-94 | 0.42948881 | 0.268 | 0.03 | 1.678E-89 | TEC_LIF | SHC4 |
| 1.549E-29 | 0.42663236 | 0.461 | 0.185 | 3.3071E-25 | TEC_LIF | ARF6 |
| 7.3759E-20 | 0.42604463 | 0.874 | 0.674 | 1.5747E-15 | TEC_LIF | YWHAZ |
| 2.7556E-70 | 0.4255567 | 0.331 | 0.058 | 5.8832E-66 | TEC_LIF | TMCC3 |
| 7.8718E-07 | 0.42529247 | 0.276 | 0.164 | 0.01680626 | TEC_LIF | CTGF |
| 5.3628E-29 | 0.4248569 | 0.469 | 0.189 | 1.145E-24 | TEC_LIF | CCNG2 |
| 8.1366E-26 | 0.4214715 | 0.35 | 0.127 | 1.7372E-21 | TEC_LIF | AL353751.1 |
| 1.652E-29 | 0.42076927 | 0.327 | 0.107 | 3.527E-25 | TEC_LIF | C3orf52 |
| 6.0676E-39 | 0.42000161 | 0.465 | 0.156 | 1.2954E-34 | TEC_LIF | HSPG2 |
| 1.2295E-29 | 0.41862651 | 0.476 | 0.191 | 2.6249E-25 | TEC_LIF | WWC1 |
| 2.753E-37 | 0.41753344 | 0.291 | 0.076 | 5.8776E-33 | TEC_LIF | TFAP2A |
| 1.8132E-25 | 0.41716991 | 0.587 | 0.283 | 3.8712E-21 | TEC_LIF | TJP1 |
| 3.3865E-21 | 0.41627606 | 0.61 | 0.329 | 7.2302E-17 | TEC_LIF | PTP4A2 |
| 1.5933E-27 | 0.41554532 | 0.236 | 0.065 | 3.4017E-23 | TEC_LIF | CX3CL1 |
| 0.0001447 | 0.41250537 | 0.366 | 0.262 | 1 | TEC_LIF | DUSP1 |
| 3.0052E-26 | 0.41039261 | 0.524 | 0.232 | 6.416E-22 | TEC_LIF | CRIM1 |
| 7.3346E-21 | 0.40990076 | 0.697 | 0.402 | 1.5659E-16 | TEC_LIF | JAK1 |
| 1.0445E-22 | 0.40813253 | 0.508 | 0.234 | 2.2299E-18 | TEC_LIF | KLF5 |
| 1.8493E-35 | 0.40772639 | 0.469 | 0.172 | 3.9483E-31 | TEC_LIF | ABI1 |

| | | | | | | |
|---|---|---|---|---|---|---|
| 2.2564E-30 | 0.40765708 | 0.307 | 0.095 | 4.8175E-26 | TEC_LIF | RAPH1 |
| 7.2927E-28 | 0.40668945 | 0.445 | 0.179 | 1.557E-23 | TEC_LIF | SLC44A2 |
| 5.5457E-42 | 0.40618413 | 0.421 | 0.129 | 1.184E-37 | TEC_LIF | EHD4 |
| 1.5431E-31 | 0.40592473 | 0.343 | 0.11 | 3.2944E-27 | TEC_LIF | RAI14 |
| 1.2084E-35 | 0.40500658 | 0.295 | 0.078 | 2.58E-31 | TEC_LIF | FSD1L |
| 9.4038E-34 | 0.40452045 | 0.343 | 0.105 | 2.0077E-29 | TEC_LIF | MALT1 |
| 8.7036E-23 | 0.40338051 | 0.39 | 0.161 | 1.8582E-18 | TEC_LIF | NTN4 |
| 3.6339E-16 | 0.40287101 | 0.358 | 0.17 | 7.7584E-12 | TEC_LIF | DSG2 |
| 1.1859E-20 | 0.40204511 | 0.547 | 0.277 | 2.5318E-16 | TEC_LIF | ASPH |
| 2.079E-16 | 0.40144526 | 0.283 | 0.115 | 4.4387E-12 | TEC_LIF | SOX9 |
| 1.5054E-22 | 0.40141905 | 0.528 | 0.26 | 3.2139E-18 | TEC_LIF | TAPBP |
| 6.6307E-32 | 0.40131252 | 0.5 | 0.199 | 1.4156E-27 | TEC_LIF | CTBP2 |
| 5.7933E-37 | 0.40055803 | 0.374 | 0.115 | 1.2369E-32 | TEC_LIF | CORO1C |
| 1.989E-37 | 0.40028213 | 0.362 | 0.108 | 4.2465E-33 | TEC_LIF | OTUD4 |
| 5.8979E-26 | 0.39911305 | 0.488 | 0.216 | 1.2592E-21 | TEC_LIF | TMOD3 |
| 4.8556E-20 | 0.39889155 | 0.445 | 0.214 | 1.0367E-15 | TEC_LIF | FNBP1L |
| 4.4351E-19 | 0.39882569 | 0.457 | 0.23 | 9.4689E-15 | TEC_LIF | TMBIM1 |
| 8.6828E-32 | 0.39859082 | 0.425 | 0.151 | 1.8538E-27 | TEC_LIF | PALLD |
| 1.7478E-24 | 0.39748771 | 0.528 | 0.25 | 3.7316E-20 | TEC_LIF | TMEM123 |
| 1.1148E-31 | 0.39715354 | 0.366 | 0.122 | 2.3801E-27 | TEC_LIF | DSE |
| 2.5598E-24 | 0.39649123 | 0.449 | 0.195 | 5.4651E-20 | TEC_LIF | DDR1 |
| 1.1886E-58 | 0.39610524 | 0.264 | 0.045 | 2.5376E-54 | TEC_LIF | PTAFR |
| 6.42E-25 | 0.39486739 | 0.425 | 0.177 | 1.3707E-20 | TEC_LIF | QSOX1 |
| 9.0293E-18 | 0.39477951 | 0.543 | 0.301 | 1.9278E-13 | TEC_LIF | WDR1 |
| 1.665E-23 | 0.39298121 | 0.535 | 0.258 | 3.5547E-19 | TEC_LIF | IL6ST |
| 4.0241E-24 | 0.39249075 | 0.453 | 0.199 | 8.5914E-20 | TEC_LIF | SUN1 |
| 6.833E-13 | 0.39235914 | 0.606 | 0.388 | 1.4588E-08 | TEC_LIF | TXNRD1 |
| 1.9871E-17 | 0.39202259 | 0.539 | 0.29 | 4.2426E-13 | TEC_LIF | MIR4435-2HG |
| 6.7605E-39 | 0.3909876 | 0.354 | 0.101 | 1.4434E-34 | TEC_LIF | FSTL1 |
| 8.418E-21 | 0.39063046 | 0.429 | 0.199 | 1.7972E-16 | TEC_LIF | IPO7 |
| 6.459E-42 | 0.39050516 | 0.236 | 0.047 | 1.379E-37 | TEC_LIF | VNN1 |
| 2.6714E-30 | 0.39043236 | 0.335 | 0.109 | 5.7035E-26 | TEC_LIF | TMEM132A |
| 1.721E-22 | 0.39041648 | 0.406 | 0.172 | 3.6743E-18 | TEC_LIF | ACTN1 |
| 4.1764E-31 | 0.38794543 | 0.386 | 0.135 | 8.9166E-27 | TEC_LIF | ERGIC1 |
| 4.5971E-27 | 0.38689528 | 0.476 | 0.199 | 9.8149E-23 | TEC_LIF | WWTR1 |
| 8.406E-14 | 0.3866262 | 0.799 | 0.597 | 1.7947E-09 | TEC_LIF | CAST |
| 5.9736E-33 | 0.38629479 | 0.441 | 0.158 | 1.2754E-28 | TEC_LIF | XIAP |
| 7.2981E-21 | 0.38556972 | 0.587 | 0.316 | 1.5581E-16 | TEC_LIF | GNB1 |
| 5.65E-22 | 0.38554105 | 0.677 | 0.378 | 1.2063E-17 | TEC_LIF | CTNNA1 |
| 1.4266E-14 | 0.38553357 | 0.453 | 0.245 | 3.0457E-10 | TEC_LIF | MGLL |
| 2.0243E-24 | 0.38449531 | 0.516 | 0.239 | 4.3218E-20 | TEC_LIF | KPNB1 |
| 8.5507E-24 | 0.38325356 | 0.346 | 0.13 | 1.8256E-19 | TEC_LIF | RUNX1 |
| 1.8738E-39 | 0.38297359 | 0.343 | 0.095 | 4.0005E-35 | TEC_LIF | PROSER2 |
| 6.1064E-30 | 0.38253699 | 0.358 | 0.121 | 1.3037E-25 | TEC_LIF | APOL6 |
| 2.672E-27 | 0.38124907 | 0.5 | 0.213 | 5.7047E-23 | TEC_LIF | CTNND1 |
| 2.9402E-24 | 0.38099313 | 0.504 | 0.229 | 6.2774E-20 | TEC_LIF | LRRFIP2 |
| 1.1328E-25 | 0.38093052 | 0.441 | 0.177 | 2.4186E-21 | TEC_LIF | CD40 |
| 3.2027E-36 | 0.37909598 | 0.366 | 0.112 | 6.8379E-32 | TEC_LIF | TMEM51 |
| 2.7847E-35 | 0.37882529 | 0.366 | 0.114 | 5.9454E-31 | TEC_LIF | MFHAS1 |
| 1.1466E-24 | 0.37875755 | 0.579 | 0.275 | 2.4481E-20 | TEC_LIF | MAP4 |

| | | | | | | |
|---|---|---|---|---|---|---|
| 1.7466E-32 | 0.37866567 | 0.465 | 0.176 | 3.729E-28 | TEC_LIF | MYO1C |
| 9.9491E-16 | 0.37831924 | 0.504 | 0.279 | 2.1241E-11 | TEC_LIF | MSN |
| 1.193E-20 | 0.37746907 | 0.311 | 0.12 | 2.5471E-16 | TEC_LIF | ARL5B |
| 3.8591E-18 | 0.37719223 | 0.469 | 0.229 | 8.2392E-14 | TEC_LIF | CDKN2AIP |
| 2.0445E-10 | 0.37712109 | 0.957 | 0.812 | 4.3651E-06 | TEC_LIF | NEAT1 |
| 5.4676E-20 | 0.37637848 | 0.346 | 0.144 | 1.1673E-15 | TEC_LIF | SGMS2 |
| 6.5796E-37 | 0.37479941 | 0.382 | 0.118 | 1.4047E-32 | TEC_LIF | CSNK1E |
| 2.2093E-29 | 0.37449533 | 0.311 | 0.097 | 4.7169E-25 | TEC_LIF | DGKH |
| 1.0254E-22 | 0.37443012 | 0.476 | 0.216 | 2.1892E-18 | TEC_LIF | RBM47 |
| 3.9797E-30 | 0.37395685 | 0.354 | 0.119 | 8.4966E-26 | TEC_LIF | SCD5 |
| 4.2013E-23 | 0.37340242 | 0.472 | 0.213 | 8.9698E-19 | TEC_LIF | PARP14 |
| 1.3571E-46 | 0.37300131 | 0.319 | 0.075 | 2.8974E-42 | TEC_LIF | ARNTL2 |
| 1.2198E-13 | 0.3724956 | 0.22 | 0.088 | 2.6042E-09 | TEC_LIF | F3 |
| 1.8994E-30 | 0.37239379 | 0.291 | 0.086 | 4.0552E-26 | TEC_LIF | FBLIM1 |
| 5.3649E-30 | 0.37237566 | 0.496 | 0.199 | 1.1454E-25 | TEC_LIF | C16orf72 |
| 4.9838E-25 | 0.37117774 | 0.445 | 0.185 | 1.064E-20 | TEC_LIF | MPRIP |
| 1.4853E-17 | 0.37046353 | 0.299 | 0.125 | 3.1712E-13 | TEC_LIF | HIVEP2 |
| 3.8628E-38 | 0.36926179 | 0.343 | 0.097 | 8.247E-34 | TEC_LIF | CGN |
| 9.6289E-39 | 0.36922301 | 0.378 | 0.111 | 2.0558E-34 | TEC_LIF | CTTNBP2NL |
| 2.6981E-17 | 0.36909381 | 0.472 | 0.238 | 5.7605E-13 | TEC_LIF | TFPI2 |
| 8.3286E-27 | 0.36710887 | 0.48 | 0.203 | 1.7781E-22 | TEC_LIF | ADAM9 |
| 2.6199E-36 | 0.36684134 | 0.315 | 0.087 | 5.5935E-32 | TEC_LIF | DAB2IP |
| 1.8917E-31 | 0.36681881 | 0.449 | 0.167 | 4.0387E-27 | TEC_LIF | LASP1 |
| 2.1936E-14 | 0.36543691 | 0.961 | 0.921 | 4.6834E-10 | TEC_LIF | ACTB |
| 1.6025E-21 | 0.36469391 | 0.61 | 0.316 | 3.4212E-17 | TEC_LIF | PAWR |
| 1.0024E-16 | 0.36412859 | 0.476 | 0.247 | 2.1401E-12 | TEC_LIF | CD47 |
| 2.9226E-27 | 0.36319221 | 0.374 | 0.136 | 6.2397E-23 | TEC_LIF | SLC44A4 |
| 2.5307E-23 | 0.36237898 | 0.453 | 0.197 | 5.4031E-19 | TEC_LIF | PLSCR1 |
| 2.3155E-27 | 0.36230641 | 0.339 | 0.117 | 4.9436E-23 | TEC_LIF | SGPP2 |
| 5.4804E-06 | 0.36116262 | 0.705 | 0.604 | 0.11700669 | TEC_LIF | GADD45A |
| 1.6588E-29 | 0.36101086 | 0.346 | 0.116 | 3.5415E-25 | TEC_LIF | SLC7A1 |
| 3.6162E-23 | 0.36092882 | 0.453 | 0.199 | 7.7206E-19 | TEC_LIF | ERO1A |
| 2.6548E-14 | 0.3606636 | 0.602 | 0.366 | 5.6681E-10 | TEC_LIF | CAMK2N1 |
| 2.2985E-19 | 0.36045728 | 0.425 | 0.198 | 4.9073E-15 | TEC_LIF | DPP4 |
| 8.0918E-38 | 0.35954755 | 0.406 | 0.126 | 1.7276E-33 | TEC_LIF | ELK4 |
| 1.9045E-15 | 0.35942455 | 0.673 | 0.427 | 4.0662E-11 | TEC_LIF | BZW1 |
| 2.6809E-29 | 0.3590358 | 0.437 | 0.167 | 5.7238E-25 | TEC_LIF | NECTIN2 |
| 4.8277E-19 | 0.35791115 | 0.555 | 0.296 | 1.0307E-14 | TEC_LIF | RAP1B |
| 1.4164E-09 | 0.35686355 | 0.476 | 0.292 | 3.0239E-05 | TEC_LIF | DST |
| 1.3653E-26 | 0.3568396 | 0.374 | 0.139 | 2.9149E-22 | TEC_LIF | GALNT3 |
| 2.3997E-22 | 0.35669285 | 0.425 | 0.183 | 5.1233E-18 | TEC_LIF | B4GALT1 |
| 1.9255E-23 | 0.35616667 | 0.421 | 0.178 | 4.1108E-19 | TEC_LIF | BTBD7 |
| 1.9767E-31 | 0.35598303 | 0.311 | 0.093 | 4.2202E-27 | TEC_LIF | C1orf116 |
| 1.5772E-31 | 0.35500327 | 0.374 | 0.124 | 3.3674E-27 | TEC_LIF | KCNK1 |
| 2.7982E-33 | 0.35216907 | 0.272 | 0.071 | 5.9743E-29 | TEC_LIF | LIPH |
| 4.2835E-38 | 0.35201253 | 0.22 | 0.045 | 9.1452E-34 | TEC_LIF | FRMD6 |
| 1.3305E-23 | 0.35189493 | 0.354 | 0.137 | 2.8407E-19 | TEC_LIF | LPGAT1 |
| 1.1235E-21 | 0.35087952 | 0.343 | 0.134 | 2.3987E-17 | TEC_LIF | YPEL2 |
| 2.07E-52 | 0.35083388 | 0.264 | 0.049 | 4.4194E-48 | TEC_LIF | RASSF3 |
| 7.7507E-21 | 0.35079687 | 0.441 | 0.199 | 1.6548E-16 | TEC_LIF | FLNB |

| | | | | | | |
|---|---|---|---|---|---|---|
| 5.4263E-43 | 0.34979153 | 0.193 | 0.032 | 1.1585E-38 | TEC_LIF | SPOCD1 |
| 1.1684E-09 | 0.34955932 | 0.185 | 0.08 | 2.4944E-05 | TEC_LIF | CXCL1 |
| 3.8355E-14 | 0.34937045 | 0.303 | 0.134 | 8.1889E-10 | TEC_LIF | TM4SF1 |
| 3.6506E-22 | 0.34835818 | 0.382 | 0.159 | 7.794E-18 | TEC_LIF | PPP2R5A |
| 1.6842E-34 | 0.3481581 | 0.339 | 0.101 | 3.5959E-30 | TEC_LIF | INAVA |
| 4.4554E-39 | 0.34809447 | 0.276 | 0.066 | 9.5122E-35 | TEC_LIF | DENND2C |
| 2.2407E-23 | 0.34795978 | 0.276 | 0.091 | 4.784E-19 | TEC_LIF | OAF |
| 2.5454E-13 | 0.34758044 | 0.622 | 0.396 | 5.4344E-09 | TEC_LIF | CDV3 |
| 3.6685E-35 | 0.34734693 | 0.323 | 0.093 | 7.8323E-31 | TEC_LIF | BMPR2 |
| 2.5711E-22 | 0.34703154 | 0.276 | 0.092 | 5.4893E-18 | TEC_LIF | ALDH1A3 |
| 1.1643E-23 | 0.34648693 | 0.449 | 0.194 | 2.4858E-19 | TEC_LIF | SEPT11 |
| 1.176E-30 | 0.3460657 | 0.378 | 0.128 | 2.5107E-26 | TEC_LIF | RALGPS2 |
| 1.5751E-26 | 0.34514587 | 0.323 | 0.11 | 3.3628E-22 | TEC_LIF | MYADM |
| 1.1123E-20 | 0.34472319 | 0.224 | 0.07 | 2.3748E-16 | TEC_LIF | FAM110C |
| 8.4142E-19 | 0.34420632 | 0.508 | 0.258 | 1.7964E-14 | TEC_LIF | HDGF |
| 5.1475E-10 | 0.34365212 | 0.232 | 0.109 | 1.099E-05 | TEC_LIF | C15orf48 |
| 5.3129E-21 | 0.34183803 | 0.35 | 0.143 | 1.1343E-16 | TEC_LIF | LNPEP |
| 2.2116E-23 | 0.34118118 | 0.374 | 0.148 | 4.7217E-19 | TEC_LIF | WWC2 |
| 8.6169E-33 | 0.34064756 | 0.35 | 0.11 | 1.8397E-28 | TEC_LIF | TNFAIP1 |
| 2.3814E-19 | 0.34010985 | 0.465 | 0.22 | 5.0844E-15 | TEC_LIF | NFAT5 |
| 9.9294E-30 | 0.3399415 | 0.213 | 0.051 | 2.1199E-25 | TEC_LIF | NFKBID |
| 4.3713E-14 | 0.33973673 | 0.445 | 0.236 | 9.3327E-10 | TEC_LIF | AAK1 |
| 5.0588E-25 | 0.33909759 | 0.39 | 0.15 | 1.0801E-20 | TEC_LIF | MARK3 |
| 5.7084E-34 | 0.33898595 | 0.343 | 0.103 | 1.2187E-29 | TEC_LIF | PXN |
| 3.1974E-25 | 0.33882521 | 0.343 | 0.124 | 6.8264E-21 | TEC_LIF | MAP7D1 |
| 5.4019E-20 | 0.33762241 | 0.37 | 0.155 | 1.1533E-15 | TEC_LIF | TRIM56 |
| 2.2836E-21 | 0.33759622 | 0.354 | 0.145 | 4.8754E-17 | TEC_LIF | GOLGA3 |
| 2.2227E-19 | 0.3370266 | 0.437 | 0.206 | 4.7455E-15 | TEC_LIF | SNX9 |
| 1.0331E-19 | 0.33691629 | 0.315 | 0.124 | 2.2057E-15 | TEC_LIF | ERBB3 |
| 1.8396E-17 | 0.33630143 | 0.37 | 0.165 | 3.9276E-13 | TEC_LIF | TYMP |
| 8.1851E-19 | 0.33621328 | 0.343 | 0.145 | 1.7475E-14 | TEC_LIF | RHOF |
| 1.3869E-25 | 0.3361009 | 0.429 | 0.172 | 2.9611E-21 | TEC_LIF | MYO10 |
| 3.2507E-23 | 0.33607 | 0.488 | 0.217 | 6.9402E-19 | TEC_LIF | LSR |
| 5.7113E-30 | 0.33476178 | 0.26 | 0.072 | 1.2194E-25 | TEC_LIF | TNFRSF21 |
| 7.7068E-25 | 0.33442192 | 0.315 | 0.11 | 1.6454E-20 | TEC_LIF | SH3RF1 |
| 2.2109E-25 | 0.3341249 | 0.232 | 0.066 | 4.7202E-21 | TEC_LIF | FOXN2 |
| 4.7922E-27 | 0.3340601 | 0.331 | 0.112 | 1.0231E-22 | TEC_LIF | MYH14 |
| 7.5921E-29 | 0.3333592 | 0.287 | 0.086 | 1.6209E-24 | TEC_LIF | TRIB1 |
| 3.8555E-14 | 0.33325977 | 0.319 | 0.149 | 8.2314E-10 | TEC_LIF | ROCK2 |
| 1.1417E-17 | 0.33282219 | 0.488 | 0.251 | 2.4375E-13 | TEC_LIF | PPP2R1A |
| 5.0091E-23 | 0.33188955 | 0.303 | 0.109 | 1.0694E-18 | TEC_LIF | OSMR |
| 2.0705E-27 | 0.33155464 | 0.37 | 0.132 | 4.4205E-23 | TEC_LIF | L1CAM |
| 2.635E-15 | 0.33125561 | 0.496 | 0.269 | 5.6258E-11 | TEC_LIF | CREBRF |
| 6.8907E-21 | 0.33062675 | 0.409 | 0.177 | 1.4712E-16 | TEC_LIF | ZFHX3 |
| 4.3666E-28 | 0.32980754 | 0.315 | 0.101 | 9.3227E-24 | TEC_LIF | PSD3 |
| 1.0837E-27 | 0.32965431 | 0.335 | 0.113 | 2.3136E-23 | TEC_LIF | GPR160 |
| 2.7168E-15 | 0.32903341 | 0.37 | 0.176 | 5.8003E-11 | TEC_LIF | SQLE |
| 2.0072E-21 | 0.32883151 | 0.425 | 0.188 | 4.2854E-17 | TEC_LIF | IDS |
| 9.4336E-19 | 0.32865179 | 0.339 | 0.142 | 2.0141E-14 | TEC_LIF | PTPN14 |
| 5.0663E-25 | 0.32829153 | 0.213 | 0.057 | 1.0817E-20 | TEC_LIF | TRAF1 |

| | | | | | | |
|---|---|---|---|---|---|---|
| 1.119E-22 | 0.32821713 | 0.358 | 0.141 | 2.3892E-18 | TEC_LIF | VASP |
| 6.3565E-44 | 0.32807427 | 0.26 | 0.054 | 1.3571E-39 | TEC_LIF | SERPINB9 |
| 5.5657E-13 | 0.32781476 | 0.307 | 0.144 | 1.1883E-08 | TEC_LIF | RETREG1 |
| 1.1127E-51 | 0.3255249 | 0.209 | 0.032 | 2.3755E-47 | TEC_LIF | NAV3 |
| 7.2097E-25 | 0.32274596 | 0.335 | 0.12 | 1.5393E-20 | TEC_LIF | KLF13 |
| 1.3898E-19 | 0.3226017 | 0.441 | 0.206 | 2.9673E-15 | TEC_LIF | GNG12 |
| 5.5683E-17 | 0.32189039 | 0.402 | 0.189 | 1.1888E-12 | TEC_LIF | GPATCH2L |
| 3.0264E-22 | 0.32187401 | 0.374 | 0.153 | 6.4613E-18 | TEC_LIF | KRAS |
| 4.0846E-09 | 0.3217704 | 0.563 | 0.388 | 8.7206E-05 | TEC_LIF | NFKBIA |
| 2.2279E-21 | 0.32175946 | 0.378 | 0.155 | 4.7566E-17 | TEC_LIF | TRAF4 |
| 5.4619E-28 | 0.32168519 | 0.283 | 0.086 | 1.1661E-23 | TEC_LIF | MTAP |
| 7.2206E-19 | 0.31970289 | 0.374 | 0.163 | 1.5416E-14 | TEC_LIF | STAT1 |
| 2.6322E-24 | 0.31890264 | 0.323 | 0.116 | 5.6197E-20 | TEC_LIF | VOPP1 |
| 2.3843E-24 | 0.31847788 | 0.386 | 0.149 | 5.0904E-20 | TEC_LIF | KIF1B |
| 6.1842E-21 | 0.31831148 | 0.394 | 0.168 | 1.3203E-16 | TEC_LIF | ERBIN |
| 3.3801E-13 | 0.31828114 | 0.492 | 0.274 | 7.2166E-09 | TEC_LIF | GTF2I |
| 2.3883E-22 | 0.31821092 | 0.402 | 0.167 | 5.099E-18 | TEC_LIF | SH3BP4 |
| 1.0307E-23 | 0.31818618 | 0.358 | 0.135 | 2.2006E-19 | TEC_LIF | VEZT |
| 5.4009E-14 | 0.31786917 | 0.386 | 0.196 | 1.1531E-09 | TEC_LIF | AGRN |
| 1.5289E-28 | 0.31745866 | 0.327 | 0.105 | 3.2642E-24 | TEC_LIF | ATP11A |
| 1.6806E-20 | 0.31630598 | 0.5 | 0.238 | 3.588E-16 | TEC_LIF | PFKP |
| 1.3693E-29 | 0.31629697 | 0.319 | 0.099 | 2.9234E-25 | TEC_LIF | MYO1B |
| 6.908E-20 | 0.31629438 | 0.378 | 0.16 | 1.4749E-15 | TEC_LIF | SEMA3B |
| 1.7503E-16 | 0.31624689 | 0.374 | 0.175 | 3.7368E-12 | TEC_LIF | USP22 |
| 9.0196E-20 | 0.31519779 | 0.488 | 0.237 | 1.9257E-15 | TEC_LIF | TOR1AIP2 |
| 1.6235E-18 | 0.31326444 | 0.264 | 0.098 | 3.4663E-14 | TEC_LIF | ECE1 |
| 1.9264E-19 | 0.31290453 | 0.421 | 0.188 | 4.1128E-15 | TEC_LIF | HIPK2 |
| 6.6933E-14 | 0.31288331 | 0.516 | 0.286 | 1.429E-09 | TEC_LIF | MAFF |
| 3.0745E-22 | 0.3126648 | 0.402 | 0.167 | 6.5641E-18 | TEC_LIF | PTPN11 |
| 2.1497E-07 | 0.31254205 | 0.311 | 0.186 | 0.00458971 | TEC_LIF | PDIA4 |
| 8.8185E-18 | 0.31250952 | 0.335 | 0.145 | 1.8828E-13 | TEC_LIF | SNX10 |
| 3.1745E-18 | 0.31245375 | 0.244 | 0.089 | 6.7775E-14 | TEC_LIF | SEMA3C |
| 9.4445E-17 | 0.31229841 | 0.354 | 0.158 | 2.0164E-12 | TEC_LIF | ITSN2 |
| 1.0526E-18 | 0.3122374 | 0.35 | 0.15 | 2.2473E-14 | TEC_LIF | BIRC2 |
| 1.7914E-09 | 0.31191635 | 0.28 | 0.145 | 3.8246E-05 | TEC_LIF | RASSF8 |
| 9.5926E-13 | 0.31159101 | 0.717 | 0.493 | 2.048E-08 | TEC_LIF | TNFRSF12A |
| 5.4812E-14 | 0.31130468 | 0.441 | 0.229 | 1.1702E-09 | TEC_LIF | PKN2 |
| 2.6478E-28 | 0.31125772 | 0.244 | 0.067 | 5.653E-24 | TEC_LIF | PPARD |
| 1.9443E-29 | 0.3111345 | 0.291 | 0.086 | 4.151E-25 | TEC_LIF | PML |
| 1.2111E-15 | 0.3108763 | 0.504 | 0.269 | 2.5857E-11 | TEC_LIF | HIF1A |
| 7.8413E-18 | 0.31025168 | 0.492 | 0.248 | 1.6741E-13 | TEC_LIF | PTP4A1 |
| 2.1117E-23 | 0.30971729 | 0.331 | 0.122 | 4.5086E-19 | TEC_LIF | MED13L |
| 4.9151E-40 | 0.30922853 | 0.256 | 0.057 | 1.0494E-35 | TEC_LIF | IRAK2 |
| 9.216E-20 | 0.30880881 | 0.366 | 0.155 | 1.9676E-15 | TEC_LIF | RIF1 |
| 3.6611E-23 | 0.3081754 | 0.287 | 0.098 | 7.8165E-19 | TEC_LIF | PLEKHG3 |
| 1.5464E-25 | 0.30813051 | 0.276 | 0.086 | 3.3015E-21 | TEC_LIF | SNTB2 |
| 2.9436E-25 | 0.30754268 | 0.291 | 0.095 | 6.2846E-21 | TEC_LIF | PTER |
| 4.0042E-14 | 0.30752632 | 0.386 | 0.197 | 8.549E-10 | TEC_LIF | ATP2A2 |
| 2.8911E-46 | 0.30728995 | 0.272 | 0.056 | 6.1725E-42 | TEC_LIF | NIPAL1 |
| 4.3661E-18 | 0.30718991 | 0.63 | 0.332 | 9.3216E-14 | TEC_LIF | PAX8 |

| | | | | | | |
|---|---|---|---|---|---|---|
| 2.9328E-20 | 0.30661261 | 0.465 | 0.217 | 6.2615E-16 | TEC_LIF | GNAI2 |
| 2.9003E-25 | 0.30655706 | 0.272 | 0.086 | 6.1921E-21 | TEC_LIF | RHPN2 |
| 1.9396E-17 | 0.30594024 | 0.445 | 0.218 | 4.1411E-13 | TEC_LIF | DIAPH1 |
| 8.3323E-13 | 0.30553211 | 0.472 | 0.264 | 1.7789E-08 | TEC_LIF | CCDC50 |
| 1.9391E-07 | 0.30489461 | 0.142 | 0.062 | 0.00413996 | TEC_LIF | SERPINE1 |
| 7.2486E-14 | 0.30469135 | 0.488 | 0.275 | 1.5476E-09 | TEC_LIF | VCP |
| 4.8665E-13 | 0.304266 | 0.449 | 0.242 | 1.039E-08 | TEC_LIF | IRS2 |
| 4.1899E-19 | 0.30421056 | 0.295 | 0.113 | 8.9455E-15 | TEC_LIF | PTPN3 |
| 1.1852E-11 | 0.30419689 | 0.244 | 0.109 | 2.5303E-07 | TEC_LIF | ARRDC4 |
| 6.9525E-35 | 0.30371417 | 0.299 | 0.081 | 1.4844E-30 | TEC_LIF | ATP8B1 |
| 8.9988E-26 | 0.30354528 | 0.276 | 0.087 | 1.9213E-21 | TEC_LIF | TTC9 |
| 3.7279E-20 | 0.30294874 | 0.524 | 0.253 | 7.959E-16 | TEC_LIF | BRD4 |
| 1.2139E-18 | 0.30294445 | 0.209 | 0.067 | 2.5917E-14 | TEC_LIF | MALL |
| 5.5717E-26 | 0.30285148 | 0.323 | 0.11 | 1.1896E-21 | TEC_LIF | AC005261.1 |
| 1.5987E-17 | 0.30251116 | 0.469 | 0.234 | 3.4131E-13 | TEC_LIF | DYNC1LI2 |
| 2.9119E-12 | 0.30237635 | 0.701 | 0.488 | 6.217E-08 | TEC_LIF | ATP1B1 |
| 4.3685E-13 | 0.30206953 | 0.646 | 0.415 | 9.3267E-09 | TEC_LIF | PTTG1IP |
| 6.166E-23 | 0.30180304 | 0.287 | 0.099 | 1.3164E-18 | TEC_LIF | FEM1B |
| 3.8103E-11 | 0.30166997 | 0.476 | 0.277 | 8.135E-07 | TEC_LIF | CLDN3 |
| 1.9307E-16 | 0.30166225 | 0.319 | 0.139 | 4.1221E-12 | TEC_LIF | ANO6 |
| 1.1308E-27 | 0.301553 | 0.236 | 0.064 | 2.4143E-23 | TEC_LIF | MYEOV |
| 8.7791E-20 | 0.3015428 | 0.319 | 0.126 | 1.8743E-15 | TEC_LIF | SLC20A2 |
| 2.5368E-12 | 0.30098403 | 0.756 | 0.531 | 5.4161E-08 | TEC_LIF | HLA-E |
| 1.1961E-20 | 0.30077659 | 0.26 | 0.087 | 2.5536E-16 | TEC_LIF | PDGFB |
| 3.2791E-17 | 0.30050987 | 0.323 | 0.138 | 7.0009E-13 | TEC_LIF | SKIL |
| 5.2551E-24 | 0.29967941 | 0.303 | 0.103 | 1.122E-19 | TEC_LIF | ITPRIPL2 |
| 7.3846E-06 | 0.29911534 | 0.311 | 0.199 | 0.15766191 | TEC_LIF | CALD1 |
| 1.744E-16 | 0.29895335 | 0.386 | 0.183 | 3.7235E-12 | TEC_LIF | PRRC2B |
| 6.5392E-15 | 0.29864177 | 0.398 | 0.197 | 1.3961E-10 | TEC_LIF | MBP |
| 7.9622E-22 | 0.29765455 | 0.343 | 0.133 | 1.6999E-17 | TEC_LIF | SPTLC2 |
| 1.5465E-21 | 0.29752183 | 0.35 | 0.139 | 3.3018E-17 | TEC_LIF | NT5C2 |
| 5.9747E-30 | 0.29678893 | 0.272 | 0.076 | 1.2756E-25 | TEC_LIF | USP31 |
| 5.6448E-13 | 0.29574821 | 0.437 | 0.24 | 1.2052E-08 | TEC_LIF | NCKAP1 |
| 2.7594E-23 | 0.29501883 | 0.256 | 0.081 | 5.8913E-19 | TEC_LIF | EFNB1 |
| 5.0675E-14 | 0.2944491 | 0.488 | 0.271 | 1.0819E-09 | TEC_LIF | CLTC |
| 2.1094E-12 | 0.2939936 | 0.177 | 0.066 | 4.5035E-08 | TEC_LIF | TRIM55 |
| 3.868E-16 | 0.29333918 | 0.504 | 0.265 | 8.2583E-12 | TEC_LIF | FARP1 |
| 9.2424E-25 | 0.29314674 | 0.303 | 0.102 | 1.9732E-20 | TEC_LIF | FRK |
| 4.3214E-16 | 0.29224025 | 0.283 | 0.116 | 9.2262E-12 | TEC_LIF | REL |
| 6.9231E-15 | 0.29093692 | 0.346 | 0.163 | 1.4781E-10 | TEC_LIF | UBE2J1 |
| 1.7835E-24 | 0.2897301 | 0.327 | 0.116 | 3.8077E-20 | TEC_LIF | CDC42EP1 |
| 1.4977E-15 | 0.28937335 | 0.39 | 0.186 | 3.1976E-11 | TEC_LIF | RNF19A |
| 1.6236E-52 | 0.28919234 | 0.224 | 0.036 | 3.4663E-48 | TEC_LIF | ADAM8 |
| 3.8149E-20 | 0.28879318 | 0.343 | 0.138 | 8.1448E-16 | TEC_LIF | USP9X |
| 3.5308E-24 | 0.28855179 | 0.248 | 0.075 | 7.5382E-20 | TEC_LIF | ATP2B4 |
| 7.7526E-22 | 0.28854002 | 0.272 | 0.092 | 1.6552E-17 | TEC_LIF | SLCO4A1 |
| 4.1434E-22 | 0.28832007 | 0.291 | 0.103 | 8.8461E-18 | TEC_LIF | TRIM25 |
| 1.6159E-13 | 0.28758334 | 0.429 | 0.229 | 3.45E-09 | TEC_LIF | CHD4 |
| 2.8164E-19 | 0.28737474 | 0.441 | 0.203 | 6.0129E-15 | TEC_LIF | SPPL2A |
| 4.1612E-12 | 0.28723798 | 0.602 | 0.387 | 8.8842E-08 | TEC_LIF | CAPN2 |

| | | | | | | |
|---|---|---|---|---|---|---|
| 1.868E-19 | 0.28706807 | 0.37 | 0.158 | 3.9881E-15 | TEC_LIF | FYTTD1 |
| 9.2872E-22 | 0.28705618 | 0.386 | 0.155 | 1.9828E-17 | TEC_LIF | BAZ1A |
| 1.0937E-11 | 0.28696745 | 0.539 | 0.341 | 2.3351E-07 | TEC_LIF | CAP1 |
| 4.2782E-25 | 0.2863917 | 0.299 | 0.1 | 9.134E-21 | TEC_LIF | SFT2D2 |
| 7.5526E-13 | 0.28636471 | 0.78 | 0.555 | 1.6125E-08 | TEC_LIF | KTN1 |
| 6.2974E-16 | 0.28584652 | 0.35 | 0.16 | 1.3445E-11 | TEC_LIF | CD82 |
| 1.8687E-15 | 0.28550664 | 0.402 | 0.199 | 3.9897E-11 | TEC_LIF | GIPC1 |
| 6.688E-18 | 0.28541715 | 0.386 | 0.175 | 1.4279E-13 | TEC_LIF | SMARCC1 |
| 3.6563E-19 | 0.28513559 | 0.276 | 0.103 | 7.8061E-15 | TEC_LIF | FBXO11 |
| 1.2263E-16 | 0.28495473 | 0.35 | 0.156 | 2.6181E-12 | TEC_LIF | ARHGAP21 |
| 5.7395E-21 | 0.28463135 | 0.311 | 0.117 | 1.2254E-16 | TEC_LIF | PANK3 |
| 2.3671E-24 | 0.28443296 | 0.224 | 0.064 | 5.0538E-20 | TEC_LIF | DOCK9 |
| 1.2532E-36 | 0.2842856 | 0.224 | 0.048 | 2.6755E-32 | TEC_LIF | ALDH1B1 |
| 2.0653E-11 | 0.28381462 | 0.543 | 0.329 | 4.4094E-07 | TEC_LIF | ANKRD12 |
| 5.4591E-24 | 0.28288853 | 0.28 | 0.092 | 1.1655E-19 | TEC_LIF | CAMSAP1 |
| 5.0388E-15 | 0.28237933 | 0.295 | 0.127 | 1.0758E-10 | TEC_LIF | OGT |
| 1.3164E-18 | 0.28162714 | 0.339 | 0.14 | 2.8106E-14 | TEC_LIF | HOOK3 |
| 3.7184E-17 | 0.2816242 | 0.311 | 0.13 | 7.9388E-13 | TEC_LIF | IGF1R |
| 2.5931E-34 | 0.28116734 | 0.213 | 0.046 | 5.5363E-30 | TEC_LIF | RASSF9 |
| 1.2828E-10 | 0.28021991 | 0.531 | 0.336 | 2.7387E-06 | TEC_LIF | EIF1AX |
| 3.3862E-39 | 0.28006735 | 0.244 | 0.053 | 7.2296E-35 | TEC_LIF | IL4R |
| 1.4324E-31 | 0.27989513 | 0.295 | 0.084 | 3.0581E-27 | TEC_LIF | CD276 |
| 4.255E-24 | 0.27968892 | 0.28 | 0.091 | 9.0843E-20 | TEC_LIF | CASZ1 |
| 3.648E-19 | 0.27954029 | 0.382 | 0.165 | 7.7884E-15 | TEC_LIF | PHC2 |
| 1.1081E-23 | 0.27933242 | 0.276 | 0.091 | 2.3657E-19 | TEC_LIF | SOAT1 |
| 1.3932E-18 | 0.27916459 | 0.268 | 0.1 | 2.9744E-14 | TEC_LIF | UBA6 |
| 1.7507E-13 | 0.27899489 | 0.555 | 0.322 | 3.7377E-09 | TEC_LIF | IL32 |
| 1.6301E-13 | 0.27880917 | 0.283 | 0.129 | 3.4802E-09 | TEC_LIF | LAMC1 |
| 6.4127E-24 | 0.27768792 | 0.366 | 0.138 | 1.3691E-19 | TEC_LIF | RNF149 |
| 5.6784E-20 | 0.2774942 | 0.35 | 0.144 | 1.2123E-15 | TEC_LIF | SAMD4B |
| 1.8791E-19 | 0.2764665 | 0.213 | 0.068 | 4.0119E-15 | TEC_LIF | GCNT3 |
| 1.0832E-20 | 0.2762083 | 0.374 | 0.154 | 2.3126E-16 | TEC_LIF | PDLIM5 |
| 1.4944E-18 | 0.27604217 | 0.343 | 0.144 | 3.1906E-14 | TEC_LIF | YAP1 |
| 1.1638E-23 | 0.27591054 | 0.283 | 0.094 | 2.4846E-19 | TEC_LIF | BCAR3 |
| 6.9064E-58 | 0.27567602 | 0.197 | 0.026 | 1.4745E-53 | TEC_LIF | AL512274.1 |
| 1.9347E-17 | 0.27546014 | 0.449 | 0.219 | 4.1306E-13 | TEC_LIF | NAMPT |
| 2.9327E-12 | 0.27536188 | 0.437 | 0.238 | 6.2613E-08 | TEC_LIF | FNBP4 |
| 2.0759E-10 | 0.27520689 | 0.413 | 0.246 | 4.4321E-06 | TEC_LIF | YPEL5 |
| 2.6698E-17 | 0.27460022 | 0.39 | 0.178 | 5.7001E-13 | TEC_LIF | FLII |
| 8.983E-15 | 0.27388295 | 0.252 | 0.103 | 1.9179E-10 | TEC_LIF | TNFAIP8 |
| 2.1547E-19 | 0.27384938 | 0.402 | 0.175 | 4.6003E-15 | TEC_LIF | FAM120A |
| 3.9266E-15 | 0.2737323 | 0.441 | 0.226 | 8.3834E-11 | TEC_LIF | SCARB2 |
| 1.2032E-17 | 0.27335135 | 0.331 | 0.139 | 2.5689E-13 | TEC_LIF | PHACTR2 |
| 1.3396E-19 | 0.27325583 | 0.28 | 0.104 | 2.8601E-15 | TEC_LIF | ABHD2 |
| 4.9651E-11 | 0.27314376 | 0.461 | 0.269 | 1.06E-06 | TEC_LIF | SMG1 |
| 1.0776E-23 | 0.27266308 | 0.307 | 0.107 | 2.3006E-19 | TEC_LIF | ZNFX1 |
| 2.4707E-20 | 0.2725338 | 0.185 | 0.053 | 5.2749E-16 | TEC_LIF | CCND2 |
| 1.1216E-07 | 0.27235155 | 0.335 | 0.201 | 0.00239472 | TEC_LIF | PIM3 |
| 1.517E-15 | 0.27148124 | 0.378 | 0.18 | 3.2388E-11 | TEC_LIF | KLF3 |
| 7.6061E-16 | 0.27139615 | 0.331 | 0.148 | 1.6239E-11 | TEC_LIF | MAPK1 |

| | | | | | | |
|---|---|---|---|---|---|---|
| 1.2738E-20 | 0.27128043 | 0.295 | 0.108 | 2.7196E-16 | TEC_LIF | SHROOM3 |
| 1.6714E-19 | 0.27063894 | 0.24 | 0.082 | 3.5685E-15 | TEC_LIF | HDAC9 |
| 3.1693E-14 | 0.27047008 | 0.382 | 0.187 | 6.7665E-10 | TEC_LIF | ZBTB43 |
| 2.4863E-20 | 0.2702517 | 0.256 | 0.088 | 5.3083E-16 | TEC_LIF | IFIT5 |
| 1.8467E-18 | 0.26937427 | 0.323 | 0.13 | 3.9427E-14 | TEC_LIF | PDLIM4 |
| 2.6724E-18 | 0.26917763 | 0.299 | 0.117 | 5.7056E-14 | TEC_LIF | SH2D4A |
| 2.8946E-14 | 0.26911421 | 0.276 | 0.12 | 6.18E-10 | TEC_LIF | TUFT1 |
| 2.3349E-29 | 0.26833415 | 0.26 | 0.071 | 4.9851E-25 | TEC_LIF | ZSWIM4 |
| 1.4556E-13 | 0.26731563 | 0.382 | 0.193 | 3.1076E-09 | TEC_LIF | IGF2R |
| 9.1619E-13 | 0.26706097 | 0.429 | 0.232 | 1.9561E-08 | TEC_LIF | NIPBL |
| 2.0031E-18 | 0.26657892 | 0.37 | 0.16 | 4.2765E-14 | TEC_LIF | PTPN12 |
| 1.3826E-18 | 0.26654619 | 0.339 | 0.14 | 2.9518E-14 | TEC_LIF | PTPRF |
| 8.6018E-18 | 0.26654038 | 0.343 | 0.147 | 1.8365E-13 | TEC_LIF | ACSL4 |
| 2.0629E-12 | 0.26594586 | 0.35 | 0.18 | 4.4043E-08 | TEC_LIF | PLEKHA1 |
| 2.5437E-17 | 0.26586132 | 0.268 | 0.104 | 5.4307E-13 | TEC_LIF | SPIN1 |
| 1.6939E-13 | 0.26560809 | 0.236 | 0.098 | 3.6164E-09 | TEC_LIF | HMGCR |
| 2.9942E-13 | 0.26555459 | 0.22 | 0.09 | 6.3925E-09 | TEC_LIF | NRP1 |
| 9.3926E-15 | 0.26526038 | 0.378 | 0.181 | 2.0053E-10 | TEC_LIF | MUC20-OT1 |
| 4.8977E-09 | 0.26523927 | 0.547 | 0.37 | 0.00010457 | TEC_LIF | HEBP2 |
| 5.8205E-23 | 0.26519202 | 0.276 | 0.092 | 1.2427E-18 | TEC_LIF | BCL9L |
| 5.8129E-15 | 0.26496194 | 0.299 | 0.13 | 1.2411E-10 | TEC_LIF | NPC1 |
| 1.2738E-20 | 0.26479461 | 0.311 | 0.117 | 2.7196E-16 | TEC_LIF | STAT6 |
| 3.5186E-15 | 0.26476616 | 0.28 | 0.119 | 7.5122E-11 | TEC_LIF | CNN2 |
| 2.3846E-16 | 0.26470031 | 0.366 | 0.166 | 5.0912E-12 | TEC_LIF | NRIP1 |
| 1.5434E-10 | 0.26431362 | 0.248 | 0.118 | 3.2953E-06 | TEC_LIF | NEURL3 |
| 4.3616E-19 | 0.26424796 | 0.358 | 0.15 | 9.3119E-15 | TEC_LIF | HGS |
| 1.2398E-24 | 0.26420504 | 0.283 | 0.092 | 2.647E-20 | TEC_LIF | PIK3CA |
| 8.1843E-12 | 0.26415031 | 0.374 | 0.203 | 1.7473E-07 | TEC_LIF | SINHCAF |
| 1.8725E-13 | 0.26403886 | 0.402 | 0.207 | 3.9978E-09 | TEC_LIF | G3BP1 |
| 8.1672E-17 | 0.26383702 | 0.213 | 0.074 | 1.7437E-12 | TEC_LIF | SEMA4B |
| 1.1021E-16 | 0.26375689 | 0.295 | 0.121 | 2.3529E-12 | TEC_LIF | NFIB |
| 4.086E-25 | 0.26353014 | 0.22 | 0.06 | 8.7236E-21 | TEC_LIF | B3GNT5 |
| 5.3198E-24 | 0.2634818 | 0.276 | 0.089 | 1.1358E-19 | TEC_LIF | ATP11B |
| 2.322E-13 | 0.26345669 | 0.303 | 0.142 | 4.9575E-09 | TEC_LIF | NR1D2 |
| 2.5277E-14 | 0.26343852 | 0.484 | 0.259 | 5.3965E-10 | TEC_LIF | RB1CC1 |
| 9.2669E-22 | 0.26275893 | 0.272 | 0.093 | 1.9785E-17 | TEC_LIF | INSIG2 |
| 3.939E-19 | 0.26253158 | 0.295 | 0.113 | 8.4098E-15 | TEC_LIF | RAB5B |
| 1.1623E-09 | 0.26249258 | 0.248 | 0.122 | 2.4815E-05 | TEC_LIF | WEE1 |
| 7.1762E-25 | 0.26247435 | 0.244 | 0.072 | 1.5321E-20 | TEC_LIF | SACS |
| 1.5423E-24 | 0.26247435 | 0.256 | 0.078 | 3.2929E-20 | TEC_LIF | MAFK |
| 3.609E-22 | 0.26199066 | 0.299 | 0.105 | 7.7052E-18 | TEC_LIF | LARP4B |
| 2.1089E-19 | 0.26188957 | 0.287 | 0.108 | 4.5024E-15 | TEC_LIF | KLHL36 |
| 4.6437E-17 | 0.26167282 | 0.382 | 0.174 | 9.9143E-13 | TEC_LIF | AGFG1 |
| 1.4805E-15 | 0.26145088 | 0.354 | 0.162 | 3.1609E-11 | TEC_LIF | KIDINS220 |
| 7.1145E-16 | 0.26056629 | 0.335 | 0.148 | 1.519E-11 | TEC_LIF | VEZF1 |
| 6.0174E-20 | 0.26055391 | 0.283 | 0.103 | 1.2847E-15 | TEC_LIF | EFR3A |
| 5.1544E-28 | 0.26022401 | 0.24 | 0.065 | 1.1005E-23 | TEC_LIF | PPP1R14C |
| 7.034E-14 | 0.26007614 | 0.429 | 0.223 | 1.5018E-09 | TEC_LIF | TRIM8 |
| 2.1529E-21 | 0.25969698 | 0.303 | 0.11 | 4.5964E-17 | TEC_LIF | EPB41L1 |
| 5.0734E-14 | 0.25960509 | 0.374 | 0.183 | 1.0832E-09 | TEC_LIF | ARHGAP18 |

| | | | | | | |
|---|---|---|---|---|---|---|
| 9.68E-14 | 0.2592963 | 0.402 | 0.202 | 2.0667E-09 | TEC_LIF | FOSL2 |
| 8.0241E-16 | 0.25927152 | 0.594 | 0.326 | 1.7131E-11 | TEC_LIF | KIF5B |
| 4.0089E-16 | 0.2586808 | 0.276 | 0.11 | 8.559E-12 | TEC_LIF | MAML2 |
| 1.0698E-14 | 0.25858929 | 0.287 | 0.126 | 2.2841E-10 | TEC_LIF | FAM49B |
| 4.2121E-22 | 0.25816941 | 0.28 | 0.096 | 8.9928E-18 | TEC_LIF | MOB1B |
| 9.6331E-35 | 0.25751893 | 0.161 | 0.028 | 2.0567E-30 | TEC_LIF | HMGA2 |
| 5.1446E-19 | 0.25734758 | 0.295 | 0.113 | 1.0984E-14 | TEC_LIF | PISD |
| 1.2362E-22 | 0.25724652 | 0.295 | 0.103 | 2.6393E-18 | TEC_LIF | SLC35E1 |
| 0.00012252 | 0.25655571 | 0.976 | 0.909 | 1 | TEC_LIF | MT-CO3 |
| 7.6881E-10 | 0.25653866 | 0.504 | 0.318 | 1.6414E-05 | TEC_LIF | AES |
| 2.5232E-28 | 0.2561104 | 0.217 | 0.054 | 5.3871E-24 | TEC_LIF | ITGB3 |
| 3.0173E-19 | 0.25588621 | 0.339 | 0.137 | 6.4419E-15 | TEC_LIF | SLMAP |
| 4.2763E-18 | 0.25560644 | 0.252 | 0.092 | 9.1299E-14 | TEC_LIF | MPZL2 |
| 8.9049E-18 | 0.25542089 | 0.307 | 0.122 | 1.9012E-13 | TEC_LIF | MICAL3 |
| 6.4238E-11 | 0.25539625 | 0.394 | 0.218 | 1.3715E-06 | TEC_LIF | GARS |
| 9.193E-05 | 0.25520677 | 0.972 | 0.926 | 1 | TEC_LIF | MT-CO1 |
| 3.7704E-18 | 0.25519362 | 0.331 | 0.138 | 8.0497E-14 | TEC_LIF | GNAQ |
| 5.3665E-10 | 0.25502309 | 0.134 | 0.049 | 1.1457E-05 | TEC_LIF | SPRR2D |
| 6.6736E-14 | 0.25495577 | 0.299 | 0.137 | 1.4248E-09 | TEC_LIF | PACSIN2 |
| 4.1081E-17 | 0.25488405 | 0.358 | 0.158 | 8.7709E-13 | TEC_LIF | CRK |
| 9.3263E-14 | 0.25459817 | 0.386 | 0.195 | 1.9912E-09 | TEC_LIF | BBX |
| 9.3583E-20 | 0.25441215 | 0.232 | 0.077 | 1.998E-15 | TEC_LIF | ARHGEF2 |
| 4.7061E-25 | 0.25405172 | 0.283 | 0.09 | 1.0048E-20 | TEC_LIF | SH2D3A |
| 1.3034E-14 | 0.25347266 | 0.303 | 0.136 | 2.7828E-10 | TEC_LIF | TMEM245 |
| 1.8622E-28 | 0.2529983 | 0.252 | 0.069 | 3.9758E-24 | TEC_LIF | IGSF3 |
| 9.5715E-27 | 0.25291905 | 0.26 | 0.076 | 2.0435E-22 | TEC_LIF | SNN |
| 8.1607E-08 | 0.25288343 | 0.201 | 0.1 | 0.00174231 | TEC_LIF | MAP3K8 |
| 1.519E-13 | 0.25248282 | 0.26 | 0.111 | 3.243E-09 | TEC_LIF | INSR |
| 7.6908E-26 | 0.25243031 | 0.244 | 0.07 | 1.642E-21 | TEC_LIF | SIPA1L3 |
| 7.3801E-21 | 0.25219421 | 0.291 | 0.106 | 1.5757E-16 | TEC_LIF | SGPL1 |
| 2.1251E-13 | 0.25201271 | 0.272 | 0.12 | 4.5372E-09 | TEC_LIF | TMEM41B |
| 1.2514E-11 | 0.25194971 | 0.437 | 0.247 | 2.6718E-07 | TEC_LIF | ARID4B |
| 1.403E-18 | 0.25188158 | 0.394 | 0.174 | 2.9955E-14 | TEC_LIF | RYBP |
| 7.3843E-21 | 0.25158244 | 0.299 | 0.11 | 1.5765E-16 | TEC_LIF | IFNGR1 |
| 1.9413E-10 | 0.25105355 | 0.327 | 0.175 | 4.1446E-06 | TEC_LIF | BACE2 |
| 1.2903E-10 | 0.25066977 | 0.563 | 0.357 | 2.7549E-06 | TEC_LIF | EIF3A |
| 1.3884E-16 | 0.25036144 | 0.303 | 0.127 | 2.9642E-12 | TEC_LIF | TOP2B |
| 6.3555E-05 | -0.250363 | 0.724 | 0.869 | 1 | TEC_LIF | RPL23 |
| 0.0069212 | -0.2507951 | 0.331 | 0.39 | 1 | TEC_LIF | HSPE1 |
| 0.00530203 | -0.2512952 | 0.291 | 0.351 | 1 | TEC_LIF | MZT2B |
| 3.899E-05 | -0.2546259 | 0.024 | 0.1 | 0.83244167 | TEC_LIF | MAFA |
| 3.1266E-05 | -0.2567316 | 0.031 | 0.113 | 0.66753914 | TEC_LIF | RARRES2 |
| 0.0016374 | -0.2585159 | 0.142 | 0.215 | 1 | TEC_LIF | CTSH |
| 0.00970398 | -0.2594022 | 0.528 | 0.561 | 1 | TEC_LIF | NCL |
| 0.00046935 | -0.2613381 | 0.248 | 0.334 | 1 | TEC_LIF | ATP6V1F |
| 0.0046903 | -0.2615457 | 0.874 | 0.936 | 1 | TEC_LIF | PTMA |
| 9.8497E-06 | -0.2626121 | 0.067 | 0.17 | 0.21029174 | TEC_LIF | TMX4 |
| 1.3346E-06 | -0.2641294 | 0.134 | 0.262 | 0.02849389 | TEC_LIF | NDUFB3 |
| 3.3033E-06 | -0.2644432 | 0.098 | 0.212 | 0.07052529 | TEC_LIF | BEX3 |
| 0.00021026 | -0.265924 | 0.213 | 0.302 | 1 | TEC_LIF | DDT |

| | | | | | | |
|---|---|---|---|---|---|---|
| 0.00018183 | -0.2662358 | 0.224 | 0.315 | 1 | TEC_LIF | PCBD1 |
| 0.00643581 | -0.2676986 | 0.701 | 0.775 | 1 | TEC_LIF | MYL12A |
| 0.0002766 | -0.2689796 | 0.528 | 0.633 | 1 | TEC_LIF | ANXA5 |
| 5.5154E-06 | -0.2707817 | 0.157 | 0.277 | 0.11775341 | TEC_LIF | NDUFS6 |
| 6.0742E-07 | -0.2743873 | 0.13 | 0.261 | 0.01296839 | TEC_LIF | NDUFA6 |
| 1.1417E-05 | -0.2744751 | 0.173 | 0.289 | 0.24375707 | TEC_LIF | RABAC1 |
| 9.2334E-07 | -0.2747184 | 0.142 | 0.271 | 0.0197133 | TEC_LIF | NDUFB8 |
| 3.9996E-05 | -0.2748382 | 0.268 | 0.374 | 0.8539179 | TEC_LIF | SNU13 |
| 9.5664E-06 | -0.2752214 | 0.224 | 0.343 | 0.20424339 | TEC_LIF | SPCS1 |
| 0.0038533 | -0.2754885 | 0.346 | 0.396 | 1 | TEC_LIF | FKBP2 |
| 5.7938E-05 | -0.2757417 | 0.031 | 0.109 | 1 | TEC_LIF | AL355075.4 |
| 0.00073058 | -0.2759615 | 0.579 | 0.628 | 1 | TEC_LIF | SLC25A6 |
| 8.1442E-07 | -0.2762573 | 0.043 | 0.154 | 0.01738795 | TEC_LIF | SNHG25 |
| 0.00022065 | -0.276604 | 0.339 | 0.427 | 1 | TEC_LIF | TMEM59 |
| 0.00241381 | -0.2770123 | 0.26 | 0.334 | 1 | TEC_LIF | SLC25A5 |
| 4.2497E-05 | -0.2770214 | 0.248 | 0.351 | 0.90731558 | TEC_LIF | COMMD6 |
| 3.811E-06 | -0.2787632 | 0.189 | 0.313 | 0.08136545 | TEC_LIF | GABARAPL2 |
| 0.00012038 | -0.2790768 | 0.555 | 0.647 | 1 | TEC_LIF | EDF1 |
| 8.402E-06 | -0.2792335 | 0.087 | 0.193 | 0.1793835 | TEC_LIF | DCXR |
| 0.0001274 | -0.2794956 | 0.063 | 0.145 | 1 | TEC_LIF | CKS2 |
| 3.6093E-06 | -0.2804726 | 0.154 | 0.276 | 0.07705767 | TEC_LIF | COX17 |
| 2.1841E-05 | -0.2809658 | 0.244 | 0.359 | 0.46630246 | TEC_LIF | EIF3K |
| 0.00037819 | -0.2819212 | 0.528 | 0.606 | 1 | TEC_LIF | PSMA7 |
| 0.00525922 | -0.2821017 | 0.728 | 0.799 | 1 | TEC_LIF | HLA-C |
| 0.00025742 | -0.28305 | 0.339 | 0.418 | 1 | TEC_LIF | DYNLRB1 |
| 0.00087319 | -0.2860246 | 0.142 | 0.223 | 1 | TEC_LIF | ADM |
| 0.00062162 | -0.2865263 | 0.075 | 0.147 | 1 | TEC_LIF | ANKRD37 |
| 0.00125687 | -0.2866269 | 0.48 | 0.567 | 1 | TEC_LIF | PPIB |
| 5.6116E-06 | -0.2877592 | 0.024 | 0.113 | 0.1198068 | TEC_LIF | HIST4H4 |
| 1.5032E-06 | -0.2880216 | 0.028 | 0.128 | 0.03209363 | TEC_LIF | AL135925.1 |
| 0.00016328 | -0.2886289 | 0.327 | 0.407 | 1 | TEC_LIF | TBCA |
| 0.00016779 | -0.289063 | 0.343 | 0.428 | 1 | TEC_LIF | ATP6V0B |
| 0.00251274 | -0.2916272 | 0.406 | 0.464 | 1 | TEC_LIF | SERP1 |
| 0.00095773 | -0.2944924 | 0.098 | 0.173 | 1 | TEC_LIF | FOSB |
| 2.4995E-06 | -0.2992645 | 0.165 | 0.288 | 0.0533646 | TEC_LIF | NDUFB11 |
| 1.5265E-05 | -0.2998084 | 0.209 | 0.324 | 0.32591554 | TEC_LIF | NPC2 |
| 2.6332E-05 | -0.3001871 | 0.693 | 0.79 | 0.56218603 | TEC_LIF | NPM1 |
| 1.5897E-05 | -0.3051521 | 0.252 | 0.364 | 0.33939734 | TEC_LIF | SNRPD2 |
| 8.6629E-06 | -0.3087107 | 0.24 | 0.364 | 0.18495335 | TEC_LIF | TRMT112 |
| 2.5953E-05 | -0.3089424 | 0.421 | 0.536 | 0.55409797 | TEC_LIF | NDUFS5 |
| 8.6157E-05 | -0.3095829 | 0.476 | 0.57 | 1 | TEC_LIF | RAN |
| 0.00034168 | -0.310247 | 0.252 | 0.337 | 1 | TEC_LIF | H2AFJ |
| 0.0029384 | -0.3110795 | 0.291 | 0.364 | 1 | TEC_LIF | HMGN2 |
| 1.351E-05 | -0.3113681 | 0.213 | 0.328 | 0.28844488 | TEC_LIF | EIF1B |
| 5.8856E-05 | -0.3124553 | 0.059 | 0.146 | 1 | TEC_LIF | ID1 |
| 2.9544E-06 | -0.3157869 | 0.106 | 0.224 | 0.06307546 | TEC_LIF | BEX2 |
| 0.0068935 | -0.3179042 | 0.055 | 0.105 | 1 | TEC_LIF | SERPINE2 |
| 1.1737E-05 | -0.3188178 | 0.205 | 0.316 | 0.25059554 | TEC_LIF | UQCR11 |
| 9.8202E-05 | -0.3210748 | 0.031 | 0.106 | 1 | TEC_LIF | ARC |
| 0.00068983 | -0.3216559 | 0.291 | 0.363 | 1 | TEC_LIF | ATP5MC3 |

| | | | | | | |
|---|---|---|---|---|---|---|
| 0.00033511 | -0.3219879 | 0.323 | 0.4 | 1 | TEC_LIF | MGST3 |
| 0.00021141 | -0.322106 | 0.094 | 0.177 | 1 | TEC_LIF | FOXC1 |
| 0.00027146 | -0.322793 | 0.165 | 0.251 | 1 | TEC_LIF | MRPS6 |
| 6.3561E-07 | -0.3257324 | 0.228 | 0.363 | 0.01357017 | TEC_LIF | NEDD8 |
| 4.8664E-06 | -0.3266632 | 0.22 | 0.342 | 0.10389726 | TEC_LIF | KRT10 |
| 4.6287E-06 | -0.3283913 | 0.256 | 0.369 | 0.09882363 | TEC_LIF | WDR83OS |
| 0.00233027 | -0.3346049 | 0.594 | 0.468 | 1 | TEC_LIF | C12orf75 |
| 1.4003E-06 | -0.3352824 | 0.028 | 0.128 | 0.02989579 | TEC_LIF | PCDH9 |
| 9.3961E-05 | -0.3366686 | 0.055 | 0.137 | 1 | TEC_LIF | AC025164.1 |
| 8.1802E-05 | -0.3368553 | 0.665 | 0.728 | 1 | TEC_LIF | CSTB |
| 1.7253E-05 | -0.3369534 | 0.496 | 0.582 | 0.36834107 | TEC_LIF | ATP6V0E1 |
| 4.253E-07 | -0.3398037 | 0.197 | 0.33 | 0.00908016 | TEC_LIF | NDUFC1 |
| 0.00707978 | -0.3398589 | 0.669 | 0.715 | 1 | TEC_LIF | HSPB1 |
| 7.3234E-07 | -0.3419687 | 0.224 | 0.352 | 0.01563543 | TEC_LIF | C19orf53 |
| 7.3015E-05 | -0.3436263 | 0.028 | 0.101 | 1 | TEC_LIF | AP001160.1 |
| 4.0396E-06 | -0.3452195 | 0.417 | 0.535 | 0.0862463 | TEC_LIF | SEC61B |
| 1.9724E-06 | -0.3453008 | 0.201 | 0.326 | 0.04211161 | TEC_LIF | PHPT1 |
| 2.9278E-06 | -0.3459005 | 0.031 | 0.128 | 0.06250943 | TEC_LIF | HIST2H2BF |
| 0.00204415 | -0.3459414 | 0.161 | 0.234 | 1 | TEC_LIF | SYNE1 |
| 1.3925E-06 | -0.3484921 | 0.303 | 0.419 | 0.0297308 | TEC_LIF | PARK7 |
| 3.6477E-07 | -0.3488233 | 0.079 | 0.203 | 0.00778792 | TEC_LIF | CHCHD7 |
| 4.8901E-07 | -0.350311 | 0.039 | 0.15 | 0.01044042 | TEC_LIF | AL118516.1 |
| 6.0104E-05 | -0.3545356 | 0.295 | 0.383 | 1 | TEC_LIF | NDUFB2 |
| 0.00132198 | -0.3591601 | 0.264 | 0.334 | 1 | TEC_LIF | CHCHD10 |
| 8.5483E-08 | -0.3599743 | 0.248 | 0.399 | 0.00182505 | TEC_LIF | DAD1 |
| 0.00035999 | -0.361101 | 0.776 | 0.854 | 1 | TEC_LIF | MYL12B |
| 1.4235E-05 | -0.3615346 | 0.213 | 0.318 | 0.30392325 | TEC_LIF | PRDX2 |
| 3.2469E-05 | -0.370237 | 0.642 | 0.73 | 0.69322281 | TEC_LIF | ENO1 |
| 0.00150994 | -0.3707405 | 0.146 | 0.218 | 1 | TEC_LIF | UBE2S |
| 5.4872E-09 | -0.3728162 | 0.374 | 0.528 | 0.00011715 | TEC_LIF | ATP5MC2 |
| 0.0020182 | -0.373448 | 0.201 | 0.266 | 1 | TEC_LIF | AMD1 |
| 4.6439E-06 | -0.3743196 | 0.291 | 0.396 | 0.09914728 | TEC_LIF | ATP5F1D |
| 6.6582E-08 | -0.3806701 | 0.181 | 0.332 | 0.00142152 | TEC_LIF | UQCRQ |
| 4.0599E-05 | -0.3847851 | 0.098 | 0.192 | 0.8667934 | TEC_LIF | TLE4 |
| 2.0448E-07 | -0.3873133 | 0.232 | 0.371 | 0.00436575 | TEC_LIF | ATP5MF |
| 9.4662E-06 | -0.3909818 | 0.091 | 0.194 | 0.20210251 | TEC_LIF | H2AFX |
| 5.8031E-05 | -0.3917103 | 0.213 | 0.305 | 1 | TEC_LIF | SNHG7 |
| 0.00453774 | -0.3935414 | 0.146 | 0.212 | 1 | TEC_LIF | ADAMTS1 |
| 2.986E-09 | -0.3943896 | 0.315 | 0.473 | 6.3752E-05 | TEC_LIF | PRDX6 |
| 1.9405E-09 | -0.396099 | 0.524 | 0.655 | 4.143E-05 | TEC_LIF | SLC25A3 |
| 2.2161E-06 | -0.3965901 | 0.307 | 0.428 | 0.04731304 | TEC_LIF | SSR4 |
| 4.951E-08 | -0.3973622 | 0.165 | 0.313 | 0.00105705 | TEC_LIF | ISCU |
| 9.8142E-08 | -0.3976503 | 0.571 | 0.694 | 0.00209532 | TEC_LIF | GPX4 |
| 1.9488E-07 | -0.397707 | 0.957 | 0.988 | 0.00416075 | TEC_LIF | EEF1A1 |
| 5.3878E-08 | -0.4008248 | 0.744 | 0.875 | 0.00115029 | TEC_LIF | RACK1 |
| 1.9894E-05 | -0.4011779 | 0.547 | 0.646 | 0.42474693 | TEC_LIF | PEBP1 |
| 9.297E-08 | -0.4096597 | 0.13 | 0.269 | 0.00198491 | TEC_LIF | GSTM3 |
| 9.024E-09 | -0.4101308 | 0.291 | 0.45 | 0.00019266 | TEC_LIF | ATP5MD |
| 0.00030287 | -0.4182892 | 0.811 | 0.84 | 1 | TEC_LIF | CD63 |
| 0.00125439 | -0.4187616 | 0.795 | 0.83 | 1 | TEC_LIF | SQSTM1 |

| | | | | | | |
|---|---|---|---|---|---|---|
| 9.8304E-08 | -0.4221109 | 0.264 | 0.399 | 0.00209878 | TEC_LIF | CYCS |
| 0.00011608 | -0.430783 | 0.728 | 0.845 | 1 | TEC_LIF | RPSA |
| 3.1215E-08 | -0.4351059 | 0.217 | 0.362 | 0.00066644 | TEC_LIF | ATP5ME |
| 0.00043557 | -0.4352299 | 0.201 | 0.285 | 1 | TEC_LIF | UCHL1 |
| 6.1045E-12 | -0.4370723 | 0.496 | 0.665 | 1.3033E-07 | TEC_LIF | UQCRB |
| 7.0965E-11 | -0.4406401 | 0.547 | 0.722 | 1.5151E-06 | TEC_LIF | RPL36AL |
| 0.0027574 | -0.4475897 | 0.587 | 0.611 | 1 | TEC_LIF | BTG1 |
| 6.1373E-06 | -0.45094 | 0.169 | 0.288 | 0.13103223 | TEC_LIF | HIST1H1C |
| 0.00327704 | -0.451819 | 0.217 | 0.277 | 1 | TEC_LIF | CITED2 |
| 3.9257E-10 | -0.4536528 | 0.209 | 0.379 | 8.3813E-06 | TEC_LIF | NDUFA1 |
| 7.4747E-11 | -0.4540931 | 0.402 | 0.583 | 1.5959E-06 | TEC_LIF | PRDX5 |
| 9.1415E-11 | -0.4556726 | 0.398 | 0.564 | 1.9517E-06 | TEC_LIF | COX6B1 |
| 1.5544E-09 | -0.4572524 | 0.224 | 0.39 | 3.3187E-05 | TEC_LIF | ATP5IF1 |
| 0.00012002 | -0.4585195 | 0.087 | 0.17 | 1 | TEC_LIF | EGLN3 |
| 2.054E-07 | -0.460896 | 0.445 | 0.567 | 0.00438534 | TEC_LIF | CST3 |
| 1.2389E-11 | -0.461181 | 0.598 | 0.751 | 2.6451E-07 | TEC_LIF | DSTN |
| 4.821E-11 | -0.4614213 | 0.272 | 0.454 | 1.0293E-06 | TEC_LIF | RPS10 |
| 3.2551E-15 | -0.4619436 | 0.528 | 0.739 | 6.9497E-11 | TEC_LIF | SKP1 |
| 9.7389E-13 | -0.4620949 | 0.677 | 0.878 | 2.0793E-08 | TEC_LIF | RPL7 |
| 8.3897E-10 | -0.4684293 | 0.299 | 0.464 | 1.7912E-05 | TEC_LIF | COX8A |
| 0.00070804 | -0.4689524 | 0.374 | 0.253 | 1 | TEC_LIF | LCN2 |
| 2.5153E-08 | -0.4721268 | 0.248 | 0.389 | 0.00053701 | TEC_LIF | POLR2L |
| 5.3149E-07 | -0.4745026 | 0.15 | 0.279 | 0.01134732 | TEC_LIF | CYB5A |
| 6.3556E-11 | -0.4746402 | 0.283 | 0.456 | 1.3569E-06 | TEC_LIF | ATP5PF |
| 4.6058E-12 | -0.474793 | 0.638 | 0.779 | 9.8334E-08 | TEC_LIF | OAZ1 |
| 1.4786E-06 | -0.4752885 | 0.492 | 0.575 | 0.03156804 | TEC_LIF | SLC3A2 |
| 2.9929E-07 | -0.4761115 | 0.067 | 0.189 | 0.00638994 | TEC_LIF | ADIRF |
| 1.3829E-12 | -0.4766487 | 0.598 | 0.785 | 2.9526E-08 | TEC_LIF | COX4I1 |
| 1.0681E-11 | -0.4769993 | 0.15 | 0.349 | 2.2803E-07 | TEC_LIF | MIF |
| 8.4084E-13 | -0.4807355 | 0.091 | 0.288 | 1.7952E-08 | TEC_LIF | C12orf57 |
| 5.4466E-08 | -0.481514 | 0.787 | 0.918 | 0.00116286 | TEC_LIF | RPLP0 |
| 0.00093149 | -0.4817491 | 0.398 | 0.468 | 1 | TEC_LIF | IFITM3 |
| 0.00033902 | -0.4818719 | 0.047 | 0.118 | 1 | TEC_LIF | NEFM |
| 6.7197E-12 | -0.482141 | 0.343 | 0.532 | 1.4347E-07 | TEC_LIF | COX6C |
| 1.0851E-07 | -0.4882928 | 0.016 | 0.125 | 0.00231677 | TEC_LIF | CXCL14 |
| 3.7044E-09 | -0.4899463 | 0.594 | 0.702 | 7.9089E-05 | TEC_LIF | MAP1LC3B |
| 0.00228337 | -0.491426 | 0.311 | 0.37 | 1 | TEC_LIF | DDIT3 |
| 7.4494E-05 | -0.493261 | 0.197 | 0.288 | 1 | TEC_LIF | CEBPD |
| 3.1712E-10 | -0.4964836 | 0.193 | 0.364 | 6.7706E-06 | TEC_LIF | COX7B |
| 0.00129492 | -0.4968689 | 0.043 | 0.103 | 1 | TEC_LIF | APOE |
| 1.1877E-11 | -0.4982118 | 0.528 | 0.693 | 2.5357E-07 | TEC_LIF | CYSTM1 |
| 1.1635E-10 | -0.4982514 | 0.366 | 0.522 | 2.484E-06 | TEC_LIF | OST4 |
| 1.7518E-06 | -0.500129 | 0.154 | 0.273 | 0.03740011 | TEC_LIF | SEPHS2 |
| 8.7337E-14 | -0.5021217 | 0.201 | 0.415 | 1.8646E-09 | TEC_LIF | UQCR10 |
| 3.9204E-17 | -0.5034755 | 0.528 | 0.755 | 8.3701E-13 | TEC_LIF | BTF3 |
| 6.1238E-15 | -0.5090075 | 0.496 | 0.669 | 1.3074E-10 | TEC_LIF | TMA7 |
| 7.2241E-11 | -0.5117342 | 0.217 | 0.401 | 1.5424E-06 | TEC_LIF | DBI |
| 1.0874E-06 | -0.5124399 | 0.008 | 0.1 | 0.02321642 | TEC_LIF | SLC12A1 |
| 2.7614E-08 | -0.5247293 | 0.094 | 0.237 | 0.00058957 | TEC_LIF | FXYD2 |
| 3.5329E-14 | -0.5277038 | 0.457 | 0.652 | 7.5427E-10 | TEC_LIF | UQCRH |

| | | | | | | |
|---|---|---|---|---|---|---|
| 1.0102E-09 | -0.5336638 | 0.732 | 0.837 | 2.1568E-05 | TEC_LIF | PPIA |
| 3.1269E-09 | -0.534909 | 0.594 | 0.718 | 6.6759E-05 | TEC_LIF | TXN |
| 1.9305E-14 | -0.5352546 | 0.724 | 0.908 | 4.1215E-10 | TEC_LIF | RPS20 |
| 8.0286E-16 | -0.5357486 | 0.579 | 0.804 | 1.7141E-11 | TEC_LIF | RPL27 |
| 8.2339E-14 | -0.5363 | 0.815 | 0.934 | 1.7579E-09 | TEC_LIF | RPL18 |
| 3.3286E-17 | -0.5370064 | 0.382 | 0.613 | 7.1066E-13 | TEC_LIF | ATP5MG |
| 2.9789E-13 | -0.5401526 | 0.508 | 0.697 | 6.3599E-09 | TEC_LIF | ZFAS1 |
| 0.0005413 | -0.5408182 | 0.165 | 0.248 | 1 | TEC_LIF | MT1X |
| 2.6436E-08 | -0.544035 | 0.059 | 0.195 | 0.00056441 | TEC_LIF | GADD45G |
| 1.4696E-13 | -0.5446414 | 0.772 | 0.901 | 3.1377E-09 | TEC_LIF | RPL5 |
| 1.0152E-13 | -0.5448316 | 0.681 | 0.897 | 2.1673E-09 | TEC_LIF | RPS11 |
| 1.7516E-18 | -0.5467382 | 0.323 | 0.576 | 3.7396E-14 | TEC_LIF | COX7A2 |
| 0.00019649 | -0.5485179 | 0.157 | 0.239 | 1 | TEC_LIF | INTS6 |
| 2.1404E-09 | -0.5489841 | 0.217 | 0.377 | 4.5698E-05 | TEC_LIF | LDHB |
| 2.916E-16 | -0.5550855 | 0.406 | 0.616 | 6.2257E-12 | TEC_LIF | SAP18 |
| 3.5091E-19 | -0.5666434 | 0.406 | 0.64 | 7.492E-15 | TEC_LIF | UBL5 |
| 4.2535E-10 | -0.5668546 | 0.807 | 0.909 | 9.0812E-06 | TEC_LIF | MYL6 |
| 0.00472555 | -0.5671028 | 0.272 | 0.328 | 1 | TEC_LIF | MT1E |
| 0.00065974 | -0.5675625 | 0.213 | 0.291 | 1 | TEC_LIF | FOS |
| 2.6229E-09 | -0.56798 | 0.059 | 0.205 | 5.5998E-05 | TEC_LIF | SGK1 |
| 1.2186E-11 | -0.5679935 | 0.827 | 0.95 | 2.6017E-07 | TEC_LIF | RPL13A |
| 7.4391E-14 | -0.5688927 | 0.768 | 0.933 | 1.5882E-09 | TEC_LIF | RPS14 |
| 0.00043271 | -0.5707352 | 0.228 | 0.303 | 1 | TEC_LIF | PLK2 |
| 9.3453E-17 | -0.5710419 | 0.661 | 0.859 | 1.9952E-12 | TEC_LIF | RPL38 |
| 2.9014E-17 | -0.5743797 | 0.26 | 0.496 | 6.1946E-13 | TEC_LIF | TOMM7 |
| 1.8298E-14 | -0.5751365 | 0.819 | 0.941 | 3.9065E-10 | TEC_LIF | RPL15 |
| 2.5026E-17 | -0.577672 | 0.484 | 0.684 | 5.343E-13 | TEC_LIF | COX6A1 |
| 6.324E-17 | -0.5782931 | 0.598 | 0.791 | 1.3502E-12 | TEC_LIF | CHCHD2 |
| 2.6937E-17 | -0.5798156 | 0.622 | 0.808 | 5.751E-13 | TEC_LIF | EEF1D |
| 3.8498E-14 | -0.5843246 | 0.866 | 0.974 | 8.2194E-10 | TEC_LIF | TPT1 |
| 0.00109307 | -0.5852217 | 0.252 | 0.313 | 1 | TEC_LIF | GLUL |
| 2.6692E-11 | -0.5864802 | 0.5 | 0.659 | 5.6987E-07 | TEC_LIF | HSPA8 |
| 4.3558E-08 | -0.5884803 | 0.059 | 0.187 | 0.00092996 | TEC_LIF | PIK3R3 |
| 1.9174E-13 | -0.5887411 | 0.803 | 0.907 | 4.0936E-09 | TEC_LIF | RPS7 |
| 2.3168E-16 | -0.5888411 | 0.622 | 0.823 | 4.9465E-12 | TEC_LIF | YBX1 |
| 9.4005E-07 | -0.5898903 | 0.276 | 0.388 | 0.02007016 | TEC_LIF | BRD2 |
| 4.6322E-17 | -0.5917484 | 0.303 | 0.532 | 9.8896E-13 | TEC_LIF | ELOB |
| 5.0865E-15 | -0.5931938 | 0.713 | 0.905 | 1.086E-10 | TEC_LIF | RPL27A |
| 2.3033E-06 | -0.5970406 | 0.811 | 0.878 | 0.04917622 | TEC_LIF | ITM2B |
| 9.9637E-14 | -0.6033586 | 0.37 | 0.573 | 2.1272E-09 | TEC_LIF | RPS27L |
| 7.3068E-19 | -0.6042629 | 0.693 | 0.891 | 1.56E-14 | TEC_LIF | RPL22 |
| 2.0738E-19 | -0.6078076 | 0.697 | 0.894 | 4.4275E-15 | TEC_LIF | NACA |
| 8.2152E-18 | -0.6083374 | 0.594 | 0.809 | 1.7539E-13 | TEC_LIF | RPL31 |
| 2.4164E-08 | -0.6111867 | 0.728 | 0.839 | 0.0005159 | TEC_LIF | PRDX1 |
| 1.5232E-18 | -0.6149087 | 0.35 | 0.587 | 3.2521E-14 | TEC_LIF | COX5B |
| 4.8683E-05 | -0.6161043 | 0.531 | 0.619 | 1 | TEC_LIF | MTRNR2L8 |
| 3.6894E-24 | -0.6168872 | 0.484 | 0.757 | 7.877E-20 | TEC_LIF | PFDN5 |
| 2.5777E-18 | -0.6173091 | 0.697 | 0.857 | 5.5034E-14 | TEC_LIF | SRP14 |
| 1.6269E-13 | -0.6199832 | 0.587 | 0.761 | 3.4734E-09 | TEC_LIF | DYNLL1 |
| 3.3995E-10 | -0.6254073 | 0.122 | 0.287 | 7.2579E-06 | TEC_LIF | POLR2A |

| | | | | | | |
|---|---|---|---|---|---|---|
| 8.074E-08 | -0.6257333 | 0.252 | 0.393 | 0.0017238 | TEC_LIF | TUBA1B |
| 5.1937E-14 | -0.6320774 | 0.213 | 0.427 | 1.1089E-09 | TEC_LIF | SNHG8 |
| 0.00242792 | -0.6333284 | 0.594 | 0.713 | 1 | TEC_LIF | MT2A |
| 9.3921E-16 | -0.6336225 | 0.815 | 0.933 | 2.0052E-11 | TEC_LIF | RPL7A |
| 3.1602E-20 | -0.6351716 | 0.504 | 0.751 | 6.747E-16 | TEC_LIF | HINT1 |
| 0.00240702 | -0.6386405 | 0.272 | 0.331 | 1 | TEC_LIF | TIMP1 |
| 4.648E-20 | -0.6387697 | 0.543 | 0.788 | 9.9235E-16 | TEC_LIF | EEF1B2 |
| 2.2793E-16 | -0.6402482 | 0.74 | 0.908 | 4.8662E-12 | TEC_LIF | RPS3 |
| 6.2567E-14 | -0.6466938 | 0.795 | 0.926 | 1.3358E-09 | TEC_LIF | RPLP2 |
| 9.0344E-08 | -0.6473102 | 0.772 | 0.875 | 0.00192885 | TEC_LIF | GSTP1 |
| 7.1225E-21 | -0.6506096 | 0.701 | 0.914 | 1.5207E-16 | TEC_LIF | RPL24 |
| 2.0943E-16 | -0.6507804 | 0.791 | 0.935 | 4.4712E-12 | TEC_LIF | RPS6 |
| 7.4485E-18 | -0.658122 | 0.685 | 0.825 | 1.5903E-13 | TEC_LIF | RPS21 |
| 0.00045229 | -0.6611265 | 0.63 | 0.639 | 1 | TEC_LIF | JUND |
| 1.0491E-19 | -0.6628555 | 0.799 | 0.936 | 2.2399E-15 | TEC_LIF | RPL6 |
| 2.6335E-09 | -0.6658973 | 0.106 | 0.258 | 5.6225E-05 | TEC_LIF | IER5L |
| 1.6161E-11 | -0.6692246 | 0.079 | 0.256 | 3.4504E-07 | TEC_LIF | ID2 |
| 2.8913E-12 | -0.6710468 | 0.063 | 0.246 | 6.1729E-08 | TEC_LIF | ID4 |
| 1.2468E-17 | -0.6724061 | 0.677 | 0.869 | 2.662E-13 | TEC_LIF | UBA52 |
| 8.6863E-14 | -0.6730896 | 0.709 | 0.869 | 1.8545E-09 | TEC_LIF | RPS4X |
| 1.6518E-19 | -0.6733374 | 0.654 | 0.873 | 3.5265E-15 | TEC_LIF | RPL10A |
| 3.0342E-23 | -0.6772501 | 0.551 | 0.773 | 6.478E-19 | TEC_LIF | COX7C |
| 0.0071659 | -0.6786302 | 0.563 | 0.564 | 1 | TEC_LIF | IER2 |
| 3.5163E-14 | -0.6814179 | 0.85 | 0.963 | 7.5072E-10 | TEC_LIF | RPL28 |
| 3.7545E-17 | -0.6858193 | 0.799 | 0.936 | 8.0158E-13 | TEC_LIF | RPL3 |
| 6.4424E-20 | -0.6905646 | 0.744 | 0.923 | 1.3755E-15 | TEC_LIF | RPL9 |
| 1.3086E-22 | -0.6931834 | 0.724 | 0.906 | 2.7939E-18 | TEC_LIF | RPS25 |
| 1.1004E-18 | -0.693631 | 0.89 | 0.975 | 2.3494E-14 | TEC_LIF | RPL13 |
| 5.3039E-17 | -0.6980157 | 0.78 | 0.937 | 1.1324E-12 | TEC_LIF | RPS27A |
| 4.8092E-05 | -0.7022222 | 0.185 | 0.279 | 1 | TEC_LIF | HES1 |
| 5.6874E-25 | -0.7038621 | 0.555 | 0.813 | 1.2143E-20 | TEC_LIF | ATP5F1E |
| 3.5666E-06 | -0.7041302 | 0.173 | 0.279 | 0.0761478 | TEC_LIF | PPP1R10 |
| 5.3514E-19 | -0.7097292 | 0.626 | 0.795 | 1.1425E-14 | TEC_LIF | PPDPF |
| 1.131E-08 | -0.7155819 | 0.283 | 0.418 | 0.00024146 | TEC_LIF | SRSF7 |
| 1.4029E-25 | -0.7166467 | 0.386 | 0.669 | 2.9953E-21 | TEC_LIF | SOD1 |
| 1.8915E-09 | -0.7264271 | 0.913 | 0.95 | 4.0383E-05 | TEC_LIF | B2M |
| 1.038E-21 | -0.728326 | 0.776 | 0.923 | 2.2161E-17 | TEC_LIF | RPL34 |
| 1.3028E-22 | -0.7291237 | 0.488 | 0.76 | 2.7814E-18 | TEC_LIF | RPS26 |
| 2.073E-05 | -0.7310678 | 0.154 | 0.263 | 0.44259067 | TEC_LIF | NUPR1 |
| 0.00162361 | -0.7367805 | 0.5 | 0.567 | 1 | TEC_LIF | GDF15 |
| 3.5592E-20 | -0.7417516 | 0.787 | 0.954 | 7.5988E-16 | TEC_LIF | RPS8 |
| 1.5759E-14 | -0.7501034 | 0.862 | 0.925 | 3.3646E-10 | TEC_LIF | RPL10 |
| 3.5911E-24 | -0.7568167 | 0.713 | 0.911 | 7.6669E-20 | TEC_LIF | RPS16 |
| 2.2312E-25 | -0.758997 | 0.898 | 0.959 | 4.7636E-21 | TEC_LIF | EIF1 |
| 4.5686E-21 | -0.7651636 | 0.752 | 0.925 | 9.7541E-17 | TEC_LIF | RPS3A |
| 2.2275E-26 | -0.7671597 | 0.638 | 0.903 | 4.7557E-22 | TEC_LIF | RPL23A |
| 2.6302E-21 | -0.7675178 | 0.78 | 0.941 | 5.6154E-17 | TEC_LIF | RPS23 |
| 9.6231E-24 | -0.7739169 | 0.756 | 0.932 | 2.0545E-19 | TEC_LIF | RPL36 |
| 3.7203E-21 | -0.7845352 | 0.677 | 0.886 | 7.9429E-17 | TEC_LIF | RPL39 |
| 1.3065E-22 | -0.7910043 | 0.795 | 0.936 | 2.7893E-18 | TEC_LIF | RPS9 |

| | | | | | | |
|---|---|---|---|---|---|---|
| 2.8128E-30 | -0.7976232 | 0.709 | 0.914 | 6.0054E-26 | TEC_LIF | FAU |
| 5.1154E-19 | -0.7978292 | 0.866 | 0.967 | 1.0921E-14 | TEC_LIF | RPS19 |
| 4.4318E-26 | -0.803633 | 0.63 | 0.901 | 9.4619E-22 | TEC_LIF | RPS13 |
| 1.8563E-19 | -0.8044662 | 0.26 | 0.515 | 3.9632E-15 | TEC_LIF | H2AFZ |
| 2.0776E-09 | -0.8106393 | 0.465 | 0.603 | 4.4357E-05 | TEC_LIF | CLU |
| 3.9526E-28 | -0.8117837 | 0.685 | 0.901 | 8.4387E-24 | TEC_LIF | RPS15A |
| 1.2623E-28 | -0.8146539 | 0.685 | 0.901 | 2.695E-24 | TEC_LIF | RPL35A |
| 5.0018E-26 | -0.8161906 | 0.736 | 0.938 | 1.0679E-21 | TEC_LIF | RPL37 |
| 5.6057E-23 | -0.8165259 | 0.815 | 0.946 | 1.1968E-18 | TEC_LIF | RPL8 |
| 5.1926E-23 | -0.8166914 | 0.815 | 0.943 | 1.1086E-18 | TEC_LIF | RPL37A |
| 2.4283E-25 | -0.8172603 | 0.768 | 0.945 | 5.1844E-21 | TEC_LIF | RPS24 |
| 6.7425E-20 | -0.8223602 | 0.748 | 0.932 | 1.4395E-15 | TEC_LIF | RPL18A |
| 1.1711E-29 | -0.8247342 | 0.413 | 0.738 | 2.5004E-25 | TEC_LIF | NDUFA4 |
| 0.000183 | -0.8287271 | 0.051 | 0.128 | 1 | TEC_LIF | GPX3 |
| 3.5268E-22 | -0.8319527 | 0.831 | 0.941 | 7.5297E-18 | TEC_LIF | RPS28 |
| 6.0906E-28 | -0.8339288 | 0.728 | 0.916 | 1.3003E-23 | TEC_LIF | RPL35 |
| 2.0884E-28 | -0.8364509 | 0.642 | 0.891 | 4.4587E-24 | TEC_LIF | RPS29 |
| 6.6087E-24 | -0.8428906 | 0.665 | 0.874 | 1.411E-19 | TEC_LIF | RPS5 |
| 1.1857E-27 | -0.856294 | 0.783 | 0.944 | 2.5315E-23 | TEC_LIF | RPL19 |
| 1.803E-30 | -0.8577011 | 0.705 | 0.926 | 3.8495E-26 | TEC_LIF | RPL14 |
| 7.9926E-10 | -0.8618449 | 0.083 | 0.237 | 1.7064E-05 | TEC_LIF | CSKMT |
| 3.5263E-33 | -0.8635926 | 0.508 | 0.824 | 7.5287E-29 | TEC_LIF | UBB |
| 1.8829E-25 | -0.8730205 | 0.74 | 0.932 | 4.02E-21 | TEC_LIF | RPL21 |
| 1.9395E-27 | -0.8825296 | 0.827 | 0.948 | 4.1408E-23 | TEC_LIF | RPL11 |
| 3.4189E-23 | -0.8885759 | 0.846 | 0.958 | 7.2993E-19 | TEC_LIF | RPS2 |
| 1.0677E-27 | -0.8899621 | 0.717 | 0.916 | 2.2796E-23 | TEC_LIF | RPL29 |
| 1.6014E-26 | -0.9098154 | 0.724 | 0.938 | 3.4189E-22 | TEC_LIF | RPL32 |
| 4.3647E-28 | -0.9119563 | 0.394 | 0.674 | 9.3186E-24 | TEC_LIF | TPI1 |
| 7.0221E-25 | -0.9227973 | 0.874 | 0.974 | 1.4992E-20 | TEC_LIF | RPLP1 |
| 4.1016E-24 | -0.9389329 | 0.799 | 0.954 | 8.757E-20 | TEC_LIF | RPS12 |
| 3.4723E-29 | -0.9412175 | 0.732 | 0.934 | 7.4134E-25 | TEC_LIF | RPL12 |
| 2.4358E-07 | -0.9488978 | 0.433 | 0.603 | 0.00520043 | TEC_LIF | SPP1 |
| 8.0174E-36 | -0.9503519 | 0.681 | 0.914 | 1.7117E-31 | TEC_LIF | RPL26 |
| 8.0725E-34 | -0.9655199 | 0.689 | 0.927 | 1.7235E-29 | TEC_LIF | RPL30 |
| 8.756E-13 | -0.9707005 | 0.445 | 0.608 | 1.8694E-08 | TEC_LIF | GADD45B |
| 6.3941E-34 | -0.9771252 | 0.76 | 0.949 | 1.3651E-29 | TEC_LIF | RPS15 |
| 9.2339E-29 | -1.006517 | 0.839 | 0.964 | 1.9714E-24 | TEC_LIF | RPS18 |
| 3.1818E-35 | -1.0083991 | 0.689 | 0.904 | 6.793E-31 | TEC_LIF | SERF2 |
| 5.5073E-09 | -1.0313639 | 0.516 | 0.682 | 0.00011758 | TEC_LIF | MTRNR2L12 |
| 4.9071E-20 | -1.0356003 | 0.724 | 0.896 | 1.0477E-15 | TEC_LIF | GAPDH |
| 9.16E-13 | -1.0640415 | 0.106 | 0.301 | 1.9557E-08 | TEC_LIF | HEXIM1 |
| 3.7926E-35 | -1.0986744 | 0.89 | 0.969 | 8.0973E-31 | TEC_LIF | RPL41 |
| 7.681E-17 | -1.1115955 | 0.913 | 0.98 | 1.6399E-12 | TEC_LIF | FTH1 |
| 4.6274E-20 | -1.1187563 | 0.657 | 0.818 | 9.8794E-16 | TEC_LIF | IGFBP7 |
| 3.8966E-47 | -1.1257898 | 0.724 | 0.96 | 8.3193E-43 | TEC_LIF | RPS27 |
| 6.3936E-15 | -1.1408143 | 0.063 | 0.274 | 1.365E-10 | TEC_LIF | HSPA2 |
| 7.7046E-11 | -1.1892307 | 0.224 | 0.38 | 1.6449E-06 | TEC_LIF | HIST1H4C |
| 1.9374E-05 | -1.219194 | 0.126 | 0.227 | 0.41362773 | TEC_LIF | DEFB1 |
| 3.6401E-15 | -1.2867671 | 0.862 | 0.963 | 7.7717E-11 | TEC_LIF | FTL |
| 5.0284E-14 | -1.2871038 | 0.083 | 0.287 | 1.0736E-09 | TEC_LIF | MAFB |

| | | | | | | |
|---|---|---|---|---|---|---|
| 1.697E-21 | -4.1142902 | 0.378 | 0.602 | 3.6231E-17 | TEC_LIF | PLCG2 |
| 0 | 3.23606752 | 0.926 | 0.595 | 0 | TEC_MMP7 | MMP7 |
| 0 | 2.69261413 | 0.795 | 0.333 | 0 | TEC_MMP7 | SLPI |
| 0 | 2.24477798 | 0.874 | 0.412 | 0 | TEC_MMP7 | C12orf75 |
| 0 | 2.16250752 | 0.629 | 0.201 | 0 | TEC_MMP7 | LCN2 |
| 7.483E-179 | 2.15690527 | 0.766 | 0.575 | 1.598E-174 | TEC_MMP7 | SPP1 |
| 0 | 2.14227655 | 0.996 | 0.908 | 0 | TEC_MMP7 | S100A11 |
| 0 | 2.08387933 | 0.632 | 0.186 | 0 | TEC_MMP7 | TFPI2 |
| 0 | 2.02559919 | 0.973 | 0.628 | 0 | TEC_MMP7 | SH3BGRL3 |
| 0 | 2.00089888 | 0.98 | 0.809 | 0 | TEC_MMP7 | ANXA2 |
| 0 | 1.90001045 | 0.993 | 0.949 | 0 | TEC_MMP7 | S100A6 |
| 0 | 1.89107149 | 0.976 | 0.853 | 0 | TEC_MMP7 | TMSB4X |
| 0 | 1.86480886 | 0.897 | 0.425 | 0 | TEC_MMP7 | TSPAN1 |
| 0 | 1.84191403 | 0.689 | 0.17 | 0 | TEC_MMP7 | CST6 |
| 0 | 1.82958838 | 0.99 | 0.853 | 0 | TEC_MMP7 | S100A10 |
| 0 | 1.68662436 | 0.919 | 0.699 | 0 | TEC_MMP7 | KRT8 |
| 0 | 1.68308064 | 0.893 | 0.609 | 0 | TEC_MMP7 | WFDC2 |
| 0 | 1.60619323 | 0.998 | 0.878 | 0 | TEC_MMP7 | GAPDH |
| 0 | 1.56123507 | 0.985 | 0.857 | 0 | TEC_MMP7 | GSTP1 |
| 0 | 1.45889035 | 0.995 | 0.913 | 0 | TEC_MMP7 | TMSB10 |
| 0 | 1.4233729 | 0.744 | 0.27 | 0 | TEC_MMP7 | TIMP1 |
| 1.828E-149 | 1.38987621 | 0.389 | 0.14 | 3.903E-145 | TEC_MMP7 | S100A4 |
| 0 | 1.32608468 | 0.994 | 0.942 | 0 | TEC_MMP7 | B2M |
| 0 | 1.32577624 | 0.978 | 0.823 | 0 | TEC_MMP7 | RPSA |
| 5.521E-163 | 1.31722103 | 0.537 | 0.24 | 1.179E-158 | TEC_MMP7 | LGALS1 |
| 6.496E-178 | 1.27586579 | 0.307 | 0.08 | 1.387E-173 | TEC_MMP7 | S100A9 |
| 0 | 1.25874066 | 0.796 | 0.387 | 0 | TEC_MMP7 | PPP1R14B |
| 1.292E-285 | 1.25737167 | 0.98 | 0.913 | 2.758E-281 | TEC_MMP7 | ACTB |
| 5.149E-195 | 1.24273305 | 0.849 | 0.62 | 1.099E-190 | TEC_MMP7 | KRT7 |
| 0 | 1.24233897 | 0.734 | 0.266 | 0 | TEC_MMP7 | CYBA |
| 2.274E-120 | 1.23425224 | 0.773 | 0.602 | 4.856E-116 | TEC_MMP7 | VIM |
| 1.083E-140 | 1.23044296 | 0.7 | 0.461 | 2.312E-136 | TEC_MMP7 | S100A2 |
| 0 | 1.2290422 | 0.983 | 0.834 | 0 | TEC_MMP7 | MYL12B |
| 0 | 1.22790922 | 0.999 | 0.944 | 0 | TEC_MMP7 | RPS12 |
| 0 | 1.22240279 | 0.999 | 0.968 | 0 | TEC_MMP7 | RPLP1 |
| 1.479E-237 | 1.20808168 | 0.966 | 0.864 | 3.158E-233 | TEC_MMP7 | ITM2B |
| 0 | 1.20074579 | 0.973 | 0.817 | 0 | TEC_MMP7 | PRDX1 |
| 0 | 1.19818366 | 0.975 | 0.82 | 0 | TEC_MMP7 | CD63 |
| 0 | 1.18676319 | 0.996 | 0.95 | 0 | TEC_MMP7 | RPS2 |
| 0 | 1.18579922 | 0.975 | 0.815 | 0 | TEC_MMP7 | PPIA |
| 0 | 1.18394929 | 0.996 | 0.918 | 0 | TEC_MMP7 | RPL18A |
| 0 | 1.16062041 | 0.997 | 0.913 | 0 | TEC_MMP7 | RPLP2 |
| 0 | 1.15734409 | 0.763 | 0.264 | 0 | TEC_MMP7 | TSPO |
| 0 | 1.15717031 | 0.968 | 0.78 | 0 | TEC_MMP7 | CFL1 |
| 4.949E-283 | 1.14652038 | 0.912 | 0.705 | 1.057E-278 | TEC_MMP7 | PFN1 |
| 0 | 1.13693817 | 0.996 | 0.903 | 0 | TEC_MMP7 | RPLP0 |
| 0 | 1.13544182 | 0.99 | 0.847 | 0 | TEC_MMP7 | UBA52 |
| 0 | 1.13146115 | 0.837 | 0.413 | 0 | TEC_MMP7 | IFITM3 |
| 2.578E-173 | 1.11159337 | 0.267 | 0.061 | 5.504E-169 | TEC_MMP7 | SAA1 |
| 0 | 1.10856646 | 0.891 | 0.538 | 0 | TEC_MMP7 | ARPC3 |

| | | | | | | |
|---|---|---|---|---|---|---|
| 0 | 1.10412905 | 0.991 | 0.867 | 0 | TEC_MMP7 | RPL39 |
| 0 | 1.09972834 | 0.998 | 0.956 | 0 | TEC_MMP7 | RPS18 |
| 6.73E-148 | 1.09640917 | 0.781 | 0.564 | 1.437E-143 | TEC_MMP7 | KRT19 |
| 2.785E-303 | 1.09496976 | 0.998 | 0.921 | 5.946E-299 | TEC_MMP7 | RPL12 |
| 0 | 1.09333442 | 0.998 | 0.955 | 0 | TEC_MMP7 | RPL28 |
| 8.545E-182 | 1.09219955 | 0.864 | 0.681 | 1.824E-177 | TEC_MMP7 | ITGB1 |
| 8.011E-301 | 1.09180536 | 0.979 | 0.897 | 1.71E-296 | TEC_MMP7 | MYL6 |
| 2.612E-246 | 1.08984617 | 0.675 | 0.281 | 5.576E-242 | TEC_MMP7 | LAMC2 |
| 0 | 1.08629208 | 0.819 | 0.368 | 0 | TEC_MMP7 | SSR4 |
| 0 | 1.07864922 | 1 | 0.962 | 0 | TEC_MMP7 | RPL41 |
| 1.1795E-80 | 1.06250667 | 0.283 | 0.114 | 2.5183E-76 | TEC_MMP7 | SERPINA1 |
| 0 | 1.05385598 | 0.999 | 0.924 | 0 | TEC_MMP7 | RPL32 |
| 0 | 1.050501 | 0.873 | 0.516 | 0 | TEC_MMP7 | GUK1 |
| 0 | 1.04839175 | 0.986 | 0.848 | 0 | TEC_MMP7 | RPS4X |
| 0 | 1.04398149 | 0.988 | 0.914 | 0 | TEC_MMP7 | RPL10 |
| 8.0143E-66 | 1.03165981 | 0.742 | 0.579 | 1.7111E-61 | TEC_MMP7 | CLU |
| 1.721E-304 | 1.0235294 | 0.935 | 0.702 | 3.673E-300 | TEC_MMP7 | NDUFA4 |
| 0 | 1.01937386 | 0.766 | 0.343 | 0 | TEC_MMP7 | RHOC |
| 0 | 1.01882786 | 0.998 | 0.935 | 0 | TEC_MMP7 | RPL8 |
| 0 | 1.0158709 | 0.6 | 0.185 | 0 | TEC_MMP7 | HIGD1A |
| 0 | 1.01205373 | 0.879 | 0.519 | 0 | TEC_MMP7 | CST3 |
| 0 | 1.00281964 | 0.631 | 0.175 | 0 | TEC_MMP7 | TGM2 |
| 0 | 0.98579896 | 0.699 | 0.245 | 0 | TEC_MMP7 | C9orf16 |
| 1.807E-286 | 0.98499382 | 0.943 | 0.739 | 3.857E-282 | TEC_MMP7 | PKM |
| 8.167E-166 | 0.97302235 | 0.605 | 0.286 | 1.744E-161 | TEC_MMP7 | IL32 |
| 6.893E-296 | 0.9683917 | 0.597 | 0.194 | 1.472E-291 | TEC_MMP7 | PDZK1IP1 |
| 0 | 0.96733263 | 0.963 | 0.761 | 0 | TEC_MMP7 | CHCHD2 |
| 0 | 0.9652924 | 0.996 | 0.9 | 0 | TEC_MMP7 | RPL29 |
| 0 | 0.95713205 | 0.996 | 0.925 | 0 | TEC_MMP7 | RPS27A |
| 3.088E-289 | 0.94927777 | 0.927 | 0.655 | 6.594E-285 | TEC_MMP7 | CYSTM1 |
| 3.25E-291 | 0.94664175 | 0.982 | 0.853 | 6.938E-287 | TEC_MMP7 | RPS5 |
| 5.8407E-83 | 0.94445867 | 0.254 | 0.095 | 1.247E-78 | TEC_MMP7 | TAGLN |
| 1.779E-263 | 0.94329683 | 0.937 | 0.75 | 3.797E-259 | TEC_MMP7 | MYL12A |
| 1.262E-270 | 0.94093724 | 0.903 | 0.633 | 2.694E-266 | TEC_MMP7 | SPINT2 |
| 4.342E-306 | 0.92780343 | 0.991 | 0.884 | 9.27E-302 | TEC_MMP7 | RPL23A |
| 1.698E-292 | 0.92617316 | 0.986 | 0.887 | 3.624E-288 | TEC_MMP7 | SERF2 |
| 0 | 0.92373227 | 0.734 | 0.306 | 0 | TEC_MMP7 | PSME1 |
| 1.514E-230 | 0.91944921 | 0.772 | 0.425 | 3.233E-226 | TEC_MMP7 | PDIA6 |
| 7.0114E-53 | 0.91879219 | 0.825 | 0.795 | 1.4969E-48 | TEC_MMP7 | KRT18 |
| 1.175E-263 | 0.9181362 | 0.73 | 0.342 | 2.508E-259 | TEC_MMP7 | CD151 |
| 4.814E-289 | 0.91535822 | 0.974 | 0.837 | 1.028E-284 | TEC_MMP7 | SRP14 |
| 1.177E-254 | 0.91147978 | 0.846 | 0.525 | 2.513E-250 | TEC_MMP7 | PPIB |
| 0 | 0.91090984 | 0.694 | 0.242 | 0 | TEC_MMP7 | CAPG |
| 1.599E-133 | 0.91002339 | 0.281 | 0.081 | 3.413E-129 | TEC_MMP7 | FXYD4 |
| 7.704E-204 | 0.90930388 | 0.947 | 0.8 | 1.645E-199 | TEC_MMP7 | YBX1 |
| 4.935E-216 | 0.90685204 | 0.899 | 0.648 | 1.054E-211 | TEC_MMP7 | EPCAM |
| 3.283E-250 | 0.90030098 | 0.993 | 0.919 | 7.009E-246 | TEC_MMP7 | RPL21 |
| 6.852E-298 | 0.89796577 | 0.994 | 0.924 | 1.463E-293 | TEC_MMP7 | RPS9 |
| 1.145E-275 | 0.89382065 | 0.996 | 0.911 | 2.444E-271 | TEC_MMP7 | RPS3A |
| 1.598E-206 | 0.89156309 | 0.806 | 0.506 | 3.412E-202 | TEC_MMP7 | CD59 |

| | | | | | | |
|---|---|---|---|---|---|---|
| 1.115E-278 | 0.88949473 | 0.83 | 0.473 | 2.381E-274 | TEC_MMP7 | OST4 |
| 3.121E-268 | 0.88840724 | 0.994 | 0.923 | 6.663E-264 | TEC_MMP7 | RPS6 |
| 4.742E-248 | 0.88575352 | 0.839 | 0.542 | 1.012E-243 | TEC_MMP7 | PRDX5 |
| 3.268E-291 | 0.87935356 | 0.998 | 0.934 | 6.977E-287 | TEC_MMP7 | RPS24 |
| 4.58E-279 | 0.8733814 | 0.978 | 0.853 | 9.778E-275 | TEC_MMP7 | RPL10A |
| 7.026E-269 | 0.86868277 | 0.999 | 0.96 | 1.5E-264 | TEC_MMP7 | RPS19 |
| 7.3467E-95 | 0.8653759 | 0.883 | 0.804 | 1.5685E-90 | TEC_MMP7 | IGFBP7 |
| 5.258E-260 | 0.86472906 | 0.985 | 0.902 | 1.123E-255 | TEC_MMP7 | RPL35 |
| 2.508E-283 | 0.86430129 | 0.998 | 0.938 | 5.355E-279 | TEC_MMP7 | RPL11 |
| 1.071E-276 | 0.86201402 | 0.998 | 0.929 | 2.287E-272 | TEC_MMP7 | RPS23 |
| 4.072E-260 | 0.85957036 | 0.994 | 0.925 | 8.693E-256 | TEC_MMP7 | RPL6 |
| 3.047E-247 | 0.85952896 | 0.703 | 0.322 | 6.504E-243 | TEC_MMP7 | TAGLN2 |
| 3.018E-231 | 0.85410397 | 0.95 | 0.759 | 6.443E-227 | TEC_MMP7 | EEF1B2 |
| 1.299E-261 | 0.85185243 | 0.994 | 0.931 | 2.772E-257 | TEC_MMP7 | RPS28 |
| 2.805E-168 | 0.84856226 | 0.95 | 0.863 | 5.989E-164 | TEC_MMP7 | HLA-A |
| 6.277E-276 | 0.84789332 | 0.998 | 0.968 | 1.34E-271 | TEC_MMP7 | TPT1 |
| 2.657E-233 | 0.843761 | 0.951 | 0.769 | 5.672E-229 | TEC_MMP7 | PPDPF |
| 2.984E-277 | 0.83939819 | 0.985 | 0.893 | 6.37E-273 | TEC_MMP7 | RPS7 |
| 5.574E-245 | 0.83693841 | 0.993 | 0.925 | 1.19E-240 | TEC_MMP7 | RPL3 |
| 5.0643E-86 | 0.83635347 | 0.109 | 0.02 | 1.0812E-81 | TEC_MMP7 | RARRES1 |
| 1.404E-262 | 0.83164198 | 0.996 | 0.908 | 2.997E-258 | TEC_MMP7 | RPL9 |
| 2.009E-299 | 0.83115715 | 0.996 | 0.933 | 4.289E-295 | TEC_MMP7 | RPL19 |
| 8.004E-264 | 0.82446569 | 0.991 | 0.882 | 1.709E-259 | TEC_MMP7 | RPS13 |
| 7.081E-206 | 0.82246743 | 0.982 | 0.88 | 1.512E-201 | TEC_MMP7 | RPS11 |
| 9.932E-207 | 0.82246042 | 0.995 | 0.932 | 2.12E-202 | TEC_MMP7 | RPL37A |
| 3.371E-261 | 0.82173914 | 0.988 | 0.892 | 7.196E-257 | TEC_MMP7 | RPS3 |
| 5.884E-269 | 0.81992056 | 0.993 | 0.919 | 1.256E-264 | TEC_MMP7 | RPL36 |
| 6.011E-235 | 0.81583937 | 0.674 | 0.303 | 1.283E-230 | TEC_MMP7 | EIF5A |
| 0 | 0.81235181 | 0.482 | 0.101 | 0 | TEC_MMP7 | FXYD5 |
| 2.151E-273 | 0.80987866 | 1 | 0.97 | 4.593E-269 | TEC_MMP7 | RPL13 |
| 7.497E-243 | 0.80807485 | 0.953 | 0.787 | 1.601E-238 | TEC_MMP7 | ATP5F1E |
| 2.364E-273 | 0.80779489 | 0.964 | 0.776 | 5.046E-269 | TEC_MMP7 | RPL27 |
| 0 | 0.80648485 | 0.629 | 0.209 | 0 | TEC_MMP7 | FKBP1A |
| 4.679E-158 | 0.80411146 | 0.778 | 0.534 | 9.989E-154 | TEC_MMP7 | MAL2 |
| 2.485E-222 | 0.80375273 | 0.99 | 0.888 | 5.306E-218 | TEC_MMP7 | RPL27A |
| 4.416E-274 | 0.80301921 | 0.553 | 0.18 | 9.428E-270 | TEC_MMP7 | IFITM2 |
| 9.251E-264 | 0.80172667 | 0.996 | 0.92 | 1.975E-259 | TEC_MMP7 | RPS14 |
| 4.578E-247 | 0.80061471 | 0.998 | 0.944 | 9.775E-243 | TEC_MMP7 | RPS8 |
| 1.543E-174 | 0.79530523 | 0.942 | 0.809 | 3.295E-170 | TEC_MMP7 | TMBIM6 |
| 9.215E-260 | 0.79273655 | 0.995 | 0.938 | 1.967E-255 | TEC_MMP7 | RPS15 |
| 5.374E-173 | 0.79269319 | 0.925 | 0.779 | 1.147E-168 | TEC_MMP7 | HLA-C |
| 1.512E-245 | 0.7903156 | 0.93 | 0.719 | 3.227E-241 | TEC_MMP7 | HINT1 |
| 1.777E-205 | 0.78897002 | 0.976 | 0.873 | 3.794E-201 | TEC_MMP7 | RPS29 |
| 6.8546E-77 | 0.78462396 | 0.33 | 0.152 | 1.4635E-72 | TEC_MMP7 | IGFBP3 |
| 5.254E-78 | 0.77793621 | 0.174 | 0.051 | 1.1217E-73 | TEC_MMP7 | MGP |
| 2.904E-134 | 0.77285168 | 0.916 | 0.814 | 6.201E-130 | TEC_MMP7 | HLA-B |
| 3.41E-256 | 0.7709248 | 0.689 | 0.295 | 7.281E-252 | TEC_MMP7 | MIF |
| 1.686E-190 | 0.77062395 | 0.992 | 0.941 | 3.6E-186 | TEC_MMP7 | RPL13A |
| 1.722E-260 | 0.77040358 | 0.912 | 0.646 | 3.676E-256 | TEC_MMP7 | COX6A1 |
| 9.279E-283 | 0.75161462 | 0.641 | 0.244 | 1.981E-278 | TEC_MMP7 | BRK1 |

| | | | | | | |
|---|---|---|---|---|---|---|
| 1.191E-243 | 0.75085422 | 0.982 | 0.874 | 2.542E-239 | TEC_MMP7 | RPL22 |
| 4.339E-242 | 0.74992842 | 0.708 | 0.328 | 9.264E-238 | TEC_MMP7 | VAMP8 |
| 8.185E-244 | 0.74650209 | 0.888 | 0.614 | 1.748E-239 | TEC_MMP7 | UQCRH |
| 8.382E-203 | 0.74390539 | 0.995 | 0.926 | 1.79E-198 | TEC_MMP7 | RPL37 |
| 1.187E-202 | 0.74187491 | 0.982 | 0.923 | 2.534E-198 | TEC_MMP7 | RPL7A |
| 4.629E-206 | 0.73886654 | 0.954 | 0.803 | 9.883E-202 | TEC_MMP7 | RPS21 |
| 3.254E-162 | 0.73875311 | 0.706 | 0.379 | 6.948E-158 | TEC_MMP7 | TPM4 |
| 3.1605E-94 | 0.73834769 | 0.794 | 0.649 | 6.7477E-90 | TEC_MMP7 | CRYAB |
| 6.55E-239 | 0.73134288 | 0.698 | 0.32 | 1.398E-234 | TEC_MMP7 | ATP5MF |
| 8.439E-185 | 0.73107249 | 0.478 | 0.175 | 1.802E-180 | TEC_MMP7 | CTSH |
| 5.15E-199 | 0.72695282 | 0.85 | 0.581 | 1.1E-194 | TEC_MMP7 | CD81 |
| 8.022E-250 | 0.72632974 | 0.71 | 0.308 | 1.713E-245 | TEC_MMP7 | TALDO1 |
| 2.436E-153 | 0.72488359 | 0.229 | 0.05 | 5.201E-149 | TEC_MMP7 | SAA2 |
| 2.845E-224 | 0.72303529 | 0.701 | 0.335 | 6.074E-220 | TEC_MMP7 | DYNLT1 |
| 1.324E-122 | 0.72147949 | 0.216 | 0.053 | 2.826E-118 | TEC_MMP7 | SPRR2A |
| 1.556E-182 | 0.7204083 | 0.655 | 0.327 | 3.322E-178 | TEC_MMP7 | SELENOW |
| 1.424E-153 | 0.71763932 | 0.492 | 0.202 | 3.041E-149 | TEC_MMP7 | ITGB6 |
| 3.123E-221 | 0.7168728 | 0.997 | 0.93 | 6.667E-217 | TEC_MMP7 | RPL15 |
| 9.905E-166 | 0.71303133 | 0.888 | 0.706 | 2.115E-161 | TEC_MMP7 | ENO1 |
| 8.054E-207 | 0.71167664 | 0.961 | 0.859 | 1.72E-202 | TEC_MMP7 | RACK1 |
| 1.862E-227 | 0.71156968 | 0.776 | 0.415 | 3.976E-223 | TEC_MMP7 | COX8A |
| 1.606E-214 | 0.71007799 | 0.998 | 0.911 | 3.43E-210 | TEC_MMP7 | RPL30 |
| 2.775E-227 | 0.70978597 | 0.991 | 0.889 | 5.924E-223 | TEC_MMP7 | RPS25 |
| 1.581E-166 | 0.70879372 | 0.802 | 0.514 | 3.375E-162 | TEC_MMP7 | CD9 |
| 1.088E-155 | 0.7053561 | 0.596 | 0.274 | 2.323E-151 | TEC_MMP7 | MAL |
| 2.781E-239 | 0.70385054 | 0.956 | 0.756 | 5.938E-235 | TEC_MMP7 | COX4I1 |
| 2.344E-224 | 0.70231867 | 0.989 | 0.884 | 5.004E-220 | TEC_MMP7 | RPS15A |
| 2.247E-194 | 0.70169676 | 0.986 | 0.898 | 4.797E-190 | TEC_MMP7 | RPL26 |
| 1.13E-190 | 0.69937722 | 0.61 | 0.273 | 2.413E-186 | TEC_MMP7 | PRDX2 |
| 3.304E-146 | 0.69842631 | 0.688 | 0.391 | 7.054E-142 | TEC_MMP7 | PDIA3 |
| 2.118E-198 | 0.69761096 | 0.992 | 0.912 | 4.521E-194 | TEC_MMP7 | RPL14 |
| 7.406E-203 | 0.69702506 | 0.98 | 0.859 | 1.581E-198 | TEC_MMP7 | RPL7 |
| 2.241E-117 | 0.6923804 | 0.963 | 0.921 | 4.784E-113 | TEC_MMP7 | ACTG1 |
| 2.819E-180 | 0.68178461 | 0.664 | 0.337 | 6.02E-176 | TEC_MMP7 | REEP5 |
| 9.315E-181 | 0.6812331 | 0.987 | 0.91 | 1.989E-176 | TEC_MMP7 | RPL34 |
| 9.873E-186 | 0.67833528 | 0.695 | 0.355 | 2.108E-181 | TEC_MMP7 | MGST3 |
| 8.571E-147 | 0.67614077 | 0.609 | 0.306 | 1.83E-142 | TEC_MMP7 | S100A16 |
| 7.734E-155 | 0.6725521 | 0.998 | 0.975 | 1.651E-150 | TEC_MMP7 | FTH1 |
| 4.787E-158 | 0.67250486 | 0.527 | 0.232 | 1.022E-153 | TEC_MMP7 | RBM3 |
| 1.515E-169 | 0.67080652 | 0.913 | 0.756 | 3.234E-165 | TEC_MMP7 | OAZ1 |
| 1.344E-184 | 0.66841486 | 0.902 | 0.701 | 2.868E-180 | TEC_MMP7 | CSTB |
| 2.782E-242 | 0.66576691 | 0.584 | 0.217 | 5.939E-238 | TEC_MMP7 | APRT |
| 1.98E-167 | 0.66263675 | 0.704 | 0.389 | 4.227E-163 | TEC_MMP7 | BSG |
| 3.592E-199 | 0.66246746 | 0.522 | 0.202 | 7.669E-195 | TEC_MMP7 | PLP2 |
| 3.119E-218 | 0.66039864 | 0.739 | 0.37 | 6.658E-214 | TEC_MMP7 | PARK7 |
| 7.133E-163 | 0.65958182 | 0.569 | 0.243 | 1.523E-158 | TEC_MMP7 | AKR1B1 |
| 1.526E-219 | 0.65863756 | 0.703 | 0.334 | 3.258E-215 | TEC_MMP7 | HSBP1 |
| 7.742E-167 | 0.65712436 | 0.726 | 0.413 | 1.653E-162 | TEC_MMP7 | LAPTM4A |
| 1.016E-253 | 0.65344753 | 0.57 | 0.2 | 2.168E-249 | TEC_MMP7 | BCAP31 |
| 1.777E-111 | 0.64796194 | 0.664 | 0.382 | 3.794E-107 | TEC_MMP7 | CCND1 |

| | | | | | | |
|---|---|---|---|---|---|---|
| 1.077E-150 | 0.64369059 | 0.475 | 0.189 | 2.3E-146 | TEC_MMP7 | CD74 |
| 3.103E-206 | 0.64250452 | 0.669 | 0.316 | 6.626E-202 | TEC_MMP7 | NEDD8 |
| 4.4608E-51 | 0.64199238 | 0.109 | 0.03 | 9.5239E-47 | TEC_MMP7 | CCL5 |
| 2.853E-100 | 0.64099371 | 0.536 | 0.274 | 6.0907E-96 | TEC_MMP7 | PLAU |
| 2.612E-171 | 0.63375787 | 0.51 | 0.21 | 5.577E-167 | TEC_MMP7 | DHRS7 |
| 2.8E-219 | 0.63190347 | 0.63 | 0.265 | 5.978E-215 | TEC_MMP7 | PGK1 |
| 2.158E-191 | 0.63153601 | 0.991 | 0.923 | 4.608E-187 | TEC_MMP7 | RPL18 |
| 2.343E-172 | 0.63132704 | 0.947 | 0.784 | 5.003E-168 | TEC_MMP7 | RPL31 |
| 2.133E-224 | 0.63021362 | 0.642 | 0.267 | 4.554E-220 | TEC_MMP7 | UQCR11 |
| 5.536E-165 | 0.62004273 | 0.727 | 0.41 | 1.182E-160 | TEC_MMP7 | RPS10 |
| 6.183E-167 | 0.61858387 | 0.674 | 0.345 | 1.32E-162 | TEC_MMP7 | POLR2L |
| 1.1759E-25 | 0.61750237 | 0.195 | 0.106 | 2.5105E-21 | TEC_MMP7 | CXCL8 |
| 1.934E-128 | 0.61635279 | 0.788 | 0.525 | 4.128E-124 | TEC_MMP7 | HMGA1 |
| 1.011E-224 | 0.61531508 | 0.586 | 0.229 | 2.158E-220 | TEC_MMP7 | CAPNS1 |
| 1.113E-173 | 0.61230474 | 0.671 | 0.34 | 2.376E-169 | TEC_MMP7 | NDUFB2 |
| 5.72E-148 | 0.61204085 | 0.543 | 0.24 | 1.221E-143 | TEC_MMP7 | SERPINB1 |
| 9.763E-257 | 0.61186213 | 0.486 | 0.145 | 2.084E-252 | TEC_MMP7 | ARPC1B |
| 1.37E-138 | 0.61158525 | 0.689 | 0.387 | 2.926E-134 | TEC_MMP7 | TMEM59 |
| 8.211E-211 | 0.60814135 | 0.593 | 0.24 | 1.753E-206 | TEC_MMP7 | ROMO1 |
| 2.714E-190 | 0.60348074 | 0.654 | 0.305 | 5.794E-186 | TEC_MMP7 | MZT2B |
| 1.026E-199 | 0.59771533 | 0.642 | 0.283 | 2.192E-195 | TEC_MMP7 | UQCRQ |
| 2.456E-132 | 0.59630918 | 0.676 | 0.382 | 5.243E-128 | TEC_MMP7 | LGALS3 |
| 8.554E-135 | 0.59630216 | 0.821 | 0.554 | 1.826E-130 | TEC_MMP7 | S100A13 |
| 1.12E-195 | 0.59554083 | 0.538 | 0.212 | 2.392E-191 | TEC_MMP7 | LAPTM4B |
| 1.551E-171 | 0.58987622 | 0.82 | 0.524 | 3.311E-167 | TEC_MMP7 | COX6B1 |
| 3.086E-140 | 0.5888679 | 0.736 | 0.443 | 6.589E-136 | TEC_MMP7 | P4HB |
| 9.899E-146 | 0.58776964 | 0.54 | 0.25 | 2.113E-141 | TEC_MMP7 | ANXA3 |
| 1.729E-222 | 0.58186985 | 0.526 | 0.182 | 3.691E-218 | TEC_MMP7 | C4orf48 |
| 5.116E-156 | 0.58165569 | 0.669 | 0.356 | 1.092E-151 | TEC_MMP7 | DAD1 |
| 3.803E-158 | 0.57956025 | 0.91 | 0.687 | 8.12E-154 | TEC_MMP7 | TXN |
| 2.859E-161 | 0.5779578 | 0.788 | 0.49 | 6.103E-157 | TEC_MMP7 | ELOB |
| 6.576E-105 | 0.57761056 | 0.968 | 0.895 | 1.404E-100 | TEC_MMP7 | RPS20 |
| 2.235E-229 | 0.57679216 | 0.564 | 0.207 | 4.772E-225 | TEC_MMP7 | ECHS1 |
| 1.297E-114 | 0.57648566 | 0.503 | 0.24 | 2.77E-110 | TEC_MMP7 | MUC1 |
| 6.595E-107 | 0.57624033 | 0.516 | 0.252 | 1.408E-102 | TEC_MMP7 | PLAUR |
| 3.062E-197 | 0.57601737 | 0.991 | 0.898 | 6.537E-193 | TEC_MMP7 | FAU |
| 2.013E-110 | 0.5758632 | 0.302 | 0.104 | 4.299E-106 | TEC_MMP7 | TNFRSF11B |
| 3.995E-149 | 0.57386607 | 0.94 | 0.785 | 8.53E-145 | TEC_MMP7 | EEF1D |
| 9.21E-100 | 0.5738221 | 0.816 | 0.637 | 1.9663E-95 | TEC_MMP7 | CALR |
| 1.349E-130 | 0.57016199 | 0.63 | 0.32 | 2.881E-126 | TEC_MMP7 | CD44 |
| 1.239E-171 | 0.56550659 | 0.621 | 0.292 | 2.646E-167 | TEC_MMP7 | PPA1 |
| 1.556E-152 | 0.56287613 | 0.607 | 0.285 | 3.322E-148 | TEC_MMP7 | CTSB |
| 2.521E-149 | 0.55907724 | 1 | 0.986 | 5.383E-145 | TEC_MMP7 | EEF1A1 |
| 1.063E-158 | 0.55588491 | 0.825 | 0.545 | 2.27E-154 | TEC_MMP7 | ATP6V0E1 |
| 3.233E-140 | 0.55562572 | 0.771 | 0.499 | 6.903E-136 | TEC_MMP7 | NDUFS5 |
| 3.811E-166 | 0.54814517 | 0.843 | 0.575 | 8.137E-162 | TEC_MMP7 | ATP5MG |
| 1.322E-165 | 0.54620351 | 0.503 | 0.209 | 2.823E-161 | TEC_MMP7 | PGAM1 |
| 2.185E-202 | 0.54016426 | 0.414 | 0.123 | 4.665E-198 | TEC_MMP7 | TMEM176B |
| 8.392E-219 | 0.53948681 | 0.435 | 0.131 | 1.792E-214 | TEC_MMP7 | PRDX4 |
| 1.477E-152 | 0.53889706 | 0.599 | 0.287 | 3.153E-148 | TEC_MMP7 | TMED9 |

| | | | | | | |
|---|---|---|---|---|---|---|
| 1.423E-150 | 0.53651668 | 0.848 | 0.604 | 3.039E-146 | TEC_MMP7 | UBL5 |
| 3.082E-182 | 0.53638627 | 0.197 | 0.03 | 6.579E-178 | TEC_MMP7 | AGR2 |
| 1.524E-132 | 0.53621696 | 0.972 | 0.888 | 3.255E-128 | TEC_MMP7 | RPL5 |
| 7.835E-107 | 0.53589561 | 0.616 | 0.352 | 1.673E-102 | TEC_MMP7 | OCIAD2 |
| 1.367E-138 | 0.53494705 | 0.431 | 0.17 | 2.919E-134 | TEC_MMP7 | DPP4 |
| 1.628E-151 | 0.53452484 | 0.577 | 0.278 | 3.475E-147 | TEC_MMP7 | SRP9 |
| 2.502E-141 | 0.53428835 | 0.706 | 0.409 | 5.342E-137 | TEC_MMP7 | ATP5MD |
| 4.943E-177 | 0.53388905 | 0.429 | 0.146 | 1.055E-172 | TEC_MMP7 | TMEM238 |
| 3.638E-141 | 0.52856059 | 0.599 | 0.304 | 7.767E-137 | TEC_MMP7 | SPCS1 |
| 6.498E-151 | 0.52791928 | 0.923 | 0.727 | 1.387E-146 | TEC_MMP7 | PFDN5 |
| 1.933E-179 | 0.5265088 | 0.575 | 0.244 | 4.128E-175 | TEC_MMP7 | NDUFB11 |
| 3.278E-132 | 0.52559863 | 0.98 | 0.896 | 6.999E-128 | TEC_MMP7 | RPS16 |
| 2.527E-103 | 0.52310505 | 0.717 | 0.487 | 5.394E-99 | TEC_MMP7 | ARPC2 |
| 8.9465E-82 | 0.52207095 | 0.846 | 0.706 | 1.9101E-77 | TEC_MMP7 | CTSD |
| 1.28E-187 | 0.52091031 | 0.44 | 0.148 | 2.732E-183 | TEC_MMP7 | TMEM54 |
| 1.083E-146 | 0.52042244 | 0.503 | 0.216 | 2.313E-142 | TEC_MMP7 | CD47 |
| 9.153E-102 | 0.51936039 | 0.495 | 0.252 | 1.9542E-97 | TEC_MMP7 | ASPH |
| 6.583E-116 | 0.51905541 | 0.162 | 0.032 | 1.406E-111 | TEC_MMP7 | PI3 |
| 1.9993E-81 | 0.51615301 | 0.855 | 0.74 | 4.2685E-77 | TEC_MMP7 | RPS26 |
| 1.544E-106 | 0.5135955 | 0.571 | 0.312 | 3.297E-102 | TEC_MMP7 | CAP1 |
| 1.663E-149 | 0.50933742 | 0.549 | 0.249 | 3.551E-145 | TEC_MMP7 | RABAC1 |
| 5.977E-172 | 0.50813836 | 0.505 | 0.2 | 1.276E-167 | TEC_MMP7 | LSM4 |
| 1.1746E-31 | 0.50584063 | 0.133 | 0.056 | 2.5077E-27 | TEC_MMP7 | CCL20 |
| 2.2558E-55 | 0.50372266 | 0.225 | 0.095 | 4.8161E-51 | TEC_MMP7 | C15orf48 |
| 3.425E-122 | 0.50279111 | 0.784 | 0.527 | 7.313E-118 | TEC_MMP7 | RHOA |
| 3.406E-124 | 0.50274842 | 0.529 | 0.26 | 7.272E-120 | TEC_MMP7 | TMED10 |
| 4.423E-136 | 0.50254351 | 0.982 | 0.899 | 9.444E-132 | TEC_MMP7 | RPL24 |
| 2.416E-117 | 0.50133942 | 0.832 | 0.618 | 5.159E-113 | TEC_MMP7 | EDF1 |
| 1.238E-174 | 0.49980508 | 0.548 | 0.227 | 2.642E-170 | TEC_MMP7 | TSTD1 |
| 8.905E-119 | 0.49914698 | 0.969 | 0.886 | 1.901E-114 | TEC_MMP7 | RPL35A |
| 1.24E-121 | 0.49902723 | 0.956 | 0.84 | 2.647E-117 | TEC_MMP7 | RPL38 |
| 3.025E-147 | 0.4987686 | 0.636 | 0.322 | 6.458E-143 | TEC_MMP7 | SNRPD2 |
| 5.094E-169 | 0.49786061 | 0.525 | 0.212 | 1.087E-164 | TEC_MMP7 | ATOX1 |
| 2.869E-215 | 0.49516283 | 0.277 | 0.053 | 6.125E-211 | TEC_MMP7 | GAL |
| 3.682E-113 | 0.49510094 | 0.458 | 0.218 | 7.862E-109 | TEC_MMP7 | RPN2 |
| 1.869E-165 | 0.49329997 | 0.518 | 0.213 | 3.991E-161 | TEC_MMP7 | TAF10 |
| 2.03E-116 | 0.49190245 | 0.897 | 0.711 | 4.334E-112 | TEC_MMP7 | SKP1 |
| 2.695E-134 | 0.48917089 | 0.628 | 0.329 | 5.753E-130 | TEC_MMP7 | WDR83OS |
| 6.027E-194 | 0.48778102 | 0.288 | 0.064 | 1.287E-189 | TEC_MMP7 | C19orf33 |
| 6.315E-146 | 0.48754081 | 0.578 | 0.275 | 1.348E-141 | TEC_MMP7 | PCBD1 |
| 6.989E-130 | 0.48661942 | 0.523 | 0.241 | 1.492E-125 | TEC_MMP7 | CYB5A |
| 1.628E-210 | 0.48645946 | 0.377 | 0.099 | 3.475E-206 | TEC_MMP7 | SMIM22 |
| 6.064E-122 | 0.48366921 | 0.652 | 0.357 | 1.295E-117 | TEC_MMP7 | ATP5F1D |
| 8.332E-167 | 0.48297886 | 0.505 | 0.203 | 1.779E-162 | TEC_MMP7 | NENF |
| 1.1301E-84 | 0.48210698 | 0.147 | 0.036 | 2.4128E-80 | TEC_MMP7 | KLK6 |
| 2.601E-141 | 0.48081807 | 0.458 | 0.193 | 5.552E-137 | TEC_MMP7 | GLO1 |
| 2.158E-121 | 0.48072197 | 0.457 | 0.2 | 4.608E-117 | TEC_MMP7 | ASAH1 |
| 7.051E-122 | 0.47796832 | 0.563 | 0.286 | 1.505E-117 | TEC_MMP7 | NPC2 |
| 6.709E-150 | 0.47513521 | 0.373 | 0.125 | 1.432E-145 | TEC_MMP7 | QPRT |
| 1.018E-136 | 0.47130069 | 0.499 | 0.222 | 2.173E-132 | TEC_MMP7 | PSMB3 |

| | | | | | | |
|---|---|---|---|---|---|---|
| 1.156E-170 | 0.46890071 | 0.356 | 0.106 | 2.469E-166 | TEC_MMP7 | CTSS |
| 1.5805E-71 | 0.46723823 | 0.79 | 0.665 | 3.3743E-67 | TEC_MMP7 | CLIC1 |
| 7.308E-135 | 0.46140278 | 0.623 | 0.318 | 1.56E-130 | TEC_MMP7 | EIF3K |
| 5.932E-148 | 0.46111767 | 0.504 | 0.215 | 1.266E-143 | TEC_MMP7 | LMAN2 |
| 5.1924E-78 | 0.46000923 | 0.307 | 0.134 | 1.1086E-73 | TEC_MMP7 | IGFBP2 |
| 1.0589E-90 | 0.45849139 | 0.745 | 0.543 | 2.2607E-86 | TEC_MMP7 | RAN |
| 3.1611E-54 | 0.45614122 | 0.875 | 0.79 | 6.749E-50 | TEC_MMP7 | CD24 |
| 4.7171E-86 | 0.45578815 | 0.633 | 0.389 | 1.0071E-81 | TEC_MMP7 | PTTG1IP |
| 3.382E-126 | 0.45421786 | 0.663 | 0.358 | 7.22E-122 | TEC_MMP7 | DBI |
| 5.4062E-61 | 0.45321367 | 0.59 | 0.387 | 1.1542E-56 | TEC_MMP7 | PHLDA2 |
| 3.964E-141 | 0.45122284 | 0.436 | 0.175 | 8.463E-137 | TEC_MMP7 | SCP2 |
| 4.752E-120 | 0.45099933 | 0.776 | 0.493 | 1.015E-115 | TEC_MMP7 | COX6C |
| 9.435E-123 | 0.4496736 | 0.531 | 0.254 | 2.014E-118 | TEC_MMP7 | PSME2 |
| 8.74E-148 | 0.44965441 | 0.452 | 0.18 | 1.866E-143 | TEC_MMP7 | NHP2 |
| 2.296E-117 | 0.4490154 | 0.573 | 0.301 | 4.901E-113 | TEC_MMP7 | KDELR1 |
| 1.971E-198 | 0.44775589 | 0.382 | 0.107 | 4.209E-194 | TEC_MMP7 | ETHE1 |
| 2.29E-136 | 0.44718639 | 0.414 | 0.163 | 4.888E-132 | TEC_MMP7 | C1QBP |
| 3.696E-119 | 0.44417126 | 0.633 | 0.338 | 7.892E-115 | TEC_MMP7 | NDUFA1 |
| 1.8187E-25 | 0.44284493 | 0.276 | 0.173 | 3.8829E-21 | TEC_MMP7 | ADIRF |
| 1.139E-105 | 0.4423963 | 0.786 | 0.54 | 2.432E-101 | TEC_MMP7 | COX7A2 |
| 1.757E-151 | 0.44234489 | 0.398 | 0.141 | 3.751E-147 | TEC_MMP7 | NME1 |
| 2.653E-142 | 0.44152254 | 0.304 | 0.089 | 5.664E-138 | TEC_MMP7 | CRABP2 |
| 6.1904E-85 | 0.43999389 | 0.54 | 0.308 | 1.3217E-80 | TEC_MMP7 | PRELID1 |
| 2.285E-112 | 0.43927953 | 0.628 | 0.337 | 4.879E-108 | TEC_MMP7 | LDHB |
| 8.971E-126 | 0.43898458 | 0.513 | 0.241 | 1.915E-121 | TEC_MMP7 | SELENOF |
| 4.362E-124 | 0.43847007 | 0.447 | 0.191 | 9.312E-120 | TEC_MMP7 | CXCL16 |
| 1.079E-113 | 0.43749173 | 0.462 | 0.212 | 2.304E-109 | TEC_MMP7 | MMP24OS |
| 2.605E-132 | 0.43715832 | 0.516 | 0.236 | 5.562E-128 | TEC_MMP7 | UFC1 |
| 1.7067E-54 | 0.4367553 | 0.635 | 0.465 | 3.6438E-50 | TEC_MMP7 | LDHA |
| 6.738E-151 | 0.43649928 | 0.478 | 0.192 | 1.439E-146 | TEC_MMP7 | PGLS |
| 2.102E-86 | 0.43559872 | 0.591 | 0.353 | 4.4877E-82 | TEC_MMP7 | MRPL33 |
| 1.7458E-51 | 0.43445444 | 0.118 | 0.035 | 3.7273E-47 | TEC_MMP7 | LTF |
| 1.861E-136 | 0.43404512 | 0.434 | 0.174 | 3.974E-132 | TEC_MMP7 | PRDX3 |
| 3.415E-136 | 0.42647644 | 0.397 | 0.149 | 7.291E-132 | TEC_MMP7 | CD99 |
| 1.528E-110 | 0.42571986 | 0.572 | 0.297 | 3.262E-106 | TEC_MMP7 | ATP6V1F |
| 8.373E-136 | 0.42521318 | 0.427 | 0.171 | 1.788E-131 | TEC_MMP7 | ARPC1A |
| 8.833E-135 | 0.42445415 | 0.379 | 0.138 | 1.886E-130 | TEC_MMP7 | PON2 |
| 3.285E-137 | 0.42387824 | 0.336 | 0.109 | 7.014E-133 | TEC_MMP7 | TMEM176A |
| 4.1261E-62 | 0.42275436 | 0.357 | 0.183 | 8.8091E-58 | TEC_MMP7 | MDK |
| 8.499E-108 | 0.42068996 | 0.907 | 0.728 | 1.815E-103 | TEC_MMP7 | BTF3 |
| 1.02E-99 | 0.4204804 | 0.635 | 0.372 | 2.1785E-95 | TEC_MMP7 | TBCA |
| 6.767E-132 | 0.41934734 | 0.437 | 0.177 | 1.445E-127 | TEC_MMP7 | MZT2A |
| 1.511E-119 | 0.41804491 | 0.435 | 0.188 | 3.227E-115 | TEC_MMP7 | GTF3C6 |
| 5.569E-103 | 0.41742765 | 0.404 | 0.18 | 1.189E-98 | TEC_MMP7 | ADAM9 |
| 6.695E-125 | 0.4150835 | 0.507 | 0.234 | 1.429E-120 | TEC_MMP7 | NDUFB8 |
| 2.2671E-89 | 0.4137373 | 0.995 | 0.95 | 4.8402E-85 | TEC_MMP7 | RPS27 |
| 8.06E-122 | 0.41363937 | 0.47 | 0.208 | 1.721E-117 | TEC_MMP7 | OSTC |
| 2.7607E-85 | 0.41249691 | 0.637 | 0.396 | 5.8941E-81 | TEC_MMP7 | ATP6V0B |
| 6.2912E-90 | 0.40999446 | 0.513 | 0.27 | 1.3432E-85 | TEC_MMP7 | MYDGF |
| 6.8739E-86 | 0.40943213 | 0.633 | 0.385 | 1.4676E-81 | TEC_MMP7 | DYNLRB1 |

| | | | | | | |
|---|---|---|---|---|---|---|
| 1.776E-141 | 0.40894582 | 0.448 | 0.179 | 3.792E-137 | TEC_MMP7 | TRAPPC1 |
| 2.162E-122 | 0.40885802 | 0.402 | 0.163 | 4.616E-118 | TEC_MMP7 | CMPK1 |
| 9.936E-107 | 0.40821165 | 0.417 | 0.187 | 2.121E-102 | TEC_MMP7 | RCN1 |
| 7.0288E-92 | 0.40809824 | 0.556 | 0.308 | 1.5006E-87 | TEC_MMP7 | ERP29 |
| 2.259E-125 | 0.40706528 | 0.513 | 0.238 | 4.823E-121 | TEC_MMP7 | ANAPC11 |
| 1.264E-102 | 0.40624321 | 0.605 | 0.331 | 2.6977E-98 | TEC_MMP7 | ATP5F1B |
| 1.0615E-82 | 0.40597649 | 0.148 | 0.037 | 2.2663E-78 | TEC_MMP7 | SPRR2D |
| 2.4344E-79 | 0.40540299 | 0.883 | 0.728 | 5.1975E-75 | TEC_MMP7 | DSTN |
| 3.6845E-66 | 0.40405256 | 0.898 | 0.814 | 7.8665E-62 | TEC_MMP7 | H3F3A |
| 4.1786E-85 | 0.40387949 | 0.529 | 0.298 | 8.9214E-81 | TEC_MMP7 | VDAC1 |
| 4.564E-131 | 0.40293944 | 0.421 | 0.169 | 9.744E-127 | TEC_MMP7 | NBDY |
| 4.014E-109 | 0.40234267 | 0.513 | 0.253 | 8.57E-105 | TEC_MMP7 | COPS9 |
| 1.8196E-59 | 0.40178765 | 0.415 | 0.232 | 3.8849E-55 | TEC_MMP7 | S100A14 |
| 6.485E-139 | 0.40162091 | 0.34 | 0.111 | 1.385E-134 | TEC_MMP7 | SLC44A4 |
| 4.1705E-69 | 0.40144057 | 0.366 | 0.176 | 8.904E-65 | TEC_MMP7 | ANXA1 |
| 3.6081E-45 | 0.39980327 | 0.645 | 0.497 | 7.7034E-41 | TEC_MMP7 | TPM3 |
| 2.4592E-86 | 0.39625308 | 0.945 | 0.855 | 5.2503E-82 | TEC_MMP7 | RPL23 |
| 8.23E-100 | 0.39600202 | 0.649 | 0.376 | 1.7572E-95 | TEC_MMP7 | UQCR10 |
| 7.1397E-83 | 0.39440787 | 0.722 | 0.497 | 1.5243E-78 | TEC_MMP7 | ATP5MC2 |
| 2.659E-122 | 0.3943163 | 0.462 | 0.203 | 5.678E-118 | TEC_MMP7 | MDH2 |
| 1.8027E-12 | 0.39407988 | 0.282 | 0.203 | 3.8487E-08 | TEC_MMP7 | GLRX |
| 1.524E-104 | 0.39295552 | 0.486 | 0.235 | 3.253E-100 | TEC_MMP7 | MINOS1 |
| 1.077E-85 | 0.39294552 | 0.548 | 0.301 | 2.2993E-81 | TEC_MMP7 | SLC25A5 |
| 1.868E-114 | 0.3927328 | 0.346 | 0.131 | 3.989E-110 | TEC_MMP7 | CAV2 |
| 9.0908E-82 | 0.39263113 | 0.597 | 0.355 | 1.9409E-77 | TEC_MMP7 | PERP |
| 5.7182E-97 | 0.39255928 | 0.554 | 0.3 | 1.2208E-92 | TEC_MMP7 | COPE |
| 1.9754E-63 | 0.39225829 | 0.771 | 0.61 | 4.2175E-59 | TEC_MMP7 | ANXA5 |
| 7.9928E-86 | 0.3913333 | 0.479 | 0.251 | 1.7065E-81 | TEC_MMP7 | GRN |
| 8.301E-113 | 0.39108764 | 0.354 | 0.134 | 1.772E-108 | TEC_MMP7 | PCSK1N |
| 9.7308E-88 | 0.39024901 | 0.497 | 0.268 | 2.0775E-83 | TEC_MMP7 | CHMP5 |
| 2.052E-73 | 0.38890832 | 0.311 | 0.137 | 4.3811E-69 | TEC_MMP7 | MGST1 |
| 2.2923E-89 | 0.38733632 | 0.455 | 0.229 | 4.8941E-85 | TEC_MMP7 | HM13 |
| 6.9584E-90 | 0.3868297 | 0.415 | 0.199 | 1.4856E-85 | TEC_MMP7 | ATP6AP2 |
| 3.855E-77 | 0.38642554 | 0.869 | 0.738 | 8.2304E-73 | TEC_MMP7 | RPL4 |
| 2.4919E-78 | 0.38532444 | 0.835 | 0.671 | 5.3203E-74 | TEC_MMP7 | GPX4 |
| 4.66E-115 | 0.38532259 | 0.451 | 0.2 | 9.949E-111 | TEC_MMP7 | DMKN |
| 9.065E-106 | 0.38397323 | 0.917 | 0.747 | 1.935E-101 | TEC_MMP7 | COX7C |
| 1.1996E-73 | 0.38326658 | 0.784 | 0.604 | 2.5612E-69 | TEC_MMP7 | SLC25A6 |
| 9.8578E-60 | 0.38297828 | 0.426 | 0.24 | 2.1046E-55 | TEC_MMP7 | SELENOM |
| 2.251E-133 | 0.38182532 | 0.406 | 0.156 | 4.807E-129 | TEC_MMP7 | TMEM147 |
| 2.241E-111 | 0.38063476 | 0.477 | 0.223 | 4.785E-107 | TEC_MMP7 | CAPZB |
| 6.267E-162 | 0.38004968 | 0.23 | 0.047 | 1.338E-157 | TEC_MMP7 | MALL |
| 6.4273E-98 | 0.37974627 | 0.53 | 0.279 | 1.3722E-93 | TEC_MMP7 | ATP5PD |
| 1.9492E-65 | 0.37570406 | 0.505 | 0.302 | 4.1616E-61 | TEC_MMP7 | TRAM1 |
| 1.6304E-91 | 0.37554796 | 0.498 | 0.258 | 3.481E-87 | TEC_MMP7 | RBX1 |
| 8.9713E-90 | 0.37409193 | 0.524 | 0.28 | 1.9154E-85 | TEC_MMP7 | TMEM50A |
| 1.4316E-77 | 0.3733523 | 0.484 | 0.267 | 3.0565E-73 | TEC_MMP7 | EIF6 |
| 3.414E-125 | 0.3725867 | 0.352 | 0.126 | 7.29E-121 | TEC_MMP7 | LGALS3BP |
| 1.0747E-79 | 0.3723336 | 0.56 | 0.319 | 2.2945E-75 | TEC_MMP7 | ANXA11 |
| 2.949E-80 | 0.37222676 | 0.59 | 0.344 | 6.2962E-76 | TEC_MMP7 | SEC61G |

| | | | | | | |
|---|---|---|---|---|---|---|
| 1.5988E-78 | 0.37150961 | 0.958 | 0.88 | 3.4134E-74 | TEC_MMP7 | NACA |
| 7.8368E-87 | 0.37020967 | 0.541 | 0.292 | 1.6732E-82 | TEC_MMP7 | PHPT1 |
| 2.3965E-79 | 0.36944679 | 0.441 | 0.227 | 5.1165E-75 | TEC_MMP7 | COTL1 |
| 1.321E-100 | 0.36889447 | 0.431 | 0.198 | 2.8202E-96 | TEC_MMP7 | COMT |
| 8.727E-57 | 0.36795235 | 0.677 | 0.502 | 1.8632E-52 | TEC_MMP7 | CDC42 |
| 5.9429E-91 | 0.36591417 | 0.215 | 0.065 | 1.2688E-86 | TEC_MMP7 | HPGD |
| 1.291E-130 | 0.36531001 | 0.343 | 0.118 | 2.756E-126 | TEC_MMP7 | EFHD2 |
| 1.8976E-98 | 0.36482271 | 0.369 | 0.155 | 4.0515E-94 | TEC_MMP7 | QSOX1 |
| 1.2606E-82 | 0.3647109 | 0.435 | 0.221 | 2.6914E-78 | TEC_MMP7 | PGRMC1 |
| 3.203E-113 | 0.36455466 | 0.291 | 0.096 | 6.839E-109 | TEC_MMP7 | GNG11 |
| 9.7582E-45 | 0.36414491 | 0.856 | 0.795 | 2.0834E-40 | TEC_MMP7 | CALM1 |
| 8.7108E-71 | 0.36359232 | 0.474 | 0.262 | 1.8597E-66 | TEC_MMP7 | FAM3C |
| 4.9016E-69 | 0.36186774 | 0.67 | 0.441 | 1.0465E-64 | TEC_MMP7 | PRDX6 |
| 1.156E-120 | 0.36180608 | 0.453 | 0.195 | 2.469E-116 | TEC_MMP7 | TMEM219 |
| 4.596E-99 | 0.36028063 | 0.449 | 0.213 | 9.8116E-95 | TEC_MMP7 | HIGD2A |
| 9.4674E-70 | 0.35864847 | 0.339 | 0.159 | 2.0213E-65 | TEC_MMP7 | CD40 |
| 1.2203E-80 | 0.35788627 | 0.552 | 0.317 | 2.6053E-76 | TEC_MMP7 | PSMD8 |
| 8.1693E-94 | 0.35659187 | 0.458 | 0.226 | 1.7442E-89 | TEC_MMP7 | RER1 |
| 7.879E-116 | 0.35550009 | 0.349 | 0.131 | 1.682E-111 | TEC_MMP7 | POLE4 |
| 2.6721E-85 | 0.3554467 | 0.462 | 0.237 | 5.705E-81 | TEC_MMP7 | TXNDC17 |
| 4.977E-99 | 0.35523014 | 0.473 | 0.229 | 1.0626E-94 | TEC_MMP7 | NDUFB3 |
| 1.2699E-96 | 0.35355895 | 0.437 | 0.206 | 2.7113E-92 | TEC_MMP7 | CALM3 |
| 2.961E-99 | 0.35261455 | 0.422 | 0.193 | 6.3216E-95 | TEC_MMP7 | ARPC4 |
| 3.6143E-72 | 0.35259774 | 0.479 | 0.261 | 7.7166E-68 | TEC_MMP7 | ITGA3 |
| 2.1232E-84 | 0.35086084 | 0.35 | 0.158 | 4.5329E-80 | TEC_MMP7 | HACD3 |
| 2.1882E-78 | 0.35009623 | 0.406 | 0.202 | 4.6718E-74 | TEC_MMP7 | PSMD2 |
| 2.1933E-87 | 0.34998644 | 0.486 | 0.253 | 4.6827E-83 | TEC_MMP7 | TMEM14C |
| 9.3149E-80 | 0.34932644 | 0.498 | 0.273 | 1.9887E-75 | TEC_MMP7 | CIB1 |
| 3.2495E-98 | 0.34723206 | 0.326 | 0.128 | 6.9376E-94 | TEC_MMP7 | TRNP1 |
| 2.1323E-67 | 0.34620388 | 0.691 | 0.462 | 4.5524E-63 | TEC_MMP7 | TOMM7 |
| 1.4392E-74 | 0.34608725 | 0.604 | 0.369 | 3.0726E-70 | TEC_MMP7 | YWHAQ |
| 4.0516E-78 | 0.3453368 | 0.525 | 0.293 | 8.6501E-74 | TEC_MMP7 | COX5A |
| 2.3009E-85 | 0.34488477 | 0.416 | 0.2 | 4.9124E-81 | TEC_MMP7 | SEC11C |
| 2.067E-95 | 0.34449019 | 0.34 | 0.139 | 4.4131E-91 | TEC_MMP7 | CD82 |
| 7.9097E-48 | 0.3440654 | 0.363 | 0.201 | 1.6887E-43 | TEC_MMP7 | SFN |
| 3.3104E-79 | 0.34311465 | 0.507 | 0.281 | 7.0677E-75 | TEC_MMP7 | NDUFC2 |
| 8.3785E-84 | 0.34274239 | 0.507 | 0.271 | 1.7888E-79 | TEC_MMP7 | DDT |
| 2.0924E-74 | 0.34266402 | 0.551 | 0.32 | 4.4672E-70 | TEC_MMP7 | COMMD6 |
| 1.4162E-93 | 0.34254072 | 0.352 | 0.15 | 3.0235E-89 | TEC_MMP7 | FAM213A |
| 5.4469E-27 | 0.34158158 | 0.39 | 0.268 | 1.1629E-22 | TEC_MMP7 | UCHL1 |
| 3.1022E-63 | 0.34035552 | 0.368 | 0.189 | 6.6231E-59 | TEC_MMP7 | SCCPDH |
| 7.7069E-89 | 0.3385028 | 0.443 | 0.217 | 1.6454E-84 | TEC_MMP7 | TUFM |
| 2.7479E-79 | 0.33554299 | 0.436 | 0.225 | 5.8669E-75 | TEC_MMP7 | PHB |
| 5.5465E-79 | 0.33533241 | 0.213 | 0.071 | 1.1842E-74 | TEC_MMP7 | LAMA3 |
| 4.1511E-78 | 0.33266711 | 0.4 | 0.199 | 8.8627E-74 | TEC_MMP7 | DAP |
| 4.1366E-69 | 0.33179638 | 0.449 | 0.243 | 8.8316E-65 | TEC_MMP7 | RPL36A |
| 2.3743E-97 | 0.3288484 | 0.382 | 0.16 | 5.0691E-93 | TEC_MMP7 | PDIA4 |
| 1.846E-115 | 0.32801339 | 0.353 | 0.133 | 3.942E-111 | TEC_MMP7 | DDOST |
| 2.054E-82 | 0.32792068 | 0.422 | 0.207 | 4.3853E-78 | TEC_MMP7 | MGAT4B |
| 1.3771E-86 | 0.32777487 | 0.435 | 0.215 | 2.94E-82 | TEC_MMP7 | PSMB5 |

| | | | | | | |
|---|---|---|---|---|---|---|
| 7.551E-85 | 0.32756686 | 0.451 | 0.228 | 1.6121E-80 | TEC_MMP7 | SMDT1 |
| 1.5636E-64 | 0.32698632 | 0.451 | 0.256 | 3.3382E-60 | TEC_MMP7 | CLTB |
| 1.484E-131 | 0.32571297 | 0.337 | 0.113 | 3.167E-127 | TEC_MMP7 | AP1M2 |
| 8.415E-94 | 0.32570872 | 0.383 | 0.168 | 1.7966E-89 | TEC_MMP7 | SLC25A39 |
| 4.6659E-78 | 0.32549572 | 0.433 | 0.223 | 9.9618E-74 | TEC_MMP7 | AP2M1 |
| 2.1843E-73 | 0.32545906 | 0.371 | 0.182 | 4.6635E-69 | TEC_MMP7 | RCN2 |
| 2.7376E-90 | 0.32526558 | 0.39 | 0.175 | 5.8449E-86 | TEC_MMP7 | NDUFAF3 |
| 4.1966E-20 | 0.32484518 | 0.395 | 0.299 | 8.9598E-16 | TEC_MMP7 | RAB13 |
| 1.9157E-72 | 0.32419126 | 0.517 | 0.294 | 4.09E-68 | TEC_MMP7 | AES |
| 2.0968E-81 | 0.32347275 | 0.453 | 0.233 | 4.4767E-77 | TEC_MMP7 | LAMTOR1 |
| 3.3447E-83 | 0.32286101 | 0.451 | 0.229 | 7.141E-79 | TEC_MMP7 | NDUFA11 |
| 5.679E-100 | 0.32282541 | 0.37 | 0.154 | 1.2124E-95 | TEC_MMP7 | CAPN1 |
| 5.1399E-43 | 0.32270489 | 0.708 | 0.569 | 1.0974E-38 | TEC_MMP7 | YWHAB |
| 4.6606E-67 | 0.32081157 | 0.722 | 0.506 | 9.9504E-63 | TEC_MMP7 | SEC61B |
| 4.3298E-41 | 0.32053274 | 0.31 | 0.177 | 9.2442E-37 | TEC_MMP7 | FDPS |
| 2.5879E-87 | 0.32006219 | 0.451 | 0.225 | 5.5252E-83 | TEC_MMP7 | EIF3I |
| 4.2445E-80 | 0.31855388 | 0.435 | 0.221 | 9.062E-76 | TEC_MMP7 | BANF1 |
| 2.049E-100 | 0.31786654 | 0.395 | 0.17 | 4.3739E-96 | TEC_MMP7 | TMEM141 |
| 2.738E-20 | 0.31773554 | 0.574 | 0.494 | 5.8456E-16 | TEC_MMP7 | CLDN7 |
| 1.767E-102 | 0.31753404 | 0.413 | 0.178 | 3.7733E-98 | TEC_MMP7 | NDUFA3 |
| 1.5357E-51 | 0.31685136 | 0.784 | 0.633 | 3.2788E-47 | TEC_MMP7 | SLC25A3 |
| 1.3632E-52 | 0.31649794 | 0.2 | 0.081 | 2.9104E-48 | TEC_MMP7 | ALDH1A3 |
| 1.1385E-69 | 0.31293872 | 0.656 | 0.423 | 2.4306E-65 | TEC_MMP7 | ATP5PF |
| 3.4288E-70 | 0.31114719 | 0.532 | 0.311 | 7.3206E-66 | TEC_MMP7 | SEC11A |
| 4.762E-99 | 0.30985224 | 0.311 | 0.118 | 1.0168E-94 | TEC_MMP7 | NBL1 |
| 1.032E-69 | 0.30838281 | 0.41 | 0.215 | 2.2033E-65 | TEC_MMP7 | KDELR2 |
| 5.298E-64 | 0.30776353 | 0.404 | 0.214 | 1.1311E-59 | TEC_MMP7 | RPS17 |
| 1.3136E-90 | 0.30682243 | 0.371 | 0.163 | 2.8045E-86 | TEC_MMP7 | CYB5R3 |
| 2.116E-85 | 0.30674033 | 0.476 | 0.241 | 4.5176E-81 | TEC_MMP7 | NDUFA2 |
| 6.5003E-32 | 0.30604769 | 0.667 | 0.535 | 1.3878E-27 | TEC_MMP7 | CANX |
| 4.0844E-67 | 0.30530892 | 0.428 | 0.232 | 8.7202E-63 | TEC_MMP7 | TMED2 |
| 6.0429E-87 | 0.30481747 | 0.417 | 0.196 | 1.2902E-82 | TEC_MMP7 | C19orf70 |
| 7.6971E-46 | 0.30172689 | 0.293 | 0.153 | 1.6433E-41 | TEC_MMP7 | SERPING1 |
| 7.0772E-62 | 0.30151449 | 0.379 | 0.201 | 1.511E-57 | TEC_MMP7 | DYNLT3 |
| 5.6383E-59 | 0.2992414 | 0.431 | 0.241 | 1.2038E-54 | TEC_MMP7 | TAPBP |
| 4.9268E-84 | 0.2983672 | 0.229 | 0.078 | 1.0519E-79 | TEC_MMP7 | HLA-DRB1 |
| 1.7178E-45 | 0.29586934 | 0.529 | 0.351 | 3.6674E-41 | TEC_MMP7 | HEBP2 |
| 5.5053E-88 | 0.29513891 | 0.355 | 0.153 | 1.1754E-83 | TEC_MMP7 | DNPH1 |
| 1.5858E-62 | 0.29415386 | 0.816 | 0.639 | 3.3856E-58 | TEC_MMP7 | UQCRB |
| 6.141E-53 | 0.29333122 | 0.514 | 0.313 | 1.3111E-48 | TEC_MMP7 | SSR3 |
| 2.2312E-78 | 0.29223445 | 0.335 | 0.149 | 4.7636E-74 | TEC_MMP7 | ARL2 |
| 2.015E-106 | 0.2921998 | 0.325 | 0.119 | 4.301E-102 | TEC_MMP7 | MBOAT7 |
| 4.4556E-47 | 0.29146379 | 0.43 | 0.262 | 9.5128E-43 | TEC_MMP7 | MSN |
| 2.5652E-62 | 0.29090642 | 0.395 | 0.211 | 5.4767E-58 | TEC_MMP7 | TMBIM1 |
| 6.0764E-59 | 0.28890029 | 0.352 | 0.183 | 1.2973E-54 | TEC_MMP7 | AP2S1 |
| 5.4058E-76 | 0.28869136 | 0.372 | 0.177 | 1.1541E-71 | TEC_MMP7 | MFSD10 |
| 2.8226E-68 | 0.2883755 | 0.352 | 0.171 | 6.0262E-64 | TEC_MMP7 | TSPAN3 |
| 3.4144E-49 | 0.28770142 | 0.988 | 0.957 | 7.2896E-45 | TEC_MMP7 | FTL |
| 3.4489E-80 | 0.28704503 | 0.372 | 0.173 | 7.3633E-76 | TEC_MMP7 | NUTF2 |
| 8.4194E-40 | 0.28616744 | 0.618 | 0.451 | 1.7975E-35 | TEC_MMP7 | SUMO2 |

| | | | | | | |
|---|---|---|---|---|---|---|
| 4.5733E-97 | 0.28600024 | 0.32 | 0.124 | 9.7641E-93 | TEC_MMP7 | CNIH1 |
| 1.5253E-68 | 0.28536857 | 0.808 | 0.62 | 3.2565E-64 | TEC_MMP7 | PEBP1 |
| 3.8536E-48 | 0.28322041 | 0.515 | 0.325 | 8.2275E-44 | TEC_MMP7 | ACADVL |
| 2.3779E-66 | 0.28231634 | 0.404 | 0.212 | 5.0768E-62 | TEC_MMP7 | PPP1CA |
| 1.4303E-58 | 0.28138145 | 0.376 | 0.202 | 3.0538E-54 | TEC_MMP7 | TMED4 |
| 2.7766E-95 | 0.28045829 | 0.209 | 0.061 | 5.928E-91 | TEC_MMP7 | RBP1 |
| 4.9228E-91 | 0.28045073 | 0.352 | 0.148 | 1.051E-86 | TEC_MMP7 | LAMTOR2 |
| 3.4827E-71 | 0.28002074 | 0.538 | 0.303 | 7.4356E-67 | TEC_MMP7 | CHCHD10 |
| 1.2556E-56 | 0.27947748 | 0.393 | 0.216 | 2.6807E-52 | TEC_MMP7 | VPS29 |
| 3.4159E-80 | 0.27927817 | 0.334 | 0.146 | 7.2928E-76 | TEC_MMP7 | POLR2E |
| 1.8435E-55 | 0.27867818 | 0.339 | 0.177 | 3.9358E-51 | TEC_MMP7 | HSD17B12 |
| 1.512E-104 | 0.27859148 | 0.291 | 0.101 | 3.227E-100 | TEC_MMP7 | CETN2 |
| 3.0607E-72 | 0.2779451 | 0.351 | 0.167 | 6.5345E-68 | TEC_MMP7 | SDHC |
| 4.2126E-82 | 0.27768909 | 0.301 | 0.122 | 8.9939E-78 | TEC_MMP7 | SH3BGRL |
| 3.4632E-76 | 0.27611555 | 0.356 | 0.165 | 7.3939E-72 | TEC_MMP7 | ATRAID |
| 1.0393E-41 | 0.27294869 | 0.419 | 0.263 | 2.2188E-37 | TEC_MMP7 | LAMP1 |
| 6.2131E-69 | 0.27226913 | 0.359 | 0.174 | 1.3265E-64 | TEC_MMP7 | STMP1 |
| 2.7742E-77 | 0.27220798 | 0.289 | 0.119 | 5.9229E-73 | TEC_MMP7 | GCSH |
| 4.1324E-59 | 0.27218454 | 0.757 | 0.557 | 8.8226E-55 | TEC_MMP7 | COX5B |
| 5.8402E-39 | 0.27160031 | 0.711 | 0.562 | 1.2469E-34 | TEC_MMP7 | ATP6V1G1 |
| 3.6183E-73 | 0.27131893 | 0.289 | 0.122 | 7.7251E-69 | TEC_MMP7 | BOLA3 |
| 1.7673E-69 | 0.2707662 | 0.462 | 0.247 | 3.7733E-65 | TEC_MMP7 | VPS28 |
| 1.8548E-66 | 0.27041469 | 0.336 | 0.161 | 3.96E-62 | TEC_MMP7 | TBCB |
| 6.3421E-63 | 0.26961115 | 0.415 | 0.223 | 1.354E-58 | TEC_MMP7 | ZNHIT1 |
| 3.4558E-65 | 0.26954627 | 0.359 | 0.179 | 7.3781E-61 | TEC_MMP7 | TMEM230 |
| 8.8841E-43 | 0.26937208 | 0.426 | 0.265 | 1.8967E-38 | TEC_MMP7 | PSMB6 |
| 5.6156E-41 | 0.2691483 | 0.301 | 0.168 | 1.1989E-36 | TEC_MMP7 | RANBP1 |
| 2.9174E-56 | 0.26886388 | 0.516 | 0.313 | 6.2287E-52 | TEC_MMP7 | NOP10 |
| 1.1945E-45 | 0.2687822 | 0.182 | 0.076 | 2.5502E-41 | TEC_MMP7 | IFI6 |
| 1.5946E-93 | 0.26718365 | 0.27 | 0.096 | 3.4046E-89 | TEC_MMP7 | PPIC |
| 7.1035E-30 | 0.26586097 | 0.434 | 0.3 | 1.5166E-25 | TEC_MMP7 | SARAF |
| 4.2532E-56 | 0.26564627 | 0.189 | 0.071 | 9.0805E-52 | TEC_MMP7 | HLA-DRA |
| 4.6089E-30 | 0.26548712 | 0.699 | 0.59 | 9.84E-26 | TEC_MMP7 | CLTA |
| 1.1391E-97 | 0.26507283 | 0.271 | 0.093 | 2.432E-93 | TEC_MMP7 | GGCT |
| 1.9994E-42 | 0.2649735 | 0.449 | 0.278 | 4.2688E-38 | TEC_MMP7 | CDH1 |
| 6.0605E-18 | 0.26363499 | 0.676 | 0.597 | 1.2939E-13 | TEC_MMP7 | APLP2 |
| 5.4745E-43 | 0.26360837 | 0.471 | 0.303 | 1.1688E-38 | TEC_MMP7 | FAM96B |
| 1.3209E-95 | 0.26308104 | 0.302 | 0.113 | 2.8202E-91 | TEC_MMP7 | TMED3 |
| 5.965E-131 | 0.26272197 | 0.215 | 0.05 | 1.274E-126 | TEC_MMP7 | QPCT |
| 9.3637E-58 | 0.26209704 | 0.548 | 0.333 | 1.9992E-53 | TEC_MMP7 | COX7B |
| 1.5025E-51 | 0.26141619 | 0.43 | 0.25 | 3.2078E-47 | TEC_MMP7 | ATP5F1C |
| 7.7635E-49 | 0.26074299 | 0.426 | 0.254 | 1.6575E-44 | TEC_MMP7 | CHMP2A |
| 9.1468E-52 | 0.26066943 | 0.444 | 0.262 | 1.9528E-47 | TEC_MMP7 | ATP5PO |
| 3.516E-55 | 0.25872982 | 0.414 | 0.233 | 7.5067E-51 | TEC_MMP7 | PPP2R1A |
| 1.4861E-43 | 0.25849452 | 0.528 | 0.349 | 3.1728E-39 | TEC_MMP7 | SNU13 |
| 6.1162E-61 | 0.25806963 | 0.389 | 0.205 | 1.3058E-56 | TEC_MMP7 | UXT |
| 4.9778E-65 | 0.25804514 | 0.274 | 0.118 | 1.0628E-60 | TEC_MMP7 | ALDH2 |
| 2.6746E-91 | 0.2551286 | 0.242 | 0.081 | 5.7103E-87 | TEC_MMP7 | TMEM45B |
| 6.2146E-53 | 0.25449197 | 0.303 | 0.152 | 1.3268E-48 | TEC_MMP7 | CENPX |
| 1.097E-105 | 0.25325803 | 0.218 | 0.06 | 2.342E-101 | TEC_MMP7 | NPDC1 |

| | | | | | | |
|---|---|---|---|---|---|---|
| 3.8042E-52 | 0.25077903 | 0.384 | 0.212 | 8.122E-48 | TEC_MMP7 | NDUFS8 |
| 1.7032E-71 | 0.2504138 | 0.26 | 0.105 | 3.6364E-67 | TEC_MMP7 | GPAA1 |
| 5.4772E-40 | -0.2500523 | 0.09 | 0.234 | 1.1694E-35 | TEC_MMP7 | GGNBP2 |
| 1.0062E-27 | -0.2503603 | 0.251 | 0.387 | 2.1482E-23 | TEC_MMP7 | UBE2B |
| 4.6632E-34 | -0.2503824 | 0.085 | 0.213 | 9.956E-30 | TEC_MMP7 | WWC1 |
| 7.6041E-39 | -0.2508732 | 0.035 | 0.153 | 1.6235E-34 | TEC_MMP7 | ZFYVE16 |
| 6.8391E-14 | -0.2510822 | 0.435 | 0.521 | 1.4601E-09 | TEC_MMP7 | H2AFZ |
| 2.5005E-48 | -0.2513015 | 0.032 | 0.171 | 5.3385E-44 | TEC_MMP7 | ANKRD17 |
| 3.99E-29 | -0.2514333 | 0.142 | 0.266 | 8.5186E-25 | TEC_MMP7 | SMARCA5 |
| 6.3798E-37 | -0.2515054 | 0.056 | 0.179 | 1.3621E-32 | TEC_MMP7 | ESF1 |
| 1.9601E-29 | -0.2517689 | 0.136 | 0.264 | 4.1849E-25 | TEC_MMP7 | TES |
| 1.114E-40 | -0.2523268 | 0.068 | 0.204 | 2.3783E-36 | TEC_MMP7 | PRRC2B |
| 3.4625E-39 | -0.2524041 | 0.083 | 0.221 | 7.3924E-35 | TEC_MMP7 | ROCK1 |
| 1.214E-14 | -0.2524051 | 0.1 | 0.175 | 2.5918E-10 | TEC_MMP7 | CTGF |
| 2.5276E-28 | -0.2528878 | 0.095 | 0.21 | 5.3964E-24 | TEC_MMP7 | NCOA7 |
| 3.3531E-37 | -0.2534216 | 0.096 | 0.233 | 7.1588E-33 | TEC_MMP7 | TRIP12 |
| 1.7011E-41 | -0.2535793 | 0.039 | 0.166 | 3.6319E-37 | TEC_MMP7 | SRSF6 |
| 2.4431E-31 | -0.2535868 | 0.083 | 0.201 | 5.2159E-27 | TEC_MMP7 | ARHGAP18 |
| 6.4762E-44 | -0.2536025 | 0.062 | 0.206 | 1.3827E-39 | TEC_MMP7 | PPP4R2 |
| 3.6718E-36 | -0.2536636 | 0.171 | 0.322 | 7.8393E-32 | TEC_MMP7 | WASF2 |
| 4.24E-27 | -0.2536711 | 0.129 | 0.245 | 9.0524E-23 | TEC_MMP7 | NFE2L1 |
| 5.1119E-39 | -0.2536783 | 0.08 | 0.217 | 1.0914E-34 | TEC_MMP7 | ARHGEF12 |
| 7.1939E-21 | -0.2539574 | 0.108 | 0.203 | 1.5359E-16 | TEC_MMP7 | BHLHE40 |
| 1.997E-28 | -0.254089 | 0.203 | 0.335 | 4.2636E-24 | TEC_MMP7 | PCBP1 |
| 2.4056E-49 | -0.2543967 | 0.022 | 0.157 | 5.1359E-45 | TEC_MMP7 | RALGAPA1 |
| 1.9769E-44 | -0.2547388 | 0.061 | 0.206 | 4.2206E-40 | TEC_MMP7 | ZNF106 |
| 6.8722E-41 | -0.2547877 | 0.054 | 0.187 | 1.4672E-36 | TEC_MMP7 | PRDM2 |
| 3.6854E-34 | -0.2549621 | 0.125 | 0.261 | 7.8684E-30 | TEC_MMP7 | NCKAP1 |
| 1.7674E-33 | -0.2551149 | 0.09 | 0.216 | 3.7735E-29 | TEC_MMP7 | MBP |
| 4.7471E-45 | -0.2552806 | 0.036 | 0.17 | 1.0135E-40 | TEC_MMP7 | BIRC6 |
| 4.7808E-43 | -0.255351 | 0.061 | 0.202 | 1.0207E-38 | TEC_MMP7 | PLEKHA1 |
| 2.5178E-43 | -0.2557232 | 0.026 | 0.151 | 5.3754E-39 | TEC_MMP7 | RSBN1L |
| 5.8088E-31 | -0.2567559 | 0.092 | 0.214 | 1.2402E-26 | TEC_MMP7 | PHLDB2 |
| 1.813E-32 | -0.2571901 | 0.179 | 0.318 | 3.8707E-28 | TEC_MMP7 | PAPOLA |
| 2.1373E-37 | -0.2574024 | 0.097 | 0.234 | 4.5632E-33 | TEC_MMP7 | THRAP3 |
| 1.1868E-49 | -0.2577327 | 0.031 | 0.173 | 2.5338E-45 | TEC_MMP7 | CWC25 |
| 5.0252E-42 | -0.2578608 | 0.079 | 0.224 | 1.0729E-37 | TEC_MMP7 | USP16 |
| 5.4973E-32 | -0.2582644 | 0.101 | 0.225 | 1.1737E-27 | TEC_MMP7 | GULP1 |
| 4.5813E-32 | -0.2584573 | 0.095 | 0.218 | 9.7811E-28 | TEC_MMP7 | IRF2BP2 |
| 9.1047E-49 | -0.2587881 | 0.033 | 0.175 | 1.9438E-44 | TEC_MMP7 | CEP350 |
| 3.9442E-38 | -0.2588052 | 0.047 | 0.169 | 8.4209E-34 | TEC_MMP7 | ASXL1 |
| 1.2808E-28 | -0.259125 | 0.124 | 0.242 | 2.7345E-24 | TEC_MMP7 | TRIM8 |
| 1.1589E-40 | -0.2605929 | 0.05 | 0.18 | 2.4743E-36 | TEC_MMP7 | OFD1 |
| 2.4023E-38 | -0.2609303 | 0.049 | 0.172 | 5.1289E-34 | TEC_MMP7 | NUDT4 |
| 3.2454E-09 | -0.2611349 | 0.342 | 0.39 | 6.9289E-05 | TEC_MMP7 | MYH9 |
| 1.7954E-29 | -0.2616191 | 0.453 | 0.591 | 3.8333E-25 | TEC_MMP7 | SET |
| 6.4802E-38 | -0.2621682 | 0.038 | 0.156 | 1.3835E-33 | TEC_MMP7 | C12orf65 |
| 8.5493E-43 | -0.2623061 | 0.044 | 0.176 | 1.8253E-38 | TEC_MMP7 | NUMA1 |
| 1.4378E-29 | -0.2628849 | 0.238 | 0.374 | 3.0697E-25 | TEC_MMP7 | UBXN4 |
| 1.1891E-48 | -0.2631897 | 0.023 | 0.158 | 2.5387E-44 | TEC_MMP7 | RNF169 |

| | | | | | | |
|---|---|---|---|---|---|---|
| 6.8299E-50 | -0.2635451 | 0.038 | 0.185 | 1.4582E-45 | TEC_MMP7 | SAFB2 |
| 2.1256E-46 | -0.2636271 | 0.06 | 0.207 | 4.5382E-42 | TEC_MMP7 | DDX6 |
| 1.2045E-27 | -0.2636403 | 0.053 | 0.151 | 2.5716E-23 | TEC_MMP7 | EFNB2 |
| 8.188E-28 | -0.2636779 | 0.135 | 0.254 | 1.7481E-23 | TEC_MMP7 | TCEAL9 |
| 5.3689E-42 | -0.2638263 | 0.054 | 0.19 | 1.1463E-37 | TEC_MMP7 | NSRP1 |
| 6.4807E-36 | -0.2639169 | 0.086 | 0.217 | 1.3836E-31 | TEC_MMP7 | FOXO3 |
| 7.5513E-21 | -0.2640344 | 0.227 | 0.335 | 1.6122E-16 | TEC_MMP7 | RRBP1 |
| 1.6414E-36 | -0.2641715 | 0.033 | 0.145 | 3.5044E-32 | TEC_MMP7 | ING1 |
| 1.4754E-48 | -0.2644048 | 0.035 | 0.177 | 3.15E-44 | TEC_MMP7 | KIAA1217 |
| 7.4024E-41 | -0.2645465 | 0.076 | 0.214 | 1.5804E-36 | TEC_MMP7 | STX3 |
| 2.9705E-31 | -0.2647434 | 0.063 | 0.175 | 6.342E-27 | TEC_MMP7 | FNIP2 |
| 1.2717E-32 | -0.2649954 | 0.081 | 0.202 | 2.7151E-28 | TEC_MMP7 | TNFRSF10B |
| 5.6663E-41 | -0.2656606 | 0.057 | 0.189 | 1.2098E-36 | TEC_MMP7 | MYEF2 |
| 9.8443E-35 | -0.2656786 | 0.114 | 0.25 | 2.1018E-30 | TEC_MMP7 | CHD4 |
| 1.6716E-29 | -0.2662325 | 0.129 | 0.252 | 3.5689E-25 | TEC_MMP7 | MKNK2 |
| 2.1062E-32 | -0.266265 | 0.215 | 0.367 | 4.4968E-28 | TEC_MMP7 | UBE2H |
| 3.5795E-37 | -0.2666965 | 0.039 | 0.156 | 7.6423E-33 | TEC_MMP7 | PCLO |
| 1.0748E-42 | -0.2666991 | 0.02 | 0.139 | 2.2947E-38 | TEC_MMP7 | KMT2E-AS1 |
| 1.3519E-12 | -0.2670347 | 0.225 | 0.297 | 2.8862E-08 | TEC_MMP7 | MT-ATP8 |
| 8.4667E-19 | -0.2674715 | 0.162 | 0.261 | 1.8076E-14 | TEC_MMP7 | MACC1 |
| 1.5466E-44 | -0.2676276 | 0.05 | 0.19 | 3.302E-40 | TEC_MMP7 | OTUD6B-AS1 |
| 1.1061E-52 | -0.2678188 | 0.03 | 0.178 | 2.3615E-48 | TEC_MMP7 | RIF1 |
| 3.8272E-33 | -0.2685064 | 0.033 | 0.137 | 8.1711E-29 | TEC_MMP7 | PTGER3 |
| 2.08E-29 | -0.2685532 | 0.068 | 0.177 | 4.4407E-25 | TEC_MMP7 | KATNBL1 |
| 3.5322E-50 | -0.2688603 | 0.026 | 0.167 | 7.5412E-46 | TEC_MMP7 | NUTM2B-AS1 |
| 2.2759E-43 | -0.269852 | 0.088 | 0.24 | 4.859E-39 | TEC_MMP7 | RBM47 |
| 3.1285E-40 | -0.2706163 | 0.027 | 0.144 | 6.6793E-36 | TEC_MMP7 | WDR60 |
| 6.4226E-37 | -0.2707372 | 0.106 | 0.244 | 1.3712E-32 | TEC_MMP7 | HNRNPA0 |
| 1.482E-32 | -0.2710222 | 0.023 | 0.12 | 3.164E-28 | TEC_MMP7 | RGS2 |
| 5.0627E-20 | -0.271156 | 0.158 | 0.261 | 1.0809E-15 | TEC_MMP7 | RAP2B |
| 1.2518E-36 | -0.2712722 | 0.088 | 0.219 | 2.6726E-32 | TEC_MMP7 | JOSD1 |
| 8.8263E-18 | -0.2716809 | 0.185 | 0.274 | 1.8844E-13 | TEC_MMP7 | PLXNB2 |
| 1.0443E-42 | -0.2717509 | 0.073 | 0.217 | 2.2295E-38 | TEC_MMP7 | SLK |
| 4.1908E-14 | -0.2740855 | 0.05 | 0.109 | 8.9473E-10 | TEC_MMP7 | TNF |
| 1.3606E-44 | -0.2740903 | 0.089 | 0.242 | 2.9049E-40 | TEC_MMP7 | RAB21 |
| 5.9044E-51 | -0.2743966 | 0.051 | 0.206 | 1.2606E-46 | TEC_MMP7 | KDM2A |
| 6.288E-45 | -0.2746757 | 0.081 | 0.233 | 1.3425E-40 | TEC_MMP7 | RSF1 |
| 4.8554E-43 | -0.2750363 | 0.042 | 0.174 | 1.0366E-38 | TEC_MMP7 | ZBTB10 |
| 3.508E-47 | -0.2754706 | 0.061 | 0.211 | 7.4895E-43 | TEC_MMP7 | MEF2A |
| 2.1309E-45 | -0.27556 | 0.074 | 0.224 | 4.5495E-41 | TEC_MMP7 | C16orf72 |
| 1.1608E-44 | -0.2756802 | 0.067 | 0.212 | 2.4784E-40 | TEC_MMP7 | ARFGEF1 |
| 1.5198E-42 | -0.276011 | 0.028 | 0.151 | 3.2449E-38 | TEC_MMP7 | HERC2 |
| 1.1721E-51 | -0.2777617 | 0.057 | 0.216 | 2.5023E-47 | TEC_MMP7 | SLC25A36 |
| 5.1512E-53 | -0.2781286 | 0.05 | 0.208 | 1.0998E-48 | TEC_MMP7 | UBE3A |
| 3.0429E-34 | -0.2782961 | 0.219 | 0.369 | 6.4965E-30 | TEC_MMP7 | VAPA |
| 7.9729E-49 | -0.2788319 | 0.057 | 0.211 | 1.7022E-44 | TEC_MMP7 | PHACTR4 |
| 2.0543E-40 | -0.2795818 | 0.039 | 0.163 | 4.386E-36 | TEC_MMP7 | EMX2 |
| 1.2573E-49 | -0.2797336 | 0.042 | 0.188 | 2.6844E-45 | TEC_MMP7 | ZNF644 |
| 2.0262E-46 | -0.2802389 | 0.042 | 0.181 | 4.3259E-42 | TEC_MMP7 | AFF1 |
| 1.0518E-35 | -0.2805015 | 0.009 | 0.104 | 2.2457E-31 | TEC_MMP7 | Z93241.1 |

| | | | | | | |
|---|---|---|---|---|---|---|
| 2.7376E-52 | -0.2806908 | 0.041 | 0.193 | 5.8449E-48 | TEC_MMP7 | MED13 |
| 1.0621E-25 | -0.2806946 | 0.177 | 0.293 | 2.2675E-21 | TEC_MMP7 | GTF2I |
| 5.4571E-36 | -0.2809786 | 0.109 | 0.245 | 1.1651E-31 | TEC_MMP7 | TERF2IP |
| 3.6839E-18 | -0.2816858 | 0.291 | 0.385 | 7.8652E-14 | TEC_MMP7 | ODC1 |
| 9.7069E-50 | -0.2817962 | 0.026 | 0.164 | 2.0724E-45 | TEC_MMP7 | CREBZF |
| 4.7799E-38 | -0.2831622 | 0.103 | 0.244 | 1.0205E-33 | TEC_MMP7 | SNRNP70 |
| 6.5311E-48 | -0.2838916 | 0.044 | 0.188 | 1.3944E-43 | TEC_MMP7 | PHIP |
| 1.9266E-43 | -0.284036 | 0.029 | 0.154 | 4.1132E-39 | TEC_MMP7 | SNX29 |
| 1.1532E-48 | -0.2843169 | 0.038 | 0.18 | 2.462E-44 | TEC_MMP7 | KIAA1109 |
| 1.497E-50 | -0.2847787 | 0.041 | 0.189 | 3.1961E-46 | TEC_MMP7 | NRIP1 |
| 2.3573E-44 | -0.2852111 | 0.068 | 0.214 | 5.0329E-40 | TEC_MMP7 | BACH1 |
| 9.3313E-49 | -0.28534 | 0.074 | 0.234 | 1.9922E-44 | TEC_MMP7 | RAD21 |
| 2.9292E-51 | -0.2863355 | 0.049 | 0.203 | 6.2539E-47 | TEC_MMP7 | KDM5A |
| 6.4677E-28 | -0.2865307 | 0.076 | 0.181 | 1.3808E-23 | TEC_MMP7 | TMX4 |
| 3.0573E-36 | -0.2867745 | 0.048 | 0.165 | 6.5273E-32 | TEC_MMP7 | MED29 |
| 1.1135E-39 | -0.2868744 | 0.075 | 0.212 | 2.3774E-35 | TEC_MMP7 | EPS8 |
| 7.1184E-50 | -0.2877093 | 0.052 | 0.203 | 1.5198E-45 | TEC_MMP7 | ARIH1 |
| 1.4938E-39 | -0.2877292 | 0.004 | 0.106 | 3.1892E-35 | TEC_MMP7 | AL137077.2 |
| 2.2619E-33 | -0.288144 | 0.096 | 0.224 | 4.8292E-29 | TEC_MMP7 | SLFN5 |
| 3.7637E-41 | -0.288629 | 0.136 | 0.292 | 8.0356E-37 | TEC_MMP7 | HNRNPD |
| 1.8768E-52 | -0.2889889 | 0.037 | 0.188 | 4.007E-48 | TEC_MMP7 | USP15 |
| 1.2084E-40 | -0.2893962 | 0.086 | 0.228 | 2.58E-36 | TEC_MMP7 | RAB3IP |
| 8.5299E-42 | -0.2907534 | 0.041 | 0.169 | 1.8211E-37 | TEC_MMP7 | ASCC3 |
| 4.2891E-42 | -0.2907751 | 0.023 | 0.143 | 9.1572E-38 | TEC_MMP7 | PKHD1 |
| 3.161E-43 | -0.2908531 | 0.021 | 0.142 | 6.7486E-39 | TEC_MMP7 | MEPCE |
| 2.0842E-38 | -0.2916912 | 0.108 | 0.249 | 4.4497E-34 | TEC_MMP7 | RBMX |
| 5.6495E-33 | -0.2927477 | 0.208 | 0.349 | 1.2062E-28 | TEC_MMP7 | KIF5B |
| 7.8211E-52 | -0.2927856 | 0.034 | 0.183 | 1.6698E-47 | TEC_MMP7 | PPP1R15B |
| 3.6617E-54 | -0.2936834 | 0.044 | 0.202 | 7.8178E-50 | TEC_MMP7 | RBM33 |
| 3.363E-22 | -0.2941555 | 0.086 | 0.18 | 7.18E-18 | TEC_MMP7 | FGF2 |
| 6.0074E-48 | -0.2955378 | 0.036 | 0.176 | 1.2826E-43 | TEC_MMP7 | HOOK2 |
| 5.6463E-40 | -0.2958717 | 0.096 | 0.241 | 1.2055E-35 | TEC_MMP7 | TEAD1 |
| 5.9251E-18 | -0.2962052 | 0.178 | 0.274 | 1.265E-13 | TEC_MMP7 | HMGB2 |
| 8.3925E-51 | -0.2962919 | 0.05 | 0.204 | 1.7918E-46 | TEC_MMP7 | XPO1 |
| 5.8835E-52 | -0.2965121 | 0.071 | 0.236 | 1.2561E-47 | TEC_MMP7 | TMF1 |
| 4.7044E-46 | -0.2968342 | 0.028 | 0.16 | 1.0044E-41 | TEC_MMP7 | GABPB1-AS1 |
| 2.4872E-48 | -0.2972738 | 0.058 | 0.209 | 5.3101E-44 | TEC_MMP7 | RNF19A |
| 7.1395E-42 | -0.2973446 | 0.073 | 0.213 | 1.5243E-37 | TEC_MMP7 | SRSF1 |
| 1.7875E-47 | -0.2976973 | 0.103 | 0.267 | 3.8162E-43 | TEC_MMP7 | PPIG |
| 3.8507E-49 | -0.2979079 | 0.01 | 0.137 | 8.2213E-45 | TEC_MMP7 | LINC00472 |
| 4.5824E-55 | -0.2981191 | 0.083 | 0.259 | 9.7835E-51 | TEC_MMP7 | PAFAH1B1 |
| 6.6111E-21 | -0.298734 | 0.381 | 0.48 | 1.4115E-16 | TEC_MMP7 | LMNA |
| 8.5774E-58 | -0.2987994 | 0.033 | 0.194 | 1.8313E-53 | TEC_MMP7 | EP300 |
| 2.7076E-42 | -0.2988742 | 0.095 | 0.244 | 5.7807E-38 | TEC_MMP7 | CLIP1 |
| 1.0983E-43 | -0.2990333 | 0.043 | 0.177 | 2.3448E-39 | TEC_MMP7 | NR2F2 |
| 6.2505E-29 | -0.2992169 | 0.167 | 0.292 | 1.3345E-24 | TEC_MMP7 | TCEA1 |
| 2.1452E-23 | -0.2995249 | 0.597 | 0.692 | 4.58E-19 | TEC_MMP7 | YBX3 |
| 7.4363E-48 | -0.3001386 | 0.069 | 0.223 | 1.5876E-43 | TEC_MMP7 | RTF1 |
| 1.6146E-43 | -0.3006274 | 0.096 | 0.249 | 3.4471E-39 | TEC_MMP7 | METAP2 |
| 8.4317E-42 | -0.3008534 | 0.01 | 0.12 | 1.8002E-37 | TEC_MMP7 | AC012306.2 |

| | | | | | | |
|---|---|---|---|---|---|---|
| 4.1743E-22 | -0.3011375 | 0.081 | 0.172 | 8.9121E-18 | TEC_MMP7 | TNFAIP3 |
| 8.6188E-35 | -0.3016478 | 0.183 | 0.328 | 1.8401E-30 | TEC_MMP7 | ARHGAP5 |
| 3.6882E-24 | -0.3016551 | 0.177 | 0.288 | 7.8742E-20 | TEC_MMP7 | SPTAN1 |
| 9.9826E-44 | -0.3017035 | 0.037 | 0.168 | 2.1313E-39 | TEC_MMP7 | DAPK1 |
| 1.6126E-45 | -0.3018892 | 0.051 | 0.191 | 3.4428E-41 | TEC_MMP7 | CCNL2 |
| 6.7463E-14 | -0.3021581 | 0.505 | 0.578 | 1.4403E-09 | TEC_MMP7 | RPS27L |
| 1.436E-43 | -0.3025588 | 0.085 | 0.233 | 3.0658E-39 | TEC_MMP7 | PATJ |
| 4.2756E-45 | -0.3028728 | 0.063 | 0.209 | 9.1284E-41 | TEC_MMP7 | ZBTB43 |
| 8.2512E-45 | -0.3029615 | 0.05 | 0.19 | 1.7616E-40 | TEC_MMP7 | TACC1 |
| 2.1415E-41 | -0.3029923 | 0.051 | 0.183 | 4.5722E-37 | TEC_MMP7 | MIR22HG |
| 6.3691E-43 | -0.3038555 | 0.041 | 0.173 | 1.3598E-38 | TEC_MMP7 | SLC20A1 |
| 5.0404E-52 | -0.3040993 | 0.047 | 0.201 | 1.0761E-47 | TEC_MMP7 | ZFHX3 |
| 4.0547E-23 | -0.3042649 | 0.034 | 0.115 | 8.6568E-19 | TEC_MMP7 | ARC |
| 3.4735E-57 | -0.3043308 | 0.051 | 0.219 | 7.416E-53 | TEC_MMP7 | SENP6 |
| 7.2716E-43 | -0.3047637 | 0.087 | 0.236 | 1.5525E-38 | TEC_MMP7 | USP53 |
| 7.7279E-56 | -0.3048272 | 0.039 | 0.199 | 1.6499E-51 | TEC_MMP7 | SETD2 |
| 3.9219E-42 | -0.3052279 | 0.086 | 0.233 | 8.3734E-38 | TEC_MMP7 | PARD6B |
| 7.4624E-52 | -0.3058249 | 0.048 | 0.202 | 1.5932E-47 | TEC_MMP7 | ZNF638 |
| 8.1565E-35 | -0.3061208 | 0.101 | 0.235 | 1.7414E-30 | TEC_MMP7 | PARP14 |
| 5.893E-58 | -0.3062088 | 0.025 | 0.181 | 1.2582E-53 | TEC_MMP7 | RBM6 |
| 3.7505E-43 | -0.3063788 | 0.089 | 0.239 | 8.0074E-39 | TEC_MMP7 | WTAP |
| 6.2751E-53 | -0.306702 | 0.05 | 0.208 | 1.3397E-48 | TEC_MMP7 | SREK1 |
| 1.7159E-39 | -0.306737 | 0.008 | 0.112 | 3.6635E-35 | TEC_MMP7 | MAFA |
| 4.5499E-34 | -0.3067729 | 0.073 | 0.196 | 9.7141E-30 | TEC_MMP7 | HMGCS1 |
| 4.6563E-49 | -0.3069073 | 0.025 | 0.162 | 9.9412E-45 | TEC_MMP7 | ILF3-DT |
| 1.8666E-46 | -0.3078264 | 0.101 | 0.262 | 3.9852E-42 | TEC_MMP7 | VAMP2 |
| 8.9165E-36 | -0.308119 | 0.191 | 0.338 | 1.9037E-31 | TEC_MMP7 | DEK |
| 2.4508E-54 | -0.3084754 | 0.05 | 0.212 | 5.2325E-50 | TEC_MMP7 | CEBPZ |
| 3.4137E-38 | -0.3089314 | 0.075 | 0.209 | 7.2882E-34 | TEC_MMP7 | SESN2 |
| 4.4301E-60 | -0.3089552 | 0.026 | 0.188 | 9.4583E-56 | TEC_MMP7 | CDK13 |
| 1.8038E-46 | -0.3090526 | 0.036 | 0.172 | 3.8511E-42 | TEC_MMP7 | IVNS1ABP |
| 4.4723E-56 | -0.3093666 | 0.067 | 0.24 | 9.5485E-52 | TEC_MMP7 | DHX36 |
| 2.4809E-41 | -0.3095081 | 0.133 | 0.287 | 5.2967E-37 | TEC_MMP7 | CCDC50 |
| 8.2748E-51 | -0.3099979 | 0.074 | 0.236 | 1.7667E-46 | TEC_MMP7 | NCOR1 |
| 2.2557E-54 | -0.3103469 | 0.05 | 0.212 | 4.8159E-50 | TEC_MMP7 | ARID1B |
| 3.4357E-54 | -0.3122786 | 0.061 | 0.226 | 7.3352E-50 | TEC_MMP7 | REST |
| 1.8164E-45 | -0.3123071 | 0.068 | 0.215 | 3.878E-41 | TEC_MMP7 | EPB41L5 |
| 1.7583E-55 | -0.3127086 | 0.037 | 0.194 | 3.7539E-51 | TEC_MMP7 | NSD1 |
| 8.8568E-39 | -0.3129202 | 0.017 | 0.125 | 1.8909E-34 | TEC_MMP7 | EGOT |
| 1.4945E-58 | -0.3129779 | 0.039 | 0.204 | 3.1907E-54 | TEC_MMP7 | FNIP1 |
| 1.5487E-15 | -0.3148475 | 0.139 | 0.219 | 3.3064E-11 | TEC_MMP7 | ADAMTS9 |
| 5.3896E-16 | -0.3154171 | 0.045 | 0.108 | 1.1507E-11 | TEC_MMP7 | CXCL3 |
| 1.3625E-44 | -0.3164583 | 0.33 | 0.51 | 2.9089E-40 | TEC_MMP7 | UBE2D3 |
| 9.6131E-27 | -0.3173984 | 0.107 | 0.214 | 2.0524E-22 | TEC_MMP7 | CHCHD7 |
| 9.9834E-46 | -0.317896 | 0.047 | 0.187 | 2.1315E-41 | TEC_MMP7 | WNK1 |
| 4.9047E-40 | -0.3185404 | 0.106 | 0.254 | 1.0472E-35 | TEC_MMP7 | MIDN |
| 2.5646E-50 | -0.3195395 | 0.052 | 0.204 | 5.4755E-46 | TEC_MMP7 | MUC20-OT1 |
| 1.3549E-56 | -0.3199448 | 0.066 | 0.241 | 2.8927E-52 | TEC_MMP7 | CCDC186 |
| 1.4261E-34 | -0.3206413 | 0.175 | 0.322 | 3.0448E-30 | TEC_MMP7 | SNHG7 |
| 9.9616E-26 | -0.3220809 | 0.213 | 0.334 | 2.1268E-21 | TEC_MMP7 | FAM107B |

| | | | | | | |
|---|---|---|---|---|---|---|
| 1.2833E-40 | -0.3225639 | 0.197 | 0.357 | 2.7398E-36 | TEC_MMP7 | SYF2 |
| 2.6081E-33 | -0.322838 | 0.108 | 0.234 | 5.5682E-29 | TEC_MMP7 | EPS8L2 |
| 9.1706E-44 | -0.322987 | 0.14 | 0.305 | 1.9579E-39 | TEC_MMP7 | XBP1 |
| 8.473E-46 | -0.3237033 | 0.111 | 0.272 | 1.809E-41 | TEC_MMP7 | PTBP3 |
| 7.0677E-47 | -0.3246106 | 0.062 | 0.21 | 1.5089E-42 | TEC_MMP7 | KLHL21 |
| 1.6333E-48 | -0.3248986 | 0.096 | 0.258 | 3.4871E-44 | TEC_MMP7 | HECTD1 |
| 2.0462E-14 | -0.3252642 | 0.613 | 0.676 | 4.3686E-10 | TEC_MMP7 | GLS |
| 6.7491E-42 | -0.3255239 | 0.06 | 0.197 | 1.4409E-37 | TEC_MMP7 | SQLE |
| 1.6548E-11 | -0.3268459 | 0.377 | 0.429 | 3.533E-07 | TEC_MMP7 | SNHG8 |
| 3.0824E-53 | -0.3270584 | 0.091 | 0.264 | 6.5808E-49 | TEC_MMP7 | TOR1AIP2 |
| 1.2096E-36 | -0.3282368 | 0.103 | 0.239 | 2.5825E-32 | TEC_MMP7 | BEX2 |
| 3.2327E-27 | -0.3290113 | 0.057 | 0.157 | 6.9018E-23 | TEC_MMP7 | XIST |
| 3.9565E-43 | -0.3298365 | 0.199 | 0.366 | 8.4472E-39 | TEC_MMP7 | MTDH |
| 1.684E-43 | -0.3304545 | 0.146 | 0.31 | 3.5954E-39 | TEC_MMP7 | TJP1 |
| 4.7407E-45 | -0.3313931 | 0.075 | 0.224 | 1.0121E-40 | TEC_MMP7 | FOSL2 |
| 8.7706E-26 | -0.332913 | 0.295 | 0.406 | 1.8725E-21 | TEC_MMP7 | CIRBP |
| 1.7472E-58 | -0.333955 | 0.06 | 0.234 | 3.7304E-54 | TEC_MMP7 | PUM1 |
| 2.4604E-39 | -0.3347081 | 0.014 | 0.121 | 5.253E-35 | TEC_MMP7 | AL355075.4 |
| 4.7782E-47 | -0.3358731 | 0.091 | 0.248 | 1.0201E-42 | TEC_MMP7 | DDX18 |
| 3.8584E-47 | -0.3361427 | 0.12 | 0.287 | 8.2376E-43 | TEC_MMP7 | SLC38A1 |
| 9.8331E-50 | -0.3365278 | 0.092 | 0.258 | 2.0994E-45 | TEC_MMP7 | AKAP13 |
| 1.3768E-54 | -0.3370574 | 0.05 | 0.211 | 2.9396E-50 | TEC_MMP7 | PRPF38B |
| 1.4576E-55 | -0.337187 | 0.09 | 0.268 | 3.112E-51 | TEC_MMP7 | SLTM |
| 6.2758E-61 | -0.3379603 | 0.039 | 0.209 | 1.3399E-56 | TEC_MMP7 | PPP4R3A |
| 2.3086E-53 | -0.3387621 | 0.066 | 0.232 | 4.9288E-49 | TEC_MMP7 | MAFG |
| 6.2851E-47 | -0.3394073 | 0.12 | 0.287 | 1.3419E-42 | TEC_MMP7 | RIOK3 |
| 1.2832E-53 | -0.3395724 | 0.068 | 0.235 | 2.7397E-49 | TEC_MMP7 | TRIP11 |
| 7.7257E-48 | -0.3402333 | 0.072 | 0.226 | 1.6494E-43 | TEC_MMP7 | DNTTIP2 |
| 1.977E-47 | -0.3413109 | 0.1 | 0.259 | 4.2209E-43 | TEC_MMP7 | ANKLE2 |
| 4.2608E-61 | -0.3413858 | 0.06 | 0.24 | 9.0969E-57 | TEC_MMP7 | QKI |
| 1.604E-64 | -0.341438 | 0.047 | 0.229 | 3.4245E-60 | TEC_MMP7 | TNRC6B |
| 3.3146E-59 | -0.3416159 | 0.042 | 0.209 | 7.0766E-55 | TEC_MMP7 | BAZ2B |
| 9.8018E-41 | -0.3416506 | 0.156 | 0.31 | 2.0927E-36 | TEC_MMP7 | HNRNPH3 |
| 4.183E-65 | -0.3423684 | 0.056 | 0.241 | 8.9306E-61 | TEC_MMP7 | PJA2 |
| 1.4651E-27 | -0.3424979 | 0.236 | 0.354 | 3.1279E-23 | TEC_MMP7 | HSPA9 |
| 2.2057E-16 | -0.3425425 | 0.82 | 0.926 | 4.7091E-12 | TEC_MMP7 | MT-ND4 |
| 7.3068E-62 | -0.344893 | 0.041 | 0.213 | 1.56E-57 | TEC_MMP7 | ATXN7L3B |
| 6.0537E-16 | -0.3450107 | 0.795 | 0.905 | 1.2925E-11 | TEC_MMP7 | MT-ND1 |
| 7.2001E-29 | -0.3472746 | 0.275 | 0.401 | 1.5372E-24 | TEC_MMP7 | MYO6 |
| 1.1337E-44 | -0.3478987 | 0.041 | 0.174 | 2.4205E-40 | TEC_MMP7 | PLEKHH2 |
| 1.6907E-46 | -0.3481694 | 0.017 | 0.142 | 3.6095E-42 | TEC_MMP7 | AL135925.1 |
| 3.3294E-46 | -0.3482874 | 0.007 | 0.126 | 7.1084E-42 | TEC_MMP7 | HIST4H4 |
| 4.5322E-39 | -0.3496503 | 0.048 | 0.171 | 9.6763E-35 | TEC_MMP7 | CDKN2B |
| 1.8846E-16 | -0.3516 | 0.817 | 0.923 | 4.0236E-12 | TEC_MMP7 | MT-ATP6 |
| 1.0068E-44 | -0.3535971 | 0.168 | 0.338 | 2.1496E-40 | TEC_MMP7 | EIF5B |
| 1.1299E-64 | -0.3540181 | 0.051 | 0.233 | 2.4123E-60 | TEC_MMP7 | PURA |
| 1.4079E-56 | -0.3552478 | 0.082 | 0.258 | 3.0059E-52 | TEC_MMP7 | NIPBL |
| 9.098E-36 | -0.3554543 | 0.006 | 0.1 | 1.9424E-31 | TEC_MMP7 | PCDH10 |
| 1.4715E-57 | -0.3560899 | 0.023 | 0.177 | 3.1415E-53 | TEC_MMP7 | RNPC3 |
| 2.2574E-43 | -0.3562322 | 0.074 | 0.217 | 4.8195E-39 | TEC_MMP7 | FAT1 |

| | | | | | | |
|---|---|---|---|---|---|---|
| 1.9009E-36 | -0.3564001 | 0.13 | 0.271 | 4.0584E-32 | TEC_MMP7 | PTP4A1 |
| 3.2213E-68 | -0.3566569 | 0.034 | 0.217 | 6.8775E-64 | TEC_MMP7 | GPATCH2L |
| 6.7539E-38 | -0.3570265 | 0.042 | 0.161 | 1.442E-33 | TEC_MMP7 | ANKRD37 |
| 2.8767E-58 | -0.3576973 | 0.045 | 0.212 | 6.1417E-54 | TEC_MMP7 | ZNF83 |
| 2.7976E-58 | -0.3593873 | 0.066 | 0.241 | 5.973E-54 | TEC_MMP7 | FAM133B |
| 3.3011E-65 | -0.3614879 | 0.019 | 0.187 | 7.0479E-61 | TEC_MMP7 | PAXBP1 |
| 2.4529E-57 | -0.3615873 | 0.115 | 0.303 | 5.2368E-53 | TEC_MMP7 | YY1 |
| 4.0691E-37 | -0.3639757 | 0.803 | 0.954 | 8.6876E-33 | TEC_MMP7 | PTMA |
| 1.5477E-60 | -0.3644796 | 0.037 | 0.205 | 3.3044E-56 | TEC_MMP7 | SPEN |
| 3.5633E-48 | -0.3644816 | 0.019 | 0.15 | 7.6077E-44 | TEC_MMP7 | CSRNP2 |
| 1.3444E-35 | -0.3653266 | 0.021 | 0.124 | 2.8703E-31 | TEC_MMP7 | AC023157.3 |
| 1.7354E-56 | -0.3659124 | 0.052 | 0.218 | 3.7051E-52 | TEC_MMP7 | UPF2 |
| 3.2903E-57 | -0.3670775 | 0.086 | 0.265 | 7.0248E-53 | TEC_MMP7 | FNBP4 |
| 1.066E-63 | -0.3685593 | 0.093 | 0.289 | 2.2759E-59 | TEC_MMP7 | SCAF11 |
| 1.4216E-66 | -0.3712181 | 0.085 | 0.283 | 3.035E-62 | TEC_MMP7 | BRD4 |
| 9.7999E-54 | -0.371306 | 0.053 | 0.212 | 2.0923E-49 | TEC_MMP7 | ERICH1 |
| 1.3305E-46 | -0.3713126 | 0.073 | 0.223 | 2.8405E-42 | TEC_MMP7 | PIM3 |
| 9.1186E-35 | -0.3714979 | 0.238 | 0.402 | 1.9468E-30 | TEC_MMP7 | HERPUD1 |
| 5.6356E-41 | -0.3718716 | 0.338 | 0.513 | 1.2032E-36 | TEC_MMP7 | DNAJA1 |
| 2.6508E-48 | -0.3719092 | 0.099 | 0.26 | 5.6594E-44 | TEC_MMP7 | KLF5 |
| 1.1704E-65 | -0.372046 | 0.037 | 0.216 | 2.4988E-61 | TEC_MMP7 | POLR2J3.1 |
| 3.9249E-46 | -0.3723298 | 0.018 | 0.143 | 8.3796E-42 | TEC_MMP7 | HOMER1 |
| 1.8484E-14 | -0.3724431 | 0.198 | 0.277 | 3.9463E-10 | TEC_MMP7 | TUBA1A |
| 6.5848E-67 | -0.3743157 | 0.048 | 0.236 | 1.4059E-62 | TEC_MMP7 | KMT2C |
| 7.1743E-61 | -0.3747378 | 0.068 | 0.25 | 1.5317E-56 | TEC_MMP7 | RNMT |
| 1.3328E-63 | -0.3765379 | 0.035 | 0.208 | 2.8455E-59 | TEC_MMP7 | PCF11 |
| 2.3876E-71 | -0.3773815 | 0.081 | 0.287 | 5.0975E-67 | TEC_MMP7 | WAC |
| 1.6893E-64 | -0.3780451 | 0.048 | 0.228 | 3.6067E-60 | TEC_MMP7 | EPC1 |
| 5.2772E-58 | -0.3792178 | 0.11 | 0.295 | 1.1267E-53 | TEC_MMP7 | OGA |
| 2.0655E-62 | -0.3797355 | 0.091 | 0.281 | 4.4099E-58 | TEC_MMP7 | SF1 |
| 2.3695E-34 | -0.380156 | 0.101 | 0.232 | 5.0589E-30 | TEC_MMP7 | GCLM |
| 3.9168E-67 | -0.3804161 | 0.049 | 0.237 | 8.3624E-63 | TEC_MMP7 | ACIN1 |
| 3.4291E-45 | -0.3804776 | 0.073 | 0.22 | 7.3211E-41 | TEC_MMP7 | ARRDC3 |
| 3.0163E-53 | -0.3807512 | 0.09 | 0.262 | 6.4399E-49 | TEC_MMP7 | LMO4 |
| 6.8941E-59 | -0.3812784 | 0.12 | 0.313 | 1.4719E-54 | TEC_MMP7 | NFE2L2 |
| 1.5261E-52 | -0.3816275 | 0.105 | 0.277 | 3.2582E-48 | TEC_MMP7 | DDX46 |
| 5.0833E-61 | -0.383099 | 0.079 | 0.263 | 1.0853E-56 | TEC_MMP7 | AAK1 |
| 4.6064E-42 | -0.383741 | 0.041 | 0.168 | 9.8346E-38 | TEC_MMP7 | PFKFB3 |
| 5.1368E-60 | -0.3844653 | 0.053 | 0.228 | 1.0967E-55 | TEC_MMP7 | THUMPD3-AS1 |
| 1.8446E-70 | -0.3851277 | 0.049 | 0.243 | 3.9381E-66 | TEC_MMP7 | USP34 |
| 4.4651E-53 | -0.3853609 | 0.105 | 0.278 | 9.533E-49 | TEC_MMP7 | NEDD4L |
| 2.2748E-63 | -0.3853623 | 0.125 | 0.329 | 4.8568E-59 | TEC_MMP7 | PNRC2 |
| 1.0014E-42 | -0.3865265 | 0.315 | 0.505 | 2.138E-38 | TEC_MMP7 | H1FX |
| 1.6043E-67 | -0.3880742 | 0.069 | 0.263 | 3.4252E-63 | TEC_MMP7 | U2SURP |
| 1.2403E-75 | -0.3881132 | 0.028 | 0.222 | 2.6481E-71 | TEC_MMP7 | CDK12 |
| 4.5098E-62 | -0.3900003 | 0.049 | 0.225 | 9.6285E-58 | TEC_MMP7 | CLK1 |
| 6.022E-13 | -0.3901595 | 0.188 | 0.262 | 1.2857E-08 | TEC_MMP7 | AKAP12 |
| 1.6529E-31 | -0.3901666 | 0.051 | 0.157 | 3.529E-27 | TEC_MMP7 | ID1 |
| 3.8545E-64 | -0.3901931 | 0.083 | 0.276 | 8.2294E-60 | TEC_MMP7 | BCLAF1 |
| 1.2791E-27 | -0.3902317 | 0.089 | 0.2 | 2.7308E-23 | TEC_MMP7 | BIRC3 |

| | | | | | | |
|---|---|---|---|---|---|---|
| 2.4576E-57 | -0.3908519 | 0.074 | 0.247 | 5.2469E-53 | TEC_MMP7 | ZNF91 |
| 3.9247E-42 | -0.3910707 | 0.593 | 0.718 | 8.3792E-38 | TEC_MMP7 | EIF4G2 |
| 3.0055E-64 | -0.3926953 | 0.199 | 0.414 | 6.4168E-60 | TEC_MMP7 | CSNK1A1 |
| 5.4365E-67 | -0.3930642 | 0.021 | 0.193 | 1.1607E-62 | TEC_MMP7 | AHI1 |
| 3.2921E-68 | -0.3935049 | 0.057 | 0.249 | 7.0287E-64 | TEC_MMP7 | NFAT5 |
| 1.2364E-58 | -0.3938676 | 0.047 | 0.215 | 2.6397E-54 | TEC_MMP7 | TOB1 |
| 4.6741E-67 | -0.3966566 | 0.063 | 0.257 | 9.9791E-63 | TEC_MMP7 | RBBP6 |
| 3.3019E-56 | -0.3967428 | 0.159 | 0.35 | 7.0495E-52 | TEC_MMP7 | EIF1B |
| 1.7551E-63 | -0.3968326 | 0.106 | 0.306 | 3.7472E-59 | TEC_MMP7 | ZNF207 |
| 2.6935E-43 | -0.3985912 | 0.151 | 0.311 | 5.7506E-39 | TEC_MMP7 | MAFF |
| 6.9603E-51 | -0.4002608 | 0.164 | 0.341 | 1.486E-46 | TEC_MMP7 | DYNC1H1 |
| 1.5807E-62 | -0.4005734 | 0.827 | 0.976 | 3.3747E-58 | TEC_MMP7 | EIF1 |
| 1.7869E-39 | -0.4016129 | 0.009 | 0.113 | 3.815E-35 | TEC_MMP7 | AP001160.1 |
| 3.5753E-56 | -0.4017683 | 0.104 | 0.287 | 7.6333E-52 | TEC_MMP7 | RB1CC1 |
| 1.6917E-16 | -0.4019359 | 0.774 | 0.881 | 3.6118E-12 | TEC_MMP7 | MT-ND3 |
| 1.1231E-61 | -0.402009 | 0.146 | 0.347 | 2.3978E-57 | TEC_MMP7 | PAWR |
| 6.9737E-54 | -0.4032255 | 0.127 | 0.309 | 1.4889E-49 | TEC_MMP7 | SRRM1 |
| 1.1313E-79 | -0.4054533 | 0.037 | 0.244 | 2.4153E-75 | TEC_MMP7 | ASH1L |
| 9.9477E-51 | -0.4059511 | 0.011 | 0.143 | 2.1238E-46 | TEC_MMP7 | PCDH9 |
| 4.4288E-68 | -0.4061743 | 0.085 | 0.286 | 9.4554E-64 | TEC_MMP7 | SRSF10 |
| 1.5453E-44 | -0.407749 | 0.09 | 0.241 | 3.2991E-40 | TEC_MMP7 | ADM |
| 5.4363E-63 | -0.4080245 | 0.047 | 0.223 | 1.1607E-58 | TEC_MMP7 | MDM4 |
| 2.4074E-70 | -0.4095884 | 0.022 | 0.202 | 5.1397E-66 | TEC_MMP7 | MYO9A |
| 9.3337E-70 | -0.410148 | 0.073 | 0.274 | 1.9928E-65 | TEC_MMP7 | TRIM44 |
| 7.8673E-66 | -0.4107984 | 0.175 | 0.39 | 1.6797E-61 | TEC_MMP7 | SRSF11 |
| 6.6388E-56 | -0.4107989 | 0.018 | 0.166 | 1.4174E-51 | TEC_MMP7 | AL118516.1 |
| 1.4392E-76 | -0.4114722 | 0.053 | 0.26 | 3.0726E-72 | TEC_MMP7 | PHF3 |
| 0.0004135 | -0.4116355 | 0.268 | 0.219 | 1 | TEC_MMP7 | DEFB1 |
| 3.0388E-45 | -0.4117561 | 0.137 | 0.293 | 6.4879E-41 | TEC_MMP7 | TCF25 |
| 2.7592E-73 | -0.4162151 | 0.026 | 0.215 | 5.891E-69 | TEC_MMP7 | TUT4 |
| 1.8688E-52 | -0.4164167 | 0.039 | 0.191 | 3.9898E-48 | TEC_MMP7 | SETX |
| 2.8701E-68 | -0.4167954 | 0.016 | 0.188 | 6.1277E-64 | TEC_MMP7 | TSPYL2 |
| 7.4239E-42 | -0.4188183 | 0.191 | 0.345 | 1.585E-37 | TEC_MMP7 | HNRNPH1 |
| 1.239E-63 | -0.4198164 | 0.052 | 0.23 | 2.6453E-59 | TEC_MMP7 | VPS13C |
| 1.7549E-24 | -0.4198255 | 0.047 | 0.134 | 3.7468E-20 | TEC_MMP7 | CXCL14 |
| 1.0697E-79 | -0.4200713 | 0.026 | 0.228 | 2.2838E-75 | TEC_MMP7 | ZFC3H1 |
| 1.64E-72 | -0.420542 | 0.057 | 0.256 | 3.5015E-68 | TEC_MMP7 | KMT2E |
| 3.5492E-54 | -0.4211642 | 0.007 | 0.143 | 7.5775E-50 | TEC_MMP7 | HIST2H2BF |
| 3.6196E-78 | -0.4228728 | 0.039 | 0.245 | 7.7278E-74 | TEC_MMP7 | NKTR |
| 8.2513E-51 | -0.4236652 | 0.047 | 0.197 | 1.7617E-46 | TEC_MMP7 | KLF4 |
| 2.6593E-69 | -0.4237324 | 0.062 | 0.258 | 5.6775E-65 | TEC_MMP7 | PKN2 |
| 4.5189E-42 | -0.4242118 | 0.163 | 0.324 | 9.6478E-38 | TEC_MMP7 | ARHGAP29 |
| 3.874E-47 | -0.424884 | 0.079 | 0.234 | 8.271E-43 | TEC_MMP7 | ZFP36 |
| 8.2135E-36 | -0.4259093 | 0.169 | 0.32 | 1.7536E-31 | TEC_MMP7 | DDIT4 |
| 1.3929E-63 | -0.4285078 | 0.025 | 0.193 | 2.9739E-59 | TEC_MMP7 | THSD7A |
| 2.147E-75 | -0.4309497 | 0.094 | 0.313 | 4.5838E-71 | TEC_MMP7 | RBM25 |
| 8.1514E-30 | -0.4311668 | 0.229 | 0.352 | 1.7403E-25 | TEC_MMP7 | HNRNPAB |
| 1.0004E-78 | -0.431581 | 0.064 | 0.278 | 2.1358E-74 | TEC_MMP7 | ARID4B |
| 3.1119E-60 | -0.4330353 | 0.084 | 0.269 | 6.6438E-56 | TEC_MMP7 | IRS2 |
| 5.6474E-20 | -0.4337356 | 0.229 | 0.341 | 1.2057E-15 | TEC_MMP7 | MT1E |

| | | | | | | |
|---|---|---|---|---|---|---|
| 3.1551E-28 | -0.4342085 | 0.17 | 0.289 | 6.7362E-24 | TEC_MMP7 | VMP1 |
| 1.1282E-39 | -0.4387879 | 0.182 | 0.337 | 2.4088E-35 | TEC_MMP7 | UGCG |
| 5.298E-62 | -0.439765 | 0.109 | 0.3 | 1.1311E-57 | TEC_MMP7 | ATRX |
| 5.4985E-39 | -0.4402601 | 0.074 | 0.209 | 1.1739E-34 | TEC_MMP7 | GADD45G |
| 7.8391E-83 | -0.4405352 | 0.054 | 0.273 | 1.6736E-78 | TEC_MMP7 | PCM1 |
| 1.0008E-76 | -0.4415217 | 0.041 | 0.244 | 2.1368E-72 | TEC_MMP7 | RERE |
| 1.7532E-68 | -0.4418662 | 0.133 | 0.34 | 3.7432E-64 | TEC_MMP7 | SF3B1 |
| 3.2365E-48 | -0.4465662 | 0.257 | 0.446 | 6.9098E-44 | TEC_MMP7 | TUBB4B |
| 3.5454E-42 | -0.44707 | 0.072 | 0.21 | 7.5694E-38 | TEC_MMP7 | H2AFX |
| 6.1599E-20 | -0.4481035 | 0.157 | 0.259 | 1.3151E-15 | TEC_MMP7 | MT1X |
| 2.1103E-86 | -0.4483756 | 0.045 | 0.267 | 4.5056E-82 | TEC_MMP7 | NSD3 |
| 3.7463E-31 | -0.4489483 | 0.111 | 0.234 | 7.9983E-27 | TEC_MMP7 | TNFAIP2 |
| 2.3534E-48 | -0.449678 | 0.108 | 0.27 | 5.0246E-44 | TEC_MMP7 | IDI1 |
| 2.0826E-78 | -0.4501498 | 0.067 | 0.283 | 4.4463E-74 | TEC_MMP7 | MPHOSPH8 |
| 3.0032E-58 | -0.4507501 | 0.04 | 0.203 | 6.4119E-54 | TEC_MMP7 | ERN1 |
| 7.4221E-80 | -0.45165 | 0.076 | 0.297 | 1.5846E-75 | TEC_MMP7 | TPR |
| 1.0369E-63 | -0.4529081 | 0.02 | 0.186 | 2.2137E-59 | TEC_MMP7 | HIST1H1E |
| 4.1934E-35 | -0.4534906 | 0.091 | 0.223 | 8.9529E-31 | TEC_MMP7 | RRAD |
| 1.6525E-71 | -0.4549346 | 0.09 | 0.3 | 3.5281E-67 | TEC_MMP7 | TAF7 |
| 3.4446E-63 | -0.4554969 | 0.052 | 0.231 | 7.3542E-59 | TEC_MMP7 | ARID5B |
| 3.8896E-64 | -0.4565768 | 0.14 | 0.341 | 8.3042E-60 | TEC_MMP7 | STAT3 |
| 9.5793E-80 | -0.4565996 | 0.045 | 0.255 | 2.0452E-75 | TEC_MMP7 | TRA2B |
| 9.0234E-31 | -0.4566852 | 0.184 | 0.313 | 1.9265E-26 | TEC_MMP7 | HSPH1 |
| 1.8371E-54 | -0.4570914 | 0.012 | 0.153 | 3.9222E-50 | TEC_MMP7 | AC025164.1 |
| 8.1574E-85 | -0.4612828 | 0.055 | 0.28 | 1.7416E-80 | TEC_MMP7 | JMJD1C |
| 3.6943E-75 | -0.4613281 | 0.086 | 0.301 | 7.8874E-71 | TEC_MMP7 | CREBRF |
| 5.3584E-61 | -0.4644574 | 0.031 | 0.196 | 1.144E-56 | TEC_MMP7 | FOXC1 |
| 2.6742E-78 | -0.4648843 | 0.073 | 0.29 | 5.7095E-74 | TEC_MMP7 | CHD2 |
| 6.5763E-35 | -0.465261 | 0.519 | 0.707 | 1.404E-30 | TEC_MMP7 | HSPA5 |
| 1.1656E-58 | -0.4661285 | 0.077 | 0.256 | 2.4886E-54 | TEC_MMP7 | CDKN2AIP |
| 6.5938E-58 | -0.4674568 | 0.018 | 0.169 | 1.4078E-53 | TEC_MMP7 | AC007952.4 |
| 3.7742E-53 | -0.4706976 | 0.126 | 0.3 | 8.0578E-49 | TEC_MMP7 | COBLL1 |
| 2.613E-37 | -0.4707392 | 0.205 | 0.37 | 5.5787E-33 | TEC_MMP7 | TXNIP |
| 7.9325E-70 | -0.4716803 | 0.171 | 0.388 | 1.6936E-65 | TEC_MMP7 | EIF3A |
| 1.7941E-58 | -0.4718506 | 0.171 | 0.37 | 3.8304E-54 | TEC_MMP7 | TSC22D1 |
| 1.6848E-81 | -0.4766541 | 0.099 | 0.33 | 3.5971E-77 | TEC_MMP7 | GPBP1 |
| 1.7713E-71 | -0.4789258 | 0.203 | 0.43 | 3.7817E-67 | TEC_MMP7 | CDV3 |
| 3.0768E-80 | -0.4796797 | 0.081 | 0.304 | 6.5689E-76 | TEC_MMP7 | BPTF |
| 3.0892E-82 | -0.4803905 | 0.207 | 0.464 | 6.5955E-78 | TEC_MMP7 | ZFAND5 |
| 8.9853E-93 | -0.4814363 | 0.669 | 0.947 | 1.9184E-88 | TEC_MMP7 | UBC |
| 4.8186E-85 | -0.4825464 | 0.068 | 0.294 | 1.0288E-80 | TEC_MMP7 | ZRANB2 |
| 8.0812E-96 | -0.4858496 | 0.042 | 0.28 | 1.7253E-91 | TEC_MMP7 | KANSL1 |
| 5.1705E-85 | -0.4943778 | 0.045 | 0.263 | 1.1039E-80 | TEC_MMP7 | CHD9 |
| 9.0087E-91 | -0.4944153 | 0.066 | 0.303 | 1.9234E-86 | TEC_MMP7 | SMG1 |
| 2.3041E-73 | -0.4959124 | 0.04 | 0.233 | 4.9193E-69 | TEC_MMP7 | MAT2A |
| 9.7404E-31 | -0.4959483 | 0.176 | 0.301 | 2.0796E-26 | TEC_MMP7 | HIST1H1C |
| 7.4491E-26 | -0.4987867 | 0.24 | 0.376 | 1.5904E-21 | TEC_MMP7 | HSPA1A |
| 1.652E-24 | -0.4996093 | 0.789 | 0.904 | 3.5271E-20 | TEC_MMP7 | MT-ND2 |
| 1.0664E-91 | -0.5015693 | 0.088 | 0.334 | 2.2768E-87 | TEC_MMP7 | AFF4 |
| 1.9457E-49 | -0.5031776 | 0.203 | 0.393 | 4.1541E-45 | TEC_MMP7 | DDIT3 |

| | | | | | | |
|---|---|---|---|---|---|---|
| 7.4626E-63 | -0.5065099 | 0.087 | 0.277 | 1.5933E-58 | TEC_MMP7 | IRF1 |
| 2.2719E-78 | -0.5068217 | 0.098 | 0.316 | 4.8504E-74 | TEC_MMP7 | HP1BP3 |
| 6.3256E-87 | -0.5141361 | 0.339 | 0.588 | 1.3505E-82 | TEC_MMP7 | LRRFIP1 |
| 1.8442E-39 | -0.5237164 | 0.028 | 0.143 | 3.9375E-35 | TEC_MMP7 | COL7A1 |
| 2.1924E-89 | -0.5247906 | 0.093 | 0.334 | 4.6808E-85 | TEC_MMP7 | YTHDC1 |
| 4.1203E-39 | -0.5262146 | 0.494 | 0.631 | 8.7969E-35 | TEC_MMP7 | MT-ND4L |
| 1.8268E-66 | -0.5309674 | 0.46 | 0.684 | 3.9002E-62 | TEC_MMP7 | HSPA8 |
| 2.6093E-32 | -0.5316249 | 0.272 | 0.41 | 5.5708E-28 | TEC_MMP7 | TXNRD1 |
| 4.5907E-75 | -0.5317984 | 0.161 | 0.387 | 9.8012E-71 | TEC_MMP7 | SPAG9 |
| 4.0974E-52 | -0.5342984 | 0.152 | 0.335 | 8.7479E-48 | TEC_MMP7 | CDKN1C |
| 9.1913E-87 | -0.5372266 | 0.106 | 0.344 | 1.9623E-82 | TEC_MMP7 | AFDN |
| 5.0143E-67 | -0.5389407 | 0.2 | 0.419 | 1.0705E-62 | TEC_MMP7 | NDRG1 |
| 3.6774E-97 | -0.5446507 | 0.052 | 0.298 | 7.8512E-93 | TEC_MMP7 | BDP1 |
| 1.2313E-27 | -0.5473092 | 0.037 | 0.129 | 2.6288E-23 | TEC_MMP7 | NEFM |
| 1.3301E-87 | -0.547983 | 0.071 | 0.303 | 2.8397E-83 | TEC_MMP7 | IFRD1 |
| 4.176E-62 | -0.5496107 | 0.304 | 0.52 | 8.9158E-58 | TEC_MMP7 | ATP1B1 |
| 9.7079E-78 | -0.5503029 | 0.02 | 0.215 | 2.0726E-73 | TEC_MMP7 | TLE4 |
| 1.6037E-47 | -0.5537029 | 0.189 | 0.363 | 3.4239E-43 | TEC_MMP7 | SNHG12 |
| 4.0481E-56 | -0.5585178 | 0.2 | 0.399 | 8.6426E-52 | TEC_MMP7 | ATF3 |
| 2.3656E-32 | -0.5599679 | 0.732 | 0.843 | 5.0506E-28 | TEC_MMP7 | MT-ND5 |
| 3.0388E-84 | -0.5653375 | 0.15 | 0.389 | 6.4878E-80 | TEC_MMP7 | GCC2 |
| 1.0258E-67 | -0.565699 | 0.02 | 0.193 | 2.19E-63 | TEC_MMP7 | FOSB |
| 9.049E-106 | -0.5659252 | 0.387 | 0.66 | 1.932E-101 | TEC_MMP7 | ATF4 |
| 3.5803E-38 | -0.5686864 | 0.009 | 0.111 | 7.644E-34 | TEC_MMP7 | SLC12A1 |
| 1.7948E-59 | -0.5709151 | 0.156 | 0.357 | 3.832E-55 | TEC_MMP7 | H1F0 |
| 9.0244E-85 | -0.5713429 | 0.113 | 0.352 | 1.9267E-80 | TEC_MMP7 | RSRC2 |
| 1.3834E-75 | -0.5713511 | 0.117 | 0.334 | 2.9536E-71 | TEC_MMP7 | YME1L1 |
| 9.9615E-80 | -0.5741776 | 0.186 | 0.424 | 2.1268E-75 | TEC_MMP7 | SRSF3 |
| 2.1409E-50 | -0.5809775 | 0.118 | 0.285 | 4.5709E-46 | TEC_MMP7 | DUSP1 |
| 7.5305E-58 | -0.5831701 | 0.124 | 0.312 | 1.6078E-53 | TEC_MMP7 | CYR61 |
| 3.184E-61 | -0.5847887 | 0.101 | 0.285 | 6.7978E-57 | TEC_MMP7 | HIST1H2AC |
| 4.249E-100 | -0.5851468 | 0.065 | 0.317 | 9.0721E-96 | TEC_MMP7 | NUFIP2 |
| 2.704E-89 | -0.5873859 | 0.094 | 0.332 | 5.773E-85 | TEC_MMP7 | C6orf62 |
| 3.3919E-53 | -0.5918923 | 0.144 | 0.329 | 7.2416E-49 | TEC_MMP7 | HSPA1B |
| 3.566E-103 | -0.5942457 | 0.113 | 0.383 | 7.613E-99 | TEC_MMP7 | DDX24 |
| 6.7521E-54 | -0.5972458 | 0.145 | 0.325 | 1.4416E-49 | TEC_MMP7 | DNAJB1 |
| 5.402E-111 | -0.5976568 | 0.076 | 0.352 | 1.153E-106 | TEC_MMP7 | BOD1L1 |
| 1.502E-106 | -0.6026151 | 0.178 | 0.471 | 3.207E-102 | TEC_MMP7 | PNRC1 |
| 3.5361E-16 | -0.6051119 | 0.528 | 0.631 | 7.5496E-12 | TEC_MMP7 | MTRNR2L8 |
| 1.581E-107 | -0.6059556 | 0.061 | 0.324 | 3.375E-103 | TEC_MMP7 | KMT2A |
| 1.778E-104 | -0.6066401 | 0.096 | 0.368 | 3.795E-100 | TEC_MMP7 | ANKRD12 |
| 2.8721E-48 | -0.6067138 | 0.197 | 0.382 | 6.132E-44 | TEC_MMP7 | KLF6 |
| 5.1462E-89 | -0.6081877 | 0.173 | 0.424 | 1.0987E-84 | TEC_MMP7 | TAF1D |
| 5.03E-107 | -0.6100329 | 0.075 | 0.346 | 1.074E-102 | TEC_MMP7 | WSB1 |
| 2.171E-104 | -0.6110392 | 0.258 | 0.535 | 4.636E-100 | TEC_MMP7 | HNRNPDL |
| 9.2159E-67 | -0.6129805 | 0.081 | 0.277 | 1.9676E-62 | TEC_MMP7 | ID2 |
| 6.555E-76 | -0.6161921 | 0.23 | 0.451 | 1.3995E-71 | TEC_MMP7 | AHNAK |
| 3.7328E-61 | -0.6196249 | 0.321 | 0.538 | 7.9696E-57 | TEC_MMP7 | CDKN1A |
| 1.907E-107 | -0.6203845 | 0.315 | 0.611 | 4.071E-103 | TEC_MMP7 | SLC3A2 |
| 1.728E-100 | -0.6217167 | 0.336 | 0.593 | 3.6884E-96 | TEC_MMP7 | NCL |

| | | | | | | |
|---|---|---|---|---|---|---|
| 1.0705E-65 | -0.6282632 | 0.022 | 0.19 | 2.2855E-61 | TEC_MMP7 | EGLN3 |
| 4.9032E-55 | -0.6289238 | 0.149 | 0.328 | 1.0468E-50 | TEC_MMP7 | PMAIP1 |
| 3.5528E-90 | -0.6293243 | 0.061 | 0.294 | 7.5851E-86 | TEC_MMP7 | AMD1 |
| 3.343E-114 | -0.6311921 | 0.045 | 0.317 | 7.136E-110 | TEC_MMP7 | PRPF4B |
| 3.824E-115 | -0.6335108 | 0.366 | 0.644 | 8.164E-111 | TEC_MMP7 | EIF4A2 |
| 1.311E-100 | -0.6414521 | 0.133 | 0.401 | 2.7992E-96 | TEC_MMP7 | GOLGA4 |
| 8.5226E-87 | -0.6468134 | 0.159 | 0.398 | 1.8196E-82 | TEC_MMP7 | MACF1 |
| 1.4119E-86 | -0.6487229 | 0.042 | 0.261 | 3.0145E-82 | TEC_MMP7 | NFKBIZ |
| 1.9326E-86 | -0.6496376 | 0.108 | 0.342 | 4.1261E-82 | TEC_MMP7 | SRSF2 |
| 3.035E-102 | -0.6498989 | 0.173 | 0.448 | 6.4798E-98 | TEC_MMP7 | NORAD |
| 4.4007E-80 | -0.6534114 | 0.079 | 0.298 | 9.3955E-76 | TEC_MMP7 | SEPHS2 |
| 3.4598E-73 | -0.6628615 | 0.111 | 0.323 | 7.3867E-69 | TEC_MMP7 | DST |
| 3.9472E-54 | -0.6635479 | 0.133 | 0.311 | 8.4272E-50 | TEC_MMP7 | RND3 |
| 4.1944E-75 | -0.6673588 | 0.06 | 0.265 | 8.9551E-71 | TEC_MMP7 | MDM2 |
| 3.2988E-37 | -0.6678582 | 0.099 | 0.227 | 7.043E-33 | TEC_MMP7 | ADAMTS1 |
| 5.2846E-83 | -0.6709119 | 0.124 | 0.356 | 1.1283E-78 | TEC_MMP7 | CEBPB |
| 9.617E-121 | -0.6788099 | 0.079 | 0.37 | 2.053E-116 | TEC_MMP7 | CPEB4 |
| 2.497E-128 | -0.6853041 | 0.071 | 0.375 | 5.331E-124 | TEC_MMP7 | LUC7L3 |
| 4.7265E-81 | -0.6885117 | 0.026 | 0.228 | 1.0091E-76 | TEC_MMP7 | SGK1 |
| 7.283E-105 | -0.6885483 | 0.627 | 0.82 | 1.555E-100 | TEC_MMP7 | HNRNPA2B1 |
| 7.595E-104 | -0.6893789 | 0.053 | 0.309 | 1.622E-99 | TEC_MMP7 | VEGFA |
| 2.9472E-76 | -0.6895796 | 0.041 | 0.239 | 6.2923E-72 | TEC_MMP7 | SOCS1 |
| 1.951E-134 | -0.6975424 | 0.152 | 0.476 | 4.166E-130 | TEC_MMP7 | PRRC2C |
| 9.9592E-89 | -0.7058159 | 0.041 | 0.265 | 2.1263E-84 | TEC_MMP7 | SLC5A3 |
| 2.056E-126 | -0.7069015 | 0.106 | 0.416 | 4.39E-122 | TEC_MMP7 | SRSF5 |
| 2.7433E-71 | -0.7073087 | 0.079 | 0.278 | 5.857E-67 | TEC_MMP7 | IER5 |
| 3.7578E-77 | -0.7080879 | 0.018 | 0.209 | 8.0228E-73 | TEC_MMP7 | PIK3R3 |
| 7.014E-126 | -0.7100834 | 0.096 | 0.4 | 1.498E-121 | TEC_MMP7 | AKAP9 |
| 1.7145E-82 | -0.7110693 | 0.14 | 0.367 | 3.6604E-78 | TEC_MMP7 | PAX8 |
| 3.986E-108 | -0.7118332 | 0.119 | 0.394 | 8.509E-104 | TEC_MMP7 | MCL1 |
| 1.054E-104 | -0.7148737 | 0.02 | 0.264 | 2.249E-100 | TEC_MMP7 | SYNE1 |
| 3.608E-120 | -0.7206667 | 0.321 | 0.648 | 7.703E-116 | TEC_MMP7 | GADD45A |
| 8.658E-100 | -0.7218443 | 0.149 | 0.411 | 1.8485E-95 | TEC_MMP7 | RSRP1 |
| 1.1059E-79 | -0.7247044 | 0.085 | 0.303 | 2.3611E-75 | TEC_MMP7 | CITED2 |
| 1.3666E-35 | -0.7251604 | 0.283 | 0.424 | 2.9177E-31 | TEC_MMP7 | RNF213 |
| 4.555E-112 | -0.7279614 | 0.154 | 0.437 | 9.725E-108 | TEC_MMP7 | FUS |
| 1.9716E-81 | -0.7309849 | 0.09 | 0.314 | 4.2094E-77 | TEC_MMP7 | CEBPD |
| 4.8671E-33 | -0.7383114 | 0.147 | 0.277 | 1.0391E-28 | TEC_MMP7 | NUPR1 |
| 1.066E-113 | -0.7420687 | 0.108 | 0.392 | 2.275E-109 | TEC_MMP7 | DDX3X |
| 3.194E-131 | -0.7499067 | 0.094 | 0.406 | 6.819E-127 | TEC_MMP7 | PNN |
| 8.4975E-98 | -0.7532783 | 0.068 | 0.315 | 1.8142E-93 | TEC_MMP7 | POLR2A |
| 5.523E-125 | -0.7651876 | 0.064 | 0.358 | 1.179E-120 | TEC_MMP7 | CCNL1 |
| 2.036E-134 | -0.76555 | 0.131 | 0.454 | 4.347E-130 | TEC_MMP7 | GOLGB1 |
| 1.604E-120 | -0.7675455 | 0.168 | 0.471 | 3.424E-116 | TEC_MMP7 | ZBTB20 |
| 7.619E-147 | -0.7696301 | 0.09 | 0.425 | 1.627E-142 | TEC_MMP7 | N4BP2L2 |
| 1.5714E-80 | -0.7723655 | 0.068 | 0.282 | 3.355E-76 | TEC_MMP7 | IER5L |
| 1.736E-142 | -0.7776461 | 0.272 | 0.592 | 3.707E-138 | TEC_MMP7 | EIF5 |
| 1.75E-112 | -0.7801979 | 0.034 | 0.3 | 3.737E-108 | TEC_MMP7 | KCNQ1OT1 |
| 1.142E-151 | -0.7825333 | 0.081 | 0.42 | 2.438E-147 | TEC_MMP7 | PNISR |
| 2.536E-96 | -0.7828284 | 0.037 | 0.272 | 5.4143E-92 | TEC_MMP7 | ID4 |

| | | | | | | |
|---|---|---|---|---|---|---|
| 2.421E-116 | -0.7838247 | 0.083 | 0.371 | 5.169E-112 | TEC_MMP7 | ZNF292 |
| 6.5064E-92 | -0.7912978 | 0.124 | 0.374 | 1.3891E-87 | TEC_MMP7 | ZFP36L2 |
| 4.424E-137 | -0.7995187 | 0.081 | 0.398 | 9.445E-133 | TEC_MMP7 | ARGLU1 |
| 5.26E-106 | -0.8031996 | 0.194 | 0.48 | 1.123E-101 | TEC_MMP7 | ZFP36L1 |
| 5.704E-154 | -0.8094633 | 0.223 | 0.562 | 1.218E-149 | TEC_MMP7 | HNRNPU |
| 3.6071E-05 | -0.8141272 | 0.165 | 0.202 | 0.77011979 | TEC_MMP7 | MT1G |
| 2.907E-130 | -0.81469 | 0.124 | 0.437 | 6.206E-126 | TEC_MMP7 | ANKRD11 |
| 2.4882E-94 | -0.8161944 | 0.036 | 0.267 | 5.3124E-90 | TEC_MMP7 | INTS6 |
| 1.707E-144 | -0.8236695 | 0.133 | 0.463 | 3.645E-140 | TEC_MMP7 | SFPQ |
| 1.4461E-69 | -0.8392308 | 0.103 | 0.302 | 3.0873E-65 | TEC_MMP7 | RASD1 |
| 3.726E-104 | -0.8454497 | 0.056 | 0.313 | 7.955E-100 | TEC_MMP7 | BTG2 |
| 8.974E-208 | -0.8670078 | 0.234 | 0.644 | 1.916E-203 | TEC_MMP7 | RBM39 |
| 3.6848E-89 | -0.8696611 | 0.104 | 0.341 | 7.8671E-85 | TEC_MMP7 | GLUL |
| 1.198E-200 | -0.8953198 | 0.509 | 0.876 | 2.558E-196 | TEC_MMP7 | SQSTM1 |
| 2.641E-166 | -0.9117671 | 0.119 | 0.479 | 5.638E-162 | TEC_MMP7 | DDX17 |
| 9.4545E-89 | -0.9121018 | 0.059 | 0.289 | 2.0185E-84 | TEC_MMP7 | EGR1 |
| 2.562E-149 | -0.9501937 | 0.095 | 0.428 | 5.469E-145 | TEC_MMP7 | BRD2 |
| 2.499E-199 | -0.9514244 | 0.164 | 0.569 | 5.335E-195 | TEC_MMP7 | SON |
| 1.1509E-86 | -0.9642136 | 0.321 | 0.601 | 2.4572E-82 | TEC_MMP7 | GDF15 |
| 4.639E-193 | -0.9679858 | 0.259 | 0.661 | 9.904E-189 | TEC_MMP7 | BTG1 |
| 4.867E-249 | -0.9722253 | 0.325 | 0.762 | 1.039E-244 | TEC_MMP7 | DDX5 |
| 1.48E-113 | -0.9751808 | 0.065 | 0.335 | 3.16E-109 | TEC_MMP7 | PLK2 |
| 6.197E-91 | -0.9763589 | 0.12 | 0.362 | 1.3231E-86 | TEC_MMP7 | RHOB |
| 6.922E-123 | -1.0073642 | 0.032 | 0.312 | 1.478E-118 | TEC_MMP7 | PPP1R10 |
| 1.506E-128 | -1.0082881 | 0.163 | 0.466 | 3.215E-124 | TEC_MMP7 | SYNE2 |
| 5.306E-105 | -1.0146589 | 0.152 | 0.42 | 1.133E-100 | TEC_MMP7 | JUNB |
| 2.7363E-79 | -1.0152573 | 0.083 | 0.299 | 5.842E-75 | TEC_MMP7 | TCIM |
| 1.676E-135 | -1.021888 | 0.143 | 0.454 | 3.578E-131 | TEC_MMP7 | SRSF7 |
| 3.73E-99 | -1.0261747 | 0.161 | 0.424 | 7.9631E-95 | TEC_MMP7 | NFKBIA |
| 1.1909E-92 | -1.0267012 | 0.037 | 0.263 | 2.5425E-88 | TEC_MMP7 | CSKMT |
| 6.685E-224 | -1.0441611 | 0.198 | 0.614 | 1.427E-219 | TEC_MMP7 | SRRM2 |
| 1.7938E-95 | -1.0759287 | 0.068 | 0.308 | 3.8297E-91 | TEC_MMP7 | HES1 |
| 2.9464E-96 | -1.0813493 | 0.076 | 0.321 | 6.2906E-92 | TEC_MMP7 | FOS |
| 5.805E-156 | -1.1180785 | 0.077 | 0.418 | 1.239E-151 | TEC_MMP7 | SLC38A2 |
| 3.757E-215 | -1.1200501 | 0.265 | 0.675 | 8.021E-211 | TEC_MMP7 | PPP1R15A |
| 1.468E-139 | -1.1831437 | 0.312 | 0.626 | 3.134E-135 | TEC_MMP7 | ELF3 |
| 2.894E-247 | -1.2455937 | 0.489 | 0.912 | 6.179E-243 | TEC_MMP7 | H3F3B |
| 3.397E-114 | -1.2633097 | 0.039 | 0.304 | 7.252E-110 | TEC_MMP7 | HSPA2 |
| 1.978E-190 | -1.3016107 | 0.274 | 0.653 | 4.223E-186 | TEC_MMP7 | GADD45B |
| 1.961E-133 | -1.3122375 | 0.037 | 0.335 | 4.187E-129 | TEC_MMP7 | HEXIM1 |
| 7.308E-236 | -1.3385179 | 0.278 | 0.691 | 1.56E-231 | TEC_MMP7 | JUND |
| 9.408E-187 | -1.3702481 | 0.243 | 0.611 | 2.009E-182 | TEC_MMP7 | IER2 |
| 1.963E-145 | -1.466138 | 0.159 | 0.48 | 4.191E-141 | TEC_MMP7 | IER3 |
| 1.276E-128 | -1.5117533 | 0.033 | 0.319 | 2.725E-124 | TEC_MMP7 | MAFB |
| 1.335E-158 | -1.5308475 | 0.08 | 0.42 | 2.851E-154 | TEC_MMP7 | HIST1H4C |
| 2.8426E-88 | -1.5797032 | 0.529 | 0.7 | 6.069E-84 | TEC_MMP7 | MTRNR2L12 |
| 9.139E-159 | -1.9029013 | 0.259 | 0.58 | 1.951E-154 | TEC_MMP7 | SOX4 |
| 4.611E-255 | -1.907152 | 0.261 | 0.688 | 9.845E-251 | TEC_MMP7 | JUN |
| 0 | -2.2857571 | 0.357 | 0.881 | 0 | TEC_MMP7 | NEAT1 |
| 0 | -2.6868513 | 0.697 | 0.976 | 0 | TEC_MMP7 | MALAT1 |

| | | | | | | |
|---|---|---|---|---|---|---|
| 1.713E-250 | -5.3514643 | 0.257 | 0.647 | 3.657E-246 | TEC_MMP7 | PLCG2 |
| 3.053E-95 | 2.54097032 | 0.955 | 0.32 | 6.5182E-91 | TEC_MT | MT1E |
| 2.5788E-71 | 2.39296561 | 1 | 0.708 | 5.5057E-67 | TEC_MT | MT2A |
| 1.704E-132 | 2.29093003 | 0.948 | 0.19 | 3.638E-128 | TEC_MT | MT1G |
| 2.6278E-96 | 2.24177992 | 0.896 | 0.24 | 5.6105E-92 | TEC_MT | MT1X |
| 1.308E-82 | 1.81200072 | 0.537 | 0.081 | 2.7926E-78 | TEC_MT | MT1H |
| 2.5827E-37 | 1.79715972 | 0.634 | 0.221 | 5.514E-33 | TEC_MT | DEFB1 |
| 3.7495E-17 | 1.5534179 | 0.731 | 0.489 | 8.0053E-13 | TEC_MT | S100A2 |
| 5.5991E-27 | 1.34062447 | 0.821 | 0.439 | 1.1954E-22 | TEC_MT | UPP1 |
| 1.6857E-75 | 1.3204011 | 0.619 | 0.116 | 3.5989E-71 | TEC_MT | MT1F |
| 1.503E-110 | 1.15489157 | 0.507 | 0.056 | 3.209E-106 | TEC_MT | MT1M |
| 8.7847E-26 | 1.04905133 | 0.657 | 0.291 | 1.8755E-21 | TEC_MT | MIR4435-2HG |
| 1.2311E-09 | 1.02233601 | 0.396 | 0.198 | 2.6284E-05 | TEC_MT | ANXA1 |
| 2.0679E-29 | 0.83254481 | 0.44 | 0.128 | 4.4149E-25 | TEC_MT | AL353751.1 |
| 2.6882E-28 | 0.83035863 | 0.716 | 0.293 | 5.7392E-24 | TEC_MT | CYTOR |
| 4.2547E-05 | 0.82512417 | 0.716 | 0.667 | 0.90837942 | TEC_MT | CRYAB |
| 9.3436E-31 | 0.75973887 | 0.403 | 0.101 | 1.9949E-26 | TEC_MT | RSAD2 |
| 1.9169E-13 | 0.74062924 | 0.701 | 0.417 | 4.0925E-09 | TEC_MT | LGALS3 |
| 3.3997E-20 | 0.73651792 | 0.851 | 0.543 | 7.2584E-16 | TEC_MT | BRI3 |
| 2.4592E-38 | 0.69648326 | 0.493 | 0.126 | 5.2503E-34 | TEC_MT | SLC30A1 |
| 1.5705E-15 | 0.69538725 | 0.336 | 0.12 | 3.3529E-11 | TEC_MT | IFIT3 |
| 4.5889E-09 | 0.63772707 | 0.873 | 0.7 | 9.7973E-05 | TEC_MT | CLDN4 |
| 1.09E-08 | 0.59237087 | 0.813 | 0.643 | 0.00023271 | TEC_MT | WFDC2 |
| 6.4917E-05 | 0.58667948 | 0.425 | 0.284 | 1 | TEC_MT | HIST1H1C |
| 4.9227E-09 | 0.58222631 | 0.425 | 0.219 | 0.0001051 | TEC_MT | SFN |
| 4.168E-10 | 0.57590791 | 0.582 | 0.329 | 8.8987E-06 | TEC_MT | H1F0 |
| 2.559E-16 | 0.57160082 | 0.881 | 0.584 | 5.4634E-12 | TEC_MT | S100A13 |
| 1.6363E-10 | 0.57006793 | 0.955 | 0.922 | 3.4935E-06 | TEC_MT | ACTB |
| 3.3314E-11 | 0.56124279 | 0.56 | 0.315 | 7.1125E-07 | TEC_MT | UGCG |
| 7.41E-13 | 0.55839211 | 0.657 | 0.378 | 1.582E-08 | TEC_MT | RHEB |
| 1.8782E-07 | 0.55396336 | 0.313 | 0.151 | 0.00400991 | TEC_MT | RCAN1 |
| 8.881E-16 | 0.54211129 | 0.418 | 0.165 | 1.8961E-11 | TEC_MT | FGF2 |
| 1.8735E-06 | 0.53647285 | 0.746 | 0.59 | 0.03999826 | TEC_MT | KRT19 |
| 3.7425E-12 | 0.5310374 | 0.709 | 0.41 | 7.9903E-08 | TEC_MT | PHLDA2 |
| 4.2414E-05 | 0.52417883 | 0.739 | 0.604 | 0.90554739 | TEC_MT | SOD2 |
| 6.9112E-10 | 0.51399163 | 0.627 | 0.368 | 1.4755E-05 | TEC_MT | CAMK2N1 |
| 1.9516E-12 | 0.50920133 | 1 | 0.987 | 4.1667E-08 | TEC_MT | EEF1A1 |
| 5.3057E-08 | 0.49994585 | 0.522 | 0.305 | 0.00113277 | TEC_MT | PLAU |
| 2.536E-11 | 0.49836115 | 0.806 | 0.565 | 5.4144E-07 | TEC_MT | HNRNPC |
| 1.6255E-14 | 0.49140032 | 0.463 | 0.203 | 3.4704E-10 | TEC_MT | CSRP1 |
| 0.00899435 | 0.49063451 | 0.172 | 0.106 | 1 | TEC_MT | AQP3 |
| 8.3738E-20 | 0.48879969 | 0.313 | 0.09 | 1.7878E-15 | TEC_MT | CAV1 |
| 4.3879E-08 | 0.48552259 | 0.388 | 0.203 | 0.00093682 | TEC_MT | MDK |
| 1.0628E-10 | 0.48480249 | 0.993 | 0.961 | 2.2691E-06 | TEC_MT | FTL |
| 4.4023E-10 | 0.48328271 | 0.799 | 0.556 | 9.399E-06 | TEC_MT | HMGA1 |
| 3.3021E-07 | 0.48270887 | 0.209 | 0.086 | 0.00705001 | TEC_MT | SAA1 |
| 4.2856E-19 | 0.47952231 | 0.216 | 0.048 | 9.1498E-15 | TEC_MT | TAC1 |
| 2.0453E-14 | 0.47698655 | 1 | 0.957 | 4.3668E-10 | TEC_MT | EIF1 |
| 1.7644E-22 | 0.47247825 | 0.246 | 0.054 | 3.7669E-18 | TEC_MT | HRK |
| 3.4778E-13 | 0.46472154 | 0.843 | 0.547 | 7.4251E-09 | TEC_MT | CD9 |

| | | | | | | |
|---|---|---|---|---|---|---|
| 2.8209E-12 | 0.46140792 | 0.836 | 0.578 | 6.0226E-08 | TEC_MT | ATP6V0E1 |
| 3.2707E-10 | 0.46127439 | 0.881 | 0.698 | 6.983E-06 | TEC_MT | MAP1LC3B |
| 1.0556E-11 | 0.459257 | 0.97 | 0.838 | 2.2536E-07 | TEC_MT | CD63 |
| 7.864E-10 | 0.45884336 | 0.209 | 0.071 | 1.679E-05 | TEC_MT | SAA2 |
| 1.4659E-10 | 0.45794299 | 0.56 | 0.316 | 3.1298E-06 | TEC_MT | NDUFB9 |
| 1.8237E-11 | 0.45313826 | 0.97 | 0.864 | 3.8937E-07 | TEC_MT | RPS4X |
| 6.7821E-14 | 0.45163649 | 0.582 | 0.298 | 1.448E-09 | TEC_MT | RAP1B |
| 1.6611E-07 | 0.45129312 | 0.619 | 0.4 | 0.00354651 | TEC_MT | LAMTOR5 |
| 3.6168E-07 | 0.45017098 | 0.828 | 0.661 | 0.00772183 | TEC_MT | TACSTD2 |
| 2.4311E-06 | 0.44846338 | 0.575 | 0.367 | 0.05190375 | TEC_MT | DDIT3 |
| 2.8804E-09 | 0.44748394 | 0.888 | 0.725 | 6.1496E-05 | TEC_MT | CSTB |
| 4.984E-08 | 0.44632647 | 0.582 | 0.368 | 0.00106409 | TEC_MT | UBE2B |
| 3.8575E-07 | 0.44577636 | 0.567 | 0.371 | 0.00823581 | TEC_MT | ODC1 |
| 0.00383557 | 0.44443788 | 0.134 | 0.072 | 1 | TEC_MT | IFIT1 |
| 3.0229E-09 | 0.44382006 | 0.642 | 0.388 | 6.454E-05 | TEC_MT | RAB11FIP1 |
| 0.000243 | 0.44331996 | 0.179 | 0.092 | 1 | TEC_MT | SNAPC1 |
| 8.1186E-07 | 0.43798009 | 0.388 | 0.218 | 0.01733331 | TEC_MT | TIMP2 |
| 1.8747E-17 | 0.4314715 | 0.381 | 0.131 | 4.0025E-13 | TEC_MT | IFIT2 |
| 1.1137E-05 | 0.42936717 | 0.44 | 0.275 | 0.23778307 | TEC_MT | SMS |
| 3.529E-08 | 0.42448255 | 0.761 | 0.563 | 0.00075343 | TEC_MT | MAL2 |
| 1.1383E-09 | 0.42362083 | 0.634 | 0.382 | 2.4302E-05 | TEC_MT | OCIAD2 |
| 1.3334E-10 | 0.4175126 | 0.493 | 0.259 | 2.8468E-06 | TEC_MT | GSTO1 |
| 2.5324E-09 | 0.41600223 | 0.276 | 0.115 | 5.4068E-05 | TEC_MT | CRABP2 |
| 3.9278E-08 | 0.41335083 | 0.94 | 0.866 | 0.00083858 | TEC_MT | RPL23 |
| 6.0984E-09 | 0.41220688 | 0.978 | 0.873 | 0.0001302 | TEC_MT | RPL7 |
| 4.4315E-12 | 0.40796782 | 0.418 | 0.187 | 9.4612E-08 | TEC_MT | EMP1 |
| 2.0984E-12 | 0.40749594 | 0.575 | 0.297 | 4.48E-08 | TEC_MT | ACTR3 |
| 6.4848E-09 | 0.40648104 | 0.978 | 0.923 | 0.00013845 | TEC_MT | RPL10 |
| 3.1895E-11 | 0.40337907 | 0.955 | 0.757 | 6.8095E-07 | TEC_MT | CALM2 |
| 0.00018481 | 0.40333073 | 0.836 | 0.748 | 1 | TEC_MT | RTN4 |
| 1.0308E-13 | 0.40293491 | 0.485 | 0.221 | 2.2009E-09 | TEC_MT | CXCL16 |
| 5.609E-06 | 0.40189299 | 0.269 | 0.136 | 0.1197532 | TEC_MT | TM4SF1 |
| 5.9654E-08 | 0.39844679 | 0.47 | 0.266 | 0.00127362 | TEC_MT | BLVRB |
| 4.003E-18 | 0.3946815 | 0.321 | 0.098 | 8.5465E-14 | TEC_MT | CD68 |
| 5.6183E-09 | 0.39299059 | 0.582 | 0.343 | 0.00011995 | TEC_MT | CNBP |
| 1.9877E-07 | 0.39217545 | 0.993 | 0.938 | 0.00424377 | TEC_MT | RPL15 |
| 1.2351E-09 | 0.39195484 | 0.873 | 0.661 | 2.637E-05 | TEC_MT | RAC1 |
| 0.00037177 | 0.39160937 | 0.224 | 0.126 | 1 | TEC_MT | HMOX1 |
| 4.0143E-09 | 0.39110264 | 1 | 0.978 | 8.5705E-05 | TEC_MT | FTH1 |
| 0.00049875 | 0.39077589 | 0.649 | 0.599 | 1 | TEC_MT | SPP1 |
| 6.6019E-14 | 0.38948506 | 0.455 | 0.195 | 1.4095E-09 | TEC_MT | MAP2K3 |
| 0.00034375 | 0.38639579 | 0.552 | 0.392 | 1 | TEC_MT | SRSF3 |
| 1.4647E-10 | 0.38320283 | 0.299 | 0.118 | 3.1271E-06 | TEC_MT | NEURL3 |
| 1.7659E-08 | 0.38265414 | 0.545 | 0.332 | 0.00037703 | TEC_MT | PTP4A2 |
| 1.206E-11 | 0.38223754 | 0.56 | 0.288 | 2.5748E-07 | TEC_MT | MAFF |
| 1.974E-08 | 0.38155489 | 0.5 | 0.299 | 0.00042145 | TEC_MT | CUTA |
| 4.2866E-08 | 0.38101926 | 0.769 | 0.583 | 0.0009152 | TEC_MT | EZR |
| 1.3219E-09 | 0.3796296 | 0.776 | 0.501 | 2.8222E-05 | TEC_MT | CLDN7 |
| 2.0481E-09 | 0.37571717 | 0.306 | 0.135 | 4.3727E-05 | TEC_MT | RMND5A |
| 5.8688E-09 | 0.37122341 | 0.381 | 0.188 | 0.0001253 | TEC_MT | RAB9A |

| | | | | | | |
|---|---|---|---|---|---|---|
| 1.136E-09 | 0.37074762 | 0.246 | 0.091 | 2.4254E-05 | TEC_MT | NR1D1 |
| 1.0721E-08 | 0.37031904 | 0.552 | 0.316 | 0.00022889 | TEC_MT | FAM107B |
| 1.036E-09 | 0.36952214 | 0.582 | 0.336 | 2.2119E-05 | TEC_MT | NOP10 |
| 6.083E-09 | 0.36801745 | 0.47 | 0.246 | 0.00012987 | TEC_MT | MACC1 |
| 2.9464E-08 | 0.36363287 | 0.545 | 0.33 | 0.00062905 | TEC_MT | ATP6V1F |
| 3.5552E-05 | 0.36243082 | 0.351 | 0.211 | 0.75902472 | TEC_MT | GLRX |
| 2.4712E-09 | 0.36017387 | 0.59 | 0.337 | 5.2761E-05 | TEC_MT | EIF1AX |
| 1.6323E-09 | 0.35879574 | 0.373 | 0.18 | 3.4849E-05 | TEC_MT | BCL10 |
| 7.2137E-10 | 0.35306082 | 0.791 | 0.513 | 1.5401E-05 | TEC_MT | ARPC2 |
| 6.4224E-05 | 0.3524938 | 0.284 | 0.162 | 1 | TEC_MT | KATNBL1 |
| 1.9941E-08 | 0.35168993 | 0.657 | 0.404 | 0.00042575 | TEC_MT | SDC4 |
| 3.0999E-08 | 0.35167137 | 0.724 | 0.468 | 0.00066184 | TEC_MT | PRDX6 |
| 8.3652E-07 | 0.35087255 | 0.537 | 0.329 | 0.01785967 | TEC_MT | TUBA1C |
| 1.0062E-06 | 0.34901716 | 0.91 | 0.749 | 0.02148157 | TEC_MT | BTF3 |
| 2.335E-08 | 0.34613842 | 0.53 | 0.308 | 0.00049853 | TEC_MT | GABARAPL2 |
| 4.0955E-09 | 0.34487397 | 0.597 | 0.341 | 8.744E-05 | TEC_MT | S100A16 |
| 4.7326E-06 | 0.34367197 | 0.515 | 0.328 | 0.10104028 | TEC_MT | TIMP1 |
| 1.2627E-06 | 0.34330476 | 0.53 | 0.335 | 0.02695799 | TEC_MT | PSMB1 |
| 2.9378E-07 | 0.34296512 | 0.194 | 0.078 | 0.00627216 | TEC_MT | AGPAT5 |
| 1.8424E-06 | 0.34107271 | 0.41 | 0.239 | 0.03933555 | TEC_MT | HES4 |
| 6.2653E-17 | 0.33770278 | 0.134 | 0.023 | 1.3376E-12 | TEC_MT | CCK |
| 1.1972E-08 | 0.33678441 | 0.351 | 0.17 | 0.0002556 | TEC_MT | SH3BP4 |
| 7.7134E-08 | 0.33442818 | 0.985 | 0.926 | 0.00164682 | TEC_MT | ACTG1 |
| 1.5831E-08 | 0.33435327 | 0.59 | 0.368 | 0.000338 | TEC_MT | SDCBP |
| 0.00215708 | 0.33214758 | 0.187 | 0.108 | 1 | TEC_MT | S100A9 |
| 3.7879E-06 | 0.33137464 | 0.754 | 0.585 | 0.08087062 | TEC_MT | YWHAB |
| 5.6985E-06 | 0.33085316 | 0.843 | 0.722 | 0.12166314 | TEC_MT | CTSD |
| 1.1139E-05 | 0.33054907 | 0.978 | 0.908 | 0.2378181 | TEC_MT | RPL26 |
| 1.5903E-06 | 0.32861782 | 0.582 | 0.381 | 0.0339525 | TEC_MT | MRPL33 |
| 7.1192E-11 | 0.32835253 | 0.306 | 0.124 | 1.52E-06 | TEC_MT | GNPDA1 |
| 4.8162E-07 | 0.32802961 | 0.299 | 0.147 | 0.01028253 | TEC_MT | SNX10 |
| 6.1742E-07 | 0.3273437 | 0.455 | 0.257 | 0.01318201 | TEC_MT | LAMB3 |
| 5.0681E-07 | 0.32727729 | 0.836 | 0.679 | 0.01082047 | TEC_MT | CLIC1 |
| 2.7537E-05 | 0.32644479 | 0.687 | 0.523 | 0.58791581 | TEC_MT | CDC42 |
| 3.4283E-07 | 0.32614176 | 0.813 | 0.629 | 0.00731936 | TEC_MT | ANXA5 |
| 2.5506E-06 | 0.32576668 | 0.493 | 0.319 | 0.05445485 | TEC_MT | NPC2 |
| 5.2268E-09 | 0.32440401 | 0.261 | 0.108 | 0.00011159 | TEC_MT | FOSL1 |
| 1.1849E-06 | 0.32242808 | 0.993 | 0.929 | 0.02529712 | TEC_MT | RPS14 |
| 3.1265E-09 | 0.32151977 | 0.515 | 0.271 | 6.6751E-05 | TEC_MT | CREBRF |
| 4.1594E-08 | 0.3213667 | 1 | 0.904 | 0.00088804 | TEC_MT | RPS7 |
| 5.4775E-09 | 0.3204328 | 0.724 | 0.45 | 0.00011695 | TEC_MT | LAPTM4A |
| 8.4304E-41 | 0.31253692 | 0.231 | 0.029 | 1.7999E-36 | TEC_MT | SLC30A2 |
| 5.7095E-07 | 0.3120285 | 0.993 | 0.919 | 0.01218988 | TEC_MT | RPL9 |
| 1.0476E-08 | 0.31188185 | 0.582 | 0.338 | 0.00022367 | TEC_MT | JUP |
| 1.146E-09 | 0.31117017 | 0.634 | 0.356 | 2.4468E-05 | TEC_MT | KLF6 |
| 4.8023E-08 | 0.31111698 | 0.597 | 0.365 | 0.0010253 | TEC_MT | SBDS |
| 9.1773E-08 | 0.31079027 | 0.515 | 0.301 | 0.00195935 | TEC_MT | WASF2 |
| 0.00563308 | 0.31078004 | 0.224 | 0.137 | 1 | TEC_MT | ISG15 |
| 2.6378E-10 | 0.31012457 | 0.418 | 0.198 | 5.6317E-06 | TEC_MT | TMEM106B |
| 1.3755E-08 | 0.30915221 | 0.336 | 0.157 | 0.00029367 | TEC_MT | FHL2 |

| | | | | | | |
|---|---|---|---|---|---|---|
| 7.8234E-07 | 0.30913222 | 0.985 | 0.898 | 0.01670305 | TEC_MT | RPL5 |
| 3.059E-06 | 0.30885636 | 0.537 | 0.346 | 0.06530888 | TEC_MT | UBE2H |
| 2.5377E-08 | 0.30833869 | 0.403 | 0.207 | 0.0005418 | TEC_MT | GULP1 |
| 6.0443E-08 | 0.30697267 | 0.41 | 0.22 | 0.00129046 | TEC_MT | TMEM167A |
| 2.5167E-10 | 0.3056488 | 0.366 | 0.163 | 5.3732E-06 | TEC_MT | HSBP1L1 |
| 1.6589E-05 | 0.30382289 | 0.94 | 0.907 | 0.35416997 | TEC_MT | MYL6 |
| 1.5435E-07 | 0.30324047 | 0.194 | 0.075 | 0.00329534 | TEC_MT | NOV |
| 7.0502E-09 | 0.3029779 | 0.493 | 0.269 | 0.00015052 | TEC_MT | UBE2I |
| 8.2915E-06 | 0.30277389 | 0.687 | 0.489 | 0.17702345 | TEC_MT | TOMM7 |
| 6.9671E-07 | 0.30027025 | 0.993 | 0.937 | 0.01487468 | TEC_MT | RPS23 |
| 2.905E-08 | 0.29945216 | 0.485 | 0.263 | 0.00062022 | TEC_MT | RIOK3 |
| 0.00014653 | 0.29871616 | 0.358 | 0.226 | 1 | TEC_MT | PRSS8 |
| 3.2059E-09 | 0.29825748 | 0.388 | 0.186 | 6.8446E-05 | TEC_MT | BEX4 |
| 0.00426289 | 0.29657304 | 0.522 | 0.42 | 1 | TEC_MT | TPM4 |
| 5.852E-06 | 0.29626525 | 0.634 | 0.431 | 0.12493943 | TEC_MT | ZBTB20 |
| 9.0437E-07 | 0.29589463 | 0.813 | 0.602 | 0.01930829 | TEC_MT | CLTA |
| 0.00041293 | 0.29568865 | 0.112 | 0.047 | 1 | TEC_MT | KRT17 |
| 1.502E-08 | 0.29565506 | 0.284 | 0.124 | 0.00032068 | TEC_MT | IFNGR2 |
| 6.5736E-05 | 0.29415349 | 0.858 | 0.8 | 1 | TEC_MT | CD24 |
| 5.6002E-08 | 0.29406161 | 0.194 | 0.073 | 0.00119565 | TEC_MT | HIST3H2A |
| 1.7308E-07 | 0.29265327 | 0.784 | 0.557 | 0.00369521 | TEC_MT | RHOA |
| 2.4288E-12 | 0.29079271 | 0.157 | 0.038 | 5.1856E-08 | TEC_MT | S100A8 |
| 5.008E-05 | 0.29039312 | 0.47 | 0.307 | 1 | TEC_MT | DSP |
| 1.8944E-06 | 0.28930567 | 0.649 | 0.46 | 0.04044465 | TEC_MT | SERP1 |
| 2.1852E-10 | 0.28922522 | 0.284 | 0.11 | 4.6654E-06 | TEC_MT | SULT1C2 |
| 1.0395E-05 | 0.2874016 | 0.418 | 0.247 | 0.22192747 | TEC_MT | MGLL |
| 4.3058E-05 | 0.28655702 | 0.328 | 0.198 | 0.91928767 | TEC_MT | CHMP1B |
| 5.8613E-06 | 0.28604338 | 0.91 | 0.772 | 0.125139 | TEC_MT | MYL12A |
| 8.1921E-07 | 0.28593085 | 0.59 | 0.363 | 0.01749006 | TEC_MT | PPP1CB |
| 4.0545E-11 | 0.28555552 | 0.985 | 0.869 | 8.6564E-07 | TEC_MT | S100A10 |
| 9.7462E-06 | 0.28459045 | 0.47 | 0.292 | 0.20808059 | TEC_MT | CD55 |
| 0.00094972 | 0.28351452 | 0.187 | 0.1 | 1 | TEC_MT | APOE |
| 2.7583E-07 | 0.28350946 | 0.41 | 0.221 | 0.00588898 | TEC_MT | GPRC5A |
| 0.00939625 | 0.28293155 | 0.716 | 0.643 | 1 | TEC_MT | PEBP1 |
| 3.8622E-08 | 0.28283448 | 0.358 | 0.179 | 0.00082457 | TEC_MT | CHP1 |
| 8.2139E-07 | 0.28260693 | 0.41 | 0.232 | 0.01753678 | TEC_MT | TMBIM1 |
| 0.00011267 | 0.28169326 | 0.769 | 0.572 | 1 | TEC_MT | SLC3A2 |
| 6.4706E-06 | 0.2809785 | 0.403 | 0.236 | 0.13814646 | TEC_MT | HECTD1 |
| 8.2044E-06 | 0.28005987 | 0.978 | 0.909 | 0.17516436 | TEC_MT | RPL24 |
| 9.0275E-06 | 0.27895114 | 0.321 | 0.176 | 0.1927362 | TEC_MT | BCL2L1 |
| 2.0393E-05 | 0.2787276 | 0.754 | 0.579 | 0.43538472 | TEC_MT | ATP6V1G1 |
| 3.2181E-09 | 0.27774728 | 0.254 | 0.1 | 6.8707E-05 | TEC_MT | MAP3K8 |
| 7.1289E-06 | 0.27715876 | 0.642 | 0.43 | 0.15220264 | TEC_MT | BZW1 |
| 3.3023E-06 | 0.27672828 | 0.53 | 0.337 | 0.07050436 | TEC_MT | SEC11A |
| 2.8773E-11 | 0.27638268 | 0.351 | 0.146 | 6.143E-07 | TEC_MT | LAMTOR3 |
| 2.8884E-06 | 0.27588206 | 0.985 | 0.913 | 0.06166682 | TEC_MT | HSP90AB1 |
| 2.5764E-05 | 0.27543933 | 0.515 | 0.334 | 0.55006055 | TEC_MT | H2AFJ |
| 1.1132E-05 | 0.27500381 | 0.806 | 0.677 | 0.23766272 | TEC_MT | YWHAZ |
| 2.1106E-10 | 0.27491723 | 0.231 | 0.082 | 4.506E-06 | TEC_MT | DHRS3 |
| 0.00048186 | 0.27468137 | 0.321 | 0.204 | 1 | TEC_MT | OSER1 |

| | | | | | | |
|---|---|---|---|---|---|---|
| 0.0002433 | 0.2732617 | 0.993 | 0.938 | 1 | TEC_MT | RPS28 |
| 6.868E-09 | 0.27323002 | 0.269 | 0.11 | 0.00014663 | TEC_MT | ISG20 |
| 3.5034E-08 | 0.27214228 | 0.993 | 0.954 | 0.00074798 | TEC_MT | S100A6 |
| 0.00010635 | 0.27208985 | 0.403 | 0.251 | 1 | TEC_MT | AKAP12 |
| 1.6325E-05 | 0.27155786 | 0.948 | 0.787 | 0.34854783 | TEC_MT | NPM1 |
| 2.9191E-06 | 0.27126137 | 0.746 | 0.533 | 0.06232353 | TEC_MT | HLA-E |
| 6.2306E-05 | 0.27001035 | 0.791 | 0.602 | 1 | TEC_MT | PSMA7 |
| 2.0241E-20 | 0.2697276 | 0.157 | 0.026 | 4.3214E-16 | TEC_MT | MT1A |
| 0.00014833 | 0.26952883 | 0.366 | 0.232 | 1 | TEC_MT | RNF181 |
| 0.00029066 | 0.26923973 | 0.41 | 0.271 | 1 | TEC_MT | NDUFV2 |
| 1.0371E-06 | 0.26781591 | 1 | 0.933 | 0.02214157 | TEC_MT | RPS27A |
| 0.00019826 | 0.26760936 | 0.164 | 0.078 | 1 | TEC_MT | LY6K |
| 5.0982E-05 | 0.26748496 | 0.425 | 0.274 | 1 | TEC_MT | ZNF706 |
| 0.00018218 | 0.26630416 | 0.231 | 0.124 | 1 | TEC_MT | GPNMB |
| 4.0868E-10 | 0.26600344 | 0.418 | 0.196 | 8.7253E-06 | TEC_MT | WBP2 |
| 0.00092018 | 0.26581851 | 0.604 | 0.471 | 1 | TEC_MT | SUMO2 |
| 3.3281E-06 | 0.26491108 | 0.41 | 0.241 | 0.07105489 | TEC_MT | PFKP |
| 6.2597E-07 | 0.26474392 | 0.44 | 0.252 | 0.01336451 | TEC_MT | ATP6V1D |
| 3.6918E-41 | 0.26460635 | 0.179 | 0.018 | 7.8819E-37 | TEC_MT | LRRTM1 |
| 6.284E-09 | 0.26453317 | 0.284 | 0.121 | 0.00013416 | TEC_MT | TUFT1 |
| 3.0959E-06 | 0.26396743 | 0.276 | 0.137 | 0.06609704 | TEC_MT | C12orf49 |
| 2.366E-09 | 0.26387244 | 0.119 | 0.03 | 5.0514E-05 | TEC_MT | VGF |
| 5.7108E-09 | 0.2630895 | 0.276 | 0.113 | 0.00012193 | TEC_MT | HSPB8 |
| 0.00013271 | 0.26280001 | 0.239 | 0.132 | 1 | TEC_MT | ABHD11 |
| 4.6343E-05 | 0.26273752 | 0.396 | 0.252 | 0.98941541 | TEC_MT | COTL1 |
| 1.1535E-10 | 0.26230752 | 0.276 | 0.105 | 2.4628E-06 | TEC_MT | ABHD2 |
| 1.1059E-11 | 0.2620042 | 0.127 | 0.028 | 2.361E-07 | TEC_MT | LINC02154 |
| 7.3876E-06 | 0.26175624 | 0.291 | 0.154 | 0.15772536 | TEC_MT | TGIF1 |
| 0.00010885 | 0.26124089 | 0.47 | 0.311 | 1 | TEC_MT | PCBD1 |
| 0.0001181 | 0.26094489 | 0.642 | 0.466 | 1 | TEC_MT | LMNA |
| 8.7344E-05 | 0.26076633 | 0.403 | 0.26 | 1 | TEC_MT | VCL |
| 0.00329951 | 0.2600782 | 0.366 | 0.264 | 1 | TEC_MT | LIMA1 |
| 1.817E-07 | 0.2594665 | 0.47 | 0.266 | 0.00387929 | TEC_MT | PAIP2 |
| 5.6875E-10 | 0.2592438 | 0.201 | 0.067 | 1.2143E-05 | TEC_MT | TEX30 |
| 0.00111058 | 0.25905492 | 0.351 | 0.238 | 1 | TEC_MT | TCEAL9 |
| 2.0212E-06 | 0.25871329 | 0.97 | 0.933 | 0.04315269 | TEC_MT | RPL3 |
| 1.613E-06 | 0.25778133 | 0.418 | 0.24 | 0.03443846 | TEC_MT | RAB11A |
| 8.5333E-07 | 0.25737188 | 0.321 | 0.166 | 0.01821852 | TEC_MT | SMIM15 |
| 2.9988E-06 | 0.25568328 | 0.41 | 0.241 | 0.06402341 | TEC_MT | PLEKHB2 |
| 7.3784E-07 | 0.25523869 | 0.366 | 0.193 | 0.01575296 | TEC_MT | CCNG2 |
| 0.00057506 | 0.25500433 | 0.306 | 0.185 | 1 | TEC_MT | ADIRF |
| 2.3951E-09 | 0.25488168 | 0.164 | 0.05 | 5.1135E-05 | TEC_MT | BEX1 |
| 0.00011164 | 0.25477899 | 0.41 | 0.264 | 1 | TEC_MT | SLC38A1 |
| 5.6807E-08 | 0.25472821 | 0.321 | 0.152 | 0.00121283 | TEC_MT | FAM200B |
| 4.5106E-05 | 0.2546838 | 0.701 | 0.495 | 0.96300339 | TEC_MT | TNFRSF12A |
| 2.0052E-06 | 0.25414637 | 0.358 | 0.2 | 0.04281088 | TEC_MT | NDUFB5 |
| 9.4747E-05 | 0.25399187 | 0.537 | 0.38 | 1 | TEC_MT | DYNLT1 |
| 8.5532E-06 | 0.25346705 | 0.612 | 0.414 | 0.18261158 | TEC_MT | DYNLRB1 |
| 0.00036366 | 0.25251251 | 0.231 | 0.134 | 1 | TEC_MT | PPIF |
| 9.1797E-06 | 0.25168483 | 0.687 | 0.485 | 0.19598639 | TEC_MT | UBE2D3 |

| | | | | | | |
|---|---|---|---|---|---|---|
| 2.7245E-08 | 0.25044226 | 0.321 | 0.147 | 0.00058168 | TEC_MT | RHOF |
| 0.00459779 | -0.250226 | 0.119 | 0.213 | 1 | TEC_MT | KMT2C |
| 0.00545458 | -0.2507554 | 0.037 | 0.112 | 1 | TEC_MT | HIST4H4 |
| 0.00460411 | -0.2524271 | 0.09 | 0.18 | 1 | TEC_MT | MYO9A |
| 0.00068826 | -0.2592284 | 0.067 | 0.174 | 1 | TEC_MT | CCNL2 |
| 0.00573293 | -0.2619232 | 0.075 | 0.158 | 1 | TEC_MT | PLEKHH2 |
| 0.00171668 | -0.2621259 | 0.037 | 0.127 | 1 | TEC_MT | AL135925.1 |
| 0.00830098 | -0.265761 | 0.194 | 0.29 | 1 | TEC_MT | HP1BP3 |
| 0.00665163 | -0.2682168 | 0.179 | 0.277 | 1 | TEC_MT | ATRX |
| 0.00014894 | -0.2691749 | 0.045 | 0.167 | 1 | TEC_MT | PAXBP1 |
| 0.00089547 | -0.2801014 | 0.067 | 0.172 | 1 | TEC_MT | AHI1 |
| 0.00012298 | -0.2863995 | 0 | 0.1 | 1 | TEC_MT | MAFA |
| 0.00519066 | -0.2868995 | 0.119 | 0.208 | 1 | TEC_MT | VPS13C |
| 0.0010772 | -0.2918926 | 0.067 | 0.169 | 1 | TEC_MT | TMX4 |
| 0.00825124 | -0.2954929 | 0.075 | 0.153 | 1 | TEC_MT | PFKFB3 |
| 0.00176256 | -0.2969039 | 0.045 | 0.134 | 1 | TEC_MT | CSRNP2 |
| 0.00108478 | -0.3002425 | 0.157 | 0.277 | 1 | TEC_MT | BPTF |
| 0.00572853 | -0.3015927 | 0.746 | 0.796 | 1 | TEC_MT | HNRNPA2B1 |
| 0.00046611 | -0.3024814 | 0.015 | 0.108 | 1 | TEC_MT | AL355075.4 |
| 0.00044911 | -0.3044486 | 0.119 | 0.24 | 1 | TEC_MT | NSD3 |
| 0.00020151 | -0.3112607 | 0.09 | 0.219 | 1 | TEC_MT | RERE |
| 0.00155702 | -0.3145548 | 0.037 | 0.127 | 1 | TEC_MT | PCDH9 |
| 0.00392023 | -0.3186647 | 0.052 | 0.136 | 1 | TEC_MT | AC025164.1 |
| 0.00642023 | -0.3195735 | 0.052 | 0.127 | 1 | TEC_MT | HIST2H2BF |
| 0.00224714 | -0.3225184 | 0.052 | 0.145 | 1 | TEC_MT | ID1 |
| 0.0022682 | -0.3261468 | 0.254 | 0.369 | 1 | TEC_MT | GOLGA4 |
| 0.00041072 | -0.3365533 | 0.149 | 0.284 | 1 | TEC_MT | PRPF4B |
| 0.00385102 | -0.3376766 | 0.03 | 0.105 | 1 | TEC_MT | ARC |
| 0.00185089 | -0.3425952 | 0.209 | 0.317 | 1 | TEC_MT | STAT3 |
| 0.00117168 | -0.359587 | 0.104 | 0.21 | 1 | TEC_MT | MAT2A |
| 0.00878532 | -0.3702506 | 0.112 | 0.191 | 1 | TEC_MT | TLE4 |
| 0.00043894 | -0.3716741 | 0.045 | 0.151 | 1 | TEC_MT | AC007952.4 |
| 0.00496316 | -0.3719263 | 0.172 | 0.265 | 1 | TEC_MT | AMD1 |
| 0.00634679 | -0.3985913 | 0.142 | 0.234 | 1 | TEC_MT | SYNE1 |
| 0.00161231 | -0.4011964 | 0.321 | 0.435 | 1 | TEC_MT | DDX17 |
| 0.00087892 | -0.4040221 | 0.067 | 0.173 | 1 | TEC_MT | FOSB |
| 0.00131525 | -0.4122293 | 0.254 | 0.363 | 1 | TEC_MT | AKAP9 |
| 0.00108826 | -0.4252761 | 0.209 | 0.328 | 1 | TEC_MT | CEBPB |
| 3.4004E-05 | -0.43105 | 0.246 | 0.415 | 0.72599259 | TEC_MT | GOLGB1 |
| 6.3485E-05 | -0.4342422 | 0.067 | 0.204 | 1 | TEC_MT | SGK1 |
| 0.00224529 | -0.4374254 | 0.284 | 0.379 | 1 | TEC_MT | RSRP1 |
| 0.00360441 | -0.447008 | 0.187 | 0.285 | 1 | TEC_MT | POLR2A |
| 0.00010244 | -0.4480973 | 0.134 | 0.278 | 1 | TEC_MT | VEGFA |
| 0.00035984 | -0.4501267 | 0.112 | 0.235 | 1 | TEC_MT | NFKBIZ |
| 0.00131984 | -0.4517661 | 0.082 | 0.186 | 1 | TEC_MT | PIK3R3 |
| 0.00170052 | -0.454083 | 0.5 | 0.562 | 1 | TEC_MT | SRRM2 |
| 0.00255418 | -0.4561699 | 0.134 | 0.238 | 1 | TEC_MT | SLC5A3 |
| 0.0004984 | -0.4628325 | 0.03 | 0.129 | 1 | TEC_MT | COL7A1 |
| 0.00362726 | -0.4664592 | 0.179 | 0.276 | 1 | TEC_MT | CITED2 |
| 0.00153902 | -0.4689142 | 0.157 | 0.268 | 1 | TEC_MT | KCNQ1OT1 |

| | | | | | | |
|---|---|---|---|---|---|---|
| 2.1659E-05 | -0.4833575 | 0.179 | 0.339 | 0.46241825 | TEC_MT | HNRNPAB |
| 0.00140782 | -0.4922634 | 0.276 | 0.392 | 1 | TEC_MT | TUBA1B |
| 9.5565E-05 | -0.4970321 | 0.045 | 0.17 | 1 | TEC_MT | EGLN3 |
| 1.6834E-05 | -0.50539 | 0.545 | 0.67 | 0.35939937 | TEC_MT | TPI1 |
| 0.00687524 | -0.5098389 | 0.045 | 0.118 | 1 | TEC_MT | NEFM |
| 0.0024145 | -0.5315152 | 0.157 | 0.254 | 1 | TEC_MT | IER5 |
| 0.0006921 | -0.5355108 | 0.187 | 0.297 | 1 | TEC_MT | DST |
| 3.3349E-05 | -0.555595 | 0.231 | 0.387 | 0.71199306 | TEC_MT | BRD2 |
| 0.00060821 | -0.5898012 | 0.985 | 0.94 | 1 | TEC_MT | MALAT1 |
| 0.00016062 | -0.5989685 | 0.127 | 0.256 | 1 | TEC_MT | IER5L |
| 0.00272401 | -0.600354 | 0.179 | 0.278 | 1 | TEC_MT | RASD1 |
| 4.7886E-05 | -0.6174006 | 0.097 | 0.239 | 1 | TEC_MT | INTS6 |
| 0.00469899 | -0.6457107 | 0.187 | 0.278 | 1 | TEC_MT | HES1 |
| 0.00014233 | -0.6478333 | 0.522 | 0.64 | 1 | TEC_MT | JUND |
| 0.00380853 | -0.7085895 | 0.179 | 0.272 | 1 | TEC_MT | TCIM |
| 2.8948E-06 | -0.7340144 | 0.052 | 0.213 | 0.0618034 | TEC_MT | ADAMTS1 |
| 1.123E-05 | -0.7413591 | 0.157 | 0.313 | 0.23976485 | TEC_MT | GLUL |
| 4.686E-06 | -0.7444457 | 0.112 | 0.278 | 0.10004578 | TEC_MT | PPP1R10 |
| 1.2401E-05 | -0.7576525 | 0.149 | 0.303 | 0.26476523 | TEC_MT | PLK2 |
| 0.00384241 | -0.7834426 | 0.56 | 0.587 | 1 | TEC_MT | ELF3 |
| 8.2569E-06 | -0.8035146 | 0.224 | 0.376 | 0.17628443 | TEC_MT | SLC38A2 |
| 0.00013484 | -0.8111455 | 0.157 | 0.291 | 1 | TEC_MT | FOS |
| 4.104E-05 | -0.8150959 | 0.194 | 0.333 | 0.87621172 | TEC_MT | RHOB |
| 1.6618E-06 | -0.8361588 | 0.276 | 0.43 | 0.03548029 | TEC_MT | SYNE2 |
| 8.1301E-05 | -0.8611342 | 0.537 | 0.634 | 1 | TEC_MT | JUN |
| 0.00133851 | -0.8807969 | 0.791 | 0.815 | 1 | TEC_MT | NEAT1 |
| 1.5466E-06 | -0.8829979 | 0.067 | 0.236 | 0.03302074 | TEC_MT | CSKMT |
| 0.00552823 | -0.9137804 | 0.373 | 0.44 | 1 | TEC_MT | IER3 |
| 0.00604361 | -0.9422808 | 0.634 | 0.679 | 1 | TEC_MT | MTRNR2L12 |
| 3.3156E-07 | -1.1090034 | 0.09 | 0.272 | 0.00707872 | TEC_MT | HSPA2 |
| 1.6104E-08 | -1.149229 | 0.09 | 0.299 | 0.00034383 | TEC_MT | HEXIM1 |
| 7.4532E-12 | -1.4234473 | 0.015 | 0.285 | 1.5913E-07 | TEC_MT | MAFB |
| 2.1405E-10 | -4.1748796 | 0.433 | 0.599 | 4.5699E-06 | TEC_MT | PLCG2 |
| 0 | 2.64695477 | 0.998 | 0.52 | 0 | TEC_prg1 | PLCG2 |
| 0 | 1.79240421 | 0.953 | 0.625 | 0 | TEC_prg1 | MTRNR2L12 |
| 0 | 1.28177477 | 0.904 | 0.469 | 0 | TEC_prg1 | SOX4 |
| 2.8864E-81 | 1.25728293 | 0.526 | 0.383 | 6.1625E-77 | TEC_prg1 | RNF213 |
| 1.231E-166 | 0.98100615 | 0.623 | 0.39 | 2.628E-162 | TEC_prg1 | SYNE2 |
| 5.5384E-81 | 0.9490548 | 0.776 | 0.587 | 1.1824E-76 | TEC_prg1 | MTRNR2L8 |
| 1.792E-227 | 0.89899954 | 0.77 | 0.521 | 3.825E-223 | TEC_prg1 | SRRM2 |
| 8.235E-193 | 0.89782981 | 0.569 | 0.317 | 1.758E-188 | TEC_prg1 | ARGLU1 |
| 7.501E-242 | 0.8649049 | 0.659 | 0.322 | 1.602E-237 | TEC_prg1 | HIST1H4C |
| 1.905E-200 | 0.85795435 | 0.322 | 0.09 | 4.066E-196 | TEC_prg1 | COL7A1 |
| 1.681E-125 | 0.8511833 | 0.968 | 0.852 | 3.59E-121 | TEC_prg1 | SAT1 |
| 5.681E-273 | 0.84358179 | 0.999 | 0.929 | 1.213E-268 | TEC_prg1 | MALAT1 |
| 7.506E-183 | 0.82515252 | 0.511 | 0.225 | 1.603E-178 | TEC_prg1 | TCIM |
| 6.672E-268 | 0.82180672 | 0.884 | 0.529 | 1.425E-263 | TEC_prg1 | ELF3 |
| 4.408E-152 | 0.79669914 | 0.606 | 0.4 | 9.412E-148 | TEC_prg1 | DDX17 |
| 8.522E-115 | 0.79318514 | 0.499 | 0.307 | 1.819E-110 | TEC_prg1 | PAX8 |
| 9.352E-118 | 0.76761018 | 0.962 | 0.803 | 1.997E-113 | TEC_prg1 | MT-ND5 |

| | | | | | | |
|---|---|---|---|---|---|---|
| 2.1753E-92 | 0.74300465 | 0.492 | 0.344 | 4.6442E-88 | TEC_prg1 | MACF1 |
| 7.116E-123 | 0.7209089 | 0.557 | 0.339 | 1.519E-118 | TEC_prg1 | SLC38A2 |
| 1.339E-112 | 0.69248532 | 0.814 | 0.595 | 2.858E-108 | TEC_prg1 | TPM1 |
| 8.137E-243 | 0.62764598 | 0.999 | 0.779 | 1.737E-238 | TEC_prg1 | NEAT1 |
| 2.254E-106 | 0.62569761 | 0.428 | 0.243 | 4.812E-102 | TEC_prg1 | TCF25 |
| 8.6622E-56 | 0.60685767 | 0.291 | 0.168 | 1.8494E-51 | TEC_prg1 | SYTL2 |
| 2.93E-144 | 0.60423648 | 0.207 | 0.05 | 6.254E-140 | TEC_prg1 | PGGHG |
| 2.5438E-63 | 0.60157201 | 0.34 | 0.195 | 5.4311E-59 | TEC_prg1 | TNFAIP2 |
| 5.977E-98 | 0.5948788 | 0.435 | 0.253 | 1.2761E-93 | TEC_prg1 | PRPF4B |
| 4.027E-106 | 0.58132721 | 0.302 | 0.129 | 8.599E-102 | TEC_prg1 | PLEKHH2 |
| 4.3229E-81 | 0.5804152 | 0.374 | 0.205 | 9.2294E-77 | TEC_prg1 | SYNE1 |
| 2.634E-193 | 0.56725369 | 0.571 | 0.244 | 5.624E-189 | TEC_prg1 | HEXIM1 |
| 3.6315E-58 | 0.56514285 | 0.475 | 0.365 | 7.7533E-54 | TEC_prg1 | N4BP2L2 |
| 4.9403E-61 | 0.56005622 | 0.467 | 0.347 | 1.0548E-56 | TEC_prg1 | PNN |
| 6.753E-144 | 0.5581689 | 0.33 | 0.124 | 1.442E-139 | TEC_prg1 | RNPC3 |
| 4.221E-136 | 0.55365032 | 0.491 | 0.236 | 9.012E-132 | TEC_prg1 | HES1 |
| 3.4649E-66 | 0.54972104 | 0.625 | 0.497 | 7.3975E-62 | TEC_prg1 | SON |
| 4.589E-186 | 0.54832696 | 0.551 | 0.231 | 9.798E-182 | TEC_prg1 | MAFB |
| 4.069E-53 | 0.54750032 | 0.452 | 0.344 | 8.6873E-49 | TEC_prg1 | AKAP9 |
| 1.0411E-97 | 0.5349837 | 0.367 | 0.189 | 2.2227E-93 | TEC_prg1 | RERE |
| 6.103E-183 | 0.53406282 | 0.484 | 0.186 | 1.303E-178 | TEC_prg1 | CSKMT |
| 6.2888E-74 | 0.53367582 | 0.399 | 0.235 | 1.3427E-69 | TEC_prg1 | HIST1H2AC |
| 1.0541E-56 | 0.53147802 | 0.437 | 0.317 | 2.2504E-52 | TEC_prg1 | LUC7L3 |
| 8.0383E-67 | 0.51898292 | 0.495 | 0.355 | 1.7162E-62 | TEC_prg1 | RSRP1 |
| 2.0381E-68 | 0.51163517 | 0.983 | 0.845 | 4.3513E-64 | TEC_prg1 | MT-ND3 |
| 2.4564E-76 | 0.50806152 | 0.367 | 0.21 | 5.2445E-72 | TEC_prg1 | CHD9 |
| 4.944E-75 | 0.5043971 | 0.334 | 0.183 | 1.0555E-70 | TEC_prg1 | VPS13C |
| 5.2953E-61 | 0.50228402 | 0.744 | 0.588 | 1.1305E-56 | TEC_prg1 | MT-ND4L |
| 6.55E-144 | 0.50009185 | 0.291 | 0.096 | 1.398E-139 | TEC_prg1 | PKHD1 |
| 1.8092E-45 | 0.49990393 | 0.458 | 0.361 | 3.8626E-41 | TEC_prg1 | PNISR |
| 7.0425E-37 | 0.49857043 | 0.359 | 0.269 | 1.5036E-32 | TEC_prg1 | COL18A1 |
| 2.492E-104 | 0.49337773 | 0.92 | 0.772 | 5.32E-100 | TEC_prg1 | HNRNPA2B1 |
| 2.87E-109 | 0.48702769 | 0.469 | 0.24 | 6.128E-105 | TEC_prg1 | RASD1 |
| 2.0348E-53 | 0.48352439 | 0.322 | 0.198 | 4.3442E-49 | TEC_prg1 | EPS8L2 |
| 1.2815E-53 | 0.48133929 | 0.394 | 0.271 | 2.7359E-49 | TEC_prg1 | KMT2A |
| 3.6491E-36 | 0.4804513 | 0.35 | 0.262 | 7.7908E-32 | TEC_prg1 | ATRX |
| 1.5445E-83 | 0.48036424 | 0.464 | 0.282 | 3.2974E-79 | TEC_prg1 | GLUL |
| 2.5777E-82 | 0.4708424 | 0.301 | 0.146 | 5.5033E-78 | TEC_prg1 | AHI1 |
| 9.847E-165 | 0.4702267 | 0.514 | 0.223 | 2.102E-160 | TEC_prg1 | HSPA2 |
| 5.6736E-66 | 0.46788744 | 0.353 | 0.21 | 1.2113E-61 | TEC_prg1 | PKN2 |
| 8.0619E-94 | 0.46654106 | 0.281 | 0.121 | 1.7212E-89 | TEC_prg1 | SLC25A29 |
| 5.2828E-52 | 0.46310309 | 0.371 | 0.246 | 1.1279E-47 | TEC_prg1 | BDP1 |
| 4.7291E-37 | 0.46112865 | 0.388 | 0.3 | 1.0097E-32 | TEC_prg1 | AFDN |
| 2.6E-99 | 0.45934534 | 0.411 | 0.204 | 5.5504E-95 | TEC_prg1 | INTS6 |
| 6.2665E-31 | 0.45651226 | 0.477 | 0.427 | 1.3379E-26 | TEC_prg1 | PRRC2C |
| 3.1085E-47 | 0.45159173 | 0.99 | 0.898 | 6.6366E-43 | TEC_prg1 | MT-ND4 |
| 5.6617E-65 | 0.44848127 | 0.308 | 0.168 | 1.2088E-60 | TEC_prg1 | TUT4 |
| 1.206E-108 | 0.44061803 | 0.279 | 0.108 | 2.575E-104 | TEC_prg1 | HERC2 |
| 2.5191E-46 | 0.43786976 | 0.993 | 0.893 | 5.3782E-42 | TEC_prg1 | MT-ATP6 |
| 1.878E-132 | 0.43583913 | 0.845 | 0.599 | 4.009E-128 | TEC_prg1 | JUND |

| | | | | | | |
|---|---|---|---|---|---|---|
| 1.9524E-60 | 0.43447114 | 0.29 | 0.157 | 4.1683E-56 | TEC_prg1 | MYO9A |
| 3.4932E-21 | 0.4304906 | 0.353 | 0.285 | 7.458E-17 | TEC_prg1 | DST |
| 1.6645E-83 | 0.4267499 | 0.334 | 0.162 | 3.5537E-79 | TEC_prg1 | TLE4 |
| 4.9584E-60 | 0.42442334 | 0.31 | 0.175 | 1.0586E-55 | TEC_prg1 | UPF2 |
| 4.2535E-44 | 0.42431803 | 0.989 | 0.881 | 9.0812E-40 | TEC_prg1 | MT-CYB |
| 7.1568E-52 | 0.42326839 | 0.305 | 0.183 | 1.528E-47 | TEC_prg1 | ZFC3H1 |
| 1.2624E-94 | 0.42154446 | 0.261 | 0.104 | 2.6952E-90 | TEC_prg1 | WDR60 |
| 3.523E-111 | 0.41479915 | 0.477 | 0.238 | 7.522E-107 | TEC_prg1 | PPP1R10 |
| 2.647E-41 | 0.41475661 | 0.365 | 0.258 | 5.6514E-37 | TEC_prg1 | BPTF |
| 4.2237E-76 | 0.4143956 | 0.178 | 0.062 | 9.0176E-72 | TEC_prg1 | AC019197.1 |
| 1.5072E-73 | 0.41245437 | 0.288 | 0.142 | 3.2179E-69 | TEC_prg1 | PAXBP1 |
| 2.8545E-59 | 0.41117851 | 0.301 | 0.171 | 6.0944E-55 | TEC_prg1 | ERICH1 |
| 8.3273E-34 | 0.41008546 | 0.549 | 0.49 | 1.7779E-29 | TEC_prg1 | HNRNPDL |
| 1.4126E-94 | 0.40777921 | 0.267 | 0.11 | 3.0158E-90 | TEC_prg1 | PLXNB1 |
| 2.0067E-67 | 0.40529568 | 0.371 | 0.207 | 4.2843E-63 | TEC_prg1 | NFKBIZ |
| 5.2687E-13 | 0.40420095 | 0.35 | 0.314 | 1.1249E-08 | TEC_prg1 | DCDC2 |
| 3.9495E-41 | 0.40356378 | 0.3 | 0.195 | 8.4322E-37 | TEC_prg1 | KMT2C |
| 1.224E-115 | 0.3988987 | 0.364 | 0.15 | 2.614E-111 | TEC_prg1 | PIK3R3 |
| 2.5619E-90 | 0.39716096 | 0.424 | 0.222 | 5.4697E-86 | TEC_prg1 | IER5L |
| 2.3797E-45 | 0.39702285 | 0.98 | 0.874 | 5.0807E-41 | TEC_prg1 | MT-ND1 |
| 3.3942E-46 | 0.39685489 | 0.269 | 0.153 | 7.2466E-42 | TEC_prg1 | THSD7A |
| 1.8127E-58 | 0.39484864 | 0.528 | 0.393 | 3.8701E-54 | TEC_prg1 | SRSF7 |
| 1.449E-91 | 0.39373333 | 0.366 | 0.171 | 3.0936E-87 | TEC_prg1 | SGK1 |
| 7.0277E-75 | 0.3902306 | 0.484 | 0.302 | 1.5004E-70 | TEC_prg1 | RHOB |
| 2.593E-66 | 0.3886354 | 0.225 | 0.1 | 5.536E-62 | TEC_prg1 | HIP1R |
| 6.576E-113 | 0.38835614 | 0.229 | 0.074 | 1.404E-108 | TEC_prg1 | GOLGA8A |
| 2.371E-32 | 0.38719146 | 0.353 | 0.25 | 5.0622E-28 | TEC_prg1 | TUBA1A |
| 2.9103E-10 | 0.38640045 | 0.25 | 0.211 | 6.2135E-06 | TEC_prg1 | PARP14 |
| 4.1732E-64 | 0.38499594 | 0.287 | 0.151 | 8.9098E-60 | TEC_prg1 | CCNL2 |
| 2.8976E-64 | 0.38473304 | 0.263 | 0.13 | 6.1865E-60 | TEC_prg1 | DAPK1 |
| 1.9105E-70 | 0.38085531 | 0.277 | 0.135 | 4.079E-66 | TEC_prg1 | HOOK2 |
| 1.9224E-29 | 0.37963274 | 0.325 | 0.241 | 4.1044E-25 | TEC_prg1 | DDX46 |
| 3.1875E-44 | 0.3781725 | 0.308 | 0.195 | 6.8052E-40 | TEC_prg1 | ACIN1 |
| 7.7194E-26 | 0.37579348 | 0.375 | 0.306 | 1.6481E-21 | TEC_prg1 | BOD1L1 |
| 1.1978E-39 | 0.37541863 | 0.589 | 0.505 | 2.5573E-35 | TEC_prg1 | HNRNPU |
| 4.5261E-52 | 0.37491476 | 0.25 | 0.134 | 9.6633E-48 | TEC_prg1 | ASCC3 |
| 4.7797E-31 | 0.37378077 | 0.339 | 0.252 | 1.0205E-26 | TEC_prg1 | ZRANB2 |
| 2.0445E-27 | 0.37284752 | 0.322 | 0.243 | 4.3651E-23 | TEC_prg1 | MPHOSPH8 |
| 1.0911E-52 | 0.37203181 | 0.427 | 0.277 | 2.3296E-48 | TEC_prg1 | PLK2 |
| 2.5129E-38 | 0.36954064 | 0.288 | 0.182 | 5.3651E-34 | TEC_prg1 | FAT1 |
| 8.0718E-21 | 0.3677455 | 0.316 | 0.253 | 1.7233E-16 | TEC_prg1 | PLXNB2 |
| 2.56E-94 | 0.36758228 | 0.232 | 0.084 | 5.4656E-90 | TEC_prg1 | KIF12 |
| 1.414E-215 | 0.36700208 | 0.917 | 0.578 | 3.019E-211 | TEC_prg1 | JUN |
| 3.1029E-37 | 0.36288849 | 0.323 | 0.223 | 6.6248E-33 | TEC_prg1 | NSD3 |
| 2.225E-122 | 0.36162258 | 0.78 | 0.522 | 4.751E-118 | TEC_prg1 | IER2 |
| 1.0613E-45 | 0.36160687 | 0.246 | 0.137 | 2.2658E-41 | TEC_prg1 | CHD3 |
| 1.4366E-69 | 0.35708084 | 0.355 | 0.186 | 3.0672E-65 | TEC_prg1 | SOCS1 |
| 5.2298E-56 | 0.3550095 | 0.265 | 0.141 | 1.1166E-51 | TEC_prg1 | RBM6 |
| 3.9831E-31 | 0.35395076 | 0.323 | 0.234 | 8.504E-27 | TEC_prg1 | TRIM44 |
| 8.5757E-54 | 0.35317484 | 0.236 | 0.12 | 1.8309E-49 | TEC_prg1 | SNX29 |

| | | | | | | |
|---|---|---|---|---|---|---|
| 1.5768E-27 | 0.35202062 | 0.295 | 0.212 | 3.3664E-23 | TEC_prg1 | ZNF91 |
| 1.3963E-33 | 0.3514592 | 0.433 | 0.325 | 2.9811E-29 | TEC_prg1 | ZFP36L2 |
| 7.7069E-80 | 0.34948256 | 0.285 | 0.124 | 1.6454E-75 | TEC_prg1 | AC007952.4 |
| 1.1504E-41 | 0.34919499 | 0.278 | 0.17 | 2.4562E-37 | TEC_prg1 | BAZ2B |
| 4.727E-75 | 0.34805078 | 0.219 | 0.085 | 1.0092E-70 | TEC_prg1 | AL355075.4 |
| 1.2797E-11 | 0.347563 | 0.374 | 0.356 | 2.7321E-07 | TEC_prg1 | GCC2 |
| 2.0624E-40 | 0.34642921 | 0.178 | 0.087 | 4.4033E-36 | TEC_prg1 | TNF |
| 9.7575E-12 | 0.34212164 | 0.278 | 0.229 | 2.0832E-07 | TEC_prg1 | SLC5A3 |
| 7.5035E-31 | 0.34100051 | 0.994 | 0.913 | 1.602E-26 | TEC_prg1 | MT-CO1 |
| 4.6427E-46 | 0.33894756 | 0.395 | 0.262 | 9.9121E-42 | TEC_prg1 | POLR2A |
| 1.4049E-88 | 0.33289426 | 0.214 | 0.077 | 2.9994E-84 | TEC_prg1 | GOLGA8B |
| 2.0252E-21 | 0.33120204 | 0.367 | 0.301 | 4.3239E-17 | TEC_prg1 | WSB1 |
| 9.6274E-26 | 0.32999167 | 0.143 | 0.075 | 2.0554E-21 | TEC_prg1 | CCDC80 |
| 2.473E-126 | 0.32950156 | 0.663 | 0.396 | 5.279E-122 | TEC_prg1 | IER3 |
| 1.7397E-21 | 0.32691027 | 0.301 | 0.234 | 3.7142E-17 | TEC_prg1 | PCM1 |
| 3.5565E-37 | 0.32291333 | 0.985 | 0.871 | 7.5931E-33 | TEC_prg1 | MT-ND2 |
| 1.0669E-05 | 0.31553823 | 0.392 | 0.399 | 0.22777624 | TEC_prg1 | ANKRD11 |
| 7.2223E-59 | 0.31320472 | 0.23 | 0.11 | 1.542E-54 | TEC_prg1 | GON4L |
| 3.6684E-71 | 0.31290364 | 0.256 | 0.111 | 7.832E-67 | TEC_prg1 | AC025164.1 |
| 5.2241E-32 | 0.31206436 | 0.147 | 0.072 | 1.1153E-27 | TEC_prg1 | ANKRD36C |
| 1.213E-127 | 0.31052589 | 0.135 | 0.022 | 2.589E-123 | TEC_prg1 | AC136475.3 |
| 7.0854E-30 | 0.3100332 | 0.274 | 0.183 | 1.5127E-25 | TEC_prg1 | CDK12 |
| 1.9075E-42 | 0.30964528 | 0.209 | 0.109 | 4.0725E-38 | TEC_prg1 | MICAL3 |
| 1.9368E-22 | 0.30661059 | 0.279 | 0.206 | 4.135E-18 | TEC_prg1 | USP34 |
| 1.1479E-30 | 0.30614434 | 0.358 | 0.26 | 2.4508E-26 | TEC_prg1 | VEGFA |
| 1.0998E-97 | 0.30529982 | 0.194 | 0.06 | 2.3481E-93 | TEC_prg1 | TRIM73 |
| 1.0126E-42 | 0.30468685 | 0.227 | 0.125 | 2.1619E-38 | TEC_prg1 | PHF14 |
| 3.2125E-23 | 0.3041068 | 0.313 | 0.237 | 6.8587E-19 | TEC_prg1 | KANSL1 |
| 1.5913E-46 | 0.30402525 | 0.267 | 0.145 | 3.3975E-42 | TEC_prg1 | HIST1H1E |
| 3.9991E-52 | 0.30378718 | 0.332 | 0.187 | 8.5382E-48 | TEC_prg1 | ADAMTS1 |
| 8.5189E-56 | 0.30359869 | 0.214 | 0.1 | 1.8188E-51 | TEC_prg1 | SHPRH |
| 3.9361E-14 | 0.3022829 | 0.308 | 0.262 | 8.4035E-10 | TEC_prg1 | TPR |
| 7.7089E-47 | 0.30199999 | 0.239 | 0.128 | 1.6458E-42 | TEC_prg1 | CREBZF |
| 3.9275E-34 | 0.30139991 | 0.224 | 0.132 | 8.3852E-30 | TEC_prg1 | EMX2 |
| 5.0076E-17 | 0.30091165 | 0.335 | 0.28 | 1.0691E-12 | TEC_prg1 | HP1BP3 |
| 1.2942E-18 | 0.29700235 | 0.267 | 0.204 | 2.7632E-14 | TEC_prg1 | PATJ |
| 8.0677E-29 | 0.29635875 | 0.244 | 0.16 | 1.7225E-24 | TEC_prg1 | NSD1 |
| 8.9184E-14 | 0.29605011 | 0.319 | 0.27 | 1.9041E-09 | TEC_prg1 | COBLL1 |
| 1.3477E-38 | 0.29551319 | 0.22 | 0.123 | 2.8773E-34 | TEC_prg1 | ZFYVE16 |
| 5.8451E-45 | 0.2953025 | 0.834 | 0.682 | 1.2479E-40 | TEC_prg1 | DDX5 |
| 5.6879E-72 | 0.29349101 | 0.304 | 0.142 | 1.2144E-67 | TEC_prg1 | EGLN3 |
| 1.2179E-52 | 0.29333238 | 0.212 | 0.102 | 2.6001E-48 | TEC_prg1 | SUGP2 |
| 6.084E-16 | 0.29239423 | 0.427 | 0.396 | 1.2989E-11 | TEC_prg1 | FUS |
| 4.3414E-18 | 0.29186011 | 0.241 | 0.18 | 9.2688E-14 | TEC_prg1 | RNF19A |
| 7.5343E-15 | 0.29084747 | 0.313 | 0.265 | 1.6086E-10 | TEC_prg1 | SMG1 |
| 1.3439E-53 | 0.28965142 | 0.208 | 0.098 | 2.8691E-49 | TEC_prg1 | HECTD4 |
| 5.3015E-28 | 0.28858494 | 0.369 | 0.27 | 1.1319E-23 | TEC_prg1 | CEBPD |
| 1.6362E-27 | 0.28690024 | 0.231 | 0.149 | 3.4934E-23 | TEC_prg1 | KIAA1109 |
| 2.0434E-69 | 0.28634246 | 0.222 | 0.09 | 4.3627E-65 | TEC_prg1 | HIST4H4 |
| 1.3622E-32 | 0.28525611 | 0.266 | 0.166 | 2.9083E-28 | TEC_prg1 | ERN1 |

| | | | | | | |
|---|---|---|---|---|---|---|
| 7.5332E-35 | 0.28336389 | 0.204 | 0.113 | 1.6083E-30 | TEC_prg1 | CCDC198 |
| 2.0516E-08 | 0.28257377 | 0.299 | 0.28 | 0.00043801 | TEC_prg1 | CDC37 |
| 2.1514E-06 | 0.28180087 | 0.412 | 0.413 | 0.04593298 | TEC_prg1 | GOLGB1 |
| 1.456E-06 | 0.28081391 | 0.36 | 0.358 | 0.03108494 | TEC_prg1 | SPAG9 |
| 1.2571E-30 | 0.2797392 | 0.233 | 0.145 | 2.6839E-26 | TEC_prg1 | NUMA1 |
| 2.3242E-58 | 0.27941368 | 0.183 | 0.076 | 4.9622E-54 | TEC_prg1 | ANKRD26 |
| 4.0555E-24 | 0.27930539 | 0.128 | 0.067 | 8.6584E-20 | TEC_prg1 | LHX1 |
| 6.1745E-34 | 0.27783537 | 0.188 | 0.102 | 1.3183E-29 | TEC_prg1 | PAX2 |
| 5.7814E-28 | 0.27466248 | 0.204 | 0.124 | 1.2343E-23 | TEC_prg1 | ITPR2 |
| 9.7965E-13 | 0.2740832 | 0.278 | 0.233 | 2.0915E-08 | TEC_prg1 | AAK1 |
| 1.392E-33 | 0.27257309 | 0.786 | 0.645 | 2.9719E-29 | TEC_prg1 | GLS |
| 2.7109E-07 | 0.27217438 | 0.396 | 0.392 | 0.00578775 | TEC_prg1 | TAF1D |
| 2.65E-11 | 0.27196241 | 0.442 | 0.417 | 5.6577E-07 | TEC_prg1 | SFPQ |
| 2.8862E-14 | 0.27076731 | 0.34 | 0.278 | 6.162E-10 | TEC_prg1 | RND3 |
| 4.4873E-12 | 0.26999186 | 0.288 | 0.244 | 9.5804E-08 | TEC_prg1 | JMJD1C |
| 1.1958E-47 | 0.26888815 | 0.193 | 0.092 | 2.553E-43 | TEC_prg1 | PPP6R2 |
| 4.6858E-27 | 0.26845111 | 0.663 | 0.578 | 1.0004E-22 | TEC_prg1 | RBM39 |
| 1.382E-35 | 0.2680328 | 0.219 | 0.125 | 2.9507E-31 | TEC_prg1 | RNF169 |
| 1.0894E-37 | 0.26591919 | 0.206 | 0.112 | 2.3258E-33 | TEC_prg1 | BRD1 |
| 3.5452E-67 | 0.26301057 | 0.161 | 0.054 | 7.5689E-63 | TEC_prg1 | HIST1H1A |
| 1.8184E-67 | 0.26215103 | 0.166 | 0.059 | 3.8822E-63 | TEC_prg1 | MAPK8IP3 |
| 7.0828E-48 | 0.25997483 | 0.213 | 0.103 | 1.5122E-43 | TEC_prg1 | LINC00472 |
| 2.4332E-29 | 0.25786715 | 0.196 | 0.115 | 5.195E-25 | TEC_prg1 | LAMA5 |
| 6.6722E-21 | 0.25727353 | 0.242 | 0.171 | 1.4245E-16 | TEC_prg1 | ZNF638 |
| 4.4201E-25 | 0.25592367 | 0.202 | 0.126 | 9.4369E-21 | TEC_prg1 | MYCBP2 |
| 2.8899E-25 | 0.25558628 | 0.281 | 0.195 | 6.1699E-21 | TEC_prg1 | MAT2A |
| 3.2066E-39 | 0.25468684 | 0.164 | 0.077 | 6.846E-35 | TEC_prg1 | PDGFB |
| 6.8492E-24 | 0.25318317 | 0.217 | 0.139 | 1.4623E-19 | TEC_prg1 | SNHG25 |
| 3.6648E-12 | 0.25289691 | 0.276 | 0.231 | 7.8244E-08 | TEC_prg1 | U2SURP |
| 1.325E-49 | 0.25259781 | 0.198 | 0.09 | 2.8289E-45 | TEC_prg1 | RGS2 |
| 2.0758E-30 | 0.25162469 | 0.207 | 0.121 | 4.4319E-26 | TEC_prg1 | RSBN1L |
| 9.1003E-07 | 0.25125914 | 0.257 | 0.231 | 0.01942916 | TEC_prg1 | SCIN |
| 1.3784E-13 | 0.25025169 | 0.236 | 0.182 | 2.9429E-09 | TEC_prg1 | ZNF83 |
| 2.9149E-22 | 0.25003034 | 0.205 | 0.133 | 6.2234E-18 | TEC_prg1 | SLC39A10 |
| 2.7084E-51 | -0.2501165 | 0.131 | 0.305 | 5.7825E-47 | TEC_prg1 | TCEA1 |
| 3.1818E-54 | -0.2502917 | 0.071 | 0.224 | 6.7931E-50 | TEC_prg1 | TINAGL1 |
| 3.8994E-69 | -0.250358 | 0.058 | 0.232 | 8.3253E-65 | TEC_prg1 | IPO7 |
| 1.4261E-76 | -0.2504801 | 0.044 | 0.223 | 3.0448E-72 | TEC_prg1 | EMC4 |
| 1.3697E-66 | -0.2506219 | 0.064 | 0.237 | 2.9243E-62 | TEC_prg1 | G3BP2 |
| 3.5777E-75 | -0.2508101 | 0.03 | 0.196 | 7.6385E-71 | TEC_prg1 | SNRPC |
| 7.4994E-33 | -0.2509391 | 0.925 | 0.929 | 1.6011E-28 | TEC_prg1 | RPL36 |
| 5.1732E-26 | -0.251037 | 0.142 | 0.258 | 1.1045E-21 | TEC_prg1 | ITGB6 |
| 1.3752E-62 | -0.2518434 | 0.063 | 0.229 | 2.9361E-58 | TEC_prg1 | PSMD12 |
| 2.8554E-38 | -0.2522781 | 0.164 | 0.321 | 6.0962E-34 | TEC_prg1 | MIR4435-2HG |
| 3.9561E-76 | -0.2523494 | 0.039 | 0.214 | 8.4463E-72 | TEC_prg1 | UFD1 |
| 6.286E-55 | -0.2523666 | 0.083 | 0.244 | 1.3421E-50 | TEC_prg1 | CCPG1 |
| 4.0245E-65 | -0.2528598 | 0.049 | 0.211 | 8.5924E-61 | TEC_prg1 | CBR1 |
| 3.9913E-78 | -0.2528929 | 0.041 | 0.22 | 8.5215E-74 | TEC_prg1 | MRPS36 |
| 2.2851E-54 | -0.2532658 | 0.119 | 0.294 | 4.8788E-50 | TEC_prg1 | RIOK3 |
| 7.0261E-13 | -0.2534059 | 0.146 | 0.218 | 1.5001E-08 | TEC_prg1 | RRAD |

| | | | | | | |
|---|---|---|---|---|---|---|
| 2.144E-73 | -0.2534605 | 0.044 | 0.217 | 4.5774E-69 | TEC_prg1 | CYB5R3 |
| 7.8517E-82 | -0.2536164 | 0.024 | 0.199 | 1.6763E-77 | TEC_prg1 | SFT2D1 |
| 5.3134E-77 | -0.2539001 | 0.032 | 0.204 | 1.1344E-72 | TEC_prg1 | ESD |
| 4.365E-78 | -0.2541113 | 0.026 | 0.194 | 9.3194E-74 | TEC_prg1 | DGUOK |
| 8.5593E-67 | -0.254115 | 0.031 | 0.184 | 1.8274E-62 | TEC_prg1 | RRAGC |
| 1.4859E-58 | -0.2544348 | 0.033 | 0.172 | 3.1723E-54 | TEC_prg1 | NUCB2 |
| 3.3619E-79 | -0.2546969 | 0.027 | 0.199 | 7.1777E-75 | TEC_prg1 | THOC7 |
| 4.4481E-81 | -0.2547592 | 0.025 | 0.198 | 9.4967E-77 | TEC_prg1 | EI24 |
| 8.5726E-73 | -0.2555138 | 0.072 | 0.261 | 1.8303E-68 | TEC_prg1 | SCARB2 |
| 2.9878E-75 | -0.2556314 | 0.056 | 0.241 | 6.3789E-71 | TEC_prg1 | PSMC1 |
| 4.5641E-82 | -0.2559027 | 0.024 | 0.198 | 9.7443E-78 | TEC_prg1 | UBE2N |
| 2.9292E-62 | -0.2559792 | 0.113 | 0.303 | 6.2538E-58 | TEC_prg1 | BAG1 |
| 2.8023E-86 | -0.2563168 | 0.014 | 0.188 | 5.9828E-82 | TEC_prg1 | APH1A |
| 1.1766E-40 | -0.256398 | 0.033 | 0.139 | 2.5121E-36 | TEC_prg1 | MT1F |
| 7.1517E-71 | -0.25643 | 0.039 | 0.204 | 1.5269E-66 | TEC_prg1 | SLC39A1 |
| 2.029E-63 | -0.2564395 | 0.048 | 0.204 | 4.3319E-59 | TEC_prg1 | DNPH1 |
| 1.527E-61 | -0.2566581 | 0.052 | 0.208 | 3.2601E-57 | TEC_prg1 | BCL10 |
| 5.0253E-38 | -0.2566842 | 0.056 | 0.166 | 1.0729E-33 | TEC_prg1 | HBEGF |
| 4.1204E-66 | -0.256702 | 0.04 | 0.198 | 8.7971E-62 | TEC_prg1 | MPC1 |
| 1.0374E-71 | -0.2568748 | 0.064 | 0.246 | 2.2149E-67 | TEC_prg1 | MED10 |
| 5.8706E-77 | -0.2570563 | 0.037 | 0.211 | 1.2534E-72 | TEC_prg1 | SAR1A |
| 8.7414E-37 | -0.2570837 | 0.059 | 0.17 | 1.8663E-32 | TEC_prg1 | CA2 |
| 7.022E-72 | -0.2573202 | 0.061 | 0.241 | 1.4992E-67 | TEC_prg1 | MIEN1 |
| 1.0845E-60 | -0.2573301 | 0.182 | 0.395 | 2.3154E-56 | TEC_prg1 | EIF3A |
| 3.694E-71 | -0.2575029 | 0.069 | 0.254 | 7.8867E-67 | TEC_prg1 | DNAJC8 |
| 7.8044E-72 | -0.2579422 | 0.073 | 0.262 | 1.6662E-67 | TEC_prg1 | DNAJC15 |
| 1.5488E-62 | -0.2580875 | 0.137 | 0.339 | 3.3067E-58 | TEC_prg1 | EID1 |
| 1.5721E-74 | -0.2581774 | 0.063 | 0.249 | 3.3563E-70 | TEC_prg1 | PSMD4 |
| 9.0283E-68 | -0.2582756 | 0.049 | 0.215 | 1.9275E-63 | TEC_prg1 | B4GALT1 |
| 9.0115E-41 | -0.2584532 | 0.044 | 0.156 | 1.924E-36 | TEC_prg1 | ISG15 |
| 5.498E-40 | -0.2584597 | 0.516 | 0.653 | 1.1738E-35 | TEC_prg1 | PABPC1 |
| 2.5121E-35 | -0.2585337 | 0.582 | 0.682 | 5.3634E-31 | TEC_prg1 | TMA7 |
| 1.0169E-74 | -0.2592589 | 0.033 | 0.202 | 2.171E-70 | TEC_prg1 | GTF3A |
| 3.015E-69 | -0.2595521 | 0.022 | 0.173 | 6.4369E-65 | TEC_prg1 | SNX10 |
| 2.0535E-64 | -0.2600018 | 0.075 | 0.25 | 4.3842E-60 | TEC_prg1 | GARS |
| 1.4084E-82 | -0.2601276 | 0.031 | 0.21 | 3.007E-78 | TEC_prg1 | CHP1 |
| 1.6129E-74 | -0.2610005 | 0.059 | 0.244 | 3.4436E-70 | TEC_prg1 | CWC15 |
| 1.2245E-47 | -0.261075 | 0.032 | 0.15 | 2.6143E-43 | TEC_prg1 | PLIN2 |
| 9.8318E-45 | -0.2613981 | 0.063 | 0.192 | 2.0991E-40 | TEC_prg1 | EMP3 |
| 1.9665E-76 | -0.2617068 | 0.119 | 0.34 | 4.1985E-72 | TEC_prg1 | DAZAP2 |
| 6.8762E-79 | -0.2617256 | 0.038 | 0.216 | 1.4681E-74 | TEC_prg1 | LYPLA1 |
| 2.8627E-76 | -0.261874 | 0.045 | 0.224 | 6.1119E-72 | TEC_prg1 | SNRPF |
| 2.6286E-41 | -0.2620271 | 0.952 | 0.944 | 5.6121E-37 | TEC_prg1 | RPS15 |
| 3.6685E-69 | -0.2624399 | 0.074 | 0.257 | 7.8323E-65 | TEC_prg1 | COMT |
| 8.5531E-77 | -0.2627716 | 0.041 | 0.217 | 1.8261E-72 | TEC_prg1 | ARL3 |
| 2.1004E-40 | -0.2629424 | 0.084 | 0.212 | 4.4844E-36 | TEC_prg1 | BHLHE40 |
| 3.6084E-67 | -0.2630188 | 0.045 | 0.209 | 7.7039E-63 | TEC_prg1 | QSOX1 |
| 3.0658E-71 | -0.263049 | 0.087 | 0.279 | 6.5455E-67 | TEC_prg1 | ZNHIT1 |
| 2.131E-53 | -0.2631887 | 0.018 | 0.136 | 4.5498E-49 | TEC_prg1 | CRABP2 |
| 3.84E-79 | -0.2632138 | 0.045 | 0.228 | 8.1984E-75 | TEC_prg1 | CCDC47 |

| | | | | | | |
|---|---|---|---|---|---|---|
| 8.6898E-85 | -0.2641752 | 0.035 | 0.223 | 1.8553E-80 | TEC_prg1 | MRPS15 |
| 1.6091E-73 | -0.2646059 | 0.046 | 0.221 | 3.4355E-69 | TEC_prg1 | BNIP3L |
| 4.793E-80 | -0.2646345 | 0.037 | 0.217 | 1.0233E-75 | TEC_prg1 | PCMT1 |
| 2.4672E-38 | -0.2646525 | 0.148 | 0.297 | 5.2674E-34 | TEC_prg1 | MUC1 |
| 3.4446E-77 | -0.2647738 | 0.045 | 0.225 | 7.3543E-73 | TEC_prg1 | SYNGR2 |
| 5.3469E-82 | -0.2649712 | 0.03 | 0.207 | 1.1416E-77 | TEC_prg1 | RPN1 |
| 1.2545E-76 | -0.2651022 | 0.037 | 0.211 | 2.6783E-72 | TEC_prg1 | SIVA1 |
| 1.2796E-61 | -0.265234 | 0.056 | 0.214 | 2.7319E-57 | TEC_prg1 | BEX4 |
| 1.8281E-20 | -0.2655418 | 0.232 | 0.351 | 3.9031E-16 | TEC_prg1 | H1F0 |
| 2.8061E-74 | -0.2660823 | 0.029 | 0.194 | 5.9911E-70 | TEC_prg1 | UBE2J1 |
| 1.7972E-52 | -0.2664695 | 0.07 | 0.218 | 3.837E-48 | TEC_prg1 | FDPS |
| 2.7605E-31 | -0.2668198 | 0.079 | 0.185 | 5.8938E-27 | TEC_prg1 | FGF2 |
| 4.3099E-75 | -0.2677838 | 0.038 | 0.21 | 9.2016E-71 | TEC_prg1 | SNRPD1 |
| 5.3106E-75 | -0.2681547 | 0.08 | 0.276 | 1.1338E-70 | TEC_prg1 | LRP10 |
| 1.9561E-73 | -0.269429 | 0.064 | 0.248 | 4.1762E-69 | TEC_prg1 | ETF1 |
| 1.3591E-79 | -0.2696725 | 0.084 | 0.293 | 2.9016E-75 | TEC_prg1 | SNRPD3 |
| 3.4383E-81 | -0.2702207 | 0.057 | 0.253 | 7.3408E-77 | TEC_prg1 | SSU72 |
| 4.0102E-39 | -0.270304 | 0.028 | 0.126 | 8.5618E-35 | TEC_prg1 | ICAM1 |
| 2.5037E-59 | -0.270506 | 0.131 | 0.321 | 5.3453E-55 | TEC_prg1 | HNRNPH3 |
| 2.4444E-76 | -0.2708213 | 0.057 | 0.242 | 5.2187E-72 | TEC_prg1 | RAB14 |
| 2.1885E-64 | -0.2710531 | 0.034 | 0.184 | 4.6724E-60 | TEC_prg1 | MTHFD2 |
| 1.3279E-65 | -0.2712956 | 0.069 | 0.243 | 2.8352E-61 | TEC_prg1 | BTG3 |
| 4.7568E-74 | -0.2724827 | 0.066 | 0.252 | 1.0156E-69 | TEC_prg1 | SPCS3 |
| 2.5041E-69 | -0.2728494 | 0.082 | 0.269 | 5.3463E-65 | TEC_prg1 | EPRS |
| 1.3169E-57 | -0.272958 | 0.063 | 0.216 | 2.8116E-53 | TEC_prg1 | SEL1L |
| 3.6824E-58 | -0.2731137 | 0.026 | 0.16 | 7.8618E-54 | TEC_prg1 | TMEM176A |
| 2.2938E-78 | -0.2736455 | 0.03 | 0.201 | 4.8973E-74 | TEC_prg1 | ARL2 |
| 3.0594E-70 | -0.2737423 | 0.061 | 0.237 | 6.5318E-66 | TEC_prg1 | ADAM9 |
| 1.1838E-06 | -0.2738633 | 0.467 | 0.519 | 0.02527327 | TEC_prg1 | CDKN1A |
| 1.0083E-79 | -0.2740152 | 0.044 | 0.229 | 2.1526E-75 | TEC_prg1 | GLRX5 |
| 3.5815E-50 | -0.2742217 | 0.144 | 0.317 | 7.6465E-46 | TEC_prg1 | TJP1 |
| 1.7727E-67 | -0.2743682 | 0.081 | 0.264 | 3.7848E-63 | TEC_prg1 | NDUFS8 |
| 4.1807E-88 | -0.2745002 | 0.022 | 0.203 | 8.9258E-84 | TEC_prg1 | SUPT4H1 |
| 1.0002E-72 | -0.2754357 | 0.124 | 0.338 | 2.1355E-68 | TEC_prg1 | WASF2 |
| 5.5178E-70 | -0.2756204 | 0.038 | 0.2 | 1.1781E-65 | TEC_prg1 | DSG2 |
| 5.5068E-48 | -0.2756727 | 0.167 | 0.348 | 1.1757E-43 | TEC_prg1 | FAM107B |
| 5.8306E-81 | -0.2759218 | 0.049 | 0.236 | 1.2448E-76 | TEC_prg1 | METTL9 |
| 1.1449E-71 | -0.2763244 | 0.024 | 0.179 | 2.4443E-67 | TEC_prg1 | VKORC1 |
| 1.2288E-68 | -0.276482 | 0.111 | 0.311 | 2.6234E-64 | TEC_prg1 | CNN3 |
| 7.2214E-70 | -0.2765735 | 0.099 | 0.295 | 1.5418E-65 | TEC_prg1 | IL6ST |
| 3.6084E-83 | -0.2767736 | 0.056 | 0.253 | 7.704E-79 | TEC_prg1 | SCOC |
| 1.6589E-62 | -0.27701 | 0.078 | 0.252 | 3.5417E-58 | TEC_prg1 | NAMPT |
| 5.0839E-85 | -0.2778677 | 0.047 | 0.242 | 1.0854E-80 | TEC_prg1 | CSNK2B |
| 5.9623E-86 | -0.2779605 | 0.033 | 0.22 | 1.273E-81 | TEC_prg1 | SDHC |
| 2.0714E-85 | -0.2780105 | 0.044 | 0.237 | 4.4225E-81 | TEC_prg1 | ZFAND6 |
| 2.0205E-73 | -0.2781897 | 0.189 | 0.435 | 4.3137E-69 | TEC_prg1 | FKBP2 |
| 1.4136E-75 | -0.27829 | 0.092 | 0.297 | 3.018E-71 | TEC_prg1 | DRAP1 |
| 1.1643E-36 | -0.2783452 | 0.505 | 0.62 | 2.4858E-32 | TEC_prg1 | CAST |
| 1.6788E-81 | -0.278375 | 0.031 | 0.209 | 3.5843E-77 | TEC_prg1 | ATP6V0D1 |
| 2.2254E-79 | -0.2783776 | 0.044 | 0.228 | 4.7513E-75 | TEC_prg1 | TMEM141 |

| | | | | | | |
|---|---|---|---|---|---|---|
| 4.8101E-81 | -0.2789878 | 0.041 | 0.224 | 1.027E-76 | TEC_prg1 | RALA |
| 6.582E-82 | -0.2793752 | 0.07 | 0.274 | 1.4053E-77 | TEC_prg1 | HNRNPF |
| 7.9386E-47 | -0.2794653 | 0.18 | 0.363 | 1.6949E-42 | TEC_prg1 | RALBP1 |
| 2.476E-77 | -0.2808563 | 0.033 | 0.206 | 5.2863E-73 | TEC_prg1 | RYBP |
| 3.0117E-60 | -0.2810478 | 0.044 | 0.193 | 6.4299E-56 | TEC_prg1 | ERRFI1 |
| 1.055E-79 | -0.2812562 | 0.183 | 0.44 | 2.2523E-75 | TEC_prg1 | YWHAQ |
| 1.3437E-76 | -0.2814865 | 0.057 | 0.243 | 2.8689E-72 | TEC_prg1 | SLIRP |
| 8.3057E-80 | -0.2818043 | 0.091 | 0.302 | 1.7733E-75 | TEC_prg1 | PAIP2 |
| 1.5771E-80 | -0.2820431 | 0.078 | 0.283 | 3.3671E-76 | TEC_prg1 | AP2M1 |
| 3.5176E-64 | -0.2820788 | 0.109 | 0.3 | 7.5101E-60 | TEC_prg1 | RPL36A |
| 1.6518E-18 | -0.2820983 | 0.1 | 0.179 | 3.5266E-14 | TEC_prg1 | CTGF |
| 8.9615E-70 | -0.2822667 | 0.135 | 0.348 | 1.9133E-65 | TEC_prg1 | DNAJB6 |
| 1.0011E-77 | -0.2827732 | 0.067 | 0.262 | 2.1374E-73 | TEC_prg1 | GNAI3 |
| 1.9328E-78 | -0.2830729 | 0.086 | 0.292 | 4.1264E-74 | TEC_prg1 | NDUFA5 |
| 1.512E-49 | -0.2831928 | 0.09 | 0.242 | 3.228E-45 | TEC_prg1 | PTGR1 |
| 7.9864E-16 | -0.283265 | 0.527 | 0.583 | 1.7051E-11 | TEC_prg1 | SLC3A2 |
| 1.0066E-75 | -0.2840987 | 0.091 | 0.296 | 2.1492E-71 | TEC_prg1 | RPS19BP1 |
| 1.3627E-80 | -0.2842963 | 0.018 | 0.185 | 2.9094E-76 | TEC_prg1 | CAV2 |
| 1.531E-69 | -0.2844329 | 0.059 | 0.233 | 3.2688E-65 | TEC_prg1 | OCLN |
| 1.2789E-61 | -0.2845962 | 0.078 | 0.247 | 2.7305E-57 | TEC_prg1 | TIMP2 |
| 3.0046E-76 | -0.2851507 | 0.05 | 0.231 | 6.4147E-72 | TEC_prg1 | SMIM14 |
| 4.5871E-81 | -0.2852842 | 0.06 | 0.256 | 9.7935E-77 | TEC_prg1 | CCT2 |
| 2.4586E-76 | -0.2853432 | 0.051 | 0.232 | 5.2492E-72 | TEC_prg1 | NDUFAF3 |
| 1.3928E-87 | -0.2855076 | 0.04 | 0.234 | 2.9735E-83 | TEC_prg1 | TMEM230 |
| 1.1164E-66 | -0.2857695 | 0.086 | 0.269 | 2.3835E-62 | TEC_prg1 | C6orf48 |
| 2.4262E-69 | -0.2859547 | 0.077 | 0.263 | 5.18E-65 | TEC_prg1 | ASAH1 |
| 3.4335E-71 | -0.2860891 | 0.063 | 0.243 | 7.3304E-67 | TEC_prg1 | RAD21 |
| 9.4489E-76 | -0.2861437 | 0.04 | 0.214 | 2.0173E-71 | TEC_prg1 | MRPL18 |
| 2.0224E-58 | -0.2862525 | 0.147 | 0.34 | 4.3179E-54 | TEC_prg1 | DSP |
| 3.7919E-81 | -0.2862669 | 0.07 | 0.273 | 8.0956E-77 | TEC_prg1 | MORF4L2 |
| 9.0183E-89 | -0.2862804 | 0.023 | 0.207 | 1.9254E-84 | TEC_prg1 | SDHD |
| 1.9405E-76 | -0.2862897 | 0.084 | 0.284 | 4.143E-72 | TEC_prg1 | TAF10 |
| 4.3191E-83 | -0.2863459 | 0.064 | 0.265 | 9.2214E-79 | TEC_prg1 | MRPL20 |
| 2.2949E-72 | -0.2863554 | 0.106 | 0.308 | 4.8995E-68 | TEC_prg1 | CLTC |
| 6.1361E-86 | -0.2865342 | 0.07 | 0.282 | 1.3101E-81 | TEC_prg1 | LAMTOR4 |
| 2.4022E-52 | -0.2868831 | 0.122 | 0.291 | 5.1287E-48 | TEC_prg1 | RB1CC1 |
| 1.2789E-71 | -0.2871133 | 0.105 | 0.308 | 2.7304E-67 | TEC_prg1 | NDUFS6 |
| 5.9812E-85 | -0.2873372 | 0.045 | 0.238 | 1.277E-80 | TEC_prg1 | TXNL4A |
| 6.4286E-88 | -0.2873842 | 0.052 | 0.256 | 1.3725E-83 | TEC_prg1 | ANXA7 |
| 7.3198E-37 | -0.2875847 | 0.088 | 0.21 | 1.5628E-32 | TEC_prg1 | THBS1 |
| 4.2974E-83 | -0.287631 | 0.032 | 0.212 | 9.1749E-79 | TEC_prg1 | HACD3 |
| 2.7889E-87 | -0.2878222 | 0.036 | 0.227 | 5.9544E-83 | TEC_prg1 | ZCRB1 |
| 1.4629E-56 | -0.2878444 | 0.024 | 0.152 | 3.1234E-52 | TEC_prg1 | AL353751.1 |
| 1.3691E-80 | -0.2881778 | 0.035 | 0.214 | 2.923E-76 | TEC_prg1 | ATP1B3 |
| 1.4784E-82 | -0.2884734 | 0.057 | 0.253 | 3.1564E-78 | TEC_prg1 | PSMA4 |
| 9.8036E-96 | -0.2888583 | 0.028 | 0.228 | 2.0931E-91 | TEC_prg1 | CCT7 |
| 1.0152E-81 | -0.2888883 | 0.073 | 0.277 | 2.1675E-77 | TEC_prg1 | KPNB1 |
| 6.755E-60 | -0.288893 | 0.025 | 0.16 | 1.4422E-55 | TEC_prg1 | TRIB3 |
| 1.281E-88 | -0.2894249 | 0.031 | 0.22 | 2.7348E-84 | TEC_prg1 | ATRAID |
| 1.2277E-88 | -0.2899309 | 0.054 | 0.258 | 2.6211E-84 | TEC_prg1 | PSMA1 |

| | | | | | | |
|---|---|---|---|---|---|---|
| 6.7276E-62 | -0.2901518 | 0.112 | 0.296 | 1.4363E-57 | TEC_prg1 | SLC38A1 |
| 9.1477E-69 | -0.2902194 | 0.064 | 0.24 | 1.953E-64 | TEC_prg1 | GFPT1 |
| 1.6684E-87 | -0.2902219 | 0.055 | 0.257 | 3.562E-83 | TEC_prg1 | STAU1 |
| 9.6256E-66 | -0.290387 | 0.147 | 0.359 | 2.0551E-61 | TEC_prg1 | ARL6IP1 |
| 7.3026E-91 | -0.2904266 | 0.087 | 0.318 | 1.5591E-86 | TEC_prg1 | MTCH1 |
| 7.2605E-60 | -0.2908306 | 0.164 | 0.369 | 1.5501E-55 | TEC_prg1 | H2AFJ |
| 2.6342E-80 | -0.2911613 | 0.099 | 0.315 | 5.6241E-76 | TEC_prg1 | RAB10 |
| 8.4798E-79 | -0.2923684 | 0.072 | 0.271 | 1.8104E-74 | TEC_prg1 | RPS17 |
| 3.9474E-37 | -0.2924687 | 0.621 | 0.706 | 8.4277E-33 | TEC_prg1 | GPX4 |
| 1.5859E-87 | -0.2929285 | 0.052 | 0.255 | 3.3858E-83 | TEC_prg1 | RWDD1 |
| 7.3629E-81 | -0.2937772 | 0.034 | 0.214 | 1.572E-76 | TEC_prg1 | TMEM54 |
| 2.5036E-89 | -0.2937881 | 0.047 | 0.249 | 5.3451E-85 | TEC_prg1 | SNRPB2 |
| 2.2513E-94 | -0.2938832 | 0.024 | 0.219 | 4.8065E-90 | TEC_prg1 | TMEM147 |
| 7.9485E-58 | -0.2939221 | 0.075 | 0.232 | 1.697E-53 | TEC_prg1 | DNTTIP2 |
| 4.1097E-80 | -0.2949346 | 0.047 | 0.231 | 8.7743E-76 | TEC_prg1 | COX14 |
| 3.1393E-68 | -0.2951823 | 0.206 | 0.447 | 6.7025E-64 | TEC_prg1 | JAK1 |
| 2.7437E-93 | -0.2953239 | 0.032 | 0.231 | 5.8577E-89 | TEC_prg1 | WBP2 |
| 8.1487E-94 | -0.2954317 | 0.028 | 0.224 | 1.7397E-89 | TEC_prg1 | FUNDC2 |
| 2.1507E-71 | -0.2959413 | 0.031 | 0.191 | 4.5917E-67 | TEC_prg1 | CAVIN1 |
| 1.0327E-92 | -0.2960746 | 0.062 | 0.278 | 2.2049E-88 | TEC_prg1 | XRCC6 |
| 1.9685E-72 | -0.296084 | 0.189 | 0.434 | 4.2027E-68 | TEC_prg1 | ATP5F1D |
| 1.3413E-82 | -0.296198 | 0.107 | 0.331 | 2.8637E-78 | TEC_prg1 | SSBP1 |
| 6.872E-88 | -0.2963889 | 0.022 | 0.204 | 1.4672E-83 | TEC_prg1 | PRNP |
| 8.0603E-83 | -0.2965182 | 0.171 | 0.426 | 1.7209E-78 | TEC_prg1 | CTNNA1 |
| 2.1497E-41 | -0.2968261 | 0.029 | 0.131 | 4.5896E-37 | TEC_prg1 | RARRES3 |
| 7.0693E-66 | -0.2971099 | 0.067 | 0.239 | 1.5093E-61 | TEC_prg1 | MAFG |
| 3.635E-86 | -0.2978496 | 0.047 | 0.243 | 7.7606E-82 | TEC_prg1 | GNG12 |
| 6.4469E-86 | -0.298062 | 0.053 | 0.253 | 1.3764E-81 | TEC_prg1 | RSL1D1 |
| 4.6696E-30 | -0.2981425 | 0.055 | 0.149 | 9.9695E-26 | TEC_prg1 | IFIT2 |
| 1.6553E-89 | -0.2985619 | 0.038 | 0.234 | 3.5341E-85 | TEC_prg1 | ILF2 |
| 9.1856E-68 | -0.2990946 | 0.133 | 0.337 | 1.9611E-63 | TEC_prg1 | PNRC2 |
| 2.0957E-73 | -0.2991236 | 0.176 | 0.412 | 4.4743E-69 | TEC_prg1 | HEBP2 |
| 2.3697E-96 | -0.2991905 | 0.027 | 0.227 | 5.0593E-92 | TEC_prg1 | EMC7 |
| 2.1298E-88 | -0.3000337 | 0.047 | 0.249 | 4.5472E-84 | TEC_prg1 | RCN1 |
| 3.0413E-93 | -0.300323 | 0.033 | 0.23 | 6.4933E-89 | TEC_prg1 | SELENOT |
| 2.5151E-92 | -0.3003299 | 0.049 | 0.257 | 5.3698E-88 | TEC_prg1 | CNIH4 |
| 8.9958E-90 | -0.3004131 | 0.037 | 0.233 | 1.9206E-85 | TEC_prg1 | NBDY |
| 8.5629E-91 | -0.3005757 | 0.049 | 0.254 | 1.8282E-86 | TEC_prg1 | GDI2 |
| 5.0359E-69 | -0.3007351 | 0.041 | 0.203 | 1.0752E-64 | TEC_prg1 | DNAJB9 |
| 7.9683E-86 | -0.3017049 | 0.052 | 0.25 | 1.7012E-81 | TEC_prg1 | SRI |
| 1.4894E-79 | -0.3018003 | 0.11 | 0.328 | 3.1798E-75 | TEC_prg1 | ACTR2 |
| 1.05E-53 | -0.3018304 | 0.055 | 0.196 | 2.2417E-49 | TEC_prg1 | PLA2G16 |
| 1.6317E-80 | -0.302188 | 0.021 | 0.189 | 3.4836E-76 | TEC_prg1 | PLS3 |
| 5.4857E-94 | -0.3023873 | 0.065 | 0.285 | 1.1712E-89 | TEC_prg1 | EIF3D |
| 2.7424E-85 | -0.3024972 | 0.033 | 0.22 | 5.8551E-81 | TEC_prg1 | RAB9A |
| 3.2993E-83 | -0.3025414 | 0.03 | 0.21 | 7.044E-79 | TEC_prg1 | CD99 |
| 1.8796E-89 | -0.302813 | 0.046 | 0.249 | 4.013E-85 | TEC_prg1 | PSMA5 |
| 6.5231E-46 | -0.3035513 | 0.1 | 0.247 | 1.3927E-41 | TEC_prg1 | GPRC5A |
| 7.1293E-90 | -0.3036537 | 0.074 | 0.293 | 1.5221E-85 | TEC_prg1 | PRKAR1A |
| 4.7537E-80 | -0.3038941 | 0.059 | 0.252 | 1.0149E-75 | TEC_prg1 | LIMS1 |

| | | | | | | |
|---|---|---|---|---|---|---|
| 9.1275E-87 | -0.3040396 | 0.07 | 0.28 | 1.9487E-82 | TEC_prg1 | TUFM |
| 1.0351E-85 | -0.3040603 | 0.023 | 0.202 | 2.2099E-81 | TEC_prg1 | NME1 |
| 1.2208E-67 | -0.3043522 | 0.032 | 0.187 | 2.6063E-63 | TEC_prg1 | SUCLG1 |
| 1.512E-86 | -0.3044771 | 0.049 | 0.246 | 3.228E-82 | TEC_prg1 | NHP2 |
| 2.1306E-77 | -0.3045531 | 0.05 | 0.234 | 4.5489E-73 | TEC_prg1 | ERO1A |
| 9.3203E-65 | -0.3047734 | 0.182 | 0.407 | 1.9899E-60 | TEC_prg1 | CAMK2N1 |
| 3.7074E-27 | -0.3049778 | 0.795 | 0.791 | 7.9152E-23 | TEC_prg1 | PPDPF |
| 6.9516E-90 | -0.3049906 | 0.017 | 0.199 | 1.4842E-85 | TEC_prg1 | PRDX4 |
| 1.1132E-88 | -0.3055231 | 0.061 | 0.271 | 2.3766E-84 | TEC_prg1 | SMIM26 |
| 8.6765E-94 | -0.3055628 | 0.029 | 0.226 | 1.8524E-89 | TEC_prg1 | TSPAN3 |
| 1.2618E-41 | -0.3059843 | 0.106 | 0.249 | 2.6939E-37 | TEC_prg1 | CD74 |
| 6.4118E-62 | -0.3061808 | 0.044 | 0.196 | 1.3689E-57 | TEC_prg1 | SERPING1 |
| 2.7762E-06 | -0.306566 | 0.3 | 0.359 | 0.0592713 | TEC_prg1 | TXNIP |
| 6.1666E-83 | -0.3073882 | 0.095 | 0.315 | 1.3166E-78 | TEC_prg1 | MRPL41 |
| 8.1413E-78 | -0.3075701 | 0.082 | 0.285 | 1.7382E-73 | TEC_prg1 | ATOX1 |
| 1.0351E-96 | -0.307694 | 0.043 | 0.254 | 2.21E-92 | TEC_prg1 | LAMP2 |
| 1.1875E-85 | -0.30772 | 0.134 | 0.377 | 2.5353E-81 | TEC_prg1 | AURKAIP1 |
| 3.0894E-54 | -0.3079926 | 0.068 | 0.216 | 6.596E-50 | TEC_prg1 | SESN2 |
| 4.806E-91 | -0.3086603 | 0.053 | 0.262 | 1.0261E-86 | TEC_prg1 | PGLS |
| 1.0241E-87 | -0.3087976 | 0.061 | 0.269 | 2.1864E-83 | TEC_prg1 | ARPC5L |
| 6.5782E-97 | -0.3093142 | 0.047 | 0.261 | 1.4045E-92 | TEC_prg1 | EIF3L |
| 2.0911E-83 | -0.3102697 | 0.1 | 0.319 | 4.4645E-79 | TEC_prg1 | FIS1 |
| 3.1956E-93 | -0.3104365 | 0.041 | 0.244 | 6.8227E-89 | TEC_prg1 | PEA15 |
| 1.9651E-92 | -0.3106178 | 0.075 | 0.297 | 4.1956E-88 | TEC_prg1 | EIF4H |
| 3.2521E-57 | -0.3106518 | 0.066 | 0.221 | 6.9432E-53 | TEC_prg1 | MSMO1 |
| 4.7451E-43 | -0.3108203 | 0.911 | 0.912 | 1.0131E-38 | TEC_prg1 | RPL35 |
| 8.3701E-87 | -0.3110991 | 0.111 | 0.344 | 1.787E-82 | TEC_prg1 | WDR1 |
| 1.2642E-65 | -0.3115381 | 0.032 | 0.181 | 2.6991E-61 | TEC_prg1 | QPRT |
| 3.2699E-74 | -0.311618 | 0.05 | 0.227 | 6.9813E-70 | TEC_prg1 | BNIP3 |
| 5.9967E-91 | -0.3116221 | 0.052 | 0.261 | 1.2803E-86 | TEC_prg1 | PSMD7 |
| 7.8311E-50 | -0.3120815 | 0.081 | 0.228 | 1.6719E-45 | TEC_prg1 | CLDN1 |
| 1.706E-103 | -0.313087 | 0.045 | 0.269 | 3.642E-99 | TEC_prg1 | ACP1 |
| 3.4626E-91 | -0.3133763 | 0.076 | 0.298 | 7.3925E-87 | TEC_prg1 | GNB2 |
| 9.325E-96 | -0.3134278 | 0.049 | 0.263 | 1.9909E-91 | TEC_prg1 | TMEM208 |
| 1.5133E-94 | -0.313507 | 0.052 | 0.265 | 3.231E-90 | TEC_prg1 | NDUFA12 |
| 9.1194E-85 | -0.3135665 | 0.034 | 0.218 | 1.947E-80 | TEC_prg1 | ARPC1B |
| 1.436E-75 | -0.3142569 | 0.053 | 0.234 | 3.0658E-71 | TEC_prg1 | ADI1 |
| 1.0102E-87 | -0.3144259 | 0.076 | 0.291 | 2.1568E-83 | TEC_prg1 | TMEM123 |
| 9.3001E-84 | -0.31443 | 0.145 | 0.389 | 1.9856E-79 | TEC_prg1 | VAPA |
| 6.8964E-95 | -0.3147392 | 0.081 | 0.314 | 1.4724E-90 | TEC_prg1 | SELENOF |
| 2.031E-100 | -0.3151304 | 0.057 | 0.283 | 4.3365E-96 | TEC_prg1 | CAMTA1 |
| 2.3018E-96 | -0.3152645 | 0.055 | 0.271 | 4.9143E-92 | TEC_prg1 | PPP1CA |
| 1.5695E-83 | -0.3159537 | 0.061 | 0.26 | 3.3509E-79 | TEC_prg1 | PSMD2 |
| 6.9882E-86 | -0.3166855 | 0.071 | 0.283 | 1.492E-81 | TEC_prg1 | PGRMC1 |
| 5.3085E-88 | -0.3168567 | 0.054 | 0.256 | 1.1334E-83 | TEC_prg1 | DYNLT3 |
| 9.5202E-53 | -0.3170812 | 0.629 | 0.747 | 2.0326E-48 | TEC_prg1 | GNAS |
| 3.939E-109 | -0.3171728 | 0.026 | 0.245 | 8.411E-105 | TEC_prg1 | RAB5C |
| 7.4294E-80 | -0.3174282 | 0.217 | 0.476 | 1.5862E-75 | TEC_prg1 | ST13 |
| 3.2575E-95 | -0.3174434 | 0.046 | 0.256 | 6.9547E-91 | TEC_prg1 | TMEM167A |
| 1.6472E-96 | -0.3177731 | 0.04 | 0.248 | 3.5168E-92 | TEC_prg1 | MCFD2 |

| | | | | | | |
|---|---|---|---|---|---|---|
| 2.5245E-82 | -0.3180284 | 0.181 | 0.435 | 5.3898E-78 | TEC_prg1 | TAX1BP1 |
| 7.0713E-82 | -0.3182513 | 0.134 | 0.369 | 1.5097E-77 | TEC_prg1 | KIF5B |
| 3.498E-75 | -0.3186754 | 0.019 | 0.178 | 7.4682E-71 | TEC_prg1 | NQO1 |
| 2.8639E-96 | -0.3193026 | 0.081 | 0.315 | 6.1145E-92 | TEC_prg1 | ARPP19 |
| 4.429E-100 | -0.3196105 | 0.029 | 0.235 | 9.4552E-96 | TEC_prg1 | NDUFB5 |
| 5.5628E-89 | -0.322793 | 0.057 | 0.264 | 1.1877E-84 | TEC_prg1 | H2AFV |
| 2.4608E-73 | -0.3231774 | 0.064 | 0.247 | 5.2539E-69 | TEC_prg1 | EPB41L4A-AS1 |
| 2.0668E-79 | -0.3234511 | 0.045 | 0.227 | 4.4126E-75 | TEC_prg1 | MAP2K3 |
| 5.0876E-86 | -0.3246692 | 0.044 | 0.237 | 1.0862E-81 | TEC_prg1 | MPC2 |
| 1.2977E-71 | -0.3249932 | 0.483 | 0.674 | 2.7706E-67 | TEC_prg1 | YWHAE |
| 1.0203E-89 | -0.3250579 | 0.108 | 0.343 | 2.1782E-85 | TEC_prg1 | RBM8A |
| 3.3221E-98 | -0.325709 | 0.033 | 0.24 | 7.0927E-94 | TEC_prg1 | CALU |
| 1.932E-83 | -0.3257469 | 0.046 | 0.237 | 4.1249E-79 | TEC_prg1 | CSRP1 |
| 5.0335E-92 | -0.3259603 | 0.088 | 0.316 | 1.0746E-87 | TEC_prg1 | SEPT7 |
| 9.9357E-56 | -0.3264068 | 0.031 | 0.166 | 2.1213E-51 | TEC_prg1 | XIST |
| 1.9674E-94 | -0.3268284 | 0.052 | 0.264 | 4.2003E-90 | TEC_prg1 | TXNL1 |
| 1.9132E-90 | -0.3271612 | 0.074 | 0.292 | 4.0848E-86 | TEC_prg1 | NDUFA11 |
| 8.6156E-95 | -0.3274187 | 0.037 | 0.241 | 1.8394E-90 | TEC_prg1 | SCP2 |
| 2.111E-108 | -0.3276736 | 0.036 | 0.262 | 4.506E-104 | TEC_prg1 | PSMB4 |
| 1.9422E-97 | -0.327678 | 0.07 | 0.3 | 4.1467E-93 | TEC_prg1 | TMCO1 |
| 1.297E-102 | -0.3276808 | 0.023 | 0.229 | 2.7698E-98 | TEC_prg1 | CMTM6 |
| 2.46E-104 | -0.3281359 | 0.04 | 0.26 | 5.252E-100 | TEC_prg1 | ARL1 |
| 1.067E-100 | -0.3291316 | 0.06 | 0.287 | 2.2785E-96 | TEC_prg1 | SNRPE |
| 3.0795E-96 | -0.3296714 | 0.034 | 0.238 | 6.5747E-92 | TEC_prg1 | AP2S1 |
| 4.3408E-88 | -0.3297381 | 0.087 | 0.309 | 9.2675E-84 | TEC_prg1 | COX17 |
| 3.301E-102 | -0.3299606 | 0.061 | 0.294 | 7.0466E-98 | TEC_prg1 | CCT8 |
| 3.917E-84 | -0.3306985 | 0.07 | 0.275 | 8.3627E-80 | TEC_prg1 | VAMP2 |
| 8.715E-107 | -0.3309492 | 0.04 | 0.264 | 1.861E-102 | TEC_prg1 | TMEM219 |
| 1.352E-35 | -0.3314512 | 0.167 | 0.311 | 2.8866E-31 | TEC_prg1 | STMN1 |
| 3.641E-106 | -0.3318012 | 0.084 | 0.335 | 7.773E-102 | TEC_prg1 | SRSF9 |
| 1.2344E-79 | -0.3320111 | 0.081 | 0.285 | 2.6354E-75 | TEC_prg1 | CD47 |
| 7.648E-99 | -0.3327896 | 0.07 | 0.301 | 1.6329E-94 | TEC_prg1 | MLEC |
| 1.6699E-93 | -0.3334639 | 0.066 | 0.285 | 3.5652E-89 | TEC_prg1 | YPEL5 |
| 5.1504E-67 | -0.3335585 | 0.185 | 0.41 | 1.0996E-62 | TEC_prg1 | ODC1 |
| 3.9184E-89 | -0.3344285 | 0.147 | 0.401 | 8.3657E-85 | TEC_prg1 | SEC62 |
| 4.1541E-29 | -0.3344721 | 0.037 | 0.12 | 8.869E-25 | TEC_prg1 | AQP3 |
| 3.3187E-78 | -0.334703 | 0.022 | 0.187 | 7.0855E-74 | TEC_prg1 | TMEM176B |
| 1.0521E-81 | -0.334836 | 0.106 | 0.327 | 2.2462E-77 | TEC_prg1 | YWHAH |
| 1.593E-105 | -0.3348426 | 0.066 | 0.304 | 3.4E-101 | TEC_prg1 | SYPL1 |
| 1.8647E-97 | -0.3348436 | 0.066 | 0.293 | 3.9811E-93 | TEC_prg1 | CCT4 |
| 8.029E-107 | -0.3348977 | 0.045 | 0.274 | 1.714E-102 | TEC_prg1 | NAA20 |
| 4.158E-103 | -0.3351378 | 0.037 | 0.255 | 8.878E-99 | TEC_prg1 | GTF3C6 |
| 2.5319E-63 | -0.3352785 | 0.128 | 0.322 | 5.4055E-59 | TEC_prg1 | MAFF |
| 2.199E-99 | -0.3354592 | 0.069 | 0.299 | 4.6944E-95 | TEC_prg1 | JTB |
| 2.654E-109 | -0.3358342 | 0.036 | 0.262 | 5.666E-105 | TEC_prg1 | C1orf43 |
| 7.106E-109 | -0.3359343 | 0.028 | 0.248 | 1.517E-104 | TEC_prg1 | TRAPPC1 |
| 3.547E-97 | -0.3363143 | 0.066 | 0.291 | 7.5728E-93 | TEC_prg1 | ARHGDIA |
| 7.5274E-61 | -0.3367636 | 0.081 | 0.249 | 1.6071E-56 | TEC_prg1 | BEX2 |
| 3.176E-106 | -0.3373308 | 0.042 | 0.265 | 6.781E-102 | TEC_prg1 | UXT |
| 1.4117E-98 | -0.337792 | 0.079 | 0.314 | 3.0141E-94 | TEC_prg1 | CCT6A |

| | | | | | | |
|---|---|---|---|---|---|---|
| 7.8881E-63 | -0.3387727 | 0.123 | 0.316 | 1.6841E-58 | TEC_prg1 | C12orf57 |
| 1.7468E-67 | -0.3388577 | 0.184 | 0.404 | 3.7294E-63 | TEC_prg1 | SELENOW |
| 7.4E-59 | -0.3389938 | 0.351 | 0.548 | 1.5799E-54 | TEC_prg1 | TPM3 |
| 2.3725E-51 | -0.3401943 | 0.589 | 0.695 | 5.0652E-47 | TEC_prg1 | YWHAZ |
| 5.3461E-56 | -0.3402386 | 0.032 | 0.165 | 1.1414E-51 | TEC_prg1 | CKS2 |
| 2.0708E-89 | -0.3412708 | 0.172 | 0.434 | 4.4211E-85 | TEC_prg1 | CAPN2 |
| 4.7833E-67 | -0.3418584 | 0.093 | 0.279 | 1.0212E-62 | TEC_prg1 | MGLL |
| 2.796E-110 | -0.3419193 | 0.042 | 0.274 | 5.97E-106 | TEC_prg1 | TMEM14B |
| 1.4115E-75 | -0.3430464 | 0.044 | 0.22 | 3.0136E-71 | TEC_prg1 | CTSL |
| 1.7813E-98 | -0.343096 | 0.074 | 0.305 | 3.803E-94 | TEC_prg1 | NUDC |
| 3.524E-106 | -0.3435951 | 0.066 | 0.308 | 7.524E-102 | TEC_prg1 | RTN3 |
| 1.554E-101 | -0.3441385 | 0.063 | 0.293 | 3.3188E-97 | TEC_prg1 | PPP2R1A |
| 1.2712E-97 | -0.3448328 | 0.081 | 0.318 | 2.714E-93 | TEC_prg1 | ANAPC16 |
| 5.057E-105 | -0.3449736 | 0.021 | 0.229 | 1.08E-100 | TEC_prg1 | C1QBP |
| 7.2121E-82 | -0.3452921 | 0.282 | 0.527 | 1.5398E-77 | TEC_prg1 | UBE2D3 |
| 5.094E-106 | -0.3457114 | 0.061 | 0.297 | 1.087E-101 | TEC_prg1 | RAB7A |
| 1.587E-106 | -0.3457502 | 0.056 | 0.288 | 3.388E-102 | TEC_prg1 | UBE2L3 |
| 1.1539E-81 | -0.3459597 | 0.012 | 0.176 | 2.4637E-77 | TEC_prg1 | FXYD5 |
| 1.501E-60 | -0.3463571 | 0.703 | 0.787 | 3.2046E-56 | TEC_prg1 | EEF2 |
| 8.263E-103 | -0.3466823 | 0.071 | 0.31 | 1.7642E-98 | TEC_prg1 | NDUFA2 |
| 1.7414E-95 | -0.3473438 | 0.08 | 0.309 | 3.7178E-91 | TEC_prg1 | ARPC5 |
| 9.5235E-68 | -0.3478031 | 0.657 | 0.77 | 2.0333E-63 | TEC_prg1 | PFDN5 |
| 6.65E-112 | -0.3484827 | 0.044 | 0.281 | 1.42E-107 | TEC_prg1 | GTF2A2 |
| 1.355E-104 | -0.348923 | 0.043 | 0.264 | 2.893E-100 | TEC_prg1 | MDH1 |
| 2.515E-105 | -0.3493413 | 0.051 | 0.278 | 5.369E-101 | TEC_prg1 | NENF |
| 4.9138E-42 | -0.3495939 | 0.494 | 0.606 | 1.0491E-37 | TEC_prg1 | S100A13 |
| 8.4656E-96 | -0.3505919 | 0.054 | 0.27 | 1.8074E-91 | TEC_prg1 | CHMP2B |
| 2.749E-119 | -0.3507802 | 0.024 | 0.257 | 5.868E-115 | TEC_prg1 | COPZ1 |
| 9.7824E-75 | -0.3511047 | 0.071 | 0.258 | 2.0885E-70 | TEC_prg1 | PRSS8 |
| 5.191E-102 | -0.3516742 | 0.055 | 0.28 | 1.1082E-97 | TEC_prg1 | PLEKHB2 |
| 2.087E-70 | -0.3520292 | 0.088 | 0.281 | 4.4558E-66 | TEC_prg1 | MRPS6 |
| 9.7836E-71 | -0.3529536 | 0.51 | 0.68 | 2.0888E-66 | TEC_prg1 | SLC25A3 |
| 8.162E-116 | -0.3537928 | 0.041 | 0.281 | 1.743E-111 | TEC_prg1 | RAB11A |
| 3.5599E-66 | -0.3545978 | 0.296 | 0.516 | 7.6005E-62 | TEC_prg1 | P4HB |
| 1.3007E-93 | -0.3546499 | 0.037 | 0.238 | 2.777E-89 | TEC_prg1 | OSER1 |
| 2.667E-121 | -0.3547889 | 0.044 | 0.294 | 5.693E-117 | TEC_prg1 | EIF3I |
| 1.636E-115 | -0.3550649 | 0.052 | 0.298 | 3.493E-111 | TEC_prg1 | ATP6V1E1 |
| 8.5038E-79 | -0.3561177 | 0.039 | 0.218 | 1.8156E-74 | TEC_prg1 | EMP1 |
| 1.2973E-94 | -0.3561969 | 0.047 | 0.257 | 2.7698E-90 | TEC_prg1 | ARL6IP5 |
| 7.165E-115 | -0.3562417 | 0.035 | 0.269 | 1.53E-110 | TEC_prg1 | STRAP |
| 8.542E-112 | -0.3564795 | 0.051 | 0.29 | 1.824E-107 | TEC_prg1 | LMAN2 |
| 1.094E-109 | -0.3567735 | 0.057 | 0.297 | 2.335E-105 | TEC_prg1 | NDUFA6 |
| 1.053E-112 | -0.3568707 | 0.042 | 0.277 | 2.249E-108 | TEC_prg1 | VPS29 |
| 3.324E-114 | -0.3569193 | 0.065 | 0.317 | 7.098E-110 | TEC_prg1 | CHMP2A |
| 1.2527E-86 | -0.3571934 | 0.076 | 0.288 | 2.6745E-82 | TEC_prg1 | COTL1 |
| 4.392E-105 | -0.3577466 | 0.125 | 0.394 | 9.376E-101 | TEC_prg1 | C19orf53 |
| 1.837E-95 | -0.3577477 | 0.088 | 0.323 | 3.9219E-91 | TEC_prg1 | ROMO1 |
| 1.434E-105 | -0.3579793 | 0.061 | 0.298 | 3.061E-101 | TEC_prg1 | NDUFB3 |
| 1.93E-115 | -0.3581519 | 0.053 | 0.299 | 4.121E-111 | TEC_prg1 | RTRAF |
| 3.31E-115 | -0.3586716 | 0.034 | 0.267 | 7.067E-111 | TEC_prg1 | ATP5PB |

| | | | | | | |
|---|---|---|---|---|---|---|
| 1.2205E-70 | -0.358683 | 0.493 | 0.663 | 2.6057E-66 | TEC_prg1 | UBL5 |
| 8.215E-61 | -0.3589447 | 0.203 | 0.418 | 1.7539E-56 | TEC_prg1 | PTMS |
| 1.099E-119 | -0.3590434 | 0.014 | 0.24 | 2.346E-115 | TEC_prg1 | ARPC1A |
| 5.2952E-98 | -0.3593213 | 0.111 | 0.358 | 1.1305E-93 | TEC_prg1 | PCBP1 |
| 1.7632E-79 | -0.359497 | 0.116 | 0.335 | 3.7645E-75 | TEC_prg1 | CDH1 |
| 4.311E-110 | -0.3595767 | 0.041 | 0.272 | 9.204E-106 | TEC_prg1 | TMBIM1 |
| 1.41E-104 | -0.3596477 | 0.045 | 0.272 | 3.01E-100 | TEC_prg1 | CALM3 |
| 1.574E-112 | -0.359967 | 0.036 | 0.266 | 3.36E-108 | TEC_prg1 | POLR2K |
| 2.4404E-47 | -0.3600572 | 0.015 | 0.121 | 5.2103E-43 | TEC_prg1 | RSAD2 |
| 8.947E-103 | -0.3610481 | 0.063 | 0.296 | 1.9102E-98 | TEC_prg1 | HM13 |
| 1.168E-110 | -0.3611708 | 0.069 | 0.318 | 2.493E-106 | TEC_prg1 | SSB |
| 1.766E-110 | -0.3612192 | 0.024 | 0.243 | 3.771E-106 | TEC_prg1 | PRDX3 |
| 3.706E-30 | -0.3619728 | 0.252 | 0.397 | 7.9123E-26 | TEC_prg1 | ATF3 |
| 1.1603E-27 | -0.3634516 | 0.038 | 0.117 | 2.4773E-23 | TEC_prg1 | SERPINE2 |
| 4.396E-114 | -0.3643939 | 0.04 | 0.274 | 9.386E-110 | TEC_prg1 | MDH2 |
| 6.676E-60 | -0.3647341 | 0.438 | 0.61 | 1.4253E-55 | TEC_prg1 | COX5B |
| 1.46E-114 | -0.3653515 | 0.067 | 0.32 | 3.117E-110 | TEC_prg1 | CCT3 |
| 1.683E-107 | -0.3656509 | 0.102 | 0.364 | 3.594E-103 | TEC_prg1 | GNB1 |
| 1.76E-107 | -0.3670023 | 0.072 | 0.317 | 3.758E-103 | TEC_prg1 | CCT5 |
| 1.868E-114 | -0.368026 | 0.05 | 0.294 | 3.989E-110 | TEC_prg1 | ATP6V1D |
| 2.03E-120 | -0.3680621 | 0.038 | 0.283 | 4.334E-116 | TEC_prg1 | PSMB5 |
| 9.988E-102 | -0.3687321 | 0.036 | 0.251 | 2.1324E-97 | TEC_prg1 | ATP5MC1 |
| 1.338E-104 | -0.3689457 | 0.09 | 0.34 | 2.857E-100 | TEC_prg1 | ACTR3 |
| 2.592E-119 | -0.3699935 | 0.052 | 0.305 | 5.534E-115 | TEC_prg1 | CAPZA2 |
| 7.123E-112 | -0.3700798 | 0.074 | 0.328 | 1.521E-107 | TEC_prg1 | RSL24D1 |
| 1.29E-116 | -0.3707046 | 0.029 | 0.26 | 2.755E-112 | TEC_prg1 | ARPC4 |
| 1.025E-113 | -0.3712292 | 0.063 | 0.314 | 2.189E-109 | TEC_prg1 | ATP5F1C |
| 8.269E-99 | -0.3712776 | 0.088 | 0.328 | 1.7655E-94 | TEC_prg1 | PSME2 |
| 1.5866E-65 | -0.371828 | 0.132 | 0.33 | 3.3875E-61 | TEC_prg1 | CYTOR |
| 9.466E-118 | -0.3722356 | 0.044 | 0.287 | 2.021E-113 | TEC_prg1 | BANF1 |
| 1.631E-109 | -0.3726207 | 0.129 | 0.407 | 3.483E-105 | TEC_prg1 | SNRPD2 |
| 4.372E-109 | -0.3726756 | 0.065 | 0.308 | 9.335E-105 | TEC_prg1 | SNRPG |
| 7.3328E-55 | -0.3733147 | 0.479 | 0.632 | 1.5656E-50 | TEC_prg1 | APLP2 |
| 5.904E-121 | -0.3735461 | 0.06 | 0.319 | 1.261E-116 | TEC_prg1 | OCIAD1 |
| 1.6976E-66 | -0.3738515 | 0.111 | 0.302 | 3.6243E-62 | TEC_prg1 | CKB |
| 1.726E-113 | -0.3738542 | 0.038 | 0.271 | 3.685E-109 | TEC_prg1 | RNF181 |
| 8.612E-103 | -0.3741175 | 0.068 | 0.302 | 1.8386E-98 | TEC_prg1 | UFM1 |
| 1.853E-113 | -0.3750898 | 0.071 | 0.324 | 3.957E-109 | TEC_prg1 | TMEM14C |
| 1.871E-115 | -0.3753851 | 0.042 | 0.281 | 3.994E-111 | TEC_prg1 | HIGD2A |
| 4.1712E-18 | -0.3756535 | 0.044 | 0.105 | 8.9056E-14 | TEC_prg1 | CCL2 |
| 7.3061E-48 | -0.375781 | 0.017 | 0.125 | 1.5599E-43 | TEC_prg1 | BST2 |
| 3.2882E-68 | -0.3760752 | 0.044 | 0.206 | 7.0204E-64 | TEC_prg1 | DCBLD2 |
| 3.985E-109 | -0.3773372 | 0.063 | 0.305 | 8.508E-105 | TEC_prg1 | TXNDC17 |
| 6.237E-124 | -0.3775529 | 0.04 | 0.291 | 1.332E-119 | TEC_prg1 | SUMO1 |
| 1.17E-106 | -0.3775549 | 0.085 | 0.335 | 2.498E-102 | TEC_prg1 | EIF6 |
| 1.851E-109 | -0.3781239 | 0.072 | 0.321 | 3.953E-105 | TEC_prg1 | CLTB |
| 1.245E-115 | -0.3783071 | 0.101 | 0.377 | 2.658E-111 | TEC_prg1 | COPE |
| 3.313E-110 | -0.3783808 | 0.057 | 0.296 | 7.073E-106 | TEC_prg1 | ATP5F1A |
| 2.4E-90 | -0.3788716 | 0.056 | 0.264 | 5.1241E-86 | TEC_prg1 | RPL22L1 |
| 1.859E-120 | -0.3790747 | 0.028 | 0.265 | 3.97E-116 | TEC_prg1 | GLO1 |

| | | | | | | |
|---|---|---|---|---|---|---|
| 2.3143E-31 | -0.3793542 | 0.038 | 0.126 | 4.941E-27 | TEC_prg1 | C15orf48 |
| 2.6783E-94 | -0.3795522 | 0.161 | 0.429 | 5.7182E-90 | TEC_prg1 | PERP |
| 1.0823E-45 | -0.3797028 | 0.05 | 0.173 | 2.3106E-41 | TEC_prg1 | RCAN1 |
| 3.438E-122 | -0.3804601 | 0.045 | 0.296 | 7.34E-118 | TEC_prg1 | CAPZB |
| 2.025E-116 | -0.3807356 | 0.058 | 0.309 | 4.324E-112 | TEC_prg1 | NDUFB8 |
| 7.195E-106 | -0.3808579 | 0.114 | 0.38 | 1.536E-101 | TEC_prg1 | EIF4B |
| 5.6299E-77 | -0.3811977 | 0.042 | 0.217 | 1.202E-72 | TEC_prg1 | PDIA4 |
| 2.5758E-13 | -0.3818542 | 0.064 | 0.117 | 5.4993E-09 | TEC_prg1 | S100A9 |
| 6.0268E-35 | -0.3828759 | 0.046 | 0.146 | 1.2867E-30 | TEC_prg1 | TNFRSF11B |
| 3.4621E-74 | -0.3829854 | 0.162 | 0.389 | 7.3917E-70 | TEC_prg1 | NUCKS1 |
| 1.468E-118 | -0.3834663 | 0.045 | 0.29 | 3.135E-114 | TEC_prg1 | RAB1A |
| 1.159E-99 | -0.3847854 | 0.073 | 0.306 | 2.4738E-95 | TEC_prg1 | BLVRB |
| 2.1829E-36 | -0.3848302 | 0.605 | 0.67 | 4.6604E-32 | TEC_prg1 | CALR |
| 3.012E-119 | -0.3850597 | 0.049 | 0.298 | 6.43E-115 | TEC_prg1 | MTPN |
| 5.867E-114 | -0.3856306 | 0.078 | 0.335 | 1.253E-109 | TEC_prg1 | SF3B5 |
| 1.723E-132 | -0.3858807 | 0.06 | 0.339 | 3.679E-128 | TEC_prg1 | RNF7 |
| 2.045E-129 | -0.3873248 | 0.026 | 0.274 | 4.366E-125 | TEC_prg1 | PSMB2 |
| 1.0727E-51 | -0.387838 | 0.556 | 0.677 | 2.2902E-47 | TEC_prg1 | PSAP |
| 2.6322E-50 | -0.3880231 | 0.879 | 0.904 | 5.6198E-46 | TEC_prg1 | SERF2 |
| 3.3405E-84 | -0.3882937 | 0.031 | 0.211 | 7.132E-80 | TEC_prg1 | CD40 |
| 7.079E-124 | -0.38843 | 0.041 | 0.293 | 1.511E-119 | TEC_prg1 | PHB |
| 4.147E-83 | -0.3887723 | 0.105 | 0.323 | 8.8539E-79 | TEC_prg1 | CDK2AP2 |
| 1.868E-128 | -0.3888792 | 0.04 | 0.297 | 3.988E-124 | TEC_prg1 | RER1 |
| 7.7569E-73 | -0.3891536 | 0.36 | 0.569 | 1.6561E-68 | TEC_prg1 | HLA-E |
| 8.102E-115 | -0.3893395 | 0.059 | 0.308 | 1.73E-110 | TEC_prg1 | TSTD1 |
| 5.803E-120 | -0.3897885 | 0.045 | 0.293 | 1.239E-115 | TEC_prg1 | PSMA3 |
| 7.8317E-67 | -0.3900301 | 0.927 | 0.943 | 1.6721E-62 | TEC_prg1 | RPL37A |
| 3.59E-107 | -0.3902924 | 0.075 | 0.32 | 7.665E-103 | TEC_prg1 | GRN |
| 1.2486E-85 | -0.3922961 | 0.139 | 0.378 | 2.6657E-81 | TEC_prg1 | HSPA9 |
| 6.2294E-79 | -0.3928503 | 0.059 | 0.249 | 1.33E-74 | TEC_prg1 | SERTAD1 |
| 6.1148E-63 | -0.3944216 | 0.523 | 0.657 | 1.3055E-58 | TEC_prg1 | CCNI |
| 8.846E-104 | -0.3952558 | 0.084 | 0.325 | 1.889E-99 | TEC_prg1 | ANP32B |
| 9.039E-121 | -0.3958741 | 0.069 | 0.331 | 1.93E-116 | TEC_prg1 | RBX1 |
| 9.03E-114 | -0.3960071 | 0.031 | 0.261 | 1.928E-109 | TEC_prg1 | CXCL16 |
| 7.054E-126 | -0.3965078 | 0.044 | 0.298 | 1.506E-121 | TEC_prg1 | TMED2 |
| 5.41E-113 | -0.3965663 | 0.072 | 0.324 | 1.155E-108 | TEC_prg1 | PDCD5 |
| 1.71E-125 | -0.3972222 | 0.054 | 0.316 | 3.652E-121 | TEC_prg1 | ANAPC11 |
| 1.692E-126 | -0.397671 | 0.032 | 0.28 | 3.613E-122 | TEC_prg1 | KDELR2 |
| 2.9985E-43 | -0.3982287 | 0.078 | 0.207 | 6.4018E-39 | TEC_prg1 | ADIRF |
| 9.123E-109 | -0.398483 | 0.069 | 0.312 | 1.948E-104 | TEC_prg1 | NDUFV2 |
| 1.016E-126 | -0.3986093 | 0.023 | 0.266 | 2.169E-122 | TEC_prg1 | ATP6AP2 |
| 2.7855E-88 | -0.3992117 | 0.125 | 0.365 | 5.9469E-84 | TEC_prg1 | CTSB |
| 1.632E-117 | -0.4000771 | 0.061 | 0.315 | 3.484E-113 | TEC_prg1 | CAPNS1 |
| 5.1028E-64 | -0.4002634 | 0.106 | 0.292 | 1.0894E-59 | TEC_prg1 | HMGB2 |
| 1.1827E-74 | -0.400821 | 0.167 | 0.396 | 2.5251E-70 | TEC_prg1 | CD44 |
| 6.37E-109 | -0.4011226 | 0.052 | 0.286 | 1.36E-104 | TEC_prg1 | DHRS7 |
| 2.082E-106 | -0.4014271 | 0.088 | 0.338 | 4.445E-102 | TEC_prg1 | SELENOS |
| 2.045E-124 | -0.4018177 | 0.033 | 0.278 | 4.366E-120 | TEC_prg1 | LSM4 |
| 2.099E-81 | -0.4023663 | 0.081 | 0.286 | 4.4814E-77 | TEC_prg1 | PTP4A1 |
| 6.943E-117 | -0.4023715 | 0.065 | 0.321 | 1.482E-112 | TEC_prg1 | LMAN1 |

| | | | | | | |
|---|---|---|---|---|---|---|
| 5.4888E-61 | -0.4025645 | 0.036 | 0.179 | 1.1719E-56 | TEC_prg1 | IGFBP2 |
| 3.8326E-78 | -0.4027799 | 0.356 | 0.568 | 8.1826E-74 | TEC_prg1 | NDUFS5 |
| 1.451E-119 | -0.4029722 | 0.042 | 0.287 | 3.097E-115 | TEC_prg1 | BCAP31 |
| 8.8359E-61 | -0.4032724 | 0.789 | 0.829 | 1.8865E-56 | TEC_prg1 | RPS21 |
| 9.774E-123 | -0.4045099 | 0.087 | 0.367 | 2.087E-118 | TEC_prg1 | ATP5MPL |
| 1.5734E-93 | -0.4056586 | 0.123 | 0.365 | 3.3592E-89 | TEC_prg1 | EIF1B |
| 6.295E-131 | -0.4058812 | 0.049 | 0.315 | 1.344E-126 | TEC_prg1 | UFC1 |
| 6.448E-135 | -0.4061692 | 0.035 | 0.3 | 1.377E-130 | TEC_prg1 | SMDT1 |
| 1.32E-99 | -0.4061797 | 0.103 | 0.351 | 2.8191E-95 | TEC_prg1 | RAB13 |
| 9.492E-121 | -0.4062357 | 0.066 | 0.325 | 2.027E-116 | TEC_prg1 | LAMP1 |
| 2.285E-121 | -0.4077375 | 0.04 | 0.286 | 4.878E-117 | TEC_prg1 | PGAM1 |
| 1.5406E-96 | -0.4078693 | 0.09 | 0.324 | 3.2893E-92 | TEC_prg1 | ANXA3 |
| 4.393E-126 | -0.4084478 | 0.037 | 0.289 | 9.38E-122 | TEC_prg1 | RPN2 |
| 4.876E-120 | -0.4087849 | 0.047 | 0.295 | 1.041E-115 | TEC_prg1 | EIF3M |
| 4.8036E-34 | -0.4092353 | 0.38 | 0.514 | 1.0256E-29 | TEC_prg1 | ATP1B1 |
| 1.892E-129 | -0.4101387 | 0.074 | 0.352 | 4.04E-125 | TEC_prg1 | TMEM258 |
| 8.3679E-86 | -0.4101678 | 0.089 | 0.303 | 1.7865E-81 | TEC_prg1 | TUBB |
| 1.499E-132 | -0.4115604 | 0.03 | 0.285 | 3.201E-128 | TEC_prg1 | LSM3 |
| 1.68E-130 | -0.4121276 | 0.095 | 0.385 | 3.588E-126 | TEC_prg1 | SEM1 |
| 2.3414E-05 | -0.4123151 | 0.227 | 0.267 | 0.49989753 | TEC_prg1 | NUPR1 |
| 1.533E-112 | -0.4126995 | 0.059 | 0.303 | 3.273E-108 | TEC_prg1 | APRT |
| 3.019E-128 | -0.4152293 | 0.038 | 0.294 | 6.446E-124 | TEC_prg1 | C4orf3 |
| 6.523E-106 | -0.4163408 | 0.051 | 0.281 | 1.393E-101 | TEC_prg1 | DNAJC3 |
| 3.2674E-44 | -0.4164526 | 0.1 | 0.239 | 6.9759E-40 | TEC_prg1 | UBE2S |
| 7.6587E-54 | -0.416545 | 0.787 | 0.824 | 1.6351E-49 | TEC_prg1 | UBB |
| 1.011E-131 | -0.4187267 | 0.042 | 0.303 | 2.159E-127 | TEC_prg1 | ERH |
| 8.579E-111 | -0.420017 | 0.095 | 0.357 | 1.832E-106 | TEC_prg1 | UQCR11 |
| 3.095E-127 | -0.4200354 | 0.061 | 0.328 | 6.608E-123 | TEC_prg1 | ATP5PO |
| 6.2051E-93 | -0.4200835 | 0.111 | 0.349 | 1.3248E-88 | TEC_prg1 | ISCU |
| 1.0601E-92 | -0.4200995 | 0.328 | 0.568 | 2.2634E-88 | TEC_prg1 | COX6C |
| 5.701E-123 | -0.4208695 | 0.027 | 0.267 | 1.217E-118 | TEC_prg1 | SEC11C |
| 4.2354E-86 | -0.4211784 | 0.378 | 0.597 | 9.0425E-82 | TEC_prg1 | COX6B1 |
| 2.8779E-92 | -0.4220623 | 0.167 | 0.429 | 6.1444E-88 | TEC_prg1 | POLR2L |
| 1.128E-119 | -0.4221087 | 0.063 | 0.322 | 2.407E-115 | TEC_prg1 | BUD31 |
| 6.8125E-91 | -0.4223886 | 0.139 | 0.385 | 1.4545E-86 | TEC_prg1 | MIF |
| 3.9236E-50 | -0.4226882 | 0.38 | 0.528 | 8.3769E-46 | TEC_prg1 | CLDN7 |
| 6.331E-131 | -0.4228802 | 0.089 | 0.378 | 1.352E-126 | TEC_prg1 | NDUFB4 |
| 1.229E-119 | -0.4242147 | 0.16 | 0.463 | 2.624E-115 | TEC_prg1 | HNRNPK |
| 5.039E-104 | -0.4243847 | 0.254 | 0.524 | 1.076E-99 | TEC_prg1 | PTGES3 |
| 2.08E-132 | -0.4248877 | 0.099 | 0.395 | 4.441E-128 | TEC_prg1 | PSMD8 |
| 4.166E-129 | -0.4252089 | 0.085 | 0.37 | 8.894E-125 | TEC_prg1 | FAM96B |
| 3.417E-143 | -0.4252337 | 0.032 | 0.305 | 7.296E-139 | TEC_prg1 | LAMTOR1 |
| 6.978E-138 | -0.4255952 | 0.032 | 0.296 | 1.49E-133 | TEC_prg1 | NDUFAB1 |
| 5.6223E-52 | -0.4258675 | 0.626 | 0.715 | 1.2004E-47 | TEC_prg1 | MAP1LC3B |
| 6.842E-138 | -0.4260792 | 0.033 | 0.301 | 1.461E-133 | TEC_prg1 | PSMB3 |
| 4.543E-130 | -0.4261336 | 0.059 | 0.329 | 9.7E-126 | TEC_prg1 | PSMB6 |
| 5.196E-104 | -0.4262898 | 0.162 | 0.442 | 1.109E-99 | TEC_prg1 | CYCS |
| 3.998E-103 | -0.4277734 | 0.028 | 0.237 | 8.535E-99 | TEC_prg1 | DPP4 |
| 1.087E-135 | -0.4286019 | 0.065 | 0.348 | 2.32E-131 | TEC_prg1 | SSR2 |
| 4.1155E-90 | -0.4286174 | 0.17 | 0.431 | 8.7865E-86 | TEC_prg1 | HSPE1 |

| | | | | | | |
|---|---|---|---|---|---|---|
| 6.787E-136 | -0.4292482 | 0.043 | 0.311 | 1.449E-131 | TEC_prg1 | PSMB7 |
| 8.044E-135 | -0.4294272 | 0.033 | 0.294 | 1.717E-130 | TEC_prg1 | UQCRFS1 |
| 1.0246E-06 | -0.4300428 | 0.086 | 0.123 | 0.02187556 | TEC_prg1 | CXCL8 |
| 2.215E-114 | -0.4305178 | 0.148 | 0.439 | 4.729E-110 | TEC_prg1 | EIF2S2 |
| 4.696E-133 | -0.4307474 | 0.028 | 0.282 | 1.003E-128 | TEC_prg1 | OSTC |
| 1.275E-146 | -0.4308401 | 0.056 | 0.348 | 2.723E-142 | TEC_prg1 | NDFIP1 |
| 1.005E-142 | -0.4311648 | 0.053 | 0.34 | 2.146E-138 | TEC_prg1 | MRPL51 |
| 9.61E-140 | -0.4317998 | 0.051 | 0.332 | 2.052E-135 | TEC_prg1 | COPS9 |
| 1.799E-137 | -0.4347445 | 0.072 | 0.361 | 3.841E-133 | TEC_prg1 | MRFAP1 |
| 1.7675E-36 | -0.435354 | 0.016 | 0.101 | 3.7736E-32 | TEC_prg1 | SAA1 |
| 9.4523E-17 | -0.43638 | 0.099 | 0.173 | 2.0181E-12 | TEC_prg1 | G0S2 |
| 6.433E-130 | -0.4365573 | 0.047 | 0.31 | 1.373E-125 | TEC_prg1 | MINOS1 |
| 6.2742E-80 | -0.4372623 | 0.919 | 0.938 | 1.3396E-75 | TEC_prg1 | RPL37 |
| 2.3151E-74 | -0.4372826 | 0.744 | 0.814 | 4.9428E-70 | TEC_prg1 | CALM1 |
| 3.966E-144 | -0.4387993 | 0.077 | 0.378 | 8.468E-140 | TEC_prg1 | EIF3H |
| 4.8122E-70 | -0.4416149 | 0.021 | 0.172 | 1.0274E-65 | TEC_prg1 | MYL9 |
| 2.994E-131 | -0.4422315 | 0.057 | 0.33 | 6.393E-127 | TEC_prg1 | NDUFB11 |
| 7.454E-122 | -0.4429203 | 0.231 | 0.525 | 1.592E-117 | TEC_prg1 | MORF4L1 |
| 3.707E-115 | -0.4429389 | 0.056 | 0.301 | 7.914E-111 | TEC_prg1 | GSTO1 |
| 3.0767E-88 | -0.4432642 | 0.916 | 0.937 | 6.5687E-84 | TEC_prg1 | RPL6 |
| 1.229E-127 | -0.4432839 | 0.071 | 0.345 | 2.623E-123 | TEC_prg1 | DDT |
| 1.4202E-66 | -0.4443328 | 0.324 | 0.523 | 3.0321E-62 | TEC_prg1 | DNAJA1 |
| 9.0818E-34 | -0.445105 | 0.959 | 0.902 | 1.939E-29 | TEC_prg1 | UBC |
| 1.511E-119 | -0.4454883 | 0.068 | 0.325 | 3.227E-115 | TEC_prg1 | MSN |
| 4.189E-86 | -0.4463693 | 0.973 | 0.974 | 8.9435E-82 | TEC_prg1 | RPL13 |
| 2.284E-124 | -0.4465808 | 0.133 | 0.432 | 4.877E-120 | TEC_prg1 | MRPL33 |
| 7.847E-132 | -0.4472946 | 0.068 | 0.346 | 1.675E-127 | TEC_prg1 | RAP1B |
| 1.524E-154 | -0.4479593 | 0.053 | 0.354 | 3.253E-150 | TEC_prg1 | EIF3F |
| 4.474E-115 | -0.4484133 | 0.118 | 0.394 | 9.551E-111 | TEC_prg1 | MZT2B |
| 6.94E-133 | -0.4485999 | 0.057 | 0.331 | 1.482E-128 | TEC_prg1 | RABAC1 |
| 2.1399E-08 | -0.4496127 | 0.611 | 0.604 | 0.00045688 | TEC_prg1 | SOD2 |
| 9.061E-132 | -0.4504218 | 0.037 | 0.295 | 1.934E-127 | TEC_prg1 | LAPTM4B |
| 3.335E-139 | -0.4510189 | 0.114 | 0.428 | 7.12E-135 | TEC_prg1 | CD164 |
| 3.494E-91 | -0.4515872 | 0.175 | 0.433 | 7.4596E-87 | TEC_prg1 | CD151 |
| 1.4042E-90 | -0.4524773 | 0.468 | 0.657 | 2.9979E-86 | TEC_prg1 | SLC25A6 |
| 3.117E-103 | -0.4530575 | 0.074 | 0.308 | 6.655E-99 | TEC_prg1 | RBM3 |
| 1.928E-135 | -0.4541206 | 0.052 | 0.327 | 4.115E-131 | TEC_prg1 | LITAF |
| 2.33E-138 | -0.4541887 | 0.086 | 0.382 | 4.975E-134 | TEC_prg1 | RAB2A |
| 6.421E-101 | -0.4549915 | 0.084 | 0.321 | 1.3708E-96 | TEC_prg1 | ASPH |
| 3.8928E-13 | -0.4553578 | 0.605 | 0.607 | 8.311E-09 | TEC_prg1 | GADD45A |
| 9.577E-101 | -0.4560123 | 0.098 | 0.343 | 2.0447E-96 | TEC_prg1 | C9orf16 |
| 1.581E-132 | -0.4563422 | 0.093 | 0.387 | 3.376E-128 | TEC_prg1 | KRT10 |
| 5.8812E-84 | -0.4564991 | 0.016 | 0.187 | 1.2556E-79 | TEC_prg1 | MGST1 |
| 1.4159E-95 | -0.4566046 | 0.028 | 0.227 | 3.0229E-91 | TEC_prg1 | SDF2L1 |
| 1.877E-130 | -0.4569323 | 0.117 | 0.419 | 4.008E-126 | TEC_prg1 | UBE2B |
| 3.4645E-11 | -0.4571674 | 0.063 | 0.109 | 7.3968E-07 | TEC_prg1 | APOE |
| 1.1093E-90 | -0.4594667 | 0.038 | 0.234 | 2.3683E-86 | TEC_prg1 | MANF |
| 4.676E-155 | -0.4609211 | 0.061 | 0.366 | 9.983E-151 | TEC_prg1 | SNX3 |
| 1.965E-109 | -0.4613866 | 0.063 | 0.303 | 4.195E-105 | TEC_prg1 | SELENOM |
| 6.671E-159 | -0.4639521 | 0.044 | 0.346 | 1.424E-154 | TEC_prg1 | CHMP5 |

| | | | | | | |
|---|---|---|---|---|---|---|
| 1.4736E-54 | -0.465762 | 0.138 | 0.311 | 3.1462E-50 | TEC_prg1 | UCHL1 |
| 1.217E-153 | -0.466064 | 0.057 | 0.36 | 2.599E-149 | TEC_prg1 | TMEM50A |
| 5.876E-60 | -0.4661881 | 0.941 | 0.928 | 1.2545E-55 | TEC_prg1 | RPL12 |
| 6.3084E-92 | -0.4676349 | 0.153 | 0.403 | 1.3468E-87 | TEC_prg1 | HMGN2 |
| 1.845E-56 | -0.4678331 | 0.145 | 0.327 | 3.9392E-52 | TEC_prg1 | HSPH1 |
| 1.035E-142 | -0.4678538 | 0.059 | 0.348 | 2.211E-138 | TEC_prg1 | CUTA |
| 4.4482E-70 | -0.4688099 | 0.961 | 0.96 | 9.497E-66 | TEC_prg1 | RPL28 |
| 1.0615E-90 | -0.4699294 | 0.618 | 0.738 | 2.2662E-86 | TEC_prg1 | RPL36AL |
| 3.727E-133 | -0.4714315 | 0.029 | 0.284 | 7.958E-129 | TEC_prg1 | PLP2 |
| 1.502E-144 | -0.4715175 | 0.075 | 0.376 | 3.207E-140 | TEC_prg1 | NDUFC1 |
| 1.027E-146 | -0.4719712 | 0.063 | 0.359 | 2.192E-142 | TEC_prg1 | ATP5PD |
| 7.258E-153 | -0.4730321 | 0.062 | 0.367 | 1.55E-148 | TEC_prg1 | SARAF |
| 1.402E-101 | -0.4731096 | 0.372 | 0.605 | 2.9937E-97 | TEC_prg1 | HNRNPC |
| 1.8421E-92 | -0.4732217 | 0.189 | 0.448 | 3.9329E-88 | TEC_prg1 | SDC4 |
| 2.963E-139 | -0.4733906 | 0.107 | 0.416 | 6.326E-135 | TEC_prg1 | PPP1CB |
| 1.202E-145 | -0.4757199 | 0.049 | 0.336 | 2.567E-141 | TEC_prg1 | SERINC1 |
| 3.329E-54 | -0.4765355 | 0.023 | 0.147 | 7.1073E-50 | TEC_prg1 | HMOX1 |
| 1.485E-111 | -0.4794539 | 0.394 | 0.624 | 3.171E-107 | TEC_prg1 | YWHAB |
| 3.7961E-65 | -0.4809471 | 0.461 | 0.609 | 8.1047E-61 | TEC_prg1 | EZR |
| 1.7108E-81 | -0.4810019 | 0.754 | 0.819 | 3.6526E-77 | TEC_prg1 | ATP5F1E |
| 5.398E-114 | -0.4815118 | 0.264 | 0.535 | 1.152E-109 | TEC_prg1 | TOMM7 |
| 2.4954E-76 | -0.482263 | 0.782 | 0.826 | 5.3277E-72 | TEC_prg1 | YBX1 |
| 3.536E-111 | -0.4845559 | 0.423 | 0.639 | 7.549E-107 | TEC_prg1 | CLTA |
| 7.205E-157 | -0.4847825 | 0.056 | 0.359 | 1.538E-152 | TEC_prg1 | GABARAPL2 |
| 1.97E-128 | -0.4857601 | 0.039 | 0.294 | 4.205E-124 | TEC_prg1 | ECHS1 |
| 3.027E-147 | -0.4857955 | 0.102 | 0.418 | 6.463E-143 | TEC_prg1 | WDR83OS |
| 4.049E-131 | -0.4858089 | 0.121 | 0.425 | 8.645E-127 | TEC_prg1 | NDUFA1 |
| 9.341E-148 | -0.4866115 | 0.031 | 0.307 | 1.994E-143 | TEC_prg1 | FKBP1A |
| 5.187E-164 | -0.4868432 | 0.053 | 0.367 | 1.107E-159 | TEC_prg1 | SRP9 |
| 7.5039E-45 | -0.4884125 | 0.067 | 0.195 | 1.6021E-40 | TEC_prg1 | IGFBP3 |
| 2.522E-128 | -0.488595 | 0.163 | 0.469 | 5.385E-124 | TEC_prg1 | PTTG1IP |
| 8.0756E-40 | -0.4886272 | 0.238 | 0.394 | 1.7241E-35 | TEC_prg1 | DDIT3 |
| 8.935E-166 | -0.4889784 | 0.037 | 0.344 | 1.908E-161 | TEC_prg1 | TMED10 |
| 4.311E-160 | -0.4890114 | 0.063 | 0.378 | 9.203E-156 | TEC_prg1 | ELOC |
| 1.064E-151 | -0.4895673 | 0.079 | 0.39 | 2.272E-147 | TEC_prg1 | ERP29 |
| 7.059E-90 | -0.491441 | 0.626 | 0.746 | 1.5071E-85 | TEC_prg1 | CSTB |
| 1.1646E-60 | -0.49278 | 0.622 | 0.707 | 2.4864E-56 | TEC_prg1 | ZFAS1 |
| 5.541E-121 | -0.4930259 | 0.37 | 0.622 | 1.183E-116 | TEC_prg1 | ATP6V1G1 |
| 5.672E-156 | -0.4931487 | 0.049 | 0.35 | 1.211E-151 | TEC_prg1 | CIB1 |
| 7.749E-149 | -0.4935339 | 0.109 | 0.433 | 1.654E-144 | TEC_prg1 | HSBP1 |
| 2.9213E-90 | -0.4941365 | 0.949 | 0.951 | 6.237E-86 | TEC_prg1 | RPS8 |
| 1.2558E-98 | -0.4942436 | 0.516 | 0.69 | 2.6812E-94 | TEC_prg1 | UQCRB |
| 9.1631E-49 | -0.4953317 | 0.378 | 0.52 | 1.9563E-44 | TEC_prg1 | TNFRSF12A |
| 8.157E-125 | -0.4956456 | 0.59 | 0.763 | 1.742E-120 | TEC_prg1 | SKP1 |
| 2.004E-163 | -0.4970926 | 0.06 | 0.379 | 4.278E-159 | TEC_prg1 | VDAC1 |
| 1.6974E-79 | -0.4974185 | 0.867 | 0.89 | 3.624E-75 | TEC_prg1 | RPS29 |
| 9.122E-157 | -0.4985149 | 0.081 | 0.401 | 1.948E-152 | TEC_prg1 | COMMD6 |
| 6.463E-129 | -0.4987669 | 0.084 | 0.365 | 1.38E-124 | TEC_prg1 | TKT |
| 1.768E-160 | -0.4990376 | 0.088 | 0.416 | 3.774E-156 | TEC_prg1 | CSDE1 |
| 2.958E-160 | -0.4994295 | 0.042 | 0.345 | 6.316E-156 | TEC_prg1 | LEPROT |

| | | | | | | |
|---|---|---|---|---|---|---|
| 1.857E-117 | -0.5008506 | 0.095 | 0.359 | 3.964E-113 | TEC_prg1 | PRDX2 |
| 9.889E-134 | -0.5014494 | 0.094 | 0.389 | 2.111E-129 | TEC_prg1 | ARF4 |
| 7.897E-156 | -0.5018546 | 0.067 | 0.378 | 1.686E-151 | TEC_prg1 | TRAM1 |
| 2.116E-114 | -0.5042685 | 0.362 | 0.608 | 4.517E-110 | TEC_prg1 | RAN |
| 1.872E-170 | -0.5067782 | 0.049 | 0.367 | 3.996E-166 | TEC_prg1 | COX7A2L |
| 6.072E-119 | -0.5068752 | 0.305 | 0.568 | 1.296E-114 | TEC_prg1 | ATP5MC2 |
| 4.3859E-61 | -0.5070861 | 0.703 | 0.764 | 9.3639E-57 | TEC_prg1 | RPS26 |
| 7.3519E-76 | -0.5072888 | 0.093 | 0.292 | 1.5696E-71 | TEC_prg1 | LAMB3 |
| 5.155E-143 | -0.5083031 | 0.267 | 0.57 | 1.1E-138 | TEC_prg1 | ARF1 |
| 9.158E-158 | -0.512096 | 0.07 | 0.386 | 1.955E-153 | TEC_prg1 | PTP4A2 |
| 5.006E-132 | -0.512276 | 0.027 | 0.278 | 1.069E-127 | TEC_prg1 | HIGD1A |
| 2.7617E-91 | -0.5134552 | 0.088 | 0.313 | 5.8962E-87 | TEC_prg1 | CYB5A |
| 1.767E-105 | -0.5147327 | 0.69 | 0.793 | 3.773E-101 | TEC_prg1 | OAZ1 |
| 6.537E-115 | -0.5159856 | 0.034 | 0.264 | 1.396E-110 | TEC_prg1 | IFITM2 |
| 5.102E-135 | -0.517729 | 0.19 | 0.497 | 1.089E-130 | TEC_prg1 | ATP5MD |
| 8.1322E-93 | -0.5185062 | 0.415 | 0.611 | 1.7362E-88 | TEC_prg1 | PRDX5 |
| 1.482E-167 | -0.5185203 | 0.053 | 0.371 | 3.164E-163 | TEC_prg1 | SNRPB |
| 4.855E-167 | -0.5185558 | 0.07 | 0.399 | 1.037E-162 | TEC_prg1 | CNBP |
| 1.098E-137 | -0.5186239 | 0.099 | 0.398 | 2.343E-133 | TEC_prg1 | ANXA11 |
| 1.98E-161 | -0.5215435 | 0.084 | 0.41 | 4.227E-157 | TEC_prg1 | EIF3K |
| 1.937E-130 | -0.5227095 | 0.064 | 0.339 | 4.135E-126 | TEC_prg1 | CD55 |
| 6.6334E-96 | -0.5228279 | 0.87 | 0.897 | 1.4162E-91 | TEC_prg1 | RPS11 |
| 3.234E-139 | -0.5239744 | 0.082 | 0.377 | 6.905E-135 | TEC_prg1 | UQCRQ |
| 2.509E-165 | -0.5279508 | 0.068 | 0.393 | 5.357E-161 | TEC_prg1 | EIF1AX |
| 9.164E-166 | -0.5280225 | 0.067 | 0.387 | 1.956E-161 | TEC_prg1 | SPCS2 |
| 2.0964E-91 | -0.5287016 | 0.306 | 0.533 | 4.4758E-87 | TEC_prg1 | HSPD1 |
| 3.447E-104 | -0.5291119 | 0.064 | 0.292 | 7.36E-100 | TEC_prg1 | S100A14 |
| 9.218E-174 | -0.5298076 | 0.043 | 0.363 | 1.968E-169 | TEC_prg1 | PGK1 |
| 5.028E-153 | -0.5305134 | 0.065 | 0.368 | 1.074E-148 | TEC_prg1 | NDUFB9 |
| 3.383E-139 | -0.5309034 | 0.062 | 0.345 | 7.223E-135 | TEC_prg1 | CAPG |
| 1.875E-133 | -0.5311855 | 0.403 | 0.648 | 4.003E-129 | TEC_prg1 | ATP5MG |
| 3.364E-163 | -0.5318928 | 0.084 | 0.415 | 7.183E-159 | TEC_prg1 | TRMT112 |
| 2.296E-117 | -0.5320996 | 0.656 | 0.79 | 4.903E-113 | TEC_prg1 | COX7C |
| 1.275E-153 | -0.5321187 | 0.044 | 0.337 | 2.721E-149 | TEC_prg1 | FAM3C |
| 2.901E-169 | -0.5322186 | 0.049 | 0.364 | 6.194E-165 | TEC_prg1 | PCBD1 |
| 1.777E-129 | -0.5326102 | 0.841 | 0.896 | 3.795E-125 | TEC_prg1 | RPL22 |
| 8.294E-141 | -0.5326834 | 0.305 | 0.588 | 1.771E-136 | TEC_prg1 | GNG5 |
| 4.184E-126 | -0.5328805 | 0.302 | 0.567 | 8.932E-122 | TEC_prg1 | CDC42 |
| 1.74E-167 | -0.5333714 | 0.113 | 0.462 | 3.716E-163 | TEC_prg1 | TBCA |
| 7.011E-127 | -0.5360854 | 0.29 | 0.56 | 1.497E-122 | TEC_prg1 | ARPC2 |
| 9.465E-161 | -0.5377212 | 0.073 | 0.392 | 2.021E-156 | TEC_prg1 | PFDN2 |
| 6.263E-187 | -0.5380423 | 0.033 | 0.363 | 1.337E-182 | TEC_prg1 | NDUFC2 |
| 8.112E-112 | -0.5384797 | 0.11 | 0.376 | 1.732E-107 | TEC_prg1 | CHCHD10 |
| 2.249E-172 | -0.5388805 | 0.052 | 0.375 | 4.802E-168 | TEC_prg1 | COX5A |
| 1.511E-169 | -0.5395101 | 0.06 | 0.385 | 3.225E-165 | TEC_prg1 | ATP6V1F |
| 7.096E-179 | -0.5418478 | 0.06 | 0.396 | 1.515E-174 | TEC_prg1 | SPCS1 |
| 3.068E-186 | -0.5423007 | 0.051 | 0.39 | 6.551E-182 | TEC_prg1 | KDELR1 |
| 7.484E-181 | -0.542463 | 0.091 | 0.446 | 1.598E-176 | TEC_prg1 | SUB1 |
| 7.7803E-96 | -0.5426157 | 0.496 | 0.672 | 1.6611E-91 | TEC_prg1 | PEBP1 |
| 8.7716E-71 | -0.5426546 | 0.015 | 0.161 | 1.8727E-66 | TEC_prg1 | TM4SF1 |

| | | | | | | |
|---|---|---|---|---|---|---|
| 3.545E-172 | -0.5436241 | 0.056 | 0.383 | 7.569E-168 | TEC_prg1 | SF3B6 |
| 2.26E-176 | -0.5451727 | 0.059 | 0.393 | 4.826E-172 | TEC_prg1 | NOP10 |
| 3.819E-157 | -0.5452378 | 0.133 | 0.465 | 8.153E-153 | TEC_prg1 | UQCR10 |
| 1.434E-125 | -0.5457084 | 0.098 | 0.376 | 3.061E-121 | TEC_prg1 | TUBA1C |
| 1.456E-134 | -0.5460259 | 0.35 | 0.614 | 3.109E-130 | TEC_prg1 | COX7A2 |
| 1.982E-106 | -0.5460588 | 0.892 | 0.907 | 4.232E-102 | TEC_prg1 | RPS20 |
| 3.484E-143 | -0.5481957 | 0.769 | 0.87 | 7.438E-139 | TEC_prg1 | SRP14 |
| 4.612E-195 | -0.5489567 | 0.046 | 0.396 | 9.847E-191 | TEC_prg1 | SEC11A |
| 2.949E-107 | -0.5494556 | 0.061 | 0.292 | 6.296E-103 | TEC_prg1 | RPS4Y1 |
| 1.248E-132 | -0.5506254 | 0.188 | 0.486 | 2.665E-128 | TEC_prg1 | NAP1L1 |
| 7.595E-182 | -0.5518883 | 0.026 | 0.346 | 1.621E-177 | TEC_prg1 | BRK1 |
| 5.432E-143 | -0.5520081 | 0.1 | 0.411 | 1.16E-138 | TEC_prg1 | COX7B |
| 7.186E-126 | -0.5522081 | 0.766 | 0.836 | 1.534E-121 | TEC_prg1 | H3F3A |
| 1.946E-140 | -0.553147 | 0.194 | 0.5 | 4.155E-136 | TEC_prg1 | RPS10 |
| 5.36E-105 | -0.5532361 | 0.945 | 0.948 | 1.144E-100 | TEC_prg1 | RPL13A |
| 1.391E-175 | -0.5533652 | 0.053 | 0.38 | 2.969E-171 | TEC_prg1 | TMED9 |
| 3.991E-120 | -0.5536739 | 0.233 | 0.507 | 8.52E-116 | TEC_prg1 | SERP1 |
| 1.34E-105 | -0.554379 | 0.678 | 0.775 | 2.86E-101 | TEC_prg1 | CALM2 |
| 7.92E-191 | -0.555905 | 0.084 | 0.448 | 1.691E-186 | TEC_prg1 | TOMM20 |
| 2.606E-93 | -0.5562641 | 0.046 | 0.249 | 5.5638E-89 | TEC_prg1 | GCLM |
| 2.265E-179 | -0.556539 | 0.074 | 0.417 | 4.837E-175 | TEC_prg1 | NEDD8 |
| 3.503E-145 | -0.5565466 | 0.323 | 0.605 | 7.479E-141 | TEC_prg1 | RHOA |
| 3.2537E-64 | -0.5569814 | 0.137 | 0.326 | 6.9466E-60 | TEC_prg1 | ITGB8 |
| 3.2846E-84 | -0.5625442 | 0.412 | 0.594 | 7.0127E-80 | TEC_prg1 | CST3 |
| 5.367E-134 | -0.5633297 | 0.14 | 0.447 | 1.146E-129 | TEC_prg1 | DBI |
| 5.329E-125 | -0.5636568 | 0.862 | 0.904 | 1.138E-120 | TEC_prg1 | RPL35A |
| 1.757E-153 | -0.5652634 | 0.21 | 0.523 | 3.751E-149 | TEC_prg1 | SUMO2 |
| 2.486E-124 | -0.5666906 | 0.296 | 0.562 | 5.307E-120 | TEC_prg1 | OST4 |
| 9.804E-159 | -0.5680677 | 0.089 | 0.413 | 2.093E-154 | TEC_prg1 | PSME1 |
| 3.5167E-27 | -0.5681292 | 0.154 | 0.265 | 7.5082E-23 | TEC_prg1 | MT1X |
| 1.053E-133 | -0.5685003 | 0.612 | 0.774 | 2.249E-129 | TEC_prg1 | DSTN |
| 1.344E-124 | -0.5689394 | 0.861 | 0.904 | 2.87E-120 | TEC_prg1 | RPS15A |
| 5.232E-125 | -0.5706833 | 0.633 | 0.771 | 1.117E-120 | TEC_prg1 | RTN4 |
| 8.67E-185 | -0.5721325 | 0.057 | 0.397 | 1.851E-180 | TEC_prg1 | GHITM |
| 2.141E-203 | -0.5728019 | 0.037 | 0.39 | 4.571E-199 | TEC_prg1 | EIF3E |
| 6.265E-128 | -0.5730222 | 0.141 | 0.432 | 1.338E-123 | TEC_prg1 | OCIAD2 |
| 4.254E-112 | -0.5733038 | 0.37 | 0.599 | 9.082E-108 | TEC_prg1 | GUK1 |
| 6.35E-106 | -0.5737094 | 0.556 | 0.705 | 1.356E-101 | TEC_prg1 | CLIC1 |
| 3.28E-195 | -0.573725 | 0.069 | 0.431 | 7.004E-191 | TEC_prg1 | SNU13 |
| 8.157E-119 | -0.574916 | 0.872 | 0.907 | 1.742E-114 | TEC_prg1 | RPL27A |
| 7.965E-175 | -0.5749631 | 0.06 | 0.387 | 1.701E-170 | TEC_prg1 | PPA1 |
| 2.9E-144 | -0.5750236 | 0.629 | 0.779 | 6.191E-140 | TEC_prg1 | RPL4 |
| 5.6684E-91 | -0.5751246 | 0.472 | 0.643 | 1.2102E-86 | TEC_prg1 | CD81 |
| 1.993E-156 | -0.5764828 | 0.856 | 0.921 | 4.255E-152 | TEC_prg1 | FAU |
| 2.9053E-96 | -0.5778141 | 0.744 | 0.808 | 6.2027E-92 | TEC_prg1 | HLA-C |
| 1.66E-150 | -0.5785593 | 0.158 | 0.481 | 3.544E-146 | TEC_prg1 | BSG |
| 1.307E-187 | -0.5803203 | 0.051 | 0.392 | 2.79E-183 | TEC_prg1 | PSMB1 |
| 2.246E-170 | -0.5807248 | 0.145 | 0.488 | 4.796E-166 | TEC_prg1 | BZW1 |
| 2.365E-183 | -0.5830822 | 0.076 | 0.424 | 5.049E-179 | TEC_prg1 | SBDS |
| 1.2786E-83 | -0.5833726 | 0.711 | 0.775 | 2.7298E-79 | TEC_prg1 | PKM |

| | | | | | | |
|---|---|---|---|---|---|---|
| 1.7872E-93 | -0.58357 | 0.644 | 0.745 | 3.8157E-89 | TEC_prg1 | ENO1 |
| 8.913E-186 | -0.5835917 | 0.041 | 0.376 | 1.903E-181 | TEC_prg1 | NPC2 |
| 4.788E-142 | -0.5881721 | 0.4 | 0.644 | 1.022E-137 | TEC_prg1 | PSMA7 |
| 3.987E-132 | -0.588312 | 0.517 | 0.712 | 8.512E-128 | TEC_prg1 | COX6A1 |
| 1.417E-117 | -0.5907308 | 0.845 | 0.873 | 3.025E-113 | TEC_prg1 | RPL10A |
| 3.067E-151 | -0.5945999 | 0.488 | 0.697 | 6.548E-147 | TEC_prg1 | RAC1 |
| 2.7543E-13 | -0.594855 | 0.884 | 0.801 | 5.8804E-09 | TEC_prg1 | IGFBP7 |
| 3.357E-25 | -0.5949972 | 0.689 | 0.657 | 7.1672E-21 | TEC_prg1 | TACSTD2 |
| 3.009E-192 | -0.5951003 | 0.077 | 0.437 | 6.423E-188 | TEC_prg1 | REEP5 |
| 3.9986E-55 | -0.5974195 | 0.204 | 0.388 | 8.5371E-51 | TEC_prg1 | KLF6 |
| 1.48E-189 | -0.5988667 | 0.111 | 0.476 | 3.161E-185 | TEC_prg1 | DYNLRB1 |
| 1.124E-180 | -0.5995372 | 0.077 | 0.422 | 2.4E-176 | TEC_prg1 | ATP5F1B |
| 7.746E-180 | -0.5996861 | 0.058 | 0.391 | 1.654E-175 | TEC_prg1 | PRELID1 |
| 2.968E-176 | -0.6009342 | 0.08 | 0.424 | 6.336E-172 | TEC_prg1 | ATP5MF |
| 7.466E-99 | -0.6014338 | 0.044 | 0.256 | 1.594E-94 | TEC_prg1 | SFN |
| 2.946E-168 | -0.6049145 | 0.171 | 0.507 | 6.29E-164 | TEC_prg1 | ATP5PF |
| 1.298E-179 | -0.6068658 | 0.159 | 0.503 | 2.771E-175 | TEC_prg1 | VDAC2 |
| 4.889E-104 | -0.6087024 | 0.113 | 0.358 | 1.044E-99 | TEC_prg1 | UGCG |
| 1.566E-174 | -0.6103254 | 0.095 | 0.437 | 3.343E-170 | TEC_prg1 | NDUFB2 |
| 4.899E-99 | -0.6105693 | 0.236 | 0.478 | 1.0459E-94 | TEC_prg1 | PPP1R14B |
| 2.195E-121 | -0.6122927 | 0.969 | 0.955 | 4.687E-117 | TEC_prg1 | EIF1 |
| 5.761E-125 | -0.613774 | 0.06 | 0.321 | 1.23E-120 | TEC_prg1 | SERPINB1 |
| 2.7179E-69 | -0.6151222 | 0.813 | 0.798 | 5.8028E-65 | TEC_prg1 | CD24 |
| 2.2111E-48 | -0.6154071 | 0.956 | 0.918 | 4.7206E-44 | TEC_prg1 | TMSB10 |
| 1.673E-18 | -0.6163609 | 0.274 | 0.375 | 3.5718E-14 | TEC_prg1 | HSPA1A |
| 1.42E-123 | -0.6173575 | 0.522 | 0.695 | 3.032E-119 | TEC_prg1 | SPINT2 |
| 3.535E-129 | -0.6185618 | 0.875 | 0.902 | 7.547E-125 | TEC_prg1 | RPL23A |
| 2.331E-189 | -0.6190708 | 0.055 | 0.401 | 4.977E-185 | TEC_prg1 | CAP1 |
| 1.107E-111 | -0.6194286 | 0.963 | 0.961 | 2.363E-107 | TEC_prg1 | RPS18 |
| 1.372E-194 | -0.6196244 | 0.068 | 0.428 | 2.929E-190 | TEC_prg1 | SDCBP |
| 9.238E-149 | -0.6196271 | 0.675 | 0.802 | 1.972E-144 | TEC_prg1 | COX4I1 |
| 8.351E-135 | -0.6205171 | 0.899 | 0.926 | 1.783E-130 | TEC_prg1 | RPL14 |
| 4.778E-133 | -0.6212223 | 0.918 | 0.936 | 1.02E-128 | TEC_prg1 | RPS9 |
| 2.701E-177 | -0.6244847 | 0.075 | 0.417 | 5.766E-173 | TEC_prg1 | ATP5MC3 |
| 4.936E-145 | -0.6251634 | 0.468 | 0.683 | 1.054E-140 | TEC_prg1 | UQCRH |
| 1.521E-186 | -0.6256502 | 0.127 | 0.484 | 3.247E-182 | TEC_prg1 | ATP6V0B |
| 0.00273061 | -0.6257477 | 0.703 | 0.608 | 1 | TEC_prg1 | VIM |
| 8.055E-152 | -0.6261449 | 0.315 | 0.598 | 1.72E-147 | TEC_prg1 | CANX |
| 7.782E-194 | -0.6278177 | 0.079 | 0.439 | 1.662E-189 | TEC_prg1 | RHEB |
| 1.4857E-60 | -0.6286268 | 0.01 | 0.135 | 3.1719E-56 | TEC_prg1 | TAGLN |
| 7.105E-140 | -0.6291914 | 0.878 | 0.913 | 1.517E-135 | TEC_prg1 | RPS16 |
| 9.208E-178 | -0.6305305 | 0.056 | 0.386 | 1.966E-173 | TEC_prg1 | SLC25A5 |
| 1.162E-197 | -0.6313555 | 0.109 | 0.476 | 2.482E-193 | TEC_prg1 | PARK7 |
| 6.479E-204 | -0.6342432 | 0.087 | 0.464 | 1.383E-199 | TEC_prg1 | LAMTOR5 |
| 5.985E-189 | -0.6343455 | 0.025 | 0.354 | 1.278E-184 | TEC_prg1 | MYDGF |
| 4.3112E-35 | -0.640384 | 0.078 | 0.19 | 9.2044E-31 | TEC_prg1 | S100A4 |
| 1.87E-111 | -0.6411596 | 0.108 | 0.367 | 3.993E-107 | TEC_prg1 | CYBA |
| 5.326E-158 | -0.6420947 | 0.101 | 0.422 | 1.137E-153 | TEC_prg1 | TAGLN2 |
| 2.391E-153 | -0.6442104 | 0.363 | 0.623 | 5.105E-149 | TEC_prg1 | ATP6V0E1 |
| 1.139E-119 | -0.644261 | 0.479 | 0.66 | 2.431E-115 | TEC_prg1 | ANXA5 |

| | | | | | | |
|---|---|---|---|---|---|---|
| 2.9279E-59 | -0.6450017 | 0.07 | 0.225 | 6.251E-55 | TEC_prg1 | ANXA1 |
| 1.812E-149 | -0.6465624 | 0.691 | 0.807 | 3.869E-145 | TEC_prg1 | NPM1 |
| 1.125E-107 | -0.6468777 | 0.289 | 0.525 | 2.401E-103 | TEC_prg1 | LDHA |
| 2.264E-183 | -0.6483019 | 0.604 | 0.779 | 4.833E-179 | TEC_prg1 | BTF3 |
| 4.924E-132 | -0.6485079 | 0.347 | 0.585 | 1.051E-127 | TEC_prg1 | BRI3 |
| 9.098E-154 | -0.6503796 | 0.879 | 0.928 | 1.942E-149 | TEC_prg1 | RPL34 |
| 4.538E-197 | -0.6526131 | 0.048 | 0.395 | 9.69E-193 | TEC_prg1 | SSR3 |
| 3.6951E-76 | -0.6529201 | 0.258 | 0.452 | 7.889E-72 | TEC_prg1 | TPM4 |
| 4.6834E-58 | -0.6529543 | 0.667 | 0.711 | 9.999E-54 | TEC_prg1 | ITGB1 |
| 5.145E-200 | -0.6536174 | 0.07 | 0.435 | 1.099E-195 | TEC_prg1 | VAMP8 |
| 4.087E-197 | -0.6548676 | 0.119 | 0.485 | 8.725E-193 | TEC_prg1 | TMEM59 |
| 2.5803E-74 | -0.6553941 | 0.062 | 0.242 | 5.509E-70 | TEC_prg1 | GLRX |
| 4.711E-188 | -0.6559141 | 0.182 | 0.526 | 1.006E-183 | TEC_prg1 | PRDX6 |
| 4.091E-122 | -0.6561263 | 0.033 | 0.271 | 8.735E-118 | TEC_prg1 | TGM2 |
| 1.295E-163 | -0.6595452 | 0.694 | 0.82 | 2.766E-159 | TEC_prg1 | RPL27 |
| 2.936E-183 | -0.6598856 | 0.837 | 0.9 | 6.269E-179 | TEC_prg1 | NACA |
| 7.361E-118 | -0.6611508 | 0.559 | 0.703 | 1.572E-113 | TEC_prg1 | EPCAM |
| 9.935E-184 | -0.6616269 | 0.064 | 0.406 | 2.121E-179 | TEC_prg1 | SELENOK |
| 4.984E-171 | -0.6694406 | 0.844 | 0.913 | 1.064E-166 | TEC_prg1 | RPS25 |
| 4.544E-189 | -0.669593 | 0.06 | 0.406 | 9.702E-185 | TEC_prg1 | EIF5A |
| 1.751E-106 | -0.6712743 | 0.921 | 0.937 | 3.738E-102 | TEC_prg1 | HSP90AA1 |
| 5.589E-149 | -0.6731626 | 0.376 | 0.623 | 1.193E-144 | TEC_prg1 | ARPC3 |
| 7.169E-156 | -0.6759255 | 0.3 | 0.578 | 1.531E-151 | TEC_prg1 | SEC61B |
| 2.41E-143 | -0.6778753 | 0.891 | 0.916 | 5.145E-139 | TEC_prg1 | RPL29 |
| 1.185E-211 | -0.6805345 | 0.067 | 0.443 | 2.529E-207 | TEC_prg1 | DYNLT1 |
| 6.623E-157 | -0.6820448 | 0.163 | 0.476 | 1.414E-152 | TEC_prg1 | SSR4 |
| 1.321E-123 | -0.6828141 | 0.09 | 0.359 | 2.82E-119 | TEC_prg1 | MAL |
| 2.744E-200 | -0.6847686 | 0.153 | 0.511 | 5.859E-196 | TEC_prg1 | LAPTM4A |
| 5.2238E-81 | -0.6870835 | 0.193 | 0.418 | 1.1153E-76 | TEC_prg1 | HERPUD1 |
| 3.584E-174 | -0.6882584 | 0.184 | 0.514 | 7.652E-170 | TEC_prg1 | COX8A |
| 3.5832E-30 | -0.6909095 | 0.036 | 0.119 | 7.6502E-26 | TEC_prg1 | FXYD4 |
| 2.1865E-90 | -0.6922012 | 0.655 | 0.737 | 4.6681E-86 | TEC_prg1 | CTSD |
| 1.824E-154 | -0.6927996 | 0.928 | 0.945 | 3.894E-150 | TEC_prg1 | RPS24 |
| 1.898E-74 | -0.6931571 | 0.147 | 0.353 | 4.0523E-70 | TEC_prg1 | ITGA2 |
| 3.662E-203 | -0.6977701 | 0.057 | 0.418 | 7.819E-199 | TEC_prg1 | TALDO1 |
| 9.592E-116 | -0.6978895 | 0.898 | 0.909 | 2.048E-111 | TEC_prg1 | MYL6 |
| 3.874E-119 | -0.6991259 | 0.071 | 0.326 | 8.27E-115 | TEC_prg1 | AKR1B1 |
| 2.054E-197 | -0.7007058 | 0.856 | 0.92 | 4.385E-193 | TEC_prg1 | RPL24 |
| 4.192E-124 | -0.7017236 | 0.065 | 0.329 | 8.949E-120 | TEC_prg1 | PLAUR |
| 5.381E-170 | -0.7026088 | 0.092 | 0.429 | 1.149E-165 | TEC_prg1 | LDHB |
| 2.273E-141 | -0.7034317 | 0.826 | 0.884 | 4.853E-137 | TEC_prg1 | HLA-A |
| 4.524E-191 | -0.7043831 | 0.043 | 0.383 | 9.658E-187 | TEC_prg1 | TSPO |
| 1.657E-155 | -0.7059706 | 0.535 | 0.72 | 3.537E-151 | TEC_prg1 | CYSTM1 |
| 2.007E-165 | -0.7064648 | 0.899 | 0.927 | 4.285E-161 | TEC_prg1 | RPL30 |
| 1.028E-234 | -0.7097687 | 0.063 | 0.46 | 2.194E-230 | TEC_prg1 | DAD1 |
| 3.309E-167 | -0.7175897 | 0.615 | 0.771 | 7.064E-163 | TEC_prg1 | HINT1 |
| 1.1485E-19 | -0.7177819 | 0.684 | 0.642 | 2.4521E-15 | TEC_prg1 | KRT7 |
| 8.741E-226 | -0.7187874 | 0.053 | 0.438 | 1.866E-221 | TEC_prg1 | SEC61G |
| 3.888E-162 | -0.7198659 | 0.81 | 0.876 | 8.3E-158 | TEC_prg1 | UBA52 |
| 7.127E-203 | -0.720584 | 0.889 | 0.94 | 1.522E-198 | TEC_prg1 | RPL18 |

| | | | | | | |
|---|---|---|---|---|---|---|
| 7.615E-151 | -0.7221988 | 0.72 | 0.82 | 1.626E-146 | TEC_prg1 | CFL1 |
| 2.48E-156 | -0.7222299 | 0.688 | 0.801 | 5.295E-152 | TEC_prg1 | EEF1B2 |
| 2.789E-175 | -0.7261003 | 0.927 | 0.95 | 5.955E-171 | TEC_prg1 | RPL11 |
| 4.244E-177 | -0.7263507 | 0.113 | 0.452 | 9.061E-173 | TEC_prg1 | RHOC |
| 6.056E-71 | -0.7280281 | 0.247 | 0.445 | 1.293E-66 | TEC_prg1 | PHLDA2 |
| 6.161E-138 | -0.7280807 | 0.374 | 0.603 | 1.315E-133 | TEC_prg1 | PPIB |
| 1.516E-141 | -0.7292761 | 0.689 | 0.79 | 3.236E-137 | TEC_prg1 | MYL12A |
| 1.117E-247 | -0.7303423 | 0.049 | 0.456 | 2.384E-243 | TEC_prg1 | POMP |
| 6.019E-122 | -0.7347919 | 0.376 | 0.584 | 1.285E-117 | TEC_prg1 | CD9 |
| 1.033E-100 | -0.7348768 | 0.416 | 0.586 | 2.2063E-96 | TEC_prg1 | HMGA1 |
| 6.573E-150 | -0.7353888 | 0.901 | 0.928 | 1.403E-145 | TEC_prg1 | RPLP2 |
| 5.225E-144 | -0.7369628 | 0.828 | 0.878 | 1.116E-139 | TEC_prg1 | RPS5 |
| 1.846E-194 | -0.7398715 | 0.134 | 0.485 | 3.941E-190 | TEC_prg1 | PDIA3 |
| 1.473E-147 | -0.743159 | 0.326 | 0.587 | 3.145E-143 | TEC_prg1 | CD59 |
| 2.663E-213 | -0.746509 | 0.087 | 0.459 | 5.686E-209 | TEC_prg1 | MGST3 |
| 3.99E-123 | -0.7498352 | 0.787 | 0.835 | 8.518E-119 | TEC_prg1 | HLA-B |
| 4.403E-177 | -0.7513252 | 0.867 | 0.917 | 9.401E-173 | TEC_prg1 | RPL26 |
| 3.068E-170 | -0.7547178 | 0.208 | 0.519 | 6.549E-166 | TEC_prg1 | PDIA6 |
| 8.919E-232 | -0.7569527 | 0.694 | 0.852 | 1.904E-227 | TEC_prg1 | TMBIM6 |
| 5.253E-145 | -0.7577137 | 0.176 | 0.467 | 1.121E-140 | TEC_prg1 | LGALS3 |
| 5.631E-120 | -0.7644689 | 0.958 | 0.956 | 1.202E-115 | TEC_prg1 | RPS2 |
| 1.479E-156 | -0.7764323 | 0.844 | 0.89 | 3.158E-152 | TEC_prg1 | RPL39 |
| 1.4497E-51 | -0.7783054 | 0.033 | 0.155 | 3.0952E-47 | TEC_prg1 | SERPINA1 |
| 5.787E-184 | -0.778646 | 0.91 | 0.934 | 1.235E-179 | TEC_prg1 | RPL7A |
| 2.063E-200 | -0.7802811 | 0.916 | 0.945 | 4.405E-196 | TEC_prg1 | RPL19 |
| 1.488E-118 | -0.7814993 | 0.634 | 0.75 | 3.176E-114 | TEC_prg1 | PFN1 |
| 7.6964E-61 | -0.7818107 | 0.173 | 0.356 | 1.6432E-56 | TEC_prg1 | MT1E |
| 2.347E-183 | -0.7862141 | 0.9 | 0.946 | 5.01E-179 | TEC_prg1 | RPS28 |
| 8.249E-162 | -0.7874267 | 0.603 | 0.756 | 1.761E-157 | TEC_prg1 | NDUFA4 |
| 7.0265E-83 | -0.7877238 | 0.145 | 0.362 | 1.5002E-78 | TEC_prg1 | IL32 |
| 2.713E-189 | -0.7883339 | 0.057 | 0.4 | 5.792E-185 | TEC_prg1 | S100A16 |
| 1.038E-171 | -0.7916584 | 0.912 | 0.938 | 2.216E-167 | TEC_prg1 | RPL32 |
| 2.623E-138 | -0.7921388 | 0.027 | 0.288 | 5.601E-134 | TEC_prg1 | PDZK1IP1 |
| 1.847E-130 | -0.8009138 | 0.389 | 0.599 | 3.944E-126 | TEC_prg1 | MAL2 |
| 2.997E-189 | -0.8028397 | 0.923 | 0.947 | 6.399E-185 | TEC_prg1 | RPL8 |
| 5.505E-204 | -0.8056303 | 0.819 | 0.882 | 1.175E-199 | TEC_prg1 | RACK1 |
| 1.1506E-73 | -0.8073244 | 0.764 | 0.774 | 2.4564E-69 | TEC_prg1 | HSP90B1 |
| 9.763E-203 | -0.8082798 | 0.8 | 0.879 | 2.084E-198 | TEC_prg1 | RPL23 |
| 2.648E-210 | -0.8119591 | 0.664 | 0.811 | 5.653E-206 | TEC_prg1 | CHCHD2 |
| 4.076E-154 | -0.8211519 | 0.92 | 0.93 | 8.703E-150 | TEC_prg1 | RPL18A |
| 4.644E-235 | -0.8270479 | 0.83 | 0.912 | 9.915E-231 | TEC_prg1 | RPL5 |
| 3.716E-202 | -0.8329529 | 0.918 | 0.937 | 7.933E-198 | TEC_prg1 | RPL3 |
| 1.69E-124 | -0.8347562 | 0.839 | 0.885 | 3.609E-120 | TEC_prg1 | ITM2B |
| 2.426E-182 | -0.8414754 | 0.888 | 0.921 | 5.18E-178 | TEC_prg1 | RPLP0 |
| 9.4555E-97 | -0.8432613 | 0.872 | 0.873 | 2.0187E-92 | TEC_prg1 | GSTP1 |
| 1.6178E-39 | -0.8436045 | 0.851 | 0.789 | 3.4541E-35 | TEC_prg1 | KRT18 |
| 9.924E-214 | -0.8495834 | 0.867 | 0.912 | 2.119E-209 | TEC_prg1 | RPS3 |
| 6.048E-203 | -0.8535667 | 0.742 | 0.853 | 1.291E-198 | TEC_prg1 | PPIA |
| 7.074E-216 | -0.8537085 | 0.851 | 0.926 | 1.51E-211 | TEC_prg1 | HSP90AB1 |
| 1.557E-122 | -0.8646549 | 0.265 | 0.506 | 3.325E-118 | TEC_prg1 | IFITM3 |

| | | | | | | |
|---|---|---|---|---|---|---|
| 1.994E-198 | -0.8658565 | 0.961 | 0.968 | 4.257E-194 | TEC_prg1 | RPL41 |
| 1.9447E-32 | -0.8796948 | 0.047 | 0.142 | 4.1519E-28 | TEC_prg1 | GPX3 |
| 1.8002E-89 | -0.8842815 | 0.971 | 0.951 | 3.8434E-85 | TEC_prg1 | S100A6 |
| 1.688E-164 | -0.8947448 | 0.597 | 0.739 | 3.603E-160 | TEC_prg1 | TXN |
| 2.3542E-57 | -0.8992107 | 0.323 | 0.466 | 5.0261E-53 | TEC_prg1 | UPP1 |
| 9.382E-219 | -0.9004824 | 0.898 | 0.934 | 2.003E-214 | TEC_prg1 | RPL21 |
| 1.948E-102 | -0.9093706 | 0.11 | 0.346 | 4.1591E-98 | TEC_prg1 | PLAU |
| 6.895E-229 | -0.9094408 | 0.915 | 0.942 | 1.472E-224 | TEC_prg1 | RPS23 |
| 8.369E-234 | -0.9109369 | 0.902 | 0.935 | 1.787E-229 | TEC_prg1 | RPS14 |
| 2.8825E-20 | -0.926476 | 0.626 | 0.585 | 6.1541E-16 | TEC_prg1 | KRT19 |
| 6.949E-264 | -0.954465 | 0.872 | 0.929 | 1.484E-259 | TEC_prg1 | RPL9 |
| 8.578E-243 | -0.9586633 | 0.849 | 0.905 | 1.831E-238 | TEC_prg1 | RPS13 |
| 4.1916E-72 | -0.9636033 | 0.24 | 0.422 | 8.9491E-68 | TEC_prg1 | TXNRD1 |
| 1.204E-266 | -0.9645554 | 0.809 | 0.887 | 2.571E-262 | TEC_prg1 | RPL7 |
| 3.161E-133 | -0.9704693 | 0.197 | 0.46 | 6.749E-129 | TEC_prg1 | CCND1 |
| 4.88E-264 | -0.9854057 | 0.974 | 0.972 | 1.042E-259 | TEC_prg1 | TPT1 |
| 1.144E-234 | -0.9881953 | 0.768 | 0.869 | 2.442E-230 | TEC_prg1 | MYL12B |
| 3.312E-132 | -0.9981957 | 0.091 | 0.376 | 7.072E-128 | TEC_prg1 | TIMP1 |
| 9.234E-229 | -1.0248891 | 0.769 | 0.857 | 1.971E-224 | TEC_prg1 | RPSA |
| 5.068E-141 | -1.025256 | 0.02 | 0.278 | 1.082E-136 | TEC_prg1 | CST6 |
| 4.177E-104 | -1.0302358 | 0.131 | 0.369 | 8.917E-100 | TEC_prg1 | LAMC2 |
| 2.137E-264 | -1.0323871 | 0.895 | 0.939 | 4.563E-260 | TEC_prg1 | RPS6 |
| 2.941E-302 | -1.0365115 | 0.835 | 0.919 | 6.278E-298 | TEC_prg1 | RPS7 |
| 3.463E-217 | -1.0460476 | 0.762 | 0.852 | 7.393E-213 | TEC_prg1 | PRDX1 |
| 4.0136E-69 | -1.065777 | 0.711 | 0.73 | 8.5689E-65 | TEC_prg1 | KRT8 |
| 3.1853E-93 | -1.0733165 | 0.067 | 0.277 | 6.8006E-89 | TEC_prg1 | TFPI2 |
| 6.541E-279 | -1.0814594 | 0.747 | 0.858 | 1.397E-274 | TEC_prg1 | CD63 |
| 6.759E-118 | -1.081988 | 0.313 | 0.518 | 1.443E-113 | TEC_prg1 | TSPAN1 |
| 2.58E-224 | -1.0873133 | 0.94 | 0.954 | 5.508E-220 | TEC_prg1 | RPS12 |
| 2.909E-283 | -1.0947117 | 0.889 | 0.928 | 6.211E-279 | TEC_prg1 | RPS3A |
| 1.8659E-51 | -1.0996745 | 0.68 | 0.684 | 3.9837E-47 | TEC_prg1 | HSPA5 |
| 1.3236E-23 | -1.1067749 | 0.692 | 0.636 | 2.826E-19 | TEC_prg1 | WFDC2 |
| 6.251E-218 | -1.1075856 | 0.91 | 0.929 | 1.335E-213 | TEC_prg1 | ACTG1 |
| 0 | -1.1158047 | 0.903 | 0.945 | 0 | TEC_prg1 | RPL15 |
| 0 | -1.1296529 | 0.984 | 0.988 | 0 | TEC_prg1 | EEF1A1 |
| 0 | -1.1340339 | 0.899 | 0.94 | 0 | TEC_prg1 | RPS27A |
| 2.0474E-75 | -1.1412203 | 0.344 | 0.495 | 4.3712E-71 | TEC_prg1 | C12orf75 |
| 4.863E-275 | -1.1517833 | 0.806 | 0.877 | 1.038E-270 | TEC_prg1 | RPS4X |
| 1.259E-105 | -1.16818 | 0.607 | 0.684 | 2.687E-101 | TEC_prg1 | SH3BGRL3 |
| 2.213E-121 | -1.1724622 | 0.067 | 0.319 | 4.724E-117 | TEC_prg1 | LGALS1 |
| 1.261E-254 | -1.1802258 | 0.911 | 0.956 | 2.693E-250 | TEC_prg1 | B2M |
| 2.156E-296 | -1.1937942 | 0.967 | 0.973 | 4.604E-292 | TEC_prg1 | RPLP1 |
| 1.667E-217 | -1.2076854 | 0.846 | 0.902 | 3.559E-213 | TEC_prg1 | GAPDH |
| 1.0444E-51 | -1.2570133 | 0.079 | 0.22 | 2.2298E-47 | TEC_prg1 | MT1G |
| 2.6022E-41 | -1.3165355 | 0.694 | 0.662 | 5.5556E-37 | TEC_prg1 | CRYAB |
| 8.493E-295 | -1.3217859 | 0.911 | 0.926 | 1.813E-290 | TEC_prg1 | RPL10 |
| 3.0164E-43 | -1.3384006 | 0.764 | 0.701 | 6.44E-39 | TEC_prg1 | MT2A |
| 3.52E-207 | -1.3464278 | 0.84 | 0.876 | 7.516E-203 | TEC_prg1 | S100A10 |
| 2.7611E-63 | -1.3525299 | 0.088 | 0.252 | 5.895E-59 | TEC_prg1 | DEFB1 |
| 9.909E-119 | -1.3673299 | 0.051 | 0.295 | 2.116E-114 | TEC_prg1 | LCN2 |

| | | | | | | |
|---|---|---|---|---|---|---|
| 7.343E-216 | -1.4429337 | 0.773 | 0.842 | 1.568E-211 | TEC_prg1 | ANXA2 |
| 5.585E-221 | -1.4638531 | 0.908 | 0.921 | 1.192E-216 | TEC_prg1 | S100A11 |
| 3.536E-272 | -1.4705032 | 0.894 | 0.927 | 7.55E-268 | TEC_prg1 | ACTB |
| 3.5E-152 | -1.5270267 | 0.889 | 0.864 | 7.473E-148 | TEC_prg1 | TMSB4X |
| 3.5628E-83 | -1.7424098 | 0.246 | 0.419 | 7.6067E-79 | TEC_prg1 | SLPI |
| 0 | -1.9040555 | 0.961 | 0.982 | 0 | TEC_prg1 | FTH1 |
| 1.641E-34 | -2.0515385 | 0.638 | 0.592 | 3.5036E-30 | TEC_prg1 | SPP1 |
| 0 | -2.1088286 | 0.933 | 0.967 | 0 | TEC_prg1 | FTL |
| 1.9108E-11 | -2.2480453 | 0.757 | 0.613 | 4.0797E-07 | TEC_prg1 | MMP7 |
| 0 | 2.46814627 | 0.869 | 0.197 | 0 | TEC_prg2 | HSPA2 |
| 0 | 2.44147749 | 0.854 | 0.212 | 0 | TEC_prg2 | MAFB |
| 0 | 2.17757831 | 0.847 | 0.23 | 0 | TEC_prg2 | HEXIM1 |
| 0 | 2.15569168 | 0.999 | 0.548 | 0 | TEC_prg2 | PLCG2 |
| 0 | 2.08577943 | 0.788 | 0.166 | 0 | TEC_prg2 | CSKMT |
| 0 | 2.03048566 | 0.982 | 0.591 | 0 | TEC_prg2 | JUN |
| 0 | 1.9764931 | 0.925 | 0.492 | 0 | TEC_prg2 | SOX4 |
| 0 | 1.97193497 | 0.827 | 0.322 | 0 | TEC_prg2 | HIST1H4C |
| 0 | 1.92953968 | 0.808 | 0.273 | 0 | TEC_prg2 | RHOB |
| 0 | 1.88653864 | 0.933 | 0.519 | 0 | TEC_prg2 | IER2 |
| 0 | 1.8037116 | 0.793 | 0.214 | 0 | TEC_prg2 | RASD1 |
| 0 | 1.75183219 | 0.74 | 0.259 | 0 | TEC_prg2 | GLUL |
| 0 | 1.73987837 | 0.952 | 0.601 | 0 | TEC_prg2 | JUND |
| 0 | 1.73753694 | 0.786 | 0.242 | 0 | TEC_prg2 | PLK2 |
| 0 | 1.71480387 | 0.71 | 0.224 | 0 | TEC_prg2 | HES1 |
| 0 | 1.7094913 | 0.788 | 0.228 | 0 | TEC_prg2 | FOS |
| 0 | 1.679923 | 0.67 | 0.125 | 0 | TEC_prg2 | PIK3R3 |
| 0 | 1.67007549 | 0.663 | 0.223 | 0 | TEC_prg2 | TCIM |
| 0 | 1.66207741 | 0.651 | 0.109 | 0 | TEC_prg2 | EGLN3 |
| 0 | 1.65647452 | 0.925 | 0.545 | 0 | TEC_prg2 | ELF3 |
| 0 | 1.64760282 | 0.875 | 0.386 | 0 | TEC_prg2 | IER3 |
| 0 | 1.63505838 | 0.718 | 0.152 | 0 | TEC_prg2 | SOCS1 |
| 0 | 1.62252639 | 0.726 | 0.197 | 0 | TEC_prg2 | IER5L |
| 0 | 1.61626267 | 0.83 | 0.332 | 0 | TEC_prg2 | JUNB |
| 0 | 1.61513359 | 0.602 | 0.163 | 0 | TEC_prg2 | ADAMTS1 |
| 0 | 1.60733333 | 0.706 | 0.205 | 0 | TEC_prg2 | EGR1 |
| 0 | 1.59742289 | 0.675 | 0.201 | 0 | TEC_prg2 | IER5 |
| 0 | 1.59326512 | 0.786 | 0.369 | 0 | TEC_prg2 | SRSF7 |
| 0 | 1.54792722 | 0.737 | 0.22 | 0 | TEC_prg2 | PPP1R10 |
| 0 | 1.54432279 | 0.679 | 0.189 | 0 | TEC_prg2 | ID4 |
| 0 | 1.52462022 | 0.956 | 0.562 | 0 | TEC_prg2 | GADD45B |
| 0 | 1.52322307 | 0.66 | 0.186 | 0 | TEC_prg2 | INTS6 |
| 0 | 1.49123641 | 0.698 | 0.142 | 0 | TEC_prg2 | SGK1 |
| 0 | 1.45622691 | 0.683 | 0.234 | 0 | TEC_prg2 | POLR2A |
| 3.538E-263 | 1.44136962 | 0.384 | 0.084 | 7.553E-259 | TEC_prg2 | NEFM |
| 1.536E-159 | 1.42811735 | 0.268 | 0.064 | 3.28E-155 | TEC_prg2 | CCDC80 |
| 0 | 1.36317836 | 0.467 | 0.055 | 0 | TEC_prg2 | AP001160.1 |
| 0 | 1.33899758 | 0.602 | 0.119 | 0 | TEC_prg2 | FOSB |
| 1.348E-291 | 1.29732592 | 0.737 | 0.33 | 2.878E-287 | TEC_prg2 | SLC38A2 |
| 0 | 1.26548573 | 0.448 | 0.044 | 0 | TEC_prg2 | PCDH10 |
| 0 | 1.25714037 | 0.547 | 0.148 | 0 | TEC_prg2 | H2AFX |

| | | | | | | |
|---|---|---|---|---|---|---|
| 1.551E-260 | 1.25562268 | 0.649 | 0.271 | 3.312E-256 | TEC_prg2 | CDKN1C |
| 1.689E-287 | 1.23417134 | 0.627 | 0.232 | 3.605E-283 | TEC_prg2 | CITED2 |
| 4.552E-231 | 1.23298013 | 0.866 | 0.644 | 9.718E-227 | TEC_prg2 | TPI1 |
| 9.851E-263 | 1.23165232 | 0.868 | 0.578 | 2.103E-258 | TEC_prg2 | BTG1 |
| 1.669E-262 | 1.2238009 | 0.688 | 0.3 | 3.564E-258 | TEC_prg2 | ZFP36L2 |
| 0 | 1.22361538 | 0.525 | 0.077 | 0 | TEC_prg2 | HIST2H2BF |
| 1.409E-186 | 1.20149471 | 0.656 | 0.358 | 3.008E-182 | TEC_prg2 | TUBA1B |
| 4.577E-250 | 1.19455556 | 0.671 | 0.3 | 9.772E-246 | TEC_prg2 | SNHG12 |
| 0 | 1.18316818 | 0.389 | 0.069 | 0 | TEC_prg2 | ARC |
| 3.742E-192 | 1.18084836 | 0.844 | 0.596 | 7.989E-188 | TEC_prg2 | PPP1R15A |
| 0 | 1.17520026 | 0.607 | 0.188 | 0 | TEC_prg2 | NFKBIZ |
| 0 | 1.16270336 | 0.499 | 0.09 | 0 | TEC_prg2 | AC025164.1 |
| 6.861E-194 | 1.1579617 | 0.598 | 0.294 | 1.465E-189 | TEC_prg2 | CEBPB |
| 2.784E-236 | 1.15768545 | 0.696 | 0.347 | 5.943E-232 | TEC_prg2 | BRD2 |
| 2.847E-218 | 1.15331044 | 0.597 | 0.242 | 6.078E-214 | TEC_prg2 | BTG2 |
| 3.865E-289 | 1.13983404 | 0.525 | 0.151 | 8.253E-285 | TEC_prg2 | GADD45G |
| 7.529E-257 | 1.13940149 | 0.587 | 0.225 | 1.607E-252 | TEC_prg2 | AMD1 |
| 1.448E-248 | 1.13913799 | 0.632 | 0.273 | 3.091E-244 | TEC_prg2 | SRSF2 |
| 0 | 1.13290692 | 0.482 | 0.084 | 0 | TEC_prg2 | HOMER1 |
| 0 | 1.12386526 | 0.536 | 0.103 | 0 | TEC_prg2 | AC007952.4 |
| 0 | 1.11636551 | 0.48 | 0.103 | 0 | TEC_prg2 | ID1 |
| 1.968E-297 | 1.08264953 | 0.621 | 0.207 | 4.201E-293 | TEC_prg2 | ID2 |
| 3.472E-207 | 1.07280227 | 0.619 | 0.302 | 7.413E-203 | TEC_prg2 | HNRNPAB |
| 0 | 1.06912027 | 0.543 | 0.138 | 0 | TEC_prg2 | ERN1 |
| 0 | 1.06749253 | 0.479 | 0.091 | 0 | TEC_prg2 | CSRNP2 |
| 3.437E-191 | 1.05024826 | 0.593 | 0.27 | 7.339E-187 | TEC_prg2 | PMAIP1 |
| 4.991E-179 | 1.04462479 | 0.857 | 0.631 | 1.066E-174 | TEC_prg2 | HSPA8 |
| 6.343E-275 | 1.04116533 | 0.534 | 0.169 | 1.354E-270 | TEC_prg2 | MAT2A |
| 3.883E-162 | 1.0382896 | 0.526 | 0.239 | 8.291E-158 | TEC_prg2 | SEPHS2 |
| 1.281E-168 | 1.03240589 | 0.605 | 0.329 | 2.734E-164 | TEC_prg2 | MCL1 |
| 8.998E-308 | 1.01901217 | 0.513 | 0.136 | 1.921E-303 | TEC_prg2 | KLF4 |
| 4.205E-134 | 1.01854782 | 0.523 | 0.26 | 8.977E-130 | TEC_prg2 | RND3 |
| 0 | 1.00383865 | 0.462 | 0.068 | 0 | TEC_prg2 | HIST4H4 |
| 0 | 0.99564114 | 0.452 | 0.055 | 0 | TEC_prg2 | MAFA |
| 1.885E-123 | 0.96203075 | 0.569 | 0.329 | 4.025E-119 | TEC_prg2 | DDX3X |
| 7.484E-180 | 0.94240777 | 0.539 | 0.227 | 1.598E-175 | TEC_prg2 | HIST1H2AC |
| 6.131E-203 | 0.94088209 | 0.612 | 0.263 | 1.309E-198 | TEC_prg2 | DDIT4 |
| 1.022E-202 | 0.94032442 | 0.978 | 0.844 | 2.182E-198 | TEC_prg2 | H3F3B |
| 1.155E-201 | 0.9347719 | 0.469 | 0.156 | 2.465E-197 | TEC_prg2 | TLE4 |
| 5.689E-192 | 0.93116601 | 0.599 | 0.266 | 1.215E-187 | TEC_prg2 | DNAJB1 |
| 0 | 0.92965204 | 0.364 | 0.052 | 0 | TEC_prg2 | AL021453.1 |
| 1.4617E-71 | 0.92841843 | 0.425 | 0.247 | 3.1206E-67 | TEC_prg2 | TUBA1A |
| 0 | 0.92696993 | 0.42 | 0.052 | 0 | TEC_prg2 | AL137077.2 |
| 2.67E-162 | 0.92585066 | 0.551 | 0.271 | 5.701E-158 | TEC_prg2 | C6orf62 |
| 2.093E-197 | 0.91790631 | 0.539 | 0.218 | 4.468E-193 | TEC_prg2 | IRF1 |
| 3.206E-161 | 0.90968559 | 0.637 | 0.346 | 6.845E-157 | TEC_prg2 | RSRP1 |
| 0 | 0.90631651 | 0.336 | 0.045 | 0 | TEC_prg2 | POU3F1 |
| 3.54E-179 | 0.90530736 | 0.685 | 0.355 | 7.558E-175 | TEC_prg2 | NFKBIA |
| 1.887E-107 | 0.90064047 | 0.612 | 0.399 | 4.029E-103 | TEC_prg2 | TUBB4B |
| 3.999E-193 | 0.8990785 | 0.616 | 0.267 | 8.538E-189 | TEC_prg2 | HSPA1B |

| | | | | | | |
|---|---|---|---|---|---|---|
| 1.479E-288 | 0.89355099 | 0.416 | 0.09 | 3.158E-284 | TEC_prg2 | AL135925.1 |
| 1.432E-224 | 0.89213369 | 0.415 | 0.113 | 3.057E-220 | TEC_prg2 | ANKRD37 |
| 3.888E-144 | 0.89115033 | 0.336 | 0.103 | 8.302E-140 | TEC_prg2 | COL7A1 |
| 3.284E-172 | 0.88865873 | 0.494 | 0.201 | 7.012E-168 | TEC_prg2 | CDKN2AIP |
| 0 | 0.88642188 | 0.39 | 0.046 | 0 | TEC_prg2 | HIST1H3A |
| 2.657E-176 | 0.88398888 | 0.543 | 0.23 | 5.672E-172 | TEC_prg2 | DUSP1 |
| 1.772E-203 | 0.87651214 | 0.475 | 0.167 | 3.783E-199 | TEC_prg2 | CHCHD7 |
| 1.534E-102 | 0.87330186 | 0.587 | 0.376 | 3.2752E-98 | TEC_prg2 | JPT1 |
| 4.05E-220 | 0.86915246 | 0.442 | 0.131 | 8.646E-216 | TEC_prg2 | HIST1H1E |
| 6.975E-148 | 0.86143513 | 0.489 | 0.22 | 1.489E-143 | TEC_prg2 | IDI1 |
| 1.677E-141 | 0.85707592 | 0.911 | 0.65 | 3.58E-137 | TEC_prg2 | MTRNR2L12 |
| 1.547E-145 | 0.85203558 | 0.544 | 0.254 | 3.304E-141 | TEC_prg2 | CEBPD |
| 7.517E-304 | 0.84928121 | 0.392 | 0.072 | 1.605E-299 | TEC_prg2 | AL355075.4 |
| 0 | 0.84620774 | 0.381 | 0.041 | 0 | TEC_prg2 | NRARP |
| 1.828E-214 | 0.84331978 | 0.438 | 0.133 | 3.902E-210 | TEC_prg2 | TSPYL2 |
| 3.24E-108 | 0.84302527 | 0.601 | 0.4 | 6.918E-104 | TEC_prg2 | SNHG8 |
| 3.755E-193 | 0.8428019 | 0.409 | 0.125 | 8.016E-189 | TEC_prg2 | CDKN2B |
| 1.039E-221 | 0.84202442 | 0.417 | 0.114 | 2.217E-217 | TEC_prg2 | AL118516.1 |
| 0 | 0.83566311 | 0.369 | 0.058 | 0 | TEC_prg2 | Z93241.1 |
| 7.727E-265 | 0.83519652 | 0.41 | 0.091 | 1.65E-260 | TEC_prg2 | PCDH9 |
| 7.269E-304 | 0.82668536 | 0.387 | 0.072 | 1.552E-299 | TEC_prg2 | AC012306.2 |
| 0 | 0.82611498 | 0.387 | 0.054 | 0 | TEC_prg2 | TAF4B |
| 6.435E-147 | 0.82377367 | 0.235 | 0.051 | 1.374E-142 | TEC_prg2 | HIST1H1A |
| 4.622E-162 | 0.82320796 | 0.398 | 0.14 | 9.867E-158 | TEC_prg2 | TMX4 |
| 2.263E-88 | 0.8224974 | 0.482 | 0.285 | 4.8315E-84 | TEC_prg2 | YME1L1 |
| 2.467E-204 | 0.82083581 | 0.253 | 0.043 | 5.266E-200 | TEC_prg2 | HIST1H2AG |
| 7.89E-145 | 0.80555985 | 0.523 | 0.235 | 1.685E-140 | TEC_prg2 | KCNQ1OT1 |
| 2.66E-262 | 0.77484903 | 0.403 | 0.093 | 5.679E-258 | TEC_prg2 | MEPCE |
| 1.7584E-67 | 0.77073911 | 0.616 | 0.464 | 3.7541E-63 | TEC_prg2 | H1FX |
| 0 | 0.76179877 | 0.396 | 0.072 | 0 | TEC_prg2 | RGS2 |
| 4.429E-198 | 0.7615638 | 0.302 | 0.065 | 9.455E-194 | TEC_prg2 | PARD6G-AS1 |
| 4.545E-124 | 0.74304675 | 0.923 | 0.78 | 9.703E-120 | TEC_prg2 | HNRNPA2B1 |
| 1.2913E-83 | 0.73266225 | 0.557 | 0.371 | 2.7569E-79 | TEC_prg2 | NDRG1 |
| 2.89E-181 | 0.72337176 | 0.39 | 0.12 | 6.171E-177 | TEC_prg2 | MED29 |
| 0 | 0.72274959 | 0.354 | 0.056 | 0 | TEC_prg2 | AC087239.1 |
| 1.8512E-77 | 0.72122452 | 0.857 | 0.697 | 3.9524E-73 | TEC_prg2 | HSPB1 |
| 4.595E-147 | 0.71577756 | 0.249 | 0.059 | 9.811E-143 | TEC_prg2 | AC093323.1 |
| 4.5444E-95 | 0.71488228 | 0.602 | 0.399 | 9.7023E-91 | TEC_prg2 | SFPQ |
| 4.574E-132 | 0.71421587 | 0.519 | 0.246 | 9.765E-128 | TEC_prg2 | VEGFA |
| 1.22E-295 | 0.70394761 | 0.271 | 0.032 | 2.605E-291 | TEC_prg2 | ADRB2 |
| 6.746E-121 | 0.70339893 | 0.385 | 0.15 | 1.44E-116 | TEC_prg2 | FOXC1 |
| 3.5332E-87 | 0.6999346 | 0.455 | 0.252 | 7.5434E-83 | TEC_prg2 | IFRD1 |
| 7.994E-151 | 0.69134076 | 0.464 | 0.184 | 1.707E-146 | TEC_prg2 | ZFP36 |
| 1.7954E-71 | 0.68910231 | 0.653 | 0.493 | 3.8332E-67 | TEC_prg2 | H2AFZ |
| 1.109E-100 | 0.68375492 | 0.5 | 0.263 | 2.3672E-96 | TEC_prg2 | CYR61 |
| 1.1E-113 | 0.68372289 | 0.284 | 0.089 | 2.35E-109 | TEC_prg2 | AC023157.3 |
| 1.361E-102 | 0.68277048 | 0.731 | 0.541 | 2.9065E-98 | TEC_prg2 | SRRM2 |
| 8.593E-203 | 0.67488 | 0.344 | 0.084 | 1.835E-198 | TEC_prg2 | DALRD3 |
| 4.184E-147 | 0.6740532 | 0.332 | 0.103 | 8.933E-143 | TEC_prg2 | IRF7 |
| 1.0934E-80 | 0.67208195 | 0.8 | 0.69 | 2.3344E-76 | TEC_prg2 | EIF4G2 |

| | | | | | | |
|---|---|---|---|---|---|---|
| 0 | 0.67158336 | 0.323 | 0.038 | 0 | TEC_prg2 | AC069224.1 |
| 1.379E-59 | 0.6629622 | 0.281 | 0.135 | 2.9442E-55 | TEC_prg2 | SLC16A3 |
| 3.493E-101 | 0.66180131 | 0.421 | 0.197 | 7.4573E-97 | TEC_prg2 | ADM |
| 1.005E-110 | 0.65192449 | 0.872 | 0.686 | 2.147E-106 | TEC_prg2 | DDX5 |
| 4.8746E-63 | 0.65092737 | 0.44 | 0.285 | 1.0407E-58 | TEC_prg2 | EIF4A1 |
| 6.642E-64 | 0.64117681 | 0.375 | 0.214 | 1.4181E-59 | TEC_prg2 | UGP2 |
| 5.947E-235 | 0.63979428 | 0.254 | 0.037 | 1.27E-230 | TEC_prg2 | C2CD4B |
| 8.001E-171 | 0.63850505 | 0.354 | 0.103 | 1.708E-166 | TEC_prg2 | ING1 |
| 3.878E-231 | 0.63169872 | 0.308 | 0.058 | 8.279E-227 | TEC_prg2 | FZD8 |
| 7.208E-155 | 0.63079492 | 0.296 | 0.079 | 1.539E-150 | TEC_prg2 | SLC12A2 |
| 2.2431E-66 | 0.63057481 | 0.638 | 0.504 | 4.7891E-62 | TEC_prg2 | HNRNPU |
| 2.0677E-79 | 0.6291674 | 0.481 | 0.277 | 4.4145E-75 | TEC_prg2 | MARCKSL1 |
| 4.0173E-67 | 0.628282 | 0.416 | 0.25 | 8.5769E-63 | TEC_prg2 | PABPC4 |
| 1.243E-177 | 0.6228754 | 0.322 | 0.082 | 2.654E-173 | TEC_prg2 | DEPP1 |
| 7.503E-262 | 0.62223082 | 0.252 | 0.031 | 1.602E-257 | TEC_prg2 | LINC02432 |
| 1.4835E-92 | 0.62175491 | 0.272 | 0.099 | 3.1673E-88 | TEC_prg2 | AKT1S1 |
| 1.5353E-91 | 0.61785676 | 0.27 | 0.097 | 3.2779E-87 | TEC_prg2 | FSCN1 |
| 4.437E-107 | 0.61730012 | 0.261 | 0.08 | 9.473E-103 | TEC_prg2 | CXCL3 |
| 1.0428E-86 | 0.61666929 | 0.985 | 0.903 | 2.2264E-82 | TEC_prg2 | UBC |
| 1.761E-65 | 0.61372252 | 0.609 | 0.424 | 3.7597E-61 | TEC_prg2 | ZFP36L1 |
| 9.5111E-97 | 0.60969134 | 0.786 | 0.577 | 2.0306E-92 | TEC_prg2 | CLU |
| 1.496E-180 | 0.60697141 | 0.349 | 0.093 | 3.194E-176 | TEC_prg2 | LINC00472 |
| 5.474E-109 | 0.60561914 | 0.284 | 0.096 | 1.169E-104 | TEC_prg2 | KPNA2 |
| 5.8872E-51 | 0.6004254 | 0.21 | 0.088 | 1.2569E-46 | TEC_prg2 | TNF |
| 1.202E-105 | 0.59748146 | 0.339 | 0.129 | 2.566E-101 | TEC_prg2 | PFKFB3 |
| 1.585E-127 | 0.58465972 | 0.318 | 0.104 | 3.383E-123 | TEC_prg2 | SOCS3 |
| 2.0885E-51 | 0.58018723 | 0.436 | 0.287 | 4.459E-47 | TEC_prg2 | SNHG7 |
| 4.513E-104 | 0.57192216 | 0.317 | 0.117 | 9.635E-100 | TEC_prg2 | EFNB2 |
| 7.359E-128 | 0.57045928 | 0.347 | 0.118 | 1.571E-123 | TEC_prg2 | ID3 |
| 3.946E-173 | 0.568068 | 0.235 | 0.043 | 8.424E-169 | TEC_prg2 | AC005837.1 |
| 2.284E-198 | 0.5660944 | 0.275 | 0.053 | 4.876E-194 | TEC_prg2 | FAM117A |
| 2.508E-175 | 0.56409531 | 0.237 | 0.044 | 5.354E-171 | TEC_prg2 | AC069185.1 |
| 7.9283E-27 | 0.56325852 | 0.471 | 0.392 | 1.6927E-22 | TEC_prg2 | FUS |
| 4.068E-153 | 0.5606054 | 0.322 | 0.092 | 8.685E-149 | TEC_prg2 | TM7SF2 |
| 3.394E-225 | 0.55901038 | 0.27 | 0.044 | 7.246E-221 | TEC_prg2 | MYCL |
| 2.781E-112 | 0.55435054 | 0.288 | 0.095 | 5.937E-108 | TEC_prg2 | KCTD12 |
| 3.445E-150 | 0.55403544 | 0.268 | 0.065 | 7.355E-146 | TEC_prg2 | HIST2H2AC |
| 9.9678E-76 | 0.55249261 | 0.276 | 0.113 | 2.1281E-71 | TEC_prg2 | SLC6A8 |
| 1.535E-137 | 0.55134978 | 0.277 | 0.076 | 3.277E-133 | TEC_prg2 | ARL4D |
| 3.807E-307 | 0.54976898 | 0.253 | 0.024 | 8.128E-303 | TEC_prg2 | HIST2H4B |
| 3.159E-112 | 0.54948614 | 0.326 | 0.118 | 6.744E-108 | TEC_prg2 | C12orf65 |
| 2.4578E-70 | 0.5470741 | 0.93 | 0.817 | 5.2473E-66 | TEC_prg2 | SQSTM1 |
| 3.755E-175 | 0.54609665 | 0.299 | 0.071 | 8.018E-171 | TEC_prg2 | C3orf58 |
| 9.5255E-38 | 0.54411067 | 0.483 | 0.364 | 2.0337E-33 | TEC_prg2 | SRSF5 |
| 6.4694E-35 | 0.53502788 | 0.935 | 0.816 | 1.3812E-30 | TEC_prg2 | MT-ND5 |
| 3.3367E-54 | 0.53120001 | 0.412 | 0.257 | 7.1239E-50 | TEC_prg2 | SPTAN1 |
| 1.335E-113 | 0.53018622 | 0.297 | 0.097 | 2.849E-109 | TEC_prg2 | SOX9 |
| 7.6572E-71 | 0.527401 | 0.246 | 0.094 | 1.6348E-66 | TEC_prg2 | EGOT |
| 6.419E-158 | 0.52434802 | 0.996 | 0.793 | 1.37E-153 | TEC_prg2 | NEAT1 |
| 4.281E-113 | 0.51998608 | 0.309 | 0.107 | 9.14E-109 | TEC_prg2 | NARF |

| | | | | | | |
|---|---|---|---|---|---|---|
| 1.268E-136 | 0.51528837 | 0.286 | 0.08 | 2.708E-132 | TEC_prg2 | PLK3 |
| 2.49E-157 | 0.51286263 | 0.285 | 0.071 | 5.317E-153 | TEC_prg2 | RND1 |
| 9.848E-271 | 0.51245169 | 0.225 | 0.022 | 2.103E-266 | TEC_prg2 | HIST1H4A |
| 1.2083E-46 | 0.50877169 | 0.469 | 0.329 | 2.5798E-42 | TEC_prg2 | TSC22D1 |
| 6.961E-131 | 0.50862251 | 0.265 | 0.073 | 1.486E-126 | TEC_prg2 | HOXD1 |
| 1.0212E-52 | 0.50676995 | 0.505 | 0.34 | 2.1803E-48 | TEC_prg2 | ARGLU1 |
| 1.5785E-93 | 0.50569161 | 0.239 | 0.076 | 3.37E-89 | TEC_prg2 | FOXQ1 |
| 1.6012E-69 | 0.50559691 | 0.3 | 0.138 | 3.4186E-65 | TEC_prg2 | KIAA1191 |
| 1.0978E-37 | 0.5055416 | 0.682 | 0.6 | 2.3437E-33 | TEC_prg2 | EIF4A2 |
| 1.4359E-91 | 0.50483754 | 0.183 | 0.047 | 3.0656E-87 | TEC_prg2 | HIST1H2BG |
| 2.501E-101 | 0.50360253 | 0.293 | 0.103 | 5.3398E-97 | TEC_prg2 | PTGER3 |
| 3.0317E-37 | 0.49292882 | 0.419 | 0.303 | 6.4726E-33 | TEC_prg2 | STAT3 |
| 3.333E-152 | 0.48379538 | 0.253 | 0.058 | 7.116E-148 | TEC_prg2 | CYB561A3 |
| 6.177E-58 | 0.47899544 | 0.797 | 0.652 | 1.3188E-53 | TEC_prg2 | GLS |
| 1.1956E-83 | 0.47582657 | 0.295 | 0.12 | 2.5526E-79 | TEC_prg2 | SLC9A1 |
| 2.5645E-81 | 0.47477951 | 0.248 | 0.09 | 5.4753E-77 | TEC_prg2 | C6orf120 |
| 6.5063E-59 | 0.47226857 | 0.244 | 0.107 | 1.3891E-54 | TEC_prg2 | FADS3 |
| 9.21E-105 | 0.47148866 | 0.34 | 0.129 | 1.966E-100 | TEC_prg2 | SNHG25 |
| 1.9976E-40 | 0.46964419 | 0.369 | 0.239 | 4.265E-36 | TEC_prg2 | PNPLA2 |
| 6.7453E-16 | 0.46342888 | 0.395 | 0.324 | 1.4401E-11 | TEC_prg2 | H1F0 |
| 1.657E-194 | 0.46340713 | 0.225 | 0.035 | 3.537E-190 | TEC_prg2 | HIST1H4B |
| 2.9181E-45 | 0.4620812 | 0.366 | 0.225 | 6.2302E-41 | TEC_prg2 | LMO4 |
| 1.1979E-67 | 0.46195287 | 0.24 | 0.094 | 2.5574E-63 | TEC_prg2 | TENT5A |
| 1.2805E-42 | 0.46103932 | 0.408 | 0.27 | 2.7339E-38 | TEC_prg2 | NUFIP2 |
| 1.141E-126 | 0.4607558 | 0.252 | 0.067 | 2.436E-122 | TEC_prg2 | FRAT2 |
| 6.415E-183 | 0.46032266 | 0.225 | 0.037 | 1.369E-178 | TEC_prg2 | AL136040.1 |
| 8.9471E-32 | 0.45713679 | 0.651 | 0.607 | 1.9102E-27 | TEC_prg2 | SAP18 |
| 1.585E-106 | 0.45684729 | 0.274 | 0.087 | 3.383E-102 | TEC_prg2 | KIF12 |
| 2.4076E-40 | 0.45656222 | 0.31 | 0.18 | 5.1403E-36 | TEC_prg2 | TOB1 |
| 5.237E-107 | 0.45631486 | 0.285 | 0.096 | 1.118E-102 | TEC_prg2 | TCTA |
| 7.9299E-34 | 0.45623943 | 0.334 | 0.222 | 1.693E-29 | TEC_prg2 | NEU1 |
| 1.865E-136 | 0.45555713 | 0.205 | 0.041 | 3.981E-132 | TEC_prg2 | IFI30 |
| 1.2658E-29 | 0.45496164 | 0.343 | 0.234 | 2.7025E-25 | TEC_prg2 | IRS2 |
| 2.1723E-86 | 0.44476472 | 0.242 | 0.082 | 4.6378E-82 | TEC_prg2 | FAM84A |
| 1.6061E-25 | 0.44458001 | 0.536 | 0.48 | 3.4291E-21 | TEC_prg2 | ATP1A1 |
| 1.241E-204 | 0.44346143 | 0.222 | 0.031 | 2.65E-200 | TEC_prg2 | NANOS1 |
| 9.998E-62 | 0.44288246 | 0.327 | 0.161 | 2.1346E-57 | TEC_prg2 | MYO9A |
| 3.93E-48 | 0.44175655 | 0.425 | 0.268 | 8.3905E-44 | TEC_prg2 | HIST1H1C |
| 1.448E-254 | 0.44095616 | 0.213 | 0.021 | 3.093E-250 | TEC_prg2 | HIST1H3C |
| 1.3023E-46 | 0.43994373 | 0.863 | 0.745 | 2.7805E-42 | TEC_prg2 | DYNLL1 |
| 5.82E-130 | 0.43965115 | 0.254 | 0.066 | 1.243E-125 | TEC_prg2 | HIST2H2BE |
| 2.3768E-40 | 0.43884938 | 0.551 | 0.413 | 5.0744E-36 | TEC_prg2 | SYNE2 |
| 1.652E-111 | 0.43815354 | 0.216 | 0.056 | 3.527E-107 | TEC_prg2 | GCH1 |
| 1.235E-186 | 0.43470661 | 0.237 | 0.04 | 2.636E-182 | TEC_prg2 | AL022069.1 |
| 4.033E-101 | 0.43454336 | 0.214 | 0.059 | 8.6106E-97 | TEC_prg2 | FAM43A |
| 8.642E-156 | 0.43185918 | 0.219 | 0.041 | 1.845E-151 | TEC_prg2 | HIST1H2BN |
| 1.31E-124 | 0.43112866 | 0.238 | 0.061 | 2.797E-120 | TEC_prg2 | KLHL15 |
| 2.1271E-89 | 0.42924969 | 0.285 | 0.104 | 4.5414E-85 | TEC_prg2 | KMT2E-AS1 |
| 5.5035E-68 | 0.42731603 | 0.207 | 0.073 | 1.175E-63 | TEC_prg2 | SLC7A8 |
| 5.0066E-72 | 0.42415554 | 0.231 | 0.084 | 1.0689E-67 | TEC_prg2 | FDXR |

| | | | | | | |
|---|---|---|---|---|---|---|
| 8.9298E-61 | 0.4234889 | 0.968 | 0.859 | 1.9065E-56 | TEC_prg2 | SAT1 |
| 3.7864E-36 | 0.42290086 | 0.303 | 0.18 | 8.084E-32 | TEC_prg2 | POLR2J3.1 |
| 7.8084E-31 | 0.41863281 | 0.288 | 0.18 | 1.6671E-26 | TEC_prg2 | KLHL21 |
| 1.508E-24 | 0.41802039 | 0.697 | 0.66 | 3.2196E-20 | TEC_prg2 | SOD1 |
| 7.4132E-79 | 0.41656152 | 0.274 | 0.108 | 1.5827E-74 | TEC_prg2 | USP38 |
| 3.9184E-47 | 0.41505488 | 0.312 | 0.17 | 8.3657E-43 | TEC_prg2 | PCF11 |
| 1.824E-196 | 0.41396978 | 0.185 | 0.022 | 3.895E-192 | TEC_prg2 | AC011498.1 |
| 9.948E-34 | 0.41234011 | 0.298 | 0.184 | 2.1239E-29 | TEC_prg2 | EPB41L5 |
| 9.1926E-50 | 0.41155171 | 0.251 | 0.118 | 1.9626E-45 | TEC_prg2 | RDH10 |
| 8.9212E-37 | 0.41046817 | 0.697 | 0.617 | 1.9047E-32 | TEC_prg2 | ATF4 |
| 8.4016E-35 | 0.41021034 | 0.54 | 0.42 | 1.7937E-30 | TEC_prg2 | DDX17 |
| 1.8229E-40 | 0.40557598 | 0.253 | 0.132 | 3.892E-36 | TEC_prg2 | SLC39A10 |
| 6.495E-162 | 0.40429626 | 0.19 | 0.029 | 1.387E-157 | TEC_prg2 | RTL5 |
| 8.0384E-49 | 0.40193563 | 0.229 | 0.105 | 1.7162E-44 | TEC_prg2 | ABTB2 |
| 1.0862E-08 | 0.40026992 | 0.589 | 0.566 | 0.00023191 | TEC_prg2 | RPS27L |
| 5.352E-172 | 0.39935525 | 0.202 | 0.032 | 1.143E-167 | TEC_prg2 | GFPT2 |
| 1.3757E-34 | 0.39443648 | 0.314 | 0.192 | 2.9372E-30 | TEC_prg2 | THUMPD3-AS1 |
| 2.563E-110 | 0.39395906 | 0.232 | 0.063 | 5.471E-106 | TEC_prg2 | ATP2B1-AS1 |
| 1.656E-253 | 0.39381124 | 0.193 | 0.016 | 3.536E-249 | TEC_prg2 | LRAT |
| 5.0443E-82 | 0.39063991 | 0.244 | 0.085 | 1.077E-77 | TEC_prg2 | SLC8B1 |
| 8.216E-242 | 0.38979064 | 0.18 | 0.014 | 1.754E-237 | TEC_prg2 | HIST1H2BB |
| 9.178E-160 | 0.389748 | 0.153 | 0.018 | 1.959E-155 | TEC_prg2 | HIST1H3H |
| 9.2733E-37 | 0.38880578 | 0.355 | 0.218 | 1.9798E-32 | TEC_prg2 | SYNE1 |
| 5.9293E-24 | 0.38755796 | 0.413 | 0.323 | 1.2659E-19 | TEC_prg2 | CPEB4 |
| 3.125E-61 | 0.3872642 | 0.212 | 0.081 | 6.6719E-57 | TEC_prg2 | ATF7IP2 |
| 1.9487E-98 | 0.3848983 | 0.213 | 0.059 | 4.1605E-94 | TEC_prg2 | HIST1H4H |
| 4.4415E-32 | 0.38314607 | 0.204 | 0.106 | 9.4825E-28 | TEC_prg2 | SERPINH1 |
| 9.787E-170 | 0.38224663 | 0.183 | 0.026 | 2.089E-165 | TEC_prg2 | CCNE2 |
| 3.0726E-62 | 0.3817663 | 0.276 | 0.121 | 6.5599E-58 | TEC_prg2 | NABP1 |
| 4.4796E-32 | 0.38078229 | 0.43 | 0.307 | 9.564E-28 | TEC_prg2 | CCNL1 |
| 4.0489E-48 | 0.37977983 | 0.245 | 0.116 | 8.6445E-44 | TEC_prg2 | LBH |
| 9.2288E-48 | 0.37956469 | 0.355 | 0.201 | 1.9703E-43 | TEC_prg2 | RERE |
| 4.9065E-75 | 0.37950608 | 0.22 | 0.075 | 1.0475E-70 | TEC_prg2 | PDGFB |
| 1.5827E-08 | 0.37911749 | 0.22 | 0.175 | 0.00033791 | TEC_prg2 | SQLE |
| 8.223E-52 | 0.37827489 | 0.286 | 0.141 | 1.7556E-47 | TEC_prg2 | SLC20A1 |
| 3.4549E-39 | 0.37789029 | 0.33 | 0.191 | 7.3762E-35 | TEC_prg2 | RRAD |
| 1.8819E-45 | 0.37771197 | 0.29 | 0.151 | 4.0179E-41 | TEC_prg2 | CTGF |
| 1.861E-129 | 0.37701073 | 0.169 | 0.029 | 3.973E-125 | TEC_prg2 | AC107959.4 |
| 1.4128E-52 | 0.37421475 | 0.27 | 0.129 | 3.0163E-48 | TEC_prg2 | NXF1 |
| 1.9029E-59 | 0.37344469 | 0.219 | 0.088 | 4.0627E-55 | TEC_prg2 | HES6 |
| 8.3876E-56 | 0.36988174 | 0.249 | 0.11 | 1.7908E-51 | TEC_prg2 | NQO2 |
| 1.741E-112 | 0.36760018 | 0.224 | 0.058 | 3.717E-108 | TEC_prg2 | DDIT4L |
| 2.4968E-67 | 0.36455143 | 0.209 | 0.074 | 5.3306E-63 | TEC_prg2 | AC091271.1 |
| 1.491E-160 | 0.3634777 | 0.182 | 0.027 | 3.184E-156 | TEC_prg2 | LINC01970 |
| 2.9266E-67 | 0.36238434 | 0.135 | 0.035 | 6.2482E-63 | TEC_prg2 | AL031777.3 |
| 6.9817E-77 | 0.3617097 | 0.206 | 0.066 | 1.4906E-72 | TEC_prg2 | MIR222HG |
| 6.6221E-19 | 0.36053671 | 0.357 | 0.275 | 1.4138E-14 | TEC_prg2 | COL18A1 |
| 7.4931E-69 | 0.35720537 | 0.193 | 0.063 | 1.5998E-64 | TEC_prg2 | CXCL2 |
| 5.155E-118 | 0.35586322 | 0.181 | 0.037 | 1.101E-113 | TEC_prg2 | AC144652.1 |
| 1.3766E-75 | 0.35250131 | 0.242 | 0.088 | 2.9391E-71 | TEC_prg2 | GPS2 |

| | | | | | | |
|---|---|---|---|---|---|---|
| 9.072E-48 | 0.35247408 | 0.258 | 0.124 | 1.9369E-43 | TEC_prg2 | SNX29 |
| 1.358E-158 | 0.35184327 | 0.165 | 0.022 | 2.899E-154 | TEC_prg2 | CABYR |
| 1.5354E-78 | 0.35097297 | 0.214 | 0.07 | 3.2781E-74 | TEC_prg2 | CCDC71L |
| 1.543E-158 | 0.34965274 | 0.167 | 0.023 | 3.293E-154 | TEC_prg2 | CEBPA |
| 1.034E-156 | 0.34759506 | 0.17 | 0.024 | 2.209E-152 | TEC_prg2 | HIST1H2AH |
| 3.6615E-63 | 0.34701643 | 0.266 | 0.111 | 7.8173E-59 | TEC_prg2 | PKHD1 |
| 5.679E-160 | 0.346803 | 0.167 | 0.022 | 1.212E-155 | TEC_prg2 | AC083973.1 |
| 1.953E-114 | 0.34372777 | 0.183 | 0.039 | 4.17E-110 | TEC_prg2 | RASL11B |
| 2.0221E-79 | 0.34315848 | 0.203 | 0.063 | 4.3171E-75 | TEC_prg2 | ZNF821 |
| 5.735E-212 | 0.3371495 | 0.186 | 0.019 | 1.224E-207 | TEC_prg2 | PRMT5-AS1 |
| 4.7741E-45 | 0.33604073 | 0.252 | 0.125 | 1.0193E-40 | TEC_prg2 | PHYH |
| 5.402E-163 | 0.33559162 | 0.171 | 0.023 | 1.153E-158 | TEC_prg2 | AL591846.2 |
| 1.168E-23 | 0.33477187 | 0.436 | 0.326 | 2.4936E-19 | TEC_prg2 | PAX8 |
| 4.3457E-06 | 0.33442242 | 0.397 | 0.397 | 0.09278096 | TEC_prg2 | CYCS |
| 7.9153E-74 | 0.33153322 | 0.157 | 0.042 | 1.6899E-69 | TEC_prg2 | PLIN5 |
| 3.6821E-66 | 0.33049481 | 0.201 | 0.071 | 7.8612E-62 | TEC_prg2 | DGAT1 |
| 2.254E-17 | 0.3300235 | 0.243 | 0.169 | 4.8123E-13 | TEC_prg2 | SLC1A5 |
| 1.4236E-28 | 0.3286875 | 0.244 | 0.143 | 3.0395E-24 | TEC_prg2 | ASXL1 |
| 7.2505E-74 | 0.3279558 | 0.195 | 0.061 | 1.548E-69 | TEC_prg2 | RGS16 |
| 3.5602E-96 | 0.32718995 | 0.187 | 0.047 | 7.6011E-92 | TEC_prg2 | FGF9 |
| 2.1133E-28 | 0.32486725 | 0.264 | 0.16 | 4.5119E-24 | TEC_prg2 | DUSP5 |
| 8.2611E-31 | 0.32477003 | 0.222 | 0.122 | 1.7638E-26 | TEC_prg2 | EPHX1 |
| 1.8891E-29 | 0.32437894 | 0.237 | 0.136 | 4.0332E-25 | TEC_prg2 | AHCY |
| 4.5551E-67 | 0.32150532 | 0.194 | 0.066 | 9.7252E-63 | TEC_prg2 | TMEM69 |
| 1.4327E-61 | 0.32052527 | 0.194 | 0.07 | 3.0587E-57 | TEC_prg2 | EPAS1 |
| 1.5336E-24 | 0.31942107 | 0.234 | 0.142 | 3.2743E-20 | TEC_prg2 | SLC7A5 |
| 4.587E-40 | 0.31823593 | 0.281 | 0.151 | 9.7931E-36 | TEC_prg2 | PAXBP1 |
| 2.169E-149 | 0.31808712 | 0.168 | 0.025 | 4.631E-145 | TEC_prg2 | AZIN2 |
| 1.0955E-30 | 0.31643197 | 0.257 | 0.149 | 2.339E-26 | TEC_prg2 | TUBA4A |
| 1.589E-21 | 0.31628859 | 0.276 | 0.185 | 3.3924E-17 | TEC_prg2 | EPS8 |
| 1.4732E-18 | 0.31542326 | 0.44 | 0.357 | 3.1454E-14 | TEC_prg2 | PNN |
| 2.967E-140 | 0.3143904 | 0.185 | 0.032 | 6.335E-136 | TEC_prg2 | TSC22D1-AS1 |
| 4.1697E-19 | 0.31285642 | 0.349 | 0.263 | 8.9024E-15 | TEC_prg2 | TCF25 |
| 9.698E-133 | 0.30964989 | 0.182 | 0.033 | 2.071E-128 | TEC_prg2 | CATSPERG |
| 1.9348E-69 | 0.3086732 | 0.198 | 0.066 | 4.1309E-65 | TEC_prg2 | CAPRIN2 |
| 1.6714E-35 | 0.30865888 | 0.239 | 0.127 | 3.5684E-31 | TEC_prg2 | TIPARP |
| 2.1215E-40 | 0.30679119 | 0.205 | 0.094 | 4.5295E-36 | TEC_prg2 | AREG |
| 1.9401E-49 | 0.30603881 | 0.19 | 0.076 | 4.1422E-45 | TEC_prg2 | PPM1D |
| 2.6062E-40 | 0.30526861 | 0.224 | 0.11 | 5.5643E-36 | TEC_prg2 | SIRT1 |
| 2.2471E-78 | 0.30025553 | 0.207 | 0.065 | 4.7975E-74 | TEC_prg2 | MST1 |
| 2.6926E-90 | 0.30024013 | 0.139 | 0.028 | 5.7487E-86 | TEC_prg2 | LINC00511 |
| 2.806E-144 | 0.29864332 | 0.147 | 0.019 | 5.991E-140 | TEC_prg2 | AP005717.1 |
| 1.38E-59 | 0.29768878 | 0.193 | 0.069 | 2.9462E-55 | TEC_prg2 | IRX2 |
| 5.2224E-12 | 0.29752183 | 0.156 | 0.103 | 1.115E-07 | TEC_prg2 | FAM53C |
| 7.4274E-43 | 0.2963047 | 0.205 | 0.093 | 1.5857E-38 | TEC_prg2 | ICE2 |
| 5.8537E-42 | 0.29462393 | 0.23 | 0.111 | 1.2498E-37 | TEC_prg2 | MEX3C |
| 8.1919E-46 | 0.29452867 | 0.21 | 0.093 | 1.749E-41 | TEC_prg2 | EXT1 |
| 6.036E-143 | 0.29305777 | 0.181 | 0.03 | 1.289E-138 | TEC_prg2 | ATRIP |
| 1.3902E-54 | 0.2920059 | 0.197 | 0.076 | 2.9681E-50 | TEC_prg2 | DNAJC3-DT |
| 1.0654E-88 | 0.29093552 | 0.153 | 0.035 | 2.2746E-84 | TEC_prg2 | LINC00663 |

| | | | | | | |
|---|---|---|---|---|---|---|
| 7.4541E-20 | 0.29051278 | 0.223 | 0.143 | 1.5914E-15 | TEC_prg2 | ZNF622 |
| 3.9298E-21 | 0.28841953 | 0.14 | 0.071 | 8.3901E-17 | TEC_prg2 | ANGPTL4 |
| 2.8366E-25 | 0.28515052 | 0.188 | 0.102 | 6.0561E-21 | TEC_prg2 | ARL4A |
| 5.0058E-13 | 0.28406844 | 0.476 | 0.429 | 1.0687E-08 | TEC_prg2 | PNRC1 |
| 2.1508E-46 | 0.28305269 | 0.171 | 0.066 | 4.592E-42 | TEC_prg2 | TMEM107 |
| 9.389E-22 | 0.28235591 | 0.234 | 0.147 | 2.0046E-17 | TEC_prg2 | NUDT4 |
| 7.344E-107 | 0.27938584 | 0.151 | 0.028 | 1.568E-102 | TEC_prg2 | PCK1 |
| 2.2161E-34 | 0.27835042 | 0.251 | 0.134 | 4.7313E-30 | TEC_prg2 | EMX2 |
| 4.4062E-17 | 0.27723792 | 0.322 | 0.24 | 9.4072E-13 | TEC_prg2 | TRIM44 |
| 2.7391E-25 | 0.27699082 | 0.233 | 0.138 | 5.8479E-21 | TEC_prg2 | SPG7 |
| 1.5465E-10 | 0.27670701 | 0.46 | 0.428 | 3.3019E-06 | TEC_prg2 | ZFAND5 |
| 3.6114E-54 | 0.27664921 | 0.163 | 0.056 | 7.7103E-50 | TEC_prg2 | AL139246.5 |
| 8.9004E-16 | 0.27364716 | 0.472 | 0.398 | 1.9002E-11 | TEC_prg2 | RNF213 |
| 3.6371E-38 | 0.27348507 | 0.256 | 0.133 | 7.7653E-34 | TEC_prg2 | CREBZF |
| 3.9327E-95 | 0.27342485 | 0.166 | 0.038 | 8.3964E-91 | TEC_prg2 | AL034549.1 |
| 2.3569E-26 | 0.27336068 | 0.299 | 0.191 | 5.032E-22 | TEC_prg2 | ZFC3H1 |
| 2.011E-133 | 0.27269445 | 0.117 | 0.012 | 4.293E-129 | TEC_prg2 | ATP1B2 |
| 1.5704E-35 | 0.27221427 | 0.222 | 0.112 | 3.3527E-31 | TEC_prg2 | EPM2AIP1 |
| 7.9494E-85 | 0.27206972 | 0.148 | 0.034 | 1.6972E-80 | TEC_prg2 | SNAI1 |
| 6.1797E-22 | 0.27195611 | 0.232 | 0.144 | 1.3194E-17 | TEC_prg2 | INSIG1 |
| 2.0901E-57 | 0.27120128 | 0.153 | 0.049 | 4.4623E-53 | TEC_prg2 | ARHGEF9 |
| 3.5126E-36 | 0.27084994 | 0.2 | 0.095 | 7.4994E-32 | TEC_prg2 | HIST1H2BD |
| 2.409E-21 | 0.269972 | 0.325 | 0.226 | 5.1431E-17 | TEC_prg2 | SLC5A3 |
| 3.2814E-10 | 0.26937942 | 0.43 | 0.39 | 7.0058E-06 | TEC_prg2 | SRSF3 |
| 9.9218E-29 | 0.26932562 | 0.188 | 0.097 | 2.1183E-24 | TEC_prg2 | RNF144B |
| 2.9617E-15 | 0.26840007 | 0.45 | 0.364 | 6.3233E-11 | TEC_prg2 | ATF3 |
| 1.0836E-46 | 0.26738688 | 0.182 | 0.073 | 2.3135E-42 | TEC_prg2 | CCNYL1 |
| 3.0549E-08 | 0.26551233 | 0.359 | 0.321 | 0.00065222 | TEC_prg2 | HNRNPH1 |
| 3.001E-12 | 0.26504543 | 0.589 | 0.557 | 6.4072E-08 | TEC_prg2 | NCL |
| 6.1559E-36 | 0.26449312 | 0.239 | 0.123 | 1.3143E-31 | TEC_prg2 | HERC2 |
| 2.0625E-18 | 0.26393748 | 0.163 | 0.095 | 4.4035E-14 | TEC_prg2 | NMRK1 |
| 1.9433E-10 | 0.26340298 | 0.202 | 0.147 | 4.149E-06 | TEC_prg2 | TSC22D3 |
| 7.7411E-53 | 0.26322322 | 0.138 | 0.043 | 1.6527E-48 | TEC_prg2 | NDUFV2-AS1 |
| 2.4596E-31 | 0.26197379 | 0.23 | 0.124 | 5.2512E-27 | TEC_prg2 | PLXNB1 |
| 2.634E-103 | 0.26183455 | 0.142 | 0.026 | 5.623E-99 | TEC_prg2 | KLLN |
| 1.7864E-54 | 0.26138527 | 0.17 | 0.059 | 3.814E-50 | TEC_prg2 | SIM2 |
| 0.00218789 | 0.26101984 | 0.366 | 0.374 | 1 | TEC_prg2 | ODC1 |
| 1.587E-19 | 0.26101649 | 0.196 | 0.119 | 3.3883E-15 | TEC_prg2 | ERBB3 |
| 1.4618E-62 | 0.26055709 | 0.155 | 0.046 | 3.121E-58 | TEC_prg2 | PRR3 |
| 2.2306E-18 | 0.25951294 | 0.303 | 0.208 | 4.7623E-14 | TEC_prg2 | TNFAIP2 |
| 2.28E-114 | 0.25907945 | 0.143 | 0.023 | 4.869E-110 | TEC_prg2 | STARD5 |
| 1.1235E-48 | 0.2590461 | 0.188 | 0.074 | 2.3986E-44 | TEC_prg2 | ERO1B |
| 4.4761E-28 | 0.25844469 | 0.214 | 0.112 | 9.5565E-24 | TEC_prg2 | CXCL14 |
| 1.8456E-18 | 0.25800966 | 0.233 | 0.151 | 3.9404E-14 | TEC_prg2 | NR2F2 |
| 4.024E-144 | 0.25768539 | 0.138 | 0.016 | 8.591E-140 | TEC_prg2 | LFNG |
| 1.834E-23 | 0.25718513 | 0.241 | 0.146 | 3.9155E-19 | TEC_prg2 | PLEKHH2 |
| 6.8637E-18 | 0.25638763 | 0.259 | 0.178 | 1.4654E-13 | TEC_prg2 | SCNN1A |
| 2.8586E-07 | 0.25388734 | 0.977 | 0.93 | 0.0061032 | TEC_prg2 | PTMA |
| 2.6302E-46 | 0.25311503 | 0.172 | 0.066 | 5.6154E-42 | TEC_prg2 | OBSL1 |
| 9.2738E-43 | 0.25310614 | 0.166 | 0.066 | 1.98E-38 | TEC_prg2 | EVC |

| | | | | | | |
|---|---|---|---|---|---|---|
| 2.0586E-17 | 0.25261987 | 0.362 | 0.272 | 4.3951E-13 | TEC_prg2 | PRPF4B |
| 2.3802E-18 | 0.25233283 | 0.274 | 0.188 | 5.0817E-14 | TEC_prg2 | UPF2 |
| 1.2391E-09 | 0.25221394 | 0.224 | 0.169 | 2.6456E-05 | TEC_prg2 | PHLDA1 |
| 7.4468E-08 | -0.2501988 | 0.978 | 0.902 | 0.00158989 | TEC_prg2 | MT-CO3 |
| 1.073E-42 | -0.2509406 | 0.05 | 0.201 | 2.2909E-38 | TEC_prg2 | RANBP1 |
| 5.4024E-42 | -0.2511112 | 0.097 | 0.271 | 1.1534E-37 | TEC_prg2 | TAF10 |
| 4.5251E-41 | -0.2513364 | 0.134 | 0.319 | 9.661E-37 | TEC_prg2 | TM9SF3 |
| 7.6675E-49 | -0.2516145 | 0.046 | 0.212 | 1.637E-44 | TEC_prg2 | EMC4 |
| 1.3261E-51 | -0.2519805 | 0.034 | 0.2 | 2.8311E-47 | TEC_prg2 | GINM1 |
| 5.6801E-35 | -0.2520593 | 0.057 | 0.191 | 1.2127E-30 | TEC_prg2 | DNAJB9 |
| 2.6646E-41 | -0.2520856 | 0.116 | 0.295 | 5.6889E-37 | TEC_prg2 | NDUFS6 |
| 9.783E-38 | -0.2521981 | 0.067 | 0.215 | 2.0887E-33 | TEC_prg2 | TINAGL1 |
| 1.0675E-43 | -0.252231 | 0.057 | 0.216 | 2.2792E-39 | TEC_prg2 | TMEM141 |
| 3.0305E-36 | -0.2525616 | 0.109 | 0.269 | 6.4701E-32 | TEC_prg2 | PTBP3 |
| 1.2535E-48 | -0.2530589 | 0.05 | 0.219 | 2.6763E-44 | TEC_prg2 | NBDY |
| 4.1175E-36 | -0.2531441 | 0.065 | 0.206 | 8.7908E-32 | TEC_prg2 | SEL1L |
| 1.9794E-48 | -0.25336 | 0.028 | 0.184 | 4.2261E-44 | TEC_prg2 | UBE2J1 |
| 1.08E-29 | -0.2534264 | 0.035 | 0.143 | 2.3057E-25 | TEC_prg2 | PLIN2 |
| 1.972E-53 | -0.2534995 | 0.031 | 0.2 | 4.2102E-49 | TEC_prg2 | CHP1 |
| 2.4297E-47 | -0.2535252 | 0.021 | 0.167 | 5.1874E-43 | TEC_prg2 | ACSL4 |
| 1.03E-49 | -0.253632 | 0.052 | 0.225 | 2.199E-45 | TEC_prg2 | TXNL4A |
| 5.2816E-44 | -0.2537122 | 0.07 | 0.238 | 1.1276E-39 | TEC_prg2 | CTNNAL1 |
| 3.845E-27 | -0.2538992 | 0.082 | 0.202 | 8.209E-23 | TEC_prg2 | EMP1 |
| 4.0373E-38 | -0.2542608 | 0.033 | 0.164 | 8.6196E-34 | TEC_prg2 | NUCB2 |
| 3.9851E-41 | -0.2543046 | 0.107 | 0.282 | 8.5081E-37 | TEC_prg2 | RPS19BP1 |
| 1.5551E-40 | -0.254572 | 0.087 | 0.252 | 3.3201E-36 | TEC_prg2 | NDUFS8 |
| 1.2783E-24 | -0.254712 | 0.263 | 0.424 | 2.7291E-20 | TEC_prg2 | SDC4 |
| 7.1473E-32 | -0.2547712 | 0.072 | 0.206 | 1.5259E-27 | TEC_prg2 | CTSL |
| 2.1829E-39 | -0.2547964 | 0.099 | 0.265 | 4.6604E-35 | TEC_prg2 | PPIG |
| 2.4434E-50 | -0.2557178 | 0.048 | 0.221 | 5.2166E-46 | TEC_prg2 | ILF2 |
| 3.3674E-46 | -0.255983 | 0.05 | 0.212 | 7.1893E-42 | TEC_prg2 | RALA |
| 1.0927E-39 | -0.2561607 | 0.123 | 0.298 | 2.333E-35 | TEC_prg2 | MRPL41 |
| 5.8321E-17 | -0.2564438 | 0.728 | 0.763 | 1.2451E-12 | TEC_prg2 | CALM2 |
| 1.2536E-46 | -0.2566511 | 0.061 | 0.228 | 2.6764E-42 | TEC_prg2 | CSNK2B |
| 1.1279E-33 | -0.2567855 | 0.097 | 0.245 | 2.4082E-29 | TEC_prg2 | ST14 |
| 6.62E-17 | -0.2569875 | 0.918 | 0.903 | 1.4134E-12 | TEC_prg2 | RPS20 |
| 3.2376E-44 | -0.2570815 | 0.044 | 0.197 | 6.9124E-40 | TEC_prg2 | CD99 |
| 7.1366E-34 | -0.2571809 | 0.039 | 0.161 | 1.5237E-29 | TEC_prg2 | RETREG1 |
| 2.4358E-52 | -0.2574322 | 0.048 | 0.225 | 5.2005E-48 | TEC_prg2 | METTL9 |
| 4.1879E-51 | -0.2574809 | 0.053 | 0.23 | 8.9412E-47 | TEC_prg2 | EMC10 |
| 1.879E-46 | -0.2575409 | 0.119 | 0.316 | 4.0117E-42 | TEC_prg2 | SSBP1 |
| 7.0711E-51 | -0.25761 | 0.038 | 0.205 | 1.5097E-46 | TEC_prg2 | LYPLA1 |
| 2.3014E-45 | -0.2576343 | 0.071 | 0.242 | 4.9134E-41 | TEC_prg2 | DIAPH1 |
| 8.9751E-39 | -0.2577017 | 0.042 | 0.18 | 1.9162E-34 | TEC_prg2 | CAVIN1 |
| 4.2914E-41 | -0.2578833 | 0.044 | 0.189 | 9.1622E-37 | TEC_prg2 | EIF4EBP1 |
| 7.4104E-45 | -0.258026 | 0.073 | 0.242 | 1.5821E-40 | TEC_prg2 | DNAJC8 |
| 1.299E-52 | -0.258138 | 0.036 | 0.206 | 2.7733E-48 | TEC_prg2 | TMEM147 |
| 4.8747E-40 | -0.258142 | 0.146 | 0.334 | 1.0408E-35 | TEC_prg2 | DNAJB6 |
| 4.0101E-44 | -0.2583361 | 0.059 | 0.22 | 8.5616E-40 | TEC_prg2 | NDUFAF3 |
| 1.0561E-48 | -0.2584655 | 0.015 | 0.162 | 2.2548E-44 | TEC_prg2 | CDCP1 |

| | | | | | | |
|---|---|---|---|---|---|---|
| 5.0353E-56 | -0.2585206 | 0.033 | 0.212 | 1.075E-51 | TEC_prg2 | MRPS15 |
| 2.5464E-47 | -0.2588537 | 0.087 | 0.271 | 5.4366E-43 | TEC_prg2 | SNRPE |
| 3.8732E-27 | -0.2589359 | 0.194 | 0.348 | 8.2693E-23 | TEC_prg2 | TUBA1C |
| 4.0333E-47 | -0.2599964 | 0.085 | 0.27 | 8.6111E-43 | TEC_prg2 | HNRNPR |
| 8.2261E-35 | -0.2604915 | 0.124 | 0.285 | 1.7563E-30 | TEC_prg2 | MET |
| 1.9063E-29 | -0.260548 | 0.137 | 0.284 | 4.0699E-25 | TEC_prg2 | TUBB |
| 6.4025E-45 | -0.2607305 | 0.046 | 0.202 | 1.3669E-40 | TEC_prg2 | ATP1B3 |
| 4.4333E-45 | -0.2608163 | 0.038 | 0.19 | 9.4651E-41 | TEC_prg2 | DSG2 |
| 2.9752E-45 | -0.2613656 | 0.071 | 0.241 | 6.3521E-41 | TEC_prg2 | SPCS3 |
| 5.5985E-35 | -0.2614189 | 0.031 | 0.151 | 1.1953E-30 | TEC_prg2 | TMEM176A |
| 8.8687E-43 | -0.2614348 | 0.119 | 0.3 | 1.8935E-38 | TEC_prg2 | RAB10 |
| 7.8664E-52 | -0.2615063 | 0.039 | 0.209 | 1.6795E-47 | TEC_prg2 | SDHC |
| 6.4767E-39 | -0.261682 | 0.097 | 0.26 | 1.3828E-34 | TEC_prg2 | VAMP2 |
| 2.6734E-35 | -0.2617652 | 0.045 | 0.176 | 5.7077E-31 | TEC_prg2 | SUCLG1 |
| 1.8144E-52 | -0.2623788 | 0.031 | 0.198 | 3.8738E-48 | TEC_prg2 | CAPZA1 |
| 5.5857E-54 | -0.2625781 | 0.037 | 0.211 | 1.1925E-49 | TEC_prg2 | FUNDC2 |
| 6.2238E-40 | -0.2628007 | 0.076 | 0.233 | 1.3288E-35 | TEC_prg2 | ATP5MC1 |
| 3.199E-43 | -0.2628854 | 0.159 | 0.36 | 6.8298E-39 | TEC_prg2 | AURKAIP1 |
| 6.6348E-49 | -0.2629358 | 0.052 | 0.223 | 1.4165E-44 | TEC_prg2 | AP2S1 |
| 8.611E-09 | -0.2629967 | 0.984 | 0.919 | 0.00018385 | TEC_prg2 | MT-CO1 |
| 5.6721E-52 | -0.2630661 | 0.066 | 0.252 | 1.211E-47 | TEC_prg2 | VPS35 |
| 4.5047E-50 | -0.263871 | 0.04 | 0.205 | 9.6175E-46 | TEC_prg2 | PPP2CA |
| 2.456E-51 | -0.2642321 | 0.019 | 0.175 | 5.2436E-47 | TEC_prg2 | CAV2 |
| 1.3018E-48 | -0.2645402 | 0.075 | 0.256 | 2.7793E-44 | TEC_prg2 | SMIM26 |
| 2.9119E-39 | -0.264773 | 0.082 | 0.241 | 6.2168E-35 | TEC_prg2 | GPRC5C |
| 2.7985E-50 | -0.2647755 | 0.052 | 0.228 | 5.9748E-46 | TEC_prg2 | ADAM9 |
| 2.3279E-41 | -0.2648459 | 0.066 | 0.223 | 4.9702E-37 | TEC_prg2 | ATP13A3 |
| 2.8088E-36 | -0.265112 | 0.012 | 0.124 | 5.9968E-32 | TEC_prg2 | ISG20 |
| 3.9087E-51 | -0.265114 | 0.063 | 0.244 | 8.3451E-47 | TEC_prg2 | STAU1 |
| 3.8229E-56 | -0.2652079 | 0.028 | 0.202 | 8.1619E-52 | TEC_prg2 | SERINC3 |
| 1.1736E-49 | -0.2652249 | 0.063 | 0.242 | 2.5056E-45 | TEC_prg2 | RWDD1 |
| 3.4775E-44 | -0.2653619 | 0.166 | 0.373 | 7.4245E-40 | TEC_prg2 | C19orf53 |
| 4.1277E-49 | -0.2655816 | 0.092 | 0.283 | 8.8125E-45 | TEC_prg2 | GNB2 |
| 5.9166E-51 | -0.2661209 | 0.04 | 0.206 | 1.2632E-46 | TEC_prg2 | ARL3 |
| 2.8284E-49 | -0.2664883 | 0.087 | 0.277 | 6.0387E-45 | TEC_prg2 | CCT4 |
| 2.7596E-42 | -0.266538 | 0.107 | 0.282 | 5.8918E-38 | TEC_prg2 | HDGF |
| 5.6046E-49 | -0.2668697 | 0.073 | 0.255 | 1.1966E-44 | TEC_prg2 | ARPC5L |
| 2.6315E-38 | -0.2670412 | 0.071 | 0.221 | 5.6182E-34 | TEC_prg2 | ADI1 |
| 9.2006E-47 | -0.2670421 | 0.041 | 0.199 | 1.9643E-42 | TEC_prg2 | QSOX1 |
| 3.9508E-24 | -0.2671958 | 0.038 | 0.133 | 8.4349E-20 | TEC_prg2 | IFIT3 |
| 4.9141E-52 | -0.2673756 | 0.04 | 0.212 | 1.0492E-47 | TEC_prg2 | CMPK1 |
| 4.2717E-49 | -0.2681829 | 0.066 | 0.243 | 9.1201E-45 | TEC_prg2 | CNIH4 |
| 8.5335E-56 | -0.2683489 | 0.055 | 0.245 | 1.8219E-51 | TEC_prg2 | ARL1 |
| 4.4999E-43 | -0.2683624 | 0.112 | 0.292 | 9.6074E-39 | TEC_prg2 | BAG1 |
| 2.117E-49 | -0.2686792 | 0.06 | 0.236 | 4.5198E-45 | TEC_prg2 | MED10 |
| 4.4989E-37 | -0.2698441 | 0.023 | 0.144 | 9.605E-33 | TEC_prg2 | AL353751.1 |
| 1.0683E-51 | -0.2699142 | 0.075 | 0.264 | 2.2809E-47 | TEC_prg2 | XRCC6 |
| 5.7915E-53 | -0.2705456 | 0.063 | 0.249 | 1.2365E-48 | TEC_prg2 | UXT |
| 2.3563E-54 | -0.2707429 | 0.016 | 0.178 | 5.0307E-50 | TEC_prg2 | F11R |
| 5.9315E-51 | -0.2707978 | 0.06 | 0.24 | 1.2664E-46 | TEC_prg2 | LAMP2 |

| | | | | | | |
|---|---|---|---|---|---|---|
| 1.4692E-42 | -0.2712013 | 0.116 | 0.3 | 3.1368E-38 | TEC_prg2 | VCP |
| 3.4778E-52 | -0.2714001 | 0.058 | 0.238 | 7.425E-48 | TEC_prg2 | PSMD4 |
| 1.1817E-06 | -0.2714768 | 0.07 | 0.113 | 0.02522948 | TEC_prg2 | S100A9 |
| 3.7439E-53 | -0.2714792 | 0.036 | 0.206 | 7.9932E-49 | TEC_prg2 | PCMT1 |
| 7.0402E-51 | -0.2716824 | 0.061 | 0.24 | 1.5031E-46 | TEC_prg2 | GDI2 |
| 8.4425E-42 | -0.2718157 | 0.109 | 0.285 | 1.8025E-37 | TEC_prg2 | TAPBP |
| 6.1231E-35 | -0.2726963 | 0.413 | 0.603 | 1.3073E-30 | TEC_prg2 | COX5B |
| 1.7385E-53 | -0.2729151 | 0.083 | 0.281 | 3.7117E-49 | TEC_prg2 | SNRPD3 |
| 3.9188E-27 | -0.2730567 | 0.582 | 0.705 | 8.3666E-23 | TEC_prg2 | GPX4 |
| 4.6061E-49 | -0.2735057 | 0.081 | 0.264 | 9.834E-45 | TEC_prg2 | LRP10 |
| 1.5189E-47 | -0.2737145 | 0.113 | 0.307 | 3.2428E-43 | TEC_prg2 | EIF3G |
| 4.5798E-50 | -0.2745045 | 0.051 | 0.222 | 9.7779E-46 | TEC_prg2 | IPO7 |
| 1.2872E-50 | -0.2746823 | 0.094 | 0.289 | 2.7481E-46 | TEC_prg2 | PAIP2 |
| 3.8788E-45 | -0.2747446 | 0.074 | 0.244 | 8.2812E-41 | TEC_prg2 | SPATS2L |
| 3.2562E-34 | -0.2749101 | 0.339 | 0.538 | 6.952E-30 | TEC_prg2 | TPM3 |
| 8.4035E-50 | -0.2754165 | 0.067 | 0.249 | 1.7941E-45 | TEC_prg2 | TXNL1 |
| 7.152E-59 | -0.2757278 | 0.033 | 0.216 | 1.5269E-54 | TEC_prg2 | CCT7 |
| 2.1419E-49 | -0.2759014 | 0.09 | 0.278 | 4.573E-45 | TEC_prg2 | PRKAR1A |
| 7.6004E-55 | -0.2762944 | 0.049 | 0.233 | 1.6227E-50 | TEC_prg2 | TRAPPC1 |
| 2.2542E-39 | -0.2763102 | 0.342 | 0.551 | 4.8128E-35 | TEC_prg2 | ELOB |
| 1.464E-45 | -0.2763493 | 0.098 | 0.28 | 3.1256E-41 | TEC_prg2 | NDUFB3 |
| 4.4749E-54 | -0.2767131 | 0.04 | 0.216 | 9.554E-50 | TEC_prg2 | ZCRB1 |
| 1.7509E-52 | -0.2774404 | 0.039 | 0.21 | 3.7383E-48 | TEC_prg2 | PSMB8 |
| 1.3757E-38 | -0.277759 | 0.158 | 0.339 | 2.9371E-34 | TEC_prg2 | DEK |
| 7.5482E-53 | -0.2782642 | 0.072 | 0.263 | 1.6116E-48 | TEC_prg2 | GTF2A2 |
| 5.9494E-52 | -0.2784278 | 0.028 | 0.191 | 1.2702E-47 | TEC_prg2 | NME1 |
| 2.2009E-53 | -0.278801 | 0.038 | 0.211 | 4.6988E-49 | TEC_prg2 | BNIP3L |
| 1.3213E-50 | -0.278905 | 0.068 | 0.252 | 2.8209E-46 | TEC_prg2 | H2AFY |
| 2.6313E-60 | -0.2791526 | 0.034 | 0.223 | 5.6178E-56 | TEC_prg2 | TMEM230 |
| 1.1275E-19 | -0.2792505 | 0.763 | 0.809 | 2.4072E-15 | TEC_prg2 | CFL1 |
| 1.1338E-55 | -0.2793789 | 0.054 | 0.242 | 2.4207E-51 | TEC_prg2 | SCOC |
| 7.4303E-58 | -0.2796589 | 0.023 | 0.196 | 1.5864E-53 | TEC_prg2 | SDHD |
| 2.2448E-48 | -0.2798052 | 0.026 | 0.178 | 4.7927E-44 | TEC_prg2 | PLS3 |
| 1.4631E-47 | -0.2800731 | 0.094 | 0.284 | 3.1237E-43 | TEC_prg2 | IL6ST |
| 7.982E-50 | -0.2802972 | 0.061 | 0.237 | 1.7042E-45 | TEC_prg2 | SEC61A1 |
| 2.7079E-55 | -0.2803289 | 0.058 | 0.245 | 5.7813E-51 | TEC_prg2 | C1orf43 |
| 1.1577E-39 | -0.2806561 | 0.078 | 0.235 | 2.4716E-35 | TEC_prg2 | CCPG1 |
| 7.0943E-51 | -0.2808178 | 0.066 | 0.248 | 1.5146E-46 | TEC_prg2 | PGLS |
| 1.7089E-37 | -0.2819951 | 0.11 | 0.272 | 3.6485E-33 | TEC_prg2 | COTL1 |
| 2.294E-50 | -0.2823285 | 0.074 | 0.258 | 4.8976E-46 | TEC_prg2 | WASL |
| 2.0578E-58 | -0.2824809 | 0.052 | 0.246 | 4.3934E-54 | TEC_prg2 | PSMA1 |
| 3.5282E-49 | -0.2825694 | 0.036 | 0.196 | 7.5328E-45 | TEC_prg2 | RYBP |
| 9.934E-55 | -0.2826076 | 0.052 | 0.236 | 2.1209E-50 | TEC_prg2 | SNRPB2 |
| 8.8034E-37 | -0.2829184 | 0.175 | 0.359 | 1.8795E-32 | TEC_prg2 | HSPA9 |
| 6.3873E-49 | -0.2833155 | 0.116 | 0.314 | 1.3637E-44 | TEC_prg2 | ACTR2 |
| 1.4784E-53 | -0.2835627 | 0.055 | 0.238 | 3.1563E-49 | TEC_prg2 | SRI |
| 1.8904E-58 | -0.2845385 | 0.035 | 0.218 | 4.0361E-54 | TEC_prg2 | SELENOT |
| 2.5976E-44 | -0.2851685 | 0.118 | 0.305 | 5.5459E-40 | TEC_prg2 | ROMO1 |
| 1.2548E-39 | -0.2851711 | 0.033 | 0.167 | 2.6789E-35 | TEC_prg2 | NQO1 |
| 6.3167E-47 | -0.2852155 | 0.133 | 0.332 | 1.3486E-42 | TEC_prg2 | NSA2 |

| | | | | | | |
|---|---|---|---|---|---|---|
| 1.7134E-43 | -0.2852409 | 0.196 | 0.41 | 3.6582E-39 | TEC_prg2 | CSNK1A1 |
| 5.2601E-36 | -0.2853825 | 0.1 | 0.253 | 1.123E-31 | TEC_prg2 | LPP |
| 2.2956E-51 | -0.285526 | 0.097 | 0.292 | 4.9012E-47 | TEC_prg2 | UBE2I |
| 8.8466E-50 | -0.2858904 | 0.097 | 0.292 | 1.8888E-45 | TEC_prg2 | ZC3H15 |
| 1.3663E-57 | -0.2862437 | 0.057 | 0.252 | 2.9171E-53 | TEC_prg2 | NDUFA12 |
| 6.9116E-48 | -0.2864972 | 0.061 | 0.233 | 1.4756E-43 | TEC_prg2 | RAD21 |
| 3.5324E-15 | -0.2865532 | 0.485 | 0.585 | 7.5417E-11 | TEC_prg2 | SLC3A2 |
| 1.326E-52 | -0.2868186 | 0.075 | 0.265 | 2.831E-48 | TEC_prg2 | KPNB1 |
| 1.7501E-51 | -0.2871741 | 0.038 | 0.207 | 3.7364E-47 | TEC_prg2 | ARPC1B |
| 2.2545E-47 | -0.2874342 | 0.071 | 0.247 | 4.8133E-43 | TEC_prg2 | PSMD2 |
| 4.3802E-58 | -0.2874581 | 0.02 | 0.194 | 9.3517E-54 | TEC_prg2 | PRNP |
| 5.3173E-47 | -0.2876219 | 0.057 | 0.224 | 1.1352E-42 | TEC_prg2 | CSRP1 |
| 6.7082E-55 | -0.2876667 | 0.173 | 0.41 | 1.4322E-50 | TEC_prg2 | CTNNA1 |
| 3.3967E-56 | -0.2879773 | 0.058 | 0.25 | 7.252E-52 | TEC_prg2 | POLR2K |
| 1.3405E-53 | -0.2887424 | 0.033 | 0.204 | 2.8619E-49 | TEC_prg2 | TMEM54 |
| 1.6684E-54 | -0.2890283 | 0.076 | 0.271 | 3.5621E-50 | TEC_prg2 | AP2M1 |
| 4.0555E-56 | -0.2890375 | 0.094 | 0.304 | 8.6585E-52 | TEC_prg2 | MTCH1 |
| 1.1479E-39 | -0.2895578 | 0.168 | 0.362 | 2.4508E-35 | TEC_prg2 | ARF4 |
| 5.6782E-47 | -0.2899543 | 0.067 | 0.24 | 1.2123E-42 | TEC_prg2 | GARS |
| 4.9131E-53 | -0.290115 | 0.077 | 0.269 | 1.0489E-48 | TEC_prg2 | BANF1 |
| 4.0641E-37 | -0.2903108 | 0.116 | 0.283 | 8.6768E-33 | TEC_prg2 | LIMA1 |
| 7.3336E-59 | -0.2904125 | 0.016 | 0.188 | 1.5657E-54 | TEC_prg2 | PRDX4 |
| 2.2084E-52 | -0.2904577 | 0.052 | 0.23 | 4.7149E-48 | TEC_prg2 | SNX9 |
| 2.0861E-56 | -0.2906267 | 0.048 | 0.235 | 4.4537E-52 | TEC_prg2 | NHP2 |
| 3.0683E-52 | -0.2906571 | 0.085 | 0.278 | 6.5509E-48 | TEC_prg2 | NDUFA11 |
| 3.8553E-37 | -0.2907966 | 0.05 | 0.186 | 8.231E-33 | TEC_prg2 | SERPING1 |
| 4.8189E-48 | -0.2911204 | 0.065 | 0.239 | 1.0288E-43 | TEC_prg2 | LIMS1 |
| 9.5652E-63 | -0.2915024 | 0.036 | 0.23 | 2.0422E-58 | TEC_prg2 | RAB5C |
| 4.9943E-57 | -0.2915802 | 0.053 | 0.245 | 1.0663E-52 | TEC_prg2 | CCT2 |
| 1.3384E-47 | -0.2923216 | 0.109 | 0.299 | 2.8575E-43 | TEC_prg2 | CNN3 |
| 2.9159E-57 | -0.2925691 | 0.052 | 0.242 | 6.2254E-53 | TEC_prg2 | PSMA4 |
| 9.8831E-49 | -0.2930029 | 0.111 | 0.305 | 2.11E-44 | TEC_prg2 | PDCD5 |
| 1.4528E-42 | -0.2938501 | 0.019 | 0.153 | 3.1018E-38 | TEC_prg2 | TRIB3 |
| 9.3418E-42 | -0.2938603 | 0.127 | 0.31 | 1.9945E-37 | TEC_prg2 | HNRNPH3 |
| 7.5722E-60 | -0.2941063 | 0.03 | 0.214 | 1.6167E-55 | TEC_prg2 | TSPAN3 |
| 1.8465E-52 | -0.2945514 | 0.038 | 0.208 | 3.9423E-48 | TEC_prg2 | RAB9A |
| 4.5889E-46 | -0.2949684 | 0.129 | 0.322 | 9.7973E-42 | TEC_prg2 | IQGAP1 |
| 1.7312E-56 | -0.295214 | 0.06 | 0.255 | 3.6962E-52 | TEC_prg2 | RNF181 |
| 3.0418E-47 | -0.295697 | 0.053 | 0.219 | 6.4942E-43 | TEC_prg2 | TMEM106B |
| 2.3492E-34 | -0.2960983 | 0.084 | 0.229 | 5.0155E-30 | TEC_prg2 | CTSH |
| 8.4529E-42 | -0.2962114 | 0.626 | 0.767 | 1.8047E-37 | TEC_prg2 | PFDN5 |
| 3.2252E-46 | -0.2964681 | 0.053 | 0.215 | 6.8859E-42 | TEC_prg2 | MAP2K3 |
| 2.7892E-44 | -0.2964713 | 0.057 | 0.215 | 5.9549E-40 | TEC_prg2 | BNIP3 |
| 4.8531E-58 | -0.2965282 | 0.043 | 0.231 | 1.0361E-53 | TEC_prg2 | PEA15 |
| 6.4464E-56 | -0.2965582 | 0.053 | 0.24 | 1.3763E-51 | TEC_prg2 | GTF3C6 |
| 1.0892E-57 | -0.2970477 | 0.058 | 0.252 | 2.3255E-53 | TEC_prg2 | SCARB2 |
| 5.8414E-34 | -0.2971088 | 0.134 | 0.291 | 1.2471E-29 | TEC_prg2 | CREBRF |
| 9.7263E-64 | -0.2974428 | 0.026 | 0.215 | 2.0766E-59 | TEC_prg2 | EMC7 |
| 1.8036E-44 | -0.2977972 | 0.051 | 0.208 | 3.8507E-40 | TEC_prg2 | ZBTB43 |
| 5.4384E-60 | -0.2983504 | 0.041 | 0.232 | 1.1611E-55 | TEC_prg2 | GNG12 |

| | | | | | | |
|---|---|---|---|---|---|---|
| 1.83E-48 | -0.2987735 | 0.165 | 0.38 | 3.907E-44 | TEC_prg2 | UBXN4 |
| 3.2836E-59 | -0.2992093 | 0.062 | 0.264 | 7.0105E-55 | TEC_prg2 | RAB11A |
| 1.2627E-38 | -0.2994887 | 0.215 | 0.416 | 2.6958E-34 | TEC_prg2 | ATP5F1D |
| 1.9615E-33 | -0.2995602 | 0.927 | 0.947 | 4.1878E-29 | TEC_prg2 | RPS15 |
| 2.4706E-41 | -0.2996365 | 0.085 | 0.248 | 5.2747E-37 | TEC_prg2 | RPL22L1 |
| 1.3512E-56 | -0.2998066 | 0.105 | 0.317 | 2.8848E-52 | TEC_prg2 | SRSF9 |
| 1.4273E-55 | -0.3003757 | 0.041 | 0.221 | 3.0473E-51 | TEC_prg2 | SMIM14 |
| 1.419E-38 | -0.300584 | 0.207 | 0.408 | 3.0295E-34 | TEC_prg2 | POLR2L |
| 6.3574E-51 | -0.3007851 | 0.095 | 0.29 | 1.3573E-46 | TEC_prg2 | SNRPG |
| 3.8534E-59 | -0.3009643 | 0.068 | 0.273 | 8.227E-55 | TEC_prg2 | UBE2L3 |
| 1.9526E-57 | -0.3010042 | 0.057 | 0.249 | 4.1688E-53 | TEC_prg2 | TMEM208 |
| 4.9128E-25 | -0.3011689 | 0.376 | 0.52 | 1.0489E-20 | TEC_prg2 | CLDN7 |
| 2.2833E-58 | -0.3017486 | 0.06 | 0.258 | 4.8748E-54 | TEC_prg2 | PPP1CA |
| 9.8387E-08 | -0.3018653 | 0.309 | 0.39 | 0.00210057 | TEC_prg2 | HERPUD1 |
| 3.9515E-30 | -0.3021037 | 0.023 | 0.126 | 8.4365E-26 | TEC_prg2 | RARRES3 |
| 5.5676E-54 | -0.3022199 | 0.051 | 0.235 | 1.1887E-49 | TEC_prg2 | BTG3 |
| 2.4466E-58 | -0.3029959 | 0.062 | 0.263 | 5.2234E-54 | TEC_prg2 | NENF |
| 1.7029E-49 | -0.3031011 | 0.179 | 0.4 | 3.6357E-45 | TEC_prg2 | C11orf58 |
| 5.431E-45 | -0.303633 | 0.03 | 0.176 | 1.1595E-40 | TEC_prg2 | TMEM176B |
| 4.3932E-40 | -0.3037694 | 0.506 | 0.685 | 9.3794E-36 | TEC_prg2 | TMA7 |
| 4.5562E-62 | -0.3038961 | 0.02 | 0.203 | 9.7275E-58 | TEC_prg2 | HACD3 |
| 3.3348E-58 | -0.3043759 | 0.065 | 0.264 | 7.1199E-54 | TEC_prg2 | HIGD2A |
| 2.1315E-46 | -0.3048536 | 0.1 | 0.286 | 4.5508E-42 | TEC_prg2 | RIOK3 |
| 1.8741E-57 | -0.3048784 | 0.065 | 0.262 | 4.0013E-53 | TEC_prg2 | HNRNPF |
| 3.5946E-53 | -0.3050078 | 0.082 | 0.276 | 7.6745E-49 | TEC_prg2 | ARHGDIA |
| 8.1836E-40 | -0.3051744 | 0.122 | 0.295 | 1.7472E-35 | TEC_prg2 | TCEA1 |
| 4.0369E-52 | -0.3055286 | 0.097 | 0.294 | 8.6187E-48 | TEC_prg2 | ARPC5 |
| 1.1047E-58 | -0.3071944 | 0.078 | 0.284 | 2.3584E-54 | TEC_prg2 | JTB |
| 2.165E-61 | -0.307222 | 0.063 | 0.269 | 4.6223E-57 | TEC_prg2 | CAMTA1 |
| 2.4272E-52 | -0.3074862 | 0.048 | 0.224 | 5.1821E-48 | TEC_prg2 | OCLN |
| 7.8231E-61 | -0.3077934 | 0.074 | 0.283 | 1.6702E-56 | TEC_prg2 | EIF4H |
| 1.0907E-53 | -0.3078485 | 0.093 | 0.295 | 2.3287E-49 | TEC_prg2 | COX17 |
| 1.1695E-54 | -0.3082192 | 0.089 | 0.29 | 2.497E-50 | TEC_prg2 | NUDC |
| 1.995E-30 | -0.3092358 | 0.102 | 0.237 | 4.2593E-26 | TEC_prg2 | BEX2 |
| 6.9888E-65 | -0.3092472 | 0.028 | 0.222 | 1.4921E-60 | TEC_prg2 | NDUFB5 |
| 2.7958E-65 | -0.3093203 | 0.048 | 0.255 | 5.9689E-61 | TEC_prg2 | ACP1 |
| 7.5315E-57 | -0.3096651 | 0.111 | 0.327 | 1.608E-52 | TEC_prg2 | WASF2 |
| 6.2324E-60 | -0.310606 | 0.094 | 0.312 | 1.3306E-55 | TEC_prg2 | RBX1 |
| 8.5701E-65 | -0.310669 | 0.035 | 0.233 | 1.8297E-60 | TEC_prg2 | PSMC1 |
| 2.119E-63 | -0.3109253 | 0.065 | 0.28 | 4.524E-59 | TEC_prg2 | CCT8 |
| 3.254E-58 | -0.3113074 | 0.075 | 0.281 | 6.9473E-54 | TEC_prg2 | NDUFA6 |
| 1.6124E-42 | -0.3113733 | 0.065 | 0.224 | 3.4424E-38 | TEC_prg2 | DNTTIP2 |
| 9.836E-59 | -0.3115361 | 0.089 | 0.3 | 2.1E-54 | TEC_prg2 | ARPP19 |
| 2.6195E-22 | -0.3120821 | 0.121 | 0.238 | 5.5926E-18 | TEC_prg2 | CD74 |
| 2.8308E-67 | -0.3128551 | 0.04 | 0.248 | 6.0438E-63 | TEC_prg2 | PSMB4 |
| 1.9098E-66 | -0.3133329 | 0.04 | 0.245 | 4.0774E-62 | TEC_prg2 | ANXA7 |
| 3.303E-58 | -0.3135488 | 0.074 | 0.277 | 7.0518E-54 | TEC_prg2 | ATP6V1D |
| 7.4335E-58 | -0.3135894 | 0.08 | 0.286 | 1.587E-53 | TEC_prg2 | MLEC |
| 2.1357E-56 | -0.3136514 | 0.142 | 0.372 | 4.5597E-52 | TEC_prg2 | PSMD8 |
| 3.5881E-62 | -0.3139625 | 0.056 | 0.259 | 7.6606E-58 | TEC_prg2 | NAA20 |

| | | | | | | |
|---|---|---|---|---|---|---|
| 4.0698E-63 | -0.3140822 | 0.039 | 0.235 | 8.689E-59 | TEC_prg2 | MCFD2 |
| 4.3393E-58 | -0.3142006 | 0.05 | 0.241 | 9.2643E-54 | TEC_prg2 | RSL1D1 |
| 2.272E-59 | -0.3148911 | 0.073 | 0.28 | 4.8506E-55 | TEC_prg2 | HM13 |
| 1.9335E-42 | -0.3152546 | 0.094 | 0.269 | 4.128E-38 | TEC_prg2 | MRPS6 |
| 1.052E-37 | -0.3155022 | 0.06 | 0.204 | 2.2461E-33 | TEC_prg2 | PDIA4 |
| 3.3237E-26 | -0.3155091 | 0.068 | 0.18 | 7.0962E-22 | TEC_prg2 | FGF2 |
| 4.5754E-64 | -0.3163033 | 0.047 | 0.248 | 9.7685E-60 | TEC_prg2 | EIF3L |
| 5.4543E-49 | -0.3163955 | 0.132 | 0.332 | 1.1645E-44 | TEC_prg2 | ARHGAP5 |
| 9.6258E-64 | -0.3169006 | 0.026 | 0.216 | 2.0551E-59 | TEC_prg2 | C1QBP |
| 8.2101E-67 | -0.3183322 | 0.022 | 0.217 | 1.7528E-62 | TEC_prg2 | CMTM6 |
| 3.5787E-52 | -0.3185127 | 0.159 | 0.382 | 7.6406E-48 | TEC_prg2 | CD46 |
| 1.1454E-49 | -0.3187359 | 0.086 | 0.272 | 2.4454E-45 | TEC_prg2 | ATOX1 |
| 5.6067E-20 | -0.3189118 | 0.096 | 0.197 | 1.197E-15 | TEC_prg2 | ADIRF |
| 2.3815E-61 | -0.3190809 | 0.043 | 0.237 | 5.0845E-57 | TEC_prg2 | RCN1 |
| 3.5606E-17 | -0.3191582 | 0.923 | 0.906 | 7.6019E-13 | TEC_prg2 | MYL6 |
| 1.6158E-63 | -0.319921 | 0.06 | 0.269 | 3.4498E-59 | TEC_prg2 | TUFM |
| 2.2949E-62 | -0.3200805 | 0.077 | 0.292 | 4.8997E-58 | TEC_prg2 | NDUFB8 |
| 2.5122E-57 | -0.3202544 | 0.04 | 0.225 | 5.3635E-53 | TEC_prg2 | OSER1 |
| 7.3289E-59 | -0.3204674 | 0.117 | 0.345 | 1.5647E-54 | TEC_prg2 | XRCC5 |
| 2.9233E-52 | -0.3205489 | 0.053 | 0.231 | 6.2413E-48 | TEC_prg2 | GFPT1 |
| 5.5376E-57 | -0.3206017 | 0.163 | 0.403 | 1.1823E-52 | TEC_prg2 | CD164 |
| 1.0255E-61 | -0.3211951 | 0.08 | 0.295 | 2.1894E-57 | TEC_prg2 | NDUFA2 |
| 1.4106E-64 | -0.3211969 | 0.042 | 0.244 | 3.0115E-60 | TEC_prg2 | TMEM167A |
| 4.3201E-45 | -0.3226141 | 0.068 | 0.236 | 9.2235E-41 | TEC_prg2 | PTGR1 |
| 1.0979E-56 | -0.3226435 | 0.08 | 0.283 | 2.3439E-52 | TEC_prg2 | PAK2 |
| 8.2237E-66 | -0.3227164 | 0.069 | 0.29 | 1.7558E-61 | TEC_prg2 | SYPL1 |
| 2.6012E-35 | -0.3227391 | 0.165 | 0.337 | 5.5536E-31 | TEC_prg2 | FAM107B |
| 1.0619E-46 | -0.3230053 | 0.062 | 0.23 | 2.2672E-42 | TEC_prg2 | MAFG |
| 3.0756E-63 | -0.3230265 | 0.048 | 0.25 | 6.5664E-59 | TEC_prg2 | MDH1 |
| 7.0299E-39 | -0.3237849 | 0.406 | 0.604 | 1.5009E-34 | TEC_prg2 | APP |
| 4.6197E-67 | -0.3238085 | 0.061 | 0.281 | 9.8631E-63 | TEC_prg2 | SMDT1 |
| 2.8803E-49 | -0.3243676 | 0.021 | 0.173 | 6.1495E-45 | TEC_prg2 | QPRT |
| 4.7605E-47 | -0.3247348 | 0.061 | 0.23 | 1.0164E-42 | TEC_prg2 | SCCPDH |
| 9.3166E-59 | -0.325522 | 0.094 | 0.306 | 1.9891E-54 | TEC_prg2 | TMEM14C |
| 4.3535E-63 | -0.3275897 | 0.067 | 0.279 | 9.2947E-59 | TEC_prg2 | TMEM123 |
| 1.5809E-65 | -0.3277032 | 0.048 | 0.257 | 3.3752E-61 | TEC_prg2 | TMBIM1 |
| 1.7E-48 | -0.3279493 | 0.462 | 0.668 | 3.6294E-44 | TEC_prg2 | EDF1 |
| 2.3839E-67 | -0.3282289 | 0.062 | 0.283 | 5.0896E-63 | TEC_prg2 | RTRAF |
| 2.7268E-55 | -0.3282638 | 0.127 | 0.346 | 5.8218E-51 | TEC_prg2 | AES |
| 2.4702E-69 | -0.32837 | 0.072 | 0.302 | 5.2739E-65 | TEC_prg2 | OCIAD1 |
| 1.2856E-53 | -0.3289776 | 0.064 | 0.253 | 2.7448E-49 | TEC_prg2 | ASAH1 |
| 5.1876E-64 | -0.3290341 | 0.077 | 0.296 | 1.1076E-59 | TEC_prg2 | UFC1 |
| 2.5212E-62 | -0.3294782 | 0.05 | 0.249 | 5.3828E-58 | TEC_prg2 | PSMD7 |
| 4.0193E-65 | -0.3296961 | 0.061 | 0.272 | 8.5811E-61 | TEC_prg2 | EIF3D |
| 1.6324E-46 | -0.3299066 | 0.119 | 0.31 | 3.4853E-42 | TEC_prg2 | TJP1 |
| 1.5966E-23 | -0.3303882 | 0.048 | 0.144 | 3.4088E-19 | TEC_prg2 | IFIT2 |
| 6.9689E-63 | -0.3306085 | 0.065 | 0.276 | 1.4879E-58 | TEC_prg2 | PSMA3 |
| 8.9632E-14 | -0.3309075 | 0.048 | 0.111 | 1.9137E-09 | TEC_prg2 | SERPINE2 |
| 7.503E-61 | -0.3310393 | 0.097 | 0.317 | 1.6019E-56 | TEC_prg2 | SF3B5 |
| 7.9452E-50 | -0.3321007 | 0.098 | 0.289 | 1.6963E-45 | TEC_prg2 | BLVRB |

| | | | | | | |
|---|---|---|---|---|---|---|
| 2.5116E-50 | -0.3321264 | 0.452 | 0.658 | 5.3623E-46 | TEC_prg2 | UBL5 |
| 1.5608E-49 | -0.3323551 | 0.598 | 0.752 | 3.3324E-45 | TEC_prg2 | SKP1 |
| 1.8958E-62 | -0.3330377 | 0.104 | 0.332 | 4.0475E-58 | TEC_prg2 | TMEM258 |
| 5.4059E-50 | -0.3331422 | 0.11 | 0.305 | 1.1542E-45 | TEC_prg2 | SPINT1 |
| 1.0431E-54 | -0.3333467 | 0.009 | 0.167 | 2.227E-50 | TEC_prg2 | FXYD5 |
| 3.5673E-68 | -0.3339429 | 0.037 | 0.246 | 7.6162E-64 | TEC_prg2 | DYNLT3 |
| 1.3261E-52 | -0.3339933 | 0.145 | 0.363 | 2.8312E-48 | TEC_prg2 | KRT10 |
| 3.1775E-46 | -0.3340924 | 0.111 | 0.297 | 6.784E-42 | TEC_prg2 | SMS |
| 2.6429E-72 | -0.3346008 | 0.041 | 0.26 | 5.6426E-68 | TEC_prg2 | TMEM14B |
| 1.4685E-71 | -0.3346951 | 0.039 | 0.255 | 3.1352E-67 | TEC_prg2 | STRAP |
| 1.4543E-64 | -0.3363209 | 0.08 | 0.3 | 3.105E-60 | TEC_prg2 | CCT6A |
| 1.198E-50 | -0.336372 | 0.156 | 0.368 | 2.5578E-46 | TEC_prg2 | MTDH |
| 1.0522E-53 | -0.3364389 | 0.167 | 0.401 | 2.2465E-49 | TEC_prg2 | NDUFA1 |
| 5.3678E-60 | -0.3367699 | 0.058 | 0.257 | 1.146E-55 | TEC_prg2 | CALM3 |
| 9.8769E-37 | -0.3384249 | 0.322 | 0.517 | 2.1087E-32 | TEC_prg2 | ACTN4 |
| 5.3408E-67 | -0.3396946 | 0.03 | 0.23 | 1.1403E-62 | TEC_prg2 | SCP2 |
| 1.5599E-66 | -0.3406047 | 0.054 | 0.267 | 3.3304E-62 | TEC_prg2 | LSM3 |
| 3.3951E-62 | -0.3417993 | 0.085 | 0.304 | 7.2486E-58 | TEC_prg2 | CLTB |
| 7.3892E-61 | -0.3418562 | 0.035 | 0.225 | 1.5776E-56 | TEC_prg2 | ERO1A |
| 4.4476E-38 | -0.3419414 | 0.955 | 0.966 | 9.4956E-34 | TEC_prg2 | RPS19 |
| 2.8897E-63 | -0.3419488 | 0.053 | 0.257 | 6.1696E-59 | TEC_prg2 | CHMP2B |
| 1.8604E-63 | -0.3420355 | 0.082 | 0.303 | 3.9719E-59 | TEC_prg2 | ANAPC16 |
| 4.3751E-66 | -0.3434624 | 0.077 | 0.3 | 9.3409E-62 | TEC_prg2 | CHMP2A |
| 3.1708E-74 | -0.3437697 | 0.053 | 0.283 | 6.7696E-70 | TEC_prg2 | ATP6V1E1 |
| 1.9166E-71 | -0.3443094 | 0.042 | 0.26 | 4.092E-67 | TEC_prg2 | MDH2 |
| 9.4007E-75 | -0.3444336 | 0.033 | 0.251 | 2.007E-70 | TEC_prg2 | TMEM219 |
| 2.9607E-59 | -0.3444516 | 0.141 | 0.375 | 6.3211E-55 | TEC_prg2 | VAPA |
| 1.1552E-72 | -0.3447942 | 0.036 | 0.253 | 2.4663E-68 | TEC_prg2 | ATP5PB |
| 2.4993E-71 | -0.3448795 | 0.056 | 0.284 | 5.3361E-67 | TEC_prg2 | RAB7A |
| 8.8952E-43 | -0.3451855 | 0.39 | 0.59 | 1.8991E-38 | TEC_prg2 | RAN |
| 2.8837E-67 | -0.3453486 | 0.086 | 0.319 | 6.1567E-63 | TEC_prg2 | MRPL51 |
| 1.1462E-54 | -0.346102 | 0.123 | 0.338 | 2.4472E-50 | TEC_prg2 | UQCR11 |
| 6.826E-75 | -0.3463261 | 0.055 | 0.289 | 1.4574E-70 | TEC_prg2 | CAPZA2 |
| 7.2047E-70 | -0.3464596 | 0.026 | 0.228 | 1.5382E-65 | TEC_prg2 | CALU |
| 5.2051E-28 | -0.3470972 | 0.129 | 0.268 | 1.1113E-23 | TEC_prg2 | AKAP12 |
| 4.8826E-49 | -0.3477593 | 0.865 | 0.916 | 1.0424E-44 | TEC_prg2 | FAU |
| 4.2276E-51 | -0.3481892 | 0.082 | 0.268 | 9.026E-47 | TEC_prg2 | TES |
| 4.4403E-68 | -0.348664 | 0.065 | 0.287 | 9.48E-64 | TEC_prg2 | TMCO1 |
| 1.0875E-36 | -0.3487728 | 0.112 | 0.273 | 2.3219E-32 | TEC_prg2 | S100A14 |
| 3.0627E-64 | -0.3492856 | 0.043 | 0.245 | 6.5389E-60 | TEC_prg2 | ARL6IP5 |
| 1.5664E-58 | -0.3495377 | 0.097 | 0.309 | 3.3442E-54 | TEC_prg2 | ANP32B |
| 1.673E-50 | -0.3496919 | 0.158 | 0.373 | 3.5718E-46 | TEC_prg2 | MZT2B |
| 5.9701E-79 | -0.3499033 | 0.012 | 0.227 | 1.2746E-74 | TEC_prg2 | ARPC1A |
| 1.7844E-70 | -0.3504271 | 0.054 | 0.276 | 3.8098E-66 | TEC_prg2 | PHB |
| 5.9937E-70 | -0.3508673 | 0.074 | 0.304 | 1.2796E-65 | TEC_prg2 | CCT3 |
| 9.4963E-71 | -0.3513115 | 0.045 | 0.262 | 2.0275E-66 | TEC_prg2 | VPS29 |
| 3.8851E-49 | -0.3515371 | 0.085 | 0.269 | 8.2946E-45 | TEC_prg2 | MGLL |
| 1.0182E-67 | -0.3515696 | 0.076 | 0.302 | 2.1738E-63 | TEC_prg2 | SSB |
| 1.365E-14 | -0.3515719 | 0.973 | 0.903 | 2.9143E-10 | TEC_prg2 | MT-CO2 |
| 8.9812E-22 | -0.3518085 | 0.947 | 0.928 | 1.9175E-17 | TEC_prg2 | RPL12 |

| | | | | | | |
|---|---|---|---|---|---|---|
| 5.2323E-70 | -0.3519712 | 0.038 | 0.25 | 1.1171E-65 | TEC_prg2 | ATP6AP2 |
| 1.7265E-51 | -0.3522547 | 0.371 | 0.598 | 3.6861E-47 | TEC_prg2 | SET |
| 4.6429E-65 | -0.3526018 | 0.087 | 0.311 | 9.9126E-61 | TEC_prg2 | RSL24D1 |
| 2.0907E-47 | -0.3526275 | 0.035 | 0.19 | 4.4636E-43 | TEC_prg2 | PLA2G16 |
| 4.0222E-75 | -0.3532027 | 0.028 | 0.246 | 8.5874E-71 | TEC_prg2 | ARPC4 |
| 6.1381E-31 | -0.3538647 | 0.015 | 0.115 | 1.3105E-26 | TEC_prg2 | RSAD2 |
| 5.7376E-65 | -0.3541283 | 0.054 | 0.262 | 1.225E-60 | TEC_prg2 | EPRS |
| 6.5902E-66 | -0.3542005 | 0.1 | 0.331 | 1.407E-61 | TEC_prg2 | WDR1 |
| 1.0448E-31 | -0.3547559 | 0.087 | 0.219 | 2.2306E-27 | TEC_prg2 | MDK |
| 6.2432E-74 | -0.3549025 | 0.045 | 0.268 | 1.3329E-69 | TEC_prg2 | PSMB5 |
| 5.1303E-74 | -0.3560609 | 0.048 | 0.275 | 1.0953E-69 | TEC_prg2 | RAB1A |
| 2.0573E-81 | -0.3570486 | 0.019 | 0.244 | 4.3923E-77 | TEC_prg2 | COPZ1 |
| 1.7942E-70 | -0.3588973 | 0.048 | 0.267 | 3.8306E-66 | TEC_prg2 | PLEKHB2 |
| 5.6291E-72 | -0.3591355 | 0.057 | 0.282 | 1.2018E-67 | TEC_prg2 | MTPN |
| 1.8161E-63 | -0.359253 | 0.116 | 0.353 | 3.8773E-59 | TEC_prg2 | NDUFC1 |
| 1.6162E-45 | -0.359526 | 0.156 | 0.351 | 3.4506E-41 | TEC_prg2 | CD2AP |
| 7.0108E-77 | -0.3600726 | 0.016 | 0.231 | 1.4968E-72 | TEC_prg2 | PRDX3 |
| 1.6354E-17 | -0.3601331 | 0.041 | 0.115 | 3.4916E-13 | TEC_prg2 | AQP3 |
| 7.7851E-33 | -0.3602308 | 0.048 | 0.169 | 1.6621E-28 | TEC_prg2 | IGFBP2 |
| 1.0296E-27 | -0.3602738 | 0.915 | 0.915 | 2.1981E-23 | TEC_prg2 | RPLP0 |
| 1.5022E-46 | -0.3625424 | 0.163 | 0.367 | 3.2071E-42 | TEC_prg2 | MIF |
| 5.479E-71 | -0.3627998 | 0.06 | 0.285 | 1.1698E-66 | TEC_prg2 | ERH |
| 1.0079E-25 | -0.3632848 | 0.756 | 0.796 | 2.1518E-21 | TEC_prg2 | PPDPF |
| 2.793E-71 | -0.3634512 | 0.072 | 0.302 | 5.9631E-67 | TEC_prg2 | CCT5 |
| 1.3451E-71 | -0.3639773 | 0.062 | 0.293 | 2.8719E-67 | TEC_prg2 | TSTD1 |
| 2.2643E-66 | -0.3640374 | 0.108 | 0.348 | 4.8342E-62 | TEC_prg2 | ATP5MPL |
| 1.4362E-77 | -0.3640968 | 0.033 | 0.259 | 3.0662E-73 | TEC_prg2 | PSMB2 |
| 1.4864E-72 | -0.364997 | 0.066 | 0.298 | 3.1734E-68 | TEC_prg2 | ANAPC11 |
| 9.2709E-69 | -0.3658604 | 0.065 | 0.288 | 1.9793E-64 | TEC_prg2 | UFM1 |
| 2.9759E-26 | -0.3659782 | 0.296 | 0.45 | 6.3535E-22 | TEC_prg2 | ZBTB20 |
| 4.9981E-62 | -0.3660441 | 0.156 | 0.4 | 1.0671E-57 | TEC_prg2 | HEBP2 |
| 2.7378E-59 | -0.3662877 | 0.07 | 0.274 | 5.8452E-55 | TEC_prg2 | CD47 |
| 3.8742E-67 | -0.3664433 | 0.083 | 0.312 | 8.2713E-63 | TEC_prg2 | RABAC1 |
| 1.986E-75 | -0.3667457 | 0.05 | 0.281 | 4.2402E-71 | TEC_prg2 | PPP2R1A |
| 5.3875E-67 | -0.3668376 | 0.138 | 0.389 | 1.1502E-62 | TEC_prg2 | SNRPD2 |
| 1.0082E-63 | -0.3669346 | 0.128 | 0.362 | 2.1525E-59 | TEC_prg2 | EIF4B |
| 1.2107E-53 | -0.3678192 | 0.063 | 0.248 | 2.5848E-49 | TEC_prg2 | PRSS8 |
| 1.9808E-65 | -0.3679289 | 0.069 | 0.287 | 4.2289E-61 | TEC_prg2 | APRT |
| 6.592E-71 | -0.3687831 | 0.058 | 0.282 | 1.4074E-66 | TEC_prg2 | ATP5F1A |
| 4.4455E-46 | -0.3690076 | 0.138 | 0.33 | 9.4911E-42 | TEC_prg2 | DSP |
| 2.6804E-65 | -0.3690654 | 0.085 | 0.31 | 5.7226E-61 | TEC_prg2 | NDUFB11 |
| 1.2745E-71 | -0.3691163 | 0.109 | 0.358 | 2.7211E-67 | TEC_prg2 | NDUFB4 |
| 1.6133E-33 | -0.3691756 | 0.013 | 0.119 | 3.4445E-29 | TEC_prg2 | BST2 |
| 1.6455E-62 | -0.3694882 | 0.093 | 0.313 | 3.5131E-58 | TEC_prg2 | PSME2 |
| 3.8281E-51 | -0.3695659 | 0.656 | 0.788 | 8.173E-47 | TEC_prg2 | EEF2 |
| 1.4369E-52 | -0.3698221 | 0.173 | 0.395 | 3.0677E-48 | TEC_prg2 | CAMK2N1 |
| 1.5181E-62 | -0.3700814 | 0.433 | 0.669 | 3.2411E-58 | TEC_prg2 | YWHAE |
| 3.5447E-80 | -0.3702111 | 0.025 | 0.252 | 7.5679E-76 | TEC_prg2 | GLO1 |
| 4.4576E-70 | -0.3702659 | 0.077 | 0.311 | 9.517E-66 | TEC_prg2 | PSMB6 |
| 3.5484E-40 | -0.3705825 | 0.026 | 0.158 | 7.5758E-36 | TEC_prg2 | XIST |

| | | | | | | |
|---|---|---|---|---|---|---|
| 9.1085E-63 | -0.3707547 | 0.123 | 0.357 | 1.9447E-58 | TEC_prg2 | KIF5B |
| 9.8678E-71 | -0.3709832 | 0.063 | 0.292 | 2.1068E-66 | TEC_prg2 | MINOS1 |
| 7.6256E-45 | -0.3720609 | 0.044 | 0.197 | 1.6281E-40 | TEC_prg2 | DCBLD2 |
| 7.1723E-75 | -0.3723214 | 0.094 | 0.343 | 1.5313E-70 | TEC_prg2 | SNX3 |
| 7.0802E-43 | -0.3727501 | 0.175 | 0.374 | 1.5116E-38 | TEC_prg2 | NUCKS1 |
| 1.0353E-79 | -0.3734298 | 0.045 | 0.281 | 2.2103E-75 | TEC_prg2 | CAPZB |
| 9.2748E-43 | -0.373585 | 0.68 | 0.788 | 1.9802E-38 | TEC_prg2 | OAZ1 |
| 1.9438E-66 | -0.3736534 | 0.094 | 0.326 | 4.1499E-62 | TEC_prg2 | DDT |
| 5.6374E-69 | -0.374246 | 0.077 | 0.304 | 1.2036E-64 | TEC_prg2 | SEPT7 |
| 1.1221E-69 | -0.3746275 | 0.07 | 0.297 | 2.3957E-65 | TEC_prg2 | NDUFV2 |
| 2.4879E-56 | -0.3746667 | 0.352 | 0.587 | 5.3116E-52 | TEC_prg2 | COX6B1 |
| 0.0023699 | -0.3750735 | 0.641 | 0.556 | 1 | TEC_prg2 | GDF15 |
| 2.301E-63 | -0.3759779 | 0.256 | 0.516 | 4.9127E-59 | TEC_prg2 | UBE2D3 |
| 6.66E-72 | -0.3763562 | 0.053 | 0.277 | 1.4219E-67 | TEC_prg2 | C4orf3 |
| 1.0912E-76 | -0.3774863 | 0.058 | 0.295 | 2.3297E-72 | TEC_prg2 | RTN3 |
| 2.6146E-56 | -0.3775654 | 0.109 | 0.323 | 5.5823E-52 | TEC_prg2 | CDH1 |
| 7.1986E-69 | -0.3777209 | 0.075 | 0.305 | 1.5369E-64 | TEC_prg2 | GRN |
| 2.968E-82 | -0.3793223 | 0.039 | 0.278 | 6.3367E-78 | TEC_prg2 | LMAN2 |
| 3.0767E-49 | -0.3794302 | 0.116 | 0.31 | 6.5688E-45 | TEC_prg2 | ITGA3 |
| 3.3079E-10 | -0.3800794 | 0.297 | 0.378 | 7.0624E-06 | TEC_prg2 | DDIT3 |
| 8.6471E-54 | -0.3805076 | 0.441 | 0.649 | 1.8462E-49 | TEC_prg2 | SLC25A6 |
| 2.5156E-72 | -0.380763 | 0.12 | 0.378 | 5.3708E-68 | TEC_prg2 | COMMD6 |
| 6.8532E-86 | -0.3819652 | 0.033 | 0.277 | 1.4632E-81 | TEC_prg2 | SUMO1 |
| 1.0379E-68 | -0.3823917 | 0.092 | 0.325 | 2.2158E-64 | TEC_prg2 | ACTR3 |
| 3.2817E-28 | -0.3847636 | 0.036 | 0.141 | 7.0064E-24 | TEC_prg2 | TNFRSF11B |
| 1.0522E-79 | -0.3851782 | 0.033 | 0.264 | 2.2464E-75 | TEC_prg2 | LSM4 |
| 2.1583E-33 | -0.3854637 | 0.668 | 0.765 | 4.6079E-29 | TEC_prg2 | RPS26 |
| 8.9797E-43 | -0.3859552 | 0.062 | 0.219 | 1.9172E-38 | TEC_prg2 | MANF |
| 1.8858E-78 | -0.3861103 | 0.084 | 0.339 | 4.0261E-74 | TEC_prg2 | TMEM50A |
| 1.7117E-57 | -0.3876318 | 0.026 | 0.201 | 3.6545E-53 | TEC_prg2 | CD40 |
| 1.9893E-42 | -0.3876872 | 0.87 | 0.918 | 4.2472E-38 | TEC_prg2 | RPL35 |
| 4.1835E-72 | -0.3877995 | 0.096 | 0.341 | 8.9318E-68 | TEC_prg2 | MRFAP1 |
| 2.5548E-45 | -0.3882984 | 0.187 | 0.392 | 5.4544E-41 | TEC_prg2 | SPTBN1 |
| 8.3933E-74 | -0.390171 | 0.099 | 0.348 | 1.792E-69 | TEC_prg2 | GNB1 |
| 3.6878E-83 | -0.3912253 | 0.097 | 0.368 | 7.8734E-79 | TEC_prg2 | SEM1 |
| 4.4709E-36 | -0.3915847 | 0.036 | 0.161 | 9.5453E-32 | TEC_prg2 | MYL9 |
| 2.3452E-48 | -0.3925295 | 0.741 | 0.832 | 5.007E-44 | TEC_prg2 | RPS21 |
| 4.0926E-77 | -0.3927004 | 0.069 | 0.313 | 8.7376E-73 | TEC_prg2 | COPS9 |
| 5.8282E-54 | -0.3930275 | 0.124 | 0.333 | 1.2443E-49 | TEC_prg2 | ISCU |
| 5.4115E-52 | -0.3931143 | 0.092 | 0.285 | 1.1553E-47 | TEC_prg2 | RB1CC1 |
| 4.7234E-77 | -0.3931591 | 0.039 | 0.266 | 1.0084E-72 | TEC_prg2 | OSTC |
| 4.5296E-51 | -0.3941894 | 0.835 | 0.892 | 9.6708E-47 | TEC_prg2 | RPS29 |
| 5.5692E-76 | -0.394888 | 0.044 | 0.272 | 1.189E-71 | TEC_prg2 | BCAP31 |
| 1.0891E-75 | -0.3955785 | 0.05 | 0.28 | 2.3253E-71 | TEC_prg2 | EIF3M |
| 5.4582E-80 | -0.396297 | 0.097 | 0.361 | 1.1653E-75 | TEC_prg2 | COPE |
| 3.9601E-60 | -0.396477 | 0.113 | 0.335 | 8.4547E-56 | TEC_prg2 | RAB13 |
| 1.9061E-88 | -0.3968217 | 0.034 | 0.284 | 4.0696E-84 | TEC_prg2 | TMED2 |
| 2.2367E-50 | -0.3973336 | 0.477 | 0.655 | 4.7753E-46 | TEC_prg2 | CCNI |
| 4.6857E-75 | -0.3985146 | 0.072 | 0.311 | 1.0004E-70 | TEC_prg2 | ATP5PO |
| 1.2089E-70 | -0.3987661 | 0.131 | 0.388 | 2.5809E-66 | TEC_prg2 | SEC62 |

| | | | | | | |
|---|---|---|---|---|---|---|
| 1.4271E-63 | -0.3991291 | 0.031 | 0.224 | 3.0468E-59 | TEC_prg2 | DPP4 |
| 4.0289E-47 | -0.3994342 | 0.381 | 0.578 | 8.6017E-43 | TEC_prg2 | LRRFIP1 |
| 1.2867E-57 | -0.3996679 | 0.159 | 0.385 | 2.7471E-53 | TEC_prg2 | EIF3A |
| 1.011E-79 | -0.3996681 | 0.047 | 0.283 | 2.1586E-75 | TEC_prg2 | PSMB3 |
| 2.2904E-91 | -0.3999626 | 0.029 | 0.281 | 4.8899E-87 | TEC_prg2 | EIF3I |
| 7.182E-78 | -0.4000625 | 0.06 | 0.3 | 1.5334E-73 | TEC_prg2 | CAPNS1 |
| 4.8413E-76 | -0.4003589 | 0.077 | 0.321 | 1.0336E-71 | TEC_prg2 | EIF6 |
| 1.6608E-23 | -0.4013831 | 0.172 | 0.297 | 3.5458E-19 | TEC_prg2 | UCHL1 |
| 1.2055E-86 | -0.4016619 | 0.07 | 0.334 | 2.5738E-82 | TEC_prg2 | EIF3F |
| 2.653E-88 | -0.4019493 | 0.034 | 0.282 | 5.6641E-84 | TEC_prg2 | RER1 |
| 1.2503E-83 | -0.4030413 | 0.037 | 0.278 | 2.6695E-79 | TEC_prg2 | UQCRFS1 |
| 6.4905E-84 | -0.4032007 | 0.05 | 0.3 | 1.3857E-79 | TEC_prg2 | ATP5F1C |
| 8.9852E-78 | -0.4045465 | 0.065 | 0.309 | 1.9183E-73 | TEC_prg2 | LITAF |
| 0.00010008 | -0.4045893 | 0.085 | 0.121 | 1 | TEC_prg2 | CXCL8 |
| 1.1111E-52 | -0.4046238 | 0.161 | 0.383 | 2.3721E-48 | TEC_prg2 | CD44 |
| 1.569E-84 | -0.4050313 | 0.033 | 0.274 | 3.3499E-80 | TEC_prg2 | RPN2 |
| 6.736E-78 | -0.4053792 | 0.026 | 0.248 | 1.4381E-73 | TEC_prg2 | CXCL16 |
| 7.1256E-37 | -0.4058451 | 0.038 | 0.167 | 1.5213E-32 | TEC_prg2 | RCAN1 |
| 3.3158E-69 | -0.4067939 | 0.051 | 0.266 | 7.0791E-65 | TEC_prg2 | PLP2 |
| 1.3592E-45 | -0.4078433 | 0.158 | 0.354 | 2.9019E-41 | TEC_prg2 | CHCHD10 |
| 1.2336E-75 | -0.4096219 | 0.066 | 0.306 | 2.6337E-71 | TEC_prg2 | BUD31 |
| 2.022E-82 | -0.4099937 | 0.089 | 0.356 | 4.3169E-78 | TEC_prg2 | ELOC |
| 1.697E-63 | -0.410545 | 0.142 | 0.387 | 3.623E-59 | TEC_prg2 | COX7B |
| 2.6794E-78 | -0.4107595 | 0.221 | 0.509 | 5.7206E-74 | TEC_prg2 | MORF4L1 |
| 2.9072E-90 | -0.4109919 | 0.021 | 0.266 | 6.2069E-86 | TEC_prg2 | KDELR2 |
| 1.3573E-89 | -0.4114557 | 0.036 | 0.288 | 2.8979E-85 | TEC_prg2 | LAMTOR1 |
| 3.0256E-54 | -0.4118569 | 0.039 | 0.214 | 6.4596E-50 | TEC_prg2 | SDF2L1 |
| 5.2665E-79 | -0.412216 | 0.087 | 0.341 | 1.1244E-74 | TEC_prg2 | PCBD1 |
| 5.6966E-65 | -0.4127746 | 0.555 | 0.739 | 1.2162E-60 | TEC_prg2 | RPL36AL |
| 2.4408E-88 | -0.412892 | 0.061 | 0.322 | 5.2112E-84 | TEC_prg2 | RNF7 |
| 0.00127355 | -0.4136268 | 0.63 | 0.623 | 1 | TEC_prg2 | VIM |
| 3.6271E-79 | -0.414741 | 0.028 | 0.252 | 7.7438E-75 | TEC_prg2 | SEC11C |
| 2.3583E-80 | -0.4147736 | 0.092 | 0.352 | 5.0349E-76 | TEC_prg2 | FAM96B |
| 1.9552E-47 | -0.4157981 | 0.153 | 0.356 | 4.1744E-43 | TEC_prg2 | RALBP1 |
| 2.8797E-79 | -0.4162544 | 0.106 | 0.37 | 6.1482E-75 | TEC_prg2 | SPCS1 |
| 8.8907E-41 | -0.4165303 | 0.759 | 0.826 | 1.8982E-36 | TEC_prg2 | YBX1 |
| 3.7031E-95 | -0.4167461 | 0.056 | 0.33 | 7.906E-91 | TEC_prg2 | NDFIP1 |
| 1.1445E-05 | -0.4170268 | 0.216 | 0.266 | 0.24435474 | TEC_prg2 | NUPR1 |
| 1.1495E-74 | -0.4174677 | 0.085 | 0.328 | 2.4541E-70 | TEC_prg2 | CUTA |
| 4.4414E-80 | -0.4175389 | 0.083 | 0.338 | 9.4823E-76 | TEC_prg2 | ATP5PD |
| 3.2636E-83 | -0.4187534 | 0.059 | 0.311 | 6.9679E-79 | TEC_prg2 | LAMP1 |
| 1.5497E-53 | -0.4189513 | 0.901 | 0.945 | 3.3085E-49 | TEC_prg2 | RPL37A |
| 5.1177E-51 | -0.4192299 | 0.199 | 0.421 | 1.0926E-46 | TEC_prg2 | DBI |
| 6.4387E-90 | -0.4192357 | 0.04 | 0.296 | 1.3747E-85 | TEC_prg2 | PSMB7 |
| 3.386E-57 | -0.4198424 | 0.413 | 0.625 | 7.229E-53 | TEC_prg2 | CAST |
| 6.0139E-69 | -0.4203721 | 0.249 | 0.508 | 1.284E-64 | TEC_prg2 | PTGES3 |
| 5.318E-67 | -0.4207678 | 0.304 | 0.562 | 1.1354E-62 | TEC_prg2 | NDUFS5 |
| 2.8096E-85 | -0.422457 | 0.032 | 0.272 | 5.9985E-81 | TEC_prg2 | PGAM1 |
| 3.5749E-49 | -0.4229582 | 0.023 | 0.176 | 7.6324E-45 | TEC_prg2 | MGST1 |
| 1.9385E-91 | -0.4231947 | 0.029 | 0.28 | 4.1387E-87 | TEC_prg2 | NDUFAB1 |

| | | | | | | |
|---|---|---|---|---|---|---|
| 3.1045E-88 | -0.42384 | 0.065 | 0.331 | 6.6281E-84 | TEC_prg2 | SSR2 |
| 1.8436E-70 | -0.4241594 | 0.066 | 0.29 | 3.9361E-66 | TEC_prg2 | SLC38A1 |
| 7.5328E-75 | -0.42456 | 0.072 | 0.309 | 1.6083E-70 | TEC_prg2 | MSN |
| 1.9371E-71 | -0.424616 | 0.06 | 0.286 | 4.1357E-67 | TEC_prg2 | GSTO1 |
| 4.5477E-78 | -0.4246626 | 0.156 | 0.436 | 9.7093E-74 | TEC_prg2 | TBCA |
| 2.9697E-71 | -0.4254486 | 0.088 | 0.323 | 6.3403E-67 | TEC_prg2 | SELENOS |
| 4.471E-57 | -0.427254 | 0.255 | 0.488 | 9.5457E-53 | TEC_prg2 | SERP1 |
| 3.776E-62 | -0.428093 | 0.115 | 0.341 | 8.0618E-58 | TEC_prg2 | PRDX2 |
| 4.7616E-48 | -0.4281085 | 0.386 | 0.586 | 1.0166E-43 | TEC_prg2 | CST3 |
| 3.0786E-59 | -0.4307231 | 0.109 | 0.327 | 6.5729E-55 | TEC_prg2 | C9orf16 |
| 4.6041E-65 | -0.4314119 | 0.619 | 0.787 | 9.8299E-61 | TEC_prg2 | COX7C |
| 4.4802E-28 | -0.4326409 | 0.036 | 0.138 | 9.5653E-24 | TEC_prg2 | HMOX1 |
| 5.2743E-45 | -0.4332734 | 0.094 | 0.268 | 1.1261E-40 | TEC_prg2 | MACC1 |
| 3.1407E-84 | -0.4336932 | 0.07 | 0.33 | 6.7054E-80 | TEC_prg2 | RAP1B |
| 6.3621E-84 | -0.4341291 | 0.035 | 0.274 | 1.3583E-79 | TEC_prg2 | DHRS7 |
| 8.5057E-70 | -0.4344198 | 0.456 | 0.687 | 1.816E-65 | TEC_prg2 | UQCRB |
| 4.9689E-85 | -0.4359011 | 0.115 | 0.398 | 1.0609E-80 | TEC_prg2 | WDR83OS |
| 2.9754E-83 | -0.4362575 | 0.04 | 0.28 | 6.3526E-79 | TEC_prg2 | LAPTM4B |
| 5.5719E-85 | -0.4392429 | 0.055 | 0.307 | 1.1896E-80 | TEC_prg2 | LMAN1 |
| 3.915E-37 | -0.4402031 | 0.229 | 0.411 | 8.3585E-33 | TEC_prg2 | RAB11FIP1 |
| 1.3592E-71 | -0.4429139 | 0.103 | 0.346 | 2.902E-67 | TEC_prg2 | TKT |
| 3.5889E-94 | -0.4437646 | 0.082 | 0.366 | 7.6623E-90 | TEC_prg2 | KDELR1 |
| 7.9992E-10 | -0.4440887 | 0.056 | 0.107 | 1.7078E-05 | TEC_prg2 | APOE |
| 4.2972E-77 | -0.4448154 | 0.131 | 0.399 | 9.1745E-73 | TEC_prg2 | UBE2B |
| 2.9111E-68 | -0.4449596 | 0.903 | 0.937 | 6.2152E-64 | TEC_prg2 | RPS9 |
| 3.1689E-71 | -0.4452151 | 0.483 | 0.702 | 6.7656E-67 | TEC_prg2 | YWHAZ |
| 5.5025E-79 | -0.4453985 | 0.102 | 0.36 | 1.1748E-74 | TEC_prg2 | ATP6V1F |
| 2.6028E-89 | -0.4471829 | 0.089 | 0.37 | 5.557E-85 | TEC_prg2 | ERP29 |
| 3.3044E-88 | -0.4473279 | 0.128 | 0.42 | 7.0548E-84 | TEC_prg2 | SUB1 |
| 7.147E-91 | -0.4475833 | 0.085 | 0.364 | 1.5259E-86 | TEC_prg2 | RAB2A |
| 2.6187E-91 | -0.4478722 | 0.074 | 0.347 | 5.5908E-87 | TEC_prg2 | SARAF |
| 5.7774E-19 | -0.4480995 | 0.82 | 0.798 | 1.2335E-14 | TEC_prg2 | CD24 |
| 1.0069E-64 | -0.448672 | 0.085 | 0.307 | 2.1497E-60 | TEC_prg2 | ASPH |
| 9.5357E-74 | -0.4499335 | 0.288 | 0.558 | 2.0359E-69 | TEC_prg2 | COX6C |
| 4.0961E-78 | -0.4512526 | 0.148 | 0.421 | 8.7453E-74 | TEC_prg2 | CAPN2 |
| 1.5626E-79 | -0.4518421 | 0.043 | 0.278 | 3.3362E-75 | TEC_prg2 | ECHS1 |
| 1.061E-105 | -0.4524131 | 0.04 | 0.329 | 2.266E-101 | TEC_prg2 | CHMP5 |
| 1.3339E-87 | -0.4525307 | 0.148 | 0.447 | 2.8478E-83 | TEC_prg2 | HNRNPK |
| 1.8239E-66 | -0.4528175 | 0.076 | 0.293 | 3.894E-62 | TEC_prg2 | RBM3 |
| 8.7626E-80 | -0.453524 | 0.146 | 0.425 | 1.8708E-75 | TEC_prg2 | TAX1BP1 |
| 7.7832E-74 | -0.4548507 | 0.076 | 0.312 | 1.6617E-69 | TEC_prg2 | ANXA3 |
| 3.3466E-56 | -0.4558122 | 0.298 | 0.521 | 7.1451E-52 | TEC_prg2 | HSPD1 |
| 5.5094E-94 | -0.4558484 | 0.058 | 0.331 | 1.1763E-89 | TEC_prg2 | CIB1 |
| 3.7842E-87 | -0.456889 | 0.125 | 0.412 | 8.0792E-83 | TEC_prg2 | HSBP1 |
| 4.868E-97 | -0.4577809 | 0.06 | 0.341 | 1.0393E-92 | TEC_prg2 | GABARAPL2 |
| 4.682E-100 | -0.4578506 | 0.06 | 0.347 | 9.9965E-96 | TEC_prg2 | COX7A2L |
| 2.2033E-58 | -0.4585091 | 0.901 | 0.939 | 4.7041E-54 | TEC_prg2 | RPL37 |
| 5.3561E-50 | -0.4588587 | 0.96 | 0.961 | 1.1435E-45 | TEC_prg2 | RPL28 |
| 1.633E-99 | -0.4590608 | 0.072 | 0.36 | 3.4861E-95 | TEC_prg2 | EIF3H |
| 1.5877E-77 | -0.4593152 | 0.126 | 0.39 | 3.3896E-73 | TEC_prg2 | TRMT112 |

| | | | | | | |
|---|---|---|---|---|---|---|
| 2.3002E-80 | -0.4629844 | 0.037 | 0.269 | 4.9108E-76 | TEC_prg2 | DNAJC3 |
| 2.1487E-13 | -0.4664328 | 0.243 | 0.337 | 4.5875E-09 | TEC_prg2 | MT1E |
| 3.2982E-65 | -0.4670819 | 0.375 | 0.605 | 7.0416E-61 | TEC_prg2 | PRDX5 |
| 1.8359E-93 | -0.4677407 | 0.084 | 0.365 | 3.9195E-89 | TEC_prg2 | SPCS2 |
| 3.5383E-85 | -0.4696093 | 0.099 | 0.37 | 7.5543E-81 | TEC_prg2 | PFDN2 |
| 9.2045E-88 | -0.4699425 | 0.126 | 0.415 | 1.9652E-83 | TEC_prg2 | MRPL33 |
| 2.5517E-96 | -0.4714496 | 0.029 | 0.291 | 5.448E-92 | TEC_prg2 | FKBP1A |
| 1.4704E-83 | -0.4715281 | 0.234 | 0.523 | 3.1393E-79 | TEC_prg2 | TOMM7 |
| 5.496E-68 | -0.4721999 | 0.043 | 0.25 | 1.1734E-63 | TEC_prg2 | IFITM2 |
| 2.0417E-97 | -0.4728317 | 0.067 | 0.354 | 4.359E-93 | TEC_prg2 | COX5A |
| 9.499E-101 | -0.473289 | 0.067 | 0.359 | 2.028E-96 | TEC_prg2 | VDAC1 |
| 6.6529E-75 | -0.474276 | 0.107 | 0.36 | 1.4204E-70 | TEC_prg2 | SLC25A5 |
| 1.3526E-73 | -0.4747404 | 0.825 | 0.895 | 2.8878E-69 | TEC_prg2 | RPL22 |
| 7.8761E-95 | -0.4749679 | 0.097 | 0.389 | 1.6816E-90 | TEC_prg2 | EIF3K |
| 9.0906E-58 | -0.4758158 | 0.705 | 0.821 | 1.9408E-53 | TEC_prg2 | ATP5F1E |
| 2.4828E-84 | -0.4784131 | 0.148 | 0.439 | 5.3008E-80 | TEC_prg2 | JAK1 |
| 8.8806E-77 | -0.4792589 | 0.194 | 0.467 | 1.896E-72 | TEC_prg2 | NAP1L1 |
| 2.265E-102 | -0.4801432 | 0.043 | 0.327 | 4.8363E-98 | TEC_prg2 | LEPROT |
| 8.0386E-79 | -0.4810698 | 0.053 | 0.29 | 1.7162E-74 | TEC_prg2 | SELENOM |
| 7.84E-105 | -0.4821482 | 0.051 | 0.341 | 1.674E-100 | TEC_prg2 | NDUFC2 |
| 7.5922E-82 | -0.4856352 | 0.132 | 0.412 | 1.6209E-77 | TEC_prg2 | NDUFB2 |
| 1.2217E-57 | -0.48587 | 0.08 | 0.276 | 2.6082E-53 | TEC_prg2 | RPS4Y1 |
| 7.045E-83 | -0.4863271 | 0.029 | 0.263 | 1.5041E-78 | TEC_prg2 | HIGD1A |
| 4.859E-60 | -0.4885929 | 0.104 | 0.319 | 1.0374E-55 | TEC_prg2 | MIR4435-2HG |
| 2.9481E-68 | -0.4890346 | 0.163 | 0.419 | 6.2942E-64 | TEC_prg2 | CD151 |
| 2.252E-101 | -0.4893673 | 0.041 | 0.319 | 4.8077E-97 | TEC_prg2 | SERINC1 |
| 6.122E-112 | -0.4899932 | 0.031 | 0.327 | 1.307E-107 | TEC_prg2 | TMED10 |
| 2.665E-103 | -0.4907841 | 0.06 | 0.351 | 5.691E-99 | TEC_prg2 | SNRPB |
| 3.2242E-81 | -0.4914117 | 0.095 | 0.358 | 6.8836E-77 | TEC_prg2 | UQCRQ |
| 2.1946E-87 | -0.4933598 | 0.172 | 0.464 | 4.6855E-83 | TEC_prg2 | BZW1 |
| 2.1203E-94 | -0.4955805 | 0.104 | 0.398 | 4.5268E-90 | TEC_prg2 | PPP1CB |
| 2.6428E-11 | -0.4958173 | 0.109 | 0.18 | 5.6424E-07 | TEC_prg2 | S100A4 |
| 5.636E-114 | -0.4978234 | 0.06 | 0.374 | 1.203E-109 | TEC_prg2 | SEC11A |
| 1.6842E-20 | -0.4988793 | 0.119 | 0.224 | 3.5958E-16 | TEC_prg2 | GLRX |
| 8.8257E-98 | -0.4999442 | 0.045 | 0.319 | 1.8843E-93 | TEC_prg2 | FAM3C |
| 3.7336E-84 | -0.5004108 | 0.276 | 0.556 | 7.9713E-80 | TEC_prg2 | ATP5MC2 |
| 7.73E-82 | -0.5012791 | 0.075 | 0.326 | 1.6504E-77 | TEC_prg2 | CAPG |
| 1.2251E-87 | -0.5032463 | 0.103 | 0.38 | 2.6156E-83 | TEC_prg2 | ANXA11 |
| 2.664E-102 | -0.504567 | 0.07 | 0.367 | 5.6873E-98 | TEC_prg2 | PTP4A2 |
| 1.131E-85 | -0.5051913 | 0.817 | 0.907 | 2.4146E-81 | TEC_prg2 | RPL35A |
| 1.711E-102 | -0.5056282 | 0.091 | 0.394 | 3.6535E-98 | TEC_prg2 | NEDD8 |
| 2.543E-100 | -0.5056991 | 0.268 | 0.576 | 5.429E-96 | TEC_prg2 | GNG5 |
| 1.198E-112 | -0.5060658 | 0.048 | 0.349 | 2.558E-108 | TEC_prg2 | SRP9 |
| 5.8325E-55 | -0.5062833 | 0.415 | 0.606 | 1.2452E-50 | TEC_prg2 | EZR |
| 6.9435E-80 | -0.5081682 | 0.072 | 0.322 | 1.4824E-75 | TEC_prg2 | CD55 |
| 2.767E-107 | -0.5085397 | 0.087 | 0.405 | 5.908E-103 | TEC_prg2 | SDCBP |
| 4.648E-107 | -0.5091739 | 0.039 | 0.326 | 9.924E-103 | TEC_prg2 | BRK1 |
| 3.2586E-86 | -0.5099437 | 0.189 | 0.478 | 6.9571E-82 | TEC_prg2 | ATP5MD |
| 5.3387E-58 | -0.5110671 | 0.822 | 0.881 | 1.1398E-53 | TEC_prg2 | HLA-A |
| 1.1863E-47 | -0.5121616 | 0.069 | 0.24 | 2.5327E-43 | TEC_prg2 | SFN |

| | | | | | | |
|---|---|---|---|---|---|---|
| 1.1446E-19 | -0.5128597 | 0.96 | 0.919 | 2.4437E-15 | TEC_prg2 | TMSB10 |
| 2.4151E-95 | -0.5135761 | 0.07 | 0.35 | 5.1562E-91 | TEC_prg2 | NDUFB9 |
| 1.3138E-70 | -0.5136024 | 0.457 | 0.682 | 2.805E-66 | TEC_prg2 | PSAP |
| 1.736E-113 | -0.5146512 | 0.085 | 0.407 | 3.707E-109 | TEC_prg2 | SNU13 |
| 3.145E-104 | -0.5157156 | 0.067 | 0.362 | 6.714E-100 | TEC_prg2 | SF3B6 |
| 1.1514E-95 | -0.5158927 | 0.139 | 0.444 | 2.4583E-91 | TEC_prg2 | UQCR10 |
| 3.1097E-27 | -0.5166014 | 0.028 | 0.126 | 6.6392E-23 | TEC_prg2 | TAGLN |
| 7.323E-100 | -0.5187759 | 0.138 | 0.451 | 1.5635E-95 | TEC_prg2 | DYNLRB1 |
| 6.5967E-06 | -0.5189296 | 0.871 | 0.807 | 0.14084051 | TEC_prg2 | IGFBP7 |
| 3.4595E-38 | -0.5191624 | 0.928 | 0.952 | 7.386E-34 | TEC_prg2 | B2M |
| 8.5967E-94 | -0.519578 | 0.185 | 0.486 | 1.8354E-89 | TEC_prg2 | ATP5PF |
| 3.2507E-63 | -0.5202509 | 0.085 | 0.302 | 6.9403E-59 | TEC_prg2 | SERPINB1 |
| 1.0336E-57 | -0.5209197 | 0.05 | 0.236 | 2.2067E-53 | TEC_prg2 | GCLM |
| 4.5174E-46 | -0.5209209 | 0.014 | 0.153 | 9.6447E-42 | TEC_prg2 | TM4SF1 |
| 6.1157E-91 | -0.5237963 | 0.114 | 0.399 | 1.3057E-86 | TEC_prg2 | ATP5MF |
| 3.8877E-89 | -0.5244059 | 0.689 | 0.843 | 8.3003E-85 | TEC_prg2 | TMBIM6 |
| 5.546E-110 | -0.525582 | 0.058 | 0.361 | 1.184E-105 | TEC_prg2 | TRAM1 |
| 1.201E-113 | -0.5282445 | 0.043 | 0.344 | 2.563E-109 | TEC_prg2 | PGK1 |
| 6.043E-82 | -0.5290293 | 0.83 | 0.905 | 1.2902E-77 | TEC_prg2 | RPS15A |
| 1.0451E-86 | -0.529156 | 0.867 | 0.929 | 2.2314E-82 | TEC_prg2 | RPL14 |
| 1.979E-111 | -0.530924 | 0.063 | 0.372 | 4.224E-107 | TEC_prg2 | NOP10 |
| 8.59E-108 | -0.5329551 | 0.07 | 0.373 | 1.834E-103 | TEC_prg2 | EIF1AX |
| 3.0331E-95 | -0.5337972 | 0.307 | 0.599 | 6.4756E-91 | TEC_prg2 | HNRNPC |
| 1.104E-108 | -0.5358266 | 0.095 | 0.414 | 2.358E-104 | TEC_prg2 | REEP5 |
| 1.773E-105 | -0.5359356 | 0.069 | 0.367 | 3.785E-101 | TEC_prg2 | PPA1 |
| 4.1195E-95 | -0.5359926 | 0.185 | 0.483 | 8.7951E-91 | TEC_prg2 | RPS10 |
| 9.849E-107 | -0.5383938 | 0.097 | 0.415 | 2.103E-102 | TEC_prg2 | RHEB |
| 3.3315E-88 | -0.5392678 | 0.836 | 0.909 | 7.1127E-84 | TEC_prg2 | RPL27A |
| 4.7112E-58 | -0.5392872 | 0.29 | 0.511 | 1.0058E-53 | TEC_prg2 | LDHA |
| 5.122E-109 | -0.5393776 | 0.054 | 0.354 | 1.094E-104 | TEC_prg2 | NPC2 |
| 1.6669E-81 | -0.539883 | 0.386 | 0.635 | 3.5588E-77 | TEC_prg2 | APLP2 |
| 2.695E-113 | -0.5404623 | 0.067 | 0.38 | 5.753E-109 | TEC_prg2 | CNBP |
| 2.8583E-76 | -0.5414634 | 0.081 | 0.325 | 6.1024E-72 | TEC_prg2 | CYTOR |
| 1.08E-116 | -0.5416043 | 0.222 | 0.558 | 2.307E-112 | TEC_prg2 | ARF1 |
| 1.3596E-52 | -0.5417592 | 0.315 | 0.52 | 2.9027E-48 | TEC_prg2 | TNFRSF12A |
| 5.842E-93 | -0.5431628 | 0.739 | 0.868 | 1.2473E-88 | TEC_prg2 | SRP14 |
| 6.1058E-80 | -0.5433597 | 0.506 | 0.702 | 1.3036E-75 | TEC_prg2 | CLIC1 |
| 1.1339E-60 | -0.54621 | 0.09 | 0.299 | 2.4209E-56 | TEC_prg2 | CYB5A |
| 1.5274E-92 | -0.5469841 | 0.104 | 0.393 | 3.2611E-88 | TEC_prg2 | ATP5MC3 |
| 8.033E-116 | -0.5506367 | 0.058 | 0.371 | 1.715E-111 | TEC_prg2 | PSMB1 |
| 2.499E-128 | -0.5516547 | 0.041 | 0.369 | 5.335E-124 | TEC_prg2 | EIF3E |
| 5.4949E-82 | -0.5522776 | 0.66 | 0.804 | 1.1732E-77 | TEC_prg2 | NPM1 |
| 3.481E-124 | -0.5524964 | 0.066 | 0.399 | 7.432E-120 | TEC_prg2 | CSDE1 |
| 8.088E-110 | -0.5534276 | 0.296 | 0.605 | 1.727E-105 | TEC_prg2 | COX7A2 |
| 3.953E-123 | -0.5560189 | 0.052 | 0.377 | 8.44E-119 | TEC_prg2 | GHITM |
| 5.0859E-80 | -0.5577117 | 0.896 | 0.938 | 1.0858E-75 | TEC_prg2 | RPL6 |
| 3.091E-114 | -0.5577943 | 0.178 | 0.509 | 6.599E-110 | TEC_prg2 | SUMO2 |
| 1.972E-110 | -0.5578567 | 0.112 | 0.438 | 4.21E-106 | TEC_prg2 | LAMTOR5 |
| 7.679E-106 | -0.5583867 | 0.109 | 0.426 | 1.639E-101 | TEC_prg2 | EIF2S2 |
| 4.2413E-87 | -0.5598843 | 0.864 | 0.912 | 9.0552E-83 | TEC_prg2 | RPS16 |

| | | | | | | |
|---|---|---|---|---|---|---|
| 2.235E-123 | -0.5598874 | 0.087 | 0.426 | 4.772E-119 | TEC_prg2 | TOMM20 |
| 1.0962E-83 | -0.5604731 | 0.828 | 0.901 | 2.3405E-79 | TEC_prg2 | RPS11 |
| 2.8093E-83 | -0.5614127 | 0.129 | 0.4 | 5.9979E-79 | TEC_prg2 | TAGLN2 |
| 7.489E-05 | -0.5617039 | 0.855 | 0.792 | 1 | TEC_prg2 | KRT18 |
| 1.215E-109 | -0.5628496 | 0.085 | 0.4 | 2.595E-105 | TEC_prg2 | ATP5F1B |
| 4.842E-112 | -0.5634412 | 0.302 | 0.615 | 1.034E-107 | TEC_prg2 | ATP6V1G1 |
| 2.0427E-82 | -0.565463 | 0.567 | 0.746 | 4.3611E-78 | TEC_prg2 | CSTB |
| 9.159E-114 | -0.5690529 | 0.339 | 0.636 | 1.955E-109 | TEC_prg2 | CLTA |
| 4.259E-121 | -0.5699412 | 0.045 | 0.361 | 9.092E-117 | TEC_prg2 | TMED9 |
| 2.357E-118 | -0.5712226 | 0.074 | 0.404 | 5.033E-114 | TEC_prg2 | SBDS |
| 1.2885E-73 | -0.5725582 | 0.675 | 0.796 | 2.751E-69 | TEC_prg2 | EEF1B2 |
| 1.9825E-95 | -0.5746855 | 0.093 | 0.382 | 4.2327E-91 | TEC_prg2 | SELENOK |
| 4.706E-101 | -0.5805692 | 0.701 | 0.84 | 1.0047E-96 | TEC_prg2 | H3F3A |
| 1.036E-102 | -0.5812184 | 0.248 | 0.552 | 2.2126E-98 | TEC_prg2 | OST4 |
| 1.179E-117 | -0.5841458 | 0.055 | 0.372 | 2.518E-113 | TEC_prg2 | PRELID1 |
| 3.763E-109 | -0.5874543 | 0.041 | 0.333 | 8.035E-105 | TEC_prg2 | MYDGF |
| 9.3709E-59 | -0.5896728 | 0.68 | 0.775 | 2.0007E-54 | TEC_prg2 | PKM |
| 5.878E-115 | -0.5902886 | 0.091 | 0.417 | 1.255E-110 | TEC_prg2 | DYNLT1 |
| 2.909E-105 | -0.5912474 | 0.148 | 0.463 | 6.212E-101 | TEC_prg2 | BSG |
| 3.801E-115 | -0.5914717 | 0.125 | 0.456 | 8.114E-111 | TEC_prg2 | PTTG1IP |
| 1.128E-117 | -0.5924097 | 0.31 | 0.62 | 2.408E-113 | TEC_prg2 | YWHAB |
| 1.22E-115 | -0.5933911 | 0.34 | 0.642 | 2.604E-111 | TEC_prg2 | ATP5MG |
| 8.5222E-75 | -0.5951296 | 0.739 | 0.847 | 1.8195E-70 | TEC_prg2 | PPIA |
| 2.461E-108 | -0.5955358 | 0.087 | 0.394 | 5.253E-104 | TEC_prg2 | PSME1 |
| 4.5183E-41 | -0.5971577 | 0.192 | 0.379 | 9.6466E-37 | TEC_prg2 | KLF6 |
| 7.9736E-80 | -0.5976239 | 0.404 | 0.641 | 1.7024E-75 | TEC_prg2 | CD81 |
| 5.0033E-69 | -0.5978361 | 0.884 | 0.917 | 1.0682E-64 | TEC_prg2 | HSP90AB1 |
| 3.7167E-67 | -0.598948 | 0.067 | 0.283 | 7.9352E-63 | TEC_prg2 | LAMB3 |
| 6.374E-124 | -0.599607 | 0.264 | 0.596 | 1.361E-119 | TEC_prg2 | RHOA |
| 7.55E-116 | -0.5998557 | 0.131 | 0.463 | 1.612E-111 | TEC_prg2 | ATP6V0B |
| 1.42E-105 | -0.605258 | 0.081 | 0.383 | 3.033E-101 | TEC_prg2 | EIF5A |
| 5.062E-105 | -0.605319 | 0.854 | 0.928 | 1.081E-100 | TEC_prg2 | RPL34 |
| 3.815E-130 | -0.6057232 | 0.131 | 0.486 | 8.145E-126 | TEC_prg2 | VDAC2 |
| 6.2065E-96 | -0.60639 | 0.876 | 0.928 | 1.3251E-91 | TEC_prg2 | RPL30 |
| 1.358E-116 | -0.6076386 | 0.087 | 0.411 | 2.898E-112 | TEC_prg2 | VAMP8 |
| 4.269E-91 | -0.6083713 | 0.841 | 0.905 | 9.1144E-87 | TEC_prg2 | RPL23A |
| 3.7795E-42 | -0.6098284 | 0.142 | 0.314 | 8.0691E-38 | TEC_prg2 | ITGB8 |
| 1.5231E-74 | -0.613462 | 0.21 | 0.467 | 3.2518E-70 | TEC_prg2 | PPP1R14B |
| 1.7425E-57 | -0.6153709 | 0.952 | 0.955 | 3.7203E-53 | TEC_prg2 | S100A6 |
| 1.802E-115 | -0.6168197 | 0.878 | 0.938 | 3.848E-111 | TEC_prg2 | RPL18 |
| 1.447E-130 | -0.6178249 | 0.104 | 0.455 | 3.09E-126 | TEC_prg2 | PARK7 |
| 1.489E-126 | -0.6223292 | 0.052 | 0.381 | 3.179E-122 | TEC_prg2 | CAP1 |
| 5.851E-125 | -0.6248354 | 0.55 | 0.78 | 1.249E-120 | TEC_prg2 | RPL4 |
| 3.431E-104 | -0.6268331 | 0.067 | 0.359 | 7.326E-100 | TEC_prg2 | TSPO |
| 1.697E-127 | -0.6290568 | 0.048 | 0.374 | 3.623E-123 | TEC_prg2 | SSR3 |
| 1.098E-99 | -0.6311763 | 0.915 | 0.951 | 2.3448E-95 | TEC_prg2 | RPL13A |
| 9.7989E-84 | -0.6312753 | 0.647 | 0.789 | 2.0921E-79 | TEC_prg2 | MYL12A |
| 5.852E-119 | -0.6317323 | 0.607 | 0.803 | 1.249E-114 | TEC_prg2 | COX4I1 |
| 9.267E-118 | -0.6331359 | 0.55 | 0.772 | 1.978E-113 | TEC_prg2 | DSTN |
| 3.503E-100 | -0.6340762 | 0.922 | 0.954 | 7.4789E-96 | TEC_prg2 | RPS8 |

| | | | | | | |
|---|---|---|---|---|---|---|
| 2.1011E-95 | -0.6342782 | 0.802 | 0.877 | 4.4857E-91 | TEC_prg2 | RPL10A |
| 5.0773E-93 | -0.6388714 | 0.864 | 0.918 | 1.084E-88 | TEC_prg2 | RPL29 |
| 6.548E-76 | -0.6404005 | 0.102 | 0.344 | 1.398E-71 | TEC_prg2 | UGCG |
| 4.833E-116 | -0.6407374 | 0.646 | 0.819 | 1.032E-111 | TEC_prg2 | RPL27 |
| 1.099E-129 | -0.6416558 | 0.068 | 0.413 | 2.347E-125 | TEC_prg2 | SEC61G |
| 5.3905E-84 | -0.6419655 | 0.68 | 0.812 | 1.1509E-79 | TEC_prg2 | HLA-C |
| 1.758E-129 | -0.6424492 | 0.22 | 0.562 | 3.754E-125 | TEC_prg2 | CDC42 |
| 2.2E-107 | -0.6434304 | 0.44 | 0.695 | 4.698E-103 | TEC_prg2 | SPINT2 |
| 2.102E-106 | -0.6438414 | 0.963 | 0.975 | 4.488E-102 | TEC_prg2 | RPL13 |
| 1.769E-114 | -0.6441772 | 0.444 | 0.709 | 3.777E-110 | TEC_prg2 | COX6A1 |
| 2.239E-133 | -0.6447302 | 0.107 | 0.464 | 4.78E-129 | TEC_prg2 | TMEM59 |
| 7.9581E-90 | -0.6472746 | 0.124 | 0.405 | 1.6991E-85 | TEC_prg2 | LDHB |
| 6.4056E-68 | -0.6472973 | 0.751 | 0.836 | 1.3676E-63 | TEC_prg2 | HLA-B |
| 8.2016E-83 | -0.6484247 | 0.027 | 0.258 | 1.751E-78 | TEC_prg2 | TGM2 |
| 1.1345E-15 | -0.6495545 | 0.642 | 0.65 | 2.4222E-11 | TEC_prg2 | KRT7 |
| 5.579E-133 | -0.6511859 | 0.156 | 0.509 | 1.191E-128 | TEC_prg2 | PRDX6 |
| 1.072E-119 | -0.6518771 | 0.817 | 0.912 | 2.288E-115 | TEC_prg2 | RPS25 |
| 3.366E-138 | -0.6548339 | 0.077 | 0.435 | 7.185E-134 | TEC_prg2 | DAD1 |
| 2.844E-135 | -0.6566918 | 0.205 | 0.554 | 6.073E-131 | TEC_prg2 | ARPC2 |
| 6.205E-146 | -0.6594808 | 0.059 | 0.43 | 1.325E-141 | TEC_prg2 | POMP |
| 7.84E-125 | -0.6600606 | 0.391 | 0.68 | 1.674E-120 | TEC_prg2 | UQCRH |
| 4.2992E-81 | -0.6626197 | 0.839 | 0.888 | 9.1788E-77 | TEC_prg2 | RPL39 |
| 3.0711E-97 | -0.6632892 | 0.416 | 0.672 | 6.5567E-93 | TEC_prg2 | PEBP1 |
| 1.1795E-96 | -0.6636506 | 0.801 | 0.878 | 2.5181E-92 | TEC_prg2 | RPS5 |
| 1.874E-123 | -0.6649313 | 0.551 | 0.773 | 4.001E-119 | TEC_prg2 | RTN4 |
| 4.717E-115 | -0.664951 | 0.287 | 0.595 | 1.007E-110 | TEC_prg2 | GUK1 |
| 8.935E-107 | -0.6655575 | 0.852 | 0.916 | 1.908E-102 | TEC_prg2 | RPL26 |
| 5.892E-139 | -0.6665616 | 0.543 | 0.776 | 1.258E-134 | TEC_prg2 | BTF3 |
| 5.5514E-74 | -0.6691227 | 0.075 | 0.312 | 1.1852E-69 | TEC_prg2 | PLAUR |
| 9.09E-73 | -0.6693715 | 0.112 | 0.351 | 1.9407E-68 | TEC_prg2 | CYBA |
| 1.0623E-46 | -0.6713536 | 0.056 | 0.217 | 2.268E-42 | TEC_prg2 | ANXA1 |
| 1.461E-131 | -0.67343 | 0.793 | 0.902 | 3.118E-127 | TEC_prg2 | NACA |
| 7.494E-110 | -0.6764825 | 0.289 | 0.578 | 1.6E-105 | TEC_prg2 | BRI3 |
| 2.162E-141 | -0.6765117 | 0.4 | 0.695 | 4.616E-137 | TEC_prg2 | RAC1 |
| 5.469E-118 | -0.6781421 | 0.888 | 0.948 | 1.168E-113 | TEC_prg2 | RPS24 |
| 7.7258E-76 | -0.681167 | 0.102 | 0.341 | 1.6495E-71 | TEC_prg2 | MAL |
| 7.26E-101 | -0.681366 | 0.563 | 0.749 | 1.55E-96 | TEC_prg2 | ENO1 |
| 1.132E-111 | -0.6830463 | 0.107 | 0.419 | 2.417E-107 | TEC_prg2 | OCIAD2 |
| 2.0652E-83 | -0.6832814 | 0.062 | 0.312 | 4.4091E-79 | TEC_prg2 | AKR1B1 |
| 2.303E-126 | -0.6839617 | 0.241 | 0.569 | 4.918E-122 | TEC_prg2 | SEC61B |
| 6.0604E-53 | -0.6911167 | 0.932 | 0.926 | 1.2939E-48 | TEC_prg2 | ACTG1 |
| 2.6553E-69 | -0.6933624 | 0.972 | 0.972 | 5.669E-65 | TEC_prg2 | RPLP1 |
| 1.079E-116 | -0.6943266 | 0.767 | 0.879 | 2.304E-112 | TEC_prg2 | RPL23 |
| 5.457E-117 | -0.6968489 | 0.136 | 0.461 | 1.165E-112 | TEC_prg2 | SSR4 |
| 1.775E-132 | -0.6977267 | 0.091 | 0.436 | 3.79E-128 | TEC_prg2 | MGST3 |
| 4.408E-139 | -0.7002411 | 0.048 | 0.398 | 9.411E-135 | TEC_prg2 | TALDO1 |
| 1.827E-101 | -0.7024942 | 0.561 | 0.752 | 3.9015E-97 | TEC_prg2 | NDUFA4 |
| 4.425E-140 | -0.7028094 | 0.825 | 0.92 | 9.447E-136 | TEC_prg2 | RPL24 |
| 1.423E-143 | -0.703145 | 0.312 | 0.64 | 3.038E-139 | TEC_prg2 | PSMA7 |
| 5.169E-134 | -0.7047019 | 0.148 | 0.499 | 1.104E-129 | TEC_prg2 | COX8A |

| | | | | | | |
|---|---|---|---|---|---|---|
| 1.528E-104 | -0.7068236 | 0.898 | 0.938 | 3.263E-100 | TEC_prg2 | RPL32 |
| 1.186E-115 | -0.7112673 | 0.388 | 0.66 | 2.533E-111 | TEC_prg2 | ANXA5 |
| 3.854E-126 | -0.7131165 | 0.558 | 0.769 | 8.228E-122 | TEC_prg2 | HINT1 |
| 2.418E-109 | -0.7175157 | 0.889 | 0.945 | 5.162E-105 | TEC_prg2 | RPS28 |
| 5.5805E-09 | -0.7178455 | 0.751 | 0.724 | 0.00011914 | TEC_prg2 | KRT8 |
| 2.409E-116 | -0.7219212 | 0.903 | 0.937 | 5.144E-112 | TEC_prg2 | RPL3 |
| 1.947E-113 | -0.7250399 | 0.857 | 0.932 | 4.157E-109 | TEC_prg2 | RPLP2 |
| 3.1513E-53 | -0.7254339 | 0.848 | 0.876 | 6.728E-49 | TEC_prg2 | GSTP1 |
| 5.6996E-88 | -0.7279814 | 0.344 | 0.585 | 1.2169E-83 | TEC_prg2 | HMGA1 |
| 1.401E-143 | -0.728779 | 0.283 | 0.617 | 2.991E-139 | TEC_prg2 | ATP6V0E1 |
| 2.0608E-29 | -0.7302219 | 0.04 | 0.147 | 4.3999E-25 | TEC_prg2 | SERPINA1 |
| 8.338E-161 | -0.7321194 | 0.108 | 0.495 | 1.78E-156 | TEC_prg2 | LAPTM4A |
| 4.879E-138 | -0.7333432 | 0.111 | 0.467 | 1.042E-133 | TEC_prg2 | PDIA3 |
| 5.101E-108 | -0.7365703 | 0.976 | 0.972 | 1.089E-103 | TEC_prg2 | TPT1 |
| 2.049E-152 | -0.7369816 | 0.227 | 0.592 | 4.374E-148 | TEC_prg2 | CANX |
| 3.957E-102 | -0.7376328 | 0.951 | 0.969 | 8.4485E-98 | TEC_prg2 | RPL41 |
| 5.709E-127 | -0.7385815 | 0.096 | 0.434 | 1.219E-122 | TEC_prg2 | RHOC |
| 3.13E-135 | -0.7476424 | 0.901 | 0.952 | 6.682E-131 | TEC_prg2 | RPL11 |
| 2.6447E-87 | -0.7500199 | 0.587 | 0.749 | 5.6465E-83 | TEC_prg2 | PFN1 |
| 1.078E-121 | -0.7505333 | 0.767 | 0.877 | 2.301E-117 | TEC_prg2 | UBA52 |
| 2.4421E-45 | -0.7507358 | 0.712 | 0.78 | 5.2139E-41 | TEC_prg2 | HSP90B1 |
| 2.577E-113 | -0.7508569 | 0.07 | 0.378 | 5.502E-109 | TEC_prg2 | S100A16 |
| 3.486E-133 | -0.75426 | 0.461 | 0.718 | 7.443E-129 | TEC_prg2 | CYSTM1 |
| 2.001E-129 | -0.7552514 | 0.879 | 0.937 | 4.271E-125 | TEC_prg2 | RPL7A |
| 5.658E-110 | -0.7560154 | 0.943 | 0.963 | 1.208E-105 | TEC_prg2 | RPS18 |
| 4.567E-146 | -0.7577718 | 0.283 | 0.619 | 9.751E-142 | TEC_prg2 | ARPC3 |
| 4.802E-125 | -0.7584065 | 0.461 | 0.706 | 1.025E-120 | TEC_prg2 | EPCAM |
| 6.5244E-31 | -0.7585609 | 0.017 | 0.117 | 1.393E-26 | TEC_prg2 | FXYD4 |
| 2.357E-110 | -0.7610922 | 0.144 | 0.453 | 5.032E-106 | TEC_prg2 | LGALS3 |
| 1.539E-126 | -0.768158 | 0.291 | 0.599 | 3.285E-122 | TEC_prg2 | PPIB |
| 5.5348E-88 | -0.775041 | 0.029 | 0.272 | 1.1817E-83 | TEC_prg2 | PDZK1IP1 |
| 1.58E-131 | -0.7762694 | 0.989 | 0.987 | 3.373E-127 | TEC_prg2 | EEF1A1 |
| 5.639E-147 | -0.7792125 | 0.884 | 0.948 | 1.204E-142 | TEC_prg2 | RPL19 |
| 2.62E-142 | -0.7811133 | 0.898 | 0.949 | 5.593E-138 | TEC_prg2 | RPL8 |
| 4.874E-135 | -0.7853887 | 0.83 | 0.914 | 1.041E-130 | TEC_prg2 | RPS3 |
| 3.171E-145 | -0.7891038 | 0.607 | 0.809 | 6.77E-141 | TEC_prg2 | CHCHD2 |
| 2.3286E-49 | -0.7894674 | 0.14 | 0.328 | 4.9716E-45 | TEC_prg2 | PLAU |
| 1.429E-80 | -0.7935395 | 0.77 | 0.845 | 3.0509E-76 | TEC_prg2 | PRDX1 |
| 8.887E-117 | -0.794976 | 0.292 | 0.582 | 1.897E-112 | TEC_prg2 | CD9 |
| 5.2181E-13 | -0.8048039 | 0.686 | 0.683 | 1.1141E-08 | TEC_prg2 | HSPA5 |
| 5.023E-130 | -0.8056046 | 0.264 | 0.579 | 1.072E-125 | TEC_prg2 | CD59 |
| 3.466E-120 | -0.8101752 | 0.883 | 0.934 | 7.401E-116 | TEC_prg2 | RPL18A |
| 7.455E-171 | -0.8121504 | 0.786 | 0.912 | 1.592E-166 | TEC_prg2 | RPL5 |
| 2.2717E-94 | -0.8155308 | 0.172 | 0.451 | 4.8501E-90 | TEC_prg2 | TPM4 |
| 5.821E-163 | -0.820653 | 0.754 | 0.887 | 1.243E-158 | TEC_prg2 | RACK1 |
| 1.981E-62 | -0.8231848 | 0.132 | 0.35 | 4.2294E-58 | TEC_prg2 | IL32 |
| 1.352E-152 | -0.8241089 | 0.141 | 0.509 | 2.887E-148 | TEC_prg2 | PDIA6 |
| 4.19E-112 | -0.8258003 | 0.539 | 0.737 | 8.946E-108 | TEC_prg2 | TXN |
| 6.922E-141 | -0.8270931 | 0.862 | 0.936 | 1.478E-136 | TEC_prg2 | RPL21 |
| 2.1312E-65 | -0.8295916 | 0.12 | 0.344 | 4.5501E-61 | TEC_prg2 | ITGA2 |

| | | | | | | |
|---|---|---|---|---|---|---|
| 1.691E-148 | -0.8355524 | 0.895 | 0.943 | 3.611E-144 | TEC_prg2 | RPS23 |
| 1.265E-69 | -0.8478289 | 0.857 | 0.897 | 2.7007E-65 | TEC_prg2 | GAPDH |
| 1.752E-161 | -0.858424 | 0.876 | 0.936 | 3.741E-157 | TEC_prg2 | RPS14 |
| 1.602E-100 | -0.861486 | 0.539 | 0.724 | 3.4204E-96 | TEC_prg2 | ITGB1 |
| 2.712E-123 | -0.8722106 | 0.54 | 0.746 | 5.79E-119 | TEC_prg2 | CTSD |
| 2.439E-133 | -0.89995 | 0.295 | 0.598 | 5.208E-129 | TEC_prg2 | MAL2 |
| 6.682E-153 | -0.9006575 | 0.821 | 0.905 | 1.427E-148 | TEC_prg2 | RPS13 |
| 1.7456E-34 | -0.9033156 | 0.115 | 0.258 | 3.7268E-30 | TEC_prg2 | TFPI2 |
| 2.947E-173 | -0.911902 | 0.774 | 0.886 | 6.291E-169 | TEC_prg2 | RPL7 |
| 4.223E-112 | -0.9385602 | 0.204 | 0.499 | 9.017E-108 | TEC_prg2 | IFITM3 |
| 6.628E-122 | -0.9482162 | 0.948 | 0.952 | 1.415E-117 | TEC_prg2 | RPS12 |
| 8.204E-161 | -0.9496247 | 0.722 | 0.869 | 1.751E-156 | TEC_prg2 | MYL12B |
| 1.292E-191 | -0.9583803 | 0.841 | 0.929 | 2.758E-187 | TEC_prg2 | RPL9 |
| 4.0667E-89 | -0.9735131 | 0.021 | 0.262 | 8.6823E-85 | TEC_prg2 | CST6 |
| 1.568E-155 | -0.9822249 | 0.724 | 0.858 | 3.348E-151 | TEC_prg2 | RPSA |
| 1.221E-202 | -0.9854985 | 0.797 | 0.918 | 2.607E-198 | TEC_prg2 | RPS7 |
| 4.2786E-85 | -0.9881703 | 0.094 | 0.359 | 9.1348E-81 | TEC_prg2 | TIMP1 |
| 1.3908E-66 | -0.9927642 | 0.25 | 0.466 | 2.9693E-62 | TEC_prg2 | UPP1 |
| 5.3668E-37 | -0.9944288 | 0.521 | 0.6 | 1.1458E-32 | TEC_prg2 | KRT19 |
| 3.591E-109 | -1.0003983 | 0.153 | 0.45 | 7.667E-105 | TEC_prg2 | CCND1 |
| 5.78E-131 | -1.0127086 | 0.789 | 0.888 | 1.234E-126 | TEC_prg2 | ITM2B |
| 4.1608E-18 | -1.0196402 | 0.651 | 0.644 | 8.8833E-14 | TEC_prg2 | WFDC2 |
| 4.917E-198 | -1.0596824 | 0.853 | 0.93 | 1.05E-193 | TEC_prg2 | RPS3A |
| 3.922E-220 | -1.0654241 | 0.869 | 0.947 | 8.374E-216 | TEC_prg2 | RPL15 |
| 1.6723E-70 | -1.0685878 | 0.079 | 0.302 | 3.5704E-66 | TEC_prg2 | LGALS1 |
| 2.993E-183 | -1.0775264 | 0.77 | 0.877 | 6.389E-179 | TEC_prg2 | RPS4X |
| 1.696E-158 | -1.0784703 | 0.898 | 0.926 | 3.621E-154 | TEC_prg2 | RPL10 |
| 1.38E-199 | -1.0807022 | 0.684 | 0.859 | 2.946E-195 | TEC_prg2 | CD63 |
| 5.548E-214 | -1.0842706 | 0.858 | 0.943 | 1.185E-209 | TEC_prg2 | RPS27A |
| 6.1131E-84 | -1.1098657 | 0.103 | 0.358 | 1.3052E-79 | TEC_prg2 | LAMC2 |
| 2.454E-205 | -1.1116378 | 0.862 | 0.94 | 5.239E-201 | TEC_prg2 | RPS6 |
| 5.8853E-87 | -1.1117445 | 0.164 | 0.42 | 1.2565E-82 | TEC_prg2 | TXNRD1 |
| 1.0687E-32 | -1.1228352 | 0.717 | 0.71 | 2.2818E-28 | TEC_prg2 | MT2A |
| 1.8798E-27 | -1.1407876 | 0.092 | 0.21 | 4.0133E-23 | TEC_prg2 | MT1G |
| 6.1013E-90 | -1.176852 | 0.54 | 0.688 | 1.3026E-85 | TEC_prg2 | SH3BGRL3 |
| 2.506E-123 | -1.1780896 | 0.223 | 0.517 | 5.35E-119 | TEC_prg2 | TSPAN1 |
| 7.806E-66 | -1.1848457 | 0.876 | 0.867 | 1.6666E-61 | TEC_prg2 | TMSB4X |
| 1.17E-128 | -1.1907734 | 0.795 | 0.879 | 2.499E-124 | TEC_prg2 | S100A10 |
| 4.728E-124 | -1.2280756 | 0.885 | 0.926 | 1.01E-119 | TEC_prg2 | ACTB |
| 7.0649E-94 | -1.2314562 | 0.245 | 0.498 | 1.5084E-89 | TEC_prg2 | C12orf75 |
| 1.5144E-30 | -1.2393416 | 0.645 | 0.67 | 3.2333E-26 | TEC_prg2 | CRYAB |
| 1.2525E-29 | -1.2716277 | 0.113 | 0.239 | 2.6741E-25 | TEC_prg2 | DEFB1 |
| 1.45E-138 | -1.2882404 | 0.874 | 0.925 | 3.096E-134 | TEC_prg2 | S100A11 |
| 5.9659E-76 | -1.3378348 | 0.052 | 0.281 | 1.2737E-71 | TEC_prg2 | LCN2 |
| 1.07E-170 | -1.4693373 | 0.714 | 0.845 | 2.285E-166 | TEC_prg2 | ANXA2 |
| 1.1464E-21 | -1.4949017 | 0.429 | 0.499 | 2.4476E-17 | TEC_prg2 | S100A2 |
| 2.035E-193 | -1.6083285 | 0.963 | 0.98 | 4.345E-189 | TEC_prg2 | FTH1 |
| 1.5096E-86 | -1.7730025 | 0.175 | 0.418 | 3.2229E-82 | TEC_prg2 | SLPI |
| 1.018E-134 | -1.7973423 | 0.949 | 0.963 | 2.174E-130 | TEC_prg2 | FTL |
| 2.2448E-26 | -1.890735 | 0.593 | 0.6 | 4.7926E-22 | TEC_prg2 | SPP1 |

| | | | | | | |
|---|---|---|---|---|---|---|
| 1.1907E-39 | -2.4784926 | 0.636 | 0.637 | 2.5422E-35 | TEC_prg2 | MMP7 |
| 2.547E-190 | 2.04971726 | 0.485 | 0.035 | 5.438E-186 | TEC_prg3 | ANKRD1 |
| 1.4731E-68 | 1.89255125 | 0.885 | 0.371 | 3.1451E-64 | TEC_prg3 | HIST1H4C |
| 6.435E-72 | 1.72790863 | 0.733 | 0.226 | 1.3739E-67 | TEC_prg3 | SYNE1 |
| 3.3669E-52 | 1.52860278 | 0.6 | 0.196 | 7.1884E-48 | TEC_prg3 | MDM4 |
| 3.812E-146 | 1.37859736 | 0.552 | 0.06 | 8.139E-142 | TEC_prg3 | HIST1H2AG |
| 1.1584E-41 | 1.27952525 | 0.933 | 0.534 | 2.4732E-37 | TEC_prg3 | SOX4 |
| 4.243E-169 | 1.25154219 | 0.515 | 0.046 | 9.058E-165 | TEC_prg3 | DIAPH3 |
| 3.2085E-38 | 1.19389454 | 0.685 | 0.272 | 6.8501E-34 | TEC_prg3 | HES1 |
| 1.2134E-31 | 1.18741076 | 0.618 | 0.281 | 2.5906E-27 | TEC_prg3 | HIST1H1C |
| 3.4811E-49 | 1.17459698 | 0.448 | 0.106 | 7.4322E-45 | TEC_prg3 | EGOT |
| 2.138E-146 | 1.16749729 | 0.503 | 0.05 | 4.565E-142 | TEC_prg3 | NUSAP1 |
| 1.817E-108 | 1.13666851 | 0.321 | 0.027 | 3.879E-104 | TEC_prg3 | C21orf58 |
| 1.1016E-22 | 1.11372818 | 0.509 | 0.237 | 2.3519E-18 | TEC_prg3 | HES4 |
| 6.7075E-36 | 1.10802281 | 0.964 | 0.629 | 1.432E-31 | TEC_prg3 | JUN |
| 1.8866E-30 | 1.09765177 | 0.739 | 0.424 | 4.0279E-26 | TEC_prg3 | SYNE2 |
| 5.252E-39 | 1.07426711 | 0.509 | 0.16 | 1.1213E-34 | TEC_prg3 | HIST1H1E |
| 3.731E-159 | 1.05008927 | 0.376 | 0.025 | 7.965E-155 | TEC_prg3 | HIST1H1D |
| 2.0931E-11 | 1.03525388 | 0.473 | 0.293 | 4.4687E-07 | TEC_prg3 | DST |
| 3.5704E-21 | 1.0252195 | 0.503 | 0.236 | 7.6228E-17 | TEC_prg3 | MDM2 |
| 7.7012E-17 | 0.981704 | 0.97 | 0.87 | 1.6442E-12 | TEC_prg3 | SAT1 |
| 6.52E-38 | 0.94910801 | 0.303 | 0.065 | 1.392E-33 | TEC_prg3 | PCNX2 |
| 2.9048E-29 | 0.94471888 | 0.612 | 0.276 | 6.2016E-25 | TEC_prg3 | BTG2 |
| 7.8283E-12 | 0.94137781 | 0.661 | 0.508 | 1.6713E-07 | TEC_prg3 | CDKN1A |
| 2.5206E-27 | 0.93041101 | 0.527 | 0.23 | 5.3814E-23 | TEC_prg3 | PKN2 |
| 9.9171E-44 | 0.91612669 | 0.436 | 0.106 | 2.1173E-39 | TEC_prg3 | AC023157.3 |
| 2.9829E-26 | 0.91208025 | 0.903 | 0.636 | 6.3685E-22 | TEC_prg3 | JUND |
| 1.5441E-25 | 0.9094089 | 0.479 | 0.196 | 3.2967E-21 | TEC_prg3 | FAT1 |
| 1.2864E-21 | 0.89068719 | 0.806 | 0.589 | 2.7464E-17 | TEC_prg3 | RBM39 |
| 1.406E-139 | 0.88115055 | 0.345 | 0.023 | 3.002E-135 | TEC_prg3 | HIST1H2BH |
| 1.2636E-12 | 0.84984331 | 0.37 | 0.184 | 2.6978E-08 | TEC_prg3 | BIRC3 |
| 2.8911E-25 | 0.83731466 | 0.582 | 0.285 | 6.1726E-21 | TEC_prg3 | HP1BP3 |
| 2.1046E-47 | 0.83299243 | 0.327 | 0.059 | 4.4932E-43 | TEC_prg3 | HIST1H2BG |
| 4.0777E-15 | 0.83089344 | 0.758 | 0.566 | 8.7058E-11 | TEC_prg3 | RPS27L |
| 1.5076E-42 | 0.82501098 | 0.982 | 0.593 | 3.2187E-38 | TEC_prg3 | PLCG2 |
| 3.9233E-18 | 0.81704822 | 0.636 | 0.371 | 8.3762E-14 | TEC_prg3 | SLC38A2 |
| 1.1763E-44 | 0.80853583 | 0.43 | 0.102 | 2.5114E-40 | TEC_prg3 | AC012306.2 |
| 7.3333E-28 | 0.7980383 | 0.576 | 0.233 | 1.5657E-23 | TEC_prg3 | INTS6 |
| 1.2023E-16 | 0.79759024 | 0.539 | 0.283 | 2.567E-12 | TEC_prg3 | CEBPD |
| 7.0574E-14 | 0.79395598 | 0.848 | 0.705 | 1.5068E-09 | TEC_prg3 | DDX5 |
| 6.0533E-24 | 0.79389697 | 0.6 | 0.272 | 1.2924E-19 | TEC_prg3 | PPP1R10 |
| 1.0767E-27 | 0.78352931 | 0.412 | 0.131 | 2.2987E-23 | TEC_prg3 | AC025164.1 |
| 2.6247E-29 | 0.74890342 | 0.448 | 0.146 | 5.6038E-25 | TEC_prg3 | AC007952.4 |
| 2.6828E-30 | 0.74819895 | 0.315 | 0.082 | 5.7279E-26 | TEC_prg3 | FAM111A |
| 2.3224E-30 | 0.7427777 | 0.303 | 0.073 | 4.9584E-26 | TEC_prg3 | FILIP1L |
| 8.4436E-20 | 0.74275436 | 0.43 | 0.188 | 1.8027E-15 | TEC_prg3 | TUT4 |
| 1.8716E-12 | 0.73724044 | 0.673 | 0.516 | 3.9958E-08 | TEC_prg3 | SON |
| 1.3291E-21 | 0.73519404 | 0.509 | 0.232 | 2.8377E-17 | TEC_prg3 | CHD9 |
| 2.8655E-15 | 0.72076733 | 0.642 | 0.43 | 6.1178E-11 | TEC_prg3 | ZBTB20 |
| 7.841E-10 | 0.72055544 | 0.436 | 0.265 | 1.674E-05 | TEC_prg3 | TUBA1A |

| | | | | | | |
|---|---|---|---|---|---|---|
| 1.9177E-20 | 0.70700606 | 0.412 | 0.168 | 4.0942E-16 | TEC_prg3 | AHI1 |
| 1.0179E-48 | 0.70382908 | 0.279 | 0.042 | 2.1733E-44 | TEC_prg3 | ARL14 |
| 3.2301E-12 | 0.69448825 | 0.467 | 0.28 | 6.8964E-08 | TEC_prg3 | PRPF4B |
| 9.1373E-14 | 0.69240423 | 0.788 | 0.678 | 1.9508E-09 | TEC_prg3 | YBX3 |
| 2.4286E-19 | 0.68399886 | 0.6 | 0.293 | 5.1852E-15 | TEC_prg3 | HEXIM1 |
| 6.603E-280 | 0.67719319 | 0.291 | 0.007 | 1.41E-275 | TEC_prg3 | HIST1H3G |
| 2.0006E-42 | 0.66412726 | 0.37 | 0.081 | 4.2712E-38 | TEC_prg3 | DDB2 |
| 6.504E-31 | 0.65549992 | 0.412 | 0.122 | 1.3886E-26 | TEC_prg3 | AL135925.1 |
| 3.3662E-08 | 0.65348989 | 0.448 | 0.31 | 0.00071868 | TEC_prg3 | WSB1 |
| 5.8466E-64 | 0.6534438 | 0.345 | 0.051 | 1.2483E-59 | TEC_prg3 | KIF20B |
| 5.3502E-35 | 0.65269825 | 0.273 | 0.055 | 1.1423E-30 | TEC_prg3 | HELLS |
| 1.1525E-15 | 0.65224959 | 0.358 | 0.15 | 2.4605E-11 | TEC_prg3 | ASCC3 |
| 4.2923E-78 | 0.64925977 | 0.248 | 0.022 | 9.164E-74 | TEC_prg3 | APOLD1 |
| 1.4031E-09 | 0.64561121 | 0.448 | 0.289 | 2.9957E-05 | TEC_prg3 | KMT2A |
| 1.0958E-13 | 0.63210465 | 0.806 | 0.608 | 2.3396E-09 | TEC_prg3 | BTG1 |
| 5.8875E-48 | 0.63102489 | 0.285 | 0.044 | 1.257E-43 | TEC_prg3 | CENPF |
| 3.3436E-12 | 0.63013931 | 0.424 | 0.233 | 7.1386E-08 | TEC_prg3 | SCIN |
| 4.4181E-12 | 0.62691943 | 0.606 | 0.412 | 9.4326E-08 | TEC_prg3 | SRSF7 |
| 2.61E-198 | 0.62499935 | 0.315 | 0.013 | 5.573E-194 | TEC_prg3 | HIST1H3B |
| 1.6681E-05 | 0.61911225 | 0.424 | 0.335 | 0.35613964 | TEC_prg3 | LUC7L3 |
| 1.4297E-28 | 0.61424372 | 0.261 | 0.058 | 3.0525E-24 | TEC_prg3 | HIST1H2BN |
| 2.3769E-11 | 0.61086112 | 0.576 | 0.399 | 5.0746E-07 | TEC_prg3 | FUS |
| 5.6068E-19 | 0.60396265 | 0.442 | 0.187 | 1.1971E-14 | TEC_prg3 | TLE4 |
| 5.5314E-22 | 0.60343444 | 0.552 | 0.23 | 1.181E-17 | TEC_prg3 | CSKMT |
| 3.782E-20 | 0.59982795 | 0.352 | 0.125 | 8.0747E-16 | TEC_prg3 | UTRN |
| 8.4779E-15 | 0.59860087 | 0.958 | 0.794 | 1.81E-10 | TEC_prg3 | HNRNPA2B1 |
| 3.3336E-11 | 0.5906041 | 0.721 | 0.559 | 7.1171E-07 | TEC_prg3 | SRRM2 |
| 4.1037E-15 | 0.58861651 | 0.388 | 0.176 | 8.7615E-11 | TEC_prg3 | MYO9A |
| 1.2612E-91 | 0.58480424 | 0.315 | 0.03 | 2.6926E-87 | TEC_prg3 | AC004264.1 |
| 2.7909E-10 | 0.58285405 | 0.515 | 0.356 | 5.9586E-06 | TEC_prg3 | SPAG9 |
| 3.5511E-17 | 0.58162112 | 0.861 | 0.583 | 7.5815E-13 | TEC_prg3 | ELF3 |
| 1.4235E-90 | 0.57829806 | 0.158 | 0.007 | 3.0391E-86 | TEC_prg3 | DNAH17 |
| 2.2039E-09 | 0.57713567 | 0.6 | 0.431 | 4.7054E-05 | TEC_prg3 | DDX17 |
| 9.4379E-08 | 0.57373421 | 0.509 | 0.381 | 0.00201499 | TEC_prg3 | N4BP2L2 |
| 7.5558E-15 | 0.56478264 | 0.364 | 0.161 | 1.6132E-10 | TEC_prg3 | OFD1 |
| 2.544E-18 | 0.56426865 | 0.836 | 0.562 | 5.4314E-14 | TEC_prg3 | GDF15 |
| 2.255E-58 | 0.5601451 | 0.255 | 0.03 | 4.8144E-54 | TEC_prg3 | MKI67 |
| 1.6664E-11 | 0.55817857 | 0.455 | 0.264 | 3.5577E-07 | TEC_prg3 | BDP1 |
| 4.2242E-10 | 0.55700968 | 0.491 | 0.318 | 9.0187E-06 | TEC_prg3 | CCNL1 |
| 1.416E-27 | 0.54909468 | 0.267 | 0.062 | 3.0231E-23 | TEC_prg3 | AC005837.1 |
| 1.8349E-17 | 0.54884332 | 0.436 | 0.181 | 3.9174E-13 | TEC_prg3 | PIK3R3 |
| 4.7615E-08 | 0.54630439 | 0.358 | 0.207 | 0.00101659 | TEC_prg3 | ARID5B |
| 2.9149E-22 | 0.54226556 | 0.279 | 0.079 | 6.2232E-18 | TEC_prg3 | MIS18BP1 |
| 4.4227E-19 | 0.54084194 | 0.285 | 0.089 | 9.4425E-15 | TEC_prg3 | Z93241.1 |
| 3.0715E-06 | 0.53873225 | 0.297 | 0.185 | 0.06557673 | TEC_prg3 | MSI2 |
| 6.3655E-80 | 0.53826284 | 0.23 | 0.018 | 1.359E-75 | TEC_prg3 | INPP5D |
| 5.4952E-16 | 0.53568891 | 0.327 | 0.127 | 1.1732E-11 | TEC_prg3 | XPC |
| 2.4208E-16 | 0.53552884 | 0.255 | 0.085 | 5.1684E-12 | TEC_prg3 | SMC4 |
| 7.7756E-08 | 0.53453456 | 0.394 | 0.259 | 0.00166008 | TEC_prg3 | SRSF10 |
| 6.256E-185 | 0.53051063 | 0.248 | 0.008 | 1.336E-180 | TEC_prg3 | ESCO2 |

| | | | | | | |
|---|---|---|---|---|---|---|
| 8.2052E-26 | 0.53032829 | 0.358 | 0.104 | 1.7518E-21 | TEC_prg3 | AL355075.4 |
| 4.2043E-27 | 0.52758189 | 0.255 | 0.057 | 8.9763E-23 | TEC_prg3 | IFI30 |
| 1.3464E-07 | 0.52233157 | 0.352 | 0.217 | 0.00287463 | TEC_prg3 | FAM133B |
| 2.0856E-16 | 0.52063586 | 0.315 | 0.118 | 4.4527E-12 | TEC_prg3 | FBXW7 |
| 5.0323E-71 | 0.51554406 | 0.188 | 0.014 | 1.0744E-66 | TEC_prg3 | TCERG1L-AS1 |
| 2.6626E-87 | 0.51391823 | 0.127 | 0.005 | 5.6847E-83 | TEC_prg3 | MUC19 |
| 5.1291E-23 | 0.50811592 | 0.291 | 0.082 | 1.0951E-18 | TEC_prg3 | ANKRD36C |
| 3.2199E-09 | 0.50608823 | 0.273 | 0.132 | 6.8745E-05 | TEC_prg3 | CREB5 |
| 1.3209E-14 | 0.49991427 | 1 | 0.94 | 2.8201E-10 | TEC_prg3 | MALAT1 |
| 3.0142E-15 | 0.4974873 | 0.982 | 0.857 | 6.4353E-11 | TEC_prg3 | H3F3B |
| 3.7977E-05 | 0.49035719 | 0.424 | 0.321 | 0.81079948 | TEC_prg3 | RSRC2 |
| 9.5523E-08 | 0.48977779 | 0.376 | 0.236 | 0.00203942 | TEC_prg3 | U2SURP |
| 8.8079E-23 | 0.48975865 | 0.267 | 0.069 | 1.8805E-18 | TEC_prg3 | HIST1H1A |
| 4.7229E-05 | 0.48854264 | 0.655 | 0.55 | 1 | TEC_prg3 | EIF5 |
| 2.8876E-07 | 0.48751154 | 0.194 | 0.085 | 0.00616506 | TEC_prg3 | CCDC80 |
| 1.8036E-06 | 0.48381408 | 0.376 | 0.253 | 0.03850716 | TEC_prg3 | DDX46 |
| 1.9331E-20 | 0.48275493 | 0.212 | 0.052 | 4.1271E-16 | TEC_prg3 | MNS1 |
| 5.5531E-08 | 0.48200062 | 0.939 | 0.828 | 0.00118558 | TEC_prg3 | SQSTM1 |
| 2.7052E-05 | 0.48052165 | 0.467 | 0.367 | 0.57755931 | TEC_prg3 | MACF1 |
| 9.1925E-09 | 0.47755724 | 0.382 | 0.217 | 0.00019626 | TEC_prg3 | NKTR |
| 1.426E-63 | 0.47299719 | 0.218 | 0.021 | 3.0445E-59 | TEC_prg3 | ASPM |
| 1.8522E-05 | 0.47048104 | 0.327 | 0.224 | 0.39544794 | TEC_prg3 | CLIP1 |
| 0.00225096 | 0.47033444 | 0.376 | 0.306 | 1 | TEC_prg3 | YME1L1 |
| 1.8128E-05 | 0.46911479 | 0.412 | 0.313 | 0.38703954 | TEC_prg3 | AFDN |
| 1.1561E-10 | 0.4683431 | 0.782 | 0.597 | 2.4683E-06 | TEC_prg3 | CLU |
| 0.00024053 | 0.46818407 | 0.479 | 0.397 | 1 | TEC_prg3 | ANKRD11 |
| 0.00018101 | 0.46344482 | 0.321 | 0.23 | 1 | TEC_prg3 | UGP2 |
| 3.0263E-07 | 0.46313332 | 0.339 | 0.201 | 0.00646108 | TEC_prg3 | CLK1 |
| 1.0078E-07 | 0.46211216 | 0.515 | 0.375 | 0.00215158 | TEC_prg3 | PNISR |
| 1.3302E-06 | 0.45816706 | 0.358 | 0.225 | 0.02840034 | TEC_prg3 | RNMT |
| 2.8689E-78 | 0.45628418 | 0.248 | 0.022 | 6.1252E-74 | TEC_prg3 | AL360012.1 |
| 3.2615E-08 | 0.4545691 | 0.297 | 0.156 | 0.00069632 | TEC_prg3 | HOOK2 |
| 4.849E-07 | 0.45116658 | 0.43 | 0.299 | 0.01035265 | TEC_prg3 | GPBP1 |
| 1.4828E-13 | 0.4510247 | 0.279 | 0.11 | 3.1657E-09 | TEC_prg3 | REV3L |
| 7.8406E-05 | 0.45089046 | 0.406 | 0.302 | 1 | TEC_prg3 | YTHDC1 |
| 7.3597E-28 | 0.44408343 | 0.23 | 0.048 | 1.5713E-23 | TEC_prg3 | ATAD2 |
| 3.368E-06 | 0.4430282 | 0.388 | 0.272 | 0.07190658 | TEC_prg3 | TAF7 |
| 2.6992E-16 | 0.44104599 | 0.333 | 0.125 | 5.7627E-12 | TEC_prg3 | PKHD1 |
| 0.00280794 | 0.44102039 | 0.455 | 0.413 | 1 | TEC_prg3 | GOLGB1 |
| 4.1309E-22 | 0.44025936 | 0.297 | 0.086 | 8.8196E-18 | TEC_prg3 | AC091271.1 |
| 1.2971E-11 | 0.43825928 | 0.303 | 0.136 | 2.7694E-07 | TEC_prg3 | ZFYVE16 |
| 1.7366E-40 | 0.4356009 | 0.194 | 0.025 | 3.7076E-36 | TEC_prg3 | ATAD5 |
| 1.7881E-05 | 0.43551802 | 0.424 | 0.302 | 0.38175694 | TEC_prg3 | ARHGAP29 |
| 5.7773E-08 | 0.43261285 | 0.558 | 0.389 | 0.00123346 | TEC_prg3 | NFKBIA |
| 1.9926E-05 | 0.43243536 | 0.23 | 0.132 | 0.42542169 | TEC_prg3 | MAST4 |
| 7.8113E-25 | 0.42866276 | 0.279 | 0.072 | 1.6677E-20 | TEC_prg3 | NSUN6 |
| 3.4796E-07 | 0.42866056 | 0.382 | 0.231 | 0.00742891 | TEC_prg3 | CDKN2AIP |
| 7.656E-29 | 0.42722691 | 0.224 | 0.043 | 1.6346E-24 | TEC_prg3 | AL031777.3 |
| 7.7546E-08 | 0.42712748 | 0.473 | 0.299 | 0.0016556 | TEC_prg3 | PLK2 |
| 1.3592E-10 | 0.42679507 | 0.509 | 0.28 | 2.9019E-06 | TEC_prg3 | MAFB |

| | | | | | | |
|---|---|---|---|---|---|---|
| 1.7894E-06 | 0.42659322 | 0.333 | 0.204 | 0.03820386 | TEC_prg3 | THUMPD3-AS1 |
| 0.00077503 | 0.42640854 | 0.418 | 0.362 | 1 | TEC_prg3 | HMGN2 |
| 2.6996E-10 | 0.42588791 | 0.291 | 0.133 | 5.7636E-06 | TEC_prg3 | RSBN1L |
| 7.8433E-09 | 0.42542115 | 0.315 | 0.168 | 0.00016745 | TEC_prg3 | CBX5 |
| 4.1601E-17 | 0.42495694 | 0.261 | 0.084 | 8.8817E-13 | TEC_prg3 | ERCC6L2 |
| 1.9055E-63 | 0.42262912 | 0.224 | 0.022 | 4.0683E-59 | TEC_prg3 | AC011447.3 |
| 2.488E-37 | 0.42239612 | 0.236 | 0.039 | 5.3119E-33 | TEC_prg3 | UBL7-AS1 |
| 1.2796E-07 | 0.42185936 | 0.352 | 0.205 | 0.00273189 | TEC_prg3 | VPS13C |
| 4.0055E-23 | 0.41425125 | 0.242 | 0.06 | 8.5518E-19 | TEC_prg3 | NRG1 |
| 2.1305E-05 | 0.4139577 | 0.248 | 0.15 | 0.45486457 | TEC_prg3 | DNM1L |
| 3.8703E-08 | 0.41298991 | 0.533 | 0.347 | 0.00082631 | TEC_prg3 | TXNIP |
| 1.6818E-11 | 0.41277211 | 0.303 | 0.135 | 3.5907E-07 | TEC_prg3 | ATF7IP |
| 9.1993E-11 | 0.41079635 | 0.261 | 0.112 | 1.9641E-06 | TEC_prg3 | MAML2 |
| 1.7377E-10 | 0.41057275 | 0.358 | 0.173 | 3.7099E-06 | TEC_prg3 | FOXC1 |
| 2.8071E-13 | 0.40911913 | 0.291 | 0.116 | 5.9931E-09 | TEC_prg3 | CENPC |
| 0.00548445 | 0.40852426 | 0.4 | 0.358 | 1 | TEC_prg3 | GCC2 |
| 2.9391E-09 | 0.40654573 | 0.442 | 0.25 | 6.275E-05 | TEC_prg3 | ID2 |
| 1.2116E-46 | 0.40632364 | 0.206 | 0.025 | 2.5869E-42 | TEC_prg3 | AC239868.2 |
| 6.8811E-05 | 0.40369037 | 0.37 | 0.271 | 1 | TEC_prg3 | SMG1 |
| 4.3456E-30 | 0.40212316 | 0.206 | 0.036 | 9.2778E-26 | TEC_prg3 | KLLN |
| 9.3464E-52 | 0.39960154 | 0.242 | 0.03 | 1.9955E-47 | TEC_prg3 | TPX2 |
| 2.0997E-07 | 0.39804169 | 0.309 | 0.17 | 0.00448278 | TEC_prg3 | THSD7A |
| 5.688E-11 | 0.39593545 | 0.491 | 0.268 | 1.2144E-06 | TEC_prg3 | TCIM |
| 5.6696E-08 | 0.39554743 | 0.297 | 0.155 | 0.00121046 | TEC_prg3 | PLEKHH2 |
| 1.7753E-08 | 0.39371966 | 0.752 | 0.562 | 0.00037903 | TEC_prg3 | IER2 |
| 1.8085E-05 | 0.39271597 | 0.224 | 0.123 | 0.38611146 | TEC_prg3 | TMPO |
| 6.5252E-05 | 0.39090353 | 0.315 | 0.217 | 1 | TEC_prg3 | DHX36 |
| 1.8397E-06 | 0.39052055 | 0.333 | 0.199 | 0.0392775 | TEC_prg3 | CALD1 |
| 8.1371E-09 | 0.38515328 | 0.17 | 0.064 | 0.00017373 | TEC_prg3 | PTPRM |
| 0.00040267 | 0.38491625 | 0.424 | 0.348 | 1 | TEC_prg3 | DDX24 |
| 8.3301E-12 | 0.38393769 | 0.212 | 0.074 | 1.7785E-07 | TEC_prg3 | HIST1H4H |
| 1.6001E-09 | 0.38261053 | 0.43 | 0.24 | 3.4162E-05 | TEC_prg3 | ID4 |
| 2.2693E-08 | 0.3813422 | 0.261 | 0.126 | 0.0004845 | TEC_prg3 | CEP290 |
| 7.7621E-47 | 0.38016656 | 0.194 | 0.022 | 1.6572E-42 | TEC_prg3 | TOP2A |
| 1.9554E-07 | 0.37750228 | 0.255 | 0.131 | 0.00417476 | TEC_prg3 | SMCHD1 |
| 6.3842E-11 | 0.37681207 | 0.152 | 0.046 | 1.363E-06 | TEC_prg3 | CEP85L |
| 0.00142832 | 0.37611159 | 0.297 | 0.218 | 1 | TEC_prg3 | CCDC186 |
| 4.9747E-07 | 0.37519222 | 0.297 | 0.166 | 0.01062103 | TEC_prg3 | TMX4 |
| 1.1133E-44 | 0.37430598 | 0.17 | 0.018 | 2.3768E-40 | TEC_prg3 | CLSPN |
| 9.2252E-13 | 0.37408391 | 0.236 | 0.084 | 1.9696E-08 | TEC_prg3 | AL450998.2 |
| 3.5833E-19 | 0.37397727 | 0.182 | 0.041 | 7.6503E-15 | TEC_prg3 | HIST1H4E |
| 7.027E-17 | 0.3724223 | 0.23 | 0.068 | 1.5003E-12 | TEC_prg3 | SRGAP3 |
| 2.8534E-12 | 0.37190755 | 0.17 | 0.051 | 6.092E-08 | TEC_prg3 | PLA2G4C |
| 3.7222E-34 | 0.36912944 | 0.224 | 0.037 | 7.9468E-30 | TEC_prg3 | HIST1H2AH |
| 4.5271E-16 | 0.36906548 | 0.248 | 0.079 | 9.6654E-12 | TEC_prg3 | INTU |
| 3.4613E-19 | 0.36858713 | 0.261 | 0.076 | 7.3898E-15 | TEC_prg3 | SRGAP2 |
| 2.3491E-10 | 0.36603406 | 0.267 | 0.118 | 5.0153E-06 | TEC_prg3 | SUGP2 |
| 5.381E-122 | 0.3649474 | 0.17 | 0.006 | 1.149E-117 | TEC_prg3 | LINC02471 |
| 2.7388E-06 | 0.36490745 | 0.218 | 0.113 | 0.05847327 | TEC_prg3 | RB1 |
| 4.4706E-09 | 0.36157062 | 0.248 | 0.114 | 9.5448E-05 | TEC_prg3 | NEDD9 |

| | | | | | | |
|---|---|---|---|---|---|---|
| 8.5597E-36 | 0.35861134 | 0.212 | 0.033 | 1.8275E-31 | TEC_prg3 | TNFSF15 |
| 2.6032E-22 | 0.35828206 | 0.242 | 0.061 | 5.5579E-18 | TEC_prg3 | SMC2 |
| 5.1717E-09 | 0.35734784 | 0.206 | 0.086 | 0.00011042 | TEC_prg3 | HMGXB4 |
| 3.689E-05 | 0.35716054 | 0.394 | 0.284 | 0.7876094 | TEC_prg3 | NUFIP2 |
| 3.7668E-20 | 0.35630693 | 0.23 | 0.059 | 8.0421E-16 | TEC_prg3 | AL022069.1 |
| 0.00092542 | 0.35533773 | 0.436 | 0.361 | 1 | TEC_prg3 | AKAP9 |
| 3.6317E-21 | 0.35270373 | 0.17 | 0.034 | 7.7537E-17 | TEC_prg3 | CCDC18 |
| 0.0004574 | 0.35252683 | 0.248 | 0.168 | 1 | TEC_prg3 | PFDN1 |
| 0.001222 | 0.35202325 | 0.358 | 0.285 | 1 | TEC_prg3 | RBM25 |
| 9.9224E-56 | 0.35149225 | 0.182 | 0.016 | 2.1184E-51 | TEC_prg3 | KNL1 |
| 7.7878E-06 | 0.35051501 | 0.267 | 0.153 | 0.16627055 | TEC_prg3 | CHD3 |
| 6.0169E-13 | 0.35024167 | 0.248 | 0.091 | 1.2846E-08 | TEC_prg3 | LINC00909 |
| 9.6259E-06 | 0.34934549 | 0.4 | 0.277 | 0.20551352 | TEC_prg3 | COBLL1 |
| 1.8132E-07 | 0.34931701 | 0.8 | 0.621 | 0.00387108 | TEC_prg3 | PPP1R15A |
| 2.6856E-15 | 0.34760655 | 0.23 | 0.073 | 5.7338E-11 | TEC_prg3 | SETBP1 |
| 0.00013718 | 0.34746779 | 0.339 | 0.239 | 1 | TEC_prg3 | LMO4 |
| 4.7837E-10 | 0.34668801 | 0.285 | 0.124 | 1.0213E-05 | TEC_prg3 | HIST2H2BF |
| 1.4344E-18 | 0.34604413 | 0.236 | 0.066 | 3.0625E-14 | TEC_prg3 | IKBIP |
| 0.00023865 | 0.34538134 | 0.321 | 0.227 | 1 | TEC_prg3 | TRA2B |
| 1.0675E-10 | 0.34483142 | 0.164 | 0.053 | 2.2791E-06 | TEC_prg3 | AL139274.2 |
| 0.00035822 | 0.34332596 | 0.321 | 0.227 | 1 | TEC_prg3 | DDX18 |
| 1.0229E-09 | 0.34292605 | 0.218 | 0.087 | 2.1839E-05 | TEC_prg3 | DNAJC3-DT |
| 3.6208E-06 | 0.34205122 | 0.327 | 0.204 | 0.07730332 | TEC_prg3 | EPC1 |
| 2.2475E-08 | 0.34192362 | 0.224 | 0.101 | 0.00047985 | TEC_prg3 | NMRK1 |
| 7.9516E-23 | 0.34110291 | 0.194 | 0.04 | 1.6977E-18 | TEC_prg3 | HIST1H3C |
| 0.00051124 | 0.3405238 | 0.358 | 0.263 | 1 | TEC_prg3 | AMD1 |
| 6.0123E-08 | 0.33795268 | 0.2 | 0.083 | 0.00128363 | TEC_prg3 | HIST1H3A |
| 1.9036E-06 | 0.33765473 | 0.23 | 0.12 | 0.04064219 | TEC_prg3 | UBAP2 |
| 3.211E-05 | 0.33355316 | 0.539 | 0.42 | 0.68555577 | TEC_prg3 | SFPQ |
| 6.3016E-38 | 0.33334409 | 0.182 | 0.024 | 1.3454E-33 | TEC_prg3 | BRCA2 |
| 8.4148E-06 | 0.33197787 | 0.224 | 0.12 | 0.17965695 | TEC_prg3 | CWF19L2 |
| 2.0481E-08 | 0.33130468 | 0.267 | 0.128 | 0.00043727 | TEC_prg3 | WDR60 |
| 1.1769E-07 | 0.33047634 | 0.291 | 0.155 | 0.00251264 | TEC_prg3 | CEP350 |
| 2.5661E-05 | 0.32955893 | 0.279 | 0.171 | 0.54786735 | TEC_prg3 | MYEF2 |
| 1.6451E-53 | 0.32625539 | 0.2 | 0.02 | 3.5123E-49 | TEC_prg3 | ANLN |
| 8.7445E-33 | 0.32599652 | 0.164 | 0.022 | 1.867E-28 | TEC_prg3 | SLFN13 |
| 9.9356E-18 | 0.32596636 | 0.176 | 0.042 | 2.1212E-13 | TEC_prg3 | NCAPD3 |
| 4.8896E-11 | 0.3250348 | 0.285 | 0.122 | 1.0439E-06 | TEC_prg3 | EPM2AIP1 |
| 0.00022817 | 0.32488465 | 0.273 | 0.181 | 1 | TEC_prg3 | ZNF638 |
| 5.231E-75 | 0.32453071 | 0.158 | 0.009 | 1.1168E-70 | TEC_prg3 | AURKB |
| 0.00066012 | 0.32280619 | 0.303 | 0.217 | 1 | TEC_prg3 | ASH1L |
| 5.2088E-07 | 0.32158381 | 0.267 | 0.139 | 0.01112088 | TEC_prg3 | RNF169 |
| 0.00391768 | 0.31796915 | 0.303 | 0.236 | 1 | TEC_prg3 | AKAP13 |
| 2.6247E-08 | 0.31708172 | 0.236 | 0.108 | 0.00056037 | TEC_prg3 | ZNHIT6 |
| 5.2499E-05 | 0.31610465 | 0.224 | 0.131 | 1 | TEC_prg3 | ZNF844 |
| 2.5043E-12 | 0.31555444 | 0.176 | 0.054 | 5.3467E-08 | TEC_prg3 | CCDC34 |
| 0.00309973 | 0.3150111 | 0.248 | 0.185 | 1 | TEC_prg3 | DNAJC1 |
| 4.5166E-15 | 0.31404713 | 0.182 | 0.049 | 9.643E-11 | TEC_prg3 | AC110769.2 |
| 0.00025659 | 0.31176702 | 0.242 | 0.153 | 1 | TEC_prg3 | ASXL1 |
| 0.00011912 | 0.31137275 | 0.776 | 0.699 | 1 | TEC_prg3 | HMGB1 |

| | | | | | | |
|---|---|---|---|---|---|---|
| 2.7464E-11 | 0.31097444 | 0.182 | 0.06 | 5.8636E-07 | TEC_prg3 | PCNT |
| 1.7558E-33 | 0.31032929 | 0.145 | 0.017 | 3.7486E-29 | TEC_prg3 | RAD51AP2 |
| 7.9274E-13 | 0.30991833 | 0.194 | 0.061 | 1.6925E-08 | TEC_prg3 | EZH2 |
| 0.00014301 | 0.30737946 | 0.2 | 0.114 | 1 | TEC_prg3 | HKDC1 |
| 3.8309E-19 | 0.30735389 | 0.121 | 0.02 | 8.1789E-15 | TEC_prg3 | HIST1H2BF |
| 1.012E-10 | 0.30715646 | 0.17 | 0.055 | 2.1607E-06 | TEC_prg3 | CRNDE |
| 0.0001838 | 0.30626788 | 0.261 | 0.168 | 1 | TEC_prg3 | USP15 |
| 0.00405714 | 0.30423893 | 0.606 | 0.559 | 1 | TEC_prg3 | KTN1 |
| 3.8809E-05 | 0.30389596 | 0.182 | 0.095 | 0.82856605 | TEC_prg3 | NET1 |
| 1.5432E-19 | 0.30306709 | 0.17 | 0.036 | 3.2948E-15 | TEC_prg3 | HIST1H2BJ |
| 2.2965E-05 | 0.30253268 | 0.291 | 0.182 | 0.49029766 | TEC_prg3 | FNIP1 |
| 1.5606E-05 | 0.30221289 | 0.224 | 0.123 | 0.33318456 | TEC_prg3 | RAB30-AS1 |
| 8.3822E-07 | 0.30207637 | 0.242 | 0.119 | 0.01789597 | TEC_prg3 | LINC00472 |
| 6.9268E-09 | 0.30198488 | 0.182 | 0.07 | 0.00014789 | TEC_prg3 | FAM13A |
| 6.8125E-08 | 0.30160469 | 0.424 | 0.251 | 0.00145448 | TEC_prg3 | IER5 |
| 0.00498423 | 0.30132953 | 0.267 | 0.196 | 1 | TEC_prg3 | UPF2 |
| 0.00152075 | 0.29988492 | 0.206 | 0.133 | 1 | TEC_prg3 | STK17A |
| 0.00365393 | 0.29977021 | 0.345 | 0.273 | 1 | TEC_prg3 | VMP1 |
| 7.4238E-06 | 0.29942573 | 0.236 | 0.126 | 0.15849798 | TEC_prg3 | HOMER1 |
| 7.6295E-06 | 0.29875126 | 0.279 | 0.162 | 0.1628896 | TEC_prg3 | ESF1 |
| 9.6201E-17 | 0.29819976 | 0.145 | 0.031 | 2.0539E-12 | TEC_prg3 | AC009812.1 |
| 3.2794E-05 | 0.29580264 | 0.261 | 0.153 | 0.70014695 | TEC_prg3 | IVNS1ABP |
| 3.6426E-07 | 0.29532442 | 0.206 | 0.094 | 0.0077769 | TEC_prg3 | ALDH1A3 |
| 0.00183221 | 0.29514508 | 0.321 | 0.241 | 1 | TEC_prg3 | TOR1AIP2 |
| 6.2074E-05 | 0.29489222 | 0.261 | 0.16 | 1 | TEC_prg3 | RBM6 |
| 3.4015E-05 | 0.29460016 | 0.212 | 0.118 | 0.72621758 | TEC_prg3 | XRN1 |
| 1.6858E-14 | 0.29439618 | 0.158 | 0.04 | 3.5992E-10 | TEC_prg3 | CEP152 |
| 8.1234E-07 | 0.29282427 | 0.224 | 0.11 | 0.0173435 | TEC_prg3 | ARL4A |
| 0.00713073 | 0.29274604 | 0.218 | 0.158 | 1 | TEC_prg3 | CDC5L |
| 2.1609E-10 | 0.29072973 | 0.164 | 0.053 | 4.6134E-06 | TEC_prg3 | RASL11B |
| 9E-09 | 0.28909605 | 0.279 | 0.132 | 0.00019215 | TEC_prg3 | PTPN13 |
| 2.4894E-05 | 0.28843392 | 0.236 | 0.134 | 0.53148498 | TEC_prg3 | OXR1 |
| 0.0005933 | 0.28584871 | 0.448 | 0.341 | 1 | TEC_prg3 | ZFP36L2 |
| 0.00518674 | 0.28511739 | 0.285 | 0.217 | 1 | TEC_prg3 | USP34 |
| 1.3459E-07 | 0.28440416 | 0.321 | 0.167 | 0.00287347 | TEC_prg3 | EGLN3 |
| 0.00600473 | 0.28342566 | 0.224 | 0.163 | 1 | TEC_prg3 | CYLD |
| 4.4551E-14 | 0.28334041 | 0.194 | 0.058 | 9.5116E-10 | TEC_prg3 | ECT2 |
| 0.00019814 | 0.282963 | 0.188 | 0.107 | 1 | TEC_prg3 | AL121944.1 |
| 7.84E-141 | 0.28259542 | 0.152 | 0.004 | 1.674E-136 | TEC_prg3 | HIST1H3F |
| 3.4839E-24 | 0.28194391 | 0.164 | 0.029 | 7.4381E-20 | TEC_prg3 | RFC3 |
| 3.9294E-33 | 0.28147378 | 0.158 | 0.02 | 8.3892E-29 | TEC_prg3 | RRM2 |
| 0.00012986 | 0.28090278 | 0.176 | 0.094 | 1 | TEC_prg3 | TSPYL4 |
| 2.9748E-08 | 0.28082034 | 0.236 | 0.106 | 0.00063513 | TEC_prg3 | ZNF701 |
| 7.1004E-06 | 0.27893094 | 0.248 | 0.137 | 0.15159332 | TEC_prg3 | PDE4DIP |
| 0.0050564 | 0.27863796 | 0.261 | 0.19 | 1 | TEC_prg3 | CFDP1 |
| 1.0008E-15 | 0.27826718 | 0.121 | 0.024 | 2.1368E-11 | TEC_prg3 | AL359513.1 |
| 2.3151E-05 | 0.27823995 | 0.412 | 0.274 | 0.49428074 | TEC_prg3 | CITED2 |
| 9.0662E-27 | 0.27822192 | 0.139 | 0.02 | 1.9356E-22 | TEC_prg3 | AL109976.1 |
| 5.0754E-18 | 0.27808242 | 0.194 | 0.047 | 1.0836E-13 | TEC_prg3 | TSC22D1-AS1 |
| 2.8782E-13 | 0.27796009 | 0.927 | 0.675 | 6.1449E-09 | TEC_prg3 | MTRNR2L12 |

| | | | | | | |
|---|---|---|---|---|---|---|
| 3.9022E-05 | 0.27680959 | 0.212 | 0.115 | 0.8331209 | TEC_prg3 | DNMBP |
| 0.00031659 | 0.27679445 | 0.315 | 0.224 | 1 | TEC_prg3 | NFAT5 |
| 0.00029866 | 0.27608344 | 0.182 | 0.099 | 1 | TEC_prg3 | CXCL3 |
| 2.196E-44 | 0.27572033 | 0.127 | 0.01 | 4.6885E-40 | TEC_prg3 | ACAD11 |
| 4.265E-194 | 0.27571583 | 0.115 | 0.001 | 9.106E-190 | TEC_prg3 | LINC01707 |
| 2.1973E-08 | 0.27478297 | 0.188 | 0.075 | 0.00046912 | TEC_prg3 | FRMD4A |
| 0.00014412 | 0.27443834 | 0.23 | 0.139 | 1 | TEC_prg3 | CCSER2 |
| 7.5372E-08 | 0.27404176 | 0.182 | 0.074 | 0.00160919 | TEC_prg3 | DDIT4L |
| 9.0738E-23 | 0.27302697 | 0.188 | 0.038 | 1.9373E-18 | TEC_prg3 | CKAP2 |
| 5.2589E-06 | 0.27295134 | 0.358 | 0.209 | 0.11227853 | TEC_prg3 | ADAMTS1 |
| 1.222E-05 | 0.27278612 | 0.206 | 0.107 | 0.26090141 | TEC_prg3 | MGA |
| 1.3144E-08 | 0.27263446 | 0.164 | 0.061 | 0.00028063 | TEC_prg3 | CEP295 |
| 4.3812E-40 | 0.2724633 | 0.115 | 0.009 | 9.3538E-36 | TEC_prg3 | CRISPLD1 |
| 1.7455E-08 | 0.27242269 | 0.2 | 0.081 | 0.00037267 | TEC_prg3 | POLH |
| 0.00345214 | 0.27213884 | 0.212 | 0.145 | 1 | TEC_prg3 | LPIN2 |
| 0.00010923 | 0.27176195 | 0.2 | 0.113 | 1 | TEC_prg3 | TRMT10C |
| 4.2773E-14 | 0.27090751 | 0.145 | 0.035 | 9.1321E-10 | TEC_prg3 | NRP2 |
| 0.00359749 | 0.27036563 | 0.23 | 0.16 | 1 | TEC_prg3 | BBC3 |
| 0.00017659 | 0.26985153 | 0.224 | 0.136 | 1 | TEC_prg3 | TAOK3 |
| 3.1785E-05 | 0.26946455 | 0.224 | 0.123 | 0.67861397 | TEC_prg3 | KMT2E-AS1 |
| 1.1798E-14 | 0.26800116 | 0.139 | 0.032 | 2.5189E-10 | TEC_prg3 | AC245014.3 |
| 1.4285E-06 | 0.26797463 | 0.17 | 0.074 | 0.0304981 | TEC_prg3 | TSGA10 |
| 0.00164257 | 0.2668235 | 0.321 | 0.238 | 1 | TEC_prg3 | NSD3 |
| 0.00500503 | 0.26679625 | 0.236 | 0.17 | 1 | TEC_prg3 | LARP7 |
| 0.0004011 | 0.26656792 | 0.224 | 0.139 | 1 | TEC_prg3 | RALGAPA1 |
| 3.0169E-12 | 0.26602495 | 0.17 | 0.05 | 6.441E-08 | TEC_prg3 | LINC02256 |
| 2.321E-165 | 0.26508141 | 0.127 | 0.002 | 4.955E-161 | TEC_prg3 | AC097478.1 |
| 4.7328E-05 | 0.26440765 | 0.273 | 0.164 | 1 | TEC_prg3 | PAXBP1 |
| 3.0081E-23 | 0.26385578 | 0.133 | 0.02 | 6.4223E-19 | TEC_prg3 | ABCA12 |
| 0.00050633 | 0.26303344 | 0.218 | 0.134 | 1 | TEC_prg3 | ZNF431 |
| 0.00955952 | 0.26279071 | 0.23 | 0.166 | 1 | TEC_prg3 | STAT1 |
| 9.071E-09 | 0.26224072 | 0.218 | 0.091 | 0.00019367 | TEC_prg3 | TRIM22 |
| 1.101E-06 | 0.26093543 | 0.188 | 0.085 | 0.02350737 | TEC_prg3 | CCDC71L |
| 0.00596425 | 0.26085498 | 0.2 | 0.137 | 1 | TEC_prg3 | NABP1 |
| 0.00115204 | 0.2603882 | 0.358 | 0.259 | 1 | TEC_prg3 | EGR1 |
| 1.7512E-09 | 0.26028985 | 0.17 | 0.06 | 3.7387E-05 | TEC_prg3 | TEX9 |
| 3.6041E-06 | 0.25977981 | 0.248 | 0.132 | 0.07694857 | TEC_prg3 | RBM5 |
| 7.7661E-07 | 0.25824 | 0.145 | 0.057 | 0.0165806 | TEC_prg3 | ADRB2 |
| 3.2866E-20 | 0.25775952 | 0.164 | 0.033 | 7.0168E-16 | TEC_prg3 | SULT1C4 |
| 4.7777E-09 | 0.25686441 | 0.127 | 0.038 | 0.000102 | TEC_prg3 | AC011498.1 |
| 0.00911427 | 0.25582776 | 0.194 | 0.137 | 1 | TEC_prg3 | EML4 |
| 4.827E-06 | 0.25580787 | 0.224 | 0.115 | 0.10305622 | TEC_prg3 | ZNF830 |
| 0.00024798 | 0.25574122 | 0.345 | 0.235 | 1 | TEC_prg3 | SLC5A3 |
| 1.4669E-07 | 0.25524055 | 0.212 | 0.095 | 0.00313184 | TEC_prg3 | FBXO34 |
| 8.4354E-10 | 0.25521892 | 0.503 | 0.287 | 1.801E-05 | TEC_prg3 | FOS |
| 0.00024016 | 0.25503228 | 0.188 | 0.106 | 1 | TEC_prg3 | C6orf120 |
| 0.00977025 | 0.25492572 | 0.376 | 0.304 | 1 | TEC_prg3 | PMAIP1 |
| 7.7672E-77 | 0.25475769 | 0.121 | 0.005 | 1.6583E-72 | TEC_prg3 | SPC25 |
| 0.00286281 | 0.25339937 | 0.236 | 0.163 | 1 | TEC_prg3 | PPP1R15B |
| 3.3576E-13 | 0.25053754 | 0.261 | 0.091 | 7.1684E-09 | TEC_prg3 | AL137077.2 |

| | | | | | | |
|---|---|---|---|---|---|---|
| 4.7233E-07 | 0.25031204 | 0.182 | 0.079 | 0.01008421 | TEC_prg3 | CAPRIN2 |
| 3.3215E-06 | -0.2503436 | 0.03 | 0.167 | 0.07091505 | TEC_prg3 | SPTSSA |
| 2.6819E-07 | -0.2504695 | 0.012 | 0.16 | 0.00572584 | TEC_prg3 | REX1BD |
| 4.1582E-07 | -0.2506145 | 0.024 | 0.176 | 0.00887771 | TEC_prg3 | PLIN3 |
| 2.295E-06 | -0.2510939 | 0.073 | 0.235 | 0.04899905 | TEC_prg3 | ASAH1 |
| 4.5871E-06 | -0.2511401 | 0.073 | 0.226 | 0.09793493 | TEC_prg3 | GPRC5C |
| 4.1154E-07 | -0.251298 | 0.036 | 0.195 | 0.00878643 | TEC_prg3 | CMPK1 |
| 1.2687E-06 | -0.2513764 | 0.048 | 0.204 | 0.02708623 | TEC_prg3 | SMIM14 |
| 2.1897E-07 | -0.251458 | 0.048 | 0.22 | 0.00467502 | TEC_prg3 | SRI |
| 8.5117E-06 | -0.2518894 | 0.006 | 0.118 | 0.18172553 | TEC_prg3 | CRABP2 |
| 6.5587E-07 | -0.2519586 | 0.073 | 0.245 | 0.01400275 | TEC_prg3 | XRCC6 |
| 3.2491E-07 | -0.2521508 | 0.036 | 0.198 | 0.00693694 | TEC_prg3 | CMTM6 |
| 0.00128016 | -0.25227 | 0.073 | 0.169 | 1 | TEC_prg3 | FGF2 |
| 2.5818E-05 | -0.252317 | 0.073 | 0.206 | 0.55122422 | TEC_prg3 | AIF1L |
| 4.8626E-05 | -0.2526122 | 0.152 | 0.313 | 1 | TEC_prg3 | RAB13 |
| 7.3121E-06 | -0.2526162 | 0.291 | 0.49 | 0.15611249 | TEC_prg3 | UBE2D3 |
| 8.0706E-05 | -0.2526746 | 0.139 | 0.29 | 1 | TEC_prg3 | CDK2AP2 |
| 8.136E-07 | -0.2526988 | 0.024 | 0.169 | 0.01737034 | TEC_prg3 | UBE2J1 |
| 2.1769E-06 | -0.2541816 | 0.061 | 0.212 | 0.04647648 | TEC_prg3 | PPM1G |
| 0.00076513 | -0.2542254 | 0.745 | 0.777 | 1 | TEC_prg3 | OAZ1 |
| 3.9951E-07 | -0.2544825 | 0.024 | 0.176 | 0.00852947 | TEC_prg3 | PRNP |
| 1.1587E-05 | -0.2551745 | 0.23 | 0.426 | 0.24737835 | TEC_prg3 | AHNAK |
| 4.5768E-07 | -0.2553795 | 0.073 | 0.249 | 0.00977149 | TEC_prg3 | CAMTA1 |
| 0.0020495 | -0.2555674 | 0.345 | 0.488 | 1 | TEC_prg3 | ATP1A1 |
| 7.3207E-07 | -0.256242 | 0.085 | 0.265 | 0.01562969 | TEC_prg3 | RPS19BP1 |
| 1.239E-06 | -0.2564652 | 0.067 | 0.231 | 0.02645163 | TEC_prg3 | POLR2K |
| 4.8901E-08 | -0.2568103 | 0.048 | 0.239 | 0.00104404 | TEC_prg3 | PPP1CA |
| 7.8191E-08 | -0.2569138 | 0.03 | 0.203 | 0.00166937 | TEC_prg3 | NBDY |
| 1.1886E-06 | -0.2569637 | 0.079 | 0.25 | 0.02537698 | TEC_prg3 | PGRMC1 |
| 1.7482E-08 | -0.2575678 | 0.036 | 0.227 | 0.00037324 | TEC_prg3 | C1orf43 |
| 2.6713E-08 | -0.2579646 | 0.012 | 0.182 | 0.00057032 | TEC_prg3 | RAP1A |
| 3.7983E-08 | -0.2579901 | 0.03 | 0.206 | 0.00081094 | TEC_prg3 | ARPC1A |
| 3.1927E-07 | -0.2580457 | 0.006 | 0.147 | 0.00681637 | TEC_prg3 | CDCP1 |
| 0.00064647 | -0.2588927 | 0.558 | 0.665 | 1 | TEC_prg3 | SOD1 |
| 4.772E-08 | -0.2591629 | 0.024 | 0.197 | 0.00101881 | TEC_prg3 | EMC7 |
| 5.7046E-06 | -0.2593415 | 0.036 | 0.169 | 0.12179282 | TEC_prg3 | ELK3 |
| 0.00709478 | -0.2593719 | 0.279 | 0.392 | 1 | TEC_prg3 | RAB11FIP1 |
| 4.478E-07 | -0.2596869 | 0.085 | 0.27 | 0.00956058 | TEC_prg3 | TSTD1 |
| 3.9949E-07 | -0.2597464 | 0.055 | 0.224 | 0.0085291 | TEC_prg3 | SPCS3 |
| 2.4916E-08 | -0.2598791 | 0.121 | 0.35 | 0.00053195 | TEC_prg3 | PSMD8 |
| 7.8814E-07 | -0.2598933 | 0.073 | 0.245 | 0.01682673 | TEC_prg3 | HIGD2A |
| 8.1155E-06 | -0.260116 | 0.061 | 0.2 | 0.17326577 | TEC_prg3 | SLK |
| 1.7872E-07 | -0.2605254 | 0.055 | 0.233 | 0.00381558 | TEC_prg3 | SCARB2 |
| 4.4586E-05 | -0.2607198 | 0.133 | 0.293 | 0.95191852 | TEC_prg3 | MAFF |
| 5.6304E-08 | -0.2611857 | 0.042 | 0.222 | 0.0012021 | TEC_prg3 | COPZ1 |
| 3.4504E-08 | -0.2616151 | 0.006 | 0.168 | 0.00073666 | TEC_prg3 | NDUFB6 |
| 4.1342E-06 | -0.2616236 | 0.085 | 0.245 | 0.08826584 | TEC_prg3 | PFKP |
| 3.1178E-05 | -0.2622792 | 0.024 | 0.136 | 0.66565006 | TEC_prg3 | ITPKC |
| 5.4313E-09 | -0.2622945 | 0.03 | 0.225 | 0.00011596 | TEC_prg3 | ARPC4 |
| 2.5701E-07 | -0.2628939 | 0.073 | 0.257 | 0.00548715 | TEC_prg3 | ATP6V1D |

| | | | | | | |
|---|---|---|---|---|---|---|
| 2.2058E-05 | -0.2629324 | 0.097 | 0.24 | 0.4709293 | TEC_prg3 | CRIM1 |
| 7.5385E-07 | -0.2634129 | 0.03 | 0.181 | 0.01609467 | TEC_prg3 | DNPH1 |
| 6.8737E-07 | -0.264284 | 0.042 | 0.197 | 0.01467539 | TEC_prg3 | SLC25A39 |
| 0.00016633 | -0.2643816 | 0.018 | 0.111 | 1 | TEC_prg3 | ICAM1 |
| 6.3141E-07 | -0.2645987 | 0.018 | 0.162 | 0.01348056 | TEC_prg3 | HYOU1 |
| 5.2645E-06 | -0.2653629 | 0.048 | 0.194 | 0.11239605 | TEC_prg3 | KLHL21 |
| 4.0624E-05 | -0.2659233 | 0.642 | 0.753 | 0.86732709 | TEC_prg3 | PFDN5 |
| 3.1097E-07 | -0.2660678 | 0.03 | 0.184 | 0.00663917 | TEC_prg3 | ALKBH7 |
| 7.5476E-06 | -0.2661591 | 0.085 | 0.238 | 0.1611406 | TEC_prg3 | LPP |
| 6.9186E-08 | -0.2664338 | 0.018 | 0.179 | 0.00147713 | TEC_prg3 | POLR2F |
| 4.9757E-08 | -0.2664349 | 0.018 | 0.185 | 0.00106232 | TEC_prg3 | TBCB |
| 7.3066E-08 | -0.266481 | 0.024 | 0.191 | 0.00155996 | TEC_prg3 | ATRAID |
| 2.4247E-08 | -0.2669907 | 0.012 | 0.181 | 0.00051767 | TEC_prg3 | RPN1 |
| 0.0004843 | -0.2674855 | 0.085 | 0.193 | 1 | TEC_prg3 | BHLHE40 |
| 2.5857E-06 | -0.267549 | 0.224 | 0.437 | 0.05520535 | TEC_prg3 | ST13 |
| 3.9267E-05 | -0.2675966 | 0.121 | 0.266 | 0.83834417 | TEC_prg3 | RB1CC1 |
| 4.1632E-06 | -0.2677755 | 0.067 | 0.217 | 0.08888415 | TEC_prg3 | SAT2 |
| 1.0193E-07 | -0.2681181 | 0.067 | 0.26 | 0.00217623 | TEC_prg3 | HM13 |
| 7.5294E-07 | -0.2685686 | 0.079 | 0.251 | 0.01607525 | TEC_prg3 | YPEL5 |
| 2.2803E-07 | -0.2693387 | 0.018 | 0.171 | 0.00486853 | TEC_prg3 | PON2 |
| 4.1376E-05 | -0.269448 | 0.455 | 0.609 | 0.88336742 | TEC_prg3 | APLP2 |
| 3.8195E-07 | -0.2701674 | 0.018 | 0.166 | 0.00815455 | TEC_prg3 | CD82 |
| 2.6051E-07 | -0.2705051 | 0.061 | 0.241 | 0.00556196 | TEC_prg3 | RPS17 |
| 1.9685E-06 | -0.2712152 | 0.073 | 0.229 | 0.04202721 | TEC_prg3 | PSMD7 |
| 1.6823E-07 | -0.2716269 | 0.042 | 0.21 | 0.00359181 | TEC_prg3 | MOB1A |
| 5.5303E-07 | -0.2720234 | 0.048 | 0.211 | 0.01180721 | TEC_prg3 | ADAM9 |
| 8.9835E-08 | -0.2721521 | 0.103 | 0.312 | 0.00191798 | TEC_prg3 | HDLBP |
| 4.6205E-08 | -0.273048 | 0.024 | 0.197 | 0.00098649 | TEC_prg3 | NUCB1 |
| 2.9473E-07 | -0.2733229 | 0.042 | 0.205 | 0.00629245 | TEC_prg3 | GIPC1 |
| 0.00236269 | -0.2733993 | 0.164 | 0.28 | 1 | TEC_prg3 | SERPINB1 |
| 3.7084E-08 | -0.2738362 | 0.103 | 0.319 | 0.00079174 | TEC_prg3 | SNX3 |
| 1.1145E-07 | -0.2738556 | 0.036 | 0.205 | 0.00237944 | TEC_prg3 | FKBP8 |
| 5.7611E-08 | -0.2738777 | 0.055 | 0.243 | 0.00122999 | TEC_prg3 | HNRNPF |
| 2.5634E-07 | -0.2739141 | 0.097 | 0.291 | 0.00547287 | TEC_prg3 | PSME2 |
| 8.761E-08 | -0.2743848 | 0.085 | 0.282 | 0.00187046 | TEC_prg3 | RAB10 |
| 9.0593E-06 | -0.2745826 | 0.012 | 0.127 | 0.19341686 | TEC_prg3 | CMYA5 |
| 9.2788E-05 | -0.2753208 | 0.073 | 0.194 | 1 | TEC_prg3 | SESN2 |
| 2.3652E-05 | -0.275467 | 0.097 | 0.248 | 0.50496062 | TEC_prg3 | IRS2 |
| 6.4946E-07 | -0.275886 | 0.103 | 0.286 | 0.01386604 | TEC_prg3 | PDCD5 |
| 1.0307E-07 | -0.2758963 | 0.073 | 0.259 | 0.00220052 | TEC_prg3 | SMDT1 |
| 5.7134E-07 | -0.2759171 | 0.036 | 0.19 | 0.01219804 | TEC_prg3 | ARPC1B |
| 4.2188E-08 | -0.2759533 | 0.073 | 0.274 | 0.00090072 | TEC_prg3 | UBE2I |
| 1.8974E-07 | -0.2767018 | 0.067 | 0.245 | 0.00405086 | TEC_prg3 | RAB11A |
| 1.6484E-08 | -0.2767151 | 0.036 | 0.229 | 0.00035193 | TEC_prg3 | ATP6AP2 |
| 2.8965E-07 | -0.2775216 | 0.073 | 0.25 | 0.00618394 | TEC_prg3 | BANF1 |
| 4.3291E-08 | -0.2775472 | 0.036 | 0.211 | 0.00092425 | TEC_prg3 | RAB5C |
| 1.777E-06 | -0.2776989 | 0.048 | 0.2 | 0.03793814 | TEC_prg3 | BNIP3 |
| 3.8683E-07 | -0.2779191 | 0.061 | 0.231 | 0.00825872 | TEC_prg3 | TXNL1 |
| 2.2405E-06 | -0.2781075 | 0.012 | 0.14 | 0.04783417 | TEC_prg3 | TMEM176A |
| 7.4443E-07 | -0.2785973 | 0.073 | 0.241 | 0.01589358 | TEC_prg3 | EPRS |

| | | | | | | |
|---|---|---|---|---|---|---|
| 5.6097E-07 | -0.2794657 | 0.042 | 0.199 | 0.01197679 | TEC_prg3 | FKBP4 |
| 1.4426E-07 | -0.2797577 | 0.242 | 0.48 | 0.00307988 | TEC_prg3 | MORF4L1 |
| 1.3107E-07 | -0.2806403 | 0.036 | 0.2 | 0.00279839 | TEC_prg3 | SELENOT |
| 5.178E-07 | -0.2810525 | 0.042 | 0.197 | 0.011055 | TEC_prg3 | C1QBP |
| 1.1379E-07 | -0.2812724 | 0.024 | 0.187 | 0.00242941 | TEC_prg3 | ATP1B3 |
| 6.1812E-07 | -0.2815432 | 0.036 | 0.187 | 0.01319684 | TEC_prg3 | TMEM54 |
| 1.4876E-07 | -0.2816473 | 0.042 | 0.213 | 0.00317598 | TEC_prg3 | SERINC2 |
| 2.5292E-07 | -0.2818635 | 0.152 | 0.37 | 0.0053999 | TEC_prg3 | WDR83OS |
| 1.5745E-05 | -0.2820396 | 0.012 | 0.123 | 0.33616305 | TEC_prg3 | MT1F |
| 9.2829E-07 | -0.2822575 | 0.055 | 0.212 | 0.01981907 | TEC_prg3 | MZT2A |
| 4.9643E-06 | -0.2824332 | 0.103 | 0.268 | 0.10598761 | TEC_prg3 | SLC38A1 |
| 1.0887E-07 | -0.2825596 | 0.055 | 0.233 | 0.00232437 | TEC_prg3 | NDUFA12 |
| 1.0463E-07 | -0.2829261 | 0.036 | 0.207 | 0.0022339 | TEC_prg3 | AP2S1 |
| 4.5379E-05 | -0.2833045 | 0.085 | 0.219 | 0.96884423 | TEC_prg3 | PTGR1 |
| 7.0143E-08 | -0.2837723 | 0.048 | 0.231 | 0.00149756 | TEC_prg3 | TMEM208 |
| 1.3563E-05 | -0.2838491 | 0.115 | 0.27 | 0.28956029 | TEC_prg3 | BLVRB |
| 1.9324E-07 | -0.2839127 | 0.055 | 0.226 | 0.00412559 | TEC_prg3 | DAP |
| 1.3904E-07 | -0.2840416 | 0.055 | 0.234 | 0.00296851 | TEC_prg3 | C1orf122 |
| 1.0529E-07 | -0.2842051 | 0.048 | 0.225 | 0.00224804 | TEC_prg3 | DYNLT3 |
| 1.4254E-07 | -0.2846373 | 0.048 | 0.225 | 0.00304317 | TEC_prg3 | ARL6IP5 |
| 7.1805E-09 | -0.284867 | 0.085 | 0.313 | 0.0001533 | TEC_prg3 | GABARAPL2 |
| 2.1012E-06 | -0.28514 | 0.03 | 0.171 | 0.04486014 | TEC_prg3 | TYMP |
| 6.121E-05 | -0.2854085 | 0.109 | 0.251 | 1 | TEC_prg3 | MRPS6 |
| 1.6785E-08 | -0.2855094 | 0.024 | 0.203 | 0.00035836 | TEC_prg3 | NDUFB5 |
| 2.0386E-08 | -0.2861667 | 0.018 | 0.19 | 0.00043525 | TEC_prg3 | TMEM147 |
| 0.00030541 | -0.2862015 | 0.018 | 0.105 | 1 | TEC_prg3 | RSAD2 |
| 2.3019E-08 | -0.2866075 | 0.024 | 0.202 | 0.00049147 | TEC_prg3 | SH3YL1 |
| 1.7022E-06 | -0.2876059 | 0.006 | 0.133 | 0.03634191 | TEC_prg3 | AL353751.1 |
| 2.8695E-08 | -0.2876975 | 0.061 | 0.253 | 0.00061263 | TEC_prg3 | UBE2L3 |
| 2.2878E-08 | -0.2877922 | 0.03 | 0.213 | 0.00048846 | TEC_prg3 | PEA15 |
| 0.00716479 | -0.2886977 | 0.939 | 0.934 | 1 | TEC_prg3 | HSP90AA1 |
| 2.9275E-05 | -0.2891781 | 0.612 | 0.72 | 0.6250198 | TEC_prg3 | RPL36AL |
| 3.388E-08 | -0.2891985 | 0.024 | 0.199 | 0.00072333 | TEC_prg3 | CYC1 |
| 6.502E-07 | -0.2908672 | 0.048 | 0.207 | 0.01388178 | TEC_prg3 | OCLN |
| 1.3895E-07 | -0.2913806 | 0.055 | 0.229 | 0.00296649 | TEC_prg3 | GLO1 |
| 1.3842E-07 | -0.2914289 | 0.061 | 0.241 | 0.0029553 | TEC_prg3 | LSM4 |
| 2.4337E-05 | -0.2918019 | 0.048 | 0.173 | 0.5195914 | TEC_prg3 | DUSP5 |
| 9.1798E-08 | -0.2918178 | 0.067 | 0.254 | 0.0019599 | TEC_prg3 | UQCRFS1 |
| 1.1249E-06 | -0.2919129 | 0.061 | 0.222 | 0.02401558 | TEC_prg3 | TIMP2 |
| 7.3478E-07 | -0.2920472 | 0.061 | 0.222 | 0.01568754 | TEC_prg3 | LIMS1 |
| 2.1305E-07 | -0.2922286 | 0.127 | 0.335 | 0.00454861 | TEC_prg3 | ATP6V1F |
| 4.1975E-07 | -0.2922392 | 0.121 | 0.315 | 0.00896163 | TEC_prg3 | NOP53 |
| 7.5834E-09 | -0.2923863 | 0.055 | 0.26 | 0.00016191 | TEC_prg3 | NDUFA11 |
| 7.0939E-07 | -0.2925218 | 0.188 | 0.408 | 0.01514546 | TEC_prg3 | TBCA |
| 3.5506E-08 | -0.2926505 | 0.067 | 0.264 | 0.00075804 | TEC_prg3 | JTB |
| 1.3532E-09 | -0.2928854 | 0.03 | 0.239 | 2.889E-05 | TEC_prg3 | SMIM26 |
| 2.1866E-08 | -0.2930253 | 0.048 | 0.239 | 0.00046683 | TEC_prg3 | TMEM14B |
| 0.00011199 | -0.2932064 | 0.055 | 0.169 | 1 | TEC_prg3 | CA12 |
| 1.3212E-08 | -0.293666 | 0.085 | 0.296 | 0.00028207 | TEC_prg3 | RNF7 |
| 9.5501E-08 | -0.29375 | 0.079 | 0.27 | 0.00203894 | TEC_prg3 | NUDC |

| | | | | | | |
|---|---|---|---|---|---|---|
| 5.7868E-08 | -0.2937541 | 0.061 | 0.25 | 0.00123548 | TEC_prg3 | RPN2 |
| 1.0095E-07 | -0.2942597 | 0.091 | 0.287 | 0.0021552 | TEC_prg3 | ATP5PO |
| 0.00648168 | -0.2946997 | 0.667 | 0.701 | 1 | TEC_prg3 | MAP1LC3B |
| 6.0611E-07 | -0.2947907 | 0.133 | 0.326 | 0.01294035 | TEC_prg3 | FAM96B |
| 2.433E-06 | -0.2948738 | 0.497 | 0.647 | 0.05194395 | TEC_prg3 | EDF1 |
| 3.0047E-07 | -0.2950655 | 0.152 | 0.362 | 0.00641512 | TEC_prg3 | ATP5ME |
| 2.3434E-06 | -0.2951986 | 0.024 | 0.159 | 0.05003128 | TEC_prg3 | PMEPA1 |
| 8.3078E-08 | -0.2952657 | 0.055 | 0.237 | 0.00177372 | TEC_prg3 | MGAT4B |
| 3.208E-09 | -0.2955005 | 0.018 | 0.208 | 6.849E-05 | TEC_prg3 | METTL9 |
| 1.1796E-08 | -0.2968987 | 0.109 | 0.341 | 0.00025184 | TEC_prg3 | SEM1 |
| 1.5797E-07 | -0.2969336 | 0.073 | 0.255 | 0.00337277 | TEC_prg3 | C4orf3 |
| 9.3803E-07 | -0.2971996 | 0.127 | 0.315 | 0.02002685 | TEC_prg3 | DNAJB6 |
| 7.6755E-08 | -0.2972567 | 0.048 | 0.224 | 0.00163872 | TEC_prg3 | TMEM167A |
| 3.0089E-06 | -0.2977829 | 0.182 | 0.386 | 0.06423942 | TEC_prg3 | PTMS |
| 4.7867E-09 | -0.2981738 | 0 | 0.173 | 0.0001022 | TEC_prg3 | CENPX |
| 1.2745E-07 | -0.2981799 | 0.073 | 0.258 | 0.00272111 | TEC_prg3 | PPP2R1A |
| 4.8855E-09 | -0.2989301 | 0.055 | 0.262 | 0.0001043 | TEC_prg3 | RAB7A |
| 8.5136E-08 | -0.2989481 | 0.055 | 0.231 | 0.00181765 | TEC_prg3 | ATP5PB |
| 3.2624E-08 | -0.2992841 | 0.055 | 0.245 | 0.00069653 | TEC_prg3 | PLEKHB2 |
| 5.8746E-06 | -0.3003676 | 0.036 | 0.17 | 0.12542173 | TEC_prg3 | FAM102A |
| 6.6102E-09 | -0.3004542 | 0.061 | 0.265 | 0.00014113 | TEC_prg3 | PRR13 |
| 1.0658E-08 | -0.3010198 | 0.073 | 0.283 | 0.00022756 | TEC_prg3 | CLTB |
| 5.1964E-07 | -0.3011086 | 0.012 | 0.154 | 0.01109431 | TEC_prg3 | NQO1 |
| 4.2223E-06 | -0.301242 | 0.042 | 0.184 | 0.09014515 | TEC_prg3 | CD40 |
| 4.9362E-08 | -0.3013448 | 0.067 | 0.257 | 0.00105389 | TEC_prg3 | ARHGDIA |
| 5.464E-09 | -0.3014726 | 0.048 | 0.253 | 0.00011666 | TEC_prg3 | RAB1A |
| 2.8694E-07 | -0.3020131 | 0.036 | 0.196 | 0.0061262 | TEC_prg3 | FAM129B |
| 1.8362E-08 | -0.3029401 | 0.048 | 0.238 | 0.00039203 | TEC_prg3 | MDH2 |
| 1.9171E-09 | -0.3031226 | 0.036 | 0.241 | 4.093E-05 | TEC_prg3 | VPS29 |
| 7.4428E-09 | -0.3034198 | 0.036 | 0.227 | 0.0001589 | TEC_prg3 | PSMB4 |
| 3.0404E-05 | -0.303454 | 0.012 | 0.116 | 0.64913489 | TEC_prg3 | RARRES3 |
| 1.152E-08 | -0.3039036 | 0.055 | 0.254 | 0.00024595 | TEC_prg3 | LMAN2 |
| 2.3794E-07 | -0.3044615 | 0.012 | 0.162 | 0.00508005 | TEC_prg3 | TMEM176B |
| 1.1713E-06 | -0.3049164 | 0.061 | 0.22 | 0.02500686 | TEC_prg3 | CCPG1 |
| 0.0018492 | -0.3055032 | 0.273 | 0.4 | 1 | TEC_prg3 | JPT1 |
| 7.8654E-09 | -0.3055543 | 0.067 | 0.282 | 0.00016793 | TEC_prg3 | MRPL41 |
| 4.4482E-09 | -0.3057816 | 0.036 | 0.234 | 9.4968E-05 | TEC_prg3 | STRAP |
| 1.1739E-08 | -0.3060661 | 0.03 | 0.213 | 0.00025063 | TEC_prg3 | GNG12 |
| 9.511E-09 | -0.3067655 | 0.024 | 0.21 | 0.00020306 | TEC_prg3 | PRDX3 |
| 1.8378E-08 | -0.3070183 | 0.03 | 0.213 | 0.00039236 | TEC_prg3 | SNX9 |
| 2.0617E-08 | -0.3073047 | 0.079 | 0.288 | 0.00044016 | TEC_prg3 | PSMB6 |
| 2.5304E-09 | -0.3077822 | 0.048 | 0.26 | 5.4024E-05 | TEC_prg3 | ATP6V1E1 |
| 2.5989E-08 | -0.3087488 | 0.03 | 0.21 | 0.00055486 | TEC_prg3 | NDUFA3 |
| 2.4686E-09 | -0.3090628 | 0.006 | 0.191 | 5.2705E-05 | TEC_prg3 | B4GALT1 |
| 8.0316E-08 | -0.3105637 | 0.079 | 0.275 | 0.00171474 | TEC_prg3 | BAG1 |
| 1.0265E-08 | -0.3107168 | 0.036 | 0.227 | 0.00021915 | TEC_prg3 | C19orf70 |
| 1.0534E-09 | -0.3107618 | 0.012 | 0.209 | 2.249E-05 | TEC_prg3 | CALU |
| 4.2331E-09 | -0.3108365 | 0.03 | 0.224 | 9.0376E-05 | TEC_prg3 | GNAI2 |
| 1.3776E-09 | -0.3109099 | 0.133 | 0.391 | 2.9412E-05 | TEC_prg3 | SUB1 |
| 7.7225E-09 | -0.3109922 | 0.024 | 0.21 | 0.00016487 | TEC_prg3 | SCP2 |

| | | | | | | |
|---|---|---|---|---|---|---|
| 3.5579E-08 | -0.31103 | 0.079 | 0.278 | 0.00075961 | TEC_prg3 | CCT6A |
| 2.0817E-09 | -0.3120647 | 0.042 | 0.252 | 4.4445E-05 | TEC_prg3 | EIF3D |
| 2.6735E-09 | -0.3122226 | 0.03 | 0.227 | 5.708E-05 | TEC_prg3 | STAU1 |
| 1.5061E-08 | -0.3124383 | 0.012 | 0.183 | 0.00032156 | TEC_prg3 | CD99 |
| 0.00020954 | -0.3125298 | 0.018 | 0.108 | 1 | TEC_prg3 | BST2 |
| 5.1485E-08 | -0.3130182 | 0.073 | 0.27 | 0.0010992 | TEC_prg3 | MINOS1 |
| 6.7321E-09 | -0.3134501 | 0.061 | 0.266 | 0.00014373 | TEC_prg3 | CAPZA2 |
| 3.3681E-10 | -0.3135127 | 0.048 | 0.279 | 7.1908E-06 | TEC_prg3 | CHMP2A |
| 1.5347E-08 | -0.3137485 | 0.048 | 0.243 | 0.00032766 | TEC_prg3 | OSTC |
| 1.9442E-08 | -0.3141709 | 0.127 | 0.353 | 0.00041509 | TEC_prg3 | VAPA |
| 5.4408E-09 | -0.3145247 | 0.036 | 0.23 | 0.00011616 | TEC_prg3 | PSMD2 |
| 3.7844E-09 | -0.3158434 | 0.048 | 0.255 | 8.0797E-05 | TEC_prg3 | NDUFAB1 |
| 1.5357E-06 | -0.315867 | 0.067 | 0.225 | 0.03278724 | TEC_prg3 | NAMPT |
| 5.5279E-08 | -0.3160251 | 0.079 | 0.275 | 0.00118021 | TEC_prg3 | NDUFV2 |
| 0.00197977 | -0.3160324 | 0.842 | 0.827 | 1 | TEC_prg3 | HLA-B |
| 7.6401E-06 | -0.3163846 | 0.382 | 0.559 | 0.16311707 | TEC_prg3 | LRRFIP1 |
| 5.364E-07 | -0.316588 | 0.024 | 0.173 | 0.01145208 | TEC_prg3 | SERPING1 |
| 2.4442E-08 | -0.3166881 | 0.048 | 0.234 | 0.00052183 | TEC_prg3 | DMKN |
| 1.4191E-07 | -0.3170821 | 0.061 | 0.237 | 0.0030298 | TEC_prg3 | CHMP2B |
| 1.933E-08 | -0.3171457 | 0.091 | 0.306 | 0.0004127 | TEC_prg3 | WASF2 |
| 1.2163E-08 | -0.3173727 | 0.085 | 0.297 | 0.00025968 | TEC_prg3 | SRSF9 |
| 7.0456E-08 | -0.3178765 | 0.073 | 0.261 | 0.00150424 | TEC_prg3 | NDUFA6 |
| 3.8064E-09 | -0.318363 | 0.055 | 0.265 | 8.1267E-05 | TEC_prg3 | GNB2 |
| 7.7257E-10 | -0.3193211 | 0.024 | 0.23 | 1.6494E-05 | TEC_prg3 | TMEM219 |
| 2.1081E-07 | -0.3197014 | 0.176 | 0.401 | 0.00450076 | TEC_prg3 | YWHAQ |
| 3.8869E-08 | -0.3200218 | 0.048 | 0.23 | 0.00082986 | TEC_prg3 | COMT |
| 1.1669E-08 | -0.320603 | 0.061 | 0.263 | 0.00024914 | TEC_prg3 | ERH |
| 6.9144E-08 | -0.3206158 | 0.127 | 0.343 | 0.00147623 | TEC_prg3 | EIF1AX |
| 3.1291E-08 | -0.3206288 | 0.03 | 0.208 | 0.00066806 | TEC_prg3 | CSRP1 |
| 7.1811E-09 | -0.3210874 | 0.067 | 0.274 | 0.00015332 | TEC_prg3 | NDUFA2 |
| 3.8049E-08 | -0.3214659 | 0.079 | 0.278 | 0.00081236 | TEC_prg3 | NDUFS6 |
| 2.3917E-10 | -0.3223727 | 0.036 | 0.259 | 5.1064E-06 | TEC_prg3 | CCT8 |
| 1.2657E-08 | -0.3225353 | 0.012 | 0.184 | 0.00027024 | TEC_prg3 | QSOX1 |
| 7.544E-09 | -0.3231249 | 0.109 | 0.341 | 0.00016106 | TEC_prg3 | AURKAIP1 |
| 9.9111E-06 | -0.3232281 | 0.048 | 0.182 | 0.2116029 | TEC_prg3 | DCBLD2 |
| 2.5186E-09 | -0.324281 | 0.024 | 0.215 | 5.3772E-05 | TEC_prg3 | TRAPPC1 |
| 4.3455E-07 | -0.3242847 | 0.091 | 0.279 | 0.00927771 | TEC_prg3 | SMS |
| 1.912E-06 | -0.3248418 | 0.115 | 0.297 | 0.04082064 | TEC_prg3 | CD55 |
| 1.3077E-08 | -0.3252942 | 0.079 | 0.289 | 0.0002792 | TEC_prg3 | COPS9 |
| 6.5402E-08 | -0.3254714 | 0 | 0.151 | 0.00139633 | TEC_prg3 | FXYD5 |
| 2.4395E-05 | -0.3260661 | 0.485 | 0.632 | 0.52082458 | TEC_prg3 | PABPC1 |
| 3.9027E-06 | -0.3269225 | 0.388 | 0.57 | 0.08332262 | TEC_prg3 | HNRNPC |
| 7.4682E-10 | -0.3281118 | 0.042 | 0.256 | 1.5945E-05 | TEC_prg3 | EIF3I |
| 4.177E-10 | -0.3292671 | 0.03 | 0.246 | 8.9179E-06 | TEC_prg3 | PSMB5 |
| 1.2171E-08 | -0.329632 | 0.073 | 0.276 | 0.00025985 | TEC_prg3 | ATP5F1C |
| 3.0465E-08 | -0.3309032 | 0.091 | 0.292 | 0.00065044 | TEC_prg3 | SERINC1 |
| 6.1701E-06 | -0.3311008 | 0.139 | 0.312 | 0.13173123 | TEC_prg3 | ISCU |
| 1.2496E-07 | -0.3313145 | 0.073 | 0.254 | 0.002668 | TEC_prg3 | ATOX1 |
| 2.1075E-05 | -0.3316243 | 0.77 | 0.809 | 0.44995208 | TEC_prg3 | ATP5F1E |
| 8.5284E-05 | -0.3323331 | 0.176 | 0.336 | 1 | TEC_prg3 | RALBP1 |

| | | | | | | |
|---|---|---|---|---|---|---|
| 6.547E-10 | -0.3331977 | 0.073 | 0.31 | 1.3978E-05 | TEC_prg3 | TMEM258 |
| 7.0139E-10 | -0.3341437 | 0.048 | 0.268 | 1.4975E-05 | TEC_prg3 | SYPL1 |
| 1.6979E-08 | -0.3341781 | 0.079 | 0.281 | 0.0003625 | TEC_prg3 | CNN3 |
| 3.6787E-09 | -0.3345703 | 0.067 | 0.279 | 7.8541E-05 | TEC_prg3 | SELENOF |
| 4.9817E-10 | -0.3361562 | 0.036 | 0.253 | 1.0636E-05 | TEC_prg3 | AP2M1 |
| 2.9192E-09 | -0.3361688 | 0.036 | 0.236 | 6.2325E-05 | TEC_prg3 | RNF181 |
| 4.2831E-10 | -0.3382297 | 0.042 | 0.26 | 9.1445E-06 | TEC_prg3 | MTPN |
| 1.1101E-09 | -0.3384153 | 0.042 | 0.254 | 2.37E-05 | TEC_prg3 | PHB |
| 5.0056E-09 | -0.3384418 | 0.055 | 0.26 | 0.00010687 | TEC_prg3 | ATP5F1A |
| 1.693E-07 | -0.3390073 | 0.03 | 0.193 | 0.00361464 | TEC_prg3 | CTSL |
| 1.2875E-07 | -0.339283 | 0.176 | 0.4 | 0.00274891 | TEC_prg3 | CYCS |
| 2.5335E-05 | -0.3394625 | 0.958 | 0.956 | 0.54090431 | TEC_prg3 | RPS27 |
| 7.406E-10 | -0.339496 | 0.036 | 0.247 | 1.5812E-05 | TEC_prg3 | LRP10 |
| 2.2374E-10 | -0.3398903 | 0.018 | 0.227 | 4.7768E-06 | TEC_prg3 | ARL1 |
| 8.2078E-08 | -0.3401165 | 0.158 | 0.385 | 0.00175237 | TEC_prg3 | NDUFB2 |
| 4.869E-10 | -0.3402542 | 0.036 | 0.257 | 1.0395E-05 | TEC_prg3 | EIF3M |
| 4.1941E-09 | -0.3408835 | 0.073 | 0.286 | 8.9544E-05 | TEC_prg3 | FIS1 |
| 1.7497E-10 | -0.340915 | 0.067 | 0.303 | 3.7357E-06 | TEC_prg3 | NDFIP1 |
| 5.0419E-10 | -0.3417896 | 0.103 | 0.353 | 1.0765E-05 | TEC_prg3 | C19orf53 |
| 5.7472E-08 | -0.3423515 | 0.055 | 0.237 | 0.00122703 | TEC_prg3 | ITGAV |
| 4.9047E-08 | -0.342832 | 0.006 | 0.164 | 0.00104715 | TEC_prg3 | PCSK1N |
| 1.7829E-07 | -0.3428354 | 0.073 | 0.254 | 0.00380639 | TEC_prg3 | CD47 |
| 3.7781E-09 | -0.3435557 | 0.055 | 0.259 | 8.0663E-05 | TEC_prg3 | CCT4 |
| 3.1229E-09 | -0.3435642 | 0.103 | 0.335 | 6.6674E-05 | TEC_prg3 | COPE |
| 2.8292E-08 | -0.3444918 | 0 | 0.159 | 0.00060404 | TEC_prg3 | QPRT |
| 1.6262E-08 | -0.3445586 | 0.067 | 0.263 | 0.00034719 | TEC_prg3 | PAK2 |
| 2.4215E-09 | -0.3463772 | 0.055 | 0.262 | 5.1698E-05 | TEC_prg3 | CTTN |
| 2.1908E-09 | -0.3469834 | 0.091 | 0.322 | 4.6774E-05 | TEC_prg3 | SNRPB |
| 4.2847E-07 | -0.347231 | 0.145 | 0.347 | 0.00914793 | TEC_prg3 | MIF |
| 7.7486E-09 | -0.3478601 | 0.085 | 0.304 | 0.00016543 | TEC_prg3 | CUTA |
| 2.4185E-10 | -0.3479299 | 0.036 | 0.26 | 5.1635E-06 | TEC_prg3 | TMED2 |
| 8.5256E-07 | -0.3479408 | 0.042 | 0.198 | 0.01820206 | TEC_prg3 | MSMO1 |
| 2.8041E-10 | -0.3484483 | 0.042 | 0.266 | 5.9867E-06 | TEC_prg3 | TMCO1 |
| 5.9059E-10 | -0.3489077 | 0.03 | 0.237 | 1.2609E-05 | TEC_prg3 | PSMB2 |
| 1.8438E-07 | -0.3491741 | 0.03 | 0.191 | 0.00393656 | TEC_prg3 | EMP1 |
| 1.5053E-06 | -0.349266 | 0.042 | 0.19 | 0.03213869 | TEC_prg3 | PDIA4 |
| 4.373E-10 | -0.3492887 | 0.055 | 0.28 | 9.3364E-06 | TEC_prg3 | CCT5 |
| 7.5559E-10 | -0.3503433 | 0.109 | 0.369 | 1.6132E-05 | TEC_prg3 | PPP1CB |
| 3.9028E-09 | -0.3510417 | 0.079 | 0.296 | 8.3325E-05 | TEC_prg3 | MRPL51 |
| 0.00022645 | -0.3512516 | 0.424 | 0.566 | 1 | TEC_prg3 | CST3 |
| 1.8363E-10 | -0.351841 | 0.091 | 0.343 | 3.9204E-06 | TEC_prg3 | ERP29 |
| 3.4532E-09 | -0.3520814 | 0.073 | 0.289 | 7.3725E-05 | TEC_prg3 | RSL24D1 |
| 1.4094E-08 | -0.3530964 | 0.091 | 0.297 | 0.0003009 | TEC_prg3 | SSBP1 |
| 7.7719E-10 | -0.354055 | 0.055 | 0.277 | 1.6593E-05 | TEC_prg3 | CAPNS1 |
| 1.1196E-09 | -0.3543905 | 0.024 | 0.228 | 2.3903E-05 | TEC_prg3 | C4orf48 |
| 5.2312E-09 | -0.3548767 | 0.097 | 0.328 | 0.00011169 | TEC_prg3 | NDUFB7 |
| 0.00063918 | -0.3551463 | 0.818 | 0.819 | 1 | TEC_prg3 | YBX1 |
| 6.295E-07 | -0.355606 | 0.539 | 0.683 | 0.01343975 | TEC_prg3 | CLIC1 |
| 1.0664E-08 | -0.356383 | 0.048 | 0.245 | 0.00022767 | TEC_prg3 | PLP2 |
| 3.6458E-09 | -0.3566008 | 0.03 | 0.224 | 7.7837E-05 | TEC_prg3 | GARS |

| | | | | | | |
|---|---|---|---|---|---|---|
| 4.1757E-10 | -0.3568523 | 0.024 | 0.231 | 8.9152E-06 | TEC_prg3 | PGLS |
| 4.0096E-10 | -0.3576178 | 0.042 | 0.258 | 8.5606E-06 | TEC_prg3 | CAPZB |
| 3.0072E-10 | -0.3599454 | 0.079 | 0.319 | 6.4203E-06 | TEC_prg3 | SRP9 |
| 1.7768E-10 | -0.3604761 | 0.036 | 0.258 | 3.7936E-06 | TEC_prg3 | RER1 |
| 9.7197E-10 | -0.3609771 | 0.055 | 0.27 | 2.0752E-05 | TEC_prg3 | PSMB7 |
| 2.2143E-05 | -0.361776 | 0.109 | 0.257 | 0.4727621 | TEC_prg3 | S100A14 |
| 1.1936E-09 | -0.3626366 | 0.024 | 0.226 | 2.5483E-05 | TEC_prg3 | CXCL16 |
| 3.0608E-09 | -0.3627785 | 0.145 | 0.398 | 6.5349E-05 | TEC_prg3 | FKBP2 |
| 8.1737E-07 | -0.3628285 | 0.509 | 0.664 | 0.01745075 | TEC_prg3 | UQCRB |
| 8.6407E-09 | -0.3633804 | 0.055 | 0.256 | 0.00018448 | TEC_prg3 | COTL1 |
| 2.4235E-08 | -0.3640159 | 0.121 | 0.344 | 0.00051742 | TEC_prg3 | JUP |
| 1.733E-09 | -0.3648387 | 0.03 | 0.23 | 3.7E-05 | TEC_prg3 | SEC11C |
| 5.0475E-09 | -0.365076 | 0.115 | 0.343 | 0.00010776 | TEC_prg3 | PFDN2 |
| 3.0573E-11 | -0.3654933 | 0.073 | 0.332 | 6.5274E-07 | TEC_prg3 | EIF3H |
| 6.6988E-10 | -0.3657031 | 0.085 | 0.324 | 1.4302E-05 | TEC_prg3 | ATP5MPL |
| 7.9416E-07 | -0.3657432 | 0.673 | 0.775 | 0.01695523 | TEC_prg3 | EEF2 |
| 1.112E-08 | -0.366331 | 0.121 | 0.351 | 0.00023742 | TEC_prg3 | UBE2H |
| 1.9572E-09 | -0.3663831 | 0.139 | 0.387 | 4.1786E-05 | TEC_prg3 | CTNNA1 |
| 0.00013082 | -0.3665092 | 0.77 | 0.798 | 1 | TEC_prg3 | HLA-C |
| 3.1879E-09 | -0.3675057 | 0.085 | 0.304 | 6.8061E-05 | TEC_prg3 | RAP1B |
| 1.5696E-09 | -0.3676733 | 0.055 | 0.268 | 3.3512E-05 | TEC_prg3 | TXNDC17 |
| 1.0133E-09 | -0.3679673 | 0.012 | 0.206 | 2.1635E-05 | TEC_prg3 | ERO1A |
| 3.0428E-10 | -0.3681198 | 0.048 | 0.272 | 6.4964E-06 | TEC_prg3 | RTN3 |
| 3.097E-09 | -0.3690745 | 0.048 | 0.249 | 6.612E-05 | TEC_prg3 | BCAP31 |
| 1.6966E-10 | -0.3692691 | 0.048 | 0.275 | 3.6222E-06 | TEC_prg3 | UFC1 |
| 8.6836E-09 | -0.3696087 | 0.091 | 0.302 | 0.00018539 | TEC_prg3 | ACTR3 |
| 4.0971E-10 | -0.369702 | 0.097 | 0.353 | 8.7474E-06 | TEC_prg3 | MZT2B |
| 2.8237E-09 | -0.3701246 | 0.097 | 0.325 | 6.0286E-05 | TEC_prg3 | AES |
| 3.3088E-06 | -0.3706943 | 0.309 | 0.498 | 0.07064387 | TEC_prg3 | ACTN4 |
| 2.9051E-11 | -0.3713033 | 0.024 | 0.253 | 6.2023E-07 | TEC_prg3 | SUMO1 |
| 1.2202E-09 | -0.3714215 | 0.2 | 0.451 | 2.6052E-05 | TEC_prg3 | VDAC2 |
| 1.3424E-08 | -0.3721009 | 0.115 | 0.333 | 0.0002866 | TEC_prg3 | SF3B6 |
| 3.7041E-05 | -0.3723514 | 0.582 | 0.693 | 0.79082413 | TEC_prg3 | GPX4 |
| 5.2377E-08 | -0.3724946 | 0.061 | 0.246 | 0.00111825 | TEC_prg3 | DNAJC3 |
| 1.7402E-10 | -0.3725196 | 0.03 | 0.249 | 3.7153E-06 | TEC_prg3 | PGAM1 |
| 1.6644E-10 | -0.37446 | 0.036 | 0.26 | 3.5535E-06 | TEC_prg3 | PSMB3 |
| 4.7949E-09 | -0.3749329 | 0.036 | 0.231 | 0.00010237 | TEC_prg3 | ST14 |
| 3.9327E-10 | -0.3751922 | 0.115 | 0.366 | 8.3963E-06 | TEC_prg3 | CSDE1 |
| 9.48E-07 | -0.3756123 | 0.048 | 0.204 | 0.02023972 | TEC_prg3 | MANF |
| 6.7629E-10 | -0.376656 | 0.097 | 0.34 | 1.4439E-05 | TEC_prg3 | EIF4B |
| 2.4999E-10 | -0.3777986 | 0.085 | 0.33 | 5.3373E-06 | TEC_prg3 | ELOC |
| 3.1182E-10 | -0.3783705 | 0.055 | 0.283 | 6.6573E-06 | TEC_prg3 | GRN |
| 2.7768E-08 | -0.3795085 | 0.315 | 0.531 | 0.00059285 | TEC_prg3 | COX6C |
| 1.8351E-09 | -0.3804197 | 0.067 | 0.282 | 3.9179E-05 | TEC_prg3 | LMAN1 |
| 3.9467E-11 | -0.3821129 | 0.036 | 0.272 | 8.4262E-07 | TEC_prg3 | NDUFB8 |
| 0.00018577 | -0.3826399 | 0.485 | 0.586 | 1 | TEC_prg3 | EZR |
| 2.9786E-10 | -0.3826595 | 0.091 | 0.337 | 6.3593E-06 | TEC_prg3 | RAB2A |
| 3.0366E-09 | -0.3861585 | 0.073 | 0.292 | 6.4832E-05 | TEC_prg3 | FAM3C |
| 7.2603E-10 | -0.3877088 | 0.158 | 0.414 | 1.5501E-05 | TEC_prg3 | UQCR10 |
| 7.5564E-05 | -0.3877341 | 0.012 | 0.108 | 1 | TEC_prg3 | AQP3 |

| | | | | | | |
|---|---|---|---|---|---|---|
| 1.1035E-10 | -0.3885603 | 0.048 | 0.277 | 2.356E-06 | TEC_prg3 | VPS28 |
| 9.0951E-07 | -0.3885805 | 0.976 | 0.965 | 0.0194181 | TEC_prg3 | RPS19 |
| 1.4786E-08 | -0.3891302 | 0.042 | 0.23 | 0.00031569 | TEC_prg3 | PRSS8 |
| 7.1092E-12 | -0.389801 | 0.055 | 0.308 | 1.5178E-07 | TEC_prg3 | EIF3F |
| 4.6834E-11 | -0.3910294 | 0.036 | 0.264 | 9.9991E-07 | TEC_prg3 | LAMTOR1 |
| 1.602E-06 | -0.3911126 | 0.012 | 0.146 | 0.03420369 | TEC_prg3 | XIST |
| 1.3641E-10 | -0.3912124 | 0.055 | 0.286 | 2.9122E-06 | TEC_prg3 | LAMP1 |
| 2.8348E-08 | -0.3912501 | 0.467 | 0.654 | 0.00060523 | TEC_prg3 | SLC25A3 |
| 5.7868E-08 | -0.3925773 | 0.376 | 0.585 | 0.00123548 | TEC_prg3 | APP |
| 3.5457E-10 | -0.3930305 | 0.158 | 0.42 | 7.5701E-06 | TEC_prg3 | DYNLRB1 |
| 9.2005E-12 | -0.3931015 | 0.012 | 0.242 | 1.9643E-07 | TEC_prg3 | KDELR2 |
| 0.00373227 | -0.3936302 | 0.945 | 0.891 | 1 | TEC_prg3 | MT-ND1 |
| 4.9308E-08 | -0.3946429 | 0.145 | 0.373 | 0.00105272 | TEC_prg3 | TAGLN2 |
| 1.5114E-07 | -0.3962524 | 0.939 | 0.945 | 0.00322682 | TEC_prg3 | RPS15 |
| 1.9957E-11 | -0.3966993 | 0.036 | 0.276 | 4.2609E-07 | TEC_prg3 | ANAPC11 |
| 8.7193E-06 | -0.397503 | 0.909 | 0.899 | 0.1861577 | TEC_prg3 | SERF2 |
| 0.0009695 | -0.3986259 | 0.588 | 0.631 | 1 | TEC_prg3 | TPM1 |
| 1.6888E-09 | -0.3987601 | 0.139 | 0.388 | 3.6056E-05 | TEC_prg3 | PERP |
| 0.00925051 | -0.399263 | 0.212 | 0.307 | 1 | TEC_prg3 | HSPA1B |
| 2.8942E-11 | -0.4000948 | 0.067 | 0.318 | 6.179E-07 | TEC_prg3 | COX7A2L |
| 4.5027E-10 | -0.4018379 | 0.036 | 0.255 | 9.6133E-06 | TEC_prg3 | ECHS1 |
| 8.8115E-10 | -0.4018861 | 0.067 | 0.288 | 1.8813E-05 | TEC_prg3 | NDUFB11 |
| 5.7836E-11 | -0.4024603 | 0.067 | 0.309 | 1.2348E-06 | TEC_prg3 | WDR1 |
| 1.9523E-07 | -0.4033175 | 0.77 | 0.826 | 0.00416826 | TEC_prg3 | H3F3A |
| 1.8292E-11 | -0.4052643 | 0.055 | 0.3 | 3.9054E-07 | TEC_prg3 | CHMP5 |
| 4.8388E-09 | -0.4058114 | 0.042 | 0.24 | 0.00010331 | TEC_prg3 | HIGD1A |
| 1.8871E-11 | -0.4071882 | 0.067 | 0.32 | 4.029E-07 | TEC_prg3 | SARAF |
| 1.6969E-10 | -0.4075705 | 0.042 | 0.265 | 3.6228E-06 | TEC_prg3 | FKBP1A |
| 9.3985E-09 | -0.4075705 | 0.133 | 0.363 | 0.00020066 | TEC_prg3 | COX7B |
| 1.4245E-10 | -0.4085184 | 0.036 | 0.264 | 3.0414E-06 | TEC_prg3 | GSTO1 |
| 8.5483E-07 | -0.408624 | 0.358 | 0.538 | 0.01825054 | TEC_prg3 | HLA-E |
| 1.1442E-09 | -0.409001 | 0.012 | 0.205 | 2.4429E-05 | TEC_prg3 | DPP4 |
| 3.3081E-10 | -0.4114372 | 0.061 | 0.285 | 7.0629E-06 | TEC_prg3 | LITAF |
| 2.1888E-06 | -0.4118579 | 0.018 | 0.149 | 0.04673003 | TEC_prg3 | MYL9 |
| 3.2241E-11 | -0.4129946 | 0.073 | 0.33 | 6.8834E-07 | TEC_prg3 | NDUFC1 |
| 2.5057E-12 | -0.4131466 | 0.079 | 0.365 | 5.3497E-08 | TEC_prg3 | SNRPD2 |
| 9.8895E-08 | -0.4168376 | 0.055 | 0.229 | 0.00211141 | TEC_prg3 | IFITM2 |
| 1.3587E-11 | -0.419699 | 0.048 | 0.297 | 2.9009E-07 | TEC_prg3 | EIF6 |
| 7.5908E-10 | -0.4219832 | 0.321 | 0.562 | 1.6206E-05 | TEC_prg3 | RHOA |
| 4.9229E-11 | -0.4243707 | 0.073 | 0.324 | 1.051E-06 | TEC_prg3 | NPC2 |
| 5.7516E-09 | -0.426503 | 0.079 | 0.291 | 0.0001228 | TEC_prg3 | ITGA3 |
| 5.2155E-10 | -0.426914 | 0.115 | 0.364 | 1.1135E-05 | TEC_prg3 | ATP5MC3 |
| 4.5538E-08 | -0.4277916 | 0.473 | 0.638 | 0.00097224 | TEC_prg3 | CCNI |
| 7.8158E-11 | -0.4278354 | 0.127 | 0.398 | 1.6687E-06 | TEC_prg3 | TAX1BP1 |
| 1.8174E-11 | -0.4280899 | 0.067 | 0.313 | 3.8802E-07 | TEC_prg3 | NDUFC2 |
| 6.8747E-11 | -0.4285226 | 0.091 | 0.342 | 1.4677E-06 | TEC_prg3 | NOP10 |
| 2.6352E-09 | -0.4290212 | 0.103 | 0.332 | 5.6262E-05 | TEC_prg3 | UQCRQ |
| 4.689E-12 | -0.4293717 | 0.042 | 0.297 | 1.0011E-07 | TEC_prg3 | BRK1 |
| 4.1608E-09 | -0.4302647 | 0.333 | 0.564 | 8.8834E-05 | TEC_prg3 | COX6B1 |
| 7.3658E-10 | -0.4305448 | 0.091 | 0.322 | 1.5726E-05 | TEC_prg3 | TKT |

| | | | | | | |
|---|---|---|---|---|---|---|
| 5.7988E-05 | -0.4307935 | 0.03 | 0.139 | 1 | TEC_prg3 | TM4SF1 |
| 6.0123E-10 | -0.4316688 | 0.073 | 0.302 | 1.2836E-05 | TEC_prg3 | CDH1 |
| 2.2656E-12 | -0.4317068 | 0.097 | 0.38 | 4.8371E-08 | TEC_prg3 | CD164 |
| 0.0005596 | -0.4319907 | 0.133 | 0.254 | 1 | TEC_prg3 | AKAP12 |
| 1.2881E-12 | -0.4328078 | 0.048 | 0.315 | 2.75E-08 | TEC_prg3 | TMEM50A |
| 2.5055E-11 | -0.4331861 | 0.079 | 0.337 | 5.3492E-07 | TEC_prg3 | PPA1 |
| 8.0662E-08 | -0.4333418 | 0.17 | 0.391 | 0.00172213 | TEC_prg3 | HSPE1 |
| 5.8076E-05 | -0.4335434 | 0.024 | 0.128 | 1 | TEC_prg3 | HMOX1 |
| 8.8105E-12 | -0.435667 | 0.018 | 0.251 | 1.881E-07 | TEC_prg3 | DHRS7 |
| 2.1835E-08 | -0.4364254 | 0 | 0.161 | 0.00046619 | TEC_prg3 | MGST1 |
| 9.0546E-13 | -0.437413 | 0.055 | 0.324 | 1.9332E-08 | TEC_prg3 | GNB1 |
| 1.0622E-11 | -0.4380178 | 0.085 | 0.344 | 2.2677E-07 | TEC_prg3 | SPCS1 |
| 9.9241E-10 | -0.4388997 | 0.127 | 0.374 | 2.1188E-05 | TEC_prg3 | CAMK2N1 |
| 4.7293E-12 | -0.4393354 | 0.055 | 0.314 | 1.0097E-07 | TEC_prg3 | ATP5PD |
| 0.00105929 | -0.4400784 | 0.255 | 0.383 | 1 | TEC_prg3 | HERPUD1 |
| 1.1558E-12 | -0.4402735 | 0.073 | 0.353 | 2.4676E-08 | TEC_prg3 | COMMD6 |
| 3.7589E-09 | -0.4404197 | 0.012 | 0.197 | 8.0251E-05 | TEC_prg3 | SDF2L1 |
| 2.5318E-09 | -0.4412856 | 0.024 | 0.216 | 5.4054E-05 | TEC_prg3 | CTSH |
| 8.9426E-08 | -0.4414447 | 0.073 | 0.257 | 0.00190925 | TEC_prg3 | RPS4Y1 |
| 1.4514E-12 | -0.4424331 | 0.085 | 0.364 | 3.0986E-08 | TEC_prg3 | NEDD8 |
| 1.2549E-12 | -0.4428552 | 0.042 | 0.306 | 2.6792E-08 | TEC_prg3 | SSR2 |
| 2.2633E-13 | -0.4432544 | 0.048 | 0.331 | 4.8321E-09 | TEC_prg3 | VDAC1 |
| 1.2454E-10 | -0.4433377 | 0.115 | 0.379 | 2.659E-06 | TEC_prg3 | NDUFA1 |
| 4.6776E-12 | -0.4449904 | 0.024 | 0.267 | 9.9867E-08 | TEC_prg3 | APRT |
| 4.2242E-10 | -0.4452689 | 0.23 | 0.483 | 9.0186E-06 | TEC_prg3 | PTGES3 |
| 9.2704E-05 | -0.4461778 | 0.358 | 0.494 | 1 | TEC_prg3 | ATP1B1 |
| 3.8565E-12 | -0.4465318 | 0.048 | 0.305 | 8.2335E-08 | TEC_prg3 | CIB1 |
| 2.6221E-11 | -0.4466251 | 0.085 | 0.334 | 5.5981E-07 | TEC_prg3 | NDUFB4 |
| 8.8665E-11 | -0.4475759 | 0.333 | 0.584 | 1.893E-06 | TEC_prg3 | ATP6V1G1 |
| 1.2771E-11 | -0.4494577 | 0.109 | 0.384 | 2.7265E-07 | TEC_prg3 | RHEB |
| 2.2049E-12 | -0.4500931 | 0.036 | 0.29 | 4.7074E-08 | TEC_prg3 | RABAC1 |
| 1.4288E-12 | -0.4506459 | 0.067 | 0.338 | 3.0505E-08 | TEC_prg3 | SPCS2 |
| 1.8942E-09 | -0.4508979 | 0.309 | 0.536 | 4.0441E-05 | TEC_prg3 | NDUFS5 |
| 4.6609E-11 | -0.4513236 | 0.127 | 0.395 | 9.951E-07 | TEC_prg3 | CAPN2 |
| 5.9913E-09 | -0.4514248 | 0.721 | 0.806 | 0.00012791 | TEC_prg3 | EEF1D |
| 3.3033E-10 | -0.4516789 | 0.048 | 0.273 | 7.0525E-06 | TEC_prg3 | RBM3 |
| 8.5637E-11 | -0.4534998 | 0.091 | 0.344 | 1.8284E-06 | TEC_prg3 | ARF4 |
| 3.6122E-13 | -0.4542401 | 0.097 | 0.393 | 7.7121E-09 | TEC_prg3 | TOMM20 |
| 1.8573E-10 | -0.454455 | 0.388 | 0.611 | 3.9653E-06 | TEC_prg3 | ATP5MG |
| 4.3765E-11 | -0.4550815 | 0.109 | 0.373 | 9.3439E-07 | TEC_prg3 | SDCBP |
| 4.8258E-11 | -0.4565398 | 0.6 | 0.756 | 1.0303E-06 | TEC_prg3 | RPL4 |
| 1.011E-10 | -0.4582062 | 0.115 | 0.371 | 2.1585E-06 | TEC_prg3 | ATP5MF |
| 4.6491E-12 | -0.4589612 | 0.048 | 0.299 | 9.9259E-08 | TEC_prg3 | LEPROT |
| 4.5936E-07 | -0.4593193 | 0.309 | 0.499 | 0.00980724 | TEC_prg3 | HSPD1 |
| 1.0988E-08 | -0.4596322 | 0.364 | 0.571 | 0.00023459 | TEC_prg3 | RAN |
| 6.2311E-10 | -0.459748 | 0.248 | 0.494 | 1.3303E-05 | TEC_prg3 | TOMM7 |
| 4.2712E-11 | -0.4597928 | 0.109 | 0.379 | 9.119E-07 | TEC_prg3 | VAMP8 |
| 3.8037E-12 | -0.4598385 | 0.248 | 0.524 | 8.1209E-08 | TEC_prg3 | ARF1 |
| 1.812E-09 | -0.4602489 | 0.309 | 0.527 | 3.8686E-05 | TEC_prg3 | CDC42 |
| 1.0448E-06 | -0.4602778 | 0.339 | 0.506 | 0.02230729 | TEC_prg3 | CLDN7 |

| | | | | | | |
|---|---|---|---|---|---|---|
| 2.3028E-07 | -0.4612019 | 0.727 | 0.805 | 0.00491649 | TEC_prg3 | RPL31 |
| 2.3843E-11 | -0.4629584 | 0.37 | 0.607 | 5.0905E-07 | TEC_prg3 | CLTA |
| 1.1689E-09 | -0.4645701 | 0.194 | 0.435 | 2.4957E-05 | TEC_prg3 | BZW1 |
| 1.1446E-06 | -0.4646181 | 0.758 | 0.789 | 0.02443719 | TEC_prg3 | NPM1 |
| 1.0605E-13 | -0.4661356 | 0.055 | 0.345 | 2.2641E-09 | TEC_prg3 | GHITM |
| 1.0844E-11 | -0.4665159 | 0.115 | 0.384 | 2.3153E-07 | TEC_prg3 | HSBP1 |
| 2.2801E-10 | -0.4673081 | 0.073 | 0.312 | 4.8679E-06 | TEC_prg3 | DSP |
| 8.528E-12 | -0.4674919 | 0.042 | 0.287 | 1.8207E-07 | TEC_prg3 | SPINT1 |
| 5.4444E-11 | -0.4676691 | 0.327 | 0.575 | 1.1624E-06 | TEC_prg3 | COX7A2 |
| 2.5374E-09 | -0.468897 | 0.097 | 0.319 | 5.4173E-05 | TEC_prg3 | PRDX2 |
| 5.4593E-12 | -0.4706008 | 0.279 | 0.545 | 1.1656E-07 | TEC_prg3 | GNG5 |
| 5.5338E-12 | -0.4713332 | 0.042 | 0.288 | 1.1815E-07 | TEC_prg3 | ROMO1 |
| 9.3321E-08 | -0.4731574 | 0.121 | 0.32 | 0.0019924 | TEC_prg3 | UGCG |
| 3.0964E-13 | -0.4735223 | 0.006 | 0.256 | 6.6109E-09 | TEC_prg3 | LAPTM4B |
| 6.1803E-12 | -0.4759929 | 0.103 | 0.371 | 1.3195E-07 | TEC_prg3 | SBDS |
| 7.5532E-09 | -0.4780898 | 0.242 | 0.464 | 0.00016126 | TEC_prg3 | COX8A |
| 5.9042E-09 | -0.4801051 | 0.642 | 0.749 | 0.00012605 | TEC_prg3 | DSTN |
| 8.3285E-14 | -0.4808096 | 0.03 | 0.298 | 1.7781E-09 | TEC_prg3 | TMED10 |
| 2.9541E-13 | -0.4828394 | 0.036 | 0.304 | 6.3069E-09 | TEC_prg3 | DDT |
| 1.2333E-06 | -0.483215 | 0.867 | 0.869 | 0.02633004 | TEC_prg3 | RPL10A |
| 1.167E-12 | -0.483882 | 0.079 | 0.349 | 2.4915E-08 | TEC_prg3 | CNBP |
| 6.2673E-11 | -0.4842611 | 0.127 | 0.401 | 1.3381E-06 | TEC_prg3 | DBI |
| 6.623E-08 | -0.4854478 | 0.606 | 0.728 | 0.00141402 | TEC_prg3 | CSTB |
| 7.7448E-14 | -0.4867636 | 0.042 | 0.326 | 1.6535E-09 | TEC_prg3 | COX5A |
| 2.7386E-14 | -0.4870277 | 0.048 | 0.339 | 5.847E-10 | TEC_prg3 | KDELR1 |
| 2.8011E-14 | -0.4878173 | 0.03 | 0.315 | 5.9804E-10 | TEC_prg3 | PGK1 |
| 4.2097E-11 | -0.4897358 | 0.048 | 0.285 | 8.9877E-07 | TEC_prg3 | ASPH |
| 1.9115E-07 | -0.4900327 | 0.885 | 0.886 | 0.00408103 | TEC_prg3 | RPS29 |
| 7.787E-13 | -0.4901061 | 0.055 | 0.322 | 1.6625E-08 | TEC_prg3 | NDUFB9 |
| 2.4242E-10 | -0.4907097 | 0.188 | 0.44 | 5.1756E-06 | TEC_prg3 | NAP1L1 |
| 7.3415E-08 | -0.4912211 | 0.085 | 0.279 | 0.0015674 | TEC_prg3 | CYB5A |
| 3.8205E-14 | -0.4915162 | 0.073 | 0.376 | 8.1567E-10 | TEC_prg3 | SNU13 |
| 8.721E-05 | -0.493174 | 0.982 | 0.909 | 1 | TEC_prg3 | MT-CO3 |
| 3.1711E-10 | -0.4931853 | 0.048 | 0.274 | 6.7704E-06 | TEC_prg3 | CKB |
| 2.9899E-11 | -0.493486 | 0.333 | 0.585 | 6.3834E-07 | TEC_prg3 | COX5B |
| 2.9856E-07 | -0.4943265 | 0.485 | 0.646 | 0.00637422 | TEC_prg3 | PEBP1 |
| 5.0632E-12 | -0.4954732 | 0.024 | 0.267 | 1.081E-07 | TEC_prg3 | SELENOM |
| 4.6854E-13 | -0.4956495 | 0.164 | 0.457 | 1.0003E-08 | TEC_prg3 | ATP5PF |
| 2.4847E-13 | -0.499299 | 0.024 | 0.286 | 5.3048E-09 | TEC_prg3 | MSN |
| 4.1951E-12 | -0.5001494 | 0.558 | 0.737 | 8.9566E-08 | TEC_prg3 | SKP1 |
| 6.7609E-09 | -0.5002964 | 0.63 | 0.747 | 0.00014435 | TEC_prg3 | HINT1 |
| 5.5347E-08 | -0.501908 | 0.042 | 0.218 | 0.00118166 | TEC_prg3 | GCLM |
| 6.6233E-10 | -0.5034899 | 0.648 | 0.77 | 1.4141E-05 | TEC_prg3 | COX7C |
| 3.0576E-10 | -0.5041165 | 0.891 | 0.897 | 6.528E-06 | TEC_prg3 | RPL35A |
| 2.0639E-07 | -0.505373 | 0.224 | 0.422 | 0.00440646 | TEC_prg3 | LGALS3 |
| 8.5375E-06 | -0.5054759 | 0.861 | 0.835 | 0.18227662 | TEC_prg3 | PPIA |
| 2.27E-12 | -0.5065793 | 0.042 | 0.302 | 4.8465E-08 | TEC_prg3 | CAPG |
| 7.137E-07 | -0.5067072 | 0.727 | 0.774 | 0.0152374 | TEC_prg3 | MYL12A |
| 9.0706E-16 | -0.5094081 | 0.036 | 0.343 | 1.9366E-11 | TEC_prg3 | SEC11A |
| 3.9768E-12 | -0.5104938 | 0.4 | 0.638 | 8.4905E-08 | TEC_prg3 | UBL5 |

| | | | | | | |
|---|---|---|---|---|---|---|
| 5.9077E-15 | -0.510572 | 0.042 | 0.337 | 1.2613E-10 | TEC_prg3 | EIF3E |
| 2.3697E-08 | -0.5113954 | 0.055 | 0.241 | 0.00050592 | TEC_prg3 | ITGB6 |
| 5.5708E-09 | -0.5115936 | 0.661 | 0.75 | 0.00011894 | TEC_prg3 | RTN4 |
| 5.2082E-14 | -0.5135874 | 0.055 | 0.34 | 1.1119E-09 | TEC_prg3 | PSMB1 |
| 1.0276E-14 | -0.5137731 | 0.03 | 0.317 | 2.1939E-10 | TEC_prg3 | PCBD1 |
| 9.6584E-09 | -0.5155081 | 0.885 | 0.898 | 0.00020621 | TEC_prg3 | RPL23A |
| 1.3974E-11 | -0.5158756 | 0.461 | 0.666 | 2.9835E-07 | TEC_prg3 | RAC1 |
| 4.7267E-13 | -0.5165049 | 0.085 | 0.365 | 1.0092E-08 | TEC_prg3 | TRMT112 |
| 1.9211E-13 | -0.5165761 | 0.061 | 0.341 | 4.1015E-09 | TEC_prg3 | PRELID1 |
| 1.1791E-11 | -0.5174239 | 0.188 | 0.454 | 2.5175E-07 | TEC_prg3 | RPS10 |
| 4.1077E-12 | -0.5178035 | 0.17 | 0.45 | 8.77E-08 | TEC_prg3 | ATP5MD |
| 2.8207E-08 | -0.5183469 | 0.085 | 0.276 | 0.00060222 | TEC_prg3 | MUC1 |
| 2.1255E-12 | -0.5198917 | 0.115 | 0.402 | 4.5379E-08 | TEC_prg3 | MGST3 |
| 3.1112E-11 | -0.5200124 | 0.061 | 0.306 | 6.6425E-07 | TEC_prg3 | C9orf16 |
| 3.3384E-06 | -0.5226397 | 0.061 | 0.215 | 0.07127382 | TEC_prg3 | GLRX |
| 5.2062E-14 | -0.5233226 | 0.073 | 0.37 | 1.1115E-09 | TEC_prg3 | ATP5F1B |
| 1.2304E-13 | -0.5234534 | 0.042 | 0.318 | 2.627E-09 | TEC_prg3 | UQCR11 |
| 1.1792E-10 | -0.5239315 | 0.091 | 0.334 | 2.5176E-06 | TEC_prg3 | TUBA1C |
| 1.6231E-08 | -0.5242371 | 0.079 | 0.287 | 0.00034654 | TEC_prg3 | AKR1B1 |
| 8.0726E-11 | -0.5252221 | 0.388 | 0.607 | 1.7235E-06 | TEC_prg3 | PSMA7 |
| 2.4092E-14 | -0.5261443 | 0.048 | 0.338 | 5.1437E-10 | TEC_prg3 | PTP4A2 |
| 9.4459E-11 | -0.5272588 | 0.224 | 0.483 | 2.0167E-06 | TEC_prg3 | P4HB |
| 1.597E-10 | -0.5315939 | 0.285 | 0.522 | 3.4096E-06 | TEC_prg3 | OST4 |
| 1.0403E-14 | -0.5322323 | 0.042 | 0.331 | 2.2211E-10 | TEC_prg3 | TRAM1 |
| 2.4185E-06 | -0.5323345 | 0.133 | 0.299 | 0.05163434 | TEC_prg3 | HSPH1 |
| 4.9513E-13 | -0.5329288 | 0.327 | 0.59 | 1.0571E-08 | TEC_prg3 | YWHAB |
| 9.6041E-12 | -0.5361269 | 0.794 | 0.855 | 2.0505E-07 | TEC_prg3 | RPL38 |
| 6.624E-10 | -0.5367396 | 0.515 | 0.692 | 1.4142E-05 | TEC_prg3 | CYSTM1 |
| 1.1929E-12 | -0.5377343 | 0.261 | 0.519 | 2.5468E-08 | TEC_prg3 | ARPC2 |
| 5.3012E-14 | -0.5382852 | 0.115 | 0.424 | 1.1318E-09 | TEC_prg3 | PTTG1IP |
| 7.0496E-08 | -0.5385166 | 0.103 | 0.302 | 0.00150509 | TEC_prg3 | MARCKSL1 |
| 2.3021E-15 | -0.538866 | 0.055 | 0.361 | 4.915E-11 | TEC_prg3 | EIF3K |
| 1.7963E-12 | -0.5389773 | 0.121 | 0.41 | 3.835E-08 | TEC_prg3 | SDC4 |
| 4.6175E-15 | -0.5407964 | 0.036 | 0.331 | 9.8583E-11 | TEC_prg3 | TMED9 |
| 6.1075E-09 | -0.5408246 | 0.109 | 0.323 | 0.00013039 | TEC_prg3 | DCDC2 |
| 2.0697E-07 | -0.5432155 | 0.055 | 0.223 | 0.00441889 | TEC_prg3 | SFN |
| 4.2767E-09 | -0.5447841 | 0.836 | 0.875 | 9.1308E-05 | TEC_prg3 | HLA-A |
| 3.4317E-12 | -0.546291 | 0.109 | 0.394 | 7.3267E-08 | TEC_prg3 | CD151 |
| 1.5112E-13 | -0.5473893 | 0.103 | 0.393 | 3.2263E-09 | TEC_prg3 | POMP |
| 6.632E-12 | -0.5486013 | 0.194 | 0.466 | 1.4159E-07 | TEC_prg3 | SERP1 |
| 4.6253E-11 | -0.5492927 | 0.715 | 0.801 | 9.8749E-07 | TEC_prg3 | RPL27 |
| 4.8447E-14 | -0.5504004 | 0.091 | 0.398 | 1.0343E-09 | TEC_prg3 | ATP5F1D |
| 3.9092E-12 | -0.5544727 | 0.279 | 0.528 | 8.3461E-08 | TEC_prg3 | ATP5MC2 |
| 2.1866E-13 | -0.5558893 | 0.4 | 0.661 | 4.6684E-09 | TEC_prg3 | PSAP |
| 1.1687E-08 | -0.5577049 | 0.727 | 0.783 | 0.00024951 | TEC_prg3 | EEF1B2 |
| 4.5971E-12 | -0.5582777 | 0.091 | 0.353 | 9.8148E-08 | TEC_prg3 | SELENOK |
| 1.0647E-05 | -0.5605896 | 0.006 | 0.116 | 0.22731557 | TEC_prg3 | TAGLN |
| 9.3738E-09 | -0.5619946 | 0.691 | 0.766 | 0.00020013 | TEC_prg3 | PKM |
| 1.3517E-12 | -0.5630709 | 0.818 | 0.888 | 2.886E-08 | TEC_prg3 | RPL22 |
| 3.1493E-13 | -0.5636843 | 0.085 | 0.377 | 6.7238E-09 | TEC_prg3 | FLNA |

| | | | | | | |
|---|---|---|---|---|---|---|
| 9.3055E-13 | -0.5668389 | 0.388 | 0.63 | 1.9867E-08 | TEC_prg3 | SLC25A6 |
| 7.0132E-16 | -0.5685907 | 0.152 | 0.477 | 1.4973E-11 | TEC_prg3 | SUMO2 |
| 1.4567E-05 | -0.5718932 | 0.067 | 0.202 | 0.31100247 | TEC_prg3 | ANXA1 |
| 1.1289E-15 | -0.5728157 | 0.048 | 0.364 | 2.4103E-11 | TEC_prg3 | PSME1 |
| 8.9305E-14 | -0.5735162 | 0.891 | 0.91 | 1.9067E-09 | TEC_prg3 | FAU |
| 1.7277E-14 | -0.5756378 | 0.024 | 0.304 | 3.6887E-10 | TEC_prg3 | MYDGF |
| 9.4657E-16 | -0.5765995 | 0.103 | 0.42 | 2.0209E-11 | TEC_prg3 | PARK7 |
| 7.75E-13 | -0.5769082 | 0.467 | 0.683 | 1.6546E-08 | TEC_prg3 | COX6A1 |
| 1.368E-10 | -0.5807572 | 0.915 | 0.934 | 2.9207E-06 | TEC_prg3 | RPL6 |
| 2.8187E-15 | -0.5815511 | 0.048 | 0.348 | 6.0179E-11 | TEC_prg3 | CAP1 |
| 2.4808E-12 | -0.5842973 | 0.867 | 0.905 | 5.2964E-08 | TEC_prg3 | RPS20 |
| 4.6011E-13 | -0.5851526 | 0.055 | 0.33 | 9.8233E-09 | TEC_prg3 | CTSB |
| 1.2613E-15 | -0.5855896 | 0.073 | 0.383 | 2.6929E-11 | TEC_prg3 | REEP5 |
| 2.0633E-11 | -0.5876333 | 0.903 | 0.929 | 4.4052E-07 | TEC_prg3 | RPL36 |
| 3.232E-12 | -0.5883923 | 0.115 | 0.388 | 6.9003E-08 | TEC_prg3 | OCIAD2 |
| 2.3387E-11 | -0.5888926 | 0.127 | 0.388 | 4.9931E-07 | TEC_prg3 | MYH9 |
| 1.3314E-12 | -0.5916165 | 0.461 | 0.67 | 2.8426E-08 | TEC_prg3 | SPINT2 |
| 8.8008E-16 | -0.5918536 | 0.036 | 0.343 | 1.879E-11 | TEC_prg3 | SSR3 |
| 5.2541E-14 | -0.5925806 | 0.267 | 0.555 | 1.1217E-09 | TEC_prg3 | CANX |
| 2.0568E-12 | -0.5940256 | 0.315 | 0.565 | 4.3913E-08 | TEC_prg3 | GUK1 |
| 3.6666E-14 | -0.5940773 | 0.061 | 0.353 | 7.8282E-10 | TEC_prg3 | EIF5A |
| 1.4764E-17 | -0.5947927 | 0.042 | 0.378 | 3.152E-13 | TEC_prg3 | HEBP2 |
| 5.7451E-15 | -0.5957388 | 0.121 | 0.43 | 1.2266E-10 | TEC_prg3 | ATP6V0B |
| 8.7983E-17 | -0.5961661 | 0.067 | 0.407 | 1.8784E-12 | TEC_prg3 | LAMTOR5 |
| 1.0252E-10 | -0.5965928 | 0.018 | 0.235 | 2.1887E-06 | TEC_prg3 | TGM2 |
| 0.00158934 | -0.596678 | 0.873 | 0.814 | 1 | TEC_prg3 | IGFBP7 |
| 1.1366E-14 | -0.5971325 | 0.079 | 0.379 | 2.4266E-10 | TEC_prg3 | SEC61G |
| 1.123E-10 | -0.5980173 | 0.448 | 0.633 | 2.3976E-06 | TEC_prg3 | ANXA5 |
| 3.1353E-15 | -0.598982 | 0.115 | 0.429 | 6.6939E-11 | TEC_prg3 | TMEM59 |
| 4.6325E-13 | -0.6001719 | 0.667 | 0.783 | 9.8904E-09 | TEC_prg3 | COX4I1 |
| 2.2531E-14 | -0.6013017 | 0.067 | 0.353 | 4.8103E-10 | TEC_prg3 | ANXA11 |
| 4.202E-16 | -0.6037852 | 0.061 | 0.386 | 8.9712E-12 | TEC_prg3 | DYNLT1 |
| 3.6949E-12 | -0.6043288 | 0.073 | 0.336 | 7.8885E-08 | TEC_prg3 | CHCHD10 |
| 2.337E-16 | -0.6121901 | 0.073 | 0.4 | 4.9895E-12 | TEC_prg3 | DAD1 |
| 2.7555E-15 | -0.6145195 | 0.036 | 0.336 | 5.8831E-11 | TEC_prg3 | SLC25A5 |
| 3.9205E-15 | -0.6155017 | 0.115 | 0.433 | 8.3703E-11 | TEC_prg3 | BSG |
| 8.0367E-10 | -0.6213076 | 0.279 | 0.489 | 1.7158E-05 | TEC_prg3 | LDHA |
| 2.7075E-14 | -0.6220393 | 0.879 | 0.907 | 5.7804E-10 | TEC_prg3 | RPS16 |
| 1.7985E-13 | -0.623919 | 0.109 | 0.401 | 3.8397E-09 | TEC_prg3 | RHOC |
| 1.4087E-15 | -0.6249684 | 0.291 | 0.584 | 3.0075E-11 | TEC_prg3 | ATP6V0E1 |
| 1.752E-10 | -0.6250879 | 0.497 | 0.662 | 3.7405E-06 | TEC_prg3 | CALR |
| 2.3855E-14 | -0.6286897 | 0.758 | 0.855 | 5.0931E-10 | TEC_prg3 | SRP14 |
| 2.9942E-10 | -0.628719 | 0.085 | 0.318 | 6.3926E-06 | TEC_prg3 | MAL |
| 3.4024E-13 | -0.630716 | 0.915 | 0.909 | 7.2642E-09 | TEC_prg3 | RPL24 |
| 8.5105E-17 | -0.6311832 | 0.564 | 0.753 | 1.817E-12 | TEC_prg3 | BTF3 |
| 2.0662E-11 | -0.6322664 | 0.958 | 0.951 | 4.4113E-07 | TEC_prg3 | RPS8 |
| 4.3978E-16 | -0.635555 | 0.382 | 0.652 | 9.3894E-12 | TEC_prg3 | UQCRH |
| 7.8977E-11 | -0.6389273 | 0.612 | 0.73 | 1.6862E-06 | TEC_prg3 | ENO1 |
| 1.5834E-11 | -0.6397114 | 0.812 | 0.866 | 3.3805E-07 | TEC_prg3 | UBA52 |
| 2.2608E-16 | -0.6456928 | 0.158 | 0.474 | 4.8269E-12 | TEC_prg3 | PRDX6 |

| | | | | | | |
|---|---|---|---|---|---|---|
| 7.5404E-07 | -0.6507823 | 0.97 | 0.91 | 0.01609874 | TEC_prg3 | MT-CO2 |
| 1.3751E-13 | -0.6531531 | 0.37 | 0.618 | 2.9359E-09 | TEC_prg3 | CD81 |
| 4.9822E-12 | -0.6537589 | 0.333 | 0.569 | 1.0637E-07 | TEC_prg3 | PPIB |
| 2.2181E-17 | -0.6564833 | 0.903 | 0.94 | 4.7356E-13 | TEC_prg3 | RPL37A |
| 1.9658E-12 | -0.657044 | 0.921 | 0.922 | 4.197E-08 | TEC_prg3 | RPL14 |
| 2.4282E-05 | -0.6583257 | 0.048 | 0.173 | 0.51842918 | TEC_prg3 | S100A4 |
| 2.0561E-06 | -0.6586776 | 0.158 | 0.329 | 0.04389734 | TEC_prg3 | IL32 |
| 4.6915E-14 | -0.6588061 | 0.145 | 0.432 | 1.0016E-09 | TEC_prg3 | PDIA3 |
| 3.6582E-15 | -0.6622098 | 0.248 | 0.537 | 7.8103E-11 | TEC_prg3 | SEC61B |
| 7.8617E-12 | -0.6626662 | 0.024 | 0.263 | 1.6785E-07 | TEC_prg3 | LAMB3 |
| 9.1344E-05 | -0.6634847 | 0.964 | 0.889 | 1 | TEC_prg3 | MT-ND2 |
| 2.359E-15 | -0.6666342 | 0.036 | 0.331 | 5.0364E-11 | TEC_prg3 | TSPO |
| 4.8518E-13 | -0.6687284 | 0.752 | 0.823 | 1.0359E-08 | TEC_prg3 | RPS21 |
| 2.9835E-14 | -0.6718746 | 0.079 | 0.378 | 6.3697E-10 | TEC_prg3 | LDHB |
| 2.5107E-14 | -0.6751619 | 0.939 | 0.947 | 5.3604E-10 | TEC_prg3 | RPL13A |
| 2.2062E-14 | -0.6787589 | 0.061 | 0.363 | 4.7103E-10 | TEC_prg3 | CD44 |
| 1.4666E-11 | -0.6794015 | 0.333 | 0.568 | 3.1312E-07 | TEC_prg3 | MAL2 |
| 4.8954E-16 | -0.6800929 | 0.848 | 0.903 | 1.0452E-11 | TEC_prg3 | RPS25 |
| 9.8788E-17 | -0.6833682 | 0.697 | 0.828 | 2.1091E-12 | TEC_prg3 | TMBIM6 |
| 6.5927E-08 | -0.6838581 | 0.618 | 0.705 | 0.00140754 | TEC_prg3 | ITGB1 |
| 5.6038E-15 | -0.6845956 | 0.273 | 0.55 | 1.1964E-10 | TEC_prg3 | BRI3 |
| 2.7809E-15 | -0.6870745 | 0.321 | 0.586 | 5.9372E-11 | TEC_prg3 | ARPC3 |
| 3.2458E-13 | -0.689404 | 0.073 | 0.348 | 6.9298E-09 | TEC_prg3 | S100A16 |
| 4.4942E-14 | -0.6911565 | 0.194 | 0.472 | 9.595E-10 | TEC_prg3 | PDIA6 |
| 1.0612E-15 | -0.693916 | 0.303 | 0.583 | 2.2657E-11 | TEC_prg3 | PRDX5 |
| 0.00010645 | -0.6985418 | 0.661 | 0.663 | 1 | TEC_prg3 | TACSTD2 |
| 3.6806E-13 | -0.7009294 | 0.503 | 0.682 | 7.8581E-09 | TEC_prg3 | EPCAM |
| 2.287E-08 | -0.7015672 | 0.945 | 0.907 | 0.00048827 | TEC_prg3 | MYL6 |
| 3.2968E-16 | -0.7018151 | 0.861 | 0.898 | 7.0387E-12 | TEC_prg3 | RPS15A |
| 2.5951E-09 | -0.7084653 | 0.085 | 0.298 | 5.5406E-05 | TEC_prg3 | ITGB8 |
| 0.00156016 | -0.708535 | 0.139 | 0.244 | 1 | TEC_prg3 | TFPI2 |
| 1.3846E-19 | -0.7089737 | 0.097 | 0.458 | 2.9561E-15 | TEC_prg3 | LAPTM4A |
| 7.5746E-05 | -0.7131065 | 0.012 | 0.107 | 1 | TEC_prg3 | FXYD4 |
| 4.3741E-13 | -0.7160177 | 0.303 | 0.547 | 9.3388E-09 | TEC_prg3 | CD59 |
| 4.2838E-14 | -0.719662 | 0.709 | 0.805 | 9.1458E-10 | TEC_prg3 | CFL1 |
| 5.6309E-17 | -0.7201109 | 0.885 | 0.923 | 1.2022E-12 | TEC_prg3 | RPL30 |
| 1.0634E-16 | -0.7240756 | 0.788 | 0.873 | 2.2703E-12 | TEC_prg3 | RACK1 |
| 2.3E-15 | -0.7243371 | 0.806 | 0.867 | 4.9104E-11 | TEC_prg3 | RPL23 |
| 6.8917E-16 | -0.7261702 | 0.879 | 0.901 | 1.4714E-11 | TEC_prg3 | RPL27A |
| 2.6983E-19 | -0.7270527 | 0.018 | 0.364 | 5.7608E-15 | TEC_prg3 | TALDO1 |
| 1.4998E-16 | -0.7284262 | 0.109 | 0.429 | 3.2022E-12 | TEC_prg3 | SSR4 |
| 8.9832E-14 | -0.7304444 | 0.582 | 0.733 | 1.9179E-09 | TEC_prg3 | NDUFA4 |
| 4.0543E-16 | -0.7317647 | 0.861 | 0.893 | 8.6559E-12 | TEC_prg3 | RPS11 |
| 6.5914E-10 | -0.7332233 | 0.206 | 0.424 | 1.4073E-05 | TEC_prg3 | TPM4 |
| 8.2839E-06 | -0.7369447 | 0.018 | 0.137 | 0.17686127 | TEC_prg3 | SERPINA1 |
| 7.0868E-15 | -0.7399314 | 0.939 | 0.943 | 1.513E-10 | TEC_prg3 | RPL8 |
| 1.3587E-11 | -0.7402413 | 0.018 | 0.248 | 2.9008E-07 | TEC_prg3 | PDZK1IP1 |
| 1.1714E-14 | -0.742807 | 0.145 | 0.442 | 2.5009E-10 | TEC_prg3 | PPP1R14B |
| 3.5086E-13 | -0.7453714 | 0.891 | 0.916 | 7.4908E-09 | TEC_prg3 | RPLP0 |
| 3.6286E-13 | -0.7454324 | 0.927 | 0.93 | 7.7471E-09 | TEC_prg3 | RPL12 |

| | | | | | | |
|---|---|---|---|---|---|---|
| 2.3309E-06 | -0.7484929 | 0.091 | 0.249 | 0.04976447 | TEC_prg3 | MT1X |
| 3.8083E-13 | -0.7502712 | 0.933 | 0.928 | 8.1308E-09 | TEC_prg3 | RPL21 |
| 8.4473E-16 | -0.7513581 | 0.921 | 0.933 | 1.8035E-11 | TEC_prg3 | RPS9 |
| 1.7321E-12 | -0.7521737 | 0.036 | 0.289 | 3.6981E-08 | TEC_prg3 | PLAUR |
| 9.9491E-18 | -0.7565484 | 0.891 | 0.935 | 2.1241E-13 | TEC_prg3 | RPL37 |
| 3.0255E-14 | -0.7607704 | 0.297 | 0.554 | 6.4594E-10 | TEC_prg3 | CD9 |
| 8.6146E-18 | -0.7641723 | 0.909 | 0.932 | 1.8392E-13 | TEC_prg3 | RPL18 |
| 8.1419E-11 | -0.7648101 | 0.758 | 0.801 | 1.7383E-06 | TEC_prg3 | CD24 |
| 4.213E-20 | -0.7700939 | 0.818 | 0.891 | 8.9948E-16 | TEC_prg3 | NACA |
| 1.4284E-12 | -0.7711985 | 0.594 | 0.717 | 3.0496E-08 | TEC_prg3 | TXN |
| 3.2777E-08 | -0.7756491 | 0.121 | 0.322 | 0.0006998 | TEC_prg3 | ITGA2 |
| 0.00974895 | -0.7759316 | 0.061 | 0.128 | 1 | TEC_prg3 | GPX3 |
| 4.3779E-14 | -0.7815824 | 0.848 | 0.87 | 9.3467E-10 | TEC_prg3 | RPS5 |
| 4.1613E-18 | -0.7838946 | 0.861 | 0.899 | 8.8844E-14 | TEC_prg3 | RPL5 |
| 4.2276E-13 | -0.7874197 | 0.927 | 0.933 | 9.0259E-09 | TEC_prg3 | RPL32 |
| 9.9328E-13 | -0.7902891 | 0.333 | 0.561 | 2.1206E-08 | TEC_prg3 | HMGA1 |
| 3.1701E-13 | -0.7980173 | 0.83 | 0.853 | 6.7682E-09 | TEC_prg3 | MYL12B |
| 2.0519E-17 | -0.8028985 | 0.982 | 0.973 | 4.3809E-13 | TEC_prg3 | RPL13 |
| 9.8546E-15 | -0.8129375 | 0.042 | 0.329 | 2.104E-10 | TEC_prg3 | CYBA |
| 5.7993E-18 | -0.8171629 | 0.921 | 0.934 | 1.2382E-13 | TEC_prg3 | RPS27A |
| 9.9376E-11 | -0.8190514 | 0.248 | 0.47 | 2.1217E-06 | TEC_prg3 | IFITM3 |
| 2.5908E-13 | -0.8230835 | 0.594 | 0.733 | 5.5313E-09 | TEC_prg3 | PFN1 |
| 1.4631E-13 | -0.8248609 | 0.594 | 0.725 | 3.1237E-09 | TEC_prg3 | CTSD |
| 6.1573E-18 | -0.8276163 | 0.842 | 0.915 | 1.3146E-13 | TEC_prg3 | HSP90AB1 |
| 1.1515E-10 | -0.830605 | 0.97 | 0.949 | 2.4584E-06 | TEC_prg3 | B2M |
| 1.03E-20 | -0.834566 | 0.612 | 0.789 | 2.1991E-16 | TEC_prg3 | CHCHD2 |
| 1.6568E-10 | -0.8355686 | 0.176 | 0.416 | 3.5372E-06 | TEC_prg3 | PHLDA2 |
| 1.5094E-16 | -0.8398035 | 0.945 | 0.942 | 3.2226E-12 | TEC_prg3 | RPS24 |
| 2.1252E-05 | -0.8428717 | 0.315 | 0.444 | 0.45372547 | TEC_prg3 | UPP1 |
| 1.6423E-18 | -0.8476289 | 0.855 | 0.913 | 3.5063E-14 | TEC_prg3 | RPL29 |
| 2.6724E-17 | -0.854942 | 0.909 | 0.939 | 5.7056E-13 | TEC_prg3 | RPS28 |
| 6.2672E-16 | -0.8552634 | 0.897 | 0.929 | 1.3381E-11 | TEC_prg3 | RPL18A |
| 2.086E-12 | -0.8559385 | 0.842 | 0.878 | 4.4535E-08 | TEC_prg3 | ITM2B |
| 3.9799E-17 | -0.8646595 | 0.873 | 0.896 | 8.4971E-13 | TEC_prg3 | RPS13 |
| 1.3149E-18 | -0.8699398 | 0.933 | 0.941 | 2.8072E-14 | TEC_prg3 | RPL19 |
| 1.7511E-19 | -0.8772199 | 0.897 | 0.905 | 3.7387E-15 | TEC_prg3 | RPS3 |
| 9.6048E-15 | -0.8773205 | 0.824 | 0.843 | 2.0506E-10 | TEC_prg3 | RPSA |
| 0.00014962 | -0.8830493 | 0.655 | 0.649 | 1 | TEC_prg3 | KRT7 |
| 1.5432E-05 | -0.8931271 | 0.558 | 0.624 | 0.3294836 | TEC_prg3 | VIM |
| 5.2188E-07 | -0.9027834 | 0.842 | 0.798 | 0.01114212 | TEC_prg3 | KRT18 |
| 1.309E-20 | -0.9040254 | 0.879 | 0.92 | 2.7948E-16 | TEC_prg3 | RPL9 |
| 3.1301E-21 | -0.9053257 | 0.794 | 0.875 | 6.6829E-17 | TEC_prg3 | RPL7 |
| 0.00097871 | -0.9077252 | 0.703 | 0.683 | 1 | TEC_prg3 | HSPA5 |
| 5.3376E-09 | -0.9139999 | 0.139 | 0.361 | 0.00011396 | TEC_prg3 | HSPA1A |
| 0.00012971 | -0.914205 | 0.758 | 0.727 | 1 | TEC_prg3 | KRT8 |
| 2.1651E-19 | -0.9189288 | 0.891 | 0.909 | 4.6225E-15 | TEC_prg3 | RPL26 |
| 1.6564E-11 | -0.9265963 | 0.012 | 0.239 | 3.5363E-07 | TEC_prg3 | CST6 |
| 1.2751E-22 | -0.9287282 | 0.836 | 0.921 | 2.7223E-18 | TEC_prg3 | RPL34 |
| 1.5066E-22 | -0.92888 | 0.891 | 0.947 | 3.2165E-18 | TEC_prg3 | RPL11 |
| 1.7925E-20 | -0.9308609 | 0.867 | 0.906 | 3.827E-16 | TEC_prg3 | RPS7 |

| | | | | | | |
|---|---|---|---|---|---|---|
| 2.2656E-21 | -0.9321432 | 0.891 | 0.931 | 4.837E-17 | TEC_prg3 | RPL7A |
| 1.054E-19 | -0.9357868 | 0.952 | 0.961 | 2.2504E-15 | TEC_prg3 | RPL28 |
| 3.0358E-22 | -0.942778 | 0.873 | 0.924 | 6.4814E-18 | TEC_prg3 | RPLP2 |
| 1.2024E-20 | -0.9427916 | 0.939 | 0.939 | 2.5672E-16 | TEC_prg3 | RPL15 |
| 3.7225E-19 | -0.9469486 | 0.758 | 0.841 | 7.9476E-15 | TEC_prg3 | CD63 |
| 1.7597E-12 | -0.9546027 | 0.17 | 0.421 | 3.7569E-08 | TEC_prg3 | CCND1 |
| 1.8371E-09 | -0.9582 | 0.109 | 0.329 | 3.9222E-05 | TEC_prg3 | MT1E |
| 5.0513E-19 | -0.9614791 | 0.921 | 0.932 | 1.0785E-14 | TEC_prg3 | RPS6 |
| 2.0551E-17 | -0.9674266 | 0.958 | 0.956 | 4.3877E-13 | TEC_prg3 | RPS2 |
| 2.1701E-19 | -0.9780964 | 0.842 | 0.883 | 4.6331E-15 | TEC_prg3 | RPL39 |
| 8.3721E-21 | -0.9840133 | 0.927 | 0.921 | 1.7874E-16 | TEC_prg3 | RPS3A |
| 7.8141E-10 | -1.0001989 | 0.976 | 0.954 | 1.6683E-05 | TEC_prg3 | S100A6 |
| 8.1715E-23 | -1.0092769 | 0.915 | 0.938 | 1.7446E-18 | TEC_prg3 | RPS23 |
| 4.4109E-26 | -1.0110827 | 0.885 | 0.934 | 9.4173E-22 | TEC_prg3 | RPL3 |
| 5.207E-21 | -1.0130095 | 0.958 | 0.961 | 1.1117E-16 | TEC_prg3 | RPS18 |
| 5.3863E-13 | -1.0154571 | 0.048 | 0.311 | 1.15E-08 | TEC_prg3 | PLAU |
| 1.338E-09 | -1.0193506 | 0.067 | 0.28 | 2.8567E-05 | TEC_prg3 | LGALS1 |
| 1.3257E-10 | -1.0341144 | 0.164 | 0.395 | 2.8303E-06 | TEC_prg3 | TXNRD1 |
| 6.083E-05 | -1.0415409 | 0.564 | 0.592 | 1 | TEC_prg3 | KRT19 |
| 1.0099E-14 | -1.0827851 | 0.042 | 0.334 | 2.1562E-10 | TEC_prg3 | TIMP1 |
| 1.7496E-10 | -1.0847349 | 0.879 | 0.87 | 3.7355E-06 | TEC_prg3 | S100A10 |
| 1.6273E-25 | -1.085722 | 0.958 | 0.929 | 3.4743E-21 | TEC_prg3 | RPS14 |
| 1.1137E-08 | -1.0947229 | 0.297 | 0.473 | 0.00023778 | TEC_prg3 | C12orf75 |
| 7.3184E-11 | -1.0977343 | 0.097 | 0.334 | 1.5625E-06 | TEC_prg3 | LAMC2 |
| 0.00243097 | -1.1017047 | 0.133 | 0.227 | 1 | TEC_prg3 | DEFB1 |
| 1.1299E-09 | -1.1087662 | 0.6 | 0.673 | 2.4123E-05 | TEC_prg3 | SH3BGRL3 |
| 1.3393E-21 | -1.1122422 | 0.752 | 0.838 | 2.8593E-17 | TEC_prg3 | PRDX1 |
| 3.6572E-22 | -1.1185561 | 0.982 | 0.972 | 7.8081E-18 | TEC_prg3 | RPLP1 |
| 2.1319E-22 | -1.1236758 | 0.812 | 0.866 | 4.5515E-18 | TEC_prg3 | RPS4X |
| 8.4936E-26 | -1.1273457 | 0.958 | 0.967 | 1.8134E-21 | TEC_prg3 | RPL41 |
| 2.2293E-15 | -1.1372436 | 0.218 | 0.488 | 4.7596E-11 | TEC_prg3 | TSPAN1 |
| 0.00014435 | -1.1512155 | 0.648 | 0.645 | 1 | TEC_prg3 | WFDC2 |
| 3.7764E-18 | -1.1600926 | 0.939 | 0.924 | 8.0626E-14 | TEC_prg3 | TMSB10 |
| 8.3452E-06 | -1.1761364 | 0.061 | 0.199 | 0.17816969 | TEC_prg3 | MT1G |
| 1.1382E-12 | -1.2255762 | 0.8 | 0.831 | 2.43E-08 | TEC_prg3 | ANXA2 |
| 7.1985E-25 | -1.2402138 | 0.939 | 0.952 | 1.5369E-20 | TEC_prg3 | RPS12 |
| 3.2755E-20 | -1.2475702 | 0.873 | 0.893 | 6.9932E-16 | TEC_prg3 | GAPDH |
| 2.2867E-32 | -1.2499973 | 0.988 | 0.972 | 4.882E-28 | TEC_prg3 | TPT1 |
| 2.0309E-16 | -1.2973762 | 0.915 | 0.919 | 4.336E-12 | TEC_prg3 | S100A11 |
| 5.5069E-30 | -1.3164534 | 0.855 | 0.927 | 1.1757E-25 | TEC_prg3 | ACTG1 |
| 2.3643E-05 | -1.3492059 | 0.776 | 0.71 | 0.50478021 | TEC_prg3 | MT2A |
| 9.5695E-11 | -1.3519917 | 0.036 | 0.258 | 2.0431E-06 | TEC_prg3 | LCN2 |
| 2.4646E-41 | -1.3849946 | 0.988 | 0.987 | 5.2619E-37 | TEC_prg3 | EEF1A1 |
| 4.2306E-28 | -1.4360713 | 0.909 | 0.924 | 9.0323E-24 | TEC_prg3 | RPL10 |
| 8.8935E-13 | -1.4741461 | 0.903 | 0.868 | 1.8988E-08 | TEC_prg3 | TMSB4X |
| 1.1206E-05 | -1.5383722 | 0.273 | 0.393 | 0.23924731 | TEC_prg3 | SLPI |
| 5.8507E-32 | -1.7156857 | 0.885 | 0.922 | 1.2491E-27 | TEC_prg3 | ACTB |
| 6.6334E-21 | -1.8708681 | 0.939 | 0.962 | 1.4162E-16 | TEC_prg3 | FTL |
| 2.52E-34 | -1.9579084 | 0.982 | 0.978 | 5.3801E-30 | TEC_prg3 | FTH1 |
| 4.244E-05 | -1.9693525 | 0.6 | 0.6 | 0.90610154 | TEC_prg3 | SPP1 |

| | | | | | | |
|---|---|---|---|---|---|---|
| 1.118E-56 | 2.13032403 | 0.719 | 0.557 | 2.3869E-52 | TEC_prg4 | GDF15 |
| 4.285E-119 | 1.87877442 | 0.731 | 0.372 | 9.15E-115 | TEC_prg4 | NFKBIA |
| 3.9865E-75 | 1.8063706 | 0.718 | 0.424 | 8.5112E-71 | TEC_prg4 | IER3 |
| 8.3178E-84 | 1.79970307 | 0.92 | 0.855 | 1.7758E-79 | TEC_prg4 | H3F3B |
| 3.7778E-52 | 1.48214447 | 0.822 | 0.623 | 8.0655E-48 | TEC_prg4 | JUN |
| 1.464E-70 | 1.42383732 | 0.817 | 0.593 | 3.1256E-66 | TEC_prg4 | GADD45B |
| 8.805E-68 | 1.40076653 | 0.747 | 0.497 | 1.8799E-63 | TEC_prg4 | CDKN1A |
| 1.553E-127 | 1.3520806 | 0.662 | 0.301 | 3.316E-123 | TEC_prg4 | CCNL1 |
| 8.2404E-39 | 1.29076621 | 0.537 | 0.339 | 1.7593E-34 | TEC_prg4 | TXNIP |
| 8.6274E-82 | 1.28974487 | 0.808 | 0.595 | 1.842E-77 | TEC_prg4 | GADD45A |
| 1.547E-144 | 1.25825309 | 0.68 | 0.302 | 3.302E-140 | TEC_prg4 | RSRC2 |
| 4.353E-116 | 1.20806148 | 0.574 | 0.235 | 9.294E-112 | TEC_prg4 | PTP4A1 |
| 3.0233E-74 | 1.15775907 | 0.495 | 0.225 | 6.4547E-70 | TEC_prg4 | MDM2 |
| 9.597E-99 | 1.0999743 | 0.311 | 0.081 | 2.049E-94 | TEC_prg4 | NR1D1 |
| 3.4147E-17 | 1.08963604 | 0.102 | 0.038 | 7.2904E-13 | TEC_prg4 | ANKRD1 |
| 5.89E-80 | 1.08928848 | 0.54 | 0.236 | 1.2575E-75 | TEC_prg4 | ID2 |
| 3.2587E-95 | 1.0874627 | 0.687 | 0.378 | 6.9573E-91 | TEC_prg4 | SRSF3 |
| 7.452E-116 | 1.07419115 | 0.838 | 0.535 | 1.591E-111 | TEC_prg4 | EIF5 |
| 1.1506E-50 | 1.07020589 | 0.22 | 0.069 | 2.4566E-46 | TEC_prg4 | CXCL2 |
| 2.3727E-65 | 1.0609259 | 0.812 | 0.613 | 5.0658E-61 | TEC_prg4 | PPP1R15A |
| 2.4919E-60 | 1.05898586 | 0.278 | 0.09 | 5.3202E-56 | TEC_prg4 | CXCL3 |
| 1.3678E-77 | 1.0345673 | 0.56 | 0.261 | 2.9202E-73 | TEC_prg4 | PPP1R10 |
| 7.9061E-87 | 1.0106719 | 0.466 | 0.187 | 1.6879E-82 | TEC_prg4 | ARRDC3 |
| 2.406E-116 | 1.00486369 | 0.738 | 0.366 | 5.136E-112 | TEC_prg4 | BRD2 |
| 4.3351E-50 | 0.99878564 | 0.614 | 0.373 | 9.2554E-46 | TEC_prg4 | JUNB |
| 1.2941E-55 | 0.99407435 | 0.425 | 0.194 | 2.7628E-51 | TEC_prg4 | RRAD |
| 4.1033E-66 | 0.98483521 | 0.54 | 0.266 | 8.7606E-62 | TEC_prg4 | BTG2 |
| 1.4375E-33 | 0.93342882 | 0.357 | 0.183 | 3.069E-29 | TEC_prg4 | GADD45G |
| 3.1877E-53 | 0.93007165 | 0.616 | 0.402 | 6.8058E-49 | TEC_prg4 | NORAD |
| 8.7758E-43 | 0.91677591 | 0.75 | 0.554 | 1.8736E-38 | TEC_prg4 | IER2 |
| 1.799E-111 | 0.89581789 | 0.544 | 0.21 | 3.84E-107 | TEC_prg4 | TRA2B |
| 2.0417E-73 | 0.88110866 | 0.799 | 0.58 | 4.359E-69 | TEC_prg4 | RBM39 |
| 2.2699E-55 | 0.86480517 | 0.546 | 0.291 | 4.8461E-51 | TEC_prg4 | PMAIP1 |
| 1.0589E-26 | 0.8489096 | 0.242 | 0.11 | 2.2608E-22 | TEC_prg4 | NEFM |
| 2.9622E-74 | 0.84499335 | 0.544 | 0.276 | 6.3243E-70 | TEC_prg4 | HNRNPH3 |
| 1.217E-112 | 0.84066621 | 0.489 | 0.173 | 2.599E-108 | TEC_prg4 | PRPF38B |
| 7.5664E-69 | 0.81075702 | 0.75 | 0.505 | 1.6154E-64 | TEC_prg4 | SON |
| 1.5951E-81 | 0.80465833 | 0.547 | 0.258 | 3.4054E-77 | TEC_prg4 | TAF7 |
| 9.9956E-73 | 0.80362905 | 0.424 | 0.173 | 2.1341E-68 | TEC_prg4 | TNFRSF10B |
| 8.3551E-95 | 0.79836006 | 0.489 | 0.19 | 1.7838E-90 | TEC_prg4 | EPC1 |
| 3.9437E-20 | 0.79287333 | 0.448 | 0.281 | 8.4198E-16 | TEC_prg4 | FOS |
| 2.4852E-43 | 0.76431071 | 0.594 | 0.356 | 5.3059E-39 | TEC_prg4 | DDIT3 |
| 4.3535E-33 | 0.75889388 | 0.946 | 0.91 | 9.2947E-29 | TEC_prg4 | UBC |
| 1.9592E-61 | 0.75718949 | 0.654 | 0.387 | 4.1829E-57 | TEC_prg4 | FUS |
| 2.6032E-88 | 0.75131473 | 0.552 | 0.247 | 5.5579E-84 | TEC_prg4 | CHD2 |
| 4.8697E-78 | 0.74592226 | 0.374 | 0.132 | 1.0397E-73 | TEC_prg4 | ILF3-DT |
| 9.36E-114 | 0.74496161 | 0.511 | 0.176 | 1.998E-109 | TEC_prg4 | TOB1 |
| 1.7003E-61 | 0.74250723 | 0.782 | 0.599 | 3.6301E-57 | TEC_prg4 | EIF4A2 |
| 9.5666E-27 | 0.74165784 | 0.336 | 0.178 | 2.0425E-22 | TEC_prg4 | BIRC3 |
| 1.468E-31 | 0.73165365 | 0.544 | 0.364 | 3.1342E-27 | TEC_prg4 | ATF3 |

| | | | | | | |
|---|---|---|---|---|---|---|
| 1.1223E-76 | 0.72924447 | 0.48 | 0.203 | 2.3962E-72 | TEC_prg4 | SERTAD1 |
| 1.4562E-38 | 0.72712913 | 0.397 | 0.204 | 3.1091E-34 | TEC_prg4 | ZFP36 |
| 2.1016E-48 | 0.724168 | 0.326 | 0.138 | 4.487E-44 | TEC_prg4 | HBEGF |
| 1.8251E-46 | 0.72268986 | 0.693 | 0.479 | 3.8967E-42 | TEC_prg4 | DNAJA1 |
| 6.7699E-19 | 0.71606076 | 0.396 | 0.259 | 1.4454E-14 | TEC_prg4 | KCNQ1OT1 |
| 3.3989E-63 | 0.70819385 | 0.837 | 0.597 | 7.2567E-59 | TEC_prg4 | BTG1 |
| 3.8469E-96 | 0.70457549 | 0.496 | 0.186 | 8.2132E-92 | TEC_prg4 | CLK1 |
| 7.964E-50 | 0.68943482 | 0.55 | 0.313 | 1.7003E-45 | TEC_prg4 | HNRNPH1 |
| 4.5592E-38 | 0.68587968 | 0.491 | 0.286 | 9.734E-34 | TEC_prg4 | HSPH1 |
| 2.757E-68 | 0.68487608 | 0.469 | 0.214 | 5.8861E-64 | TEC_prg4 | TERF2IP |
| 5.063E-48 | 0.6840657 | 0.753 | 0.564 | 1.081E-43 | TEC_prg4 | SLC3A2 |
| 5.2538E-56 | 0.6766768 | 0.579 | 0.319 | 1.1217E-51 | TEC_prg4 | CPEB4 |
| 5.1494E-31 | 0.67471879 | 0.473 | 0.266 | 1.0994E-26 | TEC_prg4 | HES1 |
| 1.2417E-40 | 0.67192974 | 0.725 | 0.551 | 2.651E-36 | TEC_prg4 | NCL |
| 1.6099E-66 | 0.66191904 | 0.448 | 0.199 | 3.4372E-62 | TEC_prg4 | PUM1 |
| 5.915E-103 | 0.65685136 | 0.488 | 0.169 | 1.2628E-98 | TEC_prg4 | PCF11 |
| 1.9482E-57 | 0.65625719 | 0.539 | 0.287 | 4.1595E-53 | TEC_prg4 | GPBP1 |
| 4.9072E-29 | 0.6549325 | 0.579 | 0.413 | 1.0477E-24 | TEC_prg4 | TUBB4B |
| 4.5578E-54 | 0.65466072 | 0.434 | 0.208 | 9.7309E-50 | TEC_prg4 | WTAP |
| 2.4433E-82 | 0.65383774 | 0.447 | 0.173 | 5.2165E-78 | TEC_prg4 | PPP4R3A |
| 4.482E-39 | 0.65242998 | 0.378 | 0.194 | 9.569E-35 | TEC_prg4 | PIM3 |
| 1.042E-134 | 0.6481815 | 0.394 | 0.097 | 2.225E-130 | TEC_prg4 | TNFRSF10D |
| 2.2999E-45 | 0.64682069 | 0.578 | 0.356 | 4.9103E-41 | TEC_prg4 | GOLGA4 |
| 1.0526E-53 | 0.64257784 | 0.447 | 0.209 | 2.2473E-49 | TEC_prg4 | ADM |
| 1.6059E-38 | 0.63885524 | 0.702 | 0.56 | 3.4286E-34 | TEC_prg4 | HNRNPC |
| 3.609E-59 | 0.63760414 | 0.459 | 0.218 | 7.7053E-55 | TEC_prg4 | KMT2E |
| 2.6132E-44 | 0.63688612 | 0.559 | 0.337 | 5.5791E-40 | TEC_prg4 | DDX24 |
| 5.7592E-44 | 0.63026767 | 0.86 | 0.698 | 1.2296E-39 | TEC_prg4 | DDX5 |
| 8.299E-28 | 0.63002003 | 0.229 | 0.102 | 1.7718E-23 | TEC_prg4 | PEG10 |
| 5.195E-40 | 0.6244154 | 0.606 | 0.361 | 1.1091E-35 | TEC_prg4 | SLC38A2 |
| 1.6239E-95 | 0.62324841 | 0.374 | 0.114 | 3.467E-91 | TEC_prg4 | MYLIP |
| 1.763E-48 | 0.62048911 | 0.702 | 0.488 | 3.764E-44 | TEC_prg4 | HNRNPDL |
| 4.6126E-67 | 0.61287675 | 0.319 | 0.109 | 9.8479E-63 | TEC_prg4 | FBXW7 |
| 3.6123E-63 | 0.6069458 | 0.534 | 0.259 | 7.7123E-59 | TEC_prg4 | IFRD1 |
| 3.8124E-05 | 0.60417386 | 0.45 | 0.411 | 0.81395781 | TEC_prg4 | PHLDA2 |
| 4.3268E-37 | 0.60052586 | 0.603 | 0.403 | 9.2377E-33 | TEC_prg4 | GOLGB1 |
| 4.6914E-87 | 0.60021278 | 0.332 | 0.098 | 1.0016E-82 | TEC_prg4 | DDX3Y |
| 6.2823E-53 | 0.59811295 | 0.46 | 0.219 | 1.3413E-48 | TEC_prg4 | RBBP6 |
| 4.7163E-61 | 0.59445767 | 0.656 | 0.378 | 1.0069E-56 | TEC_prg4 | TAF1D |
| 1.0181E-46 | 0.59130771 | 0.492 | 0.274 | 2.1735E-42 | TEC_prg4 | SRRM1 |
| 4.7249E-06 | 0.59067694 | 0.348 | 0.282 | 0.10087745 | TEC_prg4 | CEBPD |
| 2.8212E-17 | 0.58269332 | 0.21 | 0.109 | 6.0233E-13 | TEC_prg4 | NUAK2 |
| 2.5709E-05 | 0.57491897 | 0.403 | 0.356 | 0.54887995 | TEC_prg4 | KLF6 |
| 5.9283E-37 | 0.57357758 | 0.562 | 0.355 | 1.2657E-32 | TEC_prg4 | PNN |
| 1.1535E-49 | 0.57256285 | 0.246 | 0.083 | 2.4627E-45 | TEC_prg4 | GEM |
| 7.1281E-18 | 0.57053322 | 0.654 | 0.564 | 1.5218E-13 | TEC_prg4 | RPS27L |
| 1.1809E-22 | 0.56921421 | 0.588 | 0.436 | 2.5211E-18 | TEC_prg4 | ZFP36L1 |
| 2.0667E-51 | 0.56890148 | 0.447 | 0.218 | 4.4124E-47 | TEC_prg4 | RPL22L1 |
| 4.4817E-51 | 0.56687824 | 0.524 | 0.272 | 9.5684E-47 | TEC_prg4 | NUFIP2 |
| 5.3878E-16 | 0.56215819 | 0.434 | 0.295 | 1.1503E-11 | TEC_prg4 | DNAJB1 |

| | | | | | | |
|---|---|---|---|---|---|---|
| 2.1344E-49 | 0.5612569 | 0.396 | 0.181 | 4.5569E-45 | TEC_prg4 | SESN2 |
| 4.0163E-09 | 0.55798018 | 0.948 | 0.925 | 8.5748E-05 | TEC_prg4 | MT-CO1 |
| 2.6036E-40 | 0.54996571 | 0.374 | 0.181 | 5.5588E-36 | TEC_prg4 | BHLHE40 |
| 8.4181E-53 | 0.54898496 | 0.492 | 0.253 | 1.7973E-48 | TEC_prg4 | RIOK3 |
| 7.7705E-49 | 0.54151297 | 0.53 | 0.291 | 1.659E-44 | TEC_prg4 | YTHDC1 |
| 3.5713E-30 | 0.54148635 | 0.483 | 0.277 | 7.6247E-26 | TEC_prg4 | RND3 |
| 1.7053E-15 | 0.54040017 | 0.16 | 0.078 | 3.6409E-11 | TEC_prg4 | CXCL1 |
| 3.3406E-17 | 0.53703518 | 0.189 | 0.093 | 7.1321E-13 | TEC_prg4 | SLC12A1 |
| 2.0606E-50 | 0.53514545 | 0.485 | 0.248 | 4.3994E-46 | TEC_prg4 | SRSF10 |
| 2.0408E-39 | 0.53273932 | 0.64 | 0.402 | 4.3572E-35 | TEC_prg4 | SRSF7 |
| 1.9929E-44 | 0.53132505 | 0.457 | 0.24 | 4.2549E-40 | TEC_prg4 | BCLAF1 |
| 2.1946E-45 | 0.51912769 | 0.62 | 0.364 | 4.6855E-41 | TEC_prg4 | RSRP1 |
| 1.126E-18 | 0.5107973 | 0.725 | 0.634 | 2.404E-14 | TEC_prg4 | JUND |
| 7.8533E-26 | 0.50843933 | 0.983 | 0.956 | 1.6767E-21 | TEC_prg4 | EIF1 |
| 3.6636E-49 | 0.5053098 | 0.307 | 0.121 | 7.8217E-45 | TEC_prg4 | KLF10 |
| 5.4988E-82 | 0.50368696 | 0.348 | 0.11 | 1.174E-77 | TEC_prg4 | CCNT2 |
| 1.6662E-50 | 0.50281669 | 0.571 | 0.282 | 3.5574E-46 | TEC_prg4 | HEXIM1 |
| 1.6827E-18 | 0.4988759 | 0.252 | 0.137 | 3.5926E-14 | TEC_prg4 | ID3 |
| 2.4041E-16 | 0.4984287 | 0.936 | 0.912 | 5.1327E-12 | TEC_prg4 | HSP90AB1 |
| 2.0267E-78 | 0.49787337 | 0.287 | 0.08 | 4.327E-74 | TEC_prg4 | AEN |
| 1.1439E-49 | 0.49698015 | 0.384 | 0.173 | 2.4422E-45 | TEC_prg4 | XPO1 |
| 7.6255E-12 | 0.49308314 | 0.467 | 0.352 | 1.628E-07 | TEC_prg4 | HSPA1A |
| 2.9564E-53 | 0.49158991 | 0.383 | 0.167 | 6.312E-49 | TEC_prg4 | RYBP |
| 2.2674E-33 | 0.48953127 | 0.416 | 0.23 | 4.8409E-29 | TEC_prg4 | KLF5 |
| 7.3016E-10 | 0.48773907 | 0.809 | 0.818 | 1.5589E-05 | TEC_prg4 | UBB |
| 5.0492E-18 | 0.48227203 | 0.405 | 0.282 | 1.078E-13 | TEC_prg4 | NFE2L2 |
| 2.0398E-47 | 0.48110309 | 0.311 | 0.127 | 4.355E-43 | TEC_prg4 | WDR43 |
| 1.1677E-58 | 0.4806564 | 0.287 | 0.098 | 2.4931E-54 | TEC_prg4 | TRIAP1 |
| 4.5141E-17 | 0.47473493 | 0.18 | 0.088 | 9.6376E-13 | TEC_prg4 | SNAPC1 |
| 3.6213E-50 | 0.47323206 | 0.39 | 0.176 | 7.7316E-46 | TEC_prg4 | PPP4R2 |
| 2.6278E-34 | 0.47214627 | 0.326 | 0.16 | 5.6104E-30 | TEC_prg4 | BLOC1S2 |
| 9.7103E-39 | 0.47186809 | 0.451 | 0.25 | 2.0732E-34 | TEC_prg4 | WAC |
| 2.4313E-19 | 0.4699144 | 0.734 | 0.67 | 5.1907E-15 | TEC_prg4 | HNRNPA1 |
| 1.1014E-18 | 0.46740035 | 0.47 | 0.327 | 2.3514E-14 | TEC_prg4 | ZNF292 |
| 3.7118E-30 | 0.46278145 | 0.517 | 0.326 | 7.9248E-26 | TEC_prg4 | SYF2 |
| 6.9329E-18 | 0.46231793 | 0.623 | 0.513 | 1.4802E-13 | TEC_prg4 | HNRNPU |
| 3.2338E-28 | 0.46138238 | 0.279 | 0.136 | 6.9042E-24 | TEC_prg4 | CKS2 |
| 1.16E-35 | 0.46056762 | 0.518 | 0.301 | 2.4767E-31 | TEC_prg4 | SRSF2 |
| 1.482E-27 | 0.45898351 | 0.543 | 0.348 | 3.164E-23 | TEC_prg4 | MCL1 |
| 3.5256E-53 | 0.45823714 | 0.384 | 0.168 | 7.5271E-49 | TEC_prg4 | YTHDF2 |
| 2.4765E-39 | 0.45812437 | 0.53 | 0.288 | 5.2872E-35 | TEC_prg4 | PLK2 |
| 5.7456E-16 | 0.45516866 | 0.528 | 0.39 | 1.2267E-11 | TEC_prg4 | ANKRD11 |
| 3.8637E-22 | 0.45384085 | 0.732 | 0.62 | 8.249E-18 | TEC_prg4 | ATF4 |
| 3.3753E-29 | 0.4519506 | 0.444 | 0.264 | 7.2063E-25 | TEC_prg4 | CREBRF |
| 1.1034E-32 | 0.45169138 | 0.364 | 0.18 | 2.3557E-28 | TEC_prg4 | TLE4 |
| 8.2822E-17 | 0.45163989 | 0.426 | 0.281 | 1.7682E-12 | TEC_prg4 | CYR61 |
| 4.4685E-33 | 0.45142817 | 0.485 | 0.293 | 9.5403E-29 | TEC_prg4 | PNRC2 |
| 2.5707E-41 | 0.45113688 | 0.328 | 0.149 | 5.4884E-37 | TEC_prg4 | DHX9 |
| 4.8429E-22 | 0.45085475 | 0.32 | 0.185 | 1.034E-17 | TEC_prg4 | PHLDA3 |
| 8.052E-40 | 0.45021206 | 0.399 | 0.195 | 1.7191E-35 | TEC_prg4 | THUMPD3-AS1 |

| | | | | | | |
|---|---|---|---|---|---|---|
| 1.9855E-67 | 0.45000433 | 0.326 | 0.112 | 4.239E-63 | TEC_prg4 | RLF |
| 1.0636E-16 | 0.4488079 | 0.517 | 0.4 | 2.2709E-12 | TEC_prg4 | HNRNPA3 |
| 8.9218E-60 | 0.44872644 | 0.408 | 0.171 | 1.9048E-55 | TEC_prg4 | FNIP1 |
| 6.4333E-34 | 0.44740728 | 0.214 | 0.081 | 1.3735E-29 | TEC_prg4 | AC015912.3 |
| 8.6956E-40 | 0.44711976 | 0.362 | 0.177 | 1.8565E-35 | TEC_prg4 | RPL7L1 |
| 4.1742E-28 | 0.44654417 | 0.441 | 0.27 | 8.9119E-24 | TEC_prg4 | YY1 |
| 1.5858E-24 | 0.4448145 | 0.486 | 0.317 | 3.3856E-20 | TEC_prg4 | EIF1B |
| 9.7797E-36 | 0.44449457 | 0.394 | 0.209 | 2.088E-31 | TEC_prg4 | EWSR1 |
| 1.5448E-24 | 0.44438465 | 0.463 | 0.305 | 3.2981E-20 | TEC_prg4 | SF3B1 |
| 5.17E-74 | 0.44309757 | 0.358 | 0.122 | 1.1038E-69 | TEC_prg4 | KHDC4 |
| 2.7554E-09 | 0.44089697 | 0.723 | 0.699 | 5.8827E-05 | TEC_prg4 | MAP1LC3B |
| 3.3746E-64 | 0.44077812 | 0.32 | 0.109 | 7.2047E-60 | TEC_prg4 | DNAJB4 |
| 7.4398E-27 | 0.43906249 | 0.815 | 0.696 | 1.5884E-22 | TEC_prg4 | EIF4G2 |
| 4.1809E-21 | 0.43808946 | 0.288 | 0.159 | 8.9262E-17 | TEC_prg4 | MIR22HG |
| 1.1043E-28 | 0.43794288 | 0.534 | 0.351 | 2.3577E-24 | TEC_prg4 | EIF3A |
| 3.549E-30 | 0.43784587 | 0.453 | 0.271 | 7.577E-26 | TEC_prg4 | ZNF207 |
| 3.9806E-23 | 0.43690346 | 0.274 | 0.137 | 8.4985E-19 | TEC_prg4 | ID1 |
| 4.4938E-51 | 0.43617106 | 0.374 | 0.158 | 9.5942E-47 | TEC_prg4 | NOP56 |
| 5.0228E-54 | 0.43405452 | 0.288 | 0.1 | 1.0724E-49 | TEC_prg4 | TENT5A |
| 7.9499E-30 | 0.43321019 | 0.464 | 0.265 | 1.6973E-25 | TEC_prg4 | CITED2 |
| 5.7036E-78 | 0.43272696 | 0.271 | 0.073 | 1.2177E-73 | TEC_prg4 | ZBTB11 |
| 5.334E-55 | 0.43176849 | 0.373 | 0.152 | 1.1388E-50 | TEC_prg4 | PPP1R15B |
| 6.2975E-45 | 0.43125167 | 0.425 | 0.207 | 1.3445E-40 | TEC_prg4 | FAM133B |
| 8.3125E-30 | 0.43091116 | 0.348 | 0.182 | 1.7747E-25 | TEC_prg4 | ATXN7L3B |
| 1.9339E-25 | 0.4289679 | 0.55 | 0.367 | 4.1289E-21 | TEC_prg4 | PNISR |
| 1.2211E-27 | 0.42716975 | 0.876 | 0.791 | 2.607E-23 | TEC_prg4 | HNRNPA2B1 |
| 2.1466E-41 | 0.4265214 | 0.207 | 0.068 | 4.583E-37 | TEC_prg4 | RGS16 |
| 4.5786E-76 | 0.42542514 | 0.263 | 0.071 | 9.7753E-72 | TEC_prg4 | RBM4 |
| 6.1998E-31 | 0.42414817 | 0.477 | 0.295 | 1.3237E-26 | TEC_prg4 | RBM8A |
| 1.027E-59 | 0.42318491 | 0.332 | 0.121 | 2.1927E-55 | TEC_prg4 | ZNF844 |
| 1.8891E-36 | 0.42308954 | 0.399 | 0.213 | 4.0333E-32 | TEC_prg4 | RAB21 |
| 3.3089E-10 | 0.42295517 | 0.14 | 0.074 | 7.0645E-06 | TEC_prg4 | HSPA6 |
| 1.8549E-36 | 0.42234178 | 0.314 | 0.138 | 3.9603E-32 | TEC_prg4 | AL118516.1 |
| 5.634E-23 | 0.422196 | 0.137 | 0.05 | 1.2029E-18 | TEC_prg4 | CLDN14 |
| 1.6957E-06 | 0.4221562 | 0.339 | 0.26 | 0.03620345 | TEC_prg4 | DUSP1 |
| 1.2032E-28 | 0.42031111 | 0.402 | 0.238 | 2.5687E-24 | TEC_prg4 | PPIG |
| 9.7932E-12 | 0.4197209 | 0.371 | 0.269 | 2.0909E-07 | TEC_prg4 | VMP1 |
| 7.2941E-17 | 0.41940076 | 0.448 | 0.297 | 1.5573E-12 | TEC_prg4 | HSPA1B |
| 0.00048466 | 0.41909991 | 0.945 | 0.933 | 1 | TEC_prg4 | HSP90AA1 |
| 2.2162E-26 | 0.41814174 | 0.374 | 0.218 | 4.7317E-22 | TEC_prg4 | HNRNPA0 |
| 1.9236E-33 | 0.41649707 | 0.352 | 0.179 | 4.1068E-29 | TEC_prg4 | SREK1 |
| 1.1252E-48 | 0.4156024 | 0.332 | 0.136 | 2.4024E-44 | TEC_prg4 | RRM2B |
| 5.0776E-51 | 0.4155231 | 0.275 | 0.098 | 1.0841E-46 | TEC_prg4 | CBX4 |
| 3.5352E-58 | 0.41535838 | 0.17 | 0.04 | 7.5476E-54 | TEC_prg4 | CIART |
| 3.5846E-52 | 0.41230726 | 0.294 | 0.106 | 7.6531E-48 | TEC_prg4 | ZNF830 |
| 9.9208E-20 | 0.41215588 | 0.451 | 0.299 | 2.1181E-15 | TEC_prg4 | YME1L1 |
| 2.565E-36 | 0.41110745 | 0.457 | 0.242 | 5.4763E-32 | TEC_prg4 | IER5 |
| 1.632E-28 | 0.41016971 | 0.306 | 0.153 | 3.4843E-24 | TEC_prg4 | BBC3 |
| 1.4832E-39 | 0.40998898 | 0.377 | 0.18 | 3.1666E-35 | TEC_prg4 | ZBTB43 |
| 1.3932E-24 | 0.40946091 | 0.473 | 0.307 | 2.9745E-20 | TEC_prg4 | STAT3 |

| | | | | | | |
|---|---|---|---|---|---|---|
| 1.1911E-32 | 0.40922 | 0.346 | 0.175 | 2.543E-28 | TEC_prg4 | SPEN |
| 9.2609E-07 | 0.40918754 | 0.402 | 0.328 | 0.01977201 | TEC_prg4 | H1F0 |
| 4.0502E-18 | 0.4088101 | 0.16 | 0.072 | 8.6471E-14 | TEC_prg4 | FILIP1L |
| 5.4924E-48 | 0.40668172 | 0.326 | 0.135 | 1.1726E-43 | TEC_prg4 | DYRK1A |
| 1.477E-31 | 0.40536648 | 0.492 | 0.292 | 3.1534E-27 | TEC_prg4 | AFF4 |
| 9.0609E-69 | 0.40534997 | 0.275 | 0.081 | 1.9345E-64 | TEC_prg4 | SLC25A25 |
| 1.6606E-11 | 0.40513527 | 0.304 | 0.203 | 3.5455E-07 | TEC_prg4 | ARID5B |
| 6.8342E-13 | 0.40493401 | 0.6 | 0.486 | 1.4591E-08 | TEC_prg4 | ATP1B1 |
| 3.2586E-44 | 0.40304887 | 0.303 | 0.124 | 6.9571E-40 | TEC_prg4 | RBM5 |
| 8.9566E-43 | 0.40185143 | 0.301 | 0.126 | 1.9122E-38 | TEC_prg4 | YTHDF3 |
| 1.2088E-17 | 0.40173532 | 0.279 | 0.159 | 2.5807E-13 | TEC_prg4 | CTGF |
| 1.3178E-27 | 0.40114485 | 0.412 | 0.242 | 2.8135E-23 | TEC_prg4 | ARID4B |
| 2.614E-36 | 0.40068883 | 0.355 | 0.177 | 5.5809E-32 | TEC_prg4 | PPP2CA |
| 6.9674E-17 | 0.39926462 | 0.723 | 0.579 | 1.4875E-12 | TEC_prg4 | ELF3 |
| 6.698E-43 | 0.39882769 | 0.281 | 0.112 | 1.43E-38 | TEC_prg4 | DDX39B |
| 4.2711E-33 | 0.398765 | 0.326 | 0.16 | 9.1188E-29 | TEC_prg4 | ERRFI1 |
| 3.6472E-29 | 0.39734931 | 0.373 | 0.21 | 7.7867E-25 | TEC_prg4 | ETF1 |
| 5.2965E-18 | 0.39614029 | 0.501 | 0.366 | 1.1308E-13 | TEC_prg4 | ODC1 |
| 1.615E-39 | 0.39481562 | 0.332 | 0.147 | 3.4481E-35 | TEC_prg4 | SLC20A1 |
| 7.963E-22 | 0.39472452 | 0.491 | 0.331 | 1.7001E-17 | TEC_prg4 | HSPA9 |
| 1.4194E-34 | 0.39398623 | 0.319 | 0.148 | 3.0305E-30 | TEC_prg4 | ZBTB10 |
| 1.457E-22 | 0.39218025 | 0.397 | 0.234 | 3.1108E-18 | TEC_prg4 | ID4 |
| 1.4402E-36 | 0.39217082 | 0.298 | 0.134 | 3.0748E-32 | TEC_prg4 | ZNF24 |
| 1.227E-33 | 0.39196639 | 0.378 | 0.196 | 2.6196E-29 | TEC_prg4 | ABHD5 |
| 4.7284E-17 | 0.39108561 | 0.56 | 0.427 | 1.0095E-12 | TEC_prg4 | PNRC1 |
| 3.4025E-50 | 0.39087753 | 0.262 | 0.091 | 7.2642E-46 | TEC_prg4 | CDKN1B |
| 7.7435E-27 | 0.38986029 | 0.461 | 0.281 | 1.6532E-22 | TEC_prg4 | KMT2A |
| 2.382E-53 | 0.38985636 | 0.266 | 0.091 | 5.0857E-49 | TEC_prg4 | BRIX1 |
| 5.8124E-37 | 0.38942932 | 0.325 | 0.153 | 1.2409E-32 | TEC_prg4 | CRK |
| 3.0572E-13 | 0.38860685 | 0.721 | 0.612 | 6.5272E-09 | TEC_prg4 | MTRNR2L8 |
| 4.9162E-26 | 0.38837857 | 0.447 | 0.276 | 1.0496E-21 | TEC_prg4 | RBM25 |
| 2.0154E-24 | 0.38796844 | 0.265 | 0.132 | 4.3029E-20 | TEC_prg4 | EFNB2 |
| 8.077E-21 | 0.38729356 | 0.421 | 0.282 | 1.7244E-16 | TEC_prg4 | SERINC1 |
| 2.1667E-24 | 0.38699071 | 0.29 | 0.152 | 4.6258E-20 | TEC_prg4 | BAZ1A |
| 4.1619E-28 | 0.38434303 | 0.62 | 0.41 | 8.8857E-24 | TEC_prg4 | SFPQ |
| 1.0604E-07 | 0.38374272 | 0.75 | 0.68 | 0.00226392 | TEC_prg4 | HSPA5 |
| 1.3903E-08 | 0.38163896 | 0.84 | 0.828 | 0.00029682 | TEC_prg4 | MT-ND5 |
| 1.3384E-31 | 0.38134211 | 0.415 | 0.232 | 2.8574E-27 | TEC_prg4 | TOR1AIP2 |
| 3.1385E-36 | 0.38124198 | 0.322 | 0.149 | 6.7007E-32 | TEC_prg4 | IP6K2 |
| 2.5994E-34 | 0.38123851 | 0.322 | 0.152 | 5.5497E-30 | TEC_prg4 | TRA2A |
| 3.6586E-35 | 0.38116794 | 0.287 | 0.126 | 7.8111E-31 | TEC_prg4 | FAM84B |
| 3.1667E-25 | 0.380692 | 0.125 | 0.041 | 6.7608E-21 | TEC_prg4 | ARL14 |
| 1.4814E-17 | 0.37829473 | 0.325 | 0.199 | 3.1628E-13 | TEC_prg4 | FOSL2 |
| 3.2125E-33 | 0.37718045 | 0.426 | 0.235 | 6.8588E-29 | TEC_prg4 | SLTM |
| 7.7061E-44 | 0.37614148 | 0.335 | 0.146 | 1.6453E-39 | TEC_prg4 | HBP1 |
| 2.8694E-38 | 0.37587148 | 0.345 | 0.164 | 6.1261E-34 | TEC_prg4 | MED13 |
| 1.1434E-30 | 0.37561628 | 0.301 | 0.146 | 2.4412E-26 | TEC_prg4 | PURB |
| 4.511E-23 | 0.37527745 | 0.422 | 0.245 | 9.6309E-19 | TEC_prg4 | IER5L |
| 9.1465E-08 | 0.37348234 | 0.862 | 0.827 | 0.00195278 | TEC_prg4 | SQSTM1 |
| 2.6485E-25 | 0.37297976 | 0.485 | 0.302 | 5.6545E-21 | TEC_prg4 | WSB1 |

| | | | | | | |
|---|---|---|---|---|---|---|
| 9.5928E-22 | 0.37154311 | 0.488 | 0.328 | 2.0481E-17 | TEC_prg4 | LUC7L3 |
| 2.0794E-30 | 0.37110729 | 0.33 | 0.17 | 4.4396E-26 | TEC_prg4 | CCDC59 |
| 1.8302E-07 | 0.37098714 | 0.895 | 0.889 | 0.00390748 | TEC_prg4 | MT-ND2 |
| 4.3448E-69 | 0.36966488 | 0.249 | 0.068 | 9.2762E-65 | TEC_prg4 | ZC3H12A |
| 3.2922E-19 | 0.36898324 | 0.461 | 0.323 | 7.0288E-15 | TEC_prg4 | SF3B6 |
| 1.9553E-50 | 0.36879342 | 0.247 | 0.083 | 4.1747E-46 | TEC_prg4 | PER1 |
| 3.0805E-15 | 0.36779653 | 0.422 | 0.308 | 6.5768E-11 | TEC_prg4 | MRFAP1 |
| 2.4473E-12 | 0.36734305 | 0.464 | 0.334 | 5.225E-08 | TEC_prg4 | SNHG12 |
| 3.079E-34 | 0.36651984 | 0.367 | 0.185 | 6.5737E-30 | TEC_prg4 | SRSF1 |
| 1.8397E-42 | 0.36532503 | 0.284 | 0.114 | 3.9277E-38 | TEC_prg4 | FBXO22 |
| 0.0001012 | 0.36511897 | 0.533 | 0.509 | 1 | TEC_prg4 | H2AFZ |
| 3.9073E-13 | 0.36474355 | 0.665 | 0.609 | 8.3422E-09 | TEC_prg4 | SAP18 |
| 2.977E-36 | 0.36351092 | 0.281 | 0.121 | 6.356E-32 | TEC_prg4 | SLC30A1 |
| 2.1611E-44 | 0.36350535 | 0.223 | 0.074 | 4.6139E-40 | TEC_prg4 | IRX2 |
| 3.6437E-36 | 0.3632334 | 0.476 | 0.253 | 7.7792E-32 | TEC_prg4 | AMD1 |
| 9.5279E-54 | 0.36275311 | 0.304 | 0.111 | 2.0342E-49 | TEC_prg4 | C1orf52 |
| 1.8924E-31 | 0.36225135 | 0.304 | 0.148 | 4.0403E-27 | TEC_prg4 | AKIRIN1 |
| 7.9429E-20 | 0.36023141 | 0.396 | 0.241 | 1.6958E-15 | TEC_prg4 | IDI1 |
| 1.445E-33 | 0.35957682 | 0.349 | 0.168 | 3.0851E-29 | TEC_prg4 | KLF4 |
| 1.535E-13 | 0.35941917 | 0.1 | 0.042 | 3.2772E-09 | TEC_prg4 | ZNF57 |
| 6.4603E-28 | 0.35912405 | 0.307 | 0.156 | 1.3793E-23 | TEC_prg4 | CYLD |
| 1.799E-22 | 0.35825346 | 0.4 | 0.25 | 3.8409E-18 | TEC_prg4 | SYNCRIP |
| 4.6099E-26 | 0.35816878 | 0.384 | 0.216 | 9.8421E-22 | TEC_prg4 | NFAT5 |
| 4.6112E-12 | 0.35782471 | 0.691 | 0.64 | 9.8449E-08 | TEC_prg4 | YWHAE |
| 1.3237E-38 | 0.35664129 | 0.365 | 0.174 | 2.8261E-34 | TEC_prg4 | ARIH1 |
| 5.3959E-22 | 0.35656649 | 0.399 | 0.249 | 1.152E-17 | TEC_prg4 | SF1 |
| 2.6422E-36 | 0.35613533 | 0.256 | 0.104 | 5.6411E-32 | TEC_prg4 | ARRDC4 |
| 5.0933E-23 | 0.35444514 | 0.44 | 0.273 | 1.0874E-18 | TEC_prg4 | PRPF4B |
| 1.1044E-26 | 0.35297599 | 0.415 | 0.228 | 2.3579E-22 | TEC_prg4 | INTS6 |
| 3.1444E-42 | 0.35171991 | 0.306 | 0.127 | 6.7134E-38 | TEC_prg4 | CHD1 |
| 1.9909E-25 | 0.34992956 | 0.354 | 0.197 | 4.2505E-21 | TEC_prg4 | OSER1 |
| 1.1336E-09 | 0.34880351 | 0.4 | 0.321 | 2.4203E-05 | TEC_prg4 | ARL6IP1 |
| 3.6844E-67 | 0.34857617 | 0.226 | 0.059 | 7.8663E-63 | TEC_prg4 | ZNF654 |
| 2.184E-21 | 0.34821503 | 0.246 | 0.129 | 4.6628E-17 | TEC_prg4 | OAT |
| 4.8596E-32 | 0.34788155 | 0.349 | 0.18 | 1.0375E-27 | TEC_prg4 | DDX6 |
| 1.3226E-16 | 0.34782099 | 0.603 | 0.481 | 2.8237E-12 | TEC_prg4 | UBE2D3 |
| 1.4162E-13 | 0.34646517 | 0.32 | 0.21 | 3.0235E-09 | TEC_prg4 | UBE2S |
| 1.1867E-21 | 0.34602575 | 0.342 | 0.201 | 2.5337E-17 | TEC_prg4 | CSNK2A1 |
| 3.9906E-23 | 0.3457599 | 0.195 | 0.086 | 8.5199E-19 | TEC_prg4 | NR4A1 |
| 3.3659E-21 | 0.34549004 | 0.332 | 0.193 | 7.1861E-17 | TEC_prg4 | MDM4 |
| 1.3321E-44 | 0.34534829 | 0.301 | 0.12 | 2.8441E-40 | TEC_prg4 | TFRC |
| 5.4945E-27 | 0.34490412 | 0.396 | 0.227 | 1.1731E-22 | TEC_prg4 | NIPBL |
| 1.2429E-16 | 0.34447549 | 0.357 | 0.229 | 2.6536E-12 | TEC_prg4 | MIDN |
| 4.517E-13 | 0.34394839 | 0.46 | 0.338 | 9.6437E-09 | TEC_prg4 | TSC22D1 |
| 1.6278E-29 | 0.34363779 | 0.275 | 0.128 | 3.4753E-25 | TEC_prg4 | TUBB2A |
| 1.1403E-17 | 0.34227222 | 0.473 | 0.325 | 2.4346E-13 | TEC_prg4 | ANKRD12 |
| 1.0878E-20 | 0.34171823 | 0.287 | 0.153 | 2.3225E-16 | TEC_prg4 | TNFAIP3 |
| 1.5103E-20 | 0.33986939 | 0.444 | 0.292 | 3.2244E-16 | TEC_prg4 | PAPOLA |
| 3.3416E-34 | 0.33975624 | 0.272 | 0.117 | 7.1343E-30 | TEC_prg4 | CEMIP2 |
| 5.2227E-20 | 0.33860701 | 0.383 | 0.229 | 1.1151E-15 | TEC_prg4 | SLC5A3 |

| | | | | | | |
|---|---|---|---|---|---|---|
| 3.2319E-25 | 0.33818279 | 0.333 | 0.186 | 6.9002E-21 | TEC_prg4 | DNAJA2 |
| 1.0215E-37 | 0.33795234 | 0.281 | 0.115 | 2.1809E-33 | TEC_prg4 | KMT2E-AS1 |
| 4.1345E-50 | 0.33737145 | 0.281 | 0.101 | 8.8271E-46 | TEC_prg4 | ZCCHC8 |
| 3.5678E-33 | 0.33665379 | 0.314 | 0.145 | 7.6172E-29 | TEC_prg4 | IVNS1ABP |
| 1.0681E-11 | 0.33651827 | 0.213 | 0.127 | 2.2804E-07 | TEC_prg4 | CSRP2 |
| 1.4038E-15 | 0.33484998 | 0.275 | 0.167 | 2.9971E-11 | TEC_prg4 | NSRP1 |
| 7.3454E-28 | 0.33382969 | 0.373 | 0.205 | 1.5682E-23 | TEC_prg4 | RAD21 |
| 2.3721E-33 | 0.33373113 | 0.245 | 0.1 | 5.0645E-29 | TEC_prg4 | CPEB2 |
| 4.5955E-14 | 0.33337127 | 0.518 | 0.408 | 9.8113E-10 | TEC_prg4 | HNRNPK |
| 3.1467E-33 | 0.33303686 | 0.332 | 0.161 | 6.7183E-29 | TEC_prg4 | ZNF644 |
| 3.3176E-47 | 0.33253999 | 0.262 | 0.094 | 7.0832E-43 | TEC_prg4 | MRFAP1L1 |
| 1.4326E-23 | 0.33136844 | 0.39 | 0.231 | 3.0585E-19 | TEC_prg4 | C6orf48 |
| 4.7857E-52 | 0.33115702 | 0.293 | 0.106 | 1.0217E-47 | TEC_prg4 | RAF1 |
| 2.0286E-28 | 0.33015484 | 0.319 | 0.162 | 4.3311E-24 | TEC_prg4 | PRDM2 |
| 1.4918E-53 | 0.32956502 | 0.284 | 0.098 | 3.1849E-49 | TEC_prg4 | RSBN1 |
| 2.1272E-12 | 0.3292144 | 0.501 | 0.374 | 4.5417E-08 | TEC_prg4 | HERPUD1 |
| 2.8894E-19 | 0.32909296 | 0.485 | 0.342 | 6.1689E-15 | TEC_prg4 | VAPA |
| 2.5193E-29 | 0.32862946 | 0.309 | 0.154 | 5.3787E-25 | TEC_prg4 | PPP2R2A |
| 4.9764E-40 | 0.32857959 | 0.265 | 0.105 | 1.0625E-35 | TEC_prg4 | TRMT10C |
| 4.0708E-39 | 0.32811652 | 0.344 | 0.156 | 8.6911E-35 | TEC_prg4 | SAFB2 |
| 7.8295E-27 | 0.32711781 | 0.25 | 0.116 | 1.6716E-22 | TEC_prg4 | SSFA2 |
| 2.4564E-12 | 0.32498069 | 0.381 | 0.264 | 5.2445E-08 | TEC_prg4 | SEPHS2 |
| 9.6685E-24 | 0.32416583 | 0.295 | 0.156 | 2.0642E-19 | TEC_prg4 | ESF1 |
| 6.2904E-31 | 0.32307887 | 0.323 | 0.16 | 1.343E-26 | TEC_prg4 | USP15 |
| 2.7103E-37 | 0.32293686 | 0.266 | 0.109 | 5.7865E-33 | TEC_prg4 | LRIF1 |
| 2.4295E-31 | 0.32222591 | 0.339 | 0.169 | 5.1871E-27 | TEC_prg4 | SETD2 |
| 6.0449E-17 | 0.32099074 | 0.231 | 0.12 | 1.2906E-12 | TEC_prg4 | PCDH9 |
| 7.7998E-33 | 0.32093378 | 0.323 | 0.156 | 1.6653E-28 | TEC_prg4 | IST1 |
| 5.8367E-32 | 0.32070329 | 0.215 | 0.086 | 1.2461E-27 | TEC_prg4 | MITF |
| 3.6934E-33 | 0.32060525 | 0.277 | 0.122 | 7.8855E-29 | TEC_prg4 | TNRC6A |
| 2.9341E-44 | 0.31911251 | 0.291 | 0.113 | 6.2643E-40 | TEC_prg4 | SIRT1 |
| 5.1501E-46 | 0.31841123 | 0.245 | 0.085 | 1.0996E-41 | TEC_prg4 | LINC00909 |
| 1.953E-08 | 0.31829229 | 0.508 | 0.434 | 0.00041696 | TEC_prg4 | HMGN1 |
| 6.7756E-05 | 0.31821369 | 0.368 | 0.309 | 1 | TEC_prg4 | CDKN1C |
| 1.3778E-24 | 0.31782192 | 0.415 | 0.249 | 2.9417E-20 | TEC_prg4 | BRD4 |
| 6.6125E-39 | 0.31694113 | 0.245 | 0.094 | 1.4118E-34 | TEC_prg4 | CASP3 |
| 2.0529E-23 | 0.31572473 | 0.345 | 0.197 | 4.383E-19 | TEC_prg4 | C16orf72 |
| 9.0468E-27 | 0.31537717 | 0.376 | 0.208 | 1.9315E-22 | TEC_prg4 | EPB41L4A-AS1 |
| 7.9782E-13 | 0.31527435 | 0.939 | 0.808 | 1.7033E-08 | TEC_prg4 | NEAT1 |
| 1.2392E-10 | 0.3152483 | 0.737 | 0.677 | 2.6456E-06 | TEC_prg4 | YBX3 |
| 2.9635E-20 | 0.31523312 | 0.256 | 0.139 | 6.327E-16 | TEC_prg4 | MRPS18B |
| 5.2657E-44 | 0.31516377 | 0.25 | 0.091 | 1.1242E-39 | TEC_prg4 | CCNT1 |
| 9.4751E-55 | 0.31481295 | 0.252 | 0.079 | 2.0229E-50 | TEC_prg4 | PPM1D |
| 3.7264E-22 | 0.31416145 | 0.218 | 0.103 | 7.9558E-18 | TEC_prg4 | SRSF8 |
| 9.4542E-42 | 0.31348392 | 0.218 | 0.075 | 2.0185E-37 | TEC_prg4 | CMTR2 |
| 0.0024782 | 0.31335594 | 0.75 | 0.758 | 1 | TEC_prg4 | DYNLL1 |
| 1.0115E-17 | 0.31143507 | 0.377 | 0.242 | 2.1596E-13 | TEC_prg4 | KANSL1 |
| 1.8922E-34 | 0.31139696 | 0.287 | 0.128 | 4.0398E-30 | TEC_prg4 | SMNDC1 |
| 2.8342E-17 | 0.31115528 | 0.438 | 0.295 | 6.051E-13 | TEC_prg4 | EIF4A1 |
| 3.0292E-33 | 0.31025381 | 0.269 | 0.118 | 6.4674E-29 | TEC_prg4 | CTR9 |

| | | | | | | |
|---|---|---|---|---|---|---|
| 5.9111E-26 | 0.31011067 | 0.288 | 0.144 | 1.262E-21 | TEC_prg4 | BCL2L11 |
| 9.0758E-36 | 0.3091661 | 0.288 | 0.126 | 1.9377E-31 | TEC_prg4 | TFAM |
| 4.5316E-11 | 0.30898741 | 0.769 | 0.716 | 9.6749E-07 | TEC_prg4 | RPL36AL |
| 5.8885E-22 | 0.30859169 | 0.463 | 0.293 | 1.2572E-17 | TEC_prg4 | C6orf62 |
| 2.7862E-34 | 0.30776556 | 0.316 | 0.146 | 5.9486E-30 | TEC_prg4 | CWC25 |
| 4.2224E-15 | 0.30714166 | 0.326 | 0.208 | 9.0148E-11 | TEC_prg4 | PARD6B |
| 8.5551E-17 | 0.30687974 | 0.33 | 0.205 | 1.8265E-12 | TEC_prg4 | MAFG |
| 9.5249E-05 | 0.30679896 | 0.933 | 0.908 | 1 | TEC_prg4 | MT-ATP6 |
| 2.4962E-38 | 0.30645773 | 0.288 | 0.122 | 5.3295E-34 | TEC_prg4 | SPTY2D1 |
| 2.5379E-07 | 0.30616352 | 0.437 | 0.354 | 0.00541831 | TEC_prg4 | GCC2 |
| 7.2394E-39 | 0.30566274 | 0.243 | 0.092 | 1.5456E-34 | TEC_prg4 | NFIL3 |
| 2.3393E-20 | 0.30501672 | 0.387 | 0.238 | 4.9943E-16 | TEC_prg4 | IRS2 |
| 1.2679E-22 | 0.30350196 | 0.349 | 0.199 | 2.7071E-18 | TEC_prg4 | DNTTIP2 |
| 4.6685E-08 | 0.30344505 | 0.384 | 0.306 | 0.00099672 | TEC_prg4 | ISCU |
| 3.3874E-25 | 0.30257857 | 0.229 | 0.105 | 7.232E-21 | TEC_prg4 | GNAI1 |
| 1.6071E-19 | 0.30136962 | 0.416 | 0.265 | 3.4312E-15 | TEC_prg4 | SMG1 |
| 5.4658E-15 | 0.30121939 | 0.434 | 0.306 | 1.1669E-10 | TEC_prg4 | DNAJB6 |
| 1.6265E-13 | 0.30114686 | 0.368 | 0.259 | 3.4726E-09 | TEC_prg4 | HNRNPM |
| 7.2646E-39 | 0.29869725 | 0.259 | 0.103 | 1.551E-34 | TEC_prg4 | CHD8 |
| 6.9002E-10 | 0.29795171 | 0.45 | 0.35 | 1.4732E-05 | TEC_prg4 | DDX3X |
| 5.9024E-34 | 0.29751873 | 0.137 | 0.039 | 1.2602E-29 | TEC_prg4 | AC107959.4 |
| 1.1737E-21 | 0.29733419 | 0.358 | 0.211 | 2.5058E-17 | TEC_prg4 | NKTR |
| 6.9343E-33 | 0.29675268 | 0.239 | 0.097 | 1.4805E-28 | TEC_prg4 | SNHG9 |
| 4.4422E-38 | 0.2962573 | 0.247 | 0.097 | 9.4841E-34 | TEC_prg4 | POLE3 |
| 1.9416E-35 | 0.29610183 | 0.25 | 0.101 | 4.1454E-31 | TEC_prg4 | FAM53C |
| 2.0829E-40 | 0.29596462 | 0.258 | 0.099 | 4.4471E-36 | TEC_prg4 | ZMYM5 |
| 1.4188E-06 | 0.29550708 | 0.725 | 0.691 | 0.03029163 | TEC_prg4 | ZFAS1 |
| 1.751E-15 | 0.29461167 | 0.413 | 0.286 | 3.7383E-11 | TEC_prg4 | TOP1 |
| 4.3535E-49 | 0.29455002 | 0.207 | 0.061 | 9.2948E-45 | TEC_prg4 | AC132192.2 |
| 4.4567E-23 | 0.29448823 | 0.338 | 0.19 | 9.515E-19 | TEC_prg4 | SENP6 |
| 2.706E-22 | 0.29437521 | 0.3 | 0.163 | 5.7774E-18 | TEC_prg4 | NRIP1 |
| 6.9394E-37 | 0.29420683 | 0.297 | 0.128 | 1.4816E-32 | TEC_prg4 | ZNF721 |
| 6.0951E-35 | 0.29332246 | 0.215 | 0.082 | 1.3013E-30 | TEC_prg4 | SART3 |
| 1.191E-07 | 0.29206191 | 0.629 | 0.57 | 0.0025427 | TEC_prg4 | SET |
| 1.386E-37 | 0.2915162 | 0.258 | 0.102 | 2.9592E-33 | TEC_prg4 | KAT6A |
| 1.5893E-26 | 0.29148039 | 0.367 | 0.204 | 3.3932E-22 | TEC_prg4 | YWHAG |
| 1.6704E-13 | 0.29125733 | 0.278 | 0.165 | 3.5664E-09 | TEC_prg4 | FOSB |
| 1.1515E-22 | 0.29108845 | 0.262 | 0.135 | 2.4584E-18 | TEC_prg4 | EIF4E |
| 4.5768E-20 | 0.29037922 | 0.357 | 0.21 | 9.7714E-16 | TEC_prg4 | ASH1L |
| 7.0877E-49 | 0.29037552 | 0.182 | 0.051 | 1.5132E-44 | TEC_prg4 | WDR47 |
| 6.9097E-29 | 0.29010233 | 0.298 | 0.147 | 1.4752E-24 | TEC_prg4 | GTF2B |
| 3.009E-25 | 0.28978708 | 0.266 | 0.131 | 6.4241E-21 | TEC_prg4 | CAND1 |
| 2.9709E-32 | 0.28964384 | 0.243 | 0.099 | 6.343E-28 | TEC_prg4 | C6orf120 |
| 1.0156E-19 | 0.28882207 | 0.24 | 0.124 | 2.1684E-15 | TEC_prg4 | KDM6B |
| 1.4191E-24 | 0.28766262 | 0.272 | 0.137 | 3.0297E-20 | TEC_prg4 | CHTOP |
| 5.0203E-16 | 0.287218 | 0.354 | 0.228 | 1.0718E-11 | TEC_prg4 | CHMP2B |
| 5.6314E-16 | 0.28635296 | 0.201 | 0.106 | 1.2023E-11 | TEC_prg4 | ZMAT3 |
| 6.3466E-34 | 0.28586439 | 0.256 | 0.108 | 1.355E-29 | TEC_prg4 | SP3 |
| 1.0663E-29 | 0.28533336 | 0.266 | 0.123 | 2.2766E-25 | TEC_prg4 | ADNP |
| 2.4463E-29 | 0.28458278 | 0.287 | 0.137 | 5.2228E-25 | TEC_prg4 | SBNO1 |

| | | | | | | |
|---|---|---|---|---|---|---|
| 2.9016E-12 | 0.28454831 | 0.498 | 0.37 | 6.195E-08 | TEC_prg4 | SRSF5 |
| 3.4503E-20 | 0.28431407 | 0.352 | 0.211 | 7.3664E-16 | TEC_prg4 | CCDC186 |
| 2.1518E-24 | 0.28273771 | 0.354 | 0.2 | 4.5941E-20 | TEC_prg4 | G3BP2 |
| 3.0395E-17 | 0.28235666 | 0.378 | 0.225 | 6.4893E-13 | TEC_prg4 | NFKBIZ |
| 1.5759E-09 | 0.28201265 | 0.348 | 0.247 | 3.3646E-05 | TEC_prg4 | IRF1 |
| 5.6931E-36 | 0.28128364 | 0.197 | 0.069 | 1.2155E-31 | TEC_prg4 | THAP1 |
| 8.6313E-18 | 0.28104875 | 0.362 | 0.231 | 1.8428E-13 | TEC_prg4 | RNPS1 |
| 3.2996E-51 | 0.28098792 | 0.239 | 0.076 | 7.0447E-47 | TEC_prg4 | FRS2 |
| 2.6736E-29 | 0.28037247 | 0.275 | 0.128 | 5.7081E-25 | TEC_prg4 | LATS2 |
| 5.036E-34 | 0.27947681 | 0.294 | 0.13 | 1.0752E-29 | TEC_prg4 | CREBBP |
| 3.8276E-34 | 0.27882856 | 0.215 | 0.083 | 8.172E-30 | TEC_prg4 | ZNF274 |
| 1.1967E-14 | 0.27830845 | 0.127 | 0.057 | 2.555E-10 | TEC_prg4 | SCX |
| 8.0378E-47 | 0.27787154 | 0.162 | 0.042 | 1.7161E-42 | TEC_prg4 | ZNF667 |
| 2.1087E-26 | 0.27662358 | 0.319 | 0.165 | 4.5021E-22 | TEC_prg4 | EP300 |
| 1.5082E-24 | 0.27541365 | 0.285 | 0.146 | 3.22E-20 | TEC_prg4 | AP1G1 |
| 7.9266E-16 | 0.2747082 | 0.197 | 0.101 | 1.6923E-11 | TEC_prg4 | AREG |
| 8.5594E-51 | 0.27432392 | 0.179 | 0.047 | 1.8274E-46 | TEC_prg4 | TENT5C |
| 4.8051E-17 | 0.27410289 | 0.319 | 0.193 | 1.0259E-12 | TEC_prg4 | CHMP1B |
| 6.9027E-08 | 0.27408767 | 0.413 | 0.335 | 0.00147374 | TEC_prg4 | KRT10 |
| 3.7104E-14 | 0.27387478 | 0.486 | 0.356 | 7.9216E-10 | TEC_prg4 | SRSF11 |
| 3.1545E-25 | 0.2736126 | 0.265 | 0.129 | 6.7348E-21 | TEC_prg4 | TDG |
| 3.257E-28 | 0.27358301 | 0.205 | 0.084 | 6.9537E-24 | TEC_prg4 | SNRK |
| 1.1599E-12 | 0.27244164 | 0.368 | 0.256 | 2.4763E-08 | TEC_prg4 | DDX21 |
| 8.3711E-13 | 0.27229931 | 0.313 | 0.212 | 1.7872E-08 | TEC_prg4 | THRAP3 |
| 8.667E-13 | 0.27109675 | 0.556 | 0.424 | 1.8504E-08 | TEC_prg4 | ZFAND5 |
| 1.3022E-19 | 0.27081538 | 0.79 | 0.672 | 2.7801E-15 | TEC_prg4 | MTRNR2L12 |
| 2.9498E-12 | 0.27024056 | 0.298 | 0.195 | 6.2979E-08 | TEC_prg4 | EHF |
| 1.8782E-20 | 0.26998657 | 0.33 | 0.19 | 4.01E-16 | TEC_prg4 | CDK12 |
| 3.621E-15 | 0.26970612 | 0.329 | 0.208 | 7.7308E-11 | TEC_prg4 | TRIP11 |
| 6.0223E-13 | 0.26888128 | 0.393 | 0.269 | 1.2858E-08 | TEC_prg4 | BPTF |
| 1.7324E-24 | 0.26824069 | 0.261 | 0.128 | 3.6987E-20 | TEC_prg4 | GNL3 |
| 7.8767E-36 | 0.26760714 | 0.231 | 0.09 | 1.6817E-31 | TEC_prg4 | IP6K1 |
| 2.7022E-20 | 0.26742315 | 0.277 | 0.149 | 5.7692E-16 | TEC_prg4 | CHORDC1 |
| 4.034E-05 | 0.26735644 | 0.227 | 0.169 | 0.86125375 | TEC_prg4 | SETX |
| 1.183E-15 | 0.26672766 | 0.358 | 0.234 | 2.5257E-11 | TEC_prg4 | HNRNPF |
| 8.2293E-20 | 0.2662932 | 0.234 | 0.118 | 1.7569E-15 | TEC_prg4 | WEE1 |
| 4.4128E-12 | 0.26464952 | 0.437 | 0.331 | 9.4213E-08 | TEC_prg4 | EIF4B |
| 1.5515E-24 | 0.26459266 | 0.246 | 0.117 | 3.3125E-20 | TEC_prg4 | EPM2AIP1 |
| 1.3343E-13 | 0.26388183 | 0.163 | 0.081 | 2.8487E-09 | TEC_prg4 | AL021453.1 |
| 2.0852E-22 | 0.26358866 | 0.319 | 0.175 | 4.452E-18 | TEC_prg4 | RAB5A |
| 2.0782E-06 | 0.2632663 | 0.495 | 0.418 | 0.04437015 | TEC_prg4 | SNHG8 |
| 1.5208E-31 | 0.26317405 | 0.274 | 0.121 | 3.2468E-27 | TEC_prg4 | TSC22D2 |
| 4.9826E-05 | 0.26270958 | 0.352 | 0.3 | 1 | TEC_prg4 | SNHG7 |
| 4.4325E-26 | 0.26159676 | 0.262 | 0.124 | 9.4634E-22 | TEC_prg4 | UTP3 |
| 3.3593E-25 | 0.26093151 | 0.279 | 0.139 | 7.1722E-21 | TEC_prg4 | FGFR1OP2 |
| 1.8516E-17 | 0.26075894 | 0.335 | 0.204 | 3.9531E-13 | TEC_prg4 | G3BP1 |
| 9.0842E-30 | 0.26063973 | 0.255 | 0.113 | 1.9395E-25 | TEC_prg4 | NUP58 |
| 1.661E-20 | 0.2604115 | 0.258 | 0.135 | 3.5462E-16 | TEC_prg4 | SKIL |
| 3.416E-15 | 0.26015471 | 0.362 | 0.244 | 7.2933E-11 | TEC_prg4 | RAB1A |
| 1.3214E-21 | 0.25943042 | 0.183 | 0.082 | 2.8211E-17 | TEC_prg4 | BAMBI |

| | | | | | | |
|---|---|---|---|---|---|---|
| 2.5433E-26 | 0.25917612 | 0.466 | 0.273 | 5.4299E-22 | TEC_prg4 | POLR2A |
| 1.2131E-17 | 0.25863506 | 0.246 | 0.134 | 2.59E-13 | TEC_prg4 | TOB2 |
| 3.7299E-15 | 0.25803356 | 0.361 | 0.235 | 7.9633E-11 | TEC_prg4 | FNBP4 |
| 2.7298E-10 | 0.25803318 | 0.543 | 0.429 | 5.828E-06 | TEC_prg4 | PRRC2C |
| 0.00034062 | 0.25703579 | 0.929 | 0.912 | 1 | TEC_prg4 | MT-ND4 |
| 1.4006E-22 | 0.25688017 | 0.246 | 0.123 | 2.9904E-18 | TEC_prg4 | HNRNPL |
| 1.665E-13 | 0.2568193 | 0.163 | 0.083 | 3.5549E-09 | TEC_prg4 | SPOP |
| 2.2014E-35 | 0.25679597 | 0.236 | 0.092 | 4.7E-31 | TEC_prg4 | PCNA |
| 1.9448E-09 | 0.25658843 | 0.432 | 0.341 | 4.1521E-05 | TEC_prg4 | CNBP |
| 3.5539E-32 | 0.25655961 | 0.263 | 0.114 | 7.5875E-28 | TEC_prg4 | UBAP2 |
| 2.5523E-14 | 0.25617353 | 0.328 | 0.212 | 5.4491E-10 | TEC_prg4 | DHX36 |
| 6.4131E-23 | 0.25594152 | 0.277 | 0.143 | 1.3692E-18 | TEC_prg4 | SRSF6 |
| 6.2666E-06 | 0.25550244 | 0.303 | 0.24 | 0.13379175 | TEC_prg4 | DNAJC3 |
| 1.9901E-10 | 0.25489688 | 0.434 | 0.335 | 4.2488E-06 | TEC_prg4 | PFDN2 |
| 6.0749E-16 | 0.25484828 | 0.396 | 0.259 | 1.297E-11 | TEC_prg4 | BDP1 |
| 1.1025E-18 | 0.25456751 | 0.269 | 0.151 | 2.3538E-14 | TEC_prg4 | SLU7 |
| 1.6694E-09 | 0.25442524 | 0.492 | 0.391 | 3.5641E-05 | TEC_prg4 | CYCS |
| 9.9927E-28 | 0.25419811 | 0.304 | 0.148 | 2.1334E-23 | TEC_prg4 | NUDT4 |
| 3.7331E-65 | 0.25419592 | 0.118 | 0.018 | 7.9703E-61 | TEC_prg4 | AL354733.3 |
| 8.7341E-41 | 0.25376576 | 0.217 | 0.074 | 1.8647E-36 | TEC_prg4 | YOD1 |
| 9.1247E-15 | 0.2533162 | 0.239 | 0.136 | 1.9481E-10 | TEC_prg4 | EFNA1 |
| 1.0927E-12 | 0.25316454 | 0.409 | 0.297 | 2.3329E-08 | TEC_prg4 | POLR1D |
| 3.0345E-08 | 0.25314509 | 0.448 | 0.371 | 0.00064787 | TEC_prg4 | C11orf58 |
| 1.4094E-17 | 0.25313001 | 0.313 | 0.185 | 3.009E-13 | TEC_prg4 | CEBPZ |
| 7.9628E-18 | 0.25270313 | 0.31 | 0.183 | 1.7001E-13 | TEC_prg4 | PTBP1 |
| 2.0293E-36 | 0.25197437 | 0.218 | 0.08 | 4.3326E-32 | TEC_prg4 | ZBTB21 |
| 4.2201E-15 | 0.25139673 | 0.252 | 0.148 | 9.0099E-11 | TEC_prg4 | THUMPD1 |
| 6.2651E-10 | 0.25122122 | 0.344 | 0.251 | 1.3376E-05 | TEC_prg4 | CCT4 |
| 2.7863E-21 | 0.25097053 | 0.188 | 0.084 | 5.9488E-17 | TEC_prg4 | TRIB1 |
| 3.2158E-42 | 0.25086388 | 0.217 | 0.073 | 6.8657E-38 | TEC_prg4 | SESN1 |
| 4.4815E-22 | 0.25082886 | 0.265 | 0.136 | 9.5679E-18 | TEC_prg4 | GPATCH8 |
| 7.4261E-09 | 0.2507499 | 0.453 | 0.337 | 0.00015855 | TEC_prg4 | ZFP36L2 |
| 9.5559E-09 | 0.25003193 | 0.323 | 0.235 | 0.00020402 | TEC_prg4 | LMO4 |
| 3.3374E-25 | 0.25000914 | 0.316 | 0.165 | 7.1254E-21 | TEC_prg4 | CGGBP1 |
| 7.0604E-09 | -0.2519852 | 0.857 | 0.915 | 0.00015074 | TEC_prg4 | RPL35 |
| 9.2897E-14 | -0.252419 | 0.191 | 0.329 | 1.9833E-09 | TEC_prg4 | NPC2 |
| 1.8774E-14 | -0.2525161 | 0.049 | 0.155 | 4.0083E-10 | TEC_prg4 | FXYD5 |
| 7.0177E-11 | -0.2534767 | 0.16 | 0.27 | 1.4983E-06 | TEC_prg4 | SELENOM |
| 1.1069E-08 | -0.253795 | 0.191 | 0.288 | 0.00023631 | TEC_prg4 | ASPH |
| 1.2251E-09 | -0.2539376 | 0.263 | 0.379 | 2.6156E-05 | TEC_prg4 | FLNA |
| 6.6481E-07 | -0.2551471 | 0.138 | 0.212 | 0.01419365 | TEC_prg4 | ADAMTS9 |
| 2.4814E-20 | -0.2558969 | 0.105 | 0.264 | 5.2978E-16 | TEC_prg4 | CAPZB |
| 1.6907E-11 | -0.2575057 | 0.162 | 0.276 | 3.6097E-07 | TEC_prg4 | RBM3 |
| 1.7395E-11 | -0.2575399 | 0.214 | 0.329 | 3.7139E-07 | TEC_prg4 | COX5A |
| 8.3013E-13 | -0.2578 | 0.451 | 0.578 | 1.7723E-08 | TEC_prg4 | COX7A2 |
| 1.1543E-11 | -0.2580072 | 0.176 | 0.295 | 2.4645E-07 | TEC_prg4 | ITGA3 |
| 7.8427E-09 | -0.25804 | 0.137 | 0.228 | 0.00016744 | TEC_prg4 | GPRC5A |
| 1.1097E-09 | -0.2593202 | 0.873 | 0.925 | 2.3693E-05 | TEC_prg4 | RPL14 |
| 2.8971E-09 | -0.260441 | 0.888 | 0.921 | 6.1853E-05 | TEC_prg4 | RPL9 |
| 1.5603E-20 | -0.2610431 | 0.08 | 0.232 | 3.3312E-16 | TEC_prg4 | C19orf70 |

| | | | | | | |
|---|---|---|---|---|---|---|
| 2.9391E-14 | -0.2611881 | 0.587 | 0.731 | 6.2749E-10 | TEC_prg4 | CTSD |
| 5.385E-21 | -0.2613523 | 0.073 | 0.226 | 1.1497E-16 | TEC_prg4 | BLOC1S1 |
| 7.1417E-18 | -0.26145 | 0.132 | 0.282 | 1.5248E-13 | TEC_prg4 | VPS28 |
| 4.977E-10 | -0.2616893 | 0.227 | 0.338 | 1.0626E-05 | TEC_prg4 | SLC25A5 |
| 2.1045E-17 | -0.2618606 | 0.164 | 0.319 | 4.4932E-13 | TEC_prg4 | ATP5PD |
| 1.7133E-15 | -0.2618972 | 0.146 | 0.28 | 3.658E-11 | TEC_prg4 | ATP5F1C |
| 8.5149E-14 | -0.2620408 | 0.127 | 0.249 | 1.8179E-09 | TEC_prg4 | PLP2 |
| 5.7678E-13 | -0.2628556 | 0.073 | 0.179 | 1.2314E-08 | TEC_prg4 | PLA2G16 |
| 0.00016236 | -0.2637166 | 0.084 | 0.132 | 1 | TEC_prg4 | TNFRSF11B |
| 4.0955E-15 | -0.263988 | 0.1 | 0.226 | 8.7439E-11 | TEC_prg4 | TIMP2 |
| 1.2031E-11 | -0.2640626 | 0.745 | 0.831 | 2.5686E-07 | TEC_prg4 | HLA-B |
| 3.0659E-08 | -0.2665906 | 0.581 | 0.673 | 0.00065456 | TEC_prg4 | GLS |
| 1.477E-09 | -0.2669892 | 0.256 | 0.365 | 3.1535E-05 | TEC_prg4 | CD44 |
| 6.5961E-16 | -0.2670353 | 0.122 | 0.259 | 1.4083E-11 | TEC_prg4 | CD47 |
| 2.0808E-18 | -0.2674919 | 0.207 | 0.378 | 4.4426E-14 | TEC_prg4 | ARL6IP4 |
| 8.6374E-13 | -0.2688251 | 0.031 | 0.119 | 1.8441E-08 | TEC_prg4 | RARRES3 |
| 2.1853E-06 | -0.2694712 | 0.169 | 0.243 | 0.04665712 | TEC_prg4 | ITGB6 |
| 4.2584E-15 | -0.2696325 | 0.213 | 0.357 | 9.0917E-11 | TEC_prg4 | COMMD6 |
| 1.3506E-08 | -0.2697276 | 0.166 | 0.265 | 0.00028835 | TEC_prg4 | LAMB3 |
| 1.0277E-12 | -0.2706173 | 0.886 | 0.937 | 2.1941E-08 | TEC_prg4 | RPL37 |
| 1.5782E-14 | -0.2709654 | 0.831 | 0.906 | 3.3695E-10 | TEC_prg4 | RPS25 |
| 8.7575E-15 | -0.2720324 | 0.562 | 0.688 | 1.8697E-10 | TEC_prg4 | CLIC1 |
| 1.9431E-09 | -0.2723497 | 0.715 | 0.787 | 4.1486E-05 | TEC_prg4 | EEF1B2 |
| 7.5757E-06 | -0.27254 | 0.226 | 0.3 | 0.16174191 | TEC_prg4 | DST |
| 7.7217E-14 | -0.2729709 | 0.208 | 0.339 | 1.6486E-09 | TEC_prg4 | ATP6V1F |
| 1.2206E-16 | -0.2742188 | 0.119 | 0.259 | 2.6059E-12 | TEC_prg4 | ATOX1 |
| 2.3915E-13 | -0.2750292 | 0.967 | 0.972 | 5.1058E-09 | TEC_prg4 | TPT1 |
| 1.2976E-11 | -0.2751168 | 0.261 | 0.39 | 2.7704E-07 | TEC_prg4 | PTMS |
| 3.8916E-13 | -0.2755197 | 0.702 | 0.81 | 8.3086E-09 | TEC_prg4 | RPL31 |
| 1.7842E-10 | -0.2756452 | 0.547 | 0.675 | 3.8093E-06 | TEC_prg4 | TPI1 |
| 1.4901E-11 | -0.2769177 | 0.114 | 0.219 | 3.1814E-07 | TEC_prg4 | CTSH |
| 3.5554E-19 | -0.2778045 | 0.071 | 0.212 | 7.5909E-15 | TEC_prg4 | AIF1L |
| 3.2393E-18 | -0.2778841 | 0.138 | 0.294 | 6.9159E-14 | TEC_prg4 | NDUFB11 |
| 2.281E-12 | -0.2789519 | 0.338 | 0.476 | 4.8699E-08 | TEC_prg4 | PDIA6 |
| 3.5577E-13 | -0.2813642 | 0.233 | 0.366 | 7.5957E-09 | TEC_prg4 | ATP5ME |
| 7.7475E-13 | -0.2816158 | 0.4 | 0.523 | 1.6541E-08 | TEC_prg4 | ARPC2 |
| 6.2188E-09 | -0.2820018 | 0.916 | 0.935 | 0.00013277 | TEC_prg4 | RPS27A |
| 5.4575E-20 | -0.2828249 | 0.111 | 0.265 | 1.1652E-15 | TEC_prg4 | NDUFA11 |
| 1.6927E-17 | -0.28319 | 0.23 | 0.389 | 3.6139E-13 | TEC_prg4 | HSBP1 |
| 9.6812E-16 | -0.2839227 | 0.25 | 0.403 | 2.0669E-11 | TEC_prg4 | FKBP2 |
| 5.3861E-19 | -0.284562 | 0.089 | 0.233 | 1.1499E-14 | TEC_prg4 | C4orf48 |
| 3.9487E-12 | -0.2854789 | 0.121 | 0.233 | 8.4305E-08 | TEC_prg4 | IFITM2 |
| 3.753E-12 | -0.2881661 | 0.239 | 0.368 | 8.0127E-08 | TEC_prg4 | ATP5MC3 |
| 8.6898E-13 | -0.2886774 | 0.229 | 0.356 | 1.8553E-08 | TEC_prg4 | ACADVL |
| 4.5114E-12 | -0.2887202 | 0.374 | 0.502 | 9.6318E-08 | TEC_prg4 | ACTN4 |
| 4.8217E-17 | -0.289145 | 0.166 | 0.322 | 1.0294E-12 | TEC_prg4 | UQCR11 |
| 3.637E-09 | -0.2892824 | 0.866 | 0.9 | 7.7651E-05 | TEC_prg4 | RPL23A |
| 1.4817E-14 | -0.2899504 | 0.424 | 0.564 | 3.1635E-10 | TEC_prg4 | LRRFIP1 |
| 4.5717E-18 | -0.2900073 | 0.144 | 0.295 | 9.7605E-14 | TEC_prg4 | RABAC1 |
| 7.8827E-19 | -0.2913941 | 0.131 | 0.282 | 1.683E-14 | TEC_prg4 | CAPNS1 |

| | | | | | | |
|---|---|---|---|---|---|---|
| 1.1E-07 | -0.2930484 | 0.048 | 0.111 | 0.00234845 | TEC_prg4 | BST2 |
| 1.1155E-07 | -0.2935132 | 0.776 | 0.839 | 0.00238165 | TEC_prg4 | PPIA |
| 1.3736E-12 | -0.2945712 | 0.905 | 0.947 | 2.9327E-08 | TEC_prg4 | RPS15 |
| 9.7295E-15 | -0.2949311 | 0.194 | 0.333 | 2.0772E-10 | TEC_prg4 | CTSB |
| 5.7273E-19 | -0.2971867 | 0.44 | 0.61 | 1.2228E-14 | TEC_prg4 | CAST |
| 2.5476E-21 | -0.2976028 | 0.461 | 0.636 | 5.4392E-17 | TEC_prg4 | SLC25A6 |
| 2.3509E-20 | -0.2978853 | 0.144 | 0.31 | 5.0191E-16 | TEC_prg4 | CIB1 |
| 2.7662E-05 | -0.2996323 | 0.064 | 0.114 | 0.59058755 | TEC_prg4 | C15orf48 |
| 3.0892E-24 | -0.2997693 | 0.083 | 0.256 | 6.5955E-20 | TEC_prg4 | BCAP31 |
| 0.0007371 | -0.3004493 | 0.202 | 0.256 | 1 | TEC_prg4 | AKAP12 |
| 2.3145E-16 | -0.3007367 | 0.499 | 0.643 | 4.9415E-12 | TEC_prg4 | CCNI |
| 5.355E-13 | -0.3010613 | 0.716 | 0.813 | 1.1433E-08 | TEC_prg4 | ATP5F1E |
| 2.3278E-08 | -0.3037357 | 0.307 | 0.396 | 0.00049698 | TEC_prg4 | NDRG1 |
| 1.2601E-18 | -0.3048503 | 0.14 | 0.291 | 2.6904E-14 | TEC_prg4 | MSN |
| 6.1711E-13 | -0.304967 | 0.223 | 0.352 | 1.3175E-08 | TEC_prg4 | MIF |
| 1.9379E-14 | -0.3051661 | 0.218 | 0.352 | 4.1373E-10 | TEC_prg4 | CAP1 |
| 1.2407E-12 | -0.305583 | 0.338 | 0.468 | 2.6489E-08 | TEC_prg4 | COX8A |
| 4.0529E-14 | -0.3061475 | 0.266 | 0.399 | 8.653E-10 | TEC_prg4 | CAPN2 |
| 2.1255E-13 | -0.306748 | 0.127 | 0.244 | 4.5379E-09 | TEC_prg4 | HIGD1A |
| 1.3751E-20 | -0.3069388 | 0.159 | 0.334 | 2.9359E-16 | TEC_prg4 | NDUFB7 |
| 4.6929E-17 | -0.3093501 | 0.22 | 0.369 | 1.0019E-12 | TEC_prg4 | NEDD8 |
| 1.3464E-22 | -0.3094903 | 0.121 | 0.294 | 2.8745E-18 | TEC_prg4 | ROMO1 |
| 7.5965E-11 | -0.3095866 | 0.172 | 0.279 | 1.6219E-06 | TEC_prg4 | MUC1 |
| 4.2986E-13 | -0.3118459 | 0.185 | 0.309 | 9.1775E-09 | TEC_prg4 | C9orf16 |
| 4.4023E-11 | -0.3125212 | 0.272 | 0.391 | 9.3989E-07 | TEC_prg4 | MYH9 |
| 3.5904E-16 | -0.3127322 | 0.335 | 0.488 | 7.6654E-12 | TEC_prg4 | P4HB |
| 2.9217E-24 | -0.3127777 | 0.106 | 0.282 | 6.2378E-20 | TEC_prg4 | ANAPC11 |
| 1.0887E-20 | -0.31437 | 0.114 | 0.271 | 2.3243E-16 | TEC_prg4 | FKBP1A |
| 5.6767E-10 | -0.3151866 | 0.183 | 0.292 | 1.212E-05 | TEC_prg4 | PLAUR |
| 5.6476E-22 | -0.3161768 | 0.132 | 0.303 | 1.2058E-17 | TEC_prg4 | BRK1 |
| 1.2207E-17 | -0.316509 | 0.214 | 0.365 | 2.6063E-13 | TEC_prg4 | EIF3K |
| 3.8293E-20 | -0.3167606 | 0.073 | 0.219 | 8.1756E-16 | TEC_prg4 | SCCPDH |
| 2.9113E-15 | -0.3198296 | 0.546 | 0.674 | 6.2156E-11 | TEC_prg4 | SPINT2 |
| 5.4277E-25 | -0.3205612 | 0.099 | 0.277 | 1.1588E-20 | TEC_prg4 | TSTD1 |
| 2.366E-12 | -0.3224551 | 0.175 | 0.29 | 5.0514E-08 | TEC_prg4 | COL18A1 |
| 8.8136E-20 | -0.3227394 | 0.163 | 0.321 | 1.8817E-15 | TEC_prg4 | PCBD1 |
| 4.9048E-08 | -0.3229062 | 0.194 | 0.281 | 0.00104717 | TEC_prg4 | CYB5A |
| 1.4397E-14 | -0.3270828 | 0.202 | 0.336 | 3.0737E-10 | TEC_prg4 | UQCRQ |
| 3.7007E-22 | -0.3278293 | 0.148 | 0.32 | 7.9009E-18 | TEC_prg4 | PGK1 |
| 9.9643E-13 | -0.3298256 | 0.574 | 0.686 | 2.1274E-08 | TEC_prg4 | EPCAM |
| 6.546E-18 | -0.331143 | 0.671 | 0.793 | 1.3976E-13 | TEC_prg4 | CHCHD2 |
| 6.4379E-13 | -0.33197 | 0.491 | 0.622 | 1.3745E-08 | TEC_prg4 | CD81 |
| 2.6328E-17 | -0.3320095 | 0.694 | 0.806 | 5.621E-13 | TEC_prg4 | RPL27 |
| 4.1629E-16 | -0.3340192 | 0.421 | 0.574 | 8.8878E-12 | TEC_prg4 | PPIB |
| 5.2381E-12 | -0.3364607 | 0.093 | 0.194 | 1.1183E-07 | TEC_prg4 | SYTL2 |
| 1.6513E-21 | -0.3379152 | 0.108 | 0.27 | 3.5254E-17 | TEC_prg4 | GSTO1 |
| 3.5696E-14 | -0.3383307 | 0.802 | 0.891 | 7.621E-10 | TEC_prg4 | RPS29 |
| 4.0768E-19 | -0.3397465 | 0.141 | 0.295 | 8.7039E-15 | TEC_prg4 | ANXA3 |
| 6.1357E-07 | -0.3399005 | 0.345 | 0.433 | 0.01309967 | TEC_prg4 | SYNE2 |
| 1.9271E-22 | -0.3408565 | 0.354 | 0.539 | 4.1144E-18 | TEC_prg4 | COX6C |

| | | | | | | |
|---|---|---|---|---|---|---|
| 3.6522E-11 | -0.3425346 | 0.261 | 0.374 | 7.7974E-07 | TEC_prg4 | MACF1 |
| 6.694E-17 | -0.3436337 | 0.559 | 0.697 | 1.4292E-12 | TEC_prg4 | CYSTM1 |
| 4.2219E-24 | -0.3444576 | 0.1 | 0.273 | 9.0139E-20 | TEC_prg4 | APRT |
| 8.0038E-18 | -0.344554 | 0.879 | 0.933 | 1.7088E-13 | TEC_prg4 | RPL7A |
| 9.6994E-14 | -0.3446494 | 0.908 | 0.942 | 2.0708E-09 | TEC_prg4 | RPL37A |
| 7.0783E-20 | -0.3456584 | 0.109 | 0.26 | 1.5112E-15 | TEC_prg4 | ECHS1 |
| 2.6635E-19 | -0.3465422 | 0.464 | 0.617 | 5.6866E-15 | TEC_prg4 | ATP5MG |
| 6.6028E-15 | -0.3478817 | 0.112 | 0.239 | 1.4097E-10 | TEC_prg4 | TGM2 |
| 1.6498E-16 | -0.3523817 | 0.229 | 0.378 | 3.5223E-12 | TEC_prg4 | TAGLN2 |
| 6.409E-19 | -0.3531652 | 0.215 | 0.376 | 1.3683E-14 | TEC_prg4 | ATP5MF |
| 0.0001045 | -0.3549045 | 0.549 | 0.603 | 1 | TEC_prg4 | CLU |
| 5.4137E-22 | -0.3550651 | 0.25 | 0.426 | 1.1558E-17 | TEC_prg4 | PARK7 |
| 5.4484E-17 | -0.3556518 | 0.888 | 0.945 | 1.1632E-12 | TEC_prg4 | RPS24 |
| 0.00015991 | -0.3560448 | 0.067 | 0.111 | 1 | TEC_prg4 | S100A9 |
| 4.9527E-06 | -0.3574451 | 0.933 | 0.95 | 0.10573958 | TEC_prg4 | B2M |
| 1.1233E-23 | -0.3576807 | 0.197 | 0.384 | 2.3983E-19 | TEC_prg4 | HEBP2 |
| 2.0636E-19 | -0.3577169 | 0.668 | 0.805 | 4.4059E-15 | TEC_prg4 | HLA-C |
| 1.103E-20 | -0.358185 | 0.218 | 0.391 | 2.3548E-16 | TEC_prg4 | DYNLT1 |
| 1.023E-11 | -0.3595165 | 0.175 | 0.291 | 2.184E-07 | TEC_prg4 | AKR1B1 |
| 2.903E-16 | -0.3602118 | 0.381 | 0.526 | 6.1979E-12 | TEC_prg4 | OST4 |
| 8.7401E-17 | -0.3606587 | 0.793 | 0.873 | 1.866E-12 | TEC_prg4 | RPL10A |
| 5.8796E-21 | -0.3622414 | 0.358 | 0.537 | 1.2553E-16 | TEC_prg4 | ELOB |
| 4.6602E-22 | -0.3627998 | 0.419 | 0.591 | 9.9495E-18 | TEC_prg4 | COX5B |
| 1.2071E-23 | -0.3662938 | 0.387 | 0.571 | 2.5772E-19 | TEC_prg4 | COX6B1 |
| 1.1364E-16 | -0.3664488 | 0.598 | 0.739 | 2.4262E-12 | TEC_prg4 | NDUFA4 |
| 1.7277E-12 | -0.3665219 | 0.297 | 0.427 | 3.6886E-08 | TEC_prg4 | LGALS3 |
| 8.0735E-15 | -0.3675967 | 0.668 | 0.77 | 1.7237E-10 | TEC_prg4 | PKM |
| 7.9574E-23 | -0.3696326 | 0.86 | 0.937 | 1.6989E-18 | TEC_prg4 | RPS9 |
| 2.9526E-16 | -0.3700142 | 0.824 | 0.904 | 6.3039E-12 | TEC_prg4 | SERF2 |
| 1.6194E-18 | -0.3700604 | 0.169 | 0.324 | 3.4574E-14 | TEC_prg4 | PRDX2 |
| 5.4936E-17 | -0.3736628 | 0.424 | 0.569 | 1.1729E-12 | TEC_prg4 | GUK1 |
| 2.1303E-20 | -0.3752408 | 0.231 | 0.404 | 4.5483E-16 | TEC_prg4 | ATP5F1D |
| 1.099E-20 | -0.375833 | 0.789 | 0.877 | 2.3463E-16 | TEC_prg4 | RACK1 |
| 2.1055E-16 | -0.3769454 | 0.144 | 0.286 | 4.4953E-12 | TEC_prg4 | SERPINB1 |
| 4.1404E-21 | -0.3792716 | 0.282 | 0.456 | 8.8397E-17 | TEC_prg4 | ATP5MD |
| 1.4757E-22 | -0.3807756 | 0.862 | 0.938 | 3.1506E-18 | TEC_prg4 | RPL3 |
| 3.7341E-21 | -0.3812437 | 0.841 | 0.924 | 7.9723E-17 | TEC_prg4 | RPL34 |
| 1.1313E-07 | -0.3820159 | 0.066 | 0.132 | 0.00241535 | TEC_prg4 | COL7A1 |
| 6.0119E-17 | -0.382466 | 0.769 | 0.87 | 1.2835E-12 | TEC_prg4 | UBA52 |
| 1.1436E-15 | -0.3846495 | 0.199 | 0.34 | 2.4416E-11 | TEC_prg4 | CHCHD10 |
| 2.3333E-05 | -0.385614 | 0.192 | 0.265 | 0.49814901 | TEC_prg4 | NUPR1 |
| 4.3613E-17 | -0.385772 | 0.038 | 0.153 | 9.3114E-13 | TEC_prg4 | MYL9 |
| 4.6995E-05 | -0.3859826 | 0.338 | 0.393 | 1 | TEC_prg4 | TUBA1B |
| 2.1137E-18 | -0.3893422 | 0.237 | 0.393 | 4.5127E-14 | TEC_prg4 | OCIAD2 |
| 2.2518E-08 | -0.3904343 | 0.217 | 0.313 | 0.00048077 | TEC_prg4 | PLAU |
| 6.6651E-21 | -0.390952 | 0.767 | 0.875 | 1.423E-16 | TEC_prg4 | RPS5 |
| 4.2831E-17 | -0.3924525 | 0.785 | 0.879 | 9.1445E-13 | TEC_prg4 | HLA-A |
| 1.6367E-21 | -0.3970863 | 0.194 | 0.358 | 3.4943E-17 | TEC_prg4 | ANXA11 |
| 3.5079E-27 | -0.3974368 | 0.162 | 0.36 | 7.4893E-23 | TEC_prg4 | MZT2B |
| 5.7807E-22 | -0.3976168 | 0.613 | 0.753 | 1.2342E-17 | TEC_prg4 | HINT1 |

| | | | | | | |
|---|---|---|---|---|---|---|
| 2.0216E-27 | -0.3993724 | 0.536 | 0.737 | 4.3161E-23 | TEC_prg4 | CSTB |
| 2.1363E-09 | -0.399702 | 0.192 | 0.288 | 4.5611E-05 | TEC_prg4 | UCHL1 |
| 1.0343E-32 | -0.3998382 | 0.435 | 0.657 | 2.2082E-28 | TEC_prg4 | EDF1 |
| 3.289E-15 | -0.401992 | 0.905 | 0.935 | 7.022E-11 | TEC_prg4 | RPL32 |
| 6.0647E-10 | -0.4040998 | 0.042 | 0.119 | 1.2948E-05 | TEC_prg4 | TAGLN |
| 1.93E-09 | -0.4047254 | 0.223 | 0.325 | 4.1205E-05 | TEC_prg4 | ITGA2 |
| 1.5737E-22 | -0.4056031 | 0.898 | 0.949 | 3.3598E-18 | TEC_prg4 | RPL11 |
| 1.4688E-15 | -0.4062808 | 0.884 | 0.931 | 3.136E-11 | TEC_prg4 | RPL18A |
| 5.4088E-20 | -0.4079062 | 0.898 | 0.94 | 1.1548E-15 | TEC_prg4 | RPS23 |
| 1.7021E-08 | -0.4085312 | 0.118 | 0.204 | 0.00036339 | TEC_prg4 | ANXA1 |
| 4.2709E-26 | -0.4090646 | 0.124 | 0.309 | 9.1184E-22 | TEC_prg4 | CAPG |
| 5.4687E-12 | -0.4099256 | 0.574 | 0.668 | 1.1676E-07 | TEC_prg4 | TACSTD2 |
| 6.5469E-18 | -0.4099923 | 0.406 | 0.552 | 1.3978E-13 | TEC_prg4 | CD59 |
| 1.0142E-26 | -0.4131682 | 0.188 | 0.386 | 2.1652E-22 | TEC_prg4 | VAMP8 |
| 2.1653E-26 | -0.4138902 | 0.674 | 0.831 | 4.623E-22 | TEC_prg4 | RPS21 |
| 1.5404E-34 | -0.4156626 | 0.476 | 0.704 | 3.2888E-30 | TEC_prg4 | GPX4 |
| 2.4521E-23 | -0.4167382 | 0.213 | 0.391 | 5.2351E-19 | TEC_prg4 | NDUFB2 |
| 8.3149E-23 | -0.4182296 | 0.856 | 0.932 | 1.7752E-18 | TEC_prg4 | RPL36 |
| 7.1126E-21 | -0.4215398 | 0.351 | 0.513 | 1.5185E-16 | TEC_prg4 | CLDN7 |
| 1.8681E-19 | -0.4248086 | 0.838 | 0.916 | 3.9885E-15 | TEC_prg4 | RPL29 |
| 5.1446E-12 | -0.4251483 | 0.544 | 0.674 | 1.0984E-07 | TEC_prg4 | CRYAB |
| 5.7899E-18 | -0.425188 | 0.924 | 0.958 | 1.2362E-13 | TEC_prg4 | RPS2 |
| 7.0246E-20 | -0.4287329 | 0.907 | 0.954 | 1.4997E-15 | TEC_prg4 | RPS12 |
| 9.905E-11 | -0.4337371 | 0.194 | 0.301 | 2.1147E-06 | TEC_prg4 | ITGB8 |
| 1.5391E-09 | -0.4347023 | 0.627 | 0.716 | 3.2861E-05 | TEC_prg4 | MT2A |
| 1.9396E-26 | -0.4391074 | 0.365 | 0.556 | 4.141E-22 | TEC_prg4 | BRI3 |
| 1.0185E-27 | -0.4392323 | 0.648 | 0.789 | 2.1745E-23 | TEC_prg4 | COX4I1 |
| 1.3949E-18 | -0.4405334 | 0.939 | 0.963 | 2.9781E-14 | TEC_prg4 | RPS18 |
| 7.2775E-27 | -0.441733 | 0.179 | 0.371 | 1.5537E-22 | TEC_prg4 | PSME1 |
| 4.4619E-20 | -0.4460912 | 0.952 | 0.973 | 9.5262E-16 | TEC_prg4 | RPLP1 |
| 2.0511E-06 | -0.4517759 | 0.108 | 0.175 | 0.04379102 | TEC_prg4 | S100A4 |
| 3.6917E-30 | -0.4523989 | 0.335 | 0.536 | 7.8819E-26 | TEC_prg4 | ATP5MC2 |
| 5.4447E-16 | -0.4536532 | 0.208 | 0.345 | 1.1624E-11 | TEC_prg4 | PAX8 |
| 5.2536E-24 | -0.455031 | 0.677 | 0.807 | 1.1216E-19 | TEC_prg4 | CD24 |
| 8.6389E-29 | -0.45732 | 0.201 | 0.409 | 1.8444E-24 | TEC_prg4 | MGST3 |
| 7.2542E-25 | -0.4584131 | 0.94 | 0.975 | 1.5488E-20 | TEC_prg4 | RPL13 |
| 1.9698E-22 | -0.4604585 | 0.85 | 0.928 | 4.2055E-18 | TEC_prg4 | RPLP2 |
| 1.0241E-16 | -0.4636595 | 0.192 | 0.327 | 2.1865E-12 | TEC_prg4 | DCDC2 |
| 8.0667E-23 | -0.4667863 | 0.905 | 0.953 | 1.7222E-18 | TEC_prg4 | RPS8 |
| 1.4699E-19 | -0.4671283 | 0.29 | 0.447 | 3.1382E-15 | TEC_prg4 | PPP1R14B |
| 2.5281E-22 | -0.4698 | 0.838 | 0.92 | 5.3975E-18 | TEC_prg4 | RPLP0 |
| 1.4832E-18 | -0.4714645 | 0.178 | 0.333 | 3.1666E-14 | TEC_prg4 | CYBA |
| 6.2896E-34 | -0.4846354 | 0.146 | 0.371 | 1.3428E-29 | TEC_prg4 | TALDO1 |
| 2.2486E-21 | -0.4862029 | 0.189 | 0.353 | 4.8008E-17 | TEC_prg4 | S100A16 |
| 1.3284E-28 | -0.4862212 | 0.59 | 0.736 | 2.8361E-24 | TEC_prg4 | ENO1 |
| 2.5746E-19 | -0.4869452 | 0.773 | 0.877 | 5.4968E-15 | TEC_prg4 | SAT1 |
| 8.6151E-29 | -0.4900314 | 0.231 | 0.436 | 1.8393E-24 | TEC_prg4 | SSR4 |
| 1.5449E-43 | -0.495904 | 0.284 | 0.547 | 3.2983E-39 | TEC_prg4 | NDUFS5 |
| 9.9268E-27 | -0.4991195 | 0.21 | 0.401 | 2.1194E-22 | TEC_prg4 | CD151 |
| 2.718E-32 | -0.5000106 | 0.387 | 0.594 | 5.8029E-28 | TEC_prg4 | ARPC3 |

| | | | | | | |
|---|---|---|---|---|---|---|
| 9.7165E-32 | -0.5010866 | 0.802 | 0.91 | 2.0745E-27 | TEC_prg4 | RPS3 |
| 2.8746E-40 | -0.5045012 | 0.62 | 0.815 | 6.1372E-36 | TEC_prg4 | EEF1D |
| 2.1805E-32 | -0.506114 | 0.723 | 0.891 | 4.6554E-28 | TEC_prg4 | RPL39 |
| 1.6752E-25 | -0.5109965 | 0.384 | 0.56 | 3.5766E-21 | TEC_prg4 | CD9 |
| 1.0934E-28 | -0.5109991 | 0.662 | 0.812 | 2.3343E-24 | TEC_prg4 | CFL1 |
| 1.4211E-14 | -0.5188221 | 0.15 | 0.273 | 3.0341E-10 | TEC_prg4 | TUBA1A |
| 6.0571E-30 | -0.5191056 | 0.882 | 0.932 | 1.2932E-25 | TEC_prg4 | RPS14 |
| 0.00016122 | -0.5241231 | 0.189 | 0.246 | 1 | TEC_prg4 | TFPI2 |
| 6.5906E-34 | -0.5246934 | 0.384 | 0.599 | 1.4071E-29 | TEC_prg4 | S100A13 |
| 1.3575E-39 | -0.5266715 | 0.608 | 0.802 | 2.8982E-35 | TEC_prg4 | PPDPF |
| 1.0142E-26 | -0.5267035 | 0.386 | 0.575 | 2.1654E-22 | TEC_prg4 | MAL2 |
| 7.6103E-24 | -0.5349957 | 0.148 | 0.325 | 1.6248E-19 | TEC_prg4 | MAL |
| 3.0629E-41 | -0.5350176 | 0.342 | 0.593 | 6.5393E-37 | TEC_prg4 | PRDX5 |
| 1.3661E-23 | -0.5392969 | 0.729 | 0.849 | 2.9165E-19 | TEC_prg4 | RPSA |
| 1.3502E-20 | -0.5404557 | 0.576 | 0.711 | 2.8827E-16 | TEC_prg4 | ITGB1 |
| 2.4097E-35 | -0.5512264 | 0.119 | 0.339 | 5.1446E-31 | TEC_prg4 | TSPO |
| 8.285E-08 | -0.5533668 | 0.31 | 0.397 | 0.00176885 | TEC_prg4 | TXNRD1 |
| 2.3548E-18 | -0.5579187 | 0.274 | 0.426 | 5.0275E-14 | TEC_prg4 | CCND1 |
| 1.9694E-17 | -0.5609844 | 0.089 | 0.22 | 4.2046E-13 | TEC_prg4 | GLRX |
| 2.4313E-06 | -0.564673 | 0.054 | 0.109 | 0.05190777 | TEC_prg4 | FXYD4 |
| 4.0478E-26 | -0.5671788 | 0.396 | 0.568 | 8.642E-22 | TEC_prg4 | HMGA1 |
| 3.8597E-22 | -0.5724005 | 0.303 | 0.476 | 8.2405E-18 | TEC_prg4 | IFITM3 |
| 1.3803E-33 | -0.5742024 | 0.192 | 0.408 | 2.9469E-29 | TEC_prg4 | RHOC |
| 8.0466E-17 | -0.5773051 | 0.822 | 0.897 | 1.7179E-12 | TEC_prg4 | GAPDH |
| 4.9657E-42 | -0.5848585 | 0.486 | 0.728 | 1.0602E-37 | TEC_prg4 | TXN |
| 2.771E-24 | -0.6046162 | 0.086 | 0.254 | 5.9162E-20 | TEC_prg4 | PDZK1IP1 |
| 2.5377E-38 | -0.6147753 | 0.654 | 0.847 | 5.4179E-34 | TEC_prg4 | PRDX1 |
| 2.3433E-38 | -0.6162945 | 0.814 | 0.937 | 5.0029E-34 | TEC_prg4 | RPL12 |
| 1.0904E-44 | -0.6171056 | 0.691 | 0.875 | 2.3281E-40 | TEC_prg4 | RPS4X |
| 1.3575E-46 | -0.6414295 | 0.811 | 0.933 | 2.8983E-42 | TEC_prg4 | ACTG1 |
| 5.2606E-37 | -0.6494917 | 0.547 | 0.742 | 1.1231E-32 | TEC_prg4 | PFN1 |
| 1.0441E-12 | -0.6499396 | 0.211 | 0.337 | 2.2292E-08 | TEC_prg4 | TIMP1 |
| 5.5428E-14 | -0.6525915 | 0.284 | 0.413 | 1.1834E-09 | TEC_prg4 | RNF213 |
| 1.0212E-13 | -0.6572147 | 0.16 | 0.284 | 2.1803E-09 | TEC_prg4 | LGALS1 |
| 9.914E-46 | -0.6804534 | 0.776 | 0.932 | 2.1166E-41 | TEC_prg4 | RPL10 |
| 9.1E-16 | -0.6813345 | 0.115 | 0.243 | 1.9429E-11 | TEC_prg4 | CST6 |
| 2.4817E-50 | -0.7023242 | 0.789 | 0.914 | 5.2985E-46 | TEC_prg4 | MYL6 |
| 6.9065E-24 | -0.7092166 | 0.16 | 0.336 | 1.4745E-19 | TEC_prg4 | IL32 |
| 1.1907E-33 | -0.7125673 | 0.218 | 0.432 | 2.5421E-29 | TEC_prg4 | TPM4 |
| 1.6881E-47 | -0.7153346 | 0.686 | 0.848 | 3.6041E-43 | TEC_prg4 | CD63 |
| 1.5034E-09 | -0.7215022 | 0.943 | 0.962 | 3.2097E-05 | TEC_prg4 | FTL |
| 2.2119E-49 | -0.7300799 | 0.706 | 0.883 | 4.7223E-45 | TEC_prg4 | GSTP1 |
| 1.1554E-17 | -0.7551059 | 0.194 | 0.338 | 2.4668E-13 | TEC_prg4 | LAMC2 |
| 7.085E-58 | -0.7557676 | 0.873 | 0.965 | 1.5126E-53 | TEC_prg4 | RPL28 |
| 2.3429E-22 | -0.7566779 | 0.691 | 0.805 | 5.0022E-18 | TEC_prg4 | KRT18 |
| 0.00030994 | -0.773759 | 0.084 | 0.129 | 1 | TEC_prg4 | GPX3 |
| 1.637E-22 | -0.8035154 | 0.959 | 0.979 | 3.4949E-18 | TEC_prg4 | FTH1 |
| 5.2361E-12 | -0.8058988 | 0.15 | 0.261 | 1.1179E-07 | TEC_prg4 | LCN2 |
| 1.1762E-22 | -0.8077409 | 0.597 | 0.735 | 2.5113E-18 | TEC_prg4 | KRT8 |
| 9.9011E-36 | -0.8107212 | 0.735 | 0.878 | 2.1139E-31 | TEC_prg4 | S100A10 |

| | | | | | | |
|---|---|---|---|---|---|---|
| 1.7324E-55 | -0.8248232 | 0.757 | 0.933 | 3.6987E-51 | TEC_prg4 | TMSB10 |
| 4.3149E-45 | -0.8435219 | 0.406 | 0.643 | 9.2124E-41 | TEC_prg4 | TPM1 |
| 1.9671E-05 | -0.8665482 | 0.14 | 0.201 | 0.41997653 | TEC_prg4 | MT1G |
| 1.196E-36 | -0.9081023 | 0.425 | 0.635 | 2.5535E-32 | TEC_prg4 | VIM |
| 4.6458E-43 | -0.9139562 | 0.252 | 0.498 | 9.9188E-39 | TEC_prg4 | TSPAN1 |
| 5.907E-09 | -0.9196682 | 0.403 | 0.497 | 0.00012611 | TEC_prg4 | S100A2 |
| 7.601E-34 | -0.9572873 | 0.255 | 0.482 | 1.6228E-29 | TEC_prg4 | C12orf75 |
| 7.3091E-35 | -0.9964457 | 0.405 | 0.602 | 1.5605E-30 | TEC_prg4 | KRT19 |
| 3.5123E-44 | -0.9965996 | 0.789 | 0.926 | 7.4988E-40 | TEC_prg4 | S100A11 |
| 1.6984E-42 | -1.0172024 | 0.467 | 0.683 | 3.6262E-38 | TEC_prg4 | SH3BGRL3 |
| 1.5095E-56 | -1.0338649 | 0.821 | 0.928 | 3.2227E-52 | TEC_prg4 | ACTB |
| 7.9919E-57 | -1.0785411 | 0.402 | 0.663 | 1.7063E-52 | TEC_prg4 | KRT7 |
| 5.2231E-48 | -1.0830836 | 0.665 | 0.84 | 1.1151E-43 | TEC_prg4 | ANXA2 |
| 1.8245E-18 | -1.1710226 | 0.246 | 0.4 | 3.8954E-14 | TEC_prg4 | SLPI |
| 1.1016E-11 | -1.1821739 | 0.809 | 0.586 | 2.3519E-07 | TEC_prg4 | PLCG2 |
| 2.573E-65 | -1.1870556 | 0.833 | 0.962 | 5.4934E-61 | TEC_prg4 | S100A6 |
| 2.0011E-69 | -1.3401584 | 0.675 | 0.879 | 4.2723E-65 | TEC_prg4 | TMSB4X |
| 1.0046E-22 | -1.5438628 | 0.505 | 0.644 | 2.1449E-18 | TEC_prg4 | MMP7 |
| 0 | 2.68520328 | 0.793 | 0.025 | 0 | TEC_prlf | CENPF |
| 6.1E-185 | 1.94365456 | 0.752 | 0.201 | 1.302E-180 | TEC_prlf | UBE2S |
| 4.317E-216 | 1.91886801 | 0.887 | 0.27 | 9.217E-212 | TEC_prlf | STMN1 |
| 0 | 1.89746972 | 0.691 | 0.034 | 0 | TEC_prlf | CCNB1 |
| 8.548E-129 | 1.88791175 | 0.702 | 0.249 | 1.825E-124 | TEC_prlf | HMGB2 |
| 4.538E-173 | 1.88559178 | 0.992 | 0.692 | 9.689E-169 | TEC_prlf | HMGB1 |
| 0 | 1.83792666 | 0.7 | 0.058 | 0 | TEC_prlf | PTTG1 |
| 0 | 1.77609738 | 0.623 | 0.016 | 0 | TEC_prlf | MKI67 |
| 1.166E-151 | 1.73179953 | 0.835 | 0.338 | 2.489E-147 | TEC_prlf | NUCKS1 |
| 1.187E-161 | 1.70956401 | 0.862 | 0.348 | 2.535E-157 | TEC_prlf | HMGN2 |
| 3.461E-107 | 1.5842346 | 0.796 | 0.371 | 7.39E-103 | TEC_prlf | PTMS |
| 1.02E-173 | 1.55877077 | 0.614 | 0.13 | 2.179E-169 | TEC_prlf | CKS2 |
| 0 | 1.5217253 | 0.628 | 0.017 | 0 | TEC_prlf | HMMR |
| 2.8758E-86 | 1.47764468 | 0.683 | 0.266 | 6.1399E-82 | TEC_prlf | LGALS1 |
| 1.515E-114 | 1.42704621 | 0.906 | 0.499 | 3.236E-110 | TEC_prlf | H2AFZ |
| 0 | 1.4045462 | 0.565 | 0.041 | 0 | TEC_prlf | NUSAP1 |
| 0 | 1.4033166 | 0.612 | 0.016 | 0 | TEC_prlf | TPX2 |
| 0 | 1.38139737 | 0.482 | 0.011 | 0 | TEC_prlf | TOP2A |
| 0 | 1.32561509 | 0.636 | 0.016 | 0 | TEC_prlf | BIRC5 |
| 0 | 1.29239554 | 0.565 | 0.011 | 0 | TEC_prlf | CDC20 |
| 6.435E-248 | 1.25963088 | 0.556 | 0.073 | 1.374E-243 | TEC_prlf | SMC4 |
| 0 | 1.23402949 | 0.471 | 0.027 | 0 | TEC_prlf | CKAP2 |
| 0 | 1.11471378 | 0.545 | 0.016 | 0 | TEC_prlf | CDKN3 |
| 2.5549E-93 | 1.08192366 | 0.752 | 0.338 | 5.4546E-89 | TEC_prlf | MZT2B |
| 0 | 1.08141158 | 0.424 | 0.011 | 0 | TEC_prlf | ASPM |
| 2.7803E-52 | 1.0725995 | 0.708 | 0.381 | 5.9358E-48 | TEC_prlf | TUBA1B |
| 0 | 1.0594734 | 0.438 | 0.017 | 0 | TEC_prlf | AURKA |
| 7.926E-135 | 1.04209716 | 0.7 | 0.217 | 1.692E-130 | TEC_prlf | H2AFV |
| 2.2196E-58 | 1.01074365 | 0.609 | 0.28 | 4.7389E-54 | TEC_prlf | HP1BP3 |
| 0 | 1.00551547 | 0.435 | 0.01 | 0 | TEC_prlf | GTSE1 |
| 0 | 0.95588638 | 0.474 | 0.007 | 0 | TEC_prlf | CEP55 |
| 1.1915E-49 | 0.92417329 | 0.964 | 0.871 | 2.5438E-45 | TEC_prlf | GSTP1 |

| | | | | | | |
|---|---|---|---|---|---|---|
| 1.24E-111 | 0.91803788 | 0.488 | 0.114 | 2.648E-107 | TEC_prlf | TMPO |
| 2.5614E-64 | 0.90664769 | 0.986 | 0.933 | 5.4686E-60 | TEC_prlf | PTMA |
| 0 | 0.88065844 | 0.449 | 0.004 | 0 | TEC_prlf | DLGAP5 |
| 4.632E-20 | 0.87648584 | 0.915 | 0.796 | 9.8893E-16 | TEC_prlf | KRT18 |
| 0 | 0.87350305 | 0.427 | 0.012 | 0 | TEC_prlf | CCNB2 |
| 4.0909E-34 | 0.87309605 | 0.298 | 0.106 | 8.7341E-30 | TEC_prlf | RARRES2 |
| 2.2567E-54 | 0.87045195 | 0.826 | 0.561 | 4.818E-50 | TEC_prlf | RAN |
| 0 | 0.8691231 | 0.479 | 0.018 | 0 | TEC_prlf | TACC3 |
| 2.7622E-68 | 0.86379779 | 0.766 | 0.376 | 5.8973E-64 | TEC_prlf | ATP5IF1 |
| 2.809E-48 | 0.85352051 | 0.245 | 0.06 | 5.9973E-44 | TEC_prlf | SOD3 |
| 1.3057E-60 | 0.85191148 | 0.972 | 0.791 | 2.7876E-56 | TEC_prlf | HNRNPA2B1 |
| 4.4984E-51 | 0.84074047 | 0.697 | 0.388 | 9.604E-47 | TEC_prlf | DBI |
| 8.768E-110 | 0.82821106 | 0.281 | 0.039 | 1.872E-105 | TEC_prlf | THY1 |
| 1.7797E-70 | 0.82327679 | 0.584 | 0.225 | 3.7996E-66 | TEC_prlf | CALM3 |
| 6.469E-54 | 0.80850866 | 0.628 | 0.31 | 1.3811E-49 | TEC_prlf | DEK |
| 2.0444E-65 | 0.80317541 | 0.504 | 0.179 | 4.3648E-61 | TEC_prlf | HMGN3 |
| 6.0289E-67 | 0.80222279 | 0.548 | 0.204 | 1.2872E-62 | TEC_prlf | RAD21 |
| 4.634E-197 | 0.80055135 | 0.419 | 0.049 | 9.893E-193 | TEC_prlf | ECT2 |
| 1.108E-14 | 0.79789061 | 0.826 | 0.708 | 2.3656E-10 | TEC_prlf | MT2A |
| 8.4069E-38 | 0.79010669 | 0.526 | 0.261 | 1.7949E-33 | TEC_prlf | TUBB |
| 1.2871E-94 | 0.78041347 | 0.471 | 0.123 | 2.748E-90 | TEC_prlf | LSM5 |
| 2.7459E-46 | 0.77792865 | 0.928 | 0.753 | 5.8626E-42 | TEC_prlf | DYNLL1 |
| 8.9138E-90 | 0.7681776 | 0.601 | 0.199 | 1.9031E-85 | TEC_prlf | MZT2A |
| 0 | 0.76584468 | 0.369 | 0.004 | 0 | TEC_prlf | DEPDC1 |
| 8.2043E-54 | 0.76523862 | 0.601 | 0.276 | 1.7516E-49 | TEC_prlf | ANP32B |
| 1.5632E-44 | 0.76498502 | 0.931 | 0.76 | 3.3373E-40 | TEC_prlf | PKM |
| 5.8777E-16 | 0.76429754 | 0.54 | 0.365 | 1.2549E-11 | TEC_prlf | SPTBN1 |
| 0 | 0.76415464 | 0.485 | 0.039 | 0 | TEC_prlf | TK1 |
| 2.5793E-97 | 0.75908658 | 0.463 | 0.114 | 5.5068E-93 | TEC_prlf | ANP32E |
| 3.1599E-46 | 0.7539306 | 0.722 | 0.43 | 6.7465E-42 | TEC_prlf | HMGN1 |
| 1.9393E-36 | 0.74159977 | 0.545 | 0.278 | 4.1404E-32 | TEC_prlf | SRRM1 |
| 0 | 0.74118847 | 0.419 | 0.006 | 0 | TEC_prlf | PBK |
| 1.0146E-67 | 0.74091185 | 0.51 | 0.175 | 2.1662E-63 | TEC_prlf | RANBP1 |
| 4.2247E-59 | 0.73661294 | 0.956 | 0.815 | 9.0198E-55 | TEC_prlf | YBX1 |
| 4.7956E-42 | 0.73568467 | 0.755 | 0.479 | 1.0239E-37 | TEC_prlf | LDHA |
| 3.9685E-38 | 0.73562589 | 0.934 | 0.768 | 8.4729E-34 | TEC_prlf | HSP90B1 |
| 1.1773E-51 | 0.73276941 | 0.917 | 0.723 | 2.5136E-47 | TEC_prlf | ENO1 |
| 2.3331E-55 | 0.72881944 | 0.912 | 0.666 | 4.9811E-51 | TEC_prlf | HNRNPA1 |
| 0 | 0.72423637 | 0.358 | 0.013 | 0 | TEC_prlf | CENPE |
| 1.1543E-62 | 0.7166062 | 0.614 | 0.263 | 2.4645E-58 | TEC_prlf | CBX3 |
| 0 | 0.71131085 | 0.366 | 0.007 | 0 | TEC_prlf | CCNA2 |
| 6.812E-180 | 0.70960607 | 0.386 | 0.045 | 1.454E-175 | TEC_prlf | KIF20B |
| 1.0091E-43 | 0.70746594 | 0.961 | 0.814 | 2.1544E-39 | TEC_prlf | UBB |
| 0 | 0.70508155 | 0.388 | 0.018 | 0 | TEC_prlf | MAD2L1 |
| 3.8572E-50 | 0.69621829 | 0.534 | 0.234 | 8.2351E-46 | TEC_prlf | PLP2 |
| 2.5218E-45 | 0.69175329 | 0.691 | 0.398 | 5.3839E-41 | TEC_prlf | HNRNPA3 |
| 1.9422E-39 | 0.69057274 | 0.656 | 0.366 | 4.1466E-35 | TEC_prlf | LDHB |
| 0 | 0.68582625 | 0.386 | 0.012 | 0 | TEC_prlf | ANLN |
| 4.6275E-24 | 0.67777341 | 0.292 | 0.118 | 9.8798E-20 | TEC_prlf | CXCL14 |
| 0 | 0.67119929 | 0.344 | 0.013 | 0 | TEC_prlf | PRC1 |

| | | | | | | |
|---|---|---|---|---|---|---|
| 2.8865E-46 | 0.67023379 | 0.909 | 0.741 | 6.1627E-42 | TEC_prlf | HINT1 |
| 0 | 0.67014613 | 0.353 | 0.01 | 0 | TEC_prlf | PLK1 |
| 3.5757E-45 | 0.66959103 | 0.981 | 0.89 | 7.6342E-41 | TEC_prlf | GAPDH |
| 0 | 0.66920107 | 0.322 | 0.008 | 0 | TEC_prlf | CDCA8 |
| 1.8492E-27 | 0.66309153 | 0.752 | 0.555 | 3.9481E-23 | TEC_prlf | NCL |
| 0 | 0.65713842 | 0.355 | 0.004 | 0 | TEC_prlf | NUF2 |
| 2.681E-271 | 0.6544499 | 0.35 | 0.023 | 5.725E-267 | TEC_prlf | KNSTRN |
| 0 | 0.6513469 | 0.372 | 0.015 | 0 | TEC_prlf | KIF23 |
| 2.2381E-30 | 0.6479468 | 0.485 | 0.243 | 4.7784E-26 | TEC_prlf | MRPS6 |
| 9.1265E-34 | 0.64735771 | 0.587 | 0.324 | 1.9485E-29 | TEC_prlf | KIF5B |
| 3.9917E-46 | 0.64661921 | 0.587 | 0.285 | 8.5222E-42 | TEC_prlf | MRPL51 |
| 2.7642E-21 | 0.64262116 | 0.598 | 0.402 | 5.9016E-17 | TEC_prlf | JAK1 |
| 2.1516E-28 | 0.64261055 | 0.584 | 0.326 | 4.5936E-24 | TEC_prlf | RALBP1 |
| 5.6124E-48 | 0.64168814 | 0.562 | 0.255 | 1.1982E-43 | TEC_prlf | HDGF |
| 0 | 0.64078751 | 0.333 | 0.013 | 0 | TEC_prlf | ARHGAP11A |
| 9.6957E-77 | 0.63577226 | 0.383 | 0.095 | 2.07E-72 | TEC_prlf | CKS1B |
| 7.5605E-23 | 0.62893849 | 0.81 | 0.639 | 1.6142E-18 | TEC_prlf | PEBP1 |
| 1.728E-155 | 0.62795441 | 0.421 | 0.062 | 3.69E-151 | TEC_prlf | DTYMK |
| 6.8464E-21 | 0.62240789 | 0.86 | 0.728 | 1.4617E-16 | TEC_prlf | PFN1 |
| 1.1694E-28 | 0.61408999 | 0.311 | 0.124 | 2.4967E-24 | TEC_prlf | MYH10 |
| 2.9618E-35 | 0.61378309 | 0.623 | 0.354 | 6.3234E-31 | TEC_prlf | ATP5MC3 |
| 1.1733E-59 | 0.61372581 | 0.573 | 0.229 | 2.5049E-55 | TEC_prlf | LSM4 |
| 1.7875E-56 | 0.61341588 | 0.353 | 0.102 | 3.8163E-52 | TEC_prlf | BRD8 |
| 5.9038E-24 | 0.61291146 | 0.567 | 0.338 | 1.2605E-19 | TEC_prlf | MIF |
| 0 | 0.61125379 | 0.325 | 0.004 | 0 | TEC_prlf | KIF2C |
| 3.0642E-49 | 0.60997265 | 0.435 | 0.162 | 6.542E-45 | TEC_prlf | CBX5 |
| 1.2932E-24 | 0.60981603 | 0.865 | 0.724 | 2.7611E-20 | TEC_prlf | GNAS |
| 3.788E-165 | 0.6031826 | 0.317 | 0.032 | 8.087E-161 | TEC_prlf | PCLAF |
| 0 | 0.59548755 | 0.275 | 0.006 | 0 | TEC_prlf | CENPA |
| 3.1106E-35 | 0.59487312 | 0.978 | 0.896 | 6.6411E-31 | TEC_prlf | RPL23A |
| 2.9824E-42 | 0.59029137 | 0.59 | 0.282 | 6.3674E-38 | TEC_prlf | YWHAH |
| 1.1741E-08 | 0.59004624 | 0.444 | 0.315 | 0.00025067 | TEC_prlf | FAM107B |
| 6.637E-296 | 0.58907291 | 0.377 | 0.024 | 1.417E-291 | TEC_prlf | CENPW |
| 7.6391E-41 | 0.58490978 | 0.556 | 0.271 | 1.6309E-36 | TEC_prlf | SEPT7 |
| 1.385E-46 | 0.58127265 | 0.419 | 0.148 | 2.957E-42 | TEC_prlf | IGFBP2 |
| 6.5622E-34 | 0.58088353 | 0.983 | 0.91 | 1.401E-29 | TEC_prlf | RPL35 |
| 9.4248E-33 | 0.58010623 | 0.65 | 0.381 | 2.0122E-28 | TEC_prlf | SERBP1 |
| 3.393E-43 | 0.57722161 | 0.512 | 0.229 | 7.244E-39 | TEC_prlf | PDAP1 |
| 2.2494E-20 | 0.57399128 | 0.452 | 0.261 | 4.8026E-16 | TEC_prlf | GSTM3 |
| 1.3868E-35 | 0.57221737 | 0.719 | 0.453 | 2.9609E-31 | TEC_prlf | COX8A |
| 3.995E-158 | 0.570645 | 0.347 | 0.041 | 8.529E-154 | TEC_prlf | TYMS |
| 9.179E-207 | 0.56867349 | 0.394 | 0.04 | 1.96E-202 | TEC_prlf | PHF19 |
| 2.2549E-32 | 0.56516712 | 0.95 | 0.832 | 4.8142E-28 | TEC_prlf | PPIA |
| 3.9716E-28 | 0.56511745 | 0.937 | 0.796 | 8.4794E-24 | TEC_prlf | CD24 |
| 9.554E-34 | 0.56482699 | 0.727 | 0.473 | 2.0398E-29 | TEC_prlf | PTGES3 |
| 3.1E-22 | 0.56411395 | 0.961 | 0.898 | 6.6184E-18 | TEC_prlf | SERF2 |
| 8.982E-29 | 0.56218325 | 0.565 | 0.308 | 1.9177E-24 | TEC_prlf | MAL |
| 1.2063E-22 | 0.56073148 | 0.705 | 0.51 | 2.5755E-18 | TEC_prlf | TPM3 |
| 0 | 0.55972793 | 0.311 | 0.006 | 0 | TEC_prlf | TROAP |
| 1.256E-128 | 0.55787432 | 0.364 | 0.055 | 2.682E-124 | TEC_prlf | SMC2 |

| | | | | | | |
|---|---|---|---|---|---|---|
| 8.277E-249 | 0.5555449 | 0.317 | 0.02 | 1.767E-244 | TEC_prlf | UBE2T |
| 1.3336E-36 | 0.55548852 | 0.543 | 0.271 | 2.8471E-32 | TEC_prlf | CCT3 |
| 2.5462E-27 | 0.55432669 | 0.972 | 0.933 | 5.4362E-23 | TEC_prlf | HSP90AA1 |
| 4.8856E-37 | 0.55194091 | 0.854 | 0.597 | 1.0431E-32 | TEC_prlf | PSMA7 |
| 2.6737E-88 | 0.55110024 | 0.295 | 0.051 | 5.7083E-84 | TEC_prlf | METTL7A |
| 1.2981E-25 | 0.54999529 | 0.537 | 0.315 | 2.7715E-21 | TEC_prlf | NPC2 |
| 9.494E-32 | 0.54942639 | 0.587 | 0.331 | 2.027E-27 | TEC_prlf | AURKAIP1 |
| 2.8946E-43 | 0.54932035 | 0.339 | 0.113 | 6.1799E-39 | TEC_prlf | LBR |
| 6.8424E-31 | 0.54232996 | 0.612 | 0.342 | 1.4608E-26 | TEC_prlf | EIF5A |
| 0 | 0.54224552 | 0.32 | 0.006 | 0 | TEC_prlf | BUB1 |
| 1.7738E-28 | 0.53608171 | 0.926 | 0.784 | 3.7872E-24 | TEC_prlf | NPM1 |
| 0 | 0.53607059 | 0.325 | 0.003 | 0 | TEC_prlf | KIF4A |
| 4.6099E-43 | 0.5357801 | 0.57 | 0.268 | 9.8421E-39 | TEC_prlf | SMS |
| 1.0858E-42 | 0.53541042 | 0.554 | 0.256 | 2.3182E-38 | TEC_prlf | HNRNPM |
| 5.1285E-09 | 0.53251332 | 0.543 | 0.417 | 0.00010949 | TEC_prlf | TPM4 |
| 0 | 0.52965664 | 0.322 | 0.016 | 0 | TEC_prlf | RACGAP1 |
| 8.197E-303 | 0.52654478 | 0.245 | 0.008 | 1.75E-298 | TEC_prlf | UBE2C |
| 1.0113E-34 | 0.52411791 | 0.468 | 0.219 | 2.1591E-30 | TEC_prlf | PDXK |
| 3.5094E-82 | 0.52167156 | 0.394 | 0.093 | 7.4926E-78 | TEC_prlf | HMGB3 |
| 2.405E-149 | 0.51504777 | 0.353 | 0.044 | 5.134E-145 | TEC_prlf | DIAPH3 |
| 6.3055E-66 | 0.51419697 | 0.391 | 0.107 | 1.3462E-61 | TEC_prlf | DDX39A |
| 7.3438E-41 | 0.51179751 | 0.457 | 0.191 | 1.5679E-36 | TEC_prlf | GLRX5 |
| 5.6634E-51 | 0.5084193 | 0.328 | 0.093 | 1.2091E-46 | TEC_prlf | ISYNA1 |
| 9.5195E-33 | 0.50829686 | 0.521 | 0.264 | 2.0324E-28 | TEC_prlf | NDUFA2 |
| 5.739E-177 | 0.50680118 | 0.281 | 0.023 | 1.225E-172 | TEC_prlf | SGO2 |
| 8.8433E-27 | 0.50658874 | 0.694 | 0.461 | 1.8881E-22 | TEC_prlf | LMNA |
| 7.7994E-28 | 0.50624205 | 0.435 | 0.211 | 1.6652E-23 | TEC_prlf | BLOC1S1 |
| 1.1966E-25 | 0.50203923 | 0.733 | 0.49 | 2.5548E-21 | TEC_prlf | HSPD1 |
| 1.1485E-88 | 0.49637889 | 0.38 | 0.082 | 2.4521E-84 | TEC_prlf | SKA2 |
| 0 | 0.49383555 | 0.259 | 0.008 | 0 | TEC_prlf | CDK1 |
| 1.963E-38 | 0.49297778 | 0.474 | 0.204 | 4.191E-34 | TEC_prlf | UBALD2 |
| 8.7717E-23 | 0.4927143 | 0.543 | 0.326 | 1.8728E-18 | TEC_prlf | SLC25A5 |
| 0 | 0.49230532 | 0.256 | 0.004 | 0 | TEC_prlf | NEK2 |
| 2.9137E-26 | 0.48820657 | 0.595 | 0.355 | 6.2207E-22 | TEC_prlf | SNRPD2 |
| 1.0791E-36 | 0.48819481 | 0.54 | 0.261 | 2.304E-32 | TEC_prlf | SNRPG |
| 0 | 0.48724477 | 0.253 | 0.004 | 0 | TEC_prlf | HJURP |
| 7.245E-28 | 0.4869794 | 0.986 | 0.946 | 1.5468E-23 | TEC_prlf | RPL13A |
| 7.471E-22 | 0.48191547 | 0.78 | 0.579 | 1.5951E-17 | TEC_prlf | EZR |
| 6.176E-260 | 0.48182943 | 0.303 | 0.017 | 1.319E-255 | TEC_prlf | CENPM |
| 6.9196E-25 | 0.47848611 | 0.446 | 0.233 | 1.4773E-20 | TEC_prlf | COX20 |
| 9.6606E-61 | 0.47841077 | 0.306 | 0.074 | 2.0625E-56 | TEC_prlf | CAVIN3 |
| 4.4139E-28 | 0.47578737 | 0.862 | 0.661 | 9.4237E-24 | TEC_prlf | SPINT2 |
| 2.3843E-87 | 0.47552265 | 0.278 | 0.046 | 5.0905E-83 | TEC_prlf | DBF4 |
| 8.3555E-47 | 0.47462464 | 0.275 | 0.074 | 1.7839E-42 | TEC_prlf | CYS1 |
| 0 | 0.47447347 | 0.278 | 0.009 | 0 | TEC_prlf | LMNB1 |
| 4.0099E-23 | 0.47433231 | 0.576 | 0.353 | 8.5611E-19 | TEC_prlf | ATP5ME |
| 0 | 0.47374915 | 0.284 | 0.003 | 0 | TEC_prlf | NDC80 |
| 6.4152E-22 | 0.47191072 | 0.989 | 0.932 | 1.3697E-17 | TEC_prlf | RPL3 |
| 4.0344E-15 | 0.47037181 | 0.862 | 0.723 | 8.6134E-11 | TEC_prlf | KRT8 |
| 8.4228E-29 | 0.46896083 | 0.474 | 0.242 | 1.7983E-24 | TEC_prlf | GSPT1 |

| | | | | | | |
|---|---|---|---|---|---|---|
| 3.821E-31 | 0.46865704 | 0.54 | 0.287 | 8.1578E-27 | TEC_prlf | SRSF9 |
| 2.0965E-12 | 0.46796877 | 0.793 | 0.641 | 4.4761E-08 | TEC_prlf | WFDC2 |
| 1.304E-261 | 0.46773642 | 0.259 | 0.012 | 2.785E-257 | TEC_prlf | KNL1 |
| 1.6362E-34 | 0.46657765 | 0.512 | 0.252 | 3.4933E-30 | TEC_prlf | SNRPD3 |
| 8.7133E-33 | 0.46527797 | 0.408 | 0.175 | 1.8603E-28 | TEC_prlf | TMEM238 |
| 2.18E-116 | 0.46473647 | 0.333 | 0.05 | 4.655E-112 | TEC_prlf | NUDT1 |
| 3.0846E-33 | 0.46138695 | 0.534 | 0.27 | 6.5857E-29 | TEC_prlf | CCT5 |
| 6.7585E-34 | 0.46033407 | 0.397 | 0.166 | 1.4429E-29 | TEC_prlf | COA3 |
| 6.3129E-18 | 0.45821529 | 0.477 | 0.288 | 1.3478E-13 | TEC_prlf | TOP1 |
| 9.552E-214 | 0.45809082 | 0.298 | 0.021 | 2.039E-209 | TEC_prlf | ZWINT |
| 3.4096E-24 | 0.45786526 | 0.992 | 0.96 | 7.2795E-20 | TEC_prlf | RPL28 |
| 7.3946E-13 | 0.45748151 | 0.725 | 0.604 | 1.5787E-08 | TEC_prlf | APLP2 |
| 3.0658E-14 | 0.45697552 | 0.545 | 0.39 | 6.5454E-10 | TEC_prlf | FKBP2 |
| 5.0036E-50 | 0.4568035 | 0.24 | 0.055 | 1.0683E-45 | TEC_prlf | SFTA2 |
| 0 | 0.45585945 | 0.284 | 0.005 | 0 | TEC_prlf | CKAP2L |
| 6.3401E-26 | 0.45491737 | 0.311 | 0.129 | 1.3536E-21 | TEC_prlf | SUDS3 |
| 1.711E-233 | 0.451875 | 0.314 | 0.021 | 3.652E-229 | TEC_prlf | CENPU |
| 0 | 0.4516001 | 0.264 | 0.006 | 0 | TEC_prlf | KIF20A |
| 4.784E-22 | 0.45039501 | 0.606 | 0.382 | 1.0214E-17 | TEC_prlf | HSPE1 |
| 3.3775E-33 | 0.44908776 | 0.441 | 0.195 | 7.2111E-29 | TEC_prlf | ILF2 |
| 1.0103E-45 | 0.44859546 | 0.278 | 0.076 | 2.1569E-41 | TEC_prlf | MIS18BP1 |
| 1.9458E-25 | 0.4482198 | 0.493 | 0.266 | 4.1544E-21 | TEC_prlf | HNRNPD |
| 5.7437E-28 | 0.44726931 | 0.501 | 0.261 | 1.2263E-23 | TEC_prlf | NUDC |
| 1.7779E-29 | 0.44697271 | 0.421 | 0.198 | 3.7958E-25 | TEC_prlf | AP2S1 |
| 1.1661E-32 | 0.44689048 | 0.347 | 0.135 | 2.4896E-28 | TEC_prlf | CBX1 |
| 1.9404E-30 | 0.44625357 | 0.479 | 0.237 | 4.1427E-26 | TEC_prlf | LSM3 |
| 1.6403E-35 | 0.44595616 | 0.344 | 0.128 | 3.502E-31 | TEC_prlf | PSIP1 |
| 7.738E-143 | 0.4429587 | 0.289 | 0.031 | 1.652E-138 | TEC_prlf | SPDL1 |
| 6.5764E-20 | 0.44141349 | 0.433 | 0.24 | 1.4041E-15 | TEC_prlf | SLTM |
| 8.2493E-95 | 0.44114973 | 0.298 | 0.049 | 1.7612E-90 | TEC_prlf | CCDC34 |
| 5.4401E-21 | 0.44034539 | 0.512 | 0.313 | 1.1615E-16 | TEC_prlf | SNRPB |
| 1.3019E-19 | 0.44014486 | 0.983 | 0.944 | 2.7795E-15 | TEC_prlf | RPS15 |
| 9.7288E-20 | 0.43862519 | 0.565 | 0.36 | 2.0771E-15 | TEC_prlf | ATP5F1B |
| 7.2504E-31 | 0.43770768 | 0.358 | 0.149 | 1.548E-26 | TEC_prlf | TAL/DCTN3 |
| 0 | 0.43744462 | 0.248 | 0.006 | 0 | TEC_prlf | SGO1 |
| 1.178E-26 | 0.43550797 | 0.339 | 0.144 | 2.515E-22 | TEC_prlf | NUCB2 |
| 9.7724E-18 | 0.43447089 | 0.358 | 0.19 | 2.0864E-13 | TEC_prlf | BCAM |
| 5.0895E-22 | 0.43371219 | 0.725 | 0.523 | 1.0866E-17 | TEC_prlf | COX6C |
| 5.7413E-23 | 0.43284865 | 0.771 | 0.567 | 1.2258E-18 | TEC_prlf | SET |
| 1.8551E-21 | 0.43201064 | 0.388 | 0.2 | 3.9606E-17 | TEC_prlf | MPC2 |
| 5.1065E-23 | 0.4310357 | 0.857 | 0.674 | 1.0902E-18 | TEC_prlf | EPCAM |
| 9.0708E-24 | 0.4305992 | 0.369 | 0.178 | 1.9366E-19 | TEC_prlf | DUT |
| 5.7213E-17 | 0.43057509 | 0.722 | 0.522 | 1.2215E-12 | TEC_prlf | ELOB |
| 1.8729E-20 | 0.42863525 | 0.612 | 0.408 | 3.9986E-16 | TEC_prlf | HNRNPK |
| 1.313E-22 | 0.42862292 | 0.499 | 0.283 | 2.8033E-18 | TEC_prlf | PSME2 |
| 3.198E-27 | 0.42846253 | 0.826 | 0.602 | 6.8276E-23 | TEC_prlf | ATP5MG |
| 2.8186E-18 | 0.42840538 | 0.981 | 0.92 | 6.0176E-14 | TEC_prlf | ACTB |
| 2.4448E-23 | 0.42828433 | 0.854 | 0.66 | 5.2197E-19 | TEC_prlf | TMA7 |
| 4.1997E-26 | 0.42582824 | 0.562 | 0.309 | 8.9663E-22 | TEC_prlf | PRDX2 |
| 2.8274E-14 | 0.42570951 | 0.862 | 0.756 | 6.0365E-10 | TEC_prlf | CALM2 |

| | | | | | | |
|---|---|---|---|---|---|---|
| 9.11E-30 | 0.42532365 | 0.377 | 0.162 | 1.945E-25 | TEC_prlf | STIP1 |
| 8.304E-121 | 0.42465161 | 0.267 | 0.031 | 1.773E-116 | TEC_prlf | CIT |
| 9.6188E-28 | 0.42417918 | 0.73 | 0.461 | 2.0536E-23 | TEC_prlf | PDIA6 |
| 3.1862E-31 | 0.42349469 | 0.457 | 0.215 | 6.8026E-27 | TEC_prlf | PSMA4 |
| 1.7705E-32 | 0.42086823 | 0.485 | 0.227 | 3.7799E-28 | TEC_prlf | NDUFS8 |
| 1.1382E-11 | 0.4202092 | 0.435 | 0.28 | 2.4301E-07 | TEC_prlf | AKR1B1 |
| 1.2759E-40 | 0.41611735 | 0.369 | 0.132 | 2.7241E-36 | TEC_prlf | BUB3 |
| 0 | 0.4160945 | 0.259 | 0.006 | 0 | TEC_prlf | CDCA3 |
| 1.5931E-15 | 0.41536096 | 0.397 | 0.225 | 3.4012E-11 | TEC_prlf | METAP2 |
| 1.0567E-32 | 0.41389067 | 0.397 | 0.165 | 2.256E-28 | TEC_prlf | NAA38 |
| 2.0312E-33 | 0.41264668 | 0.433 | 0.188 | 4.3365E-29 | TEC_prlf | SNRPF |
| 1.887E-17 | 0.41095165 | 0.595 | 0.399 | 4.0287E-13 | TEC_prlf | TBCA |
| 1.052E-13 | 0.41031146 | 0.592 | 0.429 | 2.246E-09 | TEC_prlf | ST13 |
| 1.5685E-17 | 0.4100695 | 0.576 | 0.375 | 3.3487E-13 | TEC_prlf | RHEB |
| 1.8791E-21 | 0.40965145 | 0.556 | 0.333 | 4.0119E-17 | TEC_prlf | ERP29 |
| 1.8946E-23 | 0.40745394 | 0.485 | 0.261 | 4.0449E-19 | TEC_prlf | MINOS1 |
| 1.2117E-33 | 0.40740537 | 0.416 | 0.175 | 2.5869E-29 | TEC_prlf | SNRPD1 |
| 2.6244E-18 | 0.40714715 | 0.628 | 0.423 | 5.6032E-14 | TEC_prlf | BSG |
| 1.341E-17 | 0.40706029 | 0.562 | 0.364 | 2.8631E-13 | TEC_prlf | TAGLN2 |
| 1.2791E-17 | 0.40698491 | 0.441 | 0.262 | 2.7308E-13 | TEC_prlf | TSTD1 |
| 2.8634E-24 | 0.4065307 | 0.463 | 0.243 | 6.1133E-20 | TEC_prlf | HNRNPR |
| 5.4646E-18 | 0.40637487 | 0.559 | 0.362 | 1.1667E-13 | TEC_prlf | ATP5MF |
| 3.345E-150 | 0.40444507 | 0.273 | 0.026 | 7.141E-146 | TEC_prlf | CENPN |
| 7.8939E-40 | 0.40412006 | 0.358 | 0.128 | 1.6853E-35 | TEC_prlf | VDAC3 |
| 6.9067E-18 | 0.4038821 | 0.556 | 0.355 | 1.4746E-13 | TEC_prlf | COX7B |
| 4.0395E-56 | 0.4036849 | 0.264 | 0.059 | 8.6243E-52 | TEC_prlf | NDE1 |
| 1.513E-256 | 0.4030744 | 0.218 | 0.008 | 3.231E-252 | TEC_prlf | KIF18A |
| 2.1208E-21 | 0.40214109 | 0.537 | 0.325 | 4.5279E-17 | TEC_prlf | NDUFB4 |
| 5.601E-279 | 0.40170041 | 0.231 | 0.008 | 1.196E-274 | TEC_prlf | KIFC1 |
| 3.2192E-16 | 0.40167731 | 0.639 | 0.468 | 6.873E-12 | TEC_prlf | SUMO2 |
| 5.69E-128 | 0.40154874 | 0.292 | 0.035 | 1.215E-123 | TEC_prlf | NCAPD2 |
| 3.3501E-30 | 0.40038904 | 0.397 | 0.174 | 7.1525E-26 | TEC_prlf | TECR |
| 1.4745E-18 | 0.39875178 | 0.793 | 0.639 | 3.1481E-14 | TEC_prlf | YWHAE |
| 1.2248E-27 | 0.39756558 | 0.372 | 0.165 | 2.615E-23 | TEC_prlf | THOC7 |
| 7.0801E-23 | 0.39669399 | 0.917 | 0.772 | 1.5116E-18 | TEC_prlf | OAZ1 |
| 1.2462E-15 | 0.39493576 | 0.573 | 0.406 | 2.6606E-11 | TEC_prlf | UQCR10 |
| 3.1301E-17 | 0.39478931 | 0.915 | 0.8 | 6.6827E-13 | TEC_prlf | CALM1 |
| 1.1918E-43 | 0.39456275 | 0.35 | 0.115 | 2.5445E-39 | TEC_prlf | RPA3 |
| 5.8275E-18 | 0.39365632 | 0.554 | 0.355 | 1.2442E-13 | TEC_prlf | PSME1 |
| 3.768E-20 | 0.39343908 | 0.983 | 0.937 | 8.0446E-16 | TEC_prlf | RPL15 |
| 0 | 0.39339467 | 0.24 | 0.005 | 0 | TEC_prlf | NCAPG |
| 1.0331E-13 | 0.39301455 | 0.474 | 0.299 | 2.2057E-09 | TEC_prlf | ARHGAP29 |
| 5.7912E-17 | 0.39227443 | 0.584 | 0.393 | 1.2364E-12 | TEC_prlf | YWHAQ |
| 9.9572E-98 | 0.39213654 | 0.295 | 0.046 | 2.1259E-93 | TEC_prlf | GGH |
| 5.0091E-17 | 0.3915127 | 0.504 | 0.32 | 1.0694E-12 | TEC_prlf | NDUFB7 |
| 5.5431E-14 | 0.39079472 | 0.807 | 0.677 | 1.1834E-09 | TEC_prlf | CLIC1 |
| 1.2153E-21 | 0.39051759 | 0.981 | 0.92 | 2.5946E-17 | TEC_prlf | RPL14 |
| 0 | 0.38935366 | 0.231 | 0.006 | 0 | TEC_prlf | KIF11 |
| 2.2246E-21 | 0.38915844 | 0.474 | 0.267 | 4.7495E-17 | TEC_prlf | ANAPC11 |
| 4.54E-18 | 0.38756615 | 0.97 | 0.905 | 9.6929E-14 | TEC_prlf | RPS16 |

| | | | | | | |
|---|---|---|---|---|---|---|
| 2.2745E-10 | 0.38691939 | 0.452 | 0.321 | 4.8561E-06 | TEC_prlf | ARL6IP1 |
| 1.615E-21 | 0.38647872 | 0.444 | 0.242 | 3.448E-17 | TEC_prlf | BANF1 |
| 1.0149E-20 | 0.38612799 | 0.953 | 0.875 | 2.1668E-16 | TEC_prlf | ITM2B |
| 2.5625E-21 | 0.38565648 | 0.377 | 0.186 | 5.471E-17 | TEC_prlf | CAPRIN1 |
| 6.0642E-15 | 0.38290348 | 0.612 | 0.448 | 1.2947E-10 | TEC_prlf | ATP5PF |
| 2.0066E-17 | 0.3822582 | 0.479 | 0.288 | 4.284E-13 | TEC_prlf | NOL7 |
| 0 | 0.38178085 | 0.22 | 0.005 | 0 | TEC_prlf | IQGAP3 |
| 1.6117E-15 | 0.38166843 | 0.482 | 0.304 | 3.4409E-11 | TEC_prlf | NDUFC2 |
| 6.5732E-52 | 0.38148148 | 0.253 | 0.059 | 1.4034E-47 | TEC_prlf | DNMT1 |
| 2.7329E-54 | 0.3813276 | 0.311 | 0.082 | 5.8348E-50 | TEC_prlf | MZT1 |
| 4.8894E-14 | 0.38024757 | 0.347 | 0.194 | 1.0439E-09 | TEC_prlf | CDC42BPB |
| 2.7646E-16 | 0.37841515 | 0.499 | 0.314 | 5.9025E-12 | TEC_prlf | XRCC5 |
| 3.0256E-21 | 0.37774819 | 0.43 | 0.233 | 6.4597E-17 | TEC_prlf | VPS29 |
| 7.9928E-57 | 0.37668619 | 0.287 | 0.068 | 1.7065E-52 | TEC_prlf | EMC9 |
| 5.7384E-34 | 0.37545866 | 0.284 | 0.095 | 1.2251E-29 | TEC_prlf | SMARCA1 |
| 0 | 0.37362879 | 0.209 | 0.003 | 0 | TEC_prlf | TTK |
| 4.3561E-16 | 0.37296058 | 0.452 | 0.278 | 9.3003E-12 | TEC_prlf | PDCD5 |
| 6.7804E-05 | 0.37260805 | 0.515 | 0.426 | 1 | TEC_prlf | SYNE2 |
| 9.849E-21 | 0.37227196 | 0.347 | 0.167 | 2.1028E-16 | TEC_prlf | MYEF2 |
| 4.7444E-19 | 0.3708854 | 0.452 | 0.254 | 1.0129E-14 | TEC_prlf | NDUFB3 |
| 3.3264E-21 | 0.37061251 | 0.835 | 0.643 | 7.1019E-17 | TEC_prlf | UQCRH |
| 4.3839E-23 | 0.36998311 | 0.446 | 0.234 | 9.3596E-19 | TEC_prlf | MORF4L2 |
| 2.109E-200 | 0.36941661 | 0.264 | 0.017 | 4.502E-196 | TEC_prlf | CENPK |
| 8.4127E-21 | 0.36905784 | 0.435 | 0.236 | 1.7961E-16 | TEC_prlf | NENF |
| 3.8832E-19 | 0.36834902 | 0.512 | 0.305 | 8.2907E-15 | TEC_prlf | ATP5PD |
| 8.0777E-25 | 0.36793105 | 0.441 | 0.218 | 1.7246E-20 | TEC_prlf | DNAJC8 |
| 2.345E-269 | 0.36681789 | 0.242 | 0.009 | 5.006E-265 | TEC_prlf | KIF14 |
| 3.529E-23 | 0.3659622 | 0.287 | 0.12 | 7.5343E-19 | TEC_prlf | PKP4 |
| 1.1248E-18 | 0.36582125 | 0.978 | 0.902 | 2.4015E-14 | TEC_prlf | RPS3 |
| 3.2746E-17 | 0.36572125 | 0.956 | 0.888 | 6.9913E-13 | TEC_prlf | NACA |
| 6.3508E-14 | 0.36553934 | 0.592 | 0.417 | 1.3559E-09 | TEC_prlf | TUBB4B |
| 1.7098E-39 | 0.36497107 | 0.281 | 0.084 | 3.6505E-35 | TEC_prlf | NSD2 |
| 6.2768E-33 | 0.3648571 | 0.298 | 0.105 | 1.3401E-28 | TEC_prlf | MAZ |
| 1.7524E-21 | 0.36480876 | 0.983 | 0.942 | 3.7414E-17 | TEC_prlf | RPL8 |
| 0.00020048 | 0.36423553 | 0.463 | 0.382 | 1 | TEC_prlf | MYH9 |
| 1.2419E-13 | 0.3640476 | 0.584 | 0.411 | 2.6515E-09 | TEC_prlf | PARK7 |
| 5.1928E-17 | 0.36317587 | 0.573 | 0.378 | 1.1087E-12 | TEC_prlf | MRPL33 |
| 1.839E-12 | 0.36168792 | 0.521 | 0.359 | 3.9263E-08 | TEC_prlf | SRSF11 |
| 1.4799E-15 | 0.36144979 | 0.444 | 0.27 | 3.1595E-11 | TEC_prlf | NDUFS6 |
| 1.0376E-21 | 0.36064452 | 0.245 | 0.093 | 2.2152E-17 | TEC_prlf | SLC12A1 |
| 6.9406E-36 | 0.36060669 | 0.218 | 0.059 | 1.4818E-31 | TEC_prlf | TSPAN12 |
| 2.6696E-23 | 0.35990572 | 0.477 | 0.254 | 5.6995E-19 | TEC_prlf | ERH |
| 1.6401E-12 | 0.35961247 | 0.328 | 0.182 | 3.5016E-08 | TEC_prlf | ADIRF |
| 2.0362E-19 | 0.35847599 | 0.499 | 0.282 | 4.3473E-15 | TEC_prlf | CDK2AP2 |
| 0 | 0.35814939 | 0.19 | 0.003 | 0 | TEC_prlf | BUB1B |
| 6.1743E-23 | 0.35813124 | 0.38 | 0.184 | 1.3182E-18 | TEC_prlf | PRMT1 |
| 4.1342E-21 | 0.35752167 | 0.468 | 0.257 | 8.8266E-17 | TEC_prlf | NDUFB10 |
| 2.3459E-30 | 0.35741284 | 0.391 | 0.164 | 5.0084E-26 | TEC_prlf | CACYBP |
| 6.8839E-45 | 0.35733465 | 0.264 | 0.07 | 1.4697E-40 | TEC_prlf | CKAP5 |
| 8.5918E-15 | 0.35732682 | 0.501 | 0.33 | 1.8343E-10 | TEC_prlf | KDELR1 |

| | | | | | | |
|---|---|---|---|---|---|---|
| 9.3902E-22 | 0.35663444 | 0.317 | 0.144 | 2.0048E-17 | TEC_prlf | ITM2C |
| 2.1129E-14 | 0.35645979 | 0.672 | 0.512 | 4.511E-10 | TEC_prlf | ARPC2 |
| 2.8164E-16 | 0.3560468 | 0.543 | 0.355 | 6.013E-12 | TEC_prlf | NEDD8 |
| 1.88E-260 | 0.35582358 | 0.234 | 0.009 | 4.013E-256 | TEC_prlf | FOXM1 |
| 8.72E-18 | 0.35291917 | 0.523 | 0.316 | 1.8617E-13 | TEC_prlf | AES |
| 1.4849E-09 | 0.35015002 | 0.441 | 0.307 | 3.1702E-05 | TEC_prlf | RAB13 |
| 2.7985E-18 | 0.35009797 | 0.479 | 0.279 | 5.9748E-14 | TEC_prlf | ATP5PO |
| 1.4618E-20 | 0.34977262 | 0.38 | 0.191 | 3.121E-16 | TEC_prlf | ZCRB1 |
| 4.294E-261 | 0.34770572 | 0.201 | 0.006 | 9.167E-257 | TEC_prlf | PSRC1 |
| 5.3183E-12 | 0.34636125 | 0.769 | 0.656 | 1.1355E-07 | TEC_prlf | CALR |
| 1.1387E-19 | 0.34571796 | 0.942 | 0.84 | 2.4312E-15 | TEC_prlf | RPSA |
| 7.3039E-13 | 0.34411309 | 0.818 | 0.688 | 1.5594E-08 | TEC_prlf | GPX4 |
| 3.4986E-15 | 0.3424269 | 0.444 | 0.268 | 7.4694E-11 | TEC_prlf | COX17 |
| 5.7669E-26 | 0.34226205 | 0.273 | 0.103 | 1.2312E-21 | TEC_prlf | KMT5A |
| 1.3879E-15 | 0.34219483 | 0.579 | 0.389 | 2.9632E-11 | TEC_prlf | SRSF3 |
| 6.1538E-85 | 0.34190688 | 0.253 | 0.039 | 1.3138E-80 | TEC_prlf | DHFR |
| 1.496E-29 | 0.34050077 | 0.284 | 0.103 | 3.1939E-25 | TEC_prlf | HMG20B |
| 8.1165E-16 | 0.34043275 | 0.518 | 0.334 | 1.7329E-11 | TEC_prlf | SEC11A |
| 9.1308E-24 | 0.34016196 | 0.446 | 0.226 | 1.9494E-19 | TEC_prlf | H2AFY |
| 2.1992E-13 | 0.33928024 | 0.81 | 0.674 | 4.6954E-09 | TEC_prlf | YWHAZ |
| 7.6582E-13 | 0.3391863 | 0.882 | 0.751 | 1.635E-08 | TEC_prlf | RPS26 |
| 4.1054E-25 | 0.33910705 | 0.421 | 0.204 | 8.7649E-21 | TEC_prlf | NONO |
| 1.7545E-11 | 0.33885988 | 0.584 | 0.433 | 3.746E-07 | TEC_prlf | NAP1L1 |
| 1.1367E-12 | 0.33821574 | 0.59 | 0.416 | 2.4269E-08 | TEC_prlf | SFPQ |
| 1.6704E-26 | 0.3376168 | 0.218 | 0.072 | 3.5664E-22 | TEC_prlf | CDK5RAP2 |
| 2.4956E-17 | 0.33745326 | 0.377 | 0.198 | 5.3281E-13 | TEC_prlf | RTF1 |
| 8.3313E-16 | 0.33684531 | 0.388 | 0.223 | 1.7787E-11 | TEC_prlf | PSMD2 |
| 2.2276E-19 | 0.33677198 | 0.391 | 0.205 | 4.756E-15 | TEC_prlf | PPM1G |
| 7.0382E-17 | 0.33597673 | 0.983 | 0.903 | 1.5027E-12 | TEC_prlf | RPS7 |
| 1.1622E-27 | 0.33477158 | 0.185 | 0.053 | 2.4813E-23 | TEC_prlf | CRNDE |
| 5.9611E-17 | 0.3346221 | 0.529 | 0.319 | 1.2727E-12 | TEC_prlf | CYBA |
| 3.4881E-19 | 0.33450381 | 0.325 | 0.155 | 7.447E-15 | TEC_prlf | PNKD |
| 7.5987E-17 | 0.33411579 | 0.394 | 0.219 | 1.6223E-12 | TEC_prlf | CCT2 |
| 5.5492E-15 | 0.33399293 | 0.92 | 0.8 | 1.1848E-10 | TEC_prlf | CFL1 |
| 2.6826E-24 | 0.33391408 | 0.405 | 0.195 | 5.7274E-20 | TEC_prlf | MLF2 |
| 2.0792E-31 | 0.33368935 | 0.295 | 0.104 | 4.439E-27 | TEC_prlf | ISOC2 |
| 1.1334E-17 | 0.33316443 | 0.84 | 0.658 | 2.4198E-13 | TEC_prlf | RAC1 |
| 8.6216E-23 | 0.33304347 | 0.446 | 0.23 | 1.8407E-18 | TEC_prlf | PPP1CA |
| 7.1741E-22 | 0.33276369 | 0.353 | 0.163 | 1.5317E-17 | TEC_prlf | LMO7 |
| 4.6838E-30 | 0.33269538 | 0.273 | 0.094 | 9.9999E-26 | TEC_prlf | HDAC1 |
| 7.1892E-26 | 0.33175577 | 0.725 | 0.463 | 1.5349E-21 | TEC_prlf | C12orf75 |
| 1.5123E-20 | 0.3315707 | 0.372 | 0.187 | 3.2287E-16 | TEC_prlf | MRPS15 |
| 0.00031901 | 0.33135921 | 0.636 | 0.581 | 1 | TEC_prlf | APP |
| 2.791E-112 | 0.33124202 | 0.193 | 0.017 | 5.959E-108 | TEC_prlf | RRM2 |
| 2.3276E-26 | 0.33011707 | 0.278 | 0.106 | 4.9693E-22 | TEC_prlf | RNF5 |
| 2.1908E-06 | 0.32954773 | 0.752 | 0.646 | 0.04677312 | TEC_prlf | KRT7 |
| 1.3847E-15 | 0.32953353 | 0.983 | 0.932 | 2.9563E-11 | TEC_prlf | RPL6 |
| 1.0148E-17 | 0.32909761 | 0.416 | 0.23 | 2.1665E-13 | TEC_prlf | ARPC5L |
| 1.5709E-20 | 0.32841114 | 0.391 | 0.197 | 3.3538E-16 | TEC_prlf | NDUFAF3 |
| 4.711E-41 | 0.32804357 | 0.259 | 0.071 | 1.0058E-36 | TEC_prlf | CKLF |

| | | | | | | |
|---|---|---|---|---|---|---|
| 3.486E-236 | 0.32770751 | 0.201 | 0.007 | 7.443E-232 | TEC_prlf | PRR11 |
| 1.2403E-19 | 0.3275374 | 0.452 | 0.252 | 2.648E-15 | TEC_prlf | PSMB3 |
| 8.1539E-13 | 0.32590697 | 0.499 | 0.336 | 1.7408E-08 | TEC_prlf | SPCS1 |
| 8.2797E-23 | 0.32571334 | 0.344 | 0.157 | 1.7677E-18 | TEC_prlf | GTF2H5 |
| 7.4594E-35 | 0.32507167 | 0.212 | 0.057 | 1.5926E-30 | TEC_prlf | PCNT |
| 1.0489E-20 | 0.3248882 | 0.364 | 0.177 | 2.2394E-16 | TEC_prlf | HACD3 |
| 1.7173E-18 | 0.32485531 | 0.347 | 0.178 | 3.6665E-14 | TEC_prlf | SIVA1 |
| 8.0594E-16 | 0.32417796 | 0.405 | 0.239 | 1.7207E-11 | TEC_prlf | XRCC6 |
| 7.273E-83 | 0.32407536 | 0.253 | 0.04 | 1.5528E-78 | TEC_prlf | LMNB2 |
| 1.784E-200 | 0.32335731 | 0.24 | 0.014 | 3.81E-196 | TEC_prlf | FANCI |
| 1.1901E-20 | 0.32212637 | 0.314 | 0.145 | 2.5408E-16 | TEC_prlf | COMMD4 |
| 8.4144E-66 | 0.32103805 | 0.237 | 0.043 | 1.7965E-61 | TEC_prlf | ODF2 |
| 9.6313E-10 | 0.32061154 | 0.424 | 0.284 | 2.0563E-05 | TEC_prlf | TGOLN2 |
| 5.5828E-76 | 0.32025091 | 0.176 | 0.021 | 1.1919E-71 | TEC_prlf | GAS2L3 |
| 5.4934E-14 | 0.32016581 | 0.964 | 0.908 | 1.1728E-09 | TEC_prlf | RPL24 |
| 1.142E-15 | 0.32009469 | 0.972 | 0.902 | 2.4381E-11 | TEC_prlf | RPS20 |
| 4.7433E-23 | 0.31979203 | 0.416 | 0.204 | 1.0127E-18 | TEC_prlf | LSM7 |
| 1.8698E-48 | 0.31971538 | 0.204 | 0.042 | 3.992E-44 | TEC_prlf | HMGN5 |
| 1.223E-53 | 0.31907228 | 0.223 | 0.045 | 2.6111E-49 | TEC_prlf | ATAD2 |
| 7.4126E-20 | 0.31905799 | 0.375 | 0.19 | 1.5826E-15 | TEC_prlf | RDX |
| 1.3039E-15 | 0.31869908 | 0.967 | 0.911 | 2.7839E-11 | TEC_prlf | RPL29 |
| 1.9244E-16 | 0.31844187 | 0.452 | 0.267 | 4.1086E-12 | TEC_prlf | NDUFV2 |
| 1.1437E-16 | 0.3182211 | 0.419 | 0.237 | 2.4418E-12 | TEC_prlf | GTF2A2 |
| 1.1787E-08 | 0.31821153 | 0.256 | 0.152 | 0.00025166 | TEC_prlf | CHD3 |
| 8.8976E-26 | 0.31816332 | 0.317 | 0.13 | 1.8996E-21 | TEC_prlf | ANP32A |
| 4.011E-22 | 0.31796785 | 0.35 | 0.164 | 8.5635E-18 | TEC_prlf | SNRPC |
| 1.4295E-14 | 0.31760929 | 0.358 | 0.201 | 3.0519E-10 | TEC_prlf | IK |
| 2.8096E-15 | 0.31585427 | 0.477 | 0.3 | 5.9985E-11 | TEC_prlf | EIF3F |
| 8.0052E-11 | 0.31560601 | 0.716 | 0.567 | 1.7091E-06 | TEC_prlf | COX7A2 |
| 6.5526E-18 | 0.31503394 | 0.989 | 0.932 | 1.399E-13 | TEC_prlf | RPS9 |
| 4.0393E-17 | 0.31496623 | 0.399 | 0.223 | 8.6239E-13 | TEC_prlf | MDH1 |
| 2.4276E-37 | 0.31475949 | 0.264 | 0.079 | 5.1829E-33 | TEC_prlf | CEP70 |
| 4.6273E-21 | 0.31426561 | 0.457 | 0.244 | 9.8793E-17 | TEC_prlf | AP2M1 |
| 4.5224E-17 | 0.31414871 | 0.383 | 0.205 | 9.6554E-13 | TEC_prlf | BEX3 |
| 5.4575E-12 | 0.31379942 | 0.554 | 0.387 | 1.1652E-07 | TEC_prlf | EIF2S2 |
| 7.0616E-20 | 0.3132023 | 0.328 | 0.154 | 1.5077E-15 | TEC_prlf | TMEM160 |
| 1.6617E-12 | 0.31277315 | 0.444 | 0.29 | 3.5476E-08 | TEC_prlf | SSBP1 |
| 1.0242E-29 | 0.31251694 | 0.35 | 0.139 | 2.1867E-25 | TEC_prlf | IDH2 |
| 2.1111E-14 | 0.31239361 | 0.361 | 0.2 | 4.5071E-10 | TEC_prlf | AIF1L |
| 1.5429E-12 | 0.31151652 | 0.882 | 0.744 | 3.2941E-08 | TEC_prlf | DSTN |
| 9.1831E-18 | 0.31093274 | 0.455 | 0.255 | 1.9606E-13 | TEC_prlf | GSTO1 |
| 5.1956E-14 | 0.3107348 | 0.818 | 0.657 | 1.1093E-09 | TEC_prlf | UQCRB |
| 4.6275E-19 | 0.30979614 | 0.278 | 0.125 | 9.8798E-15 | TEC_prlf | TAF15 |
| 9.3098E-15 | 0.30947488 | 0.413 | 0.242 | 1.9876E-10 | TEC_prlf | KHDRBS1 |
| 4.719E-211 | 0.30812731 | 0.209 | 0.009 | 1.008E-206 | TEC_prlf | PARPBP |
| 1.9723E-24 | 0.30782451 | 0.981 | 0.867 | 4.2109E-20 | TEC_prlf | S100A10 |
| 3.3965E-23 | 0.30702372 | 0.295 | 0.123 | 7.2514E-19 | TEC_prlf | TIMM10 |
| 2.5718E-20 | 0.3063764 | 0.35 | 0.169 | 5.4909E-16 | TEC_prlf | GTF3A |
| 1.5857E-77 | 0.30597406 | 0.215 | 0.031 | 3.3856E-73 | TEC_prlf | CCDC18 |
| 8.1661E-10 | 0.30594301 | 0.485 | 0.346 | 1.7435E-05 | TEC_prlf | C19orf53 |

| | | | | | | |
|---|---|---|---|---|---|---|
| 9.2488E-10 | 0.30593168 | 0.408 | 0.276 | 1.9746E-05 | TEC_prlf | YY1 |
| 2.311E-230 | 0.30517477 | 0.207 | 0.008 | 4.934E-226 | TEC_prlf | CDCA2 |
| 1.9787E-20 | 0.30496881 | 0.397 | 0.202 | 4.2245E-16 | TEC_prlf | PRDX3 |
| 2.9483E-16 | 0.30466673 | 0.983 | 0.929 | 6.2946E-12 | TEC_prlf | RPL7A |
| 3.0951E-09 | 0.30461716 | 0.433 | 0.306 | 6.608E-05 | TEC_prlf | HDLBP |
| 3.785E-10 | 0.30208028 | 0.537 | 0.394 | 8.081E-06 | TEC_prlf | MGST3 |
| 7.9175E-25 | 0.30184314 | 0.212 | 0.071 | 1.6904E-20 | TEC_prlf | FRMD4B |
| 2.8233E-12 | 0.30094598 | 0.895 | 0.78 | 6.0277E-08 | TEC_prlf | EEF1B2 |
| 1.8868E-19 | 0.30051325 | 0.455 | 0.252 | 4.0283E-15 | TEC_prlf | ATP5F1A |
| 3.2326E-17 | 0.30021978 | 0.986 | 0.939 | 6.9016E-13 | TEC_prlf | RPL19 |
| 1.2882E-37 | 0.29996356 | 0.259 | 0.076 | 2.7504E-33 | TEC_prlf | DNAJC9 |
| 8.9758E-10 | 0.29933434 | 0.722 | 0.598 | 1.9163E-05 | TEC_prlf | CAST |
| 5.6693E-12 | 0.29890275 | 0.854 | 0.748 | 1.2104E-07 | TEC_prlf | BTF3 |
| 1.8434E-25 | 0.29889451 | 0.311 | 0.127 | 3.9356E-21 | TEC_prlf | RANGAP1 |
| 5.053E-233 | 0.29821945 | 0.223 | 0.009 | 1.079E-228 | TEC_prlf | SPAG5 |
| 1.8416E-23 | 0.29720174 | 0.253 | 0.097 | 3.9317E-19 | TEC_prlf | SMC1A |
| 4.533E-14 | 0.29693589 | 0.314 | 0.167 | 9.6779E-10 | TEC_prlf | BICC1 |
| 2.3059E-32 | 0.29676969 | 0.207 | 0.057 | 4.923E-28 | TEC_prlf | MAOA |
| 4.0886E-16 | 0.29665416 | 0.471 | 0.28 | 8.7291E-12 | TEC_prlf | PSMB6 |
| 1.6117E-12 | 0.29635135 | 0.391 | 0.24 | 3.4409E-08 | TEC_prlf | KPNB1 |
| 1.0486E-10 | 0.2961203 | 0.419 | 0.274 | 2.2387E-06 | TEC_prlf | GTF2I |
| 2.4084E-12 | 0.29549693 | 0.377 | 0.228 | 5.1419E-08 | TEC_prlf | CHD4 |
| 8.3561E-25 | 0.2947263 | 0.306 | 0.123 | 1.784E-20 | TEC_prlf | UQCC2 |
| 4.9657E-25 | 0.29456927 | 0.287 | 0.113 | 1.0602E-20 | TEC_prlf | TMEM106C |
| 1.487E-266 | 0.29454758 | 0.201 | 0.006 | 3.174E-262 | TEC_prlf | SKA3 |
| 4.5027E-12 | 0.29421868 | 0.424 | 0.272 | 9.6133E-08 | TEC_prlf | CCT6A |
| 1.3842E-10 | 0.29420069 | 0.711 | 0.555 | 2.9554E-06 | TEC_prlf | KTN1 |
| 1.0213E-16 | 0.29397829 | 0.992 | 0.922 | 2.1804E-12 | TEC_prlf | TMSB10 |
| 1.9053E-12 | 0.29345369 | 0.488 | 0.322 | 4.0679E-08 | TEC_prlf | VDAC1 |
| 1.008E-69 | 0.29319353 | 0.226 | 0.037 | 2.1521E-65 | TEC_prlf | ORC6 |
| 6.576E-07 | 0.29304002 | 0.314 | 0.214 | 0.01403973 | TEC_prlf | CHMP4B |
| 2.1913E-15 | 0.29289039 | 0.383 | 0.215 | 4.6784E-11 | TEC_prlf | NAP1L4 |
| 1.9427E-14 | 0.29261096 | 0.78 | 0.61 | 4.1476E-10 | TEC_prlf | CD81 |
| 2.6962E-14 | 0.29198884 | 0.725 | 0.555 | 5.7563E-10 | TEC_prlf | RHOA |
| 2.6804E-61 | 0.29198486 | 0.223 | 0.041 | 5.7226E-57 | TEC_prlf | REEP4 |
| 3.3028E-26 | 0.29144329 | 0.275 | 0.103 | 7.0516E-22 | TEC_prlf | MYL6B |
| 2.8214E-15 | 0.29138644 | 0.331 | 0.175 | 6.0237E-11 | TEC_prlf | GOT1 |
| 1.856E-39 | 0.29132634 | 0.22 | 0.056 | 3.9626E-35 | TEC_prlf | SPA17 |
| 7.1807E-12 | 0.29115852 | 0.441 | 0.276 | 1.5331E-07 | TEC_prlf | CLDN3 |
| 1.4593E-15 | 0.29110688 | 0.504 | 0.31 | 3.1156E-11 | TEC_prlf | SRP9 |
| 2.4256E-21 | 0.29072505 | 0.264 | 0.109 | 5.1787E-17 | TEC_prlf | LSM2 |
| 3.8418E-10 | 0.29053184 | 0.758 | 0.632 | 8.2023E-06 | TEC_prlf | UBL5 |
| 3.0033E-13 | 0.29046173 | 0.981 | 0.941 | 6.4121E-09 | TEC_prlf | RPS24 |
| 3.8033E-14 | 0.29017642 | 0.353 | 0.196 | 8.1201E-10 | TEC_prlf | EHF |
| 1.9136E-13 | 0.29017436 | 0.342 | 0.194 | 4.0856E-09 | TEC_prlf | STMP1 |
| 3.5256E-25 | 0.28987621 | 0.295 | 0.117 | 7.5272E-21 | TEC_prlf | NUDCD2 |
| 2.4465E-24 | 0.28951015 | 0.264 | 0.101 | 5.2232E-20 | TEC_prlf | ADD3 |
| 5.1837E-16 | 0.28912064 | 0.931 | 0.805 | 1.1067E-11 | TEC_prlf | ATP5F1E |
| 2.0393E-32 | 0.28909744 | 0.242 | 0.075 | 4.3538E-28 | TEC_prlf | RUVBL2 |
| 3.7813E-11 | 0.28841118 | 0.504 | 0.339 | 8.073E-07 | TEC_prlf | S100A16 |

| | | | | | | |
|---|---|---|---|---|---|---|
| 3.9276E-15 | 0.28838428 | 0.342 | 0.184 | 8.3855E-11 | TEC_prlf | ARL3 |
| 3.6692E-10 | 0.28721843 | 0.532 | 0.382 | 7.8336E-06 | TEC_prlf | POLR2L |
| 6.7725E-14 | 0.28672769 | 0.942 | 0.863 | 1.4459E-09 | TEC_prlf | UBA52 |
| 1.7413E-17 | 0.28622664 | 0.245 | 0.107 | 3.7176E-13 | TEC_prlf | EFHD1 |
| 7.6256E-11 | 0.28612086 | 0.466 | 0.318 | 1.6281E-06 | TEC_prlf | COX5A |
| 7.1125E-12 | 0.28590725 | 0.683 | 0.516 | 1.5185E-07 | TEC_prlf | ARF1 |
| 1.0319E-21 | 0.2839786 | 0.289 | 0.124 | 2.2032E-17 | TEC_prlf | BAD |
| 4.512E-39 | 0.28359444 | 0.237 | 0.063 | 9.6331E-35 | TEC_prlf | KIF22 |
| 2.747E-20 | 0.28310844 | 0.273 | 0.118 | 5.8648E-16 | TEC_prlf | RAB8A |
| 1.7935E-12 | 0.28279237 | 0.303 | 0.169 | 3.8292E-08 | TEC_prlf | NME1 |
| 8.9534E-18 | 0.28269966 | 0.331 | 0.165 | 1.9115E-13 | TEC_prlf | UBE2N |
| 3.076E-14 | 0.28245415 | 0.747 | 0.578 | 6.5672E-10 | TEC_prlf | ARPC3 |
| 6.9437E-11 | 0.2813998 | 0.937 | 0.87 | 1.4825E-06 | TEC_prlf | RACK1 |
| 1.1787E-08 | 0.28132712 | 0.435 | 0.311 | 0.00025165 | TEC_prlf | MRFAP1 |
| 2.8346E-12 | 0.28118654 | 0.364 | 0.219 | 6.0518E-08 | TEC_prlf | CNIH4 |
| 3.2066E-16 | 0.28105556 | 0.372 | 0.2 | 6.846E-12 | TEC_prlf | ADI1 |
| 4.0927E-15 | 0.28097605 | 0.421 | 0.247 | 8.7379E-11 | TEC_prlf | PHB |
| 3.2398E-24 | 0.28084842 | 0.339 | 0.146 | 6.917E-20 | TEC_prlf | RBBP7 |
| 1.779E-15 | 0.28081042 | 0.309 | 0.159 | 3.7983E-11 | TEC_prlf | CHCHD5 |
| 2.9889E-47 | 0.28070525 | 0.229 | 0.052 | 6.3813E-43 | TEC_prlf | RRM1 |
| 7.5422E-11 | 0.27889633 | 0.314 | 0.187 | 1.6103E-06 | TEC_prlf | ABCF1 |
| 8.0478E-18 | 0.2787579 | 0.339 | 0.169 | 1.7182E-13 | TEC_prlf | BRD7 |
| 2.6988E-09 | 0.27852765 | 0.433 | 0.3 | 5.762E-05 | TEC_prlf | PNRC2 |
| 5.9888E-14 | 0.27848636 | 0.928 | 0.851 | 1.2786E-09 | TEC_prlf | MYL12B |
| 4.818E-215 | 0.27805438 | 0.179 | 0.006 | 1.029E-210 | TEC_prlf | AURKB |
| 1.7763E-16 | 0.27610904 | 0.372 | 0.202 | 3.7924E-12 | TEC_prlf | TXNL4A |
| 8.7439E-43 | 0.27610318 | 0.231 | 0.057 | 1.8668E-38 | TEC_prlf | RPL39L |
| 4.2419E-14 | 0.27597873 | 0.3 | 0.158 | 9.0565E-10 | TEC_prlf | DDAH2 |
| 8.9138E-28 | 0.27589874 | 0.259 | 0.091 | 1.9031E-23 | TEC_prlf | ZDHHC12 |
| 1.8257E-12 | 0.27575588 | 0.793 | 0.622 | 3.8978E-08 | TEC_prlf | SLC25A6 |
| 8.8775E-13 | 0.27535199 | 0.364 | 0.214 | 1.8954E-08 | TEC_prlf | SRI |
| 2.8133E-12 | 0.2750967 | 0.97 | 0.93 | 6.0063E-08 | TEC_prlf | RPL18 |
| 2.7017E-15 | 0.27497481 | 0.402 | 0.226 | 5.7682E-11 | TEC_prlf | RBMX |
| 1.2696E-13 | 0.27351826 | 0.377 | 0.216 | 2.7105E-09 | TEC_prlf | PDCD6 |
| 4.5157E-10 | 0.27351671 | 0.661 | 0.521 | 9.641E-06 | TEC_prlf | ATP5MC2 |
| 5.7078E-12 | 0.2731824 | 0.559 | 0.39 | 1.2186E-07 | TEC_prlf | ATP5F1D |
| 9.1832E-11 | 0.27297907 | 0.545 | 0.377 | 1.9606E-06 | TEC_prlf | NDUFB2 |
| 4.2791E-20 | 0.27296168 | 0.287 | 0.125 | 9.1359E-16 | TEC_prlf | HINT2 |
| 2.1646E-09 | 0.27243736 | 0.452 | 0.306 | 4.6214E-05 | TEC_prlf | NSA2 |
| 5.3683E-20 | 0.27201026 | 0.336 | 0.159 | 1.1461E-15 | TEC_prlf | SSNA1 |
| 7.296E-10 | 0.27191988 | 0.474 | 0.333 | 1.5577E-05 | TEC_prlf | PSMB1 |
| 2.6669E-30 | 0.27155471 | 0.204 | 0.058 | 5.6938E-26 | TEC_prlf | EZH2 |
| 2.6151E-12 | 0.27152035 | 0.413 | 0.261 | 5.5832E-08 | TEC_prlf | TXNDC17 |
| 1.2615E-09 | 0.27123692 | 0.402 | 0.266 | 2.6934E-05 | TEC_prlf | FARP1 |
| 8.2345E-10 | 0.27101251 | 0.198 | 0.102 | 1.7581E-05 | TEC_prlf | NAV2 |
| 5.2597E-10 | 0.27078432 | 0.237 | 0.129 | 1.1229E-05 | TEC_prlf | CSRP2 |
| 1.5641E-10 | 0.27066267 | 0.397 | 0.258 | 3.3394E-06 | TEC_prlf | GNB2 |
| 2.4853E-12 | 0.26992876 | 0.284 | 0.155 | 5.3062E-08 | TEC_prlf | KRTCAP2 |
| 5.3371E-16 | 0.26956476 | 0.275 | 0.129 | 1.1395E-11 | TEC_prlf | CDH16 |
| 4.77E-16 | 0.26948863 | 0.306 | 0.154 | 1.0184E-11 | TEC_prlf | EBNA1BP2 |

| | | | | | | |
|---|---|---|---|---|---|---|
| 8.5789E-23 | 0.26931942 | 0.24 | 0.09 | 1.8316E-18 | TEC_prlf | KDM1A |
| 3.0313E-11 | 0.26855105 | 0.529 | 0.364 | 6.4719E-07 | TEC_prlf | ARL6IP4 |
| 4.4313E-31 | 0.2685426 | 0.19 | 0.05 | 9.4608E-27 | TEC_prlf | MNS1 |
| 3.2812E-20 | 0.26823642 | 0.116 | 0.03 | 7.0054E-16 | TEC_prlf | AKR1C2 |
| 8.0126E-12 | 0.26821028 | 0.353 | 0.21 | 1.7107E-07 | TEC_prlf | NHP2 |
| 1.959E-100 | 0.26817513 | 0.154 | 0.012 | 4.1827E-96 | TEC_prlf | INCENP |
| 6.1871E-13 | 0.26811922 | 0.86 | 0.728 | 1.321E-08 | TEC_prlf | NDUFA4 |
| 2.0889E-11 | 0.26811193 | 0.433 | 0.275 | 4.4598E-07 | TEC_prlf | ANAPC16 |
| 6.5151E-16 | 0.26705059 | 0.289 | 0.142 | 1.391E-11 | TEC_prlf | IFT57 |
| 7.5585E-06 | 0.26675772 | 0.493 | 0.391 | 0.16137324 | TEC_prlf | TAX1BP1 |
| 4.472E-19 | 0.26641119 | 0.267 | 0.116 | 9.5477E-15 | TEC_prlf | TXNDC12 |
| 1.8723E-44 | 0.26592409 | 0.209 | 0.047 | 3.9974E-40 | TEC_prlf | RNF26 |
| 3.9516E-13 | 0.26580144 | 0.474 | 0.3 | 8.4368E-09 | TEC_prlf | RBM8A |
| 1.1181E-11 | 0.26578717 | 0.702 | 0.537 | 2.3871E-07 | TEC_prlf | GNG5 |
| 7.3153E-11 | 0.26575781 | 0.736 | 0.587 | 1.5618E-06 | TEC_prlf | KRT19 |
| 6.0812E-18 | 0.26563703 | 0.331 | 0.163 | 1.2983E-13 | TEC_prlf | FKBP3 |
| 3.7957E-14 | 0.26560661 | 0.347 | 0.194 | 8.1037E-10 | TEC_prlf | POLR2J |
| 3.4926E-11 | 0.26547699 | 0.419 | 0.268 | 7.4567E-07 | TEC_prlf | ARPC5 |
| 0.00024201 | 0.26510852 | 0.562 | 0.49 | 1 | TEC_prlf | ATP1B1 |
| 1.124E-206 | 0.26504022 | 0.174 | 0.006 | 2.4E-202 | TEC_prlf | CDCA5 |
| 3.7613E-18 | 0.26494695 | 0.284 | 0.128 | 8.0305E-14 | TEC_prlf | IFI27L2 |
| 1.8221E-09 | 0.26457134 | 0.548 | 0.399 | 3.8902E-05 | TEC_prlf | PCBP2 |
| 7.884E-16 | 0.26335786 | 0.32 | 0.164 | 1.6832E-11 | TEC_prlf | PDHA1 |
| 1.5732E-45 | 0.26324822 | 0.212 | 0.047 | 3.3588E-41 | TEC_prlf | GPSM2 |
| 1.1629E-15 | 0.26324677 | 0.388 | 0.208 | 2.4828E-11 | TEC_prlf | CTSH |
| 3.2955E-13 | 0.26307031 | 0.298 | 0.16 | 7.036E-09 | TEC_prlf | SUPT16H |
| 5.0985E-14 | 0.26223993 | 0.421 | 0.254 | 1.0885E-09 | TEC_prlf | RTRAF |
| 1.099E-08 | 0.26210665 | 0.967 | 0.925 | 0.00023464 | TEC_prlf | ACTG1 |
| 3.4461E-14 | 0.26203999 | 0.95 | 0.866 | 7.3575E-10 | TEC_prlf | RPL10A |
| 2.4867E-13 | 0.26148567 | 0.408 | 0.242 | 5.309E-09 | TEC_prlf | BCAP31 |
| 1.591E-13 | 0.2613486 | 0.983 | 0.932 | 3.3968E-09 | TEC_prlf | RPS27A |
| 1.349E-12 | 0.26132084 | 0.223 | 0.108 | 2.8801E-08 | TEC_prlf | GNAI1 |
| 7.317E-11 | 0.2611352 | 0.287 | 0.162 | 1.5622E-06 | TEC_prlf | NTN4 |
| 6.8624E-18 | 0.26101825 | 0.311 | 0.147 | 1.4651E-13 | TEC_prlf | BAZ1B |
| 1.9275E-13 | 0.25964436 | 0.408 | 0.243 | 4.1152E-09 | TEC_prlf | SNX6 |
| 3.0381E-06 | 0.259268 | 0.939 | 0.913 | 0.06486419 | TEC_prlf | HSP90AB1 |
| 1.1317E-12 | 0.25917007 | 0.358 | 0.209 | 2.4162E-08 | TEC_prlf | SLIRP |
| 1.3208E-07 | 0.25891256 | 0.479 | 0.353 | 0.00281994 | TEC_prlf | UBXN4 |
| 2.9229E-18 | 0.25832064 | 0.27 | 0.119 | 6.2405E-14 | TEC_prlf | TUBB6 |
| 4.352E-148 | 0.25805601 | 0.174 | 0.01 | 9.29E-144 | TEC_prlf | TMSB15A |
| 6.3353E-09 | 0.25746786 | 0.493 | 0.353 | 0.00013526 | TEC_prlf | EIF3K |
| 2.6113E-10 | 0.25723277 | 0.35 | 0.218 | 5.5752E-06 | TEC_prlf | GNAI2 |
| 1.4397E-11 | 0.25707012 | 0.402 | 0.253 | 3.0738E-07 | TEC_prlf | NDUFA11 |
| 4.1707E-10 | 0.25683746 | 0.372 | 0.236 | 8.9045E-06 | TEC_prlf | KDELR2 |
| 7.957E-17 | 0.2564351 | 0.339 | 0.173 | 1.6988E-12 | TEC_prlf | POLR2F |
| 4.8136E-14 | 0.25611725 | 0.76 | 0.562 | 1.0277E-09 | TEC_prlf | HNRNPC |
| 1.1523E-15 | 0.25586107 | 0.149 | 0.052 | 2.4601E-11 | TEC_prlf | ARHGDIB |
| 1.2849E-12 | 0.25550129 | 0.344 | 0.196 | 2.7433E-08 | TEC_prlf | PSMC3 |
| 5.1518E-10 | 0.25448687 | 0.474 | 0.325 | 1.0999E-05 | TEC_prlf | UQCRQ |
| 1.556E-34 | 0.25374225 | 0.215 | 0.058 | 3.322E-30 | TEC_prlf | SHMT1 |

| | | | | | | |
|---|---|---|---|---|---|---|
| 3.1282E-25 | 0.25359351 | 0.24 | 0.084 | 6.6787E-21 | TEC_prlf | EXOSC8 |
| 5.9323E-10 | 0.25352026 | 0.41 | 0.267 | 1.2665E-05 | TEC_prlf | ZC3H15 |
| 1.874E-12 | 0.25287078 | 0.983 | 0.927 | 4.001E-08 | TEC_prlf | RPL18A |
| 2.796E-08 | 0.25275095 | 0.609 | 0.473 | 0.00059696 | TEC_prlf | MORF4L1 |
| 2.5827E-08 | 0.2526449 | 0.375 | 0.248 | 0.0005514 | TEC_prlf | CD47 |
| 1.8976E-60 | 0.25220475 | 0.138 | 0.016 | 4.0514E-56 | TEC_prlf | CLSPN |
| 3.8489E-10 | 0.25216409 | 0.959 | 0.895 | 8.2174E-06 | TEC_prlf | RPL35A |
| 6.9772E-12 | 0.25213888 | 0.375 | 0.23 | 1.4896E-07 | TEC_prlf | PSMB2 |
| 1.745E-10 | 0.25206959 | 0.41 | 0.269 | 3.7256E-06 | TEC_prlf | ATP5F1C |
| 6.0517E-10 | 0.25176131 | 0.468 | 0.323 | 1.292E-05 | TEC_prlf | NDUFC1 |
| 1.7986E-11 | 0.25127858 | 0.391 | 0.242 | 3.8399E-07 | TEC_prlf | PGAM1 |
| 2.2026E-12 | 0.25104961 | 0.372 | 0.218 | 4.7025E-08 | TEC_prlf | ILF3 |
| 3.8614E-11 | 0.25093961 | 0.375 | 0.232 | 8.244E-07 | TEC_prlf | ANXA4 |
| 6.5094E-10 | -0.2513612 | 0.055 | 0.18 | 1.3898E-05 | TEC_prlf | DNAJB9 |
| 8.4013E-07 | -0.2521136 | 0.096 | 0.197 | 0.01793675 | TEC_prlf | SLC25A37 |
| 7.1013E-07 | -0.2536282 | 0.116 | 0.225 | 0.0151612 | TEC_prlf | BEX2 |
| 5.8969E-05 | -0.2559633 | 0.085 | 0.163 | 1 | TEC_prlf | TNFAIP3 |
| 3.0844E-07 | -0.2571584 | 0.102 | 0.211 | 0.00658522 | TEC_prlf | SLFN5 |
| 0.00473176 | -0.2641279 | 0.245 | 0.299 | 1 | TEC_prlf | CYTOR |
| 1.0236E-05 | -0.2687106 | 0.146 | 0.242 | 0.21852986 | TEC_prlf | ITGB6 |
| 9.6265E-07 | -0.2699553 | 0.168 | 0.277 | 0.02055254 | TEC_prlf | CREBRF |
| 1.2429E-08 | -0.270946 | 0.011 | 0.101 | 0.00026536 | TEC_prlf | MAFA |
| 0.00017691 | -0.2712107 | 0.132 | 0.203 | 1 | TEC_prlf | CHCHD7 |
| 9.5654E-06 | -0.2735594 | 0.215 | 0.313 | 0.20422205 | TEC_prlf | ISCU |
| 1.5921E-06 | -0.2763701 | 0.113 | 0.217 | 0.03399096 | TEC_prlf | ZFP36 |
| 3.0394E-08 | -0.2773803 | 0.116 | 0.239 | 0.00064892 | TEC_prlf | MIDN |
| 1.5084E-05 | -0.2774248 | 0.094 | 0.177 | 0.32203941 | TEC_prlf | PHLDA1 |
| 4.754E-08 | -0.2780181 | 0.033 | 0.129 | 0.00101498 | TEC_prlf | AL135925.1 |
| 0.00197352 | -0.2787046 | 0.193 | 0.251 | 1 | TEC_prlf | IDI1 |
| 3.3159E-07 | -0.2788415 | 0.176 | 0.298 | 0.00707951 | TEC_prlf | CD55 |
| 7.1312E-05 | -0.2794792 | 0.102 | 0.178 | 1 | TEC_prlf | FOXC1 |
| 3.2302E-05 | -0.279702 | 0.091 | 0.17 | 0.68965289 | TEC_prlf | FGF2 |
| 5.9427E-10 | -0.2797988 | 0.107 | 0.246 | 1.2688E-05 | TEC_prlf | VAMP2 |
| 0.00284499 | -0.2820611 | 0.251 | 0.309 | 1 | TEC_prlf | YME1L1 |
| 0.0008976 | -0.283991 | 0.05 | 0.101 | 1 | TEC_prlf | CXCL3 |
| 5.0877E-10 | -0.2848648 | 0.066 | 0.195 | 1.0862E-05 | TEC_prlf | KLHL21 |
| 4.1121E-07 | -0.2852674 | 0.028 | 0.109 | 0.00877943 | TEC_prlf | AL355075.4 |
| 3.6062E-09 | -0.2856144 | 0.055 | 0.17 | 7.6992E-05 | TEC_prlf | MIR22HG |
| 6.39E-11 | -0.2894823 | 0.074 | 0.213 | 1.3643E-06 | TEC_prlf | GULP1 |
| 2.3774E-05 | -0.2909754 | 0.182 | 0.277 | 0.50757191 | TEC_prlf | VMP1 |
| 0.00012428 | -0.2951518 | 0.226 | 0.304 | 1 | TEC_prlf | C6orf62 |
| 8.6875E-06 | -0.2965713 | 0.052 | 0.129 | 0.18547916 | TEC_prlf | PCDH9 |
| 4.9319E-09 | -0.29799 | 0.017 | 0.114 | 0.0001053 | TEC_prlf | HIST4H4 |
| 7.989E-07 | -0.2985354 | 0.138 | 0.251 | 0.01705658 | TEC_prlf | RAP2B |
| 1.8321E-07 | -0.2989126 | 0.044 | 0.136 | 0.00391156 | TEC_prlf | CSRNP2 |
| 0.00049283 | -0.3028394 | 0.664 | 0.708 | 1 | TEC_prlf | DDX5 |
| 2.344E-09 | -0.3031228 | 0.113 | 0.25 | 5.0045E-05 | TEC_prlf | IRS2 |
| 5.3134E-06 | -0.3048604 | 0.066 | 0.148 | 0.11344031 | TEC_prlf | ANKRD37 |
| 3.5885E-09 | -0.3063443 | 0.055 | 0.17 | 7.6614E-05 | TEC_prlf | TSPYL2 |
| 1.4351E-07 | -0.3064763 | 0.039 | 0.13 | 0.00306396 | TEC_prlf | HOMER1 |

| | | | | | | |
|---|---|---|---|---|---|---|
| 8.5481E-08 | -0.3068654 | 0.149 | 0.267 | 0.00182502 | TEC_prlf | SELENOM |
| 4.5838E-07 | -0.3079124 | 0.058 | 0.15 | 0.00978646 | TEC_prlf | AL118516.1 |
| 5.608E-06 | -0.3129567 | 0.185 | 0.292 | 0.1197316 | TEC_prlf | CYR61 |
| 1.0685E-09 | -0.3141387 | 0.105 | 0.238 | 2.2812E-05 | TEC_prlf | NEU1 |
| 1.1612E-05 | -0.3160388 | 0.986 | 0.957 | 0.24792265 | TEC_prlf | EIF1 |
| 2.387E-07 | -0.3165232 | 0.174 | 0.294 | 0.00509627 | TEC_prlf | MAFF |
| 3.5285E-05 | -0.317171 | 0.105 | 0.189 | 0.75332886 | TEC_prlf | BIRC3 |
| 1.1317E-06 | -0.317334 | 0.113 | 0.212 | 0.02416109 | TEC_prlf | MAT2A |
| 0.00084731 | -0.3177411 | 0.328 | 0.392 | 1 | TEC_prlf | RAB11FIP1 |
| 7.2932E-06 | -0.3196265 | 0.281 | 0.384 | 0.1557108 | TEC_prlf | HERPUD1 |
| 9.9795E-08 | -0.3206228 | 0.025 | 0.113 | 0.00213063 | TEC_prlf | AC023157.3 |
| 5.9634E-06 | -0.3212451 | 0.314 | 0.416 | 0.12731936 | TEC_prlf | NORAD |
| 4.0666E-06 | -0.323688 | 0.567 | 0.627 | 0.08682152 | TEC_prlf | ATF4 |
| 3.5889E-07 | -0.3258254 | 0.025 | 0.107 | 0.00766229 | TEC_prlf | ARC |
| 5.1078E-08 | -0.3313777 | 0.262 | 0.396 | 0.00109052 | TEC_prlf | TAF1D |
| 2.8367E-06 | -0.3326216 | 0.19 | 0.287 | 0.06056255 | TEC_prlf | COL18A1 |
| 1.0056E-05 | -0.3352633 | 0.65 | 0.694 | 0.21469217 | TEC_prlf | ZFAS1 |
| 0.00010903 | -0.3357168 | 0.353 | 0.424 | 1 | TEC_prlf | SNHG8 |
| 1.1118E-08 | -0.3409032 | 0.127 | 0.256 | 0.00023737 | TEC_prlf | IRF1 |
| 4.6066E-08 | -0.3437204 | 0.303 | 0.435 | 0.0009835 | TEC_prlf | ZFAND5 |
| 2.6765E-05 | -0.3481988 | 0.336 | 0.415 | 0.571426 | TEC_prlf | GOLGB1 |
| 1.2992E-06 | -0.3489949 | 0.107 | 0.209 | 0.02773774 | TEC_prlf | RRAD |
| 3.3698E-11 | -0.3497048 | 0.033 | 0.159 | 7.1946E-07 | TEC_prlf | CDKN2B |
| 0.00436754 | -0.3526799 | 0.234 | 0.285 | 1 | TEC_prlf | POLR2A |
| 1.1193E-08 | -0.3551631 | 0.383 | 0.503 | 0.00023897 | TEC_prlf | HNRNPDL |
| 0.00030977 | -0.3558577 | 0.215 | 0.288 | 1 | TEC_prlf | PLAUR |
| 9.4096E-09 | -0.3573754 | 0.047 | 0.153 | 0.0002009 | TEC_prlf | AC007952.4 |
| 2.335E-05 | -0.3592248 | 0.207 | 0.304 | 0.49851656 | TEC_prlf | DDIT4 |
| 2.2649E-09 | -0.359861 | 0.025 | 0.129 | 4.8355E-05 | TEC_prlf | HIST2H2BF |
| 2.1598E-06 | -0.360867 | 0.218 | 0.314 | 0.046112 | TEC_prlf | WSB1 |
| 2.4085E-08 | -0.3622163 | 0.014 | 0.102 | 0.00051421 | TEC_prlf | AP001160.1 |
| 0.00102897 | -0.3665737 | 0.253 | 0.321 | 1 | TEC_prlf | ITGA2 |
| 2.6164E-07 | -0.3670881 | 0.198 | 0.321 | 0.00558591 | TEC_prlf | UGCG |
| 2.7503E-11 | -0.367694 | 0.047 | 0.182 | 5.872E-07 | TEC_prlf | KLF4 |
| 3.3354E-05 | -0.369208 | 0.84 | 0.891 | 0.71211812 | TEC_prlf | MT-ND2 |
| 1.8873E-06 | -0.371368 | 0.226 | 0.335 | 0.0402946 | TEC_prlf | H1F0 |
| 6.1194E-06 | -0.3719136 | 0.083 | 0.168 | 0.13065007 | TEC_prlf | CTGF |
| 3.8376E-07 | -0.3719644 | 0.234 | 0.348 | 0.00819336 | TEC_prlf | TSC22D1 |
| 0.00036877 | -0.3759645 | 0.245 | 0.329 | 1 | TEC_prlf | IL32 |
| 8.8176E-07 | -0.380492 | 0.245 | 0.362 | 0.01882555 | TEC_prlf | KLF6 |
| 4.4026E-09 | -0.3805622 | 0.033 | 0.138 | 9.3996E-05 | TEC_prlf | AC025164.1 |
| 4.7867E-07 | -0.3816065 | 0.325 | 0.436 | 0.0102197 | TEC_prlf | ZBTB20 |
| 1.541E-11 | -0.3834264 | 0.565 | 0.704 | 3.29E-07 | TEC_prlf | MAP1LC3B |
| 3.0498E-10 | -0.3871186 | 0.135 | 0.278 | 6.5114E-06 | TEC_prlf | IFRD1 |
| 7.2782E-09 | -0.3887423 | 0.168 | 0.299 | 0.00015539 | TEC_prlf | MIR4435-2HG |
| 1.3991E-06 | -0.39089 | 0.149 | 0.255 | 0.02987122 | TEC_prlf | ID2 |
| 7.8356E-05 | -0.3940951 | 0.32 | 0.416 | 1 | TEC_prlf | PHLDA2 |
| 0.00494082 | -0.3943574 | 0.253 | 0.309 | 1 | TEC_prlf | PLAU |
| 0.00784567 | -0.3953968 | 0.68 | 0.597 | 1 | TEC_prlf | SPP1 |
| 7.4669E-09 | -0.3970181 | 0.116 | 0.237 | 0.00015942 | TEC_prlf | CDKN2AIP |

| | | | | | | |
|---|---|---|---|---|---|---|
| 1.6866E-07 | -0.3994511 | 0.62 | 0.727 | 0.00360088 | TEC_prlf | CTSD |
| 1.2333E-08 | -0.4010424 | 0.314 | 0.437 | 0.00026332 | TEC_prlf | PNRC1 |
| 0.00045269 | -0.4016863 | 0.174 | 0.241 | 1 | TEC_prlf | MDM2 |
| 2.7815E-05 | -0.4042894 | 0.215 | 0.302 | 0.59384166 | TEC_prlf | MARCKSL1 |
| 8.6876E-08 | -0.4073726 | 0.185 | 0.308 | 0.0018548 | TEC_prlf | PMAIP1 |
| 9.5338E-12 | -0.4074443 | 0.05 | 0.186 | 2.0355E-07 | TEC_prlf | ERN1 |
| 4.6024E-07 | -0.4110397 | 0.223 | 0.338 | 0.00982616 | TEC_prlf | ZNF292 |
| 3.19E-08 | -0.4118656 | 0.215 | 0.346 | 0.00068107 | TEC_prlf | ZFP36L2 |
| 4.1504E-10 | -0.4227559 | 0.223 | 0.378 | 8.8611E-06 | TEC_prlf | ATF3 |
| 0.00443908 | -0.4231558 | 0.273 | 0.332 | 1 | TEC_prlf | LAMC2 |
| 3.6658E-10 | -0.4249686 | 0.493 | 0.613 | 7.8265E-06 | TEC_prlf | EIF4A2 |
| 5.0452E-06 | -0.4259172 | 0.052 | 0.13 | 0.10771419 | TEC_prlf | COL7A1 |
| 1.4557E-10 | -0.4277673 | 0.063 | 0.194 | 3.1079E-06 | TEC_prlf | TLE4 |
| 2.0058E-11 | -0.434266 | 0.083 | 0.226 | 4.2823E-07 | TEC_prlf | ADM |
| 1.9358E-07 | -0.4399497 | 0.135 | 0.246 | 0.00413293 | TEC_prlf | ID4 |
| 1.4122E-06 | -0.4443779 | 0.039 | 0.119 | 0.03015031 | TEC_prlf | NEFM |
| 1.294E-06 | -0.4493211 | 0.634 | 0.704 | 0.02762609 | TEC_prlf | CLDN4 |
| 0.00033977 | -0.4535681 | 0.212 | 0.283 | 1 | TEC_prlf | BTG2 |
| 1.9094E-10 | -0.4537992 | 0.554 | 0.671 | 4.0765E-06 | TEC_prlf | GLS |
| 2.8372E-07 | -0.4616702 | 0.116 | 0.221 | 0.00605747 | TEC_prlf | TNFAIP2 |
| 0.00513844 | -0.4618731 | 0.342 | 0.394 | 1 | TEC_prlf | TXNRD1 |
| 1.808E-09 | -0.4679569 | 0.187 | 0.324 | 3.8602E-05 | TEC_prlf | CCNL1 |
| 7.646E-13 | -0.4734355 | 0.22 | 0.387 | 1.6324E-08 | TEC_prlf | N4BP2L2 |
| 7.5525E-11 | -0.4750813 | 0.127 | 0.268 | 1.6125E-06 | TEC_prlf | AMD1 |
| 1.2518E-11 | -0.4810536 | 0.201 | 0.363 | 2.6726E-07 | TEC_prlf | MCL1 |
| 8.3717E-13 | -0.4817179 | 0.033 | 0.176 | 1.7874E-08 | TEC_prlf | FOSB |
| 3.4596E-07 | -0.4875123 | 0.187 | 0.298 | 0.00738635 | TEC_prlf | ITGB8 |
| 0.00087962 | -0.4991099 | 0.388 | 0.444 | 1 | TEC_prlf | UPP1 |
| 4.2887E-11 | -0.5053286 | 0.149 | 0.3 | 9.1563E-07 | TEC_prlf | DST |
| 2.4541E-09 | -0.5088382 | 0.058 | 0.172 | 5.2394E-05 | TEC_prlf | EGLN3 |
| 1.8525E-12 | -0.5089669 | 0.085 | 0.238 | 3.9551E-08 | TEC_prlf | NFKBIZ |
| 4.3283E-12 | -0.5163919 | 0.052 | 0.196 | 9.2409E-08 | TEC_prlf | GADD45G |
| 9.65E-09 | -0.5222244 | 0.163 | 0.29 | 0.00020603 | TEC_prlf | CEBPD |
| 8.9407E-08 | -0.5276768 | 0.193 | 0.309 | 0.00190884 | TEC_prlf | HSPA1B |
| 1.4014E-11 | -0.5292964 | 0.124 | 0.275 | 2.9921E-07 | TEC_prlf | SEPHS2 |
| 2.7051E-13 | -0.5368416 | 0.102 | 0.266 | 5.7754E-09 | TEC_prlf | HIST1H2AC |
| 2.9151E-13 | -0.5382213 | 0.273 | 0.438 | 6.2238E-09 | TEC_prlf | DDX17 |
| 9.2258E-20 | -0.5531123 | 0.116 | 0.339 | 1.9697E-15 | TEC_prlf | CPEB4 |
| 1.2955E-16 | -0.5594631 | 0.399 | 0.579 | 2.7659E-12 | TEC_prlf | SLC3A2 |
| 1.1118E-15 | -0.5607277 | 0.099 | 0.281 | 2.3738E-11 | TEC_prlf | VEGFA |
| 8.5389E-13 | -0.5635585 | 0.105 | 0.269 | 1.823E-08 | TEC_prlf | DUSP1 |
| 5.9654E-10 | -0.5673686 | 0.124 | 0.257 | 1.2736E-05 | TEC_prlf | IER5 |
| 4.6847E-12 | -0.5684887 | 0.118 | 0.271 | 1.0002E-07 | TEC_prlf | KCNQ1OT1 |
| 2.4215E-11 | -0.5770198 | 0.055 | 0.188 | 5.17E-07 | TEC_prlf | PIK3R3 |
| 0.00025767 | -0.5828088 | 0.209 | 0.279 | 1 | TEC_prlf | HES1 |
| 2.0456E-18 | -0.5836714 | 0.174 | 0.397 | 4.3673E-14 | TEC_prlf | NDRG1 |
| 1.8364E-16 | -0.5862802 | 0.146 | 0.347 | 3.9208E-12 | TEC_prlf | SNHG12 |
| 3.7813E-12 | -0.5871665 | 0.229 | 0.39 | 8.0731E-08 | TEC_prlf | BRD2 |
| 1.4049E-15 | -0.5880839 | 0.143 | 0.332 | 2.9994E-11 | TEC_prlf | CEBPB |
| 3.6396E-09 | -0.5904507 | 0.215 | 0.363 | 7.7705E-05 | TEC_prlf | HSPA1A |

| | | | | | | |
|---|---|---|---|---|---|---|
| 3.4217E-12 | -0.5986601 | 0.124 | 0.28 | 7.3053E-08 | TEC_prlf | CITED2 |
| 2.8996E-17 | -0.6092012 | 0.182 | 0.383 | 6.1907E-13 | TEC_prlf | RSRP1 |
| 6.7291E-15 | -0.6135258 | 0.052 | 0.219 | 1.4367E-10 | TEC_prlf | SOCS1 |
| 2.3078E-13 | -0.6142122 | 0.865 | 0.913 | 4.9272E-09 | TEC_prlf | UBC |
| 7.5578E-17 | -0.6162231 | 0.03 | 0.208 | 1.6136E-12 | TEC_prlf | SGK1 |
| 1.1212E-10 | -0.6176355 | 0.105 | 0.241 | 2.3938E-06 | TEC_prlf | INTS6 |
| 9.8182E-17 | -0.6278955 | 0.424 | 0.612 | 2.0962E-12 | TEC_prlf | GADD45A |
| 1.9512E-13 | -0.6329581 | 0.485 | 0.614 | 4.1657E-09 | TEC_prlf | BTG1 |
| 5.9314E-08 | -0.6356994 | 0.204 | 0.315 | 0.00126636 | TEC_prlf | GLUL |
| 2.7004E-14 | -0.6446347 | 0.264 | 0.449 | 5.7654E-10 | TEC_prlf | ZFP36L1 |
| 3.8883E-10 | -0.6500531 | 0.127 | 0.264 | 8.3015E-06 | TEC_prlf | EGR1 |
| 4.3135E-13 | -0.6590335 | 0.215 | 0.396 | 9.2093E-09 | TEC_prlf | NFKBIA |
| 6.3294E-11 | -0.6725382 | 0.08 | 0.215 | 1.3513E-06 | TEC_prlf | ADAMTS1 |
| 6.275E-09 | -0.6907952 | 0.163 | 0.28 | 0.00013397 | TEC_prlf | PPP1R10 |
| 7.4263E-21 | -0.6975098 | 0.088 | 0.319 | 1.5855E-16 | TEC_prlf | CDKN1C |
| 4.1531E-14 | -0.6994932 | 0.121 | 0.293 | 8.8668E-10 | TEC_prlf | RND3 |
| 2.1641E-15 | -0.7037224 | 0.085 | 0.26 | 4.6203E-11 | TEC_prlf | IER5L |
| 0.0019683 | -0.7220195 | 0.997 | 0.978 | 1 | TEC_prlf | FTH1 |
| 8.0112E-16 | -0.7243307 | 0.179 | 0.375 | 1.7104E-11 | TEC_prlf | DDIT3 |
| 2.1328E-14 | -0.7430187 | 0.132 | 0.306 | 4.5534E-10 | TEC_prlf | PLK2 |
| 4.0268E-19 | -0.7733821 | 0.314 | 0.516 | 8.5972E-15 | TEC_prlf | CDKN1A |
| 9.8781E-24 | -0.7758668 | 0.413 | 0.629 | 2.109E-19 | TEC_prlf | PPP1R15A |
| 4.0615E-18 | -0.7761536 | 0.077 | 0.283 | 8.6714E-14 | TEC_prlf | RASD1 |
| 0.00170586 | -0.783244 | 0.631 | 0.668 | 1 | TEC_prlf | CRYAB |
| 1.2117E-16 | -0.7911151 | 0.185 | 0.392 | 2.5869E-12 | TEC_prlf | JUNB |
| 3.1258E-20 | -0.8295305 | 0.157 | 0.381 | 6.6736E-16 | TEC_prlf | SLC38A2 |
| 4.1028E-12 | -0.840259 | 0.138 | 0.294 | 8.7594E-08 | TEC_prlf | FOS |
| 2.8321E-12 | -0.8585742 | 0.474 | 0.59 | 6.0465E-08 | TEC_prlf | ELF3 |
| 1.4248E-23 | -0.8655701 | 0.408 | 0.611 | 3.0419E-19 | TEC_prlf | SOD2 |
| 2.1783E-20 | -0.8902364 | 0.824 | 0.873 | 4.6506E-16 | TEC_prlf | SAT1 |
| 5.9052E-16 | -0.9197858 | 0.196 | 0.383 | 1.2608E-11 | TEC_prlf | HIST1H4C |
| 1.1286E-17 | -0.9231319 | 0.052 | 0.239 | 2.4096E-13 | TEC_prlf | CSKMT |
| 1.1394E-15 | -0.9441192 | 0.419 | 0.568 | 2.4326E-11 | TEC_prlf | IER2 |
| 8.9358E-16 | -0.9538729 | 0.096 | 0.276 | 1.9078E-11 | TEC_prlf | TCIM |
| 1.4222E-11 | -0.9693013 | 0.16 | 0.301 | 3.0364E-07 | TEC_prlf | HEXIM1 |
| 2.3104E-12 | -0.974251 | 0.284 | 0.444 | 4.9328E-08 | TEC_prlf | IER3 |
| 6.0494E-11 | -0.9804811 | 0.138 | 0.274 | 1.2915E-06 | TEC_prlf | HSPA2 |
| 7.2128E-19 | -1.0070018 | 0.441 | 0.61 | 1.5399E-14 | TEC_prlf | GADD45B |
| 3.2149E-42 | -1.0106658 | 0.62 | 0.835 | 6.8638E-38 | TEC_prlf | SQSTM1 |
| 1.2264E-24 | -1.0124482 | 0.091 | 0.339 | 2.6183E-20 | TEC_prlf | RHOB |
| 0.00036551 | -1.0268579 | 0.157 | 0.227 | 1 | TEC_prlf | DEFB1 |
| 8.9076E-31 | -1.1295637 | 0.43 | 0.645 | 1.9018E-26 | TEC_prlf | JUND |
| 2.9988E-19 | -1.2595982 | 0.391 | 0.571 | 6.4025E-15 | TEC_prlf | GDF15 |
| 7.0945E-20 | -1.2753559 | 0.485 | 0.638 | 1.5147E-15 | TEC_prlf | JUN |
| 7.7586E-24 | -1.3485795 | 0.052 | 0.289 | 1.6565E-19 | TEC_prlf | MAFB |
| 2.5494E-22 | -1.3638045 | 0.945 | 0.962 | 5.4429E-18 | TEC_prlf | FTL |
| 5.3846E-34 | -1.3721894 | 0.956 | 0.94 | 1.1496E-29 | TEC_prlf | MALAT1 |
| 3.4948E-19 | -1.6736165 | 0.388 | 0.544 | 7.4614E-15 | TEC_prlf | SOX4 |
| 1.9579E-45 | -1.7742678 | 0.697 | 0.818 | 4.1801E-41 | TEC_prlf | NEAT1 |
| 5.3348E-05 | -3.0373076 | 0.617 | 0.597 | 1 | TEC_prlf | PLCG2 |

| | | | | | | |
|---|---|---|---|---|---|---|
| 0 | 1.58347634 | 0.998 | 0.958 | 0 | TEC_TXNRD1 | FTL |
| 9.166E-281 | 1.34135101 | 1 | 0.976 | 1.957E-276 | TEC_TXNRD1 | FTH1 |
| 6.019E-174 | 1.24480469 | 0.724 | 0.361 | 1.285E-169 | TEC_TXNRD1 | TXNRD1 |
| 7.87E-134 | 0.74787323 | 0.493 | 0.19 | 1.68E-129 | TEC_TXNRD1 | GCLM |
| 1.982E-111 | 0.74698787 | 0.89 | 0.699 | 4.231E-107 | TEC_TXNRD1 | TXN |
| 4.567E-154 | 0.69476795 | 0.995 | 0.933 | 9.75E-150 | TEC_TXNRD1 | RPL15 |
| 5.243E-132 | 0.6815449 | 0.986 | 0.917 | 1.119E-127 | TEC_TXNRD1 | RPL10 |
| 1.333E-145 | 0.68152732 | 0.994 | 0.887 | 2.847E-141 | TEC_TXNRD1 | RPS13 |
| 1.894E-141 | 0.67359504 | 0.832 | 0.519 | 4.044E-137 | TEC_TXNRD1 | BRI3 |
| 1.411E-146 | 0.66644383 | 0.995 | 0.915 | 3.013E-142 | TEC_TXNRD1 | RPS3A |
| 8.142E-128 | 0.65849886 | 0.976 | 0.856 | 1.738E-123 | TEC_TXNRD1 | RPL23 |
| 1.3713E-65 | 0.65480222 | 0.66 | 0.422 | 2.9278E-61 | TEC_TXNRD1 | UPP1 |
| 3.937E-144 | 0.65070363 | 0.995 | 0.923 | 8.406E-140 | TEC_TXNRD1 | RPS14 |
| 2.931E-134 | 0.64537491 | 0.996 | 0.932 | 6.258E-130 | TEC_TXNRD1 | RPS23 |
| 2.516E-143 | 0.64351696 | 0.988 | 0.89 | 5.372E-139 | TEC_TXNRD1 | RPL5 |
| 2.617E-75 | 0.6416553 | 0.302 | 0.111 | 5.5873E-71 | TEC_TXNRD1 | HMOX1 |
| 5.197E-140 | 0.63310029 | 0.999 | 0.954 | 1.109E-135 | TEC_TXNRD1 | EIF1 |
| 2.285E-148 | 0.63195702 | 0.993 | 0.863 | 4.879E-144 | TEC_TXNRD1 | RPL7 |
| 2.644E-111 | 0.62438425 | 0.995 | 0.901 | 5.646E-107 | TEC_TXNRD1 | RPL26 |
| 6.797E-136 | 0.61522107 | 0.995 | 0.897 | 1.451E-131 | TEC_TXNRD1 | RPS7 |
| 8.039E-140 | 0.60858412 | 0.996 | 0.912 | 1.716E-135 | TEC_TXNRD1 | RPL9 |
| 2.969E-136 | 0.60391758 | 0.7 | 0.345 | 6.339E-132 | TEC_TXNRD1 | SEC61G |
| 1.561E-136 | 0.60247653 | 0.715 | 0.359 | 3.332E-132 | TEC_TXNRD1 | POMP |
| 7.939E-118 | 0.60219335 | 0.995 | 0.896 | 1.695E-113 | TEC_TXNRD1 | RPS3 |
| 2.06E-119 | 0.598761 | 0.998 | 0.941 | 4.399E-115 | TEC_TXNRD1 | RPL11 |
| 2.323E-120 | 0.59581309 | 0.998 | 0.928 | 4.959E-116 | TEC_TXNRD1 | RPS27A |
| 7.9519E-98 | 0.59578108 | 0.724 | 0.438 | 1.6977E-93 | TEC_TXNRD1 | SERP1 |
| 6.0502E-76 | 0.59533823 | 0.785 | 0.554 | 1.2917E-71 | TEC_TXNRD1 | SLC3A2 |
| 3.217E-115 | 0.5950593 | 0.998 | 0.915 | 6.869E-111 | TEC_TXNRD1 | RPL30 |
| 5.967E-124 | 0.59241456 | 0.982 | 0.862 | 1.274E-119 | TEC_TXNRD1 | RACK1 |
| 4.563E-142 | 0.59172817 | 0.994 | 0.88 | 9.741E-138 | TEC_TXNRD1 | NACA |
| 1.26E-128 | 0.58885971 | 0.999 | 0.986 | 2.691E-124 | TEC_TXNRD1 | EEF1A1 |
| 9.859E-125 | 0.58750142 | 0.995 | 0.926 | 2.105E-120 | TEC_TXNRD1 | RPL18 |
| 7.642E-117 | 0.57529019 | 0.941 | 0.774 | 1.632E-112 | TEC_TXNRD1 | NPM1 |
| 2.371E-110 | 0.57526329 | 0.993 | 0.899 | 5.061E-106 | TEC_TXNRD1 | RPS16 |
| 1.724E-104 | 0.57319157 | 0.998 | 0.947 | 3.681E-100 | TEC_TXNRD1 | RPS12 |
| 1.037E-122 | 0.56770456 | 0.981 | 0.855 | 2.214E-118 | TEC_TXNRD1 | RPS4X |
| 6.033E-117 | 0.56384059 | 0.991 | 0.902 | 1.288E-112 | TEC_TXNRD1 | RPL24 |
| 5.518E-99 | 0.55596322 | 0.988 | 0.925 | 1.178E-94 | TEC_TXNRD1 | RPL7A |
| 1.804E-22 | 0.55362983 | 0.321 | 0.203 | 3.8516E-18 | TEC_TXNRD1 | GLRX |
| 2.057E-101 | 0.54778094 | 0.589 | 0.294 | 4.3909E-97 | TEC_TXNRD1 | TKT |
| 3.1326E-89 | 0.54647434 | 0.952 | 0.826 | 6.688E-85 | TEC_TXNRD1 | PRDX1 |
| 7.2778E-98 | 0.54265905 | 0.993 | 0.926 | 1.5538E-93 | TEC_TXNRD1 | RPS6 |
| 4.237E-108 | 0.53804205 | 0.927 | 0.699 | 9.045E-104 | TEC_TXNRD1 | RPL36AL |
| 2.591E-99 | 0.53281678 | 0.998 | 0.958 | 5.5316E-95 | TEC_TXNRD1 | RPS18 |
| 1.171E-115 | 0.53259203 | 0.63 | 0.309 | 2.5E-111 | TEC_TXNRD1 | PSMB1 |
| 1.3037E-90 | 0.53181638 | 0.998 | 0.969 | 2.7834E-86 | TEC_TXNRD1 | RPLP1 |
| 1.017E-100 | 0.53007053 | 0.72 | 0.422 | 2.1706E-96 | TEC_TXNRD1 | VDAC2 |
| 4.416E-111 | 0.52947159 | 0.995 | 0.97 | 9.428E-107 | TEC_TXNRD1 | TPT1 |
| 9.7589E-71 | 0.52941262 | 0.857 | 0.678 | 2.0835E-66 | TEC_TXNRD1 | ZFAS1 |

| | | | | | | |
|---|---|---|---|---|---|---|
| 3.9121E-95 | 0.52789997 | 0.984 | 0.859 | 8.3523E-91 | TEC_TXNRD1 | RPS5 |
| 2.0704E-97 | 0.52570536 | 0.995 | 0.922 | 4.4204E-93 | TEC_TXNRD1 | RPL21 |
| 6.4584E-97 | 0.52557523 | 0.995 | 0.915 | 1.3789E-92 | TEC_TXNRD1 | RPL14 |
| 5.4022E-95 | 0.52467815 | 0.602 | 0.314 | 1.1534E-90 | TEC_TXNRD1 | SSR3 |
| 1.955E-117 | 0.52360949 | 0.688 | 0.361 | 4.173E-113 | TEC_TXNRD1 | TOMM20 |
| 3.0971E-89 | 0.51635895 | 0.998 | 0.971 | 6.6124E-85 | TEC_TXNRD1 | RPL13 |
| 1.467E-79 | 0.50476365 | 0.983 | 0.873 | 3.1321E-75 | TEC_TXNRD1 | RPL39 |
| 2.2251E-74 | 0.50233488 | 0.758 | 0.512 | 4.7506E-70 | TEC_TXNRD1 | SEC61B |
| 5.0722E-64 | 0.5020291 | 0.767 | 0.539 | 1.0829E-59 | TEC_TXNRD1 | HMGA1 |
| 1.2707E-86 | 0.49938707 | 0.88 | 0.683 | 2.7129E-82 | TEC_TXNRD1 | MAP1LC3B |
| 3.6468E-79 | 0.49835326 | 0.991 | 0.888 | 7.7859E-75 | TEC_TXNRD1 | RPS15A |
| 1.5845E-53 | 0.49651006 | 0.514 | 0.299 | 3.383E-49 | TEC_TXNRD1 | UGCG |
| 2.9164E-91 | 0.4945094 | 0.562 | 0.276 | 6.2266E-87 | TEC_TXNRD1 | MYDGF |
| 8.4104E-88 | 0.49350738 | 0.986 | 0.888 | 1.7956E-83 | TEC_TXNRD1 | RPL35A |
| 1.0014E-83 | 0.48751652 | 0.99 | 0.922 | 2.1379E-79 | TEC_TXNRD1 | RPL18A |
| 6.4938E-75 | 0.48682347 | 0.997 | 0.937 | 1.3864E-70 | TEC_TXNRD1 | RPS24 |
| 1.5328E-87 | 0.4853612 | 0.993 | 0.928 | 3.2725E-83 | TEC_TXNRD1 | RPL32 |
| 3.368E-106 | 0.48405221 | 0.471 | 0.193 | 7.192E-102 | TEC_TXNRD1 | EPB41L4A-AS1 |
| 2.8056E-91 | 0.47991346 | 0.609 | 0.315 | 5.99E-87 | TEC_TXNRD1 | PFDN2 |
| 2.2238E-87 | 0.47219688 | 0.917 | 0.739 | 4.7479E-83 | TEC_TXNRD1 | RPL4 |
| 8.43E-83 | 0.47110159 | 0.99 | 0.905 | 1.7998E-78 | TEC_TXNRD1 | RPL29 |
| 2.1706E-80 | 0.46950031 | 0.62 | 0.344 | 4.6343E-76 | TEC_TXNRD1 | SBDS |
| 1.1178E-72 | 0.46740765 | 0.57 | 0.309 | 2.3865E-68 | TEC_TXNRD1 | TUBA1C |
| 4.01E-108 | 0.46579187 | 0.565 | 0.26 | 8.56E-104 | TEC_TXNRD1 | RSL24D1 |
| 1.9375E-83 | 0.46550474 | 0.591 | 0.318 | 4.1365E-79 | TEC_TXNRD1 | ARF4 |
| 2.8627E-82 | 0.46415356 | 0.94 | 0.768 | 6.1119E-78 | TEC_TXNRD1 | EEF1B2 |
| 6.4175E-81 | 0.46415117 | 0.806 | 0.559 | 1.3701E-76 | TEC_TXNRD1 | ATP6V0E1 |
| 1.5139E-82 | 0.45997524 | 0.841 | 0.616 | 3.2322E-78 | TEC_TXNRD1 | CCNI |
| 1.2287E-73 | 0.45922432 | 0.995 | 0.935 | 2.6233E-69 | TEC_TXNRD1 | RPL19 |
| 5.9667E-81 | 0.45854227 | 0.831 | 0.583 | 1.2739E-76 | TEC_TXNRD1 | PSMA7 |
| 2.2505E-92 | 0.45586887 | 0.923 | 0.735 | 4.8048E-88 | TEC_TXNRD1 | BTF3 |
| 5.1225E-61 | 0.45433401 | 0.803 | 0.614 | 1.0937E-56 | TEC_TXNRD1 | ANXA5 |
| 3.7799E-68 | 0.45245132 | 0.588 | 0.338 | 8.0701E-64 | TEC_TXNRD1 | TALDO1 |
| 1.3283E-73 | 0.45238948 | 0.99 | 0.894 | 2.8359E-69 | TEC_TXNRD1 | RPS25 |
| 1.1039E-96 | 0.45143774 | 0.659 | 0.345 | 2.3567E-92 | TEC_TXNRD1 | SNU13 |
| 1.1816E-70 | 0.45130899 | 0.582 | 0.328 | 2.5227E-66 | TEC_TXNRD1 | SELENOK |
| 2.0784E-77 | 0.44261348 | 0.859 | 0.644 | 4.4374E-73 | TEC_TXNRD1 | RAC1 |
| 6.9601E-84 | 0.44148329 | 0.586 | 0.306 | 1.486E-79 | TEC_TXNRD1 | SF3B6 |
| 1.9325E-70 | 0.44028138 | 0.992 | 0.913 | 4.1258E-66 | TEC_TXNRD1 | RPL34 |
| 1.1223E-87 | 0.43972805 | 0.605 | 0.315 | 2.396E-83 | TEC_TXNRD1 | KRT10 |
| 6.6096E-76 | 0.43579211 | 0.986 | 0.893 | 1.4112E-71 | TEC_TXNRD1 | RPL27A |
| 1.3564E-77 | 0.43543795 | 0.912 | 0.76 | 2.896E-73 | TEC_TXNRD1 | EEF2 |
| 7.9984E-74 | 0.43491258 | 0.998 | 0.938 | 1.7077E-69 | TEC_TXNRD1 | RPL8 |
| 1.1395E-88 | 0.42634976 | 0.594 | 0.308 | 2.4329E-84 | TEC_TXNRD1 | EIF3E |
| 1.1962E-72 | 0.42532937 | 0.565 | 0.313 | 2.5539E-68 | TEC_TXNRD1 | SPCS2 |
| 2.2373E-70 | 0.42325395 | 0.486 | 0.243 | 4.7765E-66 | TEC_TXNRD1 | RB1CC1 |
| 1.6381E-65 | 0.4231253 | 0.995 | 0.947 | 3.4974E-61 | TEC_TXNRD1 | RPS8 |
| 1.9862E-56 | 0.41838211 | 0.814 | 0.671 | 4.2404E-52 | TEC_TXNRD1 | HSPA5 |
| 1.4025E-72 | 0.41818375 | 0.992 | 0.928 | 2.9943E-68 | TEC_TXNRD1 | RPS9 |
| 1.0739E-43 | 0.41715274 | 0.982 | 0.934 | 2.2929E-39 | TEC_TXNRD1 | RPS28 |

| | | | | | | |
|---|---|---|---|---|---|---|
| 1.0751E-80 | 0.41604546 | 0.938 | 0.773 | 2.2953E-76 | TEC_TXNRD1 | CHCHD2 |
| 6.4987E-65 | 0.41583653 | 0.668 | 0.416 | 1.3875E-60 | TEC_TXNRD1 | NAP1L1 |
| 4.3077E-69 | 0.41236716 | 0.973 | 0.898 | 9.1969E-65 | TEC_TXNRD1 | RPS20 |
| 2.5649E-86 | 0.41216559 | 0.547 | 0.269 | 5.4761E-82 | TEC_TXNRD1 | SF3B5 |
| 1.3979E-73 | 0.41131924 | 0.582 | 0.316 | 2.9846E-69 | TEC_TXNRD1 | NOP10 |
| 6.8754E-72 | 0.41110808 | 0.273 | 0.096 | 1.4679E-67 | TEC_TXNRD1 | SLC7A11 |
| 1.3522E-74 | 0.40866118 | 0.636 | 0.357 | 2.8869E-70 | TEC_TXNRD1 | RHEB |
| 5.924E-57 | 0.40733065 | 0.995 | 0.929 | 1.2648E-52 | TEC_TXNRD1 | RPL37 |
| 3.4253E-74 | 0.40623624 | 0.572 | 0.311 | 7.3131E-70 | TEC_TXNRD1 | RAB2A |
| 9.6018E-75 | 0.4047403 | 0.549 | 0.293 | 2.05E-70 | TEC_TXNRD1 | COX7A2L |
| 1.4999E-66 | 0.40193242 | 0.605 | 0.348 | 3.2024E-62 | TEC_TXNRD1 | UBE2B |
| 7.6446E-70 | 0.39997005 | 0.573 | 0.32 | 1.6321E-65 | TEC_TXNRD1 | GHITM |
| 1.8539E-48 | 0.39917147 | 0.999 | 0.964 | 3.958E-44 | TEC_TXNRD1 | RPL41 |
| 5.3934E-73 | 0.39903288 | 0.316 | 0.122 | 1.1515E-68 | TEC_TXNRD1 | TRIB3 |
| 5.3727E-64 | 0.3980032 | 0.995 | 0.902 | 1.1471E-59 | TEC_TXNRD1 | FAU |
| 1.9051E-46 | 0.39759913 | 0.38 | 0.202 | 4.0673E-42 | TEC_TXNRD1 | PTGR1 |
| 1.5578E-66 | 0.39636846 | 0.991 | 0.943 | 3.326E-62 | TEC_TXNRD1 | RPL13A |
| 9.5924E-67 | 0.39631317 | 0.805 | 0.585 | 2.048E-62 | TEC_TXNRD1 | CLTA |
| 4.6421E-63 | 0.3939205 | 0.622 | 0.37 | 9.9108E-59 | TEC_TXNRD1 | EIF2S2 |
| 9.0236E-65 | 0.39312285 | 0.627 | 0.362 | 1.9265E-60 | TEC_TXNRD1 | OCIAD2 |
| 7.5025E-82 | 0.39270834 | 0.508 | 0.245 | 1.6018E-77 | TEC_TXNRD1 | PSMB7 |
| 3.5686E-79 | 0.39116264 | 0.606 | 0.321 | 7.619E-75 | TEC_TXNRD1 | CNBP |
| 1.1281E-35 | 0.38952378 | 0.595 | 0.403 | 2.4084E-31 | TEC_TXNRD1 | LGALS3 |
| 2.158E-62 | 0.38467457 | 0.398 | 0.189 | 4.6073E-58 | TEC_TXNRD1 | DNTTIP2 |
| 9.8029E-61 | 0.38449539 | 0.7 | 0.449 | 2.0929E-56 | TEC_TXNRD1 | PRDX6 |
| 5.5193E-88 | 0.38271721 | 0.457 | 0.199 | 1.1784E-83 | TEC_TXNRD1 | RBCK1 |
| 6.5515E-66 | 0.38241916 | 0.643 | 0.373 | 1.3987E-61 | TEC_TXNRD1 | DAD1 |
| 8.3125E-52 | 0.38237999 | 0.639 | 0.406 | 1.7747E-47 | TEC_TXNRD1 | ATP6V0B |
| 1.7868E-65 | 0.38229641 | 0.521 | 0.282 | 3.8148E-61 | TEC_TXNRD1 | SSR2 |
| 1.2601E-37 | 0.38051252 | 0.451 | 0.269 | 2.6902E-33 | TEC_TXNRD1 | AKR1B1 |
| 2.2552E-68 | 0.37826588 | 0.524 | 0.276 | 4.8148E-64 | TEC_TXNRD1 | SELENOS |
| 1.6065E-62 | 0.37732017 | 0.978 | 0.879 | 3.4299E-58 | TEC_TXNRD1 | RPL22 |
| 2.0826E-48 | 0.37654342 | 0.998 | 0.957 | 4.4464E-44 | TEC_TXNRD1 | RPL28 |
| 4.8187E-56 | 0.37575731 | 0.983 | 0.929 | 1.0288E-51 | TEC_TXNRD1 | RPL3 |
| 1.4228E-53 | 0.37561797 | 0.987 | 0.925 | 3.0377E-49 | TEC_TXNRD1 | RPL12 |
| 3.5933E-69 | 0.37218825 | 0.563 | 0.305 | 7.6716E-65 | TEC_TXNRD1 | ELOC |
| 2.4869E-58 | 0.37064755 | 0.612 | 0.374 | 5.3094E-54 | TEC_TXNRD1 | TAX1BP1 |
| 2.4586E-70 | 0.36790295 | 0.562 | 0.307 | 5.2491E-66 | TEC_TXNRD1 | EIF3H |
| 4.6089E-75 | 0.3669457 | 0.547 | 0.28 | 9.84E-71 | TEC_TXNRD1 | POLR1D |
| 2.3416E-60 | 0.3667946 | 0.545 | 0.305 | 4.9993E-56 | TEC_TXNRD1 | EIF1B |
| 1.2258E-60 | 0.36536749 | 0.705 | 0.467 | 2.617E-56 | TEC_TXNRD1 | UBE2D3 |
| 1.3939E-71 | 0.36530528 | 0.466 | 0.229 | 2.9759E-67 | TEC_TXNRD1 | EIF3D |
| 1.5016E-62 | 0.3650758 | 0.865 | 0.673 | 3.2059E-58 | TEC_TXNRD1 | CYSTM1 |
| 8.4597E-63 | 0.36412112 | 0.474 | 0.244 | 1.8061E-58 | TEC_TXNRD1 | UFM1 |
| 8.283E-79 | 0.3633975 | 0.452 | 0.206 | 1.7684E-74 | TEC_TXNRD1 | PSMD7 |
| 3.8831E-57 | 0.36260379 | 0.972 | 0.859 | 8.2904E-53 | TEC_TXNRD1 | RPL10A |
| 2.2844E-47 | 0.36063332 | 0.658 | 0.431 | 4.8771E-43 | TEC_TXNRD1 | RPS10 |
| 2.6089E-65 | 0.36028584 | 0.947 | 0.786 | 5.57E-61 | TEC_TXNRD1 | RPL27 |
| 6.127E-78 | 0.36004378 | 0.421 | 0.185 | 1.3081E-73 | TEC_TXNRD1 | OSER1 |
| 1.9121E-56 | 0.35721933 | 0.76 | 0.521 | 4.0824E-52 | TEC_TXNRD1 | GNG5 |

| | | | | | | |
|---|---|---|---|---|---|---|
| 2.3738E-53 | 0.35465028 | 0.524 | 0.3 | 5.068E-49 | TEC_TXNRD1 | NDUFB9 |
| 9.1045E-46 | 0.35365143 | 0.727 | 0.506 | 1.9438E-41 | TEC_TXNRD1 | ATP5MC2 |
| 1.2878E-40 | 0.35322102 | 0.968 | 0.878 | 2.7494E-36 | TEC_TXNRD1 | RPS29 |
| 1.259E-45 | 0.35167922 | 0.996 | 0.94 | 2.6879E-41 | TEC_TXNRD1 | RPS15 |
| 3.8051E-58 | 0.34867404 | 0.974 | 0.908 | 8.1238E-54 | TEC_TXNRD1 | HSP90AB1 |
| 2.9296E-62 | 0.34832955 | 0.559 | 0.313 | 6.2548E-58 | TEC_TXNRD1 | PTP4A2 |
| 5.4306E-55 | 0.34716002 | 0.469 | 0.249 | 1.1594E-50 | TEC_TXNRD1 | BLVRB |
| 6.512E-74 | 0.34703246 | 0.964 | 0.832 | 1.3903E-69 | TEC_TXNRD1 | RPSA |
| 6.1613E-62 | 0.34613552 | 0.986 | 0.909 | 1.3154E-57 | TEC_TXNRD1 | RPLP0 |
| 3.5979E-30 | 0.34608814 | 0.429 | 0.269 | 7.6815E-26 | TEC_TXNRD1 | UCHL1 |
| 6.7579E-45 | 0.34440869 | 0.527 | 0.322 | 1.4428E-40 | TEC_TXNRD1 | HSPA9 |
| 1.8759E-51 | 0.34288347 | 0.687 | 0.461 | 4.0049E-47 | TEC_TXNRD1 | PTGES3 |
| 3.2252E-46 | 0.34147508 | 0.556 | 0.343 | 6.8858E-42 | TEC_TXNRD1 | TRMT112 |
| 2.1149E-69 | 0.34029662 | 0.4 | 0.179 | 4.5152E-65 | TEC_TXNRD1 | BNIP3 |
| 1.357E-69 | 0.3392667 | 0.447 | 0.217 | 2.8972E-65 | TEC_TXNRD1 | NAA20 |
| 4.1715E-24 | 0.33879106 | 0.829 | 0.767 | 8.9061E-20 | TEC_TXNRD1 | HSP90B1 |
| 5.5459E-56 | 0.33710993 | 0.972 | 0.855 | 1.1841E-51 | TEC_TXNRD1 | UBA52 |
| 1.0628E-33 | 0.33700885 | 0.907 | 0.814 | 2.2691E-29 | TEC_TXNRD1 | RPS21 |
| 6.0138E-63 | 0.33685653 | 0.544 | 0.298 | 1.284E-58 | TEC_TXNRD1 | SNRPB |
| 1.3847E-27 | 0.33666337 | 0.991 | 0.952 | 2.9564E-23 | TEC_TXNRD1 | RPS27 |
| 2.0203E-52 | 0.33570103 | 0.992 | 0.917 | 4.3134E-48 | TEC_TXNRD1 | RPLP2 |
| 7.6842E-54 | 0.33089334 | 0.456 | 0.243 | 1.6406E-49 | TEC_TXNRD1 | GSTO1 |
| 1.2661E-64 | 0.33042852 | 0.447 | 0.223 | 2.7032E-60 | TEC_TXNRD1 | GTF2A2 |
| 5.7354E-52 | 0.33011383 | 0.546 | 0.321 | 1.2245E-47 | TEC_TXNRD1 | EIF1AX |
| 1.9053E-47 | 0.32995634 | 0.759 | 0.554 | 4.0679E-43 | TEC_TXNRD1 | COX7A2 |
| 4.65E-55 | 0.32580687 | 0.513 | 0.288 | 9.9278E-51 | TEC_TXNRD1 | TMEM258 |
| 1.8511E-52 | 0.32574198 | 0.46 | 0.247 | 3.9521E-48 | TEC_TXNRD1 | SLC38A1 |
| 1.3715E-51 | 0.32506012 | 0.562 | 0.329 | 2.9281E-47 | TEC_TXNRD1 | COMMD6 |
| 1.2924E-47 | 0.32417248 | 0.361 | 0.179 | 2.7593E-43 | TEC_TXNRD1 | SDF2L1 |
| 2.8015E-61 | 0.32413011 | 0.508 | 0.274 | 5.9812E-57 | TEC_TXNRD1 | RNF7 |
| 9.0438E-49 | 0.3226206 | 0.699 | 0.472 | 1.9309E-44 | TEC_TXNRD1 | TOMM7 |
| 1.1278E-34 | 0.32205185 | 0.776 | 0.612 | 2.4078E-30 | TEC_TXNRD1 | SLC25A6 |
| 1.3962E-35 | 0.32091488 | 0.413 | 0.24 | 2.9808E-31 | TEC_TXNRD1 | S100A14 |
| 3.7808E-51 | 0.31896875 | 0.334 | 0.158 | 8.0721E-47 | TEC_TXNRD1 | EIF4EBP1 |
| 6.0825E-63 | 0.31765556 | 0.967 | 0.828 | 1.2986E-58 | TEC_TXNRD1 | CD63 |
| 4.8755E-53 | 0.31726914 | 0.55 | 0.317 | 1.0409E-48 | TEC_TXNRD1 | EIF4B |
| 2.4651E-55 | 0.31718507 | 0.431 | 0.221 | 5.263E-51 | TEC_TXNRD1 | EPRS |
| 1.6824E-46 | 0.31673403 | 0.913 | 0.769 | 3.5918E-42 | TEC_TXNRD1 | COX4I1 |
| 2.3103E-50 | 0.3152778 | 0.984 | 0.89 | 4.9324E-46 | TEC_TXNRD1 | RPL23A |
| 2.226E-46 | 0.3147441 | 0.621 | 0.397 | 4.7525E-42 | TEC_TXNRD1 | DYNLRB1 |
| 3.8062E-66 | 0.31394079 | 0.341 | 0.143 | 8.1262E-62 | TEC_TXNRD1 | MTHFD2 |
| 2.0405E-43 | 0.31362732 | 0.864 | 0.71 | 4.3564E-39 | TEC_TXNRD1 | CTSD |
| 1.8011E-63 | 0.31347595 | 0.419 | 0.201 | 3.8454E-59 | TEC_TXNRD1 | C8orf59 |
| 8.5539E-61 | 0.31324534 | 0.232 | 0.081 | 1.8263E-56 | TEC_TXNRD1 | AKR1C3 |
| 3.6394E-61 | 0.31185936 | 0.411 | 0.198 | 7.7702E-57 | TEC_TXNRD1 | MED10 |
| 1.1974E-48 | 0.30786137 | 0.496 | 0.287 | 2.5565E-44 | TEC_TXNRD1 | EIF3F |
| 1.2692E-67 | 0.30589976 | 0.349 | 0.148 | 2.7098E-63 | TEC_TXNRD1 | HTATIP2 |
| 3.9921E-51 | 0.30538106 | 0.489 | 0.276 | 8.5231E-47 | TEC_TXNRD1 | SSBP1 |
| 1.6433E-48 | 0.30390979 | 0.412 | 0.218 | 3.5084E-44 | TEC_TXNRD1 | CHMP2B |
| 4.9363E-44 | 0.30275138 | 0.598 | 0.379 | 1.0539E-39 | TEC_TXNRD1 | MGST3 |

| | | | | | | |
|---|---|---|---|---|---|---|
| 1.7413E-59 | 0.30268235 | 0.405 | 0.193 | 3.7178E-55 | TEC_TXNRD1 | GFPT1 |
| 9.2285E-47 | 0.30133129 | 0.55 | 0.328 | 1.9703E-42 | TEC_TXNRD1 | PSMD8 |
| 3.8518E-39 | 0.30044413 | 0.885 | 0.747 | 8.2236E-35 | TEC_TXNRD1 | CALM2 |
| 2.0067E-42 | 0.30028564 | 0.453 | 0.264 | 4.2843E-38 | TEC_TXNRD1 | BUD31 |
| 2.0119E-65 | 0.29987948 | 0.356 | 0.155 | 4.2954E-61 | TEC_TXNRD1 | IGBP1 |
| 1.067E-59 | 0.2997824 | 0.458 | 0.233 | 2.278E-55 | TEC_TXNRD1 | PSMA3 |
| 2.9321E-59 | 0.29962907 | 0.414 | 0.202 | 6.2601E-55 | TEC_TXNRD1 | RSL1D1 |
| 2.7604E-40 | 0.29892331 | 0.505 | 0.31 | 5.8935E-36 | TEC_TXNRD1 | TMED9 |
| 7.7129E-49 | 0.29833132 | 0.522 | 0.3 | 1.6467E-44 | TEC_TXNRD1 | DEK |
| 1.1189E-50 | 0.29751364 | 0.62 | 0.382 | 2.3889E-46 | TEC_TXNRD1 | LAMTOR5 |
| 9.4314E-46 | 0.29727397 | 0.991 | 0.953 | 2.0136E-41 | TEC_TXNRD1 | RPS2 |
| 4.3389E-46 | 0.29679948 | 0.721 | 0.502 | 9.2635E-42 | TEC_TXNRD1 | ARF1 |
| 7.5008E-55 | 0.29644236 | 0.411 | 0.209 | 1.6014E-50 | TEC_TXNRD1 | EIF3L |
| 3.3055E-40 | 0.29626894 | 0.764 | 0.564 | 7.0573E-36 | TEC_TXNRD1 | ATP6V1G1 |
| 2.2474E-57 | 0.29612185 | 0.427 | 0.216 | 4.7981E-53 | TEC_TXNRD1 | PSMB2 |
| 8.8749E-59 | 0.29454839 | 0.399 | 0.194 | 1.8948E-54 | TEC_TXNRD1 | PSMC1 |
| 5.1442E-46 | 0.29397772 | 0.537 | 0.32 | 1.0983E-41 | TEC_TXNRD1 | SEM1 |
| 1.0863E-61 | 0.29396003 | 0.228 | 0.079 | 2.3193E-57 | TEC_TXNRD1 | MAP2 |
| 2.7062E-51 | 0.29372086 | 0.495 | 0.274 | 5.7778E-47 | TEC_TXNRD1 | NOL7 |
| 7.3082E-37 | 0.29331655 | 0.781 | 0.616 | 1.5603E-32 | TEC_TXNRD1 | PABPC1 |
| 3.9309E-48 | 0.29323037 | 0.838 | 0.631 | 8.3925E-44 | TEC_TXNRD1 | UQCRH |
| 2.2047E-58 | 0.29280242 | 0.437 | 0.218 | 4.707E-54 | TEC_TXNRD1 | SMIM26 |
| 1.5415E-27 | 0.2923313 | 0.57 | 0.403 | 3.291E-23 | TEC_TXNRD1 | CCND1 |
| 3.1469E-33 | 0.29090692 | 0.904 | 0.795 | 6.7187E-29 | TEC_TXNRD1 | EEF1D |
| 1.1599E-55 | 0.29015565 | 0.412 | 0.203 | 2.4763E-51 | TEC_TXNRD1 | GARS |
| 3.7342E-21 | 0.2900332 | 0.606 | 0.475 | 7.9725E-17 | TEC_TXNRD1 | LDHA |
| 1.1728E-49 | 0.28975338 | 0.608 | 0.367 | 2.504E-45 | TEC_TXNRD1 | SUB1 |
| 1.1753E-44 | 0.2895686 | 0.563 | 0.342 | 2.5092E-40 | TEC_TXNRD1 | EIF3A |
| 9.3282E-49 | 0.28903158 | 0.541 | 0.316 | 1.9916E-44 | TEC_TXNRD1 | KDELR1 |
| 2.5332E-63 | 0.28890145 | 0.47 | 0.234 | 5.4084E-59 | TEC_TXNRD1 | EIF3M |
| 2.5437E-42 | 0.28844091 | 0.491 | 0.294 | 5.4308E-38 | TEC_TXNRD1 | TMEM50A |
| 3.3617E-33 | 0.2881721 | 0.717 | 0.542 | 7.1773E-29 | TEC_TXNRD1 | LRRFIP1 |
| 1.7585E-47 | 0.28779142 | 0.438 | 0.24 | 3.7544E-43 | TEC_TXNRD1 | MTPN |
| 5.9096E-34 | 0.28777616 | 0.418 | 0.249 | 1.2617E-29 | TEC_TXNRD1 | SELENOM |
| 4.7577E-54 | 0.28737788 | 0.372 | 0.18 | 1.0158E-49 | TEC_TXNRD1 | ZCRB1 |
| 5.6463E-47 | 0.2870017 | 0.505 | 0.292 | 1.2055E-42 | TEC_TXNRD1 | GABARAPL2 |
| 2.512E-56 | 0.28696958 | 0.372 | 0.176 | 5.3631E-52 | TEC_TXNRD1 | BNIP3L |
| 8.6459E-31 | 0.28682706 | 0.337 | 0.19 | 1.8459E-26 | TEC_TXNRD1 | MANF |
| 8.0092E-43 | 0.28661227 | 0.876 | 0.722 | 1.71E-38 | TEC_TXNRD1 | SKP1 |
| 7.8802E-60 | 0.2865454 | 0.365 | 0.168 | 1.6824E-55 | TEC_TXNRD1 | UFD1 |
| 2.3442E-55 | 0.28637829 | 0.455 | 0.237 | 5.0049E-51 | TEC_TXNRD1 | CCT4 |
| 8.3634E-47 | 0.28565243 | 0.443 | 0.244 | 1.7856E-42 | TEC_TXNRD1 | JTB |
| 3.8335E-50 | 0.28560804 | 0.549 | 0.317 | 8.1845E-46 | TEC_TXNRD1 | PRELID1 |
| 8.6172E-40 | 0.28520732 | 0.626 | 0.414 | 1.8398E-35 | TEC_TXNRD1 | BZW1 |
| 5.5202E-48 | 0.28486191 | 0.486 | 0.275 | 1.1786E-43 | TEC_TXNRD1 | MRPL51 |
| 7.2964E-42 | 0.28455296 | 0.505 | 0.304 | 1.5578E-37 | TEC_TXNRD1 | ATP5MPL |
| 6.3472E-13 | 0.28387911 | 0.725 | 0.657 | 1.3551E-08 | TEC_TXNRD1 | TACSTD2 |
| 8.3359E-33 | 0.28366676 | 0.274 | 0.141 | 1.7797E-28 | TEC_TXNRD1 | NQO1 |
| 2.1877E-35 | 0.28328054 | 0.73 | 0.543 | 4.6708E-31 | TEC_TXNRD1 | RHOA |
| 3.1825E-44 | 0.28277731 | 0.459 | 0.26 | 6.7947E-40 | TEC_TXNRD1 | SSB |

| | | | | | | |
|---|---|---|---|---|---|---|
| 1.8026E-48 | 0.28264684 | 0.45 | 0.245 | 3.8486E-44 | TEC_TXNRD1 | RPS19BP1 |
| 2.2493E-42 | 0.28248381 | 0.514 | 0.31 | 4.8022E-38 | TEC_TXNRD1 | TRAM1 |
| 6.7618E-39 | 0.28227042 | 0.666 | 0.459 | 1.4437E-34 | TEC_TXNRD1 | MORF4L1 |
| 2.3071E-58 | 0.2821107 | 0.33 | 0.145 | 4.9258E-54 | TEC_TXNRD1 | RNF187 |
| 9.2364E-36 | 0.28131005 | 0.404 | 0.233 | 1.972E-31 | TEC_TXNRD1 | TES |
| 6.2624E-25 | 0.28056216 | 0.979 | 0.923 | 1.337E-20 | TEC_TXNRD1 | RPL36 |
| 2.0598E-37 | 0.28033005 | 0.486 | 0.296 | 4.3977E-33 | TEC_TXNRD1 | NOP53 |
| 6.3014E-45 | 0.2784376 | 0.361 | 0.185 | 1.3453E-40 | TEC_TXNRD1 | TMEM106B |
| 1.1294E-40 | 0.27761988 | 0.558 | 0.349 | 2.4113E-36 | TEC_TXNRD1 | WDR83OS |
| 4.2443E-38 | 0.27703319 | 0.507 | 0.316 | 9.0615E-34 | TEC_TXNRD1 | COPE |
| 3.1392E-51 | 0.27691369 | 0.438 | 0.232 | 6.7022E-47 | TEC_TXNRD1 | SUMO1 |
| 2.4846E-53 | 0.27621092 | 0.425 | 0.217 | 5.3046E-49 | TEC_TXNRD1 | ARPC5L |
| 3.3291E-58 | 0.27615972 | 0.423 | 0.203 | 7.1076E-54 | TEC_TXNRD1 | BEX2 |
| 7.9098E-43 | 0.27547762 | 0.427 | 0.241 | 1.6887E-38 | TEC_TXNRD1 | TMED2 |
| 3.0247E-44 | 0.27496775 | 0.397 | 0.216 | 6.4578E-40 | TEC_TXNRD1 | STRAP |
| 1.6039E-24 | 0.27402524 | 0.952 | 0.924 | 3.4244E-20 | TEC_TXNRD1 | ACTG1 |
| 5.3303E-33 | 0.27376982 | 0.986 | 0.936 | 1.138E-28 | TEC_TXNRD1 | RPL37A |
| 1.553E-49 | 0.27346445 | 0.409 | 0.212 | 3.3157E-45 | TEC_TXNRD1 | TXNL1 |
| 5.3153E-10 | 0.2730545 | 0.342 | 0.253 | 1.1348E-05 | TEC_TXNRD1 | NUPR1 |
| 1.3296E-30 | 0.27213009 | 0.56 | 0.376 | 2.8386E-26 | TEC_TXNRD1 | CIRBP |
| 3.7843E-40 | 0.27115054 | 0.394 | 0.218 | 8.0795E-36 | TEC_TXNRD1 | RNF181 |
| 1.0555E-48 | 0.27048635 | 0.506 | 0.284 | 2.2536E-44 | TEC_TXNRD1 | WASF2 |
| 1.9064E-52 | 0.27021265 | 0.447 | 0.234 | 4.0702E-48 | TEC_TXNRD1 | NDUFAB1 |
| 2.8183E-34 | 0.27004305 | 0.408 | 0.244 | 6.0172E-30 | TEC_TXNRD1 | NDUFA6 |
| 3.3123E-48 | 0.26959042 | 0.482 | 0.267 | 7.0718E-44 | TEC_TXNRD1 | ANP32B |
| 6.4817E-42 | 0.26955365 | 0.427 | 0.238 | 1.3838E-37 | TEC_TXNRD1 | SARS |
| 5.7509E-50 | 0.26951704 | 0.441 | 0.234 | 1.2278E-45 | TEC_TXNRD1 | UQCRFS1 |
| 1.9944E-37 | 0.26932136 | 0.52 | 0.322 | 4.2581E-33 | TEC_TXNRD1 | ERP29 |
| 1.5233E-42 | 0.26813699 | 0.458 | 0.259 | 3.2523E-38 | TEC_TXNRD1 | TCEA1 |
| 2.1308E-42 | 0.26788451 | 0.478 | 0.28 | 4.5492E-38 | TEC_TXNRD1 | UBXN1 |
| 4.5762E-27 | 0.26758701 | 0.937 | 0.847 | 9.7702E-23 | TEC_TXNRD1 | RPL38 |
| 4.0605E-51 | 0.26727785 | 0.357 | 0.172 | 8.6692E-47 | TEC_TXNRD1 | CAMLG |
| 1.8546E-44 | 0.26694488 | 0.5 | 0.292 | 3.9597E-40 | TEC_TXNRD1 | NDUFC2 |
| 2.3964E-36 | 0.26623602 | 0.467 | 0.286 | 5.1164E-32 | TEC_TXNRD1 | CIB1 |
| 1.2389E-40 | 0.26472541 | 0.409 | 0.228 | 2.6451E-36 | TEC_TXNRD1 | LSM3 |
| 8.5319E-42 | 0.26384597 | 0.53 | 0.321 | 1.8216E-37 | TEC_TXNRD1 | SEC11A |
| 6.194E-29 | 0.26379701 | 0.928 | 0.82 | 1.3224E-24 | TEC_TXNRD1 | SQSTM1 |
| 5.6891E-46 | 0.26326419 | 0.409 | 0.22 | 1.2146E-41 | TEC_TXNRD1 | TMEM14B |
| 1.0343E-60 | 0.26308992 | 0.326 | 0.139 | 2.2082E-56 | TEC_TXNRD1 | RAMMET |
| 8.5769E-46 | 0.26288503 | 0.294 | 0.136 | 1.8312E-41 | TEC_TXNRD1 | PHGDH |
| 4.5574E-49 | 0.26255217 | 0.325 | 0.152 | 9.73E-45 | TEC_TXNRD1 | UBE2J1 |
| 9.0941E-57 | 0.26201813 | 0.312 | 0.134 | 1.9416E-52 | TEC_TXNRD1 | RHOD |
| 6.7621E-56 | 0.26073525 | 0.332 | 0.148 | 1.4437E-51 | TEC_TXNRD1 | EMC3 |
| 1.777E-50 | 0.26014761 | 0.225 | 0.086 | 3.794E-46 | TEC_TXNRD1 | OSGIN1 |
| 6.143E-46 | 0.25925185 | 0.269 | 0.118 | 1.3115E-41 | TEC_TXNRD1 | PLIN2 |
| 6.9542E-33 | 0.25910599 | 0.888 | 0.739 | 1.4847E-28 | TEC_TXNRD1 | PFDN5 |
| 9.2381E-50 | 0.25811175 | 0.354 | 0.172 | 1.9723E-45 | TEC_TXNRD1 | PCMT1 |
| 2.3999E-56 | 0.25779603 | 0.328 | 0.145 | 5.1239E-52 | TEC_TXNRD1 | COA6 |
| 1.6882E-37 | 0.25735847 | 0.395 | 0.225 | 3.6044E-33 | TEC_TXNRD1 | KDELR2 |
| 9.4499E-35 | 0.25698444 | 0.406 | 0.238 | 2.0176E-30 | TEC_TXNRD1 | C4orf3 |

| | | | | | | |
|---|---|---|---|---|---|---|
| 2.2566E-63 | 0.25574801 | 0.322 | 0.127 | 4.8179E-59 | TEC_TXNRD1 | XIST |
| 5.6644E-45 | 0.25567667 | 0.389 | 0.204 | 1.2094E-40 | TEC_TXNRD1 | LIMS1 |
| 8.1076E-22 | 0.25558172 | 0.83 | 0.717 | 1.731E-17 | TEC_TXNRD1 | CSTB |
| 7.3503E-63 | 0.25430537 | 0.328 | 0.136 | 1.5693E-58 | TEC_TXNRD1 | RAB32 |
| 1.4742E-40 | 0.25416588 | 0.531 | 0.325 | 3.1474E-36 | TEC_TXNRD1 | TRIR |
| 9.6503E-36 | 0.25354314 | 0.486 | 0.301 | 2.0603E-31 | TEC_TXNRD1 | SARAF |
| 5.0427E-43 | 0.25341552 | 0.425 | 0.234 | 1.0766E-38 | TEC_TXNRD1 | RAB1A |
| 8.1869E-22 | 0.25314716 | 0.694 | 0.554 | 1.7479E-17 | TEC_TXNRD1 | PPIB |
| 3.3538E-42 | 0.25274572 | 0.361 | 0.187 | 7.1603E-38 | TEC_TXNRD1 | PSMD12 |
| 1.0192E-47 | 0.25249355 | 0.345 | 0.169 | 2.1759E-43 | TEC_TXNRD1 | TTC1 |
| 8.2755E-33 | 0.25242853 | 0.305 | 0.164 | 1.7668E-28 | TEC_TXNRD1 | UBE2D1 |
| 2.0032E-36 | 0.25182462 | 0.529 | 0.333 | 4.2769E-32 | TEC_TXNRD1 | VAPA |
| 5.63E-41 | 0.25159309 | 0.377 | 0.205 | 1.202E-36 | TEC_TXNRD1 | COPZ1 |
| 4.3278E-33 | 0.25118021 | 0.553 | 0.357 | 9.2398E-29 | TEC_TXNRD1 | HEBP2 |
| 8.3959E-49 | 0.25110186 | 0.359 | 0.177 | 1.7925E-44 | TEC_TXNRD1 | FUNDC2 |
| 3.1419E-22 | 0.25052156 | 0.993 | 0.962 | 6.708E-18 | TEC_TXNRD1 | RPS19 |
| 5.5018E-39 | 0.25038776 | 0.623 | 0.407 | 1.1746E-34 | TEC_TXNRD1 | TMEM59 |
| 8.1634E-38 | 0.25030149 | 0.401 | 0.228 | 1.7429E-33 | TEC_TXNRD1 | PLEKHB2 |
| 1.5059E-15 | 0.25027968 | 0.38 | 0.273 | 3.2151E-11 | TEC_TXNRD1 | ASPH |
| 2.6577E-41 | 0.25009039 | 0.418 | 0.234 | 5.6741E-37 | TEC_TXNRD1 | UBE2L3 |
| 4.0948E-35 | 0.2500875 | 0.313 | 0.164 | 8.7425E-31 | TEC_TXNRD1 | DNAJB9 |
| 9.8728E-10 | -0.2521357 | 0.166 | 0.241 | 2.1078E-05 | TEC_TXNRD1 | FXYD2 |
| 4.7348E-05 | -0.2527169 | 0.303 | 0.336 | 1 | TEC_TXNRD1 | CPEB4 |
| 5.0525E-19 | -0.2529419 | 0.062 | 0.16 | 1.0787E-14 | TEC_TXNRD1 | DAPK1 |
| 0.00740392 | -0.2531369 | 0.666 | 0.669 | 1 | TEC_TXNRD1 | TPI1 |
| 3.3595E-06 | -0.2556783 | 0.304 | 0.348 | 0.07172612 | TEC_TXNRD1 | TSC22D1 |
| 8.4485E-25 | -0.2589787 | 0.025 | 0.13 | 1.8037E-20 | TEC_TXNRD1 | LINC00472 |
| 1.791E-21 | -0.2595944 | 0.058 | 0.164 | 3.8238E-17 | TEC_TXNRD1 | CHD3 |
| 9.6894E-23 | -0.260225 | 0.021 | 0.116 | 2.0687E-18 | TEC_TXNRD1 | RGS2 |
| 5.4549E-08 | -0.2604777 | 0.236 | 0.307 | 0.00116462 | TEC_TXNRD1 | DDIT4 |
| 1.8923E-05 | -0.2632812 | 0.3 | 0.354 | 0.40400216 | TEC_TXNRD1 | TXNIP |
| 2.0043E-09 | -0.2639583 | 0.212 | 0.285 | 4.2793E-05 | TEC_TXNRD1 | COBLL1 |
| 2.1748E-20 | -0.2641378 | 0.025 | 0.114 | 4.6432E-16 | TEC_TXNRD1 | AC012306.2 |
| 7.9413E-10 | -0.267242 | 0.173 | 0.241 | 1.6955E-05 | TEC_TXNRD1 | CHD9 |
| 1.5547E-24 | -0.2683759 | 0.031 | 0.137 | 3.3193E-20 | TEC_TXNRD1 | PKHD1 |
| 1.4267E-14 | -0.2687559 | 0.068 | 0.15 | 3.0461E-10 | TEC_TXNRD1 | ID3 |
| 2.189E-23 | -0.268788 | 0.01 | 0.101 | 4.6736E-19 | TEC_TXNRD1 | AL137077.2 |
| 4.6624E-12 | -0.2694705 | 0.09 | 0.167 | 9.9542E-08 | TEC_TXNRD1 | TNFAIP3 |
| 0.0008673 | -0.2708534 | 0.57 | 0.629 | 1 | TEC_TXNRD1 | VIM |
| 7.2294E-11 | -0.27292 | 0.251 | 0.332 | 1.5435E-06 | TEC_TXNRD1 | HNRNPH1 |
| 1.1672E-17 | -0.2766637 | 0.055 | 0.146 | 2.492E-13 | TEC_TXNRD1 | TNFSF10 |
| 2.7621E-21 | -0.2770368 | 0.018 | 0.106 | 5.897E-17 | TEC_TXNRD1 | MAFA |
| 2.2909E-08 | -0.2816144 | 0.264 | 0.319 | 0.00048911 | TEC_TXNRD1 | AFDN |
| 2.2913E-13 | -0.2844639 | 0.192 | 0.282 | 4.892E-09 | TEC_TXNRD1 | SPTAN1 |
| 2.7181E-13 | -0.2844934 | 0.267 | 0.357 | 5.8031E-09 | TEC_TXNRD1 | ACADVL |
| 2.3808E-15 | -0.2859852 | 0.123 | 0.215 | 5.083E-11 | TEC_TXNRD1 | VPS13C |
| 1.115E-18 | -0.2875943 | 0.068 | 0.165 | 2.3805E-14 | TEC_TXNRD1 | PLEKHH2 |
| 2.9455E-16 | -0.2879936 | 0.072 | 0.163 | 6.2887E-12 | TEC_TXNRD1 | IGFBP2 |
| 3.3293E-16 | -0.2886215 | 0.514 | 0.38 | 7.108E-12 | TEC_TXNRD1 | SLPI |
| 1.3475E-07 | -0.2931938 | 0.22 | 0.282 | 0.00287684 | TEC_TXNRD1 | CYB5A |

| | | | | | | |
|---|---|---|---|---|---|---|
| 1.8811E-22 | -0.2936875 | 0.024 | 0.119 | 4.0161E-18 | TEC_TXNRD1 | EGOT |
| 0.00556562 | -0.2938636 | 0.969 | 0.953 | 1 | TEC_TXNRD1 | S100A6 |
| 2.3053E-21 | -0.2974688 | 0.037 | 0.134 | 4.9217E-17 | TEC_TXNRD1 | AL135925.1 |
| 3.9919E-13 | -0.2978922 | 0.194 | 0.28 | 8.5228E-09 | TEC_TXNRD1 | TCF25 |
| 8.3699E-07 | -0.2980961 | 0.324 | 0.365 | 0.01786968 | TEC_TXNRD1 | AKAP9 |
| 1.165E-11 | -0.2987675 | 0.219 | 0.295 | 2.4874E-07 | TEC_TXNRD1 | HP1BP3 |
| 1.4967E-09 | -0.299135 | 0.111 | 0.18 | 3.1954E-05 | TEC_TXNRD1 | IGFBP3 |
| 4.2988E-05 | -0.2993441 | 0.368 | 0.384 | 0.91778805 | TEC_TXNRD1 | N4BP2L2 |
| 6.1228E-10 | -0.3026744 | 0.252 | 0.326 | 1.3072E-05 | TEC_TXNRD1 | DCDC2 |
| 5.9916E-12 | -0.3055867 | 0.245 | 0.322 | 1.2792E-07 | TEC_TXNRD1 | STAT3 |
| 2.6474E-09 | -0.3058618 | 0.279 | 0.342 | 5.6522E-05 | TEC_TXNRD1 | LUC7L3 |
| 1.4038E-15 | -0.3100747 | 0.034 | 0.108 | 2.997E-11 | TEC_TXNRD1 | TNF |
| 1.2049E-18 | -0.3126557 | 0.068 | 0.164 | 2.5725E-14 | TEC_TXNRD1 | CDKN2B |
| 1.959E-08 | -0.3136719 | 0.111 | 0.171 | 0.00041824 | TEC_TXNRD1 | CTGF |
| 2.3217E-11 | -0.3141289 | 0.217 | 0.292 | 4.9569E-07 | TEC_TXNRD1 | NUFIP2 |
| 1.5196E-05 | -0.3156851 | 0.23 | 0.268 | 0.32443295 | TEC_TXNRD1 | AMD1 |
| 9.1326E-25 | -0.3172095 | 0.016 | 0.116 | 1.9498E-20 | TEC_TXNRD1 | AL355075.4 |
| 3.5885E-06 | -0.3172388 | 0.141 | 0.19 | 0.07661403 | TEC_TXNRD1 | BIRC3 |
| 3.7061E-22 | -0.3181275 | 0.041 | 0.142 | 7.9124E-18 | TEC_TXNRD1 | CSRNP2 |
| 1.2706E-14 | -0.3188966 | 0.102 | 0.189 | 2.7128E-10 | TEC_TXNRD1 | ERN1 |
| 0.00061028 | -0.3205078 | 0.304 | 0.322 | 1 | TEC_TXNRD1 | CCNL1 |
| 4.211E-12 | -0.3245172 | 0.227 | 0.309 | 8.9904E-08 | TEC_TXNRD1 | C6orf62 |
| 1.216E-09 | -0.3251215 | 0.242 | 0.31 | 2.5962E-05 | TEC_TXNRD1 | ARHGAP29 |
| 1.4231E-08 | -0.3264515 | 0.058 | 0.111 | 0.00030382 | TEC_TXNRD1 | AQP3 |
| 4.3492E-07 | -0.3270584 | 0.333 | 0.369 | 0.0092856 | TEC_TXNRD1 | PNN |
| 0.0005294 | -0.3290025 | 0.884 | 0.87 | 1 | TEC_TXNRD1 | SAT1 |
| 8.7695E-29 | -0.329138 | 0.011 | 0.121 | 1.8723E-24 | TEC_TXNRD1 | HIST4H4 |
| 4.1528E-10 | -0.3303868 | 0.143 | 0.215 | 8.8663E-06 | TEC_TXNRD1 | ADAMTS9 |
| 8.5299E-25 | -0.3314609 | 0.102 | 0.229 | 1.8211E-20 | TEC_TXNRD1 | RERE |
| 5.9265E-19 | -0.331575 | 0.085 | 0.186 | 1.2653E-14 | TEC_TXNRD1 | KLF4 |
| 2.5549E-16 | -0.3318139 | 0.101 | 0.197 | 5.4548E-12 | TEC_TXNRD1 | SYTL2 |
| 3.2281E-13 | -0.3345649 | 0.038 | 0.106 | 6.8921E-09 | TEC_TXNRD1 | CXCL3 |
| 2.5831E-05 | -0.3391223 | 0.516 | 0.518 | 0.55148645 | TEC_TXNRD1 | SON |
| 1.9399E-19 | -0.33937 | 0.031 | 0.118 | 4.1417E-15 | TEC_TXNRD1 | AC023157.3 |
| 4.5549E-26 | -0.3395512 | 0.027 | 0.137 | 9.7246E-22 | TEC_TXNRD1 | HOMER1 |
| 3.6893E-10 | -0.341411 | 0.195 | 0.268 | 7.8766E-06 | TEC_TXNRD1 | HIST1H2AC |
| 1.3159E-20 | -0.3418974 | 0.059 | 0.161 | 2.8094E-16 | TEC_TXNRD1 | PFKFB3 |
| 6.2278E-16 | -0.3425853 | 0.203 | 0.299 | 1.3296E-11 | TEC_TXNRD1 | KMT2A |
| 4.3383E-20 | -0.3425981 | 0.091 | 0.199 | 9.2624E-16 | TEC_TXNRD1 | THBS1 |
| 1.4608E-10 | -0.3448443 | 0.042 | 0.1 | 3.1188E-06 | TEC_TXNRD1 | CCL2 |
| 8.1648E-06 | -0.345054 | 0.594 | 0.615 | 0.17431745 | TEC_TXNRD1 | MT-ND4L |
| 4.1376E-15 | -0.3466901 | 0.226 | 0.32 | 8.8337E-11 | TEC_TXNRD1 | WSB1 |
| 1.5475E-20 | -0.3493326 | 0.07 | 0.174 | 3.3039E-16 | TEC_TXNRD1 | HIST1H1E |
| 4.9067E-23 | -0.3506561 | 0.034 | 0.135 | 1.0476E-18 | TEC_TXNRD1 | PCDH9 |
| 1.1247E-11 | -0.3508466 | 0.124 | 0.198 | 2.4012E-07 | TEC_TXNRD1 | H2AFX |
| 1.6257E-28 | -0.3509136 | 0.076 | 0.211 | 3.4708E-24 | TEC_TXNRD1 | FAT1 |
| 1.1022E-15 | -0.35165 | 0.049 | 0.13 | 2.3533E-11 | TEC_TXNRD1 | CXCL14 |
| 7.4642E-17 | -0.357184 | 0.033 | 0.111 | 1.5936E-12 | TEC_TXNRD1 | ARC |
| 4.3752E-23 | -0.3590997 | 0.244 | 0.38 | 9.341E-19 | TEC_TXNRD1 | SELENOW |
| 8.136E-16 | -0.3593813 | 0.408 | 0.493 | 1.737E-11 | TEC_TXNRD1 | ATP1A1 |

| | | | | | | |
|---|---|---|---|---|---|---|
| 1.396E-21 | -0.3619058 | 0.075 | 0.185 | 2.9806E-17 | TEC_TXNRD1 | FOXC1 |
| 5.8128E-17 | -0.362117 | 0.191 | 0.291 | 1.241E-12 | TEC_TXNRD1 | PRPF4B |
| 1.4435E-08 | -0.3624312 | 0.238 | 0.301 | 0.00030818 | TEC_TXNRD1 | DST |
| 1.1397E-10 | -0.3634551 | 0.168 | 0.243 | 2.4333E-06 | TEC_TXNRD1 | SLC5A3 |
| 0.00026091 | -0.36369 | 0.918 | 0.927 | 1 | TEC_TXNRD1 | MT-CO1 |
| 2.4896E-09 | -0.3650434 | 0.322 | 0.372 | 5.3154E-05 | TEC_TXNRD1 | MACF1 |
| 3.6808E-20 | -0.3650721 | 0.11 | 0.218 | 7.8585E-16 | TEC_TXNRD1 | MAT2A |
| 6.2689E-09 | -0.368271 | 0.261 | 0.317 | 0.00013384 | TEC_TXNRD1 | SRSF2 |
| 8.5562E-13 | -0.3685973 | 0.2 | 0.283 | 1.8268E-08 | TEC_TXNRD1 | VEGFA |
| 1.6616E-17 | -0.3687366 | 0.048 | 0.137 | 3.5475E-13 | TEC_TXNRD1 | TNFRSF11B |
| 1.7275E-05 | -0.3697842 | 0.53 | 0.597 | 0.36882418 | TEC_TXNRD1 | KRT19 |
| 0.00673689 | -0.3744862 | 0.351 | 0.359 | 1 | TEC_TXNRD1 | KLF6 |
| 3.4916E-24 | -0.374715 | 0.161 | 0.296 | 7.4545E-20 | TEC_TXNRD1 | COL18A1 |
| 2.1245E-28 | -0.3770156 | 0.072 | 0.206 | 4.5358E-24 | TEC_TXNRD1 | BCAM |
| 1.2306E-21 | -0.3787004 | 0.05 | 0.153 | 2.6272E-17 | TEC_TXNRD1 | ID1 |
| 6.3555E-23 | -0.3798657 | 0.015 | 0.108 | 1.3569E-18 | TEC_TXNRD1 | AP001160.1 |
| 1.4221E-17 | -0.3799426 | 0.141 | 0.242 | 3.0363E-13 | TEC_TXNRD1 | CDKN2AIP |
| 1.2474E-13 | -0.3831694 | 0.215 | 0.308 | 2.6633E-09 | TEC_TXNRD1 | MARCKSL1 |
| 3.9278E-14 | -0.3858886 | 0.034 | 0.104 | 8.3858E-10 | TEC_TXNRD1 | SLC12A1 |
| 4.9582E-18 | -0.3868485 | 0.156 | 0.262 | 1.0586E-13 | TEC_TXNRD1 | IRF1 |
| 6.0471E-24 | -0.3873909 | 0.101 | 0.225 | 1.291E-19 | TEC_TXNRD1 | ZFP36 |
| 6.7686E-31 | -0.3913212 | 0.016 | 0.136 | 1.4451E-26 | TEC_TXNRD1 | HIST2H2BF |
| 5.6115E-11 | -0.396928 | 0.125 | 0.198 | 1.198E-06 | TEC_TXNRD1 | GADD45G |
| 1.0458E-05 | -0.4010204 | 0.973 | 0.937 | 0.22327846 | TEC_TXNRD1 | MALAT1 |
| 6.45E-10 | -0.401562 | 0.224 | 0.295 | 1.3771E-05 | TEC_TXNRD1 | CYR61 |
| 8.6847E-06 | -0.4021026 | 0.365 | 0.388 | 0.18541811 | TEC_TXNRD1 | BRD2 |
| 7.8262E-28 | -0.4090779 | 0.029 | 0.145 | 1.6709E-23 | TEC_TXNRD1 | AC025164.1 |
| 8.6393E-18 | -0.4112293 | 0.144 | 0.248 | 1.8445E-13 | TEC_TXNRD1 | ITGB6 |
| 0.00420478 | -0.4124105 | 0.092 | 0.119 | 1 | TEC_TXNRD1 | CXCL8 |
| 3.9319E-16 | -0.4149177 | 0.252 | 0.345 | 8.3945E-12 | TEC_TXNRD1 | HNRNPAB |
| 5.2569E-16 | -0.4291048 | 0.463 | 0.524 | 1.1223E-11 | TEC_TXNRD1 | HNRNPU |
| 6.163E-06 | -0.4309532 | 0.889 | 0.876 | 0.13158077 | TEC_TXNRD1 | ITM2B |
| 1.4451E-12 | -0.4336017 | 0.163 | 0.246 | 3.0852E-08 | TEC_TXNRD1 | MDM2 |
| 2.1456E-35 | -0.4411752 | 0.024 | 0.162 | 4.5809E-31 | TEC_TXNRD1 | AC007952.4 |
| 0.00310875 | -0.4414327 | 0.221 | 0.249 | 1 | TEC_TXNRD1 | MT1X |
| 8.9657E-10 | -0.4484937 | 0.254 | 0.318 | 1.9142E-05 | TEC_TXNRD1 | CDKN1C |
| 1.6615E-16 | -0.4505204 | 0.277 | 0.366 | 3.5474E-12 | TEC_TXNRD1 | MCL1 |
| 9.1936E-17 | -0.4535571 | 0.294 | 0.385 | 1.9628E-12 | TEC_TXNRD1 | RSRP1 |
| 6.7966E-16 | -0.4576055 | 0.28 | 0.365 | 1.4511E-11 | TEC_TXNRD1 | ARGLU1 |
| 2.0959E-13 | -0.4598185 | 0.226 | 0.312 | 4.4747E-09 | TEC_TXNRD1 | PMAIP1 |
| 0.00439301 | -0.4612062 | 0.1 | 0.074 | 1 | TEC_TXNRD1 | UMOD |
| 6.3273E-17 | -0.4698117 | 0.364 | 0.44 | 1.3509E-12 | TEC_TXNRD1 | DDX17 |
| 2.3374E-27 | -0.4768684 | 0.058 | 0.182 | 4.9904E-23 | TEC_TXNRD1 | FOSB |
| 5.8379E-36 | -0.4837515 | 0.053 | 0.203 | 1.2464E-31 | TEC_TXNRD1 | TLE4 |
| 3.8663E-19 | -0.4850048 | 0.163 | 0.273 | 8.2545E-15 | TEC_TXNRD1 | DUSP1 |
| 1.8212E-23 | -0.488614 | 0.221 | 0.349 | 3.8883E-19 | TEC_TXNRD1 | PAX8 |
| 1.636E-10 | -0.4906186 | 0.061 | 0.122 | 3.4928E-06 | TEC_TXNRD1 | NEFM |
| 3.3256E-18 | -0.5019889 | 0.172 | 0.275 | 7.1003E-14 | TEC_TXNRD1 | KCNQ1OT1 |
| 1.6567E-25 | -0.5035004 | 0.03 | 0.137 | 3.537E-21 | TEC_TXNRD1 | COL7A1 |
| 2.1058E-15 | -0.5062199 | 0.22 | 0.31 | 4.496E-11 | TEC_TXNRD1 | DNAJB1 |

| | | | | | | |
|---|---|---|---|---|---|---|
| 5.0665E-23 | -0.5074259 | 0.109 | 0.228 | 1.0817E-18 | TEC_TXNRD1 | TNFAIP2 |
| 1.0946E-14 | -0.509278 | 0.209 | 0.291 | 2.3369E-10 | TEC_TXNRD1 | POLR2A |
| 2.2367E-26 | -0.5167136 | 0.112 | 0.244 | 4.7753E-22 | TEC_TXNRD1 | SYNE1 |
| 1.4689E-31 | -0.525718 | 0.101 | 0.246 | 3.1361E-27 | TEC_TXNRD1 | NFKBIZ |
| 4.7745E-16 | -0.5331568 | 0.723 | 0.823 | 1.0194E-11 | TEC_TXNRD1 | IGFBP7 |
| 4.2987E-27 | -0.5340121 | 0.126 | 0.264 | 9.1777E-23 | TEC_TXNRD1 | ID2 |
| 5.4457E-43 | -0.5451722 | 0.505 | 0.683 | 1.1626E-38 | TEC_TXNRD1 | GLS |
| 1.2612E-30 | -0.5461492 | 0.048 | 0.18 | 2.6926E-26 | TEC_TXNRD1 | EGLN3 |
| 0.00195615 | -0.5539877 | 0.448 | 0.472 | 1 | TEC_TXNRD1 | C12orf75 |
| 3.3767E-17 | -0.5576419 | 0.771 | 0.834 | 7.2093E-13 | TEC_TXNRD1 | MT-ND5 |
| 5.1051E-26 | -0.5675403 | 0.155 | 0.287 | 1.0899E-21 | TEC_TXNRD1 | CITED2 |
| 4.2732E-23 | -0.5762729 | 0.186 | 0.317 | 9.1232E-19 | TEC_TXNRD1 | HSPA1B |
| 4.6124E-24 | -0.5841129 | 0.131 | 0.253 | 9.8475E-20 | TEC_TXNRD1 | ID4 |
| 1.9019E-28 | -0.5913045 | 0.155 | 0.298 | 4.0605E-24 | TEC_TXNRD1 | CEBPD |
| 7.7731E-30 | -0.59477 | 0.474 | 0.57 | 1.6595E-25 | TEC_TXNRD1 | SRRM2 |
| 1.8153E-23 | -0.601337 | 0.173 | 0.299 | 3.8756E-19 | TEC_TXNRD1 | RND3 |
| 1.7563E-24 | -0.6021805 | 0.141 | 0.264 | 3.7497E-20 | TEC_TXNRD1 | IER5 |
| 1.8202E-37 | -0.6141961 | 0.069 | 0.228 | 3.8862E-33 | TEC_TXNRD1 | SOCS1 |
| 3.5328E-40 | -0.6315814 | 0.039 | 0.198 | 7.5426E-36 | TEC_TXNRD1 | PIK3R3 |
| 6.8202E-11 | -0.6336362 | 0.05 | 0.111 | 1.4561E-06 | TEC_TXNRD1 | FXYD4 |
| 4.1559E-37 | -0.6337878 | 0.483 | 0.644 | 8.8729E-33 | TEC_TXNRD1 | TPM1 |
| 7.2311E-35 | -0.6434506 | 0.295 | 0.458 | 1.5438E-30 | TEC_TXNRD1 | ZFP36L1 |
| 1.3453E-15 | -0.6466155 | 0.226 | 0.308 | 2.8723E-11 | TEC_TXNRD1 | PLK2 |
| 1.6038E-52 | -0.6657446 | 0.028 | 0.219 | 3.4241E-48 | TEC_TXNRD1 | SGK1 |
| 6.1884E-21 | -0.670333 | 0.871 | 0.857 | 1.3212E-16 | TEC_TXNRD1 | H3F3B |
| 1.187E-61 | -0.6770768 | 0.67 | 0.808 | 2.5342E-57 | TEC_TXNRD1 | HNRNPA2B1 |
| 1.863E-17 | -0.678768 | 0.313 | 0.415 | 3.9776E-13 | TEC_TXNRD1 | RNF213 |
| 1.8517E-08 | -0.6808059 | 0.182 | 0.248 | 0.00039534 | TEC_TXNRD1 | TFPI2 |
| 1.8284E-39 | -0.6923171 | 0.1 | 0.269 | 3.9035E-35 | TEC_TXNRD1 | IER5L |
| 6.8019E-68 | -0.7128621 | 0.493 | 0.735 | 1.4522E-63 | TEC_TXNRD1 | HSPB1 |
| 0.00402752 | -0.7145014 | 0.241 | 0.193 | 1 | TEC_TXNRD1 | MT1G |
| 5.865E-33 | -0.7146658 | 0.116 | 0.273 | 1.2522E-28 | TEC_TXNRD1 | EGR1 |
| 4.6794E-09 | -0.7238283 | 0.842 | 0.812 | 9.9906E-05 | TEC_TXNRD1 | NEAT1 |
| 1.0351E-48 | -0.7321384 | 0.063 | 0.254 | 2.2098E-44 | TEC_TXNRD1 | INTS6 |
| 5.5977E-47 | -0.7429476 | 0.161 | 0.36 | 1.1951E-42 | TEC_TXNRD1 | ZFP36L2 |
| 1.2286E-43 | -0.7509925 | 0.052 | 0.226 | 2.623E-39 | TEC_TXNRD1 | ADAMTS1 |
| 1.1693E-49 | -0.7590271 | 0.099 | 0.298 | 2.4965E-45 | TEC_TXNRD1 | BTG2 |
| 1.5922E-33 | -0.7632108 | 0.174 | 0.324 | 3.3993E-29 | TEC_TXNRD1 | GLUL |
| 2.0587E-13 | -0.76975 | 0.59 | 0.607 | 4.3953E-09 | TEC_TXNRD1 | GADD45B |
| 4.4207E-37 | -0.7724849 | 0.27 | 0.429 | 9.4381E-33 | TEC_TXNRD1 | SRSF7 |
| 4.6265E-31 | -0.7879076 | 0.251 | 0.386 | 9.8777E-27 | TEC_TXNRD1 | SLC38A2 |
| 1.5315E-37 | -0.8103118 | 0.234 | 0.401 | 3.2698E-33 | TEC_TXNRD1 | JUNB |
| 5.2065E-34 | -0.8216506 | 0.245 | 0.405 | 1.1116E-29 | TEC_TXNRD1 | NFKBIA |
| 2.3105E-46 | -0.8353697 | 0.261 | 0.444 | 4.9328E-42 | TEC_TXNRD1 | SYNE2 |
| 7.6975E-39 | -0.8579066 | 0.542 | 0.648 | 1.6434E-34 | TEC_TXNRD1 | JUND |
| 8.635E-51 | -0.8581405 | 0.093 | 0.294 | 1.8436E-46 | TEC_TXNRD1 | RASD1 |
| 5.3677E-38 | -0.8643589 | 0.179 | 0.346 | 1.146E-33 | TEC_TXNRD1 | RHOB |
| 2.3916E-27 | -0.8696487 | 0.47 | 0.573 | 5.1061E-23 | TEC_TXNRD1 | IER2 |
| 2.4984E-59 | -0.8947379 | 0.075 | 0.296 | 5.3341E-55 | TEC_TXNRD1 | PPP1R10 |
| 3.9328E-36 | -0.9096634 | 0.475 | 0.661 | 8.3964E-32 | TEC_TXNRD1 | WFDC2 |

| | | | | | | |
|---|---|---|---|---|---|---|
| 3.6708E-49 | -0.9275646 | 0.06 | 0.25 | 7.8373E-45 | TEC_TXNRD1 | CSKMT |
| 1.3601E-56 | -0.962066 | 0.422 | 0.617 | 2.9038E-52 | TEC_TXNRD1 | CLU |
| 3.1708E-51 | -0.9706208 | 0.091 | 0.295 | 6.7696E-47 | TEC_TXNRD1 | HES1 |
| 1.3649E-51 | -1.0277463 | 0.105 | 0.307 | 2.914E-47 | TEC_TXNRD1 | FOS |
| 1.114E-58 | -1.043253 | 0.071 | 0.29 | 2.3784E-54 | TEC_TXNRD1 | TCIM |
| 1.0783E-40 | -1.094165 | 0.117 | 0.285 | 2.3022E-36 | TEC_TXNRD1 | HSPA2 |
| 9.3521E-37 | -1.1041721 | 0.303 | 0.453 | 1.9967E-32 | TEC_TXNRD1 | IER3 |
| 2.1171E-69 | -1.1426106 | 0.388 | 0.605 | 4.5199E-65 | TEC_TXNRD1 | ELF3 |
| 9.3517E-67 | -1.195664 | 0.079 | 0.318 | 1.9966E-62 | TEC_TXNRD1 | HEXIM1 |
| 8.1114E-50 | -1.2183079 | 0.197 | 0.394 | 1.7318E-45 | TEC_TXNRD1 | HIST1H4C |
| 3.7046E-68 | -1.3898362 | 0.064 | 0.303 | 7.9093E-64 | TEC_TXNRD1 | MAFB |
| 3.1869E-29 | -1.5048532 | 0.459 | 0.653 | 6.8039E-25 | TEC_TXNRD1 | MMP7 |
| 2.3195E-71 | -1.5088044 | 0.457 | 0.65 | 4.9522E-67 | TEC_TXNRD1 | JUN |
| 1.6765E-34 | -1.580944 | 0.33 | 0.507 | 3.5794E-30 | TEC_TXNRD1 | S100A2 |
| 5.8538E-61 | -1.6250787 | 0.532 | 0.692 | 1.2498E-56 | TEC_TXNRD1 | MTRNR2L12 |
| 9.7677E-90 | -1.8040014 | 0.293 | 0.563 | 2.0854E-85 | TEC_TXNRD1 | SOX4 |
| 1.057E-124 | -4.9163965 | 0.331 | 0.623 | 2.257E-120 | TEC_TXNRD1 | PLCG2 |
